# Nuclear actin-dependent Meg3 expression suppresses metabolic genes by affecting the chromatin architecture at sites of elevated H3K27 acetylation levels

**DOI:** 10.1101/2024.05.12.593742

**Authors:** Nadine Hosny El Said, Wael Abdrabou, Syed Raza Mahmood, Tomas Venit, Youssef Idaghdour, Piergiorgio Percipalle

## Abstract

Three-dimensional organization of the eukaryotic genome is directly affected by the nuclear β-actin pool that regulates enhancer function by affecting H3K27 acetylation levels. This actin-based mechanism, in turn, influences enhancer-dependent transcriptional regulation and plays a crucial role in driving gene expression changes observed upon compartment-switching. Using a combination of bulk RNA-seq and qPCR analyses performed on total RNA from WT mouse embryonic fibroblasts (MEFs), β-actin heterozygous (HET) MEFs, and β-actin KO MEFs, in this study we demonstrate that expression of several lncRNAs is directly affected by β-actin depletion. Among these lncRNAs, Meg3 expression increases in a β-actin dosage-dependent manner. Using ChIRP-seq, ChIRP-MS and f-RIP-qPCR, we show that β-actin depletion leads to alterations in Meg3 genomic association. It also leads to Meg3 enrichment at or close to gene regulatory sites including enhancers and promoters concomitantly with increased H3K27 acetylation levels. At these sites, specific Meg3 association with H3K27 acetylation leads to loss of promoter-enhancer interactions as revealed by the Activity by Contact (ABC) model that builds on RNA-seq, H3K27acetylation ChIP-seq, ATAC-seq and HiC-seq obtained in WT and β-actin KO MEFs. Results from metabolomics experiments in WT, HET and β-actin KO MEFs show these mechanisms contribute to the repression of genes involved in metabolic biosynthetic pathways for chondroitin, heparan, dermatan sulfate, and phospholipases, hence impacting their synthesis. We propose that at sites of actin-dependent increase in H3K27acetylation levels Meg3 interferes with promoter-enhancer interactions, potentially impairing local genome organization (or DNA looping) and negatively regulating gene expression.

## Introduction

The mammalian genome is organized hierarchically into large chromatin compartments that can be either repressed or active from a transcriptional point of view and are comprised of topologically associating domains. This complex genomic architecture comes together because of short- and long-range interactions between inter and intra chromosomal loci. These interactions include promoter-enhancer contacts that may result in the activation or repression of sets of genes or gene programs[1, 2]. Several architectural proteins are involved in these mechanisms, including CTCF and the cohesin complex[3–7]. Actin has also emerged as an essential regulator of 3D genome organization [8–11]. Using genome-wide approaches, we found that by regulating the chromatin remodeling complex SWI/SNF (BAF) through an interaction with its ATPase subunit Brg1, actin controls the 3D organization of the genome at compartment level and this impacts on transcription and key cellular functions. Upon β-actin depletion the Polycomb Repressive Complex PRC2 is recruited instead of BAF and this leads to increased methylation and chromatin compaction [12]. We further demonstrated that β-actin is important in enhancer-dependent transcriptional regulation, a mechanism that plays a crucial role in driving gene expression changes upon compartment-switching. β-actin regulates enhancer function by influencing H3K27 acetylation levels, presumably by controlling HDAC recruitment [13, 14]. Results from this study provide first mechanistic links between compartment organization, enhancer activity, and gene expression [15] and further support a role of the nuclear actin pool in the compartmentalization of gene expression. Indeed, nuclear actin is enriched at chromosomal transcription sites and it is required for transcription in complex with several factors, including heterogeneous nuclear ribonucleoproteins (hnRNPs) as part of ribonucleoprotein complexes [16, 17], chromatin remodeling complexes and epigenetic regulators. In complex with hnRNP U, actin is required to enhance RNA polymerase II-mediated transcription elongation by promoting recruitment of histone modifying enzymes [18, 19]. While these mechanisms are important at the gene level, there is also recent evidence that the actin-hnRNP U interaction promotes nuclear compartmentalization and formation of transcription hubs [20]. potentially impacting on the functional architecture of the cell nucleus. Altogether, these actin-based mechanisms have a profound effect on gene expression during neurogenesis [21], osteogenesis [22] and adipogenesis [23]. In rat peritoneal mast cells, a decrease in ATP levels leads to an elevation in nuclear actin levels, indicating a potential role for nuclear actin in cellular stress responses [24]. Similarly, an increase in nuclear actin levels has been observed in monocytic HL-60 cells following stimulation with phorbol ester, suggesting its involvement in regulating macrophage differentiation [25]. Additionally, there is evidence of physiological signals that raise cAMP levels inducing a rapid rise in nuclear actin monomer levels in vascular smooth muscle cells (VSMCs) [24]. In all these cases, cell fate commitment is affected upon β-actin deletion and this outcome is an underlying cause of major pathologies.

Chromatin modifying enzymes are regulated by long non-coding RNAs (lncRNAs) and this has a major impact on various cellular processes such as cell proliferation, differentiation, and, more generally, on the establishment of cell identity [26]. lncRNAs bind to chromatin modifying enzymes and function as guides to anchor them to specific genomic locations or as decoy to sequester them from specific genomic sites, directly affecting covalent histone modifications. lncRNAs can also affect chromatin accessibility as part of ATP-dependent chromatin remodeling complexes where they serve as scaffold to assemble the complex itself for localized chromatin modification. For instance, the Mhrt lncRNA binds to Brg1 which leads to Brg1 sequestration from genomic DNA loci and inhibits Brg1-dependent gene regulation. The lncRNA Meg3 (Maternally Expressed imprinted lncRNA Gene), on the other hand, regulates PRC2. By facilitating recruitment of the PRC2 subunits JARID2 and EZH2 - a polycomb group histone methyltransferase that targets trimethylated histone 3 lysine 27 (H3K27me3) - to the chromatin for histone H3 methylation, Meg3 is directly involved in the epigenetic control of several genes. These include genes implicated in stemness [26], Epithelial Mesenchymal Transition-EMT [27] and TGF-β pathway genes [28], where EZH2 recruitment and increased Meg3 levels mediate transcriptional repression induced by enhanced TGF-β signaling [27, 28]. Enhanced TGF-β signaling has an overall impact on chromatin accessibility with increased H3K27 acetylation at enhancers [29]. Enhanced TGF-β signaling is also downstream effect of nuclear β-actin depletion when extensive genetic reprogramming results from genome re-organization and alterations in promoter-enhancer interactions due to increased H3K27 acetylation [12, 15, 30]. However, the functional significance of a possible association of lncRNAs with increased H3K27 acetylation and their impact on genome organization upon β-actin depletion is not known.

We, therefore, set out to investigate if nuclear actin controls lncRNAs expression levels and whether this may have an impact on the spatial organization of the genome and gene expression. Using a combination of bulk RNAseq and qPCR analyses performed on total RNA isolated from wt mouse embryonic fibroblasts (MEFs), β-actin KO MEFs and heterozygous (HET) MEFs, we demonstrate that expression of several lncRNAs is directly affected by β-actin depletion. Meg3 expression is particularly upregulated in β-actin KO cells. Results from ChIRPseq, ChIRP-MS and f-RIP/qPCR experiments show alterations in Meg3 genomic association and Meg3 enrichment at or close to regulatory sites (enhancers and promoters) concomitantly with elevated H3K27 acetylation in the KO condition. At these sites, specific Meg3 enrichment correlates with loss of promoter-enhancer interactions as revealed by the Activity by Contact (ABC) model. This, in turn, contributes to repression of genes involved in chondroitin, heparan, dermatan sulfate and phospholipases biosynthetic pathways. We propose that through a chromatin sponging effect that interferes with promoter-enhancers interactions at sites of enhanced H3K27 acetylation, increased Meg3 levels impairs local genome organization (or DNA looping) and negatively impact gene expression.

## Results

### Nuclear β-actin depletion leads to increased expression of the lncRNA Meg3

To find out if β-actin depletion affects expression of lncRNAs, we analyzed RNAseq data sets obtained in fibroblasts isolated from wild type (WT), heterozygous (HET) and β-actin knockout (KO) mouse embryos, hereby referred to as WT MEFs, HET MEFs and KO MEFs [9, 10, 21]. As expected, results from principal-component analyses (PCA) performed on the normalized datasets show clear segregation among replicates of WT, KO and HET MEFs as revealed by first (PC1) and second principal components (PC2), explaining 54.5% of the variation among the samples (Figure 1a). Clustering of the transcriptomic correlation matrix based on similarity confirmed that replicates of each condition cluster together and β-actin deletion leads to changes in gene expression (Figure 1b), potentially accompanied by distinct metabolic changes in these cells. To identify gene expression alterations associated with β-actin deletion, we performed gene-by-gene analysis of covariance focusing on the comparison between β-actin KO and WT which included 46,603 genes. The analysis revealed 718 differentially expressed genes (FC ≥ |1.5|, B-H FDR < 0.05) (Figure 1c), 359 (50 %) of which are downregulated in the β-actin KO condition.

**Figure 1.**
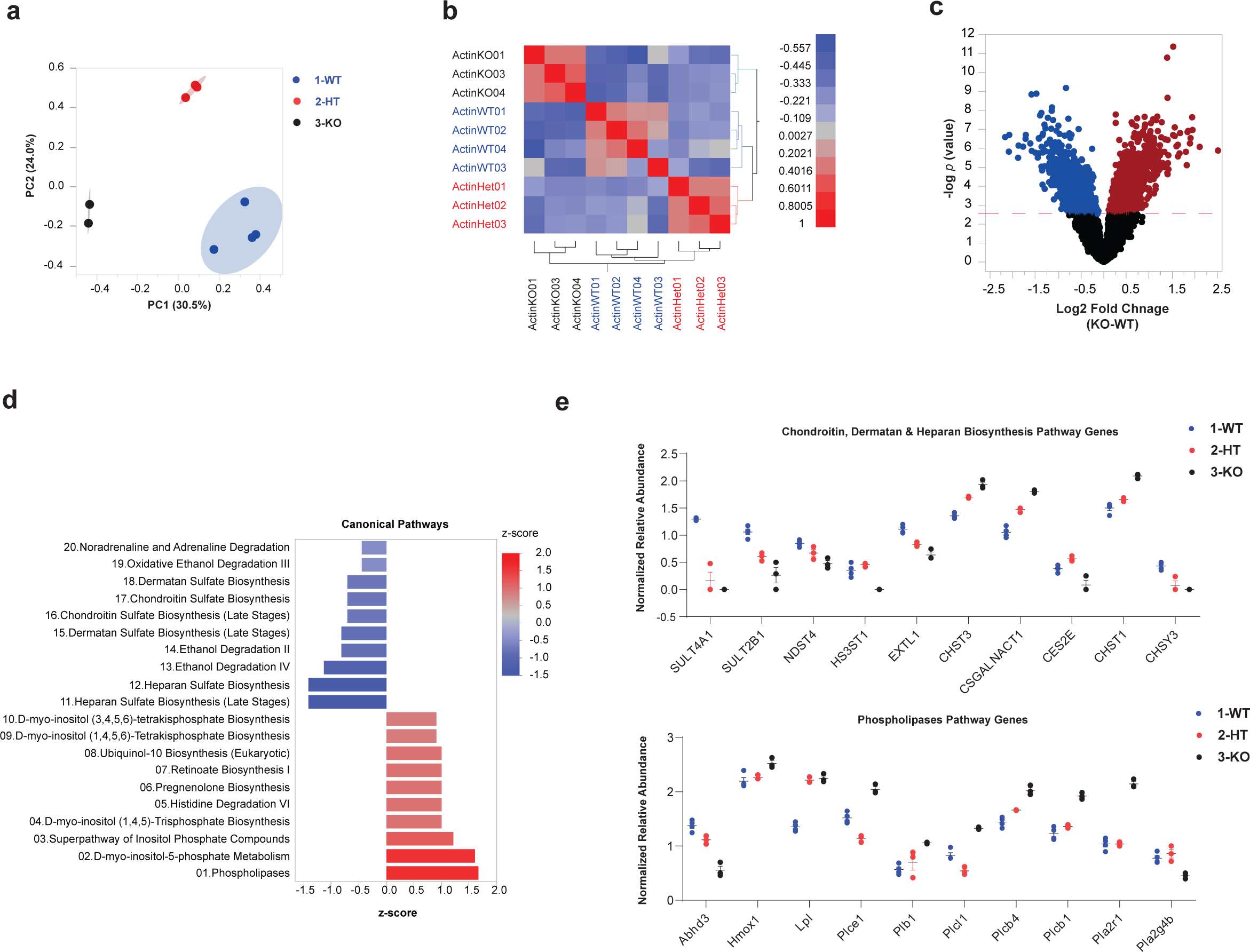
Transcriptional profiling by RNA-seq analysis shows significant differential gene expression upon β-actin depletion. **a)** Principal component analysis (PCA) showing the differential correlation of WT, HET and β-actin KO conditions; separation is highly noted between WT: black, HET: Red, KO: blue. Prin1 represents the highest common variable 1 (30.5%) and Prin2: the 2nd most common variable (24.0%). The PCA plot visualizes the relationship and variability among the wild type (WT), heterozygous (HET), and β-actin knockout (KO) samples based on their genomic RNA-seq data. Each replicate for each sample is represented by a data point in the plot, and the position of the data points reflects their similarity or dissimilarity in gene expression profiles. **b)** The heatmap represents the correlation analysis of genomic RNA-seq data from three samples: wild type (WT), heterozygous (HET), and β-actin knockout (KO) and their replicates. Each row and column in the heatmap correspond to a specific sample replicate. The color intensity represents the correlation coefficient between the expression profiles of two samples, blue lowest=-0.5 and red highest= 1. **c)** The volcano plot showing the differential expression between the transcriptomic profiles of KO and WT, significantly upregulated genes are highlighted in Red and significantly downregulated genes are highlighted in blue. **d)** pathway enrichment analysis showing the most significantly enriched Metabolic pathways in the whole RNA seq datasets. the pathways are ranked by the fold change. Positively enriched pathways are in red while downregulated pathways are in blue. **e)** Two one-way plots that show the differential expression of genes implicated in two pathways (Upper Panel; Chondroitin, heparan and dermatan Biosynthesis pathways) and (Lower Panel; Phospholipases genes). Interestingly, most of the genes implicated in these pathways show a dosage dependent effect. Wild type (WT: blue,) Heterozygous (HT: Red), Knock out (KO: Black)

Next, we examined if β-actin-dependent differential gene expression correlated with alterations in metabolic pathways. To do this we investigated the functional characteristics of the genes with significantly altered expression levels in the β-actin KO condition using Ingenuity Pathway Analysis (IPA). The analysis revealed that six of the most downregulated metabolic pathways were involved in chondroitin sulfate, heparan sulfate and dermatan sulfate biosynthesis (Heparan Sulfate Biosynthesis late stages, Heparan Sulfate Biosynthesis, Dermatan Sulfate Biosynthesis late stages, Dermatan Sulfate Biosynthesis, Chondroitin Sulfate biosynthesis late stages, and Chondroitin Sulfate Biosynthesis) (Figure 1d). Out of ten genes involved in the three aforementioned pathways, eight follow a significant β-actin dosage-dependent expression. Those that were upregulated upon the KO included CSGALNACT1, CHST3, CHST1, while SULT4A1, SULT2B1, NDST4, EXTL1, CES2E and CHSY3 were downregulated. Also, ethanol degradation pathways II, III and IV were significantly downregulated. In contrast, the phospholipases pathway was the most significantly upregulated one. Seven out of the 10 genes involved in the phospholipases pathway were β-actin dosage dependent. In particular, Abhd3 and pla2g4b were downregulated whereas Hmox1, Lpl, Plb1, Plcb4, Plcb1 were upregulated (Figure 1e). This highlights the impact of β-actin deletion on gene expression in these pathways. Notably, the transcriptomic changes that exhibited a dosage-dependent effect, with progressive alterations observed from WT to HET and KO MEFs confirm what was previously identified [12] and suggest a potential disruption of key metabolic pathways.

In metabolic diseases, expression of lncRNAs is highly dysregulated [31] and in cancer, lncRNAs can act as either oncogenes or tumor suppressors. They perform these functions by regulating transcription through direct interaction with RNA pol II, through chromatin modifiers such as Polycomb repressive complexes 1 and 2 (PRC1 and PRC2), or through interactions with Transcription factors and DNA methyltransferases [32, 33]. We, therefore, explored the potential involvement of lncRNAs in the suggested metabolic changes occurring upon β-actin depletion. Our aim was to identify lncRNAs that might be directly or indirectly associated with these alterations. Hence, we studied the impact of β-actin depletion on the expression of all identified lncRNAs in MEFs. We discovered a total of 6,622 lncRNAs, accounting for 14% of the total 46,603 generated RNAs (Figure 2a). Other noncoding RNAs, such as miRNAs, SnRNAs, SnoRNAs, and rRNAs, each comprised less than 5% of the total RNA output (Figure 2a). Among the differentially expressed lncRNAs, 6172 were downregulated whereas 450 were upregulated lncRNAs. Knowing that lncRNAs are elevated upon metabolic and cellular stress [33], in addition to playing various roles in regulation of transcription in metabolic pathways [34] with some lncRNA candidates known to be upregulated in metabolic pathways such as HOTAIR in adipogenesis [35], and SEMA 3B-AS1 and XIST in osteogenesis [36, 37], we focused on the set of upregulated lncRNAs. The ANCOVA analysis revealed Meg3 as the most upregulated lncRNA (fold change=3.09, p=0.0002) (Figure 2b,c) and upregulation appears to be β-actin dosage dependent with KO MEFs having the highest expression compared to HET and WT (Figure 2d). We also examined the expression of well-studied candidate lncRNA genes involved in transcription regulation and/or chromatin organization, such as Bvht, Malat1, and H19. Bvht exhibited increased expression in the KO compared to WT but to a lesser extent than Meg3, while Malat1, known for its involvement in speckle formation and gene silencing through interactions with Polycomb protein subunits like Ezh2 and Eed, showed no significant change [38–40]. In contrast, H19 displayed a significant decrease in the KO compared to WT (Figure 2d). Finally, the significant upregulation of Meg3 upon beta actin depletion was rescued by reintroduction of an NLS-tagged β-actin in the cell nucleus of β-actin KO MEFs (Figure 2e). Actin dependent Meg3 transcriptional regulation are dictated by changes in the epigenetic landscape that occur in the β-actin KO condition. Results from ATAC-seq show that there is a general increase in chromatin accessibility across the Meg3 gene and promoter region and this correlates with increased H3K27 acetylation levels (Figure 2f). Taken altogether, these results suggest that Meg3 expression is directly controlled by the nuclear pool of β-actin through a chromatin-based mechanism that controls H3K27 acetylation by direct HDAC recruitment.

**Figure 2.**
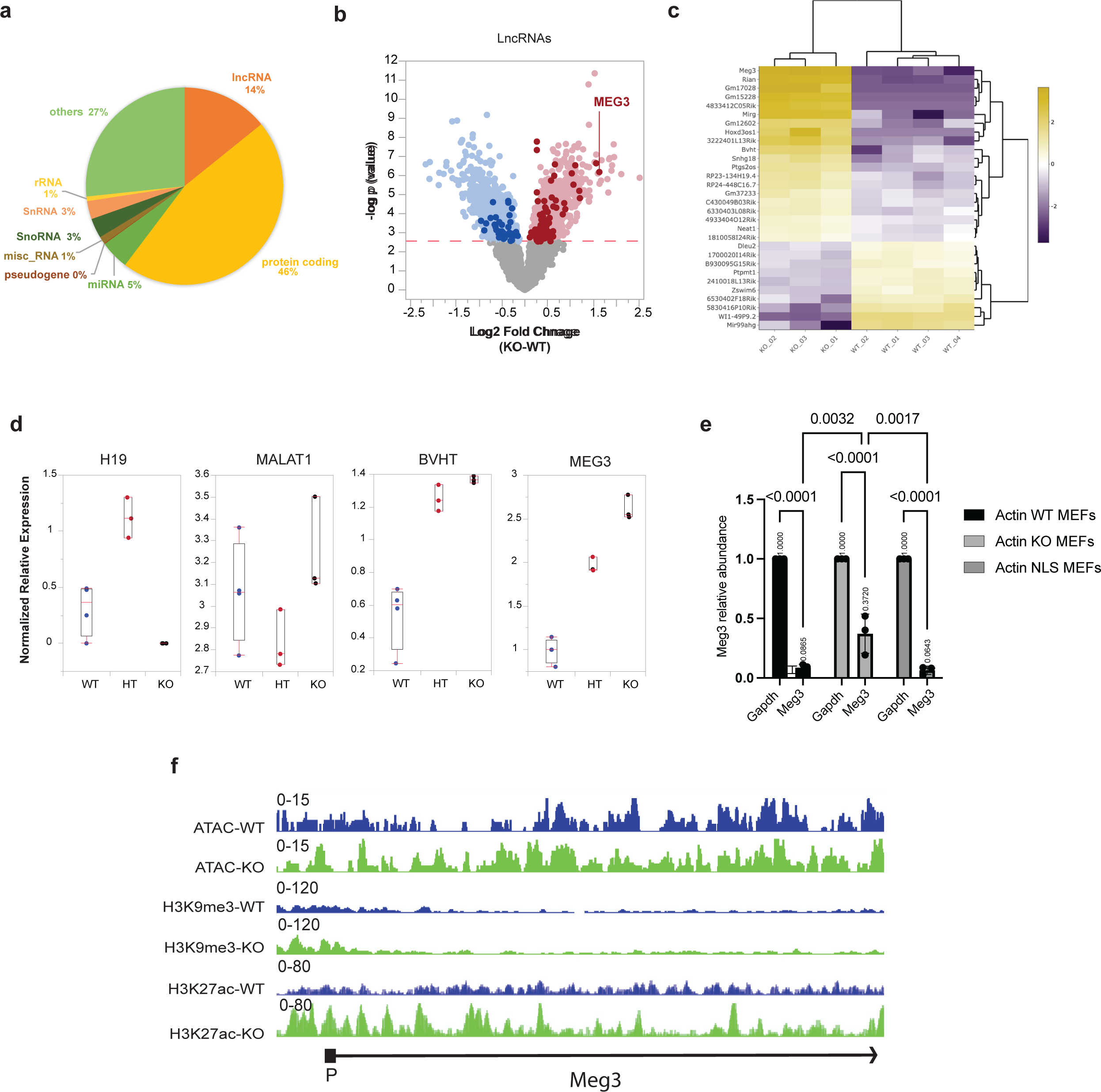
Meg3 lncRNA expression is significantly upregulated upon nuclear β-actin depletion. **a)** A pie chart showing the percentages of different RNA types differentially dysregulated upon β-actin KO where protein coding RNAs are 46%, lncRNAs are 14%, miRNAs 5%, SnoRNA 3%, SnRNA 3%, rRNA and miscRNA are 1 % each, and others are 27%. **b)** Volcano plot showing differential expression, of all genes of all lncRNAs only in WT and KO MEFs. p-values based on two-tailed Wald test corrected for multiple testing using Benjamini–Hochberg procedure. The significantly upregulated and downregulated LncRNAs are highlighted in dark red and blue, respectively. Also, the most upregulated LncRNA, *Meg3* is labeled. **c)** Heatmap of the top 30 differentially expressed candidate lncRNA purple: downregulated, yellow: upregulated. d) One-way Boxplots showing the differential expression of *Meg3*, *Malat1*, *Bvht* and *H19* in WT, HET, and KO. **e)** Rt-qPCR showing the *Meg3* relative abundance vs *Gapdh* in WT, KO and in NLS. Data are shown as mean ± SD (*n* = 3) P values indicated are based on 2-way ANOVA multiple comparisons in GraphPad Prism version 10.

### Actin dependent Meg3 enrichment at promoters/TSS of metabolic genes contributes to metabolic alterations

We next studied whether Meg3 associates with the chromatin, across the genome, and whether this potential association changes upon β-actin depletion using a combination of Chromatin Isolation by RNA Purification followed by mass spectrometry (ChIRP-MS) and Chromatin Isolation by RNA Purification followed by deep sequencing (ChIRP-seq) [41](Figure 3A). For this, we designed Meg3 specific biotinylated probes and used them to pull down chromatin-bound Meg3 from crosslinked MEFs using Streptavidin beads (Figure 3A). After mass spectrometry analysis, we identified a set of 54 proteins associated with Meg3 in the WT condition and 34 in the KO condition. 11 protein targets were common to both WT and KO condition, 43 were specific to WT and 23 were specific to KO (Figure 3b). Gene Ontology (GO) revealed that proteins associated with chromatin-bound Meg3 are involved in energy production and metabolic processes such as vacuole fusion, glycolytic processes, ATP generation from ADP, Pyruvate metabolic processes among many other metabolic terms at an enrichment of up to 300-fold (Figure 3c). On the other hand, in the β-actin KO condition, results from GO term analysis reveal that Meg3 primarily binds to proteins involved in DNA replication-dependent nucleosome assembly, nucleosome assembly, chromatin assembly, nucleosome organization and other chromatin organization and assembly-related biological processes with an enrichment of over 70 folds (Figure 3c). Compatible with this analysis, in the KO condition, the ChIRP MS revealed loss of actin association and significant high confidence enrichment of all histones, including H2B, H3 and H4, being present in the fraction of proteins associated with chromatin-bound Meg3 (Supplementary table1). These results confirm that Meg3 associates with the chromatin and that this interaction is likely to be enhanced in the β-actin KO condition when Meg3 becomes directly associated with histones. ChIRP combined with deep sequencing (ChIRP-seq) further confirmed this hypothesis, identifying a broad association of Meg3 with the WT mouse genome and this association was significantly enhanced genome-wide in β-actin KO across all chromosomes (Figure 3d). We found an overall number of 720 peaks in the KO condition in comparison to 550 in the WT condition and 288 peaks common to both (Figure 3f) with a slight increase of Meg3 association with promoters/TSS in beta actin KO MEFs (Supplementary figures 1a-b). Although the majority of Meg3 preferentially binds distal intergenic regions in both WT and KO cells (Supplementary figure 2 a-b), significant Meg3 binding at promoters was also observed in the KO cells (Supplementary Figure 3). In WT MEFs Meg3 comprise around 9.64% at promoters, the majority of which are promoters that are less than or equal to 1 Kb (5.91%). In the KO condition, 21.68% of ChIRP-seq peaks were found at promoters, mostly promoters less than or equal to 1 Kb (15.82%) (Figure 3e). Deeper analysis using upset plots showed the intersection of peaks among the different regions of the genome, where among the many intersections between genic, intergenic and distal intergenic regions, there is a noticeable increase in the promoter intersections from 9 to 13 intersections of peaks upon β-actin KO (supplementary figure 1). Metabolic pathway overrepresentation analysis by Ingenuity Pathway Analysis (IPA) for 1551 genes were identified to be bound to Meg3 in β-actin KO replicates compared to the WT replicates. Chondroitin, Heparan and Dermatan biosynthesis and phospholipases pathways were the most significantly overrepresented pathways (Figure 3g). These results suggest that in β-actin KO cells Meg3 increased binding to metabolic genes may lead to alterations of metabolic pathways due to changes in gene activity.

**Figure 3.**
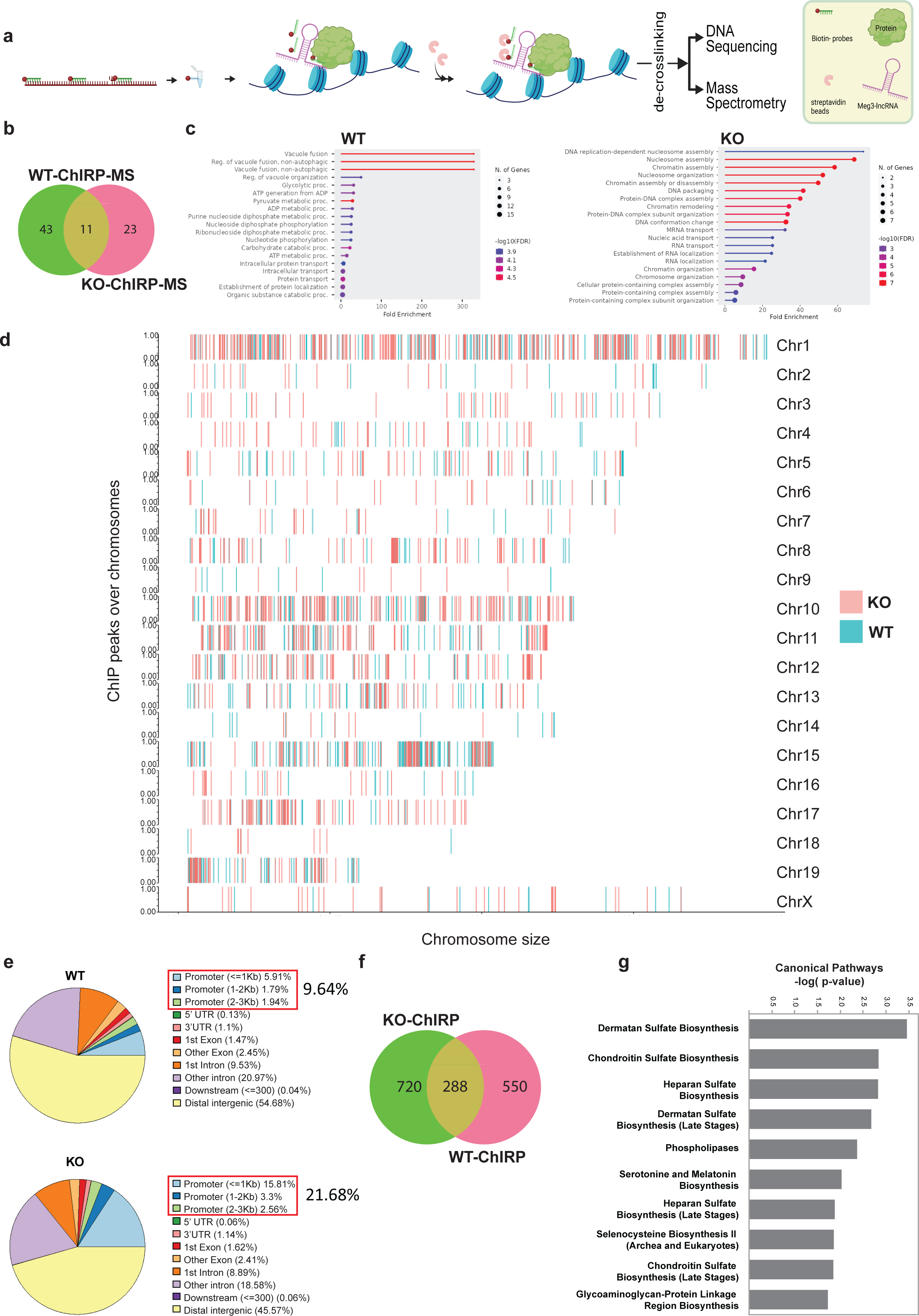
Genome-wide association of *Meg3* lncRNA is altered in the absence of β-actin. **a)** The ChIRP protocol followed to pull down *Meg3* lncRNA using biotinylated probes and streptavidin beads (adapted from[41]) . b) Merged replicates *Meg3* ChIRP peaks coverage on WT (green) and KO (red) over chromosomes, showing chromosome number and size. c) GO Analysis showing the biological processes of proteins bound to Meg3 in actin WT MEFs and actin KO MEFs. d) Global genome-wide Chromosome wise distribution/enrichment of Meg3 Peaks in actin WT MEFs (Blue) and Actin KO MEFs (Pink). e) Pie plots to visualize the genomic annotation of Peaks in WT and KO showing percentages of the annotated peaks of *Meg3* loci on the genome. f) Venn diagram showing the 550 specific peaks of *Meg3* bound genes in WT (WT-ChIRP-green) and 720 specific peaks of *Meg3* bound genes in KO (KO-ChIRP-pink) While common peaks between *Meg3*-WT-ChIRP and *Meg3*-KO-CjIRP are equal to 288. g) An overrepresentation pathway analysis using the list of genes that have been shown to bind to *Meg3* in KO relative to those in WT. Remarkably, the results show significant enrichment of Chondroitin, heparan and dermatan Biosynthesis pathways as well Phospholipases. Statistical significance is determined using right-tailed Fisher’s exact t-test and B-H FDR < 0.1. The significance of enrichment analysis represented using -log (P value) is plotted on the x axis.

To further investigate perturbations in Chondroitin, Heparan and Dermatan biosynthesis and phospholipases pathways in association with the loss of actin and, potentially, Meg3 upregulation, we performed a global metabolomic profiling of the three cell lines in three replicates using positive and negative ionization (see Methods). A total of 2,499 metabolic features were detected in all samples and replicates (n = 3). m/z values and retention time were recorded for each metabolic feature and used for further analysis (see Methods). PCA of the normalized peak dataset revealed a strong correlation structure in the metabolomic data across both conditions with the first principal component (PC1) clearly capturing the effect of partial and complete loss of function in actin and showing clear segregation between replicates of the WT, HET and KO conditions (Figure 4a). The first two principal components explain 52.9 % of the variation in the dataset (Figure 4a). Similarly, 2-way hierarchical clustering of the full metabolomic dataset including all replicates and all metabolic features (n = 2,499) displays discrete metabolic footprints for both the actin KO and actin HET from that of the WT condition (Figure 4b).

**Figure 4.**
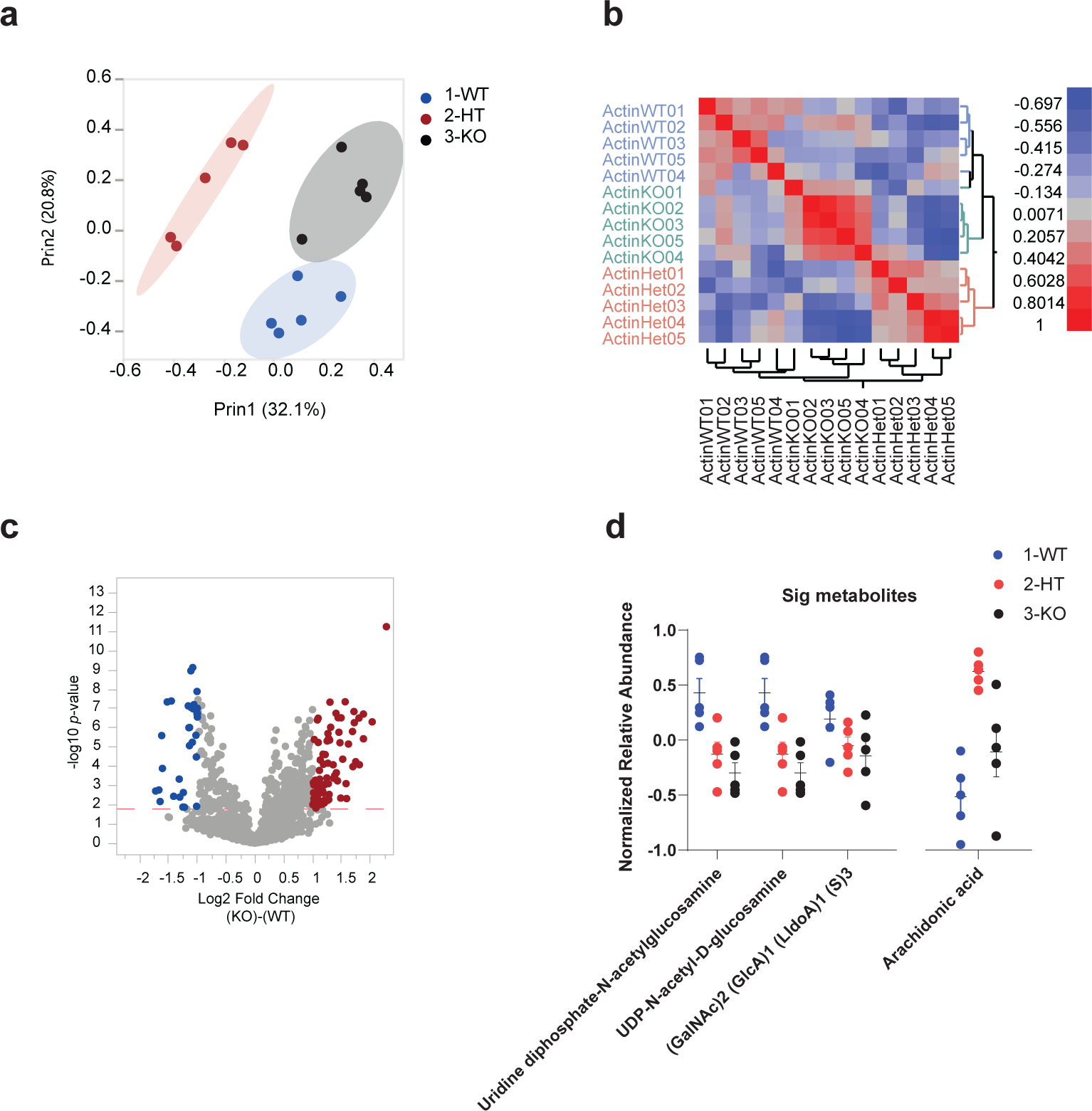
β-actin dosage dependent dysregulation of key metabolic pathways and metabolites. **a)** PCA demonstrating that WT (blue), HT (red) and KO (black) samples show distinctive clustered metabolic profiles. b) Hierarchical clustering analyses demonstrating that WT, HT, and KO samples show distinct metabolic profiles. c) shows a volcano plot demonstrating the differential expression of the normalized abundance of all the metabolic features between KO and WT. Significantly upregulated features are highlighted in red, while significantly downregulated metabolic features are highlighted in blue. d) shows a one-way plot for three different metabolites implicated in the chondroitin, heparan and dermatan pathway being downregulated in HT and KO compared to WT. It also shows another one-way plot for arachidonic acid, one of the main byproducts of phospholipase activity, being unregulated in HT and KO compared to WT.

Next, we performed metabolite-by-metabolite analysis of covariance focusing on the comparison between actin KO and WT. The analysis revealed 501 significantly differentially abundant features in actin KO compared to the WT (FC ≥ 1.5, B-H FDR < 0.1, Figure 4c). A summary of the metabolic features with statistically significant differential abundance in actin KO is shown in Supplementary Table 3. Together, these results clearly show the strong effect of β-actin loss on the cell metabolome and hint to major perturbations in metabolic resources allocation in the absence of β-actin. Next, we performed a functional analysis of curated normalized peak data using Gene Set Enrichment Analysis (GSEA) method implemented in MetaboAnalyst v5.0 to identify and annotate sets of functionally related compounds and evaluate their enrichment of potential functions defined by metabolic pathways. GSEA analysis of compounds identified using positive and negative ionization was performed separately. Putative annotation of MS peaks data was performed using m/z values and retention time dimensions to increase confidence in compounds identities and improve the accuracy of functional interpretations. In this method, a metabolite set is defined in this context as a group of metabolites with collective biological functions or common behaviors, regulations or structures. Annotated compounds were then mapped onto Mus musculus (mouse) [KEGG] (Ref: https://pubmed.ncbi.nlm.nih.gov/12466850/) for pathway activity prediction.

Of the metabolites implicated in the chondroitin, heparan and dermatan biosynthesis, we highlight UDP-GlcNAc and (GalNAc)2 (GlcA)1 (LldoA)1 (S)3. UDP-GlcNAc is extensively involved in intracellular signaling as a substrate for O-linked N-acetylglucosamine transferases (OGTs) to install the O-GlcNAc post-translational modification in a wide range of species. It is also involved in nuclear pore formation and nuclear signalling. OGTs and OG-ases play an important role in the structure of the cytoskeleton. Also, (GalNAc)2 (GlcA)1 (LldoA)1 (S)3 is an amino pentasaccharide and a galactosamine oligosaccharide. Interestingly, the levels of both metabolites showed significant dose-response between WT, Actin HET and Actin KO (Figure 4d). We also highlight arachidonic acid, one of the main intermediate metabolites of the phospholipases’ pathway and an essential fatty acid that is released from cell membrane by the activity of phospholipase A2 enzymes. Our analysis shows that the levels of arachidonic acids are significantly higher in actin HET and Actin KO than in WT (Figure 4d).

Taken together, the results of transcriptomic, ChIRP-seq and metabolomic data analyses support that the downregulation and upregulation observed in the chondroitin, heparan and dermatan biosynthesis and phospholipases pathways, respectively, are mediated by the upregulation of Meg3 upon the loss of β-actin.

### Meg3 regulates transcription of metabolic genes by mediating promoter-enhancer interactions in a β-actin dependent manner

We next investigated if changes in chromatin accessibility and epigenetic landscape accompany loss (Figure 5, top panel) or gain (Figure 5, bottom panel) of Meg3 ChIRP-seq peaks at genomic sites upon β-actin depletion. For this, we used a combination of ATAC-seq and ChIP-seq with antibodies to the BAF ATPase subunit Brg1, to the Ezh2 catalytic subunit of PRC2 which acts as the repressive H3K27 methyltransferase, and to the Suz12 RNA binding subunit of PRC2, known to be present together with Ezh2 at repressed loci [42]. Results from ATAC-seq did not show changes in chromatin accessibility between WT and KO conditions when comparing regions loss or gain of Meg3 binding (see Figure 5A, top and bottom panels) and, similarly, results from ChIP-seq experiments did not show major changes in binding of Brg1, Ezh2 and Suz12 (see Figure 5B, top and bottom panels). We next performed ChIP-seq analysis using antibodies to H3k27me3, H3K9me3 and H3K27ac, histone modifications respectively mediated by PRC2 and BRG1. Remarkably, the accumulation of H3K27me3 and H3K27ac at sites of differential Meg3 activity potentially points towards a role for Meg3 in regulation of chromatin remodeling. While we don’t see significant change in BRG1 and Ezh2 occupancy at differential ChIRP peaks, it is important that we always see distinct peak like pattern of BRG1 and Ezh2 in the heatmaps of ChIRP peaks suggesting the Meg3 usually colocalizes with these factors. Furthermore, we have shown by ChIRP-MS that Meg3 also associates with histones and nucleosome assembly and reorganization are some of the top pathways we see in Fig 3c. These observations reinforce a possible role in chromatin remodeling for Meg3 in association with BRG1 and Ezh2. Similarly, H3K27me3 and H3K27ac are marks of poised and active promoters respectively and we show increased Meg3 binding at promoter regions in the KO condition. These results indicate a potential actin-dependent functional correlation between MEG3, H3k27me3 and H3K27ac in gene expression regulation.

**Figure 5.**
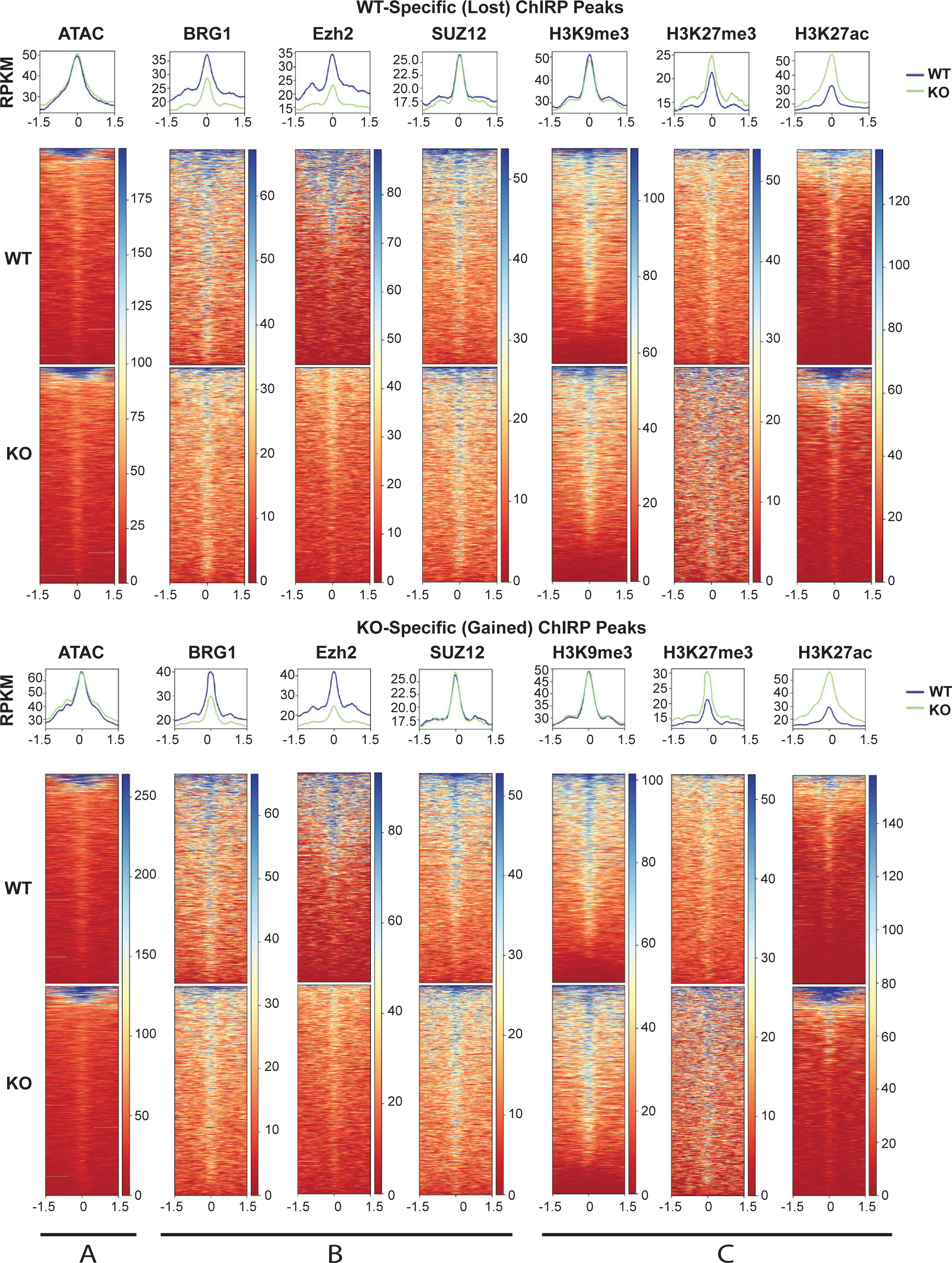
Integration of ChIP-seq, ChIRP-seq and ATAC-seq data sets shows increase of both active and repressive histone marks in KO specific Meg3 ChIRP-seq peaks. Density plots and heatmaps displaying scaled-read densities for ATAC-Seq, *Brg1*, *Ezh2*, *Suz12*, H3K9me3, H3K27me3, and H3K27ac in 3Kb regions surrounding WT-specific (top) and KO-specific (bottom) Meg3 Peaks. Scale bar shows normalized RPKM.

Nuclear actin has been shown to control acetylation levels by regulating HDAC 1 and 2 recruitment [43], which might explain a potential increase in histone acetylation levels and histone acetyl transferase activity upon β-actin depletion. Indeed, increased levels of H3K27ac were observed in β-actin KO MEFs and were shown to accompany compartment switching and enhancer-dependent transcriptional regulation [15]. To find out if these actin-dependent changes in H3K27ac levels correlate functionally with changes in Meg3 association with the genome at gene regulatory regions, either promoter or enhancer sites, we next focused on those genes that are commonly involved in the Chondroitin, Heparan and Dermatan biosynthesis and phospholipases pathways and are the most significantly overrepresented pathways in the KO condition. In particular, we focused on the gene encoding HS3ST3B1 (Heparan Sulfate-Glucosamine 3- Sulfotransferase 3B1), a type II integral membrane protein that belongs to the 3-O-sulfotransferases family, catalyzing the addition of sulfate groups at the 3-OH position of glucosamine in heparan sulfate (Figure 6a). Remarkably, in the KO condition, we found a new site of Meg3 enrichment (new ChIRP-seq peak) in intergenic regions, located between two enhancers, referred to as E1 and E2 (Figure 6z) and this binding seems to correlate with a dosage-dependent repression of the Hs3st3b1 gene (Figure 6b). Similarly, we found that the Meg3 binding site exhibits a specific increase in H3K27ac levels concomitantly with a decrease in the levels of H3K9me3 and marginal changes in ATAC signal (see Figure 6a).

**Figure 6.**
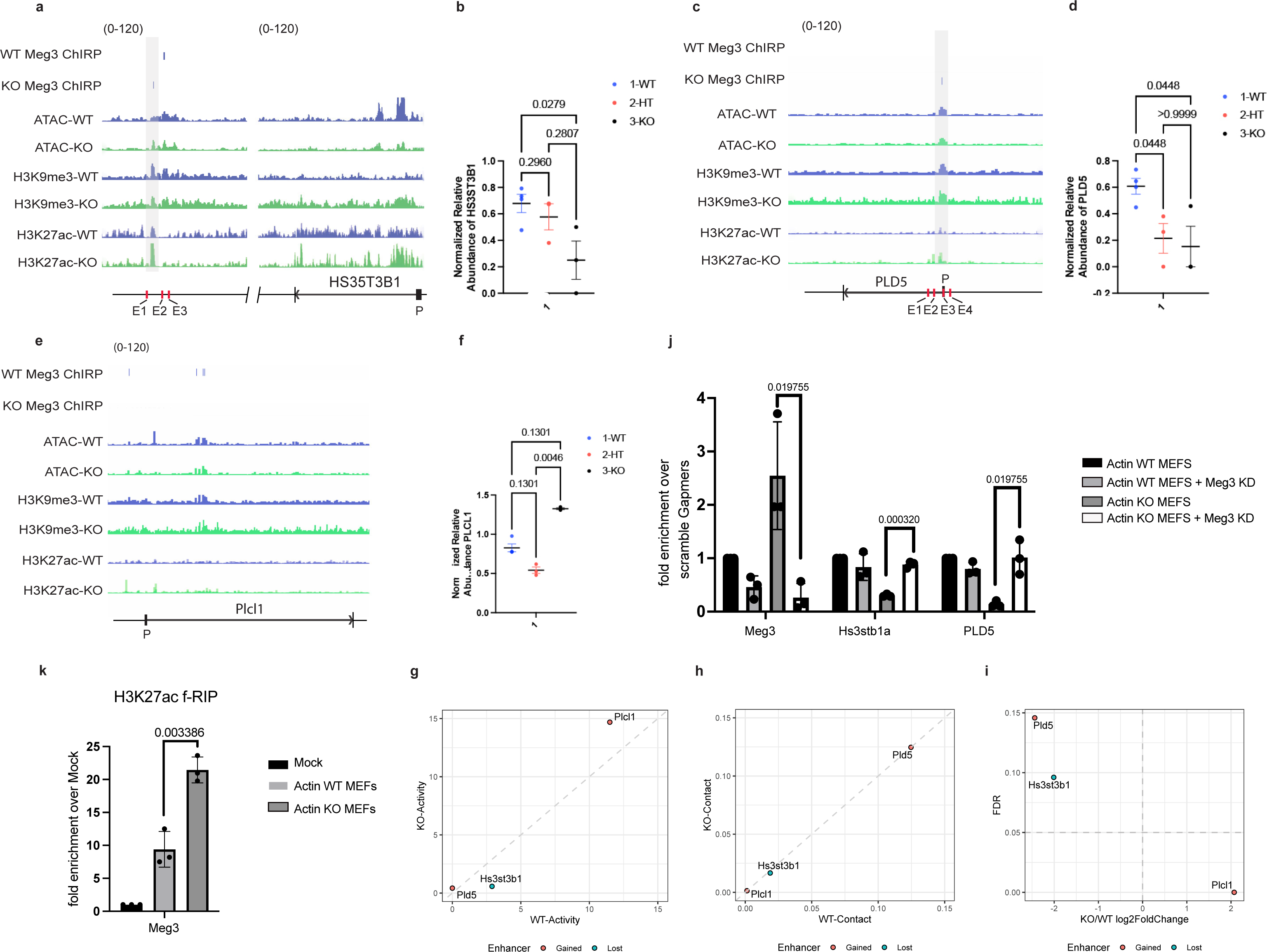
Meg3 binds to enhancer regions and affects expression of metabolic genes by interfering with promoter-enhancer contacts. IGVs (integrated genomic visualization) of Meg3 ChIRP-seq, ATAC-seq, H3K9me3 and H3K27ac ChIP-seq data in WT and β-actin KO conditions. Green: KO, Blue: WT. The reference gene body loci are shown at the bottom for Hs3st3b1 (a) Pld5 (c) and Plcl1(e). The y-axis represents RPKM (Reads Per Kilobase of sequence range per Million mapped reads) per bin. The range was set the same as in the image. shown below the tracks is the Gene body position for Hs3st3b1 (a) Pld5 (c) and Plcl1(e), (exon: box, intron: line). Plots showing the normalized relative expression of Hs3st3b1(b), Pld5 (d) and Plcl1(f) in WT (blue), Heterozygous-HT (red) and KO (black). g) Rt-qPCR analysis showing the differential enrichment of Meg3, Hs3stb1a and PLD5 , in Actin WT MEFs, Actin WT MEFS +Meg3 KD, Actin KO MEFS and Actin KO MEFS + Meg3 KD. Data are shown as mean ± SD (n = 3) P values indicated are based on multiple unpaired t-tests Holm-Šídák method in GraphPad Prism version 10. i) H3K27 acetylation f-RIP-qPCR showing Meg3 enrichment at regions of H3K27ac in WT and actin KO MEFs. Data are shown as mean ± SD (n = 3) P values indicated are based on multiple unpaired t-tests Holm-Šídák method in GraphPad Prism version 10. Results from the ABC model analysis showing enhancer activity (i) , enhancer -promoter contact (j), or fold change (k) of Hs3st3b1 , PLD5 or PLCl1 genes.

We also found changes in Meg3 enrichment peaks in the PLD5 (Phospholipase D Family Member 5) (see Figure 6c), which is part of the phospholipases biosynthesis pathway [44–46], involved in the conversion of phosphatidyl choline into 1,2-diacyl-sn-glycerol-3-phosphate (supplementary figure 4). In this case, Meg3 binds to sites within enhancers very close to the gene promoter, and upon β-actin depletion Meg3 enrichment correlates with significant increase in H3K27ac levels, no H3K9me3 changes and only marginal ones in ATAC signal (Figure 6c). Binding of Meg3 seems also to correlate with dosage dependence decrease in PLD5 mRNA levels (Figure 6d). In contrast to HS3ST3B1 and PLD5 which are both down regulated in the absence of β-actin and exhibit association of Meg3 close to or at regulatory regions such as enhancers and promoters, in the Phospholipase C like 1 enzyme gene (PLCL1), involved in converting phosphatidylinositol-4,5-bisphosphate into both the secondary messengers’ diacylglycerol and the inositol-1,4,5-triphosphate (Supplementary figure 5), Meg3 binding does not target regulatory regions (Figure 6e), it only binds far from enhancers (not shown) which seems to result in PLCL1 over expression in the KO condition (Figure 6f). To find out whether Meg3 binding is directly causing changes in expression of these genes, we next applied Meg3-specific gapmers to WT and β-actin KO cells to knock down Meg3. qPCR analysis on total RNA isolated from WT and KO cells shows that HS3ST3B1a and PLD5 mRNA levels significantly increase in the absence of Meg3 in the β-actin KO background (Figure 6g), suggesting that indeed, upon β-actin depletion Meg3 is directly involved in their transcriptional changes. We next performed f-RIP with antibody against H3K27ac from WT and β-actin KO condition. qPCR analysis with primers amplifying Meg3 show that Meg3 is specifically associated with H3K27ac and this interaction is enhanced in the β-actin KO condition (Figure 6h). Based on these results, we hypothesize that Meg3 enrichment at these specific sites close to or within gene regulatory regions is a consequence of actin dependent H3K27 acetylation.

Using HiC-Seq, ATAC-Seq, and RNA-Seq, we recently demonstrated that accessibility changes within compartment switching genes upon β-actin depletion are primarily observed in non-promoter regions such as distal regulatory elements [15]. The finding that β-actin loss induces widespread accumulation of H3K27ac [15] and that this enrichment happens at sites of Meg3 enrichment suggest that Meg3 may be involved in regulating transcription of certain metabolic genes at the level of promoter-enhancer interaction. Therefore, to find out if promoter-enhancer interactions are affected at Meg3 enrichment sites in KO condition, we applied the Activity by Contact (ABC) model of enhancer annotation based on HiC-Seq, ATAC-Seq, RNA-Seq and H3K27ac ChIP-seq in WT and KO MEFs to study if these epigenetic changes have a direct impact on enhancer activity and underlie transcriptional changes. Based on the ABC results, PLD5 and PLCl1 gained an enhancer while HS3st3b1 lost an enhancer (Figure 6 i-k and Supplementary Table 2). Looking at the Activity and Contact components of ABC score which determines enhancer gain/loss, none of the genes show a major change in Contact (see Figures 6i and 6j). So, the gain/loss of enhancer identified in these genes is primarily driven by changes in Activity, i.e. change in enhancer accessibility, particularly H3k27 acetylation (Figure 6j). Interestingly, PLCl1 upregulation can be explained by increased enhancer Activity and HS3stb1 downregulation can be explained by decreased enhancer Activity (Figure 6j). In contrast, PLD5 shows a very small increase in Activity so its downregulation cannot be entirely explained based on the ABC model. However, a close look at the IGV shows that there is an enhancer that localizes within the promoter, and this is precisely where Meg3 binding occurs concomitantly with increased H3K27ac in KO condition (Figure 6c). So, PLD5 down regulation may be due to loss of both enhancer and promoter accessibility due to Meg3 binding.

We, therefore, conclude that association of Meg3 within or close to regulatory regions of the HS3ST3B1 and PLD5 genes impacts the promoter-enhancer interaction or promoter accessibility and leads to transcriptional inhibition. In contrast, Meg3 enrichment does not affect regulation of the PLCL1 gene as its enrichment is far from the gene regulatory regions. These results also suggest that in the absence of β-actin increased Meg3 binding is guided by local increase in H3K27ac level. In addition, Meg3 primarily binds to or close to enhancers sites interfering with promoter-enhancer interaction or promoter accessibility required for gene activation, potentially contributing to alterations in the corresponding metabolic pathways.

## Discussion

In the cell nucleus β-actin plays essential functions in the 3D organization of the genome [12, 15]. We found that β-actin dependent changes in transcriptional and chromatin accessibility are highly enriched in compartment-switching regions. Furthermore, within compartment switching genes, changes in chromatin accessibility were primarily found at promoter-distal regulatory elements. We also found that loss of nuclear β-actin leads to significant accumulation of the enhancer-specific epigenetic mark H3K27ac. Using the ABC model of enhancer annotation, we showed that H3K27ac accumulation affects enhancer activity and is an underlying cause of transcriptional changes during compartment switching. From these studies we concluded that enhancer-dependent transcriptional regulation plays a crucial role in driving gene expression changes upon compartment-switching. These observations are compatible with previous studies where actin was shown to control recruitment of HDACs and HATs [13, 14]. However, the underlying mechanisms on how promoter-enhancer interactions are affected in the absence of β-actin by H3K27ac increased remained unclear.

In this study, we show that upon β-actin depletion the levels of several lncRNAs including Meg3 are significantly altered concomitantly with changes in H3K27 acetylation. H3K27ac plays an important role in the activation of several lncRNAs. For instance, CCAT1 and NEAT1 expressions are regulated by specific increase in H3K27 acetylation at their gene promoters [47, 48]. H3K27ac-dependent CCAT1 activation directly regulates SPRY4 and HOXB13 expression during cell proliferation [47]. Increased H3K27ac levels at NEAT promoter lead to increased NEAT expression, interaction with miR-212-5p followed by its suppression to regulate hepatic lipid accumulation [48]. Actin-dependent increase in H3K27ac levels at Meg3 promoter and gene coding region also seem crucial for Meg3 activation. This epigenetic landscape is accompanied by increased chromatin accessibility in the β-actin KO condition (as revealed by ATAC-seq) and might be a consequence of loss of actin-dependent HDAC recruitment that leads to alterations in the levels of H3K27 acetylation.

Importantly, the general increase in Meg3 levels observed in the KO condition has significant impact on global transcription at genomic level. ChIRP-seq using Meg3-specific probes showed that Meg3 association with the genome changes in the KO condition with respect to WT. Peaks of Meg3 enrichment are observed at both promoter and non-promoter regions. They do not specifically correlate with chromatin compaction (see ATACseq data) or chromatin remodelers such as Brg1, Ezh2 and Suz12. There is, however, significant correlation between Meg3 peaks in KO condition and increased levels of H3K27me3 and H3K27ac compared to WT. Meg3 is known to functionally associate with the PRC2 complex and to facilitate histone methyl transferase activity. So, it is not surprising that upon β-actin depletion Meg3 enrichment correlates with increased H3K27me3 levels given that PRC2 is recruited instead of the BAF complex [12]. On the other hand, the correlation between increased H3K27ac and Meg3 in KO cells is less understood, but it might be involved in regulating gene promoter and its interaction with enhancers.

So, we started by exploring if in β-actin depleted cells Meg3 binding at potential sites of H3K27 acetylation could possibly be involved in controlling gene expression. In agreement with an involvement of Meg3 in cell differentiation and metabolism [26, 49], analysis of the RNAseq data in β-actin KO cells, showed that increased Meg3 levels associate with differential expression of genes involved in metabolic pathways of chondroitin, heparan, dermatan sulfate and phospholipases biosynthesis. Metabolomic profiling confirmed perturbations in Chondroitin, Heparan and Dermatan biosynthesis and phospholipases pathways in association with Meg3 upregulation in the β-actin KO condition. Remarkably, Meg3 is enriched at intergenic sites exhibiting increased H3K27ac levels upstream the promoter of HS3ST3B1, a gene that is central in the metabolic pathway leading to heparan biosynthesis. Meg3 enrichment also peaked at an enhancer embedded within the gene promoter of the PLD5 gene (involved in phospholipase dependent metabolism). Both genes are downregulated and, in both cases, Meg3 enrichment is at sites of increased H3K27ac levels. We next applied the ABC model for enhancer annotation to find out if Meg3 could interfere with gene regulatory elements. We found that inhibition can be explained by a loss of activity-dependent (H3K27ac) promoter-enhancer interaction for HS3ST3B1 and, to a lesser extent, for the PLD5 gene. In contrast to HS3ST3B1 and PLD5, the PLCL1 gene is activated in the KO condition and Meg3 binding seems to be lost from regulatory elements, including enhancers and promoter. f-RIP qPCR experiments show that Meg3 binds directly to H3K27ac. We, therefore, propose that Meg3 contributes to repress gene expression by associating with regulatory elements, enhancers and promoters, at sites of H3K27acetylation.

Compatible with the above model, ChIRP-MS results show that in the KO condition Meg3 is primarily associated with histones in contrast to WT where Meg3 binds actin and a set of hnRNP proteins. Early studies demonstrated that actin is a component of RNPs [16, 17]. In the 40S pre-mRNP/RNP fraction isolated from rat liver extracts [50], actin binds to core hnRNPs and these interactions are important for RNP assembly and transcription[17–19]. It is, therefore, possible that actin is a component of the Meg3 RNP together with specific hnRNPs and that upon actin depletion the Meg3 RNP is disassembled and Meg3 binding to histones and thus, to the chromatin, is mediated by histone acetylation. We speculate that in the KO condition the lncRNA Meg3 is not or only partially assembled into an RNP and this leads to Meg3 collapse on the chromatin. This sponging effect on the chromatin is guided by enhanced H3K27ac and leads to a transcriptional block by affecting promoters and promoter-enhancer interactions, interfering with key metabolic pathways.

## Materials and Methods

### Cell culture

WT, HET and β-actin KO mouse embryonic fibroblasts (MEFs) are a gift of Dr. Christophe Ampe, University of Gent, Belgium. Cells were maintained and cultured with Dulbecco’s modified Eagle medium (DMEM) with high glucose, 10% fetal bovine serum (FBS) and 100 U/mL penicillin and 100 μg/mL streptomycin, in a humidified incubator with 5% CO2 at 37 °C. 2.

### ChIP-seq

WT or β-actin KO MEFs were crosslinked using 1% formaldehyde (Sigma cat. No. F8775) for 10 minutes. Then Quenching was carried out using 0.125 M Glycine for 5 minutes. Cells were then lysed using lysis buffer 1 -LB1-(50 mM Hepes KOH pH 7.5, 10 mM NaCl, 1mM EDTA, 10% glycerol, 0.5% NP–40, 0.25% Triton X-100). Nuclei were pelleted, collected then washed using lysis buffer 2 -LB2- (10 mM Tris HCl pH 8, 200 mM NaCl, 1mM EDTA, 0.5 mM EGTA). This was followed by lysis using lysis buffer 3 LB3 (10 mM Tris HCl pH 8; 100 mM NaCl, 1mM EDTA; 0.5 mM EGTA; 0.1% Na-Deoxycholate, 0.5% N-laurylsarcosine). Chromatin was then sheared using Qsonica Sonicator (4 cycles of 3 minutes at 70 % Amplitude), and then checked on 0.8% agarose gel. 100 μg of fragmented chromatin was mixed with 5μg of (*Ezh2* (D2C9) XP Rabbit mAb antibody-Cell signaling). The protein-antibody immunocomplexes were recovered by the Pierce Protein A/G Magnetic Beads (Thermo-Scientific). Beads and attached immunocomplexes were washed twice using Low salt wash buffer (LS) (0.1% SDS; 2mM EDTA, 1% Triton X-100, 20 mM Tris HCl pH8, 150 mM NaCl), and High Salt (HS) wash buffer (0.1% SDS, 2mM EDTA, 1% Triton X-100, 20 mM Tris HCl pH8, 500 mM NaCl) respectively. The beads were then re- suspended in elution buffer (50 mM Tris HCl pH8, 10 mM EDTA, 1% SDS). De-crosslinking was achieved through adding 8µl 5 M NaCl and incubating at 65°C overnight. RNase A (1 µl 10 mg/ml) was added to the tube for a 30 min incubation at 37°C. Then, 4 µl 0.5 M EDTA, 8 µl 1 M Tris-HCl, and 1 µl 20 mg/ml proteinase K (0.2 mg/ml) were added for a 2 hours incubation at 42°C to digest the chromatin. DNA was then purified by QIAquick PCR purification kit (Qiagen, Germantown, MD, USA) for qPCR analysis and sequencing. ChIP-seq library preparation was done using the TruSeq Nano DNA Library Prep Kit (Illumina, San Diego, CA, USA) and then sequenced with the HiSEq 2500 sequencing platform (performed at the NYUAD Sequencing Center).

### ChIRP-seq

Biotin labeled anti sense oligo probes were designed against Mouse *Meg3* lncRNA and purchased from IDT according to Chu et al. [51]: 1) 1 probe per 100bp of length; 2) GC% target =45; 3) oligonucleotide length=20; 4) spacing length of 60- 80bp; probes were numbered divided into even and odd numbered pools. Pools were diluted up to 100-μM concentrations. WT and β-actin KO MEFs were strongly fixed using 1% glutaraldehyde for 10 minutes. Then, Quenching was done using 1/10th volume of the 1.25 M glycine for 5 minutes. Cells were pelleted, lysed, and sonicated for 3-4 hours until clear, 30s ON, 30s OFF program by a Qsonica Sonicator (4 cycles of 3 minutes at 70 % Amplitude), and then checked on 0.8% agarose gel. Sonicated samples were centrifuged at 16100RCF for 10 min at 4 °C. aliquoted into 1 ml samples. 10 μL for RNA INPUT and 10 μL for DNA INPUT were withdrawn from the samples. Input samples were stored at -20 °C. for each 1 ml of sample 2 ml of hybridization buffer were added. 1μL of 100- pmol/μL probes per 1 mL chromatin were added to each sample. Mix well. Incubate at 37 °C for 4 hrs with shaking. C-1 magnetic beads (streptavidin conjugated magnetic beads) were washed with lysis buffer and used. 100 μL of beads per 100 pmol of probes were added to the samples and left for 30 minutes at 37°C with shaking. After 5 rounds of washing beads were resuspended in 1 ml of wash buffer. Out of each 1 ml sample, 100 μL were used for RNA isolation using Trizol. The remaining sample was used to isolate DNA. qPCR was performed to assess success of the *Meg3* lncRNA pulldown vs NONO as negative control (Miao et al., 2018).

### ChIRP-MS

The ChIRP-MS protocol was adapted from the Chang lab [52, 53]. Briefly, 20 p150 flasks of either WT or β-actin KO MEFs were cross linked for 30 minutes with 3% formaldehyde and quenched with 0.125 M glycine for 5 min. for a total of approximately 500 million cells the centrifuged at 2000 rcf. as advised 100 mg cell pellet were resuspended well in 1 mL of lysis buffer). Cells were sonicated using the Qsonica Sonicator until lysis got clear then DNA was extracted and run to check the sonication was enough to proceed.

### ATAC-seq

The ATAC-seq data were obtained from Mahmood et al [12]. Briefly, two biological replicates were used for each condition. Samples with 50,000 cells per condition were shipped in frozen medium (DMEM with 50 % FBS and 10% DMSO) on dry ice to Novogene (Beijing, China). All subsequent processing was performed by Novogene using standard DNA extraction and library preparation protocols. Cell nuclei were isolated, mixed with Tn5 Transposase with two adapters, and tagmentation was performed for 30 min at 37 °C. The fragmented DNA was purified and amplified with a limited PCR cycle using index primers. Libraries were prepared according to recommended Illumina NovaSeq6000 protocols. All ATAC-Seq processing was performed by Novogene (Beijing, China).

### HiC-seq

The HiC-seq data sets were from Mahmood et al [12]. Briefly, two biological replicates were used for each condition. Samples with 1 million cells per condition were fixed with 2% formaldehyde for 10 mins. The cell pellets were washed twice by 1× PBS and then stored at −80 °C. Frozen pellets were shipped on dry ice to Genome Technology Center at NYU Langone Health, NY. All subsequent processing was performed by Genome Technology Center at NYU Langone Health using standard DNA extraction and library preparation protocols. Hi-C was performed at Genome Technology Center at NYU Langone Health, NY from 1 million cells. Experiments were performed in duplicates following the instructions from Arima Hi-C kit (Arima Genomics, San Diego, CA). Subsequently, Illumina-compatible sequencing libraries were prepared by using a modified version of KAPA HyperPrep library kit (KAPA BioSystems, Willmington, MA). Quality check steps were performed to assess the fraction of proximally ligated DNA labeled with biotin, and the optimal number of PCR reactions needed to make libraries. The libraries were loaded into an Illumina flowcell (Illumina, San Diego, CA) on a NovaSeq instrument for paired-end 50 reads.

### ATAC-seq and ChIP-Seq (*Ezh2*) preprocessing

Raw reads were quality trimmed using Trimmomatic<u>64</u> and analyzed with FastQC (http://www.bioinformatics.babraham.ac.uk/projects/fastqc) to trim low-quality bases, systematic base calling errors, and sequencing adapter contamination. Specific parameters used were “trimmomatic_adapter.fa:2:30:10 TRAILING:3 LEADING:3 SLIDINGWINDOW:4:15 MINLEN:36”. Surviving paired reads were then aligned against the mouse reference genome (GRCm38) using Burrows-Wheeler Aligner BWA-MEM<u>65</u>. The resulting BAM alignments were cleaned, sorted, and deduplicated (PCR and Optical duplicates) with PICARD tools (http://broadinstitute.github.io/picard). Bigwig files were generated using deeptools<u>66</u> command bamCoverage -bs 10 -e --ignoreDuplicates –normalizeUsingRPKM. Encode blacklisted regions were removed and replicate bigwig files were averaged using the deeptools<u>66</u> command bigwigCompare --operation mean. Bigwig files were analyzed with computeMatrix function of deeptools<u>66</u> to plot average signal around regions of interest.

### ATAC-seq differential analysis

Differential analysis was performed using HOMER<u>60</u>. Processed bam files were converted to HOMER tag directories followed by annotation and differential analysis with the scripts annotatePeaks.pl and getDiffExpression.pl. ATAC-Seq peaks were called on cleaned, deduplicated bam files using macs2 with the parameters -q 0.05 -g mm -- keep-dup all --nomodel --shift −100 --extsize 200 -B --broad -f BAMPE<u>67</u>. Peaks common to two replicates in each condition were retained and merged using homer command mergePeaks. Differential peaks were identified and annotated using homer scripts annotatePeaks.pl and getDiffExpression.pl. Peaks showing more than twofold change were divided into 3 clusters containing 84, 391 and 539 peaks using deeptools kmeans clustering. Clusters containing 391 and 539 peaks were classified as activated and repressed and used for further analysis. Promoters showing more than twofold change in ATAC signal with FDR < 0.05 were identified using HOMER command annotatePeaks.pl tss mm10 -size -1000,100 -raw and getDiffExpression.pl. To identify enhancers showing more than twofold change in ATAC signal with FDR < 0.05, enhancer-promoter pairs for MEFs were downloaded from http://chromosome.sdsc.edu/mouse/download/Ren_supplementary_table7.xlsx18. Regions 2 kb upstream and downstream of enhancer peaks were analyzed using annotatePeaks.pl and getDiffExpression.pl. To make plots of nucleosome occupancy, bam files were filtered to remove insert sizes <30 bp and processed with the NucleoAtac<u>68</u> pipeline using default settings and previously called atac-seq peaks. The resulting bedgraph files were converted to bigwigs and nucleosomal signal was plotted in 5 kb regions surrounding TSSs of interest using deeptools.

### *Ezh2* differential analysis

Differential analysis was performed using HOMER. Processed bam files were converted to HOMER tag directories followed by annotation and differential analysis with the scripts annotatePeaks.pl and getDiffExpression.pl. Peaks were called using the macs2<u>67</u> command: macs2 callpeak -t Rep-1.bam Rep-2.bam -q 0.05 -g mm -c Input.bam (bam file for the relevant input control) --keep-dup all -B -f BAMPE --nolambda --nomodel –broad. WT and KO peaks were merged using HOMER command mergePeaks. Differential peaks were identified and annotated using homer scripts annotatePeaks.pl and getDiffExpression.pl. Overlap of TSSs with known polycomb targets for Fig. 2D was estimated using data from<u>19</u>. A transcript was regarded as a target if at least one polycomb subunit bound within 1 kb of its TSS. GC and CpG content for *Ezh2* peaks was obtained using the HOMER command annotatePeaks.pl -CpG

### *Brg1*, H3K9me3, and H3K27me3 ChIP-Seq analysis

For *Brg1*, H3K9me3, and H3K27me3 analysis, bigwig files were downloaded from GEO accession number <u>GSE100096</u>. Blacklisted regions were removed and replicates were averaged using the deeptools<u>66</u> bigwigCompare -- operation mean. Merged and cleaned bigwig files were plotted using the deeptools commands computeMatrix, plotProfile, and plotHeatmap. Heatmaps showing pairwise Spearman correlation between various epigenetic marks in Fig. 5 and <u>S4B</u> were generated with deeptools<u>66</u> using replicate merged BAM files. To generate compartment-wise heatmaps, bed files of previously identified compartments at 500 kb resolution were divided in 5 kb bins using the bedtools<u>63</u> command makewindows and supplied to deeptools command multiBamSummary and plotCorrelation with default settings.

### RNA-seq analysis

RNA seq analysis was carried out for published data of WT and KO , mouse embryonic fibroblasts (MEFs )and induced neurons from MEFs, as well as osteoblast differentiated MEFs at days 4 and 14, as described in ref. 2. DEseq was used to perform differential expression analysis between WT/KO of the different studies. Raw counts for WT, β-actin KO cells were downloaded from GSE95830 and DESeq256 was used to perform pairwise differential expression comparisons between WT/KO cells. Log2 fold change for genes overlapping different ATAC-Seq clusters was averaged. For repeat element analysis, repeat element annotation for the mouse genome was downloaded from the repeatmasker website (www.repeatmasker.org/genomes/mm10/RepeatMasker-rm405-db20140131/mm10.fa.out.gz) and filtered to exclude simple and low complexity repeats. RNA-Seq data were aligned to the genome using bowtie2 and analysis of repeat elements was performed using the Repenrich257 pipeline using default settings followed by a differential analysis of the resulting fraction_counts.txt files using edgeR58.

### Differential transcriptomic analysis

Differential transcriptomic analysis was performed using JMP Genomics 10. A Benjamini-Hochberg false discovery rate threshold of 0.1 was used to infer statistical significance.

### ChIRP-seq analysis

Adapter trimming was performed on raw fastq files using Trimmomatic. Surviving reads were aligned against the relevant reference genome (GRCm38) using Burrows-Wheeler Aligner BWA-MEM. Resulting BAM alignments were cleaned, sorted, and deduplicated (PCR and Optical duplicates) with PICARD tools. Four bam files for each condition (two replicates each for even and odd probes) were merged into a single file. Bigwig files were generated using deeptools command “bamCoverage -bs 10 -e --ignoreDuplicates –normalizeUsing RPKM”. Peaks were called on the merged files using macs2 with the following parameters: “macs2 callpeak -t merged.bam -q 0.05 -g mm -f BAMPE -c input.bam --keep-dup all”. Peaks were classified as being unique to WT, KO or shared with the HOMER command “mergePeaks" and annotated with HOMER script "AnnotatePeaks.pl". Peaks overlapping blacklisted regions were removed using bedtools. Subsequent analysis was performed using custom R scripts. ChIRP-seq Data have been deposited in the Gene Expression Omnibus with accession number GSE262113. Annotated ChIRP-seq peaks are found in (supplementary table 5)

### Integrative transcriptomic-CHIRP-seq pathway enrichment analyses

In total 1551 genes were identified to be bound to MEG3 in Actin (KO) replicates compared to WT replicates. Subsequently, we used the differential gene expression data (fold change and adjusted p value) for each of these genes between the KO and WT conditions generated by transcriptomic profiling to perform a metabolic pathway overrepresentation analysis using Ingenuity Pathway Analysis (Qiagen).

### GO term analysis

All GO term analyses were performed, and figures generated using ShinyGO

### Large-scale Ultra-Performance Liquid Chromatography High-Resolution Mass Spectrometry

5 replicates of Actin WT, HET, and KO cells were used for metabolomic profiling. Approximately 3x105 cells per sample were washed twice with ice-cold 0.9% NaCl solution and 300µl of 100% Methanol was added to each well. After 3 minutes of incubation on ice, cells were scraped using a precooled cell scraper and moved to a cold Eppendorf tube. Cell extracts were centrifuged at 15000rpm for 15 min at 4°C and 200µl of each supernatant was moved to a new tube for further processing. Next, samples were vacuum dried and reconstituted in 200µl of cyclohexane/water (1:1) solution for analysis by Ultra Performance Liquid Chromatography High-Resolution Mass Spectrometry (UPLC-HRMS) performed at the VIB Metabolomics core facility (Belgium). 10 ul of each sample were injected on a Waters Acquity UHPLC device connected to a Vion HDMS Q-TOF mass spectrometer. Chromatographic separation was carried out on an ACQUITY UPLC BEH C18 (50 × 2.1 mm, 1.7 μm) column from Watersunder under the constant temperature of 40°C. A gradient of two buffers was used for separation: buffer A (99:1:0.1 water:acetonitrile:formic acid, pH 3) and buffer B (99:1:0.1 acetonitrile:water: formic acid, pH 3), as follows: 99% A for 0.1 min decreased to 50% A in 5 min, decreased to 30% from 5 to 7 minutes, and decreased to 0% from 7 to 10 minutes. The flow rate was set to 0. 5 mL min−1. Both positive and negative Electrospray Ionization (ESI) were applied to screen for a broad array of chemical classes of metabolites present in the samples. The LockSpray ion source was operated in positive/negative electrospray ionization mode under the following specific conditions: capillary voltage, 2.5 kV; reference capillary voltage, 2.5 kV; source temperature, 120°C; desolvation gas temperature, 600°C; desolvation gas flow, 1000 L h−1; and cone gas flow, 50 L h−1. The collision energy for the full MS scan was set at 6 eV for low energy settings, for high energy settings (HDMSe) it was ramped from 28 to 70 eV. The mass range was set from 50 to 1000Da, scan time was set at 0.1s. Nitrogen (greater than 99.5%) was employed as desolvation and cone gas. Leucine-enkephalin (250 pg/μL solubilized in water:acetonitrile 1:1 [v/v], with 0.1% formic acid) was used for the lock mass calibration, with scanning every 1 min at a scan time of 0.1 s. Profile data was recorded through Unifi Workstation v2.0 (Waters). Data normalization was performed to remove potential variation resulting from instrument inter-run tuning differences. Raw MS peak data representing the abundance of each detected compound were subject to median standardization and missing values were imputed by the minimum value. Compounds with missing values in more than 50% of samples were considered missing. Standardized data that passed the quality control step were then log2 transformed and IQR normalized using JMP Genomics v8 (SAS Institute, Cary, NC) to remove potential technical artifacts and outliers.

### Metabolomic Profiling

Global metabolomic profiling of three replicates from each of the three cell lines was performed using Ultra Performance Liquid Chromatography High Resolution Mass Spectrometry (UPLC-HRMS) in positive and negative ionization modes. Overnight grown cells (approximately 3x105 per sample) were washed twice with ice-cold 0.9% NaCL solution and 300µl of 100% Methanol was added to each well. After 3 minutes incubation on ice, cells were scraped using precooled cell scraper and moved to cold Eppendorf tube. Cell extracts were spun down at 15000rpm for 15 min at 4°C and 200µl of each supernatant moved to a new tube for further processing. Next, samples were vacuum dried and reconstituted in 200µl of cyclohexane/water (1:1) solution for subsequent analysis by UPLC-HRMS performed at VIB Metabolomics core facility (Belgium). 10 ul of each sample were injected on a Waters Acquity UHPLC device connected to a Vion HDMS Q-TOF mass spectrometer. Chromatographic separation was carried out on an ACQUITY UPLC BEH C18 (50 × 2.1 mm, 1.7 μm) column from Watersunder under the constant temperature of 40°C. A gradient of two buffers was used for separation: buffer A (99:1:0.1 water:acetonitrile:formic acid, pH 3) and buffer B (99:1:0.1 acetonitrile:water:formic acid, pH 3), as follows: 99% A for 0.1 min decreased to 50% A in 5 min, decreased to 30% from 5 to 7 minutes, and decreased to 0% from 7 to 10 minutes. The flow rate was set to 0. 5 mL min−1. Both positive and negative Electrospray Ionization (ESI) were applied to screen for a broad array of chemical classes of metabolites present in the samples. The LockSpray ion source was operated in positive/negative electrospray ionization mode under the following specific conditions: capillary voltage, 2.5 kV; reference capillary voltage, 2.5 kV; source temperature, 120°C; desolvation gas temperature, 600°C; desolvation gas flow, 1000 L h−1; and cone gas flow, 50 L h−1. The collision energy for full MS scan was set at 6 eV for low energy settings, for high energy settings (HDMSe) it was ramped from 28 to 70 eV. Mass range was set from 50 to 1000Da, scan time was set at 0.1s. Nitrogen (greater than 99.5%) was employed as desolvation and cone gas. Leucine-enkephalin (250 pg/μL solubilized in water:acetonitrile 1:1 [v/v], with 0.1% formic acid) was used for the lock mass calibration, with scanning every 1 min at a scan time of 0.1 s. Profile data was recorded through Unifi Workstation v2.0 (Waters). Metabolomic data are publicly available in MendeleyData.

### Statistical analysis of the metabolomic data

Data normalization was performed to remove potential variation resulting from instrument inter-run tuning differences. Raw MS peak data representing the abundance of each detected compound were subject to median standardization and missing values were imputed by the minimum value. Compounds with missing values in more than 50% of samples were considered missing. Standardized data that passed quality control step were then log2 transformed and IQR normalized. Principal component analysis (PCA) and hierarchical clustering were done to explore the correlation structure in the data across the three conditions (WT, HT and KO). Furthermore, metabolite-by-metabolite analysis of covariance was performed to investigate differences between actin KO and WT conditions. A Benjamini-Hochberg false discovery rate threshold of 0.1 was used to infer statistical significance. Supervised and unsupervised data analyses were performed using JMP Genomics v10 (SAS Institute, Cary, NC).

### Functional and metabolic pathway enrichment analysis

Functional analysis of curated normalized peak data was performed using MetaboAnalyst v5.0 using an existing protocol [54]. Implemented Gene Set Enrichment Analysis (GSEA) method in the Functional Analysis module of MetaboAnalyst v5.0 (Accessed in 2022 from http://www.metaboanalyst.ca/) was used to identify sets of functionally related compounds and evaluate their enrichment of potential functions defined by metabolic pathways. m/z values and retention time dimensions both were used to identify and annotate compounds, and improve the accuracy of functional interpretations. Annotated compounds were then mapped onto Mus musculus (mouse) [KEGG] [55], for pathway activity prediction (Supplementary table 2). GSEA calculates Enrichment score (ES) by walking down a ranked list of metabolites, increasing a running-sum statistic when a metabolite is in the metabolite set and decreasing it when it is not. A metabolite set is defined in this context as a group of metabolites with collective biological functions or common behaviors, regulations or structures. In this method, a metabolite set are defined in this context as a group of metabolites with collective biological functions or common behaviors, regulations or structures. Annotated metabolites were then mapped onto Mus musculus (mouse) [KEGG] (Ref: https://pubmed.ncbi.nlm.nih.gov/12466850/) for pathway activity prediction.

### Formaldehyde crosslinking - RNA Immunoprecipitation (f-RIP)

Actin WT and KO MEFs were harvested by centrifugation at 500 RCF for 5 minutes, cells were rinsed with 1× phosphate-buffered saline (PBS) at room temperature. Each 5 million cells were resuspended in 1 ml of FBS and high salt free media. Subsequently cells were crosslinked for 10 minutes with 0.1% methanol free formaldehyde(Sigma) as per Hendrickson et al [40, 56]. Quenched with 125 mM of Glycine. Cells were centrifuged again at 500 RCF for 5 minutes followed by two washes with cold PBS. The resulting pellets were flash-frozen in liquid nitrogen and stored at −80°C for future use.

Upon thawing, the frozen pellets were resuspended in 1 ml of RIPA lysis buffer per tube (consisting of 50 mM Tris pH 8.0, 150 mM KCl, 0.1% SDS, 1% Triton X-100, 5 mM EDTA, and 0.5% sodium deoxycholate), supplemented with freshly prepared 0.5 mM dithiothreitol (DTT), 1× EDTA protease inhibitor cocktail (Roche), and 100 U/ml RNaseOUT (Life Technologies, 10777– 019). The cells were then incubated while rotating at 4°C for 10 minutes, followed by sonication by the Qsonica Sonicator at 10% amplitude for 1 second on and 1 second off at 30-second intervals, for a total of 90 seconds. The lysate was centrifuged at maximum speed for 10 minutes at 4°C, and the supernatant was collected. An equal volume of fRIP binding/wash buffer (150 mM KCl, 25 mM Tris pH 7.5, 5 mM EDTA, 0.5% NP-40), supplemented with freshly prepared 0.5 mM DTT, 1× EDTA protease inhibitor cocktail (Roche), and 100 U/ml RNaseOUT (Life Technologies, 10777–019), was added to dilute the samples. Fifty microliters of the lysate were set aside as the input sample and stored at −20°C for subsequent RNA purification and library preparation.

For pre-clearing, each lysate of 5 million cells was incubated with 25 μl of Pierce Protein A/G Magnetic Beads (Thermo-Scientific) on a rotor at 4°C for 30 minutes. Aliquots of 5 million cells/ml were flash-frozen and stored at −80°C for future use.

To initiate the fRIP procedure, the lysates were thawed, and 5 μg of H3K27ac antibody (ChIP grade- ab4729) were added. The lysates, along with the antibody, were left to rotate at 4°C overnight before the addition of Pierce Protein A/G Magnetic Beads (Thermo-Scientific). 50 μl of beads were added per 1 ml of lysate for 1 hour to recover immunoprecipitated complexes. Washes were performed twice using 1 ml of fRIP binding/washing buffer. After the final wash, the beads were removed and stored at −20°C. fRIP- was then followed by RNA purification, reverse transcription to cDNA (Qiagen quantitect reverse transcription kit- cat# 205311) and RT-qPCR.

### Statistical tests

Statistical significance was calculated using GraphPad Prism for macOS, GraphPad Software, La Jolla California USA, www.graphpad.com.

### Data availability

ChIRP-seq data have been deposited in the Gene Expression Omnibus with accession number GSE262113. For Ezh2 ChIP-seq analysis bigwig was downloaded from Gene Expression Omnibus with accession number GSE149987. For BRG1, H3K9me3, and H3K27me3 ChIP-seq analysis, bigwig files were downloaded from GEO accession number GSE100096.

## Acknowledgements

This work is supported by grants from New York University Abu Dhabi and the Center for Genomics and Systems Biology at New York University Abu Dhabi to PP. We thank Nizar Drou, Marc Arnoux and Mehar Sultana from the genomics core facilities at NYU Abu Dhabi, Center for Genomics and Systems Biology for technical help. We also thank Liaqat Ali from the NYU Abu Dhabi Core Technology Platform for technical help with the mass spectrometry analysis. We thank Christophe Ampe (University of Gent, Belgium) for kindly providing us with the WT MEFs, HET MEFs and β-actin KO MEFs. The authors declare no conflicts of interest.

## Author Contributions

PP conceived the research and wrote the manuscript together with NHE. NHE performed the CHIRP-seq, ChIRP-MS experiments, f-RIP and qPCR analysis. NHE and WA performed the computational analysis and made the relevant figures. TV and WA performed the metabolomics experiments and corresponding data analysis and made the relevant figures. SRM applied the ABC model to study promoter-enhancer interactions. YI worked on the manuscript and data analysis. PP supervised the research. All authors read and approved the manuscript.

## References

1. Hübner, M.R., M.A. Eckersley-Maslin, and D.L. Spector, Chromatin organization and transcriptional regulation. Curr Opin Genet Dev, 2013. 23(2): p. 89–95.

2. Pongubala, J.M.R. and C. Murre, Spatial Organization of Chromatin: Transcriptional Control of Adaptive Immune Cell Development. Frontiers in Immunology, 2021. 12.

3. Seitan, V.C., et al., Cohesin-based chromatin interactions enable regulated gene expression within preexisting architectural compartments. Genome Research, 2013. 23(12): p. 2066–2077.

4. Zuin, J., et al., Cohesin and CTCF differentially affect chromatin architecture and gene expression in human cells. Proceedings of the National Academy of Sciences, 2014. 111(3): p. 996–1001.

5. Schwarzer, W., et al., Two independent modes of chromatin organization revealed by cohesin removal. Nature, 2017. 551(7678): p. 51-56.

6. Rao, S.S.P., et al., Cohesin Loss Eliminates All Loop Domains. Cell, 2017. 171(2): p. 305–320.e24.

7. Attou, A., T. Zülske, and G. Wedemann, Cohesin and CTCF complexes mediate contacts in chromatin loops depending on nucleosome positions. Biophys J, 2022. 121(24): p. 4788–4799.

8. Visa, N. and P. Percipalle, Nuclear functions of actin. Cold Spring Harbor perspectives in biology, 2010. 2(4): p. a000620–a000620.

9. Xie, X. and P. Percipalle, An actin-based nucleoskeleton involved in gene regulation and genome organization. Biochem Biophys Res Commun, 2018. 506(2): p. 378–386.

10. Xie, X., et al., β-Actin-dependent global chromatin organization and gene expression programs control cellular identity. The FASEB Journal, 2018. 32(3): p. 1296–1314.

11. Percipalle, P. and M. Vartiainen, Cytoskeletal proteins in the cell nucleus: a special nuclear actin perspective. Molecular Biology of the Cell, 2019. 30(15): p. 1781–1785.

12. Mahmood, S.R., et al., β-actin dependent chromatin remodeling mediates compartment level changes in 3D genome architecture. Nature Communications, 2021. 12(1).

13. Serebryannyy, L.A., C.M. Cruz, and P. de Lanerolle, A Role for Nuclear Actin in HDAC 1 and 2 Regulation. Sci Rep, 2016. 6: p. 28460.

14. Viita, T., et al., Nuclear actin interactome analysis links actin to KAT14 histone acetyl transferase and mRNA splicing. J Cell Sci, 2019. 132(8).

15. Mahmood, S.R., et al., β-actin mediated H3K27ac changes demonstrate the link between compartment switching and enhancer-dependent transcriptional regulation. Genome Biology, 2023. 24(1): p. 18.

16. Percipalle, P., et al., Actin bound to the heterogeneous nuclear ribonucleoprotein hrp36 is associated with Balbiani ring mRNA from the gene to polysomes. The Journal of cell biology, 2001. 153(1): p. 229–236.

17. Percipalle, P., et al., An actin-ribonucleoprotein interaction is involved in transcription by RNA polymerase II. Proceedings of the National Academy of Sciences of the United States of America, 2003. 100(11): p. 6475–6480.

18. Obrdlik, A., et al., The histone acetyltransferase PCAF associates with actin and hnRNP U for RNA polymerase II transcription. Mol Cell Biol, 2008. 28(20): p. 6342–57.

19. Kukalev, A., et al., Actin and hnRNP U cooperate for productive transcription by RNA polymerase II. Nature Structural & Molecular Biology, 2005. 12(3): p. 238–244.

20. Wei, M., et al., Nuclear actin regulates inducible transcription by enhancing RNA polymerase II clustering. bioRxiv, 2019: p. 655043.

21. Xie, X., et al., *beta-actin regulates a heterochromatin landscape essential for optimal induction of neuronal programs during direct reprograming*. PLoS Genet, 2018. 14(12): p. e1007846.

22. Gjorgjieva, T., et al., Loss ofβ-Actin Leads to Accelerated Mineralization and Dysregulation of Osteoblast-Differentiation Genes during Osteogenic Reprogramming. Advanced Science, 2020. 7(23): p. 2002261.

23. Al-Sayegh, M.A., et al., β-actin contributes to open chromatin for activation of the adipogenic pioneer factor CEBPA during transcriptional reprograming. Mol Biol Cell, 2020. 31(23): p. 2511–2521.

24. McNeill, M.C., et al., Nuclear actin regulates cell proliferation and migration via inhibition of SRF and TEAD. Biochim Biophys Acta Mol Cell Res, 2020. 1867(7): p. 118691.

25. Xu, Y.Z., et al., Nuclear translocation of beta-actin is involved in transcriptional regulation during macrophage differentiation of HL-60 cells. Mol Biol Cell, 2010. 21(5): p. 811–20.

26. Fatica, A. and I. Bozzoni, Long non-coding RNAs: new players in cell differentiation and development. Nat Rev Genet, 2014. 15(1): p. 7–21.

27. Terashima, M., et al., MEG3 Long Noncoding RNA Contributes to the Epigenetic Regulation of Epithelial-Mesenchymal Transition in Lung Cancer Cell Lines. Journal of Biological Chemistry, 2017. 292(1): p. 82–99.

28. Mondal, T., et al., MEG3 long noncoding RNA regulates the TGF-β pathway genes through formation of RNA–DNA triplex structures. Nature Communications, 2015. 6(1): p. 7743.

29. Guerrero-Martínez, J.A., et al., TGFβ promotes widespread enhancer chromatin opening and operates on genomic regulatory domains. Nature Communications, 2020. 11(1): p. 6196.

30. Xie, X. and P. Percipalle, Elevated transforming growth factor β signaling activation in β-actin-knockout mouse embryonic fibroblasts enhances myofibroblast features. Journal of Cellular Physiology, 2018. 233(11): p. 8884–8895.

31. Daneshmoghadam, J., et al., The gene expression of long non-coding RNAs (lncRNAs): MEG3 and H19 in adipose tissues from obese women and its association with insulin resistance and obesity indices. Journal of Clinical Laboratory Analysis, 2021. 35(5).

32. Long, Y., et al., How do lncRNAs regulate transcription? Sci Adv, 2017. 3(9): p. eaao2110.

33. Connerty, P., R.B. Lock, and C.E. de Bock, Long Non-coding RNAs: Major Regulators of Cell Stress in Cancer. Front Oncol, 2020. 10: p. 285.

34. Nadhan, R. and D.N. Dhanasekaran, Regulation of Tumor Metabolome by Long Non-Coding RNAs. Journal of Molecular Signalling, 2022.

35. Kalwa, M., et al., The lncRNA HOTAIR impacts on mesenchymal stem cells via triple helix formation. Nucleic Acids Res, 2016. 44(22): p. 10631–10643.

36. Zhang, C., et al., SEMA3B-AS1-inhibited osteogenic differentiation of human mesenchymal stem cells revealed by quantitative proteomics analysis. J Cell Physiol, 2019. 234(3): p. 2491–2499.

37. Chen, X., et al., Long non-coding RNA XIST promotes osteoporosis through inhibiting bone marrow mesenchymal stem cell differentiation. Exp Ther Med, 2019. 17(1): p. 803–811.

38. Chen, X., et al., Malat1 regulates myogenic differentiation and muscle regeneration through modulating MyoD transcriptional activity. Cell Discovery, 2017. 3(1): p. 17002.

39. Li, P., et al., MALAT1 Is Associated with Poor Response to Oxaliplatin-Based Chemotherapy in Colorectal Cancer Patients and Promotes Chemoresistance through EZH2. Mol Cancer Ther, 2017. 16(4): p. 739–751.

40. El Said, N.H., et al., Malat-1-PRC2-EZH1 interaction supports adaptive oxidative stress dependent epigenome remodeling in skeletal myotubes. Cell Death & Disease, 2021. 12(10): p. 850.

41. Chu, C., J. Quinn, and H.Y. Chang, Chromatin isolation by RNA purification (ChIRP). J Vis Exp, 2012(61).

42. G Betancur, J. and Y. Tomari, Cryptic RNA-binding by PRC2 components EZH2 and SUZ12. RNA Biology, 2015. 12(9): p. 959–965.

43. Serebryannyy, L.A., et al., Persistent nuclear actin filaments inhibit transcription by RNA polymerase II. J Cell Sci, 2016. 129(18): p. 3412–25.

44. Liu, J., et al., MicroRNA miR-145-5p inhibits Phospholipase D 5 (PLD5) to downregulate cell proliferation and metastasis to mitigate prostate cancer. Bioengineered, 2021. 12(1): p. 3240–3251.

45. Dennis, E.A., Introduction to Thematic Review Series: Phospholipases: Central Role in Lipid Signaling and Disease. J Lipid Res, 2015. 56(7): p. 1245–7.

46. Kattan, R.E., et al., Interactome Analysis of Human Phospholipase D and Phosphatidic Acid-Associated Protein Network. Mol Cell Proteomics, 2022. 21(2): p. 100195.

47. Zhang, E., et al., H3K27 acetylation activated-long non-coding RNA CCAT1 affects cell proliferation and migration by regulating SPRY4 and HOXB13 expression in esophageal squamous cell carcinoma. Nucleic Acids Res, 2017. 45(6): p. 3086–3101.

48. Hu, M.J., M. Long, and R.J. Dai, Acetylation of H3K27 activated lncRNA NEAT1 and promoted hepatic lipid accumulation in non-alcoholic fatty liver disease via regulating miR-212-5p/GRIA3. Mol Cell Biochem, 2022. 477(1): p. 191–203.

49. Agostini M., M. Mancini, E. Candi. Long non-coding RNAs affecting cell metabolism in cancer. Biol Direct, 2022 17(1): p26.

50. Percipalle, P., et al., Nuclear actin is associated with a specific subset of hnRNP A/B-type proteins. Nucleic Acids Research, 2002. 30(8): p. 1725–1734.

51. Chu, C., et al., Genomic maps of long noncoding RNA occupancy reveal principles of RNA-chromatin interactions. Mol Cell, 2011. 44(4): p. 667–78.

52. Chu, C. and H.Y. Chang, ChIRP-MS: RNA-Directed Proteomic Discovery. 2018, Springer New York. p. 37-45.

53. Chu, C., et al., Systematic Discovery of Xist RNA Binding Proteins. Cell, 2015. 161(2): p. 404–416.

54. Pang, Z., et al., MetaboAnalyst 5.0: narrowing the gap between raw spectra and functional insights. Nucleic Acids Res, 2021. 49(W1): p. W388–W396.

55. Mouse Genome Sequencing, C., et al., Initial sequencing and comparative analysis of the mouse genome. Nature, 2002. 420(6915): p. 520–62.

56. Hendrickson, D., et al., Widespread RNA binding by chromatin-associated proteins. Genome Biology, 2016. 17(1): p. 1–18.

