## supplementary figure legends and supplementary figures for "Nuclear actin-dependent Meg3 expression suppresses metabolic genes by affecting the chromatin architecture at sites of elevated H3K27 acetylation levels"

**Supplementary figure 1. ChIRP peaks annotation and elevated binding at TSS in KO.** a) The Average Profile of ChIRP peaks binding to TSS region in WT and KO alone and upon overlapping, dark blue: KO; light blue: WT. b) heatmap of the *Meg3* ChIRP tags(reads), where more the tags are to falling in TSS we get redder shade. Heatmap of *Meg3* ChIRP peaks binding to TSS regions for WT and KO. c) UpSet plots for WT and KO datasets of the *Meg3* ChIRP, showing set data of *Meg3* ChIRP-seq over more than three intersecting sets. UpSet plots show intersections in a matrix, with the rows of the matrix corresponding to the sets, and the columns to the intersections between these sets. Genomic Regions, represent different regions of the genome that were bound by *Meg3* in the ChIRP-seq dataset. Each region corresponds to a specific genomic locus or set of loci. Intersection as connected dots represents the intersection of genomic regions and the Meg3 binding. It indicates the specific genomic regions where the captured *Meg3* -lncRNAs were found. The Venn diagrams within the Upset plot illustrate the overlap between different sets of genomic regions and *Meg3*. The bar shows the count of occurrences for each combination of sets. It provides a visual representation of the distribution of genomic regions and *Meg3* peaks across different subsets in WT and Actin KO MEFs.

**Supplementary Figure 2. ChIRP peaks annotation shows *Meg3* preference in binding to specific genomic regions globally in KO vs WT.** a) ChIRP-seq peak Annotation of WT-specific, KO-specific peaks and shared *Meg3* peaks in percentages of exon/intron, intergenic, Noncoding/TTS/, UTR and promoter/TSS. b) Boxplots showing average change in expression of genes overlapping or closest to (in case of intergenic peaks) *Meg3* peaks. Only genes showing differential expression with FDR <0.05 were included in the analysis. Scale bar shows normalized RPKM. WT: RED, KO: Green, Shared peaks of WT and KO: blue. p-values based on two-tailed, two-sample Wilcoxon-rank sum test.

**Supplementary Figure 3. ChIRP-seq shows Meg3 bindings to and around TSS in KO specific, WT specific peaks and Shared Meg3 peaks.**

Density plots and heatmaps displaying scaled-read densities for *Meg3* in 3Kb regions centered on WT-specific, KO-specific, and shared *Meg3* Peaks. Expression is plotted as average log2FoldChange.

**Supplementary figure 4.** Illustrations of the dermatan sulfate (a), heparan sulfate (b) and Chondroitin sulfate (c) biosynthesis generated by Ingenuity Pathway Analysis (Qiagen) demonstrating the overrepresentation of genes implicated in these pathways Predicted gene and metabolite expression and activities are highlighted. Predictions are based on the fold change in gene expression levels between KO and WT replicates (n = 42,594 genes).

**Supplementary Figure 5.** Illustrations of the phospholipases pathways generated by Ingenuity Pathway Analysis (Qiagen) demonstrating the overrepresentation of genes implicated in these pathways. Predicted gene and metabolite expression and activities are highlighted. Predictions are based on the fold change in gene expression levels between KO and WT replicates (n = 42,594 genes).

**Supplementary figure 1**

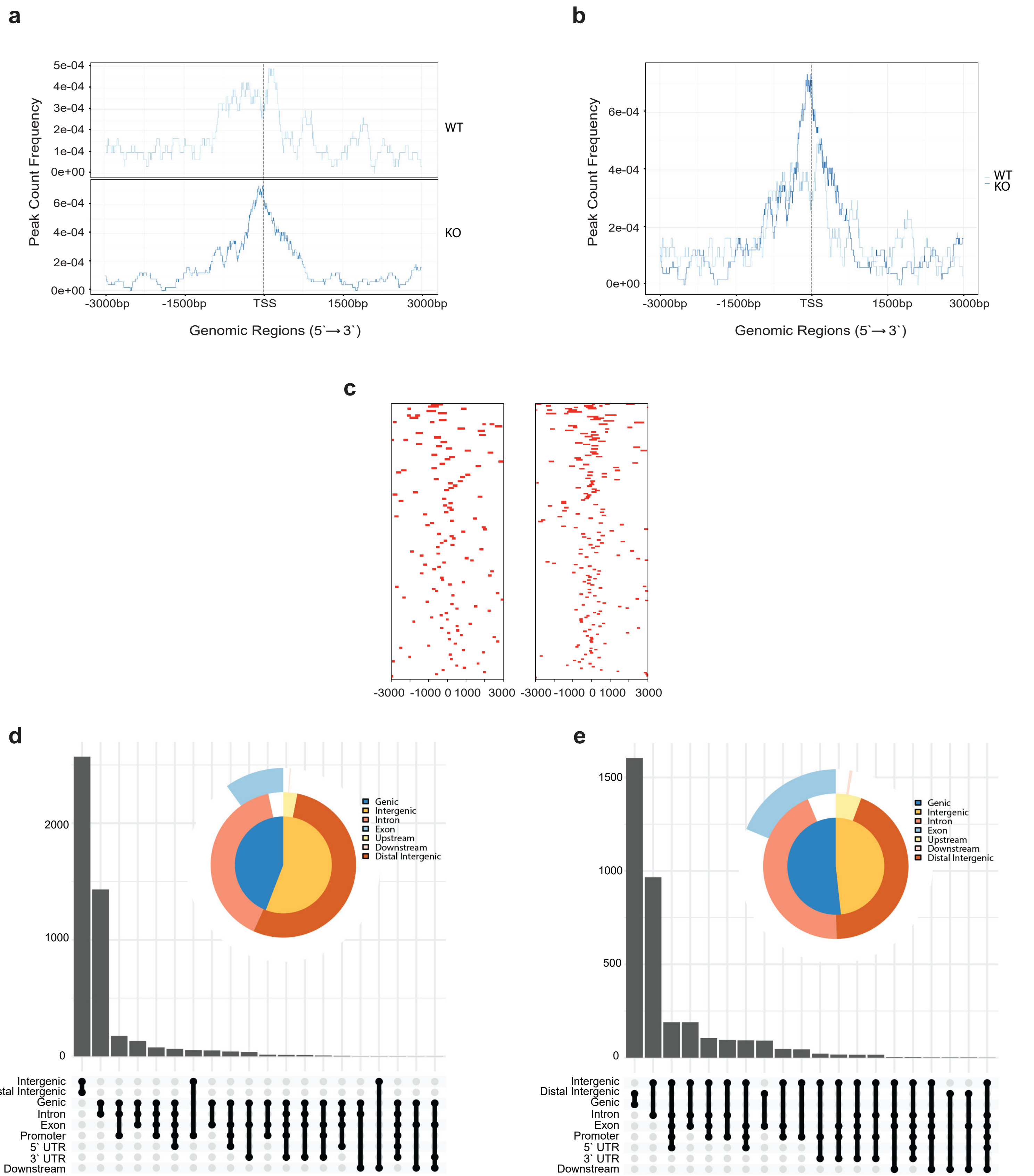

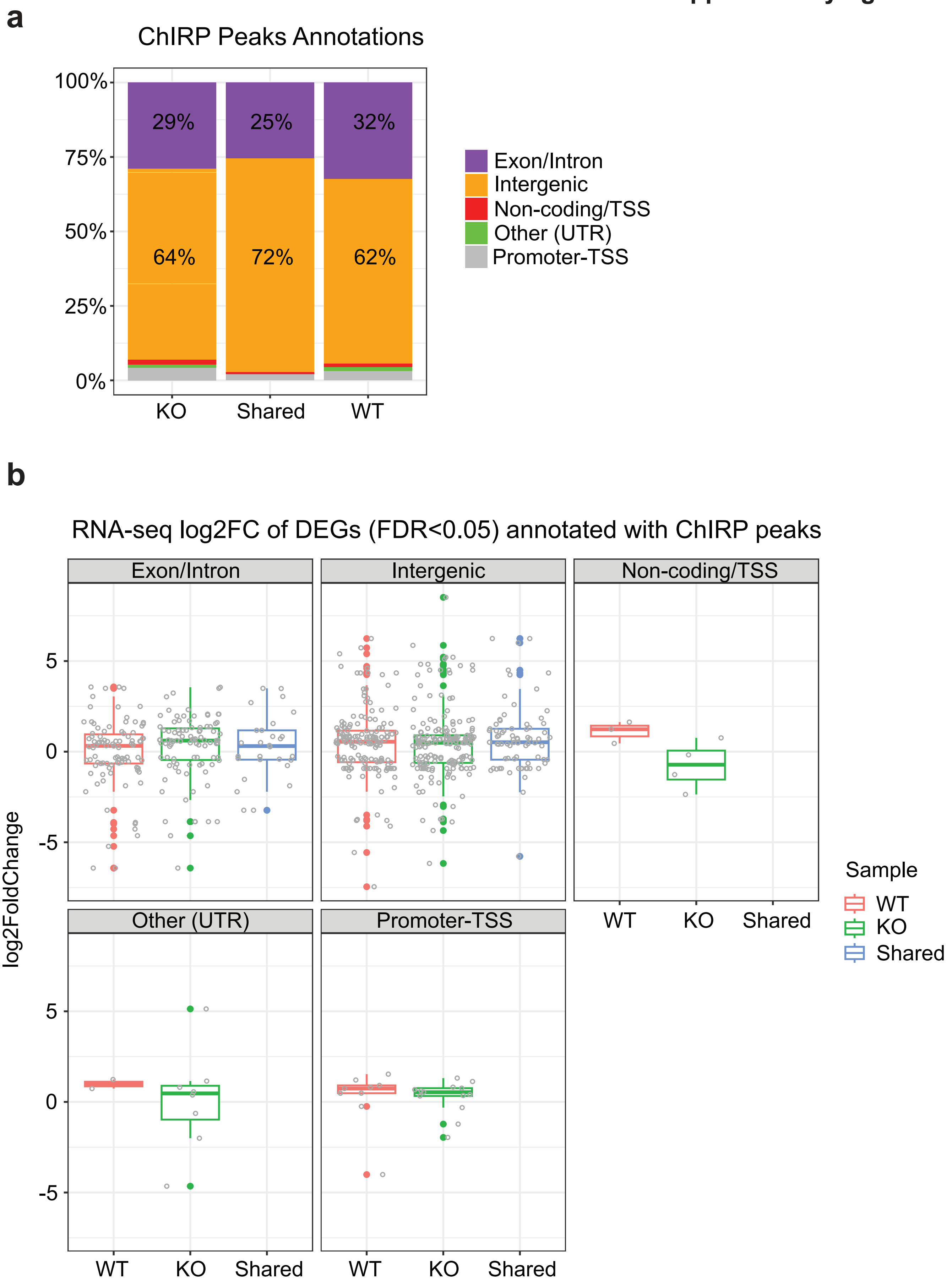
a
ChIRP Peaks Annotations
100%
75%
50%
25%
0%
29%
25%
32%
64%
72%
62%
KO
Shared
WT
Exon/Intron
Intergenic
Non-coding/TSS
Other (UTR)
Promoter-TSS
b
RNA-seq log2FC of DEGs (FDR<0.05) annotated with ChIRP peaks
Exon/Intron
Intergenic
Non-coding/TSS
Other (UTR)
Promoter-TSS
log2FoldChange
5
0
-5
WT
KO
Shared
Sample
WT
KO
Shared

**Supplementary figure 3**

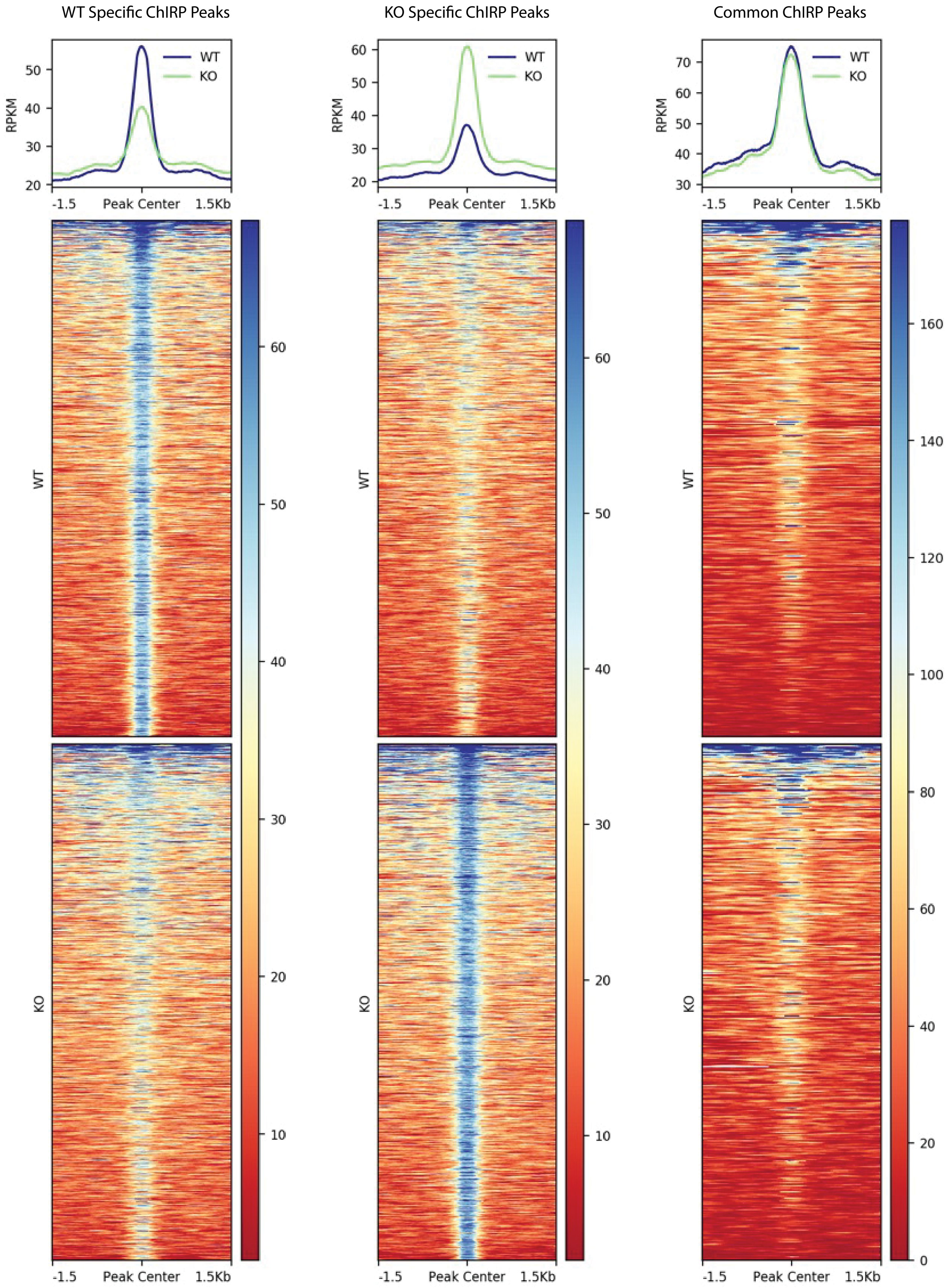

### Supplementary figure 4

**a**

#### Dermatan sulfate biosynthesis pathway

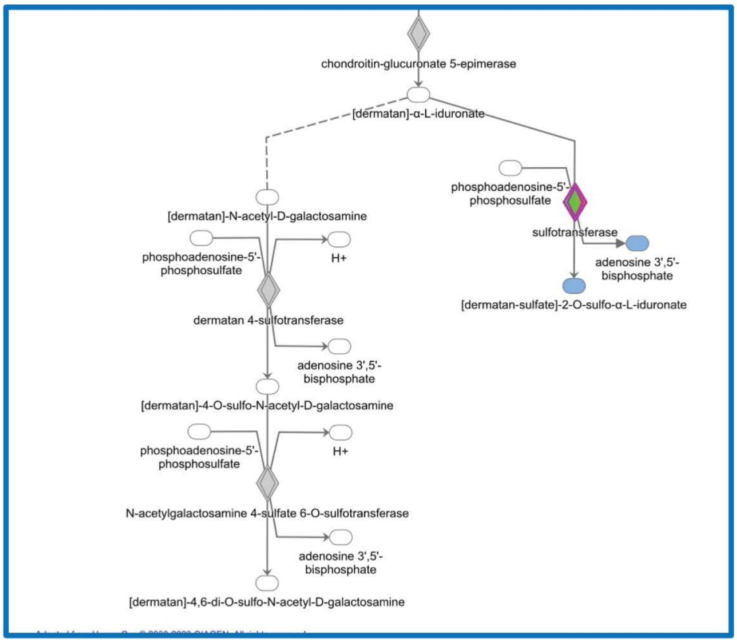


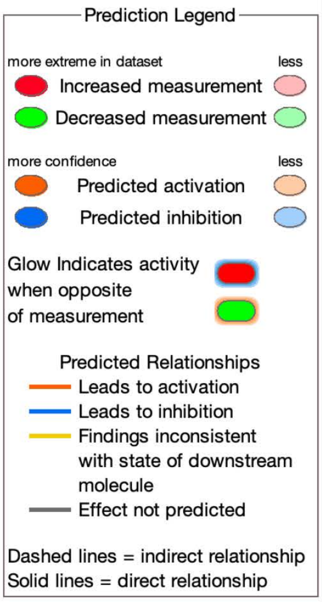


**b**

#### Heparan sulfate biosynthesis pathway

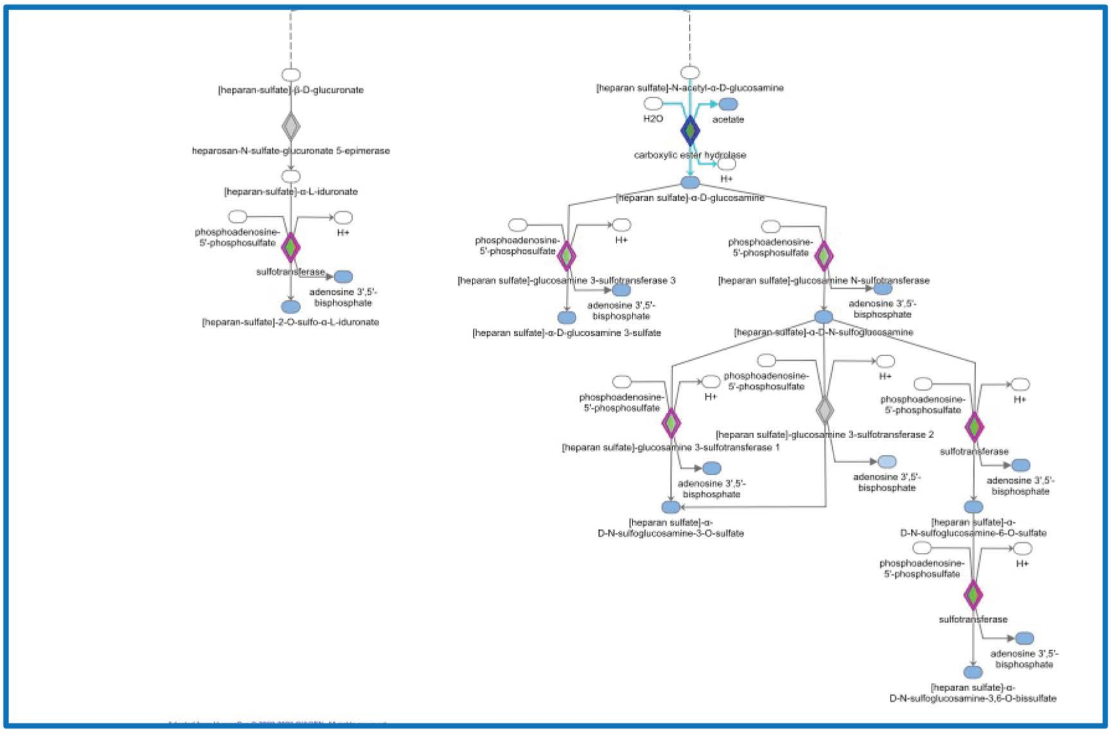


**c**

#### Chondroitin sulfate biosynthesis pathway

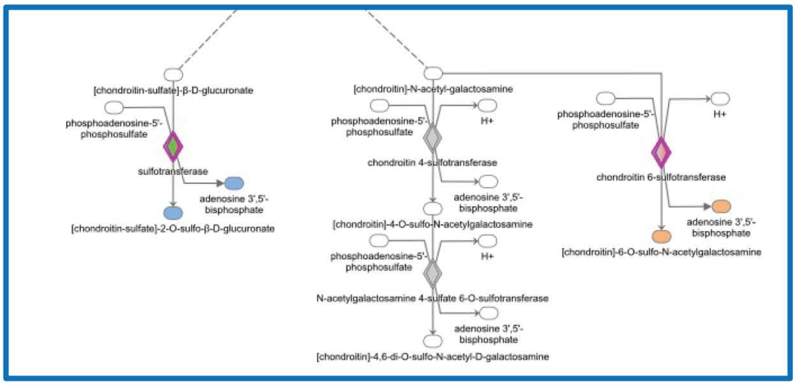

### Supplementary figure 5

#### Phospholipases biosynthesis pathways

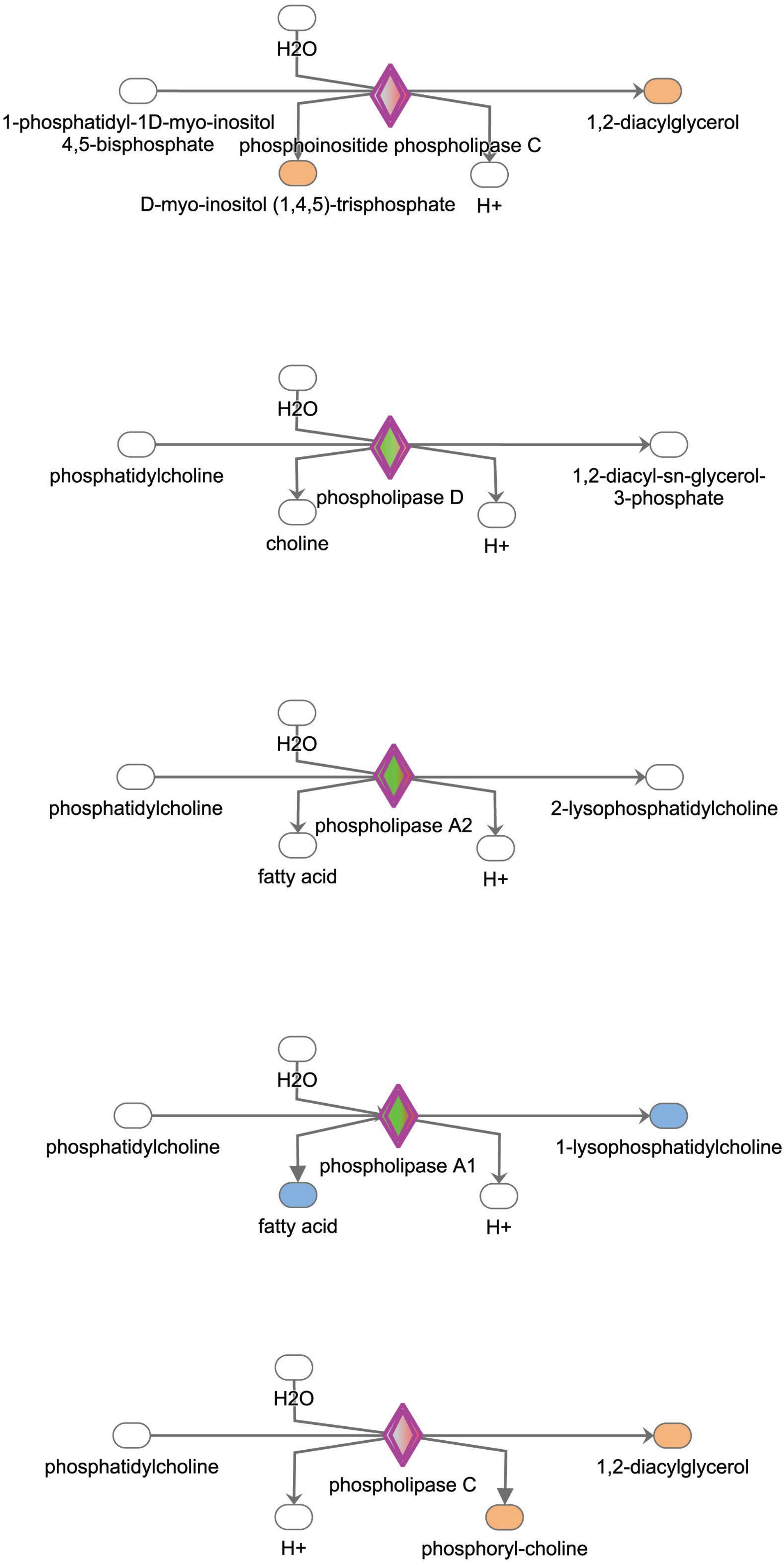


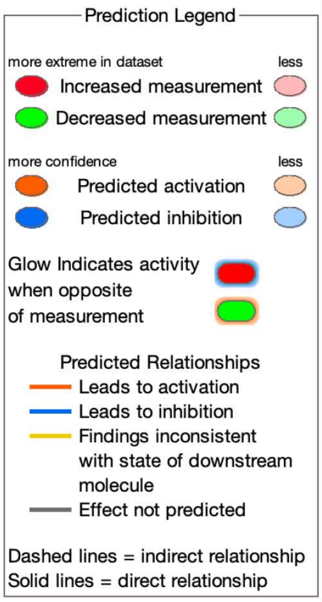
