## Supplemental table 1 for "Nuclear actin-dependent Meg3 expression suppresses metabolic genes by affecting the chromatin architecture at sites of elevated H3K27 acetylation levels"

**Supplementary Table 1**

**ChIRP-MS, WT Mouse Embryonic Fibroblasts**

| Accession | -10lgP | Coverage (% | #Peptides | #Unique | Avg. Mass | Description |
|---|---|---|---|---|---|---|
| P63260 | 239.49 | 43 | 10 | 7 | 41793 | Actin cytoplasmic 2 OS=Mus musculus OX=10090 GN=Actg1 PE=1 SV=1 |
| P60710 | 239.49 | 43 | 10 | 7 | 41737 | Actin cytoplasmic 1 OS=Mus musculus OX=10090 GN=Actb PE=1 SV=1 |
| Q8CGP6 | 36.92 | 20 | 2 | 2 | 13950 | Histone H2A type 1-H OS=Mus musculus OX=10090 GN=H2ac12 PE=1 SV=3 |
| Q64523 | 36.92 | 20 | 2 | 2 | 13988 | Histone H2A type 2-C OS=Mus musculus OX=10090 GN=H2ac20 PE=1 SV=3 |
| Q8R1M2 | 36.92 | 20 | 2 | 2 | 14045 | Histone H2A.J OS=Mus musculus OX=10090 GN=H2aj PE=1 SV=1 |
| Q8BFU2 | 36.92 | 20 | 2 | 2 | 14121 | Histone H2A type 3 OS=Mus musculus OX=10090 GN=H2ac25 PE=1 SV=3 |
| C0HKE2 | 36.92 | 20 | 2 | 2 | 14135 | Histone H2A type 1-C OS=Mus musculus OX=10090 GN=H2ac6 PE=1 SV=1 |
| C0HKE1 | 36.92 | 20 | 2 | 2 | 14135 | Histone H2A type 1-B OS=Mus musculus OX=10090 GN=H2ac4 PE=1 SV=1 |
| Q8CGP7 | 36.92 | 20 | 2 | 2 | 14150 | Histone H2A type 1-K OS=Mus musculus OX=10090 GN=H2ac15 PE=1 SV=3 |
| Q6GSS7 | 36.92 | 20 | 2 | 2 | 14095 | Histone H2A type 2-A OS=Mus musculus OX=10090 GN=Hist2h2aa2 PE=1 SV=3 |
| Q8CGP5 | 36.92 | 20 | 2 | 2 | 14162 | Histone H2A type 1-F OS=Mus musculus OX=10090 GN=Hist1h2af PE=1 SV=3 |
| P20152 | 165.27 | 18 | 7 | 5 | 53688 | Vimentin OS=Mus musculus OX=10090 GN=Vim PE=1 SV=3 |
| P52480 | 195.17 | 16 | 5 | 5 | 57845 | Pyruvate kinase PKM OS=Mus musculus OX=10090 GN=Pkm PE=1 SV=4 |
| P10126 | 176.42 | 15 | 5 | 5 | 50114 | Elongation factor 1-alpha 1 OS=Mus musculus OX=10090 GN=Eef1a1 PE=1 SV=3 |
| P16858 | 189.26 | 14 | 4 | 4 | 35810 | Glyceraldehyde-3-phosphate dehydrogenase OS=Mus musculus OX=10090 GN=Gapdh PE=1 SV=2 |
| P06151 | 75.79 | 13 | 2 | 2 | 36499 | Heat shock cognate 71 kDa protein OS=Mus musculus OX=10090 GN=Hspa8 PE=1 SV=1 |
| P06151 | 75.79 | 13 | 2 | 2 | 36499 | L-lactate dehydrogenase A chain OS=Mus musculus OX=10090 GN=Ldha PE=1 SV=3 |
| P99024 | 214.48 | 12 | 5 | 5 | 49671 | Tubulin beta-5 chain OS=Mus musculus OX=10090 GN=Tubb5 PE=1 SV=1 |
| P05213 | 178.26 | 12 | 3 | 3 | 50152 | Tubulin alpha-1B chain OS=Mus musculus OX=10090 GN=Tuba1b PE=1 SV=2 |
| P68373 | 178.26 | 12 | 3 | 3 | 49909 | Tubulin alpha-1C chain OS=Mus musculus OX=10090 GN=Tuba1c PE=1 SV=1 |
| Q9JJZ2 | 178.26 | 12 | 3 | 3 | 50052 | Tubulin alpha-8 chain OS=Mus musculus OX=10090 GN=Tuba8 PE=1 SV=1 |
| P68368 | 178.26 | 12 | 3 | 3 | 49924 | Tubulin alpha-4A chain OS=Mus musculus OX=10090 GN=Tuba4a PE=1 SV=1 |
| P68134 | 134.29 | 12 | 5 | 1 | 42051 | Actin alpha cardiac muscle 1 OS=Mus musculus OX=10090 GN=Actc1 PE=1 SV=1 |
| P45591 | 98.78 | 10 | 1 | 1 | 18710 | Annexin A2 OS=Mus musculus OX=10090 GN=Anxa2 PE=1 SV=2 |
| P07356 | 122.74 | 9 | 2 | 2 | 38676 | Actin alpha skeletal muscle OS=Mus musculus OX=10090 GN=Acta1 PE=1 SV=1 |
| P54227 | 78.18 | 9 | 1 | 1 | 17274 | Stathmin OS=Mus musculus OX=10090 GN=Stmn1 PE=1 SV=2 |
| P49312 | 47.9 | 9 | 2 | 2 | 34196 | Heterogeneous nuclear ribonucleoprotein A1 OS=Mus musculus OX=10090 GN=Hnrnpa1 PE=1 SV=2 |
| P63242 | 53 | 8 | 1 | 1 | 16832 | Eukaryotic translation initiation factor 5A-1 OS=Mus musculus OX=10090 GN=Eif5a PE=1 SV=2 |
| Q8BGY2 | 53 | 8 | 1 | 1 | 16793 | Eukaryotic translation initiation factor 5A-2 OS=Mus musculus OX=10090 GN=Eif5a2 PE=1 SV=3 |
| P62984 | 69.65 | 7 | 1 | 1 | 14728 | Ubiquitin-60S ribosomal protein L40 OS=Mus musculus OX=10090 GN=Uba52 PE=1 SV=2 |
| P62983 | 69.65 | 6 | 1 | 1 | 17951 | Ubiquitin-40S ribosomal protein S27a OS=Mus musculus OX=10090 GN=Rps27a PE=1 SV=2 |
| P17742 | 58.07 | 5 | 1 | 1 | 17971 | Peptidyl-prolyl cis-trans isomerase A OS=Mus musculus OX=10090 GN=Ppia PE=1 SV=2 |
| P17183 | 74.68 | 4 | 1 | 1 | 47297 | Gamma-enolase OS=Mus musculus OX=10090 GN=Eno2 PE=1 SV=2 |
| P17182 | 74.68 | 4 | 1 | 1 | 47141 | Alpha-enolase OS=Mus musculus OX=10090 GN=Eno1 PE=1 SV=3 |
| P21550 | 74.68 | 4 | 1 | 1 | 47025 | Beta-enolase OS=Mus musculus OX=10090 GN=Eno3 PE=1 SV=3 |
| P68254 | 69.36 | 4 | 1 | 1 | 27778 | 14-3-3 protein theta OS=Mus musculus OX=10090 GN=Ywhaq PE=1 SV=1 |
| P63101 | 69.36 | 4 | 1 | 1 | 27771 | 14-3-3 protein zeta/delta OS=Mus musculus OX=10090 GN=Ywhaz PE=1 SV=1 |
| Q9CQV8 | 69.36 | 4 | 1 | 1 | 28086 | 14-3-3 protein beta/alpha OS=Mus musculus OX=10090 GN=Ywhab PE=1 SV=3 |
| P68510 | 69.36 | 4 | 1 | 1 | 28212 | 14-3-3 protein eta OS=Mus musculus OX=10090 GN=Ywhah PE=1 SV=2 |
| P61982 | 69.36 | 4 | 1 | 1 | 28303 | 14-3-3 protein gamma OS=Mus musculus OX=10090 GN=Ywhag PE=1 SV=2 |
| O70456 | 69.36 | 4 | 1 | 1 | 27706 | 14-3-3 protein sigma OS=Mus musculus OX=10090 GN=Sfn PE=1 SV=2 |
| P62259 | 69.36 | 4 | 1 | 1 | 29174 | 14-3-3 protein epsilon OS=Mus musculus OX=10090 GN=Ywhae PE=1 SV=1 |
| P62631 | 80.94 | 4 | 2 | 2 | 50454 | Elongation factor 1-alpha 2 OS=Mus musculus OX=10090 GN=Eef1a2 PE=1 SV=1 |
| Q8VDD5 | 182.23 | 3 | 4 | 3 | 226370 | Myosin-9 OS=Mus musculus OX=10090 GN=Myh9 PE=1 SV=4 |
| P0CG49 | 69.65 | 3 | 1 | 1 | 34369 | Polyubiquitin-B OS=Mus musculus OX=10090 GN=Ubb PE=2 SV=1 |
| O08638 | 93.35 | 2 | 2 | 1 | 227026 | Cofilin-2 OS=Mus musculus OX=10090 GN=Cfl2 PE=1 SV=1 |
| P63017 | 84.58 | 2 | 1 | 1 | 70871 | Filamin-A OS=Mus musculus OX=10090 GN=Flna PE=1 SV=5 |
| P11499 | 72.24 | 2 | 2 | 2 | 83281 | Heat shock protein HSP 90-beta OS=Mus musculus OX=10090 GN=Hsp90ab1 PE=1 SV=3 |
| Q8BTM8 | 92.18 | 1 | 2 | 2 | 281219 | Myosin-11 OS=Mus musculus OX=10090 GN=Myh11 PE=1 SV=1 |
| P0CG50 | 69.65 | 1 | 1 | 1 | 82550 | Polyubiquitin-C OS=Mus musculus OX=10090 GN=Ubc PE=1 SV=2 |
| P57780 | 52.6 | 1 | 1 | 1 | 104977 | Alpha-actinin-4 OS=Mus musculus OX=10090 GN=Actn4 PE=1 SV=1 |
| P26039 | 51.54 | 1 | 1 | 1 | 269819 | Talin-1 OS=Mus musculus OX=10090 GN=Tln1 PE=1 SV=2 |
| E9Q555 | 37.18 | 0 | 2 | 2 | 584265 | E3 ubiquitin-protein ligase RNF213 OS=Mus musculus OX=10090 GN=Rnf213 PE=1 SV=3 |
| A3KFM7 | 33.99 | 0 | 1 | 1 | 305396 | Chromodomain-helicase-DNA-binding protein 6 OS=Mus musculus OX=10090 GN=Chd6 PE=1 SV=1 |

**ChIRP-MS, $\beta$-actin KO Mouse Embryonic Fibroblasts**

| Accession | -10lgP | Coverage (% | #Peptides | #Unique | Avg. Mass | filtered no keratin no rnases |
|---|---|---|---|---|---|---|
| Q64475 | 81.31 | 18 | 3 | 3 | 13952 | Histone H2B type 1-B OS=Mus musculus OX=10090 GN=H2bc3 PE=1 SV=3 |
| P10853 | 81.31 | 18 | 3 | 3 | 13936 | Histone H2B type 1-F/J/L OS=Mus musculus OX=10090 GN=H2bc15 PE=1 SV=2 |
| Q8CGP1 | 81.31 | 18 | 3 | 3 | 13920 | Histone H2B type 1-K OS=Mus musculus OX=10090 GN=H2bc12 PE=1 SV=3 |
| Q8CGP2 | 81.31 | 18 | 3 | 3 | 13992 | Histone H2B type 1-P OS=Mus musculus OX=10090 GN=Hist1h2bp PE=1 SV=3 |
| P70696 | 81.31 | 18 | 3 | 3 | 14237 | Histone H2B type 1-A OS=Mus musculus OX=10090 GN=H2bc1 PE=1 SV=3 |
| P62806 | 106.73 | 17 | 2 | 2 | 11367 | Histone H4 OS=Mus musculus OX=10090 GN=H4c16 PE=1 SV=2 |
| P07724 | 150.97 | 13 | 8 | 8 | 68693 | Albumin OS=Mus musculus OX=10090 GN=Alb PE=1 SV=3 |
| P68433 | 34.76 | 10 | 2 | 2 | 15404 | Histone H3.1 OS=Mus musculus OX=10090 GN=H3c11 PE=1 SV=2 |
| P84244 | 34.76 | 10 | 2 | 2 | 15328 | Histone H3.3 OS=Mus musculus OX=10090 GN=H3-3b PE=1 SV=2 |
| P02301 | 34.76 | 10 | 2 | 2 | 15315 | Histone H3.3C OS=Mus musculus OX=10090 GN=H3-5 PE=3 SV=3 |
| P84228 | 34.76 | 10 | 2 | 2 | 15388 | Histone H3.2 OS=Mus musculus OX=10090 GN=H3c15 PE=1 SV=2 |
| P20152 | 46.57 | 4 | 2 | 1 | 53688 | Vimentin OS=Mus musculus OX=10090 GN=Vim PE=1 SV=3 |
| P10126 | 109.27 | 4 | 3 | 3 | 50114 | Elongation factor 1-alpha 1 OS=Mus musculus OX=10090 GN=Eef1a1 PE=1 SV=3 |
| P62631 | 109.27 | 4 | 3 | 3 | 50454 | Elongation factor 1-alpha 2 OS=Mus musculus OX=10090 GN=Eef1a2 PE=1 SV=1 |
| Q9R1T2 | 31.73 | 3 | 1 | 1 | 38620 | SUMO-activating enzyme subunit 1 OS=Mus musculus OX=10090 GN=Sae1 PE=1 SV=1 |
| P63260 | 32.59 | 2 | 1 | 1 | 41793 | Actin cytoplasmic 2 OS=Mus musculus OX=10090 GN=Actg1 PE=1 SV=1 |
| P60710 | 32.59 | 2 | 1 | 1 | 41737 | Actin cytoplasmic 1 OS=Mus musculus OX=10090 GN=Actb PE=1 SV=1 |
| P68373 | 49.86 | 2 | 1 | 1 | 49909 | Tubulin alpha-1C chain OS=Mus musculus OX=10090 GN=Tuba1c PE=1 SV=1 |
| P05213 | 49.86 | 2 | 1 | 1 | 50152 | Tubulin alpha-1B chain OS=Mus musculus OX=10090 GN=Tuba1b PE=1 SV=2 |
| P68369 | 49.86 | 2 | 1 | 1 | 50136 | Tubulin alpha-1A chain OS=Mus musculus OX=10090 GN=Tuba1a PE=1 SV=1 |
| P63268 | 32.59 | 2 | 1 | 1 | 41877 | Actin gamma-enteric smooth muscle OS=Mus musculus OX=10090 GN=PE=1 SV=1 |
| Q8BFZ3 | 32.59 | 2 | 1 | 1 | 42004 | Beta-actin-like protein 2 OS=Mus musculus OX=10090 GN=Actbl2 PE=1 SV=1 |
| P68033 | 32.59 | 2 | 1 | 1 | 42019 | Actin alpha cardiac muscle 1 OS=Mus musculus OX=10090 GN=Actc1 PE=1 SV=1 |
| P62737 | 32.59 | 2 | 1 | 1 | 42009 | Actin aortic smooth muscle OS=Mus musculus OX=10090 GN=Acta2 PE=1 SV=1 |
| P68134 | 32.59 | 2 | 1 | 1 | 42051 | Actin alpha skeletal muscle OS=Mus musculus OX=10090 GN=Acta1 PE=1 SV=1 |
| Q9CX86 | 29.7 | 2 | 1 | 1 | 30530 | Heterogeneous nuclear ribonucleoprotein A0 OS=Mus musculus OX=10090 GN=Hnrnpa0 PE=1 SV=1 |
| P49312 | 29.7 | 2 | 1 | 1 | 34196 | Heterogeneous nuclear ribonucleoprotein A1 OS=Mus musculus OX=10090 GN=Hnrnpa1 PE=1 SV=2 |
| O88569 | 29.7 | 2 | 1 | 1 | 37403 | Heterogeneous nuclear ribonucleoproteins A2/B1 OS=Mus musculus OX=10090 GN=Hnrnpa2b1 PE=1 SV=2 |
| Q8BG05 | 29.7 | 2 | 1 | 1 | 39652 | Heterogeneous nuclear ribonucleoprotein A3 OS=Mus musculus OX=10090 GN=Hnrnpa3 PE=1 SV=1 |
| P11499 | 36.99 | 1 | 1 | 1 | 83281 | Heat shock protein HSP 90-beta OS=Mus musculus OX=10090 GN=Hsp90ab1 PE=1 SV=3 |
| O35709 | 28.6 | 1 | 1 | 1 | 66113 | Ectoderm-neural cortex protein 1 OS=Mus musculus OX=10090 GN=Enc1 PE=2 SV=2 |
| Q7SIG6 | 25.24 | 1 | 1 | 1 | 106805 | Arf-GAP with SH3 domain ANK repeat and PH domain-containing protein 2 OS=Mus musculus OX=10090 GN=Asap2 PE=1 SV=3 |
| Q9ERU9 | 25.02 | 1 | 1 | 1 | 341122 | E3 SUMO-protein ligase RanBP2 OS=Mus musculus OX=10090 GN=Ranbp2 PE=1 SV=2 |
| E9Q8T7 | 29.3 | 0 | 1 | 1 | 487074 | Dynein axonemal heavy chain 1 OS=Mus musculus OX=10090 GN=Dnah1 PE=1 SV=1 |

**ChIRP-MS, Venn Diagram**

| Names | total | elements | WT coverage | KO coverage |
|---|---|---|---|---|
|  |  | Actin cytopla | 43 | 2 |
|  |  | Actin cytopla | 43 | 2 |
| | | Vimentin OS= | 18 | 4 |
| KO-ChIRP-M | 11 | Elongation fa | 15 | 4 |
| | | Tubulin alpha | 12 | 2 |
| | | Tubulin alpha | 12 | 2 |
|  |  | Actin alpha c | 12 | 2 |
|  |  | Actin alpha s | 9 | 2 |
| | | Heterogeneo | 9 | 2 |
| | | Elongation fa | 4 | 4 |
| | | Heat shock p | 2 | 1 |
| WT-ChIRP-M | 43 | Histone H2A type 3 OS=Mus musculus OX=10090 GN=H2ac25 PE=1 SV=3 | | |
| | | Beta-enolase OS=Mus musculus OX=10090 GN=Eno3 PE=1 SV=3 | | |
| | | Ubiquitin-40S ribosomal protein S27a OS=Mus musculus OX=10090 GN=Rps27a PE=1 SV=2 | | |
| | | Tubulin alpha-4A chain OS=Mus musculus OX=10090 GN=Tuba4a PE=1 SV=1 | | |
| | | Stathmin OS=Mus musculus OX=10090 GN=Stmn1 PE=1 SV=2 | | |
| | | 14-3-3 protein theta OS=Mus musculus OX=10090 GN=Ywhaq PE=1 SV=1 | | |
| | | Heat shock cognate 71 kDa protein OS=Mus musculus OX=10090 GN=Hspa8 PE=1 SV=1 | | |
| | | Gamma-enolase OS=Mus musculus OX=10090 GN=Eno2 PE=1 SV=2 | | |
| | | Histone H2A type 1-C OS=Mus musculus OX=10090 GN=H2ac6 PE=1 SV=1 | | |
| | | Talin-1 OS=Mus musculus OX=10090 GN=Tln1 PE=1 SV=2 | | |
| | | 14-3-3 protein sigma OS=Mus musculus OX=10090 GN=Sfn PE=1 SV=2 | | |
| | | 14-3-3 protein eta OS=Mus musculus OX=10090 GN=Ywhah PE=1 SV=2 | | |
| | | Polyubiquitin-C OS=Mus musculus OX=10090 GN=Ubc PE=1 SV=2 | | |
| | | Polyubiquitin-B OS=Mus musculus OX=10090 GN=Ubb PE=2 SV=1 | | |
| | | Histone H2A type 1-B OS=Mus musculus OX=10090 GN=H2ac4 PE=1 SV=1 | | |
| | | 14-3-3 protein epsilon OS=Mus musculus OX=10090 GN=Ywhae PE=1 SV=1 | | |
| | | L-lactate dehydrogenase A chain OS=Mus musculus OX=10090 GN=Ldha PE=1 SV=3 | | |
| | | Cofilin-2 OS=Mus musculus OX=10090 GN=Cfl2 PE=1 SV=1 | | |
| | | Tubulin alpha-8 chain OS=Mus musculus OX=10090 GN=Tuba8 PE=1 SV=1 | | |
| | | Eukaryotic translation initiation factor 5A-1 OS=Mus musculus OX=10090 GN=Eif5a PE=1 SV=2 | | |
| | | Histone H2A.J OS=Mus musculus OX=10090 GN=H2aj PE=1 SV=1 | | |
| | | Filamin-A OS=Mus musculus OX=10090 GN=Flna PE=1 SV=5 | | |
| | | 14-3-3 protein beta/alpha OS=Mus musculus OX=10090 GN=Ywhab PE=1 SV=3 | | |
| | | Histone H2A type 2-C OS=Mus musculus OX=10090 GN=H2ac20 PE=1 SV=3 | | |
| | | Glyceraldehyde-3-phosphate dehydrogenase OS=Mus musculus OX=10090 GN=Gapdh PE=1 SV=2 | | |
| | | Pyruvate kinase PKM OS=Mus musculus OX=10090 GN=Pkm PE=1 SV=4 | | |
| | | 14-3-3 protein gamma OS=Mus musculus OX=10090 GN=Ywhag PE=1 SV=2 | | |
| | | Myosin-9 OS=Mus musculus OX=10090 GN=Myh9 PE=1 SV=4 | | |
| | | Alpha-actinin-4 OS=Mus musculus OX=10090 GN=Actn4 PE=1 SV=1 | | |
| | | Myosin-11 OS=Mus musculus OX=10090 GN=Myh11 PE=1 SV=1 | | |
| | | Alpha-enolase OS=Mus musculus OX=10090 GN=Eno1 PE=1 SV=3 | | |
| | | Histone H2A type 1-H OS=Mus musculus OX=10090 GN=H2ac12 PE=1 SV=3 | | |
| | | Tubulin beta-5 chain OS=Mus musculus OX=10090 GN=Tubb5 PE=1 SV=1 | | |
| | | Eukaryotic translation initiation factor 5A-2 OS=Mus musculus OX=10090 GN=Eif5a2 PE=1 SV=3 | | |
| | | Histone H2A type 2-A OS=Mus musculus OX=10090 GN=Hist2h2aa2 PE=1 SV=3 | | |
| | | Chromodomain-helicase-DNA-binding protein 6 OS=Mus musculus OX=10090 GN=Chd6 PE=1 SV=1 | | |
| | | 14-3-3 protein zeta/delta OS=Mus musculus OX=10090 GN=Ywhaz PE=1 SV=1 | | |
| | | E3 ubiquitin-protein ligase RNF213 OS=Mus musculus OX=10090 GN=Rnf213 PE=1 SV=3 | | |
| | | Ubiquitin-60S ribosomal protein L40 OS=Mus musculus OX=10090 GN=Uba52 PE=1 SV=2 | | |
| | | Annexin A2 OS=Mus musculus OX=10090 GN=Anxa2 PE=1 SV=2 | | |
| | | Histone H2A type 1-K OS=Mus musculus OX=10090 GN=H2ac15 PE=1 SV=3 | | |
| | | Histone H2A type 1-F OS=Mus musculus OX=10090 GN=Hist1h2af PE=1 SV=3 | | |
| | | Peptidyl-prolyl cis-trans isomerase A OS=Mus musculus OX=10090 GN=Ppia PE=1 SV=2 | | |
| KO-ChIRP-M | 23 | Actin gamma-enteric smooth muscle OS=Mus musculus OX=10090 GN=Actg2 PE=1 SV=1 |  |  |
| | | Histone H3.3 OS=Mus musculus OX=10090 GN=H3-3b PE=1 SV=2 | | |
| | | Histone H3.2 OS=Mus musculus OX=10090 GN=H3c15 PE=1 SV=2 | | |
| | | SUMO-activating enzyme subunit 1 OS=Mus musculus OX=10090 GN=Sae1 PE=1 SV=1 | | |
| | | Histone H2B type 1-K OS=Mus musculus OX=10090 GN=H2bc12 PE=1 SV=3 | | |
| | | Ectoderm-neural cortex protein 1 OS=Mus musculus OX=10090 GN=Enc1 PE=2 SV=2 | | |
| | | Heterogeneous nuclear ribonucleoprotein A0 OS=Mus musculus OX=10090 GN=Hnrnpa0 PE=1 SV=1 | | |
|  |  | Actin aortic smooth muscle OS=Mus musculus OX=10090 GN=Acta2 PE=1 SV=1 |  |  |
| | | Histone H2B type 1-A OS=Mus musculus OX=10090 GN=H2bc1 PE=1 SV=3 | | |
| | | E3 SUMO-protein ligase RanBP2 OS=Mus musculus OX=10090 GN=Ranbp2 PE=1 SV=2 | | |
| | | Beta-actin-like protein 2 OS=Mus musculus OX=10090 GN=Actbl2 PE=1 SV=1 | | |
| | | Histone H4 OS=Mus musculus OX=10090 GN=H4c16 PE=1 SV=2 | | |
| | | Tubulin alpha-1A chain OS=Mus musculus OX=10090 GN=Tuba1a PE=1 SV=1 | | |
| | | Histone H3.1 OS=Mus musculus OX=10090 GN=H3c11 PE=1 SV=2 | | |
| | | Histone H3.3C OS=Mus musculus OX=10090 GN=H3-5 PE=3 SV=3 | | |
|  |  | Arf-GAP with SH3 domain ANK repeat and PH domain-containing protein 2 OS=Mus musculus OX=10090 GN=Asap2 PE=1 SV=3 |  |  |
| | | Heterogeneous nuclear ribonucleoprotein A3 OS=Mus musculus OX=10090 GN=Hnrnpa3 PE=1 SV=1 | | |
| | | Albumin OS=Mus musculus OX=10090 GN=Alb PE=1 SV=3 | | |
| | | Histone H2B type 1-B OS=Mus musculus OX=10090 GN=H2bc3 PE=1 SV=3 | | |
| | | Dynein axonemal heavy chain 1 OS=Mus musculus OX=10090 GN=Dnah1 PE=1 SV=1 | | |
| | | Heterogeneous nuclear ribonucleoproteins A2/B1 OS=Mus musculus OX=10090 GN=Hnrnpa2b1 PE=1 SV=2 | | |
| | | Histone H2B type 1-F/J/L OS=Mus musculus OX=10090 GN=H2bc15 PE=1 SV=2 | | |
| | | Histone H2B type 1-P OS=Mus musculus OX=10090 GN=Hist1h2bp PE=1 SV=3 | | |
