## Supplemental table 2 for "Nuclear actin-dependent Meg3 expression suppresses metabolic genes by affecting the chromatin architecture at sites of elevated H3K27 acetylation levels"

**Supplementary Table 2**

| Gene | TSS_Switch | baseMean | log2FoldChar | padj | Promoter_At | Accessibility | Sample | KO | WT | Change | Enhancers |
|---|---|---|---|---|---|---|---|---|---|---|---|
| Plcl1 | TSS-B | 14.6428503 | 2.07093031 | 3.96E-06 | Decreased | No Change | KOvsWT | 2 | 1 | 1 | Gained |
| Pld5 | TSS-B | 2.3618299 | -2.4395525 | 0.14580812 | No change | No Change | KOvsWT | 4 | 3 | 1 | Gained |
| Hs3st3b1 | TSS-B | 4.20804922 | -2.0097 | 0.09608037 | Decreased | Decreased | KOvsWT | 1 | 3 | -2 | Lost |

**table showing location of Meg3 peaks and enhancers for the 3 genes**

| Gene | start | end | | Meg3 start | Meg3 end | Meg3 loci | | | | | |
|---|---|---|---|---|---|---|---|---|---|---|---|
| Plcl1 | 56972394 | 56972894 | chr1 | 55466512 | 55466700 | Intron (ENSM | -1505882 | -1506194 | -1506656 | -1506468 | Meg3 binds before |
| Plcl1 | 56973168 | 56973668 | chr1 | 55476846 | 55477259 | Intron (ENSM | -1495548 | -1495635 | -1496322 | -1496409 | Meg3 binds before |
| | | | chr1 | 55478815 | 55479009 | Intron (ENSM | -1493579 | -1493885 | -1494353 | -1494659 | Meg3 binds before |
| Pld5 | 176282263 | 176282763 | chr1 | 176284205 | 176284477 | Distal Interge | 1942 | 1714 | | | Meg3 binds after |
| Pld5 | 176282793 | 176283293 | chr1 | | | | 1412 | 1184 | | | Meg3 binds after |
| Pld5 | 176284005 | 176284505 | chr1 | | | | 200 | -28 | | | Meg3 bind On the enhancer binding site |
| Pld5 | 176288069 | 176288569 | chr1 | | | | -3864 | -4092 | | | Meg3 binds before |
| Hs3st3b1 | 63132339 | 63132839 | chr11 | 63647447 | 63647609 | intergenic | 515108 | 514770 | | | MEg3 binds after enhancer |
| Hs3st3b1 | 63914287 | 63914787 | chr11 | | | | -266840 | -267178 | | | Meg3 binds before enhancer |
| Hs3st3b1 | 63920845 | 63921345 | chr11 | | | | -273398 | -273736 | | | Meg3 binds before enhancer |

**table showing enhancers and ABC model for the 3 genes**

| id | enh_id | Gene | TSS_Switch | Compartmen | baseMean | log2FoldChar | padj | Type | Promoter_At | Accessibility | Sample | ctl_ABC | ctl_Activity | ctl_Contact | trt_ABC | trt_Activity | trt_Contact | trta_ctlc | trt_Ctrl_Activ | trt_Ctrl_Cont | trt_Ctrl_Dista | ctla_trtc | ctl_Ctrl_Activ | ctl_Ctrl_Cont | ctl_Ctrl_Dista | Dist | Enhancer | Distance | enh_switch | class |
|---|---|---|---|---|---|---|---|---|---|---|---|---|---|---|---|---|---|---|---|---|---|---|---|---|---|---|---|---|---|---|
| chr11_63132 | chr11_63132 | Hs3st3b1 | TSS-B | Stable | 4.20804922 | -2.0097 | 0.09608037 | NDE | Decreased | Decreased | KOvsWT | 0.021775 | 19.660158 | 0.002098 | 0.017316 | 18.31626 | 0.001229 | 0.028177 | 18.31626 | 0.002098 | 789695 | 0.012927 | 19.660158 | 0.001229 | 789695 | 789.695 | Lost | 500kb-5MB | Enh-A | genic |
| chr11_63914 | chr11_63914 | Hs3st3b1 | TSS-B | Stable | 4.20804922 | -2.0097 | 0.09608037 | NDE | Decreased | Decreased | KOvsWT | 0.028724 | 2.896665 | 0.018786 | 0.007339 | 0.574724 | 0.016602 | 0.007916 | 0.574724 | 0.018786 | 7747 | 0.025726 | 2.896665 | 0.016602 | 7747 | 7.747 | Lost | <10kb | Enh-B | genic |
| chr11_63920 | chr11_63920 | Hs3st3b1 | TSS-B | Stable | 4.20804922 | -2.0097 | 0.09608037 | NDE | Decreased | Decreased | KOvsWT | 0.055162 | 1.830426 | 0.057093 | 0.025736 | 0.519558 | 0.064403 | 0.021748 | 0.519558 | 0.057093 | 1189 | 0.06306 | 1.830426 | 0.064403 | 1189 | 1.189 | No Change | <10kb | Enh-B | genic |
| chr1_176282 | chr1_176282 | Pld5 | TSS-B | Stable | 2.3618299 | -2.4395525 | 0.14580812 | NDE | No change | No Change | KOvsWT | 0.240581 | 2.679446 | 0.124665 | 0.159545 | 2.25643 | 0.124665 | 0.16094 | 2.25643 | 0.124665 | 7201 | 0.236381 | 2.679446 | 0.124665 | 7201 | 7.201 | No Change | <10kb | Enh-B | intergenic |
| chr1_176282 | chr1_176282 | Pld5 | TSS-B | Stable | 2.3618299 | -2.4395525 | 0.14580812 | NDE | No change | No Change | KOvsWT | 0.141528 | 1.576259 | 0.124665 | 0.107313 | 1.517728 | 0.124665 | 0.108252 | 1.517728 | 0.124665 | 7731 | 0.139058 | 1.576259 | 0.124665 | 7731 | 7.731 | No Change | <10kb | Enh-B | intergenic |
| chr1_176284 | chr1_176284 | Pld5 | TSS-B | Stable | 2.3618299 | -2.4395525 | 0.14580812 | NDE | No change | No Change | KOvsWT | 0 | 0 | 0.124665 | 0.029902 | 0.422898 | 0.124665 | 0.030163 | 0.422898 | 0.124665 | 8943 | 0 | 0 | 0.124665 | 8943 | 8.943 | Gained | <10kb | Enh-B | intergenic |
| chr1_176288 | chr1_176288 | Pld5 | TSS-B | Stable | 2.3618299 | -2.4395525 | 0.14580812 | NDE | No change | No Change | KOvsWT | 0.059248 | 1.196163 | 0.068773 | 0.046002 | 1.179368 | 0.068773 | 0.046405 | 1.179368 | 0.068773 | 13007 | 0.058214 | 1.196163 | 0.068773 | 13007 | 13.007 | No Change | 10-100kb | Enh-B | intergenic |
| chr1_569723 | chr1_569723 | Plcl1 | TSS-B | Stable | 14.6428503 | 2.07093031 | 3.96E-06 | DE | Decreased | No Change | KOvsWT | 0.022845 | 18.743119 | 0.00133 | 0.041908 | 28.280829 | 0.001366 | 0.035036 | 28.280829 | 0.00133 | 1566699 | 0.027209 | 18.743119 | 0.001366 | 1566699 | 1566.699 | No Change | 500kb-5MB | Enh-B | intergenic |
| chr1_569731 | chr1_569731 | Plcl1 | TSS-B | Stable | 14.6428503 | 2.07093031 | 3.96E-06 | DE | Decreased | No Change | KOvsWT | 0.014006 | 11.494034 | 0.00133 | 0.021764 | 14.69084 | 0.001366 | 0.018195 | 14.69084 | 0.00133 | 1567473 | 0.016681 | 11.494034 | 0.001366 | 1567473 | 1567.473 | Gained | 500kb-5MB | Enh-B | intergenic |
