## Supplemental table 3 for "Nuclear actin-dependent Meg3 expression suppresses metabolic genes by affecting the chromatin architecture at sites of elevated H3K27 acetylation levels"

**Supplementary Table 3**

**IQR normalized expression data of ActinKO (n = 3) and ActinWT (n = 4 )replciates**

| Gene | ActinKO01 | ActinKO02 | ActinKO03 | ActinWT01 | ActinWT02 | ActinWT03 | ActinWT04 |
|---|---|---|---|---|---|---|---|
| ENSMUSG00000000001 | 3.055921842 | 2.80342795 | 2.770037827 | 2.956140257 | 2.834497071 | 3.293141483 | 2.624915771 |
| ENSMUSG00000000003 | 0 | 0 | 0 | 0 | 0 | 0 | 0 |
| ENSMUSG00000000028 | 2.506893125 | 2.289336734 | 2.307805295 | 2.412762923 | 2.376595432 | 2.731588765 | 2.245582246 |
| ENSMUSG00000000031 | 0 | 0 | 0 | 0.25 | 0.489301084 | 0.477121255 | 0 |
| ENSMUSG00000000037 | 1.558729299 | 1.438721876 | 1.403677461 | 1.527131114 | 1.462344774 | 1.477121255 | 1.278288402 |
| ENSMUSG00000000049 | 0 | 0.646240625 | 0.396240625 | 0 | 0.244650542 | 0 | 0.448487648 |
| ENSMUSG00000000056 | 2.340856238 | 2.132845365 | 2.083847589 | 2.161864607 | 2.153935427 | 2.419955748 | 1.984249047 |
| ENSMUSG00000000058 | 2.788740894 | 2.529410775 | 2.546838018 | 2.402294685 | 2.328502976 | 2.646403726 | 2.157274883 |
| ENSMUSG00000000078 | 2.869343028 | 2.646944379 | 2.664329612 | 2.715077764 | 2.646380908 | 3.035029282 | 2.430624509 |
| ENSMUSG00000000085 | 2.325351389 | 2.176839783 | 2.162762923 | 2.066696635 | 2.085294053 | 2.320146286 | 1.969165684 |
| ENSMUSG00000000088 | 1.671187741 | 1.618933358 | 1.642463902 | 1.837182039 | 1.728722551 | 1.986771734 | 1.774416565 |
| ENSMUSG00000000093 | 0 | 0.25 | 0.580482024 | 0.396240625 | 0.244650542 | 0.477121255 | 0.448487648 |
| ENSMUSG00000000094 | 0.289064826 | 0.25 | 0 | 0 | 0 | 0 | 0 |
| ENSMUSG00000000103 | 0 | 0 | 0 | 0 | 0 | 0 | 0 |
| ENSMUSG00000000120 | 0 | 0 | 0 | 0.701838731 | 0.77552387 | 0.477121255 | 0.672731473 |
| ENSMUSG00000000125 | 0.289064826 | 0 | 0 | 0.25 | 0 | 0 | 0 |
| ENSMUSG00000000126 | 1.307602651 | 1.302363341 | 1.282320754 | 1.396240625 | 1.450958105 | 1.568201724 | 1.551512352 |
| ENSMUSG00000000127 | 2.577267124 | 2.397646762 | 2.382357639 | 2.247880462 | 2.159419872 | 2.503790683 | 2.031805106 |
| ENSMUSG00000000131 | 2.533616084 | 2.362802778 | 2.33272922 | 2.320772088 | 2.336595379 | 2.614897216 | 2.20919388 |
| ENSMUSG00000000134 | 2.755750303 | 2.548996825 | 2.577937079 | 2.524178789 | 2.520546198 | 2.84135947 | 2.412651849 |
| ENSMUSG00000000142 | 1 | 1.04248125 | 0.92510993 | 1.614857905 | 1.695608647 | 1.716003344 | 1.74047993 |
| ENSMUSG00000000148 | 1.924654746 | 1.862802778 | 1.901832578 | 1.732684334 | 1.897237431 | 1.903089987 | 1.86207155 |
| ENSMUSG00000000149 | 2.496544396 | 2.414998973 | 2.424241882 | 2.044977273 | 2.232221225 | 2.394451681 | 2.203542812 |
| ENSMUSG00000000154 | 0 | 0 | 0 | 0 | 0 | 0.301029996 | 0.355418052 |
| ENSMUSG00000000157 | 0 | 0 | 0 | 0 | 0 | 0 | 0 |
| ENSMUSG00000000159 | 0.289064826 | 0 | 0 | 0 | 0 | 0 | 0 |
| ENSMUSG00000000167 | 1.205378646 | 1.04248125 | 0.976722649 | 1.27186571 | 1.212046813 | 1.397940009 | 1.193409508 |

| | | | | | | | |
|---|---|---|---|---|---|---|---|
| ENSMUSG00000000168 | 2.550525119 | 2.374961472 | 2.344844592 | 2.411864607 | 2.359374645 | 2.694605199 | 2.229218532 |
| ENSMUSG00000000171 | 2.723587515 | 2.54838134 | 2.53456795 | 2.666333979 | 2.623340676 | 3.028164419 | 2.469356413 |
| ENSMUSG00000000182 | 0.289064826 | 0.5 | 0.25 | 0 | 0 | 0 | 0 |
| ENSMUSG00000000183 | 0 | 0.25 | 0 | 0 | 0 | 0 | 0 |
| ENSMUSG00000000184 | 2.932463873 | 2.721805154 | 2.726158655 | 2.665444524 | 2.59734443 | 3.006893708 | 2.437093285 |
| ENSMUSG00000000194 | 2.435736758 | 2.268535366 | 2.286164561 | 2.245748394 | 2.281165826 | 2.485721426 | 2.117521761 |
| ENSMUSG00000000197 | 0 | 0.5 | 0.830482024 | 0.25 | 0.686820904 | 0.77815125 | 0.520678036 |
| ENSMUSG00000000202 | 0 | 0 | 0 | 0 | 0 | 0 | 0 |
| ENSMUSG00000000204 | 0 | 0 | 0.25 | 0 | 0 | 0 | 0 |
| ENSMUSG00000000214 | 0 | 0 | 0 | 0 | 0 | 0 | 0 |
| ENSMUSG00000000215 | 0 | 0 | 0 | 0 | 0 | 0 | 0 |
| ENSMUSG00000000216 | 0 | 0 | 0 | 0 | 0 | 0 | 0 |
| ENSMUSG00000000223 | 1.036286563 | 1.146240625 | 1.114857905 | 1.445339928 | 1.42077253 | 1.462397998 | 1.324583736 |
| ENSMUSG00000000244 | 0.45815691 | 0.25 | 0.5 | 0 | 0.568060967 | 0.301029996 | 0 |
| ENSMUSG00000000247 | 0 | 0 | 0 | 0.5 | 0.568060967 | 0.301029996 | 0.629532004 |
| ENSMUSG00000000248 | 0 | 0 | 0.396240625 | 0.25 | 0.387761935 | 0.477121255 | 0.224243824 |
| ENSMUSG00000000253 | 1.700465191 | 1.732684334 | 1.763820609 | 1.606566189 | 1.692703642 | 1.799340549 | 1.607811402 |
| ENSMUSG00000000263 | 0 | 0.396240625 | 0.5 | 0 | 0 | 0 | 0 |
| ENSMUSG00000000266 | 2.134369199 | 1.987091808 | 2.032320754 | 2.171625132 | 2.0510778 | 2.365487985 | 1.902443294 |
| ENSMUSG00000000275 | 2.23778581 | 2.043731421 | 2.093759858 | 2.271202097 | 2.267124351 | 2.506505032 | 2.086826862 |
| ENSMUSG00000000276 | 2.00683675 | 1.870953944 | 1.933677405 | 2.119945066 | 2.032639648 | 2.26245109 | 1.876096088 |
| ENSMUSG00000000278 | 2.961285253 | 2.696158711 | 2.71816872 | 2.694313829 | 2.65431284 | 3.0523091 | 2.484057231 |
| ENSMUSG00000000282 | 1.885732304 | 1.80479213 | 1.872963274 | 1.610735874 | 1.689774529 | 1.755874856 | 1.680001789 |
| ENSMUSG00000000290 | 1.036286563 | 0.580482024 | 0.75 | 0.75 | 0.905314583 | 0.84509804 | 0.710836105 |
| ENSMUSG00000000295 | 1.99653913 | 1.880890489 | 1.884789703 | 1.992885888 | 1.92093466 | 2.206825876 | 1.803905701 |
| ENSMUSG00000000296 | 1.87190676 | 1.843759858 | 1.841580554 | 1.52186571 | 1.319233381 | 1.322219295 | 1.329931221 |
| ENSMUSG00000000301 | 1.358731791 | 1.388647213 | 1.201838731 | 1.516522298 | 1.618292349 | 1.544068044 | 1.42167221 |
| ENSMUSG00000000303 | 0.811507563 | 1.04248125 | 0.830482024 | 0.75 | 0.77552387 | 0.477121255 | 0.710836105 |
| ENSMUSG00000000305 | 0 | 0 | 0 | 0 | 0 | 0 | 0.224243824 |
| ENSMUSG00000000308 | 0 | 0 | 0.396240625 | 0.25 | 0 | 0.477121255 | 0 |
| ENSMUSG00000000317 | 1.527823874 | 1.470660762 | 1.458222504 | 1.339388001 | 1.450958105 | 1.491361694 | 1.329931221 |
| ENSMUSG00000000318 | 0 | 0 | 0 | 0 | 0 | 0 | 0 |
| ENSMUSG00000000320 | 0.45815691 | 0.25 | 0.396240625 | 0.864857905 | 0.812711509 | 1.176091259 | 0.629532004 |
| ENSMUSG00000000325 | 1.857607105 | 1.860735874 | 1.931980114 | 1.584962501 | 1.761403742 | 1.698970004 | 1.817112247 |

| | | | | | | | |
|---|---|---|---|---|---|---|---|
| ENSMUSG00000000326 | 2.392336464 | 2.281676118 | 2.301142786 | 2.130890489 | 2.216042585 | 2.44870632 | 2.137353429 |
| ENSMUSG00000000339 | 1.952600383 | 1.870953944 | 1.92510993 | 1.862802778 | 1.87878897 | 2.103803721 | 1.879993912 |
| ENSMUSG00000000340 | 2.063793172 | 1.905512955 | 1.926839783 | 2.222185812 | 2.163285805 | 2.437750563 | 2.012457048 |
| ENSMUSG00000000346 | 3.008988783 | 2.742166698 | 2.733422664 | 2.799304173 | 2.746253085 | 3.180699201 | 2.573589458 |
| ENSMUSG00000000355 | 2.095120908 | 1.948603967 | 1.935366747 | 1.657339155 | 1.689774529 | 1.857332496 | 1.499406449 |
| ENSMUSG00000000359 | 0 | 0 | 0 | 0 | 0 | 0 | 0 |
| ENSMUSG00000000365 | 0 | 0 | 0 | 0 | 0 | 0.301029996 | 0.224243824 |
| ENSMUSG00000000374 | 2.694641526 | 2.477473271 | 2.469129237 | 2.511780978 | 2.448224646 | 2.825426118 | 2.314375784 |
| ENSMUSG00000000378 | 2.489144387 | 2.315523711 | 2.314346961 | 2.049918086 | 2.181274723 | 2.357934847 | 2.072724682 |
| ENSMUSG00000000381 | 0 | 0 | 0 | 0 | 0 | 0 | 0 |
| ENSMUSG00000000384 | 2.243591459 | 2.149047439 | 2.252107156 | 2.066696635 | 2.170892913 | 2.357934847 | 2.06186744 |
| ENSMUSG00000000385 | 0 | 0 | 0 | 0 | 0 | 0.301029996 | 0.355418052 |
| ENSMUSG00000000386 | 0 | 0 | 0.396240625 | 0 | 0.387761935 | 0 | 0 |
| ENSMUSG00000000392 | 0 | 0 | 0 | 0.25 | 0 | 0 | 0.224243824 |
| ENSMUSG00000000394 | 0 | 0 | 0 | 0 | 0 | 0 | 0 |
| ENSMUSG00000000399 | 2.587501561 | 2.383332433 | 2.387686696 | 2.46759118 | 2.399768403 | 2.728353782 | 2.270850938 |
| ENSMUSG00000000402 | 0 | 0 | 0 | 0 | 0.244650542 | 0 | 0.448487648 |
| ENSMUSG00000000409 | 1 | 1.080482024 | 1 | 1.130890489 | 0.955822902 | 0.954242509 | 1.140832123 |
| ENSMUSG00000000411 | 0 | 0 | 0 | 0 | 0 | 0 | 0 |
| ENSMUSG00000000416 | 0.578129653 | 0.646240625 | 0.864857905 | 0.5 | 0.244650542 | 0 | 0 |
| ENSMUSG00000000420 | 2.835891186 | 2.569902645 | 2.5684489 | 2.609438018 | 2.540114597 | 2.937016107 | 2.330653623 |
| ENSMUSG00000000435 | 0 | 0 | 0 | 0 | 0 | 0 | 0 |
| ENSMUSG00000000439 | 2.029919532 | 1.886723615 | 1.839388001 | 1.858657057 | 1.810629166 | 2.075546961 | 1.764880683 |
| ENSMUSG00000000440 | 2.097855556 | 2.017865591 | 1.935366747 | 1.701838731 | 1.628935049 | 1.707570176 | 1.577201671 |
| ENSMUSG00000000441 | 2.520963022 | 2.333288838 | 2.307805295 | 2.375956434 | 2.275616087 | 2.63447727 | 2.190455009 |
| ENSMUSG00000000486 | 1.432083896 | 1.339388001 | 1.438721876 | 0.701838731 | 1.091002363 | 1 | 1 |
| ENSMUSG00000000489 | 2.063793172 | 1.998588359 | 1.96759118 | 1.638647213 | 1.649303553 | 1.707570176 | 1.705896842 |
| ENSMUSG00000000530 | 0 | 0 | 0 | 0.646240625 | 0.489301084 | 0.477121255 | 0.710836105 |
| ENSMUSG00000000531 | 1.249317394 | 1.356566189 | 1.42510993 | 1.51109853 | 1.635855979 | 1.716003344 | 1.582103514 |
| ENSMUSG00000000532 | 1.971473847 | 1.858657057 | 1.910964047 | 1.966046536 | 2.022473809 | 2.167317335 | 1.990504106 |
| ENSMUSG00000000538 | 2.440557995 | 2.279735268 | 2.296218836 | 2.343759858 | 2.375332618 | 2.605305046 | 2.292939619 |
| ENSMUSG00000000544 | 0.289064826 | 0.396240625 | 0.25 | 0.396240625 | 0 | 0 | 0 |
| ENSMUSG00000000552 | 2.095120908 | 1.940387808 | 2.005591953 | 1.96293726 | 2.170892913 | 2.276461804 | 2.161359729 |
| ENSMUSG00000000555 | 3.418868765 | 3.125521476 | 3.126266798 | 2.670748646 | 2.710765025 | 3.03702788 | 2.518397386 |

| | | | | | | | |
|---|---|---|---|---|---|---|---|
| ENSMUSG00000000560 | 0.289064826 | 0.5 | 0.25 | 0.396240625 | 0.632412477 | 0.477121255 | 0.629532004 |
| ENSMUSG00000000561 | 2.626865868 | 2.355516192 | 2.333288838 | 2.53360658 | 2.486542403 | 2.829303773 | 2.265599895 |
| ENSMUSG00000000562 | 0 | 0 | 0 | 0.25 | 0 | 0 | 0.224243824 |
| ENSMUSG00000000563 | 2.824116479 | 2.58993739 | 2.581607372 | 2.804489424 | 2.709130966 | 3.151676231 | 2.549561989 |
| ENSMUSG00000000567 | 1.596667477 | 1.566696635 | 1.51109853 | 1.494319981 | 1.49445382 | 1.477121255 | 1.437266334 |
| ENSMUSG00000000568 | 3.199400276 | 2.899978211 | 2.881747051 | 2.871079856 | 2.744479432 | 3.183839037 | 2.570739083 |
| ENSMUSG00000000579 | 1.971473847 | 1.843759858 | 1.784887838 | 1.892463902 | 1.903713872 | 2.079181246 | 1.755055163 |
| ENSMUSG00000000581 | 2.434522689 | 2.225970461 | 2.221424093 | 2.409156155 | 2.318243321 | 2.693726949 | 2.155212955 |
| ENSMUSG00000000594 | 1.734388958 | 1.671625132 | 1.714495249 | 2.233422664 | 2.157861565 | 2.531478917 | 2.080634989 |
| ENSMUSG00000000600 | 2.612039956 | 2.396240625 | 2.363831805 | 2.372963274 | 2.330902415 | 2.605305046 | 2.158505762 |
| ENSMUSG00000000605 | 2.260538674 | 2.116901388 | 2.112802778 | 2.039967834 | 2.062514787 | 2.283301229 | 1.917452159 |
| ENSMUSG00000000606 | 0 | 0 | 0 | 0 | 0 | 0 | 0 |
| ENSMUSG00000000617 | 0 | 0 | 0 | 0 | 0 | 0 | 0 |
| ENSMUSG00000000627 | 0.867194479 | 0.864857905 | 0.830482024 | 1.642463902 | 1.692703642 | 1.681241237 | 1.714087544 |
| ENSMUSG00000000628 | 2.205378646 | 2.05357978 | 2.04248125 | 1.841580554 | 1.859322797 | 1.973127854 | 1.770364127 |
| ENSMUSG00000000631 | 2.711248379 | 2.544977273 | 2.561981878 | 2.505591953 | 2.634846667 | 2.84135947 | 2.534290726 |
| ENSMUSG00000000632 | 0 | 0.396240625 | 0 | 0 | 0.244650542 | 0 | 0 |
| ENSMUSG00000000673 | 0 | 0 | 0 | 0 | 0 | 0.301029996 | 0 |
| ENSMUSG00000000682 | 0 | 0 | 0 | 0 | 0 | 0 | 0 |
| ENSMUSG00000000686 | 0.747221736 | 0.864857905 | 0.701838731 | 0.830482024 | 0.686820904 | 0.84509804 | 0.629532004 |
| ENSMUSG00000000690 | 1.978790393 | 1.888647213 | 1.943696765 | 1.732684334 | 1.744182955 | 1.799340549 | 1.702561601 |
| ENSMUSG00000000693 | 2.256831695 | 2.155512955 | 2.172499493 | 1.649978211 | 1.715300549 | 1.857332496 | 1.687112305 |
| ENSMUSG00000000701 | 0 | 0 | 0 | 0 | 0 | 0 | 0 |
| ENSMUSG00000000706 | 0.811507563 | 0.25 | 0.25 | 1.26109853 | 1.106691885 | 1.176091259 | 1.054044578 |
| ENSMUSG00000000708 | 2.496544396 | 2.291854536 | 2.29745614 | 2.309601185 | 2.211376431 | 2.545307116 | 2.088864761 |
| ENSMUSG00000000711 | 2.737928151 | 2.488912477 | 2.486726569 | 2.638886943 | 2.562149491 | 2.957128198 | 2.400090259 |
| ENSMUSG00000000724 | 0.289064826 | 0.25 | 0 | 0.25 | 0.387761935 | 0.301029996 | 0.224243824 |
| ENSMUSG00000000730 | 0 | 0 | 0 | 0 | 0 | 0 | 0 |
| ENSMUSG00000000731 | 0 | 0 | 0 | 0 | 0 | 0 | 0 |
| ENSMUSG00000000732 | 1.129344651 | 0.92510993 | 1.17510993 | 1.848079356 | 1.804307395 | 2.045322979 | 1.71730663 |
| ENSMUSG00000000738 | 2.518981867 | 2.348079356 | 2.365369862 | 2.240724001 | 2.338004387 | 2.515873844 | 2.25178005 |
| ENSMUSG00000000739 | 0 | 0 | 0.25 | 0.396240625 | 0.244650542 | 0 | 0.224243824 |
| ENSMUSG00000000740 | 2.864601523 | 2.688304187 | 2.758871613 | 2.606566189 | 2.68472101 | 2.983175072 | 2.609034524 |
| ENSMUSG00000000743 | 2.523917219 | 2.392463902 | 2.417221246 | 2.319031101 | 2.382016011 | 2.654176542 | 2.283060517 |

| | | | | | | | |
|---|---|---|---|---|---|---|---|
| ENSMUSG00000000751 | 3.277146534 | 2.966626553 | 2.970565232 | 2.982499516 | 2.949497207 | 3.402605242 | 2.75609 |
| ENSMUSG00000000753 | 1.837744666 | 1.752806814 | 1.593759858 | 1.098079356 | 1 | 1.230448921 | 1.041356071 |
| ENSMUSG00000000759 | 2.805057201 | 2.608917565 | 2.593216265 | 2.441217898 | 2.398980554 | 2.710963119 | 2.275448037 |
| ENSMUSG00000000766 | 0 | 0.25 | 0 | 0 | 0 | 0 | 0 |
| ENSMUSG00000000776 | 1.99653913 | 1.896240625 | 1.955044741 | 2.009729747 | 2.0928848 | 2.269512944 | 1.996640515 |
| ENSMUSG00000000782 | 1.91631382 | 1.799918086 | 1.807204673 | 1.907339155 | 1.903713872 | 2.093421685 | 1.803905701 |
| ENSMUSG00000000787 | 3.595466952 | 3.258011432 | 3.236086509 | 3.351584473 | 3.241148929 | 3.778368343 | 2.995225229 |
| ENSMUSG00000000791 | 0.289064826 | 0.396240625 | 0.396240625 | 0.79248125 | 0.846351821 | 0.903089987 | 1.028149525 |
| ENSMUSG00000000792 | 0.747221736 | 0.896240625 | 0.75 | 0.5 | 0.489301084 | 0.477121255 | 0.74492186 |
| ENSMUSG00000000794 | 1.181541736 | 0.976722649 | 1.160964047 | 1.114857905 | 1.200473444 | 1.176091259 | 1.066254157 |
| ENSMUSG00000000804 | 2.812982046 | 2.589388001 | 2.564346961 | 2.560197833 | 2.488077665 | 2.864511081 | 2.295899033 |
| ENSMUSG00000000805 | 0 | 0 | 0 | 0 | 0 | 0 | 0 |
| ENSMUSG00000000811 | 1.944805158 | 1.782320754 | 1.779735268 | 1.67510993 | 1.66856048 | 1.799340549 | 1.562034107 |
| ENSMUSG00000000817 | 0.289064826 | 0.25 | 0.25 | 0.5 | 0.244650542 | 0 | 0.224243824 |
| ENSMUSG00000000823 | 2.146671932 | 1.959800947 | 2.041226732 | 2.192872367 | 2.252501045 | 2.421603927 | 2.107610228 |
| ENSMUSG00000000826 | 2.618346804 | 2.425543171 | 2.386723615 | 2.638647213 | 2.54899118 | 2.979092901 | 2.396970134 |
| ENSMUSG00000000827 | 2.370236862 | 2.187886015 | 2.208222504 | 2.393884812 | 2.328502976 | 2.736396502 | 2.221895853 |
| ENSMUSG00000000838 | 3.10861828 | 2.832308935 | 2.807204673 | 2.829353153 | 2.737445518 | 3.153814864 | 2.524795923 |
| ENSMUSG00000000861 | 0 | 0 | 0 | 0.25 | 0 | 0 | 0 |
| ENSMUSG00000000869 | 1.036286563 | 1.04248125 | 1 | 1.27186571 | 1.244650542 | 1.361727836 | 1.185218806 |
| ENSMUSG00000000876 | 1.418409478 | 1.451838731 | 1.494319981 | 1.598079356 | 1.603583716 | 1.86923172 | 1.605556929 |
| ENSMUSG00000000881 | 2.409468165 | 2.263142013 | 2.193696765 | 2.095926073 | 2.028157605 | 2.322219295 | 1.913982791 |
| ENSMUSG00000000884 | 1.655740331 | 1.580482024 | 1.653677461 | 1.482684334 | 1.514343934 | 1.531478917 | 1.46982607 |
| ENSMUSG00000000889 | 0 | 0 | 0.25 | 0 | 0 | 0 | 0 |
| ENSMUSG00000000901 | 2.409468165 | 2.233422664 | 2.26109853 | 2.00835575 | 2.149965125 | 2.303196057 | 2.038994634 |
| ENSMUSG00000000902 | 2.708105693 | 2.466046536 | 2.488549078 | 2.430701883 | 2.475604419 | 2.781755375 | 2.355863359 |
| ENSMUSG00000000903 | 0 | 0 | 0 | 0 | 0 | 0 | 0 |
| ENSMUSG00000000915 | 2.213719572 | 2.091580554 | 2.175975893 | 1.92510993 | 2.084333628 | 2.198657087 | 2.059651572 |
| ENSMUSG00000000916 | 1.83262764 | 1.789967834 | 1.870953944 | 1.714495249 | 1.824953989 | 1.851258349 | 1.826389311 |
| ENSMUSG00000000934 | 1.740854706 | 1.657339155 | 1.695339928 | 1.646240625 | 1.736537397 | 1.968482949 | 1.683576582 |
| ENSMUSG00000000938 | 2.113897241 | 2.009729747 | 1.976722649 | 1.681980114 | 1.739104362 | 1.819543936 | 1.574722613 |
| ENSMUSG00000000942 | 1.269664473 | 1.114857905 | 1.146240625 | 0.864857905 | 0.978602168 | 0.954242509 | 0.984950056 |
| ENSMUSG00000000948 | 1.389637216 | 1.339388001 | 1.302363341 | 1.431980114 | 1.351342427 | 1.447158031 | 1.365075947 |
| ENSMUSG00000000957 | 2.98662835 | 2.757666784 | 2.781676118 | 2.525821952 | 2.526366572 | 2.890421019 | 2.395201749 |

| | | | | | | | |
|---|---|---|---|---|---|---|---|
| ENSMUSG00000000958 | 0.960252568 | 1.226722649 | 0.896240625 | 0.976722649 | 0.877063019 | 1.146128036 | 1.100339912 |
| ENSMUSG00000000959 | 2.177057474 | 2.039967834 | 2.034887838 | 2.183677405 | 2.212046813 | 2.509202522 | 2.099849366 |
| ENSMUSG00000000976 | 2.445324132 | 2.28296424 | 2.306000419 | 2.187886015 | 2.215379757 | 2.431363764 | 2.14212705 |
| ENSMUSG00000000982 | 0 | 0 | 0.25 | 0 | 0 | 0 | 0 |
| ENSMUSG00000000983 | 0 | 0 | 0.25 | 0 | 0 | 0 | 0 |
| ENSMUSG00000001014 | 0 | 0.396240625 | 0.396240625 | 0.25 | 0.244650542 | 0 | 0 |
| ENSMUSG00000001016 | 2.844622761 | 2.593216265 | 2.556000419 | 2.667442722 | 2.616167381 | 3.009025742 | 2.437266334 |
| ENSMUSG00000001017 | 2.811999637 | 2.566403762 | 2.548689214 | 2.685998215 | 2.616167381 | 2.979548375 | 2.458199602 |
| ENSMUSG00000001018 | 2.192545868 | 2.038704527 | 2.024508021 | 2.006976499 | 1.945998439 | 2.201397124 | 1.899720848 |
| ENSMUSG00000001020 | 2.63295404 | 2.448197574 | 2.490724001 | 2.084962501 | 2.004851707 | 2.303196057 | 1.86815666 |
| ENSMUSG00000001021 | 1.249317394 | 1.02186571 | 1.130890489 | 1.160964047 | 1.27449559 | 1.230448921 | 1.168187745 |
| ENSMUSG00000001023 | 0 | 0.25 | 0 | 0 | 0.244650542 | 0.301029996 | 0 |
| ENSMUSG00000001025 | 3.699353806 | 3.340451683 | 3.404854399 | 3.280747431 | 3.217713617 | 3.752969865 | 2.976207171 |
| ENSMUSG00000001027 | 0.289064826 | 0 | 0 | 0 | 0 | 0 | 0 |
| ENSMUSG00000001029 | 0 | 0 | 0 | 0 | 0 | 0 | 0.224243824 |
| ENSMUSG00000001034 | 2.036286563 | 2.029735268 | 2.093759858 | 1.825945187 | 1.990091809 | 2.127104798 | 2.005303651 |
| ENSMUSG00000001036 | 2.610448227 | 2.430701883 | 2.387686696 | 2.445749552 | 2.440245937 | 2.713490543 | 2.299626824 |
| ENSMUSG00000001039 | 1.623015126 | 1.438721876 | 1.532320754 | 1.54745614 | 1.546663136 | 1.602059991 | 1.499406449 |
| ENSMUSG00000001052 | 2.623015126 | 2.405512955 | 2.392938161 | 2.516522298 | 2.48127199 | 2.832508913 | 2.326782038 |
| ENSMUSG00000001053 | 1.129344651 | 1.321350555 | 1.356566189 | 1.098079356 | 1.373641807 | 1.278753601 | 1.554175045 |
| ENSMUSG00000001054 | 2 | 1.961372513 | 1.945339928 | 1.981203126 | 2.046825265 | 2.187520721 | 1.995964412 |
| ENSMUSG00000001056 | 2.503815381 | 2.336628433 | 2.367918804 | 2.327084758 | 2.403292121 | 2.680335513 | 2.283617342 |
| ENSMUSG00000001062 | 1.924654746 | 1.860735874 | 1.898114259 | 1.896240625 | 1.955822902 | 2.008600172 | 1.94631981 |
| ENSMUSG00000001065 | 1.852729478 | 1.752806814 | 1.816696635 | 1.726722649 | 1.85749873 | 1.880813592 | 1.860017477 |
| ENSMUSG00000001076 | 0 | 0.25 | 0 | 0 | 0.387761935 | 0.477121255 | 0.629532004 |
| ENSMUSG00000001082 | 2.069666964 | 2.005591953 | 2.103406982 | 1.575945187 | 1.855665188 | 1.748188027 | 1.905143021 |
| ENSMUSG00000001089 | 2.952951272 | 2.701234418 | 2.674893113 | 2.713717096 | 2.6354526 | 3.052693942 | 2.455431016 |
| ENSMUSG00000001095 | 0 | 0 | 0 | 0 | 0 | 0 | 0 |
| ENSMUSG00000001098 | 2.974149285 | 2.765511534 | 2.821350555 | 2.457430684 | 2.54899118 | 2.787460475 | 2.483307832 |
| ENSMUSG00000001100 | 2.635214384 | 2.431128463 | 2.433677405 | 2.411864607 | 2.357161747 | 2.731588765 | 2.214058866 |
| ENSMUSG00000001103 | 0 | 0 | 0 | 0 | 0 | 0 | 0 |
| ENSMUSG00000001105 | 2.163307908 | 2.016522298 | 2.048689214 | 2.113831805 | 2.052132976 | 2.309630167 | 1.972821265 |
| ENSMUSG00000001119 | 3.260035113 | 3.027946934 | 3.062426514 | 2.804186465 | 2.887195467 | 3.235528447 | 2.773805813 |
| ENSMUSG00000001120 | 1.952600383 | 1.823655187 | 1.828220739 | 1.837182039 | 1.925488995 | 2.041392685 | 1.844181586 |

| | | | | | | | |
|---|---|---|---|---|---|---|---|
| ENSMUSG00000001123 | 2.591692877 | 2.360735874 | 2.370953944 | 2.406883471 | 2.354487824 | 2.741939078 | 2.212676319 |
| ENSMUSG00000001127 | 2.531694272 | 2.355516192 | 2.369945066 | 2.242885888 | 2.311739339 | 2.514547753 | 2.149789616 |
| ENSMUSG00000001128 | 1.45815691 | 1.438721876 | 1.505591953 | 1 | 1.310728002 | 1.146128036 | 1.23151414 |
| ENSMUSG00000001131 | 2.659301689 | 2.43155454 | 2.446158711 | 2.247171172 | 2.188507684 | 2.509202522 | 2.038994634 |
| ENSMUSG00000001134 | 1.686083339 | 1.584962501 | 1.678561379 | 1.695339928 | 1.635855979 | 1.880813592 | 1.514741806 |
| ENSMUSG00000001138 | 1.876566388 | 1.794977273 | 1.723704441 | 1.809601185 | 1.840643511 | 1.982271233 | 1.788855757 |
| ENSMUSG00000001143 | 2.294202867 | 2.044977273 | 2.03360658 | 2.280383379 | 2.234742351 | 2.530199698 | 2.060761403 |
| ENSMUSG00000001150 | 2.652869185 | 2.469512728 | 2.475593779 | 2.514497931 | 2.524713379 | 2.835056102 | 2.406241659 |
| ENSMUSG00000001151 | 2.499676068 | 2.388167274 | 2.423371739 | 2.406883471 | 2.473055988 | 2.71432976 | 2.413024992 |
| ENSMUSG00000001155 | 0 | 0 | 0 | 0.25 | 0 | 0 | 0 |
| ENSMUSG00000001156 | 1.827447047 | 1.67510993 | 1.671625132 | 1.711372513 | 1.695608647 | 1.886490725 | 1.603286636 |
| ENSMUSG00000001157 | 2.401648579 | 2.168989258 | 2.143411797 | 2.257666784 | 2.204601633 | 2.447158031 | 2.023210262 |
| ENSMUSG00000001158 | 2.217828299 | 2.145300145 | 2.117918804 | 2.136723615 | 2.078515538 | 2.389166084 | 2.016293244 |
| ENSMUSG00000001166 | 0.45815691 | 0.25 | 0.396240625 | 1.130890489 | 0.978602168 | 1.176091259 | 1.028149525 |
| ENSMUSG00000001168 | 0 | 0 | 0 | 0 | 0 | 0 | 0 |
| ENSMUSG00000001173 | 2.523917219 | 2.30357978 | 2.309003548 | 2.371457327 | 2.286629652 | 2.614897216 | 2.138226615 |
| ENSMUSG00000001175 | 3.212591949 | 2.939243307 | 2.94564719 | 2.96999152 | 2.870177356 | 3.366236124 | 2.701722391 |
| ENSMUSG00000001211 | 2.646335907 | 2.458222504 | 2.493244947 | 2.433677405 | 2.51377465 | 2.765668555 | 2.412465115 |
| ENSMUSG00000001225 | 0.578129653 | 0.25 | 0.5 | 0.396240625 | 0.387761935 | 0.602059991 | 0.520678036 |
| ENSMUSG00000001227 | 0.45815691 | 0.580482024 | 0.5 | 0.396240625 | 0.387761935 | 0.301029996 | 0.710836105 |
| ENSMUSG00000001228 | 2.955050433 | 2.747348625 | 2.757149194 | 2.65136988 | 2.693902717 | 3.002166062 | 2.572338386 |
| ENSMUSG00000001229 | 2.577267124 | 2.43325383 | 2.47104263 | 2.23927551 | 2.386946794 | 2.564666064 | 2.321381806 |
| ENSMUSG00000001240 | 0.289064826 | 0.580482024 | 0 | 1.160964047 | 0.978602168 | 1.113943352 | 1.066254157 |
| ENSMUSG00000001247 | 0.289064826 | 0.25 | 0.396240625 | 0.396240625 | 0.632412477 | 0.602059991 | 0.896975297 |
| ENSMUSG00000001248 | 2.20747957 | 2.060793496 | 2.078220739 | 1.794977273 | 1.90849439 | 2.017033339 | 1.86715039 |
| ENSMUSG00000001249 | 0 | 0 | 0 | 0 | 0.244650542 | 0 | 0 |
| ENSMUSG00000001260 | 0 | 0 | 0 | 0 | 0 | 0 | 0 |
| ENSMUSG00000001270 | 1.342375483 | 1.388647213 | 1.649978211 | 1.476722649 | 1.726078673 | 1.62324929 | 1.913109605 |
| ENSMUSG00000001280 | 3.039433992 | 2.762462137 | 2.737821179 | 2.854331279 | 2.777553526 | 3.197556213 | 2.557293869 |
| ENSMUSG00000001281 | 1.036286563 | 1.02186571 | 1.130890489 | 1.52186571 | 1.592135487 | 1.544068044 | 1.564611933 |
| ENSMUSG00000001285 | 2.325351389 | 2.216819685 | 2.223704441 | 2.279735268 | 2.301493158 | 2.587710965 | 2.214058866 |
| ENSMUSG00000001288 | 2.626865868 | 2.480460234 | 2.498588359 | 2.30479213 | 2.391406953 | 2.610660163 | 2.336139088 |
| ENSMUSG00000001289 | 2.623788124 | 2.434945152 | 2.445339928 | 2.505938588 | 2.466928684 | 2.81756537 | 2.331854053 |
| ENSMUSG00000001300 | 2.768497801 | 2.560495787 | 2.558404919 | 2.382357639 | 2.35849115 | 2.664641976 | 2.242122155 |

| | | | | | | | |
|---|---|---|---|---|---|---|---|
| ENSMUSG00000001305 | 2.241665209 | 2.074802005 | 2.105516192 | 2.242885888 | 2.170892913 | 2.487138375 | 2.043115906 |
| ENSMUSG00000001313 | 1.771760131 | 1.744319981 | 1.87892496 | 1.896240625 | 1.871833944 | 2.130333768 | 1.897893072 |
| ENSMUSG00000001323 | 1.771760131 | 1.695339928 | 1.602347734 | 1.868933358 | 1.770879644 | 2.056904851 | 1.657681529 |
| ENSMUSG00000001333 | 1.445324132 | 1.51109853 | 1.580482024 | 0.646240625 | 1.074582839 | 0.903089987 | 0.896975297 |
| ENSMUSG00000001334 | 0 | 0 | 0.396240625 | 0.701838731 | 0.489301084 | 0.602059991 | 0.629532004 |
| ENSMUSG00000001348 | 0 | 0 | 0.396240625 | 0.5 | 0.686820904 | 0.477121255 | 0.224243824 |
| ENSMUSG00000001349 | 3.126578257 | 2.894711098 | 2.910851319 | 1.04248125 | 1 | 1.113943352 | 0.984950056 |
| ENSMUSG00000001366 | 2.039433992 | 1.899978211 | 1.910964047 | 2.071350555 | 2.041436517 | 2.217483944 | 1.884801065 |
| ENSMUSG00000001376 | 2.820759099 | 2.598883534 | 2.597810897 | 2.51007243 | 2.433873436 | 2.748962861 | 2.244955885 |
| ENSMUSG00000001380 | 2.740271036 | 2.526477127 | 2.539336734 | 2.636482442 | 2.58568862 | 2.955206538 | 2.43220994 |
| ENSMUSG00000001383 | 2.611244851 | 2.379417347 | 2.38040011 | 2.488549078 | 2.407936347 | 2.800717078 | 2.240219117 |
| ENSMUSG00000001403 | 2.425923778 | 2.222945926 | 2.253505118 | 2.43325383 | 2.371090306 | 2.784617293 | 2.301737862 |
| ENSMUSG00000001415 | 2.871480564 | 2.661189398 | 2.703845824 | 2.649280357 | 2.666718325 | 3.003029471 | 2.551147553 |
| ENSMUSG00000001416 | 3.309396985 | 3.018033151 | 2.983330455 | 3.165667094 | 3.096510529 | 3.597914471 | 2.880433889 |
| ENSMUSG00000001418 | 2.792880208 | 2.557505109 | 2.616391601 | 2.388647213 | 2.426290456 | 2.798650645 | 2.309019868 |
| ENSMUSG00000001419 | 2.422183552 | 2.267865591 | 2.338286706 | 2.216046536 | 2.180543208 | 2.397940009 | 2.131174228 |
| ENSMUSG00000001420 | 0.811507563 | 0.92510993 | 1.061981878 | 0.896240625 | 1.057362051 | 0.84509804 | 1 |
| ENSMUSG00000001424 | 2.886966737 | 2.643884812 | 2.62521047 | 2.550530956 | 2.561403546 | 2.903089987 | 2.415439991 |
| ENSMUSG00000001435 | 3.081172036 | 2.927916743 | 3.006717301 | 2.119945066 | 2.308696595 | 2.456366033 | 2.295631111 |
| ENSMUSG00000001436 | 2.111266113 | 1.985628626 | 2.038704527 | 1.860735874 | 2.059432181 | 2.139879086 | 1.986347533 |
| ENSMUSG00000001440 | 3.704592244 | 3.338562345 | 3.318703729 | 3.397617524 | 3.305929477 | 3.849971912 | 3.058284282 |
| ENSMUSG00000001441 | 3.005911039 | 2.737456678 | 2.730088214 | 2.763820609 | 2.707653816 | 3.122870923 | 2.498262273 |
| ENSMUSG00000001444 | 0 | 0 | 0 | 0.25 | 0 | 0 | 0 |
| ENSMUSG00000001445 | 2.429630757 | 2.294977273 | 2.29248125 | 2.222185812 | 2.283908311 | 2.537819095 | 2.160953557 |
| ENSMUSG00000001467 | 2.834961347 | 2.612802778 | 2.627196291 | 2.637446167 | 2.543534996 | 2.981818607 | 2.41059183 |
| ENSMUSG00000001472 | 3.158193497 | 2.878308528 | 2.876576895 | 2.946669508 | 2.946416783 | 3.344392274 | 2.773500004 |
| ENSMUSG00000001473 | 2.863298974 | 2.727660683 | 2.814641508 | 2.452241044 | 2.592363568 | 2.804139432 | 2.552725384 |
| ENSMUSG00000001482 | 2.417143825 | 2.267194569 | 2.275821952 | 2.427270953 | 2.461024076 | 2.738780558 | 2.310046902 |
| ENSMUSG00000001493 | 0.45815691 | 0 | 0 | 0 | 0 | 0 | 0 |
| ENSMUSG00000001494 | 0 | 0 | 0 | 0 | 0 | 0 | 0.224243824 |
| ENSMUSG00000001496 | 0 | 0 | 0 | 0 | 0 | 0 | 0 |
| ENSMUSG00000001497 | 1.205378646 | 1.188721876 | 1.282320754 | 1.42510993 | 1.234113756 | 1.278753601 | 1.21680636 |
| ENSMUSG00000001504 | 1.45815691 | 1.52186571 | 1.51109853 | 1.794977273 | 1.751666404 | 1.959041392 | 1.69749337 |
| ENSMUSG00000001506 | 4.009824241 | 3.728456098 | 3.789603114 | 3.05388325 | 3.226406331 | 3.558468563 | 3.212392796 |

| | | | | | | | |
|---|---|---|---|---|---|---|---|
| ENSMUSG00000001507 | 3.05040546 | 2.847004321 | 2.848347614 | 2.524508021 | 2.644025241 | 2.866877814 | 2.496538385 |
| ENSMUSG00000001508 | 0.45815691 | 0.396240625 | 0.25 | 0 | 0.244650542 | 0.301029996 | 0.355418052 |
| ENSMUSG00000001510 | 0 | 0.25 | 0 | 0.580482024 | 0.77552387 | 0.84509804 | 1.041356071 |
| ENSMUSG00000001517 | 2.860681608 | 2.655970373 | 2.690387808 | 2.591033664 | 2.6330226 | 2.959041392 | 2.498548696 |
| ENSMUSG00000001518 | 2.010213549 | 1.87694866 | 1.919870025 | 1.938721876 | 1.966725889 | 2.164352856 | 1.86715039 |
| ENSMUSG00000001520 | 1.181541736 | 0.92510993 | 0.951838731 | 1.226722649 | 1.200473444 | 1.278753601 | 1 |
| ENSMUSG00000001521 | 2.635214384 | 2.37246199 | 2.407339155 | 2.357090043 | 2.360695755 | 2.69019608 | 2.23605706 |
| ENSMUSG00000001524 | 2.100572389 | 2.025821952 | 1.991446071 | 1.910964047 | 2.055279708 | 2.187520721 | 1.920029985 |
| ENSMUSG00000001525 | 3.844167734 | 3.498654653 | 3.528720257 | 3.387325842 | 3.409763562 | 3.910784435 | 3.234706753 |
| ENSMUSG00000001542 | 2.611244851 | 2.391037372 | 2.334962501 | 2.365369862 | 2.279510047 | 2.622214023 | 2.144273809 |
| ENSMUSG00000001552 | 2.531694272 | 2.430701883 | 2.474839231 | 2.14151351 | 2.293608479 | 2.474216264 | 2.318897036 |
| ENSMUSG00000001555 | 2.840509532 | 2.633575721 | 2.659835177 | 2.440387808 | 2.567082571 | 2.832508913 | 2.50534787 |
| ENSMUSG00000001558 | 0.811507563 | 0.830482024 | 0.830482024 | 0.646240625 | 0.632412477 | 0.77815125 | 0.853775828 |
| ENSMUSG00000001566 | 0 | 0 | 0 | 0 | 0.244650542 | 0 | 0.224243824 |
| ENSMUSG00000001569 | 2.53838222 | 2.356566189 | 2.335518667 | 2.366391601 | 2.352693888 | 2.674861141 | 2.18859465 |
| ENSMUSG00000001576 | 3.047619662 | 2.793731421 | 2.789020769 | 2.696158711 | 2.700173892 | 3.075181855 | 2.519338425 |
| ENSMUSG00000001583 | 0.289064826 | 0.25 | 0.25 | 0 | 0 | 0 | 0.224243824 |
| ENSMUSG00000001588 | 0 | 0.25 | 0 | 0.5 | 0.568060967 | 0 | 0.448487648 |
| ENSMUSG00000001604 | 1.37447073 | 1.29248125 | 1.403677461 | 1.201838731 | 1.366364104 | 1.176091259 | 1.345462945 |
| ENSMUSG00000001622 | 0 | 0 | 0 | 0 | 0 | 0 | 0.224243824 |
| ENSMUSG00000001627 | 2.858927525 | 2.624961472 | 2.595926073 | 2.603142462 | 2.49893871 | 2.876794976 | 2.318897036 |
| ENSMUSG00000001630 | 2.083999385 | 1.948603967 | 1.96293726 | 2.061981878 | 1.941702774 | 2.206825876 | 1.837623616 |
| ENSMUSG00000001632 | 2.349889783 | 2.201838731 | 2.221424093 | 2.056000419 | 2.097547432 | 2.243038049 | 2.040177507 |
| ENSMUSG00000001642 | 2.631440309 | 2.413211243 | 2.400906586 | 2.408702763 | 2.377853744 | 2.686636269 | 2.223909441 |
| ENSMUSG00000001655 | 0.289064826 | 0.79248125 | 0.79248125 | 0.75 | 0.733951626 | 0.477121255 | 0.672731473 |
| ENSMUSG00000001656 | 0.289064826 | 0 | 0 | 0.5 | 0.244650542 | 0.301029996 | 0.520678036 |
| ENSMUSG00000001657 | 0 | 0 | 0 | 0.701838731 | 0.568060967 | 0.477121255 | 0.629532004 |
| ENSMUSG00000001661 | 0.289064826 | 0.5 | 0.75 | 1.238549078 | 1.121713561 | 1.230448921 | 1.110947927 |
| ENSMUSG00000001663 | 2.185978293 | 2.117918804 | 2.090485943 | 1.905512955 | 2.023617914 | 2.10720997 | 1.949460761 |
| ENSMUSG00000001665 | 1.432083896 | 1.418106335 | 1.330482024 | 1.744319981 | 1.77552387 | 1.991226076 | 1.663407615 |
| ENSMUSG00000001666 | 1.605636262 | 1.62694866 | 1.646240625 | 1.653677461 | 1.791313678 | 1.897627091 | 1.709198048 |
| ENSMUSG00000001670 | 0 | 0 | 0 | 0 | 0 | 0 | 0 |
| ENSMUSG00000001672 | 0.289064826 | 0.25 | 0 | 0.396240625 | 0.489301084 | 0.477121255 | 0.355418052 |
| ENSMUSG00000001674 | 2.45469604 | 2.266522298 | 2.204245906 | 2.3184489 | 2.280062837 | 2.574031268 | 2.112839835 |

| | | | | | | | |
|---|---|---|---|---|---|---|---|
| ENSMUSG00000001687 | 2.097855556 | 1.96759118 | 2.016522298 | 1.995748394 | 1.925488995 | 2.201397124 | 1.838725884 |
| ENSMUSG00000001700 | 1.777675583 | 1.714495249 | 1.688721876 | 1.744319981 | 1.674752852 | 1.806179974 | 1.579661877 |
| ENSMUSG00000001707 | 2.320585253 | 2.144357207 | 2.143411797 | 2.29745614 | 2.225838481 | 2.542825427 | 2.056860222 |
| ENSMUSG00000001729 | 3.120051066 | 2.902525516 | 2.950326047 | 2.607613138 | 2.72136414 | 3.003891166 | 2.635960927 |
| ENSMUSG00000001739 | 1.249317394 | 1.42510993 | 1.388647213 | 0.646240625 | 0.978602168 | 0.903089987 | 0.876096088 |
| ENSMUSG00000001741 | 0 | 0.5 | 0.25 | 0.396240625 | 0 | 0.301029996 | 0.448487648 |
| ENSMUSG00000001750 | 2.358731791 | 2.230460234 | 2.347542239 | 2.155512955 | 2.343123155 | 2.471291711 | 2.3138695 |
| ENSMUSG00000001751 | 1.777675583 | 1.692046081 | 1.821350555 | 1.458222504 | 1.584291694 | 1.591064607 | 1.577201671 |
| ENSMUSG00000001755 | 2.188178997 | 2.009729747 | 2.027131114 | 1.763820609 | 1.974687643 | 2 | 1.869159809 |
| ENSMUSG00000001761 | 2.845984872 | 2.598615674 | 2.647646762 | 2.535526764 | 2.565117605 | 2.890979597 | 2.447651042 |
| ENSMUSG00000001763 | 1.100572389 | 1.130890489 | 1 | 1.348079356 | 1.366364104 | 1.462397998 | 1.413583902 |
| ENSMUSG00000001767 | 2.292497211 | 2.149047439 | 2.119945066 | 2.309003548 | 2.259595879 | 2.564666064 | 2.123964672 |
| ENSMUSG00000001768 | 1.771760131 | 1.610735874 | 1.618933358 | 1.602347734 | 1.514343934 | 1.602059991 | 1.520678036 |
| ENSMUSG00000001773 | 0 | 0 | 0 | 0.25 | 0 | 0 | 0 |
| ENSMUSG00000001774 | 2.545896521 | 2.411864607 | 2.391037372 | 2.581326364 | 2.49445382 | 2.8876173 | 2.318147869 |
| ENSMUSG00000001783 | 2.953652167 | 2.688304187 | 2.676624005 | 2.794510582 | 2.732574981 | 3.172310969 | 2.545378011 |
| ENSMUSG00000001785 | 2.612833548 | 2.397178659 | 2.40136988 | 2.540912419 | 2.489606278 | 2.856124444 | 2.306955966 |
| ENSMUSG00000001786 | 1.989530018 | 1.942046081 | 1.942046081 | 2.094844592 | 1.964031554 | 2.23299611 | 1.882886767 |
| ENSMUSG00000001794 | 2.794934559 | 2.561981878 | 2.599685923 | 2.591307212 | 2.570495157 | 2.937016107 | 2.399507523 |
| ENSMUSG00000001802 | 0 | 0 | 0 | 0.25 | 0.733951626 | 0.477121255 | 0.355418052 |
| ENSMUSG00000001804 | 0 | 0 | 0 | 0 | 0 | 0 | 0 |
| ENSMUSG00000001815 | 0 | 0 | 0 | 0 | 0.244650542 | 0 | 0 |
| ENSMUSG00000001819 | 0 | 0 | 0 | 0 | 0 | 0 | 0 |
| ENSMUSG00000001823 | 0 | 0 | 0 | 0 | 0 | 0 | 0 |
| ENSMUSG00000001827 | 0.289064826 | 0 | 0 | 0.25 | 0 | 0 | 0.224243824 |
| ENSMUSG00000001829 | 2.149089517 | 1.948603967 | 1.998588359 | 2.024508021 | 2.041436517 | 2.255272505 | 1.97354745 |
| ENSMUSG00000001833 | 3.210516687 | 2.915555826 | 2.923589472 | 3.026313445 | 2.930685449 | 3.391640703 | 2.703550597 |
| ENSMUSG00000001844 | 2.020182925 | 1.872963274 | 1.945339928 | 1.782320754 | 1.871833944 | 2.064457989 | 1.838725884 |
| ENSMUSG00000001847 | 3.224648428 | 2.93694335 | 2.932086429 | 3.044821776 | 2.956311085 | 3.432488256 | 2.740208752 |
| ENSMUSG00000001855 | 2.541215974 | 2.372963274 | 2.404596376 | 2.319612365 | 2.342660868 | 2.639486489 | 2.228229691 |
| ENSMUSG00000001864 | 2.643399047 | 2.425975893 | 2.420309603 | 2.300530956 | 2.289867804 | 2.575187845 | 2.159323753 |
| ENSMUSG00000001865 | 0 | 0 | 0 | 0 | 0 | 0 | 0 |
| ENSMUSG00000001870 | 3.205378646 | 2.922608639 | 2.907225288 | 2.669870025 | 2.650078855 | 3.029383778 | 2.453955645 |
| ENSMUSG00000001891 | 2.786132803 | 2.535845803 | 2.506284891 | 2.56932185 | 2.470810917 | 2.824776462 | 2.285835109 |

| | | | | | | | |
|---|---|---|---|---|---|---|---|
| ENSMUSG00000001901 | 0 | 0.25 | 0.396240625 | 0 | 0 | 0 | 0 |
| ENSMUSG00000001909 | 2.390989024 | 2.271202097 | 2.337182039 | 2.04248125 | 2.179810173 | 2.307496038 | 2.146831257 |
| ENSMUSG00000001910 | 2.926153516 | 2.699415381 | 2.732314602 | 2.636723615 | 2.70184985 | 3.062205809 | 2.597961588 |
| ENSMUSG00000001911 | 2.751223449 | 2.553276055 | 2.606041572 | 2.320192693 | 2.414035093 | 2.620136055 | 2.346838633 |
| ENSMUSG00000001918 | 2.039433992 | 1.969129237 | 1.961372513 | 2.069031101 | 2.071604288 | 2.252853031 | 1.972821265 |
| ENSMUSG00000001924 | 3.454801338 | 3.122900652 | 3.120827989 | 3.101285366 | 3.06563036 | 3.525563058 | 2.862164608 |
| ENSMUSG00000001930 | 0.867194479 | 1.04248125 | 0.830482024 | 0.5 | 0.387761935 | 0.301029996 | 0.355418052 |
| ENSMUSG00000001942 | 1.989530018 | 1.845926073 | 1.80479213 | 1.948603967 | 1.868304353 | 2.158362492 | 1.722075986 |
| ENSMUSG00000001943 | 0 | 0 | 0 | 0 | 0 | 0 | 0 |
| ENSMUSG00000001946 | 0.289064826 | 0.25 | 0.5 | 0.396240625 | 0.489301084 | 0 | 0.520678036 |
| ENSMUSG00000001948 | 0.45815691 | 0.75 | 0.25 | 0.92510993 | 0.846351821 | 0.602059991 | 0.448487648 |
| ENSMUSG00000001962 | 2.559630991 | 2.356566189 | 2.379909063 | 2.30357978 | 2.211376431 | 2.527629901 | 2.09490255 |
| ENSMUSG00000001964 | 2.641182749 | 2.414552871 | 2.462155735 | 2.277783918 | 2.277846464 | 2.544068044 | 2.16539364 |
| ENSMUSG00000001983 | 1.678702042 | 1.62694866 | 1.646240625 | 1.782320754 | 1.695608647 | 1.785329835 | 1.651852264 |
| ENSMUSG00000001985 | 0.45815691 | 0.25 | 0 | 0.25 | 0.244650542 | 0 | 0.448487648 |
| ENSMUSG00000001986 | 2.433305075 | 2.245034894 | 2.218361028 | 1.75 | 1.635855979 | 1.819543936 | 1.540636858 |
| ENSMUSG00000001988 | 0.289064826 | 0.75 | 0.580482024 | 0.396240625 | 0.733951626 | 0.477121255 | 0.853775828 |
| ENSMUSG00000001995 | 2.139333929 | 2.058404919 | 2.160061234 | 1.75835575 | 1.766173492 | 1.785329835 | 1.710836105 |
| ENSMUSG00000001998 | 2.086807695 | 1.94697564 | 1.94697564 | 2.209800947 | 2.135290471 | 2.356025857 | 1.935895857 |
| ENSMUSG00000001999 | 2.116511873 | 1.956637122 | 1.981203126 | 2.116901388 | 2.073592779 | 2.401400541 | 1.955652633 |
| ENSMUSG00000002006 | 2.057835465 | 1.982684334 | 1.959800947 | 1.991446071 | 1.98503142 | 2.264817823 | 1.907820406 |
| ENSMUSG00000002007 | 0 | 0.25 | 0 | 0.580482024 | 0.387761935 | 0 | 0.224243824 |
| ENSMUSG00000002010 | 2.556012466 | 2.366901388 | 2.375956434 | 2.335518667 | 2.341270363 | 2.627365857 | 2.191935631 |
| ENSMUSG00000002012 | 0.45815691 | 0.25 | 0.5 | 0.5 | 0.244650542 | 0.477121255 | 0.224243824 |
| ENSMUSG00000002014 | 2.422183552 | 2.219129237 | 2.256284891 | 2.098079356 | 2.124299332 | 2.367355921 | 2.02383181 |
| ENSMUSG00000002015 | 2.81249113 | 2.562278363 | 2.55357978 | 2.733791262 | 2.626252528 | 3.075911761 | 2.455594532 |
| ENSMUSG00000002017 | 2.742600832 | 2.487456678 | 2.472185812 | 2.535526764 | 2.468227512 | 2.82672252 | 2.281664246 |
| ENSMUSG00000002020 | 2.420929319 | 2.277131114 | 2.359697963 | 1.29248125 | 1.439191811 | 1.544068044 | 1.383567577 |
| ENSMUSG00000002028 | 2.905450967 | 2.654137211 | 2.646005735 | 2.692665973 | 2.668192818 | 3.038222638 | 2.500404294 |
| ENSMUSG00000002031 | 2.471720134 | 2.275165585 | 2.271202097 | 2.366391601 | 2.333760374 | 2.674861141 | 2.185972044 |
| ENSMUSG00000002032 | 0.91631382 | 0.830482024 | 0.75 | 0.92510993 | 1.074582839 | 1.204119983 | 1.100339912 |
| ENSMUSG00000002033 | 0 | 0 | 0 | 0 | 0 | 0 | 0 |
| ENSMUSG00000002043 | 1.129344651 | 1.130890489 | 1.403677461 | 0.92510993 | 1.136121934 | 1.113943352 | 1.369798885 |
| ENSMUSG00000002052 | 3.029919532 | 2.762462137 | 2.762462137 | 2.8184489 | 2.791054056 | 3.15896526 | 2.608729177 |

| | | | | | | | |
|---|---|---|---|---|---|---|---|
| ENSMUSG00000002055 | 2.841885121 | 2.571350555 | 2.589112993 | 2.741446071 | 2.699501274 | 3.072617477 | 2.51637161 |
| ENSMUSG00000002057 | 0.811507563 | 0.79248125 | 1 | 0.701838731 | 0.244650542 | 0.602059991 | 0.520678036 |
| ENSMUSG00000002058 | 2.223916471 | 2.081607372 | 2.109697963 | 2.03743678 | 2.076554666 | 2.290034611 | 2.040767326 |
| ENSMUSG00000002059 | 2.612833548 | 2.399978211 | 2.414552871 | 2.288704527 | 2.288252441 | 2.606381365 | 2.172137922 |
| ENSMUSG00000002064 | 2.355805236 | 2.173371739 | 2.179419106 | 2.208222504 | 2.175379703 | 2.456366033 | 2.062419043 |
| ENSMUSG00000002068 | 2.253091469 | 2.063166358 | 2.069031101 | 1.77186571 | 1.736537397 | 1.892094603 | 1.637827726 |
| ENSMUSG00000002076 | 0 | 0.5 | 0 | 0 | 0.733951626 | 0.602059991 | 0 |
| ENSMUSG00000002083 | 0.747221736 | 1.061981878 | 1.201838731 | 0.701838731 | 0.77552387 | 0.698970004 | 1.159323753 |
| ENSMUSG00000002100 | 0 | 0 | 0 | 0 | 0 | 0 | 0 |
| ENSMUSG00000002102 | 2.783508299 | 2.583568321 | 2.604463129 | 2.709800947 | 2.756282161 | 3.09725731 | 2.605968004 |
| ENSMUSG00000002103 | 2.199011616 | 1.990000483 | 2.006976499 | 2.21759118 | 2.196996016 | 2.481442629 | 2.040177507 |
| ENSMUSG00000002104 | 0.289064826 | 0.5 | 0.701838731 | 0.25 | 0.489301084 | 0.602059991 | 0.355418052 |
| ENSMUSG00000002105 | 2.124258828 | 2.077084758 | 2.091580554 | 1.972185812 | 2.148364414 | 2.26245109 | 2.128489437 |
| ENSMUSG00000002107 | 2.68140129 | 2.520204382 | 2.474839231 | 2.263142013 | 2.204601633 | 2.481442629 | 2.054044578 |
| ENSMUSG00000002108 | 0.578129653 | 0.396240625 | 0.5 | 0.396240625 | 0.489301084 | 0.602059991 | 0.629532004 |
| ENSMUSG00000002109 | 1.358731791 | 1.396240625 | 1.29248125 | 1.311981878 | 1.366364104 | 1.491361694 | 1.284450788 |
| ENSMUSG00000002111 | 0.289064826 | 0 | 0 | 0 | 0 | 0 | 0 |
| ENSMUSG00000002129 | 2.753493018 | 2.49929487 | 2.527131114 | 2.629171237 | 2.590305509 | 2.939019776 | 2.449821735 |
| ENSMUSG00000002147 | 2.765209749 | 2.532320754 | 2.540597832 | 2.485628626 | 2.470489023 | 2.84509804 | 2.295899033 |
| ENSMUSG00000002190 | 0.747221736 | 0.646240625 | 0.75 | 1.114857905 | 1.057362051 | 0.954242509 | 0.969165684 |
| ENSMUSG00000002204 | 0 | 0 | 0.25 | 0 | 0 | 0 | 0 |
| ENSMUSG00000002205 | 1.83262764 | 1.854463129 | 1.77186571 | 1.841580554 | 1.871833944 | 2.029383778 | 1.848480858 |
| ENSMUSG00000002210 | 1.862428343 | 1.928561379 | 1.918106335 | 1.571350555 | 1.751666404 | 1.77815125 | 1.793950594 |
| ENSMUSG00000002221 | 2.624559692 | 2.417664062 | 2.405512955 | 2.353935442 | 2.376595432 | 2.602059991 | 2.23605706 |
| ENSMUSG00000002222 | 2.525875121 | 2.301142786 | 2.26584877 | 2.374961472 | 2.260180728 | 2.64246452 | 2.089372236 |
| ENSMUSG00000002227 | 2.297593393 | 2.129909063 | 2.181980114 | 2.115881093 | 2.215379757 | 2.372912003 | 2.159731975 |
| ENSMUSG00000002228 | 0.747221736 | 0.75 | 0.864857905 | 0.580482024 | 0.733951626 | 0 | 0.629532004 |
| ENSMUSG00000002233 | 2.801542157 | 2.605778978 | 2.653907409 | 2.49929487 | 2.573875064 | 2.900913068 | 2.457225157 |
| ENSMUSG00000002240 | 0 | 0 | 0 | 0 | 0 | 0 | 0 |
| ENSMUSG00000002249 | 2.092368209 | 1.995748394 | 2.071350555 | 1.79248125 | 1.919403394 | 1.963787827 | 1.925966215 |
| ENSMUSG00000002250 | 2.039433992 | 2.025821952 | 2.117918804 | 1.910964047 | 2.102149272 | 2.164352856 | 2.112839835 |
| ENSMUSG00000002257 | 0.91631382 | 0.896240625 | 1.04248125 | 0.79248125 | 1.091002363 | 0.77815125 | 1.028149525 |
| ENSMUSG00000002265 | 0.289064826 | 0.580482024 | 0.25 | 1.476722649 | 1.427019704 | 1.447158031 | 1.355418052 |
| ENSMUSG00000002266 | 0 | 0 | 0.25 | 0 | 0 | 0 | 0.224243824 |

| | | | | | | | |
|---|---|---|---|---|---|---|---|
| ENSMUSG00000002274 | 0.578129653 | 0.580482024 | 0.79248125 | 0.951838731 | 1.091002363 | 1.278753601 | 0.876096088 |
| ENSMUSG00000002279 | 1.678702042 | 1.671625132 | 1.664552871 | 1.5 | 1.514343934 | 1.740362689 | 1.517723536 |
| ENSMUSG00000002280 | 1.993049299 | 1.884789703 | 1.985628626 | 1.807204673 | 1.875328588 | 2.029383778 | 1.82753046 |
| ENSMUSG00000002289 | 1.494443473 | 1.403677461 | 1.380890489 | 1.282320754 | 1.358933174 | 1.204119983 | 1.290497981 |
| ENSMUSG00000002297 | 2.899513054 | 2.668327269 | 2.654825739 | 2.632113528 | 2.539585427 | 2.919078092 | 2.3678751 |
| ENSMUSG00000002307 | 2.144240249 | 2.061981878 | 2.12892496 | 1.923371739 | 1.950242452 | 2.10720997 | 1.926805425 |
| ENSMUSG00000002308 | 2.105953527 | 1.94697564 | 1.92510993 | 1.955044741 | 1.955822902 | 2.212187604 | 1.89047692 |
| ENSMUSG00000002319 | 2.565888829 | 2.357613138 | 2.438721876 | 2.464883697 | 2.591450357 | 2.838219222 | 2.465732876 |
| ENSMUSG00000002320 | 2.304292853 | 2.170748646 | 2.14151351 | 2.338837774 | 2.329944595 | 2.592176757 | 2.181044355 |
| ENSMUSG00000002324 | 0.289064826 | 0 | 0 | 0 | 0 | 0 | 0.224243824 |
| ENSMUSG00000002325 | 1.587501561 | 1.527131114 | 1.54745614 | 1.874961472 | 1.934425071 | 2.089905111 | 1.801445495 |
| ENSMUSG00000002326 | 1.948720984 | 1.870953944 | 1.854463129 | 2.150906586 | 2.126012715 | 2.324282455 | 2.016928211 |
| ENSMUSG00000002329 | 1.789260561 | 1.664552871 | 1.705044741 | 1.937048212 | 1.92245931 | 2.064457989 | 1.781059827 |
| ENSMUSG00000002332 | 2.144240249 | 1.966046536 | 2.005591953 | 2.120953944 | 2.126866297 | 2.354108439 | 2.038994634 |
| ENSMUSG00000002341 | 0.289064826 | 0.25 | 0 | 0.25 | 0.244650542 | 0 | 0 |
| ENSMUSG00000002342 | 2.131864492 | 2.075945187 | 2.119945066 | 2.067865757 | 2.123439512 | 2.320146286 | 2.106650241 |
| ENSMUSG00000002343 | 2.010213549 | 2.03360658 | 2.057204673 | 1.668106335 | 1.968065382 | 1.949390007 | 1.983546516 |
| ENSMUSG00000002345 | 1.885732304 | 1.79248125 | 1.802363341 | 1.660964047 | 1.826953746 | 1.903089987 | 1.798966438 |
| ENSMUSG00000002346 | 1.639698646 | 1.649978211 | 1.732684334 | 1.571350555 | 1.718026092 | 1.851258349 | 1.665294008 |
| ENSMUSG00000002365 | 2.673935906 | 2.498234584 | 2.498234584 | 2.46293726 | 2.424460715 | 2.761175813 | 2.275733199 |
| ENSMUSG00000002372 | 2.649252229 | 2.51685859 | 2.583009137 | 2.377938163 | 2.517458689 | 2.780317312 | 2.477860296 |
| ENSMUSG00000002379 | 2.078325387 | 2.058404919 | 2.135757955 | 1.843759858 | 2.105788027 | 2.181843588 | 2.07004434 |
| ENSMUSG00000002384 | 0.45815691 | 0.396240625 | 0.396240625 | 0.646240625 | 0.489301084 | 0.698970004 | 0.629532004 |
| ENSMUSG00000002393 | 0.91631382 | 1.214495249 | 1.561981878 | 1.02186571 | 1.188507684 | 1.113943352 | 1.396774124 |
| ENSMUSG00000002395 | 1.907802659 | 1.816696635 | 1.886723615 | 1.610735874 | 1.782377602 | 1.819543936 | 1.778418869 |
| ENSMUSG00000002396 | 1.91631382 | 1.738549078 | 1.784887838 | 1.516522298 | 1.674752852 | 1.707570176 | 1.655750087 |
| ENSMUSG00000002409 | 1.358731791 | 1.356566189 | 1.388647213 | 1.17510993 | 1.366364104 | 1.397940009 | 1.40946263 |
| ENSMUSG00000002413 | 2.436947303 | 2.25835575 | 2.232684334 | 2.357613138 | 2.279510047 | 2.588831726 | 2.097385413 |
| ENSMUSG00000002416 | 1.89911417 | 1.819031101 | 1.755591953 | 2 | 1.877063019 | 2.123851641 | 1.815933683 |
| ENSMUSG00000002428 | 2.267855219 | 2.121960008 | 2.109697963 | 2.215271726 | 2.136951444 | 2.431363764 | 1.975716277 |
| ENSMUSG00000002455 | 2.453536005 | 2.284887838 | 2.298073204 | 2.468361028 | 2.444777781 | 2.748962861 | 2.307731473 |
| ENSMUSG00000002458 | 2.057835465 | 2.006976499 | 2.004202072 | 2.070192693 | 2.05001946 | 2.320146286 | 1.947894097 |
| ENSMUSG00000002459 | 0.671187741 | 0.75 | 0.79248125 | 1.080482024 | 0.931471446 | 0.954242509 | 0.916588299 |
| ENSMUSG00000002475 | 0.45815691 | 0.5 | 0.701838731 | 1.42510993 | 1.478764298 | 1.361727836 | 1.245582246 |

| | | | | | | | |
|---|---|---|---|---|---|---|---|
| ENSMUSG00000002477 | 2.644135203 | 2.448197574 | 2.413211243 | 2.563757142 | 2.456362323 | 2.848189117 | 2.301737862 |
| ENSMUSG00000002486 | 1.881174527 | 1.828220739 | 1.892463902 | 1.646240625 | 1.806427258 | 1.880813592 | 1.762103709 |
| ENSMUSG00000002489 | 2.196867486 | 2.108657057 | 2.149047439 | 2.086073977 | 2.112066955 | 2.28780173 | 2.031198704 |
| ENSMUSG00000002496 | 2.693336259 | 2.488185312 | 2.520869832 | 2.506976499 | 2.532092521 | 2.811575006 | 2.400671947 |
| ENSMUSG00000002500 | 0.289064826 | 0.25 | 0.396240625 | 0 | 0 | 0 | 0 |
| ENSMUSG00000002504 | 1.936861443 | 1.92510993 | 2.002806814 | 1.912762923 | 2.104881849 | 2.184691431 | 2.09490255 |
| ENSMUSG00000002524 | 3.01718699 | 2.777620827 | 2.825374049 | 2.725593779 | 2.783440724 | 3.12904506 | 2.66409485 |
| ENSMUSG00000002546 | 2.739101238 | 2.535845803 | 2.558704358 | 2.415889526 | 2.461354714 | 2.748962861 | 2.338495365 |
| ENSMUSG00000002550 | 2.300956576 | 2.149978211 | 2.179419106 | 2.139605178 | 2.200473444 | 2.390935107 | 2.100339912 |
| ENSMUSG00000002565 | 0.289064826 | 0 | 0 | 0.25 | 0.387761935 | 0.301029996 | 0.520678036 |
| ENSMUSG00000002578 | 1.389637216 | 1.27186571 | 1.302363341 | 1.729715809 | 1.595992969 | 1.755874856 | 1.535058868 |
| ENSMUSG00000002580 | 2.045658471 | 1.916333979 | 1.894357207 | 2.023189285 | 1.911645804 | 2.245512668 | 1.833176598 |
| ENSMUSG00000002588 | 0 | 0 | 0.25 | 0 | 0 | 0 | 0 |
| ENSMUSG00000002602 | 3.248800091 | 2.971614674 | 3.01685859 | 2.828645811 | 2.820203499 | 3.237292338 | 2.680980656 |
| ENSMUSG00000002603 | 1.890240806 | 1.811981878 | 1.837182039 | 1.606566189 | 1.662257525 | 1.792391689 | 1.753626837 |
| ENSMUSG00000002608 | 2.095120908 | 2.03360658 | 2.028435542 | 1.848079356 | 1.986303335 | 2.004321374 | 1.920884716 |
| ENSMUSG00000002617 | 1.516991255 | 1.403677461 | 1.418106335 | 1.470660762 | 1.310728002 | 1.568201724 | 1.224243824 |
| ENSMUSG00000002625 | 2.039433992 | 1.78743678 | 1.87694866 | 1.912762923 | 1.92245931 | 2.139879086 | 1.896054911 |
| ENSMUSG00000002633 | 0 | 0 | 0 | 0 | 0 | 0 | 0 |
| ENSMUSG00000002635 | 1.671187741 | 1.575945187 | 1.54745614 | 1.445339928 | 1.533172725 | 1.69019608 | 1.452143229 |
| ENSMUSG00000002658 | 2.68140129 | 2.498941788 | 2.52186571 | 2.443284802 | 2.51377465 | 2.77524626 | 2.416364058 |
| ENSMUSG00000002660 | 2.069666964 | 2.066696635 | 2.143411797 | 1.802363341 | 1.98503142 | 2.1430148 | 2.049487947 |
| ENSMUSG00000002661 | 1.227926429 | 1.226722649 | 1.302363341 | 1.214495249 | 1.327538614 | 1.278753601 | 1.365075947 |
| ENSMUSG00000002664 | 0.45815691 | 0.646240625 | 0.25 | 0.580482024 | 0.387761935 | 0.698970004 | 0.579661877 |
| ENSMUSG00000002668 | 0 | 0 | 0 | 0 | 0 | 0 | 0 |
| ENSMUSG00000002679 | 2.304292853 | 2.129909063 | 2.087182039 | 2.148114259 | 2.056322416 | 2.328379603 | 1.910475814 |
| ENSMUSG00000002688 | 2.141794304 | 1.990000483 | 1.933677405 | 1.905512955 | 1.887294348 | 2.041392685 | 1.731408809 |
| ENSMUSG00000002699 | 0.289064826 | 0.25 | 0 | 0 | 0 | 0 | 0 |
| ENSMUSG00000002718 | 3.012043979 | 2.732869058 | 2.714689525 | 2.939763985 | 2.864633702 | 3.320976677 | 2.658904651 |
| ENSMUSG00000002728 | 2.283862692 | 2.105516192 | 2.075945187 | 2.225216702 | 2.174635855 | 2.440909082 | 2.030591163 |
| ENSMUSG00000002731 | 1.967766868 | 1.848079356 | 1.825945187 | 2.059601185 | 2.048957937 | 2.206825876 | 1.896054911 |
| ENSMUSG00000002732 | 1.578129653 | 1.516522298 | 1.431980114 | 1.79745614 | 1.715300549 | 1.939519253 | 1.657681529 |
| ENSMUSG00000002733 | 2.194712275 | 2.031030328 | 2.059601185 | 2.106566189 | 2.094757251 | 2.359835482 | 1.970632877 |
| ENSMUSG00000002741 | 2.761340634 | 2.545288064 | 2.52908599 | 2.586073977 | 2.507741065 | 2.907411361 | 2.372557642 |

| | | | | | | | |
|---|---|---|---|---|---|---|---|
| ENSMUSG00000002748 | 3.355538164 | 3.057129524 | 3.041540771 | 3.060197833 | 2.99105103 | 3.444669231 | 2.776787543 |
| ENSMUSG00000002763 | 1.932832114 | 1.92510993 | 1.995748394 | 1.71759118 | 1.957204337 | 2.012837225 | 1.989124507 |
| ENSMUSG00000002767 | 2.124258828 | 2.075945187 | 2.132845365 | 1.933677405 | 2.069604532 | 2.133538908 | 2.065163042 |
| ENSMUSG00000002768 | 2.337801042 | 2.203445298 | 2.228969845 | 2.186208459 | 2.237245597 | 2.445604203 | 2.158095989 |
| ENSMUSG00000002769 | 0.45815691 | 0.396240625 | 0.5 | 0.25 | 0.568060967 | 0.602059991 | 0.355418052 |
| ENSMUSG00000002771 | 0.960252568 | 0.976722649 | 1.02186571 | 0.896240625 | 1.136121934 | 1 | 1.21680636 |
| ENSMUSG00000002778 | 2.502784398 | 2.311388177 | 2.319031101 | 2.358657057 | 2.351342427 | 2.655138435 | 2.223909441 |
| ENSMUSG00000002781 | 1.205378646 | 1.17510993 | 1.364857905 | 1.114857905 | 1.394615667 | 1.113943352 | 1.452143229 |
| ENSMUSG00000002791 | 0 | 0.396240625 | 0.25 | 0.396240625 | 0.244650542 | 0.602059991 | 0.520678036 |
| ENSMUSG00000002797 | 1.205378646 | 1.29248125 | 1.26109853 | 1.238549078 | 1.27449559 | 1.361727836 | 1.224243824 |
| ENSMUSG00000002799 | 0.867194479 | 1.080482024 | 1.160964047 | 1.04248125 | 1.380772476 | 1.204119983 | 1.486592281 |
| ENSMUSG00000002803 | 1.903481042 | 1.76109853 | 1.825945187 | 1.67510993 | 1.462344774 | 1.707570176 | 1.520678036 |
| ENSMUSG00000002804 | 1.578129653 | 1.532320754 | 1.614857905 | 1.410964047 | 1.628935049 | 1.556302501 | 1.659601507 |
| ENSMUSG00000002808 | 2.479432974 | 2.245748394 | 2.228222334 | 2.253505118 | 2.16634835 | 2.525044807 | 2.066254157 |
| ENSMUSG00000002812 | 2.74143756 | 2.543106877 | 2.591580554 | 2.55930249 | 2.632005142 | 2.937016107 | 2.506886092 |
| ENSMUSG00000002814 | 2.170239354 | 1.96293726 | 2.051142786 | 2.028435542 | 2.085294053 | 2.292256071 | 1.963977467 |
| ENSMUSG00000002820 | 1.727821383 | 1.634789703 | 1.729715809 | 1.71759118 | 1.820920161 | 1.924279286 | 1.901538355 |
| ENSMUSG00000002825 | 1.771760131 | 1.668106335 | 1.729715809 | 1.571350555 | 1.770879644 | 1.832508913 | 1.759302692 |
| ENSMUSG00000002831 | 2.708105693 | 2.512462137 | 2.533285553 | 0.646240625 | 0.955822902 | 0.698970004 | 0.672731473 |
| ENSMUSG00000002833 | 2.612039956 | 2.430274797 | 2.500703754 | 2.27053726 | 2.404070415 | 2.546542663 | 2.367231288 |
| ENSMUSG00000002835 | 2.779274431 | 2.5684489 | 2.597810897 | 2.52186571 | 2.554110285 | 2.879669206 | 2.420034153 |
| ENSMUSG00000002844 | 2.285603957 | 2.148114259 | 2.245034894 | 1.884789703 | 1.969399812 | 2.021189299 | 1.987739 |
| ENSMUSG00000002845 | 2.323768721 | 2.157339155 | 2.127938163 | 2.12694866 | 2.062514787 | 2.338456494 | 1.947894097 |
| ENSMUSG00000002846 | 2.039433992 | 1.882845365 | 1.868933358 | 1.819031101 | 1.87878897 | 2 | 1.762103709 |
| ENSMUSG00000002847 | 1.069666964 | 1.080482024 | 0.951838731 | 0.896240625 | 1 | 0.954242509 | 0.952571511 |
| ENSMUSG00000002870 | 3.038292219 | 2.835796429 | 2.855778978 | 2.654596376 | 2.740455823 | 3.042575512 | 2.637268817 |
| ENSMUSG00000002871 | 2.126809492 | 2.006976499 | 2.070192693 | 1.755591953 | 1.84445929 | 1.919078092 | 1.835407748 |
| ENSMUSG00000002881 | 2.602397159 | 2.437886015 | 2.422499493 | 2.416777886 | 2.361573756 | 2.705007959 | 2.191935631 |
| ENSMUSG00000002885 | 2.280358197 | 2.139605178 | 2.111770807 | 1.890560606 | 1.883916752 | 2.127104798 | 1.839824409 |
| ENSMUSG00000002897 | 2.188178997 | 2.039967834 | 2.058404919 | 2.136723615 | 2.181274723 | 2.397940009 | 2.012457048 |
| ENSMUSG00000002900 | 3.074935921 | 2.846735061 | 2.836905342 | 2.761610487 | 2.761686107 | 3.151982395 | 2.592440351 |
| ENSMUSG00000002908 | 0.747221736 | 0.830482024 | 0.976722649 | 1.311981878 | 1.351342427 | 1.361727836 | 1.38802993 |
| ENSMUSG00000002910 | 1.069666964 | 1.160964047 | 1.061981878 | 0.951838731 | 1.121713561 | 1.176091259 | 1.21680636 |
| ENSMUSG00000002930 | 0 | 0 | 0 | 0 | 0 | 0 | 0 |

| | | | | | | | |
|---|---|---|---|---|---|---|---|
| ENSMUSG00000002944 | 0 | 0 | 0 | 0 | 0 | 0 | 0 |
| ENSMUSG00000002948 | 2.629921064 | 2.421187155 | 2.43155454 | 2.282320754 | 2.338472809 | 2.605305046 | 2.21680636 |
| ENSMUSG00000002949 | 2.612039956 | 2.40136988 | 2.471424093 | 2.267194569 | 2.316753004 | 2.57634135 | 2.19633727 |
| ENSMUSG00000002957 | 2.896712826 | 2.646005735 | 2.644357207 | 2.677701608 | 2.683666359 | 3.021602716 | 2.53339225 |
| ENSMUSG00000002963 | 1.83262764 | 1.755591953 | 1.825945187 | 1.598079356 | 1.741652793 | 1.662757832 | 1.755055163 |
| ENSMUSG00000002968 | 1.867194479 | 1.935366747 | 1.994319981 | 1.701838731 | 1.842556557 | 1.959041392 | 1.95718226 |
| ENSMUSG00000002980 | 0.867194479 | 1.02186571 | 1.098079356 | 1.403677461 | 1.523883869 | 1.568201724 | 1.707551656 |
| ENSMUSG00000002981 | 2.536482297 | 2.356566189 | 2.376452888 | 2.410512923 | 2.434230499 | 2.736396502 | 2.294288164 |
| ENSMUSG00000002983 | 1.87190676 | 1.79745614 | 1.868933358 | 1.799918086 | 1.917865457 | 2.033423755 | 1.913109605 |
| ENSMUSG00000002984 | 2.231898196 | 2.181980114 | 2.214495249 | 1.892463902 | 2.098472613 | 2.187520721 | 2.064068234 |
| ENSMUSG00000002985 | 0.747221736 | 0.896240625 | 1.04248125 | 0.5 | 0.77552387 | 0.602059991 | 0.916588299 |
| ENSMUSG00000002992 | 0 | 0 | 0 | 0 | 0 | 0 | 0 |
| ENSMUSG00000002996 | 2.490209605 | 2.268535366 | 2.225216702 | 2.067865757 | 2.033751322 | 2.315970345 | 1.892346957 |
| ENSMUSG00000002997 | 2.788220579 | 2.557505109 | 2.559601185 | 2.393411797 | 2.353143227 | 2.667452953 | 2.150210039 |
| ENSMUSG00000003031 | 2.595015862 | 2.418548067 | 2.396709947 | 2.33272922 | 2.268839121 | 2.5132176 | 2.107610228 |
| ENSMUSG00000003032 | 1.759671389 | 1.747171172 | 1.79248125 | 1.311981878 | 1.091002363 | 1.462397998 | 1.245582246 |
| ENSMUSG00000003033 | 2.604019855 | 2.411864607 | 2.455443495 | 2.459012589 | 2.478764298 | 2.758154622 | 2.345462945 |
| ENSMUSG00000003037 | 2.467247894 | 2.358657057 | 2.366901388 | 2.352877747 | 2.331380351 | 2.606381365 | 2.205319397 |
| ENSMUSG00000003038 | 2.111266113 | 2.025821952 | 2.118933358 | 2.156427211 | 2.16097133 | 2.408239965 | 2.138662327 |
| ENSMUSG00000003039 | 2.4661223 | 2.252806814 | 2.240724001 | 2.300530956 | 2.250704826 | 2.544068044 | 2.146406417 |
| ENSMUSG00000003051 | 0 | 0.75 | 0.5 | 0.25 | 0 | 0 | 0.224243824 |
| ENSMUSG00000003053 | 0 | 0.25 | 0 | 0 | 0 | 0 | 0.224243824 |
| ENSMUSG00000003062 | 2.287337982 | 2.160061234 | 2.117918804 | 2.147178659 | 2.105788027 | 2.348304863 | 1.95411574 |
| ENSMUSG00000003068 | 2.363083473 | 2.281676118 | 2.334405475 | 2.138647213 | 2.225193812 | 2.439332694 | 2.264126018 |
| ENSMUSG00000003070 | 1.069666964 | 1.130890489 | 1.201838731 | 1.388647213 | 1.366364104 | 1.505149978 | 1.56970677 |
| ENSMUSG00000003072 | 2.585813169 | 2.448197574 | 2.554489424 | 2.243603647 | 2.420030246 | 2.650307523 | 2.420034153 |
| ENSMUSG00000003099 | 2.299278375 | 2.173371739 | 2.224461364 | 2.121960008 | 2.227124297 | 2.397940009 | 2.099358074 |
| ENSMUSG00000003119 | 2.5496035 | 2.356566189 | 2.379417347 | 2.480088214 | 2.451638175 | 2.804820679 | 2.265305657 |
| ENSMUSG00000003123 | 1.100572389 | 1.160964047 | 1.311981878 | 1.27186571 | 1.394615667 | 1.301029996 | 1.417653332 |
| ENSMUSG00000003131 | 3.054274575 | 2.767697952 | 2.758527787 | 2.925975893 | 2.818165017 | 3.274388796 | 2.613981494 |
| ENSMUSG00000003134 | 1.307602651 | 1.188721876 | 1.226722649 | 1.464495249 | 1.439191811 | 1.531478917 | 1.437266334 |
| ENSMUSG00000003135 | 2.389637216 | 2.17510993 | 2.177701608 | 2.281030328 | 2.239111448 | 2.547774705 | 2.078019652 |
| ENSMUSG00000003153 | 0.91631382 | 0.951838731 | 0.701838731 | 0.951838731 | 0.733951626 | 0.602059991 | 1.014380832 |
| ENSMUSG00000003154 | 2.078325387 | 1.972185812 | 2.013820609 | 1.918106335 | 1.900490506 | 2.127104798 | 1.819456572 |

| | | | | | | | |
|---|---|---|---|---|---|---|---|
| ENSMUSG00000003161 | 2.716227929 | 2.503156135 | 2.51348147 | 2.584684096 | 2.497151587 | 2.86569606 | 2.354078442 |
| ENSMUSG00000003166 | 2.521950082 | 2.341580554 | 2.33993739 | 2.385274154 | 2.376595432 | 2.688419822 | 2.198153757 |
| ENSMUSG00000003184 | 1.740854706 | 1.668106335 | 1.75 | 1.516522298 | 1.632412477 | 1.72427587 | 1.692344475 |
| ENSMUSG00000003190 | 1.156259305 | 1.17510993 | 1.339388001 | 1.04248125 | 1.121713561 | 0.84509804 | 1.176815335 |
| ENSMUSG00000003198 | 1.747221736 | 1.580482024 | 1.557204673 | 1.630890489 | 1.546663136 | 1.763427994 | 1.46982607 |
| ENSMUSG00000003199 | 2.349889783 | 2.263820609 | 2.289967834 | 1.858657057 | 2.135290471 | 2.193124598 | 2.076440241 |
| ENSMUSG00000003200 | 2.672564087 | 2.559601185 | 2.62019755 | 2.314346961 | 2.449252175 | 2.710117365 | 2.398923736 |
| ENSMUSG00000003206 | 0 | 0 | 0 | 0 | 0 | 0 | 0 |
| ENSMUSG00000003208 | 1.445324132 | 1.557204673 | 1.692046081 | 1.571350555 | 1.614672255 | 1.698970004 | 1.670888075 |
| ENSMUSG00000003226 | 3.301260979 | 2.989638179 | 2.975311008 | 3.088010856 | 2.994570356 | 3.458637849 | 2.737962817 |
| ENSMUSG00000003227 | 0 | 0.25 | 0 | 0.25 | 0.244650542 | 0.477121255 | 0.355418052 |
| ENSMUSG00000003228 | 1.956444026 | 1.741446071 | 1.763820609 | 1.841580554 | 1.751666404 | 1.959041392 | 1.641897157 |
| ENSMUSG00000003233 | 2.149089517 | 2.063166358 | 2.108657057 | 1.80479213 | 1.888971102 | 2.041392685 | 1.935895857 |
| ENSMUSG00000003234 | 2.363083473 | 2.204245906 | 2.186208459 | 2.263820609 | 2.273934005 | 2.496929648 | 2.10131878 |
| ENSMUSG00000003235 | 2.671187741 | 2.459406983 | 2.413211243 | 2.443284802 | 2.42262144 | 2.711807229 | 2.262051244 |
| ENSMUSG00000003269 | 2.360187403 | 2.174241882 | 2.201032755 | 2.233422664 | 2.26997769 | 2.511883361 | 2.117056606 |
| ENSMUSG00000003271 | 0.289064826 | 0 | 0.5 | 0.92510993 | 1.057362051 | 1.176091259 | 1.078019652 |
| ENSMUSG00000003273 | 0.747221736 | 0.25 | 0.92510993 | 0.5 | 0.489301084 | 0.301029996 | 0.672731473 |
| ENSMUSG00000003279 | 0.811507563 | 0.75 | 0.75 | 0.25 | 0.244650542 | 0 | 0.355418052 |
| ENSMUSG00000003282 | 1.342375483 | 1.330482024 | 1.311981878 | 1.061981878 | 0.978602168 | 1.079181246 | 0.876096088 |
| ENSMUSG00000003283 | 1.527823874 | 1.552363341 | 1.348079356 | 1.505591953 | 1.49445382 | 1.763427994 | 1.48984372 |
| ENSMUSG00000003299 | 1.924654746 | 1.80479213 | 1.87892496 | 1.912762923 | 2.09661982 | 2.243038049 | 2.074853081 |
| ENSMUSG00000003308 | 2.190368148 | 2.072504712 | 2.106566189 | 2.086073977 | 2.141887974 | 2.274157849 | 2.115189268 |
| ENSMUSG00000003309 | 0 | 0 | 0 | 0 | 0 | 0 | 0 |
| ENSMUSG00000003316 | 3.197843457 | 2.900094388 | 2.906084636 | 3.004550045 | 2.936151021 | 3.42111013 | 2.735357125 |
| ENSMUSG00000003341 | 1.470606563 | 1.388647213 | 1.458222504 | 0.5 | 0.733951626 | 0.301029996 | 0.672731473 |
| ENSMUSG00000003344 | 1.989530018 | 1.945339928 | 2.043731421 | 1.87892496 | 2.12771782 | 2.093421685 | 2.171745067 |
| ENSMUSG00000003345 | 2.331622671 | 2.309601185 | 2.421187155 | 2.069031101 | 2.267696867 | 2.357934847 | 2.317146272 |
| ENSMUSG00000003346 | 1.903481042 | 1.970660762 | 2.03743678 | 1.726722649 | 2.001219193 | 2.049218023 | 2.109044889 |
| ENSMUSG00000003348 | 1.307602651 | 1.516522298 | 1.52186571 | 1.238549078 | 1.293076518 | 1.414973348 | 1.546120303 |
| ENSMUSG00000003352 | 2.506893125 | 2.343216265 | 2.386241108 | 2.173371739 | 2.315755952 | 2.480006943 | 2.265599895 |
| ENSMUSG00000003354 | 0.671187741 | 0.25 | 0.580482024 | 0.5 | 0.632412477 | 0.301029996 | 0.224243824 |
| ENSMUSG00000003355 | 1.885732304 | 1.723704441 | 1.782320754 | 1.685366747 | 1.754126039 | 1.826074803 | 1.687112305 |
| ENSMUSG00000003360 | 2.903481042 | 2.674241882 | 2.704645544 | 2.577653195 | 2.649303553 | 2.936513742 | 2.511869741 |

| | | | | | | | |
|---|---|---|---|---|---|---|---|
| ENSMUSG00000003363 | 2.124258828 | 2.041226732 | 2.107613138 | 1.78743678 | 2.004851707 | 2.064457989 | 2.011813236 |
| ENSMUSG00000003378 | 2.013563224 | 1.919870025 | 2.039967834 | 1.79248125 | 1.978602168 | 1.954242509 | 1.952571511 |
| ENSMUSG00000003379 | 0.578129653 | 0 | 0 | 0.396240625 | 0 | 0.301029996 | 0 |
| ENSMUSG00000003380 | 1.686083339 | 1.630890489 | 1.698603967 | 1.42510993 | 1.559656853 | 1.491361694 | 1.700880998 |
| ENSMUSG00000003382 | 2.144240249 | 1.96293726 | 2.02186571 | 2.005591953 | 2.091002363 | 2.220108088 | 1.938331368 |
| ENSMUSG00000003402 | 2.752926783 | 2.566403762 | 2.601816941 | 2.510756821 | 2.603362773 | 2.892651034 | 2.518128015 |
| ENSMUSG00000003410 | 0 | 0.5 | 0.25 | 0 | 0 | 0.301029996 | 0.355418052 |
| ENSMUSG00000003411 | 1.614416215 | 1.622963274 | 1.766522298 | 1.828220739 | 1.806427258 | 1.991226076 | 1.735976162 |
| ENSMUSG00000003418 | 0 | 0 | 0 | 0.646240625 | 0.387761935 | 0.477121255 | 0.355418052 |
| ENSMUSG00000003420 | 1.129344651 | 1.201838731 | 1.26109853 | 1.25 | 1.401338844 | 1.278753601 | 1.42167221 |
| ENSMUSG00000003421 | 1.989530018 | 1.910964047 | 1.976722649 | 1.850219859 | 1.88560959 | 2.136720567 | 2.001999072 |
| ENSMUSG00000003423 | 2.192545868 | 2.100219859 | 2.152756199 | 1.903677461 | 2.086251872 | 2.187520721 | 2.043115906 |
| ENSMUSG00000003429 | 3.016886209 | 2.726158655 | 2.747525991 | 2.755938588 | 2.690120394 | 3.129367596 | 2.510631008 |
| ENSMUSG00000003435 | 2.754058486 | 2.540912419 | 2.570772088 | 2.530383379 | 2.584058335 | 2.906873535 | 2.451150342 |
| ENSMUSG00000003436 | 0.289064826 | 0 | 0 | 0.25 | 0 | 0 | 0.224243824 |
| ENSMUSG00000003437 | 2.17479713 | 2.056000419 | 2.067865757 | 2.145300145 | 2.074582839 | 2.380211242 | 1.994607951 |
| ENSMUSG00000003438 | 2.095120908 | 2.063166358 | 2.143411797 | 2.016522298 | 2.093822267 | 2.214843848 | 2.047762527 |
| ENSMUSG00000003444 | 1.655740331 | 1.638647213 | 1.732684334 | 1.714495249 | 1.704182902 | 1.84509804 | 1.69749337 |
| ENSMUSG00000003452 | 1.912079952 | 1.807204673 | 1.839388001 | 1.984159485 | 1.964031554 | 2.173186268 | 1.757893036 |
| ENSMUSG00000003458 | 2.693989403 | 2.46759118 | 2.472185812 | 2.569902645 | 2.556389881 | 2.934498451 | 2.367231288 |
| ENSMUSG00000003464 | 2.537433341 | 2.315523711 | 2.314346961 | 2.473325383 | 2.448567488 | 2.774516966 | 2.265599895 |
| ENSMUSG00000003469 | 0 | 0 | 0 | 0.396240625 | 0.244650542 | 0 | 0.448487648 |
| ENSMUSG00000003476 | 0 | 0 | 0 | 0 | 0 | 0 | 0 |
| ENSMUSG00000003477 | 0 | 0 | 0 | 0 | 0 | 0 | 0 |
| ENSMUSG00000003484 | 0 | 0 | 0 | 0 | 0 | 0 | 0 |
| ENSMUSG00000003500 | 2.333175866 | 2.236360959 | 2.275821952 | 2.153677461 | 2.265401209 | 2.397940009 | 2.211635518 |
| ENSMUSG00000003505 | 0 | 0 | 0 | 0 | 0 | 0 | 0 |
| ENSMUSG00000003518 | 2.233869984 | 2.104463129 | 2.102347734 | 2.299918086 | 2.276175004 | 2.506505032 | 2.149789616 |
| ENSMUSG00000003526 | 1.069666964 | 0.951838731 | 1.238549078 | 1.321350555 | 1.433158226 | 1.361727836 | 1.483307832 |
| ENSMUSG00000003527 | 2.075459174 | 2.006976499 | 1.995748394 | 1.987091808 | 2.057362051 | 2.113943352 | 1.934261938 |
| ENSMUSG00000003528 | 2.134369199 | 2.016522298 | 2.038704527 | 2.113831805 | 2.114724103 | 2.383815366 | 2.072190387 |
| ENSMUSG00000003531 | 1.727821383 | 1.77186571 | 1.779735268 | 1.685366747 | 1.78687392 | 1.851258349 | 1.790137008 |
| ENSMUSG00000003534 | 2.787699615 | 2.680274797 | 2.759043403 | 2.143411797 | 2.382429551 | 2.489958479 | 2.361815771 |
| ENSMUSG00000003541 | 2.23778581 | 2.156427211 | 2.31611065 | 1.916333979 | 2.039257769 | 2.152288344 | 2.02568932 |

| | | | | | | | |
|---|---|---|---|---|---|---|---|
| ENSMUSG00000003545 | 0.289064826 | 0.701838731 | 0.396240625 | 0.75 | 0.387761935 | 0.301029996 | 0.629532004 |
| ENSMUSG00000003546 | 1.686083339 | 1.646240625 | 1.819031101 | 1.532320754 | 1.686820904 | 1.643452676 | 1.762103709 |
| ENSMUSG00000003549 | 1.852729478 | 1.80479213 | 1.886723615 | 1.62694866 | 1.686820904 | 1.72427587 | 1.667169465 |
| ENSMUSG00000003555 | 0 | 0 | 0 | 0 | 0 | 0 | 0 |
| ENSMUSG00000003559 | 0.811507563 | 0.5 | 0.830482024 | 0.79248125 | 0.387761935 | 0.602059991 | 0.710836105 |
| ENSMUSG00000003573 | 2.23778581 | 2.104463129 | 2.172499493 | 1.882845365 | 2.046825265 | 2.130333768 | 2.026921761 |
| ENSMUSG00000003575 | 2.167941649 | 2.107613138 | 2.147178659 | 2.028435542 | 2.151558609 | 2.23299611 | 2.108567375 |
| ENSMUSG00000003581 | 1.707474304 | 1.695339928 | 1.777131114 | 1.657339155 | 1.848234258 | 1.792391689 | 1.729871916 |
| ENSMUSG00000003585 | 1.307602651 | 1.188721876 | 1.130890489 | 1.732684334 | 1.632412477 | 1.908485019 | 1.591689859 |
| ENSMUSG00000003604 | 1.920505136 | 1.729715809 | 1.766522298 | 2.073655187 | 1.983754904 | 2.276461804 | 1.869159809 |
| ENSMUSG00000003617 | 1.693336259 | 1.657339155 | 1.589388001 | 2.061981878 | 1.925488995 | 2.285557309 | 1.850609257 |
| ENSMUSG00000003623 | 2.399009124 | 2.220660762 | 2.192872367 | 2.290597832 | 2.207327177 | 2.521138084 | 2.054044578 |
| ENSMUSG00000003644 | 1.707474304 | 1.678561379 | 1.646240625 | 1.821350555 | 1.929985312 | 2.045322979 | 1.927642465 |
| ENSMUSG00000003657 | 0 | 0 | 0 | 0 | 0 | 0 | 0 |
| ENSMUSG00000003660 | 3.039433992 | 2.732314602 | 2.714689525 | 2.864088604 | 2.830919587 | 3.224274014 | 2.605762532 |
| ENSMUSG00000003662 | 2.078325387 | 2.025821952 | 2.028435542 | 2.204245906 | 2.201164838 | 2.469822016 | 2.07959139 |
| ENSMUSG00000003665 | 0.747221736 | 0.864857905 | 0.75 | 0.5 | 0.77552387 | 0.903089987 | 0.629532004 |
| ENSMUSG00000003680 | 2.433305075 | 2.267194569 | 2.279735268 | 2.133818844 | 2.170139537 | 2.338456494 | 2.074322292 |
| ENSMUSG00000003721 | 2.405576699 | 2.225216702 | 2.168989258 | 2.265173983 | 2.164053936 | 2.465382851 | 2.046027856 |
| ENSMUSG00000003731 | 2.853620573 | 2.637686696 | 2.63503201 | 2.801295582 | 2.724043435 | 3.13893394 | 2.544634011 |
| ENSMUSG00000003746 | 1.912079952 | 1.729715809 | 1.744319981 | 1.937048212 | 1.88560959 | 2.093421685 | 1.74196736 |
| ENSMUSG00000003752 | 1.789260561 | 1.638647213 | 1.630890489 | 1.527131114 | 1.599808748 | 1.748188027 | 1.594042709 |
| ENSMUSG00000003762 | 1.404271433 | 1.311981878 | 1.42510993 | 1.061981878 | 1.310728002 | 1.278753601 | 1.350478789 |
| ENSMUSG00000003778 | 2.788220579 | 2.480460234 | 2.497880462 | 2.53360658 | 2.507161022 | 2.840733235 | 2.337554913 |
| ENSMUSG00000003779 | 3.023157491 | 2.732869058 | 2.735994979 | 2.789967834 | 2.718823734 | 3.136403448 | 2.515150027 |
| ENSMUSG00000003808 | 2.495495257 | 2.351285366 | 2.351285366 | 2.462155735 | 2.49445382 | 2.778874472 | 2.39559556 |
| ENSMUSG00000003809 | 2.153883046 | 1.948603967 | 1.997171172 | 2.099151195 | 2.055279708 | 2.33243846 | 1.95411574 |
| ENSMUSG00000003810 | 2.857166034 | 2.622211087 | 2.629909063 | 2.53360658 | 2.536126349 | 2.861534411 | 2.386819016 |
| ENSMUSG00000003812 | 2.003432385 | 1.874961472 | 1.907339155 | 1.630890489 | 1.639266209 | 1.819543936 | 1.584526862 |
| ENSMUSG00000003813 | 2.328498818 | 2.246460484 | 2.271202097 | 2.092671852 | 2.206647761 | 2.44870632 | 2.141695983 |
| ENSMUSG00000003814 | 3.572927518 | 3.235353618 | 3.238821661 | 3.352414028 | 3.263533443 | 3.8359441 | 3.046080558 |
| ENSMUSG00000003824 | 1.639698646 | 1.505591953 | 1.593759858 | 1.614857905 | 1.584291694 | 1.755874856 | 1.584526862 |
| ENSMUSG00000003847 | 3.24254188 | 2.941528689 | 2.93123503 | 3.087112884 | 2.98846255 | 3.459693976 | 2.749105737 |
| ENSMUSG00000003848 | 2.368815967 | 2.236360959 | 2.223704441 | 2.130890489 | 2.247690665 | 2.436162647 | 2.145981019 |

| | | | | | | | |
|---|---|---|---|---|---|---|---|
| ENSMUSG00000003849 | 1.885732304 | 1.701838731 | 1.685366747 | 1.330482024 | 1.264824954 | 1.431363764 | 1.140832123 |
| ENSMUSG00000003863 | 0.747221736 | 1.04248125 | 1.130890489 | 0.92510993 | 1.020174412 | 0.77815125 | 1.20139795 |
| ENSMUSG00000003865 | 2.297593393 | 2.113831805 | 2.148114259 | 2.151832578 | 2.113840608 | 2.361727836 | 2.066254157 |
| ENSMUSG00000003868 | 2.251208701 | 2.170748646 | 2.26109853 | 1.982684334 | 2.192777367 | 2.271841607 | 2.249315158 |
| ENSMUSG00000003872 | 0.289064826 | 0.396240625 | 0.75 | 0.580482024 | 0.686820904 | 0.477121255 | 0.579661877 |
| ENSMUSG00000003873 | 2.37447073 | 2.291226732 | 2.295598588 | 1.945339928 | 2.099395374 | 2.158362492 | 1.969900112 |
| ENSMUSG00000003882 | 0 | 0 | 0 | 0 | 0 | 0 | 0 |
| ENSMUSG00000003923 | 2.575536697 | 2.327653195 | 2.299304173 | 2.415889526 | 2.321696448 | 2.698100546 | 2.175652306 |
| ENSMUSG00000003929 | 2.269664473 | 2.133818844 | 2.074802005 | 2.117918804 | 2.022473809 | 2.301029996 | 1.89141329 |
| ENSMUSG00000003934 | 1.307602651 | 1.130890489 | 1.214495249 | 1.330482024 | 1.42077253 | 1.380211242 | 1.302263476 |
| ENSMUSG00000003948 | 2.409468165 | 2.284887838 | 2.256976499 | 2.03743678 | 1.966725889 | 2.31386722 | 1.852723744 |
| ENSMUSG00000003949 | 0.747221736 | 0.580482024 | 0.864857905 | 1.04248125 | 1.057362051 | 1.301029996 | 1.054044578 |
| ENSMUSG00000003955 | 2.295901576 | 2.145300145 | 2.158248799 | 2.139605178 | 2.057362051 | 2.307496038 | 1.943148064 |
| ENSMUSG00000003970 | 3.486615732 | 3.231017546 | 3.295831306 | 3.075159638 | 3.222679761 | 3.606273853 | 3.094222085 |
| ENSMUSG00000003974 | 0 | 0.25 | 0.25 | 0.25 | 0 | 0.301029996 | 0.224243824 |
| ENSMUSG00000003992 | 2.280358197 | 2.114857905 | 2.066696635 | 2.139605178 | 2.113840608 | 2.371067862 | 1.999330888 |
| ENSMUSG00000004018 | 1.852729478 | 1.698603967 | 1.610735874 | 1.681980114 | 1.635855979 | 1.799340549 | 1.546120303 |
| ENSMUSG00000004031 | 0 | 0.25 | 0 | 0 | 0 | 0 | 0.224243824 |
| ENSMUSG00000004032 | 2.020182925 | 1.910964047 | 1.87892496 | 2.29248125 | 2.227765453 | 2.575187845 | 2.053478496 |
| ENSMUSG00000004035 | 0.867194479 | 0.396240625 | 0.830482024 | 1 | 0.978602168 | 0.954242509 | 0.829800754 |
| ENSMUSG00000004038 | 0 | 0 | 0 | 0 | 0 | 0 | 0 |
| ENSMUSG00000004040 | 2.649978134 | 2.413659007 | 2.438304187 | 2.377443751 | 2.422252432 | 2.701567985 | 2.280823582 |
| ENSMUSG00000004043 | 2.116511873 | 1.997171172 | 2.075945187 | 1.894357207 | 1.931471446 | 2.064457989 | 1.864112664 |
| ENSMUSG00000004044 | 3.305348823 | 3.013735854 | 3.061239598 | 2.998588359 | 2.934165449 | 3.374931554 | 2.813995814 |
| ENSMUSG00000004054 | 2.054824387 | 2.079353153 | 2.216046536 | 1.809601185 | 2.029283463 | 2.037426498 | 2.178359564 |
| ENSMUSG00000004056 | 2.539328946 | 2.366391601 | 2.373463862 | 2.250703754 | 2.313753359 | 2.549003262 | 2.237343405 |
| ENSMUSG00000004069 | 2.37447073 | 2.228222334 | 2.218361028 | 2.180274797 | 2.192777367 | 2.434568904 | 2.044283833 |
| ENSMUSG00000004070 | 2.105953527 | 1.972185812 | 1.997171172 | 2.168989258 | 2.112066955 | 2.383815366 | 1.975716277 |
| ENSMUSG00000004071 | 2.264212992 | 2.077084758 | 2.116901388 | 2.153677461 | 2.112954895 | 2.357934847 | 1.981429723 |
| ENSMUSG00000004085 | 2.547754129 | 2.358657057 | 2.343759858 | 2.5 | 2.452655829 | 2.78604121 | 2.291042162 |
| ENSMUSG00000004096 | 2.58665822 | 2.380890489 | 2.371457327 | 2.57991803 | 2.494152791 | 2.907411361 | 2.304878814 |
| ENSMUSG00000004098 | 1.568542279 | 1.464495249 | 1.62694866 | 1.984159485 | 2.069604532 | 2.227886705 | 2.119375723 |
| ENSMUSG00000004099 | 3.113897241 | 2.851152349 | 2.869945066 | 2.938930538 | 2.938980516 | 3.353723938 | 2.77890065 |
| ENSMUSG00000004100 | 2.016886209 | 1.990000483 | 2.065523711 | 1.735628626 | 1.968065382 | 1.982271233 | 1.988432495 |

| | | | | | | | |
|---|---|---|---|---|---|---|---|
| ENSMUSG00000004105 | 2.773381775 | 2.590759908 | 2.638167274 | 2.051142786 | 2.154724156 | 2.357934847 | 2.03780742 |
| ENSMUSG00000004110 | 0.45815691 | 0 | 0 | 0.396240625 | 0.568060967 | 0.301029996 | 0.355418052 |
| ENSMUSG00000004113 | 0.747221736 | 0.75 | 0.951838731 | 0.79248125 | 1 | 1 | 0.829800754 |
| ENSMUSG00000004127 | 2.095120908 | 1.978222334 | 1.985628626 | 2.197790722 | 2.169384551 | 2.456366033 | 2.026306127 |
| ENSMUSG00000004151 | 0.671187741 | 0.864857905 | 0.830482024 | 0.92510993 | 0.733951626 | 0.602059991 | 0.520678036 |
| ENSMUSG00000004187 | 0.867194479 | 0.92510993 | 0.830482024 | 0.646240625 | 1.039257769 | 0.77815125 | 1.185218806 |
| ENSMUSG00000004207 | 3.604534846 | 3.283084766 | 3.306791164 | 3.112093615 | 3.102418109 | 3.58726185 | 2.952887976 |
| ENSMUSG00000004221 | 2.729023206 | 2.493603647 | 2.487456678 | 2.327653195 | 2.271678687 | 2.579783597 | 2.123964672 |
| ENSMUSG00000004231 | 0 | 0 | 0 | 0.25 | 0.489301084 | 0 | 0.829800754 |
| ENSMUSG00000004233 | 1.993049299 | 1.845926073 | 1.870953944 | 1.985628626 | 1.893954095 | 2.127104798 | 1.805128823 |
| ENSMUSG00000004263 | 1.205378646 | 1.339388001 | 1.356566189 | 1.226722649 | 1.244650542 | 1.079181246 | 1.224243824 |
| ENSMUSG00000004264 | 2.720536789 | 2.512462137 | 2.519871196 | 2.42813175 | 2.472415995 | 2.790285164 | 2.347752519 |
| ENSMUSG00000004266 | 1.578129653 | 1.5 | 1.557204673 | 1.214495249 | 1.343584837 | 1.397940009 | 1.278288402 |
| ENSMUSG00000004267 | 2.095120908 | 1.894357207 | 1.988549078 | 1.678561379 | 1.773209395 | 1.826074803 | 1.710836105 |
| ENSMUSG00000004268 | 2.516991255 | 2.329353153 | 2.310197833 | 2.364857905 | 2.363759249 | 2.64246452 | 2.199599638 |
| ENSMUSG00000004270 | 2.310886388 | 2.158248799 | 2.183677405 | 2.236360959 | 2.145141048 | 2.515873844 | 2.075383 |
| ENSMUSG00000004285 | 2.339331437 | 2.233422664 | 2.285526764 | 2.017865591 | 2.150762767 | 2.303196057 | 2.108089156 |
| ENSMUSG00000004296 | 0 | 0 | 0 | 0.25 | 0 | 0 | 0 |
| ENSMUSG00000004317 | 2.708105693 | 2.483053688 | 2.464106382 | 2.528109877 | 2.448909998 | 2.827369273 | 2.281104046 |
| ENSMUSG00000004319 | 2.540273527 | 2.33993739 | 2.337182039 | 2.555094582 | 2.468875138 | 2.857935265 | 2.280823582 |
| ENSMUSG00000004328 | 0 | 0 | 0 | 0 | 0.244650542 | 0 | 0 |
| ENSMUSG00000004341 | 0 | 0 | 0 | 0 | 0 | 0 | 0 |
| ENSMUSG00000004344 | 0 | 0 | 0 | 0 | 0 | 0 | 0 |
| ENSMUSG00000004347 | 0.671187741 | 0.646240625 | 0.580482024 | 0.79248125 | 0.568060967 | 0.77815125 | 0.672731473 |
| ENSMUSG00000004356 | 2.99368599 | 2.69595419 | 2.702442032 | 2.750527944 | 2.674029951 | 3.080265627 | 2.467943337 |
| ENSMUSG00000004359 | 0.578129653 | 0.646240625 | 0.5 | 0.79248125 | 0.812711509 | 0.477121255 | 0.629532004 |
| ENSMUSG00000004360 | 1.249317394 | 1.080482024 | 1.26109853 | 0.5 | 0.632412477 | 0.301029996 | 0.355418052 |
| ENSMUSG00000004364 | 2.965050035 | 2.740724001 | 2.712351275 | 2.801142786 | 2.700341846 | 3.143639235 | 2.492917176 |
| ENSMUSG00000004366 | 0 | 0 | 0 | 0 | 0 | 0 | 0 |
| ENSMUSG00000004371 | 0.45815691 | 0.580482024 | 0.5 | 0.25 | 0.244650542 | 0 | 0.448487648 |
| ENSMUSG00000004383 | 2.371652933 | 2.232684334 | 2.210587586 | 2.069031101 | 2.053185007 | 2.374748346 | 1.909593098 |
| ENSMUSG00000004393 | 2.389637216 | 2.250703754 | 2.220660762 | 2.28743678 | 2.296256325 | 2.545307116 | 2.159731975 |
| ENSMUSG00000004394 | 2.419671301 | 2.267865591 | 2.216819685 | 2.277131114 | 2.273934005 | 2.519827994 | 2.168187745 |
| ENSMUSG00000004415 | 0 | 0.5 | 0.580482024 | 0.396240625 | 0.387761935 | 0.477121255 | 0 |

| | | | | | | | |
|---|---|---|---|---|---|---|---|
| ENSMUSG00000004446 | 2.678024491 | 2.460194481 | 2.489638179 | 2.334405475 | 2.331857641 | 2.665580991 | 2.210242554 |
| ENSMUSG00000004451 | 2.565000581 | 2.381380202 | 2.371960008 | 2.281030328 | 2.224547964 | 2.564666064 | 2.168187745 |
| ENSMUSG00000004455 | 3.154964843 | 2.875956434 | 2.851152349 | 2.900558725 | 2.811242896 | 3.27989498 | 2.644277977 |
| ENSMUSG00000004460 | 2.740271036 | 2.491806565 | 2.479715809 | 2.569031101 | 2.470489023 | 2.863917377 | 2.312345877 |
| ENSMUSG00000004462 | 2.048736215 | 1.973704441 | 1.930274797 | 1.870953944 | 1.820920161 | 1.995635195 | 1.71730663 |
| ENSMUSG00000004473 | 0.289064826 | 0 | 0 | 0.25 | 0.244650542 | 0.477121255 | 0.579661877 |
| ENSMUSG00000004500 | 1.753493018 | 1.598079356 | 1.638647213 | 1.695339928 | 1.639266209 | 1.698970004 | 1.586932193 |
| ENSMUSG00000004508 | 2 | 1.784887838 | 1.845926073 | 1.868933358 | 1.768534412 | 1.968482949 | 1.687112305 |
| ENSMUSG00000004530 | 3.116511873 | 2.879048119 | 2.88804719 | 2.5896628 | 2.618080427 | 2.932980822 | 2.487630378 |
| ENSMUSG00000004535 | 2.841427095 | 2.604726683 | 2.593759858 | 2.639605178 | 2.548733265 | 2.944975908 | 2.364643151 |
| ENSMUSG00000004540 | 0.289064826 | 0.5 | 0.396240625 | 0.396240625 | 0.568060967 | 0 | 0.629532004 |
| ENSMUSG00000004542 | 0.45815691 | 0 | 0 | 0.396240625 | 0.244650542 | 0.301029996 | 0 |
| ENSMUSG00000004552 | 0 | 0 | 0 | 0 | 0 | 0 | 0 |
| ENSMUSG00000004558 | 1.494443473 | 1.445339928 | 1.488549078 | 1.692046081 | 1.746695109 | 1.982271233 | 1.651852264 |
| ENSMUSG00000004561 | 1.655740331 | 1.571350555 | 1.571350555 | 1.606566189 | 1.695608647 | 1.857332496 | 1.683576582 |
| ENSMUSG00000004562 | 2.433305075 | 2.256976499 | 2.343759858 | 2.298073204 | 2.355381384 | 2.586587305 | 2.323355972 |
| ENSMUSG00000004565 | 2.490209605 | 2.294354884 | 2.368933358 | 2.04745614 | 2.156296347 | 2.283301229 | 2.077494038 |
| ENSMUSG00000004567 | 2.641922824 | 2.409608978 | 2.47027849 | 2.017865591 | 2.082404901 | 2.303196057 | 1.996640515 |
| ENSMUSG00000004568 | 2.800026593 | 2.570772088 | 2.609957721 | 2.32991803 | 2.344506394 | 2.62838893 | 2.236700872 |
| ENSMUSG00000004591 | 2.98662835 | 2.737638974 | 2.723893821 | 2.802363341 | 2.709785497 | 3.13289977 | 2.498977857 |
| ENSMUSG00000004609 | 0 | 0 | 0 | 0.580482024 | 0.489301084 | 0 | 0.355418052 |
| ENSMUSG00000004610 | 1.655740331 | 1.714495249 | 1.711372513 | 1.388647213 | 1.611014646 | 1.643452676 | 1.502532226 |
| ENSMUSG00000004612 | 0 | 0 | 0 | 0 | 0 | 0 | 0 |
| ENSMUSG00000004613 | 0 | 0 | 0 | 0 | 0 | 0 | 0 |
| ENSMUSG00000004626 | 2.151493169 | 1.992885888 | 2.074802005 | 1.958222504 | 2.037065488 | 2.190331698 | 1.935895857 |
| ENSMUSG00000004630 | 0 | 0.25 | 0 | 0 | 0 | 0 | 0 |
| ENSMUSG00000004631 | 2.447686928 | 2.237091808 | 2.240000483 | 2.142463902 | 2.074582839 | 2.413299764 | 1.997988498 |
| ENSMUSG00000004633 | 0.578129653 | 0.5 | 0.5 | 1.114857905 | 1.176121988 | 1.431363764 | 1.014380832 |
| ENSMUSG00000004637 | 1.445324132 | 1.5 | 1.494319981 | 1.678561379 | 1.632412477 | 1.698970004 | 1.526507301 |
| ENSMUSG00000004642 | 2.834495649 | 2.609438018 | 2.612287161 | 2.727098155 | 2.66169594 | 3.04453976 | 2.482857357 |
| ENSMUSG00000004651 | 0.671187741 | 0.79248125 | 0.580482024 | 0.580482024 | 0.77552387 | 0.602059991 | 0.74492186 |
| ENSMUSG00000004654 | 0.289064826 | 0 | 0.25 | 0 | 0 | 0 | 0 |
| ENSMUSG00000004655 | 2.278594844 | 2.167221246 | 2.17510993 | 1.396240625 | 1.445123986 | 1.505149978 | 1.502532226 |
| ENSMUSG00000004661 | 1.325351389 | 1.356566189 | 1.330482024 | 1.27186571 | 1.358933174 | 1.342422681 | 1.417653332 |

| | | | | | | | |
|---|---|---|---|---|---|---|---|
| ENSMUSG00000004665 | 3.295438978 | 3.073296058 | 3.10334087 | 2.714300868 | 2.779973806 | 3.090963077 | 2.685349273 |
| ENSMUSG00000004667 | 2.477243823 | 2.392938161 | 2.472945926 | 2.157339155 | 2.367228134 | 2.515873844 | 2.362688369 |
| ENSMUSG00000004668 | 0 | 0 | 0 | 0 | 0.244650542 | 0 | 0 |
| ENSMUSG00000004677 | 2.882006928 | 2.634061804 | 2.665444524 | 2.527131114 | 2.564131005 | 2.825426118 | 2.436400161 |
| ENSMUSG00000004698 | 0.960252568 | 0.75 | 0.79248125 | 0 | 0.387761935 | 0 | 0 |
| ENSMUSG00000004707 | 0 | 0 | 0 | 0.396240625 | 0.489301084 | 0.301029996 | 0.224243824 |
| ENSMUSG00000004709 | 0 | 0 | 0 | 0 | 0 | 0 | 0 |
| ENSMUSG00000004730 | 0 | 0 | 0 | 0 | 0 | 0 | 0 |
| ENSMUSG00000004748 | 0.811507563 | 0.896240625 | 0.951838731 | 0.92510993 | 0.905314583 | 0.84509804 | 0.853775828 |
| ENSMUSG00000004768 | 2.412042447 | 2.248588359 | 2.226722649 | 2.219129237 | 2.181274723 | 2.492760389 | 2.054044578 |
| ENSMUSG00000004771 | 2.517987749 | 2.356566189 | 2.327653195 | 2.496104615 | 2.415543455 | 2.779596491 | 2.24869599 |
| ENSMUSG00000004788 | 2.37447073 | 2.234159485 | 2.256976499 | 1.975216702 | 2.091002363 | 2.252853031 | 2.066254157 |
| ENSMUSG00000004789 | 3.018088033 | 2.751932029 | 2.762462137 | 2.672717752 | 2.625838017 | 3.008600172 | 2.497257788 |
| ENSMUSG00000004791 | 2.183765914 | 1.956637122 | 1.973704441 | 1.403677461 | 1.450958105 | 1.322219295 | 1.324583736 |
| ENSMUSG00000004798 | 2.410757293 | 2.199415381 | 2.174241882 | 2.388647213 | 2.32560195 | 2.625312451 | 2.133394659 |
| ENSMUSG00000004814 | 0 | 0 | 0 | 0 | 0 | 0 | 0 |
| ENSMUSG00000004815 | 1.881174527 | 1.794977273 | 1.83272922 | 1.602347734 | 1.810629166 | 1.716003344 | 1.707551656 |
| ENSMUSG00000004821 | 0 | 0 | 0 | 0 | 0 | 0 | 0 |
| ENSMUSG00000004837 | 0.289064826 | 0.396240625 | 0.25 | 0.25 | 0.931471446 | 0.301029996 | 0.710836105 |
| ENSMUSG00000004842 | 0.289064826 | 0.580482024 | 0.396240625 | 0.75 | 0.632412477 | 0.602059991 | 0.710836105 |
| ENSMUSG00000004843 | 2.451206209 | 2.326515422 | 2.32991803 | 2.359177886 | 2.263669614 | 2.586587305 | 2.098866035 |
| ENSMUSG00000004846 | 2.666334442 | 2.466819685 | 2.538388008 | 2.411864607 | 2.532902983 | 2.790285164 | 2.40946263 |
| ENSMUSG00000004849 | 2.710621733 | 2.53360658 | 2.590759908 | 2.580482024 | 2.579831265 | 2.933993164 | 2.473094717 |
| ENSMUSG00000004864 | 0.747221736 | 0.701838731 | 0.646240625 | 0.830482024 | 0.905314583 | 0.602059991 | 0.803905701 |
| ENSMUSG00000004865 | 2.892679599 | 2.635516136 | 2.646475362 | 2.627196291 | 2.577460832 | 2.936010796 | 2.398338893 |
| ENSMUSG00000004872 | 1.806056082 | 1.755591953 | 1.77186571 | 1.819031101 | 1.802174723 | 2.071882007 | 1.735976162 |
| ENSMUSG00000004880 | 2.572054151 | 2.389605178 | 2.399978211 | 2.45584181 | 2.464316601 | 2.776701184 | 2.3138695 |
| ENSMUSG00000004885 | 1.639698646 | 1.566696635 | 1.561981878 | 1.238549078 | 1.149965125 | 1.278753601 | 1.150210039 |
| ENSMUSG00000004891 | 3.398768344 | 3.101883333 | 3.099151195 | 3.183571558 | 3.149730181 | 3.608739919 | 2.933329884 |
| ENSMUSG00000004892 | 0 | 0 | 0.25 | 0.396240625 | 0.387761935 | 0 | 0.520678036 |
| ENSMUSG00000004894 | 0 | 0.25 | 0 | 0 | 0 | 0 | 0.224243824 |
| ENSMUSG00000004895 | 2.131864492 | 2.059601185 | 2.069031101 | 2.05357978 | 2.107593434 | 2.354108439 | 2.023210262 |
| ENSMUSG00000004896 | 1.816888701 | 1.729715809 | 1.67510993 | 1.668106335 | 1.800029087 | 1.857332496 | 1.704233519 |
| ENSMUSG00000004897 | 3.338636499 | 3.04419912 | 3.055018993 | 3.021451095 | 2.995662987 | 3.443888547 | 2.791991634 |

| | | | | | | | |
|---|---|---|---|---|---|---|---|
| ENSMUSG00000004902 | 0 | 0.396240625 | 0.5 | 0.396240625 | 0.568060967 | 0 | 0.448487648 |
| ENSMUSG00000004929 | 2.297593393 | 2.148114259 | 2.256976499 | 2.218361028 | 2.345884234 | 2.491361694 | 2.326294816 |
| ENSMUSG00000004931 | 1.847794126 | 1.866901388 | 1.998588359 | 1.584962501 | 1.870073561 | 1.886490725 | 2.068965954 |
| ENSMUSG00000004933 | 0.811507563 | 0.79248125 | 0.864857905 | 0.830482024 | 0.978602168 | 0.77815125 | 1.140832123 |
| ENSMUSG00000004934 | 2.063793172 | 2.005591953 | 1.995748394 | 1.769203899 | 1.903713872 | 2.008600172 | 1.884801065 |
| ENSMUSG00000004936 | 2.595842487 | 2.405512955 | 2.385757955 | 2.46759118 | 2.444777781 | 2.783903579 | 2.311836403 |
| ENSMUSG00000004937 | 2.553277818 | 2.439555804 | 2.534247778 | 2.334962501 | 2.488383918 | 2.604226053 | 2.456247769 |
| ENSMUSG00000004939 | 0.45815691 | 0.396240625 | 0 | 0.25 | 0.244650542 | 0.301029996 | 0.355418052 |
| ENSMUSG00000004945 | 2.299278375 | 2.160964047 | 2.174241882 | 2.147178659 | 2.187791022 | 2.451786436 | 2.120759255 |
| ENSMUSG00000004947 | 2.039433992 | 1.982684334 | 2.046218836 | 1.714495249 | 1.890639929 | 1.924279286 | 1.907820406 |
| ENSMUSG00000004948 | 0 | 0.25 | 0 | 0.25 | 0.387761935 | 0 | 0 |
| ENSMUSG00000004951 | 2.254965775 | 2.134789703 | 2.202642909 | 2.106566189 | 2.257246745 | 2.403120521 | 2.219530717 |
| ENSMUSG00000004952 | 2.167941649 | 2.043731421 | 2.154596376 | 1.866901388 | 1.958580386 | 2.017033339 | 1.960974982 |
| ENSMUSG00000004961 | 0.289064826 | 0 | 0 | 0.25 | 0.387761935 | 0 | 0.224243824 |
| ENSMUSG00000004980 | 3.747430566 | 3.39538842 | 3.393411797 | 3.445263072 | 3.34773985 | 3.906442794 | 3.106497253 |
| ENSMUSG00000004988 | 0.671187741 | 0.701838731 | 0.646240625 | 0.79248125 | 0.686820904 | 0.602059991 | 0.355418052 |
| ENSMUSG00000004994 | 1.605636262 | 1.657339155 | 1.642463902 | 1.52186571 | 1.555378544 | 1.681241237 | 1.537859885 |
| ENSMUSG00000004996 | 1.693336259 | 1.660964047 | 1.71759118 | 1.54248125 | 1.621875691 | 1.51851394 | 1.665294008 |
| ENSMUSG00000005034 | 2.721148722 | 2.50835575 | 2.487821179 | 2.546838018 | 2.46464417 | 2.862131379 | 2.319893236 |
| ENSMUSG00000005043 | 1.54867984 | 1.566696635 | 1.660964047 | 1.54745614 | 1.584291694 | 1.568201724 | 1.554175045 |
| ENSMUSG00000005045 | 1.269664473 | 1.201838731 | 1.160964047 | 0.830482024 | 1 | 0.954242509 | 0.935079929 |
| ENSMUSG00000005054 | 2.750654121 | 2.536164561 | 2.602877747 | 2.514836115 | 2.490824419 | 2.827369273 | 2.402025166 |
| ENSMUSG00000005057 | 1.289064826 | 1.356566189 | 1.51109853 | 1.188721876 | 1.351342427 | 1.230448921 | 1.425641774 |
| ENSMUSG00000005069 | 2.418409478 | 2.228969845 | 2.300530956 | 2.137686696 | 2.189936658 | 2.374748346 | 2.104721678 |
| ENSMUSG00000005078 | 2.051791411 | 1.972185812 | 1.958222504 | 1.882845365 | 1.795698282 | 2 | 1.672731473 |
| ENSMUSG00000005087 | 2.917842813 | 2.659608978 | 2.676624005 | 2.833568321 | 2.731807852 | 3.165244326 | 2.527425075 |
| ENSMUSG00000005089 | 0.45815691 | 0.396240625 | 0.396240625 | 0.5 | 0.686820904 | 0.698970004 | 0.448487648 |
| ENSMUSG00000005102 | 2.134369199 | 2 | 2.015173983 | 2.155512955 | 2.147561328 | 2.396199347 | 2.038401572 |
| ENSMUSG00000005103 | 3.562649727 | 3.239048657 | 3.229529463 | 2.964106382 | 2.94314048 | 3.355834496 | 2.736662594 |
| ENSMUSG00000005107 | 1.069666964 | 1.130890489 | 1.098079356 | 0.830482024 | 0.905314583 | 1.041392685 | 1.078019652 |
| ENSMUSG00000005124 | 3.336824221 | 3.034967766 | 2.990091002 | 2.923480622 | 2.850655544 | 3.287129621 | 2.656190065 |
| ENSMUSG00000005125 | 2.816402366 | 2.535845803 | 2.544043287 | 2.349685923 | 2.303049176 | 2.582063363 | 2.170168862 |
| ENSMUSG00000005131 | 0 | 0.25 | 0 | 0 | 0 | 0.477121255 | 0.224243824 |
| ENSMUSG00000005142 | 2.328498818 | 2.196158711 | 2.224461364 | 2.27053726 | 2.330902415 | 2.56937391 | 2.195242462 |

| | | | | | | | |
|---|---|---|---|---|---|---|---|
| ENSMUSG00000005148 | 1.325351389 | 1.388647213 | 1.388647213 | 1.54248125 | 1.546663136 | 1.62324929 | 1.493062805 |
| ENSMUSG00000005150 | 1.693336259 | 1.52186571 | 1.622963274 | 1.738549078 | 1.72341484 | 1.86332286 | 1.681794123 |
| ENSMUSG00000005161 | 2.995273477 | 2.74234663 | 2.780383379 | 2.756630861 | 2.810874786 | 3.200303183 | 2.663923178 |
| ENSMUSG00000005198 | 3.204803826 | 2.89151351 | 2.907452986 | 3.082449085 | 3.038292264 | 3.540454614 | 2.841117876 |
| ENSMUSG00000005202 | 0 | 0 | 0 | 0 | 0 | 0.301029996 | 0 |
| ENSMUSG00000005204 | 2.606442122 | 2.397646762 | 2.442046081 | 2.533285553 | 2.502486038 | 2.806179974 | 2.390438245 |
| ENSMUSG00000005220 | 0 | 0 | 0 | 0.25 | 0.387761935 | 0.301029996 | 0 |
| ENSMUSG00000005225 | 2.266038081 | 2.071350555 | 2.095926073 | 2.155512955 | 2.117361396 | 2.378397901 | 1.939139147 |
| ENSMUSG00000005232 | 0 | 0 | 0 | 0 | 0 | 0 | 0 |
| ENSMUSG00000005233 | 2.247417471 | 2.074802005 | 2.005591953 | 2.30539678 | 2.204601633 | 2.475671188 | 2.07959139 |
| ENSMUSG00000005237 | 0.671187741 | 0.580482024 | 0.396240625 | 0.25 | 0 | 0.602059991 | 0.579661877 |
| ENSMUSG00000005251 | 0 | 0 | 0.25 | 1.061981878 | 1.283908311 | 1.278753601 | 1.350478789 |
| ENSMUSG00000005262 | 2.469490009 | 2.25 | 2.278435542 | 2.397178659 | 2.367228134 | 2.67669361 | 2.19230473 |
| ENSMUSG00000005267 | 1.671187741 | 1.575945187 | 1.598079356 | 1.828220739 | 1.718026092 | 1.892094603 | 1.594042709 |
| ENSMUSG00000005268 | 0.289064826 | 0.25 | 0.580482024 | 0 | 0.244650542 | 0.301029996 | 0 |
| ENSMUSG00000005299 | 2.532656285 | 2.352347734 | 2.375459296 | 2.348615674 | 2.41479008 | 2.665580991 | 2.257861346 |
| ENSMUSG00000005312 | 2.941573724 | 2.704645544 | 2.689347502 | 2.815229883 | 2.787530338 | 3.176958981 | 2.61086058 |
| ENSMUSG00000005320 | 0 | 0 | 0 | 0.25 | 0.632412477 | 0.301029996 | 0.355418052 |
| ENSMUSG00000005338 | 0 | 0 | 0 | 0.75 | 0.733951626 | 0.602059991 | 0.74492186 |
| ENSMUSG00000005339 | 0 | 0 | 0 | 0 | 0 | 0 | 0 |
| ENSMUSG00000005354 | 2.746646916 | 2.565523711 | 2.591033664 | 2.437886015 | 2.520266629 | 2.818225894 | 2.399701885 |
| ENSMUSG00000005355 | 0 | 0.25 | 0.25 | 0.79248125 | 0 | 0.301029996 | 0 |
| ENSMUSG00000005357 | 0.578129653 | 0.580482024 | 0.25 | 1.02186571 | 0.877063019 | 0.84509804 | 1.014380832 |
| ENSMUSG00000005360 | 1.578129653 | 1.476722649 | 1.464495249 | 1 | 0.905314583 | 0.84509804 | 0.952571511 |
| ENSMUSG00000005362 | 2.156259305 | 2.02186571 | 1.953445298 | 2.009729747 | 1.882215755 | 2.178976947 | 1.796468236 |
| ENSMUSG00000005364 | 0.45815691 | 0 | 0.25 | 0.396240625 | 0 | 0.301029996 | 0.224243824 |
| ENSMUSG00000005370 | 2.880757703 | 2.611770807 | 2.592399337 | 2.654137211 | 2.533980713 | 2.925827575 | 2.380076739 |
| ENSMUSG00000005371 | 2.775534205 | 2.52053726 | 2.480460234 | 2.519203899 | 2.424827423 | 2.768638101 | 2.225910574 |
| ENSMUSG00000005373 | 0 | 0 | 0.25 | 0.75 | 0.733951626 | 0.301029996 | 0.829800754 |
| ENSMUSG00000005374 | 2.417143825 | 2.310793496 | 2.279735268 | 2.29248125 | 2.272243869 | 2.534026106 | 2.128938453 |
| ENSMUSG00000005378 | 2.20747957 | 2.114857905 | 2.154596376 | 2.039967834 | 2.195595395 | 2.326335861 | 2.121678334 |
| ENSMUSG00000005397 | 3.326355423 | 3.040912419 | 3.025739971 | 2.959111228 | 2.907281961 | 3.373279893 | 2.711948225 |
| ENSMUSG00000005410 | 3.020182925 | 2.810495787 | 2.807655233 | 2.798996825 | 2.860604721 | 3.215901813 | 2.695163117 |
| ENSMUSG00000005413 | 2.629921064 | 2.454645544 | 2.488912477 | 2.120953944 | 2.148364414 | 2.394451681 | 2.111895264 |

| | | | | | | | |
|---|---|---|---|---|---|---|---|
| ENSMUSG00000005417 | 3.313701682 | 3.042011317 | 3.049994752 | 3.120827989 | 3.124955431 | 3.562887381 | 2.940967543 |
| ENSMUSG00000005442 | 2.45469604 | 2.337182039 | 2.468361028 | 1.970660762 | 2.159419872 | 2.212187604 | 2.182568577 |
| ENSMUSG00000005447 | 2.003432385 | 1.948603967 | 1.958222504 | 1.630890489 | 1.903713872 | 1.924279286 | 1.826389311 |
| ENSMUSG00000005465 | 0.960252568 | 0.646240625 | 0.75 | 1 | 1.176121988 | 1.041392685 | 0.969165684 |
| ENSMUSG00000005469 | 2.865467634 | 2.622963274 | 2.624213444 | 2.735261902 | 2.706995316 | 3.121559844 | 2.545006225 |
| ENSMUSG00000005470 | 2.428398755 | 2.240724001 | 2.267865591 | 2.506630861 | 2.471132517 | 2.823474229 | 2.325562604 |
| ENSMUSG00000005474 | 0.289064826 | 0.5 | 0.396240625 | 0 | 0.244650542 | 0 | 0.224243824 |
| ENSMUSG00000005481 | 2.917842813 | 2.68325383 | 2.69390257 | 2.636241108 | 2.667640605 | 3.006037955 | 2.480746811 |
| ENSMUSG00000005483 | 2.388281011 | 2.220660762 | 2.290597832 | 2.324229052 | 2.305113253 | 2.653212514 | 2.187473278 |
| ENSMUSG00000005493 | 0.747221736 | 0.580482024 | 0.79248125 | 0.701838731 | 0.489301084 | 0 | 0.520678036 |
| ENSMUSG00000005501 | 2.427163103 | 2.266522298 | 2.23927551 | 2.383818844 | 2.299407739 | 2.597695186 | 2.130281771 |
| ENSMUSG00000005503 | 0 | 0 | 0 | 0 | 0 | 0 | 0 |
| ENSMUSG00000005505 | 1.631440309 | 1.52186571 | 1.571350555 | 1.834962501 | 1.756568651 | 2.029383778 | 1.649885598 |
| ENSMUSG00000005506 | 2.926900886 | 2.684311836 | 2.691010554 | 2.910625757 | 2.82753976 | 3.264581729 | 2.629532004 |
| ENSMUSG00000005510 | 2.023453784 | 1.858657057 | 1.921625132 | 2.277131114 | 2.175379703 | 2.496929648 | 2.070582188 |
| ENSMUSG00000005514 | 2.749513129 | 2.589388001 | 2.657111385 | 2.259043403 | 2.385720541 | 2.59439255 | 2.364859621 |
| ENSMUSG00000005533 | 2.160971586 | 2.105516192 | 2.056000419 | 1.799918086 | 1.906908067 | 1.995635195 | 1.786277931 |
| ENSMUSG00000005534 | 2.65645504 | 2.414998973 | 2.422499493 | 2.206637122 | 2.164053936 | 2.404833717 | 1.984950056 |
| ENSMUSG00000005540 | 0 | 0.25 | 0.396240625 | 0.25 | 0.244650542 | 0.301029996 | 0.355418052 |
| ENSMUSG00000005547 | 0 | 0 | 0 | 0 | 0 | 0 | 0 |
| ENSMUSG00000005553 | 0 | 0.5 | 0.25 | 0.25 | 0.244650542 | 0 | 0.355418052 |
| ENSMUSG00000005566 | 3.124026174 | 2.842808032 | 2.844167015 | 2.733791262 | 2.795954512 | 3.142389466 | 2.610050272 |
| ENSMUSG00000005575 | 2.448863324 | 2.255591953 | 2.277783918 | 2.207430684 | 2.21338377 | 2.541579244 | 2.075912052 |
| ENSMUSG00000005580 | 1.404271433 | 1.396240625 | 1.476722649 | 1.330482024 | 1.499532413 | 1.51851394 | 1.38802993 |
| ENSMUSG00000005583 | 1.655740331 | 1.54745614 | 1.557204673 | 1.532320754 | 1.439191811 | 1.51851394 | 1.313616061 |
| ENSMUSG00000005609 | 2.574668783 | 2.370953944 | 2.3464656 | 2.425975893 | 2.375754058 | 2.723455672 | 2.23605706 |
| ENSMUSG00000005610 | 3.727383496 | 3.357972326 | 3.328327054 | 3.406313055 | 3.3042648 | 3.859198362 | 3.037861479 |
| ENSMUSG00000005611 | 0.45815691 | 0.646240625 | 0.701838731 | 0.25 | 0 | 0.301029996 | 0.355418052 |
| ENSMUSG00000005615 | 2.585813169 | 2.437886015 | 2.417664062 | 2.519537702 | 2.435655147 | 2.783903579 | 2.285558718 |
| ENSMUSG00000005621 | 2.249317394 | 2.066696635 | 2.04745614 | 2.02053726 | 1.992595054 | 2.225309282 | 1.89141329 |
| ENSMUSG00000005625 | 2.764107946 | 2.577369088 | 2.571061437 | 2.573367912 | 2.626252528 | 2.90579588 | 2.485103482 |
| ENSMUSG00000005628 | 0.867194479 | 1.02186571 | 0.5 | 0.646240625 | 0.686820904 | 0.602059991 | 0.74492186 |
| ENSMUSG00000005640 | 0 | 0 | 0 | 0 | 0.387761935 | 0 | 0 |
| ENSMUSG00000005649 | 0 | 0 | 0 | 0 | 0 | 0.301029996 | 0 |

| | | | | | | | |
|---|---|---|---|---|---|---|---|
| ENSMUSG00000005656 | 2.81882839 | 2.58993739 | 2.58272922 | 2.668327269 | 2.550277934 | 2.959518377 | 2.38742504 |
| ENSMUSG00000005667 | 2.770132156 | 2.518535366 | 2.463717096 | 2.49929487 | 2.441646558 | 2.818225894 | 2.30669705 |
| ENSMUSG00000005672 | 1.631440309 | 1.527131114 | 1.527131114 | 0.396240625 | 0 | 0.301029996 | 0.224243824 |
| ENSMUSG00000005674 | 1.269664473 | 1.282320754 | 1.356566189 | 1.214495249 | 1.212046813 | 1.204119983 | 1.345462945 |
| ENSMUSG00000005677 | 0.289064826 | 0 | 0 | 0.25 | 0.244650542 | 0 | 0 |
| ENSMUSG00000005681 | 0 | 0 | 0 | 0.25 | 0.387761935 | 0.301029996 | 0.355418052 |
| ENSMUSG00000005682 | 2.402962068 | 2.211372513 | 2.222185812 | 2.29745614 | 2.318243321 | 2.589949601 | 2.157274883 |
| ENSMUSG00000005683 | 3.280518133 | 2.977379529 | 2.984527332 | 3.027294426 | 2.983754904 | 3.471878199 | 2.780221858 |
| ENSMUSG00000005686 | 1.876566388 | 1.845926073 | 1.828220739 | 1.561981878 | 1.642643806 | 1.77815125 | 1.556816002 |
| ENSMUSG00000005687 | 2.650702778 | 2.422499493 | 2.456637122 | 2.497525991 | 2.432082686 | 2.792391689 | 2.272582514 |
| ENSMUSG00000005696 | 0 | 0 | 0 | 0 | 0 | 0 | 0.355418052 |
| ENSMUSG00000005698 | 2.744339677 | 2.489638179 | 2.485994979 | 2.678561379 | 2.605346297 | 2.996949248 | 2.431330105 |
| ENSMUSG00000005699 | 1.482695304 | 1.494319981 | 1.53743678 | 1.188721876 | 1.588235379 | 1.361727836 | 1.511732338 |
| ENSMUSG00000005705 | 0.671187741 | 0.580482024 | 0.646240625 | 0.580482024 | 0.568060967 | 0.477121255 | 0.355418052 |
| ENSMUSG00000005716 | 0 | 0 | 0 | 0 | 0 | 0 | 0 |
| ENSMUSG00000005718 | 1.45815691 | 1.54248125 | 1.598079356 | 1.339388001 | 1.414412788 | 1.230448921 | 1.38802993 |
| ENSMUSG00000005732 | 2.886555665 | 2.671625132 | 2.669209652 | 2.653907409 | 2.630984742 | 3.010723865 | 2.531846175 |
| ENSMUSG00000005763 | 0.289064826 | 0.396240625 | 0.25 | 0 | 0 | 0 | 0 |
| ENSMUSG00000005774 | 1.907802659 | 1.714495249 | 1.774508021 | 1.909156155 | 1.964031554 | 2.075546961 | 1.898808251 |
| ENSMUSG00000005779 | 2.848245215 | 2.620953944 | 2.624712302 | 2.773519421 | 2.728257421 | 3.151982395 | 2.574722613 |
| ENSMUSG00000005800 | 0 | 0 | 0 | 0 | 0 | 0 | 0 |
| ENSMUSG00000005802 | 2.722369902 | 2.526477127 | 2.519871196 | 2.637205477 | 2.515479755 | 2.92788341 | 2.350026003 |
| ENSMUSG00000005803 | 1.693336259 | 1.54745614 | 1.494319981 | 1.71759118 | 1.674752852 | 1.919078092 | 1.62316756 |
| ENSMUSG00000005804 | 2.249317394 | 2.110735874 | 2.129909063 | 2.124961472 | 2.027028145 | 2.278753601 | 1.901538355 |
| ENSMUSG00000005813 | 2.587501561 | 2.386723615 | 2.342126615 | 2.545288064 | 2.455691304 | 2.831869774 | 2.308505126 |
| ENSMUSG00000005823 | 2.153883046 | 1.981203126 | 1.959800947 | 1.921625132 | 1.982473755 | 2.195899652 | 1.913109605 |
| ENSMUSG00000005824 | 0 | 0 | 0 | 0 | 0 | 0 | 0 |
| ENSMUSG00000005836 | 1.227926429 | 1.146240625 | 1.188721876 | 1.732684334 | 1.645989387 | 1.812913357 | 1.540636858 |
| ENSMUSG00000005846 | 2.91246663 | 2.656427211 | 2.662762923 | 2.744677615 | 2.647747821 | 3.053846427 | 2.460944696 |
| ENSMUSG00000005864 | 0 | 0 | 0 | 0 | 0.244650542 | 0 | 0 |
| ENSMUSG00000005871 | 2.888606985 | 2.656655413 | 2.642938161 | 2.736178015 | 2.65373846 | 3.038222638 | 2.448988576 |
| ENSMUSG00000005873 | 2.88366676 | 2.650906586 | 2.659382638 | 2.758699741 | 2.712715962 | 3.136720567 | 2.52716312 |
| ENSMUSG00000005881 | 2.446507203 | 2.280383379 | 2.32991803 | 2.256976499 | 2.339407793 | 2.592176757 | 2.210591357 |
| ENSMUSG00000005882 | 2.054824387 | 1.899978211 | 1.931980114 | 2.140560606 | 2.130260132 | 2.418301291 | 1.977870661 |

| | | | | | | | |
|---|---|---|---|---|---|---|---|
| ENSMUSG00000005883 | 0 | 0 | 0 | 0 | 0.489301084 | 0.301029996 | 0.355418052 |
| ENSMUSG00000005886 | 2.439357901 | 2.224461364 | 2.192872367 | 2.180274797 | 2.163285805 | 2.460897843 | 2.05853792 |
| ENSMUSG00000005891 | 0 | 0 | 0 | 0 | 0 | 0 | 0 |
| ENSMUSG00000005892 | 0 | 0 | 0 | 0 | 0 | 0 | 0 |
| ENSMUSG00000005893 | 2.457006475 | 2.28360658 | 2.243603647 | 2.231203126 | 2.126012715 | 2.499687083 | 2.031805106 |
| ENSMUSG00000005897 | 2.054824387 | 1.903677461 | 1.880890489 | 1.882845365 | 1.820920161 | 2.008600172 | 1.720494 |
| ENSMUSG00000005899 | 2.488076442 | 2.317865757 | 2.342126615 | 2.325945187 | 2.345425551 | 2.579783597 | 2.231187192 |
| ENSMUSG00000005907 | 2.045658471 | 1.919870025 | 1.896240625 | 1.899978211 | 1.925488995 | 2.029383778 | 1.743447983 |
| ENSMUSG00000005917 | 1.505869732 | 1.532320754 | 1.494319981 | 1.5 | 1.649303553 | 1.69019608 | 1.505628092 |
| ENSMUSG00000005936 | 2.773381775 | 2.530706999 | 2.517865591 | 2.614601653 | 2.565117605 | 2.952792443 | 2.372346263 |
| ENSMUSG00000005947 | 0.811507563 | 0.830482024 | 1.061981878 | 0.92510993 | 1.091002363 | 1.176091259 | 0.829800754 |
| ENSMUSG00000005949 | 2.247417471 | 2.100219859 | 2.095926073 | 1.886723615 | 1.802174723 | 2 | 1.735976162 |
| ENSMUSG00000005950 | 1.249317394 | 1.356566189 | 1.238549078 | 1.54745614 | 1.595992969 | 1.602059991 | 1.520678036 |
| ENSMUSG00000005951 | 0.960252568 | 0.896240625 | 1.04248125 | 1.238549078 | 1.264824954 | 1 | 0.984950056 |
| ENSMUSG00000005952 | 0 | 0 | 0 | 0 | 0 | 0 | 0 |
| ENSMUSG00000005958 | 2.419671301 | 2.256284891 | 2.256284891 | 1.779735268 | 1.92245931 | 2.041392685 | 1.906930404 |
| ENSMUSG00000005968 | 2.507914013 | 2.368426437 | 2.347004321 | 2.263820609 | 2.27895639 | 2.487138375 | 2.097385413 |
| ENSMUSG00000005973 | 2.713122685 | 2.503853763 | 2.480460234 | 2.631380202 | 2.547699718 | 2.942008053 | 2.377797722 |
| ENSMUSG00000005980 | 1.342375483 | 1.214495249 | 1.27186571 | 1.364857905 | 1.394615667 | 1.505149978 | 1.150210039 |
| ENSMUSG00000005981 | 2.675303226 | 2.437886015 | 2.450629091 | 2.431128463 | 2.412520257 | 2.760422483 | 2.277722321 |
| ENSMUSG00000005982 | 2.134369199 | 2.013820609 | 2.069031101 | 1.909156155 | 1.91476933 | 2.045322979 | 1.813563596 |
| ENSMUSG00000005983 | 1.227926429 | 0.92510993 | 1.130890489 | 0.830482024 | 1 | 1 | 1.20139795 |
| ENSMUSG00000005986 | 1.181541736 | 1.470660762 | 1.445339928 | 1.488549078 | 1.621875691 | 1.556302501 | 1.598697688 |
| ENSMUSG00000005994 | 0 | 0 | 0 | 0 | 0 | 0 | 0 |
| ENSMUSG00000006005 | 3.271302466 | 2.995480997 | 2.989909941 | 3.103869427 | 2.983980507 | 3.46746011 | 2.77514795 |
| ENSMUSG00000006007 | 0 | 0 | 0 | 0 | 0 | 0 | 0 |
| ENSMUSG00000006010 | 2.271465912 | 2.127938163 | 2.061981878 | 2.213717096 | 2.10031573 | 2.434568904 | 1.963977467 |
| ENSMUSG00000006014 | 0.45815691 | 0.75 | 0.701838731 | 0.896240625 | 0.846351821 | 1.113943352 | 0.829800754 |
| ENSMUSG00000006019 | 1.482695304 | 1.476722649 | 1.598079356 | 1.130890489 | 1.366364104 | 1.322219295 | 1.473249546 |
| ENSMUSG00000006021 | 1.647796617 | 1.482684334 | 1.602347734 | 1.575945187 | 1.580303447 | 1.799340549 | 1.594042709 |
| ENSMUSG00000006024 | 2.508932407 | 2.317865757 | 2.357090043 | 2.198603967 | 2.252501045 | 2.527629901 | 2.14512854 |
| ENSMUSG00000006050 | 2.389637216 | 2.175975893 | 2.196158711 | 2.154596376 | 2.153935427 | 2.414973348 | 2.052911423 |
| ENSMUSG00000006056 | 0.289064826 | 0.5 | 0.25 | 0.75 | 0.77552387 | 0.602059991 | 0.448487648 |
| ENSMUSG00000006057 | 2.4661223 | 2.345385738 | 2.327653195 | 2.201032755 | 2.327538614 | 2.552668216 | 2.197791274 |

| | | | | | | | |
|---|---|---|---|---|---|---|---|
| ENSMUSG00000006058 | 2.547754129 | 2.321350555 | 2.370953944 | 2.445749552 | 2.397795474 | 2.73239376 | 2.227237819 |
| ENSMUSG00000006095 | 2.163307908 | 2.00835575 | 2.111770807 | 1.981203126 | 2.039257769 | 2.257678575 | 2.022587518 |
| ENSMUSG00000006127 | 2.003432385 | 1.964495249 | 1.880890489 | 1.862802778 | 1.870073561 | 2.053078443 | 1.826389311 |
| ENSMUSG00000006134 | 2.569423023 | 2.341033664 | 2.345385738 | 2.381869252 | 2.354487824 | 2.675778342 | 2.207438502 |
| ENSMUSG00000006143 | 0.289064826 | 0.396240625 | 0.701838731 | 0.25 | 0.387761935 | 0 | 0.448487648 |
| ENSMUSG00000006154 | 0 | 0 | 0 | 0 | 0 | 0 | 0 |
| ENSMUSG00000006169 | 2.795957952 | 2.587182039 | 2.559601185 | 2.607090043 | 2.589846528 | 2.928395852 | 2.43361272 |
| ENSMUSG00000006179 | 0 | 0.646240625 | 0.646240625 | 1.566696635 | 1.47337555 | 1.612783857 | 1.448487648 |
| ENSMUSG00000006191 | 1.881174527 | 1.78743678 | 1.782320754 | 1.882845365 | 1.800029087 | 2.033423755 | 1.687112305 |
| ENSMUSG00000006200 | 0 | 0 | 0 | 0 | 0 | 0 | 0 |
| ENSMUSG00000006204 | 0 | 0 | 0 | 0 | 0.244650542 | 0.301029996 | 0.224243824 |
| ENSMUSG00000006205 | 2.45815691 | 2.237091808 | 2.222945926 | 2.377938163 | 2.331857641 | 2.672097858 | 2.170563634 |
| ENSMUSG00000006215 | 1.827447047 | 1.907339155 | 1.956637122 | 1.634789703 | 1.880506521 | 1.832508913 | 1.908707966 |
| ENSMUSG00000006216 | 0.289064826 | 0.396240625 | 0.396240625 | 0.25 | 0 | 0 | 0 |
| ENSMUSG00000006218 | 0.867194479 | 0.701838731 | 1.061981878 | 0.75 | 1.264824954 | 0.84509804 | 1.252393349 |
| ENSMUSG00000006219 | 2.773381775 | 2.590485943 | 2.575374049 | 2.144357207 | 2.169384551 | 2.459392488 | 2.047762527 |
| ENSMUSG00000006221 | 0.811507563 | 0.976722649 | 0.976722649 | 0 | 0 | 0.301029996 | 0.448487648 |
| ENSMUSG00000006235 | 0.91631382 | 0.896240625 | 1.061981878 | 0.830482024 | 0.978602168 | 0.903089987 | 1.110947927 |
| ENSMUSG00000006241 | 1 | 1.098079356 | 1.130890489 | 1.098079356 | 1.106691885 | 1.204119983 | 1.140832123 |
| ENSMUSG00000006262 | 2.945519866 | 2.695749552 | 2.691839214 | 2.654596376 | 2.576030893 | 2.943988875 | 2.366801367 |
| ENSMUSG00000006269 | 0 | 0 | 0 | 0 | 0.489301084 | 0 | 0.224243824 |
| ENSMUSG00000006270 | 0 | 0 | 0 | 0 | 0 | 0 | 0 |
| ENSMUSG00000006273 | 2.847342547 | 2.618933358 | 2.616136438 | 2.510414788 | 2.465625059 | 2.809559715 | 2.309019868 |
| ENSMUSG00000006276 | 2.433305075 | 2.259043403 | 2.254897682 | 2.28296424 | 2.278401864 | 2.57863921 | 2.174095069 |
| ENSMUSG00000006281 | 2.292497211 | 2.08272922 | 2.101285366 | 2.097004321 | 2.142704057 | 2.369215857 | 2.013099581 |
| ENSMUSG00000006288 | 1.806056082 | 1.741446071 | 1.744319981 | 1.930274797 | 1.970729215 | 2.127104798 | 1.837623616 |
| ENSMUSG00000006289 | 1.903481042 | 1.76109853 | 1.735628626 | 1.816696635 | 1.850106709 | 2.086359831 | 1.704233519 |
| ENSMUSG00000006299 | 2.664237041 | 2.504897682 | 2.536483036 | 2.395300145 | 2.562645913 | 2.830588669 | 2.453297759 |
| ENSMUSG00000006301 | 2.52293481 | 2.267194569 | 2.314935816 | 2.291226732 | 2.296783518 | 2.59439255 | 2.138226615 |
| ENSMUSG00000006304 | 3.126809492 | 2.8464656 | 2.866136438 | 2.88040011 | 2.798633751 | 3.266231697 | 2.612776967 |
| ENSMUSG00000006307 | 2.396352858 | 2.203445298 | 2.271202097 | 2.024508021 | 2.149165677 | 2.255272505 | 2.111421942 |
| ENSMUSG00000006310 | 0.45815691 | 0 | 0 | 0 | 0.489301084 | 0.301029996 | 0.224243824 |
| ENSMUSG00000006311 | 0.289064826 | 0.25 | 0 | 0 | 0.244650542 | 0 | 0.224243824 |
| ENSMUSG00000006313 | 0.91631382 | 0.75 | 0.830482024 | 0.396240625 | 0.489301084 | 0 | 0.672731473 |

| | | | | | | | |
|---|---|---|---|---|---|---|---|
| ENSMUSG00000006315 | 2.033115198 | 1.890560606 | 1.950224975 | 1.997171172 | 1.987570683 | 2.235528447 | 1.843097781 |
| ENSMUSG00000006333 | 3.186579638 | 3.012121718 | 3.053959077 | 2.779410775 | 2.940003825 | 3.2509077 | 2.83505382 |
| ENSMUSG00000006335 | 1.404271433 | 1.52186571 | 1.580482024 | 1.505591953 | 1.576269619 | 1.69019608 | 1.551512352 |
| ENSMUSG00000006342 | 0.91631382 | 0.92510993 | 1.061981878 | 0.79248125 | 0.955822902 | 0.602059991 | 1.078019652 |
| ENSMUSG00000006344 | 0 | 0.25 | 0 | 0 | 0 | 0 | 0 |
| ENSMUSG00000006345 | 0.45815691 | 0.5 | 0 | 0 | 0.244650542 | 0 | 0.224243824 |
| ENSMUSG00000006356 | 1.777675583 | 1.850219859 | 1.938721876 | 1.330482024 | 1.599808748 | 1.505149978 | 1.655750087 |
| ENSMUSG00000006360 | 1.734388958 | 1.688721876 | 1.71759118 | 1.364857905 | 1.478764298 | 1.568201724 | 1.483307832 |
| ENSMUSG00000006362 | 0.45815691 | 0.580482024 | 0.5 | 0.75 | 0.77552387 | 0.954242509 | 0.829800754 |
| ENSMUSG00000006369 | 2.331622671 | 2.27186571 | 2.289336734 | 2.125956434 | 2.26251048 | 2.374748346 | 2.171351735 |
| ENSMUSG00000006373 | 2.864167792 | 2.62892496 | 2.641037372 | 2.641989019 | 2.537460779 | 2.908485019 | 2.376130113 |
| ENSMUSG00000006378 | 2.20747957 | 2.131869252 | 2.201032755 | 1.984159485 | 2.179810173 | 2.320146286 | 2.139097452 |
| ENSMUSG00000006386 | 0.578129653 | 0.580482024 | 0.25 | 0 | 0 | 0 | 0.224243824 |
| ENSMUSG00000006389 | 0 | 0 | 0 | 0 | 0 | 0 | 0 |
| ENSMUSG00000006390 | 2.478339835 | 2.30357978 | 2.319612365 | 2.259043403 | 2.27449559 | 2.511883361 | 2.122594809 |
| ENSMUSG00000006392 | 2.340856238 | 2.180274797 | 2.17510993 | 2.209012589 | 2.174635855 | 2.419955748 | 2.02568932 |
| ENSMUSG00000006395 | 1.289064826 | 1.388647213 | 1.388647213 | 1.321350555 | 1.327538614 | 1.301029996 | 1.20919388 |
| ENSMUSG00000006398 | 2.683414322 | 2.471805154 | 2.516522298 | 2.469895812 | 2.459366223 | 2.778874472 | 2.334954482 |
| ENSMUSG00000006403 | 1.249317394 | 1.26109853 | 1.226722649 | 1.5 | 1.542223377 | 1.602059991 | 1.56716938 |
| ENSMUSG00000006411 | 0 | 0.25 | 0.396240625 | 0 | 0.244650542 | 0.698970004 | 0.224243824 |
| ENSMUSG00000006412 | 2.273259602 | 2.104463129 | 2.103406982 | 2.272528105 | 2.221300875 | 2.517195898 | 2.120759255 |
| ENSMUSG00000006418 | 2.179305632 | 2.009729747 | 2.059601185 | 2.21293726 | 2.204601633 | 2.484299839 | 2.06842541 |
| ENSMUSG00000006423 | 2.696591798 | 2.473704441 | 2.49539182 | 2.55479213 | 2.420401583 | 2.833147112 | 2.282502732 |
| ENSMUSG00000006435 | 1.924654746 | 1.880890489 | 1.930274797 | 1.630890489 | 1.728722551 | 1.716003344 | 1.702561601 |
| ENSMUSG00000006442 | 2.598312578 | 2.442459461 | 2.457430684 | 2.362802778 | 2.449594021 | 2.73479983 | 2.341765586 |
| ENSMUSG00000006445 | 2.544030601 | 2.359177886 | 2.425543171 | 2.366391601 | 2.498344007 | 2.717670503 | 2.411529829 |
| ENSMUSG00000006456 | 2.358731791 | 2.21759118 | 2.286801231 | 2.119945066 | 2.234742351 | 2.399673721 | 2.142987464 |
| ENSMUSG00000006457 | 1.445324132 | 1.53743678 | 1.52186571 | 0.896240625 | 0.77552387 | 0 | 0.984950056 |
| ENSMUSG00000006462 | 0.289064826 | 0.5 | 0.5 | 0 | 0.489301084 | 0.301029996 | 0.355418052 |
| ENSMUSG00000006463 | 1.777675583 | 1.701838731 | 1.79745614 | 1.735628626 | 1.741652793 | 1.929418926 | 1.692344475 |
| ENSMUSG00000006464 | 1.789260561 | 1.87892496 | 1.807204673 | 1.779735268 | 1.940259189 | 2.004321374 | 1.848480858 |
| ENSMUSG00000006469 | 0 | 0 | 0 | 0 | 0 | 0 | 0 |
| ENSMUSG00000006471 | 2.126809492 | 1.903677461 | 1.976722649 | 1.79745614 | 1.932951348 | 1.991226076 | 1.802677937 |
| ENSMUSG00000006476 | 2.282114126 | 2.114857905 | 2.214495249 | 2.145300145 | 2.254288168 | 2.413299764 | 2.192673409 |

| | | | | | | | |
|---|---|---|---|---|---|---|---|
| ENSMUSG00000006488 | 0.289064826 | 0.25 | 0.396240625 | 0 | 0 | 0.602059991 | 0.355418052 |
| ENSMUSG00000006490 | 0 | 0 | 0 | 0 | 0 | 0 | 0 |
| ENSMUSG00000006494 | 1.516991255 | 1.527131114 | 1.614857905 | 1.811981878 | 1.761403742 | 2.045322979 | 1.681794123 |
| ENSMUSG00000006498 | 3.096614757 | 2.886844141 | 2.911864607 | 2.689972046 | 2.775659544 | 3.09131516 | 2.695474789 |
| ENSMUSG00000006517 | 2.036286563 | 2.005591953 | 2.059601185 | 1.905512955 | 2.081436571 | 2.133538908 | 2.073258096 |
| ENSMUSG00000006519 | 1.156259305 | 1.098079356 | 1.098079356 | 1.985628626 | 2.265976525 | 2.334453751 | 2.242438242 |
| ENSMUSG00000006522 | 0.45815691 | 0.580482024 | 0.396240625 | 0 | 0.244650542 | 0.301029996 | 0 |
| ENSMUSG00000006526 | 1.936861443 | 1.880890489 | 1.890560606 | 1.991446071 | 2.029283463 | 2.23299611 | 1.96172822 |
| ENSMUSG00000006527 | 2.072573125 | 1.964495249 | 1.912762923 | 2.166333979 | 2.066583491 | 2.378397901 | 1.980010793 |
| ENSMUSG00000006529 | 0 | 0 | 0 | 0 | 0 | 0 | 0 |
| ENSMUSG00000006538 | 0 | 0 | 0 | 0 | 0.387761935 | 0 | 0 |
| ENSMUSG00000006542 | 1.036286563 | 0.701838731 | 0.92510993 | 1.04248125 | 1.212046813 | 1.146128036 | 1.054044578 |
| ENSMUSG00000006546 | 0 | 0 | 0 | 0 | 0 | 0 | 0 |
| ENSMUSG00000006567 | 0.289064826 | 0.25 | 0 | 0.396240625 | 0.568060967 | 0 | 0.803905701 |
| ENSMUSG00000006570 | 0 | 0 | 0 | 0 | 0 | 0 | 0 |
| ENSMUSG00000006574 | 0 | 0 | 0 | 0 | 0 | 0 | 0 |
| ENSMUSG00000006575 | 1.181541736 | 1.27186571 | 1.330482024 | 1.130890489 | 1.244650542 | 1.380211242 | 1.284450788 |
| ENSMUSG00000006576 | 1.54867984 | 1.527131114 | 1.664552871 | 1.364857905 | 1.680838453 | 1.204119983 | 1.692344475 |
| ENSMUSG00000006585 | 2.318984359 | 2.200224975 | 2.37246199 | 2.116901388 | 2.199086582 | 2.348304863 | 2.241805758 |
| ENSMUSG00000006586 | 1.156259305 | 1.098079356 | 1.04248125 | 1.061981878 | 1.057362051 | 1.204119983 | 0.853775828 |
| ENSMUSG00000006587 | 0 | 0.25 | 0 | 0.580482024 | 0.568060967 | 0.77815125 | 0.710836105 |
| ENSMUSG00000006589 | 2.575536697 | 2.43155454 | 2.54652856 | 2.254202072 | 2.522774815 | 2.675778342 | 2.442244914 |
| ENSMUSG00000006599 | 2.479432974 | 2.349685923 | 2.340485943 | 2.201032755 | 2.121713561 | 2.413299764 | 1.996640515 |
| ENSMUSG00000006611 | 0.578129653 | 0.830482024 | 0.830482024 | 1.130890489 | 1.163285805 | 0.954242509 | 1.20139795 |
| ENSMUSG00000006638 | 0.289064826 | 0.25 | 0.396240625 | 0.396240625 | 0.568060967 | 0.301029996 | 0.224243824 |
| ENSMUSG00000006641 | 1.971473847 | 1.848079356 | 1.903677461 | 1.799918086 | 1.791313678 | 1.903089987 | 1.672731473 |
| ENSMUSG00000006642 | 0.289064826 | 0 | 0 | 0 | 0 | 0 | 0 |
| ENSMUSG00000006649 | 0.289064826 | 0.396240625 | 0 | 0.396240625 | 0.244650542 | 0.301029996 | 0 |
| ENSMUSG00000006651 | 1.494443473 | 1.557204673 | 1.589388001 | 1.532320754 | 1.706995316 | 1.785329835 | 1.764880683 |
| ENSMUSG00000006673 | 2.762449763 | 2.507321807 | 2.514159429 | 2.573080408 | 2.524160589 | 2.923244019 | 2.364426535 |
| ENSMUSG00000006675 | 1.205378646 | 1.25 | 1.146240625 | 0.951838731 | 1.188507684 | 1.278753601 | 1.345462945 |
| ENSMUSG00000006676 | 2.693336259 | 2.51348147 | 2.552363341 | 2.399513125 | 2.531280194 | 2.752816431 | 2.450984564 |
| ENSMUSG00000006678 | 2.842342644 | 2.604990044 | 2.590759908 | 2.69961795 | 2.599585429 | 3.01494035 | 2.417653332 |
| ENSMUSG00000006699 | 3.345944577 | 3.029248419 | 2.999118372 | 3.12750559 | 3.024892246 | 3.548389418 | 2.806844502 |

| | | | | | | | |
|---|---|---|---|---|---|---|---|
| ENSMUSG00000006705 | 2.20747957 | 2.02186571 | 2.01109853 | 2.091580554 | 2.032639648 | 2.309630167 | 1.958704689 |
| ENSMUSG00000006711 | 0.45815691 | 0.5 | 0.75 | 0.75 | 0.632412477 | 0.77815125 | 0.579661877 |
| ENSMUSG00000006715 | 2.34539747 | 2.222945926 | 2.180274797 | 2.410964047 | 2.366364104 | 2.713490543 | 2.215778782 |
| ENSMUSG00000006717 | 2.051791411 | 1.96293726 | 2.044977273 | 1.850219859 | 1.850106709 | 2.100370545 | 1.778418869 |
| ENSMUSG00000006720 | 1.647796617 | 1.630890489 | 1.561981878 | 1.774508021 | 1.674752852 | 1.799340549 | 1.645916034 |
| ENSMUSG00000006724 | 0.45815691 | 0.580482024 | 0.396240625 | 0.580482024 | 0.812711509 | 0.602059991 | 0.448487648 |
| ENSMUSG00000006728 | 3.389144561 | 3.102679083 | 3.117283255 | 2.956934912 | 2.969634776 | 3.413969972 | 2.800321005 |
| ENSMUSG00000006731 | 1.978790393 | 1.903677461 | 1.856566189 | 2.034887838 | 2.11997913 | 2.33243846 | 2.114720744 |
| ENSMUSG00000006732 | 2.026699189 | 1.955044741 | 1.945339928 | 1.988549078 | 2.025895058 | 2.193124598 | 1.925966215 |
| ENSMUSG00000006736 | 2.380049842 | 2.222945926 | 2.20584181 | 2.360217292 | 2.31625483 | 2.638489257 | 2.197791274 |
| ENSMUSG00000006740 | 3.256154144 | 2.975970461 | 2.960194481 | 2.98654381 | 2.89972777 | 3.35564305 | 2.676964833 |
| ENSMUSG00000006763 | 2.057835465 | 1.903677461 | 1.858657057 | 1.87892496 | 1.877063019 | 2.053078443 | 1.753626837 |
| ENSMUSG00000006764 | 0 | 0.25 | 0.25 | 0 | 0 | 0 | 0.355418052 |
| ENSMUSG00000006777 | 0 | 0 | 0 | 0 | 0 | 0 | 0 |
| ENSMUSG00000006782 | 1.700465191 | 1.660964047 | 1.738549078 | 2.044977273 | 2.079491904 | 2.298853076 | 2.013740841 |
| ENSMUSG00000006784 | 1.156259305 | 0.830482024 | 0.79248125 | 1 | 0.632412477 | 0.77815125 | 0.775756176 |
| ENSMUSG00000006800 | 2.352857998 | 2.146240625 | 2.190387808 | 1.882845365 | 1.928492895 | 2.1430148 | 1.851668228 |
| ENSMUSG00000006818 | 2.331622671 | 2.208222504 | 2.232684334 | 2.338286706 | 2.28171603 | 2.622214023 | 2.14555506 |
| ENSMUSG00000006850 | 1.470606563 | 1.26109853 | 1.364857905 | 1.160964047 | 1.212046813 | 1.176091259 | 1.185218806 |
| ENSMUSG00000006906 | 2.066740409 | 1.921625132 | 1.884789703 | 1.94697564 | 1.92245931 | 2.164352856 | 1.787569411 |
| ENSMUSG00000006920 | 2.103271637 | 1.966046536 | 1.942046081 | 1.969129237 | 2.016696984 | 2.181843588 | 1.865128412 |
| ENSMUSG00000006930 | 1.655740331 | 1.618933358 | 1.692046081 | 2.192872367 | 2.311234033 | 2.465382851 | 2.254530788 |
| ENSMUSG00000006931 | 2.518981867 | 2.327653195 | 2.345385738 | 2.112802778 | 2.172394856 | 2.382017043 | 2.059651572 |
| ENSMUSG00000006932 | 3.534399734 | 3.211225469 | 3.188095162 | 3.159382638 | 3.110494531 | 3.592731766 | 2.903550452 |
| ENSMUSG00000006941 | 2.086807695 | 1.926839783 | 1.943696765 | 2.034887838 | 2 | 2.252853031 | 1.869159809 |
| ENSMUSG00000006948 | 0 | 0 | 0 | 0 | 0 | 0 | 0 |
| ENSMUSG00000006958 | 1.036286563 | 0.92510993 | 1.201838731 | 1.188721876 | 1.433158226 | 1.204119983 | 1.365075947 |
| ENSMUSG00000006998 | 3.335705046 | 3.020121114 | 3.005331758 | 3.110411848 | 3.05281406 | 3.521530341 | 2.829286175 |
| ENSMUSG00000007021 | 1 | 0.92510993 | 0.864857905 | 1.52186571 | 1.504538967 | 1.69019608 | 1.537859885 |
| ENSMUSG00000007029 | 2.946945619 | 2.769036882 | 2.839112993 | 2.52219706 | 2.673124238 | 2.919078092 | 2.604423774 |
| ENSMUSG00000007030 | 0.671187741 | 0.75 | 0.830482024 | 0.5 | 0.812711509 | 0.84509804 | 0.775756176 |
| ENSMUSG00000007033 | 1.129344651 | 0.92510993 | 0.864857905 | 0.896240625 | 0.978602168 | 0.903089987 | 0.829800754 |
| ENSMUSG00000007034 | 0 | 0 | 0 | 0.396240625 | 0 | 0 | 0.224243824 |
| ENSMUSG00000007035 | 1.227926429 | 1.098079356 | 1.146240625 | 0.976722649 | 1.234113756 | 0.954242509 | 1.078019652 |

| | | | | | | | |
|---|---|---|---|---|---|---|---|
| ENSMUSG00000007036 | 2.144240249 | 2.024508021 | 2.113831805 | 1.928561379 | 2.116484486 | 2.230448921 | 2.13996595 |
| ENSMUSG00000007038 | 2.340856238 | 2.237091808 | 2.30539678 | 1.937048212 | 2.088159748 | 2.23299611 | 2.03601839 |
| ENSMUSG00000007039 | 2.514991096 | 2.453044326 | 2.547147211 | 2.015173983 | 2.273934005 | 2.309630167 | 2.310303152 |
| ENSMUSG00000007041 | 3.237076571 | 2.961960089 | 2.985078399 | 2.801295582 | 2.802677689 | 3.218010043 | 2.637175571 |
| ENSMUSG00000007050 | 2.410757293 | 2.267865591 | 2.256284891 | 2.21759118 | 2.192777367 | 2.506505032 | 2.085803078 |
| ENSMUSG00000007080 | 2.985332211 | 2.670529191 | 2.69595419 | 2.720469677 | 2.68472101 | 3.070776463 | 2.498691813 |
| ENSMUSG00000007097 | 0.289064826 | 0 | 0 | 0.25 | 0 | 0 | 0.355418052 |
| ENSMUSG00000007107 | 0 | 0 | 0 | 0 | 0 | 0 | 0 |
| ENSMUSG00000007122 | 0.747221736 | 0.5 | 0 | 0.701838731 | 0.632412477 | 0.698970004 | 0.829800754 |
| ENSMUSG00000007207 | 1.614416215 | 1.622963274 | 1.701838731 | 1.571350555 | 1.709785497 | 1.826074803 | 1.71890424 |
| ENSMUSG00000007209 | 0 | 0 | 0 | 0 | 0 | 0 | 0 |
| ENSMUSG00000007216 | 1.156259305 | 1.160964047 | 1.160964047 | 1 | 0.978602168 | 1.113943352 | 1.121219121 |
| ENSMUSG00000007279 | 0 | 0 | 0 | 0.396240625 | 0.489301084 | 0.301029996 | 0.629532004 |
| ENSMUSG00000007338 | 2.635965117 | 2.413659007 | 2.425543171 | 2.546838018 | 2.461024076 | 2.850033258 | 2.345462945 |
| ENSMUSG00000007379 | 1.91631382 | 1.789967834 | 1.802363341 | 1.843759858 | 1.868304353 | 2.068185862 | 1.714087544 |
| ENSMUSG00000007411 | 2.688735382 | 2.482684334 | 2.490362424 | 2.478596283 | 2.445469852 | 2.768638101 | 2.291042162 |
| ENSMUSG00000007415 | 2.669114668 | 2.449009902 | 2.450629091 | 2.485994979 | 2.382842606 | 2.761175813 | 2.230204359 |
| ENSMUSG00000007440 | 0 | 0 | 0 | 0 | 0 | 0 | 0 |
| ENSMUSG00000007457 | 0 | 0 | 0 | 0 | 0 | 0 | 0 |
| ENSMUSG00000007458 | 2.800026593 | 2.605516192 | 2.568157446 | 2.620701989 | 2.548733265 | 2.948412966 | 2.389035574 |
| ENSMUSG00000007476 | 1.876566388 | 1.695339928 | 1.664552871 | 1.52186571 | 1.635855979 | 1.612783857 | 1.517723536 |
| ENSMUSG00000007480 | 0.45815691 | 0 | 0 | 0 | 0 | 0 | 0 |
| ENSMUSG00000007564 | 3.245334908 | 2.946158711 | 2.945851886 | 2.899629453 | 2.896467625 | 3.320354033 | 2.716066686 |
| ENSMUSG00000007570 | 1.693336259 | 1.779735268 | 1.914552871 | 1.494319981 | 1.674752852 | 1.602059991 | 1.694069894 |
| ENSMUSG00000007589 | 1.794934559 | 1.807204673 | 1.77186571 | 1.975216702 | 2.020174412 | 2.155336037 | 1.898808251 |
| ENSMUSG00000007591 | 0.671187741 | 0.79248125 | 0.75 | 0.701838731 | 0.568060967 | 0.77815125 | 0.775756176 |
| ENSMUSG00000007594 | 1.678702042 | 1.575945187 | 1.505591953 | 1.438721876 | 1.366364104 | 1.544068044 | 1.365075947 |
| ENSMUSG00000007603 | 1.783508299 | 1.87694866 | 1.959800947 | 1.668106335 | 1.931471446 | 1.875061263 | 1.989124507 |
| ENSMUSG00000007610 | 1.811507563 | 1.825945187 | 1.886723615 | 1.664552871 | 1.87878897 | 1.908485019 | 1.837623616 |
| ENSMUSG00000007613 | 2.611244851 | 2.3843046 | 2.359697963 | 2.196158711 | 2.138604637 | 2.457881897 | 1.962479708 |
| ENSMUSG00000007617 | 2.003432385 | 1.845926073 | 1.87694866 | 2.081607372 | 2.03596422 | 2.267171728 | 1.864112664 |
| ENSMUSG00000007646 | 2.042557845 | 1.928561379 | 1.856566189 | 2.061981878 | 1.987570683 | 2.294466226 | 1.835407748 |
| ENSMUSG00000007653 | 0.289064826 | 0 | 0 | 0 | 0 | 0 | 0 |
| ENSMUSG00000007655 | 3.233691115 | 2.923915824 | 2.935050597 | 2.917664062 | 2.822349135 | 3.302114377 | 2.643364349 |

| | | | | | | | |
|---|---|---|---|---|---|---|---|
| ENSMUSG00000007656 | 2.892679599 | 2.640560606 | 2.63015467 | 2.794043287 | 2.703018303 | 3.143639235 | 2.508833311 |
| ENSMUSG00000007659 | 2.126809492 | 2.015173983 | 2.006976499 | 1.921625132 | 1.970729215 | 2.120573931 | 1.91917299 |
| ENSMUSG00000007670 | 2.880757703 | 2.667442722 | 2.739819376 | 2.517865591 | 2.58731141 | 2.859138297 | 2.522677108 |
| ENSMUSG00000007682 | 0 | 0 | 0 | 0 | 0 | 0.301029996 | 0 |
| ENSMUSG00000007721 | 1.822201287 | 1.870953944 | 2.04745614 | 1.630890489 | 1.789100779 | 1.819543936 | 2.034820183 |
| ENSMUSG00000007739 | 3.189970974 | 2.89541784 | 2.869818758 | 3.104265337 | 3.005064224 | 3.484442208 | 2.789497017 |
| ENSMUSG00000007777 | 1.83262764 | 1.657339155 | 1.695339928 | 1.75 | 1.763796674 | 1.826074803 | 1.610050272 |
| ENSMUSG00000007783 | 1.944805158 | 1.892463902 | 1.992885888 | 1.701838731 | 1.880506521 | 1.86923172 | 1.873141588 |
| ENSMUSG00000007805 | 1.596667477 | 1.54248125 | 1.593759858 | 1.552363341 | 1.584291694 | 1.544068044 | 1.591689859 |
| ENSMUSG00000007812 | 2.446507203 | 2.222185812 | 2.225216702 | 2.274508021 | 2.165585202 | 2.471291711 | 1.988432495 |
| ENSMUSG00000007815 | 3.126115402 | 2.824085671 | 2.807204673 | 2.906883471 | 2.833231787 | 3.295347148 | 2.646369581 |
| ENSMUSG00000007817 | 2.630681378 | 2.466046536 | 2.482314602 | 2.304186465 | 2.397003204 | 2.639486489 | 2.327511498 |
| ENSMUSG00000007827 | 2.418409478 | 2.227473271 | 2.237091808 | 2.247171172 | 2.17760189 | 2.492760389 | 2.042530357 |
| ENSMUSG00000007833 | 2.023453784 | 2.023189285 | 2.166333979 | 1.708222504 | 1.938809675 | 1.86332286 | 2.096394553 |
| ENSMUSG00000007836 | 2.947656672 | 2.675326615 | 2.653447366 | 2.81742778 | 2.745958094 | 3.1532049 | 2.553571809 |
| ENSMUSG00000007837 | 1.445324132 | 1.476722649 | 1.51109853 | 1.114857905 | 1.327538614 | 1.204119983 | 1.444790289 |
| ENSMUSG00000007850 | 3.476244943 | 3.149687602 | 3.136964627 | 3.238003291 | 3.175117347 | 3.66029616 | 2.930216069 |
| ENSMUSG00000007867 | 2.060824957 | 1.905512955 | 1.979715809 | 1.779735268 | 1.751666404 | 1.949390007 | 1.707551656 |
| ENSMUSG00000007872 | 2.578990401 | 2.527131114 | 2.656198864 | 1.854463129 | 2.068600388 | 2.152288344 | 2.146406417 |
| ENSMUSG00000007877 | 0.289064826 | 0.396240625 | 0.396240625 | 0 | 0.244650542 | 0 | 0.224243824 |
| ENSMUSG00000007880 | 2.818344313 | 2.626204747 | 2.614857905 | 2.414998973 | 2.46464417 | 2.748962861 | 2.343157052 |
| ENSMUSG00000007888 | 2.057835465 | 2.004202072 | 2.061981878 | 1.610735874 | 1.751666404 | 1.903089987 | 1.777090262 |
| ENSMUSG00000007891 | 3.142685399 | 2.849418658 | 2.894475209 | 2.655741736 | 2.632005142 | 3.01745073 | 2.486443709 |
| ENSMUSG00000007892 | 2.928019436 | 2.864473459 | 3.022528105 | 2.448197574 | 2.7475775 | 2.843855423 | 2.835306665 |
| ENSMUSG00000007907 | 0 | 0 | 0 | 0 | 0 | 0 | 0 |
| ENSMUSG00000007908 | 1.952600383 | 1.87892496 | 1.854463129 | 0.92510993 | 1.188507684 | 1.255272505 | 0.969165684 |
| ENSMUSG00000007944 | 0 | 0 | 0 | 0 | 0 | 0 | 0 |
| ENSMUSG00000007946 | 0.289064826 | 0 | 0 | 0 | 0.244650542 | 0 | 0 |
| ENSMUSG00000007950 | 2.020182925 | 2.028435542 | 2.116901388 | 1.653677461 | 1.890639929 | 1.819543936 | 2.073258096 |
| ENSMUSG00000007987 | 2.029919532 | 1.973704441 | 1.928561379 | 2.006976499 | 1.977302144 | 2.164352856 | 1.907820406 |
| ENSMUSG00000007989 | 1.631440309 | 1.685366747 | 1.575945187 | 1.909156155 | 1.782377602 | 1.934498451 | 1.63986884 |
| ENSMUSG00000008028 | 0 | 0 | 0 | 0.25 | 0 | 0 | 0.224243824 |
| ENSMUSG00000008035 | 2.060824957 | 2.015173983 | 2.044977273 | 1.494319981 | 1.572189156 | 1.51851394 | 1.605556929 |
| ENSMUSG00000008036 | 2.160971586 | 2.137686696 | 2.172499493 | 2.034887838 | 2.105788027 | 2.311753861 | 2.198515833 |

| | | | | | | | |
|---|---|---|---|---|---|---|---|
| ENSMUSG00000008090 | 2.203267085 | 2.073655187 | 2.102347734 | 1.799918086 | 1.937354184 | 2.056904851 | 1.871156825 |
| ENSMUSG00000008129 | 0.578129653 | 0.75 | 0.951838731 | 1.02186571 | 1.057362051 | 0.84509804 | 1 |
| ENSMUSG00000008136 | 2.668421347 | 2.467976309 | 2.426839783 | 2.027131114 | 2.06759338 | 2.257678575 | 1.935079929 |
| ENSMUSG00000008140 | 2.045658471 | 2.034887838 | 2.140560606 | 1.726722649 | 1.853822071 | 1.924279286 | 1.981429723 |
| ENSMUSG00000008153 | 0.289064826 | 0.646240625 | 0 | 0.701838731 | 0.77552387 | 0.903089987 | 0.520678036 |
| ENSMUSG00000008167 | 1.578129653 | 1.646240625 | 1.729715809 | 1.5 | 1.639266209 | 1.591064607 | 1.709198048 |
| ENSMUSG00000008193 | 0.289064826 | 0.25 | 0.75 | 0.580482024 | 0.686820904 | 0.698970004 | 0.710836105 |
| ENSMUSG00000008200 | 2.343889214 | 2.201032755 | 2.160061234 | 2.407339155 | 2.318243321 | 2.652246341 | 2.170563634 |
| ENSMUSG00000008206 | 2.078325387 | 1.956637122 | 1.898114259 | 1.622963274 | 1.611014646 | 1.755874856 | 1.514741806 |
| ENSMUSG00000008226 | 2.063793172 | 1.903677461 | 1.896240625 | 2.088286706 | 2.046825265 | 2.307496038 | 1.929310074 |
| ENSMUSG00000008301 | 2.278594844 | 2.140560606 | 2.148114259 | 2.150906586 | 2.097547432 | 2.374748346 | 2.008574816 |
| ENSMUSG00000008305 | 1.91631382 | 1.834962501 | 1.868933358 | 1.692046081 | 1.66856048 | 1.77815125 | 1.607811402 |
| ENSMUSG00000008307 | 1.269664473 | 1.330482024 | 1.201838731 | 1.418106335 | 1.264824954 | 1.255272505 | 1.224243824 |
| ENSMUSG00000008318 | 1.663535556 | 1.653677461 | 1.784887838 | 1.464495249 | 1.504538967 | 1.531478917 | 1.559435575 |
| ENSMUSG00000008333 | 2.35433422 | 2.148114259 | 2.177701608 | 2.352347734 | 2.299407739 | 2.618048097 | 2.137353429 |
| ENSMUSG00000008348 | 2.797488347 | 2.540282971 | 2.543731421 | 2.439972046 | 2.503073808 | 2.806179974 | 2.331614323 |
| ENSMUSG00000008373 | 2.378662046 | 2.234159485 | 2.196158711 | 2.234894804 | 2.239731213 | 2.465382851 | 2.113781657 |
| ENSMUSG00000008384 | 1.816888701 | 1.744319981 | 1.862802778 | 1.584962501 | 1.712553795 | 1.73239376 | 1.726776051 |
| ENSMUSG00000008393 | 2.709994145 | 2.478222334 | 2.469895812 | 2.550837001 | 2.517176663 | 2.845718018 | 2.364643151 |
| ENSMUSG00000008398 | 2.671187741 | 2.504897682 | 2.478596283 | 2.31611065 | 2.344045917 | 2.650307523 | 2.239900853 |
| ENSMUSG00000008429 | 2.033115198 | 1.935366747 | 1.901832578 | 1.948603967 | 1.917865457 | 2.130333768 | 1.803905701 |
| ENSMUSG00000008435 | 2.081172036 | 1.882845365 | 1.880890489 | 1.777131114 | 1.777823267 | 1.924279286 | 1.729871916 |
| ENSMUSG00000008438 | 0.289064826 | 0.830482024 | 0.396240625 | 0.580482024 | 0.387761935 | 0 | 0.224243824 |
| ENSMUSG00000008450 | 2.360187403 | 2.161864607 | 2.192872367 | 2.27053726 | 2.235996193 | 2.537819095 | 2.086826862 |
| ENSMUSG00000008461 | 0 | 0.396240625 | 0.25 | 0 | 0 | 0 | 0 |
| ENSMUSG00000008475 | 3.104980289 | 2.833707982 | 2.831466895 | 2.646709947 | 2.581245921 | 2.986771734 | 2.425462392 |
| ENSMUSG00000008482 | 0.747221736 | 1.114857905 | 1.061981878 | 0.830482024 | 0.955822902 | 1.146128036 | 1 |
| ENSMUSG00000008489 | 1.069666964 | 0.976722649 | 0.896240625 | 1.04248125 | 0.568060967 | 0.84509804 | 0.853775828 |
| ENSMUSG00000008496 | 0.289064826 | 0.580482024 | 0 | 0.5 | 0.387761935 | 0.477121255 | 0.520678036 |
| ENSMUSG00000008540 | 1.907802659 | 1.738549078 | 1.809601185 | 1 | 0.978602168 | 1.176091259 | 0.969165684 |
| ENSMUSG00000008575 | 2.838668326 | 2.598615674 | 2.53360658 | 2.58272922 | 2.522774815 | 2.855519156 | 2.323601898 |
| ENSMUSG00000008590 | 0.289064826 | 0 | 0 | 0 | 0 | 0 | 0 |
| ENSMUSG00000008601 | 0 | 0.5 | 0.25 | 0.396240625 | 0.387761935 | 0 | 0.224243824 |
| ENSMUSG00000008604 | 2.65645504 | 2.463717096 | 2.503156135 | 2.490362424 | 2.495655374 | 2.786751422 | 2.380695513 |

| | | | | | | | |
|---|---|---|---|---|---|---|---|
| ENSMUSG00000008658 | 1.516991255 | 1.356566189 | 1.380890489 | 0.646240625 | 0.568060967 | 0.477121255 | 0.448487648 |
| ENSMUSG00000008668 | 3.129803938 | 2.889605178 | 2.916777886 | 2.753679482 | 2.840643511 | 3.181271772 | 2.688229252 |
| ENSMUSG00000008682 | 3.059740347 | 2.825802487 | 2.840759908 | 2.724839231 | 2.745662857 | 3.087426457 | 2.579550454 |
| ENSMUSG00000008683 | 2.945162665 | 2.673371739 | 2.675759597 | 2.775001306 | 2.669661176 | 3.151676231 | 2.483007585 |
| ENSMUSG00000008690 | 3.088331566 | 2.834405475 | 2.845791065 | 2.783124932 | 2.807172415 | 3.175511813 | 2.649616488 |
| ENSMUSG00000008730 | 2.769588082 | 2.529410775 | 2.511439915 | 2.513820609 | 2.457700547 | 2.794488047 | 2.316895388 |
| ENSMUSG00000008734 | 1.932832114 | 1.858657057 | 1.834962501 | 1.821350555 | 1.85749873 | 1.959041392 | 1.764880683 |
| ENSMUSG00000008763 | 2.759113486 | 2.543731421 | 2.525493918 | 2.6343046 | 2.540642974 | 2.92788341 | 2.341765586 |
| ENSMUSG00000008789 | 0 | 0 | 0 | 0 | 0 | 0 | 0 |
| ENSMUSG00000008813 | 0 | 0 | 0 | 0 | 0 | 0 | 0 |
| ENSMUSG00000008822 | 1.596667477 | 1.580482024 | 1.681980114 | 1.470660762 | 1.546663136 | 1.662757832 | 1.508694612 |
| ENSMUSG00000008843 | 0 | 0 | 0 | 0 | 0 | 0 | 0 |
| ENSMUSG00000008845 | 0.289064826 | 0.396240625 | 0.25 | 0 | 0 | 0.301029996 | 0 |
| ENSMUSG00000008855 | 2.163307908 | 2.137686696 | 2.240000483 | 1.708222504 | 1.959951091 | 1.977723605 | 2.028761664 |
| ENSMUSG00000008859 | 2.631440309 | 2.435366747 | 2.411414608 | 2.552667836 | 2.441296923 | 2.82672252 | 2.276587178 |
| ENSMUSG00000008874 | 0 | 0 | 0 | 0 | 0 | 0 | 0 |
| ENSMUSG00000008892 | 2.730819472 | 2.510756821 | 2.475216702 | 2.623213655 | 2.516611933 | 2.932473765 | 2.328723622 |
| ENSMUSG00000008932 | 0.45815691 | 0.25 | 0 | 0.25 | 0 | 0.602059991 | 0.224243824 |
| ENSMUSG00000008958 | 2.124258828 | 2.130890489 | 2.174241882 | 1.96293726 | 2.108492685 | 2.23299611 | 2.090889903 |
| ENSMUSG00000008976 | 2.750654121 | 2.539967834 | 2.53743678 | 2.579635702 | 2.479392893 | 2.857332496 | 2.330413001 |
| ENSMUSG00000008999 | 0 | 0 | 0.396240625 | 0 | 0.387761935 | 0 | 0.355418052 |
| ENSMUSG00000009013 | 2.897514811 | 2.678776002 | 2.664775991 | 2.716239979 | 2.676553652 | 3.100370545 | 2.540133724 |
| ENSMUSG00000009030 | 2.414600936 | 2.24929487 | 2.214495249 | 2.259043403 | 2.142704057 | 2.469822016 | 2.018194418 |
| ENSMUSG00000009035 | 2.716846213 | 2.595656007 | 2.654825739 | 2.38040011 | 2.516611933 | 2.758154622 | 2.442585457 |
| ENSMUSG00000009047 | 0 | 0 | 0 | 0 | 0 | 0 | 0 |
| ENSMUSG00000009070 | 0 | 0 | 0 | 0 | 0 | 0 | 0.355418052 |
| ENSMUSG00000009073 | 2.659301689 | 2.453044326 | 2.415889526 | 2.518200634 | 2.480646735 | 2.834420704 | 2.352733261 |
| ENSMUSG00000009075 | 0 | 0.25 | 0.396240625 | 0 | 0.244650542 | 0 | 0 |
| ENSMUSG00000009076 | 1.432083896 | 1.418106335 | 1.418106335 | 1.451838731 | 1.509475496 | 1.591064607 | 1.520678036 |
| ENSMUSG00000009079 | 3.110545643 | 2.861383055 | 2.882235604 | 2.778272746 | 2.775252365 | 3.129689892 | 2.647365154 |
| ENSMUSG00000009090 | 2.551444705 | 2.341580554 | 2.409608978 | 2.422499493 | 2.459366223 | 2.737192643 | 2.311070686 |
| ENSMUSG00000009092 | 0.289064826 | 0.396240625 | 0.5 | 0.396240625 | 0.632412477 | 0 | 0.355418052 |
| ENSMUSG00000009093 | 0 | 0 | 0.396240625 | 0.5 | 0.387761935 | 0 | 0.355418052 |
| ENSMUSG00000009097 | 0 | 0 | 0 | 0 | 0.244650542 | 0 | 0.224243824 |

| | | | | | | | |
|---|---|---|---|---|---|---|---|
| ENSMUSG00000009108 | 1.269664473 | 1.25 | 1.238549078 | 1.080482024 | 1 | 1 | 0.969165684 |
| ENSMUSG00000009112 | 2.533616084 | 2.31728167 | 2.319031101 | 2.47104263 | 2.451638175 | 2.788875116 | 2.33233298 |
| ENSMUSG00000009114 | 0 | 0.396240625 | 0 | 0 | 0.244650542 | 0 | 0 |
| ENSMUSG00000009115 | 0 | 0 | 0 | 0 | 0 | 0 | 0 |
| ENSMUSG00000009145 | 1 | 0.976722649 | 1.04248125 | 1.02186571 | 1.136121934 | 0.84509804 | 0.952571511 |
| ENSMUSG00000009185 | 0 | 0 | 0.25 | 0 | 0 | 0 | 0.355418052 |
| ENSMUSG00000009207 | 2.269664473 | 2.112802778 | 2.101285366 | 2.247880462 | 2.21338377 | 2.46834733 | 2.027536225 |
| ENSMUSG00000009210 | 0 | 0 | 0 | 0 | 0 | 0 | 0 |
| ENSMUSG00000009214 | 0 | 0 | 0 | 0 | 0 | 0 | 0 |
| ENSMUSG00000009216 | 0 | 0 | 0 | 0 | 0 | 0 | 0 |
| ENSMUSG00000009246 | 0 | 0.396240625 | 0.5 | 0 | 0 | 0 | 0.224243824 |
| ENSMUSG00000009248 | 0 | 0 | 0 | 0 | 0 | 0 | 0 |
| ENSMUSG00000009281 | 0 | 0.396240625 | 0 | 0 | 0 | 0 | 0 |
| ENSMUSG00000009291 | 2.884907314 | 2.629417347 | 2.633575721 | 2.823798739 | 2.811365514 | 3.215637563 | 2.624237034 |
| ENSMUSG00000009292 | 0.960252568 | 1 | 1.098079356 | 0.92510993 | 0.905314583 | 0.77815125 | 0.853775828 |
| ENSMUSG00000009293 | 2.476144923 | 2.347542239 | 2.309003548 | 2.314935816 | 2.368949874 | 2.638489257 | 2.236700872 |
| ENSMUSG00000009350 | 0 | 0 | 0 | 0.25 | 0 | 0 | 0.224243824 |
| ENSMUSG00000009356 | 0 | 0.25 | 0 | 0.25 | 0.568060967 | 0.301029996 | 0.448487648 |
| ENSMUSG00000009376 | 2.951545939 | 2.702039943 | 2.67619206 | 2.654366866 | 2.531280194 | 2.954724791 | 2.363558619 |
| ENSMUSG00000009378 | 0.289064826 | 0 | 0 | 0 | 0 | 0 | 0 |
| ENSMUSG00000009394 | 0.289064826 | 0.5 | 0.79248125 | 0.951838731 | 0.905314583 | 1.041392685 | 0.829800754 |
| ENSMUSG00000009406 | 2.440557995 | 2.233422664 | 2.250703754 | 2.046218836 | 2.022473809 | 2.269512944 | 1.972093446 |
| ENSMUSG00000009418 | 2.743181252 | 2.51109853 | 2.519203899 | 2.512462137 | 2.476871767 | 2.803457116 | 2.353854633 |
| ENSMUSG00000009470 | 3.322324702 | 3.018702615 | 2.998941788 | 3.146768569 | 3.055218277 | 3.54567815 | 2.81155704 |
| ENSMUSG00000009471 | 0 | 0 | 0 | 0 | 0 | 0 | 0 |
| ENSMUSG00000009487 | 0 | 0.25 | 0 | 0 | 0 | 0 | 0 |
| ENSMUSG00000009535 | 2.390989024 | 2.198603967 | 2.188721876 | 2.232684334 | 2.165585202 | 2.5132176 | 1.991877847 |
| ENSMUSG00000009545 | 0 | 0 | 0 | 0.25 | 0 | 0 | 0.224243824 |
| ENSMUSG00000009549 | 2.273259602 | 2.097004321 | 2.145300145 | 2.279735268 | 2.237245597 | 2.564666064 | 2.134278581 |
| ENSMUSG00000009551 | 0.289064826 | 0 | 0 | 0.25 | 0 | 0 | 0.355418052 |
| ENSMUSG00000009555 | 2.495495257 | 2.344844592 | 2.381869252 | 2.182829758 | 2.275056283 | 2.392696953 | 2.13295179 |
| ENSMUSG00000009563 | 1.852729478 | 1.755591953 | 1.850219859 | 1.685366747 | 1.866526233 | 1.968482949 | 1.853775828 |
| ENSMUSG00000009566 | 1.885732304 | 1.868933358 | 1.992885888 | 1.858657057 | 1.958580386 | 2.017033339 | 1.97354745 |
| ENSMUSG00000009569 | 2.503815381 | 2.269203899 | 2.274508021 | 2.162762923 | 2.136121934 | 2.414973348 | 1.993927582 |

| | | | | | | | |
|---|---|---|---|---|---|---|---|
| ENSMUSG00000009575 | 3.652672703 | 3.291305267 | 3.273106704 | 3.350453208 | 3.245260665 | 3.769820258 | 3.002632559 |
| ENSMUSG00000009580 | 0 | 0 | 0 | 0 | 0 | 0 | 0 |
| ENSMUSG00000009585 | 2.310886388 | 2.098079356 | 2.111770807 | 2.254897682 | 2.16174451 | 2.45484486 | 2.057979654 |
| ENSMUSG00000009588 | 0 | 0 | 0 | 0 | 0 | 0 | 0 |
| ENSMUSG00000009596 | 0 | 0 | 0 | 0.25 | 0.387761935 | 0 | 0.224243824 |
| ENSMUSG00000009614 | 1.847794126 | 1.76109853 | 1.864857905 | 1.27186571 | 1.319233381 | 1.342422681 | 1.369798885 |
| ENSMUSG00000009621 | 2.258689303 | 2.130890489 | 2.103406982 | 2.041226732 | 2.06149024 | 2.212187604 | 1.972093446 |
| ENSMUSG00000009628 | 0.91631382 | 0.864857905 | 0.951838731 | 0.75 | 0.905314583 | 0.84509804 | 0.775756176 |
| ENSMUSG00000009630 | 2.690713405 | 2.487091808 | 2.427701608 | 2.543731421 | 2.461024076 | 2.820201459 | 2.269981661 |
| ENSMUSG00000009633 | 0.45815691 | 0 | 0 | 0 | 0 | 0 | 0.355418052 |
| ENSMUSG00000009640 | 2.066740409 | 1.935366747 | 1.964495249 | 1.990000483 | 2.069604532 | 2.227886705 | 1.912234055 |
| ENSMUSG00000009646 | 0 | 0 | 0 | 0 | 0 | 0 | 0 |
| ENSMUSG00000009647 | 2.3855553 | 2.179419106 | 2.190387808 | 2.159156155 | 2.163285805 | 2.378397901 | 2.028761664 |
| ENSMUSG00000009654 | 0.811507563 | 0.79248125 | 0.92510993 | 0.580482024 | 0.632412477 | 0.602059991 | 0.74492186 |
| ENSMUSG00000009670 | 0 | 0 | 0 | 0 | 0 | 0 | 0.224243824 |
| ENSMUSG00000009681 | 2.457006475 | 2.277131114 | 2.345926073 | 2.289967834 | 2.343584837 | 2.572871602 | 2.245894972 |
| ENSMUSG00000009687 | 1.912079952 | 1.799918086 | 1.884789703 | 1.841580554 | 1.895599581 | 2.068185862 | 1.743447983 |
| ENSMUSG00000009731 | 1.714367558 | 1.660964047 | 1.646240625 | 1.42510993 | 1.467903253 | 1.505149978 | 1.369798885 |
| ENSMUSG00000009733 | 2.153883046 | 1.997171172 | 2.012462137 | 1.970660762 | 1.988833497 | 2.220108088 | 1.899720848 |
| ENSMUSG00000009734 | 0 | 0 | 0 | 0 | 0 | 0 | 0 |
| ENSMUSG00000009739 | 2 | 1.870953944 | 1.866901388 | 1.979715809 | 1.957204337 | 2.161368002 | 1.884801065 |
| ENSMUSG00000009741 | 2.578990401 | 2.364857905 | 2.332168732 | 2.353406982 | 2.383255179 | 2.656098202 | 2.213368331 |
| ENSMUSG00000009772 | 2.179305632 | 2.032320754 | 2.098079356 | 1.964495249 | 2.033751322 | 2.113943352 | 1.941550454 |
| ENSMUSG00000009828 | 2.131864492 | 1.958222504 | 1.933677405 | 1.979715809 | 1.950242452 | 2.195899652 | 1.879993912 |
| ENSMUSG00000009863 | 2.558729299 | 2.396240625 | 2.439139081 | 2.365369862 | 2.393415815 | 2.683947131 | 2.286387183 |
| ENSMUSG00000009876 | 0 | 0.25 | 0 | 0 | 0 | 0 | 0 |
| ENSMUSG00000009894 | 2.544030601 | 2.357613138 | 2.348079356 | 2.457826811 | 2.480646735 | 2.818885415 | 2.359843777 |
| ENSMUSG00000009900 | 0 | 0.396240625 | 0 | 0 | 0.244650542 | 0 | 0 |
| ENSMUSG00000009905 | 2.243591459 | 2.099151195 | 2.092671852 | 1.898114259 | 1.871833944 | 2.025305865 | 1.749303626 |
| ENSMUSG00000009907 | 2.699822119 | 2.469512728 | 2.437048212 | 2.54652856 | 2.450276722 | 2.802773725 | 2.288037776 |
| ENSMUSG00000009927 | 2.930617777 | 2.663659007 | 2.661864607 | 2.813314145 | 2.711253771 | 3.174059808 | 2.513923814 |
| ENSMUSG00000009941 | 0.289064826 | 0 | 0 | 0.79248125 | 0.387761935 | 0.301029996 | 0.355418052 |
| ENSMUSG00000009995 | 2.036286563 | 1.92510993 | 1.926839783 | 1.848079356 | 1.910073614 | 2.1430148 | 1.782372264 |
| ENSMUSG00000010021 | 0.578129653 | 0 | 0.396240625 | 0.830482024 | 1 | 0.84509804 | 1.089372236 |

| | | | | | | | |
|---|---|---|---|---|---|---|---|
| ENSMUSG00000010025 | 2.410757293 | 2.225216702 | 2.229715809 | 2.438721876 | 2.363323232 | 2.683047038 | 2.230859913 |
| ENSMUSG00000010044 | 0.45815691 | 0.25 | 0.79248125 | 0.646240625 | 0.568060967 | 0.602059991 | 0.355418052 |
| ENSMUSG00000010045 | 2.033115198 | 1.888647213 | 2.00835575 | 1.886723615 | 1.969399812 | 2.10720997 | 1.979298987 |
| ENSMUSG00000010047 | 2.100572389 | 2.029735268 | 2.094844592 | 1.784887838 | 1.966725889 | 1.991226076 | 2.005303651 |
| ENSMUSG00000010048 | 1.967766868 | 1.926839783 | 2 | 1.872963274 | 2 | 2.117271296 | 2.001334086 |
| ENSMUSG00000010051 | 1.129344651 | 1.02186571 | 1.130890489 | 1.02186571 | 1.200473444 | 1.113943352 | 0.916588299 |
| ENSMUSG00000010054 | 1.894701088 | 1.794977273 | 1.809601185 | 1.848079356 | 1.830919587 | 2.025305865 | 1.71890424 |
| ENSMUSG00000010057 | 1.783508299 | 1.747171172 | 1.720660762 | 1.799918086 | 1.851969278 | 1.991226076 | 1.757893036 |
| ENSMUSG00000010064 | 0.289064826 | 0.25 | 0.5 | 1.17510993 | 1.283908311 | 1.079181246 | 1.20919388 |
| ENSMUSG00000010066 | 0 | 0.25 | 0.25 | 0.75 | 0.387761935 | 0.301029996 | 0.629532004 |
| ENSMUSG00000010067 | 1.890240806 | 1.896240625 | 1.901832578 | 1.78743678 | 1.818885826 | 2.033423755 | 1.851668228 |
| ENSMUSG00000010080 | 1.505869732 | 1.451838731 | 1.445339928 | 1.410964047 | 1.484072009 | 1.531478917 | 1.594042709 |
| ENSMUSG00000010086 | 0 | 0 | 0 | 0 | 0.686820904 | 0.301029996 | 0 |
| ENSMUSG00000010095 | 2.857166034 | 2.598347614 | 2.583288838 | 2.494319981 | 2.523883869 | 2.82672252 | 2.390638131 |
| ENSMUSG00000010097 | 2.945519866 | 2.687467357 | 2.690803092 | 2.742706225 | 2.710765025 | 3.092720645 | 2.549194983 |
| ENSMUSG00000010110 | 2.611244851 | 2.396709947 | 2.428990498 | 2.421187155 | 2.404458919 | 2.658964843 | 2.253005488 |
| ENSMUSG00000010122 | 0 | 0 | 0 | 0 | 0.387761935 | 0.602059991 | 0 |
| ENSMUSG00000010136 | 0 | 0 | 0 | 0 | 0 | 0 | 0 |
| ENSMUSG00000010142 | 0 | 0 | 0 | 0 | 0 | 0 | 0 |
| ENSMUSG00000010154 | 1.129344651 | 1.146240625 | 1.321350555 | 1.561981878 | 1.671670246 | 1.672097858 | 1.73898563 |
| ENSMUSG00000010175 | 0 | 0.25 | 0 | 0.75 | 0.489301084 | 0.602059991 | 0.896975297 |
| ENSMUSG00000010205 | 2.593357679 | 2.484894804 | 2.51348147 | 2.464883697 | 2.574115252 | 2.857935265 | 2.544882201 |
| ENSMUSG00000010277 | 3.192150764 | 2.907794264 | 2.896827182 | 3.096600356 | 3.024020832 | 3.50623436 | 2.809920916 |
| ENSMUSG00000010290 | 2.235832493 | 2.095926073 | 2.041226732 | 2.129909063 | 2.063536368 | 2.346352974 | 1.937521567 |
| ENSMUSG00000010307 | 1.558729299 | 1.54248125 | 1.445339928 | 1.438721876 | 1.380772476 | 1.531478917 | 1.455757965 |
| ENSMUSG00000010311 | 1.181541736 | 1.282320754 | 1.061981878 | 0.396240625 | 0 | 0 | 0.224243824 |
| ENSMUSG00000010342 | 0 | 0.396240625 | 0.25 | 0.396240625 | 0.568060967 | 0.698970004 | 0.448487648 |
| ENSMUSG00000010358 | 1.794934559 | 1.807204673 | 1.843759858 | 1.950224975 | 1.991345651 | 2.274157849 | 1.886704102 |
| ENSMUSG00000010362 | 1.53838222 | 1.51109853 | 1.516522298 | 1.622963274 | 1.618292349 | 1.77815125 | 1.572224412 |
| ENSMUSG00000010376 | 2.514991096 | 2.284887838 | 2.322504712 | 2.540912419 | 2.488995627 | 2.895974732 | 2.34130043 |
| ENSMUSG00000010392 | 2.583267694 | 2.372963274 | 2.391989019 | 2.488912477 | 2.430282804 | 2.816903839 | 2.29374942 |
| ENSMUSG00000010406 | 1.342375483 | 1.29248125 | 1.557204673 | 1.470660762 | 1.433158226 | 1.62324929 | 1.67819947 |
| ENSMUSG00000010435 | 0 | 0 | 0 | 0 | 0 | 0 | 0 |
| ENSMUSG00000010453 | 2.655740331 | 2.415444524 | 2.449009902 | 2.459012589 | 2.462015062 | 2.782472624 | 2.312854549 |

| | | | | | | | |
|---|---|---|---|---|---|---|---|
| ENSMUSG00000010461 | 2.149089517 | 1.979715809 | 1.950224975 | 2.106566189 | 1.983754904 | 2.184691431 | 1.836517579 |
| ENSMUSG00000010476 | 1.940852214 | 1.884789703 | 1.845926073 | 2.04745614 | 1.947418788 | 2.290034611 | 1.87902385 |
| ENSMUSG00000010492 | 0.671187741 | 0.396240625 | 0.25 | 0.701838731 | 0.77552387 | 0.698970004 | 0.672731473 |
| ENSMUSG00000010505 | 0 | 0 | 0 | 0 | 0.244650542 | 0 | 0 |
| ENSMUSG00000010517 | 2.709994145 | 2.484527332 | 2.444929839 | 2.485628626 | 2.374910674 | 2.736396502 | 2.225244904 |
| ENSMUSG00000010529 | 0 | 0 | 0 | 0 | 0 | 0 | 0 |
| ENSMUSG00000010538 | 0.747221736 | 0.701838731 | 0.701838731 | 0.646240625 | 0.387761935 | 0.84509804 | 0.672731473 |
| ENSMUSG00000010554 | 2.644135203 | 2.472566069 | 2.414552871 | 2.491085217 | 2.458034312 | 2.771587481 | 2.281104046 |
| ENSMUSG00000010592 | 0 | 0 | 0 | 0.25 | 0 | 0 | 0 |
| ENSMUSG00000010601 | 1.129344651 | 1.130890489 | 1.080482024 | 0.951838731 | 1.020174412 | 0.77815125 | 0.935079929 |
| ENSMUSG00000010607 | 1.307602651 | 1.364857905 | 1.488549078 | 1 | 1.223252711 | 1.278753601 | 1.455757965 |
| ENSMUSG00000010608 | 3.088838288 | 2.798073204 | 2.811388177 | 2.791226732 | 2.68436981 | 3.093771781 | 2.500688828 |
| ENSMUSG00000010609 | 1.587501561 | 1.339388001 | 1.445339928 | 1.54248125 | 1.628935049 | 1.69019608 | 1.559435575 |
| ENSMUSG00000010651 | 0.91631382 | 0.5 | 0.92510993 | 0.646240625 | 0.812711509 | 0.477121255 | 0.853775828 |
| ENSMUSG00000010660 | 1.903481042 | 1.825945187 | 1.923371739 | 1.830482024 | 1.940259189 | 1.949390007 | 1.928477344 |
| ENSMUSG00000010663 | 2.921262688 | 2.647646762 | 2.632845365 | 2.704645544 | 2.690465922 | 3.096562438 | 2.51637161 |
| ENSMUSG00000010721 | 1.596667477 | 1.561981878 | 1.505591953 | 1.741446071 | 1.645989387 | 1.903089987 | 1.572224412 |
| ENSMUSG00000010751 | 1.036286563 | 0.92510993 | 0.646240625 | 1.634789703 | 1.628935049 | 1.698970004 | 1.502532226 |
| ENSMUSG00000010755 | 2.575536697 | 2.347004321 | 2.360735874 | 2.271202097 | 2.245869735 | 2.57054294 | 2.136039214 |
| ENSMUSG00000010760 | 0 | 0.25 | 0 | 0 | 0 | 0 | 0.224243824 |
| ENSMUSG00000010796 | 0 | 0 | 0 | 0 | 0 | 0 | 0 |
| ENSMUSG00000010797 | 0 | 0 | 0.5 | 0.580482024 | 0.489301084 | 0.477121255 | 0.224243824 |
| ENSMUSG00000010803 | 0 | 0 | 0.25 | 0 | 0.244650542 | 0 | 0.224243824 |
| ENSMUSG00000010825 | 0 | 0 | 0.25 | 0.580482024 | 0.489301084 | 0.602059991 | 0.448487648 |
| ENSMUSG00000010830 | 2.705574381 | 2.457826811 | 2.483053688 | 2.421187155 | 2.366364104 | 2.703291378 | 2.209543816 |
| ENSMUSG00000010841 | 0 | 0 | 0 | 0 | 0 | 0 | 0 |
| ENSMUSG00000010911 | 1.700465191 | 1.575945187 | 1.584962501 | 1.905512955 | 1.877063019 | 2.117271296 | 1.763495175 |
| ENSMUSG00000010914 | 2.020182925 | 1.961372513 | 1.907339155 | 2.151832578 | 2.111176775 | 2.383815366 | 1.996640515 |
| ENSMUSG00000010936 | 2.525875121 | 2.38040011 | 2.397178659 | 2.201838731 | 2.282265378 | 2.484299839 | 2.15104925 |
| ENSMUSG00000011008 | 1 | 0.79248125 | 1.080482024 | 0.25 | 0.244650542 | 0.301029996 | 0 |
| ENSMUSG00000011034 | 0 | 0 | 0 | 0.25 | 0 | 0 | 0 |
| ENSMUSG00000011052 | 0 | 0.25 | 0 | 0 | 0 | 0 | 0 |
| ENSMUSG00000011096 | 1.890240806 | 1.87694866 | 1.976722649 | 1.660964047 | 1.761403742 | 1.857332496 | 1.885753983 |
| ENSMUSG00000011114 | 2.603209296 | 2.403217124 | 2.402294685 | 2.52053726 | 2.51576314 | 2.833147112 | 2.356752139 |

| | | | | | | | |
|---|---|---|---|---|---|---|---|
| ENSMUSG00000011118 | 0.45815691 | 0 | 0 | 0 | 0 | 0 | 0 |
| ENSMUSG00000011148 | 2.146671932 | 2.057204673 | 2.161864607 | 1.214495249 | 1.223252711 | 1.278753601 | 1.284450788 |
| ENSMUSG00000011154 | 0 | 0 | 0 | 0 | 0 | 0 | 0 |
| ENSMUSG00000011158 | 2.23973002 | 2.119945066 | 2.111770807 | 2.00835575 | 2.053185007 | 2.161368002 | 1.963977467 |
| ENSMUSG00000011171 | 0 | 0 | 0 | 0.396240625 | 0.244650542 | 0.301029996 | 0.224243824 |
| ENSMUSG00000011179 | 2.712498852 | 2.531676118 | 2.488549078 | 2.566696635 | 2.500716831 | 2.848804701 | 2.331614323 |
| ENSMUSG00000011254 | 1.956444026 | 1.890560606 | 1.903677461 | 2.099151195 | 2.025895058 | 2.292256071 | 1.910475814 |
| ENSMUSG00000011256 | 2.435736758 | 2.252806814 | 2.236360959 | 2.032320754 | 2.024758322 | 2.264817823 | 1.977870661 |
| ENSMUSG00000011257 | 2.623015126 | 2.433677405 | 2.464495249 | 2.314346961 | 2.292543754 | 2.575187845 | 2.187847501 |
| ENSMUSG00000011263 | 0 | 0 | 0 | 0 | 0 | 0 | 0.448487648 |
| ENSMUSG00000011267 | 0.91631382 | 0.646240625 | 0.830482024 | 0.580482024 | 0.632412477 | 0.477121255 | 0.935079929 |
| ENSMUSG00000011305 | 0 | 0.5 | 0.396240625 | 0.25 | 0.489301084 | 0 | 0.579661877 |
| ENSMUSG00000011306 | 2.201144779 | 2.066696635 | 2.072504712 | 2.101285366 | 2.128567294 | 2.346352974 | 2.044866219 |
| ENSMUSG00000011349 | 0 | 0 | 0 | 0 | 0 | 0 | 0 |
| ENSMUSG00000011350 | 0 | 0 | 0 | 0 | 0 | 0 | 0 |
| ENSMUSG00000011382 | 0.578129653 | 0.5 | 0.396240625 | 0.896240625 | 0.632412477 | 0.84509804 | 0.672731473 |
| ENSMUSG00000011427 | 1.686083339 | 1.668106335 | 1.571350555 | 1.738549078 | 1.680838453 | 1.832508913 | 1.631625962 |
| ENSMUSG00000011463 | 0 | 0 | 0 | 0 | 0 | 0 | 0 |
| ENSMUSG00000011486 | 0 | 0 | 0 | 0 | 0.244650542 | 0.301029996 | 0.355418052 |
| ENSMUSG00000011589 | 1.587501561 | 1.476722649 | 1.571350555 | 1.302363341 | 1.462344774 | 1.431363764 | 1.437266334 |
| ENSMUSG00000011632 | 0 | 0 | 0.25 | 0.396240625 | 0 | 0 | 0 |
| ENSMUSG00000011658 | 1.325351389 | 1.226722649 | 1.364857905 | 1.238549078 | 1.254881871 | 1.204119983 | 1.176815335 |
| ENSMUSG00000011751 | 0.578129653 | 0.701838731 | 0.25 | 0 | 0.387761935 | 0.301029996 | 0.672731473 |
| ENSMUSG00000011752 | 3.397803828 | 3.113960227 | 3.115817229 | 3.132174571 | 3.090835783 | 3.571592383 | 2.904286458 |
| ENSMUSG00000011831 | 2.769043298 | 2.56728167 | 2.522858847 | 2.503505118 | 2.417419887 | 2.753583059 | 2.235411964 |
| ENSMUSG00000011832 | 2.016886209 | 1.843759858 | 1.884789703 | 1.774508021 | 1.820920161 | 1.982271233 | 1.74047993 |
| ENSMUSG00000011837 | 2.267855219 | 2.198603967 | 2.187886015 | 1.807204673 | 1.964031554 | 2.012837225 | 1.882886767 |
| ENSMUSG00000011877 | 2.694641526 | 2.519203899 | 2.581607372 | 2.098079356 | 2.25190332 | 2.392696953 | 2.282223478 |
| ENSMUSG00000011884 | 2.53552908 | 2.381380202 | 2.424241882 | 2.389126514 | 2.435299524 | 2.737987326 | 2.332093605 |
| ENSMUSG00000011958 | 2.928763473 | 2.668548067 | 2.641989019 | 2.735628626 | 2.665423072 | 3.069668097 | 2.470451221 |
| ENSMUSG00000011960 | 2.781925631 | 2.544354884 | 2.542794199 | 2.600753006 | 2.534786856 | 2.889861721 | 2.330894065 |
| ENSMUSG00000012017 | 2.361637952 | 2.187048212 | 2.32991803 | 1.769203899 | 1.961316494 | 1.919078092 | 2.074322292 |
| ENSMUSG00000012042 | 0 | 0 | 0 | 0 | 0 | 0 | 0 |
| ENSMUSG00000012076 | 2.537433341 | 2.323080408 | 2.320192693 | 2.336628433 | 2.245869735 | 2.593286067 | 2.138226615 |

| | | | | | | | |
|---|---|---|---|---|---|---|---|
| ENSMUSG00000012114 | 2.452372734 | 2.28743678 | 2.311981878 | 2.156427211 | 2.221952687 | 2.408239965 | 2.166993225 |
| ENSMUSG00000012117 | 2.317377295 | 2.134789703 | 2.159156155 | 2.266522298 | 2.222603297 | 2.545307116 | 2.128938453 |
| ENSMUSG00000012123 | 0.289064826 | 0.646240625 | 0.830482024 | 0.896240625 | 0.846351821 | 0.698970004 | 0.579661877 |
| ENSMUSG00000012126 | 1.156259305 | 0.75 | 0.951838731 | 0.646240625 | 0.812711509 | 1 | 1.089372236 |
| ENSMUSG00000012187 | 0 | 0 | 0 | 0 | 0 | 0 | 0.224243824 |
| ENSMUSG00000012211 | 0 | 0.25 | 0 | 0 | 0 | 0 | 0 |
| ENSMUSG00000012282 | 0 | 0 | 0 | 0 | 0 | 0 | 0 |
| ENSMUSG00000012296 | 2.476144923 | 2.344302633 | 2.399978211 | 2.105516192 | 2.292010185 | 2.359835482 | 2.243069492 |
| ENSMUSG00000012350 | 0 | 0 | 0 | 0 | 0 | 0 | 0 |
| ENSMUSG00000012396 | 1.100572389 | 1.114857905 | 1.214495249 | 1.061981878 | 0.905314583 | 1.079181246 | 0.853775828 |
| ENSMUSG00000012405 | 2.833095427 | 2.652525516 | 2.623463862 | 2.775985857 | 2.807048332 | 3.174059808 | 2.643181314 |
| ENSMUSG00000012422 | 2.752926783 | 2.492166698 | 2.473325383 | 2.601816941 | 2.49745007 | 2.857332496 | 2.285005227 |
| ENSMUSG00000012428 | 0.289064826 | 0.646240625 | 0.646240625 | 0.79248125 | 0.812711509 | 1.079181246 | 0.803905701 |
| ENSMUSG00000012429 | 1.655740331 | 1.642463902 | 1.62694866 | 1.794977273 | 1.614672255 | 1.880813592 | 1.56970677 |
| ENSMUSG00000012443 | 3.098350844 | 2.819612365 | 2.787119146 | 2.969800079 | 2.860604721 | 3.312388949 | 2.661769559 |
| ENSMUSG00000012483 | 2.124258828 | 2.013820609 | 1.943696765 | 1.931980114 | 1.846351821 | 2.056904851 | 1.774416565 |
| ENSMUSG00000012519 | 2.389637216 | 2.162762923 | 2.181128463 | 2.125956434 | 2.107593434 | 2.322219295 | 1.955652633 |
| ENSMUSG00000012520 | 0 | 0 | 0 | 0 | 0 | 0 | 0 |
| ENSMUSG00000012535 | 2.800026593 | 2.617156011 | 2.584684096 | 2.625459296 | 2.563636668 | 2.967547976 | 2.390238236 |
| ENSMUSG00000012609 | 2.243591459 | 2.120953944 | 2.036164561 | 2.023189285 | 2.015530186 | 2.278753601 | 1.933441873 |
| ENSMUSG00000012640 | 1.876566388 | 1.80479213 | 1.811981878 | 1.789967834 | 1.674752852 | 1.919078092 | 1.610050272 |
| ENSMUSG00000012705 | 0 | 0 | 0 | 0.25 | 0.244650542 | 0 | 0 |
| ENSMUSG00000012777 | 0 | 0 | 0 | 0 | 0 | 0 | 0 |
| ENSMUSG00000012819 | 1 | 1.17510993 | 1.080482024 | 0.646240625 | 0.568060967 | 0.84509804 | 0.803905701 |
| ENSMUSG00000012848 | 3.112942387 | 2.911189398 | 2.940179987 | 2.678561379 | 2.814903034 | 3.136086097 | 2.713498798 |
| ENSMUSG00000012889 | 1.881174527 | 1.841580554 | 1.841580554 | 1.660964047 | 1.709785497 | 1.755874856 | 1.694069894 |
| ENSMUSG00000013033 | 2.183765914 | 2.123963757 | 2.081607372 | 1.950224975 | 2.025895058 | 2.178976947 | 1.95718226 |
| ENSMUSG00000013076 | 2.803052234 | 2.55357978 | 2.556903986 | 2.54745614 | 2.522774815 | 2.860338007 | 2.372979986 |
| ENSMUSG00000013083 | 0 | 0 | 0 | 0 | 0 | 0 | 0 |
| ENSMUSG00000013089 | 1.358731791 | 1.29248125 | 1.348079356 | 1.711372513 | 1.689774529 | 2 | 1.610050272 |
| ENSMUSG00000013091 | 0.289064826 | 0.396240625 | 0.25 | 0.396240625 | 0.387761935 | 0.477121255 | 0 |
| ENSMUSG00000013150 | 1.89911417 | 1.958222504 | 2.01109853 | 1.864857905 | 1.931471446 | 2.037426498 | 1.865128412 |
| ENSMUSG00000013155 | 1.516991255 | 1.62694866 | 1.807204673 | 1.566696635 | 1.680838453 | 1.740362689 | 1.793950594 |
| ENSMUSG00000013158 | 0 | 0 | 0.396240625 | 0.396240625 | 0.489301084 | 0 | 0.355418052 |

| | | | | | | | |
|---|---|---|---|---|---|---|---|
| ENSMUSG00000013160 | 2.791849225 | 2.544666213 | 2.609438018 | 2.505938588 | 2.497151587 | 2.851258349 | 2.371499364 |
| ENSMUSG00000013236 | 2.583267694 | 2.446567407 | 2.528435542 | 2.254897682 | 2.438487308 | 2.555094449 | 2.412278274 |
| ENSMUSG00000013275 | 2.179305632 | 2.114857905 | 2.121960008 | 2.125956434 | 2.109389652 | 2.350248018 | 2.026921761 |
| ENSMUSG00000013338 | 0 | 0 | 0 | 0 | 0 | 0.301029996 | 0.224243824 |
| ENSMUSG00000013353 | 0 | 0 | 0 | 0 | 0 | 0 | 0 |
| ENSMUSG00000013367 | 1.822201287 | 1.634789703 | 1.766522298 | 1.458222504 | 1.584291694 | 1.491361694 | 1.532233387 |
| ENSMUSG00000013415 | 2.66563648 | 2.42813175 | 2.416333979 | 2.362287161 | 2.388168801 | 2.632457292 | 2.252086845 |
| ENSMUSG00000013418 | 0 | 0.25 | 0.25 | 0 | 0.244650542 | 0 | 0.448487648 |
| ENSMUSG00000013419 | 2.144240249 | 2.112802778 | 2.191217898 | 1.606566189 | 1.736537397 | 1.799340549 | 1.762103709 |
| ENSMUSG00000013465 | 2.555102908 | 2.360735874 | 2.357090043 | 2.30175358 | 2.363323232 | 2.627365857 | 2.202829437 |
| ENSMUSG00000013483 | 0 | 0 | 0.25 | 0.25 | 0 | 0 | 0.224243824 |
| ENSMUSG00000013495 | 2.042557845 | 1.973704441 | 1.940387808 | 1.821350555 | 1.816839698 | 2.033423755 | 1.806347338 |
| ENSMUSG00000013523 | 0 | 0 | 0 | 0 | 0.244650542 | 0 | 0 |
| ENSMUSG00000013539 | 1.928763473 | 1.852347734 | 1.854463129 | 1.821350555 | 1.862942891 | 1.977723605 | 1.757893036 |
| ENSMUSG00000013562 | 0 | 0.25 | 0 | 0 | 0 | 0 | 0 |
| ENSMUSG00000013584 | 0 | 0 | 0 | 0.25 | 0.489301084 | 0 | 0 |
| ENSMUSG00000013593 | 2.911305519 | 2.650442697 | 2.663211243 | 2.793106877 | 2.787923604 | 3.182414652 | 2.613680783 |
| ENSMUSG00000013611 | 0.289064826 | 0.25 | 0 | 0 | 0.489301084 | 0.301029996 | 0 |
| ENSMUSG00000013622 | 2.163307908 | 1.930274797 | 1.943696765 | 1.940387808 | 1.888971102 | 2.198657087 | 1.849546807 |
| ENSMUSG00000013629 | 2.943013009 | 2.745213402 | 2.801906117 | 2.538388008 | 2.690811111 | 2.902546779 | 2.580773997 |
| ENSMUSG00000013643 | 0 | 0 | 0 | 0 | 0 | 0 | 0 |
| ENSMUSG00000013646 | 2.179305632 | 2.088286706 | 2.12892496 | 2.113831805 | 2.242813819 | 2.413299764 | 2.164992504 |
| ENSMUSG00000013653 | 0 | 0 | 0 | 0.25 | 0.244650542 | 0.301029996 | 0.224243824 |
| ENSMUSG00000013662 | 2.925029944 | 2.694108258 | 2.661414608 | 2.680915236 | 2.580774999 | 2.992553518 | 2.393027179 |
| ENSMUSG00000013663 | 3.158622101 | 2.867156011 | 2.839800121 | 2.824945101 | 2.705674618 | 3.120244796 | 2.494225486 |
| ENSMUSG00000013668 | 0 | 0 | 0 | 0 | 0 | 0 | 0 |
| ENSMUSG00000013698 | 2.883252421 | 2.608657057 | 2.61434522 | 2.768535366 | 2.681901575 | 3.137986733 | 2.533777617 |
| ENSMUSG00000013701 | 1.936861443 | 1.868933358 | 1.839388001 | 2.05479213 | 2.009637626 | 2.328379603 | 1.958704689 |
| ENSMUSG00000013707 | 0 | 0 | 0.396240625 | 0.701838731 | 0.244650542 | 0 | 0.355418052 |
| ENSMUSG00000013736 | 2.457006475 | 2.28743678 | 2.272528105 | 2.312574604 | 2.197694248 | 2.507855872 | 2.043115906 |
| ENSMUSG00000013766 | 0 | 0 | 0 | 0 | 0 | 0 | 0 |
| ENSMUSG00000013787 | 2.69268209 | 2.501756817 | 2.619692405 | 2.351285366 | 2.505998068 | 2.725094521 | 2.479838059 |
| ENSMUSG00000013822 | 2.318984359 | 2.198603967 | 2.164552871 | 2.230460234 | 2.280614762 | 2.511883361 | 2.133394659 |
| ENSMUSG00000013833 | 2.051791411 | 2 | 2.125956434 | 1.714495249 | 1.961316494 | 1.934498451 | 2.018825668 |

| | | | | | | | |
|---|---|---|---|---|---|---|---|
| ENSMUSG00000013846 | 1.989530018 | 1.839388001 | 1.811981878 | 1.201838731 | 1.200473444 | 1.397940009 | 1.21680636 |
| ENSMUSG00000013858 | 2.283862692 | 2.209800947 | 2.308404919 | 1.995748394 | 2.192777367 | 2.257678575 | 2.220546465 |
| ENSMUSG00000013878 | 2.119110214 | 1.959800947 | 1.966046536 | 2.071350555 | 1.947418788 | 2.193124598 | 1.845261772 |
| ENSMUSG00000013921 | 2.310886388 | 2.160061234 | 2.213717096 | 2.131869252 | 2.126866297 | 2.342422681 | 2.103267674 |
| ENSMUSG00000013928 | 0 | 0.25 | 0.25 | 0 | 0.387761935 | 0 | 0.520678036 |
| ENSMUSG00000013936 | 0 | 0 | 0 | 0 | 0.387761935 | 0.301029996 | 0 |
| ENSMUSG00000013973 | 1.445324132 | 1.464495249 | 1.418106335 | 1.356566189 | 1.450958105 | 1.544068044 | 1.369798885 |
| ENSMUSG00000013974 | 0 | 0.25 | 0 | 0 | 0 | 0 | 0 |
| ENSMUSG00000013997 | 1.794934559 | 1.646240625 | 1.688721876 | 1.809601185 | 1.704182902 | 2.053078443 | 1.705896842 |
| ENSMUSG00000014030 | 0 | 0 | 0 | 0 | 0 | 0 | 0 |
| ENSMUSG00000014039 | 1.989530018 | 1.905512955 | 1.953445298 | 1.821350555 | 1.868304353 | 1.982271233 | 1.769001954 |
| ENSMUSG00000014074 | 2.518981867 | 2.294977273 | 2.273849256 | 2.333288838 | 2.227765453 | 2.571708832 | 2.126687118 |
| ENSMUSG00000014075 | 1.771760131 | 1.821350555 | 1.821350555 | 1.807204673 | 1.66856048 | 2.025305865 | 1.603286636 |
| ENSMUSG00000014077 | 2.619128496 | 2.373463862 | 2.359177886 | 2.682405186 | 2.604906476 | 3.040206628 | 2.429740345 |
| ENSMUSG00000014104 | 0 | 0 | 0 | 0 | 0 | 0 | 0 |
| ENSMUSG00000014158 | 1.765759561 | 1.701838731 | 1.837182039 | 1.638647213 | 1.836786029 | 1.929418926 | 1.809975587 |
| ENSMUSG00000014164 | 0 | 0.25 | 0 | 0.701838731 | 0.489301084 | 0.301029996 | 0.829800754 |
| ENSMUSG00000014177 | 2.095120908 | 1.976722649 | 1.950224975 | 2.095926073 | 1.992595054 | 2.303196057 | 1.845261772 |
| ENSMUSG00000014195 | 2.857166034 | 2.599151195 | 2.609438018 | 2.70881523 | 2.622504286 | 3.049218023 | 2.441391986 |
| ENSMUSG00000014198 | 0 | 0.25 | 0 | 0.396240625 | 0.489301084 | 0.301029996 | 0.224243824 |
| ENSMUSG00000014226 | 2.830751226 | 2.625707951 | 2.572216519 | 2.703044326 | 2.590305509 | 3.011147361 | 2.424743868 |
| ENSMUSG00000014232 | 1.631440309 | 1.458222504 | 1.516522298 | 1.642463902 | 1.628935049 | 1.826074803 | 1.546120303 |
| ENSMUSG00000014243 | 1.249317394 | 1.282320754 | 1.25 | 1.364857905 | 1.176121988 | 1.477121255 | 1.259064007 |
| ENSMUSG00000014245 | 1.655740331 | 1.54248125 | 1.51109853 | 1.54745614 | 1.680838453 | 1.698970004 | 1.511732338 |
| ENSMUSG00000014294 | 2.097855556 | 2.095926073 | 2.182829758 | 1.882845365 | 2.034859505 | 2.100370545 | 2.072724682 |
| ENSMUSG00000014301 | 1.482695304 | 1.464495249 | 1.348079356 | 1.302363341 | 1.407936347 | 1.278753601 | 1.479989695 |
| ENSMUSG00000014303 | 2.428398755 | 2.267194569 | 2.288071211 | 2.204245906 | 2.317250476 | 2.471291711 | 2.204609933 |
| ENSMUSG00000014313 | 2.69788693 | 2.503156135 | 2.511439915 | 2.40136988 | 2.429560273 | 2.767155866 | 2.274018449 |
| ENSMUSG00000014329 | 1.982401085 | 1.850219859 | 1.763820609 | 1.87892496 | 1.806427258 | 1.934498451 | 1.612273755 |
| ENSMUSG00000014349 | 3.00745275 | 2.753505118 | 2.747171172 | 2.827937079 | 2.771566462 | 3.199755177 | 2.596590088 |
| ENSMUSG00000014351 | 0 | 0.396240625 | 0 | 0 | 0 | 0 | 0 |
| ENSMUSG00000014353 | 2.016886209 | 1.868933358 | 1.938721876 | 2.199415381 | 2.126012715 | 2.423245874 | 1.970632877 |
| ENSMUSG00000014355 | 3.109583075 | 2.822216519 | 2.803731547 | 3.009472521 | 2.928316901 | 3.395326393 | 2.710092562 |
| ENSMUSG00000014361 | 1.527823874 | 1.396240625 | 1.488549078 | 1.470660762 | 1.394615667 | 1.51851394 | 1.340368108 |

| | | | | | | | |
|---|---|---|---|---|---|---|---|
| ENSMUSG00000014402 | 2.418409478 | 2.218361028 | 2.152756199 | 2.197790722 | 2.19206933 | 2.477121255 | 2.034820183 |
| ENSMUSG00000014418 | 2.233869984 | 2.08272922 | 2.038704527 | 2.171625132 | 2.112066955 | 2.340444115 | 1.986347533 |
| ENSMUSG00000014426 | 2.423434025 | 2.236360959 | 2.244319981 | 2.133818844 | 2.171644683 | 2.374748346 | 2.072190387 |
| ENSMUSG00000014444 | 2.877408028 | 2.695134942 | 2.769203899 | 2.484159485 | 2.594182934 | 2.836956737 | 2.552846438 |
| ENSMUSG00000014453 | 0.289064826 | 0 | 0 | 0 | 0 | 0 | 0 |
| ENSMUSG00000014470 | 1.894701088 | 1.834962501 | 1.92510993 | 1.598079356 | 1.814781639 | 1.919078092 | 1.783679399 |
| ENSMUSG00000014496 | 2.481610694 | 2.261780978 | 2.27053726 | 2.544666213 | 2.433873436 | 2.790285164 | 2.2551389 |
| ENSMUSG00000014498 | 2.880757703 | 2.662538553 | 2.695544798 | 2.70881523 | 2.732574981 | 3.08170727 | 2.605659747 |
| ENSMUSG00000014503 | 1.771760131 | 1.688721876 | 1.580482024 | 1.516522298 | 1.572189156 | 1.568201724 | 1.40946263 |
| ENSMUSG00000014504 | 2.392336464 | 2.168106335 | 2.213717096 | 2.257666784 | 2.204601633 | 2.465382851 | 2.085803078 |
| ENSMUSG00000014529 | 0 | 0 | 0 | 0 | 0 | 0 | 0 |
| ENSMUSG00000014542 | 0 | 0 | 0 | 0 | 0 | 0 | 0 |
| ENSMUSG00000014543 | 0 | 0 | 0 | 0 | 0 | 0 | 0 |
| ENSMUSG00000014547 | 1.470606563 | 1.464495249 | 1.364857905 | 1.476722649 | 1.533172725 | 1.643452676 | 1.433437704 |
| ENSMUSG00000014550 | 2.508932407 | 2.329353153 | 2.29745614 | 2.347542239 | 2.291475807 | 2.598790507 | 2.162170545 |
| ENSMUSG00000014551 | 2.231898196 | 2.073655187 | 2.061981878 | 2.090485943 | 2.13194489 | 2.298853076 | 2.002662694 |
| ENSMUSG00000014554 | 1.734388958 | 1.649978211 | 1.664552871 | 1.747171172 | 1.655839964 | 1.880813592 | 1.637827726 |
| ENSMUSG00000014592 | 1.516991255 | 1.505591953 | 1.516522298 | 1.494319981 | 1.546663136 | 1.643452676 | 1.483307832 |
| ENSMUSG00000014599 | 3.396595037 | 3.101949713 | 3.105975941 | 2.977004315 | 3.001147593 | 3.43264866 | 2.798740124 |
| ENSMUSG00000014601 | 2.484856099 | 2.324229052 | 2.3464656 | 2.358135475 | 2.41591954 | 2.662757832 | 2.242754021 |
| ENSMUSG00000014602 | 2.619128496 | 2.441217898 | 2.523189285 | 1.418106335 | 1.509475496 | 1.568201724 | 1.511732338 |
| ENSMUSG00000014603 | 0.289064826 | 0.396240625 | 0.5 | 1 | 1.074582839 | 1 | 0.876096088 |
| ENSMUSG00000014606 | 2.582415738 | 2.381869252 | 2.387686696 | 2.411414608 | 2.368520226 | 2.695481676 | 2.256654198 |
| ENSMUSG00000014609 | 0 | 0 | 0 | 0 | 0 | 0 | 0.224243824 |
| ENSMUSG00000014633 | 1.903481042 | 1.78743678 | 1.860735874 | 1.858657057 | 1.871833944 | 2.071882007 | 1.863093717 |
| ENSMUSG00000014668 | 2.530730035 | 2.333288838 | 2.328787391 | 2.307204673 | 2.278401864 | 2.568201724 | 2.181807364 |
| ENSMUSG00000014686 | 0 | 0 | 0 | 0 | 0 | 0 | 0 |
| ENSMUSG00000014704 | 1.036286563 | 1.330482024 | 1.201838731 | 1.04248125 | 1.106691885 | 1.079181246 | 1.054044578 |
| ENSMUSG00000014725 | 0.45815691 | 0.646240625 | 0.5 | 0.580482024 | 0.686820904 | 0.301029996 | 0.710836105 |
| ENSMUSG00000014747 | 0.45815691 | 0.396240625 | 0.580482024 | 0.75 | 0.568060967 | 0.602059991 | 0.672731473 |
| ENSMUSG00000014748 | 2.610448227 | 2.354463129 | 2.393411797 | 2.42293588 | 2.439894911 | 2.788168371 | 2.312854549 |
| ENSMUSG00000014763 | 2.35727108 | 2.171625132 | 2.188721876 | 2.209012589 | 2.158641579 | 2.450249108 | 2.043115906 |
| ENSMUSG00000014767 | 2.413323654 | 2.212155735 | 2.211372513 | 2.27908599 | 2.164820399 | 2.505149978 | 2.021338422 |
| ENSMUSG00000014769 | 3.022860988 | 2.757494337 | 2.760927716 | 2.797764804 | 2.771429206 | 3.232233521 | 2.599432116 |

| | | | | | | | |
|---|---|---|---|---|---|---|---|
| ENSMUSG00000014771 | 0 | 0 | 0 | 0 | 0 | 0 | 0 |
| ENSMUSG00000014773 | 0.289064826 | 0.396240625 | 0.396240625 | 0.396240625 | 0.489301084 | 0 | 0.448487648 |
| ENSMUSG00000014776 | 1 | 0.830482024 | 1.02186571 | 1.130890489 | 1 | 1.041392685 | 1.193409508 |
| ENSMUSG00000014778 | 2.22189694 | 2.150906586 | 2.184523065 | 1.992885888 | 2.12771782 | 2.230448921 | 2.096890362 |
| ENSMUSG00000014782 | 1.876566388 | 1.784887838 | 1.910964047 | 1.741446071 | 1.895599581 | 2.029383778 | 1.817112247 |
| ENSMUSG00000014786 | 2.039433992 | 1.903677461 | 1.950224975 | 1.854463129 | 1.959951091 | 1.963787827 | 1.864112664 |
| ENSMUSG00000014791 | 1.249317394 | 1.146240625 | 1.098079356 | 0.896240625 | 1.074582839 | 1.113943352 | 1.159323753 |
| ENSMUSG00000014813 | 0.45815691 | 0.396240625 | 0.25 | 0 | 0.387761935 | 0 | 0.448487648 |
| ENSMUSG00000014837 | 1.857607105 | 1.779735268 | 1.821350555 | 1.79745614 | 1.941702774 | 1.991226076 | 1.82753046 |
| ENSMUSG00000014846 | 1.069666964 | 1.080482024 | 1.17510993 | 1.160964047 | 1.439191811 | 1.230448921 | 1.505628092 |
| ENSMUSG00000014850 | 2.194712275 | 2.02186571 | 1.958222504 | 2.269203899 | 2.21736452 | 2.531478917 | 2.05853792 |
| ENSMUSG00000014852 | 0 | 0 | 0 | 0 | 0.244650542 | 0 | 0 |
| ENSMUSG00000014856 | 2.205378646 | 2.077084758 | 2.100219859 | 1.914552871 | 2.058398634 | 2.136720567 | 1.978585612 |
| ENSMUSG00000014859 | 2.414600936 | 2.246460484 | 2.266522298 | 2.24929487 | 2.289867804 | 2.609594409 | 2.166594069 |
| ENSMUSG00000014867 | 3.167522521 | 2.890679857 | 2.876576895 | 2.818303202 | 2.829874135 | 3.230704314 | 2.64373011 |
| ENSMUSG00000014873 | 2.119110214 | 1.984159485 | 1.940387808 | 1.880890489 | 1.85749873 | 2.136720567 | 1.812372009 |
| ENSMUSG00000014905 | 1.876566388 | 1.741446071 | 1.75 | 1.75835575 | 1.576269619 | 1.892094603 | 1.466365979 |
| ENSMUSG00000014907 | 2.131864492 | 1.990000483 | 1.992885888 | 2.024508021 | 1.957204337 | 2.195899652 | 1.86207155 |
| ENSMUSG00000014932 | 2.470606563 | 2.310793496 | 2.30539678 | 2.419870025 | 2.317250476 | 2.627365857 | 2.125328759 |
| ENSMUSG00000014956 | 3.223183227 | 2.927163209 | 2.895888233 | 2.926083993 | 2.832077581 | 3.28057837 | 2.609644355 |
| ENSMUSG00000014959 | 2.654307229 | 2.442872367 | 2.430274797 | 2.517194569 | 2.512061267 | 2.846955325 | 2.387828426 |
| ENSMUSG00000014980 | 1.993049299 | 1.907339155 | 1.896240625 | 1.991446071 | 1.92093466 | 2.201397124 | 1.786277931 |
| ENSMUSG00000015001 | 0 | 0 | 0 | 0 | 0 | 0.301029996 | 0 |
| ENSMUSG00000015002 | 2.93940544 | 2.697790722 | 2.697383411 | 2.807505109 | 2.711416536 | 3.128399269 | 2.502956138 |
| ENSMUSG00000015013 | 2.063793172 | 1.858657057 | 1.919870025 | 2.032320754 | 1.996316946 | 2.243038049 | 1.874129423 |
| ENSMUSG00000015016 | 1.777675583 | 1.646240625 | 1.714495249 | 1.584962501 | 1.726078673 | 1.857332496 | 1.750751125 |
| ENSMUSG00000015023 | 2.412042447 | 2.245748394 | 2.27186571 | 2.327084758 | 2.297309926 | 2.610660163 | 2.160139682 |
| ENSMUSG00000015027 | 1.944805158 | 1.896240625 | 1.921625132 | 1.987091808 | 2.015530186 | 2.167317335 | 1.912234055 |
| ENSMUSG00000015053 | 1.956444026 | 2 | 2.03743678 | 1.681980114 | 1.880506521 | 1.929418926 | 1.89047692 |
| ENSMUSG00000015083 | 0.960252568 | 1.080482024 | 1.04248125 | 0.896240625 | 0.905314583 | 1.113943352 | 0.853775828 |
| ENSMUSG00000015085 | 0.289064826 | 0 | 0 | 0.25 | 0.489301084 | 0 | 0.448487648 |
| ENSMUSG00000015087 | 2.643399047 | 2.428561379 | 2.464883697 | 2.517865591 | 2.534518346 | 2.803457116 | 2.394412686 |
| ENSMUSG00000015090 | 0.289064826 | 0.5 | 0.580482024 | 0.25 | 0.568060967 | 0.477121255 | 0.672731473 |
| ENSMUSG00000015092 | 2.289064826 | 2.272528105 | 2.314935816 | 1.96759118 | 2.208005287 | 2.195899652 | 2.20139795 |

| | | | | | | | |
|---|---|---|---|---|---|---|---|
| ENSMUSG00000015093 | 0.811507563 | 0.580482024 | 0.701838731 | 0.646240625 | 0.905314583 | 0.698970004 | 0.896975297 |
| ENSMUSG00000015094 | 1.847794126 | 1.860735874 | 1.96759118 | 1.821350555 | 2.042520869 | 2.041392685 | 2.080113611 |
| ENSMUSG00000015095 | 2.235832493 | 2.105516192 | 2.154596376 | 2.064346961 | 2.149165677 | 2.290034611 | 2.030591163 |
| ENSMUSG00000015112 | 2.539328946 | 2.341033664 | 2.328220739 | 2.159156155 | 2.062514787 | 2.432969291 | 1.975716277 |
| ENSMUSG00000015120 | 3.066473338 | 2.791854536 | 2.796063874 | 2.767194569 | 2.704182902 | 3.106870544 | 2.526638571 |
| ENSMUSG00000015126 | 1.816888701 | 1.843759858 | 1.905512955 | 1.723704441 | 1.873585591 | 1.929418926 | 1.822941512 |
| ENSMUSG00000015127 | 1.747221736 | 1.71759118 | 1.784887838 | 1.692046081 | 1.744182955 | 1.707570176 | 1.764880683 |
| ENSMUSG00000015133 | 2.721148722 | 2.533285553 | 2.541854536 | 2.299918086 | 2.383255179 | 2.633468456 | 2.229547475 |
| ENSMUSG00000015134 | 1.249317394 | 1.17510993 | 1.29248125 | 0.864857905 | 0.905314583 | 0.84509804 | 0.916588299 |
| ENSMUSG00000015143 | 3.644669774 | 3.326729028 | 3.311870634 | 2.821928095 | 2.823772347 | 3.225825991 | 2.629818343 |
| ENSMUSG00000015149 | 2.211649928 | 2.147178659 | 2.153677461 | 1.981203126 | 2.080465577 | 2.260071388 | 2.035419841 |
| ENSMUSG00000015165 | 3.124491352 | 2.823798739 | 2.825374049 | 2.848347614 | 2.793770609 | 3.238297068 | 2.618755249 |
| ENSMUSG00000015176 | 3.01386641 | 2.772528105 | 2.767697952 | 2.744498842 | 2.743440671 | 3.122215878 | 2.566821826 |
| ENSMUSG00000015189 | 2.481610694 | 2.275165585 | 2.235628626 | 2.102347734 | 1.992595054 | 2.344392274 | 1.920029985 |
| ENSMUSG00000015202 | 2.083999385 | 1.970660762 | 1.976722649 | 1.870953944 | 1.866526233 | 2.029383778 | 1.811176017 |
| ENSMUSG00000015214 | 2.17479713 | 2.024508021 | 1.978222334 | 2.256284891 | 2.171644683 | 2.459392488 | 2.01501956 |
| ENSMUSG00000015217 | 2.495495257 | 2.252107156 | 2.27186571 | 2.192046081 | 2.126866297 | 2.421603927 | 1.971363985 |
| ENSMUSG00000015222 | 2.578990401 | 2.379909063 | 2.320772088 | 1.907339155 | 1.902105869 | 2.096910013 | 1.74492186 |
| ENSMUSG00000015224 | 0.289064826 | 0.580482024 | 0.5 | 0.396240625 | 0 | 0 | 0.355418052 |
| ENSMUSG00000015242 | 0 | 0 | 0 | 0 | 0 | 0 | 0 |
| ENSMUSG00000015243 | 1.631440309 | 1.476722649 | 1.488549078 | 1.575945187 | 1.559656853 | 1.748188027 | 1.508694612 |
| ENSMUSG00000015247 | 2.310886388 | 2.14151351 | 2.121960008 | 2.116901388 | 2.099395374 | 2.378397901 | 1.943148064 |
| ENSMUSG00000015289 | 2.048736215 | 1.970660762 | 1.935366747 | 2.023189285 | 2.003645018 | 2.322219295 | 1.879993912 |
| ENSMUSG00000015290 | 2.282114126 | 2.129909063 | 2.159156155 | 2.328220739 | 2.282265378 | 2.58546073 | 2.112367894 |
| ENSMUSG00000015291 | 2.867194479 | 2.566989271 | 2.583847589 | 2.73654381 | 2.69373167 | 3.083144143 | 2.495961709 |
| ENSMUSG00000015305 | 1.903481042 | 1.79248125 | 1.914552871 | 2.12694866 | 2.080465577 | 2.315970345 | 1.972093446 |
| ENSMUSG00000015312 | 2.10861828 | 2.004202072 | 2.135757955 | 2.093759858 | 2.219338185 | 2.399673721 | 2.264126018 |
| ENSMUSG00000015314 | 0 | 0 | 0 | 0 | 0.387761935 | 0 | 0 |
| ENSMUSG00000015316 | 0.45815691 | 0.25 | 0.5 | 0.5 | 0.244650542 | 0 | 0.224243824 |
| ENSMUSG00000015335 | 1.700465191 | 1.726722649 | 1.856566189 | 1.403677461 | 1.692703642 | 1.591064607 | 1.734460864 |
| ENSMUSG00000015337 | 0.45815691 | 0.5 | 0.646240625 | 0 | 0.489301084 | 0.301029996 | 0.672731473 |
| ENSMUSG00000015340 | 0 | 0 | 0 | 0 | 0 | 0 | 0 |
| ENSMUSG00000015341 | 2.544030601 | 2.367410454 | 2.351285366 | 2.372963274 | 2.318738698 | 2.673020907 | 2.185218806 |
| ENSMUSG00000015342 | 1.87190676 | 1.769203899 | 1.814346961 | 1.752806814 | 1.706995316 | 1.838849091 | 1.62316756 |

| | | | | | | | |
|---|---|---|---|---|---|---|---|
| ENSMUSG00000015354 | 2.124258828 | 1.96759118 | 2.046218836 | 1.926839783 | 1.944572352 | 2.240549248 | 1.840919217 |
| ENSMUSG00000015355 | 0 | 0 | 0 | 0 | 0 | 0 | 0 |
| ENSMUSG00000015357 | 2.302628051 | 2.129909063 | 2.152756199 | 2.353935442 | 2.305627392 | 2.618048097 | 2.123508695 |
| ENSMUSG00000015363 | 2.340856238 | 2.219895812 | 2.26584877 | 2.012462137 | 2.206647761 | 2.354108439 | 2.178744473 |
| ENSMUSG00000015365 | 0 | 0 | 0 | 0.25 | 0 | 0 | 0 |
| ENSMUSG00000015377 | 1.700465191 | 1.532320754 | 1.668106335 | 1.561981878 | 1.65906333 | 1.612783857 | 1.502532226 |
| ENSMUSG00000015396 | 0.289064826 | 0.25 | 0.25 | 1.04248125 | 0.686820904 | 0.602059991 | 0.74492186 |
| ENSMUSG00000015401 | 0 | 0 | 0 | 0 | 0 | 0 | 0 |
| ENSMUSG00000015405 | 0 | 0 | 0 | 0 | 0 | 0 | 0 |
| ENSMUSG00000015437 | 0 | 0 | 0 | 0 | 0 | 0 | 0.224243824 |
| ENSMUSG00000015441 | 0 | 0 | 0 | 0 | 0 | 0 | 0 |
| ENSMUSG00000015443 | 0 | 0 | 0 | 0 | 0 | 0 | 0 |
| ENSMUSG00000015451 | 0.747221736 | 0.646240625 | 0.75 | 0.864857905 | 1.136121934 | 0.77815125 | 1.014380832 |
| ENSMUSG00000015452 | 0.45815691 | 0.75 | 0.951838731 | 0.951838731 | 1.020174412 | 0.903089987 | 1.078019652 |
| ENSMUSG00000015461 | 2.183765914 | 2.121960008 | 2.160061234 | 1.995748394 | 2.1962964 | 2.318063335 | 2.255442528 |
| ENSMUSG00000015467 | 0 | 0.75 | 0.646240625 | 0.25 | 0.568060967 | 0 | 0.520678036 |
| ENSMUSG00000015468 | 0.289064826 | 0.75 | 0.79248125 | 0.646240625 | 0.978602168 | 0.602059991 | 1 |
| ENSMUSG00000015474 | 1.605636262 | 1.464495249 | 1.488549078 | 1.657339155 | 1.665423072 | 1.897627091 | 1.591689859 |
| ENSMUSG00000015476 | 0.45815691 | 0.25 | 0 | 0.25 | 0.244650542 | 0 | 0.224243824 |
| ENSMUSG00000015478 | 1.614416215 | 1.598079356 | 1.668106335 | 1.54248125 | 1.66856048 | 1.716003344 | 1.655750087 |
| ENSMUSG00000015484 | 0 | 0 | 0 | 0.25 | 0 | 0 | 0 |
| ENSMUSG00000015488 | 2.310886388 | 2.185366747 | 2.185366747 | 2.202642909 | 2.173890435 | 2.489958479 | 2.080634989 |
| ENSMUSG00000015501 | 1.993049299 | 1.862802778 | 1.802363341 | 1.995748394 | 1.940259189 | 2.187520721 | 1.860017477 |
| ENSMUSG00000015519 | 0 | 0 | 0 | 0 | 0 | 0 | 0 |
| ENSMUSG00000015522 | 2.559630991 | 2.354990044 | 2.36434522 | 2.500703754 | 2.450617578 | 2.762678564 | 2.282223478 |
| ENSMUSG00000015533 | 1.289064826 | 1.238549078 | 1.238549078 | 1.061981878 | 1 | 1.113943352 | 0.853775828 |
| ENSMUSG00000015536 | 2.213719572 | 2.074802005 | 2.077084758 | 2.352877747 | 2.256066288 | 2.620136055 | 2.079068325 |
| ENSMUSG00000015542 | 1.527823874 | 1.566696635 | 1.580482024 | 1.403677461 | 1.514343934 | 1.568201724 | 1.483307832 |
| ENSMUSG00000015568 | 2.333175866 | 2.189555804 | 2.219895812 | 1.388647213 | 1.27449559 | 1.447158031 | 1.307989563 |
| ENSMUSG00000015575 | 2.577267124 | 2.414552871 | 2.411864607 | 2.511439915 | 2.432082686 | 2.85308953 | 2.318147869 |
| ENSMUSG00000015579 | 1.269664473 | 1.27186571 | 1.494319981 | 1 | 1.414412788 | 1.322219295 | 1.564611933 |
| ENSMUSG00000015597 | 2.340856238 | 2.168989258 | 2.127938163 | 2.242166698 | 2.182733222 | 2.515873844 | 2.025071334 |
| ENSMUSG00000015599 | 0.747221736 | 0.75 | 1.02186571 | 1.130890489 | 1.373641807 | 1.079181246 | 1.448487648 |
| ENSMUSG00000015605 | 2.563218389 | 2.431980114 | 2.506630861 | 2.075945187 | 2.205967036 | 2.324282455 | 2.123052075 |

| | | | | | | | |
|---|---|---|---|---|---|---|---|
| ENSMUSG00000015619 | 1.975148165 | 1.909156155 | 2 | 1.784887838 | 1.814781639 | 1.963787827 | 1.779742043 |
| ENSMUSG00000015627 | 0 | 0 | 0 | 0.25 | 0.244650542 | 0 | 0 |
| ENSMUSG00000015647 | 2.824116479 | 2.683889006 | 2.747525991 | 2.057204673 | 2.345425551 | 2.44870632 | 2.318397784 |
| ENSMUSG00000015652 | 2.069666964 | 1.92510993 | 1.910964047 | 1.950224975 | 1.932951348 | 2.170261715 | 1.822941512 |
| ENSMUSG00000015653 | 2.841885121 | 2.601551251 | 2.570482507 | 2.678561379 | 2.60092323 | 3.004321374 | 2.42167221 |
| ENSMUSG00000015656 | 3.433305075 | 3.119439567 | 3.09801226 | 3.253810201 | 3.156111746 | 3.695131298 | 2.931720444 |
| ENSMUSG00000015659 | 2.066740409 | 1.848079356 | 1.919870025 | 1.701838731 | 1.665423072 | 1.755874856 | 1.612273755 |
| ENSMUSG00000015665 | 0 | 0 | 0 | 0 | 0 | 0 | 0 |
| ENSMUSG00000015668 | 2.480523256 | 2.280383379 | 2.232684334 | 2.242166698 | 2.187791022 | 2.540329475 | 2.059651572 |
| ENSMUSG00000015671 | 2.966750129 | 2.708222504 | 2.688930538 | 2.90136988 | 2.809151845 | 3.277150614 | 2.588995228 |
| ENSMUSG00000015672 | 2.039433992 | 1.945339928 | 1.935366747 | 2.099151195 | 2.021325983 | 2.352182518 | 1.930969131 |
| ENSMUSG00000015697 | 2.626865868 | 2.406883471 | 2.387686696 | 2.469512728 | 2.444777781 | 2.774516966 | 2.253311124 |
| ENSMUSG00000015702 | 1.389637216 | 1.396240625 | 1.464495249 | 1.238549078 | 1.27449559 | 1.380211242 | 1.168187745 |
| ENSMUSG00000015709 | 2.531694272 | 2.253505118 | 2.31728167 | 2.132845365 | 2.086251872 | 2.357934847 | 2.001334086 |
| ENSMUSG00000015711 | 2.326928073 | 2.122963274 | 2.120953944 | 2.250703754 | 2.227765453 | 2.481442629 | 2.066254157 |
| ENSMUSG00000015714 | 2.82886633 | 2.570772088 | 2.573655187 | 2.740362424 | 2.666163799 | 3.061829307 | 2.503520492 |
| ENSMUSG00000015721 | 0 | 0 | 0 | 0.580482024 | 0.244650542 | 0 | 0.355418052 |
| ENSMUSG00000015733 | 3.163942822 | 2.899047439 | 2.87233656 | 2.812574604 | 2.728257421 | 3.183554534 | 2.519069836 |
| ENSMUSG00000015745 | 2.455852858 | 2.352347734 | 2.383332433 | 1.843759858 | 2.013184954 | 2.045322979 | 1.982842456 |
| ENSMUSG00000015747 | 2.22592627 | 1.991446071 | 2.031030328 | 2.145300145 | 2.056322416 | 2.330413773 | 1.938331368 |
| ENSMUSG00000015748 | 2.547754129 | 2.328220739 | 2.327653195 | 2.370953944 | 2.344045917 | 2.620136055 | 2.190455009 |
| ENSMUSG00000015749 | 3.070726095 | 2.802972074 | 2.764328719 | 2.957925775 | 2.886700645 | 3.353531559 | 2.705670528 |
| ENSMUSG00000015750 | 2.271465912 | 2.179419106 | 2.245748394 | 2.132845365 | 2.252501045 | 2.423245874 | 2.179128924 |
| ENSMUSG00000015755 | 2.945519866 | 2.682617534 | 2.64127552 | 2.633575721 | 2.546403514 | 2.905256049 | 2.366586192 |
| ENSMUSG00000015757 | 2.396352858 | 2.237821179 | 2.178561379 | 2.352347734 | 2.27729019 | 2.597695186 | 2.123508695 |
| ENSMUSG00000015759 | 2.489144387 | 2.339388001 | 2.325945187 | 2.561090959 | 2.458367761 | 2.867467488 | 2.330894065 |
| ENSMUSG00000015766 | 2.282114126 | 2.175975893 | 2.182829758 | 2.202642909 | 2.160196454 | 2.40654018 | 2.073258096 |
| ENSMUSG00000015776 | 2.592526109 | 2.432829758 | 2.415444524 | 2.398114259 | 2.410998892 | 2.649334859 | 2.256351705 |
| ENSMUSG00000015787 | 0 | 0 | 0 | 0 | 0 | 0 | 0 |
| ENSMUSG00000015790 | 2.116511873 | 1.969129237 | 2.00835575 | 1.942046081 | 1.932951348 | 2.113943352 | 1.834294096 |
| ENSMUSG00000015804 | 2.511972754 | 2.335518667 | 2.347004321 | 2.245748394 | 2.268839121 | 2.547774705 | 2.191196167 |
| ENSMUSG00000015806 | 2.051791411 | 1.880890489 | 1.898114259 | 1.987091808 | 2.00243419 | 2.176091259 | 1.910475814 |
| ENSMUSG00000015812 | 0.811507563 | 0.5 | 0.646240625 | 0.896240625 | 0.812711509 | 0.954242509 | 0.74492186 |
| ENSMUSG00000015829 | 0 | 0.25 | 0.25 | 0.396240625 | 0 | 0.301029996 | 0 |

| | | | | | | | |
|---|---|---|---|---|---|---|---|
| ENSMUSG00000015837 | 3.267525419 | 2.967976309 | 2.975311008 | 2.736360959 | 2.757140935 | 3.116939647 | 2.641161057 |
| ENSMUSG00000015839 | 2.266038081 | 2.122963274 | 2.127938163 | 2.349151195 | 2.238490593 | 2.539076099 | 2.090889903 |
| ENSMUSG00000015843 | 0 | 0 | 0 | 0.25 | 0 | 0 | 0.224243824 |
| ENSMUSG00000015846 | 2.435736758 | 2.358135475 | 2.367410454 | 2.05357978 | 2.136121934 | 2.301029996 | 2.065163042 |
| ENSMUSG00000015850 | 2.066740409 | 2.017865591 | 2.111770807 | 1.916333979 | 2.111176775 | 2.071882007 | 2.117986249 |
| ENSMUSG00000015852 | 0 | 0 | 0 | 0 | 0 | 0 | 0 |
| ENSMUSG00000015854 | 0 | 0 | 0 | 0 | 0 | 0 | 0 |
| ENSMUSG00000015869 | 2.507914013 | 2.312574604 | 2.308404919 | 2.362287161 | 2.384078881 | 2.636487896 | 2.267652107 |
| ENSMUSG00000015879 | 0.289064826 | 0.396240625 | 0.5 | 0.396240625 | 0.387761935 | 0.301029996 | 0.224243824 |
| ENSMUSG00000015880 | 2.763003223 | 2.531353368 | 2.545598588 | 2.616646584 | 2.5461437 | 2.916980047 | 2.337319372 |
| ENSMUSG00000015882 | 2.320585253 | 2.131869252 | 2.187886015 | 2.134789703 | 2.004851707 | 2.28780173 | 1.856911705 |
| ENSMUSG00000015889 | 2.636714502 | 2.428561379 | 2.392938161 | 2.534247778 | 2.449252175 | 2.876217841 | 2.315638039 |
| ENSMUSG00000015890 | 0.289064826 | 0.25 | 0 | 0.25 | 0 | 0.301029996 | 0 |
| ENSMUSG00000015932 | 3.433194207 | 3.103076301 | 3.068303202 | 2.829353153 | 2.739855812 | 3.1775365 | 2.54076252 |
| ENSMUSG00000015937 | 2.597490839 | 2.420309603 | 2.357090043 | 2.626452888 | 2.621875691 | 3.005180513 | 2.443604939 |
| ENSMUSG00000015942 | 1.83262764 | 1.794977273 | 1.782320754 | 1.593759858 | 1.686820904 | 1.770852012 | 1.63986884 |
| ENSMUSG00000015943 | 0.747221736 | 1.098079356 | 1.188721876 | 0.830482024 | 0.931471446 | 0.698970004 | 1.278288402 |
| ENSMUSG00000015944 | 2.455852858 | 2.27186571 | 2.282320754 | 2.294354884 | 2.331857641 | 2.564666064 | 2.198153757 |
| ENSMUSG00000015947 | 0 | 0.25 | 0 | 0 | 0 | 0 | 0 |
| ENSMUSG00000015950 | 0.45815691 | 0.396240625 | 0.830482024 | 0.5 | 0.489301084 | 0.301029996 | 0.224243824 |
| ENSMUSG00000015957 | 0.45815691 | 0.580482024 | 0.25 | 0.646240625 | 0.905314583 | 0.602059991 | 1 |
| ENSMUSG00000015961 | 2.8302808 | 2.591580554 | 2.581326364 | 2.703445298 | 2.629756355 | 3.015359755 | 2.450984564 |
| ENSMUSG00000015962 | 0 | 0.25 | 0.25 | 0 | 0.387761935 | 0 | 0.520678036 |
| ENSMUSG00000015966 | 0.45815691 | 0.25 | 0.25 | 0 | 0 | 0.301029996 | 0 |
| ENSMUSG00000015968 | 0 | 0 | 0 | 0 | 0.244650542 | 0 | 0.224243824 |
| ENSMUSG00000015970 | 0 | 0 | 0 | 0 | 0 | 0 | 0 |
| ENSMUSG00000015971 | 2.039433992 | 1.981203126 | 1.994319981 | 2.17510993 | 2.126866297 | 2.421603927 | 2.015657029 |
| ENSMUSG00000015980 | 0.91631382 | 0.864857905 | 0.951838731 | 1.098079356 | 0.877063019 | 0.954242509 | 1.041356071 |
| ENSMUSG00000015981 | 0 | 0 | 0.25 | 0.830482024 | 0.733951626 | 0.77815125 | 0.672731473 |
| ENSMUSG00000015994 | 2.647066901 | 2.407794264 | 2.400906586 | 2.574515642 | 2.488995627 | 2.857935265 | 2.303835214 |
| ENSMUSG00000016018 | 2.68005385 | 2.465271726 | 2.461764337 | 2.702039943 | 2.60666248 | 2.987666265 | 2.43115385 |
| ENSMUSG00000016024 | 0.811507563 | 0.580482024 | 0.75 | 0.79248125 | 0.77552387 | 0.903089987 | 0.672731473 |
| ENSMUSG00000016028 | 1.156259305 | 1.356566189 | 1.188721876 | 1.364857905 | 1.499532413 | 1.505149978 | 1.562034107 |
| ENSMUSG00000016087 | 0.289064826 | 0 | 0.396240625 | 1.302363341 | 1.343584837 | 1.278753601 | 1.176815335 |

| | | | | | | | |
|---|---|---|---|---|---|---|---|
| ENSMUSG00000016128 | 2.508932407 | 2.343759858 | 2.328220739 | 2.248588359 | 2.299407739 | 2.530199698 | 2.151886289 |
| ENSMUSG00000016150 | 0.289064826 | 0.25 | 0 | 0 | 0 | 0 | 0 |
| ENSMUSG00000016179 | 0 | 0.25 | 0 | 0 | 0 | 0 | 0 |
| ENSMUSG00000016181 | 2.116511873 | 1.972185812 | 1.969129237 | 1.992885888 | 2.020174412 | 2.195899652 | 1.830929954 |
| ENSMUSG00000016194 | 0 | 0 | 0 | 0 | 0 | 0 | 0 |
| ENSMUSG00000016200 | 0.289064826 | 0.25 | 0 | 1.794977273 | 1.754126039 | 1.968482949 | 1.647906904 |
| ENSMUSG00000016206 | 1.960252568 | 1.858657057 | 1.896240625 | 1.580482024 | 1.563883923 | 1.633468456 | 1.526507301 |
| ENSMUSG00000016239 | 1.631440309 | 1.614857905 | 1.747171172 | 1.311981878 | 1.234113756 | 1.301029996 | 1.238624656 |
| ENSMUSG00000016252 | 2.105953527 | 1.96293726 | 2.02053726 | 1.973704441 | 2 | 2.133538908 | 1.953344547 |
| ENSMUSG00000016253 | 2.181541736 | 2.090485943 | 2.163659007 | 2.180274797 | 2.233484039 | 2.447158031 | 2.140399326 |
| ENSMUSG00000016255 | 0 | 0 | 0 | 0 | 0 | 0 | 0 |
| ENSMUSG00000016256 | 2.17479713 | 2.074802005 | 2.122963274 | 2.234894804 | 2.275056283 | 2.530199698 | 2.176815335 |
| ENSMUSG00000016257 | 2.410757293 | 2.212155735 | 2.197790722 | 2.344844592 | 2.28171603 | 2.597695186 | 2.156863548 |
| ENSMUSG00000016262 | 2.778209214 | 2.537119146 | 2.485628626 | 2.331045137 | 2.28771234 | 2.624282096 | 2.137353429 |
| ENSMUSG00000016283 | 0 | 0.25 | 0.5 | 0.25 | 0.244650542 | 0.477121255 | 0.448487648 |
| ENSMUSG00000016308 | 2.399009124 | 2.227473271 | 2.248588359 | 2.345385738 | 2.276175004 | 2.619093331 | 2.133394659 |
| ENSMUSG00000016319 | 3.046780281 | 2.818594538 | 2.802819987 | 2.794043287 | 2.79183235 | 3.174931594 | 2.61307852 |
| ENSMUSG00000016327 | 0 | 0.25 | 0 | 0 | 0 | 0 | 0 |
| ENSMUSG00000016344 | 1.494443473 | 1.445339928 | 1.431980114 | 1.372963274 | 1.456697355 | 1.462397998 | 1.499406449 |
| ENSMUSG00000016346 | 0.747221736 | 0.5 | 0.580482024 | 0.396240625 | 0.632412477 | 0.301029996 | 0.355418052 |
| ENSMUSG00000016349 | 0 | 0 | 0 | 0.396240625 | 0.244650542 | 0 | 0.355418052 |
| ENSMUSG00000016356 | 0.91631382 | 1 | 1.04248125 | 0.864857905 | 1.091002363 | 0.77815125 | 1.066254157 |
| ENSMUSG00000016382 | 3.346217862 | 3.045520982 | 3.010414788 | 2.727847997 | 2.614886232 | 3.035429738 | 2.40946263 |
| ENSMUSG00000016386 | 1.639698646 | 1.51109853 | 1.527131114 | 1.54248125 | 1.509475496 | 1.662757832 | 1.459332757 |
| ENSMUSG00000016409 | 2.287337982 | 2.137686696 | 2.125956434 | 2.274508021 | 2.215379757 | 2.496929648 | 2.040177507 |
| ENSMUSG00000016427 | 2.083999385 | 1.984159485 | 2.087182039 | 1.87892496 | 1.905314583 | 2.1430148 | 1.853775828 |
| ENSMUSG00000016458 | 0.289064826 | 0.25 | 0 | 0.5 | 0.244650542 | 0.602059991 | 0.355418052 |
| ENSMUSG00000016477 | 2.42468076 | 2.222185812 | 2.19697564 | 2.263820609 | 2.235369829 | 2.509202522 | 2.076440241 |
| ENSMUSG00000016481 | 2.269664473 | 2.080482024 | 2.070192693 | 2.189555804 | 2.113840608 | 2.361727836 | 1.944737824 |
| ENSMUSG00000016487 | 3.097855556 | 2.834405475 | 2.804186465 | 2.752632026 | 2.681901575 | 3.090610708 | 2.479230803 |
| ENSMUSG00000016493 | 0.671187741 | 0.830482024 | 0.646240625 | 1 | 0.846351821 | 0.698970004 | 0.520678036 |
| ENSMUSG00000016494 | 1.036286563 | 1.080482024 | 1.146240625 | 2.164552871 | 2.115605392 | 2.424881637 | 1.992562536 |
| ENSMUSG00000016495 | 2.388281011 | 2.229715809 | 2.178561379 | 2.27908599 | 2.1962964 | 2.531478917 | 2.057979654 |
| ENSMUSG00000016496 | 1.100572389 | 1.130890489 | 1.214495249 | 1.282320754 | 1.310728002 | 1.380211242 | 1.131174228 |

| | | | | | | | |
|---|---|---|---|---|---|---|---|
| ENSMUSG00000016498 | 0 | 0 | 0.25 | 0 | 0 | 0 | 0 |
| ENSMUSG00000016503 | 1.289064826 | 1.311981878 | 1.26109853 | 1.302363341 | 1.149965125 | 1.176091259 | 1.238624656 |
| ENSMUSG00000016510 | 1.87190676 | 1.705044741 | 1.705044741 | 1.811981878 | 1.761403742 | 1.959041392 | 1.674564426 |
| ENSMUSG00000016520 | 1.971473847 | 1.87892496 | 1.901832578 | 1.610735874 | 1.611014646 | 1.740362689 | 1.694069894 |
| ENSMUSG00000016524 | 0 | 0.25 | 0 | 0 | 0 | 0 | 0 |
| ENSMUSG00000016526 | 1.269664473 | 1.282320754 | 1.330482024 | 1.388647213 | 1.49445382 | 1.602059991 | 1.355418052 |
| ENSMUSG00000016528 | 2.86633195 | 2.620449858 | 2.645300145 | 2.521534056 | 2.512633319 | 2.855519156 | 2.386616756 |
| ENSMUSG00000016529 | 0 | 0 | 0 | 0 | 0 | 0 | 0 |
| ENSMUSG00000016534 | 3.402604253 | 3.102612838 | 3.075588332 | 2.944108258 | 2.817563216 | 3.30427505 | 2.623362273 |
| ENSMUSG00000016541 | 3.142017256 | 2.825802487 | 2.824372375 | 2.920967967 | 2.824245479 | 3.276691529 | 2.640976771 |
| ENSMUSG00000016552 | 1.734388958 | 1.642463902 | 1.657339155 | 2.070192693 | 2.030405741 | 2.176091259 | 1.984249047 |
| ENSMUSG00000016554 | 2.938317055 | 2.701032755 | 2.68155454 | 2.634789703 | 2.687518101 | 3.049605613 | 2.517993245 |
| ENSMUSG00000016559 | 3.503347069 | 3.185366747 | 3.187676747 | 3.277906187 | 3.199984585 | 3.710540448 | 2.999604783 |
| ENSMUSG00000016619 | 2.929506184 | 2.672499493 | 2.658248799 | 2.692046081 | 2.651624369 | 3.022428371 | 2.468415048 |
| ENSMUSG00000016624 | 0.671187741 | 0.646240625 | 0.79248125 | 0.580482024 | 0.568060967 | 0.477121255 | 0.803905701 |
| ENSMUSG00000016626 | 0 | 0 | 0 | 0 | 0 | 0 | 0 |
| ENSMUSG00000016637 | 2.026699189 | 1.916333979 | 1.959800947 | 1.825945187 | 1.962676635 | 2.103803721 | 1.866140981 |
| ENSMUSG00000016664 | 2.890648263 | 2.682192712 | 2.671187155 | 2.623963757 | 2.632615969 | 2.978636948 | 2.486592281 |
| ENSMUSG00000016756 | 3.085022778 | 2.838424552 | 2.819902645 | 2.768702615 | 2.664866505 | 3.033021445 | 2.430977499 |
| ENSMUSG00000016757 | 2.834961347 | 2.64151351 | 2.656655413 | 2.404596376 | 2.493248162 | 2.776701184 | 2.38742504 |
| ENSMUSG00000016758 | 1.404271433 | 1.532320754 | 1.516522298 | 0.580482024 | 0.877063019 | 0.301029996 | 0.896975297 |
| ENSMUSG00000016763 | 0.289064826 | 0 | 0 | 0.75 | 0.77552387 | 0.602059991 | 0.710836105 |
| ENSMUSG00000016831 | 2.241665209 | 2.099151195 | 2.086073977 | 2.404596376 | 2.375754058 | 2.705007959 | 2.20389891 |
| ENSMUSG00000016833 | 2.134369199 | 2.027131114 | 2.002806814 | 2.089388001 | 2.066583491 | 2.361727836 | 1.884801065 |
| ENSMUSG00000016918 | 1.623015126 | 1.51109853 | 1.5 | 0.701838731 | 0.489301084 | 0.602059991 | 0.520678036 |
| ENSMUSG00000016921 | 3.013563224 | 2.748588359 | 2.729156481 | 2.9645924 | 2.881412441 | 3.33243846 | 2.689262976 |
| ENSMUSG00000016933 | 2.644135203 | 2.459012589 | 2.426408097 | 2.389126514 | 2.37406527 | 2.678518379 | 2.23151414 |
| ENSMUSG00000016940 | 1.989530018 | 1.892463902 | 1.914552871 | 1.782320754 | 1.804307395 | 1.880813592 | 1.747849621 |
| ENSMUSG00000016942 | 0 | 0 | 0 | 0.396240625 | 0.244650542 | 0.477121255 | 0.355418052 |
| ENSMUSG00000016946 | 2.837282068 | 2.625956434 | 2.580200137 | 2.393411797 | 2.405621873 | 2.729974286 | 2.294288164 |
| ENSMUSG00000016982 | 0 | 0.25 | 0.396240625 | 0.25 | 0 | 0.301029996 | 0 |
| ENSMUSG00000016984 | 2.113897241 | 2.056000419 | 1.96759118 | 2.167221246 | 2.099395374 | 2.29666519 | 1.950241257 |
| ENSMUSG00000016995 | 0 | 0 | 0 | 0 | 0 | 0.301029996 | 0 |
| ENSMUSG00000016998 | 0 | 0 | 0 | 0 | 0 | 0 | 0 |

| | | | | | | | |
|---|---|---|---|---|---|---|---|
| ENSMUSG00000017000 | 0 | 0 | 0 | 0 | 0 | 0 | 0 |
| ENSMUSG00000017002 | 0 | 0 | 0.25 | 0 | 0.244650542 | 0 | 0.448487648 |
| ENSMUSG00000017003 | 0 | 0 | 0 | 0 | 0 | 0 | 0 |
| ENSMUSG00000017004 | 0 | 0 | 0 | 0 | 0 | 0 | 0 |
| ENSMUSG00000017007 | 0 | 0 | 0 | 0 | 0.244650542 | 0 | 0.224243824 |
| ENSMUSG00000017009 | 2.604828841 | 2.426408097 | 2.399513125 | 2.483422664 | 2.500125119 | 2.808210973 | 2.343157052 |
| ENSMUSG00000017049 | 0 | 0 | 0 | 0 | 0 | 0 | 0 |
| ENSMUSG00000017057 | 2.414600936 | 2.206637122 | 2.20584181 | 2.278435542 | 2.193483986 | 2.552668216 | 2.040767326 |
| ENSMUSG00000017064 | 0 | 0 | 0 | 0 | 0 | 0 | 0 |
| ENSMUSG00000017119 | 2.777141269 | 2.536164561 | 2.503505118 | 2.643411797 | 2.571705985 | 2.963787827 | 2.409839468 |
| ENSMUSG00000017132 | 1.907802659 | 1.860735874 | 1.914552871 | 1.956637122 | 1.996316946 | 2.161368002 | 1.93179548 |
| ENSMUSG00000017144 | 2.897114011 | 2.663211243 | 2.602612838 | 2.716819685 | 2.650078855 | 3.043755127 | 2.449322097 |
| ENSMUSG00000017146 | 2.788220579 | 2.566989271 | 2.535207443 | 2.715271726 | 2.630371083 | 3.034628457 | 2.448153266 |
| ENSMUSG00000017165 | 0 | 0.396240625 | 0 | 0 | 0 | 0 | 0 |
| ENSMUSG00000017167 | 1.759671389 | 1.678561379 | 1.701838731 | 1.494319981 | 1.645989387 | 1.662757832 | 1.543390198 |
| ENSMUSG00000017176 | 2.477243823 | 2.307805295 | 2.319031101 | 2.336073977 | 2.375332618 | 2.666517981 | 2.258763761 |
| ENSMUSG00000017188 | 2.280358197 | 2.137686696 | 2.164552871 | 2.02053726 | 2.045754077 | 2.309630167 | 1.970632877 |
| ENSMUSG00000017195 | 0 | 0 | 0.25 | 0.5 | 0.244650542 | 0.301029996 | 0.355418052 |
| ENSMUSG00000017204 | 0 | 0.25 | 0.25 | 0 | 0.244650542 | 0.301029996 | 0.520678036 |
| ENSMUSG00000017210 | 2.592526109 | 2.371457327 | 2.410964047 | 2.288071211 | 2.372368362 | 2.593286067 | 2.257861346 |
| ENSMUSG00000017211 | 0 | 0 | 0 | 0.25 | 0 | 0 | 0 |
| ENSMUSG00000017221 | 2.95990777 | 2.755071375 | 2.773024104 | 2.637927065 | 2.790143872 | 3.044931546 | 2.679429412 |
| ENSMUSG00000017264 | 2.754058486 | 2.540282971 | 2.552972074 | 2.606303976 | 2.56338924 | 2.910090546 | 2.422397581 |
| ENSMUSG00000017286 | 2.217828299 | 2.057204673 | 2.039967834 | 2.208222504 | 2.103973339 | 2.440909082 | 2.009225098 |
| ENSMUSG00000017288 | 2.396352858 | 2.185366747 | 2.20584181 | 2.261780978 | 2.257835496 | 2.562292864 | 2.121678334 |
| ENSMUSG00000017291 | 3.100326135 | 2.828787391 | 2.791383785 | 2.884547233 | 2.781844838 | 3.203304916 | 2.565194949 |
| ENSMUSG00000017299 | 1.867194479 | 1.802363341 | 1.705044741 | 1.864857905 | 1.948833444 | 2.056904851 | 1.847411384 |
| ENSMUSG00000017300 | 0 | 0 | 0 | 0 | 0 | 0.301029996 | 0 |
| ENSMUSG00000017307 | 1.205378646 | 1.17510993 | 1.321350555 | 1.321350555 | 1.42077253 | 1.505149978 | 1.462868481 |
| ENSMUSG00000017309 | 0 | 0 | 0 | 0 | 0 | 0 | 0.224243824 |
| ENSMUSG00000017310 | 0 | 0 | 0 | 0 | 0 | 0 | 0 |
| ENSMUSG00000017311 | 0 | 0 | 0 | 0 | 0 | 0 | 0 |
| ENSMUSG00000017314 | 1.99653913 | 1.888647213 | 1.899978211 | 1.951838731 | 2.043601899 | 2.155336037 | 1.970632877 |
| ENSMUSG00000017316 | 0 | 0 | 0 | 0 | 0 | 0 | 0 |

| | | | | | | | |
|---|---|---|---|---|---|---|---|
| ENSMUSG00000017344 | 0 | 0 | 0 | 0 | 0.244650542 | 0 | 0 |
| ENSMUSG00000017376 | 2.402962068 | 2.210587586 | 2.201032755 | 2.231203126 | 2.176121988 | 2.491361694 | 2.026921761 |
| ENSMUSG00000017386 | 2.540273527 | 2.418989258 | 2.458617762 | 2.041226732 | 2.267124351 | 2.411619706 | 2.197065089 |
| ENSMUSG00000017390 | 0.867194479 | 0.79248125 | 0.646240625 | 0.951838731 | 1.039257769 | 0.77815125 | 0.853775828 |
| ENSMUSG00000017400 | 1.789260561 | 1.77186571 | 1.79745614 | 1.634789703 | 1.599808748 | 1.799340549 | 1.594042709 |
| ENSMUSG00000017404 | 3.026699189 | 2.781191884 | 2.812278363 | 2.717783796 | 2.73699175 | 3.147367108 | 2.604423774 |
| ENSMUSG00000017405 | 1.721148722 | 1.618933358 | 1.67510993 | 1.505591953 | 1.607318736 | 1.716003344 | 1.616675393 |
| ENSMUSG00000017412 | 0.960252568 | 0.646240625 | 0.79248125 | 0.580482024 | 0.568060967 | 0.602059991 | 0.629532004 |
| ENSMUSG00000017417 | 0 | 0.396240625 | 0.5 | 0.5 | 0.632412477 | 0.301029996 | 0.629532004 |
| ENSMUSG00000017418 | 2.425923778 | 2.250703754 | 2.199415381 | 2.320772088 | 2.240967488 | 2.555094449 | 2.052911423 |
| ENSMUSG00000017421 | 3.181338949 | 2.887927065 | 2.845926073 | 2.919209652 | 2.87317422 | 3.295347148 | 2.660817406 |
| ENSMUSG00000017428 | 2.868914203 | 2.638167274 | 2.614088604 | 2.689972046 | 2.638266616 | 3.038620162 | 2.488222085 |
| ENSMUSG00000017446 | 2.29078455 | 2.070192693 | 2.079353153 | 1.87694866 | 1.902105869 | 2.123851641 | 1.903345709 |
| ENSMUSG00000017453 | 0.289064826 | 0.25 | 0.396240625 | 0 | 0 | 0.301029996 | 0 |
| ENSMUSG00000017466 | 3.76027916 | 3.398989186 | 3.40705442 | 3.341751287 | 3.283232735 | 3.834357113 | 3.064964261 |
| ENSMUSG00000017478 | 2.615992899 | 2.450224975 | 2.501055117 | 2.37892496 | 2.470166836 | 2.728353782 | 2.403180579 |
| ENSMUSG00000017485 | 2.874454827 | 2.634061804 | 2.613574823 | 2.832168732 | 2.757712293 | 3.169380495 | 2.532491271 |
| ENSMUSG00000017491 | 1.036286563 | 0.976722649 | 0.951838731 | 1.04248125 | 1.074582839 | 1.146128036 | 1.041356071 |
| ENSMUSG00000017493 | 1.759671389 | 1.634789703 | 1.657339155 | 1.431980114 | 1.445123986 | 1.447158031 | 1.505628092 |
| ENSMUSG00000017499 | 2.579849376 | 2.396709947 | 2.378431899 | 2.517865591 | 2.460693128 | 2.824776462 | 2.308247447 |
| ENSMUSG00000017548 | 2.995590251 | 2.760927716 | 2.748941788 | 2.885395165 | 2.790534238 | 3.224791956 | 2.593936131 |
| ENSMUSG00000017550 | 2.658591846 | 2.437048212 | 2.414106216 | 2.606828211 | 2.524713379 | 2.895974732 | 2.320390187 |
| ENSMUSG00000017561 | 2.5496035 | 2.373463862 | 2.369945066 | 2.428561379 | 2.377853744 | 2.730782276 | 2.222568443 |
| ENSMUSG00000017588 | 0 | 0 | 0 | 0 | 0 | 0 | 0 |
| ENSMUSG00000017607 | 0.289064826 | 0 | 0.25 | 0.5 | 0.244650542 | 0.301029996 | 0.448487648 |
| ENSMUSG00000017615 | 3.007144864 | 2.747880462 | 2.761951389 | 2.798073204 | 2.744923682 | 3.159266331 | 2.568901575 |
| ENSMUSG00000017631 | 2.864601523 | 2.61306031 | 2.617918804 | 2.556603048 | 2.546663136 | 2.88536122 | 2.369798885 |
| ENSMUSG00000017639 | 1 | 1 | 1.201838731 | 1.5 | 1.445123986 | 1.591064607 | 1.551512352 |
| ENSMUSG00000017652 | 0.45815691 | 0.701838731 | 0.79248125 | 1.061981878 | 1.200473444 | 0.903089987 | 1.089372236 |
| ENSMUSG00000017664 | 1.862428343 | 1.916333979 | 1.964495249 | 1.777131114 | 1.859322797 | 1.897627091 | 1.832055226 |
| ENSMUSG00000017667 | 2.119110214 | 1.991446071 | 1.985628626 | 2.070192693 | 2.021325983 | 2.212187604 | 1.902443294 |
| ENSMUSG00000017670 | 2.427163103 | 2.277131114 | 2.289967834 | 2.293106877 | 2.348168747 | 2.593286067 | 2.230859913 |
| ENSMUSG00000017677 | 3.035999251 | 2.715465585 | 2.742526473 | 2.803731547 | 2.763656362 | 3.126780577 | 2.551390798 |
| ENSMUSG00000017679 | 2.292497211 | 2.123963757 | 2.172499493 | 2.353406982 | 2.305113253 | 2.610660163 | 2.126234965 |

| | | | | | | | |
|---|---|---|---|---|---|---|---|
| ENSMUSG00000017686 | 2.700465191 | 2.506630861 | 2.490724001 | 2.617156011 | 2.533980713 | 2.949877704 | 2.361378588 |
| ENSMUSG00000017688 | 0 | 0 | 0 | 0 | 0 | 0 | 0 |
| ENSMUSG00000017692 | 1.890240806 | 1.75 | 1.705044741 | 1.94697564 | 1.916320788 | 2.103803721 | 1.837623616 |
| ENSMUSG00000017697 | 1.289064826 | 1.188721876 | 1.380890489 | 1.622963274 | 1.662257525 | 1.826074803 | 1.574722613 |
| ENSMUSG00000017707 | 2.558729299 | 2.333288838 | 2.314935816 | 2.556000419 | 2.434943542 | 2.846337112 | 2.302263476 |
| ENSMUSG00000017713 | 0.578129653 | 0.701838731 | 1.04248125 | 0.580482024 | 0.812711509 | 0.77815125 | 0.952571511 |
| ENSMUSG00000017715 | 2.484856099 | 2.361253711 | 2.393884812 | 2.192872367 | 2.292010185 | 2.492760389 | 2.198153757 |
| ENSMUSG00000017716 | 2.804556862 | 2.5684489 | 2.590485943 | 2.710194481 | 2.650078855 | 3.044147621 | 2.513103749 |
| ENSMUSG00000017718 | 1.342375483 | 1.302363341 | 1.160964047 | 0.896240625 | 1.057362051 | 1.146128036 | 1.054044578 |
| ENSMUSG00000017720 | 0 | 0.25 | 0.25 | 0.25 | 0.244650542 | 0.602059991 | 0.224243824 |
| ENSMUSG00000017721 | 2.373064211 | 2.234159485 | 2.27186571 | 2.226722649 | 2.308696595 | 2.56937391 | 2.252393349 |
| ENSMUSG00000017723 | 0 | 0.5 | 0 | 0 | 0 | 0.477121255 | 0 |
| ENSMUSG00000017724 | 1.663535556 | 1.557204673 | 1.584962501 | 1.747171172 | 1.824953989 | 2.06069784 | 1.709198048 |
| ENSMUSG00000017733 | 0 | 0 | 0 | 0 | 0 | 0 | 0 |
| ENSMUSG00000017734 | 0.811507563 | 0.646240625 | 0.92510993 | 0.830482024 | 1.039257769 | 0.602059991 | 0.629532004 |
| ENSMUSG00000017737 | 1.129344651 | 1.282320754 | 1.26109853 | 1.54745614 | 1.559656853 | 1.653212514 | 1.556816002 |
| ENSMUSG00000017740 | 0.867194479 | 0.75 | 0.646240625 | 1.17510993 | 1.27449559 | 1.176091259 | 1.110947927 |
| ENSMUSG00000017747 | 1.765759561 | 1.769203899 | 1.892463902 | 1.698603967 | 1.853822071 | 1.982271233 | 1.795211864 |
| ENSMUSG00000017754 | 1.816888701 | 1.777131114 | 1.799918086 | 1.593759858 | 1.639266209 | 1.755874856 | 1.633706455 |
| ENSMUSG00000017756 | 2.635214384 | 2.488912477 | 2.481573999 | 2.41942991 | 2.509763742 | 2.741939078 | 2.389436958 |
| ENSMUSG00000017760 | 2.779806021 | 2.596735061 | 2.61817271 | 2.392463902 | 2.419658517 | 2.742725131 | 2.306955966 |
| ENSMUSG00000017764 | 1.445324132 | 1.431980114 | 1.470660762 | 1.54248125 | 1.628935049 | 1.681241237 | 1.601000298 |
| ENSMUSG00000017765 | 2.432083896 | 2.364857905 | 2.391037372 | 2.222945926 | 2.342660868 | 2.487138375 | 2.284450788 |
| ENSMUSG00000017767 | 0 | 0 | 0.25 | 0 | 0.244650542 | 0 | 0 |
| ENSMUSG00000017774 | 3.262045719 | 3.02401406 | 3.035047676 | 2.976910451 | 3.028157605 | 3.413299764 | 2.845016592 |
| ENSMUSG00000017776 | 3.152581175 | 2.877320042 | 2.862673943 | 2.900326632 | 2.81924568 | 3.252124553 | 2.607913505 |
| ENSMUSG00000017778 | 2.00683675 | 1.964495249 | 2.04745614 | 1.896240625 | 1.982473755 | 2.155336037 | 1.863093717 |
| ENSMUSG00000017781 | 3.069666964 | 2.794821776 | 2.794821776 | 2.843488164 | 2.763375569 | 3.199480915 | 2.581660949 |
| ENSMUSG00000017801 | 2.349889783 | 2.219895812 | 2.196158711 | 2.223704441 | 2.178339521 | 2.481442629 | 2.064616101 |
| ENSMUSG00000017802 | 2.54867984 | 2.345385738 | 2.344302633 | 2.350219859 | 2.300972958 | 2.621176282 | 2.18859465 |
| ENSMUSG00000017817 | 2.078325387 | 1.964495249 | 1.997171172 | 0.25 | 0 | 0 | 0 |
| ENSMUSG00000017830 | 0.91631382 | 0.830482024 | 0.701838731 | 1 | 1.020174412 | 1 | 1.140832123 |
| ENSMUSG00000017831 | 2.662129038 | 2.472566069 | 2.435366747 | 2.462546709 | 2.35043857 | 2.702430536 | 2.166194421 |
| ENSMUSG00000017832 | 0.289064826 | 0.396240625 | 0.25 | 0.5 | 0.568060967 | 0.477121255 | 0.579661877 |

| | | | | | | | |
|---|---|---|---|---|---|---|---|
| ENSMUSG00000017837 | 2.121692466 | 1.972185812 | 2.051142786 | 1.918106335 | 1.955822902 | 2.056904851 | 1.89141329 |
| ENSMUSG00000017843 | 2.68005385 | 2.509729747 | 2.496460484 | 2.43155454 | 2.364194729 | 2.754348336 | 2.254834987 |
| ENSMUSG00000017858 | 2.060824957 | 1.862802778 | 1.856566189 | 2.04745614 | 1.997548906 | 2.303196057 | 1.901538355 |
| ENSMUSG00000017861 | 2.320585253 | 2.19697564 | 2.216046536 | 2.242166698 | 2.349985771 | 2.527629901 | 2.231840759 |
| ENSMUSG00000017868 | 0 | 0.25 | 0 | 0.25 | 0 | 0 | 0.355418052 |
| ENSMUSG00000017897 | 0 | 0 | 0 | 0 | 0.244650542 | 0.477121255 | 0.355418052 |
| ENSMUSG00000017922 | 0 | 0 | 0 | 0 | 0 | 0 | 0 |
| ENSMUSG00000017929 | 2.144240249 | 2.081607372 | 2.136723615 | 1.994319981 | 2.099395374 | 2.195899652 | 1.989815042 |
| ENSMUSG00000017943 | 0.289064826 | 0 | 0 | 0.580482024 | 0.686820904 | 0.477121255 | 0.629532004 |
| ENSMUSG00000017950 | 0 | 0 | 0 | 0 | 0 | 0 | 0 |
| ENSMUSG00000017969 | 1.771760131 | 1.552363341 | 1.698603967 | 1.410964047 | 1.655839964 | 1.672097858 | 1.607811402 |
| ENSMUSG00000017978 | 0 | 0 | 0 | 0 | 0 | 0 | 0.355418052 |
| ENSMUSG00000017999 | 2.471720134 | 2.32991803 | 2.342126615 | 2.349685923 | 2.390196125 | 2.664641976 | 2.322370395 |
| ENSMUSG00000018001 | 2.551444705 | 2.329353153 | 2.347004321 | 2.361253711 | 2.315256368 | 2.613841822 | 2.200320163 |
| ENSMUSG00000018008 | 0.289064826 | 0.5 | 0 | 0.75 | 0.387761935 | 0.301029996 | 0.74492186 |
| ENSMUSG00000018012 | 1.837744666 | 1.848079356 | 1.872963274 | 0.92510993 | 1.212046813 | 1.041392685 | 1.121219121 |
| ENSMUSG00000018040 | 2.43085913 | 2.254897682 | 2.336628433 | 2.294354884 | 2.367659357 | 2.619093331 | 2.245269217 |
| ENSMUSG00000018042 | 3.240434765 | 2.97540529 | 3.01685859 | 2.841307212 | 2.877977809 | 3.270212855 | 2.760260311 |
| ENSMUSG00000018068 | 2.600768124 | 2.381869252 | 2.362802778 | 2.465271726 | 2.403292121 | 2.753583059 | 2.222232323 |
| ENSMUSG00000018076 | 2.69268209 | 2.478969845 | 2.464106382 | 2.246460484 | 2.284999367 | 2.521138084 | 2.14212705 |
| ENSMUSG00000018102 | 0.45815691 | 0.830482024 | 0.830482024 | 1.188721876 | 1.163285805 | 1.301029996 | 1.272006351 |
| ENSMUSG00000018126 | 0.45815691 | 0.25 | 0.701838731 | 0.701838731 | 0.812711509 | 0.698970004 | 1.121219121 |
| ENSMUSG00000018143 | 1.936861443 | 1.886723615 | 2.027131114 | 1.816696635 | 1.966725889 | 2.08278537 | 1.943148064 |
| ENSMUSG00000018160 | 2.726616086 | 2.47934302 | 2.487821179 | 2.526149688 | 2.49625462 | 2.784617293 | 2.302525964 |
| ENSMUSG00000018166 | 1.156259305 | 1.238549078 | 1.17510993 | 1.668106335 | 1.632412477 | 1.755874856 | 1.610050272 |
| ENSMUSG00000018167 | 2.20747957 | 2.036164561 | 2.080482024 | 2.034887838 | 1.998776581 | 2.264817823 | 1.971363985 |
| ENSMUSG00000018168 | 0 | 0 | 0 | 0 | 0 | 0 | 0.224243824 |
| ENSMUSG00000018169 | 0 | 0.396240625 | 0 | 0 | 0 | 0 | 0.355418052 |
| ENSMUSG00000018171 | 3.113420088 | 2.826942508 | 2.798843052 | 2.581326364 | 2.501897287 | 2.873320602 | 2.324828731 |
| ENSMUSG00000018189 | 2.445324132 | 2.25835575 | 2.259729747 | 2.342671852 | 2.252501045 | 2.580924976 | 2.076440241 |
| ENSMUSG00000018196 | 2.172524468 | 1.985628626 | 1.940387808 | 2.001406137 | 1.950242452 | 2.164352856 | 1.844181586 |
| ENSMUSG00000018199 | 2.652148299 | 2.464883697 | 2.459406983 | 2.680701883 | 2.587542629 | 2.950851459 | 2.369372364 |
| ENSMUSG00000018209 | 2.43085913 | 2.234894804 | 2.247880462 | 2.416333979 | 2.361135028 | 2.699837726 | 2.220208236 |
| ENSMUSG00000018211 | 0 | 0 | 0 | 0 | 0 | 0 | 0 |

| | | | | | | | |
|---|---|---|---|---|---|---|---|
| ENSMUSG00000018217 | 2.452372734 | 2.28296424 | 2.306603048 | 2.30357978 | 2.267696867 | 2.596597096 | 2.152721168 |
| ENSMUSG00000018238 | 0.289064826 | 0.646240625 | 0.25 | 0.646240625 | 0.733951626 | 0.602059991 | 0.520678036 |
| ENSMUSG00000018239 | 1.771760131 | 1.763820609 | 1.695339928 | 1.966046536 | 1.887294348 | 2.133538908 | 1.793950594 |
| ENSMUSG00000018259 | 0 | 0 | 0 | 0 | 0 | 0 | 0 |
| ENSMUSG00000018263 | 0 | 0 | 0.396240625 | 0 | 0.244650542 | 0 | 0 |
| ENSMUSG00000018286 | 2.829809843 | 2.637927065 | 2.69059551 | 2.68346568 | 2.739555424 | 3.113943352 | 2.60016488 |
| ENSMUSG00000018287 | 2.432083896 | 2.295598588 | 2.289967834 | 2.230460234 | 2.289867804 | 2.530199698 | 2.14212705 |
| ENSMUSG00000018293 | 3.302324644 | 2.999559455 | 3.019203899 | 2.867156011 | 2.817081036 | 3.28645647 | 2.704990635 |
| ENSMUSG00000018322 | 1.971473847 | 1.834962501 | 1.821350555 | 1.959800947 | 1.944572352 | 2.178976947 | 1.881925353 |
| ENSMUSG00000018326 | 3.097359679 | 2.804489424 | 2.799304173 | 2.997259909 | 2.899345783 | 3.36398783 | 2.68518852 |
| ENSMUSG00000018334 | 2.163307908 | 1.997171172 | 2.025821952 | 1.76109853 | 1.911645804 | 2.056904851 | 1.839824409 |
| ENSMUSG00000018339 | 0 | 0.580482024 | 0.25 | 0.25 | 0.489301084 | 0.477121255 | 0.74492186 |
| ENSMUSG00000018340 | 3.339886555 | 3.035925519 | 2.998234584 | 2.867791783 | 2.840530656 | 3.24723655 | 2.6634936 |
| ENSMUSG00000018341 | 0 | 0.25 | 0 | 0.701838731 | 0.812711509 | 0.77815125 | 0.448487648 |
| ENSMUSG00000018347 | 1.54867984 | 1.53743678 | 1.53743678 | 1.668106335 | 1.655839964 | 1.968482949 | 1.574722613 |
| ENSMUSG00000018362 | 2.891055323 | 2.639126514 | 2.611512352 | 2.745926548 | 2.63504876 | 3.080626487 | 2.48540179 |
| ENSMUSG00000018363 | 2.736751754 | 2.492885888 | 2.481944491 | 2.561388177 | 2.494152791 | 2.868056362 | 2.311581365 |
| ENSMUSG00000018372 | 2.020182925 | 1.87694866 | 1.909156155 | 1.910964047 | 1.898867717 | 2.053078443 | 1.802677937 |
| ENSMUSG00000018377 | 2.703665763 | 2.467205639 | 2.453445298 | 2.518869787 | 2.434230499 | 2.778874472 | 2.272006351 |
| ENSMUSG00000018378 | 2.205378646 | 2.089388001 | 2.106566189 | 1.828220739 | 2.039257769 | 2.075546961 | 2.008574816 |
| ENSMUSG00000018379 | 3.340579416 | 3.020204382 | 3.002107156 | 3.126514897 | 3.042775527 | 3.513350799 | 2.802174324 |
| ENSMUSG00000018381 | 0.289064826 | 0 | 0.25 | 0 | 0 | 0 | 0 |
| ENSMUSG00000018387 | 1.325351389 | 1.321350555 | 1.348079356 | 1.575945187 | 1.677808768 | 1.672097858 | 1.665294008 |
| ENSMUSG00000018395 | 2.325351389 | 2.162762923 | 2.152756199 | 2.345385738 | 2.27449559 | 2.644438589 | 2.152303998 |
| ENSMUSG00000018398 | 2.824593906 | 2.604199382 | 2.607090043 | 2.759901129 | 2.757283861 | 3.128722284 | 2.596378571 |
| ENSMUSG00000018401 | 2.830751226 | 2.571061437 | 2.562870603 | 2.608135475 | 2.576746587 | 2.918554531 | 2.402218022 |
| ENSMUSG00000018405 | 1.765759561 | 1.668106335 | 1.614857905 | 1.678561379 | 1.580303447 | 1.748188027 | 1.582103514 |
| ENSMUSG00000018411 | 0.289064826 | 0 | 0.396240625 | 0.646240625 | 0.877063019 | 0.84509804 | 0.710836105 |
| ENSMUSG00000018412 | 2.5496035 | 2.370449858 | 2.352877747 | 2.377443751 | 2.336595379 | 2.665580991 | 2.193041668 |
| ENSMUSG00000018415 | 2.211649928 | 2.01109853 | 1.942046081 | 2.202642909 | 2.153935427 | 2.45484486 | 2.009225098 |
| ENSMUSG00000018417 | 2.760228547 | 2.528435542 | 2.505938588 | 2.403217124 | 2.296256325 | 2.613841822 | 2.171745067 |
| ENSMUSG00000018425 | 2.623015126 | 2.405970373 | 2.390560606 | 2.373463862 | 2.296783518 | 2.653212514 | 2.149789616 |
| ENSMUSG00000018427 | 1.505869732 | 1.54745614 | 1.5 | 1.464495249 | 1.445123986 | 1.51851394 | 1.448487648 |
| ENSMUSG00000018428 | 2.662129038 | 2.458222504 | 2.481573999 | 2.353935442 | 2.389791591 | 2.662757832 | 2.233793574 |

| | | | | | | | |
|---|---|---|---|---|---|---|---|
| ENSMUSG00000018433 | 2.68140129 | 2.465659339 | 2.459800947 | 2.613831805 | 2.523883869 | 2.923761961 | 2.364643151 |
| ENSMUSG00000018442 | 2.45930418 | 2.280383379 | 2.28743678 | 2.342126615 | 2.275616087 | 2.545307116 | 2.120298735 |
| ENSMUSG00000018446 | 2.947656672 | 2.669209652 | 2.690387808 | 2.766690483 | 2.685247156 | 3.099680641 | 2.485848738 |
| ENSMUSG00000018449 | 2.003432385 | 1.816696635 | 1.83272922 | 1.856566189 | 1.804307395 | 2.071882007 | 1.71890424 |
| ENSMUSG00000018451 | 0.91631382 | 0.92510993 | 0.830482024 | 0.864857905 | 0.686820904 | 0.698970004 | 0.935079929 |
| ENSMUSG00000018459 | 0.289064826 | 0 | 0 | 0.25 | 0 | 0 | 0 |
| ENSMUSG00000018470 | 1.289064826 | 1.356566189 | 1.380890489 | 1.598079356 | 1.576269619 | 1.748188027 | 1.548827561 |
| ENSMUSG00000018474 | 3.005292755 | 2.781837385 | 2.785367139 | 2.80930249 | 2.807544405 | 3.185542155 | 2.675062526 |
| ENSMUSG00000018476 | 1.944805158 | 1.850219859 | 1.896240625 | 1.580482024 | 1.751666404 | 1.73239376 | 1.805128823 |
| ENSMUSG00000018479 | 0 | 0.580482024 | 0 | 0 | 0 | 0 | 0.579661877 |
| ENSMUSG00000018481 | 2.917842813 | 2.668548067 | 2.611253711 | 2.748411515 | 2.662070429 | 3.070037867 | 2.466207819 |
| ENSMUSG00000018486 | 0 | 0 | 0 | 0 | 0 | 0 | 0 |
| ENSMUSG00000018500 | 1.37447073 | 1.26109853 | 1.188721876 | 1 | 0.686820904 | 0.954242509 | 0.969165684 |
| ENSMUSG00000018501 | 2.940129454 | 2.702843673 | 2.671843921 | 2.882723495 | 2.806924204 | 3.249442961 | 2.616973335 |
| ENSMUSG00000018507 | 1.53838222 | 1.589388001 | 1.622963274 | 1.201838731 | 1.163285805 | 1.230448921 | 1.159323753 |
| ENSMUSG00000018509 | 0.867194479 | 0.75 | 0.830482024 | 1.17510993 | 1.302012594 | 1.278753601 | 1.193409508 |
| ENSMUSG00000018537 | 2.105953527 | 1.994319981 | 2.024508021 | 1.896240625 | 1.979897422 | 2.093421685 | 1.912234055 |
| ENSMUSG00000018541 | 2.217828299 | 2.071350555 | 2.093759858 | 2.136723615 | 2.148364414 | 2.372912003 | 2.045447559 |
| ENSMUSG00000018543 | 0.671187741 | 0.701838731 | 0.25 | 0 | 0.686820904 | 0.698970004 | 0.448487648 |
| ENSMUSG00000018547 | 2.624559692 | 2.399513125 | 2.406883471 | 2.490000483 | 2.496553863 | 2.832508913 | 2.321381806 |
| ENSMUSG00000018548 | 2.881174527 | 2.645300145 | 2.621205724 | 2.660964047 | 2.551560013 | 2.954242509 | 2.37424372 |
| ENSMUSG00000018554 | 0 | 0 | 0 | 0.396240625 | 0.244650542 | 0 | 0.672731473 |
| ENSMUSG00000018559 | 2.740271036 | 2.523849256 | 2.538071211 | 2.440387808 | 2.454008153 | 2.797267541 | 2.326782038 |
| ENSMUSG00000018565 | 2.53552908 | 2.325374049 | 2.324802005 | 2.359177886 | 2.319233381 | 2.701567985 | 2.193409508 |
| ENSMUSG00000018566 | 0.867194479 | 0.646240625 | 0.896240625 | 0.79248125 | 0.905314583 | 1 | 0.896975297 |
| ENSMUSG00000018567 | 3.196867486 | 2.883697303 | 2.90079067 | 2.780545225 | 2.668927761 | 3.126780577 | 2.502532226 |
| ENSMUSG00000018569 | 0 | 0 | 0 | 0 | 0.244650542 | 0 | 0.224243824 |
| ENSMUSG00000018570 | 0.960252568 | 1.26109853 | 1.201838731 | 0.864857905 | 1.039257769 | 1.230448921 | 1.121219121 |
| ENSMUSG00000018572 | 2.63370885 | 2.434100483 | 2.468745337 | 2.227473271 | 2.235369829 | 2.489958479 | 2.111421942 |
| ENSMUSG00000018574 | 2.442947872 | 2.28743678 | 2.301142786 | 2.360735874 | 2.387761935 | 2.706717782 | 2.260561074 |
| ENSMUSG00000018581 | 0.289064826 | 0.5 | 0.396240625 | 0 | 0.244650542 | 0.477121255 | 0.224243824 |
| ENSMUSG00000018583 | 3.150620716 | 2.863960227 | 2.87694866 | 2.967205639 | 2.945998439 | 3.361538971 | 2.760515198 |
| ENSMUSG00000018585 | 2.163307908 | 1.973704441 | 2.072504712 | 2.087182039 | 2.169384551 | 2.413299764 | 2.119375723 |
| ENSMUSG00000018589 | 0 | 0 | 0 | 0 | 0 | 0 | 0 |

| | | | | | | | |
|---|---|---|---|---|---|---|---|
| ENSMUSG00000018593 | 3.829552733 | 3.459530143 | 3.405798909 | 3.497747576 | 3.443348642 | 4.04289039 | 3.206139789 |
| ENSMUSG00000018595 | 0 | 0.25 | 0.25 | 0.580482024 | 0.244650542 | 0.301029996 | 0.448487648 |
| ENSMUSG00000018599 | 1.614416215 | 1.634789703 | 1.698603967 | 1.614857905 | 1.665423072 | 1.556302501 | 1.681794123 |
| ENSMUSG00000018604 | 0 | 0 | 0 | 0.5 | 0.489301084 | 0 | 0.579661877 |
| ENSMUSG00000018620 | 0 | 0 | 0 | 0 | 0 | 0 | 0 |
| ENSMUSG00000018623 | 0 | 0 | 0 | 0 | 0 | 0 | 0 |
| ENSMUSG00000018634 | 0 | 0 | 0 | 0 | 0 | 0 | 0.224243824 |
| ENSMUSG00000018648 | 1.964026642 | 1.938721876 | 1.916333979 | 2.113831805 | 2.165585202 | 2.320146286 | 2.044283833 |
| ENSMUSG00000018651 | 2.278594844 | 2.114857905 | 2.05357978 | 2.133818844 | 2.076554666 | 2.389166084 | 1.95718226 |
| ENSMUSG00000018654 | 0 | 0 | 0 | 0.25 | 0 | 0 | 0.224243824 |
| ENSMUSG00000018656 | 0 | 0 | 0 | 0 | 0 | 0.301029996 | 0 |
| ENSMUSG00000018659 | 1.777675583 | 1.71759118 | 1.714495249 | 1.899978211 | 1.86473911 | 2.152288344 | 1.782372264 |
| ENSMUSG00000018661 | 2.185978293 | 2.077084758 | 2.091580554 | 1.997171172 | 2.094757251 | 2.281033367 | 2.007923223 |
| ENSMUSG00000018666 | 2.845077293 | 2.587458523 | 2.571928095 | 2.655284028 | 2.592135487 | 2.986771734 | 2.392033886 |
| ENSMUSG00000018669 | 2.275045611 | 2.137686696 | 2.159156155 | 2.025821952 | 2.137779008 | 2.320146286 | 2.034219411 |
| ENSMUSG00000018672 | 2.231898196 | 2.066696635 | 2.049918086 | 2.015173983 | 1.978602168 | 2.161368002 | 1.872150729 |
| ENSMUSG00000018677 | 2.638209243 | 2.512121718 | 2.569902645 | 2.233422664 | 2.465951417 | 2.610660163 | 2.396185378 |
| ENSMUSG00000018678 | 2 | 1.890560606 | 1.928561379 | 1.80479213 | 1.877063019 | 1.954242509 | 1.952571511 |
| ENSMUSG00000018697 | 2.52293481 | 2.30539678 | 2.324229052 | 2.282320754 | 2.266550905 | 2.544068044 | 2.110473216 |
| ENSMUSG00000018698 | 0 | 0 | 0.396240625 | 0.79248125 | 0.846351821 | 0.903089987 | 0.853775828 |
| ENSMUSG00000018707 | 3.471214334 | 3.152525516 | 3.165778327 | 3.014667063 | 3.035314806 | 3.418301291 | 2.842307272 |
| ENSMUSG00000018727 | 1 | 1 | 1 | 0.976722649 | 0.846351821 | 0.903089987 | 0.853775828 |
| ENSMUSG00000018733 | 1.794934559 | 1.732684334 | 1.78743678 | 1.67510993 | 1.758994476 | 1.838849091 | 1.594042709 |
| ENSMUSG00000018736 | 2.510961766 | 2.291226732 | 2.330482024 | 2.320192693 | 2.265976525 | 2.605305046 | 2.190083792 |
| ENSMUSG00000018740 | 0.867194479 | 1.02186571 | 0.646240625 | 1.02186571 | 1 | 0.903089987 | 1.014380832 |
| ENSMUSG00000018750 | 1.881174527 | 1.860735874 | 1.905512955 | 2.071350555 | 2.093822267 | 2.276461804 | 1.991191705 |
| ENSMUSG00000018752 | 0.289064826 | 0 | 0.396240625 | 0.5 | 0.244650542 | 0.477121255 | 0.448487648 |
| ENSMUSG00000018761 | 2.188178997 | 2.120953944 | 2.103406982 | 2.081607372 | 2.125157063 | 2.315970345 | 1.974272009 |
| ENSMUSG00000018765 | 2.710621733 | 2.530383379 | 2.556903986 | 2.518200634 | 2.576984829 | 2.863917377 | 2.462068294 |
| ENSMUSG00000018770 | 2.787699615 | 2.561388177 | 2.600753006 | 2.726158655 | 2.691672611 | 3.067070856 | 2.53274895 |
| ENSMUSG00000018774 | 1.036286563 | 0.951838731 | 1.302363341 | 0.896240625 | 1.106691885 | 0.84509804 | 1.066254157 |
| ENSMUSG00000018776 | 0.289064826 | 0 | 0.5 | 0.25 | 0.387761935 | 0 | 0.579661877 |
| ENSMUSG00000018796 | 2.470606563 | 2.263820609 | 2.271202097 | 2.205044741 | 2.16097133 | 2.482873584 | 2.056860222 |
| ENSMUSG00000018800 | 2.322180025 | 2.113831805 | 2.172499493 | 2.3464656 | 2.224547964 | 2.550228353 | 2.071655208 |

| | | | | | | | |
|---|---|---|---|---|---|---|---|
| ENSMUSG00000018809 | 2.119110214 | 1.982684334 | 1.966046536 | 1.862802778 | 1.84445929 | 2.100370545 | 1.752192176 |
| ENSMUSG00000018819 | 2.203267085 | 1.987091808 | 1.940387808 | 1.311981878 | 1.293076518 | 1.447158031 | 1.369798885 |
| ENSMUSG00000018820 | 2.203267085 | 2.120953944 | 2.12892496 | 2.149047439 | 2.201854879 | 2.414973348 | 2.087337541 |
| ENSMUSG00000018821 | 2.136858952 | 2.073655187 | 2.275165585 | 1.928561379 | 2.110284345 | 2.247973266 | 2.175263699 |
| ENSMUSG00000018822 | 0 | 0 | 0 | 0.25 | 0 | 0.301029996 | 0.355418052 |
| ENSMUSG00000018830 | 0.578129653 | 0.646240625 | 0.830482024 | 0.396240625 | 0.568060967 | 0.477121255 | 0.448487648 |
| ENSMUSG00000018841 | 1.985980784 | 1.907339155 | 1.886723615 | 1.688721876 | 1.768534412 | 1.963787827 | 1.722075986 |
| ENSMUSG00000018844 | 0.289064826 | 0.396240625 | 0.396240625 | 0 | 0.568060967 | 0.477121255 | 0.448487648 |
| ENSMUSG00000018845 | 0 | 0 | 0 | 0 | 0 | 0 | 0.224243824 |
| ENSMUSG00000018846 | 2.841427095 | 2.617918804 | 2.617410454 | 2.785047676 | 2.702017009 | 3.128076013 | 2.484804898 |
| ENSMUSG00000018848 | 2.901898374 | 2.666111821 | 2.60839636 | 2.817573832 | 2.722627519 | 3.165837625 | 2.522145228 |
| ENSMUSG00000018849 | 2.388281011 | 2.268535366 | 2.316696635 | 2.452241044 | 2.480333692 | 2.762678564 | 2.382339813 |
| ENSMUSG00000018858 | 2.078325387 | 1.860735874 | 1.894357207 | 2.036164561 | 2.040348824 | 2.271841607 | 1.907820406 |
| ENSMUSG00000018861 | 1.445324132 | 1.552363341 | 1.630890489 | 1.364857905 | 1.555378544 | 1.301029996 | 1.577201671 |
| ENSMUSG00000018862 | 0 | 0 | 0 | 0 | 0 | 0 | 0 |
| ENSMUSG00000018865 | 0 | 0 | 0 | 1.27186571 | 1.283908311 | 1.322219295 | 1.313616061 |
| ENSMUSG00000018868 | 0 | 0 | 0 | 0 | 0 | 0 | 0 |
| ENSMUSG00000018882 | 2.628396264 | 2.41942991 | 2.450224975 | 2.481573999 | 2.44788147 | 2.812913357 | 2.30409643 |
| ENSMUSG00000018893 | 0 | 0.25 | 0 | 0 | 0 | 0 | 0 |
| ENSMUSG00000018899 | 2.402962068 | 2.214495249 | 2.200224975 | 2.608657057 | 2.559656853 | 2.946452265 | 2.413769991 |
| ENSMUSG00000018900 | 1.470606563 | 1.410964047 | 1.557204673 | 1.726722649 | 1.718026092 | 1.924279286 | 1.635773653 |
| ENSMUSG00000018906 | 2.457006475 | 2.266522298 | 2.28743678 | 2.24929487 | 2.288252441 | 2.553883027 | 2.13206423 |
| ENSMUSG00000018907 | 0 | 0 | 0 | 0 | 0.244650542 | 0 | 0.355418052 |
| ENSMUSG00000018909 | 2.292497211 | 2.12694866 | 2.135757955 | 2.260414788 | 2.195595395 | 2.457881897 | 2.103753068 |
| ENSMUSG00000018914 | 0 | 0 | 0 | 0 | 0 | 0 | 0 |
| ENSMUSG00000018916 | 0 | 0 | 0 | 0.396240625 | 0 | 0.477121255 | 0.355418052 |
| ENSMUSG00000018919 | 0.289064826 | 0 | 0.25 | 0 | 0 | 0 | 0 |
| ENSMUSG00000018920 | 1.156259305 | 1.080482024 | 1.214495249 | 1.723704441 | 1.715300549 | 1.812913357 | 1.643912836 |
| ENSMUSG00000018921 | 2.457006475 | 2.3184489 | 2.329353153 | 2.167221246 | 2.259595879 | 2.431363764 | 2.171351735 |
| ENSMUSG00000018923 | 1.777675583 | 1.852347734 | 1.901832578 | 1.830482024 | 1.848234258 | 2.053078443 | 1.829800754 |
| ENSMUSG00000018924 | 0.289064826 | 0.25 | 0 | 0.396240625 | 0 | 0.602059991 | 0.448487648 |
| ENSMUSG00000018925 | 0 | 0.25 | 0 | 0.25 | 0.244650542 | 0 | 0.448487648 |
| ENSMUSG00000018927 | 0.45815691 | 0.396240625 | 0.5 | 0.396240625 | 0 | 0.301029996 | 0 |
| ENSMUSG00000018930 | 0 | 0.25 | 0 | 0 | 0 | 0 | 0 |

| | | | | | | | |
|---|---|---|---|---|---|---|---|
| ENSMUSG00000018931 | 2.223916471 | 2.081607372 | 2.017865591 | 2.166333979 | 2.195595395 | 2.457881897 | 2.093904044 |
| ENSMUSG00000018932 | 2.534573679 | 2.345385738 | 2.331045137 | 2.468361028 | 2.452655829 | 2.806179974 | 2.307473178 |
| ENSMUSG00000018965 | 2.967766868 | 2.71816872 | 2.680488403 | 2.697179583 | 2.617018904 | 3.021602716 | 2.43361272 |
| ENSMUSG00000018973 | 0 | 0.25 | 0 | 0 | 0 | 0 | 0 |
| ENSMUSG00000018974 | 2.698532991 | 2.53199858 | 2.563757142 | 2.338286706 | 2.438487308 | 2.650307523 | 2.333050044 |
| ENSMUSG00000018983 | 1.678702042 | 1.571350555 | 1.634789703 | 1.738549078 | 1.791313678 | 1.86332286 | 1.674564426 |
| ENSMUSG00000018986 | 0 | 0 | 0 | 0 | 0 | 0 | 0 |
| ENSMUSG00000018995 | 1.714367558 | 1.552363341 | 1.532320754 | 1.630890489 | 1.584291694 | 1.662757832 | 1.455757965 |
| ENSMUSG00000018999 | 2.524897319 | 2.334962501 | 2.306000419 | 2.291226732 | 2.196996016 | 2.46686762 | 2.041356071 |
| ENSMUSG00000019027 | 0.45815691 | 0.396240625 | 0.396240625 | 0.646240625 | 0.632412477 | 0.477121255 | 0.74492186 |
| ENSMUSG00000019039 | 1.568542279 | 1.476722649 | 1.527131114 | 1.339388001 | 1.674752852 | 1.531478917 | 1.657681529 |
| ENSMUSG00000019054 | 2.600768124 | 2.440803092 | 2.447790722 | 2.316696635 | 2.369379 | 2.595496222 | 2.26794422 |
| ENSMUSG00000019055 | 2.475043119 | 2.342126615 | 2.390083209 | 2.333288838 | 2.42077253 | 2.716837723 | 2.322123531 |
| ENSMUSG00000019066 | 1.647796617 | 1.610735874 | 1.606566189 | 1.868933358 | 1.900490506 | 2.053078443 | 1.824094866 |
| ENSMUSG00000019080 | 0.91631382 | 1.114857905 | 1.146240625 | 0.896240625 | 1.234113756 | 0.84509804 | 1.245582246 |
| ENSMUSG00000019082 | 1.867194479 | 1.763820609 | 1.834962501 | 1.552363341 | 1.773209395 | 1.681241237 | 1.752192176 |
| ENSMUSG00000019087 | 3.104736624 | 2.865497738 | 2.874836908 | 2.831466895 | 2.790924173 | 3.237794993 | 2.622094539 |
| ENSMUSG00000019088 | 1.881174527 | 1.660964047 | 1.794977273 | 1.614857905 | 1.53772706 | 1.72427587 | 1.496250175 |
| ENSMUSG00000019102 | 1.036286563 | 1.114857905 | 1.061981878 | 0 | 0.244650542 | 0 | 0 |
| ENSMUSG00000019122 | 2.406877894 | 2.273849256 | 2.173371739 | 1.330482024 | 1.254881871 | 1.431363764 | 1.238624656 |
| ENSMUSG00000019124 | 2.540273527 | 2.338286706 | 2.379417347 | 2.446158711 | 2.383255179 | 2.707570176 | 2.25604893 |
| ENSMUSG00000019132 | 2.896712826 | 2.643175057 | 2.633575721 | 2.791697688 | 2.710765025 | 3.131297797 | 2.517588597 |
| ENSMUSG00000019139 | 2.260538674 | 2.173371739 | 2.275821952 | 2.064346961 | 2.304598364 | 2.361727836 | 2.263238458 |
| ENSMUSG00000019143 | 2.29078455 | 2.088286706 | 2.077084758 | 2.144357207 | 2.12771782 | 2.33243846 | 1.98213686 |
| ENSMUSG00000019146 | 0 | 0.25 | 0 | 0 | 0 | 0 | 0 |
| ENSMUSG00000019158 | 0.91631382 | 1.201838731 | 1.282320754 | 0.976722649 | 1.163285805 | 1.079181246 | 1.38802993 |
| ENSMUSG00000019173 | 2.733796161 | 2.573367912 | 2.612029076 | 2.510414788 | 2.581481146 | 2.867467488 | 2.485699824 |
| ENSMUSG00000019178 | 0.45815691 | 0 | 0 | 0.396240625 | 0.244650542 | 0.301029996 | 0.520678036 |
| ENSMUSG00000019179 | 3.22592627 | 2.960587586 | 2.992436563 | 2.878185116 | 2.928492895 | 3.351603072 | 2.778358596 |
| ENSMUSG00000019188 | 2.365959569 | 2.203445298 | 2.23927551 | 2.240000483 | 2.236621448 | 2.495544338 | 2.135599953 |
| ENSMUSG00000019189 | 2.667031238 | 2.430274797 | 2.431980114 | 2.546218836 | 2.520266629 | 2.829303773 | 2.321876478 |
| ENSMUSG00000019194 | 0.960252568 | 0.864857905 | 1.04248125 | 0.701838731 | 0.568060967 | 0.903089987 | 0.579661877 |
| ENSMUSG00000019210 | 2.635214384 | 2.443284802 | 2.42510993 | 2.325374049 | 2.359374645 | 2.680335513 | 2.194876701 |
| ENSMUSG00000019214 | 2.156259305 | 2.124961472 | 2.198603967 | 1.959800947 | 2.212715924 | 2.28780173 | 2.194876701 |

| | | | | | | | |
|---|---|---|---|---|---|---|---|
| ENSMUSG00000019230 | 1.928763473 | 1.953445298 | 1.886723615 | 1.302363341 | 1.223252711 | 1.342422681 | 1.20919388 |
| ENSMUSG00000019232 | 0 | 0.396240625 | 0 | 0 | 0.244650542 | 0 | 0 |
| ENSMUSG00000019235 | 0.867194479 | 0.25 | 0.5 | 0.646240625 | 0.489301084 | 0.477121255 | 0.710836105 |
| ENSMUSG00000019254 | 2.072573125 | 2.036164561 | 2.118933358 | 1.886723615 | 2.064555002 | 2.06069784 | 2.043115906 |
| ENSMUSG00000019256 | 2.245508853 | 2.057204673 | 1.972185812 | 2.074802005 | 1.987570683 | 2.212187604 | 1.883845332 |
| ENSMUSG00000019261 | 1.639698646 | 1.738549078 | 2 | 1.29248125 | 1.649303553 | 1.505149978 | 1.935079929 |
| ENSMUSG00000019278 | 0 | 0 | 0 | 0 | 0 | 0 | 0 |
| ENSMUSG00000019295 | 2.231898196 | 2.063166358 | 2.073655187 | 2.025821952 | 2.055279708 | 2.290034611 | 1.972821265 |
| ENSMUSG00000019297 | 2.211649928 | 2.058404919 | 2.056000419 | 2.296838018 | 2.294670002 | 2.544068044 | 2.159323753 |
| ENSMUSG00000019301 | 0.578129653 | 0.25 | 0.396240625 | 0.25 | 0 | 0 | 0.224243824 |
| ENSMUSG00000019302 | 2.664937347 | 2.478969845 | 2.528109877 | 2.393411797 | 2.409087918 | 2.673941999 | 2.258763761 |
| ENSMUSG00000019303 | 2.033115198 | 1.923371739 | 1.972185812 | 1.843759858 | 1.919403394 | 2 | 1.850609257 |
| ENSMUSG00000019312 | 0 | 0 | 0.25 | 0.25 | 0.387761935 | 0.301029996 | 0.224243824 |
| ENSMUSG00000019320 | 0.289064826 | 0.701838731 | 0.580482024 | 0.5 | 0.568060967 | 0.301029996 | 0.829800754 |
| ENSMUSG00000019326 | 2.048736215 | 1.888647213 | 1.862802778 | 0.896240625 | 0.877063019 | 0.84509804 | 0.710836105 |
| ENSMUSG00000019338 | 2.402962068 | 2.200224975 | 2.21293726 | 2.125956434 | 2.207327177 | 2.401400541 | 2.123508695 |
| ENSMUSG00000019359 | 0.45815691 | 0.25 | 0.396240625 | 0.396240625 | 0.244650542 | 0 | 0 |
| ENSMUSG00000019362 | 2.363083473 | 2.281030328 | 2.352347734 | 2.081607372 | 2.184185718 | 2.342422681 | 2.227237819 |
| ENSMUSG00000019368 | 0 | 0.25 | 0.25 | 0.396240625 | 0 | 0 | 0 |
| ENSMUSG00000019370 | 2.808046693 | 2.591853688 | 2.623963757 | 2.575945187 | 2.605126455 | 2.952792443 | 2.518397386 |
| ENSMUSG00000019373 | 2.593357679 | 2.392463902 | 2.387686696 | 2.550224651 | 2.467578695 | 2.881954971 | 2.347067347 |
| ENSMUSG00000019428 | 2.563218389 | 2.483053688 | 2.61306031 | 2.231203126 | 2.488995627 | 2.617000341 | 2.54858238 |
| ENSMUSG00000019429 | 0 | 0 | 0 | 0 | 0 | 0 | 0 |
| ENSMUSG00000019432 | 3.035424034 | 2.780221461 | 2.766354034 | 2.755938588 | 2.781844838 | 3.169674434 | 2.616078685 |
| ENSMUSG00000019433 | 2.215778996 | 2.193696765 | 2.319031101 | 2.090485943 | 2.241584006 | 2.31386722 | 2.325806859 |
| ENSMUSG00000019437 | 1.794934559 | 1.593759858 | 1.668106335 | 1.54248125 | 1.584291694 | 1.740362689 | 1.437266334 |
| ENSMUSG00000019461 | 2.29078455 | 2.171625132 | 2.223704441 | 2.051142786 | 2.165585202 | 2.409933123 | 2.122594809 |
| ENSMUSG00000019464 | 1.940852214 | 1.898114259 | 1.886723615 | 1.610735874 | 1.782377602 | 1.819543936 | 1.795211864 |
| ENSMUSG00000019467 | 2.508932407 | 2.313166358 | 2.363831805 | 2.187886015 | 2.200473444 | 2.365487985 | 2.111895264 |
| ENSMUSG00000019470 | 2.472830741 | 2.353406982 | 2.362287161 | 2.039967834 | 2.176862714 | 2.260071388 | 2.107610228 |
| ENSMUSG00000019471 | 2.734980913 | 2.548073204 | 2.628678514 | 2.593216265 | 2.686122326 | 2.95568775 | 2.600791646 |
| ENSMUSG00000019478 | 1.647796617 | 1.598079356 | 1.657339155 | 1.80479213 | 1.782377602 | 1.986771734 | 1.82753046 |
| ENSMUSG00000019487 | 2.330063669 | 2.202642909 | 2.229715809 | 2.095926073 | 2.160196454 | 2.33243846 | 2.060206963 |
| ENSMUSG00000019489 | 0 | 0 | 0 | 0 | 0 | 0.301029996 | 0.224243824 |

| | | | | | | | |
|---|---|---|---|---|---|---|---|
| ENSMUSG00000019494 | 2.787177998 | 2.57508814 | 2.568157446 | 2.663659007 | 2.614458147 | 3.025715384 | 2.472008833 |
| ENSMUSG00000019505 | 2.564110437 | 2.341580554 | 2.370449858 | 2.559899632 | 2.62166591 | 2.953759692 | 2.474485519 |
| ENSMUSG00000019518 | 2.066740409 | 1.959800947 | 1.950224975 | 1.923371739 | 1.944572352 | 2.11058971 | 1.842010333 |
| ENSMUSG00000019528 | 2.331622671 | 2.163659007 | 2.146240625 | 2.286164561 | 2.208682096 | 2.579783597 | 2.053478496 |
| ENSMUSG00000019539 | 2.455852858 | 2.359697963 | 2.386241108 | 1.933677405 | 2.149965125 | 2.198657087 | 2.182188194 |
| ENSMUSG00000019558 | 2.752926783 | 2.534887838 | 2.597004321 | 2.203445298 | 2.249502249 | 2.499687083 | 2.114251541 |
| ENSMUSG00000019564 | 1.771760131 | 1.67510993 | 1.747171172 | 1.606566189 | 1.628935049 | 1.740362689 | 1.621017948 |
| ENSMUSG00000019577 | 2.793394745 | 2.537119146 | 2.471424093 | 2.049918086 | 1.974687643 | 2.152288344 | 1.795211864 |
| ENSMUSG00000019578 | 2.435736758 | 2.264497931 | 2.309601185 | 2.176839783 | 2.197694248 | 2.397940009 | 2.158915016 |
| ENSMUSG00000019579 | 2.414600936 | 2.25 | 2.366391601 | 2.174241882 | 2.221952687 | 2.46834733 | 2.204254617 |
| ENSMUSG00000019590 | 0.811507563 | 0.896240625 | 0.79248125 | 1.214495249 | 1.188507684 | 1.041392685 | 1.150210039 |
| ENSMUSG00000019647 | 0 | 0 | 0 | 0.5 | 0.387761935 | 0.477121255 | 0.520678036 |
| ENSMUSG00000019659 | 1.960252568 | 2.038704527 | 2.046218836 | 1.843759858 | 2.107593434 | 2.164352856 | 2.041356071 |
| ENSMUSG00000019689 | 1.342375483 | 1.372963274 | 1.464495249 | 1.188721876 | 1.223252711 | 1.380211242 | 1.355418052 |
| ENSMUSG00000019699 | 2.551444705 | 2.362802778 | 2.310197833 | 2.112802778 | 2.022473809 | 2.303196057 | 1.893277937 |
| ENSMUSG00000019710 | 2.160971586 | 1.970660762 | 1.985628626 | 1.994319981 | 2.012006468 | 2.201397124 | 1.840919217 |
| ENSMUSG00000019715 | 2.498634788 | 2.302363341 | 2.294354884 | 2.349151195 | 2.310221245 | 2.618048097 | 2.180662175 |
| ENSMUSG00000019718 | 1.494443473 | 1.52186571 | 1.488549078 | 0.79248125 | 1 | 0.84509804 | 0.984950056 |
| ENSMUSG00000019726 | 2.367390214 | 2.192046081 | 2.20584181 | 2.067865757 | 2.039257769 | 2.320146286 | 1.861046144 |
| ENSMUSG00000019731 | 2.693989403 | 2.472945926 | 2.477847997 | 2.563757142 | 2.531009002 | 2.859138297 | 2.383567577 |
| ENSMUSG00000019732 | 1.249317394 | 1.238549078 | 1.160964047 | 1.311981878 | 1.319233381 | 1.322219295 | 1.329931221 |
| ENSMUSG00000019734 | 0.867194479 | 0.830482024 | 0.79248125 | 0.5 | 0.686820904 | 0.477121255 | 0.520678036 |
| ENSMUSG00000019737 | 0.45815691 | 0.580482024 | 0.5 | 0.580482024 | 0.489301084 | 0 | 0.710836105 |
| ENSMUSG00000019738 | 1.631440309 | 1.532320754 | 1.726722649 | 1.380890489 | 1.642643806 | 1.51851394 | 1.702561601 |
| ENSMUSG00000019756 | 0.45815691 | 0.25 | 0.396240625 | 0.396240625 | 0.568060967 | 0 | 0.629532004 |
| ENSMUSG00000019761 | 1.181541736 | 1.311981878 | 1.356566189 | 1.130890489 | 1.039257769 | 1.204119983 | 1.159323753 |
| ENSMUSG00000019762 | 0 | 0 | 0 | 0 | 0 | 0 | 0 |
| ENSMUSG00000019763 | 2.185978293 | 2.04745614 | 2.032320754 | 2.004202072 | 1.951645858 | 2.206825876 | 1.877074955 |
| ENSMUSG00000019767 | 0.289064826 | 0.396240625 | 0 | 0.25 | 0.387761935 | 0 | 0.224243824 |
| ENSMUSG00000019768 | 0.578129653 | 0.580482024 | 0.396240625 | 0.701838731 | 0.632412477 | 0.477121255 | 0.520678036 |
| ENSMUSG00000019772 | 0 | 0 | 0 | 0 | 0 | 0 | 0 |
| ENSMUSG00000019773 | 2.355805236 | 2.20584181 | 2.19697564 | 2.20584181 | 2.16097133 | 2.463892989 | 2.013099581 |
| ENSMUSG00000019774 | 1.596667477 | 1.53743678 | 1.54248125 | 1.741446071 | 1.692703642 | 1.86923172 | 1.551512352 |
| ENSMUSG00000019775 | 1.100572389 | 1.160964047 | 1.114857905 | 0.646240625 | 0.632412477 | 0.84509804 | 0.74492186 |

| | | | | | | | |
|---|---|---|---|---|---|---|---|
| ENSMUSG00000019777 | 2.954002173 | 2.699820405 | 2.687676747 | 2.718553234 | 2.623340676 | 3.046104787 | 2.456247769 |
| ENSMUSG00000019779 | 2.444137694 | 2.271202097 | 2.253505118 | 1.282320754 | 1.136121934 | 1.342422681 | 1.089372236 |
| ENSMUSG00000019782 | 2.40817504 | 2.219129237 | 2.215271726 | 2.37246199 | 2.269408865 | 2.656098202 | 2.130281771 |
| ENSMUSG00000019785 | 0 | 0 | 0 | 0 | 0.244650542 | 0 | 0 |
| ENSMUSG00000019787 | 0 | 0 | 0 | 0 | 0 | 0 | 0 |
| ENSMUSG00000019789 | 1.358731791 | 1.438721876 | 1.348079356 | 0.396240625 | 0.905314583 | 0.477121255 | 0.579661877 |
| ENSMUSG00000019790 | 2.472830741 | 2.306603048 | 2.304186465 | 2.388647213 | 2.363323232 | 2.650307523 | 2.180662175 |
| ENSMUSG00000019791 | 1.325351389 | 1.27186571 | 1.29248125 | 1.160964047 | 1.302012594 | 1.361727836 | 1.20919388 |
| ENSMUSG00000019792 | 2.103271637 | 1.910964047 | 1.843759858 | 1.994319981 | 1.974687643 | 2.136720567 | 1.812372009 |
| ENSMUSG00000019794 | 2.299278375 | 2.113831805 | 2.094844592 | 2.212155735 | 2.121713561 | 2.428134794 | 2.023210262 |
| ENSMUSG00000019795 | 2.658591846 | 2.43662858 | 2.451435968 | 2.528435542 | 2.461024076 | 2.835690571 | 2.301211393 |
| ENSMUSG00000019796 | 1.587501561 | 1.598079356 | 1.476722649 | 1.923371739 | 1.859322797 | 2.06069784 | 1.732938436 |
| ENSMUSG00000019797 | 1.948720984 | 1.858657057 | 1.862802778 | 1.894357207 | 1.855665188 | 2.008600172 | 1.773071385 |
| ENSMUSG00000019802 | 2.916696594 | 2.675326615 | 2.645300145 | 2.77053726 | 2.680838453 | 3.090963077 | 2.478774612 |
| ENSMUSG00000019803 | 0 | 0 | 0 | 0 | 0 | 0.301029996 | 0 |
| ENSMUSG00000019804 | 2.779274431 | 2.563166358 | 2.538388008 | 2.59995299 | 2.557901472 | 2.955206538 | 2.415994748 |
| ENSMUSG00000019806 | 1.069666964 | 1.061981878 | 1 | 1.04248125 | 0.955822902 | 1.301029996 | 1.159323753 |
| ENSMUSG00000019808 | 1.558729299 | 1.5 | 1.532320754 | 1.664552871 | 1.731346772 | 1.838849091 | 1.554175045 |
| ENSMUSG00000019809 | 2.163307908 | 1.96293726 | 1.948603967 | 2.099151195 | 1.969399812 | 2.290034611 | 1.86715039 |
| ENSMUSG00000019810 | 2.505869732 | 2.272528105 | 2.332168732 | 2.27053726 | 2.157861565 | 2.472756449 | 2.037212175 |
| ENSMUSG00000019813 | 2.092368209 | 1.982684334 | 1.951838731 | 2.02053726 | 1.926994141 | 2.155336037 | 1.836517579 |
| ENSMUSG00000019814 | 2.507914013 | 2.277131114 | 2.281030328 | 2.445749552 | 2.355827317 | 2.698970004 | 2.224577862 |
| ENSMUSG00000019815 | 0 | 0 | 0 | 0 | 0.244650542 | 0 | 0 |
| ENSMUSG00000019817 | 2.020182925 | 1.990000483 | 1.958222504 | 1.821350555 | 1.791313678 | 2.053078443 | 1.753626837 |
| ENSMUSG00000019818 | 3.223733281 | 2.932617534 | 2.927916743 | 3.006371414 | 2.897141297 | 3.408409578 | 2.692580302 |
| ENSMUSG00000019820 | 3.057562627 | 2.791226732 | 2.800530956 | 2.725970461 | 2.6656084 | 3.042969073 | 2.48059553 |
| ENSMUSG00000019822 | 1.975148165 | 1.905512955 | 1.843759858 | 1.903677461 | 1.937354184 | 2.173186268 | 1.863093717 |
| ENSMUSG00000019823 | 2.245508853 | 2.131869252 | 2.184523065 | 1.75835575 | 1.900490506 | 1.924279286 | 1.896054911 |
| ENSMUSG00000019826 | 1.876566388 | 1.75835575 | 1.816696635 | 1.819031101 | 1.834841362 | 1.959041392 | 1.752192176 |
| ENSMUSG00000019828 | 0.578129653 | 0 | 0.25 | 0 | 0 | 0 | 0 |
| ENSMUSG00000019831 | 1.956444026 | 1.843759858 | 1.825945187 | 1.940387808 | 1.953043706 | 2.184691431 | 1.833176598 |
| ENSMUSG00000019832 | 2.072573125 | 1.94697564 | 1.970660762 | 2.017865591 | 1.977302144 | 2.164352856 | 1.846338363 |
| ENSMUSG00000019834 | 0 | 0 | 0 | 0 | 0 | 0 | 0 |
| ENSMUSG00000019836 | 0.289064826 | 0 | 0 | 0 | 0 | 0.301029996 | 0.224243824 |

| | | | | | | | |
|---|---|---|---|---|---|---|---|
| ENSMUSG00000019837 | 2.190368148 | 2.074802005 | 2.056000419 | 2.260414788 | 2.174635855 | 2.522444234 | 2.064068234 |
| ENSMUSG00000019838 | 2.039433992 | 1.837182039 | 1.896240625 | 1.890560606 | 1.85749873 | 2.103803721 | 1.762103709 |
| ENSMUSG00000019841 | 2.686747932 | 2.467976309 | 2.433677405 | 2.564935816 | 2.505123331 | 2.854913022 | 2.288586107 |
| ENSMUSG00000019842 | 1.714367558 | 1.741446071 | 1.752806814 | 1.618933358 | 1.72073075 | 1.84509804 | 1.722075986 |
| ENSMUSG00000019843 | 2.603209296 | 2.375459296 | 2.363831805 | 2.109697963 | 2.093822267 | 2.281033367 | 1.927642465 |
| ENSMUSG00000019845 | 1.867194479 | 1.75 | 1.752806814 | 1.828220739 | 1.836786029 | 2.017033339 | 1.737484395 |
| ENSMUSG00000019846 | 1.700465191 | 1.54745614 | 1.584962501 | 0.79248125 | 0.846351821 | 0.602059991 | 0.969165684 |
| ENSMUSG00000019848 | 0 | 0.25 | 0 | 0 | 0.244650542 | 0 | 0.224243824 |
| ENSMUSG00000019849 | 2.840049992 | 2.610735874 | 2.611253711 | 2.585796429 | 2.567082571 | 2.931457871 | 2.429208684 |
| ENSMUSG00000019850 | 2.584966402 | 2.418106335 | 2.385274154 | 2.215271726 | 2.208682096 | 2.46834733 | 2.044866219 |
| ENSMUSG00000019851 | 2.302628051 | 2.106566189 | 2.168106335 | 2.32991803 | 2.219338185 | 2.565847819 | 2.105204898 |
| ENSMUSG00000019852 | 0.671187741 | 1.04248125 | 0.79248125 | 1.04248125 | 1.149965125 | 1 | 1.100339912 |
| ENSMUSG00000019853 | 1.936861443 | 1.777131114 | 1.779735268 | 1.901832578 | 1.828942237 | 2.158362492 | 1.773071385 |
| ENSMUSG00000019854 | 2.519973621 | 2.245034894 | 2.252806814 | 2.278435542 | 2.237245597 | 2.51054501 | 2.115189268 |
| ENSMUSG00000019856 | 0.811507563 | 0.75 | 0.830482024 | 1.080482024 | 1.106691885 | 1.278753601 | 0.984950056 |
| ENSMUSG00000019857 | 2.17479713 | 2.063166358 | 2.028435542 | 2.17510993 | 2.06759338 | 2.383815366 | 1.912234055 |
| ENSMUSG00000019861 | 2.377269616 | 2.213717096 | 2.139605178 | 2.231203126 | 2.176121988 | 2.471291711 | 2.017561934 |
| ENSMUSG00000019863 | 2 | 1.899978211 | 1.882845365 | 1.933677405 | 1.873585591 | 2.1430148 | 1.819456572 |
| ENSMUSG00000019864 | 1.771760131 | 1.685366747 | 1.711372513 | 1.789967834 | 1.72073075 | 1.886490725 | 1.548827561 |
| ENSMUSG00000019865 | 0.45815691 | 0 | 0.396240625 | 0.396240625 | 0.387761935 | 0 | 0.224243824 |
| ENSMUSG00000019866 | 2.461589295 | 2.275165585 | 2.267865591 | 2.412314044 | 2.364629672 | 2.707570176 | 2.178359564 |
| ENSMUSG00000019867 | 0.289064826 | 0.396240625 | 0 | 0 | 0.244650542 | 0.301029996 | 0 |
| ENSMUSG00000019868 | 2.361637952 | 2.179419106 | 2.145300145 | 2.170748646 | 2.150762767 | 2.450249108 | 2.012457048 |
| ENSMUSG00000019872 | 1.765759561 | 1.688721876 | 1.557204673 | 1.146240625 | 1.163285805 | 1.301029996 | 1.078019652 |
| ENSMUSG00000019873 | 2.816402366 | 2.599685923 | 2.599151195 | 2.633088981 | 2.578648039 | 2.996073654 | 2.366586192 |
| ENSMUSG00000019874 | 1.156259305 | 1.114857905 | 0.864857905 | 0.5 | 0.387761935 | 0.602059991 | 0.448487648 |
| ENSMUSG00000019876 | 1.036286563 | 1.130890489 | 1.114857905 | 0.951838731 | 1.091002363 | 0.602059991 | 1 |
| ENSMUSG00000019877 | 3.206907639 | 2.893411797 | 2.890560606 | 2.996282593 | 2.882015098 | 3.387567779 | 2.693129897 |
| ENSMUSG00000019878 | 2.365959569 | 2.181128463 | 2.144357207 | 2.228969845 | 2.16174451 | 2.44870632 | 1.991191705 |
| ENSMUSG00000019880 | 0.867194479 | 1.02186571 | 0.92510993 | 0.75 | 0.846351821 | 0.77815125 | 0.829800754 |
| ENSMUSG00000019883 | 1.777675583 | 1.638647213 | 1.642463902 | 1.649978211 | 1.592135487 | 1.69019608 | 1.459332757 |
| ENSMUSG00000019888 | 0.289064826 | 0.25 | 0 | 0 | 0 | 0 | 0 |
| ENSMUSG00000019889 | 2.732608034 | 2.471424093 | 2.449415381 | 2.494319981 | 2.356717497 | 2.785329835 | 2.234117901 |
| ENSMUSG00000019890 | 0 | 0 | 0 | 0 | 0.244650542 | 0 | 0 |

| | | | | | | | |
|---|---|---|---|---|---|---|---|
| ENSMUSG00000019891 | 1.936861443 | 1.807204673 | 1.83272922 | 1.918106335 | 1.897237431 | 2.120573931 | 1.811176017 |
| ENSMUSG00000019892 | 0 | 0.396240625 | 0 | 0.25 | 0.387761935 | 0 | 0.224243824 |
| ENSMUSG00000019893 | 0 | 0 | 0 | 0 | 0 | 0.301029996 | 0 |
| ENSMUSG00000019894 | 2.158622101 | 2.073655187 | 2.05479213 | 1.403677461 | 1.212046813 | 1.380211242 | 1.176815335 |
| ENSMUSG00000019897 | 2.330063669 | 2.201032755 | 2.188721876 | 2.233422664 | 2.211376431 | 2.51054501 | 2.077494038 |
| ENSMUSG00000019899 | 0 | 0.25 | 0 | 0.396240625 | 0.846351821 | 0.84509804 | 0.520678036 |
| ENSMUSG00000019900 | 0 | 0 | 0 | 0 | 0.244650542 | 0.301029996 | 0 |
| ENSMUSG00000019905 | 0 | 0 | 0 | 0 | 0 | 0 | 0 |
| ENSMUSG00000019906 | 0 | 0.25 | 0.25 | 0.79248125 | 0.812711509 | 0.698970004 | 0.710836105 |
| ENSMUSG00000019907 | 2.755187126 | 2.541226732 | 2.536483036 | 2.505938588 | 2.420030246 | 2.795880017 | 2.244955885 |
| ENSMUSG00000019909 | 0 | 0 | 0 | 0 | 0 | 0 | 0 |
| ENSMUSG00000019913 | 0 | 0.396240625 | 0.5 | 0.25 | 0 | 0 | 0.355418052 |
| ENSMUSG00000019916 | 2.91285295 | 2.650674716 | 2.638167274 | 2.639126514 | 2.55104774 | 2.983626287 | 2.394807458 |
| ENSMUSG00000019917 | 2.429630757 | 2.253505118 | 2.224461364 | 2.252806814 | 2.106691885 | 2.426511261 | 2.01501956 |
| ENSMUSG00000019920 | 3.121692466 | 2.868806694 | 2.846869716 | 2.832589179 | 2.738652723 | 3.182414652 | 2.537606247 |
| ENSMUSG00000019923 | 3.100572389 | 2.837596686 | 2.810793496 | 2.887686696 | 2.82151627 | 3.264581729 | 2.62307016 |
| ENSMUSG00000019927 | 2.336265009 | 2.148114259 | 2.091580554 | 2.209012589 | 2.145141048 | 2.424881637 | 1.95718226 |
| ENSMUSG00000019929 | 1.129344651 | 1.098079356 | 0.976722649 | 0.92510993 | 0.846351821 | 1.113943352 | 0.896975297 |
| ENSMUSG00000019932 | 0 | 0 | 0 | 0 | 0 | 0 | 0 |
| ENSMUSG00000019933 | 0 | 0 | 0 | 0.25 | 0 | 0 | 0 |
| ENSMUSG00000019935 | 0.671187741 | 0.79248125 | 0.701838731 | 0.25 | 0.387761935 | 0.301029996 | 0.579661877 |
| ENSMUSG00000019936 | 0 | 0 | 0 | 0 | 0 | 0 | 0 |
| ENSMUSG00000019942 | 3.097855556 | 2.83593523 | 2.816989271 | 2.942769185 | 2.881914726 | 3.356025857 | 2.686312118 |
| ENSMUSG00000019943 | 2.927646919 | 2.65892953 | 2.649745743 | 2.684100483 | 2.600031926 | 2.990338855 | 2.382339813 |
| ENSMUSG00000019944 | 1.647796617 | 1.642463902 | 1.602347734 | 1.380890489 | 1.283908311 | 1.591064607 | 1.417653332 |
| ENSMUSG00000019945 | 0 | 0 | 0 | 0 | 0 | 0 | 0 |
| ENSMUSG00000019947 | 2.390989024 | 2.264497931 | 2.228222334 | 2.038704527 | 2.014359518 | 2.206825876 | 1.924281235 |
| ENSMUSG00000019948 | 1.876566388 | 1.834962501 | 1.747171172 | 1.981203126 | 1.868304353 | 2.158362492 | 1.737484395 |
| ENSMUSG00000019951 | 2.607246427 | 2.418548067 | 2.450224975 | 2.478222334 | 2.40985355 | 2.781755375 | 2.237984665 |
| ENSMUSG00000019952 | 2.086807695 | 1.964495249 | 1.935366747 | 2.113831805 | 2.029283463 | 2.367355921 | 1.936709733 |
| ENSMUSG00000019960 | 1.482695304 | 1.470660762 | 1.505591953 | 1.52186571 | 1.611014646 | 1.785329835 | 1.572224412 |
| ENSMUSG00000019961 | 3.270320442 | 2.962742038 | 2.942769185 | 3.059899632 | 2.945411925 | 3.431846046 | 2.735769948 |
| ENSMUSG00000019966 | 2.10861828 | 1.978222334 | 2.064346961 | 1.959800947 | 1.862942891 | 2.021189299 | 1.710836105 |
| ENSMUSG00000019969 | 2.627631768 | 2.397178659 | 2.379417347 | 2.393411797 | 2.310728002 | 2.650307523 | 2.177588371 |

| | | | | | | | |
|---|---|---|---|---|---|---|---|
| ENSMUSG00000019970 | 2.247417471 | 2.140560606 | 2.135757955 | 2.148114259 | 2.115605392 | 2.315970345 | 1.984249047 |
| ENSMUSG00000019971 | 2.060824957 | 2.006976499 | 2.029735268 | 2.109697963 | 2.033751322 | 2.294466226 | 1.86715039 |
| ENSMUSG00000019975 | 2.564110437 | 2.392938161 | 2.353406982 | 2.368426437 | 2.283908311 | 2.62324929 | 2.142557543 |
| ENSMUSG00000019977 | 2.629921064 | 2.403677461 | 2.405054956 | 2.527457664 | 2.447194116 | 2.803457116 | 2.295899033 |
| ENSMUSG00000019978 | 3.19881715 | 2.915444524 | 2.904366866 | 2.939347502 | 2.893371499 | 3.344981414 | 2.692894471 |
| ENSMUSG00000019979 | 2.616779011 | 2.368933358 | 2.357613138 | 2.413211243 | 2.357605438 | 2.6599162 | 2.181807364 |
| ENSMUSG00000019982 | 1 | 0.92510993 | 1.02186571 | 1.061981878 | 1.039257769 | 1 | 1.014380832 |
| ENSMUSG00000019984 | 2.617563644 | 2.387205477 | 2.367918804 | 2.408702763 | 2.368949874 | 2.672097858 | 2.242122155 |
| ENSMUSG00000019986 | 2.131864492 | 2.012462137 | 1.966046536 | 2.192046081 | 2.086251872 | 2.411619706 | 1.965468324 |
| ENSMUSG00000019987 | 0.578129653 | 0.25 | 0.75 | 0.396240625 | 0.244650542 | 0.301029996 | 0.520678036 |
| ENSMUSG00000019988 | 2.588343201 | 2.354463129 | 2.27908599 | 2.456239685 | 2.352693888 | 2.728353782 | 2.21543553 |
| ENSMUSG00000019989 | 0 | 0.396240625 | 0 | 0 | 0 | 0 | 0 |
| ENSMUSG00000019990 | 0.289064826 | 0 | 0.25 | 0 | 0 | 0 | 0.224243824 |
| ENSMUSG00000019992 | 2.124258828 | 1.992885888 | 2.024508021 | 2.089388001 | 2.048957937 | 2.338456494 | 1.943148064 |
| ENSMUSG00000019996 | 0.578129653 | 0.396240625 | 0 | 0.75 | 0.877063019 | 1.146128036 | 0.984950056 |
| ENSMUSG00000019997 | 3.009295312 | 2.755591953 | 2.72104263 | 2.504897682 | 2.489606278 | 2.81756537 | 2.255745871 |
| ENSMUSG00000019998 | 2.618346804 | 2.417221246 | 2.387205477 | 2.501055117 | 2.421513257 | 2.791690649 | 2.236379126 |
| ENSMUSG00000020000 | 0.289064826 | 0.396240625 | 0.396240625 | 0.976722649 | 0.955822902 | 0.698970004 | 0.710836105 |
| ENSMUSG00000020003 | 1.605636262 | 1.660964047 | 1.571350555 | 1.856566189 | 1.77552387 | 1.959041392 | 1.655750087 |
| ENSMUSG00000020007 | 0 | 0 | 0 | 0 | 0.244650542 | 0 | 0 |
| ENSMUSG00000020009 | 2.371652933 | 2.210587586 | 2.183677405 | 2.286164561 | 2.253097759 | 2.525044807 | 2.084261301 |
| ENSMUSG00000020010 | 0 | 0 | 0 | 0 | 0 | 0 | 0 |
| ENSMUSG00000020014 | 0.747221736 | 0.701838731 | 0.701838731 | 0.75 | 0.387761935 | 0.477121255 | 0 |
| ENSMUSG00000020015 | 2.450036412 | 2.25 | 2.209800947 | 2.233422664 | 2.220647857 | 2.459392488 | 2.020084484 |
| ENSMUSG00000020017 | 0.867194479 | 0.830482024 | 0.79248125 | 0.580482024 | 0.632412477 | 0.602059991 | 0.629532004 |
| ENSMUSG00000020018 | 0.671187741 | 0.5 | 0.5 | 0.580482024 | 0.686820904 | 0.477121255 | 0.876096088 |
| ENSMUSG00000020019 | 1.527823874 | 1.589388001 | 1.505591953 | 1.723704441 | 1.677808768 | 1.785329835 | 1.616675393 |
| ENSMUSG00000020020 | 0.867194479 | 0.830482024 | 0.864857905 | 1.17510993 | 1.136121934 | 1 | 1.100339912 |
| ENSMUSG00000020021 | 1.978790393 | 1.828220739 | 1.811981878 | 2.052363341 | 1.964031554 | 2.195899652 | 1.822941512 |
| ENSMUSG00000020022 | 2.042557845 | 1.940387808 | 1.942046081 | 2.168989258 | 2.112954895 | 2.414973348 | 2.012457048 |
| ENSMUSG00000020023 | 2.124258828 | 1.931980114 | 1.912762923 | 2.099151195 | 1.996316946 | 2.285557309 | 1.908707966 |
| ENSMUSG00000020024 | 2.295901576 | 2.182829758 | 2.201838731 | 2.220660762 | 2.132784263 | 2.416640507 | 1.997988498 |
| ENSMUSG00000020027 | 2.013563224 | 1.862802778 | 1.852347734 | 1.67510993 | 1.652586889 | 1.770852012 | 1.540636858 |
| ENSMUSG00000020029 | 3.05206805 | 2.788546302 | 2.778760913 | 2.621457327 | 2.539320544 | 2.932980822 | 2.356308054 |

| | | | | | | | |
|---|---|---|---|---|---|---|---|
| ENSMUSG00000020032 | 2.447686928 | 2.252107156 | 2.225970461 | 2.262462137 | 2.25190332 | 2.437750563 | 2.125328759 |
| ENSMUSG00000020033 | 0 | 0 | 0 | 0.25 | 0.244650542 | 0 | 0 |
| ENSMUSG00000020034 | 1.289064826 | 1.214495249 | 1.302363341 | 1.396240625 | 1.366364104 | 1.255272505 | 1.307989563 |
| ENSMUSG00000020037 | 0 | 0 | 0 | 0 | 0 | 0 | 0 |
| ENSMUSG00000020038 | 2.086807695 | 1.868933358 | 1.988549078 | 1.953445298 | 1.973373093 | 2.10720997 | 1.869159809 |
| ENSMUSG00000020042 | 1.686083339 | 1.571350555 | 1.729715809 | 1.580482024 | 1.625423019 | 1.73239376 | 1.526507301 |
| ENSMUSG00000020044 | 3.292497211 | 3.007062857 | 3.023354391 | 2.658702763 | 2.593274423 | 2.995635195 | 2.414513278 |
| ENSMUSG00000020048 | 3.635896925 | 3.31168515 | 3.280909113 | 3.373620154 | 3.274891466 | 3.835436895 | 3.034492628 |
| ENSMUSG00000020051 | 0 | 0 | 0 | 0 | 0 | 0 | 0 |
| ENSMUSG00000020052 | 0 | 0 | 0 | 0 | 0 | 0 | 0 |
| ENSMUSG00000020053 | 1.800532393 | 1.752806814 | 1.62694866 | 2.142463902 | 2.029283463 | 2.356025857 | 1.882886767 |
| ENSMUSG00000020056 | 2.181541736 | 2.051142786 | 2.061981878 | 2.095926073 | 2.069604532 | 2.346352974 | 1.963229454 |
| ENSMUSG00000020057 | 1.852729478 | 1.769203899 | 1.79745614 | 1.794977273 | 1.797870327 | 1.897627091 | 1.704233519 |
| ENSMUSG00000020059 | 0 | 0.701838731 | 0.25 | 0.396240625 | 0.489301084 | 0.477121255 | 0.629532004 |
| ENSMUSG00000020061 | 0 | 0.25 | 0 | 0 | 0.387761935 | 0 | 0 |
| ENSMUSG00000020062 | 0.747221736 | 0.646240625 | 0.25 | 0.646240625 | 0.568060967 | 0.602059991 | 0.448487648 |
| ENSMUSG00000020063 | 2.309247751 | 2.137686696 | 2.121960008 | 2.350753006 | 2.310728002 | 2.62838893 | 2.101807105 |
| ENSMUSG00000020064 | 2.591692877 | 2.365881093 | 2.354463129 | 2.424676167 | 2.329944595 | 2.685741739 | 2.169773608 |
| ENSMUSG00000020067 | 0.45815691 | 0.25 | 0.396240625 | 0 | 0 | 0 | 0 |
| ENSMUSG00000020069 | 2.29078455 | 2.106566189 | 2.132845365 | 2.240724001 | 2.202543574 | 2.495544338 | 2.044866219 |
| ENSMUSG00000020070 | 2.219867581 | 2.041226732 | 2.079353153 | 2.21759118 | 2.175379703 | 2.432969291 | 2.029372647 |
| ENSMUSG00000020072 | 1.205378646 | 1.160964047 | 1.160964047 | 1 | 0.931471446 | 0.602059991 | 0.876096088 |
| ENSMUSG00000020074 | 2.935766391 | 2.698197574 | 2.714106382 | 2.788071211 | 2.687866184 | 3.093771781 | 2.487630378 |
| ENSMUSG00000020075 | 3.157120054 | 2.886361795 | 2.887927065 | 3.00158152 | 2.924244626 | 3.368286885 | 2.714087544 |
| ENSMUSG00000020076 | 2.733202521 | 2.52219706 | 2.498588359 | 2.559899632 | 2.48127199 | 2.854306042 | 2.300155876 |
| ENSMUSG00000020077 | 0 | 0.25 | 0.25 | 0.25 | 0.387761935 | 0.301029996 | 0.355418052 |
| ENSMUSG00000020078 | 2.944089222 | 2.705443495 | 2.712351275 | 2.692872367 | 2.620404601 | 3.023663918 | 2.459978497 |
| ENSMUSG00000020079 | 2.217828299 | 2.027131114 | 2.031030328 | 2.17510993 | 2.135290471 | 2.416640507 | 1.990504106 |
| ENSMUSG00000020080 | 0.289064826 | 0.396240625 | 0 | 0.396240625 | 0 | 0.301029996 | 0.224243824 |
| ENSMUSG00000020081 | 0 | 0 | 0.25 | 0 | 0 | 0 | 0.355418052 |
| ENSMUSG00000020083 | 0.578129653 | 0.830482024 | 0.830482024 | 1.04248125 | 1.136121934 | 1.255272505 | 1.176815335 |
| ENSMUSG00000020085 | 2 | 1.843759858 | 1.899978211 | 2.031030328 | 2.103062484 | 2.315970345 | 1.995964412 |
| ENSMUSG00000020086 | 2.267855219 | 2.05357978 | 2.14151351 | 2.184523065 | 2.230318492 | 2.436162647 | 2.123508695 |
| ENSMUSG00000020087 | 1.578129653 | 1.418106335 | 1.516522298 | 1.593759858 | 1.599808748 | 1.653212514 | 1.67819947 |

| | | | | | | | |
|---|---|---|---|---|---|---|---|
| ENSMUSG00000020088 | 3.151275227 | 2.876204747 | 2.85035322 | 2.908475852 | 2.840417764 | 3.282395505 | 2.645461849 |
| ENSMUSG00000020089 | 2.742019602 | 2.518535366 | 2.500703754 | 2.704845198 | 2.646576505 | 3.063708559 | 2.472164183 |
| ENSMUSG00000020090 | 0 | 0 | 0.25 | 0.396240625 | 0.244650542 | 0 | 0.448487648 |
| ENSMUSG00000020091 | 2.422183552 | 2.252107156 | 2.26584877 | 2.272528105 | 2.227765453 | 2.532754379 | 2.126687118 |
| ENSMUSG00000020092 | 2.048736215 | 1.87892496 | 2.01109853 | 1.955044741 | 2.032639648 | 2.225309282 | 1.994607951 |
| ENSMUSG00000020096 | 0 | 0.25 | 0.5 | 0 | 0.387761935 | 0.301029996 | 0.355418052 |
| ENSMUSG00000020097 | 3.059740347 | 2.766522298 | 2.772858847 | 2.784247778 | 2.718823734 | 3.141449773 | 2.520945293 |
| ENSMUSG00000020098 | 0 | 0 | 0 | 0.5 | 0.387761935 | 0.477121255 | 0.672731473 |
| ENSMUSG00000020099 | 3.029919532 | 2.836628433 | 2.877567418 | 2.540282971 | 2.648720957 | 2.868056362 | 2.528862053 |
| ENSMUSG00000020100 | 2.119110214 | 2.00835575 | 1.950224975 | 2.066696635 | 1.962676635 | 2.260071388 | 1.910475814 |
| ENSMUSG00000020101 | 1.678702042 | 1.527131114 | 1.458222504 | 1.188721876 | 1.176121988 | 1.146128036 | 1.140832123 |
| ENSMUSG00000020102 | 1.289064826 | 1.29248125 | 1.302363341 | 0.951838731 | 0.77552387 | 0.602059991 | 0.579661877 |
| ENSMUSG00000020105 | 2.292497211 | 2.137686696 | 2.185366747 | 2.069031101 | 2.033751322 | 2.278753601 | 1.972093446 |
| ENSMUSG00000020107 | 2.361637952 | 2.234894804 | 2.277131114 | 2.250703754 | 2.232853197 | 2.53529412 | 2.123964672 |
| ENSMUSG00000020108 | 2.326928073 | 2.132845365 | 2.202642909 | 1.807204673 | 1.789100779 | 1.954242509 | 1.811176017 |
| ENSMUSG00000020109 | 2.334723298 | 2.25 | 2.243603647 | 2.161864607 | 2.238490593 | 2.434568904 | 2.153969461 |
| ENSMUSG00000020111 | 2.803052234 | 2.570192693 | 2.580200137 | 2.530706999 | 2.517458689 | 2.867467488 | 2.37466387 |
| ENSMUSG00000020114 | 3.157334963 | 2.851551251 | 2.826230417 | 2.907566781 | 2.818885826 | 3.274619619 | 2.614981852 |
| ENSMUSG00000020115 | 2.6088504 | 2.40136988 | 2.414552871 | 2.409608978 | 2.338004387 | 2.666517981 | 2.228889254 |
| ENSMUSG00000020116 | 2.499676068 | 2.3184489 | 2.342126615 | 2.488912477 | 2.392211871 | 2.733197265 | 2.249624299 |
| ENSMUSG00000020120 | 0.289064826 | 0.646240625 | 0.864857905 | 0.864857905 | 0.846351821 | 0.84509804 | 0.775756176 |
| ENSMUSG00000020121 | 2.163307908 | 2.069031101 | 2.065523711 | 1.25 | 1.149965125 | 1.230448921 | 1.176815335 |
| ENSMUSG00000020122 | 2.304292853 | 2.051142786 | 2.051142786 | 1.79745614 | 1.808534465 | 1.939519253 | 1.725216937 |
| ENSMUSG00000020123 | 0.45815691 | 0 | 0.25 | 0.701838731 | 0.632412477 | 0 | 0.224243824 |
| ENSMUSG00000020124 | 2.745494892 | 2.510756821 | 2.494677615 | 2.661414608 | 2.54143406 | 2.967547976 | 2.364426535 |
| ENSMUSG00000020125 | 0 | 0 | 0 | 0 | 0 | 0 | 0 |
| ENSMUSG00000020128 | 2.462727174 | 2.264497931 | 2.224461364 | 2.431128463 | 2.345425551 | 2.699837726 | 2.13295179 |
| ENSMUSG00000020130 | 2.787699615 | 2.547147211 | 2.529735268 | 2.6343046 | 2.5461437 | 2.95230801 | 2.37466387 |
| ENSMUSG00000020131 | 0.811507563 | 1.04248125 | 1.214495249 | 0.5 | 0.77552387 | 0.602059991 | 1.176815335 |
| ENSMUSG00000020132 | 2.469490009 | 2.279735268 | 2.256976499 | 2.26109853 | 2.240349892 | 2.561101384 | 2.138226615 |
| ENSMUSG00000020133 | 2.20747957 | 2.079353153 | 2.131869252 | 1.969129237 | 2.101233692 | 2.243038049 | 2.083229353 |
| ENSMUSG00000020134 | 2.20747957 | 2.02186571 | 1.979715809 | 2.182829758 | 2.139428339 | 2.390935107 | 1.936709733 |
| ENSMUSG00000020135 | 1.816888701 | 1.695339928 | 1.843759858 | 1.403677461 | 1.632412477 | 1.531478917 | 1.759302692 |
| ENSMUSG00000020137 | 2.196867486 | 2.004202072 | 2.013820609 | 1.976722649 | 1.929985312 | 2.130333768 | 1.767634022 |

| | | | | | | | |
|---|---|---|---|---|---|---|---|
| ENSMUSG00000020140 | 0.867194479 | 0.646240625 | 0.646240625 | 1.51109853 | 1.358933174 | 1.612783857 | 1.462868481 |
| ENSMUSG00000020142 | 2.386920383 | 2.224461364 | 2.187886015 | 2.187048212 | 2.159419872 | 2.385606274 | 2.028149525 |
| ENSMUSG00000020143 | 0 | 0 | 0 | 0 | 0 | 0 | 0 |
| ENSMUSG00000020149 | 2.997170519 | 2.735811849 | 2.725782169 | 2.883697303 | 2.778227492 | 3.228656958 | 2.57910438 |
| ENSMUSG00000020150 | 1.432083896 | 1.470660762 | 1.557204673 | 1.226722649 | 1.380772476 | 1.414973348 | 1.259064007 |
| ENSMUSG00000020151 | 0.289064826 | 0.396240625 | 0.25 | 0 | 0 | 0.301029996 | 0.355418052 |
| ENSMUSG00000020152 | 3.31414447 | 3.026067782 | 3.023519421 | 3.111899965 | 3.014773138 | 3.496652939 | 2.779381714 |
| ENSMUSG00000020153 | 1.936861443 | 1.914552871 | 2.087182039 | 1.681980114 | 1.977302144 | 1.982271233 | 2.126234965 |
| ENSMUSG00000020154 | 2.275045611 | 2.073655187 | 2.013820609 | 1.723704441 | 1.662257525 | 1.903089987 | 1.574722613 |
| ENSMUSG00000020155 | 0.289064826 | 0.701838731 | 0.25 | 0 | 0.244650542 | 0 | 0 |
| ENSMUSG00000020156 | 2.480523256 | 2.309601185 | 2.337182039 | 2.250703754 | 2.342660868 | 2.478566496 | 2.243069492 |
| ENSMUSG00000020159 | 0 | 0 | 0 | 0 | 0 | 0 | 0 |
| ENSMUSG00000020160 | 2.363083473 | 2.118933358 | 2.111770807 | 1.866901388 | 1.795698282 | 1.995635195 | 1.67819947 |
| ENSMUSG00000020163 | 2.054824387 | 2.051142786 | 2.187886015 | 1.779735268 | 2.044679629 | 2.08278537 | 2.151886289 |
| ENSMUSG00000020164 | 0 | 0 | 0 | 0 | 0 | 0 | 0 |
| ENSMUSG00000020166 | 2.638209243 | 2.421187155 | 2.415444524 | 2.582449085 | 2.489301084 | 2.84260924 | 2.338025481 |
| ENSMUSG00000020167 | 2.539328946 | 2.474461364 | 2.597542239 | 2.167221246 | 2.35849115 | 2.44870632 | 2.421308913 |
| ENSMUSG00000020168 | 0 | 0 | 0 | 0 | 0 | 0 | 0 |
| ENSMUSG00000020169 | 0 | 0.25 | 0 | 0 | 0 | 0.301029996 | 0.520678036 |
| ENSMUSG00000020170 | 2.811999637 | 2.562870603 | 2.535526764 | 2.540597832 | 2.501307552 | 2.84509804 | 2.324338556 |
| ENSMUSG00000020171 | 2.165631214 | 2.04745614 | 2.065523711 | 2.160061234 | 2.085294053 | 2.380211242 | 1.997315209 |
| ENSMUSG00000020173 | 1.578129653 | 1.52186571 | 1.52186571 | 1.188721876 | 1.020174412 | 1.113943352 | 1.041356071 |
| ENSMUSG00000020175 | 1.623015126 | 1.464495249 | 1.482684334 | 1.505591953 | 1.563883923 | 1.579783597 | 1.486592281 |
| ENSMUSG00000020176 | 3.444677402 | 3.11817271 | 3.083009137 | 2.762121718 | 2.697136965 | 3.10754913 | 2.524795923 |
| ENSMUSG00000020177 | 0 | 0.25 | 0 | 0 | 0 | 0 | 0 |
| ENSMUSG00000020178 | 0.867194479 | 1.02186571 | 1.114857905 | 0.646240625 | 0.387761935 | 0.477121255 | 0.876096088 |
| ENSMUSG00000020180 | 2.804055922 | 2.583009137 | 2.55205859 | 2.659156155 | 2.588004614 | 3.024074987 | 2.439163791 |
| ENSMUSG00000020181 | 1.663535556 | 1.589388001 | 1.5 | 1.282320754 | 1.254881871 | 1.113943352 | 1.168187745 |
| ENSMUSG00000020182 | 0 | 0.25 | 0 | 0 | 0.489301084 | 0 | 0.448487648 |
| ENSMUSG00000020183 | 0.289064826 | 0.5 | 0.25 | 1.114857905 | 1 | 1.255272505 | 1.089372236 |
| ENSMUSG00000020184 | 2.768497801 | 2.522528105 | 2.530706999 | 2.418106335 | 2.389791591 | 2.73479983 | 2.20496486 |
| ENSMUSG00000020185 | 2.521950082 | 2.328220739 | 2.33272922 | 2.341580554 | 2.320220671 | 2.656098202 | 2.159731975 |
| ENSMUSG00000020186 | 2.158622101 | 2.038704527 | 2.027131114 | 2.051142786 | 2.047893212 | 2.225309282 | 1.972821265 |
| ENSMUSG00000020189 | 2.552362269 | 2.364857905 | 2.334962501 | 2.484894804 | 2.404458919 | 2.764922985 | 2.22088434 |

| | | | | | | | |
|---|---|---|---|---|---|---|---|
| ENSMUSG00000020190 | 2.782453855 | 2.629417347 | 2.68662858 | 2.494319981 | 2.626045333 | 2.875061263 | 2.583647744 |
| ENSMUSG00000020191 | 0 | 0 | 0.25 | 0 | 0 | 0 | 0.224243824 |
| ENSMUSG00000020193 | 0.45815691 | 0 | 0 | 0 | 0 | 0 | 0.224243824 |
| ENSMUSG00000020196 | 2.657168526 | 2.475593779 | 2.53743678 | 2.370953944 | 2.480020371 | 2.752048448 | 2.367231288 |
| ENSMUSG00000020198 | 2.856724497 | 2.634789703 | 2.652986735 | 2.539336734 | 2.583123351 | 2.888740961 | 2.470763344 |
| ENSMUSG00000020205 | 1.912079952 | 1.926839783 | 1.898114259 | 1.62694866 | 1.55104774 | 1.770852012 | 1.559435575 |
| ENSMUSG00000020211 | 2.289064826 | 2.160061234 | 2.218361028 | 1.956637122 | 2.157861565 | 2.307496038 | 2.119375723 |
| ENSMUSG00000020212 | 2.069666964 | 2.03743678 | 1.990000483 | 2.05357978 | 2.016696984 | 2.214843848 | 1.905143021 |
| ENSMUSG00000020213 | 0 | 0 | 0 | 0 | 0.244650542 | 0 | 0 |
| ENSMUSG00000020214 | 0 | 0 | 0 | 0 | 0 | 0 | 0 |
| ENSMUSG00000020216 | 0 | 0.25 | 0.25 | 0 | 0 | 0 | 0.224243824 |
| ENSMUSG00000020218 | 0.578129653 | 0 | 0 | 0.951838731 | 1.039257769 | 1.113943352 | 1.168187745 |
| ENSMUSG00000020219 | 2.205378646 | 2.153677461 | 2.267194569 | 1.83272922 | 2.167869715 | 2.214843848 | 2.217148162 |
| ENSMUSG00000020220 | 2.552362269 | 2.331607372 | 2.360735874 | 2.373963757 | 2.336595379 | 2.617000341 | 2.158095989 |
| ENSMUSG00000020224 | 2.163307908 | 2.004202072 | 2.027131114 | 2.124961472 | 2.122577592 | 2.389166084 | 1.984950056 |
| ENSMUSG00000020225 | 2.136858952 | 1.976722649 | 1.96293726 | 2.228222334 | 2.11997913 | 2.44870632 | 2.037212175 |
| ENSMUSG00000020226 | 0.45815691 | 0.25 | 0.75 | 0.701838731 | 0.244650542 | 0 | 0.448487648 |
| ENSMUSG00000020227 | 1.418409478 | 1.311981878 | 1.27186571 | 1.282320754 | 1.319233381 | 1.568201724 | 1.329931221 |
| ENSMUSG00000020228 | 2.158622101 | 2.034887838 | 2.038704527 | 2.125956434 | 2.078515538 | 2.276461804 | 1.988432495 |
| ENSMUSG00000020229 | 0 | 0 | 0 | 0 | 0 | 0.301029996 | 0 |
| ENSMUSG00000020230 | 2.381433035 | 2.237821179 | 2.285526764 | 2.159156155 | 2.283908311 | 2.484299839 | 2.205673547 |
| ENSMUSG00000020231 | 2.600768124 | 2.436208459 | 2.464106382 | 2.430274797 | 2.532633036 | 2.807535028 | 2.411529829 |
| ENSMUSG00000020232 | 2.033115198 | 1.933677405 | 2.052363341 | 1.789967834 | 1.910073614 | 1.934498451 | 1.964723754 |
| ENSMUSG00000020234 | 0.45815691 | 0.580482024 | 0.396240625 | 0.396240625 | 0.387761935 | 0.301029996 | 0.629532004 |
| ENSMUSG00000020235 | 2.43085913 | 2.325945187 | 2.362802778 | 2.079353153 | 2.328502976 | 2.40654018 | 2.30565931 |
| ENSMUSG00000020238 | 2.512981297 | 2.403677461 | 2.498588359 | 2.146240625 | 2.410998892 | 2.514547753 | 2.410027723 |
| ENSMUSG00000020241 | 2.857166034 | 2.683889006 | 2.76601727 | 2.209012589 | 2.311739339 | 2.5289167 | 2.323109859 |
| ENSMUSG00000020246 | 2.479432974 | 2.244319981 | 2.279735268 | 2.30479213 | 2.253693465 | 2.57634135 | 2.109044889 |
| ENSMUSG00000020248 | 2.355805236 | 2.143411797 | 2.135757955 | 2.21293726 | 2.109389652 | 2.414973348 | 2.000667731 |
| ENSMUSG00000020250 | 3.226290649 | 2.943181737 | 2.94961795 | 2.896944379 | 2.861243942 | 3.309417226 | 2.687591469 |
| ENSMUSG00000020251 | 1.753493018 | 1.67510993 | 1.685366747 | 1.527131114 | 1.499532413 | 1.653212514 | 1.417653332 |
| ENSMUSG00000020253 | 2.215778996 | 2.092671852 | 2.143411797 | 2.109697963 | 2.164820399 | 2.389166084 | 2.077494038 |
| ENSMUSG00000020255 | 2.577267124 | 2.365881093 | 2.337182039 | 2.461764337 | 2.416670511 | 2.761927838 | 2.249315158 |
| ENSMUSG00000020256 | 2.484856099 | 2.228222334 | 2.121960008 | 1.894357207 | 1.911645804 | 2.045322979 | 1.784981274 |

| | | | | | | | |
|---|---|---|---|---|---|---|---|
| ENSMUSG00000020257 | 2.817859673 | 2.57163944 | 2.544666213 | 2.730831871 | 2.653163144 | 3.043755127 | 2.464144677 |
| ENSMUSG00000020258 | 1.205378646 | 1.114857905 | 1.080482024 | 0.951838731 | 1.136121934 | 1.041392685 | 1.100339912 |
| ENSMUSG00000020260 | 2.629159361 | 2.478969845 | 2.525165585 | 2.168106335 | 2.354487824 | 2.540329475 | 2.293209778 |
| ENSMUSG00000020261 | 2.203267085 | 2.013820609 | 2.104463129 | 1.96293726 | 1.947418788 | 2.139879086 | 1.833176598 |
| ENSMUSG00000020262 | 1.842799666 | 1.76109853 | 1.789967834 | 1.819031101 | 1.887294348 | 2.004321374 | 1.839824409 |
| ENSMUSG00000020263 | 2.063793172 | 1.903677461 | 1.892463902 | 2.061981878 | 1.998776581 | 2.152288344 | 1.888596011 |
| ENSMUSG00000020264 | 0 | 0 | 0 | 0 | 0 | 0 | 0 |
| ENSMUSG00000020265 | 2.868914203 | 2.603935442 | 2.621708755 | 2.764667063 | 2.68436981 | 3.126456113 | 2.522809941 |
| ENSMUSG00000020267 | 2.89911417 | 2.643411797 | 2.672499493 | 2.760585845 | 2.687866184 | 3.089551883 | 2.546860895 |
| ENSMUSG00000020268 | 1.269664473 | 1.114857905 | 1.214495249 | 1.214495249 | 1.163285805 | 1.361727836 | 1.150210039 |
| ENSMUSG00000020270 | 0 | 0 | 0 | 0 | 0 | 0 | 0 |
| ENSMUSG00000020271 | 2.768497801 | 2.552972074 | 2.528760913 | 2.62246199 | 2.539850111 | 2.944482672 | 2.389035574 |
| ENSMUSG00000020272 | 2.312518611 | 2.216046536 | 2.219895812 | 2.027131114 | 2.126012715 | 2.307496038 | 2.082194103 |
| ENSMUSG00000020273 | 2.185978293 | 1.984159485 | 2.01109853 | 2.146240625 | 2.068600388 | 2.354108439 | 1.924281235 |
| ENSMUSG00000020275 | 2.188178997 | 1.984159485 | 2.029735268 | 2.243603647 | 2.184185718 | 2.487138375 | 2.040177507 |
| ENSMUSG00000020277 | 2.737340367 | 2.588010856 | 2.666777886 | 2.521202097 | 2.656791073 | 2.925312091 | 2.608321598 |
| ENSMUSG00000020279 | 0 | 0.25 | 0.396240625 | 0.5 | 0 | 0 | 0.224243824 |
| ENSMUSG00000020280 | 2.253091469 | 2.043731421 | 2.065523711 | 2.355516192 | 2.297309926 | 2.620136055 | 2.07959139 |
| ENSMUSG00000020282 | 2.144240249 | 2.072504712 | 2.140560606 | 1.852347734 | 2.108492685 | 2.127104798 | 2.124420007 |
| ENSMUSG00000020283 | 2.243591459 | 2.064346961 | 2.074802005 | 2.29248125 | 2.221952687 | 2.488550717 | 2.050061048 |
| ENSMUSG00000020284 | 1.87190676 | 1.807204673 | 1.899978211 | 1.823655187 | 1.931471446 | 1.959041392 | 1.915722126 |
| ENSMUSG00000020286 | 0 | 0 | 0 | 0 | 0.244650542 | 0 | 0 |
| ENSMUSG00000020287 | 1.358731791 | 1.438721876 | 1.451838731 | 1.5 | 1.559656853 | 1.633468456 | 1.508694612 |
| ENSMUSG00000020288 | 2.100572389 | 1.959800947 | 1.969129237 | 2.158248799 | 2.141887974 | 2.374748346 | 2.003324957 |
| ENSMUSG00000020289 | 2.194712275 | 2.039967834 | 2.049918086 | 1.953445298 | 2.06149024 | 2.206825876 | 1.930969131 |
| ENSMUSG00000020290 | 3.211461269 | 2.916999634 | 2.904366866 | 3.063757142 | 2.963314892 | 3.415307292 | 2.735563602 |
| ENSMUSG00000020295 | 0 | 0 | 0 | 0 | 0 | 0 | 0 |
| ENSMUSG00000020297 | 0 | 0 | 0 | 0.25 | 0 | 0 | 0 |
| ENSMUSG00000020299 | 0 | 0 | 0 | 0 | 0 | 0 | 0 |
| ENSMUSG00000020300 | 2.256831695 | 2.107613138 | 2.111770807 | 2.314346961 | 2.205284995 | 2.498310554 | 2.059095225 |
| ENSMUSG00000020303 | 1 | 0.79248125 | 0.79248125 | 0.580482024 | 0.632412477 | 0 | 0.629532004 |
| ENSMUSG00000020305 | 2.060824957 | 1.858657057 | 1.732684334 | 1.948603967 | 1.855665188 | 2.117271296 | 1.687112305 |
| ENSMUSG00000020307 | 2.34539747 | 2.243603647 | 2.300530956 | 2.075945187 | 2.173890435 | 2.340444115 | 2.177202084 |
| ENSMUSG00000020308 | 0.91631382 | 1.02186571 | 1.348079356 | 0.75 | 1.283908311 | 1.041392685 | 1.586932193 |

| | | | | | | | |
|---|---|---|---|---|---|---|---|
| ENSMUSG00000020309 | 1.069666964 | 1.188721876 | 1.302363341 | 1.664552871 | 1.621875691 | 1.886490725 | 1.476637173 |
| ENSMUSG00000020310 | 0.578129653 | 0.396240625 | 0.396240625 | 0.25 | 0.387761935 | 0.477121255 | 0.710836105 |
| ENSMUSG00000020311 | 2.219867581 | 2.038704527 | 2.048689214 | 2.198603967 | 2.097547432 | 2.414973348 | 1.947894097 |
| ENSMUSG00000020312 | 0.91631382 | 1.04248125 | 1.214495249 | 1.214495249 | 1.343584837 | 1.176091259 | 1.345462945 |
| ENSMUSG00000020315 | 3.433858977 | 3.119692405 | 3.095926073 | 3.138227302 | 3.08916564 | 3.528531061 | 2.866783697 |
| ENSMUSG00000020317 | 0 | 0 | 0 | 0 | 0 | 0 | 0 |
| ENSMUSG00000020319 | 1.747221736 | 1.695339928 | 1.54248125 | 1.752806814 | 1.628935049 | 1.944482672 | 1.582103514 |
| ENSMUSG00000020321 | 2.941213125 | 2.677270953 | 2.665667094 | 2.826515422 | 2.76519674 | 3.211921084 | 2.572907655 |
| ENSMUSG00000020323 | 0.289064826 | 0.25 | 0 | 0 | 0 | 0 | 0 |
| ENSMUSG00000020325 | 1.156259305 | 1.214495249 | 1.282320754 | 0.701838731 | 1.039257769 | 0.77815125 | 1.392431569 |
| ENSMUSG00000020326 | 3.058380605 | 2.790597832 | 2.776477127 | 2.674241882 | 2.582654929 | 2.979092901 | 2.389035574 |
| ENSMUSG00000020327 | 0.289064826 | 0.25 | 0 | 0 | 0 | 0 | 0.355418052 |
| ENSMUSG00000020328 | 2.427163103 | 2.267865591 | 2.298073204 | 2.372963274 | 2.278401864 | 2.633468456 | 2.156863548 |
| ENSMUSG00000020329 | 1.857607105 | 1.92510993 | 1.981203126 | 1.54745614 | 1.84445929 | 1.886490725 | 1.974272009 |
| ENSMUSG00000020330 | 2.715608727 | 2.519537702 | 2.500352049 | 2.789336734 | 2.687518101 | 3.118595365 | 2.477249313 |
| ENSMUSG00000020331 | 1.389637216 | 1.29248125 | 1.388647213 | 1.532320754 | 1.580303447 | 1.740362689 | 1.603286636 |
| ENSMUSG00000020332 | 0 | 0 | 0 | 0 | 0 | 0 | 0 |
| ENSMUSG00000020333 | 0.960252568 | 1.02186571 | 1.098079356 | 1.779735268 | 1.680838453 | 1.919078092 | 1.56716938 |
| ENSMUSG00000020334 | 0.91631382 | 0.701838731 | 0.896240625 | 1.160964047 | 1.121713561 | 1.176091259 | 1.089372236 |
| ENSMUSG00000020335 | 0.578129653 | 0.701838731 | 0.25 | 0.701838731 | 0.877063019 | 0.301029996 | 0.629532004 |
| ENSMUSG00000020340 | 0.91631382 | 0.830482024 | 0.830482024 | 1.726722649 | 1.751666404 | 1.934498451 | 1.657681529 |
| ENSMUSG00000020346 | 2.360187403 | 2.240724001 | 2.288071211 | 2.101285366 | 2.212046813 | 2.369215857 | 2.129386845 |
| ENSMUSG00000020349 | 3.122160254 | 2.84727338 | 2.797610505 | 3.034967766 | 2.931296931 | 3.431041945 | 2.707701675 |
| ENSMUSG00000020354 | 0 | 0 | 0 | 0.396240625 | 0.244650542 | 0 | 0.355418052 |
| ENSMUSG00000020357 | 1.181541736 | 1.26109853 | 1.214495249 | 1.657339155 | 1.754126039 | 1.86923172 | 1.774416565 |
| ENSMUSG00000020358 | 3.054824387 | 2.817719824 | 2.813904687 | 2.993334655 | 2.90979544 | 3.401400541 | 2.745055516 |
| ENSMUSG00000020359 | 1.920505136 | 1.732684334 | 1.774508021 | 2.081607372 | 2.004851707 | 2.139879086 | 1.86815666 |
| ENSMUSG00000020361 | 3.295130293 | 3.00402796 | 2.973514962 | 3.180221376 | 3.082404901 | 3.59505509 | 2.855537562 |
| ENSMUSG00000020362 | 3.064330585 | 2.819612365 | 2.78456795 | 3.063461871 | 2.96696264 | 3.437592032 | 2.723007172 |
| ENSMUSG00000020363 | 0.811507563 | 0.951838731 | 1.04248125 | 1.794977273 | 1.832885921 | 2 | 1.653807046 |
| ENSMUSG00000020364 | 1.432083896 | 1.418106335 | 1.282320754 | 1.561981878 | 1.489301084 | 1.591064607 | 1.38802993 |
| ENSMUSG00000020366 | 2.380049842 | 2.121960008 | 2.12694866 | 2.385274154 | 2.359374645 | 2.687528961 | 2.184841529 |
| ENSMUSG00000020368 | 3.487005755 | 3.141335032 | 3.108657057 | 3.252937849 | 3.169606773 | 3.668199484 | 2.93078103 |
| ENSMUSG00000020372 | 3.516809818 | 3.245302622 | 3.256241621 | 3.209308425 | 3.248863928 | 3.69818757 | 3.071265431 |

| | | | | | | | |
|---|---|---|---|---|---|---|---|
| ENSMUSG00000020374 | 0 | 0 | 0 | 0 | 0.244650542 | 0 | 0 |
| ENSMUSG00000020375 | 2.086807695 | 1.976722649 | 1.978222334 | 2.094844592 | 2.043601899 | 2.357934847 | 1.977870661 |
| ENSMUSG00000020376 | 1.358731791 | 1.388647213 | 1.330482024 | 2.02186571 | 1.957204337 | 2.217483944 | 1.848480858 |
| ENSMUSG00000020377 | 0 | 0.5 | 0.25 | 0.25 | 0.387761935 | 0.301029996 | 0.355418052 |
| ENSMUSG00000020380 | 2.492331912 | 2.323655187 | 2.311388177 | 2.493603647 | 2.401730365 | 2.718501689 | 2.223239637 |
| ENSMUSG00000020381 | 1.036286563 | 0.951838731 | 0.951838731 | 1.130890489 | 1.121713561 | 1.176091259 | 1.20139795 |
| ENSMUSG00000020383 | 0.289064826 | 0 | 0 | 0 | 0.244650542 | 0 | 0 |
| ENSMUSG00000020385 | 2.370236862 | 2.140560606 | 2.154596376 | 2.277131114 | 2.164820399 | 2.462397998 | 2.029372647 |
| ENSMUSG00000020386 | 2.331622671 | 2.129909063 | 2.114857905 | 2.272528105 | 2.245260665 | 2.487138375 | 2.072190387 |
| ENSMUSG00000020387 | 2.227926429 | 2.064346961 | 2.097004321 | 2 | 2.107593434 | 2.206825876 | 2.001999072 |
| ENSMUSG00000020388 | 1.227926429 | 1.348079356 | 1.29248125 | 1.598079356 | 1.78687392 | 1.832508913 | 1.87511425 |
| ENSMUSG00000020389 | 1.404271433 | 1.339388001 | 1.201838731 | 1.410964047 | 1.445123986 | 1.51851394 | 1.307989563 |
| ENSMUSG00000020390 | 2.678024491 | 2.481944491 | 2.477473271 | 2.592126615 | 2.507741065 | 2.893761762 | 2.341765586 |
| ENSMUSG00000020392 | 2.260538674 | 2.140560606 | 2.131869252 | 2.256284891 | 2.24889942 | 2.555094449 | 2.128489437 |
| ENSMUSG00000020393 | 2.201144779 | 2.05357978 | 2.104463129 | 2.034887838 | 2.136951444 | 2.264817823 | 2.043115906 |
| ENSMUSG00000020395 | 0 | 0 | 0 | 0.25 | 0 | 0 | 0 |
| ENSMUSG00000020396 | 0.289064826 | 0.25 | 0 | 0 | 0.568060967 | 0 | 0.579661877 |
| ENSMUSG00000020397 | 1.639698646 | 1.54745614 | 1.561981878 | 1.688721876 | 1.655839964 | 1.770852012 | 1.537859885 |
| ENSMUSG00000020399 | 0 | 0.25 | 0.25 | 0.25 | 0.387761935 | 0.301029996 | 0.224243824 |
| ENSMUSG00000020400 | 2.307602651 | 2.137686696 | 2.202642909 | 2.175975893 | 2.328021124 | 2.51054501 | 2.242438242 |
| ENSMUSG00000020401 | 0 | 0.396240625 | 0 | 0 | 0 | 0 | 0 |
| ENSMUSG00000020402 | 3.018088033 | 2.746816003 | 2.759386738 | 2.990091002 | 2.938296657 | 3.372359583 | 2.731478495 |
| ENSMUSG00000020405 | 0 | 0 | 0 | 0 | 0 | 0 | 0 |
| ENSMUSG00000020407 | 1.100572389 | 0.951838731 | 0.92510993 | 0.92510993 | 0.978602168 | 1 | 0.935079929 |
| ENSMUSG00000020409 | 2.365959569 | 2.162762923 | 2.180274797 | 2.365369862 | 2.289867804 | 2.63748973 | 2.152721168 |
| ENSMUSG00000020411 | 1.205378646 | 1.080482024 | 1.114857905 | 0.79248125 | 0.77552387 | 0.84509804 | 0.629532004 |
| ENSMUSG00000020412 | 2.488076442 | 2.310197833 | 2.281030328 | 2.349685923 | 2.338472809 | 2.673941999 | 2.223909441 |
| ENSMUSG00000020413 | 2.203267085 | 2.049918086 | 2.027131114 | 2.113831805 | 2.022473809 | 2.28780173 | 1.906930404 |
| ENSMUSG00000020415 | 2.487005755 | 2.253505118 | 2.254897682 | 2.448197574 | 2.386129765 | 2.765668555 | 2.209893373 |
| ENSMUSG00000020419 | 0.45815691 | 0.396240625 | 0.396240625 | 0.396240625 | 0 | 0.301029996 | 0.224243824 |
| ENSMUSG00000020420 | 1.307602651 | 1.201838731 | 1.214495249 | 1.356566189 | 1.27449559 | 1.380211242 | 1.121219121 |
| ENSMUSG00000020422 | 2.949075151 | 2.685998215 | 2.699820405 | 2.580200137 | 2.58731141 | 2.916980047 | 2.426894671 |
| ENSMUSG00000020423 | 2.063793172 | 1.930274797 | 1.858657057 | 1.828220739 | 1.789100779 | 2.037426498 | 1.753626837 |
| ENSMUSG00000020424 | 1.418409478 | 1.388647213 | 1.42510993 | 1.226722649 | 1.358933174 | 1.380211242 | 1.396774124 |

| | | | | | | | |
|---|---|---|---|---|---|---|---|
| ENSMUSG00000020427 | 2.201144779 | 1.975216702 | 2.032320754 | 0.79248125 | 0.568060967 | 1 | 0.74492186 |
| ENSMUSG00000020428 | 0 | 0 | 0 | 0 | 0 | 0 | 0 |
| ENSMUSG00000020429 | 0 | 0 | 0 | 0 | 0 | 0 | 0 |
| ENSMUSG00000020430 | 2.697239867 | 2.485261902 | 2.451838731 | 2.504202072 | 2.448909998 | 2.80140371 | 2.282502732 |
| ENSMUSG00000020431 | 2.016886209 | 1.943696765 | 1.905512955 | 1.445339928 | 1.484072009 | 1.462397998 | 1.313616061 |
| ENSMUSG00000020432 | 2.170239354 | 2.02053726 | 2.066696635 | 1.729715809 | 1.741652793 | 1.880813592 | 1.665294008 |
| ENSMUSG00000020434 | 0.671187741 | 1.02186571 | 0.896240625 | 0.976722649 | 0.632412477 | 0.84509804 | 0.74492186 |
| ENSMUSG00000020435 | 0 | 0.25 | 0.396240625 | 0.701838731 | 0.877063019 | 1 | 0.876096088 |
| ENSMUSG00000020436 | 0 | 0 | 0 | 0 | 0 | 0 | 0 |
| ENSMUSG00000020437 | 0.289064826 | 0.25 | 0.396240625 | 0 | 0.387761935 | 0.301029996 | 0.224243824 |
| ENSMUSG00000020439 | 2.134369199 | 2.066696635 | 2.186208459 | 1.71759118 | 1.953043706 | 1.995635195 | 1.951796623 |
| ENSMUSG00000020440 | 2.149089517 | 2.110735874 | 2.226722649 | 1.938721876 | 2.056322416 | 2.206825876 | 2.128938453 |
| ENSMUSG00000020441 | 1.516991255 | 1.372963274 | 1.482684334 | 1.618933358 | 1.614672255 | 1.785329835 | 1.579661877 |
| ENSMUSG00000020444 | 1.837744666 | 1.870953944 | 1.882845365 | 1.868933358 | 1.974687643 | 2.167317335 | 1.865128412 |
| ENSMUSG00000020447 | 0 | 0 | 0.25 | 0 | 0.244650542 | 0 | 0 |
| ENSMUSG00000020448 | 2.349889783 | 2.192872367 | 2.184523065 | 2.173371739 | 2.086251872 | 2.361727836 | 2.001334086 |
| ENSMUSG00000020451 | 2.448863324 | 2.273849256 | 2.31611065 | 2.17510993 | 2.197694248 | 2.46834733 | 2.109044889 |
| ENSMUSG00000020453 | 2.188178997 | 2.079353153 | 2.117918804 | 1.825945187 | 1.86473911 | 2.037426498 | 1.820622395 |
| ENSMUSG00000020454 | 2.687411469 | 2.486726569 | 2.466433318 | 2.482314602 | 2.404846996 | 2.778874472 | 2.256654198 |
| ENSMUSG00000020455 | 2.036286563 | 1.903677461 | 1.931980114 | 1.905512955 | 1.931471446 | 2.071882007 | 1.903345709 |
| ENSMUSG00000020456 | 2.827920678 | 2.606303976 | 2.612029076 | 2.696363117 | 2.692875188 | 3.080265627 | 2.52519166 |
| ENSMUSG00000020457 | 2.754058486 | 2.481203126 | 2.494677615 | 2.620701989 | 2.513204446 | 2.95568775 | 2.364643151 |
| ENSMUSG00000020458 | 3.223916471 | 2.901138307 | 2.906655413 | 2.967109189 | 2.854473682 | 3.325515663 | 2.657243576 |
| ENSMUSG00000020459 | 2.029919532 | 1.926839783 | 1.930274797 | 2.059601185 | 1.98503142 | 2.214843848 | 1.808770687 |
| ENSMUSG00000020460 | 3.007144864 | 2.75 | 2.775657972 | 2.936523596 | 2.844570897 | 3.290034611 | 2.631151251 |
| ENSMUSG00000020461 | 0.45815691 | 0.25 | 0.580482024 | 0.5 | 0 | 0.698970004 | 0.710836105 |
| ENSMUSG00000020462 | 2.051791411 | 2.029735268 | 2.017865591 | 2.264497931 | 2.169384551 | 2.451786436 | 2.060761403 |
| ENSMUSG00000020463 | 2.728422728 | 2.506630861 | 2.521534056 | 2.737274289 | 2.658496636 | 3.084576278 | 2.45362687 |
| ENSMUSG00000020464 | 2.337801042 | 2.219895812 | 2.196158711 | 2.422062577 | 2.349985771 | 2.656098202 | 2.162979334 |
| ENSMUSG00000020467 | 0 | 0.25 | 0.25 | 1.54745614 | 1.628935049 | 1.69019608 | 1.486592281 |
| ENSMUSG00000020469 | 0 | 0 | 0.396240625 | 0.25 | 0 | 0 | 0 |
| ENSMUSG00000020471 | 2.325351389 | 2.228222334 | 2.25835575 | 2.058404919 | 2.178339521 | 2.369215857 | 2.148525059 |
| ENSMUSG00000020472 | 2.267855219 | 2.198603967 | 2.184523065 | 2.159156155 | 2.254881871 | 2.46686762 | 2.158095989 |
| ENSMUSG00000020473 | 2.910529644 | 2.714689525 | 2.827937079 | 2.140560606 | 2.314255074 | 2.472756449 | 2.237984665 |

| | | | | | | | |
|---|---|---|---|---|---|---|---|
| ENSMUSG00000020474 | 1.639698646 | 1.606566189 | 1.52186571 | 1.431980114 | 1.628935049 | 1.633468456 | 1.572224412 |
| ENSMUSG00000020475 | 0.578129653 | 0.830482024 | 0.701838731 | 0.580482024 | 0.77552387 | 0.77815125 | 0.952571511 |
| ENSMUSG00000020476 | 2.472830741 | 2.329353153 | 2.408248799 | 2.247171172 | 2.342660868 | 2.589949601 | 2.292939619 |
| ENSMUSG00000020477 | 1.771760131 | 1.708222504 | 1.75 | 1.711372513 | 1.766173492 | 1.826074803 | 1.709198048 |
| ENSMUSG00000020481 | 0 | 0 | 0 | 0 | 0 | 0 | 0 |
| ENSMUSG00000020482 | 2.192545868 | 2.044977273 | 2.03743678 | 2.163659007 | 2.091002363 | 2.40654018 | 1.968429585 |
| ENSMUSG00000020483 | 2.817859673 | 2.576515422 | 2.585518667 | 2.522528105 | 2.469198506 | 2.822168079 | 2.335428845 |
| ENSMUSG00000020484 | 2.3855553 | 2.204245906 | 2.173371739 | 2.12694866 | 2.11997913 | 2.397940009 | 1.970632877 |
| ENSMUSG00000020485 | 2.450036412 | 2.255591953 | 2.25 | 2.172499493 | 2.085294053 | 2.432969291 | 1.997315209 |
| ENSMUSG00000020486 | 0.289064826 | 0.580482024 | 0.646240625 | 0.79248125 | 1.057362051 | 0.84509804 | 1.121219121 |
| ENSMUSG00000020490 | 0 | 0 | 0 | 0.25 | 0 | 0 | 0 |
| ENSMUSG00000020491 | 1.686083339 | 1.610735874 | 1.527131114 | 1.921625132 | 1.818885826 | 2.049218023 | 1.705896842 |
| ENSMUSG00000020492 | 2.622240692 | 2.419870025 | 2.367918804 | 2.368933358 | 2.325116119 | 2.683947131 | 2.183707049 |
| ENSMUSG00000020493 | 2.487005755 | 2.284887838 | 2.236360959 | 2.476346751 | 2.390196125 | 2.737987326 | 2.221221862 |
| ENSMUSG00000020495 | 2.395018353 | 2.202642909 | 2.185366747 | 2.254897682 | 2.218023635 | 2.531478917 | 2.087847416 |
| ENSMUSG00000020496 | 2.650702778 | 2.50835575 | 2.55357978 | 2.598079356 | 2.628729423 | 2.943494516 | 2.522677108 |
| ENSMUSG00000020513 | 1.783508299 | 1.763820609 | 1.660964047 | 1.79745614 | 1.736537397 | 1.929418926 | 1.62316756 |
| ENSMUSG00000020514 | 2.026699189 | 1.919870025 | 1.96759118 | 2.088286706 | 2.033751322 | 2.305351369 | 1.926805425 |
| ENSMUSG00000020515 | 2.529763563 | 2.332168732 | 2.301142786 | 2.402756199 | 2.356717497 | 2.711807229 | 2.203186321 |
| ENSMUSG00000020516 | 2.733796161 | 2.515173983 | 2.526149688 | 2.630645383 | 2.531821953 | 2.908485019 | 2.320886377 |
| ENSMUSG00000020519 | 1.734388958 | 1.54248125 | 1.634789703 | 1.638647213 | 1.712553795 | 1.851258349 | 1.700880998 |
| ENSMUSG00000020520 | 2.283862692 | 2.088286706 | 2.122963274 | 2.117918804 | 2.214715681 | 2.465382851 | 2.123508695 |
| ENSMUSG00000020521 | 2.574668783 | 2.381380202 | 2.365369862 | 2.385274154 | 2.277846464 | 2.622214023 | 2.125328759 |
| ENSMUSG00000020522 | 2.45815691 | 2.263820609 | 2.265173983 | 2.452642909 | 2.381601986 | 2.694605199 | 2.203186321 |
| ENSMUSG00000020523 | 2.410757293 | 2.240724001 | 2.228969845 | 2.388647213 | 2.280614762 | 2.649334859 | 2.139097452 |
| ENSMUSG00000020524 | 0.289064826 | 0 | 0 | 0 | 0 | 0 | 0 |
| ENSMUSG00000020525 | 2.095120908 | 1.953445298 | 1.928561379 | 1.991446071 | 1.923977404 | 2.187520721 | 1.819456572 |
| ENSMUSG00000020526 | 1.948720984 | 1.809601185 | 1.858657057 | 1.854463129 | 1.814781639 | 2.08278537 | 1.695786161 |
| ENSMUSG00000020527 | 2.312518611 | 2.175975893 | 2.240000483 | 2.120953944 | 2.189222894 | 2.444044796 | 2.088356488 |
| ENSMUSG00000020528 | 1.867194479 | 1.79745614 | 1.766522298 | 2.181980114 | 2.16174451 | 2.416640507 | 2.016928211 |
| ENSMUSG00000020530 | 2.81249113 | 2.564346961 | 2.539652422 | 2.593488164 | 2.512633319 | 2.888740961 | 2.341067602 |
| ENSMUSG00000020532 | 2.947301297 | 2.674024543 | 2.66942991 | 2.813609537 | 2.758709949 | 3.141449773 | 2.553451027 |
| ENSMUSG00000020534 | 2.333175866 | 2.191217898 | 2.157339155 | 2.27908599 | 2.216042585 | 2.46686762 | 2.100829716 |
| ENSMUSG00000020536 | 2.330063669 | 2.238549078 | 2.28296424 | 2.080482024 | 2.236621448 | 2.418301291 | 2.188221291 |

| | | | | | | | |
|---|---|---|---|---|---|---|---|
| ENSMUSG00000020537 | 2.556920045 | 2.31728167 | 2.331607372 | 2.506976499 | 2.545363112 | 2.875639937 | 2.412651849 |
| ENSMUSG00000020538 | 2.266038081 | 2.203445298 | 2.33272922 | 2.105516192 | 2.329944595 | 2.416640507 | 2.367660638 |
| ENSMUSG00000020541 | 2.172524468 | 2.031030328 | 2.023189285 | 1.997171172 | 1.928492895 | 2.217483944 | 1.801445495 |
| ENSMUSG00000020542 | 0.747221736 | 0.896240625 | 1.02186571 | 1.04248125 | 0.905314583 | 0.903089987 | 0.896975297 |
| ENSMUSG00000020544 | 0.91631382 | 0.75 | 0.701838731 | 0.580482024 | 0.686820904 | 0.602059991 | 0.952571511 |
| ENSMUSG00000020545 | 0 | 0 | 0 | 0 | 0 | 0 | 0 |
| ENSMUSG00000020546 | 2.331622671 | 2.164552871 | 2.161864607 | 2.23927551 | 2.19135987 | 2.499687083 | 2.03780742 |
| ENSMUSG00000020547 | 2.637462542 | 2.42293588 | 2.426408097 | 2.28360658 | 2.24829556 | 2.532754379 | 2.096890362 |
| ENSMUSG00000020548 | 0 | 0 | 0 | 0 | 0 | 0 | 0 |
| ENSMUSG00000020549 | 2.251208701 | 2.02053726 | 2.034887838 | 2.143411797 | 2.164053936 | 2.383815366 | 2.027536225 |
| ENSMUSG00000020553 | 1.249317394 | 0.92510993 | 0.92510993 | 1.403677461 | 1.373641807 | 1.531478917 | 1.259064007 |
| ENSMUSG00000020561 | 2.201144779 | 2.03360658 | 2.061981878 | 2.046218836 | 1.992595054 | 2.267171728 | 1.853775828 |
| ENSMUSG00000020562 | 0 | 0 | 0 | 0 | 0 | 0.301029996 | 0 |
| ENSMUSG00000020564 | 2.124258828 | 2.059601185 | 2.069031101 | 1.961372513 | 1.969399812 | 2.176091259 | 1.825244123 |
| ENSMUSG00000020566 | 0.578129653 | 0.396240625 | 0.396240625 | 1.311981878 | 1.509475496 | 1.51851394 | 1.252393349 |
| ENSMUSG00000020570 | 2.85804771 | 2.606566189 | 2.603406982 | 2.532320754 | 2.431003859 | 2.819543936 | 2.276018109 |
| ENSMUSG00000020571 | 3.112703332 | 2.838700086 | 2.799764706 | 2.835518667 | 2.828475363 | 3.25139485 | 2.657769048 |
| ENSMUSG00000020572 | 2.794421919 | 2.539967834 | 2.545288064 | 2.480088214 | 2.376595432 | 2.738780558 | 2.20708628 |
| ENSMUSG00000020573 | 0 | 0 | 0 | 0 | 0 | 0 | 0 |
| ENSMUSG00000020576 | 2.323768721 | 2.234159485 | 2.166333979 | 2.221424093 | 2.221300875 | 2.50242712 | 2.060206963 |
| ENSMUSG00000020577 | 1 | 1.226722649 | 1.188721876 | 1.311981878 | 1.335652905 | 1.361727836 | 1.265599895 |
| ENSMUSG00000020580 | 3.021077525 | 2.771699921 | 2.739638179 | 2.739094039 | 2.645205039 | 3.040997692 | 2.42021657 |
| ENSMUSG00000020581 | 0 | 0 | 0 | 0 | 0 | 0 | 0 |
| ENSMUSG00000020583 | 1.100572389 | 1.201838731 | 0.951838731 | 0.976722649 | 1.121713561 | 1.113943352 | 1.066254157 |
| ENSMUSG00000020585 | 3.132777036 | 2.878555227 | 2.890799068 | 2.824802005 | 2.747283614 | 3.183839037 | 2.607913505 |
| ENSMUSG00000020589 | 0.289064826 | 0 | 0.25 | 1.238549078 | 1.091002363 | 1.255272505 | 1.028149525 |
| ENSMUSG00000020590 | 2.605636262 | 2.399047439 | 2.411864607 | 2.337734795 | 2.275056283 | 2.588831726 | 2.081675233 |
| ENSMUSG00000020591 | 0 | 0 | 0 | 0 | 0 | 0 | 0 |
| ENSMUSG00000020592 | 2.63895461 | 2.451435968 | 2.483791262 | 2.108657057 | 2.170892913 | 2.29666519 | 2.125782179 |
| ENSMUSG00000020593 | 1.516991255 | 1.372963274 | 1.431980114 | 1.403677461 | 1.456697355 | 1.447158031 | 1.452143229 |
| ENSMUSG00000020594 | 3.098103274 | 2.837320307 | 2.825374049 | 2.822216519 | 2.744775661 | 3.173768823 | 2.528731682 |
| ENSMUSG00000020598 | 0.578129653 | 0.5 | 0.646240625 | 0 | 0.632412477 | 0.602059991 | 0.224243824 |
| ENSMUSG00000020599 | 1.418409478 | 1.311981878 | 1.451838731 | 1.622963274 | 1.603583716 | 1.716003344 | 1.537859885 |
| ENSMUSG00000020600 | 0 | 0 | 0.25 | 0 | 0 | 0 | 0 |

| | | | | | | | |
|---|---|---|---|---|---|---|---|
| ENSMUSG00000020601 | 1.655740331 | 1.494319981 | 1.532320754 | 0.701838731 | 0.686820904 | 0.698970004 | 0.710836105 |
| ENSMUSG00000020604 | 1.181541736 | 1.311981878 | 1.364857905 | 1.080482024 | 1.074582839 | 1.079181246 | 1.041356071 |
| ENSMUSG00000020605 | 2.256831695 | 2.131869252 | 2.156427211 | 1.874961472 | 1.828942237 | 2.100370545 | 1.805128823 |
| ENSMUSG00000020607 | 0.45815691 | 0.580482024 | 0.646240625 | 0.396240625 | 0.244650542 | 0.477121255 | 0 |
| ENSMUSG00000020608 | 2.869771412 | 2.668548067 | 2.650442697 | 2.692046081 | 2.599361968 | 2.976808337 | 2.376547822 |
| ENSMUSG00000020609 | 1.37447073 | 1.311981878 | 1.238549078 | 0.646240625 | 0.846351821 | 0.698970004 | 0.520678036 |
| ENSMUSG00000020610 | 2.348397716 | 2.192872367 | 2.181980114 | 2.247171172 | 2.145141048 | 2.457881897 | 2.034820183 |
| ENSMUSG00000020611 | 2.867625075 | 2.624462959 | 2.62521047 | 2.645064642 | 2.580303447 | 2.973589623 | 2.401445908 |
| ENSMUSG00000020612 | 3.305951036 | 3.009729747 | 3.003330668 | 3.059750439 | 2.999928151 | 3.443262987 | 2.795954856 |
| ENSMUSG00000020614 | 1.759671389 | 1.708222504 | 1.634789703 | 0.701838731 | 0.978602168 | 0.903089987 | 0.896975297 |
| ENSMUSG00000020617 | 0 | 0 | 0.25 | 0 | 0 | 0 | 0 |
| ENSMUSG00000020620 | 1.342375483 | 1.321350555 | 1.282320754 | 1 | 0.978602168 | 0.84509804 | 0.952571511 |
| ENSMUSG00000020621 | 1.505869732 | 1.476722649 | 1.380890489 | 1.431980114 | 1.427019704 | 1.579783597 | 1.345462945 |
| ENSMUSG00000020622 | 0 | 0.25 | 0 | 0.25 | 0 | 0 | 0.224243824 |
| ENSMUSG00000020623 | 1.727821383 | 1.589388001 | 1.557204673 | 1.51109853 | 1.576269619 | 1.707570176 | 1.383567577 |
| ENSMUSG00000020624 | 0 | 0 | 0 | 0 | 0 | 0 | 0 |
| ENSMUSG00000020627 | 0 | 0.25 | 0.396240625 | 0.5 | 0.489301084 | 0.477121255 | 0.672731473 |
| ENSMUSG00000020628 | 2.083999385 | 2.02053726 | 2.024508021 | 1.918106335 | 1.975997314 | 2.155336037 | 1.88765144 |
| ENSMUSG00000020629 | 2.105953527 | 1.864857905 | 1.948603967 | 2.131869252 | 2.044679629 | 2.298853076 | 1.89047692 |
| ENSMUSG00000020630 | 1.753493018 | 1.67510993 | 1.723704441 | 1.470660762 | 1.484072009 | 1.544068044 | 1.523605797 |
| ENSMUSG00000020633 | 1.100572389 | 0.896240625 | 0.864857905 | 0.75 | 0.77552387 | 0.84509804 | 0.74492186 |
| ENSMUSG00000020634 | 1.952600383 | 1.866901388 | 1.837182039 | 2.094844592 | 2.040348824 | 2.31386722 | 1.897893072 |
| ENSMUSG00000020635 | 1.069666964 | 0.896240625 | 1 | 1.201838731 | 1.223252711 | 1.230448921 | 1.21680636 |
| ENSMUSG00000020636 | 0 | 0 | 0 | 0 | 0 | 0 | 0 |
| ENSMUSG00000020638 | 0.671187741 | 0.5 | 0.396240625 | 0.896240625 | 0.931471446 | 0.698970004 | 0.672731473 |
| ENSMUSG00000020639 | 0 | 0 | 0.25 | 0.25 | 0.387761935 | 0 | 0.520678036 |
| ENSMUSG00000020640 | 2.45815691 | 2.252107156 | 2.263142013 | 2.33993739 | 2.303566328 | 2.640481437 | 2.132508314 |
| ENSMUSG00000020641 | 0 | 0 | 0 | 0.25 | 0 | 0.602059991 | 0 |
| ENSMUSG00000020642 | 0.960252568 | 1.04248125 | 0.92510993 | 1 | 0.905314583 | 1 | 1.014380832 |
| ENSMUSG00000020644 | 2.423434025 | 2.26109853 | 2.311388177 | 1.956637122 | 1.935892666 | 2.227886705 | 1.849546807 |
| ENSMUSG00000020646 | 1.614416215 | 1.396240625 | 1.339388001 | 1.67510993 | 1.519146132 | 1.799340549 | 1.462868481 |
| ENSMUSG00000020647 | 2.029919532 | 1.886723615 | 1.923371739 | 1.964495249 | 1.905314583 | 2.127104798 | 1.801445495 |
| ENSMUSG00000020648 | 1.587501561 | 1.527131114 | 1.494319981 | 1.458222504 | 1.327538614 | 1.462397998 | 1.313616061 |
| ENSMUSG00000020649 | 3.033404503 | 2.776967729 | 2.774508021 | 2.864216935 | 2.778900174 | 3.247973266 | 2.612776967 |

| | | | | | | | |
|---|---|---|---|---|---|---|---|
| ENSMUSG00000020650 | 2.129344651 | 2.002806814 | 1.948603967 | 1.909156155 | 1.768534412 | 2.004321374 | 1.647906904 |
| ENSMUSG00000020651 | 0 | 0 | 0 | 0 | 0 | 0 | 0 |
| ENSMUSG00000020652 | 1.967766868 | 1.874961472 | 1.850219859 | 2.002806814 | 1.978602168 | 2.230448921 | 1.826389311 |
| ENSMUSG00000020653 | 1.527823874 | 1.532320754 | 1.482684334 | 1.356566189 | 1.234113756 | 1.322219295 | 1.313616061 |
| ENSMUSG00000020654 | 1.920505136 | 1.77186571 | 1.809601185 | 1.884789703 | 1.919403394 | 2.045322979 | 1.86715039 |
| ENSMUSG00000020656 | 0.578129653 | 0.646240625 | 0.79248125 | 0.701838731 | 0.812711509 | 0.602059991 | 0.803905701 |
| ENSMUSG00000020657 | 1.960252568 | 1.858657057 | 1.854463129 | 1.860735874 | 1.741652793 | 2.033423755 | 1.700880998 |
| ENSMUSG00000020658 | 1.342375483 | 1.160964047 | 1.25 | 1.602347734 | 1.528558853 | 1.51851394 | 1.476637173 |
| ENSMUSG00000020659 | 2.016886209 | 1.872963274 | 1.75835575 | 1.845926073 | 1.746695109 | 1.963787827 | 1.631625962 |
| ENSMUSG00000020660 | 0 | 0 | 0 | 0 | 0.244650542 | 0 | 0.224243824 |
| ENSMUSG00000020661 | 2.352857998 | 2.174241882 | 2.174241882 | 2.133818844 | 2.152352662 | 2.334453751 | 2.055173778 |
| ENSMUSG00000020664 | 2.930617777 | 2.676839783 | 2.646240625 | 2.661864607 | 2.580067435 | 2.984977126 | 2.399313044 |
| ENSMUSG00000020668 | 2.302628051 | 2.130890489 | 2.153677461 | 1.966046536 | 1.92245931 | 2.117271296 | 1.853775828 |
| ENSMUSG00000020669 | 1.418409478 | 1.25 | 1.330482024 | 1.160964047 | 1.212046813 | 1.204119983 | 1.168187745 |
| ENSMUSG00000020671 | 2.929134994 | 2.684523065 | 2.684523065 | 2.636241108 | 2.539585427 | 2.937016107 | 2.371711297 |
| ENSMUSG00000020672 | 0 | 0 | 0 | 0 | 0 | 0 | 0 |
| ENSMUSG00000020673 | 0 | 0 | 0 | 0 | 0 | 0 | 0 |
| ENSMUSG00000020674 | 3.125652033 | 2.855647608 | 2.833288838 | 2.773684376 | 2.782111321 | 3.183269844 | 2.604939337 |
| ENSMUSG00000020676 | 0.289064826 | 0 | 0 | 0 | 0 | 0 | 0 |
| ENSMUSG00000020677 | 2.664937347 | 2.461764337 | 2.411864607 | 2.539336734 | 2.465951417 | 2.846955325 | 2.285835109 |
| ENSMUSG00000020679 | 0.289064826 | 0.396240625 | 0.864857905 | 0.701838731 | 0.632412477 | 0.602059991 | 0.775756176 |
| ENSMUSG00000020680 | 3.012652342 | 2.706239685 | 2.722756048 | 2.56168515 | 2.542223377 | 2.880813592 | 2.372768883 |
| ENSMUSG00000020681 | 0 | 0.25 | 0.25 | 0.396240625 | 0.568060967 | 0 | 0.520678036 |
| ENSMUSG00000020682 | 1.89911417 | 1.729715809 | 1.825945187 | 1.364857905 | 1.439191811 | 1.414973348 | 1.493062805 |
| ENSMUSG00000020684 | 0.867194479 | 0.646240625 | 0.864857905 | 0.92510993 | 1.020174412 | 0.954242509 | 1.140832123 |
| ENSMUSG00000020686 | 0 | 0 | 0 | 0 | 0 | 0 | 0 |
| ENSMUSG00000020687 | 2.936131728 | 2.69059551 | 2.672717752 | 2.790125437 | 2.706170459 | 3.102090526 | 2.502249309 |
| ENSMUSG00000020689 | 2.219867581 | 2.130890489 | 2.091580554 | 1.188721876 | 1.293076518 | 1.301029996 | 1.20919388 |
| ENSMUSG00000020690 | 0 | 0 | 0 | 0 | 0 | 0 | 0 |
| ENSMUSG00000020691 | 2.179305632 | 2.027131114 | 2.034887838 | 2.113831805 | 2.042520869 | 2.334453751 | 1.949460761 |
| ENSMUSG00000020692 | 1.678702042 | 1.75 | 1.874961472 | 1.348079356 | 1.645989387 | 1.431363764 | 1.645916034 |
| ENSMUSG00000020694 | 2.641182749 | 2.443284802 | 2.443284802 | 2.480460234 | 2.380772476 | 2.746634199 | 2.22557791 |
| ENSMUSG00000020695 | 2.753493018 | 2.571928095 | 2.63015467 | 2.335518667 | 2.449252175 | 2.63748973 | 2.361815771 |
| ENSMUSG00000020696 | 1.811507563 | 1.752806814 | 1.678561379 | 1.714495249 | 1.66856048 | 1.806179974 | 1.601000298 |

| | | | | | | | |
|---|---|---|---|---|---|---|---|
| ENSMUSG00000020697 | 2.485932312 | 2.263142013 | 2.255591953 | 2.307805295 | 2.273371525 | 2.517195898 | 2.119837558 |
| ENSMUSG00000020698 | 0.289064826 | 0.396240625 | 0.25 | 0.396240625 | 0.568060967 | 0 | 0.355418052 |
| ENSMUSG00000020700 | 2.74143756 | 2.508011432 | 2.498588359 | 2.458222504 | 2.431723443 | 2.790285164 | 2.259363975 |
| ENSMUSG00000020701 | 0 | 0 | 0 | 0 | 0 | 0 | 0 |
| ENSMUSG00000020702 | 0 | 0 | 0 | 0 | 0 | 0 | 0 |
| ENSMUSG00000020703 | 0 | 0 | 0 | 0 | 0 | 0.301029996 | 0.224243824 |
| ENSMUSG00000020704 | 0 | 0 | 0 | 0 | 0 | 0 | 0 |
| ENSMUSG00000020705 | 2.893489371 | 2.630645383 | 2.613831805 | 2.718937338 | 2.635250738 | 3.036229544 | 2.444790289 |
| ENSMUSG00000020706 | 2.945876762 | 2.695749552 | 2.709800947 | 2.800684011 | 2.748896963 | 3.115943177 | 2.529513121 |
| ENSMUSG00000020707 | 1.727821383 | 1.62694866 | 1.571350555 | 1.602347734 | 1.55104774 | 1.591064607 | 1.546120303 |
| ENSMUSG00000020708 | 2.969792941 | 2.723514962 | 2.729715809 | 2.748765117 | 2.746695109 | 3.108565024 | 2.5875851 |
| ENSMUSG00000020709 | 0.289064826 | 0.396240625 | 0 | 0.701838731 | 0.568060967 | 0.477121255 | 0.710836105 |
| ENSMUSG00000020712 | 0.578129653 | 0.830482024 | 0.5 | 0.646240625 | 0.632412477 | 0.698970004 | 0.579661877 |
| ENSMUSG00000020713 | 0 | 0 | 0 | 0 | 0 | 0 | 0 |
| ENSMUSG00000020715 | 2.337801042 | 2.120953944 | 2.109697963 | 2.366391601 | 2.311739339 | 2.640481437 | 2.197065089 |
| ENSMUSG00000020716 | 2.906628471 | 2.662762923 | 2.638647213 | 2.654366866 | 2.604025187 | 2.934498451 | 2.402603388 |
| ENSMUSG00000020717 | 0.289064826 | 0.580482024 | 0.5 | 0.396240625 | 0.489301084 | 0 | 0.579661877 |
| ENSMUSG00000020718 | 1.631440309 | 1.532320754 | 1.52186571 | 1.53743678 | 1.523883869 | 1.544068044 | 1.473249546 |
| ENSMUSG00000020719 | 3.626935553 | 3.253417904 | 3.257408082 | 3.370544428 | 3.277322936 | 3.815046176 | 3.03114352 |
| ENSMUSG00000020720 | 2.926153516 | 2.690179987 | 2.680061065 | 2.798073204 | 2.734409318 | 3.155639634 | 2.544634011 |
| ENSMUSG00000020721 | 2.607246427 | 2.375956434 | 2.398581151 | 2.502457154 | 2.465625059 | 2.81756537 | 2.312345877 |
| ENSMUSG00000020722 | 0 | 0 | 0 | 0 | 0 | 0 | 0 |
| ENSMUSG00000020723 | 0 | 0.5 | 0.25 | 0 | 0.387761935 | 0 | 0 |
| ENSMUSG00000020728 | 1.753493018 | 1.634789703 | 1.701838731 | 1.752806814 | 1.607318736 | 1.86332286 | 1.572224412 |
| ENSMUSG00000020732 | 0.289064826 | 0.580482024 | 0.75 | 0.5 | 0.387761935 | 0.301029996 | 0.224243824 |
| ENSMUSG00000020733 | 2.368815967 | 2.231203126 | 2.31728167 | 2.08272922 | 2.167109853 | 2.365487985 | 2.14212705 |
| ENSMUSG00000020734 | 0 | 0 | 0 | 0 | 0 | 0 | 0 |
| ENSMUSG00000020736 | 1.989530018 | 1.888647213 | 1.964495249 | 1.948603967 | 1.995080671 | 2.161368002 | 1.954885099 |
| ENSMUSG00000020737 | 2.971473847 | 2.76685859 | 2.788071211 | 2.691632227 | 2.793770609 | 3.063333359 | 2.694851144 |
| ENSMUSG00000020738 | 3.178489522 | 2.899513125 | 2.879294313 | 3.006371414 | 2.867886777 | 3.353723938 | 2.691083819 |
| ENSMUSG00000020739 | 2.933934888 | 2.667442722 | 2.675975893 | 2.650674716 | 2.669111257 | 3.035429738 | 2.48540179 |
| ENSMUSG00000020740 | 2.336265009 | 2.194519282 | 2.191217898 | 2.179419106 | 2.203230928 | 2.42975228 | 2.073790632 |
| ENSMUSG00000020741 | 2.751792 | 2.527131114 | 2.577084758 | 2.405512955 | 2.483451699 | 2.710117365 | 2.342230073 |
| ENSMUSG00000020743 | 1.885732304 | 1.777131114 | 1.854463129 | 1.854463129 | 1.90849439 | 1.913813852 | 1.876096088 |

| | | | | | | | |
|---|---|---|---|---|---|---|---|
| ENSMUSG00000020744 | 1.852729478 | 1.799918086 | 1.854463129 | 1.79248125 | 1.87878897 | 1.944482672 | 1.793950594 |
| ENSMUSG00000020745 | 3.344026554 | 3.017110604 | 2.99770327 | 3.106828211 | 3.009147943 | 3.492900011 | 2.770672913 |
| ENSMUSG00000020747 | 2.241665209 | 2.194519282 | 2.231203126 | 1.907339155 | 2.201164838 | 2.225309282 | 2.141264342 |
| ENSMUSG00000020752 | 1.985980784 | 1.837182039 | 1.933677405 | 1.784887838 | 1.828942237 | 1.929418926 | 1.798966438 |
| ENSMUSG00000020755 | 2.533616084 | 2.326515422 | 2.334405475 | 2.336628433 | 2.308696595 | 2.599883072 | 2.185972044 |
| ENSMUSG00000020758 | 0.811507563 | 0.92510993 | 0.75 | 0.646240625 | 0.733951626 | 0.77815125 | 0.74492186 |
| ENSMUSG00000020766 | 2.116511873 | 2.118933358 | 2.265173983 | 1.816696635 | 2.075570129 | 2.053078443 | 2.202472158 |
| ENSMUSG00000020770 | 2.177057474 | 2.095926073 | 2.158248799 | 1.852347734 | 2.072599934 | 2.127104798 | 2.07004434 |
| ENSMUSG00000020773 | 2.172524468 | 2.137686696 | 2.266522298 | 1.75835575 | 1.965381292 | 2.004321374 | 2.026306127 |
| ENSMUSG00000020774 | 0.289064826 | 0.5 | 0.396240625 | 0.646240625 | 0.489301084 | 0 | 0.520678036 |
| ENSMUSG00000020775 | 2.17479713 | 2.056000419 | 2.104463129 | 2.114857905 | 2.124299332 | 2.394451681 | 2.099358074 |
| ENSMUSG00000020776 | 2.35727108 | 2.252806814 | 2.307805295 | 2.064346961 | 2.180543208 | 2.311753861 | 2.123508695 |
| ENSMUSG00000020777 | 2.849145934 | 2.601551251 | 2.620953944 | 2.591853688 | 2.56338924 | 2.926856709 | 2.395398714 |
| ENSMUSG00000020778 | 2.003432385 | 1.896240625 | 1.935366747 | 1.755591953 | 1.846351821 | 2.056904851 | 1.787569411 |
| ENSMUSG00000020780 | 2.731416512 | 2.54961126 | 2.544666213 | 2.520204382 | 2.567816622 | 2.916453949 | 2.44122113 |
| ENSMUSG00000020781 | 1.89911417 | 1.843759858 | 1.880890489 | 1.77186571 | 1.890639929 | 2.068185862 | 1.811176017 |
| ENSMUSG00000020782 | 1.527823874 | 1.575945187 | 1.774508021 | 1.695339928 | 1.986303335 | 1.892094603 | 2.073258096 |
| ENSMUSG00000020783 | 2.496544396 | 2.336628433 | 2.327653195 | 2.405970373 | 2.354487824 | 2.670245853 | 2.159731975 |
| ENSMUSG00000020785 | 1.806056082 | 1.777131114 | 1.837182039 | 1.678561379 | 1.715300549 | 1.875061263 | 1.651852264 |
| ENSMUSG00000020787 | 0 | 0 | 0.25 | 0 | 0.244650542 | 0 | 0 |
| ENSMUSG00000020788 | 0.747221736 | 0.864857905 | 0.830482024 | 1.29248125 | 1.264824954 | 1.041392685 | 1.20919388 |
| ENSMUSG00000020790 | 3.00093891 | 2.745926548 | 2.697994205 | 2.753679482 | 2.704016765 | 3.113274692 | 2.500404294 |
| ENSMUSG00000020792 | 2.428398755 | 2.288704527 | 2.312574604 | 2.234159485 | 2.304082723 | 2.495544338 | 2.22088434 |
| ENSMUSG00000020793 | 0.289064826 | 0.5 | 0.396240625 | 0.396240625 | 0.632412477 | 0.477121255 | 0.629532004 |
| ENSMUSG00000020794 | 2.66563648 | 2.481203126 | 2.422499493 | 2.393411797 | 2.302531265 | 2.657055853 | 2.121678334 |
| ENSMUSG00000020798 | 0 | 0.396240625 | 0.25 | 0.396240625 | 0.632412477 | 0.602059991 | 0.629532004 |
| ENSMUSG00000020799 | 0 | 0 | 0 | 0 | 0 | 0 | 0 |
| ENSMUSG00000020801 | 1.894701088 | 1.782320754 | 1.660964047 | 1.646240625 | 1.595992969 | 1.968482949 | 1.667169465 |
| ENSMUSG00000020802 | 2.435736758 | 2.280383379 | 2.30539678 | 2.247171172 | 2.333760374 | 2.56937391 | 2.268818981 |
| ENSMUSG00000020803 | 2.546826359 | 2.359177886 | 2.387686696 | 2.418106335 | 2.377015368 | 2.71432976 | 2.22557791 |
| ENSMUSG00000020804 | 0.289064826 | 0.25 | 0.396240625 | 0 | 0.244650542 | 0 | 0 |
| ENSMUSG00000020805 | 0 | 0 | 0 | 0.5 | 0.568060967 | 0.698970004 | 0.803905701 |
| ENSMUSG00000020806 | 1.83262764 | 1.784887838 | 1.870953944 | 1.678561379 | 1.92245931 | 1.812913357 | 1.872150729 |
| ENSMUSG00000020807 | 2.141794304 | 2 | 1.998588359 | 1.96293726 | 1.958580386 | 2.164352856 | 1.805128823 |

| | | | | | | | |
|---|---|---|---|---|---|---|---|
| ENSMUSG00000020808 | 2.330063669 | 2.205044741 | 2.275821952 | 2.213717096 | 2.227765453 | 2.46834733 | 2.175263699 |
| ENSMUSG00000020810 | 0.289064826 | 0.396240625 | 0 | 1.146240625 | 1.176121988 | 1.146128036 | 1.335191751 |
| ENSMUSG00000020811 | 0 | 0 | 0 | 0 | 0 | 0 | 0 |
| ENSMUSG00000020812 | 1.663535556 | 1.476722649 | 1.593759858 | 1.488549078 | 1.692703642 | 1.556302501 | 1.621017948 |
| ENSMUSG00000020814 | 2.325351389 | 2.101285366 | 2.161864607 | 2.212155735 | 2.195595395 | 2.494154594 | 2.072724682 |
| ENSMUSG00000020817 | 2.923527123 | 2.685577359 | 2.666999634 | 2.717783796 | 2.622922729 | 3.009450896 | 2.411529829 |
| ENSMUSG00000020818 | 2.58665822 | 2.360735874 | 2.363317659 | 2.228969845 | 2.229044281 | 2.472756449 | 2.055173778 |
| ENSMUSG00000020821 | 3.059196982 | 2.832589179 | 2.900674716 | 2.62694866 | 2.708967143 | 3.011570444 | 2.611466982 |
| ENSMUSG00000020823 | 2.809533441 | 2.608917565 | 2.623213655 | 2.572216519 | 2.583591153 | 2.932980822 | 2.490137691 |
| ENSMUSG00000020826 | 1.470606563 | 1.27186571 | 1.364857905 | 0.951838731 | 0.931471446 | 1.041392685 | 1.028149525 |
| ENSMUSG00000020827 | 2.591692877 | 2.38040011 | 2.438304187 | 2.309601185 | 2.357161747 | 2.612783857 | 2.301737862 |
| ENSMUSG00000020828 | 2.294202867 | 2.124961472 | 2.181128463 | 2.009729747 | 2.09661982 | 2.271841607 | 1.981429723 |
| ENSMUSG00000020829 | 1.205378646 | 1.25 | 1.311981878 | 1.610735874 | 1.599808748 | 1.857332496 | 1.669034113 |
| ENSMUSG00000020830 | 0 | 0 | 0.25 | 0 | 0 | 0 | 0 |
| ENSMUSG00000020831 | 0.671187741 | 0.580482024 | 0.896240625 | 0.580482024 | 0.686820904 | 0.602059991 | 0.896975297 |
| ENSMUSG00000020832 | 2.039433992 | 1.901832578 | 1.914552871 | 1.951838731 | 2 | 2.149219113 | 1.87511425 |
| ENSMUSG00000020834 | 1.920505136 | 1.816696635 | 1.841580554 | 1.914552871 | 1.888971102 | 2.100370545 | 1.854824501 |
| ENSMUSG00000020836 | 2.016886209 | 1.841580554 | 1.874961472 | 1.114857905 | 1.176121988 | 1.278753601 | 1.150210039 |
| ENSMUSG00000020838 | 0.578129653 | 0 | 0.396240625 | 0.646240625 | 0.733951626 | 0.698970004 | 0.520678036 |
| ENSMUSG00000020839 | 0 | 0 | 0 | 0 | 0 | 0 | 0 |
| ENSMUSG00000020840 | 2.652148299 | 2.453445298 | 2.371457327 | 2.511439915 | 2.462015062 | 2.81756537 | 2.291856721 |
| ENSMUSG00000020841 | 3.096117401 | 2.825659731 | 2.79652856 | 3.033927322 | 2.928932497 | 3.397766256 | 2.698729343 |
| ENSMUSG00000020843 | 2.342375483 | 2.184523065 | 2.271202097 | 2.252806814 | 2.259595879 | 2.521138084 | 2.122136896 |
| ENSMUSG00000020844 | 2.609650079 | 2.402294685 | 2.418548067 | 2.320192693 | 2.317250476 | 2.565847819 | 2.238944178 |
| ENSMUSG00000020846 | 2.302628051 | 2.107613138 | 2.087182039 | 2.04248125 | 2.021325983 | 2.274157849 | 1.94631981 |
| ENSMUSG00000020847 | 0.960252568 | 0.92510993 | 0.976722649 | 1.080482024 | 1.149965125 | 0.954242509 | 1.224243824 |
| ENSMUSG00000020848 | 1.036286563 | 0.951838731 | 1 | 1.17510993 | 1.106691885 | 1.230448921 | 1.054044578 |
| ENSMUSG00000020849 | 3.571258591 | 3.22938964 | 3.222850999 | 3.357939687 | 3.249608524 | 3.811440944 | 3.039210024 |
| ENSMUSG00000020850 | 3.501562669 | 3.150558725 | 3.136723615 | 3.165889526 | 3.120745371 | 3.611510887 | 2.89743451 |
| ENSMUSG00000020857 | 0 | 0.396240625 | 0.580482024 | 0.25 | 0.568060967 | 0.301029996 | 0.579661877 |
| ENSMUSG00000020859 | 3.034848022 | 2.785207443 | 2.784087585 | 2.800684011 | 2.689947504 | 3.097951071 | 2.513514042 |
| ENSMUSG00000020863 | 2.89911417 | 2.662538553 | 2.644357207 | 2.729715809 | 2.671852323 | 3.038620162 | 2.475563135 |
| ENSMUSG00000020864 | 2.719923957 | 2.492885888 | 2.474839231 | 2.490724001 | 2.446160569 | 2.805500858 | 2.311581365 |
| ENSMUSG00000020865 | 1.069666964 | 0.92510993 | 1.098079356 | 1.080482024 | 1.039257769 | 1 | 1.110947927 |

| | | | | | | | |
|---|---|---|---|---|---|---|---|
| ENSMUSG00000020866 | 2.348397716 | 2.228222334 | 2.209800947 | 1.701838731 | 1.741652793 | 1.963787827 | 1.82753046 |
| ENSMUSG00000020867 | 0.289064826 | 0.5 | 0.5 | 0 | 0 | 0 | 0 |
| ENSMUSG00000020868 | 1.952600383 | 1.942046081 | 2.044977273 | 1.910964047 | 2.048957937 | 2.158362492 | 2.074322292 |
| ENSMUSG00000020869 | 3.300956576 | 3.016690483 | 3.012377063 | 2.982684334 | 2.945411925 | 3.373095987 | 2.796923892 |
| ENSMUSG00000020870 | 0.578129653 | 0.25 | 0 | 0 | 0.244650542 | 0.301029996 | 0.448487648 |
| ENSMUSG00000020871 | 0.289064826 | 0.75 | 0.646240625 | 1.214495249 | 1.136121934 | 1.230448921 | 1.278288402 |
| ENSMUSG00000020872 | 0 | 0 | 0 | 0 | 0.489301084 | 0.301029996 | 0 |
| ENSMUSG00000020873 | 2.519973621 | 2.284247778 | 2.336628433 | 2.396709947 | 2.32316608 | 2.664641976 | 2.209893373 |
| ENSMUSG00000020875 | 2.762449763 | 2.552972074 | 2.538388008 | 2.549304173 | 2.475604419 | 2.877371346 | 2.312600313 |
| ENSMUSG00000020876 | 2.029919532 | 1.964495249 | 2.03360658 | 1.816696635 | 1.92245931 | 2.068185862 | 1.86715039 |
| ENSMUSG00000020877 | 1.707474304 | 1.747171172 | 1.845926073 | 1.642463902 | 1.832885921 | 1.785329835 | 1.896975297 |
| ENSMUSG00000020878 | 0.867194479 | 1 | 0.896240625 | 1.080482024 | 1.020174412 | 1 | 1.014380832 |
| ENSMUSG00000020882 | 1.249317394 | 1.282320754 | 1.410964047 | 1.130890489 | 1.445123986 | 1.278753601 | 1.365075947 |
| ENSMUSG00000020883 | 2.373064211 | 2.136723615 | 2.133818844 | 2.078220739 | 2.063536368 | 2.318063335 | 1.942350245 |
| ENSMUSG00000020884 | 0 | 0.396240625 | 0.396240625 | 0.25 | 0.244650542 | 0 | 0.224243824 |
| ENSMUSG00000020886 | 2.160971586 | 2.117918804 | 2.098079356 | 1.94697564 | 2.040348824 | 2.178976947 | 1.981429723 |
| ENSMUSG00000020887 | 0.289064826 | 0 | 0 | 0.5 | 0.387761935 | 0 | 0 |
| ENSMUSG00000020888 | 2.401648579 | 2.28296424 | 2.405512955 | 1.948603967 | 2.085294053 | 2.139879086 | 2.113781657 |
| ENSMUSG00000020889 | 2.013563224 | 1.928561379 | 1.916333979 | 1.388647213 | 1.611014646 | 1.653212514 | 1.508694612 |
| ENSMUSG00000020890 | 0 | 0 | 0 | 0.396240625 | 0.489301084 | 0.301029996 | 0.448487648 |
| ENSMUSG00000020891 | 0.45815691 | 0.25 | 0.5 | 0.79248125 | 0.632412477 | 0.301029996 | 0.74492186 |
| ENSMUSG00000020892 | 0.91631382 | 0.701838731 | 0.701838731 | 1.04248125 | 1.057362051 | 0.903089987 | 1.028149525 |
| ENSMUSG00000020893 | 1.707474304 | 1.708222504 | 1.76109853 | 1.685366747 | 1.877063019 | 1.934498451 | 1.845261772 |
| ENSMUSG00000020894 | 2.201144779 | 1.976722649 | 1.984159485 | 2.191217898 | 2.171644683 | 2.477121255 | 2.024452165 |
| ENSMUSG00000020895 | 1.596667477 | 1.561981878 | 1.668106335 | 1.593759858 | 1.649303553 | 1.72427587 | 1.633706455 |
| ENSMUSG00000020897 | 2.441754646 | 2.245748394 | 2.299304173 | 2.424241882 | 2.412899576 | 2.767155866 | 2.316895388 |
| ENSMUSG00000020898 | 2.483777101 | 2.293106877 | 2.310793496 | 2.25 | 2.301493158 | 2.598790507 | 2.198877505 |
| ENSMUSG00000020899 | 2.462727174 | 2.311388177 | 2.338837774 | 2.341033664 | 2.395014713 | 2.56937391 | 2.276871337 |
| ENSMUSG00000020900 | 3.464788124 | 3.164217931 | 3.135757955 | 3.007666784 | 3.012630862 | 3.40534636 | 2.825139813 |
| ENSMUSG00000020901 | 1.806056082 | 1.723704441 | 1.774508021 | 1.653677461 | 1.746695109 | 1.851258349 | 1.665294008 |
| ENSMUSG00000020902 | 2.76355595 | 2.551142786 | 2.605253214 | 2.452241044 | 2.488689905 | 2.731588765 | 2.39814371 |
| ENSMUSG00000020903 | 2.075459174 | 1.899978211 | 1.848079356 | 2.074802005 | 2.012006468 | 2.225309282 | 1.86715039 |
| ENSMUSG00000020904 | 0 | 0 | 0 | 0.830482024 | 0.632412477 | 0.602059991 | 0.629532004 |
| ENSMUSG00000020905 | 1.482695304 | 1.53743678 | 1.445339928 | 1.403677461 | 1.47337555 | 1.462397998 | 1.42167221 |

| | | | | | | | |
|---|---|---|---|---|---|---|---|
| ENSMUSG00000020907 | 0 | 0 | 0 | 0 | 0 | 0 | 0 |
| ENSMUSG00000020908 | 0 | 0 | 0 | 0.25 | 0 | 0 | 0 |
| ENSMUSG00000020910 | 1.605636262 | 1.634789703 | 1.54248125 | 1.711372513 | 1.671670246 | 1.949390007 | 1.625302983 |
| ENSMUSG00000020911 | 2.235832493 | 2.201838731 | 2.26584877 | 2.077084758 | 2.278401864 | 2.31386722 | 2.250549951 |
| ENSMUSG00000020912 | 0 | 0 | 0 | 0 | 0 | 0 | 0 |
| ENSMUSG00000020913 | 0 | 0 | 0 | 0 | 0 | 0 | 0 |
| ENSMUSG00000020914 | 3.622804059 | 3.267404398 | 3.259300935 | 3.44510931 | 3.329492956 | 3.898560645 | 3.086175738 |
| ENSMUSG00000020916 | 0 | 0 | 0 | 0 | 0 | 0 | 0 |
| ENSMUSG00000020917 | 3.222080938 | 2.95871651 | 2.961470509 | 3.021285115 | 3.034077619 | 3.469085299 | 2.844623916 |
| ENSMUSG00000020918 | 2.521950082 | 2.388167274 | 2.407339155 | 2.365881093 | 2.436010412 | 2.67669361 | 2.319893236 |
| ENSMUSG00000020919 | 2.444137694 | 2.240724001 | 2.319031101 | 2.081607372 | 2.211376431 | 2.354108439 | 2.10131878 |
| ENSMUSG00000020921 | 1.907802659 | 1.816696635 | 1.744319981 | 1.888647213 | 1.957204337 | 2.113943352 | 1.906037947 |
| ENSMUSG00000020922 | 2.4661223 | 2.275821952 | 2.268535366 | 2.320192693 | 2.265976525 | 2.592176757 | 2.142557543 |
| ENSMUSG00000020923 | 2.876566388 | 2.662314044 | 2.67619206 | 2.556000419 | 2.607973774 | 2.893206753 | 2.463188002 |
| ENSMUSG00000020925 | 2.479432974 | 2.321928095 | 2.311981878 | 2.52053726 | 2.395413309 | 2.769377326 | 2.253311124 |
| ENSMUSG00000020926 | 0.811507563 | 0.864857905 | 1.04248125 | 0.79248125 | 0.877063019 | 0.301029996 | 1.028149525 |
| ENSMUSG00000020928 | 0.289064826 | 0.25 | 0 | 0 | 0.387761935 | 0 | 0.224243824 |
| ENSMUSG00000020929 | 3.118167235 | 2.855778978 | 2.864601653 | 2.887806901 | 2.889561001 | 3.27989498 | 2.709272686 |
| ENSMUSG00000020930 | 0.91631382 | 1.04248125 | 0.896240625 | 0.79248125 | 0.686820904 | 0.698970004 | 0.876096088 |
| ENSMUSG00000020932 | 0 | 0.396240625 | 0.646240625 | 0.701838731 | 0.489301084 | 0.477121255 | 0 |
| ENSMUSG00000020935 | 2.364524 | 2.311388177 | 2.392938161 | 2.161864607 | 2.302531265 | 2.496929648 | 2.337554913 |
| ENSMUSG00000020936 | 3.052896866 | 2.792950572 | 2.763311781 | 2.853935442 | 2.807544405 | 3.260071388 | 2.621703466 |
| ENSMUSG00000020937 | 2.072573125 | 1.997171172 | 2.029735268 | 1.695339928 | 1.947418788 | 1.973127854 | 1.975716277 |
| ENSMUSG00000020940 | 0.289064826 | 0.396240625 | 0.396240625 | 0 | 0.387761935 | 0.301029996 | 0.224243824 |
| ENSMUSG00000020941 | 2.192545868 | 2.145300145 | 2.113831805 | 1.912762923 | 2.170892913 | 2.220108088 | 2.082194103 |
| ENSMUSG00000020945 | 0 | 0 | 0 | 0 | 0 | 0 | 0 |
| ENSMUSG00000020946 | 2.808542865 | 2.560197833 | 2.536483036 | 2.629417347 | 2.595767222 | 2.996073654 | 2.428320637 |
| ENSMUSG00000020948 | 1.647796617 | 1.630890489 | 1.52186571 | 1.614857905 | 1.559656853 | 1.568201724 | 1.452143229 |
| ENSMUSG00000020949 | 2.340856238 | 2.208222504 | 2.215271726 | 2.129909063 | 2.118236133 | 2.350248018 | 1.969165684 |
| ENSMUSG00000020950 | 2.542156295 | 2.310793496 | 2.333847589 | 2.078220739 | 2.030405741 | 2.298853076 | 1.902443294 |
| ENSMUSG00000020952 | 2.593357679 | 2.36434522 | 2.358657057 | 2.385757955 | 2.291475807 | 2.685741739 | 2.180662175 |
| ENSMUSG00000020953 | 1.100572389 | 1.302363341 | 1.160964047 | 1.664552871 | 1.607318736 | 1.799340549 | 1.505628092 |
| ENSMUSG00000020954 | 2.853620573 | 2.642226539 | 2.58993739 | 2.59995299 | 2.531009002 | 2.853698212 | 2.308505126 |
| ENSMUSG00000020955 | 1.568542279 | 1.396240625 | 1.482684334 | 1.302363341 | 1.200473444 | 1.322219295 | 1.159323753 |

| | | | | | | | |
|---|---|---|---|---|---|---|---|
| ENSMUSG00000020956 | 1.671187741 | 1.657339155 | 1.618933358 | 1.634789703 | 1.555378544 | 1.707570176 | 1.444790289 |
| ENSMUSG00000020961 | 0.578129653 | 0.830482024 | 0.830482024 | 1.114857905 | 1.200473444 | 1.146128036 | 1.193409508 |
| ENSMUSG00000020962 | 2.35433422 | 2.207430684 | 2.179419106 | 2.051142786 | 1.97205363 | 2.123851641 | 1.811176017 |
| ENSMUSG00000020963 | 0.289064826 | 0.396240625 | 0.396240625 | 0 | 0 | 0 | 0.224243824 |
| ENSMUSG00000020964 | 2.804055922 | 2.574515642 | 2.540597832 | 2.540597832 | 2.480959501 | 2.873901598 | 2.335902514 |
| ENSMUSG00000020973 | 2.141794304 | 2.013820609 | 2.049918086 | 2.038704527 | 1.966725889 | 2.187520721 | 1.869159809 |
| ENSMUSG00000020974 | 2.267855219 | 2.070192693 | 2.024508021 | 2.05479213 | 2.024758322 | 2.352182518 | 1.902443294 |
| ENSMUSG00000020978 | 2.683414322 | 2.439555804 | 2.433677405 | 2.404596376 | 2.313753359 | 2.64246452 | 2.173704585 |
| ENSMUSG00000020982 | 2.643399047 | 2.443696765 | 2.450629091 | 2.435366747 | 2.357161747 | 2.683047038 | 2.201756416 |
| ENSMUSG00000020986 | 2.814451333 | 2.560495787 | 2.538704527 | 2.484159485 | 2.418913884 | 2.77815125 | 2.237664194 |
| ENSMUSG00000020988 | 2.172524468 | 2.032320754 | 2.04745614 | 2.125956434 | 2.057362051 | 2.320146286 | 1.913982791 |
| ENSMUSG00000020990 | 0.91631382 | 0.976722649 | 0.976722649 | 0.396240625 | 0.244650542 | 0.477121255 | 0.579661877 |
| ENSMUSG00000020992 | 0 | 0.25 | 0.25 | 0.25 | 0.244650542 | 0 | 0.224243824 |
| ENSMUSG00000020993 | 2.243591459 | 2.12892496 | 2.104463129 | 2.075945187 | 1.958580386 | 2.235528447 | 1.86715039 |
| ENSMUSG00000020994 | 2.952249197 | 2.710783978 | 2.707430684 | 2.69595419 | 2.616167381 | 2.958563883 | 2.445465687 |
| ENSMUSG00000021000 | 2.565888829 | 2.358657057 | 2.320192693 | 2.285526764 | 2.196996016 | 2.523746467 | 2.064068234 |
| ENSMUSG00000021003 | 2.337801042 | 2.171625132 | 2.182829758 | 2.198603967 | 2.12771782 | 2.416640507 | 1.966211184 |
| ENSMUSG00000021007 | 1.655740331 | 1.527131114 | 1.649978211 | 1.660964047 | 1.584291694 | 1.799340549 | 1.448487648 |
| ENSMUSG00000021009 | 2.645603628 | 2.46293726 | 2.476722649 | 2.400906586 | 2.387761935 | 2.661812686 | 2.235734673 |
| ENSMUSG00000021010 | 1 | 0.951838731 | 0.646240625 | 0.396240625 | 0.489301084 | 0 | 0.355418052 |
| ENSMUSG00000021012 | 2.799520178 | 2.552972074 | 2.517530236 | 2.578504176 | 2.49982889 | 2.857935265 | 2.311836403 |
| ENSMUSG00000021013 | 2.172524468 | 1.973704441 | 2.031030328 | 1.899978211 | 1.880506521 | 2.133538908 | 1.788855757 |
| ENSMUSG00000021018 | 2.17479713 | 2.043731421 | 2.089388001 | 2.028435542 | 1.998776581 | 2.245512668 | 1.877074955 |
| ENSMUSG00000021022 | 2.22592627 | 2.094844592 | 2.05479213 | 2.078220739 | 2 | 2.283301229 | 1.923435442 |
| ENSMUSG00000021023 | 1.885732304 | 1.811981878 | 1.782320754 | 1.784887838 | 1.709785497 | 1.903089987 | 1.546120303 |
| ENSMUSG00000021024 | 2.98109172 | 2.705642708 | 2.704845198 | 2.742166698 | 2.622085347 | 3.059563418 | 2.451977959 |
| ENSMUSG00000021025 | 2.415874319 | 2.314935816 | 2.368426437 | 2.162762923 | 2.210031835 | 2.457881897 | 2.170563634 |
| ENSMUSG00000021027 | 2.578129653 | 2.361770807 | 2.360217292 | 2.389605178 | 2.322187006 | 2.64246452 | 2.126234965 |
| ENSMUSG00000021028 | 1.940852214 | 1.80479213 | 1.807204673 | 1.78743678 | 1.655839964 | 1.977723605 | 1.62316756 |
| ENSMUSG00000021032 | 0.289064826 | 0.25 | 0.25 | 0.25 | 0.489301084 | 0.301029996 | 0.355418052 |
| ENSMUSG00000021033 | 2.247417471 | 2.032320754 | 2.025821952 | 2.029735268 | 1.97205363 | 2.255272505 | 1.86815666 |
| ENSMUSG00000021036 | 2.688073951 | 2.461372513 | 2.468361028 | 2.442459461 | 2.369807605 | 2.72427587 | 2.198877505 |
| ENSMUSG00000021037 | 2.893084681 | 2.647646762 | 2.646709947 | 2.591853688 | 2.604025187 | 2.92788341 | 2.454448184 |
| ENSMUSG00000021038 | 2.34539747 | 2.117918804 | 2.131869252 | 2.140560606 | 2.064555002 | 2.328379603 | 1.94631981 |

| | | | | | | | |
|---|---|---|---|---|---|---|---|
| ENSMUSG00000021039 | 2.427163103 | 2.273189285 | 2.21293726 | 2.334405475 | 2.192777367 | 2.496929648 | 2.066798338 |
| ENSMUSG00000021040 | 2.057835465 | 1.874961472 | 1.870953944 | 1.860735874 | 1.806427258 | 2.025305865 | 1.729871916 |
| ENSMUSG00000021044 | 2.045658471 | 1.945339928 | 1.901832578 | 1.552363341 | 1.652586889 | 1.740362689 | 1.554175045 |
| ENSMUSG00000021047 | 1.249317394 | 1.02186571 | 1.160964047 | 0.646240625 | 0.733951626 | 0.77815125 | 0.672731473 |
| ENSMUSG00000021048 | 2.792365035 | 2.591853688 | 2.615369862 | 2.478596283 | 2.537993141 | 2.813580989 | 2.399313044 |
| ENSMUSG00000021051 | 2.622240692 | 2.395770692 | 2.385757955 | 2.341033664 | 2.300451989 | 2.553883027 | 2.109521699 |
| ENSMUSG00000021054 | 2.388281011 | 2.228969845 | 2.250703754 | 2.293731421 | 2.253693465 | 2.571708832 | 2.082194103 |
| ENSMUSG00000021055 | 0.578129653 | 0.701838731 | 1.114857905 | 0.75 | 0.568060967 | 0.602059991 | 0.629532004 |
| ENSMUSG00000021056 | 0 | 0 | 0 | 0 | 0 | 0 | 0 |
| ENSMUSG00000021057 | 1 | 0.976722649 | 1.04248125 | 1.79745614 | 1.763796674 | 1.903089987 | 1.616675393 |
| ENSMUSG00000021061 | 0.671187741 | 0.896240625 | 0.896240625 | 1.02186571 | 0.955822902 | 0.903089987 | 0.876096088 |
| ENSMUSG00000021062 | 0.578129653 | 0.396240625 | 0.646240625 | 1.729715809 | 1.706995316 | 1.875061263 | 1.584526862 |
| ENSMUSG00000021065 | 2.417143825 | 2.23927551 | 2.23927551 | 1.973704441 | 1.917865457 | 2.113943352 | 1.786277931 |
| ENSMUSG00000021066 | 1.482695304 | 1.364857905 | 1.282320754 | 1.214495249 | 1.091002363 | 1.146128036 | 1.028149525 |
| ENSMUSG00000021067 | 2.615992899 | 2.427270953 | 2.398581151 | 2.356041572 | 2.253693465 | 2.604226053 | 2.138662327 |
| ENSMUSG00000021068 | 2.640441358 | 2.436208459 | 2.42293588 | 2.358657057 | 2.334234459 | 2.624282096 | 2.167391888 |
| ENSMUSG00000021069 | 0.578129653 | 0 | 0 | 1.04248125 | 0.978602168 | 1.146128036 | 1 |
| ENSMUSG00000021070 | 0 | 0.25 | 0.5 | 0.79248125 | 1.039257769 | 0.84509804 | 1.159323753 |
| ENSMUSG00000021071 | 0 | 0 | 0 | 0 | 0 | 0 | 0 |
| ENSMUSG00000021072 | 2.897915226 | 2.651601303 | 2.654596376 | 2.627691042 | 2.567572108 | 2.937517892 | 2.355863359 |
| ENSMUSG00000021076 | 2.7985055 | 2.572792675 | 2.539336734 | 2.619439567 | 2.529104803 | 2.914343157 | 2.348891258 |
| ENSMUSG00000021078 | 0 | 0 | 0 | 0 | 0 | 0 | 0.224243824 |
| ENSMUSG00000021079 | 2.095120908 | 1.938721876 | 1.956637122 | 1.868933358 | 1.851969278 | 2.037426498 | 1.767634022 |
| ENSMUSG00000021081 | 0 | 0 | 0 | 0 | 0 | 0 | 0 |
| ENSMUSG00000021086 | 0 | 0 | 0 | 0 | 0 | 0 | 0 |
| ENSMUSG00000021087 | 0.289064826 | 0.25 | 0.25 | 0.75 | 0.568060967 | 0.301029996 | 0.629532004 |
| ENSMUSG00000021090 | 0 | 0 | 0 | 0 | 0 | 0 | 0.355418052 |
| ENSMUSG00000021091 | 0.289064826 | 0.580482024 | 0.701838731 | 0.25 | 0 | 0 | 0.355418052 |
| ENSMUSG00000021094 | 1.631440309 | 1.557204673 | 1.646240625 | 1.618933358 | 1.484072009 | 1.716003344 | 1.476637173 |
| ENSMUSG00000021095 | 0 | 0 | 0 | 0 | 0 | 0 | 0.355418052 |
| ENSMUSG00000021096 | 2.66283289 | 2.460980263 | 2.453445298 | 2.336073977 | 2.25665701 | 2.505149978 | 2.085803078 |
| ENSMUSG00000021097 | 1 | 0.830482024 | 0.951838731 | 1.54745614 | 1.499532413 | 1.505149978 | 1.369798885 |
| ENSMUSG00000021098 | 0 | 0 | 0 | 0 | 0.568060967 | 0 | 0 |
| ENSMUSG00000021099 | 0 | 0 | 0 | 0 | 0 | 0 | 0 |

| | | | | | | | |
|---|---|---|---|---|---|---|---|
| ENSMUSG00000021101 | 0 | 0 | 0 | 0 | 0 | 0 | 0 |
| ENSMUSG00000021102 | 2.051791411 | 2.006976499 | 2.058404919 | 1.903677461 | 1.941702774 | 2.093421685 | 1.975716277 |
| ENSMUSG00000021103 | 2.264212992 | 2.121960008 | 2.125956434 | 2.065523711 | 2.032639648 | 2.305351369 | 1.922587433 |
| ENSMUSG00000021108 | 0.91631382 | 0.864857905 | 0.701838731 | 0.864857905 | 0.877063019 | 1.255272505 | 0.935079929 |
| ENSMUSG00000021109 | 3.166262603 | 2.891751343 | 2.862287161 | 2.706438458 | 2.619138767 | 3.029789471 | 2.436226648 |
| ENSMUSG00000021111 | 3.118167235 | 2.853935442 | 2.822648722 | 2.858917565 | 2.774164253 | 3.190891717 | 2.567516562 |
| ENSMUSG00000021112 | 2.637462542 | 2.417221246 | 2.394828984 | 2.315523711 | 2.223900931 | 2.531478917 | 2.051204213 |
| ENSMUSG00000021113 | 2.165631214 | 2.027131114 | 1.96759118 | 1.890560606 | 1.808534465 | 2.079181246 | 1.732938436 |
| ENSMUSG00000021114 | 2.65645504 | 2.436208459 | 2.454645544 | 2.267865591 | 2.227124297 | 2.591064607 | 2.114251541 |
| ENSMUSG00000021115 | 2.396352858 | 2.247880462 | 2.231203126 | 2.211372513 | 2.204601633 | 2.523746467 | 2.073258096 |
| ENSMUSG00000021116 | 2.977138777 | 2.732314602 | 2.703845824 | 2.702442032 | 2.59304693 | 2.997386384 | 2.427073261 |
| ENSMUSG00000021118 | 0.578129653 | 0.5 | 0.580482024 | 0.830482024 | 0.931471446 | 1 | 0.775756176 |
| ENSMUSG00000021120 | 1.307602651 | 1.29248125 | 1.27186571 | 1.388647213 | 1.394615667 | 1.568201724 | 1.319146374 |
| ENSMUSG00000021123 | 0.960252568 | 0.396240625 | 0.976722649 | 1.061981878 | 0.978602168 | 1.041392685 | 1.089372236 |
| ENSMUSG00000021124 | 2.70302761 | 2.480831871 | 2.493603647 | 2.281030328 | 2.170139537 | 2.522444234 | 2.056860222 |
| ENSMUSG00000021125 | 1.100572389 | 1.04248125 | 1.080482024 | 1.061981878 | 0.931471446 | 1.041392685 | 0.853775828 |
| ENSMUSG00000021127 | 1.777675583 | 1.777131114 | 1.738549078 | 1.29248125 | 1.499532413 | 1.602059991 | 1.455757965 |
| ENSMUSG00000021130 | 0.289064826 | 0.25 | 0.5 | 0.580482024 | 0.733951626 | 0.602059991 | 0.579661877 |
| ENSMUSG00000021131 | 2.68408318 | 2.464883697 | 2.482684334 | 2.42510993 | 2.351342427 | 2.727541257 | 2.182948513 |
| ENSMUSG00000021133 | 2.683414322 | 2.459406983 | 2.472566069 | 2.252806814 | 2.251304582 | 2.51054501 | 2.132508314 |
| ENSMUSG00000021134 | 2.976476295 | 2.682192712 | 2.738185312 | 2.752981517 | 2.70732472 | 3.103461622 | 2.521479147 |
| ENSMUSG00000021135 | 0.289064826 | 0.25 | 0.580482024 | 0.396240625 | 0.387761935 | 0.301029996 | 0.355418052 |
| ENSMUSG00000021136 | 2.22189694 | 2.067865757 | 2.065523711 | 0.5 | 0.489301084 | 0.602059991 | 0.224243824 |
| ENSMUSG00000021139 | 0.578129653 | 0.396240625 | 0.580482024 | 0.25 | 0.387761935 | 0.301029996 | 0.355418052 |
| ENSMUSG00000021140 | 2.432083896 | 2.250703754 | 2.228222334 | 2.211372513 | 2.196996016 | 2.437750563 | 2.056860222 |
| ENSMUSG00000021143 | 2.649978134 | 2.496816003 | 2.525821952 | 2.263142013 | 2.325116119 | 2.532754379 | 2.237343405 |
| ENSMUSG00000021144 | 2.504843822 | 2.37246199 | 2.42813175 | 2.119945066 | 2.245869735 | 2.445604203 | 2.206027309 |
| ENSMUSG00000021147 | 2.264212992 | 2.113831805 | 2.105516192 | 2.221424093 | 2.133621644 | 2.459392488 | 1.982842456 |
| ENSMUSG00000021148 | 0 | 0 | 0 | 0 | 0 | 0 | 0 |
| ENSMUSG00000021149 | 2.669806838 | 2.430274797 | 2.429847205 | 2.559899632 | 2.470166836 | 2.85308953 | 2.281664246 |
| ENSMUSG00000021156 | 2.711874084 | 2.528760913 | 2.474461364 | 2.584684096 | 2.528012058 | 2.921166051 | 2.391237049 |
| ENSMUSG00000021171 | 3.056195755 | 2.777294426 | 2.777294426 | 2.619439567 | 2.54143406 | 2.88592634 | 2.345922158 |
| ENSMUSG00000021175 | 2.163307908 | 2.059601185 | 2.052363341 | 1.930274797 | 1.913211023 | 2.064457989 | 1.771720588 |
| ENSMUSG00000021176 | 1.753493018 | 1.557204673 | 1.638647213 | 1.566696635 | 1.439191811 | 1.602059991 | 1.413583902 |

| | | | | | | | |
|---|---|---|---|---|---|---|---|
| ENSMUSG00000021177 | 2.23973002 | 2.013820609 | 2.04248125 | 1.94697564 | 2.019019072 | 2.120573931 | 1.911356129 |
| ENSMUSG00000021178 | 2.974149285 | 2.69961795 | 2.689139081 | 2.740000483 | 2.660757975 | 3.065206128 | 2.487630378 |
| ENSMUSG00000021179 | 2.072573125 | 1.938721876 | 1.981203126 | 1.848079356 | 1.892300902 | 2.068185862 | 1.857950279 |
| ENSMUSG00000021180 | 1.721148722 | 1.681980114 | 1.571350555 | 1.598079356 | 1.607318736 | 1.72427587 | 1.441050184 |
| ENSMUSG00000021182 | 1.771760131 | 1.720660762 | 1.735628626 | 1.201838731 | 1.433158226 | 1.278753601 | 1.486592281 |
| ENSMUSG00000021185 | 1.663535556 | 1.561981878 | 1.584962501 | 0.5 | 0.686820904 | 0.477121255 | 0.775756176 |
| ENSMUSG00000021186 | 2.858927525 | 2.591033664 | 2.575374049 | 2.219895812 | 2.157079823 | 2.471291711 | 2.020712061 |
| ENSMUSG00000021187 | 0.289064826 | 0.580482024 | 0.25 | 0.75 | 0.244650542 | 0.301029996 | 0.448487648 |
| ENSMUSG00000021188 | 2.572927518 | 2.409608978 | 2.407794264 | 2.405054956 | 2.371516839 | 2.678518379 | 2.181044355 |
| ENSMUSG00000021189 | 2.254965775 | 2.05357978 | 2.083847589 | 1.984159485 | 1.906908067 | 2.130333768 | 1.762103709 |
| ENSMUSG00000021190 | 2.759671389 | 2.56728167 | 2.5658173 | 2.366391601 | 2.358933174 | 2.678518379 | 2.231840759 |
| ENSMUSG00000021192 | 2.310886388 | 2.185366747 | 2.178561379 | 2.098079356 | 2.076554666 | 2.322219295 | 1.916588299 |
| ENSMUSG00000021193 | 3.045377547 | 2.778923488 | 2.788229644 | 2.77053726 | 2.689428323 | 3.133219457 | 2.495961709 |
| ENSMUSG00000021194 | 0 | 0 | 0 | 0 | 0.489301084 | 0 | 0.224243824 |
| ENSMUSG00000021196 | 2.361637952 | 2.153677461 | 2.183677405 | 2.331607372 | 2.309205544 | 2.596597096 | 2.235088933 |
| ENSMUSG00000021198 | 0.289064826 | 0.25 | 0 | 0 | 0.387761935 | 0.477121255 | 0 |
| ENSMUSG00000021200 | 0 | 0.25 | 0 | 0 | 0.244650542 | 0 | 0.224243824 |
| ENSMUSG00000021203 | 1.837744666 | 1.630890489 | 1.614857905 | 1.638647213 | 1.628935049 | 1.812913357 | 1.526507301 |
| ENSMUSG00000021207 | 0 | 0 | 0 | 0 | 0 | 0 | 0 |
| ENSMUSG00000021208 | 0 | 0 | 0 | 0 | 0 | 0 | 0.224243824 |
| ENSMUSG00000021209 | 0 | 0.646240625 | 0.25 | 1.114857905 | 1.039257769 | 1.255272505 | 1.028149525 |
| ENSMUSG00000021210 | 0 | 0 | 0 | 0 | 0 | 0 | 0 |
| ENSMUSG00000021211 | 0 | 0 | 0 | 0 | 0 | 0 | 0 |
| ENSMUSG00000021213 | 0.45815691 | 0.580482024 | 0.396240625 | 0.5 | 0 | 0.301029996 | 0.355418052 |
| ENSMUSG00000021214 | 0 | 0 | 0 | 0 | 0 | 0 | 0 |
| ENSMUSG00000021215 | 2.663535556 | 2.454245906 | 2.445749552 | 2.441632227 | 2.407166541 | 2.744292983 | 2.271140178 |
| ENSMUSG00000021216 | 0 | 0.25 | 0.25 | 0 | 0 | 0 | 0 |
| ENSMUSG00000021217 | 1.777675583 | 1.729715809 | 1.705044741 | 1.505591953 | 1.489301084 | 1.672097858 | 1.476637173 |
| ENSMUSG00000021218 | 3.341409331 | 3.029410775 | 3.019287379 | 3.182723662 | 3.071545633 | 3.594834356 | 2.845163722 |
| ENSMUSG00000021219 | 0 | 0.25 | 0.396240625 | 0 | 0 | 0 | 0 |
| ENSMUSG00000021221 | 0 | 0 | 0 | 0.25 | 0.244650542 | 0 | 0.224243824 |
| ENSMUSG00000021222 | 1.806056082 | 1.723704441 | 1.789967834 | 1.714495249 | 1.768534412 | 1.903089987 | 1.720494 |
| ENSMUSG00000021223 | 0 | 0 | 0.25 | 0 | 0.244650542 | 0 | 0 |
| ENSMUSG00000021224 | 2.273259602 | 2.101285366 | 2.108657057 | 2.061981878 | 2.003645018 | 2.247973266 | 1.933441873 |

| | | | | | | | |
|---|---|---|---|---|---|---|---|
| ENSMUSG00000021226 | 1.289064826 | 1.238549078 | 1.226722649 | 1 | 1.039257769 | 0.954242509 | 1.066254157 |
| ENSMUSG00000021228 | 0 | 0 | 0 | 0 | 0 | 0 | 0 |
| ENSMUSG00000021234 | 1.227926429 | 1.321350555 | 1.25 | 1.201838731 | 1.188507684 | 1.361727836 | 1.252393349 |
| ENSMUSG00000021235 | 2.134369199 | 1.926839783 | 1.94697564 | 1.794977273 | 1.887294348 | 2.029383778 | 1.788855757 |
| ENSMUSG00000021236 | 2.199011616 | 2.056000419 | 2.034887838 | 2.245748394 | 2.169384551 | 2.434568904 | 2.022587518 |
| ENSMUSG00000021238 | 2.00683675 | 1.777131114 | 1.825945187 | 1.870953944 | 1.793512788 | 1.995635195 | 1.712465909 |
| ENSMUSG00000021239 | 0 | 0 | 0 | 0.25 | 0 | 0 | 0 |
| ENSMUSG00000021240 | 2.089597219 | 1.982684334 | 2.060793496 | 1.735628626 | 1.72073075 | 1.944482672 | 1.71730663 |
| ENSMUSG00000021241 | 1.777675583 | 1.738549078 | 1.811981878 | 1.75835575 | 1.768534412 | 1.913813852 | 1.629532004 |
| ENSMUSG00000021242 | 3.079880513 | 2.807204673 | 2.849017389 | 2.601551251 | 2.540642974 | 2.940018155 | 2.415625016 |
| ENSMUSG00000021243 | 2.20747957 | 1.987091808 | 2.034887838 | 2.102347734 | 1.975997314 | 2.334453751 | 1.861046144 |
| ENSMUSG00000021244 | 2.689395766 | 2.478222334 | 2.454645544 | 2.462546709 | 2.391406953 | 2.750508395 | 2.223574712 |
| ENSMUSG00000021245 | 1.885732304 | 1.685366747 | 1.642463902 | 1.814346961 | 1.706995316 | 1.944482672 | 1.651852264 |
| ENSMUSG00000021248 | 2.953652167 | 2.691632227 | 2.701435968 | 2.680274797 | 2.61998316 | 3.02489596 | 2.445634316 |
| ENSMUSG00000021250 | 1.227926429 | 1.146240625 | 1.364857905 | 1.02186571 | 1.254881871 | 1.301029996 | 1.121219121 |
| ENSMUSG00000021252 | 2.343889214 | 2.17510993 | 2.145300145 | 2.136723615 | 2.111176775 | 2.334453751 | 1.966211184 |
| ENSMUSG00000021253 | 2.638209243 | 2.453044326 | 2.404596376 | 2.049918086 | 2.121713561 | 2.338456494 | 2.02383181 |
| ENSMUSG00000021254 | 2.089597219 | 1.976722649 | 1.966046536 | 1.886723615 | 1.898867717 | 2.053078443 | 1.815933683 |
| ENSMUSG00000021255 | 0 | 0.701838731 | 0.580482024 | 1.160964047 | 1.212046813 | 1.255272505 | 1.110947927 |
| ENSMUSG00000021256 | 0.867194479 | 1.130890489 | 1.201838731 | 0.896240625 | 1.057362051 | 0.903089987 | 1.193409508 |
| ENSMUSG00000021257 | 1.342375483 | 1.226722649 | 1.339388001 | 1.488549078 | 1.628935049 | 1.73239376 | 1.496250175 |
| ENSMUSG00000021258 | 2.315764015 | 2.153677461 | 2.132845365 | 2.093759858 | 2.075570129 | 2.322219295 | 1.975716277 |
| ENSMUSG00000021259 | 0.578129653 | 0.701838731 | 0.646240625 | 0.580482024 | 0.686820904 | 0.698970004 | 0.520678036 |
| ENSMUSG00000021260 | 0.91631382 | 0.976722649 | 0.864857905 | 0.830482024 | 0.632412477 | 0.698970004 | 0.876096088 |
| ENSMUSG00000021262 | 2.276824003 | 2.088286706 | 2.120953944 | 1.982684334 | 2.033751322 | 2.222716471 | 1.941550454 |
| ENSMUSG00000021263 | 1.494443473 | 1.42510993 | 1.388647213 | 1.201838731 | 1.302012594 | 1 | 1.159323753 |
| ENSMUSG00000021264 | 2.649978134 | 2.438304187 | 2.430701883 | 2.413659007 | 2.343123155 | 2.725911632 | 2.166993225 |
| ENSMUSG00000021265 | 1 | 1.238549078 | 1.214495249 | 1.114857905 | 1.293076518 | 1.278753601 | 1.296434211 |
| ENSMUSG00000021266 | 2.690713405 | 2.455044741 | 2.477473271 | 2.291854536 | 2.318243321 | 2.555094449 | 2.21474793 |
| ENSMUSG00000021268 | 2.776070581 | 2.520204382 | 2.549918086 | 1 | 1 | 1.146128036 | 0.803905701 |
| ENSMUSG00000021270 | 3.504189653 | 3.190906838 | 3.167442722 | 3.227847997 | 3.138895575 | 3.651181062 | 2.885840473 |
| ENSMUSG00000021271 | 1.714367558 | 1.671625132 | 1.714495249 | 1.557204673 | 1.519146132 | 1.73239376 | 1.537859885 |
| ENSMUSG00000021273 | 2.429630757 | 2.203445298 | 2.179419106 | 2.382845365 | 2.32560195 | 2.686636269 | 2.186723532 |
| ENSMUSG00000021275 | 2.23778581 | 2.105516192 | 2.110735874 | 1.981203126 | 2.073592779 | 2.274157849 | 1.934261938 |

| | | | | | | | |
|---|---|---|---|---|---|---|---|
| ENSMUSG00000021276 | 2.373064211 | 2.172499493 | 2.164552871 | 2.130890489 | 2.118236133 | 2.416640507 | 2.012457048 |
| ENSMUSG00000021277 | 2.590021402 | 2.418989258 | 2.439972046 | 2.272528105 | 2.336124456 | 2.584331224 | 2.214403581 |
| ENSMUSG00000021278 | 0 | 0 | 0 | 0.25 | 0.244650542 | 0 | 0 |
| ENSMUSG00000021279 | 2.786655728 | 2.601285366 | 2.652063705 | 2.424676167 | 2.462344774 | 2.699837726 | 2.387828426 |
| ENSMUSG00000021280 | 1.358731791 | 1.282320754 | 1.557204673 | 0.646240625 | 1.020174412 | 0.903089987 | 1.168187745 |
| ENSMUSG00000021281 | 2.557825653 | 2.402294685 | 2.437467357 | 2.162762923 | 2.218681522 | 2.480006943 | 2.126687118 |
| ENSMUSG00000021282 | 3.304745739 | 2.994498842 | 2.971519396 | 3.009901129 | 2.903147178 | 3.371990911 | 2.681143513 |
| ENSMUSG00000021285 | 2.151493169 | 2.032320754 | 1.953445298 | 1.83272922 | 1.903713872 | 2.117271296 | 1.832055226 |
| ENSMUSG00000021286 | 1.655740331 | 1.67510993 | 1.738549078 | 1.396240625 | 1.42077253 | 1.51851394 | 1.517723536 |
| ENSMUSG00000021287 | 1.928763473 | 1.777131114 | 1.845926073 | 1.610735874 | 1.701347897 | 1.612783857 | 1.633706455 |
| ENSMUSG00000021288 | 2.673935906 | 2.514159429 | 2.561981878 | 2.418106335 | 2.493248162 | 2.757396029 | 2.36247044 |
| ENSMUSG00000021290 | 2.420929319 | 2.144357207 | 2.213717096 | 2.164552871 | 2.109389652 | 2.45484486 | 1.98213686 |
| ENSMUSG00000021294 | 0 | 0 | 0 | 0.5 | 0.387761935 | 0.301029996 | 0.984950056 |
| ENSMUSG00000021298 | 0 | 0.25 | 0 | 0 | 0 | 0 | 0 |
| ENSMUSG00000021301 | 2.190368148 | 2.004202072 | 2.046218836 | 1.25 | 1.121713561 | 1.146128036 | 1.176815335 |
| ENSMUSG00000021302 | 2.139333929 | 2.061981878 | 2.032320754 | 2.167221246 | 2.029283463 | 2.338456494 | 1.913109605 |
| ENSMUSG00000021303 | 0 | 0 | 0 | 0.5 | 0.489301084 | 0 | 0.629532004 |
| ENSMUSG00000021306 | 2.473938397 | 2.253505118 | 2.274508021 | 2.115881093 | 2.079491904 | 2.305351369 | 1.927642465 |
| ENSMUSG00000021311 | 2.129344651 | 2.002806814 | 1.982684334 | 2.142463902 | 2.110284345 | 2.33243846 | 1.965468324 |
| ENSMUSG00000021313 | 0.289064826 | 0.25 | 0 | 0 | 0.244650542 | 0 | 0 |
| ENSMUSG00000021314 | 0 | 0.25 | 0.396240625 | 1.04248125 | 0.846351821 | 1.041392685 | 0.876096088 |
| ENSMUSG00000021318 | 2.574668783 | 2.363831805 | 2.362802778 | 2.47934302 | 2.4345872 | 2.748188027 | 2.284450788 |
| ENSMUSG00000021319 | 1.129344651 | 1.29248125 | 1.282320754 | 0.5 | 0.877063019 | 0.602059991 | 0.775756176 |
| ENSMUSG00000021322 | 0 | 0 | 0.25 | 0 | 0 | 0 | 0 |
| ENSMUSG00000021326 | 2.527823874 | 2.312574604 | 2.343759858 | 2.416777886 | 2.372368362 | 2.696356389 | 2.199238773 |
| ENSMUSG00000021327 | 2.627631768 | 2.434100483 | 2.416333979 | 2.310793496 | 2.24889942 | 2.526339277 | 2.062969708 |
| ENSMUSG00000021335 | 0 | 0 | 0 | 0 | 0 | 0 | 0 |
| ENSMUSG00000021336 | 0 | 0 | 0 | 0 | 0 | 0 | 0 |
| ENSMUSG00000021337 | 0 | 0 | 0 | 0 | 0 | 0 | 0 |
| ENSMUSG00000021338 | 1.985980784 | 1.874961472 | 1.866901388 | 1.953445298 | 1.968065382 | 2.222716471 | 1.904245613 |
| ENSMUSG00000021339 | 2.051791411 | 1.953445298 | 1.87694866 | 2.013820609 | 1.958580386 | 2.227886705 | 1.826389311 |
| ENSMUSG00000021340 | 1.647796617 | 1.557204673 | 1.5 | 1.747171172 | 1.763796674 | 1.86923172 | 1.653807046 |
| ENSMUSG00000021342 | 0 | 0 | 0.25 | 0 | 0 | 0 | 0 |
| ENSMUSG00000021345 | 0 | 0 | 0 | 0 | 0 | 0 | 0 |

| | | | | | | | |
|---|---|---|---|---|---|---|---|
| ENSMUSG00000021346 | 0 | 0 | 0 | 0 | 0 | 0 | 0 |
| ENSMUSG00000021347 | 0 | 0 | 0 | 0 | 0 | 0 | 0 |
| ENSMUSG00000021348 | 0 | 0 | 0 | 0 | 0 | 0 | 0 |
| ENSMUSG00000021356 | 0 | 0.25 | 0.396240625 | 0.25 | 0.244650542 | 0.301029996 | 0.355418052 |
| ENSMUSG00000021357 | 2.467247894 | 2.272528105 | 2.293731421 | 2.362287161 | 2.306140782 | 2.649334859 | 2.194510526 |
| ENSMUSG00000021359 | 0.289064826 | 0.25 | 0.25 | 0.580482024 | 0 | 0 | 0.448487648 |
| ENSMUSG00000021360 | 2.126809492 | 1.970660762 | 1.914552871 | 1.618933358 | 1.504538967 | 1.568201724 | 1.455757965 |
| ENSMUSG00000021361 | 2.438154344 | 2.30357978 | 2.345926073 | 2.360735874 | 2.408704478 | 2.700703717 | 2.26501115 |
| ENSMUSG00000021362 | 0 | 0 | 0 | 0 | 0 | 0 | 0 |
| ENSMUSG00000021363 | 0 | 0 | 0 | 0.5 | 0.489301084 | 0.301029996 | 0.355418052 |
| ENSMUSG00000021364 | 0 | 0 | 0.396240625 | 0.864857905 | 0.489301084 | 0.602059991 | 0.629532004 |
| ENSMUSG00000021365 | 2.43085913 | 2.241446071 | 2.187048212 | 2.251406137 | 2.183460217 | 2.505149978 | 2.081675233 |
| ENSMUSG00000021366 | 2.249317394 | 2.134789703 | 2.119945066 | 1.955044741 | 1.964031554 | 2.100370545 | 1.832055226 |
| ENSMUSG00000021367 | 1.693336259 | 1.598079356 | 1.62694866 | 1.188721876 | 1.176121988 | 1.230448921 | 0.952571511 |
| ENSMUSG00000021368 | 1.952600383 | 1.763820609 | 1.752806814 | 1.92510993 | 1.906908067 | 2.1430148 | 1.791413205 |
| ENSMUSG00000021371 | 2.066740409 | 1.912762923 | 1.942046081 | 2.138647213 | 2.097547432 | 2.378397901 | 1.932619724 |
| ENSMUSG00000021373 | 1.655740331 | 1.571350555 | 1.630890489 | 1.26109853 | 1.293076518 | 1.301029996 | 1.259064007 |
| ENSMUSG00000021374 | 2.982727785 | 2.703445298 | 2.700224975 | 2.820772088 | 2.746400488 | 3.163757524 | 2.53390597 |
| ENSMUSG00000021375 | 2.365959569 | 2.221424093 | 2.224461364 | 2.201032755 | 2.230318492 | 2.489958479 | 2.101807105 |
| ENSMUSG00000021376 | 1.614416215 | 1.552363341 | 1.571350555 | 1.634789703 | 1.546663136 | 1.799340549 | 1.425641774 |
| ENSMUSG00000021377 | 3.07519763 | 2.862673943 | 2.836628433 | 3.03199858 | 2.945495772 | 3.428782511 | 2.719049087 |
| ENSMUSG00000021379 | 0.747221736 | 1 | 0.830482024 | 0.864857905 | 0.568060967 | 0.954242509 | 0.629532004 |
| ENSMUSG00000021381 | 0 | 0 | 0 | 0.396240625 | 0.568060967 | 0.301029996 | 0.448487648 |
| ENSMUSG00000021384 | 0 | 0 | 0.396240625 | 0 | 0.387761935 | 0.698970004 | 0.629532004 |
| ENSMUSG00000021385 | 2.020182925 | 1.950224975 | 1.92510993 | 2.015173983 | 1.968065382 | 2.245512668 | 1.832055226 |
| ENSMUSG00000021388 | 0.747221736 | 0.92510993 | 0.701838731 | 0.25 | 0.568060967 | 0.602059991 | 0.355418052 |
| ENSMUSG00000021390 | 2.620687497 | 2.392463902 | 2.23927551 | 1.858657057 | 1.797870327 | 1.929418926 | 1.676387053 |
| ENSMUSG00000021391 | 1.890240806 | 1.802363341 | 1.802363341 | 1.870953944 | 1.850106709 | 2.181843588 | 1.731408809 |
| ENSMUSG00000021392 | 2.351376531 | 2.201838731 | 2.207430684 | 2.313166358 | 2.243427123 | 2.537819095 | 2.033014511 |
| ENSMUSG00000021395 | 2.748369006 | 2.523189285 | 2.461764337 | 2.524178789 | 2.477187895 | 2.822168079 | 2.295362967 |
| ENSMUSG00000021396 | 0 | 0 | 0 | 0 | 0 | 0 | 0 |
| ENSMUSG00000021400 | 2.496544396 | 2.311388177 | 2.299304173 | 2.337182039 | 2.352243976 | 2.64738297 | 2.233468922 |
| ENSMUSG00000021403 | 2.163307908 | 2.00835575 | 2.110735874 | 1.79248125 | 1.706995316 | 1.913813852 | 1.618853957 |
| ENSMUSG00000021404 | 0.91631382 | 0.75 | 0.896240625 | 0.25 | 0.489301084 | 0 | 0.224243824 |

| | | | | | | | |
|---|---|---|---|---|---|---|---|
| ENSMUSG00000021408 | 2.287337982 | 2.084962501 | 2.110735874 | 2.267865591 | 2.180543208 | 2.5132176 | 2.02568932 |
| ENSMUSG00000021411 | 1.867194479 | 1.67510993 | 1.782320754 | 1.769203899 | 1.804307395 | 1.851258349 | 1.720494 |
| ENSMUSG00000021413 | 2.796468708 | 2.556000419 | 2.533285553 | 2.664552871 | 2.563636668 | 2.910624405 | 2.379663565 |
| ENSMUSG00000021414 | 0 | 0 | 0 | 0.25 | 0 | 0 | 0.224243824 |
| ENSMUSG00000021415 | 0 | 0.25 | 0 | 0 | 0 | 0 | 0 |
| ENSMUSG00000021416 | 0.91631382 | 0.580482024 | 0.864857905 | 0.951838731 | 0.846351821 | 1.113943352 | 0.775756176 |
| ENSMUSG00000021417 | 2.346900291 | 2.183677405 | 2.158248799 | 2.300530956 | 2.275056283 | 2.58546073 | 2.123964672 |
| ENSMUSG00000021418 | 1.87190676 | 1.649978211 | 1.741446071 | 1.735628626 | 1.671670246 | 1.812913357 | 1.655750087 |
| ENSMUSG00000021420 | 1.678702042 | 1.593759858 | 1.660964047 | 1.606566189 | 1.718026092 | 1.919078092 | 1.594042709 |
| ENSMUSG00000021423 | 0 | 0 | 0 | 0.25 | 0 | 0 | 0 |
| ENSMUSG00000021427 | 3.198427946 | 2.904825739 | 2.896592674 | 3.06589066 | 2.97205363 | 3.457579147 | 2.75968608 |
| ENSMUSG00000021428 | 2.260538674 | 2.098079356 | 2.056000419 | 2.259729747 | 2.176862714 | 2.480006943 | 2.005303651 |
| ENSMUSG00000021431 | 2.013563224 | 1.880890489 | 1.880890489 | 2.059601185 | 1.998776581 | 2.133538908 | 1.86815666 |
| ENSMUSG00000021432 | 1.777675583 | 1.738549078 | 1.735628626 | 1.845926073 | 1.84445929 | 2.08278537 | 1.69749337 |
| ENSMUSG00000021439 | 0.289064826 | 0 | 0.25 | 0 | 0 | 0 | 0 |
| ENSMUSG00000021440 | 0 | 0 | 0 | 0 | 0 | 0 | 0 |
| ENSMUSG00000021441 | 0 | 0 | 0 | 0 | 0 | 0 | 0 |
| ENSMUSG00000021448 | 0 | 0 | 0 | 0.25 | 0.387761935 | 0.477121255 | 0 |
| ENSMUSG00000021451 | 1.205378646 | 1.339388001 | 1.26109853 | 1.589388001 | 1.625423019 | 1.698970004 | 1.496250175 |
| ENSMUSG00000021453 | 1.920505136 | 1.953445298 | 2.100219859 | 1.602347734 | 1.818885826 | 1.740362689 | 2.017561934 |
| ENSMUSG00000021456 | 0 | 0 | 0 | 0 | 0 | 0 | 0 |
| ENSMUSG00000021457 | 2.495495257 | 2.313757142 | 2.338837774 | 2.114857905 | 2.146756411 | 2.390935107 | 2.020712061 |
| ENSMUSG00000021458 | 1.982401085 | 1.87694866 | 1.834962501 | 1.94697564 | 2.012006468 | 2.230448921 | 1.838725884 |
| ENSMUSG00000021460 | 1.952600383 | 1.870953944 | 1.839388001 | 1.923371739 | 1.900490506 | 2.130333768 | 1.784981274 |
| ENSMUSG00000021461 | 1.876566388 | 1.668106335 | 1.726722649 | 1.872963274 | 1.782377602 | 2.056904851 | 1.704233519 |
| ENSMUSG00000021464 | 1.342375483 | 1.302363341 | 1.494319981 | 1.638647213 | 1.78010778 | 1.857332496 | 1.82753046 |
| ENSMUSG00000021466 | 2.260538674 | 2.182829758 | 2.155512955 | 2.086073977 | 2.070605826 | 2.322219295 | 1.977870661 |
| ENSMUSG00000021468 | 2.37447073 | 2.211372513 | 2.162762923 | 2.337734795 | 2.275616087 | 2.602059991 | 2.119837558 |
| ENSMUSG00000021469 | 2.172524468 | 2.019203899 | 2.02053726 | 1.114857905 | 1.057362051 | 1.278753601 | 1.168187745 |
| ENSMUSG00000021470 | 2.235832493 | 2.083847589 | 2.052363341 | 2.317865757 | 2.216704172 | 2.544068044 | 2.059095225 |
| ENSMUSG00000021474 | 2.800532393 | 2.539336734 | 2.546838018 | 2.796992648 | 2.736385816 | 3.205475037 | 2.557890211 |
| ENSMUSG00000021476 | 2.136858952 | 1.985628626 | 2.028435542 | 2.301142786 | 2.25190332 | 2.5289167 | 2.130281771 |
| ENSMUSG00000021477 | 3.039433992 | 2.755071375 | 2.774178789 | 2.746282593 | 2.672579693 | 3.135450699 | 2.497401477 |
| ENSMUSG00000021478 | 0 | 0 | 0 | 0.25 | 0.489301084 | 0 | 0.224243824 |

| | | | | | | | |
|---|---|---|---|---|---|---|---|
| ENSMUSG00000021481 | 2.199011616 | 2.01109853 | 2.02186571 | 2.149978211 | 2.11997913 | 2.389166084 | 2.005960535 |
| ENSMUSG00000021482 | 1.693336259 | 1.552363341 | 1.584962501 | 1.657339155 | 1.628935049 | 1.716003344 | 1.562034107 |
| ENSMUSG00000021483 | 1.227926429 | 1.26109853 | 1.311981878 | 1.04248125 | 1.149965125 | 1.204119983 | 1.252393349 |
| ENSMUSG00000021484 | 2.774459379 | 2.612029076 | 2.652294685 | 2.760927716 | 2.803054443 | 3.166430114 | 2.654338125 |
| ENSMUSG00000021485 | 1.605636262 | 1.692046081 | 1.816696635 | 1.505591953 | 1.614672255 | 1.556302501 | 1.732938436 |
| ENSMUSG00000021486 | 2.84689048 | 2.676839783 | 2.717205639 | 2.713911792 | 2.781444737 | 3.133858125 | 2.663665499 |
| ENSMUSG00000021488 | 2.932463873 | 2.671187155 | 2.663211243 | 2.860606298 | 2.778227492 | 3.194514342 | 2.560619314 |
| ENSMUSG00000021490 | 0 | 0 | 0 | 0 | 0 | 0 | 0 |
| ENSMUSG00000021492 | 0 | 0 | 0 | 0 | 0 | 0 | 0 |
| ENSMUSG00000021493 | 2.723587515 | 2.548689214 | 2.595926073 | 2.129909063 | 2.271678687 | 2.414973348 | 2.228559641 |
| ENSMUSG00000021494 | 2.381433035 | 2.237091808 | 2.251406137 | 2.312574604 | 2.338472809 | 2.610660163 | 2.229218532 |
| ENSMUSG00000021495 | 2.203267085 | 2.072504712 | 2.162762923 | 2.025821952 | 2.212046813 | 2.281033367 | 2.15146804 |
| ENSMUSG00000021496 | 1.527823874 | 1.403677461 | 1.330482024 | 1.396240625 | 1.53772706 | 1.579783597 | 1.405288179 |
| ENSMUSG00000021497 | 2.336265009 | 2.149978211 | 2.179419106 | 2.412314044 | 2.324629618 | 2.688419822 | 2.257861346 |
| ENSMUSG00000021499 | 0.578129653 | 0.580482024 | 0.396240625 | 0.701838731 | 0.733951626 | 0.698970004 | 0.579661877 |
| ENSMUSG00000021500 | 2.860681608 | 2.620953944 | 2.627938163 | 2.79745614 | 2.718345365 | 3.116939647 | 2.537606247 |
| ENSMUSG00000021501 | 1.932832114 | 1.884789703 | 1.931980114 | 2.077084758 | 2.121713561 | 2.260071388 | 2.006616089 |
| ENSMUSG00000021504 | 1.727821383 | 1.76109853 | 1.802363341 | 1.738549078 | 1.810629166 | 1.977723605 | 1.793950594 |
| ENSMUSG00000021506 | 0 | 0 | 0 | 0 | 0 | 0 | 0 |
| ENSMUSG00000021508 | 0 | 0 | 0 | 0.396240625 | 0.244650542 | 0 | 0.520678036 |
| ENSMUSG00000021509 | 0.289064826 | 0.701838731 | 0.646240625 | 1.372963274 | 1.42077253 | 1.568201724 | 1.401059159 |
| ENSMUSG00000021510 | 1.811507563 | 1.708222504 | 1.668106335 | 1.862802778 | 1.812711509 | 1.959041392 | 1.635773653 |
| ENSMUSG00000021514 | 1.982401085 | 1.858657057 | 1.834962501 | 1.956637122 | 1.887294348 | 2.133538908 | 1.81475081 |
| ENSMUSG00000021518 | 2.573799059 | 2.362802778 | 2.3464656 | 2.590485943 | 2.48127199 | 2.893206753 | 2.336611718 |
| ENSMUSG00000021519 | 2.22189694 | 2.05479213 | 2.072504712 | 2.211372513 | 2.139428339 | 2.390935107 | 1.95718226 |
| ENSMUSG00000021520 | 2.266038081 | 2.113831805 | 2.101285366 | 2.399513125 | 2.318243321 | 2.63748973 | 2.150210039 |
| ENSMUSG00000021532 | 2.029919532 | 1.864857905 | 1.956637122 | 1.937048212 | 1.954436039 | 2.133538908 | 1.795211864 |
| ENSMUSG00000021534 | 0 | 0 | 0 | 0.25 | 0.244650542 | 0.477121255 | 0.355418052 |
| ENSMUSG00000021536 | 0 | 0 | 0 | 0 | 0 | 0 | 0 |
| ENSMUSG00000021537 | 2.556920045 | 2.338286706 | 2.369439567 | 2.537119146 | 2.402902329 | 2.816903839 | 2.266774181 |
| ENSMUSG00000021538 | 0 | 0 | 0 | 0 | 0 | 0 | 0 |
| ENSMUSG00000021539 | 0 | 0 | 0 | 0 | 0 | 0 | 0 |
| ENSMUSG00000021540 | 2.900309677 | 2.62019755 | 2.613317659 | 2.7543761 | 2.691155963 | 3.077004327 | 2.479989695 |
| ENSMUSG00000021541 | 0 | 0 | 0 | 0 | 0 | 0 | 0 |

| | | | | | | | |
|---|---|---|---|---|---|---|---|
| ENSMUSG00000021545 | 0 | 0 | 0 | 0 | 0 | 0 | 0 |
| ENSMUSG00000021546 | 3.537433341 | 3.207034121 | 3.195851886 | 3.277906187 | 3.183844492 | 3.722222464 | 2.982842456 |
| ENSMUSG00000021548 | 2.196867486 | 2.081607372 | 2.079353153 | 2.136723615 | 2.040348824 | 2.365487985 | 1.925966215 |
| ENSMUSG00000021549 | 2.790815687 | 2.524836953 | 2.526149688 | 2.479715809 | 2.393415815 | 2.73479983 | 2.182568577 |
| ENSMUSG00000021550 | 0.747221736 | 0.646240625 | 0.701838731 | 2.039967834 | 1.969399812 | 2.235528447 | 1.874129423 |
| ENSMUSG00000021552 | 2.151493169 | 2.009729747 | 1.973704441 | 2.338286706 | 2.219338185 | 2.547774705 | 2.080634989 |
| ENSMUSG00000021553 | 0.45815691 | 0.25 | 0.25 | 0 | 0.244650542 | 0.301029996 | 0.448487648 |
| ENSMUSG00000021555 | 2.612039956 | 2.399047439 | 2.350219859 | 2.540597832 | 2.452316937 | 2.848804701 | 2.301737862 |
| ENSMUSG00000021556 | 2.589183146 | 2.373463862 | 2.392463902 | 2.720087202 | 2.677271408 | 3.094121596 | 2.499691862 |
| ENSMUSG00000021557 | 2.390989024 | 2.245748394 | 2.201838731 | 2.402756199 | 2.314756076 | 2.672097858 | 2.170168862 |
| ENSMUSG00000021559 | 0 | 0 | 0 | 0.75 | 0.489301084 | 0.698970004 | 0.74492186 |
| ENSMUSG00000021565 | 0 | 0 | 0 | 0.25 | 0 | 0.477121255 | 0.224243824 |
| ENSMUSG00000021566 | 0 | 0 | 0 | 0 | 0 | 0 | 0 |
| ENSMUSG00000021567 | 1 | 1.04248125 | 0.976722649 | 1.53743678 | 1.467903253 | 1.716003344 | 1.48984372 |
| ENSMUSG00000021569 | 2.460448303 | 2.240000483 | 2.228222334 | 2.445339928 | 2.379941013 | 2.719331287 | 2.182188194 |
| ENSMUSG00000021572 | 1.516991255 | 1.580482024 | 1.482684334 | 1.698603967 | 1.698489937 | 1.897627091 | 1.663407615 |
| ENSMUSG00000021573 | 0.45815691 | 0.864857905 | 0.701838731 | 0.830482024 | 0.632412477 | 0.77815125 | 0.876096088 |
| ENSMUSG00000021575 | 0.747221736 | 0.92510993 | 0.864857905 | 1.769203899 | 1.72341484 | 1.924279286 | 1.574722613 |
| ENSMUSG00000021576 | 2.4661223 | 2.304186465 | 2.316696635 | 2.501055117 | 2.426655268 | 2.814247596 | 2.311070686 |
| ENSMUSG00000021577 | 3.033693607 | 2.762291968 | 2.750879478 | 2.942872367 | 2.885112539 | 3.361538971 | 2.687272105 |
| ENSMUSG00000021578 | 2.539328946 | 2.365881093 | 2.340485943 | 2.501055117 | 2.442693389 | 2.77524626 | 2.237022298 |
| ENSMUSG00000021579 | 0 | 0 | 0 | 0 | 0 | 0 | 0 |
| ENSMUSG00000021583 | 2.134369199 | 1.966046536 | 1.990000483 | 2.43325383 | 2.309205544 | 2.711807229 | 2.130728307 |
| ENSMUSG00000021585 | 2.771760131 | 2.526804269 | 2.540912419 | 2.696158711 | 2.611878676 | 3.004321374 | 2.410967356 |
| ENSMUSG00000021587 | 0 | 0 | 0 | 0 | 0.244650542 | 0 | 0.448487648 |
| ENSMUSG00000021589 | 2.787699615 | 2.5684489 | 2.519537702 | 2.648581151 | 2.541170562 | 2.919601024 | 2.348663831 |
| ENSMUSG00000021590 | 0.578129653 | 1.04248125 | 0.896240625 | 1 | 1.020174412 | 1.278753601 | 0.916588299 |
| ENSMUSG00000021591 | 2.199011616 | 2.116901388 | 2.051142786 | 1.992885888 | 1.893954095 | 2.220108088 | 1.813563596 |
| ENSMUSG00000021592 | 1.721148722 | 1.566696635 | 1.527131114 | 1.692046081 | 1.642643806 | 1.903089987 | 1.499406449 |
| ENSMUSG00000021594 | 1.53838222 | 1.598079356 | 1.575945187 | 1.774508021 | 1.77552387 | 1.934498451 | 1.68886578 |
| ENSMUSG00000021595 | 2.95990777 | 2.724650347 | 2.712351275 | 2.818011631 | 2.779034557 | 3.168202747 | 2.564144762 |
| ENSMUSG00000021596 | 0.671187741 | 0.5 | 0.646240625 | 0.396240625 | 0.489301084 | 0 | 0.355418052 |
| ENSMUSG00000021597 | 2.299278375 | 2.139605178 | 2.118933358 | 2.341033664 | 2.304598364 | 2.586587305 | 2.096394553 |
| ENSMUSG00000021598 | 0 | 0 | 0 | 0 | 0 | 0 | 0 |

| | | | | | | | |
|---|---|---|---|---|---|---|---|
| ENSMUSG00000021600 | 0 | 0 | 0 | 0 | 0 | 0 | 0 |
| ENSMUSG00000021604 | 0 | 0 | 0 | 0 | 0 | 0 | 0 |
| ENSMUSG00000021606 | 1.993049299 | 1.898114259 | 1.942046081 | 1.935366747 | 1.998776581 | 2.209515015 | 1.943943921 |
| ENSMUSG00000021607 | 1.800532393 | 1.75 | 1.807204673 | 1.896240625 | 1.846351821 | 2.089905111 | 1.855869786 |
| ENSMUSG00000021608 | 2.423434025 | 2.271202097 | 2.256976499 | 2.464495249 | 2.447194116 | 2.777426822 | 2.302000776 |
| ENSMUSG00000021609 | 0 | 0 | 0 | 0.25 | 0.387761935 | 0.602059991 | 0.448487648 |
| ENSMUSG00000021610 | 2.793908648 | 2.581607372 | 2.555698726 | 2.718553234 | 2.658874533 | 3.084576278 | 2.484506038 |
| ENSMUSG00000021611 | 0.671187741 | 0.79248125 | 0.25 | 1.410964047 | 1.380772476 | 1.397940009 | 1.329931221 |
| ENSMUSG00000021612 | 0.867194479 | 0.701838731 | 0.830482024 | 0.830482024 | 0.812711509 | 0.77815125 | 0.629532004 |
| ENSMUSG00000021613 | 0 | 0 | 0 | 0 | 0 | 0 | 0 |
| ENSMUSG00000021614 | 2.742600832 | 2.497880462 | 2.524178789 | 2.286164561 | 2.181274723 | 2.489958479 | 2.015657029 |
| ENSMUSG00000021615 | 1.707474304 | 1.630890489 | 1.571350555 | 1.784887838 | 1.639266209 | 1.944482672 | 1.579661877 |
| ENSMUSG00000021619 | 1.181541736 | 1.130890489 | 1.214495249 | 1.238549078 | 1.188507684 | 0.954242509 | 1.110947927 |
| ENSMUSG00000021620 | 0 | 0 | 0 | 0 | 0.244650542 | 0 | 0 |
| ENSMUSG00000021621 | 2.136858952 | 1.96293726 | 1.991446071 | 2.074802005 | 2.094757251 | 2.269512944 | 1.927642465 |
| ENSMUSG00000021622 | 0 | 0 | 0 | 0 | 0 | 0 | 0 |
| ENSMUSG00000021624 | 0 | 0.25 | 0 | 0.396240625 | 0 | 0 | 0 |
| ENSMUSG00000021629 | 2.5496035 | 2.37246199 | 2.345926073 | 2.518200634 | 2.454345426 | 2.802773725 | 2.272294561 |
| ENSMUSG00000021631 | 0 | 0 | 0 | 0 | 0 | 0 | 0 |
| ENSMUSG00000021635 | 2.302628051 | 2.115881093 | 2.078220739 | 2.174241882 | 2.103973339 | 2.414973348 | 2.003324957 |
| ENSMUSG00000021636 | 0 | 0 | 0.396240625 | 0 | 0 | 0 | 0 |
| ENSMUSG00000021638 | 0 | 0.25 | 0 | 0.79248125 | 0.905314583 | 0.77815125 | 0.672731473 |
| ENSMUSG00000021639 | 2.081172036 | 1.969129237 | 1.943696765 | 2.179419106 | 2.153935427 | 2.482873584 | 2.006616089 |
| ENSMUSG00000021640 | 0 | 0 | 0 | 0 | 0 | 0 | 0 |
| ENSMUSG00000021643 | 1.558729299 | 1.557204673 | 1.614857905 | 1.494319981 | 1.49445382 | 1.707570176 | 1.483307832 |
| ENSMUSG00000021645 | 2.144240249 | 1.994319981 | 1.953445298 | 2.184523065 | 2.10031573 | 2.390935107 | 2.050061048 |
| ENSMUSG00000021646 | 1.924654746 | 1.882845365 | 1.850219859 | 2.080482024 | 1.998776581 | 2.230448921 | 1.907820406 |
| ENSMUSG00000021647 | 0 | 0 | 0 | 0 | 0 | 0 | 0 |
| ENSMUSG00000021650 | 2.086807695 | 1.973704441 | 1.955044741 | 2.129909063 | 2.075570129 | 2.330413773 | 1.920884716 |
| ENSMUSG00000021660 | 3.133460136 | 2.847676593 | 2.828504176 | 3.070844447 | 2.974224243 | 3.486855355 | 2.770611179 |
| ENSMUSG00000021661 | 1.837744666 | 1.816696635 | 1.79248125 | 1.850219859 | 1.826953746 | 2.017033339 | 1.702561601 |
| ENSMUSG00000021662 | 2.330063669 | 2.155512955 | 2.165444524 | 2.155512955 | 2.128567294 | 2.376576957 | 1.98213686 |
| ENSMUSG00000021665 | 2.219867581 | 2.049918086 | 2.04248125 | 2.123963757 | 2.004851707 | 2.340444115 | 1.915722126 |
| ENSMUSG00000021666 | 2.177057474 | 2.016522298 | 2.015173983 | 2.130890489 | 2.147561328 | 2.396199347 | 2.008574816 |

| | | | | | | | |
|---|---|---|---|---|---|---|---|
| ENSMUSG00000021668 | 2.235832493 | 2.110735874 | 2.103406982 | 2.174241882 | 2.09661982 | 2.330413773 | 1.935895857 |
| ENSMUSG00000021669 | 2.703665763 | 2.48927551 | 2.457034121 | 2.686418581 | 2.601368039 | 2.992995098 | 2.38702115 |
| ENSMUSG00000021670 | 2.682073381 | 2.442046081 | 2.442459461 | 2.614088604 | 2.596218572 | 2.936010796 | 2.411717103 |
| ENSMUSG00000021671 | 2.095120908 | 1.916333979 | 1.903677461 | 2.124961472 | 2.075570129 | 2.318063335 | 1.96172822 |
| ENSMUSG00000021675 | 0.289064826 | 0.25 | 0.25 | 0.5 | 0.632412477 | 0.301029996 | 0.629532004 |
| ENSMUSG00000021676 | 0.578129653 | 0.646240625 | 0.701838731 | 1.42510993 | 1.489301084 | 1.602059991 | 1.360283038 |
| ENSMUSG00000021678 | 0.45815691 | 0.25 | 0.5 | 1.732684334 | 1.695608647 | 1.740362689 | 1.540636858 |
| ENSMUSG00000021679 | 0 | 0 | 0 | 0 | 0 | 0 | 0 |
| ENSMUSG00000021680 | 0 | 0 | 0 | 0.79248125 | 0.568060967 | 0.301029996 | 0.74492186 |
| ENSMUSG00000021681 | 2.427163103 | 2.259043403 | 2.289967834 | 2.422499493 | 2.371090306 | 2.723455672 | 2.240219117 |
| ENSMUSG00000021684 | 1.325351389 | 1.302363341 | 1.403677461 | 1.214495249 | 1.106691885 | 1 | 1.131174228 |
| ENSMUSG00000021685 | 0 | 0.25 | 0 | 0 | 0.244650542 | 0 | 0 |
| ENSMUSG00000021686 | 2.619128496 | 2.430274797 | 2.413659007 | 2.503853763 | 2.452994396 | 2.809559715 | 2.280261923 |
| ENSMUSG00000021687 | 2.386920383 | 2.256976499 | 2.26109853 | 2.462155735 | 2.386129765 | 2.7355989 | 2.193776931 |
| ENSMUSG00000021690 | 2.267855219 | 2.136723615 | 2.139605178 | 2.324802005 | 2.203230928 | 2.527629901 | 2.062419043 |
| ENSMUSG00000021692 | 2.023453784 | 1.874961472 | 1.845926073 | 2.066696635 | 1.975997314 | 2.250420002 | 1.86815666 |
| ENSMUSG00000021693 | 2.655024396 | 2.405054956 | 2.390083209 | 2.585240691 | 2.491735274 | 2.840106094 | 2.282223478 |
| ENSMUSG00000021694 | 1.765759561 | 1.714495249 | 1.688721876 | 1.841580554 | 1.791313678 | 2.021189299 | 1.71890424 |
| ENSMUSG00000021696 | 1.596667477 | 1.42510993 | 1.527131114 | 1.720660762 | 1.695608647 | 1.84509804 | 1.526507301 |
| ENSMUSG00000021697 | 1.948720984 | 1.789967834 | 1.802363341 | 1.825945187 | 1.806427258 | 1.963787827 | 1.67819947 |
| ENSMUSG00000021699 | 2.243591459 | 2.106566189 | 2.134789703 | 1.580482024 | 1.592135487 | 1.643452676 | 1.517723536 |
| ENSMUSG00000021700 | 0.289064826 | 0.25 | 0.646240625 | 0.25 | 0 | 0.301029996 | 0 |
| ENSMUSG00000021701 | 2.940490991 | 2.698603967 | 2.69225283 | 2.947688938 | 2.88560959 | 3.316808752 | 2.674730545 |
| ENSMUSG00000021702 | 0 | 0 | 0 | 0.580482024 | 0.686820904 | 0 | 0 |
| ENSMUSG00000021703 | 2.188178997 | 2.005591953 | 1.991446071 | 2.343216265 | 2.298884454 | 2.605305046 | 2.13206423 |
| ENSMUSG00000021704 | 2.121692466 | 1.991446071 | 1.96759118 | 2.211372513 | 2.185632262 | 2.382017043 | 1.997988498 |
| ENSMUSG00000021706 | 2.412042447 | 2.226722649 | 2.238549078 | 2.387205477 | 2.302531265 | 2.63447727 | 2.113781657 |
| ENSMUSG00000021707 | 2.559630991 | 2.386723615 | 2.348079356 | 2.648114259 | 2.552582334 | 2.967547976 | 2.370437612 |
| ENSMUSG00000021708 | 0.45815691 | 0.75 | 0.25 | 0 | 0.387761935 | 0.602059991 | 0.355418052 |
| ENSMUSG00000021709 | 3.114849914 | 2.837043717 | 2.80175358 | 2.87007133 | 2.763375569 | 3.19893187 | 2.537479353 |
| ENSMUSG00000021710 | 2.677345838 | 2.462155735 | 2.472566069 | 2.541226732 | 2.493248162 | 2.857935265 | 2.332093605 |
| ENSMUSG00000021711 | 2.626865868 | 2.438721876 | 2.399513125 | 2.605516192 | 2.520825546 | 2.906873535 | 2.356752139 |
| ENSMUSG00000021712 | 2.380049842 | 2.162762923 | 2.175975893 | 2.37694866 | 2.294670002 | 2.57634135 | 2.122136896 |
| ENSMUSG00000021713 | 2.29078455 | 2.079353153 | 2.038704527 | 2.284247778 | 2.212046813 | 2.5289167 | 2.07004434 |

| | | | | | | | |
|---|---|---|---|---|---|---|---|
| ENSMUSG00000021714 | 1.827447047 | 1.80479213 | 1.799918086 | 2.038704527 | 1.968065382 | 2.292256071 | 1.865128412 |
| ENSMUSG00000021715 | 2.042557845 | 1.942046081 | 1.937048212 | 2.023189285 | 1.923977404 | 2.164352856 | 1.836517579 |
| ENSMUSG00000021716 | 2.092368209 | 2.001406137 | 1.964495249 | 2.064346961 | 2.078515538 | 2.309630167 | 1.91917299 |
| ENSMUSG00000021718 | 0 | 0 | 0 | 0 | 0 | 0 | 0 |
| ENSMUSG00000021719 | 0 | 0 | 0 | 0 | 0 | 0.301029996 | 0 |
| ENSMUSG00000021720 | 0.289064826 | 0.25 | 0 | 0 | 0.568060967 | 0.301029996 | 0.224243824 |
| ENSMUSG00000021721 | 0 | 0 | 0 | 0 | 0 | 0 | 0 |
| ENSMUSG00000021725 | 2.333175866 | 2.160964047 | 2.127938163 | 2.178561379 | 2.12771782 | 2.404833717 | 1.993927582 |
| ENSMUSG00000021728 | 2.4661223 | 2.257666784 | 2.220660762 | 2.228969845 | 2.10031573 | 2.439332694 | 2.036615833 |
| ENSMUSG00000021730 | 0 | 0.396240625 | 0.5 | 0.25 | 0.387761935 | 0.301029996 | 0 |
| ENSMUSG00000021731 | 2.179305632 | 2.056000419 | 2.029735268 | 2.181128463 | 2.141887974 | 2.45484486 | 1.987044015 |
| ENSMUSG00000021732 | 1.655740331 | 1.614857905 | 1.671625132 | 1.482684334 | 1.49445382 | 1.477121255 | 1.350478789 |
| ENSMUSG00000021733 | 2.716846213 | 2.488185312 | 2.463717096 | 2.5658173 | 2.466603229 | 2.831229694 | 2.281104046 |
| ENSMUSG00000021737 | 2.517987749 | 2.347542239 | 2.37246199 | 2.678990498 | 2.58731141 | 2.967079734 | 2.419851634 |
| ENSMUSG00000021738 | 2.100572389 | 1.942046081 | 1.901832578 | 2.101285366 | 2.05001946 | 2.346352974 | 1.950241257 |
| ENSMUSG00000021741 | 0 | 0 | 0.25 | 0 | 0 | 0 | 0.224243824 |
| ENSMUSG00000021743 | 0 | 0 | 0 | 0 | 0 | 0 | 0 |
| ENSMUSG00000021745 | 2.048736215 | 1.894357207 | 1.981203126 | 2.036164561 | 1.997548906 | 2.184691431 | 1.854824501 |
| ENSMUSG00000021747 | 1.45815691 | 1.52186571 | 1.431980114 | 1.451838731 | 1.358933174 | 1.612783857 | 1.379042812 |
| ENSMUSG00000021748 | 2.37447073 | 2.185366747 | 2.192046081 | 2.482314602 | 2.449252175 | 2.780317312 | 2.274304872 |
| ENSMUSG00000021749 | 0 | 0 | 0 | 0 | 0 | 0 | 0 |
| ENSMUSG00000021750 | 0 | 0.25 | 0.25 | 0.5 | 0.387761935 | 0.602059991 | 0.629532004 |
| ENSMUSG00000021751 | 0.45815691 | 0.25 | 0 | 0.5 | 0.387761935 | 0 | 0.355418052 |
| ENSMUSG00000021752 | 1.655740331 | 1.494319981 | 1.516522298 | 1.79745614 | 1.782377602 | 1.986771734 | 1.67819947 |
| ENSMUSG00000021754 | 1.671187741 | 1.681980114 | 1.642463902 | 1.899978211 | 1.832885921 | 2.056904851 | 1.812372009 |
| ENSMUSG00000021756 | 2.991772994 | 2.707430684 | 2.694108258 | 3.051371955 | 2.97368284 | 3.452706227 | 2.768691568 |
| ENSMUSG00000021758 | 0.289064826 | 0.25 | 0 | 0.396240625 | 0.244650542 | 0 | 0.355418052 |
| ENSMUSG00000021759 | 2.179305632 | 2.057204673 | 2.039967834 | 2.129909063 | 2.072599934 | 2.365487985 | 1.97354745 |
| ENSMUSG00000021760 | 2.511972754 | 2.324229052 | 2.326515422 | 2.532642641 | 2.458367761 | 2.80685803 | 2.292127786 |
| ENSMUSG00000021763 | 0.578129653 | 0.25 | 0.5 | 1.598079356 | 1.580303447 | 1.763427994 | 1.296434211 |
| ENSMUSG00000021764 | 2.478339835 | 2.254897682 | 2.286801231 | 2.466046536 | 2.405621873 | 2.748188027 | 2.265599895 |
| ENSMUSG00000021765 | 1.783508299 | 1.698603967 | 1.711372513 | 1.364857905 | 1.335652905 | 1.431363764 | 1.259064007 |
| ENSMUSG00000021767 | 2.063793172 | 1.903677461 | 1.912762923 | 1.990000483 | 1.929985312 | 2.079181246 | 1.842010333 |
| ENSMUSG00000021768 | 0.289064826 | 0 | 0.396240625 | 0.396240625 | 0 | 0 | 0 |

| | | | | | | | |
|---|---|---|---|---|---|---|---|
| ENSMUSG00000021770 | 2.343889214 | 2.176839783 | 2.205044741 | 2.424676167 | 2.368520226 | 2.674861141 | 2.167790061 |
| ENSMUSG00000021771 | 2.943372055 | 2.68346568 | 2.683677405 | 2.847945152 | 2.754414284 | 3.199755177 | 2.555498219 |
| ENSMUSG00000021772 | 1.527823874 | 1.372963274 | 1.438721876 | 1.723704441 | 1.66856048 | 1.832508913 | 1.502532226 |
| ENSMUSG00000021773 | 0.811507563 | 1 | 1 | 0.75 | 0.931471446 | 0.301029996 | 0.969165684 |
| ENSMUSG00000021774 | 1.516991255 | 1.566696635 | 1.593759858 | 1.874961472 | 1.756568651 | 1.929418926 | 1.680001789 |
| ENSMUSG00000021775 | 2.45815691 | 2.256976499 | 2.215271726 | 2.387205477 | 2.297835549 | 2.591064607 | 2.108567375 |
| ENSMUSG00000021779 | 0.45815691 | 0.646240625 | 0.396240625 | 0.5 | 0.244650542 | 0.301029996 | 0.520678036 |
| ENSMUSG00000021782 | 2.487005755 | 2.333847589 | 2.348615674 | 2.483053688 | 2.49804628 | 2.835056102 | 2.403948569 |
| ENSMUSG00000021785 | 1.885732304 | 1.720660762 | 1.634789703 | 2.03743678 | 1.870073561 | 2.206825876 | 1.778418869 |
| ENSMUSG00000021786 | 1.249317394 | 1.04248125 | 1.080482024 | 1.356566189 | 1.343584837 | 1.491361694 | 1.290497981 |
| ENSMUSG00000021789 | 0 | 0 | 0 | 0 | 0 | 0 | 0 |
| ENSMUSG00000021790 | 0 | 0 | 0 | 0 | 0 | 0 | 0 |
| ENSMUSG00000021791 | 0 | 0 | 0 | 0 | 0 | 0 | 0 |
| ENSMUSG00000021792 | 1.389637216 | 1.282320754 | 1.372963274 | 1.348079356 | 1.414412788 | 1.51851394 | 1.272006351 |
| ENSMUSG00000021794 | 2.823638505 | 2.566696635 | 2.599418658 | 2.861253711 | 2.785029426 | 3.214843848 | 2.589319772 |
| ENSMUSG00000021795 | 0 | 0 | 0 | 0 | 0.244650542 | 0 | 0 |
| ENSMUSG00000021796 | 2.733796161 | 2.475216702 | 2.440387808 | 2.725970461 | 2.601368039 | 3.011147361 | 2.408329475 |
| ENSMUSG00000021797 | 0 | 0 | 0 | 0 | 0 | 0 | 0 |
| ENSMUSG00000021798 | 1.885732304 | 1.777131114 | 1.79745614 | 1.04248125 | 1.335652905 | 1.278753601 | 1.066254157 |
| ENSMUSG00000021799 | 0 | 0 | 0 | 0 | 0 | 0 | 0 |
| ENSMUSG00000021803 | 2.626098559 | 2.388167274 | 2.350753006 | 2.263820609 | 2.192777367 | 2.496929648 | 2.06186744 |
| ENSMUSG00000021804 | 0 | 0 | 0 | 0 | 0 | 0 | 0 |
| ENSMUSG00000021806 | 2.492331912 | 2.324802005 | 2.360217292 | 2.219129237 | 2.302531265 | 2.545307116 | 2.269981661 |
| ENSMUSG00000021807 | 2.45930418 | 2.310197833 | 2.282320754 | 2.519203899 | 2.444777781 | 2.796574333 | 2.277722321 |
| ENSMUSG00000021809 | 1.269664473 | 1.238549078 | 1.330482024 | 1.552363341 | 1.588235379 | 1.643452676 | 1.459332757 |
| ENSMUSG00000021810 | 2.217828299 | 2.134789703 | 2.077084758 | 2.366391601 | 2.330902415 | 2.674861141 | 2.208492871 |
| ENSMUSG00000021811 | 2.402962068 | 2.191217898 | 2.222185812 | 2.556000419 | 2.462344774 | 2.854306042 | 2.31789776 |
| ENSMUSG00000021814 | 2.556012466 | 2.325945187 | 2.342671852 | 2.561388177 | 2.542486091 | 2.897077003 | 2.381929521 |
| ENSMUSG00000021815 | 0.960252568 | 1.061981878 | 0.701838731 | 1.302363341 | 1.327538614 | 1.204119983 | 1.100339912 |
| ENSMUSG00000021816 | 2.635965117 | 2.450224975 | 2.400442697 | 2.726722649 | 2.669844292 | 3.033423755 | 2.427430145 |
| ENSMUSG00000021819 | 2.66283289 | 2.446158711 | 2.547147211 | 2.418106335 | 2.517458689 | 2.739572344 | 2.479382724 |
| ENSMUSG00000021820 | 2.617563644 | 2.415444524 | 2.405970373 | 2.411864607 | 2.420401583 | 2.698970004 | 2.237984665 |
| ENSMUSG00000021822 | 2.614416215 | 2.415889526 | 2.378431899 | 2.094844592 | 2.084333628 | 2.350248018 | 1.939944915 |
| ENSMUSG00000021823 | 3.203844027 | 2.959702496 | 2.940387808 | 2.883088981 | 2.886403419 | 3.286680969 | 2.701187212 |

| | | | | | | | |
|---|---|---|---|---|---|---|---|
| ENSMUSG00000021824 | 2.323768721 | 2.149047439 | 2.123963757 | 2.385274154 | 2.31625483 | 2.663700925 | 2.162575192 |
| ENSMUSG00000021830 | 1.87190676 | 1.76109853 | 1.75 | 1.839388001 | 1.784632921 | 2.021189299 | 1.645916034 |
| ENSMUSG00000021831 | 2.512981297 | 2.302363341 | 2.252107156 | 2.559899632 | 2.453670557 | 2.880241776 | 2.273157653 |
| ENSMUSG00000021832 | 2.789779581 | 2.554186465 | 2.532320754 | 2.879048119 | 2.795056894 | 3.23146959 | 2.580107184 |
| ENSMUSG00000021835 | 1.765759561 | 1.763820609 | 1.744319981 | 1.561981878 | 1.689774529 | 1.785329835 | 1.627424404 |
| ENSMUSG00000021838 | 2.262379879 | 2.106566189 | 2.131869252 | 2.223704441 | 2.219993628 | 2.436162647 | 2.066254157 |
| ENSMUSG00000021840 | 2.853620573 | 2.582168732 | 2.560197833 | 2.780545225 | 2.710275601 | 3.120244796 | 2.512829932 |
| ENSMUSG00000021843 | 2.842342644 | 2.61434522 | 2.594031347 | 2.863960227 | 2.765336442 | 3.201397124 | 2.553813239 |
| ENSMUSG00000021846 | 1.470606563 | 1.488549078 | 1.557204673 | 1.348079356 | 1.223252711 | 1.255272505 | 1.259064007 |
| ENSMUSG00000021848 | 0 | 0 | 0 | 0.25 | 0 | 0 | 0 |
| ENSMUSG00000021850 | 0 | 0 | 0 | 0 | 0 | 0 | 0 |
| ENSMUSG00000021852 | 0 | 0 | 0 | 0 | 0 | 0 | 0 |
| ENSMUSG00000021866 | 1.771760131 | 1.711372513 | 1.816696635 | 1.933677405 | 2.028157605 | 2.195899652 | 2.043115906 |
| ENSMUSG00000021867 | 0 | 0 | 0 | 0 | 0 | 0 | 0 |
| ENSMUSG00000021868 | 1.936861443 | 1.83272922 | 1.890560606 | 2.005591953 | 2.080465577 | 2.260071388 | 1.987739 |
| ENSMUSG00000021870 | 2.583267694 | 2.396709947 | 2.412762923 | 2.618933358 | 2.533172725 | 2.913283902 | 2.363123786 |
| ENSMUSG00000021871 | 2.227926429 | 2.032320754 | 2.065523711 | 2.204245906 | 2.152352662 | 2.437750563 | 2.028149525 |
| ENSMUSG00000021872 | 0 | 0 | 0 | 0 | 0 | 0 | 0 |
| ENSMUSG00000021874 | 0.45815691 | 0.5 | 0.25 | 0.5 | 0.77552387 | 0.602059991 | 0.520678036 |
| ENSMUSG00000021876 | 1.727821383 | 1.671625132 | 1.698603967 | 1.841580554 | 1.820920161 | 2.079181246 | 1.71890424 |
| ENSMUSG00000021877 | 3.009907694 | 2.748057566 | 2.710391087 | 2.914441259 | 2.831730581 | 3.267640982 | 2.619740976 |
| ENSMUSG00000021879 | 0 | 0.396240625 | 0.396240625 | 0.396240625 | 0 | 0 | 0 |
| ENSMUSG00000021880 | 0 | 0 | 0 | 0 | 0 | 0 | 0 |
| ENSMUSG00000021884 | 1.358731791 | 1.25 | 1.282320754 | 1.464495249 | 1.445123986 | 1.612783857 | 1.360283038 |
| ENSMUSG00000021886 | 0 | 0.396240625 | 0 | 0 | 0 | 0 | 0.355418052 |
| ENSMUSG00000021890 | 2.468370459 | 2.27186571 | 2.27053726 | 2.544977273 | 2.474332504 | 2.850646235 | 2.321629237 |
| ENSMUSG00000021891 | 2.045658471 | 1.882845365 | 1.899978211 | 2.120953944 | 2.020174412 | 2.340444115 | 1.945529784 |
| ENSMUSG00000021892 | 1.944805158 | 1.874961472 | 1.899978211 | 2.321350555 | 2.313753359 | 2.59439255 | 2.214058866 |
| ENSMUSG00000021893 | 2.251208701 | 2.146240625 | 2.14151351 | 2.491085217 | 2.422990063 | 2.80140371 | 2.249624299 |
| ENSMUSG00000021895 | 1.789260561 | 1.618933358 | 1.610735874 | 2.027131114 | 1.940259189 | 2.198657087 | 1.847411384 |
| ENSMUSG00000021898 | 0 | 0.25 | 0.396240625 | 0.951838731 | 0.489301084 | 0.477121255 | 0.224243824 |
| ENSMUSG00000021900 | 1.924654746 | 1.821350555 | 1.777131114 | 1.976722649 | 2.009637626 | 2.155336037 | 1.930969131 |
| ENSMUSG00000021901 | 2.442947872 | 2.30357978 | 2.28296424 | 2.435366747 | 2.47528687 | 2.789580712 | 2.358080746 |
| ENSMUSG00000021902 | 1.227926429 | 1.114857905 | 1.226722649 | 1.348079356 | 1.380772476 | 1.568201724 | 1.252393349 |

| | | | | | | | |
|---|---|---|---|---|---|---|---|
| ENSMUSG00000021903 | 0.578129653 | 0.646240625 | 0.580482024 | 0.580482024 | 0.568060967 | 0.602059991 | 0.520678036 |
| ENSMUSG00000021904 | 1.129344651 | 0.896240625 | 0.75 | 1.080482024 | 1.254881871 | 1.397940009 | 1.272006351 |
| ENSMUSG00000021905 | 2.144240249 | 1.94697564 | 1.955044741 | 2.194519282 | 2.176862714 | 2.374748346 | 2.019455688 |
| ENSMUSG00000021906 | 1.811507563 | 1.692046081 | 1.610735874 | 1.935366747 | 1.888971102 | 2.152288344 | 1.798966438 |
| ENSMUSG00000021907 | 0 | 0 | 0 | 0 | 0 | 0 | 0 |
| ENSMUSG00000021908 | 0 | 0 | 0 | 0.25 | 0.244650542 | 0 | 0 |
| ENSMUSG00000021910 | 3.050960393 | 2.807354922 | 2.87019755 | 2.952039943 | 2.983453875 | 3.351603072 | 2.875917793 |
| ENSMUSG00000021911 | 2.258689303 | 2.079353153 | 2.078220739 | 2.31728167 | 2.254881871 | 2.604226053 | 2.105687397 |
| ENSMUSG00000021913 | 1.686083339 | 1.705044741 | 1.763820609 | 1.061981878 | 1.121713561 | 1.113943352 | 1.121219121 |
| ENSMUSG00000021916 | 2.060824957 | 1.884789703 | 1.880890489 | 2.075945187 | 1.988833497 | 2.238046103 | 1.842010333 |
| ENSMUSG00000021917 | 2.450036412 | 2.222945926 | 2.21293726 | 2.552667836 | 2.452655829 | 2.84509804 | 2.295094601 |
| ENSMUSG00000021918 | 1.765759561 | 1.657339155 | 1.584962501 | 1.872963274 | 1.808534465 | 2.079181246 | 1.725216937 |
| ENSMUSG00000021919 | 0 | 0 | 0 | 0 | 0 | 0.301029996 | 0 |
| ENSMUSG00000021922 | 0.45815691 | 0.580482024 | 0.25 | 0.5 | 0.632412477 | 0 | 0.448487648 |
| ENSMUSG00000021927 | 0.578129653 | 0.79248125 | 0.701838731 | 0.701838731 | 0.812711509 | 0.84509804 | 0.579661877 |
| ENSMUSG00000021928 | 1.587501561 | 1.51109853 | 1.516522298 | 1.794977273 | 1.836786029 | 2 | 1.663407615 |
| ENSMUSG00000021929 | 2.776606269 | 2.530383379 | 2.507321807 | 2.701838731 | 2.611446925 | 3.01745073 | 2.416548555 |
| ENSMUSG00000021930 | 1.890240806 | 1.744319981 | 1.720660762 | 2.025821952 | 1.911645804 | 2.206825876 | 1.793950594 |
| ENSMUSG00000021932 | 2.010213549 | 1.870953944 | 1.862802778 | 2.188721876 | 2.091944837 | 2.414973348 | 1.95411574 |
| ENSMUSG00000021933 | 0 | 0 | 0.25 | 0 | 0 | 0 | 0 |
| ENSMUSG00000021936 | 2.209569962 | 2.071350555 | 1.990000483 | 2.293106877 | 2.21338377 | 2.545307116 | 2.06842541 |
| ENSMUSG00000021938 | 2.386920383 | 2.173371739 | 2.160964047 | 2.514159429 | 2.446849937 | 2.796574333 | 2.285005227 |
| ENSMUSG00000021939 | 3.134142119 | 2.833847589 | 2.879909063 | 2.862158142 | 2.791573109 | 3.25358029 | 2.625689732 |
| ENSMUSG00000021940 | 0 | 0 | 0 | 0 | 0 | 0 | 0 |
| ENSMUSG00000021943 | 0 | 0 | 0 | 0 | 0 | 0 | 0 |
| ENSMUSG00000021944 | 1.721148722 | 1.685366747 | 1.708222504 | 1.681980114 | 1.72073075 | 1.792391689 | 1.594042709 |
| ENSMUSG00000021945 | 2.405576699 | 2.209012589 | 2.199415381 | 2.408702763 | 2.340340306 | 2.655138435 | 2.136039214 |
| ENSMUSG00000021947 | 1.249317394 | 1.226722649 | 1.403677461 | 1.476722649 | 1.618292349 | 1.579783597 | 1.526507301 |
| ENSMUSG00000021948 | 2.523917219 | 2.349685923 | 2.328220739 | 2.376452888 | 2.3873546 | 2.682145076 | 2.261455999 |
| ENSMUSG00000021950 | 1.920505136 | 1.708222504 | 1.71759118 | 1.330482024 | 1.327538614 | 1.568201724 | 1.340368108 |
| ENSMUSG00000021951 | 1.765759561 | 1.646240625 | 1.657339155 | 2.025821952 | 2.012006468 | 2.303196057 | 1.886704102 |
| ENSMUSG00000021952 | 2.395018353 | 2.168106335 | 2.182829758 | 2.512802235 | 2.419658517 | 2.751279104 | 2.21680636 |
| ENSMUSG00000021953 | 0 | 0 | 0 | 0 | 0 | 0 | 0 |
| ENSMUSG00000021957 | 2.520963022 | 2.428990498 | 2.469129237 | 2.434523065 | 2.614672255 | 2.8162413 | 2.538872454 |

| | | | | | | | |
|---|---|---|---|---|---|---|---|
| ENSMUSG00000021958 | 1.678702042 | 1.646240625 | 1.54745614 | 1.837182039 | 1.77552387 | 2.017033339 | 1.694069894 |
| ENSMUSG00000021959 | 2.414600936 | 2.240000483 | 2.240000483 | 2.503505118 | 2.457366467 | 2.750508395 | 2.326782038 |
| ENSMUSG00000021961 | 0 | 0 | 0 | 0 | 0 | 0 | 0 |
| ENSMUSG00000021962 | 2.165631214 | 2.063166358 | 1.997171172 | 2.23927551 | 2.204601633 | 2.498310554 | 2.079068325 |
| ENSMUSG00000021963 | 2.322180025 | 2.14151351 | 2.105516192 | 2.424241882 | 2.344506394 | 2.721810615 | 2.242754021 |
| ENSMUSG00000021965 | 2.233869984 | 2.04248125 | 2.064346961 | 2.30539678 | 2.234113756 | 2.557507202 | 2.110473216 |
| ENSMUSG00000021966 | 0 | 0 | 0 | 0 | 0 | 0 | 0 |
| ENSMUSG00000021967 | 1.432083896 | 1.505591953 | 1.630890489 | 1.618933358 | 1.55104774 | 1.857332496 | 1.683576582 |
| ENSMUSG00000021969 | 2.557825653 | 2.344844592 | 2.354990044 | 2.688095162 | 2.59734443 | 2.98721923 | 2.395004664 |
| ENSMUSG00000021972 | 2.139333929 | 2.006976499 | 1.961372513 | 2.199415381 | 2.172394856 | 2.471291711 | 2.02568932 |
| ENSMUSG00000021973 | 2.414600936 | 2.209800947 | 2.181980114 | 2.477098155 | 2.407551654 | 2.790988475 | 2.221559033 |
| ENSMUSG00000021974 | 0 | 0.25 | 0 | 0 | 0 | 0 | 0.224243824 |
| ENSMUSG00000021975 | 2.051791411 | 1.862802778 | 1.862802778 | 2.130890489 | 2.053185007 | 2.403120521 | 1.98213686 |
| ENSMUSG00000021977 | 0 | 0 | 0 | 0 | 0 | 0 | 0 |
| ENSMUSG00000021978 | 2.432083896 | 2.293106877 | 2.366391601 | 2.390560606 | 2.42519375 | 2.710117365 | 2.309790449 |
| ENSMUSG00000021981 | 2.297593393 | 2.109697963 | 2.122963274 | 2.406427211 | 2.326087113 | 2.688419822 | 2.161765391 |
| ENSMUSG00000021982 | 1.623015126 | 1.527131114 | 1.516522298 | 1.841580554 | 1.744182955 | 2 | 1.702561601 |
| ENSMUSG00000021983 | 0 | 0 | 0 | 0.701838731 | 0.387761935 | 0.301029996 | 0.448487648 |
| ENSMUSG00000021986 | 0 | 0.25 | 0.25 | 0.25 | 0.244650542 | 0 | 0.448487648 |
| ENSMUSG00000021987 | 2.35433422 | 2.208222504 | 2.231944491 | 2.471424093 | 2.381601986 | 2.707570176 | 2.222232323 |
| ENSMUSG00000021990 | 2.097855556 | 2 | 1.92510993 | 1.774508021 | 1.741652793 | 1.838849091 | 1.729871916 |
| ENSMUSG00000021991 | 0.289064826 | 0 | 0 | 0.396240625 | 0 | 0 | 0.224243824 |
| ENSMUSG00000021993 | 1.837744666 | 1.774508021 | 1.708222504 | 2.063166358 | 1.970729215 | 2.245512668 | 1.877074955 |
| ENSMUSG00000021994 | 2.177057474 | 2.051142786 | 2.036164561 | 1.62694866 | 1.639266209 | 1.698970004 | 1.493062805 |
| ENSMUSG00000021996 | 2.789260561 | 2.51584877 | 2.50835575 | 2.726346751 | 2.662070429 | 3.058426024 | 2.450652753 |
| ENSMUSG00000021997 | 0 | 0 | 0 | 0 | 0 | 0 | 0 |
| ENSMUSG00000021998 | 0.671187741 | 0.976722649 | 0.830482024 | 0.951838731 | 1 | 1 | 1.028149525 |
| ENSMUSG00000021999 | 0.289064826 | 0 | 0 | 0 | 0 | 0 | 0 |
| ENSMUSG00000022000 | 2.3855553 | 2.146240625 | 2.150906586 | 2.378431899 | 2.338472809 | 2.615950052 | 2.199960101 |
| ENSMUSG00000022002 | 0 | 0 | 0 | 0 | 0 | 0 | 0 |
| ENSMUSG00000022003 | 1.971473847 | 1.872963274 | 1.839388001 | 1.890560606 | 1.902105869 | 2.167317335 | 1.815933683 |
| ENSMUSG00000022008 | 2.269664473 | 2.097004321 | 2.059601185 | 2.428990498 | 2.32560195 | 2.629409599 | 2.164992504 |
| ENSMUSG00000022009 | 2.057835465 | 1.905512955 | 1.933677405 | 2.168989258 | 2.098472613 | 2.431363764 | 1.993927582 |
| ENSMUSG00000022010 | 2.926527369 | 2.70444578 | 2.731759292 | 2.634789703 | 2.555378544 | 2.948412966 | 2.438130195 |

| | | | | | | | |
|---|---|---|---|---|---|---|---|
| ENSMUSG00000022012 | 0.289064826 | 0.25 | 0.25 | 0.75 | 0.632412477 | 0.602059991 | 0.829800754 |
| ENSMUSG00000022013 | 1.800532393 | 1.634789703 | 1.681980114 | 1.76109853 | 1.712553795 | 1.929418926 | 1.627424404 |
| ENSMUSG00000022014 | 0 | 0 | 0 | 0.25 | 0 | 0.477121255 | 0.355418052 |
| ENSMUSG00000022015 | 0.289064826 | 0 | 0 | 0.5 | 0.489301084 | 0.301029996 | 0.629532004 |
| ENSMUSG00000022016 | 2.690055106 | 2.474461364 | 2.424241882 | 2.725970461 | 2.650465868 | 3.019531685 | 2.42903127 |
| ENSMUSG00000022018 | 0.289064826 | 0 | 0 | 0.896240625 | 0.905314583 | 0.698970004 | 0.803905701 |
| ENSMUSG00000022019 | 2.039433992 | 1.94697564 | 1.94697564 | 2.038704527 | 2.058398634 | 2.240549248 | 1.86715039 |
| ENSMUSG00000022020 | 1.847794126 | 1.845926073 | 1.779735268 | 2.095926073 | 2.008447216 | 2.243038049 | 1.869159809 |
| ENSMUSG00000022021 | 2.730819472 | 2.514497931 | 2.522528105 | 2.81508288 | 2.71658578 | 3.138302698 | 2.512007086 |
| ENSMUSG00000022022 | 0.45815691 | 0.951838731 | 0.75 | 1.372963274 | 1.27449559 | 1.301029996 | 1.20919388 |
| ENSMUSG00000022023 | 2.269664473 | 2.160061234 | 2.195339928 | 2.319031101 | 2.247690665 | 2.537819095 | 2.121219121 |
| ENSMUSG00000022024 | 2.500714755 | 2.271202097 | 2.240000483 | 2.549304173 | 2.448909998 | 2.835690571 | 2.299096905 |
| ENSMUSG00000022025 | 0 | 0 | 0 | 0 | 0 | 0 | 0 |
| ENSMUSG00000022026 | 0 | 0 | 0 | 0 | 0 | 0 | 0 |
| ENSMUSG00000022031 | 2.410757293 | 2.198603967 | 2.23927551 | 2.398114259 | 2.339407793 | 2.707570176 | 2.187847501 |
| ENSMUSG00000022032 | 0 | 0.396240625 | 0 | 0.646240625 | 0.632412477 | 0.602059991 | 0.579661877 |
| ENSMUSG00000022033 | 2.423434025 | 2.256284891 | 2.244319981 | 2.487091808 | 2.422252432 | 2.758154622 | 2.255745871 |
| ENSMUSG00000022034 | 2 | 1.940387808 | 1.880890489 | 2.274508021 | 2.187072901 | 2.575187845 | 2.026921761 |
| ENSMUSG00000022035 | 2.066740409 | 2 | 1.976722649 | 2.259043403 | 2.194892994 | 2.482873584 | 2.048913829 |
| ENSMUSG00000022037 | 1.249317394 | 1.282320754 | 1.339388001 | 1.372963274 | 1.319233381 | 1.591064607 | 1.284450788 |
| ENSMUSG00000022039 | 0 | 0 | 0 | 0.396240625 | 0 | 0 | 0 |
| ENSMUSG00000022040 | 0 | 0.25 | 0 | 1.527131114 | 1.509475496 | 1.62324929 | 1.369798885 |
| ENSMUSG00000022041 | 0 | 0 | 0 | 0.25 | 0.244650542 | 0 | 0 |
| ENSMUSG00000022043 | 2.724194991 | 2.474839231 | 2.484159485 | 2.584405475 | 2.57935845 | 2.955206538 | 2.427964734 |
| ENSMUSG00000022044 | 0.747221736 | 0.25 | 0.5 | 0.951838731 | 1.020174412 | 0.698970004 | 0.829800754 |
| ENSMUSG00000022048 | 2.23973002 | 2.049918086 | 2.044977273 | 1.945339928 | 1.992595054 | 2.238046103 | 1.87805087 |
| ENSMUSG00000022051 | 2.393679564 | 2.230460234 | 2.215271726 | 2.51584877 | 2.427019704 | 2.805500858 | 2.268527657 |
| ENSMUSG00000022052 | 2.615992899 | 2.38040011 | 2.373463862 | 2.652525516 | 2.545363112 | 2.932980822 | 2.3678751 |
| ENSMUSG00000022053 | 0.45815691 | 0.5 | 0.25 | 1.356566189 | 1.351342427 | 1.361727836 | 1.224243824 |
| ENSMUSG00000022054 | 0.289064826 | 0.25 | 0 | 0.580482024 | 0.244650542 | 0.477121255 | 0.355418052 |
| ENSMUSG00000022055 | 0 | 0 | 0 | 0.25 | 0.244650542 | 0 | 0 |
| ENSMUSG00000022056 | 0 | 0 | 0 | 0 | 0 | 0 | 0 |
| ENSMUSG00000022057 | 0 | 0 | 0 | 0 | 0 | 0 | 0 |
| ENSMUSG00000022061 | 0 | 0 | 0 | 0.5 | 0 | 0 | 0.224243824 |

| | | | | | | | |
|---|---|---|---|---|---|---|---|
| ENSMUSG00000022064 | 1.700465191 | 1.618933358 | 1.678561379 | 1.845926073 | 1.756568651 | 2.004321374 | 1.653807046 |
| ENSMUSG00000022066 | 0 | 0 | 0 | 0 | 0 | 0 | 0 |
| ENSMUSG00000022070 | 2.016886209 | 1.860735874 | 1.886723615 | 2.114857905 | 2.126012715 | 2.367355921 | 1.995964412 |
| ENSMUSG00000022074 | 2.196867486 | 2.048689214 | 2.05357978 | 2.26109853 | 2.200473444 | 2.505149978 | 2.057979654 |
| ENSMUSG00000022075 | 2.00683675 | 1.83272922 | 1.841580554 | 1.96293726 | 1.941702774 | 2.238046103 | 1.903345709 |
| ENSMUSG00000022076 | 0 | 0.25 | 0 | 0 | 0 | 0 | 0 |
| ENSMUSG00000022085 | 0 | 0 | 0.25 | 0 | 0 | 0.301029996 | 0 |
| ENSMUSG00000022089 | 1.827447047 | 1.747171172 | 1.819031101 | 1.992885888 | 1.959951091 | 2.252853031 | 1.899720848 |
| ENSMUSG00000022090 | 2.072573125 | 1.988549078 | 2.038704527 | 1.938721876 | 2.066583491 | 2.23299611 | 2.096890362 |
| ENSMUSG00000022091 | 2.348397716 | 2.140560606 | 2.240000483 | 2.30479213 | 2.407166541 | 2.633468456 | 2.305919057 |
| ENSMUSG00000022092 | 1.432083896 | 1.380890489 | 1.464495249 | 1.618933358 | 1.563883923 | 1.908485019 | 1.429563222 |
| ENSMUSG00000022094 | 2.310886388 | 2.194519282 | 2.147178659 | 2.299918086 | 2.304082723 | 2.612783857 | 2.164992504 |
| ENSMUSG00000022095 | 1.857607105 | 1.834962501 | 1.825945187 | 1.854463129 | 1.957204337 | 2.093421685 | 1.893277937 |
| ENSMUSG00000022096 | 1.129344651 | 0.79248125 | 0.92510993 | 1.67510993 | 1.704182902 | 1.73239376 | 1.731408809 |
| ENSMUSG00000022097 | 0 | 0 | 0 | 0 | 0 | 0 | 0 |
| ENSMUSG00000022098 | 2.574668783 | 2.410512923 | 2.389126514 | 2.360735874 | 2.48376199 | 2.752048448 | 2.379663565 |
| ENSMUSG00000022099 | 1.307602651 | 1.02186571 | 1.080482024 | 1.17510993 | 1.310728002 | 1.361727836 | 1.340368108 |
| ENSMUSG00000022100 | 2.706208651 | 2.461372513 | 2.451838731 | 2.711960089 | 2.636258897 | 3.062205809 | 2.457225157 |
| ENSMUSG00000022101 | 0 | 0 | 0 | 0 | 0.244650542 | 0 | 0 |
| ENSMUSG00000022102 | 0.578129653 | 0.5 | 0.396240625 | 0.646240625 | 0.244650542 | 0.477121255 | 0.629532004 |
| ENSMUSG00000022103 | 0.671187741 | 0.5 | 0.580482024 | 0.75 | 0.846351821 | 0.698970004 | 0.853775828 |
| ENSMUSG00000022105 | 1.993049299 | 1.903677461 | 1.862802778 | 2.034887838 | 1.890639929 | 2.204119983 | 1.775756176 |
| ENSMUSG00000022106 | 2.342375483 | 2.174241882 | 2.163659007 | 2.238549078 | 2.21338377 | 2.44870632 | 2.081155529 |
| ENSMUSG00000022108 | 2.793394745 | 2.564935816 | 2.560495787 | 2.694108258 | 2.605566002 | 3.065206128 | 2.459009404 |
| ENSMUSG00000022109 | 2.167941649 | 2.006976499 | 2.031030328 | 2.261780978 | 2.246477757 | 2.489958479 | 2.018194418 |
| ENSMUSG00000022110 | 2.524897319 | 2.357090043 | 2.345926073 | 2.583288838 | 2.483451699 | 2.869818208 | 2.317146272 |
| ENSMUSG00000022111 | 2.095120908 | 1.956637122 | 1.933677405 | 2.228969845 | 2.147561328 | 2.457881897 | 2.025071334 |
| ENSMUSG00000022112 | 0 | 0 | 0.25 | 0 | 0 | 0 | 0.224243824 |
| ENSMUSG00000022113 | 0.671187741 | 0.396240625 | 0.25 | 0.79248125 | 0.489301084 | 0.698970004 | 0.448487648 |
| ENSMUSG00000022114 | 1.37447073 | 1.339388001 | 1.42510993 | 1.505591953 | 1.433158226 | 1.792391689 | 1.496250175 |
| ENSMUSG00000022116 | 0 | 0 | 0 | 0 | 0 | 0 | 0 |
| ENSMUSG00000022119 | 2.481610694 | 2.311981878 | 2.284887838 | 2.519537702 | 2.46464417 | 2.769377326 | 2.229876084 |
| ENSMUSG00000022120 | 2 | 1.921625132 | 1.888647213 | 2.046218836 | 1.977302144 | 2.283301229 | 1.865128412 |
| ENSMUSG00000022122 | 0 | 0 | 0 | 0 | 0.244650542 | 0 | 0 |

| | | | | | | | |
|---|---|---|---|---|---|---|---|
| ENSMUSG00000022123 | 0.289064826 | 0.5 | 0.396240625 | 0.951838731 | 0.632412477 | 0.954242509 | 0.520678036 |
| ENSMUSG00000022124 | 2.37447073 | 2.168989258 | 2.184523065 | 2.449415381 | 2.369807605 | 2.730782276 | 2.200320163 |
| ENSMUSG00000022125 | 2.211649928 | 2.105516192 | 2.101285366 | 2.191217898 | 2.145141048 | 2.444044796 | 1.99866039 |
| ENSMUSG00000022126 | 0 | 0 | 0 | 0 | 0 | 0 | 0 |
| ENSMUSG00000022129 | 0 | 0 | 0 | 0 | 0 | 0 | 0 |
| ENSMUSG00000022130 | 1.912079952 | 1.789967834 | 1.726722649 | 1.909156155 | 1.85749873 | 2.093421685 | 1.760706232 |
| ENSMUSG00000022131 | 1.847794126 | 1.848079356 | 1.799918086 | 1.845926073 | 1.875328588 | 2.089905111 | 1.821784031 |
| ENSMUSG00000022132 | 0 | 0 | 0 | 0.5 | 0 | 0 | 0.355418052 |
| ENSMUSG00000022136 | 2.559630991 | 2.392938161 | 2.353935442 | 2.727473271 | 2.635654347 | 3.045714059 | 2.434311835 |
| ENSMUSG00000022139 | 2.720536789 | 2.521534056 | 2.507666784 | 2.638647213 | 2.55990691 | 2.953276337 | 2.365939808 |
| ENSMUSG00000022141 | 3.021375299 | 2.758871613 | 2.768535366 | 2.80930249 | 2.727014069 | 3.145196406 | 2.510355088 |
| ENSMUSG00000022142 | 3.140901301 | 2.822648722 | 2.823798739 | 2.900210529 | 2.820203499 | 3.244277121 | 2.601521341 |
| ENSMUSG00000022144 | 2.315764015 | 2.155512955 | 2.120953944 | 2.285526764 | 2.257246745 | 2.583198774 | 2.122594809 |
| ENSMUSG00000022146 | 2.388281011 | 2.21759118 | 2.195339928 | 2.178561379 | 2.112954895 | 2.409933123 | 2.001999072 |
| ENSMUSG00000022148 | 0 | 0.5 | 0 | 0 | 0.387761935 | 0.698970004 | 0.355418052 |
| ENSMUSG00000022149 | 0 | 0 | 0 | 0 | 0 | 0 | 0 |
| ENSMUSG00000022150 | 2.601583437 | 2.355516192 | 2.367918804 | 2.735261902 | 2.637063356 | 3.08278537 | 2.481502159 |
| ENSMUSG00000022151 | 2.183765914 | 2.038704527 | 2.038704527 | 2.087182039 | 2.031524462 | 2.274157849 | 1.846338363 |
| ENSMUSG00000022155 | 0 | 0 | 0 | 0 | 0 | 0 | 0 |
| ENSMUSG00000022156 | 0 | 0 | 0 | 0 | 0 | 0 | 0.224243824 |
| ENSMUSG00000022157 | 0.45815691 | 0.396240625 | 0.396240625 | 0 | 0.244650542 | 0.301029996 | 0.355418052 |
| ENSMUSG00000022159 | 1 | 1 | 1.146240625 | 1.321350555 | 1.335652905 | 1.414973348 | 1.110947927 |
| ENSMUSG00000022160 | 1.740854706 | 1.732684334 | 1.685366747 | 1.841580554 | 1.820920161 | 2.08278537 | 1.710836105 |
| ENSMUSG00000022174 | 2.20747957 | 2.058404919 | 2.099151195 | 2.235628626 | 2.254288168 | 2.57863921 | 2.13691595 |
| ENSMUSG00000022175 | 2.373064211 | 2.228222334 | 2.289967834 | 2.248588359 | 2.373217836 | 2.618048097 | 2.302000776 |
| ENSMUSG00000022176 | 1.129344651 | 1.04248125 | 1.27186571 | 0.75 | 0.568060967 | 0.602059991 | 0.448487648 |
| ENSMUSG00000022177 | 1.99653913 | 1.975216702 | 1.918106335 | 1.914552871 | 2.056322416 | 2.198657087 | 1.95718226 |
| ENSMUSG00000022178 | 2.509948321 | 2.324802005 | 2.336628433 | 2.604463129 | 2.610148495 | 2.921166051 | 2.43361272 |
| ENSMUSG00000022179 | 1.432083896 | 1.458222504 | 1.54248125 | 1.482684334 | 1.489301084 | 1.591064607 | 1.448487648 |
| ENSMUSG00000022180 | 0 | 0 | 0 | 1.238549078 | 1.136121934 | 1 | 1.089372236 |
| ENSMUSG00000022181 | 0 | 0 | 0 | 0.396240625 | 0 | 0 | 0 |
| ENSMUSG00000022184 | 1.944805158 | 1.890560606 | 1.78743678 | 1.87694866 | 1.704182902 | 2.008600172 | 1.669034113 |
| ENSMUSG00000022185 | 2.876566388 | 2.663659007 | 2.687676747 | 2.701435968 | 2.745367373 | 3.067814511 | 2.578657689 |
| ENSMUSG00000022186 | 3.306852728 | 3.002194687 | 3.002019603 | 2.936313535 | 2.848234258 | 3.29710365 | 2.642631585 |

| | | | | | | | |
|---|---|---|---|---|---|---|---|
| ENSMUSG00000022187 | 0.289064826 | 0.396240625 | 0 | 0.646240625 | 0.489301084 | 0 | 0.520678036 |
| ENSMUSG00000022191 | 2.933200029 | 2.670967967 | 2.661864607 | 2.666777886 | 2.594182934 | 2.985426474 | 2.436226648 |
| ENSMUSG00000022193 | 2.054824387 | 1.979715809 | 2.006976499 | 2.135757955 | 2.199086582 | 2.440909082 | 2.089878917 |
| ENSMUSG00000022194 | 1.663535556 | 1.516522298 | 1.67510993 | 1.657339155 | 1.746695109 | 1.785329835 | 1.627424404 |
| ENSMUSG00000022195 | 2.468370459 | 2.272528105 | 2.250703754 | 2.222945926 | 2.202543574 | 2.482873584 | 2.088864761 |
| ENSMUSG00000022197 | 0.747221736 | 1 | 1.04248125 | 1.27186571 | 1.358933174 | 1.544068044 | 1.296434211 |
| ENSMUSG00000022199 | 0.671187741 | 0.830482024 | 0.92510993 | 1.356566189 | 1.546663136 | 1.380211242 | 1.543390198 |
| ENSMUSG00000022200 | 2.856282491 | 2.635516136 | 2.592399337 | 2.697587124 | 2.618715812 | 3.035429738 | 2.42366308 |
| ENSMUSG00000022201 | 2.989851182 | 2.744856299 | 2.715271726 | 2.731759292 | 2.671123451 | 3.085290578 | 2.485997583 |
| ENSMUSG00000022203 | 0.671187741 | 1.061981878 | 1.080482024 | 0.25 | 0.489301084 | 0.477121255 | 0.672731473 |
| ENSMUSG00000022204 | 2.213719572 | 1.992885888 | 2.064346961 | 2.282320754 | 2.293608479 | 2.602059991 | 2.160546876 |
| ENSMUSG00000022205 | 2.692026894 | 2.474461364 | 2.468361028 | 2.678776002 | 2.575792006 | 2.982723388 | 2.406051188 |
| ENSMUSG00000022206 | 2.50175086 | 2.319031101 | 2.127938163 | 1.950224975 | 1.895599581 | 2.093421685 | 1.712465909 |
| ENSMUSG00000022208 | 0 | 0.25 | 0.25 | 0.79248125 | 0.877063019 | 0.903089987 | 0.916588299 |
| ENSMUSG00000022209 | 0 | 0 | 0 | 0 | 0 | 0 | 0 |
| ENSMUSG00000022210 | 1.686083339 | 1.653677461 | 1.685366747 | 1.914552871 | 1.928492895 | 2.064457989 | 1.913982791 |
| ENSMUSG00000022211 | 0 | 0.580482024 | 0.580482024 | 0.75 | 0.905314583 | 0.301029996 | 1.054044578 |
| ENSMUSG00000022212 | 0 | 0 | 0 | 0 | 0 | 0 | 0 |
| ENSMUSG00000022214 | 1.944805158 | 1.901832578 | 1.892463902 | 2.200224975 | 2.179810173 | 2.418301291 | 2.028149525 |
| ENSMUSG00000022215 | 0 | 0 | 0 | 0 | 0 | 0 | 0 |
| ENSMUSG00000022216 | 2.029919532 | 1.892463902 | 2.002806814 | 2.383332433 | 2.361573756 | 2.591064607 | 2.280542874 |
| ENSMUSG00000022217 | 1.069666964 | 1.188721876 | 1.146240625 | 1.26109853 | 1.149965125 | 1.204119983 | 1.350478789 |
| ENSMUSG00000022218 | 0 | 0.25 | 0.396240625 | 0.5 | 0.244650542 | 0.602059991 | 0.520678036 |
| ENSMUSG00000022219 | 0.289064826 | 0 | 0.25 | 0.25 | 0.387761935 | 0.477121255 | 0.629532004 |
| ENSMUSG00000022220 | 0 | 0.25 | 0 | 0.646240625 | 0.568060967 | 0.301029996 | 0.224243824 |
| ENSMUSG00000022221 | 1.83262764 | 1.638647213 | 1.807204673 | 2.199415381 | 2.189936658 | 2.496929648 | 2.071119143 |
| ENSMUSG00000022223 | 1.404271433 | 1.25 | 1.282320754 | 1.418106335 | 1.514343934 | 1.431363764 | 1.425641774 |
| ENSMUSG00000022225 | 0 | 0 | 0 | 0 | 0 | 0 | 0 |
| ENSMUSG00000022226 | 0 | 0 | 0 | 0 | 0 | 0 | 0 |
| ENSMUSG00000022227 | 0 | 0 | 0 | 0 | 0 | 0 | 0 |
| ENSMUSG00000022228 | 2.163307908 | 2.039967834 | 2.03360658 | 2.097004321 | 2.071604288 | 2.342422681 | 1.914853627 |
| ENSMUSG00000022229 | 0 | 0 | 0 | 0 | 0 | 0 | 0 |
| ENSMUSG00000022231 | 1.37447073 | 1.396240625 | 1.575945187 | 1.816696635 | 1.683842353 | 1.959041392 | 1.631625962 |
| ENSMUSG00000022234 | 3.510685616 | 3.171515687 | 3.163211243 | 3.200326047 | 3.143709563 | 3.670153045 | 2.9163523 |

| | | | | | | | |
|---|---|---|---|---|---|---|---|
| ENSMUSG00000022235 | 0.289064826 | 0.396240625 | 0.396240625 | 0.5 | 0.846351821 | 0.477121255 | 0.355418052 |
| ENSMUSG00000022236 | 1.069666964 | 1.226722649 | 0.976722649 | 0.701838731 | 0.77552387 | 0.84509804 | 0.952571511 |
| ENSMUSG00000022237 | 0.960252568 | 0.951838731 | 1.080482024 | 1.584962501 | 1.47337555 | 1.672097858 | 1.444790289 |
| ENSMUSG00000022240 | 1.181541736 | 1.214495249 | 1.114857905 | 0.75 | 0.244650542 | 0.477121255 | 0.355418052 |
| ENSMUSG00000022241 | 3.079103675 | 2.804943388 | 2.77401406 | 2.731573999 | 2.701515294 | 3.07809415 | 2.562973876 |
| ENSMUSG00000022243 | 0 | 0 | 0 | 0 | 0 | 0 | 0 |
| ENSMUSG00000022244 | 1.91631382 | 1.837182039 | 1.830482024 | 1.589388001 | 1.680838453 | 1.785329835 | 1.674564426 |
| ENSMUSG00000022245 | 0 | 0 | 0 | 0.25 | 0 | 0 | 0.224243824 |
| ENSMUSG00000022246 | 2.930247575 | 2.705443495 | 2.704245906 | 2.495748394 | 2.417419887 | 2.793790385 | 2.303835214 |
| ENSMUSG00000022247 | 2.753493018 | 2.535845803 | 2.491806565 | 2.535526764 | 2.466928684 | 2.866287339 | 2.314628629 |
| ENSMUSG00000022248 | 2.149089517 | 2.005591953 | 1.958222504 | 1.943696765 | 1.866526233 | 2.045322979 | 1.728327686 |
| ENSMUSG00000022249 | 0 | 0 | 0 | 0 | 0 | 0 | 0 |
| ENSMUSG00000022253 | 2.40817504 | 2.265173983 | 2.240724001 | 2.277783918 | 2.219338185 | 2.542825427 | 2.080113611 |
| ENSMUSG00000022255 | 3.093371294 | 2.817865757 | 2.801295582 | 2.884425937 | 2.787399152 | 3.214578954 | 2.577089398 |
| ENSMUSG00000022257 | 2.483777101 | 2.252806814 | 2.263820609 | 2.185366747 | 2.16174451 | 2.447158031 | 2.03780742 |
| ENSMUSG00000022261 | 3.141794304 | 2.860865403 | 2.839800121 | 2.77236262 | 2.712229237 | 3.112269768 | 2.538998802 |
| ENSMUSG00000022262 | 0 | 0.396240625 | 0.25 | 0.5 | 0.244650542 | 0.301029996 | 0 |
| ENSMUSG00000022263 | 2.9901721 | 2.760927716 | 2.777294426 | 2.54248125 | 2.553092387 | 2.875639937 | 2.424384006 |
| ENSMUSG00000022265 | 3.016886209 | 2.809003548 | 2.772031423 | 2.251406137 | 2.260180728 | 2.51851394 | 2.155212955 |
| ENSMUSG00000022269 | 0 | 0 | 0 | 0 | 0 | 0 | 0 |
| ENSMUSG00000022270 | 2.328498818 | 2.151832578 | 2.218361028 | 2.535526764 | 2.452655829 | 2.831229694 | 2.282223478 |
| ENSMUSG00000022272 | 2.929134994 | 2.667664062 | 2.701838731 | 2.574229052 | 2.588235379 | 2.894316063 | 2.458199602 |
| ENSMUSG00000022280 | 2.649978134 | 2.431980114 | 2.409608978 | 2.459012589 | 2.382842606 | 2.689308859 | 2.222232323 |
| ENSMUSG00000022283 | 3.70470791 | 3.360509086 | 3.346162216 | 3.395770692 | 3.287298763 | 3.856849679 | 3.052188252 |
| ENSMUSG00000022285 | 3.498445185 | 3.147880587 | 3.119313081 | 3.221757543 | 3.125459295 | 3.638089722 | 2.901620727 |
| ENSMUSG00000022286 | 0.289064826 | 0.396240625 | 0.396240625 | 0 | 0.387761935 | 0.301029996 | 0 |
| ENSMUSG00000022288 | 0 | 0 | 0 | 0 | 0 | 0 | 0 |
| ENSMUSG00000022292 | 2.328498818 | 2.12892496 | 2.131869252 | 2.262462137 | 2.210704773 | 2.514547753 | 2.013099581 |
| ENSMUSG00000022295 | 2.938317055 | 2.686208459 | 2.696771581 | 2.610476676 | 2.513204446 | 2.921166051 | 2.368517634 |
| ENSMUSG00000022296 | 1.129344651 | 1.226722649 | 1.146240625 | 1.098079356 | 1.091002363 | 1.230448921 | 1.131174228 |
| ENSMUSG00000022297 | 2.883252421 | 2.604199382 | 2.591307212 | 2.716626553 | 2.644615633 | 3.041392685 | 2.454119908 |
| ENSMUSG00000022299 | 2.305951036 | 2.168989258 | 2.168989258 | 2.146240625 | 2.05001946 | 2.322219295 | 1.920029985 |
| ENSMUSG00000022300 | 2.763003223 | 2.518869787 | 2.518200634 | 2.539652422 | 2.489301084 | 2.856124444 | 2.31789776 |
| ENSMUSG00000022303 | 0 | 0 | 0 | 0 | 0 | 0 | 0 |

| | | | | | | | |
|---|---|---|---|---|---|---|---|
| ENSMUSG00000022304 | 0.289064826 | 0 | 0 | 0.25 | 0.244650542 | 0 | 0 |
| ENSMUSG00000022305 | 2.579849376 | 2.345926073 | 2.324229052 | 2.298073204 | 2.257246745 | 2.564666064 | 2.127589533 |
| ENSMUSG00000022306 | 2.349889783 | 2.148114259 | 2.181128463 | 2.237091808 | 2.176121988 | 2.45484486 | 2.013740841 |
| ENSMUSG00000022307 | 2.886144188 | 2.645770692 | 2.638167274 | 2.675543171 | 2.584058335 | 2.974511693 | 2.402218022 |
| ENSMUSG00000022309 | 1.482695304 | 1.53743678 | 1.418106335 | 1.114857905 | 1.163285805 | 0.84509804 | 1.110947927 |
| ENSMUSG00000022311 | 0 | 0 | 0 | 0 | 0.244650542 | 0 | 0.355418052 |
| ENSMUSG00000022312 | 3.254114862 | 2.956835676 | 2.947179583 | 2.988003291 | 2.919583888 | 3.411451342 | 2.746189372 |
| ENSMUSG00000022313 | 2.205378646 | 2.084962501 | 2.094844592 | 1.998588359 | 1.965381292 | 2.217483944 | 1.873141588 |
| ENSMUSG00000022314 | 3.501938967 | 3.158248799 | 3.144946832 | 3.212497861 | 3.116122771 | 3.645716939 | 2.900961164 |
| ENSMUSG00000022315 | 0 | 0.25 | 0 | 0 | 0.244650542 | 0 | 0 |
| ENSMUSG00000022321 | 1.944805158 | 1.782320754 | 1.777131114 | 1.130890489 | 1.091002363 | 1.041392685 | 1.041356071 |
| ENSMUSG00000022322 | 2.765209749 | 2.588562345 | 2.571061437 | 2.607351685 | 2.522774815 | 2.921166051 | 2.344542559 |
| ENSMUSG00000022323 | 2.075459174 | 2.001406137 | 1.991446071 | 1.96293726 | 1.902105869 | 2.152288344 | 1.771720588 |
| ENSMUSG00000022324 | 2.34539747 | 2.142463902 | 2.166333979 | 1.54745614 | 1.467903253 | 1.69019608 | 1.340368108 |
| ENSMUSG00000022325 | 2.542156295 | 2.343216265 | 2.328787391 | 2.383332433 | 2.355827317 | 2.632457292 | 2.187847501 |
| ENSMUSG00000022329 | 2.537433341 | 2.321928095 | 2.293731421 | 2.457034121 | 2.337535342 | 2.706717782 | 2.204254617 |
| ENSMUSG00000022330 | 0 | 0 | 0 | 0.5 | 0.632412477 | 0.477121255 | 0.224243824 |
| ENSMUSG00000022332 | 2.337801042 | 2.131869252 | 2.138647213 | 2.102347734 | 2.004851707 | 2.227886705 | 1.89047692 |
| ENSMUSG00000022335 | 1.721148722 | 1.752806814 | 1.660964047 | 1.698603967 | 1.804307395 | 1.875061263 | 1.694069894 |
| ENSMUSG00000022336 | 3.204803826 | 2.910400054 | 2.887686696 | 2.947179583 | 2.851859987 | 3.325310372 | 2.638385672 |
| ENSMUSG00000022337 | 2.544030601 | 2.379909063 | 2.369439567 | 2.42813175 | 2.32657161 | 2.662757832 | 2.156451689 |
| ENSMUSG00000022338 | 2.751792 | 2.543419284 | 2.525821952 | 2.555698726 | 2.474332504 | 2.816903839 | 2.319146374 |
| ENSMUSG00000022339 | 2.179305632 | 2.05357978 | 2.024508021 | 2.080482024 | 1.993840051 | 2.342422681 | 1.872150729 |
| ENSMUSG00000022340 | 0.671187741 | 0.75 | 0.896240625 | 0.896240625 | 1 | 0.698970004 | 0.803905701 |
| ENSMUSG00000022342 | 0 | 0 | 0 | 0 | 0 | 0 | 0 |
| ENSMUSG00000022346 | 2.251208701 | 2.157339155 | 2.202642909 | 2.079353153 | 2.128567294 | 2.340444115 | 2.084261301 |
| ENSMUSG00000022347 | 0 | 0 | 0 | 0 | 0 | 0 | 0 |
| ENSMUSG00000022350 | 3.272119056 | 2.986452396 | 2.988185312 | 2.79575375 | 2.735322923 | 3.152288344 | 2.55053864 |
| ENSMUSG00000022351 | 2.811014908 | 2.573655187 | 2.608135475 | 2.659608978 | 2.636460183 | 3.025715384 | 2.460300884 |
| ENSMUSG00000022353 | 2.129344651 | 1.955044741 | 2.004202072 | 2.267865591 | 2.264824954 | 2.489958479 | 2.189712149 |
| ENSMUSG00000022354 | 2.651426165 | 2.437467357 | 2.476346751 | 2.514497931 | 2.433158226 | 2.783903579 | 2.333765523 |
| ENSMUSG00000022357 | 0 | 0 | 0 | 0 | 0 | 0 | 0 |
| ENSMUSG00000022358 | 1.727821383 | 1.610735874 | 1.488549078 | 1.470660762 | 1.42077253 | 1.505149978 | 1.302263476 |
| ENSMUSG00000022359 | 2.086807695 | 1.945339928 | 1.864857905 | 1.698603967 | 1.746695109 | 1.86923172 | 1.610050272 |

| | | | | | | | |
|---|---|---|---|---|---|---|---|
| ENSMUSG00000022360 | 3.326928073 | 3.027294426 | 3.01610149 | 3.045831306 | 2.960031555 | 3.417803723 | 2.725926557 |
| ENSMUSG00000022361 | 1.944805158 | 1.937048212 | 1.945339928 | 2.472566069 | 2.356272688 | 2.758154622 | 2.183707049 |
| ENSMUSG00000022362 | 0.289064826 | 0.25 | 0 | 0 | 0 | 0 | 0 |
| ENSMUSG00000022364 | 2.594187595 | 2.356566189 | 2.347004321 | 2.441217898 | 2.363759249 | 2.688419822 | 2.19230473 |
| ENSMUSG00000022365 | 2.874454827 | 2.628431899 | 2.642226539 | 2.526477127 | 2.536660727 | 2.88422877 | 2.41820431 |
| ENSMUSG00000022366 | 0 | 0 | 0 | 0 | 0 | 0 | 0 |
| ENSMUSG00000022367 | 2.172524468 | 2.029735268 | 2.002806814 | 1.729715809 | 1.625423019 | 1.929418926 | 1.540636858 |
| ENSMUSG00000022369 | 2.626865868 | 2.374961472 | 2.337734795 | 2.499647607 | 2.433873436 | 2.779596491 | 2.275733199 |
| ENSMUSG00000022370 | 2.582415738 | 2.396240625 | 2.400442697 | 2.480460234 | 2.378690133 | 2.77524626 | 2.252086845 |
| ENSMUSG00000022371 | 0.747221736 | 0.701838731 | 0.951838731 | 0.396240625 | 0.489301084 | 0.477121255 | 0.355418052 |
| ENSMUSG00000022372 | 1.470606563 | 1.410964047 | 1.339388001 | 1.098079356 | 1.020174412 | 1.079181246 | 1 |
| ENSMUSG00000022375 | 0 | 0 | 0 | 0 | 0 | 0 | 0 |
| ENSMUSG00000022376 | 0.45815691 | 0.25 | 0.25 | 0 | 0 | 0 | 0.224243824 |
| ENSMUSG00000022377 | 3.18537608 | 2.918106335 | 2.889126514 | 2.705642708 | 2.66579363 | 3.06669855 | 2.492188045 |
| ENSMUSG00000022378 | 2.769043298 | 2.55357978 | 2.540282971 | 2.54248125 | 2.437074068 | 2.848189117 | 2.297768298 |
| ENSMUSG00000022382 | 0.45815691 | 0 | 0.396240625 | 1.098079356 | 1.149965125 | 1.278753601 | 1.193409508 |
| ENSMUSG00000022383 | 0.747221736 | 0.396240625 | 0.646240625 | 0.896240625 | 1.039257769 | 0.84509804 | 0.969165684 |
| ENSMUSG00000022385 | 2.565000581 | 2.428561379 | 2.445339928 | 2.204245906 | 2.339407793 | 2.511883361 | 2.283060517 |
| ENSMUSG00000022386 | 1.734388958 | 1.575945187 | 1.678561379 | 1.711372513 | 1.728722551 | 1.857332496 | 1.670888075 |
| ENSMUSG00000022387 | 2.69594272 | 2.517194569 | 2.532642641 | 2.319031101 | 2.379941013 | 2.638489257 | 2.307989563 |
| ENSMUSG00000022388 | 0 | 0 | 0 | 0 | 0 | 0 | 0 |
| ENSMUSG00000022389 | 2.773920925 | 2.519203899 | 2.484159485 | 2.357090043 | 2.329944595 | 2.630427875 | 2.170957925 |
| ENSMUSG00000022390 | 2.87190676 | 2.651832578 | 2.686838457 | 2.722375991 | 2.738351308 | 3.087781418 | 2.629532004 |
| ENSMUSG00000022391 | 3.085278234 | 2.851285366 | 2.871583063 | 2.829635702 | 2.901631529 | 3.264109156 | 2.725074826 |
| ENSMUSG00000022394 | 2.514991096 | 2.344844592 | 2.408702763 | 2.349151195 | 2.418913884 | 2.693726949 | 2.299361973 |
| ENSMUSG00000022400 | 2.902690459 | 2.640321986 | 2.64881437 | 2.704245906 | 2.651045594 | 3.053846427 | 2.465891268 |
| ENSMUSG00000022401 | 2.342375483 | 2.158248799 | 2.158248799 | 2.205044741 | 2.154724156 | 2.437750563 | 1.996640515 |
| ENSMUSG00000022403 | 3.367779539 | 3.055170155 | 3.056678306 | 3.024425741 | 2.943899253 | 3.407390904 | 2.726563886 |
| ENSMUSG00000022404 | 2.686083339 | 2.457430684 | 2.454245906 | 2.528109877 | 2.439894911 | 2.828015064 | 2.293479712 |
| ENSMUSG00000022407 | 2.624559692 | 2.407794264 | 2.447790722 | 2.263142013 | 2.362449578 | 2.591064607 | 2.249933144 |
| ENSMUSG00000022408 | 0 | 0 | 0 | 0 | 0 | 0 | 0 |
| ENSMUSG00000022412 | 2.446507203 | 2.277131114 | 2.274508021 | 2.216819685 | 2.219993628 | 2.495544338 | 2.07959139 |
| ENSMUSG00000022414 | 2.331622671 | 2.182829758 | 2.26584877 | 2.05357978 | 2.211376431 | 2.318063335 | 2.161765391 |
| ENSMUSG00000022415 | 1.289064826 | 1.396240625 | 1.5 | 1.580482024 | 1.72341484 | 1.755874856 | 1.732938436 |

| | | | | | | | |
|---|---|---|---|---|---|---|---|
| ENSMUSG00000022416 | 0 | 0.5 | 0.5 | 0 | 0.568060967 | 0.698970004 | 0.672731473 |
| ENSMUSG00000022419 | 2.312518611 | 2.164552871 | 2.147178659 | 1.930274797 | 1.887294348 | 2.037426498 | 1.757893036 |
| ENSMUSG00000022420 | 2.121692466 | 2.070192693 | 2.078220739 | 1.828220739 | 1.944572352 | 2.103803721 | 1.924281235 |
| ENSMUSG00000022421 | 2.4661223 | 2.268535366 | 2.328787391 | 2.526477127 | 2.523329778 | 2.8876173 | 2.385603553 |
| ENSMUSG00000022422 | 1.978790393 | 1.866901388 | 1.890560606 | 2.025821952 | 1.925488995 | 2.136720567 | 1.846338363 |
| ENSMUSG00000022425 | 1.289064826 | 1.356566189 | 1.42510993 | 1.858657057 | 1.746695109 | 1.982271233 | 1.663407615 |
| ENSMUSG00000022426 | 2.756312721 | 2.513142013 | 2.505591953 | 2.586628433 | 2.528012058 | 2.932980822 | 2.365508166 |
| ENSMUSG00000022427 | 2.840968566 | 2.599685923 | 2.626452888 | 2.678346629 | 2.649885189 | 3.022428371 | 2.464780893 |
| ENSMUSG00000022428 | 2.271465912 | 2.060793496 | 2.079353153 | 2.065523711 | 2.125157063 | 2.36361198 | 2.013099581 |
| ENSMUSG00000022429 | 0 | 0 | 0 | 0 | 0 | 0 | 0 |
| ENSMUSG00000022431 | 0 | 0 | 0 | 0.25 | 0 | 0.477121255 | 0.355418052 |
| ENSMUSG00000022432 | 0.45815691 | 0.5 | 0.75 | 0.396240625 | 0.812711509 | 0 | 0.448487648 |
| ENSMUSG00000022433 | 2.740854706 | 2.573367912 | 2.562574604 | 2.362287161 | 2.412899576 | 2.707570176 | 2.334479423 |
| ENSMUSG00000022434 | 2.572927518 | 2.306000419 | 2.348615674 | 2.207430684 | 2.156296347 | 2.51851394 | 2.055736903 |
| ENSMUSG00000022435 | 0 | 0 | 0 | 0 | 0 | 0 | 0 |
| ENSMUSG00000022436 | 2.317377295 | 2.242885888 | 2.319612365 | 2.215271726 | 2.342197974 | 2.519827994 | 2.315133728 |
| ENSMUSG00000022437 | 2.709994145 | 2.506976499 | 2.550224651 | 2.465659339 | 2.49445382 | 2.762678564 | 2.34500308 |
| ENSMUSG00000022438 | 2.606442122 | 2.412314044 | 2.446567407 | 2.224461364 | 2.254288168 | 2.58546073 | 2.146406417 |
| ENSMUSG00000022439 | 0.867194479 | 0.5 | 0.646240625 | 0.396240625 | 0.568060967 | 0.301029996 | 0.355418052 |
| ENSMUSG00000022440 | 2.201144779 | 2.077084758 | 2.113831805 | 2.142463902 | 2.165585202 | 2.421603927 | 2.095400649 |
| ENSMUSG00000022441 | 0 | 0 | 0 | 0 | 0 | 0 | 0 |
| ENSMUSG00000022442 | 2.253091469 | 2.099151195 | 2.121960008 | 2.075945187 | 2.077536464 | 2.324282455 | 1.979298987 |
| ENSMUSG00000022443 | 3.91746109 | 3.600852883 | 3.646578012 | 3.187990603 | 3.326713983 | 3.679609572 | 3.193776931 |
| ENSMUSG00000022445 | 0 | 0 | 0 | 0 | 0 | 0 | 0 |
| ENSMUSG00000022449 | 0 | 0 | 0 | 0.580482024 | 0.568060967 | 0.698970004 | 0.775756176 |
| ENSMUSG00000022450 | 2.229917041 | 2.107613138 | 2.194519282 | 2.113831805 | 2.068600388 | 2.26245109 | 2.104721678 |
| ENSMUSG00000022451 | 2.875300735 | 2.621960008 | 2.604990044 | 2.717205639 | 2.651817083 | 3.046495164 | 2.454939973 |
| ENSMUSG00000022452 | 2.343889214 | 2.234894804 | 2.336628433 | 2.161864607 | 2.183460217 | 2.469822016 | 2.187847501 |
| ENSMUSG00000022453 | 2.188178997 | 2.090485943 | 2.119945066 | 1.898114259 | 2.054233912 | 2.158362492 | 1.966952343 |
| ENSMUSG00000022454 | 0.289064826 | 0.396240625 | 0.646240625 | 0.396240625 | 0.733951626 | 0.84509804 | 0.520678036 |
| ENSMUSG00000022455 | 0 | 0 | 0 | 0 | 0 | 0 | 0 |
| ENSMUSG00000022456 | 1.156259305 | 1.02186571 | 0.92510993 | 1.27186571 | 1.264824954 | 1.176091259 | 1.265599895 |
| ENSMUSG00000022462 | 3.398406911 | 3.077866129 | 3.058105232 | 3.06442062 | 3.006477747 | 3.457881897 | 2.786042561 |
| ENSMUSG00000022463 | 2.877408028 | 2.702843673 | 2.763990059 | 2.632113528 | 2.728412533 | 3.02489596 | 2.687431826 |

| | | | | | | | |
|---|---|---|---|---|---|---|---|
| ENSMUSG00000022464 | 1.759671389 | 1.732684334 | 1.698603967 | 2.684100483 | 2.6165934 | 3.031004281 | 2.444790289 |
| ENSMUSG00000022466 | 2.310886388 | 2.162762923 | 2.181980114 | 2.248588359 | 2.21736452 | 2.522444234 | 2.123508695 |
| ENSMUSG00000022468 | 0 | 0.25 | 0 | 0 | 0 | 0 | 0 |
| ENSMUSG00000022469 | 1.700465191 | 1.67510993 | 1.858657057 | 1.438721876 | 1.625423019 | 1.77815125 | 1.723650273 |
| ENSMUSG00000022471 | 2.730819472 | 2.460194481 | 2.477098155 | 2.593216265 | 2.535323264 | 2.926342447 | 2.376130113 |
| ENSMUSG00000022472 | 2.708736128 | 2.467976309 | 2.461764337 | 2.632357639 | 2.603804521 | 2.970346876 | 2.406241659 |
| ENSMUSG00000022474 | 2.292497211 | 2.142463902 | 2.20584181 | 2.073655187 | 2.208682096 | 2.387389826 | 2.109521699 |
| ENSMUSG00000022475 | 2.777675583 | 2.612545062 | 2.661864607 | 2.376452888 | 2.549763795 | 2.729974286 | 2.447818537 |
| ENSMUSG00000022476 | 2.31414447 | 2.150906586 | 2.185366747 | 2.19697564 | 2.240967488 | 2.418301291 | 2.112839835 |
| ENSMUSG00000022477 | 3.196867486 | 2.935682619 | 2.945544798 | 2.904137211 | 2.920214892 | 3.320146286 | 2.768505194 |
| ENSMUSG00000022479 | 2.307602651 | 2.132845365 | 2.148114259 | 2.340485943 | 2.315755952 | 2.618048097 | 2.20139795 |
| ENSMUSG00000022483 | 2.67325056 | 2.561981878 | 2.629417347 | 1.646240625 | 1.78687392 | 1.939519253 | 1.873141588 |
| ENSMUSG00000022484 | 0 | 0 | 0.396240625 | 0.580482024 | 0.489301084 | 0.84509804 | 0.775756176 |
| ENSMUSG00000022485 | 0 | 0.396240625 | 0.25 | 1 | 0.733951626 | 0.954242509 | 0.876096088 |
| ENSMUSG00000022487 | 0.671187741 | 0.75 | 0.25 | 0 | 0 | 0 | 0.224243824 |
| ENSMUSG00000022488 | 0.289064826 | 0.951838731 | 0.5 | 0.396240625 | 0 | 0 | 0 |
| ENSMUSG00000022489 | 0.960252568 | 0.976722649 | 1.098079356 | 0.5 | 0.568060967 | 0.77815125 | 0.579661877 |
| ENSMUSG00000022490 | 0.289064826 | 0.25 | 0.25 | 0 | 0 | 0 | 0.355418052 |
| ENSMUSG00000022491 | 0 | 0 | 0 | 0 | 0 | 0 | 0 |
| ENSMUSG00000022494 | 0 | 0 | 0 | 0 | 0 | 0 | 0 |
| ENSMUSG00000022496 | 0 | 0 | 0 | 0 | 0 | 0 | 0 |
| ENSMUSG00000022498 | 2.149089517 | 2.038704527 | 2.019203899 | 2.066696635 | 2.065570703 | 2.315970345 | 1.963229454 |
| ENSMUSG00000022500 | 2.655740331 | 2.450224975 | 2.43662858 | 2.310197833 | 2.29413964 | 2.622214023 | 2.163786106 |
| ENSMUSG00000022501 | 0 | 0.25 | 0.396240625 | 0 | 0 | 0 | 0 |
| ENSMUSG00000022503 | 2.083999385 | 1.892463902 | 1.975216702 | 1.868933358 | 1.997548906 | 2.08278537 | 1.882886767 |
| ENSMUSG00000022504 | 0.45815691 | 0.396240625 | 0.25 | 0.5 | 0.244650542 | 0 | 0.224243824 |
| ENSMUSG00000022505 | 2.668421347 | 2.437886015 | 2.43325383 | 2.257666784 | 2.212046813 | 2.550228353 | 2.071655208 |
| ENSMUSG00000022507 | 2.469490009 | 2.27053726 | 2.286801231 | 2.238549078 | 2.117361396 | 2.440909082 | 1.984950056 |
| ENSMUSG00000022508 | 1.482695304 | 1.634789703 | 1.54745614 | 1.494319981 | 1.489301084 | 1.568201724 | 1.392431569 |
| ENSMUSG00000022510 | 0.45815691 | 0.580482024 | 0.396240625 | 0 | 0 | 0 | 0 |
| ENSMUSG00000022512 | 1 | 0.92510993 | 0.92510993 | 1.339388001 | 1.302012594 | 1.342422681 | 1.21680636 |
| ENSMUSG00000022514 | 1.936861443 | 1.802363341 | 1.87694866 | 1.973704441 | 1.897237431 | 2.096910013 | 1.790137008 |
| ENSMUSG00000022515 | 2.078325387 | 1.969129237 | 1.981203126 | 1.928561379 | 2.008447216 | 2.164352856 | 1.881925353 |
| ENSMUSG00000022516 | 2.023453784 | 1.870953944 | 1.943696765 | 1.890560606 | 1.986303335 | 2.139879086 | 1.913109605 |

| | | | | | | | |
|---|---|---|---|---|---|---|---|
| ENSMUSG00000022517 | 2.537433341 | 2.350753006 | 2.408702763 | 2.173371739 | 2.254288168 | 2.501059262 | 2.176428123 |
| ENSMUSG00000022518 | 0.289064826 | 0.5 | 0.701838731 | 0 | 0 | 0.301029996 | 0.448487648 |
| ENSMUSG00000022519 | 1.789260561 | 1.685366747 | 1.552363341 | 1.532320754 | 1.499532413 | 1.763427994 | 1.345462945 |
| ENSMUSG00000022521 | 2.40033094 | 2.247880462 | 2.25 | 2.196158711 | 2.222603297 | 2.440909082 | 2.118913228 |
| ENSMUSG00000022523 | 0 | 0 | 0 | 0 | 0.244650542 | 0 | 0 |
| ENSMUSG00000022525 | 0.289064826 | 0 | 0 | 0 | 0 | 0 | 0 |
| ENSMUSG00000022526 | 2.307602651 | 2.181128463 | 2.181128463 | 2.170748646 | 2.165585202 | 2.376576957 | 2.060761403 |
| ENSMUSG00000022528 | 1.975148165 | 1.866901388 | 1.845926073 | 1.431980114 | 1.47337555 | 1.556302501 | 1.462868481 |
| ENSMUSG00000022529 | 2.054824387 | 1.959800947 | 1.923371739 | 2.012462137 | 1.964031554 | 2.222716471 | 1.837623616 |
| ENSMUSG00000022533 | 3.212403715 | 2.942872367 | 2.923698289 | 2.833009137 | 2.755851989 | 3.198106999 | 2.553692546 |
| ENSMUSG00000022534 | 0 | 0 | 0 | 0 | 0.244650542 | 0 | 0 |
| ENSMUSG00000022536 | 2.877828211 | 2.629909063 | 2.643648382 | 2.661189398 | 2.651624369 | 3.02325246 | 2.475563135 |
| ENSMUSG00000022537 | 1.862428343 | 1.825945187 | 1.766522298 | 1.321350555 | 1.27449559 | 1.447158031 | 1.345462945 |
| ENSMUSG00000022538 | 2.476144923 | 2.255591953 | 2.290597832 | 2.315523711 | 2.269408865 | 2.579783597 | 2.14555506 |
| ENSMUSG00000022540 | 1.693336259 | 1.62694866 | 1.705044741 | 1.649978211 | 1.816839698 | 1.934498451 | 1.833176598 |
| ENSMUSG00000022542 | 0 | 0 | 0 | 0.25 | 0 | 0 | 0 |
| ENSMUSG00000022543 | 0.45815691 | 0.396240625 | 0.25 | 0.5 | 0.568060967 | 0.477121255 | 0.355418052 |
| ENSMUSG00000022544 | 1.568542279 | 1.396240625 | 1.482684334 | 1.403677461 | 1.414412788 | 1.579783597 | 1.479989695 |
| ENSMUSG00000022545 | 1.842799666 | 1.735628626 | 1.763820609 | 1.807204673 | 1.804307395 | 1.944482672 | 1.728327686 |
| ENSMUSG00000022546 | 1.470606563 | 1.53743678 | 1.54745614 | 1.42510993 | 1.649303553 | 1.447158031 | 1.616675393 |
| ENSMUSG00000022548 | 0 | 0 | 0 | 0 | 0 | 0.477121255 | 0.224243824 |
| ENSMUSG00000022550 | 1.806056082 | 1.850219859 | 1.948603967 | 1.630890489 | 1.923977404 | 1.913813852 | 1.87805087 |
| ENSMUSG00000022551 | 2.904664113 | 2.649047439 | 2.699820405 | 2.695134942 | 2.726234744 | 3.115277591 | 2.582214061 |
| ENSMUSG00000022552 | 2.146671932 | 1.958222504 | 1.956637122 | 1.969129237 | 2.06149024 | 2.212187604 | 1.972821265 |
| ENSMUSG00000022553 | 2.460448303 | 2.259043403 | 2.246460484 | 2.189555804 | 2.246477757 | 2.463892989 | 2.119837558 |
| ENSMUSG00000022554 | 1.967766868 | 1.850219859 | 1.882845365 | 1.811981878 | 1.929985312 | 2.045322979 | 1.822941512 |
| ENSMUSG00000022555 | 1.99653913 | 1.910964047 | 1.916333979 | 1.985628626 | 2.025895058 | 2.178976947 | 1.889537832 |
| ENSMUSG00000022556 | 2.414600936 | 2.273849256 | 2.333847589 | 2.188721876 | 2.308696595 | 2.494154594 | 2.202114485 |
| ENSMUSG00000022557 | 2.636714502 | 2.499647607 | 2.563461871 | 2.43155454 | 2.548733265 | 2.77815125 | 2.487926367 |
| ENSMUSG00000022558 | 2.67325056 | 2.538388008 | 2.658702763 | 2.121960008 | 2.322187006 | 2.411619706 | 2.288859925 |
| ENSMUSG00000022559 | 1.89911417 | 1.774508021 | 1.896240625 | 1.516522298 | 1.834841362 | 1.832508913 | 1.793950594 |
| ENSMUSG00000022560 | 1.862428343 | 1.821350555 | 1.956637122 | 1.566696635 | 1.746695109 | 1.86332286 | 1.791413205 |
| ENSMUSG00000022561 | 2.183765914 | 2.124961472 | 2.280383379 | 1.862802778 | 2.121713561 | 2.193124598 | 2.153137801 |
| ENSMUSG00000022562 | 1.445324132 | 1.482684334 | 1.5 | 1.130890489 | 1.319233381 | 1.176091259 | 1.350478789 |

| | | | | | | | |
|---|---|---|---|---|---|---|---|
| ENSMUSG00000022564 | 2.729622822 | 2.53456795 | 2.605778978 | 2.399513125 | 2.532902983 | 2.80685803 | 2.397557454 |
| ENSMUSG00000022565 | 3.181744425 | 2.990633641 | 3.097743752 | 2.664998973 | 2.883916752 | 3.075546961 | 2.85281953 |
| ENSMUSG00000022568 | 2.619908725 | 2.539967834 | 2.631135429 | 2.170748646 | 2.467903253 | 2.586587305 | 2.472474658 |
| ENSMUSG00000022570 | 2.336265009 | 2.172499493 | 2.206637122 | 2.129909063 | 2.229681962 | 2.389166084 | 2.104237736 |
| ENSMUSG00000022571 | 2.34539747 | 2.282320754 | 2.309601185 | 2.177701608 | 2.307165329 | 2.525044807 | 2.272006351 |
| ENSMUSG00000022574 | 1 | 1.114857905 | 1.04248125 | 0.79248125 | 0.905314583 | 0.84509804 | 1.014380832 |
| ENSMUSG00000022575 | 2.042557845 | 1.950224975 | 1.966046536 | 2.339388001 | 2.356272688 | 2.608526034 | 2.247765009 |
| ENSMUSG00000022577 | 0 | 0 | 0 | 0.25 | 0 | 0.301029996 | 0.355418052 |
| ENSMUSG00000022579 | 0 | 0 | 0 | 0 | 0 | 0 | 0 |
| ENSMUSG00000022580 | 0.960252568 | 0.92510993 | 0.951838731 | 1.080482024 | 1.302012594 | 1.255272505 | 1.572224412 |
| ENSMUSG00000022582 | 0 | 0 | 0 | 0 | 0 | 0 | 0 |
| ENSMUSG00000022583 | 0 | 0.5 | 0.25 | 0.396240625 | 0.387761935 | 0 | 0.448487648 |
| ENSMUSG00000022584 | 0.671187741 | 0.25 | 0.396240625 | 0.25 | 0.387761935 | 0 | 0 |
| ENSMUSG00000022586 | 0.289064826 | 0.396240625 | 0.646240625 | 0 | 0 | 0.301029996 | 0.224243824 |
| ENSMUSG00000022587 | 2.188178997 | 1.950224975 | 1.979715809 | 1.75 | 1.683842353 | 1.954242509 | 1.676387053 |
| ENSMUSG00000022589 | 0 | 0 | 0 | 0 | 0 | 0 | 0 |
| ENSMUSG00000022591 | 0 | 0 | 0 | 0 | 0 | 0 | 0 |
| ENSMUSG00000022594 | 1.783508299 | 1.681980114 | 1.708222504 | 1.868933358 | 1.911645804 | 2.100370545 | 1.782372264 |
| ENSMUSG00000022595 | 0 | 0.396240625 | 0.25 | 0.25 | 0 | 0 | 0.224243824 |
| ENSMUSG00000022596 | 0 | 0 | 0.25 | 0 | 0 | 0 | 0 |
| ENSMUSG00000022598 | 0 | 0 | 0.25 | 0 | 0 | 0.477121255 | 0.224243824 |
| ENSMUSG00000022601 | 2.151493169 | 2.013820609 | 1.973704441 | 2.159156155 | 2.090057366 | 2.352182518 | 1.927642465 |
| ENSMUSG00000022602 | 1.83262764 | 1.816696635 | 1.858657057 | 1.04248125 | 1.234113756 | 1.113943352 | 1.23151414 |
| ENSMUSG00000022603 | 0 | 0 | 0.25 | 0 | 0 | 0 | 0 |
| ENSMUSG00000022604 | 2.092368209 | 1.910964047 | 1.938721876 | 1.814346961 | 1.784632921 | 1.991226076 | 1.663407615 |
| ENSMUSG00000022607 | 3.010824585 | 2.757149194 | 2.711764337 | 2.700224975 | 2.672034306 | 3.056142262 | 2.480292754 |
| ENSMUSG00000022610 | 2.213719572 | 2.029735268 | 2.093759858 | 1.92510993 | 1.910073614 | 2.120573931 | 1.769001954 |
| ENSMUSG00000022613 | 0 | 0 | 0 | 0 | 0 | 0 | 0 |
| ENSMUSG00000022614 | 2.608049185 | 2.492166698 | 2.54216803 | 2.186208459 | 2.373217836 | 2.507855872 | 2.324828731 |
| ENSMUSG00000022615 | 0.91631382 | 0.646240625 | 0.830482024 | 0.5 | 0.877063019 | 0.301029996 | 0.803905701 |
| ENSMUSG00000022617 | 2.190368148 | 2.044977273 | 2.046218836 | 2.005591953 | 2.042520869 | 2.206825876 | 1.923435442 |
| ENSMUSG00000022619 | 0 | 0.25 | 0 | 0 | 0.244650542 | 0 | 0.224243824 |
| ENSMUSG00000022620 | 2.22189694 | 2.131869252 | 2.215271726 | 1.898114259 | 1.981187939 | 2.113943352 | 1.983546516 |
| ENSMUSG00000022621 | 1.967766868 | 1.816696635 | 1.819031101 | 1.982684334 | 1.934425071 | 2.181843588 | 1.822941512 |

| | | | | | | | |
|---|---|---|---|---|---|---|---|
| ENSMUSG00000022622 | 0 | 0 | 0 | 0 | 0 | 0 | 0 |
| ENSMUSG00000022623 | 0 | 0 | 0 | 0.646240625 | 0.632412477 | 0.602059991 | 1.028149525 |
| ENSMUSG00000022629 | 2.395018353 | 2.201032755 | 2.26584877 | 2.058404919 | 2.016696984 | 2.301029996 | 1.888596011 |
| ENSMUSG00000022634 | 2.360187403 | 2.237821179 | 2.266522298 | 2.169870025 | 2.133621644 | 2.411619706 | 2.047762527 |
| ENSMUSG00000022635 | 2.518981867 | 2.319031101 | 2.343759858 | 2.363317659 | 2.28445426 | 2.627365857 | 2.168981648 |
| ENSMUSG00000022636 | 2.016886209 | 1.870953944 | 1.874961472 | 2.043731421 | 1.986303335 | 2.212187604 | 1.835407748 |
| ENSMUSG00000022637 | 2.163307908 | 2.025821952 | 1.956637122 | 1.735628626 | 1.639266209 | 1.944482672 | 1.562034107 |
| ENSMUSG00000022639 | 2.029919532 | 1.892463902 | 1.928561379 | 1.850219859 | 1.777823267 | 1.924279286 | 1.699191618 |
| ENSMUSG00000022641 | 2.503815381 | 2.333288838 | 2.313757142 | 2.406883471 | 2.333760374 | 2.629409599 | 2.171745067 |
| ENSMUSG00000022650 | 0 | 0 | 0 | 0 | 0.244650542 | 0 | 0 |
| ENSMUSG00000022651 | 0 | 0 | 0 | 0 | 0 | 0 | 0 |
| ENSMUSG00000022652 | 0 | 0 | 0 | 0 | 0 | 0 | 0 |
| ENSMUSG00000022656 | 2.476144923 | 2.294354884 | 2.281030328 | 2.107613138 | 2.05001946 | 2.342422681 | 1.879993912 |
| ENSMUSG00000022657 | 0 | 0 | 0 | 0 | 0 | 0 | 0 |
| ENSMUSG00000022658 | 0.747221736 | 0.396240625 | 0.646240625 | 0 | 0.387761935 | 0.301029996 | 0.224243824 |
| ENSMUSG00000022659 | 0 | 0 | 0 | 0 | 0 | 0 | 0 |
| ENSMUSG00000022661 | 0.289064826 | 0.396240625 | 0 | 0.396240625 | 0 | 0.301029996 | 0.224243824 |
| ENSMUSG00000022663 | 2.417143825 | 2.237821179 | 2.215271726 | 2.311981878 | 2.26997769 | 2.558708571 | 2.099849366 |
| ENSMUSG00000022664 | 2.179305632 | 1.958222504 | 1.978222334 | 2.078220739 | 2.031524462 | 2.301029996 | 1.911356129 |
| ENSMUSG00000022665 | 2.91746109 | 2.643175057 | 2.603671309 | 2.703845824 | 2.641256943 | 3.04060234 | 2.484057231 |
| ENSMUSG00000022667 | 0 | 0 | 0.25 | 0 | 0 | 0 | 0.224243824 |
| ENSMUSG00000022668 | 2.119110214 | 1.933677405 | 1.884789703 | 1.937048212 | 1.870073561 | 2.167317335 | 1.806347338 |
| ENSMUSG00000022671 | 1.568542279 | 1.580482024 | 1.54745614 | 1.339388001 | 1.47337555 | 1.568201724 | 1.473249546 |
| ENSMUSG00000022672 | 2.156259305 | 1.970660762 | 1.970660762 | 2.002806814 | 1.923977404 | 2.167317335 | 1.821784031 |
| ENSMUSG00000022673 | 3.130949947 | 2.863317659 | 2.828928915 | 2.848749629 | 2.806924204 | 3.238297068 | 2.598592633 |
| ENSMUSG00000022674 | 2.371652933 | 2.212155735 | 2.238549078 | 2.325374049 | 2.254881871 | 2.523746467 | 2.057979654 |
| ENSMUSG00000022676 | 1.99653913 | 1.951838731 | 1.858657057 | 1.823655187 | 1.782377602 | 1.963787827 | 1.629532004 |
| ENSMUSG00000022677 | 2.209569962 | 2.102347734 | 2.034887838 | 2.105516192 | 2.071604288 | 2.399673721 | 1.950241257 |
| ENSMUSG00000022678 | 2.662129038 | 2.453044326 | 2.493244947 | 2.437048212 | 2.425559697 | 2.725911632 | 2.272870211 |
| ENSMUSG00000022679 | 1.445324132 | 1.42510993 | 1.451838731 | 1.410964047 | 1.264824954 | 1.447158031 | 1.302263476 |
| ENSMUSG00000022680 | 2.809038447 | 2.577084758 | 2.5684489 | 2.573080408 | 2.471132517 | 2.824125834 | 2.316393034 |
| ENSMUSG00000022681 | 2.045658471 | 1.825945187 | 1.870953944 | 2.006976499 | 1.965381292 | 2.235528447 | 1.837623616 |
| ENSMUSG00000022682 | 2.756312721 | 2.511780978 | 2.482684334 | 2.539652422 | 2.468227512 | 2.815577748 | 2.296434211 |
| ENSMUSG00000022683 | 0.289064826 | 0 | 0 | 0.396240625 | 0.244650542 | 0 | 0.224243824 |

| | | | | | | | |
|---|---|---|---|---|---|---|---|
| ENSMUSG00000022684 | 2.139333929 | 2.066696635 | 2.028435542 | 2.090485943 | 2.031524462 | 2.356025857 | 1.930140666 |
| ENSMUSG00000022685 | 2.267855219 | 2.065523711 | 2.044977273 | 2.113831805 | 2.057362051 | 2.31386722 | 1.913982791 |
| ENSMUSG00000022686 | 0 | 0 | 0 | 0.92510993 | 1.057362051 | 0.602059991 | 0.74492186 |
| ENSMUSG00000022687 | 1.325351389 | 1.348079356 | 1.348079356 | 1.410964047 | 1.519146132 | 1.462397998 | 1.520678036 |
| ENSMUSG00000022696 | 0 | 0 | 0 | 0 | 0.244650542 | 0 | 0 |
| ENSMUSG00000022698 | 3.089597219 | 2.847138876 | 2.826515422 | 2.859957721 | 2.745071641 | 3.178113252 | 2.544012703 |
| ENSMUSG00000022701 | 1.53838222 | 1.566696635 | 1.51109853 | 1.54248125 | 1.576269619 | 1.763427994 | 1.429563222 |
| ENSMUSG00000022702 | 1.924654746 | 1.83272922 | 1.87892496 | 1.698603967 | 1.810629166 | 1.886490725 | 1.726776051 |
| ENSMUSG00000022704 | 2.227926429 | 2.089388001 | 2.03743678 | 2.072504712 | 2.015530186 | 2.346352974 | 1.911356129 |
| ENSMUSG00000022705 | 0 | 0 | 0 | 0 | 0 | 0 | 0 |
| ENSMUSG00000022706 | 2.177057474 | 1.988549078 | 2.027131114 | 2.120953944 | 2.052132976 | 2.322219295 | 1.939139147 |
| ENSMUSG00000022707 | 1.842799666 | 1.769203899 | 1.782320754 | 1.985628626 | 1.888971102 | 2.123851641 | 1.777090262 |
| ENSMUSG00000022708 | 2.660010325 | 2.425975893 | 2.406427211 | 2.255591953 | 2.157861565 | 2.444044796 | 2.019455688 |
| ENSMUSG00000022710 | 2.870626864 | 2.633332433 | 2.610735874 | 2.698400827 | 2.6165934 | 3.001300933 | 2.435879342 |
| ENSMUSG00000022711 | 2.375872521 | 2.17510993 | 2.166333979 | 2.21293726 | 2.209357611 | 2.428134794 | 2.049487947 |
| ENSMUSG00000022715 | 0 | 0 | 0 | 0 | 0 | 0 | 0 |
| ENSMUSG00000022718 | 2.3855553 | 2.228969845 | 2.216046536 | 2.273849256 | 2.222603297 | 2.478566496 | 2.076967568 |
| ENSMUSG00000022721 | 2.300956576 | 2.12892496 | 2.182829758 | 2.057204673 | 2.079491904 | 2.303196057 | 2.025071334 |
| ENSMUSG00000022722 | 1.956444026 | 1.894357207 | 1.901832578 | 1.92510993 | 1.830919587 | 2.068185862 | 1.702561601 |
| ENSMUSG00000022723 | 2.083999385 | 1.888647213 | 1.973704441 | 2.032320754 | 1.964031554 | 2.243038049 | 1.812372009 |
| ENSMUSG00000022724 | 1.99653913 | 1.779735268 | 1.811981878 | 1.850219859 | 1.84445929 | 2.037426498 | 1.692344475 |
| ENSMUSG00000022738 | 0 | 0 | 0 | 0 | 0 | 0 | 0 |
| ENSMUSG00000022742 | 2.389637216 | 2.182829758 | 2.206637122 | 2.259043403 | 2.221300875 | 2.519827994 | 2.073258096 |
| ENSMUSG00000022744 | 2.58665822 | 2.379909063 | 2.382357639 | 2.497880462 | 2.444084351 | 2.788168371 | 2.294288164 |
| ENSMUSG00000022747 | 0.747221736 | 0.701838731 | 0.646240625 | 1.080482024 | 1.212046813 | 1.176091259 | 1.014380832 |
| ENSMUSG00000022748 | 1.269664473 | 1.348079356 | 1.476722649 | 1.527131114 | 1.614672255 | 1.612783857 | 1.543390198 |
| ENSMUSG00000022749 | 2.45469604 | 2.227473271 | 2.209800947 | 2.304186465 | 2.227124297 | 2.514547753 | 2.063519437 |
| ENSMUSG00000022750 | 2.653588827 | 2.449415381 | 2.442872367 | 2.274508021 | 2.283361516 | 2.545307116 | 2.187473278 |
| ENSMUSG00000022751 | 1.978790393 | 1.852347734 | 1.864857905 | 1.933677405 | 1.880506521 | 2.100370545 | 1.737484395 |
| ENSMUSG00000022752 | 2.992730589 | 2.760756821 | 2.729715809 | 2.816403762 | 2.721205899 | 3.155336037 | 2.510355088 |
| ENSMUSG00000022753 | 0.578129653 | 0.646240625 | 0.79248125 | 0.580482024 | 0.244650542 | 0.477121255 | 0.520678036 |
| ENSMUSG00000022754 | 0.747221736 | 0.864857905 | 0.864857905 | 0.896240625 | 0.686820904 | 0.602059991 | 0.579661877 |
| ENSMUSG00000022755 | 0 | 0 | 0 | 0 | 0 | 0 | 0 |
| ENSMUSG00000022756 | 0.747221736 | 0.896240625 | 0.976722649 | 0.25 | 0.846351821 | 0.698970004 | 0.916588299 |

| | | | | | | | |
|---|---|---|---|---|---|---|---|
| ENSMUSG00000022757 | 2.608049185 | 2.439555804 | 2.389126514 | 2.324229052 | 2.295199569 | 2.662757832 | 2.179896458 |
| ENSMUSG00000022758 | 1.036286563 | 1.098079356 | 1.080482024 | 0.5 | 0.733951626 | 0.602059991 | 0.520678036 |
| ENSMUSG00000022759 | 0 | 0 | 0.580482024 | 0.25 | 0.387761935 | 0 | 0 |
| ENSMUSG00000022760 | 1.734388958 | 1.681980114 | 1.794977273 | 1.339388001 | 1.514343934 | 1.414973348 | 1.659601507 |
| ENSMUSG00000022761 | 2.69788693 | 2.480088214 | 2.493961992 | 2.368933358 | 2.423358301 | 2.655138435 | 2.293479712 |
| ENSMUSG00000022762 | 0 | 0 | 0 | 0 | 0.244650542 | 0 | 0 |
| ENSMUSG00000022763 | 0.45815691 | 0.580482024 | 0.396240625 | 0.701838731 | 0.568060967 | 0.301029996 | 0.579661877 |
| ENSMUSG00000022765 | 2.251208701 | 2.112802778 | 2.079353153 | 2.252107156 | 2.153144932 | 2.440909082 | 2.044866219 |
| ENSMUSG00000022766 | 0 | 0 | 0 | 0.396240625 | 0.244650542 | 0 | 0 |
| ENSMUSG00000022768 | 0.671187741 | 0.646240625 | 0.5 | 0.646240625 | 0.632412477 | 0.84509804 | 0.775756176 |
| ENSMUSG00000022769 | 1.867194479 | 1.80479213 | 1.79745614 | 1.784887838 | 1.850106709 | 2.149219113 | 1.846338363 |
| ENSMUSG00000022770 | 2.740271036 | 2.556000419 | 2.52053726 | 2.564935816 | 2.493851506 | 2.88592634 | 2.292398623 |
| ENSMUSG00000022771 | 2.422183552 | 2.232684334 | 2.255591953 | 2.333288838 | 2.338004387 | 2.638489257 | 2.191935631 |
| ENSMUSG00000022772 | 2.635965117 | 2.401832578 | 2.394357207 | 2.369439567 | 2.28445426 | 2.599883072 | 2.086826862 |
| ENSMUSG00000022773 | 1.205378646 | 1.130890489 | 1.160964047 | 1.146240625 | 1.212046813 | 1.230448921 | 1.100339912 |
| ENSMUSG00000022774 | 2.699178053 | 2.459012589 | 2.453044326 | 2.518200634 | 2.471453824 | 2.84135947 | 2.300684065 |
| ENSMUSG00000022779 | 2.393679564 | 2.181980114 | 2.201838731 | 2.166333979 | 2.146756411 | 2.447158031 | 2.043115906 |
| ENSMUSG00000022780 | 0 | 0 | 0 | 0 | 0 | 0 | 0.224243824 |
| ENSMUSG00000022781 | 2.946589638 | 2.693284802 | 2.666111821 | 2.724461364 | 2.659440622 | 3.056904851 | 2.449322097 |
| ENSMUSG00000022783 | 0 | 0 | 0 | 0 | 0.244650542 | 0 | 0 |
| ENSMUSG00000022787 | 1.614416215 | 1.614857905 | 1.602347734 | 1.610735874 | 1.595992969 | 1.748188027 | 1.496250175 |
| ENSMUSG00000022788 | 1.686083339 | 1.649978211 | 1.589388001 | 1.77186571 | 1.706995316 | 1.892094603 | 1.548827561 |
| ENSMUSG00000022789 | 2.840968566 | 2.587182039 | 2.569902645 | 2.668989258 | 2.598018225 | 2.997386384 | 2.385603553 |
| ENSMUSG00000022790 | 1.205378646 | 1.080482024 | 1.188721876 | 0.25 | 0.632412477 | 0 | 0.355418052 |
| ENSMUSG00000022791 | 2.5496035 | 2.435787849 | 2.513142013 | 2.154596376 | 2.338004387 | 2.444044796 | 2.338964569 |
| ENSMUSG00000022792 | 2.029919532 | 1.926839783 | 1.899978211 | 2.001406137 | 1.929985312 | 2.093421685 | 1.773071385 |
| ENSMUSG00000022793 | 1.721148722 | 1.752806814 | 1.71759118 | 1.96759118 | 1.955822902 | 2.146128036 | 1.774416565 |
| ENSMUSG00000022797 | 3.181744425 | 2.897412787 | 2.887927065 | 3.099819481 | 2.988314062 | 3.467756051 | 2.746920932 |
| ENSMUSG00000022798 | 0 | 0 | 0 | 0 | 0 | 0 | 0 |
| ENSMUSG00000022799 | 2.678702042 | 2.439972046 | 2.471424093 | 2.408702763 | 2.349985771 | 2.684845362 | 2.20919388 |
| ENSMUSG00000022800 | 2.825547124 | 2.58272922 | 2.556000419 | 2.68473417 | 2.588465994 | 2.930949031 | 2.388633692 |
| ENSMUSG00000022801 | 2.546826359 | 2.319612365 | 2.275165585 | 2.359697963 | 2.305627392 | 2.631443769 | 2.167790061 |
| ENSMUSG00000022802 | 1.794934559 | 1.809601185 | 1.779735268 | 1.75835575 | 1.674752852 | 1.819543936 | 1.562034107 |
| ENSMUSG00000022803 | 0.671187741 | 0.701838731 | 0.580482024 | 0 | 0.489301084 | 0.301029996 | 0.448487648 |

| | | | | | | | |
|---|---|---|---|---|---|---|---|
| ENSMUSG00000022805 | 0.671187741 | 0.25 | 0.701838731 | 0.25 | 0 | 0.301029996 | 0.520678036 |
| ENSMUSG00000022807 | 2.406877894 | 2.263820609 | 2.269203899 | 2.266522298 | 2.244650542 | 2.494154594 | 2.100829716 |
| ENSMUSG00000022808 | 2.634462296 | 2.396709947 | 2.394828984 | 2.528760913 | 2.424460715 | 2.818225894 | 2.296968506 |
| ENSMUSG00000022809 | 0 | 0 | 0 | 0 | 0 | 0 | 0 |
| ENSMUSG00000022811 | 2.742019602 | 2.551142786 | 2.514836115 | 2.639126514 | 2.529377461 | 2.908485019 | 2.333527206 |
| ENSMUSG00000022812 | 2.915930694 | 2.656883471 | 2.627938163 | 2.530706999 | 2.490215874 | 2.854913022 | 2.295631111 |
| ENSMUSG00000022814 | 2.682073381 | 2.469895812 | 2.475593779 | 2.497525991 | 2.468227512 | 2.80140371 | 2.316644308 |
| ENSMUSG00000022816 | 3.682805338 | 3.335205929 | 3.283165094 | 3.251713 | 3.163647487 | 3.694166296 | 2.946570778 |
| ENSMUSG00000022817 | 2.91285295 | 2.737091808 | 2.735078399 | 2.382845365 | 2.454682377 | 2.710963119 | 2.366155413 |
| ENSMUSG00000022818 | 0 | 0 | 0 | 0 | 0 | 0 | 0 |
| ENSMUSG00000022820 | 1.837744666 | 1.794977273 | 1.821350555 | 1.630890489 | 1.692703642 | 1.707570176 | 1.647906904 |
| ENSMUSG00000022821 | 0 | 0.25 | 0.5 | 0.25 | 0 | 0.301029996 | 0 |
| ENSMUSG00000022822 | 2.716846213 | 2.518200634 | 2.514497931 | 2.479715809 | 2.477187895 | 2.766412847 | 2.307731473 |
| ENSMUSG00000022824 | 0 | 0.25 | 0.25 | 0 | 0 | 0 | 0 |
| ENSMUSG00000022827 | 1.907802659 | 1.735628626 | 1.752806814 | 1.83272922 | 1.758994476 | 1.973127854 | 1.690609803 |
| ENSMUSG00000022828 | 2.267855219 | 2.100219859 | 2.103406982 | 2.064346961 | 2.027028145 | 2.271841607 | 1.863093717 |
| ENSMUSG00000022829 | 0 | 0 | 0 | 0 | 0 | 0 | 0.355418052 |
| ENSMUSG00000022831 | 0 | 0 | 0 | 0 | 0 | 0 | 0 |
| ENSMUSG00000022832 | 0 | 0 | 0 | 0 | 0 | 0 | 0 |
| ENSMUSG00000022833 | 2.072573125 | 1.888647213 | 1.874961472 | 1.898114259 | 1.78010778 | 2.033423755 | 1.71730663 |
| ENSMUSG00000022836 | 1.721148722 | 1.705044741 | 1.598079356 | 1.364857905 | 1.366364104 | 1.397940009 | 1.413583902 |
| ENSMUSG00000022837 | 2.196867486 | 2.01109853 | 1.972185812 | 2.100219859 | 2.020174412 | 2.264817823 | 1.850609257 |
| ENSMUSG00000022838 | 0.960252568 | 0.92510993 | 0.92510993 | 1.098079356 | 1.149965125 | 1.255272505 | 1.168187745 |
| ENSMUSG00000022840 | 0.289064826 | 0.701838731 | 0.5 | 0.75 | 0.686820904 | 0.301029996 | 0.448487648 |
| ENSMUSG00000022841 | 3.066473338 | 2.767362442 | 2.760927716 | 2.880890489 | 2.851203528 | 3.309630167 | 2.68663243 |
| ENSMUSG00000022842 | 1.881174527 | 1.744319981 | 1.75 | 1.593759858 | 1.677808768 | 1.924279286 | 1.540636858 |
| ENSMUSG00000022843 | 1.505869732 | 1.51109853 | 1.51109853 | 1.642463902 | 1.701347897 | 1.72427587 | 1.631625962 |
| ENSMUSG00000022844 | 2.373064211 | 2.255591953 | 2.220660762 | 1.935366747 | 1.87878897 | 2.158362492 | 1.857950279 |
| ENSMUSG00000022847 | 0.45815691 | 0.25 | 0.5 | 0.396240625 | 0.387761935 | 0.301029996 | 0.803905701 |
| ENSMUSG00000022848 | 2.103271637 | 1.976722649 | 1.948603967 | 2 | 1.962676635 | 2.130333768 | 1.835407748 |
| ENSMUSG00000022849 | 1.948720984 | 1.860735874 | 1.862802778 | 1.919870025 | 1.86473911 | 2.10720997 | 1.842010333 |
| ENSMUSG00000022853 | 0.671187741 | 0.830482024 | 0.646240625 | 0.896240625 | 0.632412477 | 0.698970004 | 0.579661877 |
| ENSMUSG00000022855 | 2.671876482 | 2.43155454 | 2.364857905 | 2.523519421 | 2.439191811 | 2.784617293 | 2.257559981 |
| ENSMUSG00000022856 | 1.765759561 | 1.602347734 | 1.566696635 | 1.445339928 | 1.572189156 | 1.662757832 | 1.40946263 |

| | | | | | | | |
|---|---|---|---|---|---|---|---|
| ENSMUSG00000022857 | 0 | 0 | 0 | 0 | 0 | 0 | 0 |
| ENSMUSG00000022858 | 2.908583619 | 2.669209652 | 2.646475362 | 2.755071375 | 2.674572266 | 3.114944416 | 2.478165355 |
| ENSMUSG00000022860 | 0.578129653 | 0.396240625 | 0.580482024 | 0.396240625 | 0.387761935 | 0.602059991 | 0.448487648 |
| ENSMUSG00000022861 | 0.289064826 | 0 | 0 | 0.25 | 0.489301084 | 0.477121255 | 0.355418052 |
| ENSMUSG00000022863 | 1.605636262 | 1.52186571 | 1.532320754 | 1.470660762 | 1.523883869 | 1.707570176 | 1.448487648 |
| ENSMUSG00000022864 | 1.631440309 | 1.552363341 | 1.606566189 | 1.610735874 | 1.595992969 | 1.77815125 | 1.505628092 |
| ENSMUSG00000022865 | 2.271465912 | 2.131869252 | 2.066696635 | 2.570192693 | 2.502780045 | 2.823474229 | 2.323601898 |
| ENSMUSG00000022867 | 2.37447073 | 2.231944491 | 2.209800947 | 2.391037372 | 2.305627392 | 2.657055853 | 2.141695983 |
| ENSMUSG00000022868 | 0 | 0 | 0 | 0 | 0 | 0 | 0 |
| ENSMUSG00000022871 | 0 | 0.5 | 0.396240625 | 0.25 | 0 | 0 | 0.224243824 |
| ENSMUSG00000022875 | 0 | 0 | 0 | 0 | 0 | 0 | 0 |
| ENSMUSG00000022876 | 0 | 0 | 0 | 0 | 0 | 0.301029996 | 0 |
| ENSMUSG00000022877 | 0 | 0 | 0 | 0 | 0 | 0 | 0 |
| ENSMUSG00000022878 | 0 | 0 | 0.25 | 0 | 0 | 0 | 0 |
| ENSMUSG00000022881 | 2.340856238 | 2.152756199 | 2.149978211 | 2.256284891 | 2.187791022 | 2.496929648 | 2.038994634 |
| ENSMUSG00000022883 | 1.912079952 | 1.83272922 | 1.671625132 | 2.015173983 | 2.040348824 | 2.176091259 | 1.918313719 |
| ENSMUSG00000022884 | 3.275693177 | 2.945032405 | 2.922390314 | 2.996727156 | 2.939577806 | 3.373279893 | 2.718106421 |
| ENSMUSG00000022885 | 2.092368209 | 1.940387808 | 1.903677461 | 1.494319981 | 1.366364104 | 1.544068044 | 1.355418052 |
| ENSMUSG00000022886 | 0 | 0 | 0 | 0 | 0 | 0 | 0 |
| ENSMUSG00000022887 | 2.036286563 | 1.79248125 | 1.79745614 | 1.985628626 | 1.92093466 | 2.176091259 | 1.792684387 |
| ENSMUSG00000022889 | 2.124258828 | 2.017865591 | 2.002806814 | 2.146240625 | 2.059432181 | 2.328379603 | 1.972093446 |
| ENSMUSG00000022890 | 2.615205302 | 2.393884812 | 2.420309603 | 2.431128463 | 2.395014713 | 2.716003344 | 2.236379126 |
| ENSMUSG00000022892 | 3.210516687 | 2.973230556 | 2.95584181 | 2.79544336 | 2.805431261 | 3.223236273 | 2.655044876 |
| ENSMUSG00000022893 | 2.682073381 | 2.461764337 | 2.445339928 | 2.259043403 | 2.207327177 | 2.445604203 | 2.02383181 |
| ENSMUSG00000022894 | 2.838206752 | 2.621960008 | 2.578504176 | 1.930274797 | 1.948833444 | 2.113943352 | 1.786277931 |
| ENSMUSG00000022895 | 2.777141269 | 2.548996825 | 2.545598588 | 2.163659007 | 2.172394856 | 2.42975228 | 2.065163042 |
| ENSMUSG00000022897 | 2.574668783 | 2.372963274 | 2.321928095 | 2.27053726 | 2.235369829 | 2.534026106 | 2.084776044 |
| ENSMUSG00000022898 | 2.590021402 | 2.394357207 | 2.389126514 | 2.29745614 | 2.237868644 | 2.547774705 | 2.120759255 |
| ENSMUSG00000022899 | 0.45815691 | 0 | 0 | 0 | 0 | 0 | 0 |
| ENSMUSG00000022900 | 0 | 0.25 | 0 | 0 | 0 | 0 | 0 |
| ENSMUSG00000022901 | 0 | 0 | 0 | 0 | 0 | 0 | 0 |
| ENSMUSG00000022902 | 0 | 0 | 0 | 0 | 0 | 0 | 0 |
| ENSMUSG00000022905 | 2.643399047 | 2.42293588 | 2.409156155 | 2.501756817 | 2.430643515 | 2.773054693 | 2.224911555 |
| ENSMUSG00000022906 | 1.747221736 | 1.52186571 | 1.571350555 | 2.102347734 | 2.003645018 | 2.367355921 | 1.888596011 |

| | | | | | | | |
|---|---|---|---|---|---|---|---|
| ENSMUSG00000022911 | 2.241665209 | 2.088286706 | 2.025821952 | 2.133818844 | 2.064555002 | 2.348304863 | 1.888596011 |
| ENSMUSG00000022912 | 2.23973002 | 2.044977273 | 2.061981878 | 2.213717096 | 2.170892913 | 2.450249108 | 2.01501956 |
| ENSMUSG00000022913 | 2.075459174 | 1.992885888 | 1.973704441 | 1.981203126 | 1.947418788 | 2.123851641 | 1.850609257 |
| ENSMUSG00000022914 | 2.729622822 | 2.476346751 | 2.456239685 | 2.524508021 | 2.48127199 | 2.825426118 | 2.297768298 |
| ENSMUSG00000022915 | 0 | 0 | 0 | 0 | 0 | 0 | 0 |
| ENSMUSG00000022931 | 0 | 0 | 0 | 0 | 0 | 0 | 0 |
| ENSMUSG00000022935 | 0 | 0 | 0 | 0 | 0.387761935 | 0 | 0 |
| ENSMUSG00000022938 | 0 | 0 | 0 | 0 | 0 | 0 | 0 |
| ENSMUSG00000022940 | 1.912079952 | 1.837182039 | 1.799918086 | 1.741446071 | 1.718026092 | 1.77815125 | 1.631625962 |
| ENSMUSG00000022941 | 1.156259305 | 1.188721876 | 0.79248125 | 1.160964047 | 1.163285805 | 0.903089987 | 1.121219121 |
| ENSMUSG00000022945 | 2.572054151 | 2.373463862 | 2.39151351 | 2.267865591 | 2.244650542 | 2.491361694 | 2.13996595 |
| ENSMUSG00000022946 | 1.558729299 | 1.494319981 | 1.53743678 | 1.671625132 | 1.797870327 | 1.963787827 | 1.769001954 |
| ENSMUSG00000022947 | 1.036286563 | 0.896240625 | 0.896240625 | 1.598079356 | 1.572189156 | 1.748188027 | 1.505628092 |
| ENSMUSG00000022948 | 1.205378646 | 1.348079356 | 1.388647213 | 1.214495249 | 1.264824954 | 1.342422681 | 1.302263476 |
| ENSMUSG00000022949 | 0 | 0 | 0 | 0.25 | 0.244650542 | 0.301029996 | 0.224243824 |
| ENSMUSG00000022951 | 1.876566388 | 1.890560606 | 1.909156155 | 1.930274797 | 1.818885826 | 2.096910013 | 1.801445495 |
| ENSMUSG00000022952 | 2.039433992 | 2.027131114 | 1.945339928 | 2.153677461 | 2.187791022 | 2.411619706 | 2.055173778 |
| ENSMUSG00000022956 | 2.719310222 | 2.530059469 | 2.541854536 | 2.577937079 | 2.580774999 | 2.915927212 | 2.403180579 |
| ENSMUSG00000022957 | 2.489144387 | 2.341033664 | 2.349151195 | 2.377443751 | 2.382016011 | 2.722633923 | 2.252086845 |
| ENSMUSG00000022960 | 2.113897241 | 1.92510993 | 1.930274797 | 2.119945066 | 2.047893212 | 2.238046103 | 1.953344547 |
| ENSMUSG00000022961 | 3.303385603 | 2.984619235 | 2.984159485 | 3.04861227 | 2.988388314 | 3.417305583 | 2.764188671 |
| ENSMUSG00000022962 | 2.673935906 | 2.527783918 | 2.497880462 | 2.451838731 | 2.441995847 | 2.781755375 | 2.295899033 |
| ENSMUSG00000022964 | 1.967766868 | 1.843759858 | 1.910964047 | 2.016522298 | 2.013184954 | 2.255272505 | 1.91917299 |
| ENSMUSG00000022965 | 2.322180025 | 2.190387808 | 2.200224975 | 2.288704527 | 2.26251048 | 2.552668216 | 2.133394659 |
| ENSMUSG00000022967 | 2.580706586 | 2.380890489 | 2.389605178 | 2.37892496 | 2.334234459 | 2.649334859 | 2.199238773 |
| ENSMUSG00000022969 | 2.023453784 | 1.848079356 | 1.903677461 | 1.845926073 | 1.816839698 | 2.064457989 | 1.681794123 |
| ENSMUSG00000022971 | 1.876566388 | 1.726722649 | 1.664552871 | 1.799918086 | 1.770879644 | 1.963787827 | 1.69749337 |
| ENSMUSG00000022972 | 1.993049299 | 1.862802778 | 1.866901388 | 1.905512955 | 1.861137485 | 2.06069784 | 1.778418869 |
| ENSMUSG00000022973 | 2.310886388 | 2.148114259 | 2.159156155 | 2.232684334 | 2.164053936 | 2.46834733 | 2.027536225 |
| ENSMUSG00000022974 | 2.471720134 | 2.27186571 | 2.247171172 | 2.244319981 | 2.205284995 | 2.46686762 | 2.071119143 |
| ENSMUSG00000022978 | 1.985980784 | 1.899978211 | 1.860735874 | 1.938721876 | 1.848234258 | 2.1430148 | 1.842010333 |
| ENSMUSG00000022982 | 2.742600832 | 2.525821952 | 2.566989271 | 2.617918804 | 2.548475162 | 2.935003151 | 2.472474658 |
| ENSMUSG00000022983 | 2.275045611 | 2.115881093 | 2.167221246 | 2.043731421 | 2.065570703 | 2.28780173 | 1.997988498 |
| ENSMUSG00000022986 | 0 | 0.25 | 0 | 0 | 0 | 0 | 0.224243824 |

| | | | | | | | |
|---|---|---|---|---|---|---|---|
| ENSMUSG00000022987 | 1.389637216 | 1.27186571 | 1.17510993 | 1.516522298 | 1.478764298 | 1.579783597 | 1.462868481 |
| ENSMUSG00000022991 | 0 | 0 | 0 | 0 | 0 | 0 | 0 |
| ENSMUSG00000022992 | 2.755187126 | 2.524836953 | 2.526804269 | 2.525821952 | 2.474013807 | 2.838849091 | 2.327511498 |
| ENSMUSG00000022993 | 0.671187741 | 0.580482024 | 0.75 | 0.646240625 | 0.387761935 | 0.477121255 | 0.448487648 |
| ENSMUSG00000022994 | 2.701748369 | 2.496104615 | 2.533285553 | 2.365369862 | 2.409087918 | 2.62838893 | 2.316644308 |
| ENSMUSG00000022995 | 2.769043298 | 2.557204673 | 2.517530236 | 2.718745337 | 2.612740596 | 3.001300933 | 2.444282813 |
| ENSMUSG00000022996 | 2.267855219 | 2.094844592 | 2.198603967 | 2.157339155 | 2.165585202 | 2.365487985 | 2.134278581 |
| ENSMUSG00000022997 | 0 | 0 | 0.646240625 | 0 | 0 | 0 | 0.224243824 |
| ENSMUSG00000022999 | 1.816888701 | 1.711372513 | 1.695339928 | 1.688721876 | 1.816839698 | 1.763427994 | 1.743447983 |
| ENSMUSG00000023000 | 0 | 0 | 0 | 0.396240625 | 0.686820904 | 0.301029996 | 0.672731473 |
| ENSMUSG00000023004 | 2.998115814 | 2.740181499 | 2.755765312 | 2.697994205 | 2.721838437 | 3.067814511 | 2.504505747 |
| ENSMUSG00000023007 | 2.016886209 | 1.923371739 | 1.969129237 | 1.769203899 | 1.866526233 | 1.913813852 | 1.845261772 |
| ENSMUSG00000023008 | 2.858927525 | 2.659835177 | 2.68346568 | 2.482314602 | 2.507741065 | 2.82672252 | 2.367660638 |
| ENSMUSG00000023009 | 1.721148722 | 1.774508021 | 1.888647213 | 1.418106335 | 1.728722551 | 1.51851394 | 1.874129423 |
| ENSMUSG00000023010 | 3.086043666 | 2.881502527 | 2.919650035 | 2.771036003 | 2.887788338 | 3.206825876 | 2.791181543 |
| ENSMUSG00000023011 | 0.671187741 | 0.830482024 | 0.79248125 | 0 | 0 | 0 | 0 |
| ENSMUSG00000023013 | 0.289064826 | 0 | 0 | 0 | 0.244650542 | 0 | 0 |
| ENSMUSG00000023015 | 3.239200679 | 2.942872367 | 2.949009902 | 3.005938588 | 2.945160263 | 3.393048466 | 2.77178211 |
| ENSMUSG00000023017 | 1.811507563 | 1.763820609 | 1.714495249 | 1.464495249 | 1.628935049 | 1.681241237 | 1.579661877 |
| ENSMUSG00000023018 | 2.463861956 | 2.276477127 | 2.289967834 | 2.252107156 | 2.288252441 | 2.591064607 | 2.200320163 |
| ENSMUSG00000023019 | 0.960252568 | 0.646240625 | 0.580482024 | 0.396240625 | 0.387761935 | 0.477121255 | 0.520678036 |
| ENSMUSG00000023020 | 1.936861443 | 1.969129237 | 2.025821952 | 1.618933358 | 1.810629166 | 1.755874856 | 1.853775828 |
| ENSMUSG00000023021 | 2.946589638 | 2.681980114 | 2.656198864 | 2.85035322 | 2.77687827 | 3.21031852 | 2.574156532 |
| ENSMUSG00000023022 | 3.29280785 | 3.010414788 | 3.027457664 | 2.723514962 | 2.686122326 | 3.047664195 | 2.526375977 |
| ENSMUSG00000023025 | 3.09186576 | 2.842944161 | 2.823798739 | 2.918989258 | 2.856205454 | 3.252853031 | 2.594894066 |
| ENSMUSG00000023026 | 2.696591798 | 2.449009902 | 2.440803092 | 2.347004321 | 2.319233381 | 2.62324929 | 2.226242896 |
| ENSMUSG00000023027 | 2.584117913 | 2.370449858 | 2.411864607 | 2.447383411 | 2.359815565 | 2.692846919 | 2.202829437 |
| ENSMUSG00000023030 | 2.744339677 | 2.510756821 | 2.536801231 | 2.531030328 | 2.515479755 | 2.813580989 | 2.34638072 |
| ENSMUSG00000023031 | 1.069666964 | 1.146240625 | 1.098079356 | 1.238549078 | 1.176121988 | 1.322219295 | 1.041356071 |
| ENSMUSG00000023032 | 2.029919532 | 1.825945187 | 1.882845365 | 1.784887838 | 1.898867717 | 2.025305865 | 1.822941512 |
| ENSMUSG00000023033 | 0.578129653 | 0.25 | 0.25 | 0.896240625 | 1 | 0.84509804 | 0.74492186 |
| ENSMUSG00000023034 | 1.307602651 | 1.458222504 | 1.451838731 | 1.214495249 | 1.254881871 | 1.301029996 | 1.392431569 |
| ENSMUSG00000023036 | 1.227926429 | 1.27186571 | 1.311981878 | 1.17510993 | 1.176121988 | 1.204119983 | 1.140832123 |
| ENSMUSG00000023039 | 1.783508299 | 1.814346961 | 1.769203899 | 0.580482024 | 0.686820904 | 0.84509804 | 0.520678036 |

| | | | | | | | |
|---|---|---|---|---|---|---|---|
| ENSMUSG00000023041 | 0 | 0 | 0 | 0 | 0 | 0 | 0 |
| ENSMUSG00000023043 | 0.289064826 | 0.5 | 0.5 | 1.146240625 | 1.091002363 | 1.079181246 | 1.100339912 |
| ENSMUSG00000023044 | 2.131864492 | 1.988549078 | 2.036164561 | 1.994319981 | 2.064555002 | 2.193124598 | 1.983546516 |
| ENSMUSG00000023045 | 0.45815691 | 0.580482024 | 1.02186571 | 0.646240625 | 0.489301084 | 0.602059991 | 0.355418052 |
| ENSMUSG00000023046 | 0.45815691 | 0.976722649 | 0.701838731 | 0.701838731 | 1.039257769 | 0.698970004 | 1.224243824 |
| ENSMUSG00000023047 | 0.289064826 | 0.580482024 | 0.396240625 | 0 | 0.244650542 | 0.301029996 | 0.579661877 |
| ENSMUSG00000023048 | 2.342375483 | 2.171625132 | 2.225216702 | 2.367918804 | 2.399374698 | 2.700703717 | 2.283617342 |
| ENSMUSG00000023050 | 2.29078455 | 2.137686696 | 2.215271726 | 2.063166358 | 2.088159748 | 2.240549248 | 2.001999072 |
| ENSMUSG00000023051 | 2.358731791 | 2.198603967 | 2.210587586 | 2.093759858 | 2.198391101 | 2.324282455 | 2.097385413 |
| ENSMUSG00000023052 | 0.747221736 | 0.646240625 | 0.75 | 0.896240625 | 0.955822902 | 1.079181246 | 0.803905701 |
| ENSMUSG00000023055 | 2.245508853 | 2.107613138 | 2.17510993 | 2.015173983 | 2.145949654 | 2.271841607 | 2.098866035 |
| ENSMUSG00000023057 | 0 | 0 | 0 | 0 | 0 | 0 | 0 |
| ENSMUSG00000023064 | 0.289064826 | 0.396240625 | 0.25 | 0 | 0 | 0 | 0 |
| ENSMUSG00000023066 | 2.285603957 | 2.064346961 | 2.081607372 | 2.101285366 | 2.040348824 | 2.320146286 | 1.91917299 |
| ENSMUSG00000023067 | 2.625329835 | 2.449820405 | 2.43155454 | 1.726722649 | 1.683842353 | 1.77815125 | 1.641897157 |
| ENSMUSG00000023068 | 2.792365035 | 2.575374049 | 2.539967834 | 2.666333979 | 2.568060967 | 2.963787827 | 2.372134746 |
| ENSMUSG00000023070 | 0 | 0 | 0 | 0 | 0 | 0 | 0 |
| ENSMUSG00000023072 | 1.852729478 | 1.77186571 | 1.837182039 | 1.802363341 | 1.816839698 | 2.008600172 | 1.746389053 |
| ENSMUSG00000023073 | 0 | 0 | 0.25 | 0.25 | 0 | 0 | 0 |
| ENSMUSG00000023074 | 2.172524468 | 2.097004321 | 2.075945187 | 2.013820609 | 1.938809675 | 2.198657087 | 1.803905701 |
| ENSMUSG00000023075 | 2.405576699 | 2.269203899 | 2.231203126 | 2.338286706 | 2.298884454 | 2.62838893 | 2.140399326 |
| ENSMUSG00000023078 | 0 | 0.25 | 0 | 0 | 0 | 0 | 0 |
| ENSMUSG00000023079 | 2.573799059 | 2.399978211 | 2.410061234 | 2.24929487 | 2.298884454 | 2.544068044 | 2.178359564 |
| ENSMUSG00000023083 | 0 | 0.5 | 0 | 0.396240625 | 0.568060967 | 0.301029996 | 0.448487648 |
| ENSMUSG00000023084 | 0.45815691 | 0 | 0.25 | 0.5 | 0.244650542 | 0.301029996 | 0.224243824 |
| ENSMUSG00000023087 | 2.674620126 | 2.467205639 | 2.466433318 | 2.497525991 | 2.468227512 | 2.808210973 | 2.282781745 |
| ENSMUSG00000023088 | 2.83912939 | 2.575374049 | 2.574515642 | 2.467976309 | 2.465951417 | 2.790285164 | 2.283060517 |
| ENSMUSG00000023089 | 2.22189694 | 2.03743678 | 2.057204673 | 2.104463129 | 2.047893212 | 2.352182518 | 1.939139147 |
| ENSMUSG00000023092 | 3.351241591 | 3.057430023 | 3.029491926 | 2.256976499 | 2.212715924 | 2.51054501 | 2.097385413 |
| ENSMUSG00000023093 | 0 | 0 | 0 | 0 | 0 | 0 | 0 |
| ENSMUSG00000023094 | 1.069666964 | 1.188721876 | 1.282320754 | 1.146240625 | 1.176121988 | 1.204119983 | 0.984950056 |
| ENSMUSG00000023104 | 2.652148299 | 2.445339928 | 2.455443495 | 2.435366747 | 2.514912301 | 2.768638101 | 2.365508166 |
| ENSMUSG00000023106 | 2.888197527 | 2.656427211 | 2.666333979 | 2.670309603 | 2.556642263 | 2.962369336 | 2.387223158 |
| ENSMUSG00000023110 | 2.519973621 | 2.308404919 | 2.254897682 | 2.579635702 | 2.537194296 | 2.911157609 | 2.379870218 |

| | | | | | | | |
|---|---|---|---|---|---|---|---|
| ENSMUSG00000023118 | 2.661423995 | 2.513820609 | 2.563166358 | 2.352347734 | 2.486234548 | 2.667452953 | 2.396774124 |
| ENSMUSG00000023120 | 0 | 0 | 0 | 0 | 0 | 0 | 0 |
| ENSMUSG00000023122 | 0 | 0 | 0 | 0 | 0 | 0 | 0 |
| ENSMUSG00000023132 | 0 | 0 | 0 | 0 | 0 | 0 | 0 |
| ENSMUSG00000023140 | 0 | 0 | 0 | 0 | 0.387761935 | 0 | 0.355418052 |
| ENSMUSG00000023143 | 1.753493018 | 1.695339928 | 1.828220739 | 1.396240625 | 1.576269619 | 1.579783597 | 1.62316756 |
| ENSMUSG00000023147 | 2.448863324 | 2.294354884 | 2.298073204 | 1.964495249 | 1.945998439 | 2.176091259 | 1.781059827 |
| ENSMUSG00000023150 | 3 | 2.735445311 | 2.700629091 | 2.778435542 | 2.691328263 | 3.084576278 | 2.497688664 |
| ENSMUSG00000023151 | 0 | 0.25 | 0 | 0 | 0 | 0 | 0 |
| ENSMUSG00000023153 | 0 | 0 | 0 | 0 | 0.244650542 | 0 | 0.224243824 |
| ENSMUSG00000023156 | 2.075459174 | 1.896240625 | 1.880890489 | 2.097004321 | 2.031524462 | 2.348304863 | 1.885753983 |
| ENSMUSG00000023159 | 0 | 0 | 0 | 0 | 0 | 0 | 0 |
| ENSMUSG00000023165 | 0 | 0 | 0 | 0 | 0 | 0 | 0 |
| ENSMUSG00000023169 | 3.097359679 | 2.808853984 | 2.785207443 | 2.681980114 | 2.626045333 | 3.03702788 | 2.467470936 |
| ENSMUSG00000023170 | 2.075459174 | 1.907339155 | 1.985628626 | 1.886723615 | 1.964031554 | 2.053078443 | 1.881925353 |
| ENSMUSG00000023175 | 3.237253994 | 2.957826811 | 2.986635201 | 2.933359771 | 2.95517094 | 3.349471799 | 2.821784031 |
| ENSMUSG00000023176 | 0 | 0 | 0 | 0 | 0 | 0 | 0 |
| ENSMUSG00000023185 | 0 | 0 | 0 | 0 | 0 | 0 | 0 |
| ENSMUSG00000023186 | 2.29078455 | 2.097004321 | 2.066696635 | 2.365369862 | 2.287171411 | 2.595496222 | 2.167790061 |
| ENSMUSG00000023191 | 1.827447047 | 1.671625132 | 1.774508021 | 1.527131114 | 1.677808768 | 1.633468456 | 1.645916034 |
| ENSMUSG00000023192 | 0 | 0 | 0 | 0.25 | 0.244650542 | 0 | 0 |
| ENSMUSG00000023206 | 0.811507563 | 0.896240625 | 0.951838731 | 1.418106335 | 1.439191811 | 1.477121255 | 1.290497981 |
| ENSMUSG00000023210 | 0 | 0 | 0 | 0 | 0 | 0 | 0 |
| ENSMUSG00000023216 | 0.45815691 | 0.396240625 | 0.396240625 | 0 | 0.632412477 | 0 | 0 |
| ENSMUSG00000023224 | 0.867194479 | 0.646240625 | 0.896240625 | 0.580482024 | 0.686820904 | 0 | 0.579661877 |
| ENSMUSG00000023232 | 1.811507563 | 1.720660762 | 1.79745614 | 1.53743678 | 1.639266209 | 1.716003344 | 1.56970677 |
| ENSMUSG00000023235 | 1.83262764 | 1.653677461 | 1.747171172 | 1.494319981 | 1.467903253 | 1.491361694 | 1.374453863 |
| ENSMUSG00000023236 | 0.45815691 | 0 | 0 | 0 | 0.244650542 | 0.301029996 | 0.355418052 |
| ENSMUSG00000023243 | 0 | 0 | 0.396240625 | 1.561981878 | 1.652586889 | 1.748188027 | 1.574722613 |
| ENSMUSG00000023247 | 0 | 0 | 0 | 0 | 0 | 0 | 0 |
| ENSMUSG00000023249 | 2.194712275 | 1.991446071 | 2.090485943 | 1.888647213 | 2.042520869 | 2.155336037 | 1.94631981 |
| ENSMUSG00000023257 | 0 | 0 | 0 | 0 | 0 | 0 | 0 |
| ENSMUSG00000023259 | 1.54867984 | 1.42510993 | 1.445339928 | 1.464495249 | 1.572189156 | 1.544068044 | 1.523605797 |
| ENSMUSG00000023262 | 1.920505136 | 1.821350555 | 1.860735874 | 1.747171172 | 1.84445929 | 1.991226076 | 1.811176017 |

| | | | | | | | |
|---|---|---|---|---|---|---|---|
| ENSMUSG00000023263 | 0 | 0 | 0 | 0 | 0 | 0 | 0 |
| ENSMUSG00000023266 | 1.100572389 | 0.92510993 | 1 | 0.396240625 | 0.905314583 | 0.477121255 | 1.089372236 |
| ENSMUSG00000023267 | 0 | 0 | 0 | 0 | 0.244650542 | 0.301029996 | 0 |
| ENSMUSG00000023272 | 2.330063669 | 2.262462137 | 2.267194569 | 2.226722649 | 2.265401209 | 2.51851394 | 2.165794278 |
| ENSMUSG00000023274 | 0 | 0 | 0 | 0 | 0 | 0 | 0 |
| ENSMUSG00000023277 | 1.993049299 | 1.984159485 | 1.988549078 | 1.959800947 | 2.075570129 | 2.243038049 | 2.074322292 |
| ENSMUSG00000023279 | 0 | 0 | 0 | 0 | 0 | 0 | 0 |
| ENSMUSG00000023284 | 1.881174527 | 1.766522298 | 1.75835575 | 1.657339155 | 1.625423019 | 1.770852012 | 1.56716938 |
| ENSMUSG00000023286 | 2.158622101 | 1.990000483 | 2.027131114 | 2.066696635 | 2.065570703 | 2.274157849 | 1.987739 |
| ENSMUSG00000023289 | 0 | 0 | 0 | 0 | 0 | 0 | 0 |
| ENSMUSG00000023307 | 2.872757846 | 2.623963757 | 2.567573832 | 2.709997768 | 2.589846528 | 2.995196292 | 2.423302014 |
| ENSMUSG00000023328 | 0 | 0.25 | 0.25 | 0.25 | 0 | 0 | 0.224243824 |
| ENSMUSG00000023330 | 2.129344651 | 2.02186571 | 1.956637122 | 1.685366747 | 1.645989387 | 1.886490725 | 1.520678036 |
| ENSMUSG00000023333 | 0 | 0 | 0 | 0 | 0 | 0 | 0 |
| ENSMUSG00000023336 | 0 | 0 | 0 | 0 | 0 | 0 | 0 |
| ENSMUSG00000023341 | 0.289064826 | 0 | 0 | 0.5 | 0.387761935 | 0.77815125 | 0.224243824 |
| ENSMUSG00000023345 | 1.881174527 | 1.848079356 | 1.899978211 | 1.905512955 | 2.032639648 | 2.152288344 | 1.918313719 |
| ENSMUSG00000023348 | 2.505869732 | 2.407339155 | 2.486726569 | 2.157339155 | 2.358048572 | 2.477121255 | 2.298300395 |
| ENSMUSG00000023349 | 0 | 0 | 0 | 0 | 0 | 0 | 0 |
| ENSMUSG00000023350 | 0 | 0 | 0 | 0 | 0 | 0 | 0.224243824 |
| ENSMUSG00000023353 | 2.37447073 | 2.331607372 | 2.413659007 | 1.961372513 | 2.170892913 | 2.247973266 | 2.185972044 |
| ENSMUSG00000023367 | 2.190368148 | 2.044977273 | 2.101285366 | 2.111770807 | 2.114724103 | 2.426511261 | 2.011813236 |
| ENSMUSG00000023387 | 0 | 0 | 0 | 0 | 0 | 0 | 0 |
| ENSMUSG00000023391 | 1.156259305 | 1.27186571 | 1.282320754 | 0.79248125 | 1.200473444 | 1.113943352 | 1.21680636 |
| ENSMUSG00000023393 | 0.578129653 | 0.5 | 0.75 | 0.25 | 0.387761935 | 0.301029996 | 0.629532004 |
| ENSMUSG00000023403 | 0 | 0.25 | 0 | 0 | 0 | 0 | 0 |
| ENSMUSG00000023411 | 1.847794126 | 1.692046081 | 1.75835575 | 1.769203899 | 1.861137485 | 1.986771734 | 1.806347338 |
| ENSMUSG00000023433 | 0 | 0 | 0 | 0 | 0 | 0 | 0 |
| ENSMUSG00000023439 | 0.289064826 | 0 | 0.396240625 | 0.5 | 0 | 0.301029996 | 0.224243824 |
| ENSMUSG00000023443 | 0.45815691 | 0.75 | 0.75 | 1.29248125 | 1.136121934 | 1.342422681 | 1.340368108 |
| ENSMUSG00000023452 | 2.17479713 | 1.956637122 | 1.959800947 | 1.841580554 | 1.887294348 | 1.973127854 | 1.769001954 |
| ENSMUSG00000023456 | 3.326068802 | 3.031272635 | 3.07401396 | 3.033285553 | 3.054787955 | 3.504878459 | 2.880961074 |
| ENSMUSG00000023460 | 2.615205302 | 2.367918804 | 2.315523711 | 2.450629091 | 2.354487824 | 2.694605199 | 2.207438502 |
| ENSMUSG00000023467 | 0 | 0.396240625 | 0.25 | 0 | 0.387761935 | 0 | 0 |

| | | | | | | | |
|---|---|---|---|---|---|---|---|
| ENSMUSG00000023473 | 0.45815691 | 0.25 | 0.5 | 0.396240625 | 0.489301084 | 0.301029996 | 0.224243824 |
| ENSMUSG00000023484 | 0.811507563 | 0.864857905 | 0.25 | 1.061981878 | 1.223252711 | 1.322219295 | 1.365075947 |
| ENSMUSG00000023495 | 2.65645504 | 2.487456678 | 2.577653195 | 2.267865591 | 2.466277473 | 2.592176757 | 2.395201749 |
| ENSMUSG00000023505 | 2.596667477 | 2.379417347 | 2.431980114 | 2.465659339 | 2.412899576 | 2.736396502 | 2.296166733 |
| ENSMUSG00000023571 | 0.91631382 | 0.830482024 | 1.098079356 | 0.701838731 | 1 | 0.477121255 | 1.089372236 |
| ENSMUSG00000023572 | 2.124258828 | 2.017865591 | 2.01109853 | 2.127938163 | 2.045754077 | 2.303196057 | 1.953344547 |
| ENSMUSG00000023577 | 0 | 0 | 0 | 0 | 0 | 0 | 0 |
| ENSMUSG00000023707 | 1.678702042 | 1.552363341 | 1.741446071 | 1.372963274 | 1.555378544 | 1.477121255 | 1.562034107 |
| ENSMUSG00000023723 | 2.243591459 | 2.113831805 | 2.133818844 | 2.056000419 | 2.046825265 | 2.357934847 | 1.988432495 |
| ENSMUSG00000023737 | 0.578129653 | 0.580482024 | 0.396240625 | 0.701838731 | 0.489301084 | 0.301029996 | 0.520678036 |
| ENSMUSG00000023755 | 1.37447073 | 1.42510993 | 1.476722649 | 1.51109853 | 1.462344774 | 1.69019608 | 1.473249546 |
| ENSMUSG00000023764 | 2.048736215 | 1.933677405 | 1.951838731 | 1.94697564 | 2 | 2.120573931 | 1.89047692 |
| ENSMUSG00000023781 | 0.289064826 | 0.396240625 | 0.25 | 0 | 0.244650542 | 0 | 0 |
| ENSMUSG00000023791 | 1.862428343 | 1.819031101 | 1.714495249 | 1.819031101 | 1.698489937 | 1.968482949 | 1.651852264 |
| ENSMUSG00000023795 | 1.37447073 | 1.29248125 | 1.130890489 | 1.27186571 | 1.264824954 | 1.255272505 | 1.140832123 |
| ENSMUSG00000023800 | 2.564110437 | 2.348079356 | 2.3464656 | 2.269871196 | 2.230318492 | 2.51851394 | 2.105204898 |
| ENSMUSG00000023802 | 0 | 0 | 0 | 0 | 0 | 0 | 0 |
| ENSMUSG00000023805 | 2.527823874 | 2.326515422 | 2.320772088 | 1.864857905 | 1.882215755 | 1.991226076 | 1.812372009 |
| ENSMUSG00000023806 | 0 | 0.396240625 | 0.5 | 0 | 0.244650542 | 0.301029996 | 0.803905701 |
| ENSMUSG00000023809 | 2.418409478 | 2.21759118 | 2.231203126 | 2.088286706 | 2.091944837 | 2.423245874 | 1.985649549 |
| ENSMUSG00000023826 | 0.289064826 | 0.25 | 0.5 | 0.5 | 0.568060967 | 0.698970004 | 0.224243824 |
| ENSMUSG00000023827 | 2.22189694 | 2.08272922 | 2.078220739 | 1.894357207 | 1.931471446 | 2.173186268 | 1.864112664 |
| ENSMUSG00000023828 | 0 | 0 | 0 | 0 | 0 | 0 | 0 |
| ENSMUSG00000023829 | 0 | 0 | 0 | 0 | 0 | 0 | 0 |
| ENSMUSG00000023830 | 2.72721917 | 2.488912477 | 2.500352049 | 2.499647607 | 2.509475496 | 2.834420704 | 2.347752519 |
| ENSMUSG00000023832 | 2.499676068 | 2.27186571 | 2.246460484 | 2.463327389 | 2.410235744 | 2.728353782 | 2.249624299 |
| ENSMUSG00000023845 | 2.826972877 | 2.607613138 | 2.5658173 | 2.554186465 | 2.460030299 | 2.808210973 | 2.273444839 |
| ENSMUSG00000023852 | 2.865034804 | 2.646005735 | 2.623213655 | 2.68176739 | 2.604245716 | 2.982271233 | 2.407002425 |
| ENSMUSG00000023861 | 1.857607105 | 1.794977273 | 1.744319981 | 1.92510993 | 1.888971102 | 2.093421685 | 1.715701091 |
| ENSMUSG00000023868 | 0.671187741 | 0.830482024 | 0.864857905 | 0.864857905 | 0.733951626 | 0.477121255 | 0.629532004 |
| ENSMUSG00000023873 | 0.45815691 | 0 | 0.25 | 0.396240625 | 0 | 0 | 0 |
| ENSMUSG00000023882 | 1.494443473 | 1.470660762 | 1.482684334 | 1.348079356 | 1.27449559 | 1.462397998 | 1.245582246 |
| ENSMUSG00000023883 | 2.555102908 | 2.319031101 | 2.291226732 | 2.440803092 | 2.338472809 | 2.67669361 | 2.201756416 |
| ENSMUSG00000023885 | 1.558729299 | 1.438721876 | 1.388647213 | 1.470660762 | 1.414412788 | 1.544068044 | 1.369798885 |

| | | | | | | | |
|---|---|---|---|---|---|---|---|
| ENSMUSG00000023886 | 2.334723298 | 2.173371739 | 2.133818844 | 1.201838731 | 1.223252711 | 1.255272505 | 1.150210039 |
| ENSMUSG00000023892 | 1.975148165 | 1.828220739 | 1.850219859 | 1.909156155 | 1.802174723 | 2.029383778 | 1.670888075 |
| ENSMUSG00000023902 | 0 | 0 | 0 | 0 | 0 | 0 | 0 |
| ENSMUSG00000023903 | 0 | 0.580482024 | 0.396240625 | 1.04248125 | 1.121713561 | 0.954242509 | 1.140832123 |
| ENSMUSG00000023904 | 2.512981297 | 2.448603967 | 2.457430684 | 1.978222334 | 2.115605392 | 2.250420002 | 2.098866035 |
| ENSMUSG00000023905 | 2.68140129 | 2.545598588 | 2.581888161 | 2.484894804 | 2.529922146 | 2.841984805 | 2.423482597 |
| ENSMUSG00000023906 | 0 | 0 | 0 | 0 | 0 | 0 | 0 |
| ENSMUSG00000023908 | 1.811507563 | 1.823655187 | 1.964495249 | 1.54248125 | 1.818885826 | 1.763427994 | 1.892346957 |
| ENSMUSG00000023909 | 1.623015126 | 1.580482024 | 1.606566189 | 1.610735874 | 1.662257525 | 1.819543936 | 1.631625962 |
| ENSMUSG00000023911 | 0.671187741 | 0.79248125 | 0.976722649 | 0.701838731 | 0.77552387 | 0.698970004 | 0.829800754 |
| ENSMUSG00000023912 | 1.596667477 | 1.532320754 | 1.53743678 | 1.380890489 | 1.366364104 | 1.414973348 | 1.355418052 |
| ENSMUSG00000023913 | 1.432083896 | 1.348079356 | 1.282320754 | 1.130890489 | 1.057362051 | 1.278753601 | 1.176815335 |
| ENSMUSG00000023914 | 0 | 0.396240625 | 0.75 | 0 | 0 | 0 | 0 |
| ENSMUSG00000023915 | 1.587501561 | 1.614857905 | 1.584962501 | 1.735628626 | 1.686820904 | 1.77815125 | 1.594042709 |
| ENSMUSG00000023918 | 0.289064826 | 0 | 0 | 0 | 0 | 0 | 0.355418052 |
| ENSMUSG00000023919 | 2.307602651 | 2.178561379 | 2.155512955 | 2.294977273 | 2.197694248 | 2.523746467 | 2.063519437 |
| ENSMUSG00000023921 | 2.435736758 | 2.211372513 | 2.234894804 | 2.223704441 | 2.163285805 | 2.492760389 | 2.017561934 |
| ENSMUSG00000023923 | 2.433305075 | 2.210587586 | 2.216046536 | 2.207430684 | 2.182004726 | 2.478566496 | 2.048913829 |
| ENSMUSG00000023926 | 0 | 0 | 0.5 | 0.25 | 0.244650542 | 0 | 0.224243824 |
| ENSMUSG00000023927 | 1.54867984 | 1.622963274 | 1.516522298 | 1.321350555 | 1.27449559 | 1.380211242 | 1.159323753 |
| ENSMUSG00000023930 | 0 | 0 | 0 | 0 | 0 | 0 | 0 |
| ENSMUSG00000023931 | 0.289064826 | 0.25 | 0.5 | 0.580482024 | 0.686820904 | 0.301029996 | 0.520678036 |
| ENSMUSG00000023932 | 2.814939948 | 2.564346961 | 2.55479213 | 2.708420187 | 2.612740596 | 3.003029471 | 2.433962466 |
| ENSMUSG00000023935 | 0.578129653 | 0.580482024 | 0.580482024 | 0.580482024 | 0 | 0.301029996 | 0.579661877 |
| ENSMUSG00000023938 | 1.596667477 | 1.642463902 | 1.763820609 | 1.584962501 | 1.746695109 | 1.86332286 | 1.792684387 |
| ENSMUSG00000023939 | 2.016886209 | 1.96293726 | 2.108657057 | 1.75 | 1.990091809 | 2.017033339 | 1.982842456 |
| ENSMUSG00000023940 | 2.599132701 | 2.374462959 | 2.333847589 | 2.427270953 | 2.335652905 | 2.69019608 | 2.19230473 |
| ENSMUSG00000023942 | 3.041991609 | 2.808255107 | 2.82991803 | 2.300530956 | 2.379941013 | 2.626340367 | 2.269981661 |
| ENSMUSG00000023943 | 0 | 0 | 0 | 0 | 0 | 0 | 0 |
| ENSMUSG00000023944 | 4.175078004 | 3.748378347 | 3.725782169 | 3.793877642 | 3.727718564 | 4.37451006 | 3.463746406 |
| ENSMUSG00000023945 | 0.578129653 | 0.830482024 | 0.75 | 0.25 | 0.686820904 | 0 | 0.224243824 |
| ENSMUSG00000023947 | 1.747221736 | 1.720660762 | 1.688721876 | 2.165444524 | 2.187072901 | 2.372912003 | 2.10278155 |
| ENSMUSG00000023949 | 0 | 0 | 0 | 0 | 0 | 0 | 0 |
| ENSMUSG00000023951 | 2.556012466 | 2.39151351 | 2.367918804 | 1.766522298 | 1.882215755 | 2.008600172 | 1.811176017 |

| | | | | | | | |
|---|---|---|---|---|---|---|---|
| ENSMUSG00000023952 | 2.258689303 | 2.063166358 | 2.151832578 | 1.991446071 | 2.151558609 | 2.269512944 | 2.047762527 |
| ENSMUSG00000023953 | 2.418409478 | 2.277131114 | 2.269871196 | 2.316696635 | 2.255474576 | 2.5289167 | 2.105204898 |
| ENSMUSG00000023959 | 0.671187741 | 0.646240625 | 0.646240625 | 0.951838731 | 1 | 0.698970004 | 0.672731473 |
| ENSMUSG00000023960 | 2.587501561 | 2.341580554 | 2.371960008 | 2.221424093 | 2.10031573 | 2.437750563 | 2.001334086 |
| ENSMUSG00000023961 | 2.425923778 | 2.263820609 | 2.23927551 | 2.275821952 | 2.212046813 | 2.531478917 | 2.05629905 |
| ENSMUSG00000023963 | 2.013563224 | 1.799918086 | 1.903677461 | 1.464495249 | 1.366364104 | 1.477121255 | 1.131174228 |
| ENSMUSG00000023964 | 0 | 0 | 0.25 | 0 | 0.244650542 | 0 | 0 |
| ENSMUSG00000023965 | 2.273259602 | 2.101285366 | 2.069031101 | 2.127938163 | 2.024758322 | 2.29666519 | 1.889537832 |
| ENSMUSG00000023966 | 1.249317394 | 1.17510993 | 1.080482024 | 1.282320754 | 1.467903253 | 1.380211242 | 1.40946263 |
| ENSMUSG00000023967 | 2.365959569 | 2.182829758 | 2.236360959 | 2.197790722 | 2.196996016 | 2.447158031 | 2.11845007 |
| ENSMUSG00000023968 | 0 | 0 | 0.25 | 0.5 | 0.387761935 | 0 | 0.448487648 |
| ENSMUSG00000023971 | 2.042557845 | 1.958222504 | 2.004202072 | 1.79745614 | 1.875328588 | 2.008600172 | 1.854824501 |
| ENSMUSG00000023972 | 2.518981867 | 2.424241882 | 2.449009902 | 2.233422664 | 2.409087918 | 2.541579244 | 2.390438245 |
| ENSMUSG00000023973 | 2.40033094 | 2.294977273 | 2.324802005 | 2.194519282 | 2.260764609 | 2.462397998 | 2.208141797 |
| ENSMUSG00000023977 | 2.885320013 | 2.644828984 | 2.604199382 | 2.624712302 | 2.598242538 | 2.921686475 | 2.42167221 |
| ENSMUSG00000023978 | 0 | 0 | 0 | 0 | 0 | 0 | 0 |
| ENSMUSG00000023979 | 0 | 0 | 0.25 | 0.25 | 0.387761935 | 0.301029996 | 0.224243824 |
| ENSMUSG00000023980 | 2.051791411 | 1.901832578 | 1.921625132 | 1.919870025 | 1.986303335 | 2.190331698 | 1.871156825 |
| ENSMUSG00000023982 | 0.747221736 | 0.25 | 0.646240625 | 0.79248125 | 0.877063019 | 0.84509804 | 0.853775828 |
| ENSMUSG00000023984 | 0 | 0 | 0.5 | 0.5 | 0.632412477 | 0.301029996 | 0.629532004 |
| ENSMUSG00000023987 | 0 | 0 | 0 | 0 | 0 | 0 | 0 |
| ENSMUSG00000023988 | 2.399009124 | 2.263142013 | 2.331607372 | 2.202642909 | 2.275056283 | 2.505149978 | 2.182948513 |
| ENSMUSG00000023990 | 1.418409478 | 1.494319981 | 1.584962501 | 1.080482024 | 1.439191811 | 1.204119983 | 1.574722613 |
| ENSMUSG00000023991 | 2.156259305 | 2.131869252 | 2.209800947 | 1.828220739 | 1.992595054 | 2.096910013 | 2.041943747 |
| ENSMUSG00000023992 | 0 | 0.25 | 0 | 0 | 0 | 0 | 0 |
| ENSMUSG00000023993 | 0 | 0 | 0 | 0 | 0 | 0 | 0 |
| ENSMUSG00000023994 | 2.698532991 | 2.494319981 | 2.484527332 | 2.508011432 | 2.461685042 | 2.785329835 | 2.324338556 |
| ENSMUSG00000023995 | 0 | 0 | 0 | 0 | 0 | 0 | 0 |
| ENSMUSG00000023999 | 0.960252568 | 0.976722649 | 1.04248125 | 0.25 | 0.387761935 | 0.602059991 | 0.355418052 |
| ENSMUSG00000024002 | 2.80555694 | 2.586905342 | 2.619692405 | 2.527783918 | 2.552327032 | 2.880813592 | 2.441904013 |
| ENSMUSG00000024006 | 2.838668326 | 2.575374049 | 2.58076369 | 2.607874401 | 2.549248906 | 2.897627091 | 2.365724059 |
| ENSMUSG00000024007 | 2.54867984 | 2.408702763 | 2.354990044 | 2.306000419 | 2.342197974 | 2.660865478 | 2.241805758 |
| ENSMUSG00000024008 | 0.578129653 | 0.646240625 | 0.646240625 | 0.580482024 | 0.812711509 | 0.698970004 | 0.672731473 |
| ENSMUSG00000024011 | 0.578129653 | 0.25 | 0.25 | 0.5 | 0.812711509 | 0.477121255 | 0.579661877 |

| | | | | | | | |
|---|---|---|---|---|---|---|---|
| ENSMUSG00000024012 | 3.120755314 | 2.873713896 | 2.899513125 | 2.75123067 | 2.816235631 | 3.16790781 | 2.665464954 |
| ENSMUSG00000024013 | 0 | 0.396240625 | 0.25 | 0 | 0.568060967 | 0.477121255 | 0.579661877 |
| ENSMUSG00000024014 | 1.822201287 | 1.726722649 | 1.735628626 | 1.630890489 | 1.645989387 | 1.799340549 | 1.562034107 |
| ENSMUSG00000024018 | 1.342375483 | 1.214495249 | 1.26109853 | 1.589388001 | 1.55104774 | 1.748188027 | 1.523605797 |
| ENSMUSG00000024019 | 1.482695304 | 1.403677461 | 1.458222504 | 1.482684334 | 1.55104774 | 1.643452676 | 1.429563222 |
| ENSMUSG00000024026 | 2.69788693 | 2.469895812 | 2.455443495 | 2.792324674 | 2.718823734 | 3.158060794 | 2.519338425 |
| ENSMUSG00000024027 | 0 | 0.25 | 0 | 0 | 0 | 0 | 0 |
| ENSMUSG00000024028 | 0 | 0 | 0 | 0 | 0 | 0 | 0 |
| ENSMUSG00000024029 | 0 | 0 | 0 | 0 | 0 | 0 | 0 |
| ENSMUSG00000024030 | 0.289064826 | 0.25 | 0 | 1.02186571 | 0.905314583 | 1.041392685 | 0.916588299 |
| ENSMUSG00000024032 | 0 | 0 | 0 | 0 | 0 | 0 | 0 |
| ENSMUSG00000024033 | 0 | 0.25 | 0 | 1.04248125 | 1.149965125 | 1.113943352 | 1.131174228 |
| ENSMUSG00000024034 | 0 | 0 | 0.25 | 0 | 0 | 0 | 0 |
| ENSMUSG00000024036 | 0 | 0.396240625 | 0 | 1.04248125 | 1.212046813 | 1.204119983 | 1.140832123 |
| ENSMUSG00000024037 | 2.294202867 | 2.155512955 | 2.205044741 | 1.998588359 | 2.143518259 | 2.227886705 | 2.051774285 |
| ENSMUSG00000024038 | 2.083999385 | 1.985628626 | 1.990000483 | 1.87892496 | 1.974687643 | 2.120573931 | 1.898808251 |
| ENSMUSG00000024039 | 0.289064826 | 0.701838731 | 0.976722649 | 1.061981878 | 1.188507684 | 1.041392685 | 1.23151414 |
| ENSMUSG00000024041 | 0 | 0 | 0 | 0 | 0 | 0 | 0 |
| ENSMUSG00000024042 | 1.89911417 | 1.837182039 | 1.823655187 | 1.723704441 | 1.78687392 | 1.963787827 | 1.836517579 |
| ENSMUSG00000024043 | 2.295901576 | 2.087182039 | 2.110735874 | 2.331607372 | 2.25190332 | 2.579783597 | 2.100829716 |
| ENSMUSG00000024044 | 3.194908664 | 2.924784656 | 2.914998973 | 2.825374049 | 2.791183891 | 3.206825876 | 2.596166915 |
| ENSMUSG00000024045 | 2.766308649 | 2.535526764 | 2.561388177 | 2.564641508 | 2.574834836 | 2.862727528 | 2.386819016 |
| ENSMUSG00000024048 | 3.520513585 | 3.220994918 | 3.201536701 | 2.994141031 | 2.969086282 | 3.43964843 | 2.831441926 |
| ENSMUSG00000024049 | 1.249317394 | 1.372963274 | 1.282320754 | 0.396240625 | 0.489301084 | 0.301029996 | 0.775756176 |
| ENSMUSG00000024050 | 2.492331912 | 2.385757955 | 2.424676167 | 2.207430684 | 2.340805641 | 2.509202522 | 2.288586107 |
| ENSMUSG00000024052 | 2.682073381 | 2.44697564 | 2.421625132 | 2.257666784 | 2.244650542 | 2.495544338 | 2.096890362 |
| ENSMUSG00000024053 | 2.37447073 | 2.198603967 | 2.293731421 | 1.557204673 | 1.467903253 | 1.662757832 | 1.526507301 |
| ENSMUSG00000024054 | 3.08984989 | 2.820772088 | 2.780545225 | 2.908475852 | 2.789100779 | 3.255031163 | 2.583207287 |
| ENSMUSG00000024055 | 1.432083896 | 1.188721876 | 1.403677461 | 1.146240625 | 1.27449559 | 1.414973348 | 1.272006351 |
| ENSMUSG00000024056 | 2.637462542 | 2.45584181 | 2.440803092 | 2.467205639 | 2.365931294 | 2.743509765 | 2.221559033 |
| ENSMUSG00000024059 | 1.734388958 | 1.711372513 | 1.711372513 | 1.79745614 | 1.802174723 | 1.963787827 | 1.729871916 |
| ENSMUSG00000024063 | 2.249317394 | 2.113831805 | 2.051142786 | 1.903677461 | 1.897237431 | 2.149219113 | 1.784981274 |
| ENSMUSG00000024064 | 0 | 0 | 0 | 0.646240625 | 0.733951626 | 0.77815125 | 0.448487648 |
| ENSMUSG00000024065 | 2.105953527 | 1.923371739 | 1.937048212 | 1.931980114 | 1.882215755 | 2.155336037 | 1.757893036 |

| | | | | | | | |
|---|---|---|---|---|---|---|---|
| ENSMUSG00000024066 | 1.53838222 | 1.356566189 | 1.593759858 | 0 | 0.244650542 | 0.477121255 | 0.355418052 |
| ENSMUSG00000024067 | 2.320585253 | 2.074802005 | 2.119945066 | 2.193696765 | 2.089109832 | 2.484299839 | 2.022587518 |
| ENSMUSG00000024068 | 2.513987407 | 2.304186465 | 2.290597832 | 2.451032755 | 2.365064079 | 2.691965103 | 2.191935631 |
| ENSMUSG00000024069 | 2.134369199 | 1.945339928 | 1.935366747 | 1.982684334 | 1.945998439 | 2.230448921 | 1.818286533 |
| ENSMUSG00000024070 | 2.91631382 | 2.664998973 | 2.647880587 | 2.64151351 | 2.557398328 | 2.951823035 | 2.372557642 |
| ENSMUSG00000024072 | 2.605636262 | 2.369439567 | 2.362802778 | 2.453445298 | 2.382016011 | 2.767897616 | 2.227568781 |
| ENSMUSG00000024073 | 3.287180641 | 2.988549078 | 2.96816872 | 2.998146086 | 2.944404198 | 3.364175633 | 2.725429986 |
| ENSMUSG00000024074 | 3.293893266 | 3.033766986 | 3.021285115 | 2.845926073 | 2.829990449 | 3.215901813 | 2.640146179 |
| ENSMUSG00000024076 | 0.289064826 | 0 | 0.25 | 1.160964047 | 1.163285805 | 1.342422681 | 1.176815335 |
| ENSMUSG00000024077 | 2.708105693 | 2.491806565 | 2.478969845 | 2.616901388 | 2.524437092 | 2.894316063 | 2.355418052 |
| ENSMUSG00000024078 | 2.300956576 | 2.176839783 | 2.151832578 | 2.252107156 | 2.212046813 | 2.472756449 | 2.026306127 |
| ENSMUSG00000024079 | 2.629921064 | 2.415889526 | 2.422062577 | 2.667221246 | 2.545883696 | 2.970346876 | 2.367016399 |
| ENSMUSG00000024081 | 2.697239867 | 2.504202072 | 2.496816003 | 2.600753006 | 2.489301084 | 2.895422546 | 2.303573788 |
| ENSMUSG00000024082 | 2.209569962 | 2.024508021 | 1.998588359 | 2.03360658 | 2.008447216 | 2.267171728 | 1.847411384 |
| ENSMUSG00000024083 | 3.0249321 | 2.762291968 | 2.747171172 | 2.791540771 | 2.678166555 | 3.118264726 | 2.48807426 |
| ENSMUSG00000024084 | 2.23973002 | 2.12892496 | 2.204245906 | 1.618933358 | 1.439191811 | 1.662757832 | 1.374453863 |
| ENSMUSG00000024085 | 3.091362705 | 2.838424552 | 2.8464656 | 2.763990059 | 2.682255238 | 3.095169351 | 2.489990739 |
| ENSMUSG00000024087 | 3.617135846 | 3.276518036 | 3.321531135 | 3.041305267 | 2.970104237 | 3.42357352 | 2.758726757 |
| ENSMUSG00000024088 | 0 | 0 | 0 | 0 | 0 | 0 | 0 |
| ENSMUSG00000024091 | 2.884080688 | 2.646240625 | 2.592671852 | 2.592399337 | 2.505415151 | 2.923761961 | 2.338964569 |
| ENSMUSG00000024095 | 2.82886633 | 2.586351312 | 2.574229052 | 2.638886943 | 2.576508184 | 2.953759692 | 2.368945281 |
| ENSMUSG00000024096 | 2.632197861 | 2.466046536 | 2.491085217 | 2.399978211 | 2.426290456 | 2.717670503 | 2.311326126 |
| ENSMUSG00000024097 | 3.002810412 | 2.736178015 | 2.726722649 | 2.805094582 | 2.748896963 | 3.137670537 | 2.547599797 |
| ENSMUSG00000024098 | 2.811999637 | 2.606828211 | 2.603935442 | 2.673371739 | 2.632005142 | 2.999565488 | 2.472164183 |
| ENSMUSG00000024099 | 2.611244851 | 2.389605178 | 2.360217292 | 2.522858847 | 2.416295225 | 2.822168079 | 2.290770186 |
| ENSMUSG00000024101 | 2.339331437 | 2.208222504 | 2.221424093 | 2.051142786 | 2.11997913 | 2.29666519 | 2.066798338 |
| ENSMUSG00000024104 | 2.887787667 | 2.630645383 | 2.636964627 | 2.500352049 | 2.439543536 | 2.792391689 | 2.300947836 |
| ENSMUSG00000024105 | 0 | 0 | 0 | 0 | 0 | 0 | 0 |
| ENSMUSG00000024107 | 0 | 0 | 0 | 0 | 0 | 0 | 0 |
| ENSMUSG00000024109 | 0 | 0 | 0 | 0 | 0 | 0 | 0 |
| ENSMUSG00000024112 | 1.342375483 | 1.348079356 | 1.364857905 | 0.5 | 0.733951626 | 0.698970004 | 0.896975297 |
| ENSMUSG00000024114 | 1.342375483 | 1.364857905 | 1.226722649 | 0 | 0.387761935 | 0.301029996 | 0 |
| ENSMUSG00000024116 | 0 | 0 | 0 | 0 | 0 | 0 | 0 |
| ENSMUSG00000024118 | 2.069666964 | 1.937048212 | 1.931980114 | 1.923371739 | 1.938809675 | 2.130333768 | 1.858985529 |

| | | | | | | | |
|---|---|---|---|---|---|---|---|
| ENSMUSG00000024120 | 2.970129664 | 2.692046081 | 2.69390257 | 2.639126514 | 2.55990691 | 2.958563883 | 2.379456779 |
| ENSMUSG00000024121 | 1.806056082 | 1.726722649 | 1.802363341 | 1.649978211 | 1.65906333 | 1.579783597 | 1.653807046 |
| ENSMUSG00000024122 | 2.7985055 | 2.56932185 | 2.536483036 | 2.599418658 | 2.557146486 | 2.901458321 | 2.344312053 |
| ENSMUSG00000024124 | 0 | 0 | 0.5 | 0 | 0 | 0 | 0 |
| ENSMUSG00000024125 | 0 | 0 | 0 | 0 | 0 | 0 | 0 |
| ENSMUSG00000024127 | 2.699178053 | 2.445749552 | 2.387686696 | 2.285526764 | 2.218023635 | 2.567026366 | 2.086315375 |
| ENSMUSG00000024128 | 0 | 0 | 0 | 0 | 0 | 0 | 0 |
| ENSMUSG00000024130 | 2.544964605 | 2.415444524 | 2.417664062 | 2.446567407 | 2.514628232 | 2.794488047 | 2.338495365 |
| ENSMUSG00000024131 | 0.671187741 | 0 | 0.5 | 0.701838731 | 0.632412477 | 0.84509804 | 0.520678036 |
| ENSMUSG00000024132 | 2.029919532 | 2 | 2.001406137 | 2.04248125 | 2.048957937 | 2.257678575 | 1.941550454 |
| ENSMUSG00000024134 | 0.960252568 | 1 | 1.146240625 | 1.17510993 | 1.091002363 | 0.903089987 | 0.952571511 |
| ENSMUSG00000024135 | 2.392336464 | 2.155512955 | 2.137686696 | 2.30539678 | 2.190648982 | 2.517195898 | 2.063519437 |
| ENSMUSG00000024136 | 0.867194479 | 0.951838731 | 0.830482024 | 0.79248125 | 0.978602168 | 0.602059991 | 0.853775828 |
| ENSMUSG00000024137 | 1.811507563 | 1.741446071 | 1.75 | 1.552363341 | 1.733951626 | 1.556302501 | 1.766260282 |
| ENSMUSG00000024140 | 0.45815691 | 0.646240625 | 0.75 | 1.356566189 | 1.467903253 | 1.612783857 | 1.483307832 |
| ENSMUSG00000024142 | 2.231898196 | 2.051142786 | 2.048689214 | 2.221424093 | 2.237868644 | 2.496929648 | 2.061314894 |
| ENSMUSG00000024143 | 2.871053932 | 2.62892496 | 2.621708755 | 2.317865757 | 2.275616087 | 2.5289167 | 2.094403682 |
| ENSMUSG00000024145 | 1.837744666 | 1.779735268 | 1.766522298 | 1.914552871 | 1.791313678 | 1.991226076 | 1.704233519 |
| ENSMUSG00000024146 | 2.396352858 | 2.193696765 | 2.208222504 | 2.348079356 | 2.287171411 | 2.617000341 | 2.061314894 |
| ENSMUSG00000024150 | 2.608049185 | 2.418106335 | 2.396240625 | 2.391989019 | 2.368090054 | 2.678518379 | 2.224577862 |
| ENSMUSG00000024151 | 2.682744391 | 2.458222504 | 2.443284802 | 2.406883471 | 2.33281029 | 2.663700925 | 2.173704585 |
| ENSMUSG00000024154 | 0 | 0 | 0 | 0 | 0.387761935 | 0 | 0 |
| ENSMUSG00000024155 | 0.289064826 | 0.396240625 | 0 | 0.75 | 0.489301084 | 0.477121255 | 0.710836105 |
| ENSMUSG00000024158 | 2.029919532 | 1.899978211 | 1.888647213 | 2.052363341 | 2.094757251 | 2.367355921 | 2.005960535 |
| ENSMUSG00000024160 | 1.036286563 | 1.02186571 | 1.188721876 | 0.951838731 | 0.812711509 | 0.954242509 | 1.028149525 |
| ENSMUSG00000024163 | 2.794934559 | 2.564346961 | 2.584684096 | 2.424241882 | 2.520266629 | 2.713490543 | 2.392431569 |
| ENSMUSG00000024164 | 1.671187741 | 1.505591953 | 1.606566189 | 1.29248125 | 1.234113756 | 1.301029996 | 1.20139795 |
| ENSMUSG00000024165 | 3.069401761 | 2.787754135 | 2.761610487 | 2.912875055 | 2.825189844 | 3.284205068 | 2.626654587 |
| ENSMUSG00000024168 | 0.671187741 | 0.646240625 | 0.580482024 | 0.580482024 | 0.846351821 | 0.477121255 | 0.629532004 |
| ENSMUSG00000024169 | 2.289064826 | 2.107613138 | 2.107613138 | 2.038704527 | 2.033751322 | 2.252853031 | 1.856911705 |
| ENSMUSG00000024170 | 2.072573125 | 1.998588359 | 2.013820609 | 1.870953944 | 2.05001946 | 2.146128036 | 1.942350245 |
| ENSMUSG00000024171 | 0 | 0 | 0 | 0 | 0 | 0 | 0 |
| ENSMUSG00000024172 | 0 | 0 | 0 | 0 | 0 | 0 | 0 |
| ENSMUSG00000024173 | 0.671187741 | 1 | 0.864857905 | 0.25 | 0 | 0 | 0.520678036 |

| | | | | | | | |
|---|---|---|---|---|---|---|---|
| ENSMUSG00000024174 | 1.936861443 | 1.83272922 | 1.732684334 | 1.777131114 | 1.736537397 | 1.924279286 | 1.641897157 |
| ENSMUSG00000024175 | 0 | 0 | 0 | 0 | 0 | 0 | 0 |
| ENSMUSG00000024176 | 0 | 0 | 0.25 | 0.75 | 0.733951626 | 0.301029996 | 0.710836105 |
| ENSMUSG00000024177 | 1.747221736 | 1.660964047 | 1.695339928 | 1.71759118 | 1.795698282 | 1.84509804 | 1.670888075 |
| ENSMUSG00000024180 | 1.307602651 | 1.372963274 | 1.311981878 | 1.557204673 | 1.599808748 | 1.716003344 | 1.56716938 |
| ENSMUSG00000024181 | 2.419671301 | 2.323080408 | 2.377443751 | 2.26109853 | 2.349985771 | 2.572871602 | 2.286111264 |
| ENSMUSG00000024182 | 2.722979153 | 2.504897682 | 2.504202072 | 2.469512728 | 2.488077665 | 2.774516966 | 2.321381806 |
| ENSMUSG00000024184 | 0 | 0 | 0 | 0 | 0 | 0 | 0 |
| ENSMUSG00000024186 | 0 | 0.5 | 0.25 | 0.396240625 | 0.568060967 | 0 | 0.520678036 |
| ENSMUSG00000024187 | 2.439357901 | 2.231944491 | 2.265173983 | 2.295598588 | 2.366796383 | 2.627365857 | 2.25178005 |
| ENSMUSG00000024188 | 2.77230138 | 2.501756817 | 2.502806814 | 2.56611065 | 2.523329778 | 2.832508913 | 2.345462945 |
| ENSMUSG00000024190 | 2.20747957 | 2.048689214 | 2.056000419 | 2.181128463 | 2.17760189 | 2.450249108 | 2.086315375 |
| ENSMUSG00000024191 | 1.806056082 | 1.782320754 | 1.811981878 | 1.726722649 | 1.692703642 | 1.886490725 | 1.694069894 |
| ENSMUSG00000024193 | 1.894701088 | 1.782320754 | 1.868933358 | 1.714495249 | 1.806427258 | 1.838849091 | 1.800208341 |
| ENSMUSG00000024194 | 2.188178997 | 2.103406982 | 2.219895812 | 2.036164561 | 2.062514787 | 2.278753601 | 2.049487947 |
| ENSMUSG00000024197 | 2.463861956 | 2.306603048 | 2.37892496 | 2.189555804 | 2.205284995 | 2.356025857 | 2.19156611 |
| ENSMUSG00000024201 | 2.023453784 | 2.002806814 | 2.080482024 | 1.779735268 | 1.961316494 | 2.004321374 | 1.98213686 |
| ENSMUSG00000024205 | 0 | 0 | 0 | 0 | 0 | 0 | 0 |
| ENSMUSG00000024206 | 1.269664473 | 1.321350555 | 1.27186571 | 1.238549078 | 1.343584837 | 1.361727836 | 1.444790289 |
| ENSMUSG00000024207 | 0 | 0 | 0 | 0 | 0 | 0 | 0 |
| ENSMUSG00000024208 | 2.262379879 | 2.155512955 | 2.235628626 | 2.093759858 | 2.154724156 | 2.371067862 | 2.158505762 |
| ENSMUSG00000024209 | 0.289064826 | 0 | 0 | 0.646240625 | 0.632412477 | 0.698970004 | 0.579661877 |
| ENSMUSG00000024210 | 0 | 0.646240625 | 0.396240625 | 0 | 0 | 0.301029996 | 0.224243824 |
| ENSMUSG00000024211 | 0 | 0 | 0 | 0 | 0 | 0 | 0 |
| ENSMUSG00000024212 | 2.528794847 | 2.44697564 | 2.508699741 | 2.226722649 | 2.415543455 | 2.5289167 | 2.391037533 |
| ENSMUSG00000024213 | 2.604828841 | 2.428561379 | 2.432405186 | 2.27908599 | 2.373217836 | 2.627365857 | 2.267652107 |
| ENSMUSG00000024215 | 0 | 0 | 0 | 0 | 0 | 0 | 0.520678036 |
| ENSMUSG00000024217 | 2.294202867 | 2.14151351 | 2.262462137 | 2.155512955 | 2.212715924 | 2.399673721 | 2.206380685 |
| ENSMUSG00000024218 | 2.336265009 | 2.161864607 | 2.132845365 | 2.232684334 | 2.225193812 | 2.519827994 | 2.120298735 |
| ENSMUSG00000024219 | 1.975148165 | 1.970660762 | 2.02186571 | 1.789967834 | 1.944572352 | 2.004321374 | 1.968429585 |
| ENSMUSG00000024220 | 1.971473847 | 1.948603967 | 2.057204673 | 1.685366747 | 1.981187939 | 1.963787827 | 2.030591163 |
| ENSMUSG00000024222 | 2.491272108 | 2.323080408 | 2.33272922 | 2.464495249 | 2.47844958 | 2.795880017 | 2.311326126 |
| ENSMUSG00000024223 | 0 | 0 | 0 | 0 | 0 | 0 | 0.224243824 |
| ENSMUSG00000024224 | 0 | 0 | 0 | 0 | 0 | 0 | 0 |

| | | | | | | | |
|---|---|---|---|---|---|---|---|
| ENSMUSG00000024225 | 0 | 0 | 0 | 0 | 0 | 0 | 0 |
| ENSMUSG00000024227 | 0 | 0 | 0 | 0 | 0 | 0 | 0 |
| ENSMUSG00000024228 | 1.800532393 | 1.75 | 1.782320754 | 1.755591953 | 1.756568651 | 1.977723605 | 1.532233387 |
| ENSMUSG00000024231 | 2.599951214 | 2.389126514 | 2.342126615 | 2.444929839 | 2.365497953 | 2.686636269 | 2.196701384 |
| ENSMUSG00000024232 | 1.181541736 | 1.160964047 | 1.02186571 | 0.75 | 0.568060967 | 0.477121255 | 0.829800754 |
| ENSMUSG00000024233 | 0 | 0 | 0 | 0 | 0 | 0 | 0 |
| ENSMUSG00000024234 | 2.215778996 | 2.005591953 | 2.006976499 | 2.03360658 | 2.006054284 | 2.281033367 | 1.845261772 |
| ENSMUSG00000024235 | 1.358731791 | 0.976722649 | 1.098079356 | 0.951838731 | 0.846351821 | 0.903089987 | 0.876096088 |
| ENSMUSG00000024236 | 2.610448227 | 2.403677461 | 2.362287161 | 2.005591953 | 2.024758322 | 2.278753601 | 1.86815666 |
| ENSMUSG00000024238 | 2.578990401 | 2.352877747 | 2.31728167 | 2.271202097 | 2.211376431 | 2.503790683 | 2.03601839 |
| ENSMUSG00000024240 | 2.320585253 | 2.107613138 | 2.081607372 | 2.142463902 | 2.121713561 | 2.33243846 | 1.991877847 |
| ENSMUSG00000024241 | 2.719310222 | 2.52053726 | 2.491085217 | 2.51348147 | 2.438487308 | 2.797267541 | 2.241805758 |
| ENSMUSG00000024242 | 2.397683106 | 2.241446071 | 2.219129237 | 2.229715809 | 2.198391101 | 2.484299839 | 2.034219411 |
| ENSMUSG00000024245 | 0 | 0 | 0 | 0 | 0 | 0 | 0 |
| ENSMUSG00000024246 | 1.639698646 | 1.52186571 | 1.557204673 | 1.282320754 | 1.121713561 | 1.278753601 | 1.20139795 |
| ENSMUSG00000024247 | 1.894701088 | 1.899978211 | 1.937048212 | 1.464495249 | 1.65906333 | 1.662757832 | 1.647906904 |
| ENSMUSG00000024248 | 2.81249113 | 2.548689214 | 2.558105232 | 2.434100483 | 2.412140531 | 2.720159303 | 2.280542874 |
| ENSMUSG00000024251 | 2.448863324 | 2.201838731 | 2.215271726 | 2.296218836 | 2.247084732 | 2.546542663 | 2.106650241 |
| ENSMUSG00000024253 | 2.048736215 | 1.898114259 | 1.909156155 | 1.928561379 | 1.88560959 | 2.120573931 | 1.790137008 |
| ENSMUSG00000024254 | 0 | 0.25 | 0 | 0.25 | 0 | 0 | 0 |
| ENSMUSG00000024256 | 0 | 0 | 0 | 0 | 0 | 0 | 0 |
| ENSMUSG00000024258 | 2.337801042 | 2.149047439 | 2.110735874 | 2.212155735 | 2.164053936 | 2.481442629 | 2.018194418 |
| ENSMUSG00000024259 | 2.413323654 | 2.226722649 | 2.257666784 | 2.314935816 | 2.21736452 | 2.526339277 | 2.065709059 |
| ENSMUSG00000024260 | 2.561428547 | 2.353935442 | 2.37892496 | 2.291226732 | 2.25665701 | 2.51851394 | 2.141264342 |
| ENSMUSG00000024261 | 0 | 0 | 0 | 0 | 0 | 0 | 0 |
| ENSMUSG00000024266 | 0 | 0 | 0 | 0 | 0 | 0 | 0 |
| ENSMUSG00000024268 | 1.45815691 | 1.566696635 | 1.622963274 | 1.610735874 | 1.572189156 | 1.672097858 | 1.523605797 |
| ENSMUSG00000024269 | 2.235832493 | 2.04745614 | 2.063166358 | 2.136723615 | 2.014359518 | 2.336459734 | 1.883845332 |
| ENSMUSG00000024270 | 2.846437921 | 2.629909063 | 2.645064642 | 2.501055117 | 2.438839735 | 2.762678564 | 2.279699287 |
| ENSMUSG00000024271 | 2.752359777 | 2.532320754 | 2.502806814 | 2.576800202 | 2.489301084 | 2.860338007 | 2.29131391 |
| ENSMUSG00000024273 | 1.647796617 | 1.527131114 | 1.52186571 | 1.53743678 | 1.603583716 | 1.72427587 | 1.517723536 |
| ENSMUSG00000024274 | 0.671187741 | 0.830482024 | 0.701838731 | 0.396240625 | 0.489301084 | 0.602059991 | 0.629532004 |
| ENSMUSG00000024276 | 2.432083896 | 2.247880462 | 2.206637122 | 2.205044741 | 2.12771782 | 2.436162647 | 1.943943921 |
| ENSMUSG00000024277 | 2.395018353 | 2.191217898 | 2.254202072 | 2.309003548 | 2.255474576 | 2.56937391 | 2.109997807 |

| | | | | | | | |
|---|---|---|---|---|---|---|---|
| ENSMUSG00000024283 | 2.801542157 | 2.576800202 | 2.526477127 | 2.58076369 | 2.48561803 | 2.858537198 | 2.29883162 |
| ENSMUSG00000024286 | 2.571178952 | 2.388167274 | 2.37246199 | 2.405512955 | 2.314255074 | 2.633468456 | 2.153137801 |
| ENSMUSG00000024287 | 2.573799059 | 2.405054956 | 2.37246199 | 2.442872367 | 2.395811454 | 2.722633923 | 2.223239637 |
| ENSMUSG00000024290 | 2.671187741 | 2.535845803 | 2.531676118 | 2.5684489 | 2.482830298 | 2.805500858 | 2.26794422 |
| ENSMUSG00000024292 | 0 | 0.25 | 0 | 0 | 0 | 0 | 0 |
| ENSMUSG00000024293 | 2.153883046 | 1.988549078 | 1.990000483 | 2.044977273 | 2.029283463 | 2.170261715 | 1.835407748 |
| ENSMUSG00000024294 | 2.91898589 | 2.69059551 | 2.652294685 | 2.706438458 | 2.638666793 | 3.042575512 | 2.416917233 |
| ENSMUSG00000024295 | 0.45815691 | 0.5 | 0.396240625 | 0.701838731 | 0.568060967 | 0 | 0.672731473 |
| ENSMUSG00000024298 | 2.540273527 | 2.33272922 | 2.335518667 | 2.405970373 | 2.35043857 | 2.702430536 | 2.15104925 |
| ENSMUSG00000024299 | 2.165631214 | 2.05479213 | 2.078220739 | 1.589388001 | 1.812711509 | 1.880813592 | 1.809975587 |
| ENSMUSG00000024300 | 0 | 0 | 0 | 0 | 0.244650542 | 0 | 0 |
| ENSMUSG00000024301 | 1.789260561 | 1.843759858 | 1.882845365 | 1.5 | 1.800029087 | 1.69019608 | 1.778418869 |
| ENSMUSG00000024302 | 2.217828299 | 2.08272922 | 2.071350555 | 1.388647213 | 1.283908311 | 1.342422681 | 1.360283038 |
| ENSMUSG00000024304 | 2.685417684 | 2.474461364 | 2.484894804 | 2.444108258 | 2.407936347 | 2.748188027 | 2.226242896 |
| ENSMUSG00000024306 | 0 | 0 | 0 | 0 | 0 | 0 | 0 |
| ENSMUSG00000024308 | 2.565000581 | 2.394828984 | 2.460980263 | 2.667442722 | 2.689947504 | 3.043362278 | 2.635586271 |
| ENSMUSG00000024309 | 2.253091469 | 2.184523065 | 2.212155735 | 2.065523711 | 2.126866297 | 2.326335861 | 2.020712061 |
| ENSMUSG00000024312 | 2.331622671 | 2.160964047 | 2.215271726 | 2.143411797 | 2.230953877 | 2.411619706 | 2.176815335 |
| ENSMUSG00000024313 | 0 | 0 | 0 | 0 | 0 | 0 | 0 |
| ENSMUSG00000024317 | 2.10861828 | 1.998588359 | 1.995748394 | 2.041226732 | 1.973373093 | 2.209515015 | 1.837623616 |
| ENSMUSG00000024319 | 2.318984359 | 2.171625132 | 2.227473271 | 2.149978211 | 2.235369829 | 2.475671188 | 2.191196167 |
| ENSMUSG00000024325 | 1.53838222 | 1.575945187 | 1.618933358 | 1.330482024 | 1.642643806 | 1.612783857 | 1.631625962 |
| ENSMUSG00000024327 | 1.358731791 | 1.42510993 | 1.494319981 | 1.282320754 | 1.489301084 | 1.505149978 | 1.559435575 |
| ENSMUSG00000024330 | 1.37447073 | 1.396240625 | 1.494319981 | 1.02186571 | 1.223252711 | 0.84509804 | 1.340368108 |
| ENSMUSG00000024331 | 1.404271433 | 1.27186571 | 1.348079356 | 1.330482024 | 1.244650542 | 1.322219295 | 1.252393349 |
| ENSMUSG00000024334 | 0 | 0 | 0 | 0 | 0 | 0 | 0 |
| ENSMUSG00000024335 | 3.133460136 | 2.857743793 | 2.857482435 | 2.914106216 | 2.867468707 | 3.306639441 | 2.709347306 |
| ENSMUSG00000024338 | 0.671187741 | 0.646240625 | 0.701838731 | 1.948603967 | 1.955822902 | 2.11058971 | 1.885753983 |
| ENSMUSG00000024339 | 1 | 1.080482024 | 0.951838731 | 1.653677461 | 1.810629166 | 1.939519253 | 1.824094866 |
| ENSMUSG00000024340 | 0 | 0 | 0 | 0 | 0 | 0 | 0 |
| ENSMUSG00000024346 | 2.375872521 | 2.243603647 | 2.286164561 | 2.390083209 | 2.297835549 | 2.673020907 | 2.20919388 |
| ENSMUSG00000024347 | 0 | 0.25 | 0 | 0.396240625 | 0 | 0 | 0.448487648 |
| ENSMUSG00000024349 | 0.867194479 | 1 | 0.701838731 | 1.403677461 | 1.302012594 | 1.544068044 | 1.369798885 |
| ENSMUSG00000024350 | 2.205378646 | 2.041226732 | 2.023189285 | 2.205044741 | 2.149165677 | 2.42975228 | 1.986347533 |

| | | | | | | | |
|---|---|---|---|---|---|---|---|
| ENSMUSG00000024352 | 1.37447073 | 1.410964047 | 1.388647213 | 1.26109853 | 1.188507684 | 1.230448921 | 1.168187745 |
| ENSMUSG00000024353 | 0 | 0 | 0 | 0 | 0 | 0 | 0 |
| ENSMUSG00000024354 | 0.578129653 | 1.02186571 | 0.646240625 | 0.701838731 | 0.846351821 | 0.477121255 | 0.74492186 |
| ENSMUSG00000024357 | 1.862428343 | 1.779735268 | 1.864857905 | 1.681980114 | 1.770879644 | 1.924279286 | 1.710836105 |
| ENSMUSG00000024359 | 3.53993029 | 3.183200849 | 3.159552407 | 3.149338563 | 3.036482892 | 3.554004321 | 2.825296265 |
| ENSMUSG00000024360 | 3.21857101 | 2.919099472 | 2.895770692 | 2.936103353 | 2.855124093 | 3.282395505 | 2.641345239 |
| ENSMUSG00000024365 | 0 | 0 | 0 | 0 | 0 | 0 | 0 |
| ENSMUSG00000024366 | 0 | 0 | 0 | 0 | 0 | 0 | 0.224243824 |
| ENSMUSG00000024369 | 2.124258828 | 2.02053726 | 2.091580554 | 2.048689214 | 2.078515538 | 2.257678575 | 2.037212175 |
| ENSMUSG00000024370 | 2.590021402 | 2.333847589 | 2.30357978 | 2.486360959 | 2.375754058 | 2.752048448 | 2.243699513 |
| ENSMUSG00000024371 | 0 | 0.5 | 0.396240625 | 0.864857905 | 0.686820904 | 1.079181246 | 0.775756176 |
| ENSMUSG00000024376 | 0.811507563 | 0.976722649 | 0.75 | 1.438721876 | 1.373641807 | 1.505149978 | 1.302263476 |
| ENSMUSG00000024378 | 2.58665822 | 2.38040011 | 2.337182039 | 2.454645544 | 2.382842606 | 2.7355989 | 2.222904214 |
| ENSMUSG00000024379 | 1.100572389 | 0.896240625 | 0.92510993 | 0.646240625 | 0.568060967 | 0.698970004 | 0.829800754 |
| ENSMUSG00000024381 | 2.544964605 | 2.319031101 | 2.370449858 | 2.218361028 | 2.249502249 | 2.472756449 | 2.139097452 |
| ENSMUSG00000024382 | 2.557825653 | 2.357090043 | 2.321350555 | 2.358657057 | 2.368520226 | 2.64246452 | 2.213368331 |
| ENSMUSG00000024383 | 2.684750966 | 2.466433318 | 2.452241044 | 2.428561379 | 2.361573756 | 2.674861141 | 2.142557543 |
| ENSMUSG00000024384 | 2.711248379 | 2.469512728 | 2.465271726 | 2.568740118 | 2.477187895 | 2.854913022 | 2.30565931 |
| ENSMUSG00000024386 | 0 | 0.25 | 0 | 0 | 0 | 0 | 0 |
| ENSMUSG00000024387 | 2.670497861 | 2.428990498 | 2.428990498 | 2.468361028 | 2.431363834 | 2.825426118 | 2.312600313 |
| ENSMUSG00000024388 | 0 | 0 | 0 | 0 | 0 | 0 | 0 |
| ENSMUSG00000024391 | 0.747221736 | 0.701838731 | 0.646240625 | 0.396240625 | 0.387761935 | 0.301029996 | 0.74492186 |
| ENSMUSG00000024392 | 2.983380418 | 2.775657972 | 2.846195938 | 2.568157446 | 2.708967143 | 2.974971994 | 2.634177434 |
| ENSMUSG00000024393 | 3.289534546 | 3.087389422 | 3.151890374 | 2.806753548 | 2.997042149 | 3.262925469 | 2.921427462 |
| ENSMUSG00000024395 | 1.156259305 | 1.330482024 | 1.488549078 | 0.25 | 0 | 0 | 0.355418052 |
| ENSMUSG00000024397 | 0.289064826 | 0.580482024 | 0.396240625 | 0.75 | 0.632412477 | 0.602059991 | 0.629532004 |
| ENSMUSG00000024399 | 0 | 0.701838731 | 0.25 | 0.5 | 0.244650542 | 0.301029996 | 0.448487648 |
| ENSMUSG00000024400 | 2.470606563 | 2.337182039 | 2.359177886 | 2.327084758 | 2.305113253 | 2.57054294 | 2.214058866 |
| ENSMUSG00000024401 | 0 | 0 | 0 | 0 | 0 | 0.301029996 | 0 |
| ENSMUSG00000024402 | 0.289064826 | 0 | 0 | 0.25 | 0 | 0 | 0.224243824 |
| ENSMUSG00000024403 | 1.156259305 | 1.098079356 | 1.188721876 | 1 | 1.091002363 | 1 | 1.110947927 |
| ENSMUSG00000024404 | 2.592526109 | 2.408248799 | 2.390560606 | 2.435787849 | 2.383667271 | 2.702430536 | 2.217830684 |
| ENSMUSG00000024406 | 0.289064826 | 0 | 0 | 0.25 | 0 | 0 | 0 |
| ENSMUSG00000024409 | 0.45815691 | 0.396240625 | 0.580482024 | 0.25 | 0.812711509 | 0.301029996 | 0.710836105 |

| | | | | | | | |
|---|---|---|---|---|---|---|---|
| ENSMUSG00000024410 | 2.17479713 | 1.948603967 | 1.998588359 | 2.001406137 | 2.022473809 | 2.222716471 | 1.865128412 |
| ENSMUSG00000024411 | 0 | 0 | 0 | 0 | 0 | 0 | 0 |
| ENSMUSG00000024413 | 2.323768721 | 2.139605178 | 2.195339928 | 2.08272922 | 2.071604288 | 2.283301229 | 1.899720848 |
| ENSMUSG00000024414 | 2.209569962 | 2.098079356 | 2.105516192 | 2.081607372 | 2.070605826 | 2.348304863 | 2.008574816 |
| ENSMUSG00000024420 | 1.99653913 | 1.930274797 | 1.870953944 | 0.5 | 0.568060967 | 0.477121255 | 0.579661877 |
| ENSMUSG00000024421 | 0.811507563 | 0.79248125 | 0.864857905 | 0.646240625 | 0.568060967 | 0.477121255 | 0.74492186 |
| ENSMUSG00000024422 | 2.420929319 | 2.28360658 | 2.342126615 | 2.201032755 | 2.285543632 | 2.51054501 | 2.223909441 |
| ENSMUSG00000024423 | 2.631440309 | 2.399513125 | 2.408702763 | 2.30357978 | 2.229681962 | 2.546542663 | 2.096890362 |
| ENSMUSG00000024424 | 1.924654746 | 1.681980114 | 1.708222504 | 1.741446071 | 1.674752852 | 1.903089987 | 1.598697688 |
| ENSMUSG00000024425 | 2.395018353 | 2.25 | 2.19697564 | 2.277131114 | 2.206647761 | 2.545307116 | 2.062969708 |
| ENSMUSG00000024426 | 2.063793172 | 1.896240625 | 1.898114259 | 1.898114259 | 1.883916752 | 1.973127854 | 1.830929954 |
| ENSMUSG00000024427 | 1.100572389 | 1.061981878 | 0.864857905 | 0.864857905 | 1.039257769 | 1 | 0.935079929 |
| ENSMUSG00000024429 | 2.609650079 | 2.422062577 | 2.513820609 | 2.360735874 | 2.44234479 | 2.683047038 | 2.385806448 |
| ENSMUSG00000024430 | 0.671187741 | 0.92510993 | 0.75 | 1.02186571 | 0.686820904 | 0.903089987 | 0.876096088 |
| ENSMUSG00000024431 | 2.74143756 | 2.539020769 | 2.555094582 | 2.46759118 | 2.404846996 | 2.687528961 | 2.210939786 |
| ENSMUSG00000024436 | 1.944805158 | 1.708222504 | 1.784887838 | 1.708222504 | 1.749189509 | 1.875061263 | 1.627424404 |
| ENSMUSG00000024437 | 0.45815691 | 0.25 | 0 | 0 | 0 | 0 | 0 |
| ENSMUSG00000024440 | 0.289064826 | 0.396240625 | 0 | 0.396240625 | 0 | 0 | 0 |
| ENSMUSG00000024442 | 2.153883046 | 2.039967834 | 2.079353153 | 2.017865591 | 2.083370582 | 2.260071388 | 2.030591163 |
| ENSMUSG00000024446 | 1.445324132 | 1.54745614 | 1.516522298 | 1.160964047 | 1.351342427 | 1.255272505 | 1.496250175 |
| ENSMUSG00000024448 | 0 | 0 | 0 | 0 | 0 | 0 | 0 |
| ENSMUSG00000024451 | 1.269664473 | 1.146240625 | 1.130890489 | 0.5 | 0.489301084 | 0.301029996 | 0.355418052 |
| ENSMUSG00000024454 | 2.630681378 | 2.42510993 | 2.441632227 | 2.51109853 | 2.466277473 | 2.818885415 | 2.280261923 |
| ENSMUSG00000024456 | 2.610448227 | 2.403217124 | 2.408702763 | 2.437886015 | 2.405621873 | 2.681241237 | 2.238304819 |
| ENSMUSG00000024457 | 2.348397716 | 2.229715809 | 2.233422664 | 2.252107156 | 2.278401864 | 2.521138084 | 2.160546876 |
| ENSMUSG00000024459 | 1.036286563 | 1.080482024 | 1.061981878 | 0.5 | 0.733951626 | 0 | 0.672731473 |
| ENSMUSG00000024462 | 2.203267085 | 2.023189285 | 2.086073977 | 1.850219859 | 1.882215755 | 2.025305865 | 1.767634022 |
| ENSMUSG00000024471 | 0 | 0 | 0 | 0 | 0 | 0 | 0 |
| ENSMUSG00000024472 | 2.707474304 | 2.486726569 | 2.456239685 | 2.614857905 | 2.542486091 | 2.86332286 | 2.331854053 |
| ENSMUSG00000024474 | 2.77230138 | 2.502457154 | 2.503156135 | 2.673807073 | 2.549763795 | 3.000434077 | 2.405097146 |
| ENSMUSG00000024477 | 2.149089517 | 2.069031101 | 2.060793496 | 2.122963274 | 2.0510778 | 2.318063335 | 1.896975297 |
| ENSMUSG00000024479 | 0.578129653 | 0.396240625 | 0 | 0.79248125 | 0.568060967 | 0.698970004 | 0.355418052 |
| ENSMUSG00000024480 | 2.409468165 | 2.230460234 | 2.230460234 | 2.30539678 | 2.227765453 | 2.567026366 | 2.044283833 |
| ENSMUSG00000024481 | 0.867194479 | 0.580482024 | 0.5 | 0.25 | 0.387761935 | 0.301029996 | 0.224243824 |

| | | | | | | | |
|---|---|---|---|---|---|---|---|
| ENSMUSG00000024483 | 2.825070787 | 2.578787391 | 2.531676118 | 2.564641508 | 2.469198506 | 2.837588438 | 2.300684065 |
| ENSMUSG00000024485 | 0 | 0 | 0 | 0.25 | 0 | 0 | 0 |
| ENSMUSG00000024486 | 2.111266113 | 2.002806814 | 1.992885888 | 2.049918086 | 2.038163331 | 2.31386722 | 1.908707966 |
| ENSMUSG00000024487 | 2.569423023 | 2.39151351 | 2.391989019 | 2.356041572 | 2.322676882 | 2.626340367 | 2.176040447 |
| ENSMUSG00000024491 | 2.581562037 | 2.373463862 | 2.400906586 | 2.437886015 | 2.356272688 | 2.675778342 | 2.199238773 |
| ENSMUSG00000024493 | 3.045658471 | 2.745213402 | 2.690387808 | 2.668106335 | 2.605346297 | 2.99343623 | 2.424203925 |
| ENSMUSG00000024497 | 0 | 0 | 0 | 0.25 | 0 | 0 | 0 |
| ENSMUSG00000024498 | 2.786655728 | 2.561981878 | 2.54652856 | 2.608657057 | 2.532092521 | 2.892651034 | 2.366370874 |
| ENSMUSG00000024500 | 0.867194479 | 1.04248125 | 1 | 0.25 | 0.632412477 | 0 | 0.224243824 |
| ENSMUSG00000024501 | 3.188378486 | 2.926083993 | 2.883818844 | 2.665889526 | 2.591221684 | 2.971275849 | 2.421127112 |
| ENSMUSG00000024502 | 1.227926429 | 1.214495249 | 1.26109853 | 0.830482024 | 0.733951626 | 0.954242509 | 0.803905701 |
| ENSMUSG00000024503 | 0 | 0 | 0 | 0 | 0 | 0 | 0 |
| ENSMUSG00000024505 | 1.418409478 | 1.356566189 | 1.372963274 | 1.618933358 | 1.489301084 | 1.568201724 | 1.429563222 |
| ENSMUSG00000024507 | 2.63370885 | 2.375459296 | 2.349151195 | 2.400906586 | 2.346799814 | 2.653212514 | 2.199599638 |
| ENSMUSG00000024510 | 0 | 0 | 0 | 0 | 0 | 0 | 0 |
| ENSMUSG00000024511 | 1.967766868 | 1.784887838 | 1.884789703 | 1.755591953 | 1.66856048 | 1.908485019 | 1.56716938 |
| ENSMUSG00000024512 | 0 | 0.580482024 | 0.25 | 0.79248125 | 0.489301084 | 0.477121255 | 0.520678036 |
| ENSMUSG00000024513 | 2.429630757 | 2.236360959 | 2.206637122 | 2.231944491 | 2.167869715 | 2.469822016 | 2.021338422 |
| ENSMUSG00000024515 | 2.678702042 | 2.472945926 | 2.42510993 | 2.369945066 | 2.355827317 | 2.658964843 | 2.172922202 |
| ENSMUSG00000024516 | 1.876566388 | 1.809601185 | 1.830482024 | 1.92510993 | 1.85749873 | 2.127104798 | 1.771720588 |
| ENSMUSG00000024517 | 0 | 0 | 0 | 0 | 0 | 0 | 0 |
| ENSMUSG00000024518 | 0 | 0 | 0 | 0 | 0 | 0 | 0.355418052 |
| ENSMUSG00000024519 | 0 | 0 | 0 | 0 | 0 | 0 | 0 |
| ENSMUSG00000024521 | 2.439357901 | 2.274508021 | 2.254897682 | 1.926839783 | 1.887294348 | 2.071882007 | 1.746389053 |
| ENSMUSG00000024524 | 1.307602651 | 1.188721876 | 1.114857905 | 1.29248125 | 1.254881871 | 1.176091259 | 1.089372236 |
| ENSMUSG00000024525 | 1.686083339 | 1.657339155 | 1.711372513 | 1.5 | 1.621875691 | 1.698970004 | 1.645916034 |
| ENSMUSG00000024526 | 0 | 0 | 0 | 0 | 0.244650542 | 0 | 0.355418052 |
| ENSMUSG00000024527 | 2.520963022 | 2.345385738 | 2.357090043 | 2.343216265 | 2.280062837 | 2.544068044 | 2.170957925 |
| ENSMUSG00000024528 | 1.978790393 | 1.959800947 | 2.005591953 | 1.814346961 | 1.828942237 | 2.025305865 | 1.702561601 |
| ENSMUSG00000024529 | 3.354869736 | 3.038783614 | 2.969895812 | 2.192872367 | 2.12771782 | 2.399673721 | 1.960974982 |
| ENSMUSG00000024530 | 1.289064826 | 1.29248125 | 1.27186571 | 1.080482024 | 1.074582839 | 1.176091259 | 1 |
| ENSMUSG00000024532 | 0 | 0 | 0.25 | 0 | 0 | 0 | 0 |
| ENSMUSG00000024533 | 2.35433422 | 2.168989258 | 2.160061234 | 2.12892496 | 2.081436571 | 2.298853076 | 1.921737195 |
| ENSMUSG00000024534 | 0 | 0 | 0 | 0.25 | 0 | 0.301029996 | 0.224243824 |

| | | | | | | | |
|---|---|---|---|---|---|---|---|
| ENSMUSG00000024535 | 1.993049299 | 1.866901388 | 1.923371739 | 1.777131114 | 1.611014646 | 1.913813852 | 1.572224412 |
| ENSMUSG00000024537 | 2.121692466 | 1.985628626 | 1.994319981 | 1.953445298 | 1.979897422 | 2.161368002 | 1.86207155 |
| ENSMUSG00000024538 | 2.814451333 | 2.567865757 | 2.553276055 | 2.602347734 | 2.542486091 | 2.953276337 | 2.393027179 |
| ENSMUSG00000024539 | 2.339331437 | 2.231944491 | 2.206637122 | 2.296838018 | 2.212046813 | 2.586587305 | 2.094403682 |
| ENSMUSG00000024542 | 2.807052575 | 2.584405475 | 2.541854536 | 2.588286706 | 2.495955124 | 2.869818208 | 2.339198915 |
| ENSMUSG00000024544 | 2.042557845 | 1.921625132 | 1.937048212 | 1.410964047 | 1.42077253 | 1.633468456 | 1.441050184 |
| ENSMUSG00000024546 | 0 | 0 | 0 | 0 | 0 | 0 | 0 |
| ENSMUSG00000024548 | 2.312518611 | 2.163659007 | 2.125956434 | 1.668106335 | 1.621875691 | 1.77815125 | 1.577201671 |
| ENSMUSG00000024552 | 0 | 0.396240625 | 0 | 0 | 0 | 0 | 0.224243824 |
| ENSMUSG00000024553 | 0 | 0 | 0 | 0 | 0 | 0 | 0 |
| ENSMUSG00000024556 | 2.579849376 | 2.392938161 | 2.38040011 | 2.487821179 | 2.404070415 | 2.818225894 | 2.245894972 |
| ENSMUSG00000024558 | 0.867194479 | 0.5 | 0.830482024 | 0.5 | 0.244650542 | 0.602059991 | 0.224243824 |
| ENSMUSG00000024560 | 2.165631214 | 2.048689214 | 2.119945066 | 1.819031101 | 2.039257769 | 1.977723605 | 1.994607951 |
| ENSMUSG00000024561 | 2.190368148 | 2.067865757 | 2.102347734 | 1.898114259 | 1.947418788 | 2.100370545 | 1.921737195 |
| ENSMUSG00000024563 | 2.540273527 | 2.310793496 | 2.252107156 | 2.277131114 | 2.205967036 | 2.492760389 | 2.059651572 |
| ENSMUSG00000024565 | 0 | 0 | 0 | 0 | 0.244650542 | 0 | 0 |
| ENSMUSG00000024566 | 2.382811656 | 2.215271726 | 2.228969845 | 2.184523065 | 2.179075613 | 2.416640507 | 2.048913829 |
| ENSMUSG00000024570 | 1.912079952 | 1.816696635 | 1.910964047 | 1.726722649 | 1.770879644 | 1.785329835 | 1.774416565 |
| ENSMUSG00000024571 | 2.612039956 | 2.488185312 | 2.492166698 | 2.234894804 | 2.29094062 | 2.563481085 | 2.225910574 |
| ENSMUSG00000024575 | 0.45815691 | 0.396240625 | 0.396240625 | 0.25 | 0.244650542 | 0 | 0.224243824 |
| ENSMUSG00000024576 | 3.150620716 | 2.869692405 | 2.841580554 | 2.852612838 | 2.761403742 | 3.196176185 | 2.555498219 |
| ENSMUSG00000024578 | 0 | 0 | 0 | 0 | 0 | 0 | 0 |
| ENSMUSG00000024579 | 1.54867984 | 1.660964047 | 1.726722649 | 1.238549078 | 1.49445382 | 1.51851394 | 1.452143229 |
| ENSMUSG00000024580 | 2.23973002 | 2.110735874 | 2.086073977 | 2.132845365 | 2.05001946 | 2.401400541 | 1.941550454 |
| ENSMUSG00000024581 | 2.34539747 | 2.200224975 | 2.216046536 | 2.091580554 | 2.138604637 | 2.396199347 | 1.986347533 |
| ENSMUSG00000024583 | 2.937589884 | 2.671187155 | 2.651138307 | 2.732869058 | 2.651624369 | 3.044931546 | 2.463666693 |
| ENSMUSG00000024587 | 3.198233207 | 2.901022465 | 2.849552315 | 2.888287319 | 2.807172415 | 3.241546481 | 2.617865519 |
| ENSMUSG00000024588 | 2.489144387 | 2.277131114 | 2.302972074 | 2.408248799 | 2.342197974 | 2.668385917 | 2.191935631 |
| ENSMUSG00000024589 | 2.231898196 | 2.005591953 | 2.05479213 | 1.907339155 | 1.897237431 | 2.113943352 | 1.822941512 |
| ENSMUSG00000024590 | 2.903875772 | 2.658702763 | 2.612287161 | 2.62521047 | 2.602699114 | 2.985426474 | 2.449155379 |
| ENSMUSG00000024592 | 1.578129653 | 1.52186571 | 1.54745614 | 1.330482024 | 1.373641807 | 1.602059991 | 1.252393349 |
| ENSMUSG00000024593 | 2.645603628 | 2.446567407 | 2.435366747 | 0.701838731 | 0.812711509 | 0.477121255 | 0.629532004 |
| ENSMUSG00000024594 | 2.35727108 | 2.196158711 | 2.232684334 | 2.057204673 | 2.071604288 | 2.303196057 | 2.005960535 |
| ENSMUSG00000024597 | 2.946233352 | 2.686208459 | 2.68325383 | 2.459406983 | 2.351342427 | 2.691965103 | 2.172530299 |

| | | | | | | | |
|---|---|---|---|---|---|---|---|
| ENSMUSG00000024598 | 0.960252568 | 0.976722649 | 1.098079356 | 0.976722649 | 1.106691885 | 1.113943352 | 1.066254157 |
| ENSMUSG00000024600 | 1.036286563 | 0.701838731 | 0.896240625 | 0.396240625 | 0.244650542 | 0 | 0.520678036 |
| ENSMUSG00000024601 | 2.243591459 | 2.104463129 | 2.065523711 | 2.145300145 | 2.113840608 | 2.361727836 | 2.02383181 |
| ENSMUSG00000024603 | 2.692026894 | 2.472185812 | 2.449415381 | 2.451032755 | 2.375332618 | 2.748962861 | 2.229876084 |
| ENSMUSG00000024604 | 2.513987407 | 2.293106877 | 2.291854536 | 2.321350555 | 2.310728002 | 2.615950052 | 2.151886289 |
| ENSMUSG00000024608 | 3.253262209 | 3.018033151 | 3.027865435 | 2.863317659 | 2.968144018 | 3.300595484 | 2.775999146 |
| ENSMUSG00000024610 | 1.822201287 | 1.681980114 | 1.505591953 | 1.79248125 | 1.903713872 | 2.049218023 | 1.833176598 |
| ENSMUSG00000024613 | 2.703665763 | 2.556000419 | 2.608657057 | 2.29248125 | 2.474332504 | 2.622214023 | 2.423302014 |
| ENSMUSG00000024614 | 2.904664113 | 2.672281101 | 2.663211243 | 2.736726569 | 2.64026298 | 3.058426024 | 2.43291209 |
| ENSMUSG00000024617 | 0.671187741 | 0.864857905 | 0.976722649 | 0.580482024 | 0.632412477 | 0.698970004 | 0.775756176 |
| ENSMUSG00000024619 | 0 | 0.25 | 0.25 | 0 | 0 | 0 | 0 |
| ENSMUSG00000024620 | 3.171071754 | 2.929097697 | 2.925867762 | 2.399978211 | 2.458367761 | 2.720159303 | 2.33637549 |
| ENSMUSG00000024621 | 0.289064826 | 0.5 | 0.396240625 | 0.396240625 | 0.686820904 | 0.77815125 | 0.520678036 |
| ENSMUSG00000024622 | 2.405576699 | 2.254202072 | 2.27908599 | 2.168106335 | 2.181274723 | 2.451786436 | 2.082194103 |
| ENSMUSG00000024639 | 2.756312721 | 2.510756821 | 2.47104263 | 2.464883697 | 2.369379 | 2.681241237 | 2.189340078 |
| ENSMUSG00000024640 | 3.294512239 | 2.969704321 | 2.891394534 | 3.028923488 | 2.954844506 | 3.422425676 | 2.731687462 |
| ENSMUSG00000024642 | 1.956444026 | 1.843759858 | 1.850219859 | 1.505591953 | 1.595992969 | 1.748188027 | 1.48984372 |
| ENSMUSG00000024644 | 2.339331437 | 2.148114259 | 2.185366747 | 2.163659007 | 2.215379757 | 2.445604203 | 2.078019652 |
| ENSMUSG00000024645 | 1.903481042 | 1.763820609 | 1.77186571 | 1.777131114 | 1.680838453 | 1.968482949 | 1.683576582 |
| ENSMUSG00000024646 | 2.352857998 | 2.153677461 | 2.168106335 | 2.140560606 | 2.009637626 | 2.322219295 | 1.87902385 |
| ENSMUSG00000024647 | 0 | 0 | 0 | 0 | 0 | 0 | 0 |
| ENSMUSG00000024650 | 0 | 0 | 0 | 0 | 0 | 0 | 0 |
| ENSMUSG00000024653 | 0 | 0 | 0.25 | 0 | 0 | 0 | 0 |
| ENSMUSG00000024654 | 1.907802659 | 1.75835575 | 1.789967834 | 1.930274797 | 1.873585591 | 2.071882007 | 1.826389311 |
| ENSMUSG00000024658 | 0.91631382 | 0.951838731 | 0.701838731 | 0.646240625 | 0.733951626 | 0.301029996 | 0.710836105 |
| ENSMUSG00000024659 | 3.861238181 | 3.485811849 | 3.474768411 | 3.199415381 | 3.096018299 | 3.640978057 | 2.875471627 |
| ENSMUSG00000024660 | 2.856724497 | 2.635274154 | 2.654825739 | 2.653907409 | 2.663191519 | 3.050379756 | 2.547722783 |
| ENSMUSG00000024661 | 3.569183005 | 3.207430684 | 3.262036564 | 3.149222142 | 3.041372628 | 3.585235063 | 2.883061263 |
| ENSMUSG00000024663 | 2.026699189 | 1.948603967 | 2.048689214 | 1.606566189 | 1.665423072 | 1.799340549 | 1.596378571 |
| ENSMUSG00000024664 | 2.901898374 | 2.680488403 | 2.687257846 | 2.643175057 | 2.645205039 | 2.968949681 | 2.502390799 |
| ENSMUSG00000024665 | 2.582415738 | 2.392463902 | 2.433677405 | 2.438721876 | 2.410235744 | 2.757396029 | 2.310815043 |
| ENSMUSG00000024666 | 2.177057474 | 2.01109853 | 2.02186571 | 2.036164561 | 1.969399812 | 2.255272505 | 1.87902385 |
| ENSMUSG00000024667 | 1.862428343 | 1.711372513 | 1.671625132 | 1.75835575 | 1.768534412 | 1.986771734 | 1.61448206 |
| ENSMUSG00000024668 | 2.502784398 | 2.299304173 | 2.269871196 | 2.390083209 | 2.346799814 | 2.685741739 | 2.161359729 |

| | | | | | | | |
|---|---|---|---|---|---|---|---|
| ENSMUSG00000024669 | 0 | 0 | 0.396240625 | 0 | 0 | 0 | 0 |
| ENSMUSG00000024670 | 0.289064826 | 0.396240625 | 0 | 0 | 0 | 0 | 0 |
| ENSMUSG00000024672 | 0 | 0.25 | 0.25 | 0.25 | 0.489301084 | 0 | 0.224243824 |
| ENSMUSG00000024673 | 0.289064826 | 0.5 | 0.646240625 | 0.396240625 | 0.686820904 | 0.84509804 | 0.448487648 |
| ENSMUSG00000024675 | 0 | 0 | 0 | 0 | 0 | 0 | 0.224243824 |
| ENSMUSG00000024677 | 0 | 0 | 0 | 0 | 0 | 0 | 0 |
| ENSMUSG00000024678 | 0.45815691 | 0.646240625 | 0.580482024 | 0.25 | 0.387761935 | 0 | 0.224243824 |
| ENSMUSG00000024679 | 0 | 0 | 0 | 0 | 0.244650542 | 0 | 0.224243824 |
| ENSMUSG00000024680 | 0 | 0 | 0 | 0 | 0.244650542 | 0 | 0.224243824 |
| ENSMUSG00000024681 | 0 | 0 | 0 | 0 | 0 | 0 | 0 |
| ENSMUSG00000024682 | 0 | 0 | 0.25 | 0.25 | 0.387761935 | 0 | 0.224243824 |
| ENSMUSG00000024683 | 2.333175866 | 2.170748646 | 2.200224975 | 2.237821179 | 2.180543208 | 2.45484486 | 2.055736903 |
| ENSMUSG00000024687 | 2.863298974 | 2.623713896 | 2.586351312 | 2.633818844 | 2.566101456 | 2.946943271 | 2.40471474 |
| ENSMUSG00000024691 | 2.735572029 | 2.531676118 | 2.503156135 | 1.061981878 | 1.091002363 | 1.176091259 | 0.935079929 |
| ENSMUSG00000024694 | 0 | 0 | 0 | 0 | 0 | 0 | 0 |
| ENSMUSG00000024695 | 3.178080867 | 2.896709947 | 2.870449858 | 2.94961795 | 2.861775743 | 3.306425028 | 2.635304994 |
| ENSMUSG00000024696 | 0.811507563 | 0.951838731 | 0.701838731 | 1.080482024 | 1.136121934 | 1.230448921 | 1.028149525 |
| ENSMUSG00000024697 | 0 | 0.25 | 0.25 | 0.5 | 0.632412477 | 0.477121255 | 0.520678036 |
| ENSMUSG00000024712 | 2.509948321 | 2.273189285 | 2.312574604 | 2.377443751 | 2.309205544 | 2.667452953 | 2.161765391 |
| ENSMUSG00000024713 | 0 | 0 | 0 | 1.188721876 | 1.091002363 | 1.176091259 | 1.110947927 |
| ENSMUSG00000024725 | 2.540273527 | 2.325945187 | 2.377443751 | 2.320772088 | 2.337535342 | 2.614897216 | 2.21302251 |
| ENSMUSG00000024726 | 2.315764015 | 2.181128463 | 2.094844592 | 2.220660762 | 2.138604637 | 2.471291711 | 1.978585612 |
| ENSMUSG00000024727 | 0.867194479 | 0.830482024 | 0.25 | 0.976722649 | 0.686820904 | 0.698970004 | 0.672731473 |
| ENSMUSG00000024728 | 0 | 0 | 0 | 0 | 0 | 0 | 0 |
| ENSMUSG00000024729 | 0 | 0 | 0 | 0.25 | 0 | 0 | 0 |
| ENSMUSG00000024730 | 0 | 0 | 0 | 0 | 0 | 0 | 0 |
| ENSMUSG00000024731 | 0 | 0 | 0 | 0 | 0 | 0 | 0 |
| ENSMUSG00000024732 | 2.095120908 | 2.03743678 | 2.084962501 | 2.029735268 | 2.053185007 | 2.257678575 | 2.031198704 |
| ENSMUSG00000024734 | 0 | 0 | 0 | 0 | 0 | 0 | 0.224243824 |
| ENSMUSG00000024735 | 3.132777036 | 2.846869716 | 2.838837774 | 2.844979954 | 2.807792181 | 3.256477206 | 2.631341219 |
| ENSMUSG00000024736 | 2.287337982 | 2.293731421 | 2.436208459 | 1.802363341 | 2.101233692 | 2.120573931 | 2.268527657 |
| ENSMUSG00000024737 | 0.45815691 | 0.75 | 0.896240625 | 0.396240625 | 0.244650542 | 0 | 0.629532004 |
| ENSMUSG00000024738 | 0 | 0 | 0.25 | 0 | 0 | 0 | 0 |
| ENSMUSG00000024740 | 3.916244187 | 3.503810201 | 3.50250088 | 3.489819376 | 3.426848252 | 3.99206726 | 3.181426084 |

| | | | | | | | |
|---|---|---|---|---|---|---|---|
| ENSMUSG00000024742 | 0.811507563 | 1.061981878 | 1.17510993 | 0.951838731 | 1.091002363 | 0.77815125 | 1.089372236 |
| ENSMUSG00000024743 | 0.671187741 | 1.02186571 | 0.92510993 | 1.214495249 | 1.327538614 | 1.322219295 | 1.23151414 |
| ENSMUSG00000024747 | 1.156259305 | 1.188721876 | 1.238549078 | 0.25 | 0 | 0 | 0 |
| ENSMUSG00000024749 | 0 | 0 | 0 | 0.25 | 0 | 0 | 0 |
| ENSMUSG00000024750 | 2.811507563 | 2.592671852 | 2.554186465 | 2.53296424 | 2.416670511 | 2.783903579 | 2.24838596 |
| ENSMUSG00000024754 | 2.630681378 | 2.410964047 | 2.404137211 | 2.33993739 | 2.307676489 | 2.575187845 | 2.166993225 |
| ENSMUSG00000024757 | 0.289064826 | 0.25 | 0 | 0.25 | 0 | 0 | 0 |
| ENSMUSG00000024758 | 3.173765637 | 2.909269414 | 2.882723495 | 2.99832306 | 2.895503 | 3.367355921 | 2.691714761 |
| ENSMUSG00000024759 | 3.204036163 | 2.918658415 | 2.87694866 | 3.016690483 | 2.913670063 | 3.397592434 | 2.690055906 |
| ENSMUSG00000024761 | 0 | 0.25 | 0 | 0 | 0 | 0 | 0 |
| ENSMUSG00000024764 | 2.694641526 | 2.485994979 | 2.483422664 | 2.42293588 | 2.461024076 | 2.750508395 | 2.297768298 |
| ENSMUSG00000024766 | 0 | 0 | 0 | 1.418106335 | 1.283908311 | 1.477121255 | 1.20919388 |
| ENSMUSG00000024767 | 2.856724497 | 2.588286706 | 2.638167274 | 2.585240691 | 2.603804521 | 2.948901761 | 2.490137691 |
| ENSMUSG00000024768 | 0 | 0 | 0 | 0 | 0 | 0 | 0 |
| ENSMUSG00000024769 | 1.952600383 | 1.972185812 | 2.02186571 | 1.638647213 | 1.898867717 | 1.851258349 | 1.874129423 |
| ENSMUSG00000024770 | 0 | 0 | 0 | 0 | 0 | 0 | 0 |
| ENSMUSG00000024771 | 0 | 0 | 0 | 0 | 0.244650542 | 0.301029996 | 0 |
| ENSMUSG00000024772 | 2.85848785 | 2.679204865 | 2.700629091 | 2.662987153 | 2.754990069 | 3.055760465 | 2.647906904 |
| ENSMUSG00000024773 | 2.258689303 | 2.161864607 | 2.241446071 | 2.05479213 | 2.208682096 | 2.326335861 | 2.222904214 |
| ENSMUSG00000024774 | 0 | 0 | 0 | 0 | 0 | 0 | 0 |
| ENSMUSG00000024776 | 1.985980784 | 1.888647213 | 1.830482024 | 1.852347734 | 1.804307395 | 2.033423755 | 1.694069894 |
| ENSMUSG00000024777 | 2.302628051 | 2.136723615 | 2.221424093 | 2.260414788 | 2.318738698 | 2.519827994 | 2.260561074 |
| ENSMUSG00000024778 | 2.591692877 | 2.379417347 | 2.338837774 | 1.994319981 | 1.88560959 | 2.089905111 | 1.798966438 |
| ENSMUSG00000024780 | 2.508932407 | 2.276477127 | 2.254897682 | 2.288071211 | 2.228405446 | 2.471291711 | 2.066798338 |
| ENSMUSG00000024781 | 2.796468708 | 2.557505109 | 2.550837001 | 2.542794199 | 2.467578695 | 2.863917377 | 2.296701469 |
| ENSMUSG00000024782 | 2.817374469 | 2.585240691 | 2.53296424 | 2.579635702 | 2.496553863 | 2.881954971 | 2.315638039 |
| ENSMUSG00000024784 | 0 | 0.25 | 0.5 | 0 | 0.244650542 | 0.477121255 | 0.775756176 |
| ENSMUSG00000024785 | 2.455852858 | 2.269203899 | 2.246460484 | 2.223704441 | 2.299407739 | 2.539076099 | 2.139531992 |
| ENSMUSG00000024786 | 0 | 0 | 0 | 0 | 0 | 0 | 0 |
| ENSMUSG00000024787 | 1.989530018 | 1.994319981 | 2.03743678 | 1.723704441 | 1.92093466 | 1.838849091 | 1.909593098 |
| ENSMUSG00000024789 | 2.679378494 | 2.437048212 | 2.418548067 | 2.535845803 | 2.434943542 | 2.824125834 | 2.2945572 |
| ENSMUSG00000024790 | 1.912079952 | 1.80479213 | 1.894357207 | 1.657339155 | 1.741652793 | 1.908485019 | 1.797719748 |
| ENSMUSG00000024791 | 2.427163103 | 2.241446071 | 2.257666784 | 2.362287161 | 2.322187006 | 2.638489257 | 2.16539364 |
| ENSMUSG00000024792 | 1.827447047 | 1.735628626 | 1.729715809 | 1.458222504 | 1.652586889 | 1.748188027 | 1.69749337 |

| | | | | | | | |
|---|---|---|---|---|---|---|---|
| ENSMUSG00000024793 | 0.671187741 | 0.580482024 | 0.75 | 0.580482024 | 0.77552387 | 0.477121255 | 0.710836105 |
| ENSMUSG00000024795 | 2.984682628 | 2.745926548 | 2.754202072 | 2.768535366 | 2.698658695 | 3.121231455 | 2.48614636 |
| ENSMUSG00000024797 | 2.146671932 | 2.03360658 | 2.151832578 | 1.926839783 | 2.043601899 | 2.173186268 | 2.052343353 |
| ENSMUSG00000024798 | 0.289064826 | 0.25 | 0.396240625 | 0.25 | 0.387761935 | 0 | 0.448487648 |
| ENSMUSG00000024799 | 1.358731791 | 1.418106335 | 1.464495249 | 1.380890489 | 1.514343934 | 1.681241237 | 1.508694612 |
| ENSMUSG00000024800 | 2.370236862 | 2.21759118 | 2.172499493 | 2.354463129 | 2.260764609 | 2.640481437 | 2.109521699 |
| ENSMUSG00000024803 | 3.369333216 | 3.098883534 | 3.096937025 | 2.745034894 | 2.711253771 | 3.08170727 | 2.517048279 |
| ENSMUSG00000024805 | 2.304292853 | 2.107613138 | 2.113831805 | 2.03360658 | 1.997548906 | 2.252853031 | 1.874129423 |
| ENSMUSG00000024806 | 0 | 0.25 | 0 | 0 | 0 | 0 | 0 |
| ENSMUSG00000024807 | 2.339331437 | 2.248588359 | 2.285526764 | 2.099151195 | 2.19135987 | 2.338456494 | 2.125782179 |
| ENSMUSG00000024810 | 1.129344651 | 1.061981878 | 0.951838731 | 1.130890489 | 0.905314583 | 0.84509804 | 0.952571511 |
| ENSMUSG00000024811 | 3.30504739 | 3.008785687 | 2.963522295 | 3.026231576 | 2.941024176 | 3.40277707 | 2.717452193 |
| ENSMUSG00000024812 | 2.53838222 | 2.385757955 | 2.428561379 | 2.401832578 | 2.4345872 | 2.705007959 | 2.325562604 |
| ENSMUSG00000024815 | 0 | 0 | 0 | 0 | 0 | 0 | 0 |
| ENSMUSG00000024816 | 2.428398755 | 2.332168732 | 2.372963274 | 2.136723615 | 2.278401864 | 2.390935107 | 2.257258335 |
| ENSMUSG00000024817 | 3.08984989 | 2.792794199 | 2.76109853 | 2.680488403 | 2.618927353 | 2.996073654 | 2.422035099 |
| ENSMUSG00000024818 | 0.747221736 | 0.79248125 | 0.701838731 | 0.75 | 0.568060967 | 0.301029996 | 0.896975297 |
| ENSMUSG00000024824 | 2.507914013 | 2.331607372 | 2.359697963 | 2.123963757 | 2.149165677 | 2.281033367 | 2.042530357 |
| ENSMUSG00000024826 | 2.667726872 | 2.493244947 | 2.448603967 | 2.562574604 | 2.568792988 | 2.888740961 | 2.425462392 |
| ENSMUSG00000024827 | 2.149089517 | 2.001406137 | 2.041226732 | 2.313166358 | 2.305627392 | 2.605305046 | 2.168981648 |
| ENSMUSG00000024829 | 2.351376531 | 2.209012589 | 2.195339928 | 2.160964047 | 2.182733222 | 2.423245874 | 2.110947927 |
| ENSMUSG00000024830 | 2.078325387 | 1.972185812 | 2.031030328 | 1.992885888 | 2.093822267 | 2.281033367 | 1.989815042 |
| ENSMUSG00000024831 | 1.87190676 | 1.860735874 | 1.923371739 | 1.646240625 | 1.84445929 | 1.832508913 | 1.832055226 |
| ENSMUSG00000024833 | 2.659301689 | 2.506630861 | 2.54961126 | 2.500703754 | 2.519146132 | 2.86332286 | 2.415439991 |
| ENSMUSG00000024835 | 2.949075151 | 2.735811849 | 2.794666213 | 2.524508021 | 2.641653746 | 2.921166051 | 2.553692546 |
| ENSMUSG00000024837 | 0 | 0 | 0 | 0 | 0 | 0.301029996 | 0 |
| ENSMUSG00000024841 | 2.56677519 | 2.348615674 | 2.389126514 | 2.277131114 | 2.243427123 | 2.592176757 | 2.150210039 |
| ENSMUSG00000024842 | 0.578129653 | 0.5 | 0.701838731 | 0.580482024 | 0.489301084 | 0.602059991 | 0.629532004 |
| ENSMUSG00000024843 | 2.377269616 | 2.240000483 | 2.216819685 | 2.357613138 | 2.346799814 | 2.599883072 | 2.180279543 |
| ENSMUSG00000024844 | 2.958871665 | 2.768033151 | 2.762802235 | 2.70920984 | 2.71352568 | 3.102090526 | 2.550294753 |
| ENSMUSG00000024845 | 2.406877894 | 2.279735268 | 2.260414788 | 2.140560606 | 2.198391101 | 2.463892989 | 2.120298735 |
| ENSMUSG00000024846 | 1.129344651 | 1.17510993 | 1.339388001 | 0.864857905 | 0.632412477 | 0.77815125 | 0.952571511 |
| ENSMUSG00000024847 | 2.560530738 | 2.449820405 | 2.483791262 | 2.324229052 | 2.427019704 | 2.688419822 | 2.362688369 |
| ENSMUSG00000024851 | 2.399009124 | 2.289967834 | 2.415889526 | 2.077084758 | 2.255474576 | 2.340444115 | 2.286111264 |

| | | | | | | | |
|---|---|---|---|---|---|---|---|
| ENSMUSG00000024853 | 3.285129787 | 3.000791627 | 3.03783343 | 2.976628711 | 3.006759774 | 3.402605242 | 2.844967533 |
| ENSMUSG00000024854 | 1.647796617 | 1.747171172 | 1.799918086 | 1.552363341 | 1.588235379 | 1.653212514 | 1.559435575 |
| ENSMUSG00000024855 | 2.503815381 | 2.321350555 | 2.338837774 | 2.370449858 | 2.407551654 | 2.687528961 | 2.276018109 |
| ENSMUSG00000024856 | 1.975148165 | 2.03360658 | 2.160061234 | 1.747171172 | 1.98503142 | 2.037426498 | 2.027536225 |
| ENSMUSG00000024857 | 0 | 0 | 0 | 0 | 0 | 0 | 0 |
| ENSMUSG00000024858 | 2.716846213 | 2.539652422 | 2.547147211 | 2.485628626 | 2.562893863 | 2.850033258 | 2.432385621 |
| ENSMUSG00000024862 | 2.227926429 | 2.094844592 | 2.205044741 | 1.991446071 | 2.148364414 | 2.301029996 | 2.098373247 |
| ENSMUSG00000024863 | 0 | 0 | 0 | 0 | 0 | 0 | 0 |
| ENSMUSG00000024866 | 1 | 1.356566189 | 1.27186571 | 1.356566189 | 1.427019704 | 1.462397998 | 1.383567577 |
| ENSMUSG00000024867 | 1.227926429 | 1.238549078 | 1.339388001 | 1.114857905 | 0.978602168 | 1.301029996 | 1.014380832 |
| ENSMUSG00000024868 | 0.289064826 | 0.396240625 | 0.25 | 0 | 0 | 0 | 0.355418052 |
| ENSMUSG00000024869 | 1.418409478 | 1.330482024 | 1.52186571 | 1.372963274 | 1.542223377 | 1.681241237 | 1.459332757 |
| ENSMUSG00000024870 | 2.774459379 | 2.564641508 | 2.635516136 | 2.602877747 | 2.65981751 | 2.998259338 | 2.595212748 |
| ENSMUSG00000024871 | 0.578129653 | 0.5 | 0.79248125 | 0.5 | 0.877063019 | 0.602059991 | 0.853775828 |
| ENSMUSG00000024873 | 0.867194479 | 1 | 1.061981878 | 1.04248125 | 1.074582839 | 1.146128036 | 1.20139795 |
| ENSMUSG00000024875 | 2.310886388 | 2.115881093 | 2.130890489 | 1.972185812 | 2.029283463 | 2.238046103 | 1.947107911 |
| ENSMUSG00000024878 | 2.163307908 | 2.059601185 | 2.01109853 | 2.154596376 | 2.041436517 | 2.274157849 | 1.889537832 |
| ENSMUSG00000024883 | 2.435736758 | 2.280383379 | 2.309601185 | 2.20584181 | 2.219993628 | 2.426511261 | 2.168187745 |
| ENSMUSG00000024885 | 0.91631382 | 0.864857905 | 1 | 1.321350555 | 1.343584837 | 1.462397998 | 1.479989695 |
| ENSMUSG00000024887 | 2.229917041 | 2.013820609 | 2.041226732 | 2.159156155 | 2.054233912 | 2.334453751 | 1.936709733 |
| ENSMUSG00000024889 | 1.876566388 | 1.950224975 | 1.919870025 | 1.903677461 | 1.910073614 | 2.113943352 | 1.895131899 |
| ENSMUSG00000024891 | 1.53838222 | 1.589388001 | 1.575945187 | 1.380890489 | 1.433158226 | 1.477121255 | 1.466365979 |
| ENSMUSG00000024892 | 2.081172036 | 2.019203899 | 2.103406982 | 2.016522298 | 2.176862714 | 2.326335861 | 2.158095989 |
| ENSMUSG00000024896 | 2.519973621 | 2.327653195 | 2.331607372 | 2.31611065 | 2.264824954 | 2.568201724 | 2.112367894 |
| ENSMUSG00000024897 | 1.432083896 | 1.321350555 | 1.470660762 | 1.226722649 | 1.136121934 | 1.255272505 | 1.168187745 |
| ENSMUSG00000024899 | 2.209569962 | 2.073655187 | 2.009729747 | 2.056000419 | 2 | 2.267171728 | 1.888596011 |
| ENSMUSG00000024900 | 2.879922803 | 2.645770692 | 2.696158711 | 2.412762923 | 2.371090306 | 2.72916479 | 2.243069492 |
| ENSMUSG00000024901 | 1.036286563 | 1.061981878 | 1.330482024 | 1.04248125 | 1.264824954 | 1.146128036 | 1.20139795 |
| ENSMUSG00000024902 | 2.292497211 | 2.159156155 | 2.182829758 | 2.215271726 | 2.16634835 | 2.44870632 | 2.091394214 |
| ENSMUSG00000024903 | 0 | 0 | 0 | 0 | 0 | 0 | 0 |
| ENSMUSG00000024905 | 0.289064826 | 0.5 | 0.396240625 | 0.25 | 0.489301084 | 0.477121255 | 0.579661877 |
| ENSMUSG00000024906 | 2.095120908 | 1.96759118 | 1.984159485 | 1.606566189 | 1.784632921 | 1.857332496 | 1.822941512 |
| ENSMUSG00000024907 | 0 | 0 | 0 | 0 | 0 | 0.77815125 | 0.355418052 |
| ENSMUSG00000024908 | 3.033404503 | 2.769370839 | 2.749471282 | 2.805094582 | 2.733951626 | 3.190051418 | 2.560382912 |

| | | | | | | | |
|---|---|---|---|---|---|---|---|
| ENSMUSG00000024909 | 2.904664113 | 2.718745337 | 2.722375991 | 2.481573999 | 2.612309899 | 2.868644438 | 2.482255748 |
| ENSMUSG00000024910 | 1.289064826 | 1.098079356 | 1.201838731 | 0.75 | 1 | 0.77815125 | 0.710836105 |
| ENSMUSG00000024911 | 2.177057474 | 1.998588359 | 2.05357978 | 2.028435542 | 2.077536464 | 2.276461804 | 1.980010793 |
| ENSMUSG00000024912 | 1.948720984 | 1.898114259 | 1.931980114 | 1.809601185 | 1.97205363 | 2.127104798 | 1.915722126 |
| ENSMUSG00000024913 | 2.328498818 | 2.198603967 | 2.252806814 | 2.342671852 | 2.439191811 | 2.662757832 | 2.383771753 |
| ENSMUSG00000024914 | 2.613625632 | 2.459012589 | 2.54961126 | 2.334962501 | 2.437781395 | 2.662757832 | 2.369585695 |
| ENSMUSG00000024921 | 2.312518611 | 2.097004321 | 2.119945066 | 2.341033664 | 2.273371525 | 2.602059991 | 2.166993225 |
| ENSMUSG00000024922 | 0.578129653 | 0.396240625 | 0.580482024 | 0.5 | 0 | 0.698970004 | 0.448487648 |
| ENSMUSG00000024924 | 2.687411469 | 2.422499493 | 2.3843046 | 1.29248125 | 1.254881871 | 1.322219295 | 1.185218806 |
| ENSMUSG00000024925 | 1.558729299 | 1.52186571 | 1.642463902 | 1.431980114 | 1.47337555 | 1.531478917 | 1.594042709 |
| ENSMUSG00000024926 | 2.269664473 | 2.063166358 | 2.122963274 | 2.059601185 | 2.084333628 | 2.264817823 | 1.974994949 |
| ENSMUSG00000024927 | 2.611244851 | 2.420309603 | 2.474461364 | 2.322504712 | 2.475921683 | 2.703291378 | 2.368945281 |
| ENSMUSG00000024935 | 0 | 0.25 | 0.25 | 0.396240625 | 0.244650542 | 0 | 0.520678036 |
| ENSMUSG00000024936 | 0 | 0 | 0.25 | 0.25 | 0.387761935 | 0 | 0 |
| ENSMUSG00000024937 | 2.800026593 | 2.652063705 | 2.714689525 | 2.255591953 | 2.357605438 | 2.553883027 | 2.313616061 |
| ENSMUSG00000024939 | 0 | 0.25 | 0 | 0 | 0 | 0 | 0 |
| ENSMUSG00000024940 | 2.720536789 | 2.584962501 | 2.643175057 | 2.237821179 | 2.375332618 | 2.526339277 | 2.38702115 |
| ENSMUSG00000024941 | 2.714367558 | 2.523519421 | 2.559899632 | 2.394828984 | 2.539055463 | 2.768638101 | 2.434660827 |
| ENSMUSG00000024942 | 2.448863324 | 2.251406137 | 2.3184489 | 2.289336734 | 2.382016011 | 2.608526034 | 2.261753759 |
| ENSMUSG00000024943 | 3.027873102 | 2.787912707 | 2.756457917 | 2.791383785 | 2.688387666 | 3.07773118 | 2.48345785 |
| ENSMUSG00000024944 | 2.307602651 | 2.274508021 | 2.338286706 | 2.019203899 | 2.164053936 | 2.342422681 | 2.143416814 |
| ENSMUSG00000024947 | 2.446507203 | 2.331045137 | 2.406427211 | 2.271202097 | 2.361573756 | 2.539076099 | 2.32409319 |
| ENSMUSG00000024948 | 2.177057474 | 2.089388001 | 2.164552871 | 2.193696765 | 2.269408865 | 2.482873584 | 2.175263699 |
| ENSMUSG00000024949 | 3.104736624 | 2.839112993 | 2.849552315 | 2.861899965 | 2.831846286 | 3.255754787 | 2.690530733 |
| ENSMUSG00000024952 | 2.451206209 | 2.347004321 | 2.43155454 | 2.165444524 | 2.285543632 | 2.40654018 | 2.269110043 |
| ENSMUSG00000024953 | 2.602397159 | 2.460194481 | 2.513820609 | 2.223704441 | 2.382429551 | 2.591064607 | 2.345233094 |
| ENSMUSG00000024955 | 1.700465191 | 1.76109853 | 1.874961472 | 1.330482024 | 1.649303553 | 1.643452676 | 1.670888075 |
| ENSMUSG00000024957 | 0 | 0.25 | 0 | 0.25 | 0 | 0 | 0 |
| ENSMUSG00000024958 | 1.721148722 | 1.738549078 | 1.807204673 | 1.614857905 | 1.766173492 | 1.792391689 | 1.825244123 |
| ENSMUSG00000024959 | 2.023453784 | 2.01109853 | 2.112802778 | 1.852347734 | 2.017859938 | 2.103803721 | 2.034219411 |
| ENSMUSG00000024960 | 2.742600832 | 2.53199858 | 2.57991803 | 2.430701883 | 2.469844355 | 2.766412847 | 2.384994105 |
| ENSMUSG00000024962 | 2.010213549 | 1.87694866 | 1.948603967 | 1.575945187 | 1.655839964 | 1.643452676 | 1.680001789 |
| ENSMUSG00000024963 | 1.482695304 | 1.451838731 | 1.527131114 | 1.17510993 | 1.433158226 | 1.342422681 | 1.466365979 |
| ENSMUSG00000024965 | 0.960252568 | 0.896240625 | 0.896240625 | 0.79248125 | 0.877063019 | 0.602059991 | 1.041356071 |

| | | | | | | | |
|---|---|---|---|---|---|---|---|
| ENSMUSG00000024966 | 3.342237598 | 3.0308687 | 2.998588359 | 3.192562732 | 3.116536129 | 3.646893624 | 2.912711921 |
| ENSMUSG00000024968 | 2.22189694 | 2.094844592 | 2.086073977 | 1.850219859 | 2.006054284 | 2.086359831 | 1.895131899 |
| ENSMUSG00000024969 | 2.461589295 | 2.314935816 | 2.369945066 | 2.206637122 | 2.267124351 | 2.509202522 | 2.241172035 |
| ENSMUSG00000024970 | 2.217828299 | 2.074802005 | 2.08272922 | 1.87694866 | 2.014359518 | 2.184691431 | 1.972093446 |
| ENSMUSG00000024972 | 0 | 0 | 0 | 0 | 0 | 0 | 0 |
| ENSMUSG00000024973 | 0 | 0 | 0 | 0 | 0 | 0 | 0 |
| ENSMUSG00000024974 | 3.188777179 | 2.914664448 | 2.880890489 | 2.97104263 | 2.87685941 | 3.35755372 | 2.672480718 |
| ENSMUSG00000024975 | 2.223916471 | 2.027131114 | 1.966046536 | 2.075945187 | 1.977302144 | 2.243038049 | 1.899720848 |
| ENSMUSG00000024976 | 2.844167734 | 2.646944379 | 2.605253214 | 2.640083209 | 2.521662266 | 2.95616843 | 2.346609758 |
| ENSMUSG00000024978 | 2.141794304 | 2.025821952 | 1.961372513 | 1.975216702 | 1.986303335 | 2.204119983 | 1.884801065 |
| ENSMUSG00000024979 | 0 | 0 | 0 | 0 | 0 | 0 | 0 |
| ENSMUSG00000024981 | 2.914394662 | 2.672281101 | 2.679847205 | 2.844573714 | 2.754270191 | 3.165837625 | 2.5314585 |
| ENSMUSG00000024982 | 2.69529263 | 2.421187155 | 2.434945152 | 2.493961992 | 2.398980554 | 2.744292983 | 2.264126018 |
| ENSMUSG00000024983 | 2.160971586 | 1.982684334 | 2.036164561 | 2.075945187 | 1.974687643 | 2.235528447 | 1.93179548 |
| ENSMUSG00000024985 | 2.045658471 | 1.848079356 | 1.860735874 | 1.841580554 | 1.784632921 | 2.008600172 | 1.74196736 |
| ENSMUSG00000024986 | 1.482695304 | 1.282320754 | 1.388647213 | 1.388647213 | 1.373641807 | 1.491361694 | 1.345462945 |
| ENSMUSG00000024987 | 0.289064826 | 0 | 0.396240625 | 0 | 0 | 0 | 0.224243824 |
| ENSMUSG00000024989 | 2.606442122 | 2.403677461 | 2.390560606 | 2.517530236 | 2.418913884 | 2.750508395 | 2.248075634 |
| ENSMUSG00000024990 | 0.867194479 | 0.75 | 0.396240625 | 0.396240625 | 0.632412477 | 0.77815125 | 0.775756176 |
| ENSMUSG00000024991 | 3.683596844 | 3.33780383 | 3.328716608 | 3.336178002 | 3.24151153 | 3.77524626 | 3.005691971 |
| ENSMUSG00000024992 | 0 | 0 | 0 | 0 | 0 | 0 | 0 |
| ENSMUSG00000024993 | 2.340856238 | 2.174241882 | 2.123963757 | 2.267194569 | 2.138604637 | 2.462397998 | 2.019455688 |
| ENSMUSG00000024997 | 2.648525058 | 2.43662858 | 2.433677405 | 2.586905342 | 2.509475496 | 2.935507266 | 2.32848156 |
| ENSMUSG00000024998 | 2.141794304 | 1.984159485 | 2.005591953 | 1.458222504 | 1.55104774 | 1.643452676 | 1.425641774 |
| ENSMUSG00000024999 | 2.511972754 | 2.343216265 | 2.331607372 | 2.369439567 | 2.272243869 | 2.597695186 | 2.116590781 |
| ENSMUSG00000025001 | 3.224465559 | 2.940699316 | 2.92510993 | 3.051371955 | 2.951728238 | 3.427486109 | 2.709869165 |
| ENSMUSG00000025002 | 0.578129653 | 0.646240625 | 0.646240625 | 0.5 | 0.812711509 | 0.84509804 | 0.629532004 |
| ENSMUSG00000025003 | 0 | 0 | 0 | 0 | 0 | 0 | 0 |
| ENSMUSG00000025004 | 0 | 0 | 0 | 0 | 0 | 0 | 0 |
| ENSMUSG00000025006 | 2.283862692 | 2.137686696 | 2.133818844 | 1.584962501 | 1.592135487 | 1.755874856 | 1.520678036 |
| ENSMUSG00000025007 | 2.931357196 | 2.664552871 | 2.661414608 | 2.472945926 | 2.520825546 | 2.822168079 | 2.375921056 |
| ENSMUSG00000025008 | 1.99653913 | 1.886723615 | 1.811981878 | 1.784887838 | 1.733951626 | 1.954242509 | 1.685349273 |
| ENSMUSG00000025010 | 2.010213549 | 1.882845365 | 1.830482024 | 1.894357207 | 1.84445929 | 2.021189299 | 1.71890424 |
| ENSMUSG00000025013 | 0 | 0.25 | 0 | 0.25 | 0 | 0 | 0.355418052 |

| | | | | | | | |
|---|---|---|---|---|---|---|---|
| ENSMUSG00000025014 | 0 | 0 | 0 | 0 | 0 | 0 | 0 |
| ENSMUSG00000025016 | 3.404746548 | 3.104924222 | 3.073870493 | 3.19209778 | 3.079147613 | 3.600646236 | 2.846093998 |
| ENSMUSG00000025017 | 0.671187741 | 0.580482024 | 0.701838731 | 1.04248125 | 1.074582839 | 0.954242509 | 1.150210039 |
| ENSMUSG00000025019 | 1.985980784 | 1.866901388 | 1.860735874 | 1.990000483 | 1.964031554 | 2.161368002 | 1.769001954 |
| ENSMUSG00000025020 | 0 | 0 | 0.580482024 | 0.701838731 | 0.812711509 | 0.477121255 | 0.710836105 |
| ENSMUSG00000025024 | 2.657168526 | 2.396240625 | 2.351285366 | 2.464883697 | 2.391003805 | 2.718501689 | 2.200679824 |
| ENSMUSG00000025025 | 1.806056082 | 1.705044741 | 1.705044741 | 1.912762923 | 1.810629166 | 2.064457989 | 1.753626837 |
| ENSMUSG00000025026 | 2.081172036 | 1.958222504 | 1.969129237 | 2.158248799 | 2.160196454 | 2.459392488 | 2.079068325 |
| ENSMUSG00000025027 | 2.888606985 | 2.696771581 | 2.722566069 | 2.667885267 | 2.731192945 | 3.05804623 | 2.606173345 |
| ENSMUSG00000025034 | 2.188178997 | 2.122963274 | 2.208222504 | 1.892463902 | 2.037065488 | 2.158362492 | 2.044866219 |
| ENSMUSG00000025035 | 2.317377295 | 2.176839783 | 2.208222504 | 2.225216702 | 2.256066288 | 2.485721426 | 2.167790061 |
| ENSMUSG00000025036 | 1.87190676 | 1.752806814 | 1.714495249 | 1.905512955 | 1.877063019 | 1.986771734 | 1.782372264 |
| ENSMUSG00000025037 | 2.985656624 | 2.70881523 | 2.736726569 | 2.642938161 | 2.547699718 | 2.951823035 | 2.361815771 |
| ENSMUSG00000025038 | 0.289064826 | 0.25 | 0 | 0 | 0 | 0 | 0.224243824 |
| ENSMUSG00000025040 | 2.406877894 | 2.205044741 | 2.210587586 | 2.218361028 | 2.149165677 | 2.409933123 | 1.976436 |
| ENSMUSG00000025041 | 2.540273527 | 2.322504712 | 2.293106877 | 2.374961472 | 2.305113253 | 2.630427875 | 2.143416814 |
| ENSMUSG00000025043 | 0 | 0 | 0 | 0 | 0 | 0 | 0 |
| ENSMUSG00000025044 | 0 | 0.25 | 0 | 0 | 0 | 0 | 0 |
| ENSMUSG00000025047 | 2.800026593 | 2.570772088 | 2.588010856 | 2.545598588 | 2.569036659 | 2.884795364 | 2.427786636 |
| ENSMUSG00000025049 | 1.881174527 | 1.802363341 | 1.79248125 | 1.998588359 | 1.962676635 | 2.089905111 | 1.829800754 |
| ENSMUSG00000025050 | 2.045658471 | 1.858657057 | 1.839388001 | 2.091580554 | 2.031524462 | 2.250420002 | 1.861046144 |
| ENSMUSG00000025051 | 0 | 0 | 0 | 0 | 0 | 0 | 0 |
| ENSMUSG00000025056 | 0 | 0 | 0 | 0 | 0 | 0 | 0 |
| ENSMUSG00000025058 | 0 | 0 | 0.25 | 0.5 | 0.244650542 | 0 | 0.224243824 |
| ENSMUSG00000025059 | 2.351376531 | 2.197790722 | 2.214495249 | 2.202642909 | 2.132784263 | 2.413299764 | 1.95794437 |
| ENSMUSG00000025060 | 3.018088033 | 2.781514779 | 2.758183632 | 2.879171237 | 2.809398495 | 3.237292338 | 2.586714264 |
| ENSMUSG00000025064 | 0.45815691 | 0.25 | 0.5 | 0 | 0 | 0.301029996 | 0.224243824 |
| ENSMUSG00000025066 | 2.974482511 | 2.739819376 | 2.705443495 | 2.725782169 | 2.663004918 | 3.089905111 | 2.492479894 |
| ENSMUSG00000025068 | 2.615205302 | 2.442872367 | 2.466433318 | 2.344302633 | 2.29413964 | 2.62838893 | 2.170563634 |
| ENSMUSG00000025069 | 0.91631382 | 0.864857905 | 0.646240625 | 0.580482024 | 0.489301084 | 0.301029996 | 0.224243824 |
| ENSMUSG00000025075 | 0 | 0.25 | 0 | 0 | 0 | 0 | 0 |
| ENSMUSG00000025076 | 2.141794304 | 1.995748394 | 1.945339928 | 2.25 | 2.206647761 | 2.5289167 | 2.072724682 |
| ENSMUSG00000025077 | 2.267855219 | 2.029735268 | 2.077084758 | 2.009729747 | 1.979897422 | 2.149219113 | 1.851668228 |
| ENSMUSG00000025078 | 2.395018353 | 2.256284891 | 2.248588359 | 2.225216702 | 2.156296347 | 2.444044796 | 2.043700396 |

| | | | | | | | |
|---|---|---|---|---|---|---|---|
| ENSMUSG00000025081 | 0 | 0 | 0 | 0 | 0 | 0 | 0 |
| ENSMUSG00000025082 | 0.289064826 | 0 | 0 | 0.864857905 | 0.733951626 | 0.77815125 | 0.803905701 |
| ENSMUSG00000025083 | 1.800532393 | 1.738549078 | 1.634789703 | 2.067865757 | 2.159419872 | 2.354108439 | 2.077494038 |
| ENSMUSG00000025085 | 2.728422728 | 2.505244985 | 2.494319981 | 2.475593779 | 2.439543536 | 2.742725131 | 2.289406866 |
| ENSMUSG00000025086 | 1.940852214 | 1.79745614 | 1.852347734 | 1.802363341 | 1.793512788 | 1.977723605 | 1.710836105 |
| ENSMUSG00000025089 | 1.342375483 | 1.364857905 | 1.410964047 | 0.396240625 | 0.686820904 | 0.77815125 | 0.803905701 |
| ENSMUSG00000025090 | 0 | 0 | 0 | 0 | 0 | 0 | 0 |
| ENSMUSG00000025091 | 0 | 0 | 0 | 0 | 0 | 0 | 0 |
| ENSMUSG00000025092 | 1.890240806 | 1.837182039 | 1.850219859 | 1.51109853 | 1.519146132 | 1.698970004 | 1.532233387 |
| ENSMUSG00000025094 | 1.671187741 | 1.505591953 | 1.458222504 | 1.566696635 | 1.519146132 | 1.653212514 | 1.296434211 |
| ENSMUSG00000025102 | 1.505869732 | 1.494319981 | 1.470660762 | 1.282320754 | 1.394615667 | 1.380211242 | 1.379042812 |
| ENSMUSG00000025103 | 2.254965775 | 2.064346961 | 2.098079356 | 2.200224975 | 2.122577592 | 2.408239965 | 1.963977467 |
| ENSMUSG00000025104 | 2.275045611 | 2.139605178 | 2.152756199 | 2.251406137 | 2.16634835 | 2.45484486 | 1.969165684 |
| ENSMUSG00000025105 | 0 | 0.396240625 | 0.25 | 0.5 | 0.568060967 | 0.477121255 | 0.448487648 |
| ENSMUSG00000025127 | 0.45815691 | 0 | 0.25 | 0.396240625 | 0 | 0 | 0.355418052 |
| ENSMUSG00000025128 | 0 | 0 | 0 | 0 | 0 | 0 | 0 |
| ENSMUSG00000025129 | 0 | 0 | 0.396240625 | 0 | 0 | 0 | 0.355418052 |
| ENSMUSG00000025130 | 3.708678855 | 3.358852456 | 3.354133415 | 3.342160717 | 3.270211646 | 3.819280647 | 3.070801959 |
| ENSMUSG00000025132 | 3.437606122 | 3.182617534 | 3.23129588 | 3.01593303 | 3.091501633 | 3.500236475 | 2.962035858 |
| ENSMUSG00000025133 | 2.280358197 | 2.109697963 | 2.063166358 | 2.119945066 | 2.077536464 | 2.401400541 | 1.916588299 |
| ENSMUSG00000025134 | 2.777675583 | 2.561090959 | 2.58272922 | 2.632845365 | 2.617231464 | 2.954724791 | 2.457550299 |
| ENSMUSG00000025135 | 2.177057474 | 2.084962501 | 2.140560606 | 1.83272922 | 1.951645858 | 2.136720567 | 1.952571511 |
| ENSMUSG00000025137 | 2.054824387 | 1.918106335 | 2.03360658 | 1.94697564 | 2.176862714 | 2.247973266 | 2.121678334 |
| ENSMUSG00000025138 | 1.944805158 | 1.866901388 | 1.845926073 | 1.79248125 | 1.875328588 | 1.963787827 | 1.853775828 |
| ENSMUSG00000025139 | 2.514991096 | 2.315523711 | 2.302363341 | 2.291226732 | 2.254881871 | 2.5774918 | 2.122136896 |
| ENSMUSG00000025140 | 1.989530018 | 1.916333979 | 1.755591953 | 1.488549078 | 1.563883923 | 1.672097858 | 1.559435575 |
| ENSMUSG00000025141 | 0 | 0 | 0 | 0 | 0 | 0 | 0 |
| ENSMUSG00000025142 | 2.231898196 | 2.134789703 | 2.181980114 | 1.985628626 | 2.126012715 | 2.267171728 | 2.096890362 |
| ENSMUSG00000025144 | 1.671187741 | 1.698603967 | 1.685366747 | 1.705044741 | 1.78687392 | 1.886490725 | 1.707551656 |
| ENSMUSG00000025145 | 2.057835465 | 1.933677405 | 2.056000419 | 1.752806814 | 2.027028145 | 1.963787827 | 1.949460761 |
| ENSMUSG00000025147 | 1.87190676 | 1.729715809 | 1.723704441 | 1.618933358 | 1.741652793 | 1.748188027 | 1.612273755 |
| ENSMUSG00000025150 | 2.751223449 | 2.550837001 | 2.61306031 | 1.29248125 | 1.450958105 | 1.431363764 | 1.324583736 |
| ENSMUSG00000025151 | 3.495304221 | 3.181075168 | 3.180755233 | 2.991265689 | 3.010684664 | 3.447313109 | 2.859642604 |
| ENSMUSG00000025153 | 3.256154144 | 3.029654172 | 3.095115266 | 2.79652856 | 2.991492869 | 3.230704314 | 2.885277875 |

| | | | | | | | |
|---|---|---|---|---|---|---|---|
| ENSMUSG00000025154 | 2.53552908 | 2.290597832 | 2.281030328 | 2.584962501 | 2.524713379 | 2.911157609 | 2.354302096 |
| ENSMUSG00000025155 | 2.269664473 | 2.097004321 | 2.186208459 | 2.150906586 | 2.265976525 | 2.421603927 | 2.130728307 |
| ENSMUSG00000025156 | 2.729023206 | 2.534887838 | 2.55175358 | 2.439555804 | 2.526641353 | 2.819543936 | 2.37361247 |
| ENSMUSG00000025157 | 2.177057474 | 2.002806814 | 2.097004321 | 2.081607372 | 2.094757251 | 2.255272505 | 1.98213686 |
| ENSMUSG00000025158 | 1.862428343 | 1.892463902 | 1.898114259 | 1.814346961 | 2.013184954 | 2.056904851 | 1.939139147 |
| ENSMUSG00000025159 | 2.644870062 | 2.450629091 | 2.439972046 | 2.399513125 | 2.379941013 | 2.697229343 | 2.266481009 |
| ENSMUSG00000025161 | 0.867194479 | 0.79248125 | 0.864857905 | 1.794977273 | 1.988833497 | 2.103803721 | 1.994607951 |
| ENSMUSG00000025162 | 3.022564273 | 2.786801231 | 2.796063874 | 2.723893821 | 2.717866348 | 3.069668097 | 2.579438993 |
| ENSMUSG00000025163 | 0 | 0 | 0 | 0 | 0 | 0 | 0 |
| ENSMUSG00000025165 | 0 | 0 | 0 | 0 | 0.244650542 | 0 | 0.224243824 |
| ENSMUSG00000025169 | 1.494443473 | 1.438721876 | 1.476722649 | 1.610735874 | 1.55104774 | 1.812913357 | 1.532233387 |
| ENSMUSG00000025170 | 0.91631382 | 0.5 | 0.896240625 | 1.17510993 | 1.188507684 | 1.113943352 | 1.21680636 |
| ENSMUSG00000025171 | 2.039433992 | 2.005591953 | 2.134789703 | 1.527131114 | 1.828942237 | 1.812913357 | 1.876096088 |
| ENSMUSG00000025172 | 0.289064826 | 0.25 | 0.25 | 0 | 0 | 0 | 0 |
| ENSMUSG00000025173 | 2.70430294 | 2.47027849 | 2.463327389 | 2.489638179 | 2.452655829 | 2.827369273 | 2.351382464 |
| ENSMUSG00000025175 | 0.289064826 | 0 | 0.25 | 0.580482024 | 0.387761935 | 0.477121255 | 0.803905701 |
| ENSMUSG00000025176 | 0.960252568 | 1.146240625 | 1.160964047 | 1.25 | 1.212046813 | 1.146128036 | 1.20919388 |
| ENSMUSG00000025178 | 2.641922824 | 2.450224975 | 2.444519282 | 2.461764337 | 2.489911207 | 2.804139432 | 2.349118526 |
| ENSMUSG00000025184 | 2.35433422 | 2.143411797 | 2.134789703 | 2.159156155 | 2.110284345 | 2.440909082 | 2.013099581 |
| ENSMUSG00000025185 | 2.791849225 | 2.605253214 | 2.626452888 | 2.666556001 | 2.683314108 | 3.031812271 | 2.576189806 |
| ENSMUSG00000025188 | 1.83262764 | 1.789967834 | 1.87892496 | 1.701838731 | 1.861137485 | 1.944482672 | 1.88765144 |
| ENSMUSG00000025189 | 0 | 0.25 | 0.25 | 0.5 | 0.387761935 | 0.477121255 | 0.355418052 |
| ENSMUSG00000025190 | 2.5496035 | 2.311981878 | 2.382357639 | 2.344844592 | 2.35043857 | 2.681241237 | 2.217830684 |
| ENSMUSG00000025192 | 2.095120908 | 1.973704441 | 2.02186571 | 1.931980114 | 1.978602168 | 2.164352856 | 1.838725884 |
| ENSMUSG00000025193 | 1.432083896 | 1.42510993 | 1.410964047 | 1.458222504 | 1.358933174 | 1.51851394 | 1.437266334 |
| ENSMUSG00000025194 | 0 | 0 | 0.25 | 0 | 0 | 0 | 0.355418052 |
| ENSMUSG00000025195 | 2.339331437 | 2.242885888 | 2.30479213 | 2.133818844 | 2.179075613 | 2.380211242 | 2.090384805 |
| ENSMUSG00000025196 | 0 | 0 | 0 | 0.25 | 0 | 0 | 0 |
| ENSMUSG00000025197 | 0 | 0.25 | 0 | 0 | 0.244650542 | 0 | 0 |
| ENSMUSG00000025198 | 2.701107274 | 2.489638179 | 2.491806565 | 2.691839214 | 2.591221684 | 3.011147361 | 2.39794841 |
| ENSMUSG00000025199 | 2.827447047 | 2.590211771 | 2.56932185 | 2.571061437 | 2.48127199 | 2.85672889 | 2.319395519 |
| ENSMUSG00000025200 | 2.201144779 | 2.016522298 | 2.049918086 | 2.118933358 | 2.042520869 | 2.31386722 | 1.94631981 |
| ENSMUSG00000025201 | 0 | 0 | 0 | 0 | 0 | 0 | 0 |
| ENSMUSG00000025202 | 0.289064826 | 0.580482024 | 0.396240625 | 0.5 | 0.489301084 | 0.301029996 | 0.355418052 |

| | | | | | | | |
|---|---|---|---|---|---|---|---|
| ENSMUSG00000025203 | 3.753081287 | 3.399571294 | 3.38199141 | 3.423671088 | 3.369404227 | 3.932727367 | 3.164627404 |
| ENSMUSG00000025204 | 2.520963022 | 2.360217292 | 2.410061234 | 2.402756199 | 2.415543455 | 2.729974286 | 2.355640783 |
| ENSMUSG00000025207 | 1.516991255 | 1.418106335 | 1.451838731 | 1.646240625 | 1.674752852 | 1.826074803 | 1.591689859 |
| ENSMUSG00000025208 | 2.129344651 | 2.057204673 | 2.129909063 | 1.992885888 | 2.088159748 | 2.283301229 | 2.051774285 |
| ENSMUSG00000025209 | 2.413323654 | 2.271202097 | 2.254897682 | 2.132845365 | 2.107593434 | 2.326335861 | 2.018825668 |
| ENSMUSG00000025212 | 2.674620126 | 2.481573999 | 2.470660762 | 2.383332433 | 2.398190943 | 2.694605199 | 2.263238458 |
| ENSMUSG00000025213 | 1 | 1.098079356 | 1.02186571 | 1.214495249 | 1.234113756 | 1.146128036 | 1.224243824 |
| ENSMUSG00000025215 | 0 | 0 | 0 | 0.580482024 | 0.568060967 | 0.477121255 | 0.579661877 |
| ENSMUSG00000025216 | 0 | 0 | 0 | 0 | 0.244650542 | 0 | 0.355418052 |
| ENSMUSG00000025217 | 2.217828299 | 2.05479213 | 2.078220739 | 2.182829758 | 2.103062484 | 2.376576957 | 2.017561934 |
| ENSMUSG00000025218 | 1.936861443 | 1.907339155 | 1.969129237 | 1.872963274 | 1.977302144 | 2.113943352 | 1.952571511 |
| ENSMUSG00000025219 | 0 | 0 | 0 | 0 | 0 | 0 | 0.224243824 |
| ENSMUSG00000025220 | 2.93904296 | 2.64151351 | 2.636241108 | 2.655512955 | 2.56462465 | 2.934498451 | 2.387626796 |
| ENSMUSG00000025221 | 0.289064826 | 0.25 | 0.25 | 0.396240625 | 0 | 0 | 0.224243824 |
| ENSMUSG00000025223 | 2.766857015 | 2.57508814 | 2.629171237 | 2.284247778 | 2.414035093 | 2.57634135 | 2.357195614 |
| ENSMUSG00000025224 | 2.879922803 | 2.648347781 | 2.654596376 | 2.670309603 | 2.677450619 | 3.021602716 | 2.51365069 |
| ENSMUSG00000025225 | 2.676666078 | 2.519203899 | 2.578220739 | 2.586073977 | 2.679594084 | 2.973127854 | 2.576864734 |
| ENSMUSG00000025226 | 0.960252568 | 1.098079356 | 1.226722649 | 0.976722649 | 1.163285805 | 0.954242509 | 1.245582246 |
| ENSMUSG00000025227 | 1.069666964 | 1.130890489 | 1.214495249 | 1.061981878 | 1.149965125 | 1.255272505 | 1.473249546 |
| ENSMUSG00000025228 | 3.111026095 | 2.834684096 | 2.837872852 | 2.820772088 | 2.841770088 | 3.225050696 | 2.657681529 |
| ENSMUSG00000025229 | 0 | 0 | 0 | 0 | 0.244650542 | 0 | 0.224243824 |
| ENSMUSG00000025231 | 2.517987749 | 2.343216265 | 2.357090043 | 2.307805295 | 2.322676882 | 2.571708832 | 2.198877505 |
| ENSMUSG00000025232 | 2.834961347 | 2.578220739 | 2.618426437 | 2.238549078 | 2.168627946 | 2.46686762 | 2.077494038 |
| ENSMUSG00000025234 | 2.907802659 | 2.660287149 | 2.633332433 | 2.718361028 | 2.64540129 | 3.015359755 | 2.43830269 |
| ENSMUSG00000025235 | 1.771760131 | 1.62694866 | 1.575945187 | 1.78743678 | 1.741652793 | 1.892094603 | 1.627424404 |
| ENSMUSG00000025236 | 2.312518611 | 2.135757955 | 2.14151351 | 2.178561379 | 2.162515999 | 2.45331834 | 2.053478496 |
| ENSMUSG00000025237 | 1.912079952 | 1.766522298 | 1.777131114 | 1.769203899 | 1.726078673 | 1.968482949 | 1.659601507 |
| ENSMUSG00000025239 | 2.671876482 | 2.47104263 | 2.481203126 | 2.503156135 | 2.480333692 | 2.809559715 | 2.317146272 |
| ENSMUSG00000025240 | 2.598312578 | 2.423371739 | 2.400442697 | 2.575374049 | 2.484072009 | 2.891537458 | 2.307989563 |
| ENSMUSG00000025241 | 2.183765914 | 2.072504712 | 2.106566189 | 2.025821952 | 2.068600388 | 2.301029996 | 1.947894097 |
| ENSMUSG00000025243 | 0 | 0 | 0.25 | 0.396240625 | 0.244650542 | 0.301029996 | 0 |
| ENSMUSG00000025245 | 2.029919532 | 1.892463902 | 1.870953944 | 2.029735268 | 1.944572352 | 2.130333768 | 1.797719748 |
| ENSMUSG00000025246 | 2.980106992 | 2.733053688 | 2.699820405 | 2.671625132 | 2.577698593 | 2.959518377 | 2.382339813 |
| ENSMUSG00000025255 | 1.53838222 | 1.557204673 | 1.476722649 | 1.678561379 | 1.645989387 | 1.740362689 | 1.483307832 |

| | | | | | | | |
|---|---|---|---|---|---|---|---|
| ENSMUSG00000025257 | 0.289064826 | 0.79248125 | 0.92510993 | 0.864857905 | 0.77552387 | 0.84509804 | 0.876096088 |
| ENSMUSG00000025260 | 2.203267085 | 2.071350555 | 2.034887838 | 2.193696765 | 2.184185718 | 2.496929648 | 2.053478496 |
| ENSMUSG00000025261 | 3.354468164 | 3.043731421 | 3.031111115 | 3.224224996 | 3.157815628 | 3.648164779 | 2.921582365 |
| ENSMUSG00000025262 | 1.663535556 | 1.532320754 | 1.464495249 | 1.92510993 | 1.782377602 | 2.029383778 | 1.692344475 |
| ENSMUSG00000025264 | 2.097855556 | 1.942046081 | 1.950224975 | 2.215271726 | 2.182004726 | 2.462397998 | 2.02383181 |
| ENSMUSG00000025265 | 2.057835465 | 2.002806814 | 1.882845365 | 1.94697564 | 1.973373093 | 2.120573931 | 1.895131899 |
| ENSMUSG00000025266 | 2.559630991 | 2.343216265 | 2.359697963 | 2.643884812 | 2.53344226 | 2.913283902 | 2.341533092 |
| ENSMUSG00000025268 | 2.797997234 | 2.577369088 | 2.608135475 | 2.267194569 | 2.314756076 | 2.575187845 | 2.234765579 |
| ENSMUSG00000025269 | 2.086807695 | 2 | 1.938721876 | 2.078220739 | 2.074582839 | 2.348304863 | 1.951796623 |
| ENSMUSG00000025270 | 0.811507563 | 0.701838731 | 0.5 | 0.79248125 | 0.387761935 | 0.602059991 | 0.672731473 |
| ENSMUSG00000025271 | 1.568542279 | 1.445339928 | 1.505591953 | 1.618933358 | 1.580303447 | 1.770852012 | 1.540636858 |
| ENSMUSG00000025272 | 1.129344651 | 1.214495249 | 1.238549078 | 1.660964047 | 1.665423072 | 1.857332496 | 1.540636858 |
| ENSMUSG00000025277 | 1.777675583 | 1.67510993 | 1.653677461 | 1.951838731 | 1.87878897 | 2.167317335 | 1.760706232 |
| ENSMUSG00000025278 | 3.526762047 | 3.218072528 | 3.226299736 | 3.147880587 | 3.179767006 | 3.605412798 | 2.971496737 |
| ENSMUSG00000025279 | 0.289064826 | 0.25 | 0.25 | 0 | 0 | 0 | 0.224243824 |
| ENSMUSG00000025280 | 2.213719572 | 2.034887838 | 2.024508021 | 2.219895812 | 2.208682096 | 2.432969291 | 2.039586611 |
| ENSMUSG00000025283 | 2.433305075 | 2.237821179 | 2.264497931 | 2.231944491 | 2.156296347 | 2.460897843 | 2.009225098 |
| ENSMUSG00000025287 | 2.606442122 | 2.393884812 | 2.393411797 | 2.651832578 | 2.576984829 | 2.989449818 | 2.406812401 |
| ENSMUSG00000025288 | 0 | 0 | 0 | 0 | 0 | 0 | 0 |
| ENSMUSG00000025289 | 2.651426165 | 2.403677461 | 2.456239685 | 2.589388001 | 2.471453824 | 2.86332286 | 2.362906151 |
| ENSMUSG00000025290 | 2.932463873 | 2.680274797 | 2.67619206 | 2.956040802 | 2.843677054 | 3.308777774 | 2.637175571 |
| ENSMUSG00000025314 | 1.903481042 | 1.83272922 | 1.75835575 | 1.982684334 | 1.883916752 | 2.146128036 | 1.801445495 |
| ENSMUSG00000025316 | 2.243591459 | 2.134789703 | 2.149978211 | 2.012462137 | 2.068600388 | 2.173186268 | 1.976436 |
| ENSMUSG00000025317 | 0 | 0 | 0.396240625 | 0 | 0 | 0.477121255 | 0.224243824 |
| ENSMUSG00000025318 | 0 | 0 | 0.396240625 | 0.396240625 | 0.387761935 | 0 | 0.355418052 |
| ENSMUSG00000025321 | 2.072573125 | 2.029735268 | 1.948603967 | 1.52186571 | 1.394615667 | 1.477121255 | 1.272006351 |
| ENSMUSG00000025323 | 1.707474304 | 1.610735874 | 1.614857905 | 1.614857905 | 1.47337555 | 1.653212514 | 1.38802993 |
| ENSMUSG00000025324 | 1.885732304 | 1.809601185 | 1.850219859 | 1.671625132 | 1.655839964 | 1.806179974 | 1.651852264 |
| ENSMUSG00000025326 | 2.698532991 | 2.511780978 | 2.491806565 | 2.58272922 | 2.488689905 | 2.843855423 | 2.260262214 |
| ENSMUSG00000025328 | 0.578129653 | 0.396240625 | 0.25 | 0.25 | 0.489301084 | 0.477121255 | 0.520678036 |
| ENSMUSG00000025329 | 0 | 0 | 0 | 0 | 0 | 0 | 0 |
| ENSMUSG00000025330 | 0 | 0 | 0 | 0 | 0 | 0 | 0 |
| ENSMUSG00000025332 | 2.708736128 | 2.483422664 | 2.505244985 | 2.631380202 | 2.614886232 | 2.923244019 | 2.474176968 |
| ENSMUSG00000025333 | 0 | 0 | 0 | 0 | 0.244650542 | 0 | 0 |

| | | | | | | | |
|---|---|---|---|---|---|---|---|
| ENSMUSG00000025337 | 2.544964605 | 2.342671852 | 2.371960008 | 2.263820609 | 2.242199448 | 2.509202522 | 2.142557543 |
| ENSMUSG00000025340 | 2.493389029 | 2.299304173 | 2.293106877 | 2.130890489 | 2.136951444 | 2.382017043 | 2.013099581 |
| ENSMUSG00000025347 | 0 | 0 | 0 | 0 | 0.244650542 | 0 | 0 |
| ENSMUSG00000025348 | 1.639698646 | 1.566696635 | 1.671625132 | 1.532320754 | 1.72073075 | 1.838849091 | 1.71890424 |
| ENSMUSG00000025350 | 1.227926429 | 1.17510993 | 1.214495249 | 0.864857905 | 0.955822902 | 0.77815125 | 0.935079929 |
| ENSMUSG00000025351 | 3.191953071 | 2.905627364 | 2.915667094 | 2.771036003 | 2.731807852 | 3.136403448 | 2.542516675 |
| ENSMUSG00000025352 | 2.249317394 | 2.043731421 | 2.106566189 | 2.257666784 | 2.215379757 | 2.499687083 | 2.075383 |
| ENSMUSG00000025353 | 1.727821383 | 1.566696635 | 1.678561379 | 1.711372513 | 1.731346772 | 1.924279286 | 1.61448206 |
| ENSMUSG00000025354 | 2.604828841 | 2.422499493 | 2.438304187 | 2.457034121 | 2.441296923 | 2.764176132 | 2.28967999 |
| ENSMUSG00000025355 | 1.734388958 | 1.54745614 | 1.646240625 | 1.061981878 | 1 | 1.230448921 | 1 |
| ENSMUSG00000025357 | 1.765759561 | 1.602347734 | 1.622963274 | 1.856566189 | 1.808534465 | 1.995635195 | 1.731408809 |
| ENSMUSG00000025358 | 2.675985209 | 2.413211243 | 2.429419106 | 2.598079356 | 2.531009002 | 2.925312091 | 2.392033886 |
| ENSMUSG00000025359 | 0.671187741 | 0.701838731 | 0.92510993 | 1.02186571 | 1.057362051 | 0.84509804 | 0.935079929 |
| ENSMUSG00000025362 | 2.899911556 | 2.750352049 | 2.819612365 | 2.490362424 | 2.703684258 | 2.902546779 | 2.677952919 |
| ENSMUSG00000025364 | 3.105467192 | 2.832168732 | 2.821350555 | 2.987638974 | 2.907188525 | 3.387567779 | 2.706198347 |
| ENSMUSG00000025366 | 3.002810412 | 2.760585845 | 2.805245713 | 2.71293726 | 2.810874786 | 3.099680641 | 2.664009025 |
| ENSMUSG00000025369 | 2.800532393 | 2.560197833 | 2.614088604 | 2.635274154 | 2.643433859 | 3.030599722 | 2.528209673 |
| ENSMUSG00000025370 | 0 | 0 | 0 | 0 | 0 | 0 | 0 |
| ENSMUSG00000025371 | 2.231898196 | 2.142463902 | 2.199415381 | 1.997171172 | 2.071604288 | 2.222716471 | 2.043115906 |
| ENSMUSG00000025372 | 2.482695304 | 2.365369862 | 2.426839783 | 2.067865757 | 2.282813873 | 2.352182518 | 2.243069492 |
| ENSMUSG00000025373 | 2.260538674 | 2.193696765 | 2.201838731 | 2 | 2.112954895 | 2.225309282 | 2.046027856 |
| ENSMUSG00000025374 | 2.100572389 | 1.973704441 | 1.921625132 | 2.039967834 | 2.145141048 | 2.367355921 | 2.024452165 |
| ENSMUSG00000025375 | 1.269664473 | 1.380890489 | 1.505591953 | 1.160964047 | 1.467903253 | 1.255272505 | 1.61448206 |
| ENSMUSG00000025377 | 1.800532393 | 1.744319981 | 1.841580554 | 1.584962501 | 1.770879644 | 1.785329835 | 1.74196736 |
| ENSMUSG00000025380 | 0.671187741 | 0 | 0.5 | 0 | 0.387761935 | 0 | 0.224243824 |
| ENSMUSG00000025381 | 2.488076442 | 2.30357978 | 2.300530956 | 2.27908599 | 2.247690665 | 2.575187845 | 2.149368646 |
| ENSMUSG00000025383 | 0.289064826 | 0.25 | 0 | 0.25 | 0.244650542 | 0.477121255 | 0.579661877 |
| ENSMUSG00000025384 | 1.890240806 | 1.926839783 | 2.046218836 | 1.527131114 | 1.72341484 | 1.579783597 | 1.824094866 |
| ENSMUSG00000025386 | 1.100572389 | 0.864857905 | 0.976722649 | 0.864857905 | 0.812711509 | 0.903089987 | 0.829800754 |
| ENSMUSG00000025389 | 0 | 0.25 | 0 | 0 | 0 | 0 | 0.224243824 |
| ENSMUSG00000025393 | 3.846273233 | 3.456090533 | 3.457480223 | 3.576533228 | 3.496307446 | 4.106870544 | 3.260411678 |
| ENSMUSG00000025395 | 2.748369006 | 2.544043287 | 2.520869832 | 2.734159485 | 2.658118333 | 3.030599722 | 2.428676148 |
| ENSMUSG00000025396 | 0 | 0 | 0 | 0 | 0 | 0 | 0 |
| ENSMUSG00000025400 | 0 | 0 | 0 | 0 | 0 | 0 | 0 |

| | | | | | | | |
|---|---|---|---|---|---|---|---|
| ENSMUSG00000025401 | 1.100572389 | 0.830482024 | 1.061981878 | 0.92510993 | 0.905314583 | 0.602059991 | 0.853775828 |
| ENSMUSG00000025402 | 2.199011616 | 2.097004321 | 2.134789703 | 1.688721876 | 1.784632921 | 1.897627091 | 1.784981274 |
| ENSMUSG00000025403 | 2.845531329 | 2.674241882 | 2.69059551 | 2.432405186 | 2.623549464 | 2.773054693 | 2.554656826 |
| ENSMUSG00000025404 | 2.635214384 | 2.422499493 | 2.438721876 | 2.456637122 | 2.465951417 | 2.770115295 | 2.376964992 |
| ENSMUSG00000025405 | 0 | 0 | 0 | 0 | 0 | 0 | 0 |
| ENSMUSG00000025407 | 0.45815691 | 0 | 0.396240625 | 0.830482024 | 0.733951626 | 0.84509804 | 0.876096088 |
| ENSMUSG00000025408 | 2.31414447 | 2.178561379 | 2.225216702 | 1.821350555 | 1.761403742 | 1.944482672 | 1.803905701 |
| ENSMUSG00000025409 | 2.129344651 | 2.02186571 | 2.178561379 | 1.834962501 | 1.948833444 | 2 | 2.034820183 |
| ENSMUSG00000025410 | 2.985656624 | 2.740543258 | 2.750879478 | 2.729156481 | 2.7726626 | 3.128722284 | 2.610354375 |
| ENSMUSG00000025413 | 2.31414447 | 2.098079356 | 2.107613138 | 2.259729747 | 2.214050354 | 2.540329475 | 2.087337541 |
| ENSMUSG00000025417 | 2.631440309 | 2.379909063 | 2.388647213 | 2.370449858 | 2.347256716 | 2.621176282 | 2.201756416 |
| ENSMUSG00000025418 | 0 | 0 | 0 | 0 | 0 | 0 | 0 |
| ENSMUSG00000025420 | 1.516991255 | 1.42510993 | 1.396240625 | 1.418106335 | 1.42077253 | 1.579783597 | 1.345462945 |
| ENSMUSG00000025421 | 2.343889214 | 2.156427211 | 2.143411797 | 2.138647213 | 2.043601899 | 2.354108439 | 1.947107911 |
| ENSMUSG00000025422 | 0 | 0.25 | 0.580482024 | 0.976722649 | 0.931471446 | 0.602059991 | 1.014380832 |
| ENSMUSG00000025423 | 2.636714502 | 2.480831871 | 2.444929839 | 2.407339155 | 2.340805641 | 2.684845362 | 2.142987464 |
| ENSMUSG00000025425 | 0 | 0 | 0 | 0 | 0 | 0 | 0 |
| ENSMUSG00000025427 | 0 | 0 | 0 | 0.701838731 | 0.632412477 | 0.602059991 | 0.520678036 |
| ENSMUSG00000025428 | 3.455011854 | 3.155226774 | 3.168216819 | 3.165444524 | 3.167065104 | 3.624178926 | 2.972159679 |
| ENSMUSG00000025429 | 0.91631382 | 0.896240625 | 0.976722649 | 0.864857905 | 0.812711509 | 1.041392685 | 0.896975297 |
| ENSMUSG00000025431 | 0 | 0 | 0 | 0 | 0 | 0 | 0 |
| ENSMUSG00000025432 | 0.45815691 | 0.75 | 0.75 | 0.580482024 | 0.686820904 | 0.477121255 | 0.520678036 |
| ENSMUSG00000025433 | 0 | 0 | 0 | 0 | 0 | 0 | 0 |
| ENSMUSG00000025436 | 1.389637216 | 1.214495249 | 1.29248125 | 1.372963274 | 1.387761935 | 1.447158031 | 1.374453863 |
| ENSMUSG00000025437 | 2.340856238 | 2.178561379 | 2.195339928 | 2.344844592 | 2.273371525 | 2.57054294 | 2.103753068 |
| ENSMUSG00000025439 | 2.405576699 | 2.193696765 | 2.160061234 | 2.230460234 | 2.160196454 | 2.511883361 | 2.062969708 |
| ENSMUSG00000025450 | 0.747221736 | 0.75 | 0.896240625 | 0.79248125 | 1.106691885 | 0.698970004 | 0.803905701 |
| ENSMUSG00000025451 | 2.647796617 | 2.452241044 | 2.413659007 | 2.675326615 | 2.554871788 | 2.949877704 | 2.356085783 |
| ENSMUSG00000025453 | 2.891055323 | 2.677270953 | 2.66942991 | 2.68346568 | 2.670940998 | 3.039017322 | 2.507582874 |
| ENSMUSG00000025461 | 0 | 0 | 0 | 0 | 0 | 0 | 0 |
| ENSMUSG00000025464 | 1 | 1.17510993 | 1.04248125 | 0.951838731 | 1 | 0.84509804 | 0.829800754 |
| ENSMUSG00000025465 | 2.328498818 | 2.177701608 | 2.108657057 | 2.04248125 | 2.019019072 | 2.178976947 | 1.926805425 |
| ENSMUSG00000025466 | 1.249317394 | 1.201838731 | 1.27186571 | 0.5 | 0.733951626 | 0.77815125 | 0.629532004 |
| ENSMUSG00000025467 | 0 | 0 | 0 | 0 | 0 | 0 | 0 |

| | | | | | | | |
|---|---|---|---|---|---|---|---|
| ENSMUSG00000025468 | 0 | 0 | 0 | 0 | 0 | 0 | 0 |
| ENSMUSG00000025469 | 0 | 0 | 0 | 0 | 0 | 0.477121255 | 0.224243824 |
| ENSMUSG00000025470 | 1.527823874 | 1.606566189 | 1.557204673 | 1.557204673 | 1.652586889 | 1.755874856 | 1.629532004 |
| ENSMUSG00000025473 | 1.205378646 | 1.27186571 | 1.26109853 | 1.302363341 | 1.484072009 | 1.51851394 | 1.329931221 |
| ENSMUSG00000025474 | 2.386920383 | 2.216046536 | 2.174241882 | 2.057204673 | 2.082404901 | 2.374748346 | 1.999330888 |
| ENSMUSG00000025475 | 0.578129653 | 0.580482024 | 1.080482024 | 0.396240625 | 0.244650542 | 0 | 0.355418052 |
| ENSMUSG00000025477 | 2.100572389 | 2.031030328 | 2.025821952 | 1.882845365 | 1.855665188 | 2.064457989 | 1.791413205 |
| ENSMUSG00000025478 | 0 | 0 | 0 | 0.864857905 | 1.091002363 | 0.77815125 | 0.969165684 |
| ENSMUSG00000025479 | 0.45815691 | 0.580482024 | 0.5 | 0.5 | 0.686820904 | 0.301029996 | 0.520678036 |
| ENSMUSG00000025480 | 0 | 0 | 0 | 0 | 0 | 0 | 0 |
| ENSMUSG00000025481 | 0 | 0 | 0 | 0.25 | 0 | 0 | 0 |
| ENSMUSG00000025482 | 0 | 0 | 0 | 0 | 0 | 0 | 0 |
| ENSMUSG00000025484 | 1.890240806 | 1.789967834 | 1.777131114 | 1.67510993 | 1.761403742 | 1.851258349 | 1.647906904 |
| ENSMUSG00000025485 | 2.051791411 | 1.959800947 | 2.016522298 | 1.854463129 | 1.957204337 | 2.093421685 | 1.889537832 |
| ENSMUSG00000025486 | 1.37447073 | 1.451838731 | 1.29248125 | 1.330482024 | 1.462344774 | 1.342422681 | 1.307989563 |
| ENSMUSG00000025487 | 2.667726872 | 2.463327389 | 2.452241044 | 2.511780978 | 2.463332064 | 2.814913181 | 2.307214675 |
| ENSMUSG00000025488 | 0 | 0 | 0 | 0 | 0 | 0 | 0 |
| ENSMUSG00000025489 | 0.289064826 | 0.5 | 0.5 | 0 | 0.686820904 | 0.477121255 | 0.355418052 |
| ENSMUSG00000025491 | 0.578129653 | 0.646240625 | 0.5 | 1.098079356 | 1.039257769 | 1.255272505 | 1.185218806 |
| ENSMUSG00000025492 | 2.262379879 | 2.084962501 | 2.097004321 | 2.121960008 | 2.097547432 | 2.36361198 | 1.977154126 |
| ENSMUSG00000025494 | 0.45815691 | 0.396240625 | 0.75 | 0.5 | 0.686820904 | 0.477121255 | 0.853775828 |
| ENSMUSG00000025495 | 1.920505136 | 1.83272922 | 1.839388001 | 1.918106335 | 1.926994141 | 2.158362492 | 1.873141588 |
| ENSMUSG00000025496 | 0 | 0 | 0 | 0 | 0.244650542 | 0 | 0.224243824 |
| ENSMUSG00000025497 | 0 | 0 | 0 | 0 | 0.244650542 | 0 | 0 |
| ENSMUSG00000025498 | 0.578129653 | 0.580482024 | 0.75 | 0.92510993 | 1.057362051 | 0.602059991 | 1.121219121 |
| ENSMUSG00000025499 | 2.158622101 | 2.097004321 | 2.093759858 | 2.05479213 | 2.150762767 | 2.374748346 | 2.070582188 |
| ENSMUSG00000025500 | 0.578129653 | 0.396240625 | 0.396240625 | 0.25 | 0.77552387 | 0.698970004 | 0.896975297 |
| ENSMUSG00000025503 | 2.45930418 | 2.377938163 | 2.394828984 | 2.262462137 | 2.394615667 | 2.545307116 | 2.301737862 |
| ENSMUSG00000025504 | 0 | 0 | 0 | 0 | 0 | 0.301029996 | 0.355418052 |
| ENSMUSG00000025505 | 1.249317394 | 1.238549078 | 1.226722649 | 1.27186571 | 1.310728002 | 1.361727836 | 1.272006351 |
| ENSMUSG00000025507 | 1.596667477 | 1.729715809 | 1.630890489 | 1.26109853 | 1.533172725 | 1.51851394 | 1.526507301 |
| ENSMUSG00000025508 | 2.52293481 | 2.439139081 | 2.588562345 | 2.158248799 | 2.348168747 | 2.445604203 | 2.441733427 |
| ENSMUSG00000025509 | 1.87190676 | 1.769203899 | 1.852347734 | 1.802363341 | 1.883916752 | 2.100370545 | 1.889537832 |
| ENSMUSG00000025510 | 2.682744391 | 2.489638179 | 2.488912477 | 2.319031101 | 2.341270363 | 2.600972896 | 2.229547475 |

| | | | | | | | |
|---|---|---|---|---|---|---|---|
| ENSMUSG00000025511 | 2.40033094 | 2.24929487 | 2.229715809 | 2.083847589 | 2.151558609 | 2.320146286 | 2.062969708 |
| ENSMUSG00000025512 | 2.160971586 | 1.998588359 | 1.995748394 | 2.027131114 | 2.032639648 | 2.220108088 | 1.912234055 |
| ENSMUSG00000025515 | 0.45815691 | 0 | 0.25 | 0.75 | 0.568060967 | 0.301029996 | 0.672731473 |
| ENSMUSG00000025519 | 0 | 0 | 0 | 0 | 0 | 0 | 0 |
| ENSMUSG00000025521 | 1.978790393 | 1.872963274 | 1.958222504 | 1.825945187 | 1.888971102 | 2.152288344 | 1.779742043 |
| ENSMUSG00000025525 | 1.867194479 | 1.698603967 | 1.695339928 | 1.848079356 | 1.784632921 | 2.025305865 | 1.685349273 |
| ENSMUSG00000025527 | 0 | 0 | 0 | 0.25 | 0.387761935 | 0.301029996 | 0 |
| ENSMUSG00000025528 | 0 | 0 | 0 | 0 | 0 | 0 | 0 |
| ENSMUSG00000025529 | 0 | 0 | 0 | 0 | 0 | 0 | 0 |
| ENSMUSG00000025531 | 2.215778996 | 2.04745614 | 2.031030328 | 2.267194569 | 2.155511127 | 2.480006943 | 1.990504106 |
| ENSMUSG00000025532 | 2.365959569 | 2.215271726 | 2.222945926 | 2.225216702 | 2.21338377 | 2.45331834 | 2.066798338 |
| ENSMUSG00000025533 | 2.163307908 | 1.992885888 | 2.066696635 | 1.995748394 | 2.135290471 | 2.336459734 | 2.084776044 |
| ENSMUSG00000025534 | 2.819311906 | 2.564641508 | 2.575374049 | 2.426839783 | 2.412520257 | 2.758911892 | 2.244014063 |
| ENSMUSG00000025537 | 0.578129653 | 0.701838731 | 0.701838731 | 0.646240625 | 0.812711509 | 0.602059991 | 0.829800754 |
| ENSMUSG00000025538 | 2.185978293 | 2.061981878 | 2.121960008 | 1.819031101 | 1.88560959 | 1.986771734 | 1.86815666 |
| ENSMUSG00000025544 | 2.85804771 | 2.607351685 | 2.573942234 | 2.890441316 | 2.791183891 | 3.251881455 | 2.579438993 |
| ENSMUSG00000025545 | 1.227926429 | 1.114857905 | 1.238549078 | 1.458222504 | 1.358933174 | 1.579783597 | 1.307989563 |
| ENSMUSG00000025551 | 0 | 0 | 0 | 0 | 0 | 0 | 0 |
| ENSMUSG00000025555 | 2.826022917 | 2.623213655 | 2.612802778 | 2.702241044 | 2.722785123 | 3.054613055 | 2.561916443 |
| ENSMUSG00000025557 | 0 | 0.396240625 | 0.25 | 0.646240625 | 0.387761935 | 0.477121255 | 0.520678036 |
| ENSMUSG00000025558 | 2 | 1.970660762 | 2.00835575 | 2.174241882 | 2.173143437 | 2.44870632 | 2.03780742 |
| ENSMUSG00000025571 | 2.492331912 | 2.363831805 | 2.368933358 | 2.324229052 | 2.297835549 | 2.593286067 | 2.22557791 |
| ENSMUSG00000025572 | 1.249317394 | 1.130890489 | 1.17510993 | 1.29248125 | 1.42077253 | 1.176091259 | 1.476637173 |
| ENSMUSG00000025573 | 0 | 0 | 0.25 | 0 | 0 | 0 | 0 |
| ENSMUSG00000025574 | 2.784560084 | 2.598347614 | 2.631380202 | 2.43662858 | 2.564131005 | 2.859738566 | 2.434486378 |
| ENSMUSG00000025575 | 2.382811656 | 2.269203899 | 2.310793496 | 2.145300145 | 2.201164838 | 2.436162647 | 2.166194421 |
| ENSMUSG00000025576 | 0 | 0 | 0 | 0 | 0 | 0 | 0 |
| ENSMUSG00000025577 | 2.17479713 | 2.049918086 | 2.075945187 | 1.935366747 | 1.948833444 | 2.164352856 | 1.94074868 |
| ENSMUSG00000025578 | 1.623015126 | 1.630890489 | 1.720660762 | 1.403677461 | 1.563883923 | 1.462397998 | 1.589319772 |
| ENSMUSG00000025579 | 2.740854706 | 2.642463902 | 2.749118372 | 2.205044741 | 2.42077253 | 2.592176757 | 2.402410762 |
| ENSMUSG00000025580 | 2.815915463 | 2.580200137 | 2.606566189 | 2.586905342 | 2.612309899 | 2.945960704 | 2.493353867 |
| ENSMUSG00000025582 | 0.289064826 | 0.25 | 0 | 0.646240625 | 0.846351821 | 0.477121255 | 0.876096088 |
| ENSMUSG00000025583 | 2.578129653 | 2.386241108 | 2.399978211 | 2.275821952 | 2.343584837 | 2.57863921 | 2.224243824 |
| ENSMUSG00000025584 | 1.734388958 | 1.698603967 | 1.653677461 | 1.708222504 | 1.706995316 | 1.880813592 | 1.612273755 |

| | | | | | | | |
|---|---|---|---|---|---|---|---|
| ENSMUSG00000025586 | 1.700465191 | 1.488549078 | 1.618933358 | 1.114857905 | 1.106691885 | 1.079181246 | 1.110947927 |
| ENSMUSG00000025588 | 0 | 0 | 0 | 0.25 | 0 | 0 | 0 |
| ENSMUSG00000025591 | 2.051791411 | 1.956637122 | 1.935366747 | 1.919870025 | 1.917865457 | 2.120573931 | 1.821784031 |
| ENSMUSG00000025592 | 0.289064826 | 0 | 0.25 | 0 | 0 | 0 | 0 |
| ENSMUSG00000025597 | 0 | 0 | 0 | 0 | 0 | 0 | 0 |
| ENSMUSG00000025602 | 1.505869732 | 1.54745614 | 1.516522298 | 1.54745614 | 1.542223377 | 1.544068044 | 1.42167221 |
| ENSMUSG00000025607 | 2.339331437 | 2.181980114 | 2.150906586 | 2.224461364 | 2.090057366 | 2.460897843 | 1.978585612 |
| ENSMUSG00000025608 | 1.948720984 | 1.87892496 | 1.856566189 | 1.814346961 | 1.715300549 | 1.913813852 | 1.627424404 |
| ENSMUSG00000025609 | 2.717463581 | 2.422062577 | 2.445339928 | 2.560495787 | 2.49745007 | 2.860338007 | 2.306178596 |
| ENSMUSG00000025610 | 0 | 0 | 0 | 0.396240625 | 0.489301084 | 0 | 0 |
| ENSMUSG00000025612 | 2.333175866 | 2.171625132 | 2.161864607 | 2.05479213 | 1.990091809 | 2.28780173 | 1.896975297 |
| ENSMUSG00000025613 | 3.378409222 | 3.048919946 | 3.025001306 | 3.144062032 | 3.034664196 | 3.544316142 | 2.810194179 |
| ENSMUSG00000025616 | 2.377269616 | 2.211372513 | 2.199415381 | 2.275821952 | 2.198391101 | 2.460897843 | 2.047185336 |
| ENSMUSG00000025626 | 2.750084015 | 2.53743678 | 2.504550045 | 2.593759858 | 2.505415151 | 2.892094603 | 2.329690062 |
| ENSMUSG00000025630 | 2.719310222 | 2.517530236 | 2.516185691 | 2.60839636 | 2.507741065 | 2.935507266 | 2.334479423 |
| ENSMUSG00000025644 | 0.671187741 | 0.25 | 0.646240625 | 0.646240625 | 0.632412477 | 0.602059991 | 0.579661877 |
| ENSMUSG00000025645 | 1.655740331 | 1.62694866 | 1.62694866 | 1.552363341 | 1.572189156 | 1.672097858 | 1.586932193 |
| ENSMUSG00000025646 | 0.578129653 | 0.646240625 | 0.701838731 | 0.79248125 | 0.77552387 | 0.477121255 | 0.710836105 |
| ENSMUSG00000025647 | 2.413323654 | 2.263142013 | 2.251406137 | 2.27053726 | 2.283361516 | 2.606381365 | 2.202472158 |
| ENSMUSG00000025648 | 2.17479713 | 2.03360658 | 2.039967834 | 2.009729747 | 2.040348824 | 2.238046103 | 1.947894097 |
| ENSMUSG00000025650 | 2.612833548 | 2.481203126 | 2.545908845 | 2.313757142 | 2.526091577 | 2.572871602 | 2.478774612 |
| ENSMUSG00000025651 | 2.800026593 | 2.629417347 | 2.686838457 | 2.602612838 | 2.686122326 | 2.982723388 | 2.583867747 |
| ENSMUSG00000025652 | 0 | 0 | 0 | 0 | 0 | 0 | 0 |
| ENSMUSG00000025656 | 1.811507563 | 1.747171172 | 1.660964047 | 1.777131114 | 1.611014646 | 1.785329835 | 1.529383012 |
| ENSMUSG00000025658 | 0.578129653 | 0.396240625 | 0.701838731 | 0 | 0 | 0 | 0.224243824 |
| ENSMUSG00000025665 | 1.920505136 | 1.784887838 | 1.732684334 | 1.981203126 | 1.937354184 | 2.212187604 | 1.767634022 |
| ENSMUSG00000025666 | 2.442947872 | 2.27053726 | 2.236360959 | 1.5 | 1.439191811 | 1.612783857 | 1.324583736 |
| ENSMUSG00000025701 | 0 | 0.25 | 0 | 0.25 | 0 | 0.477121255 | 0.520678036 |
| ENSMUSG00000025702 | 2.748369006 | 2.530059469 | 2.549918086 | 2.374961472 | 2.308696595 | 2.621176282 | 2.183328003 |
| ENSMUSG00000025716 | 0 | 0 | 0 | 0 | 0 | 0 | 0 |
| ENSMUSG00000025722 | 1.876566388 | 1.711372513 | 1.726722649 | 1.732684334 | 1.736537397 | 1.934498451 | 1.667169465 |
| ENSMUSG00000025723 | 0.45815691 | 0.646240625 | 0.580482024 | 0.646240625 | 0.955822902 | 0.903089987 | 0.876096088 |
| ENSMUSG00000025724 | 2.292497211 | 2.130890489 | 2.100219859 | 2.19697564 | 2.048957937 | 2.374748346 | 1.96172822 |
| ENSMUSG00000025726 | 0 | 0 | 0.25 | 0 | 0 | 0 | 0 |

| | | | | | | | |
|---|---|---|---|---|---|---|---|
| ENSMUSG00000025727 | 0 | 0.25 | 0 | 0 | 0.387761935 | 0.301029996 | 0.448487648 |
| ENSMUSG00000025728 | 2.506893125 | 2.310197833 | 2.357613138 | 2.364857905 | 2.395811454 | 2.668385917 | 2.239582276 |
| ENSMUSG00000025730 | 2.203267085 | 2.029735268 | 2.064346961 | 1.83272922 | 2.003645018 | 1.991226076 | 1.896975297 |
| ENSMUSG00000025731 | 1.91631382 | 1.76109853 | 1.819031101 | 1.782320754 | 1.871833944 | 1.977723605 | 1.911356129 |
| ENSMUSG00000025732 | 1.389637216 | 1.348079356 | 1.339388001 | 1.214495249 | 1.373641807 | 1.146128036 | 1.441050184 |
| ENSMUSG00000025733 | 1.952600383 | 1.784887838 | 1.907339155 | 1.886723615 | 1.988833497 | 2.041392685 | 1.895131899 |
| ENSMUSG00000025735 | 0.578129653 | 0.396240625 | 0.5 | 0 | 0.387761935 | 0 | 0.448487648 |
| ENSMUSG00000025736 | 1.325351389 | 1.364857905 | 1.302363341 | 1.410964047 | 1.414412788 | 1.531478917 | 1.365075947 |
| ENSMUSG00000025737 | 2.00683675 | 1.819031101 | 1.935366747 | 1.79248125 | 1.931471446 | 1.934498451 | 1.888596011 |
| ENSMUSG00000025738 | 1.587501561 | 1.638647213 | 1.505591953 | 1.339388001 | 1.528558853 | 1.591064607 | 1.591689859 |
| ENSMUSG00000025739 | 0.289064826 | 0 | 0.25 | 0.25 | 0 | 0 | 0.448487648 |
| ENSMUSG00000025742 | 2.330063669 | 2.181980114 | 2.189555804 | 2.394828984 | 2.307165329 | 2.684845362 | 2.141264342 |
| ENSMUSG00000025743 | 2.974815471 | 2.747348625 | 2.829635702 | 2.232684334 | 2.283361516 | 2.489958479 | 2.240219117 |
| ENSMUSG00000025745 | 2.963342983 | 2.689347502 | 2.686418581 | 2.70083098 | 2.653163144 | 3.022840611 | 2.501682732 |
| ENSMUSG00000025746 | 1.156259305 | 0.896240625 | 0.646240625 | 0.25 | 0.387761935 | 0.477121255 | 0 |
| ENSMUSG00000025747 | 2.821240383 | 2.61817271 | 2.609177886 | 2.650906586 | 2.560656022 | 2.940018155 | 2.400671947 |
| ENSMUSG00000025754 | 0 | 0 | 0 | 0 | 0 | 0 | 0 |
| ENSMUSG00000025757 | 2.899911556 | 2.67140621 | 2.703845824 | 2.818303202 | 2.730114275 | 3.14332713 | 2.515150027 |
| ENSMUSG00000025758 | 2.69788693 | 2.475593779 | 2.466046536 | 2.598347614 | 2.482519186 | 2.874481818 | 2.326050929 |
| ENSMUSG00000025759 | 1.789260561 | 1.67510993 | 1.580482024 | 1.75835575 | 1.712553795 | 1.880813592 | 1.535058868 |
| ENSMUSG00000025762 | 1.912079952 | 1.839388001 | 1.80479213 | 1.823655187 | 1.766173492 | 1.897627091 | 1.723650273 |
| ENSMUSG00000025764 | 2.878667311 | 2.681341564 | 2.641037372 | 2.264497931 | 2.219338185 | 2.477121255 | 2.065163042 |
| ENSMUSG00000025766 | 1.505869732 | 1.476722649 | 1.388647213 | 1.54745614 | 1.499532413 | 1.672097858 | 1.462868481 |
| ENSMUSG00000025774 | 0 | 0 | 0 | 0 | 0 | 0 | 0 |
| ENSMUSG00000025776 | 0 | 0 | 0 | 0 | 0 | 0 | 0.224243824 |
| ENSMUSG00000025777 | 0.289064826 | 0.25 | 0.25 | 0.396240625 | 0.244650542 | 0.301029996 | 0.355418052 |
| ENSMUSG00000025779 | 1.45815691 | 1.29248125 | 1.25 | 1.188721876 | 1.057362051 | 1.361727836 | 1.041356071 |
| ENSMUSG00000025780 | 0.578129653 | 0.580482024 | 0.580482024 | 0.896240625 | 0.733951626 | 0.84509804 | 0.775756176 |
| ENSMUSG00000025781 | 3.02167286 | 2.740543258 | 2.743603647 | 2.815229883 | 2.736537397 | 3.160168293 | 2.558960864 |
| ENSMUSG00000025782 | 2.036286563 | 1.948603967 | 1.961372513 | 1.997171172 | 1.944572352 | 2.245512668 | 1.857950279 |
| ENSMUSG00000025783 | 0.289064826 | 0 | 0.396240625 | 0.25 | 0 | 0 | 0 |
| ENSMUSG00000025784 | 0 | 0.25 | 0 | 0.396240625 | 0 | 0 | 0 |
| ENSMUSG00000025785 | 2.078325387 | 1.973704441 | 1.937048212 | 1.998588359 | 1.996316946 | 2.276461804 | 1.870159858 |
| ENSMUSG00000025786 | 2.817374469 | 2.560495787 | 2.546218836 | 2.684311836 | 2.61424391 | 3.004321374 | 2.437957607 |

| | | | | | | | |
|---|---|---|---|---|---|---|---|
| ENSMUSG00000025787 | 0.289064826 | 0.5 | 0 | 0.25 | 0.387761935 | 0.301029996 | 0 |
| ENSMUSG00000025789 | 1 | 0.896240625 | 0.79248125 | 1.330482024 | 1.212046813 | 1.491361694 | 1.168187745 |
| ENSMUSG00000025790 | 2.116511873 | 2.064346961 | 2.031030328 | 1.720660762 | 1.77552387 | 1.903089987 | 1.712465909 |
| ENSMUSG00000025791 | 2.440557995 | 2.28360658 | 2.294977273 | 2.259729747 | 2.263669614 | 2.545307116 | 2.105687397 |
| ENSMUSG00000025792 | 2.256831695 | 2.166333979 | 2.259729747 | 1.990000483 | 2.235369829 | 2.281033367 | 2.233143945 |
| ENSMUSG00000025793 | 2.404271433 | 2.25 | 2.290597832 | 2.150906586 | 2.247084732 | 2.457881897 | 2.21474793 |
| ENSMUSG00000025794 | 3.173765637 | 2.929419106 | 2.919539989 | 2.978596283 | 2.933992261 | 3.376394442 | 2.737962817 |
| ENSMUSG00000025795 | 2.063793172 | 1.955044741 | 1.956637122 | 2.056000419 | 2.048957937 | 2.324282455 | 1.951019875 |
| ENSMUSG00000025804 | 0 | 0 | 0 | 0 | 0 | 0 | 0 |
| ENSMUSG00000025808 | 0 | 0 | 0 | 0 | 0 | 0 | 0 |
| ENSMUSG00000025809 | 3.793114167 | 3.416694695 | 3.375054866 | 3.388946851 | 3.265638218 | 3.832445041 | 3.022955648 |
| ENSMUSG00000025810 | 2.871053932 | 2.644828984 | 2.619439567 | 2.294977273 | 2.261347526 | 2.515873844 | 2.073258096 |
| ENSMUSG00000025812 | 2.435736758 | 2.247171172 | 2.25835575 | 2.348079356 | 2.339407793 | 2.660865478 | 2.197791274 |
| ENSMUSG00000025813 | 0.289064826 | 0 | 0.25 | 1 | 0.877063019 | 0.84509804 | 0.969165684 |
| ENSMUSG00000025815 | 1.671187741 | 1.598079356 | 1.571350555 | 1.476722649 | 1.445123986 | 1.579783597 | 1.238624656 |
| ENSMUSG00000025816 | 2.363083473 | 2.121960008 | 2.133818844 | 2.256976499 | 2.149965125 | 2.492760389 | 2.038401572 |
| ENSMUSG00000025817 | 2.513987407 | 2.342671852 | 2.28743678 | 2.372963274 | 2.31325093 | 2.599883072 | 2.164992504 |
| ENSMUSG00000025821 | 2.026699189 | 1.903677461 | 1.935366747 | 1.841580554 | 1.977302144 | 2.053078443 | 1.886704102 |
| ENSMUSG00000025823 | 2.897915226 | 2.62246199 | 2.608657057 | 2.692665973 | 2.632208868 | 3.01911629 | 2.456410872 |
| ENSMUSG00000025825 | 2.146671932 | 2.117918804 | 2.12694866 | 1.945339928 | 2 | 2.195899652 | 1.901538355 |
| ENSMUSG00000025838 | 0.289064826 | 0 | 0 | 0 | 0 | 0 | 0 |
| ENSMUSG00000025839 | 0 | 0.25 | 0 | 0 | 0 | 0 | 0 |
| ENSMUSG00000025854 | 1.952600383 | 1.819031101 | 1.914552871 | 1.527131114 | 1.628935049 | 1.633468456 | 1.631625962 |
| ENSMUSG00000025855 | 2.036286563 | 1.992885888 | 1.948603967 | 2.060793496 | 2.111176775 | 2.338456494 | 2.030591163 |
| ENSMUSG00000025856 | 2.217828299 | 2.155512955 | 2.150906586 | 1.708222504 | 1.800029087 | 1.886490725 | 1.784981274 |
| ENSMUSG00000025857 | 2.153883046 | 2.107613138 | 2.091580554 | 1.951838731 | 2.037065488 | 2.136720567 | 1.965468324 |
| ENSMUSG00000025858 | 2.56677519 | 2.434945152 | 2.477473271 | 2.295598588 | 2.359815565 | 2.605305046 | 2.319893236 |
| ENSMUSG00000025860 | 2.956791698 | 2.688304187 | 2.665667094 | 2.755765312 | 2.67130581 | 3.061452479 | 2.451647167 |
| ENSMUSG00000025862 | 3.007144864 | 2.761610487 | 2.728222334 | 2.806301859 | 2.681193183 | 3.127104798 | 2.472474658 |
| ENSMUSG00000025867 | 1.967766868 | 1.890560606 | 1.905512955 | 1.940387808 | 1.977302144 | 2.127104798 | 1.898808251 |
| ENSMUSG00000025868 | 2.078325387 | 1.976722649 | 2.070192693 | 2.039967834 | 2.153935427 | 2.324282455 | 2.162170545 |
| ENSMUSG00000025869 | 2.304292853 | 2.191217898 | 2.177701608 | 2.293731421 | 2.32898417 | 2.559906625 | 2.177974197 |
| ENSMUSG00000025870 | 1.54867984 | 1.488549078 | 1.476722649 | 1.311981878 | 1.572189156 | 1.431363764 | 1.473249546 |
| ENSMUSG00000025871 | 2.575536697 | 2.382357639 | 2.373963757 | 2.548073204 | 2.495055108 | 2.900367129 | 2.357638483 |

| | | | | | | | |
|---|---|---|---|---|---|---|---|
| ENSMUSG00000025872 | 2.447686928 | 2.254897682 | 2.243603647 | 2.503156135 | 2.424827423 | 2.766412847 | 2.282781745 |
| ENSMUSG00000025873 | 2.716227929 | 2.537119146 | 2.476722649 | 2.693490842 | 2.615740848 | 3.029789471 | 2.473713589 |
| ENSMUSG00000025875 | 1.903481042 | 1.735628626 | 1.852347734 | 1.618933358 | 1.686820904 | 1.707570176 | 1.665294008 |
| ENSMUSG00000025876 | 0 | 0 | 0 | 0 | 0 | 0 | 0.448487648 |
| ENSMUSG00000025877 | 0 | 0.25 | 0 | 0 | 0 | 0 | 0 |
| ENSMUSG00000025878 | 2.310886388 | 2.140560606 | 2.120953944 | 2.323655187 | 2.228405446 | 2.555094449 | 2.09490255 |
| ENSMUSG00000025880 | 2.153883046 | 1.959800947 | 1.975216702 | 1.692046081 | 1.709785497 | 1.806179974 | 1.612273755 |
| ENSMUSG00000025885 | 0.671187741 | 0.79248125 | 0.580482024 | 0.646240625 | 0.812711509 | 0.602059991 | 0.710836105 |
| ENSMUSG00000025887 | 2.151493169 | 2.012462137 | 1.984159485 | 2.213717096 | 2.101233692 | 2.501059262 | 1.974994949 |
| ENSMUSG00000025888 | 0 | 0 | 0 | 0 | 0 | 0.301029996 | 0 |
| ENSMUSG00000025889 | 1.432083896 | 1.311981878 | 1.321350555 | 0.25 | 0.387761935 | 0.477121255 | 0.520678036 |
| ENSMUSG00000025892 | 0 | 0.25 | 0.396240625 | 0 | 0 | 0.477121255 | 0.224243824 |
| ENSMUSG00000025893 | 1.129344651 | 1.061981878 | 1.201838731 | 1.114857905 | 1 | 1.342422681 | 1.110947927 |
| ENSMUSG00000025894 | 2.223916471 | 2.075945187 | 2.046218836 | 2.12694866 | 2.098472613 | 2.346352974 | 1.95718226 |
| ENSMUSG00000025898 | 2.141794304 | 1.959800947 | 1.976722649 | 2.201838731 | 2.089109832 | 2.33243846 | 1.930969131 |
| ENSMUSG00000025899 | 2.192545868 | 2.077084758 | 2.046218836 | 2.222945926 | 2.06759338 | 2.403120521 | 1.935079929 |
| ENSMUSG00000025900 | 0 | 0 | 0 | 0 | 0 | 0 | 0 |
| ENSMUSG00000025902 | 0 | 0 | 0 | 0 | 0 | 0 | 0 |
| ENSMUSG00000025903 | 2.494443473 | 2.30357978 | 2.288071211 | 2.525165585 | 2.429198453 | 2.821513528 | 2.230532302 |
| ENSMUSG00000025905 | 0 | 0 | 0 | 0 | 0 | 0 | 0 |
| ENSMUSG00000025907 | 2.937589884 | 2.705642708 | 2.676624005 | 2.589388001 | 2.500421099 | 2.828659897 | 2.289133511 |
| ENSMUSG00000025909 | 0 | 0 | 0 | 0 | 0 | 0 | 0 |
| ENSMUSG00000025911 | 0.671187741 | 0.5 | 0.646240625 | 0.396240625 | 0.812711509 | 0.698970004 | 0.710836105 |
| ENSMUSG00000025912 | 1.99653913 | 1.96293726 | 1.866901388 | 2.04248125 | 2.017859938 | 2.11058971 | 1.788855757 |
| ENSMUSG00000025915 | 1.794934559 | 1.720660762 | 1.701838731 | 1.79745614 | 1.621875691 | 1.934498451 | 1.594042709 |
| ENSMUSG00000025916 | 0 | 0 | 0 | 0 | 0 | 0.602059991 | 0.224243824 |
| ENSMUSG00000025917 | 2.682744391 | 2.444519282 | 2.463327389 | 2.627938163 | 2.529104803 | 2.944975908 | 2.354078442 |
| ENSMUSG00000025920 | 2.201144779 | 2.03360658 | 2.027131114 | 2.156427211 | 2.085294053 | 2.36361198 | 1.955652633 |
| ENSMUSG00000025921 | 1.800532393 | 1.732684334 | 1.642463902 | 1.839388001 | 1.795698282 | 2 | 1.651852264 |
| ENSMUSG00000025922 | 0.289064826 | 0.25 | 0.25 | 0 | 0.489301084 | 0 | 0.355418052 |
| ENSMUSG00000025925 | 2.10861828 | 1.943696765 | 1.961372513 | 2.027131114 | 1.995080671 | 2.227886705 | 1.885753983 |
| ENSMUSG00000025927 | 0.289064826 | 0.25 | 0 | 0 | 0 | 0 | 0 |
| ENSMUSG00000025929 | 0 | 0 | 0 | 0 | 0 | 0 | 0 |
| ENSMUSG00000025930 | 0 | 0.25 | 0 | 0.25 | 0 | 0 | 0.448487648 |

| | | | | | | | |
|---|---|---|---|---|---|---|---|
| ENSMUSG00000025931 | 1.89911417 | 1.784887838 | 1.83272922 | 1.571350555 | 1.652586889 | 1.826074803 | 1.582103514 |
| ENSMUSG00000025932 | 1.771760131 | 1.79745614 | 1.688721876 | 1.711372513 | 1.718026092 | 1.939519253 | 1.61448206 |
| ENSMUSG00000025933 | 1.227926429 | 1.26109853 | 1.201838731 | 1.431980114 | 1.55104774 | 1.672097858 | 1.340368108 |
| ENSMUSG00000025934 | 0.289064826 | 0.396240625 | 0.25 | 0.25 | 0 | 0 | 0 |
| ENSMUSG00000025935 | 2.786132803 | 2.550530956 | 2.528760913 | 2.67140621 | 2.5845249 | 2.963315511 | 2.398533958 |
| ENSMUSG00000025936 | 0 | 0 | 0 | 0 | 0 | 0 | 0 |
| ENSMUSG00000025937 | 2.23778581 | 2.123963757 | 2.127938163 | 2.116901388 | 2.070605826 | 2.318063335 | 1.966952343 |
| ENSMUSG00000025938 | 0.45815691 | 0.580482024 | 0.896240625 | 1.061981878 | 1.212046813 | 1.079181246 | 0.935079929 |
| ENSMUSG00000025939 | 2.355805236 | 2.182829758 | 2.168106335 | 2.219895812 | 2.148364414 | 2.463892989 | 1.988432495 |
| ENSMUSG00000025940 | 2.23778581 | 2.036164561 | 2.039967834 | 2.108657057 | 1.998776581 | 2.356025857 | 1.837623616 |
| ENSMUSG00000025945 | 0 | 0 | 0 | 0 | 0 | 0 | 0 |
| ENSMUSG00000025946 | 0.289064826 | 0.25 | 0.25 | 0 | 0.244650542 | 0.301029996 | 0.224243824 |
| ENSMUSG00000025949 | 2.477243823 | 2.211372513 | 2.273189285 | 2.353406982 | 2.27729019 | 2.588831726 | 2.089878917 |
| ENSMUSG00000025950 | 2.528794847 | 2.310197833 | 2.310793496 | 2.365369862 | 2.271112598 | 2.608526034 | 2.135599953 |
| ENSMUSG00000025952 | 0 | 0 | 0 | 0 | 0 | 0 | 0 |
| ENSMUSG00000025955 | 0 | 0 | 0 | 0.5 | 0 | 0 | 0.224243824 |
| ENSMUSG00000025956 | 2.136858952 | 1.945339928 | 1.96759118 | 1.856566189 | 1.822942838 | 2.068185862 | 1.74196736 |
| ENSMUSG00000025958 | 2.624559692 | 2.418989258 | 2.406427211 | 2.514497931 | 2.423358301 | 2.788168371 | 2.228559641 |
| ENSMUSG00000025959 | 2.233869984 | 2.025821952 | 2.080482024 | 1.668106335 | 1.72341484 | 1.785329835 | 1.605556929 |
| ENSMUSG00000025961 | 0.289064826 | 0.396240625 | 0.25 | 0.5 | 0.489301084 | 0.477121255 | 0 |
| ENSMUSG00000025962 | 2.266038081 | 2.115881093 | 2.097004321 | 2.120953944 | 2.068600388 | 2.399673721 | 1.968429585 |
| ENSMUSG00000025963 | 0.289064826 | 0 | 0.25 | 0 | 0 | 0 | 0 |
| ENSMUSG00000025964 | 0.289064826 | 0.646240625 | 0.701838731 | 0.646240625 | 0.568060967 | 0.477121255 | 0.520678036 |
| ENSMUSG00000025967 | 3.278755457 | 2.998941788 | 3.018368038 | 3.021285115 | 2.974224243 | 3.435366507 | 2.770364127 |
| ENSMUSG00000025968 | 2.759671389 | 2.541540771 | 2.508699741 | 2.660061234 | 2.570979987 | 3.002166062 | 2.39814371 |
| ENSMUSG00000025969 | 3.225196554 | 2.92845402 | 2.931448068 | 2.732314602 | 2.664494972 | 3.072249898 | 2.534290726 |
| ENSMUSG00000025971 | 1.87190676 | 1.774508021 | 1.807204673 | 1.992885888 | 1.913211023 | 2.130333768 | 1.792684387 |
| ENSMUSG00000025977 | 0 | 0 | 0.396240625 | 0.25 | 0 | 0 | 0.355418052 |
| ENSMUSG00000025978 | 1.647796617 | 1.606566189 | 1.649978211 | 1.5 | 1.489301084 | 1.643452676 | 1.483307832 |
| ENSMUSG00000025979 | 2.472830741 | 2.352877747 | 2.332168732 | 2.424241882 | 2.379941013 | 2.72427587 | 2.162170545 |
| ENSMUSG00000025980 | 3.39997086 | 3.084196369 | 3.072648722 | 3.195339928 | 3.094592434 | 3.614264287 | 2.853058869 |
| ENSMUSG00000025981 | 2.328498818 | 2.133818844 | 2.160061234 | 2.296218836 | 2.240349892 | 2.522444234 | 2.092400484 |
| ENSMUSG00000025982 | 3.365307651 | 3.055774173 | 3.028516912 | 3.115369862 | 3.020716796 | 3.519696767 | 2.786630664 |
| ENSMUSG00000025983 | 0.91631382 | 0.92510993 | 0.75 | 0.79248125 | 1 | 0.698970004 | 0.672731473 |

| | | | | | | | |
|---|---|---|---|---|---|---|---|
| ENSMUSG00000025986 | 2.406877894 | 2.199415381 | 2.198603967 | 2.232684334 | 2.176121988 | 2.525044807 | 2.026306127 |
| ENSMUSG00000025991 | 0 | 0 | 0 | 0.25 | 0 | 0 | 0.224243824 |
| ENSMUSG00000025993 | 1.857607105 | 1.671625132 | 1.557204673 | 1.130890489 | 1 | 0.77815125 | 0.853775828 |
| ENSMUSG00000025995 | 2.723587515 | 2.486360959 | 2.461764337 | 2.476346751 | 2.420401583 | 2.769377326 | 2.230204359 |
| ENSMUSG00000025997 | 2.063793172 | 1.937048212 | 1.935366747 | 1.940387808 | 1.842556557 | 2.089905111 | 1.723650273 |
| ENSMUSG00000026000 | 2.245508853 | 2.063166358 | 1.987091808 | 1.976722649 | 1.932951348 | 2.198657087 | 1.858985529 |
| ENSMUSG00000026003 | 2.68005385 | 2.447383411 | 2.448603967 | 2.428561379 | 2.365931294 | 2.713490543 | 2.198877505 |
| ENSMUSG00000026004 | 2.199011616 | 2.049918086 | 2.070192693 | 1.799918086 | 1.773209395 | 1.968482949 | 1.663407615 |
| ENSMUSG00000026005 | 2.348397716 | 2.165444524 | 2.12892496 | 2.23927551 | 2.173143437 | 2.496929648 | 2.049487947 |
| ENSMUSG00000026009 | 0.289064826 | 0.25 | 0.396240625 | 0.396240625 | 0 | 0.301029996 | 0.224243824 |
| ENSMUSG00000026011 | 0 | 0 | 0 | 0 | 0 | 0.301029996 | 0 |
| ENSMUSG00000026012 | 0.45815691 | 0.25 | 0.25 | 0.25 | 0 | 0.477121255 | 0.355418052 |
| ENSMUSG00000026014 | 2.450036412 | 2.240724001 | 2.308404919 | 2.363317659 | 2.298360391 | 2.64738297 | 2.148525059 |
| ENSMUSG00000026017 | 1.707474304 | 1.488549078 | 1.575945187 | 1.729715809 | 1.706995316 | 1.838849091 | 1.551512352 |
| ENSMUSG00000026018 | 0.960252568 | 1.061981878 | 1 | 1.396240625 | 1.450958105 | 1.462397998 | 1.340368108 |
| ENSMUSG00000026019 | 2.444137694 | 2.250703754 | 2.254897682 | 2.337734795 | 2.259595879 | 2.552668216 | 2.086826862 |
| ENSMUSG00000026020 | 3.018987133 | 2.76024365 | 2.768702615 | 2.88015467 | 2.77983978 | 3.237040791 | 2.587149975 |
| ENSMUSG00000026021 | 2.794421919 | 2.559601185 | 2.56168515 | 2.699415381 | 2.628111827 | 3.029789471 | 2.454284088 |
| ENSMUSG00000026023 | 0.578129653 | 0.396240625 | 0.396240625 | 0.25 | 0 | 0 | 0 |
| ENSMUSG00000026024 | 2.699822119 | 2.493961992 | 2.512462137 | 2.444108258 | 2.400946889 | 2.701567985 | 2.254530788 |
| ENSMUSG00000026027 | 1.822201287 | 1.763820609 | 1.75 | 1.823655187 | 1.797870327 | 1.982271233 | 1.714087544 |
| ENSMUSG00000026028 | 2.718080037 | 2.544666213 | 2.492526473 | 2.661639678 | 2.596669346 | 2.949390007 | 2.419120525 |
| ENSMUSG00000026029 | 2.336265009 | 2.186208459 | 2.186208459 | 2.293731421 | 2.210031835 | 2.547774705 | 2.096394553 |
| ENSMUSG00000026031 | 2.420929319 | 2.253505118 | 2.250703754 | 2.281676118 | 2.179810173 | 2.572871602 | 2.062419043 |
| ENSMUSG00000026032 | 2.153883046 | 2.064346961 | 2.03743678 | 2.149047439 | 2.141887974 | 2.411619706 | 1.994607951 |
| ENSMUSG00000026034 | 2.506893125 | 2.320772088 | 2.26584877 | 2.420309603 | 2.367228134 | 2.657055853 | 2.210242554 |
| ENSMUSG00000026035 | 2.217828299 | 2.048689214 | 1.997171172 | 2.097004321 | 2.072599934 | 2.315970345 | 1.939139147 |
| ENSMUSG00000026036 | 2.188178997 | 2.067865757 | 2.052363341 | 2.23927551 | 2.175379703 | 2.480006943 | 2.017561934 |
| ENSMUSG00000026037 | 2.545896521 | 2.350219859 | 2.311388177 | 2.476346751 | 2.371516839 | 2.742725131 | 2.211982824 |
| ENSMUSG00000026039 | 2.063793172 | 2.04248125 | 2.03360658 | 2.168989258 | 2.083370582 | 2.378397901 | 1.963977467 |
| ENSMUSG00000026042 | 3.258689303 | 2.998765117 | 2.995659283 | 3.115817229 | 3.151277928 | 3.57205799 | 2.954535617 |
| ENSMUSG00000026043 | 3.024341402 | 2.80357978 | 2.70584181 | 1.557204673 | 1.614672255 | 1.748188027 | 1.523605797 |
| ENSMUSG00000026047 | 2.370236862 | 2.140560606 | 2.112802778 | 2.244319981 | 2.173143437 | 2.387389826 | 2.019455688 |
| ENSMUSG00000026048 | 2.192545868 | 2.064346961 | 2.049918086 | 2.117918804 | 2.065570703 | 2.356025857 | 1.882886767 |

| | | | | | | | |
|---|---|---|---|---|---|---|---|
| ENSMUSG00000026049 | 2.262379879 | 2.069031101 | 2.071350555 | 2.176839783 | 2.126866297 | 2.390935107 | 1.945529784 |
| ENSMUSG00000026051 | 0 | 0.25 | 0 | 0.5 | 0.387761935 | 0 | 0 |
| ENSMUSG00000026058 | 0 | 0.5 | 0.5 | 0.25 | 0.387761935 | 0 | 0.355418052 |
| ENSMUSG00000026062 | 0.289064826 | 0.25 | 0.396240625 | 0.92510993 | 0.905314583 | 0.954242509 | 0.74492186 |
| ENSMUSG00000026063 | 0 | 0 | 0 | 0 | 0 | 0 | 0 |
| ENSMUSG00000026064 | 2.957832978 | 2.711960089 | 2.686418581 | 2.624961472 | 2.561403546 | 2.935003151 | 2.347752519 |
| ENSMUSG00000026065 | 0 | 0 | 0 | 0 | 0 | 0 | 0 |
| ENSMUSG00000026068 | 1.249317394 | 1.29248125 | 1.188721876 | 1.061981878 | 0.955822902 | 0.903089987 | 0.896975297 |
| ENSMUSG00000026069 | 1.527823874 | 1.458222504 | 1.321350555 | 0.92510993 | 0.812711509 | 0.477121255 | 0.355418052 |
| ENSMUSG00000026070 | 0 | 0 | 0.25 | 0.25 | 0.568060967 | 0.301029996 | 0 |
| ENSMUSG00000026072 | 2.294202867 | 2.116901388 | 2.121960008 | 2.144357207 | 2.087207099 | 2.324282455 | 1.929310074 |
| ENSMUSG00000026073 | 0.289064826 | 0 | 0.396240625 | 0 | 0.387761935 | 0.477121255 | 0.224243824 |
| ENSMUSG00000026074 | 3.154964843 | 2.87007133 | 2.872838018 | 2.925326615 | 2.874406897 | 3.282622113 | 2.709794665 |
| ENSMUSG00000026077 | 0.45815691 | 0.396240625 | 0 | 0.646240625 | 0.77552387 | 0.301029996 | 0.629532004 |
| ENSMUSG00000026078 | 2.706841958 | 2.518869787 | 2.472566069 | 2.502107156 | 2.456026973 | 2.830588669 | 2.288859925 |
| ENSMUSG00000026080 | 1.83262764 | 1.630890489 | 1.678561379 | 1.823655187 | 1.789100779 | 1.973127854 | 1.700880998 |
| ENSMUSG00000026082 | 2.269664473 | 2.115881093 | 2.099151195 | 2.084962501 | 2.020174412 | 2.326335861 | 1.86207155 |
| ENSMUSG00000026083 | 2.965050035 | 2.733422664 | 2.738549078 | 2.863960227 | 2.766591274 | 3.18582536 | 2.56238684 |
| ENSMUSG00000026085 | 0 | 0 | 0 | 0 | 0 | 0 | 0 |
| ENSMUSG00000026087 | 2.451206209 | 2.200224975 | 2.197790722 | 2.299304173 | 2.234113756 | 2.57054294 | 2.135599953 |
| ENSMUSG00000026088 | 1.89911417 | 1.75835575 | 1.799918086 | 2.05357978 | 1.934425071 | 2.214843848 | 1.892346957 |
| ENSMUSG00000026090 | 0.578129653 | 0.396240625 | 0.25 | 0.75 | 0.846351821 | 0.698970004 | 0.803905701 |
| ENSMUSG00000026094 | 2.655740331 | 2.394828984 | 2.395300145 | 2.324229052 | 2.227124297 | 2.568201724 | 2.073790632 |
| ENSMUSG00000026095 | 1.867194479 | 1.660964047 | 1.685366747 | 1.794977273 | 1.715300549 | 1.826074803 | 1.520678036 |
| ENSMUSG00000026096 | 1.714367558 | 1.505591953 | 1.580482024 | 1.726722649 | 1.576269619 | 1.770852012 | 1.526507301 |
| ENSMUSG00000026097 | 2.105953527 | 2.02186571 | 1.991446071 | 2.08272922 | 2.033751322 | 2.336459734 | 1.896054911 |
| ENSMUSG00000026098 | 1.967766868 | 1.870953944 | 1.828220739 | 1.928561379 | 1.862942891 | 2.130333768 | 1.732938436 |
| ENSMUSG00000026100 | 0 | 0 | 0 | 0 | 0 | 0 | 0 |
| ENSMUSG00000026102 | 1.940852214 | 1.802363341 | 1.777131114 | 2 | 1.928492895 | 2.204119983 | 1.812372009 |
| ENSMUSG00000026103 | 3.070726095 | 2.802667836 | 2.790440436 | 2.867918804 | 2.778900174 | 3.173186268 | 2.535058868 |
| ENSMUSG00000026104 | 1.852729478 | 1.76109853 | 1.630890489 | 1.837182039 | 1.770879644 | 2.045322979 | 1.685349273 |
| ENSMUSG00000026107 | 2.231898196 | 2.070192693 | 2.084962501 | 1.943696765 | 1.875328588 | 2.152288344 | 1.773071385 |
| ENSMUSG00000026109 | 2.213719572 | 2.001406137 | 1.998588359 | 1.75 | 1.674752852 | 1.892094603 | 1.572224412 |
| ENSMUSG00000026110 | 1.181541736 | 1.201838731 | 1.114857905 | 1.799918086 | 1.758994476 | 1.86332286 | 1.627424404 |

| | | | | | | | |
|---|---|---|---|---|---|---|---|
| ENSMUSG00000026111 | 2.100572389 | 1.943696765 | 1.931980114 | 2.016522298 | 2.001219193 | 2.220108088 | 1.86715039 |
| ENSMUSG00000026112 | 2.590857977 | 2.414998973 | 2.402756199 | 2.501055117 | 2.388981128 | 2.774516966 | 2.246831342 |
| ENSMUSG00000026113 | 1.989530018 | 1.854463129 | 1.894357207 | 1.918106335 | 1.97205363 | 2.123851641 | 1.861046144 |
| ENSMUSG00000026114 | 0.289064826 | 0 | 0.396240625 | 0.396240625 | 0 | 0 | 0 |
| ENSMUSG00000026116 | 2.685417684 | 2.47027849 | 2.484894804 | 2.583009137 | 2.535055162 | 2.930949031 | 2.39183486 |
| ENSMUSG00000026117 | 0 | 0 | 0 | 0.25 | 0 | 0 | 0.355418052 |
| ENSMUSG00000026121 | 1.982401085 | 1.903677461 | 1.92510993 | 1.705044741 | 1.824953989 | 1.982271233 | 1.770364127 |
| ENSMUSG00000026123 | 2.726616086 | 2.501055117 | 2.47104263 | 2.342671852 | 2.346799814 | 2.658011397 | 2.216121671 |
| ENSMUSG00000026124 | 0 | 0 | 0 | 0 | 0 | 0 | 0 |
| ENSMUSG00000026125 | 0 | 0 | 0 | 0 | 0 | 0 | 0 |
| ENSMUSG00000026126 | 0.289064826 | 0.5 | 0.701838731 | 0.701838731 | 0.877063019 | 0.77815125 | 0.952571511 |
| ENSMUSG00000026127 | 2.446507203 | 2.261780978 | 2.266522298 | 2.281030328 | 2.286629652 | 2.537819095 | 2.164992504 |
| ENSMUSG00000026131 | 2.126809492 | 1.951838731 | 1.964495249 | 2.074802005 | 2.043601899 | 2.158362492 | 1.928477344 |
| ENSMUSG00000026134 | 2.50175086 | 2.28743678 | 2.301142786 | 2.474461364 | 2.41591954 | 2.767897616 | 2.272006351 |
| ENSMUSG00000026135 | 2.386920383 | 2.221424093 | 2.222945926 | 2.180274797 | 2.196996016 | 2.408239965 | 2.073790632 |
| ENSMUSG00000026141 | 0 | 0 | 0.25 | 0.25 | 0.244650542 | 0.301029996 | 0 |
| ENSMUSG00000026142 | 2.22592627 | 2.017865591 | 1.995748394 | 2.132845365 | 2.131103516 | 2.401400541 | 2.007270316 |
| ENSMUSG00000026147 | 0 | 0 | 0 | 0 | 0 | 0 | 0 |
| ENSMUSG00000026149 | 0 | 0.25 | 0.396240625 | 0.25 | 0 | 0.477121255 | 0.224243824 |
| ENSMUSG00000026150 | 2.625329835 | 2.429419106 | 2.411414608 | 2.472945926 | 2.417419887 | 2.807535028 | 2.237984665 |
| ENSMUSG00000026153 | 2.312518611 | 2.104463129 | 2.024508021 | 2.336628433 | 2.276175004 | 2.562292864 | 2.074853081 |
| ENSMUSG00000026154 | 1.87190676 | 1.741446071 | 1.732684334 | 1.723704441 | 1.655839964 | 1.919078092 | 1.582103514 |
| ENSMUSG00000026155 | 1.967766868 | 1.845926073 | 1.814346961 | 1.870953944 | 1.868304353 | 2.08278537 | 1.773071385 |
| ENSMUSG00000026156 | 0.960252568 | 0.79248125 | 0.701838731 | 0.951838731 | 1.020174412 | 0.698970004 | 0.775756176 |
| ENSMUSG00000026158 | 1.342375483 | 1.302363341 | 1.080482024 | 1.610735874 | 1.414412788 | 1.681241237 | 1.441050184 |
| ENSMUSG00000026159 | 2.809533441 | 2.573080408 | 2.533927322 | 2.632113528 | 2.571464152 | 2.945960704 | 2.375083475 |
| ENSMUSG00000026162 | 1.227926429 | 1.311981878 | 1.27186571 | 1.339388001 | 1.456697355 | 1.62324929 | 1.329931221 |
| ENSMUSG00000026163 | 0 | 0.25 | 0 | 0 | 0 | 0 | 0 |
| ENSMUSG00000026166 | 0 | 0.5 | 0.25 | 0.701838731 | 0.489301084 | 0.698970004 | 0.853775828 |
| ENSMUSG00000026167 | 1.693336259 | 1.634789703 | 1.685366747 | 1.431980114 | 1.380772476 | 1.612783857 | 1.579661877 |
| ENSMUSG00000026170 | 0.578129653 | 0.79248125 | 0.864857905 | 1.25 | 1.188507684 | 1.204119983 | 1.176815335 |
| ENSMUSG00000026171 | 2.00683675 | 1.901832578 | 1.958222504 | 1.96759118 | 2.041436517 | 2.250420002 | 1.928477344 |
| ENSMUSG00000026172 | 1.876566388 | 1.755591953 | 1.708222504 | 1.755591953 | 1.701347897 | 1.880813592 | 1.589319772 |
| ENSMUSG00000026173 | 0.91631382 | 0.830482024 | 0.864857905 | 0.830482024 | 0.812711509 | 0.602059991 | 0.579661877 |

| | | | | | | | |
|---|---|---|---|---|---|---|---|
| ENSMUSG00000026174 | 2.787177998 | 2.556000419 | 2.537754135 | 2.647646762 | 2.554618136 | 2.959994838 | 2.398533958 |
| ENSMUSG00000026175 | 0 | 0.5 | 0.396240625 | 0.396240625 | 0.387761935 | 0.477121255 | 0 |
| ENSMUSG00000026176 | 2.660010325 | 2.514159429 | 2.558704358 | 2.340485943 | 2.434230499 | 2.656098202 | 2.349799373 |
| ENSMUSG00000026177 | 0.91631382 | 1.04248125 | 1.04248125 | 0.396240625 | 0.568060967 | 0.477121255 | 0.448487648 |
| ENSMUSG00000026179 | 2.196867486 | 2.059601185 | 2.155512955 | 2.155512955 | 2.198391101 | 2.413299764 | 2.165794278 |
| ENSMUSG00000026180 | 0.671187741 | 0.701838731 | 0.701838731 | 0.75 | 0.387761935 | 0.477121255 | 0.672731473 |
| ENSMUSG00000026181 | 2.676666078 | 2.403217124 | 2.432405186 | 2.43325383 | 2.422990063 | 2.746634199 | 2.269981661 |
| ENSMUSG00000026182 | 0 | 0 | 0 | 0 | 0 | 0 | 0 |
| ENSMUSG00000026185 | 0.289064826 | 0 | 0 | 0.396240625 | 0.244650542 | 0 | 0 |
| ENSMUSG00000026187 | 2.439357901 | 2.25 | 2.231944491 | 2.225970461 | 2.094757251 | 2.401400541 | 1.962479708 |
| ENSMUSG00000026188 | 1 | 0.396240625 | 0.5 | 1.201838731 | 1.091002363 | 1.230448921 | 0.969165684 |
| ENSMUSG00000026189 | 1.358731791 | 1.482684334 | 1.311981878 | 1.527131114 | 1.542223377 | 1.612783857 | 1.466365979 |
| ENSMUSG00000026192 | 2.89911417 | 2.649047439 | 2.632601584 | 2.702039943 | 2.685597484 | 3.094471129 | 2.520276735 |
| ENSMUSG00000026193 | 4.363429177 | 3.932862906 | 3.918920358 | 3.452617805 | 3.434062514 | 3.983400738 | 3.207150349 |
| ENSMUSG00000026196 | 2.083999385 | 1.931980114 | 1.953445298 | 2.03743678 | 1.98503142 | 2.209515015 | 1.809975587 |
| ENSMUSG00000026197 | 1.53838222 | 1.671625132 | 1.668106335 | 1.610735874 | 1.706995316 | 1.819543936 | 1.67819947 |
| ENSMUSG00000026198 | 1.100572389 | 1.02186571 | 1.098079356 | 1 | 1.223252711 | 1.278753601 | 1.028149525 |
| ENSMUSG00000026199 | 1.822201287 | 1.642463902 | 1.642463902 | 1.494319981 | 1.689774529 | 1.69019608 | 1.496250175 |
| ENSMUSG00000026200 | 1.325351389 | 1.302363341 | 1.282320754 | 1.516522298 | 1.580303447 | 1.643452676 | 1.413583902 |
| ENSMUSG00000026201 | 2.273259602 | 2.084962501 | 2.098079356 | 2.220660762 | 2.225193812 | 2.51054501 | 2.099358074 |
| ENSMUSG00000026202 | 2.273259602 | 2.102347734 | 2.127938163 | 2.036164561 | 2.038163331 | 2.322219295 | 1.959463225 |
| ENSMUSG00000026203 | 2.057835465 | 2.005591953 | 2.001406137 | 1.898114259 | 1.951645858 | 1.968482949 | 1.89047692 |
| ENSMUSG00000026204 | 1.671187741 | 1.71759118 | 1.766522298 | 1.080482024 | 1.264824954 | 1 | 1.340368108 |
| ENSMUSG00000026205 | 0.45815691 | 0.25 | 0.396240625 | 0.396240625 | 0.489301084 | 0.301029996 | 0.355418052 |
| ENSMUSG00000026207 | 2.42468076 | 2.31611065 | 2.399047439 | 1.928561379 | 2.014359518 | 2.045322979 | 2.050061048 |
| ENSMUSG00000026208 | 1.181541736 | 1.339388001 | 1.356566189 | 0.976722649 | 0.905314583 | 0.954242509 | 0.935079929 |
| ENSMUSG00000026209 | 2.395018353 | 2.266522298 | 2.274508021 | 2.061981878 | 2.195595395 | 2.372912003 | 2.139531992 |
| ENSMUSG00000026211 | 2.243591459 | 2.117918804 | 2.181128463 | 2.174241882 | 2.297835549 | 2.45331834 | 2.249315158 |
| ENSMUSG00000026213 | 2.045658471 | 1.972185812 | 2.060793496 | 1.886723615 | 1.986303335 | 2.133538908 | 1.941550454 |
| ENSMUSG00000026219 | 3.311480663 | 2.999911934 | 2.976910451 | 3.127443751 | 3.028223932 | 3.503790683 | 2.799418591 |
| ENSMUSG00000026220 | 0 | 0 | 0 | 0 | 0 | 0 | 0.224243824 |
| ENSMUSG00000026222 | 0.578129653 | 0.646240625 | 0 | 0.580482024 | 0.733951626 | 1.041392685 | 0.775756176 |
| ENSMUSG00000026223 | 2.476144923 | 2.308404919 | 2.336073977 | 2.352877747 | 2.389386592 | 2.729974286 | 2.318897036 |
| ENSMUSG00000026224 | 0 | 0 | 0 | 0 | 0.387761935 | 0 | 0 |

| | | | | | | | |
|---|---|---|---|---|---|---|---|
| ENSMUSG00000026226 | 0 | 0 | 0 | 0 | 0 | 0 | 0 |
| ENSMUSG00000026227 | 0 | 0 | 0 | 0 | 0.244650542 | 0 | 0 |
| ENSMUSG00000026228 | 0 | 0.5 | 0.396240625 | 0.580482024 | 0.387761935 | 0.698970004 | 0.520678036 |
| ENSMUSG00000026229 | 3.227200208 | 2.929204865 | 2.897529794 | 3.068667336 | 2.960995697 | 3.440121603 | 2.748114472 |
| ENSMUSG00000026234 | 3.736537507 | 3.391335032 | 3.389186382 | 3.54753333 | 3.431046228 | 4.009918085 | 3.177921612 |
| ENSMUSG00000026235 | 2.310886388 | 2.130890489 | 2.066696635 | 1.789967834 | 1.665423072 | 1.838849091 | 1.586932193 |
| ENSMUSG00000026237 | 0 | 0 | 0 | 0 | 0.244650542 | 0 | 0 |
| ENSMUSG00000026238 | 3.364654711 | 3.075374049 | 3.117918804 | 3.080482024 | 2.955660024 | 3.476106717 | 2.842257801 |
| ENSMUSG00000026239 | 2.342375483 | 2.169870025 | 2.12892496 | 2.242885888 | 2.194892994 | 2.489958479 | 2.081675233 |
| ENSMUSG00000026240 | 2.318984359 | 2.107613138 | 2.121960008 | 2.233422664 | 2.19206933 | 2.482873584 | 2.055173778 |
| ENSMUSG00000026241 | 0 | 0 | 0 | 0 | 0 | 0 | 0 |
| ENSMUSG00000026245 | 2.485932312 | 2.281030328 | 2.251406137 | 2.459406983 | 2.378690133 | 2.73479983 | 2.227237819 |
| ENSMUSG00000026246 | 0.289064826 | 0.396240625 | 0.396240625 | 0 | 0 | 0 | 0 |
| ENSMUSG00000026247 | 0 | 0 | 0.25 | 0.580482024 | 0.489301084 | 0.301029996 | 0.520678036 |
| ENSMUSG00000026248 | 2.00683675 | 1.890560606 | 1.856566189 | 1.901832578 | 1.944572352 | 2.161368002 | 1.82753046 |
| ENSMUSG00000026249 | 1.971473847 | 1.830482024 | 1.784887838 | 1.04248125 | 0.877063019 | 1 | 0.952571511 |
| ENSMUSG00000026251 | 0 | 0 | 0 | 0 | 0 | 0 | 0 |
| ENSMUSG00000026253 | 0 | 0 | 0 | 0 | 0 | 0 | 0 |
| ENSMUSG00000026254 | 2.670497861 | 2.444108258 | 2.439555804 | 2.602082435 | 2.511201447 | 2.926342447 | 2.354078442 |
| ENSMUSG00000026255 | 0 | 0.396240625 | 0.5 | 0.25 | 0.632412477 | 0.84509804 | 0.629532004 |
| ENSMUSG00000026258 | 0 | 0 | 0.25 | 0 | 0 | 0 | 0 |
| ENSMUSG00000026259 | 1.181541736 | 1.160964047 | 1.311981878 | 1.238549078 | 1.407936347 | 1.380211242 | 1.355418052 |
| ENSMUSG00000026260 | 2.671876482 | 2.452642909 | 2.451435968 | 2.505591953 | 2.457032069 | 2.844477176 | 2.335191751 |
| ENSMUSG00000026269 | 2.153883046 | 2.025821952 | 2.088286706 | 1.910964047 | 2.046825265 | 2.146128036 | 2.082194103 |
| ENSMUSG00000026270 | 1.978790393 | 1.874961472 | 1.950224975 | 1.789967834 | 1.948833444 | 2.033423755 | 1.853775828 |
| ENSMUSG00000026271 | 0.671187741 | 0.580482024 | 0.25 | 0.396240625 | 0.244650542 | 0.301029996 | 0.355418052 |
| ENSMUSG00000026272 | 0 | 0 | 0 | 0 | 0 | 0 | 0 |
| ENSMUSG00000026273 | 1.765759561 | 1.688721876 | 1.705044741 | 1.782320754 | 1.761403742 | 1.897627091 | 1.657681529 |
| ENSMUSG00000026274 | 2.300956576 | 2.111770807 | 2.016522298 | 2.087182039 | 2.089109832 | 2.290034611 | 1.944737824 |
| ENSMUSG00000026275 | 2.342375483 | 2.170748646 | 2.225216702 | 2.397646762 | 2.310221245 | 2.666517981 | 2.176040447 |
| ENSMUSG00000026276 | 3.323191714 | 3.020786718 | 2.995124159 | 3.006284891 | 2.916320788 | 3.39707055 | 2.718759328 |
| ENSMUSG00000026277 | 2.568542279 | 2.348079356 | 2.381869252 | 2.406883471 | 2.386946794 | 2.746634199 | 2.238624656 |
| ENSMUSG00000026278 | 2.469490009 | 2.344302633 | 2.416333979 | 1.695339928 | 1.814781639 | 1.880813592 | 1.763495175 |
| ENSMUSG00000026279 | 1.948720984 | 1.79248125 | 1.87892496 | 1.819031101 | 1.850106709 | 2.021189299 | 1.858985529 |

| | | | | | | | |
|---|---|---|---|---|---|---|---|
| ENSMUSG00000026280 | 2.45469604 | 2.240724001 | 2.254202072 | 2.30479213 | 2.272808147 | 2.62324929 | 2.130281771 |
| ENSMUSG00000026281 | 2.435736758 | 2.279735268 | 2.30357978 | 2.225216702 | 2.179075613 | 2.503790683 | 2.056860222 |
| ENSMUSG00000026283 | 2.205378646 | 2.05357978 | 2.098079356 | 2.100219859 | 2.078515538 | 2.359835482 | 1.924281235 |
| ENSMUSG00000026285 | 0 | 0 | 0 | 0 | 0 | 0 | 0.224243824 |
| ENSMUSG00000026288 | 0 | 0 | 0 | 0.25 | 0.489301084 | 0.301029996 | 0.520678036 |
| ENSMUSG00000026289 | 2.280358197 | 2.168106335 | 2.142463902 | 2.170748646 | 2.167109853 | 2.357934847 | 2.038401572 |
| ENSMUSG00000026295 | 0 | 0 | 0 | 0 | 0 | 0 | 0 |
| ENSMUSG00000026301 | 0.289064826 | 0 | 0.25 | 0.396240625 | 0.244650542 | 0 | 0 |
| ENSMUSG00000026303 | 0.747221736 | 0.5 | 0.25 | 0.580482024 | 0.244650542 | 0.698970004 | 0.672731473 |
| ENSMUSG00000026304 | 0 | 0.25 | 0 | 0 | 0 | 0 | 0.224243824 |
| ENSMUSG00000026305 | 2.91246663 | 2.678346629 | 2.684311836 | 2.629663289 | 2.565609873 | 2.933993164 | 2.388231309 |
| ENSMUSG00000026307 | 1.759671389 | 1.678561379 | 1.777131114 | 1.557204673 | 1.665423072 | 1.812913357 | 1.750751125 |
| ENSMUSG00000026308 | 1 | 0.75 | 0.830482024 | 0.701838731 | 0.733951626 | 0.84509804 | 0.952571511 |
| ENSMUSG00000026309 | 2.512981297 | 2.344844592 | 2.310793496 | 2.289336734 | 2.268839121 | 2.545307116 | 2.116124284 |
| ENSMUSG00000026311 | 2.078325387 | 2.005591953 | 1.964495249 | 2.123963757 | 2.091002363 | 2.294466226 | 1.948678378 |
| ENSMUSG00000026312 | 0 | 0 | 0 | 0.25 | 0 | 0 | 0 |
| ENSMUSG00000026313 | 2.134369199 | 2.012462137 | 2.013820609 | 1.789967834 | 1.859322797 | 2.025305865 | 1.766260282 |
| ENSMUSG00000026315 | 2.111266113 | 1.933677405 | 2.025821952 | 0.75 | 0.77552387 | 0.698970004 | 0.775756176 |
| ENSMUSG00000026317 | 2.163307908 | 2.083847589 | 2.091580554 | 1.898114259 | 1.957204337 | 2.164352856 | 1.913109605 |
| ENSMUSG00000026319 | 2.37447073 | 2.169870025 | 2.212155735 | 2.334405475 | 2.264247757 | 2.592176757 | 2.123052075 |
| ENSMUSG00000026321 | 1.342375483 | 1.403677461 | 1.372963274 | 1.238549078 | 1.176121988 | 1.342422681 | 1.23151414 |
| ENSMUSG00000026322 | 0 | 0 | 0.25 | 0 | 0.244650542 | 0 | 0 |
| ENSMUSG00000026327 | 0 | 0 | 0.25 | 0 | 0 | 0 | 0 |
| ENSMUSG00000026331 | 0.289064826 | 0.5 | 0.701838731 | 0.580482024 | 0.244650542 | 0.602059991 | 0.448487648 |
| ENSMUSG00000026333 | 1.783508299 | 1.708222504 | 1.664552871 | 1.821350555 | 1.726078673 | 1.944482672 | 1.577201671 |
| ENSMUSG00000026335 | 2.56677519 | 2.328787391 | 2.392463902 | 2.412314044 | 2.339407793 | 2.698970004 | 2.187473278 |
| ENSMUSG00000026336 | 0 | 0 | 0 | 0 | 0 | 0 | 0 |
| ENSMUSG00000026339 | 2.596667477 | 2.412314044 | 2.416777886 | 2.397178659 | 2.323654601 | 2.683947131 | 2.190083792 |
| ENSMUSG00000026341 | 3.203074594 | 2.923153875 | 2.897529794 | 2.862545062 | 2.745958094 | 3.18920949 | 2.5494397 |
| ENSMUSG00000026342 | 2.287337982 | 2.137686696 | 2.100219859 | 2.133818844 | 2.068600388 | 2.326335861 | 1.91917299 |
| ENSMUSG00000026343 | 0.289064826 | 0.25 | 0 | 0.25 | 0.244650542 | 0.602059991 | 0.355418052 |
| ENSMUSG00000026344 | 0.45815691 | 0.396240625 | 0.646240625 | 0.396240625 | 0.387761935 | 0.301029996 | 0.355418052 |
| ENSMUSG00000026347 | 0 | 0 | 0 | 0 | 0 | 0 | 0 |
| ENSMUSG00000026348 | 0 | 0.25 | 0.25 | 0.5 | 0.489301084 | 0.301029996 | 0.520678036 |

| | | | | | | | |
|---|---|---|---|---|---|---|---|
| ENSMUSG00000026349 | 2.53552908 | 2.307204673 | 2.304186465 | 2.382357639 | 2.325116119 | 2.617000341 | 2.121219121 |
| ENSMUSG00000026353 | 2.755187126 | 2.518869787 | 2.52908599 | 2.615625568 | 2.54899118 | 2.947433722 | 2.397557454 |
| ENSMUSG00000026354 | 0 | 0.25 | 0.25 | 0.580482024 | 0.244650542 | 0.301029996 | 0.355418052 |
| ENSMUSG00000026355 | 3.173559032 | 2.870449858 | 2.851684121 | 2.87521047 | 2.808410859 | 3.259593879 | 2.625012618 |
| ENSMUSG00000026356 | 2.726616086 | 2.500352049 | 2.501406137 | 2.645770692 | 2.560905373 | 2.964730921 | 2.376964992 |
| ENSMUSG00000026357 | 0 | 0 | 0 | 0 | 0 | 0 | 0 |
| ENSMUSG00000026358 | 0 | 0 | 0 | 0 | 0 | 0 | 0 |
| ENSMUSG00000026360 | 1.069666964 | 1.160964047 | 1.061981878 | 0.79248125 | 0.812711509 | 0.903089987 | 0.448487648 |
| ENSMUSG00000026361 | 2.653588827 | 2.429419106 | 2.417221246 | 2.575659731 | 2.480959501 | 2.872738827 | 2.278571072 |
| ENSMUSG00000026365 | 1.156259305 | 1.080482024 | 0.864857905 | 0.864857905 | 0.812711509 | 0.698970004 | 0.710836105 |
| ENSMUSG00000026368 | 0 | 0.25 | 0.25 | 0 | 0 | 0 | 0 |
| ENSMUSG00000026374 | 2.839589945 | 2.628678514 | 2.61817271 | 2.701234418 | 2.625630579 | 3.051152522 | 2.439507591 |
| ENSMUSG00000026377 | 2.497590902 | 2.244319981 | 2.306603048 | 2.451435968 | 2.37023569 | 2.713490543 | 2.201756416 |
| ENSMUSG00000026380 | 0.960252568 | 1.114857905 | 0.75 | 1.561981878 | 1.489301084 | 1.568201724 | 1.462868481 |
| ENSMUSG00000026383 | 1.876566388 | 1.744319981 | 1.77186571 | 2.004202072 | 1.938809675 | 2.245512668 | 1.822941512 |
| ENSMUSG00000026384 | 2.256831695 | 2.127938163 | 2.150906586 | 2.397646762 | 2.303566328 | 2.627365857 | 2.090384805 |
| ENSMUSG00000026385 | 2.557825653 | 2.352877747 | 2.399047439 | 2.401832578 | 2.350890788 | 2.737192643 | 2.223239637 |
| ENSMUSG00000026387 | 0 | 0 | 0 | 0 | 0 | 0 | 0 |
| ENSMUSG00000026388 | 1.181541736 | 1.214495249 | 1.130890489 | 1.29248125 | 1.343584837 | 1.477121255 | 1.168187745 |
| ENSMUSG00000026389 | 1.894701088 | 1.923371739 | 1.988549078 | 1.657339155 | 1.671670246 | 1.740362689 | 1.670888075 |
| ENSMUSG00000026390 | 0 | 0 | 0 | 0 | 0 | 0 | 0 |
| ENSMUSG00000026393 | 2.839589945 | 2.585796429 | 2.563166358 | 2.506284891 | 2.433873436 | 2.812244697 | 2.251472963 |
| ENSMUSG00000026394 | 0 | 0 | 0 | 0 | 0 | 0 | 0 |
| ENSMUSG00000026395 | 0 | 0 | 0 | 0 | 0 | 0 | 0.355418052 |
| ENSMUSG00000026398 | 0 | 0 | 0 | 0.25 | 0.244650542 | 0.698970004 | 0 |
| ENSMUSG00000026399 | 0.671187741 | 0.396240625 | 0.580482024 | 1.04248125 | 1.091002363 | 1.278753601 | 0.896975297 |
| ENSMUSG00000026401 | 0 | 0 | 0.25 | 0 | 0 | 0 | 0 |
| ENSMUSG00000026404 | 1.903481042 | 1.735628626 | 1.71759118 | 1.828220739 | 1.810629166 | 2.089905111 | 1.755055163 |
| ENSMUSG00000026405 | 0 | 0.25 | 0.25 | 0.25 | 0 | 0.301029996 | 0.629532004 |
| ENSMUSG00000026407 | 0 | 0 | 0 | 0 | 0 | 0 | 0 |
| ENSMUSG00000026409 | 1.87190676 | 1.701838731 | 1.664552871 | 1.664552871 | 1.635855979 | 1.72427587 | 1.56970677 |
| ENSMUSG00000026411 | 2.397683106 | 2.247880462 | 2.275165585 | 2.164552871 | 2.21736452 | 2.44870632 | 2.217489603 |
| ENSMUSG00000026413 | 2.278594844 | 2.108657057 | 2.12694866 | 2.072504712 | 2.0928848 | 2.328379603 | 1.993927582 |
| ENSMUSG00000026414 | 2.072573125 | 1.991446071 | 1.935366747 | 0.864857905 | 0.877063019 | 1.041392685 | 0.984950056 |

| | | | | | | | |
|---|---|---|---|---|---|---|---|
| ENSMUSG00000026415 | 0 | 0 | 0 | 0 | 0 | 0 | 0 |
| ENSMUSG00000026416 | 0 | 0 | 0 | 0 | 0 | 0 | 0 |
| ENSMUSG00000026417 | 0 | 0 | 0 | 0 | 0 | 0 | 0 |
| ENSMUSG00000026418 | 0.45815691 | 0.701838731 | 0.646240625 | 0 | 0 | 0 | 0.224243824 |
| ENSMUSG00000026420 | 0 | 0 | 0 | 0.396240625 | 0.489301084 | 0.477121255 | 0.579661877 |
| ENSMUSG00000026421 | 3.314291961 | 3.031030328 | 3.021948576 | 2.682829758 | 2.674572266 | 3.035829825 | 2.510631008 |
| ENSMUSG00000026424 | 0 | 0 | 0 | 0 | 0 | 0 | 0 |
| ENSMUSG00000026425 | 2.607246427 | 2.407794264 | 2.38040011 | 2.374961472 | 2.29094062 | 2.643452676 | 2.165794278 |
| ENSMUSG00000026426 | 2.53552908 | 2.333288838 | 2.327084758 | 2.168989258 | 2.220647857 | 2.494154594 | 2.093904044 |
| ENSMUSG00000026427 | 2.340856238 | 2.172499493 | 2.187048212 | 2.144357207 | 2.209357611 | 2.396199347 | 2.083745739 |
| ENSMUSG00000026429 | 1.944805158 | 1.843759858 | 1.868933358 | 1.837182039 | 1.777823267 | 1.973127854 | 1.702561601 |
| ENSMUSG00000026430 | 1.596667477 | 1.505591953 | 1.532320754 | 1.860735874 | 1.851969278 | 1.982271233 | 1.694069894 |
| ENSMUSG00000026432 | 0 | 0 | 0 | 0 | 0.244650542 | 0 | 0 |
| ENSMUSG00000026433 | 1.568542279 | 1.51109853 | 1.610735874 | 1.71759118 | 1.715300549 | 1.929418926 | 1.610050272 |
| ENSMUSG00000026434 | 3.144018482 | 2.808255107 | 2.820047698 | 2.989909941 | 2.872453165 | 3.356599436 | 2.636148092 |
| ENSMUSG00000026435 | 1.605636262 | 1.464495249 | 1.571350555 | 1.51109853 | 1.576269619 | 1.568201724 | 1.479989695 |
| ENSMUSG00000026436 | 2.280358197 | 2.088286706 | 2.089388001 | 2.124961472 | 2.102149272 | 2.322219295 | 1.944737824 |
| ENSMUSG00000026437 | 2.37447073 | 2.269871196 | 2.26584877 | 2.154596376 | 2.242813819 | 2.477121255 | 2.13996595 |
| ENSMUSG00000026439 | 2.425923778 | 2.252107156 | 2.240724001 | 2.337182039 | 2.260180728 | 2.544068044 | 2.092400484 |
| ENSMUSG00000026442 | 0 | 0 | 0 | 0.25 | 0 | 0.301029996 | 0.629532004 |
| ENSMUSG00000026443 | 0 | 0 | 0 | 0 | 0.244650542 | 0 | 0.355418052 |
| ENSMUSG00000026447 | 1.771760131 | 1.695339928 | 1.688721876 | 1.482684334 | 1.584291694 | 1.698970004 | 1.473249546 |
| ENSMUSG00000026450 | 0 | 0 | 0 | 0 | 0 | 0 | 0.224243824 |
| ENSMUSG00000026452 | 0.747221736 | 0.75 | 0.896240625 | 0.396240625 | 0.632412477 | 0.477121255 | 0.629532004 |
| ENSMUSG00000026455 | 2.315764015 | 2.090485943 | 2.079353153 | 2.251406137 | 2.19206933 | 2.51054501 | 2.077494038 |
| ENSMUSG00000026456 | 2.432083896 | 2.213717096 | 2.212155735 | 2.147178659 | 2.147561328 | 2.392696953 | 2.014380832 |
| ENSMUSG00000026457 | 2.786655728 | 2.551448312 | 2.54652856 | 2.58993739 | 2.573153518 | 2.948412966 | 2.454284088 |
| ENSMUSG00000026458 | 0.45815691 | 0.646240625 | 0.75 | 0.646240625 | 0.733951626 | 0.301029996 | 0.579661877 |
| ENSMUSG00000026459 | 0 | 0 | 0 | 0 | 0 | 0 | 0 |
| ENSMUSG00000026463 | 2.811014908 | 2.583847589 | 2.536164561 | 2.186208459 | 2.110284345 | 2.428134794 | 2.006616089 |
| ENSMUSG00000026466 | 2.660010325 | 2.467976309 | 2.460980263 | 2.554186465 | 2.510051751 | 2.889301703 | 2.32409319 |
| ENSMUSG00000026468 | 0 | 0 | 0.25 | 0.75 | 0.846351821 | 0.77815125 | 0.672731473 |
| ENSMUSG00000026469 | 2.476144923 | 2.286164561 | 2.269871196 | 2.500352049 | 2.432800077 | 2.788168371 | 2.253005488 |
| ENSMUSG00000026470 | 2.396352858 | 2.266522298 | 2.259729747 | 2.092671852 | 2.125157063 | 2.309630167 | 2.059651572 |

| | | | | | | | |
|---|---|---|---|---|---|---|---|
| ENSMUSG00000026471 | 0.671187741 | 0.896240625 | 0.701838731 | 1.403677461 | 1.310728002 | 1.397940009 | 1.185218806 |
| ENSMUSG00000026473 | 2.190368148 | 1.998588359 | 2.059601185 | 2.25835575 | 2.208682096 | 2.514547753 | 2.095400649 |
| ENSMUSG00000026475 | 1.418409478 | 1.445339928 | 1.410964047 | 1.516522298 | 1.584291694 | 1.662757832 | 1.514741806 |
| ENSMUSG00000026478 | 3.117222119 | 2.860347007 | 2.866264042 | 2.816989271 | 2.818285254 | 3.211387553 | 2.651138495 |
| ENSMUSG00000026479 | 1.678702042 | 1.614857905 | 1.527131114 | 1.732684334 | 1.692703642 | 1.84509804 | 1.633706455 |
| ENSMUSG00000026480 | 0.45815691 | 0.396240625 | 0.25 | 0.580482024 | 0.686820904 | 0.77815125 | 0.579661877 |
| ENSMUSG00000026482 | 2.17479713 | 2.059601185 | 2.013820609 | 1.864857905 | 1.877063019 | 2.10720997 | 1.764880683 |
| ENSMUSG00000026483 | 2.785609222 | 2.52053726 | 2.523849256 | 2.440803092 | 2.379107586 | 2.73239376 | 2.240219117 |
| ENSMUSG00000026484 | 2.565888829 | 2.334405475 | 2.331607372 | 2.493603647 | 2.392211871 | 2.750508395 | 2.228559641 |
| ENSMUSG00000026489 | 0.747221736 | 0.830482024 | 0.75 | 1.214495249 | 1.223252711 | 1.113943352 | 1.20139795 |
| ENSMUSG00000026490 | 2.887787667 | 2.668327269 | 2.670529191 | 2.637686696 | 2.565363825 | 2.957607287 | 2.395988891 |
| ENSMUSG00000026491 | 3.056743041 | 2.785686319 | 2.783285553 | 2.864857905 | 2.780509399 | 3.178976947 | 2.575400606 |
| ENSMUSG00000026492 | 1.614416215 | 1.575945187 | 1.532320754 | 1.723704441 | 1.704182902 | 1.886490725 | 1.591689859 |
| ENSMUSG00000026494 | 1.129344651 | 1.188721876 | 1.311981878 | 1.061981878 | 1.234113756 | 1.146128036 | 1.131174228 |
| ENSMUSG00000026495 | 1.37447073 | 1.311981878 | 1.25 | 1.445339928 | 1.310728002 | 1.462397998 | 1.150210039 |
| ENSMUSG00000026496 | 2.921640949 | 2.717398461 | 2.716239979 | 2.755765312 | 2.791313678 | 3.114944416 | 2.656277989 |
| ENSMUSG00000026497 | 0.289064826 | 0.396240625 | 0 | 0.25 | 0 | 0 | 0 |
| ENSMUSG00000026499 | 2.615992899 | 2.441632227 | 2.401832578 | 2.457034121 | 2.368090054 | 2.760422483 | 2.216121671 |
| ENSMUSG00000026500 | 0 | 0 | 0 | 0 | 0 | 0 | 0.355418052 |
| ENSMUSG00000026502 | 2.721148722 | 2.488912477 | 2.476722649 | 2.439555804 | 2.349985771 | 2.69019608 | 2.155212955 |
| ENSMUSG00000026504 | 1.794934559 | 1.692046081 | 1.782320754 | 1.763820609 | 1.692703642 | 1.924279286 | 1.647906904 |
| ENSMUSG00000026509 | 3.216711746 | 2.92140621 | 2.91175216 | 2.911864607 | 2.835643405 | 3.290034611 | 2.648717833 |
| ENSMUSG00000026510 | 2.205378646 | 2.081607372 | 2.074802005 | 2.069031101 | 2.090057366 | 2.378397901 | 1.984950056 |
| ENSMUSG00000026511 | 2.520963022 | 2.28360658 | 2.308404919 | 2.388167274 | 2.321205208 | 2.657055853 | 2.222904214 |
| ENSMUSG00000026514 | 0.289064826 | 0 | 0.25 | 0.25 | 0 | 0 | 0 |
| ENSMUSG00000026516 | 2.724801584 | 2.51109853 | 2.496104615 | 2.613574823 | 2.550791325 | 2.925827575 | 2.358301651 |
| ENSMUSG00000026519 | 2.448863324 | 2.24929487 | 2.279735268 | 2.130890489 | 2.198391101 | 2.372912003 | 2.066254157 |
| ENSMUSG00000026520 | 2.258689303 | 2.213717096 | 2.288071211 | 2.089388001 | 2.230318492 | 2.340444115 | 2.16539364 |
| ENSMUSG00000026523 | 0 | 0 | 0 | 0 | 0 | 0 | 0 |
| ENSMUSG00000026525 | 0.811507563 | 0.79248125 | 0.864857905 | 0.75 | 0.686820904 | 0.77815125 | 0.896975297 |
| ENSMUSG00000026526 | 2.580706586 | 2.379909063 | 2.377938163 | 2.44697564 | 2.393014957 | 2.767897616 | 2.222904214 |
| ENSMUSG00000026527 | 0 | 0 | 0 | 0 | 0 | 0 | 0 |
| ENSMUSG00000026531 | 0 | 0 | 0 | 0 | 0 | 0 | 0 |
| ENSMUSG00000026532 | 0 | 0 | 0.25 | 1.784887838 | 1.808534465 | 1.944482672 | 1.641897157 |

| | | | | | | | |
|---|---|---|---|---|---|---|---|
| ENSMUSG00000026535 | 2.280358197 | 2.149978211 | 2.157339155 | 1.732684334 | 1.662257525 | 1.799340549 | 1.517723536 |
| ENSMUSG00000026536 | 0.747221736 | 1.061981878 | 1.114857905 | 1.214495249 | 1.212046813 | 1.113943352 | 1.131174228 |
| ENSMUSG00000026542 | 0 | 0 | 0 | 0 | 0 | 0 | 0 |
| ENSMUSG00000026544 | 0.578129653 | 0.580482024 | 0.701838731 | 0.75 | 0.877063019 | 0.903089987 | 0.853775828 |
| ENSMUSG00000026546 | 0.289064826 | 0.75 | 0.5 | 0.396240625 | 0.632412477 | 0.698970004 | 0.74492186 |
| ENSMUSG00000026547 | 3.261878538 | 3.009129268 | 3.004289097 | 2.75 | 2.791054056 | 3.170555059 | 2.627808626 |
| ENSMUSG00000026548 | 0 | 0 | 0 | 0 | 0 | 0 | 0 |
| ENSMUSG00000026553 | 3.323047337 | 2.984711114 | 2.971233412 | 3.101351856 | 3.023079973 | 3.521661015 | 2.817326071 |
| ENSMUSG00000026554 | 2.626865868 | 2.402294685 | 2.444929839 | 2.481573999 | 2.468875138 | 2.764922985 | 2.339433092 |
| ENSMUSG00000026556 | 2.048736215 | 1.910964047 | 1.907339155 | 1.720660762 | 1.754126039 | 1.954242509 | 1.71730663 |
| ENSMUSG00000026558 | 2.7985055 | 2.572792675 | 2.589388001 | 2.625956434 | 2.608627598 | 2.961421094 | 2.465415859 |
| ENSMUSG00000026560 | 0 | 0 | 0 | 0 | 0 | 0 | 0 |
| ENSMUSG00000026563 | 1.389637216 | 1.321350555 | 1.42510993 | 1.54248125 | 1.504538967 | 1.716003344 | 1.511732338 |
| ENSMUSG00000026564 | 1.036286563 | 0.830482024 | 0.864857905 | 0.25 | 0.244650542 | 0.301029996 | 0 |
| ENSMUSG00000026565 | 2.276824003 | 2.091580554 | 2.095926073 | 2.228969845 | 2.17760189 | 2.459392488 | 1.997988498 |
| ENSMUSG00000026566 | 2.492331912 | 2.27908599 | 2.273189285 | 2.29248125 | 2.242813819 | 2.50242712 | 2.119375723 |
| ENSMUSG00000026567 | 0.578129653 | 0 | 0 | 0 | 0.244650542 | 0 | 0 |
| ENSMUSG00000026568 | 1.948720984 | 1.862802778 | 1.848079356 | 2.038704527 | 1.990091809 | 2.217483944 | 1.848480858 |
| ENSMUSG00000026571 | 2.262379879 | 2.114857905 | 2.101285366 | 2.097004321 | 1.991345651 | 2.326335861 | 1.947894097 |
| ENSMUSG00000026572 | 0 | 0 | 0 | 0.25 | 0 | 0 | 0 |
| ENSMUSG00000026573 | 0 | 0 | 0 | 0 | 0 | 0 | 0 |
| ENSMUSG00000026574 | 0.91631382 | 0.5 | 0.580482024 | 0.396240625 | 0.244650542 | 0.477121255 | 0 |
| ENSMUSG00000026575 | 1.91631382 | 1.79248125 | 1.80479213 | 1.982684334 | 1.882215755 | 2.158362492 | 1.746389053 |
| ENSMUSG00000026576 | 2.718080037 | 2.528109877 | 2.514497931 | 2.155512955 | 2.103062484 | 2.397940009 | 1.944737824 |
| ENSMUSG00000026577 | 2.033115198 | 1.945339928 | 1.886723615 | 1.998588359 | 1.97205363 | 2.240549248 | 1.847411384 |
| ENSMUSG00000026578 | 1.777675583 | 1.692046081 | 1.566696635 | 1.688721876 | 1.649303553 | 1.785329835 | 1.607811402 |
| ENSMUSG00000026579 | 0 | 0 | 0.25 | 0.396240625 | 0.244650542 | 0.84509804 | 0.579661877 |
| ENSMUSG00000026580 | 0 | 0.646240625 | 0.396240625 | 0.25 | 0 | 0.698970004 | 0.520678036 |
| ENSMUSG00000026581 | 0 | 0 | 0 | 0 | 0 | 0 | 0 |
| ENSMUSG00000026582 | 0 | 0 | 0.25 | 0 | 0 | 0 | 0 |
| ENSMUSG00000026584 | 2.020182925 | 1.988549078 | 1.882845365 | 2.04745614 | 1.988833497 | 2.167317335 | 1.853775828 |
| ENSMUSG00000026585 | 2.471720134 | 2.294977273 | 2.285526764 | 2.319612365 | 2.224547964 | 2.568201724 | 2.075383 |
| ENSMUSG00000026586 | 2.820759099 | 2.584684096 | 2.547764804 | 2.313757142 | 2.192777367 | 2.485721426 | 2.049487947 |
| ENSMUSG00000026587 | 0 | 0 | 0.25 | 0 | 0.244650542 | 0.477121255 | 0 |

| | | | | | | | |
|---|---|---|---|---|---|---|---|
| ENSMUSG00000026589 | 2.282114126 | 2.061981878 | 2.025821952 | 2.004202072 | 1.962676635 | 2.176091259 | 1.834294096 |
| ENSMUSG00000026592 | 0.289064826 | 0.25 | 0 | 0 | 0.244650542 | 0.301029996 | 0 |
| ENSMUSG00000026594 | 1.975148165 | 1.951838731 | 1.894357207 | 1.919870025 | 1.848234258 | 2.053078443 | 1.707551656 |
| ENSMUSG00000026596 | 2.774459379 | 2.519203899 | 2.551448312 | 2.579635702 | 2.547958389 | 2.866287339 | 2.379456779 |
| ENSMUSG00000026600 | 2.834495649 | 2.564641508 | 2.588010856 | 2.506284891 | 2.418540977 | 2.790988475 | 2.26501115 |
| ENSMUSG00000026601 | 0.45815691 | 0.864857905 | 0.396240625 | 0.580482024 | 0.846351821 | 0.477121255 | 0.579661877 |
| ENSMUSG00000026602 | 0 | 0.25 | 0.25 | 0.92510993 | 0.978602168 | 1.041392685 | 1.028149525 |
| ENSMUSG00000026603 | 2.163307908 | 2 | 2.04745614 | 2.204245906 | 2.129414728 | 2.356025857 | 1.984249047 |
| ENSMUSG00000026604 | 2.757995436 | 2.574515642 | 2.5684489 | 2.578787391 | 2.571464152 | 2.910090546 | 2.428142734 |
| ENSMUSG00000026605 | 2.893893668 | 2.656655413 | 2.67813175 | 2.797147211 | 2.72514079 | 3.092720645 | 2.492479894 |
| ENSMUSG00000026608 | 2.305951036 | 2.158248799 | 2.168989258 | 2.154596376 | 2.173143437 | 2.383815366 | 2.059095225 |
| ENSMUSG00000026609 | 0 | 0 | 0 | 0 | 0 | 0 | 0 |
| ENSMUSG00000026610 | 0 | 0.25 | 0 | 0.580482024 | 0.489301084 | 0.477121255 | 0 |
| ENSMUSG00000026611 | 0.45815691 | 0.396240625 | 0.396240625 | 0.25 | 0.244650542 | 0.301029996 | 0.448487648 |
| ENSMUSG00000026614 | 0 | 0 | 0 | 0 | 0.244650542 | 0.301029996 | 0.224243824 |
| ENSMUSG00000026615 | 3.183160496 | 2.912650756 | 2.853935442 | 3.009472521 | 2.917593359 | 3.37821615 | 2.710613221 |
| ENSMUSG00000026616 | 0.747221736 | 0.646240625 | 0.896240625 | 0.646240625 | 0.632412477 | 0.477121255 | 0.448487648 |
| ENSMUSG00000026617 | 2.339331437 | 2.168106335 | 2.156427211 | 2.323080408 | 2.28771234 | 2.631443769 | 2.153969461 |
| ENSMUSG00000026618 | 2.640441358 | 2.452642909 | 2.469129237 | 2.565229883 | 2.485926424 | 2.868056362 | 2.355195169 |
| ENSMUSG00000026620 | 1.894701088 | 1.744319981 | 1.811981878 | 1.874961472 | 1.873585591 | 2.089905111 | 1.82753046 |
| ENSMUSG00000026621 | 0 | 0 | 0 | 0 | 0 | 0 | 0 |
| ENSMUSG00000026622 | 2.708736128 | 2.458617762 | 2.451435968 | 2.465659339 | 2.417045398 | 2.748188027 | 2.290497981 |
| ENSMUSG00000026623 | 2.862428343 | 2.631380202 | 2.609957721 | 2.765005089 | 2.68401826 | 3.089551883 | 2.476483958 |
| ENSMUSG00000026626 | 2.45469604 | 2.277783918 | 2.264497931 | 2.398581151 | 2.335180723 | 2.688419822 | 2.222904214 |
| ENSMUSG00000026627 | 1.944805158 | 1.819031101 | 1.809601185 | 1.852347734 | 1.868304353 | 2.103803721 | 1.774416565 |
| ENSMUSG00000026628 | 1.494443473 | 1.431980114 | 1.356566189 | 1.755591953 | 1.649303553 | 1.886490725 | 1.601000298 |
| ENSMUSG00000026630 | 0.289064826 | 0.580482024 | 0.75 | 0.701838731 | 0.632412477 | 0.698970004 | 0.672731473 |
| ENSMUSG00000026632 | 1.527823874 | 1.431980114 | 1.52186571 | 1.723704441 | 1.692703642 | 1.716003344 | 1.540636858 |
| ENSMUSG00000026634 | 2.493389029 | 2.296838018 | 2.280383379 | 2.32991803 | 2.276733037 | 2.580924976 | 2.117056606 |
| ENSMUSG00000026637 | 1.894701088 | 1.841580554 | 1.848079356 | 1.458222504 | 1.427019704 | 1.579783597 | 1.437266334 |
| ENSMUSG00000026638 | 0.45815691 | 0.5 | 0.646240625 | 1.061981878 | 1.091002363 | 0.903089987 | 1.041356071 |
| ENSMUSG00000026639 | 0.671187741 | 0.830482024 | 0.79248125 | 1.17510993 | 1.234113756 | 1 | 1.272006351 |
| ENSMUSG00000026640 | 2.457006475 | 2.27186571 | 2.306603048 | 2.231203126 | 2.174635855 | 2.389166084 | 2.032410373 |
| ENSMUSG00000026641 | 2.245508853 | 2.044977273 | 2.12892496 | 1.956637122 | 2.074582839 | 2.146128036 | 1.972821265 |

| | | | | | | | |
|---|---|---|---|---|---|---|---|
| ENSMUSG00000026643 | 2.714367558 | 2.475216702 | 2.449820405 | 2.609177886 | 2.530737602 | 2.915927212 | 2.331614323 |
| ENSMUSG00000026644 | 0.289064826 | 0.25 | 0.25 | 0 | 0.387761935 | 0.602059991 | 0.520678036 |
| ENSMUSG00000026645 | 0 | 0 | 0 | 0 | 0 | 0 | 0 |
| ENSMUSG00000026646 | 2.078325387 | 1.990000483 | 1.953445298 | 2.015173983 | 1.955822902 | 2.240549248 | 1.828667599 |
| ENSMUSG00000026648 | 1.87190676 | 1.779735268 | 1.802363341 | 1.901832578 | 1.739104362 | 2.049218023 | 1.637827726 |
| ENSMUSG00000026649 | 1.036286563 | 1.02186571 | 0.896240625 | 0.896240625 | 1.057362051 | 0.954242509 | 0.876096088 |
| ENSMUSG00000026650 | 0 | 0 | 0 | 0 | 0 | 0.301029996 | 0.355418052 |
| ENSMUSG00000026655 | 2.582415738 | 2.336628433 | 2.320772088 | 2.539967834 | 2.453670557 | 2.825426118 | 2.259064007 |
| ENSMUSG00000026656 | 0.45815691 | 0 | 0 | 0 | 0 | 0 | 0 |
| ENSMUSG00000026657 | 2.70238848 | 2.486726569 | 2.488912477 | 2.259043403 | 2.234742351 | 2.549003262 | 2.119837558 |
| ENSMUSG00000026659 | 1.816888701 | 1.698603967 | 1.705044741 | 1.685366747 | 1.695608647 | 1.86332286 | 1.546120303 |
| ENSMUSG00000026662 | 2.782453855 | 2.519537702 | 2.514159429 | 2.520869832 | 2.463660548 | 2.857935265 | 2.302000776 |
| ENSMUSG00000026663 | 2.623788124 | 2.398114259 | 2.398114259 | 2.509729747 | 2.42883626 | 2.796574333 | 2.275162625 |
| ENSMUSG00000026664 | 2.203267085 | 2.006976499 | 1.992885888 | 1.888647213 | 1.744182955 | 1.982271233 | 1.712465909 |
| ENSMUSG00000026667 | 2.879086229 | 2.6343046 | 2.609957721 | 2.730646101 | 2.638666793 | 3.006893708 | 2.457062464 |
| ENSMUSG00000026668 | 0 | 0 | 0 | 0 | 0 | 0 | 0 |
| ENSMUSG00000026669 | 2.556920045 | 2.355516192 | 2.386241108 | 2.446567407 | 2.398585969 | 2.718501689 | 2.228559641 |
| ENSMUSG00000026670 | 2.638209243 | 2.432405186 | 2.394828984 | 2.391037372 | 2.340805641 | 2.680335513 | 2.202114485 |
| ENSMUSG00000026672 | 2.203267085 | 2.019203899 | 2.087182039 | 2.032320754 | 1.990091809 | 2.173186268 | 1.852723744 |
| ENSMUSG00000026674 | 2.325351389 | 2.117918804 | 2.005591953 | 2.990904654 | 2.90324169 | 3.359076226 | 2.689818231 |
| ENSMUSG00000026675 | 2.273259602 | 2.12892496 | 2.142463902 | 2.078220739 | 2.042520869 | 2.29666519 | 1.882886767 |
| ENSMUSG00000026676 | 0.578129653 | 0 | 0 | 0 | 0 | 0 | 0 |
| ENSMUSG00000026678 | 0.671187741 | 0.580482024 | 0.701838731 | 0.396240625 | 0 | 0 | 0 |
| ENSMUSG00000026679 | 0.91631382 | 0.396240625 | 0.75 | 0.79248125 | 0.877063019 | 0.954242509 | 0.803905701 |
| ENSMUSG00000026683 | 2.693989403 | 2.467976309 | 2.461764337 | 2.621708755 | 2.517458689 | 2.95568775 | 2.358522405 |
| ENSMUSG00000026686 | 0 | 0 | 0 | 0 | 0 | 0 | 0 |
| ENSMUSG00000026687 | 2.315764015 | 2.146240625 | 2.164552871 | 2.206637122 | 2.212715924 | 2.507855872 | 2.069505596 |
| ENSMUSG00000026688 | 2.051791411 | 1.87892496 | 2.013820609 | 1.51109853 | 1.603583716 | 1.707570176 | 1.532233387 |
| ENSMUSG00000026691 | 0.45815691 | 0 | 0.25 | 0 | 0 | 0 | 0 |
| ENSMUSG00000026692 | 0 | 0 | 0 | 0 | 0 | 0 | 0 |
| ENSMUSG00000026694 | 2.285603957 | 2.111770807 | 2.12694866 | 2.001406137 | 2.017859938 | 2.23299611 | 1.934261938 |
| ENSMUSG00000026696 | 2.10861828 | 1.990000483 | 2.005591953 | 2.074802005 | 1.950242452 | 2.238046103 | 1.852723744 |
| ENSMUSG00000026697 | 0 | 0 | 0 | 0 | 0 | 0 | 0 |
| ENSMUSG00000026698 | 1.99653913 | 1.901832578 | 1.843759858 | 1.841580554 | 1.736537397 | 1.908485019 | 1.653807046 |

| | | | | | | | |
|---|---|---|---|---|---|---|---|
| ENSMUSG00000026700 | 0 | 0 | 0 | 0 | 0 | 0 | 0.224243824 |
| ENSMUSG00000026701 | 2.8736072 | 2.637205477 | 2.623213655 | 2.505938588 | 2.478134581 | 2.844477176 | 2.290770186 |
| ENSMUSG00000026705 | 2.177057474 | 2.031030328 | 2.088286706 | 2.180274797 | 2.104881849 | 2.397940009 | 1.944737824 |
| ENSMUSG00000026707 | 1.727821383 | 1.622963274 | 1.681980114 | 1.711372513 | 1.635855979 | 1.792391689 | 1.514741806 |
| ENSMUSG00000026708 | 2.097855556 | 2.009729747 | 1.964495249 | 2.023189285 | 1.983754904 | 2.255272505 | 1.908707966 |
| ENSMUSG00000026709 | 2.136858952 | 1.938721876 | 1.950224975 | 2.084962501 | 2.00243419 | 2.311753861 | 1.880961074 |
| ENSMUSG00000026712 | 0 | 0 | 0 | 0.25 | 0.244650542 | 0 | 0.520678036 |
| ENSMUSG00000026715 | 0.578129653 | 0.5 | 0.5 | 0.5 | 0.632412477 | 0.698970004 | 0.74492186 |
| ENSMUSG00000026718 | 2.644135203 | 2.417664062 | 2.401832578 | 2.43662858 | 2.364194729 | 2.685741739 | 2.208492871 |
| ENSMUSG00000026721 | 2.318984359 | 2.140560606 | 2.131869252 | 2.267194569 | 2.189222894 | 2.469822016 | 2.038994634 |
| ENSMUSG00000026723 | 1.445324132 | 1.470660762 | 1.52186571 | 1.784887838 | 1.692703642 | 1.944482672 | 1.577201671 |
| ENSMUSG00000026725 | 0.45815691 | 0.396240625 | 0.396240625 | 0 | 0.244650542 | 0 | 0.224243824 |
| ENSMUSG00000026726 | 1.842799666 | 1.657339155 | 1.75835575 | 1.67510993 | 1.55104774 | 1.69019608 | 1.455757965 |
| ENSMUSG00000026727 | 2.804055922 | 2.544043287 | 2.560793496 | 2.374462959 | 2.298360391 | 2.651278014 | 2.14512854 |
| ENSMUSG00000026728 | 3.990142936 | 3.620796493 | 3.680101149 | 3.423698289 | 3.406099624 | 3.91603261 | 3.243971188 |
| ENSMUSG00000026729 | 0 | 0 | 0 | 0 | 0 | 0 | 0 |
| ENSMUSG00000026730 | 2.097855556 | 1.951838731 | 1.978222334 | 1.870953944 | 1.728722551 | 1.919078092 | 1.647906904 |
| ENSMUSG00000026734 | 0 | 0 | 0 | 0 | 0 | 0 | 0 |
| ENSMUSG00000026735 | 0 | 0 | 0 | 0 | 0 | 0 | 0 |
| ENSMUSG00000026736 | 0.45815691 | 0 | 0.580482024 | 0.25 | 0.244650542 | 0.301029996 | 0 |
| ENSMUSG00000026737 | 1.940852214 | 1.87694866 | 1.80479213 | 1.864857905 | 1.645989387 | 2.017033339 | 1.625302983 |
| ENSMUSG00000026739 | 2.653588827 | 2.473704441 | 2.42293588 | 2.410061234 | 2.306140782 | 2.621176282 | 2.110473216 |
| ENSMUSG00000026740 | 2.315764015 | 2.186208459 | 2.135757955 | 2.147178659 | 2.031524462 | 2.281033367 | 1.904245613 |
| ENSMUSG00000026743 | 2.415874319 | 2.247171172 | 2.240724001 | 2.125956434 | 2.06149024 | 2.225309282 | 1.908707966 |
| ENSMUSG00000026748 | 0.289064826 | 0.580482024 | 0.701838731 | 0.396240625 | 0.387761935 | 0.301029996 | 0.224243824 |
| ENSMUSG00000026749 | 2.655024396 | 2.435366747 | 2.437048212 | 2.30175358 | 2.282813873 | 2.614897216 | 2.160546876 |
| ENSMUSG00000026750 | 2.590857977 | 2.358657057 | 2.360217292 | 2.552667836 | 2.453670557 | 2.853698212 | 2.283060517 |
| ENSMUSG00000026751 | 0 | 0.25 | 0 | 0 | 0 | 0 | 0 |
| ENSMUSG00000026753 | 2.561428547 | 2.367410454 | 2.321928095 | 2.503156135 | 2.41591954 | 2.777426822 | 2.23281864 |
| ENSMUSG00000026754 | 2.436947303 | 2.214495249 | 2.230460234 | 2.271202097 | 2.234742351 | 2.463892989 | 2.064068234 |
| ENSMUSG00000026755 | 2.482695304 | 2.254202072 | 2.286801231 | 2.296218836 | 2.218681522 | 2.579783597 | 2.105687397 |
| ENSMUSG00000026761 | 2.544030601 | 2.361253711 | 2.337182039 | 2.33272922 | 2.281165826 | 2.571708832 | 2.098866035 |
| ENSMUSG00000026764 | 2.294202867 | 2.123963757 | 2.149047439 | 1.769203899 | 1.680838453 | 1.740362689 | 1.56970677 |
| ENSMUSG00000026765 | 0 | 0 | 0.25 | 0.396240625 | 0.387761935 | 0.301029996 | 0.224243824 |

| | | | | | | | |
|---|---|---|---|---|---|---|---|
| ENSMUSG00000026766 | 2.704939145 | 2.476722649 | 2.453445298 | 2.545598588 | 2.444084351 | 2.844477176 | 2.307731473 |
| ENSMUSG00000026767 | 2.339331437 | 2.143411797 | 2.147178659 | 2.236360959 | 2.094757251 | 2.404833717 | 1.989815042 |
| ENSMUSG00000026768 | 1.100572389 | 1.02186571 | 0.976722649 | 0.5 | 0.387761935 | 0.301029996 | 0.448487648 |
| ENSMUSG00000026770 | 0 | 0.25 | 0.25 | 0.25 | 0.244650542 | 0 | 0 |
| ENSMUSG00000026771 | 2.199011616 | 2.142463902 | 2.099151195 | 2.132845365 | 2.028157605 | 2.274157849 | 1.869159809 |
| ENSMUSG00000026773 | 2.203267085 | 2.05479213 | 2.070192693 | 1.830482024 | 1.840643511 | 2.017033339 | 1.74492186 |
| ENSMUSG00000026774 | 0 | 0 | 0 | 0 | 0 | 0 | 0 |
| ENSMUSG00000026775 | 2.987274912 | 2.738912477 | 2.71759118 | 2.758527787 | 2.655268067 | 3.093071306 | 2.461747664 |
| ENSMUSG00000026778 | 0.671187741 | 0.5 | 0.396240625 | 0.5 | 0.733951626 | 0.301029996 | 0.579661877 |
| ENSMUSG00000026779 | 2.422183552 | 2.259043403 | 2.241446071 | 2.337182039 | 2.201854879 | 2.503790683 | 2.049487947 |
| ENSMUSG00000026781 | 2.390989024 | 2.233422664 | 2.235628626 | 2.274508021 | 2.216042585 | 2.526339277 | 2.048913829 |
| ENSMUSG00000026782 | 2.388281011 | 2.250703754 | 2.220660762 | 2.317865757 | 2.254288168 | 2.565847819 | 2.097879707 |
| ENSMUSG00000026784 | 1.83262764 | 1.708222504 | 1.698603967 | 1.83272922 | 1.741652793 | 1.963787827 | 1.627424404 |
| ENSMUSG00000026785 | 1.568542279 | 1.532320754 | 1.630890489 | 1.380890489 | 1.546663136 | 1.380211242 | 1.508694612 |
| ENSMUSG00000026786 | 2.35727108 | 2.134789703 | 2.130890489 | 1.96759118 | 1.913211023 | 2.127104798 | 1.829800754 |
| ENSMUSG00000026787 | 0.45815691 | 0.25 | 0.25 | 0.396240625 | 0.387761935 | 0.477121255 | 0.355418052 |
| ENSMUSG00000026788 | 2.126809492 | 2.017865591 | 1.956637122 | 2.027131114 | 1.974687643 | 2.271841607 | 1.871156825 |
| ENSMUSG00000026790 | 2.496544396 | 2.367918804 | 2.359697963 | 2.340485943 | 2.312243922 | 2.630427875 | 2.185972044 |
| ENSMUSG00000026791 | 1.227926429 | 1.403677461 | 1.470660762 | 1.080482024 | 1.407936347 | 1.361727836 | 1.483307832 |
| ENSMUSG00000026792 | 1.912079952 | 1.823655187 | 1.834962501 | 1.642463902 | 1.655839964 | 1.799340549 | 1.695786161 |
| ENSMUSG00000026796 | 3.106196482 | 2.886120381 | 2.952843673 | 2.51007243 | 2.655458802 | 2.877371346 | 2.564261618 |
| ENSMUSG00000026797 | 2.581562037 | 2.410964047 | 2.439139081 | 2.510756821 | 2.532092521 | 2.876217841 | 2.395201749 |
| ENSMUSG00000026798 | 1.631440309 | 1.552363341 | 1.606566189 | 1.532320754 | 1.588235379 | 1.602059991 | 1.574722613 |
| ENSMUSG00000026799 | 1.852729478 | 1.777131114 | 1.747171172 | 1.649978211 | 1.842556557 | 1.86923172 | 1.728327686 |
| ENSMUSG00000026803 | 2.033115198 | 1.928561379 | 1.948603967 | 1.987091808 | 1.944572352 | 2.149219113 | 1.832055226 |
| ENSMUSG00000026805 | 0 | 0 | 0.25 | 0 | 0 | 0 | 0 |
| ENSMUSG00000026806 | 2.013563224 | 1.940387808 | 1.940387808 | 1.953445298 | 1.880506521 | 2.06069784 | 1.797719748 |
| ENSMUSG00000026807 | 0.289064826 | 0.25 | 0 | 0.25 | 0 | 0 | 0 |
| ENSMUSG00000026809 | 0 | 0.25 | 0.25 | 0 | 0.387761935 | 0.602059991 | 0.579661877 |
| ENSMUSG00000026810 | 2.181541736 | 2.043731421 | 2.038704527 | 1.976722649 | 2.046825265 | 2.181843588 | 1.912234055 |
| ENSMUSG00000026811 | 2.227926429 | 2.08272922 | 2.146240625 | 2.087182039 | 2.115605392 | 2.376576957 | 1.988432495 |
| ENSMUSG00000026812 | 2.552362269 | 2.385757955 | 2.352877747 | 2.289967834 | 2.302012594 | 2.615950052 | 2.141695983 |
| ENSMUSG00000026814 | 0.578129653 | 0.580482024 | 0.951838731 | 2.024508021 | 2.156296347 | 2.320146286 | 2.106169177 |
| ENSMUSG00000026815 | 0 | 0 | 0 | 0.25 | 0 | 0 | 0 |

| | | | | | | | |
|---|---|---|---|---|---|---|---|
| ENSMUSG00000026816 | 2.315764015 | 2.158248799 | 2.161864607 | 2.222185812 | 2.225838481 | 2.509202522 | 2.05853792 |
| ENSMUSG00000026817 | 2.351376531 | 2.168989258 | 2.203445298 | 1.802363341 | 1.84445929 | 2.093421685 | 1.835407748 |
| ENSMUSG00000026818 | 0.289064826 | 0 | 0 | 0 | 0.244650542 | 0.301029996 | 0.224243824 |
| ENSMUSG00000026819 | 2.095120908 | 2.004202072 | 2.074802005 | 2.058404919 | 2.128567294 | 2.257678575 | 1.999330888 |
| ENSMUSG00000026820 | 1.903481042 | 1.864857905 | 1.905512955 | 1.649978211 | 1.840643511 | 1.929418926 | 1.774416565 |
| ENSMUSG00000026821 | 2.144240249 | 2.098079356 | 2.111770807 | 1.848079356 | 2.012006468 | 2.056904851 | 1.96172822 |
| ENSMUSG00000026822 | 0.747221736 | 0.396240625 | 0.580482024 | 0 | 0.387761935 | 0 | 0.355418052 |
| ENSMUSG00000026824 | 0 | 0 | 0.25 | 0.25 | 0 | 0 | 0 |
| ENSMUSG00000026825 | 2.086807695 | 1.966046536 | 2.015173983 | 1.988549078 | 2.13194489 | 2.267171728 | 2.034820183 |
| ENSMUSG00000026826 | 1.418409478 | 1.214495249 | 1.282320754 | 1.226722649 | 1.188507684 | 1.301029996 | 1.014380832 |
| ENSMUSG00000026827 | 2.67325056 | 2.468745337 | 2.468361028 | 2.439555804 | 2.363323232 | 2.710963119 | 2.211287839 |
| ENSMUSG00000026828 | 0 | 0 | 0 | 0 | 0 | 0 | 0 |
| ENSMUSG00000026829 | 0.289064826 | 0 | 0.25 | 0 | 0 | 0.301029996 | 0.224243824 |
| ENSMUSG00000026830 | 0 | 0 | 0 | 0 | 0 | 0 | 0 |
| ENSMUSG00000026831 | 0 | 0 | 0 | 0.25 | 0 | 0.301029996 | 0.448487648 |
| ENSMUSG00000026832 | 0.747221736 | 0.896240625 | 0.580482024 | 0 | 0 | 0 | 0.224243824 |
| ENSMUSG00000026833 | 1.596667477 | 1.593759858 | 1.668106335 | 1.54745614 | 1.47337555 | 1.653212514 | 1.493062805 |
| ENSMUSG00000026834 | 0.289064826 | 0.701838731 | 0.396240625 | 0.396240625 | 0.568060967 | 0.602059991 | 0.224243824 |
| ENSMUSG00000026835 | 0 | 0 | 0 | 0 | 0 | 0 | 0 |
| ENSMUSG00000026836 | 2.539328946 | 2.344844592 | 2.325374049 | 2.086073977 | 2.029283463 | 2.260071388 | 1.885753983 |
| ENSMUSG00000026837 | 3.034559718 | 2.857743793 | 2.899861995 | 2.597004321 | 2.689428323 | 2.949877704 | 2.662201431 |
| ENSMUSG00000026839 | 0.747221736 | 0.951838731 | 0.701838731 | 0.830482024 | 0.489301084 | 0.903089987 | 0.710836105 |
| ENSMUSG00000026840 | 0.289064826 | 0.5 | 0.5 | 0.396240625 | 0.877063019 | 0.602059991 | 0.896975297 |
| ENSMUSG00000026841 | 0 | 0.25 | 0.25 | 0.25 | 0.244650542 | 0.301029996 | 0.629532004 |
| ENSMUSG00000026842 | 2.811014908 | 2.638886943 | 2.645300145 | 2.392463902 | 2.460030299 | 2.693726949 | 2.383567577 |
| ENSMUSG00000026843 | 2.666334442 | 2.456637122 | 2.494319981 | 2.426408097 | 2.408704478 | 2.741939078 | 2.264126018 |
| ENSMUSG00000026848 | 2.674620126 | 2.438304187 | 2.452241044 | 2.470660762 | 2.463332064 | 2.763427994 | 2.305399353 |
| ENSMUSG00000026849 | 2.264212992 | 2.114857905 | 2.133818844 | 2.069031101 | 2.043601899 | 2.294466226 | 1.953344547 |
| ENSMUSG00000026851 | 2.146671932 | 2.004202072 | 2.056000419 | 2.147178659 | 2.113840608 | 2.357934847 | 1.970632877 |
| ENSMUSG00000026853 | 2.423434025 | 2.254897682 | 2.281676118 | 2.27908599 | 2.268839121 | 2.53529412 | 2.191196167 |
| ENSMUSG00000026854 | 2.247417471 | 2.146240625 | 2.209800947 | 1.997171172 | 2.079491904 | 2.276461804 | 2.009874076 |
| ENSMUSG00000026856 | 1.876566388 | 1.830482024 | 1.899978211 | 1.782320754 | 1.88560959 | 1.982271233 | 1.849546807 |
| ENSMUSG00000026857 | 1.982401085 | 1.979715809 | 2.006976499 | 1.646240625 | 1.83872004 | 1.924279286 | 1.900630878 |
| ENSMUSG00000026858 | 2.17479713 | 2.05479213 | 2.089388001 | 1.956637122 | 2.091002363 | 2.247973266 | 2.008574816 |

| | | | | | | | |
|---|---|---|---|---|---|---|---|
| ENSMUSG00000026860 | 2.194712275 | 2.140560606 | 2.21293726 | 1.96759118 | 2.167869715 | 2.230448921 | 2.116590781 |
| ENSMUSG00000026864 | 3.515901449 | 3.21293726 | 3.165667094 | 3.305736451 | 3.211257996 | 3.753506457 | 2.987233704 |
| ENSMUSG00000026866 | 0 | 0 | 0 | 0 | 0 | 0 | 0 |
| ENSMUSG00000026867 | 2.867194479 | 2.612287161 | 2.592944161 | 2.664998973 | 2.606881368 | 2.983175072 | 2.396577994 |
| ENSMUSG00000026869 | 2.734388958 | 2.476346751 | 2.454645544 | 2.576230417 | 2.495955124 | 2.86569606 | 2.321381806 |
| ENSMUSG00000026870 | 0.289064826 | 0.25 | 0.580482024 | 0.25 | 0.244650542 | 0.301029996 | 0.448487648 |
| ENSMUSG00000026872 | 2.307602651 | 2.209800947 | 2.174241882 | 2.150906586 | 2.098472613 | 2.264817823 | 1.932619724 |
| ENSMUSG00000026873 | 2.124258828 | 1.978222334 | 2.03360658 | 1.976722649 | 2.066583491 | 2.240549248 | 1.943148064 |
| ENSMUSG00000026874 | 0 | 0.396240625 | 0 | 0 | 0 | 0 | 0.355418052 |
| ENSMUSG00000026875 | 1.227926429 | 1.282320754 | 1.311981878 | 1.634789703 | 1.773209395 | 1.826074803 | 1.710836105 |
| ENSMUSG00000026878 | 3.015982563 | 2.739638179 | 2.702843673 | 2.779248419 | 2.690983579 | 3.128076013 | 2.49378997 |
| ENSMUSG00000026879 | 3.381558553 | 3.128123393 | 3.166000691 | 2.747348625 | 2.855232378 | 3.154119526 | 2.736525424 |
| ENSMUSG00000026880 | 2.381433035 | 2.205044741 | 2.195339928 | 2.21293726 | 2.163285805 | 2.491361694 | 2.041356071 |
| ENSMUSG00000026882 | 0 | 0 | 0 | 0 | 0 | 0 | 0 |
| ENSMUSG00000026883 | 2.438154344 | 2.332168732 | 2.359697963 | 2.004202072 | 2.175379703 | 2.290034611 | 2.117986249 |
| ENSMUSG00000026885 | 1 | 1.02186571 | 1.04248125 | 0.580482024 | 0.568060967 | 0.477121255 | 0.579661877 |
| ENSMUSG00000026887 | 1.903481042 | 1.928561379 | 1.884789703 | 1.852347734 | 1.887294348 | 2.021189299 | 1.734460864 |
| ENSMUSG00000026888 | 1.800532393 | 1.76109853 | 1.799918086 | 0.864857905 | 0.931471446 | 1 | 0.803905701 |
| ENSMUSG00000026889 | 2.446507203 | 2.261780978 | 2.21293726 | 2.278435542 | 2.214715681 | 2.532754379 | 2.041356071 |
| ENSMUSG00000026890 | 1.505869732 | 1.602347734 | 1.488549078 | 1.445339928 | 1.433158226 | 1.662757832 | 1.502532226 |
| ENSMUSG00000026893 | 1.53838222 | 1.464495249 | 1.557204673 | 1.464495249 | 1.49445382 | 1.707570176 | 1.345462945 |
| ENSMUSG00000026894 | 0.289064826 | 0.5 | 0.5 | 0.646240625 | 0.489301084 | 0.477121255 | 0.579661877 |
| ENSMUSG00000026895 | 2.399009124 | 2.269203899 | 2.289336734 | 2.179419106 | 2.297309926 | 2.506505032 | 2.177974197 |
| ENSMUSG00000026896 | 1.505869732 | 1.532320754 | 1.464495249 | 1.807204673 | 1.836786029 | 2.041392685 | 1.669034113 |
| ENSMUSG00000026904 | 0 | 0 | 0 | 0 | 0 | 0 | 0 |
| ENSMUSG00000026914 | 2.944447344 | 2.682192712 | 2.689347502 | 2.70083098 | 2.598690736 | 3.006893708 | 2.434835182 |
| ENSMUSG00000026915 | 2.415874319 | 2.206637122 | 2.185366747 | 2.124961472 | 2.080465577 | 2.260071388 | 1.910475814 |
| ENSMUSG00000026917 | 2.69788693 | 2.476346751 | 2.453845824 | 2.505591953 | 2.436719872 | 2.812913357 | 2.302525964 |
| ENSMUSG00000026918 | 2.631440309 | 2.478969845 | 2.457826811 | 2.28743678 | 2.364629672 | 2.546542663 | 2.236700872 |
| ENSMUSG00000026919 | 0 | 0 | 0 | 0 | 0 | 0 | 0 |
| ENSMUSG00000026921 | 1.325351389 | 1.339388001 | 1.410964047 | 1.27186571 | 1.407936347 | 1.301029996 | 1.483307832 |
| ENSMUSG00000026922 | 2.026699189 | 1.938721876 | 2.031030328 | 1.899978211 | 1.982473755 | 2.08278537 | 1.899720848 |
| ENSMUSG00000026923 | 2.119110214 | 2.013820609 | 2.132845365 | 1.614857905 | 1.836786029 | 1.851258349 | 1.911356129 |
| ENSMUSG00000026924 | 2.683414322 | 2.474083101 | 2.46759118 | 2.442872367 | 2.449594021 | 2.748962861 | 2.310303152 |

| | | | | | | | |
|---|---|---|---|---|---|---|---|
| ENSMUSG00000026925 | 2.113897241 | 1.888647213 | 1.942046081 | 1.837182039 | 1.814781639 | 2.021189299 | 1.777090262 |
| ENSMUSG00000026926 | 2.790815687 | 2.552972074 | 2.543419284 | 2.595115266 | 2.545883696 | 2.919078092 | 2.380283128 |
| ENSMUSG00000026927 | 2.336265009 | 2.120953944 | 2.134789703 | 2.077084758 | 2.133621644 | 2.326335861 | 2.001334086 |
| ENSMUSG00000026928 | 0.289064826 | 0.396240625 | 0 | 0.25 | 0 | 0.301029996 | 0.224243824 |
| ENSMUSG00000026930 | 2.141794304 | 2.051142786 | 2.060793496 | 1.816696635 | 1.970729215 | 1.949390007 | 2.028761664 |
| ENSMUSG00000026931 | 0.289064826 | 0 | 0 | 0 | 0 | 0 | 0 |
| ENSMUSG00000026932 | 2.829338353 | 2.63040011 | 2.643411797 | 2.060793496 | 2.167109853 | 2.334453751 | 2.068965954 |
| ENSMUSG00000026933 | 2.651426165 | 2.441217898 | 2.429419106 | 2.319031101 | 2.32898417 | 2.583198774 | 2.171745067 |
| ENSMUSG00000026934 | 0 | 0 | 0.25 | 0 | 0 | 0 | 0 |
| ENSMUSG00000026936 | 0 | 0 | 0 | 0 | 0 | 0 | 0 |
| ENSMUSG00000026937 | 0 | 0 | 0 | 0 | 0 | 0 | 0 |
| ENSMUSG00000026938 | 0.289064826 | 0 | 0.580482024 | 0.25 | 0 | 0 | 0 |
| ENSMUSG00000026939 | 0 | 0.25 | 0 | 1.188721876 | 1.121713561 | 1.322219295 | 1.185218806 |
| ENSMUSG00000026940 | 0.960252568 | 0.75 | 0.951838731 | 0.701838731 | 0.77552387 | 0.77815125 | 0.775756176 |
| ENSMUSG00000026941 | 1.342375483 | 1.282320754 | 1.339388001 | 1.080482024 | 1.293076518 | 1.146128036 | 1.193409508 |
| ENSMUSG00000026942 | 2.530730035 | 2.372963274 | 2.30175358 | 2.036164561 | 2.147561328 | 2.376576957 | 2.025071334 |
| ENSMUSG00000026943 | 0 | 0 | 0 | 0 | 0 | 0 | 0 |
| ENSMUSG00000026944 | 2.219867581 | 2.181980114 | 2.24929487 | 2.228222334 | 2.362449578 | 2.561101384 | 2.31789776 |
| ENSMUSG00000026946 | 1.418409478 | 1.226722649 | 1.330482024 | 1.681980114 | 1.559656853 | 1.86332286 | 1.473249546 |
| ENSMUSG00000026950 | 0.811507563 | 1.080482024 | 0.830482024 | 0.896240625 | 0.931471446 | 0.903089987 | 0.853775828 |
| ENSMUSG00000026955 | 1.707474304 | 1.614857905 | 1.657339155 | 1.51109853 | 1.655839964 | 1.653212514 | 1.596378571 |
| ENSMUSG00000026956 | 2.649252229 | 2.450224975 | 2.564052172 | 2.001406137 | 2.09661982 | 2.271841607 | 2.036615833 |
| ENSMUSG00000026958 | 2.477243823 | 2.262462137 | 2.352877747 | 1.860735874 | 1.926994141 | 2.113943352 | 1.830929954 |
| ENSMUSG00000026959 | 0 | 0.25 | 0 | 0.25 | 0.244650542 | 0 | 0 |
| ENSMUSG00000026960 | 2.442947872 | 2.284887838 | 2.286164561 | 2.25 | 2.16634835 | 2.484299839 | 2.020712061 |
| ENSMUSG00000026961 | 0 | 0 | 0 | 0 | 0 | 0 | 0 |
| ENSMUSG00000026963 | 0 | 0 | 0 | 0 | 0 | 0 | 0 |
| ENSMUSG00000026965 | 2.799013148 | 2.561981878 | 2.572792675 | 2.491446071 | 2.54143406 | 2.870403905 | 2.384383506 |
| ENSMUSG00000026966 | 2.156259305 | 2.084962501 | 2.104463129 | 1.96759118 | 1.948833444 | 2.130333768 | 1.924281235 |
| ENSMUSG00000026969 | 0 | 0 | 0.25 | 0 | 0.244650542 | 0 | 0 |
| ENSMUSG00000026970 | 2.926153516 | 2.719512728 | 2.702039943 | 2.566989271 | 2.506289166 | 2.892094603 | 2.326050929 |
| ENSMUSG00000026971 | 0 | 0.5 | 0.25 | 0.5 | 0.244650542 | 0.602059991 | 0.448487648 |
| ENSMUSG00000026972 | 1.721148722 | 1.71759118 | 1.769203899 | 1.726722649 | 1.895599581 | 1.968482949 | 1.753626837 |
| ENSMUSG00000026974 | 2.442947872 | 2.28743678 | 2.268535366 | 2.215271726 | 2.25665701 | 2.498310554 | 2.108567375 |

| | | | | | | | |
|---|---|---|---|---|---|---|---|
| ENSMUSG00000026975 | 1.596667477 | 1.51109853 | 1.5 | 1.580482024 | 1.528558853 | 1.556302501 | 1.417653332 |
| ENSMUSG00000026976 | 0.747221736 | 0.396240625 | 0 | 0.25 | 0.568060967 | 0.301029996 | 0.520678036 |
| ENSMUSG00000026977 | 2.762449763 | 2.563461871 | 2.521202097 | 2.54745614 | 2.454008153 | 2.810232518 | 2.259963078 |
| ENSMUSG00000026979 | 0.45815691 | 0 | 0.25 | 0.5 | 0.568060967 | 0 | 0.579661877 |
| ENSMUSG00000026980 | 3.042274823 | 2.78472793 | 2.776640735 | 2.688930538 | 2.610798307 | 3.012837225 | 2.400671947 |
| ENSMUSG00000026981 | 0.289064826 | 0 | 0 | 0.701838731 | 0.244650542 | 0 | 0.448487648 |
| ENSMUSG00000026983 | 0 | 0 | 0 | 0 | 0 | 0 | 0 |
| ENSMUSG00000026984 | 0 | 0 | 0 | 0 | 0 | 0 | 0 |
| ENSMUSG00000026985 | 0 | 0 | 0 | 0 | 0 | 0 | 0 |
| ENSMUSG00000026986 | 0 | 0 | 0 | 0 | 0 | 0 | 0 |
| ENSMUSG00000026987 | 2.423434025 | 2.186208459 | 2.228222334 | 2.168989258 | 2.124299332 | 2.416640507 | 1.986347533 |
| ENSMUSG00000026988 | 2.121692466 | 1.981203126 | 2.029735268 | 1.681980114 | 1.706995316 | 1.826074803 | 1.635773653 |
| ENSMUSG00000026989 | 0 | 0 | 0 | 0 | 0 | 0.301029996 | 0 |
| ENSMUSG00000026991 | 2.648525058 | 2.436208459 | 2.451032755 | 2.439555804 | 2.385310841 | 2.698970004 | 2.267652107 |
| ENSMUSG00000026994 | 1 | 1.17510993 | 0.864857905 | 0.580482024 | 0.568060967 | 0.477121255 | 0.672731473 |
| ENSMUSG00000026999 | 1.867194479 | 1.799918086 | 1.76109853 | 2.006976499 | 1.961316494 | 2.181843588 | 1.818286533 |
| ENSMUSG00000027001 | 1.45815691 | 1.482684334 | 1.356566189 | 1.638647213 | 1.674752852 | 1.86332286 | 1.523605797 |
| ENSMUSG00000027002 | 3.014472122 | 2.744498842 | 2.716819685 | 2.901601303 | 2.791443417 | 3.2219356 | 2.58791106 |
| ENSMUSG00000027004 | 0 | 0 | 0 | 0.75 | 0.846351821 | 0.477121255 | 0.579661877 |
| ENSMUSG00000027006 | 2.961285253 | 2.708222504 | 2.67813175 | 2.791854536 | 2.709785497 | 3.145507171 | 2.520276735 |
| ENSMUSG00000027007 | 2.317377295 | 2.168106335 | 2.187886015 | 2.172499493 | 2.124299332 | 2.374748346 | 1.997988498 |
| ENSMUSG00000027009 | 0.578129653 | 0.701838731 | 0.5 | 0.75 | 0.686820904 | 0.698970004 | 0.355418052 |
| ENSMUSG00000027010 | 2.167941649 | 2.049918086 | 2.04248125 | 2.222945926 | 2.143518259 | 2.424881637 | 2.060206963 |
| ENSMUSG00000027011 | 2.340856238 | 2.125956434 | 2.142463902 | 2.320772088 | 2.25010405 | 2.597695186 | 2.05853792 |
| ENSMUSG00000027012 | 2.820759099 | 2.547147211 | 2.53199858 | 2.727473271 | 2.624383383 | 3.05422991 | 2.460461957 |
| ENSMUSG00000027014 | 2.192545868 | 2.04745614 | 2.001406137 | 2.237091808 | 2.176862714 | 2.51054501 | 2.035419841 |
| ENSMUSG00000027015 | 0 | 0.5 | 0.396240625 | 0.701838731 | 0.733951626 | 0.84509804 | 0.629532004 |
| ENSMUSG00000027016 | 0 | 0 | 0 | 0 | 0.387761935 | 0 | 0 |
| ENSMUSG00000027018 | 2.735572029 | 2.540912419 | 2.494319981 | 2.724083101 | 2.624175088 | 3.075911761 | 2.435009443 |
| ENSMUSG00000027022 | 0 | 0 | 0 | 0 | 0 | 0 | 0 |
| ENSMUSG00000027030 | 1.847794126 | 1.671625132 | 1.692046081 | 2.002806814 | 1.932951348 | 2.204119983 | 1.860017477 |
| ENSMUSG00000027035 | 2.632197861 | 2.390083209 | 2.37246199 | 2.296838018 | 2.227765453 | 2.525044807 | 2.07959139 |
| ENSMUSG00000027048 | 0 | 0 | 0.5 | 0 | 0 | 0 | 0 |
| ENSMUSG00000027067 | 3.22592627 | 2.910625757 | 2.912987153 | 3.125583645 | 3.028820309 | 3.537944959 | 2.822521089 |

| | | | | | | | |
|---|---|---|---|---|---|---|---|
| ENSMUSG00000027068 | 1.734388958 | 1.708222504 | 1.646240625 | 1.557204673 | 1.456697355 | 1.662757832 | 1.278288402 |
| ENSMUSG00000027070 | 1.876566388 | 1.741446071 | 1.741446071 | 2.662987153 | 2.629140555 | 2.994756945 | 2.410967356 |
| ENSMUSG00000027071 | 0 | 0.701838731 | 0.5 | 0.25 | 0.568060967 | 0.301029996 | 0.520678036 |
| ENSMUSG00000027072 | 0 | 0 | 0 | 0 | 0 | 0 | 0 |
| ENSMUSG00000027073 | 0 | 0 | 0 | 0 | 0 | 0 | 0 |
| ENSMUSG00000027074 | 0 | 0 | 0 | 0.5 | 0.77552387 | 0.477121255 | 0.448487648 |
| ENSMUSG00000027075 | 1.54867984 | 1.52186571 | 1.464495249 | 0.830482024 | 0.686820904 | 0.602059991 | 0.710836105 |
| ENSMUSG00000027076 | 1.663535556 | 1.584962501 | 1.580482024 | 1.54248125 | 1.689774529 | 1.875061263 | 1.641897157 |
| ENSMUSG00000027077 | 0 | 0.25 | 0.25 | 0 | 0 | 0 | 0.224243824 |
| ENSMUSG00000027078 | 1.639698646 | 1.51109853 | 1.54248125 | 1.79745614 | 1.72341484 | 2.033423755 | 1.702561601 |
| ENSMUSG00000027079 | 1.89911417 | 1.779735268 | 1.71759118 | 1.901832578 | 1.855665188 | 2.120573931 | 1.74047993 |
| ENSMUSG00000027080 | 2 | 1.935366747 | 1.870953944 | 2.143411797 | 2.057362051 | 2.411619706 | 1.953344547 |
| ENSMUSG00000027082 | 1.100572389 | 1.146240625 | 1.061981878 | 2.195339928 | 2.107593434 | 2.444044796 | 1.95794437 |
| ENSMUSG00000027086 | 1.837744666 | 1.610735874 | 1.701838731 | 1.75835575 | 1.736537397 | 1.963787827 | 1.594042709 |
| ENSMUSG00000027087 | 3.105467192 | 2.823080408 | 2.825231123 | 2.905970373 | 2.815993715 | 3.247482261 | 2.574043197 |
| ENSMUSG00000027088 | 1.907802659 | 1.76109853 | 1.732684334 | 1.950224975 | 1.880506521 | 2.079181246 | 1.746389053 |
| ENSMUSG00000027091 | 2.528794847 | 2.359697963 | 2.345385738 | 2.51109853 | 2.412520257 | 2.831869774 | 2.233793574 |
| ENSMUSG00000027099 | 2.352857998 | 2.200224975 | 2.111770807 | 2.250703754 | 2.136121934 | 2.462397998 | 2.021963573 |
| ENSMUSG00000027102 | 1.783508299 | 1.811981878 | 1.807204673 | 0.25 | 0.244650542 | 0 | 0 |
| ENSMUSG00000027104 | 2.496544396 | 2.28296424 | 2.27053726 | 2.360217292 | 2.301493158 | 2.604226053 | 2.123508695 |
| ENSMUSG00000027107 | 0 | 0.25 | 0.25 | 0.25 | 0.387761935 | 0 | 0 |
| ENSMUSG00000027108 | 2.550525119 | 2.361770807 | 2.312574604 | 2.606303976 | 2.505123331 | 2.902002891 | 2.351607989 |
| ENSMUSG00000027109 | 2.569423023 | 2.385757955 | 2.36434522 | 2.551142786 | 2.44788147 | 2.846955325 | 2.253616472 |
| ENSMUSG00000027111 | 2.271465912 | 2.099151195 | 2.120953944 | 2.291854536 | 2.24889942 | 2.558708571 | 2.082194103 |
| ENSMUSG00000027115 | 2.066740409 | 1.984159485 | 1.948603967 | 2.132845365 | 2.057362051 | 2.340444115 | 1.886704102 |
| ENSMUSG00000027120 | 0 | 0 | 0 | 0 | 0 | 0 | 0 |
| ENSMUSG00000027122 | 2.40033094 | 2.164552871 | 2.12694866 | 2.095926073 | 2.097547432 | 2.346352974 | 1.922587433 |
| ENSMUSG00000027130 | 2.033115198 | 1.907339155 | 1.931980114 | 2.05479213 | 1.950242452 | 2.227886705 | 1.824094866 |
| ENSMUSG00000027131 | 2.029919532 | 1.926839783 | 1.96759118 | 2.131869252 | 2.158641579 | 2.424881637 | 2.032410373 |
| ENSMUSG00000027132 | 2.097855556 | 1.931980114 | 1.972185812 | 2.152756199 | 2.085294053 | 2.33243846 | 1.892346957 |
| ENSMUSG00000027133 | 2.026699189 | 1.868933358 | 1.862802778 | 2.112802778 | 2.022473809 | 2.382017043 | 1.922587433 |
| ENSMUSG00000027134 | 1.482695304 | 1.54248125 | 1.708222504 | 1.75835575 | 1.905314583 | 1.924279286 | 1.935895857 |
| ENSMUSG00000027157 | 0 | 0 | 0 | 0 | 0 | 0 | 0 |
| ENSMUSG00000027160 | 2.188178997 | 2.074802005 | 2.114857905 | 2.061981878 | 2.007252778 | 2.301029996 | 1.909593098 |

| | | | | | | | |
|---|---|---|---|---|---|---|---|
| ENSMUSG00000027162 | 2.623788124 | 2.451435968 | 2.442872367 | 2.710194481 | 2.617231464 | 3.001300933 | 2.406622266 |
| ENSMUSG00000027163 | 1.956444026 | 1.852347734 | 1.839388001 | 1.825945187 | 1.948833444 | 2 | 1.910475814 |
| ENSMUSG00000027164 | 2.160971586 | 2 | 1.938721876 | 2.203445298 | 2.143518259 | 2.413299764 | 1.980010793 |
| ENSMUSG00000027165 | 1.759671389 | 1.714495249 | 1.668106335 | 1.956637122 | 1.861137485 | 2.152288344 | 1.764880683 |
| ENSMUSG00000027166 | 1.45815691 | 1.476722649 | 1.380890489 | 1.557204673 | 1.53772706 | 1.716003344 | 1.523605797 |
| ENSMUSG00000027167 | 1.83262764 | 1.763820609 | 1.752806814 | 1.888647213 | 1.840643511 | 2.041392685 | 1.71730663 |
| ENSMUSG00000027168 | 0.867194479 | 0.75 | 0.79248125 | 1.614857905 | 1.546663136 | 1.579783597 | 1.496250175 |
| ENSMUSG00000027170 | 2.42468076 | 2.267865591 | 2.281030328 | 2.464495249 | 2.386946794 | 2.767155866 | 2.22088434 |
| ENSMUSG00000027171 | 0.578129653 | 0.896240625 | 0.896240625 | 1.27186571 | 1.223252711 | 1.322219295 | 1.168187745 |
| ENSMUSG00000027173 | 1.837744666 | 1.695339928 | 1.766522298 | 1.978222334 | 1.98503142 | 2.152288344 | 1.821784031 |
| ENSMUSG00000027175 | 1.837744666 | 1.671625132 | 1.638647213 | 1.923371739 | 1.848234258 | 2.086359831 | 1.757893036 |
| ENSMUSG00000027176 | 2.578990401 | 2.37246199 | 2.326515422 | 2.374462959 | 2.333285652 | 2.617000341 | 2.16858494 |
| ENSMUSG00000027177 | 2.612039956 | 2.399047439 | 2.382845365 | 2.559003548 | 2.476238662 | 2.818225894 | 2.304357435 |
| ENSMUSG00000027180 | 2.395018353 | 2.235628626 | 2.228222334 | 2.512121718 | 2.42519375 | 2.783903579 | 2.247765009 |
| ENSMUSG00000027184 | 3.373320295 | 3.069394501 | 3.033766986 | 3.216964466 | 3.128667098 | 3.621176282 | 2.891115648 |
| ENSMUSG00000027185 | 2.251208701 | 2.110735874 | 2.162762923 | 2.184523065 | 2.229681962 | 2.469822016 | 2.074322292 |
| ENSMUSG00000027186 | 0.289064826 | 0 | 0 | 0.396240625 | 0 | 0 | 0.355418052 |
| ENSMUSG00000027187 | 2.256831695 | 2.169870025 | 2.122963274 | 2.275821952 | 2.288791717 | 2.572871602 | 2.176428123 |
| ENSMUSG00000027188 | 0.289064826 | 0.396240625 | 0 | 0.646240625 | 0.846351821 | 0.84509804 | 0.969165684 |
| ENSMUSG00000027189 | 2.765759561 | 2.519203899 | 2.497525991 | 2.711764337 | 2.614672255 | 3.020361283 | 2.439163791 |
| ENSMUSG00000027193 | 2.854509768 | 2.627938163 | 2.601816941 | 2.798073204 | 2.703517887 | 3.131618664 | 2.524002992 |
| ENSMUSG00000027194 | 2.103271637 | 1.990000483 | 1.982684334 | 2.075945187 | 2.025895058 | 2.31386722 | 1.914853627 |
| ENSMUSG00000027195 | 2.506893125 | 2.313166358 | 2.331607372 | 2.495034894 | 2.395014713 | 2.789580712 | 2.256351705 |
| ENSMUSG00000027196 | 0.45815691 | 0.396240625 | 0.396240625 | 0.646240625 | 0.632412477 | 0.698970004 | 0.448487648 |
| ENSMUSG00000027198 | 2.450036412 | 2.289336734 | 2.325374049 | 2.451435968 | 2.512061267 | 2.820857989 | 2.382954278 |
| ENSMUSG00000027199 | 0 | 0.25 | 0 | 0.864857905 | 1.074582839 | 1.146128036 | 0.935079929 |
| ENSMUSG00000027200 | 0.91631382 | 1.02186571 | 0.79248125 | 0.25 | 0.244650542 | 0 | 0.224243824 |
| ENSMUSG00000027201 | 2.523917219 | 2.3184489 | 2.311981878 | 2.441632227 | 2.404846996 | 2.779596491 | 2.232493008 |
| ENSMUSG00000027202 | 0.45815691 | 0.396240625 | 0.396240625 | 0.396240625 | 0.244650542 | 0 | 0.224243824 |
| ENSMUSG00000027203 | 2.336265009 | 2.147178659 | 2.144357207 | 2.399978211 | 2.268268456 | 2.663700925 | 2.178744473 |
| ENSMUSG00000027204 | 2.899911556 | 2.655741736 | 2.642226539 | 2.839800121 | 2.811855561 | 3.203304916 | 2.60266469 |
| ENSMUSG00000027206 | 2.924654746 | 2.703244868 | 2.691217898 | 2.797610505 | 2.690120394 | 3.111598525 | 2.489696634 |
| ENSMUSG00000027207 | 2.336265009 | 2.178561379 | 2.192872367 | 2.05479213 | 2 | 2.260071388 | 1.913982791 |
| ENSMUSG00000027208 | 1.129344651 | 0.830482024 | 0.976722649 | 1.566696635 | 1.450958105 | 1.612783857 | 1.360283038 |

| | | | | | | | |
|---|---|---|---|---|---|---|---|
| ENSMUSG00000027209 | 0.289064826 | 0.396240625 | 0.25 | 0.25 | 0 | 0 | 0 |
| ENSMUSG00000027210 | 1.700465191 | 1.735628626 | 1.732684334 | 1.78743678 | 1.749189509 | 1.949390007 | 1.670888075 |
| ENSMUSG00000027215 | 2.095120908 | 1.982684334 | 1.982684334 | 2.124961472 | 2.173143437 | 2.404833717 | 2.101807105 |
| ENSMUSG00000027217 | 1.181541736 | 1.372963274 | 1.25 | 0.951838731 | 0.931471446 | 0.77815125 | 0.916588299 |
| ENSMUSG00000027219 | 0 | 0 | 0.25 | 0 | 0 | 0 | 0 |
| ENSMUSG00000027220 | 1.389637216 | 1.356566189 | 1.356566189 | 1.905512955 | 1.938809675 | 2.127104798 | 1.872150729 |
| ENSMUSG00000027221 | 2.121692466 | 2.044977273 | 2.107613138 | 1.364857905 | 1.542223377 | 1.568201724 | 1.526507301 |
| ENSMUSG00000027222 | 1.227926429 | 1.27186571 | 1.403677461 | 1.339388001 | 1.504538967 | 1.431363764 | 1.554175045 |
| ENSMUSG00000027223 | 1.482695304 | 1.42510993 | 1.464495249 | 1.825945187 | 1.86473911 | 2.004321374 | 1.830929954 |
| ENSMUSG00000027224 | 0.867194479 | 0.976722649 | 1.04248125 | 0.5 | 0.733951626 | 0.301029996 | 0 |
| ENSMUSG00000027225 | 0 | 0 | 0.25 | 0 | 0 | 0 | 0 |
| ENSMUSG00000027227 | 2.526850636 | 2.28743678 | 2.367918804 | 1.953445298 | 1.929985312 | 2.146128036 | 1.792684387 |
| ENSMUSG00000027229 | 0 | 0 | 0 | 0 | 0 | 0 | 0 |
| ENSMUSG00000027230 | 2.52293481 | 2.379417347 | 2.424676167 | 2.075945187 | 2.154724156 | 2.328379603 | 2.182568577 |
| ENSMUSG00000027233 | 0 | 0.25 | 0 | 0 | 0 | 0 | 0.224243824 |
| ENSMUSG00000027236 | 2.051791411 | 1.923371739 | 1.87892496 | 2.044977273 | 1.964031554 | 2.240549248 | 1.854824501 |
| ENSMUSG00000027238 | 0.747221736 | 0.25 | 0.79248125 | 1.130890489 | 1.074582839 | 1 | 1.140832123 |
| ENSMUSG00000027239 | 0.289064826 | 0.396240625 | 0.25 | 0.976722649 | 1.149965125 | 1.230448921 | 1.290497981 |
| ENSMUSG00000027242 | 2.330063669 | 2.190387808 | 2.137686696 | 2.269871196 | 2.212715924 | 2.469822016 | 2.027536225 |
| ENSMUSG00000027243 | 1.505869732 | 1.321350555 | 1.410964047 | 1.348079356 | 1.394615667 | 1.591064607 | 1.290497981 |
| ENSMUSG00000027244 | 2.029919532 | 1.890560606 | 1.880890489 | 1.972185812 | 2 | 2.204119983 | 1.927642465 |
| ENSMUSG00000027245 | 1.587501561 | 1.488549078 | 1.561981878 | 1.494319981 | 1.49445382 | 1.672097858 | 1.429563222 |
| ENSMUSG00000027246 | 0.289064826 | 0.25 | 0 | 0.396240625 | 0 | 0 | 0 |
| ENSMUSG00000027247 | 2.783508299 | 2.529735268 | 2.53743678 | 2.523189285 | 2.49893871 | 2.849419414 | 2.357638483 |
| ENSMUSG00000027248 | 3.526052658 | 3.199466034 | 3.172281101 | 3.262164288 | 3.140250123 | 3.686189234 | 2.913069858 |
| ENSMUSG00000027249 | 0 | 0 | 0 | 0.25 | 0 | 0 | 0.224243824 |
| ENSMUSG00000027253 | 1.686083339 | 1.649978211 | 1.642463902 | 1.755591953 | 1.848234258 | 2.123851641 | 1.790137008 |
| ENSMUSG00000027254 | 2.634462296 | 2.459012589 | 2.501055117 | 2.319612365 | 2.360255934 | 2.640481437 | 2.240537069 |
| ENSMUSG00000027255 | 2.013563224 | 1.862802778 | 1.862802778 | 2.067865757 | 2.079491904 | 2.301029996 | 1.951796623 |
| ENSMUSG00000027257 | 1.816888701 | 1.799918086 | 1.899978211 | 1.698603967 | 1.855665188 | 1.875061263 | 1.910475814 |
| ENSMUSG00000027259 | 1.912079952 | 1.856566189 | 1.79248125 | 1.823655187 | 1.761403742 | 1.924279286 | 1.649885598 |
| ENSMUSG00000027261 | 0.289064826 | 0.25 | 0.25 | 0 | 0 | 0 | 0 |
| ENSMUSG00000027263 | 2.351376531 | 2.151832578 | 2.153677461 | 2.210587586 | 2.208682096 | 2.484299839 | 2.04833869 |
| ENSMUSG00000027270 | 0 | 0 | 0 | 0 | 0 | 0 | 0 |

| | | | | | | | |
|---|---|---|---|---|---|---|---|
| ENSMUSG00000027272 | 2.524897319 | 2.310197833 | 2.288704527 | 2.449820405 | 2.356272688 | 2.702430536 | 2.190083792 |
| ENSMUSG00000027273 | 0 | 0 | 0 | 0.25 | 0 | 0 | 0 |
| ENSMUSG00000027274 | 2.00683675 | 1.921625132 | 1.888647213 | 1.856566189 | 1.848234258 | 2.086359831 | 1.645916034 |
| ENSMUSG00000027276 | 1.978790393 | 1.866901388 | 1.799918086 | 1.364857905 | 1.283908311 | 1.380211242 | 1.313616061 |
| ENSMUSG00000027281 | 1.783508299 | 1.649978211 | 1.668106335 | 1.837182039 | 1.758994476 | 2 | 1.616675393 |
| ENSMUSG00000027282 | 2.745494892 | 2.519871196 | 2.525165585 | 2.796373731 | 2.706005256 | 3.144262774 | 2.538493112 |
| ENSMUSG00000027284 | 2.119110214 | 2.009729747 | 2.024508021 | 1.955044741 | 2.001219193 | 2.164352856 | 1.861046144 |
| ENSMUSG00000027285 | 2.196867486 | 2.03360658 | 2.025821952 | 2.212155735 | 2.099395374 | 2.334453751 | 1.972821265 |
| ENSMUSG00000027286 | 1.852729478 | 1.807204673 | 1.825945187 | 1.850219859 | 1.84445929 | 1.973127854 | 1.723650273 |
| ENSMUSG00000027287 | 2.470606563 | 2.27186571 | 2.259729747 | 2.353935442 | 2.260764609 | 2.575187845 | 2.109521699 |
| ENSMUSG00000027288 | 3.119580905 | 2.820627327 | 2.822648722 | 2.870701989 | 2.810506292 | 3.230448921 | 2.595425028 |
| ENSMUSG00000027291 | 2.330063669 | 2.145300145 | 2.139605178 | 2.218361028 | 2.187072901 | 2.480006943 | 2.060761403 |
| ENSMUSG00000027293 | 2.158622101 | 2.028435542 | 2.095926073 | 2.464495249 | 2.411760395 | 2.759667845 | 2.285558718 |
| ENSMUSG00000027296 | 0.578129653 | 0.396240625 | 0.646240625 | 0.701838731 | 0.931471446 | 0.477121255 | 0.935079929 |
| ENSMUSG00000027297 | 0.289064826 | 0.25 | 0 | 1.02186571 | 1.163285805 | 1.079181246 | 0.935079929 |
| ENSMUSG00000027298 | 2.066740409 | 1.961372513 | 1.96293726 | 2.069031101 | 2.16634835 | 2.394451681 | 2.123964672 |
| ENSMUSG00000027300 | 1.655740331 | 1.575945187 | 1.602347734 | 1.602347734 | 1.695608647 | 1.707570176 | 1.589319772 |
| ENSMUSG00000027301 | 0 | 0 | 0 | 0 | 0 | 0.301029996 | 0 |
| ENSMUSG00000027303 | 2.575536697 | 2.375459296 | 2.367918804 | 2.539967834 | 2.48561803 | 2.848804701 | 2.334003665 |
| ENSMUSG00000027304 | 2.46499366 | 2.278435542 | 2.284247778 | 2.512121718 | 2.44788147 | 2.810232518 | 2.302000776 |
| ENSMUSG00000027305 | 1.800532393 | 1.711372513 | 1.744319981 | 1.991446071 | 1.902105869 | 2.158362492 | 1.801445495 |
| ENSMUSG00000027306 | 2.541215974 | 2.339388001 | 2.348079356 | 2.649047439 | 2.571222152 | 2.986771734 | 2.4027959 |
| ENSMUSG00000027309 | 2.213719572 | 2.069031101 | 2.070192693 | 2.222185812 | 2.212046813 | 2.488550717 | 2.026921761 |
| ENSMUSG00000027312 | 2.475043119 | 2.319031101 | 2.31728167 | 2.483053688 | 2.417419887 | 2.726727209 | 2.252086845 |
| ENSMUSG00000027313 | 1.940852214 | 1.76109853 | 1.720660762 | 1.214495249 | 1.358933174 | 1.301029996 | 1.379042812 |
| ENSMUSG00000027314 | 0 | 0 | 0 | 0.5 | 0.489301084 | 0.698970004 | 0.224243824 |
| ENSMUSG00000027315 | 0.811507563 | 0.830482024 | 0.864857905 | 1.828220739 | 1.773209395 | 1.973127854 | 1.722075986 |
| ENSMUSG00000027316 | 0.289064826 | 0.75 | 0.646240625 | 0.701838731 | 0.846351821 | 0.602059991 | 0.853775828 |
| ENSMUSG00000027317 | 0 | 0 | 0 | 0 | 0.244650542 | 0 | 0 |
| ENSMUSG00000027318 | 0 | 0.25 | 0.25 | 0 | 0.244650542 | 0 | 0.355418052 |
| ENSMUSG00000027322 | 0.671187741 | 0.976722649 | 0.79248125 | 0.701838731 | 0.931471446 | 0.84509804 | 0.853775828 |
| ENSMUSG00000027323 | 2.404271433 | 2.235628626 | 2.204245906 | 2.460194481 | 2.425559697 | 2.72916479 | 2.262645396 |
| ENSMUSG00000027324 | 1.404271433 | 1.372963274 | 1.42510993 | 1.458222504 | 1.49445382 | 1.556302501 | 1.448487648 |
| ENSMUSG00000027326 | 2.445324132 | 2.296218836 | 2.321350555 | 2.588837774 | 2.500716831 | 2.815577748 | 2.279980727 |

| | | | | | | | |
|---|---|---|---|---|---|---|---|
| ENSMUSG00000027327 | 1.944805158 | 1.850219859 | 1.894357207 | 1.79248125 | 1.902105869 | 2.045322979 | 1.889537832 |
| ENSMUSG00000027329 | 0.811507563 | 0.830482024 | 0.896240625 | 0.79248125 | 0.905314583 | 0.903089987 | 1.028149525 |
| ENSMUSG00000027330 | 2.273259602 | 2.121960008 | 2.188721876 | 2.265173983 | 2.258423266 | 2.546542663 | 2.149789616 |
| ENSMUSG00000027331 | 2.43085913 | 2.173371739 | 2.228222334 | 2.462155735 | 2.3873546 | 2.737192643 | 2.249315158 |
| ENSMUSG00000027332 | 2.317377295 | 2.142463902 | 2.192872367 | 2.338286706 | 2.359374645 | 2.6599162 | 2.241489052 |
| ENSMUSG00000027333 | 1.342375483 | 1.464495249 | 1.494319981 | 1.364857905 | 1.572189156 | 1.579783597 | 1.637827726 |
| ENSMUSG00000027335 | 0.747221736 | 1.04248125 | 1.114857905 | 0.396240625 | 0.244650542 | 0 | 0.520678036 |
| ENSMUSG00000027338 | 0 | 0 | 0 | 0 | 0 | 0 | 0 |
| ENSMUSG00000027339 | 2.188178997 | 2.029735268 | 2.001406137 | 1.802363341 | 1.72341484 | 1.84509804 | 1.655750087 |
| ENSMUSG00000027340 | 2.192545868 | 2.060793496 | 2.109697963 | 2.245748394 | 2.245869735 | 2.534026106 | 2.07959139 |
| ENSMUSG00000027341 | 1.852729478 | 1.729715809 | 1.71759118 | 1.856566189 | 1.850106709 | 1.995635195 | 1.68886578 |
| ENSMUSG00000027342 | 2.685417684 | 2.520204382 | 2.539020769 | 2.5684489 | 2.534249632 | 2.888740961 | 2.40946263 |
| ENSMUSG00000027344 | 0.45815691 | 0.580482024 | 0.25 | 0.396240625 | 0.632412477 | 0.602059991 | 0.803905701 |
| ENSMUSG00000027345 | 0 | 0 | 0 | 0 | 0 | 0 | 0 |
| ENSMUSG00000027346 | 1.989530018 | 1.837182039 | 1.852347734 | 2.132845365 | 2.045754077 | 2.255272505 | 1.865128412 |
| ENSMUSG00000027347 | 0.289064826 | 0.25 | 0.396240625 | 0 | 0.244650542 | 0.477121255 | 0.224243824 |
| ENSMUSG00000027349 | 2.283862692 | 2.118933358 | 2.089388001 | 2.350753006 | 2.260180728 | 2.613841822 | 2.140399326 |
| ENSMUSG00000027350 | 0 | 0.396240625 | 0 | 0 | 0.489301084 | 0 | 0 |
| ENSMUSG00000027351 | 2.53552908 | 2.329353153 | 2.309601185 | 2.381380202 | 2.283908311 | 2.617000341 | 2.119837558 |
| ENSMUSG00000027353 | 1.867194479 | 1.854463129 | 1.78743678 | 2.031030328 | 1.987570683 | 2.260071388 | 1.833176598 |
| ENSMUSG00000027355 | 0 | 0 | 0 | 0 | 0 | 0 | 0 |
| ENSMUSG00000027356 | 0.578129653 | 0 | 0.25 | 0 | 0 | 0 | 0 |
| ENSMUSG00000027357 | 1.944805158 | 1.896240625 | 1.744319981 | 2.083847589 | 1.92093466 | 2.198657087 | 1.818286533 |
| ENSMUSG00000027358 | 0.91631382 | 0.896240625 | 0.701838731 | 1.720660762 | 1.639266209 | 1.86923172 | 1.594042709 |
| ENSMUSG00000027359 | 0.45815691 | 0 | 0 | 0 | 0 | 0 | 0.224243824 |
| ENSMUSG00000027360 | 0 | 0 | 0 | 0 | 0.244650542 | 0.301029996 | 0.224243824 |
| ENSMUSG00000027361 | 2.235832493 | 2.157339155 | 2.097004321 | 2.174241882 | 2.093822267 | 2.434568904 | 1.987739 |
| ENSMUSG00000027363 | 2.45815691 | 2.289967834 | 2.311981878 | 2.479715809 | 2.398190943 | 2.760422483 | 2.266481009 |
| ENSMUSG00000027364 | 0 | 0.25 | 0 | 0.25 | 0.387761935 | 0.301029996 | 0.448487648 |
| ENSMUSG00000027365 | 2.784560084 | 2.5964656 | 2.58076369 | 2.732499516 | 2.629345941 | 3.02325246 | 2.420398883 |
| ENSMUSG00000027366 | 2.589183146 | 2.348615674 | 2.360217292 | 2.58412664 | 2.498344007 | 2.880813592 | 2.310303152 |
| ENSMUSG00000027367 | 2.489144387 | 2.295598588 | 2.332168732 | 2.581607372 | 2.498641484 | 2.835690571 | 2.313108585 |
| ENSMUSG00000027368 | 0.578129653 | 0.830482024 | 0.830482024 | 0.864857905 | 0.812711509 | 0.77815125 | 0.710836105 |
| ENSMUSG00000027371 | 0 | 0 | 0 | 1.061981878 | 1.188507684 | 1.146128036 | 1.150210039 |

| | | | | | | | |
|---|---|---|---|---|---|---|---|
| ENSMUSG00000027374 | 2.086807695 | 1.921625132 | 1.894357207 | 2.041226732 | 2.105788027 | 2.230448921 | 1.947894097 |
| ENSMUSG00000027375 | 0 | 0 | 0 | 1 | 0.931471446 | 0.903089987 | 0.775756176 |
| ENSMUSG00000027376 | 0 | 0 | 0 | 0 | 0 | 0 | 0 |
| ENSMUSG00000027377 | 0 | 0 | 0 | 0.5 | 0.387761935 | 0.301029996 | 0.224243824 |
| ENSMUSG00000027378 | 1.932832114 | 1.834962501 | 1.823655187 | 2.041226732 | 1.990091809 | 2.240549248 | 1.857950279 |
| ENSMUSG00000027379 | 2.629921064 | 2.44697564 | 2.431128463 | 2.706040802 | 2.618715812 | 3.046495164 | 2.415994748 |
| ENSMUSG00000027380 | 0.811507563 | 0.5 | 0.5 | 0 | 0 | 0 | 0 |
| ENSMUSG00000027381 | 2 | 1.852347734 | 1.852347734 | 1.92510993 | 1.900490506 | 2.096910013 | 1.808770687 |
| ENSMUSG00000027384 | 1.623015126 | 1.598079356 | 1.62694866 | 1.794977273 | 1.649303553 | 1.977723605 | 1.637827726 |
| ENSMUSG00000027386 | 0.289064826 | 0.580482024 | 0.646240625 | 0.396240625 | 0.931471446 | 0.477121255 | 0.829800754 |
| ENSMUSG00000027387 | 1.389637216 | 1.348079356 | 1.364857905 | 1.5 | 1.504538967 | 1.62324929 | 1.355418052 |
| ENSMUSG00000027394 | 2.196867486 | 2.056000419 | 2.05479213 | 2.05479213 | 2.075570129 | 2.31386722 | 2.005960535 |
| ENSMUSG00000027395 | 2.042557845 | 1.890560606 | 1.880890489 | 2.052363341 | 2.023617914 | 2.276461804 | 1.951796623 |
| ENSMUSG00000027397 | 2.594187595 | 2.334962501 | 2.375956434 | 2.388167274 | 2.336595379 | 2.698100546 | 2.202472158 |
| ENSMUSG00000027398 | 0 | 0 | 0 | 0 | 0 | 0 | 0 |
| ENSMUSG00000027399 | 0 | 0.25 | 0.396240625 | 0.25 | 0.244650542 | 0.301029996 | 0 |
| ENSMUSG00000027400 | 0 | 0 | 0 | 0 | 0 | 0 | 0 |
| ENSMUSG00000027401 | 0.289064826 | 0 | 0.25 | 0 | 0.244650542 | 0 | 0 |
| ENSMUSG00000027403 | 0 | 0.25 | 0 | 0 | 0 | 0 | 0 |
| ENSMUSG00000027404 | 2.686083339 | 2.579070383 | 2.636241108 | 2.490000483 | 2.619350055 | 2.875061263 | 2.608015577 |
| ENSMUSG00000027405 | 2.848695818 | 2.599685923 | 2.615369862 | 2.714689525 | 2.692360299 | 3.045714059 | 2.495239414 |
| ENSMUSG00000027406 | 2.528794847 | 2.319612365 | 2.329353153 | 2.489638179 | 2.458367761 | 2.777426822 | 2.335902514 |
| ENSMUSG00000027408 | 0.289064826 | 0.646240625 | 0.830482024 | 0.951838731 | 1.020174412 | 1.176091259 | 1.089372236 |
| ENSMUSG00000027409 | 0.289064826 | 0 | 0.25 | 0 | 0.244650542 | 0.301029996 | 0 |
| ENSMUSG00000027411 | 2.3855553 | 2.215271726 | 2.26584877 | 2.28360658 | 2.29993025 | 2.59439255 | 2.174095069 |
| ENSMUSG00000027412 | 1.37447073 | 1.494319981 | 1.418106335 | 0.79248125 | 1.057362051 | 1.041392685 | 1.028149525 |
| ENSMUSG00000027416 | 0 | 0 | 0 | 0 | 0 | 0 | 0 |
| ENSMUSG00000027419 | 0.671187741 | 0.396240625 | 0.5 | 0.396240625 | 0 | 0.301029996 | 0.448487648 |
| ENSMUSG00000027420 | 0.289064826 | 0 | 0.5 | 0 | 0 | 0 | 0.355418052 |
| ENSMUSG00000027422 | 2.904664113 | 2.714883697 | 2.690179987 | 2.606303976 | 2.656791073 | 2.975891136 | 2.53582519 |
| ENSMUSG00000027423 | 2.829809843 | 2.590485943 | 2.576800202 | 2.710194481 | 2.625007532 | 3.021189299 | 2.421853705 |
| ENSMUSG00000027424 | 1.89911417 | 1.698603967 | 1.668106335 | 1.747171172 | 1.749189509 | 1.913813852 | 1.596378571 |
| ENSMUSG00000027425 | 2.309247751 | 2.127938163 | 2.175975893 | 2.120953944 | 2.118236133 | 2.350248018 | 2.019455688 |
| ENSMUSG00000027427 | 2.215778996 | 2.041226732 | 2.114857905 | 2.316696635 | 2.24829556 | 2.57863921 | 2.064616101 |

| | | | | | | | |
|---|---|---|---|---|---|---|---|
| ENSMUSG00000027428 | 2.010213549 | 1.809601185 | 1.802363341 | 2.044977273 | 1.993840051 | 2.230448921 | 1.826389311 |
| ENSMUSG00000027429 | 2.507914013 | 2.30357978 | 2.277131114 | 2.541854536 | 2.481584202 | 2.85308953 | 2.334241631 |
| ENSMUSG00000027430 | 1.45815691 | 1.403677461 | 1.302363341 | 1.681980114 | 1.709785497 | 1.880813592 | 1.69749337 |
| ENSMUSG00000027431 | 0 | 0 | 0 | 0 | 0 | 0 | 0 |
| ENSMUSG00000027433 | 2.840049992 | 2.603671309 | 2.57163944 | 2.715659339 | 2.618504144 | 3.01494035 | 2.428142734 |
| ENSMUSG00000027434 | 0 | 0 | 0 | 0 | 0 | 0.301029996 | 0 |
| ENSMUSG00000027435 | 0.91631382 | 0.896240625 | 0.75 | 0.5 | 0.244650542 | 0.602059991 | 0.629532004 |
| ENSMUSG00000027438 | 1.432083896 | 1.26109853 | 1.410964047 | 1.464495249 | 1.450958105 | 1.462397998 | 1.452143229 |
| ENSMUSG00000027439 | 2.201144779 | 2.005591953 | 1.969129237 | 2.147178659 | 2.106691885 | 2.369215857 | 1.981429723 |
| ENSMUSG00000027442 | 0 | 0 | 0 | 0 | 0 | 0 | 0 |
| ENSMUSG00000027443 | 0 | 0 | 0 | 0 | 0 | 0 | 0 |
| ENSMUSG00000027444 | 0 | 0 | 0 | 0 | 0 | 0 | 0 |
| ENSMUSG00000027445 | 0 | 0 | 0 | 0 | 0 | 0 | 0 |
| ENSMUSG00000027446 | 0 | 0.25 | 0 | 0 | 0 | 0 | 0 |
| ENSMUSG00000027447 | 2.37447073 | 2.256284891 | 2.328787391 | 2.152756199 | 2.163285805 | 2.488550717 | 2.200320163 |
| ENSMUSG00000027452 | 0 | 0 | 0 | 0.25 | 0 | 0.301029996 | 0.224243824 |
| ENSMUSG00000027454 | 1.789260561 | 1.755591953 | 1.79745614 | 1.92510993 | 1.897237431 | 2.113943352 | 1.817112247 |
| ENSMUSG00000027455 | 2.434522689 | 2.253505118 | 2.235628626 | 2.369439567 | 2.408704478 | 2.727541257 | 2.245894972 |
| ENSMUSG00000027456 | 0.747221736 | 0.396240625 | 0.580482024 | 0.5 | 0.387761935 | 0 | 0.355418052 |
| ENSMUSG00000027457 | 0.45815691 | 0.396240625 | 0.396240625 | 0.79248125 | 0.955822902 | 1.041392685 | 0.935079929 |
| ENSMUSG00000027459 | 1.37447073 | 1.396240625 | 1.53743678 | 1.372963274 | 1.504538967 | 1.505149978 | 1.586932193 |
| ENSMUSG00000027460 | 0.578129653 | 0.396240625 | 0.580482024 | 0.580482024 | 0.733951626 | 0.301029996 | 0.710836105 |
| ENSMUSG00000027463 | 0 | 0 | 0.646240625 | 0.25 | 0.686820904 | 0.602059991 | 0.579661877 |
| ENSMUSG00000027465 | 2.592526109 | 2.370449858 | 2.347004321 | 2.433677405 | 2.439894911 | 2.741151599 | 2.275733199 |
| ENSMUSG00000027466 | 2.105953527 | 2.038704527 | 2.098079356 | 1.928561379 | 2.097547432 | 2.212187604 | 2.112367894 |
| ENSMUSG00000027468 | 0 | 0 | 0 | 0 | 0 | 0 | 0 |
| ENSMUSG00000027469 | 3.002499078 | 2.707430684 | 2.724839231 | 2.87246199 | 2.800408677 | 3.226599905 | 2.608117616 |
| ENSMUSG00000027470 | 1 | 0.79248125 | 0.976722649 | 0.25 | 0 | 0 | 0 |
| ENSMUSG00000027472 | 2.253091469 | 2.04745614 | 2.057204673 | 2.129909063 | 2.140250123 | 2.423245874 | 1.988432495 |
| ENSMUSG00000027474 | 0 | 0 | 0 | 0 | 0 | 0 | 0 |
| ENSMUSG00000027475 | 2.40817504 | 2.187048212 | 2.181128463 | 2.388167274 | 2.319233381 | 2.681241237 | 2.197065089 |
| ENSMUSG00000027478 | 1.205378646 | 1.330482024 | 1.160964047 | 1.527131114 | 1.514343934 | 1.568201724 | 1.537859885 |
| ENSMUSG00000027479 | 3.095619452 | 2.826515422 | 2.788704527 | 3.063683347 | 2.957771595 | 3.447778009 | 2.743044848 |
| ENSMUSG00000027480 | 0 | 0 | 0 | 0 | 0 | 0 | 0 |

| | | | | | | | |
|---|---|---|---|---|---|---|---|
| ENSMUSG00000027481 | 0 | 0 | 0 | 0 | 0 | 0 | 0 |
| ENSMUSG00000027482 | 0 | 0 | 0 | 0 | 0 | 0 | 0 |
| ENSMUSG00000027483 | 0 | 0 | 0 | 0 | 0 | 0 | 0 |
| ENSMUSG00000027484 | 0 | 0 | 0 | 0 | 0 | 0 | 0 |
| ENSMUSG00000027485 | 0 | 0 | 0 | 0 | 0 | 0 | 0 |
| ENSMUSG00000027487 | 1.727821383 | 1.566696635 | 1.54745614 | 1.79248125 | 1.78687392 | 2.017033339 | 1.657681529 |
| ENSMUSG00000027488 | 1.940852214 | 1.819031101 | 1.916333979 | 1.681980114 | 1.804307395 | 1.799340549 | 1.923435442 |
| ENSMUSG00000027489 | 0.867194479 | 0.646240625 | 0.830482024 | 0.701838731 | 0.733951626 | 0.477121255 | 0.896975297 |
| ENSMUSG00000027490 | 2.22592627 | 2.060793496 | 2.098079356 | 2.061981878 | 2.159419872 | 2.328379603 | 2.062419043 |
| ENSMUSG00000027495 | 1.482695304 | 1.29248125 | 1.438721876 | 1.618933358 | 1.704182902 | 1.913813852 | 1.577201671 |
| ENSMUSG00000027496 | 2.521950082 | 2.363317659 | 2.331607372 | 2.453845824 | 2.477503739 | 2.790285164 | 2.308505126 |
| ENSMUSG00000027498 | 2.227926429 | 2.05357978 | 2.074802005 | 2.029735268 | 2.103973339 | 2.340444115 | 2.015657029 |
| ENSMUSG00000027499 | 2.334723298 | 2.143411797 | 2.130890489 | 2.162762923 | 2.119108707 | 2.437750563 | 1.935895857 |
| ENSMUSG00000027500 | 1.269664473 | 1.238549078 | 1.311981878 | 1.837182039 | 1.728722551 | 1.886490725 | 1.670888075 |
| ENSMUSG00000027502 | 2.442947872 | 2.244319981 | 2.213717096 | 2.319612365 | 2.342660868 | 2.617000341 | 2.214403581 |
| ENSMUSG00000027505 | 0.289064826 | 0.396240625 | 0.25 | 0.25 | 0 | 0.301029996 | 0.355418052 |
| ENSMUSG00000027506 | 2.181541736 | 2.051142786 | 2.051142786 | 2.201032755 | 2.135290471 | 2.394451681 | 2 |
| ENSMUSG00000027508 | 1.87190676 | 1.714495249 | 1.720660762 | 2.188721876 | 2.079491904 | 2.330413773 | 1.992562536 |
| ENSMUSG00000027509 | 2.267855219 | 2.150906586 | 2.143411797 | 2.242885888 | 2.196996016 | 2.495544338 | 2.068965954 |
| ENSMUSG00000027510 | 1.45815691 | 1.431980114 | 1.634789703 | 1.380890489 | 1.603583716 | 1.556302501 | 1.69749337 |
| ENSMUSG00000027513 | 0 | 0.25 | 0 | 0 | 0 | 0 | 0 |
| ENSMUSG00000027514 | 0 | 0 | 0 | 0.25 | 0 | 0.301029996 | 0.448487648 |
| ENSMUSG00000027517 | 0 | 0 | 0 | 0 | 0 | 0 | 0 |
| ENSMUSG00000027518 | 0 | 0 | 0 | 0 | 0 | 0 | 0 |
| ENSMUSG00000027519 | 2.572927518 | 2.37892496 | 2.365881093 | 2.442046081 | 2.393415815 | 2.723455672 | 2.229218532 |
| ENSMUSG00000027520 | 0 | 0 | 0 | 1.188721876 | 1.244650542 | 1.079181246 | 1 |
| ENSMUSG00000027522 | 2.205378646 | 2.075945187 | 2.038704527 | 2.120953944 | 2.145141048 | 2.326335861 | 2.014380832 |
| ENSMUSG00000027523 | 1.269664473 | 0.92510993 | 0.896240625 | 0.976722649 | 0.931471446 | 1.113943352 | 1.078019652 |
| ENSMUSG00000027524 | 0 | 0 | 0 | 0 | 0 | 0 | 0 |
| ENSMUSG00000027525 | 0 | 0 | 0 | 0 | 0.387761935 | 0 | 0 |
| ENSMUSG00000027528 | 0 | 0 | 0 | 0 | 0 | 0 | 0 |
| ENSMUSG00000027530 | 0 | 0 | 0 | 0 | 0 | 0 | 0 |
| ENSMUSG00000027531 | 2.524897319 | 2.351285366 | 2.328787391 | 2.490000483 | 2.402121453 | 2.764922985 | 2.281664246 |
| ENSMUSG00000027533 | 2.737928151 | 2.490000483 | 2.475970461 | 2.387686696 | 2.31625483 | 2.679427897 | 2.190455009 |

| | | | | | | | |
|---|---|---|---|---|---|---|---|
| ENSMUSG00000027534 | 1.932832114 | 1.766522298 | 1.83272922 | 1.755591953 | 1.674752852 | 1.875061263 | 1.548827561 |
| ENSMUSG00000027536 | 2.17479713 | 2.066696635 | 2.078220739 | 1.988549078 | 1.948833444 | 2.184691431 | 1.820622395 |
| ENSMUSG00000027540 | 2.223916471 | 2.147178659 | 2.219129237 | 2.230460234 | 2.337535342 | 2.551449998 | 2.270850938 |
| ENSMUSG00000027544 | 0.578129653 | 0.896240625 | 0.896240625 | 1.080482024 | 0.955822902 | 1.041392685 | 1.140832123 |
| ENSMUSG00000027546 | 0 | 0 | 0 | 1.396240625 | 1.484072009 | 1.556302501 | 1.483307832 |
| ENSMUSG00000027547 | 0 | 0 | 0 | 0 | 0 | 0 | 0 |
| ENSMUSG00000027550 | 2.360187403 | 2.157339155 | 2.201032755 | 2.204245906 | 2.095689764 | 2.309630167 | 1.925966215 |
| ENSMUSG00000027551 | 1.663535556 | 1.688721876 | 1.821350555 | 1.598079356 | 1.712553795 | 1.770852012 | 1.842010333 |
| ENSMUSG00000027552 | 2 | 1.856566189 | 1.866901388 | 1.953445298 | 1.871833944 | 2.100370545 | 1.726776051 |
| ENSMUSG00000027555 | 2.405576699 | 2.23927551 | 2.216819685 | 2.079353153 | 1.995080671 | 2.298853076 | 1.89141329 |
| ENSMUSG00000027556 | 0 | 0 | 0 | 0 | 0 | 0 | 0 |
| ENSMUSG00000027559 | 0.45815691 | 0 | 0 | 0.5 | 0.733951626 | 0.602059991 | 0.710836105 |
| ENSMUSG00000027560 | 0.671187741 | 0.75 | 0.951838731 | 0 | 0.244650542 | 0 | 0.224243824 |
| ENSMUSG00000027562 | 0 | 0.701838731 | 0 | 0.701838731 | 0.387761935 | 0.77815125 | 0.710836105 |
| ENSMUSG00000027564 | 0 | 0 | 0 | 0 | 0 | 0 | 0 |
| ENSMUSG00000027566 | 2.711874084 | 2.490724001 | 2.496460484 | 2.619186551 | 2.566592354 | 2.976808337 | 2.429563222 |
| ENSMUSG00000027568 | 0 | 0 | 0 | 0 | 0.244650542 | 0.477121255 | 0 |
| ENSMUSG00000027569 | 1.800532393 | 1.688721876 | 1.747171172 | 1.782320754 | 1.804307395 | 1.84509804 | 1.680001789 |
| ENSMUSG00000027570 | 0.45815691 | 0.396240625 | 0 | 0.25 | 0.244650542 | 0 | 0.520678036 |
| ENSMUSG00000027573 | 2.188178997 | 2.073655187 | 2.075945187 | 2.205044741 | 2.145141048 | 2.397940009 | 2.007270316 |
| ENSMUSG00000027574 | 0 | 0 | 0 | 0 | 0.568060967 | 0.602059991 | 0.355418052 |
| ENSMUSG00000027575 | 2.480523256 | 2.257666784 | 2.273189285 | 2.192872367 | 2.255474576 | 2.45331834 | 2.13206423 |
| ENSMUSG00000027577 | 0 | 0 | 0 | 0.580482024 | 0.489301084 | 0.477121255 | 0.224243824 |
| ENSMUSG00000027579 | 0 | 0 | 0 | 0 | 0 | 0.301029996 | 0.224243824 |
| ENSMUSG00000027580 | 1.936861443 | 1.852347734 | 1.882845365 | 2.106566189 | 2.235996193 | 2.348304863 | 2.12713864 |
| ENSMUSG00000027581 | 0 | 0 | 0 | 0 | 0 | 0 | 0 |
| ENSMUSG00000027582 | 1.759671389 | 1.622963274 | 1.75 | 1.692046081 | 1.842556557 | 1.838849091 | 1.747849621 |
| ENSMUSG00000027583 | 1.269664473 | 1.321350555 | 1.282320754 | 1.598079356 | 1.614672255 | 1.770852012 | 1.48984372 |
| ENSMUSG00000027584 | 0.45815691 | 0 | 0 | 0.396240625 | 0 | 0 | 0 |
| ENSMUSG00000027589 | 2.051791411 | 1.966046536 | 1.898114259 | 2.107613138 | 2.032639648 | 2.255272505 | 1.896975297 |
| ENSMUSG00000027593 | 2.879922803 | 2.657794264 | 2.664775991 | 2.65892953 | 2.674933346 | 3.031812271 | 2.552119436 |
| ENSMUSG00000027596 | 0 | 0 | 0 | 0 | 0 | 0 | 0 |
| ENSMUSG00000027597 | 2.223916471 | 2.136723615 | 2.12892496 | 2.101285366 | 2.240967488 | 2.431363764 | 2.125328759 |
| ENSMUSG00000027598 | 2.577267124 | 2.381380202 | 2.341580554 | 2.560197833 | 2.484381755 | 2.821513528 | 2.286662868 |

| | | | | | | | |
|---|---|---|---|---|---|---|---|
| ENSMUSG00000027599 | 2.572927518 | 2.347542239 | 2.348079356 | 2.460587586 | 2.397399561 | 2.723455672 | 2.223239637 |
| ENSMUSG00000027601 | 2.56677519 | 2.358135475 | 2.315523711 | 2.390560606 | 2.329464709 | 2.652246341 | 2.14725554 |
| ENSMUSG00000027602 | 1.740854706 | 1.642463902 | 1.708222504 | 1.552363341 | 1.744182955 | 1.740362689 | 1.704233519 |
| ENSMUSG00000027603 | 0.289064826 | 0.92510993 | 0.864857905 | 0.646240625 | 0.812711509 | 0.77815125 | 1.100339912 |
| ENSMUSG00000027605 | 1.289064826 | 1.114857905 | 1.321350555 | 1.79248125 | 1.791313678 | 1.908485019 | 1.603286636 |
| ENSMUSG00000027606 | 0 | 0 | 0 | 0 | 0 | 0 | 0 |
| ENSMUSG00000027610 | 2.092368209 | 1.942046081 | 1.970660762 | 2.072504712 | 2.124299332 | 2.307496038 | 2.051774285 |
| ENSMUSG00000027611 | 1.45815691 | 1.356566189 | 1.311981878 | 1.678561379 | 1.718026092 | 1.886490725 | 1.625302983 |
| ENSMUSG00000027612 | 0 | 0.25 | 0.25 | 0.580482024 | 0.77552387 | 0.602059991 | 0.74492186 |
| ENSMUSG00000027613 | 2.472830741 | 2.308404919 | 2.383818844 | 2.361770807 | 2.462015062 | 2.702430536 | 2.401639109 |
| ENSMUSG00000027615 | 2.057835465 | 1.899978211 | 1.870953944 | 2.031030328 | 1.941702774 | 2.086359831 | 1.856911705 |
| ENSMUSG00000027618 | 2.060824957 | 1.956637122 | 1.938721876 | 1.976722649 | 1.997548906 | 2.235528447 | 1.906037947 |
| ENSMUSG00000027620 | 3.060011764 | 2.773024104 | 2.757149194 | 2.936103353 | 2.84445929 | 3.265289626 | 2.641897157 |
| ENSMUSG00000027624 | 2.596667477 | 2.400906586 | 2.430701883 | 2.449009902 | 2.495355369 | 2.748188027 | 2.3847907 |
| ENSMUSG00000027628 | 2.081172036 | 1.937048212 | 1.892463902 | 2.032320754 | 2.041436517 | 2.267171728 | 1.956418351 |
| ENSMUSG00000027630 | 2.507914013 | 2.265173983 | 2.255591953 | 2.273849256 | 2.204601633 | 2.506505032 | 2.031805106 |
| ENSMUSG00000027634 | 2.404271433 | 2.155512955 | 2.145300145 | 2.445339928 | 2.355827317 | 2.758911892 | 2.211635518 |
| ENSMUSG00000027635 | 2.295901576 | 2.098079356 | 2.108657057 | 2.207430684 | 2.152352662 | 2.484299839 | 2.041943747 |
| ENSMUSG00000027636 | 0 | 0 | 0 | 0.396240625 | 0.489301084 | 0.602059991 | 0.629532004 |
| ENSMUSG00000027637 | 1.342375483 | 1.451838731 | 1.494319981 | 1.571350555 | 1.489301084 | 1.568201724 | 1.448487648 |
| ENSMUSG00000027639 | 2.804055922 | 2.549304173 | 2.502457154 | 2.863703337 | 2.790274042 | 3.236033147 | 2.628192393 |
| ENSMUSG00000027641 | 2.527823874 | 2.336073977 | 2.324802005 | 2.450629091 | 2.356717497 | 2.720985744 | 2.217830684 |
| ENSMUSG00000027642 | 3.060011764 | 2.786642169 | 2.77236262 | 2.891632446 | 2.862836946 | 3.293141483 | 2.687751032 |
| ENSMUSG00000027643 | 0 | 0 | 0 | 0 | 0 | 0.301029996 | 0 |
| ENSMUSG00000027646 | 2.370236862 | 2.27053726 | 2.30479213 | 2.159156155 | 2.308186911 | 2.440909082 | 2.243384656 |
| ENSMUSG00000027649 | 2.075459174 | 1.976722649 | 2.002806814 | 2.073655187 | 2.188507684 | 2.344392274 | 2.044283833 |
| ENSMUSG00000027650 | 2.048736215 | 1.896240625 | 1.858657057 | 2.03743678 | 2.042520869 | 2.320146286 | 1.93179548 |
| ENSMUSG00000027651 | 2.211649928 | 2.104463129 | 2.102347734 | 2.20584181 | 2.221952687 | 2.498310554 | 2.053478496 |
| ENSMUSG00000027652 | 2.642661589 | 2.42293588 | 2.383332433 | 2.568740118 | 2.488077665 | 2.862131379 | 2.304357435 |
| ENSMUSG00000027654 | 2.100572389 | 1.958222504 | 1.94697564 | 2.059601185 | 2.104881849 | 2.292256071 | 2.018825668 |
| ENSMUSG00000027655 | 1.867194479 | 1.711372513 | 1.729715809 | 1.79745614 | 1.78010778 | 1.949390007 | 1.659601507 |
| ENSMUSG00000027656 | 1.249317394 | 1.348079356 | 1.410964047 | 1.02186571 | 0.931471446 | 1.041392685 | 1.078019652 |
| ENSMUSG00000027660 | 2.578990401 | 2.412314044 | 2.352877747 | 2.344302633 | 2.302531265 | 2.588831726 | 2.132508314 |
| ENSMUSG00000027661 | 1.777675583 | 1.723704441 | 1.705044741 | 1.678561379 | 1.744182955 | 1.908485019 | 1.667169465 |

| | | | | | | | |
|---|---|---|---|---|---|---|---|
| ENSMUSG00000027663 | 2.534573679 | 2.314935816 | 2.319031101 | 2.150906586 | 2.090057366 | 2.292256071 | 1.979298987 |
| ENSMUSG00000027665 | 2.643399047 | 2.453445298 | 2.395300145 | 2.455044741 | 2.366796383 | 2.704150517 | 2.192673409 |
| ENSMUSG00000027667 | 2.243591459 | 2.044977273 | 2.059601185 | 2.071350555 | 2.043601899 | 2.255272505 | 1.917452159 |
| ENSMUSG00000027668 | 2.505869732 | 2.328220739 | 2.300530956 | 2.507321807 | 2.446505421 | 2.781755375 | 2.270561438 |
| ENSMUSG00000027669 | 2.43085913 | 2.244319981 | 2.252107156 | 2.359697963 | 2.282813873 | 2.605305046 | 2.14555506 |
| ENSMUSG00000027670 | 0 | 0.5 | 0.396240625 | 0.396240625 | 0.387761935 | 0 | 0.224243824 |
| ENSMUSG00000027671 | 2.552362269 | 2.341033664 | 2.344302633 | 2.453044326 | 2.384078881 | 2.746634199 | 2.282223478 |
| ENSMUSG00000027673 | 2.380049842 | 2.134789703 | 2.168106335 | 2.222185812 | 2.11997913 | 2.460897843 | 2.019455688 |
| ENSMUSG00000027674 | 0 | 0 | 0 | 0 | 0 | 0 | 0 |
| ENSMUSG00000027676 | 1.342375483 | 1.17510993 | 1.061981878 | 1.42510993 | 1.319233381 | 1.447158031 | 1.272006351 |
| ENSMUSG00000027677 | 2.504843822 | 2.332168732 | 2.314935816 | 2.439139081 | 2.381187475 | 2.617000341 | 2.182568577 |
| ENSMUSG00000027678 | 2.199011616 | 2.079353153 | 2.034887838 | 2.145300145 | 2.174635855 | 2.432969291 | 2.033014511 |
| ENSMUSG00000027679 | 1.771760131 | 1.720660762 | 1.688721876 | 1.774508021 | 1.706995316 | 1.86332286 | 1.616675393 |
| ENSMUSG00000027680 | 2.794934559 | 2.581888161 | 2.527131114 | 2.673807073 | 2.591907258 | 2.983175072 | 2.386819016 |
| ENSMUSG00000027684 | 2.31414447 | 2.168989258 | 2.163659007 | 1.938721876 | 1.882215755 | 2.120573931 | 1.775756176 |
| ENSMUSG00000027690 | 0 | 0 | 0 | 0 | 0 | 0 | 0 |
| ENSMUSG00000027692 | 1 | 1.302363341 | 1.356566189 | 0.896240625 | 0.733951626 | 0.477121255 | 0.672731473 |
| ENSMUSG00000027694 | 0 | 0 | 0 | 0 | 0 | 0 | 0 |
| ENSMUSG00000027695 | 1.765759561 | 1.720660762 | 1.71759118 | 1.79248125 | 1.726078673 | 1.851258349 | 1.649885598 |
| ENSMUSG00000027698 | 2.223916471 | 2.061981878 | 2.063166358 | 2.137686696 | 2.075570129 | 2.352182518 | 1.930140666 |
| ENSMUSG00000027699 | 2.939767604 | 2.685366747 | 2.680488403 | 2.904596376 | 2.804307395 | 3.259115844 | 2.599641647 |
| ENSMUSG00000027702 | 0.289064826 | 0 | 0.25 | 0 | 0 | 0.301029996 | 0.224243824 |
| ENSMUSG00000027703 | 0 | 0 | 0 | 0 | 0 | 0 | 0 |
| ENSMUSG00000027706 | 2.797488347 | 2.609438018 | 2.581326364 | 2.663211243 | 2.590076093 | 3.006893708 | 2.434660827 |
| ENSMUSG00000027708 | 2.373064211 | 2.149978211 | 2.177701608 | 2.278435542 | 2.158641579 | 2.472756449 | 2.01116814 |
| ENSMUSG00000027709 | 2.045658471 | 1.930274797 | 2.009729747 | 1.926839783 | 1.898867717 | 2.170261715 | 1.802677937 |
| ENSMUSG00000027710 | 2.289064826 | 2.156427211 | 2.161864607 | 2.214495249 | 2.180543208 | 2.481442629 | 2.044283833 |
| ENSMUSG00000027712 | 3.363869832 | 3.044821776 | 3.039494612 | 2.947077626 | 2.878180774 | 3.349083169 | 2.712096219 |
| ENSMUSG00000027713 | 1 | 1.02186571 | 0.896240625 | 0.896240625 | 0.632412477 | 0.698970004 | 0.896975297 |
| ENSMUSG00000027714 | 2.310886388 | 2.162762923 | 2.156427211 | 2.273849256 | 2.209357611 | 2.498310554 | 2.042530357 |
| ENSMUSG00000027715 | 3.014472122 | 2.714883697 | 2.713132377 | 2.8843046 | 2.783971086 | 3.238547888 | 2.598487544 |
| ENSMUSG00000027716 | 0 | 0.25 | 0.25 | 0.79248125 | 0.77552387 | 0.602059991 | 0.629532004 |
| ENSMUSG00000027718 | 0 | 0 | 0 | 0 | 0 | 0 | 0 |
| ENSMUSG00000027719 | 0 | 0 | 0 | 0 | 0 | 0 | 0 |

| | | | | | | | |
|---|---|---|---|---|---|---|---|
| ENSMUSG00000027720 | 0 | 0.25 | 0 | 0 | 0.244650542 | 0 | 0 |
| ENSMUSG00000027722 | 2.287337982 | 2.119945066 | 2.094844592 | 2.169870025 | 2.034859505 | 2.334453751 | 1.921737195 |
| ENSMUSG00000027737 | 1.623015126 | 1.438721876 | 1.302363341 | 1.25 | 1.302012594 | 1.342422681 | 1.20919388 |
| ENSMUSG00000027739 | 2.089597219 | 1.995748394 | 1.940387808 | 2.012462137 | 1.981187939 | 2.247973266 | 1.825244123 |
| ENSMUSG00000027742 | 2.333175866 | 2.143411797 | 2.148114259 | 2.268535366 | 2.196996016 | 2.519827994 | 2.046607113 |
| ENSMUSG00000027744 | 0.289064826 | 0.396240625 | 0 | 0 | 0 | 0 | 0 |
| ENSMUSG00000027746 | 2.435736758 | 2.252806814 | 2.246460484 | 2.215271726 | 2.154724156 | 2.418301291 | 1.989815042 |
| ENSMUSG00000027748 | 0 | 0 | 0 | 0 | 0 | 0.301029996 | 0 |
| ENSMUSG00000027750 | 2.480523256 | 2.28296424 | 2.146240625 | 0.580482024 | 0.632412477 | 0.954242509 | 0.520678036 |
| ENSMUSG00000027751 | 2.550525119 | 2.324229052 | 2.296838018 | 2.284887838 | 2.285543632 | 2.526339277 | 2.115657114 |
| ENSMUSG00000027752 | 2.485932312 | 2.255591953 | 2.25 | 2.375956434 | 2.359374645 | 2.664641976 | 2.19156611 |
| ENSMUSG00000027761 | 0 | 0 | 0 | 0.25 | 0 | 0 | 0 |
| ENSMUSG00000027762 | 0 | 0 | 0 | 0 | 0 | 0 | 0 |
| ENSMUSG00000027763 | 3.065671095 | 2.815964005 | 2.818740118 | 2.685156011 | 2.652779079 | 3.007320953 | 2.415994748 |
| ENSMUSG00000027765 | 0.867194479 | 0.864857905 | 0.951838731 | 1.04248125 | 0.77552387 | 1.079181246 | 0.916588299 |
| ENSMUSG00000027770 | 2.747795766 | 2.505938588 | 2.468745337 | 2.64127552 | 2.524160589 | 2.953276337 | 2.335665766 |
| ENSMUSG00000027774 | 2.641182749 | 2.438304187 | 2.426839783 | 2.483791262 | 2.437427909 | 2.789580712 | 2.27487696 |
| ENSMUSG00000027775 | 2.985980784 | 2.717398461 | 2.727285762 | 2.607090043 | 2.540906867 | 2.916980047 | 2.34638072 |
| ENSMUSG00000027776 | 0 | 0 | 0 | 0 | 0 | 0 | 0 |
| ENSMUSG00000027777 | 0.747221736 | 0.830482024 | 0.79248125 | 0.646240625 | 0.489301084 | 0.477121255 | 0.224243824 |
| ENSMUSG00000027778 | 2.536482297 | 2.351285366 | 2.333288838 | 2.481944491 | 2.388168801 | 2.740362689 | 2.233468922 |
| ENSMUSG00000027782 | 2.967766868 | 2.689555804 | 2.659382638 | 2.7808687 | 2.669294659 | 3.078456818 | 2.474022582 |
| ENSMUSG00000027784 | 2.275045611 | 2.039967834 | 2.117918804 | 2.219895812 | 2.13194489 | 2.426511261 | 2.007270316 |
| ENSMUSG00000027787 | 2.540273527 | 2.327084758 | 2.326515422 | 2.451032755 | 2.369379 | 2.721810615 | 2.211635518 |
| ENSMUSG00000027788 | 0.289064826 | 0 | 0.25 | 0 | 0 | 0 | 0 |
| ENSMUSG00000027790 | 0 | 0 | 0 | 0 | 0 | 0.301029996 | 0 |
| ENSMUSG00000027792 | 1.181541736 | 1.29248125 | 1.25 | 1.282320754 | 1.188507684 | 1.255272505 | 1.089372236 |
| ENSMUSG00000027793 | 0.45815691 | 0.396240625 | 0.396240625 | 0.5 | 0 | 0 | 0 |
| ENSMUSG00000027794 | 0 | 0.5 | 0.25 | 0.25 | 0.244650542 | 0 | 0 |
| ENSMUSG00000027796 | 1.827447047 | 1.744319981 | 1.741446071 | 1.816696635 | 1.671670246 | 1.977723605 | 1.645916034 |
| ENSMUSG00000027797 | 2.862428343 | 2.613831805 | 2.587182039 | 2.55357978 | 2.480020371 | 2.85672889 | 2.290497981 |
| ENSMUSG00000027799 | 2.409468165 | 2.257666784 | 2.197790722 | 2.232684334 | 2.168627946 | 2.492760389 | 2.002662694 |
| ENSMUSG00000027800 | 2.026699189 | 1.928561379 | 1.862802778 | 2.061981878 | 2 | 2.315970345 | 1.843097781 |
| ENSMUSG00000027801 | 1.069666964 | 1.02186571 | 1.114857905 | 0.701838731 | 0.387761935 | 0.602059991 | 0.520678036 |

| | | | | | | | |
|---|---|---|---|---|---|---|---|
| ENSMUSG00000027803 | 2.824116479 | 2.60839636 | 2.579353153 | 2.548689214 | 2.502486038 | 2.865103975 | 2.338730052 |
| ENSMUSG00000027804 | 2.756874381 | 2.558105232 | 2.536164561 | 2.691217898 | 2.626045333 | 3.026533265 | 2.427251752 |
| ENSMUSG00000027805 | 2.278594844 | 2.132845365 | 2.110735874 | 2.360735874 | 2.276733037 | 2.622214023 | 2.124420007 |
| ENSMUSG00000027806 | 2.647066901 | 2.451838731 | 2.442872367 | 2.422499493 | 2.367659357 | 2.686636269 | 2.194510526 |
| ENSMUSG00000027808 | 2.96059708 | 2.683677405 | 2.645300145 | 2.741986676 | 2.632412477 | 3.067814511 | 2.448821686 |
| ENSMUSG00000027809 | 2.576402808 | 2.386723615 | 2.3464656 | 2.490724001 | 2.398585969 | 2.741939078 | 2.244328309 |
| ENSMUSG00000027810 | 2.53838222 | 2.336628433 | 2.323080408 | 2.461372513 | 2.377434805 | 2.750508395 | 2.20496486 |
| ENSMUSG00000027811 | 2.105953527 | 2.027131114 | 2.066696635 | 2.083847589 | 2.038163331 | 2.307496038 | 1.963977467 |
| ENSMUSG00000027820 | 1.663535556 | 1.681980114 | 1.67510993 | 1.431980114 | 1.394615667 | 1.462397998 | 1.265599895 |
| ENSMUSG00000027822 | 2.181541736 | 2.04248125 | 2.03360658 | 2.110735874 | 2.030405741 | 2.264817823 | 1.86207155 |
| ENSMUSG00000027823 | 2.868484936 | 2.602877747 | 2.567573832 | 2.686838457 | 2.602699114 | 2.981818607 | 2.39183486 |
| ENSMUSG00000027824 | 0 | 0 | 0 | 0 | 0 | 0.301029996 | 0 |
| ENSMUSG00000027827 | 1.342375483 | 1.364857905 | 1.410964047 | 0.5 | 0.733951626 | 0.602059991 | 0.629532004 |
| ENSMUSG00000027828 | 3.315175817 | 3.026640735 | 2.991446071 | 3.065156389 | 2.969243082 | 3.444981112 | 2.757764581 |
| ENSMUSG00000027829 | 2.492331912 | 2.265173983 | 2.247171172 | 2.365369862 | 2.327538614 | 2.592176757 | 2.151886289 |
| ENSMUSG00000027831 | 1.527823874 | 1.410964047 | 1.388647213 | 1.130890489 | 1.136121934 | 1.113943352 | 1.078019652 |
| ENSMUSG00000027832 | 2.96059708 | 2.689972046 | 2.69595419 | 2.650674716 | 2.61424391 | 2.986771734 | 2.424023744 |
| ENSMUSG00000027833 | 1.794934559 | 1.732684334 | 1.777131114 | 1.52186571 | 1.484072009 | 1.447158031 | 1.355418052 |
| ENSMUSG00000027834 | 1.181541736 | 1.348079356 | 1.098079356 | 1.557204673 | 1.387761935 | 1.69019608 | 1.46982607 |
| ENSMUSG00000027835 | 2.364524 | 2.228222334 | 2.201838731 | 2.265173983 | 2.197694248 | 2.496929648 | 2.033617522 |
| ENSMUSG00000027840 | 0.811507563 | 0 | 0 | 0.951838731 | 0.905314583 | 0 | 0.579661877 |
| ENSMUSG00000027843 | 0.671187741 | 0.396240625 | 0.5 | 0.646240625 | 0.77552387 | 0.301029996 | 0.710836105 |
| ENSMUSG00000027845 | 1.857607105 | 1.76109853 | 1.729715809 | 1.907339155 | 1.86473911 | 2.033423755 | 1.734460864 |
| ENSMUSG00000027848 | 2.060824957 | 1.901832578 | 1.945339928 | 0.951838731 | 1.074582839 | 1.113943352 | 0.984950056 |
| ENSMUSG00000027849 | 0.671187741 | 0.25 | 0.79248125 | 0 | 0 | 0 | 0 |
| ENSMUSG00000027852 | 2.910141165 | 2.680274797 | 2.641751343 | 2.721614674 | 2.636057495 | 3.024074987 | 2.444621219 |
| ENSMUSG00000027854 | 2.342375483 | 2.177701608 | 2.242166698 | 2.222185812 | 2.183460217 | 2.42975228 | 2.040767326 |
| ENSMUSG00000027855 | 1.505869732 | 1.494319981 | 1.372963274 | 1.618933358 | 1.467903253 | 1.672097858 | 1.360283038 |
| ENSMUSG00000027857 | 0.91631382 | 0.976722649 | 0.92510993 | 0.79248125 | 0.877063019 | 0.301029996 | 0.448487648 |
| ENSMUSG00000027858 | 2.692026894 | 2.501055117 | 2.434945152 | 2.103406982 | 2.008447216 | 2.334453751 | 1.851668228 |
| ENSMUSG00000027859 | 1.83262764 | 1.807204673 | 1.819031101 | 1.614857905 | 1.733951626 | 1.875061263 | 1.700880998 |
| ENSMUSG00000027860 | 2.194712275 | 1.961372513 | 1.988549078 | 1.995748394 | 1.970729215 | 2.214843848 | 1.877074955 |
| ENSMUSG00000027861 | 0 | 0.25 | 0 | 0.25 | 0.387761935 | 0 | 0 |
| ENSMUSG00000027863 | 0 | 0 | 0 | 0 | 0 | 0 | 0 |

| | | | | | | | |
|---|---|---|---|---|---|---|---|
| ENSMUSG00000027864 | 2.913238913 | 2.691632227 | 2.720087202 | 2.586073977 | 2.593274423 | 2.919601024 | 2.475101739 |
| ENSMUSG00000027865 | 2.111266113 | 1.919870025 | 1.953445298 | 2.052363341 | 2.015530186 | 2.283301229 | 1.848480858 |
| ENSMUSG00000027867 | 0 | 0 | 0 | 0 | 0 | 0 | 0 |
| ENSMUSG00000027868 | 2.205378646 | 2.043731421 | 2.03360658 | 2.228969845 | 2.182733222 | 2.477121255 | 2.03780742 |
| ENSMUSG00000027869 | 0 | 0 | 0 | 0 | 0 | 0 | 0 |
| ENSMUSG00000027870 | 0 | 0 | 0 | 0 | 0 | 0 | 0 |
| ENSMUSG00000027871 | 0 | 0 | 0 | 0 | 0 | 0 | 0 |
| ENSMUSG00000027875 | 0 | 0 | 0 | 0 | 0 | 0 | 0 |
| ENSMUSG00000027876 | 0 | 0 | 0.25 | 0 | 0 | 0 | 0 |
| ENSMUSG00000027878 | 2.791849225 | 2.58076369 | 2.599418658 | 2.460194481 | 2.523052406 | 2.758154622 | 2.384587167 |
| ENSMUSG00000027879 | 2.537433341 | 2.338286706 | 2.3464656 | 2.360217292 | 2.322187006 | 2.660865478 | 2.170957925 |
| ENSMUSG00000027880 | 0.289064826 | 0.25 | 0.396240625 | 0 | 0 | 0 | 0 |
| ENSMUSG00000027881 | 2.795957952 | 2.546218836 | 2.539652422 | 2.637927065 | 2.576984829 | 2.921686475 | 2.384383506 |
| ENSMUSG00000027882 | 2.299278375 | 2.17510993 | 2.166333979 | 2.225970461 | 2.11997913 | 2.445604203 | 1.996640515 |
| ENSMUSG00000027883 | 2.565888829 | 2.358135475 | 2.33272922 | 2.379417347 | 2.33042383 | 2.679427897 | 2.141695983 |
| ENSMUSG00000027884 | 2.247417471 | 2.066696635 | 2.058404919 | 2.252107156 | 2.203916946 | 2.507855872 | 2.071119143 |
| ENSMUSG00000027886 | 0 | 0 | 0 | 0.396240625 | 0.632412477 | 0.301029996 | 0.710836105 |
| ENSMUSG00000027887 | 0 | 0 | 0 | 0 | 0.387761935 | 0.301029996 | 0.579661877 |
| ENSMUSG00000027889 | 2.455852858 | 2.324229052 | 2.3464656 | 2.230460234 | 2.338004387 | 2.584331224 | 2.263238458 |
| ENSMUSG00000027890 | 0.578129653 | 0.396240625 | 0.5 | 0.896240625 | 0.955822902 | 0.77815125 | 1 |
| ENSMUSG00000027893 | 3.2972863 | 2.979156481 | 2.96166642 | 3.01593303 | 2.919042129 | 3.399327532 | 2.705670528 |
| ENSMUSG00000027894 | 2.743760867 | 2.525493918 | 2.5896628 | 2.315523711 | 2.346342321 | 2.607455023 | 2.206027309 |
| ENSMUSG00000027895 | 0 | 0 | 0 | 0 | 0 | 0 | 0 |
| ENSMUSG00000027896 | 1.342375483 | 1.330482024 | 1.26109853 | 1.380890489 | 1.351342427 | 1.255272505 | 1.252393349 |
| ENSMUSG00000027900 | 1.956444026 | 1.938721876 | 1.868933358 | 2.034887838 | 1.979897422 | 2.250420002 | 1.857950279 |
| ENSMUSG00000027901 | 0.91631382 | 0.396240625 | 0.5 | 0.75 | 0.632412477 | 0.477121255 | 0.876096088 |
| ENSMUSG00000027902 | 0 | 0 | 0 | 0 | 0 | 0 | 0 |
| ENSMUSG00000027905 | 2.365959569 | 2.191217898 | 2.159156155 | 2.372963274 | 2.305113253 | 2.611723308 | 2.160953557 |
| ENSMUSG00000027907 | 3.137985763 | 2.823942234 | 2.834405475 | 2.998765117 | 2.911184122 | 3.416141031 | 2.72214771 |
| ENSMUSG00000027908 | 0 | 0 | 0 | 0 | 0 | 0 | 0 |
| ENSMUSG00000027912 | 0 | 0 | 0 | 0 | 0.244650542 | 0 | 0.224243824 |
| ENSMUSG00000027913 | 0 | 0 | 0.25 | 0.25 | 0 | 0 | 0.224243824 |
| ENSMUSG00000027919 | 0 | 0 | 0 | 0 | 0 | 0 | 0 |
| ENSMUSG00000027923 | 0 | 0 | 0 | 0 | 0 | 0 | 0 |

| | | | | | | | |
|---|---|---|---|---|---|---|---|
| ENSMUSG00000027925 | 0 | 0.396240625 | 0.25 | 0 | 0 | 0 | 0 |
| ENSMUSG00000027927 | 0.289064826 | 0.25 | 0.25 | 0 | 0 | 0 | 0 |
| ENSMUSG00000027931 | 1.53838222 | 1.410964047 | 1.403677461 | 1.188721876 | 0.931471446 | 1.146128036 | 1.185218806 |
| ENSMUSG00000027932 | 1.289064826 | 1.130890489 | 1.146240625 | 1.445339928 | 1.614672255 | 1.681241237 | 1.667169465 |
| ENSMUSG00000027933 | 2.746646916 | 2.528109877 | 2.52053726 | 2.618679987 | 2.625630579 | 2.961421094 | 2.460139731 |
| ENSMUSG00000027935 | 1.948720984 | 1.862802778 | 1.931980114 | 1.926839783 | 1.90849439 | 2.158362492 | 1.82753046 |
| ENSMUSG00000027936 | 2.183765914 | 2.036164561 | 2.065523711 | 2.063166358 | 2.112066955 | 2.31386722 | 2.021338422 |
| ENSMUSG00000027937 | 2.063793172 | 1.966046536 | 2.004202072 | 2.02186571 | 1.965381292 | 2.190331698 | 1.86815666 |
| ENSMUSG00000027938 | 0.671187741 | 0.830482024 | 0.580482024 | 0.92510993 | 0.686820904 | 0.954242509 | 0.775756176 |
| ENSMUSG00000027939 | 0.960252568 | 1.02186571 | 1.061981878 | 1.26109853 | 1.234113756 | 1.278753601 | 1.284450788 |
| ENSMUSG00000027940 | 2.662129038 | 2.428561379 | 2.506976499 | 2.693696765 | 2.664494972 | 3.059941888 | 2.524927889 |
| ENSMUSG00000027942 | 2.53552908 | 2.317865757 | 2.307805295 | 2.299918086 | 2.234113756 | 2.549003262 | 2.116590781 |
| ENSMUSG00000027944 | 2.309247751 | 2.113831805 | 2.091580554 | 2.207430684 | 2.121713561 | 2.521138084 | 2.032410373 |
| ENSMUSG00000027947 | 1.944805158 | 1.708222504 | 1.741446071 | 1.701838731 | 1.689774529 | 1.812913357 | 1.621017948 |
| ENSMUSG00000027950 | 0.671187741 | 0.701838731 | 0.701838731 | 0.79248125 | 0.733951626 | 0.301029996 | 0.629532004 |
| ENSMUSG00000027951 | 2.262379879 | 2.078220739 | 2.092671852 | 2.158248799 | 2.179810173 | 2.431363764 | 2.080634989 |
| ENSMUSG00000027952 | 1.936861443 | 1.87892496 | 1.905512955 | 2.115881093 | 2.174635855 | 2.365487985 | 2.077494038 |
| ENSMUSG00000027953 | 1.83262764 | 1.653677461 | 1.671625132 | 1.688721876 | 1.706995316 | 1.851258349 | 1.643912836 |
| ENSMUSG00000027954 | 1.857607105 | 1.642463902 | 1.649978211 | 1.814346961 | 1.898867717 | 1.949390007 | 1.71890424 |
| ENSMUSG00000027955 | 3.030793509 | 2.768535366 | 2.749647607 | 2.501406137 | 2.425559697 | 2.804139432 | 2.221221862 |
| ENSMUSG00000027956 | 2.029919532 | 1.938721876 | 1.970660762 | 1.566696635 | 1.563883923 | 1.755874856 | 1.505628092 |
| ENSMUSG00000027957 | 2.481610694 | 2.325374049 | 2.277783918 | 2.574515642 | 2.482519186 | 2.820857989 | 2.26735973 |
| ENSMUSG00000027959 | 1.924654746 | 1.837182039 | 1.784887838 | 2.069031101 | 2.00243419 | 2.294466226 | 1.869159809 |
| ENSMUSG00000027961 | 0.289064826 | 0.580482024 | 0.25 | 0.864857905 | 0.489301084 | 0.698970004 | 0.579661877 |
| ENSMUSG00000027962 | 2.256831695 | 2.177701608 | 2.063166358 | 2.507321807 | 2.391809641 | 2.765668555 | 2.235734673 |
| ENSMUSG00000027963 | 2.280358197 | 2.063166358 | 2.102347734 | 2.200224975 | 2.16097133 | 2.440909082 | 2.008574816 |
| ENSMUSG00000027965 | 0 | 0 | 0 | 0 | 0 | 0 | 0 |
| ENSMUSG00000027966 | 0.289064826 | 0.25 | 0 | 1.602347734 | 1.563883923 | 1.716003344 | 1.540636858 |
| ENSMUSG00000027967 | 0 | 0 | 0 | 0 | 0 | 0 | 0 |
| ENSMUSG00000027968 | 2.521950082 | 2.337734795 | 2.344844592 | 2.601019284 | 2.515479755 | 2.905256049 | 2.37340178 |
| ENSMUSG00000027971 | 0.45815691 | 0.396240625 | 0.580482024 | 0.830482024 | 0.877063019 | 0.77815125 | 0.916588299 |
| ENSMUSG00000027973 | 0 | 0 | 0 | 0 | 0 | 0 | 0 |
| ENSMUSG00000027977 | 0.747221736 | 0.896240625 | 0.79248125 | 0.951838731 | 1.039257769 | 0.84509804 | 0.74492186 |
| ENSMUSG00000027978 | 0.289064826 | 0.25 | 0.396240625 | 0 | 0 | 0.477121255 | 0.355418052 |

| | | | | | | | |
|---|---|---|---|---|---|---|---|
| ENSMUSG00000027981 | 1.920505136 | 1.87694866 | 1.830482024 | 1.973704441 | 1.94314048 | 2.201397124 | 1.774416565 |
| ENSMUSG00000027983 | 1.129344651 | 1.160964047 | 1.130890489 | 1.214495249 | 1.188507684 | 1.301029996 | 1.176815335 |
| ENSMUSG00000027984 | 2.42468076 | 2.234159485 | 2.282320754 | 2.357613138 | 2.378690133 | 2.6599162 | 2.310815043 |
| ENSMUSG00000027985 | 1.307602651 | 1.321350555 | 1.25 | 1.302363341 | 1.163285805 | 1.278753601 | 1.21680636 |
| ENSMUSG00000027993 | 2.405576699 | 2.201838731 | 2.231203126 | 2.306603048 | 2.25190332 | 2.57863921 | 2.093904044 |
| ENSMUSG00000027994 | 2.170239354 | 2.031030328 | 2.02053726 | 1.984159485 | 1.981187939 | 2.136720567 | 1.87902385 |
| ENSMUSG00000027995 | 2.033115198 | 1.909156155 | 1.950224975 | 1.763820609 | 1.731346772 | 1.995635195 | 1.633706455 |
| ENSMUSG00000027996 | 1.527823874 | 1.53743678 | 1.470660762 | 1.214495249 | 1.302012594 | 1.204119983 | 1.324583736 |
| ENSMUSG00000027997 | 2.111266113 | 1.958222504 | 1.928561379 | 2.046218836 | 2.041436517 | 2.281033367 | 1.952571511 |
| ENSMUSG00000027998 | 2.627631768 | 2.37694866 | 2.365881093 | 2.47104263 | 2.415166968 | 2.807535028 | 2.278571072 |
| ENSMUSG00000027999 | 1.975148165 | 1.907339155 | 1.862802778 | 1.905512955 | 1.86473911 | 2.113943352 | 1.791413205 |
| ENSMUSG00000028001 | 0 | 0 | 0 | 0 | 0 | 0 | 0 |
| ENSMUSG00000028003 | 0 | 0 | 0 | 0.25 | 0 | 0 | 0.224243824 |
| ENSMUSG00000028004 | 0 | 0 | 0 | 0 | 0 | 0 | 0 |
| ENSMUSG00000028005 | 1.205378646 | 1.188721876 | 1.17510993 | 1.188721876 | 1.039257769 | 1.301029996 | 1.078019652 |
| ENSMUSG00000028007 | 2.231898196 | 2.105516192 | 2.079353153 | 2.172499493 | 2.110284345 | 2.359835482 | 1.980721036 |
| ENSMUSG00000028008 | 0 | 0 | 0 | 0 | 0 | 0 | 0 |
| ENSMUSG00000028009 | 0.289064826 | 0.396240625 | 0.25 | 0 | 0 | 0 | 0 |
| ENSMUSG00000028010 | 2.185978293 | 2.04248125 | 2.02186571 | 2.113831805 | 2.079491904 | 2.29666519 | 1.947107911 |
| ENSMUSG00000028011 | 0 | 0.580482024 | 0 | 0.580482024 | 0 | 0 | 0.355418052 |
| ENSMUSG00000028012 | 0.578129653 | 0 | 0.396240625 | 0.5 | 0.244650542 | 0.301029996 | 0.448487648 |
| ENSMUSG00000028013 | 2.241665209 | 2.108657057 | 2.027131114 | 2.163659007 | 2.065570703 | 2.376576957 | 1.938331368 |
| ENSMUSG00000028015 | 1.227926429 | 1.282320754 | 1.321350555 | 1.488549078 | 1.499532413 | 1.662757832 | 1.473249546 |
| ENSMUSG00000028016 | 2.233869984 | 2.084962501 | 2.043731421 | 2.193696765 | 2.117361396 | 2.421603927 | 1.987044015 |
| ENSMUSG00000028017 | 0.867194479 | 0.830482024 | 0.830482024 | 1.130890489 | 1.074582839 | 1.041392685 | 1.078019652 |
| ENSMUSG00000028018 | 2.315764015 | 2.145300145 | 2.093759858 | 2.232684334 | 2.195595395 | 2.51054501 | 2.051774285 |
| ENSMUSG00000028019 | 2.103271637 | 1.894357207 | 1.825945187 | 1.741446071 | 1.632412477 | 1.857332496 | 1.535058868 |
| ENSMUSG00000028020 | 1.227926429 | 1.098079356 | 0.951838731 | 1.29248125 | 1.200473444 | 1.591064607 | 1.21680636 |
| ENSMUSG00000028023 | 1.721148722 | 1.598079356 | 1.711372513 | 1.87892496 | 1.903713872 | 2.113943352 | 1.967691807 |
| ENSMUSG00000028024 | 0 | 0.5 | 0.25 | 0 | 0 | 0 | 0 |
| ENSMUSG00000028028 | 1.700465191 | 1.606566189 | 1.653677461 | 1.752806814 | 1.751666404 | 1.838849091 | 1.564611933 |
| ENSMUSG00000028029 | 2.45469604 | 2.294977273 | 2.27908599 | 2.409608978 | 2.357161747 | 2.725911632 | 2.187098622 |
| ENSMUSG00000028030 | 1.800532393 | 1.896240625 | 1.828220739 | 1.982684334 | 1.90849439 | 2.206825876 | 1.846338363 |
| ENSMUSG00000028031 | 1.100572389 | 1.25 | 1.25 | 0.830482024 | 0.686820904 | 0.77815125 | 0.448487648 |

| | | | | | | | |
|---|---|---|---|---|---|---|---|
| ENSMUSG00000028032 | 2.583267694 | 2.439139081 | 2.455044741 | 2.463717096 | 2.508609346 | 2.804820679 | 2.361378588 |
| ENSMUSG00000028033 | 1.181541736 | 1.214495249 | 1.061981878 | 1.27186571 | 1.176121988 | 1.041392685 | 1.054044578 |
| ENSMUSG00000028034 | 2.934668455 | 2.702843673 | 2.633332433 | 2.698400827 | 2.637264184 | 2.981365509 | 2.436573582 |
| ENSMUSG00000028035 | 2.53552908 | 2.331607372 | 2.349685923 | 2.286164561 | 2.198391101 | 2.51054501 | 2.033014511 |
| ENSMUSG00000028036 | 0.578129653 | 0.701838731 | 0.580482024 | 0.25 | 0 | 0 | 0 |
| ENSMUSG00000028037 | 0 | 0.25 | 0 | 0.25 | 0.244650542 | 0.301029996 | 0 |
| ENSMUSG00000028039 | 1.100572389 | 1.226722649 | 1.214495249 | 1.311981878 | 1.106691885 | 1.146128036 | 1.272006351 |
| ENSMUSG00000028040 | 1.707474304 | 1.62694866 | 1.708222504 | 1.42510993 | 1.563883923 | 1.612783857 | 1.554175045 |
| ENSMUSG00000028041 | 2.211649928 | 2.134789703 | 2.254202072 | 2.071350555 | 2.27729019 | 2.359835482 | 2.277155247 |
| ENSMUSG00000028042 | 2.116511873 | 2.058404919 | 2.124961472 | 1.695339928 | 1.808534465 | 1.826074803 | 1.884801065 |
| ENSMUSG00000028044 | 2.597490839 | 2.409156155 | 2.426408097 | 2.366391601 | 2.386538516 | 2.660865478 | 2.296701469 |
| ENSMUSG00000028047 | 2.026699189 | 1.880890489 | 1.96759118 | 1.886723615 | 1.998776581 | 2.149219113 | 1.947894097 |
| ENSMUSG00000028048 | 2.595842487 | 2.391037372 | 2.434100483 | 2.274508021 | 2.230953877 | 2.514547753 | 2.119837558 |
| ENSMUSG00000028049 | 2.048736215 | 2.101285366 | 2.138647213 | 1.856566189 | 2.090057366 | 2.093421685 | 2.105687397 |
| ENSMUSG00000028051 | 0.45815691 | 0.5 | 0.75 | 0.5 | 0.77552387 | 0.698970004 | 0.952571511 |
| ENSMUSG00000028053 | 2.877828211 | 2.622211087 | 2.620953944 | 2.733422664 | 2.660193997 | 3.055378331 | 2.467470936 |
| ENSMUSG00000028057 | 2.194712275 | 2.024508021 | 2.060793496 | 2.093759858 | 2.063536368 | 2.322219295 | 1.958704689 |
| ENSMUSG00000028059 | 2.787177998 | 2.526149688 | 2.52219706 | 2.432829758 | 2.456026973 | 2.763427994 | 2.317146272 |
| ENSMUSG00000028060 | 2.562324428 | 2.351816941 | 2.363317659 | 2.414552871 | 2.363759249 | 2.664641976 | 2.20779034 |
| ENSMUSG00000028062 | 2.010213549 | 1.888647213 | 1.991446071 | 1.899978211 | 1.988833497 | 2.096910013 | 1.872150729 |
| ENSMUSG00000028063 | 2.978790393 | 2.855121653 | 2.972375991 | 2.471805154 | 2.706665604 | 2.829946696 | 2.730081923 |
| ENSMUSG00000028064 | 0.671187741 | 0.701838731 | 0.701838731 | 0.580482024 | 0.632412477 | 0.301029996 | 0.672731473 |
| ENSMUSG00000028066 | 2.256831695 | 2.135757955 | 2.254202072 | 2.121960008 | 2.185632262 | 2.392696953 | 2.193776931 |
| ENSMUSG00000028068 | 2.627631768 | 2.443696765 | 2.482314602 | 2.491806565 | 2.565363825 | 2.808210973 | 2.445634316 |
| ENSMUSG00000028069 | 2.209569962 | 2.106566189 | 2.044977273 | 2.171625132 | 2.16174451 | 2.401400541 | 2.009874076 |
| ENSMUSG00000028070 | 2.245508853 | 2.092671852 | 2.144357207 | 2.222185812 | 2.212715924 | 2.456366033 | 2.116124284 |
| ENSMUSG00000028071 | 0 | 0 | 0 | 0 | 0 | 0 | 0 |
| ENSMUSG00000028072 | 0 | 0 | 0 | 0.396240625 | 0 | 0.301029996 | 0.579661877 |
| ENSMUSG00000028073 | 1.37447073 | 1.438721876 | 1.488549078 | 1.774508021 | 1.926994141 | 2.053078443 | 1.949460761 |
| ENSMUSG00000028076 | 0.747221736 | 0.646240625 | 0.580482024 | 1.114857905 | 1.074582839 | 1.230448921 | 1.066254157 |
| ENSMUSG00000028078 | 1.924654746 | 1.782320754 | 1.735628626 | 1.695339928 | 1.741652793 | 1.681241237 | 1.62316756 |
| ENSMUSG00000028080 | 2.194712275 | 2.057204673 | 2.070192693 | 2.255591953 | 2.188507684 | 2.51851394 | 2.057420422 |
| ENSMUSG00000028081 | 3.140901301 | 2.845115266 | 2.84253562 | 2.967398461 | 2.888971102 | 3.370698093 | 2.684302948 |
| ENSMUSG00000028082 | 2.245508853 | 2.051142786 | 2.009729747 | 2.306000419 | 2.267124351 | 2.580924976 | 2.15146804 |

| | | | | | | | |
|---|---|---|---|---|---|---|---|
| ENSMUSG00000028085 | 1.700465191 | 1.53743678 | 1.62694866 | 1.580482024 | 1.559656853 | 1.698970004 | 1.529383012 |
| ENSMUSG00000028086 | 2.042557845 | 1.930274797 | 1.860735874 | 1.892463902 | 1.800029087 | 1.986771734 | 1.714087544 |
| ENSMUSG00000028088 | 0.960252568 | 0.79248125 | 0.75 | 1.061981878 | 1.091002363 | 0.954242509 | 0.916588299 |
| ENSMUSG00000028089 | 2.209569962 | 2.027131114 | 1.991446071 | 2.113831805 | 2.113840608 | 2.36361198 | 1.978585612 |
| ENSMUSG00000028093 | 1.45815691 | 1.494319981 | 1.527131114 | 1.671625132 | 1.751666404 | 1.892094603 | 1.635773653 |
| ENSMUSG00000028096 | 2.069666964 | 1.912762923 | 1.823655187 | 2.100219859 | 2.029283463 | 2.28780173 | 1.953344547 |
| ENSMUSG00000028098 | 2.482695304 | 2.273849256 | 2.254202072 | 2.399047439 | 2.31625483 | 2.701567985 | 2.175263699 |
| ENSMUSG00000028099 | 2.300956576 | 2.12892496 | 2.103406982 | 2.242885888 | 2.198391101 | 2.496929648 | 2.054044578 |
| ENSMUSG00000028100 | 0.289064826 | 0.79248125 | 0.976722649 | 0.79248125 | 0.77552387 | 1.041392685 | 0.853775828 |
| ENSMUSG00000028101 | 2.194712275 | 2.049918086 | 2.074802005 | 2.006976499 | 2.112954895 | 2.309630167 | 2.008574816 |
| ENSMUSG00000028102 | 1.847794126 | 1.584962501 | 1.720660762 | 1.807204673 | 1.754126039 | 1.963787827 | 1.645916034 |
| ENSMUSG00000028104 | 1.418409478 | 1.410964047 | 1.464495249 | 1.695339928 | 1.618292349 | 1.838849091 | 1.579661877 |
| ENSMUSG00000028106 | 2.43085913 | 2.271202097 | 2.24929487 | 2.321928095 | 2.268268456 | 2.631443769 | 2.110947927 |
| ENSMUSG00000028107 | 2.256831695 | 2.116901388 | 2.142463902 | 2.152756199 | 2.17760189 | 2.346352974 | 2.072190387 |
| ENSMUSG00000028108 | 2.617563644 | 2.481944491 | 2.557204673 | 2.193696765 | 2.301493158 | 2.539076099 | 2.234765579 |
| ENSMUSG00000028109 | 0 | 0 | 0 | 0 | 0 | 0 | 0 |
| ENSMUSG00000028111 | 0.45815691 | 0.646240625 | 0.25 | 0.79248125 | 0.812711509 | 0.602059991 | 0.775756176 |
| ENSMUSG00000028114 | 2.256831695 | 2.173371739 | 2.142463902 | 2.277783918 | 2.244650542 | 2.480006943 | 2.092400484 |
| ENSMUSG00000028115 | 0 | 0.701838731 | 0.25 | 0.396240625 | 0.568060967 | 0.602059991 | 0.355418052 |
| ENSMUSG00000028116 | 0.289064826 | 0 | 0.396240625 | 0 | 0.244650542 | 0 | 0.448487648 |
| ENSMUSG00000028121 | 2.048736215 | 1.938721876 | 1.886723615 | 1.777131114 | 1.78687392 | 1.875061263 | 1.762103709 |
| ENSMUSG00000028124 | 2.559630991 | 2.353935442 | 2.3184489 | 2.522528105 | 2.436719872 | 2.797959644 | 2.269110043 |
| ENSMUSG00000028125 | 0.578129653 | 0.580482024 | 0.396240625 | 0.396240625 | 0.489301084 | 0.602059991 | 0.629532004 |
| ENSMUSG00000028126 | 2.605636262 | 2.402294685 | 2.429847205 | 2.398114259 | 2.407936347 | 2.710963119 | 2.287488514 |
| ENSMUSG00000028127 | 2.626865868 | 2.431128463 | 2.418989258 | 2.564641508 | 2.485926424 | 2.852479994 | 2.309276932 |
| ENSMUSG00000028128 | 2.792365035 | 2.592671852 | 2.615881093 | 1.75 | 1.733951626 | 1.886490725 | 1.637827726 |
| ENSMUSG00000028132 | 1.482695304 | 1.53743678 | 1.52186571 | 1.618933358 | 1.576269619 | 1.763427994 | 1.379042812 |
| ENSMUSG00000028133 | 1.325351389 | 1.188721876 | 1.238549078 | 1.226722649 | 1.358933174 | 1.204119983 | 1.168187745 |
| ENSMUSG00000028134 | 2.557825653 | 2.311388177 | 2.26584877 | 2.354463129 | 2.308696595 | 2.607455023 | 2.113781657 |
| ENSMUSG00000028136 | 2.736751754 | 2.509043403 | 2.505244985 | 2.705044741 | 2.656410937 | 3.042181595 | 2.44495927 |
| ENSMUSG00000028137 | 0.671187741 | 0.75 | 0.75 | 0.75 | 0.632412477 | 0.477121255 | 0.710836105 |
| ENSMUSG00000028138 | 2.620687497 | 2.420309603 | 2.403677461 | 2.569612365 | 2.518303419 | 2.895974732 | 2.337790282 |
| ENSMUSG00000028139 | 0.45815691 | 0.25 | 0.396240625 | 0.396240625 | 0.632412477 | 0.301029996 | 0.355418052 |
| ENSMUSG00000028140 | 2.512981297 | 2.325374049 | 2.322504712 | 2.394828984 | 2.372793355 | 2.674861141 | 2.222568443 |

| | | | | | | | |
|---|---|---|---|---|---|---|---|
| ENSMUSG00000028141 | 0.671187741 | 0.79248125 | 0.396240625 | 0 | 0.686820904 | 0 | 0.224243824 |
| ENSMUSG00000028145 | 1.777675583 | 1.476722649 | 1.52186571 | 1.630890489 | 1.614672255 | 1.740362689 | 1.548827561 |
| ENSMUSG00000028148 | 0 | 0 | 0 | 0 | 0 | 0 | 0 |
| ENSMUSG00000028149 | 2.706208651 | 2.482684334 | 2.439972046 | 2.601285366 | 2.538790182 | 2.910090546 | 2.337790282 |
| ENSMUSG00000028150 | 1.445324132 | 1.403677461 | 1.388647213 | 0.580482024 | 0.632412477 | 0.698970004 | 0.74492186 |
| ENSMUSG00000028152 | 2.436947303 | 2.242885888 | 2.230460234 | 2.294354884 | 2.27895639 | 2.555094449 | 2.145981019 |
| ENSMUSG00000028156 | 2.780867173 | 2.5684489 | 2.535526764 | 2.807805295 | 2.695097733 | 3.139564266 | 2.505768111 |
| ENSMUSG00000028158 | 1.505869732 | 1.53743678 | 1.470660762 | 1.566696635 | 1.555378544 | 1.672097858 | 1.425641774 |
| ENSMUSG00000028159 | 1.811507563 | 1.755591953 | 1.729715809 | 1.874961472 | 1.861137485 | 2.053078443 | 1.787569411 |
| ENSMUSG00000028161 | 2.592526109 | 2.392463902 | 2.370449858 | 2.370449858 | 2.292543754 | 2.574031268 | 2.115189268 |
| ENSMUSG00000028163 | 2.736751754 | 2.531353368 | 2.572216519 | 2.526477127 | 2.549506445 | 2.881954971 | 2.400284271 |
| ENSMUSG00000028164 | 2.185978293 | 2.03360658 | 2.118933358 | 1.899978211 | 1.832885921 | 2.170261715 | 1.760706232 |
| ENSMUSG00000028165 | 2.544964605 | 2.381380202 | 2.349151195 | 2.491446071 | 2.432800077 | 2.757396029 | 2.259064007 |
| ENSMUSG00000028167 | 1.249317394 | 1.302363341 | 1.17510993 | 0.864857905 | 0.733951626 | 0.698970004 | 0.876096088 |
| ENSMUSG00000028172 | 0 | 0 | 0 | 0 | 0 | 0 | 0.224243824 |
| ENSMUSG00000028173 | 3.366089831 | 3.040361712 | 3.014497931 | 3.304867767 | 3.194272066 | 3.733598461 | 2.970466485 |
| ENSMUSG00000028174 | 0 | 0 | 0 | 0 | 0 | 0 | 0 |
| ENSMUSG00000028175 | 2.323768721 | 2.216819685 | 2.21759118 | 2.391037372 | 2.318738698 | 2.680335513 | 2.156039304 |
| ENSMUSG00000028176 | 1.156259305 | 1.146240625 | 1.146240625 | 0.25 | 0 | 0 | 0 |
| ENSMUSG00000028177 | 0 | 0.396240625 | 0.396240625 | 0.25 | 0.387761935 | 0.602059991 | 0.448487648 |
| ENSMUSG00000028179 | 1.862428343 | 1.580482024 | 1.464495249 | 1.602347734 | 1.588235379 | 1.716003344 | 1.433437704 |
| ENSMUSG00000028180 | 2.865900016 | 2.649745743 | 2.633818844 | 2.73654381 | 2.660570082 | 3.056142262 | 2.458037398 |
| ENSMUSG00000028182 | 0.747221736 | 0.580482024 | 0.79248125 | 0.830482024 | 0.733951626 | 0.77815125 | 0.355418052 |
| ENSMUSG00000028184 | 2.578129653 | 2.38040011 | 2.319612365 | 2.077084758 | 2.103062484 | 2.342422681 | 1.976436 |
| ENSMUSG00000028185 | 0 | 0 | 0 | 0 | 0 | 0 | 0 |
| ENSMUSG00000028186 | 0 | 0 | 0 | 0 | 0.244650542 | 0 | 0 |
| ENSMUSG00000028187 | 2.153883046 | 2.019203899 | 1.987091808 | 2.168106335 | 2.111176775 | 2.382017043 | 1.984249047 |
| ENSMUSG00000028188 | 1.527823874 | 1.339388001 | 1.27186571 | 1.396240625 | 1.212046813 | 1.255272505 | 1.110947927 |
| ENSMUSG00000028189 | 1.721148722 | 1.566696635 | 1.606566189 | 1.646240625 | 1.642643806 | 1.77815125 | 1.496250175 |
| ENSMUSG00000028191 | 2.349889783 | 2.184523065 | 2.135757955 | 2.316696635 | 2.279510047 | 2.547774705 | 2.138662327 |
| ENSMUSG00000028194 | 2.654307229 | 2.434100483 | 2.392463902 | 2.528435542 | 2.450276722 | 2.829946696 | 2.296968506 |
| ENSMUSG00000028195 | 3.189573422 | 2.908816164 | 2.927701608 | 2.878431899 | 2.895309758 | 3.277150614 | 2.693286753 |
| ENSMUSG00000028197 | 0.811507563 | 0.79248125 | 0.79248125 | 0.701838731 | 0.686820904 | 0.477121255 | 0.853775828 |
| ENSMUSG00000028199 | 2.20747957 | 2.004202072 | 2.065523711 | 1.981203126 | 1.986303335 | 2.214843848 | 1.879993912 |

| | | | | | | | |
|---|---|---|---|---|---|---|---|
| ENSMUSG00000028207 | 2.585813169 | 2.343216265 | 2.321928095 | 2.449415381 | 2.401730365 | 2.754348336 | 2.210591357 |
| ENSMUSG00000028211 | 2.396352858 | 2.181128463 | 2.190387808 | 1.78743678 | 1.677808768 | 1.949390007 | 1.551512352 |
| ENSMUSG00000028212 | 2.517987749 | 2.309003548 | 2.338286706 | 2.387686696 | 2.292010185 | 2.672097858 | 2.108089156 |
| ENSMUSG00000028214 | 1.857607105 | 1.726722649 | 1.809601185 | 1.348079356 | 1.254881871 | 1.491361694 | 1.319146374 |
| ENSMUSG00000028217 | 0 | 0.25 | 0.25 | 0.396240625 | 0 | 0 | 0.579661877 |
| ENSMUSG00000028218 | 2.415874319 | 2.242166698 | 2.256284891 | 2.367410454 | 2.249502249 | 2.559906625 | 2.093904044 |
| ENSMUSG00000028221 | 2.524897319 | 2.348079356 | 2.351816941 | 2.325945187 | 2.241584006 | 2.557507202 | 2.07004434 |
| ENSMUSG00000028222 | 0.289064826 | 0 | 0 | 0.25 | 0.244650542 | 0.301029996 | 0 |
| ENSMUSG00000028223 | 2.294202867 | 2.143411797 | 2.08272922 | 2.079353153 | 2.003645018 | 2.257678575 | 1.861046144 |
| ENSMUSG00000028224 | 2.392336464 | 2.271202097 | 2.206637122 | 2.31728167 | 2.244039363 | 2.575187845 | 2.09490255 |
| ENSMUSG00000028226 | 2.013563224 | 1.828220739 | 1.837182039 | 1.557204673 | 1.433158226 | 1.568201724 | 1.413583902 |
| ENSMUSG00000028228 | 2.734980913 | 2.528435542 | 2.496104615 | 2.6343046 | 2.538259021 | 2.902002891 | 2.327996893 |
| ENSMUSG00000028229 | 1.249317394 | 1.201838731 | 1.25 | 1.372963274 | 1.327538614 | 1.431363764 | 1.224243824 |
| ENSMUSG00000028232 | 2.163307908 | 2.046218836 | 2.001406137 | 2.063166358 | 2.037065488 | 2.230448921 | 1.851668228 |
| ENSMUSG00000028233 | 2.337801042 | 2.187886015 | 2.207430684 | 2.271202097 | 2.222603297 | 2.51054501 | 2.060761403 |
| ENSMUSG00000028234 | 2.91285295 | 2.701234418 | 2.734159485 | 2.584405475 | 2.578885001 | 2.944975908 | 2.43291209 |
| ENSMUSG00000028236 | 0 | 0 | 0 | 0 | 0 | 0 | 0 |
| ENSMUSG00000028238 | 0 | 0 | 0.25 | 0 | 0 | 0 | 0 |
| ENSMUSG00000028240 | 0 | 0 | 0 | 0 | 0 | 0 | 0 |
| ENSMUSG00000028243 | 2.131864492 | 1.988549078 | 1.956637122 | 2.158248799 | 2.085294053 | 2.320146286 | 1.936709733 |
| ENSMUSG00000028244 | 0 | 0 | 0 | 0 | 0 | 0 | 0 |
| ENSMUSG00000028245 | 2.382811656 | 2.163659007 | 2.205044741 | 2.262462137 | 2.19135987 | 2.45484486 | 2.026306127 |
| ENSMUSG00000028246 | 2.370236862 | 2.252806814 | 2.225216702 | 2.132845365 | 2.031524462 | 2.322219295 | 1.901538355 |
| ENSMUSG00000028247 | 2.020182925 | 1.948603967 | 1.914552871 | 1.991446071 | 1.938809675 | 2.204119983 | 1.808770687 |
| ENSMUSG00000028248 | 2.727821383 | 2.501055117 | 2.506630861 | 2.508699741 | 2.454682377 | 2.769377326 | 2.269691383 |
| ENSMUSG00000028249 | 3.066206095 | 2.805547785 | 2.777294426 | 2.861899965 | 2.759562844 | 3.206556044 | 2.56472862 |
| ENSMUSG00000028251 | 1.903481042 | 1.774508021 | 1.80479213 | 1.789967834 | 1.751666404 | 1.986771734 | 1.723650273 |
| ENSMUSG00000028252 | 2.269664473 | 2.08272922 | 2.099151195 | 2.245748394 | 2.172394856 | 2.46686762 | 2.000667731 |
| ENSMUSG00000028255 | 0 | 0 | 0 | 0.25 | 0.387761935 | 0 | 0 |
| ENSMUSG00000028256 | 2.249317394 | 2.125956434 | 2.118933358 | 2.279735268 | 2.169384551 | 2.46834733 | 2.006616089 |
| ENSMUSG00000028259 | 0 | 0 | 0 | 0 | 0 | 0 | 0 |
| ENSMUSG00000028261 | 2.181541736 | 2.004202072 | 2.009729747 | 2.093759858 | 1.974687643 | 2.28780173 | 1.860017477 |
| ENSMUSG00000028262 | 0 | 0.701838731 | 0.396240625 | 0 | 0 | 0 | 0 |
| ENSMUSG00000028264 | 0 | 0 | 0 | 0 | 0 | 0 | 0 |

| | | | | | | | |
|---|---|---|---|---|---|---|---|
| ENSMUSG00000028266 | 2.352857998 | 2.189555804 | 2.163659007 | 2.027131114 | 1.977302144 | 2.238046103 | 1.876096088 |
| ENSMUSG00000028268 | 1.432083896 | 1.321350555 | 1.26109853 | 1.914552871 | 1.86473911 | 2.103803721 | 1.714087544 |
| ENSMUSG00000028270 | 1.445324132 | 1.380890489 | 1.458222504 | 2.065523711 | 1.964031554 | 2.220108088 | 1.863093717 |
| ENSMUSG00000028271 | 2.386920383 | 2.245748394 | 2.242166698 | 2.347004321 | 2.330902415 | 2.64246452 | 2.193409508 |
| ENSMUSG00000028273 | 3.156474659 | 2.886844141 | 2.863703337 | 2.482314602 | 2.418540977 | 2.743509765 | 2.25178005 |
| ENSMUSG00000028274 | 2.285603957 | 2.116901388 | 2.087182039 | 2.218361028 | 2.126866297 | 2.409933123 | 2.009874076 |
| ENSMUSG00000028277 | 2.675303226 | 2.44697564 | 2.46293726 | 2.55388325 | 2.49804628 | 2.860936621 | 2.341067602 |
| ENSMUSG00000028278 | 2.089597219 | 1.912762923 | 1.997171172 | 1.76109853 | 1.645989387 | 1.819543936 | 1.546120303 |
| ENSMUSG00000028280 | 0 | 0 | 0 | 0 | 0 | 0 | 0.224243824 |
| ENSMUSG00000028282 | 2.688073951 | 2.49539182 | 2.472566069 | 2.551142786 | 2.448224646 | 2.783188691 | 2.264126018 |
| ENSMUSG00000028284 | 2.812982046 | 2.583847589 | 2.556603048 | 2.724272282 | 2.60468636 | 2.996949248 | 2.434486378 |
| ENSMUSG00000028287 | 0.289064826 | 0.75 | 0.701838731 | 0.25 | 0.387761935 | 0.301029996 | 0.224243824 |
| ENSMUSG00000028289 | 1.568542279 | 1.321350555 | 1.364857905 | 1.445339928 | 1.074582839 | 1.397940009 | 1.110947927 |
| ENSMUSG00000028291 | 2.410757293 | 2.191217898 | 2.175975893 | 2.234894804 | 2.172394856 | 2.456366033 | 2.014380832 |
| ENSMUSG00000028292 | 1.960252568 | 1.914552871 | 1.899978211 | 1.994319981 | 1.861137485 | 2.093421685 | 1.762103709 |
| ENSMUSG00000028293 | 2.194712275 | 2.031030328 | 2.048689214 | 2.087182039 | 2.046825265 | 2.344392274 | 1.885753983 |
| ENSMUSG00000028294 | 0.45815691 | 0.25 | 0.25 | 0.580482024 | 0.733951626 | 0.602059991 | 0.710836105 |
| ENSMUSG00000028295 | 1.516991255 | 1.445339928 | 1.552363341 | 1.54745614 | 1.559656853 | 1.544068044 | 1.441050184 |
| ENSMUSG00000028298 | 0 | 0 | 0 | 0 | 0 | 0 | 0 |
| ENSMUSG00000028300 | 2.069666964 | 1.87892496 | 1.816696635 | 1.955044741 | 1.964031554 | 2.178976947 | 1.788855757 |
| ENSMUSG00000028307 | 0 | 0 | 0 | 0 | 0 | 0 | 0.224243824 |
| ENSMUSG00000028309 | 2.670497861 | 2.435366747 | 2.436208459 | 2.5658173 | 2.496553863 | 2.828659897 | 2.307473178 |
| ENSMUSG00000028310 | 0 | 0 | 0 | 0 | 0 | 0 | 0 |
| ENSMUSG00000028312 | 3.101555957 | 2.830059111 | 2.828078937 | 2.943696765 | 2.836786029 | 3.277379975 | 2.62297273 |
| ENSMUSG00000028314 | 0 | 0 | 0 | 0 | 0 | 0 | 0 |
| ENSMUSG00000028318 | 1.967766868 | 1.834962501 | 1.868933358 | 2.038704527 | 2.019019072 | 2.23299611 | 1.871156825 |
| ENSMUSG00000028322 | 1.903481042 | 1.87694866 | 1.834962501 | 1.837182039 | 1.832885921 | 2.037426498 | 1.755055163 |
| ENSMUSG00000028327 | 0 | 0 | 0 | 0 | 0 | 0.301029996 | 0.355418052 |
| ENSMUSG00000028328 | 0.578129653 | 0.896240625 | 0.79248125 | 0.5 | 0.568060967 | 0.301029996 | 0.520678036 |
| ENSMUSG00000028329 | 1.527823874 | 1.410964047 | 1.42510993 | 1.657339155 | 1.665423072 | 1.69019608 | 1.559435575 |
| ENSMUSG00000028330 | 2.842342644 | 2.630645383 | 2.600486531 | 2.710587586 | 2.624591555 | 3.045714059 | 2.461426716 |
| ENSMUSG00000028331 | 1.289064826 | 1.201838731 | 1.160964047 | 1.188721876 | 1.212046813 | 1.204119983 | 1.121219121 |
| ENSMUSG00000028332 | 0 | 0 | 0 | 0 | 0 | 0 | 0 |
| ENSMUSG00000028333 | 3.220052477 | 2.903101948 | 2.905054956 | 2.932723662 | 2.844682468 | 3.314077992 | 2.657155915 |

| | | | | | | | |
|---|---|---|---|---|---|---|---|
| ENSMUSG00000028334 | 2.638209243 | 2.430274797 | 2.501406137 | 2.385274154 | 2.428473696 | 2.740362689 | 2.382544765 |
| ENSMUSG00000028337 | 0.671187741 | 0.701838731 | 0.75 | 1.67510993 | 1.662257525 | 1.875061263 | 1.63986884 |
| ENSMUSG00000028339 | 1.389637216 | 1.364857905 | 1.482684334 | 0 | 0.387761935 | 0.301029996 | 0.224243824 |
| ENSMUSG00000028341 | 0.45815691 | 0.5 | 0.25 | 0.580482024 | 0.846351821 | 1 | 0.775756176 |
| ENSMUSG00000028343 | 2.485932312 | 2.28743678 | 2.295598588 | 2.351285366 | 2.263090523 | 2.624282096 | 2.125782179 |
| ENSMUSG00000028344 | 2.033115198 | 1.821350555 | 1.850219859 | 1.909156155 | 1.822942838 | 2.045322979 | 1.700880998 |
| ENSMUSG00000028345 | 2.537433341 | 2.323080408 | 2.333288838 | 2.363317659 | 2.277846464 | 2.608526034 | 2.095400649 |
| ENSMUSG00000028347 | 1.862428343 | 1.779735268 | 1.720660762 | 1.950224975 | 1.92245931 | 2.146128036 | 1.757893036 |
| ENSMUSG00000028348 | 0.747221736 | 0.580482024 | 0.951838731 | 1.188721876 | 1.149965125 | 1 | 1.041356071 |
| ENSMUSG00000028351 | 0 | 0 | 0.25 | 0.25 | 0 | 0 | 0 |
| ENSMUSG00000028354 | 0 | 0 | 0 | 1.098079356 | 1.091002363 | 1 | 1.078019652 |
| ENSMUSG00000028356 | 0 | 0 | 0 | 0 | 0 | 0 | 0 |
| ENSMUSG00000028357 | 0 | 0 | 0 | 0 | 0 | 0 | 0 |
| ENSMUSG00000028358 | 2.203267085 | 2.130890489 | 2.097004321 | 1.685366747 | 1.72073075 | 1.832508913 | 1.655750087 |
| ENSMUSG00000028359 | 0.289064826 | 0 | 0.396240625 | 0 | 0 | 0 | 0 |
| ENSMUSG00000028360 | 0 | 0 | 0 | 0.25 | 0 | 0 | 0 |
| ENSMUSG00000028362 | 0 | 0 | 0 | 0.25 | 0 | 0 | 0 |
| ENSMUSG00000028364 | 3.065938681 | 2.822648722 | 2.8158173 | 2.702843673 | 2.699837743 | 3.038620162 | 2.525718559 |
| ENSMUSG00000028367 | 3.051791411 | 2.800684011 | 2.803731547 | 2.786005217 | 2.794928476 | 3.218535505 | 2.60699341 |
| ENSMUSG00000028369 | 0.960252568 | 1 | 0.75 | 0.830482024 | 0.931471446 | 0.602059991 | 0.775756176 |
| ENSMUSG00000028370 | 1.940852214 | 1.823655187 | 1.882845365 | 0.701838731 | 0.77552387 | 0.954242509 | 0.629532004 |
| ENSMUSG00000028373 | 0 | 0 | 0 | 0 | 0 | 0.301029996 | 0 |
| ENSMUSG00000028378 | 2.194712275 | 2.052363341 | 2.079353153 | 2.218361028 | 2.184185718 | 2.460897843 | 2.05853792 |
| ENSMUSG00000028381 | 2.638209243 | 2.434945152 | 2.42813175 | 2.593216265 | 2.503367327 | 2.88536122 | 2.327268527 |
| ENSMUSG00000028382 | 3.006528408 | 2.745926548 | 2.714300868 | 2.847676593 | 2.771840816 | 3.183839037 | 2.529383012 |
| ENSMUSG00000028383 | 2.036286563 | 1.926839783 | 1.918106335 | 2.060793496 | 1.997548906 | 2.320146286 | 1.927642465 |
| ENSMUSG00000028385 | 2.294202867 | 2.175975893 | 2.190387808 | 2.080482024 | 2.056322416 | 2.318063335 | 1.947894097 |
| ENSMUSG00000028386 | 0 | 0 | 0 | 0 | 0 | 0 | 0 |
| ENSMUSG00000028389 | 1.794934559 | 1.755591953 | 1.688721876 | 2.013820609 | 1.954436039 | 2.204119983 | 1.764880683 |
| ENSMUSG00000028391 | 1.358731791 | 1.388647213 | 1.445339928 | 1.561981878 | 1.504538967 | 1.568201724 | 1.48984372 |
| ENSMUSG00000028392 | 0.867194479 | 0.701838731 | 0.646240625 | 0.896240625 | 0.568060967 | 1 | 0.935079929 |
| ENSMUSG00000028393 | 2.552362269 | 2.408702763 | 2.452642909 | 2.132845365 | 2.365931294 | 2.487138375 | 2.289952884 |
| ENSMUSG00000028394 | 2.530730035 | 2.337734795 | 2.344844592 | 2.407339155 | 2.331380351 | 2.691965103 | 2.230204359 |
| ENSMUSG00000028396 | 0 | 0 | 0 | 0 | 0 | 0 | 0 |

| | | | | | | | |
|---|---|---|---|---|---|---|---|
| ENSMUSG00000028397 | 2.35727108 | 2.181128463 | 2.151832578 | 2.255591953 | 2.1962964 | 2.477121255 | 2.041356071 |
| ENSMUSG00000028398 | 1.693336259 | 1.678561379 | 1.660964047 | 1.580482024 | 1.568060967 | 1.72427587 | 1.605556929 |
| ENSMUSG00000028399 | 2.048736215 | 1.912762923 | 1.828220739 | 1.973704441 | 1.917865457 | 2.193124598 | 1.731408809 |
| ENSMUSG00000028402 | 2.830751226 | 2.600753006 | 2.575374049 | 2.548689214 | 2.462015062 | 2.830588669 | 2.293209778 |
| ENSMUSG00000028403 | 2.436947303 | 2.28743678 | 2.269871196 | 2.333288838 | 2.24829556 | 2.575187845 | 2.07004434 |
| ENSMUSG00000028405 | 2.66283289 | 2.46293726 | 2.43662858 | 2.476346751 | 2.447194116 | 2.820857989 | 2.292939619 |
| ENSMUSG00000028407 | 0.289064826 | 0.75 | 0.701838731 | 0.580482024 | 0.387761935 | 0.477121255 | 0.629532004 |
| ENSMUSG00000028409 | 2.871480564 | 2.604199382 | 2.588562345 | 2.731203126 | 2.650465868 | 3.096910013 | 2.458037398 |
| ENSMUSG00000028410 | 3.000313205 | 2.723325383 | 2.679204865 | 2.863446264 | 2.769225805 | 3.185258765 | 2.574383083 |
| ENSMUSG00000028411 | 2.045658471 | 1.926839783 | 1.884789703 | 1.955044741 | 1.91476933 | 2.178976947 | 1.778418869 |
| ENSMUSG00000028412 | 2.555102908 | 2.327653195 | 2.320192693 | 2.447790722 | 2.365497953 | 2.711807229 | 2.175652306 |
| ENSMUSG00000028413 | 2.87190676 | 2.623463862 | 2.644593173 | 2.512121718 | 2.483451699 | 2.815577748 | 2.3030503 |
| ENSMUSG00000028414 | 2.462727174 | 2.273849256 | 2.23927551 | 2.276477127 | 2.241584006 | 2.559906625 | 2.062969708 |
| ENSMUSG00000028415 | 0.289064826 | 0.25 | 0.25 | 0 | 0 | 0 | 0 |
| ENSMUSG00000028416 | 2.743760867 | 2.501055117 | 2.52186571 | 2.628678514 | 2.523606932 | 2.954242509 | 2.356308054 |
| ENSMUSG00000028417 | 0 | 0 | 0 | 0.25 | 0.244650542 | 0 | 0 |
| ENSMUSG00000028419 | 2.497590902 | 2.314346961 | 2.311981878 | 2.449820405 | 2.385310841 | 2.767155866 | 2.237984665 |
| ENSMUSG00000028420 | 1.876566388 | 1.794977273 | 1.769203899 | 1.705044741 | 1.642643806 | 1.812913357 | 1.546120303 |
| ENSMUSG00000028423 | 2.764659211 | 2.539967834 | 2.524178789 | 2.561090959 | 2.506289166 | 2.850646235 | 2.339900938 |
| ENSMUSG00000028426 | 3.144461898 | 2.871079856 | 2.859438018 | 2.884789703 | 2.788185538 | 3.250663919 | 2.61187062 |
| ENSMUSG00000028427 | 0.289064826 | 0.25 | 0.396240625 | 0.25 | 0.568060967 | 0 | 0 |
| ENSMUSG00000028430 | 2.326928073 | 2.27053726 | 2.273189285 | 2.170748646 | 2.261347526 | 2.462397998 | 2.17331363 |
| ENSMUSG00000028431 | 2.737340367 | 2.524178789 | 2.510756821 | 2.477847997 | 2.48684999 | 2.781036939 | 2.2945572 |
| ENSMUSG00000028433 | 2.786132803 | 2.607090043 | 2.616391601 | 2.512121718 | 2.52691592 | 2.840733235 | 2.435357683 |
| ENSMUSG00000028434 | 0.289064826 | 0.580482024 | 0.79248125 | 0.896240625 | 0.931471446 | 1 | 0.829800754 |
| ENSMUSG00000028435 | 0.45815691 | 0.580482024 | 0.25 | 0.25 | 0.77552387 | 0.698970004 | 0.579661877 |
| ENSMUSG00000028436 | 2.488076442 | 2.289336734 | 2.332168732 | 2.438304187 | 2.35849115 | 2.757396029 | 2.208492871 |
| ENSMUSG00000028437 | 2.461589295 | 2.271202097 | 2.281030328 | 2.324229052 | 2.27895639 | 2.588831726 | 2.13691595 |
| ENSMUSG00000028438 | 1.967766868 | 1.843759858 | 1.821350555 | 2.00835575 | 1.950242452 | 2.123851641 | 1.812372009 |
| ENSMUSG00000028439 | 1.505869732 | 1.445339928 | 1.505591953 | 1.606566189 | 1.625423019 | 1.785329835 | 1.647906904 |
| ENSMUSG00000028441 | 0 | 0.25 | 0.25 | 0.580482024 | 0.568060967 | 0.477121255 | 0.224243824 |
| ENSMUSG00000028443 | 1.967766868 | 1.903677461 | 1.892463902 | 1.87694866 | 1.834841362 | 2.045322979 | 1.735976162 |
| ENSMUSG00000028444 | 0.289064826 | 0.5 | 0.79248125 | 1.17510993 | 1.310728002 | 1 | 1.313616061 |
| ENSMUSG00000028445 | 0.578129653 | 0.396240625 | 0.5 | 0.580482024 | 0.812711509 | 0.698970004 | 1.140832123 |

| | | | | | | | |
|---|---|---|---|---|---|---|---|
| ENSMUSG00000028447 | 2.100572389 | 1.973704441 | 2.028435542 | 1.92510993 | 2.014359518 | 2.136720567 | 2.009874076 |
| ENSMUSG00000028451 | 0.45815691 | 0.396240625 | 0.25 | 0.25 | 0.387761935 | 0.301029996 | 0.224243824 |
| ENSMUSG00000028452 | 3.162035173 | 2.889006748 | 2.871457327 | 2.956934912 | 2.908028564 | 3.335658452 | 2.727270561 |
| ENSMUSG00000028453 | 1.847794126 | 1.735628626 | 1.71759118 | 1.735628626 | 1.820920161 | 1.968482949 | 1.747849621 |
| ENSMUSG00000028454 | 1.623015126 | 1.584962501 | 1.657339155 | 1.505591953 | 1.704182902 | 1.662757832 | 1.625302983 |
| ENSMUSG00000028455 | 2.428398755 | 2.23927551 | 2.286801231 | 2.363831805 | 2.362449578 | 2.666517981 | 2.226242896 |
| ENSMUSG00000028456 | 1.678702042 | 1.589388001 | 1.646240625 | 2.017865591 | 1.991345651 | 2.176091259 | 1.861046144 |
| ENSMUSG00000028457 | 0.45815691 | 0.830482024 | 0.79248125 | 0.646240625 | 0.489301084 | 0.477121255 | 0.672731473 |
| ENSMUSG00000028458 | 1.789260561 | 1.685366747 | 1.752806814 | 1.571350555 | 1.652586889 | 1.806179974 | 1.647906904 |
| ENSMUSG00000028459 | 0.289064826 | 0.25 | 0.396240625 | 0.75 | 0.568060967 | 0.301029996 | 0.224243824 |
| ENSMUSG00000028460 | 0 | 0 | 0 | 0 | 0 | 0 | 0 |
| ENSMUSG00000028461 | 1.87190676 | 1.777131114 | 1.856566189 | 1.880890489 | 1.814781639 | 2.075546961 | 1.74492186 |
| ENSMUSG00000028463 | 1.342375483 | 1.445339928 | 1.418106335 | 1.02186571 | 0.978602168 | 0.77815125 | 1.014380832 |
| ENSMUSG00000028464 | 3.437386631 | 3.151254112 | 3.18086191 | 2.511780978 | 2.602034204 | 2.891537458 | 2.470451221 |
| ENSMUSG00000028465 | 3.358333922 | 3.10353917 | 3.129355835 | 2.868553234 | 2.985705357 | 3.288919606 | 2.837974745 |
| ENSMUSG00000028466 | 2.304292853 | 2.118933358 | 2.131869252 | 2.266522298 | 2.235369829 | 2.46686762 | 2.099358074 |
| ENSMUSG00000028467 | 1.87190676 | 1.774508021 | 1.79745614 | 1.92510993 | 1.979897422 | 2.127104798 | 1.869159809 |
| ENSMUSG00000028468 | 2.377269616 | 2.181128463 | 2.245748394 | 2.153677461 | 2.234113756 | 2.459392488 | 2.092400484 |
| ENSMUSG00000028469 | 2.231898196 | 2.118933358 | 2.165444524 | 2.198603967 | 2.198391101 | 2.416640507 | 2.119837558 |
| ENSMUSG00000028470 | 1.37447073 | 1.238549078 | 1.26109853 | 1.451838731 | 1.42077253 | 1.579783597 | 1.499406449 |
| ENSMUSG00000028475 | 0.45815691 | 0.25 | 0.396240625 | 0 | 0.568060967 | 0 | 0.355418052 |
| ENSMUSG00000028476 | 1.655740331 | 1.54248125 | 1.418106335 | 1.685366747 | 1.665423072 | 1.832508913 | 1.56716938 |
| ENSMUSG00000028478 | 2.819794862 | 2.59727338 | 2.617664719 | 2.581326364 | 2.571222152 | 2.932473765 | 2.451316035 |
| ENSMUSG00000028479 | 1.964026642 | 1.916333979 | 1.94697564 | 1.799918086 | 1.861137485 | 1.986771734 | 1.797719748 |
| ENSMUSG00000028480 | 3.016283996 | 2.765680192 | 2.749471282 | 2.641989019 | 2.621875691 | 2.999565488 | 2.463666693 |
| ENSMUSG00000028483 | 2.029919532 | 1.921625132 | 1.96293726 | 2.019203899 | 1.925488995 | 2.146128036 | 1.802677937 |
| ENSMUSG00000028484 | 3.013259818 | 2.756976499 | 2.722375991 | 2.835657575 | 2.744479432 | 3.185542155 | 2.54076252 |
| ENSMUSG00000028487 | 1.53838222 | 1.54248125 | 1.527131114 | 1.744319981 | 1.746695109 | 1.903089987 | 1.63986884 |
| ENSMUSG00000028488 | 0.289064826 | 0 | 0.5 | 0.896240625 | 0.77552387 | 0.903089987 | 0.803905701 |
| ENSMUSG00000028492 | 0 | 0 | 0 | 0 | 0 | 0 | 0.224243824 |
| ENSMUSG00000028494 | 2.711248379 | 2.487821179 | 2.514836115 | 2.345926073 | 2.362449578 | 2.64738297 | 2.204254617 |
| ENSMUSG00000028495 | 2.921640949 | 2.693284802 | 2.670748646 | 2.760756821 | 2.697306371 | 3.103461622 | 2.525981686 |
| ENSMUSG00000028496 | 2.497590902 | 2.328787391 | 2.293106877 | 2.137686696 | 2.093822267 | 2.340444115 | 1.929310074 |
| ENSMUSG00000028497 | 2.457006475 | 2.255591953 | 2.298689214 | 2.237091808 | 2.157861565 | 2.474216264 | 2.013099581 |

| | | | | | | | |
|---|---|---|---|---|---|---|---|
| ENSMUSG00000028514 | 2.893084681 | 2.637205477 | 2.613317659 | 2.717012713 | 2.663004918 | 3.038620162 | 2.449821735 |
| ENSMUSG00000028517 | 1.993049299 | 1.928561379 | 1.87694866 | 1.339388001 | 1.106691885 | 1.278753601 | 1.168187745 |
| ENSMUSG00000028518 | 1.811507563 | 1.695339928 | 1.630890489 | 1.67510993 | 1.655839964 | 1.826074803 | 1.562034107 |
| ENSMUSG00000028519 | 0 | 0 | 0 | 0 | 0 | 0 | 0 |
| ENSMUSG00000028520 | 0 | 0 | 0 | 0 | 0 | 0 | 0 |
| ENSMUSG00000028521 | 1.940852214 | 1.892463902 | 1.843759858 | 1.948603967 | 1.917865457 | 2.178976947 | 1.786277931 |
| ENSMUSG00000028522 | 2.631440309 | 2.428990498 | 2.405512955 | 2.505591953 | 2.383667271 | 2.777426822 | 2.20779034 |
| ENSMUSG00000028523 | 0 | 0 | 0 | 0 | 0 | 0 | 0 |
| ENSMUSG00000028524 | 1.578129653 | 1.372963274 | 1.488549078 | 1.431980114 | 1.351342427 | 1.544068044 | 1.259064007 |
| ENSMUSG00000028525 | 1.885732304 | 1.78743678 | 1.685366747 | 1.571350555 | 1.53772706 | 1.591064607 | 1.444790289 |
| ENSMUSG00000028527 | 1.707474304 | 1.566696635 | 1.602347734 | 1.238549078 | 1.234113756 | 1.255272505 | 1.252393349 |
| ENSMUSG00000028528 | 0 | 0 | 0 | 0.25 | 0.387761935 | 0 | 0.224243824 |
| ENSMUSG00000028530 | 3.133460136 | 2.842671852 | 2.833428607 | 2.904366866 | 2.810260415 | 3.261976191 | 2.612877516 |
| ENSMUSG00000028532 | 2.192545868 | 2.08272922 | 2.092671852 | 2.004202072 | 2.044679629 | 2.29666519 | 1.941550454 |
| ENSMUSG00000028533 | 0 | 0 | 0 | 0 | 0 | 0 | 0.224243824 |
| ENSMUSG00000028536 | 0 | 0 | 0 | 0 | 0 | 0 | 0 |
| ENSMUSG00000028538 | 1.894701088 | 1.839388001 | 1.926839783 | 1.76109853 | 1.893954095 | 1.959041392 | 1.812372009 |
| ENSMUSG00000028539 | 1 | 1.04248125 | 1.114857905 | 1.130890489 | 1.039257769 | 1.146128036 | 1.193409508 |
| ENSMUSG00000028540 | 1.847794126 | 1.811981878 | 1.828220739 | 1.638647213 | 1.814781639 | 1.913813852 | 1.753626837 |
| ENSMUSG00000028541 | 2.23778581 | 2.129909063 | 2.161864607 | 1.930274797 | 1.997548906 | 2.053078443 | 1.945529784 |
| ENSMUSG00000028542 | 1.87190676 | 1.720660762 | 1.664552871 | 1.130890489 | 1.283908311 | 1.255272505 | 1.238624656 |
| ENSMUSG00000028544 | 0 | 0 | 0 | 0 | 0 | 0 | 0 |
| ENSMUSG00000028545 | 1.587501561 | 1.584962501 | 1.505591953 | 1.557204673 | 1.580303447 | 1.591064607 | 1.543390198 |
| ENSMUSG00000028546 | 0.747221736 | 1.04248125 | 1.04248125 | 0.646240625 | 0.387761935 | 0.301029996 | 0.224243824 |
| ENSMUSG00000028549 | 1.678702042 | 1.755591953 | 1.729715809 | 1.766522298 | 1.695608647 | 1.963787827 | 1.556816002 |
| ENSMUSG00000028550 | 1.777675583 | 1.701838731 | 1.729715809 | 1.698603967 | 1.671670246 | 1.799340549 | 1.56970677 |
| ENSMUSG00000028551 | 2.355805236 | 2.184523065 | 2.21759118 | 2.133818844 | 2.094757251 | 2.303196057 | 1.987739 |
| ENSMUSG00000028552 | 2.731416512 | 2.513820609 | 2.493603647 | 2.650674716 | 2.55764999 | 2.949877704 | 2.357638483 |
| ENSMUSG00000028553 | 0 | 0 | 0 | 0.25 | 0.387761935 | 0 | 0 |
| ENSMUSG00000028555 | 1.614416215 | 1.464495249 | 1.593759858 | 1.339388001 | 1.351342427 | 1.361727836 | 1.278288402 |
| ENSMUSG00000028556 | 2.944805158 | 2.674024543 | 2.639844273 | 2.691217898 | 2.621875691 | 3.028571253 | 2.447651042 |
| ENSMUSG00000028557 | 2.886555665 | 2.640083209 | 2.630890489 | 2.595115266 | 2.517458689 | 2.914871818 | 2.349799373 |
| ENSMUSG00000028558 | 0.578129653 | 0.25 | 0.396240625 | 0.25 | 0 | 0 | 0.355418052 |
| ENSMUSG00000028559 | 2.734388958 | 2.494677615 | 2.492885888 | 2.583288838 | 2.460693128 | 2.856124444 | 2.315638039 |

| | | | | | | | |
|---|---|---|---|---|---|---|---|
| ENSMUSG00000028560 | 2.873182739 | 2.629663289 | 2.613831805 | 2.662538553 | 2.553856086 | 2.954242509 | 2.376756474 |
| ENSMUSG00000028563 | 1.822201287 | 1.732684334 | 1.714495249 | 1.794977273 | 1.777823267 | 1.913813852 | 1.562034107 |
| ENSMUSG00000028565 | 1.614416215 | 1.593759858 | 1.638647213 | 1.42510993 | 1.380772476 | 1.602059991 | 1.340368108 |
| ENSMUSG00000028567 | 2.436947303 | 2.230460234 | 2.276477127 | 2.289967834 | 2.184909731 | 2.568201724 | 2.080113611 |
| ENSMUSG00000028568 | 2.587501561 | 2.370449858 | 2.354990044 | 2.46759118 | 2.351342427 | 2.678518379 | 2.189340078 |
| ENSMUSG00000028571 | 0 | 0.396240625 | 0 | 0 | 0.387761935 | 0.301029996 | 0 |
| ENSMUSG00000028572 | 0 | 0 | 0.25 | 1.54248125 | 1.427019704 | 1.531478917 | 1.290497981 |
| ENSMUSG00000028573 | 0 | 0 | 0 | 0 | 0 | 0 | 0 |
| ENSMUSG00000028575 | 0 | 0.5 | 0 | 0.25 | 0 | 0.301029996 | 0.355418052 |
| ENSMUSG00000028576 | 2.381433035 | 2.227473271 | 2.250703754 | 2.329353153 | 2.227765453 | 2.582063363 | 2.074853081 |
| ENSMUSG00000028577 | 2.906236339 | 2.661414608 | 2.642226539 | 2.723325383 | 2.603804521 | 3.009025742 | 2.418387761 |
| ENSMUSG00000028578 | 1.956444026 | 1.898114259 | 1.811981878 | 1.898114259 | 1.87878897 | 2.11058971 | 1.734460864 |
| ENSMUSG00000028580 | 2.760228547 | 2.551142786 | 2.572216519 | 2.51007243 | 2.491128299 | 2.805500858 | 2.340368108 |
| ENSMUSG00000028581 | 0.289064826 | 0 | 0.25 | 0 | 0 | 0 | 0 |
| ENSMUSG00000028582 | 2.299278375 | 2.146240625 | 2.188721876 | 2.087182039 | 2.201164838 | 2.340444115 | 2.06186744 |
| ENSMUSG00000028583 | 0.289064826 | 0.25 | 0.25 | 2.368933358 | 2.253097759 | 2.599883072 | 2.13206423 |
| ENSMUSG00000028584 | 0 | 0 | 0 | 0 | 0 | 0 | 0 |
| ENSMUSG00000028587 | 2.283862692 | 2.073655187 | 2.117918804 | 2.002806814 | 1.94314048 | 2.161368002 | 1.87511425 |
| ENSMUSG00000028589 | 0 | 0 | 0 | 0 | 0 | 0 | 0 |
| ENSMUSG00000028590 | 0 | 0 | 0 | 0 | 0 | 0 | 0 |
| ENSMUSG00000028591 | 0 | 0 | 0 | 0 | 0 | 0 | 0 |
| ENSMUSG00000028593 | 0 | 0 | 0 | 0 | 0 | 0 | 0 |
| ENSMUSG00000028597 | 0 | 0 | 0 | 1.311981878 | 1.351342427 | 1.505149978 | 1.307989563 |
| ENSMUSG00000028599 | 2.739101238 | 2.501756817 | 2.536483036 | 2.306000419 | 2.328502976 | 2.63447727 | 2.195972745 |
| ENSMUSG00000028600 | 0.867194479 | 0.75 | 0.92510993 | 0.580482024 | 0.489301084 | 0 | 0.355418052 |
| ENSMUSG00000028601 | 1.783508299 | 1.688721876 | 1.738549078 | 1.789967834 | 1.836786029 | 1.908485019 | 1.877074955 |
| ENSMUSG00000028602 | 0 | 0 | 0.25 | 0 | 0.244650542 | 0 | 0.355418052 |
| ENSMUSG00000028603 | 2.580706586 | 2.422499493 | 2.426408097 | 2.461764337 | 2.412520257 | 2.803457116 | 2.263534582 |
| ENSMUSG00000028607 | 2.151493169 | 2.012462137 | 2.029735268 | 2.028435542 | 1.975997314 | 2.285557309 | 1.923435442 |
| ENSMUSG00000028608 | 2.160971586 | 2.019203899 | 1.990000483 | 1.970660762 | 2.047893212 | 2.23299611 | 1.89047692 |
| ENSMUSG00000028609 | 2.492331912 | 2.32991803 | 2.343216265 | 2.415889526 | 2.304598364 | 2.715167358 | 2.203186321 |
| ENSMUSG00000028610 | 0 | 0 | 0 | 0 | 0 | 0 | 0 |
| ENSMUSG00000028612 | 0 | 0 | 0 | 0 | 0 | 0 | 0 |
| ENSMUSG00000028613 | 2.196867486 | 2.073655187 | 2.094844592 | 2.234894804 | 2.261347526 | 2.480006943 | 2.110947927 |

| | | | | | | | |
|---|---|---|---|---|---|---|---|
| ENSMUSG00000028614 | 2.776606269 | 2.514497931 | 2.495034894 | 2.706438458 | 2.60906281 | 3.024074987 | 2.434486378 |
| ENSMUSG00000028617 | 2.564110437 | 2.352877747 | 2.348615674 | 2.323080408 | 2.261929482 | 2.565847819 | 2.163382972 |
| ENSMUSG00000028618 | 2.864601523 | 2.607351685 | 2.604463129 | 2.691632227 | 2.629961383 | 3.06595298 | 2.446980196 |
| ENSMUSG00000028619 | 1.978790393 | 1.823655187 | 1.910964047 | 1.850219859 | 1.866526233 | 1.991226076 | 1.771720588 |
| ENSMUSG00000028621 | 1.404271433 | 1.494319981 | 1.431980114 | 1.602347734 | 1.509475496 | 1.431363764 | 1.46982607 |
| ENSMUSG00000028622 | 2.346900291 | 2.155512955 | 2.197790722 | 2.245748394 | 2.220647857 | 2.503790683 | 2.106650241 |
| ENSMUSG00000028626 | 0.289064826 | 0 | 0 | 0 | 0 | 0 | 0.224243824 |
| ENSMUSG00000028629 | 2.153883046 | 2.006976499 | 2 | 1.933677405 | 1.88560959 | 2.025305865 | 1.778418869 |
| ENSMUSG00000028630 | 1.759671389 | 1.668106335 | 1.777131114 | 1.494319981 | 1.499532413 | 1.716003344 | 1.502532226 |
| ENSMUSG00000028631 | 0 | 0 | 0 | 0 | 0 | 0 | 0.224243824 |
| ENSMUSG00000028633 | 3.047060262 | 2.764497931 | 2.74539182 | 2.868679987 | 2.829291987 | 3.245265839 | 2.606378556 |
| ENSMUSG00000028634 | 1.753493018 | 1.811981878 | 1.852347734 | 1.67510993 | 1.773209395 | 1.838849091 | 1.661510159 |
| ENSMUSG00000028635 | 0.289064826 | 0 | 0 | 0 | 0 | 0 | 0.355418052 |
| ENSMUSG00000028636 | 1.470606563 | 1.54745614 | 1.438721876 | 1.438721876 | 1.53772706 | 1.602059991 | 1.392431569 |
| ENSMUSG00000028637 | 1.181541736 | 1.098079356 | 1.238549078 | 1.130890489 | 1.121713561 | 1.301029996 | 1.131174228 |
| ENSMUSG00000028639 | 3.330205638 | 3.071855953 | 3.074658852 | 2.999647607 | 2.976458394 | 3.419955748 | 2.783679399 |
| ENSMUSG00000028640 | 0 | 0 | 0.25 | 0 | 0 | 0 | 0 |
| ENSMUSG00000028641 | 2.339331437 | 2.175975893 | 2.181980114 | 2.178561379 | 2.209357611 | 2.488550717 | 2.137790317 |
| ENSMUSG00000028642 | 0 | 0 | 0 | 0.396240625 | 0 | 0 | 0 |
| ENSMUSG00000028643 | 1.647796617 | 1.681980114 | 1.678561379 | 1.685366747 | 1.739104362 | 1.819543936 | 1.637827726 |
| ENSMUSG00000028644 | 0.289064826 | 0.25 | 0.25 | 0.5 | 0 | 0 | 0 |
| ENSMUSG00000028645 | 2.471720134 | 2.345385738 | 2.399513125 | 1.912762923 | 2.084333628 | 2.252853031 | 2.083229353 |
| ENSMUSG00000028646 | 2.891461985 | 2.646944379 | 2.659835177 | 2.451032755 | 2.418540977 | 2.745074792 | 2.24869599 |
| ENSMUSG00000028647 | 1.714367558 | 1.618933358 | 1.580482024 | 1.841580554 | 1.777823267 | 1.982271233 | 1.680001789 |
| ENSMUSG00000028648 | 2.013563224 | 1.928561379 | 2.002806814 | 2.001406137 | 1.919403394 | 2.08278537 | 1.853775828 |
| ENSMUSG00000028649 | 3.376889054 | 3.067792798 | 3.060495787 | 3.125956434 | 3.072599934 | 3.532244644 | 2.837020792 |
| ENSMUSG00000028651 | 2.066740409 | 1.990000483 | 1.953445298 | 2.02053726 | 1.973373093 | 2.264817823 | 1.874129423 |
| ENSMUSG00000028653 | 1.894701088 | 1.744319981 | 1.75 | 1.774508021 | 1.816839698 | 1.954242509 | 1.653807046 |
| ENSMUSG00000028654 | 1.700465191 | 1.618933358 | 1.53743678 | 1.311981878 | 1.394615667 | 1.342422681 | 1.193409508 |
| ENSMUSG00000028655 | 0.671187741 | 0.830482024 | 0.976722649 | 1.146240625 | 1.310728002 | 1.342422681 | 1.20139795 |
| ENSMUSG00000028656 | 3.284338304 | 2.983053688 | 2.982499516 | 2.816989271 | 2.765894696 | 3.174059808 | 2.585294136 |
| ENSMUSG00000028657 | 2.595842487 | 2.387205477 | 2.403217124 | 2.350219859 | 2.253693465 | 2.57634135 | 2.123964672 |
| ENSMUSG00000028661 | 0 | 0 | 0 | 0 | 0.489301084 | 0.301029996 | 0.355418052 |
| ENSMUSG00000028664 | 1.639698646 | 1.653677461 | 1.622963274 | 1.598079356 | 1.731346772 | 1.806179974 | 1.692344475 |

| | | | | | | | |
|---|---|---|---|---|---|---|---|
| ENSMUSG00000028668 | 2.626865868 | 2.406883471 | 2.369945066 | 2.535207443 | 2.453332638 | 2.783903579 | 2.295899033 |
| ENSMUSG00000028669 | 2.146671932 | 1.998588359 | 2.02186571 | 2.101285366 | 2.107593434 | 2.36361198 | 1.994607951 |
| ENSMUSG00000028670 | 2.153883046 | 2.08272922 | 2.187048212 | 1.912762923 | 2.137779008 | 2.220108088 | 2.133394659 |
| ENSMUSG00000028671 | 1.655740331 | 1.54248125 | 1.602347734 | 1.646240625 | 1.870073561 | 1.806179974 | 1.894206246 |
| ENSMUSG00000028672 | 2.131864492 | 1.981203126 | 1.995748394 | 2.027131114 | 2.083370582 | 2.309630167 | 1.980721036 |
| ENSMUSG00000028673 | 2.282114126 | 2.066696635 | 2.112802778 | 2.234894804 | 2.211376431 | 2.539076099 | 2.14512854 |
| ENSMUSG00000028675 | 2.556920045 | 2.329353153 | 2.261780978 | 2.439139081 | 2.366796383 | 2.715167358 | 2.21680636 |
| ENSMUSG00000028676 | 3.009601615 | 2.757321807 | 2.728409357 | 2.79838134 | 2.716906358 | 3.133858125 | 2.525059802 |
| ENSMUSG00000028677 | 2.337801042 | 2.160964047 | 2.190387808 | 2.019203899 | 2.130260132 | 2.309630167 | 2.038401572 |
| ENSMUSG00000028678 | 2.719923957 | 2.46293726 | 2.500703754 | 2.58272922 | 2.529377461 | 2.897627091 | 2.370437612 |
| ENSMUSG00000028680 | 1.578129653 | 1.593759858 | 1.602347734 | 1.54745614 | 1.741652793 | 1.826074803 | 1.651852264 |
| ENSMUSG00000028681 | 0.45815691 | 0.396240625 | 0.5 | 0.5 | 0.812711509 | 0.477121255 | 0.74492186 |
| ENSMUSG00000028683 | 2.269664473 | 2.112802778 | 2.083847589 | 2.221424093 | 2.190648982 | 2.426511261 | 2.023210262 |
| ENSMUSG00000028684 | 2.072573125 | 1.937048212 | 2.009729747 | 1.899978211 | 1.98503142 | 2.167317335 | 1.906930404 |
| ENSMUSG00000028687 | 1.37447073 | 1.410964047 | 1.431980114 | 1.364857905 | 1.484072009 | 1.361727836 | 1.505628092 |
| ENSMUSG00000028688 | 2.003432385 | 1.87892496 | 1.83272922 | 1.828220739 | 1.859322797 | 2.068185862 | 1.753626837 |
| ENSMUSG00000028689 | 1.404271433 | 1.380890489 | 1.302363341 | 1.356566189 | 1.528558853 | 1.544068044 | 1.401059159 |
| ENSMUSG00000028690 | 1.734388958 | 1.705044741 | 1.723704441 | 1.653677461 | 1.580303447 | 1.812913357 | 1.508694612 |
| ENSMUSG00000028691 | 3.418179645 | 3.100686406 | 3.084266085 | 3.154079774 | 3.036936105 | 3.571475904 | 2.829903571 |
| ENSMUSG00000028692 | 3.22428261 | 2.949111315 | 2.953545471 | 2.928883266 | 2.917230235 | 3.360971884 | 2.764880683 |
| ENSMUSG00000028693 | 3.081429861 | 2.833847589 | 2.800071401 | 2.935261394 | 2.844905505 | 3.277150614 | 2.622192234 |
| ENSMUSG00000028696 | 2.141794304 | 1.991446071 | 1.938721876 | 2.074802005 | 2.023617914 | 2.326335861 | 1.881925353 |
| ENSMUSG00000028698 | 2.555102908 | 2.314346961 | 2.321928095 | 2.449415381 | 2.376174996 | 2.700703717 | 2.191196167 |
| ENSMUSG00000028699 | 0.45815691 | 0 | 0.25 | 0 | 0.244650542 | 0 | 0.224243824 |
| ENSMUSG00000028700 | 2.524897319 | 2.360217292 | 2.405054956 | 2.357613138 | 2.426290456 | 2.692846919 | 2.252699564 |
| ENSMUSG00000028701 | 1.614416215 | 1.53743678 | 1.593759858 | 1.732684334 | 1.731346772 | 1.826074803 | 1.594042709 |
| ENSMUSG00000028702 | 2.37447073 | 2.191217898 | 2.192046081 | 2.371960008 | 2.325116119 | 2.643452676 | 2.157685697 |
| ENSMUSG00000028703 | 2.266038081 | 2.152756199 | 2.149047439 | 2.081607372 | 2.151558609 | 2.352182518 | 2.075383 |
| ENSMUSG00000028706 | 1.967766868 | 1.769203899 | 1.819031101 | 1.970660762 | 1.893954095 | 2.127104798 | 1.830929954 |
| ENSMUSG00000028707 | 0.811507563 | 0.951838731 | 0.75 | 1.25 | 1.234113756 | 1.380211242 | 1.176815335 |
| ENSMUSG00000028708 | 1.940852214 | 1.910964047 | 1.898114259 | 1.979715809 | 1.965381292 | 2.170261715 | 1.856911705 |
| ENSMUSG00000028709 | 1.418409478 | 1.458222504 | 1.396240625 | 1.646240625 | 1.533172725 | 1.748188027 | 1.46982607 |
| ENSMUSG00000028710 | 1.967766868 | 1.79248125 | 1.814346961 | 1.982684334 | 1.94314048 | 2.117271296 | 1.783679399 |
| ENSMUSG00000028712 | 0 | 0.396240625 | 0 | 0 | 0 | 0 | 0.224243824 |

| | | | | | | | |
|---|---|---|---|---|---|---|---|
| ENSMUSG00000028713 | 0 | 0 | 0 | 0 | 0 | 0 | 0 |
| ENSMUSG00000028715 | 0 | 0 | 0 | 0 | 0 | 0 | 0 |
| ENSMUSG00000028716 | 0 | 0 | 0 | 0 | 0.244650542 | 0 | 0 |
| ENSMUSG00000028717 | 0 | 0 | 0 | 0 | 0 | 0 | 0 |
| ENSMUSG00000028718 | 2.370236862 | 2.192046081 | 2.174241882 | 2.243603647 | 2.226481975 | 2.505149978 | 2.061314894 |
| ENSMUSG00000028719 | 2.63370885 | 2.412762923 | 2.383332433 | 2.54652856 | 2.468227512 | 2.82672252 | 2.273444839 |
| ENSMUSG00000028729 | 2.671876482 | 2.460587586 | 2.473704441 | 2.456637122 | 2.38035699 | 2.773786445 | 2.265305657 |
| ENSMUSG00000028730 | 0.289064826 | 0 | 0 | 0.25 | 0 | 0.301029996 | 0 |
| ENSMUSG00000028736 | 0 | 0 | 0 | 0 | 0 | 0 | 0 |
| ENSMUSG00000028737 | 1.777675583 | 1.701838731 | 1.726722649 | 1.856566189 | 1.834841362 | 1.944482672 | 1.732938436 |
| ENSMUSG00000028738 | 0 | 0 | 0.25 | 0.396240625 | 0.387761935 | 0 | 0.579661877 |
| ENSMUSG00000028741 | 2.43085913 | 2.237091808 | 2.240000483 | 2.309003548 | 2.300451989 | 2.626340367 | 2.143416814 |
| ENSMUSG00000028743 | 1.578129653 | 1.649978211 | 1.769203899 | 1.464495249 | 1.77552387 | 1.69019608 | 1.802677937 |
| ENSMUSG00000028744 | 1.714367558 | 1.681980114 | 1.708222504 | 1.53743678 | 1.572189156 | 1.633468456 | 1.543390198 |
| ENSMUSG00000028745 | 3.033115198 | 2.803124096 | 2.804943388 | 2.677055433 | 2.64026298 | 3.009875634 | 2.470138797 |
| ENSMUSG00000028747 | 0 | 0 | 0 | 0 | 0.244650542 | 0 | 0 |
| ENSMUSG00000028749 | 0 | 0 | 0 | 0 | 0 | 0 | 0 |
| ENSMUSG00000028750 | 0 | 0 | 0 | 0 | 0 | 0 | 0 |
| ENSMUSG00000028751 | 0 | 0 | 0.396240625 | 0 | 0 | 0 | 0 |
| ENSMUSG00000028753 | 0 | 0 | 0 | 0 | 0 | 0 | 0 |
| ENSMUSG00000028755 | 1.156259305 | 1.282320754 | 1.380890489 | 1.160964047 | 1.27449559 | 1.079181246 | 1.21680636 |
| ENSMUSG00000028756 | 1.920505136 | 1.78743678 | 1.825945187 | 1.580482024 | 1.751666404 | 1.77815125 | 1.694069894 |
| ENSMUSG00000028757 | 3.034271214 | 2.767865591 | 2.763311781 | 2.762972164 | 2.753837558 | 3.14113609 | 2.58287655 |
| ENSMUSG00000028758 | 0 | 0 | 0.25 | 0.5 | 0.244650542 | 0 | 0.355418052 |
| ENSMUSG00000028759 | 2.857607105 | 2.629171237 | 2.637686696 | 2.629417347 | 2.572430494 | 2.937016107 | 2.424563987 |
| ENSMUSG00000028760 | 2.54867984 | 2.341580554 | 2.324802005 | 2.233422664 | 2.249502249 | 2.460897843 | 2.096890362 |
| ENSMUSG00000028763 | 3.409468165 | 3.221900356 | 3.310533014 | 2.727473271 | 2.973218118 | 3.196728723 | 2.946032751 |
| ENSMUSG00000028766 | 1.100572389 | 1.098079356 | 0.951838731 | 0.646240625 | 0.733951626 | 0.77815125 | 0.775756176 |
| ENSMUSG00000028771 | 2.709365611 | 2.505938588 | 2.480460234 | 2.368426437 | 2.28771234 | 2.640481437 | 2.153137801 |
| ENSMUSG00000028772 | 2.307602651 | 2.090485943 | 2.095926073 | 2.244319981 | 2.186353316 | 2.522444234 | 2.072724682 |
| ENSMUSG00000028773 | 1.54867984 | 1.476722649 | 1.618933358 | 1.02186571 | 0.955822902 | 1.204119983 | 1.041356071 |
| ENSMUSG00000028776 | 2.533616084 | 2.441632227 | 2.505591953 | 2.219895812 | 2.362886675 | 2.575187845 | 2.376547822 |
| ENSMUSG00000028777 | 0 | 0 | 0 | 0 | 0 | 0 | 0 |
| ENSMUSG00000028778 | 0 | 0.5 | 0.5 | 0.25 | 0.489301084 | 0 | 0.520678036 |

| | | | | | | | |
|---|---|---|---|---|---|---|---|
| ENSMUSG00000028779 | 2.380049842 | 2.193696765 | 2.223704441 | 2.209012589 | 2.263669614 | 2.50242712 | 2.148102439 |
| ENSMUSG00000028780 | 2.527823874 | 2.3464656 | 2.267194569 | 2.168989258 | 2.081436571 | 2.390935107 | 1.94631981 |
| ENSMUSG00000028782 | 0.811507563 | 0.864857905 | 0.79248125 | 1.160964047 | 1.223252711 | 1.255272505 | 1.444790289 |
| ENSMUSG00000028784 | 0.289064826 | 0.25 | 0 | 0 | 0 | 0.301029996 | 0.224243824 |
| ENSMUSG00000028785 | 0.45815691 | 0.396240625 | 0.580482024 | 0 | 0.244650542 | 0.301029996 | 0 |
| ENSMUSG00000028786 | 0.289064826 | 0 | 0 | 0 | 0 | 0 | 0.355418052 |
| ENSMUSG00000028788 | 3.116748756 | 2.821928095 | 2.833707982 | 2.883940344 | 2.787923604 | 3.219846386 | 2.557055024 |
| ENSMUSG00000028789 | 1.686083339 | 1.71759118 | 1.819031101 | 1.575945187 | 1.797870327 | 1.819543936 | 1.805128823 |
| ENSMUSG00000028790 | 3.147552676 | 2.849151195 | 2.836905342 | 2.885878804 | 2.81148809 | 3.259115844 | 2.606686129 |
| ENSMUSG00000028792 | 2.644135203 | 2.486360959 | 2.505591953 | 2.512802235 | 2.537194296 | 2.86569606 | 2.427964734 |
| ENSMUSG00000028793 | 2.429630757 | 2.247880462 | 2.264497931 | 2.337182039 | 2.345884234 | 2.625312451 | 2.223574712 |
| ENSMUSG00000028794 | 0.289064826 | 0 | 0.396240625 | 0.396240625 | 0.489301084 | 0 | 0.579661877 |
| ENSMUSG00000028795 | 1.307602651 | 1.226722649 | 1.302363341 | 1.42510993 | 1.484072009 | 1.556302501 | 1.554175045 |
| ENSMUSG00000028796 | 2.551444705 | 2.442872367 | 2.492526473 | 2.179419106 | 2.344966272 | 2.514547753 | 2.272006351 |
| ENSMUSG00000028797 | 2.417143825 | 2.28743678 | 2.330482024 | 2.296218836 | 2.278401864 | 2.598790507 | 2.16937787 |
| ENSMUSG00000028798 | 2.835891186 | 2.615113974 | 2.630645383 | 2.610735874 | 2.642643806 | 3.006893708 | 2.496970219 |
| ENSMUSG00000028799 | 1.639698646 | 1.78743678 | 1.79248125 | 1.54745614 | 1.603583716 | 1.51851394 | 1.715701091 |
| ENSMUSG00000028800 | 2.621464818 | 2.400906586 | 2.418548067 | 2.552363341 | 2.506580024 | 2.896526217 | 2.344312053 |
| ENSMUSG00000028801 | 0 | 0.25 | 0 | 0 | 0.244650542 | 0 | 0.224243824 |
| ENSMUSG00000028803 | 1.578129653 | 1.516522298 | 1.54745614 | 1.614857905 | 1.66856048 | 1.792391689 | 1.582103514 |
| ENSMUSG00000028804 | 0.289064826 | 0.25 | 0.396240625 | 0.25 | 0.387761935 | 0.477121255 | 0.448487648 |
| ENSMUSG00000028807 | 1.482695304 | 1.431980114 | 1.396240625 | 1.380890489 | 1.373641807 | 1.591064607 | 1.425641774 |
| ENSMUSG00000028809 | 3.028166065 | 2.734343455 | 2.741986676 | 2.734527332 | 2.71352568 | 3.095518042 | 2.551269198 |
| ENSMUSG00000028811 | 2.838206752 | 2.577369088 | 2.516522298 | 2.549918086 | 2.516894411 | 2.854913022 | 2.335902514 |
| ENSMUSG00000028813 | 0 | 0 | 0.25 | 0 | 0 | 0 | 0 |
| ENSMUSG00000028820 | 3.265541121 | 2.966819685 | 2.947994205 | 3.008613774 | 2.91976429 | 3.37051309 | 2.702332941 |
| ENSMUSG00000028821 | 2.22189694 | 2.012462137 | 2.072504712 | 2.129909063 | 2.056322416 | 2.307496038 | 1.982842456 |
| ENSMUSG00000028822 | 2.390989024 | 2.210587586 | 2.200224975 | 2.191217898 | 2.165585202 | 2.463892989 | 2.074853081 |
| ENSMUSG00000028825 | 0.671187741 | 0.580482024 | 1 | 1 | 0.846351821 | 0.698970004 | 1.066254157 |
| ENSMUSG00000028826 | 2.413323654 | 2.24929487 | 2.171625132 | 2.241446071 | 2.195595395 | 2.506505032 | 2.050061048 |
| ENSMUSG00000028830 | 2.343889214 | 2.172499493 | 2.188721876 | 2.135757955 | 2.189936658 | 2.440909082 | 2.089372236 |
| ENSMUSG00000028832 | 2.897915226 | 2.645535495 | 2.657339155 | 2.637686696 | 2.576984829 | 2.946452265 | 2.418020755 |
| ENSMUSG00000028833 | 2.020182925 | 1.992885888 | 2.138647213 | 1.747171172 | 1.948833444 | 2.068185862 | 2.00464543 |
| ENSMUSG00000028834 | 0 | 0.25 | 0 | 0 | 0 | 0 | 0 |

| | | | | | | | |
|---|---|---|---|---|---|---|---|
| ENSMUSG00000028836 | 0.578129653 | 0.396240625 | 0.5 | 0.646240625 | 0.632412477 | 0.301029996 | 0.629532004 |
| ENSMUSG00000028837 | 2.732608034 | 2.519871196 | 2.501406137 | 2.545908845 | 2.540642974 | 2.894316063 | 2.383771753 |
| ENSMUSG00000028838 | 0.747221736 | 0.580482024 | 0.580482024 | 1.146240625 | 1.057362051 | 1.041392685 | 1.20139795 |
| ENSMUSG00000028840 | 0.289064826 | 0.896240625 | 0.830482024 | 0.646240625 | 1 | 0.602059991 | 1 |
| ENSMUSG00000028841 | 0 | 0.5 | 0.79248125 | 0.701838731 | 0.686820904 | 0.301029996 | 0.876096088 |
| ENSMUSG00000028842 | 2.595842487 | 2.411864607 | 2.418106335 | 2.511780978 | 2.428473696 | 2.795880017 | 2.212676319 |
| ENSMUSG00000028843 | 2.063793172 | 1.87694866 | 2.013820609 | 1.834962501 | 1.959951091 | 2.064457989 | 1.883845332 |
| ENSMUSG00000028845 | 0.91631382 | 0.864857905 | 0.79248125 | 0.701838731 | 0.877063019 | 0.84509804 | 0.775756176 |
| ENSMUSG00000028847 | 2.519973621 | 2.354990044 | 2.36434522 | 2.377938163 | 2.331857641 | 2.654176542 | 2.22088434 |
| ENSMUSG00000028848 | 1.558729299 | 1.571350555 | 1.646240625 | 1.488549078 | 1.635855979 | 1.602059991 | 1.535058868 |
| ENSMUSG00000028849 | 2.718695584 | 2.537754135 | 2.621708755 | 2.262462137 | 2.357605438 | 2.540329475 | 2.35250852 |
| ENSMUSG00000028850 | 1.655740331 | 1.657339155 | 1.638647213 | 1.388647213 | 1.595992969 | 1.579783597 | 1.594042709 |
| ENSMUSG00000028851 | 2.740271036 | 2.597810897 | 2.646709947 | 2.595926073 | 2.629345941 | 2.941511433 | 2.527425075 |
| ENSMUSG00000028854 | 2.253091469 | 2.168106335 | 2.25 | 2.066696635 | 2.21736452 | 2.342422681 | 2.178744473 |
| ENSMUSG00000028857 | 2.285603957 | 2.172499493 | 2.114857905 | 2.144357207 | 2.175379703 | 2.376576957 | 2.013740841 |
| ENSMUSG00000028859 | 0.289064826 | 0.25 | 0.580482024 | 0.25 | 0.387761935 | 0 | 0 |
| ENSMUSG00000028860 | 0.671187741 | 0.396240625 | 0.646240625 | 0.701838731 | 0.877063019 | 0.602059991 | 0.775756176 |
| ENSMUSG00000028861 | 2.165631214 | 2.061981878 | 2.03743678 | 2.125956434 | 2.044679629 | 2.372912003 | 1.943148064 |
| ENSMUSG00000028862 | 0.867194479 | 0.864857905 | 1.04248125 | 0.79248125 | 0.905314583 | 1.079181246 | 0.952571511 |
| ENSMUSG00000028863 | 2.124258828 | 1.972185812 | 1.961372513 | 1.948603967 | 1.988833497 | 2.217483944 | 1.926805425 |
| ENSMUSG00000028864 | 1.036286563 | 1.02186571 | 0.92510993 | 0 | 0 | 0 | 0 |
| ENSMUSG00000028865 | 0 | 0 | 0.25 | 0 | 0 | 0 | 0 |
| ENSMUSG00000028868 | 2.74143756 | 2.564346961 | 2.543731421 | 2.410964047 | 2.422990063 | 2.716003344 | 2.321629237 |
| ENSMUSG00000028869 | 2.515992375 | 2.284887838 | 2.312574604 | 2.418989258 | 2.354040194 | 2.656098202 | 2.190083792 |
| ENSMUSG00000028871 | 0 | 0 | 0 | 0.25 | 0.387761935 | 0.301029996 | 0.672731473 |
| ENSMUSG00000028873 | 2.595015862 | 2.402756199 | 2.431128463 | 2.457430684 | 2.429921724 | 2.789580712 | 2.341533092 |
| ENSMUSG00000028874 | 0 | 0 | 0 | 0 | 0 | 0 | 0 |
| ENSMUSG00000028876 | 0.289064826 | 0 | 0 | 0 | 0 | 0 | 0 |
| ENSMUSG00000028878 | 2.322180025 | 2.149978211 | 2.137686696 | 2.146240625 | 2.034859505 | 2.359835482 | 1.969900112 |
| ENSMUSG00000028879 | 2.709994145 | 2.505591953 | 2.469129237 | 2.556000419 | 2.491128299 | 2.8876173 | 2.311836403 |
| ENSMUSG00000028882 | 2.669114668 | 2.425975893 | 2.460587586 | 2.467205639 | 2.441995847 | 2.781036939 | 2.301737862 |
| ENSMUSG00000028883 | 2.906628471 | 2.680701883 | 2.631135429 | 2.089388001 | 2.019019072 | 2.267171728 | 1.877074955 |
| ENSMUSG00000028884 | 2.518981867 | 2.333847589 | 2.351816941 | 2.390560606 | 2.393415815 | 2.716837723 | 2.236700872 |
| ENSMUSG00000028885 | 1.740854706 | 1.657339155 | 1.71759118 | 1.657339155 | 1.689774529 | 1.875061263 | 1.621017948 |

| | | | | | | | |
|---|---|---|---|---|---|---|---|
| ENSMUSG00000028886 | 2.578990401 | 2.358657057 | 2.358135475 | 2.43155454 | 2.362449578 | 2.741151599 | 2.218171406 |
| ENSMUSG00000028889 | 2.151493169 | 2.019203899 | 2 | 2.039967834 | 2.013184954 | 2.245512668 | 1.846338363 |
| ENSMUSG00000028890 | 2.116511873 | 1.984159485 | 2.017865591 | 1.984159485 | 1.92245931 | 2.146128036 | 1.853775828 |
| ENSMUSG00000028893 | 2.172524468 | 2.066696635 | 2.029735268 | 1.789967834 | 1.744182955 | 1.86923172 | 1.74196736 |
| ENSMUSG00000028894 | 2.326928073 | 2.187886015 | 2.143411797 | 2.344302633 | 2.300972958 | 2.643452676 | 2.183328003 |
| ENSMUSG00000028896 | 2.504843822 | 2.349685923 | 2.38040011 | 2.242885888 | 2.28608706 | 2.536558443 | 2.197428385 |
| ENSMUSG00000028898 | 2.121692466 | 1.964495249 | 2.013820609 | 2.006976499 | 1.988833497 | 2.222716471 | 1.865128412 |
| ENSMUSG00000028899 | 2.266038081 | 2.115881093 | 2.031030328 | 2.197790722 | 2.132784263 | 2.45484486 | 2.001999072 |
| ENSMUSG00000028901 | 2.371652933 | 2.209800947 | 2.197790722 | 2.192046081 | 2.136121934 | 2.403120521 | 2.021338422 |
| ENSMUSG00000028902 | 2.761340634 | 2.502806814 | 2.512462137 | 2.596735061 | 2.567816622 | 2.894869657 | 2.37466387 |
| ENSMUSG00000028906 | 2.528794847 | 2.317865757 | 2.30479213 | 2.407339155 | 2.335652905 | 2.643452676 | 2.178744473 |
| ENSMUSG00000028907 | 2.358731791 | 2.235628626 | 2.232684334 | 2.237091808 | 2.199086582 | 2.503790683 | 2.07004434 |
| ENSMUSG00000028909 | 0.289064826 | 0.25 | 0.646240625 | 0 | 0.387761935 | 0.602059991 | 0.629532004 |
| ENSMUSG00000028910 | 2.083999385 | 2.009729747 | 2.016522298 | 1.969129237 | 2.016696984 | 2.225309282 | 1.938331368 |
| ENSMUSG00000028911 | 2.010213549 | 1.92510993 | 1.914552871 | 1.802363341 | 1.87878897 | 1.959041392 | 1.808770687 |
| ENSMUSG00000028914 | 2.042557845 | 1.919870025 | 1.961372513 | 1.868933358 | 1.883916752 | 2.041392685 | 1.833176598 |
| ENSMUSG00000028917 | 2.060824957 | 2.028435542 | 2.028435542 | 1.769203899 | 1.977302144 | 2.029383778 | 1.958704689 |
| ENSMUSG00000028919 | 1.928763473 | 1.898114259 | 1.984159485 | 1.5 | 1.746695109 | 1.556302501 | 1.899720848 |
| ENSMUSG00000028920 | 2.355805236 | 2.241446071 | 2.189555804 | 2.205044741 | 2.222603297 | 2.474216264 | 2.106650241 |
| ENSMUSG00000028923 | 2.163307908 | 2.104463129 | 2.187886015 | 1.991446071 | 2.146756411 | 2.250420002 | 2.14555506 |
| ENSMUSG00000028926 | 2.346900291 | 2.160964047 | 2.173371739 | 2.061981878 | 1.913211023 | 2.26245109 | 1.809975587 |
| ENSMUSG00000028927 | 0.578129653 | 0.580482024 | 0.646240625 | 0.580482024 | 0.632412477 | 0.301029996 | 0.710836105 |
| ENSMUSG00000028931 | 1.470606563 | 1.505591953 | 1.552363341 | 1.52186571 | 1.701347897 | 1.633468456 | 1.705896842 |
| ENSMUSG00000028932 | 3.143130234 | 2.867537609 | 2.841853688 | 2.928024263 | 2.863366354 | 3.345177617 | 2.655750087 |
| ENSMUSG00000028933 | 2.213719572 | 2.061981878 | 2.060793496 | 2.051142786 | 2.008447216 | 2.264817823 | 1.902443294 |
| ENSMUSG00000028936 | 2.932463873 | 2.70042709 | 2.684523065 | 2.745213402 | 2.648720957 | 3.103803721 | 2.487037588 |
| ENSMUSG00000028937 | 2.589183146 | 2.434523065 | 2.504897682 | 2.405970373 | 2.560156791 | 2.761927838 | 2.484655503 |
| ENSMUSG00000028938 | 0 | 0 | 0 | 0 | 0 | 0 | 0 |
| ENSMUSG00000028940 | 0 | 0 | 0 | 0 | 0 | 0 | 0 |
| ENSMUSG00000028943 | 0.671187741 | 0.396240625 | 0.646240625 | 0.976722649 | 1.039257769 | 0.84509804 | 1.278288402 |
| ENSMUSG00000028944 | 2.505869732 | 2.306603048 | 2.26584877 | 2.03360658 | 1.997548906 | 2.245512668 | 1.892346957 |
| ENSMUSG00000028945 | 2.804556862 | 2.58076369 | 2.563757142 | 2.417221246 | 2.326087113 | 2.667452953 | 2.186723532 |
| ENSMUSG00000028946 | 0 | 0.25 | 0 | 0 | 0 | 0 | 0 |
| ENSMUSG00000028948 | 2.455852858 | 2.277783918 | 2.291854536 | 2.320192693 | 2.296256325 | 2.567026366 | 2.147679267 |

| | | | | | | | |
|---|---|---|---|---|---|---|---|
| ENSMUSG00000028949 | 1.771760131 | 1.741446071 | 1.726722649 | 1.646240625 | 1.635855979 | 1.748188027 | 1.543390198 |
| ENSMUSG00000028950 | 0.289064826 | 0.79248125 | 0.701838731 | 0.580482024 | 0.733951626 | 0.84509804 | 0.853775828 |
| ENSMUSG00000028952 | 1.54867984 | 1.614857905 | 1.630890489 | 1.321350555 | 1.677808768 | 1.556302501 | 1.657681529 |
| ENSMUSG00000028953 | 2.984357457 | 2.748765117 | 2.731203126 | 2.748588359 | 2.731346772 | 3.114944416 | 2.580329607 |
| ENSMUSG00000028954 | 2.513987407 | 2.281030328 | 2.293106877 | 2.339388001 | 2.329944595 | 2.658011397 | 2.212329757 |
| ENSMUSG00000028955 | 2.801542157 | 2.563166358 | 2.564641508 | 2.675326615 | 2.581481146 | 2.98811284 | 2.426715983 |
| ENSMUSG00000028957 | 2.089597219 | 1.830482024 | 1.898114259 | 1.622963274 | 1.576269619 | 1.698970004 | 1.582103514 |
| ENSMUSG00000028958 | 1.269664473 | 1.451838731 | 1.516522298 | 1.080482024 | 1.293076518 | 1.113943352 | 1.365075947 |
| ENSMUSG00000028959 | 2.231898196 | 2.107613138 | 2.230460234 | 1.882845365 | 2.086251872 | 2.10720997 | 2.076440241 |
| ENSMUSG00000028960 | 2.957832978 | 2.712742038 | 2.695544798 | 2.591853688 | 2.573875064 | 2.92890769 | 2.425821057 |
| ENSMUSG00000028961 | 2.757995436 | 2.547764804 | 2.576230417 | 2.552363341 | 2.613385658 | 2.918030337 | 2.502249309 |
| ENSMUSG00000028962 | 2.629921064 | 2.491806565 | 2.559601185 | 2.175975893 | 2.399374698 | 2.509202522 | 2.365724059 |
| ENSMUSG00000028963 | 0 | 0 | 0 | 0 | 0 | 0 | 0 |
| ENSMUSG00000028964 | 2.684750966 | 2.459406983 | 2.498941788 | 2.401832578 | 2.397795474 | 2.73239376 | 2.285558718 |
| ENSMUSG00000028965 | 1.558729299 | 1.458222504 | 1.356566189 | 1.388647213 | 1.319233381 | 1.431363764 | 1.396774124 |
| ENSMUSG00000028967 | 2.457006475 | 2.269203899 | 2.200224975 | 2.110735874 | 2.057362051 | 2.352182518 | 1.939139147 |
| ENSMUSG00000028969 | 2.289064826 | 2.168989258 | 2.173371739 | 2.005591953 | 2.031524462 | 2.267171728 | 1.939944915 |
| ENSMUSG00000028970 | 2.299278375 | 2.151832578 | 2.189555804 | 2.159156155 | 2.090057366 | 2.390935107 | 1.950241257 |
| ENSMUSG00000028971 | 0.45815691 | 0.396240625 | 0.701838731 | 0.580482024 | 0.846351821 | 0.301029996 | 0.629532004 |
| ENSMUSG00000028972 | 0 | 0.25 | 0 | 0 | 0.244650542 | 0 | 0 |
| ENSMUSG00000028973 | 1.89911417 | 1.839388001 | 1.899978211 | 1.695339928 | 1.868304353 | 1.908485019 | 1.888596011 |
| ENSMUSG00000028974 | 1.881174527 | 1.688721876 | 1.729715809 | 1.926839783 | 1.917865457 | 2.123851641 | 1.846338363 |
| ENSMUSG00000028975 | 1.700465191 | 1.726722649 | 1.858657057 | 1.431980114 | 1.66856048 | 1.662757832 | 1.817112247 |
| ENSMUSG00000028976 | 0 | 0.25 | 0.25 | 0.25 | 0 | 0.301029996 | 0.224243824 |
| ENSMUSG00000028977 | 0.811507563 | 1 | 0.951838731 | 1.188721876 | 1.244650542 | 1.301029996 | 1.335191751 |
| ENSMUSG00000028978 | 0 | 0.396240625 | 0 | 0.25 | 0.244650542 | 0 | 0.448487648 |
| ENSMUSG00000028979 | 0.91631382 | 0.830482024 | 0.79248125 | 0.580482024 | 0.955822902 | 0.301029996 | 0.775756176 |
| ENSMUSG00000028980 | 2.35727108 | 2.195339928 | 2.234894804 | 2.148114259 | 2.229044281 | 2.445604203 | 2.121219121 |
| ENSMUSG00000028982 | 1.623015126 | 1.54248125 | 1.602347734 | 1.589388001 | 1.66856048 | 1.851258349 | 1.572224412 |
| ENSMUSG00000028986 | 2.312518611 | 2.094844592 | 2.115881093 | 2.259043403 | 2.158641579 | 2.445604203 | 2.008574816 |
| ENSMUSG00000028988 | 2.003432385 | 1.864857905 | 1.910964047 | 2.072504712 | 2.058398634 | 2.274157849 | 2.029982479 |
| ENSMUSG00000028989 | 0.289064826 | 0.396240625 | 0.25 | 0 | 0.387761935 | 0.301029996 | 0.448487648 |
| ENSMUSG00000028990 | 2.307602651 | 2.136723615 | 2.103406982 | 2.060793496 | 2.009637626 | 2.255272505 | 1.918313719 |
| ENSMUSG00000028991 | 2.518981867 | 2.392463902 | 2.388167274 | 2.30479213 | 2.343123155 | 2.588831726 | 2.234765579 |

| | | | | | | | |
|---|---|---|---|---|---|---|---|
| ENSMUSG00000028992 | 1.842799666 | 1.657339155 | 1.664552871 | 1.752806814 | 1.715300549 | 1.995635195 | 1.680001789 |
| ENSMUSG00000028995 | 2.545896521 | 2.338837774 | 2.319031101 | 2.395770692 | 2.348623881 | 2.662757832 | 2.179896458 |
| ENSMUSG00000028996 | 0 | 0 | 0 | 0.25 | 0 | 0 | 0 |
| ENSMUSG00000028998 | 2.297593393 | 2.172499493 | 2.187886015 | 2.153677461 | 2.189222894 | 2.482873584 | 2.029372647 |
| ENSMUSG00000028999 | 2.315764015 | 2.149047439 | 2.136723615 | 2.095926073 | 2.071604288 | 2.344392274 | 1.97354745 |
| ENSMUSG00000029001 | 1.505869732 | 1.557204673 | 1.614857905 | 1.54248125 | 1.66856048 | 1.740362689 | 1.661510159 |
| ENSMUSG00000029003 | 2.063793172 | 2.005591953 | 2.005591953 | 1.994319981 | 1.979897422 | 2.23299611 | 1.977154126 |
| ENSMUSG00000029004 | 2.659301689 | 2.474461364 | 2.467976309 | 2.442046081 | 2.371090306 | 2.728353782 | 2.219869654 |
| ENSMUSG00000029005 | 0.45815691 | 0.580482024 | 0.75 | 0.896240625 | 0.846351821 | 1.041392685 | 0.829800754 |
| ENSMUSG00000029007 | 2.042557845 | 1.930274797 | 1.923371739 | 2.078220739 | 2.089109832 | 2.336459734 | 2.043700396 |
| ENSMUSG00000029009 | 2.348397716 | 2.192872367 | 2.264497931 | 2.016522298 | 2.093822267 | 2.209515015 | 2.007270316 |
| ENSMUSG00000029012 | 2.194712275 | 2.03743678 | 2.005591953 | 2.064346961 | 1.986303335 | 2.187520721 | 1.842010333 |
| ENSMUSG00000029014 | 2.680728114 | 2.469129237 | 2.433677405 | 2.47934302 | 2.384900665 | 2.760422483 | 2.222232323 |
| ENSMUSG00000029015 | 0 | 0 | 0 | 0 | 0 | 0 | 0 |
| ENSMUSG00000029016 | 2.151493169 | 2.006976499 | 2.079353153 | 1.987091808 | 2.033751322 | 2.167317335 | 1.921737195 |
| ENSMUSG00000029017 | 2.787177998 | 2.527783918 | 2.529410775 | 2.646944379 | 2.561154547 | 2.940018155 | 2.403948569 |
| ENSMUSG00000029019 | 0.45815691 | 0.646240625 | 0 | 0 | 0.568060967 | 0 | 0.224243824 |
| ENSMUSG00000029020 | 2.789260561 | 2.539967834 | 2.56728167 | 2.51007243 | 2.533172725 | 2.872156273 | 2.392232788 |
| ENSMUSG00000029022 | 1.885732304 | 1.705044741 | 1.858657057 | 1.593759858 | 1.655839964 | 1.69019608 | 1.663407615 |
| ENSMUSG00000029026 | 0 | 0 | 0 | 0 | 0 | 0 | 0 |
| ENSMUSG00000029027 | 1.482695304 | 1.214495249 | 1.356566189 | 1.364857905 | 1.310728002 | 1.342422681 | 1.272006351 |
| ENSMUSG00000029028 | 2.215778996 | 2.107613138 | 2.075945187 | 2.057204673 | 2.115605392 | 2.315970345 | 2.032410373 |
| ENSMUSG00000029029 | 1.847794126 | 1.882845365 | 1.858657057 | 1.634789703 | 1.715300549 | 1.755874856 | 1.710836105 |
| ENSMUSG00000029030 | 2.194712275 | 2.057204673 | 2.073655187 | 2.058404919 | 2.072599934 | 2.326335861 | 1.958704689 |
| ENSMUSG00000029032 | 0 | 0 | 0 | 0.25 | 0.812711509 | 0 | 0.775756176 |
| ENSMUSG00000029033 | 2.183765914 | 2.044977273 | 2.144357207 | 1.978222334 | 2.084333628 | 2.139879086 | 2.085289968 |
| ENSMUSG00000029034 | 2.348397716 | 2.200224975 | 2.189555804 | 2.290597832 | 2.271112598 | 2.633468456 | 2.112839835 |
| ENSMUSG00000029036 | 2.455852858 | 2.28743678 | 2.291226732 | 2.20584181 | 2.297309926 | 2.534026106 | 2.190083792 |
| ENSMUSG00000029038 | 2.124258828 | 2.002806814 | 2.02053726 | 2.028435542 | 1.951645858 | 2.220108088 | 1.970632877 |
| ENSMUSG00000029044 | 0 | 0 | 0 | 0 | 0 | 0 | 0 |
| ENSMUSG00000029047 | 1.129344651 | 1.356566189 | 1.27186571 | 1.321350555 | 1.462344774 | 1.361727836 | 1.479989695 |
| ENSMUSG00000029048 | 2.669114668 | 2.467205639 | 2.454245906 | 2.519537702 | 2.436719872 | 2.841984805 | 2.309019868 |
| ENSMUSG00000029049 | 0.91631382 | 1.02186571 | 0.75 | 0.396240625 | 0.846351821 | 0.903089987 | 0.853775828 |
| ENSMUSG00000029050 | 2.582415738 | 2.396240625 | 2.462155735 | 2.263142013 | 2.311234033 | 2.587710965 | 2.296166733 |

| | | | | | | | |
|---|---|---|---|---|---|---|---|
| ENSMUSG00000029053 | 0.960252568 | 1.114857905 | 1.114857905 | 1.681980114 | 1.736537397 | 1.897627091 | 1.676387053 |
| ENSMUSG00000029054 | 0 | 0 | 0 | 0 | 0 | 0 | 0 |
| ENSMUSG00000029055 | 0 | 0 | 0 | 0 | 0.387761935 | 0 | 0.355418052 |
| ENSMUSG00000029056 | 2.181541736 | 2.015173983 | 2.073655187 | 2.036164561 | 2.135290471 | 2.260071388 | 2.07004434 |
| ENSMUSG00000029059 | 0 | 0 | 0 | 1.25 | 1.244650542 | 1.462397998 | 1.383567577 |
| ENSMUSG00000029060 | 1.816888701 | 1.75835575 | 1.880890489 | 1.482684334 | 1.751666404 | 1.602059991 | 1.821784031 |
| ENSMUSG00000029061 | 1.940852214 | 1.868933358 | 1.969129237 | 0.79248125 | 1 | 1.146128036 | 1.041356071 |
| ENSMUSG00000029062 | 2.721759759 | 2.510756821 | 2.550224651 | 2.538071211 | 2.492946102 | 2.84260924 | 2.338964569 |
| ENSMUSG00000029063 | 2.729622822 | 2.529410775 | 2.50835575 | 2.505938588 | 2.509475496 | 2.848804701 | 2.371075081 |
| ENSMUSG00000029064 | 3.428623026 | 3.10839636 | 3.096532984 | 3.118743346 | 3.043601899 | 3.553154548 | 2.828873922 |
| ENSMUSG00000029066 | 2.415874319 | 2.291226732 | 2.273849256 | 2.322504712 | 2.334707908 | 2.592176757 | 2.232493008 |
| ENSMUSG00000029068 | 2.623015126 | 2.390560606 | 2.410964047 | 2.488912477 | 2.522497007 | 2.763427994 | 2.366370874 |
| ENSMUSG00000029070 | 2.534573679 | 2.30175358 | 2.362802778 | 2.119945066 | 2.145141048 | 2.397940009 | 2.118913228 |
| ENSMUSG00000029071 | 2.23778581 | 2.142463902 | 2.256976499 | 2.031030328 | 2.16097133 | 2.294466226 | 2.222232323 |
| ENSMUSG00000029072 | 0.45815691 | 0 | 0.396240625 | 0.701838731 | 0.733951626 | 0.602059991 | 0.829800754 |
| ENSMUSG00000029073 | 2.042557845 | 1.966046536 | 1.955044741 | 1.981203126 | 1.991345651 | 2.204119983 | 1.865128412 |
| ENSMUSG00000029074 | 0 | 0 | 0 | 0 | 0 | 0 | 0 |
| ENSMUSG00000029075 | 0.45815691 | 0.701838731 | 0.79248125 | 0.646240625 | 0.978602168 | 0 | 0.775756176 |
| ENSMUSG00000029076 | 2.833562691 | 2.608657057 | 2.591580554 | 2.689139081 | 2.640660901 | 3.050379756 | 2.470138797 |
| ENSMUSG00000029082 | 0 | 0 | 0.25 | 0.25 | 0.387761935 | 0.301029996 | 0.355418052 |
| ENSMUSG00000029084 | 0.671187741 | 0.92510993 | 1.130890489 | 1.339388001 | 1.188507684 | 1.342422681 | 1.252393349 |
| ENSMUSG00000029086 | 1.129344651 | 1.201838731 | 1.061981878 | 1.26109853 | 1.057362051 | 1.176091259 | 1.066254157 |
| ENSMUSG00000029088 | 0.747221736 | 0.92510993 | 0.701838731 | 1.146240625 | 1.223252711 | 1.176091259 | 1.028149525 |
| ENSMUSG00000029089 | 1.989530018 | 1.766522298 | 1.809601185 | 1.868933358 | 1.795698282 | 2.029383778 | 1.676387053 |
| ENSMUSG00000029090 | 2.644870062 | 2.427701608 | 2.385757955 | 2.342671852 | 2.315755952 | 2.612783857 | 2.179128924 |
| ENSMUSG00000029092 | 0.811507563 | 0.951838731 | 0.701838731 | 0.951838731 | 0.77552387 | 0.477121255 | 0.629532004 |
| ENSMUSG00000029093 | 2.478339835 | 2.333847589 | 2.307204673 | 1.71759118 | 1.758994476 | 1.819543936 | 1.734460864 |
| ENSMUSG00000029094 | 2.603209296 | 2.379909063 | 2.380890489 | 2.247171172 | 2.276733037 | 2.471291711 | 2.145981019 |
| ENSMUSG00000029095 | 0 | 0 | 0 | 0.25 | 0.77552387 | 0.77815125 | 0.853775828 |
| ENSMUSG00000029096 | 0.867194479 | 1.114857905 | 1.080482024 | 0.5 | 0.812711509 | 0.477121255 | 0.74492186 |
| ENSMUSG00000029097 | 1.721148722 | 1.671625132 | 1.789967834 | 1.557204673 | 1.677808768 | 1.591064607 | 1.601000298 |
| ENSMUSG00000029098 | 2.057835465 | 1.884789703 | 1.894357207 | 1.901832578 | 1.940259189 | 2.136720567 | 1.834294096 |
| ENSMUSG00000029101 | 2.326928073 | 2.121960008 | 2.152756199 | 1.896240625 | 1.998776581 | 2.225309282 | 1.98213686 |
| ENSMUSG00000029102 | 0 | 0 | 0 | 0.396240625 | 0 | 0.301029996 | 0 |

| | | | | | | | |
|---|---|---|---|---|---|---|---|
| ENSMUSG00000029103 | 2.307602651 | 2.136723615 | 2.160964047 | 2.354990044 | 2.383667271 | 2.72427587 | 2.260262214 |
| ENSMUSG00000029104 | 2.895104213 | 2.637686696 | 2.637205477 | 2.673589472 | 2.627493148 | 2.957607287 | 2.460300884 |
| ENSMUSG00000029106 | 3.13140747 | 2.868426437 | 2.89541784 | 2.848749629 | 2.811733113 | 3.25139485 | 2.668780475 |
| ENSMUSG00000029108 | 2.334723298 | 2.174241882 | 2.159156155 | 2.078220739 | 2.057362051 | 2.346352974 | 1.886704102 |
| ENSMUSG00000029110 | 3.053999397 | 2.763142013 | 2.756630861 | 2.752981517 | 2.702851618 | 3.088844563 | 2.507582874 |
| ENSMUSG00000029111 | 2.401648579 | 2.207430684 | 2.203445298 | 2.134789703 | 2.151558609 | 2.457881897 | 2.066798338 |
| ENSMUSG00000029112 | 0 | 0 | 0 | 0 | 0 | 0 | 0 |
| ENSMUSG00000029119 | 2.172524468 | 2.149978211 | 2.216819685 | 1.926839783 | 2.112954895 | 2.214843848 | 2.117521761 |
| ENSMUSG00000029120 | 0 | 0 | 0.396240625 | 0.580482024 | 0.244650542 | 0.301029996 | 0.355418052 |
| ENSMUSG00000029121 | 0.811507563 | 1.061981878 | 1.080482024 | 0.976722649 | 1.020174412 | 1.204119983 | 1.078019652 |
| ENSMUSG00000029122 | 2.427163103 | 2.241446071 | 2.293106877 | 2.155512955 | 2.151558609 | 2.305351369 | 2.075912052 |
| ENSMUSG00000029123 | 0.747221736 | 0.79248125 | 0.79248125 | 0 | 0.489301084 | 0.477121255 | 0.224243824 |
| ENSMUSG00000029125 | 1.876566388 | 1.856566189 | 1.933677405 | 1.890560606 | 1.90849439 | 2.161368002 | 1.871156825 |
| ENSMUSG00000029126 | 2.201144779 | 2.059601185 | 2.03360658 | 2.201838731 | 2.144330586 | 2.411619706 | 2.028149525 |
| ENSMUSG00000029127 | 1.714367558 | 1.552363341 | 1.685366747 | 1.302363341 | 1.514343934 | 1.579783597 | 1.505628092 |
| ENSMUSG00000029128 | 2.361637952 | 2.187048212 | 2.146240625 | 2.240724001 | 2.172394856 | 2.499687083 | 2.031805106 |
| ENSMUSG00000029130 | 0.811507563 | 0.75 | 0.864857905 | 1.02186571 | 1.188507684 | 1.146128036 | 1.159323753 |
| ENSMUSG00000029131 | 2.607246427 | 2.399047439 | 2.390560606 | 2.381869252 | 2.325116119 | 2.613841822 | 2.176428123 |
| ENSMUSG00000029134 | 1.069666964 | 1.061981878 | 1.04248125 | 0.580482024 | 0.686820904 | 0.477121255 | 0.520678036 |
| ENSMUSG00000029135 | 2.699822119 | 2.491085217 | 2.514497931 | 2.319031101 | 2.333285652 | 2.582063363 | 2.179128924 |
| ENSMUSG00000029136 | 0.91631382 | 0.580482024 | 0.951838731 | 1.061981878 | 0.955822902 | 1.079181246 | 0.952571511 |
| ENSMUSG00000029138 | 0 | 0 | 0.25 | 0 | 0 | 0 | 0 |
| ENSMUSG00000029141 | 2.367390214 | 2.185366747 | 2.137686696 | 2.269203899 | 2.201854879 | 2.477121255 | 2.067883961 |
| ENSMUSG00000029145 | 2.126809492 | 2.072504712 | 2.031030328 | 2.012462137 | 2.136951444 | 2.344392274 | 2.026306127 |
| ENSMUSG00000029146 | 2.673935906 | 2.474083101 | 2.50835575 | 2.459406983 | 2.528558853 | 2.7930916 | 2.375293074 |
| ENSMUSG00000029147 | 3.16054539 | 2.878801758 | 2.887205477 | 2.884789703 | 2.859215759 | 3.288696261 | 2.656277989 |
| ENSMUSG00000029148 | 2.811507563 | 2.595385738 | 2.56168515 | 2.603142462 | 2.57935845 | 2.912753304 | 2.411342446 |
| ENSMUSG00000029149 | 1.100572389 | 1.04248125 | 1.02186571 | 0.79248125 | 0.905314583 | 0.954242509 | 0.935079929 |
| ENSMUSG00000029151 | 0.671187741 | 0.5 | 0.79248125 | 0.75 | 1.149965125 | 0.84509804 | 1.21680636 |
| ENSMUSG00000029152 | 2.897915226 | 2.640560606 | 2.611253711 | 2.635274154 | 2.545102531 | 2.940516485 | 2.39989613 |
| ENSMUSG00000029153 | 1.936861443 | 1.71759118 | 1.839388001 | 1.660964047 | 1.588235379 | 1.748188027 | 1.444790289 |
| ENSMUSG00000029154 | 0 | 0 | 0 | 0 | 0 | 0 | 0 |
| ENSMUSG00000029155 | 0 | 0 | 0 | 0 | 0 | 0 | 0 |
| ENSMUSG00000029156 | 1.960252568 | 1.901832578 | 1.907339155 | 2.166333979 | 2.085294053 | 2.401400541 | 1.987044015 |

| | | | | | | | |
|---|---|---|---|---|---|---|---|
| ENSMUSG00000029158 | 0 | 0 | 0 | 0 | 0 | 0 | 0 |
| ENSMUSG00000029161 | 1.993049299 | 1.890560606 | 1.943696765 | 1.931980114 | 1.931471446 | 2.167317335 | 1.769001954 |
| ENSMUSG00000029162 | 1.445324132 | 1.602347734 | 1.622963274 | 1.396240625 | 1.632412477 | 1.612783857 | 1.589319772 |
| ENSMUSG00000029163 | 2.502784398 | 2.421187155 | 2.468745337 | 1.723704441 | 1.988833497 | 1.982271233 | 2.066798338 |
| ENSMUSG00000029165 | 1.940852214 | 1.825945187 | 1.839388001 | 1.708222504 | 1.746695109 | 1.84509804 | 1.690609803 |
| ENSMUSG00000029166 | 1.857607105 | 1.866901388 | 1.850219859 | 1.868933358 | 1.905314583 | 1.991226076 | 1.888596011 |
| ENSMUSG00000029167 | 1.794934559 | 1.62694866 | 1.708222504 | 1.098079356 | 0.877063019 | 0.84509804 | 0.672731473 |
| ENSMUSG00000029168 | 0.45815691 | 0.5 | 0.580482024 | 1.160964047 | 1.223252711 | 1.301029996 | 1.296434211 |
| ENSMUSG00000029169 | 3.273096858 | 2.965659339 | 2.937990603 | 3.034967766 | 2.951316146 | 3.410777233 | 2.719193869 |
| ENSMUSG00000029171 | 2.333175866 | 2.174241882 | 2.134789703 | 2.30357978 | 2.244650542 | 2.605305046 | 2.116124284 |
| ENSMUSG00000029173 | 1.985980784 | 1.896240625 | 1.918106335 | 1.919870025 | 1.882215755 | 2.089905111 | 1.762103709 |
| ENSMUSG00000029174 | 2.584966402 | 2.411414608 | 2.434100483 | 2.467205639 | 2.442693389 | 2.770115295 | 2.325562604 |
| ENSMUSG00000029175 | 2.384185734 | 2.252806814 | 2.286801231 | 2.060793496 | 2.095689764 | 2.271841607 | 2.005303651 |
| ENSMUSG00000029176 | 2.955747812 | 2.678776002 | 2.655512955 | 2.697994205 | 2.588696459 | 2.968482949 | 2.421127112 |
| ENSMUSG00000029177 | 2.711874084 | 2.465659339 | 2.517530236 | 2.570482507 | 2.524437092 | 2.859738566 | 2.374453863 |
| ENSMUSG00000029178 | 2.8302808 | 2.606303976 | 2.593216265 | 2.468361028 | 2.397399561 | 2.7355989 | 2.240219117 |
| ENSMUSG00000029179 | 2.089597219 | 1.96759118 | 1.926839783 | 1.945339928 | 1.905314583 | 2.146128036 | 1.808770687 |
| ENSMUSG00000029182 | 0 | 0 | 0 | 0 | 0 | 0 | 0 |
| ENSMUSG00000029184 | 0 | 0 | 0 | 0 | 0 | 0 | 0 |
| ENSMUSG00000029185 | 2.802549482 | 2.630645383 | 2.603142462 | 2.366391601 | 2.289867804 | 2.626340367 | 2.167790061 |
| ENSMUSG00000029186 | 2.346900291 | 2.19697564 | 2.235628626 | 2.257666784 | 2.162515999 | 2.44870632 | 2.021338422 |
| ENSMUSG00000029188 | 0 | 0 | 0 | 0 | 0 | 0 | 0.224243824 |
| ENSMUSG00000029189 | 0.45815691 | 0 | 0.25 | 0.25 | 0.387761935 | 0 | 0.224243824 |
| ENSMUSG00000029190 | 2.724194991 | 2.506630861 | 2.475593779 | 2.549918086 | 2.488995627 | 2.86332286 | 2.326294816 |
| ENSMUSG00000029191 | 2.945876762 | 2.720469677 | 2.717783796 | 2.804337976 | 2.705674618 | 3.148602655 | 2.516100545 |
| ENSMUSG00000029192 | 2.534573679 | 2.302363341 | 2.284247778 | 2.382357639 | 2.314756076 | 2.631443769 | 2.14725554 |
| ENSMUSG00000029193 | 0 | 0 | 0 | 0 | 0 | 0 | 0 |
| ENSMUSG00000029195 | 0.960252568 | 0.79248125 | 0.701838731 | 0.75 | 0.632412477 | 0.77815125 | 0.74492186 |
| ENSMUSG00000029196 | 2.29078455 | 2.073655187 | 2.137686696 | 2.142463902 | 2.039257769 | 2.367355921 | 1.980721036 |
| ENSMUSG00000029198 | 2.594187595 | 2.437048212 | 2.408702763 | 2.37892496 | 2.368520226 | 2.698970004 | 2.252393349 |
| ENSMUSG00000029199 | 2.364524 | 2.166333979 | 2.178561379 | 2.247171172 | 2.129414728 | 2.436162647 | 2 |
| ENSMUSG00000029201 | 2.997485856 | 2.759043403 | 2.757666784 | 2.757494337 | 2.670393071 | 3.114610984 | 2.51637161 |
| ENSMUSG00000029202 | 3.118167235 | 2.853142462 | 2.822504712 | 2.862029076 | 2.784103553 | 3.216429831 | 2.56705357 |
| ENSMUSG00000029203 | 2.793908648 | 2.586628433 | 2.542794199 | 2.626452888 | 2.538790182 | 2.963315511 | 2.356752139 |

| | | | | | | | |
|---|---|---|---|---|---|---|---|
| ENSMUSG00000029204 | 0.289064826 | 0 | 0 | 0.25 | 0 | 0.301029996 | 0 |
| ENSMUSG00000029205 | 0 | 0 | 0 | 0 | 0 | 0 | 0 |
| ENSMUSG00000029206 | 1.445324132 | 1.25 | 1.27186571 | 0.976722649 | 1.020174412 | 1.079181246 | 0.916588299 |
| ENSMUSG00000029207 | 2.851836474 | 2.630890489 | 2.58993739 | 2.3464656 | 2.306140782 | 2.600972896 | 2.162575192 |
| ENSMUSG00000029208 | 2.231898196 | 2.05357978 | 2.086073977 | 2.159156155 | 2.10031573 | 2.374748346 | 1.960974982 |
| ENSMUSG00000029209 | 1.91631382 | 1.769203899 | 1.735628626 | 1.634789703 | 1.467903253 | 1.681241237 | 1.486592281 |
| ENSMUSG00000029211 | 0.289064826 | 0 | 0 | 0 | 0.244650542 | 0 | 0 |
| ENSMUSG00000029212 | 0.45815691 | 0.646240625 | 0.25 | 0 | 0.387761935 | 0 | 0.629532004 |
| ENSMUSG00000029213 | 2.545896521 | 2.335518667 | 2.352347734 | 2.280383379 | 2.210031835 | 2.445604203 | 2.029982479 |
| ENSMUSG00000029217 | 1.777675583 | 1.685366747 | 1.614857905 | 1.747171172 | 1.677808768 | 1.880813592 | 1.584526862 |
| ENSMUSG00000029219 | 0 | 0 | 0 | 0.396240625 | 0 | 0 | 0.224243824 |
| ENSMUSG00000029221 | 2.676666078 | 2.468745337 | 2.448197574 | 2.513142013 | 2.429560273 | 2.783188691 | 2.265305657 |
| ENSMUSG00000029223 | 2.103271637 | 2.086073977 | 2.058404919 | 1.931980114 | 2.059432181 | 2.181843588 | 2.018194418 |
| ENSMUSG00000029227 | 2.766857015 | 2.534247778 | 2.536801231 | 2.5896628 | 2.505998068 | 2.898725182 | 2.368945281 |
| ENSMUSG00000029228 | 0.289064826 | 0.396240625 | 0.646240625 | 0 | 0.244650542 | 0.477121255 | 0.355418052 |
| ENSMUSG00000029229 | 2.116511873 | 1.950224975 | 1.96293726 | 2.023189285 | 1.950242452 | 2.201397124 | 1.846338363 |
| ENSMUSG00000029231 | 1.45815691 | 1.464495249 | 1.557204673 | 1.02186571 | 0.905314583 | 1 | 0.984950056 |
| ENSMUSG00000029233 | 2.190368148 | 2.072504712 | 2.067865757 | 1.938721876 | 1.897237431 | 2.10720997 | 1.74492186 |
| ENSMUSG00000029234 | 2.696591798 | 2.451435968 | 2.427270953 | 2.385757955 | 2.320713283 | 2.63447727 | 2.16858494 |
| ENSMUSG00000029235 | 0 | 0 | 0 | 0 | 0 | 0 | 0.224243824 |
| ENSMUSG00000029236 | 0 | 0.25 | 0 | 0 | 0 | 0 | 0.224243824 |
| ENSMUSG00000029238 | 2.832159327 | 2.59995299 | 2.586905342 | 2.622963274 | 2.531280194 | 2.891537458 | 2.344772901 |
| ENSMUSG00000029245 | 0 | 0.25 | 0 | 0.25 | 0 | 0.301029996 | 0.224243824 |
| ENSMUSG00000029246 | 2.629921064 | 2.388167274 | 2.380890489 | 2.394357207 | 2.338940611 | 2.668385917 | 2.145981019 |
| ENSMUSG00000029247 | 3.239024083 | 2.951032755 | 2.90904286 | 2.977004315 | 2.870695876 | 3.348304863 | 2.65820629 |
| ENSMUSG00000029248 | 0 | 0 | 0 | 0 | 0 | 0 | 0.224243824 |
| ENSMUSG00000029249 | 2.642661589 | 2.42510993 | 2.428990498 | 2.3843046 | 2.33281029 | 2.618048097 | 2.141264342 |
| ENSMUSG00000029250 | 3.01386641 | 2.74234663 | 2.710391087 | 2.841853688 | 2.722627519 | 3.165837625 | 2.533263692 |
| ENSMUSG00000029253 | 2.494443473 | 2.327084758 | 2.325374049 | 2.276477127 | 2.21338377 | 2.492760389 | 1.996640515 |
| ENSMUSG00000029254 | 0 | 0.580482024 | 0 | 0.5 | 0.244650542 | 0.477121255 | 0 |
| ENSMUSG00000029255 | 0 | 0 | 0 | 0 | 0 | 0 | 0 |
| ENSMUSG00000029260 | 0 | 0 | 0 | 0 | 0 | 0 | 0 |
| ENSMUSG00000029263 | 2.185978293 | 2.017865591 | 1.987091808 | 1.96759118 | 1.92093466 | 2.103803721 | 1.815933683 |
| ENSMUSG00000029265 | 2.711874084 | 2.476722649 | 2.477847997 | 2.471805154 | 2.37406527 | 2.767897616 | 2.189340078 |

| | | | | | | | |
|---|---|---|---|---|---|---|---|
| ENSMUSG00000029267 | 2.724194991 | 2.467205639 | 2.469129237 | 2.430274797 | 2.349985771 | 2.669316881 | 2.196701384 |
| ENSMUSG00000029268 | 0 | 0 | 0 | 0 | 0 | 0 | 0 |
| ENSMUSG00000029269 | 0 | 0 | 0 | 0 | 0 | 0 | 0 |
| ENSMUSG00000029270 | 2.158622101 | 2.00835575 | 1.966046536 | 1.821350555 | 1.814781639 | 2.004321374 | 1.653807046 |
| ENSMUSG00000029272 | 0 | 0 | 0 | 0 | 0 | 0 | 0 |
| ENSMUSG00000029273 | 0 | 0 | 0 | 0 | 0 | 0 | 0 |
| ENSMUSG00000029275 | 0 | 0 | 0.25 | 0.25 | 0 | 0.301029996 | 0.224243824 |
| ENSMUSG00000029276 | 2.020182925 | 1.888647213 | 1.864857905 | 1.894357207 | 1.808534465 | 2.056904851 | 1.676387053 |
| ENSMUSG00000029279 | 1 | 1.226722649 | 1.080482024 | 1.226722649 | 1.264824954 | 1.230448921 | 1.193409508 |
| ENSMUSG00000029280 | 0 | 0 | 0 | 0 | 0 | 0 | 0 |
| ENSMUSG00000029281 | 0 | 0 | 0 | 0 | 0 | 0 | 0 |
| ENSMUSG00000029282 | 0 | 0 | 0 | 0 | 0 | 0 | 0 |
| ENSMUSG00000029283 | 2.534573679 | 2.351816941 | 2.357090043 | 2.306603048 | 2.245869735 | 2.551449998 | 2.109521699 |
| ENSMUSG00000029286 | 0 | 0 | 0 | 0 | 0 | 0 | 0 |
| ENSMUSG00000029287 | 2.036286563 | 1.821350555 | 1.914552871 | 1.42510993 | 1.335652905 | 1.462397998 | 1.307989563 |
| ENSMUSG00000029288 | 0 | 0 | 0 | 0 | 0 | 0 | 0 |
| ENSMUSG00000029290 | 2.557825653 | 2.353406982 | 2.360735874 | 2.399513125 | 2.330902415 | 2.645422269 | 2.186348006 |
| ENSMUSG00000029291 | 2.149089517 | 1.987091808 | 2.02053726 | 1.982684334 | 1.94314048 | 2.227886705 | 1.855869786 |
| ENSMUSG00000029298 | 1 | 0.75 | 0.951838731 | 1.25 | 1.351342427 | 1.380211242 | 1.252393349 |
| ENSMUSG00000029299 | 0 | 0 | 0 | 0 | 0 | 0 | 0 |
| ENSMUSG00000029304 | 2.68005385 | 2.458617762 | 2.39151351 | 1.668106335 | 1.595992969 | 1.748188027 | 1.56970677 |
| ENSMUSG00000029306 | 0 | 0 | 0 | 0 | 0 | 0 | 0 |
| ENSMUSG00000029307 | 0 | 0 | 0 | 0 | 0 | 0 | 0 |
| ENSMUSG00000029309 | 0 | 0.25 | 0.396240625 | 0.25 | 0 | 0 | 0.224243824 |
| ENSMUSG00000029310 | 2.441754646 | 2.261780978 | 2.322504712 | 2.140560606 | 2.140250123 | 2.324282455 | 2.049487947 |
| ENSMUSG00000029311 | 2.326928073 | 2.150906586 | 2.156427211 | 1.898114259 | 1.78687392 | 2.033423755 | 1.702561601 |
| ENSMUSG00000029312 | 1.811507563 | 1.763820609 | 1.695339928 | 1.685366747 | 1.698489937 | 1.832508913 | 1.647906904 |
| ENSMUSG00000029313 | 3.065938681 | 2.796992648 | 2.808404919 | 2.557805295 | 2.532902983 | 2.894869657 | 2.386211856 |
| ENSMUSG00000029314 | 1.671187741 | 1.593759858 | 1.464495249 | 1.935366747 | 1.880506521 | 2.158362492 | 1.771720588 |
| ENSMUSG00000029319 | 2.144240249 | 1.96293726 | 1.994319981 | 1.926839783 | 1.898867717 | 2.123851641 | 1.802677937 |
| ENSMUSG00000029320 | 0 | 0 | 0 | 0 | 0 | 0 | 0 |
| ENSMUSG00000029321 | 0 | 0 | 0 | 0 | 0 | 0 | 0 |
| ENSMUSG00000029322 | 0.578129653 | 0.701838731 | 0.396240625 | 0.396240625 | 0 | 0 | 0.224243824 |
| ENSMUSG00000029326 | 2.253091469 | 2.086073977 | 2.12892496 | 1.909156155 | 1.961316494 | 2.158362492 | 1.913982791 |

| | | | | | | | |
|---|---|---|---|---|---|---|---|
| ENSMUSG00000029328 | 3.153883046 | 2.842671852 | 2.84253562 | 2.827795165 | 2.764077132 | 3.140193679 | 2.571196834 |
| ENSMUSG00000029330 | 0.45815691 | 0.396240625 | 0.5 | 0.951838731 | 1.176121988 | 1.041392685 | 0.853775828 |
| ENSMUSG00000029331 | 0 | 0 | 0 | 0 | 0 | 0 | 0 |
| ENSMUSG00000029333 | 1.325351389 | 1.25 | 1.226722649 | 1.634789703 | 1.614672255 | 1.716003344 | 1.529383012 |
| ENSMUSG00000029334 | 2.733202521 | 2.557505109 | 2.503853763 | 1.921625132 | 1.842556557 | 2.025305865 | 1.663407615 |
| ENSMUSG00000029335 | 0.45815691 | 0.25 | 0.396240625 | 0.646240625 | 0.387761935 | 0.301029996 | 0.448487648 |
| ENSMUSG00000029337 | 0 | 0 | 0.25 | 1.130890489 | 1.149965125 | 1.342422681 | 1.054044578 |
| ENSMUSG00000029338 | 2.516991255 | 2.310197833 | 2.307805295 | 2.223704441 | 2.148364414 | 2.431363764 | 2.018825668 |
| ENSMUSG00000029343 | 0.289064826 | 0 | 0.25 | 0 | 0 | 0 | 0 |
| ENSMUSG00000029344 | 2.355805236 | 2.281030328 | 2.350753006 | 1.945339928 | 2.091002363 | 2.158362492 | 2.090889903 |
| ENSMUSG00000029345 | 2.388281011 | 2.30539678 | 2.390560606 | 2.073655187 | 2.1962964 | 2.322219295 | 2.20389891 |
| ENSMUSG00000029346 | 1.418409478 | 1.321350555 | 1.396240625 | 1.339388001 | 1.351342427 | 1.204119983 | 1.319146374 |
| ENSMUSG00000029348 | 0.747221736 | 0.701838731 | 0.5 | 1.080482024 | 1.387761935 | 1.322219295 | 1.335191751 |
| ENSMUSG00000029352 | 0.45815691 | 0.25 | 0 | 0.25 | 0.632412477 | 0.477121255 | 0.74492186 |
| ENSMUSG00000029359 | 0.811507563 | 0.830482024 | 0.976722649 | 0.396240625 | 0.812711509 | 0.301029996 | 0.935079929 |
| ENSMUSG00000029360 | 0.747221736 | 0.896240625 | 0.701838731 | 0.396240625 | 0.733951626 | 0 | 0.803905701 |
| ENSMUSG00000029361 | 0 | 0 | 0.25 | 0.25 | 0.244650542 | 0.301029996 | 0.224243824 |
| ENSMUSG00000029363 | 2.737340367 | 2.495748394 | 2.489638179 | 2.564346961 | 2.495055108 | 2.862727528 | 2.34130043 |
| ENSMUSG00000029364 | 2.787699615 | 2.617664719 | 2.638886943 | 2.40136988 | 2.438839735 | 2.751279104 | 2.31789776 |
| ENSMUSG00000029366 | 2.926900886 | 2.689972046 | 2.689763985 | 2.629909063 | 2.54143406 | 2.937016107 | 2.352283622 |
| ENSMUSG00000029368 | 0 | 0 | 0.25 | 0 | 0 | 0 | 0 |
| ENSMUSG00000029369 | 0.289064826 | 0 | 0 | 0 | 0 | 0 | 0 |
| ENSMUSG00000029370 | 1.289064826 | 1.396240625 | 1.160964047 | 0.25 | 0.812711509 | 0.477121255 | 0.224243824 |
| ENSMUSG00000029371 | 2.036286563 | 1.910964047 | 1.972185812 | 1.584962501 | 1.576269619 | 1.602059991 | 1.441050184 |
| ENSMUSG00000029372 | 0.289064826 | 0 | 0.25 | 0.396240625 | 0.244650542 | 0.301029996 | 0.224243824 |
| ENSMUSG00000029373 | 0 | 0 | 0 | 0.25 | 0 | 0 | 0 |
| ENSMUSG00000029375 | 0.578129653 | 0.701838731 | 0.646240625 | 0 | 0 | 0 | 0 |
| ENSMUSG00000029376 | 2.348397716 | 2.216046536 | 2.140560606 | 2.154596376 | 2.057362051 | 2.324282455 | 1.880961074 |
| ENSMUSG00000029377 | 1.87190676 | 1.830482024 | 1.834962501 | 1.532320754 | 1.467903253 | 1.579783597 | 1.369798885 |
| ENSMUSG00000029378 | 0.91631382 | 0.830482024 | 1 | 0.396240625 | 0.568060967 | 0.477121255 | 0.520678036 |
| ENSMUSG00000029379 | 0.45815691 | 0 | 0.25 | 0.580482024 | 0.244650542 | 0.954242509 | 0.74492186 |
| ENSMUSG00000029380 | 2.445324132 | 2.245034894 | 2.364857905 | 2.309601185 | 2.236621448 | 2.57634135 | 2.212676319 |
| ENSMUSG00000029381 | 1.753493018 | 1.735628626 | 1.755591953 | 1.774508021 | 1.94314048 | 2.056904851 | 1.95718226 |
| ENSMUSG00000029384 | 0 | 0 | 0.25 | 0 | 0.244650542 | 0 | 0.355418052 |

| | | | | | | | |
|---|---|---|---|---|---|---|---|
| ENSMUSG00000029385 | 2.471720134 | 2.253505118 | 2.263820609 | 1.975216702 | 1.897237431 | 2.173186268 | 1.824094866 |
| ENSMUSG00000029386 | 1.952600383 | 1.845926073 | 1.882845365 | 1.870953944 | 1.932951348 | 2.117271296 | 1.838725884 |
| ENSMUSG00000029387 | 2.305951036 | 2.160061234 | 2.135757955 | 2.164552871 | 2.118236133 | 2.397940009 | 2.000667731 |
| ENSMUSG00000029388 | 2.531694272 | 2.386723615 | 2.366901388 | 2.420309603 | 2.372793355 | 2.72427587 | 2.262348456 |
| ENSMUSG00000029389 | 2.103271637 | 1.991446071 | 2.058404919 | 1.918106335 | 1.928492895 | 2.037426498 | 1.869159809 |
| ENSMUSG00000029390 | 3.351646281 | 3.040597832 | 3.021699921 | 3.017781781 | 2.922817097 | 3.413969972 | 2.709794665 |
| ENSMUSG00000029392 | 2.089597219 | 2.029735268 | 2.080482024 | 1.866901388 | 1.90849439 | 2.049218023 | 1.913109605 |
| ENSMUSG00000029394 | 2.111266113 | 1.958222504 | 1.976722649 | 2.015173983 | 1.965381292 | 2.176091259 | 1.87511425 |
| ENSMUSG00000029397 | 2.485932312 | 2.291854536 | 2.28296424 | 2.27908599 | 2.194189193 | 2.477121255 | 2.020084484 |
| ENSMUSG00000029401 | 2.054824387 | 1.872963274 | 1.860735874 | 1.821350555 | 1.944572352 | 2.10720997 | 1.896054911 |
| ENSMUSG00000029402 | 1.862428343 | 1.71759118 | 1.729715809 | 1.653677461 | 1.763796674 | 1.886490725 | 1.667169465 |
| ENSMUSG00000029403 | 2.153883046 | 1.956637122 | 2.001406137 | 2.136723615 | 2.065570703 | 2.294466226 | 1.902443294 |
| ENSMUSG00000029404 | 2.185978293 | 2.138647213 | 2.201032755 | 1.918106335 | 2.062514787 | 2.235528447 | 2.07004434 |
| ENSMUSG00000029405 | 2.928763473 | 2.686418581 | 2.680701883 | 2.719895812 | 2.648526544 | 3.056142262 | 2.468257888 |
| ENSMUSG00000029406 | 2.095120908 | 2.095926073 | 2.14151351 | 1.653677461 | 1.902105869 | 1.86332286 | 1.885753983 |
| ENSMUSG00000029407 | 3.038863496 | 2.776149688 | 2.753505118 | 2.821061437 | 2.759704793 | 3.142076461 | 2.54858238 |
| ENSMUSG00000029408 | 0.811507563 | 0.864857905 | 0.864857905 | 0.951838731 | 1.310728002 | 1.255272505 | 1.21680636 |
| ENSMUSG00000029409 | 0.289064826 | 0.25 | 0.25 | 1 | 0.733951626 | 1 | 0.853775828 |
| ENSMUSG00000029410 | 0.811507563 | 0.79248125 | 0.92510993 | 0.396240625 | 0.733951626 | 0.602059991 | 0.853775828 |
| ENSMUSG00000029413 | 2.278594844 | 2.109697963 | 2.145300145 | 2.204245906 | 2.197694248 | 2.550228353 | 2.096394553 |
| ENSMUSG00000029414 | 2.845984872 | 2.569612365 | 2.557505109 | 2.580200137 | 2.530737602 | 2.875639937 | 2.351607989 |
| ENSMUSG00000029415 | 2.630681378 | 2.40136988 | 2.396709947 | 2.388647213 | 2.317250476 | 2.600972896 | 2.13691595 |
| ENSMUSG00000029416 | 2.247417471 | 2.130890489 | 2.064346961 | 1.866901388 | 1.905314583 | 2.068185862 | 1.763495175 |
| ENSMUSG00000029417 | 0 | 0 | 0.25 | 0.396240625 | 0.387761935 | 0 | 0.224243824 |
| ENSMUSG00000029419 | 0 | 0 | 0 | 0 | 0 | 0 | 0 |
| ENSMUSG00000029420 | 0.289064826 | 0 | 0 | 0 | 0 | 0 | 0.224243824 |
| ENSMUSG00000029422 | 2.834029431 | 2.573942234 | 2.56932185 | 2.601816941 | 2.539585427 | 2.902546779 | 2.357417124 |
| ENSMUSG00000029423 | 0 | 0 | 0 | 0 | 0 | 0 | 0 |
| ENSMUSG00000029426 | 2.793908648 | 2.531030328 | 2.535207443 | 2.562574604 | 2.474969034 | 2.85308953 | 2.298300395 |
| ENSMUSG00000029427 | 2.479432974 | 2.323655187 | 2.373963757 | 2.317865757 | 2.319727371 | 2.574031268 | 2.211287839 |
| ENSMUSG00000029428 | 2.752926783 | 2.512462137 | 2.506976499 | 2.353406982 | 2.35043857 | 2.709269961 | 2.209893373 |
| ENSMUSG00000029430 | 3.515901449 | 3.192459461 | 3.1925111 | 3.126886726 | 3.066047671 | 3.54740546 | 2.867973935 |
| ENSMUSG00000029431 | 0.289064826 | 0 | 0 | 0 | 0 | 0 | 0 |
| ENSMUSG00000029432 | 2.593357679 | 2.366901388 | 2.371457327 | 2.418548067 | 2.352693888 | 2.741151599 | 2.228559641 |

| | | | | | | | |
|---|---|---|---|---|---|---|---|
| ENSMUSG00000029433 | 2.346900291 | 2.172499493 | 2.157339155 | 2.211372513 | 2.162515999 | 2.480006943 | 2.078019652 |
| ENSMUSG00000029434 | 2.78298141 | 2.574515642 | 2.566989271 | 2.52219706 | 2.48684999 | 2.84135947 | 2.355418052 |
| ENSMUSG00000029436 | 1.069666964 | 0.951838731 | 1.098079356 | 1.098079356 | 1.039257769 | 0.903089987 | 1.23151414 |
| ENSMUSG00000029437 | 0 | 0 | 0 | 0 | 0 | 0 | 0 |
| ENSMUSG00000029438 | 2.144240249 | 2.012462137 | 2.03360658 | 1.79248125 | 1.78687392 | 2.012837225 | 1.734460864 |
| ENSMUSG00000029439 | 2.786132803 | 2.55539678 | 2.576515422 | 2.407339155 | 2.437427909 | 2.671172843 | 2.327754286 |
| ENSMUSG00000029440 | 2.318984359 | 2.203445298 | 2.221424093 | 2.135757955 | 2.221300875 | 2.423245874 | 2.181044355 |
| ENSMUSG00000029442 | 0.578129653 | 0.646240625 | 0.864857905 | 0.701838731 | 0.733951626 | 0.301029996 | 0.74492186 |
| ENSMUSG00000029445 | 0 | 0 | 0 | 0 | 0 | 0 | 0 |
| ENSMUSG00000029446 | 2.457006475 | 2.256284891 | 2.227473271 | 2.129909063 | 2.203916946 | 2.45484486 | 2.082194103 |
| ENSMUSG00000029447 | 3.489435171 | 3.168161582 | 3.144180131 | 3.227145066 | 3.117515919 | 3.642563437 | 2.898558913 |
| ENSMUSG00000029449 | 1.516991255 | 1.42510993 | 1.431980114 | 1.282320754 | 1.433158226 | 1.447158031 | 1.360283038 |
| ENSMUSG00000029451 | 0 | 0 | 0 | 0 | 0 | 0 | 0 |
| ENSMUSG00000029452 | 0.747221736 | 0.396240625 | 0.896240625 | 0.580482024 | 0.632412477 | 0.477121255 | 0.520678036 |
| ENSMUSG00000029454 | 1.100572389 | 1 | 0.976722649 | 0.646240625 | 0.955822902 | 0.602059991 | 0.876096088 |
| ENSMUSG00000029455 | 3.045658471 | 2.856566189 | 2.958518988 | 2.574229052 | 2.684194078 | 2.969881644 | 2.656629445 |
| ENSMUSG00000029456 | 1.418409478 | 1.53743678 | 1.602347734 | 1.130890489 | 1.335652905 | 1.361727836 | 1.265599895 |
| ENSMUSG00000029458 | 2.66283289 | 2.421625132 | 2.430274797 | 2.37246199 | 2.395413309 | 2.6599162 | 2.242438242 |
| ENSMUSG00000029461 | 2.674620126 | 2.441217898 | 2.435366747 | 2.366391601 | 2.310728002 | 2.662757832 | 2.157274883 |
| ENSMUSG00000029462 | 2.845984872 | 2.603935442 | 2.612545062 | 2.517530236 | 2.467578695 | 2.855519156 | 2.29131391 |
| ENSMUSG00000029463 | 2.183765914 | 2.065523711 | 2.036164561 | 2.107613138 | 2.027028145 | 2.322219295 | 1.980010793 |
| ENSMUSG00000029464 | 2.215778996 | 2.185366747 | 2.203445298 | 2.038704527 | 2.112066955 | 2.352182518 | 2.041356071 |
| ENSMUSG00000029465 | 2.830751226 | 2.597542239 | 2.62521047 | 2.563461871 | 2.527190274 | 2.962369336 | 2.437612155 |
| ENSMUSG00000029466 | 2.679378494 | 2.491806565 | 2.469512728 | 2.3843046 | 2.386129765 | 2.687528961 | 2.250241695 |
| ENSMUSG00000029467 | 3.49910842 | 3.160964047 | 3.137866988 | 3.082168732 | 3.024624348 | 3.496929648 | 2.807340903 |
| ENSMUSG00000029468 | 1.069666964 | 0.951838731 | 0.830482024 | 0.701838731 | 0.812711509 | 0.602059991 | 0.672731473 |
| ENSMUSG00000029469 | 2.300956576 | 2.161864607 | 2.156427211 | 2.160061234 | 2.072599934 | 2.350248018 | 1.918313719 |
| ENSMUSG00000029470 | 2.126809492 | 1.987091808 | 2.03743678 | 1.953445298 | 1.928492895 | 2.133538908 | 1.809975587 |
| ENSMUSG00000029471 | 2.507914013 | 2.323655187 | 2.355516192 | 2.260414788 | 2.279510047 | 2.527629901 | 2.187847501 |
| ENSMUSG00000029472 | 3.386424506 | 3.087458523 | 3.091990177 | 3.065523711 | 3.042520869 | 3.499136995 | 2.860017477 |
| ENSMUSG00000029474 | 2.339331437 | 2.178561379 | 2.133818844 | 2.118933358 | 2.085294053 | 2.428134794 | 1.967691807 |
| ENSMUSG00000029475 | 2.302628051 | 2.177701608 | 2.226722649 | 2.222945926 | 2.223900931 | 2.45484486 | 2.159731975 |
| ENSMUSG00000029477 | 0 | 0.25 | 0 | 0.25 | 0.244650542 | 0.301029996 | 0 |
| ENSMUSG00000029478 | 2.645603628 | 2.565229883 | 2.693490842 | 2.065523711 | 2.316753004 | 2.311753861 | 2.381313105 |

| | | | | | | | |
|---|---|---|---|---|---|---|---|
| ENSMUSG00000029480 | 2.397683106 | 2.208222504 | 2.333288838 | 1.985628626 | 2.141069998 | 2.240549248 | 2.118913228 |
| ENSMUSG00000029482 | 2.517987749 | 2.397646762 | 2.470660762 | 2.154596376 | 2.356717497 | 2.488550717 | 2.293479712 |
| ENSMUSG00000029484 | 3.426149384 | 3.131196638 | 3.121708755 | 3.047069938 | 2.947418788 | 3.447468131 | 2.750882395 |
| ENSMUSG00000029486 | 2.317377295 | 2.175975893 | 2.143411797 | 2.21759118 | 2.13194489 | 2.392696953 | 1.989815042 |
| ENSMUSG00000029490 | 1.205378646 | 1.080482024 | 1.02186571 | 0.396240625 | 0.387761935 | 0.602059991 | 0.448487648 |
| ENSMUSG00000029491 | 1 | 0.580482024 | 0.646240625 | 0 | 0.568060967 | 0.301029996 | 0.448487648 |
| ENSMUSG00000029499 | 1.307602651 | 1.146240625 | 0.951838731 | 1.330482024 | 1.387761935 | 1.397940009 | 1.278288402 |
| ENSMUSG00000029500 | 2.083999385 | 2.03360658 | 2.023189285 | 1.903677461 | 1.964031554 | 2.187520721 | 1.845261772 |
| ENSMUSG00000029501 | 2.742019602 | 2.514497931 | 2.510756821 | 2.573942234 | 2.485000434 | 2.856124444 | 2.288312058 |
| ENSMUSG00000029502 | 2.954701306 | 2.672281101 | 2.67140621 | 2.597542239 | 2.540642974 | 2.926342447 | 2.37800557 |
| ENSMUSG00000029503 | 0.91631382 | 0.830482024 | 0.701838731 | 0.864857905 | 0.955822902 | 0.698970004 | 0.775756176 |
| ENSMUSG00000029504 | 2.37447073 | 2.168989258 | 2.208222504 | 2.161864607 | 2.170139537 | 2.361727836 | 2.003324957 |
| ENSMUSG00000029505 | 3.109342085 | 2.859177886 | 2.855384727 | 2.76024365 | 2.755708482 | 3.117602692 | 2.593189105 |
| ENSMUSG00000029507 | 2.275045611 | 2.136723615 | 2.219895812 | 2.002806814 | 2.063536368 | 2.26245109 | 1.95794437 |
| ENSMUSG00000029510 | 0.811507563 | 0.79248125 | 0.646240625 | 1.02186571 | 0.931471446 | 0.84509804 | 1 |
| ENSMUSG00000029512 | 2.527823874 | 2.365881093 | 2.407794264 | 2.122963274 | 2.234742351 | 2.311753861 | 2.17331363 |
| ENSMUSG00000029513 | 2.331622671 | 2.187048212 | 2.176839783 | 2.074802005 | 2.064555002 | 2.298853076 | 1.986347533 |
| ENSMUSG00000029516 | 2.420929319 | 2.256976499 | 2.274508021 | 2.04248125 | 2.091944837 | 2.348304863 | 2.028761664 |
| ENSMUSG00000029517 | 0 | 0.5 | 0.25 | 0.25 | 0 | 0 | 0.224243824 |
| ENSMUSG00000029518 | 2.77230138 | 2.583568321 | 2.589388001 | 2.462155735 | 2.491431917 | 2.788168371 | 2.355863359 |
| ENSMUSG00000029521 | 2.235832493 | 2.101285366 | 2.080482024 | 1.979715809 | 1.893954095 | 2.08278537 | 1.775756176 |
| ENSMUSG00000029522 | 0 | 0 | 0 | 0 | 0.244650542 | 0 | 0 |
| ENSMUSG00000029524 | 1.671187741 | 1.646240625 | 1.598079356 | 1.53743678 | 1.572189156 | 1.556302501 | 1.540636858 |
| ENSMUSG00000029526 | 0 | 0 | 0 | 0 | 0 | 0 | 0.448487648 |
| ENSMUSG00000029528 | 2.635214384 | 2.465659339 | 2.513820609 | 2.386241108 | 2.460693128 | 2.742725131 | 2.376964992 |
| ENSMUSG00000029530 | 0 | 0.25 | 0 | 0 | 0 | 0 | 0.355418052 |
| ENSMUSG00000029534 | 2.057835465 | 1.890560606 | 1.903677461 | 1.809601185 | 1.704182902 | 1.982271233 | 1.627424404 |
| ENSMUSG00000029535 | 1.989530018 | 1.898114259 | 1.916333979 | 1.914552871 | 1.868304353 | 2.045322979 | 1.770364127 |
| ENSMUSG00000029536 | 2.192545868 | 2.04745614 | 2.03360658 | 2.08272922 | 2.033751322 | 2.309630167 | 1.920884716 |
| ENSMUSG00000029538 | 2.604828841 | 2.475216702 | 2.509729747 | 2.341033664 | 2.401338844 | 2.645422269 | 2.291856721 |
| ENSMUSG00000029541 | 0 | 0.25 | 0 | 0 | 0 | 0 | 0 |
| ENSMUSG00000029544 | 0.289064826 | 0.646240625 | 0.25 | 0 | 0 | 0 | 0.355418052 |
| ENSMUSG00000029545 | 2.392336464 | 2.30539678 | 2.349151195 | 1.921625132 | 2.124299332 | 2.136720567 | 2.149789616 |
| ENSMUSG00000029546 | 0 | 0.25 | 0 | 0 | 0 | 0 | 0 |

| | | | | | | | |
|---|---|---|---|---|---|---|---|
| ENSMUSG00000029547 | 2.506893125 | 2.438721876 | 2.555698726 | 2.184523065 | 2.462015062 | 2.496929648 | 2.501257147 |
| ENSMUSG00000029550 | 2.53838222 | 2.406427211 | 2.37892496 | 2.046218836 | 2.145141048 | 2.361727836 | 2.064068234 |
| ENSMUSG00000029551 | 1.912079952 | 1.828220739 | 1.864857905 | 1.747171172 | 1.92093466 | 1.995635195 | 1.802677937 |
| ENSMUSG00000029552 | 2.693989403 | 2.485628626 | 2.475970461 | 2.417664062 | 2.410998892 | 2.742725131 | 2.262051244 |
| ENSMUSG00000029553 | 0 | 0 | 0 | 0 | 0 | 0 | 0 |
| ENSMUSG00000029554 | 2.494443473 | 2.299918086 | 2.324229052 | 2.231203126 | 2.337065673 | 2.540329475 | 2.24651952 |
| ENSMUSG00000029556 | 0.289064826 | 0.25 | 0.25 | 0 | 0.244650542 | 0 | 0.355418052 |
| ENSMUSG00000029557 | 1.647796617 | 1.649978211 | 1.527131114 | 1.5 | 1.49445382 | 1.568201724 | 1.584526862 |
| ENSMUSG00000029559 | 2.188178997 | 2.03360658 | 2.110735874 | 1.907339155 | 2.001219193 | 2.071882007 | 1.966952343 |
| ENSMUSG00000029560 | 2.409468165 | 2.259043403 | 2.382845365 | 2.066696635 | 2.221300875 | 2.322219295 | 2.21302251 |
| ENSMUSG00000029561 | 1.036286563 | 0.79248125 | 0.701838731 | 1.188721876 | 0.978602168 | 1.322219295 | 1.041356071 |
| ENSMUSG00000029563 | 1.605636262 | 1.51109853 | 1.557204673 | 0.92510993 | 0.77552387 | 0.698970004 | 0.775756176 |
| ENSMUSG00000029564 | 0 | 0 | 0.25 | 0 | 0.387761935 | 0.301029996 | 0.224243824 |
| ENSMUSG00000029569 | 2.481610694 | 2.23927551 | 2.240000483 | 2.177701608 | 2.103062484 | 2.404833717 | 1.947894097 |
| ENSMUSG00000029570 | 2.086807695 | 1.978222334 | 2.02186571 | 1.657339155 | 1.84445929 | 1.929418926 | 1.846338363 |
| ENSMUSG00000029571 | 2.983054229 | 2.747171172 | 2.742706225 | 2.619439567 | 2.518022067 | 2.912753304 | 2.31789776 |
| ENSMUSG00000029575 | 2.054824387 | 1.970660762 | 1.992885888 | 1.894357207 | 1.929985312 | 2.155336037 | 1.874129423 |
| ENSMUSG00000029576 | 1.129344651 | 1.410964047 | 1.445339928 | 1.602347734 | 1.848234258 | 1.838849091 | 1.865128412 |
| ENSMUSG00000029577 | 2.921262688 | 2.74539182 | 2.746993631 | 2.548073204 | 2.640461996 | 2.913813852 | 2.507861167 |
| ENSMUSG00000029578 | 2.72721917 | 2.504202072 | 2.504897682 | 2.525821952 | 2.537194296 | 2.815577748 | 2.364426535 |
| ENSMUSG00000029580 | 2.438154344 | 2.23927551 | 2.275165585 | 3.499162505 | 3.55048942 | 4.072874597 | 3.336837049 |
| ENSMUSG00000029581 | 3.287180641 | 3.063018511 | 3.13954538 | 2.83076369 | 2.97197586 | 3.29136885 | 2.900548253 |
| ENSMUSG00000029586 | 0 | 0 | 0.396240625 | 0 | 0 | 0.301029996 | 0 |
| ENSMUSG00000029587 | 2.136858952 | 1.961372513 | 1.997171172 | 1.937048212 | 1.950242452 | 2.155336037 | 1.842010333 |
| ENSMUSG00000029591 | 1.948720984 | 1.87892496 | 1.943696765 | 1.769203899 | 1.793512788 | 1.977723605 | 1.774416565 |
| ENSMUSG00000029592 | 2.312518611 | 2.132845365 | 2.180274797 | 2.090485943 | 2.104881849 | 2.338456494 | 2.015657029 |
| ENSMUSG00000029594 | 2.415874319 | 2.26109853 | 2.293106877 | 2.048689214 | 2.19206933 | 2.33243846 | 2.10278155 |
| ENSMUSG00000029595 | 0 | 0 | 0 | 0 | 0 | 0 | 0 |
| ENSMUSG00000029596 | 0 | 0 | 0.396240625 | 0 | 0.387761935 | 0 | 0.520678036 |
| ENSMUSG00000029597 | 0.45815691 | 0.5 | 0.580482024 | 0.25 | 0.244650542 | 0 | 0.448487648 |
| ENSMUSG00000029598 | 2.660010325 | 2.528109877 | 2.583847589 | 2.168106335 | 2.291475807 | 2.5132176 | 2.227237819 |
| ENSMUSG00000029599 | 2.381433035 | 2.328787391 | 2.420309603 | 1.96293726 | 2.199086582 | 2.187520721 | 2.269981661 |
| ENSMUSG00000029600 | 1.53838222 | 1.575945187 | 1.705044741 | 1.321350555 | 1.489301084 | 1.230448921 | 1.537859885 |
| ENSMUSG00000029601 | 0.747221736 | 0.830482024 | 0.79248125 | 0.5 | 0.489301084 | 0 | 0.74492186 |

| | | | | | | | |
|---|---|---|---|---|---|---|---|
| ENSMUSG00000029602 | 0.289064826 | 0 | 0.5 | 0 | 0 | 0.301029996 | 0.224243824 |
| ENSMUSG00000029603 | 0.289064826 | 0.25 | 0.25 | 0 | 0.244650542 | 0.301029996 | 0.355418052 |
| ENSMUSG00000029605 | 0.289064826 | 0.396240625 | 0 | 0.976722649 | 0.568060967 | 0.954242509 | 0.672731473 |
| ENSMUSG00000029607 | 0.867194479 | 1.04248125 | 1.04248125 | 1 | 1.074582839 | 0.77815125 | 1.054044578 |
| ENSMUSG00000029608 | 0 | 0 | 0 | 0.25 | 0 | 0.301029996 | 0.224243824 |
| ENSMUSG00000029610 | 2.057835465 | 1.948603967 | 1.992885888 | 1.928561379 | 1.935892666 | 2.120573931 | 1.885753983 |
| ENSMUSG00000029613 | 2.485932312 | 2.314935816 | 2.291226732 | 2.457430684 | 2.373641807 | 2.718501689 | 2.229547475 |
| ENSMUSG00000029614 | 3.463655862 | 3.169925001 | 3.157282226 | 3.062204265 | 3.067296654 | 3.506640306 | 2.882581172 |
| ENSMUSG00000029616 | 2.721148722 | 2.546838018 | 2.599151195 | 2.42510993 | 2.48561803 | 2.745074792 | 2.413397706 |
| ENSMUSG00000029617 | 2.66283289 | 2.462155735 | 2.463327389 | 2.564935816 | 2.488995627 | 2.859138297 | 2.326782038 |
| ENSMUSG00000029618 | 0 | 0 | 0 | 0 | 0 | 0 | 0 |
| ENSMUSG00000029620 | 0 | 0 | 0 | 0 | 0 | 0 | 0 |
| ENSMUSG00000029621 | 2.715608727 | 2.531676118 | 2.515173983 | 2.512121718 | 2.555884575 | 2.883661435 | 2.410967356 |
| ENSMUSG00000029622 | 2.826022917 | 2.714300868 | 2.78743678 | 2.450224975 | 2.60468636 | 2.792391689 | 2.564378432 |
| ENSMUSG00000029623 | 2.822201287 | 2.583568321 | 2.624213444 | 2.587734795 | 2.55990691 | 2.906873535 | 2.462548643 |
| ENSMUSG00000029624 | 2.033115198 | 1.988549078 | 1.988549078 | 1.864857905 | 1.992595054 | 2.096910013 | 1.914853627 |
| ENSMUSG00000029625 | 2.368815967 | 2.197790722 | 2.231203126 | 2.095926073 | 2.130260132 | 2.365487985 | 2.100339912 |
| ENSMUSG00000029627 | 1.205378646 | 1.25 | 1.238549078 | 1.339388001 | 1.149965125 | 1.041392685 | 1.150210039 |
| ENSMUSG00000029629 | 2.488076442 | 2.307805295 | 2.269871196 | 2.149047439 | 2.126866297 | 2.36361198 | 1.969900112 |
| ENSMUSG00000029630 | 0 | 0 | 0 | 0 | 0 | 0 | 0 |
| ENSMUSG00000029632 | 2.968105231 | 2.714689525 | 2.703645616 | 2.587458523 | 2.506580024 | 2.890421019 | 2.349572583 |
| ENSMUSG00000029633 | 0 | 0 | 0 | 0.396240625 | 0 | 0 | 0 |
| ENSMUSG00000029634 | 2.673935906 | 2.485994979 | 2.521202097 | 2.464495249 | 2.441646558 | 2.782472624 | 2.282223478 |
| ENSMUSG00000029635 | 2.660717759 | 2.433677405 | 2.437886015 | 2.333847589 | 2.2705456 | 2.591064607 | 2.152303998 |
| ENSMUSG00000029636 | 1.307602651 | 1.372963274 | 1.29248125 | 2.102347734 | 2.047893212 | 2.305351369 | 1.943148064 |
| ENSMUSG00000029638 | 0.45815691 | 0.580482024 | 0.25 | 0.79248125 | 0.387761935 | 0.477121255 | 0.448487648 |
| ENSMUSG00000029640 | 2.509948321 | 2.377443751 | 2.333847589 | 2.31728167 | 2.260764609 | 2.567026366 | 2.102294694 |
| ENSMUSG00000029641 | 0.960252568 | 0.951838731 | 1.04248125 | 0.25 | 0.568060967 | 0.698970004 | 0.916588299 |
| ENSMUSG00000029642 | 2.425923778 | 2.298073204 | 2.349151195 | 2.353406982 | 2.304082723 | 2.648360011 | 2.199238773 |
| ENSMUSG00000029644 | 0 | 0 | 0 | 0 | 0 | 0 | 0 |
| ENSMUSG00000029646 | 0 | 0 | 0 | 0 | 0.244650542 | 0 | 0 |
| ENSMUSG00000029647 | 2.577267124 | 2.31611065 | 2.310793496 | 2.28360658 | 2.261929482 | 2.471291711 | 2.063519437 |
| ENSMUSG00000029648 | 1.269664473 | 1.396240625 | 1.348079356 | 1.763820609 | 1.677808768 | 1.838849091 | 1.517723536 |
| ENSMUSG00000029649 | 2.689395766 | 2.5 | 2.468745337 | 2.628185116 | 2.555884575 | 2.970346876 | 2.378420866 |

| | | | | | | | |
|---|---|---|---|---|---|---|---|
| ENSMUSG00000029650 | 1 | 0.896240625 | 0.896240625 | 1.146240625 | 1.020174412 | 1.176091259 | 1.028149525 |
| ENSMUSG00000029651 | 0.578129653 | 0.5 | 0.25 | 0 | 0 | 0.477121255 | 0 |
| ENSMUSG00000029655 | 2.513987407 | 2.360735874 | 2.367918804 | 2.484894804 | 2.417419887 | 2.720985744 | 2.206733675 |
| ENSMUSG00000029656 | 0 | 0 | 0 | 0 | 0 | 0 | 0 |
| ENSMUSG00000029657 | 3.055099022 | 2.785207443 | 2.727660683 | 2.979622648 | 2.912106884 | 3.355259906 | 2.692737424 |
| ENSMUSG00000029658 | 0 | 0.25 | 0 | 0 | 0 | 0 | 0 |
| ENSMUSG00000029659 | 1.907802659 | 1.75 | 1.71759118 | 1.061981878 | 1.039257769 | 1 | 0.876096088 |
| ENSMUSG00000029660 | 0.867194479 | 0.92510993 | 0.79248125 | 0.79248125 | 0.686820904 | 1 | 0.829800754 |
| ENSMUSG00000029661 | 2.444137694 | 2.349685923 | 2.268535366 | 2.049918086 | 2.094757251 | 2.238046103 | 2.090384805 |
| ENSMUSG00000029663 | 0 | 0.25 | 0 | 0 | 0 | 0 | 0 |
| ENSMUSG00000029664 | 0 | 0 | 0 | 0.25 | 0.244650542 | 0 | 0.355418052 |
| ENSMUSG00000029669 | 2.10861828 | 1.935366747 | 1.987091808 | 1.584962501 | 1.427019704 | 1.672097858 | 1.437266334 |
| ENSMUSG00000029670 | 2.095120908 | 1.973704441 | 1.928561379 | 2.029735268 | 1.964031554 | 2.178976947 | 1.864112664 |
| ENSMUSG00000029671 | 0.811507563 | 0.75 | 0.701838731 | 0 | 0.244650542 | 0 | 0 |
| ENSMUSG00000029672 | 2.360187403 | 2.160964047 | 2.157339155 | 2.067865757 | 1.973373093 | 2.243038049 | 1.861046144 |
| ENSMUSG00000029673 | 0.289064826 | 0.25 | 0.25 | 0.976722649 | 1.039257769 | 1.041392685 | 1.100339912 |
| ENSMUSG00000029674 | 2.051791411 | 1.916333979 | 2.039967834 | 2.043731421 | 2.117361396 | 2.301029996 | 2.085803078 |
| ENSMUSG00000029675 | 0.289064826 | 0.25 | 0 | 0.396240625 | 0.244650542 | 0 | 0.520678036 |
| ENSMUSG00000029676 | 2.229917041 | 2.073655187 | 2.02053726 | 2.029735268 | 2.013184954 | 2.281033367 | 1.830929954 |
| ENSMUSG00000029678 | 0 | 0 | 0 | 0 | 0 | 0 | 0 |
| ENSMUSG00000029679 | 0 | 0 | 0 | 0 | 0 | 0 | 0 |
| ENSMUSG00000029680 | 0 | 0 | 0 | 0 | 0 | 0 | 0 |
| ENSMUSG00000029681 | 1.989530018 | 1.955044741 | 1.997171172 | 1.685366747 | 1.836786029 | 2.017033339 | 1.815933683 |
| ENSMUSG00000029682 | 0 | 0 | 0 | 0 | 0 | 0 | 0 |
| ENSMUSG00000029683 | 0 | 0 | 0 | 0 | 0 | 0 | 0 |
| ENSMUSG00000029684 | 2.621464818 | 2.439972046 | 2.416333979 | 2.42813175 | 2.343123155 | 2.682145076 | 2.176040447 |
| ENSMUSG00000029685 | 0.45815691 | 0.864857905 | 0.396240625 | 0.830482024 | 0.733951626 | 0.477121255 | 0.629532004 |
| ENSMUSG00000029686 | 2.942293987 | 2.68155454 | 2.65136988 | 2.653907409 | 2.591678881 | 2.954242509 | 2.388834696 |
| ENSMUSG00000029687 | 2.817374469 | 2.576515422 | 2.547147211 | 2.560793496 | 2.451298303 | 2.788875116 | 2.285835109 |
| ENSMUSG00000029695 | 0.289064826 | 0.396240625 | 0.396240625 | 0.830482024 | 0.568060967 | 0.477121255 | 0.629532004 |
| ENSMUSG00000029697 | 0 | 0 | 0 | 0 | 0 | 0 | 0 |
| ENSMUSG00000029699 | 0.960252568 | 1.098079356 | 0.79248125 | 1.02186571 | 1.039257769 | 1.079181246 | 0.969165684 |
| ENSMUSG00000029700 | 0.289064826 | 0.79248125 | 0.580482024 | 0.396240625 | 0.244650542 | 0.301029996 | 0.520678036 |
| ENSMUSG00000029701 | 2.767404661 | 2.509043403 | 2.535207443 | 2.577084758 | 2.540114597 | 2.893761762 | 2.379456779 |

| | | | | | | | |
|---|---|---|---|---|---|---|---|
| ENSMUSG00000029703 | 2.235832493 | 2.137686696 | 2.212155735 | 2.088286706 | 2.207327177 | 2.385606274 | 2.187847501 |
| ENSMUSG00000029705 | 2.900707419 | 2.677701608 | 2.666556001 | 2.519537702 | 2.488689905 | 2.80685803 | 2.342925556 |
| ENSMUSG00000029706 | 0 | 0 | 0 | 0 | 0 | 0 | 0 |
| ENSMUSG00000029707 | 0.289064826 | 0 | 0.25 | 0.25 | 0.244650542 | 0.301029996 | 0 |
| ENSMUSG00000029708 | 1.876566388 | 1.854463129 | 1.901832578 | 1.714495249 | 1.770879644 | 1.973127854 | 1.687112305 |
| ENSMUSG00000029710 | 2.380049842 | 2.266522298 | 2.356566189 | 2.181980114 | 2.302531265 | 2.514547753 | 2.262645396 |
| ENSMUSG00000029711 | 0 | 0 | 0 | 0 | 0 | 0 | 0 |
| ENSMUSG00000029712 | 0 | 0 | 0 | 0 | 0.244650542 | 0 | 0 |
| ENSMUSG00000029713 | 3.067806989 | 2.888167274 | 2.963034832 | 2.528435542 | 2.773755345 | 2.958563883 | 2.717961151 |
| ENSMUSG00000029714 | 2.231898196 | 2.072504712 | 2.20584181 | 1.747171172 | 1.851969278 | 1.939519253 | 1.937521567 |
| ENSMUSG00000029715 | 1.811507563 | 1.841580554 | 1.860735874 | 1.451838731 | 1.662257525 | 1.819543936 | 1.71890424 |
| ENSMUSG00000029716 | 0.289064826 | 0.25 | 0.396240625 | 0 | 0.387761935 | 0.477121255 | 0.448487648 |
| ENSMUSG00000029718 | 3.320294638 | 3.055547785 | 3.114601653 | 2.7543761 | 2.909238434 | 3.20167018 | 2.792972597 |
| ENSMUSG00000029720 | 1 | 0.830482024 | 0.896240625 | 0.896240625 | 0.905314583 | 0.77815125 | 0.829800754 |
| ENSMUSG00000029722 | 2.330063669 | 2.164552871 | 2.243603647 | 2.061981878 | 2.123439512 | 2.318063335 | 2.060761403 |
| ENSMUSG00000029723 | 2.163307908 | 2.157339155 | 2.322504712 | 1.868933358 | 1.945998439 | 2 | 2.065163042 |
| ENSMUSG00000029725 | 0.811507563 | 0.864857905 | 1.160964047 | 0.646240625 | 0.955822902 | 0.698970004 | 1.193409508 |
| ENSMUSG00000029726 | 2.382811656 | 2.242885888 | 2.269203899 | 2.160964047 | 2.225193812 | 2.387389826 | 2.125328759 |
| ENSMUSG00000029727 | 0 | 0 | 0 | 0.25 | 0.387761935 | 0 | 0.579661877 |
| ENSMUSG00000029729 | 2.425923778 | 2.227473271 | 2.208222504 | 2.315523711 | 2.232853197 | 2.561101384 | 2.06186744 |
| ENSMUSG00000029730 | 3.173145515 | 2.891394534 | 2.877567418 | 2.8918702 | 2.873688359 | 3.298853076 | 2.68220009 |
| ENSMUSG00000029735 | 1.342375483 | 1.451838731 | 1.321350555 | 1.561981878 | 1.568060967 | 1.672097858 | 1.345462945 |
| ENSMUSG00000029736 | 0 | 0 | 0 | 0 | 0 | 0 | 0 |
| ENSMUSG00000029752 | 2.873182739 | 2.58076369 | 2.472185812 | 2.385757955 | 2.327055443 | 2.606381365 | 2.157274883 |
| ENSMUSG00000029754 | 0.867194479 | 0.976722649 | 0.75 | 0.79248125 | 0.77552387 | 0.477121255 | 0.710836105 |
| ENSMUSG00000029755 | 0.289064826 | 0.830482024 | 0.92510993 | 0.25 | 0.387761935 | 0 | 0.579661877 |
| ENSMUSG00000029757 | 0.960252568 | 1.201838731 | 1.04248125 | 1.188721876 | 1 | 1.041392685 | 1.066254157 |
| ENSMUSG00000029759 | 2.373064211 | 2.225970461 | 2.183677405 | 1.51109853 | 1.212046813 | 1.342422681 | 1.238624656 |
| ENSMUSG00000029761 | 3.871945483 | 3.497060219 | 3.467374364 | 3.196822607 | 3.109705685 | 3.60031933 | 2.897100601 |
| ENSMUSG00000029762 | 2.468370459 | 2.296218836 | 2.28743678 | 2.083847589 | 2.0510778 | 2.374748346 | 1.93179548 |
| ENSMUSG00000029763 | 2.625329835 | 2.42293588 | 2.415889526 | 2.410512923 | 2.314756076 | 2.693726949 | 2.1908258 |
| ENSMUSG00000029765 | 1.307602651 | 1.080482024 | 1.311981878 | 0.5 | 0.244650542 | 0 | 0.224243824 |
| ENSMUSG00000029766 | 0 | 0 | 0 | 0 | 0 | 0 | 0 |
| ENSMUSG00000029767 | 3.554771667 | 3.210980263 | 3.17255407 | 3.246015592 | 3.125207453 | 3.672374979 | 2.905956692 |

| | | | | | | | |
|---|---|---|---|---|---|---|---|
| ENSMUSG00000029769 | 2.151493169 | 2.006976499 | 2.080482024 | 1.918106335 | 1.931471446 | 2.133538908 | 1.861046144 |
| ENSMUSG00000029771 | 1.623015126 | 1.720660762 | 1.653677461 | 1.732684334 | 1.754126039 | 1.875061263 | 1.728327686 |
| ENSMUSG00000029772 | 2.266038081 | 2.081607372 | 2.048689214 | 2.242166698 | 2.160196454 | 2.428134794 | 1.982842456 |
| ENSMUSG00000029775 | 2.749513129 | 2.526477127 | 2.487091808 | 2.512802235 | 2.445123986 | 2.794488047 | 2.260561074 |
| ENSMUSG00000029776 | 2.219867581 | 2.039967834 | 2.048689214 | 2.130890489 | 2.075570129 | 2.378397901 | 1.932619724 |
| ENSMUSG00000029777 | 3.166262603 | 2.866008788 | 2.788229644 | 2.82163944 | 2.740455823 | 3.165244326 | 2.543140861 |
| ENSMUSG00000029778 | 2.172524468 | 2.043731421 | 2.029735268 | 0.396240625 | 0.489301084 | 0 | 0.224243824 |
| ENSMUSG00000029780 | 2.121692466 | 1.916333979 | 1.942046081 | 2.070192693 | 1.981187939 | 2.217483944 | 1.82753046 |
| ENSMUSG00000029781 | 3.175209014 | 2.860476676 | 2.859438018 | 2.77053726 | 2.701682612 | 3.133538908 | 2.545501844 |
| ENSMUSG00000029782 | 2.585813169 | 2.398114259 | 2.37892496 | 2.471805154 | 2.374910674 | 2.741939078 | 2.217489603 |
| ENSMUSG00000029784 | 0 | 0.396240625 | 0.25 | 0.396240625 | 0 | 0.477121255 | 0.355418052 |
| ENSMUSG00000029787 | 2.42468076 | 2.263820609 | 2.222945926 | 2.241446071 | 2.199780694 | 2.445604203 | 2.003324957 |
| ENSMUSG00000029788 | 0 | 0 | 0 | 0 | 0 | 0 | 0 |
| ENSMUSG00000029790 | 1.881174527 | 1.782320754 | 1.76109853 | 1.79248125 | 1.749189509 | 1.86923172 | 1.655750087 |
| ENSMUSG00000029797 | 0.289064826 | 0 | 0 | 0 | 0 | 0 | 0.224243824 |
| ENSMUSG00000029798 | 1.800532393 | 1.678561379 | 1.653677461 | 1.880890489 | 1.866526233 | 2.041392685 | 1.676387053 |
| ENSMUSG00000029802 | 2.276824003 | 2.154596376 | 2.084962501 | 2.342671852 | 2.2705456 | 2.57863921 | 2.088864761 |
| ENSMUSG00000029804 | 2.003432385 | 1.886723615 | 1.892463902 | 2.110735874 | 2.021325983 | 2.338456494 | 1.938331368 |
| ENSMUSG00000029810 | 2.6088504 | 2.418548067 | 2.395770692 | 2.494319981 | 2.47528687 | 2.855519156 | 2.334479423 |
| ENSMUSG00000029811 | 0 | 0 | 0 | 0 | 0 | 0 | 0 |
| ENSMUSG00000029813 | 0 | 0 | 0 | 0 | 0 | 0 | 0 |
| ENSMUSG00000029814 | 2.410757293 | 2.192872367 | 2.202642909 | 2.009729747 | 1.94314048 | 2.184691431 | 1.811176017 |
| ENSMUSG00000029815 | 1.568542279 | 1.464495249 | 1.476722649 | 1.606566189 | 1.427019704 | 1.698970004 | 1.455757965 |
| ENSMUSG00000029816 | 0.289064826 | 0.396240625 | 0.580482024 | 0 | 0.489301084 | 0.301029996 | 0.579661877 |
| ENSMUSG00000029817 | 2.891461985 | 2.644828984 | 2.645535495 | 2.716046536 | 2.628935049 | 3.050379756 | 2.465732876 |
| ENSMUSG00000029819 | 0 | 0 | 0 | 0 | 0.244650542 | 0.301029996 | 0 |
| ENSMUSG00000029821 | 1.907802659 | 1.75835575 | 1.821350555 | 1.638647213 | 1.704182902 | 1.770852012 | 1.505628092 |
| ENSMUSG00000029822 | 2.451206209 | 2.310197833 | 2.304186465 | 2.32991803 | 2.322676882 | 2.546542663 | 2.175652306 |
| ENSMUSG00000029823 | 2.980763736 | 2.707430684 | 2.694108258 | 2.746282593 | 2.66653358 | 3.050379756 | 2.477554948 |
| ENSMUSG00000029826 | 2.590857977 | 2.356041572 | 2.372963274 | 2.573367912 | 2.52691592 | 2.896526217 | 2.344312053 |
| ENSMUSG00000029828 | 0.45815691 | 0.25 | 0.25 | 0.701838731 | 0.568060967 | 0.301029996 | 0.629532004 |
| ENSMUSG00000029829 | 0 | 0 | 0 | 0 | 0 | 0 | 0 |
| ENSMUSG00000029830 | 0 | 0.25 | 0 | 0.25 | 0 | 0 | 0.224243824 |
| ENSMUSG00000029831 | 0 | 0 | 0 | 0 | 0 | 0 | 0 |

| | | | | | | | |
|---|---|---|---|---|---|---|---|
| ENSMUSG00000029832 | 1.418409478 | 1.410964047 | 1.403677461 | 1.872963274 | 1.768534412 | 2.045322979 | 1.680001789 |
| ENSMUSG00000029833 | 2.317377295 | 2.174241882 | 2.088286706 | 2.101285366 | 2.068600388 | 2.305351369 | 1.920884716 |
| ENSMUSG00000029836 | 3.079103675 | 2.810048763 | 2.772528105 | 2.764497931 | 2.700173892 | 3.096214585 | 2.466997845 |
| ENSMUSG00000029837 | 0 | 0 | 0 | 0 | 0 | 0 | 0 |
| ENSMUSG00000029838 | 2.909752324 | 2.69390257 | 2.660287149 | 1.698603967 | 1.683842353 | 1.875061263 | 1.535058868 |
| ENSMUSG00000029840 | 3.194515793 | 2.91997997 | 2.912987153 | 2.940387808 | 2.832539716 | 3.291590826 | 2.610253039 |
| ENSMUSG00000029843 | 0 | 0.396240625 | 0.25 | 0.25 | 0 | 0 | 0 |
| ENSMUSG00000029844 | 1.100572389 | 1.25 | 1.282320754 | 1.114857905 | 1.091002363 | 1.176091259 | 1.066254157 |
| ENSMUSG00000029847 | 0 | 0 | 0 | 0 | 0.244650542 | 0 | 0.355418052 |
| ENSMUSG00000029848 | 0.289064826 | 0 | 0 | 0 | 0.244650542 | 0 | 0 |
| ENSMUSG00000029851 | 1.816888701 | 1.766522298 | 1.688721876 | 1.988549078 | 1.897237431 | 2.155336037 | 1.832055226 |
| ENSMUSG00000029859 | 0.91631382 | 0.951838731 | 0.951838731 | 1.226722649 | 1.373641807 | 1.380211242 | 1.20919388 |
| ENSMUSG00000029860 | 3.052896866 | 2.883940344 | 2.90654133 | 2.514497931 | 2.750794189 | 2.953759692 | 2.646550823 |
| ENSMUSG00000029861 | 2.016886209 | 1.837182039 | 1.856566189 | 1.841580554 | 1.818885826 | 1.949390007 | 1.723650273 |
| ENSMUSG00000029862 | 0.91631382 | 0.864857905 | 0.864857905 | 0.75 | 0.568060967 | 0.301029996 | 0 |
| ENSMUSG00000029863 | 2.440557995 | 2.242885888 | 2.205044741 | 2.175975893 | 2.160196454 | 2.480006943 | 2.020084484 |
| ENSMUSG00000029864 | 0 | 0 | 0 | 0 | 0.632412477 | 0 | 0.224243824 |
| ENSMUSG00000029865 | 0 | 0 | 0 | 0 | 0 | 0 | 0 |
| ENSMUSG00000029866 | 0 | 0 | 0 | 0 | 0 | 0 | 0 |
| ENSMUSG00000029867 | 0 | 0 | 0 | 0 | 0 | 0 | 0 |
| ENSMUSG00000029868 | 0.45815691 | 0.25 | 0.25 | 0.396240625 | 0 | 0.477121255 | 0 |
| ENSMUSG00000029869 | 1.759671389 | 1.646240625 | 1.634789703 | 1.061981878 | 1.302012594 | 1.113943352 | 1.324583736 |
| ENSMUSG00000029875 | 0.747221736 | 1 | 0.79248125 | 0.25 | 0.568060967 | 0 | 0.448487648 |
| ENSMUSG00000029878 | 0.289064826 | 0 | 0.25 | 0 | 0.387761935 | 0.477121255 | 0.224243824 |
| ENSMUSG00000029882 | 0 | 0 | 0 | 0 | 0 | 0 | 0 |
| ENSMUSG00000029883 | 0 | 0 | 0 | 0 | 0 | 0 | 0 |
| ENSMUSG00000029885 | 0 | 0 | 0 | 0 | 0 | 0 | 0 |
| ENSMUSG00000029909 | 0 | 0 | 0 | 0 | 0 | 0 | 0 |
| ENSMUSG00000029910 | 2.782453855 | 2.559601185 | 2.545288064 | 2.639605178 | 2.538524702 | 2.931966115 | 2.380901508 |
| ENSMUSG00000029911 | 2.227926429 | 2.072504712 | 2.072504712 | 2.075945187 | 1.969399812 | 2.212187604 | 1.832055226 |
| ENSMUSG00000029913 | 1.978790393 | 1.892463902 | 1.823655187 | 1.76109853 | 1.777823267 | 1.908485019 | 1.637827726 |
| ENSMUSG00000029915 | 0 | 0 | 0 | 0 | 0 | 0 | 0 |
| ENSMUSG00000029916 | 1.907802659 | 1.830482024 | 1.830482024 | 1.860735874 | 1.797870327 | 2.021189299 | 1.737484395 |
| ENSMUSG00000029917 | 0 | 0 | 0 | 0 | 0 | 0 | 0 |

| | | | | | | | |
|---|---|---|---|---|---|---|---|
| ENSMUSG00000029918 | 2.233869984 | 2.00835575 | 2.02186571 | 2.075945187 | 1.988833497 | 2.28780173 | 1.889537832 |
| ENSMUSG00000029919 | 0.45815691 | 0.396240625 | 0.5 | 0.5 | 0.812711509 | 0.477121255 | 0.355418052 |
| ENSMUSG00000029920 | 2.641182749 | 2.422499493 | 2.415444524 | 2.369439567 | 2.29040462 | 2.617000341 | 2.127589533 |
| ENSMUSG00000029922 | 2.057835465 | 1.888647213 | 1.938721876 | 2.060793496 | 2.044679629 | 2.283301229 | 1.904245613 |
| ENSMUSG00000029923 | 0 | 0 | 0 | 0 | 0 | 0 | 0.355418052 |
| ENSMUSG00000029924 | 2.517987749 | 2.328787391 | 2.291226732 | 2.383818844 | 2.298360391 | 2.619093331 | 2.166993225 |
| ENSMUSG00000029925 | 0.671187741 | 0 | 0.580482024 | 0.396240625 | 0.387761935 | 0.301029996 | 0.448487648 |
| ENSMUSG00000029992 | 2.91746109 | 2.642226539 | 2.625459296 | 2.698806992 | 2.643236511 | 3.035029282 | 2.458847605 |
| ENSMUSG00000029993 | 2.089597219 | 2 | 1.984159485 | 1.972185812 | 1.91476933 | 2.123851641 | 1.874129423 |
| ENSMUSG00000029994 | 2.900707419 | 2.63503201 | 2.636241108 | 2.617664719 | 2.570979987 | 2.954724791 | 2.416364058 |
| ENSMUSG00000029998 | 2.68408318 | 2.484894804 | 2.472566069 | 2.59727338 | 2.542223377 | 2.932473765 | 2.365292129 |
| ENSMUSG00000029999 | 1.587501561 | 1.575945187 | 1.51109853 | 1.726722649 | 1.739104362 | 1.838849091 | 1.612273755 |
| ENSMUSG00000030000 | 0.867194479 | 0.92510993 | 0.951838731 | 1.505591953 | 1.607318736 | 1.653212514 | 1.514741806 |
| ENSMUSG00000030001 | 0 | 0 | 0 | 0 | 0 | 0 | 0 |
| ENSMUSG00000030002 | 2.857607105 | 2.631869252 | 2.5964656 | 2.710587586 | 2.605566002 | 3.030194785 | 2.44122113 |
| ENSMUSG00000030004 | 0 | 0 | 0 | 0 | 0 | 0 | 0 |
| ENSMUSG00000030007 | 3.298972521 | 3.006544399 | 3.001406137 | 3.088355636 | 3.024289189 | 3.506369717 | 2.804462239 |
| ENSMUSG00000030008 | 1.783508299 | 1.584962501 | 1.698603967 | 1.698603967 | 1.66856048 | 1.838849091 | 1.548827561 |
| ENSMUSG00000030016 | 2.893489371 | 2.658021603 | 2.657339155 | 2.707430684 | 2.607537217 | 3.014520539 | 2.402988297 |
| ENSMUSG00000030017 | 0 | 0 | 0 | 0 | 0 | 0 | 0 |
| ENSMUSG00000030019 | 1.881174527 | 1.83272922 | 1.937048212 | 1.777131114 | 1.741652793 | 2.029383778 | 1.81475081 |
| ENSMUSG00000030020 | 1.936861443 | 1.839388001 | 1.819031101 | 1.516522298 | 1.546663136 | 1.602059991 | 1.425641774 |
| ENSMUSG00000030022 | 1.358731791 | 1.476722649 | 1.311981878 | 1.226722649 | 1.121713561 | 1.342422681 | 1.252393349 |
| ENSMUSG00000030029 | 2.853175263 | 2.655970373 | 2.635999613 | 2.355516192 | 2.406008674 | 2.685741739 | 2.268527657 |
| ENSMUSG00000030030 | 0.91631382 | 0.646240625 | 0.830482024 | 0.92510993 | 1.091002363 | 0.77815125 | 1.041356071 |
| ENSMUSG00000030031 | 1.678702042 | 1.532320754 | 1.464495249 | 1.62694866 | 1.632412477 | 1.662757832 | 1.452143229 |
| ENSMUSG00000030032 | 1.614416215 | 1.54248125 | 1.552363341 | 1.348079356 | 1.427019704 | 1.491361694 | 1.429563222 |
| ENSMUSG00000030034 | 1.205378646 | 1.396240625 | 1.431980114 | 1.26109853 | 1.343584837 | 1.361727836 | 1.392431569 |
| ENSMUSG00000030035 | 2.181541736 | 2.101285366 | 2.099151195 | 1.819031101 | 1.970729215 | 2.133538908 | 1.910475814 |
| ENSMUSG00000030036 | 2.283862692 | 2.108657057 | 2.162762923 | 1.970660762 | 2.094757251 | 2.245512668 | 2.056860222 |
| ENSMUSG00000030037 | 0 | 0 | 0 | 0 | 0 | 0 | 0.355418052 |
| ENSMUSG00000030041 | 0.811507563 | 0.830482024 | 0.92510993 | 0.646240625 | 0 | 0.301029996 | 0.579661877 |
| ENSMUSG00000030042 | 2.40033094 | 2.216819685 | 2.193696765 | 2.301142786 | 2.234113756 | 2.564666064 | 2.097879707 |
| ENSMUSG00000030043 | 0.289064826 | 0 | 0 | 0.25 | 0 | 0 | 0.448487648 |

| | | | | | | | |
|---|---|---|---|---|---|---|---|
| ENSMUSG00000030045 | 2.163307908 | 2.048689214 | 2.029735268 | 2.133818844 | 2.046825265 | 2.354108439 | 1.977154126 |
| ENSMUSG00000030046 | 0 | 0 | 0 | 0 | 0 | 0 | 0 |
| ENSMUSG00000030047 | 0 | 0 | 0 | 0.25 | 0 | 0 | 0.224243824 |
| ENSMUSG00000030048 | 0 | 0 | 0 | 0 | 0 | 0 | 0 |
| ENSMUSG00000030049 | 0 | 0 | 0 | 0 | 0 | 0 | 0 |
| ENSMUSG00000030050 | 0 | 0 | 0 | 0 | 0 | 0 | 0.224243824 |
| ENSMUSG00000030051 | 1.827447047 | 1.711372513 | 1.692046081 | 1.828220739 | 1.741652793 | 1.959041392 | 1.692344475 |
| ENSMUSG00000030054 | 0 | 0 | 0 | 0 | 0 | 0 | 0 |
| ENSMUSG00000030055 | 0.289064826 | 0.25 | 0.5 | 0.25 | 0.244650542 | 0 | 0.629532004 |
| ENSMUSG00000030056 | 2.097855556 | 2.009729747 | 2.066696635 | 2.172499493 | 2.132784263 | 2.371067862 | 2.013740841 |
| ENSMUSG00000030057 | 3.251208701 | 2.961176441 | 2.963522295 | 2.990543258 | 2.930860266 | 3.400365273 | 2.722506097 |
| ENSMUSG00000030058 | 3.139782351 | 2.859177886 | 2.865369862 | 2.771202097 | 2.807668315 | 3.179838928 | 2.617370165 |
| ENSMUSG00000030059 | 2.564110437 | 2.369439567 | 2.369945066 | 2.362287161 | 2.302531265 | 2.605305046 | 2.123508695 |
| ENSMUSG00000030060 | 2.192545868 | 2.017865591 | 1.961372513 | 2.163659007 | 2.117361396 | 2.357934847 | 1.993245779 |
| ENSMUSG00000030061 | 2.761895567 | 2.57163944 | 2.52219706 | 2.645770692 | 2.547181807 | 2.940516485 | 2.359623923 |
| ENSMUSG00000030062 | 3.0249321 | 2.780221461 | 2.756803721 | 2.829776894 | 2.783042429 | 3.23299611 | 2.635492539 |
| ENSMUSG00000030064 | 0.578129653 | 0.396240625 | 0.396240625 | 1.238549078 | 1.176121988 | 1.301029996 | 1.041356071 |
| ENSMUSG00000030067 | 2.649978134 | 2.453845824 | 2.439139081 | 2.333288838 | 2.297309926 | 2.609594409 | 2.187847501 |
| ENSMUSG00000030068 | 0 | 0.25 | 0.396240625 | 0 | 0 | 0 | 0 |
| ENSMUSG00000030069 | 0 | 0 | 0 | 0 | 0 | 0 | 0 |
| ENSMUSG00000030074 | 2.397683106 | 2.199415381 | 2.254897682 | 2.634547233 | 2.552071544 | 2.950364854 | 2.365724059 |
| ENSMUSG00000030075 | 0 | 0.396240625 | 0.25 | 0.25 | 0.489301084 | 0 | 0.224243824 |
| ENSMUSG00000030077 | 0 | 0 | 0 | 0 | 0 | 0 | 0 |
| ENSMUSG00000030079 | 2.479432974 | 2.290597832 | 2.320192693 | 2.359177886 | 2.30971376 | 2.565847819 | 2.186723532 |
| ENSMUSG00000030082 | 2.99432171 | 2.77908599 | 2.760414788 | 2.684311836 | 2.716746105 | 3.04060234 | 2.574269828 |
| ENSMUSG00000030083 | 1.418409478 | 1.330482024 | 1.445339928 | 1.282320754 | 1.394615667 | 1.255272505 | 1.379042812 |
| ENSMUSG00000030084 | 3.018088033 | 2.85839636 | 2.88503201 | 2.480088214 | 2.612309899 | 2.828015064 | 2.527294124 |
| ENSMUSG00000030086 | 1.789260561 | 1.671625132 | 1.726722649 | 1.566696635 | 1.680838453 | 1.681241237 | 1.61448206 |
| ENSMUSG00000030087 | 0 | 0 | 0.396240625 | 0 | 0.244650542 | 0.301029996 | 0.448487648 |
| ENSMUSG00000030088 | 0.578129653 | 0.396240625 | 0.79248125 | 0.25 | 0 | 0 | 0.224243824 |
| ENSMUSG00000030089 | 1.482695304 | 1.476722649 | 1.494319981 | 1.238549078 | 1.293076518 | 1.380211242 | 1.473249546 |
| ENSMUSG00000030091 | 1.181541736 | 1.25 | 1.348079356 | 1.080482024 | 1.223252711 | 1.041392685 | 1.168187745 |
| ENSMUSG00000030092 | 0 | 0 | 0 | 0 | 0 | 0 | 0 |
| ENSMUSG00000030093 | 0.45815691 | 0 | 0 | 0.25 | 0.244650542 | 0.301029996 | 0.520678036 |

| | | | | | | | |
|---|---|---|---|---|---|---|---|
| ENSMUSG00000030094 | 1.967766868 | 1.868933358 | 1.852347734 | 1.752806814 | 1.816839698 | 1.982271233 | 1.73898563 |
| ENSMUSG00000030095 | 2.654307229 | 2.453445298 | 2.444519282 | 2.384789703 | 2.400946889 | 2.719331287 | 2.226242896 |
| ENSMUSG00000030096 | 3.289378031 | 3.008699741 | 2.986726569 | 2.776477127 | 2.769225805 | 3.180412633 | 2.579884607 |
| ENSMUSG00000030098 | 0 | 0 | 0 | 0.396240625 | 0.387761935 | 0 | 0.224243824 |
| ENSMUSG00000030101 | 2.343889214 | 2.231944491 | 2.234894804 | 2.218361028 | 2.123439512 | 2.46834733 | 2.018194418 |
| ENSMUSG00000030102 | 2.599951214 | 2.391037372 | 2.388647213 | 2.313166358 | 2.255474576 | 2.562292864 | 2.135160095 |
| ENSMUSG00000030103 | 2.271465912 | 2.140560606 | 2.14151351 | 1.896240625 | 1.808534465 | 2.049218023 | 1.786277931 |
| ENSMUSG00000030104 | 2.579849376 | 2.373463862 | 2.343216265 | 2.458617762 | 2.390600196 | 2.740362689 | 2.254530788 |
| ENSMUSG00000030105 | 2.903085937 | 2.633818844 | 2.615881093 | 2.572216519 | 2.48684999 | 2.860338007 | 2.315385981 |
| ENSMUSG00000030107 | 0.45815691 | 0.5 | 0.5 | 0.864857905 | 0.846351821 | 1.079181246 | 0.969165684 |
| ENSMUSG00000030108 | 0 | 0 | 0 | 0 | 0.387761935 | 0 | 0 |
| ENSMUSG00000030109 | 0.289064826 | 0 | 0 | 0.25 | 0.244650542 | 0 | 0.224243824 |
| ENSMUSG00000030110 | 1.482695304 | 1.372963274 | 1.282320754 | 1.04248125 | 1.074582839 | 0.84509804 | 0.896975297 |
| ENSMUSG00000030111 | 0 | 0 | 0 | 0 | 0 | 0 | 0 |
| ENSMUSG00000030114 | 0 | 0 | 0 | 0 | 0 | 0 | 0 |
| ENSMUSG00000030116 | 0 | 0 | 0 | 0 | 0 | 0 | 0 |
| ENSMUSG00000030117 | 0.289064826 | 0.25 | 0 | 0 | 0 | 0 | 0 |
| ENSMUSG00000030120 | 2.825070787 | 2.632845365 | 2.687257846 | 2.453044326 | 2.56462465 | 2.808885867 | 2.503097319 |
| ENSMUSG00000030122 | 2.903481042 | 2.745748394 | 2.834684096 | 2.350753006 | 2.429560273 | 2.693726949 | 2.447148038 |
| ENSMUSG00000030123 | 2.499676068 | 2.347004321 | 2.458222504 | 1.802363341 | 1.929985312 | 2.053078443 | 1.953344547 |
| ENSMUSG00000030124 | 0.747221736 | 0.92510993 | 0.896240625 | 0.396240625 | 0.632412477 | 0.301029996 | 0.74492186 |
| ENSMUSG00000030125 | 0.45815691 | 0 | 0.25 | 0.25 | 0.244650542 | 0.301029996 | 0 |
| ENSMUSG00000030126 | 2.271465912 | 2.064346961 | 2.090485943 | 2.140560606 | 2.09661982 | 2.354108439 | 1.933441873 |
| ENSMUSG00000030127 | 2.553277818 | 2.417664062 | 2.442872367 | 2.228969845 | 2.331857641 | 2.603144373 | 2.291042162 |
| ENSMUSG00000030131 | 0 | 0 | 0 | 0 | 0 | 0 | 0 |
| ENSMUSG00000030134 | 1.249317394 | 1.29248125 | 1.348079356 | 0.79248125 | 0.846351821 | 0.903089987 | 0.853775828 |
| ENSMUSG00000030137 | 0.45815691 | 0.396240625 | 0.396240625 | 0.580482024 | 0.387761935 | 0.301029996 | 0.710836105 |
| ENSMUSG00000030138 | 2.732012699 | 2.519537702 | 2.539020769 | 2.541226732 | 2.513204446 | 2.833147112 | 2.326538518 |
| ENSMUSG00000030142 | 0 | 0 | 0 | 0 | 0 | 0 | 0 |
| ENSMUSG00000030143 | 0 | 0 | 0 | 0 | 0 | 0 | 0 |
| ENSMUSG00000030144 | 0 | 0 | 0 | 0 | 0 | 0 | 0 |
| ENSMUSG00000030145 | 1.83262764 | 1.726722649 | 1.735628626 | 1.735628626 | 1.645989387 | 1.819543936 | 1.554175045 |
| ENSMUSG00000030147 | 0 | 0 | 0 | 0 | 0 | 0 | 0 |
| ENSMUSG00000030148 | 0 | 0 | 0.5 | 0.5 | 0.489301084 | 0.301029996 | 0.224243824 |

| | | | | | | | |
|---|---|---|---|---|---|---|---|
| ENSMUSG00000030149 | 0 | 0 | 0 | 0 | 0 | 0 | 0 |
| ENSMUSG00000030154 | 0 | 0 | 0 | 0 | 0.244650542 | 0.301029996 | 0.224243824 |
| ENSMUSG00000030155 | 0 | 0 | 0 | 0 | 0 | 0 | 0.520678036 |
| ENSMUSG00000030156 | 0.289064826 | 0 | 0.396240625 | 0.25 | 0 | 0.477121255 | 0 |
| ENSMUSG00000030157 | 1.307602651 | 1.130890489 | 0.976722649 | 0.25 | 0.387761935 | 0.477121255 | 0.224243824 |
| ENSMUSG00000030158 | 0 | 0 | 0 | 0 | 0 | 0 | 0 |
| ENSMUSG00000030159 | 0 | 0 | 0 | 0 | 0 | 0 | 0 |
| ENSMUSG00000030160 | 0 | 0 | 0 | 0 | 0 | 0 | 0 |
| ENSMUSG00000030161 | 2.404271433 | 2.209012589 | 2.216819685 | 2.03743678 | 1.977302144 | 2.212187604 | 1.902443294 |
| ENSMUSG00000030162 | 0 | 0 | 0 | 0 | 0.244650542 | 0 | 0 |
| ENSMUSG00000030165 | 0 | 0 | 0 | 0 | 0 | 0 | 0 |
| ENSMUSG00000030166 | 1.822201287 | 1.695339928 | 1.705044741 | 1.54248125 | 1.584291694 | 1.643452676 | 1.466365979 |
| ENSMUSG00000030167 | 0 | 0 | 0 | 0 | 0 | 0 | 0 |
| ENSMUSG00000030168 | 2.325351389 | 2.139605178 | 2.143411797 | 2.281676118 | 2.279510047 | 2.62838893 | 2.111895264 |
| ENSMUSG00000030170 | 0.289064826 | 0 | 0.580482024 | 1.380890489 | 1.394615667 | 1.397940009 | 1.425641774 |
| ENSMUSG00000030172 | 2.502784398 | 2.289336734 | 2.27053726 | 2.387686696 | 2.327055443 | 2.654176542 | 2.174485081 |
| ENSMUSG00000030173 | 0 | 0 | 0 | 0 | 0 | 0 | 0 |
| ENSMUSG00000030177 | 2.010213549 | 1.839388001 | 1.868933358 | 1.921625132 | 1.890639929 | 2.146128036 | 1.735976162 |
| ENSMUSG00000030178 | 0 | 0 | 0 | 0 | 0 | 0 | 0 |
| ENSMUSG00000030180 | 2.778742163 | 2.519871196 | 2.481573999 | 2.583568321 | 2.515196142 | 2.878521796 | 2.320390187 |
| ENSMUSG00000030187 | 0.289064826 | 0.25 | 0.580482024 | 0.5 | 0.387761935 | 0.301029996 | 0.448487648 |
| ENSMUSG00000030188 | 2.146671932 | 1.961372513 | 1.937048212 | 1.991446071 | 1.888971102 | 2.1430148 | 1.732938436 |
| ENSMUSG00000030189 | 2.932463873 | 2.760927716 | 2.771036003 | 2.524836953 | 2.605346297 | 2.868056362 | 2.508278158 |
| ENSMUSG00000030194 | 0 | 0 | 0 | 0 | 0 | 0 | 0 |
| ENSMUSG00000030196 | 0 | 0 | 0 | 0 | 0 | 0 | 0 |
| ENSMUSG00000030199 | 2.455852858 | 2.341033664 | 2.29745614 | 2.240724001 | 2.206647761 | 2.474216264 | 2.080634989 |
| ENSMUSG00000030200 | 0 | 0 | 0 | 0 | 0 | 0 | 0 |
| ENSMUSG00000030201 | 2.878247972 | 2.639844273 | 2.582168732 | 2.612802778 | 2.526091577 | 2.894869657 | 2.346838633 |
| ENSMUSG00000030203 | 2.20747957 | 2.064346961 | 2.03360658 | 2.212155735 | 2.134457044 | 2.436162647 | 2 |
| ENSMUSG00000030204 | 2.533616084 | 2.348079356 | 2.333288838 | 2.385274154 | 2.382016011 | 2.687528961 | 2.213368331 |
| ENSMUSG00000030205 | 0 | 0 | 0 | 0.25 | 0 | 0 | 0 |
| ENSMUSG00000030206 | 0.45815691 | 0.25 | 0 | 0 | 0 | 0 | 0.355418052 |
| ENSMUSG00000030207 | 1.978790393 | 1.854463129 | 1.839388001 | 1.961372513 | 1.925488995 | 2.1430148 | 1.820622395 |
| ENSMUSG00000030208 | 3.391969412 | 3.079847469 | 3.053124096 | 3.139784514 | 3.031524462 | 3.560026249 | 2.809537961 |

| | | | | | | | |
|---|---|---|---|---|---|---|---|
| ENSMUSG00000030209 | 0 | 0 | 0 | 0.25 | 0.244650542 | 0 | 0 |
| ENSMUSG00000030213 | 2.457006475 | 2.282320754 | 2.307805295 | 2.231944491 | 2.173890435 | 2.451786436 | 2.040177507 |
| ENSMUSG00000030214 | 0 | 0 | 0 | 0.25 | 0.244650542 | 0.477121255 | 0.224243824 |
| ENSMUSG00000030216 | 2.610448227 | 2.445749552 | 2.428990498 | 2.400906586 | 2.400946889 | 2.706717782 | 2.296166733 |
| ENSMUSG00000030217 | 0 | 0 | 0 | 0 | 0 | 0 | 0 |
| ENSMUSG00000030218 | 1.867194479 | 1.698603967 | 1.779735268 | 0 | 0.489301084 | 0.602059991 | 0.448487648 |
| ENSMUSG00000030219 | 0 | 0.25 | 0 | 0 | 0 | 0 | 0 |
| ENSMUSG00000030220 | 0.960252568 | 1 | 0.864857905 | 1.470660762 | 1.327538614 | 1.462397998 | 1.278288402 |
| ENSMUSG00000030222 | 1.100572389 | 1.061981878 | 1.160964047 | 0.701838731 | 0.77552387 | 0.602059991 | 0.775756176 |
| ENSMUSG00000030223 | 0 | 0 | 0 | 0 | 0 | 0 | 0 |
| ENSMUSG00000030224 | 3.076764441 | 2.806000419 | 2.783285553 | 2.864985962 | 2.766452068 | 3.19893187 | 2.553692546 |
| ENSMUSG00000030225 | 1.867194479 | 1.809601185 | 1.850219859 | 1.864857905 | 1.897237431 | 2.127104798 | 1.817112247 |
| ENSMUSG00000030226 | 0 | 0 | 0 | 0 | 0 | 0.301029996 | 0 |
| ENSMUSG00000030228 | 0 | 0 | 0 | 0 | 0 | 0 | 0 |
| ENSMUSG00000030230 | 0 | 0 | 0 | 0 | 0 | 0 | 0 |
| ENSMUSG00000030231 | 2.181541736 | 2.029735268 | 2.009729747 | 1.926839783 | 1.851969278 | 2.086359831 | 1.787569411 |
| ENSMUSG00000030232 | 2.856724497 | 2.617410454 | 2.598347614 | 2.594844592 | 2.49625462 | 2.877371346 | 2.318147869 |
| ENSMUSG00000030235 | 0 | 0 | 0 | 0 | 0 | 0 | 0 |
| ENSMUSG00000030236 | 0 | 0 | 0 | 0 | 0 | 0 | 0 |
| ENSMUSG00000030237 | 0 | 0 | 0 | 0 | 0 | 0 | 0.224243824 |
| ENSMUSG00000030243 | 2.276824003 | 2.112802778 | 2.066696635 | 2.155512955 | 2.070605826 | 2.390935107 | 1.927642465 |
| ENSMUSG00000030244 | 0 | 0 | 0 | 0 | 0 | 0 | 0 |
| ENSMUSG00000030245 | 2.63895461 | 2.402294685 | 2.397178659 | 2.379909063 | 2.330902415 | 2.640481437 | 2.184841529 |
| ENSMUSG00000030246 | 1.867194479 | 1.811981878 | 1.823655187 | 2.066696635 | 2.012006468 | 2.354108439 | 1.913109605 |
| ENSMUSG00000030247 | 0 | 0 | 0 | 0.25 | 0 | 0.477121255 | 0.355418052 |
| ENSMUSG00000030249 | 0.289064826 | 0.25 | 0.25 | 0.580482024 | 0.244650542 | 0.301029996 | 0.355418052 |
| ENSMUSG00000030254 | 2.392336464 | 2.185366747 | 2.186208459 | 2.373963757 | 2.2705456 | 2.624282096 | 2.116124284 |
| ENSMUSG00000030255 | 2.492331912 | 2.255591953 | 2.264497931 | 1.92510993 | 1.880506521 | 2.113943352 | 1.752192176 |
| ENSMUSG00000030256 | 1.747221736 | 1.54745614 | 1.53743678 | 0.701838731 | 0.77552387 | 0.602059991 | 0.916588299 |
| ENSMUSG00000030257 | 2.22592627 | 2.090485943 | 2.113831805 | 1.741446071 | 1.662257525 | 1.838849091 | 1.62316756 |
| ENSMUSG00000030259 | 2.215778996 | 2.095926073 | 2.135757955 | 1.96293726 | 1.982473755 | 2.184691431 | 1.897893072 |
| ENSMUSG00000030263 | 0.289064826 | 0.396240625 | 0.25 | 0 | 0 | 0 | 0 |
| ENSMUSG00000030264 | 2.258689303 | 2.081607372 | 2.108657057 | 2.122963274 | 2.038163331 | 2.342422681 | 1.888596011 |
| ENSMUSG00000030265 | 2.750654121 | 2.539967834 | 2.51685859 | 2.573367912 | 2.462674177 | 2.858537198 | 2.290497981 |

| | | | | | | | |
|---|---|---|---|---|---|---|---|
| ENSMUSG00000030268 | 2.804055922 | 2.558105232 | 2.502806814 | 2.469512728 | 2.443041643 | 2.797267541 | 2.294288164 |
| ENSMUSG00000030269 | 2.17479713 | 2.00835575 | 2.024508021 | 2.001406137 | 1.935892666 | 2.139879086 | 1.852723744 |
| ENSMUSG00000030270 | 0 | 0.580482024 | 0.396240625 | 0 | 0.387761935 | 0.602059991 | 0.224243824 |
| ENSMUSG00000030271 | 1.876566388 | 1.80479213 | 1.896240625 | 1.634789703 | 1.744182955 | 1.86332286 | 1.749303626 |
| ENSMUSG00000030272 | 2.386920383 | 2.225216702 | 2.30175358 | 2.266522298 | 2.225193812 | 2.557507202 | 2.139097452 |
| ENSMUSG00000030275 | 2.998430437 | 2.731017546 | 2.704245906 | 2.78296424 | 2.70184985 | 3.076276255 | 2.500119509 |
| ENSMUSG00000030276 | 1.700465191 | 1.622963274 | 1.62694866 | 1.238549078 | 1.358933174 | 1.146128036 | 1.224243824 |
| ENSMUSG00000030278 | 0 | 0 | 0 | 0 | 0 | 0 | 0 |
| ENSMUSG00000030279 | 2.054824387 | 1.903677461 | 1.961372513 | 2.109697963 | 2.041436517 | 2.320146286 | 1.912234055 |
| ENSMUSG00000030281 | 2.036286563 | 2.03743678 | 2.114857905 | 1.874961472 | 2.082404901 | 2.139879086 | 2.055173778 |
| ENSMUSG00000030282 | 2.141794304 | 2.012462137 | 2.01109853 | 2.105516192 | 2.05001946 | 2.326335861 | 1.970632877 |
| ENSMUSG00000030283 | 0 | 0.25 | 0.396240625 | 0.5 | 0.244650542 | 0 | 0.355418052 |
| ENSMUSG00000030284 | 1.993049299 | 1.966046536 | 1.964495249 | 1.843759858 | 1.991345651 | 2.11058971 | 1.960219986 |
| ENSMUSG00000030286 | 2.578129653 | 2.357090043 | 2.352347734 | 2.49539182 | 2.42262144 | 2.813580989 | 2.304357435 |
| ENSMUSG00000030287 | 1.867194479 | 1.71759118 | 1.732684334 | 1.698603967 | 1.706995316 | 1.792391689 | 1.594042709 |
| ENSMUSG00000030291 | 2.209569962 | 2.071350555 | 2.006976499 | 2.064346961 | 2.012006468 | 2.276461804 | 1.902443294 |
| ENSMUSG00000030292 | 0.747221736 | 1.114857905 | 1.061981878 | 0.25 | 0 | 0 | 0 |
| ENSMUSG00000030298 | 2.867194479 | 2.652525516 | 2.650442697 | 2.589388001 | 2.616806216 | 2.972665592 | 2.475255611 |
| ENSMUSG00000030301 | 2.097855556 | 1.940387808 | 1.935366747 | 1.899978211 | 1.919403394 | 2.068185862 | 1.807561282 |
| ENSMUSG00000030302 | 0 | 0.25 | 0 | 0 | 0.244650542 | 0.301029996 | 0.224243824 |
| ENSMUSG00000030303 | 1.069666964 | 1.188721876 | 1.214495249 | 1.27186571 | 1.200473444 | 1.278753601 | 1.066254157 |
| ENSMUSG00000030304 | 2.560530738 | 2.369945066 | 2.350753006 | 2.43662858 | 2.363323232 | 2.680335513 | 2.193409508 |
| ENSMUSG00000030306 | 0.578129653 | 0.580482024 | 0 | 1.188721876 | 1.188507684 | 1.041392685 | 1.028149525 |
| ENSMUSG00000030307 | 0 | 0 | 0 | 0.25 | 0 | 0 | 0 |
| ENSMUSG00000030309 | 1.956444026 | 1.816696635 | 1.747171172 | 1.872963274 | 1.770879644 | 2.053078443 | 1.641897157 |
| ENSMUSG00000030310 | 0.289064826 | 0.25 | 0 | 0 | 0 | 0.301029996 | 0.448487648 |
| ENSMUSG00000030313 | 2.081172036 | 1.901832578 | 1.945339928 | 2.095926073 | 2.045754077 | 2.28780173 | 1.900630878 |
| ENSMUSG00000030314 | 2.075459174 | 1.982684334 | 2.02053726 | 1.905512955 | 1.977302144 | 2.139879086 | 1.825244123 |
| ENSMUSG00000030315 | 2.256831695 | 2.162762923 | 2.192046081 | 1.931980114 | 2.009637626 | 2.184691431 | 1.980721036 |
| ENSMUSG00000030316 | 2.051791411 | 1.916333979 | 1.890560606 | 1.937048212 | 1.846351821 | 2.123851641 | 1.763495175 |
| ENSMUSG00000030317 | 0 | 0 | 0 | 0 | 0.244650542 | 0 | 0 |
| ENSMUSG00000030319 | 2.156259305 | 2.064346961 | 2.166333979 | 1.955044741 | 2.055279708 | 2.167317335 | 2.074322292 |
| ENSMUSG00000030321 | 0.45815691 | 0.646240625 | 0.25 | 0.5 | 0.686820904 | 0 | 0.672731473 |
| ENSMUSG00000030322 | 2.020182925 | 1.843759858 | 1.854463129 | 1.94697564 | 1.86473911 | 2.045322979 | 1.774416565 |

| | | | | | | | |
|---|---|---|---|---|---|---|---|
| ENSMUSG00000030323 | 2.158622101 | 2.046218836 | 2.006976499 | 1.964495249 | 1.947418788 | 2.1430148 | 1.913982791 |
| ENSMUSG00000030324 | 0.960252568 | 0.396240625 | 0.79248125 | 0 | 0.387761935 | 0.301029996 | 0 |
| ENSMUSG00000030325 | 0 | 0 | 0 | 0 | 0 | 0 | 0 |
| ENSMUSG00000030327 | 2.461589295 | 2.269871196 | 2.191217898 | 2.300530956 | 2.276175004 | 2.561101384 | 2.117986249 |
| ENSMUSG00000030329 | 1.885732304 | 1.744319981 | 1.886723615 | 0.580482024 | 0.877063019 | 0.84509804 | 0.896975297 |
| ENSMUSG00000030330 | 0.747221736 | 0.5 | 0.646240625 | 0.92510993 | 0.905314583 | 0.77815125 | 0.710836105 |
| ENSMUSG00000030335 | 2.247417471 | 2.150906586 | 2.202642909 | 2.152756199 | 2.155511127 | 2.40654018 | 2.089878917 |
| ENSMUSG00000030336 | 0.45815691 | 0.5 | 0.5 | 0.830482024 | 0.77552387 | 0.84509804 | 0.853775828 |
| ENSMUSG00000030337 | 1.596667477 | 1.575945187 | 1.505591953 | 1.766522298 | 1.625423019 | 1.897627091 | 1.631625962 |
| ENSMUSG00000030339 | 2.561428547 | 2.399047439 | 2.460194481 | 2.167221246 | 2.322187006 | 2.494154594 | 2.23281864 |
| ENSMUSG00000030340 | 1.307602651 | 1.25 | 1.214495249 | 1.080482024 | 1.212046813 | 1.230448921 | 1.054044578 |
| ENSMUSG00000030341 | 2.603209296 | 2.402756199 | 2.463717096 | 2.327653195 | 2.362449578 | 2.602059991 | 2.229876084 |
| ENSMUSG00000030342 | 2.812982046 | 2.62019755 | 2.583009137 | 2.55479213 | 2.501012315 | 2.911690159 | 2.344081383 |
| ENSMUSG00000030344 | 0.289064826 | 0 | 0 | 0 | 0 | 0 | 0 |
| ENSMUSG00000030345 | 0 | 0 | 0 | 0 | 0 | 0 | 0 |
| ENSMUSG00000030346 | 2.34539747 | 2.171625132 | 2.184523065 | 2.187886015 | 2.163285805 | 2.411619706 | 2.051774285 |
| ENSMUSG00000030347 | 2.134369199 | 1.901832578 | 2.001406137 | 1.894357207 | 1.888971102 | 2.086359831 | 1.74196736 |
| ENSMUSG00000030350 | 0 | 0 | 0 | 0 | 0.244650542 | 0 | 0 |
| ENSMUSG00000030351 | 0.289064826 | 0 | 0.396240625 | 0 | 0 | 0 | 0 |
| ENSMUSG00000030352 | 2.595015862 | 2.388647213 | 2.424241882 | 2.430701883 | 2.473055988 | 2.781036939 | 2.373190952 |
| ENSMUSG00000030353 | 1.470606563 | 1.431980114 | 1.52186571 | 1.752806814 | 1.754126039 | 1.913813852 | 1.699191618 |
| ENSMUSG00000030357 | 2.941573724 | 2.728409357 | 2.756284891 | 2.70042709 | 2.72152231 | 3.106190897 | 2.586387095 |
| ENSMUSG00000030359 | 0 | 0 | 0 | 0 | 0 | 0 | 0.355418052 |
| ENSMUSG00000030361 | 0 | 0 | 0 | 0.25 | 0 | 0 | 0 |
| ENSMUSG00000030363 | 0 | 0 | 0 | 0 | 0 | 0 | 0 |
| ENSMUSG00000030364 | 0.289064826 | 0 | 0.396240625 | 0.25 | 0.244650542 | 0.698970004 | 0.224243824 |
| ENSMUSG00000030365 | 0 | 0.25 | 0 | 0 | 0.244650542 | 0 | 0 |
| ENSMUSG00000030366 | 0 | 0 | 0 | 0 | 0 | 0 | 0 |
| ENSMUSG00000030368 | 0 | 0 | 0 | 0 | 0 | 0 | 0 |
| ENSMUSG00000030373 | 0 | 0 | 0 | 0 | 0 | 0 | 0 |
| ENSMUSG00000030374 | 2.40817504 | 2.268535366 | 2.321928095 | 2.193696765 | 2.279510047 | 2.477121255 | 2.218171406 |
| ENSMUSG00000030376 | 0 | 0 | 0 | 0 | 0.489301084 | 0.602059991 | 0.629532004 |
| ENSMUSG00000030378 | 0 | 0 | 0 | 0 | 0 | 0 | 0 |
| ENSMUSG00000030380 | 1 | 1.080482024 | 1.061981878 | 0.864857905 | 1 | 0.698970004 | 0.916588299 |

| | | | | | | | |
|---|---|---|---|---|---|---|---|
| ENSMUSG00000030382 | 0.45815691 | 0 | 0 | 0 | 0.244650542 | 0 | 0 |
| ENSMUSG00000030385 | 0 | 0 | 0 | 0 | 0 | 0 | 0 |
| ENSMUSG00000030386 | 1.747221736 | 1.618933358 | 1.698603967 | 1.614857905 | 1.514343934 | 1.707570176 | 1.508694612 |
| ENSMUSG00000030393 | 2.026699189 | 1.888647213 | 1.850219859 | 1.843759858 | 1.739104362 | 1.986771734 | 1.629532004 |
| ENSMUSG00000030397 | 1.714367558 | 1.638647213 | 1.794977273 | 1.610735874 | 1.77552387 | 1.755874856 | 1.710836105 |
| ENSMUSG00000030399 | 0 | 0 | 0 | 0 | 0 | 0 | 0 |
| ENSMUSG00000030400 | 1.678702042 | 1.738549078 | 1.779735268 | 1.552363341 | 1.692703642 | 1.770852012 | 1.695786161 |
| ENSMUSG00000030401 | 1.036286563 | 1.17510993 | 1.130890489 | 1.04248125 | 1.091002363 | 1.176091259 | 1.150210039 |
| ENSMUSG00000030402 | 0 | 0 | 0 | 0 | 0 | 0 | 0 |
| ENSMUSG00000030403 | 2.577267124 | 2.423371739 | 2.468361028 | 1.979715809 | 2.041436517 | 2.278753601 | 1.983546516 |
| ENSMUSG00000030406 | 0 | 0.25 | 0 | 0 | 0 | 0 | 0 |
| ENSMUSG00000030407 | 1.432083896 | 1.418106335 | 1.494319981 | 1.438721876 | 1.611014646 | 1.672097858 | 1.582103514 |
| ENSMUSG00000030409 | 1.734388958 | 1.784887838 | 1.834962501 | 1.26109853 | 1.427019704 | 1.431363764 | 1.493062805 |
| ENSMUSG00000030410 | 1.940852214 | 1.953445298 | 2.015173983 | 1.747171172 | 1.937354184 | 1.986771734 | 1.943148064 |
| ENSMUSG00000030411 | 0.45815691 | 0.25 | 0.25 | 0 | 0.387761935 | 0.77815125 | 0.224243824 |
| ENSMUSG00000030413 | 0.289064826 | 0 | 0 | 0.25 | 0.244650542 | 0 | 0 |
| ENSMUSG00000030417 | 2.029919532 | 1.903677461 | 1.928561379 | 2.028435542 | 1.929985312 | 2.247973266 | 1.87902385 |
| ENSMUSG00000030421 | 2.254965775 | 2.078220739 | 2.125956434 | 2.058404919 | 2.013184954 | 2.230448921 | 1.87902385 |
| ENSMUSG00000030423 | 1.740854706 | 1.584962501 | 1.598079356 | 1.561981878 | 1.595992969 | 1.662757832 | 1.444790289 |
| ENSMUSG00000030424 | 0.960252568 | 0.864857905 | 0.864857905 | 1.114857905 | 0.905314583 | 0.903089987 | 0.935079929 |
| ENSMUSG00000030427 | 0 | 0 | 0 | 0 | 0 | 0 | 0 |
| ENSMUSG00000030428 | 0.671187741 | 0.5 | 0.701838731 | 0.646240625 | 0.733951626 | 0.301029996 | 0.579661877 |
| ENSMUSG00000030431 | 0.747221736 | 0.79248125 | 0.864857905 | 0.75 | 0.77552387 | 0.477121255 | 0.775756176 |
| ENSMUSG00000030432 | 2.593357679 | 2.458222504 | 2.495034894 | 2.306000419 | 2.443041643 | 2.653212514 | 2.343850547 |
| ENSMUSG00000030433 | 0 | 0 | 0 | 0 | 0 | 0.301029996 | 0 |
| ENSMUSG00000030435 | 2.87233252 | 2.646005735 | 2.684523065 | 2.569031101 | 2.656410937 | 2.930439595 | 2.507164985 |
| ENSMUSG00000030443 | 0.747221736 | 0.896240625 | 0.92510993 | 0.976722649 | 0.846351821 | 1.079181246 | 0.952571511 |
| ENSMUSG00000030446 | 0.91631382 | 0.951838731 | 0.896240625 | 1.201838731 | 1.057362051 | 1.113943352 | 0.853775828 |
| ENSMUSG00000030447 | 2.81052167 | 2.563757142 | 2.564346961 | 2.623963757 | 2.594636314 | 2.968015714 | 2.399701885 |
| ENSMUSG00000030450 | 0 | 0 | 0 | 0 | 0 | 0 | 0 |
| ENSMUSG00000030451 | 2.780867173 | 2.54248125 | 2.548996825 | 2.525493918 | 2.514628232 | 2.822168079 | 2.331614323 |
| ENSMUSG00000030452 | 2.438154344 | 2.232684334 | 2.232684334 | 2.290597832 | 2.179075613 | 2.514547753 | 2.051204213 |
| ENSMUSG00000030463 | 0 | 0 | 0 | 0 | 0 | 0 | 0.224243824 |
| ENSMUSG00000030465 | 2.089597219 | 1.96293726 | 1.958222504 | 1.784887838 | 1.777823267 | 1.995635195 | 1.669034113 |

| | | | | | | | |
|---|---|---|---|---|---|---|---|
| ENSMUSG00000030468 | 1.811507563 | 1.782320754 | 1.825945187 | 1.747171172 | 1.795698282 | 1.897627091 | 1.700880998 |
| ENSMUSG00000030469 | 1.816888701 | 1.755591953 | 1.741446071 | 1.834962501 | 1.761403742 | 1.973127854 | 1.635773653 |
| ENSMUSG00000030470 | 0 | 0 | 0 | 0 | 0 | 0 | 0 |
| ENSMUSG00000030471 | 2.089597219 | 1.928561379 | 1.910964047 | 1.995748394 | 1.957204337 | 2.225309282 | 1.797719748 |
| ENSMUSG00000030472 | 0 | 0 | 0 | 0 | 0 | 0 | 0 |
| ENSMUSG00000030474 | 0 | 0 | 0.25 | 0 | 0 | 0 | 0 |
| ENSMUSG00000030483 | 0 | 0 | 0 | 0 | 0 | 0 | 0 |
| ENSMUSG00000030484 | 0 | 0 | 0 | 0 | 0 | 0 | 0 |
| ENSMUSG00000030486 | 0.45815691 | 0.79248125 | 0.75 | 0.646240625 | 0.489301084 | 0.698970004 | 0.355418052 |
| ENSMUSG00000030491 | 0 | 0 | 0 | 0 | 0 | 0 | 0 |
| ENSMUSG00000030492 | 0 | 0 | 0 | 0 | 0 | 0 | 0 |
| ENSMUSG00000030493 | 1.578129653 | 1.488549078 | 1.464495249 | 1.552363341 | 1.625423019 | 1.740362689 | 1.476637173 |
| ENSMUSG00000030494 | 1.249317394 | 1.114857905 | 1.25 | 1.464495249 | 1.559656853 | 1.612783857 | 1.417653332 |
| ENSMUSG00000030495 | 0 | 0 | 0 | 0 | 0 | 0 | 0 |
| ENSMUSG00000030498 | 1.783508299 | 1.634789703 | 1.580482024 | 1.238549078 | 1.212046813 | 1.230448921 | 1.20919388 |
| ENSMUSG00000030499 | 0 | 0.25 | 0 | 0.25 | 0 | 0 | 0 |
| ENSMUSG00000030500 | 0 | 0 | 0 | 0 | 0 | 0 | 0 |
| ENSMUSG00000030505 | 2.192545868 | 2.028435542 | 2.015173983 | 2.049918086 | 2.066583491 | 2.245512668 | 1.894206246 |
| ENSMUSG00000030507 | 0 | 0 | 0 | 0 | 0 | 0 | 0 |
| ENSMUSG00000030509 | 1.847794126 | 1.825945187 | 1.769203899 | 1.769203899 | 1.744182955 | 1.892094603 | 1.598697688 |
| ENSMUSG00000030510 | 0.45815691 | 0.5 | 0.396240625 | 0.25 | 0.568060967 | 0 | 0.448487648 |
| ENSMUSG00000030512 | 2.452372734 | 2.31611065 | 2.313757142 | 2.331607372 | 2.286629652 | 2.593286067 | 2.15438449 |
| ENSMUSG00000030513 | 1.227926429 | 1.226722649 | 1.302363341 | 0.396240625 | 0.568060967 | 0.602059991 | 0.710836105 |
| ENSMUSG00000030515 | 2.075459174 | 1.894357207 | 1.894357207 | 1.966046536 | 1.94314048 | 2.127104798 | 1.793950594 |
| ENSMUSG00000030516 | 2.757995436 | 2.518200634 | 2.487456678 | 2.693696765 | 2.590763894 | 3.022428371 | 2.443265468 |
| ENSMUSG00000030518 | 0 | 0 | 0.25 | 0.25 | 0.568060967 | 0 | 0.355418052 |
| ENSMUSG00000030519 | 0 | 0 | 0 | 0 | 0 | 0 | 0.224243824 |
| ENSMUSG00000030521 | 2.278594844 | 2.144357207 | 2.139605178 | 2.219895812 | 2.076554666 | 2.423245874 | 1.968429585 |
| ENSMUSG00000030522 | 2.060824957 | 1.991446071 | 1.961372513 | 2.057204673 | 2.020174412 | 2.240549248 | 1.880961074 |
| ENSMUSG00000030523 | 0 | 0 | 0 | 0 | 0.244650542 | 0 | 0 |
| ENSMUSG00000030525 | 0 | 0 | 0 | 0.396240625 | 0 | 0 | 0 |
| ENSMUSG00000030527 | 2.023453784 | 1.918106335 | 1.896240625 | 1.698603967 | 1.850106709 | 1.949390007 | 1.812372009 |
| ENSMUSG00000030528 | 2.31414447 | 2.169870025 | 2.186208459 | 2.28296424 | 2.203230928 | 2.505149978 | 2.033014511 |
| ENSMUSG00000030530 | 2.381433035 | 2.286164561 | 2.382357639 | 1.937048212 | 2.104881849 | 2.190331698 | 2.133836922 |

| | | | | | | | |
|---|---|---|---|---|---|---|---|
| ENSMUSG00000030532 | 1.605636262 | 1.505591953 | 1.593759858 | 1.380890489 | 1.358933174 | 1.62324929 | 1.40946263 |
| ENSMUSG00000030533 | 2.22189694 | 2.12892496 | 2.223704441 | 1.972185812 | 2.128567294 | 2.214843848 | 2.137790317 |
| ENSMUSG00000030534 | 1.655740331 | 1.664552871 | 1.653677461 | 1.580482024 | 1.632412477 | 1.740362689 | 1.520678036 |
| ENSMUSG00000030536 | 3.317377295 | 3.01109853 | 2.989819376 | 2.885878804 | 2.834611872 | 3.252124553 | 2.651406343 |
| ENSMUSG00000030538 | 1.936861443 | 1.79745614 | 1.892463902 | 1.87694866 | 1.871833944 | 2.075546961 | 1.87902385 |
| ENSMUSG00000030539 | 1.924654746 | 1.78743678 | 1.899978211 | 1.606566189 | 1.793512788 | 1.86923172 | 1.821784031 |
| ENSMUSG00000030541 | 2.93904296 | 2.694108258 | 2.70404592 | 2.441632227 | 2.501602543 | 2.775974331 | 2.359183766 |
| ENSMUSG00000030543 | 0 | 0.25 | 0.25 | 0.396240625 | 0.568060967 | 0.477121255 | 0.629532004 |
| ENSMUSG00000030544 | 0 | 0 | 0 | 0 | 0 | 0 | 0 |
| ENSMUSG00000030545 | 1.45815691 | 1.488549078 | 1.505591953 | 1.566696635 | 1.592135487 | 1.662757832 | 1.499406449 |
| ENSMUSG00000030546 | 0.289064826 | 0 | 0 | 0 | 0 | 0 | 0.224243824 |
| ENSMUSG00000030549 | 0 | 0 | 0.25 | 0 | 0.244650542 | 0.301029996 | 0.355418052 |
| ENSMUSG00000030551 | 2.151493169 | 2.024508021 | 2.01109853 | 1.933677405 | 1.880506521 | 2.053078443 | 1.798966438 |
| ENSMUSG00000030553 | 0 | 0 | 0 | 0 | 0 | 0 | 0 |
| ENSMUSG00000030554 | 2.060824957 | 1.872963274 | 1.864857905 | 1.26109853 | 1.358933174 | 1.531478917 | 1.324583736 |
| ENSMUSG00000030555 | 1.940852214 | 1.828220739 | 1.78743678 | 1.79745614 | 1.66856048 | 1.838849091 | 1.653807046 |
| ENSMUSG00000030556 | 1.700465191 | 1.580482024 | 1.638647213 | 1.598079356 | 1.533172725 | 1.681241237 | 1.486592281 |
| ENSMUSG00000030557 | 2.667031238 | 2.461764337 | 2.454645544 | 2.302972074 | 2.241584006 | 2.542825427 | 2.082194103 |
| ENSMUSG00000030559 | 0 | 0.5 | 0.25 | 1.130890489 | 0.955822902 | 1.113943352 | 0.984950056 |
| ENSMUSG00000030560 | 1.912079952 | 1.834962501 | 1.720660762 | 2.016522298 | 1.937354184 | 2.285557309 | 1.842010333 |
| ENSMUSG00000030562 | 1.847794126 | 1.660964047 | 1.729715809 | 1.388647213 | 1.42077253 | 1.568201724 | 1.296434211 |
| ENSMUSG00000030577 | 0 | 0.25 | 0 | 0.25 | 0.244650542 | 0 | 0 |
| ENSMUSG00000030579 | 0 | 0 | 0 | 0 | 0 | 0 | 0 |
| ENSMUSG00000030583 | 2.256831695 | 2.139605178 | 2.228222334 | 1.894357207 | 2.080465577 | 2.161368002 | 2.093904044 |
| ENSMUSG00000030584 | 1.494443473 | 1.396240625 | 1.476722649 | 1.42510993 | 1.621875691 | 1.477121255 | 1.437266334 |
| ENSMUSG00000030587 | 1.205378646 | 1.214495249 | 1.380890489 | 0 | 0.489301084 | 0.477121255 | 0.74492186 |
| ENSMUSG00000030588 | 1.920505136 | 1.868933358 | 1.940387808 | 1.575945187 | 1.808534465 | 1.716003344 | 1.811176017 |
| ENSMUSG00000030589 | 0 | 0.396240625 | 0.5 | 0 | 0 | 0.301029996 | 0.224243824 |
| ENSMUSG00000030590 | 0.578129653 | 0.580482024 | 0.701838731 | 0.79248125 | 0.77552387 | 0.77815125 | 0.829800754 |
| ENSMUSG00000030591 | 2.590857977 | 2.40136988 | 2.420748646 | 2.496816003 | 2.415166968 | 2.79518459 | 2.27142916 |
| ENSMUSG00000030592 | 1.069666964 | 0.896240625 | 0.75 | 0.830482024 | 0.905314583 | 1.079181246 | 1 |
| ENSMUSG00000030595 | 1.307602651 | 1.388647213 | 1.494319981 | 1.372963274 | 1.456697355 | 1.397940009 | 1.649885598 |
| ENSMUSG00000030598 | 1.358731791 | 1.238549078 | 1.364857905 | 1.27186571 | 1.366364104 | 1.255272505 | 1.324583736 |
| ENSMUSG00000030600 | 0 | 0 | 0 | 0.830482024 | 0.905314583 | 0.477121255 | 0.896975297 |

| | | | | | | | |
|---|---|---|---|---|---|---|---|
| ENSMUSG00000030602 | 2.511972754 | 2.402294685 | 2.47027849 | 2.166333979 | 2.327538614 | 2.506505032 | 2.302263476 |
| ENSMUSG00000030603 | 2.54867984 | 2.374462959 | 2.398114259 | 2.450224975 | 2.449935537 | 2.760422483 | 2.315133728 |
| ENSMUSG00000030604 | 1.181541736 | 1.061981878 | 1.04248125 | 1.04248125 | 1 | 1.204119983 | 0.916588299 |
| ENSMUSG00000030605 | 2.667726872 | 2.499647607 | 2.544043287 | 2.343216265 | 2.478134581 | 2.731588765 | 2.381518708 |
| ENSMUSG00000030606 | 0 | 0 | 0 | 0 | 0.244650542 | 0 | 0.355418052 |
| ENSMUSG00000030607 | 0.578129653 | 0.25 | 0.25 | 0.25 | 0.244650542 | 0.301029996 | 0.224243824 |
| ENSMUSG00000030609 | 2.307602651 | 2.152756199 | 2.214495249 | 1.995748394 | 2.006054284 | 2.198657087 | 1.932619724 |
| ENSMUSG00000030610 | 1.45815691 | 1.589388001 | 1.5 | 1.29248125 | 1.427019704 | 1.462397998 | 1.345462945 |
| ENSMUSG00000030611 | 1.837744666 | 1.819031101 | 1.819031101 | 1.698603967 | 1.758994476 | 1.903089987 | 1.807561282 |
| ENSMUSG00000030612 | 1.928763473 | 1.685366747 | 1.692046081 | 1.799918086 | 1.850106709 | 2.033423755 | 1.756477211 |
| ENSMUSG00000030613 | 1.678702042 | 1.708222504 | 1.705044741 | 1.622963274 | 1.439191811 | 1.806179974 | 1.425641774 |
| ENSMUSG00000030614 | 1.932832114 | 1.78743678 | 1.779735268 | 1.741446071 | 1.677808768 | 1.977723605 | 1.559435575 |
| ENSMUSG00000030615 | 2.136858952 | 2.02053726 | 2.04745614 | 1.997171172 | 1.892300902 | 2.130333768 | 1.825244123 |
| ENSMUSG00000030616 | 2.428398755 | 2.20584181 | 2.185366747 | 1.602347734 | 1.618292349 | 1.792391689 | 1.46982607 |
| ENSMUSG00000030617 | 0 | 0 | 0 | 0 | 0 | 0 | 0 |
| ENSMUSG00000030619 | 2.37447073 | 2.114857905 | 2.152756199 | 2.137686696 | 2.11997913 | 2.401400541 | 1.994607951 |
| ENSMUSG00000030621 | 1.753493018 | 1.664552871 | 1.606566189 | 1.445339928 | 1.414412788 | 1.414973348 | 1.252393349 |
| ENSMUSG00000030623 | 0.45815691 | 0.646240625 | 0.646240625 | 0.5 | 0.568060967 | 0.301029996 | 0.579661877 |
| ENSMUSG00000030629 | 2.258689303 | 2.153677461 | 2.157339155 | 2.229715809 | 2.140250123 | 2.499687083 | 2.033014511 |
| ENSMUSG00000030630 | 2.003432385 | 1.744319981 | 1.858657057 | 1.566696635 | 1.584291694 | 1.633468456 | 1.508694612 |
| ENSMUSG00000030636 | 0 | 0 | 0 | 0 | 0 | 0 | 0 |
| ENSMUSG00000030638 | 1 | 1.17510993 | 0.830482024 | 1.02186571 | 1.091002363 | 1.041392685 | 1.100339912 |
| ENSMUSG00000030641 | 2.119110214 | 2.013820609 | 1.984159485 | 1.953445298 | 1.900490506 | 2.089905111 | 1.722075986 |
| ENSMUSG00000030643 | 1.605636262 | 1.593759858 | 1.438721876 | 1.71759118 | 1.689774529 | 1.77815125 | 1.496250175 |
| ENSMUSG00000030647 | 1.952600383 | 1.916333979 | 1.979715809 | 1.868933358 | 1.941702774 | 2.136720567 | 1.959463225 |
| ENSMUSG00000030649 | 1.655740331 | 1.532320754 | 1.52186571 | 1.571350555 | 1.572189156 | 1.785329835 | 1.459332757 |
| ENSMUSG00000030650 | 0.91631382 | 0.79248125 | 0.951838731 | 0.951838731 | 0.978602168 | 1 | 0.829800754 |
| ENSMUSG00000030651 | 0 | 0 | 0 | 0 | 0 | 0 | 0 |
| ENSMUSG00000030652 | 1.894701088 | 1.843759858 | 1.910964047 | 1.819031101 | 1.890639929 | 2.103803721 | 1.857950279 |
| ENSMUSG00000030653 | 1.45815691 | 1.27186571 | 1.238549078 | 0.864857905 | 1 | 0.84509804 | 1 |
| ENSMUSG00000030654 | 2.902294604 | 2.65136988 | 2.607351685 | 2.617918804 | 2.538259021 | 2.91750551 | 2.363775816 |
| ENSMUSG00000030655 | 3.070726095 | 2.803731547 | 2.762462137 | 2.790912419 | 2.715943749 | 3.118264726 | 2.500404294 |
| ENSMUSG00000030657 | 0 | 0 | 0 | 0.5 | 0.387761935 | 0.301029996 | 0.520678036 |
| ENSMUSG00000030659 | 1.806056082 | 1.681980114 | 1.735628626 | 1.671625132 | 1.674752852 | 1.832508913 | 1.574722613 |

| | | | | | | | |
|---|---|---|---|---|---|---|---|
| ENSMUSG00000030660 | 2.769043298 | 2.509043403 | 2.514497931 | 2.530706999 | 2.448567488 | 2.787460475 | 2.246207397 |
| ENSMUSG00000030662 | 3.193728939 | 2.897178659 | 2.894121087 | 3.036084898 | 2.960112001 | 3.400710637 | 2.748312968 |
| ENSMUSG00000030663 | 2.512981297 | 2.266522298 | 2.245748394 | 2.40136988 | 2.29993025 | 2.638489257 | 2.182948513 |
| ENSMUSG00000030664 | 0 | 0 | 0 | 0 | 0 | 0 | 0 |
| ENSMUSG00000030666 | 0 | 0 | 0 | 0 | 0 | 0 | 0 |
| ENSMUSG00000030669 | 0 | 0.25 | 0.25 | 0.25 | 0 | 0.301029996 | 0 |
| ENSMUSG00000030670 | 0 | 0 | 0 | 0 | 0 | 0 | 0 |
| ENSMUSG00000030671 | 1.989530018 | 1.79745614 | 1.729715809 | 1.695339928 | 1.662257525 | 1.903089987 | 1.535058868 |
| ENSMUSG00000030672 | 0.289064826 | 0.580482024 | 0.580482024 | 0.5 | 0.632412477 | 0.954242509 | 0.672731473 |
| ENSMUSG00000030674 | 0 | 0 | 0 | 0 | 0.244650542 | 0 | 0 |
| ENSMUSG00000030677 | 2.480523256 | 2.284887838 | 2.293106877 | 2.370449858 | 2.305627392 | 2.631443769 | 2.220546465 |
| ENSMUSG00000030678 | 2.455852858 | 2.316696635 | 2.334405475 | 2.164552871 | 2.203230928 | 2.401400541 | 2.093904044 |
| ENSMUSG00000030680 | 0 | 0 | 0 | 0 | 0 | 0 | 0 |
| ENSMUSG00000030681 | 2.160971586 | 2.160964047 | 2.247171172 | 1.959800947 | 2.188507684 | 2.212187604 | 2.175652306 |
| ENSMUSG00000030682 | 2.249317394 | 2.165444524 | 2.17510993 | 2.088286706 | 2.169384551 | 2.336459734 | 2.074853081 |
| ENSMUSG00000030683 | 0.289064826 | 0.25 | 0.646240625 | 0 | 0.387761935 | 0 | 0 |
| ENSMUSG00000030685 | 1.404271433 | 1.380890489 | 1.438721876 | 1.188721876 | 1.335652905 | 1.204119983 | 1.369798885 |
| ENSMUSG00000030688 | 1.181541736 | 1.214495249 | 1.146240625 | 1.532320754 | 1.662257525 | 1.69019608 | 1.647906904 |
| ENSMUSG00000030689 | 1.862428343 | 1.80479213 | 1.807204673 | 1.720660762 | 1.789100779 | 1.924279286 | 1.847411384 |
| ENSMUSG00000030691 | 2.020182925 | 1.937048212 | 1.916333979 | 1.870953944 | 1.836786029 | 1.944482672 | 1.746389053 |
| ENSMUSG00000030693 | 0.578129653 | 0.25 | 0.580482024 | 0.25 | 0.387761935 | 0.301029996 | 0.355418052 |
| ENSMUSG00000030695 | 3.150402317 | 2.914998973 | 2.957331584 | 2.64881437 | 2.745810507 | 3.091666958 | 2.636895673 |
| ENSMUSG00000030697 | 2.485932312 | 2.320192693 | 2.321350555 | 2.225970461 | 2.230953877 | 2.50242712 | 2.191935631 |
| ENSMUSG00000030699 | 0 | 0.5 | 0.25 | 0.25 | 0 | 0 | 0 |
| ENSMUSG00000030701 | 0 | 0.25 | 0.25 | 0.580482024 | 0.489301084 | 0.301029996 | 0.520678036 |
| ENSMUSG00000030703 | 0.811507563 | 0.701838731 | 0.75 | 0.25 | 0.244650542 | 0 | 0.355418052 |
| ENSMUSG00000030704 | 2.712498852 | 2.508011432 | 2.480088214 | 2.552667836 | 2.466603229 | 2.870988814 | 2.273731771 |
| ENSMUSG00000030706 | 1.960252568 | 1.825945187 | 1.782320754 | 2.016522298 | 1.902105869 | 2.250420002 | 1.833176598 |
| ENSMUSG00000030707 | 0.578129653 | 0.830482024 | 1.080482024 | 1.130890489 | 1.433158226 | 1.322219295 | 1.437266334 |
| ENSMUSG00000030708 | 0 | 0 | 0.396240625 | 0.396240625 | 0.244650542 | 0.602059991 | 0 |
| ENSMUSG00000030711 | 0.747221736 | 0.580482024 | 0.580482024 | 0.75 | 0.733951626 | 0.77815125 | 0.579661877 |
| ENSMUSG00000030713 | 0 | 0 | 0.25 | 0.25 | 0.489301084 | 0 | 0 |
| ENSMUSG00000030714 | 1.721148722 | 1.695339928 | 1.681980114 | 1.532320754 | 1.746695109 | 1.707570176 | 1.657681529 |
| ENSMUSG00000030717 | 2.378662046 | 2.209012589 | 2.177701608 | 1.816696635 | 1.870073561 | 2.004321374 | 1.777090262 |

| | | | | | | | |
|---|---|---|---|---|---|---|---|
| ENSMUSG00000030718 | 2.371652933 | 2.225216702 | 2.228969845 | 2.231944491 | 2.250704826 | 2.46686762 | 2.15104925 |
| ENSMUSG00000030720 | 1.747221736 | 1.692046081 | 1.741446071 | 1.214495249 | 1.283908311 | 1.397940009 | 1.168187745 |
| ENSMUSG00000030722 | 1.686083339 | 1.698603967 | 1.653677461 | 1.884789703 | 1.836786029 | 2.017033339 | 1.68886578 |
| ENSMUSG00000030724 | 0 | 0 | 0 | 0 | 0 | 0 | 0 |
| ENSMUSG00000030725 | 0.747221736 | 1.114857905 | 0.976722649 | 0.92510993 | 0.812711509 | 0.477121255 | 0.969165684 |
| ENSMUSG00000030726 | 2.119110214 | 2.031030328 | 2.013820609 | 2.107613138 | 2.053185007 | 2.326335861 | 1.933441873 |
| ENSMUSG00000030727 | 1.89911417 | 1.894357207 | 1.964495249 | 1.552363341 | 1.84445929 | 1.799340549 | 1.857950279 |
| ENSMUSG00000030729 | 1.912079952 | 1.735628626 | 1.769203899 | 1.951838731 | 1.851969278 | 2.127104798 | 1.743447983 |
| ENSMUSG00000030730 | 0.289064826 | 0.5 | 0.396240625 | 0.25 | 0.387761935 | 0 | 0.672731473 |
| ENSMUSG00000030731 | 0 | 0 | 0 | 0.5 | 0.244650542 | 0.301029996 | 0.520678036 |
| ENSMUSG00000030732 | 0 | 0 | 0 | 0 | 0 | 0.301029996 | 0 |
| ENSMUSG00000030733 | 2.100572389 | 1.994319981 | 2.074802005 | 1.864857905 | 2.032639648 | 2.100370545 | 2.033617522 |
| ENSMUSG00000030735 | 0 | 0.25 | 0.25 | 0 | 0 | 0 | 0.355418052 |
| ENSMUSG00000030737 | 0 | 0 | 0 | 0 | 0 | 0 | 0 |
| ENSMUSG00000030738 | 3.208240925 | 2.920967967 | 2.898347781 | 2.961372513 | 2.881311898 | 3.340047318 | 2.71526183 |
| ENSMUSG00000030739 | 0.578129653 | 0.646240625 | 0.79248125 | 0.646240625 | 0.877063019 | 1 | 1.150210039 |
| ENSMUSG00000030741 | 2.089597219 | 2.004202072 | 2.135757955 | 1.681980114 | 1.90849439 | 1.86332286 | 2.003985867 |
| ENSMUSG00000030742 | 0.45815691 | 0.5 | 0.701838731 | 0.580482024 | 0.489301084 | 0 | 0.520678036 |
| ENSMUSG00000030744 | 3.360055284 | 3.046838018 | 3.060719092 | 3.063904687 | 3.028555377 | 3.514813295 | 2.81561151 |
| ENSMUSG00000030745 | 0 | 0.396240625 | 0 | 0.25 | 0 | 0 | 0 |
| ENSMUSG00000030747 | 1.494443473 | 1.53743678 | 1.54745614 | 1.403677461 | 1.568060967 | 1.662757832 | 1.48984372 |
| ENSMUSG00000030748 | 1.952600383 | 1.892463902 | 1.914552871 | 1.532320754 | 1.645989387 | 1.84509804 | 1.661510159 |
| ENSMUSG00000030750 | 2.165631214 | 1.998588359 | 2.036164561 | 1.997171172 | 2.029283463 | 2.204119983 | 1.857950279 |
| ENSMUSG00000030751 | 2.755750303 | 2.513142013 | 2.492166698 | 2.685998215 | 2.599361968 | 3.007747778 | 2.417285492 |
| ENSMUSG00000030752 | 1.587501561 | 1.516522298 | 1.532320754 | 1.438721876 | 1.394615667 | 1.62324929 | 1.425641774 |
| ENSMUSG00000030753 | 2.641182749 | 2.448197574 | 2.416777886 | 2.432829758 | 2.355827317 | 2.699837726 | 2.176815335 |
| ENSMUSG00000030754 | 2.938317055 | 2.682192712 | 2.656427211 | 2.787119146 | 2.706665604 | 3.130333768 | 2.491164492 |
| ENSMUSG00000030757 | 0 | 0 | 0 | 0.646240625 | 0.489301084 | 0.301029996 | 0.355418052 |
| ENSMUSG00000030759 | 2.855840016 | 2.606566189 | 2.592399337 | 2.575659731 | 2.476871767 | 2.873901598 | 2.279417602 |
| ENSMUSG00000030760 | 2.026699189 | 1.964495249 | 1.966046536 | 2.129909063 | 2.006054284 | 2.326335861 | 1.913109605 |
| ENSMUSG00000030761 | 1.907802659 | 1.723704441 | 1.747171172 | 0.864857905 | 0.955822902 | 0.903089987 | 0.952571511 |
| ENSMUSG00000030762 | 0 | 0 | 0 | 0 | 0 | 0 | 0 |
| ENSMUSG00000030763 | 1.794934559 | 1.744319981 | 1.802363341 | 1.843759858 | 1.808534465 | 2.021189299 | 1.728327686 |
| ENSMUSG00000030766 | 2.160971586 | 2.001406137 | 2.05357978 | 1.955044741 | 1.987570683 | 2.139879086 | 1.904245613 |

| | | | | | | | |
|---|---|---|---|---|---|---|---|
| ENSMUSG00000030768 | 2.361637952 | 2.233422664 | 2.273189285 | 2.170748646 | 2.232853197 | 2.437750563 | 2.163786106 |
| ENSMUSG00000030769 | 0 | 0 | 0 | 0 | 0 | 0 | 0 |
| ENSMUSG00000030770 | 3.10253721 | 2.823511578 | 2.813018511 | 2.831466895 | 2.753981828 | 3.194514342 | 2.560027986 |
| ENSMUSG00000030771 | 0.45815691 | 0.79248125 | 0.646240625 | 0.5 | 0.387761935 | 0.477121255 | 0.224243824 |
| ENSMUSG00000030772 | 0.91631382 | 0.701838731 | 0.92510993 | 0.92510993 | 0.905314583 | 1 | 1 |
| ENSMUSG00000030774 | 2.460448303 | 2.219129237 | 2.225970461 | 2.263820609 | 2.152352662 | 2.519827994 | 2.038401572 |
| ENSMUSG00000030775 | 0 | 0 | 0 | 0 | 0 | 0 | 0 |
| ENSMUSG00000030779 | 2.644135203 | 2.445749552 | 2.42813175 | 2.540597832 | 2.466277473 | 2.826074803 | 2.307731473 |
| ENSMUSG00000030780 | 1.663535556 | 1.606566189 | 1.638647213 | 1.356566189 | 1.519146132 | 1.447158031 | 1.459332757 |
| ENSMUSG00000030781 | 0 | 0.5 | 0.396240625 | 0 | 0.244650542 | 0 | 0 |
| ENSMUSG00000030782 | 2.553277818 | 2.396240625 | 2.436208459 | 1.907339155 | 2.009637626 | 2.176091259 | 1.991877847 |
| ENSMUSG00000030785 | 0 | 0 | 0.25 | 0 | 0.244650542 | 0 | 0 |
| ENSMUSG00000030786 | 0.45815691 | 0 | 0 | 0 | 0.387761935 | 0 | 0.224243824 |
| ENSMUSG00000030787 | 0 | 0 | 0 | 0 | 0 | 0 | 0 |
| ENSMUSG00000030788 | 1.989530018 | 1.905512955 | 1.975216702 | 2.107613138 | 2.043601899 | 2.290034611 | 1.871156825 |
| ENSMUSG00000030789 | 0 | 0 | 0 | 0 | 0 | 0 | 0 |
| ENSMUSG00000030790 | 1.404271433 | 1.25 | 1.238549078 | 0.646240625 | 0.244650542 | 0 | 0.629532004 |
| ENSMUSG00000030792 | 0 | 0.25 | 0.25 | 0.79248125 | 0.77552387 | 0.602059991 | 0.775756176 |
| ENSMUSG00000030793 | 0 | 0.25 | 0.396240625 | 0.25 | 0.244650542 | 0 | 0.224243824 |
| ENSMUSG00000030795 | 3.158193497 | 2.849017389 | 2.879909063 | 2.868045779 | 2.837697472 | 3.23578087 | 2.66503742 |
| ENSMUSG00000030796 | 2.65645504 | 2.465659339 | 2.49539182 | 2.368426437 | 2.406395052 | 2.709269961 | 2.33637549 |
| ENSMUSG00000030798 | 0 | 0.5 | 0.580482024 | 0.396240625 | 0.489301084 | 0.602059991 | 0.520678036 |
| ENSMUSG00000030800 | 0 | 0 | 0 | 0 | 0 | 0 | 0 |
| ENSMUSG00000030801 | 1.794934559 | 1.67510993 | 1.671625132 | 1.53743678 | 1.588235379 | 1.755874856 | 1.46982607 |
| ENSMUSG00000030802 | 2.307602651 | 2.137686696 | 2.189555804 | 2.142463902 | 2.188507684 | 2.450249108 | 2.112839835 |
| ENSMUSG00000030804 | 0 | 0 | 0 | 0 | 0 | 0 | 0 |
| ENSMUSG00000030805 | 2.172524468 | 2.03743678 | 2.137686696 | 1.94697564 | 2.052132976 | 2.181843588 | 1.947894097 |
| ENSMUSG00000030806 | 0.578129653 | 0 | 0.5 | 1.26109853 | 1.136121934 | 1.146128036 | 1.066254157 |
| ENSMUSG00000030811 | 1.99653913 | 1.880890489 | 1.942046081 | 1.802363341 | 1.846351821 | 1.903089987 | 1.849546807 |
| ENSMUSG00000030814 | 2.042557845 | 1.92510993 | 1.912762923 | 1.868933358 | 1.955822902 | 2.08278537 | 1.864112664 |
| ENSMUSG00000030815 | 1.505869732 | 1.451838731 | 1.482684334 | 1.516522298 | 1.572189156 | 1.662757832 | 1.499406449 |
| ENSMUSG00000030816 | 2.346900291 | 2.216819685 | 2.26109853 | 2.136723615 | 2.254881871 | 2.419955748 | 2.179512918 |
| ENSMUSG00000030822 | 2.081172036 | 1.955044741 | 1.992885888 | 1.850219859 | 1.903713872 | 2.008600172 | 1.796468236 |
| ENSMUSG00000030823 | 0.289064826 | 0.580482024 | 0.396240625 | 0.25 | 0.244650542 | 0.77815125 | 0.672731473 |

| | | | | | | | |
|---|---|---|---|---|---|---|---|
| ENSMUSG00000030824 | 2.66283289 | 2.5 | 2.551448312 | 2.443696765 | 2.495655374 | 2.803457116 | 2.435009443 |
| ENSMUSG00000030825 | 0 | 0.25 | 0 | 0 | 0.387761935 | 0.477121255 | 0 |
| ENSMUSG00000030826 | 1.89911417 | 1.916333979 | 1.937048212 | 1.752806814 | 1.887294348 | 1.995635195 | 1.872150729 |
| ENSMUSG00000030827 | 0.671187741 | 0.896240625 | 0.5 | 0 | 0 | 0.301029996 | 0 |
| ENSMUSG00000030830 | 0 | 0 | 0 | 0 | 0.244650542 | 0 | 0.224243824 |
| ENSMUSG00000030834 | 0.289064826 | 0 | 0.25 | 0 | 0.489301084 | 0 | 0 |
| ENSMUSG00000030835 | 2.718080037 | 2.559003548 | 2.567573832 | 2.51685859 | 2.618504144 | 2.918030337 | 2.462228489 |
| ENSMUSG00000030838 | 0 | 0 | 0 | 0.25 | 0 | 0 | 0 |
| ENSMUSG00000030839 | 1.205378646 | 1.339388001 | 1.364857905 | 1.364857905 | 1.47337555 | 1.505149978 | 1.417653332 |
| ENSMUSG00000030842 | 2.463861956 | 2.319031101 | 2.33993739 | 2.071350555 | 2.159419872 | 2.330413773 | 2.043115906 |
| ENSMUSG00000030844 | 1.587501561 | 1.464495249 | 1.464495249 | 1.839388001 | 1.728722551 | 1.977723605 | 1.676387053 |
| ENSMUSG00000030846 | 2.518981867 | 2.352877747 | 2.30357978 | 2.411414608 | 2.357161747 | 2.729974286 | 2.203542812 |
| ENSMUSG00000030847 | 2.185978293 | 2.104463129 | 2.134789703 | 1.903677461 | 2.003645018 | 2.206825876 | 1.950241257 |
| ENSMUSG00000030849 | 2.305951036 | 2.177701608 | 2.146240625 | 1.614857905 | 1.715300549 | 1.72427587 | 1.529383012 |
| ENSMUSG00000030850 | 2.572927518 | 2.368933358 | 2.370449858 | 2.261780978 | 2.201854879 | 2.437750563 | 2.017561934 |
| ENSMUSG00000030851 | 0 | 0 | 0 | 0 | 0 | 0 | 0 |
| ENSMUSG00000030852 | 2.266038081 | 2.117918804 | 2.139605178 | 1.970660762 | 2.010824034 | 2.214843848 | 1.94074868 |
| ENSMUSG00000030854 | 0.289064826 | 0.580482024 | 0 | 0.25 | 0.489301084 | 0.301029996 | 0.579661877 |
| ENSMUSG00000030858 | 0 | 0 | 0.25 | 0 | 0.244650542 | 0 | 0 |
| ENSMUSG00000030859 | 0 | 0 | 0 | 0 | 0.387761935 | 0 | 0 |
| ENSMUSG00000030861 | 2.170239354 | 2.001406137 | 1.955044741 | 1.995748394 | 1.951645858 | 2.230448921 | 1.833176598 |
| ENSMUSG00000030862 | 0 | 0 | 0 | 0 | 0 | 0 | 0 |
| ENSMUSG00000030865 | 0 | 0 | 0 | 0 | 0 | 0 | 0 |
| ENSMUSG00000030866 | 0 | 0 | 0 | 0 | 0 | 0 | 0 |
| ENSMUSG00000030867 | 2.427163103 | 2.25 | 2.328787391 | 2.211372513 | 2.265401209 | 2.426511261 | 2.267652107 |
| ENSMUSG00000030868 | 2.337801042 | 2.142463902 | 2.091580554 | 2.238549078 | 2.147561328 | 2.478566496 | 2.032410373 |
| ENSMUSG00000030869 | 2.163307908 | 1.969129237 | 1.978222334 | 2.02053726 | 1.941702774 | 2.235528447 | 1.850609257 |
| ENSMUSG00000030870 | 2.473938397 | 2.279735268 | 2.235628626 | 2.30175358 | 2.289330171 | 2.542825427 | 2.121678334 |
| ENSMUSG00000030871 | 1.494443473 | 1.54248125 | 1.458222504 | 1.458222504 | 1.450958105 | 1.740362689 | 1.429563222 |
| ENSMUSG00000030872 | 2.6088504 | 2.395770692 | 2.414106216 | 2.321350555 | 2.307676489 | 2.591064607 | 2.158095989 |
| ENSMUSG00000030873 | 0 | 0 | 0 | 0 | 0 | 0 | 0.224243824 |
| ENSMUSG00000030876 | 2.577267124 | 2.345926073 | 2.337182039 | 2.406427211 | 2.307165329 | 2.704150517 | 2.162575192 |
| ENSMUSG00000030877 | 0 | 0.25 | 0 | 0 | 0 | 0.301029996 | 0 |
| ENSMUSG00000030878 | 2.260538674 | 2.094844592 | 2.097004321 | 1.998588359 | 1.940259189 | 2.285557309 | 1.851668228 |

| | | | | | | | |
|---|---|---|---|---|---|---|---|
| ENSMUSG00000030879 | 2.151493169 | 2.090485943 | 2.073655187 | 1.969129237 | 1.969399812 | 2.230448921 | 1.939944915 |
| ENSMUSG00000030880 | 1.985980784 | 1.852347734 | 1.852347734 | 1.701838731 | 1.820920161 | 1.886490725 | 1.743447983 |
| ENSMUSG00000030881 | 2.275045611 | 2.138647213 | 2.154596376 | 2.135757955 | 2.214050354 | 2.378397901 | 2.064068234 |
| ENSMUSG00000030882 | 1.470606563 | 1.396240625 | 1.321350555 | 1.364857905 | 1.264824954 | 1.278753601 | 1.150210039 |
| ENSMUSG00000030884 | 2.713122685 | 2.465659339 | 2.5 | 2.612029076 | 2.506289166 | 2.893761762 | 2.325806859 |
| ENSMUSG00000030887 | 0.289064826 | 0.580482024 | 0.701838731 | 0.25 | 0 | 0 | 0.355418052 |
| ENSMUSG00000030888 | 2.297593393 | 2.110735874 | 2.111770807 | 2.031030328 | 2.058398634 | 2.230448921 | 1.866140981 |
| ENSMUSG00000030889 | 0 | 0.25 | 0.5 | 0.5 | 0.568060967 | 0 | 0 |
| ENSMUSG00000030890 | 0.747221736 | 0.830482024 | 0.79248125 | 0 | 0.244650542 | 0 | 0.520678036 |
| ENSMUSG00000030894 | 2.785084983 | 2.556603048 | 2.567865757 | 2.275821952 | 2.261347526 | 2.589949601 | 2.115189268 |
| ENSMUSG00000030895 | 0.867194479 | 0.646240625 | 1.04248125 | 0.701838731 | 0.733951626 | 0.698970004 | 0.710836105 |
| ENSMUSG00000030897 | 0 | 0.25 | 0 | 0 | 0 | 0 | 0 |
| ENSMUSG00000030898 | 0 | 0 | 0 | 0 | 0 | 0 | 0 |
| ENSMUSG00000030905 | 0 | 0 | 0 | 0 | 0.387761935 | 0 | 0 |
| ENSMUSG00000030909 | 0 | 0 | 0 | 0 | 0 | 0 | 0 |
| ENSMUSG00000030911 | 0 | 0 | 0 | 0 | 0 | 0 | 0 |
| ENSMUSG00000030917 | 1.269664473 | 1.380890489 | 1.356566189 | 1.321350555 | 1.387761935 | 1.380211242 | 1.313616061 |
| ENSMUSG00000030921 | 0 | 0 | 0 | 0.396240625 | 0.489301084 | 0 | 0.224243824 |
| ENSMUSG00000030922 | 1.100572389 | 0.830482024 | 1 | 1.080482024 | 0.877063019 | 0.954242509 | 0.853775828 |
| ENSMUSG00000030924 | 1.53838222 | 1.561981878 | 1.494319981 | 1.610735874 | 1.504538967 | 1.568201724 | 1.473249546 |
| ENSMUSG00000030929 | 1.822201287 | 1.614857905 | 1.649978211 | 1.769203899 | 1.689774529 | 1.880813592 | 1.56716938 |
| ENSMUSG00000030930 | 1.881174527 | 1.819031101 | 1.794977273 | 1.653677461 | 1.645989387 | 1.770852012 | 1.548827561 |
| ENSMUSG00000030934 | 2.845077293 | 2.595385738 | 2.590211771 | 2.58272922 | 2.540114597 | 2.90579588 | 2.367231288 |
| ENSMUSG00000030935 | 0.91631382 | 0.5 | 0 | 0.701838731 | 0.244650542 | 0 | 0.520678036 |
| ENSMUSG00000030942 | 2.267855219 | 2.098079356 | 2.102347734 | 2.147178659 | 2.078515538 | 2.359835482 | 1.935079929 |
| ENSMUSG00000030945 | 0 | 0.25 | 0 | 0.5 | 0.244650542 | 0 | 0 |
| ENSMUSG00000030946 | 1.516991255 | 1.54248125 | 1.618933358 | 1.692046081 | 1.72341484 | 1.944482672 | 1.659601507 |
| ENSMUSG00000030954 | 0 | 0 | 0 | 0 | 0 | 0 | 0 |
| ENSMUSG00000030956 | 1.418409478 | 1.488549078 | 1.321350555 | 1.201838731 | 1.234113756 | 1.462397998 | 1.329931221 |
| ENSMUSG00000030960 | 1.655740331 | 1.688721876 | 1.642463902 | 1.664552871 | 1.665423072 | 1.792391689 | 1.616675393 |
| ENSMUSG00000030963 | 0 | 0 | 0 | 0 | 0 | 0 | 0 |
| ENSMUSG00000030965 | 2.167941649 | 1.987091808 | 1.972185812 | 2.031030328 | 1.945998439 | 2.222716471 | 1.839824409 |
| ENSMUSG00000030966 | 1.205378646 | 1.214495249 | 1.04248125 | 1.482684334 | 1.401338844 | 1.579783597 | 1.290497981 |
| ENSMUSG00000030967 | 2.266038081 | 2.132845365 | 2.121960008 | 2.155512955 | 2.099395374 | 2.303196057 | 1.954885099 |

| | | | | | | | |
|---|---|---|---|---|---|---|---|
| ENSMUSG00000030968 | 0 | 0 | 0 | 0 | 0 | 0 | 0 |
| ENSMUSG00000030970 | 2.149089517 | 2.135757955 | 2.090485943 | 1.943696765 | 2.007252778 | 2.127104798 | 1.87902385 |
| ENSMUSG00000030972 | 0 | 0 | 0 | 0 | 0 | 0 | 0 |
| ENSMUSG00000030976 | 0 | 0 | 0 | 0 | 0 | 0 | 0 |
| ENSMUSG00000030978 | 3.262546859 | 2.96614327 | 2.931980114 | 3.048843052 | 2.955415567 | 3.450557009 | 2.744721271 |
| ENSMUSG00000030979 | 1.596667477 | 1.445339928 | 1.51109853 | 1.5 | 1.528558853 | 1.681241237 | 1.40946263 |
| ENSMUSG00000030980 | 2.494443473 | 2.295598588 | 2.306000419 | 2.353935442 | 2.28771234 | 2.622214023 | 2.123052075 |
| ENSMUSG00000030981 | 0 | 0.25 | 0 | 0 | 0 | 0.301029996 | 0 |
| ENSMUSG00000030982 | 2.558729299 | 2.331045137 | 2.366901388 | 2.186208459 | 2.235369829 | 2.501059262 | 2.095400649 |
| ENSMUSG00000030983 | 2.20747957 | 2.066696635 | 2.073655187 | 2.101285366 | 2.060462711 | 2.322219295 | 1.912234055 |
| ENSMUSG00000030986 | 2.083999385 | 1.950224975 | 1.959800947 | 2.089388001 | 2.008447216 | 2.276461804 | 1.886704102 |
| ENSMUSG00000030987 | 2.016886209 | 1.899978211 | 1.862802778 | 2.029735268 | 2.031524462 | 2.222716471 | 1.921737195 |
| ENSMUSG00000030990 | 1.964026642 | 1.828220739 | 1.856566189 | 1.858657057 | 1.826953746 | 1.913813852 | 1.687112305 |
| ENSMUSG00000030994 | 0 | 0 | 0 | 0 | 0 | 0 | 0 |
| ENSMUSG00000030996 | 0 | 0 | 0 | 0 | 0 | 0 | 0 |
| ENSMUSG00000031004 | 3.225379103 | 2.953144611 | 2.929204865 | 3.08897541 | 2.990682407 | 3.458335626 | 2.766947881 |
| ENSMUSG00000031007 | 2.968781135 | 2.738003291 | 2.760756821 | 2.596735061 | 2.530194174 | 2.933487288 | 2.336847775 |
| ENSMUSG00000031010 | 3.309098465 | 3.019203899 | 2.996104615 | 3.044899533 | 2.962197183 | 3.407560849 | 2.712392005 |
| ENSMUSG00000031012 | 2.337801042 | 2.180274797 | 2.147178659 | 2.20584181 | 2.134457044 | 2.424881637 | 1.995964412 |
| ENSMUSG00000031015 | 2.410757293 | 2.267865591 | 2.225216702 | 2.092671852 | 2.06759338 | 2.322219295 | 1.966211184 |
| ENSMUSG00000031016 | 2.066740409 | 1.907339155 | 1.930274797 | 1.828220739 | 1.72073075 | 1.929418926 | 1.661510159 |
| ENSMUSG00000031021 | 2.131864492 | 1.899978211 | 1.862802778 | 2.099151195 | 2.028157605 | 2.283301229 | 1.918313719 |
| ENSMUSG00000031022 | 0.578129653 | 0 | 0.25 | 0.25 | 0.244650542 | 0 | 0.448487648 |
| ENSMUSG00000031023 | 1.714367558 | 1.695339928 | 1.714495249 | 1.584962501 | 1.504538967 | 1.763427994 | 1.526507301 |
| ENSMUSG00000031024 | 2.520963022 | 2.348615674 | 2.32991803 | 2.103406982 | 2.125157063 | 2.309630167 | 2.052911423 |
| ENSMUSG00000031026 | 0.960252568 | 1.04248125 | 1.061981878 | 0.396240625 | 0.244650542 | 0 | 0.448487648 |
| ENSMUSG00000031027 | 0 | 0 | 0.396240625 | 0 | 0.387761935 | 0 | 0 |
| ENSMUSG00000031028 | 0.289064826 | 0 | 0 | 0.896240625 | 0.686820904 | 0.602059991 | 0.672731473 |
| ENSMUSG00000031029 | 2.658591846 | 2.399513125 | 2.426839783 | 2.427701608 | 2.398190943 | 2.748188027 | 2.239582276 |
| ENSMUSG00000031059 | 2.363083473 | 2.242166698 | 2.298689214 | 2.193696765 | 2.186353316 | 2.46834733 | 2.15104925 |
| ENSMUSG00000031060 | 2.409468165 | 2.259043403 | 2.279735268 | 2.101285366 | 2.251304582 | 2.416640507 | 2.163786106 |
| ENSMUSG00000031065 | 2.958871665 | 2.701032755 | 2.703845824 | 2.656655413 | 2.642841485 | 3.018284308 | 2.489990739 |
| ENSMUSG00000031066 | 2.254965775 | 2.072504712 | 2.059601185 | 2.149047439 | 2.078515538 | 2.403120521 | 1.971363985 |
| ENSMUSG00000031068 | 2.528794847 | 2.310197833 | 2.280383379 | 2.374462959 | 2.289867804 | 2.57634135 | 2.146406417 |

| | | | | | | | |
|---|---|---|---|---|---|---|---|
| ENSMUSG00000031070 | 1.418409478 | 1.431980114 | 1.482684334 | 0.951838731 | 0.733951626 | 0.954242509 | 1 |
| ENSMUSG00000031072 | 1.789260561 | 1.692046081 | 1.657339155 | 1.744319981 | 1.741652793 | 1.982271233 | 1.647906904 |
| ENSMUSG00000031073 | 0 | 0 | 0 | 0 | 0 | 0 | 0 |
| ENSMUSG00000031074 | 0 | 0 | 0 | 0 | 0 | 0 | 0 |
| ENSMUSG00000031075 | 2.17479713 | 1.950224975 | 1.981203126 | 1.610735874 | 1.632412477 | 1.770852012 | 1.554175045 |
| ENSMUSG00000031077 | 1.587501561 | 1.464495249 | 1.451838731 | 1.610735874 | 1.607318736 | 1.748188027 | 1.554175045 |
| ENSMUSG00000031078 | 3.103760538 | 2.850086449 | 2.853671309 | 2.719704321 | 2.726078673 | 3.071513805 | 2.568555877 |
| ENSMUSG00000031079 | 1.069666964 | 1.061981878 | 1.201838731 | 1.356566189 | 1.358933174 | 1.462397998 | 1.23151414 |
| ENSMUSG00000031085 | 0 | 0 | 0 | 0 | 0 | 0 | 0 |
| ENSMUSG00000031089 | 0 | 0 | 0 | 0 | 0 | 0 | 0 |
| ENSMUSG00000031090 | 1.342375483 | 1.160964047 | 1.17510993 | 0.864857905 | 1.327538614 | 1.342422681 | 1.176815335 |
| ENSMUSG00000031093 | 2.685417684 | 2.483422664 | 2.496104615 | 2.42813175 | 2.368090054 | 2.7084209 | 2.161765391 |
| ENSMUSG00000031095 | 2.917079017 | 2.67510993 | 2.652294685 | 2.733053688 | 2.637464898 | 3.012415375 | 2.441562752 |
| ENSMUSG00000031097 | 0 | 0 | 0.25 | 0 | 0 | 0 | 0 |
| ENSMUSG00000031098 | 0 | 0 | 0 | 0 | 0.244650542 | 0 | 0.224243824 |
| ENSMUSG00000031099 | 1.181541736 | 1.238549078 | 1.04248125 | 1.692046081 | 1.427019704 | 1.707570176 | 1.452143229 |
| ENSMUSG00000031101 | 0 | 0 | 0 | 0 | 0 | 0 | 0 |
| ENSMUSG00000031103 | 2.638209243 | 2.407794264 | 2.402294685 | 2.490000483 | 2.426290456 | 2.781036939 | 2.278288402 |
| ENSMUSG00000031104 | 0.45815691 | 0.25 | 0 | 0.5 | 0.632412477 | 0 | 0.672731473 |
| ENSMUSG00000031105 | 1.827447047 | 1.695339928 | 1.695339928 | 1.774508021 | 1.665423072 | 1.826074803 | 1.572224412 |
| ENSMUSG00000031107 | 2.060824957 | 1.959800947 | 1.976722649 | 1.950224975 | 1.957204337 | 2.181843588 | 1.819456572 |
| ENSMUSG00000031109 | 2.042557845 | 1.903677461 | 1.903677461 | 1.921625132 | 1.846351821 | 2.089905111 | 1.692344475 |
| ENSMUSG00000031111 | 0.867194479 | 0.92510993 | 1.04248125 | 0.830482024 | 0.955822902 | 0.954242509 | 0.803905701 |
| ENSMUSG00000031112 | 1.227926429 | 1.114857905 | 1.160964047 | 0.976722649 | 0.77552387 | 1.204119983 | 0.876096088 |
| ENSMUSG00000031118 | 0 | 0 | 0 | 0 | 0 | 0 | 0 |
| ENSMUSG00000031119 | 1.924654746 | 1.848079356 | 1.79248125 | 1.870953944 | 1.810629166 | 2.071882007 | 1.723650273 |
| ENSMUSG00000031125 | 0 | 0 | 0 | 0 | 0 | 0 | 0 |
| ENSMUSG00000031129 | 1.269664473 | 1.214495249 | 1.27186571 | 1.372963274 | 1.264824954 | 1.505149978 | 1.185218806 |
| ENSMUSG00000031130 | 0 | 0 | 0 | 0 | 0 | 0 | 0 |
| ENSMUSG00000031131 | 0 | 0 | 0 | 0 | 0 | 0 | 0 |
| ENSMUSG00000031132 | 0 | 0 | 0 | 0 | 0 | 0 | 0 |
| ENSMUSG00000031133 | 1.100572389 | 0.830482024 | 1.114857905 | 1.146240625 | 1.057362051 | 1.278753601 | 0.969165684 |
| ENSMUSG00000031134 | 2.381433035 | 2.212155735 | 2.201032755 | 2.322504712 | 2.241584006 | 2.530199698 | 2.111895264 |
| ENSMUSG00000031137 | 0 | 0 | 0 | 0.25 | 0.244650542 | 0 | 0 |

| | | | | | | | |
|---|---|---|---|---|---|---|---|
| ENSMUSG00000031138 | 0 | 0 | 0 | 0 | 0 | 0 | 0 |
| ENSMUSG00000031139 | 0 | 0 | 0 | 0 | 0 | 0.301029996 | 0 |
| ENSMUSG00000031142 | 0 | 0 | 0 | 0 | 0 | 0 | 0 |
| ENSMUSG00000031143 | 0 | 0 | 0 | 0 | 0 | 0 | 0 |
| ENSMUSG00000031144 | 0.45815691 | 0 | 0.25 | 0.580482024 | 0.489301084 | 0.477121255 | 0.448487648 |
| ENSMUSG00000031145 | 0.578129653 | 0.25 | 0 | 0.951838731 | 0.812711509 | 0.698970004 | 0.520678036 |
| ENSMUSG00000031146 | 2.299278375 | 2.176839783 | 2.220660762 | 2.120953944 | 2.116484486 | 2.376576957 | 2.02568932 |
| ENSMUSG00000031147 | 0.45815691 | 0.396240625 | 0.5 | 0.396240625 | 0.244650542 | 0 | 0.448487648 |
| ENSMUSG00000031148 | 2.531694272 | 2.323655187 | 2.337182039 | 2.407794264 | 2.347256716 | 2.686636269 | 2.209543816 |
| ENSMUSG00000031149 | 2.412042447 | 2.203445298 | 2.235628626 | 2.254897682 | 2.205284995 | 2.567026366 | 2.086315375 |
| ENSMUSG00000031150 | 1.816888701 | 1.741446071 | 1.76109853 | 1.735628626 | 1.802174723 | 1.959041392 | 1.796468236 |
| ENSMUSG00000031153 | 2.35433422 | 2.17510993 | 2.178561379 | 2.200224975 | 2.258423266 | 2.475671188 | 2.13295179 |
| ENSMUSG00000031154 | 2.499676068 | 2.307204673 | 2.324802005 | 2.31611065 | 2.319727371 | 2.553883027 | 2.189340078 |
| ENSMUSG00000031155 | 1.800532393 | 1.701838731 | 1.606566189 | 1.784887838 | 1.78010778 | 1.954242509 | 1.705896842 |
| ENSMUSG00000031156 | 2.054824387 | 1.987091808 | 2.01109853 | 1.769203899 | 1.875328588 | 2.004321374 | 1.771720588 |
| ENSMUSG00000031157 | 2.337801042 | 2.180274797 | 2.205044741 | 2.259729747 | 2.230318492 | 2.519827994 | 2.143416814 |
| ENSMUSG00000031158 | 2.078325387 | 2.058404919 | 2.133818844 | 1.888647213 | 1.990091809 | 2.127104798 | 1.941550454 |
| ENSMUSG00000031160 | 0 | 0 | 0 | 0 | 0 | 0 | 0 |
| ENSMUSG00000031161 | 2.45930418 | 2.323655187 | 2.334405475 | 2.359177886 | 2.390196125 | 2.600972896 | 2.254834987 |
| ENSMUSG00000031162 | 0 | 0 | 0 | 0.396240625 | 0 | 0 | 0 |
| ENSMUSG00000031163 | 0 | 0 | 0 | 0 | 0 | 0 | 0 |
| ENSMUSG00000031165 | 0 | 0.25 | 0.396240625 | 0.396240625 | 0.244650542 | 0.301029996 | 0.629532004 |
| ENSMUSG00000031166 | 2.299278375 | 2.20584181 | 2.210587586 | 2.100219859 | 2.088159748 | 2.324282455 | 1.95718226 |
| ENSMUSG00000031167 | 3.519883559 | 3.17510993 | 3.173970188 | 3.247082412 | 3.139670239 | 3.667266119 | 2.896640914 |
| ENSMUSG00000031168 | 2.305951036 | 2.098079356 | 2.12694866 | 2.203445298 | 2.198391101 | 2.437750563 | 2.039586611 |
| ENSMUSG00000031169 | 1.516991255 | 1.451838731 | 1.339388001 | 1.29248125 | 1.478764298 | 1.431363764 | 1.329931221 |
| ENSMUSG00000031170 | 0 | 0 | 0 | 0 | 0 | 0.301029996 | 0.224243824 |
| ENSMUSG00000031171 | 2.103271637 | 2.038704527 | 2.044977273 | 2.038704527 | 2.06759338 | 2.252853031 | 1.915722126 |
| ENSMUSG00000031173 | 0 | 0 | 0 | 0 | 0 | 0 | 0 |
| ENSMUSG00000031174 | 1.578129653 | 1.418106335 | 1.321350555 | 1.634789703 | 1.499532413 | 1.602059991 | 1.401059159 |
| ENSMUSG00000031176 | 2.675303226 | 2.467976309 | 2.482684334 | 2.434100483 | 2.347256716 | 2.668385917 | 2.158915016 |
| ENSMUSG00000031179 | 0 | 0 | 0 | 0 | 0 | 0 | 0 |
| ENSMUSG00000031181 | 0 | 0 | 0 | 0 | 0 | 0 | 0 |
| ENSMUSG00000031182 | 0 | 0 | 0 | 0 | 0 | 0 | 0 |

| | | | | | | | |
|---|---|---|---|---|---|---|---|
| ENSMUSG00000031189 | 1.54867984 | 1.418106335 | 1.42510993 | 1.130890489 | 1.057362051 | 1 | 0.935079929 |
| ENSMUSG00000031194 | 0 | 0 | 0 | 0 | 0 | 0 | 0 |
| ENSMUSG00000031196 | 1.505869732 | 1.476722649 | 1.464495249 | 1.388647213 | 1.254881871 | 1.431363764 | 1.089372236 |
| ENSMUSG00000031197 | 2.65645504 | 2.459012589 | 2.457430684 | 2.643175057 | 2.5461437 | 2.922206277 | 2.363123786 |
| ENSMUSG00000031198 | 2.513987407 | 2.33993739 | 2.354990044 | 2.435366747 | 2.297835549 | 2.687528961 | 2.14725554 |
| ENSMUSG00000031200 | 1.783508299 | 1.614857905 | 1.664552871 | 1.76109853 | 1.733951626 | 1.806179974 | 1.554175045 |
| ENSMUSG00000031201 | 2.364524 | 2.147178659 | 2.170748646 | 2.28296424 | 2.234113756 | 2.484299839 | 2.03601839 |
| ENSMUSG00000031202 | 1.671187741 | 1.606566189 | 1.532320754 | 1.566696635 | 1.489301084 | 1.633468456 | 1.369798885 |
| ENSMUSG00000031204 | 0 | 0 | 0 | 0 | 0 | 0 | 0 |
| ENSMUSG00000031207 | 3.846931597 | 3.475051607 | 3.463692751 | 3.193181737 | 3.14542665 | 3.633973156 | 2.916941975 |
| ENSMUSG00000031209 | 0.45815691 | 0 | 0 | 0 | 0 | 0 | 0 |
| ENSMUSG00000031210 | 0 | 0.5 | 0 | 0 | 0.244650542 | 0.602059991 | 0 |
| ENSMUSG00000031212 | 0.45815691 | 0.5 | 0.580482024 | 0.25 | 0 | 0 | 0 |
| ENSMUSG00000031214 | 1.822201287 | 1.71759118 | 1.741446071 | 1.695339928 | 1.645989387 | 1.892094603 | 1.591689859 |
| ENSMUSG00000031216 | 1.156259305 | 1.02186571 | 1.238549078 | 1.458222504 | 1.373641807 | 1.477121255 | 1.444790289 |
| ENSMUSG00000031217 | 1.558729299 | 1.681980114 | 1.642463902 | 1.464495249 | 1.618292349 | 1.672097858 | 1.635773653 |
| ENSMUSG00000031220 | 0.578129653 | 0.92510993 | 0.646240625 | 0.701838731 | 0.632412477 | 0.602059991 | 0.579661877 |
| ENSMUSG00000031221 | 2.483777101 | 2.284247778 | 2.27053726 | 2.310793496 | 2.280614762 | 2.588831726 | 2.089372236 |
| ENSMUSG00000031224 | 0.289064826 | 0 | 0.25 | 0.396240625 | 0.244650542 | 0.477121255 | 0.579661877 |
| ENSMUSG00000031226 | 2.508932407 | 2.31611065 | 2.310793496 | 2.338286706 | 2.269408865 | 2.621176282 | 2.098373247 |
| ENSMUSG00000031227 | 2.241665209 | 2.137686696 | 2.167221246 | 2.181128463 | 2.112954895 | 2.390935107 | 2.013099581 |
| ENSMUSG00000031229 | 3.182149508 | 2.887566452 | 2.893293446 | 2.984343455 | 2.890346003 | 3.31993844 | 2.641621316 |
| ENSMUSG00000031230 | 0 | 0 | 0 | 0 | 0 | 0 | 0 |
| ENSMUSG00000031231 | 2.658591846 | 2.461764337 | 2.465271726 | 2.478596283 | 2.432800077 | 2.77815125 | 2.289406866 |
| ENSMUSG00000031232 | 2.896712826 | 2.642463902 | 2.621708755 | 2.644828984 | 2.547699718 | 2.944975908 | 2.350026003 |
| ENSMUSG00000031233 | 0 | 0 | 0 | 0 | 0 | 0 | 0 |
| ENSMUSG00000031239 | 1.822201287 | 1.695339928 | 1.685366747 | 1.720660762 | 1.72341484 | 1.982271233 | 1.633706455 |
| ENSMUSG00000031241 | 0 | 0 | 0 | 0 | 0 | 0 | 0 |
| ENSMUSG00000031242 | 2.330063669 | 2.143411797 | 2.124961472 | 2.200224975 | 2.122577592 | 2.421603927 | 1.975716277 |
| ENSMUSG00000031245 | 2.45930418 | 2.277783918 | 2.359697963 | 2.365881093 | 2.282813873 | 2.611723308 | 2.16858494 |
| ENSMUSG00000031246 | 3.400810563 | 3.117219638 | 3.073152306 | 2.745926548 | 2.657550118 | 3.074084689 | 2.462548643 |
| ENSMUSG00000031250 | 0 | 0 | 0 | 0 | 0 | 0 | 0 |
| ENSMUSG00000031253 | 0 | 0 | 0.25 | 0 | 0.244650542 | 0.477121255 | 0.224243824 |
| ENSMUSG00000031255 | 1.482695304 | 1.356566189 | 1.214495249 | 1.78743678 | 1.674752852 | 1.86923172 | 1.559435575 |

| | | | | | | | |
|---|---|---|---|---|---|---|---|
| ENSMUSG00000031256 | 2.496544396 | 2.333847589 | 2.30357978 | 2.550530956 | 2.506580024 | 2.848189117 | 2.33471704 |
| ENSMUSG00000031257 | 0 | 0 | 0.396240625 | 0.396240625 | 0.387761935 | 0 | 0.224243824 |
| ENSMUSG00000031258 | 0.578129653 | 0.25 | 0.646240625 | 0.646240625 | 0.632412477 | 0 | 0.672731473 |
| ENSMUSG00000031262 | 2.396352858 | 2.228969845 | 2.193696765 | 2.495748394 | 2.387761935 | 2.755112266 | 2.223909441 |
| ENSMUSG00000031264 | 0 | 0.25 | 0.25 | 0 | 0 | 0 | 0.355418052 |
| ENSMUSG00000031266 | 1.852729478 | 1.784887838 | 1.819031101 | 1.814346961 | 1.744182955 | 2.064457989 | 1.700880998 |
| ENSMUSG00000031270 | 0 | 0 | 0 | 0 | 0 | 0 | 0 |
| ENSMUSG00000031271 | 0 | 0 | 0 | 0 | 0 | 0 | 0 |
| ENSMUSG00000031273 | 1.129344651 | 1.17510993 | 1.282320754 | 0.951838731 | 0.978602168 | 0.84509804 | 1.028149525 |
| ENSMUSG00000031274 | 3.015379043 | 2.785526764 | 2.761269263 | 2.762802235 | 2.763375569 | 3.147676324 | 2.556576804 |
| ENSMUSG00000031278 | 2.292497211 | 2.103406982 | 2.118933358 | 2.483422664 | 2.384490012 | 2.748962861 | 2.214058866 |
| ENSMUSG00000031283 | 1.482695304 | 1.438721876 | 1.445339928 | 1.451838731 | 1.27449559 | 1.414973348 | 1.21680636 |
| ENSMUSG00000031284 | 2.515992375 | 2.386723615 | 2.338837774 | 2.383818844 | 2.267124351 | 2.622214023 | 2.109521699 |
| ENSMUSG00000031285 | 0 | 0 | 0 | 0.25 | 0 | 0.301029996 | 0 |
| ENSMUSG00000031286 | 1.036286563 | 1.02186571 | 1.080482024 | 1.201838731 | 1.136121934 | 1 | 1.110947927 |
| ENSMUSG00000031289 | 0 | 0.25 | 0.396240625 | 0 | 0 | 0 | 0 |
| ENSMUSG00000031290 | 2.53552908 | 2.313757142 | 2.27053726 | 2.349151195 | 2.27895639 | 2.568201724 | 2.054044578 |
| ENSMUSG00000031292 | 1.734388958 | 1.606566189 | 1.668106335 | 2.05357978 | 1.978602168 | 2.269512944 | 1.821784031 |
| ENSMUSG00000031293 | 0 | 0 | 0 | 0.25 | 0.387761935 | 0 | 0 |
| ENSMUSG00000031294 | 0 | 0.25 | 0 | 0.25 | 0.489301084 | 0.301029996 | 0.224243824 |
| ENSMUSG00000031295 | 2.119110214 | 1.886723615 | 1.942046081 | 2.08272922 | 2.056322416 | 2.31386722 | 1.925124822 |
| ENSMUSG00000031297 | 1.45815691 | 1.348079356 | 1.302363341 | 1.160964047 | 1.176121988 | 0.954242509 | 1.078019652 |
| ENSMUSG00000031298 | 1.389637216 | 1.42510993 | 1.311981878 | 1.52186571 | 1.533172725 | 1.672097858 | 1.374453863 |
| ENSMUSG00000031299 | 2.719310222 | 2.490000483 | 2.490000483 | 2.624961472 | 2.56239779 | 2.938019097 | 2.39559556 |
| ENSMUSG00000031302 | 0.289064826 | 0.25 | 0.5 | 0.5 | 0.489301084 | 0 | 0.448487648 |
| ENSMUSG00000031303 | 0.671187741 | 0.396240625 | 0.580482024 | 1.238549078 | 1.176121988 | 1.176091259 | 1.100339912 |
| ENSMUSG00000031304 | 0 | 0 | 0 | 0.580482024 | 0.733951626 | 0.602059991 | 0.520678036 |
| ENSMUSG00000031309 | 2.726616086 | 2.497880462 | 2.486726569 | 2.632845365 | 2.546403514 | 2.913813852 | 2.352283622 |
| ENSMUSG00000031310 | 2.507914013 | 2.336628433 | 2.314346961 | 2.263820609 | 2.323654601 | 2.545307116 | 2.137790317 |
| ENSMUSG00000031311 | 3.407114039 | 3.074515642 | 3.04745614 | 3.114281083 | 3.032770615 | 3.525692525 | 2.837824308 |
| ENSMUSG00000031312 | 0.960252568 | 0.580482024 | 0.75 | 0.396240625 | 0.846351821 | 0.301029996 | 0.710836105 |
| ENSMUSG00000031314 | 2.917079017 | 2.632113528 | 2.631135429 | 2.780545225 | 2.66113346 | 3.08350262 | 2.479382724 |
| ENSMUSG00000031320 | 3.559712867 | 3.206140257 | 3.188669691 | 3.259686889 | 3.159785535 | 3.718667735 | 2.923435442 |
| ENSMUSG00000031323 | 0 | 0 | 0 | 0 | 0 | 0.301029996 | 0.224243824 |

| | | | | | | | |
|---|---|---|---|---|---|---|---|
| ENSMUSG00000031325 | 0 | 0.646240625 | 0.646240625 | 0.25 | 0 | 0 | 0.224243824 |
| ENSMUSG00000031326 | 0 | 0 | 0 | 0 | 0 | 0 | 0 |
| ENSMUSG00000031327 | 2.051791411 | 1.921625132 | 1.880890489 | 1.852347734 | 1.832885921 | 2 | 1.655750087 |
| ENSMUSG00000031328 | 4.309329158 | 3.86977138 | 3.859982064 | 3.458148344 | 3.467635992 | 3.975753389 | 3.211493331 |
| ENSMUSG00000031329 | 0.45815691 | 0.25 | 0.646240625 | 0.25 | 0.387761935 | 0 | 0.224243824 |
| ENSMUSG00000031330 | 0 | 0 | 0 | 0 | 0 | 0 | 0 |
| ENSMUSG00000031333 | 2.414600936 | 2.206637122 | 2.222945926 | 2.386723615 | 2.333760374 | 2.64738297 | 2.162170545 |
| ENSMUSG00000031337 | 1.693336259 | 1.52186571 | 1.575945187 | 1.685366747 | 1.674752852 | 1.755874856 | 1.508694612 |
| ENSMUSG00000031340 | 2.060824957 | 1.864857905 | 1.83272922 | 1.802363341 | 1.784632921 | 1.832508913 | 1.612273755 |
| ENSMUSG00000031342 | 0 | 0 | 0.25 | 0.79248125 | 0.632412477 | 0.477121255 | 0.579661877 |
| ENSMUSG00000031343 | 1.156259305 | 1.17510993 | 1.061981878 | 0.25 | 0.632412477 | 0 | 0.355418052 |
| ENSMUSG00000031344 | 0 | 0 | 0 | 0 | 0.244650542 | 0.301029996 | 0 |
| ENSMUSG00000031347 | 2.205378646 | 2.092671852 | 2.070192693 | 2.294977273 | 2.193483986 | 2.489958479 | 2.050633136 |
| ENSMUSG00000031349 | 2.364524 | 2.201032755 | 2.237091808 | 2.389605178 | 2.324629618 | 2.6599162 | 2.179128924 |
| ENSMUSG00000031351 | 1.623015126 | 1.464495249 | 1.431980114 | 1.83272922 | 1.751666404 | 1.944482672 | 1.633706455 |
| ENSMUSG00000031352 | 2.23778581 | 2.109697963 | 2.095926073 | 2.214495249 | 2.189222894 | 2.436162647 | 1.997315209 |
| ENSMUSG00000031353 | 3.168360356 | 2.897295742 | 2.893648382 | 2.99929487 | 2.902295427 | 3.38309693 | 2.721573465 |
| ENSMUSG00000031354 | 0 | 0 | 0 | 0 | 0 | 0 | 0 |
| ENSMUSG00000031355 | 0.289064826 | 0.396240625 | 0.5 | 0.75 | 0.846351821 | 0.903089987 | 0.853775828 |
| ENSMUSG00000031357 | 2.339331437 | 2.153677461 | 2.172499493 | 2.451435968 | 2.349985771 | 2.706717782 | 2.202829437 |
| ENSMUSG00000031358 | 2.285603957 | 2.118933358 | 2.113831805 | 2.286164561 | 2.193483986 | 2.514547753 | 2.064068234 |
| ENSMUSG00000031360 | 2.282114126 | 2.117918804 | 2.14151351 | 2.211372513 | 2.073592779 | 2.404833717 | 1.962479708 |
| ENSMUSG00000031362 | 0 | 0 | 0 | 0 | 0 | 0 | 0 |
| ENSMUSG00000031364 | 0 | 0.25 | 0 | 0 | 0 | 0.301029996 | 0 |
| ENSMUSG00000031365 | 2.747795766 | 2.512462137 | 2.492166698 | 2.457430684 | 2.420030246 | 2.710963119 | 2.240219117 |
| ENSMUSG00000031367 | 2.299278375 | 2.107613138 | 2.092671852 | 2.226722649 | 2.176121988 | 2.369215857 | 1.963229454 |
| ENSMUSG00000031370 | 2.337801042 | 2.150906586 | 2.123963757 | 2.325374049 | 2.25190332 | 2.539076099 | 2.11845007 |
| ENSMUSG00000031371 | 2.675985209 | 2.465271726 | 2.45584181 | 2.263142013 | 2.186353316 | 2.531478917 | 2.064616101 |
| ENSMUSG00000031372 | 0 | 0 | 0 | 0 | 0 | 0 | 0 |
| ENSMUSG00000031373 | 2.153883046 | 2.029735268 | 1.984159485 | 2.099151195 | 2.06759338 | 2.303196057 | 1.914853627 |
| ENSMUSG00000031374 | 0 | 0 | 0 | 0 | 0.244650542 | 0 | 0 |
| ENSMUSG00000031375 | 2.124258828 | 1.955044741 | 1.854463129 | 2.928239205 | 2.891129262 | 3.350248018 | 2.693521895 |
| ENSMUSG00000031376 | 0.289064826 | 0.580482024 | 0.580482024 | 0.25 | 0.489301084 | 0.477121255 | 0.629532004 |
| ENSMUSG00000031377 | 0 | 0 | 0 | 0 | 0 | 0 | 0 |

| | | | | | | | |
|---|---|---|---|---|---|---|---|
| ENSMUSG00000031378 | 1.971473847 | 1.837182039 | 1.886723615 | 1.80479213 | 1.932951348 | 2.056904851 | 1.86715039 |
| ENSMUSG00000031379 | 1.596667477 | 1.62694866 | 1.566696635 | 1.678561379 | 1.53772706 | 1.681241237 | 1.437266334 |
| ENSMUSG00000031380 | 1.269664473 | 1.29248125 | 1.302363341 | 1.664552871 | 1.555378544 | 1.792391689 | 1.476637173 |
| ENSMUSG00000031381 | 2.078325387 | 1.882845365 | 1.909156155 | 2.046218836 | 1.997548906 | 2.274157849 | 1.847411384 |
| ENSMUSG00000031382 | 0 | 0 | 0 | 0 | 0 | 0 | 0 |
| ENSMUSG00000031383 | 2.183765914 | 2.08272922 | 2.072504712 | 1.994319981 | 2.120847411 | 2.146128036 | 2.041943747 |
| ENSMUSG00000031384 | 0 | 0 | 0 | 0 | 0 | 0 | 0 |
| ENSMUSG00000031385 | 0 | 0.580482024 | 0.5 | 0 | 0.387761935 | 0 | 0.224243824 |
| ENSMUSG00000031386 | 3.155396777 | 2.908589325 | 2.934100483 | 2.859827865 | 2.881311898 | 3.254064453 | 2.743783545 |
| ENSMUSG00000031387 | 2.253091469 | 2.110735874 | 2.21759118 | 1.188721876 | 1.283908311 | 1.322219295 | 1.150210039 |
| ENSMUSG00000031388 | 2.201144779 | 2.064346961 | 2.051142786 | 1.866901388 | 2.048957937 | 2.167317335 | 1.982842456 |
| ENSMUSG00000031389 | 0.671187741 | 0.580482024 | 0.396240625 | 0.646240625 | 0.733951626 | 0.602059991 | 0.672731473 |
| ENSMUSG00000031390 | 0 | 0 | 0 | 0.25 | 0.244650542 | 0.301029996 | 0 |
| ENSMUSG00000031391 | 0.960252568 | 1.080482024 | 1.02186571 | 1.51109853 | 1.584291694 | 1.544068044 | 1.401059159 |
| ENSMUSG00000031392 | 2.635214384 | 2.469512728 | 2.448603967 | 2.434523065 | 2.419286397 | 2.741151599 | 2.298300395 |
| ENSMUSG00000031393 | 2.788740894 | 2.548073204 | 2.50835575 | 2.55357978 | 2.493851506 | 2.84135947 | 2.312345877 |
| ENSMUSG00000031394 | 0 | 0 | 0 | 0.25 | 0.489301084 | 0 | 0.224243824 |
| ENSMUSG00000031397 | 0.45815691 | 0.25 | 0.5 | 0.580482024 | 0.387761935 | 0.477121255 | 0 |
| ENSMUSG00000031398 | 2.496544396 | 2.343216265 | 2.352877747 | 2.148114259 | 2.250704826 | 2.387389826 | 2.085289968 |
| ENSMUSG00000031399 | 1.952600383 | 1.854463129 | 1.852347734 | 1.890560606 | 1.919403394 | 2.064457989 | 1.832055226 |
| ENSMUSG00000031400 | 2.684750966 | 2.437048212 | 2.478596283 | 2.606566189 | 2.573394197 | 2.974971994 | 2.39183486 |
| ENSMUSG00000031402 | 2.545896521 | 2.342671852 | 2.342671852 | 2.492166698 | 2.406008674 | 2.760422483 | 2.233468922 |
| ENSMUSG00000031403 | 2.755750303 | 2.55357978 | 2.541226732 | 2.667885267 | 2.573153518 | 2.971739591 | 2.367016399 |
| ENSMUSG00000031409 | 0 | 0 | 0 | 0 | 0 | 0 | 0 |
| ENSMUSG00000031410 | 0 | 0 | 0 | 0 | 0 | 0 | 0 |
| ENSMUSG00000031411 | 0 | 0 | 0 | 0 | 0 | 0 | 0 |
| ENSMUSG00000031422 | 3.224648428 | 2.935577359 | 2.904252057 | 3.023354391 | 2.953207795 | 3.427810573 | 2.762926662 |
| ENSMUSG00000031424 | 0 | 0 | 0 | 0 | 0 | 0 | 0 |
| ENSMUSG00000031425 | 0.811507563 | 0.79248125 | 0.75 | 1.02186571 | 1.121713561 | 1 | 0.876096088 |
| ENSMUSG00000031428 | 0.960252568 | 0.92510993 | 0.864857905 | 1.226722649 | 1.310728002 | 1.322219295 | 1.252393349 |
| ENSMUSG00000031429 | 2.042557845 | 1.856566189 | 1.856566189 | 2.202642909 | 2.124299332 | 2.450249108 | 1.984249047 |
| ENSMUSG00000031430 | 0 | 0 | 0 | 0 | 0 | 0 | 0 |
| ENSMUSG00000031431 | 1.753493018 | 1.660964047 | 1.618933358 | 1.505591953 | 1.595992969 | 1.740362689 | 1.473249546 |
| ENSMUSG00000031432 | 2.405576699 | 2.190387808 | 2.192872367 | 2.473325383 | 2.399768403 | 2.719331287 | 2.213368331 |

| | | | | | | | |
|---|---|---|---|---|---|---|---|
| ENSMUSG00000031433 | 1.759671389 | 1.714495249 | 1.668106335 | 2.009729747 | 1.90849439 | 2.193124598 | 1.755055163 |
| ENSMUSG00000031434 | 2.016886209 | 1.864857905 | 1.845926073 | 2.006976499 | 1.917865457 | 2.190331698 | 1.821784031 |
| ENSMUSG00000031438 | 1.789260561 | 1.630890489 | 1.732684334 | 1.75835575 | 1.635855979 | 1.838849091 | 1.562034107 |
| ENSMUSG00000031441 | 2.910529644 | 2.713911792 | 2.691010554 | 2.738549078 | 2.737898704 | 3.101747074 | 2.58287655 |
| ENSMUSG00000031442 | 1.714367558 | 1.720660762 | 1.75 | 1.238549078 | 1.467903253 | 1.477121255 | 1.505628092 |
| ENSMUSG00000031443 | 0 | 0 | 0 | 0 | 0 | 0 | 0 |
| ENSMUSG00000031444 | 0 | 0.396240625 | 0 | 0.396240625 | 0 | 0.301029996 | 0.224243824 |
| ENSMUSG00000031445 | 0.747221736 | 0.830482024 | 0.5 | 1.04248125 | 0.733951626 | 0.903089987 | 0.935079929 |
| ENSMUSG00000031446 | 3.068605137 | 2.804034889 | 2.80205859 | 2.708024712 | 2.687518101 | 3.062205809 | 2.484954224 |
| ENSMUSG00000031447 | 3.90168209 | 3.541952532 | 3.555566656 | 3.195134942 | 3.140925453 | 3.661244609 | 2.945745437 |
| ENSMUSG00000031448 | 0 | 0 | 0 | 0 | 0 | 0 | 0 |
| ENSMUSG00000031449 | 0.289064826 | 0 | 0 | 0 | 0 | 0 | 0.224243824 |
| ENSMUSG00000031450 | 0.578129653 | 0.701838731 | 0.646240625 | 0.580482024 | 0.877063019 | 0.77815125 | 0.803905701 |
| ENSMUSG00000031451 | 1.623015126 | 1.602347734 | 1.678561379 | 1.664552871 | 1.744182955 | 1.812913357 | 1.659601507 |
| ENSMUSG00000031452 | 0 | 0 | 0 | 0 | 0.489301084 | 0 | 0 |
| ENSMUSG00000031453 | 2.886966737 | 2.64881437 | 2.657339155 | 2.366391601 | 2.415166968 | 2.709269961 | 2.281943983 |
| ENSMUSG00000031458 | 2.192545868 | 2.048689214 | 2.153677461 | 1.83272922 | 1.846351821 | 2.075546961 | 1.726776051 |
| ENSMUSG00000031461 | 0.867194479 | 0.79248125 | 0.701838731 | 0.79248125 | 0.978602168 | 0.954242509 | 0.803905701 |
| ENSMUSG00000031465 | 1.389637216 | 1.17510993 | 1.238549078 | 0.830482024 | 0.978602168 | 1 | 0.935079929 |
| ENSMUSG00000031467 | 2.22592627 | 2.073655187 | 2.069031101 | 2.188721876 | 2.133621644 | 2.342422681 | 1.968429585 |
| ENSMUSG00000031471 | 0 | 0 | 0 | 0 | 0 | 0 | 0 |
| ENSMUSG00000031478 | 1.587501561 | 1.580482024 | 1.488549078 | 1.52186571 | 1.563883923 | 1.69019608 | 1.505628092 |
| ENSMUSG00000031479 | 2.496544396 | 2.334962501 | 2.324229052 | 2.449009902 | 2.364629672 | 2.733999287 | 2.249624299 |
| ENSMUSG00000031480 | 1.100572389 | 1.061981878 | 1.17510993 | 0.896240625 | 1.074582839 | 1.041392685 | 1.028149525 |
| ENSMUSG00000031481 | 0.289064826 | 0 | 0.396240625 | 0 | 0 | 0.301029996 | 0 |
| ENSMUSG00000031482 | 2.072573125 | 1.903677461 | 1.918106335 | 1.966046536 | 1.953043706 | 2.130333768 | 1.787569411 |
| ENSMUSG00000031483 | 2.779274431 | 2.547764804 | 2.523849256 | 2.47934302 | 2.434230499 | 2.813580989 | 2.291856721 |
| ENSMUSG00000031485 | 2.599951214 | 2.439972046 | 2.429419106 | 2.30175358 | 2.27729019 | 2.59439255 | 2.148102439 |
| ENSMUSG00000031486 | 2.497590902 | 2.358657057 | 2.343216265 | 2.142463902 | 2.201164838 | 2.383815366 | 2.122594809 |
| ENSMUSG00000031487 | 1.806056082 | 1.741446071 | 1.657339155 | 1.602347734 | 1.655839964 | 1.785329835 | 1.69749337 |
| ENSMUSG00000031488 | 1.894701088 | 1.769203899 | 1.78743678 | 1.970660762 | 1.938809675 | 2.209515015 | 1.879993912 |
| ENSMUSG00000031489 | 0 | 0.396240625 | 0.25 | 0.646240625 | 0.733951626 | 0.301029996 | 0.448487648 |
| ENSMUSG00000031490 | 2.370236862 | 2.206637122 | 2.181980114 | 1.843759858 | 1.996316946 | 2.100370545 | 1.984950056 |
| ENSMUSG00000031491 | 0 | 0 | 0 | 0 | 0 | 0 | 0 |

| | | | | | | | |
|---|---|---|---|---|---|---|---|
| ENSMUSG00000031492 | 0 | 0 | 0 | 0 | 0 | 0 | 0 |
| ENSMUSG00000031493 | 0.578129653 | 0.75 | 0.864857905 | 0.25 | 0.632412477 | 0.477121255 | 0.672731473 |
| ENSMUSG00000031494 | 0.289064826 | 0 | 0 | 0 | 0 | 0 | 0 |
| ENSMUSG00000031495 | 0 | 0 | 0 | 0 | 0 | 0 | 0 |
| ENSMUSG00000031497 | 0.289064826 | 0.396240625 | 0 | 0.25 | 0.387761935 | 0 | 0 |
| ENSMUSG00000031502 | 3.751481977 | 3.48900327 | 3.514455635 | 3.008441779 | 3.154538731 | 3.514547753 | 3.006764893 |
| ENSMUSG00000031503 | 3.38243612 | 3.181820582 | 3.24293079 | 2.747525991 | 2.885808215 | 3.180699201 | 2.797606174 |
| ENSMUSG00000031504 | 1.686083339 | 1.505591953 | 1.598079356 | 1.29248125 | 1.302012594 | 1.278753601 | 1.365075947 |
| ENSMUSG00000031505 | 2.280358197 | 2.170748646 | 2.197790722 | 1.923371739 | 1.981187939 | 2.100370545 | 1.914853627 |
| ENSMUSG00000031506 | 0.45815691 | 0 | 0.580482024 | 0.580482024 | 0.387761935 | 0.477121255 | 0.629532004 |
| ENSMUSG00000031508 | 2.652148299 | 2.427701608 | 2.426839783 | 2.263142013 | 2.271112598 | 2.460897843 | 2.121219121 |
| ENSMUSG00000031509 | 0 | 0 | 0.25 | 0.25 | 0 | 0 | 0 |
| ENSMUSG00000031510 | 0 | 0 | 0 | 0 | 0 | 0 | 0 |
| ENSMUSG00000031511 | 2.613625632 | 2.418106335 | 2.414998973 | 2.219895812 | 2.237868644 | 2.481442629 | 2.135599953 |
| ENSMUSG00000031512 | 0 | 0 | 0.25 | 0 | 0 | 0 | 0 |
| ENSMUSG00000031513 | 2.373064211 | 2.229715809 | 2.232684334 | 2.229715809 | 2.187072901 | 2.523746467 | 2.077494038 |
| ENSMUSG00000031516 | 2.131864492 | 1.998588359 | 2.00835575 | 2.038704527 | 2.076554666 | 2.301029996 | 2.009225098 |
| ENSMUSG00000031517 | 0 | 0 | 0 | 0.25 | 0 | 0.477121255 | 0.355418052 |
| ENSMUSG00000031518 | 0.289064826 | 0 | 0 | 0 | 0 | 0 | 0 |
| ENSMUSG00000031519 | 0.671187741 | 0.5 | 0.701838731 | 0.580482024 | 0.632412477 | 0.477121255 | 0.629532004 |
| ENSMUSG00000031520 | 1.747221736 | 1.657339155 | 1.708222504 | 1.681980114 | 1.625423019 | 1.681241237 | 1.535058868 |
| ENSMUSG00000031521 | 2.323768721 | 2.166333979 | 2.201838731 | 1.899978211 | 1.806427258 | 2.096910013 | 1.778418869 |
| ENSMUSG00000031523 | 2.644135203 | 2.454645544 | 2.439972046 | 2.462155735 | 2.455355315 | 2.769377326 | 2.325318165 |
| ENSMUSG00000031527 | 2.766308649 | 2.517530236 | 2.492166698 | 2.640083209 | 2.560656022 | 2.923244019 | 2.368303597 |
| ENSMUSG00000031529 | 2.81052167 | 2.602612838 | 2.556903986 | 2.568740118 | 2.506870642 | 2.889301703 | 2.343157052 |
| ENSMUSG00000031530 | 1.867194479 | 1.747171172 | 1.837182039 | 1.860735874 | 1.941702774 | 2.033423755 | 1.893277937 |
| ENSMUSG00000031532 | 2.472830741 | 2.313166358 | 2.285526764 | 2.342126615 | 2.329464709 | 2.591064607 | 2.190083792 |
| ENSMUSG00000031533 | 2.179305632 | 2.057204673 | 2.031030328 | 2.08272922 | 1.978602168 | 2.301029996 | 1.93179548 |
| ENSMUSG00000031534 | 1.707474304 | 1.593759858 | 1.54745614 | 1.482684334 | 1.439191811 | 1.643452676 | 1.355418052 |
| ENSMUSG00000031535 | 0 | 0 | 0 | 0 | 0 | 0 | 0 |
| ENSMUSG00000031536 | 2.039433992 | 1.918106335 | 1.898114259 | 2.025821952 | 2.015530186 | 2.235528447 | 1.873141588 |
| ENSMUSG00000031537 | 2.481610694 | 2.242885888 | 2.269871196 | 2.293731421 | 2.317250476 | 2.568201724 | 2.163786106 |
| ENSMUSG00000031538 | 2.514991096 | 2.336073977 | 2.33993739 | 2.337734795 | 2.317747248 | 2.636487896 | 2.183707049 |
| ENSMUSG00000031539 | 2.036286563 | 1.779735268 | 1.763820609 | 1.899978211 | 1.826953746 | 2.06069784 | 1.729871916 |

| | | | | | | | |
|---|---|---|---|---|---|---|---|
| ENSMUSG00000031540 | 2.870199357 | 2.643411797 | 2.627196291 | 2.700022747 | 2.628935049 | 3.053078443 | 2.454776126 |
| ENSMUSG00000031543 | 1.54867984 | 1.584962501 | 1.584962501 | 1.201838731 | 1.351342427 | 1.230448921 | 1.379042812 |
| ENSMUSG00000031545 | 2.893084681 | 2.651832578 | 2.665889526 | 2.616391601 | 2.58568862 | 2.921166051 | 2.413955973 |
| ENSMUSG00000031546 | 2.435736758 | 2.229715809 | 2.26109853 | 2.26584877 | 2.233484039 | 2.557507202 | 2.117056606 |
| ENSMUSG00000031548 | 1.418409478 | 1.201838731 | 1.238549078 | 1.96759118 | 1.855665188 | 2.170261715 | 1.787569411 |
| ENSMUSG00000031549 | 0.578129653 | 0.25 | 0 | 0.25 | 0 | 0.301029996 | 0 |
| ENSMUSG00000031551 | 0 | 0 | 0.396240625 | 0 | 0 | 0 | 0.224243824 |
| ENSMUSG00000031552 | 0.289064826 | 0 | 0 | 0 | 0 | 0 | 0 |
| ENSMUSG00000031553 | 0.45815691 | 0 | 0 | 0.25 | 0 | 0 | 0.224243824 |
| ENSMUSG00000031554 | 0.811507563 | 0.830482024 | 0.79248125 | 0.396240625 | 0.387761935 | 0.477121255 | 0.579661877 |
| ENSMUSG00000031555 | 3.152363801 | 2.900558725 | 2.916777886 | 2.84821351 | 2.789753073 | 3.238297068 | 2.601625449 |
| ENSMUSG00000031556 | 2.266038081 | 2.104463129 | 2.133818844 | 2.131869252 | 2.116484486 | 2.440909082 | 1.968429585 |
| ENSMUSG00000031557 | 2.800532393 | 2.545288064 | 2.539336734 | 2.49539182 | 2.47528687 | 2.835056102 | 2.344542559 |
| ENSMUSG00000031558 | 2.619908725 | 2.453845824 | 2.434523065 | 2.216819685 | 2.201164838 | 2.457881897 | 2.034219411 |
| ENSMUSG00000031559 | 0.289064826 | 0.25 | 0.396240625 | 0.5 | 0.387761935 | 0 | 0 |
| ENSMUSG00000031561 | 0.45815691 | 0 | 0.396240625 | 2.335518667 | 2.309205544 | 2.631443769 | 2.199599638 |
| ENSMUSG00000031562 | 2.149089517 | 2.039967834 | 2.005591953 | 1.896240625 | 1.820920161 | 2.029383778 | 1.790137008 |
| ENSMUSG00000031563 | 2.989851182 | 2.786164561 | 2.76685859 | 2.686838457 | 2.709785497 | 3.053078443 | 2.566937719 |
| ENSMUSG00000031565 | 2.838206752 | 2.653677461 | 2.701637405 | 2.366391601 | 2.417419887 | 2.677606953 | 2.314375784 |
| ENSMUSG00000031568 | 2.435736758 | 2.259043403 | 2.265173983 | 2.282320754 | 2.214050354 | 2.531478917 | 2.090384805 |
| ENSMUSG00000031570 | 2.34539747 | 2.159156155 | 2.158248799 | 2.148114259 | 2.160196454 | 2.394451681 | 2.040177507 |
| ENSMUSG00000031574 | 1.404271433 | 1.282320754 | 1.188721876 | 1.42510993 | 1.319233381 | 1.431363764 | 1.20919388 |
| ENSMUSG00000031575 | 2.759113486 | 2.518869787 | 2.505591953 | 2.5896628 | 2.54143406 | 2.924795996 | 2.364643151 |
| ENSMUSG00000031576 | 0 | 0.25 | 0 | 0 | 0 | 0 | 0 |
| ENSMUSG00000031577 | 1.816888701 | 1.726722649 | 1.77186571 | 1.695339928 | 1.726078673 | 1.929418926 | 1.598697688 |
| ENSMUSG00000031578 | 2.601583437 | 2.379417347 | 2.385274154 | 2.460587586 | 2.378690133 | 2.720985744 | 2.218171406 |
| ENSMUSG00000031583 | 2.29078455 | 2.154596376 | 2.08272922 | 2.212155735 | 2.134457044 | 2.463892989 | 1.966952343 |
| ENSMUSG00000031584 | 2.475043119 | 2.308404919 | 2.276477127 | 2.340485943 | 2.312243922 | 2.609594409 | 2.184463811 |
| ENSMUSG00000031585 | 2.483777101 | 2.269203899 | 2.242166698 | 2.394828984 | 2.28171603 | 2.646403726 | 2.150629917 |
| ENSMUSG00000031586 | 1.069666964 | 1.02186571 | 1.098079356 | 1.27186571 | 1.319233381 | 1.301029996 | 1.324583736 |
| ENSMUSG00000031590 | 2.289064826 | 2.163659007 | 2.135757955 | 2.231203126 | 2.12771782 | 2.436162647 | 2.018194418 |
| ENSMUSG00000031591 | 2.922773676 | 2.685366747 | 2.703445298 | 2.626700859 | 2.546663136 | 2.971739591 | 2.388432563 |
| ENSMUSG00000031592 | 3.076503714 | 2.78199858 | 2.808105232 | 2.920858323 | 2.814295641 | 3.265053789 | 2.603183062 |
| ENSMUSG00000031594 | 0 | 0.5 | 0.830482024 | 0.396240625 | 0.244650542 | 0.301029996 | 0.520678036 |

| | | | | | | | |
|---|---|---|---|---|---|---|---|
| ENSMUSG00000031595 | 2.334723298 | 2.08272922 | 2.124961472 | 1.723704441 | 1.744182955 | 1.919078092 | 1.651852264 |
| ENSMUSG00000031596 | 2.333175866 | 2.147178659 | 2.108657057 | 1.843759858 | 1.83872004 | 2.021189299 | 1.687112305 |
| ENSMUSG00000031600 | 2.5496035 | 2.365369862 | 2.358135475 | 2.476722649 | 2.416670511 | 2.72916479 | 2.222568443 |
| ENSMUSG00000031601 | 2.707474304 | 2.496460484 | 2.46293726 | 2.57991803 | 2.501307552 | 2.866877814 | 2.292127786 |
| ENSMUSG00000031603 | 0 | 0 | 0 | 0.25 | 0 | 0 | 0 |
| ENSMUSG00000031604 | 2.641922824 | 2.463327389 | 2.488912477 | 2.519871196 | 2.427383764 | 2.825426118 | 2.263534582 |
| ENSMUSG00000031605 | 2.429630757 | 2.263142013 | 2.237091808 | 2.160061234 | 2.122577592 | 2.444044796 | 2.018194418 |
| ENSMUSG00000031608 | 2.249317394 | 2.091580554 | 2.063166358 | 2.230460234 | 2.153144932 | 2.399673721 | 1.986347533 |
| ENSMUSG00000031609 | 1.847794126 | 1.664552871 | 1.79745614 | 1.67510993 | 1.621875691 | 1.86923172 | 1.589319772 |
| ENSMUSG00000031610 | 0 | 0 | 0 | 0 | 0 | 0 | 0 |
| ENSMUSG00000031613 | 0 | 0.25 | 0 | 0 | 0 | 0 | 0 |
| ENSMUSG00000031616 | 0.289064826 | 0.25 | 0 | 0 | 0 | 0 | 0.224243824 |
| ENSMUSG00000031617 | 2.348397716 | 2.231944491 | 2.191217898 | 2.353406982 | 2.25010405 | 2.599883072 | 2.094403682 |
| ENSMUSG00000031618 | 0.45815691 | 0 | 0.701838731 | 0.5 | 0.733951626 | 0.477121255 | 0.629532004 |
| ENSMUSG00000031620 | 0 | 0 | 0 | 0 | 0 | 0 | 0 |
| ENSMUSG00000031621 | 0 | 0 | 0 | 0 | 0 | 0 | 0 |
| ENSMUSG00000031622 | 2.597490839 | 2.432829758 | 2.443696765 | 2.487091808 | 2.495355369 | 2.836324116 | 2.380076739 |
| ENSMUSG00000031626 | 2.450036412 | 2.281030328 | 2.177701608 | 1.638647213 | 1.635855979 | 1.770852012 | 1.529383012 |
| ENSMUSG00000031627 | 1.978790393 | 1.837182039 | 1.901832578 | 1.916333979 | 1.917865457 | 2.170261715 | 1.792684387 |
| ENSMUSG00000031628 | 2.54867984 | 2.39151351 | 2.32991803 | 2.428561379 | 2.363323232 | 2.713490543 | 2.18859465 |
| ENSMUSG00000031629 | 1.759671389 | 1.692046081 | 1.692046081 | 1.741446071 | 1.715300549 | 1.84509804 | 1.574722613 |
| ENSMUSG00000031631 | 2.315764015 | 2.184523065 | 2.160061234 | 2.229715809 | 2.164820399 | 2.463892989 | 2.021963573 |
| ENSMUSG00000031633 | 3.332470585 | 3.095385738 | 3.136663337 | 3.013311781 | 3.018405887 | 3.460145817 | 2.889623317 |
| ENSMUSG00000031634 | 2.381433035 | 2.181980114 | 2.188721876 | 2.29745614 | 2.240967488 | 2.556302501 | 2.073790632 |
| ENSMUSG00000031635 | 0 | 0.25 | 0 | 0 | 0 | 0 | 0.224243824 |
| ENSMUSG00000031636 | 0.578129653 | 0.5 | 0.396240625 | 0.25 | 0 | 0 | 0.224243824 |
| ENSMUSG00000031637 | 0.45815691 | 0.580482024 | 0.646240625 | 0.5 | 0.632412477 | 0.301029996 | 0.579661877 |
| ENSMUSG00000031639 | 1.678702042 | 1.571350555 | 1.494319981 | 1.735628626 | 1.704182902 | 1.995635195 | 1.596378571 |
| ENSMUSG00000031640 | 0 | 0 | 0 | 0 | 0 | 0 | 0 |
| ENSMUSG00000031641 | 1.663535556 | 1.602347734 | 1.54248125 | 1.622963274 | 1.55104774 | 1.792391689 | 1.466365979 |
| ENSMUSG00000031642 | 2.183765914 | 2.052363341 | 2.04745614 | 2.032320754 | 2.075570129 | 2.294466226 | 1.994607951 |
| ENSMUSG00000031644 | 2.167941649 | 2.025821952 | 2.00835575 | 2.04745614 | 2 | 2.240549248 | 1.819456572 |
| ENSMUSG00000031645 | 0 | 0 | 0 | 0 | 0 | 0 | 0 |
| ENSMUSG00000031647 | 2.167941649 | 2.057204673 | 2.107613138 | 1.973704441 | 1.954436039 | 2.103803721 | 1.722075986 |

| | | | | | | | |
|---|---|---|---|---|---|---|---|
| ENSMUSG00000031651 | 0 | 0 | 0 | 0 | 0 | 0 | 0 |
| ENSMUSG00000031652 | 2.571178952 | 2.416333979 | 2.385274154 | 2.405054956 | 2.356272688 | 2.674861141 | 2.173704585 |
| ENSMUSG00000031654 | 0.578129653 | 0.25 | 0.25 | 0.830482024 | 0.632412477 | 0.77815125 | 0.916588299 |
| ENSMUSG00000031657 | 2.751223449 | 2.507666784 | 2.5 | 2.738185312 | 2.644025241 | 3.067814511 | 2.460944696 |
| ENSMUSG00000031659 | 1.156259305 | 1.188721876 | 1.226722649 | 1.841580554 | 1.848234258 | 1.986771734 | 1.790137008 |
| ENSMUSG00000031660 | 2.590857977 | 2.356566189 | 2.387205477 | 2.54216803 | 2.448224646 | 2.809559715 | 2.29374942 |
| ENSMUSG00000031661 | 0 | 0 | 0 | 0.79248125 | 1.091002363 | 1.230448921 | 1.168187745 |
| ENSMUSG00000031662 | 0 | 0 | 0 | 0 | 0 | 0 | 0 |
| ENSMUSG00000031665 | 1.389637216 | 1.431980114 | 1.356566189 | 0 | 0 | 0 | 0 |
| ENSMUSG00000031666 | 2.295901576 | 2.150906586 | 2.12694866 | 2.246460484 | 2.200473444 | 2.463892989 | 2.074322292 |
| ENSMUSG00000031667 | 2.129344651 | 1.981203126 | 1.987091808 | 2.149978211 | 2.078515538 | 2.338456494 | 1.981429723 |
| ENSMUSG00000031668 | 2.042557845 | 1.96759118 | 1.928561379 | 1.909156155 | 1.928492895 | 2.225309282 | 1.826389311 |
| ENSMUSG00000031669 | 2.165631214 | 2.019203899 | 2.006976499 | 2.148114259 | 2.074582839 | 2.365487985 | 1.97354745 |
| ENSMUSG00000031671 | 1.765759561 | 1.642463902 | 1.766522298 | 1.837182039 | 1.793512788 | 2.017033339 | 1.704233519 |
| ENSMUSG00000031672 | 2.948011745 | 2.677270953 | 2.703244868 | 2.705642708 | 2.690120394 | 3.062957834 | 2.515829254 |
| ENSMUSG00000031673 | 2.485932312 | 2.28743678 | 2.25835575 | 2.370953944 | 2.259595879 | 2.58546073 | 2.116124284 |
| ENSMUSG00000031681 | 2.213719572 | 2.064346961 | 2.104463129 | 1.956637122 | 1.961316494 | 2.167317335 | 1.863093717 |
| ENSMUSG00000031682 | 0 | 0 | 0 | 0 | 0 | 0 | 0 |
| ENSMUSG00000031683 | 2.506893125 | 2.344302633 | 2.301142786 | 2.450224975 | 2.400161669 | 2.767897616 | 2.229876084 |
| ENSMUSG00000031684 | 2.165631214 | 2.02053726 | 1.984159485 | 1.96759118 | 1.903713872 | 2.155336037 | 1.798966438 |
| ENSMUSG00000031688 | 0 | 0 | 0 | 0 | 0 | 0 | 0 |
| ENSMUSG00000031691 | 2.944447344 | 2.678561379 | 2.671187155 | 2.59727338 | 2.617868378 | 2.941511433 | 2.438991754 |
| ENSMUSG00000031696 | 3.37100986 | 3.050224651 | 3.031837385 | 3.041540771 | 2.931820217 | 3.409425869 | 2.712982766 |
| ENSMUSG00000031697 | 2.66283289 | 2.464495249 | 2.464495249 | 2.526804269 | 2.434943542 | 2.838849091 | 2.302525964 |
| ENSMUSG00000031698 | 0.45815691 | 0.396240625 | 0.396240625 | 0.75 | 0.632412477 | 0 | 0.520678036 |
| ENSMUSG00000031700 | 2.35727108 | 2.167221246 | 2.031030328 | 1.921625132 | 1.932951348 | 2.136720567 | 1.834294096 |
| ENSMUSG00000031701 | 3.090859043 | 2.808255107 | 2.787912707 | 2.894711098 | 2.80204887 | 3.265760917 | 2.594787769 |
| ENSMUSG00000031703 | 2.640441358 | 2.446158711 | 2.406883471 | 2.527131114 | 2.42262144 | 2.80140371 | 2.254834987 |
| ENSMUSG00000031706 | 1.707474304 | 1.708222504 | 1.845926073 | 1.403677461 | 1.715300549 | 1.505149978 | 1.811176017 |
| ENSMUSG00000031708 | 2.732012699 | 2.495748394 | 2.504897682 | 2.471805154 | 2.528831933 | 2.858537198 | 2.37403344 |
| ENSMUSG00000031709 | 1.956444026 | 1.856566189 | 1.890560606 | 2.058404919 | 2.0510778 | 2.31386722 | 1.969165684 |
| ENSMUSG00000031710 | 0 | 0 | 0 | 0 | 0 | 0 | 0 |
| ENSMUSG00000031711 | 2.262379879 | 2.075945187 | 2.084962501 | 2.179419106 | 2.148364414 | 2.390935107 | 1.991191705 |
| ENSMUSG00000031712 | 1.100572389 | 1.080482024 | 0.92510993 | 1.282320754 | 1.188507684 | 1.079181246 | 1.121219121 |

| | | | | | | | |
|---|---|---|---|---|---|---|---|
| ENSMUSG00000031714 | 2.307602651 | 2.117918804 | 2.117918804 | 2.070192693 | 2.007252778 | 2.225309282 | 1.870159858 |
| ENSMUSG00000031715 | 3.001563678 | 2.779897405 | 2.746460484 | 2.888167274 | 2.773482476 | 3.198106999 | 2.565311426 |
| ENSMUSG00000031722 | 0.45815691 | 0.25 | 0.396240625 | 0 | 0 | 0 | 0 |
| ENSMUSG00000031723 | 1.759671389 | 1.634789703 | 1.602347734 | 1.711372513 | 1.628935049 | 1.73239376 | 1.582103514 |
| ENSMUSG00000031725 | 0 | 0 | 0 | 0 | 0 | 0 | 0 |
| ENSMUSG00000031727 | 0 | 0 | 0 | 0 | 0 | 0 | 0 |
| ENSMUSG00000031728 | 1.924654746 | 1.779735268 | 1.821350555 | 1.71759118 | 1.754126039 | 1.826074803 | 1.735976162 |
| ENSMUSG00000031729 | 2.682744391 | 2.429419106 | 2.437467357 | 2.54652856 | 2.482519186 | 2.843232778 | 2.331854053 |
| ENSMUSG00000031730 | 1.806056082 | 1.80479213 | 1.858657057 | 1.685366747 | 1.789100779 | 1.934498451 | 1.773071385 |
| ENSMUSG00000031731 | 2.891868252 | 2.616391601 | 2.581607372 | 2.773024104 | 2.671488075 | 3.114277297 | 2.490724833 |
| ENSMUSG00000031732 | 2.46499366 | 2.233422664 | 2.241446071 | 2.369945066 | 2.311739339 | 2.641474111 | 2.147679267 |
| ENSMUSG00000031734 | 0.45815691 | 0.5 | 0.92510993 | 0.5 | 0.632412477 | 0.698970004 | 0.853775828 |
| ENSMUSG00000031736 | 1.181541736 | 1.04248125 | 1.04248125 | 1.02186571 | 1.020174412 | 0.954242509 | 0.916588299 |
| ENSMUSG00000031737 | 1.181541736 | 1.17510993 | 1.098079356 | 1.27186571 | 1.407936347 | 1.113943352 | 1.319146374 |
| ENSMUSG00000031738 | 0 | 0 | 0 | 0 | 0 | 0 | 0 |
| ENSMUSG00000031740 | 2.992411635 | 2.784087585 | 2.738003291 | 2.687048212 | 2.660945767 | 3.069298012 | 2.505488011 |
| ENSMUSG00000031748 | 2.167941649 | 2.019203899 | 2.081607372 | 1.458222504 | 1.478764298 | 1.447158031 | 1.360283038 |
| ENSMUSG00000031749 | 2.619908725 | 2.447383411 | 2.517194569 | 2.327653195 | 2.421143088 | 2.641474111 | 2.342462067 |
| ENSMUSG00000031750 | 1.100572389 | 1.201838731 | 1.080482024 | 1.226722649 | 1.335652905 | 1.301029996 | 1.392431569 |
| ENSMUSG00000031751 | 2.996854944 | 2.740181499 | 2.677055433 | 2.823942234 | 2.767008562 | 3.167021796 | 2.568671151 |
| ENSMUSG00000031753 | 2.608049185 | 2.390560606 | 2.37892496 | 2.426408097 | 2.427019704 | 2.7355989 | 2.25604893 |
| ENSMUSG00000031754 | 2.670497861 | 2.414106216 | 2.392463902 | 2.56168515 | 2.465951417 | 2.856124444 | 2.271717885 |
| ENSMUSG00000031755 | 1.707474304 | 1.646240625 | 1.464495249 | 1.752806814 | 1.698489937 | 1.908485019 | 1.618853957 |
| ENSMUSG00000031756 | 2.285603957 | 2.100219859 | 2.090485943 | 2.242166698 | 2.210704773 | 2.485721426 | 2.054609671 |
| ENSMUSG00000031757 | 0 | 0 | 0 | 0 | 0 | 0 | 0 |
| ENSMUSG00000031758 | 1.527823874 | 1.451838731 | 1.431980114 | 1.42510993 | 1.49445382 | 1.544068044 | 1.444790289 |
| ENSMUSG00000031760 | 0 | 0.25 | 0.25 | 0 | 0.244650542 | 0 | 0.355418052 |
| ENSMUSG00000031762 | 2.051791411 | 2.041226732 | 2.118933358 | 0.951838731 | 1.456697355 | 1.278753601 | 1.508694612 |
| ENSMUSG00000031765 | 2.20747957 | 2.214495249 | 2.396240625 | 1.302363341 | 1.523883869 | 1.477121255 | 1.731408809 |
| ENSMUSG00000031766 | 0 | 0 | 0 | 0 | 0.244650542 | 0 | 0 |
| ENSMUSG00000031767 | 1.470606563 | 1.470660762 | 1.438721876 | 1.146240625 | 1.254881871 | 1.176091259 | 1.224243824 |
| ENSMUSG00000031770 | 2.384185734 | 2.188721876 | 2.177701608 | 2.03743678 | 2.056322416 | 2.209515015 | 1.932619724 |
| ENSMUSG00000031772 | 0.45815691 | 0 | 0 | 0 | 0 | 0 | 0 |
| ENSMUSG00000031774 | 2.368815967 | 2.177701608 | 2.19697564 | 2.186208459 | 2.089109832 | 2.439332694 | 2.03601839 |

| | | | | | | | |
|---|---|---|---|---|---|---|---|
| ENSMUSG00000031775 | 0.45815691 | 0.25 | 0.580482024 | 0.25 | 0.387761935 | 0.602059991 | 0.579661877 |
| ENSMUSG00000031776 | 2.615992899 | 2.408248799 | 2.418548067 | 2.598615674 | 2.584291694 | 2.953276337 | 2.420034153 |
| ENSMUSG00000031778 | 2.247417471 | 2.112802778 | 2.089388001 | 2 | 2.029283463 | 2.190331698 | 1.948678378 |
| ENSMUSG00000031779 | 0 | 0 | 0 | 0 | 0 | 0 | 0 |
| ENSMUSG00000031780 | 0 | 0.580482024 | 0.396240625 | 0.646240625 | 0.77552387 | 0.602059991 | 0.710836105 |
| ENSMUSG00000031781 | 2.504843822 | 2.342126615 | 2.336628433 | 2.508699741 | 2.51377465 | 2.812244697 | 2.369585695 |
| ENSMUSG00000031782 | 2.095120908 | 1.973704441 | 1.909156155 | 2.173371739 | 2.1962964 | 2.450249108 | 2.075383 |
| ENSMUSG00000031783 | 2.444137694 | 2.284247778 | 2.300530956 | 2.356041572 | 2.362011938 | 2.653212514 | 2.205319397 |
| ENSMUSG00000031785 | 2.167941649 | 2.070192693 | 2.078220739 | 1.938721876 | 1.97205363 | 2.173186268 | 1.993927582 |
| ENSMUSG00000031786 | 0 | 0 | 0 | 0 | 0 | 0.301029996 | 0 |
| ENSMUSG00000031787 | 2.066740409 | 2.05357978 | 2.146240625 | 1.910964047 | 2.078515538 | 2.214843848 | 2.078019652 |
| ENSMUSG00000031788 | 2.086807695 | 2.067865757 | 2.144357207 | 1.823655187 | 2.137779008 | 2.170261715 | 2.184085651 |
| ENSMUSG00000031789 | 0.867194479 | 0.646240625 | 0.25 | 1 | 0.931471446 | 1.041392685 | 1.028149525 |
| ENSMUSG00000031790 | 1.156259305 | 1.080482024 | 1.372963274 | 1.25 | 1.401338844 | 1.51851394 | 1.42167221 |
| ENSMUSG00000031791 | 1.948720984 | 1.825945187 | 1.850219859 | 1.77186571 | 1.873585591 | 2.075546961 | 1.747849621 |
| ENSMUSG00000031792 | 2.153883046 | 1.964495249 | 1.970660762 | 1.970660762 | 1.982473755 | 2.139879086 | 1.883845332 |
| ENSMUSG00000031796 | 2.320585253 | 2.136723615 | 2.194519282 | 2.144357207 | 2.045754077 | 2.31386722 | 1.96172822 |
| ENSMUSG00000031799 | 3.704245055 | 3.361512352 | 3.327866129 | 3.254810804 | 3.139815299 | 3.682956926 | 2.922355771 |
| ENSMUSG00000031802 | 0 | 0 | 0 | 0 | 0 | 0 | 0 |
| ENSMUSG00000031803 | 0 | 0.25 | 0 | 0.5 | 0 | 0 | 0.224243824 |
| ENSMUSG00000031805 | 1.527823874 | 1.26109853 | 1.54248125 | 1.160964047 | 1.244650542 | 1.301029996 | 1.417653332 |
| ENSMUSG00000031807 | 1.822201287 | 1.807204673 | 1.958222504 | 1.571350555 | 1.751666404 | 1.77815125 | 1.840919217 |
| ENSMUSG00000031808 | 2.029919532 | 1.987091808 | 2.159156155 | 1.311981878 | 1.450958105 | 1.397940009 | 1.607811402 |
| ENSMUSG00000031809 | 0 | 0 | 0 | 0 | 0 | 0 | 0.355418052 |
| ENSMUSG00000031812 | 2.786655728 | 2.570192693 | 2.558704358 | 2.459800947 | 2.483141135 | 2.827369273 | 2.3396671 |
| ENSMUSG00000031813 | 1.700465191 | 1.701838731 | 1.854463129 | 1.593759858 | 1.733951626 | 1.806179974 | 1.844181586 |
| ENSMUSG00000031816 | 2.227926429 | 2.110735874 | 2.134789703 | 2.106566189 | 2.212046813 | 2.401400541 | 2.129834618 |
| ENSMUSG00000031818 | 3.07519763 | 2.854858388 | 2.90136988 | 2.760756821 | 2.855989447 | 3.186673867 | 2.702485399 |
| ENSMUSG00000031819 | 2.506893125 | 2.334405475 | 2.350753006 | 2.46293726 | 2.462015062 | 2.764922985 | 2.333050044 |
| ENSMUSG00000031820 | 2.300956576 | 2.177701608 | 2.253505118 | 2.070192693 | 2.182004726 | 2.298853076 | 2.181426084 |
| ENSMUSG00000031821 | 2.428398755 | 2.322504712 | 2.363317659 | 2.223704441 | 2.334707908 | 2.514547753 | 2.253005488 |
| ENSMUSG00000031822 | 1.777675583 | 1.819031101 | 1.943696765 | 1.451838731 | 1.78687392 | 1.698970004 | 1.879993912 |
| ENSMUSG00000031823 | 2.215778996 | 2.143411797 | 2.174241882 | 2.080482024 | 2.179075613 | 2.336459734 | 2.147679267 |
| ENSMUSG00000031824 | 1.867194479 | 1.698603967 | 1.735628626 | 1.912762923 | 2.017859938 | 2.252853031 | 1.917452159 |

| | | | | | | | |
|---|---|---|---|---|---|---|---|
| ENSMUSG00000031825 | 1.325351389 | 1.146240625 | 1.130890489 | 0.646240625 | 0.632412477 | 0.477121255 | 0.355418052 |
| ENSMUSG00000031826 | 2.529763563 | 2.374961472 | 2.422499493 | 2.391989019 | 2.414412788 | 2.666517981 | 2.359403919 |
| ENSMUSG00000031827 | 2.299278375 | 2.243603647 | 2.267194569 | 2.075945187 | 2.239111448 | 2.342422681 | 2.272870211 |
| ENSMUSG00000031828 | 0.289064826 | 0.830482024 | 0.396240625 | 0.79248125 | 1.188507684 | 1.113943352 | 1.290497981 |
| ENSMUSG00000031831 | 0 | 0 | 0 | 0 | 0 | 0.301029996 | 0.448487648 |
| ENSMUSG00000031832 | 1.516991255 | 1.54248125 | 1.657339155 | 1.302363341 | 1.555378544 | 1.397940009 | 1.643912836 |
| ENSMUSG00000031833 | 2.016886209 | 1.94697564 | 2.070192693 | 1.969129237 | 2.056322416 | 2.161368002 | 2.027536225 |
| ENSMUSG00000031834 | 2.343889214 | 2.224461364 | 2.363317659 | 2.144357207 | 2.277846464 | 2.440909082 | 2.32507354 |
| ENSMUSG00000031835 | 3.023749867 | 2.759043403 | 2.774178789 | 2.80479213 | 2.788316432 | 3.196728723 | 2.639035395 |
| ENSMUSG00000031837 | 0 | 0.5 | 0.25 | 0 | 0.244650542 | 0.477121255 | 0.224243824 |
| ENSMUSG00000031838 | 2.167941649 | 2.063166358 | 2.052363341 | 2.032320754 | 2.102149272 | 2.338456494 | 2.046607113 |
| ENSMUSG00000031839 | 2.565000581 | 2.369945066 | 2.394357207 | 2.48927551 | 2.478134581 | 2.836956737 | 2.335428845 |
| ENSMUSG00000031840 | 1.358731791 | 1.410964047 | 1.451838731 | 1.380890489 | 1.489301084 | 1.662757832 | 1.523605797 |
| ENSMUSG00000031841 | 2.156259305 | 2.059601185 | 2.078220739 | 2.029735268 | 2.075570129 | 2.276461804 | 1.936709733 |
| ENSMUSG00000031842 | 0.289064826 | 0.396240625 | 0.25 | 0.701838731 | 0.77552387 | 0.84509804 | 0.672731473 |
| ENSMUSG00000031843 | 2.066740409 | 1.978222334 | 1.959800947 | 2.152756199 | 2.085294053 | 2.336459734 | 1.94074868 |
| ENSMUSG00000031844 | 0 | 0 | 0 | 0 | 0 | 0 | 0 |
| ENSMUSG00000031845 | 0 | 0 | 0 | 0 | 0 | 0 | 0 |
| ENSMUSG00000031847 | 1.069666964 | 0.830482024 | 0.896240625 | 1 | 1.039257769 | 1.204119983 | 1.150210039 |
| ENSMUSG00000031848 | 2.289064826 | 2.169870025 | 2.266522298 | 2.064346961 | 2.189222894 | 2.394451681 | 2.1908258 |
| ENSMUSG00000031849 | 0.45815691 | 0.25 | 0 | 0.5 | 0.77552387 | 0.301029996 | 0.520678036 |
| ENSMUSG00000031851 | 1.307602651 | 1.26109853 | 1.238549078 | 1.54248125 | 1.528558853 | 1.643452676 | 1.551512352 |
| ENSMUSG00000031853 | 0 | 0 | 0 | 0 | 0.489301084 | 0 | 0 |
| ENSMUSG00000031858 | 2.562324428 | 2.329353153 | 2.301142786 | 2.403217124 | 2.399768403 | 2.681241237 | 2.238304819 |
| ENSMUSG00000031860 | 0.960252568 | 1.201838731 | 1.27186571 | 1.201838731 | 1.200473444 | 1.113943352 | 1.345462945 |
| ENSMUSG00000031861 | 1.912079952 | 1.78743678 | 1.905512955 | 1.811981878 | 1.78010778 | 1.995635195 | 1.68886578 |
| ENSMUSG00000031862 | 2.509948321 | 2.370953944 | 2.378431899 | 2.232684334 | 2.402902329 | 2.57863921 | 2.256956407 |
| ENSMUSG00000031864 | 2.463861956 | 2.233422664 | 2.243603647 | 2.273849256 | 2.238490593 | 2.494154594 | 2.084261301 |
| ENSMUSG00000031865 | 2.63370885 | 2.528109877 | 2.59727338 | 2.250703754 | 2.423358301 | 2.563481085 | 2.438130195 |
| ENSMUSG00000031870 | 0 | 0 | 0 | 0 | 0 | 0 | 0 |
| ENSMUSG00000031871 | 0 | 0 | 0 | 0.396240625 | 0.244650542 | 0.301029996 | 0 |
| ENSMUSG00000031872 | 0 | 0.25 | 0 | 0.701838731 | 0.568060967 | 0.602059991 | 0.74492186 |
| ENSMUSG00000031875 | 2.556012466 | 2.344302633 | 2.395300145 | 2.098079356 | 2.13194489 | 2.367355921 | 2.032410373 |
| ENSMUSG00000031876 | 0.289064826 | 0.5 | 0.396240625 | 0.25 | 0.387761935 | 0 | 0.224243824 |

| | | | | | | | |
|---|---|---|---|---|---|---|---|
| ENSMUSG00000031877 | 0.578129653 | 0 | 0.25 | 0.5 | 0.632412477 | 0.77815125 | 0.629532004 |
| ENSMUSG00000031878 | 2.640441358 | 2.416333979 | 2.387686696 | 2.533927322 | 2.448224646 | 2.790285164 | 2.239263384 |
| ENSMUSG00000031879 | 2.029919532 | 1.969129237 | 2.002806814 | 1.866901388 | 1.990091809 | 2.117271296 | 1.970632877 |
| ENSMUSG00000031880 | 1.794934559 | 1.726722649 | 1.755591953 | 1.505591953 | 1.683842353 | 1.770852012 | 1.74492186 |
| ENSMUSG00000031881 | 0 | 0 | 0 | 0 | 0.244650542 | 0 | 0 |
| ENSMUSG00000031883 | 0.289064826 | 0.580482024 | 0.580482024 | 0.396240625 | 0.568060967 | 0.477121255 | 0.672731473 |
| ENSMUSG00000031884 | 0 | 0 | 0 | 0 | 0 | 0 | 0.224243824 |
| ENSMUSG00000031885 | 2.936496745 | 2.666333979 | 2.647646762 | 2.854858388 | 2.751230566 | 3.183554534 | 2.54475813 |
| ENSMUSG00000031886 | 0 | 0 | 0.25 | 0.396240625 | 0.387761935 | 0.301029996 | 0.448487648 |
| ENSMUSG00000031887 | 1.558729299 | 1.53743678 | 1.698603967 | 1.571350555 | 1.689774529 | 1.633468456 | 1.749303626 |
| ENSMUSG00000031889 | 2.129344651 | 1.976722649 | 1.964495249 | 2.038704527 | 1.983754904 | 2.294466226 | 1.888596011 |
| ENSMUSG00000031891 | 0 | 0.580482024 | 0 | 0.646240625 | 0.733951626 | 0.77815125 | 0.579661877 |
| ENSMUSG00000031893 | 0.578129653 | 0 | 0.396240625 | 0.5 | 0.568060967 | 0 | 0.579661877 |
| ENSMUSG00000031896 | 0.671187741 | 0.396240625 | 0.580482024 | 0.396240625 | 0.846351821 | 0.301029996 | 0.579661877 |
| ENSMUSG00000031897 | 1.578129653 | 1.77186571 | 1.714495249 | 1.910964047 | 2.107593434 | 2.195899652 | 2.121219121 |
| ENSMUSG00000031898 | 0 | 0 | 0 | 0 | 0 | 0 | 0 |
| ENSMUSG00000031901 | 1.87190676 | 1.726722649 | 1.726722649 | 1.681980114 | 1.715300549 | 1.838849091 | 1.645916034 |
| ENSMUSG00000031902 | 2.903085937 | 2.636482442 | 2.608135475 | 2.702442032 | 2.653930024 | 3.069298012 | 2.446476147 |
| ENSMUSG00000031903 | 2.029919532 | 1.901832578 | 1.866901388 | 1.862802778 | 1.990091809 | 2.11058971 | 1.856911705 |
| ENSMUSG00000031904 | 2.654307229 | 2.383818844 | 2.396240625 | 2.555094582 | 2.477819301 | 2.856124444 | 2.298300395 |
| ENSMUSG00000031906 | 2.144240249 | 2.083847589 | 2.019203899 | 1.482684334 | 1.603583716 | 1.62324929 | 1.596378571 |
| ENSMUSG00000031907 | 0.578129653 | 1.02186571 | 1.080482024 | 0.396240625 | 0.733951626 | 0.477121255 | 1.054044578 |
| ENSMUSG00000031910 | 1.289064826 | 1.226722649 | 1.188721876 | 1.302363341 | 1.234113756 | 1 | 1.193409508 |
| ENSMUSG00000031913 | 2.476144923 | 2.231944491 | 2.25835575 | 2.381380202 | 2.317747248 | 2.673941999 | 2.187847501 |
| ENSMUSG00000031916 | 2.092368209 | 1.880890489 | 1.973704441 | 1.938721876 | 1.926994141 | 2.214843848 | 1.861046144 |
| ENSMUSG00000031917 | 2.497590902 | 2.309601185 | 2.296838018 | 2.403677461 | 2.325116119 | 2.707570176 | 2.172530299 |
| ENSMUSG00000031918 | 2.785609222 | 2.53743678 | 2.524508021 | 2.730831871 | 2.626666553 | 3.040206628 | 2.435705549 |
| ENSMUSG00000031919 | 0 | 0.25 | 0 | 0 | 0 | 0 | 0 |
| ENSMUSG00000031921 | 2.315764015 | 2.122963274 | 2.104463129 | 2.191217898 | 2.196996016 | 2.447158031 | 2.039586611 |
| ENSMUSG00000031922 | 2.541215974 | 2.323080408 | 2.342126615 | 2.451435968 | 2.356272688 | 2.671172843 | 2.20389891 |
| ENSMUSG00000031924 | 3.10861828 | 2.817719824 | 2.812870603 | 2.922390314 | 2.855124093 | 3.312600439 | 2.655750087 |
| ENSMUSG00000031925 | 2.203267085 | 2.041226732 | 2.02053726 | 1.930274797 | 1.877063019 | 2.100370545 | 1.808770687 |
| ENSMUSG00000031927 | 0.45815691 | 0 | 0.396240625 | 0 | 0 | 0 | 0 |
| ENSMUSG00000031928 | 2.623015126 | 2.399047439 | 2.418548067 | 2.549304173 | 2.474013807 | 2.834420704 | 2.304878814 |

| | | | | | | | |
|---|---|---|---|---|---|---|---|
| ENSMUSG00000031930 | 2.596667477 | 2.408702763 | 2.415889526 | 2.343216265 | 2.360255934 | 2.643452676 | 2.255745871 |
| ENSMUSG00000031931 | 1.903481042 | 1.837182039 | 1.76109853 | 1.916333979 | 1.868304353 | 2.123851641 | 1.71730663 |
| ENSMUSG00000031932 | 0 | 0 | 0.25 | 0 | 0 | 0 | 0.448487648 |
| ENSMUSG00000031933 | 0 | 0 | 0 | 0 | 0 | 0 | 0 |
| ENSMUSG00000031934 | 1.358731791 | 1.418106335 | 1.42510993 | 1.92510993 | 1.795698282 | 2.049218023 | 1.74196736 |
| ENSMUSG00000031935 | 2.333175866 | 2.215271726 | 2.170748646 | 2.29248125 | 2.230318492 | 2.521138084 | 2.082712142 |
| ENSMUSG00000031936 | 0 | 0 | 0 | 0 | 0 | 0 | 0 |
| ENSMUSG00000031937 | 0 | 0 | 0 | 0.864857905 | 0.877063019 | 0.84509804 | 0.803905701 |
| ENSMUSG00000031938 | 2.116511873 | 1.988549078 | 1.988549078 | 2.016522298 | 1.897237431 | 2.195899652 | 1.817112247 |
| ENSMUSG00000031939 | 2.22189694 | 2.071350555 | 2.091580554 | 2.202642909 | 2.158641579 | 2.477121255 | 2.006616089 |
| ENSMUSG00000031948 | 3.029627799 | 2.766354034 | 2.745926548 | 2.90654133 | 2.857928771 | 3.317227349 | 2.680817717 |
| ENSMUSG00000031949 | 1.777675583 | 1.668106335 | 1.688721876 | 1.75835575 | 1.749189509 | 1.934498451 | 1.690609803 |
| ENSMUSG00000031950 | 2.414600936 | 2.192046081 | 2.199415381 | 2.402294685 | 2.333285652 | 2.645422269 | 2.197791274 |
| ENSMUSG00000031951 | 1.714367558 | 1.614857905 | 1.584962501 | 1.930274797 | 1.875328588 | 2.100370545 | 1.813563596 |
| ENSMUSG00000031952 | 0 | 0 | 0.25 | 0.396240625 | 0.244650542 | 0 | 0.448487648 |
| ENSMUSG00000031953 | 1.445324132 | 1.311981878 | 1.380890489 | 1.557204673 | 1.514343934 | 1.643452676 | 1.459332757 |
| ENSMUSG00000031954 | 2.904270129 | 2.674241882 | 2.661414608 | 2.780059469 | 2.683314108 | 3.106870544 | 2.515693522 |
| ENSMUSG00000031955 | 2.670497861 | 2.594844592 | 2.676839783 | 2.222945926 | 2.436719872 | 2.592176757 | 2.445971311 |
| ENSMUSG00000031957 | 0.45815691 | 0 | 0 | 0 | 0 | 0 | 0 |
| ENSMUSG00000031958 | 0 | 0 | 0.25 | 0 | 0.244650542 | 0.301029996 | 0.224243824 |
| ENSMUSG00000031959 | 2.22189694 | 2.093759858 | 2.117918804 | 2.077084758 | 2.095689764 | 2.318063335 | 1.984249047 |
| ENSMUSG00000031960 | 3.327499938 | 3.031595458 | 3.006111781 | 2.903332264 | 2.928316901 | 3.332842267 | 2.77398916 |
| ENSMUSG00000031962 | 0 | 0 | 0 | 0 | 0 | 0 | 0 |
| ENSMUSG00000031963 | 1.989530018 | 1.918106335 | 1.918106335 | 1.903677461 | 1.855665188 | 2.041392685 | 1.714087544 |
| ENSMUSG00000031965 | 1.822201287 | 1.708222504 | 1.732684334 | 2.29248125 | 2.229681962 | 2.532754379 | 2.062969708 |
| ENSMUSG00000031966 | 0 | 0 | 0 | 0 | 0 | 0 | 0 |
| ENSMUSG00000031967 | 2.563218389 | 2.384789703 | 2.36434522 | 2.385757955 | 2.432082686 | 2.721810615 | 2.279980727 |
| ENSMUSG00000031969 | 2.113897241 | 1.940387808 | 1.979715809 | 2.036164561 | 2.009637626 | 2.198657087 | 1.901538355 |
| ENSMUSG00000031970 | 1.307602651 | 1.146240625 | 1.25 | 1.418106335 | 1.433158226 | 1.544068044 | 1.441050184 |
| ENSMUSG00000031971 | 1.623015126 | 1.445339928 | 1.27186571 | 1.664552871 | 1.639266209 | 1.748188027 | 1.574722613 |
| ENSMUSG00000031972 | 1.156259305 | 1.388647213 | 1.321350555 | 0 | 0.244650542 | 0.301029996 | 0.224243824 |
| ENSMUSG00000031974 | 2.294202867 | 2.157339155 | 2.12694866 | 2.262462137 | 2.247690665 | 2.488550717 | 2.100829716 |
| ENSMUSG00000031976 | 2.58665822 | 2.343759858 | 2.342671852 | 2.476346751 | 2.449594021 | 2.781755375 | 2.255745871 |
| ENSMUSG00000031979 | 2.367390214 | 2.185366747 | 2.17510993 | 2.168989258 | 2.186353316 | 2.354108439 | 2.072724682 |

| | | | | | | | |
|---|---|---|---|---|---|---|---|
| ENSMUSG00000031980 | 0 | 0 | 0 | 0 | 0 | 0.301029996 | 0 |
| ENSMUSG00000031981 | 0.289064826 | 0 | 0.25 | 0 | 0 | 0 | 0 |
| ENSMUSG00000031982 | 1.623015126 | 1.53743678 | 1.482684334 | 1.602347734 | 1.607318736 | 1.755874856 | 1.529383012 |
| ENSMUSG00000031983 | 2.855397071 | 2.657339155 | 2.67510993 | 2.561388177 | 2.567082571 | 2.858537198 | 2.461266122 |
| ENSMUSG00000031984 | 2.092368209 | 2.028435542 | 2.025821952 | 2.025821952 | 1.983754904 | 2.225309282 | 1.880961074 |
| ENSMUSG00000031985 | 2.587501561 | 2.378431899 | 2.368426437 | 2.417664062 | 2.422252432 | 2.765668555 | 2.226242896 |
| ENSMUSG00000031986 | 1.881174527 | 1.809601185 | 1.77186571 | 1.825945187 | 1.850106709 | 2.103803721 | 1.777090262 |
| ENSMUSG00000031987 | 2.420929319 | 2.314935816 | 2.302972074 | 2.259043403 | 2.276733037 | 2.540329475 | 2.130281771 |
| ENSMUSG00000031988 | 2.54867984 | 2.336628433 | 2.306000419 | 2.477473271 | 2.382016011 | 2.736396502 | 2.242438242 |
| ENSMUSG00000031990 | 2.185978293 | 2.01109853 | 2.031030328 | 2.192046081 | 2.138604637 | 2.423245874 | 2.046607113 |
| ENSMUSG00000031991 | 0 | 0 | 0 | 0 | 0 | 0 | 0 |
| ENSMUSG00000031993 | 2.409468165 | 2.245034894 | 2.201838731 | 2.349151195 | 2.289330171 | 2.604226053 | 2.161359729 |
| ENSMUSG00000031994 | 0 | 0 | 0 | 0.25 | 0.244650542 | 0 | 0 |
| ENSMUSG00000031995 | 0 | 0.25 | 0 | 0.25 | 0 | 0 | 0.355418052 |
| ENSMUSG00000031996 | 3.307752474 | 2.992166698 | 2.961176441 | 3.017614104 | 2.939748275 | 3.425697213 | 2.737279141 |
| ENSMUSG00000031997 | 0.289064826 | 0.25 | 0.5 | 0 | 0 | 0 | 0 |
| ENSMUSG00000031998 | 0.289064826 | 0 | 0.25 | 0.396240625 | 0.244650542 | 0 | 0.224243824 |
| ENSMUSG00000032000 | 1.83262764 | 1.660964047 | 1.695339928 | 2.117918804 | 2.038163331 | 2.260071388 | 1.900630878 |
| ENSMUSG00000032002 | 2.513987407 | 2.359697963 | 2.31728167 | 2.508011432 | 2.368520226 | 2.754348336 | 2.235734673 |
| ENSMUSG00000032006 | 0.747221736 | 0.396240625 | 0.5 | 1.080482024 | 1 | 1.176091259 | 1.028149525 |
| ENSMUSG00000032009 | 1.928763473 | 1.870953944 | 1.845926073 | 2.060793496 | 1.992595054 | 2.298853076 | 1.903345709 |
| ENSMUSG00000032010 | 0.960252568 | 1.201838731 | 0.896240625 | 1.188721876 | 1.200473444 | 1.255272505 | 1.110947927 |
| ENSMUSG00000032011 | 1.307602651 | 1.214495249 | 1.311981878 | 0 | 0 | 0 | 0.355418052 |
| ENSMUSG00000032012 | 1.614416215 | 1.552363341 | 1.664552871 | 0.976722649 | 1.401338844 | 1.176091259 | 1.313616061 |
| ENSMUSG00000032013 | 0 | 0 | 0 | 0 | 0 | 0 | 0 |
| ENSMUSG00000032014 | 2.245508853 | 2.064346961 | 2.080482024 | 1.688721876 | 1.846351821 | 1.880813592 | 1.729871916 |
| ENSMUSG00000032015 | 0 | 0 | 0 | 0 | 0 | 0 | 0 |
| ENSMUSG00000032017 | 0.45815691 | 0.25 | 0.75 | 0.25 | 0.568060967 | 0 | 0.224243824 |
| ENSMUSG00000032018 | 2.389637216 | 2.234159485 | 2.231203126 | 2.267194569 | 2.153144932 | 2.539076099 | 2.045447559 |
| ENSMUSG00000032020 | 1.37447073 | 1.458222504 | 1.364857905 | 1.685366747 | 1.611014646 | 1.785329835 | 1.514741806 |
| ENSMUSG00000032021 | 0 | 0 | 0 | 0 | 0 | 0 | 0.224243824 |
| ENSMUSG00000032023 | 0 | 0 | 0 | 0 | 0 | 0 | 0 |
| ENSMUSG00000032024 | 2.020182925 | 1.870953944 | 1.926839783 | 1.593759858 | 1.509475496 | 1.633468456 | 1.46982607 |
| ENSMUSG00000032026 | 2.590857977 | 2.377443751 | 2.385274154 | 2.304186465 | 2.267696867 | 2.549003262 | 2.119375723 |

| | | | | | | | |
|---|---|---|---|---|---|---|---|
| ENSMUSG00000032028 | 0.289064826 | 0 | 0.25 | 0 | 0.632412477 | 0 | 0.355418052 |
| ENSMUSG00000032030 | 2.657880793 | 2.455044741 | 2.457826811 | 2.593488164 | 2.50483127 | 2.873320602 | 2.316141565 |
| ENSMUSG00000032033 | 0 | 0 | 0 | 0 | 0 | 0 | 0 |
| ENSMUSG00000032034 | 0 | 0 | 0 | 0 | 0 | 0 | 0 |
| ENSMUSG00000032035 | 2.824593906 | 2.597810897 | 2.554186465 | 2.723704441 | 2.683842353 | 3.059563418 | 2.489402261 |
| ENSMUSG00000032036 | 1.516991255 | 1.464495249 | 1.458222504 | 1.396240625 | 1.335652905 | 1.255272505 | 1.307989563 |
| ENSMUSG00000032038 | 2.451206209 | 2.275165585 | 2.280383379 | 2.036164561 | 1.951645858 | 2.276461804 | 1.826389311 |
| ENSMUSG00000032040 | 2.295901576 | 2.116901388 | 2.122963274 | 2.161864607 | 2.180543208 | 2.434568904 | 2.065163042 |
| ENSMUSG00000032041 | 2.215778996 | 2.073655187 | 2.097004321 | 2.207430684 | 2.160196454 | 2.451786436 | 2.006616089 |
| ENSMUSG00000032042 | 2.778742163 | 2.555698726 | 2.511439915 | 2.700224975 | 2.652394595 | 3.056523724 | 2.477860296 |
| ENSMUSG00000032044 | 1.928763473 | 1.884789703 | 1.872963274 | 1.910964047 | 1.968065382 | 2.146128036 | 1.837623616 |
| ENSMUSG00000032046 | 2.213719572 | 2.081607372 | 2.086073977 | 2.153677461 | 2.113840608 | 2.428134794 | 2.027536225 |
| ENSMUSG00000032047 | 2.572927518 | 2.382357639 | 2.421187155 | 2.580482024 | 2.486542403 | 2.890979597 | 2.335191751 |
| ENSMUSG00000032048 | 0 | 0 | 0 | 0 | 0 | 0 | 0 |
| ENSMUSG00000032050 | 3.045096433 | 2.787595492 | 2.776149688 | 2.929526178 | 2.81792442 | 3.286231854 | 2.62847992 |
| ENSMUSG00000032051 | 1.614416215 | 1.464495249 | 1.52186571 | 1.71759118 | 1.665423072 | 1.851258349 | 1.559435575 |
| ENSMUSG00000032053 | 0 | 0 | 0 | 0 | 0 | 0 | 0 |
| ENSMUSG00000032056 | 0.289064826 | 0 | 0.25 | 0 | 0 | 0 | 0 |
| ENSMUSG00000032057 | 0.91631382 | 1 | 0.92510993 | 0.396240625 | 0.489301084 | 0.301029996 | 0.224243824 |
| ENSMUSG00000032058 | 2.652869185 | 2.448197574 | 2.415444524 | 2.497525991 | 2.454008153 | 2.767897616 | 2.304618229 |
| ENSMUSG00000032059 | 2.039433992 | 1.982684334 | 1.953445298 | 2.015173983 | 1.934425071 | 2.227886705 | 1.869159809 |
| ENSMUSG00000032060 | 2.531694272 | 2.323655187 | 2.310197833 | 1.933677405 | 1.964031554 | 2.190331698 | 1.839824409 |
| ENSMUSG00000032062 | 1.707474304 | 1.488549078 | 1.54248125 | 1.618933358 | 1.592135487 | 1.707570176 | 1.433437704 |
| ENSMUSG00000032064 | 2.413323654 | 2.216819685 | 2.228969845 | 1.899978211 | 1.86473911 | 2.117271296 | 1.710836105 |
| ENSMUSG00000032065 | 0 | 0 | 0 | 0 | 0 | 0 | 0 |
| ENSMUSG00000032066 | 0 | 0 | 0.25 | 0.25 | 0.244650542 | 0.477121255 | 0.448487648 |
| ENSMUSG00000032067 | 1.418409478 | 1.330482024 | 1.494319981 | 1.848079356 | 1.704182902 | 1.826074803 | 1.645916034 |
| ENSMUSG00000032068 | 0.289064826 | 0 | 0 | 0 | 0.244650542 | 0 | 0.224243824 |
| ENSMUSG00000032076 | 2.273259602 | 2.088286706 | 2.110735874 | 2.406883471 | 2.322676882 | 2.626340367 | 2.124874702 |
| ENSMUSG00000032077 | 1.890240806 | 1.862802778 | 1.848079356 | 2.013820609 | 1.991345651 | 2.252853031 | 1.895131899 |
| ENSMUSG00000032078 | 2.402962068 | 2.183677405 | 2.192872367 | 2.299918086 | 2.294670002 | 2.621176282 | 2.158915016 |
| ENSMUSG00000032079 | 0 | 0 | 0 | 0 | 0 | 0 | 0.224243824 |
| ENSMUSG00000032080 | 0 | 0 | 0 | 0 | 0 | 0 | 0 |
| ENSMUSG00000032081 | 0 | 0 | 0 | 0 | 0 | 0 | 0 |

| | | | | | | | |
|---|---|---|---|---|---|---|---|
| ENSMUSG00000032083 | 0.289064826 | 0 | 0 | 0 | 0 | 0 | 0.355418052 |
| ENSMUSG00000032085 | 3.694937608 | 3.376514897 | 3.36514597 | 2.288704527 | 2.306140782 | 2.5774918 | 2.170563634 |
| ENSMUSG00000032086 | 2.612833548 | 2.43155454 | 2.391989019 | 2.512462137 | 2.491431917 | 2.848189117 | 2.316393034 |
| ENSMUSG00000032087 | 0 | 0 | 0 | 0 | 0 | 0 | 0 |
| ENSMUSG00000032089 | 0 | 0 | 0 | 0 | 0 | 0 | 0 |
| ENSMUSG00000032091 | 0 | 0 | 0 | 0 | 0 | 0 | 0 |
| ENSMUSG00000032092 | 0.578129653 | 0.701838731 | 0.75 | 1.516522298 | 1.47337555 | 1.579783597 | 1.345462945 |
| ENSMUSG00000032093 | 0 | 0 | 0 | 0 | 0 | 0 | 0 |
| ENSMUSG00000032094 | 0 | 0 | 0 | 0 | 0.387761935 | 0 | 0 |
| ENSMUSG00000032096 | 3.216711746 | 2.927270953 | 2.899861995 | 3.085796429 | 2.982775621 | 3.488409689 | 2.767447037 |
| ENSMUSG00000032097 | 3.152798436 | 2.866774008 | 2.851418333 | 2.994141031 | 2.918499537 | 3.371621927 | 2.690293407 |
| ENSMUSG00000032098 | 0 | 0 | 0 | 0 | 0 | 0 | 0 |
| ENSMUSG00000032099 | 0 | 0 | 0 | 0 | 0 | 0 | 0 |
| ENSMUSG00000032101 | 0 | 0 | 0 | 1.302363341 | 1.091002363 | 1.230448921 | 1.100339912 |
| ENSMUSG00000032103 | 0 | 0 | 0 | 0 | 0 | 0 | 0 |
| ENSMUSG00000032105 | 0 | 0 | 0 | 0 | 0 | 0 | 0 |
| ENSMUSG00000032108 | 0 | 0 | 0 | 0 | 0 | 0 | 0 |
| ENSMUSG00000032109 | 0 | 0 | 0 | 0.5 | 0.846351821 | 0.698970004 | 1.041356071 |
| ENSMUSG00000032110 | 0.578129653 | 0.396240625 | 0.396240625 | 0.396240625 | 0.244650542 | 0.301029996 | 0.355418052 |
| ENSMUSG00000032112 | 2.326928073 | 2.168989258 | 2.172499493 | 2.220660762 | 2.180543208 | 2.460897843 | 2.059651572 |
| ENSMUSG00000032113 | 2.358731791 | 2.223704441 | 2.155512955 | 2.343759858 | 2.247084732 | 2.614897216 | 2.109521699 |
| ENSMUSG00000032114 | 1.639698646 | 1.614857905 | 1.584962501 | 1.695339928 | 1.677808768 | 1.84509804 | 1.657681529 |
| ENSMUSG00000032115 | 2.905843838 | 2.679419106 | 2.647412787 | 2.849418658 | 2.847017357 | 3.247727833 | 2.659688509 |
| ENSMUSG00000032116 | 3.137760644 | 2.859827865 | 2.826942508 | 3.036721709 | 2.922011567 | 3.420120848 | 2.691162754 |
| ENSMUSG00000032118 | 0 | 0.580482024 | 0.79248125 | 1.226722649 | 1.074582839 | 1 | 1.066254157 |
| ENSMUSG00000032119 | 1.903481042 | 1.825945187 | 1.769203899 | 1.951838731 | 1.944572352 | 2.075546961 | 1.808770687 |
| ENSMUSG00000032120 | 1.890240806 | 1.825945187 | 1.830482024 | 1.634789703 | 1.866526233 | 1.991226076 | 1.86715039 |
| ENSMUSG00000032121 | 1.404271433 | 1.348079356 | 1.321350555 | 1.630890489 | 1.559656853 | 1.662757832 | 1.511732338 |
| ENSMUSG00000032122 | 1.794934559 | 1.708222504 | 1.726722649 | 1.602347734 | 1.621875691 | 1.792391689 | 1.523605797 |
| ENSMUSG00000032123 | 2.177057474 | 2.041226732 | 2.034887838 | 2.167221246 | 2.125157063 | 2.44870632 | 2.024452165 |
| ENSMUSG00000032125 | 0 | 0 | 0 | 0 | 0 | 0 | 0 |
| ENSMUSG00000032126 | 2.273259602 | 2.174241882 | 2.178561379 | 2.220660762 | 2.19206933 | 2.471291711 | 2.078019652 |
| ENSMUSG00000032127 | 2.585813169 | 2.391989019 | 2.385757955 | 2.357090043 | 2.369807605 | 2.658011397 | 2.24651952 |
| ENSMUSG00000032128 | 0.45815691 | 0.646240625 | 0.79248125 | 0 | 0 | 0.477121255 | 0.224243824 |

| | | | | | | | |
|---|---|---|---|---|---|---|---|
| ENSMUSG00000032131 | 0.867194479 | 0.701838731 | 0.864857905 | 1 | 1.188507684 | 0.903089987 | 1.140832123 |
| ENSMUSG00000032134 | 0 | 0 | 0 | 0 | 0 | 0 | 0 |
| ENSMUSG00000032135 | 2.933934888 | 2.697179583 | 2.688721876 | 2.356041572 | 2.32560195 | 2.64246452 | 2.222232323 |
| ENSMUSG00000032171 | 2.323768721 | 2.224461364 | 2.25835575 | 2.125956434 | 2.275616087 | 2.374748346 | 2.257861346 |
| ENSMUSG00000032172 | 0 | 0.396240625 | 0.701838731 | 0.701838731 | 0.812711509 | 0.698970004 | 0.853775828 |
| ENSMUSG00000032174 | 0 | 0.396240625 | 0.25 | 0 | 0.632412477 | 0.698970004 | 0.853775828 |
| ENSMUSG00000032175 | 1.816888701 | 1.845926073 | 1.802363341 | 1.67510993 | 1.706995316 | 1.785329835 | 1.787569411 |
| ENSMUSG00000032177 | 2.194712275 | 2.070192693 | 2.139605178 | 1.937048212 | 2.028157605 | 2.214843848 | 1.965468324 |
| ENSMUSG00000032178 | 2.919366221 | 2.702642909 | 2.684311836 | 2.679847205 | 2.695608647 | 3.013679697 | 2.522012122 |
| ENSMUSG00000032179 | 0 | 0 | 0 | 0 | 0 | 0 | 0 |
| ENSMUSG00000032180 | 2.153883046 | 2.081607372 | 2.098079356 | 1.985628626 | 2.132784263 | 2.281033367 | 2.034219411 |
| ENSMUSG00000032181 | 0 | 0.25 | 0.25 | 0 | 0 | 0.477121255 | 0 |
| ENSMUSG00000032182 | 1.623015126 | 1.505591953 | 1.593759858 | 1.54745614 | 1.514343934 | 1.579783597 | 1.466365979 |
| ENSMUSG00000032184 | 1.445324132 | 1.380890489 | 1.356566189 | 1.53743678 | 1.528558853 | 1.62324929 | 1.502532226 |
| ENSMUSG00000032185 | 2.601583437 | 2.418106335 | 2.442459461 | 2.414998973 | 2.463660548 | 2.758911892 | 2.341997913 |
| ENSMUSG00000032186 | 1.876566388 | 1.825945187 | 1.766522298 | 2.158248799 | 2.091944837 | 2.348304863 | 1.93179548 |
| ENSMUSG00000032187 | 3.07519763 | 2.840622952 | 2.875334905 | 2.784887838 | 2.846906522 | 3.19893187 | 2.735494791 |
| ENSMUSG00000032191 | 0 | 0.25 | 0 | 0 | 0 | 0 | 0 |
| ENSMUSG00000032192 | 1.445324132 | 1.482684334 | 1.552363341 | 1.410964047 | 1.555378544 | 1.447158031 | 1.564611933 |
| ENSMUSG00000032193 | 2.608049185 | 2.45584181 | 2.506630861 | 2.580200137 | 2.643828223 | 2.965201701 | 2.529773182 |
| ENSMUSG00000032194 | 2.543094501 | 2.343216265 | 2.442459461 | 2.315523711 | 2.349985771 | 2.598790507 | 2.287763261 |
| ENSMUSG00000032198 | 1.940852214 | 1.845926073 | 1.912762923 | 1.741446071 | 1.951645858 | 2.008600172 | 1.954885099 |
| ENSMUSG00000032199 | 2.992092437 | 2.740724001 | 2.729156481 | 2.89127552 | 2.80330539 | 3.26245109 | 2.606378556 |
| ENSMUSG00000032202 | 1.759671389 | 1.566696635 | 1.62694866 | 1.852347734 | 1.78687392 | 1.944482672 | 1.667169465 |
| ENSMUSG00000032204 | 0.45815691 | 0 | 0 | 0 | 0.244650542 | 0 | 0 |
| ENSMUSG00000032207 | 0 | 0 | 0 | 0 | 0 | 0 | 0 |
| ENSMUSG00000032212 | 2.557825653 | 2.361770807 | 2.409608978 | 2.487091808 | 2.423358301 | 2.761927838 | 2.249933144 |
| ENSMUSG00000032215 | 2.20747957 | 2.061981878 | 1.988549078 | 2.136723615 | 2.126866297 | 2.389166084 | 1.989815042 |
| ENSMUSG00000032216 | 3.902654487 | 3.520412467 | 3.501822531 | 3.521326617 | 3.421839554 | 4.007021926 | 3.168747288 |
| ENSMUSG00000032217 | 2.493389029 | 2.334962501 | 2.264497931 | 2.30175358 | 2.277846464 | 2.611723308 | 2.126234965 |
| ENSMUSG00000032218 | 2.53838222 | 2.343216265 | 2.370953944 | 2.533927322 | 2.483141135 | 2.873901598 | 2.361597253 |
| ENSMUSG00000032220 | 2.76355595 | 2.52219706 | 2.531676118 | 2.355516192 | 2.339407793 | 2.574031268 | 2.210242554 |
| ENSMUSG00000032221 | 1.91631382 | 1.752806814 | 1.799918086 | 2.049918086 | 1.978602168 | 2.260071388 | 1.877074955 |
| ENSMUSG00000032224 | 0.671187741 | 0.92510993 | 0.951838731 | 1.061981878 | 1.091002363 | 1.230448921 | 1.028149525 |

| | | | | | | | |
|---|---|---|---|---|---|---|---|
| ENSMUSG00000032226 | 0.45815691 | 0.5 | 0 | 0.396240625 | 0.387761935 | 0 | 0.448487648 |
| ENSMUSG00000032228 | 2.971809217 | 2.702642909 | 2.65892953 | 2.711960089 | 2.641653746 | 3.050766311 | 2.465891268 |
| ENSMUSG00000032231 | 3.601287144 | 3.247437319 | 3.253068836 | 3.433624485 | 3.365395913 | 3.941113727 | 3.128346438 |
| ENSMUSG00000032232 | 2.378662046 | 2.222185812 | 2.23927551 | 2.508011432 | 2.517458689 | 2.819543936 | 2.362906151 |
| ENSMUSG00000032235 | 2.054824387 | 1.979715809 | 1.914552871 | 2.05357978 | 2.027028145 | 2.276461804 | 1.849546807 |
| ENSMUSG00000032238 | 0.45815691 | 0.896240625 | 0.580482024 | 1.146240625 | 1.106691885 | 1.361727836 | 1.054044578 |
| ENSMUSG00000032239 | 2.042557845 | 1.959800947 | 2.017865591 | 2.079353153 | 2.090057366 | 2.356025857 | 2.007923223 |
| ENSMUSG00000032243 | 2.373064211 | 2.215271726 | 2.163659007 | 0.830482024 | 0.905314583 | 0.84509804 | 0.74492186 |
| ENSMUSG00000032244 | 2.789779581 | 2.571061437 | 2.537119146 | 2.666999634 | 2.581010538 | 2.985875357 | 2.382749586 |
| ENSMUSG00000032245 | 1.978790393 | 1.837182039 | 1.868933358 | 1.945339928 | 1.988833497 | 2.195899652 | 1.899720848 |
| ENSMUSG00000032246 | 0.45815691 | 0.864857905 | 0.830482024 | 0.396240625 | 0.244650542 | 0 | 0.224243824 |
| ENSMUSG00000032249 | 2.971473847 | 2.72027849 | 2.675543171 | 2.725593779 | 2.639864609 | 3.071513805 | 2.496105974 |
| ENSMUSG00000032251 | 2.010213549 | 1.905512955 | 1.886723615 | 1.839388001 | 1.818885826 | 2.025305865 | 1.685349273 |
| ENSMUSG00000032252 | 2.674620126 | 2.459012589 | 2.441217898 | 2.538388008 | 2.4345872 | 2.845718018 | 2.274591042 |
| ENSMUSG00000032253 | 2.99653913 | 2.74929487 | 2.714106382 | 2.780545225 | 2.687169675 | 3.046104787 | 2.452968314 |
| ENSMUSG00000032254 | 2.786655728 | 2.533285553 | 2.541226732 | 2.79248125 | 2.717546641 | 3.1430148 | 2.505628092 |
| ENSMUSG00000032257 | 0 | 0 | 0.25 | 0 | 0.244650542 | 0.301029996 | 0 |
| ENSMUSG00000032258 | 1.948720984 | 1.890560606 | 1.880890489 | 2 | 1.919403394 | 2.187520721 | 1.860017477 |
| ENSMUSG00000032259 | 0 | 0 | 0 | 0 | 0 | 0 | 0 |
| ENSMUSG00000032261 | 1.867194479 | 1.80479213 | 1.777131114 | 1.766522298 | 1.718026092 | 1.84509804 | 1.607811402 |
| ENSMUSG00000032262 | 1.989530018 | 1.943696765 | 1.926839783 | 2.041226732 | 1.968065382 | 2.206825876 | 1.801445495 |
| ENSMUSG00000032263 | 2.103271637 | 1.972185812 | 1.948603967 | 2.099151195 | 2.03596422 | 2.352182518 | 1.939139147 |
| ENSMUSG00000032264 | 2.493389029 | 2.265173983 | 2.276477127 | 2.391037372 | 2.318243321 | 2.665580991 | 2.153553898 |
| ENSMUSG00000032265 | 1.956444026 | 1.809601185 | 1.75 | 1.160964047 | 1 | 1.113943352 | 1.089372236 |
| ENSMUSG00000032267 | 2.360187403 | 2.14151351 | 2.110735874 | 2.229715809 | 2.199086582 | 2.515873844 | 2.013099581 |
| ENSMUSG00000032268 | 0 | 0 | 0 | 0 | 0 | 0 | 0 |
| ENSMUSG00000032269 | 0 | 0 | 0 | 0 | 0 | 0 | 0 |
| ENSMUSG00000032271 | 0 | 0 | 0 | 0 | 0 | 0 | 0 |
| ENSMUSG00000032274 | 0 | 0 | 0 | 0 | 0.244650542 | 0 | 0 |
| ENSMUSG00000032278 | 0.811507563 | 0.580482024 | 0.580482024 | 1 | 0.905314583 | 0.903089987 | 0.803905701 |
| ENSMUSG00000032279 | 2.806056082 | 2.560495787 | 2.57508814 | 2.748941788 | 2.669111257 | 3.076640444 | 2.492771481 |
| ENSMUSG00000032280 | 2.349889783 | 2.203445298 | 2.274508021 | 2.118933358 | 2.16174451 | 2.357934847 | 2.124420007 |
| ENSMUSG00000032281 | 1.811507563 | 1.83272922 | 1.821350555 | 0.951838731 | 1.121713561 | 1 | 1.110947927 |
| ENSMUSG00000032285 | 1.53838222 | 1.494319981 | 1.584962501 | 1.698603967 | 1.72073075 | 1.851258349 | 1.629532004 |

| | | | | | | | |
|---|---|---|---|---|---|---|---|
| ENSMUSG00000032288 | 1.867194479 | 1.845926073 | 1.850219859 | 1.909156155 | 1.851969278 | 2.123851641 | 1.756477211 |
| ENSMUSG00000032289 | 0.747221736 | 1.04248125 | 1.160964047 | 0.25 | 0.632412477 | 0 | 0.74492186 |
| ENSMUSG00000032290 | 2.589183146 | 2.373463862 | 2.401832578 | 2.561981878 | 2.499532413 | 2.85672889 | 2.337554913 |
| ENSMUSG00000032291 | 0.671187741 | 0.864857905 | 0.701838731 | 1.098079356 | 1.020174412 | 1.079181246 | 1.014380832 |
| ENSMUSG00000032292 | 0 | 0 | 0 | 0 | 0 | 0 | 0 |
| ENSMUSG00000032293 | 2.983054229 | 2.708222504 | 2.67140621 | 2.869439567 | 2.76055529 | 3.21031852 | 2.549317365 |
| ENSMUSG00000032294 | 3.753750144 | 3.424866002 | 3.464616684 | 3.331396636 | 3.405894953 | 3.879726497 | 3.21228247 |
| ENSMUSG00000032295 | 2.075459174 | 1.998588359 | 2.044977273 | 1.809601185 | 2.032639648 | 2.093421685 | 1.999330888 |
| ENSMUSG00000032297 | 0.45815691 | 0.396240625 | 0 | 0.396240625 | 0 | 0 | 0 |
| ENSMUSG00000032298 | 1.54867984 | 1.431980114 | 1.380890489 | 1.339388001 | 1.427019704 | 1.414973348 | 1.401059159 |
| ENSMUSG00000032299 | 2.165631214 | 2.05479213 | 2.180274797 | 1.874961472 | 1.983754904 | 2.161368002 | 1.958704689 |
| ENSMUSG00000032300 | 2.203267085 | 2.137686696 | 2.238549078 | 1.821350555 | 1.975997314 | 2.06069784 | 1.991877847 |
| ENSMUSG00000032301 | 2.84371221 | 2.599685923 | 2.602877747 | 2.822648722 | 2.6978141 | 3.177824972 | 2.529643178 |
| ENSMUSG00000032303 | 0 | 0 | 0 | 0 | 0.244650542 | 0 | 0 |
| ENSMUSG00000032305 | 2.042557845 | 1.894357207 | 1.892463902 | 2.015173983 | 1.951645858 | 2.193124598 | 1.865128412 |
| ENSMUSG00000032306 | 2.219867581 | 2.038704527 | 2.027131114 | 2.237091808 | 2.182733222 | 2.51054501 | 2.048913829 |
| ENSMUSG00000032307 | 2.17479713 | 2.074802005 | 2.02186571 | 2.092671852 | 2.077536464 | 2.294466226 | 1.929310074 |
| ENSMUSG00000032308 | 1.631440309 | 1.53743678 | 1.557204673 | 1.580482024 | 1.614672255 | 1.633468456 | 1.584526862 |
| ENSMUSG00000032309 | 2.185978293 | 2.072504712 | 2.058404919 | 2.109697963 | 2.084333628 | 2.383815366 | 1.984950056 |
| ENSMUSG00000032310 | 0 | 0 | 0 | 0 | 0 | 0 | 0 |
| ENSMUSG00000032311 | 1.069666964 | 0.646240625 | 0.646240625 | 0.830482024 | 1.039257769 | 0.602059991 | 0.896975297 |
| ENSMUSG00000032312 | 2.472830741 | 2.309003548 | 2.368933358 | 2.189555804 | 2.317747248 | 2.485721426 | 2.283339049 |
| ENSMUSG00000032313 | 0.45815691 | 0.79248125 | 0.5 | 0 | 0.387761935 | 0.477121255 | 0.355418052 |
| ENSMUSG00000032314 | 2.786132803 | 2.562278363 | 2.561090959 | 2.701637405 | 2.610148495 | 3.011993115 | 2.401059159 |
| ENSMUSG00000032315 | 0 | 0.25 | 0.396240625 | 0.646240625 | 0.387761935 | 0.477121255 | 0.520678036 |
| ENSMUSG00000032316 | 1.993049299 | 1.903677461 | 1.950224975 | 1.931980114 | 2.004851707 | 2.152288344 | 1.871156825 |
| ENSMUSG00000032318 | 0.45815691 | 0.5 | 0.5 | 0.864857905 | 0.846351821 | 0.602059991 | 1.224243824 |
| ENSMUSG00000032320 | 2.922773676 | 2.672499493 | 2.652294685 | 2.78296424 | 2.675113748 | 3.128399269 | 2.477402167 |
| ENSMUSG00000032322 | 0.45815691 | 0.701838731 | 0.75 | 0.25 | 0.77552387 | 0.602059991 | 0.74492186 |
| ENSMUSG00000032323 | 0 | 0 | 0.396240625 | 0.25 | 0 | 0 | 0 |
| ENSMUSG00000032324 | 2.703665763 | 2.509043403 | 2.485628626 | 2.592671852 | 2.546922567 | 2.95616843 | 2.359403919 |
| ENSMUSG00000032327 | 0.747221736 | 0.79248125 | 0.701838731 | 1.62694866 | 1.731346772 | 1.86923172 | 1.699191618 |
| ENSMUSG00000032328 | 2.983706352 | 2.742526473 | 2.741265689 | 2.87246199 | 2.777148527 | 3.200029267 | 2.554416025 |
| ENSMUSG00000032329 | 2.188178997 | 2.120953944 | 2.113831805 | 2.111770807 | 2.112954895 | 2.397940009 | 2.007923223 |

| | | | | | | | |
|---|---|---|---|---|---|---|---|
| ENSMUSG00000032330 | 2.744917684 | 2.506284891 | 2.546838018 | 2.606566189 | 2.543273062 | 2.942999593 | 2.362906151 |
| ENSMUSG00000032332 | 3.200564088 | 2.927486345 | 2.900442697 | 2.820047698 | 2.758709949 | 3.159266331 | 2.561798737 |
| ENSMUSG00000032333 | 1.647796617 | 1.606566189 | 1.67510993 | 1.476722649 | 1.632412477 | 1.799340549 | 1.690609803 |
| ENSMUSG00000032334 | 2.678024491 | 2.551448312 | 2.611770807 | 2.219895812 | 2.352243976 | 2.527629901 | 2.375711864 |
| ENSMUSG00000032336 | 3.071782542 | 2.809003548 | 2.786642169 | 2.886361795 | 2.795570098 | 3.241795431 | 2.591582503 |
| ENSMUSG00000032338 | 0 | 0 | 0 | 0 | 0 | 0.602059991 | 0.448487648 |
| ENSMUSG00000032340 | 3.146230861 | 2.868553234 | 2.875583645 | 2.673589472 | 2.59554133 | 2.992111488 | 2.40471474 |
| ENSMUSG00000032342 | 2.020182925 | 1.83272922 | 1.848079356 | 2.015173983 | 1.947418788 | 2.173186268 | 1.822941512 |
| ENSMUSG00000032343 | 0 | 0 | 0 | 0 | 0 | 0 | 0 |
| ENSMUSG00000032344 | 2.042557845 | 1.979715809 | 1.994319981 | 2.254202072 | 2.167109853 | 2.492760389 | 2.005303651 |
| ENSMUSG00000032346 | 0.45815691 | 0.580482024 | 0.79248125 | 0.580482024 | 0.489301084 | 0.84509804 | 0.672731473 |
| ENSMUSG00000032348 | 2.026699189 | 1.87892496 | 1.919870025 | 1.896240625 | 1.86473911 | 2.096910013 | 1.726776051 |
| ENSMUSG00000032349 | 2.595015862 | 2.431980114 | 2.429847205 | 2.524178789 | 2.458034312 | 2.84509804 | 2.285835109 |
| ENSMUSG00000032350 | 2.35727108 | 2.172499493 | 2.207430684 | 2.321928095 | 2.242199448 | 2.522444234 | 2.073258096 |
| ENSMUSG00000032352 | 2.045658471 | 1.935366747 | 1.864857905 | 1.809601185 | 1.83872004 | 1.949390007 | 1.74196736 |
| ENSMUSG00000032353 | 2.358731791 | 2.142463902 | 2.167221246 | 2.036164561 | 2.045754077 | 2.283301229 | 1.95794437 |
| ENSMUSG00000032355 | 0 | 0.25 | 0 | 0.25 | 0.387761935 | 0 | 0 |
| ENSMUSG00000032356 | 1.069666964 | 0.79248125 | 1.098079356 | 0.25 | 0 | 0.301029996 | 0.355418052 |
| ENSMUSG00000032357 | 0 | 0 | 0 | 0 | 0 | 0 | 0 |
| ENSMUSG00000032358 | 0 | 0 | 0 | 0.396240625 | 0.387761935 | 0.477121255 | 0.224243824 |
| ENSMUSG00000032359 | 1.822201287 | 1.777131114 | 1.837182039 | 2.04745614 | 2.028157605 | 2.290034611 | 1.849546807 |
| ENSMUSG00000032360 | 0 | 0 | 0 | 0 | 0 | 0 | 0 |
| ENSMUSG00000032363 | 2.105953527 | 1.979715809 | 1.981203126 | 1.649978211 | 1.741652793 | 1.672097858 | 1.759302692 |
| ENSMUSG00000032366 | 3.876949162 | 3.52947164 | 3.499515371 | 3.373588901 | 3.331380351 | 3.873611197 | 3.125823368 |
| ENSMUSG00000032368 | 0 | 0 | 0.25 | 1.951838731 | 1.91476933 | 2.053078443 | 1.89141329 |
| ENSMUSG00000032369 | 2.482695304 | 2.260414788 | 2.293731421 | 2.343759858 | 2.337535342 | 2.652246341 | 2.178359564 |
| ENSMUSG00000032370 | 1.862428343 | 1.809601185 | 1.752806814 | 2.031030328 | 1.965381292 | 2.28780173 | 1.893277937 |
| ENSMUSG00000032372 | 2.386920383 | 2.184523065 | 2.201838731 | 2.083847589 | 2.022473809 | 2.271841607 | 1.866140981 |
| ENSMUSG00000032373 | 0.45815691 | 0.896240625 | 0.896240625 | 1.238549078 | 1.149965125 | 1.414973348 | 1.159323753 |
| ENSMUSG00000032374 | 2.770132156 | 2.54838134 | 2.516185691 | 2.719895812 | 2.633631671 | 3.010299957 | 2.422940544 |
| ENSMUSG00000032375 | 2.045658471 | 1.890560606 | 1.940387808 | 2.070192693 | 1.955822902 | 2.23299611 | 1.800208341 |
| ENSMUSG00000032376 | 2.295901576 | 2.060793496 | 2.059601185 | 2.246460484 | 2.124299332 | 2.465382851 | 2.017561934 |
| ENSMUSG00000032377 | 1.639698646 | 1.571350555 | 1.552363341 | 1.339388001 | 1.407936347 | 1.531478917 | 1.20919388 |
| ENSMUSG00000032380 | 0.811507563 | 0.646240625 | 0.864857905 | 1.114857905 | 1.039257769 | 0.602059991 | 0.896975297 |

| | | | | | | | |
|---|---|---|---|---|---|---|---|
| ENSMUSG00000032381 | 2.334723298 | 2.121960008 | 2.103406982 | 2.138647213 | 2.113840608 | 2.350248018 | 1.977870661 |
| ENSMUSG00000032382 | 2.631440309 | 2.430701883 | 2.433677405 | 2.628678514 | 2.542486091 | 2.929418926 | 2.415625016 |
| ENSMUSG00000032383 | 2.23973002 | 2.077084758 | 2.182829758 | 2.127938163 | 2.119108707 | 2.382017043 | 2.123052075 |
| ENSMUSG00000032384 | 2.577267124 | 2.376452888 | 2.363831805 | 2.479715809 | 2.388575198 | 2.746634199 | 2.231187192 |
| ENSMUSG00000032386 | 2.45930418 | 2.251406137 | 2.259729747 | 2.528109877 | 2.445469852 | 2.8162413 | 2.266187571 |
| ENSMUSG00000032387 | 1.903481042 | 1.864857905 | 1.79248125 | 1.671625132 | 1.814781639 | 1.875061263 | 1.766260282 |
| ENSMUSG00000032388 | 2.713745586 | 2.486360959 | 2.499647607 | 2.514159429 | 2.422252432 | 2.779596491 | 2.290770186 |
| ENSMUSG00000032392 | 1.671187741 | 1.552363341 | 1.557204673 | 1.598079356 | 1.555378544 | 1.707570176 | 1.520678036 |
| ENSMUSG00000032393 | 2.722369902 | 2.504897682 | 2.502806814 | 2.684945152 | 2.579121805 | 2.962369336 | 2.387626796 |
| ENSMUSG00000032394 | 0 | 0 | 0 | 0.396240625 | 0.568060967 | 0.301029996 | 0.629532004 |
| ENSMUSG00000032396 | 2.183765914 | 2.078220739 | 2.052363341 | 2.192046081 | 2.172394856 | 2.439332694 | 2.044866219 |
| ENSMUSG00000032397 | 2.636714502 | 2.40136988 | 2.388647213 | 2.586351312 | 2.518022067 | 2.920123326 | 2.330413001 |
| ENSMUSG00000032398 | 1.806056082 | 1.678561379 | 1.71759118 | 1.905512955 | 1.800029087 | 2.11058971 | 1.722075986 |
| ENSMUSG00000032399 | 3.789685262 | 3.423943007 | 3.419154566 | 3.497126795 | 3.436031299 | 4.021396057 | 3.18923853 |
| ENSMUSG00000032400 | 2.470606563 | 2.282320754 | 2.261780978 | 2.428990498 | 2.364194729 | 2.710117365 | 2.190455009 |
| ENSMUSG00000032401 | 1.482695304 | 1.51109853 | 1.445339928 | 1.098079356 | 1.020174412 | 0.84509804 | 0.829800754 |
| ENSMUSG00000032402 | 2.621464818 | 2.406427211 | 2.365881093 | 2.431980114 | 2.396209151 | 2.72427587 | 2.275162625 |
| ENSMUSG00000032403 | 1.156259305 | 1.214495249 | 1.188721876 | 1.114857905 | 1.293076518 | 1.414973348 | 1.168187745 |
| ENSMUSG00000032405 | 2.352857998 | 2.109697963 | 2.114857905 | 2.280383379 | 2.240349892 | 2.506505032 | 2.062969708 |
| ENSMUSG00000032407 | 2.800532393 | 2.576515422 | 2.581326364 | 2.695544798 | 2.619350055 | 3.017033339 | 2.413769991 |
| ENSMUSG00000032409 | 2.481610694 | 2.269871196 | 2.237091808 | 2.386241108 | 2.27895639 | 2.57054294 | 2.105204898 |
| ENSMUSG00000032410 | 2.63895461 | 2.413211243 | 2.388167274 | 2.618933358 | 2.524713379 | 2.888740961 | 2.336847775 |
| ENSMUSG00000032411 | 2.43085913 | 2.236360959 | 2.231944491 | 2.397178659 | 2.362011938 | 2.723455672 | 2.202829437 |
| ENSMUSG00000032412 | 2.875300735 | 2.645770692 | 2.619186551 | 2.719704321 | 2.625630579 | 3.05422991 | 2.464462941 |
| ENSMUSG00000032413 | 2.151493169 | 1.985628626 | 1.997171172 | 2.093759858 | 1.987570683 | 2.173186268 | 1.845261772 |
| ENSMUSG00000032415 | 1.765759561 | 1.566696635 | 1.598079356 | 1.698603967 | 1.628935049 | 1.832508913 | 1.548827561 |
| ENSMUSG00000032417 | 1.036286563 | 0.92510993 | 0.701838731 | 1.29248125 | 1.106691885 | 1.342422681 | 1.110947927 |
| ENSMUSG00000032418 | 2.856724497 | 2.594031347 | 2.572792675 | 2.721233412 | 2.620193943 | 3.044931546 | 2.454612197 |
| ENSMUSG00000032419 | 2.029919532 | 1.982684334 | 1.966046536 | 1.910964047 | 1.871833944 | 2.06069784 | 1.763495175 |
| ENSMUSG00000032420 | 0.811507563 | 0.976722649 | 1.061981878 | 1 | 0.931471446 | 1 | 0.74492186 |
| ENSMUSG00000032422 | 2.365959569 | 2.118933358 | 2.087182039 | 2.193696765 | 2.105788027 | 2.440909082 | 1.977154126 |
| ENSMUSG00000032423 | 3.149308605 | 2.888527288 | 2.855910299 | 2.93166098 | 2.845239797 | 3.289588953 | 2.605042351 |
| ENSMUSG00000032425 | 1.614416215 | 1.470660762 | 1.42510993 | 1.53743678 | 1.42077253 | 1.799340549 | 1.42167221 |
| ENSMUSG00000032431 | 2.478339835 | 2.273189285 | 2.294354884 | 2.265173983 | 2.220647857 | 2.547774705 | 2.158505762 |

| | | | | | | | |
|---|---|---|---|---|---|---|---|
| ENSMUSG00000032434 | 2.241665209 | 2.059601185 | 2.061981878 | 2.233422664 | 2.203230928 | 2.472756449 | 2.080634989 |
| ENSMUSG00000032435 | 2.554191361 | 2.31611065 | 2.281676118 | 2.468745337 | 2.463003274 | 2.793790385 | 2.297235323 |
| ENSMUSG00000032436 | 2.262379879 | 2.134789703 | 2.146240625 | 1.978222334 | 2.074582839 | 2.31386722 | 2.000667731 |
| ENSMUSG00000032437 | 2.851389255 | 2.638407323 | 2.586351312 | 2.743244947 | 2.675474275 | 3.072249898 | 2.481049162 |
| ENSMUSG00000032440 | 2.652148299 | 2.443284802 | 2.455044741 | 2.785047676 | 2.76069684 | 3.160768562 | 2.619051282 |
| ENSMUSG00000032443 | 0 | 0.25 | 0 | 0.25 | 0.244650542 | 0 | 0.355418052 |
| ENSMUSG00000032446 | 0.747221736 | 0.701838731 | 0.396240625 | 1.201838731 | 1.106691885 | 1.079181246 | 1.089372236 |
| ENSMUSG00000032449 | 2.572054151 | 2.314935816 | 2.311981878 | 2.458222504 | 2.389791591 | 2.699837726 | 2.200320163 |
| ENSMUSG00000032451 | 0 | 0 | 0 | 0 | 0 | 0 | 0 |
| ENSMUSG00000032452 | 0 | 0 | 0 | 0.896240625 | 0.931471446 | 0.77815125 | 0.935079929 |
| ENSMUSG00000032454 | 0 | 0 | 0 | 0 | 0 | 0 | 0.355418052 |
| ENSMUSG00000032456 | 1.53838222 | 1.42510993 | 1.396240625 | 1.418106335 | 1.27449559 | 1.322219295 | 1.290497981 |
| ENSMUSG00000032458 | 3.162672026 | 2.88040011 | 2.863446264 | 2.989819376 | 2.896178514 | 3.376029182 | 2.68309143 |
| ENSMUSG00000032459 | 2.075459174 | 1.961372513 | 1.948603967 | 2.122963274 | 2.047893212 | 2.31386722 | 1.945529784 |
| ENSMUSG00000032462 | 2.527823874 | 2.339388001 | 2.269871196 | 2.566403762 | 2.513489663 | 2.880241776 | 2.327268527 |
| ENSMUSG00000032463 | 2.016886209 | 1.856566189 | 1.905512955 | 2.080482024 | 1.897237431 | 2.190331698 | 1.82753046 |
| ENSMUSG00000032468 | 2.35433422 | 2.179419106 | 2.142463902 | 2.327084758 | 2.202543574 | 2.534026106 | 2.054044578 |
| ENSMUSG00000032469 | 2.183765914 | 2.034887838 | 2.044977273 | 2.181980114 | 2.093822267 | 2.378397901 | 1.980721036 |
| ENSMUSG00000032470 | 2.444137694 | 2.294977273 | 2.245034894 | 2.376452888 | 2.369807605 | 2.682145076 | 2.223239637 |
| ENSMUSG00000032473 | 0 | 0 | 0 | 0 | 0 | 0 | 0.224243824 |
| ENSMUSG00000032475 | 2.278594844 | 2.118933358 | 2.079353153 | 2.179419106 | 2.086251872 | 2.365487985 | 1.905143021 |
| ENSMUSG00000032477 | 2.482695304 | 2.262462137 | 2.247171172 | 2.36434522 | 2.344045917 | 2.641474111 | 2.188221291 |
| ENSMUSG00000032478 | 2.131864492 | 1.987091808 | 1.96293726 | 1.964495249 | 1.917865457 | 2.209515015 | 1.836517579 |
| ENSMUSG00000032479 | 3.240082541 | 2.934417466 | 2.936523596 | 2.977191971 | 2.926286637 | 3.369772289 | 2.751406944 |
| ENSMUSG00000032480 | 2.630681378 | 2.437048212 | 2.457430684 | 2.398581151 | 2.490824419 | 2.713490543 | 2.345462945 |
| ENSMUSG00000032481 | 3.140230291 | 2.843488164 | 2.824945101 | 2.935682619 | 2.861882007 | 3.281487888 | 2.656014146 |
| ENSMUSG00000032482 | 0.671187741 | 0.75 | 0.79248125 | 0.830482024 | 0.877063019 | 0.77815125 | 0.803905701 |
| ENSMUSG00000032484 | 0 | 0 | 0 | 0 | 0 | 0 | 0 |
| ENSMUSG00000032485 | 2.305951036 | 2.122963274 | 2.229715809 | 2.071350555 | 2.199780694 | 2.324282455 | 2.177202084 |
| ENSMUSG00000032487 | 2.808542865 | 2.623463862 | 2.640321986 | 1.784887838 | 1.754126039 | 1.968482949 | 1.601000298 |
| ENSMUSG00000032489 | 0.867194479 | 0.951838731 | 1 | 0.92510993 | 1.212046813 | 1.342422681 | 1.054044578 |
| ENSMUSG00000032491 | 2.22592627 | 2.113831805 | 2.174241882 | 1.830482024 | 1.950242452 | 2.120573931 | 1.909593098 |
| ENSMUSG00000032492 | 0.578129653 | 0.25 | 0.701838731 | 0.864857905 | 1.074582839 | 0.698970004 | 0.952571511 |
| ENSMUSG00000032493 | 0 | 0 | 0 | 0 | 0 | 0 | 0 |

| | | | | | | | |
|---|---|---|---|---|---|---|---|
| ENSMUSG00000032494 | 0 | 0 | 0 | 0 | 0 | 0 | 0 |
| ENSMUSG00000032495 | 0.289064826 | 0.25 | 0 | 0.25 | 0.244650542 | 0.477121255 | 0.355418052 |
| ENSMUSG00000032496 | 0 | 0 | 0 | 0 | 0 | 0 | 0 |
| ENSMUSG00000032497 | 2.351376531 | 2.188721876 | 2.17510993 | 2.313757142 | 2.242199448 | 2.564666064 | 2.096394553 |
| ENSMUSG00000032498 | 2.177057474 | 1.931980114 | 1.942046081 | 2.02053726 | 1.947418788 | 2.238046103 | 1.87902385 |
| ENSMUSG00000032500 | 0 | 0 | 0.25 | 0 | 0.244650542 | 0.477121255 | 0.224243824 |
| ENSMUSG00000032501 | 2.167941649 | 1.990000483 | 2.05357978 | 1.642463902 | 1.603583716 | 1.826074803 | 1.589319772 |
| ENSMUSG00000032502 | 0 | 0 | 0 | 0 | 0.244650542 | 0 | 0 |
| ENSMUSG00000032503 | 0 | 0 | 0 | 0 | 0 | 0 | 0.224243824 |
| ENSMUSG00000032504 | 3.025521962 | 2.770204382 | 2.760414788 | 2.871331547 | 2.804682415 | 3.237292338 | 2.632194697 |
| ENSMUSG00000032507 | 1.568542279 | 1.488549078 | 1.52186571 | 1.653677461 | 1.649303553 | 1.86923172 | 1.607811402 |
| ENSMUSG00000032508 | 1.912079952 | 1.830482024 | 1.821350555 | 2.046218836 | 2.039257769 | 2.28780173 | 1.926805425 |
| ENSMUSG00000032511 | 0 | 0.25 | 0 | 0.580482024 | 0.568060967 | 0.602059991 | 0.876096088 |
| ENSMUSG00000032512 | 2.516991255 | 2.296838018 | 2.31728167 | 2.413211243 | 2.335180723 | 2.677606953 | 2.184841529 |
| ENSMUSG00000032513 | 1.857607105 | 1.843759858 | 1.860735874 | 1.606566189 | 1.83872004 | 1.73239376 | 1.767634022 |
| ENSMUSG00000032514 | 0 | 0.5 | 0.5 | 0.25 | 0.632412477 | 0.301029996 | 0.629532004 |
| ENSMUSG00000032515 | 1.505869732 | 1.42510993 | 1.557204673 | 1.42510993 | 1.576269619 | 1.477121255 | 1.610050272 |
| ENSMUSG00000032517 | 0 | 0 | 0 | 0 | 0 | 0 | 0 |
| ENSMUSG00000032518 | 3.505310458 | 3.23106395 | 3.246593845 | 3.160456493 | 3.163059566 | 3.622214023 | 2.991285356 |
| ENSMUSG00000032519 | 1.83262764 | 1.711372513 | 1.819031101 | 1.671625132 | 1.820920161 | 1.892094603 | 1.806347338 |
| ENSMUSG00000032523 | 0.289064826 | 0.5 | 0.646240625 | 0 | 0 | 0 | 0 |
| ENSMUSG00000032525 | 3.034559718 | 2.765680192 | 2.756111781 | 2.827653195 | 2.775930736 | 3.165244326 | 2.558009347 |
| ENSMUSG00000032526 | 1.389637216 | 1.488549078 | 1.482684334 | 1.634789703 | 1.559656853 | 1.740362689 | 1.425641774 |
| ENSMUSG00000032527 | 1.985980784 | 1.850219859 | 1.868933358 | 1.928561379 | 2 | 2.093421685 | 1.914853627 |
| ENSMUSG00000032528 | 0.578129653 | 0.396240625 | 0.5 | 0.646240625 | 0.568060967 | 0.698970004 | 0.224243824 |
| ENSMUSG00000032530 | 0 | 0 | 0 | 0 | 0 | 0 | 0 |
| ENSMUSG00000032531 | 3.196867486 | 2.950123875 | 2.985170162 | 2.681980114 | 2.73425682 | 3.034227261 | 2.642631585 |
| ENSMUSG00000032532 | 0 | 0 | 0 | 0 | 0 | 0 | 0 |
| ENSMUSG00000032534 | 2.188178997 | 2.015173983 | 2.039967834 | 2.111770807 | 2.068600388 | 2.267171728 | 1.920884716 |
| ENSMUSG00000032536 | 2.134369199 | 2.081607372 | 2.124961472 | 2.048689214 | 2.121713561 | 2.271841607 | 2.075912052 |
| ENSMUSG00000032537 | 0 | 0 | 0 | 0 | 0 | 0 | 0 |
| ENSMUSG00000032540 | 2.211649928 | 2.078220739 | 2.071350555 | 2.119945066 | 2.080465577 | 2.359835482 | 1.984249047 |
| ENSMUSG00000032547 | 2.917079017 | 2.694929839 | 2.702039943 | 2.77908599 | 2.727014069 | 3.146748014 | 2.556217675 |
| ENSMUSG00000032548 | 2.425923778 | 2.228222334 | 2.26109853 | 2.103406982 | 2.104881849 | 2.303196057 | 1.980721036 |

| | | | | | | | |
|---|---|---|---|---|---|---|---|
| ENSMUSG00000032549 | 1.903481042 | 1.841580554 | 1.852347734 | 1.527131114 | 1.607318736 | 1.72427587 | 1.520678036 |
| ENSMUSG00000032551 | 1.83262764 | 1.692046081 | 1.638647213 | 1.87892496 | 1.770879644 | 1.995635195 | 1.62316756 |
| ENSMUSG00000032553 | 2.273259602 | 2.157339155 | 2.146240625 | 2.165444524 | 2.194892994 | 2.456366033 | 2.057420422 |
| ENSMUSG00000032554 | 0.289064826 | 0.5 | 0.25 | 0.25 | 0 | 0.698970004 | 0.224243824 |
| ENSMUSG00000032555 | 3.118874669 | 2.840622952 | 2.840759908 | 2.965949776 | 2.884216078 | 3.33243846 | 2.699499435 |
| ENSMUSG00000032556 | 0.289064826 | 0 | 0 | 0 | 0 | 0 | 0 |
| ENSMUSG00000032557 | 2.479432974 | 2.272528105 | 2.257666784 | 2.295598588 | 2.253693465 | 2.572871602 | 2.117521761 |
| ENSMUSG00000032558 | 1.614416215 | 1.403677461 | 1.5 | 1.54248125 | 1.618292349 | 1.69019608 | 1.529383012 |
| ENSMUSG00000032560 | 3.120285948 | 2.852745317 | 2.852612838 | 2.87521047 | 2.7738917 | 3.229681842 | 2.569936457 |
| ENSMUSG00000032561 | 0.289064826 | 0.25 | 0.396240625 | 0.701838731 | 0.686820904 | 0.698970004 | 0.829800754 |
| ENSMUSG00000032562 | 3.181338949 | 2.924133229 | 2.956537804 | 2.92140621 | 2.975535632 | 3.376394442 | 2.84022295 |
| ENSMUSG00000032563 | 2.479432974 | 2.252107156 | 2.241446071 | 2.410512923 | 2.307165329 | 2.656098202 | 2.163382972 |
| ENSMUSG00000032564 | 0 | 0 | 0 | 0 | 0 | 0 | 0 |
| ENSMUSG00000032565 | 1.129344651 | 1.17510993 | 1.188721876 | 1.282320754 | 1.302012594 | 1.255272505 | 1.176815335 |
| ENSMUSG00000032566 | 0 | 0 | 0 | 0 | 0 | 0 | 0 |
| ENSMUSG00000032567 | 1.647796617 | 1.720660762 | 1.54248125 | 1.630890489 | 1.499532413 | 1.612783857 | 1.466365979 |
| ENSMUSG00000032570 | 2.569423023 | 2.36434522 | 2.345385738 | 2.467205639 | 2.398190943 | 2.733999287 | 2.203542812 |
| ENSMUSG00000032571 | 2.199011616 | 2.05357978 | 2.052363341 | 2.156427211 | 2.143518259 | 2.356025857 | 2.008574816 |
| ENSMUSG00000032572 | 0 | 0.25 | 0.5 | 0.92510993 | 0.877063019 | 0.77815125 | 0.896975297 |
| ENSMUSG00000032575 | 2.575536697 | 2.39151351 | 2.383332433 | 2.505591953 | 2.446849937 | 2.835690571 | 2.335428845 |
| ENSMUSG00000032577 | 1.605636262 | 1.5 | 1.52186571 | 1.396240625 | 1.401338844 | 1.579783597 | 1.290497981 |
| ENSMUSG00000032578 | 1.639698646 | 1.571350555 | 1.606566189 | 1.282320754 | 1.283908311 | 1.255272505 | 1.252393349 |
| ENSMUSG00000032579 | 1.605636262 | 1.557204673 | 1.638647213 | 1.598079356 | 1.72341484 | 1.716003344 | 1.651852264 |
| ENSMUSG00000032580 | 2.809533441 | 2.589388001 | 2.562574604 | 2.697994205 | 2.653546792 | 3.013679697 | 2.432034165 |
| ENSMUSG00000032582 | 2.651426165 | 2.43155454 | 2.414552871 | 2.550530956 | 2.439543536 | 2.791690649 | 2.253616472 |
| ENSMUSG00000032583 | 1.445324132 | 1.51109853 | 1.516522298 | 1.356566189 | 1.603583716 | 1.51851394 | 1.661510159 |
| ENSMUSG00000032584 | 0.289064826 | 0 | 0.25 | 0.646240625 | 0.632412477 | 0 | 0.710836105 |
| ENSMUSG00000032586 | 2.167941649 | 2.006976499 | 2.004202072 | 2.166333979 | 2.136951444 | 2.424881637 | 2.053478496 |
| ENSMUSG00000032589 | 0.578129653 | 0.75 | 0.976722649 | 0.646240625 | 0.931471446 | 0.477121255 | 1 |
| ENSMUSG00000032590 | 2.295901576 | 2.162762923 | 2.201032755 | 2.25835575 | 2.308186911 | 2.547774705 | 2.206380685 |
| ENSMUSG00000032591 | 0.289064826 | 0.25 | 0 | 0.5 | 0.568060967 | 0 | 0.520678036 |
| ENSMUSG00000032593 | 1.036286563 | 1.080482024 | 1.188721876 | 1.114857905 | 1.380772476 | 1.255272505 | 1.302263476 |
| ENSMUSG00000032594 | 2.593357679 | 2.390083209 | 2.401832578 | 2.347004321 | 2.396209151 | 2.64246452 | 2.264126018 |
| ENSMUSG00000032595 | 0 | 0 | 0 | 0 | 0.244650542 | 0 | 0 |

| | | | | | | | |
|---|---|---|---|---|---|---|---|
| ENSMUSG00000032596 | 1.181541736 | 1.098079356 | 1.26109853 | 1.54745614 | 1.715300549 | 1.716003344 | 1.627424404 |
| ENSMUSG00000032598 | 1.894701088 | 1.823655187 | 1.866901388 | 1.614857905 | 1.859322797 | 1.86332286 | 1.805128823 |
| ENSMUSG00000032599 | 2.253091469 | 2.116901388 | 2.145300145 | 2.092671852 | 2.126012715 | 2.307496038 | 2.066254157 |
| ENSMUSG00000032601 | 2.910917762 | 2.68473417 | 2.690387808 | 2.748234584 | 2.668744168 | 3.062957834 | 2.476177308 |
| ENSMUSG00000032602 | 2.089597219 | 1.914552871 | 1.92510993 | 2.074802005 | 2.017859938 | 2.340444115 | 1.950241257 |
| ENSMUSG00000032604 | 2.795446569 | 2.521534056 | 2.54248125 | 2.66942991 | 2.611014646 | 2.989449818 | 2.444621219 |
| ENSMUSG00000032606 | 2.144240249 | 2.001406137 | 2.024508021 | 2.090485943 | 2.081436571 | 2.344392274 | 1.96172822 |
| ENSMUSG00000032607 | 1.482695304 | 1.445339928 | 1.482684334 | 1.494319981 | 1.478764298 | 1.51851394 | 1.42167221 |
| ENSMUSG00000032609 | 1.494443473 | 1.54745614 | 1.606566189 | 1.516522298 | 1.744182955 | 1.792391689 | 1.694069894 |
| ENSMUSG00000032611 | 0.289064826 | 0.25 | 0.646240625 | 0.396240625 | 0.489301084 | 0.301029996 | 0.579661877 |
| ENSMUSG00000032612 | 2.845531329 | 2.567865757 | 2.568157446 | 2.714495249 | 2.662444522 | 3.059184618 | 2.478012861 |
| ENSMUSG00000032615 | 1.734388958 | 1.678561379 | 1.688721876 | 1.711372513 | 1.736537397 | 1.799340549 | 1.674564426 |
| ENSMUSG00000032621 | 2.593357679 | 2.393884812 | 2.397646762 | 2.49539182 | 2.42883626 | 2.709269961 | 2.262348456 |
| ENSMUSG00000032623 | 0 | 0 | 0.396240625 | 0 | 0 | 0 | 0 |
| ENSMUSG00000032624 | 2.677345838 | 2.483422664 | 2.423371739 | 2.457826811 | 2.388575198 | 2.713490543 | 2.222232323 |
| ENSMUSG00000032625 | 2.003432385 | 1.919870025 | 1.839388001 | 1.886723615 | 1.880506521 | 2.041392685 | 1.712465909 |
| ENSMUSG00000032633 | 2.473938397 | 2.307204673 | 2.352347734 | 2.149978211 | 2.194189193 | 2.367355921 | 2.109521699 |
| ENSMUSG00000032637 | 2.641922824 | 2.481573999 | 2.52219706 | 2.331045137 | 2.405234647 | 2.633468456 | 2.3030503 |
| ENSMUSG00000032640 | 2.048736215 | 1.884789703 | 1.901832578 | 1.720660762 | 1.766173492 | 1.929418926 | 1.612273755 |
| ENSMUSG00000032641 | 1.205378646 | 1.26109853 | 1.26109853 | 0.830482024 | 1.057362051 | 0.84509804 | 1 |
| ENSMUSG00000032643 | 2.22189694 | 2.191217898 | 2.281676118 | 1.735628626 | 1.916320788 | 1.949390007 | 1.865128412 |
| ENSMUSG00000032648 | 0.747221736 | 0.75 | 0.896240625 | 0.396240625 | 0.632412477 | 0.602059991 | 0.803905701 |
| ENSMUSG00000032649 | 0 | 0 | 0 | 0.396240625 | 0.244650542 | 0.301029996 | 0.448487648 |
| ENSMUSG00000032652 | 1.663535556 | 1.62694866 | 1.54248125 | 1.54745614 | 1.542223377 | 1.72427587 | 1.441050184 |
| ENSMUSG00000032656 | 1.249317394 | 1.26109853 | 1.25 | 1.04248125 | 1.366364104 | 1.278753601 | 1.290497981 |
| ENSMUSG00000032657 | 0.578129653 | 1.04248125 | 1.226722649 | 1.575945187 | 1.77552387 | 1.770852012 | 1.783679399 |
| ENSMUSG00000032661 | 0.578129653 | 0.396240625 | 0 | 0.646240625 | 0.77552387 | 0.301029996 | 0.710836105 |
| ENSMUSG00000032666 | 2.365959569 | 2.206637122 | 2.160061234 | 2.295598588 | 2.184185718 | 2.451786436 | 2.023210262 |
| ENSMUSG00000032667 | 2.427163103 | 2.224461364 | 2.234159485 | 2.208222504 | 2.176862714 | 2.434568904 | 2.01501956 |
| ENSMUSG00000032671 | 0 | 0 | 0 | 0 | 0 | 0 | 0 |
| ENSMUSG00000032673 | 1.269664473 | 1.311981878 | 1.27186571 | 1.692046081 | 1.563883923 | 1.826074803 | 1.517723536 |
| ENSMUSG00000032679 | 0 | 0 | 0 | 1.130890489 | 1 | 1.322219295 | 1.089372236 |
| ENSMUSG00000032680 | 0.671187741 | 0.25 | 0 | 0.5 | 0.244650542 | 0.301029996 | 0.355418052 |
| ENSMUSG00000032688 | 1.837744666 | 1.642463902 | 1.664552871 | 1.955044741 | 1.830919587 | 2.123851641 | 1.759302692 |

| | | | | | | | |
|---|---|---|---|---|---|---|---|
| ENSMUSG00000032690 | 0 | 0 | 0 | 0.396240625 | 0.244650542 | 0.301029996 | 0 |
| ENSMUSG00000032691 | 0 | 0 | 0 | 0 | 0.244650542 | 0 | 0 |
| ENSMUSG00000032698 | 0 | 0.396240625 | 0.25 | 0.396240625 | 0.244650542 | 0.477121255 | 0.448487648 |
| ENSMUSG00000032702 | 2.124258828 | 1.992885888 | 2.028435542 | 1.774508021 | 1.840643511 | 2.029383778 | 1.811176017 |
| ENSMUSG00000032705 | 2.036286563 | 1.898114259 | 1.823655187 | 1.714495249 | 1.715300549 | 1.851258349 | 1.621017948 |
| ENSMUSG00000032712 | 2.814451333 | 2.5896628 | 2.545908845 | 2.467205639 | 2.393816219 | 2.749736316 | 2.221895853 |
| ENSMUSG00000032714 | 2.495495257 | 2.361770807 | 2.393884812 | 2.312574604 | 2.376174996 | 2.64246452 | 2.341997913 |
| ENSMUSG00000032715 | 1.783508299 | 1.668106335 | 1.516522298 | 1.080482024 | 1.234113756 | 1.176091259 | 1.272006351 |
| ENSMUSG00000032717 | 0 | 0 | 0 | 0.396240625 | 0.632412477 | 0.602059991 | 1.041356071 |
| ENSMUSG00000032718 | 0 | 0 | 0 | 0.646240625 | 0.686820904 | 0.477121255 | 0.355418052 |
| ENSMUSG00000032719 | 0 | 0 | 0 | 0 | 0 | 0 | 0 |
| ENSMUSG00000032724 | 1.655740331 | 1.71759118 | 1.75835575 | 1.698603967 | 1.836786029 | 1.86332286 | 1.801445495 |
| ENSMUSG00000032725 | 0.289064826 | 0.25 | 0.396240625 | 0.396240625 | 0.244650542 | 0.477121255 | 0 |
| ENSMUSG00000032726 | 0.747221736 | 0.396240625 | 0.396240625 | 0.864857905 | 0.877063019 | 0.77815125 | 0.853775828 |
| ENSMUSG00000032727 | 2.401648579 | 2.181980114 | 2.155512955 | 2.402294685 | 2.33281029 | 2.621176282 | 2.158095989 |
| ENSMUSG00000032733 | 2.260538674 | 2.100219859 | 2.133818844 | 1.998588359 | 2.03596422 | 2.201397124 | 1.938331368 |
| ENSMUSG00000032735 | 0.811507563 | 0.864857905 | 1.04248125 | 0.646240625 | 0.632412477 | 0.602059991 | 0.672731473 |
| ENSMUSG00000032737 | 2.663535556 | 2.511780978 | 2.524836953 | 2.24929487 | 2.33281029 | 2.505149978 | 2.217489603 |
| ENSMUSG00000032739 | 0.289064826 | 0 | 0.5 | 0 | 0.244650542 | 0 | 0.224243824 |
| ENSMUSG00000032740 | 2.66283289 | 2.484527332 | 2.459406983 | 2.608657057 | 2.527738343 | 2.892651034 | 2.307731473 |
| ENSMUSG00000032741 | 2.267855219 | 2.173371739 | 2.175975893 | 2.081607372 | 2.167109853 | 2.324282455 | 2.104237736 |
| ENSMUSG00000032743 | 1.964026642 | 1.898114259 | 1.802363341 | 1.698603967 | 1.820920161 | 1.959041392 | 1.710836105 |
| ENSMUSG00000032744 | 0 | 0 | 0 | 0 | 0.244650542 | 0 | 0.224243824 |
| ENSMUSG00000032745 | 2.777675583 | 2.58272922 | 2.566696635 | 2.814494265 | 2.718983046 | 3.146438135 | 2.486443709 |
| ENSMUSG00000032750 | 0.289064826 | 0.701838731 | 0.951838731 | 0.830482024 | 0.978602168 | 0.954242509 | 0.672731473 |
| ENSMUSG00000032754 | 2.113897241 | 1.959800947 | 2.059601185 | 1.76109853 | 1.84445929 | 2 | 1.853775828 |
| ENSMUSG00000032757 | 2.318984359 | 2.146240625 | 2.14151351 | 2.036164561 | 1.98503142 | 2.240549248 | 1.833176598 |
| ENSMUSG00000032758 | 0 | 0 | 0 | 0 | 0 | 0 | 0 |
| ENSMUSG00000032763 | 2.146671932 | 2.075945187 | 2.136723615 | 1.850219859 | 1.962676635 | 2.012837225 | 1.87805087 |
| ENSMUSG00000032766 | 2.153883046 | 2.049918086 | 2.032320754 | 1.610735874 | 1.625423019 | 1.77815125 | 1.511732338 |
| ENSMUSG00000032769 | 0 | 0.25 | 0.25 | 0 | 0 | 0.301029996 | 0.224243824 |
| ENSMUSG00000032773 | 0.578129653 | 0 | 0 | 0 | 0.244650542 | 0.698970004 | 0.355418052 |
| ENSMUSG00000032776 | 0.578129653 | 0.396240625 | 0.5 | 0.5 | 0.686820904 | 0.602059991 | 0.672731473 |
| ENSMUSG00000032777 | 2.68005385 | 2.471424093 | 2.491446071 | 2.427701608 | 2.485000434 | 2.738780558 | 2.350026003 |

| | | | | | | | |
|---|---|---|---|---|---|---|---|
| ENSMUSG00000032782 | 2.081172036 | 1.943696765 | 2.043731421 | 2.02186571 | 2.11997913 | 2.283301229 | 2.050633136 |
| ENSMUSG00000032783 | 2.330063669 | 2.178561379 | 2.240000483 | 2.127938163 | 2.223252711 | 2.426511261 | 2.160546876 |
| ENSMUSG00000032786 | 2.29078455 | 2.215271726 | 2.250703754 | 2.099151195 | 2.183460217 | 2.428134794 | 2.113781657 |
| ENSMUSG00000032788 | 2.158622101 | 2.052363341 | 2.123963757 | 2.118933358 | 2.187791022 | 2.419955748 | 2.074322292 |
| ENSMUSG00000032796 | 0.289064826 | 0 | 0 | 0.830482024 | 1.121713561 | 0.84509804 | 1.089372236 |
| ENSMUSG00000032802 | 2.229917041 | 2.015173983 | 2.059601185 | 2.198603967 | 2.141069998 | 2.444044796 | 1.991877847 |
| ENSMUSG00000032803 | 3.188976382 | 2.886723615 | 2.850753006 | 2.923153875 | 2.829058858 | 3.281260687 | 2.618557742 |
| ENSMUSG00000032806 | 2.072573125 | 1.928561379 | 1.890560606 | 1.903677461 | 1.893954095 | 2.155336037 | 1.786277931 |
| ENSMUSG00000032807 | 0 | 0 | 0 | 0.25 | 0.686820904 | 0.301029996 | 0.355418052 |
| ENSMUSG00000032808 | 0 | 0 | 0 | 0 | 0 | 0 | 0 |
| ENSMUSG00000032812 | 1.978790393 | 1.953445298 | 1.988549078 | 1.708222504 | 1.873585591 | 1.897627091 | 1.844181586 |
| ENSMUSG00000032815 | 2.432083896 | 2.260414788 | 2.269203899 | 2.376452888 | 2.433516012 | 2.649334859 | 2.299626824 |
| ENSMUSG00000032816 | 1 | 0.830482024 | 0.896240625 | 1.311981878 | 1.223252711 | 1.278753601 | 1.193409508 |
| ENSMUSG00000032818 | 0 | 0 | 0 | 0 | 0 | 0 | 0 |
| ENSMUSG00000032826 | 1.822201287 | 1.755591953 | 1.79248125 | 1.83272922 | 1.78687392 | 1.929418926 | 1.690609803 |
| ENSMUSG00000032827 | 2.097855556 | 1.916333979 | 1.938721876 | 1.660964047 | 1.639266209 | 1.653212514 | 1.505628092 |
| ENSMUSG00000032834 | 2.516991255 | 2.313166358 | 2.309601185 | 2.31728167 | 2.388168801 | 2.666517981 | 2.231187192 |
| ENSMUSG00000032839 | 1.623015126 | 1.476722649 | 1.482684334 | 1.610735874 | 1.628935049 | 1.819543936 | 1.38802993 |
| ENSMUSG00000032840 | 2.360187403 | 2.185366747 | 2.221424093 | 2.107613138 | 2.097547432 | 2.352182518 | 1.976436 |
| ENSMUSG00000032841 | 1.753493018 | 1.720660762 | 1.705044741 | 1.53743678 | 1.55104774 | 1.672097858 | 1.493062805 |
| ENSMUSG00000032842 | 1.389637216 | 1.54745614 | 1.476722649 | 1.356566189 | 1.559656853 | 1.51851394 | 1.548827561 |
| ENSMUSG00000032845 | 0.867194479 | 0.864857905 | 0.896240625 | 0.92510993 | 0.733951626 | 0.698970004 | 0.672731473 |
| ENSMUSG00000032846 | 1.558729299 | 1.458222504 | 1.42510993 | 1.75 | 1.812711509 | 2.053078443 | 1.692344475 |
| ENSMUSG00000032849 | 2.033115198 | 1.926839783 | 1.966046536 | 2.044977273 | 2.001219193 | 2.161368002 | 1.902443294 |
| ENSMUSG00000032850 | 1.418409478 | 1.464495249 | 1.451838731 | 1.410964047 | 1.47337555 | 1.612783857 | 1.502532226 |
| ENSMUSG00000032852 | 0 | 0 | 0.25 | 0 | 0 | 0.301029996 | 0 |
| ENSMUSG00000032854 | 0.811507563 | 1.061981878 | 1.080482024 | 1.692046081 | 1.731346772 | 1.913813852 | 1.546120303 |
| ENSMUSG00000032855 | 3.038863496 | 2.825945187 | 2.854199382 | 2.5684489 | 2.687866184 | 2.965671971 | 2.555498219 |
| ENSMUSG00000032860 | 0.578129653 | 0.864857905 | 0.896240625 | 1.17510993 | 1.254881871 | 1.301029996 | 1.21680636 |
| ENSMUSG00000032864 | 0 | 0 | 0 | 0 | 0 | 0.301029996 | 0 |
| ENSMUSG00000032867 | 2.887377404 | 2.636723615 | 2.665221818 | 2.577084758 | 2.618292349 | 2.956648579 | 2.461908018 |
| ENSMUSG00000032869 | 2.083999385 | 1.969129237 | 1.918106335 | 2.116901388 | 2.115605392 | 2.357934847 | 2.031805106 |
| ENSMUSG00000032870 | 2.390989024 | 2.184523065 | 2.201032755 | 2.214495249 | 2.153935427 | 2.401400541 | 2.031198704 |
| ENSMUSG00000032872 | 2.235832493 | 2.108657057 | 2.107613138 | 2.338286706 | 2.219993628 | 2.53529412 | 2.081155529 |

| | | | | | | | |
|---|---|---|---|---|---|---|---|
| ENSMUSG00000032875 | 2.705574381 | 2.601816941 | 2.647178659 | 2.209012589 | 2.371942857 | 2.544068044 | 2.344312053 |
| ENSMUSG00000032878 | 0 | 0.25 | 0.396240625 | 0 | 0 | 0 | 0 |
| ENSMUSG00000032883 | 2.484856099 | 2.273849256 | 2.277131114 | 2.356041572 | 2.271112598 | 2.613841822 | 2.136039214 |
| ENSMUSG00000032889 | 0.578129653 | 0.5 | 0.25 | 0.830482024 | 0.77552387 | 0.602059991 | 0.579661877 |
| ENSMUSG00000032890 | 0.578129653 | 0.701838731 | 0.646240625 | 1.380890489 | 1.42077253 | 1.505149978 | 1.340368108 |
| ENSMUSG00000032892 | 0 | 0 | 0 | 0 | 0.244650542 | 0 | 0 |
| ENSMUSG00000032894 | 0 | 0 | 0 | 0 | 0 | 0 | 0 |
| ENSMUSG00000032897 | 2.336265009 | 2.187048212 | 2.190387808 | 2.21293726 | 2.256066288 | 2.498310554 | 2.126234965 |
| ENSMUSG00000032898 | 2.348397716 | 2.204245906 | 2.214495249 | 2.144357207 | 2.162515999 | 2.354108439 | 2.05629905 |
| ENSMUSG00000032899 | 2.081172036 | 1.975216702 | 1.910964047 | 1.566696635 | 1.523883869 | 1.72427587 | 1.437266334 |
| ENSMUSG00000032900 | 0 | 0 | 0 | 0 | 0 | 0 | 0 |
| ENSMUSG00000032902 | 2.603209296 | 2.398581151 | 2.377938163 | 2.490362424 | 2.402902329 | 2.741939078 | 2.24869599 |
| ENSMUSG00000032905 | 2.199011616 | 2.013820609 | 1.982684334 | 2.058404919 | 2.016696984 | 2.271841607 | 1.933441873 |
| ENSMUSG00000032908 | 0.289064826 | 0.580482024 | 0.701838731 | 0.92510993 | 0.955822902 | 0.954242509 | 0.775756176 |
| ENSMUSG00000032911 | 0 | 0 | 0.5 | 1.862802778 | 2.077536464 | 2.164352856 | 2.085803078 |
| ENSMUSG00000032913 | 2.057835465 | 1.958222504 | 1.975216702 | 2.088286706 | 2.013184954 | 2.29666519 | 1.881925353 |
| ENSMUSG00000032915 | 0 | 0 | 0 | 0.25 | 0 | 0 | 0 |
| ENSMUSG00000032921 | 0 | 0.25 | 0.25 | 0 | 0 | 0 | 0.224243824 |
| ENSMUSG00000032925 | 0.289064826 | 0.580482024 | 0.5 | 0.5 | 0.244650542 | 0.477121255 | 0.520678036 |
| ENSMUSG00000032932 | 2.764107946 | 2.517865591 | 2.512462137 | 2.597004321 | 2.535323264 | 2.914343157 | 2.310046902 |
| ENSMUSG00000032936 | 0.289064826 | 0.25 | 0 | 0.951838731 | 1.091002363 | 0.84509804 | 1.014380832 |
| ENSMUSG00000032937 | 0 | 0 | 0 | 0 | 0 | 0 | 0 |
| ENSMUSG00000032939 | 2.901104782 | 2.645770692 | 2.627443751 | 2.761780978 | 2.698489937 | 3.109915863 | 2.514605617 |
| ENSMUSG00000032940 | 0.45815691 | 0.396240625 | 0 | 0.580482024 | 0.846351821 | 0.477121255 | 0.579661877 |
| ENSMUSG00000032942 | 0 | 0 | 0 | 0 | 0 | 0 | 0 |
| ENSMUSG00000032946 | 0.289064826 | 0.75 | 0.864857905 | 0.75 | 0.489301084 | 0.477121255 | 0.579661877 |
| ENSMUSG00000032948 | 0 | 0.396240625 | 0 | 0 | 0 | 0 | 0 |
| ENSMUSG00000032952 | 1.45815691 | 1.25 | 1.238549078 | 1.27186571 | 1.212046813 | 1.531478917 | 1.168187745 |
| ENSMUSG00000032959 | 3.002187511 | 2.77401406 | 2.817719824 | 2.715271726 | 2.708967143 | 3.065579715 | 2.569131835 |
| ENSMUSG00000032965 | 1.952600383 | 1.870953944 | 1.882845365 | 1.953445298 | 1.86473911 | 2.133538908 | 1.74047993 |
| ENSMUSG00000032966 | 2.956791698 | 2.70920984 | 2.676408097 | 2.796838018 | 2.728412533 | 3.128076013 | 2.520945293 |
| ENSMUSG00000032968 | 0.289064826 | 0.864857905 | 0.830482024 | 0.830482024 | 0.733951626 | 0 | 0.896975297 |
| ENSMUSG00000032977 | 2.054824387 | 1.992885888 | 2.060793496 | 1.850219859 | 1.996316946 | 2.096910013 | 2 |
| ENSMUSG00000032978 | 0 | 0 | 0 | 0 | 0 | 0 | 0.224243824 |

| | | | | | | | |
|---|---|---|---|---|---|---|---|
| ENSMUSG00000032985 | 0 | 0 | 0 | 0 | 0 | 0 | 0 |
| ENSMUSG00000032987 | 0 | 0 | 0 | 0 | 0 | 0 | 0 |
| ENSMUSG00000032988 | 0.289064826 | 0.580482024 | 0.5 | 0.25 | 0.387761935 | 0 | 0.520678036 |
| ENSMUSG00000032997 | 1.940852214 | 2.031030328 | 2.176839783 | 1.634789703 | 1.938809675 | 1.763427994 | 2.130281771 |
| ENSMUSG00000032998 | 2.543094501 | 2.331607372 | 2.28360658 | 2.302972074 | 2.221952687 | 2.57054294 | 2.087337541 |
| ENSMUSG00000032999 | 0 | 0 | 0 | 0 | 0.244650542 | 0 | 0.224243824 |
| ENSMUSG00000033004 | 2.68005385 | 2.466046536 | 2.439139081 | 2.583009137 | 2.536126349 | 2.859738566 | 2.346609758 |
| ENSMUSG00000033006 | 0 | 0 | 0 | 0 | 0 | 0 | 0 |
| ENSMUSG00000033007 | 0 | 0.25 | 0 | 0 | 0.244650542 | 0.301029996 | 0 |
| ENSMUSG00000033009 | 2.45469604 | 2.256976499 | 2.208222504 | 2.414998973 | 2.376595432 | 2.701567985 | 2.190455009 |
| ENSMUSG00000033014 | 2.562324428 | 2.380890489 | 2.339388001 | 2.445749552 | 2.359374645 | 2.694605199 | 2.199599638 |
| ENSMUSG00000033015 | 0 | 0 | 0 | 0 | 0 | 0 | 0 |
| ENSMUSG00000033016 | 2.144240249 | 2.048689214 | 2.089388001 | 1.77186571 | 1.905314583 | 2.012837225 | 1.869159809 |
| ENSMUSG00000033020 | 2.307602651 | 2.219129237 | 2.33272922 | 2.03743678 | 2.242813819 | 2.29666519 | 2.233468922 |
| ENSMUSG00000033021 | 2.215778996 | 2.067865757 | 2.063166358 | 1.959800947 | 2.114724103 | 2.269512944 | 2.040177507 |
| ENSMUSG00000033022 | 0.747221736 | 0.5 | 0.580482024 | 1.061981878 | 1.039257769 | 1.176091259 | 0.969165684 |
| ENSMUSG00000033024 | 0 | 0 | 0 | 0 | 0 | 0 | 0 |
| ENSMUSG00000033027 | 0 | 0 | 0 | 0 | 0 | 0 | 0 |
| ENSMUSG00000033029 | 1.036286563 | 0.701838731 | 0.864857905 | 0.646240625 | 0.733951626 | 0.77815125 | 0.896975297 |
| ENSMUSG00000033031 | 2.872757846 | 2.660287149 | 2.633332433 | 2.833009137 | 2.730576966 | 3.167021796 | 2.522012122 |
| ENSMUSG00000033032 | 1.289064826 | 1.372963274 | 1.302363341 | 1.42510993 | 1.407936347 | 1.643452676 | 1.429563222 |
| ENSMUSG00000033033 | 1.631440309 | 1.630890489 | 1.744319981 | 1.380890489 | 1.635855979 | 1.672097858 | 1.579661877 |
| ENSMUSG00000033036 | 1.578129653 | 1.364857905 | 1.458222504 | 1.598079356 | 1.514343934 | 1.681241237 | 1.374453863 |
| ENSMUSG00000033039 | 2.211649928 | 2.088286706 | 2.156427211 | 2.05479213 | 2.095689764 | 2.309630167 | 2.081675233 |
| ENSMUSG00000033044 | 0 | 0 | 0 | 0 | 0 | 0 | 0 |
| ENSMUSG00000033047 | 3.124026174 | 2.843216265 | 2.860865403 | 2.891632446 | 2.863577895 | 3.317436497 | 2.696252658 |
| ENSMUSG00000033053 | 0 | 0 | 0 | 0 | 0 | 0 | 0 |
| ENSMUSG00000033054 | 2.647066901 | 2.414552871 | 2.420309603 | 2.530059469 | 2.460361869 | 2.859138297 | 2.253921531 |
| ENSMUSG00000033055 | 2.129344651 | 1.981203126 | 2.064346961 | 1.988549078 | 2.09661982 | 2.255272505 | 2.05853792 |
| ENSMUSG00000033059 | 2.502784398 | 2.304186465 | 2.311981878 | 2.414998973 | 2.439191811 | 2.7355989 | 2.304357435 |
| ENSMUSG00000033060 | 1.789260561 | 1.692046081 | 1.692046081 | 1.848079356 | 1.804307395 | 2.064457989 | 1.687112305 |
| ENSMUSG00000033061 | 0 | 0 | 0 | 0 | 0 | 0 | 0 |
| ENSMUSG00000033063 | 0 | 0 | 0 | 0 | 0 | 0 | 0 |
| ENSMUSG00000033065 | 2.575536697 | 2.400442697 | 2.445749552 | 2.473325383 | 2.539850111 | 2.815577748 | 2.407002425 |

| | | | | | | | |
|---|---|---|---|---|---|---|---|
| ENSMUSG00000033066 | 0 | 0 | 0.25 | 0.75 | 0.931471446 | 0.954242509 | 0.829800754 |
| ENSMUSG00000033068 | 1.678702042 | 1.527131114 | 1.532320754 | 1.692046081 | 1.733951626 | 1.792391689 | 1.637827726 |
| ENSMUSG00000033075 | 2.842799666 | 2.592399337 | 2.572216519 | 2.63503201 | 2.554618136 | 2.95230801 | 2.39421512 |
| ENSMUSG00000033080 | 0 | 0 | 0 | 0 | 0 | 0.301029996 | 0 |
| ENSMUSG00000033082 | 0.289064826 | 0 | 0 | 0 | 0.244650542 | 0 | 0.355418052 |
| ENSMUSG00000033083 | 1.822201287 | 1.653677461 | 1.649978211 | 1.950224975 | 1.919403394 | 2.086359831 | 1.81475081 |
| ENSMUSG00000033088 | 2.339331437 | 2.27908599 | 2.397646762 | 2.119945066 | 2.28608706 | 2.380211242 | 2.37424372 |
| ENSMUSG00000033096 | 2.103271637 | 1.945339928 | 1.834962501 | 2.214495249 | 2.156296347 | 2.484299839 | 2.031198704 |
| ENSMUSG00000033099 | 2.315764015 | 2.170748646 | 2.234159485 | 2.181128463 | 2.198391101 | 2.424881637 | 2.116124284 |
| ENSMUSG00000033102 | 2.342375483 | 2.175975893 | 2.182829758 | 2.299918086 | 2.232221225 | 2.550228353 | 2.052343353 |
| ENSMUSG00000033105 | 2.384185734 | 2.181128463 | 2.200224975 | 2.321350555 | 2.359815565 | 2.618048097 | 2.194143936 |
| ENSMUSG00000033106 | 2.285603957 | 2.134789703 | 2.119945066 | 2.224461364 | 2.178339521 | 2.40654018 | 2.021338422 |
| ENSMUSG00000033107 | 0.747221736 | 0.580482024 | 0.5 | 0.75 | 0.931471446 | 1.113943352 | 0.829800754 |
| ENSMUSG00000033111 | 1.753493018 | 1.622963274 | 1.642463902 | 1.828220739 | 1.739104362 | 1.934498451 | 1.661510159 |
| ENSMUSG00000033114 | 1.482695304 | 1.516522298 | 1.458222504 | 1.566696635 | 1.439191811 | 1.633468456 | 1.329931221 |
| ENSMUSG00000033122 | 0 | 0 | 0 | 0 | 0 | 0 | 0 |
| ENSMUSG00000033124 | 2.116511873 | 1.950224975 | 2.019203899 | 2.059601185 | 2.126866297 | 2.340444115 | 2.031805106 |
| ENSMUSG00000033126 | 1.671187741 | 1.51109853 | 1.532320754 | 1.657339155 | 1.514343934 | 1.698970004 | 1.523605797 |
| ENSMUSG00000033128 | 2.461589295 | 2.336628433 | 2.389605178 | 2.084962501 | 2.264824954 | 2.426511261 | 2.245894972 |
| ENSMUSG00000033147 | 1.389637216 | 1.226722649 | 1.42510993 | 1.505591953 | 1.53772706 | 1.556302501 | 1.462868481 |
| ENSMUSG00000033149 | 2.91285295 | 2.689555804 | 2.673371739 | 2.799304173 | 2.718185765 | 3.121231455 | 2.53531451 |
| ENSMUSG00000033152 | 0.671187741 | 0.79248125 | 1.04248125 | 1.201838731 | 1.358933174 | 1.176091259 | 1.392431569 |
| ENSMUSG00000033156 | 0 | 0 | 0 | 0 | 0 | 0 | 0 |
| ENSMUSG00000033157 | 2.045658471 | 1.852347734 | 1.872963274 | 1.914552871 | 1.777823267 | 2.017033339 | 1.714087544 |
| ENSMUSG00000033159 | 2.201144779 | 2.03360658 | 2.089388001 | 1.892463902 | 1.966725889 | 2.139879086 | 1.906037947 |
| ENSMUSG00000033161 | 3.329495312 | 3.0352873 | 3.052896038 | 2.959604019 | 2.987719484 | 3.392345155 | 2.774843407 |
| ENSMUSG00000033166 | 2.597490839 | 2.417664062 | 2.403217124 | 2.700629091 | 2.60666248 | 2.968482949 | 2.417101415 |
| ENSMUSG00000033170 | 1.852729478 | 1.814346961 | 1.912762923 | 1.802363341 | 2.071604288 | 2.012837225 | 2.082194103 |
| ENSMUSG00000033174 | 1.325351389 | 1.17510993 | 1.388647213 | 0.864857905 | 0.877063019 | 0.77815125 | 0.935079929 |
| ENSMUSG00000033177 | 0 | 0 | 0.25 | 0 | 0.244650542 | 0 | 0 |
| ENSMUSG00000033182 | 0 | 0 | 0 | 0 | 0 | 0 | 0 |
| ENSMUSG00000033184 | 2.942653653 | 2.700629091 | 2.675543171 | 2.746993631 | 2.66856048 | 3.067442843 | 2.457875113 |
| ENSMUSG00000033186 | 2.352857998 | 2.229715809 | 2.181128463 | 2.351285366 | 2.29413964 | 2.621176282 | 2.134278581 |
| ENSMUSG00000033187 | 0 | 0 | 0 | 0 | 0 | 0 | 0 |

| | | | | | | | |
|---|---|---|---|---|---|---|---|
| ENSMUSG00000033191 | 0 | 0.25 | 0 | 0.25 | 0.489301084 | 0 | 0.224243824 |
| ENSMUSG00000033192 | 1.783508299 | 1.657339155 | 1.671625132 | 1.42510993 | 1.394615667 | 1.531478917 | 1.302263476 |
| ENSMUSG00000033196 | 0 | 0 | 0 | 0 | 0 | 0 | 0 |
| ENSMUSG00000033200 | 0 | 0 | 0 | 0 | 0 | 0 | 0 |
| ENSMUSG00000033207 | 1.249317394 | 0.976722649 | 1.214495249 | 0.92510993 | 0.955822902 | 0.477121255 | 0.916588299 |
| ENSMUSG00000033208 | 0 | 0.25 | 0.25 | 0.580482024 | 0.489301084 | 0.301029996 | 0.224243824 |
| ENSMUSG00000033209 | 2.651426165 | 2.510414788 | 2.56728167 | 2.089388001 | 2.197694248 | 2.389166084 | 2.197065089 |
| ENSMUSG00000033210 | 0 | 0 | 0 | 0 | 0 | 0 | 0 |
| ENSMUSG00000033213 | 0 | 0.25 | 0 | 0 | 0 | 0 | 0.224243824 |
| ENSMUSG00000033214 | 0.45815691 | 0.25 | 0.701838731 | 0.646240625 | 0.244650542 | 0 | 0.520678036 |
| ENSMUSG00000033216 | 1.948720984 | 1.884789703 | 1.860735874 | 1.916333979 | 1.974687643 | 2.017033339 | 1.87902385 |
| ENSMUSG00000033219 | 0 | 0 | 0 | 0 | 0 | 0 | 0 |
| ENSMUSG00000033220 | 0.747221736 | 0.79248125 | 0.646240625 | 0.951838731 | 0.812711509 | 0.77815125 | 0.74492186 |
| ENSMUSG00000033222 | 2.333175866 | 2.172499493 | 2.196158711 | 2.302363341 | 2.245869735 | 2.547774705 | 2.100339912 |
| ENSMUSG00000033227 | 0.289064826 | 0 | 0.5 | 0.580482024 | 0.812711509 | 0.77815125 | 1.078019652 |
| ENSMUSG00000033228 | 3.277146534 | 2.981110348 | 2.962546709 | 3.02655894 | 2.935634121 | 3.396373728 | 2.723864356 |
| ENSMUSG00000033233 | 1.655740331 | 1.622963274 | 1.552363341 | 1.403677461 | 1.478764298 | 1.579783597 | 1.396774124 |
| ENSMUSG00000033237 | 2.834961347 | 2.615113974 | 2.579635702 | 2.58412664 | 2.550791325 | 2.88592634 | 2.386009215 |
| ENSMUSG00000033249 | 1.249317394 | 1.282320754 | 1.396240625 | 1.146240625 | 1.244650542 | 1.255272505 | 1.335191751 |
| ENSMUSG00000033253 | 2.280358197 | 2.145300145 | 2.202642909 | 1.933677405 | 2.119108707 | 2.193124598 | 2.060761403 |
| ENSMUSG00000033255 | 0 | 0 | 0 | 0 | 0 | 0 | 0 |
| ENSMUSG00000033256 | 1.678702042 | 1.54248125 | 1.634789703 | 1.214495249 | 1.462344774 | 1.380211242 | 1.392431569 |
| ENSMUSG00000033257 | 2.771218178 | 2.522528105 | 2.502806814 | 2.674241882 | 2.631597336 | 2.980457892 | 2.436226648 |
| ENSMUSG00000033268 | 1.069666964 | 1.061981878 | 1.27186571 | 0.580482024 | 0.568060967 | 0.301029996 | 0.448487648 |
| ENSMUSG00000033272 | 2.573799059 | 2.377938163 | 2.40136988 | 2.405970373 | 2.417045398 | 2.715167358 | 2.299096905 |
| ENSMUSG00000033276 | 1.53838222 | 1.418106335 | 1.330482024 | 1.610735874 | 1.599808748 | 1.681241237 | 1.582103514 |
| ENSMUSG00000033278 | 1.847794126 | 1.843759858 | 1.880890489 | 1.79745614 | 1.797870327 | 1.913813852 | 1.68886578 |
| ENSMUSG00000033282 | 2.129344651 | 2.046218836 | 1.942046081 | 2.134789703 | 2.066583491 | 2.352182518 | 1.955652633 |
| ENSMUSG00000033285 | 2.583267694 | 2.405512955 | 2.392463902 | 2.469895812 | 2.408320622 | 2.693726949 | 2.222232323 |
| ENSMUSG00000033287 | 2.251208701 | 2.206637122 | 2.323655187 | 2.175975893 | 2.26997769 | 2.485721426 | 2.259963078 |
| ENSMUSG00000033294 | 2.371652933 | 2.251406137 | 2.299304173 | 2.168106335 | 2.221300875 | 2.383815366 | 2.124420007 |
| ENSMUSG00000033295 | 2.842799666 | 2.702642909 | 2.783285553 | 2.367918804 | 2.498344007 | 2.701567985 | 2.474947794 |
| ENSMUSG00000033306 | 3.435074977 | 3.12233656 | 3.096532984 | 2.767194569 | 2.69356054 | 3.098297536 | 2.471386688 |
| ENSMUSG00000033307 | 2.340856238 | 2.296218836 | 2.452241044 | 1.969129237 | 2.1962964 | 2.278753601 | 2.296968506 |

| | | | | | | | |
|---|---|---|---|---|---|---|---|
| ENSMUSG00000033308 | 0.45815691 | 0.701838731 | 0.396240625 | 0.864857905 | 0.812711509 | 0.903089987 | 0.672731473 |
| ENSMUSG00000033313 | 1.181541736 | 1.396240625 | 1.431980114 | 1.130890489 | 1.264824954 | 1.255272505 | 1.40946263 |
| ENSMUSG00000033316 | 0 | 0 | 0 | 0 | 0 | 0 | 0 |
| ENSMUSG00000033318 | 0.289064826 | 0.5 | 0.5 | 0.5 | 0.244650542 | 0.477121255 | 0.579661877 |
| ENSMUSG00000033319 | 2.489144387 | 2.248588359 | 2.23927551 | 2.296218836 | 2.194892994 | 2.542825427 | 2.043115906 |
| ENSMUSG00000033323 | 2.406877894 | 2.289336734 | 2.3184489 | 1.926839783 | 2.048957937 | 2.164352856 | 1.97354745 |
| ENSMUSG00000033326 | 2.706208651 | 2.505244985 | 2.492526473 | 2.42510993 | 2.445469852 | 2.737192643 | 2.309790449 |
| ENSMUSG00000033327 | 0.811507563 | 0.896240625 | 1.02186571 | 0.896240625 | 0.877063019 | 0.84509804 | 0.916588299 |
| ENSMUSG00000033335 | 2.644135203 | 2.460980263 | 2.47934302 | 2.524836953 | 2.553092387 | 2.888179494 | 2.476023874 |
| ENSMUSG00000033342 | 0 | 0 | 0 | 0 | 0 | 0 | 0.355418052 |
| ENSMUSG00000033343 | 0 | 0 | 0 | 0 | 0 | 0 | 0 |
| ENSMUSG00000033350 | 1.876566388 | 1.802363341 | 1.848079356 | 1.52186571 | 1.65906333 | 1.681241237 | 1.629532004 |
| ENSMUSG00000033352 | 2.661423995 | 2.429847205 | 2.378431899 | 2.548996825 | 2.458367761 | 2.85672889 | 2.280261923 |
| ENSMUSG00000033355 | 0.578129653 | 0.25 | 0.25 | 0.92510993 | 0.846351821 | 0.84509804 | 0.803905701 |
| ENSMUSG00000033356 | 1.903481042 | 1.874961472 | 1.807204673 | 1.819031101 | 1.812711509 | 2.008600172 | 1.699191618 |
| ENSMUSG00000033361 | 0 | 0 | 0 | 0.951838731 | 0.905314583 | 0.698970004 | 0.896975297 |
| ENSMUSG00000033364 | 2.541215974 | 2.328220739 | 2.320772088 | 2.455044741 | 2.3873546 | 2.722633923 | 2.216121671 |
| ENSMUSG00000033365 | 2.554191361 | 2.373463862 | 2.411414608 | 2.214495249 | 2.325116119 | 2.515873844 | 2.244955885 |
| ENSMUSG00000033368 | 0.867194479 | 0.830482024 | 0.646240625 | 0.646240625 | 0.846351821 | 0.77815125 | 0.448487648 |
| ENSMUSG00000033373 | 2.146671932 | 2.019203899 | 2.038704527 | 1.799918086 | 1.87878897 | 2.017033339 | 1.854824501 |
| ENSMUSG00000033377 | 0 | 0 | 0 | 0.25 | 0.244650542 | 0.301029996 | 0 |
| ENSMUSG00000033379 | 2.625329835 | 2.410061234 | 2.501406137 | 2.188721876 | 2.284999367 | 2.439332694 | 2.160139682 |
| ENSMUSG00000033382 | 2.69594272 | 2.490724001 | 2.466046536 | 2.476722649 | 2.382016011 | 2.720985744 | 2.204609933 |
| ENSMUSG00000033383 | 0 | 0 | 0 | 0 | 0 | 0 | 0 |
| ENSMUSG00000033386 | 2.282114126 | 2.091580554 | 2.166333979 | 2.158248799 | 2.146756411 | 2.320146286 | 1.980721036 |
| ENSMUSG00000033389 | 1.505869732 | 1.42510993 | 1.51109853 | 1.557204673 | 1.568060967 | 1.544068044 | 1.551512352 |
| ENSMUSG00000033392 | 2.475043119 | 2.286801231 | 2.302972074 | 2.359697963 | 2.282265378 | 2.627365857 | 2.165794278 |
| ENSMUSG00000033396 | 2.50175086 | 2.260414788 | 2.247171172 | 2.242166698 | 2.240967488 | 2.491361694 | 2.07004434 |
| ENSMUSG00000033400 | 2.648525058 | 2.391989019 | 2.372963274 | 2.449415381 | 2.376174996 | 2.710963119 | 2.184841529 |
| ENSMUSG00000033405 | 1.205378646 | 1.098079356 | 1.098079356 | 1.42510993 | 1.427019704 | 1.707570176 | 1.350478789 |
| ENSMUSG00000033409 | 0 | 0 | 0 | 0 | 0 | 0 | 0 |
| ENSMUSG00000033411 | 2.536482297 | 2.370953944 | 2.322504712 | 2.381869252 | 2.314756076 | 2.598790507 | 2.129386845 |
| ENSMUSG00000033416 | 2.304292853 | 2.052363341 | 2.115881093 | 2.074802005 | 2.070605826 | 2.311753861 | 1.969165684 |
| ENSMUSG00000033417 | 2.992411635 | 2.725593779 | 2.725216702 | 2.666333979 | 2.549248906 | 2.969881644 | 2.382749586 |

| | | | | | | | |
|---|---|---|---|---|---|---|---|
| ENSMUSG00000033419 | 1.129344651 | 1.061981878 | 1 | 0.896240625 | 0.812711509 | 1.041392685 | 0.876096088 |
| ENSMUSG00000033420 | 2.584966402 | 2.333847589 | 2.30539678 | 2.125956434 | 2.020174412 | 2.307496038 | 1.926805425 |
| ENSMUSG00000033423 | 2.455852858 | 2.277783918 | 2.295598588 | 2.104463129 | 2.168627946 | 2.432969291 | 2.110473216 |
| ENSMUSG00000033427 | 0.45815691 | 0 | 0.25 | 0.396240625 | 0.244650542 | 0.301029996 | 0.355418052 |
| ENSMUSG00000033429 | 1.342375483 | 1.098079356 | 1.226722649 | 1.160964047 | 1.136121934 | 1.255272505 | 1.041356071 |
| ENSMUSG00000033430 | 2.00683675 | 1.921625132 | 1.914552871 | 2.081607372 | 2.056322416 | 2.357934847 | 1.94631981 |
| ENSMUSG00000033434 | 1.156259305 | 1.130890489 | 1.17510993 | 1.02186571 | 1.091002363 | 0.77815125 | 1.20139795 |
| ENSMUSG00000033436 | 2.834495649 | 2.584962501 | 2.585796429 | 2.740543258 | 2.698827372 | 3.111262514 | 2.51406029 |
| ENSMUSG00000033439 | 1.558729299 | 1.445339928 | 1.380890489 | 1.54745614 | 1.542223377 | 1.643452676 | 1.369798885 |
| ENSMUSG00000033444 | 2.808046693 | 2.558404919 | 2.528760913 | 2.59995299 | 2.580539302 | 2.941014244 | 2.403756742 |
| ENSMUSG00000033446 | 1.527823874 | 1.488549078 | 1.445339928 | 1.78743678 | 1.698489937 | 1.908485019 | 1.586932193 |
| ENSMUSG00000033450 | 1.156259305 | 0.92510993 | 0.864857905 | 0.701838731 | 0.77552387 | 0.954242509 | 0.952571511 |
| ENSMUSG00000033453 | 0 | 0.396240625 | 0.396240625 | 0.5 | 0.244650542 | 0.301029996 | 0.448487648 |
| ENSMUSG00000033454 | 2.52293481 | 2.349685923 | 2.331607372 | 2.289967834 | 2.197694248 | 2.447158031 | 2.035419841 |
| ENSMUSG00000033458 | 1.578129653 | 1.42510993 | 1.470660762 | 1.51109853 | 1.445123986 | 1.477121255 | 1.396774124 |
| ENSMUSG00000033460 | 2.016886209 | 1.892463902 | 1.87892496 | 1.794977273 | 1.642643806 | 1.954242509 | 1.676387053 |
| ENSMUSG00000033467 | 1.036286563 | 1.061981878 | 1.214495249 | 0.5 | 0.846351821 | 0.477121255 | 1.20919388 |
| ENSMUSG00000033470 | 0 | 0 | 0 | 0 | 0 | 0 | 0 |
| ENSMUSG00000033475 | 0 | 0.580482024 | 0.830482024 | 0.396240625 | 0.489301084 | 0.602059991 | 0.224243824 |
| ENSMUSG00000033478 | 2.485932312 | 2.343216265 | 2.333288838 | 2.242166698 | 2.212715924 | 2.491361694 | 2.117056606 |
| ENSMUSG00000033486 | 1.129344651 | 1.061981878 | 1.146240625 | 1.02186571 | 1 | 1.176091259 | 0.969165684 |
| ENSMUSG00000033487 | 2.597490839 | 2.394357207 | 2.383818844 | 2.472185812 | 2.388575198 | 2.703291378 | 2.187098622 |
| ENSMUSG00000033488 | 0.671187741 | 0.896240625 | 0.79248125 | 0.646240625 | 0.877063019 | 0.77815125 | 0.672731473 |
| ENSMUSG00000033491 | 0.671187741 | 0.5 | 0.830482024 | 0.79248125 | 0.877063019 | 0.77815125 | 0.710836105 |
| ENSMUSG00000033498 | 0 | 0 | 0.25 | 0 | 0.244650542 | 0 | 0 |
| ENSMUSG00000033499 | 2.823159983 | 2.603406982 | 2.563757142 | 2.613317659 | 2.546663136 | 2.962842681 | 2.386211856 |
| ENSMUSG00000033501 | 0.289064826 | 0 | 0 | 0 | 0 | 0 | 0.224243824 |
| ENSMUSG00000033502 | 1.759671389 | 1.589388001 | 1.660964047 | 1.681980114 | 1.677808768 | 1.851258349 | 1.621017948 |
| ENSMUSG00000033508 | 0 | 0.25 | 0.25 | 0 | 0 | 0 | 0 |
| ENSMUSG00000033510 | 0 | 0 | 0 | 0 | 0.489301084 | 0.602059991 | 0.629532004 |
| ENSMUSG00000033520 | 0 | 0 | 0 | 0 | 0.244650542 | 0 | 0.224243824 |
| ENSMUSG00000033526 | 2.217828299 | 2.074802005 | 2.115881093 | 2.162762923 | 2.150762767 | 2.340444115 | 2.012457048 |
| ENSMUSG00000033530 | 1.568542279 | 1.532320754 | 1.557204673 | 1.708222504 | 1.739104362 | 1.812913357 | 1.67819947 |
| ENSMUSG00000033533 | 0 | 0 | 0 | 0 | 0 | 0 | 0 |

| | | | | | | | |
|---|---|---|---|---|---|---|---|
| ENSMUSG00000033538 | 0.960252568 | 1.238549078 | 1.098079356 | 1.464495249 | 1.401338844 | 1.662757832 | 1.345462945 |
| ENSMUSG00000033540 | 1.686083339 | 1.54248125 | 1.602347734 | 1.372963274 | 1.467903253 | 1.447158031 | 1.360283038 |
| ENSMUSG00000033542 | 2.677345838 | 2.454245906 | 2.43155454 | 2.443284802 | 2.373217836 | 2.683947131 | 2.20139795 |
| ENSMUSG00000033543 | 2.35727108 | 2.231944491 | 2.221424093 | 2.329353153 | 2.261929482 | 2.540329475 | 2.112839835 |
| ENSMUSG00000033544 | 0 | 0 | 0 | 0.25 | 0.244650542 | 0 | 0.224243824 |
| ENSMUSG00000033545 | 2.275045611 | 2.173371739 | 2.20584181 | 1.948603967 | 2.052132976 | 2.28780173 | 2.006616089 |
| ENSMUSG00000033554 | 1.952600383 | 1.80479213 | 1.80479213 | 1.899978211 | 1.877063019 | 2.012837225 | 1.74492186 |
| ENSMUSG00000033557 | 2.589183146 | 2.394828984 | 2.388167274 | 2.511780978 | 2.448909998 | 2.789580712 | 2.243384656 |
| ENSMUSG00000033565 | 3.223183227 | 2.94992159 | 2.930915236 | 2.868933358 | 2.828007872 | 3.239799818 | 2.66563581 |
| ENSMUSG00000033569 | 0.747221736 | 1.04248125 | 1 | 1 | 0.846351821 | 1.041392685 | 0.775756176 |
| ENSMUSG00000033576 | 1.069666964 | 1.146240625 | 1.17510993 | 0.5 | 0.632412477 | 0.602059991 | 0.710836105 |
| ENSMUSG00000033577 | 2.121692466 | 2.017865591 | 2 | 2.355516192 | 2.265401209 | 2.605305046 | 2.097879707 |
| ENSMUSG00000033578 | 0 | 0 | 0 | 0 | 0 | 0 | 0.224243824 |
| ENSMUSG00000033579 | 0 | 0 | 0.25 | 0 | 0.489301084 | 0.301029996 | 0.355418052 |
| ENSMUSG00000033581 | 2.556012466 | 2.360735874 | 2.345385738 | 2.00835575 | 2.016696984 | 2.230448921 | 1.897893072 |
| ENSMUSG00000033585 | 1.822201287 | 1.708222504 | 1.681980114 | 1.388647213 | 1.445123986 | 1.491361694 | 1.46982607 |
| ENSMUSG00000033589 | 1.967766868 | 1.926839783 | 1.961372513 | 1.905512955 | 2.083370582 | 2.315970345 | 2.026306127 |
| ENSMUSG00000033590 | 1.944805158 | 1.807204673 | 1.894357207 | 0.976722649 | 1.136121934 | 1.176091259 | 0.935079929 |
| ENSMUSG00000033594 | 0.811507563 | 1.02186571 | 0.864857905 | 1.372963274 | 1.414412788 | 1.397940009 | 1.425641774 |
| ENSMUSG00000033595 | 0 | 0 | 0 | 0 | 0.489301084 | 0.301029996 | 0.355418052 |
| ENSMUSG00000033596 | 2.817859673 | 2.559003548 | 2.544977273 | 2.760756821 | 2.666163799 | 3.061075324 | 2.468572133 |
| ENSMUSG00000033597 | 0.811507563 | 0.896240625 | 0.896240625 | 0.864857905 | 1.074582839 | 0.954242509 | 1.302263476 |
| ENSMUSG00000033610 | 2.095120908 | 1.955044741 | 1.940387808 | 1.784887838 | 1.761403742 | 1.929418926 | 1.63986884 |
| ENSMUSG00000033615 | 0 | 0.25 | 0 | 0 | 0.244650542 | 0 | 0 |
| ENSMUSG00000033618 | 1.505869732 | 1.364857905 | 1.226722649 | 1.339388001 | 1.380772476 | 1.462397998 | 1.307989563 |
| ENSMUSG00000033623 | 2.970466116 | 2.718361028 | 2.68155454 | 2.609697963 | 2.554618136 | 2.933487288 | 2.389637464 |
| ENSMUSG00000033624 | 2.292497211 | 2.079353153 | 2.064346961 | 2.391037372 | 2.273371525 | 2.604226053 | 2.156039304 |
| ENSMUSG00000033628 | 2.201144779 | 1.997171172 | 2.064346961 | 2.090485943 | 2.012006468 | 2.267171728 | 1.896975297 |
| ENSMUSG00000033629 | 2.619128496 | 2.390560606 | 2.386241108 | 2.670089881 | 2.585920905 | 2.989004616 | 2.417653332 |
| ENSMUSG00000033632 | 1.862428343 | 1.763820609 | 1.732684334 | 1.96759118 | 1.832885921 | 2.071882007 | 1.73898563 |
| ENSMUSG00000033633 | 0.289064826 | 0 | 0 | 0.25 | 0 | 0 | 0.224243824 |
| ENSMUSG00000033634 | 0 | 0 | 0 | 0 | 0 | 0 | 0 |
| ENSMUSG00000033644 | 0.289064826 | 0.25 | 0.396240625 | 0.25 | 0.489301084 | 0 | 0.355418052 |
| ENSMUSG00000033653 | 2.271465912 | 2.083847589 | 2.091580554 | 2.051142786 | 1.958580386 | 2.267171728 | 1.886704102 |

| | | | | | | | |
|---|---|---|---|---|---|---|---|
| ENSMUSG00000033658 | 2.151493169 | 1.959800947 | 1.928561379 | 2.111770807 | 2.075570129 | 2.292256071 | 1.978585612 |
| ENSMUSG00000033669 | 2.229917041 | 2.052363341 | 2.083847589 | 2.211372513 | 2.134457044 | 2.411619706 | 2.043700396 |
| ENSMUSG00000033671 | 2.53838222 | 2.360217292 | 2.333288838 | 2.474839231 | 2.403292121 | 2.698100546 | 2.232493008 |
| ENSMUSG00000033676 | 0 | 0 | 0 | 0.396240625 | 0 | 0 | 0.224243824 |
| ENSMUSG00000033684 | 2.831690488 | 2.603406982 | 2.611253711 | 2.522528105 | 2.547699718 | 2.859738566 | 2.414513278 |
| ENSMUSG00000033685 | 2.718080037 | 2.539967834 | 2.595656007 | 2.246460484 | 2.321205208 | 2.549003262 | 2.280823582 |
| ENSMUSG00000033688 | 0 | 0.25 | 0.25 | 0 | 0 | 0.301029996 | 0 |
| ENSMUSG00000033697 | 1.753493018 | 1.664552871 | 1.705044741 | 1.561981878 | 1.77552387 | 1.86332286 | 1.760706232 |
| ENSMUSG00000033701 | 2.092368209 | 1.896240625 | 2.025821952 | 1.981203126 | 2.021325983 | 2.247973266 | 1.999330888 |
| ENSMUSG00000033703 | 1.800532393 | 1.75 | 1.866901388 | 1.622963274 | 1.86473911 | 1.968482949 | 1.832055226 |
| ENSMUSG00000033705 | 2.395018353 | 2.161864607 | 2.180274797 | 2.089388001 | 2.0510778 | 2.23299611 | 1.882886767 |
| ENSMUSG00000033706 | 2.422183552 | 2.260414788 | 2.248588359 | 2.234894804 | 2.271678687 | 2.5132176 | 2.103267674 |
| ENSMUSG00000033707 | 1.307602651 | 1.311981878 | 1.27186571 | 1.061981878 | 1.264824954 | 1.278753601 | 1.302263476 |
| ENSMUSG00000033712 | 2.235832493 | 2.105516192 | 2.121960008 | 2.235628626 | 2.29993025 | 2.522444234 | 2.193776931 |
| ENSMUSG00000033713 | 1.822201287 | 1.75 | 1.809601185 | 1.580482024 | 1.528558853 | 1.69019608 | 1.396774124 |
| ENSMUSG00000033715 | 1.83262764 | 1.807204673 | 1.825945187 | 1.080482024 | 1.188507684 | 1.204119983 | 1.100339912 |
| ENSMUSG00000033717 | 0.578129653 | 0.5 | 0.864857905 | 0 | 0 | 0 | 0.224243824 |
| ENSMUSG00000033720 | 1.663535556 | 1.571350555 | 1.561981878 | 1.445339928 | 1.439191811 | 1.505149978 | 1.459332757 |
| ENSMUSG00000033721 | 1.53838222 | 1.431980114 | 1.470660762 | 1.505591953 | 1.380772476 | 1.397940009 | 1.324583736 |
| ENSMUSG00000033722 | 2.126809492 | 2.025821952 | 2.036164561 | 1.888647213 | 1.987570683 | 2.136720567 | 1.97354745 |
| ENSMUSG00000033726 | 0 | 0 | 0 | 0 | 0 | 0 | 0 |
| ENSMUSG00000033728 | 2.103271637 | 2.032320754 | 2.048689214 | 1.882845365 | 1.955822902 | 2 | 1.898808251 |
| ENSMUSG00000033730 | 0 | 0.5 | 0.580482024 | 0 | 0.387761935 | 0.301029996 | 0.355418052 |
| ENSMUSG00000033731 | 0 | 0 | 0 | 0 | 0 | 0 | 0 |
| ENSMUSG00000033732 | 3.256154144 | 2.944724619 | 2.914106216 | 3.027131114 | 2.953453786 | 3.42975228 | 2.749039747 |
| ENSMUSG00000033735 | 1.842799666 | 1.814346961 | 1.901832578 | 1.692046081 | 1.820920161 | 1.851258349 | 1.759302692 |
| ENSMUSG00000033737 | 0 | 0 | 0 | 0.5 | 0 | 0 | 0.224243824 |
| ENSMUSG00000033739 | 1.036286563 | 1.061981878 | 1.160964047 | 1.431980114 | 1.414412788 | 1.431363764 | 1.360283038 |
| ENSMUSG00000033740 | 0 | 0 | 0 | 0.396240625 | 0 | 0 | 0 |
| ENSMUSG00000033751 | 1.482695304 | 1.516522298 | 1.54745614 | 1.5 | 1.592135487 | 1.662757832 | 1.627424404 |
| ENSMUSG00000033752 | 1.404271433 | 1.27186571 | 1.321350555 | 1.51109853 | 1.467903253 | 1.612783857 | 1.417653332 |
| ENSMUSG00000033760 | 2.22189694 | 2.108657057 | 2.038704527 | 2.124961472 | 2.069604532 | 2.338456494 | 1.918313719 |
| ENSMUSG00000033762 | 2.072573125 | 1.973704441 | 2.015173983 | 2.023189285 | 2.138604637 | 2.250420002 | 2.079068325 |
| ENSMUSG00000033763 | 2.440557995 | 2.296218836 | 2.306603048 | 1.622963274 | 1.768534412 | 1.740362689 | 1.752192176 |

| | | | | | | | |
|---|---|---|---|---|---|---|---|
| ENSMUSG00000033765 | 0 | 0 | 0 | 0 | 0 | 0 | 0 |
| ENSMUSG00000033767 | 2.267855219 | 2.097004321 | 2.098079356 | 2.070192693 | 2.039257769 | 2.214843848 | 1.928477344 |
| ENSMUSG00000033768 | 0 | 0 | 0 | 0.25 | 0.733951626 | 0.477121255 | 0.520678036 |
| ENSMUSG00000033769 | 2.487005755 | 2.260414788 | 2.263142013 | 2.329353153 | 2.259595879 | 2.571708832 | 2.118913228 |
| ENSMUSG00000033770 | 0 | 0 | 0.25 | 0 | 0 | 0 | 0 |
| ENSMUSG00000033773 | 2.097855556 | 1.991446071 | 1.985628626 | 1.991446071 | 1.928492895 | 2.201397124 | 1.784981274 |
| ENSMUSG00000033774 | 0.45815691 | 0.25 | 0 | 0.5 | 0.244650542 | 0.301029996 | 0.224243824 |
| ENSMUSG00000033777 | 0 | 0 | 0.25 | 0 | 0.244650542 | 0 | 0.224243824 |
| ENSMUSG00000033781 | 1.944805158 | 1.823655187 | 1.782320754 | 1.898114259 | 1.814781639 | 2.103803721 | 1.792684387 |
| ENSMUSG00000033788 | 2.476144923 | 2.25 | 2.302363341 | 1.638647213 | 1.683842353 | 1.69019608 | 1.543390198 |
| ENSMUSG00000033790 | 2.199011616 | 2.038704527 | 2.065523711 | 2.078220739 | 1.998776581 | 2.283301229 | 1.907820406 |
| ENSMUSG00000033792 | 2.343889214 | 2.189555804 | 2.179419106 | 2.277783918 | 2.21338377 | 2.501059262 | 2.009225098 |
| ENSMUSG00000033793 | 2.729023206 | 2.481203126 | 2.492526473 | 2.492166698 | 2.395014713 | 2.723455672 | 2.210242554 |
| ENSMUSG00000033794 | 0 | 0 | 0 | 0 | 0 | 0 | 0 |
| ENSMUSG00000033799 | 2.675303226 | 2.491806565 | 2.455443495 | 2.663659007 | 2.600700615 | 2.926342447 | 2.356530173 |
| ENSMUSG00000033805 | 0.45815691 | 0 | 0 | 0.580482024 | 0.244650542 | 0 | 0.224243824 |
| ENSMUSG00000033808 | 2.495495257 | 2.307805295 | 2.261780978 | 2.357613138 | 2.296783518 | 2.600972896 | 2.104721678 |
| ENSMUSG00000033809 | 1.794934559 | 1.732684334 | 1.723704441 | 1.681980114 | 1.733951626 | 1.982271233 | 1.69749337 |
| ENSMUSG00000033813 | 2.897514811 | 2.651601303 | 2.64151351 | 2.80357978 | 2.708639269 | 3.150449409 | 2.487334119 |
| ENSMUSG00000033819 | 2.00683675 | 1.978222334 | 2.072504712 | 1.714495249 | 2.019019072 | 1.929418926 | 2.044283833 |
| ENSMUSG00000033825 | 0 | 0 | 0 | 0 | 0 | 0 | 0 |
| ENSMUSG00000033826 | 0 | 0 | 0.25 | 1 | 0.812711509 | 1 | 0.829800754 |
| ENSMUSG00000033831 | 0 | 0 | 0 | 0 | 0 | 0 | 0 |
| ENSMUSG00000033834 | 0 | 0 | 0 | 0 | 0 | 0 | 0 |
| ENSMUSG00000033837 | 0.289064826 | 0 | 0 | 0 | 0.387761935 | 0 | 0.448487648 |
| ENSMUSG00000033845 | 2.317377295 | 2.12694866 | 2.144357207 | 2.269871196 | 2.219338185 | 2.561101384 | 2.093403634 |
| ENSMUSG00000033847 | 1.069666964 | 1.188721876 | 1.080482024 | 1.282320754 | 1.106691885 | 1.176091259 | 1.110947927 |
| ENSMUSG00000033849 | 0.45815691 | 0 | 0 | 0.25 | 0.387761935 | 0 | 0.448487648 |
| ENSMUSG00000033850 | 0 | 0 | 0 | 0 | 0 | 0 | 0 |
| ENSMUSG00000033852 | 0 | 0 | 0 | 0 | 0 | 0 | 0 |
| ENSMUSG00000033854 | 0.289064826 | 0.5 | 0.396240625 | 0.396240625 | 0.387761935 | 0.301029996 | 0.355418052 |
| ENSMUSG00000033855 | 2.506893125 | 2.307204673 | 2.273849256 | 2.342126615 | 2.298360391 | 2.633468456 | 2.14555506 |
| ENSMUSG00000033857 | 0.960252568 | 1.160964047 | 1.201838731 | 1.188721876 | 1.310728002 | 1.255272505 | 1.369798885 |
| ENSMUSG00000033860 | 0 | 0 | 0 | 0 | 0 | 0 | 0 |

| | | | | | | | |
|---|---|---|---|---|---|---|---|
| ENSMUSG00000033862 | 2.029919532 | 1.821350555 | 1.858657057 | 1.964495249 | 1.94314048 | 2.181843588 | 1.892346957 |
| ENSMUSG00000033863 | 1.903481042 | 1.858657057 | 1.872963274 | 1.566696635 | 1.542223377 | 1.672097858 | 1.42167221 |
| ENSMUSG00000033871 | 0.671187741 | 1.02186571 | 1.04248125 | 0.25 | 0.812711509 | 0.301029996 | 0.629532004 |
| ENSMUSG00000033872 | 0 | 0.25 | 0.25 | 0 | 0 | 0 | 0 |
| ENSMUSG00000033880 | 0.578129653 | 0.701838731 | 0.830482024 | 1.618933358 | 1.635855979 | 1.698970004 | 1.603286636 |
| ENSMUSG00000033882 | 0 | 0 | 0 | 0 | 0.244650542 | 0 | 0 |
| ENSMUSG00000033883 | 2.026699189 | 1.866901388 | 1.921625132 | 2.04745614 | 1.944572352 | 2.217483944 | 1.815933683 |
| ENSMUSG00000033885 | 2.023453784 | 1.834962501 | 1.860735874 | 2.066696635 | 2.040348824 | 2.40654018 | 1.949460761 |
| ENSMUSG00000033898 | 0 | 0 | 0 | 0 | 0 | 0 | 0 |
| ENSMUSG00000033900 | 2.010213549 | 1.872963274 | 1.848079356 | 1.872963274 | 1.800029087 | 1.963787827 | 1.695786161 |
| ENSMUSG00000033902 | 1.960252568 | 1.958222504 | 1.854463129 | 2.00835575 | 2.064555002 | 2.26245109 | 1.943148064 |
| ENSMUSG00000033904 | 2.363083473 | 2.157339155 | 2.148114259 | 2.289967834 | 2.246477757 | 2.521138084 | 2.071655208 |
| ENSMUSG00000033906 | 2.010213549 | 1.972185812 | 2.009729747 | 1.912762923 | 1.83872004 | 2.139879086 | 1.674564426 |
| ENSMUSG00000033909 | 2.402962068 | 2.237821179 | 2.228969845 | 2.075945187 | 2.221300875 | 2.459392488 | 2.096394553 |
| ENSMUSG00000033910 | 0 | 0 | 0.25 | 0.25 | 0 | 0 | 0 |
| ENSMUSG00000033916 | 2.390989024 | 2.209012589 | 2.203445298 | 2.187048212 | 2.184185718 | 2.46686762 | 2.035419841 |
| ENSMUSG00000033917 | 2.092368209 | 1.992885888 | 1.985628626 | 2.127938163 | 2.10031573 | 2.361727836 | 1.987739 |
| ENSMUSG00000033918 | 2.136858952 | 1.988549078 | 2.00835575 | 2.122963274 | 2.033751322 | 2.357934847 | 1.907820406 |
| ENSMUSG00000033931 | 2.223916471 | 2.113831805 | 2.115881093 | 2.254202072 | 2.187791022 | 2.509202522 | 2.02568932 |
| ENSMUSG00000033933 | 2.287337982 | 2.052363341 | 2.031030328 | 2.254202072 | 2.21338377 | 2.45331834 | 2.060206963 |
| ENSMUSG00000033938 | 2.124258828 | 2.145300145 | 2.247171172 | 1.868933358 | 2.187072901 | 2.230448921 | 2.222232323 |
| ENSMUSG00000033940 | 2.725407296 | 2.452642909 | 2.477098155 | 2.463717096 | 2.421513257 | 2.804820679 | 2.264421331 |
| ENSMUSG00000033943 | 2.865034804 | 2.61817271 | 2.617918804 | 2.713717096 | 2.637464898 | 3.027349608 | 2.423302014 |
| ENSMUSG00000033948 | 0.45815691 | 0.701838731 | 0.701838731 | 1.160964047 | 1.188507684 | 1.301029996 | 1.193409508 |
| ENSMUSG00000033949 | 0.91631382 | 1.061981878 | 1.02186571 | 1.445339928 | 1.327538614 | 1.505149978 | 1.355418052 |
| ENSMUSG00000033952 | 2.745494892 | 2.492526473 | 2.490724001 | 2.616136438 | 2.539320544 | 2.902546779 | 2.351607989 |
| ENSMUSG00000033953 | 2.81249113 | 2.577369088 | 2.533927322 | 2.666556001 | 2.612525313 | 2.979548375 | 2.36808942 |
| ENSMUSG00000033955 | 2.731416512 | 2.52908599 | 2.587182039 | 2.444519282 | 2.573634712 | 2.770852012 | 2.453133078 |
| ENSMUSG00000033960 | 2.676666078 | 2.416777886 | 2.405054956 | 2.298073204 | 2.258423266 | 2.608526034 | 2.119837558 |
| ENSMUSG00000033961 | 1.342375483 | 1.311981878 | 1.201838731 | 1.114857905 | 1.310728002 | 1.278753601 | 1.245582246 |
| ENSMUSG00000033963 | 0.289064826 | 0.25 | 0 | 0.25 | 0 | 0 | 0.224243824 |
| ENSMUSG00000033964 | 2.682073381 | 2.459406983 | 2.452241044 | 2.588837774 | 2.491735274 | 2.796574333 | 2.259663665 |
| ENSMUSG00000033965 | 1.249317394 | 1.348079356 | 1.29248125 | 1.380890489 | 1.462344774 | 1.591064607 | 1.417653332 |
| ENSMUSG00000033966 | 0.45815691 | 0.25 | 0.396240625 | 0 | 0 | 0 | 0 |

| | | | | | | | |
|---|---|---|---|---|---|---|---|
| ENSMUSG00000033967 | 1.100572389 | 0.951838731 | 1.061981878 | 0.951838731 | 0.846351821 | 0.84509804 | 1.041356071 |
| ENSMUSG00000033970 | 2.583267694 | 2.379417347 | 2.33993739 | 2.378431899 | 2.320713283 | 2.688419822 | 2.189340078 |
| ENSMUSG00000033972 | 1.881174527 | 1.80479213 | 1.83272922 | 1.763820609 | 1.72341484 | 1.924279286 | 1.601000298 |
| ENSMUSG00000033981 | 0 | 0 | 0 | 0.25 | 0 | 0 | 0.355418052 |
| ENSMUSG00000033983 | 1.816888701 | 1.688721876 | 1.714495249 | 1.83272922 | 1.820920161 | 2.068185862 | 1.769001954 |
| ENSMUSG00000033985 | 1.89911417 | 1.664552871 | 1.698603967 | 1.874961472 | 1.842556557 | 2 | 1.669034113 |
| ENSMUSG00000033987 | 1.269664473 | 1.160964047 | 1.238549078 | 0.830482024 | 1.223252711 | 1.146128036 | 1.159323753 |
| ENSMUSG00000033991 | 2.50175086 | 2.336073977 | 2.289336734 | 2.496460484 | 2.434943542 | 2.773054693 | 2.239582276 |
| ENSMUSG00000033998 | 0 | 0 | 0 | 0.5 | 0.244650542 | 0.301029996 | 0 |
| ENSMUSG00000034000 | 0 | 0.25 | 0 | 0 | 0 | 0 | 0 |
| ENSMUSG00000034006 | 1.783508299 | 1.809601185 | 1.735628626 | 1.388647213 | 1.588235379 | 1.707570176 | 1.551512352 |
| ENSMUSG00000034007 | 1.993049299 | 1.872963274 | 1.87694866 | 1.991446071 | 1.840643511 | 2.08278537 | 1.715701091 |
| ENSMUSG00000034009 | 0.289064826 | 0 | 0.25 | 0 | 0 | 0 | 0.224243824 |
| ENSMUSG00000034021 | 2.905843838 | 2.645064642 | 2.618933358 | 2.644828984 | 2.583123351 | 2.924795996 | 2.372134746 |
| ENSMUSG00000034022 | 2.800026593 | 2.617156011 | 2.646475362 | 2.515511534 | 2.623340676 | 2.895422546 | 2.520678036 |
| ENSMUSG00000034023 | 2.251208701 | 2.080482024 | 2.04745614 | 2.242166698 | 2.210031835 | 2.515873844 | 2.07004434 |
| ENSMUSG00000034024 | 3.430412868 | 3.127258172 | 3.083917372 | 3.254332601 | 3.169206672 | 3.705692697 | 2.943111842 |
| ENSMUSG00000034028 | 0 | 0 | 0 | 0 | 0 | 0 | 0 |
| ENSMUSG00000034031 | 0 | 0 | 0 | 0 | 0 | 0 | 0 |
| ENSMUSG00000034032 | 2 | 1.916333979 | 1.978222334 | 1.961372513 | 2.103062484 | 2.217483944 | 2.01116814 |
| ENSMUSG00000034035 | 0 | 0.580482024 | 0.25 | 0.5 | 0.686820904 | 0 | 0.672731473 |
| ENSMUSG00000034037 | 0 | 0.25 | 0 | 0.25 | 0.244650542 | 0.301029996 | 0 |
| ENSMUSG00000034039 | 0 | 0 | 0 | 0 | 0 | 0 | 0 |
| ENSMUSG00000034040 | 2.057835465 | 1.966046536 | 2.048689214 | 1.622963274 | 1.618292349 | 1.806179974 | 1.579661877 |
| ENSMUSG00000034041 | 0 | 0 | 0.25 | 0.25 | 0.387761935 | 0.301029996 | 0.224243824 |
| ENSMUSG00000034042 | 2.245508853 | 2.105516192 | 2.056000419 | 2.134789703 | 2.043601899 | 2.322219295 | 1.920884716 |
| ENSMUSG00000034055 | 2.380049842 | 2.181980114 | 2.211372513 | 2.24929487 | 2.192777367 | 2.469822016 | 2.043115906 |
| ENSMUSG00000034057 | 0.289064826 | 0.396240625 | 0.396240625 | 0.5 | 0.489301084 | 0.477121255 | 0.629532004 |
| ENSMUSG00000034059 | 1.036286563 | 0.864857905 | 0.92510993 | 0.75 | 0.77552387 | 0.602059991 | 0.74492186 |
| ENSMUSG00000034063 | 0 | 0 | 0 | 0 | 0 | 0 | 0 |
| ENSMUSG00000034064 | 2.496544396 | 2.302972074 | 2.278435542 | 2.25835575 | 2.187072901 | 2.495544338 | 2.055173778 |
| ENSMUSG00000034066 | 2.060824957 | 1.956637122 | 1.910964047 | 1.994319981 | 1.954436039 | 2.178976947 | 1.857950279 |
| ENSMUSG00000034071 | 1.342375483 | 1.214495249 | 1.27186571 | 1.238549078 | 1.149965125 | 1.146128036 | 1.041356071 |
| ENSMUSG00000034075 | 2.641182749 | 2.433677405 | 2.414998973 | 2.633818844 | 2.617656201 | 2.997386384 | 2.437439291 |

| | | | | | | | |
|---|---|---|---|---|---|---|---|
| ENSMUSG00000034083 | 1.89911417 | 1.856566189 | 1.79745614 | 1.970660762 | 1.90849439 | 2.130333768 | 1.74047993 |
| ENSMUSG00000034087 | 0 | 0 | 0 | 0 | 0 | 0 | 0 |
| ENSMUSG00000034088 | 3.635555794 | 3.280989928 | 3.26007243 | 3.37694866 | 3.286246732 | 3.839792184 | 3.055839184 |
| ENSMUSG00000034098 | 0 | 0.25 | 0 | 0 | 0 | 0 | 0 |
| ENSMUSG00000034101 | 2.253091469 | 2.067865757 | 2.058404919 | 2.089388001 | 1.978602168 | 2.357934847 | 1.896054911 |
| ENSMUSG00000034105 | 1.445324132 | 1.52186571 | 1.464495249 | 1.516522298 | 1.611014646 | 1.707570176 | 1.603286636 |
| ENSMUSG00000034107 | 0.289064826 | 0 | 0 | 0 | 0.244650542 | 0 | 0 |
| ENSMUSG00000034108 | 2.075459174 | 1.937048212 | 1.950224975 | 1.997171172 | 2.006054284 | 2.198657087 | 1.896975297 |
| ENSMUSG00000034109 | 2.688735382 | 2.483422664 | 2.47934302 | 2.475970461 | 2.42077253 | 2.760422483 | 2.258162431 |
| ENSMUSG00000034110 | 1.852729478 | 1.701838731 | 1.726722649 | 1.714495249 | 1.770879644 | 1.838849091 | 1.695786161 |
| ENSMUSG00000034111 | 2.358731791 | 2.206637122 | 2.203445298 | 2.228969845 | 2.196996016 | 2.488550717 | 2.031198704 |
| ENSMUSG00000034112 | 0 | 0 | 0 | 0 | 0.244650542 | 0 | 0 |
| ENSMUSG00000034115 | 0 | 0 | 0 | 0 | 0 | 0.301029996 | 0 |
| ENSMUSG00000034116 | 0 | 0 | 0 | 0 | 0 | 0 | 0 |
| ENSMUSG00000034117 | 0.289064826 | 0.396240625 | 0.25 | 0.25 | 0 | 0 | 0.520678036 |
| ENSMUSG00000034118 | 2.511972754 | 2.31611065 | 2.302972074 | 2.225970461 | 2.205284995 | 2.491361694 | 2.107130591 |
| ENSMUSG00000034120 | 2.995590251 | 2.722185812 | 2.733238223 | 2.771202097 | 2.717066537 | 3.09482038 | 2.554897447 |
| ENSMUSG00000034121 | 1.181541736 | 1.061981878 | 1.188721876 | 1.26109853 | 1.319233381 | 1.322219295 | 1.383567577 |
| ENSMUSG00000034122 | 0 | 0 | 0 | 0 | 0 | 0 | 0 |
| ENSMUSG00000034126 | 2.227926429 | 2.083847589 | 2.078220739 | 1.96759118 | 1.98503142 | 2.201397124 | 1.853775828 |
| ENSMUSG00000034127 | 0.289064826 | 0 | 0 | 0 | 0 | 0 | 0.224243824 |
| ENSMUSG00000034135 | 2.482695304 | 2.30175358 | 2.349685923 | 2.362802778 | 2.429921724 | 2.721810615 | 2.286662868 |
| ENSMUSG00000034139 | 0 | 0 | 0 | 0 | 0 | 0 | 0 |
| ENSMUSG00000034145 | 0.289064826 | 0.25 | 0 | 0.25 | 0.489301084 | 0 | 0.224243824 |
| ENSMUSG00000034151 | 0 | 0 | 0 | 0 | 0 | 0 | 0 |
| ENSMUSG00000034152 | 2.530730035 | 2.295598588 | 2.28360658 | 2.552363341 | 2.474332504 | 2.850033258 | 2.30669705 |
| ENSMUSG00000034154 | 2.312518611 | 2.103406982 | 2.05357978 | 2.28296424 | 2.257835496 | 2.5132176 | 2.086826862 |
| ENSMUSG00000034156 | 2.181541736 | 2.006976499 | 2.165444524 | 0.79248125 | 1.057362051 | 0.84509804 | 1.066254157 |
| ENSMUSG00000034157 | 2.34539747 | 2.170748646 | 2.199415381 | 2.05357978 | 2.068600388 | 2.33243846 | 1.930140666 |
| ENSMUSG00000034158 | 2.954701306 | 2.72104263 | 2.696771581 | 2.533285553 | 2.468551474 | 2.8162413 | 2.279980727 |
| ENSMUSG00000034159 | 0 | 0 | 0 | 0 | 0.387761935 | 0 | 0.224243824 |
| ENSMUSG00000034160 | 2.987597818 | 2.721995533 | 2.695339928 | 2.78296424 | 2.70184985 | 3.074084689 | 2.503661426 |
| ENSMUSG00000034161 | 1.857607105 | 1.886723615 | 1.951838731 | 1.589388001 | 1.709785497 | 1.819543936 | 1.792684387 |
| ENSMUSG00000034163 | 2.505869732 | 2.248588359 | 2.248588359 | 2.327653195 | 2.266550905 | 2.494154594 | 2.071655208 |

| | | | | | | | |
|---|---|---|---|---|---|---|---|
| ENSMUSG00000034164 | 0.289064826 | 0.646240625 | 0.25 | 0.580482024 | 0.568060967 | 0.301029996 | 0.74492186 |
| ENSMUSG00000034165 | 2.266038081 | 2.196158711 | 2.284887838 | 2.136723615 | 2.304598364 | 2.469822016 | 2.297235323 |
| ENSMUSG00000034168 | 2.219867581 | 2.079353153 | 2.161864607 | 1.819031101 | 1.85749873 | 2.06069784 | 1.916588299 |
| ENSMUSG00000034171 | 0.867194479 | 1 | 0.951838731 | 1.330482024 | 1.387761935 | 1.477121255 | 1.360283038 |
| ENSMUSG00000034173 | 1.587501561 | 1.575945187 | 1.53743678 | 1.527131114 | 1.380772476 | 1.579783597 | 1.345462945 |
| ENSMUSG00000034175 | 1.671187741 | 1.584962501 | 1.630890489 | 1.25 | 1.450958105 | 1.361727836 | 1.511732338 |
| ENSMUSG00000034177 | 1.876566388 | 1.720660762 | 1.752806814 | 0.646240625 | 0.877063019 | 0.84509804 | 0.853775828 |
| ENSMUSG00000034185 | 0 | 0 | 0 | 0 | 0 | 0 | 0 |
| ENSMUSG00000034187 | 2.638209243 | 2.393411797 | 2.404596376 | 2.617410454 | 2.582889218 | 2.95230801 | 2.413397706 |
| ENSMUSG00000034189 | 2.205378646 | 2.080482024 | 2.069031101 | 2.027131114 | 2.062514787 | 2.209515015 | 1.953344547 |
| ENSMUSG00000034190 | 2.251208701 | 2.14151351 | 2.089388001 | 2.324229052 | 2.28608706 | 2.561101384 | 2.160546876 |
| ENSMUSG00000034192 | 2.299278375 | 2.137686696 | 2.137686696 | 2.284887838 | 2.216042585 | 2.556302501 | 2.113781657 |
| ENSMUSG00000034194 | 1.920505136 | 1.837182039 | 1.907339155 | 1.890560606 | 1.938809675 | 2.100370545 | 1.88765144 |
| ENSMUSG00000034201 | 2.086807695 | 2.05479213 | 2.177701608 | 1.839388001 | 2.022473809 | 2.06069784 | 2.106650241 |
| ENSMUSG00000034203 | 1.948720984 | 1.909156155 | 1.896240625 | 1.930274797 | 1.966725889 | 2.152288344 | 1.832055226 |
| ENSMUSG00000034205 | 3.058925034 | 2.778760913 | 2.727473271 | 3.033686792 | 2.941448453 | 3.445448514 | 2.745322664 |
| ENSMUSG00000034206 | 2.343889214 | 2.143411797 | 2.181980114 | 2.368933358 | 2.298360391 | 2.593286067 | 2.158095989 |
| ENSMUSG00000034209 | 0.671187741 | 0.896240625 | 0.5 | 0.75 | 0.812711509 | 0.602059991 | 0.803905701 |
| ENSMUSG00000034210 | 2.565000581 | 2.315523711 | 2.321928095 | 2.425543171 | 2.386538516 | 2.720159303 | 2.218171406 |
| ENSMUSG00000034211 | 2.368815967 | 2.242885888 | 2.252806814 | 2.139605178 | 2.173890435 | 2.439332694 | 2.041356071 |
| ENSMUSG00000034212 | 0.289064826 | 0.580482024 | 0.396240625 | 0.25 | 0.489301084 | 0.477121255 | 0.629532004 |
| ENSMUSG00000034216 | 1.83262764 | 1.912762923 | 1.928561379 | 1.678561379 | 1.871833944 | 1.982271233 | 1.928477344 |
| ENSMUSG00000034218 | 2.781396738 | 2.55479213 | 2.533927322 | 2.654596376 | 2.565363825 | 2.930439595 | 2.363558619 |
| ENSMUSG00000034219 | 0.45815691 | 0 | 0.25 | 0 | 0 | 0 | 0 |
| ENSMUSG00000034220 | 2.146671932 | 2.169870025 | 2.208222504 | 1.958222504 | 2.105788027 | 2.204119983 | 2.069505596 |
| ENSMUSG00000034224 | 0.289064826 | 0 | 0 | 0 | 0 | 0 | 0 |
| ENSMUSG00000034226 | 0.578129653 | 0.75 | 0.701838731 | 0.830482024 | 0.931471446 | 1.079181246 | 1.078019652 |
| ENSMUSG00000034227 | 0.45815691 | 0.896240625 | 0.701838731 | 0 | 0.632412477 | 0.602059991 | 0.448487648 |
| ENSMUSG00000034235 | 2.124258828 | 1.930274797 | 1.931980114 | 2.157339155 | 2.071604288 | 2.380211242 | 1.934261938 |
| ENSMUSG00000034239 | 0 | 0 | 0 | 0 | 0 | 0 | 0 |
| ENSMUSG00000034243 | 2.86633195 | 2.619945066 | 2.599685923 | 2.575945187 | 2.523329778 | 2.860338007 | 2.336139088 |
| ENSMUSG00000034245 | 1.404271433 | 1.418106335 | 1.516522298 | 1.188721876 | 1.27449559 | 1.079181246 | 1.238624656 |
| ENSMUSG00000034247 | 2.139333929 | 2.03743678 | 2.080482024 | 1.969129237 | 2.060462711 | 2.209515015 | 2.003985867 |
| ENSMUSG00000034248 | 2.382811656 | 2.19697564 | 2.233422664 | 2.343759858 | 2.319233381 | 2.519827994 | 2.162170545 |

| | | | | | | | |
|---|---|---|---|---|---|---|---|
| ENSMUSG00000034252 | 2.628396264 | 2.430274797 | 2.414106216 | 2.470660762 | 2.398585969 | 2.710963119 | 2.219869654 |
| ENSMUSG00000034254 | 2.435736758 | 2.289967834 | 2.352347734 | 2.194519282 | 2.337065673 | 2.526339277 | 2.298300395 |
| ENSMUSG00000034255 | 1.505869732 | 1.557204673 | 1.630890489 | 0.864857905 | 1.310728002 | 1.041392685 | 1.284450788 |
| ENSMUSG00000034258 | 1.156259305 | 1.080482024 | 1.146240625 | 0.92510993 | 1 | 0.903089987 | 0.984950056 |
| ENSMUSG00000034259 | 1.842799666 | 1.845926073 | 1.87892496 | 1.705044741 | 1.824953989 | 1.954242509 | 1.843097781 |
| ENSMUSG00000034263 | 2.343889214 | 2.186208459 | 2.202642909 | 2.379909063 | 2.319727371 | 2.620136055 | 2.197791274 |
| ENSMUSG00000034265 | 0 | 0.25 | 0 | 0.646240625 | 0.568060967 | 0.477121255 | 0.629532004 |
| ENSMUSG00000034266 | 0 | 0 | 0.25 | 0 | 0 | 0.301029996 | 0 |
| ENSMUSG00000034269 | 3.010519179 | 2.721805154 | 2.672717752 | 2.664552871 | 2.588235379 | 2.985426474 | 2.403564802 |
| ENSMUSG00000034271 | 2.381433035 | 2.252107156 | 2.269871196 | 1.945339928 | 1.992595054 | 2.103803721 | 2.031198704 |
| ENSMUSG00000034274 | 2.395018353 | 2.187886015 | 2.160964047 | 2.216046536 | 2.202543574 | 2.475671188 | 2.057420422 |
| ENSMUSG00000034275 | 1.249317394 | 1.29248125 | 1.348079356 | 1.17510993 | 1.358933174 | 1 | 1.396774124 |
| ENSMUSG00000034278 | 1.358731791 | 1.348079356 | 1.311981878 | 1.638647213 | 1.611014646 | 1.792391689 | 1.601000298 |
| ENSMUSG00000034282 | 0.45815691 | 0.396240625 | 0.701838731 | 0.25 | 0.632412477 | 0.602059991 | 0.520678036 |
| ENSMUSG00000034285 | 2.013563224 | 1.903677461 | 1.933677405 | 2.065523711 | 2.014359518 | 2.201397124 | 1.905143021 |
| ENSMUSG00000034290 | 2.797997234 | 2.533285553 | 2.537119146 | 2.548073204 | 2.495055108 | 2.834420704 | 2.291042162 |
| ENSMUSG00000034292 | 2.042557845 | 1.899978211 | 1.975216702 | 1.912762923 | 1.979897422 | 2.170261715 | 1.811176017 |
| ENSMUSG00000034295 | 2.013563224 | 1.880890489 | 1.848079356 | 1.96759118 | 2.006054284 | 2.235528447 | 1.86815666 |
| ENSMUSG00000034297 | 2.969118676 | 2.723325383 | 2.692665973 | 2.769203899 | 2.677092106 | 3.072984745 | 2.462068294 |
| ENSMUSG00000034300 | 2.530730035 | 2.404137211 | 2.394828984 | 2.423807073 | 2.420401583 | 2.721810615 | 2.277722321 |
| ENSMUSG00000034303 | 1.482695304 | 1.494319981 | 1.470660762 | 1.614857905 | 1.592135487 | 1.69019608 | 1.483307832 |
| ENSMUSG00000034308 | 0 | 0 | 0.396240625 | 0 | 0 | 0 | 0 |
| ENSMUSG00000034310 | 0 | 0 | 0 | 0 | 0 | 0 | 0 |
| ENSMUSG00000034311 | 2.781925631 | 2.556301859 | 2.550530956 | 2.740181499 | 2.638066357 | 3.014520539 | 2.445465687 |
| ENSMUSG00000034312 | 2.401648579 | 2.205044741 | 2.26584877 | 2.063166358 | 2.135290471 | 2.372912003 | 2.080634989 |
| ENSMUSG00000034317 | 2.83262764 | 2.585240691 | 2.575374049 | 2.670748646 | 2.592363568 | 2.970346876 | 2.369585695 |
| ENSMUSG00000034320 | 2.215778996 | 2.048689214 | 2.023189285 | 2.228969845 | 2.167109853 | 2.409933123 | 2.050633136 |
| ENSMUSG00000034321 | 2.264212992 | 2.135757955 | 2.112802778 | 2.083847589 | 2.037065488 | 2.298853076 | 1.935895857 |
| ENSMUSG00000034324 | 0 | 0 | 0 | 0 | 0 | 0 | 0 |
| ENSMUSG00000034327 | 2.269664473 | 2.116901388 | 2.104463129 | 2.518869787 | 2.449594021 | 2.781036939 | 2.289406866 |
| ENSMUSG00000034329 | 2.475043119 | 2.27908599 | 2.264497931 | 2.382357639 | 2.315755952 | 2.619093331 | 2.109521699 |
| ENSMUSG00000034330 | 2.139333929 | 2.002806814 | 2.100219859 | 1.890560606 | 1.937354184 | 2.06069784 | 1.846338363 |
| ENSMUSG00000034333 | 2.20747957 | 2.012462137 | 2.012462137 | 2.043731421 | 2.014359518 | 2.278753601 | 1.830929954 |
| ENSMUSG00000034334 | 0.671187741 | 0.830482024 | 0.75 | 0.79248125 | 0.955822902 | 1 | 0.803905701 |

| | | | | | | | |
|---|---|---|---|---|---|---|---|
| ENSMUSG00000034336 | 0 | 0 | 0.25 | 0.92510993 | 0.978602168 | 1.079181246 | 0.935079929 |
| ENSMUSG00000034339 | 0 | 0 | 0 | 0 | 0 | 0 | 0 |
| ENSMUSG00000034341 | 2.669114668 | 2.455443495 | 2.497171172 | 2.392463902 | 2.449935537 | 2.780317312 | 2.338495365 |
| ENSMUSG00000034342 | 2.543094501 | 2.323655187 | 2.298689214 | 2.418548067 | 2.358048572 | 2.737987326 | 2.194143936 |
| ENSMUSG00000034343 | 2.445324132 | 2.218361028 | 2.206637122 | 2.294354884 | 2.225193812 | 2.517195898 | 2.082194103 |
| ENSMUSG00000034345 | 2.473938397 | 2.298073204 | 2.320192693 | 2.349151195 | 2.295728342 | 2.641474111 | 2.181044355 |
| ENSMUSG00000034349 | 3.231718479 | 2.959702496 | 2.969129237 | 3.129355835 | 3.029944058 | 3.521007252 | 2.792453634 |
| ENSMUSG00000034353 | 0 | 0.5 | 0.646240625 | 0.646240625 | 0.489301084 | 0 | 0.710836105 |
| ENSMUSG00000034354 | 2.666334442 | 2.431128463 | 2.408248799 | 2.375956434 | 2.374910674 | 2.656098202 | 2.224577862 |
| ENSMUSG00000034359 | 0 | 0 | 0 | 0 | 0 | 0 | 0 |
| ENSMUSG00000034361 | 2.654307229 | 2.46759118 | 2.524508021 | 2.198603967 | 2.221952687 | 2.488550717 | 2.134278581 |
| ENSMUSG00000034362 | 0 | 0.25 | 0.25 | 0 | 0 | 0 | 0 |
| ENSMUSG00000034371 | 1.903481042 | 1.794977273 | 1.819031101 | 1.557204673 | 1.689774529 | 1.698970004 | 1.643912836 |
| ENSMUSG00000034377 | 3.246377489 | 2.956736413 | 2.9645924 | 2.914552871 | 2.865686361 | 3.290702243 | 2.662374019 |
| ENSMUSG00000034379 | 1.418409478 | 1.380890489 | 1.438721876 | 1.431980114 | 1.373641807 | 1.380211242 | 1.193409508 |
| ENSMUSG00000034382 | 1.069666964 | 1.04248125 | 1.114857905 | 1.330482024 | 1.351342427 | 1.477121255 | 1.392431569 |
| ENSMUSG00000034384 | 0 | 0 | 0 | 0 | 0.387761935 | 0 | 0 |
| ENSMUSG00000034387 | 0 | 0 | 0 | 0 | 0 | 0 | 0 |
| ENSMUSG00000034390 | 2.365959569 | 2.237091808 | 2.278435542 | 2.174241882 | 2.16097133 | 2.472756449 | 2.114720744 |
| ENSMUSG00000034391 | 0 | 0.25 | 0.25 | 0 | 0.244650542 | 0.477121255 | 0 |
| ENSMUSG00000034394 | 1.890240806 | 1.763820609 | 1.777131114 | 1.505591953 | 1.588235379 | 1.602059991 | 1.586932193 |
| ENSMUSG00000034401 | 2.179305632 | 2.03360658 | 2.051142786 | 2.05479213 | 1.974687643 | 2.28780173 | 1.81475081 |
| ENSMUSG00000034402 | 0.45815691 | 0.580482024 | 0.25 | 0 | 0 | 0 | 0 |
| ENSMUSG00000034403 | 2.417143825 | 2.29248125 | 2.289967834 | 2.205044741 | 2.25190332 | 2.489958479 | 2.179896458 |
| ENSMUSG00000034412 | 1.928763473 | 1.938721876 | 2.029735268 | 1.723704441 | 1.937354184 | 2.041392685 | 1.920029985 |
| ENSMUSG00000034413 | 1.707474304 | 1.516522298 | 1.557204673 | 1.339388001 | 1.467903253 | 1.531478917 | 1.374453863 |
| ENSMUSG00000034416 | 0 | 0 | 0 | 0 | 0 | 0 | 0 |
| ENSMUSG00000034422 | 0.867194479 | 0.951838731 | 1 | 2.044977273 | 1.957204337 | 2.23299611 | 1.857950279 |
| ENSMUSG00000034424 | 2.536482297 | 2.360735874 | 2.344844592 | 2.496460484 | 2.440245937 | 2.782472624 | 2.26501115 |
| ENSMUSG00000034427 | 0 | 0.25 | 0.25 | 0.396240625 | 0.489301084 | 0 | 0.520678036 |
| ENSMUSG00000034429 | 1.596667477 | 1.580482024 | 1.708222504 | 1.482684334 | 1.53772706 | 1.568201724 | 1.687112305 |
| ENSMUSG00000034430 | 2.119110214 | 1.961372513 | 1.981203126 | 1.816696635 | 1.846351821 | 2.064457989 | 1.73898563 |
| ENSMUSG00000034432 | 2.731416512 | 2.510414788 | 2.501406137 | 2.498234584 | 2.472415995 | 2.84260924 | 2.301474734 |
| ENSMUSG00000034435 | 0 | 0 | 0 | 0.701838731 | 0.489301084 | 0.77815125 | 0.520678036 |

| | | | | | | | |
|---|---|---|---|---|---|---|---|
| ENSMUSG00000034437 | 0.289064826 | 0.396240625 | 0 | 0 | 0.489301084 | 0 | 0 |
| ENSMUSG00000034438 | 0 | 0 | 0 | 0 | 0 | 0 | 0 |
| ENSMUSG00000034442 | 1.827447047 | 1.80479213 | 1.744319981 | 1.784887838 | 1.686820904 | 1.875061263 | 1.562034107 |
| ENSMUSG00000034445 | 1.693336259 | 1.738549078 | 1.668106335 | 1.403677461 | 1.484072009 | 1.653212514 | 1.452143229 |
| ENSMUSG00000034449 | 1.631440309 | 1.396240625 | 1.51109853 | 1.418106335 | 1.555378544 | 1.361727836 | 1.523605797 |
| ENSMUSG00000034450 | 0 | 0 | 0 | 0 | 0 | 0 | 0.224243824 |
| ENSMUSG00000034452 | 0 | 0 | 0 | 0 | 0 | 0 | 0 |
| ENSMUSG00000034453 | 2.523917219 | 2.277783918 | 2.311981878 | 2.403217124 | 2.333760374 | 2.668385917 | 2.198515833 |
| ENSMUSG00000034456 | 0 | 0.25 | 0 | 0 | 0 | 0 | 0 |
| ENSMUSG00000034457 | 2.243591459 | 2.095926073 | 2.03743678 | 1.282320754 | 0.978602168 | 1.397940009 | 1.159323753 |
| ENSMUSG00000034459 | 0.671187741 | 0.396240625 | 0 | 0.92510993 | 0.877063019 | 0.77815125 | 0.74492186 |
| ENSMUSG00000034460 | 2.241665209 | 2.086073977 | 2.02186571 | 1.966046536 | 1.928492895 | 2.079181246 | 1.809975587 |
| ENSMUSG00000034462 | 2.845531329 | 2.611253711 | 2.614088604 | 2.474839231 | 2.421513257 | 2.783903579 | 2.276587178 |
| ENSMUSG00000034463 | 2.243591459 | 2.071350555 | 2.179419106 | 2.110735874 | 2.056322416 | 2.354108439 | 1.984950056 |
| ENSMUSG00000034467 | 0 | 0 | 0 | 0 | 0 | 0 | 0.224243824 |
| ENSMUSG00000034471 | 2.045658471 | 2.038704527 | 2.137686696 | 1.75835575 | 1.981187939 | 2.017033339 | 2.066798338 |
| ENSMUSG00000034472 | 0.289064826 | 0 | 0 | 0 | 0 | 0 | 0 |
| ENSMUSG00000034473 | 2.016886209 | 1.919870025 | 1.852347734 | 1.899978211 | 1.842556557 | 2.127104798 | 1.753626837 |
| ENSMUSG00000034480 | 2.063793172 | 1.907339155 | 1.905512955 | 2.061981878 | 1.954436039 | 2.230448921 | 1.840919217 |
| ENSMUSG00000034482 | 0 | 0 | 0 | 0 | 0 | 0 | 0 |
| ENSMUSG00000034484 | 2.616779011 | 2.420309603 | 2.441217898 | 2.347004321 | 2.26251048 | 2.615950052 | 2.131619535 |
| ENSMUSG00000034485 | 2.287337982 | 2.183677405 | 2.193696765 | 2.324229052 | 2.35043857 | 2.609594409 | 2.23605706 |
| ENSMUSG00000034486 | 0 | 0.396240625 | 0.25 | 0.701838731 | 0.632412477 | 0.602059991 | 0.579661877 |
| ENSMUSG00000034487 | 2.336265009 | 2.216046536 | 2.160061234 | 2.149047439 | 2.10031573 | 2.376576957 | 1.960219986 |
| ENSMUSG00000034488 | 1.714367558 | 1.692046081 | 1.657339155 | 1.04248125 | 1.136121934 | 1.146128036 | 1.028149525 |
| ENSMUSG00000034493 | 0 | 0 | 0 | 0.25 | 0.244650542 | 0 | 0 |
| ENSMUSG00000034501 | 2.287337982 | 2.12892496 | 2.136723615 | 2.041226732 | 1.982473755 | 2.250420002 | 1.886704102 |
| ENSMUSG00000034509 | 2.126809492 | 1.979715809 | 2.023189285 | 1.938721876 | 2.033751322 | 2.227886705 | 1.95411574 |
| ENSMUSG00000034518 | 2.399009124 | 2.216046536 | 2.183677405 | 2.25835575 | 2.236621448 | 2.472756449 | 2.065709059 |
| ENSMUSG00000034520 | 2.441754646 | 2.259729747 | 2.257666784 | 2.319031101 | 2.254881871 | 2.547774705 | 2.087847416 |
| ENSMUSG00000034522 | 2.023453784 | 1.959800947 | 1.979715809 | 1.823655187 | 1.868304353 | 1.995635195 | 1.813563596 |
| ENSMUSG00000034525 | 2.804556862 | 2.57991803 | 2.574802005 | 2.54838134 | 2.535323264 | 2.851258349 | 2.377173376 |
| ENSMUSG00000034528 | 0.45815691 | 0 | 0.396240625 | 0 | 0 | 0.301029996 | 0.224243824 |
| ENSMUSG00000034532 | 0.45815691 | 0.5 | 0.25 | 0.75 | 0.955822902 | 0.77815125 | 0.74492186 |

| | | | | | | | |
|---|---|---|---|---|---|---|---|
| ENSMUSG00000034533 | 0 | 0 | 0 | 0 | 0 | 0.301029996 | 0 |
| ENSMUSG00000034538 | 1 | 1.214495249 | 1.130890489 | 1.226722649 | 1.106691885 | 0.954242509 | 1.028149525 |
| ENSMUSG00000034543 | 2.69529263 | 2.465659339 | 2.472185812 | 2.504202072 | 2.480020371 | 2.765668555 | 2.292127786 |
| ENSMUSG00000034544 | 2.339331437 | 2.173371739 | 2.182829758 | 2.30357978 | 2.1962964 | 2.527629901 | 2.068965954 |
| ENSMUSG00000034551 | 0.811507563 | 1.02186571 | 0.79248125 | 1.388647213 | 1.234113756 | 1.322219295 | 1.159323753 |
| ENSMUSG00000034552 | 0 | 0 | 0 | 0 | 0 | 0 | 0 |
| ENSMUSG00000034555 | 0 | 0 | 0 | 0 | 0 | 0 | 0 |
| ENSMUSG00000034557 | 2.299278375 | 2.161864607 | 2.145300145 | 2.333847589 | 2.254288168 | 2.596597096 | 2.083229353 |
| ENSMUSG00000034560 | 2.750084015 | 2.537119146 | 2.538704527 | 2.520869832 | 2.470810917 | 2.805500858 | 2.294826012 |
| ENSMUSG00000034563 | 0 | 0.25 | 0 | 0 | 0 | 0 | 0 |
| ENSMUSG00000034566 | 2.751792 | 2.518535366 | 2.517194569 | 2.627938163 | 2.549506445 | 2.961895474 | 2.351382464 |
| ENSMUSG00000034570 | 1.036286563 | 1.238549078 | 1.061981878 | 1.061981878 | 1.136121934 | 1.230448921 | 1.110947927 |
| ENSMUSG00000034573 | 2.418409478 | 2.201838731 | 2.205044741 | 1.928561379 | 1.913211023 | 2.198657087 | 1.801445495 |
| ENSMUSG00000034574 | 2.389637216 | 2.187048212 | 2.181128463 | 2.168989258 | 2.129414728 | 2.403120521 | 1.976436 |
| ENSMUSG00000034575 | 2.389637216 | 2.231203126 | 2.222185812 | 2.21293726 | 2.223252711 | 2.442479769 | 2.083229353 |
| ENSMUSG00000034579 | 0.578129653 | 0.75 | 0.75 | 0.5 | 0.568060967 | 0 | 0 |
| ENSMUSG00000034583 | 0 | 0 | 0 | 0 | 0 | 0 | 0 |
| ENSMUSG00000034584 | 0.811507563 | 0.830482024 | 0.580482024 | 0.5 | 0.632412477 | 0.602059991 | 0.520678036 |
| ENSMUSG00000034586 | 2.013563224 | 1.892463902 | 1.964495249 | 1.821350555 | 1.945998439 | 2.064457989 | 1.895131899 |
| ENSMUSG00000034587 | 1.404271433 | 1.403677461 | 1.356566189 | 1.282320754 | 1.401338844 | 1.447158031 | 1.296434211 |
| ENSMUSG00000034591 | 2.089597219 | 1.907339155 | 1.916333979 | 2.071350555 | 2.014359518 | 2.281033367 | 1.850609257 |
| ENSMUSG00000034593 | 3.162247566 | 2.87246199 | 2.882479632 | 2.866264042 | 2.792996582 | 3.232487866 | 2.593082247 |
| ENSMUSG00000034595 | 2.679378494 | 2.467976309 | 2.533285553 | 2.310793496 | 2.335652905 | 2.557507202 | 2.225244904 |
| ENSMUSG00000034601 | 2.254965775 | 2.063166358 | 2.031030328 | 1.997171172 | 1.948833444 | 2.113943352 | 1.813563596 |
| ENSMUSG00000034602 | 2.767951589 | 2.563461871 | 2.547764804 | 2.549918086 | 2.541960468 | 2.865103975 | 2.366370874 |
| ENSMUSG00000034607 | 0 | 0 | 0 | 0 | 0 | 0 | 0 |
| ENSMUSG00000034610 | 2.511972754 | 2.328220739 | 2.284887838 | 2.289336734 | 2.216042585 | 2.503790683 | 2.034820183 |
| ENSMUSG00000034612 | 2.556920045 | 2.36434522 | 2.326515422 | 2.326515422 | 2.303566328 | 2.592176757 | 2.156039304 |
| ENSMUSG00000034613 | 1.568542279 | 1.488549078 | 1.692046081 | 1.321350555 | 1.293076518 | 1.397940009 | 1.238624656 |
| ENSMUSG00000034614 | 1.771760131 | 1.584962501 | 1.668106335 | 1.201838731 | 1.319233381 | 1.397940009 | 1.278288402 |
| ENSMUSG00000034616 | 1.907802659 | 1.819031101 | 1.919870025 | 1.476722649 | 1.639266209 | 1.662757832 | 1.667169465 |
| ENSMUSG00000034617 | 2.280358197 | 2.174241882 | 2.122963274 | 2.156427211 | 2.120847411 | 2.390935107 | 2.00464543 |
| ENSMUSG00000034620 | 2.355805236 | 2.169870025 | 2.185366747 | 2.254202072 | 2.111176775 | 2.40654018 | 1.959463225 |
| ENSMUSG00000034621 | 2.678024491 | 2.445339928 | 2.459406983 | 2.506284891 | 2.468875138 | 2.807535028 | 2.339198915 |

| | | | | | | | |
|---|---|---|---|---|---|---|---|
| ENSMUSG00000034623 | 0 | 0 | 0 | 0 | 0 | 0 | 0 |
| ENSMUSG00000034634 | 0 | 0 | 0 | 0 | 0 | 0 | 0 |
| ENSMUSG00000034636 | 2.644870062 | 2.42813175 | 2.391989019 | 2.573655187 | 2.455019006 | 2.861534411 | 2.287488514 |
| ENSMUSG00000034639 | 1.418409478 | 1.330482024 | 1.238549078 | 1.364857905 | 1.283908311 | 1.462397998 | 1.238624656 |
| ENSMUSG00000034640 | 2.20747957 | 2.03360658 | 2.02186571 | 2.252806814 | 2.143518259 | 2.428134794 | 1.955652633 |
| ENSMUSG00000034641 | 0.45815691 | 0 | 0.25 | 0 | 0 | 0 | 0 |
| ENSMUSG00000034645 | 0.811507563 | 0.580482024 | 0.396240625 | 0.75 | 0.77552387 | 0.477121255 | 0.672731473 |
| ENSMUSG00000034647 | 2.619128496 | 2.469895812 | 2.518869787 | 2.362802778 | 2.250704826 | 2.544068044 | 2.092400484 |
| ENSMUSG00000034648 | 0 | 0 | 0 | 0 | 0 | 0 | 0 |
| ENSMUSG00000034652 | 0.867194479 | 0.79248125 | 0.79248125 | 0.830482024 | 0.387761935 | 0.477121255 | 0 |
| ENSMUSG00000034653 | 2.116511873 | 1.970660762 | 2.015173983 | 2.088286706 | 2.043601899 | 2.257678575 | 1.837623616 |
| ENSMUSG00000034656 | 2.016886209 | 1.860735874 | 1.958222504 | 1.27186571 | 1.514343934 | 1.633468456 | 1.56716938 |
| ENSMUSG00000034659 | 2.682073381 | 2.505938588 | 2.526804269 | 2.359697963 | 2.467253839 | 2.713490543 | 2.334479423 |
| ENSMUSG00000034660 | 0.578129653 | 0 | 0 | 0 | 0 | 0 | 0 |
| ENSMUSG00000034663 | 2.550525119 | 2.299918086 | 2.298073204 | 2.181128463 | 2.117361396 | 2.356025857 | 1.94074868 |
| ENSMUSG00000034664 | 1.614416215 | 1.5 | 1.571350555 | 1.17510993 | 1.283908311 | 1.176091259 | 1.245582246 |
| ENSMUSG00000034667 | 2.903085937 | 2.646709947 | 2.626204747 | 2.599685923 | 2.562149491 | 2.91750551 | 2.385603553 |
| ENSMUSG00000034673 | 2.095120908 | 2.004202072 | 2.032320754 | 2.03360658 | 2.071604288 | 2.247973266 | 2 |
| ENSMUSG00000034674 | 2.312518611 | 2.187048212 | 2.127938163 | 2.38040011 | 2.357605438 | 2.663700925 | 2.201039086 |
| ENSMUSG00000034675 | 2.641182749 | 2.47934302 | 2.518869787 | 2.430701883 | 2.56585575 | 2.783188691 | 2.54475813 |
| ENSMUSG00000034677 | 0 | 0 | 0 | 0 | 0 | 0 | 0 |
| ENSMUSG00000034681 | 2.742600832 | 2.543419284 | 2.56932185 | 2.488912477 | 2.510627067 | 2.820201459 | 2.431506263 |
| ENSMUSG00000034683 | 0 | 0.396240625 | 0 | 0.25 | 0.244650542 | 0 | 0.355418052 |
| ENSMUSG00000034684 | 2.547754129 | 2.444519282 | 2.508011432 | 2.036164561 | 2.155511127 | 2.276461804 | 2.149789616 |
| ENSMUSG00000034685 | 1.693336259 | 1.698603967 | 1.850219859 | 1.311981878 | 1.649303553 | 1.414973348 | 1.726776051 |
| ENSMUSG00000034686 | 0 | 0.701838731 | 0.646240625 | 0 | 0.387761935 | 0.602059991 | 0.952571511 |
| ENSMUSG00000034687 | 2.131864492 | 1.950224975 | 1.930274797 | 2.120953944 | 2.138604637 | 2.372912003 | 1.993245779 |
| ENSMUSG00000034689 | 0 | 0 | 0 | 0 | 0 | 0 | 0 |
| ENSMUSG00000034690 | 0 | 0.25 | 0 | 0.25 | 0 | 0 | 0 |
| ENSMUSG00000034701 | 0 | 0 | 0 | 0 | 0 | 0 | 0 |
| ENSMUSG00000034706 | 0.867194479 | 0.92510993 | 0.951838731 | 0.701838731 | 0.686820904 | 1.041392685 | 0.775756176 |
| ENSMUSG00000034707 | 3.057562627 | 2.799764706 | 2.796838018 | 2.674024543 | 2.621035819 | 3.061829307 | 2.473713589 |
| ENSMUSG00000034708 | 3.15969169 | 2.890679857 | 2.958123621 | 2.484159485 | 2.496852851 | 2.843232778 | 2.398728906 |
| ENSMUSG00000034709 | 2.436947303 | 2.268535366 | 2.210587586 | 2.226722649 | 2.208005287 | 2.475671188 | 2.121678334 |

| | | | | | | | |
|---|---|---|---|---|---|---|---|
| ENSMUSG00000034714 | 2.231898196 | 2.106566189 | 2.093759858 | 2.097004321 | 2.151558609 | 2.328379603 | 2.066798338 |
| ENSMUSG00000034723 | 2.438154344 | 2.209012589 | 2.185366747 | 2.302972074 | 2.21736452 | 2.534026106 | 2.054044578 |
| ENSMUSG00000034724 | 2.749513129 | 2.524836953 | 2.522528105 | 2.506976499 | 2.414035093 | 2.771587481 | 2.228229691 |
| ENSMUSG00000034729 | 2.192545868 | 2.070192693 | 2.041226732 | 2.092671852 | 2.059432181 | 2.336459734 | 1.999330888 |
| ENSMUSG00000034730 | 0 | 0 | 0 | 0 | 0 | 0 | 0 |
| ENSMUSG00000034731 | 1.794934559 | 1.769203899 | 1.701838731 | 1.571350555 | 1.504538967 | 1.633468456 | 1.396774124 |
| ENSMUSG00000034732 | 0 | 0 | 0 | 0 | 0 | 0 | 0 |
| ENSMUSG00000034738 | 0.289064826 | 0.5 | 0.5 | 0.5 | 0.686820904 | 0.477121255 | 0.579661877 |
| ENSMUSG00000034739 | 0.671187741 | 0.646240625 | 0.646240625 | 0.580482024 | 0.387761935 | 0.301029996 | 0.775756176 |
| ENSMUSG00000034744 | 2.149089517 | 2.059601185 | 2.115881093 | 1.698603967 | 1.763796674 | 1.968482949 | 1.704233519 |
| ENSMUSG00000034748 | 1.623015126 | 1.598079356 | 1.708222504 | 1.388647213 | 1.584291694 | 1.491361694 | 1.598697688 |
| ENSMUSG00000034751 | 2.541215974 | 2.358135475 | 2.356566189 | 2.123963757 | 2.12771782 | 2.389166084 | 2.020712061 |
| ENSMUSG00000034755 | 0.289064826 | 0.396240625 | 0 | 0 | 0.244650542 | 0 | 0 |
| ENSMUSG00000034757 | 2.235832493 | 2.079353153 | 2.125956434 | 1.991446071 | 2.107593434 | 2.301029996 | 2.007270316 |
| ENSMUSG00000034758 | 1.783508299 | 1.747171172 | 1.802363341 | 1.130890489 | 1.27449559 | 1.255272505 | 1.369798885 |
| ENSMUSG00000034761 | 2.768497801 | 2.523849256 | 2.486360959 | 2.412314044 | 2.343584837 | 2.668385917 | 2.168981648 |
| ENSMUSG00000034762 | 0.747221736 | 0.951838731 | 1.238549078 | 0.5 | 0.733951626 | 0.954242509 | 1.054044578 |
| ENSMUSG00000034764 | 0.811507563 | 1.02186571 | 0.896240625 | 0.25 | 0.387761935 | 0.698970004 | 0.224243824 |
| ENSMUSG00000034765 | 2.215778996 | 2.067865757 | 2.063166358 | 1.811981878 | 1.836786029 | 1.913813852 | 1.749303626 |
| ENSMUSG00000034768 | 0.289064826 | 0.25 | 0.25 | 0 | 0 | 0 | 0 |
| ENSMUSG00000034771 | 1.857607105 | 1.692046081 | 1.848079356 | 1.584962501 | 1.709785497 | 1.477121255 | 1.720494 |
| ENSMUSG00000034773 | 2.095120908 | 1.92510993 | 1.933677405 | 1.882845365 | 1.94314048 | 2.008600172 | 1.852723744 |
| ENSMUSG00000034774 | 0 | 0 | 0 | 0 | 0 | 0 | 0 |
| ENSMUSG00000034777 | 0.671187741 | 0.701838731 | 1.114857905 | 1.201838731 | 1.351342427 | 1.531478917 | 1.514741806 |
| ENSMUSG00000034780 | 1.693336259 | 1.580482024 | 1.5 | 1.505591953 | 1.47337555 | 1.653212514 | 1.401059159 |
| ENSMUSG00000034781 | 2.590857977 | 2.428990498 | 2.482314602 | 2.459406983 | 2.560406495 | 2.790285164 | 2.445802857 |
| ENSMUSG00000034783 | 0 | 0 | 0 | 0 | 0 | 0 | 0 |
| ENSMUSG00000034785 | 0 | 0 | 0 | 0 | 0 | 0 | 0 |
| ENSMUSG00000034786 | 0.45815691 | 0.396240625 | 0.79248125 | 0.580482024 | 0.905314583 | 0.77815125 | 0.829800754 |
| ENSMUSG00000034789 | 2.192545868 | 2.009729747 | 2.059601185 | 2.00835575 | 2.031524462 | 2.247973266 | 1.939944915 |
| ENSMUSG00000034792 | 0 | 0 | 0 | 0.25 | 0 | 0 | 0 |
| ENSMUSG00000034793 | 2.097855556 | 2.029735268 | 2.043731421 | 1.561981878 | 1.754126039 | 1.792391689 | 1.653807046 |
| ENSMUSG00000034795 | 1.205378646 | 1.27186571 | 1.282320754 | 1.29248125 | 1.188507684 | 1.397940009 | 1.159323753 |
| ENSMUSG00000034796 | 0 | 0 | 0 | 0.25 | 0 | 0 | 0.224243824 |

| | | | | | | | |
|---|---|---|---|---|---|---|---|
| ENSMUSG00000034799 | 1.54867984 | 1.438721876 | 1.330482024 | 1.602347734 | 1.621875691 | 1.653212514 | 1.508694612 |
| ENSMUSG00000034800 | 1.100572389 | 1.27186571 | 1.02186571 | 1.214495249 | 1.319233381 | 1.322219295 | 1.23151414 |
| ENSMUSG00000034801 | 2.349889783 | 2.132845365 | 2.127938163 | 2.08272922 | 1.987570683 | 2.278753601 | 1.898808251 |
| ENSMUSG00000034807 | 2.639698646 | 2.469512728 | 2.459406983 | 2.450629091 | 2.47337555 | 2.782472624 | 2.34500308 |
| ENSMUSG00000034810 | 0.45815691 | 0.25 | 0.5 | 0 | 0 | 0 | 0 |
| ENSMUSG00000034813 | 0.289064826 | 0.5 | 0.580482024 | 0.79248125 | 0.632412477 | 0.903089987 | 0.775756176 |
| ENSMUSG00000034818 | 0.747221736 | 0.75 | 0.830482024 | 1.080482024 | 1.039257769 | 1.113943352 | 1.028149525 |
| ENSMUSG00000034820 | 2.76355595 | 2.508699741 | 2.517530236 | 2.640321986 | 2.588926773 | 2.967079734 | 2.406432018 |
| ENSMUSG00000034825 | 1.129344651 | 1.080482024 | 1.238549078 | 1.614857905 | 1.592135487 | 1.698970004 | 1.479989695 |
| ENSMUSG00000034826 | 2.614416215 | 2.382845365 | 2.405512955 | 2.337734795 | 2.280062837 | 2.605305046 | 2.160953557 |
| ENSMUSG00000034829 | 0 | 0 | 0 | 0.580482024 | 0.387761935 | 0.698970004 | 0.224243824 |
| ENSMUSG00000034832 | 2.528794847 | 2.366901388 | 2.343759858 | 2.291226732 | 2.35849115 | 2.574031268 | 2.261157966 |
| ENSMUSG00000034833 | 0 | 0 | 0 | 0 | 0 | 0 | 0 |
| ENSMUSG00000034837 | 0.578129653 | 0.646240625 | 0.75 | 0.580482024 | 0.489301084 | 0 | 0.520678036 |
| ENSMUSG00000034839 | 1.129344651 | 1.226722649 | 1.238549078 | 0.92510993 | 1 | 1.041392685 | 0.935079929 |
| ENSMUSG00000034842 | 0 | 0.396240625 | 0 | 0.25 | 0.244650542 | 0 | 0.355418052 |
| ENSMUSG00000034845 | 0.578129653 | 0.75 | 1.061981878 | 0.580482024 | 0.877063019 | 0.698970004 | 0.876096088 |
| ENSMUSG00000034848 | 2.380049842 | 2.180274797 | 2.168989258 | 2.091580554 | 2.029283463 | 2.324282455 | 1.896975297 |
| ENSMUSG00000034850 | 2.479432974 | 2.279735268 | 2.254897682 | 2.422499493 | 2.373641807 | 2.693726949 | 2.238624656 |
| ENSMUSG00000034853 | 1.227926429 | 1.02186571 | 0.976722649 | 0.396240625 | 0.632412477 | 0.602059991 | 0.775756176 |
| ENSMUSG00000034854 | 1.45815691 | 1.458222504 | 1.642463902 | 1.593759858 | 1.806427258 | 1.716003344 | 1.755055163 |
| ENSMUSG00000034855 | 1.036286563 | 1.098079356 | 1.04248125 | 1.964495249 | 1.98503142 | 2.190331698 | 1.925124822 |
| ENSMUSG00000034858 | 1.87190676 | 1.653677461 | 1.76109853 | 1.850219859 | 1.824953989 | 1.991226076 | 1.787569411 |
| ENSMUSG00000034863 | 1.885732304 | 1.862802778 | 1.969129237 | 1.766522298 | 1.892300902 | 2.008600172 | 1.922587433 |
| ENSMUSG00000034867 | 2.215778996 | 2.059601185 | 2.086073977 | 1.979715809 | 1.934425071 | 2.187520721 | 1.815933683 |
| ENSMUSG00000034868 | 2.477243823 | 2.327084758 | 2.361770807 | 2.512462137 | 2.535858857 | 2.889301703 | 2.417653332 |
| ENSMUSG00000034871 | 0 | 0 | 0 | 0 | 0 | 0 | 0 |
| ENSMUSG00000034872 | 0 | 0 | 0 | 0.25 | 0 | 0 | 0 |
| ENSMUSG00000034875 | 1.816888701 | 1.738549078 | 1.735628626 | 1.811981878 | 1.768534412 | 1.968482949 | 1.710836105 |
| ENSMUSG00000034880 | 1.432083896 | 1.494319981 | 1.649978211 | 1.348079356 | 1.514343934 | 1.568201724 | 1.723650273 |
| ENSMUSG00000034881 | 0.289064826 | 0.646240625 | 0 | 0.25 | 0.244650542 | 0 | 0.448487648 |
| ENSMUSG00000034883 | 1.721148722 | 1.618933358 | 1.714495249 | 1.630890489 | 1.639266209 | 1.73239376 | 1.548827561 |
| ENSMUSG00000034889 | 2.285603957 | 2.234159485 | 2.33993739 | 1.898114259 | 2.107593434 | 2.158362492 | 2.191196167 |
| ENSMUSG00000034891 | 0 | 0 | 0 | 0 | 0 | 0 | 0.448487648 |

| | | | | | | | |
|---|---|---|---|---|---|---|---|
| ENSMUSG00000034892 | 2.592526109 | 2.493244947 | 2.58993739 | 2.136723615 | 2.312243922 | 2.482873584 | 2.288586107 |
| ENSMUSG00000034893 | 2.326928073 | 2.158248799 | 2.17510993 | 2.321928095 | 2.264247757 | 2.584331224 | 2.128489437 |
| ENSMUSG00000034898 | 0 | 0.25 | 0 | 0 | 0 | 0 | 0 |
| ENSMUSG00000034902 | 2.519973621 | 2.374961472 | 2.437886015 | 2.320772088 | 2.464971436 | 2.681241237 | 2.403564802 |
| ENSMUSG00000034903 | 2.609650079 | 2.403677461 | 2.445339928 | 2.288704527 | 2.233484039 | 2.534026106 | 2.092902448 |
| ENSMUSG00000034906 | 2.572054151 | 2.36434522 | 2.33272922 | 2.53743678 | 2.472095563 | 2.847572659 | 2.352957847 |
| ENSMUSG00000034908 | 2.630681378 | 2.430274797 | 2.437467357 | 2.451838731 | 2.467578695 | 2.777426822 | 2.341067602 |
| ENSMUSG00000034910 | 1.445324132 | 1.388647213 | 1.431980114 | 1.96293726 | 1.871833944 | 2.11058971 | 1.792684387 |
| ENSMUSG00000034911 | 0.289064826 | 0.25 | 0 | 0.25 | 0.387761935 | 0 | 0 |
| ENSMUSG00000034912 | 0.289064826 | 0.75 | 0.5 | 0.25 | 0.568060967 | 0.477121255 | 0.448487648 |
| ENSMUSG00000034913 | 0 | 0 | 0 | 0 | 0 | 0 | 0 |
| ENSMUSG00000034917 | 0 | 0 | 0 | 0.830482024 | 0.489301084 | 0.301029996 | 0.876096088 |
| ENSMUSG00000034918 | 0 | 0 | 0 | 0.25 | 0 | 0 | 0 |
| ENSMUSG00000034919 | 0 | 0 | 0 | 0 | 0 | 0 | 0 |
| ENSMUSG00000034923 | 0 | 0 | 0 | 0 | 0 | 0 | 0 |
| ENSMUSG00000034926 | 2.42468076 | 2.316696635 | 2.353406982 | 2.343759858 | 2.435655147 | 2.694605199 | 2.354748941 |
| ENSMUSG00000034928 | 2.419671301 | 2.254202072 | 2.29745614 | 2.158248799 | 2.25190332 | 2.390935107 | 2.185595644 |
| ENSMUSG00000034930 | 1.389637216 | 1.339388001 | 1.372963274 | 1.17510993 | 1.351342427 | 1.204119983 | 1.437266334 |
| ENSMUSG00000034931 | 2.649978134 | 2.43325383 | 2.429847205 | 2.418548067 | 2.393014957 | 2.764176132 | 2.265599895 |
| ENSMUSG00000034932 | 2.10861828 | 2.02186571 | 2.12892496 | 2.04745614 | 2.13194489 | 2.336459734 | 2.133836922 |
| ENSMUSG00000034936 | 1.663535556 | 1.62694866 | 1.738549078 | 1.27186571 | 1.394615667 | 1.462397998 | 1.313616061 |
| ENSMUSG00000034940 | 2.440557995 | 2.28360658 | 2.291226732 | 2.315523711 | 2.318738698 | 2.596597096 | 2.154798988 |
| ENSMUSG00000034947 | 1.777675583 | 1.692046081 | 1.708222504 | 1.961372513 | 1.979897422 | 2.271841607 | 1.851668228 |
| ENSMUSG00000034949 | 1.069666964 | 1.080482024 | 1.160964047 | 1.061981878 | 1.163285805 | 1.079181246 | 1.350478789 |
| ENSMUSG00000034951 | 1.89911417 | 1.834962501 | 1.848079356 | 1.799918086 | 1.850106709 | 1.977723605 | 1.828667599 |
| ENSMUSG00000034957 | 0.45815691 | 0.25 | 0.396240625 | 0.646240625 | 0.77552387 | 0.301029996 | 0.579661877 |
| ENSMUSG00000034958 | 0 | 0 | 0 | 0 | 0 | 0 | 0 |
| ENSMUSG00000034959 | 0.960252568 | 0.830482024 | 0.830482024 | 1.02186571 | 0.812711509 | 0.954242509 | 0.896975297 |
| ENSMUSG00000034968 | 0 | 0.25 | 0 | 0 | 0.244650542 | 0 | 0 |
| ENSMUSG00000034973 | 2.337801042 | 2.156427211 | 2.123963757 | 2.299304173 | 2.240967488 | 2.532754379 | 2.085289968 |
| ENSMUSG00000034974 | 2.078325387 | 1.990000483 | 2.088286706 | 1.62694866 | 1.824953989 | 1.875061263 | 1.897893072 |
| ENSMUSG00000034981 | 2.151493169 | 2.016522298 | 2.034887838 | 1.580482024 | 1.49445382 | 1.544068044 | 1.433437704 |
| ENSMUSG00000034987 | 0 | 0 | 0 | 0 | 0 | 0 | 0 |
| ENSMUSG00000034990 | 0 | 0 | 0 | 0 | 0 | 0 | 0.224243824 |

| | | | | | | | |
|---|---|---|---|---|---|---|---|
| ENSMUSG00000034993 | 3.117694944 | 2.853935442 | 2.947281512 | 2.631869252 | 2.664680788 | 2.995196292 | 2.584636584 |
| ENSMUSG00000034994 | 3.87023824 | 3.591324302 | 3.640456229 | 3.358982663 | 3.530082188 | 3.915663604 | 3.396577994 |
| ENSMUSG00000034997 | 1.036286563 | 1.061981878 | 0.951838731 | 0 | 0 | 0 | 0 |
| ENSMUSG00000034998 | 2.392336464 | 2.203445298 | 2.218361028 | 2.209800947 | 2.167109853 | 2.426511261 | 2.006616089 |
| ENSMUSG00000035000 | 0 | 0 | 0 | 0 | 0 | 0 | 0.224243824 |
| ENSMUSG00000035004 | 0.811507563 | 0.580482024 | 0.951838731 | 0.79248125 | 0.905314583 | 0.77815125 | 0.710836105 |
| ENSMUSG00000035007 | 1.920505136 | 1.850219859 | 1.937048212 | 1.930274797 | 1.932951348 | 2.133538908 | 1.872150729 |
| ENSMUSG00000035011 | 1.99653913 | 1.926839783 | 2.094844592 | 1.868933358 | 1.906908067 | 2.075546961 | 1.985649549 |
| ENSMUSG00000035020 | 0.45815691 | 0.396240625 | 0.396240625 | 0.79248125 | 0.387761935 | 0.301029996 | 0.224243824 |
| ENSMUSG00000035021 | 2.853620573 | 2.613574823 | 2.612545062 | 2.62246199 | 2.528558853 | 2.91750551 | 2.369158893 |
| ENSMUSG00000035024 | 2.615205302 | 2.388167274 | 2.387686696 | 2.536164561 | 2.489301084 | 2.808210973 | 2.309276932 |
| ENSMUSG00000035027 | 2.542156295 | 2.475216702 | 2.591033664 | 2.150906586 | 2.42519375 | 2.474216264 | 2.456084583 |
| ENSMUSG00000035031 | 0 | 0 | 0 | 0 | 0 | 0 | 0 |
| ENSMUSG00000035032 | 0 | 0.396240625 | 0.25 | 0 | 0 | 0.301029996 | 0 |
| ENSMUSG00000035033 | 0.289064826 | 0.5 | 0.25 | 0.25 | 0 | 0 | 0 |
| ENSMUSG00000035041 | 0.289064826 | 0 | 0.25 | 0.580482024 | 0.489301084 | 0 | 0.355418052 |
| ENSMUSG00000035042 | 0.811507563 | 0.646240625 | 0.75 | 1.201838731 | 1.091002363 | 1.278753601 | 1.150210039 |
| ENSMUSG00000035045 | 1.777675583 | 1.552363341 | 1.642463902 | 1.708222504 | 1.509475496 | 1.653212514 | 1.502532226 |
| ENSMUSG00000035047 | 1.753493018 | 1.660964047 | 1.732684334 | 1.811981878 | 1.751666404 | 1.959041392 | 1.792684387 |
| ENSMUSG00000035048 | 1.975148165 | 1.799918086 | 1.802363341 | 1.896240625 | 1.937354184 | 2.149219113 | 1.871156825 |
| ENSMUSG00000035049 | 2.4661223 | 2.298689214 | 2.344844592 | 2.186208459 | 2.318738698 | 2.444044796 | 2.202472158 |
| ENSMUSG00000035051 | 2.418409478 | 2.262462137 | 2.245748394 | 2.226722649 | 2.215379757 | 2.53529412 | 2.085803078 |
| ENSMUSG00000035057 | 0 | 0.25 | 0 | 0 | 0 | 0 | 0.224243824 |
| ENSMUSG00000035062 | 2.057835465 | 1.931980114 | 1.886723615 | 1.894357207 | 1.797870327 | 1.995635195 | 1.734460864 |
| ENSMUSG00000035064 | 2.276824003 | 2.158248799 | 2.190387808 | 1.860735874 | 1.871833944 | 2.053078443 | 1.770364127 |
| ENSMUSG00000035067 | 1.596667477 | 1.51109853 | 1.53743678 | 1.630890489 | 1.504538967 | 1.707570176 | 1.493062805 |
| ENSMUSG00000035069 | 1.596667477 | 1.356566189 | 1.431980114 | 1.5 | 1.563883923 | 1.643452676 | 1.479989695 |
| ENSMUSG00000035078 | 2.131864492 | 1.912762923 | 2.013820609 | 2.209012589 | 2.164820399 | 2.445604203 | 2.006616089 |
| ENSMUSG00000035085 | 0.289064826 | 0.701838731 | 0.25 | 0.396240625 | 0 | 0 | 0.629532004 |
| ENSMUSG00000035086 | 2.552362269 | 2.320772088 | 2.324802005 | 2.439555804 | 2.385310841 | 2.730782276 | 2.252699564 |
| ENSMUSG00000035093 | 2.531694272 | 2.361253711 | 2.33272922 | 2.32991803 | 2.28771234 | 2.575187845 | 2.151886289 |
| ENSMUSG00000035095 | 1.100572389 | 1.130890489 | 1.130890489 | 1.080482024 | 1.020174412 | 1.361727836 | 1.159323753 |
| ENSMUSG00000035104 | 1.156259305 | 1.146240625 | 1.098079356 | 0.92510993 | 0.931471446 | 1 | 0.803905701 |
| ENSMUSG00000035105 | 2.054824387 | 1.918106335 | 1.888647213 | 1.823655187 | 1.689774529 | 1.934498451 | 1.643912836 |

| | | | | | | | |
|---|---|---|---|---|---|---|---|
| ENSMUSG00000035107 | 2.834961347 | 2.621457327 | 2.591580554 | 2.726534749 | 2.674572266 | 3.089551883 | 2.490284576 |
| ENSMUSG00000035109 | 1.37447073 | 1.431980114 | 1.476722649 | 1.02186571 | 0.978602168 | 1.146128036 | 0.952571511 |
| ENSMUSG00000035112 | 1.721148722 | 1.657339155 | 1.705044741 | 1.51109853 | 1.686820904 | 1.672097858 | 1.582103514 |
| ENSMUSG00000035121 | 0 | 0.25 | 0.25 | 0.25 | 0 | 0.602059991 | 0.355418052 |
| ENSMUSG00000035125 | 1.881174527 | 1.744319981 | 1.701838731 | 1.799918086 | 1.736537397 | 1.851258349 | 1.551512352 |
| ENSMUSG00000035126 | 1.289064826 | 1.061981878 | 1.226722649 | 1.114857905 | 1.091002363 | 1.079181246 | 1 |
| ENSMUSG00000035129 | 0 | 0 | 0 | 0 | 0.244650542 | 0.477121255 | 0 |
| ENSMUSG00000035131 | 0 | 0 | 0 | 0 | 0 | 0 | 0.224243824 |
| ENSMUSG00000035133 | 2.767951589 | 2.580200137 | 2.552972074 | 2.51109853 | 2.412520257 | 2.810904281 | 2.201039086 |
| ENSMUSG00000035139 | 1.903481042 | 1.76109853 | 1.779735268 | 1.870953944 | 1.851969278 | 2.064457989 | 1.781059827 |
| ENSMUSG00000035142 | 1.605636262 | 1.42510993 | 1.54745614 | 1.470660762 | 1.401338844 | 1.491361694 | 1.313616061 |
| ENSMUSG00000035148 | 0 | 0 | 0 | 0 | 0 | 0 | 0 |
| ENSMUSG00000035150 | 3.130491921 | 2.849151195 | 2.832168732 | 2.928990498 | 2.83940011 | 3.289588953 | 2.612575776 |
| ENSMUSG00000035151 | 2.233869984 | 2.075945187 | 2.04745614 | 2.138647213 | 2.062514787 | 2.359835482 | 1.886704102 |
| ENSMUSG00000035152 | 3.072046237 | 2.788862683 | 2.786005217 | 2.936523596 | 2.853930756 | 3.296226287 | 2.65592615 |
| ENSMUSG00000035158 | 1.445324132 | 1.348079356 | 1.380890489 | 1.311981878 | 1.366364104 | 1.414973348 | 1.176815335 |
| ENSMUSG00000035161 | 2.126809492 | 1.981203126 | 1.969129237 | 2.067865757 | 2.020174412 | 2.26245109 | 1.874129423 |
| ENSMUSG00000035164 | 2.081172036 | 2.001406137 | 2.002806814 | 2.061981878 | 2.048957937 | 2.23299611 | 1.87511425 |
| ENSMUSG00000035165 | 0 | 0 | 0 | 0.396240625 | 0 | 0.477121255 | 0.224243824 |
| ENSMUSG00000035168 | 2.777141269 | 2.55539678 | 2.562278363 | 2.320772088 | 2.31325093 | 2.586587305 | 2.216121671 |
| ENSMUSG00000035171 | 2.447686928 | 2.257666784 | 2.233422664 | 2.357613138 | 2.250704826 | 2.605305046 | 2.117521761 |
| ENSMUSG00000035172 | 1.975148165 | 1.858657057 | 2.001406137 | 1.458222504 | 1.621875691 | 1.531478917 | 1.770364127 |
| ENSMUSG00000035173 | 2.612833548 | 2.416333979 | 2.391037372 | 2.496460484 | 2.449252175 | 2.80140371 | 2.279417602 |
| ENSMUSG00000035177 | 0 | 0.396240625 | 0 | 0.25 | 0 | 0.301029996 | 0.448487648 |
| ENSMUSG00000035179 | 0.578129653 | 0.25 | 0 | 0.25 | 0 | 0.301029996 | 0.710836105 |
| ENSMUSG00000035181 | 2.842342644 | 2.609697963 | 2.580200137 | 2.420748646 | 2.402121453 | 2.720159303 | 2.241489052 |
| ENSMUSG00000035183 | 0.811507563 | 0.580482024 | 0.75 | 1 | 1.149965125 | 1.041392685 | 1.100339912 |
| ENSMUSG00000035184 | 1.181541736 | 1.17510993 | 1.04248125 | 0.646240625 | 0.686820904 | 0.301029996 | 0.520678036 |
| ENSMUSG00000035186 | 0 | 0.396240625 | 0 | 0.580482024 | 0.244650542 | 0.477121255 | 0.355418052 |
| ENSMUSG00000035187 | 0 | 0 | 0 | 0.976722649 | 0.978602168 | 0.84509804 | 1.078019652 |
| ENSMUSG00000035189 | 0.45815691 | 0.25 | 0.25 | 0.396240625 | 0.387761935 | 0 | 0 |
| ENSMUSG00000035191 | 0 | 0 | 0.25 | 0 | 0 | 0 | 0 |
| ENSMUSG00000035198 | 2.592526109 | 2.415889526 | 2.483053688 | 2.344302633 | 2.453332638 | 2.695481676 | 2.330653623 |
| ENSMUSG00000035199 | 2.795957952 | 2.556000419 | 2.584684096 | 2.715659339 | 2.642248115 | 3.044147621 | 2.455103737 |

| | | | | | | | |
|---|---|---|---|---|---|---|---|
| ENSMUSG00000035200 | 0 | 0.396240625 | 0 | 0.396240625 | 0.244650542 | 0 | 0.224243824 |
| ENSMUSG00000035201 | 0 | 0.396240625 | 0 | 0 | 0 | 0 | 0 |
| ENSMUSG00000035202 | 3.154532461 | 2.817573832 | 2.812722634 | 2.873463862 | 2.74845769 | 3.09131516 | 2.594681436 |
| ENSMUSG00000035203 | 2.149089517 | 2.075945187 | 2.172499493 | 2.00835575 | 2.104881849 | 2.274157849 | 2.117056606 |
| ENSMUSG00000035206 | 1.721148722 | 1.738549078 | 1.807204673 | 1.638647213 | 1.903713872 | 1.826074803 | 1.956418351 |
| ENSMUSG00000035208 | 0 | 0.25 | 0 | 0.5 | 0.387761935 | 0 | 0.355418052 |
| ENSMUSG00000035211 | 0.289064826 | 0.580482024 | 0.5 | 0.396240625 | 0.387761935 | 0.84509804 | 0.629532004 |
| ENSMUSG00000035212 | 2.370236862 | 2.203445298 | 2.220660762 | 2.152756199 | 2.057362051 | 2.33243846 | 1.945529784 |
| ENSMUSG00000035215 | 1.358731791 | 1.25 | 1.42510993 | 1.380890489 | 1.47337555 | 1.681241237 | 1.46982607 |
| ENSMUSG00000035226 | 0 | 0 | 0 | 0 | 0 | 0 | 0.355418052 |
| ENSMUSG00000035227 | 2.516991255 | 2.331607372 | 2.334405475 | 2.481203126 | 2.420401583 | 2.783188691 | 2.271140178 |
| ENSMUSG00000035228 | 0.747221736 | 0.896240625 | 0.864857905 | 0.5 | 0.686820904 | 0.698970004 | 0.829800754 |
| ENSMUSG00000035232 | 2.578129653 | 2.38040011 | 2.379909063 | 2.413659007 | 2.341270363 | 2.689308859 | 2.164188739 |
| ENSMUSG00000035234 | 2.066740409 | 1.866901388 | 1.858657057 | 1.899978211 | 1.797870327 | 1.959041392 | 1.705896842 |
| ENSMUSG00000035235 | 1.418409478 | 1.438721876 | 1.410964047 | 1.678561379 | 1.55104774 | 1.69019608 | 1.392431569 |
| ENSMUSG00000035236 | 2.500714755 | 2.312574604 | 2.308404919 | 2.42510993 | 2.345425551 | 2.620136055 | 2.137353429 |
| ENSMUSG00000035237 | 1.249317394 | 1.214495249 | 1.302363341 | 1.17510993 | 1.200473444 | 1.342422681 | 1.252393349 |
| ENSMUSG00000035238 | 0 | 0 | 0 | 0 | 0 | 0 | 0 |
| ENSMUSG00000035239 | 1.505869732 | 1.396240625 | 1.214495249 | 1.396240625 | 1.319233381 | 1.397940009 | 1.340368108 |
| ENSMUSG00000035242 | 2.850493374 | 2.642701109 | 2.696567407 | 2.620953944 | 2.699164483 | 3.007747778 | 2.625302983 |
| ENSMUSG00000035245 | 2.326928073 | 2.144357207 | 2.100219859 | 2.201032755 | 2.126866297 | 2.447158031 | 1.95718226 |
| ENSMUSG00000035246 | 2.285603957 | 2.174241882 | 2.112802778 | 2.144357207 | 2.077536464 | 2.322219295 | 1.939944915 |
| ENSMUSG00000035247 | 3.180120149 | 2.897763694 | 2.878555227 | 2.876452888 | 2.807792181 | 3.23325001 | 2.593189105 |
| ENSMUSG00000035248 | 2.517987749 | 2.306603048 | 2.269203899 | 2.435787849 | 2.372368362 | 2.683047038 | 2.203186321 |
| ENSMUSG00000035258 | 1.404271433 | 1.214495249 | 1.17510993 | 1.080482024 | 1.039257769 | 0.903089987 | 0.853775828 |
| ENSMUSG00000035262 | 0.45815691 | 0.25 | 0.5 | 0.25 | 0 | 0 | 0.448487648 |
| ENSMUSG00000035266 | 2.361637952 | 2.131869252 | 2.21293726 | 2.112802778 | 2.040348824 | 2.340444115 | 1.995286892 |
| ENSMUSG00000035268 | 1.623015126 | 1.62694866 | 1.71759118 | 1.470660762 | 1.712553795 | 1.832508913 | 1.767634022 |
| ENSMUSG00000035270 | 0.578129653 | 0.25 | 0.5 | 0.5 | 0.387761935 | 0.698970004 | 0.355418052 |
| ENSMUSG00000035273 | 0.811507563 | 0.75 | 0.580482024 | 2.095926073 | 2.012006468 | 2.240549248 | 1.870159858 |
| ENSMUSG00000035274 | 2.295901576 | 2.119945066 | 2.152756199 | 2.125956434 | 2.016696984 | 2.330413773 | 1.923435442 |
| ENSMUSG00000035275 | 0.289064826 | 0.646240625 | 0.830482024 | 1.061981878 | 1.223252711 | 1.255272505 | 1.159323753 |
| ENSMUSG00000035277 | 0 | 0 | 0 | 0 | 0 | 0 | 0 |
| ENSMUSG00000035278 | 1.989530018 | 1.888647213 | 1.926839783 | 1.834962501 | 1.900490506 | 2.037426498 | 1.872150729 |

| | | | | | | | |
|---|---|---|---|---|---|---|---|
| ENSMUSG00000035279 | 1.99653913 | 1.874961472 | 2.012462137 | 1.114857905 | 1.319233381 | 1.397940009 | 1.224243824 |
| ENSMUSG00000035283 | 0 | 0 | 0 | 0 | 0.244650542 | 0 | 0.579661877 |
| ENSMUSG00000035284 | 2.612833548 | 2.364857905 | 2.355516192 | 2.414106216 | 2.347256716 | 2.674861141 | 2.182948513 |
| ENSMUSG00000035285 | 0.960252568 | 1.130890489 | 1.188721876 | 0.75 | 0.733951626 | 0.301029996 | 0.710836105 |
| ENSMUSG00000035293 | 2.730221576 | 2.473325383 | 2.443696765 | 2.45584181 | 2.397795474 | 2.72916479 | 2.175263699 |
| ENSMUSG00000035295 | 0.45815691 | 0.396240625 | 0.396240625 | 0.5 | 0.244650542 | 0.602059991 | 0 |
| ENSMUSG00000035296 | 0 | 0 | 0 | 0 | 0 | 0 | 0 |
| ENSMUSG00000035297 | 2.823159983 | 2.593759858 | 2.557805295 | 2.663659007 | 2.573153518 | 2.960946196 | 2.429740345 |
| ENSMUSG00000035298 | 0.289064826 | 0 | 0.5 | 0.25 | 0.244650542 | 0 | 0.224243824 |
| ENSMUSG00000035299 | 1.967766868 | 1.79745614 | 1.868933358 | 1.705044741 | 1.736537397 | 1.77815125 | 1.670888075 |
| ENSMUSG00000035305 | 0.960252568 | 1.04248125 | 0.976722649 | 1.302363341 | 1.310728002 | 1.397940009 | 1.20919388 |
| ENSMUSG00000035310 | 2.562324428 | 2.396709947 | 2.417221246 | 2.391037372 | 2.31325093 | 2.648360011 | 2.152721168 |
| ENSMUSG00000035311 | 2.572054151 | 2.307204673 | 2.352347734 | 2.360217292 | 2.303566328 | 2.631443769 | 2.117521761 |
| ENSMUSG00000035314 | 1.852729478 | 1.868933358 | 1.848079356 | 1.671625132 | 1.926994141 | 1.949390007 | 1.907820406 |
| ENSMUSG00000035325 | 3.308500783 | 3.009644026 | 3.005851961 | 2.768869787 | 2.74713658 | 3.148294097 | 2.597012708 |
| ENSMUSG00000035329 | 2.116511873 | 2.013820609 | 2.02186571 | 2.113831805 | 2.001219193 | 2.260071388 | 1.836517579 |
| ENSMUSG00000035337 | 0 | 0 | 0.25 | 0.25 | 0.244650542 | 0 | 0 |
| ENSMUSG00000035342 | 2.156259305 | 2.137686696 | 2.300530956 | 1.598079356 | 1.92093466 | 1.892094603 | 2.087847416 |
| ENSMUSG00000035349 | 0.747221736 | 0.646240625 | 0.580482024 | 0.396240625 | 0.244650542 | 0.602059991 | 0 |
| ENSMUSG00000035351 | 2.254965775 | 2.149047439 | 2.117918804 | 2.147178659 | 2.111176775 | 2.42975228 | 1.956418351 |
| ENSMUSG00000035352 | 0 | 0 | 0 | 0 | 0 | 0 | 0 |
| ENSMUSG00000035354 | 2.282114126 | 2.147178659 | 2.173371739 | 2.089388001 | 2.063536368 | 2.330413773 | 1.972821265 |
| ENSMUSG00000035355 | 0.289064826 | 0.25 | 0.25 | 0.25 | 0.489301084 | 0 | 0.579661877 |
| ENSMUSG00000035356 | 2.469490009 | 2.313757142 | 2.28360658 | 1.928561379 | 1.897237431 | 2.041392685 | 1.766260282 |
| ENSMUSG00000035357 | 2.010213549 | 1.992885888 | 1.943696765 | 1.494319981 | 1.533172725 | 1.568201724 | 1.473249546 |
| ENSMUSG00000035364 | 0.289064826 | 0.75 | 0.646240625 | 0.396240625 | 0.387761935 | 0.301029996 | 0.355418052 |
| ENSMUSG00000035365 | 2.172524468 | 2.039967834 | 2.025821952 | 2.119945066 | 2.045754077 | 2.367355921 | 1.925124822 |
| ENSMUSG00000035367 | 1.614416215 | 1.610735874 | 1.557204673 | 1.571350555 | 1.603583716 | 1.681241237 | 1.459332757 |
| ENSMUSG00000035370 | 1.325351389 | 1.356566189 | 1.42510993 | 1.04248125 | 1.319233381 | 1.146128036 | 1.42167221 |
| ENSMUSG00000035371 | 0 | 0 | 0 | 0 | 0 | 0 | 0 |
| ENSMUSG00000035372 | 2.40817504 | 2.269871196 | 2.30479213 | 2.192046081 | 2.296783518 | 2.532754379 | 2.20919388 |
| ENSMUSG00000035373 | 2.900309677 | 2.64881437 | 2.677701608 | 1.811981878 | 1.758994476 | 1.963787827 | 1.643912836 |
| ENSMUSG00000035376 | 2.199011616 | 2.01109853 | 2.05357978 | 2.131869252 | 2.062514787 | 2.342422681 | 1.941550454 |
| ENSMUSG00000035378 | 1.852729478 | 1.75 | 1.726722649 | 1.610735874 | 1.642643806 | 1.792391689 | 1.56716938 |

| | | | | | | | |
|---|---|---|---|---|---|---|---|
| ENSMUSG00000035382 | 2.323768721 | 2.146240625 | 2.178561379 | 2.191217898 | 2.256066288 | 2.5132176 | 2.090889903 |
| ENSMUSG00000035383 | 0.45815691 | 0.75 | 0.646240625 | 0.701838731 | 0.733951626 | 0.84509804 | 0.803905701 |
| ENSMUSG00000035385 | 3.123560476 | 2.866391601 | 2.921296699 | 2.53743678 | 2.562893863 | 2.887054378 | 2.404331881 |
| ENSMUSG00000035387 | 0 | 0 | 0 | 0 | 0 | 0 | 0 |
| ENSMUSG00000035390 | 1.671187741 | 1.571350555 | 1.610735874 | 1.482684334 | 1.42077253 | 1.579783597 | 1.520678036 |
| ENSMUSG00000035392 | 1.857607105 | 1.726722649 | 1.755591953 | 1.681980114 | 1.692703642 | 1.959041392 | 1.643912836 |
| ENSMUSG00000035394 | 0.289064826 | 0 | 0.25 | 0 | 0.568060967 | 0 | 0.224243824 |
| ENSMUSG00000035395 | 0 | 0 | 0 | 0 | 0.244650542 | 0 | 0 |
| ENSMUSG00000035397 | 2.063793172 | 1.905512955 | 1.979715809 | 1.982684334 | 1.988833497 | 2.26245109 | 1.95411574 |
| ENSMUSG00000035399 | 1.91631382 | 1.87892496 | 1.834962501 | 1.928561379 | 1.846351821 | 2.113943352 | 1.709198048 |
| ENSMUSG00000035401 | 2.31414447 | 2.171625132 | 2.166333979 | 2.160061234 | 2.098472613 | 2.376576957 | 1.974272009 |
| ENSMUSG00000035403 | 0.45815691 | 0.25 | 0.396240625 | 0 | 0.244650542 | 0 | 0 |
| ENSMUSG00000035407 | 0.811507563 | 0.646240625 | 0.25 | 0.396240625 | 0 | 0.301029996 | 0.579661877 |
| ENSMUSG00000035413 | 2.584117913 | 2.412314044 | 2.424241882 | 2.168106335 | 2.267696867 | 2.521138084 | 2.179896458 |
| ENSMUSG00000035420 | 0 | 0 | 0 | 0 | 0 | 0 | 0 |
| ENSMUSG00000035427 | 0 | 0 | 0 | 0 | 0 | 0 | 0 |
| ENSMUSG00000035429 | 0 | 0.25 | 0.25 | 0 | 0 | 0 | 0 |
| ENSMUSG00000035431 | 0.289064826 | 0 | 0 | 0 | 0 | 0 | 0 |
| ENSMUSG00000035435 | 0.289064826 | 0.396240625 | 0 | 0 | 0.244650542 | 0.301029996 | 0.448487648 |
| ENSMUSG00000035437 | 2.789779581 | 2.547764804 | 2.538071211 | 2.520869832 | 2.422990063 | 2.808210973 | 2.259363975 |
| ENSMUSG00000035439 | 2.325351389 | 2.155512955 | 2.17510993 | 2.122963274 | 2.148364414 | 2.404833717 | 2.060761403 |
| ENSMUSG00000035441 | 2.131864492 | 2.024508021 | 2.027131114 | 1.916333979 | 1.947418788 | 2.155336037 | 1.86715039 |
| ENSMUSG00000035443 | 1.639698646 | 1.52186571 | 1.458222504 | 1.634789703 | 1.584291694 | 1.857332496 | 1.499406449 |
| ENSMUSG00000035448 | 0 | 0 | 0 | 0 | 0 | 0 | 0 |
| ENSMUSG00000035451 | 1.978790393 | 1.907339155 | 1.821350555 | 0.5 | 0.846351821 | 0.602059991 | 0.710836105 |
| ENSMUSG00000035454 | 0 | 0 | 0 | 0 | 0 | 0 | 0 |
| ENSMUSG00000035455 | 2.541215974 | 2.319031101 | 2.325374049 | 2.357090043 | 2.295199569 | 2.64246452 | 2.124874702 |
| ENSMUSG00000035456 | 0.289064826 | 0 | 0.25 | 0.580482024 | 0.387761935 | 0 | 0.355418052 |
| ENSMUSG00000035458 | 0.671187741 | 0.5 | 0 | 0.5 | 0.568060967 | 0.602059991 | 0.629532004 |
| ENSMUSG00000035459 | 0.811507563 | 0.5 | 0.701838731 | 0.646240625 | 0.489301084 | 0.477121255 | 0.448487648 |
| ENSMUSG00000035469 | 2.337801042 | 2.172499493 | 2.114857905 | 2.242885888 | 2.187072901 | 2.428134794 | 2.038401572 |
| ENSMUSG00000035472 | 0.289064826 | 0 | 0 | 0.396240625 | 0 | 0 | 0.224243824 |
| ENSMUSG00000035473 | 1.964026642 | 1.845926073 | 1.884789703 | 1.959800947 | 1.846351821 | 2.214843848 | 1.774416565 |
| ENSMUSG00000035476 | 2.368815967 | 2.176839783 | 2.125956434 | 2.166333979 | 2.108492685 | 2.342422681 | 1.949460761 |

| | | | | | | | |
|---|---|---|---|---|---|---|---|
| ENSMUSG00000035478 | 2.779806021 | 2.636482442 | 2.709406983 | 2.37892496 | 2.625423019 | 2.750508395 | 2.559791152 |
| ENSMUSG00000035486 | 0 | 0 | 0 | 0.25 | 0 | 0 | 0.448487648 |
| ENSMUSG00000035491 | 0 | 0 | 0 | 0 | 0 | 0 | 0 |
| ENSMUSG00000035493 | 2.209569962 | 2.001406137 | 2.006976499 | 2.116901388 | 2.095689764 | 2.396199347 | 2.013740841 |
| ENSMUSG00000035495 | 2.295901576 | 2.131869252 | 2.123963757 | 2.123963757 | 2.070605826 | 2.274157849 | 1.963977467 |
| ENSMUSG00000035498 | 0 | 0 | 0 | 0 | 0 | 0.301029996 | 0.224243824 |
| ENSMUSG00000035504 | 1.129344651 | 1.238549078 | 1.146240625 | 1.494319981 | 1.621875691 | 1.681241237 | 1.535058868 |
| ENSMUSG00000035505 | 2.051791411 | 1.899978211 | 1.928561379 | 1.701838731 | 1.751666404 | 1.908485019 | 1.695786161 |
| ENSMUSG00000035506 | 0.45815691 | 0.25 | 0.75 | 0.396240625 | 0.244650542 | 0.698970004 | 0.710836105 |
| ENSMUSG00000035509 | 0.45815691 | 0.580482024 | 0.701838731 | 1.201838731 | 1.074582839 | 1.041392685 | 1.150210039 |
| ENSMUSG00000035513 | 0.289064826 | 0 | 0.25 | 0 | 0.846351821 | 0.301029996 | 0.984950056 |
| ENSMUSG00000035517 | 2.340856238 | 2.207430684 | 2.161864607 | 2.300530956 | 2.25901006 | 2.531478917 | 2.103753068 |
| ENSMUSG00000035521 | 1.794934559 | 1.802363341 | 1.901832578 | 1.653677461 | 1.698489937 | 1.84509804 | 1.661510159 |
| ENSMUSG00000035522 | 0 | 0 | 0 | 0 | 0 | 0 | 0 |
| ENSMUSG00000035523 | 0 | 0 | 0 | 0 | 0 | 0 | 0 |
| ENSMUSG00000035528 | 0 | 0 | 0 | 0 | 0 | 0 | 0 |
| ENSMUSG00000035529 | 2.657168526 | 2.460587586 | 2.401832578 | 2.359177886 | 2.304082723 | 2.63748973 | 2.164992504 |
| ENSMUSG00000035530 | 3.253262209 | 2.924024543 | 2.931448068 | 2.902756199 | 2.835872226 | 3.270445908 | 2.630199734 |
| ENSMUSG00000035539 | 0.45815691 | 0.396240625 | 0.830482024 | 0.701838731 | 0.632412477 | 0.301029996 | 0.579661877 |
| ENSMUSG00000035540 | 0 | 0 | 0 | 0 | 0 | 0 | 0 |
| ENSMUSG00000035545 | 2.35727108 | 2.207430684 | 2.282320754 | 2.063166358 | 2.210031835 | 2.283301229 | 2.117986249 |
| ENSMUSG00000035547 | 1.647796617 | 1.584962501 | 1.671625132 | 1.598079356 | 1.603583716 | 1.653212514 | 1.643912836 |
| ENSMUSG00000035551 | 0.45815691 | 0.25 | 0.25 | 0.25 | 0 | 0 | 0.224243824 |
| ENSMUSG00000035557 | 0 | 0 | 0 | 0 | 0 | 0 | 0.224243824 |
| ENSMUSG00000035559 | 1.956444026 | 1.816696635 | 1.83272922 | 1.774508021 | 1.859322797 | 1.991226076 | 1.791413205 |
| ENSMUSG00000035560 | 0 | 0.25 | 0 | 0 | 0 | 0 | 0.224243824 |
| ENSMUSG00000035561 | 0.578129653 | 0.896240625 | 0.580482024 | 1.130890489 | 1.283908311 | 1.322219295 | 1.335191751 |
| ENSMUSG00000035566 | 0.45815691 | 0.580482024 | 0.646240625 | 0.396240625 | 0.244650542 | 0 | 0.224243824 |
| ENSMUSG00000035569 | 3.014472122 | 2.778760913 | 2.785047676 | 2.650674716 | 2.686471788 | 2.987666265 | 2.545873061 |
| ENSMUSG00000035572 | 2.45930418 | 2.259729747 | 2.230460234 | 2.385274154 | 2.28445426 | 2.59439255 | 2.130728307 |
| ENSMUSG00000035575 | 2.795446569 | 2.562574604 | 2.579070383 | 2.75402796 | 2.662631419 | 3.077367905 | 2.462068294 |
| ENSMUSG00000035576 | 0.289064826 | 0 | 0.25 | 0.25 | 0.244650542 | 0 | 0 |
| ENSMUSG00000035578 | 0.289064826 | 0.701838731 | 0.75 | 0.830482024 | 0.931471446 | 0.698970004 | 0.672731473 |
| ENSMUSG00000035580 | 0 | 0 | 0 | 0 | 0 | 0 | 0 |

| | | | | | | | |
|---|---|---|---|---|---|---|---|
| ENSMUSG00000035582 | 0 | 0 | 0 | 0 | 0 | 0 | 0 |
| ENSMUSG00000035585 | 1.928763473 | 1.868933358 | 1.991446071 | 1.598079356 | 1.851969278 | 1.897627091 | 1.830929954 |
| ENSMUSG00000035592 | 0 | 0 | 0 | 0 | 0 | 0 | 0 |
| ENSMUSG00000035594 | 0.45815691 | 0.25 | 0.25 | 0.976722649 | 0.978602168 | 0.84509804 | 0.629532004 |
| ENSMUSG00000035595 | 1.470606563 | 1.566696635 | 1.494319981 | 1.388647213 | 1.555378544 | 1.544068044 | 1.479989695 |
| ENSMUSG00000035596 | 2.196867486 | 2.05357978 | 2.05479213 | 1.995748394 | 2.076554666 | 2.257678575 | 1.989124507 |
| ENSMUSG00000035597 | 2.640441358 | 2.399047439 | 2.369439567 | 2.371457327 | 2.322676882 | 2.559906625 | 2.116590781 |
| ENSMUSG00000035601 | 1.876566388 | 1.649978211 | 1.714495249 | 1.79745614 | 1.77552387 | 1.949390007 | 1.669034113 |
| ENSMUSG00000035606 | 0 | 0 | 0 | 0 | 0 | 0 | 0 |
| ENSMUSG00000035614 | 2.508932407 | 2.358135475 | 2.334962501 | 2.214495249 | 2.138604637 | 2.382017043 | 1.990504106 |
| ENSMUSG00000035615 | 0 | 0 | 0 | 0.92510993 | 0.931471446 | 0.903089987 | 0.829800754 |
| ENSMUSG00000035620 | 2.213719572 | 2.091580554 | 2.066696635 | 2.225970461 | 2.112066955 | 2.352182518 | 1.964723754 |
| ENSMUSG00000035621 | 2.336265009 | 2.272528105 | 2.348615674 | 1.985628626 | 2.105788027 | 2.269512944 | 2.135599953 |
| ENSMUSG00000035623 | 2.325351389 | 2.172499493 | 2.168989258 | 2.247880462 | 2.1962964 | 2.457881897 | 2.039586611 |
| ENSMUSG00000035626 | 0 | 0 | 0 | 0 | 0 | 0 | 0 |
| ENSMUSG00000035629 | 2.565000581 | 2.386241108 | 2.434945152 | 2.352877747 | 2.371090306 | 2.602059991 | 2.248075634 |
| ENSMUSG00000035632 | 2.17479713 | 2.039967834 | 2.110735874 | 2.04248125 | 2.044679629 | 2.23299611 | 2.016928211 |
| ENSMUSG00000035637 | 1.655740331 | 1.698603967 | 1.774508021 | 1.602347734 | 1.715300549 | 1.73239376 | 1.715701091 |
| ENSMUSG00000035638 | 0 | 0 | 0 | 0 | 0 | 0.301029996 | 0 |
| ENSMUSG00000035640 | 1.289064826 | 1.356566189 | 1.488549078 | 0.896240625 | 1.188507684 | 0.602059991 | 1.224243824 |
| ENSMUSG00000035642 | 1.307602651 | 1.302363341 | 1.302363341 | 1.566696635 | 1.47337555 | 1.69019608 | 1.46982607 |
| ENSMUSG00000035649 | 2.179305632 | 2.012462137 | 2.036164561 | 2.070192693 | 2.003645018 | 2.303196057 | 1.832055226 |
| ENSMUSG00000035651 | 0 | 0 | 0 | 0 | 0 | 0 | 0 |
| ENSMUSG00000035653 | 0 | 0 | 0 | 0 | 0 | 0 | 0 |
| ENSMUSG00000035666 | 2.793394745 | 2.515173983 | 2.506284891 | 2.558704358 | 2.491431917 | 2.829946696 | 2.292669234 |
| ENSMUSG00000035671 | 1.842799666 | 1.779735268 | 1.852347734 | 1.80479213 | 1.954436039 | 2.075546961 | 1.947107911 |
| ENSMUSG00000035673 | 2.390989024 | 2.282320754 | 2.352347734 | 2.083847589 | 2.25901006 | 2.372912003 | 2.271140178 |
| ENSMUSG00000035674 | 1.852729478 | 1.823655187 | 1.834962501 | 1.848079356 | 1.851969278 | 2.1430148 | 1.766260282 |
| ENSMUSG00000035678 | 1.470606563 | 1.476722649 | 1.464495249 | 1.431980114 | 1.366364104 | 1.491361694 | 1.433437704 |
| ENSMUSG00000035681 | 0.578129653 | 0.646240625 | 0.25 | 0.701838731 | 0.686820904 | 0.602059991 | 0.710836105 |
| ENSMUSG00000035683 | 2.6088504 | 2.372963274 | 2.368933358 | 2.573655187 | 2.514628232 | 2.880241776 | 2.338260508 |
| ENSMUSG00000035686 | 0 | 0 | 0 | 0.396240625 | 0 | 0 | 0.224243824 |
| ENSMUSG00000035692 | 0.289064826 | 0 | 0.5 | 1.160964047 | 1.212046813 | 1.278753601 | 1.193409508 |
| ENSMUSG00000035694 | 0.45815691 | 0.5 | 0.396240625 | 0.580482024 | 0.244650542 | 0.477121255 | 0.448487648 |

| | | | | | | | |
|---|---|---|---|---|---|---|---|
| ENSMUSG00000035696 | 2.595842487 | 2.397178659 | 2.333847589 | 2.293106877 | 2.285543632 | 2.556302501 | 2.09490255 |
| ENSMUSG00000035697 | 0.91631382 | 1.04248125 | 0.951838731 | 0.646240625 | 1.106691885 | 1 | 1.21680636 |
| ENSMUSG00000035699 | 0.289064826 | 0 | 0 | 0 | 0.244650542 | 0 | 0 |
| ENSMUSG00000035704 | 1.783508299 | 1.589388001 | 1.634789703 | 1.678561379 | 1.642643806 | 1.812913357 | 1.607811402 |
| ENSMUSG00000035711 | 0.45815691 | 0.976722649 | 1.080482024 | 0.580482024 | 0.931471446 | 0.477121255 | 0.984950056 |
| ENSMUSG00000035713 | 1.404271433 | 1.388647213 | 1.29248125 | 1.146240625 | 1.380772476 | 1.380211242 | 1.302263476 |
| ENSMUSG00000035722 | 2.310886388 | 2.225970461 | 2.306000419 | 1.931980114 | 2.129414728 | 2.155336037 | 2.070582188 |
| ENSMUSG00000035725 | 2.22189694 | 2.061981878 | 2.127938163 | 2.30479213 | 2.243427123 | 2.542825427 | 2.053478496 |
| ENSMUSG00000035726 | 2.845077293 | 2.601551251 | 2.588010856 | 2.851816941 | 2.749627873 | 3.18920949 | 2.55041672 |
| ENSMUSG00000035735 | 1.920505136 | 1.901832578 | 1.955044741 | 1.575945187 | 1.814781639 | 1.826074803 | 1.86207155 |
| ENSMUSG00000035745 | 0.747221736 | 0.396240625 | 0.896240625 | 0 | 0.387761935 | 0.477121255 | 0.672731473 |
| ENSMUSG00000035754 | 2.292497211 | 2.220660762 | 2.304186465 | 2.046218836 | 2.234113756 | 2.315970345 | 2.198153757 |
| ENSMUSG00000035757 | 1.631440309 | 1.593759858 | 1.552363341 | 1.618933358 | 1.645989387 | 1.84509804 | 1.535058868 |
| ENSMUSG00000035759 | 1.307602651 | 1.330482024 | 1.403677461 | 1.476722649 | 1.407936347 | 1.681241237 | 1.350478789 |
| ENSMUSG00000035762 | 2.016886209 | 1.845926073 | 1.816696635 | 2.004202072 | 1.964031554 | 2.195899652 | 1.811176017 |
| ENSMUSG00000035764 | 2.37447073 | 2.234894804 | 2.156427211 | 2.246460484 | 2.16174451 | 2.439332694 | 1.993927582 |
| ENSMUSG00000035765 | 2.179305632 | 1.970660762 | 1.997171172 | 2.005591953 | 2.044679629 | 2.281033367 | 1.907820406 |
| ENSMUSG00000035769 | 1.445324132 | 1.470660762 | 1.505591953 | 1.571350555 | 1.645989387 | 1.707570176 | 1.603286636 |
| ENSMUSG00000035770 | 2.918605213 | 2.66942991 | 2.652294685 | 2.675543171 | 2.63322574 | 3.016197354 | 2.463028281 |
| ENSMUSG00000035772 | 2.20747957 | 2.109697963 | 2.122963274 | 2.192046081 | 2.131103516 | 2.383815366 | 1.995964412 |
| ENSMUSG00000035773 | 0.747221736 | 0.25 | 0.396240625 | 0.396240625 | 0.489301084 | 0.477121255 | 0.710836105 |
| ENSMUSG00000035775 | 0.289064826 | 0 | 0 | 0.25 | 0.244650542 | 0 | 0 |
| ENSMUSG00000035776 | 2.572927518 | 2.374961472 | 2.394828984 | 2.465271726 | 2.405621873 | 2.72427587 | 2.263534582 |
| ENSMUSG00000035778 | 2.604019855 | 2.394828984 | 2.379909063 | 2.196158711 | 2.113840608 | 2.421603927 | 2.003324957 |
| ENSMUSG00000035780 | 0 | 0 | 0 | 0 | 0 | 0 | 0 |
| ENSMUSG00000035781 | 2.235832493 | 2.091580554 | 2.107613138 | 2.095926073 | 2.102149272 | 2.372912003 | 2.053478496 |
| ENSMUSG00000035783 | 4.63007198 | 4.158720484 | 4.118830447 | 2.433677405 | 2.419286397 | 2.758911892 | 2.303573788 |
| ENSMUSG00000035785 | 0 | 0 | 0 | 0 | 0 | 0 | 0 |
| ENSMUSG00000035790 | 1.964026642 | 1.79248125 | 1.807204673 | 1.910964047 | 1.818885826 | 2.064457989 | 1.775756176 |
| ENSMUSG00000035798 | 2.317377295 | 2.124961472 | 2.124961472 | 2.299918086 | 2.252501045 | 2.58546073 | 2.082712142 |
| ENSMUSG00000035799 | 1.852729478 | 1.777131114 | 1.837182039 | 1.75835575 | 1.749189509 | 1.977723605 | 1.687112305 |
| ENSMUSG00000035804 | 0 | 0 | 0 | 0 | 0 | 0 | 0 |
| ENSMUSG00000035805 | 0 | 0.25 | 0 | 0 | 0 | 0 | 0 |
| ENSMUSG00000035811 | 0 | 0 | 0 | 0 | 0 | 0 | 0 |

| | | | | | | | |
|---|---|---|---|---|---|---|---|
| ENSMUSG00000035818 | 0 | 0 | 0 | 0.5 | 0.489301084 | 0 | 0.448487648 |
| ENSMUSG00000035824 | 1.783508299 | 1.705044741 | 1.784887838 | 1.77186571 | 1.689774529 | 1.903089987 | 1.670888075 |
| ENSMUSG00000035828 | 2.092368209 | 2.002806814 | 2.078220739 | 1.814346961 | 1.995080671 | 2.079181246 | 1.943943921 |
| ENSMUSG00000035829 | 1.558729299 | 1.438721876 | 1.311981878 | 1.516522298 | 1.655839964 | 1.681241237 | 1.610050272 |
| ENSMUSG00000035831 | 0 | 0 | 0 | 0 | 0 | 0 | 0 |
| ENSMUSG00000035834 | 2.003432385 | 1.884789703 | 1.841580554 | 1.976722649 | 1.968065382 | 2.238046103 | 1.847411384 |
| ENSMUSG00000035835 | 1.432083896 | 1.431980114 | 1.561981878 | 0.896240625 | 1.200473444 | 0.84509804 | 1.259064007 |
| ENSMUSG00000035836 | 0 | 0 | 0 | 0 | 0 | 0 | 0 |
| ENSMUSG00000035840 | 2.205378646 | 2.065523711 | 2.102347734 | 2.142463902 | 2.076554666 | 2.369215857 | 1.909593098 |
| ENSMUSG00000035842 | 2.346900291 | 2.165444524 | 2.161864607 | 2.213717096 | 2.239111448 | 2.478566496 | 2.082194103 |
| ENSMUSG00000035845 | 1.759671389 | 1.769203899 | 1.714495249 | 1.744319981 | 1.706995316 | 1.832508913 | 1.649885598 |
| ENSMUSG00000035847 | 2.740854706 | 2.51007243 | 2.502107156 | 2.628185116 | 2.556137318 | 2.929418926 | 2.343157052 |
| ENSMUSG00000035849 | 0 | 0.396240625 | 0 | 0.580482024 | 0 | 0.477121255 | 0.224243824 |
| ENSMUSG00000035851 | 2.716227929 | 2.478222334 | 2.462155735 | 2.448603967 | 2.384490012 | 2.741151599 | 2.230859913 |
| ENSMUSG00000035852 | 0.289064826 | 0 | 0.396240625 | 0.396240625 | 0.387761935 | 0 | 0 |
| ENSMUSG00000035860 | 0 | 0 | 0 | 0 | 0 | 0 | 0 |
| ENSMUSG00000035861 | 0.45815691 | 0 | 0.646240625 | 0.5 | 0.733951626 | 0.477121255 | 0.520678036 |
| ENSMUSG00000035863 | 2.297593393 | 2.243603647 | 2.359697963 | 1.979715809 | 2.176862714 | 2.257678575 | 2.223239637 |
| ENSMUSG00000035864 | 1 | 1.04248125 | 0.976722649 | 1.130890489 | 1.020174412 | 1.079181246 | 0.984950056 |
| ENSMUSG00000035868 | 1.587501561 | 1.470660762 | 1.42510993 | 1.476722649 | 1.394615667 | 1.716003344 | 1.437266334 |
| ENSMUSG00000035873 | 2.282114126 | 2.143411797 | 2.098079356 | 2.104463129 | 1.991345651 | 2.264817823 | 1.912234055 |
| ENSMUSG00000035875 | 0.578129653 | 0.580482024 | 0.5 | 0.5 | 0.489301084 | 0 | 0.355418052 |
| ENSMUSG00000035877 | 2 | 1.880890489 | 1.834962501 | 1.953445298 | 1.92245931 | 2.10720997 | 1.787569411 |
| ENSMUSG00000035878 | 1.678702042 | 1.488549078 | 1.464495249 | 1.75 | 1.662257525 | 1.84509804 | 1.618853957 |
| ENSMUSG00000035885 | 2.439357901 | 2.284887838 | 2.410061234 | 2.279735268 | 2.205967036 | 2.511883361 | 2.249933144 |
| ENSMUSG00000035890 | 1.944805158 | 1.882845365 | 2.009729747 | 1.653677461 | 1.870073561 | 1.892094603 | 2.059095225 |
| ENSMUSG00000035891 | 2.219867581 | 2.156427211 | 2.159156155 | 2.046218836 | 2.056322416 | 2.271841607 | 1.986347533 |
| ENSMUSG00000035896 | 0 | 0 | 0 | 0 | 0 | 0 | 0 |
| ENSMUSG00000035898 | 2.842799666 | 2.612545062 | 2.600753006 | 2.551142786 | 2.451638175 | 2.846955325 | 2.259363975 |
| ENSMUSG00000035900 | 2.63295404 | 2.42813175 | 2.499647607 | 2.442459461 | 2.530194174 | 2.768638101 | 2.412838474 |
| ENSMUSG00000035901 | 2.841885121 | 2.598883534 | 2.572792675 | 2.659835177 | 2.591450357 | 2.970811611 | 2.423843462 |
| ENSMUSG00000035910 | 0 | 0 | 0 | 0 | 0 | 0 | 0 |
| ENSMUSG00000035914 | 2.333175866 | 2.209012589 | 2.229715809 | 2.264497931 | 2.279510047 | 2.591064607 | 2.210939786 |
| ENSMUSG00000035916 | 0 | 0.580482024 | 0.646240625 | 0.896240625 | 0.733951626 | 0.602059991 | 0.579661877 |

| | | | | | | | |
|---|---|---|---|---|---|---|---|
| ENSMUSG00000035919 | 1.881174527 | 1.729715809 | 1.752806814 | 1.991446071 | 1.970729215 | 2.181843588 | 1.798966438 |
| ENSMUSG00000035923 | 0 | 0 | 0 | 0 | 0 | 0 | 0 |
| ENSMUSG00000035929 | 1.418409478 | 1.388647213 | 1.458222504 | 2.167221246 | 2.19206933 | 2.45331834 | 2.149789616 |
| ENSMUSG00000035930 | 0 | 0 | 0.396240625 | 0 | 0.387761935 | 0 | 0 |
| ENSMUSG00000035932 | 0 | 0 | 0 | 0 | 0 | 0 | 0 |
| ENSMUSG00000035933 | 2.045658471 | 1.845926073 | 1.841580554 | 1.926839783 | 1.83872004 | 2.012837225 | 1.709198048 |
| ENSMUSG00000035934 | 1.45815691 | 1.348079356 | 1.403677461 | 1.744319981 | 1.746695109 | 1.986771734 | 1.687112305 |
| ENSMUSG00000035936 | 1.759671389 | 1.708222504 | 1.688721876 | 2.01109853 | 1.966725889 | 2.250420002 | 1.838725884 |
| ENSMUSG00000035941 | 2.605636262 | 2.380890489 | 2.371457327 | 2.410512923 | 2.336595379 | 2.691081492 | 2.168981648 |
| ENSMUSG00000035944 | 1.358731791 | 1.188721876 | 1.29248125 | 1.571350555 | 1.533172725 | 1.633468456 | 1.517723536 |
| ENSMUSG00000035946 | 0 | 0 | 0 | 0 | 0 | 0 | 0 |
| ENSMUSG00000035948 | 0.289064826 | 0.25 | 0 | 0.5 | 0.244650542 | 0.301029996 | 0.355418052 |
| ENSMUSG00000035949 | 2.52293481 | 2.329353153 | 2.337182039 | 2.399978211 | 2.315256368 | 2.607455023 | 2.175263699 |
| ENSMUSG00000035951 | 0 | 0.25 | 0.25 | 0 | 0.489301084 | 0 | 0 |
| ENSMUSG00000035953 | 2.355805236 | 2.19697564 | 2.205044741 | 2.17510993 | 2.150762767 | 2.361727836 | 2.033014511 |
| ENSMUSG00000035954 | 1.932832114 | 1.726722649 | 1.839388001 | 1.777131114 | 1.712553795 | 1.924279286 | 1.603286636 |
| ENSMUSG00000035958 | 2.40033094 | 2.247171172 | 2.209012589 | 2.321350555 | 2.227765453 | 2.537819095 | 2.091897741 |
| ENSMUSG00000035960 | 1.671187741 | 1.505591953 | 1.575945187 | 1.516522298 | 1.692703642 | 1.698970004 | 1.672731473 |
| ENSMUSG00000035963 | 0 | 0 | 0 | 0 | 0 | 0 | 0 |
| ENSMUSG00000035964 | 0 | 0 | 0 | 0 | 0 | 0.477121255 | 0.520678036 |
| ENSMUSG00000035967 | 1.932832114 | 1.862802778 | 1.78743678 | 1.930274797 | 1.900490506 | 2.06069784 | 1.781059827 |
| ENSMUSG00000035969 | 2.377269616 | 2.273849256 | 2.344844592 | 1.657339155 | 1.855665188 | 1.792391689 | 1.909593098 |
| ENSMUSG00000035983 | 0.45815691 | 0.5 | 0.5 | 0.25 | 0.387761935 | 0.301029996 | 0.224243824 |
| ENSMUSG00000035984 | 0 | 0.396240625 | 0.25 | 0.25 | 0.489301084 | 0.301029996 | 0.448487648 |
| ENSMUSG00000035992 | 2.708736128 | 2.485261902 | 2.502107156 | 2.468745337 | 2.360255934 | 2.701567985 | 2.176428123 |
| ENSMUSG00000036002 | 1.857607105 | 1.828220739 | 1.905512955 | 1.741446071 | 1.768534412 | 1.897627091 | 1.779742043 |
| ENSMUSG00000036006 | 2.134369199 | 2.016522298 | 2.057204673 | 2.254202072 | 2.220647857 | 2.485721426 | 2.079068325 |
| ENSMUSG00000036009 | 1.777675583 | 1.62694866 | 1.649978211 | 1.747171172 | 1.671670246 | 1.812913357 | 1.629532004 |
| ENSMUSG00000036013 | 0 | 0 | 0 | 0 | 0 | 0 | 0 |
| ENSMUSG00000036019 | 0.867194479 | 0.79248125 | 0.396240625 | 1.080482024 | 0.812711509 | 0.698970004 | 0.74492186 |
| ENSMUSG00000036022 | 1.747221736 | 1.744319981 | 1.769203899 | 1.956637122 | 1.861137485 | 2.103803721 | 1.715701091 |
| ENSMUSG00000036023 | 2.069666964 | 1.969129237 | 1.937048212 | 2.288071211 | 2.240967488 | 2.572871602 | 2.094403682 |
| ENSMUSG00000036026 | 2.578990401 | 2.358135475 | 2.396240625 | 2.199415381 | 2.255474576 | 2.492760389 | 2.18859465 |
| ENSMUSG00000036027 | 0 | 0 | 0 | 0 | 0 | 0 | 0 |

| | | | | | | | |
|---|---|---|---|---|---|---|---|
| ENSMUSG00000036030 | 1.631440309 | 1.438721876 | 1.482684334 | 1.561981878 | 1.387761935 | 1.698970004 | 1.374453863 |
| ENSMUSG00000036036 | 2.247417471 | 2.149047439 | 2.089388001 | 2.031030328 | 2.009637626 | 2.181843588 | 1.880961074 |
| ENSMUSG00000036040 | 0.289064826 | 0.896240625 | 0.976722649 | 0.25 | 0.568060967 | 0.477121255 | 0.355418052 |
| ENSMUSG00000036045 | 0 | 0.25 | 0.25 | 0 | 0 | 0 | 0.355418052 |
| ENSMUSG00000036046 | 2.756312721 | 2.593759858 | 2.623963757 | 2.670089881 | 2.666348738 | 3.02201574 | 2.540007818 |
| ENSMUSG00000036052 | 2.063793172 | 1.907339155 | 1.981203126 | 1.695339928 | 1.686820904 | 1.770852012 | 1.68886578 |
| ENSMUSG00000036053 | 2.569423023 | 2.350753006 | 2.331607372 | 2.414106216 | 2.320220671 | 2.672097858 | 2.182948513 |
| ENSMUSG00000036054 | 2.054824387 | 1.894357207 | 1.938721876 | 1.935366747 | 1.978602168 | 2.068185862 | 1.849546807 |
| ENSMUSG00000036057 | 2.641922824 | 2.465271726 | 2.528435542 | 2.240724001 | 2.392211871 | 2.630427875 | 2.318397784 |
| ENSMUSG00000036061 | 2.078325387 | 1.972185812 | 1.985628626 | 2.002806814 | 2.007252778 | 2.209515015 | 1.98213686 |
| ENSMUSG00000036062 | 0 | 0 | 0 | 0.701838731 | 0.733951626 | 0.477121255 | 0.520678036 |
| ENSMUSG00000036067 | 1.249317394 | 1.364857905 | 1.494319981 | 1.552363341 | 1.665423072 | 1.716003344 | 1.725216937 |
| ENSMUSG00000036073 | 1.358731791 | 1.330482024 | 1.388647213 | 1.42510993 | 1.514343934 | 1.477121255 | 1.523605797 |
| ENSMUSG00000036078 | 2.358731791 | 2.187886015 | 2.233422664 | 2.266522298 | 2.235996193 | 2.499687083 | 2.163786106 |
| ENSMUSG00000036083 | 0 | 0 | 0 | 0 | 0 | 0 | 0 |
| ENSMUSG00000036086 | 2.131864492 | 1.982684334 | 1.959800947 | 2.051142786 | 2.009637626 | 2.28780173 | 1.906037947 |
| ENSMUSG00000036087 | 2.69529263 | 2.474083101 | 2.428561379 | 2.495034894 | 2.443737124 | 2.745855195 | 2.252393349 |
| ENSMUSG00000036091 | 0.578129653 | 0.580482024 | 0.25 | 0.396240625 | 0.77552387 | 0.77815125 | 0.520678036 |
| ENSMUSG00000036093 | 2.974149285 | 2.723704441 | 2.659835177 | 2.604463129 | 2.506870642 | 2.875061263 | 2.327268527 |
| ENSMUSG00000036095 | 0 | 0 | 0 | 0 | 0.244650542 | 0 | 0 |
| ENSMUSG00000036097 | 2.947301297 | 2.679847205 | 2.686838457 | 2.748941788 | 2.656980987 | 3.029789471 | 2.465098532 |
| ENSMUSG00000036098 | 0.811507563 | 1.146240625 | 1.25 | 1.714495249 | 1.770879644 | 1.857332496 | 1.712465909 |
| ENSMUSG00000036099 | 2.333175866 | 2.114857905 | 2.129909063 | 2.189555804 | 2.159419872 | 2.385606274 | 2.022587518 |
| ENSMUSG00000036103 | 2.804055922 | 2.588286706 | 2.600219859 | 2.353935442 | 2.267696867 | 2.62324929 | 2.164590871 |
| ENSMUSG00000036104 | 2.612833548 | 2.370953944 | 2.3843046 | 2.52186571 | 2.460693128 | 2.802773725 | 2.30409643 |
| ENSMUSG00000036106 | 1.614416215 | 1.53743678 | 1.589388001 | 1.488549078 | 1.607318736 | 1.653212514 | 1.692344475 |
| ENSMUSG00000036109 | 2.036286563 | 1.955044741 | 1.918106335 | 1.935366747 | 1.848234258 | 2.089905111 | 1.71730663 |
| ENSMUSG00000036110 | 0 | 0 | 0 | 0 | 0 | 0 | 0 |
| ENSMUSG00000036111 | 0 | 0 | 0 | 0.396240625 | 0.489301084 | 0.602059991 | 0.629532004 |
| ENSMUSG00000036112 | 3.101064463 | 2.835518667 | 2.82665784 | 2.981759292 | 2.882616728 | 3.36078269 | 2.695163117 |
| ENSMUSG00000036114 | 1.578129653 | 1.593759858 | 1.67510993 | 1.356566189 | 1.607318736 | 1.544068044 | 1.647906904 |
| ENSMUSG00000036117 | 0 | 0 | 0 | 0 | 0 | 0 | 0.448487648 |
| ENSMUSG00000036120 | 1.693336259 | 1.561981878 | 1.618933358 | 1.557204673 | 1.655839964 | 1.86332286 | 1.607811402 |
| ENSMUSG00000036123 | 0 | 0 | 0 | 0 | 0 | 0 | 0.224243824 |

| | | | | | | | |
|---|---|---|---|---|---|---|---|
| ENSMUSG00000036131 | 0 | 0 | 0 | 0 | 0 | 0 | 0 |
| ENSMUSG00000036136 | 0.960252568 | 0.830482024 | 0.79248125 | 1.04248125 | 1.091002363 | 0.84509804 | 0.672731473 |
| ENSMUSG00000036138 | 1.783508299 | 1.811981878 | 1.834962501 | 1.685366747 | 1.940259189 | 1.982271233 | 1.850609257 |
| ENSMUSG00000036139 | 0 | 0 | 0.5 | 1.27186571 | 1.091002363 | 1.278753601 | 1.176815335 |
| ENSMUSG00000036144 | 0 | 0 | 0 | 0 | 0 | 0 | 0 |
| ENSMUSG00000036151 | 0.747221736 | 0.79248125 | 0.896240625 | 0.864857905 | 0.489301084 | 0.602059991 | 0.876096088 |
| ENSMUSG00000036155 | 2.235832493 | 2.109697963 | 2.077084758 | 2.149047439 | 2.133621644 | 2.330413773 | 1.969165684 |
| ENSMUSG00000036158 | 1.227926429 | 1.130890489 | 1.214495249 | 1.580482024 | 1.671670246 | 1.77815125 | 1.610050272 |
| ENSMUSG00000036160 | 2.217828299 | 1.985628626 | 2.077084758 | 2.02053726 | 2.028157605 | 2.209515015 | 1.927642465 |
| ENSMUSG00000036162 | 0 | 0 | 0 | 0 | 0.244650542 | 0 | 0 |
| ENSMUSG00000036167 | 2.819794862 | 2.603406982 | 2.578504176 | 2.711176441 | 2.611014646 | 3.003891166 | 2.44445206 |
| ENSMUSG00000036168 | 0.578129653 | 0.864857905 | 0.646240625 | 0.75 | 0.877063019 | 0.77815125 | 0.710836105 |
| ENSMUSG00000036169 | 0 | 0 | 0 | 0 | 0 | 0 | 0 |
| ENSMUSG00000036172 | 0 | 0 | 0 | 0.25 | 0 | 0 | 0.224243824 |
| ENSMUSG00000036180 | 2.755750303 | 2.58076369 | 2.612802778 | 2.460194481 | 2.543534996 | 2.761175813 | 2.449322097 |
| ENSMUSG00000036181 | 1.307602651 | 1.372963274 | 1.372963274 | 1.843759858 | 1.87878897 | 2.025305865 | 1.843097781 |
| ENSMUSG00000036185 | 0.45815691 | 0.25 | 0.396240625 | 0.396240625 | 0.244650542 | 0 | 0 |
| ENSMUSG00000036186 | 1.639698646 | 1.642463902 | 1.681980114 | 1.201838731 | 1.467903253 | 1.431363764 | 1.610050272 |
| ENSMUSG00000036188 | 2.149089517 | 1.979715809 | 1.990000483 | 1.981203126 | 1.893954095 | 2.184691431 | 1.784981274 |
| ENSMUSG00000036192 | 1.249317394 | 1.160964047 | 1.160964047 | 1.201838731 | 1.283908311 | 1.278753601 | 1.078019652 |
| ENSMUSG00000036196 | 0.45815691 | 0.25 | 0.5 | 0.396240625 | 0.489301084 | 0 | 0.355418052 |
| ENSMUSG00000036197 | 2.655740331 | 2.442459461 | 2.40136988 | 2.676624005 | 2.604906476 | 2.984077034 | 2.426358309 |
| ENSMUSG00000036198 | 1.249317394 | 1.146240625 | 1.080482024 | 1.678561379 | 1.618292349 | 1.755874856 | 1.559435575 |
| ENSMUSG00000036199 | 2.22189694 | 2.176839783 | 2.284887838 | 1.985628626 | 2.203230928 | 2.276461804 | 2.206027309 |
| ENSMUSG00000036202 | 2.852729478 | 2.591580554 | 2.559899632 | 2.675759597 | 2.583123351 | 2.915927212 | 2.368731528 |
| ENSMUSG00000036206 | 2.750084015 | 2.545288064 | 2.54838134 | 2.333847589 | 2.376595432 | 2.597695186 | 2.310303152 |
| ENSMUSG00000036208 | 2.156259305 | 2.005591953 | 1.984159485 | 1.950224975 | 1.974687643 | 2.164352856 | 1.792684387 |
| ENSMUSG00000036211 | 0 | 0 | 0 | 0 | 0 | 0.301029996 | 0 |
| ENSMUSG00000036214 | 1.205378646 | 1.27186571 | 1.302363341 | 1.238549078 | 1.176121988 | 1.146128036 | 1.168187745 |
| ENSMUSG00000036216 | 0 | 0 | 0 | 0.25 | 0 | 0.477121255 | 0.224243824 |
| ENSMUSG00000036218 | 0 | 0 | 0 | 0 | 0 | 0 | 0 |
| ENSMUSG00000036223 | 2.072573125 | 1.918106335 | 1.935366747 | 1.959800947 | 1.950242452 | 2.187520721 | 1.849546807 |
| ENSMUSG00000036225 | 1.516991255 | 1.52186571 | 1.532320754 | 1.302363341 | 1.27449559 | 1.491361694 | 1.335191751 |
| ENSMUSG00000036231 | 0 | 0 | 0 | 0 | 0 | 0 | 0 |

| | | | | | | | |
|---|---|---|---|---|---|---|---|
| ENSMUSG00000036241 | 2.708105693 | 2.535207443 | 2.501406137 | 2.475593779 | 2.400554497 | 2.796574333 | 2.255745871 |
| ENSMUSG00000036242 | 0 | 0 | 0 | 0 | 0 | 0 | 0 |
| ENSMUSG00000036244 | 0 | 0 | 0.25 | 0 | 0 | 0 | 0 |
| ENSMUSG00000036246 | 2.060824957 | 1.945339928 | 2.027131114 | 1.866901388 | 2.025895058 | 2.152288344 | 1.98213686 |
| ENSMUSG00000036249 | 1.700465191 | 1.630890489 | 1.54248125 | 1.75835575 | 1.683842353 | 1.880813592 | 1.523605797 |
| ENSMUSG00000036251 | 0 | 0 | 0.25 | 0.5 | 0.387761935 | 0.477121255 | 0.448487648 |
| ENSMUSG00000036256 | 3.146892292 | 2.882601584 | 2.873963757 | 2.284247778 | 2.25010405 | 2.5132176 | 2.11845007 |
| ENSMUSG00000036257 | 2.312518611 | 2.12694866 | 2.134789703 | 2.071350555 | 1.998776581 | 2.238046103 | 1.882886767 |
| ENSMUSG00000036264 | 0 | 0 | 0 | 0 | 0 | 0 | 0 |
| ENSMUSG00000036270 | 2.624559692 | 2.439555804 | 2.47934302 | 2.382845365 | 2.490215874 | 2.716003344 | 2.388231309 |
| ENSMUSG00000036273 | 2.647066901 | 2.417664062 | 2.416333979 | 2.43662858 | 2.363759249 | 2.686636269 | 2.16858494 |
| ENSMUSG00000036275 | 2.997800954 | 2.723893821 | 2.726158655 | 2.888767098 | 2.833231787 | 3.268577972 | 2.666659054 |
| ENSMUSG00000036278 | 1.036286563 | 1.130890489 | 1.188721876 | 0.92510993 | 1.163285805 | 1.176091259 | 1.089372236 |
| ENSMUSG00000036281 | 2.039433992 | 1.950224975 | 2.001406137 | 1.856566189 | 1.935892666 | 2.086359831 | 1.830929954 |
| ENSMUSG00000036282 | 2.172524468 | 2.038704527 | 2.031030328 | 2.319612365 | 2.2705456 | 2.553883027 | 2.075383 |
| ENSMUSG00000036285 | 2.205378646 | 2.039967834 | 2.041226732 | 2.065523711 | 2.054233912 | 2.330413773 | 1.985649549 |
| ENSMUSG00000036291 | 1.800532393 | 1.726722649 | 1.701838731 | 1.970660762 | 1.90849439 | 2.184691431 | 1.786277931 |
| ENSMUSG00000036292 | 0.45815691 | 0.864857905 | 1.061981878 | 1.214495249 | 1.188507684 | 1.462397998 | 1 |
| ENSMUSG00000036295 | 0.45815691 | 0 | 0.25 | 0 | 0 | 0 | 0 |
| ENSMUSG00000036298 | 1.129344651 | 1.17510993 | 1.201838731 | 1.146240625 | 1.057362051 | 1.322219295 | 1.159323753 |
| ENSMUSG00000036299 | 2.231898196 | 2.012462137 | 2.044977273 | 2.031030328 | 1.945998439 | 2.29666519 | 1.929310074 |
| ENSMUSG00000036304 | 0 | 0.396240625 | 0 | 0.25 | 0.489301084 | 0.301029996 | 0.629532004 |
| ENSMUSG00000036305 | 1.227926429 | 1.160964047 | 1.114857905 | 1.282320754 | 1.264824954 | 1.431363764 | 1.20919388 |
| ENSMUSG00000036306 | 0.578129653 | 0.25 | 0.396240625 | 0.75 | 0.77552387 | 0.84509804 | 0.803905701 |
| ENSMUSG00000036309 | 2.933200029 | 2.68325383 | 2.670089881 | 2.892463902 | 2.813199692 | 3.246252312 | 2.589859954 |
| ENSMUSG00000036315 | 2.092368209 | 1.930274797 | 1.94697564 | 1.987091808 | 1.955822902 | 2.217483944 | 1.845261772 |
| ENSMUSG00000036322 | 0 | 0 | 0 | 0 | 0 | 0 | 0 |
| ENSMUSG00000036323 | 2.983706352 | 2.736909234 | 2.71293726 | 2.82608783 | 2.750939708 | 3.186391216 | 2.563676916 |
| ENSMUSG00000036327 | 2.181541736 | 2.074802005 | 2.034887838 | 2.113831805 | 2.102149272 | 2.326335861 | 1.992562536 |
| ENSMUSG00000036330 | 0.289064826 | 0 | 0.396240625 | 0.396240625 | 0.489301084 | 0.698970004 | 0.579661877 |
| ENSMUSG00000036333 | 2.762449763 | 2.50835575 | 2.517194569 | 2.540912419 | 2.496852851 | 2.85672889 | 2.314628629 |
| ENSMUSG00000036334 | 2.072573125 | 1.884789703 | 1.92510993 | 1.711372513 | 1.645989387 | 1.73239376 | 1.520678036 |
| ENSMUSG00000036339 | 1.671187741 | 1.54745614 | 1.552363341 | 1.735628626 | 1.718026092 | 1.892094603 | 1.579661877 |
| ENSMUSG00000036352 | 2.273259602 | 2.127938163 | 2.072504712 | 2.157339155 | 2.146756411 | 2.475671188 | 2.009874076 |

| | | | | | | | |
|---|---|---|---|---|---|---|---|
| ENSMUSG00000036353 | 0 | 0.25 | 0 | 0 | 0 | 0 | 0.224243824 |
| ENSMUSG00000036356 | 1.83262764 | 1.779735268 | 1.799918086 | 1.080482024 | 0.955822902 | 1.176091259 | 1 |
| ENSMUSG00000036357 | 0 | 0 | 0 | 0 | 0 | 0 | 0 |
| ENSMUSG00000036362 | 0 | 0 | 0 | 0 | 0 | 0 | 0 |
| ENSMUSG00000036368 | 1.857607105 | 1.78743678 | 1.843759858 | 1.646240625 | 1.572189156 | 1.875061263 | 1.499406449 |
| ENSMUSG00000036371 | 3.286077588 | 3.018535366 | 3.044588406 | 3.017949381 | 2.963872422 | 3.408239965 | 2.817058769 |
| ENSMUSG00000036372 | 2.504843822 | 2.243603647 | 2.289967834 | 2.314346961 | 2.225838481 | 2.608526034 | 2.141695983 |
| ENSMUSG00000036376 | 1.789260561 | 1.850219859 | 1.79248125 | 1.872963274 | 1.906908067 | 2.004321374 | 1.880961074 |
| ENSMUSG00000036377 | 2.196867486 | 2.108657057 | 2.095926073 | 1.410964047 | 1.462344774 | 1.51851394 | 1.413583902 |
| ENSMUSG00000036381 | 0.578129653 | 0.5 | 0.25 | 0.580482024 | 0.244650542 | 0.477121255 | 0.355418052 |
| ENSMUSG00000036390 | 1.596667477 | 1.482684334 | 1.476722649 | 1.431980114 | 1.427019704 | 1.579783597 | 1.413583902 |
| ENSMUSG00000036391 | 2.440557995 | 2.290597832 | 2.275821952 | 2.375956434 | 2.306653428 | 2.622214023 | 2.109521699 |
| ENSMUSG00000036395 | 0 | 0 | 0 | 0.92510993 | 0.846351821 | 0.477121255 | 0.74492186 |
| ENSMUSG00000036398 | 2.530730035 | 2.286164561 | 2.277131114 | 2.418989258 | 2.361573756 | 2.673941999 | 2.214058866 |
| ENSMUSG00000036401 | 0 | 0 | 0 | 0 | 0 | 0 | 0 |
| ENSMUSG00000036402 | 2.820759099 | 2.588010856 | 2.559601185 | 2.614088604 | 2.502486038 | 2.883661435 | 2.329690062 |
| ENSMUSG00000036403 | 2.203267085 | 2.025821952 | 2.072504712 | 2.168989258 | 2.132784263 | 2.348304863 | 1.964723754 |
| ENSMUSG00000036411 | 1.516991255 | 1.396240625 | 1.470660762 | 1.561981878 | 1.499532413 | 1.568201724 | 1.476637173 |
| ENSMUSG00000036412 | 0.671187741 | 0.830482024 | 0.830482024 | 0.5 | 0.632412477 | 0.602059991 | 0.876096088 |
| ENSMUSG00000036422 | 0 | 0 | 0 | 0 | 0 | 0 | 0 |
| ENSMUSG00000036427 | 2.870199357 | 2.637446167 | 2.648347781 | 2.569031101 | 2.639066517 | 2.959518377 | 2.445802857 |
| ENSMUSG00000036430 | 1.37447073 | 1.51109853 | 1.458222504 | 1.339388001 | 1.373641807 | 1.51851394 | 1.520678036 |
| ENSMUSG00000036432 | 1.837744666 | 1.752806814 | 1.701838731 | 1.766522298 | 1.580303447 | 1.959041392 | 1.598697688 |
| ENSMUSG00000036435 | 2.629921064 | 2.397178659 | 2.394828984 | 2.353935442 | 2.308186911 | 2.640481437 | 2.131174228 |
| ENSMUSG00000036437 | 1.036286563 | 0.976722649 | 1.02186571 | 0.25 | 0.244650542 | 0.602059991 | 0.355418052 |
| ENSMUSG00000036438 | 3.447258323 | 3.106369547 | 3.083568321 | 3.118489841 | 3.017449921 | 3.536179532 | 2.799870113 |
| ENSMUSG00000036442 | 2.126809492 | 2.041226732 | 2.063166358 | 2.058404919 | 2.070605826 | 2.29666519 | 2 |
| ENSMUSG00000036446 | 0 | 0 | 0 | 0 | 0 | 0 | 0 |
| ENSMUSG00000036449 | 0 | 0 | 0 | 0.25 | 0 | 0 | 0 |
| ENSMUSG00000036450 | 2.643399047 | 2.452642909 | 2.462546709 | 2.574802005 | 2.54143406 | 2.881954971 | 2.360063482 |
| ENSMUSG00000036452 | 1.129344651 | 1.080482024 | 1.201838731 | 1.311981878 | 1.264824954 | 1.414973348 | 1.21680636 |
| ENSMUSG00000036459 | 2.003432385 | 1.77186571 | 1.890560606 | 1.660964047 | 1.662257525 | 1.785329835 | 1.612273755 |
| ENSMUSG00000036461 | 2.217828299 | 2.063166358 | 2.090485943 | 2.218361028 | 2.184185718 | 2.424881637 | 2.007270316 |
| ENSMUSG00000036463 | 0 | 0 | 0 | 0 | 0 | 0 | 0 |

| | | | | | | | |
|---|---|---|---|---|---|---|---|
| ENSMUSG00000036466 | 0.960252568 | 0.976722649 | 0.976722649 | 1.17510993 | 1.074582839 | 1.176091259 | 1 |
| ENSMUSG00000036469 | 0.45815691 | 0.396240625 | 0.25 | 0.5 | 0.387761935 | 0.698970004 | 0.520678036 |
| ENSMUSG00000036473 | 2.278594844 | 2.097004321 | 2.145300145 | 2.083847589 | 2.113840608 | 2.348304863 | 1.989815042 |
| ENSMUSG00000036478 | 2.611244851 | 2.375459296 | 2.391037372 | 2.219129237 | 2.150762767 | 2.475671188 | 1.997315209 |
| ENSMUSG00000036480 | 0 | 0.25 | 0 | 0 | 0 | 0 | 0 |
| ENSMUSG00000036492 | 0.578129653 | 0.396240625 | 0.75 | 0.580482024 | 0 | 0.698970004 | 0.448487648 |
| ENSMUSG00000036499 | 2.752926783 | 2.548073204 | 2.55930249 | 2.633088981 | 2.535591162 | 2.908485019 | 2.340601441 |
| ENSMUSG00000036500 | 0.578129653 | 0.75 | 1 | 0.25 | 0 | 0 | 0 |
| ENSMUSG00000036501 | 2.584966402 | 2.352347734 | 2.337734795 | 2.340485943 | 2.273371525 | 2.563481085 | 2.098373247 |
| ENSMUSG00000036502 | 0 | 0.5 | 0.25 | 0.396240625 | 0.244650542 | 0.301029996 | 0.520678036 |
| ENSMUSG00000036503 | 2.565000581 | 2.358135475 | 2.347542239 | 2.384789703 | 2.29040462 | 2.693726949 | 2.14725554 |
| ENSMUSG00000036504 | 1.445324132 | 1.431980114 | 1.356566189 | 1.330482024 | 1.319233381 | 1.380211242 | 1.100339912 |
| ENSMUSG00000036510 | 0 | 0 | 0 | 0 | 0 | 0 | 0 |
| ENSMUSG00000036513 | 2.260538674 | 2.113831805 | 2.122963274 | 2.143411797 | 2.078515538 | 2.385606274 | 2.001999072 |
| ENSMUSG00000036521 | 0 | 0 | 0 | 0 | 0 | 0 | 0 |
| ENSMUSG00000036523 | 0.671187741 | 0.396240625 | 0.580482024 | 0.396240625 | 0.244650542 | 0.477121255 | 0.520678036 |
| ENSMUSG00000036526 | 0 | 0 | 0 | 0 | 0 | 0 | 0 |
| ENSMUSG00000036528 | 1.181541736 | 1.201838731 | 1.26109853 | 0.701838731 | 0.489301084 | 0.477121255 | 0.579661877 |
| ENSMUSG00000036529 | 2.859805489 | 2.680274797 | 2.7543761 | 2.525821952 | 2.665423072 | 2.876217841 | 2.586823247 |
| ENSMUSG00000036533 | 2.336265009 | 2.197790722 | 2.136723615 | 1.814346961 | 1.793512788 | 2.004321374 | 1.704233519 |
| ENSMUSG00000036534 | 2.283862692 | 2.160964047 | 2.180274797 | 1.729715809 | 1.853822071 | 1.991226076 | 1.749303626 |
| ENSMUSG00000036537 | 1 | 1.214495249 | 1.17510993 | 0.92510993 | 1.163285805 | 1.146128036 | 1.176815335 |
| ENSMUSG00000036545 | 2.294202867 | 2.201838731 | 2.223704441 | 1.160964047 | 1.039257769 | 1.146128036 | 1.014380832 |
| ENSMUSG00000036550 | 3.303082747 | 2.964203638 | 2.952039943 | 3.09110207 | 3.004284367 | 3.469085299 | 2.780281785 |
| ENSMUSG00000036551 | 0 | 0 | 0.25 | 0 | 0.244650542 | 0 | 0.224243824 |
| ENSMUSG00000036552 | 1.890240806 | 1.828220739 | 1.789967834 | 1.714495249 | 1.773209395 | 1.913813852 | 1.56716938 |
| ENSMUSG00000036553 | 0.811507563 | 0.701838731 | 0.580482024 | 0.646240625 | 0.877063019 | 0.301029996 | 0.916588299 |
| ENSMUSG00000036555 | 2.139333929 | 2 | 1.987091808 | 1.997171172 | 2.05001946 | 2.247973266 | 1.914853627 |
| ENSMUSG00000036557 | 0 | 0 | 0 | 0.25 | 0 | 0 | 0 |
| ENSMUSG00000036560 | 0 | 0 | 0.396240625 | 0 | 0 | 0 | 0 |
| ENSMUSG00000036561 | 2.095120908 | 1.961372513 | 2.023189285 | 2.115881093 | 2.136951444 | 2.318063335 | 2.017561934 |
| ENSMUSG00000036564 | 2.086807695 | 2.002806814 | 2.034887838 | 1.848079356 | 1.961316494 | 2.139879086 | 1.828667599 |
| ENSMUSG00000036565 | 2.726012129 | 2.58076369 | 2.6343046 | 2.232684334 | 2.42519375 | 2.583198774 | 2.424203925 |
| ENSMUSG00000036568 | 2.075459174 | 1.892463902 | 1.958222504 | 1.894357207 | 1.810629166 | 1.982271233 | 1.769001954 |

| | | | | | | | |
|---|---|---|---|---|---|---|---|
| ENSMUSG00000036570 | 0 | 0 | 0 | 0 | 0 | 0 | 0 |
| ENSMUSG00000036572 | 2.371652933 | 2.180274797 | 2.256284891 | 2.299304173 | 2.260180728 | 2.503790683 | 2.110473216 |
| ENSMUSG00000036574 | 0 | 0 | 0 | 0 | 0 | 0 | 0 |
| ENSMUSG00000036578 | 0 | 0 | 0.25 | 0 | 0 | 0 | 0 |
| ENSMUSG00000036580 | 2.589183146 | 2.421625132 | 2.43155454 | 2.42293588 | 2.379941013 | 2.692846919 | 2.253005488 |
| ENSMUSG00000036585 | 0.578129653 | 0.25 | 0.396240625 | 0.25 | 0.244650542 | 0 | 0 |
| ENSMUSG00000036586 | 0 | 0 | 0 | 0 | 0 | 0 | 0 |
| ENSMUSG00000036587 | 0 | 0.25 | 0.25 | 0 | 0 | 0.301029996 | 0.355418052 |
| ENSMUSG00000036591 | 2.672564087 | 2.474461364 | 2.461372513 | 2.419870025 | 2.344045917 | 2.651278014 | 2.200679824 |
| ENSMUSG00000036594 | 0 | 0 | 0 | 0.580482024 | 0.244650542 | 0.477121255 | 0.448487648 |
| ENSMUSG00000036596 | 0 | 0 | 0 | 0.25 | 0 | 0 | 0.355418052 |
| ENSMUSG00000036598 | 0 | 0 | 0 | 0 | 0.244650542 | 0.301029996 | 0.355418052 |
| ENSMUSG00000036599 | 2.039433992 | 2.04745614 | 2.080482024 | 1.62694866 | 1.888971102 | 1.897627091 | 1.923435442 |
| ENSMUSG00000036602 | 2.153883046 | 1.979715809 | 2.04248125 | 0.25 | 0 | 0 | 0.355418052 |
| ENSMUSG00000036606 | 3.274883562 | 3.024261125 | 3.074802005 | 2.931980114 | 3.032115293 | 3.39497672 | 2.907011414 |
| ENSMUSG00000036611 | 1.269664473 | 1.302363341 | 1.238549078 | 1.160964047 | 1.121713561 | 1.079181246 | 1.20139795 |
| ENSMUSG00000036613 | 1.967766868 | 1.899978211 | 1.959800947 | 1.723704441 | 1.768534412 | 1.991226076 | 1.683576582 |
| ENSMUSG00000036615 | 1.993049299 | 1.799918086 | 1.75835575 | 1.882845365 | 1.873585591 | 2.133538908 | 1.782372264 |
| ENSMUSG00000036617 | 1.269664473 | 1.356566189 | 1.348079356 | 1.410964047 | 1.519146132 | 1.643452676 | 1.444790289 |
| ENSMUSG00000036620 | 2.742600832 | 2.523519421 | 2.554186465 | 2.336628433 | 2.37406527 | 2.643452676 | 2.257559981 |
| ENSMUSG00000036622 | 1.494443473 | 1.657339155 | 1.75835575 | 1.527131114 | 1.784632921 | 1.77815125 | 1.966211184 |
| ENSMUSG00000036632 | 2.167941649 | 1.975216702 | 1.991446071 | 2.089388001 | 1.992595054 | 2.281033367 | 1.874129423 |
| ENSMUSG00000036634 | 0.289064826 | 0 | 0.5 | 0.25 | 0.244650542 | 0 | 0.355418052 |
| ENSMUSG00000036636 | 2.694641526 | 2.466433318 | 2.481944491 | 2.26584877 | 2.351793489 | 2.617000341 | 2.239263384 |
| ENSMUSG00000036639 | 1.982401085 | 1.870953944 | 1.841580554 | 1.87892496 | 1.848234258 | 2.071882007 | 1.737484395 |
| ENSMUSG00000036641 | 1.100572389 | 1 | 0.92510993 | 0.92510993 | 0.955822902 | 1.113943352 | 1.078019652 |
| ENSMUSG00000036644 | 2.435736758 | 2.257666784 | 2.281676118 | 2.322504712 | 2.41779398 | 2.670245853 | 2.313108585 |
| ENSMUSG00000036646 | 2.742600832 | 2.496816003 | 2.460980263 | 2.535845803 | 2.532092521 | 2.879669206 | 2.34638072 |
| ENSMUSG00000036647 | 0 | 0 | 0 | 0 | 0 | 0 | 0 |
| ENSMUSG00000036655 | 0.45815691 | 0.25 | 0.5 | 0.5 | 0 | 0.301029996 | 0 |
| ENSMUSG00000036658 | 0 | 0 | 0 | 0 | 0 | 0 | 0 |
| ENSMUSG00000036661 | 1.759671389 | 1.75835575 | 1.823655187 | 1.622963274 | 1.797870327 | 1.897627091 | 1.702561601 |
| ENSMUSG00000036667 | 2.769588082 | 2.529410775 | 2.525165585 | 2.567865757 | 2.520825546 | 2.857332496 | 2.330653623 |
| ENSMUSG00000036672 | 2.295901576 | 2.120953944 | 2.160964047 | 2.094844592 | 2.182733222 | 2.369215857 | 2.080634989 |

| | | | | | | | |
|---|---|---|---|---|---|---|---|
| ENSMUSG00000036676 | 2.788740894 | 2.576515422 | 2.596735061 | 2.77037086 | 2.676194227 | 3.093421685 | 2.441733427 |
| ENSMUSG00000036678 | 2.686747932 | 2.485628626 | 2.492885888 | 2.460587586 | 2.494754592 | 2.80685803 | 2.343388382 |
| ENSMUSG00000036686 | 2.262379879 | 2.172499493 | 2.244319981 | 2.091580554 | 2.26251048 | 2.29666519 | 2.243069492 |
| ENSMUSG00000036687 | 0 | 0.25 | 0.25 | 0 | 0.387761935 | 0 | 0.448487648 |
| ENSMUSG00000036693 | 2.613625632 | 2.377443751 | 2.416777886 | 2.395300145 | 2.349532392 | 2.677606953 | 2.214058866 |
| ENSMUSG00000036698 | 3.144240249 | 2.893766616 | 2.890083209 | 2.740362424 | 2.725610043 | 3.104145551 | 2.552483141 |
| ENSMUSG00000036699 | 0.578129653 | 0.25 | 0.25 | 0.25 | 0.244650542 | 0.301029996 | 0 |
| ENSMUSG00000036707 | 2.807052575 | 2.589388001 | 2.563757142 | 2.654137211 | 2.570495157 | 2.981365509 | 2.39794841 |
| ENSMUSG00000036712 | 2.699822119 | 2.475970461 | 2.488185312 | 2.810346841 | 2.699837743 | 3.117933835 | 2.495094762 |
| ENSMUSG00000036718 | 2.637462542 | 2.43662858 | 2.494677615 | 2.14151351 | 2.263669614 | 2.478566496 | 2.197428385 |
| ENSMUSG00000036721 | 1.54867984 | 1.494319981 | 1.593759858 | 1.580482024 | 1.584291694 | 1.799340549 | 1.486592281 |
| ENSMUSG00000036731 | 0 | 0 | 0 | 0 | 0 | 0 | 0 |
| ENSMUSG00000036733 | 0 | 0 | 0.25 | 0 | 0 | 0.301029996 | 0 |
| ENSMUSG00000036737 | 2.576402808 | 2.347542239 | 2.351816941 | 2.468745337 | 2.437074068 | 2.774516966 | 2.250857915 |
| ENSMUSG00000036743 | 0 | 0 | 0 | 0 | 0 | 0 | 0 |
| ENSMUSG00000036744 | 0 | 0 | 0 | 0 | 0 | 0 | 0 |
| ENSMUSG00000036745 | 2.579849376 | 2.374961472 | 2.37246199 | 2.538704527 | 2.475604419 | 2.816903839 | 2.272582514 |
| ENSMUSG00000036748 | 2.576402808 | 2.466819685 | 2.521202097 | 2.247880462 | 2.33042383 | 2.5774918 | 2.271140178 |
| ENSMUSG00000036749 | 0 | 0.5 | 0 | 0 | 0.387761935 | 0.301029996 | 0 |
| ENSMUSG00000036751 | 2.530730035 | 2.323655187 | 2.340485943 | 2.411864607 | 2.392211871 | 2.739572344 | 2.254226303 |
| ENSMUSG00000036752 | 2.667031238 | 2.436208459 | 2.504202072 | 2.58412664 | 2.621875691 | 2.95616843 | 2.518128015 |
| ENSMUSG00000036760 | 0 | 0 | 0 | 0 | 0 | 0 | 0 |
| ENSMUSG00000036764 | 1.663535556 | 1.602347734 | 1.685366747 | 1.843759858 | 1.78687392 | 2.06069784 | 1.651852264 |
| ENSMUSG00000036766 | 0 | 0.25 | 0.25 | 0 | 0 | 0.301029996 | 0.520678036 |
| ENSMUSG00000036768 | 2.523917219 | 2.325945187 | 2.381869252 | 2.544354884 | 2.474013807 | 2.826074803 | 2.296166733 |
| ENSMUSG00000036769 | 2.051791411 | 1.959800947 | 1.96759118 | 1.96759118 | 1.893954095 | 2.130333768 | 1.769001954 |
| ENSMUSG00000036770 | 0 | 0 | 0.396240625 | 0 | 0.244650542 | 0 | 0.224243824 |
| ENSMUSG00000036775 | 1.940852214 | 1.841580554 | 1.850219859 | 1.814346961 | 1.840643511 | 2.053078443 | 1.753626837 |
| ENSMUSG00000036777 | 3.144240249 | 2.885395165 | 2.871205724 | 3.021782825 | 2.946918141 | 3.408748606 | 2.72372165 |
| ENSMUSG00000036779 | 2.484856099 | 2.260414788 | 2.259729747 | 2.453845824 | 2.380772476 | 2.72427587 | 2.205319397 |
| ENSMUSG00000036781 | 2.659301689 | 2.443284802 | 2.446158711 | 2.502107156 | 2.434943542 | 2.865103975 | 2.314881277 |
| ENSMUSG00000036782 | 2.131864492 | 1.92510993 | 1.935366747 | 1.777131114 | 1.662257525 | 1.880813592 | 1.520678036 |
| ENSMUSG00000036790 | 0.45815691 | 0.396240625 | 0.25 | 0.396240625 | 0.387761935 | 0 | 0 |
| ENSMUSG00000036792 | 2.181541736 | 2.100219859 | 2.05357978 | 2.077084758 | 2.001219193 | 2.235528447 | 1.881925353 |

| | | | | | | | |
|---|---|---|---|---|---|---|---|
| ENSMUSG00000036800 | 0.289064826 | 0 | 0 | 0 | 0 | 0 | 0 |
| ENSMUSG00000036805 | 0 | 0 | 0 | 0 | 0 | 0 | 0 |
| ENSMUSG00000036810 | 2.552362269 | 2.340485943 | 2.330482024 | 2.33993739 | 2.238490593 | 2.622214023 | 2.086826862 |
| ENSMUSG00000036813 | 0 | 0.25 | 0 | 0 | 0 | 0 | 0 |
| ENSMUSG00000036814 | 0 | 0 | 0 | 0.25 | 0 | 0 | 0 |
| ENSMUSG00000036815 | 1.249317394 | 0.976722649 | 1.114857905 | 1.321350555 | 1.163285805 | 1.146128036 | 0.935079929 |
| ENSMUSG00000036816 | 0 | 0 | 0 | 0 | 0 | 0 | 0 |
| ENSMUSG00000036817 | 2.795446569 | 2.569031101 | 2.576230417 | 2.551142786 | 2.537460779 | 2.895974732 | 2.376756474 |
| ENSMUSG00000036819 | 2.026699189 | 1.940387808 | 1.931980114 | 2.074802005 | 2.048957937 | 2.303196057 | 1.942350245 |
| ENSMUSG00000036820 | 2.010213549 | 1.950224975 | 2.131869252 | 1.214495249 | 1.49445382 | 1.204119983 | 1.594042709 |
| ENSMUSG00000036822 | 2.778742163 | 2.55175358 | 2.541226732 | 2.580200137 | 2.476238662 | 2.840106094 | 2.312854549 |
| ENSMUSG00000036825 | 2.330063669 | 2.19697564 | 2.134789703 | 2.3843046 | 2.305113253 | 2.617000341 | 2.134719638 |
| ENSMUSG00000036826 | 0 | 0 | 0 | 0 | 0 | 0 | 0 |
| ENSMUSG00000036832 | 0.867194479 | 0.951838731 | 1.04248125 | 0.976722649 | 1.020174412 | 1.079181246 | 0.829800754 |
| ENSMUSG00000036833 | 1.800532393 | 1.630890489 | 1.769203899 | 1.51109853 | 1.618292349 | 1.707570176 | 1.559435575 |
| ENSMUSG00000036834 | 1.516991255 | 1.321350555 | 1.438721876 | 1.201838731 | 1.106691885 | 1.255272505 | 1.110947927 |
| ENSMUSG00000036835 | 1.956444026 | 1.87892496 | 1.845926073 | 1.807204673 | 1.733951626 | 1.929418926 | 1.653807046 |
| ENSMUSG00000036840 | 2.013563224 | 1.919870025 | 1.848079356 | 1.866901388 | 1.816839698 | 2.012837225 | 1.672731473 |
| ENSMUSG00000036845 | 1.527823874 | 1.302363341 | 1.29248125 | 1.226722649 | 1.302012594 | 1.342422681 | 1.365075947 |
| ENSMUSG00000036850 | 1.747221736 | 1.782320754 | 1.79248125 | 1.735628626 | 1.677808768 | 1.897627091 | 1.643912836 |
| ENSMUSG00000036853 | 0.91631382 | 0.79248125 | 0.830482024 | 0.25 | 0 | 0.301029996 | 0.224243824 |
| ENSMUSG00000036854 | 2.144240249 | 1.988549078 | 2.073655187 | 1.410964047 | 1.450958105 | 1.707570176 | 1.499406449 |
| ENSMUSG00000036855 | 0 | 0 | 0 | 0 | 0 | 0 | 0 |
| ENSMUSG00000036856 | 1.932832114 | 1.858657057 | 2.025821952 | 1.890560606 | 2.038163331 | 2.075546961 | 1.985649549 |
| ENSMUSG00000036858 | 0.671187741 | 0 | 0.25 | 0.25 | 0.244650542 | 0 | 0.448487648 |
| ENSMUSG00000036860 | 1.445324132 | 1.516522298 | 1.561981878 | 1.438721876 | 1.55104774 | 1.591064607 | 1.659601507 |
| ENSMUSG00000036862 | 1.686083339 | 1.571350555 | 1.618933358 | 1.62694866 | 1.756568651 | 1.880813592 | 1.663407615 |
| ENSMUSG00000036863 | 1.740854706 | 1.711372513 | 1.711372513 | 1.774508021 | 1.756568651 | 1.973127854 | 1.61448206 |
| ENSMUSG00000036864 | 1.54867984 | 1.476722649 | 1.464495249 | 1.214495249 | 1.335652905 | 1.322219295 | 1.417653332 |
| ENSMUSG00000036867 | 1.605636262 | 1.622963274 | 1.738549078 | 1.488549078 | 1.407936347 | 1.579783597 | 1.616675393 |
| ENSMUSG00000036872 | 0 | 0 | 0 | 0.25 | 0 | 0.301029996 | 0.224243824 |
| ENSMUSG00000036873 | 1.912079952 | 1.732684334 | 1.794977273 | 2.039967834 | 1.953043706 | 2.225309282 | 1.871156825 |
| ENSMUSG00000036875 | 2.278594844 | 2.124961472 | 2.160964047 | 2.151832578 | 2.157861565 | 2.357934847 | 2.085803078 |
| ENSMUSG00000036879 | 2.35727108 | 2.197790722 | 2.172499493 | 2.263142013 | 2.175379703 | 2.471291711 | 2.01116814 |

| | | | | | | | |
|---|---|---|---|---|---|---|---|
| ENSMUSG00000036880 | 2.307602651 | 2.139605178 | 2.151832578 | 1.987091808 | 2.043601899 | 2.305351369 | 1.927642465 |
| ENSMUSG00000036882 | 0.671187741 | 1.160964047 | 1.080482024 | 0.701838731 | 0.978602168 | 1.041392685 | 1.159323753 |
| ENSMUSG00000036885 | 1.269664473 | 1.302363341 | 1.114857905 | 0.976722649 | 0.877063019 | 0.903089987 | 0.935079929 |
| ENSMUSG00000036887 | 0 | 0 | 0 | 0 | 0 | 0 | 0 |
| ENSMUSG00000036890 | 1.789260561 | 1.71759118 | 1.653677461 | 1.671625132 | 1.584291694 | 1.73239376 | 1.523605797 |
| ENSMUSG00000036892 | 0.45815691 | 0 | 0.25 | 0 | 0 | 0.301029996 | 0.224243824 |
| ENSMUSG00000036893 | 2.519973621 | 2.370449858 | 2.338837774 | 2.273849256 | 2.292543754 | 2.503790683 | 2.132508314 |
| ENSMUSG00000036894 | 1.907802659 | 1.79745614 | 1.807204673 | 1.79248125 | 1.770879644 | 1.86332286 | 1.63986884 |
| ENSMUSG00000036896 | 0 | 0 | 0 | 0 | 0 | 0 | 0 |
| ENSMUSG00000036898 | 2.111266113 | 1.972185812 | 1.96759118 | 1.982684334 | 1.905314583 | 2.139879086 | 1.746389053 |
| ENSMUSG00000036899 | 0 | 0 | 0 | 0 | 0 | 0 | 0 |
| ENSMUSG00000036902 | 0.960252568 | 1.080482024 | 0.976722649 | 2.234894804 | 2.144330586 | 2.419955748 | 1.980010793 |
| ENSMUSG00000036904 | 1.663535556 | 1.505591953 | 1.458222504 | 1.02186571 | 1.057362051 | 0.698970004 | 1.168187745 |
| ENSMUSG00000036905 | 0 | 0 | 0 | 0 | 0 | 0 | 0 |
| ENSMUSG00000036907 | 0 | 0 | 0 | 0 | 0 | 0 | 0 |
| ENSMUSG00000036908 | 1.445324132 | 1.348079356 | 1.418106335 | 1.061981878 | 1.254881871 | 1 | 1.365075947 |
| ENSMUSG00000036912 | 0 | 0.25 | 0.25 | 0 | 0 | 0.301029996 | 0 |
| ENSMUSG00000036913 | 0 | 0 | 0.25 | 0.25 | 0 | 0 | 0.224243824 |
| ENSMUSG00000036915 | 0 | 0 | 0.25 | 0 | 0 | 0.301029996 | 0 |
| ENSMUSG00000036916 | 2.475043119 | 2.256976499 | 2.244319981 | 2.245748394 | 2.225193812 | 2.487138375 | 2.051774285 |
| ENSMUSG00000036918 | 1.948720984 | 1.909156155 | 1.978222334 | 1.75835575 | 1.855665188 | 1.908485019 | 1.813563596 |
| ENSMUSG00000036921 | 0 | 0 | 0 | 0 | 0 | 0 | 0 |
| ENSMUSG00000036923 | 0 | 0.25 | 0.5 | 0.75 | 0.568060967 | 0 | 0.579661877 |
| ENSMUSG00000036924 | 0 | 0 | 0 | 0 | 0 | 0 | 0 |
| ENSMUSG00000036925 | 0 | 0 | 0 | 0 | 0 | 0 | 0 |
| ENSMUSG00000036928 | 0.811507563 | 0.396240625 | 0.5 | 1 | 1.149965125 | 0.698970004 | 0.803905701 |
| ENSMUSG00000036931 | 0 | 0 | 0 | 0.396240625 | 0.812711509 | 0.477121255 | 0.710836105 |
| ENSMUSG00000036932 | 2.850493374 | 2.613574823 | 2.619186551 | 2.596195938 | 2.528831933 | 2.922206277 | 2.362906151 |
| ENSMUSG00000036934 | 2.086807695 | 1.912762923 | 1.868933358 | 1.981203126 | 1.897237431 | 2.201397124 | 1.846338363 |
| ENSMUSG00000036938 | 0 | 0 | 0 | 0 | 0 | 0 | 0.224243824 |
| ENSMUSG00000036940 | 2.95713908 | 2.66942991 | 2.641751343 | 2.665889526 | 2.571222152 | 2.976808337 | 2.412278274 |
| ENSMUSG00000036941 | 1.771760131 | 1.598079356 | 1.598079356 | 1.653677461 | 1.592135487 | 1.770852012 | 1.493062805 |
| ENSMUSG00000036943 | 2.544030601 | 2.369945066 | 2.321928095 | 2.636723615 | 2.552582334 | 2.947433722 | 2.373823024 |
| ENSMUSG00000036944 | 0.289064826 | 0 | 0 | 0.396240625 | 0 | 0.301029996 | 0.224243824 |

| | | | | | | | |
|---|---|---|---|---|---|---|---|
| ENSMUSG00000036948 | 1.842799666 | 1.807204673 | 1.870953944 | 1.634789703 | 1.793512788 | 1.799340549 | 1.781059827 |
| ENSMUSG00000036949 | 0 | 0 | 0 | 0 | 0 | 0 | 0 |
| ENSMUSG00000036951 | 0 | 0 | 0 | 0 | 0 | 0 | 0 |
| ENSMUSG00000036955 | 2.404271433 | 2.230460234 | 2.182829758 | 2.389126514 | 2.348168747 | 2.691081492 | 2.20496486 |
| ENSMUSG00000036957 | 1.156259305 | 1.214495249 | 1.372963274 | 0.976722649 | 1 | 0.77815125 | 1.21680636 |
| ENSMUSG00000036958 | 0 | 0 | 0 | 0 | 0 | 0 | 0 |
| ENSMUSG00000036959 | 2.136858952 | 2.01109853 | 2.03743678 | 1.964495249 | 1.950242452 | 2.167317335 | 1.86815666 |
| ENSMUSG00000036960 | 0.671187741 | 0.701838731 | 0.79248125 | 0 | 0 | 0 | 0 |
| ENSMUSG00000036961 | 0 | 0 | 0 | 0 | 0 | 0 | 0 |
| ENSMUSG00000036962 | 0 | 0 | 0 | 0 | 0 | 0 | 0 |
| ENSMUSG00000036964 | 0 | 0.5 | 0.396240625 | 0.25 | 0.568060967 | 0.301029996 | 0.224243824 |
| ENSMUSG00000036966 | 2.547754129 | 2.359177886 | 2.416333979 | 2.131869252 | 2.24829556 | 2.460897843 | 2.155212955 |
| ENSMUSG00000036968 | 2.477243823 | 2.282320754 | 2.290597832 | 2.428561379 | 2.369807605 | 2.705007959 | 2.228889254 |
| ENSMUSG00000036972 | 0.289064826 | 0 | 0 | 1.146240625 | 1.176121988 | 1.204119983 | 1.168187745 |
| ENSMUSG00000036975 | 1.881174527 | 1.714495249 | 1.769203899 | 1.75 | 1.689774529 | 1.949390007 | 1.672731473 |
| ENSMUSG00000036977 | 2.020182925 | 1.923371739 | 1.905512955 | 2.00835575 | 1.903713872 | 2.227886705 | 1.755055163 |
| ENSMUSG00000036980 | 2.380049842 | 2.218361028 | 2.226722649 | 2.19697564 | 2.233484039 | 2.474216264 | 2.059651572 |
| ENSMUSG00000036983 | 1.852729478 | 1.692046081 | 1.705044741 | 1.622963274 | 1.614672255 | 1.86332286 | 1.551512352 |
| ENSMUSG00000036985 | 2.22592627 | 2.094844592 | 2.073655187 | 2.110735874 | 2.033751322 | 2.40654018 | 1.923435442 |
| ENSMUSG00000036986 | 2.390989024 | 2.275821952 | 2.281030328 | 2.211372513 | 2.28608706 | 2.475671188 | 2.179128924 |
| ENSMUSG00000036989 | 2.312518611 | 2.181980114 | 2.234894804 | 1.901832578 | 1.987570683 | 2.127104798 | 1.966211184 |
| ENSMUSG00000036990 | 2.977800208 | 2.709604019 | 2.687467357 | 2.77037086 | 2.678523979 | 3.067070856 | 2.486740784 |
| ENSMUSG00000036992 | 1.989530018 | 1.92510993 | 1.933677405 | 1.938721876 | 1.925488995 | 2.113943352 | 1.843097781 |
| ENSMUSG00000036995 | 1.663535556 | 1.532320754 | 1.575945187 | 1.67510993 | 1.766173492 | 1.812913357 | 1.690609803 |
| ENSMUSG00000037001 | 1.069666964 | 0.951838731 | 1.17510993 | 1.388647213 | 1.450958105 | 1.51851394 | 1.265599895 |
| ENSMUSG00000037003 | 2.595842487 | 2.474083101 | 2.529410775 | 2.174241882 | 2.372368362 | 2.568201724 | 2.333050044 |
| ENSMUSG00000037005 | 1.036286563 | 1.02186571 | 1.04248125 | 0.75 | 0.686820904 | 0.77815125 | 0.74492186 |
| ENSMUSG00000037007 | 1.876566388 | 1.862802778 | 1.794977273 | 1.976722649 | 1.855665188 | 2.1430148 | 1.731408809 |
| ENSMUSG00000037010 | 1.932832114 | 1.774508021 | 1.882845365 | 1.841580554 | 1.791313678 | 1.897627091 | 1.61448206 |
| ENSMUSG00000037012 | 2.819311906 | 2.596735061 | 2.654825739 | 2.550837001 | 2.546922567 | 2.878521796 | 2.4638261 |
| ENSMUSG00000037013 | 2.649252229 | 2.444108258 | 2.444929839 | 2.393884812 | 2.367659357 | 2.658964843 | 2.208492871 |
| ENSMUSG00000037014 | 0 | 0 | 0 | 0 | 0 | 0 | 0 |
| ENSMUSG00000037016 | 0 | 0 | 0 | 0.79248125 | 0.686820904 | 0.698970004 | 0.355418052 |
| ENSMUSG00000037017 | 2.241665209 | 2.061981878 | 2.052363341 | 2.077084758 | 2.027028145 | 2.309630167 | 1.930140666 |

| | | | | | | | |
|---|---|---|---|---|---|---|---|
| ENSMUSG00000037020 | 2.003432385 | 1.852347734 | 1.956637122 | 1.819031101 | 1.928492895 | 2.06069784 | 1.874129423 |
| ENSMUSG00000037022 | 1.686083339 | 1.566696635 | 1.516522298 | 1.584962501 | 1.523883869 | 1.643452676 | 1.417653332 |
| ENSMUSG00000037025 | 0 | 0 | 0 | 0.396240625 | 0.244650542 | 0.477121255 | 0.710836105 |
| ENSMUSG00000037028 | 0 | 0 | 0 | 0 | 0 | 0 | 0 |
| ENSMUSG00000037029 | 2.285603957 | 2.186208459 | 2.132845365 | 2.115881093 | 2.102149272 | 2.348304863 | 1.936709733 |
| ENSMUSG00000037031 | 2.089597219 | 1.978222334 | 1.985628626 | 2.034887838 | 2.065570703 | 2.307496038 | 1.963977467 |
| ENSMUSG00000037032 | 2.039433992 | 1.987091808 | 2.016522298 | 1.75835575 | 1.964031554 | 1.949390007 | 1.917452159 |
| ENSMUSG00000037033 | 0 | 0 | 0 | 0 | 0 | 0 | 0 |
| ENSMUSG00000037034 | 0 | 0 | 0 | 0 | 0 | 0 | 0 |
| ENSMUSG00000037035 | 1.156259305 | 1.25 | 1.201838731 | 0.951838731 | 1.074582839 | 0.301029996 | 1.185218806 |
| ENSMUSG00000037049 | 2.192545868 | 2.105516192 | 2.111770807 | 2.08272922 | 2.105788027 | 2.330413773 | 2.040177507 |
| ENSMUSG00000037053 | 0 | 0 | 0 | 0 | 0 | 0 | 0 |
| ENSMUSG00000037058 | 2.723587515 | 2.524508021 | 2.501055117 | 2.523849256 | 2.432441564 | 2.825426118 | 2.295362967 |
| ENSMUSG00000037060 | 1.920505136 | 1.828220739 | 1.899978211 | 1.321350555 | 1.414412788 | 1.380211242 | 1.459332757 |
| ENSMUSG00000037062 | 3.074935921 | 2.805547785 | 2.786164561 | 2.908589325 | 2.816114694 | 3.270212855 | 2.595000328 |
| ENSMUSG00000037064 | 0 | 0 | 0 | 0 | 0 | 0 | 0 |
| ENSMUSG00000037070 | 2.340856238 | 2.155512955 | 2.150906586 | 2.105516192 | 2.094757251 | 2.371067862 | 1.958704689 |
| ENSMUSG00000037071 | 2.898714905 | 2.637205477 | 2.659382638 | 2.759386738 | 2.705178079 | 3.131618664 | 2.544509845 |
| ENSMUSG00000037072 | 3.136181401 | 2.875334905 | 2.862416135 | 2.98129588 | 2.867991218 | 3.367355921 | 2.684464142 |
| ENSMUSG00000037075 | 2.541215974 | 2.32991803 | 2.30539678 | 2.368933358 | 2.323654601 | 2.611723308 | 2.132508314 |
| ENSMUSG00000037085 | 1.639698646 | 1.634789703 | 1.653677461 | 1.638647213 | 1.592135487 | 1.826074803 | 1.62316756 |
| ENSMUSG00000037086 | 0 | 0 | 0 | 0 | 0 | 0 | 0 |
| ENSMUSG00000037089 | 2.521950082 | 2.333847589 | 2.374462959 | 2.193696765 | 2.283361516 | 2.564666064 | 2.156863548 |
| ENSMUSG00000037095 | 0 | 0 | 0 | 0.25 | 0 | 0 | 0 |
| ENSMUSG00000037096 | 0 | 0 | 0 | 0 | 0 | 0 | 0 |
| ENSMUSG00000037098 | 2.418409478 | 2.273849256 | 2.321928095 | 2.154596376 | 2.282813873 | 2.492760389 | 2.225244904 |
| ENSMUSG00000037101 | 0 | 0 | 0 | 0 | 0 | 0 | 0 |
| ENSMUSG00000037103 | 2.34539747 | 2.225216702 | 2.300530956 | 2.157339155 | 2.303566328 | 2.451786436 | 2.22557791 |
| ENSMUSG00000037104 | 2.712498852 | 2.523189285 | 2.524508021 | 2.382845365 | 2.329944595 | 2.630427875 | 2.182948513 |
| ENSMUSG00000037106 | 1.54867984 | 1.364857905 | 1.470660762 | 1.04248125 | 1 | 0.954242509 | 0.935079929 |
| ENSMUSG00000037108 | 1.663535556 | 1.482684334 | 1.54248125 | 1.62694866 | 1.563883923 | 1.643452676 | 1.433437704 |
| ENSMUSG00000037110 | 2.057835465 | 1.868933358 | 1.87892496 | 2.089388001 | 2.057362051 | 2.292256071 | 1.916588299 |
| ENSMUSG00000037111 | 2.910141165 | 2.657111385 | 2.644593173 | 2.777294426 | 2.718185765 | 3.138302698 | 2.518801024 |
| ENSMUSG00000037112 | 2.156259305 | 2.039967834 | 1.998588359 | 1.997171172 | 1.91476933 | 2.214843848 | 1.86715039 |

| | | | | | | | |
|---|---|---|---|---|---|---|---|
| ENSMUSG00000037119 | 3.171487332 | 2.892226539 | 2.858135475 | 2.951838731 | 2.862200607 | 3.31386722 | 2.641989051 |
| ENSMUSG00000037124 | 0 | 0 | 0 | 0 | 0 | 0 | 0 |
| ENSMUSG00000037126 | 1.759671389 | 1.735628626 | 1.811981878 | 1.809601185 | 1.945998439 | 1.982271233 | 1.926805425 |
| ENSMUSG00000037129 | 0.289064826 | 0 | 0.25 | 0 | 0 | 0 | 0.224243824 |
| ENSMUSG00000037130 | 0 | 0 | 0 | 0 | 0 | 0 | 0 |
| ENSMUSG00000037134 | 2.188178997 | 1.990000483 | 1.942046081 | 2.046218836 | 2.016696984 | 2.225309282 | 1.872150729 |
| ENSMUSG00000037138 | 1.036286563 | 1 | 1.146240625 | 1.330482024 | 1.234113756 | 1.380211242 | 1.238624656 |
| ENSMUSG00000037139 | 0 | 0 | 0.25 | 0.25 | 0.244650542 | 0 | 0.224243824 |
| ENSMUSG00000037140 | 0 | 0 | 0 | 0 | 0.244650542 | 0 | 0 |
| ENSMUSG00000037143 | 0 | 0 | 0.25 | 0.396240625 | 0.244650542 | 0.698970004 | 0.520678036 |
| ENSMUSG00000037145 | 0 | 0 | 0 | 0 | 0 | 0 | 0 |
| ENSMUSG00000037148 | 2.320585253 | 2.157339155 | 2.159156155 | 2.277131114 | 2.223900931 | 2.514547753 | 2.114720744 |
| ENSMUSG00000037149 | 2.953301867 | 2.679419106 | 2.661414608 | 2.845791065 | 2.719301455 | 3.149834697 | 2.536844137 |
| ENSMUSG00000037151 | 2.042557845 | 1.914552871 | 1.953445298 | 1.943696765 | 1.970729215 | 2.158362492 | 1.882886767 |
| ENSMUSG00000037152 | 1.894701088 | 1.841580554 | 1.894357207 | 1.903677461 | 1.895599581 | 2.089905111 | 1.86815666 |
| ENSMUSG00000037157 | 0 | 0 | 0 | 0 | 0 | 0 | 0 |
| ENSMUSG00000037159 | 0 | 0 | 0 | 0 | 0 | 0.301029996 | 0 |
| ENSMUSG00000037161 | 0.578129653 | 0.701838731 | 0.396240625 | 1.061981878 | 1.335652905 | 1.204119983 | 1.392431569 |
| ENSMUSG00000037166 | 0.867194479 | 0.951838731 | 1.04248125 | 0.25 | 0.387761935 | 0 | 0.224243824 |
| ENSMUSG00000037167 | 0 | 0 | 0.25 | 0 | 0 | 0 | 0.224243824 |
| ENSMUSG00000037169 | 0.289064826 | 0 | 0 | 0.5 | 0.387761935 | 0.301029996 | 0.579661877 |
| ENSMUSG00000037171 | 0 | 0 | 0 | 0.25 | 0 | 0.301029996 | 0 |
| ENSMUSG00000037172 | 1.693336259 | 1.54248125 | 1.494319981 | 1.814346961 | 1.802174723 | 1.959041392 | 1.667169465 |
| ENSMUSG00000037174 | 2.450036412 | 2.281030328 | 2.234894804 | 2.3843046 | 2.337535342 | 2.674861141 | 2.164992504 |
| ENSMUSG00000037185 | 1.771760131 | 1.766522298 | 1.830482024 | 1.958222504 | 2.058398634 | 2.23299611 | 2.03601839 |
| ENSMUSG00000037188 | 0 | 0.396240625 | 0 | 0.25 | 0.244650542 | 0 | 0.448487648 |
| ENSMUSG00000037190 | 1.678702042 | 1.711372513 | 1.671625132 | 1.671625132 | 1.802174723 | 1.633468456 | 1.720494 |
| ENSMUSG00000037196 | 0.45815691 | 0.396240625 | 0.646240625 | 0.396240625 | 0.632412477 | 0.301029996 | 0.629532004 |
| ENSMUSG00000037197 | 2.662129038 | 2.472185812 | 2.483053688 | 2.51109853 | 2.441995847 | 2.808210973 | 2.295362967 |
| ENSMUSG00000037202 | 0 | 0 | 0 | 0 | 0 | 0 | 0 |
| ENSMUSG00000037204 | 1.99653913 | 1.969129237 | 2.064346961 | 1.816696635 | 1.974687643 | 2.071882007 | 1.952571511 |
| ENSMUSG00000037206 | 1 | 1.188721876 | 0.951838731 | 1.061981878 | 0.931471446 | 1.041392685 | 1.159323753 |
| ENSMUSG00000037210 | 2.544030601 | 2.408248799 | 2.369945066 | 2.286801231 | 2.277846464 | 2.5774918 | 2.185218806 |
| ENSMUSG00000037211 | 1.269664473 | 1.372963274 | 1.372963274 | 0.701838731 | 1.020174412 | 0.954242509 | 0.952571511 |

| | | | | | | | |
|---|---|---|---|---|---|---|---|
| ENSMUSG00000037214 | 1.470606563 | 1.438721876 | 1.51109853 | 1.418106335 | 1.462344774 | 1.62324929 | 1.466365979 |
| ENSMUSG00000037216 | 1.418409478 | 1.42510993 | 1.282320754 | 1.458222504 | 1.478764298 | 1.643452676 | 1.437266334 |
| ENSMUSG00000037217 | 1.734388958 | 1.622963274 | 1.602347734 | 1.451838731 | 1.373641807 | 1.568201724 | 1.379042812 |
| ENSMUSG00000037221 | 1.587501561 | 1.695339928 | 1.77186571 | 1.458222504 | 1.698489937 | 1.556302501 | 1.63986884 |
| ENSMUSG00000037224 | 0 | 0.25 | 0.25 | 0.5 | 0.686820904 | 0.602059991 | 0.803905701 |
| ENSMUSG00000037225 | 1.663535556 | 1.606566189 | 1.54745614 | 1.348079356 | 1.343584837 | 1.361727836 | 1.20919388 |
| ENSMUSG00000037234 | 2.947656672 | 2.706239685 | 2.665444524 | 2.748411515 | 2.645793463 | 3.026941628 | 2.443774541 |
| ENSMUSG00000037235 | 2.569423023 | 2.384789703 | 2.407339155 | 2.315523711 | 2.29094062 | 2.621176282 | 2.248075634 |
| ENSMUSG00000037236 | 3.127733152 | 2.83897541 | 2.83412664 | 2.855121653 | 2.765615679 | 3.198657087 | 2.532362355 |
| ENSMUSG00000037239 | 1.482695304 | 1.445339928 | 1.580482024 | 1.593759858 | 1.542223377 | 1.653212514 | 1.459332757 |
| ENSMUSG00000037242 | 3.310886388 | 3.011525211 | 2.971614674 | 2.936208459 | 2.882215755 | 3.345373731 | 2.695163117 |
| ENSMUSG00000037243 | 1.789260561 | 1.67510993 | 1.638647213 | 1.741446071 | 1.789100779 | 1.806179974 | 1.735976162 |
| ENSMUSG00000037246 | 0 | 0 | 0 | 0 | 0 | 0 | 0 |
| ENSMUSG00000037247 | 0 | 0.25 | 0.5 | 0.25 | 0 | 0 | 0 |
| ENSMUSG00000037251 | 2.066740409 | 1.953445298 | 1.958222504 | 1.958222504 | 1.898867717 | 2.08278537 | 1.769001954 |
| ENSMUSG00000037253 | 2.751223449 | 2.516522298 | 2.488912477 | 2.43325383 | 2.335180723 | 2.666517981 | 2.154798988 |
| ENSMUSG00000037254 | 1.505869732 | 1.410964047 | 1.238549078 | 0.5 | 0.632412477 | 0 | 0.448487648 |
| ENSMUSG00000037257 | 2.307602651 | 2.164552871 | 2.188721876 | 2.263820609 | 2.259595879 | 2.588831726 | 2.13206423 |
| ENSMUSG00000037259 | 0.671187741 | 0.896240625 | 0.75 | 1.438721876 | 1.394615667 | 1.477121255 | 1.278288402 |
| ENSMUSG00000037260 | 2.254965775 | 2.039967834 | 1.990000483 | 2.158248799 | 2.054233912 | 2.365487985 | 1.945529784 |
| ENSMUSG00000037262 | 2.010213549 | 1.874961472 | 1.916333979 | 1.92510993 | 1.893954095 | 2.127104798 | 1.792684387 |
| ENSMUSG00000037263 | 0 | 0 | 0 | 0 | 0.244650542 | 0 | 0 |
| ENSMUSG00000037266 | 2.840968566 | 2.600219859 | 2.616136438 | 2.584405475 | 2.587542629 | 2.851258349 | 2.413024992 |
| ENSMUSG00000037270 | 2.703665763 | 2.479715809 | 2.445749552 | 2.689139081 | 2.606443457 | 2.94792362 | 2.372346263 |
| ENSMUSG00000037275 | 2.588343201 | 2.391037372 | 2.373963757 | 2.555094582 | 2.555125257 | 2.866877814 | 2.364209774 |
| ENSMUSG00000037278 | 2.40817504 | 2.225970461 | 2.214495249 | 2.325945187 | 2.234742351 | 2.59439255 | 2.147679267 |
| ENSMUSG00000037279 | 0 | 0 | 0.25 | 0 | 0 | 0 | 0 |
| ENSMUSG00000037280 | 0 | 0 | 0 | 0.5 | 0.632412477 | 0.477121255 | 0.448487648 |
| ENSMUSG00000037286 | 2.716227929 | 2.493961992 | 2.484159485 | 2.645300145 | 2.532092521 | 2.942999593 | 2.330894065 |
| ENSMUSG00000037287 | 2.337801042 | 2.193696765 | 2.160061234 | 2.228222334 | 2.21736452 | 2.457881897 | 2.084261301 |
| ENSMUSG00000037295 | 2.072573125 | 2.005591953 | 2.016522298 | 2.122963274 | 2.238490593 | 2.397940009 | 2.140832123 |
| ENSMUSG00000037296 | 2.129344651 | 1.976722649 | 2.038704527 | 2.110735874 | 2.074582839 | 2.361727836 | 1.928477344 |
| ENSMUSG00000037300 | 2.537433341 | 2.355516192 | 2.374462959 | 2.460587586 | 2.439894911 | 2.757396029 | 2.294288164 |
| ENSMUSG00000037306 | 2.081172036 | 1.919870025 | 2.001406137 | 1.80479213 | 1.795698282 | 1.892094603 | 1.762103709 |

| | | | | | | | |
|---|---|---|---|---|---|---|---|
| ENSMUSG00000037307 | 0 | 0 | 0 | 0 | 0 | 0 | 0 |
| ENSMUSG00000037313 | 2.938680166 | 2.680061065 | 2.689972046 | 2.7844079 | 2.715783059 | 3.13289977 | 2.522411277 |
| ENSMUSG00000037315 | 2.315764015 | 2.118933358 | 2.103406982 | 2.087182039 | 1.983754904 | 2.217483944 | 1.845261772 |
| ENSMUSG00000037316 | 2.163307908 | 1.984159485 | 1.982684334 | 2.152756199 | 2.056322416 | 2.374748346 | 1.918313719 |
| ENSMUSG00000037318 | 0 | 0.396240625 | 0 | 0.396240625 | 0.244650542 | 0 | 0.448487648 |
| ENSMUSG00000037321 | 1.069666964 | 1.17510993 | 1.114857905 | 1.649978211 | 1.828942237 | 1.77815125 | 1.792684387 |
| ENSMUSG00000037325 | 2.158622101 | 1.950224975 | 2.005591953 | 2.058404919 | 2.032639648 | 2.269512944 | 1.860017477 |
| ENSMUSG00000037326 | 1.777675583 | 1.741446071 | 1.811981878 | 1.302363341 | 1.736537397 | 1.431363764 | 1.783679399 |
| ENSMUSG00000037331 | 2.994003971 | 2.763651079 | 2.745926548 | 2.781353368 | 2.766591274 | 3.147676324 | 2.581439439 |
| ENSMUSG00000037334 | 0 | 0 | 0 | 0 | 0 | 0 | 0 |
| ENSMUSG00000037335 | 0 | 0 | 0 | 0 | 0 | 0 | 0 |
| ENSMUSG00000037336 | 0.45815691 | 1 | 1.02186571 | 1.061981878 | 1 | 1.079181246 | 1.110947927 |
| ENSMUSG00000037337 | 1.307602651 | 1.26109853 | 1.201838731 | 1 | 1.200473444 | 1.079181246 | 1.131174228 |
| ENSMUSG00000037339 | 2.141794304 | 2.083847589 | 2.156427211 | 1.854463129 | 1.950242452 | 2.021189299 | 1.97354745 |
| ENSMUSG00000037341 | 1.605636262 | 1.488549078 | 1.516522298 | 1.418106335 | 1.49445382 | 1.591064607 | 1.459332757 |
| ENSMUSG00000037343 | 2.834961347 | 2.591580554 | 2.58272922 | 2.653677461 | 2.576746587 | 2.927370363 | 2.390438245 |
| ENSMUSG00000037344 | 1.890240806 | 1.880890489 | 1.921625132 | 1.589388001 | 1.822942838 | 1.819543936 | 1.860017477 |
| ENSMUSG00000037346 | 0 | 0 | 0 | 0 | 0 | 0 | 0 |
| ENSMUSG00000037347 | 0.960252568 | 1.080482024 | 1.114857905 | 0.79248125 | 0.77552387 | 1.146128036 | 0.935079929 |
| ENSMUSG00000037348 | 1.516991255 | 1.464495249 | 1.516522298 | 1.388647213 | 1.456697355 | 1.447158031 | 1.511732338 |
| ENSMUSG00000037349 | 1.647796617 | 1.602347734 | 1.752806814 | 1.593759858 | 1.808534465 | 1.755874856 | 1.818286533 |
| ENSMUSG00000037351 | 2.786655728 | 2.62246199 | 2.64151351 | 2.680274797 | 2.717866348 | 3.049992857 | 2.56238684 |
| ENSMUSG00000037353 | 2.116511873 | 1.928561379 | 1.988549078 | 1.789967834 | 1.85749873 | 2.086359831 | 1.815933683 |
| ENSMUSG00000037355 | 2.095120908 | 1.943696765 | 1.964495249 | 1.896240625 | 1.911645804 | 1.991226076 | 1.732938436 |
| ENSMUSG00000037358 | 0 | 0 | 0 | 0 | 0 | 0 | 0 |
| ENSMUSG00000037361 | 2.445324132 | 2.254202072 | 2.242166698 | 2.311981878 | 2.222603297 | 2.527629901 | 2.086315375 |
| ENSMUSG00000037362 | 1.960252568 | 1.825945187 | 1.985628626 | 1.464495249 | 1.380772476 | 1.491361694 | 1.40946263 |
| ENSMUSG00000037363 | 1.205378646 | 1.04248125 | 1.04248125 | 1.130890489 | 0.931471446 | 1.041392685 | 0.952571511 |
| ENSMUSG00000037364 | 2.810027849 | 2.623713896 | 2.646240625 | 2.589388001 | 2.654886286 | 2.908485019 | 2.530292677 |
| ENSMUSG00000037366 | 2.089597219 | 1.969129237 | 1.964495249 | 1.828220739 | 1.824953989 | 2.041392685 | 1.764880683 |
| ENSMUSG00000037369 | 2.386920383 | 2.189555804 | 2.180274797 | 2.160964047 | 2.105788027 | 2.367355921 | 1.949460761 |
| ENSMUSG00000037370 | 1.596667477 | 1.602347734 | 1.598079356 | 1.410964047 | 1.49445382 | 1.612783857 | 1.40946263 |
| ENSMUSG00000037373 | 2.864167792 | 2.651832578 | 2.722375991 | 2.495748394 | 2.615527388 | 2.821513528 | 2.552361952 |
| ENSMUSG00000037375 | 1.53838222 | 1.53743678 | 1.532320754 | 1.552363341 | 1.49445382 | 1.662757832 | 1.46982607 |

| | | | | | | | |
|---|---|---|---|---|---|---|---|
| ENSMUSG00000037376 | 2.264212992 | 2.107613138 | 2.05357978 | 2.252806814 | 2.208682096 | 2.50242712 | 2.060206963 |
| ENSMUSG00000037379 | 0.671187741 | 0.5 | 0.701838731 | 0.396240625 | 0.244650542 | 0 | 0.224243824 |
| ENSMUSG00000037386 | 1.205378646 | 1.098079356 | 1.098079356 | 1.364857905 | 1.335652905 | 1.556302501 | 1.284450788 |
| ENSMUSG00000037390 | 0 | 0 | 0.25 | 0.25 | 0 | 0 | 0 |
| ENSMUSG00000037393 | 0 | 0.25 | 0 | 0 | 0 | 0 | 0 |
| ENSMUSG00000037395 | 1.936861443 | 1.839388001 | 1.884789703 | 1.916333979 | 1.808534465 | 2.021189299 | 1.722075986 |
| ENSMUSG00000037400 | 2.635214384 | 2.420748646 | 2.415889526 | 2.649745743 | 2.521940732 | 2.912753304 | 2.344312053 |
| ENSMUSG00000037405 | 0.45815691 | 0.79248125 | 0.580482024 | 1.226722649 | 1.319233381 | 1.447158031 | 1.345462945 |
| ENSMUSG00000037406 | 0 | 0.580482024 | 0.25 | 0 | 0.244650542 | 0 | 0.224243824 |
| ENSMUSG00000037408 | 1.924654746 | 1.841580554 | 1.926839783 | 1.854463129 | 1.950242452 | 2.120573931 | 1.845261772 |
| ENSMUSG00000037410 | 2.215778996 | 2.143411797 | 2.198603967 | 2.066696635 | 2.132784263 | 2.394451681 | 2.015657029 |
| ENSMUSG00000037411 | 3.100818499 | 2.836766914 | 2.837872852 | 2.43325383 | 2.444777781 | 2.769377326 | 2.276587178 |
| ENSMUSG00000037415 | 2.550525119 | 2.344302633 | 2.362802778 | 2.399978211 | 2.412899576 | 2.67669361 | 2.287488514 |
| ENSMUSG00000037416 | 2.707474304 | 2.491085217 | 2.485994979 | 2.531030328 | 2.446849937 | 2.751279104 | 2.264421331 |
| ENSMUSG00000037418 | 0.289064826 | 0.25 | 0.25 | 0 | 0 | 0 | 0.224243824 |
| ENSMUSG00000037419 | 1.663535556 | 1.618933358 | 1.606566189 | 1.979715809 | 1.923977404 | 2.152288344 | 1.801445495 |
| ENSMUSG00000037424 | 0 | 0 | 0 | 0 | 0 | 0 | 0 |
| ENSMUSG00000037426 | 2.436947303 | 2.237091808 | 2.227473271 | 2.212155735 | 2.16097133 | 2.445604203 | 2.047762527 |
| ENSMUSG00000037428 | 0.289064826 | 0 | 0 | 0 | 0 | 0 | 0.224243824 |
| ENSMUSG00000037432 | 1.494443473 | 1.431980114 | 1.380890489 | 1.598079356 | 1.563883923 | 1.77815125 | 1.508694612 |
| ENSMUSG00000037434 | 1.920505136 | 1.789967834 | 1.75 | 1.763820609 | 1.770879644 | 1.897627091 | 1.591689859 |
| ENSMUSG00000037437 | 0.289064826 | 0.580482024 | 0.396240625 | 0 | 0.568060967 | 0.602059991 | 0.224243824 |
| ENSMUSG00000037438 | 0 | 0 | 0 | 0 | 0 | 0 | 0 |
| ENSMUSG00000037440 | 0.671187741 | 0.646240625 | 0.79248125 | 1.226722649 | 1.020174412 | 1.176091259 | 0.984950056 |
| ENSMUSG00000037443 | 2.325351389 | 2.207430684 | 2.176839783 | 2.200224975 | 2.181274723 | 2.462397998 | 2.103753068 |
| ENSMUSG00000037446 | 0.289064826 | 0.5 | 0.580482024 | 0.5 | 0.489301084 | 0 | 0.448487648 |
| ENSMUSG00000037447 | 1.37447073 | 1.458222504 | 1.403677461 | 1.678561379 | 1.77552387 | 1.919078092 | 1.746389053 |
| ENSMUSG00000037451 | 0 | 0 | 0 | 0 | 0 | 0 | 0 |
| ENSMUSG00000037455 | 1.975148165 | 1.903677461 | 1.856566189 | 2.058404919 | 1.992595054 | 2.274157849 | 1.845261772 |
| ENSMUSG00000037458 | 3.248109354 | 2.95594132 | 2.944519282 | 3.039336734 | 2.936753115 | 3.411114419 | 2.70506625 |
| ENSMUSG00000037461 | 2.661423995 | 2.414998973 | 2.457826811 | 2.509386738 | 2.474650913 | 2.771587481 | 2.353406551 |
| ENSMUSG00000037463 | 0 | 0.25 | 0 | 0 | 0 | 0 | 0.224243824 |
| ENSMUSG00000037465 | 2.396352858 | 2.243603647 | 2.228222334 | 2.025821952 | 2.030405741 | 2.307496038 | 1.900630878 |
| ENSMUSG00000037466 | 2.299278375 | 2.167221246 | 2.226722649 | 2.012462137 | 2.09661982 | 2.255272505 | 1.994607951 |

| | | | | | | | |
|---|---|---|---|---|---|---|---|
| ENSMUSG00000037469 | 0 | 0 | 0 | 0 | 0 | 0 | 0 |
| ENSMUSG00000037470 | 3.015076955 | 2.727847997 | 2.707430684 | 2.912314044 | 2.843677054 | 3.27600199 | 2.634835654 |
| ENSMUSG00000037474 | 2.673935906 | 2.478222334 | 2.444929839 | 2.644357207 | 2.531009002 | 2.923244019 | 2.366370874 |
| ENSMUSG00000037475 | 2.874031229 | 2.62019755 | 2.629417347 | 2.687257846 | 2.61424391 | 2.974971994 | 2.37800557 |
| ENSMUSG00000037477 | 0 | 0 | 0 | 0 | 0 | 0 | 0 |
| ENSMUSG00000037482 | 0 | 0 | 0 | 0 | 0 | 0 | 0 |
| ENSMUSG00000037486 | 2.521950082 | 2.367918804 | 2.332168732 | 2.348615674 | 2.278401864 | 2.574031268 | 2.075912052 |
| ENSMUSG00000037487 | 3.398406911 | 3.078858159 | 3.055698726 | 3.058928774 | 2.981112157 | 3.442793226 | 2.766947881 |
| ENSMUSG00000037490 | 0.289064826 | 0.396240625 | 0 | 0.75 | 0.733951626 | 0.477121255 | 0.672731473 |
| ENSMUSG00000037492 | 0 | 0 | 0.25 | 0.25 | 0.244650542 | 0 | 0.355418052 |
| ENSMUSG00000037493 | 1.794934559 | 1.732684334 | 1.80479213 | 2.05479213 | 2.041436517 | 2.301029996 | 1.892346957 |
| ENSMUSG00000037499 | 1.885732304 | 1.766522298 | 1.880890489 | 1.723704441 | 1.744182955 | 1.886490725 | 1.747849621 |
| ENSMUSG00000037503 | 3.233154048 | 2.943078643 | 2.925218289 | 2.899280357 | 2.854690619 | 3.272073788 | 2.65899184 |
| ENSMUSG00000037509 | 1.156259305 | 1.282320754 | 1.226722649 | 1.25 | 1.343584837 | 1.447158031 | 1.335191751 |
| ENSMUSG00000037514 | 2.217828299 | 2.013820609 | 1.995748394 | 2.178561379 | 2.122577592 | 2.404833717 | 1.966211184 |
| ENSMUSG00000037519 | 2.553277818 | 2.391037372 | 2.3843046 | 2.345926073 | 2.326087113 | 2.613841822 | 2.19230473 |
| ENSMUSG00000037523 | 2.111266113 | 1.935366747 | 1.953445298 | 2.048689214 | 2.0510778 | 2.350248018 | 1.987739 |
| ENSMUSG00000037525 | 1.647796617 | 1.575945187 | 1.580482024 | 1.356566189 | 1.394615667 | 1.505149978 | 1.441050184 |
| ENSMUSG00000037526 | 1.707474304 | 1.532320754 | 1.695339928 | 1.789967834 | 1.726078673 | 2.056904851 | 1.707551656 |
| ENSMUSG00000037529 | 0 | 0 | 0 | 0 | 0 | 0 | 0 |
| ENSMUSG00000037531 | 2.029919532 | 1.858657057 | 1.816696635 | 1.921625132 | 1.875328588 | 2.079181246 | 1.759302692 |
| ENSMUSG00000037533 | 2.590021402 | 2.399513125 | 2.345385738 | 2.561388177 | 2.489301084 | 2.835690571 | 2.307473178 |
| ENSMUSG00000037535 | 0 | 0 | 0 | 0 | 0.244650542 | 0 | 0 |
| ENSMUSG00000037536 | 1.822201287 | 1.705044741 | 1.738549078 | 1.92510993 | 1.916320788 | 2.120573931 | 1.774416565 |
| ENSMUSG00000037537 | 0 | 0 | 0 | 0 | 0 | 0 | 0 |
| ENSMUSG00000037541 | 1.325351389 | 1.226722649 | 1.445339928 | 1.388647213 | 1.351342427 | 1.462397998 | 1.324583736 |
| ENSMUSG00000037542 | 0 | 0.25 | 0 | 0 | 0 | 0 | 0 |
| ENSMUSG00000037544 | 2.479432974 | 2.308404919 | 2.298689214 | 2.569612365 | 2.499235686 | 2.912753304 | 2.363123786 |
| ENSMUSG00000037548 | 0.45815691 | 0.25 | 0.5 | 0.830482024 | 0.686820904 | 0.602059991 | 0.520678036 |
| ENSMUSG00000037552 | 2.410757293 | 2.267194569 | 2.325945187 | 1.953445298 | 2.027028145 | 2.222716471 | 2.019455688 |
| ENSMUSG00000037553 | 2.199011616 | 2.095926073 | 2.174241882 | 2.015173983 | 2.160196454 | 2.269512944 | 2.112367894 |
| ENSMUSG00000037563 | 2.792365035 | 2.619186551 | 2.630890489 | 2.504202072 | 2.54274861 | 2.875639937 | 2.451647167 |
| ENSMUSG00000037568 | 2.381433035 | 2.165444524 | 2.166333979 | 2.086073977 | 2.093822267 | 2.361727836 | 1.978585612 |
| ENSMUSG00000037570 | 2.506893125 | 2.421625132 | 2.435366747 | 2.219129237 | 2.389791591 | 2.629409599 | 2.331134329 |

| | | | | | | | |
|---|---|---|---|---|---|---|---|
| ENSMUSG00000037572 | 2.53552908 | 2.350753006 | 2.304186465 | 2.572504712 | 2.478764298 | 2.833147112 | 2.314375784 |
| ENSMUSG00000037573 | 2.348397716 | 2.173371739 | 2.220660762 | 2.002806814 | 1.961316494 | 2.269512944 | 1.913982791 |
| ENSMUSG00000037577 | 0.289064826 | 0.25 | 0.25 | 0 | 0.387761935 | 0.301029996 | 0.448487648 |
| ENSMUSG00000037578 | 0.289064826 | 0 | 0 | 0.25 | 0 | 0 | 0 |
| ENSMUSG00000037579 | 0 | 0 | 0 | 0.701838731 | 0.733951626 | 0.301029996 | 0.710836105 |
| ENSMUSG00000037580 | 1.181541736 | 1.130890489 | 1.146240625 | 1.403677461 | 1.335652905 | 1.380211242 | 1.401059159 |
| ENSMUSG00000037583 | 0 | 0.25 | 0 | 0 | 0 | 0 | 0 |
| ENSMUSG00000037593 | 0.578129653 | 0.25 | 0.646240625 | 0.830482024 | 0.846351821 | 0.602059991 | 0.672731473 |
| ENSMUSG00000037594 | 1.753493018 | 1.664552871 | 1.752806814 | 1.566696635 | 1.614672255 | 1.763427994 | 1.537859885 |
| ENSMUSG00000037600 | 0.45815691 | 0.25 | 0 | 0.75 | 0.632412477 | 0.77815125 | 0.775756176 |
| ENSMUSG00000037601 | 2.789779581 | 2.578504176 | 2.577937079 | 2.633818844 | 2.570737655 | 2.976349979 | 2.390038102 |
| ENSMUSG00000037603 | 0 | 0 | 0 | 0 | 0 | 0 | 0 |
| ENSMUSG00000037605 | 1.494443473 | 1.26109853 | 1.27186571 | 0.864857905 | 0.812711509 | 0.84509804 | 0.935079929 |
| ENSMUSG00000037606 | 2.051791411 | 2.04745614 | 2.04745614 | 1.975216702 | 2.063536368 | 2.173186268 | 2.01501956 |
| ENSMUSG00000037608 | 3.038006286 | 2.7844079 | 2.772858847 | 2.950022747 | 2.847571014 | 3.304705898 | 2.645279997 |
| ENSMUSG00000037610 | 0 | 0 | 0 | 0 | 0 | 0 | 0 |
| ENSMUSG00000037613 | 2.196867486 | 2.032320754 | 2.038704527 | 1.896240625 | 1.855665188 | 2.149219113 | 1.729871916 |
| ENSMUSG00000037617 | 1.54867984 | 1.53743678 | 1.348079356 | 1.561981878 | 1.546663136 | 1.568201724 | 1.466365979 |
| ENSMUSG00000037621 | 2.060824957 | 1.992885888 | 2.028435542 | 1.438721876 | 1.542223377 | 1.556302501 | 1.601000298 |
| ENSMUSG00000037622 | 2.227926429 | 2.157339155 | 2.225216702 | 2.01109853 | 2.207327177 | 2.376576957 | 2.182568577 |
| ENSMUSG00000037624 | 1.307602651 | 1.321350555 | 1.25 | 1.160964047 | 1.234113756 | 1.113943352 | 1.100339912 |
| ENSMUSG00000037625 | 0 | 0 | 0 | 0 | 0 | 0 | 0 |
| ENSMUSG00000037627 | 0 | 0 | 0 | 0.25 | 0.244650542 | 0 | 0.224243824 |
| ENSMUSG00000037628 | 1.470606563 | 1.482684334 | 1.470660762 | 1.685366747 | 1.649303553 | 1.857332496 | 1.591689859 |
| ENSMUSG00000037636 | 1.129344651 | 1.25 | 1.201838731 | 0.864857905 | 0.812711509 | 0.301029996 | 0.775756176 |
| ENSMUSG00000037638 | 1.418409478 | 1.418106335 | 1.476722649 | 1.098079356 | 1.343584837 | 1.230448921 | 1.340368108 |
| ENSMUSG00000037640 | 2 | 1.843759858 | 1.852347734 | 1.945339928 | 1.861137485 | 2.096910013 | 1.755055163 |
| ENSMUSG00000037643 | 2.884080688 | 2.651832578 | 2.614601653 | 2.859177886 | 2.754702294 | 3.190611798 | 2.58276623 |
| ENSMUSG00000037646 | 2.541215974 | 2.331045137 | 2.351816941 | 2.443284802 | 2.372368362 | 2.717670503 | 2.213713783 |
| ENSMUSG00000037649 | 1.205378646 | 1.311981878 | 1.226722649 | 1.214495249 | 1.27449559 | 1.431363764 | 1.272006351 |
| ENSMUSG00000037652 | 2.688735382 | 2.472566069 | 2.453445298 | 2.577084758 | 2.463003274 | 2.835690571 | 2.292939619 |
| ENSMUSG00000037653 | 0 | 0 | 0 | 0 | 0 | 0 | 0 |
| ENSMUSG00000037656 | 2.435736758 | 2.291854536 | 2.331607372 | 2.069031101 | 2.141887974 | 2.298853076 | 2.065709059 |
| ENSMUSG00000037660 | 0 | 0 | 0 | 0 | 0 | 0 | 0 |

| | | | | | | | |
|---|---|---|---|---|---|---|---|
| ENSMUSG00000037661 | 0.747221736 | 0.580482024 | 0.5 | 0.25 | 0.686820904 | 0.602059991 | 0.672731473 |
| ENSMUSG00000037664 | 0 | 0 | 0.25 | 1.061981878 | 0.978602168 | 1.041392685 | 1.110947927 |
| ENSMUSG00000037669 | 2.445324132 | 2.259729747 | 2.234894804 | 2.201032755 | 2.16174451 | 2.445604203 | 2.01116814 |
| ENSMUSG00000037674 | 2.436947303 | 2.243603647 | 2.21293726 | 2.462155735 | 2.371090306 | 2.72427587 | 2.209893373 |
| ENSMUSG00000037679 | 2.811999637 | 2.641989019 | 2.735994979 | 2.176839783 | 2.371942857 | 2.530199698 | 2.376130113 |
| ENSMUSG00000037681 | 0 | 0.25 | 0.25 | 0.79248125 | 0.568060967 | 0.602059991 | 0.579661877 |
| ENSMUSG00000037683 | 0 | 0 | 0 | 0 | 0 | 0 | 0 |
| ENSMUSG00000037685 | 1.721148722 | 1.575945187 | 1.54248125 | 2.174241882 | 2.110284345 | 2.403120521 | 1.964723754 |
| ENSMUSG00000037686 | 0 | 0 | 0 | 0.5 | 0.733951626 | 0.77815125 | 0.74492186 |
| ENSMUSG00000037689 | 0 | 0 | 0 | 0 | 0 | 0 | 0 |
| ENSMUSG00000037692 | 1.989530018 | 2.017865591 | 2.164552871 | 1.418106335 | 1.671670246 | 1.633468456 | 1.845261772 |
| ENSMUSG00000037697 | 1.765759561 | 1.566696635 | 1.638647213 | 1.96759118 | 1.919403394 | 2.049218023 | 1.757893036 |
| ENSMUSG00000037703 | 1.881174527 | 1.854463129 | 1.87892496 | 1.930274797 | 2.063536368 | 2.173186268 | 2.043700396 |
| ENSMUSG00000037705 | 0 | 0.25 | 0.701838731 | 0 | 0 | 0.301029996 | 0 |
| ENSMUSG00000037706 | 2.777675583 | 2.547147211 | 2.532642641 | 2.223704441 | 2.25010405 | 2.557507202 | 2.174874624 |
| ENSMUSG00000037708 | 0.45815691 | 0.5 | 0.396240625 | 0 | 0.244650542 | 0.301029996 | 0.520678036 |
| ENSMUSG00000037709 | 0.578129653 | 0.701838731 | 0.701838731 | 0.701838731 | 0.686820904 | 0.602059991 | 0.579661877 |
| ENSMUSG00000037710 | 2.256831695 | 2.067865757 | 2.116901388 | 2.193696765 | 2.176121988 | 2.432969291 | 2.059651572 |
| ENSMUSG00000037712 | 2.983054229 | 2.758527787 | 2.735078399 | 2.794821776 | 2.704182902 | 3.099335278 | 2.512144372 |
| ENSMUSG00000037716 | 0 | 0 | 0 | 0.396240625 | 0.489301084 | 0.301029996 | 0.629532004 |
| ENSMUSG00000037720 | 2.729023206 | 2.543731421 | 2.538704527 | 2.599418658 | 2.533172725 | 2.92992956 | 2.344312053 |
| ENSMUSG00000037722 | 2.111266113 | 1.926839783 | 1.860735874 | 2.195339928 | 2.085294053 | 2.367355921 | 1.927642465 |
| ENSMUSG00000037725 | 2.686083339 | 2.473704441 | 2.476722649 | 2.418106335 | 2.349985771 | 2.697229343 | 2.190083792 |
| ENSMUSG00000037727 | 0 | 0 | 0 | 0 | 0 | 0 | 0 |
| ENSMUSG00000037730 | 2.219867581 | 2.02053726 | 2.041226732 | 2.224461364 | 2.130260132 | 2.426511261 | 2.002662694 |
| ENSMUSG00000037731 | 0 | 0.25 | 0 | 0 | 0.244650542 | 0 | 0.710836105 |
| ENSMUSG00000037735 | 0 | 0 | 0 | 0.25 | 0 | 0 | 0 |
| ENSMUSG00000037736 | 1.700465191 | 1.653677461 | 1.653677461 | 1 | 0.812711509 | 1.041392685 | 0.896975297 |
| ENSMUSG00000037737 | 0 | 0 | 0 | 0.25 | 0.387761935 | 0.477121255 | 0.448487648 |
| ENSMUSG00000037738 | 0 | 0 | 0.396240625 | 0 | 0 | 0.301029996 | 0 |
| ENSMUSG00000037740 | 2.016886209 | 1.87892496 | 2.02186571 | 2.001406137 | 2.033751322 | 2.269512944 | 2.054609671 |
| ENSMUSG00000037742 | 4.334250725 | 3.878038508 | 3.860428056 | 3.936267568 | 3.834051018 | 4.540379535 | 3.553060921 |
| ENSMUSG00000037747 | 0.45815691 | 0.580482024 | 0.5 | 0.896240625 | 0.686820904 | 0.602059991 | 0.672731473 |
| ENSMUSG00000037750 | 2.179305632 | 2.02053726 | 2.092671852 | 2.032320754 | 2.027028145 | 2.193124598 | 1.976436 |

| | | | | | | | |
|---|---|---|---|---|---|---|---|
| ENSMUSG00000037752 | 1.707474304 | 1.708222504 | 1.614857905 | 1.638647213 | 1.645989387 | 1.851258349 | 1.605556929 |
| ENSMUSG00000037754 | 0.45815691 | 0.396240625 | 0.25 | 0.5 | 0.387761935 | 0.477121255 | 0.355418052 |
| ENSMUSG00000037759 | 0 | 0 | 0 | 0.25 | 0 | 0 | 0 |
| ENSMUSG00000037761 | 1.249317394 | 1.27186571 | 1.410964047 | 1.26109853 | 1.514343934 | 1.662757832 | 1.582103514 |
| ENSMUSG00000037762 | 0.867194479 | 0.896240625 | 0.830482024 | 1.02186571 | 1.039257769 | 0.903089987 | 0.829800754 |
| ENSMUSG00000037771 | 0 | 0 | 0 | 0 | 0 | 0 | 0 |
| ENSMUSG00000037772 | 1.358731791 | 1.311981878 | 1.410964047 | 1.201838731 | 1.188507684 | 1.414973348 | 1.20919388 |
| ENSMUSG00000037773 | 1.890240806 | 1.747171172 | 1.819031101 | 1.807204673 | 1.802174723 | 1.919078092 | 1.764880683 |
| ENSMUSG00000037780 | 0 | 0 | 0 | 0 | 0 | 0 | 0 |
| ENSMUSG00000037784 | 2.489144387 | 2.328787391 | 2.339388001 | 2.269203899 | 2.26997769 | 2.450249108 | 2.139097452 |
| ENSMUSG00000037787 | 1.777675583 | 1.630890489 | 1.664552871 | 1.445339928 | 1.514343934 | 1.556302501 | 1.455757965 |
| ENSMUSG00000037788 | 2.384185734 | 2.174241882 | 2.177701608 | 2.092671852 | 2.058398634 | 2.326335861 | 1.896975297 |
| ENSMUSG00000037790 | 0 | 0 | 0 | 0 | 0 | 0 | 0 |
| ENSMUSG00000037791 | 2.495495257 | 2.311388177 | 2.365369862 | 2.196158711 | 2.247084732 | 2.444044796 | 2.164590871 |
| ENSMUSG00000037795 | 2.419671301 | 2.28360658 | 2.226722649 | 2.28296424 | 2.187072901 | 2.50242712 | 2.017561934 |
| ENSMUSG00000037797 | 0 | 0 | 0 | 0 | 0 | 0 | 0 |
| ENSMUSG00000037798 | 0 | 0 | 0 | 0 | 0 | 0 | 0 |
| ENSMUSG00000037801 | 0.45815691 | 0.5 | 0.25 | 0.25 | 0 | 0.477121255 | 0.579661877 |
| ENSMUSG00000037805 | 2.799013148 | 2.568157446 | 2.579070383 | 2.503156135 | 2.538790182 | 2.836324116 | 2.424384006 |
| ENSMUSG00000037808 | 2.410757293 | 2.21759118 | 2.19697564 | 2.307805295 | 2.261929482 | 2.603144373 | 2.074853081 |
| ENSMUSG00000037813 | 0.671187741 | 0.396240625 | 1.04248125 | 0.75 | 0.877063019 | 0.77815125 | 0.829800754 |
| ENSMUSG00000037815 | 3.297132669 | 3.002019603 | 3.006284891 | 2.932935823 | 2.899632312 | 3.329397879 | 2.727552799 |
| ENSMUSG00000037816 | 1.952600383 | 1.874961472 | 1.854463129 | 1.821350555 | 1.866526233 | 2.096910013 | 1.762103709 |
| ENSMUSG00000037818 | 1.307602651 | 1.302363341 | 1.29248125 | 1.476722649 | 1.47337555 | 1.544068044 | 1.405288179 |
| ENSMUSG00000037820 | 2.703665763 | 2.541226732 | 2.531030328 | 1.839388001 | 1.919403394 | 2.033423755 | 1.885753983 |
| ENSMUSG00000037822 | 2.393679564 | 2.227473271 | 2.19697564 | 2.187886015 | 2.138604637 | 2.432969291 | 2.019455688 |
| ENSMUSG00000037824 | 2.134369199 | 1.942046081 | 1.994319981 | 2.094844592 | 2.122577592 | 2.356025857 | 2.01116814 |
| ENSMUSG00000037826 | 1.358731791 | 1.380890489 | 1.214495249 | 1.527131114 | 1.533172725 | 1.643452676 | 1.444790289 |
| ENSMUSG00000037827 | 0 | 0 | 0 | 0 | 0 | 0 | 0 |
| ENSMUSG00000037833 | 0 | 0.25 | 0 | 0.25 | 0.244650542 | 0.301029996 | 0 |
| ENSMUSG00000037843 | 0 | 0 | 0 | 0 | 0 | 0 | 0 |
| ENSMUSG00000037845 | 1.269664473 | 1.282320754 | 1.330482024 | 1.364857905 | 1.358933174 | 1.544068044 | 1.307989563 |
| ENSMUSG00000037846 | 1.605636262 | 1.575945187 | 1.614857905 | 1.692046081 | 1.709785497 | 1.832508913 | 1.556816002 |
| ENSMUSG00000037847 | 1.794934559 | 1.614857905 | 1.668106335 | 1.476722649 | 1.499532413 | 1.602059991 | 1.345462945 |

| | | | | | | | |
|---|---|---|---|---|---|---|---|
| ENSMUSG00000037849 | 0 | 0 | 0 | 0 | 0 | 0 | 0 |
| ENSMUSG00000037851 | 3.120755314 | 2.828928915 | 2.768368038 | 2.91997997 | 2.867782306 | 3.319314304 | 2.659079006 |
| ENSMUSG00000037852 | 3.411809077 | 3.085865836 | 3.06728167 | 2.896827182 | 2.835299896 | 3.271841607 | 2.653452508 |
| ENSMUSG00000037855 | 2.095120908 | 2.013820609 | 2.023189285 | 1.828220739 | 1.814781639 | 1.934498451 | 1.68886578 |
| ENSMUSG00000037857 | 2.669806838 | 2.469129237 | 2.464883697 | 2.523519421 | 2.410235744 | 2.755112266 | 2.224243824 |
| ENSMUSG00000037860 | 0 | 0 | 0.25 | 0.5 | 0 | 0 | 0 |
| ENSMUSG00000037868 | 0.578129653 | 0.976722649 | 1 | 0.396240625 | 0.244650542 | 0.477121255 | 0.74492186 |
| ENSMUSG00000037872 | 0 | 0 | 0 | 0.396240625 | 0 | 0 | 0 |
| ENSMUSG00000037876 | 2.747795766 | 2.535845803 | 2.496460484 | 2.594844592 | 2.527190274 | 2.900367129 | 2.340834605 |
| ENSMUSG00000037884 | 0 | 0 | 0 | 0 | 0 | 0 | 0 |
| ENSMUSG00000037885 | 2.219867581 | 2.110735874 | 2.111770807 | 2.187048212 | 2.186353316 | 2.440909082 | 2.06842541 |
| ENSMUSG00000037887 | 1.596667477 | 1.634789703 | 1.54248125 | 1.769203899 | 1.842556557 | 1.924279286 | 1.732938436 |
| ENSMUSG00000037890 | 2.165631214 | 2 | 2.002806814 | 2.004202072 | 1.929985312 | 2.204119983 | 1.874129423 |
| ENSMUSG00000037892 | 2.508932407 | 2.32991803 | 2.315523711 | 2.388647213 | 2.359815565 | 2.654176542 | 2.157274883 |
| ENSMUSG00000037894 | 2.913624518 | 2.688304187 | 2.70584181 | 2.792794199 | 2.744182955 | 3.159266331 | 2.6191499 |
| ENSMUSG00000037896 | 2.760228547 | 2.56932185 | 2.51685859 | 2.492166698 | 2.462674177 | 2.838219222 | 2.276018109 |
| ENSMUSG00000037902 | 1.663535556 | 1.438721876 | 1.566696635 | 2.174241882 | 2.192777367 | 2.42975228 | 2.042530357 |
| ENSMUSG00000037904 | 1.181541736 | 1.311981878 | 1.396240625 | 1.02186571 | 1.136121934 | 1.146128036 | 1.340368108 |
| ENSMUSG00000037905 | 2.439357901 | 2.222945926 | 2.275821952 | 2.149047439 | 2.171644683 | 2.40654018 | 2.038994634 |
| ENSMUSG00000037907 | 2.126809492 | 2.107613138 | 2.194519282 | 1.610735874 | 1.840643511 | 1.72427587 | 1.830929954 |
| ENSMUSG00000037910 | 0 | 0 | 0 | 0 | 0 | 0 | 0 |
| ENSMUSG00000037913 | 0 | 0 | 0 | 0 | 0 | 0 | 0 |
| ENSMUSG00000037916 | 2.748369006 | 2.546838018 | 2.586351312 | 2.429847205 | 2.554871788 | 2.814913181 | 2.442925641 |
| ENSMUSG00000037921 | 0.289064826 | 0 | 0 | 0.646240625 | 0.812711509 | 0.602059991 | 0.710836105 |
| ENSMUSG00000037922 | 0.289064826 | 0 | 0 | 0 | 0.244650542 | 0 | 0 |
| ENSMUSG00000037924 | 0 | 0 | 0 | 0 | 0 | 0 | 0 |
| ENSMUSG00000037926 | 2.392336464 | 2.195339928 | 2.191217898 | 2.296838018 | 2.198391101 | 2.531478917 | 2.10131878 |
| ENSMUSG00000037933 | 2.77230138 | 2.558105232 | 2.604463129 | 2.600753006 | 2.603141692 | 2.939519253 | 2.497832161 |
| ENSMUSG00000037935 | 2.721148722 | 2.503505118 | 2.486726569 | 2.569031101 | 2.49625462 | 2.839478047 | 2.355640783 |
| ENSMUSG00000037936 | 2.744917684 | 2.525493918 | 2.5684489 | 2.529410775 | 2.561403546 | 2.918030337 | 2.460139731 |
| ENSMUSG00000037938 | 1.671187741 | 1.527131114 | 1.584962501 | 1.527131114 | 1.603583716 | 1.792391689 | 1.548827561 |
| ENSMUSG00000037940 | 0.45815691 | 0.580482024 | 0 | 0.25 | 0.387761935 | 0.477121255 | 0.224243824 |
| ENSMUSG00000037942 | 0 | 0 | 0 | 0 | 0.244650542 | 0 | 0 |
| ENSMUSG00000037944 | 0 | 0 | 0 | 0.25 | 0 | 0 | 0 |

| | | | | | | | |
|---|---|---|---|---|---|---|---|
| ENSMUSG00000037946 | 2.813472384 | 2.601019284 | 2.623963757 | 2.42813175 | 2.455355315 | 2.778874472 | 2.312345877 |
| ENSMUSG00000037949 | 2.460448303 | 2.273189285 | 2.237821179 | 2.314346961 | 2.317747248 | 2.666517981 | 2.180662175 |
| ENSMUSG00000037953 | 0 | 0.25 | 0.25 | 0 | 0 | 0.301029996 | 0.224243824 |
| ENSMUSG00000037957 | 1.993049299 | 1.907339155 | 1.882845365 | 1.860735874 | 1.816839698 | 1.982271233 | 1.725216937 |
| ENSMUSG00000037958 | 2.22592627 | 2.051142786 | 2.094844592 | 2.185366747 | 2.11997913 | 2.457881897 | 1.977870661 |
| ENSMUSG00000037960 | 1.920505136 | 1.830482024 | 2.006976499 | 1.77186571 | 1.911645804 | 2 | 1.975716277 |
| ENSMUSG00000037962 | 0.747221736 | 0.580482024 | 0.580482024 | 0.5 | 0.632412477 | 0.698970004 | 0.520678036 |
| ENSMUSG00000037965 | 2.452372734 | 2.28296424 | 2.259729747 | 2.369439567 | 2.327538614 | 2.588831726 | 2.158505762 |
| ENSMUSG00000037966 | 1.747221736 | 1.794977273 | 1.942046081 | 1.848079356 | 1.969399812 | 2.056904851 | 2.044283833 |
| ENSMUSG00000037971 | 1.432083896 | 1.396240625 | 1.339388001 | 1.532320754 | 1.467903253 | 1.556302501 | 1.38802993 |
| ENSMUSG00000037972 | 2.551444705 | 2.333847589 | 2.370449858 | 2.230460234 | 2.180543208 | 2.437750563 | 2.044866219 |
| ENSMUSG00000037973 | 0 | 0 | 0 | 0 | 0 | 0 | 0 |
| ENSMUSG00000037974 | 0 | 0 | 0 | 0 | 0 | 0 | 0.224243824 |
| ENSMUSG00000037977 | 0.747221736 | 0.701838731 | 0.79248125 | 0.25 | 0.686820904 | 0.477121255 | 0.876096088 |
| ENSMUSG00000037979 | 1.727821383 | 1.714495249 | 1.777131114 | 1.52186571 | 1.652586889 | 1.806179974 | 1.726776051 |
| ENSMUSG00000037982 | 0 | 0 | 0 | 0 | 0 | 0 | 0 |
| ENSMUSG00000037984 | 0 | 0 | 0 | 0 | 0 | 0 | 0 |
| ENSMUSG00000037989 | 0 | 0 | 0.25 | 0.896240625 | 1.302012594 | 0.77815125 | 1.168187745 |
| ENSMUSG00000037990 | 1.358731791 | 1.418106335 | 1.482684334 | 1 | 0.955822902 | 1.041392685 | 1.041356071 |
| ENSMUSG00000037991 | 1.494443473 | 1.505591953 | 1.464495249 | 1.51109853 | 1.47337555 | 1.602059991 | 1.413583902 |
| ENSMUSG00000037992 | 2.203267085 | 2.122963274 | 2.205044741 | 1.561981878 | 1.77552387 | 1.763427994 | 1.826389311 |
| ENSMUSG00000037993 | 2.671187741 | 2.483791262 | 2.53360658 | 2.374462959 | 2.522218979 | 2.719331287 | 2.384383506 |
| ENSMUSG00000037994 | 0.45815691 | 0.580482024 | 0.580482024 | 0.5 | 0.244650542 | 0.301029996 | 0.448487648 |
| ENSMUSG00000037995 | 1.852729478 | 1.766522298 | 1.823655187 | 0.92510993 | 1.176121988 | 1.079181246 | 1.185218806 |
| ENSMUSG00000037996 | 0 | 0 | 0 | 0 | 0 | 0 | 0 |
| ENSMUSG00000037997 | 2.016886209 | 1.868933358 | 1.834962501 | 1.955044741 | 1.855665188 | 2.133538908 | 1.778418869 |
| ENSMUSG00000037999 | 1.45815691 | 1.348079356 | 1.431980114 | 1.698603967 | 1.592135487 | 1.770852012 | 1.56970677 |
| ENSMUSG00000038000 | 2.22189694 | 2.104463129 | 2.155512955 | 2.060793496 | 2.09661982 | 2.344392274 | 2.041356071 |
| ENSMUSG00000038002 | 2.45469604 | 2.246460484 | 2.231944491 | 2.266522298 | 2.210704773 | 2.432969291 | 2.089878917 |
| ENSMUSG00000038005 | 2.3855553 | 2.206637122 | 2.201838731 | 2.275821952 | 2.145141048 | 2.496929648 | 2.046027856 |
| ENSMUSG00000038007 | 0.811507563 | 0.701838731 | 0.701838731 | 1.04248125 | 1 | 1.146128036 | 1 |
| ENSMUSG00000038009 | 0.867194479 | 0.896240625 | 0.864857905 | 0.701838731 | 0.877063019 | 0.77815125 | 0.896975297 |
| ENSMUSG00000038010 | 1.605636262 | 1.403677461 | 1.380890489 | 1.470660762 | 1.319233381 | 1.301029996 | 1.252393349 |
| ENSMUSG00000038011 | 0.45815691 | 0.396240625 | 0.79248125 | 0.5 | 0.632412477 | 0.477121255 | 0.520678036 |

| | | | | | | | |
|---|---|---|---|---|---|---|---|
| ENSMUSG00000038013 | 2.317377295 | 2.132845365 | 2.14151351 | 2.186208459 | 2.157079823 | 2.421603927 | 2.051204213 |
| ENSMUSG00000038014 | 3.160118759 | 2.919870025 | 2.933041856 | 2.914106216 | 2.887195467 | 3.290702243 | 2.720782209 |
| ENSMUSG00000038015 | 0 | 0 | 0 | 0 | 0 | 0 | 0.224243824 |
| ENSMUSG00000038020 | 1.631440309 | 1.482684334 | 1.482684334 | 1.52186571 | 1.607318736 | 1.62324929 | 1.610050272 |
| ENSMUSG00000038022 | 1.578129653 | 1.575945187 | 1.653677461 | 1.5 | 1.489301084 | 1.643452676 | 1.466365979 |
| ENSMUSG00000038023 | 2.723587515 | 2.510414788 | 2.51584877 | 2.447383411 | 2.424827423 | 2.741939078 | 2.276587178 |
| ENSMUSG00000038024 | 2.56765967 | 2.377938163 | 2.373463862 | 2.446158711 | 2.347713026 | 2.683947131 | 2.189712149 |
| ENSMUSG00000038025 | 2.483777101 | 2.370953944 | 2.385757955 | 2.284887838 | 2.356272688 | 2.620136055 | 2.285835109 |
| ENSMUSG00000038026 | 0 | 0 | 0 | 0 | 0 | 0 | 0 |
| ENSMUSG00000038028 | 2.026699189 | 1.870953944 | 1.961372513 | 1.901832578 | 1.890639929 | 2.064457989 | 1.770364127 |
| ENSMUSG00000038034 | 1.867194479 | 1.784887838 | 1.935366747 | 1.410964047 | 1.652586889 | 1.568201724 | 1.774416565 |
| ENSMUSG00000038037 | 0.45815691 | 0.75 | 0.864857905 | 0.646240625 | 0.877063019 | 0.77815125 | 1.066254157 |
| ENSMUSG00000038039 | 2.53838222 | 2.360735874 | 2.333847589 | 2.527457664 | 2.424460715 | 2.7930916 | 2.243069492 |
| ENSMUSG00000038042 | 1.985980784 | 1.884789703 | 1.937048212 | 1.75 | 1.746695109 | 1.986771734 | 1.74047993 |
| ENSMUSG00000038044 | 0 | 0 | 0 | 0 | 0 | 0 | 0 |
| ENSMUSG00000038045 | 1.036286563 | 0.701838731 | 0.79248125 | 0.701838731 | 0.568060967 | 0.698970004 | 0.579661877 |
| ENSMUSG00000038046 | 1.734388958 | 1.688721876 | 1.668106335 | 1.752806814 | 1.731346772 | 1.892094603 | 1.67819947 |
| ENSMUSG00000038047 | 2.682073381 | 2.484159485 | 2.469512728 | 2.543731421 | 2.506289166 | 2.831229694 | 2.310815043 |
| ENSMUSG00000038048 | 0 | 0 | 0 | 0 | 0 | 0 | 0 |
| ENSMUSG00000038055 | 1.45815691 | 1.505591953 | 1.566696635 | 1.410964047 | 1.559656853 | 1.447158031 | 1.526507301 |
| ENSMUSG00000038056 | 2.477243823 | 2.309601185 | 2.301142786 | 2.30479213 | 2.224547964 | 2.527629901 | 2.097879707 |
| ENSMUSG00000038057 | 0.289064826 | 0.396240625 | 0.396240625 | 0 | 0.387761935 | 0.301029996 | 0.579661877 |
| ENSMUSG00000038058 | 1.678702042 | 1.575945187 | 1.681980114 | 1.339388001 | 1.302012594 | 1.255272505 | 1.313616061 |
| ENSMUSG00000038059 | 1.89911417 | 1.80479213 | 1.741446071 | 1.870953944 | 1.903713872 | 2.139879086 | 1.777090262 |
| ENSMUSG00000038060 | 0 | 0 | 0.396240625 | 0.5 | 0.568060967 | 0.301029996 | 0.579661877 |
| ENSMUSG00000038064 | 0 | 0.580482024 | 0 | 0.25 | 0.568060967 | 0 | 0.448487648 |
| ENSMUSG00000038065 | 1.505869732 | 1.464495249 | 1.388647213 | 1.146240625 | 1.319233381 | 1.230448921 | 1.238624656 |
| ENSMUSG00000038067 | 0 | 0.25 | 0.396240625 | 0 | 0 | 0 | 0.448487648 |
| ENSMUSG00000038068 | 1.418409478 | 1.27186571 | 1.302363341 | 1.668106335 | 1.686820904 | 1.819543936 | 1.572224412 |
| ENSMUSG00000038069 | 2.266038081 | 2.057204673 | 2.02186571 | 2.05479213 | 1.983754904 | 2.217483944 | 1.856911705 |
| ENSMUSG00000038070 | 2.299278375 | 2.161864607 | 2.159156155 | 2.222945926 | 2.130260132 | 2.399673721 | 1.991191705 |
| ENSMUSG00000038071 | 0 | 0 | 0 | 0 | 0 | 0 | 0 |
| ENSMUSG00000038072 | 2.036286563 | 1.948603967 | 1.892463902 | 2.08272922 | 1.968065382 | 2.301029996 | 1.914853627 |
| ENSMUSG00000038074 | 2.305951036 | 2.149978211 | 2.14151351 | 2.134789703 | 2.111176775 | 2.408239965 | 1.917452159 |

| | | | | | | | |
|---|---|---|---|---|---|---|---|
| ENSMUSG00000038077 | 0 | 0 | 0 | 0.5 | 0.387761935 | 0.602059991 | 0.355418052 |
| ENSMUSG00000038079 | 2.136858952 | 2.031030328 | 1.981203126 | 2.121960008 | 2.103062484 | 2.399673721 | 1.96172822 |
| ENSMUSG00000038080 | 1.99653913 | 1.914552871 | 1.870953944 | 2.189555804 | 2.143518259 | 2.387389826 | 1.974994949 |
| ENSMUSG00000038084 | 2.746071301 | 2.50835575 | 2.493244947 | 2.541226732 | 2.507451162 | 2.839478047 | 2.304357435 |
| ENSMUSG00000038085 | 1.129344651 | 1.114857905 | 0.92510993 | 1.114857905 | 0.905314583 | 1.113943352 | 0.935079929 |
| ENSMUSG00000038086 | 1.358731791 | 1.188721876 | 1.330482024 | 0.396240625 | 0 | 0 | 0.355418052 |
| ENSMUSG00000038092 | 0 | 0 | 0 | 0 | 0 | 0 | 0 |
| ENSMUSG00000038094 | 0 | 0 | 0 | 0 | 0 | 0 | 0 |
| ENSMUSG00000038095 | 3.189772246 | 2.909835177 | 2.885999613 | 2.872085569 | 2.790143872 | 3.216693599 | 2.584965527 |
| ENSMUSG00000038102 | 2.550525119 | 2.348079356 | 2.299304173 | 2.387205477 | 2.360255934 | 2.69019608 | 2.210242554 |
| ENSMUSG00000038112 | 0 | 0 | 0 | 0 | 0.244650542 | 0 | 0 |
| ENSMUSG00000038115 | 0 | 0 | 0 | 0 | 0.244650542 | 0.301029996 | 0.224243824 |
| ENSMUSG00000038116 | 2.557825653 | 2.409156155 | 2.388167274 | 2.493244947 | 2.432800077 | 2.758911892 | 2.260561074 |
| ENSMUSG00000038119 | 2.755750303 | 2.485261902 | 2.451032755 | 2.636241108 | 2.57935845 | 2.982271233 | 2.405288179 |
| ENSMUSG00000038121 | 2.641922824 | 2.403217124 | 2.43155454 | 2.441217898 | 2.395413309 | 2.702430536 | 2.212676319 |
| ENSMUSG00000038122 | 1.971473847 | 1.79745614 | 1.814346961 | 1.943696765 | 1.855665188 | 2.079181246 | 1.712465909 |
| ENSMUSG00000038126 | 2.65645504 | 2.435366747 | 2.434523065 | 2.457430684 | 2.384078881 | 2.716003344 | 2.239900853 |
| ENSMUSG00000038127 | 2.854065408 | 2.633575721 | 2.602082435 | 2.731944491 | 2.61998316 | 3.005609445 | 2.408707635 |
| ENSMUSG00000038128 | 1.100572389 | 1.130890489 | 1.160964047 | 1.282320754 | 1.373641807 | 1.447158031 | 1.224243824 |
| ENSMUSG00000038132 | 0.45815691 | 0.75 | 0.896240625 | 0.396240625 | 0.733951626 | 0 | 0.355418052 |
| ENSMUSG00000038135 | 0 | 0 | 0.25 | 0 | 0.244650542 | 0 | 0 |
| ENSMUSG00000038141 | 2.31414447 | 2.169870025 | 2.154596376 | 2.255591953 | 2.206647761 | 2.499687083 | 2.079068325 |
| ENSMUSG00000038143 | 0 | 0.25 | 0.25 | 1.321350555 | 1.264824954 | 1.447158031 | 1.185218806 |
| ENSMUSG00000038145 | 2.054824387 | 1.899978211 | 1.940387808 | 1.92510993 | 1.919403394 | 2.008600172 | 1.892346957 |
| ENSMUSG00000038146 | 2.223916471 | 2.131869252 | 2.194519282 | 1.098079356 | 1.234113756 | 0.903089987 | 1.21680636 |
| ENSMUSG00000038147 | 0.91631382 | 0.5 | 0.646240625 | 0.580482024 | 0.632412477 | 0.602059991 | 0.579661877 |
| ENSMUSG00000038148 | 0 | 0 | 0 | 0 | 0 | 0 | 0 |
| ENSMUSG00000038150 | 2.351376531 | 2.223704441 | 2.194519282 | 2.058404919 | 2.164820399 | 2.401400541 | 2.113781657 |
| ENSMUSG00000038151 | 0 | 0 | 0 | 0.25 | 0 | 0 | 0 |
| ENSMUSG00000038152 | 1.100572389 | 1.226722649 | 1.403677461 | 0.646240625 | 0.955822902 | 1.079181246 | 0.969165684 |
| ENSMUSG00000038155 | 0 | 0 | 0 | 0 | 0 | 0 | 0 |
| ENSMUSG00000038156 | 0.811507563 | 0.5 | 0.396240625 | 0.92510993 | 0.955822902 | 1 | 1.014380832 |
| ENSMUSG00000038160 | 2.382811656 | 2.245748394 | 2.222945926 | 2.242885888 | 2.173890435 | 2.481442629 | 2.043115906 |
| ENSMUSG00000038165 | 0 | 0 | 0 | 0 | 0 | 0 | 0 |

| | | | | | | | |
|---|---|---|---|---|---|---|---|
| ENSMUSG00000038167 | 1.205378646 | 1.146240625 | 1.114857905 | 0.580482024 | 0.978602168 | 0.77815125 | 0.629532004 |
| ENSMUSG00000038168 | 1.623015126 | 1.593759858 | 1.54248125 | 1.622963274 | 1.645989387 | 1.62324929 | 1.523605797 |
| ENSMUSG00000038170 | 2.80555694 | 2.56728167 | 2.5658173 | 2.371960008 | 2.359815565 | 2.646403726 | 2.202114485 |
| ENSMUSG00000038172 | 2.45930418 | 2.284247778 | 2.302972074 | 2.244319981 | 2.199780694 | 2.432969291 | 2.00464543 |
| ENSMUSG00000038173 | 0 | 0 | 0 | 0 | 0 | 0 | 0 |
| ENSMUSG00000038174 | 2.124258828 | 1.948603967 | 1.988549078 | 2.102347734 | 2.058398634 | 2.326335861 | 1.882886767 |
| ENSMUSG00000038175 | 1.129344651 | 1.04248125 | 1.061981878 | 1.602347734 | 1.528558853 | 1.716003344 | 1.520678036 |
| ENSMUSG00000038178 | 1.928763473 | 1.755591953 | 1.860735874 | 1.642463902 | 1.603583716 | 1.812913357 | 1.586932193 |
| ENSMUSG00000038179 | 0 | 0 | 0 | 0 | 0 | 0 | 0 |
| ENSMUSG00000038180 | 0.45815691 | 0.75 | 0.75 | 0.701838731 | 0.77552387 | 0.77815125 | 0.829800754 |
| ENSMUSG00000038181 | 1.932832114 | 1.868933358 | 1.994319981 | 1.720660762 | 1.861137485 | 1.924279286 | 1.845261772 |
| ENSMUSG00000038187 | 2.254965775 | 2.05479213 | 2.078220739 | 1.988549078 | 1.883916752 | 2.155336037 | 1.792684387 |
| ENSMUSG00000038188 | 0.671187741 | 0.701838731 | 0.646240625 | 0.396240625 | 0.387761935 | 0.602059991 | 0.520678036 |
| ENSMUSG00000038192 | 0 | 0 | 0 | 0 | 0 | 0 | 0.224243824 |
| ENSMUSG00000038193 | 0 | 0 | 0 | 0.25 | 0 | 0 | 0 |
| ENSMUSG00000038194 | 0.289064826 | 0 | 0 | 0 | 0 | 0 | 0.224243824 |
| ENSMUSG00000038195 | 0.578129653 | 0.646240625 | 0.951838731 | 0.951838731 | 1.136121934 | 0.477121255 | 0.775756176 |
| ENSMUSG00000038199 | 0 | 0 | 0 | 0 | 0 | 0 | 0 |
| ENSMUSG00000038201 | 0 | 0 | 0 | 0.396240625 | 0 | 0 | 0.224243824 |
| ENSMUSG00000038203 | 0 | 0 | 0.396240625 | 0 | 0 | 0 | 0.355418052 |
| ENSMUSG00000038204 | 0 | 0 | 0 | 0 | 0 | 0 | 0 |
| ENSMUSG00000038205 | 2.247417471 | 2.043731421 | 2.092671852 | 1.938721876 | 1.862942891 | 2.041392685 | 1.704233519 |
| ENSMUSG00000038206 | 1.928763473 | 1.87892496 | 1.862802778 | 1.916333979 | 1.793512788 | 2.133538908 | 1.757893036 |
| ENSMUSG00000038208 | 1.527823874 | 1.438721876 | 1.566696635 | 1.431980114 | 1.55104774 | 1.579783597 | 1.591689859 |
| ENSMUSG00000038209 | 0 | 0 | 0 | 0 | 0 | 0 | 0 |
| ENSMUSG00000038210 | 0.671187741 | 0.580482024 | 0.396240625 | 0.701838731 | 0.632412477 | 0.602059991 | 0.74492186 |
| ENSMUSG00000038212 | 2.565888829 | 2.370449858 | 2.32991803 | 2.570482507 | 2.492946102 | 2.848804701 | 2.30331215 |
| ENSMUSG00000038213 | 1.614416215 | 1.580482024 | 1.584962501 | 1.784887838 | 1.736537397 | 1.949390007 | 1.743447983 |
| ENSMUSG00000038214 | 2.075459174 | 2.064346961 | 2.002806814 | 1.87892496 | 1.910073614 | 2.089905111 | 1.838725884 |
| ENSMUSG00000038215 | 1.936861443 | 1.837182039 | 1.76109853 | 1.80479213 | 1.728722551 | 2.025305865 | 1.674564426 |
| ENSMUSG00000038216 | 0 | 0 | 0 | 0 | 0 | 0 | 0 |
| ENSMUSG00000038217 | 0.960252568 | 0.75 | 0.896240625 | 0.976722649 | 1.074582839 | 0.84509804 | 0.829800754 |
| ENSMUSG00000038218 | 0 | 0 | 0 | 0 | 0 | 0 | 0 |
| ENSMUSG00000038224 | 0.45815691 | 0.646240625 | 0.25 | 0.25 | 0 | 0 | 0.224243824 |

| | | | | | | | |
|---|---|---|---|---|---|---|---|
| ENSMUSG00000038225 | 1.771760131 | 1.681980114 | 1.622963274 | 1.602347734 | 1.572189156 | 1.62324929 | 1.496250175 |
| ENSMUSG00000038227 | 2.083999385 | 1.942046081 | 1.905512955 | 1.364857905 | 1.283908311 | 1.544068044 | 1.340368108 |
| ENSMUSG00000038233 | 0.578129653 | 0.25 | 0.396240625 | 0.5 | 0.632412477 | 0.477121255 | 0.710836105 |
| ENSMUSG00000038235 | 2.181541736 | 1.992885888 | 1.961372513 | 2.352877747 | 2.321205208 | 2.687528961 | 2.178359564 |
| ENSMUSG00000038236 | 1.759671389 | 1.755591953 | 1.782320754 | 1.403677461 | 1.499532413 | 1.544068044 | 1.466365979 |
| ENSMUSG00000038239 | 0.289064826 | 0.5 | 0 | 0 | 0.489301084 | 0.301029996 | 0 |
| ENSMUSG00000038240 | 1.734388958 | 1.606566189 | 1.701838731 | 1.777131114 | 1.680838453 | 1.913813852 | 1.661510159 |
| ENSMUSG00000038241 | 2.388281011 | 2.288071211 | 2.323655187 | 2.224461364 | 2.356272688 | 2.509202522 | 2.272294561 |
| ENSMUSG00000038242 | 0 | 0.25 | 0 | 0 | 0.244650542 | 0 | 0.224243824 |
| ENSMUSG00000038244 | 3.018088033 | 2.762802235 | 2.761439915 | 2.215271726 | 2.190648982 | 2.450249108 | 2.075912052 |
| ENSMUSG00000038246 | 0 | 0 | 0 | 0 | 0 | 0 | 0 |
| ENSMUSG00000038248 | 0 | 0.646240625 | 0.701838731 | 1.311981878 | 1.394615667 | 1.568201724 | 1.245582246 |
| ENSMUSG00000038250 | 2.352857998 | 2.143411797 | 2.088286706 | 2.199415381 | 2.170139537 | 2.383815366 | 1.989815042 |
| ENSMUSG00000038252 | 3.059740347 | 2.790597832 | 2.788862683 | 2.65892953 | 2.671488075 | 3.02489596 | 2.489549481 |
| ENSMUSG00000038253 | 1.777675583 | 1.692046081 | 1.614857905 | 1.445339928 | 1.200473444 | 1.447158031 | 1.383567577 |
| ENSMUSG00000038255 | 0 | 0 | 0 | 0 | 0 | 0 | 0 |
| ENSMUSG00000038256 | 2.196867486 | 2.120953944 | 2.099151195 | 1.978222334 | 2.047893212 | 2.214843848 | 1.912234055 |
| ENSMUSG00000038257 | 0 | 0 | 0 | 0.25 | 0 | 0 | 0 |
| ENSMUSG00000038259 | 0.289064826 | 0.396240625 | 0 | 0.25 | 0.244650542 | 0.301029996 | 0.520678036 |
| ENSMUSG00000038260 | 1.631440309 | 1.614857905 | 1.671625132 | 1.25 | 1.559656853 | 1.342422681 | 1.618853957 |
| ENSMUSG00000038264 | 1.156259305 | 1.330482024 | 1.330482024 | 1.5 | 1.599808748 | 1.612783857 | 1.496250175 |
| ENSMUSG00000038267 | 2.013563224 | 1.821350555 | 1.850219859 | 2.001406137 | 1.986303335 | 2.167317335 | 1.902443294 |
| ENSMUSG00000038268 | 1.289064826 | 1.339388001 | 1.25 | 1.552363341 | 1.514343934 | 1.698970004 | 1.396774124 |
| ENSMUSG00000038271 | 1.663535556 | 1.62694866 | 1.698603967 | 1.622963274 | 1.715300549 | 1.838849091 | 1.746389053 |
| ENSMUSG00000038274 | 2.54867984 | 2.491806565 | 2.621205724 | 2.078220739 | 2.371090306 | 2.397940009 | 2.455103737 |
| ENSMUSG00000038276 | 0.578129653 | 0.5 | 0.646240625 | 0.25 | 0.489301084 | 0.477121255 | 0.224243824 |
| ENSMUSG00000038279 | 2.765759561 | 2.572504712 | 2.600486531 | 2.480088214 | 2.549763795 | 2.810232518 | 2.401639109 |
| ENSMUSG00000038280 | 2.461589295 | 2.278435542 | 2.278435542 | 2.355516192 | 2.289330171 | 2.62838893 | 2.137790317 |
| ENSMUSG00000038286 | 1.771760131 | 1.671625132 | 1.688721876 | 1.664552871 | 1.66856048 | 1.886490725 | 1.633706455 |
| ENSMUSG00000038290 | 2.409468165 | 2.225970461 | 2.240000483 | 2.293106877 | 2.268268456 | 2.507855872 | 2.134719638 |
| ENSMUSG00000038291 | 2.390989024 | 2.215271726 | 2.219895812 | 2.368933358 | 2.28445426 | 2.681241237 | 2.124874702 |
| ENSMUSG00000038292 | 0 | 0.396240625 | 0.396240625 | 0.25 | 0.632412477 | 0.301029996 | 0.579661877 |
| ENSMUSG00000038295 | 0.578129653 | 0.580482024 | 0.580482024 | 0.580482024 | 0.387761935 | 0.477121255 | 0.448487648 |
| ENSMUSG00000038296 | 1.249317394 | 1.488549078 | 1.302363341 | 0.25 | 0.632412477 | 0.477121255 | 0.520678036 |

| | | | | | | | |
|---|---|---|---|---|---|---|---|
| ENSMUSG00000038298 | 0 | 0 | 0 | 0 | 0 | 0.301029996 | 0 |
| ENSMUSG00000038299 | 2.715608727 | 2.493244947 | 2.491806565 | 2.509729747 | 2.452316937 | 2.802089258 | 2.261157966 |
| ENSMUSG00000038300 | 0 | 0 | 0 | 0 | 0 | 0 | 0 |
| ENSMUSG00000038301 | 2.196867486 | 2.087182039 | 2.064346961 | 2.134789703 | 2.019019072 | 2.336459734 | 1.886704102 |
| ENSMUSG00000038302 | 1.707474304 | 1.638647213 | 1.602347734 | 1.671625132 | 1.671670246 | 1.897627091 | 1.543390198 |
| ENSMUSG00000038304 | 0 | 0.25 | 0.25 | 0.5 | 0.489301084 | 0.698970004 | 0.629532004 |
| ENSMUSG00000038305 | 0.45815691 | 0 | 0 | 1.130890489 | 0.905314583 | 1.255272505 | 1.066254157 |
| ENSMUSG00000038311 | 0.289064826 | 0 | 0.25 | 0 | 0 | 0 | 0 |
| ENSMUSG00000038312 | 2.036286563 | 1.899978211 | 1.901832578 | 2.031030328 | 2.064555002 | 2.303196057 | 1.990504106 |
| ENSMUSG00000038319 | 2.003432385 | 1.882845365 | 1.990000483 | 0.79248125 | 0.812711509 | 0.477121255 | 1.20139795 |
| ENSMUSG00000038323 | 1.944805158 | 1.841580554 | 1.828220739 | 1.975216702 | 1.892300902 | 2.170261715 | 1.757893036 |
| ENSMUSG00000038324 | 2.698532991 | 2.530383379 | 2.514836115 | 2.658248799 | 2.652586889 | 3.039810554 | 2.518801024 |
| ENSMUSG00000038327 | 0.45815691 | 0.25 | 0 | 0 | 0 | 0 | 0 |
| ENSMUSG00000038330 | 0 | 0 | 0 | 0 | 0 | 0 | 0 |
| ENSMUSG00000038331 | 1.54867984 | 1.711372513 | 1.638647213 | 1.75 | 1.726078673 | 1.897627091 | 1.612273755 |
| ENSMUSG00000038332 | 2.229917041 | 2.100219859 | 2.032320754 | 2.238549078 | 2.143518259 | 2.495544338 | 2.001999072 |
| ENSMUSG00000038335 | 2.611244851 | 2.425975893 | 2.418106335 | 2.527131114 | 2.474013807 | 2.808210973 | 2.274304872 |
| ENSMUSG00000038342 | 2.575536697 | 2.3464656 | 2.310197833 | 2.341580554 | 2.312243922 | 2.59439255 | 2.179512918 |
| ENSMUSG00000038344 | 1.847794126 | 1.744319981 | 1.634789703 | 2.001406137 | 1.938809675 | 2.204119983 | 1.770364127 |
| ENSMUSG00000038346 | 2.434522689 | 2.203445298 | 2.205044741 | 2.220660762 | 2.199780694 | 2.484299839 | 2.024452165 |
| ENSMUSG00000038347 | 1.37447073 | 1.348079356 | 1.214495249 | 1.339388001 | 1.223252711 | 1.230448921 | 1.265599895 |
| ENSMUSG00000038349 | 1.307602651 | 1.321350555 | 1.348079356 | 0.976722649 | 0.77552387 | 0.77815125 | 0.775756176 |
| ENSMUSG00000038351 | 2.119110214 | 2.048689214 | 2.083847589 | 1.823655187 | 1.873585591 | 1.973127854 | 1.856911705 |
| ENSMUSG00000038352 | 1.069666964 | 0.75 | 0.951838731 | 0.864857905 | 0.877063019 | 0.84509804 | 0.935079929 |
| ENSMUSG00000038354 | 0.45815691 | 0 | 0.396240625 | 0.864857905 | 1.188507684 | 1.079181246 | 1.20139795 |
| ENSMUSG00000038357 | 0 | 0 | 0.396240625 | 0 | 0 | 0.301029996 | 0 |
| ENSMUSG00000038365 | 2.331622671 | 2.215271726 | 2.145300145 | 2.027131114 | 2.008447216 | 2.278753601 | 1.958704689 |
| ENSMUSG00000038366 | 3.267360421 | 2.997880462 | 3.023766825 | 2.841717147 | 2.856852685 | 3.278296208 | 2.698806435 |
| ENSMUSG00000038368 | 2.211649928 | 2.095926073 | 2.041226732 | 1.988549078 | 2.014359518 | 2.195899652 | 1.913982791 |
| ENSMUSG00000038369 | 2.484856099 | 2.360217292 | 2.31611065 | 2.431980114 | 2.384078881 | 2.700703717 | 2.244955885 |
| ENSMUSG00000038370 | 0.811507563 | 0.701838731 | 0.396240625 | 1.488549078 | 1.351342427 | 1.431363764 | 1.374453863 |
| ENSMUSG00000038371 | 2.43085913 | 2.254202072 | 2.244319981 | 2.105516192 | 2.090057366 | 2.346352974 | 1.920884716 |
| ENSMUSG00000038372 | 2.196867486 | 2.071350555 | 2.05479213 | 2.173371739 | 2.131103516 | 2.434568904 | 2.044866219 |
| ENSMUSG00000038374 | 2.496544396 | 2.30539678 | 2.328787391 | 2.511780978 | 2.466277473 | 2.826074803 | 2.380695513 |

| | | | | | | | |
|---|---|---|---|---|---|---|---|
| ENSMUSG00000038375 | 2.364524 | 2.21293726 | 2.192046081 | 2.331045137 | 2.25901006 | 2.568201724 | 2.14725554 |
| ENSMUSG00000038379 | 2.439357901 | 2.269203899 | 2.291226732 | 2.454245906 | 2.403681482 | 2.760422483 | 2.21543553 |
| ENSMUSG00000038383 | 2.063793172 | 1.995748394 | 1.964495249 | 2.078220739 | 2.062514787 | 2.324282455 | 1.953344547 |
| ENSMUSG00000038384 | 2.086807695 | 2.044977273 | 2.073655187 | 1.834962501 | 1.84445929 | 2.008600172 | 1.901538355 |
| ENSMUSG00000038387 | 2.078325387 | 1.975216702 | 1.96759118 | 1.958222504 | 1.959951091 | 2.198657087 | 1.869159809 |
| ENSMUSG00000038388 | 2.364524 | 2.220660762 | 2.225216702 | 2.186208459 | 2.077536464 | 2.380211242 | 1.924281235 |
| ENSMUSG00000038390 | 1.227926429 | 1.098079356 | 1.311981878 | 1.17510993 | 1.327538614 | 1.278753601 | 1.335191751 |
| ENSMUSG00000038393 | 2.251208701 | 2.079353153 | 2.044977273 | 1.955044741 | 1.926994141 | 2.201397124 | 1.844181586 |
| ENSMUSG00000038398 | 2.427163103 | 2.262462137 | 2.320772088 | 2.231203126 | 2.225193812 | 2.491361694 | 2.114251541 |
| ENSMUSG00000038400 | 2.722979153 | 2.547764804 | 2.561388177 | 2.222185812 | 2.223900931 | 2.501059262 | 2.156863548 |
| ENSMUSG00000038402 | 1.100572389 | 1.130890489 | 1.17510993 | 1.302363341 | 1.264824954 | 1.255272505 | 1.290497981 |
| ENSMUSG00000038403 | 0 | 0 | 0 | 0 | 0.387761935 | 0.477121255 | 0 |
| ENSMUSG00000038406 | 2.266038081 | 2.180274797 | 2.360735874 | 1.94697564 | 2.124299332 | 2.193124598 | 2.178359564 |
| ENSMUSG00000038408 | 0.289064826 | 0.25 | 0.25 | 0.5 | 0.244650542 | 0.301029996 | 0.224243824 |
| ENSMUSG00000038412 | 2.013563224 | 1.884789703 | 1.898114259 | 2.080482024 | 1.996316946 | 2.276461804 | 1.869159809 |
| ENSMUSG00000038415 | 0.45815691 | 0.5 | 0.646240625 | 0.580482024 | 0 | 0.301029996 | 0.520678036 |
| ENSMUSG00000038416 | 2.969118676 | 2.684523065 | 2.684945152 | 2.593488164 | 2.522497007 | 2.87909588 | 2.324338556 |
| ENSMUSG00000038417 | 2.50175086 | 2.309003548 | 2.335518667 | 2.428561379 | 2.386129765 | 2.766412847 | 2.253005488 |
| ENSMUSG00000038418 | 2.144240249 | 2.00835575 | 2.052363341 | 1.843759858 | 1.959951091 | 2.037426498 | 1.844181586 |
| ENSMUSG00000038421 | 0 | 0 | 0 | 0 | 0 | 0 | 0 |
| ENSMUSG00000038422 | 1.596667477 | 1.54745614 | 1.54248125 | 1.388647213 | 1.599808748 | 1.462397998 | 1.448487648 |
| ENSMUSG00000038425 | 1.663535556 | 1.614857905 | 1.598079356 | 1.464495249 | 1.519146132 | 1.602059991 | 1.473249546 |
| ENSMUSG00000038429 | 2.923903336 | 2.713717096 | 2.741986676 | 2.643175057 | 2.71417212 | 3.01745073 | 2.556097877 |
| ENSMUSG00000038437 | 2.213719572 | 2.106566189 | 2.107613138 | 1.894357207 | 2.030405741 | 2.08278537 | 1.952571511 |
| ENSMUSG00000038446 | 2.582415738 | 2.368426437 | 2.324229052 | 2.467205639 | 2.398190943 | 2.739572344 | 2.228229691 |
| ENSMUSG00000038451 | 1.527823874 | 1.476722649 | 1.54248125 | 1.17510993 | 1.264824954 | 1.278753601 | 1.396774124 |
| ENSMUSG00000038453 | 0 | 0 | 0 | 0.396240625 | 0.387761935 | 0.301029996 | 0.579661877 |
| ENSMUSG00000038456 | 2.569423023 | 2.410964047 | 2.399978211 | 2.133818844 | 2.170139537 | 2.436162647 | 2.076440241 |
| ENSMUSG00000038457 | 0 | 0 | 0 | 0 | 0 | 0 | 0 |
| ENSMUSG00000038459 | 2.305951036 | 2.08272922 | 2.088286706 | 2.116901388 | 2.109389652 | 2.336459734 | 1.920029985 |
| ENSMUSG00000038462 | 2.666334442 | 2.469129237 | 2.489638179 | 2.614601653 | 2.547699718 | 2.953759692 | 2.398533958 |
| ENSMUSG00000038463 | 2.889424695 | 2.670529191 | 2.685787849 | 1.901832578 | 1.931471446 | 2.10720997 | 1.885753983 |
| ENSMUSG00000038467 | 2.520963022 | 2.359177886 | 2.379417347 | 2.492166698 | 2.498344007 | 2.822168079 | 2.386819016 |
| ENSMUSG00000038473 | 0 | 0.580482024 | 0.830482024 | 0.5 | 0.812711509 | 0.698970004 | 0.579661877 |

| | | | | | | | |
|---|---|---|---|---|---|---|---|
| ENSMUSG00000038481 | 2.402962068 | 2.194519282 | 2.194519282 | 2.271202097 | 2.182004726 | 2.525044807 | 2.044866219 |
| ENSMUSG00000038482 | 3.137084558 | 2.864473459 | 2.869818758 | 2.78360658 | 2.743737772 | 3.149219113 | 2.557413225 |
| ENSMUSG00000038485 | 2.427163103 | 2.247880462 | 2.246460484 | 2.367410454 | 2.319727371 | 2.621176282 | 2.181807364 |
| ENSMUSG00000038486 | 1.693336259 | 1.649978211 | 1.678561379 | 1.653677461 | 1.763796674 | 1.770852012 | 1.670888075 |
| ENSMUSG00000038489 | 1.494443473 | 1.330482024 | 1.396240625 | 1.5 | 1.445123986 | 1.556302501 | 1.40946263 |
| ENSMUSG00000038491 | 0 | 0 | 0 | 0 | 0 | 0 | 0 |
| ENSMUSG00000038495 | 2.65645504 | 2.457430684 | 2.457034121 | 2.557505109 | 2.519146132 | 2.894869657 | 2.354078442 |
| ENSMUSG00000038496 | 0 | 0 | 0 | 0 | 0 | 0 | 0 |
| ENSMUSG00000038497 | 2.441754646 | 2.291854536 | 2.311981878 | 2.218361028 | 2.208682096 | 2.460897843 | 2.098373247 |
| ENSMUSG00000038498 | 0 | 0 | 0 | 0 | 0 | 0 | 0 |
| ENSMUSG00000038500 | 1.975148165 | 1.896240625 | 1.856566189 | 1.937048212 | 1.983754904 | 2.103803721 | 1.861046144 |
| ENSMUSG00000038502 | 2.390989024 | 2.271202097 | 2.338286706 | 1.919870025 | 2.190648982 | 2.164352856 | 2.143845596 |
| ENSMUSG00000038503 | 2.572927518 | 2.353406982 | 2.363831805 | 2.348615674 | 2.305113253 | 2.597695186 | 2.160953557 |
| ENSMUSG00000038506 | 2.348397716 | 2.187048212 | 2.201032755 | 2.134789703 | 2.141887974 | 2.309630167 | 1.974272009 |
| ENSMUSG00000038507 | 2.160971586 | 2.004202072 | 1.979715809 | 2.240724001 | 2.210704773 | 2.506505032 | 2.092902448 |
| ENSMUSG00000038508 | 0.45815691 | 0.701838731 | 0.646240625 | 0.25 | 0 | 0 | 0.224243824 |
| ENSMUSG00000038510 | 2.315764015 | 2.124961472 | 2.069031101 | 2.263142013 | 2.185632262 | 2.511883361 | 1.999330888 |
| ENSMUSG00000038515 | 1.129344651 | 1.130890489 | 1.061981878 | 1.282320754 | 1.327538614 | 1.462397998 | 1.40946263 |
| ENSMUSG00000038517 | 0.91631382 | 0.864857905 | 1.114857905 | 0.830482024 | 0.978602168 | 0.477121255 | 1.014380832 |
| ENSMUSG00000038518 | 2.111266113 | 1.942046081 | 1.978222334 | 1.809601185 | 1.818885826 | 2.123851641 | 1.759302692 |
| ENSMUSG00000038520 | 2.023453784 | 1.982684334 | 1.945339928 | 1.575945187 | 1.804307395 | 1.806179974 | 1.773071385 |
| ENSMUSG00000038521 | 1.527823874 | 1.445339928 | 1.606566189 | 0.864857905 | 0.733951626 | 1 | 0.853775828 |
| ENSMUSG00000038522 | 0 | 0 | 0 | 0 | 0 | 0 | 0 |
| ENSMUSG00000038523 | 0.747221736 | 0.951838731 | 1.17510993 | 0.5 | 0.489301084 | 0.602059991 | 0.520678036 |
| ENSMUSG00000038524 | 1.418409478 | 1.54745614 | 1.62694866 | 1.282320754 | 1.528558853 | 1.204119983 | 1.496250175 |
| ENSMUSG00000038525 | 2.402962068 | 2.201838731 | 2.166333979 | 2.188721876 | 2.16634835 | 2.419955748 | 2.021338422 |
| ENSMUSG00000038526 | 0.960252568 | 0.75 | 0.646240625 | 0.646240625 | 0.489301084 | 0.477121255 | 0.672731473 |
| ENSMUSG00000038527 | 1.578129653 | 1.445339928 | 1.488549078 | 1.130890489 | 1.074582839 | 1.113943352 | 1.185218806 |
| ENSMUSG00000038528 | 0 | 0.396240625 | 0.25 | 0 | 0.244650542 | 0 | 0.224243824 |
| ENSMUSG00000038530 | 0.671187741 | 0.79248125 | 0.864857905 | 0.92510993 | 0.931471446 | 1.079181246 | 0.853775828 |
| ENSMUSG00000038533 | 2.300956576 | 2.110735874 | 2.095926073 | 2.232684334 | 2.16634835 | 2.499687083 | 2.047185336 |
| ENSMUSG00000038534 | 1.516991255 | 1.575945187 | 1.747171172 | 1.606566189 | 1.832885921 | 1.707570176 | 1.784981274 |
| ENSMUSG00000038535 | 2.235832493 | 2.106566189 | 2.038704527 | 2.248588359 | 2.143518259 | 2.42975228 | 2.02383181 |
| ENSMUSG00000038537 | 0 | 0 | 0 | 0 | 0 | 0 | 0 |

| | | | | | | | |
|---|---|---|---|---|---|---|---|
| ENSMUSG00000038538 | 2.615205302 | 2.360217292 | 2.341580554 | 2.405054956 | 2.367659357 | 2.691081492 | 2.170957925 |
| ENSMUSG00000038539 | 2.134369199 | 1.964495249 | 1.984159485 | 1.571350555 | 1.509475496 | 1.707570176 | 1.551512352 |
| ENSMUSG00000038540 | 0.747221736 | 0.864857905 | 0.92510993 | 1.02186571 | 0.632412477 | 0.903089987 | 0.829800754 |
| ENSMUSG00000038541 | 0 | 0 | 0 | 0 | 0.244650542 | 0 | 0 |
| ENSMUSG00000038542 | 2.445324132 | 2.263142013 | 2.289967834 | 2.212155735 | 2.173890435 | 2.463892989 | 2.093403634 |
| ENSMUSG00000038543 | 1.269664473 | 1.348079356 | 1.321350555 | 1.201838731 | 1.136121934 | 1.361727836 | 1.078019652 |
| ENSMUSG00000038544 | 2.23973002 | 2.019203899 | 2.02053726 | 2.041226732 | 2.007252778 | 2.214843848 | 1.888596011 |
| ENSMUSG00000038545 | 2.602397159 | 2.459800947 | 2.523849256 | 2.278435542 | 2.440245937 | 2.582063363 | 2.392033886 |
| ENSMUSG00000038546 | 2.497590902 | 2.323655187 | 2.288071211 | 2.432829758 | 2.386538516 | 2.710117365 | 2.194510526 |
| ENSMUSG00000038550 | 1.37447073 | 1.29248125 | 1.04248125 | 1.17510993 | 1.039257769 | 1.204119983 | 1.131174228 |
| ENSMUSG00000038552 | 2.113897241 | 2.023189285 | 2.052363341 | 2.05479213 | 2.082404901 | 2.267171728 | 1.971363985 |
| ENSMUSG00000038555 | 1 | 1.160964047 | 1.321350555 | 0.396240625 | 0.244650542 | 0.301029996 | 0.448487648 |
| ENSMUSG00000038560 | 0 | 0 | 0 | 0 | 0 | 0 | 0 |
| ENSMUSG00000038563 | 2.016886209 | 1.870953944 | 1.845926073 | 1.896240625 | 1.814781639 | 1.919078092 | 1.670888075 |
| ENSMUSG00000038564 | 2.278594844 | 2.121960008 | 2.118933358 | 2.041226732 | 2.111176775 | 2.290034611 | 1.969900112 |
| ENSMUSG00000038567 | 0 | 0.396240625 | 0 | 0 | 0 | 0 | 0 |
| ENSMUSG00000038569 | 0.747221736 | 1.080482024 | 0.976722649 | 1.04248125 | 0.877063019 | 1.204119983 | 0.935079929 |
| ENSMUSG00000038570 | 0.578129653 | 0.5 | 0.25 | 0.79248125 | 0.489301084 | 0.77815125 | 0.579661877 |
| ENSMUSG00000038572 | 0 | 0 | 0 | 0 | 0 | 0 | 0 |
| ENSMUSG00000038576 | 0.671187741 | 0.5 | 0.580482024 | 0.864857905 | 1 | 1.041392685 | 0.916588299 |
| ENSMUSG00000038578 | 0.671187741 | 0.701838731 | 0.580482024 | 0.951838731 | 1.020174412 | 0.698970004 | 0.829800754 |
| ENSMUSG00000038580 | 0 | 0 | 0 | 0 | 0.244650542 | 0 | 0 |
| ENSMUSG00000038582 | 2.360187403 | 2.269871196 | 2.245748394 | 2.124961472 | 2.123439512 | 2.369215857 | 1.987739 |
| ENSMUSG00000038583 | 0 | 0 | 0 | 0.396240625 | 0 | 0 | 0 |
| ENSMUSG00000038587 | 3.075981771 | 2.810048763 | 2.844709178 | 2.718745337 | 2.710438817 | 3.045714059 | 2.543639344 |
| ENSMUSG00000038591 | 0.578129653 | 0.5 | 0.5 | 0 | 0.387761935 | 0 | 0.520678036 |
| ENSMUSG00000038593 | 2 | 1.799918086 | 1.860735874 | 1.695339928 | 1.744182955 | 1.897627091 | 1.699191618 |
| ENSMUSG00000038594 | 1.325351389 | 1.27186571 | 1.214495249 | 1.42510993 | 1.380772476 | 1.591064607 | 1.307989563 |
| ENSMUSG00000038598 | 0.289064826 | 0 | 0.25 | 0 | 0 | 0.301029996 | 0.579661877 |
| ENSMUSG00000038599 | 0 | 0 | 0 | 0 | 0 | 0 | 0 |
| ENSMUSG00000038600 | 0 | 0 | 0.396240625 | 0.396240625 | 0.387761935 | 0.477121255 | 0.520678036 |
| ENSMUSG00000038602 | 0 | 0.396240625 | 0.25 | 0.701838731 | 0.387761935 | 0.698970004 | 0.579661877 |
| ENSMUSG00000038604 | 2.557825653 | 2.403677461 | 2.446567407 | 2.312574604 | 2.424827423 | 2.679427897 | 2.323601898 |
| ENSMUSG00000038605 | 1.671187741 | 1.575945187 | 1.692046081 | 1.598079356 | 1.559656853 | 1.799340549 | 1.529383012 |

| | | | | | | | |
|---|---|---|---|---|---|---|---|
| ENSMUSG00000038607 | 0.289064826 | 0.25 | 0 | 0.396240625 | 0 | 0.301029996 | 0.579661877 |
| ENSMUSG00000038608 | 0.867194479 | 0.646240625 | 0.830482024 | 0.976722649 | 1.057362051 | 0.84509804 | 0.775756176 |
| ENSMUSG00000038611 | 2.523917219 | 2.32991803 | 2.386723615 | 2.221424093 | 2.343584837 | 2.527629901 | 2.206027309 |
| ENSMUSG00000038612 | 2.910141165 | 2.602347734 | 2.611770807 | 2.850753006 | 2.730731061 | 3.172894698 | 2.55053864 |
| ENSMUSG00000038615 | 3.192940598 | 2.931873767 | 2.920638935 | 2.736726569 | 2.778092802 | 3.128399269 | 2.641897157 |
| ENSMUSG00000038618 | 1.747221736 | 1.664552871 | 1.816696635 | 1.53743678 | 1.72073075 | 1.643452676 | 1.746389053 |
| ENSMUSG00000038619 | 2.718695584 | 2.490724001 | 2.453044326 | 2.584405475 | 2.541960468 | 2.864511081 | 2.380901508 |
| ENSMUSG00000038622 | 2.183765914 | 1.998588359 | 1.96293726 | 2.091580554 | 2.017859938 | 2.338456494 | 1.909593098 |
| ENSMUSG00000038623 | 0.747221736 | 0.580482024 | 0.79248125 | 0.830482024 | 1.057362051 | 0.954242509 | 1.078019652 |
| ENSMUSG00000038624 | 0 | 0 | 0.25 | 0 | 0 | 0 | 0 |
| ENSMUSG00000038628 | 2.060824957 | 2.009729747 | 2.017865591 | 2.119945066 | 2.046825265 | 2.344392274 | 1.94074868 |
| ENSMUSG00000038630 | 0.960252568 | 0.896240625 | 1.02186571 | 0.951838731 | 1 | 1.113943352 | 0.952571511 |
| ENSMUSG00000038633 | 2.894701088 | 2.655741736 | 2.672499493 | 2.755591953 | 2.696797911 | 3.11991541 | 2.517588597 |
| ENSMUSG00000038637 | 0.91631382 | 0.896240625 | 0.976722649 | 0.951838731 | 1.200473444 | 0.602059991 | 1.131174228 |
| ENSMUSG00000038641 | 0.289064826 | 0 | 0.25 | 0 | 0 | 0 | 0 |
| ENSMUSG00000038642 | 0 | 0 | 0 | 0.25 | 0 | 0 | 0 |
| ENSMUSG00000038644 | 2.17479713 | 2.103406982 | 2.23927551 | 1.726722649 | 2.05001946 | 1.982271233 | 2.111421942 |
| ENSMUSG00000038646 | 2.045658471 | 1.882845365 | 1.837182039 | 1.830482024 | 1.756568651 | 2.025305865 | 1.621017948 |
| ENSMUSG00000038648 | 2.54867984 | 2.355516192 | 2.316696635 | 2.219129237 | 2.167869715 | 2.494154594 | 2.065709059 |
| ENSMUSG00000038650 | 2.796468708 | 2.574515642 | 2.636482442 | 2.426839783 | 2.507451162 | 2.804139432 | 2.37340178 |
| ENSMUSG00000038651 | 0 | 0 | 0 | 0 | 0 | 0 | 0 |
| ENSMUSG00000038656 | 0 | 0 | 0 | 0 | 0 | 0 | 0 |
| ENSMUSG00000038658 | 2.604828841 | 2.382845365 | 2.3464656 | 2.446567407 | 2.398980554 | 2.726727209 | 2.222568443 |
| ENSMUSG00000038663 | 0 | 0.25 | 0 | 0 | 0 | 0 | 0 |
| ENSMUSG00000038664 | 2.927646919 | 2.679847205 | 2.659608978 | 2.796992648 | 2.732268329 | 3.140508043 | 2.538366565 |
| ENSMUSG00000038665 | 0.45815691 | 0.580482024 | 0.25 | 0 | 0 | 0 | 0 |
| ENSMUSG00000038668 | 2.089597219 | 1.94697564 | 1.916333979 | 1.856566189 | 1.814781639 | 2.056904851 | 1.777090262 |
| ENSMUSG00000038670 | 0 | 0.25 | 0 | 0 | 0 | 0.602059991 | 0.448487648 |
| ENSMUSG00000038671 | 2.095120908 | 1.886723615 | 1.874961472 | 2.002806814 | 2.033751322 | 2.212187604 | 1.877074955 |
| ENSMUSG00000038676 | 0 | 0 | 0 | 0 | 0 | 0 | 0 |
| ENSMUSG00000038677 | 2.029919532 | 1.905512955 | 1.75835575 | 1.973704441 | 2.056322416 | 2.190331698 | 1.950241257 |
| ENSMUSG00000038679 | 2.477243823 | 2.333288838 | 2.248588359 | 2.246460484 | 2.168627946 | 2.418301291 | 1.977870661 |
| ENSMUSG00000038683 | 2.452372734 | 2.309601185 | 2.28296424 | 2.394357207 | 2.33281029 | 2.615950052 | 2.16937787 |
| ENSMUSG00000038685 | 2.450036412 | 2.236360959 | 2.264497931 | 2.426408097 | 2.423726155 | 2.673020907 | 2.244642249 |

| | | | | | | | |
|---|---|---|---|---|---|---|---|
| ENSMUSG00000038690 | 2.23778581 | 2.063166358 | 2.104463129 | 2.161864607 | 2.097547432 | 2.434568904 | 2.002662694 |
| ENSMUSG00000038691 | 0 | 0 | 0 | 0 | 0 | 0 | 0 |
| ENSMUSG00000038692 | 1.700465191 | 1.714495249 | 1.726722649 | 1.388647213 | 1.439191811 | 1.397940009 | 1.444790289 |
| ENSMUSG00000038695 | 1.505869732 | 1.445339928 | 1.705044741 | 1.311981878 | 1.509475496 | 1.204119983 | 1.68886578 |
| ENSMUSG00000038696 | 2.448863324 | 2.291226732 | 2.295598588 | 2.166333979 | 2.200473444 | 2.462397998 | 2.121219121 |
| ENSMUSG00000038697 | 2.348397716 | 2.174241882 | 2.181980114 | 2.167221246 | 2.180543208 | 2.434568904 | 2.041943747 |
| ENSMUSG00000038700 | 1.558729299 | 1.356566189 | 1.458222504 | 1.339388001 | 1.394615667 | 1.447158031 | 1.350478789 |
| ENSMUSG00000038702 | 2.026699189 | 1.951838731 | 1.921625132 | 1.931980114 | 1.870073561 | 2.071882007 | 1.705896842 |
| ENSMUSG00000038704 | 0 | 0 | 0 | 0 | 0 | 0 | 0 |
| ENSMUSG00000038705 | 2.188178997 | 2.027131114 | 2.057204673 | 2.153677461 | 2.130260132 | 2.357934847 | 2.029982479 |
| ENSMUSG00000038708 | 2.715608727 | 2.54216803 | 2.538071211 | 2.521534056 | 2.502191785 | 2.838219222 | 2.359843777 |
| ENSMUSG00000038709 | 0 | 0 | 0 | 0 | 0 | 0 | 0 |
| ENSMUSG00000038712 | 2.131864492 | 2.016522298 | 1.988549078 | 2.178561379 | 2.196996016 | 2.423245874 | 2.096394553 |
| ENSMUSG00000038717 | 2.275045611 | 2.102347734 | 2.111770807 | 2.08272922 | 2.077536464 | 2.352182518 | 1.951796623 |
| ENSMUSG00000038718 | 2.075459174 | 2 | 1.942046081 | 2.05357978 | 2.007252778 | 2.269512944 | 1.87805087 |
| ENSMUSG00000038721 | 1.944805158 | 1.916333979 | 1.982684334 | 1.681980114 | 1.686820904 | 1.903089987 | 1.669034113 |
| ENSMUSG00000038722 | 2.512981297 | 2.325374049 | 2.331045137 | 2.513820609 | 2.368520226 | 2.798650645 | 2.249624299 |
| ENSMUSG00000038725 | 0.45815691 | 0.580482024 | 0.75 | 0.25 | 0 | 0.301029996 | 0.448487648 |
| ENSMUSG00000038729 | 0.289064826 | 0.25 | 0 | 0 | 0.632412477 | 0 | 0 |
| ENSMUSG00000038732 | 2.40033094 | 2.186208459 | 2.170748646 | 2.198603967 | 2.121713561 | 2.457881897 | 1.983546516 |
| ENSMUSG00000038733 | 3.001563678 | 2.752107156 | 2.718361028 | 2.918989258 | 2.840869114 | 3.267640982 | 2.602872139 |
| ENSMUSG00000038736 | 2.704939145 | 2.508699741 | 2.469895812 | 2.536801231 | 2.446505421 | 2.771587481 | 2.230859913 |
| ENSMUSG00000038738 | 0 | 0 | 0.25 | 0 | 0 | 0 | 0.448487648 |
| ENSMUSG00000038740 | 2.213719572 | 2.060793496 | 2.125956434 | 1.937048212 | 1.981187939 | 2.198657087 | 1.906930404 |
| ENSMUSG00000038742 | 0.747221736 | 0.580482024 | 0.951838731 | 0.5 | 0.686820904 | 0 | 0.935079929 |
| ENSMUSG00000038745 | 0 | 0 | 0 | 0 | 0 | 0 | 0 |
| ENSMUSG00000038750 | 0 | 0 | 0 | 0 | 0 | 0 | 0 |
| ENSMUSG00000038751 | 0 | 0.25 | 0 | 0 | 0 | 0 | 0.224243824 |
| ENSMUSG00000038754 | 0 | 0.25 | 0.25 | 0.5 | 0.244650542 | 0.301029996 | 0.224243824 |
| ENSMUSG00000038756 | 0.960252568 | 0.75 | 0.79248125 | 0.646240625 | 0.686820904 | 0.602059991 | 0.579661877 |
| ENSMUSG00000038759 | 3.155828264 | 2.870575946 | 2.836212671 | 2.873463862 | 2.80828721 | 3.241048151 | 2.608423541 |
| ENSMUSG00000038760 | 0 | 0 | 0 | 0 | 0 | 0 | 0 |
| ENSMUSG00000038762 | 2.898714905 | 2.683677405 | 2.731573999 | 2.697790722 | 2.715622296 | 3.04453976 | 2.61458208 |
| ENSMUSG00000038763 | 0.289064826 | 0 | 0.25 | 0.25 | 0.244650542 | 0 | 0 |

| | | | | | | | |
|---|---|---|---|---|---|---|---|
| ENSMUSG00000038764 | 1.747221736 | 1.589388001 | 1.634789703 | 1.729715809 | 1.588235379 | 1.799340549 | 1.562034107 |
| ENSMUSG00000038765 | 0 | 0 | 0 | 0.896240625 | 0.955822902 | 0.77815125 | 1.121219121 |
| ENSMUSG00000038766 | 2.81052167 | 2.564346961 | 2.565229883 | 2.747348625 | 2.659629116 | 3.07809415 | 2.4836078 |
| ENSMUSG00000038768 | 0 | 0.25 | 0 | 0 | 0.244650542 | 0 | 0.224243824 |
| ENSMUSG00000038770 | 0 | 0 | 0 | 0 | 0 | 0 | 0 |
| ENSMUSG00000038773 | 2.662129038 | 2.451435968 | 2.438721876 | 2.435787849 | 2.41479008 | 2.743509765 | 2.283895396 |
| ENSMUSG00000038774 | 2.743181252 | 2.517530236 | 2.514497931 | 2.532642641 | 2.420030246 | 2.818225894 | 2.251165585 |
| ENSMUSG00000038775 | 1.857607105 | 1.744319981 | 1.834962501 | 1.29248125 | 1.358933174 | 1.531478917 | 1.42167221 |
| ENSMUSG00000038776 | 2.669806838 | 2.417664062 | 2.466819685 | 2.097004321 | 2.087207099 | 2.372912003 | 1.987044015 |
| ENSMUSG00000038777 | 1.227926429 | 1.17510993 | 1.188721876 | 1.438721876 | 1.546663136 | 1.69019608 | 1.520678036 |
| ENSMUSG00000038780 | 2.745494892 | 2.550530956 | 2.55175358 | 2.441217898 | 2.440596613 | 2.758154622 | 2.327754286 |
| ENSMUSG00000038781 | 1.036286563 | 0.951838731 | 1.130890489 | 1.226722649 | 1.293076518 | 1.146128036 | 1.319146374 |
| ENSMUSG00000038782 | 0.45815691 | 0.25 | 0.646240625 | 0.396240625 | 0.489301084 | 0.602059991 | 0.672731473 |
| ENSMUSG00000038784 | 2.192545868 | 2.013820609 | 1.976722649 | 2.098079356 | 2.021325983 | 2.356025857 | 1.921737195 |
| ENSMUSG00000038791 | 0 | 0 | 0 | 0 | 0 | 0 | 0 |
| ENSMUSG00000038792 | 0 | 0 | 0 | 0 | 0 | 0 | 0 |
| ENSMUSG00000038793 | 0.960252568 | 0.701838731 | 1.04248125 | 0.830482024 | 0.931471446 | 1 | 0.984950056 |
| ENSMUSG00000038797 | 1.069666964 | 1.17510993 | 1.27186571 | 0.75 | 1.121713561 | 0.602059991 | 1.176815335 |
| ENSMUSG00000038801 | 0 | 0 | 0 | 0 | 0 | 0 | 0 |
| ENSMUSG00000038803 | 1.596667477 | 1.532320754 | 1.668106335 | 1.214495249 | 1.401338844 | 1.342422681 | 1.40946263 |
| ENSMUSG00000038805 | 0 | 0.25 | 0 | 1.238549078 | 1.358933174 | 1.380211242 | 1.340368108 |
| ENSMUSG00000038806 | 2.425923778 | 2.27053726 | 2.294977273 | 2.312574604 | 2.289867804 | 2.559906625 | 2.158915016 |
| ENSMUSG00000038807 | 2.190368148 | 2.006976499 | 2.099151195 | 1.839388001 | 1.800029087 | 1.959041392 | 1.806347338 |
| ENSMUSG00000038811 | 0.45815691 | 0.646240625 | 0.5 | 0 | 0.244650542 | 0 | 0 |
| ENSMUSG00000038812 | 1.867194479 | 1.854463129 | 1.938721876 | 1.78743678 | 1.795698282 | 1.963787827 | 1.871156825 |
| ENSMUSG00000038816 | 2.158622101 | 2.05357978 | 2.071350555 | 2.343759858 | 2.268268456 | 2.592176757 | 2.131619535 |
| ENSMUSG00000038822 | 2.241665209 | 2.118933358 | 2.048689214 | 2.122963274 | 2.059432181 | 2.389166084 | 1.939139147 |
| ENSMUSG00000038827 | 2.273259602 | 2.113831805 | 2.088286706 | 2.209800947 | 2.157079823 | 2.447158031 | 2.010521755 |
| ENSMUSG00000038828 | 2.54867984 | 2.380890489 | 2.409156155 | 2.226722649 | 2.377434805 | 2.532754379 | 2.250857915 |
| ENSMUSG00000038831 | 1.671187741 | 1.54745614 | 1.602347734 | 1.75835575 | 1.766173492 | 2.053078443 | 1.690609803 |
| ENSMUSG00000038836 | 1.432083896 | 1.431980114 | 1.451838731 | 1.348079356 | 1.310728002 | 1.278753601 | 1.224243824 |
| ENSMUSG00000038838 | 1.989530018 | 1.839388001 | 1.901832578 | 1.527131114 | 1.836786029 | 1.716003344 | 1.803905701 |
| ENSMUSG00000038840 | 0 | 0 | 0 | 0 | 0 | 0 | 0 |
| ENSMUSG00000038843 | 1.740854706 | 1.622963274 | 1.685366747 | 1.188721876 | 1.106691885 | 1.176091259 | 0.984950056 |

| | | | | | | | |
|---|---|---|---|---|---|---|---|
| ENSMUSG00000038844 | 2.205378646 | 2.02186571 | 2.092671852 | 2.083847589 | 2.029283463 | 2.260071388 | 1.826389311 |
| ENSMUSG00000038845 | 2.514991096 | 2.30175358 | 2.361770807 | 2.373463862 | 2.418913884 | 2.719331287 | 2.286938317 |
| ENSMUSG00000038848 | 2.542156295 | 2.325945187 | 2.30539678 | 2.425543171 | 2.408704478 | 2.740362689 | 2.237984665 |
| ENSMUSG00000038855 | 2.192545868 | 2.071350555 | 2.153677461 | 2.027131114 | 2.134457044 | 2.359835482 | 2.00464543 |
| ENSMUSG00000038859 | 1.753493018 | 1.614857905 | 1.610735874 | 1.642463902 | 1.689774529 | 1.832508913 | 1.610050272 |
| ENSMUSG00000038860 | 1.432083896 | 1.339388001 | 1.330482024 | 1.602347734 | 1.572189156 | 1.602059991 | 1.405288179 |
| ENSMUSG00000038861 | 2.367390214 | 2.171625132 | 2.176839783 | 2.248588359 | 2.233484039 | 2.540329475 | 2.07959139 |
| ENSMUSG00000038866 | 2.229917041 | 2.083847589 | 2.077084758 | 2.104463129 | 2.048957937 | 2.250420002 | 1.87902385 |
| ENSMUSG00000038871 | 2.121692466 | 1.997171172 | 2.005591953 | 1.990000483 | 2.00243419 | 2.264817823 | 1.87902385 |
| ENSMUSG00000038872 | 2.352857998 | 2.260414788 | 2.296838018 | 2.02053726 | 2.114724103 | 2.173186268 | 1.978585612 |
| ENSMUSG00000038876 | 2.278594844 | 2.146240625 | 2.149047439 | 2.136723615 | 2.025895058 | 2.36361198 | 1.943148064 |
| ENSMUSG00000038879 | 0.747221736 | 0.5 | 0.580482024 | 0.580482024 | 0.733951626 | 0.77815125 | 0.710836105 |
| ENSMUSG00000038880 | 0 | 0.396240625 | 0.580482024 | 0 | 0.244650542 | 0.477121255 | 0.876096088 |
| ENSMUSG00000038883 | 0 | 0 | 0 | 0 | 0 | 0 | 0 |
| ENSMUSG00000038884 | 0.671187741 | 0.25 | 0.25 | 0.976722649 | 0.812711509 | 0.954242509 | 1.054044578 |
| ENSMUSG00000038886 | 2.595015862 | 2.447383411 | 2.485994979 | 2.212155735 | 2.279510047 | 2.509202522 | 2.179128924 |
| ENSMUSG00000038888 | 1.470606563 | 1.418106335 | 1.516522298 | 1.356566189 | 1.373641807 | 1.380211242 | 1.401059159 |
| ENSMUSG00000038891 | 0 | 0 | 0 | 0 | 0 | 0 | 0 |
| ENSMUSG00000038893 | 1.445324132 | 1.451838731 | 1.606566189 | 1.589388001 | 1.671670246 | 1.812913357 | 1.71730663 |
| ENSMUSG00000038894 | 1.907802659 | 2.00835575 | 2.090485943 | 1.445339928 | 1.66856048 | 1.491361694 | 1.73898563 |
| ENSMUSG00000038895 | 1.494443473 | 1.42510993 | 1.557204673 | 1.25 | 1.427019704 | 1.301029996 | 1.493062805 |
| ENSMUSG00000038900 | 3.132777036 | 2.900558725 | 2.964980743 | 2.832449085 | 2.788185538 | 3.214578954 | 2.71386689 |
| ENSMUSG00000038902 | 2.500714755 | 2.327653195 | 2.319612365 | 2.326515422 | 2.303049176 | 2.617000341 | 2.159731975 |
| ENSMUSG00000038903 | 0.45815691 | 0.79248125 | 0.864857905 | 0.5 | 0 | 0.602059991 | 0.355418052 |
| ENSMUSG00000038909 | 2.858927525 | 2.591307212 | 2.587458523 | 2.69225283 | 2.618504144 | 3.027349608 | 2.44445206 |
| ENSMUSG00000038910 | 1.418409478 | 1.410964047 | 1.396240625 | 1.5 | 1.407936347 | 1.505149978 | 1.345462945 |
| ENSMUSG00000038914 | 2.604828841 | 2.436208459 | 2.444108258 | 2.497880462 | 2.468227512 | 2.772321707 | 2.313108585 |
| ENSMUSG00000038916 | 1.269664473 | 1.27186571 | 1.364857905 | 0.5 | 0.568060967 | 0.301029996 | 0.775756176 |
| ENSMUSG00000038917 | 0.811507563 | 0.79248125 | 0.896240625 | 0.5 | 0.686820904 | 0.301029996 | 0.579661877 |
| ENSMUSG00000038925 | 0.289064826 | 0.25 | 0.25 | 0.25 | 0.387761935 | 0.301029996 | 0.224243824 |
| ENSMUSG00000038930 | 1.482695304 | 1.464495249 | 1.54248125 | 1.356566189 | 1.327538614 | 1.278753601 | 1.224243824 |
| ENSMUSG00000038932 | 0 | 0 | 0 | 0.580482024 | 0.489301084 | 0.301029996 | 0.829800754 |
| ENSMUSG00000038936 | 2.361637952 | 2.188721876 | 2.167221246 | 2.381380202 | 2.320220671 | 2.645422269 | 2.160139682 |
| ENSMUSG00000038943 | 2.998430437 | 2.73945689 | 2.723514962 | 2.839800121 | 2.739104362 | 3.172602931 | 2.549317365 |

| | | | | | | | |
|---|---|---|---|---|---|---|---|
| ENSMUSG00000038949 | 2.219867581 | 2.107613138 | 2.117918804 | 2.086073977 | 2.093822267 | 2.315970345 | 1.966952343 |
| ENSMUSG00000038954 | 2.026699189 | 1.919870025 | 1.856566189 | 1.821350555 | 1.770879644 | 1.954242509 | 1.74047993 |
| ENSMUSG00000038957 | 2.35727108 | 2.180274797 | 2.171625132 | 2.280383379 | 2.257835496 | 2.534026106 | 2.095897983 |
| ENSMUSG00000038963 | 0.867194479 | 1.114857905 | 1.348079356 | 1.561981878 | 1.625423019 | 1.69019608 | 1.605556929 |
| ENSMUSG00000038965 | 2.622240692 | 2.412762923 | 2.390083209 | 2.544354884 | 2.476238662 | 2.866877814 | 2.331854053 |
| ENSMUSG00000038967 | 1.325351389 | 1.438721876 | 1.396240625 | 1.42510993 | 1.439191811 | 1.62324929 | 1.433437704 |
| ENSMUSG00000038968 | 0 | 0 | 0 | 0 | 0 | 0 | 0 |
| ENSMUSG00000038970 | 2.481610694 | 2.3464656 | 2.353935442 | 2.341033664 | 2.352243976 | 2.618048097 | 2.222232323 |
| ENSMUSG00000038973 | 0 | 0 | 0 | 0 | 0 | 0 | 0 |
| ENSMUSG00000038975 | 2.675985209 | 2.434945152 | 2.440803092 | 2.514836115 | 2.465951417 | 2.799340549 | 2.291042162 |
| ENSMUSG00000038976 | 2.603209296 | 2.444929839 | 2.474839231 | 2.186208459 | 2.317747248 | 2.517195898 | 2.27142916 |
| ENSMUSG00000038980 | 0 | 0 | 0 | 0 | 0 | 0 | 0 |
| ENSMUSG00000038982 | 1.707474304 | 1.688721876 | 1.695339928 | 1.77186571 | 1.777823267 | 1.929418926 | 1.699191618 |
| ENSMUSG00000038984 | 1.99653913 | 1.860735874 | 1.821350555 | 1.580482024 | 1.584291694 | 1.672097858 | 1.448487648 |
| ENSMUSG00000038987 | 1.389637216 | 1.302363341 | 1.464495249 | 1.26109853 | 1.293076518 | 1.176091259 | 1.238624656 |
| ENSMUSG00000038990 | 2.003432385 | 1.874961472 | 1.896240625 | 2.041226732 | 2.038163331 | 2.264817823 | 1.900630878 |
| ENSMUSG00000038991 | 3.139109537 | 2.8918702 | 2.859438018 | 2.869313081 | 2.835070703 | 3.266466895 | 2.661077362 |
| ENSMUSG00000038994 | 0.289064826 | 0 | 0 | 0 | 0 | 0.301029996 | 0.224243824 |
| ENSMUSG00000038997 | 0 | 0.25 | 0.5 | 0.25 | 0 | 0.477121255 | 0.355418052 |
| ENSMUSG00000039000 | 2.966070923 | 2.678561379 | 2.652756199 | 2.714495249 | 2.626045333 | 3.026941628 | 2.446812267 |
| ENSMUSG00000039001 | 2.599951214 | 2.49539182 | 2.610476676 | 2.302972074 | 2.49625462 | 2.667452953 | 2.479078811 |
| ENSMUSG00000039004 | 0.289064826 | 0 | 0 | 0.646240625 | 0.905314583 | 0.698970004 | 0.853775828 |
| ENSMUSG00000039005 | 1.482695304 | 1.431980114 | 1.52186571 | 1.403677461 | 1.188507684 | 1.544068044 | 1.313616061 |
| ENSMUSG00000039007 | 2.269664473 | 2.072504712 | 2.05357978 | 1.741446071 | 1.726078673 | 1.929418926 | 1.591689859 |
| ENSMUSG00000039013 | 0 | 0.396240625 | 0.396240625 | 0 | 0 | 0 | 0 |
| ENSMUSG00000039016 | 2.089597219 | 2.032320754 | 2.012462137 | 2.061981878 | 2.015530186 | 2.311753861 | 1.999330888 |
| ENSMUSG00000039018 | 1.558729299 | 1.51109853 | 1.557204673 | 1.356566189 | 1.445123986 | 1.633468456 | 1.329931221 |
| ENSMUSG00000039021 | 0.289064826 | 0.396240625 | 0.79248125 | 0.580482024 | 0.632412477 | 0.602059991 | 0.672731473 |
| ENSMUSG00000039031 | 2.22189694 | 1.997171172 | 1.991446071 | 2.127938163 | 2.057362051 | 2.371067862 | 1.954885099 |
| ENSMUSG00000039032 | 0.289064826 | 0 | 0.25 | 0 | 0.244650542 | 0 | 0.224243824 |
| ENSMUSG00000039033 | 1.747221736 | 1.54745614 | 1.618933358 | 1.664552871 | 1.655839964 | 1.812913357 | 1.520678036 |
| ENSMUSG00000039037 | 0 | 0 | 0 | 0 | 0.244650542 | 0.477121255 | 0 |
| ENSMUSG00000039041 | 2.020182925 | 1.990000483 | 2.00835575 | 1.909156155 | 2.038163331 | 2.11058971 | 2.021963573 |
| ENSMUSG00000039043 | 2.273259602 | 2.119945066 | 2.112802778 | 2.170748646 | 2.155511127 | 2.485721426 | 2.030591163 |

| | | | | | | | |
|---|---|---|---|---|---|---|---|
| ENSMUSG00000039046 | 2.442947872 | 2.255591953 | 2.214495249 | 2.27908599 | 2.232853197 | 2.541579244 | 2.065709059 |
| ENSMUSG00000039047 | 2.556920045 | 2.381869252 | 2.375956434 | 2.454245906 | 2.417419887 | 2.764176132 | 2.252393349 |
| ENSMUSG00000039048 | 1.975148165 | 1.802363341 | 1.843759858 | 1.933677405 | 1.992595054 | 2.161368002 | 1.805128823 |
| ENSMUSG00000039050 | 2.181541736 | 1.975216702 | 1.951838731 | 2.098079356 | 2.052132976 | 2.283301229 | 1.950241257 |
| ENSMUSG00000039055 | 2.141794304 | 1.96293726 | 1.964495249 | 2.019203899 | 2 | 2.252853031 | 1.924281235 |
| ENSMUSG00000039057 | 0.45815691 | 0.25 | 0 | 0.396240625 | 0.489301084 | 0.301029996 | 0.448487648 |
| ENSMUSG00000039058 | 1.783508299 | 1.618933358 | 1.634789703 | 1.602347734 | 1.478764298 | 1.62324929 | 1.429563222 |
| ENSMUSG00000039059 | 0 | 0 | 0 | 0 | 0 | 0 | 0 |
| ENSMUSG00000039062 | 1.928763473 | 1.744319981 | 1.802363341 | 1.732684334 | 1.777823267 | 2.004321374 | 1.692344475 |
| ENSMUSG00000039063 | 1.578129653 | 1.552363341 | 1.516522298 | 1.42510993 | 1.373641807 | 1.477121255 | 1.40946263 |
| ENSMUSG00000039065 | 1.800532393 | 1.814346961 | 1.779735268 | 1.738549078 | 1.704182902 | 1.924279286 | 1.67819947 |
| ENSMUSG00000039067 | 2.987597818 | 2.753330668 | 2.742166698 | 2.869818758 | 2.799395527 | 3.226084116 | 2.625496416 |
| ENSMUSG00000039068 | 2.790815687 | 2.554489424 | 2.546218836 | 2.689763985 | 2.582185884 | 2.967079734 | 2.398728906 |
| ENSMUSG00000039069 | 1.714367558 | 1.698603967 | 1.671625132 | 1.646240625 | 1.695608647 | 1.62324929 | 1.605556929 |
| ENSMUSG00000039070 | 0 | 0 | 0 | 0 | 0 | 0 | 0 |
| ENSMUSG00000039079 | 0 | 0 | 0.396240625 | 0 | 0 | 0 | 0.355418052 |
| ENSMUSG00000039081 | 2.033115198 | 1.914552871 | 1.995748394 | 1.755591953 | 1.718026092 | 1.913813852 | 1.764880683 |
| ENSMUSG00000039084 | 0.960252568 | 0.646240625 | 0.951838731 | 0.75 | 0.686820904 | 0.477121255 | 0.829800754 |
| ENSMUSG00000039086 | 1.432083896 | 1.372963274 | 1.372963274 | 1.396240625 | 1.343584837 | 1.491361694 | 1.307989563 |
| ENSMUSG00000039087 | 2.163307908 | 2.027131114 | 2.064346961 | 1.870953944 | 1.974687643 | 2.100370545 | 1.903345709 |
| ENSMUSG00000039089 | 2.003432385 | 1.870953944 | 1.777131114 | 1.774508021 | 1.715300549 | 1.954242509 | 1.603286636 |
| ENSMUSG00000039092 | 0 | 0 | 0 | 0 | 0 | 0 | 0 |
| ENSMUSG00000039095 | 0 | 0 | 0 | 0.79248125 | 0.489301084 | 0.477121255 | 0.672731473 |
| ENSMUSG00000039096 | 1.671187741 | 1.711372513 | 1.729715809 | 1.602347734 | 1.621875691 | 1.716003344 | 1.532233387 |
| ENSMUSG00000039097 | 0.45815691 | 0 | 0 | 0 | 0.244650542 | 0.301029996 | 0.355418052 |
| ENSMUSG00000039098 | 0 | 0 | 0 | 0 | 0 | 0 | 0 |
| ENSMUSG00000039099 | 0 | 0 | 0.25 | 0.701838731 | 0.489301084 | 0.301029996 | 0.224243824 |
| ENSMUSG00000039100 | 2.948366515 | 2.700224975 | 2.663435195 | 2.673807073 | 2.612740596 | 2.995196292 | 2.4027959 |
| ENSMUSG00000039103 | 2.512981297 | 2.350753006 | 2.366901388 | 2.171625132 | 2.131103516 | 2.408239965 | 1.953344547 |
| ENSMUSG00000039105 | 2.740271036 | 2.54652856 | 2.548689214 | 2.393884812 | 2.340340306 | 2.643452676 | 2.217489603 |
| ENSMUSG00000039106 | 0 | 0 | 0 | 0 | 0 | 0 | 0 |
| ENSMUSG00000039108 | 2.199011616 | 2.058404919 | 2.109697963 | 2.016522298 | 2.033751322 | 2.193124598 | 1.985649549 |
| ENSMUSG00000039109 | 0 | 0 | 0 | 0 | 0 | 0 | 0 |
| ENSMUSG00000039110 | 0.289064826 | 0.5 | 0.79248125 | 1.130890489 | 1.091002363 | 1.041392685 | 1.168187745 |

| | | | | | | | |
|---|---|---|---|---|---|---|---|
| ENSMUSG00000039114 | 0.960252568 | 1.160964047 | 1.302363341 | 0.25 | 0.489301084 | 0.477121255 | 0.579661877 |
| ENSMUSG00000039115 | 1.036286563 | 1.02186571 | 1.04248125 | 0.92510993 | 1.121713561 | 1.079181246 | 0.896975297 |
| ENSMUSG00000039116 | 2.591692877 | 2.369945066 | 2.37246199 | 2.086073977 | 1.996316946 | 2.269512944 | 1.856911705 |
| ENSMUSG00000039117 | 1.967766868 | 1.819031101 | 1.821350555 | 1.839388001 | 1.883916752 | 2.113943352 | 1.844181586 |
| ENSMUSG00000039126 | 2.970129664 | 2.713911792 | 2.699009902 | 1.994319981 | 1.959951091 | 2.243038049 | 1.833176598 |
| ENSMUSG00000039128 | 2.664937347 | 2.486360959 | 2.474083101 | 2.572792675 | 2.485926424 | 2.884795364 | 2.316141565 |
| ENSMUSG00000039130 | 2.35433422 | 2.151832578 | 2.149047439 | 2.209012589 | 2.225193812 | 2.496929648 | 2.102294694 |
| ENSMUSG00000039131 | 0.960252568 | 0.864857905 | 0.92510993 | 0.25 | 0.489301084 | 0.477121255 | 0.579661877 |
| ENSMUSG00000039133 | 0.289064826 | 0 | 0 | 0.25 | 0.387761935 | 0.602059991 | 0 |
| ENSMUSG00000039137 | 1.527823874 | 1.527131114 | 1.614857905 | 1.080482024 | 1.188507684 | 0.954242509 | 1.23151414 |
| ENSMUSG00000039138 | 0 | 0 | 0 | 0 | 0 | 0 | 0 |
| ENSMUSG00000039145 | 2.479432974 | 2.281676118 | 2.250703754 | 2.104463129 | 2.087207099 | 2.344392274 | 1.95794437 |
| ENSMUSG00000039146 | 0 | 0 | 0 | 0.25 | 0 | 0 | 0 |
| ENSMUSG00000039148 | 2.647066901 | 2.479715809 | 2.54838134 | 2.418548067 | 2.502780045 | 2.736396502 | 2.409274047 |
| ENSMUSG00000039153 | 2.415874319 | 2.219129237 | 2.223704441 | 2.115881093 | 2.10031573 | 2.320146286 | 1.910475814 |
| ENSMUSG00000039154 | 0 | 0 | 0 | 0.396240625 | 0.387761935 | 0.477121255 | 0.224243824 |
| ENSMUSG00000039155 | 0 | 0 | 0 | 0.5 | 0.489301084 | 0.477121255 | 0.629532004 |
| ENSMUSG00000039156 | 2.283862692 | 2.169870025 | 2.178561379 | 2.12892496 | 2.086251872 | 2.348304863 | 1.984950056 |
| ENSMUSG00000039157 | 1.862428343 | 1.732684334 | 1.752806814 | 1.657339155 | 1.800029087 | 1.954242509 | 1.722075986 |
| ENSMUSG00000039158 | 1.614416215 | 1.54248125 | 1.67510993 | 1.25 | 1.136121934 | 0.903089987 | 1.224243824 |
| ENSMUSG00000039159 | 2.448863324 | 2.275821952 | 2.250703754 | 2.132845365 | 2.169384551 | 2.396199347 | 2.018825668 |
| ENSMUSG00000039163 | 1.734388958 | 1.678561379 | 1.708222504 | 1.755591953 | 1.695608647 | 1.913813852 | 1.532233387 |
| ENSMUSG00000039164 | 1.714367558 | 1.642463902 | 1.571350555 | 1.532320754 | 1.635855979 | 1.662757832 | 1.556816002 |
| ENSMUSG00000039166 | 1.837744666 | 1.692046081 | 1.668106335 | 1.678561379 | 1.611014646 | 1.799340549 | 1.493062805 |
| ENSMUSG00000039167 | 0 | 0.25 | 0 | 0.396240625 | 0.568060967 | 0 | 0.355418052 |
| ENSMUSG00000039168 | 2.969118676 | 2.766522298 | 2.831185778 | 2.411414608 | 2.432800077 | 2.746634199 | 2.375083475 |
| ENSMUSG00000039174 | 0 | 0 | 0 | 0 | 0 | 0 | 0 |
| ENSMUSG00000039176 | 2.213719572 | 2.129909063 | 2.180274797 | 2.14151351 | 2.221300875 | 2.376576957 | 2.110473216 |
| ENSMUSG00000039178 | 2.312518611 | 2.144357207 | 2.169870025 | 2.195339928 | 2.122577592 | 2.481442629 | 1.978585612 |
| ENSMUSG00000039179 | 0 | 0 | 0 | 0 | 0 | 0 | 0 |
| ENSMUSG00000039182 | 1.867194479 | 1.858657057 | 1.845926073 | 1.726722649 | 1.632412477 | 1.886490725 | 1.598697688 |
| ENSMUSG00000039183 | 2.205378646 | 2.091580554 | 2.112802778 | 1.970660762 | 2.091002363 | 2.230448921 | 1.995286892 |
| ENSMUSG00000039187 | 2.201144779 | 1.948603967 | 2.02053726 | 2.166333979 | 2.133621644 | 2.413299764 | 1.987044015 |
| ENSMUSG00000039191 | 2.774459379 | 2.551448312 | 2.494677615 | 2.5684489 | 2.512918998 | 2.893206753 | 2.332572178 |

| | | | | | | | |
|---|---|---|---|---|---|---|---|
| ENSMUSG00000039193 | 0 | 0.25 | 0.396240625 | 0 | 0 | 0 | 0.224243824 |
| ENSMUSG00000039194 | 0 | 0 | 0 | 0.25 | 0.244650542 | 0 | 0.448487648 |
| ENSMUSG00000039195 | 1.129344651 | 1.464495249 | 1.54248125 | 1.51109853 | 1.563883923 | 1.51851394 | 1.709198048 |
| ENSMUSG00000039196 | 0 | 0 | 0 | 0 | 0 | 0 | 0 |
| ENSMUSG00000039197 | 2.533616084 | 2.375956434 | 2.340485943 | 2.662762923 | 2.598914622 | 2.997386384 | 2.414141848 |
| ENSMUSG00000039198 | 0 | 0 | 0 | 0 | 0 | 0 | 0 |
| ENSMUSG00000039199 | 1.647796617 | 1.614857905 | 1.678561379 | 1.516522298 | 1.683842353 | 1.681241237 | 1.694069894 |
| ENSMUSG00000039200 | 1 | 0.75 | 0.830482024 | 1.27186571 | 1.244650542 | 1.477121255 | 1.089372236 |
| ENSMUSG00000039201 | 1.876566388 | 1.823655187 | 1.811981878 | 1.782320754 | 1.751666404 | 1.973127854 | 1.747849621 |
| ENSMUSG00000039202 | 2.611244851 | 2.382845365 | 2.410061234 | 2.363831805 | 2.364629672 | 2.62324929 | 2.210939786 |
| ENSMUSG00000039205 | 2.139333929 | 2.048689214 | 2.114857905 | 1.898114259 | 2.093822267 | 2.198657087 | 2.006616089 |
| ENSMUSG00000039206 | 1.907802659 | 1.811981878 | 1.898114259 | 1.841580554 | 1.906908067 | 2.071882007 | 1.86715039 |
| ENSMUSG00000039208 | 2.170239354 | 2.088286706 | 2.025821952 | 1.834962501 | 1.92093466 | 2.079181246 | 1.853775828 |
| ENSMUSG00000039209 | 0 | 0 | 0 | 0 | 0 | 0 | 0 |
| ENSMUSG00000039210 | 2.003432385 | 1.984159485 | 2.004202072 | 1.843759858 | 1.92093466 | 2.136720567 | 1.800208341 |
| ENSMUSG00000039215 | 0 | 0 | 0 | 0 | 0 | 0 | 0 |
| ENSMUSG00000039217 | 0.45815691 | 0.5 | 0.701838731 | 1.29248125 | 1.343584837 | 1.397940009 | 1.259064007 |
| ENSMUSG00000039218 | 3.583577065 | 3.225546666 | 3.235903425 | 3.232453296 | 3.204118463 | 3.652439748 | 2.982425699 |
| ENSMUSG00000039219 | 2.570301912 | 2.363317659 | 2.345385738 | 2.531676118 | 2.410235744 | 2.787460475 | 2.244642249 |
| ENSMUSG00000039220 | 2.614416215 | 2.427701608 | 2.483791262 | 2.350219859 | 2.366796383 | 2.704150517 | 2.265305657 |
| ENSMUSG00000039221 | 1.771760131 | 1.653677461 | 1.602347734 | 1.848079356 | 1.793512788 | 1.919078092 | 1.598697688 |
| ENSMUSG00000039224 | 0 | 0 | 0 | 0.25 | 0 | 0 | 0 |
| ENSMUSG00000039230 | 2.660717759 | 2.432405186 | 2.453445298 | 2.397646762 | 2.404846996 | 2.695481676 | 2.281664246 |
| ENSMUSG00000039231 | 2.647796617 | 2.402294685 | 2.393411797 | 2.485628626 | 2.432800077 | 2.808885867 | 2.271140178 |
| ENSMUSG00000039232 | 1.505869732 | 1.445339928 | 1.527131114 | 1.438721876 | 1.555378544 | 1.698970004 | 1.455757965 |
| ENSMUSG00000039233 | 2.103271637 | 1.979715809 | 1.96759118 | 2.119945066 | 2.086251872 | 2.269512944 | 1.94074868 |
| ENSMUSG00000039234 | 2.911692915 | 2.655284028 | 2.640083209 | 2.589388001 | 2.571222152 | 2.890979597 | 2.418387761 |
| ENSMUSG00000039236 | 1.249317394 | 1.188721876 | 1.26109853 | 1.330482024 | 1.47337555 | 1.491361694 | 1.459332757 |
| ENSMUSG00000039238 | 0.45815691 | 0.396240625 | 0.25 | 0 | 0.244650542 | 0 | 0 |
| ENSMUSG00000039239 | 2.413323654 | 2.235628626 | 2.233422664 | 2.209012589 | 2.149165677 | 2.428134794 | 1.993927582 |
| ENSMUSG00000039242 | 2.35433422 | 2.160964047 | 2.117918804 | 2.219895812 | 2.165585202 | 2.492760389 | 1.989815042 |
| ENSMUSG00000039244 | 2.048736215 | 1.888647213 | 2.023189285 | 1.769203899 | 1.934425071 | 2.045322979 | 1.980010793 |
| ENSMUSG00000039246 | 1.227926429 | 1.146240625 | 1.080482024 | 1.311981878 | 1.407936347 | 1.361727836 | 1.20919388 |
| ENSMUSG00000039252 | 0.811507563 | 0.701838731 | 0.580482024 | 0.79248125 | 0.686820904 | 0.477121255 | 0.803905701 |

| | | | | | | | |
|---|---|---|---|---|---|---|---|
| ENSMUSG00000039253 | 1.978790393 | 1.809601185 | 1.769203899 | 1.931980114 | 1.892300902 | 2.123851641 | 1.774416565 |
| ENSMUSG00000039254 | 2.054824387 | 1.921625132 | 1.926839783 | 1.845926073 | 1.938809675 | 2.130333768 | 1.819456572 |
| ENSMUSG00000039257 | 0 | 0 | 0 | 0 | 0 | 0 | 0 |
| ENSMUSG00000039262 | 3.076242824 | 2.863960227 | 2.89869778 | 2.711960089 | 2.775252365 | 3.09482038 | 2.666744179 |
| ENSMUSG00000039263 | 1.985980784 | 1.96293726 | 2.012462137 | 1.593759858 | 1.808534465 | 1.832508913 | 1.828667599 |
| ENSMUSG00000039264 | 0 | 0.25 | 0 | 0 | 0 | 0 | 0 |
| ENSMUSG00000039269 | 0 | 0 | 0 | 0 | 0 | 0 | 0 |
| ENSMUSG00000039270 | 2.245508853 | 2.025821952 | 2.02186571 | 1.975216702 | 1.94314048 | 2.170261715 | 1.766260282 |
| ENSMUSG00000039275 | 2.900309677 | 2.663882681 | 2.658475852 | 2.563166358 | 2.54899118 | 2.898176483 | 2.403948569 |
| ENSMUSG00000039278 | 0 | 0 | 0 | 0 | 0 | 0 | 0 |
| ENSMUSG00000039285 | 2.40033094 | 2.262462137 | 2.21293726 | 2.359697963 | 2.276175004 | 2.56937391 | 2.100829716 |
| ENSMUSG00000039286 | 2.883252421 | 2.661639678 | 2.640560606 | 2.643648382 | 2.60710012 | 2.986771734 | 2.426715983 |
| ENSMUSG00000039294 | 2.40033094 | 2.173371739 | 2.215271726 | 2.28743678 | 2.286629652 | 2.580924976 | 2.132508314 |
| ENSMUSG00000039296 | 0 | 0.25 | 0 | 0 | 0 | 0 | 0.224243824 |
| ENSMUSG00000039298 | 2.765759561 | 2.532320754 | 2.524836953 | 2.575374049 | 2.505123331 | 2.872156273 | 2.340601441 |
| ENSMUSG00000039304 | 0.671187741 | 0.25 | 0 | 0 | 0.489301084 | 0.301029996 | 0 |
| ENSMUSG00000039307 | 1.956444026 | 1.830482024 | 1.823655187 | 1.777131114 | 1.840643511 | 1.934498451 | 1.790137008 |
| ENSMUSG00000039308 | 2.026699189 | 1.940387808 | 1.96293726 | 2.089388001 | 2.137779008 | 2.392696953 | 2.048913829 |
| ENSMUSG00000039313 | 0 | 0 | 0 | 0 | 0.568060967 | 0 | 0.224243824 |
| ENSMUSG00000039315 | 0 | 0 | 0 | 0 | 0 | 0 | 0 |
| ENSMUSG00000039316 | 1.37447073 | 1.214495249 | 1.27186571 | 1.488549078 | 1.595992969 | 1.505149978 | 1.496250175 |
| ENSMUSG00000039318 | 2.688073951 | 2.467205639 | 2.487456678 | 2.42813175 | 2.393415815 | 2.717670503 | 2.279417602 |
| ENSMUSG00000039321 | 0 | 0 | 0 | 0 | 0 | 0 | 0 |
| ENSMUSG00000039323 | 1.740854706 | 1.755591953 | 1.79745614 | 1.945339928 | 2.069604532 | 2.184691431 | 2.151886289 |
| ENSMUSG00000039328 | 1.940852214 | 1.830482024 | 1.794977273 | 1.438721876 | 1.489301084 | 1.602059991 | 1.546120303 |
| ENSMUSG00000039329 | 0 | 0 | 0 | 0 | 0.244650542 | 0 | 0.224243824 |
| ENSMUSG00000039330 | 0 | 0 | 0.25 | 0.396240625 | 0.387761935 | 0.477121255 | 0.224243824 |
| ENSMUSG00000039335 | 0 | 0 | 0 | 0 | 0 | 0 | 0 |
| ENSMUSG00000039337 | 0 | 0 | 0 | 0.25 | 0 | 0 | 0.224243824 |
| ENSMUSG00000039339 | 0 | 0 | 0 | 0 | 0 | 0 | 0 |
| ENSMUSG00000039342 | 0 | 0 | 0 | 0 | 0 | 0 | 0 |
| ENSMUSG00000039345 | 1.404271433 | 1.348079356 | 1.330482024 | 1.17510993 | 1.387761935 | 1.322219295 | 1.383567577 |
| ENSMUSG00000039347 | 1.920505136 | 1.789967834 | 1.816696635 | 1.938721876 | 1.895599581 | 2.209515015 | 1.828667599 |
| ENSMUSG00000039349 | 0.45815691 | 0 | 0.25 | 0.951838731 | 0.812711509 | 0.602059991 | 0.935079929 |

| | | | | | | | |
|---|---|---|---|---|---|---|---|
| ENSMUSG00000039354 | 2.103271637 | 1.923371739 | 2.002806814 | 1.882845365 | 1.926994141 | 2.037426498 | 1.834294096 |
| ENSMUSG00000039356 | 2.433305075 | 2.243603647 | 2.267865591 | 2.400442697 | 2.364194729 | 2.673941999 | 2.204609933 |
| ENSMUSG00000039357 | 1.700465191 | 1.610735874 | 1.614857905 | 2.048689214 | 2.001219193 | 2.23299611 | 1.850609257 |
| ENSMUSG00000039358 | 0 | 0 | 0 | 0 | 0 | 0 | 0 |
| ENSMUSG00000039361 | 3.058925034 | 2.808704358 | 2.774508021 | 2.687886015 | 2.591450357 | 3.003460532 | 2.406812401 |
| ENSMUSG00000039364 | 0 | 0 | 0 | 0 | 0 | 0 | 0 |
| ENSMUSG00000039367 | 2.672564087 | 2.473325383 | 2.457034121 | 2.661639678 | 2.651624369 | 3.008174184 | 2.474176968 |
| ENSMUSG00000039372 | 0.289064826 | 0.396240625 | 0 | 0.396240625 | 0.244650542 | 0 | 0.355418052 |
| ENSMUSG00000039375 | 0.960252568 | 1.160964047 | 1.214495249 | 1.201838731 | 1.163285805 | 0.954242509 | 1.100339912 |
| ENSMUSG00000039376 | 0.45815691 | 0.25 | 0.25 | 0.25 | 0.812711509 | 0 | 0.579661877 |
| ENSMUSG00000039377 | 1.100572389 | 1.061981878 | 1.160964047 | 1.04248125 | 1.121713561 | 1.041392685 | 1.319146374 |
| ENSMUSG00000039382 | 2.060824957 | 1.940387808 | 1.987091808 | 1.816696635 | 1.850106709 | 1.963787827 | 1.821784031 |
| ENSMUSG00000039384 | 1.903481042 | 1.809601185 | 1.789967834 | 1.843759858 | 1.903713872 | 2.079181246 | 1.833176598 |
| ENSMUSG00000039385 | 0 | 0 | 0 | 0 | 0 | 0.301029996 | 0.448487648 |
| ENSMUSG00000039391 | 0.289064826 | 0.5 | 0.396240625 | 0.25 | 0.244650542 | 0 | 0.224243824 |
| ENSMUSG00000039395 | 1.445324132 | 1.282320754 | 1.302363341 | 1.571350555 | 1.639266209 | 1.851258349 | 1.546120303 |
| ENSMUSG00000039396 | 2.111266113 | 2.004202072 | 1.994319981 | 2.112802778 | 2.0510778 | 2.28780173 | 1.966952343 |
| ENSMUSG00000039405 | 2.888197527 | 2.666333979 | 2.619692405 | 2.477847997 | 2.406395052 | 2.79518459 | 2.246831342 |
| ENSMUSG00000039410 | 0 | 0 | 0 | 0 | 0 | 0 | 0.224243824 |
| ENSMUSG00000039414 | 2.617563644 | 2.37892496 | 2.373963757 | 2.443696765 | 2.42262144 | 2.740362689 | 2.293479712 |
| ENSMUSG00000039419 | 0 | 0 | 0 | 0 | 0 | 0 | 0.224243824 |
| ENSMUSG00000039427 | 1.91631382 | 1.802363341 | 1.799918086 | 1.527131114 | 1.618292349 | 1.740362689 | 1.61448206 |
| ENSMUSG00000039428 | 1.964026642 | 1.837182039 | 1.79745614 | 1.930274797 | 1.816839698 | 1.944482672 | 1.692344475 |
| ENSMUSG00000039431 | 1.631440309 | 1.42510993 | 1.372963274 | 1.657339155 | 1.652586889 | 1.748188027 | 1.535058868 |
| ENSMUSG00000039438 | 0 | 0 | 0 | 0.25 | 0 | 0 | 0 |
| ENSMUSG00000039449 | 2.463861956 | 2.267865591 | 2.284247778 | 2.313166358 | 2.212046813 | 2.558708571 | 2.092400484 |
| ENSMUSG00000039450 | 1.568542279 | 1.54248125 | 1.622963274 | 1.098079356 | 1.176121988 | 1.255272505 | 1.272006351 |
| ENSMUSG00000039452 | 0.578129653 | 0.580482024 | 0.896240625 | 0.896240625 | 1 | 0.698970004 | 0.896975297 |
| ENSMUSG00000039456 | 2.401648579 | 2.262462137 | 2.218361028 | 2.290597832 | 2.245260665 | 2.550228353 | 2.10131878 |
| ENSMUSG00000039457 | 1.686083339 | 1.67510993 | 1.903677461 | 1.809601185 | 1.916320788 | 1.973127854 | 1.837623616 |
| ENSMUSG00000039458 | 2.396352858 | 2.236360959 | 2.200224975 | 2.345385738 | 2.264824954 | 2.57634135 | 2.12713864 |
| ENSMUSG00000039461 | 1.806056082 | 1.671625132 | 1.701838731 | 1.845926073 | 1.766173492 | 1.903089987 | 1.669034113 |
| ENSMUSG00000039462 | 0.578129653 | 0.864857905 | 0.580482024 | 0.396240625 | 0 | 0 | 0.224243824 |
| ENSMUSG00000039463 | 1.99653913 | 1.926839783 | 1.966046536 | 1.970660762 | 2.059432181 | 2.152288344 | 1.944737824 |

| | | | | | | | |
|---|---|---|---|---|---|---|---|
| ENSMUSG00000039470 | 2.304292853 | 2.079353153 | 2.14151351 | 2.389126514 | 2.334234459 | 2.654176542 | 2.139097452 |
| ENSMUSG00000039473 | 2.787177998 | 2.520869832 | 2.535845803 | 2.488912477 | 2.44788147 | 2.775974331 | 2.285005227 |
| ENSMUSG00000039474 | 1.789260561 | 1.901832578 | 1.896240625 | 1.646240625 | 1.846351821 | 1.934498451 | 1.863093717 |
| ENSMUSG00000039476 | 1.816888701 | 1.821350555 | 1.83272922 | 1.610735874 | 1.873585591 | 1.959041392 | 1.819456572 |
| ENSMUSG00000039477 | 2.768497801 | 2.635516136 | 2.773849256 | 2.130890489 | 2.300451989 | 2.419955748 | 2.359623923 |
| ENSMUSG00000039478 | 1.907802659 | 1.741446071 | 1.75835575 | 1.856566189 | 1.846351821 | 2.012837225 | 1.665294008 |
| ENSMUSG00000039480 | 1.99653913 | 1.852347734 | 1.856566189 | 1.933677405 | 1.890639929 | 2.075546961 | 1.704233519 |
| ENSMUSG00000039481 | 0 | 0 | 0 | 0 | 0.244650542 | 0 | 0.224243824 |
| ENSMUSG00000039483 | 2.057835465 | 2.02053726 | 2.127938163 | 1.634789703 | 1.92245931 | 1.880813592 | 1.966952343 |
| ENSMUSG00000039485 | 1.747221736 | 1.688721876 | 1.606566189 | 1.903677461 | 1.903713872 | 2.079181246 | 1.73898563 |
| ENSMUSG00000039488 | 0 | 0 | 0 | 0 | 0 | 0 | 0 |
| ENSMUSG00000039492 | 0.289064826 | 0.25 | 0 | 0 | 0.387761935 | 0.301029996 | 0.224243824 |
| ENSMUSG00000039496 | 0.91631382 | 0.79248125 | 0.830482024 | 0.79248125 | 0.489301084 | 0.477121255 | 0.672731473 |
| ENSMUSG00000039497 | 2.610448227 | 2.409156155 | 2.391989019 | 2.396240625 | 2.314756076 | 2.67669361 | 2.170563634 |
| ENSMUSG00000039501 | 2.141794304 | 2.016522298 | 2.028435542 | 2.237091808 | 2.241584006 | 2.506505032 | 2.146406417 |
| ENSMUSG00000039508 | 0.671187741 | 0.5 | 0.25 | 0.25 | 0.632412477 | 0 | 0 |
| ENSMUSG00000039509 | 2.743181252 | 2.509043403 | 2.498234584 | 2.606566189 | 2.557146486 | 2.913283902 | 2.38172418 |
| ENSMUSG00000039512 | 2.467247894 | 2.295598588 | 2.317865757 | 2.30357978 | 2.328021124 | 2.550228353 | 2.203542812 |
| ENSMUSG00000039515 | 2.867194479 | 2.645535495 | 2.660738556 | 2.560793496 | 2.594862785 | 2.8876173 | 2.46982607 |
| ENSMUSG00000039518 | 2.167941649 | 2.049918086 | 2.01109853 | 1.02186571 | 1.074582839 | 0.903089987 | 0.916588299 |
| ENSMUSG00000039519 | 0 | 0 | 0 | 0.25 | 0.387761935 | 0.477121255 | 0.355418052 |
| ENSMUSG00000039521 | 0.671187741 | 0.580482024 | 0.5 | 0 | 0.632412477 | 0.477121255 | 0 |
| ENSMUSG00000039523 | 2.342375483 | 2.201838731 | 2.193696765 | 2.19697564 | 2.26251048 | 2.51054501 | 2.11845007 |
| ENSMUSG00000039529 | 0.45815691 | 0.396240625 | 0 | 0.75 | 0.489301084 | 0.84509804 | 0.775756176 |
| ENSMUSG00000039530 | 2.652148299 | 2.424241882 | 2.411414608 | 2.475216702 | 2.366364104 | 2.776701184 | 2.228889254 |
| ENSMUSG00000039531 | 1.87190676 | 1.701838731 | 1.685366747 | 1.843759858 | 1.763796674 | 2.025305865 | 1.618853957 |
| ENSMUSG00000039533 | 0 | 0 | 0 | 0.25 | 0 | 0.301029996 | 0.224243824 |
| ENSMUSG00000039536 | 2.545896521 | 2.327653195 | 2.302972074 | 2.481573999 | 2.444777781 | 2.8162413 | 2.294288164 |
| ENSMUSG00000039539 | 0 | 0 | 0 | 0 | 0 | 0 | 0 |
| ENSMUSG00000039540 | 0 | 0 | 0 | 0 | 0 | 0 | 0 |
| ENSMUSG00000039542 | 2.556012466 | 2.353935442 | 2.334962501 | 1.956637122 | 1.900490506 | 2.1430148 | 1.888596011 |
| ENSMUSG00000039543 | 0 | 0 | 0.25 | 0 | 0 | 0.602059991 | 0.224243824 |
| ENSMUSG00000039545 | 0.671187741 | 0.5 | 0.701838731 | 0.580482024 | 0.632412477 | 0.477121255 | 0.448487648 |
| ENSMUSG00000039546 | 0 | 0 | 0 | 0 | 0 | 0 | 0 |

| | | | | | | | |
|---|---|---|---|---|---|---|---|
| ENSMUSG00000039552 | 0.578129653 | 0 | 0 | 0 | 0 | 0.301029996 | 0.355418052 |
| ENSMUSG00000039555 | 0 | 0 | 0 | 0 | 0 | 0 | 0 |
| ENSMUSG00000039556 | 0.289064826 | 0.396240625 | 0 | 0.580482024 | 0.632412477 | 0.301029996 | 0.448487648 |
| ENSMUSG00000039563 | 0.811507563 | 0.79248125 | 0.75 | 0.396240625 | 0.686820904 | 0.477121255 | 0.520678036 |
| ENSMUSG00000039568 | 1.956444026 | 1.910964047 | 2.002806814 | 1.476722649 | 1.433158226 | 1.491361694 | 1.514741806 |
| ENSMUSG00000039577 | 1.53838222 | 1.464495249 | 1.657339155 | 1.27186571 | 1.572189156 | 1.591064607 | 1.676387053 |
| ENSMUSG00000039578 | 1.885732304 | 1.75 | 1.777131114 | 1.598079356 | 1.462344774 | 1.681241237 | 1.444790289 |
| ENSMUSG00000039579 | 0.671187741 | 0.580482024 | 0.75 | 0.864857905 | 0.846351821 | 0.84509804 | 0.710836105 |
| ENSMUSG00000039585 | 2.785084983 | 2.55357978 | 2.541540771 | 2.632113528 | 2.504246421 | 2.872156273 | 2.321629237 |
| ENSMUSG00000039599 | 1.936861443 | 1.872963274 | 1.834962501 | 2.019203899 | 1.990091809 | 2.235528447 | 1.899720848 |
| ENSMUSG00000039601 | 2.199011616 | 2.03360658 | 2.081607372 | 1.25 | 1 | 1.041392685 | 1 |
| ENSMUSG00000039607 | 2.194712275 | 1.988549078 | 1.933677405 | 2.080482024 | 1.977302144 | 2.298853076 | 1.864112664 |
| ENSMUSG00000039608 | 0 | 0 | 0 | 0 | 0 | 0 | 0 |
| ENSMUSG00000039611 | 1.445324132 | 1.348079356 | 1.445339928 | 1.62694866 | 1.706995316 | 1.886490725 | 1.616675393 |
| ENSMUSG00000039615 | 1.876566388 | 1.890560606 | 2.056000419 | 1.610735874 | 1.814781639 | 1.770852012 | 1.914853627 |
| ENSMUSG00000039616 | 1.269664473 | 1.26109853 | 1.321350555 | 1.51109853 | 1.467903253 | 1.681241237 | 1.379042812 |
| ENSMUSG00000039617 | 0 | 0 | 0 | 0 | 0 | 0 | 0 |
| ENSMUSG00000039620 | 0.289064826 | 0 | 0 | 0.25 | 0.632412477 | 0.301029996 | 0.355418052 |
| ENSMUSG00000039621 | 0.578129653 | 0.580482024 | 0.75 | 1.396240625 | 1.599808748 | 1.505149978 | 1.589319772 |
| ENSMUSG00000039623 | 1.971473847 | 1.870953944 | 1.978222334 | 1.580482024 | 1.680838453 | 1.716003344 | 1.766260282 |
| ENSMUSG00000039628 | 0.289064826 | 0 | 0 | 0 | 0.387761935 | 0 | 0 |
| ENSMUSG00000039629 | 1.129344651 | 1.160964047 | 1.146240625 | 1.634789703 | 1.611014646 | 1.86332286 | 1.577201671 |
| ENSMUSG00000039630 | 2.964709183 | 2.67813175 | 2.645535495 | 2.748234584 | 2.682961504 | 3.030194785 | 2.466365979 |
| ENSMUSG00000039632 | 0.578129653 | 0.646240625 | 0.646240625 | 0.701838731 | 0.77552387 | 0.602059991 | 0.876096088 |
| ENSMUSG00000039633 | 1.842799666 | 1.705044741 | 1.67510993 | 1.348079356 | 1.433158226 | 1.477121255 | 1.278288402 |
| ENSMUSG00000039634 | 1.759671389 | 1.584962501 | 1.561981878 | 1.720660762 | 1.674752852 | 1.832508913 | 1.476637173 |
| ENSMUSG00000039637 | 1.765759561 | 1.763820609 | 1.894357207 | 1.71759118 | 1.842556557 | 1.963787827 | 1.817112247 |
| ENSMUSG00000039639 | 0 | 0 | 0 | 0.25 | 0.387761935 | 0 | 0 |
| ENSMUSG00000039640 | 2.297593393 | 2.139605178 | 2.26109853 | 2.078220739 | 2.140250123 | 2.372912003 | 2.133836922 |
| ENSMUSG00000039646 | 2.361637952 | 2.313757142 | 2.382357639 | 1.825945187 | 2.019019072 | 2.056904851 | 2.025071334 |
| ENSMUSG00000039648 | 1.289064826 | 1.214495249 | 1.330482024 | 1.146240625 | 1.343584837 | 1.301029996 | 1.329931221 |
| ENSMUSG00000039652 | 1.693336259 | 1.638647213 | 1.575945187 | 1.839388001 | 1.715300549 | 1.886490725 | 1.655750087 |
| ENSMUSG00000039653 | 0.289064826 | 0.396240625 | 0.25 | 0 | 0.387761935 | 0 | 0.520678036 |
| ENSMUSG00000039656 | 2.241665209 | 2.131869252 | 2.208222504 | 1.909156155 | 2.047893212 | 2.117271296 | 2.021963573 |

| | | | | | | | |
|---|---|---|---|---|---|---|---|
| ENSMUSG00000039660 | 1.87190676 | 1.766522298 | 1.821350555 | 1.708222504 | 1.87878897 | 1.897627091 | 1.820622395 |
| ENSMUSG00000039661 | 0 | 0 | 0 | 0 | 0 | 0 | 0 |
| ENSMUSG00000039662 | 2.512981297 | 2.344844592 | 2.347004321 | 2.385757955 | 2.430282804 | 2.705007959 | 2.297235323 |
| ENSMUSG00000039670 | 1.069666964 | 0.976722649 | 1.080482024 | 0.92510993 | 1.057362051 | 1.041392685 | 1.140832123 |
| ENSMUSG00000039671 | 2.297593393 | 2.074802005 | 2.150906586 | 2.266522298 | 2.282813873 | 2.549003262 | 2.189712149 |
| ENSMUSG00000039672 | 0 | 0.25 | 0 | 0 | 0 | 0 | 0.224243824 |
| ENSMUSG00000039676 | 0 | 0 | 0 | 0 | 0 | 0 | 0.224243824 |
| ENSMUSG00000039678 | 2.141794304 | 2.034887838 | 2.049918086 | 2.049918086 | 2.057362051 | 2.243038049 | 1.941550454 |
| ENSMUSG00000039680 | 2.428398755 | 2.259729747 | 2.298689214 | 2.225970461 | 2.144330586 | 2.421603927 | 2.034219411 |
| ENSMUSG00000039682 | 3.006528408 | 2.781191884 | 2.762972164 | 2.688095162 | 2.659440622 | 3.042575512 | 2.492771481 |
| ENSMUSG00000039683 | 0 | 0 | 0 | 0 | 0 | 0 | 0.224243824 |
| ENSMUSG00000039684 | 0.45815691 | 0.25 | 0.396240625 | 0.25 | 0 | 0.301029996 | 0.224243824 |
| ENSMUSG00000039686 | 2.149089517 | 1.978222334 | 2.00835575 | 1.823655187 | 2.025895058 | 2.045322979 | 1.920884716 |
| ENSMUSG00000039691 | 0 | 0 | 0 | 0 | 0 | 0 | 0 |
| ENSMUSG00000039693 | 1.87190676 | 1.799918086 | 1.777131114 | 1.841580554 | 1.806427258 | 1.986771734 | 1.757893036 |
| ENSMUSG00000039697 | 2.063793172 | 1.978222334 | 1.969129237 | 2.071350555 | 1.990091809 | 2.243038049 | 1.840919217 |
| ENSMUSG00000039699 | 0 | 0 | 0 | 0 | 0 | 0.301029996 | 0.224243824 |
| ENSMUSG00000039701 | 2.141794304 | 1.969129237 | 2.00835575 | 2.04745614 | 1.995080671 | 2.225309282 | 1.815933683 |
| ENSMUSG00000039703 | 3.015982563 | 2.76109853 | 2.745748394 | 2.858265941 | 2.820562012 | 3.232487866 | 2.645007027 |
| ENSMUSG00000039704 | 2.488076442 | 2.333288838 | 2.316696635 | 2.497525991 | 2.414412788 | 2.745074792 | 2.225244904 |
| ENSMUSG00000039706 | 0 | 0 | 0 | 0 | 0 | 0 | 0 |
| ENSMUSG00000039710 | 0 | 0 | 0.25 | 0 | 0 | 0 | 0.224243824 |
| ENSMUSG00000039713 | 1.765759561 | 1.688721876 | 1.769203899 | 1.552363341 | 1.751666404 | 1.748188027 | 1.747849621 |
| ENSMUSG00000039714 | 0.289064826 | 0.25 | 0 | 0 | 0 | 0 | 0 |
| ENSMUSG00000039715 | 1.91631382 | 1.78743678 | 1.823655187 | 1.51109853 | 1.718026092 | 1.62324929 | 1.653807046 |
| ENSMUSG00000039716 | 0.671187741 | 0.830482024 | 0.864857905 | 1.201838731 | 1.327538614 | 1.51851394 | 1.150210039 |
| ENSMUSG00000039717 | 0 | 0 | 0 | 0 | 0 | 0 | 0 |
| ENSMUSG00000039720 | 0 | 0 | 0 | 0 | 0 | 0 | 0 |
| ENSMUSG00000039725 | 1.568542279 | 1.445339928 | 1.431980114 | 1.52186571 | 1.580303447 | 1.62324929 | 1.543390198 |
| ENSMUSG00000039728 | 0.289064826 | 0.25 | 0.25 | 0.25 | 0 | 0 | 0 |
| ENSMUSG00000039735 | 2.883252421 | 2.637686696 | 2.599685923 | 2.716819685 | 2.618504144 | 3.012837225 | 2.420763202 |
| ENSMUSG00000039737 | 2.201144779 | 2.063166358 | 2.071350555 | 2.091580554 | 2.126012715 | 2.322219295 | 2.01501956 |
| ENSMUSG00000039738 | 2.406877894 | 2.294977273 | 2.27186571 | 2.246460484 | 2.25665701 | 2.539076099 | 2.120298735 |
| ENSMUSG00000039740 | 2.258689303 | 2.107613138 | 2.061981878 | 2.131869252 | 2.079491904 | 2.354108439 | 1.93179548 |

| | | | | | | | |
|---|---|---|---|---|---|---|---|
| ENSMUSG00000039741 | 2.328498818 | 2.252806814 | 2.387205477 | 2.012462137 | 2.226481975 | 2.340444115 | 2.226906518 |
| ENSMUSG00000039742 | 0.289064826 | 0.580482024 | 0.5 | 0.580482024 | 0.568060967 | 0.698970004 | 0.803905701 |
| ENSMUSG00000039745 | 1.404271433 | 1.438721876 | 1.356566189 | 1.52186571 | 1.533172725 | 1.69019608 | 1.379042812 |
| ENSMUSG00000039747 | 2.105953527 | 1.982684334 | 2.036164561 | 1.926839783 | 1.926994141 | 2.120573931 | 1.858985529 |
| ENSMUSG00000039748 | 2.337801042 | 2.166333979 | 2.201838731 | 2.299918086 | 2.296256325 | 2.557507202 | 2.141264342 |
| ENSMUSG00000039753 | 2.610448227 | 2.373463862 | 2.324802005 | 2.296218836 | 2.280614762 | 2.559906625 | 2.07959139 |
| ENSMUSG00000039754 | 1.445324132 | 1.445339928 | 1.464495249 | 1.431980114 | 1.580303447 | 1.544068044 | 1.511732338 |
| ENSMUSG00000039756 | 2.610448227 | 2.416333979 | 2.418548067 | 2.600219859 | 2.520825546 | 2.907411361 | 2.333765523 |
| ENSMUSG00000039759 | 1.418409478 | 1.516522298 | 1.62694866 | 1.29248125 | 1.467903253 | 1.397940009 | 1.612273755 |
| ENSMUSG00000039760 | 0 | 0 | 0 | 0 | 0 | 0 | 0 |
| ENSMUSG00000039763 | 1.45815691 | 1.380890489 | 1.388647213 | 1.438721876 | 1.244650542 | 1.278753601 | 1.168187745 |
| ENSMUSG00000039765 | 2.209569962 | 2.04745614 | 2.090485943 | 1.657339155 | 1.671670246 | 1.799340549 | 1.56716938 |
| ENSMUSG00000039768 | 2.54867984 | 2.313166358 | 2.373463862 | 2.327653195 | 2.395811454 | 2.674861141 | 2.260262214 |
| ENSMUSG00000039770 | 2.363083473 | 2.107613138 | 2.087182039 | 2.136723615 | 2.037065488 | 2.307496038 | 1.956418351 |
| ENSMUSG00000039771 | 2.103271637 | 1.942046081 | 2.025821952 | 2.004202072 | 2.00243419 | 2.326335861 | 1.952571511 |
| ENSMUSG00000039774 | 1.181541736 | 1.380890489 | 1.5 | 1.54248125 | 1.595992969 | 1.51851394 | 1.493062805 |
| ENSMUSG00000039775 | 0 | 0 | 0 | 0 | 0 | 0 | 0 |
| ENSMUSG00000039781 | 1.53838222 | 1.610735874 | 1.741446071 | 1.410964047 | 1.674752852 | 1.633468456 | 1.790137008 |
| ENSMUSG00000039782 | 1.789260561 | 1.642463902 | 1.692046081 | 1.311981878 | 1.212046813 | 1.079181246 | 1 |
| ENSMUSG00000039783 | 0.747221736 | 0 | 0.25 | 0.25 | 0.387761935 | 0.301029996 | 0 |
| ENSMUSG00000039785 | 0 | 0 | 0 | 0 | 0 | 0 | 0 |
| ENSMUSG00000039787 | 1.647796617 | 1.610735874 | 1.668106335 | 1.685366747 | 1.695608647 | 1.812913357 | 1.672731473 |
| ENSMUSG00000039789 | 1.912079952 | 1.769203899 | 1.809601185 | 1.866901388 | 1.816839698 | 2.017033339 | 1.676387053 |
| ENSMUSG00000039795 | 1.771760131 | 1.580482024 | 1.664552871 | 1.744319981 | 1.766173492 | 1.908485019 | 1.657681529 |
| ENSMUSG00000039798 | 0.289064826 | 0.646240625 | 0.79248125 | 0.75 | 0.489301084 | 0.477121255 | 0.775756176 |
| ENSMUSG00000039801 | 2.88283767 | 2.620953944 | 2.617918804 | 2.673589472 | 2.619561216 | 2.985875357 | 2.430801052 |
| ENSMUSG00000039804 | 2.258689303 | 2.101285366 | 2.102347734 | 2.193696765 | 2.234113756 | 2.459392488 | 2.093403634 |
| ENSMUSG00000039809 | 0 | 0 | 0 | 0 | 0.244650542 | 0.301029996 | 0.355418052 |
| ENSMUSG00000039810 | 1.655740331 | 1.630890489 | 1.738549078 | 1.476722649 | 1.528558853 | 1.633468456 | 1.543390198 |
| ENSMUSG00000039813 | 0.45815691 | 0.75 | 0.646240625 | 0.976722649 | 0.978602168 | 0.77815125 | 1.21680636 |
| ENSMUSG00000039814 | 1.205378646 | 1.080482024 | 1.201838731 | 0.75 | 1.020174412 | 0.77815125 | 0.829800754 |
| ENSMUSG00000039824 | 1.655740331 | 1.671625132 | 1.729715809 | 1.282320754 | 1.504538967 | 1.544068044 | 1.496250175 |
| ENSMUSG00000039826 | 2.283862692 | 2.168106335 | 2.165444524 | 2.080482024 | 2.06759338 | 2.382017043 | 2.007270316 |
| ENSMUSG00000039828 | 2.390989024 | 2.225970461 | 2.198603967 | 2.199415381 | 2.153935427 | 2.440909082 | 2.055173778 |

| | | | | | | | |
|---|---|---|---|---|---|---|---|
| ENSMUSG00000039830 | 0 | 0 | 0 | 0 | 0 | 0 | 0 |
| ENSMUSG00000039831 | 2.616779011 | 2.404596376 | 2.349151195 | 2.719512728 | 2.649303553 | 3.02201574 | 2.456573893 |
| ENSMUSG00000039834 | 1.867194479 | 1.841580554 | 1.892463902 | 1.653677461 | 1.851969278 | 1.826074803 | 1.916588299 |
| ENSMUSG00000039835 | 2.033115198 | 1.979715809 | 1.979715809 | 2.087182039 | 2.024758322 | 2.338456494 | 1.951796623 |
| ENSMUSG00000039838 | 0.45815691 | 0 | 0.396240625 | 0.5 | 0.489301084 | 0.477121255 | 0.355418052 |
| ENSMUSG00000039840 | 2.312518611 | 2.216819685 | 2.154596376 | 2.095926073 | 2.094757251 | 2.328379603 | 1.977154126 |
| ENSMUSG00000039841 | 2.00683675 | 1.880890489 | 1.894357207 | 1.990000483 | 1.92245931 | 2.161368002 | 1.764880683 |
| ENSMUSG00000039842 | 2.299278375 | 2.181128463 | 2.172499493 | 2.262462137 | 2.185632262 | 2.475671188 | 2.043700396 |
| ENSMUSG00000039844 | 2.717463581 | 2.564346961 | 2.564641508 | 2.411864607 | 2.473055988 | 2.711807229 | 2.345692633 |
| ENSMUSG00000039849 | 2.163307908 | 2.08272922 | 2.094844592 | 1.907339155 | 2 | 2.146128036 | 1.922587433 |
| ENSMUSG00000039850 | 2.066740409 | 1.933677405 | 1.914552871 | 1.850219859 | 1.947418788 | 2.068185862 | 1.913982791 |
| ENSMUSG00000039851 | 0 | 0 | 0 | 0 | 0 | 0 | 0.224243824 |
| ENSMUSG00000039852 | 2.833562691 | 2.617918804 | 2.648581151 | 2.484894804 | 2.493248162 | 2.789580712 | 2.390837894 |
| ENSMUSG00000039853 | 0.289064826 | 0 | 0 | 0.701838731 | 0.955822902 | 1.079181246 | 1.041356071 |
| ENSMUSG00000039860 | 0.45815691 | 0.25 | 0.5 | 0.75 | 0.846351821 | 0.698970004 | 0.829800754 |
| ENSMUSG00000039865 | 0.289064826 | 0.396240625 | 0 | 0 | 0.244650542 | 0.301029996 | 0 |
| ENSMUSG00000039873 | 1.069666964 | 1 | 1.098079356 | 0.396240625 | 0.686820904 | 0.477121255 | 0.829800754 |
| ENSMUSG00000039878 | 0.91631382 | 0.75 | 0.75 | 0.75 | 0.905314583 | 0.903089987 | 0.896975297 |
| ENSMUSG00000039879 | 1.971473847 | 2.002806814 | 1.942046081 | 1.845926073 | 1.78010778 | 1.973127854 | 1.672731473 |
| ENSMUSG00000039883 | 0.289064826 | 0.25 | 0.5 | 0.396240625 | 0.489301084 | 0.698970004 | 0.629532004 |
| ENSMUSG00000039886 | 2.410757293 | 2.228222334 | 2.281030328 | 2.114857905 | 2.113840608 | 2.369215857 | 2.031805106 |
| ENSMUSG00000039887 | 2.146671932 | 2.001406137 | 2.002806814 | 2.150906586 | 2.095689764 | 2.324282455 | 1.930140666 |
| ENSMUSG00000039891 | 0.578129653 | 0.646240625 | 0.701838731 | 0 | 0.489301084 | 0.477121255 | 0 |
| ENSMUSG00000039899 | 0.960252568 | 0.701838731 | 0.896240625 | 0.896240625 | 0.77552387 | 1.079181246 | 0.876096088 |
| ENSMUSG00000039901 | 2.196867486 | 2.002806814 | 2.012462137 | 2.208222504 | 2.16097133 | 2.385606274 | 2.035419841 |
| ENSMUSG00000039903 | 1.470606563 | 1.516522298 | 1.516522298 | 1.114857905 | 1.106691885 | 1.301029996 | 1.168187745 |
| ENSMUSG00000039904 | 0.289064826 | 0.25 | 0 | 0 | 0 | 0.301029996 | 0 |
| ENSMUSG00000039908 | 1.614416215 | 1.431980114 | 1.54248125 | 1.396240625 | 1.467903253 | 1.431363764 | 1.360283038 |
| ENSMUSG00000039910 | 2.690713405 | 2.454245906 | 2.510414788 | 2.321928095 | 2.260764609 | 2.515873844 | 2.112839835 |
| ENSMUSG00000039911 | 1.614416215 | 1.575945187 | 1.557204673 | 1.671625132 | 1.618292349 | 1.72427587 | 1.62316756 |
| ENSMUSG00000039913 | 0 | 0 | 0 | 0 | 0.244650542 | 0 | 0 |
| ENSMUSG00000039914 | 1.822201287 | 1.681980114 | 1.660964047 | 1.814346961 | 1.749189509 | 1.939519253 | 1.601000298 |
| ENSMUSG00000039917 | 2.026699189 | 1.901832578 | 1.984159485 | 1.930274797 | 1.973373093 | 2.100370545 | 1.925966215 |
| ENSMUSG00000039929 | 2.282114126 | 2.107613138 | 2.131869252 | 2.092671852 | 2.107593434 | 2.328379603 | 2.008574816 |

| | | | | | | | |
|---|---|---|---|---|---|---|---|
| ENSMUSG00000039934 | 1.862428343 | 1.75 | 1.714495249 | 1.638647213 | 1.584291694 | 1.826074803 | 1.476637173 |
| ENSMUSG00000039936 | 1.639698646 | 1.557204673 | 1.664552871 | 1.646240625 | 1.768534412 | 1.949390007 | 1.818286533 |
| ENSMUSG00000039942 | 1.307602651 | 1.080482024 | 1.130890489 | 0.646240625 | 0.733951626 | 0.477121255 | 0.520678036 |
| ENSMUSG00000039943 | 2.121692466 | 2.00835575 | 1.950224975 | 1.488549078 | 1.414412788 | 1.531478917 | 1.329931221 |
| ENSMUSG00000039952 | 3.038577955 | 2.783124932 | 2.800071401 | 2.869692405 | 2.87990128 | 3.267640982 | 2.702561601 |
| ENSMUSG00000039953 | 2.852283215 | 2.658021603 | 2.678990498 | 2.603671309 | 2.726858342 | 2.981818607 | 2.595849172 |
| ENSMUSG00000039954 | 0.578129653 | 0.5 | 0.701838731 | 0.646240625 | 0.568060967 | 0.602059991 | 0.629532004 |
| ENSMUSG00000039956 | 0 | 0 | 0 | 0 | 0 | 0 | 0 |
| ENSMUSG00000039958 | 0.867194479 | 0.976722649 | 1.146240625 | 1.04248125 | 1.302012594 | 1.361727836 | 1.066254157 |
| ENSMUSG00000039959 | 2.902690459 | 2.656655413 | 2.681128463 | 2.564641508 | 2.56165237 | 2.868644438 | 2.408896549 |
| ENSMUSG00000039960 | 2.413323654 | 2.190387808 | 2.21293726 | 2.264497931 | 2.212046813 | 2.539076099 | 2.124874702 |
| ENSMUSG00000039962 | 0 | 0 | 0 | 0 | 0 | 0 | 0 |
| ENSMUSG00000039963 | 0.289064826 | 0.580482024 | 0.25 | 0.646240625 | 0.812711509 | 0.477121255 | 0.579661877 |
| ENSMUSG00000039967 | 2.569423023 | 2.381869252 | 2.365881093 | 2.413211243 | 2.327055443 | 2.638489257 | 2.163382972 |
| ENSMUSG00000039968 | 2.209569962 | 2.084962501 | 2.074802005 | 2.154596376 | 2.052132976 | 2.301029996 | 1.896975297 |
| ENSMUSG00000039976 | 2.051791411 | 2.046218836 | 2.01109853 | 2.08272922 | 2.151558609 | 2.356025857 | 2.076967568 |
| ENSMUSG00000039977 | 0 | 0 | 0 | 0 | 0 | 0 | 0 |
| ENSMUSG00000039981 | 0 | 0.25 | 0.25 | 0 | 0.387761935 | 0 | 0.224243824 |
| ENSMUSG00000039982 | 2.600768124 | 2.393884812 | 2.377938163 | 1.741446071 | 1.66856048 | 1.875061263 | 1.667169465 |
| ENSMUSG00000039983 | 1.789260561 | 1.622963274 | 1.723704441 | 1.799918086 | 1.802174723 | 2.021189299 | 1.732938436 |
| ENSMUSG00000039985 | 2.346900291 | 2.19697564 | 2.123963757 | 2.090485943 | 2.025895058 | 2.294466226 | 1.885753983 |
| ENSMUSG00000039987 | 2.078325387 | 1.926839783 | 1.937048212 | 2.124961472 | 2.044679629 | 2.222716471 | 1.869159809 |
| ENSMUSG00000039988 | 2.571178952 | 2.353935442 | 2.345926073 | 2.530706999 | 2.429921724 | 2.794488047 | 2.25604893 |
| ENSMUSG00000039989 | 1.647796617 | 1.598079356 | 1.692046081 | 1.51109853 | 1.632412477 | 1.62324929 | 1.645916034 |
| ENSMUSG00000039990 | 2.081172036 | 1.979715809 | 1.935366747 | 2.070192693 | 2.048957937 | 2.267171728 | 1.874129423 |
| ENSMUSG00000039994 | 2.66563648 | 2.464495249 | 2.485994979 | 2.473704441 | 2.543273062 | 2.828659897 | 2.452143229 |
| ENSMUSG00000039997 | 1.249317394 | 1.26109853 | 1.201838731 | 1.114857905 | 1.057362051 | 1.146128036 | 1.066254157 |
| ENSMUSG00000040003 | 1.205378646 | 1.25 | 1.396240625 | 0.896240625 | 0.877063019 | 1.230448921 | 1.089372236 |
| ENSMUSG00000040006 | 1.975148165 | 1.892463902 | 1.898114259 | 2.149978211 | 2.046825265 | 2.359835482 | 1.947107911 |
| ENSMUSG00000040007 | 1.912079952 | 1.789967834 | 1.845926073 | 1.841580554 | 1.964031554 | 2.068185862 | 1.916588299 |
| ENSMUSG00000040009 | 1.036286563 | 0.92510993 | 1.188721876 | 1.02186571 | 1.176121988 | 0.954242509 | 1.245582246 |
| ENSMUSG00000040010 | 2.836355329 | 2.547764804 | 2.517865591 | 2.524836953 | 2.532092521 | 2.874481818 | 2.39989613 |
| ENSMUSG00000040013 | 0 | 0 | 0 | 0 | 0.244650542 | 0 | 0 |
| ENSMUSG00000040016 | 0 | 0 | 0 | 0 | 0 | 0.301029996 | 0.355418052 |

| | | | | | | | |
|---|---|---|---|---|---|---|---|
| ENSMUSG00000040017 | 0.45815691 | 0 | 0 | 0 | 0 | 0 | 0.224243824 |
| ENSMUSG00000040018 | 2.463861956 | 2.275165585 | 2.29745614 | 2.313757142 | 2.291475807 | 2.589949601 | 2.16858494 |
| ENSMUSG00000040021 | 2.784560084 | 2.552972074 | 2.524178789 | 2.612029076 | 2.556389881 | 2.897077003 | 2.327754286 |
| ENSMUSG00000040022 | 2.388281011 | 2.240724001 | 2.216819685 | 2.211372513 | 2.118236133 | 2.346352974 | 1.963229454 |
| ENSMUSG00000040025 | 2.488076442 | 2.286801231 | 2.277131114 | 2.31611065 | 2.28171603 | 2.592176757 | 2.109521699 |
| ENSMUSG00000040026 | 0.45815691 | 0 | 0 | 0.25 | 0 | 0 | 0.224243824 |
| ENSMUSG00000040028 | 2.926900886 | 2.666111821 | 2.621960008 | 2.588837774 | 2.504538967 | 2.868056362 | 2.343850547 |
| ENSMUSG00000040029 | 2.615992899 | 2.43155454 | 2.417221246 | 2.510414788 | 2.41591954 | 2.773054693 | 2.262645396 |
| ENSMUSG00000040033 | 2.141794304 | 2.009729747 | 1.995748394 | 2.029735268 | 2.020174412 | 2.285557309 | 1.965468324 |
| ENSMUSG00000040034 | 2.370236862 | 2.137686696 | 2.140560606 | 2.334405475 | 2.281165826 | 2.558708571 | 2.109997807 |
| ENSMUSG00000040035 | 0 | 0 | 0.25 | 0 | 0.686820904 | 0 | 0.629532004 |
| ENSMUSG00000040037 | 0.960252568 | 0.701838731 | 0.951838731 | 0.79248125 | 0.686820904 | 0.602059991 | 0.629532004 |
| ENSMUSG00000040040 | 1.800532393 | 1.571350555 | 1.566696635 | 1.903677461 | 1.871833944 | 2.004321374 | 1.715701091 |
| ENSMUSG00000040043 | 2.786132803 | 2.52053726 | 2.501055117 | 2.656655413 | 2.598466708 | 3.003891166 | 2.419303457 |
| ENSMUSG00000040044 | 2.358731791 | 2.169870025 | 2.168989258 | 2.334405475 | 2.23158812 | 2.559906625 | 2.083745739 |
| ENSMUSG00000040046 | 0 | 0 | 0.396240625 | 0.396240625 | 0 | 0 | 0.355418052 |
| ENSMUSG00000040048 | 2.299278375 | 2.188721876 | 2.225216702 | 2.202642909 | 2.225838481 | 2.492760389 | 2.156451689 |
| ENSMUSG00000040054 | 2.734980913 | 2.514159429 | 2.495034894 | 2.584962501 | 2.570979987 | 2.939019776 | 2.403372748 |
| ENSMUSG00000040055 | 0 | 0 | 0 | 0 | 0.244650542 | 0 | 0 |
| ENSMUSG00000040061 | 0.289064826 | 0 | 0 | 0 | 0.244650542 | 0 | 0.224243824 |
| ENSMUSG00000040065 | 0 | 0 | 0 | 0 | 0 | 0 | 0 |
| ENSMUSG00000040078 | 0 | 0 | 0 | 0 | 0 | 0 | 0 |
| ENSMUSG00000040084 | 2.693989403 | 2.452241044 | 2.466046536 | 2.680488403 | 2.624799605 | 3.014100322 | 2.460139731 |
| ENSMUSG00000040086 | 0.671187741 | 0.25 | 0 | 0 | 0 | 0 | 0 |
| ENSMUSG00000040093 | 1.445324132 | 1.482684334 | 1.445339928 | 1.080482024 | 1.223252711 | 1.255272505 | 1.193409508 |
| ENSMUSG00000040097 | 2.309247751 | 2.187048212 | 2.275165585 | 1.987091808 | 2.09661982 | 2.173186268 | 2.150210039 |
| ENSMUSG00000040102 | 1.985980784 | 1.807204673 | 1.843759858 | 1.870953944 | 1.789100779 | 1.991226076 | 1.687112305 |
| ENSMUSG00000040105 | 1.881174527 | 1.774508021 | 1.738549078 | 1.839388001 | 1.820920161 | 2.037426498 | 1.67819947 |
| ENSMUSG00000040111 | 1.964026642 | 1.79248125 | 1.830482024 | 1.695339928 | 1.614672255 | 1.740362689 | 1.554175045 |
| ENSMUSG00000040112 | 2.103271637 | 1.982684334 | 1.969129237 | 1.976722649 | 1.945998439 | 2.247973266 | 1.837623616 |
| ENSMUSG00000040113 | 0 | 0 | 0 | 0 | 0 | 0 | 0 |
| ENSMUSG00000040118 | 2.550525119 | 2.330482024 | 2.342671852 | 2.216046536 | 2.142704057 | 2.444044796 | 1.981429723 |
| ENSMUSG00000040121 | 0.91631382 | 0.92510993 | 0.951838731 | 0.92510993 | 0.978602168 | 0.84509804 | 0.829800754 |
| ENSMUSG00000040123 | 2.113897241 | 1.930274797 | 1.945339928 | 2.198603967 | 2.114724103 | 2.367355921 | 1.934261938 |

| | | | | | | | |
|---|---|---|---|---|---|---|---|
| ENSMUSG00000040124 | 2.253091469 | 2.081607372 | 2.066696635 | 1.985628626 | 1.910073614 | 2.120573931 | 1.784981274 |
| ENSMUSG00000040125 | 0.45815691 | 0.396240625 | 0.25 | 0.5 | 0.489301084 | 0 | 0.224243824 |
| ENSMUSG00000040127 | 0 | 0 | 0 | 0 | 0 | 0 | 0 |
| ENSMUSG00000040128 | 1.964026642 | 1.80479213 | 1.830482024 | 1.741446071 | 1.739104362 | 1.954242509 | 1.672731473 |
| ENSMUSG00000040132 | 0 | 0 | 0 | 0 | 0 | 0 | 0 |
| ENSMUSG00000040133 | 2.095120908 | 1.956637122 | 1.96293726 | 1.76109853 | 1.662257525 | 1.851258349 | 1.574722613 |
| ENSMUSG00000040134 | 0 | 0 | 0 | 0 | 0 | 0 | 0 |
| ENSMUSG00000040136 | 0.289064826 | 0 | 0.25 | 0.701838731 | 0.733951626 | 0.602059991 | 0.829800754 |
| ENSMUSG00000040138 | 0 | 0 | 0 | 0 | 0 | 0 | 0 |
| ENSMUSG00000040139 | 1.205378646 | 1.098079356 | 1.114857905 | 1.380890489 | 1.293076518 | 1.397940009 | 1.296434211 |
| ENSMUSG00000040140 | 0.289064826 | 0.5 | 0.25 | 0 | 0 | 0.301029996 | 0 |
| ENSMUSG00000040146 | 1.470606563 | 1.53743678 | 1.557204673 | 1.606566189 | 1.665423072 | 1.73239376 | 1.71890424 |
| ENSMUSG00000040147 | 0 | 0 | 0 | 0 | 0.387761935 | 0.301029996 | 0.224243824 |
| ENSMUSG00000040148 | 0 | 0 | 0 | 0 | 0 | 0 | 0 |
| ENSMUSG00000040151 | 2.803052234 | 2.543419284 | 2.512462137 | 2.626204747 | 2.541170562 | 2.930439595 | 2.389837845 |
| ENSMUSG00000040152 | 3.765909384 | 3.423671088 | 3.363446264 | 3.11093015 | 3.106160484 | 3.536810866 | 2.872736603 |
| ENSMUSG00000040154 | 0 | 0 | 0 | 0 | 0 | 0 | 0 |
| ENSMUSG00000040158 | 2.81052167 | 2.588010856 | 2.598347614 | 2.54216803 | 2.536927612 | 2.854913022 | 2.389236329 |
| ENSMUSG00000040163 | 0.578129653 | 0 | 0 | 0.25 | 0.244650542 | 0 | 0 |
| ENSMUSG00000040164 | 0 | 0 | 0 | 0 | 0 | 0 | 0 |
| ENSMUSG00000040165 | 1.404271433 | 1.321350555 | 1.226722649 | 1.431980114 | 1.343584837 | 1.204119983 | 1.21680636 |
| ENSMUSG00000040167 | 2.057835465 | 1.979715809 | 1.940387808 | 2.024508021 | 1.970729215 | 2.178976947 | 1.813563596 |
| ENSMUSG00000040170 | 0 | 0 | 0 | 0 | 0 | 0 | 0 |
| ENSMUSG00000040174 | 1.928763473 | 1.671625132 | 1.80479213 | 1.823655187 | 1.712553795 | 1.892094603 | 1.653807046 |
| ENSMUSG00000040177 | 1.753493018 | 1.642463902 | 1.614857905 | 1.685366747 | 1.655839964 | 1.86332286 | 1.61448206 |
| ENSMUSG00000040181 | 1.325351389 | 1.146240625 | 1.17510993 | 1.25 | 1.223252711 | 1.342422681 | 1.110947927 |
| ENSMUSG00000040183 | 1.269664473 | 1.321350555 | 1.488549078 | 1.330482024 | 1.407936347 | 1.447158031 | 1.335191751 |
| ENSMUSG00000040187 | 1.639698646 | 1.54248125 | 1.527131114 | 1.701838731 | 1.683842353 | 1.72427587 | 1.548827561 |
| ENSMUSG00000040188 | 2.342375483 | 2.17510993 | 2.219895812 | 2.301142786 | 2.329464709 | 2.62838893 | 2.218171406 |
| ENSMUSG00000040189 | 0.45815691 | 0 | 0.396240625 | 0.701838731 | 0.846351821 | 0.301029996 | 0.74492186 |
| ENSMUSG00000040195 | 2.300956576 | 2.090485943 | 2.12694866 | 2.281030328 | 2.234742351 | 2.534026106 | 2.073790632 |
| ENSMUSG00000040197 | 0 | 0 | 0 | 0 | 0 | 0 | 0 |
| ENSMUSG00000040204 | 2.856724497 | 2.621457327 | 2.622963274 | 2.864601653 | 2.755564916 | 3.222716471 | 2.556457139 |
| ENSMUSG00000040205 | 0.289064826 | 0 | 0 | 0 | 0 | 0 | 0 |

| | | | | | | | |
|---|---|---|---|---|---|---|---|
| ENSMUSG00000040209 | 2.415874319 | 2.254897682 | 2.263820609 | 2.314346961 | 2.269408865 | 2.496929648 | 2.107610228 |
| ENSMUSG00000040212 | 2.042557845 | 1.843759858 | 1.923371739 | 1.916333979 | 2.029283463 | 2.250420002 | 1.935895857 |
| ENSMUSG00000040213 | 1.527823874 | 1.54248125 | 1.51109853 | 1.898114259 | 1.749189509 | 1.880813592 | 1.694069894 |
| ENSMUSG00000040217 | 0 | 0 | 0 | 0 | 0 | 0 | 0 |
| ENSMUSG00000040219 | 1.54867984 | 1.482684334 | 1.42510993 | 1.482684334 | 1.358933174 | 1.51851394 | 1.307989563 |
| ENSMUSG00000040220 | 2.048736215 | 1.992885888 | 1.998588359 | 1.860735874 | 1.968065382 | 2.158362492 | 1.955652633 |
| ENSMUSG00000040225 | 3.244812638 | 2.968649298 | 2.954645544 | 3.034007462 | 2.964111094 | 3.416307587 | 2.745055516 |
| ENSMUSG00000040229 | 0 | 0.25 | 0.25 | 0 | 0 | 0 | 0 |
| ENSMUSG00000040231 | 0.45815691 | 0.25 | 0.396240625 | 0.25 | 0.568060967 | 0.954242509 | 0.710836105 |
| ENSMUSG00000040234 | 2.346900291 | 2.219895812 | 2.246460484 | 2.121960008 | 2.060462711 | 2.382017043 | 2.005303651 |
| ENSMUSG00000040236 | 2.280358197 | 2.134789703 | 2.154596376 | 1.969129237 | 1.988833497 | 2.181843588 | 1.917452159 |
| ENSMUSG00000040242 | 2.482695304 | 2.277131114 | 2.31728167 | 2.388167274 | 2.297309926 | 2.622214023 | 2.150629917 |
| ENSMUSG00000040247 | 0.45815691 | 0.646240625 | 0.646240625 | 0.701838731 | 0.632412477 | 0.301029996 | 0.984950056 |
| ENSMUSG00000040249 | 3.563705197 | 3.322036281 | 3.355253214 | 2.774508021 | 2.971119265 | 3.226599905 | 2.86897765 |
| ENSMUSG00000040250 | 2.442947872 | 2.269203899 | 2.216046536 | 2.405970373 | 2.315256368 | 2.646403726 | 2.146406417 |
| ENSMUSG00000040253 | 0.811507563 | 0.864857905 | 0.830482024 | 1.839388001 | 1.698489937 | 1.973127854 | 1.616675393 |
| ENSMUSG00000040254 | 1.765759561 | 1.732684334 | 1.660964047 | 1.380890489 | 1.293076518 | 1.544068044 | 1.193409508 |
| ENSMUSG00000040258 | 0.289064826 | 0.396240625 | 0.646240625 | 0.830482024 | 0.931471446 | 0.698970004 | 0.916588299 |
| ENSMUSG00000040260 | 1.721148722 | 1.711372513 | 1.630890489 | 1.188721876 | 1.106691885 | 1.041392685 | 1.028149525 |
| ENSMUSG00000040263 | 2.315764015 | 2.214495249 | 2.218361028 | 2.139605178 | 2.268839121 | 2.40654018 | 2.162979334 |
| ENSMUSG00000040264 | 1.568542279 | 1.25 | 1.403677461 | 1.816696635 | 1.744182955 | 1.924279286 | 1.633706455 |
| ENSMUSG00000040265 | 1 | 1.02186571 | 0.79248125 | 0.92510993 | 1.074582839 | 1.204119983 | 0.952571511 |
| ENSMUSG00000040268 | 2.334723298 | 2.148114259 | 2.167221246 | 2.300530956 | 2.234113756 | 2.514547753 | 2.117986249 |
| ENSMUSG00000040269 | 1.721148722 | 1.557204673 | 1.711372513 | 1.681980114 | 1.628935049 | 1.857332496 | 1.694069894 |
| ENSMUSG00000040270 | 1.971473847 | 1.912762923 | 1.868933358 | 1.858657057 | 1.848234258 | 2.041392685 | 1.694069894 |
| ENSMUSG00000040272 | 1.647796617 | 1.566696635 | 1.552363341 | 1.649978211 | 1.680838453 | 1.86332286 | 1.591689859 |
| ENSMUSG00000040274 | 2.620687497 | 2.403677461 | 2.410512923 | 2.62892496 | 2.582654929 | 2.908485019 | 2.398338893 |
| ENSMUSG00000040276 | 0.289064826 | 0 | 0 | 0.79248125 | 0.812711509 | 0.602059991 | 0.876096088 |
| ENSMUSG00000040280 | 0.289064826 | 0 | 0 | 0 | 0.387761935 | 0.301029996 | 0 |
| ENSMUSG00000040282 | 1.727821383 | 1.726722649 | 1.610735874 | 1.642463902 | 1.618292349 | 1.880813592 | 1.543390198 |
| ENSMUSG00000040283 | 0 | 0 | 0.396240625 | 0.396240625 | 0 | 0.301029996 | 0 |
| ENSMUSG00000040284 | 0 | 0 | 0 | 0 | 0 | 0 | 0 |
| ENSMUSG00000040287 | 1.069666964 | 0.830482024 | 0.92510993 | 0.896240625 | 0.978602168 | 0.77815125 | 0.876096088 |
| ENSMUSG00000040289 | 1.036286563 | 0.976722649 | 0.79248125 | 0.396240625 | 0.568060967 | 0.602059991 | 0.355418052 |

| | | | | | | | |
|---|---|---|---|---|---|---|---|
| ENSMUSG00000040296 | 2.144240249 | 1.970660762 | 1.982684334 | 2.429847205 | 2.364194729 | 2.719331287 | 2.192673409 |
| ENSMUSG00000040297 | 2.56677519 | 2.365881093 | 2.345385738 | 2.466819685 | 2.351342427 | 2.706717782 | 2.1908258 |
| ENSMUSG00000040298 | 0 | 0 | 0.396240625 | 0.580482024 | 0.568060967 | 0.301029996 | 0.579661877 |
| ENSMUSG00000040302 | 1.800532393 | 1.660964047 | 1.552363341 | 1.688721876 | 1.599808748 | 1.73239376 | 1.479989695 |
| ENSMUSG00000040310 | 0.289064826 | 0.25 | 0.25 | 0.92510993 | 0.905314583 | 1.041392685 | 0.896975297 |
| ENSMUSG00000040312 | 2.256831695 | 2.177701608 | 2.243603647 | 1.926839783 | 2.040348824 | 2.120573931 | 2.039586611 |
| ENSMUSG00000040314 | 0 | 0 | 0 | 0 | 0 | 0 | 0 |
| ENSMUSG00000040321 | 1.907802659 | 1.866901388 | 1.75 | 2.05357978 | 1.888971102 | 2.167317335 | 1.779742043 |
| ENSMUSG00000040322 | 2.838206752 | 2.611770807 | 2.599418658 | 2.732314602 | 2.640461996 | 3.056523724 | 2.47169791 |
| ENSMUSG00000040323 | 0 | 0 | 0 | 0 | 0.244650542 | 0 | 0 |
| ENSMUSG00000040325 | 2.590021402 | 2.366901388 | 2.338286706 | 2.433677405 | 2.402121453 | 2.715167358 | 2.226906518 |
| ENSMUSG00000040327 | 0.747221736 | 0.646240625 | 0.79248125 | 0.646240625 | 0.77552387 | 0.301029996 | 0.74492186 |
| ENSMUSG00000040328 | 0 | 0 | 0 | 0.25 | 0.387761935 | 0.477121255 | 0.224243824 |
| ENSMUSG00000040329 | 1.269664473 | 1.080482024 | 1.188721876 | 1.226722649 | 1.200473444 | 1.255272505 | 1.100339912 |
| ENSMUSG00000040331 | 2.37447073 | 2.187886015 | 2.187048212 | 2.300530956 | 2.212715924 | 2.559906625 | 2.112367894 |
| ENSMUSG00000040339 | 2.727821383 | 2.504550045 | 2.48927551 | 2.61434522 | 2.538790182 | 2.887054378 | 2.341765586 |
| ENSMUSG00000040340 | 0 | 0.25 | 0 | 0 | 0 | 0 | 0 |
| ENSMUSG00000040345 | 0 | 0.25 | 0 | 0.25 | 0.568060967 | 0.301029996 | 0.520678036 |
| ENSMUSG00000040350 | 1.389637216 | 1.25 | 1.321350555 | 1.130890489 | 1.136121934 | 1.255272505 | 1.110947927 |
| ENSMUSG00000040351 | 2.833095427 | 2.593216265 | 2.58993739 | 2.545598588 | 2.48684999 | 2.848804701 | 2.301737862 |
| ENSMUSG00000040354 | 2.829809843 | 2.613574823 | 2.575945187 | 2.643648382 | 2.671488075 | 3.01911629 | 2.510768879 |
| ENSMUSG00000040356 | 2.199011616 | 2.145300145 | 2.185366747 | 2.150906586 | 2.283908311 | 2.411619706 | 2.226906518 |
| ENSMUSG00000040359 | 2.072573125 | 1.892463902 | 1.956637122 | 1.985628626 | 1.893954095 | 2.227886705 | 1.847411384 |
| ENSMUSG00000040363 | 2.480523256 | 2.299304173 | 2.296838018 | 2.098079356 | 2.158641579 | 2.40654018 | 2.046027856 |
| ENSMUSG00000040364 | 0.289064826 | 0.396240625 | 0.25 | 0 | 0.244650542 | 0 | 0.355418052 |
| ENSMUSG00000040365 | 2.134369199 | 2.04248125 | 2.072504712 | 2.041226732 | 2.11997913 | 2.252853031 | 2.042530357 |
| ENSMUSG00000040367 | 1.156259305 | 1.080482024 | 1.130890489 | 0.580482024 | 0.489301084 | 0.477121255 | 0.672731473 |
| ENSMUSG00000040370 | 1.852729478 | 1.67510993 | 1.642463902 | 1.5 | 1.576269619 | 1.69019608 | 1.486592281 |
| ENSMUSG00000040372 | 1.445324132 | 1.25 | 1.188721876 | 1.29248125 | 1.176121988 | 1.278753601 | 1.066254157 |
| ENSMUSG00000040373 | 0 | 0 | 0 | 0 | 0 | 0 | 0 |
| ENSMUSG00000040374 | 2.194712275 | 2.032320754 | 2.048689214 | 2.091580554 | 2.013184954 | 2.28780173 | 1.865128412 |
| ENSMUSG00000040380 | 0.671187741 | 0.830482024 | 0.646240625 | 0.5 | 0.489301084 | 0.477121255 | 0.629532004 |
| ENSMUSG00000040383 | 2.598312578 | 2.374462959 | 2.383332433 | 2.528109877 | 2.446849937 | 2.850033258 | 2.315638039 |
| ENSMUSG00000040385 | 3.350295786 | 3.062426514 | 3.067062393 | 3.084056897 | 3.070899785 | 3.543198586 | 2.885624205 |

| | | | | | | | |
|---|---|---|---|---|---|---|---|
| ENSMUSG00000040387 | 1 | 0.864857905 | 1 | 1.146240625 | 1.020174412 | 1.113943352 | 1.014380832 |
| ENSMUSG00000040389 | 2.069666964 | 1.995748394 | 1.991446071 | 2.226722649 | 2.179810173 | 2.426511261 | 1.989124507 |
| ENSMUSG00000040390 | 1.568542279 | 1.642463902 | 1.698603967 | 1.451838731 | 1.603583716 | 1.612783857 | 1.69749337 |
| ENSMUSG00000040396 | 2.63295404 | 2.425975893 | 2.405054956 | 2.324802005 | 2.223252711 | 2.536558443 | 2.084776044 |
| ENSMUSG00000040405 | 0 | 0 | 0 | 0 | 0 | 0 | 0 |
| ENSMUSG00000040407 | 2.686747932 | 2.467205639 | 2.475970461 | 2.536801231 | 2.469844355 | 2.779596491 | 2.239263384 |
| ENSMUSG00000040410 | 1.857607105 | 1.622963274 | 1.75 | 1.668106335 | 1.665423072 | 1.86923172 | 1.556816002 |
| ENSMUSG00000040412 | 0 | 0 | 0 | 0.5 | 0.489301084 | 0.301029996 | 0.448487648 |
| ENSMUSG00000040413 | 0 | 0 | 0 | 0 | 0 | 0 | 0 |
| ENSMUSG00000040414 | 1.852729478 | 1.83272922 | 1.75 | 1.584962501 | 1.824953989 | 1.785329835 | 1.749303626 |
| ENSMUSG00000040415 | 2.247417471 | 2.101285366 | 2.169870025 | 1.907339155 | 2.06149024 | 2.103803721 | 2.003985867 |
| ENSMUSG00000040420 | 1.181541736 | 1.214495249 | 1.26109853 | 0.830482024 | 1 | 0.903089987 | 0.829800754 |
| ENSMUSG00000040423 | 2.618346804 | 2.389126514 | 2.379417347 | 2.415889526 | 2.349078429 | 2.662757832 | 2.150210039 |
| ENSMUSG00000040424 | 0.91631382 | 1.080482024 | 1.25 | 1.04248125 | 0.877063019 | 0.602059991 | 1.078019652 |
| ENSMUSG00000040428 | 0.45815691 | 0.25 | 0.701838731 | 0.5 | 0.568060967 | 0 | 0.829800754 |
| ENSMUSG00000040429 | 1.227926429 | 1.098079356 | 1.146240625 | 1.160964047 | 0.905314583 | 0.954242509 | 0.775756176 |
| ENSMUSG00000040430 | 1.721148722 | 1.634789703 | 1.458222504 | 1.720660762 | 1.686820904 | 1.897627091 | 1.589319772 |
| ENSMUSG00000040432 | 0 | 0 | 0 | 0 | 0 | 0 | 0 |
| ENSMUSG00000040433 | 2.722979153 | 2.528760913 | 2.484527332 | 2.356566189 | 2.320220671 | 2.583198774 | 2.134278581 |
| ENSMUSG00000040434 | 0.289064826 | 0.79248125 | 0.5 | 0.25 | 0.568060967 | 0.301029996 | 0.853775828 |
| ENSMUSG00000040435 | 1.862428343 | 1.828220739 | 1.80479213 | 1.811981878 | 1.797870327 | 2.086359831 | 1.793950594 |
| ENSMUSG00000040441 | 0.671187741 | 0.25 | 0.396240625 | 0.5 | 0.489301084 | 0.477121255 | 0.520678036 |
| ENSMUSG00000040446 | 2.546826359 | 2.324229052 | 2.28360658 | 2.265173983 | 2.239731213 | 2.499687083 | 2.059095225 |
| ENSMUSG00000040447 | 1.227926429 | 1.238549078 | 1.098079356 | 1.54248125 | 1.450958105 | 1.544068044 | 1.448487648 |
| ENSMUSG00000040451 | 2.572054151 | 2.400906586 | 2.37246199 | 2.277131114 | 2.195595395 | 2.514547753 | 2.055173778 |
| ENSMUSG00000040452 | 0 | 0 | 0 | 0 | 0 | 0 | 0 |
| ENSMUSG00000040455 | 2.448863324 | 2.261780978 | 2.25 | 2.475970461 | 2.358048572 | 2.73479983 | 2.191935631 |
| ENSMUSG00000040456 | 0 | 0 | 0.396240625 | 0 | 0 | 0 | 0 |
| ENSMUSG00000040459 | 2.745494892 | 2.512462137 | 2.48927551 | 2.404137211 | 2.381601986 | 2.635483747 | 2.17331363 |
| ENSMUSG00000040462 | 2.399009124 | 2.293731421 | 2.279735268 | 2.301142786 | 2.367659357 | 2.592176757 | 2.254530788 |
| ENSMUSG00000040463 | 3.312666678 | 3.025575954 | 3.021036003 | 2.971138033 | 2.998127167 | 3.402261382 | 2.790311331 |
| ENSMUSG00000040464 | 2.095120908 | 1.937048212 | 1.907339155 | 1.943696765 | 1.853822071 | 2.130333768 | 1.692344475 |
| ENSMUSG00000040466 | 1.100572389 | 1.282320754 | 1.527131114 | 1.26109853 | 1.244650542 | 1 | 1.466365979 |
| ENSMUSG00000040467 | 0 | 0.396240625 | 0.25 | 0 | 0.244650542 | 0.301029996 | 0.224243824 |

| | | | | | | | |
|---|---|---|---|---|---|---|---|
| ENSMUSG00000040471 | 0 | 0 | 0.5 | 0 | 0.244650542 | 0 | 0 |
| ENSMUSG00000040472 | 1.614416215 | 1.566696635 | 1.618933358 | 1.681980114 | 1.828942237 | 1.792391689 | 1.764880683 |
| ENSMUSG00000040473 | 1.639698646 | 1.561981878 | 1.516522298 | 1.649978211 | 1.572189156 | 1.716003344 | 1.405288179 |
| ENSMUSG00000040478 | 0 | 0 | 0 | 0 | 0 | 0 | 0 |
| ENSMUSG00000040479 | 2.276824003 | 2.134789703 | 2.185366747 | 2.123963757 | 2.26251048 | 2.396199347 | 2.21302251 |
| ENSMUSG00000040481 | 3.058380605 | 2.810048763 | 2.792011317 | 2.871331547 | 2.810506292 | 3.242541428 | 2.626461846 |
| ENSMUSG00000040482 | 1.721148722 | 1.701838731 | 1.744319981 | 1.53743678 | 1.72341484 | 1.69019608 | 1.790137008 |
| ENSMUSG00000040483 | 0.671187741 | 0.864857905 | 0.79248125 | 1.42510993 | 1.407936347 | 1.568201724 | 1.350478789 |
| ENSMUSG00000040485 | 0 | 0 | 0 | 0 | 0 | 0 | 0 |
| ENSMUSG00000040488 | 1.227926429 | 1.364857905 | 1.438721876 | 1.098079356 | 1.283908311 | 0.84509804 | 1.452143229 |
| ENSMUSG00000040489 | 0 | 0 | 0 | 0 | 0 | 0 | 0 |
| ENSMUSG00000040490 | 0 | 0 | 0 | 0 | 0 | 0 | 0 |
| ENSMUSG00000040495 | 0 | 0 | 0 | 0.580482024 | 0.244650542 | 0.301029996 | 0.579661877 |
| ENSMUSG00000040498 | 0 | 0 | 0.25 | 0 | 0 | 0 | 0 |
| ENSMUSG00000040502 | 1.036286563 | 1.04248125 | 0.864857905 | 1.098079356 | 1.106691885 | 1.113943352 | 1.238624656 |
| ENSMUSG00000040505 | 0 | 0 | 0 | 0 | 0 | 0 | 0.224243824 |
| ENSMUSG00000040506 | 2.243591459 | 2.079353153 | 2.067865757 | 2.183677405 | 2.19206933 | 2.440909082 | 2.066798338 |
| ENSMUSG00000040511 | 2.275045611 | 2.094844592 | 2.087182039 | 1.953445298 | 1.938809675 | 2.184691431 | 1.848480858 |
| ENSMUSG00000040514 | 0 | 0 | 0 | 0 | 0 | 0 | 0.224243824 |
| ENSMUSG00000040520 | 2.310886388 | 2.106566189 | 2.092671852 | 2.164552871 | 2.066583491 | 2.344392274 | 1.89141329 |
| ENSMUSG00000040521 | 1.960252568 | 1.886723615 | 1.978222334 | 1.856566189 | 1.92245931 | 1.986771734 | 1.920029985 |
| ENSMUSG00000040522 | 0 | 0 | 0 | 0 | 0 | 0 | 0 |
| ENSMUSG00000040524 | 2.604828841 | 2.420748646 | 2.403217124 | 2.448197574 | 2.42883626 | 2.740362689 | 2.296434211 |
| ENSMUSG00000040525 | 0 | 0 | 0 | 0 | 0.244650542 | 0 | 0.224243824 |
| ENSMUSG00000040528 | 0.45815691 | 0.79248125 | 0.396240625 | 0.396240625 | 0.387761935 | 0.477121255 | 0.629532004 |
| ENSMUSG00000040532 | 1.596667477 | 1.589388001 | 1.685366747 | 1.527131114 | 1.572189156 | 1.763427994 | 1.546120303 |
| ENSMUSG00000040533 | 0 | 0 | 0 | 0 | 0 | 0 | 0 |
| ENSMUSG00000040536 | 1.069666964 | 0.976722649 | 1.080482024 | 0.580482024 | 0.387761935 | 0 | 0.224243824 |
| ENSMUSG00000040537 | 1.663535556 | 1.54745614 | 1.51109853 | 2.04745614 | 1.983754904 | 2.187520721 | 1.853775828 |
| ENSMUSG00000040540 | 0 | 0 | 0.25 | 0.396240625 | 0 | 0 | 0.224243824 |
| ENSMUSG00000040541 | 0 | 0 | 0 | 0 | 0 | 0 | 0 |
| ENSMUSG00000040543 | 0 | 0.396240625 | 0.396240625 | 0.5 | 0.387761935 | 0.301029996 | 0.579661877 |
| ENSMUSG00000040548 | 2.315764015 | 2.152756199 | 2.199415381 | 2.327084758 | 2.310728002 | 2.591064607 | 2.156451689 |
| ENSMUSG00000040549 | 2.998115814 | 2.705044741 | 2.708222504 | 2.986909234 | 2.886502522 | 3.366236124 | 2.668103133 |

| | | | | | | | |
|---|---|---|---|---|---|---|---|
| ENSMUSG00000040550 | 2.54867984 | 2.362802778 | 2.347004321 | 2.434523065 | 2.385310841 | 2.737192643 | 2.193776931 |
| ENSMUSG00000040552 | 1.249317394 | 1.282320754 | 1.146240625 | 1.201838731 | 1.091002363 | 1.146128036 | 1.028149525 |
| ENSMUSG00000040554 | 0.578129653 | 0.75 | 0.864857905 | 0.25 | 0.387761935 | 0.77815125 | 0.224243824 |
| ENSMUSG00000040557 | 1.470606563 | 1.188721876 | 1.51109853 | 1.27186571 | 1.387761935 | 1.414973348 | 1.313616061 |
| ENSMUSG00000040560 | 2.409468165 | 2.224461364 | 2.259043403 | 2.214495249 | 2.153144932 | 2.416640507 | 2.026921761 |
| ENSMUSG00000040562 | 1.881174527 | 1.692046081 | 1.630890489 | 1.714495249 | 1.686820904 | 1.913813852 | 1.63986884 |
| ENSMUSG00000040563 | 1.53838222 | 1.589388001 | 1.711372513 | 1.642463902 | 1.773209395 | 1.73239376 | 1.809975587 |
| ENSMUSG00000040564 | 0 | 0 | 0 | 0 | 0 | 0 | 0 |
| ENSMUSG00000040565 | 2.945876762 | 2.689347502 | 2.655970373 | 2.713132377 | 2.6330226 | 3.049992857 | 2.432561206 |
| ENSMUSG00000040569 | 0 | 0.25 | 0 | 0 | 0 | 0 | 0 |
| ENSMUSG00000040570 | 1.325351389 | 1.302363341 | 1.29248125 | 1.356566189 | 1.373641807 | 1.361727836 | 1.265599895 |
| ENSMUSG00000040576 | 0 | 0 | 0 | 0.25 | 0 | 0 | 0 |
| ENSMUSG00000040583 | 0 | 0.25 | 0.25 | 0 | 0 | 0 | 0 |
| ENSMUSG00000040584 | 0.671187741 | 0.5 | 0.25 | 0.25 | 0 | 0.301029996 | 0 |
| ENSMUSG00000040586 | 1.960252568 | 1.755591953 | 1.799918086 | 2.005591953 | 1.926994141 | 2.23299611 | 1.809975587 |
| ENSMUSG00000040591 | 1.881174527 | 1.741446071 | 1.774508021 | 1.982684334 | 1.982473755 | 2.214843848 | 1.852723744 |
| ENSMUSG00000040592 | 0 | 0 | 0 | 0 | 0 | 0 | 0 |
| ENSMUSG00000040594 | 1.852729478 | 1.744319981 | 1.766522298 | 1.671625132 | 1.686820904 | 1.929418926 | 1.554175045 |
| ENSMUSG00000040596 | 2.414600936 | 2.203445298 | 2.205044741 | 2.154596376 | 2.113840608 | 2.383815366 | 1.942350245 |
| ENSMUSG00000040599 | 2.631440309 | 2.40136988 | 2.386241108 | 2.451435968 | 2.333760374 | 2.727541257 | 2.213368331 |
| ENSMUSG00000040600 | 0 | 0 | 0 | 0.25 | 0 | 0 | 0 |
| ENSMUSG00000040601 | 0 | 0 | 0 | 0 | 0 | 0 | 0 |
| ENSMUSG00000040605 | 0 | 0 | 0 | 1.321350555 | 1.223252711 | 1.230448921 | 1.131174228 |
| ENSMUSG00000040606 | 0 | 0.580482024 | 0.396240625 | 1.146240625 | 1.176121988 | 1.204119983 | 1.089372236 |
| ENSMUSG00000040610 | 0 | 0 | 0 | 0 | 0 | 0 | 0.224243824 |
| ENSMUSG00000040612 | 2.289064826 | 2.119945066 | 2.190387808 | 2.457034121 | 2.375332618 | 2.718501689 | 2.178359564 |
| ENSMUSG00000040613 | 0.867194479 | 0.896240625 | 0.79248125 | 1.080482024 | 1.106691885 | 1.146128036 | 1.066254157 |
| ENSMUSG00000040614 | 0 | 0 | 0 | 0 | 0 | 0 | 0 |
| ENSMUSG00000040616 | 0 | 0 | 0 | 0.830482024 | 0.905314583 | 0.954242509 | 1.140832123 |
| ENSMUSG00000040618 | 2.451206209 | 2.275165585 | 2.26109853 | 2.049918086 | 2.202543574 | 2.390935107 | 2.116590781 |
| ENSMUSG00000040620 | 2.68140129 | 2.447790722 | 2.448197574 | 2.5 | 2.448567488 | 2.812913357 | 2.274591042 |
| ENSMUSG00000040621 | 1.269664473 | 1.238549078 | 1.372963274 | 1.42510993 | 1.414412788 | 1.579783597 | 1.355418052 |
| ENSMUSG00000040624 | 0.578129653 | 0 | 0.864857905 | 0.79248125 | 0.955822902 | 0.903089987 | 0.829800754 |
| ENSMUSG00000040627 | 0.289064826 | 0.25 | 0 | 0 | 0 | 0.301029996 | 0.355418052 |

| | | | | | | | |
|---|---|---|---|---|---|---|---|
| ENSMUSG00000040629 | 0 | 0 | 0 | 0 | 0 | 0 | 0 |
| ENSMUSG00000040631 | 1.789260561 | 1.660964047 | 1.816696635 | 1.823655187 | 1.998776581 | 2.06069784 | 1.916588299 |
| ENSMUSG00000040632 | 0 | 0 | 0 | 0 | 0 | 0 | 0 |
| ENSMUSG00000040640 | 1.129344651 | 1.146240625 | 1.114857905 | 1.388647213 | 1.212046813 | 1.431363764 | 1.150210039 |
| ENSMUSG00000040648 | 2.650702778 | 2.433677405 | 2.425543171 | 2.51685859 | 2.41779398 | 2.761175813 | 2.232167048 |
| ENSMUSG00000040649 | 0 | 0 | 0 | 0.25 | 0 | 0 | 0 |
| ENSMUSG00000040650 | 0 | 0 | 0 | 0 | 0 | 0 | 0 |
| ENSMUSG00000040651 | 2.305951036 | 2.101285366 | 2.109697963 | 2.302972074 | 2.216704172 | 2.558708571 | 2.031198704 |
| ENSMUSG00000040652 | 2.414600936 | 2.260414788 | 2.252806814 | 2.434523065 | 2.424460715 | 2.758911892 | 2.292398623 |
| ENSMUSG00000040653 | 0.747221736 | 0.75 | 0.864857905 | 1.380890489 | 1.223252711 | 1.278753601 | 1.21680636 |
| ENSMUSG00000040657 | 0 | 0 | 0 | 0 | 0 | 0 | 0 |
| ENSMUSG00000040658 | 2.156259305 | 1.985628626 | 2.115881093 | 2.175975893 | 2.119108707 | 2.371067862 | 2.082712142 |
| ENSMUSG00000040659 | 2.241665209 | 2.149047439 | 2.187886015 | 2.00835575 | 2.12771782 | 2.212187604 | 2.076967568 |
| ENSMUSG00000040660 | 0 | 0 | 0 | 0 | 0 | 0 | 0 |
| ENSMUSG00000040661 | 2.693989403 | 2.474839231 | 2.449820405 | 2.53296424 | 2.508320156 | 2.804820679 | 2.342230073 |
| ENSMUSG00000040663 | 2.213719572 | 2.064346961 | 2.105516192 | 1.87694866 | 1.987570683 | 2.103803721 | 1.913109605 |
| ENSMUSG00000040666 | 1.432083896 | 1.372963274 | 1.388647213 | 0.396240625 | 0.244650542 | 0.301029996 | 0 |
| ENSMUSG00000040667 | 2.814939948 | 2.567573832 | 2.559003548 | 2.676408097 | 2.628523678 | 3.006893708 | 2.475255611 |
| ENSMUSG00000040669 | 2.22189694 | 2.084962501 | 2.105516192 | 1.970660762 | 1.986303335 | 2.209515015 | 1.922587433 |
| ENSMUSG00000040675 | 2.769043298 | 2.516522298 | 2.469512728 | 2.418106335 | 2.373641807 | 2.703291378 | 2.240219117 |
| ENSMUSG00000040680 | 0 | 0 | 0.396240625 | 0.25 | 0 | 0 | 0 |
| ENSMUSG00000040681 | 3.242716994 | 2.939347502 | 2.936733534 | 3.039336734 | 2.933125047 | 3.427648371 | 2.731269393 |
| ENSMUSG00000040687 | 1.948720984 | 1.809601185 | 1.799918086 | 2.013820609 | 2.057362051 | 2.198657087 | 1.965468324 |
| ENSMUSG00000040688 | 2.478339835 | 2.30539678 | 2.306603048 | 2.228222334 | 2.391809641 | 2.563481085 | 2.276871337 |
| ENSMUSG00000040690 | 2.58665822 | 2.471424093 | 2.59995299 | 1.984159485 | 2.237245597 | 2.247973266 | 2.226906518 |
| ENSMUSG00000040693 | 0 | 0 | 0 | 0 | 0 | 0 | 0 |
| ENSMUSG00000040694 | 0.45815691 | 0 | 0.396240625 | 0.25 | 0 | 0.477121255 | 0.355418052 |
| ENSMUSG00000040697 | 2.054824387 | 1.884789703 | 1.930274797 | 1.995748394 | 1.935892666 | 2.173186268 | 1.879993912 |
| ENSMUSG00000040699 | 2.499676068 | 2.383818844 | 2.361253711 | 2.138647213 | 2.242813819 | 2.459392488 | 2.136477879 |
| ENSMUSG00000040701 | 1.614416215 | 1.482684334 | 1.571350555 | 1.698603967 | 1.741652793 | 1.826074803 | 1.694069894 |
| ENSMUSG00000040703 | 0.811507563 | 0.701838731 | 0.79248125 | 0.864857905 | 0.77552387 | 0.903089987 | 0.803905701 |
| ENSMUSG00000040705 | 0 | 0 | 0 | 0 | 0 | 0.477121255 | 0.224243824 |
| ENSMUSG00000040706 | 0 | 0.396240625 | 0.25 | 0 | 0 | 0.477121255 | 0 |
| ENSMUSG00000040710 | 0.45815691 | 0.25 | 0 | 0.25 | 0.489301084 | 0.77815125 | 0.355418052 |

| | | | | | | | |
|---|---|---|---|---|---|---|---|
| ENSMUSG00000040711 | 3.033404503 | 2.761780978 | 2.756284891 | 2.603935442 | 2.573634712 | 2.925827575 | 2.43378764 |
| ENSMUSG00000040712 | 2.075459174 | 1.975216702 | 2.08272922 | 1.769203899 | 1.850106709 | 2.021189299 | 1.906930404 |
| ENSMUSG00000040713 | 1.721148722 | 1.688721876 | 1.825945187 | 1.77186571 | 1.816839698 | 2.079181246 | 1.769001954 |
| ENSMUSG00000040714 | 0.867194479 | 0.976722649 | 1.17510993 | 0.75 | 1.163285805 | 0.954242509 | 1.437266334 |
| ENSMUSG00000040715 | 0 | 0 | 0 | 0 | 0 | 0 | 0 |
| ENSMUSG00000040717 | 1.249317394 | 1.25 | 1.098079356 | 1.938721876 | 1.900490506 | 2.120573931 | 1.808770687 |
| ENSMUSG00000040720 | 2.797997234 | 2.562870603 | 2.532320754 | 2.617664719 | 2.545363112 | 2.938019097 | 2.362688369 |
| ENSMUSG00000040721 | 1.470606563 | 1.348079356 | 1.445339928 | 1.610735874 | 1.766173492 | 1.785329835 | 1.783679399 |
| ENSMUSG00000040722 | 2.163307908 | 2.016522298 | 2.04745614 | 2.227473271 | 2.24829556 | 2.465382851 | 2.091897741 |
| ENSMUSG00000040723 | 0.747221736 | 0.79248125 | 0.25 | 0.75 | 0.877063019 | 0.77815125 | 0.672731473 |
| ENSMUSG00000040724 | 0.289064826 | 0.25 | 0 | 0 | 0.244650542 | 0 | 0 |
| ENSMUSG00000040725 | 2.737928151 | 2.543419284 | 2.52908599 | 2.483422664 | 2.488689905 | 2.824776462 | 2.359623923 |
| ENSMUSG00000040726 | 0 | 0 | 0 | 0 | 0 | 0 | 0 |
| ENSMUSG00000040728 | 0 | 0 | 0 | 0.25 | 0.244650542 | 0 | 0 |
| ENSMUSG00000040729 | 1.358731791 | 1.431980114 | 1.418106335 | 1.634789703 | 1.445123986 | 1.653212514 | 1.417653332 |
| ENSMUSG00000040731 | 3.248109354 | 3.015427176 | 3.051677287 | 2.929419106 | 2.964826143 | 3.346157302 | 2.839025852 |
| ENSMUSG00000040732 | 0 | 0 | 0 | 0 | 0 | 0 | 0 |
| ENSMUSG00000040734 | 1.53838222 | 1.561981878 | 1.649978211 | 1.130890489 | 1.264824954 | 1.322219295 | 1.405288179 |
| ENSMUSG00000040738 | 2.584966402 | 2.418989258 | 2.392463902 | 2.544043287 | 2.449252175 | 2.846337112 | 2.26794422 |
| ENSMUSG00000040740 | 0 | 0.25 | 0.396240625 | 0.25 | 0.387761935 | 0 | 0 |
| ENSMUSG00000040746 | 1.907802659 | 1.779735268 | 1.811981878 | 1.80479213 | 1.873585591 | 2 | 1.790137008 |
| ENSMUSG00000040747 | 0 | 0.25 | 0 | 0 | 0 | 0 | 0.224243824 |
| ENSMUSG00000040749 | 1.827447047 | 1.630890489 | 1.642463902 | 1.907339155 | 1.859322797 | 2.06069784 | 1.723650273 |
| ENSMUSG00000040751 | 0.811507563 | 0.864857905 | 0.92510993 | 0.5 | 1 | 0.602059991 | 0.672731473 |
| ENSMUSG00000040752 | 0 | 0 | 0.25 | 0 | 0 | 0 | 0 |
| ENSMUSG00000040759 | 0 | 0 | 0 | 0 | 0 | 0 | 0 |
| ENSMUSG00000040760 | 2.446507203 | 2.269203899 | 2.261780978 | 2.559899632 | 2.472415995 | 2.822821645 | 2.260859657 |
| ENSMUSG00000040761 | 2.540273527 | 2.422062577 | 2.432405186 | 2.263820609 | 2.267696867 | 2.489958479 | 2.242754021 |
| ENSMUSG00000040767 | 1.83262764 | 1.837182039 | 1.843759858 | 1.907339155 | 2.019019072 | 2.193124598 | 1.911356129 |
| ENSMUSG00000040770 | 0 | 0 | 0 | 0 | 0 | 0 | 0 |
| ENSMUSG00000040771 | 2.060824957 | 1.899978211 | 1.892463902 | 1.92510993 | 1.840643511 | 2.08278537 | 1.709198048 |
| ENSMUSG00000040774 | 2.223916471 | 2.024508021 | 2.017865591 | 2.129909063 | 2.037065488 | 2.33243846 | 1.883845332 |
| ENSMUSG00000040775 | 0.289064826 | 0 | 0.25 | 0.396240625 | 0.244650542 | 0 | 0 |
| ENSMUSG00000040782 | 2.545896521 | 2.349685923 | 2.304186465 | 2.412314044 | 2.294670002 | 2.664641976 | 2.147679267 |

| | | | | | | | |
|---|---|---|---|---|---|---|---|
| ENSMUSG00000040785 | 3.151275227 | 2.883332433 | 2.835379706 | 2.842399337 | 2.734104256 | 3.179838928 | 2.528340254 |
| ENSMUSG00000040794 | 0 | 0 | 0 | 0.25 | 0 | 0 | 0.224243824 |
| ENSMUSG00000040795 | 1.740854706 | 1.688721876 | 1.638647213 | 1.598079356 | 1.639266209 | 1.832508913 | 1.551512352 |
| ENSMUSG00000040797 | 0 | 0 | 0 | 0 | 0 | 0 | 0 |
| ENSMUSG00000040808 | 0 | 0 | 0 | 0 | 0 | 0 | 0 |
| ENSMUSG00000040809 | 0 | 0 | 0 | 0 | 0 | 0 | 0 |
| ENSMUSG00000040811 | 1.325351389 | 1.396240625 | 1.494319981 | 0.701838731 | 1 | 0.84509804 | 1.078019652 |
| ENSMUSG00000040812 | 0.578129653 | 0.396240625 | 0.396240625 | 0.75 | 0.632412477 | 0.698970004 | 0.74492186 |
| ENSMUSG00000040813 | 2.269664473 | 2.133818844 | 2.219129237 | 1.991446071 | 2.097547432 | 2.252853031 | 2.044866219 |
| ENSMUSG00000040818 | 2.22189694 | 2.028435542 | 2.067865757 | 2.382845365 | 2.297309926 | 2.59439255 | 2.122594809 |
| ENSMUSG00000040820 | 1.432083896 | 1.527131114 | 1.606566189 | 1.372963274 | 1.504538967 | 1.579783597 | 1.46982607 |
| ENSMUSG00000040822 | 2.010213549 | 1.868933358 | 1.919870025 | 2.05479213 | 2.007252778 | 2.264817823 | 1.900630878 |
| ENSMUSG00000040824 | 2.31414447 | 2.199415381 | 2.24929487 | 2.233422664 | 2.227765453 | 2.558708571 | 2.156451689 |
| ENSMUSG00000040828 | 0.578129653 | 0.396240625 | 0.396240625 | 0.5 | 0.489301084 | 0.301029996 | 0.224243824 |
| ENSMUSG00000040829 | 0.289064826 | 0.25 | 0.701838731 | 0.701838731 | 0.812711509 | 0.698970004 | 0.853775828 |
| ENSMUSG00000040836 | 2.349889783 | 2.183677405 | 2.171625132 | 2.074802005 | 2.058398634 | 2.326335861 | 1.983546516 |
| ENSMUSG00000040838 | 0 | 0 | 0 | 0 | 0 | 0 | 0 |
| ENSMUSG00000040840 | 1.432083896 | 1.146240625 | 1.330482024 | 1.584962501 | 1.489301084 | 1.633468456 | 1.493062805 |
| ENSMUSG00000040841 | 1.307602651 | 1.418106335 | 1.348079356 | 1.02186571 | 1.234113756 | 1.204119983 | 1.193409508 |
| ENSMUSG00000040842 | 2.651426165 | 2.499647607 | 2.512121718 | 2.44697564 | 2.490824419 | 2.769377326 | 2.388834696 |
| ENSMUSG00000040843 | 2.427163103 | 2.23927551 | 2.204245906 | 2.275165585 | 2.204601633 | 2.551449998 | 2.083745739 |
| ENSMUSG00000040848 | 2.378662046 | 2.198603967 | 2.171625132 | 2.390083209 | 2.353143227 | 2.672097858 | 2.156451689 |
| ENSMUSG00000040850 | 2.980106992 | 2.710194481 | 2.685366747 | 2.891632446 | 2.794285685 | 3.236789099 | 2.581328627 |
| ENSMUSG00000040852 | 2.434522689 | 2.281030328 | 2.254897682 | 2.413211243 | 2.374910674 | 2.736396502 | 2.257258335 |
| ENSMUSG00000040855 | 0.671187741 | 0.25 | 0.830482024 | 1.04248125 | 0.955822902 | 1 | 1 |
| ENSMUSG00000040856 | 1.936861443 | 1.926839783 | 2.013820609 | 0.701838731 | 0.877063019 | 0.77815125 | 0.775756176 |
| ENSMUSG00000040857 | 1.91631382 | 1.907339155 | 2.027131114 | 1.657339155 | 1.72073075 | 1.838849091 | 1.81475081 |
| ENSMUSG00000040859 | 2.103271637 | 2.028435542 | 1.979715809 | 1.998588359 | 2.028157605 | 2.181843588 | 1.916588299 |
| ENSMUSG00000040860 | 1.37447073 | 1.482684334 | 1.681980114 | 1.25 | 1.588235379 | 1.431363764 | 1.737484395 |
| ENSMUSG00000040865 | 2.584117913 | 2.400906586 | 2.404137211 | 2.538071211 | 2.489911207 | 2.807535028 | 2.304618229 |
| ENSMUSG00000040866 | 0 | 0 | 0 | 0 | 0 | 0 | 0 |
| ENSMUSG00000040867 | 0.45815691 | 0.580482024 | 0.5 | 0 | 0.489301084 | 0 | 0.629532004 |
| ENSMUSG00000040875 | 1.37447073 | 1.403677461 | 1.52186571 | 1 | 1.149965125 | 1.079181246 | 1.365075947 |
| ENSMUSG00000040877 | 1.45815691 | 1.396240625 | 1.527131114 | 1.114857905 | 1.310728002 | 1.397940009 | 1.121219121 |

| | | | | | | | |
|---|---|---|---|---|---|---|---|
| ENSMUSG00000040883 | 1.269664473 | 1.130890489 | 1.348079356 | 1.160964047 | 1.351342427 | 1.230448921 | 1.355418052 |
| ENSMUSG00000040888 | 2.183765914 | 2.017865591 | 2.03743678 | 2.091580554 | 2.10031573 | 2.359835482 | 1.997315209 |
| ENSMUSG00000040891 | 0.671187741 | 0.580482024 | 0.396240625 | 0.5 | 0.387761935 | 0 | 0.629532004 |
| ENSMUSG00000040896 | 0 | 0 | 0 | 0 | 0 | 0 | 0 |
| ENSMUSG00000040899 | 0 | 0 | 0 | 0 | 0 | 0 | 0 |
| ENSMUSG00000040901 | 0 | 0 | 0 | 0 | 0 | 0 | 0 |
| ENSMUSG00000040904 | 0.289064826 | 0.396240625 | 0.25 | 0.25 | 0.244650542 | 0 | 0.448487648 |
| ENSMUSG00000040907 | 0 | 0.396240625 | 0 | 0.25 | 0.244650542 | 0.301029996 | 0.672731473 |
| ENSMUSG00000040913 | 1.806056082 | 1.67510993 | 1.741446071 | 1.380890489 | 1.504538967 | 1.51851394 | 1.473249546 |
| ENSMUSG00000040918 | 2.144240249 | 2.041226732 | 2.001406137 | 1.94697564 | 1.86473911 | 2.130333768 | 1.759302692 |
| ENSMUSG00000040919 | 0 | 0 | 0 | 0 | 0 | 0 | 0 |
| ENSMUSG00000040928 | 2.40033094 | 2.26109853 | 2.241446071 | 2.252107156 | 2.187072901 | 2.460897843 | 2.026921761 |
| ENSMUSG00000040929 | 2.048736215 | 1.975216702 | 1.945339928 | 1.969129237 | 1.902105869 | 2.021189299 | 1.687112305 |
| ENSMUSG00000040935 | 0 | 0 | 0 | 0 | 0 | 0 | 0 |
| ENSMUSG00000040936 | 1.614416215 | 1.505591953 | 1.622963274 | 1.403677461 | 1.212046813 | 1.361727836 | 1.313616061 |
| ENSMUSG00000040938 | 0 | 0.396240625 | 0.5 | 0.5 | 0 | 0 | 0.224243824 |
| ENSMUSG00000040940 | 2.149089517 | 2.075945187 | 2.176839783 | 1.843759858 | 2.062514787 | 2.093421685 | 2.082194103 |
| ENSMUSG00000040943 | 2.057835465 | 1.956637122 | 1.899978211 | 2.064346961 | 2.010824034 | 2.164352856 | 1.844181586 |
| ENSMUSG00000040945 | 3.016283996 | 2.825659731 | 2.859697963 | 2.503853763 | 2.66856048 | 2.921166051 | 2.592118924 |
| ENSMUSG00000040950 | 0 | 0 | 0 | 0 | 0 | 0 | 0 |
| ENSMUSG00000040952 | 2.7985055 | 2.655512955 | 2.68662858 | 2.404596376 | 2.590305509 | 2.795880017 | 2.448988576 |
| ENSMUSG00000040957 | 1 | 0.976722649 | 1.080482024 | 1.114857905 | 1.212046813 | 1.397940009 | 1.296434211 |
| ENSMUSG00000040963 | 0.289064826 | 0 | 0 | 0 | 0 | 0 | 0 |
| ENSMUSG00000040964 | 2.307602651 | 2.231944491 | 2.31611065 | 1.794977273 | 2.039257769 | 2.053078443 | 2.041943747 |
| ENSMUSG00000040966 | 0 | 0 | 0 | 0 | 0 | 0 | 0 |
| ENSMUSG00000040969 | 0 | 0 | 0 | 0.396240625 | 0 | 0 | 0.355418052 |
| ENSMUSG00000040972 | 0 | 0 | 0 | 0 | 0 | 0 | 0 |
| ENSMUSG00000040978 | 0.289064826 | 0.25 | 0 | 0 | 0.686820904 | 0.477121255 | 0.448487648 |
| ENSMUSG00000040985 | 0 | 0 | 0.25 | 0 | 0 | 0 | 0 |
| ENSMUSG00000040987 | 0 | 0 | 0.396240625 | 0 | 0 | 0 | 0.224243824 |
| ENSMUSG00000040990 | 2.81052167 | 2.583568321 | 2.559003548 | 2.558404919 | 2.466928684 | 2.86569606 | 2.298566117 |
| ENSMUSG00000040997 | 2.346900291 | 2.210587586 | 2.210587586 | 2.280383379 | 2.230318492 | 2.592176757 | 2.138662327 |
| ENSMUSG00000040998 | 2.616779011 | 2.423371739 | 2.41942991 | 2.540282971 | 2.482519186 | 2.837588438 | 2.308505126 |
| ENSMUSG00000041000 | 0.91631382 | 0.976722649 | 0.951838731 | 1.188721876 | 1.27449559 | 1.462397998 | 1.355418052 |

| | | | | | | | |
|---|---|---|---|---|---|---|---|
| ENSMUSG00000041009 | 0 | 0 | 0 | 0 | 0 | 0 | 0 |
| ENSMUSG00000041012 | 0.289064826 | 0.25 | 0 | 0.830482024 | 0.846351821 | 0.602059991 | 0.916588299 |
| ENSMUSG00000041014 | 0 | 0 | 0 | 0 | 0 | 0 | 0 |
| ENSMUSG00000041020 | 1.036286563 | 0.951838731 | 0.896240625 | 1.201838731 | 1.176121988 | 1.301029996 | 1.054044578 |
| ENSMUSG00000041025 | 2.390989024 | 2.181980114 | 2.218361028 | 2.168989258 | 2.141069998 | 2.404833717 | 2.025071334 |
| ENSMUSG00000041028 | 2.943730793 | 2.677916743 | 2.656883471 | 2.831326364 | 2.751085167 | 3.206556044 | 2.568325207 |
| ENSMUSG00000041035 | 2.026699189 | 1.872963274 | 1.910964047 | 1.981203126 | 2.045754077 | 2.230448921 | 1.91917299 |
| ENSMUSG00000041037 | 2.036286563 | 1.830482024 | 1.909156155 | 1.811981878 | 1.820920161 | 1.959041392 | 1.763495175 |
| ENSMUSG00000041040 | 2.103271637 | 2.015173983 | 1.961372513 | 1.935366747 | 1.919403394 | 2.086359831 | 1.74047993 |
| ENSMUSG00000041044 | 0 | 0 | 0 | 0 | 0 | 0 | 0 |
| ENSMUSG00000041046 | 0 | 0.25 | 0.396240625 | 0.5 | 0.77552387 | 0.903089987 | 0.876096088 |
| ENSMUSG00000041052 | 0 | 0 | 0.25 | 0 | 0 | 0 | 0 |
| ENSMUSG00000041057 | 2.867194479 | 2.678346629 | 2.643648382 | 2.731759292 | 2.66579363 | 3.0523091 | 2.46062295 |
| ENSMUSG00000041058 | 2.381433035 | 2.168989258 | 2.148114259 | 2.351816941 | 2.273371525 | 2.605305046 | 2.082194103 |
| ENSMUSG00000041062 | 0.289064826 | 0.5 | 0.646240625 | 0.396240625 | 0.632412477 | 0.301029996 | 0.448487648 |
| ENSMUSG00000041064 | 2.039433992 | 1.998588359 | 1.958222504 | 1.923371739 | 1.995080671 | 2.10720997 | 1.917452159 |
| ENSMUSG00000041068 | 0 | 0 | 0 | 0 | 0 | 0 | 0 |
| ENSMUSG00000041073 | 1.623015126 | 1.494319981 | 1.464495249 | 1.282320754 | 1.478764298 | 1.431363764 | 1.508694612 |
| ENSMUSG00000041075 | 2.233869984 | 2.073655187 | 2.059601185 | 2.013820609 | 1.968065382 | 2.146128036 | 1.853775828 |
| ENSMUSG00000041078 | 0 | 0 | 0 | 0 | 0 | 0 | 0.448487648 |
| ENSMUSG00000041079 | 1.734388958 | 1.610735874 | 1.653677461 | 1.744319981 | 1.751666404 | 1.959041392 | 1.633706455 |
| ENSMUSG00000041084 | 2.578990401 | 2.414998973 | 2.381380202 | 2.570482507 | 2.501307552 | 2.914871818 | 2.339198915 |
| ENSMUSG00000041096 | 2.136858952 | 1.969129237 | 2.019203899 | 2.154596376 | 2.093822267 | 2.383815366 | 2.01116814 |
| ENSMUSG00000041112 | 1.982401085 | 1.784887838 | 1.77186571 | 1.766522298 | 1.751666404 | 1.963787827 | 1.653807046 |
| ENSMUSG00000041115 | 1.37447073 | 1.527131114 | 1.622963274 | 1.410964047 | 1.514343934 | 1.556302501 | 1.502532226 |
| ENSMUSG00000041117 | 0.578129653 | 0.5 | 0.75 | 0.701838731 | 0.244650542 | 0.698970004 | 0.448487648 |
| ENSMUSG00000041119 | 0.91631382 | 0.701838731 | 0.75 | 1.27186571 | 1.200473444 | 1.361727836 | 1.140832123 |
| ENSMUSG00000041120 | 2.199011616 | 2.12892496 | 2.150906586 | 1.649978211 | 1.758994476 | 1.886490725 | 1.67819947 |
| ENSMUSG00000041124 | 2.545896521 | 2.308404919 | 2.314346961 | 2.372963274 | 2.293608479 | 2.62324929 | 2.152303998 |
| ENSMUSG00000041126 | 2.386920383 | 2.178561379 | 2.209012589 | 2.032320754 | 2.062514787 | 2.252853031 | 1.967691807 |
| ENSMUSG00000041130 | 2.351376531 | 2.219895812 | 2.332168732 | 2.056000419 | 2.246477757 | 2.338456494 | 2.197065089 |
| ENSMUSG00000041132 | 1.129344651 | 0.951838731 | 0.864857905 | 1.080482024 | 0.931471446 | 1.041392685 | 0.829800754 |
| ENSMUSG00000041133 | 3.080397602 | 2.787119146 | 2.785845803 | 3.030545225 | 2.930335554 | 3.41480628 | 2.725926557 |
| ENSMUSG00000041134 | 0.578129653 | 0.396240625 | 0.25 | 0.25 | 0 | 0 | 0 |

| | | | | | | | |
|---|---|---|---|---|---|---|---|
| ENSMUSG00000041135 | 2.401648579 | 2.225216702 | 2.214495249 | 2.105516192 | 1.962676635 | 2.201397124 | 1.854824501 |
| ENSMUSG00000041138 | 0 | 0 | 0 | 0 | 0 | 0 | 0 |
| ENSMUSG00000041141 | 0 | 0 | 0 | 0 | 0 | 0 | 0 |
| ENSMUSG00000041143 | 1.596667477 | 1.589388001 | 1.634789703 | 0.5 | 0.244650542 | 0 | 0 |
| ENSMUSG00000041144 | 1.069666964 | 1.098079356 | 0.951838731 | 1.160964047 | 0.812711509 | 0.698970004 | 0.672731473 |
| ENSMUSG00000041147 | 2.432083896 | 2.234159485 | 2.245748394 | 2.304186465 | 2.235369829 | 2.564666064 | 2.079068325 |
| ENSMUSG00000041153 | 2.010213549 | 1.919870025 | 1.886723615 | 1.916333979 | 1.887294348 | 1.973127854 | 1.674564426 |
| ENSMUSG00000041161 | 1.578129653 | 1.53743678 | 1.642463902 | 1.445339928 | 1.484072009 | 1.51851394 | 1.473249546 |
| ENSMUSG00000041164 | 2.156259305 | 2.155512955 | 2.256976499 | 1.914552871 | 2.134457044 | 2.133538908 | 2.168981648 |
| ENSMUSG00000041165 | 0 | 0.396240625 | 0.25 | 0 | 0.244650542 | 0 | 0 |
| ENSMUSG00000041168 | 2.881174527 | 2.644121087 | 2.677270953 | 2.512802235 | 2.607537217 | 2.871572936 | 2.474176968 |
| ENSMUSG00000041180 | 1.952600383 | 1.789967834 | 1.769203899 | 1.933677405 | 1.913211023 | 2.06069784 | 1.667169465 |
| ENSMUSG00000041187 | 1.614416215 | 1.678561379 | 1.695339928 | 1.738549078 | 1.850106709 | 1.991226076 | 1.89141329 |
| ENSMUSG00000041189 | 1.269664473 | 1.302363341 | 1.372963274 | 1.890560606 | 1.900490506 | 2.049218023 | 1.850609257 |
| ENSMUSG00000041193 | 0 | 0 | 0 | 0 | 0 | 0 | 0 |
| ENSMUSG00000041199 | 1.885732304 | 1.882845365 | 1.862802778 | 1.735628626 | 1.916320788 | 1.959041392 | 1.908707966 |
| ENSMUSG00000041202 | 0 | 0.25 | 0.25 | 0.25 | 0.244650542 | 0.301029996 | 0 |
| ENSMUSG00000041203 | 2.346900291 | 2.180274797 | 2.199415381 | 2.247880462 | 2.19135987 | 2.527629901 | 2.146406417 |
| ENSMUSG00000041205 | 0.45815691 | 0.580482024 | 0.580482024 | 0.701838731 | 0.568060967 | 0.602059991 | 0.710836105 |
| ENSMUSG00000041215 | 2.580706586 | 2.352877747 | 2.354990044 | 2.335518667 | 2.309205544 | 2.586587305 | 2.146831257 |
| ENSMUSG00000041216 | 0.45815691 | 0 | 0.79248125 | 0 | 0 | 0 | 0 |
| ENSMUSG00000041219 | 2.588343201 | 2.400906586 | 2.40136988 | 2.584684096 | 2.535858857 | 2.912222057 | 2.365939808 |
| ENSMUSG00000041220 | 2.37447073 | 2.218361028 | 2.228222334 | 2.549304173 | 2.472736137 | 2.848804701 | 2.282223478 |
| ENSMUSG00000041225 | 2.418409478 | 2.193696765 | 2.267865591 | 2.103406982 | 2.046825265 | 2.294466226 | 1.902443294 |
| ENSMUSG00000041229 | 2.105953527 | 1.969129237 | 1.96759118 | 2.187048212 | 2.124299332 | 2.394451681 | 1.991877847 |
| ENSMUSG00000041231 | 2.31414447 | 2.108657057 | 2.110735874 | 2.302972074 | 2.202543574 | 2.532754379 | 2.075912052 |
| ENSMUSG00000041235 | 1.568542279 | 1.649978211 | 1.606566189 | 1.494319981 | 1.439191811 | 1.579783597 | 1.396774124 |
| ENSMUSG00000041236 | 2.673935906 | 2.443284802 | 2.418989258 | 2.483053688 | 2.398190943 | 2.768638101 | 2.2551389 |
| ENSMUSG00000041237 | 0 | 0 | 0 | 0 | 0 | 0 | 0 |
| ENSMUSG00000041238 | 2.428398755 | 2.242885888 | 2.263142013 | 2.333288838 | 2.23158812 | 2.582063363 | 2.108089156 |
| ENSMUSG00000041241 | 1.881174527 | 1.886723615 | 1.868933358 | 1.657339155 | 1.832885921 | 1.875061263 | 1.760706232 |
| ENSMUSG00000041245 | 1.771760131 | 1.714495249 | 1.692046081 | 1.943696765 | 1.91476933 | 2.155336037 | 1.783679399 |
| ENSMUSG00000041247 | 0 | 0 | 0 | 0.25 | 0 | 0.301029996 | 0 |
| ENSMUSG00000041248 | 0 | 0 | 0 | 0 | 0 | 0 | 0 |

| | | | | | | | |
|---|---|---|---|---|---|---|---|
| ENSMUSG00000041255 | 0 | 0 | 0 | 0 | 0 | 0 | 0 |
| ENSMUSG00000041258 | 2.479432974 | 2.285526764 | 2.281030328 | 2.160964047 | 2.195595395 | 2.44870632 | 2.081675233 |
| ENSMUSG00000041261 | 0.289064826 | 0 | 0.396240625 | 0 | 0.387761935 | 0 | 0 |
| ENSMUSG00000041263 | 1.036286563 | 0.951838731 | 1.02186571 | 1.330482024 | 1.327538614 | 1.414973348 | 1.374453863 |
| ENSMUSG00000041264 | 2.556012466 | 2.366391601 | 2.272528105 | 2.300530956 | 2.275056283 | 2.545307116 | 2.109521699 |
| ENSMUSG00000041268 | 2.188178997 | 2.048689214 | 1.979715809 | 2.240724001 | 2.17760189 | 2.436162647 | 1.993927582 |
| ENSMUSG00000041272 | 0.747221736 | 0.25 | 0.396240625 | 0.25 | 0.244650542 | 0.301029996 | 0.224243824 |
| ENSMUSG00000041278 | 2.35433422 | 2.173371739 | 2.159156155 | 2.353406982 | 2.28171603 | 2.564666064 | 2.120298735 |
| ENSMUSG00000041287 | 0.289064826 | 0 | 0 | 0 | 0.244650542 | 0 | 0.224243824 |
| ENSMUSG00000041293 | 0 | 0 | 0 | 0 | 0 | 0.301029996 | 0 |
| ENSMUSG00000041297 | 2.533616084 | 2.340485943 | 2.325374049 | 2.456637122 | 2.402512107 | 2.711807229 | 2.210591357 |
| ENSMUSG00000041298 | 2.271465912 | 2.067865757 | 2.060793496 | 1.886723615 | 1.850106709 | 2.123851641 | 1.74047993 |
| ENSMUSG00000041301 | 0.45815691 | 0.5 | 0.25 | 0.864857905 | 0.568060967 | 0.477121255 | 0.579661877 |
| ENSMUSG00000041303 | 2.483777101 | 2.286164561 | 2.257666784 | 2.377938163 | 2.252501045 | 2.652246341 | 2.153553898 |
| ENSMUSG00000041308 | 2.432083896 | 2.254202072 | 2.253505118 | 2.328220739 | 2.244650542 | 2.553883027 | 2.087847416 |
| ENSMUSG00000041309 | 0 | 0 | 0 | 0 | 0 | 0 | 0 |
| ENSMUSG00000041313 | 2.736751754 | 2.505244985 | 2.47104263 | 2.491085217 | 2.488383918 | 2.808885867 | 2.330653623 |
| ENSMUSG00000041319 | 2.119110214 | 1.979715809 | 2.075945187 | 1.618933358 | 1.880506521 | 1.959041392 | 1.863093717 |
| ENSMUSG00000041323 | 0.45815691 | 0.25 | 0.25 | 0 | 0.244650542 | 0 | 0 |
| ENSMUSG00000041324 | 2.146671932 | 2.083847589 | 2.084962501 | 1.777131114 | 1.806427258 | 2.008600172 | 1.728327686 |
| ENSMUSG00000041328 | 2.40033094 | 2.241446071 | 2.244319981 | 2.293106877 | 2.197694248 | 2.545307116 | 2.073790632 |
| ENSMUSG00000041329 | 0.747221736 | 0.830482024 | 0.830482024 | 1.27186571 | 1.27449559 | 1.361727836 | 1.302263476 |
| ENSMUSG00000041333 | 0 | 0 | 0 | 0 | 0 | 0 | 0 |
| ENSMUSG00000041341 | 2.713745586 | 2.488912477 | 2.452642909 | 2.41942991 | 2.354934887 | 2.69019608 | 2.193041668 |
| ENSMUSG00000041343 | 1.227926429 | 1.130890489 | 1.282320754 | 1.146240625 | 1.057362051 | 1.041392685 | 1 |
| ENSMUSG00000041346 | 1.924654746 | 1.769203899 | 1.752806814 | 1.926839783 | 2.028157605 | 2.127104798 | 1.926805425 |
| ENSMUSG00000041347 | 0 | 0 | 0.25 | 0.25 | 0.244650542 | 0.301029996 | 0.224243824 |
| ENSMUSG00000041351 | 1.623015126 | 1.593759858 | 1.701838731 | 1.431980114 | 1.614672255 | 1.556302501 | 1.584526862 |
| ENSMUSG00000041353 | 1.558729299 | 1.51109853 | 1.516522298 | 1.868933358 | 1.808534465 | 2.021189299 | 1.687112305 |
| ENSMUSG00000041354 | 2.089597219 | 2.02053726 | 2.134789703 | 1.884789703 | 2.045754077 | 2.06069784 | 2.075383 |
| ENSMUSG00000041355 | 2.811999637 | 2.578787391 | 2.562574604 | 2.702642909 | 2.714656174 | 3.104828404 | 2.504786699 |
| ENSMUSG00000041358 | 0 | 0 | 0 | 0.75 | 0.632412477 | 0.698970004 | 0.672731473 |
| ENSMUSG00000041359 | 0 | 0 | 0 | 0 | 0 | 0 | 0 |
| ENSMUSG00000041360 | 2.819311906 | 2.622712719 | 2.604726683 | 2.543419284 | 2.522497007 | 2.859138297 | 2.350704945 |

| | | | | | | | |
|---|---|---|---|---|---|---|---|
| ENSMUSG00000041361 | 1.129344651 | 1.061981878 | 1 | 1.25 | 1.223252711 | 1.204119983 | 1.066254157 |
| ENSMUSG00000041362 | 0.289064826 | 0.580482024 | 0.5 | 1.62694866 | 1.645989387 | 1.681241237 | 1.48984372 |
| ENSMUSG00000041372 | 0 | 0.25 | 0 | 0.830482024 | 0.931471446 | 0.77815125 | 0.710836105 |
| ENSMUSG00000041375 | 1.881174527 | 1.755591953 | 1.854463129 | 1.688721876 | 1.812711509 | 1.86332286 | 1.801445495 |
| ENSMUSG00000041377 | 0 | 0.396240625 | 0.25 | 0.25 | 0 | 0.602059991 | 0.224243824 |
| ENSMUSG00000041378 | 0 | 0 | 0 | 0 | 0 | 0 | 0 |
| ENSMUSG00000041380 | 0 | 0.25 | 0 | 0 | 0 | 0 | 0 |
| ENSMUSG00000041390 | 2.602397159 | 2.399047439 | 2.401832578 | 2.165444524 | 1.983754904 | 2.311753861 | 1.896975297 |
| ENSMUSG00000041396 | 1.568542279 | 1.372963274 | 1.364857905 | 1.5 | 1.433158226 | 1.568201724 | 1.383567577 |
| ENSMUSG00000041399 | 0 | 0 | 0 | 0 | 0 | 0 | 0 |
| ENSMUSG00000041406 | 2.364524 | 2.231203126 | 2.228222334 | 2.320772088 | 2.224547964 | 2.50242712 | 2.089878917 |
| ENSMUSG00000041408 | 2.669806838 | 2.461372513 | 2.463717096 | 2.710783978 | 2.624799605 | 3.020361283 | 2.397752991 |
| ENSMUSG00000041415 | 2.840968566 | 2.588010856 | 2.575659731 | 2.492885888 | 2.449252175 | 2.804139432 | 2.280261923 |
| ENSMUSG00000041417 | 2.540273527 | 2.335518667 | 2.331045137 | 2.363317659 | 2.324629618 | 2.622214023 | 2.167391888 |
| ENSMUSG00000041420 | 1.623015126 | 1.458222504 | 1.54248125 | 1.5 | 1.680838453 | 1.672097858 | 1.605556929 |
| ENSMUSG00000041423 | 0.289064826 | 0 | 0 | 0 | 0.632412477 | 0 | 0.355418052 |
| ENSMUSG00000041426 | 1.932832114 | 1.87694866 | 1.779735268 | 2.025821952 | 1.94314048 | 2.247973266 | 1.813563596 |
| ENSMUSG00000041429 | 1.587501561 | 1.593759858 | 1.54248125 | 1.311981878 | 1.509475496 | 1.505149978 | 1.564611933 |
| ENSMUSG00000041431 | 2.862428343 | 2.601019284 | 2.592671852 | 2.850219859 | 2.745662857 | 3.20709554 | 2.558723247 |
| ENSMUSG00000041438 | 2.732608034 | 2.501406137 | 2.511780978 | 2.523189285 | 2.441296923 | 2.799340549 | 2.27142916 |
| ENSMUSG00000041439 | 1.693336259 | 1.54248125 | 1.431980114 | 1.695339928 | 1.66856048 | 1.86923172 | 1.643912836 |
| ENSMUSG00000041440 | 1.753493018 | 1.593759858 | 1.54745614 | 1.850219859 | 1.828942237 | 2.004321374 | 1.69749337 |
| ENSMUSG00000041444 | 2.177057474 | 2.097004321 | 2.08272922 | 2.224461364 | 2.145141048 | 2.404833717 | 1.962479708 |
| ENSMUSG00000041445 | 0 | 0 | 0.25 | 0.396240625 | 0.568060967 | 0 | 0.355418052 |
| ENSMUSG00000041449 | 2.315764015 | 2.046218836 | 2.087182039 | 1.330482024 | 1.283908311 | 1.556302501 | 1.284450788 |
| ENSMUSG00000041453 | 3.144461898 | 2.861124322 | 2.882357639 | 2.971328765 | 2.862306743 | 3.362482475 | 2.699499435 |
| ENSMUSG00000041459 | 3.276662644 | 2.984986613 | 2.949415381 | 3.078149845 | 2.990165688 | 3.454997217 | 2.758982855 |
| ENSMUSG00000041460 | 0.289064826 | 0.5 | 0.5 | 0.5 | 0.489301084 | 0.301029996 | 0.448487648 |
| ENSMUSG00000041468 | 0 | 0 | 0 | 0 | 0 | 0 | 0 |
| ENSMUSG00000041471 | 1.578129653 | 1.54248125 | 1.54248125 | 1.705044741 | 1.662257525 | 1.886490725 | 1.476637173 |
| ENSMUSG00000041476 | 0 | 0 | 0 | 0 | 0 | 0 | 0 |
| ENSMUSG00000041477 | 1.069666964 | 1.214495249 | 1.098079356 | 1.188721876 | 1.293076518 | 1.230448921 | 1.278288402 |
| ENSMUSG00000041479 | 0.289064826 | 0.396240625 | 0.396240625 | 0 | 0.244650542 | 0 | 0.520678036 |
| ENSMUSG00000041481 | 2.436947303 | 2.206637122 | 2.223704441 | 1.992885888 | 1.910073614 | 2.164352856 | 1.852723744 |

| | | | | | | | |
|---|---|---|---|---|---|---|---|
| ENSMUSG00000041482 | 1.777675583 | 1.634789703 | 1.561981878 | 1.701838731 | 1.65906333 | 1.806179974 | 1.548827561 |
| ENSMUSG00000041483 | 2.479432974 | 2.296838018 | 2.261780978 | 2.333288838 | 2.223252711 | 2.545307116 | 2.085803078 |
| ENSMUSG00000041488 | 2.233869984 | 2.066696635 | 2.102347734 | 1.961372513 | 1.83872004 | 2.139879086 | 1.766260282 |
| ENSMUSG00000041491 | 2.016886209 | 1.825945187 | 1.87694866 | 1.79745614 | 1.78687392 | 1.934498451 | 1.705896842 |
| ENSMUSG00000041498 | 2.404271433 | 2.229715809 | 2.222185812 | 2.405054956 | 2.349532392 | 2.649334859 | 2.20139795 |
| ENSMUSG00000041505 | 0 | 0 | 0 | 0 | 0 | 0 | 0 |
| ENSMUSG00000041506 | 1.932832114 | 1.921625132 | 2.056000419 | 1.75835575 | 2.001219193 | 2.004321374 | 2.052911423 |
| ENSMUSG00000041512 | 0 | 0 | 0 | 0 | 0 | 0 | 0 |
| ENSMUSG00000041515 | 0 | 0 | 0.25 | 1.04248125 | 0.931471446 | 0.77815125 | 1 |
| ENSMUSG00000041523 | 0 | 0 | 0 | 0 | 0 | 0 | 0 |
| ENSMUSG00000041528 | 2.054824387 | 1.956637122 | 2.005591953 | 1.898114259 | 2.025895058 | 2.152288344 | 1.988432495 |
| ENSMUSG00000041530 | 2.703665763 | 2.481944491 | 2.454645544 | 2.528109877 | 2.464971436 | 2.835690571 | 2.329931221 |
| ENSMUSG00000041534 | 0 | 0 | 0 | 0 | 0 | 0 | 0 |
| ENSMUSG00000041536 | 0 | 0 | 0 | 0 | 0 | 0 | 0 |
| ENSMUSG00000041538 | 0 | 0 | 0 | 0 | 0 | 0 | 0 |
| ENSMUSG00000041540 | 1.418409478 | 1.238549078 | 1.160964047 | 0.396240625 | 0.632412477 | 0.301029996 | 0.520678036 |
| ENSMUSG00000041544 | 0 | 0 | 0 | 0.396240625 | 0 | 0 | 0.224243824 |
| ENSMUSG00000041548 | 2.748369006 | 2.566403762 | 2.591307212 | 2.400906586 | 2.397399561 | 2.698970004 | 2.271140178 |
| ENSMUSG00000041550 | 0.289064826 | 0 | 0 | 0 | 0 | 0 | 0 |
| ENSMUSG00000041552 | 0.747221736 | 0 | 0.75 | 0 | 0 | 0 | 0 |
| ENSMUSG00000041556 | 1 | 1.25 | 1.311981878 | 1.482684334 | 1.741652793 | 1.72427587 | 1.777090262 |
| ENSMUSG00000041559 | 2.163307908 | 1.940387808 | 1.848079356 | 0.646240625 | 0.568060967 | 0.477121255 | 0.629532004 |
| ENSMUSG00000041560 | 1.956444026 | 1.890560606 | 2.044977273 | 1.807204673 | 1.961316494 | 2.008600172 | 2.016928211 |
| ENSMUSG00000041565 | 0 | 0 | 0 | 0 | 0 | 0 | 0 |
| ENSMUSG00000041566 | 0.578129653 | 0 | 0.396240625 | 0.25 | 0.632412477 | 0.301029996 | 0.520678036 |
| ENSMUSG00000041567 | 0 | 0 | 0.396240625 | 0 | 0 | 0 | 0.224243824 |
| ENSMUSG00000041570 | 2.767404661 | 2.556603048 | 2.54248125 | 2.602877747 | 2.529377461 | 2.90579588 | 2.345233094 |
| ENSMUSG00000041571 | 2.666334442 | 2.480460234 | 2.55479213 | 2.223704441 | 2.32657161 | 2.523746467 | 2.21474793 |
| ENSMUSG00000041577 | 0.578129653 | 0.79248125 | 0.701838731 | 0 | 0.244650542 | 0 | 0 |
| ENSMUSG00000041578 | 0.45815691 | 0.5 | 0.396240625 | 0.5 | 0.244650542 | 0.301029996 | 0.224243824 |
| ENSMUSG00000041583 | 0 | 0 | 0 | 0 | 0 | 0 | 0 |
| ENSMUSG00000041592 | 0.45815691 | 0.25 | 0.25 | 0.25 | 0.632412477 | 0 | 0.224243824 |
| ENSMUSG00000041594 | 2.163307908 | 2.024508021 | 1.94697564 | 1.92510993 | 1.873585591 | 2.086359831 | 1.743447983 |
| ENSMUSG00000041596 | 0 | 0 | 0 | 0.951838731 | 0.931471446 | 0.77815125 | 0.579661877 |

| | | | | | | | |
|---|---|---|---|---|---|---|---|
| ENSMUSG00000041598 | 2.141794304 | 2 | 2.099151195 | 1.830482024 | 1.970729215 | 2.133538908 | 1.916588299 |
| ENSMUSG00000041605 | 1.614416215 | 1.598079356 | 1.755591953 | 1.516522298 | 1.580303447 | 1.672097858 | 1.690609803 |
| ENSMUSG00000041607 | 1.663535556 | 1.561981878 | 1.598079356 | 1.516522298 | 1.55104774 | 1.579783597 | 1.429563222 |
| ENSMUSG00000041608 | 0 | 0.25 | 0 | 1.04248125 | 1.057362051 | 1.079181246 | 0.969165684 |
| ENSMUSG00000041609 | 1.205378646 | 1.364857905 | 1.418106335 | 0.830482024 | 1.244650542 | 0.903089987 | 1.365075947 |
| ENSMUSG00000041616 | 0 | 0.25 | 0 | 0 | 0 | 0 | 0 |
| ENSMUSG00000041617 | 1.181541736 | 1.302363341 | 1.438721876 | 1.238549078 | 1.254881871 | 1.041392685 | 1.245582246 |
| ENSMUSG00000041620 | 0 | 0 | 0 | 0 | 0 | 0 | 0 |
| ENSMUSG00000041623 | 1.993049299 | 1.809601185 | 1.839388001 | 1.794977273 | 1.763796674 | 1.929418926 | 1.657681529 |
| ENSMUSG00000041624 | 1.771760131 | 1.646240625 | 1.729715809 | 1.458222504 | 1.528558853 | 1.556302501 | 1.224243824 |
| ENSMUSG00000041625 | 1.558729299 | 1.614857905 | 1.438721876 | 1.552363341 | 1.72341484 | 1.826074803 | 1.633706455 |
| ENSMUSG00000041629 | 1.989530018 | 1.910964047 | 1.940387808 | 1.741446071 | 1.822942838 | 1.963787827 | 1.846338363 |
| ENSMUSG00000041632 | 1.924654746 | 1.874961472 | 1.880890489 | 1.890560606 | 2.03596422 | 2.173186268 | 1.913982791 |
| ENSMUSG00000041633 | 0.747221736 | 0.5 | 0.701838731 | 1 | 0.955822902 | 1.041392685 | 0.916588299 |
| ENSMUSG00000041638 | 2.975148165 | 2.788071211 | 2.858917565 | 2.439972046 | 2.638266616 | 2.812913357 | 2.58342759 |
| ENSMUSG00000041642 | 2.50175086 | 2.311981878 | 2.31611065 | 2.013820609 | 2.141887974 | 2.285557309 | 2.05853792 |
| ENSMUSG00000041644 | 0 | 0 | 0 | 0 | 0 | 0 | 0 |
| ENSMUSG00000041645 | 2.740271036 | 2.503156135 | 2.496460484 | 2.450224975 | 2.443737124 | 2.786751422 | 2.29131391 |
| ENSMUSG00000041649 | 0.91631382 | 1.061981878 | 1.17510993 | 1.516522298 | 1.373641807 | 1.653212514 | 1.383567577 |
| ENSMUSG00000041650 | 1.806056082 | 1.634789703 | 1.705044741 | 2.009729747 | 1.935892666 | 2.164352856 | 1.797719748 |
| ENSMUSG00000041653 | 0.578129653 | 0.580482024 | 0 | 0.580482024 | 0.387761935 | 0.84509804 | 0.803905701 |
| ENSMUSG00000041654 | 1.907802659 | 1.882845365 | 1.845926073 | 1.910964047 | 1.887294348 | 2.139879086 | 1.797719748 |
| ENSMUSG00000041658 | 1.920505136 | 1.784887838 | 1.755591953 | 1.618933358 | 1.595992969 | 1.832508913 | 1.584526862 |
| ENSMUSG00000041660 | 0 | 0 | 0 | 0 | 0 | 0 | 0 |
| ENSMUSG00000041669 | 0 | 0 | 0 | 0 | 0.244650542 | 0 | 0 |
| ENSMUSG00000041670 | 0 | 0 | 0 | 0.25 | 0 | 0 | 0 |
| ENSMUSG00000041671 | 1.605636262 | 1.470660762 | 1.516522298 | 1.5 | 1.380772476 | 1.477121255 | 1.455757965 |
| ENSMUSG00000041673 | 0 | 0 | 0 | 0 | 0 | 0 | 0 |
| ENSMUSG00000041674 | 0.289064826 | 0 | 0.25 | 0.396240625 | 0 | 0 | 0.355418052 |
| ENSMUSG00000041679 | 0.747221736 | 0.896240625 | 0.92510993 | 0.951838731 | 1.057362051 | 0.698970004 | 1.121219121 |
| ENSMUSG00000041681 | 0 | 0 | 0 | 0 | 0 | 0.301029996 | 0 |
| ENSMUSG00000041684 | 2.262379879 | 2.089388001 | 2.056000419 | 2.19697564 | 2.088159748 | 2.326335861 | 1.933441873 |
| ENSMUSG00000041685 | 2.572927518 | 2.371960008 | 2.351285366 | 2.505591953 | 2.454008153 | 2.742725131 | 2.226906518 |
| ENSMUSG00000041688 | 2.364524 | 2.171625132 | 2.145300145 | 2.584962501 | 2.538790182 | 2.920645001 | 2.360721703 |

| | | | | | | | |
|---|---|---|---|---|---|---|---|
| ENSMUSG00000041695 | 0.960252568 | 1.188721876 | 1.02186571 | 1.339388001 | 1.42077253 | 1.230448921 | 1.176815335 |
| ENSMUSG00000041696 | 1.700465191 | 1.532320754 | 1.571350555 | 0.92510993 | 0.812711509 | 0.698970004 | 0.74492186 |
| ENSMUSG00000041697 | 2.478339835 | 2.364857905 | 2.502107156 | 2.229715809 | 2.296256325 | 2.521138084 | 2.352058568 |
| ENSMUSG00000041698 | 0 | 0 | 0.25 | 0 | 0 | 0 | 0 |
| ENSMUSG00000041700 | 0 | 0 | 0 | 0 | 0 | 0 | 0 |
| ENSMUSG00000041702 | 2.614416215 | 2.463717096 | 2.421187155 | 2.328787391 | 2.271112598 | 2.571708832 | 2.089372236 |
| ENSMUSG00000041703 | 0.811507563 | 1.098079356 | 1.04248125 | 1.614857905 | 1.568060967 | 1.77815125 | 1.499406449 |
| ENSMUSG00000041707 | 0 | 0 | 0 | 0 | 0 | 0 | 0 |
| ENSMUSG00000041708 | 0 | 0 | 0 | 0 | 0 | 0 | 0 |
| ENSMUSG00000041710 | 0 | 0 | 0 | 0.25 | 0.244650542 | 0 | 0 |
| ENSMUSG00000041712 | 2.621464818 | 2.400442697 | 2.378431899 | 2.277783918 | 2.240967488 | 2.580924976 | 2.130728307 |
| ENSMUSG00000041716 | 1.45815691 | 1.51109853 | 1.566696635 | 1.505591953 | 1.467903253 | 1.477121255 | 1.374453863 |
| ENSMUSG00000041718 | 2.075459174 | 2.016522298 | 2.006976499 | 2.334405475 | 2.253097759 | 2.534026106 | 2.092400484 |
| ENSMUSG00000041720 | 2.588343201 | 2.394828984 | 2.397178659 | 2.485628626 | 2.440245937 | 2.720985744 | 2.269400844 |
| ENSMUSG00000041722 | 0 | 0 | 0 | 0 | 0 | 0 | 0 |
| ENSMUSG00000041729 | 1.890240806 | 1.75 | 1.825945187 | 1.557204673 | 1.683842353 | 1.716003344 | 1.596378571 |
| ENSMUSG00000041730 | 0 | 0 | 0 | 0 | 0 | 0 | 0 |
| ENSMUSG00000041731 | 1.852729478 | 1.774508021 | 1.910964047 | 0.75 | 0.955822902 | 1.146128036 | 0.916588299 |
| ENSMUSG00000041733 | 2.23778581 | 2.129909063 | 2.121960008 | 2.072504712 | 2.03596422 | 2.274157849 | 1.920029985 |
| ENSMUSG00000041734 | 3.246030252 | 2.954345857 | 2.953144611 | 2.818157446 | 2.782909563 | 3.174931594 | 2.600582859 |
| ENSMUSG00000041735 | 0 | 0 | 0 | 0 | 0 | 0 | 0 |
| ENSMUSG00000041736 | 2.555102908 | 2.445339928 | 2.609177886 | 1.834962501 | 2.107593434 | 2.089905111 | 2.174485081 |
| ENSMUSG00000041737 | 0 | 0 | 0 | 0.25 | 0.568060967 | 0 | 0 |
| ENSMUSG00000041740 | 3.143130234 | 2.880645383 | 2.871205724 | 2.830482024 | 2.83940011 | 3.241795431 | 2.66503742 |
| ENSMUSG00000041741 | 0 | 0.25 | 0.396240625 | 0 | 0 | 0 | 0 |
| ENSMUSG00000041747 | 2.484856099 | 2.269203899 | 2.277783918 | 2.392938161 | 2.357605438 | 2.683047038 | 2.174874624 |
| ENSMUSG00000041750 | 0 | 0 | 0 | 0 | 0 | 0 | 0 |
| ENSMUSG00000041754 | 0 | 0 | 0 | 0 | 0 | 0 | 0 |
| ENSMUSG00000041757 | 1.205378646 | 1.146240625 | 1.04248125 | 1.464495249 | 1.542223377 | 1.72427587 | 1.535058868 |
| ENSMUSG00000041762 | 1.389637216 | 1.098079356 | 1.201838731 | 1.575945187 | 1.559656853 | 1.69019608 | 1.476637173 |
| ENSMUSG00000041763 | 2.91746109 | 2.664329612 | 2.632845365 | 2.763990059 | 2.673667945 | 3.069668097 | 2.452473516 |
| ENSMUSG00000041765 | 2.029919532 | 1.953445298 | 1.907339155 | 2.065523711 | 2.106691885 | 2.36361198 | 1.993245779 |
| ENSMUSG00000041769 | 2.317377295 | 2.14151351 | 2.118933358 | 2.256976499 | 2.182004726 | 2.489958479 | 2.050061048 |
| ENSMUSG00000041771 | 0 | 0 | 0 | 0.25 | 0 | 0 | 0 |

| | | | | | | | |
|---|---|---|---|---|---|---|---|
| ENSMUSG00000041773 | 2.773381775 | 2.538071211 | 2.538388008 | 2.337182039 | 2.312747785 | 2.595496222 | 2.176040447 |
| ENSMUSG00000041774 | 0.811507563 | 0.864857905 | 1.098079356 | 1.311981878 | 1.302012594 | 1.079181246 | 1.374453863 |
| ENSMUSG00000041775 | 1.358731791 | 1.17510993 | 1.17510993 | 1.098079356 | 1.200473444 | 1.204119983 | 1.245582246 |
| ENSMUSG00000041777 | 1.707474304 | 1.62694866 | 1.589388001 | 1.799918086 | 1.739104362 | 1.908485019 | 1.653807046 |
| ENSMUSG00000041779 | 2.436947303 | 2.234894804 | 2.227473271 | 2.242885888 | 2.251304582 | 2.550228353 | 2.086826862 |
| ENSMUSG00000041781 | 2.934668455 | 2.657794264 | 2.672062577 | 2.664106216 | 2.586616842 | 2.988558957 | 2.392630228 |
| ENSMUSG00000041782 | 0.578129653 | 0.396240625 | 0.396240625 | 0.864857905 | 0.905314583 | 0.698970004 | 0.952571511 |
| ENSMUSG00000041789 | 0 | 0 | 0 | 0 | 0.244650542 | 0 | 0 |
| ENSMUSG00000041791 | 0 | 0 | 0 | 0 | 0 | 0 | 0 |
| ENSMUSG00000041794 | 0.289064826 | 0.25 | 0 | 0 | 0 | 0.301029996 | 0.224243824 |
| ENSMUSG00000041797 | 0 | 0.396240625 | 0.25 | 1.67510993 | 1.499532413 | 1.707570176 | 1.448487648 |
| ENSMUSG00000041798 | 0 | 0 | 0.396240625 | 0 | 0.244650542 | 0 | 0.448487648 |
| ENSMUSG00000041801 | 2.333175866 | 2.250703754 | 2.25 | 1.860735874 | 1.947418788 | 2.06069784 | 1.858985529 |
| ENSMUSG00000041805 | 0 | 0 | 0 | 0 | 0 | 0 | 0 |
| ENSMUSG00000041809 | 0.747221736 | 0.951838731 | 0.830482024 | 0.976722649 | 0.955822902 | 1.113943352 | 0.710836105 |
| ENSMUSG00000041815 | 3.004363607 | 2.756630861 | 2.734159485 | 2.711764337 | 2.737898704 | 3.114277297 | 2.572793882 |
| ENSMUSG00000041817 | 1.129344651 | 1.17510993 | 1.146240625 | 1.403677461 | 1.358933174 | 1.591064607 | 1.319146374 |
| ENSMUSG00000041827 | 0 | 0 | 0 | 0.25 | 0 | 0 | 0 |
| ENSMUSG00000041828 | 0 | 0 | 0 | 0 | 0 | 0 | 0 |
| ENSMUSG00000041831 | 0 | 0 | 0.396240625 | 0.25 | 0 | 0 | 0 |
| ENSMUSG00000041836 | 1.227926429 | 1.396240625 | 1.339388001 | 1.364857905 | 1.380772476 | 1.397940009 | 1.245582246 |
| ENSMUSG00000041837 | 1.924654746 | 1.794977273 | 1.766522298 | 1.823655187 | 1.789100779 | 2.075546961 | 1.702561601 |
| ENSMUSG00000041840 | 2.103271637 | 1.910964047 | 1.994319981 | 2.052363341 | 2.031524462 | 2.318063335 | 1.87805087 |
| ENSMUSG00000041841 | 2.881174527 | 2.641037372 | 2.732314602 | 2.486726569 | 2.58335733 | 2.859138297 | 2.517183443 |
| ENSMUSG00000041842 | 2.013563224 | 2.009729747 | 2.001406137 | 1.914552871 | 1.94314048 | 2.198657087 | 1.853775828 |
| ENSMUSG00000041845 | 2.269664473 | 2.109697963 | 2.105516192 | 2.183677405 | 2.153144932 | 2.447158031 | 2.092902448 |
| ENSMUSG00000041846 | 2.761895567 | 2.521202097 | 2.509729747 | 2.391037372 | 2.32316608 | 2.650307523 | 2.143416814 |
| ENSMUSG00000041849 | 1.100572389 | 1.160964047 | 1.098079356 | 1.146240625 | 0.931471446 | 1.113943352 | 1.028149525 |
| ENSMUSG00000041852 | 2.862863886 | 2.653447366 | 2.617156011 | 2.636723615 | 2.572912674 | 2.954724791 | 2.397361799 |
| ENSMUSG00000041857 | 0 | 0.25 | 0 | 0 | 0 | 0 | 0 |
| ENSMUSG00000041859 | 3.127040599 | 2.859177886 | 2.853935442 | 2.895064642 | 2.88381692 | 3.297541668 | 2.715554737 |
| ENSMUSG00000041870 | 2.842342644 | 2.658702763 | 2.668106335 | 2.725028016 | 2.731346772 | 3.114944416 | 2.587149975 |
| ENSMUSG00000041872 | 0.671187741 | 0.646240625 | 0.25 | 0.396240625 | 0.632412477 | 0.301029996 | 0.579661877 |
| ENSMUSG00000041878 | 0.45815691 | 0 | 0 | 0.25 | 0.387761935 | 0.301029996 | 0.579661877 |

| | | | | | | | |
|---|---|---|---|---|---|---|---|
| ENSMUSG00000041879 | 3.279076498 | 2.960685795 | 2.938826222 | 3.059003548 | 2.983227935 | 3.457579147 | 2.77091973 |
| ENSMUSG00000041881 | 2.280358197 | 2.198603967 | 2.284247778 | 1.976722649 | 2.109389652 | 2.23299611 | 2.131174228 |
| ENSMUSG00000041885 | 0 | 0 | 0 | 0 | 0 | 0 | 0 |
| ENSMUSG00000041886 | 1.740854706 | 1.614857905 | 1.642463902 | 0.646240625 | 0.387761935 | 0.602059991 | 0.672731473 |
| ENSMUSG00000041889 | 2.163307908 | 2.060793496 | 2.046218836 | 2.013820609 | 2.046825265 | 2.212187604 | 1.977154126 |
| ENSMUSG00000041890 | 2.727821383 | 2.531030328 | 2.526149688 | 2.439555804 | 2.438134528 | 2.752816431 | 2.307731473 |
| ENSMUSG00000041891 | 2.942293987 | 2.697383411 | 2.658702763 | 2.713911792 | 2.646380908 | 3.042181595 | 2.456899689 |
| ENSMUSG00000041895 | 2.23778581 | 2.12694866 | 2.104463129 | 2.046218836 | 2.029283463 | 2.274157849 | 1.960219986 |
| ENSMUSG00000041907 | 0 | 0.25 | 0 | 0.5 | 0.387761935 | 0.301029996 | 0.224243824 |
| ENSMUSG00000041911 | 1.227926429 | 1.238549078 | 1.302363341 | 1.160964047 | 1.106691885 | 1.204119983 | 1.168187745 |
| ENSMUSG00000041912 | 2.351376531 | 2.188721876 | 2.140560606 | 1.959800947 | 1.940259189 | 2.204119983 | 1.760706232 |
| ENSMUSG00000041915 | 2.531694272 | 2.328220739 | 2.289336734 | 2.347004321 | 2.25190332 | 2.561101384 | 2.111895264 |
| ENSMUSG00000041920 | 1.289064826 | 1.26109853 | 1.114857905 | 1.114857905 | 0.955822902 | 1.255272505 | 1.066254157 |
| ENSMUSG00000041921 | 1.940852214 | 1.888647213 | 1.926839783 | 2.034887838 | 2.037065488 | 2.267171728 | 1.877074955 |
| ENSMUSG00000041923 | 1.325351389 | 1.238549078 | 1.372963274 | 1 | 0.812711509 | 0.954242509 | 0.876096088 |
| ENSMUSG00000041926 | 2.406877894 | 2.241446071 | 2.281676118 | 2.264497931 | 2.276175004 | 2.586587305 | 2.148525059 |
| ENSMUSG00000041930 | 0.289064826 | 0.5 | 0.396240625 | 0.580482024 | 0.733951626 | 0.301029996 | 0.984950056 |
| ENSMUSG00000041935 | 2.751223449 | 2.483053688 | 2.475593779 | 2.463327389 | 2.3873546 | 2.755112266 | 2.219869654 |
| ENSMUSG00000041936 | 2.312518611 | 2.211372513 | 2.306603048 | 2.26109853 | 2.368949874 | 2.562292864 | 2.302525964 |
| ENSMUSG00000041939 | 2.179305632 | 2.101285366 | 2.216046536 | 1.864857905 | 2.101233692 | 2.053078443 | 2.121678334 |
| ENSMUSG00000041945 | 0.811507563 | 0.864857905 | 0.830482024 | 0.864857905 | 0.812711509 | 0.77815125 | 1.041356071 |
| ENSMUSG00000041949 | 1.956444026 | 1.858657057 | 1.903677461 | 1.973704441 | 1.977302144 | 2.079181246 | 1.828667599 |
| ENSMUSG00000041954 | 0.867194479 | 0.79248125 | 0.896240625 | 0 | 0 | 0.301029996 | 0.224243824 |
| ENSMUSG00000041957 | 2.542156295 | 2.344844592 | 2.351285366 | 2.338837774 | 2.247084732 | 2.580924976 | 2.115189268 |
| ENSMUSG00000041958 | 2.453536005 | 2.316696635 | 2.3464656 | 2.273849256 | 2.371516839 | 2.583198774 | 2.321876478 |
| ENSMUSG00000041959 | 2.963000732 | 2.684311836 | 2.69697564 | 2.912875055 | 2.829291987 | 3.323870607 | 2.639591263 |
| ENSMUSG00000041961 | 2.039433992 | 1.901832578 | 1.938721876 | 1.80479213 | 1.834841362 | 2.012837225 | 1.714087544 |
| ENSMUSG00000041966 | 1.623015126 | 1.71759118 | 1.678561379 | 1.937048212 | 1.850106709 | 2.012837225 | 1.692344475 |
| ENSMUSG00000041974 | 2.342375483 | 2.144357207 | 2.131869252 | 2.087182039 | 2.003645018 | 2.336459734 | 1.858985529 |
| ENSMUSG00000041975 | 1.734388958 | 1.593759858 | 1.566696635 | 1.747171172 | 1.728722551 | 1.977723605 | 1.577201671 |
| ENSMUSG00000041977 | 2.429630757 | 2.187886015 | 2.20584181 | 2.260414788 | 2.261347526 | 2.552668216 | 2.128039798 |
| ENSMUSG00000041984 | 0 | 0 | 0 | 0 | 0 | 0 | 0 |
| ENSMUSG00000041986 | 0.289064826 | 0 | 0.25 | 0.896240625 | 1.091002363 | 0.903089987 | 0.935079929 |
| ENSMUSG00000041991 | 0 | 0 | 0 | 0 | 0 | 0 | 0 |

| | | | | | | | |
|---|---|---|---|---|---|---|---|
| ENSMUSG00000041992 | 0.45815691 | 0 | 0.25 | 0.75 | 0.877063019 | 0.84509804 | 0.520678036 |
| ENSMUSG00000041995 | 2.00683675 | 1.841580554 | 1.935366747 | 1.96293726 | 1.940259189 | 2.173186268 | 1.89047692 |
| ENSMUSG00000041997 | 2.470606563 | 2.338837774 | 2.263820609 | 2.43155454 | 2.348623881 | 2.706717782 | 2.167790061 |
| ENSMUSG00000042002 | 0 | 0 | 0 | 0 | 0 | 0 | 0 |
| ENSMUSG00000042010 | 1.587501561 | 1.396240625 | 1.598079356 | 1.130890489 | 1.358933174 | 1 | 1.355418052 |
| ENSMUSG00000042015 | 2.036286563 | 1.894357207 | 1.984159485 | 2.143411797 | 2.117361396 | 2.416640507 | 2.008574816 |
| ENSMUSG00000042029 | 3.011739465 | 2.749647607 | 2.757839149 | 2.752981517 | 2.666348738 | 3.060320029 | 2.462388606 |
| ENSMUSG00000042031 | 0 | 0 | 0 | 0 | 0 | 0 | 0 |
| ENSMUSG00000042032 | 2.537433341 | 2.355516192 | 2.328787391 | 2.376452888 | 2.31625483 | 2.635483747 | 2.135160095 |
| ENSMUSG00000042035 | 2.100572389 | 1.969129237 | 2.034887838 | 1.77186571 | 1.868304353 | 1.995635195 | 1.817112247 |
| ENSMUSG00000042041 | 0 | 0 | 0 | 0 | 0 | 0 | 0 |
| ENSMUSG00000042042 | 2.476144923 | 2.293731421 | 2.288071211 | 2.377938163 | 2.305113253 | 2.567026366 | 2.082712142 |
| ENSMUSG00000042043 | 2.590021402 | 2.340485943 | 2.391037372 | 2.589388001 | 2.51576314 | 2.888179494 | 2.333765523 |
| ENSMUSG00000042045 | 0 | 0 | 0 | 0 | 0 | 0 | 0 |
| ENSMUSG00000042046 | 2.371652933 | 2.206637122 | 2.174241882 | 2.263820609 | 2.208005287 | 2.526339277 | 2.092902448 |
| ENSMUSG00000042050 | 2.146671932 | 2.023189285 | 1.992885888 | 1.96759118 | 1.892300902 | 2.146128036 | 1.809975587 |
| ENSMUSG00000042055 | 2.292497211 | 2.175975893 | 2.138647213 | 2.158248799 | 2.162515999 | 2.390935107 | 2.044866219 |
| ENSMUSG00000042063 | 2.196867486 | 2.051142786 | 2.027131114 | 2.015173983 | 1.983754904 | 2.222716471 | 1.777090262 |
| ENSMUSG00000042064 | 0 | 0 | 0 | 0 | 0 | 0 | 0 |
| ENSMUSG00000042066 | 2.247417471 | 2.108657057 | 2.206637122 | 1.87694866 | 1.945998439 | 2.068185862 | 1.991191705 |
| ENSMUSG00000042073 | 1.714367558 | 1.614857905 | 1.723704441 | 1.482684334 | 1.635855979 | 1.568201724 | 1.505628092 |
| ENSMUSG00000042078 | 0.578129653 | 0.5 | 0.580482024 | 0.25 | 0 | 0.477121255 | 0.579661877 |
| ENSMUSG00000042079 | 3.107651248 | 2.84995299 | 2.844979954 | 2.736360959 | 2.673305567 | 3.068556895 | 2.50897195 |
| ENSMUSG00000042082 | 2.121692466 | 2.012462137 | 2.044977273 | 2.102347734 | 2.056322416 | 2.324282455 | 1.948678378 |
| ENSMUSG00000042087 | 0.289064826 | 0.25 | 0.75 | 0.25 | 0.244650542 | 0 | 0.448487648 |
| ENSMUSG00000042092 | 0 | 0 | 0 | 0 | 0 | 0 | 0 |
| ENSMUSG00000042096 | 0 | 0 | 0 | 0 | 0.387761935 | 0 | 0.355418052 |
| ENSMUSG00000042097 | 1.100572389 | 1.214495249 | 1.27186571 | 1.451838731 | 1.394615667 | 1.431363764 | 1.245582246 |
| ENSMUSG00000042099 | 1.734388958 | 1.62694866 | 1.660964047 | 1.388647213 | 1.592135487 | 1.62324929 | 1.546120303 |
| ENSMUSG00000042102 | 0 | 0 | 0 | 0 | 0 | 0 | 0 |
| ENSMUSG00000042104 | 2.278594844 | 2.137686696 | 2.113831805 | 2.307204673 | 2.219338185 | 2.567026366 | 2.029372647 |
| ENSMUSG00000042105 | 2.190368148 | 1.975216702 | 2.016522298 | 2.105516192 | 2.017859938 | 2.340444115 | 1.898808251 |
| ENSMUSG00000042106 | 0.45815691 | 0.92510993 | 1 | 1.02186571 | 1.223252711 | 1.113943352 | 1.168187745 |
| ENSMUSG00000042109 | 1.605636262 | 1.614857905 | 1.584962501 | 1.735628626 | 1.800029087 | 1.86923172 | 1.759302692 |

| | | | | | | | |
|---|---|---|---|---|---|---|---|
| ENSMUSG00000042111 | 1.842799666 | 1.75 | 1.809601185 | 1.784887838 | 1.78010778 | 1.934498451 | 1.71890424 |
| ENSMUSG00000042115 | 2.026699189 | 1.886723615 | 1.912762923 | 1.856566189 | 1.940259189 | 2.025305865 | 1.872150729 |
| ENSMUSG00000042116 | 0.811507563 | 1.02186571 | 0.646240625 | 1.080482024 | 1.244650542 | 1.301029996 | 1.150210039 |
| ENSMUSG00000042118 | 0.671187741 | 0.646240625 | 0.701838731 | 0.5 | 0.568060967 | 0.477121255 | 0.629532004 |
| ENSMUSG00000042121 | 2.477243823 | 2.322504712 | 2.280383379 | 2.03360658 | 2.084333628 | 2.290034611 | 1.989124507 |
| ENSMUSG00000042124 | 0 | 0 | 0 | 0 | 0 | 0 | 0 |
| ENSMUSG00000042129 | 1.53838222 | 1.482684334 | 1.396240625 | 0.92510993 | 0.905314583 | 0.903089987 | 0.853775828 |
| ENSMUSG00000042133 | 2.546826359 | 2.368426437 | 2.393411797 | 2.599418658 | 2.50570673 | 2.872156273 | 2.327511498 |
| ENSMUSG00000042138 | 1.985980784 | 1.894357207 | 1.852347734 | 1.907339155 | 1.893954095 | 2.049218023 | 1.726776051 |
| ENSMUSG00000042148 | 1.91631382 | 1.799918086 | 1.819031101 | 1.814346961 | 1.932951348 | 2.053078443 | 1.882886767 |
| ENSMUSG00000042155 | 2.229917041 | 2.027131114 | 2.028435542 | 2.263820609 | 2.210704773 | 2.51054501 | 2.011813236 |
| ENSMUSG00000042156 | 2.116511873 | 1.985628626 | 1.997171172 | 2.201838731 | 2.107593434 | 2.411619706 | 2.029372647 |
| ENSMUSG00000042157 | 0 | 0 | 0 | 0 | 0 | 0 | 0 |
| ENSMUSG00000042165 | 0.289064826 | 0.25 | 0 | 0.25 | 0 | 0.301029996 | 0 |
| ENSMUSG00000042167 | 2.355805236 | 2.183677405 | 2.182829758 | 2.234894804 | 2.146756411 | 2.477121255 | 1.982842456 |
| ENSMUSG00000042178 | 1.505869732 | 1.593759858 | 1.714495249 | 1.302363341 | 1.542223377 | 1.531478917 | 1.68886578 |
| ENSMUSG00000042179 | 0 | 0 | 0.25 | 0 | 0 | 0 | 0 |
| ENSMUSG00000042182 | 1.631440309 | 1.575945187 | 1.584962501 | 1.622963274 | 1.576269619 | 1.763427994 | 1.486592281 |
| ENSMUSG00000042184 | 0 | 0 | 0 | 0 | 0 | 0 | 0 |
| ENSMUSG00000042185 | 2.269664473 | 2.107613138 | 2.091580554 | 2.143411797 | 2.175379703 | 2.359835482 | 2.043700396 |
| ENSMUSG00000042189 | 0 | 0 | 0 | 0 | 0 | 0 | 0 |
| ENSMUSG00000042190 | 1.358731791 | 1.372963274 | 1.438721876 | 0.896240625 | 1.020174412 | 1.301029996 | 1 |
| ENSMUSG00000042195 | 0.289064826 | 0.396240625 | 0.580482024 | 1.5 | 1.523883869 | 1.748188027 | 1.417653332 |
| ENSMUSG00000042197 | 2.65645504 | 2.429419106 | 2.400442697 | 2.515511534 | 2.402121453 | 2.762678564 | 2.226574877 |
| ENSMUSG00000042198 | 1.596667477 | 1.458222504 | 1.470660762 | 1.52186571 | 1.528558853 | 1.653212514 | 1.355418052 |
| ENSMUSG00000042200 | 0 | 0 | 0 | 0 | 0 | 0.301029996 | 0 |
| ENSMUSG00000042202 | 2.227926429 | 2.081607372 | 2.065523711 | 2.171625132 | 2.103973339 | 2.424881637 | 1.947107911 |
| ENSMUSG00000042203 | 2.054824387 | 2.04248125 | 2.027131114 | 2.02053726 | 2.041436517 | 2.206825876 | 1.95718226 |
| ENSMUSG00000042207 | 2.412042447 | 2.234159485 | 2.160061234 | 2.161864607 | 2.079491904 | 2.392696953 | 1.967691807 |
| ENSMUSG00000042208 | 2.351376531 | 2.183677405 | 2.199415381 | 2.452241044 | 2.385310841 | 2.698100546 | 2.195242462 |
| ENSMUSG00000042210 | 1.578129653 | 1.714495249 | 1.589388001 | 1.610735874 | 1.793512788 | 1.903089987 | 1.670888075 |
| ENSMUSG00000042211 | 2.573799059 | 2.290597832 | 2.327653195 | 2.347542239 | 2.308696595 | 2.646403726 | 2.174485081 |
| ENSMUSG00000042212 | 0 | 0 | 0 | 0 | 0 | 0 | 0 |
| ENSMUSG00000042213 | 1.389637216 | 1.364857905 | 1.238549078 | 1.188721876 | 1.121713561 | 1.230448921 | 1.078019652 |

| | | | | | | | |
|---|---|---|---|---|---|---|---|
| ENSMUSG00000042215 | 2.518981867 | 2.296218836 | 2.315523711 | 2.218361028 | 2.225193812 | 2.515873844 | 2.043115906 |
| ENSMUSG00000042216 | 0.747221736 | 0.79248125 | 0.864857905 | 1.464495249 | 1.603583716 | 1.672097858 | 1.596378571 |
| ENSMUSG00000042219 | 0 | 0 | 0 | 0 | 0 | 0 | 0 |
| ENSMUSG00000042225 | 2.294202867 | 2.136723615 | 2.134789703 | 2.415444524 | 2.339407793 | 2.662757832 | 2.166194421 |
| ENSMUSG00000042228 | 1.389637216 | 1.339388001 | 1.5 | 1.642463902 | 1.642643806 | 1.698970004 | 1.535058868 |
| ENSMUSG00000042229 | 2.358731791 | 2.153677461 | 2.152756199 | 2.202642909 | 2.172394856 | 2.411619706 | 2.02568932 |
| ENSMUSG00000042240 | 0 | 0 | 0 | 0 | 0 | 0 | 0 |
| ENSMUSG00000042243 | 0 | 0 | 0 | 0 | 0 | 0 | 0 |
| ENSMUSG00000042244 | 0 | 0 | 0 | 0 | 0 | 0 | 0 |
| ENSMUSG00000042246 | 0.867194479 | 0.701838731 | 0.396240625 | 1 | 1.223252711 | 1.255272505 | 1.014380832 |
| ENSMUSG00000042248 | 0 | 0 | 0 | 0 | 0 | 0 | 0 |
| ENSMUSG00000042249 | 0 | 0 | 0.396240625 | 0.25 | 0.387761935 | 0.301029996 | 0.224243824 |
| ENSMUSG00000042250 | 0 | 0 | 0 | 0 | 0 | 0 | 0 |
| ENSMUSG00000042251 | 0 | 0.396240625 | 0.396240625 | 0.5 | 0.387761935 | 0.698970004 | 0.579661877 |
| ENSMUSG00000042254 | 0.289064826 | 0.25 | 0.25 | 0 | 0.244650542 | 0.301029996 | 0.224243824 |
| ENSMUSG00000042256 | 1.989530018 | 1.872963274 | 1.919870025 | 1.905512955 | 1.793512788 | 2.045322979 | 1.651852264 |
| ENSMUSG00000042258 | 0 | 0 | 0 | 0 | 0 | 0 | 0 |
| ENSMUSG00000042262 | 0 | 0 | 0 | 0 | 0 | 0 | 0 |
| ENSMUSG00000042265 | 0 | 0 | 0 | 0 | 0 | 0 | 0 |
| ENSMUSG00000042268 | 0 | 0 | 0.25 | 0 | 0.244650542 | 0 | 0 |
| ENSMUSG00000042269 | 0 | 0 | 0 | 0 | 0.244650542 | 0 | 0.224243824 |
| ENSMUSG00000042271 | 1.975148165 | 1.888647213 | 1.821350555 | 2.088286706 | 1.978602168 | 2.285557309 | 1.882886767 |
| ENSMUSG00000042272 | 2.322180025 | 2.122963274 | 2.098079356 | 2.262462137 | 2.199780694 | 2.436162647 | 2.030591163 |
| ENSMUSG00000042275 | 1.811507563 | 1.80479213 | 1.83272922 | 1.735628626 | 1.766173492 | 1.949390007 | 1.764880683 |
| ENSMUSG00000042279 | 0 | 0 | 0 | 0 | 0 | 0 | 0 |
| ENSMUSG00000042282 | 0 | 0 | 0 | 0 | 0 | 0 | 0 |
| ENSMUSG00000042284 | 1.960252568 | 1.872963274 | 1.830482024 | 1.671625132 | 1.621875691 | 1.77815125 | 1.441050184 |
| ENSMUSG00000042286 | 0.45815691 | 0.580482024 | 0.396240625 | 0.646240625 | 0.846351821 | 0.903089987 | 0.876096088 |
| ENSMUSG00000042289 | 1.404271433 | 1.431980114 | 1.445339928 | 1.348079356 | 1.401338844 | 1.204119983 | 1.405288179 |
| ENSMUSG00000042292 | 2.544964605 | 2.449820405 | 2.513142013 | 2.104463129 | 2.305627392 | 2.44870632 | 2.24869599 |
| ENSMUSG00000042293 | 0.289064826 | 0.646240625 | 0.75 | 0.25 | 0.489301084 | 0.301029996 | 0.74492186 |
| ENSMUSG00000042298 | 2.170239354 | 1.976722649 | 1.997171172 | 2.193696765 | 2.111176775 | 2.369215857 | 1.991877847 |
| ENSMUSG00000042302 | 2.209569962 | 2.012462137 | 2.044977273 | 2.115881093 | 2.091002363 | 2.392696953 | 1.960974982 |
| ENSMUSG00000042303 | 1.358731791 | 1.445339928 | 1.566696635 | 1.27186571 | 1.523883869 | 1.491361694 | 1.413583902 |

| | | | | | | | |
|---|---|---|---|---|---|---|---|
| ENSMUSG00000042305 | 2.680728114 | 2.468745337 | 2.442459461 | 2.582168732 | 2.519986838 | 2.889861721 | 2.326538518 |
| ENSMUSG00000042306 | 0 | 0 | 0.396240625 | 0.5 | 0.568060967 | 0.477121255 | 0.224243824 |
| ENSMUSG00000042308 | 2.086807695 | 1.933677405 | 1.953445298 | 1.864857905 | 1.944572352 | 2.037426498 | 1.87805087 |
| ENSMUSG00000042312 | 1.876566388 | 1.720660762 | 1.845926073 | 2.004202072 | 1.86473911 | 2.181843588 | 1.806347338 |
| ENSMUSG00000042320 | 1.249317394 | 1.25 | 1.282320754 | 1.146240625 | 1.163285805 | 1.301029996 | 1.185218806 |
| ENSMUSG00000042323 | 2.908973551 | 2.67813175 | 2.633088981 | 2.914552871 | 2.833116536 | 3.263872677 | 2.602457109 |
| ENSMUSG00000042328 | 1.948720984 | 1.830482024 | 1.921625132 | 1.71759118 | 1.806427258 | 1.792391689 | 1.725216937 |
| ENSMUSG00000042331 | 2.976144659 | 2.710194481 | 2.688513092 | 2.735994979 | 2.694415363 | 3.064457989 | 2.499263648 |
| ENSMUSG00000042333 | 0.289064826 | 0 | 0 | 0.25 | 0.686820904 | 0.301029996 | 0.629532004 |
| ENSMUSG00000042340 | 1.269664473 | 1.160964047 | 1.17510993 | 1.282320754 | 1.149965125 | 1.278753601 | 1.054044578 |
| ENSMUSG00000042345 | 0 | 0 | 0 | 0 | 0 | 0 | 0 |
| ENSMUSG00000042348 | 1.989530018 | 1.825945187 | 1.807204673 | 2.03360658 | 1.992595054 | 2.204119983 | 1.871156825 |
| ENSMUSG00000042349 | 1.753493018 | 1.561981878 | 1.62694866 | 1.688721876 | 1.756568651 | 1.857332496 | 1.637827726 |
| ENSMUSG00000042350 | 2.494443473 | 2.30479213 | 2.29745614 | 2.12694866 | 2.118236133 | 2.399673721 | 1.981429723 |
| ENSMUSG00000042351 | 0 | 0 | 0 | 0 | 0 | 0 | 0.224243824 |
| ENSMUSG00000042353 | 0 | 0 | 0 | 0 | 0 | 0 | 0 |
| ENSMUSG00000042354 | 2.53838222 | 2.352347734 | 2.39151351 | 2.556301859 | 2.487464361 | 2.814913181 | 2.30409643 |
| ENSMUSG00000042357 | 1.069666964 | 1.146240625 | 1.02186571 | 0.5 | 0.686820904 | 0.698970004 | 0.448487648 |
| ENSMUSG00000042359 | 1.342375483 | 1.214495249 | 1.29248125 | 1.741446071 | 1.741652793 | 1.924279286 | 1.61448206 |
| ENSMUSG00000042360 | 0 | 0 | 0 | 0.25 | 0 | 0 | 0 |
| ENSMUSG00000042363 | 2.392336464 | 2.167221246 | 2.169870025 | 2.143411797 | 2.104881849 | 2.356025857 | 1.911356129 |
| ENSMUSG00000042364 | 2.615205302 | 2.395300145 | 2.386241108 | 2.387205477 | 2.366364104 | 2.648360011 | 2.204609933 |
| ENSMUSG00000042367 | 2.010213549 | 1.950224975 | 2.001406137 | 1.575945187 | 1.65906333 | 1.812913357 | 1.649885598 |
| ENSMUSG00000042369 | 2.141794304 | 1.943696765 | 1.997171172 | 2.091580554 | 2.062514787 | 2.344392274 | 1.937521567 |
| ENSMUSG00000042371 | 0 | 0 | 0 | 0 | 0 | 0 | 0 |
| ENSMUSG00000042372 | 0 | 0 | 0 | 0 | 0 | 0 | 0 |
| ENSMUSG00000042377 | 0 | 0.25 | 0.25 | 0.396240625 | 0.77552387 | 0 | 0.876096088 |
| ENSMUSG00000042379 | 0 | 0 | 0.25 | 0 | 0.244650542 | 0 | 0 |
| ENSMUSG00000042380 | 1.727821383 | 1.774508021 | 1.779735268 | 1.532320754 | 1.677808768 | 1.748188027 | 1.612273755 |
| ENSMUSG00000042385 | 0 | 0 | 0 | 0 | 0 | 0 | 0 |
| ENSMUSG00000042386 | 0.289064826 | 0.25 | 0.25 | 0.396240625 | 0.244650542 | 0 | 0 |
| ENSMUSG00000042388 | 0 | 0.25 | 0.5 | 0.396240625 | 0.568060967 | 0.477121255 | 0.984950056 |
| ENSMUSG00000042389 | 1.655740331 | 1.571350555 | 1.614857905 | 1.598079356 | 1.649303553 | 1.72427587 | 1.526507301 |
| ENSMUSG00000042390 | 2.683414322 | 2.445749552 | 2.42293588 | 2.53743678 | 2.452994396 | 2.811575006 | 2.284173211 |

| | | | | | | | |
|---|---|---|---|---|---|---|---|
| ENSMUSG00000042396 | 2.509948321 | 2.330482024 | 2.308404919 | 2.42813175 | 2.369807605 | 2.7084209 | 2.200679824 |
| ENSMUSG00000042401 | 0 | 0 | 0 | 0.25 | 0 | 0 | 0.224243824 |
| ENSMUSG00000042402 | 0 | 0 | 0 | 0 | 0 | 0 | 0 |
| ENSMUSG00000042404 | 1.389637216 | 1.388647213 | 1.431980114 | 1.130890489 | 1.327538614 | 1.322219295 | 1.290497981 |
| ENSMUSG00000042406 | 3.163731291 | 2.91152716 | 2.902525516 | 2.768033151 | 2.77795806 | 3.161068385 | 2.605659747 |
| ENSMUSG00000042408 | 1.971473847 | 1.814346961 | 1.774508021 | 1.912762923 | 1.848234258 | 2.103803721 | 1.726776051 |
| ENSMUSG00000042410 | 2.759113486 | 2.540282971 | 2.530059469 | 2.747171172 | 2.667456342 | 3.066325925 | 2.440536803 |
| ENSMUSG00000042414 | 0 | 0 | 0 | 0 | 0 | 0 | 0 |
| ENSMUSG00000042417 | 0.289064826 | 0.396240625 | 0.25 | 0.79248125 | 0.733951626 | 0.477121255 | 0.829800754 |
| ENSMUSG00000042419 | 1.325351389 | 1.26109853 | 1.552363341 | 1.160964047 | 1.343584837 | 1.176091259 | 1.526507301 |
| ENSMUSG00000042423 | 1.852729478 | 1.854463129 | 1.912762923 | 1.614857905 | 1.773209395 | 1.73239376 | 1.802677937 |
| ENSMUSG00000042425 | 0 | 0 | 0 | 0.25 | 0.244650542 | 0 | 0.224243824 |
| ENSMUSG00000042426 | 2.45469604 | 2.265173983 | 2.296218836 | 2.507321807 | 2.419658517 | 2.7930916 | 2.238624656 |
| ENSMUSG00000042428 | 2.425923778 | 2.269203899 | 2.267194569 | 2.240000483 | 2.230318492 | 2.503790683 | 2.124420007 |
| ENSMUSG00000042429 | 0.811507563 | 0.75 | 0.701838731 | 0.580482024 | 0.387761935 | 0.301029996 | 0.448487648 |
| ENSMUSG00000042433 | 0 | 0 | 0 | 0 | 0 | 0 | 0 |
| ENSMUSG00000042436 | 0.289064826 | 0.396240625 | 0.25 | 0 | 0 | 0 | 0.224243824 |
| ENSMUSG00000042439 | 2.245508853 | 2.157339155 | 2.175975893 | 1.695339928 | 1.773209395 | 1.86923172 | 1.712465909 |
| ENSMUSG00000042444 | 2.131864492 | 2.056000419 | 2.028435542 | 2.148114259 | 2.055279708 | 2.278753601 | 1.913982791 |
| ENSMUSG00000042446 | 2.740271036 | 2.46293726 | 2.454245906 | 2.491806565 | 2.427019704 | 2.799340549 | 2.244328309 |
| ENSMUSG00000042447 | 2.201144779 | 2.029735268 | 2.016522298 | 2.012462137 | 1.974687643 | 2.161368002 | 1.798966438 |
| ENSMUSG00000042448 | 0 | 0 | 0.25 | 0 | 0 | 0 | 0 |
| ENSMUSG00000042451 | 0 | 0 | 0 | 0 | 0 | 0 | 0 |
| ENSMUSG00000042453 | 0 | 0 | 0.25 | 0.25 | 0.244650542 | 0 | 0 |
| ENSMUSG00000042459 | 0 | 0 | 0 | 0 | 0 | 0 | 0 |
| ENSMUSG00000042460 | 2.608049185 | 2.3843046 | 2.345385738 | 2.368426437 | 2.266550905 | 2.586587305 | 2.075912052 |
| ENSMUSG00000042462 | 2.010213549 | 1.894357207 | 1.972185812 | 1.79248125 | 1.911645804 | 2.064457989 | 1.892346957 |
| ENSMUSG00000042472 | 2.506893125 | 2.289967834 | 2.274508021 | 2.170748646 | 2.143518259 | 2.416640507 | 1.986347533 |
| ENSMUSG00000042473 | 2.427163103 | 2.234894804 | 2.200224975 | 2.338837774 | 2.242813819 | 2.580924976 | 2.074853081 |
| ENSMUSG00000042474 | 0 | 0 | 0 | 0 | 0 | 0 | 0 |
| ENSMUSG00000042476 | 0.45815691 | 0.646240625 | 0.79248125 | 0.976722649 | 0.733951626 | 0.698970004 | 0.710836105 |
| ENSMUSG00000042477 | 0 | 0.25 | 0 | 1.04248125 | 1.106691885 | 1.041392685 | 1.041356071 |
| ENSMUSG00000042485 | 1.740854706 | 1.678561379 | 1.705044741 | 1.29248125 | 1.310728002 | 1.322219295 | 1.252393349 |
| ENSMUSG00000042487 | 2.302628051 | 2.179419106 | 2.202642909 | 2.30539678 | 2.276733037 | 2.562292864 | 2.15146804 |

| | | | | | | | |
|---|---|---|---|---|---|---|---|
| ENSMUSG00000042489 | 2.648525058 | 2.423371739 | 2.440387808 | 2.606041572 | 2.536927612 | 2.907411361 | 2.354525595 |
| ENSMUSG00000042492 | 2.227926429 | 2.105516192 | 2.189555804 | 2.08272922 | 2.137779008 | 2.328379603 | 2.114720744 |
| ENSMUSG00000042496 | 1.789260561 | 1.602347734 | 1.688721876 | 1.671625132 | 1.642643806 | 1.819543936 | 1.616675393 |
| ENSMUSG00000042498 | 1.205378646 | 1.25 | 1.418106335 | 1.226722649 | 1.188507684 | 1.230448921 | 1.054044578 |
| ENSMUSG00000042499 | 0 | 0 | 0 | 0 | 0.244650542 | 0 | 0 |
| ENSMUSG00000042500 | 1.470606563 | 1.330482024 | 1.364857905 | 1.634789703 | 1.618292349 | 1.72427587 | 1.42167221 |
| ENSMUSG00000042501 | 0.747221736 | 0.701838731 | 1.080482024 | 0.92510993 | 1.074582839 | 0.954242509 | 0.775756176 |
| ENSMUSG00000042502 | 2.415874319 | 2.247171172 | 2.264497931 | 2.254897682 | 2.237245597 | 2.472756449 | 2.129834618 |
| ENSMUSG00000042505 | 1.721148722 | 1.464495249 | 1.488549078 | 1.410964047 | 1.234113756 | 1.491361694 | 1.245582246 |
| ENSMUSG00000042506 | 2.701107274 | 2.480831871 | 2.472566069 | 2.626204747 | 2.598690736 | 2.964730921 | 2.425462392 |
| ENSMUSG00000042507 | 2.340856238 | 2.206637122 | 2.214495249 | 2.074802005 | 2.126866297 | 2.294466226 | 2.051204213 |
| ENSMUSG00000042508 | 2.490209605 | 2.284247778 | 2.277783918 | 2.312574604 | 2.24829556 | 2.542825427 | 2.03780742 |
| ENSMUSG00000042510 | 1.325351389 | 1.146240625 | 1.130890489 | 0.976722649 | 0.846351821 | 0.77815125 | 0.853775828 |
| ENSMUSG00000042514 | 0 | 0 | 0 | 0 | 0 | 0.301029996 | 0 |
| ENSMUSG00000042515 | 2.34539747 | 2.168106335 | 2.179419106 | 2.492166698 | 2.401730365 | 2.761927838 | 2.208492871 |
| ENSMUSG00000042520 | 3.115087743 | 2.822792675 | 2.81168515 | 2.849284952 | 2.805929611 | 3.20709554 | 2.623556869 |
| ENSMUSG00000042523 | 2.267855219 | 2.105516192 | 2.080482024 | 2.098079356 | 2.057362051 | 2.342422681 | 1.896975297 |
| ENSMUSG00000042524 | 2.721148722 | 2.549918086 | 2.595115266 | 2.441217898 | 2.537460779 | 2.756636108 | 2.445128164 |
| ENSMUSG00000042525 | 0 | 0 | 0 | 0 | 0 | 0 | 0 |
| ENSMUSG00000042529 | 0 | 0 | 0.396240625 | 0.25 | 0 | 0 | 0.224243824 |
| ENSMUSG00000042532 | 0.45815691 | 0.25 | 0.396240625 | 0.396240625 | 0.387761935 | 0 | 0.355418052 |
| ENSMUSG00000042535 | 2.571178952 | 2.355516192 | 2.42510993 | 2.433677405 | 2.492341207 | 2.785329835 | 2.370011934 |
| ENSMUSG00000042540 | 0 | 0 | 0 | 0.25 | 0 | 0 | 0 |
| ENSMUSG00000042541 | 2.688073951 | 2.515511534 | 2.564052172 | 2.368933358 | 2.355827317 | 2.669316881 | 2.256654198 |
| ENSMUSG00000042548 | 2.642661589 | 2.371457327 | 2.386723615 | 2.520869832 | 2.476238662 | 2.820201459 | 2.338260508 |
| ENSMUSG00000042549 | 1.036286563 | 1.04248125 | 0.976722649 | 1.214495249 | 1.163285805 | 1.255272505 | 1.20139795 |
| ENSMUSG00000042554 | 0 | 0 | 0 | 0 | 0 | 0 | 0 |
| ENSMUSG00000042557 | 2.622240692 | 2.409156155 | 2.396240625 | 2.495748394 | 2.414035093 | 2.787460475 | 2.26794422 |
| ENSMUSG00000042558 | 1.985980784 | 1.894357207 | 1.975216702 | 1.54745614 | 1.751666404 | 1.851258349 | 1.775756176 |
| ENSMUSG00000042564 | 1.069666964 | 1.17510993 | 1.188721876 | 1.061981878 | 1.163285805 | 1.255272505 | 1.176815335 |
| ENSMUSG00000042567 | 0.289064826 | 0 | 0 | 0.25 | 0 | 0 | 0.224243824 |
| ENSMUSG00000042569 | 1.985980784 | 1.886723615 | 1.909156155 | 2.058404919 | 2.111176775 | 2.367355921 | 1.933441873 |
| ENSMUSG00000042570 | 2.069666964 | 2.025821952 | 2.034887838 | 1.819031101 | 1.981187939 | 2.053078443 | 1.997315209 |
| ENSMUSG00000042572 | 2.613625632 | 2.356566189 | 2.363831805 | 2.410964047 | 2.354487824 | 2.692846919 | 2.218511769 |

| | | | | | | | |
|---|---|---|---|---|---|---|---|
| ENSMUSG00000042579 | 1.325351389 | 1.17510993 | 1.098079356 | 1.080482024 | 0.905314583 | 1.342422681 | 0.984950056 |
| ENSMUSG00000042581 | 0.45815691 | 0.25 | 0.580482024 | 0.25 | 0.387761935 | 0.301029996 | 0 |
| ENSMUSG00000042589 | 0.671187741 | 0.75 | 0.646240625 | 0.896240625 | 1 | 0.954242509 | 1.028149525 |
| ENSMUSG00000042590 | 2.597490839 | 2.38040011 | 2.365369862 | 2.601285366 | 2.511488286 | 2.904174368 | 2.326294816 |
| ENSMUSG00000042594 | 2.339331437 | 2.215271726 | 2.246460484 | 2.116901388 | 2.21736452 | 2.382017043 | 2.098373247 |
| ENSMUSG00000042595 | 2.060824957 | 1.956637122 | 1.959800947 | 2.12694866 | 2.079491904 | 2.290034611 | 1.900630878 |
| ENSMUSG00000042596 | 0 | 0 | 0 | 0 | 0 | 0 | 0 |
| ENSMUSG00000042599 | 1.920505136 | 1.80479213 | 1.738549078 | 1.671625132 | 1.628935049 | 1.908485019 | 1.441050184 |
| ENSMUSG00000042604 | 0.671187741 | 0.646240625 | 0.5 | 0.580482024 | 0.387761935 | 0.602059991 | 0.579661877 |
| ENSMUSG00000042605 | 2.777675583 | 2.533285553 | 2.529410775 | 2.367410454 | 2.377015368 | 2.645422269 | 2.210591357 |
| ENSMUSG00000042606 | 2.179305632 | 2.039967834 | 2.058404919 | 2.086073977 | 2.090057366 | 2.334453751 | 1.94631981 |
| ENSMUSG00000042607 | 1.181541736 | 0.976722649 | 1.080482024 | 1.238549078 | 1.176121988 | 1.041392685 | 1.054044578 |
| ENSMUSG00000042608 | 1.993049299 | 1.837182039 | 1.907339155 | 1.692046081 | 1.826953746 | 1.851258349 | 1.778418869 |
| ENSMUSG00000042613 | 2.45469604 | 2.359697963 | 2.417221246 | 2.024508021 | 2.0928848 | 2.344392274 | 2.021338422 |
| ENSMUSG00000042616 | 1.811507563 | 1.77186571 | 1.774508021 | 1.921625132 | 1.875328588 | 2.120573931 | 1.822941512 |
| ENSMUSG00000042622 | 1.631440309 | 1.505591953 | 1.482684334 | 1.646240625 | 1.65906333 | 1.851258349 | 1.591689859 |
| ENSMUSG00000042625 | 2.395018353 | 2.263820609 | 2.232684334 | 2.107613138 | 2.207327177 | 2.396199347 | 2.13206423 |
| ENSMUSG00000042626 | 2.72721917 | 2.523519421 | 2.560793496 | 2.533285553 | 2.519706825 | 2.84260924 | 2.370650241 |
| ENSMUSG00000042628 | 2.340856238 | 2.208222504 | 2.263142013 | 1.979715809 | 1.923977404 | 2.252853031 | 1.913109605 |
| ENSMUSG00000042631 | 0 | 0 | 0 | 0 | 0 | 0 | 0 |
| ENSMUSG00000042632 | 2.273259602 | 2.166333979 | 2.207430684 | 1.958222504 | 2.16097133 | 2.305351369 | 2.123508695 |
| ENSMUSG00000042638 | 0 | 0 | 0 | 0 | 0.244650542 | 0 | 0 |
| ENSMUSG00000042641 | 0 | 0 | 0 | 0.396240625 | 0 | 0.301029996 | 0 |
| ENSMUSG00000042642 | 1.822201287 | 1.695339928 | 1.634789703 | 1.571350555 | 1.744182955 | 1.755874856 | 1.67819947 |
| ENSMUSG00000042644 | 2.869771412 | 2.667664062 | 2.741626363 | 2.454645544 | 2.525265304 | 2.752048448 | 2.472164183 |
| ENSMUSG00000042647 | 1.418409478 | 1.42510993 | 1.410964047 | 1.321350555 | 1.27449559 | 1.544068044 | 1.313616061 |
| ENSMUSG00000042650 | 2.747795766 | 2.533285553 | 2.521202097 | 2.592399337 | 2.557901472 | 2.884795364 | 2.393225472 |
| ENSMUSG00000042655 | 0 | 0 | 0 | 0.25 | 0.244650542 | 0 | 0 |
| ENSMUSG00000042659 | 1.639698646 | 1.403677461 | 1.431980114 | 1.42510993 | 1.49445382 | 1.612783857 | 1.345462945 |
| ENSMUSG00000042660 | 2.167941649 | 2.051142786 | 2.072504712 | 2.02053726 | 2.058398634 | 2.257678575 | 2.052343353 |
| ENSMUSG00000042662 | 0.747221736 | 0.580482024 | 0.580482024 | 0.25 | 0.244650542 | 0.301029996 | 0.520678036 |
| ENSMUSG00000042670 | 1.890240806 | 1.77186571 | 1.747171172 | 1.921625132 | 1.836786029 | 2.130333768 | 1.731408809 |
| ENSMUSG00000042671 | 0 | 0 | 0 | 0 | 0.244650542 | 0 | 0.224243824 |
| ENSMUSG00000042672 | 0.747221736 | 0.5 | 0.79248125 | 1.061981878 | 0.931471446 | 1.113943352 | 0.935079929 |

| | | | | | | | |
|---|---|---|---|---|---|---|---|
| ENSMUSG00000042675 | 1.325351389 | 1.380890489 | 1.451838731 | 1.130890489 | 1.223252711 | 1.278753601 | 1.345462945 |
| ENSMUSG00000042677 | 1.936861443 | 1.919870025 | 2.027131114 | 1.339388001 | 1.607318736 | 1.633468456 | 1.584526862 |
| ENSMUSG00000042678 | 0 | 0 | 0 | 0.5 | 0.244650542 | 0 | 0.448487648 |
| ENSMUSG00000042680 | 1.516991255 | 1.5 | 1.403677461 | 1.27186571 | 1.42077253 | 1.447158031 | 1.324583736 |
| ENSMUSG00000042682 | 2.312518611 | 2.120953944 | 2.120953944 | 2.235628626 | 2.208005287 | 2.530199698 | 2.053478496 |
| ENSMUSG00000042684 | 0 | 0 | 0 | 0.701838731 | 0.733951626 | 0.602059991 | 0.520678036 |
| ENSMUSG00000042686 | 1.156259305 | 1.226722649 | 1.226722649 | 0.951838731 | 1.039257769 | 1.079181246 | 0.935079929 |
| ENSMUSG00000042688 | 3.020182925 | 2.762121718 | 2.734527332 | 2.814788692 | 2.754702294 | 3.186391216 | 2.552967446 |
| ENSMUSG00000042694 | 2.156259305 | 2.015173983 | 1.998588359 | 2.166333979 | 2.078515538 | 2.357934847 | 2.002662694 |
| ENSMUSG00000042699 | 3.37280797 | 3.04753333 | 3.022610819 | 3.159156155 | 3.052504639 | 3.561816664 | 2.824147193 |
| ENSMUSG00000042700 | 2.791849225 | 2.579635702 | 2.585240691 | 2.295598588 | 2.306140782 | 2.57863921 | 2.16858494 |
| ENSMUSG00000042705 | 2.054824387 | 1.92510993 | 1.919870025 | 1.979715809 | 1.87878897 | 2.217483944 | 1.855869786 |
| ENSMUSG00000042707 | 0.289064826 | 0 | 0 | 0.25 | 0.244650542 | 0 | 0.520678036 |
| ENSMUSG00000042708 | 0.747221736 | 0.396240625 | 0.75 | 1.061981878 | 0.812711509 | 0.698970004 | 0.829800754 |
| ENSMUSG00000042709 | 2.069666964 | 1.973704441 | 1.931980114 | 2.102347734 | 2.032639648 | 2.252853031 | 1.95718226 |
| ENSMUSG00000042712 | 2.988565037 | 2.728222334 | 2.730088214 | 2.705443495 | 2.606005002 | 3.049605613 | 2.447148038 |
| ENSMUSG00000042717 | 0 | 0 | 0 | 0 | 0 | 0 | 0 |
| ENSMUSG00000042719 | 2.728422728 | 2.51007243 | 2.503853763 | 2.49929487 | 2.456362323 | 2.806179974 | 2.290497981 |
| ENSMUSG00000042724 | 0.578129653 | 0.75 | 0.830482024 | 0.396240625 | 0.387761935 | 0.477121255 | 0.710836105 |
| ENSMUSG00000042726 | 2.285603957 | 2.117918804 | 2.156427211 | 2.015173983 | 2.136951444 | 2.298853076 | 2.025071334 |
| ENSMUSG00000042729 | 2.409468165 | 2.185366747 | 2.216046536 | 2.205044741 | 2.221952687 | 2.480006943 | 2.080634989 |
| ENSMUSG00000042734 | 0.747221736 | 0.5 | 0.830482024 | 0.75 | 0.77552387 | 0.602059991 | 0.672731473 |
| ENSMUSG00000042737 | 1.249317394 | 1.410964047 | 1.494319981 | 0.79248125 | 1.149965125 | 0.903089987 | 1.496250175 |
| ENSMUSG00000042742 | 2.40033094 | 2.245034894 | 2.183677405 | 2.148114259 | 2.066583491 | 2.338456494 | 1.914853627 |
| ENSMUSG00000042743 | 1.707474304 | 1.618933358 | 1.646240625 | 1.990000483 | 1.888971102 | 2.187520721 | 1.826389311 |
| ENSMUSG00000042744 | 2.591692877 | 2.416777886 | 2.431980114 | 2.240000483 | 2.360255934 | 2.540329475 | 2.26794422 |
| ENSMUSG00000042745 | 1.45815691 | 1.575945187 | 1.62694866 | 1.410964047 | 1.42077253 | 1.505149978 | 1.483307832 |
| ENSMUSG00000042747 | 2.126809492 | 2.08272922 | 2.145300145 | 1.866901388 | 1.925488995 | 2.086359831 | 1.965468324 |
| ENSMUSG00000042750 | 0 | 0 | 0 | 0.701838731 | 0.846351821 | 0.477121255 | 0.355418052 |
| ENSMUSG00000042751 | 0.747221736 | 0.701838731 | 0.646240625 | 1 | 1.136121934 | 1.176091259 | 1.041356071 |
| ENSMUSG00000042757 | 0 | 0 | 0 | 0.25 | 0 | 0.301029996 | 0.448487648 |
| ENSMUSG00000042759 | 1.740854706 | 1.726722649 | 1.671625132 | 1.634789703 | 1.655839964 | 1.826074803 | 1.56970677 |
| ENSMUSG00000042761 | 0 | 0.25 | 0 | 0.25 | 0.244650542 | 0 | 0 |
| ENSMUSG00000042763 | 0.45815691 | 0.25 | 0.5 | 1.04248125 | 1.27449559 | 1.278753601 | 1.176815335 |

| | | | | | | | |
|---|---|---|---|---|---|---|---|
| ENSMUSG00000042766 | 1.89911417 | 1.77186571 | 1.886723615 | 1.410964047 | 1.568060967 | 1.579783597 | 1.616675393 |
| ENSMUSG00000042770 | 1.753493018 | 1.649978211 | 1.741446071 | 1.606566189 | 1.592135487 | 1.819543936 | 1.564611933 |
| ENSMUSG00000042772 | 2.771760131 | 2.505938588 | 2.478596283 | 2.51584877 | 2.477503739 | 2.797959644 | 2.320390187 |
| ENSMUSG00000042774 | 0 | 0 | 0 | 0 | 0 | 0 | 0 |
| ENSMUSG00000042784 | 0.671187741 | 0.646240625 | 0.951838731 | 1.160964047 | 1 | 0.903089987 | 1.054044578 |
| ENSMUSG00000042787 | 2.095120908 | 2.013820609 | 1.985628626 | 2.029735268 | 2.025895058 | 2.167317335 | 1.918313719 |
| ENSMUSG00000042788 | 0 | 0 | 0.25 | 0 | 0.244650542 | 0 | 0 |
| ENSMUSG00000042790 | 2.163307908 | 2.044977273 | 2.046218836 | 2.043731421 | 1.978602168 | 2.260071388 | 1.86815666 |
| ENSMUSG00000042793 | 1.205378646 | 1.339388001 | 1.476722649 | 1.130890489 | 1.163285805 | 1.301029996 | 1.121219121 |
| ENSMUSG00000042796 | 0 | 0 | 0 | 0 | 0 | 0 | 0 |
| ENSMUSG00000042797 | 0.45815691 | 0.701838731 | 0 | 0.580482024 | 0.568060967 | 0.77815125 | 0.629532004 |
| ENSMUSG00000042800 | 0 | 0 | 0 | 0 | 0 | 0.301029996 | 0.224243824 |
| ENSMUSG00000042801 | 0 | 0 | 0 | 0 | 0 | 0 | 0 |
| ENSMUSG00000042804 | 1.686083339 | 1.634789703 | 1.729715809 | 1.657339155 | 1.628935049 | 1.748188027 | 1.683576582 |
| ENSMUSG00000042807 | 0.811507563 | 1.146240625 | 0.864857905 | 1 | 1.020174412 | 1.255272505 | 1.078019652 |
| ENSMUSG00000042808 | 0 | 0 | 0 | 0 | 0 | 0 | 0 |
| ENSMUSG00000042810 | 2.095120908 | 1.907339155 | 1.955044741 | 1.837182039 | 1.954436039 | 2.041392685 | 1.849546807 |
| ENSMUSG00000042812 | 0.867194479 | 0.5 | 0.580482024 | 1.664552871 | 1.572189156 | 1.69019608 | 1.586932193 |
| ENSMUSG00000042814 | 1.714367558 | 1.54248125 | 1.622963274 | 1.77186571 | 1.692703642 | 1.86923172 | 1.596378571 |
| ENSMUSG00000042816 | 0.289064826 | 0.25 | 0 | 0 | 0.244650542 | 0 | 0.224243824 |
| ENSMUSG00000042817 | 0 | 0.25 | 0.25 | 0.580482024 | 0.733951626 | 0.477121255 | 0.579661877 |
| ENSMUSG00000042821 | 1.527823874 | 1.622963274 | 1.766522298 | 0.830482024 | 1.264824954 | 1.204119983 | 1.383567577 |
| ENSMUSG00000042826 | 0.578129653 | 0.646240625 | 0.580482024 | 1.214495249 | 1.149965125 | 1 | 1.100339912 |
| ENSMUSG00000042828 | 0.289064826 | 0.25 | 0 | 0.5 | 0.244650542 | 0.477121255 | 0.520678036 |
| ENSMUSG00000042831 | 1.358731791 | 1.27186571 | 1.348079356 | 1.330482024 | 1.335652905 | 1.633468456 | 1.452143229 |
| ENSMUSG00000042834 | 2.495495257 | 2.272528105 | 2.220660762 | 1.943696765 | 1.834841362 | 2.198657087 | 1.753626837 |
| ENSMUSG00000042842 | 2.368815967 | 2.169870025 | 2.198603967 | 1.330482024 | 1.200473444 | 1.255272505 | 1.23151414 |
| ENSMUSG00000042845 | 0 | 0 | 0 | 0 | 0 | 0 | 0 |
| ENSMUSG00000042846 | 0 | 0.25 | 0 | 0.25 | 0 | 0 | 0.224243824 |
| ENSMUSG00000042848 | 0 | 0 | 0 | 0 | 0 | 0 | 0 |
| ENSMUSG00000042849 | 0 | 0 | 0 | 0 | 0 | 0 | 0 |
| ENSMUSG00000042851 | 1.494443473 | 1.380890489 | 1.438721876 | 1.557204673 | 1.49445382 | 1.681241237 | 1.38802993 |
| ENSMUSG00000042854 | 0.747221736 | 1 | 1.146240625 | 1.238549078 | 1.091002363 | 1.301029996 | 1.110947927 |
| ENSMUSG00000042857 | 1.129344651 | 1.114857905 | 1.26109853 | 1.201838731 | 1.212046813 | 1.255272505 | 1.131174228 |

| | | | | | | | |
|---|---|---|---|---|---|---|---|
| ENSMUSG00000042861 | 0 | 0 | 0 | 0 | 0 | 0 | 0 |
| ENSMUSG00000042863 | 0 | 0 | 0 | 0 | 0 | 0 | 0 |
| ENSMUSG00000042869 | 0 | 0 | 0 | 0 | 0 | 0 | 0 |
| ENSMUSG00000042870 | 2.243591459 | 2.123963757 | 2.181980114 | 1.874961472 | 1.964031554 | 2.086359831 | 1.877074955 |
| ENSMUSG00000042873 | 0.811507563 | 0.951838731 | 0.864857905 | 1.642463902 | 1.652586889 | 1.72427587 | 1.564611933 |
| ENSMUSG00000042874 | 0 | 0 | 0 | 0 | 0 | 0 | 0 |
| ENSMUSG00000042888 | 0 | 0 | 0 | 0 | 0 | 0 | 0 |
| ENSMUSG00000042894 | 0 | 0 | 0 | 0 | 0 | 0 | 0 |
| ENSMUSG00000042895 | 0 | 0 | 0 | 0.25 | 0 | 0 | 0 |
| ENSMUSG00000042901 | 2.662129038 | 2.442872367 | 2.444519282 | 2.46293726 | 2.39421617 | 2.746634199 | 2.219869654 |
| ENSMUSG00000042903 | 2.003432385 | 1.933677405 | 2.017865591 | 1.961372513 | 2.041436517 | 2.190331698 | 1.966952343 |
| ENSMUSG00000042909 | 0 | 0 | 0 | 0 | 0 | 0 | 0 |
| ENSMUSG00000042918 | 0 | 0.25 | 0.396240625 | 0 | 0.632412477 | 0 | 0.355418052 |
| ENSMUSG00000042938 | 0 | 0 | 0 | 0 | 0 | 0 | 0 |
| ENSMUSG00000042940 | 0 | 0 | 0 | 0 | 0 | 0 | 0 |
| ENSMUSG00000042942 | 1.714367558 | 1.598079356 | 1.580482024 | 1.961372513 | 1.893954095 | 2.08278537 | 1.808770687 |
| ENSMUSG00000042961 | 0 | 0 | 0 | 0.5 | 0.489301084 | 0 | 0.355418052 |
| ENSMUSG00000042962 | 0.578129653 | 0 | 0.25 | 0.25 | 0.489301084 | 0.602059991 | 0.629532004 |
| ENSMUSG00000042976 | 0 | 0 | 0 | 0 | 0 | 0 | 0 |
| ENSMUSG00000042978 | 1.655740331 | 1.618933358 | 1.606566189 | 1.527131114 | 1.576269619 | 1.556302501 | 1.579661877 |
| ENSMUSG00000042985 | 0.289064826 | 0.396240625 | 0.396240625 | 0.646240625 | 0.489301084 | 0.301029996 | 0.520678036 |
| ENSMUSG00000042988 | 0.811507563 | 0.75 | 1.04248125 | 0.79248125 | 0.812711509 | 0.698970004 | 0.952571511 |
| ENSMUSG00000042992 | 1.249317394 | 1.380890489 | 1.396240625 | 1.282320754 | 1.039257769 | 1.397940009 | 1.307989563 |
| ENSMUSG00000042993 | 0 | 0 | 0 | 0 | 0 | 0 | 0 |
| ENSMUSG00000042997 | 2.116511873 | 1.978222334 | 1.990000483 | 1.80479213 | 1.826953746 | 1.954242509 | 1.681794123 |
| ENSMUSG00000043003 | 0 | 0 | 0 | 0 | 0 | 0 | 0 |
| ENSMUSG00000043004 | 1.827447047 | 1.799918086 | 1.735628626 | 1.720660762 | 1.652586889 | 1.903089987 | 1.520678036 |
| ENSMUSG00000043008 | 0 | 0 | 0 | 0 | 0 | 0 | 0 |
| ENSMUSG00000043013 | 0 | 0 | 0 | 0 | 0 | 0 | 0 |
| ENSMUSG00000043015 | 1.765759561 | 1.598079356 | 1.602347734 | 1.705044741 | 1.639266209 | 1.770852012 | 1.535058868 |
| ENSMUSG00000043017 | 0.747221736 | 0.396240625 | 0.5 | 0.5 | 0.387761935 | 0.477121255 | 0.579661877 |
| ENSMUSG00000043019 | 2.666334442 | 2.469512728 | 2.430701883 | 2.568740118 | 2.490520278 | 2.851869601 | 2.323109859 |
| ENSMUSG00000043020 | 0 | 0 | 0 | 0.25 | 0 | 0 | 0 |
| ENSMUSG00000043024 | 0 | 0 | 0 | 0.396240625 | 0.244650542 | 0.477121255 | 0.224243824 |

| | | | | | | | |
|---|---|---|---|---|---|---|---|
| ENSMUSG00000043029 | 0 | 0 | 0 | 0 | 0.244650542 | 0 | 0.224243824 |
| ENSMUSG00000043036 | 0.671187741 | 0 | 0.396240625 | 0.396240625 | 0.387761935 | 0.602059991 | 0.520678036 |
| ENSMUSG00000043050 | 0 | 0.25 | 0 | 0 | 0 | 0 | 0 |
| ENSMUSG00000043051 | 0 | 0.25 | 0.25 | 0.580482024 | 0.686820904 | 0.477121255 | 0.448487648 |
| ENSMUSG00000043059 | 1.924654746 | 1.83272922 | 1.880890489 | 1.571350555 | 1.692703642 | 1.716003344 | 1.818286533 |
| ENSMUSG00000043060 | 0 | 0 | 0 | 0 | 0 | 0 | 0 |
| ENSMUSG00000043061 | 2.267855219 | 2.155512955 | 2.08272922 | 2.080482024 | 2.112066955 | 2.330413773 | 1.920884716 |
| ENSMUSG00000043065 | 2.078325387 | 1.938721876 | 1.845926073 | 1.899978211 | 1.866526233 | 2.093421685 | 1.73898563 |
| ENSMUSG00000043066 | 0 | 0 | 0 | 0 | 0 | 0 | 0 |
| ENSMUSG00000043067 | 2.842342644 | 2.599685923 | 2.614088604 | 2.509386738 | 2.458700896 | 2.803457116 | 2.257861346 |
| ENSMUSG00000043068 | 0.289064826 | 0.580482024 | 0.580482024 | 0.79248125 | 0.632412477 | 0.77815125 | 0.775756176 |
| ENSMUSG00000043073 | 0 | 0 | 0 | 0 | 0 | 0 | 0 |
| ENSMUSG00000043079 | 2.251208701 | 2.117918804 | 2.05479213 | 2.071350555 | 2.055279708 | 2.28780173 | 2.029982479 |
| ENSMUSG00000043085 | 0.289064826 | 0.25 | 0 | 0 | 0.387761935 | 0.301029996 | 0.224243824 |
| ENSMUSG00000043087 | 0 | 0 | 0 | 0 | 0 | 0 | 0 |
| ENSMUSG00000043088 | 0 | 0.25 | 0.25 | 0 | 0.387761935 | 0 | 0.224243824 |
| ENSMUSG00000043089 | 0 | 0 | 0 | 0 | 0 | 0 | 0 |
| ENSMUSG00000043090 | 2.223916471 | 2.027131114 | 2.027131114 | 2.170748646 | 2.114724103 | 2.346352974 | 1.95411574 |
| ENSMUSG00000043091 | 2.405576699 | 2.267194569 | 2.234894804 | 2.306000419 | 2.319233381 | 2.613841822 | 2.171745067 |
| ENSMUSG00000043099 | 0.289064826 | 0 | 0.580482024 | 0.25 | 0.244650542 | 0 | 0.224243824 |
| ENSMUSG00000043102 | 0 | 0 | 0 | 0.25 | 0.489301084 | 0 | 0.448487648 |
| ENSMUSG00000043110 | 0 | 0 | 0 | 0 | 0 | 0 | 0 |
| ENSMUSG00000043119 | 0 | 0 | 0 | 0 | 0 | 0 | 0 |
| ENSMUSG00000043122 | 0 | 0 | 0 | 0.25 | 0 | 0 | 0 |
| ENSMUSG00000043126 | 0 | 0.25 | 0 | 0.25 | 0.244650542 | 0 | 0.355418052 |
| ENSMUSG00000043131 | 3.134822988 | 2.854463129 | 2.825802487 | 2.876824781 | 2.76798032 | 3.19893187 | 2.56970677 |
| ENSMUSG00000043140 | 1.932832114 | 1.83272922 | 1.799918086 | 1.825945187 | 1.78010778 | 1.929418926 | 1.670888075 |
| ENSMUSG00000043144 | 0 | 0 | 0.25 | 0 | 0 | 0 | 0 |
| ENSMUSG00000043145 | 0 | 0 | 0 | 0 | 0 | 0 | 0 |
| ENSMUSG00000043153 | 1.53838222 | 1.418106335 | 1.445339928 | 1.17510993 | 1.020174412 | 1.176091259 | 1.054044578 |
| ENSMUSG00000043154 | 2.508932407 | 2.343759858 | 2.350753006 | 2.313166358 | 2.197694248 | 2.5289167 | 2.051774285 |
| ENSMUSG00000043155 | 0.578129653 | 0.976722649 | 1 | 0.75 | 0.812711509 | 0.954242509 | 1.066254157 |
| ENSMUSG00000043157 | 0 | 0.25 | 0.5 | 0 | 0 | 0 | 0 |
| ENSMUSG00000043162 | 1.45815691 | 1.380890489 | 1.431980114 | 1.27186571 | 1.121713561 | 1.380211242 | 1.284450788 |

| | | | | | | | |
|---|---|---|---|---|---|---|---|
| ENSMUSG00000043164 | 0 | 0 | 0 | 0 | 0 | 0 | 0 |
| ENSMUSG00000043165 | 0 | 0 | 0 | 0.396240625 | 0 | 0 | 0.448487648 |
| ENSMUSG00000043168 | 0 | 0 | 0 | 0 | 0 | 0 | 0 |
| ENSMUSG00000043183 | 2.309247751 | 2.116901388 | 2.101285366 | 2.298073204 | 2.225193812 | 2.522444234 | 2.060206963 |
| ENSMUSG00000043190 | 1.516991255 | 1.54248125 | 1.470660762 | 1.657339155 | 1.639266209 | 1.740362689 | 1.508694612 |
| ENSMUSG00000043192 | 1.269664473 | 1.330482024 | 1.380890489 | 1.25 | 1.188507684 | 1.079181246 | 1.350478789 |
| ENSMUSG00000043207 | 2.737340367 | 2.529410775 | 2.506630861 | 2.653907409 | 2.560905373 | 2.975431809 | 2.37800557 |
| ENSMUSG00000043219 | 1.129344651 | 1.061981878 | 1.146240625 | 0.646240625 | 0.877063019 | 0.602059991 | 0.984950056 |
| ENSMUSG00000043223 | 0 | 0 | 0.580482024 | 0.5 | 0.489301084 | 0.477121255 | 0.672731473 |
| ENSMUSG00000043226 | 0 | 0 | 0 | 0 | 0 | 0 | 0 |
| ENSMUSG00000043230 | 0 | 0 | 0 | 0 | 0 | 0 | 0 |
| ENSMUSG00000043241 | 2.490209605 | 2.307204673 | 2.281676118 | 2.45584181 | 2.357161747 | 2.701567985 | 2.20779034 |
| ENSMUSG00000043243 | 1.307602651 | 1.146240625 | 1.321350555 | 0.896240625 | 0.905314583 | 0.602059991 | 0.969165684 |
| ENSMUSG00000043251 | 0.867194479 | 0.896240625 | 0.976722649 | 0.75 | 1.121713561 | 1 | 1.140832123 |
| ENSMUSG00000043252 | 2.854065408 | 2.619186551 | 2.579070383 | 2.397646762 | 2.292543754 | 2.656098202 | 2.13996595 |
| ENSMUSG00000043257 | 1.432083896 | 1.380890489 | 1.410964047 | 1.339388001 | 1.387761935 | 1.431363764 | 1.383567577 |
| ENSMUSG00000043259 | 1.944805158 | 1.839388001 | 1.874961472 | 1.282320754 | 1.057362051 | 1.361727836 | 1.21680636 |
| ENSMUSG00000043262 | 2 | 1.860735874 | 1.87694866 | 1.958222504 | 1.897237431 | 2.117271296 | 1.694069894 |
| ENSMUSG00000043263 | 0 | 0 | 0 | 0 | 0 | 0 | 0 |
| ENSMUSG00000043267 | 0 | 0 | 0 | 0 | 0 | 0 | 0 |
| ENSMUSG00000043274 | 0 | 0 | 0 | 0 | 0 | 0 | 0 |
| ENSMUSG00000043279 | 2.565888829 | 2.37694866 | 2.367918804 | 2.420748646 | 2.381601986 | 2.681241237 | 2.262942063 |
| ENSMUSG00000043282 | 0.45815691 | 0.646240625 | 0.5 | 0.580482024 | 0.489301084 | 0.77815125 | 0.520678036 |
| ENSMUSG00000043284 | 2.054824387 | 1.882845365 | 1.907339155 | 1.899978211 | 1.969399812 | 2.149219113 | 1.89047692 |
| ENSMUSG00000043286 | 0.289064826 | 0.25 | 0.25 | 0.396240625 | 0.387761935 | 0 | 0.355418052 |
| ENSMUSG00000043289 | 0.811507563 | 0.396240625 | 0.396240625 | 0 | 0.489301084 | 0.301029996 | 0.355418052 |
| ENSMUSG00000043290 | 0.578129653 | 0.75 | 0.864857905 | 0.646240625 | 0.931471446 | 0.698970004 | 0.803905701 |
| ENSMUSG00000043295 | 0 | 0 | 0 | 0 | 0.244650542 | 0 | 0 |
| ENSMUSG00000043298 | 0 | 0 | 0 | 0 | 0 | 0 | 0 |
| ENSMUSG00000043300 | 2.310886388 | 2.183677405 | 2.171625132 | 2.284247778 | 2.164820399 | 2.469822016 | 2.037212175 |
| ENSMUSG00000043301 | 0 | 0 | 0 | 0 | 0 | 0 | 0 |
| ENSMUSG00000043308 | 0 | 0 | 0 | 0 | 0 | 0 | 0 |
| ENSMUSG00000043310 | 0 | 0 | 0 | 0 | 0 | 0 | 0 |
| ENSMUSG00000043311 | 1.806056082 | 1.755591953 | 1.705044741 | 1.75 | 1.842556557 | 1.991226076 | 1.709198048 |

| | | | | | | | |
|---|---|---|---|---|---|---|---|
| ENSMUSG00000043312 | 0 | 0 | 0 | 0 | 0 | 0 | 0 |
| ENSMUSG00000043313 | 0.867194479 | 0.92510993 | 0.896240625 | 0.5 | 0.632412477 | 0.602059991 | 0.224243824 |
| ENSMUSG00000043314 | 0 | 0 | 0 | 0 | 0 | 0 | 0 |
| ENSMUSG00000043319 | 0 | 0 | 0 | 0 | 0 | 0 | 0 |
| ENSMUSG00000043323 | 2.092368209 | 1.976722649 | 2.058404919 | 1.809601185 | 1.866526233 | 2.008600172 | 1.889537832 |
| ENSMUSG00000043329 | 0 | 0 | 0 | 0 | 0 | 0 | 0 |
| ENSMUSG00000043331 | 0 | 0 | 0 | 0 | 0 | 0 | 0 |
| ENSMUSG00000043333 | 0.811507563 | 0.396240625 | 0.5 | 0.5 | 0.489301084 | 0.477121255 | 0.672731473 |
| ENSMUSG00000043336 | 2.945519866 | 2.721614674 | 2.685156011 | 2.632113528 | 2.502780045 | 2.914343157 | 2.34615152 |
| ENSMUSG00000043340 | 0.747221736 | 0.646240625 | 0.75 | 0.75 | 0.733951626 | 0.602059991 | 0.672731473 |
| ENSMUSG00000043342 | 1.205378646 | 1.330482024 | 1.226722649 | 0.25 | 0 | 0 | 0 |
| ENSMUSG00000043346 | 0 | 0 | 0 | 0.25 | 0 | 0 | 0 |
| ENSMUSG00000043354 | 0 | 0 | 0 | 0 | 0 | 0 | 0 |
| ENSMUSG00000043357 | 0 | 0 | 0 | 0 | 0 | 0 | 0 |
| ENSMUSG00000043366 | 0 | 0 | 0 | 0 | 0 | 0 | 0 |
| ENSMUSG00000043372 | 1.249317394 | 1.339388001 | 1.410964047 | 1.130890489 | 1.343584837 | 1.230448921 | 1.324583736 |
| ENSMUSG00000043383 | 0 | 0 | 0 | 0 | 0 | 0 | 0 |
| ENSMUSG00000043384 | 2.116511873 | 1.92510993 | 1.923371739 | 2.246460484 | 2.215379757 | 2.450249108 | 2.006616089 |
| ENSMUSG00000043385 | 0 | 0 | 0 | 0.25 | 0 | 0 | 0.224243824 |
| ENSMUSG00000043388 | 0.578129653 | 0.396240625 | 0.580482024 | 1.061981878 | 1.163285805 | 1.301029996 | 1.100339912 |
| ENSMUSG00000043391 | 1.37447073 | 1.464495249 | 1.364857905 | 1.958222504 | 1.830919587 | 2.071882007 | 1.781059827 |
| ENSMUSG00000043398 | 0 | 0 | 0 | 0 | 0 | 0 | 0.224243824 |
| ENSMUSG00000043410 | 0 | 0 | 0 | 0 | 0 | 0 | 0 |
| ENSMUSG00000043411 | 2.719923957 | 2.458222504 | 2.451435968 | 2.539652422 | 2.465951417 | 2.788875116 | 2.27142916 |
| ENSMUSG00000043415 | 1.404271433 | 1.311981878 | 1.372963274 | 1.668106335 | 1.628935049 | 1.672097858 | 1.511732338 |
| ENSMUSG00000043418 | 0.289064826 | 0 | 0 | 0 | 0 | 0 | 0 |
| ENSMUSG00000043419 | 0.671187741 | 0.580482024 | 0.830482024 | 1.566696635 | 1.509475496 | 1.591064607 | 1.433437704 |
| ENSMUSG00000043421 | 1.789260561 | 1.602347734 | 1.580482024 | 1.445339928 | 1.484072009 | 1.579783597 | 1.413583902 |
| ENSMUSG00000043424 | 1.614416215 | 1.445339928 | 1.5 | 1.711372513 | 1.635855979 | 1.77815125 | 1.459332757 |
| ENSMUSG00000043429 | 0 | 0 | 0 | 0 | 0 | 0 | 0 |
| ENSMUSG00000043430 | 0 | 0.25 | 0.25 | 0 | 0 | 0 | 0 |
| ENSMUSG00000043432 | 0.91631382 | 0.830482024 | 0.830482024 | 0.701838731 | 0.877063019 | 1 | 0.935079929 |
| ENSMUSG00000043439 | 1.358731791 | 1.458222504 | 1.410964047 | 1.571350555 | 1.621875691 | 1.799340549 | 1.582103514 |
| ENSMUSG00000043441 | 1.205378646 | 1.061981878 | 1.080482024 | 0.75 | 0.77552387 | 0.602059991 | 0.448487648 |

| | | | | | | | |
|---|---|---|---|---|---|---|---|
| ENSMUSG00000043445 | 1.527823874 | 1.638647213 | 1.695339928 | 1.482684334 | 1.599808748 | 1.612783857 | 1.795211864 |
| ENSMUSG00000043448 | 0 | 0 | 0 | 0 | 0 | 0 | 0 |
| ENSMUSG00000043453 | 0 | 0 | 0 | 0 | 0 | 0 | 0 |
| ENSMUSG00000043456 | 0 | 0.25 | 0 | 0.5 | 0.686820904 | 0.698970004 | 0.672731473 |
| ENSMUSG00000043458 | 0 | 0 | 0.25 | 0 | 0 | 0 | 0 |
| ENSMUSG00000043460 | 0 | 0 | 0 | 0 | 0 | 0 | 0 |
| ENSMUSG00000043461 | 0 | 0 | 0 | 0 | 0 | 0 | 0 |
| ENSMUSG00000043463 | 0.578129653 | 0 | 0 | 0.396240625 | 0.387761935 | 0.301029996 | 0.355418052 |
| ENSMUSG00000043467 | 2.072573125 | 1.912762923 | 1.858657057 | 1.988549078 | 1.898867717 | 2.1430148 | 1.783679399 |
| ENSMUSG00000043468 | 0 | 0 | 0 | 0 | 0 | 0 | 0 |
| ENSMUSG00000043472 | 0 | 0 | 0 | 0 | 0 | 0 | 0 |
| ENSMUSG00000043483 | 0 | 0.25 | 0.25 | 0.580482024 | 0 | 0 | 0 |
| ENSMUSG00000043484 | 0 | 0 | 0 | 0 | 0 | 0 | 0 |
| ENSMUSG00000043485 | 0 | 0 | 0 | 0 | 0 | 0 | 0 |
| ENSMUSG00000043487 | 1 | 1.114857905 | 1.114857905 | 0.864857905 | 0.812711509 | 0.84509804 | 0.74492186 |
| ENSMUSG00000043488 | 0 | 0 | 0 | 0 | 0 | 0 | 0 |
| ENSMUSG00000043496 | 0.289064826 | 0.580482024 | 0.25 | 0 | 0 | 0 | 0.355418052 |
| ENSMUSG00000043498 | 0.289064826 | 0.646240625 | 0.701838731 | 0 | 0.632412477 | 0.301029996 | 0 |
| ENSMUSG00000043501 | 1.100572389 | 0.896240625 | 1.160964047 | 0.75 | 1.091002363 | 0.698970004 | 0.896975297 |
| ENSMUSG00000043505 | 0 | 0 | 0 | 0 | 0 | 0 | 0 |
| ENSMUSG00000043510 | 1.655740331 | 1.53743678 | 1.516522298 | 1.431980114 | 1.445123986 | 1.579783597 | 1.401059159 |
| ENSMUSG00000043518 | 0.45815691 | 0.25 | 0.25 | 0.25 | 0.244650542 | 0 | 0.224243824 |
| ENSMUSG00000043522 | 0 | 0 | 0 | 0 | 0.244650542 | 0 | 0 |
| ENSMUSG00000043529 | 0 | 0 | 0 | 0 | 0 | 0 | 0 |
| ENSMUSG00000043531 | 1.794934559 | 1.784887838 | 1.688721876 | 1.080482024 | 1.163285805 | 1.301029996 | 1.014380832 |
| ENSMUSG00000043535 | 2.83912939 | 2.594031347 | 2.592671852 | 2.674024543 | 2.604466107 | 2.958085849 | 2.4054791 |
| ENSMUSG00000043537 | 0 | 0 | 0 | 0 | 0 | 0 | 0 |
| ENSMUSG00000043541 | 0.289064826 | 0.951838731 | 0.75 | 0.580482024 | 0.931471446 | 0.77815125 | 0.579661877 |
| ENSMUSG00000043542 | 2.363083473 | 2.23927551 | 2.162762923 | 2.203445298 | 2.172394856 | 2.462397998 | 2.033617522 |
| ENSMUSG00000043549 | 0 | 0 | 0 | 0.5 | 0.244650542 | 0.301029996 | 0.355418052 |
| ENSMUSG00000043556 | 1.156259305 | 1.26109853 | 1.26109853 | 0.580482024 | 0.905314583 | 0.602059991 | 0.916588299 |
| ENSMUSG00000043557 | 1.227926429 | 1.282320754 | 1.282320754 | 1.51109853 | 1.618292349 | 1.643452676 | 1.637827726 |
| ENSMUSG00000043569 | 0 | 0 | 0 | 0 | 0 | 0 | 0 |
| ENSMUSG00000043572 | 1.639698646 | 1.593759858 | 1.646240625 | 1.488549078 | 1.559656853 | 1.612783857 | 1.417653332 |

| | | | | | | | |
|---|---|---|---|---|---|---|---|
| ENSMUSG00000043583 | 0 | 0 | 0 | 0 | 0 | 0 | 0 |
| ENSMUSG00000043587 | 2.29078455 | 2.164552871 | 2.228222334 | 1.841580554 | 1.802174723 | 2.117271296 | 1.757893036 |
| ENSMUSG00000043592 | 0 | 0 | 0 | 0 | 0 | 0 | 0 |
| ENSMUSG00000043602 | 1.205378646 | 1.238549078 | 1.238549078 | 1.396240625 | 1.335652905 | 1.447158031 | 1.335191751 |
| ENSMUSG00000043605 | 0 | 0 | 0 | 0 | 0 | 0 | 0 |
| ENSMUSG00000043613 | 0.811507563 | 0.75 | 0.701838731 | 0.25 | 0 | 0 | 0.224243824 |
| ENSMUSG00000043614 | 0.747221736 | 0.976722649 | 1.061981878 | 0.864857905 | 0.978602168 | 1.041392685 | 1.20919388 |
| ENSMUSG00000043618 | 0.289064826 | 0.5 | 0.5 | 0.396240625 | 0.686820904 | 0 | 0.520678036 |
| ENSMUSG00000043621 | 0 | 0 | 0 | 0 | 0 | 0 | 0 |
| ENSMUSG00000043623 | 0 | 0 | 0 | 0 | 0 | 0 | 0 |
| ENSMUSG00000043629 | 0 | 0.396240625 | 0 | 0.580482024 | 0.387761935 | 0.301029996 | 0.520678036 |
| ENSMUSG00000043631 | 0.578129653 | 0.79248125 | 0.5 | 0.646240625 | 0.686820904 | 0.77815125 | 0.803905701 |
| ENSMUSG00000043633 | 0.91631382 | 0.75 | 0.701838731 | 0.5 | 0.568060967 | 0.602059991 | 0.710836105 |
| ENSMUSG00000043635 | 1.100572389 | 1.02186571 | 1.04248125 | 0.580482024 | 0.877063019 | 0.602059991 | 0.710836105 |
| ENSMUSG00000043639 | 1.53838222 | 1.488549078 | 1.431980114 | 1.02186571 | 1.057362051 | 1.146128036 | 1 |
| ENSMUSG00000043644 | 0.578129653 | 0.646240625 | 0.5 | 0.396240625 | 0.568060967 | 0.301029996 | 0.355418052 |
| ENSMUSG00000043648 | 0.289064826 | 0.396240625 | 0.396240625 | 0.396240625 | 0.387761935 | 0.301029996 | 0.448487648 |
| ENSMUSG00000043659 | 0 | 0 | 0 | 0 | 0 | 0 | 0 |
| ENSMUSG00000043661 | 0.289064826 | 0.396240625 | 0 | 0.25 | 0.387761935 | 0 | 0 |
| ENSMUSG00000043664 | 0.289064826 | 0.396240625 | 0.25 | 0.580482024 | 0.489301084 | 0.477121255 | 0.629532004 |
| ENSMUSG00000043668 | 0 | 0 | 0 | 0 | 0 | 0 | 0 |
| ENSMUSG00000043670 | 0.289064826 | 0.25 | 0 | 0.396240625 | 0.733951626 | 0.602059991 | 0.224243824 |
| ENSMUSG00000043671 | 2.170239354 | 2.029735268 | 2.059601185 | 2.048689214 | 2.030405741 | 2.305351369 | 1.901538355 |
| ENSMUSG00000043673 | 0.289064826 | 0 | 0 | 1.188721876 | 1.074582839 | 1.041392685 | 0.969165684 |
| ENSMUSG00000043681 | 0 | 0 | 0 | 0 | 0 | 0 | 0 |
| ENSMUSG00000043683 | 2.312518611 | 2.14151351 | 2.142463902 | 2.088286706 | 2.119108707 | 2.326335861 | 2.020712061 |
| ENSMUSG00000043687 | 0.578129653 | 0.701838731 | 0.864857905 | 0.25 | 0.244650542 | 0.301029996 | 0.853775828 |
| ENSMUSG00000043692 | 0 | 0 | 0 | 0 | 0 | 0 | 0 |
| ENSMUSG00000043698 | 0 | 0 | 0 | 0 | 0 | 0 | 0 |
| ENSMUSG00000043702 | 1.940852214 | 1.821350555 | 1.938721876 | 2.070192693 | 2.03596422 | 2.281033367 | 1.927642465 |
| ENSMUSG00000043705 | 0 | 0 | 0 | 0 | 0 | 0 | 0 |
| ENSMUSG00000043715 | 0 | 0 | 0 | 0 | 0 | 0 | 0 |
| ENSMUSG00000043716 | 3.319130149 | 3.00835575 | 3.005418511 | 3.142878913 | 3.036806675 | 3.556181847 | 2.840670723 |
| ENSMUSG00000043719 | 0 | 0 | 0 | 0.25 | 0 | 0 | 0.224243824 |

| | | | | | | | |
|---|---|---|---|---|---|---|---|
| ENSMUSG00000043727 | 0 | 0 | 0 | 0 | 0 | 0 | 0 |
| ENSMUSG00000043733 | 3.30700282 | 3.020703584 | 3.006371414 | 2.992076699 | 2.953043706 | 3.396373728 | 2.779862063 |
| ENSMUSG00000043740 | 0 | 0.701838731 | 0.25 | 0.5 | 0.387761935 | 0 | 0.224243824 |
| ENSMUSG00000043747 | 0.289064826 | 0 | 0.5 | 0 | 0.244650542 | 0 | 0 |
| ENSMUSG00000043753 | 0.45815691 | 0.75 | 0.580482024 | 0.580482024 | 0.733951626 | 0.698970004 | 0.803905701 |
| ENSMUSG00000043760 | 0 | 0 | 0.25 | 0 | 0 | 0 | 0 |
| ENSMUSG00000043770 | 0 | 0 | 0 | 0 | 0 | 0 | 0 |
| ENSMUSG00000043773 | 1.181541736 | 1 | 1.098079356 | 1.476722649 | 1.343584837 | 1.397940009 | 1.313616061 |
| ENSMUSG00000043782 | 0 | 0.25 | 0.396240625 | 0.396240625 | 0.244650542 | 0.301029996 | 0.629532004 |
| ENSMUSG00000043787 | 0 | 0 | 0 | 0 | 0 | 0 | 0 |
| ENSMUSG00000043789 | 0.671187741 | 0.951838731 | 0.580482024 | 0.396240625 | 0.387761935 | 0.477121255 | 0.520678036 |
| ENSMUSG00000043794 | 0 | 0 | 0.25 | 0 | 0 | 0 | 0 |
| ENSMUSG00000043795 | 1.156259305 | 0.75 | 0.92510993 | 0.396240625 | 0.387761935 | 0.698970004 | 0.224243824 |
| ENSMUSG00000043800 | 0.289064826 | 0.5 | 0.79248125 | 0.5 | 0.244650542 | 0.698970004 | 0.629532004 |
| ENSMUSG00000043801 | 0 | 0 | 0 | 0 | 0 | 0 | 0.224243824 |
| ENSMUSG00000043807 | 0.747221736 | 0.701838731 | 0.646240625 | 0.5 | 0.978602168 | 0.903089987 | 0.853775828 |
| ENSMUSG00000043811 | 1.100572389 | 1.160964047 | 1.311981878 | 0.5 | 0.686820904 | 0.301029996 | 0.876096088 |
| ENSMUSG00000043822 | 1.806056082 | 1.79248125 | 1.982684334 | 1.505591953 | 1.706995316 | 1.755874856 | 1.722075986 |
| ENSMUSG00000043827 | 0 | 0 | 0 | 0 | 0 | 0 | 0 |
| ENSMUSG00000043831 | 1.596667477 | 1.505591953 | 1.482684334 | 1.410964047 | 1.533172725 | 1.414973348 | 1.413583902 |
| ENSMUSG00000043832 | 0 | 0 | 0 | 0 | 0 | 0 | 0 |
| ENSMUSG00000043833 | 0.289064826 | 0.79248125 | 0.580482024 | 0.646240625 | 0.568060967 | 0 | 0.224243824 |
| ENSMUSG00000043843 | 0 | 0 | 0 | 0 | 0 | 0 | 0 |
| ENSMUSG00000043850 | 0.671187741 | 0.5 | 0 | 0.396240625 | 0.244650542 | 0.301029996 | 0.355418052 |
| ENSMUSG00000043855 | 0 | 0 | 0 | 0 | 0 | 0 | 0 |
| ENSMUSG00000043857 | 0.289064826 | 0 | 0 | 0.396240625 | 0.489301084 | 0 | 0.74492186 |
| ENSMUSG00000043858 | 2.497590902 | 2.329353153 | 2.350753006 | 2.075945187 | 2.221300875 | 2.369215857 | 2.108089156 |
| ENSMUSG00000043859 | 0 | 0 | 0 | 0 | 0 | 0 | 0 |
| ENSMUSG00000043865 | 0 | 0 | 0 | 0 | 0 | 0 | 0 |
| ENSMUSG00000043866 | 1.940852214 | 1.807204673 | 1.816696635 | 1.814346961 | 1.733951626 | 1.944482672 | 1.700880998 |
| ENSMUSG00000043870 | 0 | 0 | 0 | 0 | 0 | 0 | 0 |
| ENSMUSG00000043872 | 1.932832114 | 1.866901388 | 1.807204673 | 1.916333979 | 1.806427258 | 2.021189299 | 1.720494 |
| ENSMUSG00000043873 | 0 | 0 | 0 | 0 | 0 | 0 | 0 |
| ENSMUSG00000043880 | 0 | 0 | 0 | 0 | 0 | 0 | 0 |

| | | | | | | | |
|---|---|---|---|---|---|---|---|
| ENSMUSG00000043881 | 1.614416215 | 1.552363341 | 1.618933358 | 1.862802778 | 1.797870327 | 1.949390007 | 1.63986884 |
| ENSMUSG00000043885 | 2.3855553 | 2.164552871 | 2.176839783 | 2.408702763 | 2.331857641 | 2.641474111 | 2.17331363 |
| ENSMUSG00000043889 | 1.269664473 | 1.25 | 1.238549078 | 0.701838731 | 0.931471446 | 0.477121255 | 0.876096088 |
| ENSMUSG00000043892 | 0 | 0 | 0 | 0 | 0 | 0 | 0 |
| ENSMUSG00000043895 | 2.381433035 | 2.180274797 | 2.256976499 | 2.267194569 | 2.314756076 | 2.509202522 | 2.180279543 |
| ENSMUSG00000043897 | 0 | 0 | 0 | 0 | 0 | 0 | 0 |
| ENSMUSG00000043903 | 1.920505136 | 1.858657057 | 1.892463902 | 1.380890489 | 1.599808748 | 1.431363764 | 1.572224412 |
| ENSMUSG00000043909 | 2.653588827 | 2.423371739 | 2.42510993 | 2.509386738 | 2.474332504 | 2.804139432 | 2.322370395 |
| ENSMUSG00000043911 | 0 | 0 | 0 | 0 | 0 | 0 | 0 |
| ENSMUSG00000043913 | 0 | 0 | 0.25 | 0.25 | 0 | 0.477121255 | 0.355418052 |
| ENSMUSG00000043923 | 1.289064826 | 1.372963274 | 1.26109853 | 1.5 | 1.572189156 | 1.544068044 | 1.448487648 |
| ENSMUSG00000043924 | 0 | 0 | 0.580482024 | 0.396240625 | 0.733951626 | 0.602059991 | 0.629532004 |
| ENSMUSG00000043925 | 0 | 0.25 | 0 | 0 | 0 | 0 | 0 |
| ENSMUSG00000043929 | 1.678702042 | 1.610735874 | 1.464495249 | 1.660964047 | 1.642643806 | 1.819543936 | 1.46982607 |
| ENSMUSG00000043931 | 0 | 0 | 0 | 0 | 0 | 0 | 0 |
| ENSMUSG00000043932 | 0 | 0 | 0 | 0 | 0 | 0 | 0 |
| ENSMUSG00000043939 | 0.289064826 | 0.396240625 | 0.25 | 0.396240625 | 0.244650542 | 0.301029996 | 0.224243824 |
| ENSMUSG00000043940 | 2.954351886 | 2.713911792 | 2.707232457 | 2.595656007 | 2.601145705 | 2.912753304 | 2.435531663 |
| ENSMUSG00000043943 | 0 | 0.25 | 0.5 | 0.864857905 | 0.568060967 | 0.477121255 | 0.629532004 |
| ENSMUSG00000043945 | 0 | 0 | 0 | 0 | 0 | 0 | 0 |
| ENSMUSG00000043948 | 0 | 0 | 0 | 0 | 0 | 0 | 0 |
| ENSMUSG00000043953 | 0 | 0.396240625 | 0.25 | 0.25 | 0 | 0.301029996 | 0.448487648 |
| ENSMUSG00000043962 | 2.896311254 | 2.635274154 | 2.666333979 | 2.651601303 | 2.620825538 | 2.992995098 | 2.457062464 |
| ENSMUSG00000043964 | 1.806056082 | 1.726722649 | 1.698603967 | 1.62694866 | 1.65906333 | 1.812913357 | 1.532233387 |
| ENSMUSG00000043969 | 2.023453784 | 1.919870025 | 1.909156155 | 1.799918086 | 1.72341484 | 1.903089987 | 1.577201671 |
| ENSMUSG00000043972 | 0 | 0 | 0 | 0.25 | 0.244650542 | 0.301029996 | 0 |
| ENSMUSG00000043982 | 0 | 0 | 0 | 0 | 0 | 0 | 0 |
| ENSMUSG00000043986 | 0 | 0 | 0 | 0.25 | 0 | 0 | 0.224243824 |
| ENSMUSG00000043987 | 2.209569962 | 2.119945066 | 2.17510993 | 2.08272922 | 2.164820399 | 2.350248018 | 2.077494038 |
| ENSMUSG00000043991 | 2.631440309 | 2.42813175 | 2.400906586 | 2.491806565 | 2.431723443 | 2.804139432 | 2.234441902 |
| ENSMUSG00000043993 | 0.578129653 | 0.646240625 | 0.5 | 0.701838731 | 0.905314583 | 0.698970004 | 0.74492186 |
| ENSMUSG00000043998 | 2.636714502 | 2.431128463 | 2.434945152 | 2.38040011 | 2.300451989 | 2.608526034 | 2.163786106 |
| ENSMUSG00000043999 | 0.289064826 | 0 | 0.5 | 0.5 | 0.686820904 | 0.602059991 | 0.355418052 |
| ENSMUSG00000044005 | 1.289064826 | 1.321350555 | 1.356566189 | 1.614857905 | 1.645989387 | 1.681241237 | 1.554175045 |

| | | | | | | | |
|---|---|---|---|---|---|---|---|
| ENSMUSG00000044006 | 0.578129653 | 0.396240625 | 0 | 0.25 | 0.733951626 | 0.301029996 | 0.224243824 |
| ENSMUSG00000044014 | 0 | 0 | 0.25 | 0.25 | 0 | 0 | 0 |
| ENSMUSG00000044017 | 0 | 0 | 0 | 0 | 0 | 0 | 0 |
| ENSMUSG00000044018 | 2.401648579 | 2.197790722 | 2.170748646 | 2.262462137 | 2.134457044 | 2.463892989 | 2.000667731 |
| ENSMUSG00000044021 | 0.289064826 | 0 | 0.25 | 0 | 0 | 0 | 0 |
| ENSMUSG00000044022 | 1.325351389 | 1.080482024 | 0.92510993 | 0.896240625 | 0.686820904 | 0.77815125 | 0.710836105 |
| ENSMUSG00000044024 | 0.289064826 | 0.396240625 | 0.25 | 0.25 | 0 | 0.301029996 | 0.629532004 |
| ENSMUSG00000044025 | 0 | 0 | 0 | 0 | 0 | 0 | 0 |
| ENSMUSG00000044026 | 2.083999385 | 2.012462137 | 1.959800947 | 2.061981878 | 2.024758322 | 2.274157849 | 1.837623616 |
| ENSMUSG00000044029 | 0 | 0 | 0 | 0 | 0 | 0 | 0 |
| ENSMUSG00000044030 | 1.605636262 | 1.668106335 | 1.774508021 | 1.458222504 | 1.621875691 | 1.740362689 | 1.712465909 |
| ENSMUSG00000044033 | 1.568542279 | 1.51109853 | 1.431980114 | 1.926839783 | 1.866526233 | 2.161368002 | 1.687112305 |
| ENSMUSG00000044034 | 0 | 0 | 0.396240625 | 0 | 0.244650542 | 0 | 0.224243824 |
| ENSMUSG00000044037 | 1.753493018 | 1.532320754 | 1.714495249 | 2.097004321 | 2.179075613 | 2.413299764 | 2.127589533 |
| ENSMUSG00000044039 | 0 | 0 | 0 | 0 | 0 | 0 | 0 |
| ENSMUSG00000044040 | 0 | 0 | 0 | 0 | 0 | 0 | 0 |
| ENSMUSG00000044041 | 1.227926429 | 1.27186571 | 1.330482024 | 0.580482024 | 0.387761935 | 0.477121255 | 0.520678036 |
| ENSMUSG00000044042 | 2.378662046 | 2.240724001 | 2.212155735 | 2.231203126 | 2.13194489 | 2.348304863 | 1.978585612 |
| ENSMUSG00000044043 | 1 | 0.580482024 | 0.646240625 | 0.25 | 0.489301084 | 0 | 0.355418052 |
| ENSMUSG00000044052 | 0.289064826 | 0.646240625 | 0.5 | 0.25 | 0.387761935 | 0 | 0.448487648 |
| ENSMUSG00000044055 | 0.671187741 | 0 | 0 | 0 | 0.387761935 | 0.301029996 | 0.224243824 |
| ENSMUSG00000044056 | 0.45815691 | 0.396240625 | 0.646240625 | 0.580482024 | 0.489301084 | 0.84509804 | 0.710836105 |
| ENSMUSG00000044058 | 0 | 0 | 0 | 0 | 0 | 0 | 0 |
| ENSMUSG00000044060 | 1.707474304 | 1.701838731 | 1.630890489 | 1.589388001 | 1.467903253 | 1.612783857 | 1.448487648 |
| ENSMUSG00000044061 | 0 | 0 | 0 | 0 | 0 | 0 | 0 |
| ENSMUSG00000044062 | 0 | 0 | 0 | 0 | 0 | 0 | 0 |
| ENSMUSG00000044065 | 0 | 0 | 0 | 0 | 0 | 0 | 0 |
| ENSMUSG00000044066 | 2.307602651 | 2.199415381 | 2.140560606 | 2.246460484 | 2.192777367 | 2.465382851 | 2.051774285 |
| ENSMUSG00000044067 | 0.289064826 | 0.25 | 0 | 0.25 | 0 | 0 | 0 |
| ENSMUSG00000044068 | 1.482695304 | 1.410964047 | 1.42510993 | 1.732684334 | 1.603583716 | 1.86923172 | 1.572224412 |
| ENSMUSG00000044071 | 0.289064826 | 0 | 0.396240625 | 0 | 0 | 0 | 0.224243824 |
| ENSMUSG00000044072 | 0.747221736 | 0.701838731 | 0.830482024 | 0.92510993 | 0.812711509 | 0.84509804 | 0.935079929 |
| ENSMUSG00000044080 | 1.671187741 | 1.642463902 | 1.723704441 | 1.580482024 | 1.528558853 | 1.698970004 | 1.505628092 |
| ENSMUSG00000044081 | 0.289064826 | 0.5 | 0 | 0.646240625 | 0.568060967 | 0.84509804 | 0.710836105 |

| | | | | | | | |
|---|---|---|---|---|---|---|---|
| ENSMUSG00000044083 | 0.578129653 | 0 | 0.701838731 | 0.396240625 | 0.244650542 | 0 | 0.355418052 |
| ENSMUSG00000044084 | 0 | 0 | 0 | 0 | 0 | 0 | 0 |
| ENSMUSG00000044086 | 0 | 0 | 0 | 0 | 0 | 0 | 0 |
| ENSMUSG00000044092 | 0.289064826 | 0.580482024 | 0.5 | 0.396240625 | 0.489301084 | 0.602059991 | 0.520678036 |
| ENSMUSG00000044098 | 2.029919532 | 1.935366747 | 1.882845365 | 1.903677461 | 1.800029087 | 2.089905111 | 1.704233519 |
| ENSMUSG00000044103 | 0.747221736 | 0.580482024 | 0.701838731 | 0.896240625 | 0.489301084 | 0.301029996 | 0.775756176 |
| ENSMUSG00000044106 | 0 | 0 | 0 | 0 | 0 | 0 | 0 |
| ENSMUSG00000044107 | 0 | 0 | 0 | 0 | 0 | 0 | 0 |
| ENSMUSG00000044117 | 0.45815691 | 0.5 | 0.79248125 | 0.646240625 | 0.244650542 | 0.301029996 | 0.520678036 |
| ENSMUSG00000044120 | 0 | 0 | 0 | 0 | 0 | 0 | 0 |
| ENSMUSG00000044121 | 0 | 0 | 0 | 0 | 0 | 0 | 0 |
| ENSMUSG00000044122 | 0.671187741 | 0.79248125 | 0.92510993 | 0.75 | 0.733951626 | 0.698970004 | 0.629532004 |
| ENSMUSG00000044125 | 0 | 0 | 0 | 0 | 0 | 0 | 0 |
| ENSMUSG00000044134 | 1.789260561 | 1.784887838 | 1.862802778 | 1.403677461 | 1.65906333 | 1.612783857 | 1.720494 |
| ENSMUSG00000044139 | 0.45815691 | 0.396240625 | 0.580482024 | 0 | 0 | 0.301029996 | 0.520678036 |
| ENSMUSG00000044141 | 0 | 0 | 0.396240625 | 0 | 0.244650542 | 0 | 0 |
| ENSMUSG00000044145 | 0.578129653 | 0.5 | 0.5 | 0.25 | 0.489301084 | 0.602059991 | 0.448487648 |
| ENSMUSG00000044147 | 2.903481042 | 2.681128463 | 2.679419106 | 2.503156135 | 2.451638175 | 2.753583059 | 2.299891458 |
| ENSMUSG00000044148 | 2.447686928 | 2.21759118 | 2.232684334 | 2.24929487 | 2.179075613 | 2.46686762 | 2.084261301 |
| ENSMUSG00000044149 | 2.069666964 | 1.998588359 | 1.96293726 | 1.995748394 | 1.98503142 | 2.23299611 | 1.839824409 |
| ENSMUSG00000044150 | 2.089597219 | 1.943696765 | 1.981203126 | 2.179419106 | 2.118236133 | 2.376576957 | 1.944737824 |
| ENSMUSG00000044155 | 2.594187595 | 2.405970373 | 2.399513125 | 2.327653195 | 2.218681522 | 2.539076099 | 2.103753068 |
| ENSMUSG00000044156 | 0 | 0 | 0.25 | 0.5 | 0.387761935 | 0 | 0.224243824 |
| ENSMUSG00000044162 | 0.289064826 | 0.396240625 | 0.396240625 | 0.25 | 0.244650542 | 0.301029996 | 0.448487648 |
| ENSMUSG00000044164 | 0.671187741 | 0.701838731 | 0.951838731 | 0.864857905 | 0.955822902 | 1.204119983 | 0.896975297 |
| ENSMUSG00000044165 | 0 | 0.25 | 0.396240625 | 0 | 0 | 0.301029996 | 0 |
| ENSMUSG00000044167 | 2.280358197 | 2.129909063 | 2.04248125 | 1.933677405 | 1.883916752 | 2.133538908 | 1.767634022 |
| ENSMUSG00000044170 | 0 | 0 | 0 | 0 | 0 | 0.301029996 | 0 |
| ENSMUSG00000044172 | 0 | 0 | 0 | 0 | 0 | 0 | 0 |
| ENSMUSG00000044176 | 0.747221736 | 0.5 | 0.5 | 0.976722649 | 1.020174412 | 0.954242509 | 0.829800754 |
| ENSMUSG00000044177 | 0.289064826 | 0 | 0.646240625 | 0 | 0.244650542 | 0.301029996 | 0.355418052 |
| ENSMUSG00000044186 | 0 | 0 | 0 | 0 | 0 | 0 | 0 |
| ENSMUSG00000044197 | 1.516991255 | 1.532320754 | 1.561981878 | 1.339388001 | 1.358933174 | 1.505149978 | 1.425641774 |
| ENSMUSG00000044199 | 0 | 0.25 | 0 | 0.396240625 | 0.244650542 | 0.301029996 | 0.224243824 |

| | | | | | | | |
|---|---|---|---|---|---|---|---|
| ENSMUSG00000044201 | 2.136858952 | 1.951838731 | 1.961372513 | 2.156427211 | 2.112066955 | 2.404833717 | 1.995286892 |
| ENSMUSG00000044205 | 0 | 0 | 0 | 0 | 0 | 0 | 0 |
| ENSMUSG00000044206 | 0 | 0 | 0 | 0 | 0 | 0 | 0 |
| ENSMUSG00000044211 | 0 | 0 | 0 | 0.396240625 | 0 | 0 | 0 |
| ENSMUSG00000044213 | 0 | 0 | 0 | 0 | 0 | 0 | 0 |
| ENSMUSG00000044216 | 1.129344651 | 1.146240625 | 1.311981878 | 0.580482024 | 0.931471446 | 0.903089987 | 1.185218806 |
| ENSMUSG00000044217 | 2.10861828 | 2.034887838 | 2.016522298 | 1.04248125 | 1.293076518 | 1 | 1.369798885 |
| ENSMUSG00000044220 | 0 | 0 | 0 | 0.646240625 | 0.632412477 | 0 | 0.629532004 |
| ENSMUSG00000044221 | 2.85804771 | 2.624712302 | 2.566403762 | 2.633818844 | 2.531551177 | 2.928395852 | 2.365724059 |
| ENSMUSG00000044222 | 0 | 0 | 0 | 0 | 0 | 0 | 0 |
| ENSMUSG00000044224 | 2.35727108 | 2.196158711 | 2.166333979 | 2.277783918 | 2.219993628 | 2.537819095 | 2.074853081 |
| ENSMUSG00000044227 | 0 | 0 | 0 | 0 | 0 | 0 | 0 |
| ENSMUSG00000044229 | 1.181541736 | 1.356566189 | 1.25 | 1.532320754 | 1.484072009 | 1.431363764 | 1.425641774 |
| ENSMUSG00000044231 | 1.069666964 | 1.098079356 | 1.061981878 | 0.646240625 | 0.846351821 | 0.602059991 | 0.916588299 |
| ENSMUSG00000044243 | 0 | 0 | 0 | 0 | 0 | 0 | 0 |
| ENSMUSG00000044244 | 0.91631382 | 0.580482024 | 0.92510993 | 0.951838731 | 0.955822902 | 0.698970004 | 0.803905701 |
| ENSMUSG00000044248 | 0 | 0 | 0 | 0 | 0 | 0 | 0 |
| ENSMUSG00000044249 | 0 | 0 | 0 | 0 | 0 | 0 | 0 |
| ENSMUSG00000044250 | 0.45815691 | 0.701838731 | 0.75 | 1.302363341 | 1.439191811 | 1.431363764 | 1.355418052 |
| ENSMUSG00000044252 | 2.417143825 | 2.21293726 | 2.226722649 | 2.213717096 | 2.108492685 | 2.365487985 | 1.986347533 |
| ENSMUSG00000044254 | 1.342375483 | 1.364857905 | 1.557204673 | 0.79248125 | 1.057362051 | 0.602059991 | 0.969165684 |
| ENSMUSG00000044258 | 0.811507563 | 0.646240625 | 0.92510993 | 0.580482024 | 0.387761935 | 0.84509804 | 0.672731473 |
| ENSMUSG00000044265 | 0 | 0 | 0 | 0 | 0 | 0 | 0 |
| ENSMUSG00000044268 | 0 | 0 | 0 | 0 | 0 | 0 | 0 |
| ENSMUSG00000044276 | 0 | 0.25 | 0.25 | 0.5 | 0.387761935 | 0.301029996 | 0.520678036 |
| ENSMUSG00000044279 | 0 | 0 | 0 | 0 | 0 | 0 | 0.224243824 |
| ENSMUSG00000044285 | 0.289064826 | 0.25 | 0.25 | 0.5 | 0.387761935 | 0 | 0.448487648 |
| ENSMUSG00000044286 | 0 | 0 | 0 | 0.25 | 0 | 0 | 0 |
| ENSMUSG00000044287 | 0 | 0 | 0 | 0 | 0 | 0 | 0 |
| ENSMUSG00000044288 | 0.811507563 | 0.580482024 | 0.580482024 | 0.25 | 0.568060967 | 0.301029996 | 0.629532004 |
| ENSMUSG00000044292 | 0 | 0 | 0 | 0 | 0 | 0 | 0 |
| ENSMUSG00000044293 | 0 | 0 | 0 | 0 | 0 | 0 | 0 |
| ENSMUSG00000044294 | 0 | 0 | 0 | 0 | 0 | 0 | 0 |
| ENSMUSG00000044296 | 0 | 0 | 0 | 0.580482024 | 0.489301084 | 0.477121255 | 0.355418052 |

| | | | | | | | |
|---|---|---|---|---|---|---|---|
| ENSMUSG00000044303 | 2.612039956 | 2.51685859 | 2.601285366 | 2.178561379 | 2.367228134 | 2.462397998 | 2.402025166 |
| ENSMUSG00000044306 | 0 | 0 | 0 | 0 | 0 | 0 | 0 |
| ENSMUSG00000044308 | 2.558729299 | 2.375956434 | 2.321928095 | 2.640083209 | 2.539585427 | 2.919601024 | 2.367446034 |
| ENSMUSG00000044309 | 0 | 0 | 0 | 0 | 0 | 0 | 0 |
| ENSMUSG00000044312 | 0 | 0 | 0 | 0 | 0 | 0 | 0 |
| ENSMUSG00000044313 | 0.960252568 | 0.92510993 | 0.976722649 | 0.5 | 0 | 0.301029996 | 0.448487648 |
| ENSMUSG00000044317 | 0 | 0.396240625 | 0.396240625 | 0.396240625 | 0.568060967 | 0.477121255 | 0.355418052 |
| ENSMUSG00000044320 | 0.45815691 | 0.25 | 0.25 | 0 | 0.568060967 | 0.301029996 | 0 |
| ENSMUSG00000044322 | 0.289064826 | 0 | 0 | 0 | 0 | 0 | 0 |
| ENSMUSG00000044328 | 1.54867984 | 1.606566189 | 1.732684334 | 1.348079356 | 1.580303447 | 1.431363764 | 1.694069894 |
| ENSMUSG00000044330 | 0 | 0 | 0 | 0 | 0 | 0 | 0 |
| ENSMUSG00000044337 | 1.342375483 | 1.130890489 | 0.951838731 | 1.17510993 | 1.335652905 | 1.431363764 | 1.185218806 |
| ENSMUSG00000044338 | 0 | 0 | 0.25 | 0 | 0 | 0.301029996 | 0 |
| ENSMUSG00000044339 | 1.181541736 | 1.061981878 | 1.04248125 | 0.75 | 0.733951626 | 0.698970004 | 1 |
| ENSMUSG00000044340 | 2.471720134 | 2.262462137 | 2.281030328 | 2.092671852 | 1.993840051 | 2.225309282 | 1.909593098 |
| ENSMUSG00000044345 | 2.593357679 | 2.374961472 | 2.405970373 | 2.326515422 | 2.300451989 | 2.575187845 | 2.198153757 |
| ENSMUSG00000044348 | 1.37447073 | 1.238549078 | 1.339388001 | 1.634789703 | 1.572189156 | 1.755874856 | 1.496250175 |
| ENSMUSG00000044349 | 0.45815691 | 0.646240625 | 0.396240625 | 0.580482024 | 1.020174412 | 0.903089987 | 0.803905701 |
| ENSMUSG00000044350 | 1.890240806 | 1.811981878 | 1.78743678 | 2.009729747 | 1.916320788 | 2.181843588 | 1.757893036 |
| ENSMUSG00000044352 | 0 | 0.25 | 0 | 0.5 | 0.244650542 | 0 | 0 |
| ENSMUSG00000044359 | 0 | 0.25 | 0.25 | 0 | 0 | 0.301029996 | 0.355418052 |
| ENSMUSG00000044361 | 0.578129653 | 0.396240625 | 0.396240625 | 0.25 | 0.387761935 | 0 | 0.224243824 |
| ENSMUSG00000044362 | 0.289064826 | 0.5 | 0.5 | 0.646240625 | 0.244650542 | 0 | 0.448487648 |
| ENSMUSG00000044364 | 0 | 0.25 | 0 | 0.25 | 0 | 0.477121255 | 0.448487648 |
| ENSMUSG00000044365 | 1.325351389 | 1.201838731 | 1.146240625 | 0.580482024 | 0.846351821 | 0.602059991 | 0.520678036 |
| ENSMUSG00000044367 | 1.53838222 | 1.470660762 | 1.54745614 | 1.348079356 | 1.394615667 | 1.361727836 | 1.350478789 |
| ENSMUSG00000044375 | 0 | 0 | 0 | 0 | 0 | 0 | 0 |
| ENSMUSG00000044377 | 0 | 0 | 0.25 | 0 | 0 | 0 | 0 |
| ENSMUSG00000044378 | 0 | 0 | 0 | 0.25 | 0 | 0 | 0 |
| ENSMUSG00000044387 | 0 | 0 | 0 | 0 | 0 | 0 | 0 |
| ENSMUSG00000044390 | 1.249317394 | 1.348079356 | 1.25 | 1.348079356 | 1.310728002 | 1.397940009 | 1.302263476 |
| ENSMUSG00000044393 | 0.578129653 | 0.25 | 0 | 1.080482024 | 0.877063019 | 1.079181246 | 0.969165684 |
| ENSMUSG00000044400 | 0 | 0 | 0 | 0 | 0.387761935 | 0 | 0.224243824 |
| ENSMUSG00000044405 | 0 | 0.25 | 0 | 0 | 0.244650542 | 0 | 0 |

| | | | | | | | |
|---|---|---|---|---|---|---|---|
| ENSMUSG00000044408 | 2.553277818 | 2.33272922 | 2.382357639 | 2.359177886 | 2.301493158 | 2.613841822 | 2.143416814 |
| ENSMUSG00000044424 | 0.45815691 | 0.646240625 | 0.75 | 0.580482024 | 0.387761935 | 0 | 0.710836105 |
| ENSMUSG00000044429 | 0 | 0 | 0 | 0 | 0 | 0 | 0 |
| ENSMUSG00000044430 | 0 | 0 | 0 | 0 | 0.244650542 | 0 | 0 |
| ENSMUSG00000044433 | 0.289064826 | 0.580482024 | 0.646240625 | 0.646240625 | 0.905314583 | 0.77815125 | 0.829800754 |
| ENSMUSG00000044434 | 0 | 0 | 0 | 0 | 0 | 0 | 0 |
| ENSMUSG00000044441 | 0 | 0 | 0 | 0 | 0 | 0 | 0 |
| ENSMUSG00000044442 | 1.967766868 | 1.882845365 | 1.880890489 | 1.903677461 | 1.812711509 | 2.130333768 | 1.74047993 |
| ENSMUSG00000044444 | 0 | 0 | 0 | 0 | 0 | 0 | 0 |
| ENSMUSG00000044447 | 2.384185734 | 2.192046081 | 2.198603967 | 2.466819685 | 2.397003204 | 2.723455672 | 2.24869599 |
| ENSMUSG00000044452 | 2.170239354 | 2.017865591 | 2.004202072 | 2.04745614 | 2.003645018 | 2.193124598 | 1.896054911 |
| ENSMUSG00000044453 | 0 | 0 | 0 | 0 | 0 | 0 | 0.224243824 |
| ENSMUSG00000044454 | 0 | 0 | 0 | 0 | 0 | 0 | 0 |
| ENSMUSG00000044456 | 1.771760131 | 1.782320754 | 1.903677461 | 1.77186571 | 1.938809675 | 2.033423755 | 1.906930404 |
| ENSMUSG00000044457 | 0 | 0 | 0 | 0 | 0 | 0 | 0 |
| ENSMUSG00000044461 | 0.289064826 | 0 | 0 | 0.75 | 0.733951626 | 0.477121255 | 0.74492186 |
| ENSMUSG00000044465 | 2.213719572 | 2.048689214 | 2.063166358 | 2.019203899 | 1.962676635 | 2.089905111 | 1.86715039 |
| ENSMUSG00000044468 | 1.156259305 | 1 | 1.188721876 | 1.642463902 | 1.450958105 | 1.716003344 | 1.429563222 |
| ENSMUSG00000044469 | 1.494443473 | 1.445339928 | 1.571350555 | 1.410964047 | 1.335652905 | 1.342422681 | 1.401059159 |
| ENSMUSG00000044471 | 0 | 0 | 0 | 0.25 | 0 | 0 | 0 |
| ENSMUSG00000044475 | 1.978790393 | 1.896240625 | 1.886723615 | 1.747171172 | 1.888971102 | 2.017033339 | 1.757893036 |
| ENSMUSG00000044477 | 2.460448303 | 2.307204673 | 2.368426437 | 2.112802778 | 2.158641579 | 2.399673721 | 2.201756416 |
| ENSMUSG00000044485 | 0 | 0 | 0 | 0 | 0 | 0 | 0 |
| ENSMUSG00000044487 | 0 | 0 | 0 | 0 | 0 | 0 | 0 |
| ENSMUSG00000044496 | 2.223916471 | 2.149047439 | 2.157339155 | 1.92510993 | 2.135290471 | 2.243038049 | 2.117056606 |
| ENSMUSG00000044499 | 0 | 0 | 0 | 0 | 0 | 0 | 0.224243824 |
| ENSMUSG00000044501 | 1.614416215 | 1.610735874 | 1.610735874 | 1.747171172 | 1.698489937 | 1.857332496 | 1.473249546 |
| ENSMUSG00000044502 | 2.307602651 | 2.158248799 | 2.147178659 | 2.244319981 | 2.215379757 | 2.501059262 | 2.065163042 |
| ENSMUSG00000044505 | 0.960252568 | 0.976722649 | 0.864857905 | 0.5 | 0.632412477 | 0.477121255 | 0.579661877 |
| ENSMUSG00000044515 | 0 | 0 | 0 | 0 | 0 | 0 | 0 |
| ENSMUSG00000044518 | 0 | 0 | 0 | 0 | 0 | 0 | 0 |
| ENSMUSG00000044519 | 1.647796617 | 1.53743678 | 1.5 | 1.505591953 | 1.478764298 | 1.612783857 | 1.335191751 |
| ENSMUSG00000044522 | 0.45815691 | 0.701838731 | 0.5 | 0 | 0.244650542 | 0 | 0 |
| ENSMUSG00000044526 | 0 | 0 | 0 | 0 | 0 | 0 | 0 |

| | | | | | | | |
|---|---|---|---|---|---|---|---|
| ENSMUSG00000044528 | 1.907802659 | 1.77186571 | 1.862802778 | 1.660964047 | 1.576269619 | 1.913813852 | 1.546120303 |
| ENSMUSG00000044533 | 3.2769853 | 3.064125892 | 3.081747794 | 2.864473459 | 2.962596771 | 3.326949994 | 2.823308934 |
| ENSMUSG00000044534 | 0.289064826 | 0.25 | 0.25 | 0.25 | 0 | 0 | 0 |
| ENSMUSG00000044537 | 0 | 0 | 0 | 0 | 0 | 0 | 0 |
| ENSMUSG00000044538 | 0 | 0 | 0 | 0 | 0 | 0 | 0 |
| ENSMUSG00000044542 | 0.289064826 | 0 | 0 | 0 | 0 | 0 | 0 |
| ENSMUSG00000044544 | 0 | 0.5 | 0 | 0.396240625 | 0 | 0.301029996 | 0.224243824 |
| ENSMUSG00000044548 | 0.671187741 | 0.646240625 | 0.25 | 1.061981878 | 0.955822902 | 0.84509804 | 0.984950056 |
| ENSMUSG00000044550 | 0.45815691 | 0.646240625 | 0.396240625 | 0.396240625 | 0.632412477 | 0.301029996 | 0.629532004 |
| ENSMUSG00000044551 | 1.527823874 | 1.488549078 | 1.388647213 | 1.445339928 | 1.588235379 | 1.698970004 | 1.535058868 |
| ENSMUSG00000044556 | 0 | 0 | 0.25 | 0 | 0 | 0 | 0.355418052 |
| ENSMUSG00000044560 | 0 | 0 | 0 | 0 | 0 | 0 | 0 |
| ENSMUSG00000044562 | 0.45815691 | 0.701838731 | 1 | 0.701838731 | 0.733951626 | 1 | 0.952571511 |
| ENSMUSG00000044566 | 1.389637216 | 1.330482024 | 1.080482024 | 1.114857905 | 1 | 0.903089987 | 0.952571511 |
| ENSMUSG00000044573 | 2.736751754 | 2.524178789 | 2.503156135 | 2.464495249 | 2.454008153 | 2.800029359 | 2.276871337 |
| ENSMUSG00000044574 | 0 | 0 | 0.25 | 0 | 0.244650542 | 0 | 0 |
| ENSMUSG00000044576 | 1.714367558 | 1.610735874 | 1.741446071 | 0.79248125 | 0.877063019 | 0.903089987 | 0.952571511 |
| ENSMUSG00000044581 | 0 | 0.25 | 0.5 | 0.25 | 0.244650542 | 0 | 0.224243824 |
| ENSMUSG00000044583 | 0.45815691 | 0.396240625 | 0.25 | 0.580482024 | 0.489301084 | 0 | 0.355418052 |
| ENSMUSG00000044594 | 0 | 0 | 0 | 0 | 0 | 0 | 0 |
| ENSMUSG00000044595 | 0.91631382 | 1.02186571 | 0.864857905 | 0.701838731 | 0.733951626 | 0.477121255 | 1 |
| ENSMUSG00000044597 | 0 | 0 | 0 | 0 | 0 | 0 | 0 |
| ENSMUSG00000044600 | 2.561428547 | 2.399978211 | 2.363317659 | 2.544354884 | 2.434230499 | 2.803457116 | 2.249933144 |
| ENSMUSG00000044609 | 0 | 0.25 | 0.25 | 0.580482024 | 0 | 0.301029996 | 0 |
| ENSMUSG00000044617 | 2.048736215 | 1.933677405 | 1.858657057 | 1.882845365 | 1.828942237 | 2.071882007 | 1.82753046 |
| ENSMUSG00000044624 | 0 | 0 | 0 | 0 | 0 | 0 | 0 |
| ENSMUSG00000044626 | 1 | 0.701838731 | 1.04248125 | 1.061981878 | 0.568060967 | 0.602059991 | 0.775756176 |
| ENSMUSG00000044627 | 2.920505136 | 2.661864607 | 2.705244173 | 2.684945152 | 2.633834461 | 3.036628895 | 2.462868481 |
| ENSMUSG00000044628 | 0 | 0 | 0.25 | 0.25 | 0.568060967 | 0.477121255 | 0.448487648 |
| ENSMUSG00000044629 | 1.389637216 | 1.311981878 | 1.356566189 | 1.348079356 | 1.264824954 | 1.176091259 | 1.176815335 |
| ENSMUSG00000044633 | 0 | 0 | 0 | 0 | 0 | 0 | 0 |
| ENSMUSG00000044636 | 2.300956576 | 2.138647213 | 2.190387808 | 2.125956434 | 2.133621644 | 2.383815366 | 2.016928211 |
| ENSMUSG00000044639 | 0 | 0 | 0 | 0 | 0 | 0 | 0 |
| ENSMUSG00000044641 | 1.558729299 | 1.571350555 | 1.476722649 | 1.51109853 | 1.592135487 | 1.491361694 | 1.48984372 |

| | | | | | | | |
|---|---|---|---|---|---|---|---|
| ENSMUSG00000044645 | 0 | 0 | 0 | 0 | 0 | 0 | 0 |
| ENSMUSG00000044646 | 1.37447073 | 1.403677461 | 1.339388001 | 1.42510993 | 1.450958105 | 1.568201724 | 1.529383012 |
| ENSMUSG00000044647 | 0 | 0.25 | 0.25 | 0.5 | 0.489301084 | 0.477121255 | 0.355418052 |
| ENSMUSG00000044649 | 0 | 0.25 | 0 | 0 | 0 | 0 | 0 |
| ENSMUSG00000044664 | 0 | 0 | 0 | 0 | 0.244650542 | 0.301029996 | 0 |
| ENSMUSG00000044667 | 0.45815691 | 0.646240625 | 0.396240625 | 0.25 | 0.244650542 | 0 | 0.224243824 |
| ENSMUSG00000044674 | 2.312518611 | 2.169870025 | 2.142463902 | 1.966046536 | 1.977302144 | 2.193124598 | 1.933441873 |
| ENSMUSG00000044676 | 0.671187741 | 0.580482024 | 0.976722649 | 1.372963274 | 1.335652905 | 1.342422681 | 1.313616061 |
| ENSMUSG00000044678 | 0 | 0 | 0 | 0 | 0 | 0 | 0 |
| ENSMUSG00000044681 | 0 | 0 | 0 | 0.5 | 0 | 0 | 0.355418052 |
| ENSMUSG00000044689 | 0 | 0 | 0 | 0 | 0 | 0 | 0 |
| ENSMUSG00000044694 | 0.289064826 | 0.25 | 0.5 | 0.396240625 | 0.387761935 | 0.602059991 | 0.448487648 |
| ENSMUSG00000044697 | 0 | 0 | 0 | 0 | 0 | 0 | 0 |
| ENSMUSG00000044700 | 2.172524468 | 2.158248799 | 2.225216702 | 2.00835575 | 2.141887974 | 2.29666519 | 2.155626394 |
| ENSMUSG00000044701 | 0 | 0.25 | 0 | 0 | 0 | 0 | 0 |
| ENSMUSG00000044702 | 1.837744666 | 1.695339928 | 1.62694866 | 1.714495249 | 1.655839964 | 1.72427587 | 1.564611933 |
| ENSMUSG00000044703 | 0 | 0 | 0.25 | 0.25 | 0 | 0.301029996 | 0 |
| ENSMUSG00000044705 | 0 | 0 | 0 | 0 | 0 | 0 | 0 |
| ENSMUSG00000044707 | 0.45815691 | 0.580482024 | 0.701838731 | 0.951838731 | 0.978602168 | 1.146128036 | 1 |
| ENSMUSG00000044708 | 0.45815691 | 0 | 0.396240625 | 0.580482024 | 0.877063019 | 0.602059991 | 0.579661877 |
| ENSMUSG00000044709 | 1.960252568 | 1.894357207 | 1.964495249 | 1.845926073 | 1.890639929 | 2.071882007 | 1.824094866 |
| ENSMUSG00000044712 | 2.211649928 | 2.095926073 | 2.079353153 | 2.031030328 | 2.019019072 | 2.173186268 | 1.828667599 |
| ENSMUSG00000044715 | 2.190368148 | 2.043731421 | 2.05357978 | 2 | 1.928492895 | 2.198657087 | 1.812372009 |
| ENSMUSG00000044716 | 0.289064826 | 0 | 0.25 | 1.130890489 | 1.366364104 | 1.301029996 | 1.335191751 |
| ENSMUSG00000044719 | 0.45815691 | 0.580482024 | 0.5 | 0 | 0.244650542 | 0 | 0 |
| ENSMUSG00000044724 | 0 | 0 | 0.396240625 | 0 | 0 | 0 | 0 |
| ENSMUSG00000044726 | 0.45815691 | 0.396240625 | 0.25 | 0 | 0 | 0 | 0.224243824 |
| ENSMUSG00000044730 | 2.029919532 | 1.933677405 | 1.92510993 | 1.83272922 | 1.870073561 | 2.086359831 | 1.770364127 |
| ENSMUSG00000044734 | 2.382811656 | 2.155512955 | 2.208222504 | 1.668106335 | 1.478764298 | 1.812913357 | 1.493062805 |
| ENSMUSG00000044737 | 0 | 0 | 0 | 0 | 0.244650542 | 0 | 0 |
| ENSMUSG00000044743 | 0 | 0 | 0 | 0 | 0 | 0 | 0 |
| ENSMUSG00000044748 | 0 | 0 | 0 | 0 | 0 | 0 | 0 |
| ENSMUSG00000044749 | 0 | 0 | 0 | 0.396240625 | 0.387761935 | 0 | 0.224243824 |
| ENSMUSG00000044751 | 0 | 0 | 0.25 | 0.25 | 0.244650542 | 0.301029996 | 0.355418052 |

| | | | | | | | |
|---|---|---|---|---|---|---|---|
| ENSMUSG00000044757 | 0.45815691 | 0 | 0.25 | 0.396240625 | 0.244650542 | 0 | 0.224243824 |
| ENSMUSG00000044763 | 2.149089517 | 1.997171172 | 2.04745614 | 2.066696635 | 1.965381292 | 2.201397124 | 1.828667599 |
| ENSMUSG00000044768 | 1.912079952 | 1.843759858 | 1.766522298 | 1.918106335 | 1.818885826 | 2.068185862 | 1.661510159 |
| ENSMUSG00000044770 | 0.45815691 | 0 | 0.25 | 0.5 | 0.244650542 | 0.602059991 | 0.448487648 |
| ENSMUSG00000044772 | 0 | 0 | 0 | 0 | 0 | 0 | 0 |
| ENSMUSG00000044783 | 3.296671435 | 2.965174758 | 2.9645924 | 3.025165585 | 2.98927812 | 3.434728542 | 2.782372264 |
| ENSMUSG00000044786 | 1.527823874 | 1.557204673 | 1.602347734 | 1.388647213 | 1.27449559 | 1.462397998 | 1.508694612 |
| ENSMUSG00000044787 | 0 | 0 | 0 | 0 | 0 | 0.301029996 | 0 |
| ENSMUSG00000044788 | 0.867194479 | 0.396240625 | 0.580482024 | 0.864857905 | 0.955822902 | 0.903089987 | 0.969165684 |
| ENSMUSG00000044791 | 2.905843838 | 2.650210529 | 2.636482442 | 2.728969845 | 2.628111827 | 3.036229544 | 2.439507591 |
| ENSMUSG00000044792 | 1.867194479 | 1.71759118 | 1.779735268 | 1.886723615 | 1.902105869 | 2.133538908 | 1.798966438 |
| ENSMUSG00000044795 | 1.734388958 | 1.695339928 | 1.668106335 | 1.890560606 | 1.828942237 | 2.064457989 | 1.720494 |
| ENSMUSG00000044798 | 0 | 0 | 0 | 0 | 0 | 0 | 0 |
| ENSMUSG00000044801 | 0 | 0 | 0 | 0 | 0 | 0 | 0 |
| ENSMUSG00000044807 | 2.029919532 | 1.862802778 | 1.794977273 | 2.016522298 | 1.944572352 | 2.187520721 | 1.833176598 |
| ENSMUSG00000044811 | 0 | 0 | 0 | 0 | 0 | 0 | 0 |
| ENSMUSG00000044813 | 1.881174527 | 1.819031101 | 1.933677405 | 1.777131114 | 1.913211023 | 2.071882007 | 1.909593098 |
| ENSMUSG00000044814 | 0.289064826 | 0.396240625 | 0 | 0.5 | 0 | 0.477121255 | 0.355418052 |
| ENSMUSG00000044816 | 0 | 0 | 0.25 | 0.25 | 0.387761935 | 0 | 0 |
| ENSMUSG00000044819 | 0 | 0 | 0 | 0 | 0 | 0 | 0 |
| ENSMUSG00000044820 | 0.289064826 | 0 | 0 | 0.396240625 | 0.387761935 | 0.301029996 | 0 |
| ENSMUSG00000044824 | 0 | 0 | 0 | 0 | 0 | 0 | 0 |
| ENSMUSG00000044827 | 1.944805158 | 1.799918086 | 1.825945187 | 1.26109853 | 1.200473444 | 1.255272505 | 1.041356071 |
| ENSMUSG00000044835 | 0.747221736 | 0.701838731 | 0.75 | 0.864857905 | 0.686820904 | 0.77815125 | 0.876096088 |
| ENSMUSG00000044847 | 2.075459174 | 1.990000483 | 1.937048212 | 1.973704441 | 1.926994141 | 2.217483944 | 1.86815666 |
| ENSMUSG00000044854 | 0.578129653 | 0.580482024 | 0.5 | 0.396240625 | 0.387761935 | 0 | 0.355418052 |
| ENSMUSG00000044857 | 2.199011616 | 2.103406982 | 2.155512955 | 1.80479213 | 1.998776581 | 1.959041392 | 1.985649549 |
| ENSMUSG00000044860 | 0 | 0.25 | 0 | 0.25 | 0.387761935 | 0.698970004 | 0 |
| ENSMUSG00000044863 | 0 | 0 | 0 | 0 | 0 | 0 | 0 |
| ENSMUSG00000044864 | 2.841885121 | 2.615113974 | 2.624961472 | 2.420309603 | 2.413278488 | 2.727541257 | 2.26794422 |
| ENSMUSG00000044867 | 0 | 0 | 0 | 0 | 0 | 0 | 0 |
| ENSMUSG00000044876 | 1.358731791 | 1.26109853 | 1.372963274 | 1.238549078 | 1.254881871 | 1.255272505 | 1.369798885 |
| ENSMUSG00000044881 | 1.289064826 | 1.17510993 | 1.282320754 | 1.238549078 | 1.310728002 | 1.380211242 | 1.259064007 |
| ENSMUSG00000044894 | 1.783508299 | 1.96293726 | 2.143411797 | 1.77186571 | 1.898867717 | 1.903089987 | 2.115189268 |

| | | | | | | | |
|---|---|---|---|---|---|---|---|
| ENSMUSG00000044897 | 0 | 0 | 0 | 0 | 0 | 0 | 0 |
| ENSMUSG00000044899 | 0 | 0 | 0 | 0 | 0 | 0 | 0 |
| ENSMUSG00000044903 | 0 | 0 | 0 | 0 | 0 | 0 | 0 |
| ENSMUSG00000044906 | 2.003432385 | 1.956637122 | 1.909156155 | 2.028435542 | 2 | 2.193124598 | 1.853775828 |
| ENSMUSG00000044912 | 0 | 0 | 0 | 0 | 0 | 0 | 0 |
| ENSMUSG00000044916 | 0.45815691 | 0.79248125 | 0.25 | 0.864857905 | 1 | 0.84509804 | 0.935079929 |
| ENSMUSG00000044921 | 0.671187741 | 0.5 | 0.5 | 0 | 0 | 0 | 0 |
| ENSMUSG00000044923 | 0 | 0 | 0 | 0 | 0 | 0 | 0 |
| ENSMUSG00000044927 | 0.811507563 | 0.951838731 | 1.160964047 | 0.830482024 | 1.039257769 | 1 | 1.245582246 |
| ENSMUSG00000044933 | 0 | 0 | 0 | 0 | 0 | 0 | 0 |
| ENSMUSG00000044934 | 2.254965775 | 2.159156155 | 2.127938163 | 2.333288838 | 2.242813819 | 2.58546073 | 2.092400484 |
| ENSMUSG00000044937 | 1.227926429 | 1.26109853 | 1.160964047 | 1.130890489 | 1.200473444 | 1.255272505 | 1 |
| ENSMUSG00000044938 | 0 | 0 | 0 | 0 | 0 | 0 | 0 |
| ENSMUSG00000044948 | 2.583267694 | 2.440803092 | 2.442459461 | 1.978222334 | 2.039257769 | 2.264817823 | 1.925124822 |
| ENSMUSG00000044949 | 1.837744666 | 1.825945187 | 1.823655187 | 1.782320754 | 1.818885826 | 1.995635195 | 1.750751125 |
| ENSMUSG00000044950 | 2.045658471 | 1.928561379 | 1.959800947 | 2.108657057 | 1.993840051 | 2.269512944 | 1.896975297 |
| ENSMUSG00000044951 | 0 | 0.25 | 0 | 0.396240625 | 0 | 0 | 0.224243824 |
| ENSMUSG00000044952 | 1.418409478 | 1.380890489 | 1.226722649 | 1.42510993 | 1.439191811 | 1.278753601 | 1.284450788 |
| ENSMUSG00000044957 | 0.289064826 | 0 | 0 | 0.25 | 0.244650542 | 0 | 0.224243824 |
| ENSMUSG00000044966 | 1.482695304 | 1.380890489 | 1.388647213 | 1.5 | 1.555378544 | 1.51851394 | 1.296434211 |
| ENSMUSG00000044968 | 1.494443473 | 1.54745614 | 1.584962501 | 1.589388001 | 1.499532413 | 1.698970004 | 1.459332757 |
| ENSMUSG00000044976 | 0 | 0 | 0 | 0 | 0 | 0 | 0 |
| ENSMUSG00000044982 | 1.269664473 | 1.130890489 | 1.080482024 | 1.431980114 | 1.254881871 | 1.380211242 | 1.21680636 |
| ENSMUSG00000044985 | 0 | 0 | 0 | 0 | 0 | 0 | 0 |
| ENSMUSG00000044986 | 1.678702042 | 1.642463902 | 1.766522298 | 1.451838731 | 1.665423072 | 1.612783857 | 1.699191618 |
| ENSMUSG00000044988 | 0 | 0 | 0 | 0 | 0 | 0 | 0 |
| ENSMUSG00000044991 | 1 | 1 | 1.061981878 | 1.080482024 | 0.978602168 | 0.903089987 | 0.952571511 |
| ENSMUSG00000044994 | 0 | 0 | 0 | 0 | 0 | 0 | 0 |
| ENSMUSG00000044997 | 0 | 0 | 0 | 0 | 0 | 0 | 0 |
| ENSMUSG00000045004 | 0 | 0 | 0 | 0.25 | 0.244650542 | 0 | 0.224243824 |
| ENSMUSG00000045005 | 1.857607105 | 1.711372513 | 1.774508021 | 1.445339928 | 1.450958105 | 1.477121255 | 1.355418052 |
| ENSMUSG00000045007 | 1.289064826 | 1.302363341 | 1.356566189 | 0.701838731 | 0.812711509 | 0.77815125 | 0.853775828 |
| ENSMUSG00000045008 | 0 | 0.25 | 0 | 0.25 | 0.387761935 | 0.602059991 | 0.448487648 |
| ENSMUSG00000045009 | 0 | 0.25 | 0.396240625 | 0.25 | 0.489301084 | 0 | 0.672731473 |

| | | | | | | | |
|---|---|---|---|---|---|---|---|
| ENSMUSG00000045010 | 0 | 0 | 0 | 0 | 0.387761935 | 0 | 0.224243824 |
| ENSMUSG00000045013 | 0 | 0 | 0 | 0 | 0 | 0 | 0 |
| ENSMUSG00000045019 | 0 | 0 | 0 | 0.25 | 0.244650542 | 0 | 0 |
| ENSMUSG00000045022 | 0 | 0 | 0 | 0 | 0 | 0 | 0 |
| ENSMUSG00000045027 | 1.389637216 | 1.27186571 | 1.339388001 | 0.701838731 | 0.77552387 | 0.602059991 | 0.672731473 |
| ENSMUSG00000045030 | 0 | 0 | 0 | 0 | 0 | 0 | 0 |
| ENSMUSG00000045031 | 1.325351389 | 1.02186571 | 1.146240625 | 1.330482024 | 1.212046813 | 1.380211242 | 1.245582246 |
| ENSMUSG00000045034 | 0 | 0 | 0 | 0.396240625 | 0.244650542 | 0.301029996 | 0.224243824 |
| ENSMUSG00000045036 | 0.289064826 | 0.396240625 | 0 | 0.25 | 0.244650542 | 0 | 0 |
| ENSMUSG00000045038 | 1.789260561 | 1.75 | 1.729715809 | 1.729715809 | 1.618292349 | 1.86923172 | 1.659601507 |
| ENSMUSG00000045039 | 2.451206209 | 2.306603048 | 2.438721876 | 2.023189285 | 2.185632262 | 2.227886705 | 2.212329757 |
| ENSMUSG00000045045 | 2.092368209 | 2.148114259 | 2.313757142 | 1.418106335 | 1.773209395 | 1.672097858 | 1.936709733 |
| ENSMUSG00000045048 | 0 | 0 | 0 | 0 | 0 | 0 | 0 |
| ENSMUSG00000045052 | 0 | 0 | 0 | 0 | 0 | 0 | 0 |
| ENSMUSG00000045053 | 0 | 0 | 0 | 0 | 0 | 0 | 0 |
| ENSMUSG00000045055 | 0.671187741 | 0.92510993 | 0.951838731 | 0.646240625 | 0.77552387 | 0.602059991 | 0.74492186 |
| ENSMUSG00000045062 | 0 | 0.25 | 0.25 | 0 | 0.244650542 | 0 | 0.355418052 |
| ENSMUSG00000045064 | 1.404271433 | 1.27186571 | 1.29248125 | 1.146240625 | 1.234113756 | 1.146128036 | 1.252393349 |
| ENSMUSG00000045065 | 0 | 0 | 0 | 0 | 0 | 0.301029996 | 0 |
| ENSMUSG00000045071 | 1.857607105 | 1.701838731 | 1.668106335 | 1.741446071 | 1.72341484 | 1.799340549 | 1.667169465 |
| ENSMUSG00000045075 | 0.811507563 | 0.5 | 0.646240625 | 0.5 | 0.244650542 | 0 | 0.355418052 |
| ENSMUSG00000045078 | 2.560530738 | 2.373463862 | 2.358135475 | 2.348079356 | 2.32316608 | 2.67669361 | 2.219530717 |
| ENSMUSG00000045083 | 0 | 0 | 0 | 0 | 0 | 0 | 0 |
| ENSMUSG00000045087 | 0.289064826 | 0 | 0 | 0.396240625 | 0 | 0.301029996 | 0.224243824 |
| ENSMUSG00000045091 | 0 | 0 | 0 | 0 | 0 | 0 | 0 |
| ENSMUSG00000045092 | 2.045658471 | 1.994319981 | 1.982684334 | 1.92510993 | 1.919403394 | 2.127104798 | 1.786277931 |
| ENSMUSG00000045094 | 0.578129653 | 0.396240625 | 0.701838731 | 0.79248125 | 1 | 1.079181246 | 0.853775828 |
| ENSMUSG00000045095 | 2.103271637 | 1.975216702 | 1.951838731 | 1.476722649 | 1.467903253 | 1.51851394 | 1.455757965 |
| ENSMUSG00000045098 | 2.599132701 | 2.398114259 | 2.366901388 | 2.471805154 | 2.427019704 | 2.778874472 | 2.254834987 |
| ENSMUSG00000045100 | 1.505869732 | 1.238549078 | 1.302363341 | 1.04248125 | 1.27449559 | 1.380211242 | 1.278288402 |
| ENSMUSG00000045102 | 0.578129653 | 0.701838731 | 0.701838731 | 0.701838731 | 0.489301084 | 0.477121255 | 0.672731473 |
| ENSMUSG00000045103 | 1.639698646 | 1.580482024 | 1.527131114 | 1.188721876 | 1.106691885 | 1.278753601 | 1.121219121 |
| ENSMUSG00000045104 | 0 | 0 | 0 | 0 | 0 | 0 | 0 |
| ENSMUSG00000045106 | 1.249317394 | 1.04248125 | 1.080482024 | 1.438721876 | 1.366364104 | 1.431363764 | 1.23151414 |

| | | | | | | | |
|---|---|---|---|---|---|---|---|
| ENSMUSG00000045107 | 1.325351389 | 1.330482024 | 1.302363341 | 1.17510993 | 1.136121934 | 1.278753601 | 1.23151414 |
| ENSMUSG00000045109 | 0 | 0 | 0 | 0 | 0 | 0 | 0 |
| ENSMUSG00000045111 | 0 | 0 | 0 | 0 | 0 | 0 | 0 |
| ENSMUSG00000045114 | 0 | 0 | 0.25 | 0 | 0 | 0 | 0.224243824 |
| ENSMUSG00000045126 | 0 | 0 | 0 | 0 | 0 | 0 | 0 |
| ENSMUSG00000045128 | 3.072309764 | 2.874836908 | 2.887686696 | 2.666777886 | 2.834267356 | 3.123524981 | 2.725287969 |
| ENSMUSG00000045132 | 0 | 0 | 0 | 0 | 0 | 0 | 0 |
| ENSMUSG00000045136 | 2.43085913 | 2.21759118 | 2.253505118 | 2.223704441 | 2.237868644 | 2.492760389 | 2.130728307 |
| ENSMUSG00000045140 | 1.881174527 | 1.784887838 | 1.814346961 | 1.83272922 | 1.763796674 | 1.963787827 | 1.584526862 |
| ENSMUSG00000045148 | 0 | 0 | 0 | 0 | 0 | 0 | 0 |
| ENSMUSG00000045150 | 0 | 0 | 0 | 0 | 0 | 0 | 0 |
| ENSMUSG00000045160 | 1.558729299 | 1.593759858 | 1.53743678 | 1.688721876 | 1.645989387 | 1.698970004 | 1.508694612 |
| ENSMUSG00000045165 | 0 | 0.25 | 0.396240625 | 0.646240625 | 0.489301084 | 0.698970004 | 0.579661877 |
| ENSMUSG00000045174 | 0 | 0 | 0 | 0 | 0 | 0 | 0 |
| ENSMUSG00000045176 | 1.289064826 | 1.25 | 1.282320754 | 1.302363341 | 1.394615667 | 1.51851394 | 1.433437704 |
| ENSMUSG00000045178 | 0 | 0 | 0 | 0 | 0 | 0 | 0 |
| ENSMUSG00000045179 | 0 | 0 | 0 | 0 | 0 | 0 | 0 |
| ENSMUSG00000045180 | 1.470606563 | 1.451838731 | 1.372963274 | 2.201032755 | 2.19135987 | 2.459392488 | 1.985649549 |
| ENSMUSG00000045193 | 1.885732304 | 1.802363341 | 1.852347734 | 1.711372513 | 1.749189509 | 1.799340549 | 1.710836105 |
| ENSMUSG00000045201 | 0 | 0 | 0 | 0 | 0 | 0 | 0 |
| ENSMUSG00000045202 | 0 | 0 | 0 | 0 | 0 | 0 | 0 |
| ENSMUSG00000045204 | 0 | 0 | 0 | 0 | 0 | 0 | 0 |
| ENSMUSG00000045205 | 2.623788124 | 2.392463902 | 2.367918804 | 2.500352049 | 2.426290456 | 2.78604121 | 2.224243824 |
| ENSMUSG00000045210 | 2.502784398 | 2.329353153 | 2.329353153 | 2.457826811 | 2.38035699 | 2.705863712 | 2.184085651 |
| ENSMUSG00000045211 | 1.614416215 | 1.571350555 | 1.614857905 | 1.777131114 | 1.862942891 | 1.986771734 | 1.756477211 |
| ENSMUSG00000045215 | 0.867194479 | 0.75 | 0.646240625 | 1.130890489 | 1.057362051 | 1.113943352 | 1.089372236 |
| ENSMUSG00000045216 | 2.440557995 | 2.281676118 | 2.301142786 | 2.100219859 | 2.012006468 | 2.245512668 | 1.979298987 |
| ENSMUSG00000045217 | 0 | 0 | 0 | 0 | 0 | 0 | 0 |
| ENSMUSG00000045225 | 0 | 0 | 0 | 0 | 0 | 0 | 0 |
| ENSMUSG00000045231 | 0.45815691 | 0.79248125 | 0.75 | 0.5 | 0.387761935 | 0.301029996 | 0.224243824 |
| ENSMUSG00000045232 | 0 | 0 | 0 | 0 | 0.244650542 | 0 | 0 |
| ENSMUSG00000045236 | 0 | 0 | 0 | 0 | 0 | 0 | 0 |
| ENSMUSG00000045237 | 2.126809492 | 1.928561379 | 1.903677461 | 1.894357207 | 1.804307395 | 2.053078443 | 1.695786161 |
| ENSMUSG00000045238 | 0.289064826 | 0 | 0 | 0.396240625 | 0 | 0 | 0.448487648 |

| | | | | | | | |
|---|---|---|---|---|---|---|---|
| ENSMUSG00000045246 | 0 | 0 | 0 | 0 | 0 | 0 | 0 |
| ENSMUSG00000045248 | 1.721148722 | 1.708222504 | 1.79248125 | 1.575945187 | 1.635855979 | 1.662757832 | 1.683576582 |
| ENSMUSG00000045251 | 1.289064826 | 1.26109853 | 1.160964047 | 1.02186571 | 1.039257769 | 1 | 1.150210039 |
| ENSMUSG00000045252 | 1.568542279 | 1.660964047 | 1.708222504 | 1.321350555 | 1.542223377 | 1.544068044 | 1.709198048 |
| ENSMUSG00000045257 | 1.249317394 | 1.146240625 | 1.146240625 | 1.160964047 | 1.212046813 | 1.301029996 | 1.259064007 |
| ENSMUSG00000045259 | 0 | 0 | 0 | 0 | 0 | 0 | 0 |
| ENSMUSG00000045267 | 0 | 0 | 0 | 0 | 0 | 0 | 0 |
| ENSMUSG00000045268 | 1.432083896 | 1.29248125 | 1.282320754 | 1.17510993 | 1.27449559 | 1.414973348 | 1.21680636 |
| ENSMUSG00000045273 | 2.209569962 | 2.106566189 | 2.092671852 | 2.252107156 | 2.189936658 | 2.437750563 | 2.086826862 |
| ENSMUSG00000045275 | 0.811507563 | 0.896240625 | 0.75 | 0.830482024 | 1 | 0.698970004 | 0.710836105 |
| ENSMUSG00000045281 | 0 | 0 | 0 | 0 | 0 | 0 | 0 |
| ENSMUSG00000045282 | 1.100572389 | 0.976722649 | 0.92510993 | 1.098079356 | 0.955822902 | 0.698970004 | 1.041356071 |
| ENSMUSG00000045284 | 0 | 0 | 0 | 0 | 0 | 0 | 0 |
| ENSMUSG00000045287 | 0.578129653 | 0.580482024 | 1.02186571 | 1 | 1 | 0.903089987 | 1.100339912 |
| ENSMUSG00000045288 | 0 | 0.25 | 0 | 0 | 0 | 0.301029996 | 0.224243824 |
| ENSMUSG00000045291 | 0.45815691 | 0.396240625 | 0 | 0.25 | 0.244650542 | 0.301029996 | 0.224243824 |
| ENSMUSG00000045294 | 2.31414447 | 2.151832578 | 2.188721876 | 2.325945187 | 2.267696867 | 2.608526034 | 2.134278581 |
| ENSMUSG00000045302 | 2.604828841 | 2.37694866 | 2.413659007 | 2.356041572 | 2.406395052 | 2.655138435 | 2.257258335 |
| ENSMUSG00000045306 | 0 | 0 | 0 | 0 | 0 | 0 | 0 |
| ENSMUSG00000045312 | 2.371652933 | 2.269871196 | 2.251406137 | 1.997171172 | 1.973373093 | 2.225309282 | 1.840919217 |
| ENSMUSG00000045314 | 0 | 0.5 | 0.5 | 0 | 0.632412477 | 0.301029996 | 0.520678036 |
| ENSMUSG00000045316 | 1.605636262 | 1.552363341 | 1.598079356 | 1.516522298 | 1.53772706 | 1.653212514 | 1.433437704 |
| ENSMUSG00000045318 | 0 | 0 | 0 | 0 | 0 | 0 | 0 |
| ENSMUSG00000045319 | 1.129344651 | 0.896240625 | 1.098079356 | 1.098079356 | 1.091002363 | 0.84509804 | 1.089372236 |
| ENSMUSG00000045322 | 0 | 0 | 0 | 0 | 0 | 0 | 0 |
| ENSMUSG00000045326 | 0.578129653 | 0.75 | 0.646240625 | 0.79248125 | 0.568060967 | 0.477121255 | 0.672731473 |
| ENSMUSG00000045327 | 0.578129653 | 0.646240625 | 0.25 | 1.061981878 | 0.846351821 | 1.041392685 | 0.896975297 |
| ENSMUSG00000045328 | 2.915163385 | 2.700224975 | 2.693696765 | 2.875334905 | 2.78661101 | 3.206825876 | 2.576977086 |
| ENSMUSG00000045330 | 0 | 0.25 | 0 | 0 | 0 | 0 | 0 |
| ENSMUSG00000045331 | 0 | 0 | 0 | 0 | 0 | 0 | 0 |
| ENSMUSG00000045333 | 0 | 0 | 0 | 0.25 | 0 | 0.602059991 | 0.672731473 |
| ENSMUSG00000045336 | 0 | 0 | 0 | 0 | 0 | 0 | 0 |
| ENSMUSG00000045337 | 0 | 0 | 0 | 0 | 0 | 0 | 0 |
| ENSMUSG00000045339 | 0 | 0 | 0 | 0 | 0 | 0 | 0 |

| | | | | | | | |
|---|---|---|---|---|---|---|---|
| ENSMUSG00000045340 | 0 | 0 | 0 | 0 | 0 | 0 | 0 |
| ENSMUSG00000045341 | 0 | 0 | 0 | 0 | 0 | 0 | 0 |
| ENSMUSG00000045345 | 0.811507563 | 0.396240625 | 0.5 | 0.580482024 | 0.77552387 | 0.903089987 | 0.629532004 |
| ENSMUSG00000045348 | 1.721148722 | 1.76109853 | 1.83272922 | 1.410964047 | 1.542223377 | 1.612783857 | 1.694069894 |
| ENSMUSG00000045349 | 0.867194479 | 0.646240625 | 1.02186571 | 0.580482024 | 0.489301084 | 0.602059991 | 0.803905701 |
| ENSMUSG00000045350 | 0.289064826 | 0 | 0.25 | 0 | 0 | 0.602059991 | 0.355418052 |
| ENSMUSG00000045362 | 0.671187741 | 0.646240625 | 0.396240625 | 0.5 | 0.244650542 | 0 | 0.224243824 |
| ENSMUSG00000045364 | 0.289064826 | 0 | 0.25 | 0 | 0 | 0 | 0 |
| ENSMUSG00000045374 | 2.476144923 | 2.367918804 | 2.449415381 | 1.948603967 | 2.063536368 | 2.195899652 | 2.076440241 |
| ENSMUSG00000045377 | 1.227926429 | 1.201838731 | 1.364857905 | 1.54248125 | 1.698489937 | 1.832508913 | 1.661510159 |
| ENSMUSG00000045378 | 0 | 0 | 0 | 0.25 | 0 | 0 | 0 |
| ENSMUSG00000045381 | 0.45815691 | 0 | 0 | 0 | 0 | 0 | 0 |
| ENSMUSG00000045382 | 0.91631382 | 0.864857905 | 0.864857905 | 0.5 | 0.244650542 | 0.301029996 | 0 |
| ENSMUSG00000045391 | 0 | 0 | 0 | 0.25 | 0 | 0 | 0 |
| ENSMUSG00000045392 | 0 | 0 | 0 | 0 | 0 | 0 | 0 |
| ENSMUSG00000045394 | 0.289064826 | 0 | 0 | 0.896240625 | 0.733951626 | 0.77815125 | 0.896975297 |
| ENSMUSG00000045395 | 0 | 0 | 0 | 0 | 0 | 0 | 0 |
| ENSMUSG00000045404 | 0 | 0 | 0 | 0 | 0 | 0 | 0 |
| ENSMUSG00000045409 | 2.337801042 | 2.116901388 | 2.133818844 | 2.191217898 | 2.159419872 | 2.380211242 | 1.997315209 |
| ENSMUSG00000045410 | 2.003432385 | 1.79745614 | 1.777131114 | 1.937048212 | 1.832885921 | 2.170261715 | 1.760706232 |
| ENSMUSG00000045411 | 2.17479713 | 2.072504712 | 2.08272922 | 1.938721876 | 2.027028145 | 2.158362492 | 1.939944915 |
| ENSMUSG00000045414 | 2.395018353 | 2.253505118 | 2.216819685 | 2.425975893 | 2.358048572 | 2.672097858 | 2.154798988 |
| ENSMUSG00000045417 | 0 | 0 | 0 | 0 | 0 | 0 | 0 |
| ENSMUSG00000045421 | 0 | 0 | 0 | 0 | 0 | 0 | 0 |
| ENSMUSG00000045427 | 2.604828841 | 2.363317659 | 2.359697963 | 2.582168732 | 2.506289166 | 2.896526217 | 2.322617071 |
| ENSMUSG00000045435 | 2.010213549 | 1.856566189 | 1.850219859 | 1.782320754 | 1.763796674 | 1.995635195 | 1.603286636 |
| ENSMUSG00000045438 | 1.971473847 | 1.931980114 | 1.938721876 | 1.959800947 | 1.997548906 | 2.173186268 | 1.952571511 |
| ENSMUSG00000045440 | 0 | 0 | 0 | 0 | 0 | 0 | 0 |
| ENSMUSG00000045441 | 1.389637216 | 1.364857905 | 1.160964047 | 1.302363341 | 1.244650542 | 1.146128036 | 1.193409508 |
| ENSMUSG00000045455 | 0.289064826 | 0.25 | 0 | 0 | 0.387761935 | 0.301029996 | 0.224243824 |
| ENSMUSG00000045463 | 0 | 0 | 0 | 0 | 0 | 0 | 0 |
| ENSMUSG00000045464 | 1.445324132 | 1.388647213 | 1.27186571 | 1.364857905 | 1.335652905 | 1.397940009 | 1.245582246 |
| ENSMUSG00000045466 | 1.605636262 | 1.451838731 | 1.532320754 | 1.51109853 | 1.563883923 | 1.591064607 | 1.511732338 |
| ENSMUSG00000045467 | 0 | 0.25 | 0.25 | 0 | 0.387761935 | 0.602059991 | 0.355418052 |

| | | | | | | | |
|---|---|---|---|---|---|---|---|
| ENSMUSG00000045471 | 0 | 0 | 0 | 0 | 0 | 0 | 0 |
| ENSMUSG00000045474 | 0 | 0 | 0 | 0 | 0 | 0 | 0 |
| ENSMUSG00000045475 | 0 | 0 | 0 | 0 | 0 | 0 | 0 |
| ENSMUSG00000045479 | 0 | 0 | 0 | 0 | 0 | 0 | 0 |
| ENSMUSG00000045482 | 3.054549572 | 2.820916792 | 2.824945101 | 2.80388325 | 2.871420525 | 3.201397124 | 2.713130288 |
| ENSMUSG00000045493 | 0 | 0 | 0 | 0 | 0 | 0 | 0 |
| ENSMUSG00000045498 | 0 | 0.25 | 0 | 0 | 0 | 0.301029996 | 0 |
| ENSMUSG00000045502 | 0 | 0 | 0 | 0 | 0 | 0 | 0 |
| ENSMUSG00000045503 | 1.771760131 | 1.695339928 | 1.729715809 | 1.769203899 | 1.822942838 | 1.913813852 | 1.786277931 |
| ENSMUSG00000045506 | 0.289064826 | 0.580482024 | 0.5 | 0 | 0 | 0 | 0.448487648 |
| ENSMUSG00000045508 | 0 | 0 | 0 | 0 | 0 | 0 | 0 |
| ENSMUSG00000045509 | 0 | 0 | 0 | 0 | 0 | 0 | 0 |
| ENSMUSG00000045514 | 0 | 0 | 0 | 0 | 0 | 0 | 0 |
| ENSMUSG00000045515 | 0.289064826 | 0 | 0 | 0 | 0 | 0 | 0 |
| ENSMUSG00000045518 | 0.289064826 | 0 | 0.25 | 0.25 | 0 | 0 | 0.224243824 |
| ENSMUSG00000045519 | 1.806056082 | 1.755591953 | 1.738549078 | 1.763820609 | 1.754126039 | 1.924279286 | 1.594042709 |
| ENSMUSG00000045521 | 0 | 0 | 0.25 | 0.396240625 | 0.244650542 | 0 | 0.448487648 |
| ENSMUSG00000045528 | 0 | 0 | 0 | 0 | 0 | 0 | 0 |
| ENSMUSG00000045532 | 0.289064826 | 0.396240625 | 0.580482024 | 1.188721876 | 1.343584837 | 1.380211242 | 1.417653332 |
| ENSMUSG00000045534 | 0 | 0 | 0 | 0 | 0 | 0 | 0 |
| ENSMUSG00000045538 | 1.811507563 | 1.630890489 | 1.732684334 | 1.711372513 | 1.680838453 | 1.908485019 | 1.663407615 |
| ENSMUSG00000045539 | 0 | 0 | 0 | 0 | 0 | 0 | 0 |
| ENSMUSG00000045540 | 0 | 0 | 0 | 0 | 0 | 0 | 0 |
| ENSMUSG00000045545 | 0 | 0 | 0.25 | 1 | 1.319233381 | 0.954242509 | 1.365075947 |
| ENSMUSG00000045551 | 0.747221736 | 0.5 | 0.701838731 | 0.75 | 0.632412477 | 0 | 0.224243824 |
| ENSMUSG00000045555 | 0.289064826 | 0.396240625 | 0.396240625 | 0 | 0.387761935 | 0 | 0.224243824 |
| ENSMUSG00000045559 | 0 | 0 | 0 | 0 | 0 | 0 | 0 |
| ENSMUSG00000045566 | 0 | 0 | 0 | 0 | 0 | 0 | 0 |
| ENSMUSG00000045569 | 0 | 0 | 0 | 0 | 0 | 0 | 0 |
| ENSMUSG00000045573 | 1.307602651 | 1.372963274 | 1.380890489 | 0.646240625 | 0.812711509 | 0.477121255 | 0.672731473 |
| ENSMUSG00000045575 | 0 | 0 | 0 | 0 | 0 | 0 | 0 |
| ENSMUSG00000045576 | 1.686083339 | 1.593759858 | 1.708222504 | 1.888647213 | 1.877063019 | 2.033423755 | 1.726776051 |
| ENSMUSG00000045581 | 0 | 0 | 0 | 0 | 0 | 0 | 0 |
| ENSMUSG00000045584 | 0 | 0 | 0 | 0 | 0 | 0 | 0 |

| | | | | | | | |
|---|---|---|---|---|---|---|---|
| ENSMUSG00000045587 | 0 | 0 | 0 | 0 | 0 | 0 | 0 |
| ENSMUSG00000045589 | 0 | 0.25 | 0 | 0.580482024 | 0.489301084 | 0.602059991 | 0.672731473 |
| ENSMUSG00000045591 | 0 | 0.396240625 | 0 | 0 | 0 | 0 | 0 |
| ENSMUSG00000045594 | 2.404271433 | 2.190387808 | 2.206637122 | 2.267865591 | 2.233484039 | 2.551449998 | 2.103753068 |
| ENSMUSG00000045598 | 1.89911417 | 1.794977273 | 1.874961472 | 1.566696635 | 1.683842353 | 1.886490725 | 1.705896842 |
| ENSMUSG00000045608 | 0 | 0 | 0 | 0 | 0 | 0 | 0 |
| ENSMUSG00000045609 | 0 | 0 | 0 | 0 | 0 | 0 | 0 |
| ENSMUSG00000045613 | 0.289064826 | 0 | 0.25 | 0 | 0.244650542 | 0 | 0.224243824 |
| ENSMUSG00000045620 | 0 | 0 | 0 | 0 | 0 | 0 | 0 |
| ENSMUSG00000045624 | 2.543094501 | 2.368426437 | 2.38040011 | 2.403217124 | 2.32657161 | 2.631443769 | 2.113781657 |
| ENSMUSG00000045625 | 0.289064826 | 0.5 | 0.25 | 0.396240625 | 0.244650542 | 0 | 0.355418052 |
| ENSMUSG00000045629 | 1.227926429 | 1.17510993 | 1.330482024 | 0.5 | 0.387761935 | 0.301029996 | 0.224243824 |
| ENSMUSG00000045631 | 0 | 0 | 0.25 | 0 | 0 | 0 | 0 |
| ENSMUSG00000045636 | 2.036286563 | 1.864857905 | 1.892463902 | 1.552363341 | 1.504538967 | 1.707570176 | 1.329931221 |
| ENSMUSG00000045639 | 1.852729478 | 1.660964047 | 1.638647213 | 1.580482024 | 1.635855979 | 1.826074803 | 1.663407615 |
| ENSMUSG00000045648 | 0 | 0 | 0 | 0 | 0 | 0 | 0 |
| ENSMUSG00000045655 | 0 | 0 | 0 | 0 | 0 | 0 | 0 |
| ENSMUSG00000045657 | 0 | 0.396240625 | 0.25 | 0.25 | 0 | 0.301029996 | 0 |
| ENSMUSG00000045658 | 0 | 0.25 | 0 | 0.580482024 | 0.77552387 | 0.698970004 | 0.74492186 |
| ENSMUSG00000045659 | 0.747221736 | 0.5 | 0.75 | 0.701838731 | 0.632412477 | 0.477121255 | 0.916588299 |
| ENSMUSG00000045662 | 0.289064826 | 0.396240625 | 0.396240625 | 0 | 0 | 0 | 0.224243824 |
| ENSMUSG00000045664 | 1.985980784 | 1.890560606 | 1.972185812 | 1.747171172 | 1.773209395 | 1.982271233 | 1.712465909 |
| ENSMUSG00000045665 | 2.441754646 | 2.30539678 | 2.367410454 | 2.095926073 | 2.131103516 | 2.359835482 | 2.164590871 |
| ENSMUSG00000045667 | 0.578129653 | 0.830482024 | 0.646240625 | 1.098079356 | 1.176121988 | 1.079181246 | 1.185218806 |
| ENSMUSG00000045671 | 2.081172036 | 2.009729747 | 1.973704441 | 2.060793496 | 2.043601899 | 2.342422681 | 1.927642465 |
| ENSMUSG00000045672 | 2.023453784 | 1.990000483 | 1.994319981 | 1.146240625 | 1.293076518 | 1.204119983 | 1.245582246 |
| ENSMUSG00000045678 | 0 | 0 | 0 | 0 | 0 | 0 | 0 |
| ENSMUSG00000045679 | 1.940852214 | 1.726722649 | 1.807204673 | 1.848079356 | 1.756568651 | 1.986771734 | 1.651852264 |
| ENSMUSG00000045680 | 0 | 0 | 0 | 0 | 0 | 0 | 0.224243824 |
| ENSMUSG00000045684 | 0 | 0 | 0 | 0 | 0 | 0 | 0 |
| ENSMUSG00000045689 | 0 | 0 | 0 | 0 | 0 | 0 | 0 |
| ENSMUSG00000045690 | 1.269664473 | 0.976722649 | 1.061981878 | 0.92510993 | 1.020174412 | 0.903089987 | 0.952571511 |
| ENSMUSG00000045691 | 1.623015126 | 1.505591953 | 1.451838731 | 1.841580554 | 1.873585591 | 2.045322979 | 1.732938436 |
| ENSMUSG00000045693 | 0 | 0 | 0 | 0 | 0 | 0 | 0 |

| | | | | | | | |
|---|---|---|---|---|---|---|---|
| ENSMUSG00000045694 | 0 | 0.25 | 0.25 | 0 | 0.244650542 | 0 | 0 |
| ENSMUSG00000045699 | 0.289064826 | 0.25 | 0 | 0.25 | 0.244650542 | 0 | 0.355418052 |
| ENSMUSG00000045708 | 0 | 0 | 0 | 0 | 0 | 0 | 0 |
| ENSMUSG00000045709 | 0 | 0.25 | 0 | 0.25 | 0.244650542 | 0 | 0.355418052 |
| ENSMUSG00000045713 | 0 | 0 | 0 | 0 | 0 | 0 | 0 |
| ENSMUSG00000045725 | 0 | 0 | 0 | 0.25 | 0 | 0 | 0 |
| ENSMUSG00000045730 | 1.470606563 | 1.388647213 | 1.302363341 | 1.02186571 | 0.877063019 | 0.77815125 | 0.876096088 |
| ENSMUSG00000045731 | 0 | 0 | 0 | 0 | 0 | 0 | 0.224243824 |
| ENSMUSG00000045733 | 0 | 0 | 0 | 0 | 0 | 0 | 0 |
| ENSMUSG00000045744 | 0 | 0 | 0 | 0 | 0 | 0 | 0 |
| ENSMUSG00000045746 | 0.289064826 | 0 | 0.25 | 0.396240625 | 0.489301084 | 0.477121255 | 0.520678036 |
| ENSMUSG00000045751 | 2.432083896 | 2.237821179 | 2.273849256 | 2.319031101 | 2.256066288 | 2.523746467 | 2.071119143 |
| ENSMUSG00000045752 | 1.960252568 | 1.923371739 | 2.01109853 | 1.62694866 | 1.802174723 | 1.913813852 | 1.790137008 |
| ENSMUSG00000045757 | 0.289064826 | 0.75 | 0.646240625 | 0.896240625 | 1.020174412 | 1.079181246 | 0.935079929 |
| ENSMUSG00000045761 | 0 | 0.5 | 0.5 | 0.646240625 | 0.905314583 | 0.77815125 | 0.829800754 |
| ENSMUSG00000045763 | 2.724194991 | 2.508011432 | 2.498941788 | 2.498234584 | 2.403681482 | 2.804820679 | 2.277722321 |
| ENSMUSG00000045767 | 3.022564273 | 2.768535366 | 2.740181499 | 2.863703337 | 2.782244487 | 3.223495941 | 2.566705891 |
| ENSMUSG00000045775 | 1.181541736 | 0.75 | 0.896240625 | 0.701838731 | 0.568060967 | 0.84509804 | 0.876096088 |
| ENSMUSG00000045776 | 0.289064826 | 0.396240625 | 0 | 0.25 | 0 | 0 | 0.224243824 |
| ENSMUSG00000045777 | 0.45815691 | 0.5 | 0.580482024 | 0.25 | 0.244650542 | 0.301029996 | 0.579661877 |
| ENSMUSG00000045780 | 0 | 0 | 0 | 0 | 0 | 0 | 0 |
| ENSMUSG00000045790 | 1.418409478 | 1.214495249 | 1.29248125 | 1.451838731 | 1.462344774 | 1.380211242 | 1.313616061 |
| ENSMUSG00000045792 | 0 | 0 | 0 | 0 | 0 | 0 | 0 |
| ENSMUSG00000045794 | 0 | 0 | 0 | 0 | 0 | 0 | 0 |
| ENSMUSG00000045795 | 1.432083896 | 1.201838731 | 1.410964047 | 1.380890489 | 1.407936347 | 1.431363764 | 1.42167221 |
| ENSMUSG00000045797 | 0 | 0 | 0 | 0 | 0 | 0 | 0 |
| ENSMUSG00000045799 | 0 | 0 | 0 | 0 | 0 | 0 | 0.224243824 |
| ENSMUSG00000045802 | 0 | 0 | 0 | 0 | 0 | 0 | 0 |
| ENSMUSG00000045812 | 0 | 0 | 0 | 0 | 0 | 0 | 0 |
| ENSMUSG00000045813 | 0 | 0 | 0 | 0 | 0 | 0 | 0 |
| ENSMUSG00000045815 | 0 | 0 | 0 | 0 | 0 | 0 | 0 |
| ENSMUSG00000045817 | 3.023157491 | 2.745213402 | 2.765173983 | 2.575374049 | 2.49445382 | 2.85672889 | 2.31209124 |
| ENSMUSG00000045822 | 1.578129653 | 1.410964047 | 1.482684334 | 1.580482024 | 1.588235379 | 1.662757832 | 1.452143229 |
| ENSMUSG00000045824 | 0 | 0 | 0 | 0 | 0 | 0 | 0 |

| | | | | | | | |
|---|---|---|---|---|---|---|---|
| ENSMUSG00000045826 | 0 | 0.25 | 0.25 | 0.5 | 0.244650542 | 0.477121255 | 0.448487648 |
| ENSMUSG00000045827 | 2.399009124 | 2.203445298 | 2.140560606 | 2.299304173 | 2.189936658 | 2.53529412 | 2.064068234 |
| ENSMUSG00000045835 | 0 | 0 | 0 | 0 | 0 | 0 | 0 |
| ENSMUSG00000045838 | 2.134369199 | 1.956637122 | 2.005591953 | 2.105516192 | 2.099395374 | 2.264817823 | 1.995964412 |
| ENSMUSG00000045842 | 0 | 0 | 0 | 0 | 0 | 0 | 0 |
| ENSMUSG00000045854 | 1.432083896 | 1.396240625 | 1.482684334 | 1.431980114 | 1.358933174 | 1.568201724 | 1.38802993 |
| ENSMUSG00000045867 | 1.100572389 | 0.976722649 | 1.17510993 | 1 | 1.057362051 | 1.176091259 | 1.159323753 |
| ENSMUSG00000045868 | 0 | 0 | 0 | 0 | 0 | 0 | 0 |
| ENSMUSG00000045871 | 0 | 0 | 0 | 0 | 0 | 0 | 0 |
| ENSMUSG00000045875 | 0 | 0.25 | 0 | 0 | 0 | 0 | 0 |
| ENSMUSG00000045876 | 0.289064826 | 0.25 | 0 | 0.396240625 | 0 | 0 | 0.355418052 |
| ENSMUSG00000045877 | 0 | 0 | 0.25 | 0 | 0 | 0.301029996 | 0.520678036 |
| ENSMUSG00000045883 | 0 | 0 | 0 | 0 | 0 | 0 | 0 |
| ENSMUSG00000045886 | 0 | 0 | 0 | 0 | 0 | 0 | 0 |
| ENSMUSG00000045896 | 2.583267694 | 2.376452888 | 2.341033664 | 2.489638179 | 2.40985355 | 2.767155866 | 2.256654198 |
| ENSMUSG00000045903 | 1.890240806 | 1.884789703 | 1.905512955 | 0.646240625 | 0.632412477 | 0.301029996 | 0.355418052 |
| ENSMUSG00000045912 | 0 | 0 | 0 | 0.580482024 | 0.632412477 | 0.301029996 | 0.775756176 |
| ENSMUSG00000045915 | 0 | 0 | 0 | 0 | 0 | 0 | 0 |
| ENSMUSG00000045917 | 1.91631382 | 1.732684334 | 1.729715809 | 1.330482024 | 1.394615667 | 1.431363764 | 1.444790289 |
| ENSMUSG00000045928 | 0 | 0 | 0 | 0 | 0 | 0 | 0 |
| ENSMUSG00000045929 | 0 | 0 | 0 | 0 | 0.244650542 | 0 | 0 |
| ENSMUSG00000045930 | 0.578129653 | 0.830482024 | 0.701838731 | 0 | 0 | 0 | 0 |
| ENSMUSG00000045932 | 2.614416215 | 2.388647213 | 2.385757955 | 2.225970461 | 2.154724156 | 2.477121255 | 1.991191705 |
| ENSMUSG00000045934 | 1.822201287 | 1.723704441 | 1.738549078 | 1.685366747 | 1.715300549 | 1.959041392 | 1.651852264 |
| ENSMUSG00000045942 | 0 | 0.25 | 0 | 0.5 | 0 | 0 | 0.224243824 |
| ENSMUSG00000045946 | 0 | 0 | 0 | 0 | 0 | 0 | 0.224243824 |
| ENSMUSG00000045948 | 1.227926429 | 1.29248125 | 1.552363341 | 1.02186571 | 1.223252711 | 1.113943352 | 1.523605797 |
| ENSMUSG00000045952 | 0 | 0 | 0 | 0 | 0 | 0 | 0.355418052 |
| ENSMUSG00000045954 | 3.054274575 | 2.842671852 | 2.833987141 | 2.371960008 | 2.334234459 | 2.607455023 | 2.151886289 |
| ENSMUSG00000045962 | 3.421613914 | 3.09921808 | 3.067938701 | 3.121142795 | 3.029944058 | 3.512684396 | 2.813509528 |
| ENSMUSG00000045967 | 0 | 0 | 0 | 0 | 0 | 0 | 0 |
| ENSMUSG00000045968 | 0.289064826 | 0.396240625 | 0.5 | 0.5 | 0.632412477 | 0.301029996 | 0.579661877 |
| ENSMUSG00000045969 | 2.526850636 | 2.357613138 | 2.368933358 | 2.159156155 | 2.170892913 | 2.426511261 | 2.053478496 |
| ENSMUSG00000045973 | 2.473938397 | 2.302363341 | 2.325945187 | 2.30479213 | 2.254881871 | 2.523746467 | 2.159323753 |

| | | | | | | | |
|---|---|---|---|---|---|---|---|
| ENSMUSG00000045975 | 2.086807695 | 1.866901388 | 1.83272922 | 1.854463129 | 1.802174723 | 1.977723605 | 1.705896842 |
| ENSMUSG00000045980 | 1.753493018 | 1.816696635 | 1.852347734 | 1.470660762 | 1.746695109 | 1.681241237 | 1.657681529 |
| ENSMUSG00000045983 | 3.589030555 | 3.264878366 | 3.28001896 | 3.248234584 | 3.213736824 | 3.693287157 | 3.017820827 |
| ENSMUSG00000045989 | 0 | 0 | 0 | 0 | 0 | 0 | 0 |
| ENSMUSG00000045991 | 1.505869732 | 1.339388001 | 1.238549078 | 1.04248125 | 0.931471446 | 1.204119983 | 0.952571511 |
| ENSMUSG00000045994 | 0 | 0 | 0 | 0 | 0 | 0 | 0 |
| ENSMUSG00000045996 | 2.066740409 | 1.882845365 | 1.884789703 | 2.025821952 | 1.955822902 | 2.161368002 | 1.812372009 |
| ENSMUSG00000046000 | 0 | 0 | 0 | 0 | 0 | 0 | 0 |
| ENSMUSG00000046005 | 0 | 0 | 0 | 0.396240625 | 0.489301084 | 0 | 0.355418052 |
| ENSMUSG00000046006 | 0 | 0 | 0 | 0 | 0 | 0 | 0 |
| ENSMUSG00000046008 | 0 | 0.25 | 0 | 0 | 0 | 0 | 0 |
| ENSMUSG00000046010 | 1.862428343 | 1.825945187 | 1.79745614 | 1.839388001 | 1.842556557 | 2.021189299 | 1.737484395 |
| ENSMUSG00000046016 | 0 | 0 | 0 | 0 | 0.244650542 | 0 | 0 |
| ENSMUSG00000046020 | 2.352857998 | 2.180274797 | 2.174241882 | 2.164552871 | 2.186353316 | 2.396199347 | 2.03601839 |
| ENSMUSG00000046027 | 1.54867984 | 1.482684334 | 1.532320754 | 1.396240625 | 1.427019704 | 1.505149978 | 1.365075947 |
| ENSMUSG00000046031 | 0 | 0 | 0 | 0 | 0 | 0 | 0 |
| ENSMUSG00000046032 | 2.647066901 | 2.441632227 | 2.465659339 | 2.469129237 | 2.391406953 | 2.736396502 | 2.229876084 |
| ENSMUSG00000046034 | 2.358731791 | 2.190387808 | 2.176839783 | 2.247880462 | 2.212046813 | 2.489958479 | 2.01501956 |
| ENSMUSG00000046041 | 0 | 0 | 0 | 0 | 0 | 0 | 0 |
| ENSMUSG00000046049 | 0 | 0 | 0 | 0 | 0 | 0 | 0.224243824 |
| ENSMUSG00000046056 | 0.867194479 | 0.646240625 | 0.75 | 1.114857905 | 1.121713561 | 1.146128036 | 1.110947927 |
| ENSMUSG00000046057 | 0 | 0 | 0 | 0 | 0 | 0 | 0 |
| ENSMUSG00000046058 | 1.227926429 | 1.282320754 | 1.27186571 | 1.339388001 | 1.254881871 | 1.301029996 | 1.238624656 |
| ENSMUSG00000046062 | 2.85848785 | 2.558105232 | 2.54248125 | 2.626452888 | 2.565117605 | 2.938019097 | 2.378628314 |
| ENSMUSG00000046070 | 0 | 0 | 0 | 0 | 0 | 0 | 0 |
| ENSMUSG00000046079 | 2.026699189 | 1.886723615 | 1.987091808 | 1.87694866 | 1.882215755 | 2.037426498 | 1.779742043 |
| ENSMUSG00000046080 | 0 | 0 | 0 | 0 | 0 | 0 | 0.224243824 |
| ENSMUSG00000046082 | 0 | 0 | 0 | 0 | 0 | 0 | 0 |
| ENSMUSG00000046085 | 0 | 0 | 0 | 0 | 0 | 0 | 0.224243824 |
| ENSMUSG00000046088 | 0 | 0 | 0.25 | 0 | 0 | 0 | 0 |
| ENSMUSG00000046093 | 0 | 0.25 | 0 | 0.646240625 | 0.568060967 | 0.77815125 | 0.74492186 |
| ENSMUSG00000046095 | 0 | 0 | 0 | 0 | 0 | 0 | 0 |
| ENSMUSG00000046096 | 2.177057474 | 2.058404919 | 2.038704527 | 2.091580554 | 2.040348824 | 2.303196057 | 1.880961074 |
| ENSMUSG00000046101 | 0.671187741 | 0.25 | 0.5 | 0.75 | 0.244650542 | 0.301029996 | 0.520678036 |

| | | | | | | | |
|---|---|---|---|---|---|---|---|
| ENSMUSG00000046108 | 0 | 0 | 0 | 0 | 0.244650542 | 0.301029996 | 0.355418052 |
| ENSMUSG00000046109 | 0 | 0 | 0 | 0 | 0 | 0 | 0 |
| ENSMUSG00000046110 | 0 | 0 | 0 | 0 | 0 | 0 | 0 |
| ENSMUSG00000046111 | 2.320585253 | 2.157339155 | 2.138647213 | 2.300530956 | 2.196996016 | 2.517195898 | 2.064616101 |
| ENSMUSG00000046119 | 0 | 0 | 0 | 0 | 0 | 0 | 0 |
| ENSMUSG00000046130 | 0 | 0 | 0 | 0 | 0 | 0 | 0 |
| ENSMUSG00000046133 | 0 | 0 | 0 | 0.25 | 0.244650542 | 0.301029996 | 0 |
| ENSMUSG00000046138 | 2.526850636 | 2.334962501 | 2.354463129 | 2.302972074 | 2.212046813 | 2.552668216 | 2.053478496 |
| ENSMUSG00000046139 | 2.74143756 | 2.528435542 | 2.544666213 | 2.53456795 | 2.476871767 | 2.858537198 | 2.296701469 |
| ENSMUSG00000046145 | 0 | 0 | 0 | 0 | 0 | 0 | 0 |
| ENSMUSG00000046150 | 0 | 0 | 0 | 0 | 0 | 0 | 0 |
| ENSMUSG00000046152 | 2.078325387 | 1.994319981 | 1.921625132 | 1.933677405 | 1.911645804 | 2.123851641 | 1.788855757 |
| ENSMUSG00000046157 | 0.867194479 | 0.646240625 | 0.5 | 1.226722649 | 1.212046813 | 1.301029996 | 1.20139795 |
| ENSMUSG00000046159 | 0 | 0 | 0 | 0 | 0 | 0 | 0 |
| ENSMUSG00000046160 | 0 | 0 | 0 | 0 | 0 | 0 | 0 |
| ENSMUSG00000046167 | 0 | 0.25 | 0.25 | 0 | 0 | 0 | 0 |
| ENSMUSG00000046168 | 0 | 0.25 | 0.25 | 0.5 | 0.489301084 | 0.301029996 | 0.629532004 |
| ENSMUSG00000046169 | 2.023453784 | 1.845926073 | 1.752806814 | 1.726722649 | 1.746695109 | 1.897627091 | 1.591689859 |
| ENSMUSG00000046173 | 0 | 0 | 0 | 0 | 0 | 0 | 0 |
| ENSMUSG00000046178 | 0 | 0 | 0 | 0 | 0 | 0.301029996 | 0 |
| ENSMUSG00000046179 | 2.069666964 | 1.948603967 | 1.975216702 | 1.970660762 | 1.973373093 | 2.178976947 | 1.893277937 |
| ENSMUSG00000046180 | 0 | 0 | 0 | 0 | 0 | 0 | 0 |
| ENSMUSG00000046182 | 0 | 0.25 | 0 | 0.25 | 0 | 0 | 0 |
| ENSMUSG00000046185 | 1.678702042 | 1.516522298 | 1.561981878 | 1.580482024 | 1.652586889 | 1.748188027 | 1.586932193 |
| ENSMUSG00000046186 | 2.849595564 | 2.596195938 | 2.603671309 | 2.596195938 | 2.482519186 | 2.868644438 | 2.319395519 |
| ENSMUSG00000046191 | 0.867194479 | 1.160964047 | 0.830482024 | 0 | 0.387761935 | 0 | 0.224243824 |
| ENSMUSG00000046192 | 0.289064826 | 0 | 0 | 0 | 0.387761935 | 0.301029996 | 0.224243824 |
| ENSMUSG00000046196 | 0.289064826 | 0.25 | 0.396240625 | 0 | 0.387761935 | 0.477121255 | 0.355418052 |
| ENSMUSG00000046201 | 2.8302808 | 2.560197833 | 2.526149688 | 2.563166358 | 2.487771146 | 2.877946952 | 2.316393034 |
| ENSMUSG00000046203 | 0 | 0 | 0 | 0 | 0 | 0 | 0 |
| ENSMUSG00000046204 | 0.45815691 | 0.396240625 | 0.646240625 | 0.976722649 | 1.223252711 | 1.113943352 | 1.100339912 |
| ENSMUSG00000046207 | 0 | 0 | 0.25 | 0.396240625 | 0.387761935 | 0 | 0 |
| ENSMUSG00000046210 | 0 | 0 | 0 | 0 | 0 | 0 | 0 |
| ENSMUSG00000046213 | 0 | 0 | 0 | 0 | 0 | 0 | 0 |

| | | | | | | | |
|---|---|---|---|---|---|---|---|
| ENSMUSG00000046215 | 0 | 0.25 | 0 | 0 | 0 | 0 | 0 |
| ENSMUSG00000046219 | 0 | 0.580482024 | 0.396240625 | 0.5 | 0.489301084 | 0 | 0.224243824 |
| ENSMUSG00000046223 | 2.251208701 | 2.149978211 | 2.237821179 | 1.642463902 | 1.728722551 | 1.799340549 | 1.74196736 |
| ENSMUSG00000046229 | 0.578129653 | 0.5 | 0.646240625 | 0 | 0 | 0.301029996 | 0.710836105 |
| ENSMUSG00000046230 | 2.730819472 | 2.518869787 | 2.501406137 | 2.616391601 | 2.528285562 | 2.883093359 | 2.331614323 |
| ENSMUSG00000046240 | 0.289064826 | 0 | 0 | 0 | 0 | 0 | 0 |
| ENSMUSG00000046242 | 0 | 0 | 0 | 0 | 0 | 0 | 0 |
| ENSMUSG00000046245 | 0.289064826 | 0.646240625 | 0.75 | 0.25 | 0 | 0.301029996 | 0 |
| ENSMUSG00000046248 | 0 | 0 | 0 | 0 | 0 | 0 | 0 |
| ENSMUSG00000046258 | 0 | 0 | 0 | 0 | 0 | 0 | 0 |
| ENSMUSG00000046259 | 0 | 0 | 0 | 0 | 0 | 0 | 0 |
| ENSMUSG00000046262 | 0 | 0 | 0 | 0 | 0 | 0 | 0 |
| ENSMUSG00000046269 | 1.568542279 | 1.364857905 | 1.505591953 | 1.321350555 | 1.414412788 | 1.477121255 | 1.245582246 |
| ENSMUSG00000046272 | 0 | 0 | 0 | 0 | 0 | 0 | 0 |
| ENSMUSG00000046275 | 0 | 0 | 0.25 | 0.25 | 0 | 0 | 0 |
| ENSMUSG00000046280 | 0.289064826 | 0.5 | 0.5 | 1.646240625 | 1.649303553 | 1.579783597 | 1.603286636 |
| ENSMUSG00000046282 | 0 | 0 | 0 | 0 | 0 | 0 | 0 |
| ENSMUSG00000046287 | 0 | 0 | 0 | 0 | 0 | 0 | 0 |
| ENSMUSG00000046295 | 1.418409478 | 1.364857905 | 1.557204673 | 1.364857905 | 1.504538967 | 1.301029996 | 1.508694612 |
| ENSMUSG00000046300 | 0 | 0 | 0 | 0 | 0 | 0 | 0 |
| ENSMUSG00000046311 | 2.331622671 | 2.163659007 | 2.149978211 | 2.294354884 | 2.199780694 | 2.488550717 | 2.032410373 |
| ENSMUSG00000046312 | 1.578129653 | 1.552363341 | 1.552363341 | 1.71759118 | 1.645989387 | 1.924279286 | 1.700880998 |
| ENSMUSG00000046314 | 1.876566388 | 1.75 | 1.747171172 | 1.494319981 | 1.439191811 | 1.662757832 | 1.313616061 |
| ENSMUSG00000046317 | 0 | 0 | 0 | 0 | 0 | 0.477121255 | 0.224243824 |
| ENSMUSG00000046318 | 1.777675583 | 1.75835575 | 1.741446071 | 0.864857905 | 0.686820904 | 0.954242509 | 0.520678036 |
| ENSMUSG00000046321 | 0 | 0 | 0 | 0 | 0 | 0 | 0 |
| ENSMUSG00000046323 | 0 | 0 | 0 | 0 | 0 | 0 | 0 |
| ENSMUSG00000046324 | 2.614416215 | 2.359697963 | 2.32991803 | 2.537754135 | 2.448224646 | 2.843232778 | 2.282781745 |
| ENSMUSG00000046329 | 1.87190676 | 1.708222504 | 1.766522298 | 1.814346961 | 1.806427258 | 2.06069784 | 1.753626837 |
| ENSMUSG00000046330 | 2.773381775 | 2.527457664 | 2.571061437 | 2.494677615 | 2.534518346 | 2.878521796 | 2.402025166 |
| ENSMUSG00000046334 | 0 | 0 | 0 | 0 | 0 | 0 | 0.224243824 |
| ENSMUSG00000046337 | 0 | 0 | 0 | 0.25 | 0.244650542 | 0.301029996 | 0 |
| ENSMUSG00000046338 | 0 | 0 | 0 | 0 | 0.244650542 | 0 | 0.355418052 |
| ENSMUSG00000046341 | 0 | 0 | 0.25 | 0 | 0 | 0 | 0.224243824 |

| | | | | | | | |
|---|---|---|---|---|---|---|---|
| ENSMUSG00000046345 | 0.578129653 | 0.396240625 | 0.396240625 | 0.396240625 | 0.568060967 | 0 | 0.579661877 |
| ENSMUSG00000046351 | 2.167941649 | 2.090485943 | 2.029735268 | 2.176839783 | 2.089109832 | 2.346352974 | 1.939139147 |
| ENSMUSG00000046352 | 0.91631382 | 0.75 | 0.580482024 | 0.5 | 0.244650542 | 0.301029996 | 0.355418052 |
| ENSMUSG00000046354 | 0 | 0 | 0 | 0 | 0 | 0 | 0 |
| ENSMUSG00000046364 | 2.837282068 | 2.609697963 | 2.670967967 | 2.645300145 | 2.58335733 | 2.953759692 | 2.462228489 |
| ENSMUSG00000046367 | 0 | 0 | 0 | 0 | 0 | 0 | 0 |
| ENSMUSG00000046372 | 0.45815691 | 0.580482024 | 0.396240625 | 0.396240625 | 0.489301084 | 0 | 0.448487648 |
| ENSMUSG00000046378 | 0.45815691 | 0.25 | 0.396240625 | 0 | 0.387761935 | 0.698970004 | 0.224243824 |
| ENSMUSG00000046380 | 1.960252568 | 1.848079356 | 1.852347734 | 1.76109853 | 1.806427258 | 1.959041392 | 1.746389053 |
| ENSMUSG00000046387 | 1.53838222 | 1.51109853 | 1.516522298 | 0.646240625 | 0.387761935 | 0.602059991 | 0.803905701 |
| ENSMUSG00000046388 | 0 | 0 | 0 | 0 | 0 | 0 | 0 |
| ENSMUSG00000046396 | 0 | 0 | 0 | 0 | 0 | 0 | 0 |
| ENSMUSG00000046402 | 1.578129653 | 1.458222504 | 1.681980114 | 1.752806814 | 1.758994476 | 1.84509804 | 1.725216937 |
| ENSMUSG00000046404 | 1.944805158 | 1.769203899 | 1.735628626 | 1.87694866 | 1.826953746 | 2.117271296 | 1.669034113 |
| ENSMUSG00000046408 | 0 | 0.396240625 | 0.5 | 0.5 | 0.244650542 | 0.301029996 | 0.579661877 |
| ENSMUSG00000046410 | 0.578129653 | 0.25 | 0.5 | 1.146240625 | 1.212046813 | 1.113943352 | 1.131174228 |
| ENSMUSG00000046411 | 0 | 0 | 0 | 0 | 0 | 0 | 0 |
| ENSMUSG00000046413 | 0 | 0.396240625 | 0.396240625 | 0 | 0.387761935 | 0 | 0.224243824 |
| ENSMUSG00000046415 | 0.289064826 | 0.646240625 | 0.396240625 | 0.646240625 | 0.244650542 | 0.477121255 | 0.448487648 |
| ENSMUSG00000046417 | 1.862428343 | 1.741446071 | 1.735628626 | 1.685366747 | 1.701347897 | 1.86332286 | 1.594042709 |
| ENSMUSG00000046431 | 0 | 0 | 0 | 0 | 0 | 0 | 0 |
| ENSMUSG00000046432 | 2.619128496 | 2.444108258 | 2.48927551 | 2.54961126 | 2.510339527 | 2.862727528 | 2.414327616 |
| ENSMUSG00000046434 | 3.699002226 | 3.319576063 | 3.296141363 | 3.370827989 | 3.280776929 | 3.839540893 | 3.034738325 |
| ENSMUSG00000046435 | 0 | 0.25 | 0 | 0 | 0 | 0 | 0.224243824 |
| ENSMUSG00000046438 | 0 | 0 | 0 | 0 | 0 | 0 | 0 |
| ENSMUSG00000046440 | 0.45815691 | 0.396240625 | 0.25 | 0.396240625 | 0.387761935 | 0 | 0.448487648 |
| ENSMUSG00000046441 | 2.103271637 | 1.910964047 | 1.914552871 | 1.87892496 | 1.86473911 | 2.06069784 | 1.728327686 |
| ENSMUSG00000046442 | 1.967766868 | 1.899978211 | 1.905512955 | 2.049918086 | 2.033751322 | 2.264817823 | 1.848480858 |
| ENSMUSG00000046447 | 0.91631382 | 1.04248125 | 0.951838731 | 0.951838731 | 0.733951626 | 0.954242509 | 0.935079929 |
| ENSMUSG00000046449 | 0.811507563 | 1.04248125 | 1.130890489 | 1.188721876 | 1.200473444 | 1.230448921 | 0.969165684 |
| ENSMUSG00000046450 | 0 | 0 | 0 | 0 | 0 | 0 | 0 |
| ENSMUSG00000046456 | 0 | 0 | 0 | 0 | 0 | 0 | 0 |
| ENSMUSG00000046460 | 0 | 0 | 0 | 0 | 0 | 0 | 0 |
| ENSMUSG00000046463 | 1.249317394 | 1.27186571 | 1.226722649 | 0.896240625 | 0.931471446 | 0.602059991 | 0.853775828 |

| | | | | | | | |
|---|---|---|---|---|---|---|---|
| ENSMUSG00000046470 | 0 | 0 | 0.25 | 0.25 | 0.244650542 | 0.477121255 | 0 |
| ENSMUSG00000046474 | 0 | 0 | 0 | 0 | 0 | 0 | 0 |
| ENSMUSG00000046480 | 0 | 0.25 | 0 | 0.396240625 | 0 | 0 | 0 |
| ENSMUSG00000046486 | 0 | 0 | 0 | 0 | 0 | 0 | 0 |
| ENSMUSG00000046487 | 0 | 0.25 | 0 | 0.25 | 0.244650542 | 0 | 0 |
| ENSMUSG00000046490 | 0 | 0 | 0 | 0 | 0 | 0 | 0 |
| ENSMUSG00000046491 | 0 | 0 | 0 | 0 | 0.387761935 | 0 | 0.355418052 |
| ENSMUSG00000046493 | 0 | 0 | 0 | 0 | 0 | 0 | 0 |
| ENSMUSG00000046500 | 0 | 0 | 0 | 0 | 0 | 0 | 0 |
| ENSMUSG00000046516 | 1.671187741 | 1.598079356 | 1.638647213 | 1.42510993 | 1.53772706 | 1.568201724 | 1.401059159 |
| ENSMUSG00000046518 | 0 | 0 | 0 | 0 | 0 | 0 | 0 |
| ENSMUSG00000046519 | 2.139333929 | 2.019203899 | 1.994319981 | 2.192046081 | 2.131103516 | 2.42975228 | 1.971363985 |
| ENSMUSG00000046523 | 0 | 0 | 0 | 0.25 | 0 | 0.698970004 | 0 |
| ENSMUSG00000046532 | 2.023453784 | 1.862802778 | 1.87694866 | 1.431980114 | 1.254881871 | 1.477121255 | 1.140832123 |
| ENSMUSG00000046541 | 1.445324132 | 1.445339928 | 1.470660762 | 1.330482024 | 1.499532413 | 1.278753601 | 1.476637173 |
| ENSMUSG00000046546 | 0.91631382 | 0.864857905 | 1 | 1.27186571 | 1.467903253 | 1.505149978 | 1.532233387 |
| ENSMUSG00000046550 | 0.747221736 | 0.830482024 | 0.701838731 | 1.226722649 | 1.106691885 | 1.176091259 | 1.078019652 |
| ENSMUSG00000046556 | 1.876566388 | 1.839388001 | 1.868933358 | 1.837182039 | 1.87878897 | 1.954242509 | 1.847411384 |
| ENSMUSG00000046561 | 2.013563224 | 1.87694866 | 1.894357207 | 1.789967834 | 1.683842353 | 1.86923172 | 1.618853957 |
| ENSMUSG00000046562 | 2.529763563 | 2.39151351 | 2.40136988 | 2.394828984 | 2.453670557 | 2.743509765 | 2.326538518 |
| ENSMUSG00000046567 | 1.740854706 | 1.688721876 | 1.67510993 | 1.763820609 | 1.698489937 | 1.908485019 | 1.562034107 |
| ENSMUSG00000046572 | 2.295901576 | 2.148114259 | 2.143411797 | 2.08272922 | 1.983754904 | 2.303196057 | 1.857950279 |
| ENSMUSG00000046573 | 1.358731791 | 1.372963274 | 1.388647213 | 1.566696635 | 1.433158226 | 1.612783857 | 1.307989563 |
| ENSMUSG00000046574 | 1.952600383 | 1.976722649 | 2.129909063 | 1.602347734 | 1.800029087 | 1.633468456 | 1.908707966 |
| ENSMUSG00000046580 | 0.289064826 | 0 | 0.25 | 0 | 0.244650542 | 0 | 0.224243824 |
| ENSMUSG00000046585 | 0 | 0 | 0 | 0 | 0 | 0 | 0 |
| ENSMUSG00000046589 | 1.894701088 | 1.872963274 | 1.837182039 | 1.858657057 | 1.812711509 | 1.949390007 | 1.775756176 |
| ENSMUSG00000046590 | 0 | 0 | 0 | 0 | 0 | 0 | 0 |
| ENSMUSG00000046591 | 2.194712275 | 2.013820609 | 2.012462137 | 2.214495249 | 2.108492685 | 2.387389826 | 1.98213686 |
| ENSMUSG00000046593 | 0 | 0 | 0 | 0 | 0 | 0 | 0 |
| ENSMUSG00000046598 | 0.45815691 | 0.396240625 | 0.5 | 1.330482024 | 1.489301084 | 1.361727836 | 1.329931221 |
| ENSMUSG00000046603 | 1.740854706 | 1.714495249 | 1.646240625 | 1.695339928 | 1.674752852 | 1.84509804 | 1.562034107 |
| ENSMUSG00000046605 | 1.596667477 | 1.610735874 | 1.516522298 | 1.660964047 | 1.645989387 | 1.770852012 | 1.526507301 |
| ENSMUSG00000046607 | 0 | 0 | 0.396240625 | 0 | 0 | 0 | 0.224243824 |

| | | | | | | | |
|---|---|---|---|---|---|---|---|
| ENSMUSG00000046610 | 0.671187741 | 0.701838731 | 0.396240625 | 0.25 | 0.387761935 | 0.301029996 | 0.520678036 |
| ENSMUSG00000046613 | 0 | 0.5 | 0.25 | 0.396240625 | 0.77552387 | 0 | 0.710836105 |
| ENSMUSG00000046615 | 0 | 0 | 0 | 0 | 0 | 0 | 0 |
| ENSMUSG00000046618 | 0.45815691 | 0.646240625 | 0.646240625 | 0.79248125 | 0.686820904 | 0.477121255 | 0.629532004 |
| ENSMUSG00000046620 | 0 | 0 | 0 | 0 | 0 | 0 | 0 |
| ENSMUSG00000046623 | 1.358731791 | 1.321350555 | 1.330482024 | 1.431980114 | 1.462344774 | 1.414973348 | 1.319146374 |
| ENSMUSG00000046634 | 0.289064826 | 0 | 0 | 0 | 0.244650542 | 0.477121255 | 0 |
| ENSMUSG00000046636 | 1.129344651 | 0.92510993 | 0.830482024 | 1.061981878 | 0.77552387 | 0.903089987 | 0.896975297 |
| ENSMUSG00000046637 | 0 | 0 | 0 | 0 | 0 | 0 | 0.355418052 |
| ENSMUSG00000046643 | 0 | 0 | 0 | 0 | 0 | 0 | 0 |
| ENSMUSG00000046650 | 0 | 0 | 0 | 0 | 0 | 0 | 0 |
| ENSMUSG00000046652 | 0 | 0 | 0 | 0 | 0 | 0 | 0 |
| ENSMUSG00000046658 | 1.811507563 | 1.755591953 | 1.752806814 | 1.622963274 | 1.66856048 | 1.681241237 | 1.61448206 |
| ENSMUSG00000046667 | 0.45815691 | 0.79248125 | 0.396240625 | 1.080482024 | 0.978602168 | 1.079181246 | 0.896975297 |
| ENSMUSG00000046668 | 1.890240806 | 1.852347734 | 1.823655187 | 1.705044741 | 1.773209395 | 1.929418926 | 1.770364127 |
| ENSMUSG00000046671 | 2.397683106 | 2.222185812 | 2.234894804 | 2.247880462 | 2.179810173 | 2.436162647 | 2.078544414 |
| ENSMUSG00000046675 | 1.971473847 | 1.886723615 | 1.866901388 | 1.75835575 | 1.72073075 | 1.826074803 | 1.584526862 |
| ENSMUSG00000046676 | 0 | 0 | 0 | 0 | 0 | 0 | 0 |
| ENSMUSG00000046678 | 0 | 0 | 0 | 0 | 0 | 0 | 0 |
| ENSMUSG00000046679 | 1.827447047 | 1.738549078 | 1.848079356 | 1.634789703 | 1.635855979 | 1.77815125 | 1.612273755 |
| ENSMUSG00000046683 | 0 | 0 | 0 | 0 | 0 | 0 | 0 |
| ENSMUSG00000046687 | 0 | 0 | 0 | 0 | 0.244650542 | 0 | 0 |
| ENSMUSG00000046688 | 1.827447047 | 1.708222504 | 1.67510993 | 1.494319981 | 1.533172725 | 1.69019608 | 1.413583902 |
| ENSMUSG00000046691 | 2.904664113 | 2.698806992 | 2.704645544 | 2.530706999 | 2.595992969 | 2.910090546 | 2.462388606 |
| ENSMUSG00000046694 | 2.026699189 | 1.811981878 | 1.959800947 | 1.720660762 | 1.793512788 | 1.819543936 | 1.792684387 |
| ENSMUSG00000046697 | 0 | 0 | 0 | 0 | 0 | 0 | 0 |
| ENSMUSG00000046699 | 0 | 0 | 0 | 0 | 0.387761935 | 0 | 0 |
| ENSMUSG00000046707 | 2.497590902 | 2.322504712 | 2.309601185 | 2.379909063 | 2.27729019 | 2.664641976 | 2.164590871 |
| ENSMUSG00000046709 | 0.45815691 | 0.396240625 | 0.396240625 | 0.25 | 0.568060967 | 0 | 0.224243824 |
| ENSMUSG00000046711 | 1.982401085 | 1.972185812 | 2.039967834 | 1.646240625 | 1.888971102 | 1.707570176 | 1.922587433 |
| ENSMUSG00000046714 | 1.655740331 | 1.610735874 | 1.695339928 | 1.348079356 | 1.401338844 | 1.204119983 | 1.396774124 |
| ENSMUSG00000046716 | 0 | 0 | 0 | 0 | 0 | 0 | 0 |
| ENSMUSG00000046717 | 0 | 0 | 0 | 0 | 0 | 0 | 0 |
| ENSMUSG00000046718 | 1.753493018 | 1.741446071 | 1.779735268 | 1.884789703 | 1.866526233 | 2.021189299 | 1.947107911 |

| | | | | | | | |
|---|---|---|---|---|---|---|---|
| ENSMUSG00000046719 | 0.578129653 | 0.646240625 | 0.5 | 0.25 | 0 | 0.477121255 | 0.520678036 |
| ENSMUSG00000046721 | 0.289064826 | 0.580482024 | 0.701838731 | 0.701838731 | 0.812711509 | 0.477121255 | 0.775756176 |
| ENSMUSG00000046722 | 2.693989403 | 2.503853763 | 2.518535366 | 2.46759118 | 2.485926424 | 2.812244697 | 2.354302096 |
| ENSMUSG00000046723 | 0 | 0 | 0 | 0 | 0 | 0 | 0 |
| ENSMUSG00000046727 | 1.714367558 | 1.698603967 | 1.811981878 | 1.642463902 | 1.782377602 | 1.86332286 | 1.74492186 |
| ENSMUSG00000046731 | 1.960252568 | 1.843759858 | 1.79745614 | 1.755591953 | 1.814781639 | 2.029383778 | 1.685349273 |
| ENSMUSG00000046733 | 2.020182925 | 1.937048212 | 1.935366747 | 1.994319981 | 2.044679629 | 2.276461804 | 1.990504106 |
| ENSMUSG00000046743 | 2.63295404 | 2.416333979 | 2.357613138 | 1.747171172 | 1.731346772 | 1.944482672 | 1.647906904 |
| ENSMUSG00000046747 | 0.289064826 | 0 | 0.25 | 0 | 0.244650542 | 0.301029996 | 0 |
| ENSMUSG00000046748 | 0 | 0 | 0 | 0 | 0.244650542 | 0.301029996 | 0.355418052 |
| ENSMUSG00000046750 | 0 | 0 | 0 | 0 | 0 | 0 | 0 |
| ENSMUSG00000046753 | 1.876566388 | 1.784887838 | 1.79248125 | 2.036164561 | 1.957204337 | 2.201397124 | 1.802677937 |
| ENSMUSG00000046755 | 0 | 0 | 0 | 0 | 0 | 0 | 0 |
| ENSMUSG00000046756 | 2.4661223 | 2.333288838 | 2.361770807 | 2.381869252 | 2.429198453 | 2.698100546 | 2.314628629 |
| ENSMUSG00000046761 | 1.596667477 | 1.634789703 | 1.738549078 | 1.53743678 | 1.731346772 | 1.77815125 | 1.854824501 |
| ENSMUSG00000046764 | 0 | 0 | 0 | 0 | 0 | 0 | 0.224243824 |
| ENSMUSG00000046768 | 2.020182925 | 1.864857905 | 1.907339155 | 1.660964047 | 1.592135487 | 1.748188027 | 1.601000298 |
| ENSMUSG00000046774 | 0 | 0 | 0 | 0 | 0 | 0 | 0 |
| ENSMUSG00000046782 | 1 | 1.080482024 | 1.02186571 | 0 | 0 | 0 | 0 |
| ENSMUSG00000046785 | 2.640441358 | 2.460587586 | 2.442046081 | 2.522528105 | 2.469521578 | 2.781036939 | 2.264716375 |
| ENSMUSG00000046790 | 0 | 0 | 0 | 0 | 0 | 0 | 0 |
| ENSMUSG00000046791 | 1.876566388 | 1.766522298 | 1.76109853 | 1.638647213 | 1.639266209 | 1.880813592 | 1.586932193 |
| ENSMUSG00000046792 | 1.418409478 | 1.42510993 | 1.5 | 1.080482024 | 1.293076518 | 0.903089987 | 1.313616061 |
| ENSMUSG00000046793 | 0 | 0 | 0 | 0 | 0 | 0 | 0.355418052 |
| ENSMUSG00000046794 | 0.578129653 | 0.580482024 | 0 | 0.79248125 | 1 | 1.113943352 | 0.916588299 |
| ENSMUSG00000046798 | 2.6088504 | 2.388167274 | 2.36434522 | 2.444929839 | 2.377853744 | 2.709269961 | 2.235411964 |
| ENSMUSG00000046804 | 0 | 0 | 0 | 0 | 0 | 0 | 0 |
| ENSMUSG00000046805 | 0.747221736 | 0.396240625 | 0.646240625 | 0 | 0 | 0.301029996 | 0.224243824 |
| ENSMUSG00000046806 | 1 | 1.160964047 | 1.238549078 | 0.976722649 | 1.074582839 | 1 | 1.014380832 |
| ENSMUSG00000046807 | 2.339331437 | 2.115881093 | 2.132845365 | 1.784887838 | 1.701347897 | 1.908485019 | 1.680001789 |
| ENSMUSG00000046808 | 2.023453784 | 1.816696635 | 1.864857905 | 1.828220739 | 1.814781639 | 1.963787827 | 1.645916034 |
| ENSMUSG00000046811 | 0 | 0 | 0.25 | 0 | 0 | 0 | 0 |
| ENSMUSG00000046814 | 1.527823874 | 1.54248125 | 1.618933358 | 1.575945187 | 1.782377602 | 1.72427587 | 1.762103709 |
| ENSMUSG00000046818 | 2.045658471 | 1.843759858 | 1.83272922 | 1.04248125 | 1.020174412 | 1.041392685 | 0.853775828 |

| | | | | | | | |
|---|---|---|---|---|---|---|---|
| ENSMUSG00000046822 | 1.912079952 | 1.83272922 | 1.882845365 | 1.618933358 | 1.87878897 | 1.792391689 | 1.830929954 |
| ENSMUSG00000046826 | 0 | 0 | 0 | 0.25 | 0.244650542 | 0 | 0.448487648 |
| ENSMUSG00000046828 | 0 | 0.5 | 0 | 0.396240625 | 0 | 0 | 0.224243824 |
| ENSMUSG00000046834 | 0 | 0 | 0 | 0 | 0 | 0 | 0 |
| ENSMUSG00000046836 | 2.646335907 | 2.400906586 | 2.384789703 | 2.441632227 | 2.376595432 | 2.672097858 | 2.207438502 |
| ENSMUSG00000046840 | 0 | 0 | 0 | 0 | 0 | 0 | 0 |
| ENSMUSG00000046841 | 2.974482511 | 2.768535366 | 2.781837385 | 2.599151195 | 2.639864609 | 2.938019097 | 2.525981686 |
| ENSMUSG00000046844 | 0 | 0 | 0 | 0.25 | 0.244650542 | 0 | 0.355418052 |
| ENSMUSG00000046845 | 0 | 0 | 0 | 0 | 0 | 0 | 0 |
| ENSMUSG00000046846 | 0 | 0 | 0 | 0 | 0 | 0 | 0 |
| ENSMUSG00000046854 | 0 | 0.5 | 0.396240625 | 0.830482024 | 0.905314583 | 0.903089987 | 0.952571511 |
| ENSMUSG00000046856 | 0 | 0.25 | 0 | 0 | 0 | 0 | 0.224243824 |
| ENSMUSG00000046861 | 1.881174527 | 1.841580554 | 1.884789703 | 1.830482024 | 1.842556557 | 1.949390007 | 1.760706232 |
| ENSMUSG00000046862 | 2.192545868 | 2.046218836 | 1.992885888 | 1.990000483 | 1.945998439 | 2.161368002 | 1.843097781 |
| ENSMUSG00000046865 | 2.205378646 | 2.122963274 | 2.151832578 | 2.05357978 | 2.174635855 | 2.348304863 | 2.109521699 |
| ENSMUSG00000046873 | 2.251208701 | 2.079353153 | 2.080482024 | 2.344302633 | 2.272808147 | 2.605305046 | 2.069505596 |
| ENSMUSG00000046876 | 2.136858952 | 1.979715809 | 2.004202072 | 1.918106335 | 1.88560959 | 2.071882007 | 1.775756176 |
| ENSMUSG00000046879 | 2.420929319 | 2.240000483 | 2.166333979 | 2.221424093 | 2.184185718 | 2.42975228 | 2.025071334 |
| ENSMUSG00000046881 | 0 | 0 | 0 | 0 | 0 | 0 | 0 |
| ENSMUSG00000046886 | 0 | 0 | 0 | 0.25 | 0 | 0 | 0 |
| ENSMUSG00000046897 | 2.933200029 | 2.649513125 | 2.659156155 | 2.682192712 | 2.630371083 | 2.997386384 | 2.468415048 |
| ENSMUSG00000046899 | 0 | 0 | 0 | 0 | 0 | 0 | 0.224243824 |
| ENSMUSG00000046908 | 0 | 0 | 0 | 0 | 0 | 0 | 0.224243824 |
| ENSMUSG00000046909 | 1.857607105 | 1.695339928 | 1.62694866 | 1.726722649 | 1.625423019 | 1.806179974 | 1.473249546 |
| ENSMUSG00000046913 | 0 | 0 | 0 | 0 | 0 | 0 | 0 |
| ENSMUSG00000046916 | 0 | 0 | 0 | 0 | 0 | 0 | 0 |
| ENSMUSG00000046922 | 0 | 0 | 0 | 0 | 0 | 0 | 0 |
| ENSMUSG00000046924 | 0 | 0 | 0 | 0 | 0 | 0 | 0 |
| ENSMUSG00000046932 | 0 | 0 | 0 | 0 | 0 | 0 | 0 |
| ENSMUSG00000046934 | 0.289064826 | 0 | 0.580482024 | 0.25 | 0.489301084 | 0 | 0.224243824 |
| ENSMUSG00000046942 | 0 | 0 | 0 | 0 | 0 | 0 | 0 |
| ENSMUSG00000046947 | 1.45815691 | 1.488549078 | 1.532320754 | 1.532320754 | 1.414412788 | 1.505149978 | 1.405288179 |
| ENSMUSG00000046949 | 2.177057474 | 2.034887838 | 2.043731421 | 2.205044741 | 2.080465577 | 2.439332694 | 2.002662694 |
| ENSMUSG00000046952 | 0 | 0 | 0 | 0 | 0 | 0 | 0 |

| | | | | | | | |
|---|---|---|---|---|---|---|---|
| ENSMUSG00000046957 | 0 | 0 | 0 | 0 | 0 | 0 | 0 |
| ENSMUSG00000046958 | 0 | 0 | 0 | 0.646240625 | 0.387761935 | 0.477121255 | 0.224243824 |
| ENSMUSG00000046959 | 0 | 0.396240625 | 0.5 | 0.580482024 | 0.489301084 | 0.301029996 | 0.520678036 |
| ENSMUSG00000046961 | 0.578129653 | 0.75 | 0.79248125 | 0.79248125 | 0.77552387 | 0.903089987 | 0.520678036 |
| ENSMUSG00000046962 | 1.932832114 | 1.87694866 | 1.905512955 | 1.850219859 | 1.818885826 | 1.886490725 | 1.695786161 |
| ENSMUSG00000046971 | 0 | 0.396240625 | 0.25 | 0 | 0.244650542 | 0 | 0.448487648 |
| ENSMUSG00000046974 | 0 | 0 | 0 | 0 | 0 | 0 | 0 |
| ENSMUSG00000046975 | 0 | 0 | 0 | 0 | 0 | 0 | 0 |
| ENSMUSG00000046982 | 1.876566388 | 1.779735268 | 1.834962501 | 1.769203899 | 1.741652793 | 1.84509804 | 1.725216937 |
| ENSMUSG00000046985 | 2.446507203 | 2.27053726 | 2.21293726 | 2.317865757 | 2.237245597 | 2.568201724 | 2.104237736 |
| ENSMUSG00000046991 | 0 | 0.396240625 | 0 | 0 | 0 | 0 | 0.224243824 |
| ENSMUSG00000046993 | 0 | 0.396240625 | 0.25 | 0 | 0.244650542 | 0 | 0.355418052 |
| ENSMUSG00000046994 | 2.013563224 | 1.943696765 | 1.884789703 | 2.034887838 | 1.973373093 | 2.227886705 | 1.829800754 |
| ENSMUSG00000046995 | 0 | 0 | 0 | 0 | 0 | 0 | 0 |
| ENSMUSG00000046997 | 1.227926429 | 1.114857905 | 1.146240625 | 1.29248125 | 1.310728002 | 1.255272505 | 1.307989563 |
| ENSMUSG00000046999 | 0 | 0 | 0.25 | 0.25 | 0.387761935 | 0 | 0 |
| ENSMUSG00000047002 | 0 | 0 | 0 | 0 | 0 | 0 | 0 |
| ENSMUSG00000047003 | 2.227926429 | 2.048689214 | 1.991446071 | 1.910964047 | 1.928492895 | 2.056904851 | 1.819456572 |
| ENSMUSG00000047013 | 0 | 0.5 | 0.25 | 0.25 | 0.632412477 | 0.301029996 | 0.520678036 |
| ENSMUSG00000047014 | 0 | 0 | 0 | 0 | 0 | 0 | 0 |
| ENSMUSG00000047021 | 0 | 0 | 0 | 0 | 0 | 0 | 0 |
| ENSMUSG00000047022 | 1.54867984 | 1.618933358 | 1.571350555 | 1.505591953 | 1.319233381 | 1.531478917 | 1.313616061 |
| ENSMUSG00000047025 | 0 | 0 | 0 | 0 | 0 | 0 | 0 |
| ENSMUSG00000047026 | 0 | 0 | 0 | 0 | 0 | 0 | 0 |
| ENSMUSG00000047030 | 1.89911417 | 1.874961472 | 1.882845365 | 1.848079356 | 1.882215755 | 2.117271296 | 1.869159809 |
| ENSMUSG00000047031 | 0 | 0 | 0 | 0 | 0 | 0 | 0 |
| ENSMUSG00000047033 | 0.671187741 | 0.580482024 | 0.396240625 | 0.25 | 0 | 0 | 0 |
| ENSMUSG00000047034 | 0 | 0 | 0 | 0.25 | 0 | 0 | 0 |
| ENSMUSG00000047036 | 2.3855553 | 2.194519282 | 2.19697564 | 2.3464656 | 2.295728342 | 2.537819095 | 2.066254157 |
| ENSMUSG00000047037 | 1.671187741 | 1.622963274 | 1.62694866 | 1.657339155 | 1.621875691 | 1.698970004 | 1.511732338 |
| ENSMUSG00000047039 | 0 | 0 | 0 | 0 | 0 | 0 | 0 |
| ENSMUSG00000047040 | 0.45815691 | 0.25 | 0 | 0.396240625 | 0 | 0 | 0.224243824 |
| ENSMUSG00000047044 | 2.419671301 | 2.21293726 | 2.170748646 | 2.240000483 | 2.163285805 | 2.480006943 | 2.029372647 |
| ENSMUSG00000047045 | 2.417143825 | 2.210587586 | 2.211372513 | 2.385274154 | 2.344966272 | 2.701567985 | 2.190083792 |

| | | | | | | | |
|---|---|---|---|---|---|---|---|
| ENSMUSG00000047046 | 0 | 0 | 0 | 0 | 0 | 0 | 0 |
| ENSMUSG00000047048 | 0 | 0 | 0 | 0 | 0 | 0 | 0 |
| ENSMUSG00000047050 | 0 | 0 | 0 | 0 | 0 | 0 | 0 |
| ENSMUSG00000047053 | 0 | 0 | 0 | 0 | 0 | 0 | 0 |
| ENSMUSG00000047061 | 0 | 0 | 0 | 0.5 | 0 | 0 | 0 |
| ENSMUSG00000047067 | 1.129344651 | 1.080482024 | 1.080482024 | 1.04248125 | 0.955822902 | 0.903089987 | 1.028149525 |
| ENSMUSG00000047079 | 0 | 0 | 0 | 0 | 0 | 0 | 0 |
| ENSMUSG00000047084 | 1.885732304 | 1.77186571 | 1.744319981 | 1.80479213 | 1.814781639 | 1.959041392 | 1.725216937 |
| ENSMUSG00000047085 | 0 | 0 | 0.25 | 0 | 0 | 0 | 0.224243824 |
| ENSMUSG00000047090 | 2.160971586 | 2.017865591 | 2.043731421 | 1.80479213 | 1.893954095 | 2.004321374 | 1.773071385 |
| ENSMUSG00000047094 | 0 | 0 | 0 | 0 | 0 | 0 | 0 |
| ENSMUSG00000047098 | 1.663535556 | 1.671625132 | 1.646240625 | 1.584962501 | 1.746695109 | 1.806179974 | 1.752192176 |
| ENSMUSG00000047102 | 0 | 0 | 0 | 0 | 0 | 0 | 0 |
| ENSMUSG00000047104 | 0.747221736 | 0.75 | 0.75 | 0.75 | 0.489301084 | 0.602059991 | 0.775756176 |
| ENSMUSG00000047108 | 0 | 0 | 0.25 | 0.5 | 0 | 0 | 0.224243824 |
| ENSMUSG00000047109 | 0 | 0 | 0 | 0 | 0 | 0 | 0 |
| ENSMUSG00000047115 | 1.249317394 | 0.830482024 | 0.92510993 | 1.25 | 1.254881871 | 1.278753601 | 1.121219121 |
| ENSMUSG00000047117 | 1.036286563 | 1.04248125 | 1.061981878 | 1.445339928 | 1.380772476 | 1.51851394 | 1.302263476 |
| ENSMUSG00000047123 | 2.033115198 | 1.87694866 | 1.921625132 | 1.794977273 | 1.826953746 | 1.995635195 | 1.793950594 |
| ENSMUSG00000047126 | 3.721093129 | 3.355022951 | 3.3203014 | 3.29224636 | 3.195099728 | 3.703978825 | 2.946248069 |
| ENSMUSG00000047129 | 0 | 0 | 0 | 0 | 0 | 0 | 0 |
| ENSMUSG00000047139 | 3.384310426 | 3.078928915 | 3.056979181 | 3.001493839 | 2.895889166 | 3.390758529 | 2.694538871 |
| ENSMUSG00000047141 | 2.092368209 | 1.953445298 | 1.912762923 | 2.023189285 | 1.979897422 | 2.209515015 | 1.753626837 |
| ENSMUSG00000047143 | 0.811507563 | 1.080482024 | 1.061981878 | 1.146240625 | 1.200473444 | 1.230448921 | 1.319146374 |
| ENSMUSG00000047146 | 1.842799666 | 1.688721876 | 1.575945187 | 1.890560606 | 1.855665188 | 2.056904851 | 1.674564426 |
| ENSMUSG00000047149 | 0 | 0 | 0 | 0 | 0 | 0 | 0 |
| ENSMUSG00000047150 | 0.45815691 | 0 | 0 | 0.5 | 0.387761935 | 0.477121255 | 0 |
| ENSMUSG00000047153 | 1.89911417 | 1.777131114 | 1.83272922 | 1.921625132 | 1.951645858 | 2.117271296 | 1.882886767 |
| ENSMUSG00000047155 | 0.578129653 | 0.25 | 0.396240625 | 0.5 | 0.244650542 | 0.301029996 | 0.355418052 |
| ENSMUSG00000047161 | 0 | 0 | 0 | 0 | 0 | 0 | 0 |
| ENSMUSG00000047168 | 0 | 0 | 0 | 0 | 0 | 0 | 0 |
| ENSMUSG00000047171 | 0 | 0 | 0 | 0 | 0 | 0.301029996 | 0 |
| ENSMUSG00000047180 | 0.960252568 | 1 | 1.098079356 | 1.668106335 | 1.614672255 | 1.740362689 | 1.505628092 |
| ENSMUSG00000047181 | 2.412042447 | 2.234894804 | 2.348079356 | 2.043731421 | 2.145949654 | 2.322219295 | 2.177588371 |

| | | | | | | | |
|---|---|---|---|---|---|---|---|
| ENSMUSG00000047182 | 0 | 0 | 0.25 | 0 | 0 | 0 | 0 |
| ENSMUSG00000047187 | 2.965390609 | 2.701838731 | 2.67619206 | 2.700629091 | 2.595767222 | 2.995196292 | 2.429208684 |
| ENSMUSG00000047189 | 0 | 0 | 0 | 0 | 0 | 0 | 0 |
| ENSMUSG00000047193 | 2.231898196 | 2.048689214 | 2.065523711 | 2.161864607 | 2.0928848 | 2.387389826 | 1.95411574 |
| ENSMUSG00000047197 | 0 | 0 | 0 | 0 | 0 | 0 | 0 |
| ENSMUSG00000047203 | 0 | 0 | 0 | 0 | 0 | 0 | 0 |
| ENSMUSG00000047205 | 2.282114126 | 2.14151351 | 2.143411797 | 1.919870025 | 1.895599581 | 2.103803721 | 1.775756176 |
| ENSMUSG00000047207 | 0 | 0 | 0 | 0 | 0 | 0 | 0 |
| ENSMUSG00000047213 | 2.789260561 | 2.564052172 | 2.53743678 | 2.666777886 | 2.600700615 | 2.961421094 | 2.384994105 |
| ENSMUSG00000047215 | 3.243591459 | 2.928776002 | 2.938617499 | 3.001756817 | 2.904562212 | 3.418798291 | 2.706725307 |
| ENSMUSG00000047216 | 0 | 0 | 0 | 0 | 0 | 0 | 0 |
| ENSMUSG00000047220 | 0.289064826 | 0.25 | 0.25 | 0.396240625 | 0.387761935 | 0.301029996 | 0 |
| ENSMUSG00000047221 | 1.936861443 | 1.75 | 1.779735268 | 1.882845365 | 1.773209395 | 1.886490725 | 1.651852264 |
| ENSMUSG00000047222 | 0 | 0 | 0 | 0 | 0 | 0 | 0 |
| ENSMUSG00000047225 | 0 | 0 | 0 | 0 | 0 | 0 | 0 |
| ENSMUSG00000047227 | 1.129344651 | 1.04248125 | 1.114857905 | 1.02186571 | 0.978602168 | 1.113943352 | 1.041356071 |
| ENSMUSG00000047228 | 0 | 0 | 0 | 0.25 | 0 | 0 | 0 |
| ENSMUSG00000047230 | 0 | 0 | 0 | 0.25 | 0.244650542 | 0 | 0 |
| ENSMUSG00000047237 | 0 | 0 | 0 | 0 | 0 | 0 | 0 |
| ENSMUSG00000047238 | 1.771760131 | 1.779735268 | 1.735628626 | 1.494319981 | 1.49445382 | 1.602059991 | 1.466365979 |
| ENSMUSG00000047242 | 1.837744666 | 1.638647213 | 1.752806814 | 1.552363341 | 1.568060967 | 1.643452676 | 1.401059159 |
| ENSMUSG00000047246 | 0.45815691 | 0.75 | 0.25 | 0.396240625 | 0.77552387 | 0.477121255 | 0.629532004 |
| ENSMUSG00000047248 | 2.251208701 | 2.046218836 | 2.036164561 | 2.142463902 | 2.11997913 | 2.423245874 | 2.002662694 |
| ENSMUSG00000047250 | 2.497590902 | 2.302363341 | 2.265173983 | 1.53743678 | 1.533172725 | 1.62324929 | 1.396774124 |
| ENSMUSG00000047253 | 0.45815691 | 0.5 | 0.5 | 0.396240625 | 0.244650542 | 0.477121255 | 0.579661877 |
| ENSMUSG00000047257 | 0 | 0 | 0 | 0 | 0 | 0 | 0 |
| ENSMUSG00000047259 | 0.289064826 | 0.396240625 | 0.396240625 | 0 | 0 | 0 | 0 |
| ENSMUSG00000047260 | 2.158622101 | 2.02186571 | 2.069031101 | 2.125956434 | 2.086251872 | 2.356025857 | 1.990504106 |
| ENSMUSG00000047261 | 0.671187741 | 1 | 0.830482024 | 1.146240625 | 0.978602168 | 1.146128036 | 1.20919388 |
| ENSMUSG00000047264 | 2.231898196 | 2.070192693 | 2.112802778 | 1.714495249 | 1.704182902 | 1.838849091 | 1.764880683 |
| ENSMUSG00000047281 | 0.289064826 | 0.75 | 0.75 | 0.396240625 | 0.568060967 | 0 | 0.672731473 |
| ENSMUSG00000047284 | 2.294202867 | 2.17510993 | 2.20584181 | 2.038704527 | 2.193483986 | 2.307496038 | 2.096394553 |
| ENSMUSG00000047286 | 0 | 0 | 0 | 0 | 0 | 0 | 0 |
| ENSMUSG00000047293 | 0 | 0 | 0.25 | 0.25 | 0.387761935 | 0 | 0.224243824 |

| | | | | | | | |
|---|---|---|---|---|---|---|---|
| ENSMUSG00000047295 | 0 | 0 | 0 | 0 | 0 | 0 | 0 |
| ENSMUSG00000047298 | 0 | 0.25 | 0 | 0 | 0 | 0 | 0 |
| ENSMUSG00000047307 | 0 | 0 | 0 | 0 | 0 | 0 | 0 |
| ENSMUSG00000047324 | 0 | 0 | 0 | 0 | 0 | 0 | 0 |
| ENSMUSG00000047330 | 0 | 0 | 0 | 0 | 0 | 0 | 0 |
| ENSMUSG00000047342 | 1.100572389 | 1.098079356 | 1.146240625 | 1.671625132 | 1.625423019 | 1.908485019 | 1.546120303 |
| ENSMUSG00000047343 | 0.289064826 | 0.25 | 0 | 0 | 0 | 0.477121255 | 0.448487648 |
| ENSMUSG00000047344 | 1.558729299 | 1.396240625 | 1.505591953 | 1.664552871 | 1.555378544 | 1.716003344 | 1.392431569 |
| ENSMUSG00000047347 | 1.623015126 | 1.67510993 | 1.726722649 | 1.54248125 | 1.542223377 | 1.579783597 | 1.551512352 |
| ENSMUSG00000047352 | 0 | 0 | 0 | 0 | 0 | 0 | 0 |
| ENSMUSG00000047356 | 0 | 0 | 0 | 0 | 0 | 0 | 0 |
| ENSMUSG00000047361 | 0 | 0 | 0 | 0 | 0 | 0 | 0 |
| ENSMUSG00000047363 | 0.45815691 | 0.25 | 0.396240625 | 0 | 0 | 0 | 0 |
| ENSMUSG00000047368 | 2.491272108 | 2.266522298 | 2.27053726 | 2.319031101 | 2.247084732 | 2.572871602 | 2.084261301 |
| ENSMUSG00000047369 | 0 | 0 | 0 | 0 | 0 | 0 | 0 |
| ENSMUSG00000047370 | 0 | 0 | 0 | 0 | 0 | 0 | 0.224243824 |
| ENSMUSG00000047371 | 1.671187741 | 1.649978211 | 1.688721876 | 1.42510993 | 1.489301084 | 1.591064607 | 1.543390198 |
| ENSMUSG00000047379 | 2.003432385 | 1.938721876 | 1.956637122 | 2.136723615 | 2.107593434 | 2.311753861 | 2.031198704 |
| ENSMUSG00000047383 | 0 | 0 | 0 | 0 | 0 | 0 | 0 |
| ENSMUSG00000047384 | 0 | 0 | 0 | 0 | 0 | 0 | 0.224243824 |
| ENSMUSG00000047388 | 2.136858952 | 1.997171172 | 2.00835575 | 2.069031101 | 2.040348824 | 2.257678575 | 1.952571511 |
| ENSMUSG00000047390 | 0 | 0 | 0 | 0 | 0 | 0 | 0 |
| ENSMUSG00000047394 | 0 | 0 | 0 | 0 | 0 | 0 | 0.448487648 |
| ENSMUSG00000047407 | 2.481610694 | 2.274508021 | 2.203445298 | 1.96759118 | 1.925488995 | 2.139879086 | 1.774416565 |
| ENSMUSG00000047409 | 2.256831695 | 2.139605178 | 2.099151195 | 2.028435542 | 2.00243419 | 2.250420002 | 1.922587433 |
| ENSMUSG00000047412 | 2.219867581 | 2.048689214 | 2.015173983 | 2.024508021 | 1.977302144 | 2.152288344 | 1.828667599 |
| ENSMUSG00000047414 | 2.605636262 | 2.394357207 | 2.351816941 | 2.242166698 | 2.121713561 | 2.495544338 | 1.995286892 |
| ENSMUSG00000047415 | 0.289064826 | 0 | 0 | 0.396240625 | 0.244650542 | 0 | 0.224243824 |
| ENSMUSG00000047417 | 2.153883046 | 2.163659007 | 2.269871196 | 1.955044741 | 2.184185718 | 2.235528447 | 2.229547475 |
| ENSMUSG00000047419 | 0 | 0 | 0 | 0.25 | 0.244650542 | 0 | 0 |
| ENSMUSG00000047420 | 1.389637216 | 1.201838731 | 1.188721876 | 1.080482024 | 0.812711509 | 1.113943352 | 0.935079929 |
| ENSMUSG00000047423 | 2.269664473 | 2.124961472 | 2.26584877 | 1.87892496 | 2.062514787 | 2.113943352 | 2.077494038 |
| ENSMUSG00000047428 | 1.181541736 | 1.311981878 | 1.505591953 | 0.951838731 | 1.335652905 | 0.84509804 | 1.523605797 |
| ENSMUSG00000047434 | 2.023453784 | 1.926839783 | 1.951838731 | 2.023189285 | 2.058398634 | 2.309630167 | 1.897893072 |

| | | | | | | | |
|---|---|---|---|---|---|---|---|
| ENSMUSG00000047441 | 0 | 0 | 0 | 0 | 0 | 0 | 0 |
| ENSMUSG00000047443 | 1.907802659 | 1.854463129 | 1.79248125 | 1.843759858 | 1.855665188 | 1.991226076 | 1.757893036 |
| ENSMUSG00000047444 | 0 | 0 | 0 | 0 | 0 | 0 | 0 |
| ENSMUSG00000047446 | 1.558729299 | 1.566696635 | 1.642463902 | 1.53743678 | 1.445123986 | 1.505149978 | 1.369798885 |
| ENSMUSG00000047454 | 2.124258828 | 1.976722649 | 1.96293726 | 2.04745614 | 1.990091809 | 2.238046103 | 1.847411384 |
| ENSMUSG00000047459 | 2.373064211 | 2.161864607 | 2.193696765 | 2.362287161 | 2.29993025 | 2.656098202 | 2.228229691 |
| ENSMUSG00000047462 | 0 | 0 | 0 | 0 | 0 | 0 | 0 |
| ENSMUSG00000047466 | 2.086807695 | 1.923371739 | 1.916333979 | 1.882845365 | 1.846351821 | 1.986771734 | 1.702561601 |
| ENSMUSG00000047473 | 1.389637216 | 1.458222504 | 1.532320754 | 1.372963274 | 1.433158226 | 1.51851394 | 1.383567577 |
| ENSMUSG00000047485 | 0 | 0.396240625 | 0 | 0 | 0 | 0 | 0 |
| ENSMUSG00000047490 | 0 | 0 | 0 | 0 | 0 | 0 | 0 |
| ENSMUSG00000047492 | 0 | 0 | 0 | 0 | 0 | 0 | 0 |
| ENSMUSG00000047495 | 0.45815691 | 0.580482024 | 0.25 | 0 | 0 | 0 | 0 |
| ENSMUSG00000047496 | 0.289064826 | 0.25 | 0 | 0.396240625 | 0.568060967 | 0.301029996 | 0.672731473 |
| ENSMUSG00000047497 | 1.876566388 | 1.77186571 | 1.738549078 | 1.282320754 | 1.310728002 | 1.431363764 | 1.150210039 |
| ENSMUSG00000047501 | 0.289064826 | 0 | 0 | 0 | 0.244650542 | 0 | 0.224243824 |
| ENSMUSG00000047502 | 0 | 0 | 0 | 0 | 0 | 0 | 0 |
| ENSMUSG00000047507 | 0.289064826 | 0.25 | 0.5 | 0 | 0 | 0 | 0.355418052 |
| ENSMUSG00000047508 | 0 | 0 | 0 | 0 | 0 | 0 | 0 |
| ENSMUSG00000047509 | 0 | 0 | 0 | 0 | 0 | 0 | 0 |
| ENSMUSG00000047511 | 0 | 0 | 0 | 0 | 0 | 0 | 0 |
| ENSMUSG00000047514 | 2.452372734 | 2.254897682 | 2.26109853 | 2.382845365 | 2.301493158 | 2.643452676 | 2.200320163 |
| ENSMUSG00000047515 | 0.289064826 | 0.396240625 | 0.580482024 | 0.75 | 0.489301084 | 0.477121255 | 0.520678036 |
| ENSMUSG00000047517 | 0 | 0 | 0 | 0 | 0 | 0 | 0 |
| ENSMUSG00000047518 | 0 | 0 | 0 | 0 | 0 | 0 | 0 |
| ENSMUSG00000047528 | 0.289064826 | 0.580482024 | 0.5 | 0.25 | 0.244650542 | 0 | 0.448487648 |
| ENSMUSG00000047531 | 0 | 0 | 0 | 0 | 0 | 0 | 0 |
| ENSMUSG00000047534 | 2.388281011 | 2.218361028 | 2.238549078 | 2.191217898 | 2.098472613 | 2.338456494 | 1.94631981 |
| ENSMUSG00000047535 | 0 | 0 | 0 | 0 | 0 | 0 | 0 |
| ENSMUSG00000047539 | 2.352857998 | 2.195339928 | 2.171625132 | 2.370449858 | 2.299407739 | 2.605305046 | 2.141264342 |
| ENSMUSG00000047544 | 0 | 0 | 0 | 0 | 0 | 0 | 0 |
| ENSMUSG00000047545 | 0 | 0 | 0 | 0 | 0 | 0 | 0 |
| ENSMUSG00000047547 | 2.144240249 | 1.958222504 | 2.104463129 | 2.254202072 | 2.320220671 | 2.531478917 | 2.277722321 |
| ENSMUSG00000047554 | 2.294202867 | 2.099151195 | 2.116901388 | 2.241446071 | 2.130260132 | 2.439332694 | 1.979298987 |

| | | | | | | | |
|---|---|---|---|---|---|---|---|
| ENSMUSG00000047557 | 2.448863324 | 2.274508021 | 2.242885888 | 2.111770807 | 2.045754077 | 2.292256071 | 1.930969131 |
| ENSMUSG00000047562 | 0 | 0 | 0 | 0.25 | 0 | 0 | 0.224243824 |
| ENSMUSG00000047564 | 0 | 0 | 0 | 0 | 0 | 0 | 0 |
| ENSMUSG00000047565 | 0 | 0 | 0.580482024 | 0 | 0 | 0.477121255 | 0 |
| ENSMUSG00000047583 | 1.53838222 | 1.339388001 | 1.339388001 | 1.396240625 | 1.401338844 | 1.672097858 | 1.379042812 |
| ENSMUSG00000047586 | 0 | 0 | 0 | 0 | 0 | 0 | 0 |
| ENSMUSG00000047591 | 0 | 0.25 | 0 | 0 | 0 | 0 | 0.224243824 |
| ENSMUSG00000047592 | 0 | 0 | 0.25 | 0 | 0.244650542 | 0 | 0 |
| ENSMUSG00000047594 | 0 | 0 | 0 | 0 | 0 | 0 | 0 |
| ENSMUSG00000047603 | 1.156259305 | 1.26109853 | 1.302363341 | 1.214495249 | 1.244650542 | 1.176091259 | 1.185218806 |
| ENSMUSG00000047604 | 0.960252568 | 1.04248125 | 1 | 0.951838731 | 1.188507684 | 1.146128036 | 1.168187745 |
| ENSMUSG00000047606 | 0.289064826 | 0.25 | 0 | 0 | 0 | 0 | 0 |
| ENSMUSG00000047613 | 1.740854706 | 1.618933358 | 1.602347734 | 1.52186571 | 1.66856048 | 1.681241237 | 1.670888075 |
| ENSMUSG00000047617 | 1.069666964 | 1.321350555 | 1.464495249 | 1.380890489 | 1.588235379 | 1.579783597 | 1.540636858 |
| ENSMUSG00000047619 | 0 | 0 | 0 | 0 | 0 | 0 | 0 |
| ENSMUSG00000047626 | 0 | 0 | 0 | 0 | 0 | 0 | 0 |
| ENSMUSG00000047631 | 0.289064826 | 0.25 | 0 | 0.701838731 | 0.489301084 | 0.477121255 | 0.629532004 |
| ENSMUSG00000047632 | 1.100572389 | 1.098079356 | 1.098079356 | 1.302363341 | 1.343584837 | 1.462397998 | 1.272006351 |
| ENSMUSG00000047635 | 1.967766868 | 1.872963274 | 1.882845365 | 1.80479213 | 1.761403742 | 1.929418926 | 1.633706455 |
| ENSMUSG00000047636 | 0 | 0 | 0 | 0 | 0 | 0 | 0 |
| ENSMUSG00000047638 | 0 | 0 | 0.5 | 0.5 | 0 | 0.301029996 | 0.224243824 |
| ENSMUSG00000047641 | 0 | 0.75 | 0.646240625 | 0 | 0 | 0 | 0 |
| ENSMUSG00000047642 | 0 | 0 | 0 | 0 | 0 | 0.301029996 | 0.355418052 |
| ENSMUSG00000047643 | 0 | 0 | 0 | 0 | 0 | 0 | 0 |
| ENSMUSG00000047648 | 2.578990401 | 2.359177886 | 2.349151195 | 2.394828984 | 2.302012594 | 2.641474111 | 2.117521761 |
| ENSMUSG00000047649 | 1.822201287 | 1.777131114 | 1.87694866 | 1.870953944 | 1.868304353 | 2.021189299 | 1.764880683 |
| ENSMUSG00000047652 | 0 | 0 | 0 | 0 | 0 | 0 | 0 |
| ENSMUSG00000047654 | 0 | 0 | 0 | 0 | 0 | 0.301029996 | 0.224243824 |
| ENSMUSG00000047655 | 0 | 0 | 0 | 0 | 0 | 0 | 0 |
| ENSMUSG00000047656 | 1.358731791 | 1.226722649 | 1.372963274 | 0.951838731 | 1.244650542 | 1.041392685 | 1.401059159 |
| ENSMUSG00000047657 | 0 | 0 | 0 | 0 | 0 | 0 | 0 |
| ENSMUSG00000047658 | 0 | 0.580482024 | 0.5 | 0.396240625 | 0.686820904 | 0.602059991 | 0.775756176 |
| ENSMUSG00000047661 | 0 | 0 | 0 | 0 | 0 | 0 | 0 |
| ENSMUSG00000047667 | 0 | 0 | 0 | 0 | 0 | 0 | 0 |

| | | | | | | | |
|---|---|---|---|---|---|---|---|
| ENSMUSG00000047669 | 1.37447073 | 1.27186571 | 1.388647213 | 1.226722649 | 1.136121934 | 1.079181246 | 1.089372236 |
| ENSMUSG00000047671 | 0 | 0.25 | 0 | 0 | 0 | 0.301029996 | 0.224243824 |
| ENSMUSG00000047674 | 0 | 0 | 0 | 0 | 0 | 0 | 0 |
| ENSMUSG00000047675 | 3.060283005 | 2.859957721 | 2.851684121 | 2.769537702 | 2.821992434 | 3.20871002 | 2.661077362 |
| ENSMUSG00000047676 | 0.671187741 | 0.79248125 | 0.79248125 | 0 | 0.77552387 | 0.477121255 | 0.672731473 |
| ENSMUSG00000047678 | 0 | 0 | 0 | 0.25 | 0 | 0 | 0 |
| ENSMUSG00000047686 | 0 | 0 | 0 | 0 | 0 | 0 | 0 |
| ENSMUSG00000047692 | 0 | 0 | 0 | 0 | 0 | 0 | 0 |
| ENSMUSG00000047694 | 2.188178997 | 2.016522298 | 2.064346961 | 2.311981878 | 2.216042585 | 2.562292864 | 2.050633136 |
| ENSMUSG00000047696 | 0.45815691 | 0 | 0 | 0.646240625 | 0.733951626 | 0.698970004 | 0.520678036 |
| ENSMUSG00000047702 | 0 | 0 | 0 | 0 | 0 | 0 | 0 |
| ENSMUSG00000047710 | 2.604828841 | 2.410061234 | 2.395770692 | 2.248588359 | 2.252501045 | 2.498310554 | 2.100339912 |
| ENSMUSG00000047712 | 1.789260561 | 1.723704441 | 1.552363341 | 1.660964047 | 1.555378544 | 1.785329835 | 1.559435575 |
| ENSMUSG00000047714 | 2.70430294 | 2.500703754 | 2.444108258 | 2.613574823 | 2.547440858 | 2.94792362 | 2.331134329 |
| ENSMUSG00000047716 | 0 | 0 | 0 | 0 | 0 | 0 | 0 |
| ENSMUSG00000047719 | 2.139333929 | 2.057204673 | 2.041226732 | 2.123963757 | 2.094757251 | 2.374748346 | 2.052911423 |
| ENSMUSG00000047720 | 0 | 0 | 0 | 0 | 0 | 0 | 0 |
| ENSMUSG00000047721 | 1.494443473 | 1.634789703 | 1.698603967 | 1.26109853 | 1.53772706 | 1.322219295 | 1.596378571 |
| ENSMUSG00000047728 | 0 | 0.396240625 | 0.5 | 0 | 0 | 0 | 0 |
| ENSMUSG00000047730 | 0 | 0 | 0.25 | 0 | 0.244650542 | 0 | 0.224243824 |
| ENSMUSG00000047731 | 2.649252229 | 2.446158711 | 2.496460484 | 2.33272922 | 2.411379849 | 2.665580991 | 2.3030503 |
| ENSMUSG00000047733 | 0.289064826 | 0.25 | 0.396240625 | 0 | 0 | 0 | 0 |
| ENSMUSG00000047735 | 1.806056082 | 1.738549078 | 1.692046081 | 1.910964047 | 1.789100779 | 2.056904851 | 1.670888075 |
| ENSMUSG00000047746 | 0 | 0 | 0 | 0 | 0 | 0 | 0 |
| ENSMUSG00000047747 | 2.048736215 | 1.914552871 | 1.912762923 | 1.356566189 | 1.163285805 | 1.146128036 | 1.185218806 |
| ENSMUSG00000047749 | 2.192545868 | 2.01109853 | 2.02053726 | 1.823655187 | 1.828942237 | 2 | 1.699191618 |
| ENSMUSG00000047751 | 0 | 0 | 0 | 0 | 0 | 0 | 0 |
| ENSMUSG00000047757 | 1.982401085 | 1.821350555 | 1.856566189 | 2.04745614 | 1.982473755 | 2.250420002 | 1.87511425 |
| ENSMUSG00000047759 | 0 | 0.5 | 0.5 | 0.396240625 | 0.489301084 | 0.77815125 | 0.448487648 |
| ENSMUSG00000047766 | 2.156259305 | 1.933677405 | 1.930274797 | 2.160964047 | 2.0928848 | 2.394451681 | 1.936709733 |
| ENSMUSG00000047767 | 1.647796617 | 1.488549078 | 1.488549078 | 1.516522298 | 1.665423072 | 1.653212514 | 1.598697688 |
| ENSMUSG00000047773 | 0 | 0 | 0 | 0 | 0 | 0 | 0 |
| ENSMUSG00000047777 | 2.249317394 | 2.069031101 | 2.136723615 | 2.063166358 | 2.103062484 | 2.303196057 | 2.009225098 |
| ENSMUSG00000047786 | 0 | 0.396240625 | 0 | 0.25 | 0 | 0.301029996 | 0.355418052 |

| | | | | | | | |
|---|---|---|---|---|---|---|---|
| ENSMUSG00000047787 | 1.036286563 | 1.080482024 | 0.92510993 | 1.214495249 | 1.335652905 | 1.322219295 | 1.38802993 |
| ENSMUSG00000047789 | 1.952600383 | 1.726722649 | 1.80479213 | 1.907339155 | 1.897237431 | 2.161368002 | 1.774416565 |
| ENSMUSG00000047793 | 1.932832114 | 1.799918086 | 1.769203899 | 1.593759858 | 1.546663136 | 1.698970004 | 1.579661877 |
| ENSMUSG00000047794 | 0 | 0 | 0 | 0 | 0 | 0 | 0 |
| ENSMUSG00000047797 | 0 | 0 | 0 | 0 | 0 | 0 | 0 |
| ENSMUSG00000047798 | 0 | 0 | 0.25 | 0 | 0 | 0 | 0 |
| ENSMUSG00000047799 | 0 | 0 | 0 | 0 | 0 | 0 | 0 |
| ENSMUSG00000047804 | 2.203267085 | 2.038704527 | 2.046218836 | 2.184523065 | 2.069604532 | 2.367355921 | 1.951019875 |
| ENSMUSG00000047807 | 0.867194479 | 0.5 | 0.92510993 | 1.146240625 | 1.039257769 | 1.255272505 | 0.710836105 |
| ENSMUSG00000047810 | 0.289064826 | 0.580482024 | 0.580482024 | 0 | 0.77552387 | 0.301029996 | 0.74492186 |
| ENSMUSG00000047819 | 0 | 0.396240625 | 0 | 0.25 | 0.387761935 | 0.477121255 | 0.355418052 |
| ENSMUSG00000047821 | 2.185978293 | 2.081607372 | 2.046218836 | 2.178561379 | 2.193483986 | 2.482873584 | 2.059651572 |
| ENSMUSG00000047822 | 0 | 0 | 0.25 | 0 | 0 | 0 | 0.224243824 |
| ENSMUSG00000047824 | 2.445324132 | 2.29745614 | 2.348615674 | 2.255591953 | 2.298884454 | 2.517195898 | 2.211635518 |
| ENSMUSG00000047828 | 0 | 0 | 0 | 0 | 0 | 0 | 0 |
| ENSMUSG00000047832 | 2.571178952 | 2.366901388 | 2.375956434 | 2.201838731 | 2.208005287 | 2.457881897 | 2.075383 |
| ENSMUSG00000047841 | 0 | 0 | 0 | 0 | 0 | 0 | 0 |
| ENSMUSG00000047842 | 1.389637216 | 1.27186571 | 1.396240625 | 0.701838731 | 0.568060967 | 0.698970004 | 0.74492186 |
| ENSMUSG00000047843 | 2.213719572 | 2.016522298 | 2.093759858 | 1.809601185 | 1.818885826 | 2.029383778 | 1.74047993 |
| ENSMUSG00000047844 | 0 | 0 | 0 | 1.098079356 | 0.812711509 | 1 | 0.74492186 |
| ENSMUSG00000047854 | 0.747221736 | 0.5 | 0.701838731 | 0.5 | 0.387761935 | 0.602059991 | 0.579661877 |
| ENSMUSG00000047861 | 0 | 0 | 0 | 0 | 0 | 0 | 0 |
| ENSMUSG00000047866 | 2.516991255 | 2.356566189 | 2.320772088 | 2.399978211 | 2.331380351 | 2.667452953 | 2.197065089 |
| ENSMUSG00000047867 | 0 | 0.396240625 | 0.396240625 | 0.25 | 0.244650542 | 0 | 0 |
| ENSMUSG00000047868 | 0 | 0 | 0 | 0 | 0 | 0 | 0 |
| ENSMUSG00000047875 | 1.036286563 | 0.976722649 | 1.080482024 | 1.403677461 | 1.310728002 | 1.255272505 | 1.224243824 |
| ENSMUSG00000047878 | 0.578129653 | 1 | 0.976722649 | 1.532320754 | 1.572189156 | 1.602059991 | 1.425641774 |
| ENSMUSG00000047879 | 2.943013009 | 2.695544798 | 2.662762923 | 2.734894804 | 2.653163144 | 3.059184618 | 2.448988576 |
| ENSMUSG00000047880 | 0.91631382 | 0.896240625 | 0.79248125 | 0.396240625 | 0.77552387 | 0.477121255 | 0.355418052 |
| ENSMUSG00000047881 | 2.396352858 | 2.241446071 | 2.269871196 | 2.172499493 | 2.167869715 | 2.421603927 | 1.991191705 |
| ENSMUSG00000047884 | 0 | 0 | 0 | 0 | 0 | 0 | 0 |
| ENSMUSG00000047888 | 2.87614493 | 2.629171237 | 2.598347614 | 2.702642909 | 2.650078855 | 3.005180513 | 2.454448184 |
| ENSMUSG00000047889 | 0 | 0 | 0 | 0 | 0 | 0 | 0 |
| ENSMUSG00000047894 | 0.45815691 | 0.25 | 0.396240625 | 0.580482024 | 0.489301084 | 0.477121255 | 0.448487648 |

| | | | | | | | |
|---|---|---|---|---|---|---|---|
| ENSMUSG00000047897 | 0 | 0 | 0 | 0 | 0 | 0 | 0 |
| ENSMUSG00000047898 | 0 | 0 | 0.25 | 0.396240625 | 0 | 0 | 0 |
| ENSMUSG00000047904 | 0 | 0 | 0.396240625 | 0.5 | 0.632412477 | 0.301029996 | 0.355418052 |
| ENSMUSG00000047905 | 0 | 0.25 | 0.25 | 0.5 | 0.244650542 | 0.602059991 | 0.355418052 |
| ENSMUSG00000047907 | 0 | 0 | 0 | 0 | 0.244650542 | 0 | 0 |
| ENSMUSG00000047909 | 2.146671932 | 1.948603967 | 1.998588359 | 1.987091808 | 1.92093466 | 2.096910013 | 1.833176598 |
| ENSMUSG00000047910 | 1.389637216 | 1.114857905 | 1.238549078 | 0.25 | 0.244650542 | 0 | 0.224243824 |
| ENSMUSG00000047911 | 0.867194479 | 0.580482024 | 0.701838731 | 0.830482024 | 0.244650542 | 0.698970004 | 0.853775828 |
| ENSMUSG00000047921 | 2.172524468 | 2.080482024 | 2.144357207 | 1.948603967 | 2.024758322 | 2.195899652 | 2.00464543 |
| ENSMUSG00000047935 | 0.289064826 | 0.5 | 0.396240625 | 0.396240625 | 0.686820904 | 0 | 0.224243824 |
| ENSMUSG00000047938 | 0.289064826 | 0.25 | 0 | 0.396240625 | 0 | 0 | 0.224243824 |
| ENSMUSG00000047940 | 0 | 0 | 0 | 0 | 0 | 0.301029996 | 0 |
| ENSMUSG00000047945 | 2.922396442 | 2.687467357 | 2.707826811 | 2.538704527 | 2.570979987 | 2.914871818 | 2.499691862 |
| ENSMUSG00000047953 | 0 | 0 | 0.25 | 0 | 0.244650542 | 0 | 0 |
| ENSMUSG00000047959 | 0 | 0 | 0 | 0 | 0 | 0 | 0 |
| ENSMUSG00000047960 | 0 | 0 | 0 | 0 | 0 | 0 | 0 |
| ENSMUSG00000047963 | 2.349889783 | 2.189555804 | 2.130890489 | 1.943696765 | 1.905314583 | 2.136720567 | 1.74047993 |
| ENSMUSG00000047965 | 0.289064826 | 0 | 0.25 | 0.25 | 0.387761935 | 0 | 0 |
| ENSMUSG00000047969 | 0 | 0 | 0 | 0 | 0 | 0 | 0 |
| ENSMUSG00000047976 | 0 | 0 | 0 | 0.25 | 0.244650542 | 0 | 0 |
| ENSMUSG00000047977 | 0.747221736 | 0.646240625 | 0.830482024 | 0.580482024 | 0.877063019 | 0.477121255 | 0.520678036 |
| ENSMUSG00000047980 | 0 | 0 | 0 | 0 | 0 | 0 | 0 |
| ENSMUSG00000047986 | 0 | 0.5 | 0 | 0.864857905 | 0.489301084 | 0.301029996 | 0.629532004 |
| ENSMUSG00000047988 | 0.747221736 | 0.701838731 | 0.646240625 | 0.580482024 | 0.632412477 | 0.77815125 | 0.448487648 |
| ENSMUSG00000047989 | 2.134369199 | 1.995748394 | 2.041226732 | 2.198603967 | 2.141069998 | 2.418301291 | 2.030591163 |
| ENSMUSG00000047990 | 0 | 0 | 0 | 0 | 0 | 0 | 0 |
| ENSMUSG00000047992 | 0 | 0 | 0.25 | 0.25 | 0.387761935 | 0.477121255 | 0.710836105 |
| ENSMUSG00000047995 | 0 | 0 | 0 | 0 | 0 | 0 | 0 |
| ENSMUSG00000047996 | 2.151493169 | 2.00835575 | 1.991446071 | 1.884789703 | 1.808534465 | 2.053078443 | 1.669034113 |
| ENSMUSG00000048000 | 2.737340367 | 2.512802235 | 2.502107156 | 2.569612365 | 2.512633319 | 2.871572936 | 2.357195614 |
| ENSMUSG00000048001 | 0 | 0 | 0 | 0 | 0 | 0 | 0 |
| ENSMUSG00000048003 | 0 | 0 | 0 | 0 | 0 | 0 | 0 |
| ENSMUSG00000048004 | 0 | 0 | 0 | 0 | 0 | 0 | 0 |
| ENSMUSG00000048007 | 1.249317394 | 1.25 | 1.356566189 | 1.25 | 1.380772476 | 1.380211242 | 1.340368108 |

| | | | | | | | |
|---|---|---|---|---|---|---|---|
| ENSMUSG00000048012 | 0.960252568 | 0.864857905 | 0.79248125 | 0.830482024 | 0.568060967 | 1 | 0.803905701 |
| ENSMUSG00000048013 | 0 | 0 | 0.25 | 0 | 0 | 0 | 0 |
| ENSMUSG00000048015 | 0 | 0 | 0 | 0 | 0 | 0 | 0 |
| ENSMUSG00000048022 | 0 | 0 | 0 | 0 | 0 | 0 | 0 |
| ENSMUSG00000048027 | 1.857607105 | 1.763820609 | 1.819031101 | 1.678561379 | 1.632412477 | 1.875061263 | 1.627424404 |
| ENSMUSG00000048029 | 1 | 1.146240625 | 0.976722649 | 0.92510993 | 1.039257769 | 1.204119983 | 1.014380832 |
| ENSMUSG00000048031 | 0 | 0 | 0 | 0 | 0 | 0 | 0 |
| ENSMUSG00000048038 | 0 | 0 | 0 | 0 | 0 | 0 | 0 |
| ENSMUSG00000048039 | 2.368815967 | 2.195339928 | 2.166333979 | 2.288071211 | 2.242813819 | 2.534026106 | 2.078544414 |
| ENSMUSG00000048040 | 1.307602651 | 1.410964047 | 1.410964047 | 1.653677461 | 1.576269619 | 1.748188027 | 1.455757965 |
| ENSMUSG00000048047 | 2.457006475 | 2.307805295 | 2.294977273 | 2.339388001 | 2.237868644 | 2.542825427 | 2.048913829 |
| ENSMUSG00000048058 | 2.069666964 | 1.87892496 | 1.912762923 | 1.87892496 | 1.870073561 | 2.017033339 | 1.712465909 |
| ENSMUSG00000048062 | 0 | 0 | 0 | 0 | 0 | 0 | 0 |
| ENSMUSG00000048065 | 0 | 0 | 0 | 0 | 0.244650542 | 0 | 0 |
| ENSMUSG00000048067 | 0 | 0 | 0 | 0 | 0 | 0 | 0 |
| ENSMUSG00000048070 | 0 | 0 | 0 | 0 | 0 | 0 | 0 |
| ENSMUSG00000048076 | 3.221344457 | 2.972185812 | 2.94441657 | 3.033847163 | 3.027892175 | 3.502017215 | 2.852867412 |
| ENSMUSG00000048077 | 0 | 0 | 0 | 0 | 0 | 0 | 0 |
| ENSMUSG00000048078 | 1.389637216 | 1.238549078 | 1.226722649 | 0.864857905 | 0.905314583 | 0.77815125 | 1.014380832 |
| ENSMUSG00000048080 | 0 | 0 | 0 | 0 | 0 | 0 | 0 |
| ENSMUSG00000048087 | 1.631440309 | 1.54745614 | 1.593759858 | 1.692046081 | 1.728722551 | 1.707570176 | 1.618853957 |
| ENSMUSG00000048096 | 2.516991255 | 2.323080408 | 2.348079356 | 0.396240625 | 0.733951626 | 0.301029996 | 0.579661877 |
| ENSMUSG00000048100 | 2.069666964 | 1.937048212 | 1.905512955 | 2.071350555 | 2.075570129 | 2.307496038 | 1.89141329 |
| ENSMUSG00000048101 | 0 | 0 | 0 | 0 | 0 | 0 | 0 |
| ENSMUSG00000048106 | 1.876566388 | 1.774508021 | 1.738549078 | 1.852347734 | 1.733951626 | 2.068185862 | 1.69749337 |
| ENSMUSG00000048108 | 2.804556862 | 2.544043287 | 2.480460234 | 2.559899632 | 2.488077665 | 2.755874856 | 2.279980727 |
| ENSMUSG00000048109 | 2.183765914 | 2.05357978 | 2.043731421 | 2.113831805 | 2.060462711 | 2.374748346 | 1.904245613 |
| ENSMUSG00000048118 | 2.256831695 | 2.105516192 | 2.093759858 | 2.004202072 | 1.962676635 | 2.217483944 | 1.863093717 |
| ENSMUSG00000048120 | 0.578129653 | 0.25 | 0.580482024 | 0.5 | 0.244650542 | 0 | 0.224243824 |
| ENSMUSG00000048126 | 2.556012466 | 2.391037372 | 2.333288838 | 0.396240625 | 0 | 0.301029996 | 0.579661877 |
| ENSMUSG00000048138 | 0 | 0.25 | 0.25 | 0.646240625 | 0.905314583 | 0.602059991 | 0.672731473 |
| ENSMUSG00000048142 | 0.671187741 | 0.864857905 | 0.864857905 | 0.864857905 | 1 | 0.77815125 | 0.969165684 |
| ENSMUSG00000048148 | 0 | 0.5 | 0.25 | 0.25 | 0 | 0 | 0.579661877 |
| ENSMUSG00000048153 | 0 | 0 | 0 | 0 | 0 | 0 | 0 |

| | | | | | | | |
|---|---|---|---|---|---|---|---|
| ENSMUSG00000048154 | 2.722369902 | 2.55930249 | 2.633332433 | 2.310793496 | 2.514059407 | 2.59439255 | 2.430271133 |
| ENSMUSG00000048155 | 0 | 0 | 0 | 0 | 0 | 0 | 0 |
| ENSMUSG00000048163 | 0.578129653 | 0.396240625 | 0.25 | 0.25 | 0.387761935 | 0.301029996 | 0.224243824 |
| ENSMUSG00000048164 | 0.45815691 | 0.580482024 | 0.701838731 | 0.396240625 | 0.632412477 | 0.301029996 | 0.355418052 |
| ENSMUSG00000048170 | 2.53552908 | 2.368933358 | 2.372963274 | 2.403217124 | 2.372368362 | 2.672097858 | 2.200679824 |
| ENSMUSG00000048173 | 0 | 0 | 0 | 0 | 0 | 0 | 0 |
| ENSMUSG00000048174 | 1.389637216 | 1.372963274 | 1.445339928 | 1.482684334 | 1.467903253 | 1.591064607 | 1.433437704 |
| ENSMUSG00000048175 | 2.302628051 | 2.156427211 | 2.151832578 | 2.089388001 | 2.10031573 | 2.320146286 | 1.999330888 |
| ENSMUSG00000048185 | 0 | 0 | 0.25 | 0 | 0 | 0 | 0 |
| ENSMUSG00000048186 | 0.811507563 | 0.830482024 | 0.5 | 0.951838731 | 1.057362051 | 1.146128036 | 0.952571511 |
| ENSMUSG00000048188 | 0 | 0 | 0 | 0 | 0 | 0 | 0 |
| ENSMUSG00000048191 | 0.45815691 | 0.646240625 | 0.5 | 0.5 | 0.568060967 | 0.698970004 | 0.579661877 |
| ENSMUSG00000048197 | 0 | 0 | 0 | 0 | 0 | 0 | 0 |
| ENSMUSG00000048200 | 1.100572389 | 1.061981878 | 1.04248125 | 0.951838731 | 0.955822902 | 1 | 0.984950056 |
| ENSMUSG00000048206 | 0 | 0 | 0 | 0 | 0 | 0 | 0 |
| ENSMUSG00000048215 | 0 | 0 | 0 | 0 | 0 | 0 | 0 |
| ENSMUSG00000048216 | 1.578129653 | 1.54745614 | 1.580482024 | 1.114857905 | 1.176121988 | 1 | 1.078019652 |
| ENSMUSG00000048217 | 0 | 0 | 0 | 0.25 | 0 | 0 | 0 |
| ENSMUSG00000048218 | 0.91631382 | 1.061981878 | 1.130890489 | 2.056000419 | 2.030405741 | 2.206825876 | 1.854824501 |
| ENSMUSG00000048222 | 1.700465191 | 1.532320754 | 1.580482024 | 1.602347734 | 1.635855979 | 1.62324929 | 1.523605797 |
| ENSMUSG00000048226 | 0 | 0 | 0 | 0 | 0 | 0 | 0 |
| ENSMUSG00000048230 | 0 | 0.25 | 0 | 0.5 | 0.387761935 | 0 | 0.355418052 |
| ENSMUSG00000048231 | 0 | 0 | 0 | 0 | 0 | 0 | 0 |
| ENSMUSG00000048232 | 2.139333929 | 1.892463902 | 1.950224975 | 1.888647213 | 1.897237431 | 2.133538908 | 1.880961074 |
| ENSMUSG00000048234 | 1.985980784 | 1.884789703 | 1.898114259 | 2.084962501 | 1.983754904 | 2.220108088 | 1.811176017 |
| ENSMUSG00000048236 | 0 | 0 | 0 | 0 | 0 | 0 | 0 |
| ENSMUSG00000048240 | 1.558729299 | 1.311981878 | 1.403677461 | 1.972185812 | 2.00243419 | 2.238046103 | 1.901538355 |
| ENSMUSG00000048249 | 2.144240249 | 1.979715809 | 2.00835575 | 1.930274797 | 1.875328588 | 2.08278537 | 1.731408809 |
| ENSMUSG00000048251 | 0 | 0 | 0.25 | 0.25 | 0.489301084 | 0 | 0.520678036 |
| ENSMUSG00000048261 | 0 | 0 | 0 | 0 | 0 | 0 | 0.224243824 |
| ENSMUSG00000048264 | 2.323768721 | 2.135757955 | 2.146240625 | 2.176839783 | 2.129414728 | 2.411619706 | 1.988432495 |
| ENSMUSG00000048271 | 2.705574381 | 2.452642909 | 2.466046536 | 2.419870025 | 2.397399561 | 2.698100546 | 2.245582246 |
| ENSMUSG00000048277 | 2.328498818 | 2.195339928 | 2.257666784 | 2.302363341 | 2.358048572 | 2.614897216 | 2.268818981 |
| ENSMUSG00000048279 | 2.718080037 | 2.492166698 | 2.474461364 | 2.604199382 | 2.487157309 | 2.830588669 | 2.296701469 |

| | | | | | | | |
|---|---|---|---|---|---|---|---|
| ENSMUSG00000048280 | 1.605636262 | 1.52186571 | 1.5 | 1.705044741 | 1.680838453 | 1.86332286 | 1.483307832 |
| ENSMUSG00000048281 | 0 | 0 | 0 | 0 | 0 | 0 | 0 |
| ENSMUSG00000048284 | 0 | 0 | 0 | 0 | 0 | 0 | 0 |
| ENSMUSG00000048285 | 2.701748369 | 2.494677615 | 2.492885888 | 2.500352049 | 2.48127199 | 2.811575006 | 2.347980588 |
| ENSMUSG00000048292 | 0.289064826 | 0.396240625 | 0.25 | 0 | 0 | 0 | 0 |
| ENSMUSG00000048294 | 0 | 0 | 0 | 0 | 0 | 0 | 0 |
| ENSMUSG00000048299 | 0 | 0 | 0 | 0 | 0 | 0 | 0 |
| ENSMUSG00000048304 | 0 | 0 | 0 | 0 | 0 | 0 | 0.224243824 |
| ENSMUSG00000048307 | 2.583267694 | 2.306603048 | 2.298689214 | 2.354463129 | 2.360695755 | 2.671172843 | 2.184841529 |
| ENSMUSG00000048310 | 2.183765914 | 2.089388001 | 2.097004321 | 2.149978211 | 2.164053936 | 2.440909082 | 2.066798338 |
| ENSMUSG00000048312 | 0 | 0 | 0 | 0 | 0 | 0 | 0 |
| ENSMUSG00000048316 | 0 | 0 | 0 | 0 | 0 | 0 | 0 |
| ENSMUSG00000048327 | 2.446507203 | 2.275165585 | 2.255591953 | 2.510414788 | 2.429560273 | 2.78604121 | 2.271140178 |
| ENSMUSG00000048329 | 0 | 0 | 0 | 0 | 0 | 0 | 0 |
| ENSMUSG00000048330 | 0 | 0 | 0.25 | 0.396240625 | 0.244650542 | 0 | 0.224243824 |
| ENSMUSG00000048332 | 2.660010325 | 2.478596283 | 2.513820609 | 2.300530956 | 2.288791717 | 2.542825427 | 2.157274883 |
| ENSMUSG00000048334 | 0.289064826 | 0 | 0 | 0 | 0 | 0 | 0 |
| ENSMUSG00000048337 | 0.671187741 | 0.5 | 0.396240625 | 0.25 | 0.244650542 | 0.477121255 | 0.448487648 |
| ENSMUSG00000048347 | 1.249317394 | 1.114857905 | 1.080482024 | 0.396240625 | 0.489301084 | 0.602059991 | 0.224243824 |
| ENSMUSG00000048349 | 0 | 0 | 0.25 | 0.396240625 | 0.387761935 | 0.301029996 | 0.448487648 |
| ENSMUSG00000048351 | 1.857607105 | 1.794977273 | 1.825945187 | 1.823655187 | 1.836786029 | 2.071882007 | 1.743447983 |
| ENSMUSG00000048355 | 0.45815691 | 0.5 | 0 | 0.580482024 | 0.812711509 | 0.698970004 | 0.520678036 |
| ENSMUSG00000048356 | 0 | 0 | 0 | 0 | 0 | 0 | 0 |
| ENSMUSG00000048368 | 1.631440309 | 1.557204673 | 1.464495249 | 1.282320754 | 1.234113756 | 1.397940009 | 1.193409508 |
| ENSMUSG00000048371 | 1.693336259 | 1.618933358 | 1.618933358 | 1.720660762 | 1.726078673 | 1.897627091 | 1.591689859 |
| ENSMUSG00000048373 | 0.289064826 | 0.5 | 0 | 0.25 | 0.489301084 | 0.477121255 | 0.224243824 |
| ENSMUSG00000048376 | 3.046219753 | 2.83076369 | 2.8158173 | 2.694519282 | 2.666348738 | 3.012837225 | 2.488369842 |
| ENSMUSG00000048377 | 0 | 0 | 0 | 0 | 0 | 0 | 0 |
| ENSMUSG00000048378 | 0 | 0 | 0 | 0 | 0 | 0 | 0 |
| ENSMUSG00000048379 | 2.389637216 | 2.245748394 | 2.242885888 | 2.497171172 | 2.413656993 | 2.713490543 | 2.204609933 |
| ENSMUSG00000048385 | 0 | 0 | 0 | 0 | 0 | 0 | 0.224243824 |
| ENSMUSG00000048387 | 2.010213549 | 1.979715809 | 2.077084758 | 1.372963274 | 1.509475496 | 1.491361694 | 1.681794123 |
| ENSMUSG00000048388 | 2.227926429 | 2.032320754 | 2.073655187 | 2.057204673 | 2.022473809 | 2.206825876 | 1.844181586 |
| ENSMUSG00000048391 | 0 | 0 | 0 | 0 | 0 | 0 | 0 |

| | | | | | | | |
|---|---|---|---|---|---|---|---|
| ENSMUSG00000048398 | 0 | 0 | 0 | 0 | 0 | 0 | 0 |
| ENSMUSG00000048399 | 0 | 0 | 0.25 | 0 | 0 | 0 | 0 |
| ENSMUSG00000048400 | 0 | 0 | 0 | 0 | 0 | 0 | 0 |
| ENSMUSG00000048402 | 1.985980784 | 1.976722649 | 2.025821952 | 1.602347734 | 1.78010778 | 1.86332286 | 1.712465909 |
| ENSMUSG00000048406 | 0.747221736 | 0.5 | 0.896240625 | 0.646240625 | 0.812711509 | 0.602059991 | 0.853775828 |
| ENSMUSG00000048410 | 2.269664473 | 2.086073977 | 2.066696635 | 2.142463902 | 2.10031573 | 2.298853076 | 1.936709733 |
| ENSMUSG00000048411 | 0 | 0 | 0 | 0 | 0 | 0 | 0 |
| ENSMUSG00000048416 | 1.800532393 | 1.634789703 | 1.701838731 | 1.668106335 | 1.519146132 | 1.838849091 | 1.502532226 |
| ENSMUSG00000048424 | 0 | 0.25 | 0.396240625 | 0 | 0 | 0.301029996 | 0 |
| ENSMUSG00000048425 | 0 | 0 | 0 | 0 | 0 | 0 | 0 |
| ENSMUSG00000048429 | 2.377269616 | 2.145300145 | 2.158248799 | 2.316696635 | 2.27449559 | 2.558708571 | 2.095400649 |
| ENSMUSG00000048439 | 1.83262764 | 1.653677461 | 1.695339928 | 1.755591953 | 1.665423072 | 1.903089987 | 1.612273755 |
| ENSMUSG00000048440 | 1.940852214 | 1.75 | 1.777131114 | 1.610735874 | 1.65906333 | 1.740362689 | 1.598697688 |
| ENSMUSG00000048442 | 0.867194479 | 0.396240625 | 0.580482024 | 0.701838731 | 0.632412477 | 0 | 0.629532004 |
| ENSMUSG00000048445 | 1.527823874 | 1.470660762 | 1.494319981 | 1.396240625 | 1.462344774 | 1.380211242 | 1.508694612 |
| ENSMUSG00000048450 | 0 | 0 | 0 | 1.388647213 | 1.467903253 | 1.301029996 | 1.433437704 |
| ENSMUSG00000048455 | 0 | 0 | 0 | 0 | 0 | 0 | 0 |
| ENSMUSG00000048456 | 0 | 0 | 0 | 0 | 0 | 0 | 0 |
| ENSMUSG00000048458 | 1.663535556 | 1.54745614 | 1.618933358 | 1.160964047 | 1.212046813 | 1.361727836 | 1.150210039 |
| ENSMUSG00000048469 | 0 | 0 | 0 | 0 | 0 | 0 | 0 |
| ENSMUSG00000048473 | 0 | 0 | 0 | 0 | 0 | 0 | 0 |
| ENSMUSG00000048478 | 0.289064826 | 0.25 | 0.25 | 0.646240625 | 0.387761935 | 0.602059991 | 0.448487648 |
| ENSMUSG00000048480 | 0 | 0 | 0 | 0 | 0 | 0 | 0 |
| ENSMUSG00000048481 | 1.069666964 | 1 | 1.098079356 | 0.976722649 | 1.149965125 | 1.176091259 | 1.066254157 |
| ENSMUSG00000048482 | 2.185978293 | 2.03743678 | 2 | 1.744319981 | 1.645989387 | 1.892094603 | 1.579661877 |
| ENSMUSG00000048483 | 0 | 0 | 0 | 0 | 0 | 0 | 0 |
| ENSMUSG00000048484 | 0.289064826 | 0.396240625 | 0 | 0.25 | 0.244650542 | 0.301029996 | 0 |
| ENSMUSG00000048485 | 0.289064826 | 0 | 0.25 | 0.396240625 | 0.489301084 | 0 | 0.355418052 |
| ENSMUSG00000048486 | 1.842799666 | 1.729715809 | 1.671625132 | 1.914552871 | 1.928492895 | 2.089905111 | 1.779742043 |
| ENSMUSG00000048489 | 0 | 0.5 | 0 | 0 | 0 | 0 | 0 |
| ENSMUSG00000048490 | 2.136858952 | 2.04248125 | 1.96759118 | 2.006976499 | 1.925488995 | 2.117271296 | 1.811176017 |
| ENSMUSG00000048495 | 1.862428343 | 1.789967834 | 1.76109853 | 1.918106335 | 1.877063019 | 2.041392685 | 1.709198048 |
| ENSMUSG00000048497 | 1.867194479 | 1.811981878 | 1.845926073 | 1.901832578 | 1.85749873 | 2.089905111 | 1.74492186 |
| ENSMUSG00000048498 | 0 | 0 | 0 | 0 | 0 | 0 | 0 |

| | | | | | | | |
|---|---|---|---|---|---|---|---|
| ENSMUSG00000048500 | 0 | 0 | 0 | 0 | 0 | 0 | 0 |
| ENSMUSG00000048501 | 0 | 0 | 0 | 0 | 0 | 0 | 0 |
| ENSMUSG00000048502 | 0 | 0 | 0 | 0 | 0 | 0 | 0 |
| ENSMUSG00000048503 | 1.205378646 | 1.25 | 1.311981878 | 1.584962501 | 1.523883869 | 1.698970004 | 1.462868481 |
| ENSMUSG00000048516 | 0 | 0 | 0 | 0 | 0 | 0 | 0 |
| ENSMUSG00000048520 | 0.747221736 | 0.701838731 | 0.25 | 0.396240625 | 0.387761935 | 0.477121255 | 0.224243824 |
| ENSMUSG00000048521 | 0.91631382 | 0.75 | 0.646240625 | 0.976722649 | 0.955822902 | 1.113943352 | 1.054044578 |
| ENSMUSG00000048528 | 0 | 0 | 0 | 0.25 | 0.244650542 | 0 | 0.224243824 |
| ENSMUSG00000048534 | 0 | 0 | 0 | 0 | 0 | 0 | 0.355418052 |
| ENSMUSG00000048537 | 2.598312578 | 2.462546709 | 2.536801231 | 2.364857905 | 2.581481146 | 2.689308859 | 2.527817612 |
| ENSMUSG00000048538 | 0 | 0 | 0 | 0 | 0 | 0 | 0 |
| ENSMUSG00000048540 | 0 | 0 | 0 | 0 | 0 | 0 | 0 |
| ENSMUSG00000048544 | 0 | 0 | 0.25 | 0 | 0 | 0 | 0 |
| ENSMUSG00000048546 | 2.425923778 | 2.264497931 | 2.332168732 | 2.172499493 | 2.200473444 | 2.465382851 | 2.148102439 |
| ENSMUSG00000048550 | 1.89911417 | 1.830482024 | 1.726722649 | 1.678561379 | 1.639266209 | 1.913813852 | 1.514741806 |
| ENSMUSG00000048559 | 0 | 0 | 0 | 0 | 0 | 0 | 0 |
| ENSMUSG00000048562 | 1.956444026 | 1.854463129 | 1.809601185 | 0.976722649 | 1 | 1.041392685 | 0.803905701 |
| ENSMUSG00000048572 | 0.45815691 | 0.976722649 | 0.580482024 | 0.25 | 0.244650542 | 0 | 0 |
| ENSMUSG00000048573 | 0 | 0 | 0 | 0 | 0 | 0 | 0 |
| ENSMUSG00000048574 | 0 | 0 | 0 | 0 | 0 | 0 | 0 |
| ENSMUSG00000048578 | 3.3771428 | 3.062870603 | 3.044510582 | 3.076586638 | 3.025561104 | 3.518908574 | 2.807065219 |
| ENSMUSG00000048581 | 1.432083896 | 1.330482024 | 1.431980114 | 1.348079356 | 1.27449559 | 1.431363764 | 1.193409508 |
| ENSMUSG00000048582 | 0 | 0.396240625 | 0.701838731 | 0.396240625 | 0.632412477 | 0.602059991 | 0.775756176 |
| ENSMUSG00000048583 | 0.289064826 | 0 | 0 | 0.92510993 | 0.733951626 | 0.903089987 | 0.952571511 |
| ENSMUSG00000048592 | 0 | 0 | 0 | 0 | 0 | 0.301029996 | 0.224243824 |
| ENSMUSG00000048600 | 0 | 0 | 0 | 0 | 0 | 0 | 0 |
| ENSMUSG00000048602 | 0 | 0 | 0.396240625 | 0 | 0.244650542 | 0 | 0 |
| ENSMUSG00000048603 | 0.671187741 | 0.79248125 | 0.92510993 | 0.646240625 | 0.686820904 | 0.477121255 | 0.775756176 |
| ENSMUSG00000048612 | 3.622099732 | 3.266059383 | 3.269370839 | 3.179900682 | 3.107646394 | 3.60948769 | 2.897225857 |
| ENSMUSG00000048616 | 0.289064826 | 0.701838731 | 0.396240625 | 1.04248125 | 1.020174412 | 1.255272505 | 1.140832123 |
| ENSMUSG00000048617 | 0.671187741 | 0.830482024 | 0.75 | 0.580482024 | 0.244650542 | 0.301029996 | 0.520678036 |
| ENSMUSG00000048620 | 0 | 0 | 0 | 0 | 0 | 0 | 0 |
| ENSMUSG00000048621 | 0.578129653 | 0.864857905 | 1.02186571 | 0.580482024 | 0.244650542 | 0.301029996 | 0.520678036 |
| ENSMUSG00000048626 | 0 | 0 | 0 | 0 | 0 | 0 | 0 |

| | | | | | | | |
|---|---|---|---|---|---|---|---|
| ENSMUSG00000048628 | 0 | 0 | 0 | 0 | 0 | 0 | 0 |
| ENSMUSG00000048636 | 0 | 0.25 | 0.25 | 0 | 0 | 0 | 0 |
| ENSMUSG00000048643 | 0 | 0 | 0 | 0 | 0 | 0 | 0 |
| ENSMUSG00000048644 | 2.361637952 | 2.159156155 | 2.195339928 | 1.649978211 | 1.683842353 | 1.908485019 | 1.651852264 |
| ENSMUSG00000048647 | 0.747221736 | 0.646240625 | 0.79248125 | 0.75 | 0.846351821 | 1.041392685 | 0.896975297 |
| ENSMUSG00000048652 | 0 | 0 | 0 | 0 | 0 | 0 | 0 |
| ENSMUSG00000048655 | 0.289064826 | 0.5 | 0.580482024 | 0.646240625 | 0.244650542 | 0.698970004 | 0.355418052 |
| ENSMUSG00000048661 | 2.139333929 | 2.038704527 | 2.015173983 | 2.181980114 | 2.117361396 | 2.394451681 | 1.980010793 |
| ENSMUSG00000048668 | 2.160971586 | 1.950224975 | 1.966046536 | 1.87694866 | 1.883916752 | 2.158362492 | 1.779742043 |
| ENSMUSG00000048677 | 1.783508299 | 1.557204673 | 1.668106335 | 1.589388001 | 1.588235379 | 1.763427994 | 1.596378571 |
| ENSMUSG00000048686 | 0 | 0 | 0 | 0 | 0 | 0 | 0 |
| ENSMUSG00000048693 | 0 | 0 | 0 | 0 | 0 | 0 | 0 |
| ENSMUSG00000048696 | 2.194712275 | 2.112802778 | 2.200224975 | 1.766522298 | 1.862942891 | 1.991226076 | 1.904245613 |
| ENSMUSG00000048697 | 0 | 0 | 0 | 0 | 0 | 0 | 0 |
| ENSMUSG00000048699 | 0 | 0 | 0 | 0 | 0 | 0 | 0 |
| ENSMUSG00000048701 | 2.719310222 | 2.495034894 | 2.474461364 | 2.406883471 | 2.318738698 | 2.63447727 | 2.19156611 |
| ENSMUSG00000048703 | 0 | 0 | 0 | 0 | 0 | 0 | 0 |
| ENSMUSG00000048706 | 2.170239354 | 2.044977273 | 2.01109853 | 2.192872367 | 2.160196454 | 2.419955748 | 2.00464543 |
| ENSMUSG00000048707 | 1.707474304 | 1.692046081 | 1.774508021 | 1.67510993 | 1.739104362 | 1.851258349 | 1.690609803 |
| ENSMUSG00000048709 | 0 | 0 | 0.396240625 | 0 | 0.244650542 | 0 | 0 |
| ENSMUSG00000048720 | 2.227926429 | 2.099151195 | 2.114857905 | 2.061981878 | 1.979897422 | 2.176091259 | 1.837623616 |
| ENSMUSG00000048721 | 0 | 0.25 | 0.396240625 | 0.830482024 | 0.489301084 | 0.84509804 | 0.876096088 |
| ENSMUSG00000048728 | 0.578129653 | 0.646240625 | 0.580482024 | 0.951838731 | 0.846351821 | 0.954242509 | 0.775756176 |
| ENSMUSG00000048731 | 1.036286563 | 1 | 1 | 0.864857905 | 0.812711509 | 0.301029996 | 0.629532004 |
| ENSMUSG00000048732 | 2.097855556 | 1.894357207 | 1.903677461 | 2.028435542 | 1.938809675 | 2.155336037 | 1.791413205 |
| ENSMUSG00000048745 | 0 | 0 | 0 | 0 | 0 | 0 | 0 |
| ENSMUSG00000048747 | 0 | 0 | 0 | 0 | 0 | 0 | 0 |
| ENSMUSG00000048752 | 0 | 0 | 0 | 0 | 0 | 0 | 0 |
| ENSMUSG00000048755 | 2.039433992 | 1.958222504 | 1.903677461 | 1.744319981 | 1.806427258 | 1.973127854 | 1.746389053 |
| ENSMUSG00000048756 | 2.457006475 | 2.247171172 | 2.263142013 | 2.069031101 | 2.008447216 | 2.307496038 | 1.969900112 |
| ENSMUSG00000048758 | 1.734388958 | 1.649978211 | 1.741446071 | 1.854463129 | 1.816839698 | 2.004321374 | 1.803905701 |
| ENSMUSG00000048763 | 1.482695304 | 1.431980114 | 1.418106335 | 1.238549078 | 1.293076518 | 1.380211242 | 1.272006351 |
| ENSMUSG00000048764 | 0 | 0 | 0.5 | 0.580482024 | 0.632412477 | 0.602059991 | 0.629532004 |
| ENSMUSG00000048766 | 0 | 0 | 0 | 0 | 0 | 0 | 0 |

| | | | | | | | |
|---|---|---|---|---|---|---|---|
| ENSMUSG00000048772 | 1.205378646 | 1.146240625 | 1.282320754 | 0.92510993 | 1.200473444 | 1 | 1.278288402 |
| ENSMUSG00000048775 | 0 | 0 | 0 | 0 | 0.244650542 | 0 | 0 |
| ENSMUSG00000048776 | 0 | 0.25 | 0.646240625 | 0.396240625 | 0.387761935 | 0 | 0.355418052 |
| ENSMUSG00000048779 | 0 | 0 | 0.396240625 | 0 | 0 | 0.301029996 | 0 |
| ENSMUSG00000048782 | 0.867194479 | 0.396240625 | 0.701838731 | 0.25 | 0.489301084 | 0.77815125 | 0.355418052 |
| ENSMUSG00000048787 | 1.857607105 | 1.777131114 | 1.816696635 | 1.685366747 | 1.618292349 | 1.792391689 | 1.493062805 |
| ENSMUSG00000048794 | 0 | 0 | 0.830482024 | 0 | 0.387761935 | 0.477121255 | 0.520678036 |
| ENSMUSG00000048796 | 1.578129653 | 1.438721876 | 1.438721876 | 1.681980114 | 1.677808768 | 1.748188027 | 1.554175045 |
| ENSMUSG00000048799 | 2.471720134 | 2.192046081 | 2.179419106 | 2.185366747 | 2.123439512 | 2.419955748 | 2.003324957 |
| ENSMUSG00000048806 | 0 | 0 | 0 | 0 | 0 | 0 | 0 |
| ENSMUSG00000048807 | 2.033115198 | 2.01109853 | 2.099151195 | 1.598079356 | 1.791313678 | 1.826074803 | 1.854824501 |
| ENSMUSG00000048810 | 0 | 0 | 0 | 0 | 0 | 0 | 0 |
| ENSMUSG00000048814 | 0 | 0 | 0.25 | 0.864857905 | 0.812711509 | 0.84509804 | 0.803905701 |
| ENSMUSG00000048824 | 0 | 0 | 0 | 0 | 0 | 0 | 0 |
| ENSMUSG00000048826 | 1.867194479 | 1.802363341 | 1.890560606 | 1.04248125 | 1.234113756 | 1.113943352 | 1.374453863 |
| ENSMUSG00000048827 | 0.91631382 | 0.79248125 | 0.580482024 | 0.951838731 | 1 | 1.041392685 | 0.969165684 |
| ENSMUSG00000048830 | 0 | 0 | 0 | 0 | 0 | 0 | 0 |
| ENSMUSG00000048832 | 2.258689303 | 2.140560606 | 2.164552871 | 2.027131114 | 2.090057366 | 2.206825876 | 2.007923223 |
| ENSMUSG00000048833 | 2.560530738 | 2.405054956 | 2.373963757 | 2.429847205 | 2.373641807 | 2.688419822 | 2.183328003 |
| ENSMUSG00000048834 | 0 | 0 | 0 | 0 | 0 | 0 | 0 |
| ENSMUSG00000048852 | 0.960252568 | 0.580482024 | 0.5 | 0.701838731 | 0.489301084 | 0.477121255 | 0.448487648 |
| ENSMUSG00000048856 | 1.269664473 | 1.214495249 | 1.146240625 | 0.864857905 | 1.074582839 | 0.77815125 | 1.054044578 |
| ENSMUSG00000048865 | 0.289064826 | 0 | 0.25 | 0.5 | 0.489301084 | 0.301029996 | 0 |
| ENSMUSG00000048874 | 2.818344313 | 2.617410454 | 2.611253711 | 2.606566189 | 2.536393639 | 2.911690159 | 2.361597253 |
| ENSMUSG00000048875 | 0 | 0 | 0 | 0 | 0 | 0 | 0 |
| ENSMUSG00000048878 | 2.231898196 | 2.147178659 | 2.220660762 | 2.079353153 | 2.181274723 | 2.374748346 | 2.183707049 |
| ENSMUSG00000048895 | 1.587501561 | 1.54248125 | 1.664552871 | 1.238549078 | 1.121713561 | 1.278753601 | 1.23151414 |
| ENSMUSG00000048897 | 1.614416215 | 1.708222504 | 1.681980114 | 1.482684334 | 1.611014646 | 1.568201724 | 1.627424404 |
| ENSMUSG00000048899 | 0.289064826 | 0.79248125 | 0 | 0.75 | 0.846351821 | 0.903089987 | 0.876096088 |
| ENSMUSG00000048904 | 0 | 0 | 0 | 0 | 0 | 0 | 0 |
| ENSMUSG00000048905 | 0.289064826 | 0.5 | 0.5 | 0.25 | 0.387761935 | 0.477121255 | 0.224243824 |
| ENSMUSG00000048911 | 2.355805236 | 2.179419106 | 2.151832578 | 2.14151351 | 2.088159748 | 2.36361198 | 1.930969131 |
| ENSMUSG00000048915 | 1.852729478 | 1.784887838 | 1.809601185 | 1.852347734 | 1.816839698 | 1.959041392 | 1.723650273 |
| ENSMUSG00000048916 | 0 | 0 | 0.25 | 0 | 0 | 0 | 0 |

| | | | | | | | |
|---|---|---|---|---|---|---|---|
| ENSMUSG00000048919 | 0 | 0 | 0 | 0 | 0 | 0 | 0 |
| ENSMUSG00000048920 | 1.568542279 | 1.610735874 | 1.622963274 | 1.606566189 | 1.65906333 | 1.633468456 | 1.591689859 |
| ENSMUSG00000048921 | 0.91631382 | 1.098079356 | 1.146240625 | 0.864857905 | 0.877063019 | 0.77815125 | 1 |
| ENSMUSG00000048922 | 2.337801042 | 2.173371739 | 2.161864607 | 2.413659007 | 2.361573756 | 2.691965103 | 2.251165585 |
| ENSMUSG00000048924 | 1.325351389 | 1.061981878 | 1.282320754 | 1.464495249 | 1.310728002 | 1.491361694 | 1.329931221 |
| ENSMUSG00000048930 | 2.083999385 | 2.059601185 | 2.05479213 | 1.777131114 | 1.903713872 | 1.973127854 | 1.915722126 |
| ENSMUSG00000048933 | 0 | 0 | 0 | 0 | 0 | 0 | 0 |
| ENSMUSG00000048938 | 0 | 0.396240625 | 0.25 | 0.25 | 0 | 0 | 0 |
| ENSMUSG00000048939 | 0 | 0 | 0 | 0 | 0 | 0 | 0 |
| ENSMUSG00000048949 | 1.227926429 | 1.238549078 | 1.146240625 | 1.311981878 | 1.121713561 | 1.322219295 | 1.089372236 |
| ENSMUSG00000048960 | 2.936496745 | 2.68473417 | 2.668548067 | 2.244319981 | 2.209357611 | 2.485721426 | 2.03601839 |
| ENSMUSG00000048965 | 1.069666964 | 0.896240625 | 0.976722649 | 1.098079356 | 1.149965125 | 1.204119983 | 1.121219121 |
| ENSMUSG00000048967 | 0.289064826 | 0.5 | 0.79248125 | 0 | 0.244650542 | 0.301029996 | 0.829800754 |
| ENSMUSG00000048970 | 2.22592627 | 2.104463129 | 2.039967834 | 2.04745614 | 2.030405741 | 2.31386722 | 1.901538355 |
| ENSMUSG00000048978 | 0 | 0 | 0 | 0.5 | 0.568060967 | 0.602059991 | 0.710836105 |
| ENSMUSG00000048981 | 0 | 0 | 0 | 0 | 0 | 0 | 0 |
| ENSMUSG00000048982 | 0 | 0 | 0.25 | 0 | 0 | 0 | 0 |
| ENSMUSG00000048988 | 0.671187741 | 0.951838731 | 1.098079356 | 0 | 0 | 0.301029996 | 0.448487648 |
| ENSMUSG00000048992 | 0 | 0.5 | 0.5 | 0.396240625 | 0 | 0 | 0 |
| ENSMUSG00000048994 | 0 | 0 | 0 | 0 | 0 | 0 | 0 |
| ENSMUSG00000048996 | 0 | 0 | 0 | 0 | 0.244650542 | 0 | 0 |
| ENSMUSG00000048997 | 1.800532393 | 1.711372513 | 1.777131114 | 1.505591953 | 1.812711509 | 1.69019608 | 1.759302692 |
| ENSMUSG00000049001 | 2.060824957 | 1.845926073 | 1.685366747 | 1.779735268 | 1.726078673 | 1.968482949 | 1.633706455 |
| ENSMUSG00000049008 | 0 | 0 | 0 | 0 | 0 | 0 | 0 |
| ENSMUSG00000049010 | 0 | 0 | 0 | 0 | 0 | 0 | 0 |
| ENSMUSG00000049011 | 0 | 0 | 0 | 0 | 0 | 0 | 0 |
| ENSMUSG00000049013 | 0 | 0 | 0 | 0 | 0 | 0 | 0.224243824 |
| ENSMUSG00000049015 | 0 | 0 | 0 | 0 | 0 | 0 | 0 |
| ENSMUSG00000049018 | 0 | 0 | 0 | 0 | 0 | 0 | 0 |
| ENSMUSG00000049028 | 0 | 0 | 0 | 0 | 0 | 0 | 0 |
| ENSMUSG00000049036 | 1.036286563 | 1.29248125 | 1.494319981 | 0.896240625 | 1.163285805 | 0.602059991 | 1.313616061 |
| ENSMUSG00000049037 | 0 | 0 | 0 | 0 | 0 | 0 | 0 |
| ENSMUSG00000049038 | 1.83262764 | 1.79745614 | 1.698603967 | 1.794977273 | 1.718026092 | 1.908485019 | 1.591689859 |
| ENSMUSG00000049041 | 0 | 0 | 0 | 0 | 0 | 0 | 0 |

| | | | | | | | |
|---|---|---|---|---|---|---|---|
| ENSMUSG00000049044 | 0 | 0 | 0.25 | 0 | 0.244650542 | 0 | 0.224243824 |
| ENSMUSG00000049047 | 2.569423023 | 2.381869252 | 2.379417347 | 2.409608978 | 2.364194729 | 2.705863712 | 2.172922202 |
| ENSMUSG00000049052 | 0 | 0 | 0 | 0 | 0 | 0 | 0 |
| ENSMUSG00000049057 | 0 | 0 | 0 | 0 | 0 | 0 | 0 |
| ENSMUSG00000049062 | 0 | 0 | 0 | 0 | 0 | 0 | 0 |
| ENSMUSG00000049073 | 0 | 0 | 0 | 0 | 0 | 0 | 0 |
| ENSMUSG00000049076 | 2.771218178 | 2.574229052 | 2.572216519 | 2.533285553 | 2.480020371 | 2.836956737 | 2.311326126 |
| ENSMUSG00000049086 | 0.289064826 | 0.580482024 | 0.25 | 0.646240625 | 0.686820904 | 0.602059991 | 1.041356071 |
| ENSMUSG00000049090 | 2.167941649 | 2.05357978 | 2.063166358 | 1.988549078 | 1.934425071 | 2.146128036 | 1.813563596 |
| ENSMUSG00000049091 | 1.847794126 | 1.794977273 | 1.821350555 | 1.799918086 | 1.818885826 | 1.924279286 | 1.707551656 |
| ENSMUSG00000049092 | 0.960252568 | 0.701838731 | 0.75 | 1.02186571 | 1.149965125 | 0.954242509 | 0.935079929 |
| ENSMUSG00000049093 | 0.578129653 | 0.396240625 | 0.25 | 0.79248125 | 0.877063019 | 0.77815125 | 0.74492186 |
| ENSMUSG00000049094 | 0 | 0 | 0 | 0 | 0 | 0 | 0 |
| ENSMUSG00000049097 | 0.45815691 | 0.92510993 | 0.896240625 | 0.79248125 | 0.812711509 | 1.041392685 | 0.969165684 |
| ENSMUSG00000049098 | 0 | 0 | 0 | 0 | 0 | 0 | 0 |
| ENSMUSG00000049100 | 0.289064826 | 0.580482024 | 0.396240625 | 0.25 | 0.686820904 | 0.301029996 | 0 |
| ENSMUSG00000049103 | 0.289064826 | 0 | 0.25 | 0 | 0 | 0 | 0 |
| ENSMUSG00000049106 | 2.406877894 | 2.216819685 | 2.234894804 | 2.105516192 | 2.117361396 | 2.330413773 | 2.01501956 |
| ENSMUSG00000049107 | 0 | 0 | 0.25 | 0.5 | 0.489301084 | 0.602059991 | 0.579661877 |
| ENSMUSG00000049109 | 0 | 0 | 0 | 0 | 0.244650542 | 0.301029996 | 0.224243824 |
| ENSMUSG00000049112 | 0.289064826 | 0 | 0 | 0 | 0.489301084 | 0 | 0.224243824 |
| ENSMUSG00000049115 | 0 | 0.25 | 0 | 0.396240625 | 0.568060967 | 0.301029996 | 0.355418052 |
| ENSMUSG00000049119 | 1.505869732 | 1.54745614 | 1.671625132 | 1.238549078 | 1.244650542 | 1.462397998 | 1.307989563 |
| ENSMUSG00000049122 | 0 | 0 | 0 | 0 | 0.244650542 | 0 | 0.224243824 |
| ENSMUSG00000049123 | 0 | 0.25 | 0.5 | 0.396240625 | 0.244650542 | 0 | 0.224243824 |
| ENSMUSG00000049124 | 0 | 0 | 0 | 0 | 0 | 0 | 0 |
| ENSMUSG00000049128 | 0 | 0 | 0 | 0 | 0 | 0 | 0 |
| ENSMUSG00000049130 | 0 | 0 | 0 | 0 | 0 | 0 | 0 |
| ENSMUSG00000049133 | 0 | 0 | 0 | 0 | 0 | 0 | 0 |
| ENSMUSG00000049134 | 0.289064826 | 0.25 | 0 | 0 | 0 | 0.301029996 | 0.672731473 |
| ENSMUSG00000049148 | 0 | 0 | 0 | 0 | 0 | 0 | 0 |
| ENSMUSG00000049149 | 0 | 0 | 0 | 0 | 0 | 0 | 0 |
| ENSMUSG00000049152 | 0 | 0 | 0 | 0 | 0 | 0 | 0.224243824 |
| ENSMUSG00000049154 | 0 | 0 | 0 | 0 | 0 | 0 | 0 |

| | | | | | | | |
|---|---|---|---|---|---|---|---|
| ENSMUSG00000049160 | 0 | 0 | 0 | 0.396240625 | 0 | 0 | 0 |
| ENSMUSG00000049164 | 2.310886388 | 2.151832578 | 2.14151351 | 2.369439567 | 2.254881871 | 2.530199698 | 2.076967568 |
| ENSMUSG00000049168 | 0 | 0 | 0 | 0 | 0 | 0 | 0 |
| ENSMUSG00000049173 | 0 | 0 | 0 | 0 | 0 | 0 | 0 |
| ENSMUSG00000049176 | 0 | 0 | 0 | 0.25 | 0 | 0 | 0 |
| ENSMUSG00000049184 | 1.505869732 | 1.431980114 | 1.51109853 | 1.54248125 | 1.427019704 | 1.414973348 | 1.417653332 |
| ENSMUSG00000049191 | 1.852729478 | 1.692046081 | 1.642463902 | 1.311981878 | 1.351342427 | 1.322219295 | 1.265599895 |
| ENSMUSG00000049202 | 0 | 0 | 0 | 0 | 0 | 0 | 0.224243824 |
| ENSMUSG00000049214 | 0 | 0 | 0 | 0 | 0.244650542 | 0 | 0.355418052 |
| ENSMUSG00000049217 | 0 | 0 | 0 | 0 | 0 | 0 | 0 |
| ENSMUSG00000049225 | 2.045658471 | 1.814346961 | 1.880890489 | 1.660964047 | 1.595992969 | 1.806179974 | 1.511732338 |
| ENSMUSG00000049229 | 0 | 0 | 0 | 0 | 0 | 0 | 0 |
| ENSMUSG00000049230 | 0 | 0.25 | 0.25 | 0 | 0 | 0.301029996 | 0 |
| ENSMUSG00000049231 | 0 | 0 | 0 | 0 | 0 | 0 | 0 |
| ENSMUSG00000049232 | 1.960252568 | 1.78743678 | 1.723704441 | 1.720660762 | 1.65906333 | 1.792391689 | 1.514741806 |
| ENSMUSG00000049233 | 0.289064826 | 0 | 0 | 0.25 | 0.244650542 | 0.477121255 | 0.224243824 |
| ENSMUSG00000049235 | 0 | 0 | 0 | 0 | 0 | 0 | 0 |
| ENSMUSG00000049241 | 0 | 0 | 0 | 0 | 0 | 0 | 0 |
| ENSMUSG00000049246 | 0 | 0 | 0 | 0.25 | 0.244650542 | 0.477121255 | 0 |
| ENSMUSG00000049249 | 0.289064826 | 0.580482024 | 0.396240625 | 0.25 | 0.489301084 | 0.301029996 | 0.448487648 |
| ENSMUSG00000049252 | 0 | 0.25 | 0 | 0.25 | 0.387761935 | 0.301029996 | 0.224243824 |
| ENSMUSG00000049265 | 0.747221736 | 1 | 0.830482024 | 0.25 | 0.244650542 | 0.301029996 | 0 |
| ENSMUSG00000049280 | 0 | 0 | 0 | 0 | 0 | 0 | 0 |
| ENSMUSG00000049281 | 0 | 0.396240625 | 0.396240625 | 0 | 0.387761935 | 0.301029996 | 0 |
| ENSMUSG00000049285 | 0.747221736 | 0.864857905 | 0.701838731 | 0.896240625 | 0.387761935 | 1 | 0.853775828 |
| ENSMUSG00000049287 | 1.527823874 | 1.372963274 | 1.488549078 | 1.451838731 | 1.523883869 | 1.633468456 | 1.537859885 |
| ENSMUSG00000049288 | 2.643399047 | 2.425543171 | 2.425975893 | 2.461372513 | 2.38035699 | 2.722633923 | 2.251165585 |
| ENSMUSG00000049291 | 0.289064826 | 0 | 0 | 0 | 0 | 0 | 0 |
| ENSMUSG00000049295 | 1.389637216 | 1.51109853 | 1.62694866 | 1.321350555 | 1.49445382 | 1.397940009 | 1.694069894 |
| ENSMUSG00000049299 | 2.262379879 | 2.08272922 | 2.077084758 | 2.269203899 | 2.173143437 | 2.542825427 | 2.050633136 |
| ENSMUSG00000049300 | 1.693336259 | 1.657339155 | 1.62694866 | 1.809601185 | 1.751666404 | 1.886490725 | 1.546120303 |
| ENSMUSG00000049303 | 1.631440309 | 1.494319981 | 1.552363341 | 1.606566189 | 1.770879644 | 1.681241237 | 1.596378571 |
| ENSMUSG00000049305 | 2.136858952 | 2.031030328 | 2.106566189 | 2.060793496 | 2.109389652 | 2.292256071 | 2.04833869 |
| ENSMUSG00000049307 | 1.623015126 | 1.634789703 | 1.575945187 | 1.451838731 | 1.394615667 | 1.579783597 | 1.355418052 |

| | | | | | | | |
|---|---|---|---|---|---|---|---|
| ENSMUSG00000049313 | 2.103271637 | 2.031030328 | 2.048689214 | 1.811981878 | 1.800029087 | 1.954242509 | 1.712465909 |
| ENSMUSG00000049314 | 0 | 0 | 0 | 0 | 0 | 0 | 0 |
| ENSMUSG00000049315 | 0 | 0 | 0 | 0 | 0 | 0 | 0 |
| ENSMUSG00000049321 | 1.596667477 | 1.54745614 | 1.482684334 | 1.62694866 | 1.572189156 | 1.633468456 | 1.476637173 |
| ENSMUSG00000049323 | 2.715608727 | 2.506976499 | 2.492885888 | 2.508699741 | 2.4345872 | 2.803457116 | 2.278005485 |
| ENSMUSG00000049327 | 2.883252421 | 2.680701883 | 2.707430684 | 2.642701109 | 2.605346297 | 2.951337519 | 2.461105449 |
| ENSMUSG00000049334 | 0 | 0 | 0 | 0 | 0 | 0 | 0 |
| ENSMUSG00000049336 | 0 | 0 | 0 | 0.25 | 0 | 0 | 0.224243824 |
| ENSMUSG00000049339 | 2.488076442 | 2.343216265 | 2.373963757 | 2.265173983 | 2.295199569 | 2.525044807 | 2.184841529 |
| ENSMUSG00000049349 | 0 | 0 | 0 | 0 | 0 | 0 | 0 |
| ENSMUSG00000049350 | 0 | 0 | 0 | 0 | 0 | 0 | 0 |
| ENSMUSG00000049353 | 0 | 0 | 0 | 0 | 0 | 0 | 0 |
| ENSMUSG00000049354 | 3.041424604 | 2.756630861 | 2.74929487 | 2.821350555 | 2.784765139 | 3.205745541 | 2.620919898 |
| ENSMUSG00000049357 | 0.811507563 | 0.79248125 | 0.580482024 | 0.92510993 | 0.905314583 | 0.84509804 | 0.952571511 |
| ENSMUSG00000049362 | 0 | 0 | 0 | 0 | 0 | 0 | 0 |
| ENSMUSG00000049372 | 0 | 0 | 0 | 0 | 0 | 0 | 0 |
| ENSMUSG00000049382 | 0.747221736 | 0.580482024 | 0.79248125 | 0 | 0 | 0.301029996 | 0.224243824 |
| ENSMUSG00000049387 | 0 | 0 | 0 | 0.25 | 0 | 0 | 0 |
| ENSMUSG00000049396 | 0.289064826 | 0.580482024 | 0.580482024 | 0 | 0.244650542 | 0.301029996 | 0.520678036 |
| ENSMUSG00000049401 | 2.229917041 | 2.167221246 | 2.193696765 | 2.295598588 | 2.405234647 | 2.617000341 | 2.364209774 |
| ENSMUSG00000049404 | 1.129344651 | 1.114857905 | 1.080482024 | 0.646240625 | 0.244650542 | 0.698970004 | 0.579661877 |
| ENSMUSG00000049409 | 1.325351389 | 1.188721876 | 1.226722649 | 1.339388001 | 1.293076518 | 1.447158031 | 1.252393349 |
| ENSMUSG00000049410 | 0 | 0 | 0 | 0 | 0 | 0 | 0 |
| ENSMUSG00000049411 | 1.389637216 | 1.282320754 | 1.160964047 | 1.403677461 | 1.351342427 | 1.602059991 | 1.319146374 |
| ENSMUSG00000049414 | 0 | 0 | 0 | 0 | 0 | 0 | 0 |
| ENSMUSG00000049420 | 1.227926429 | 1 | 0.951838731 | 0.896240625 | 1.020174412 | 1.041392685 | 0.672731473 |
| ENSMUSG00000049421 | 2.422183552 | 2.191217898 | 2.185366747 | 2.216819685 | 2.103062484 | 2.414973348 | 1.956418351 |
| ENSMUSG00000049422 | 0.578129653 | 0.75 | 1.114857905 | 0.580482024 | 0.733951626 | 0.602059991 | 0.710836105 |
| ENSMUSG00000049427 | 0 | 0 | 0 | 0 | 0 | 0 | 0.355418052 |
| ENSMUSG00000049436 | 0 | 0 | 0 | 0.25 | 0 | 0 | 0 |
| ENSMUSG00000049439 | 2.495495257 | 2.286164561 | 2.277131114 | 2.320772088 | 2.229044281 | 2.583198774 | 2.116124284 |
| ENSMUSG00000049456 | 0 | 0 | 0 | 0 | 0 | 0 | 0 |
| ENSMUSG00000049460 | 0 | 0.25 | 0 | 0.25 | 0 | 0 | 0 |
| ENSMUSG00000049470 | 3.139109537 | 2.861383055 | 2.840485943 | 3.006025195 | 2.935806505 | 3.399327532 | 2.715188562 |

| | | | | | | | |
|---|---|---|---|---|---|---|---|
| ENSMUSG00000049476 | 0 | 0 | 0 | 0 | 0 | 0 | 0 |
| ENSMUSG00000049477 | 0 | 0 | 0 | 0 | 0 | 0 | 0 |
| ENSMUSG00000049482 | 1.83262764 | 1.769203899 | 1.884789703 | 1.614857905 | 1.814781639 | 1.851258349 | 1.898808251 |
| ENSMUSG00000049488 | 1.806056082 | 1.610735874 | 1.630890489 | 1.67510993 | 1.595992969 | 1.73239376 | 1.444790289 |
| ENSMUSG00000049489 | 1.857607105 | 1.807204673 | 1.78743678 | 1.685366747 | 1.712553795 | 1.908485019 | 1.699191618 |
| ENSMUSG00000049491 | 0 | 0 | 0 | 0 | 0 | 0 | 0 |
| ENSMUSG00000049493 | 0.671187741 | 0.580482024 | 0.580482024 | 1.04248125 | 1.057362051 | 0.84509804 | 0.969165684 |
| ENSMUSG00000049498 | 0 | 0 | 0 | 0 | 0 | 0 | 0 |
| ENSMUSG00000049502 | 1.482695304 | 1.26109853 | 1.372963274 | 1.96293726 | 1.895599581 | 2.133538908 | 1.800208341 |
| ENSMUSG00000049504 | 2.063793172 | 1.940387808 | 1.921625132 | 1.823655187 | 1.895599581 | 2.100370545 | 1.818286533 |
| ENSMUSG00000049506 | 0 | 0 | 0 | 0 | 0 | 0 | 0 |
| ENSMUSG00000049511 | 1.156259305 | 1.098079356 | 1.27186571 | 0.92510993 | 0.846351821 | 0.903089987 | 0.74492186 |
| ENSMUSG00000049515 | 0 | 0.396240625 | 0 | 0 | 0.244650542 | 0 | 0 |
| ENSMUSG00000049516 | 2.146671932 | 2.04248125 | 1.976722649 | 2.072504712 | 1.973373093 | 2.23299611 | 1.903345709 |
| ENSMUSG00000049517 | 2.542156295 | 2.40136988 | 2.439139081 | 2.42293588 | 2.426655268 | 2.765668555 | 2.302788238 |
| ENSMUSG00000049521 | 2.584966402 | 2.46293726 | 2.461764337 | 2.231203126 | 2.397795474 | 2.617000341 | 2.338495365 |
| ENSMUSG00000049526 | 0.289064826 | 0.79248125 | 0.864857905 | 0.396240625 | 0.686820904 | 0.602059991 | 0.520678036 |
| ENSMUSG00000049528 | 0 | 0 | 0 | 0 | 0 | 0 | 0 |
| ENSMUSG00000049530 | 0 | 0 | 0 | 0 | 0 | 0 | 0 |
| ENSMUSG00000049532 | 2.105953527 | 1.959800947 | 2.015173983 | 2.02186571 | 2.066583491 | 2.290034611 | 1.95718226 |
| ENSMUSG00000049536 | 1.707474304 | 1.494319981 | 1.5 | 1.311981878 | 1.188507684 | 1.397940009 | 1.140832123 |
| ENSMUSG00000049537 | 0 | 0 | 0 | 0 | 0 | 0 | 0 |
| ENSMUSG00000049538 | 1.069666964 | 1.080482024 | 1.04248125 | 0.580482024 | 0.489301084 | 0.954242509 | 0.853775828 |
| ENSMUSG00000049539 | 0.289064826 | 0.25 | 0 | 0.396240625 | 0.568060967 | 0 | 0.224243824 |
| ENSMUSG00000049548 | 0 | 0 | 0 | 0 | 0 | 0 | 0 |
| ENSMUSG00000049550 | 2.837282068 | 2.627443751 | 2.621205724 | 2.679204865 | 2.59304693 | 2.974050903 | 2.463507208 |
| ENSMUSG00000049551 | 0.45815691 | 0.646240625 | 0.79248125 | 0.580482024 | 0.733951626 | 0.698970004 | 0.984950056 |
| ENSMUSG00000049553 | 2.811999637 | 2.598615674 | 2.591307212 | 2.539652422 | 2.543534996 | 2.881954971 | 2.394412686 |
| ENSMUSG00000049555 | 0.45815691 | 0 | 0.701838731 | 0.79248125 | 0.733951626 | 0.698970004 | 0.74492186 |
| ENSMUSG00000049556 | 0 | 0.5 | 0 | 0 | 0.489301084 | 0.301029996 | 0.672731473 |
| ENSMUSG00000049560 | 0 | 0 | 0 | 0 | 0 | 0 | 0 |
| ENSMUSG00000049561 | 0 | 0 | 0 | 0 | 0 | 0 | 0 |
| ENSMUSG00000049562 | 1.205378646 | 1.17510993 | 1.339388001 | 0.864857905 | 1.188507684 | 1.041392685 | 1.307989563 |
| ENSMUSG00000049565 | 0.578129653 | 0.5 | 0.5 | 0 | 0.244650542 | 0 | 0.224243824 |

| | | | | | | | |
|---|---|---|---|---|---|---|---|
| ENSMUSG00000049571 | 0.45815691 | 0 | 0 | 0.396240625 | 0 | 0 | 0.224243824 |
| ENSMUSG00000049573 | 0 | 0 | 0 | 0 | 0 | 0 | 0 |
| ENSMUSG00000049576 | 0.578129653 | 0 | 0.25 | 0 | 0.244650542 | 0.477121255 | 0.520678036 |
| ENSMUSG00000049577 | 1.482695304 | 1.701838731 | 1.807204673 | 1.302363341 | 1.584291694 | 1.591064607 | 1.807561282 |
| ENSMUSG00000049580 | 1.89911417 | 1.821350555 | 1.80479213 | 1.561981878 | 1.741652793 | 1.740362689 | 1.676387053 |
| ENSMUSG00000049583 | 0.289064826 | 0.25 | 0 | 0 | 0 | 0 | 0 |
| ENSMUSG00000049588 | 0.960252568 | 0.646240625 | 0.580482024 | 1.114857905 | 1.136121934 | 1.176091259 | 1.054044578 |
| ENSMUSG00000049593 | 0 | 0 | 0 | 0 | 0 | 0 | 0 |
| ENSMUSG00000049598 | 0 | 0 | 0 | 0 | 0 | 0 | 0 |
| ENSMUSG00000049600 | 1.036286563 | 1.380890489 | 1.464495249 | 1.080482024 | 1.310728002 | 1.322219295 | 1.413583902 |
| ENSMUSG00000049604 | 1.156259305 | 1.27186571 | 1.339388001 | 1.482684334 | 1.768534412 | 1.653212514 | 1.759302692 |
| ENSMUSG00000049605 | 0 | 0 | 0 | 0 | 0 | 0 | 0 |
| ENSMUSG00000049606 | 2.760228547 | 2.555094582 | 2.51348147 | 2.457034121 | 2.349532392 | 2.658964843 | 2.150210039 |
| ENSMUSG00000049608 | 0 | 0.25 | 0.25 | 0 | 0 | 0 | 0 |
| ENSMUSG00000049612 | 0 | 0 | 0 | 0.25 | 0.244650542 | 0 | 0 |
| ENSMUSG00000049618 | 0 | 0 | 0 | 0 | 0 | 0 | 0 |
| ENSMUSG00000049620 | 0.289064826 | 0 | 0.396240625 | 0 | 0 | 0 | 0 |
| ENSMUSG00000049624 | 1.932832114 | 1.816696635 | 1.864857905 | 1.814346961 | 1.820920161 | 2.049218023 | 1.757893036 |
| ENSMUSG00000049625 | 0 | 0.25 | 0 | 0 | 0 | 0 | 0 |
| ENSMUSG00000049630 | 1.558729299 | 1.339388001 | 1.380890489 | 0.79248125 | 0.77552387 | 0.602059991 | 0.775756176 |
| ENSMUSG00000049641 | 0 | 0 | 0 | 0.25 | 0.489301084 | 0 | 0 |
| ENSMUSG00000049643 | 1.952600383 | 1.874961472 | 1.894357207 | 1.593759858 | 1.791313678 | 1.851258349 | 1.747849621 |
| ENSMUSG00000049648 | 0 | 0 | 0 | 0 | 0 | 0 | 0 |
| ENSMUSG00000049649 | 0 | 0 | 0.25 | 0.25 | 0 | 0 | 0.224243824 |
| ENSMUSG00000049653 | 0.289064826 | 0 | 0 | 0 | 0.244650542 | 0 | 0.355418052 |
| ENSMUSG00000049657 | 2.063793172 | 1.961372513 | 1.905512955 | 1.92510993 | 1.948833444 | 2.100370545 | 1.864112664 |
| ENSMUSG00000049658 | 2.585813169 | 2.366901388 | 2.308404919 | 2.544354884 | 2.48684999 | 2.819543936 | 2.283060517 |
| ENSMUSG00000049659 | 2.471720134 | 2.310793496 | 2.310793496 | 2.474083101 | 2.395811454 | 2.730782276 | 2.226906518 |
| ENSMUSG00000049670 | 2.165631214 | 2.00835575 | 2.048689214 | 2.081607372 | 2.112954895 | 2.361727836 | 2.01501956 |
| ENSMUSG00000049672 | 2.144240249 | 1.995748394 | 2.013820609 | 1.926839783 | 1.893954095 | 2.170261715 | 1.782372264 |
| ENSMUSG00000049674 | 0 | 0 | 0 | 0 | 0 | 0 | 0 |
| ENSMUSG00000049676 | 0.671187741 | 1 | 1 | 0.701838731 | 0.686820904 | 0.698970004 | 0.710836105 |
| ENSMUSG00000049680 | 2.134369199 | 1.987091808 | 2.00835575 | 1.890560606 | 1.986303335 | 2.033423755 | 1.908707966 |
| ENSMUSG00000049685 | 0 | 0 | 0 | 0 | 0 | 0 | 0 |

| | | | | | | | |
|---|---|---|---|---|---|---|---|
| ENSMUSG00000049686 | 1.663535556 | 1.598079356 | 1.708222504 | 1.451838731 | 1.555378544 | 1.51851394 | 1.712465909 |
| ENSMUSG00000049687 | 0.45815691 | 0.396240625 | 0 | 0.646240625 | 0.489301084 | 0.301029996 | 0.355418052 |
| ENSMUSG00000049690 | 1.558729299 | 1.494319981 | 1.470660762 | 1.214495249 | 1.223252711 | 1.278753601 | 1.150210039 |
| ENSMUSG00000049691 | 0 | 0 | 0 | 0 | 0 | 0 | 0.224243824 |
| ENSMUSG00000049692 | 0 | 0 | 0 | 0 | 0 | 0 | 0 |
| ENSMUSG00000049694 | 0 | 0 | 0 | 0 | 0 | 0 | 0 |
| ENSMUSG00000049699 | 1.249317394 | 1 | 1.080482024 | 1.330482024 | 1.319233381 | 1.041392685 | 1.131174228 |
| ENSMUSG00000049708 | 0 | 0 | 0 | 0 | 0 | 0 | 0 |
| ENSMUSG00000049709 | 0 | 0.25 | 0.396240625 | 0.396240625 | 0.244650542 | 0 | 0.224243824 |
| ENSMUSG00000049717 | 2.360187403 | 2.242885888 | 2.180274797 | 2.03360658 | 1.951645858 | 2.225309282 | 1.826389311 |
| ENSMUSG00000049719 | 0.289064826 | 0.701838731 | 0.5 | 0.25 | 0.244650542 | 0.301029996 | 0.224243824 |
| ENSMUSG00000049721 | 0 | 0 | 0 | 0 | 0 | 0 | 0 |
| ENSMUSG00000049723 | 0.811507563 | 0.75 | 0.75 | 0.701838731 | 0.568060967 | 0.698970004 | 0.672731473 |
| ENSMUSG00000049728 | 1.53838222 | 1.418106335 | 1.571350555 | 1.372963274 | 1.439191811 | 1.361727836 | 1.496250175 |
| ENSMUSG00000049734 | 0 | 0 | 0 | 0 | 0 | 0 | 0 |
| ENSMUSG00000049737 | 0 | 0 | 0 | 0 | 0 | 0 | 0 |
| ENSMUSG00000049739 | 2.201144779 | 2.08272922 | 2.064346961 | 1.990000483 | 2.06759338 | 2.195899652 | 1.913982791 |
| ENSMUSG00000049740 | 0 | 0 | 0 | 0 | 0 | 0 | 0 |
| ENSMUSG00000049741 | 0 | 0 | 0.25 | 0.25 | 0 | 0 | 0.224243824 |
| ENSMUSG00000049744 | 0 | 0 | 0.25 | 0 | 0 | 0 | 0 |
| ENSMUSG00000049751 | 2.273259602 | 2.130890489 | 2.137686696 | 2.100219859 | 2.031524462 | 2.274157849 | 1.944737824 |
| ENSMUSG00000049755 | 1.847794126 | 1.726722649 | 1.777131114 | 1.985628626 | 1.925488995 | 2.176091259 | 1.854824501 |
| ENSMUSG00000049758 | 0 | 0 | 0 | 0 | 0 | 0 | 0 |
| ENSMUSG00000049760 | 2.194712275 | 2.044977273 | 2.059601185 | 1.938721876 | 2.00243419 | 2.212187604 | 1.900630878 |
| ENSMUSG00000049761 | 0 | 0 | 0 | 0 | 0 | 0 | 0 |
| ENSMUSG00000049764 | 2.351376531 | 2.211372513 | 2.180274797 | 2.147178659 | 2.148364414 | 2.396199347 | 2.01116814 |
| ENSMUSG00000049775 | 3.332046844 | 3.02028763 | 2.997082412 | 2.484527332 | 2.407936347 | 2.795880017 | 2.262942063 |
| ENSMUSG00000049791 | 1.37447073 | 1.330482024 | 1.410964047 | 1.080482024 | 1.136121934 | 1.255272505 | 1.054044578 |
| ENSMUSG00000049792 | 2.158622101 | 2.048689214 | 2.044977273 | 1.955044741 | 1.926994141 | 2.117271296 | 1.834294096 |
| ENSMUSG00000049796 | 0 | 0 | 0 | 0 | 0 | 0 | 0 |
| ENSMUSG00000049797 | 0 | 0 | 0 | 0 | 0 | 0 | 0 |
| ENSMUSG00000049799 | 0.671187741 | 0.701838731 | 0.646240625 | 0.646240625 | 0.733951626 | 0 | 0.710836105 |
| ENSMUSG00000049800 | 2.585813169 | 2.39151351 | 2.37694866 | 2.400906586 | 2.392211871 | 2.673941999 | 2.219191424 |
| ENSMUSG00000049804 | 2.655740331 | 2.456637122 | 2.480831871 | 2.484527332 | 2.458700896 | 2.775974331 | 2.286938317 |

| | | | | | | | |
|---|---|---|---|---|---|---|---|
| ENSMUSG00000049806 | 0 | 0 | 0 | 0 | 0 | 0 | 0 |
| ENSMUSG00000049807 | 2.427163103 | 2.288071211 | 2.367410454 | 2.153677461 | 2.259595879 | 2.394451681 | 2.215091912 |
| ENSMUSG00000049809 | 0 | 0 | 0 | 0 | 0 | 0 | 0 |
| ENSMUSG00000049811 | 1.036286563 | 1.238549078 | 1.061981878 | 1.080482024 | 1.163285805 | 1.230448921 | 1.150210039 |
| ENSMUSG00000049815 | 0 | 0 | 0 | 0 | 0 | 0 | 0 |
| ENSMUSG00000049823 | 1.721148722 | 1.809601185 | 1.866901388 | 1.516522298 | 1.65906333 | 1.531478917 | 1.705896842 |
| ENSMUSG00000049832 | 0.289064826 | 0 | 0.396240625 | 0.396240625 | 0.387761935 | 0.301029996 | 0.448487648 |
| ENSMUSG00000049843 | 0 | 0 | 0 | 0 | 0 | 0 | 0 |
| ENSMUSG00000049848 | 0 | 0 | 0 | 0 | 0 | 0 | 0 |
| ENSMUSG00000049858 | 2.066740409 | 1.910964047 | 1.937048212 | 2.110735874 | 2.143518259 | 2.411619706 | 1.986347533 |
| ENSMUSG00000049864 | 0 | 0 | 0 | 0 | 0 | 0 | 0 |
| ENSMUSG00000049866 | 2.847342547 | 2.603671309 | 2.589388001 | 2.566696635 | 2.533980713 | 2.893761762 | 2.39989613 |
| ENSMUSG00000049871 | 1.269664473 | 1.130890489 | 1.080482024 | 0.830482024 | 0.905314583 | 0.84509804 | 0.74492186 |
| ENSMUSG00000049872 | 0.671187741 | 0.75 | 0.75 | 0.646240625 | 0.489301084 | 0.477121255 | 0.629532004 |
| ENSMUSG00000049878 | 2.53552908 | 2.361770807 | 2.29248125 | 2.275821952 | 2.216042585 | 2.561101384 | 2.055736903 |
| ENSMUSG00000049881 | 2.320585253 | 2.158248799 | 2.172499493 | 2.213717096 | 2.241584006 | 2.545307116 | 2.092902448 |
| ENSMUSG00000049882 | 1.890240806 | 1.747171172 | 1.649978211 | 1.561981878 | 1.559656853 | 1.662757832 | 1.42167221 |
| ENSMUSG00000049891 | 0.289064826 | 0.5 | 0.5 | 0.396240625 | 0.244650542 | 0 | 0.355418052 |
| ENSMUSG00000049892 | 0.45815691 | 0 | 0.5 | 0.830482024 | 0.568060967 | 0 | 0.775756176 |
| ENSMUSG00000049894 | 0 | 0 | 0 | 0 | 0 | 0 | 0 |
| ENSMUSG00000049897 | 0.289064826 | 0.5 | 0.5 | 0.396240625 | 0.489301084 | 0.301029996 | 0.355418052 |
| ENSMUSG00000049902 | 0 | 0 | 0.396240625 | 0 | 0 | 0 | 0 |
| ENSMUSG00000049904 | 1.227926429 | 1.238549078 | 1.330482024 | 1.25 | 1.302012594 | 1.414973348 | 1.193409508 |
| ENSMUSG00000049907 | 1.269664473 | 1.130890489 | 1.130890489 | 0.951838731 | 1.121713561 | 1.079181246 | 1.054044578 |
| ENSMUSG00000049908 | 0 | 0 | 0 | 0 | 0.244650542 | 0 | 0.224243824 |
| ENSMUSG00000049916 | 0 | 0 | 0 | 1.146240625 | 1.283908311 | 1.380211242 | 1.307989563 |
| ENSMUSG00000049922 | 1.806056082 | 1.766522298 | 1.708222504 | 1.828220739 | 1.855665188 | 2.093421685 | 1.793950594 |
| ENSMUSG00000049926 | 0 | 0 | 0 | 0 | 0 | 0 | 0 |
| ENSMUSG00000049928 | 0 | 0 | 0 | 0 | 0 | 0 | 0 |
| ENSMUSG00000049929 | 1.940852214 | 1.825945187 | 1.779735268 | 1.816696635 | 1.563883923 | 1.84509804 | 1.523605797 |
| ENSMUSG00000049932 | 1.920505136 | 1.79248125 | 1.799918086 | 2.027131114 | 2.066583491 | 2.274157849 | 2.025071334 |
| ENSMUSG00000049939 | 0 | 0 | 0 | 0 | 0.244650542 | 0 | 0.355418052 |
| ENSMUSG00000049940 | 2.652869185 | 2.422499493 | 2.412762923 | 2.394828984 | 2.342197974 | 2.727541257 | 2.158505762 |
| ENSMUSG00000049946 | 0 | 0 | 0 | 0 | 0 | 0 | 0 |

| | | | | | | | |
|---|---|---|---|---|---|---|---|
| ENSMUSG00000049950 | 1.707474304 | 1.580482024 | 1.610735874 | 1.593759858 | 1.519146132 | 1.707570176 | 1.483307832 |
| ENSMUSG00000049957 | 2.23973002 | 2.064346961 | 2.031030328 | 2.131869252 | 2.143518259 | 2.348304863 | 2.034219411 |
| ENSMUSG00000049960 | 1.907802659 | 1.825945187 | 1.811981878 | 1.998588359 | 1.983754904 | 2.247973266 | 1.926805425 |
| ENSMUSG00000049969 | 1.928763473 | 1.830482024 | 1.821350555 | 1.839388001 | 1.85749873 | 2.053078443 | 1.74047993 |
| ENSMUSG00000049971 | 1.578129653 | 1.418106335 | 1.410964047 | 1.348079356 | 1.176121988 | 1.113943352 | 1.066254157 |
| ENSMUSG00000049972 | 0 | 0 | 0 | 0 | 0 | 0 | 0 |
| ENSMUSG00000049985 | 0.289064826 | 0 | 0.5 | 0.646240625 | 0.733951626 | 0 | 0.520678036 |
| ENSMUSG00000049988 | 0 | 0 | 0 | 0 | 0 | 0 | 0 |
| ENSMUSG00000049999 | 0.671187741 | 0.580482024 | 0.646240625 | 0.580482024 | 0.686820904 | 0.301029996 | 0.224243824 |
| ENSMUSG00000050002 | 1.45815691 | 1.302363341 | 1.348079356 | 1.660964047 | 1.686820904 | 1.799340549 | 1.621017948 |
| ENSMUSG00000050010 | 0 | 0 | 0.25 | 0.25 | 0.244650542 | 0 | 0 |
| ENSMUSG00000050014 | 1.568542279 | 1.580482024 | 1.418106335 | 0.646240625 | 0.568060967 | 1 | 0.579661877 |
| ENSMUSG00000050015 | 0 | 0 | 0 | 0 | 0 | 0 | 0 |
| ENSMUSG00000050017 | 2.914394662 | 2.661189398 | 2.632113528 | 2.655970373 | 2.571222152 | 2.944482672 | 2.416179455 |
| ENSMUSG00000050022 | 0.747221736 | 0.79248125 | 0.79248125 | 0.25 | 0.568060967 | 0 | 0.579661877 |
| ENSMUSG00000050023 | 0 | 0 | 0 | 0 | 0 | 0 | 0 |
| ENSMUSG00000050028 | 0 | 0 | 0 | 0 | 0 | 0 | 0 |
| ENSMUSG00000050029 | 2.690713405 | 2.445749552 | 2.425543171 | 2.472566069 | 2.380772476 | 2.772321707 | 2.207438502 |
| ENSMUSG00000050030 | 0 | 0 | 0 | 0 | 0 | 0 | 0 |
| ENSMUSG00000050035 | 0.960252568 | 0.951838731 | 0.976722649 | 1.114857905 | 1 | 1.041392685 | 0.952571511 |
| ENSMUSG00000050043 | 2.283862692 | 2.024508021 | 2.028435542 | 2.330482024 | 2.273371525 | 2.612783857 | 2.115189268 |
| ENSMUSG00000050050 | 0 | 0.646240625 | 0.396240625 | 0.701838731 | 0.846351821 | 0.903089987 | 0.829800754 |
| ENSMUSG00000050052 | 1.505869732 | 1.388647213 | 1.302363341 | 1.593759858 | 1.568060967 | 1.62324929 | 1.505628092 |
| ENSMUSG00000050058 | 0.289064826 | 0 | 0 | 0 | 0 | 0 | 0 |
| ENSMUSG00000050063 | 0 | 0 | 0 | 0 | 0.244650542 | 0 | 0.224243824 |
| ENSMUSG00000050064 | 1.655740331 | 1.67510993 | 1.660964047 | 1.657339155 | 1.739104362 | 1.763427994 | 1.607811402 |
| ENSMUSG00000050066 | 0 | 0 | 0.25 | 0 | 0 | 0 | 0 |
| ENSMUSG00000050069 | 0 | 0.396240625 | 0 | 0 | 0.733951626 | 0.698970004 | 0.224243824 |
| ENSMUSG00000050071 | 0.45815691 | 0.79248125 | 0.830482024 | 1.201838731 | 1.310728002 | 1.255272505 | 1.355418052 |
| ENSMUSG00000050074 | 0 | 0 | 0 | 0 | 0 | 0 | 0 |
| ENSMUSG00000050075 | 0 | 0 | 0 | 0 | 0 | 0 | 0 |
| ENSMUSG00000050079 | 2.336265009 | 2.194519282 | 2.190387808 | 2.253505118 | 2.167869715 | 2.521138084 | 2.026306127 |
| ENSMUSG00000050085 | 0 | 0 | 0 | 0 | 0 | 0 | 0 |
| ENSMUSG00000050087 | 0 | 0 | 0 | 0 | 0 | 0 | 0 |

| | | | | | | | |
|---|---|---|---|---|---|---|---|
| ENSMUSG00000050088 | 2.304292853 | 2.168989258 | 2.163659007 | 2.067865757 | 2.048957937 | 2.322219295 | 1.960219986 |
| ENSMUSG00000050089 | 0 | 0 | 0 | 0 | 0 | 0 | 0 |
| ENSMUSG00000050092 | 0 | 0.25 | 0 | 0 | 0 | 0 | 0 |
| ENSMUSG00000050097 | 0.811507563 | 0.5 | 0.830482024 | 0.830482024 | 0.77552387 | 0.77815125 | 0.579661877 |
| ENSMUSG00000050100 | 0 | 0 | 0 | 0.25 | 0 | 0 | 0 |
| ENSMUSG00000050102 | 0 | 0 | 0 | 0 | 0 | 0 | 0 |
| ENSMUSG00000050103 | 0 | 0 | 0 | 0.25 | 0 | 0 | 0.224243824 |
| ENSMUSG00000050105 | 0 | 0 | 0 | 0 | 0 | 0 | 0 |
| ENSMUSG00000050106 | 0.289064826 | 0.25 | 0.580482024 | 0 | 0 | 0 | 0.224243824 |
| ENSMUSG00000050107 | 2.042557845 | 1.899978211 | 1.886723615 | 2.098079356 | 1.919403394 | 2.283301229 | 1.881925353 |
| ENSMUSG00000050108 | 0 | 0.25 | 0 | 0.25 | 0 | 0 | 0.355418052 |
| ENSMUSG00000050114 | 0 | 0 | 0 | 0 | 0 | 0 | 0 |
| ENSMUSG00000050121 | 0 | 0 | 0 | 0 | 0 | 0 | 0 |
| ENSMUSG00000050122 | 0 | 0 | 0 | 0 | 0 | 0 | 0 |
| ENSMUSG00000050128 | 0 | 0 | 0 | 0 | 0 | 0 | 0 |
| ENSMUSG00000050132 | 0.867194479 | 0.864857905 | 0.701838731 | 1.364857905 | 1.504538967 | 1.531478917 | 1.455757965 |
| ENSMUSG00000050134 | 0 | 0 | 0 | 0 | 0 | 0 | 0 |
| ENSMUSG00000050138 | 0 | 0.25 | 0.25 | 0 | 0 | 0 | 0 |
| ENSMUSG00000050141 | 0 | 0 | 0 | 0 | 0 | 0 | 0 |
| ENSMUSG00000050144 | 2.348397716 | 2.160061234 | 2.204245906 | 2.139605178 | 2.155511127 | 2.414973348 | 2.070582188 |
| ENSMUSG00000050147 | 0 | 0.25 | 0.25 | 0.25 | 0.244650542 | 0 | 0 |
| ENSMUSG00000050148 | 2.406877894 | 2.296838018 | 2.289336734 | 2.425975893 | 2.421143088 | 2.669316881 | 2.276018109 |
| ENSMUSG00000050150 | 0.671187741 | 0 | 0 | 0.396240625 | 0.632412477 | 0.602059991 | 0.579661877 |
| ENSMUSG00000050157 | 0.289064826 | 0 | 0 | 0 | 0 | 0.301029996 | 0 |
| ENSMUSG00000050158 | 0 | 0 | 0 | 0 | 0 | 0 | 0 |
| ENSMUSG00000050164 | 0 | 0.25 | 0.25 | 0.5 | 0.244650542 | 0.301029996 | 0.224243824 |
| ENSMUSG00000050174 | 0.578129653 | 0.25 | 0.396240625 | 0.25 | 0.244650542 | 0.301029996 | 0.355418052 |
| ENSMUSG00000050179 | 0 | 0 | 0 | 0 | 0 | 0 | 0 |
| ENSMUSG00000050188 | 1.727821383 | 1.723704441 | 1.789967834 | 1.593759858 | 1.607318736 | 1.662757832 | 1.584526862 |
| ENSMUSG00000050189 | 0 | 0 | 0 | 0 | 0 | 0 | 0 |
| ENSMUSG00000050190 | 0 | 0 | 0 | 0 | 0 | 0 | 0 |
| ENSMUSG00000050192 | 2.211649928 | 2.111770807 | 2.072504712 | 2.160964047 | 2.080465577 | 2.439332694 | 1.95411574 |
| ENSMUSG00000050195 | 0.578129653 | 0.75 | 0.580482024 | 0.580482024 | 0.686820904 | 0.477121255 | 0.74492186 |
| ENSMUSG00000050197 | 0 | 0 | 0 | 0 | 0 | 0 | 0 |

| | | | | | | | |
|---|---|---|---|---|---|---|---|
| ENSMUSG00000050198 | 0 | 0 | 0 | 0 | 0 | 0 | 0 |
| ENSMUSG00000050199 | 2.529763563 | 2.299304173 | 2.302363341 | 1.995748394 | 1.90849439 | 2.117271296 | 1.790137008 |
| ENSMUSG00000050201 | 0 | 0 | 0 | 0 | 0 | 0.301029996 | 0 |
| ENSMUSG00000050211 | 0.289064826 | 0 | 0 | 0.25 | 0.387761935 | 0.301029996 | 0.448487648 |
| ENSMUSG00000050212 | 0.91631382 | 1.098079356 | 1.380890489 | 0.864857905 | 1.234113756 | 1.113943352 | 1.586932193 |
| ENSMUSG00000050213 | 2.23973002 | 2.017865591 | 1.979715809 | 2.031030328 | 1.990091809 | 2.222716471 | 1.898808251 |
| ENSMUSG00000050215 | 0 | 0 | 0 | 0 | 0 | 0 | 0 |
| ENSMUSG00000050217 | 0 | 0 | 0 | 0 | 0 | 0 | 0 |
| ENSMUSG00000050222 | 0 | 0.5 | 0.396240625 | 0 | 0.244650542 | 0.77815125 | 0.629532004 |
| ENSMUSG00000050224 | 0 | 0 | 0 | 0 | 0 | 0 | 0 |
| ENSMUSG00000050229 | 2.209569962 | 2.04248125 | 2.063166358 | 2.165444524 | 2.071604288 | 2.396199347 | 1.958704689 |
| ENSMUSG00000050232 | 0 | 0 | 0 | 0 | 0 | 0 | 0 |
| ENSMUSG00000050234 | 0.289064826 | 0.396240625 | 0.580482024 | 0.396240625 | 0.77552387 | 0.698970004 | 0.629532004 |
| ENSMUSG00000050239 | 0 | 0 | 0 | 0 | 0 | 0 | 0 |
| ENSMUSG00000050240 | 1.37447073 | 1.302363341 | 1.321350555 | 1.330482024 | 1.264824954 | 1.397940009 | 1.131174228 |
| ENSMUSG00000050241 | 0 | 0.396240625 | 0.25 | 0.25 | 0.387761935 | 0 | 0 |
| ENSMUSG00000050243 | 0 | 0 | 0 | 0 | 0 | 0 | 0 |
| ENSMUSG00000050244 | 2.701748369 | 2.487456678 | 2.466046536 | 2.58272922 | 2.51377465 | 2.851869601 | 2.353406551 |
| ENSMUSG00000050248 | 2.410757293 | 2.243603647 | 2.289967834 | 2.061981878 | 2.159419872 | 2.348304863 | 2.076967568 |
| ENSMUSG00000050251 | 0 | 0 | 0 | 0 | 0 | 0 | 0 |
| ENSMUSG00000050266 | 0 | 0 | 0 | 0 | 0 | 0 | 0 |
| ENSMUSG00000050270 | 0.671187741 | 0.75 | 0.864857905 | 0.976722649 | 0.733951626 | 1.079181246 | 0.579661877 |
| ENSMUSG00000050271 | 1.494443473 | 1.445339928 | 1.470660762 | 1.17510993 | 1.310728002 | 1.342422681 | 1.335191751 |
| ENSMUSG00000050272 | 1.342375483 | 1.302363341 | 1.226722649 | 0.79248125 | 0.931471446 | 0.77815125 | 0.829800754 |
| ENSMUSG00000050276 | 0 | 0 | 0 | 0 | 0 | 0 | 0 |
| ENSMUSG00000050281 | 0 | 0 | 0 | 0 | 0 | 0 | 0 |
| ENSMUSG00000050288 | 2.497590902 | 2.344302633 | 2.37892496 | 2.123963757 | 2.164053936 | 2.392696953 | 2.189712149 |
| ENSMUSG00000050295 | 2.105953527 | 1.923371739 | 1.907339155 | 1.979715809 | 1.968065382 | 2.225309282 | 1.896975297 |
| ENSMUSG00000050296 | 0.289064826 | 0.25 | 0 | 0.25 | 0 | 0.477121255 | 0.224243824 |
| ENSMUSG00000050299 | 1.129344651 | 1 | 0.951838731 | 0.976722649 | 1.039257769 | 0.903089987 | 0.984950056 |
| ENSMUSG00000050304 | 0 | 0.25 | 0.396240625 | 0.5 | 0.489301084 | 0.301029996 | 0.355418052 |
| ENSMUSG00000050310 | 2.721759759 | 2.528760913 | 2.477098155 | 2.530059469 | 2.46464417 | 2.803457116 | 2.285835109 |
| ENSMUSG00000050312 | 1.614416215 | 1.54248125 | 1.580482024 | 1.557204673 | 1.542223377 | 1.740362689 | 1.496250175 |
| ENSMUSG00000050315 | 2.003432385 | 1.910964047 | 1.907339155 | 0.864857905 | 0.877063019 | 1.113943352 | 0.969165684 |

| | | | | | | | |
|---|---|---|---|---|---|---|---|
| ENSMUSG00000050321 | 0 | 0 | 0 | 0 | 0 | 0 | 0 |
| ENSMUSG00000050323 | 1.482695304 | 1.516522298 | 1.330482024 | 1.62694866 | 1.484072009 | 1.716003344 | 1.459332757 |
| ENSMUSG00000050328 | 0.45815691 | 0 | 0 | 0 | 0.244650542 | 0 | 0.579661877 |
| ENSMUSG00000050332 | 2.211649928 | 2.092671852 | 2.005591953 | 2.044977273 | 1.974687643 | 2.173186268 | 1.855869786 |
| ENSMUSG00000050334 | 1.771760131 | 1.75 | 1.708222504 | 1.769203899 | 1.709785497 | 1.875061263 | 1.629532004 |
| ENSMUSG00000050335 | 2.432083896 | 2.254202072 | 2.416333979 | 2.595656007 | 2.606443457 | 2.937016107 | 2.559079607 |
| ENSMUSG00000050343 | 0 | 0 | 0 | 0 | 0 | 0 | 0 |
| ENSMUSG00000050345 | 0.578129653 | 0.79248125 | 0.75 | 0.5 | 0.244650542 | 0 | 0.224243824 |
| ENSMUSG00000050347 | 0 | 0 | 0 | 0 | 0 | 0 | 0 |
| ENSMUSG00000050350 | 0 | 0.396240625 | 0 | 0.25 | 0 | 0 | 0.224243824 |
| ENSMUSG00000050357 | 0.578129653 | 0.864857905 | 0.79248125 | 0.580482024 | 0.877063019 | 0.77815125 | 0.952571511 |
| ENSMUSG00000050359 | 0.578129653 | 0.646240625 | 0.646240625 | 1 | 0.846351821 | 0.84509804 | 1 |
| ENSMUSG00000050366 | 0 | 0 | 0 | 0 | 0 | 0 | 0 |
| ENSMUSG00000050368 | 0 | 0 | 0 | 0 | 0 | 0 | 0 |
| ENSMUSG00000050370 | 1.358731791 | 1.356566189 | 1.348079356 | 0.75 | 0.955822902 | 0.77815125 | 1.014380832 |
| ENSMUSG00000050373 | 1.707474304 | 1.561981878 | 1.695339928 | 1.566696635 | 1.677808768 | 1.672097858 | 1.676387053 |
| ENSMUSG00000050377 | 0 | 0.25 | 0.25 | 0 | 0 | 0 | 0 |
| ENSMUSG00000050379 | 2.323768721 | 2.185366747 | 2.133818844 | 2.256976499 | 2.199780694 | 2.521138084 | 2.065709059 |
| ENSMUSG00000050382 | 1.614416215 | 1.610735874 | 1.614857905 | 1.372963274 | 1.628935049 | 1.681241237 | 1.653807046 |
| ENSMUSG00000050383 | 0 | 0 | 0 | 0 | 0 | 0 | 0 |
| ENSMUSG00000050390 | 2.35727108 | 2.225970461 | 2.288071211 | 2.066696635 | 2.186353316 | 2.315970345 | 2.138662327 |
| ENSMUSG00000050394 | 1.727821383 | 1.606566189 | 1.681980114 | 1.403677461 | 1.433158226 | 1.477121255 | 1.307989563 |
| ENSMUSG00000050395 | 1.227926429 | 1.17510993 | 1.04248125 | 1.380890489 | 1.380772476 | 1.255272505 | 1.355418052 |
| ENSMUSG00000050397 | 0 | 0.25 | 0 | 0 | 0 | 0 | 0.355418052 |
| ENSMUSG00000050410 | 2.605636262 | 2.472566069 | 2.53296424 | 2.389126514 | 2.452316937 | 2.716837723 | 2.369158893 |
| ENSMUSG00000050423 | 0 | 0 | 0 | 0 | 0 | 0 | 0 |
| ENSMUSG00000050424 | 0 | 0.25 | 0.25 | 0.25 | 0.244650542 | 0.301029996 | 0.224243824 |
| ENSMUSG00000050425 | 0 | 0 | 0 | 0 | 0 | 0 | 0 |
| ENSMUSG00000050428 | 1.470606563 | 1.5 | 1.593759858 | 1.188721876 | 1.42077253 | 1.447158031 | 1.496250175 |
| ENSMUSG00000050431 | 0 | 0 | 0 | 0 | 0 | 0 | 0 |
| ENSMUSG00000050435 | 0 | 0 | 0 | 0 | 0 | 0 | 0 |
| ENSMUSG00000050439 | 0 | 0 | 0.25 | 0.396240625 | 0.244650542 | 0.477121255 | 0.520678036 |
| ENSMUSG00000050440 | 0 | 0 | 0 | 0 | 0 | 0 | 0 |
| ENSMUSG00000050445 | 0 | 0 | 0 | 0.25 | 0 | 0 | 0 |

| | | | | | | | |
|---|---|---|---|---|---|---|---|
| ENSMUSG00000050447 | 2.151493169 | 1.988549078 | 1.972185812 | 1.5 | 1.53772706 | 1.681241237 | 1.417653332 |
| ENSMUSG00000050463 | 0.289064826 | 0 | 0.25 | 0.25 | 0.489301084 | 0.301029996 | 0.355418052 |
| ENSMUSG00000050468 | 0 | 0 | 0 | 0 | 0 | 0 | 0 |
| ENSMUSG00000050471 | 2.317377295 | 2.131869252 | 2.121960008 | 2.259043403 | 2.167869715 | 2.45331834 | 2.005303651 |
| ENSMUSG00000050473 | 0 | 0 | 0 | 0 | 0.244650542 | 0 | 0.355418052 |
| ENSMUSG00000050478 | 0 | 0 | 0 | 0 | 0 | 0 | 0 |
| ENSMUSG00000050490 | 0 | 0 | 0 | 0.396240625 | 0.568060967 | 0 | 0.629532004 |
| ENSMUSG00000050493 | 0 | 0 | 0 | 0 | 0 | 0 | 0 |
| ENSMUSG00000050503 | 0.671187741 | 0.75 | 0.580482024 | 0.646240625 | 0.733951626 | 0.77815125 | 0.896975297 |
| ENSMUSG00000050504 | 0 | 0 | 0 | 0 | 0 | 0 | 0 |
| ENSMUSG00000050505 | 0.578129653 | 0.25 | 0 | 0 | 0 | 0.301029996 | 0.448487648 |
| ENSMUSG00000050511 | 0 | 0 | 0 | 0.396240625 | 0 | 0.301029996 | 0.355418052 |
| ENSMUSG00000050520 | 0 | 0 | 0 | 0 | 0 | 0 | 0 |
| ENSMUSG00000050526 | 0.289064826 | 0 | 0 | 0 | 0.244650542 | 0 | 0 |
| ENSMUSG00000050530 | 2.23778581 | 2.14151351 | 2.119945066 | 2.043731421 | 2.079491904 | 2.28780173 | 1.95411574 |
| ENSMUSG00000050533 | 0 | 0 | 0 | 0 | 0 | 0 | 0 |
| ENSMUSG00000050534 | 0 | 0 | 0 | 0 | 0 | 0 | 0.520678036 |
| ENSMUSG00000050538 | 1.418409478 | 1.302363341 | 1.226722649 | 1.146240625 | 1.121713561 | 1.041392685 | 1.121219121 |
| ENSMUSG00000050541 | 1.389637216 | 1.364857905 | 1.516522298 | 0.92510993 | 0.733951626 | 0.84509804 | 0.896975297 |
| ENSMUSG00000050545 | 0 | 0 | 0.396240625 | 0.25 | 0.387761935 | 0.301029996 | 0 |
| ENSMUSG00000050549 | 1.036286563 | 0.976722649 | 1.146240625 | 1.438721876 | 1.445123986 | 1.397940009 | 1.252393349 |
| ENSMUSG00000050550 | 1.527823874 | 1.571350555 | 1.464495249 | 1.720660762 | 1.628935049 | 1.763427994 | 1.496250175 |
| ENSMUSG00000050552 | 1.989530018 | 1.981203126 | 2.057204673 | 1.735628626 | 1.910073614 | 1.954242509 | 1.87511425 |
| ENSMUSG00000050553 | 0 | 0 | 0 | 0 | 0 | 0 | 0 |
| ENSMUSG00000050555 | 1.91631382 | 1.882845365 | 1.830482024 | 1.890560606 | 1.871833944 | 2 | 1.728327686 |
| ENSMUSG00000050556 | 0 | 0 | 0 | 0.25 | 0.387761935 | 0.602059991 | 0.579661877 |
| ENSMUSG00000050558 | 0 | 0.25 | 0.25 | 0 | 0.568060967 | 0 | 0.448487648 |
| ENSMUSG00000050565 | 2.914009768 | 2.654366866 | 2.639126514 | 2.742885888 | 2.673124238 | 3.07114529 | 2.47169791 |
| ENSMUSG00000050567 | 2.483777101 | 2.316696635 | 2.325374049 | 2.255591953 | 2.273934005 | 2.547774705 | 2.207438502 |
| ENSMUSG00000050577 | 0 | 0 | 0 | 0 | 0 | 0 | 0 |
| ENSMUSG00000050578 | 0.289064826 | 0.396240625 | 0.646240625 | 0 | 0 | 0 | 0.355418052 |
| ENSMUSG00000050587 | 0.671187741 | 0.896240625 | 0.896240625 | 0.25 | 0 | 0.301029996 | 0 |
| ENSMUSG00000050592 | 0.289064826 | 0 | 0.25 | 0.25 | 0.489301084 | 0.301029996 | 0.629532004 |
| ENSMUSG00000050600 | 0 | 0 | 0 | 0 | 0 | 0 | 0 |

| | | | | | | | |
|---|---|---|---|---|---|---|---|
| ENSMUSG00000050603 | 0 | 0 | 0 | 0 | 0 | 0 | 0 |
| ENSMUSG00000050605 | 1.404271433 | 1.445339928 | 1.388647213 | 1.557204673 | 1.514343934 | 1.602059991 | 1.476637173 |
| ENSMUSG00000050608 | 2.539328946 | 2.325374049 | 2.341580554 | 2.362802778 | 2.35043857 | 2.677606953 | 2.153137801 |
| ENSMUSG00000050612 | 0.289064826 | 0.396240625 | 0.25 | 0.25 | 0.244650542 | 0.301029996 | 0.355418052 |
| ENSMUSG00000050613 | 0 | 0 | 0 | 0 | 0 | 0 | 0 |
| ENSMUSG00000050619 | 2.069666964 | 1.953445298 | 1.921625132 | 2.046218836 | 1.997548906 | 2.260071388 | 1.873141588 |
| ENSMUSG00000050621 | 0.91631382 | 0.92510993 | 0.75 | 1.080482024 | 0.931471446 | 0.84509804 | 1.100339912 |
| ENSMUSG00000050623 | 0.45815691 | 0.25 | 0.25 | 0.396240625 | 0.387761935 | 0.301029996 | 0 |
| ENSMUSG00000050625 | 0 | 0 | 0 | 0 | 0 | 0 | 0 |
| ENSMUSG00000050627 | 2.305951036 | 2.186208459 | 2.179419106 | 2.314935816 | 2.275616087 | 2.580924976 | 2.182188194 |
| ENSMUSG00000050628 | 1.714367558 | 1.741446071 | 1.774508021 | 1.532320754 | 1.642643806 | 1.832508913 | 1.63986884 |
| ENSMUSG00000050635 | 0 | 0 | 0 | 0 | 0 | 0 | 0 |
| ENSMUSG00000050640 | 0 | 0.5 | 0 | 0.5 | 0.244650542 | 0.301029996 | 0.355418052 |
| ENSMUSG00000050641 | 0 | 0 | 0 | 0 | 0 | 0 | 0 |
| ENSMUSG00000050645 | 0 | 0 | 0 | 0 | 0 | 0 | 0 |
| ENSMUSG00000050650 | 0 | 0.25 | 0 | 0 | 0 | 0 | 0 |
| ENSMUSG00000050654 | 0 | 0 | 0 | 0 | 0 | 0 | 0 |
| ENSMUSG00000050663 | 0 | 0 | 0 | 0 | 0 | 0 | 0 |
| ENSMUSG00000050666 | 0 | 0 | 0 | 0 | 0 | 0.301029996 | 0 |
| ENSMUSG00000050668 | 2.083999385 | 2.024508021 | 1.956637122 | 2.180274797 | 2.103062484 | 2.369215857 | 2.000667731 |
| ENSMUSG00000050671 | 0 | 0.580482024 | 0.79248125 | 0.701838731 | 0.877063019 | 0.954242509 | 0.916588299 |
| ENSMUSG00000050675 | 0.960252568 | 0.79248125 | 0.580482024 | 0.646240625 | 0.812711509 | 0.77815125 | 0.579661877 |
| ENSMUSG00000050677 | 0.671187741 | 1.061981878 | 1.04248125 | 1.080482024 | 0.877063019 | 1.113943352 | 0.853775828 |
| ENSMUSG00000050685 | 0 | 0 | 0 | 0 | 0 | 0 | 0 |
| ENSMUSG00000050697 | 2.774997138 | 2.554186465 | 2.481944491 | 2.652063705 | 2.576269619 | 2.967547976 | 2.3678751 |
| ENSMUSG00000050700 | 0 | 0 | 0 | 0 | 0 | 0 | 0 |
| ENSMUSG00000050702 | 0 | 0.25 | 0.396240625 | 0.5 | 0 | 0 | 0.224243824 |
| ENSMUSG00000050704 | 0 | 0 | 0 | 0 | 0 | 0 | 0 |
| ENSMUSG00000050705 | 2.029919532 | 1.935366747 | 1.935366747 | 1.828220739 | 1.877063019 | 2.049218023 | 1.798966438 |
| ENSMUSG00000050708 | 3.157979029 | 2.938721876 | 3.053503873 | 2.51007243 | 2.60092323 | 2.918554531 | 2.565194949 |
| ENSMUSG00000050711 | 0 | 0 | 0 | 0 | 0 | 0 | 0 |
| ENSMUSG00000050714 | 2 | 1.935366747 | 1.888647213 | 1.903677461 | 1.800029087 | 2.017033339 | 1.625302983 |
| ENSMUSG00000050721 | 2.513987407 | 2.353935442 | 2.383332433 | 1.987091808 | 2.105788027 | 2.252853031 | 2.002662694 |
| ENSMUSG00000050730 | 2.541215974 | 2.307204673 | 2.273189285 | 2.28296424 | 2.25901006 | 2.545307116 | 2.069505596 |

| | | | | | | | |
|---|---|---|---|---|---|---|---|
| ENSMUSG00000050732 | 2.35433422 | 2.185366747 | 2.199415381 | 2.300530956 | 2.271112598 | 2.58546073 | 2.167790061 |
| ENSMUSG00000050737 | 0 | 0.396240625 | 0.5 | 1.114857905 | 1.057362051 | 1.301029996 | 1.159323753 |
| ENSMUSG00000050742 | 0 | 0 | 0 | 0 | 0 | 0 | 0 |
| ENSMUSG00000050747 | 0.578129653 | 0.396240625 | 0.396240625 | 0.5 | 0.568060967 | 0.477121255 | 0.355418052 |
| ENSMUSG00000050751 | 0 | 0 | 0 | 0.25 | 0.387761935 | 0 | 0.448487648 |
| ENSMUSG00000050756 | 0 | 0 | 0 | 0 | 0 | 0 | 0 |
| ENSMUSG00000050761 | 0.578129653 | 0.25 | 0.5 | 0.396240625 | 0.387761935 | 0 | 0.672731473 |
| ENSMUSG00000050762 | 0.811507563 | 1.04248125 | 1.061981878 | 0.25 | 0.77552387 | 0 | 0.520678036 |
| ENSMUSG00000050763 | 0 | 0 | 0 | 0 | 0 | 0 | 0 |
| ENSMUSG00000050765 | 0 | 0 | 0.25 | 0 | 0 | 0 | 0 |
| ENSMUSG00000050766 | 0 | 0 | 0 | 0 | 0 | 0 | 0 |
| ENSMUSG00000050772 | 0 | 0 | 0 | 0 | 0 | 0 | 0 |
| ENSMUSG00000050776 | 0 | 0 | 0 | 0 | 0 | 0 | 0 |
| ENSMUSG00000050777 | 1.837744666 | 1.678561379 | 1.868933358 | 1.532320754 | 1.614672255 | 1.707570176 | 1.61448206 |
| ENSMUSG00000050781 | 0 | 0 | 0 | 0 | 0 | 0 | 0 |
| ENSMUSG00000050783 | 0 | 0 | 0 | 0 | 0 | 0 | 0 |
| ENSMUSG00000050786 | 1.578129653 | 1.606566189 | 1.630890489 | 1.668106335 | 1.53772706 | 1.72427587 | 1.452143229 |
| ENSMUSG00000050788 | 0 | 0 | 0 | 0 | 0 | 0 | 0 |
| ENSMUSG00000050796 | 1.227926429 | 1.388647213 | 1.380890489 | 1.26109853 | 1.327538614 | 1.397940009 | 1.238624656 |
| ENSMUSG00000050799 | 0 | 0 | 0 | 0 | 0.244650542 | 0 | 0.224243824 |
| ENSMUSG00000050803 | 0 | 0 | 0 | 0 | 0 | 0 | 0 |
| ENSMUSG00000050808 | 0 | 0 | 0 | 0 | 0 | 0 | 0 |
| ENSMUSG00000050810 | 0 | 0 | 0 | 0 | 0 | 0 | 0 |
| ENSMUSG00000050812 | 2.945876762 | 2.664775991 | 2.643411797 | 2.721614674 | 2.637464898 | 3.020361283 | 2.440194096 |
| ENSMUSG00000050813 | 0 | 0 | 0 | 0 | 0 | 0 | 0 |
| ENSMUSG00000050815 | 0 | 0 | 0 | 0 | 0 | 0 | 0 |
| ENSMUSG00000050818 | 0 | 0 | 0 | 0 | 0 | 0 | 0 |
| ENSMUSG00000050821 | 1.227926429 | 1.372963274 | 1.482684334 | 1.339388001 | 1.373641807 | 1.414973348 | 1.529383012 |
| ENSMUSG00000050822 | 0.578129653 | 0.701838731 | 0.580482024 | 1 | 1.091002363 | 1.204119983 | 1.176815335 |
| ENSMUSG00000050824 | 0 | 0 | 0 | 0 | 0.244650542 | 0 | 0.224243824 |
| ENSMUSG00000050830 | 0 | 0 | 0 | 0 | 0 | 0 | 0 |
| ENSMUSG00000050840 | 0 | 0 | 0 | 0 | 0 | 0 | 0 |
| ENSMUSG00000050844 | 0 | 0 | 0 | 0 | 0 | 0 | 0 |
| ENSMUSG00000050846 | 2.131864492 | 1.92510993 | 1.854463129 | 1.928561379 | 1.919403394 | 2.130333768 | 1.806347338 |

| | | | | | | | |
|---|---|---|---|---|---|---|---|
| ENSMUSG00000050853 | 0 | 0 | 0 | 0 | 0 | 0 | 0 |
| ENSMUSG00000050854 | 0 | 0.25 | 0 | 0 | 0 | 0 | 0 |
| ENSMUSG00000050855 | 1.389637216 | 1.282320754 | 1.26109853 | 1.410964047 | 1.433158226 | 1.491361694 | 1.278288402 |
| ENSMUSG00000050856 | 2.235832493 | 2.103406982 | 2.123963757 | 2.079353153 | 2.083370582 | 2.357934847 | 2.006616089 |
| ENSMUSG00000050860 | 0 | 0 | 0 | 0.25 | 0.244650542 | 0 | 0 |
| ENSMUSG00000050865 | 0 | 0 | 0 | 0 | 0 | 0 | 0 |
| ENSMUSG00000050866 | 0 | 0 | 0 | 0 | 0 | 0 | 0 |
| ENSMUSG00000050870 | 0 | 0 | 0 | 0 | 0 | 0 | 0 |
| ENSMUSG00000050875 | 0 | 0 | 0 | 0 | 0 | 0 | 0 |
| ENSMUSG00000050876 | 0 | 0 | 0 | 0 | 0 | 0 | 0 |
| ENSMUSG00000050890 | 2.083999385 | 1.905512955 | 1.892463902 | 1.992885888 | 1.85749873 | 2.170261715 | 1.792684387 |
| ENSMUSG00000050891 | 1.907802659 | 1.782320754 | 1.789967834 | 1.868933358 | 1.830919587 | 2.089905111 | 1.676387053 |
| ENSMUSG00000050896 | 0 | 0 | 0 | 0 | 0 | 0 | 0 |
| ENSMUSG00000050900 | 0 | 0 | 0 | 0 | 0.244650542 | 0 | 0 |
| ENSMUSG00000050901 | 0 | 0 | 0 | 0 | 0 | 0 | 0 |
| ENSMUSG00000050908 | 1.867194479 | 1.692046081 | 1.685366747 | 1.302363341 | 1.264824954 | 1.301029996 | 1.290497981 |
| ENSMUSG00000050910 | 2.282114126 | 2.147178659 | 2.158248799 | 2.113831805 | 2.157861565 | 2.372912003 | 2.090889903 |
| ENSMUSG00000050912 | 2.903085937 | 2.639844273 | 2.615369862 | 2.754202072 | 2.655839964 | 3.084576278 | 2.453791299 |
| ENSMUSG00000050914 | 1.227926429 | 1.114857905 | 1.080482024 | 0.92510993 | 1 | 0.954242509 | 1.078019652 |
| ENSMUSG00000050917 | 0 | 0 | 0 | 0 | 0 | 0 | 0 |
| ENSMUSG00000050919 | 0 | 0 | 0.25 | 0 | 0 | 0 | 0 |
| ENSMUSG00000050921 | 0 | 0 | 0 | 0 | 0 | 0 | 0 |
| ENSMUSG00000050926 | 0 | 0 | 0 | 0 | 0 | 0 | 0 |
| ENSMUSG00000050930 | 0.289064826 | 0 | 0.25 | 0.396240625 | 0.387761935 | 0 | 0 |
| ENSMUSG00000050931 | 2.158622101 | 1.992885888 | 1.998588359 | 2.255591953 | 2.223900931 | 2.495544338 | 2.065709059 |
| ENSMUSG00000050933 | 0 | 0 | 0 | 0 | 0 | 0 | 0 |
| ENSMUSG00000050936 | 0 | 0.396240625 | 0.580482024 | 0.25 | 0.387761935 | 0 | 0.355418052 |
| ENSMUSG00000050944 | 1.289064826 | 1.26109853 | 1.321350555 | 1.17510993 | 1.057362051 | 1 | 1.041356071 |
| ENSMUSG00000050945 | 1.307602651 | 1.42510993 | 1.330482024 | 1.410964047 | 1.254881871 | 1.447158031 | 1.20919388 |
| ENSMUSG00000050947 | 1.505869732 | 1.438721876 | 1.552363341 | 1.87892496 | 1.866526233 | 2.100370545 | 1.788855757 |
| ENSMUSG00000050953 | 2.821721112 | 2.598615674 | 2.5964656 | 2.522528105 | 2.499235686 | 2.822168079 | 2.306178596 |
| ENSMUSG00000050954 | 1.587501561 | 1.51109853 | 1.516522298 | 1.54248125 | 1.499532413 | 1.653212514 | 1.479989695 |
| ENSMUSG00000050957 | 1.389637216 | 1.282320754 | 1.348079356 | 1.29248125 | 1.234113756 | 1.414973348 | 1.20139795 |
| ENSMUSG00000050961 | 0 | 0 | 0 | 0 | 0 | 0 | 0 |

| | | | | | | | |
|---|---|---|---|---|---|---|---|
| ENSMUSG00000050963 | 0 | 0 | 0 | 0 | 0 | 0 | 0 |
| ENSMUSG00000050965 | 2.836818956 | 2.597004321 | 2.593488164 | 2.487821179 | 2.402512107 | 2.749736316 | 2.246207397 |
| ENSMUSG00000050966 | 0 | 0.25 | 0 | 0 | 0 | 0 | 0.224243824 |
| ENSMUSG00000050967 | 0 | 0 | 0 | 0.25 | 0.387761935 | 0 | 0 |
| ENSMUSG00000050973 | 1.269664473 | 1.311981878 | 1.396240625 | 1.348079356 | 1.327538614 | 1.380211242 | 1.193409508 |
| ENSMUSG00000050974 | 0 | 0 | 0 | 0 | 0 | 0 | 0 |
| ENSMUSG00000050978 | 0 | 0 | 0 | 0 | 0 | 0 | 0 |
| ENSMUSG00000050982 | 0 | 0 | 0 | 0 | 0 | 0 | 0 |
| ENSMUSG00000050989 | 2.644135203 | 2.425543171 | 2.474083101 | 2.341580554 | 2.400946889 | 2.661812686 | 2.271717885 |
| ENSMUSG00000050994 | 0 | 0.396240625 | 0 | 0.646240625 | 0.387761935 | 0 | 0.355418052 |
| ENSMUSG00000050996 | 0 | 0 | 0 | 0 | 0 | 0 | 0 |
| ENSMUSG00000051000 | 1.765759561 | 1.614857905 | 1.705044741 | 1.809601185 | 1.797870327 | 1.919078092 | 1.667169465 |
| ENSMUSG00000051002 | 0 | 0 | 0 | 0 | 0 | 0 | 0 |
| ENSMUSG00000051003 | 0 | 0 | 0 | 0 | 0 | 0 | 0 |
| ENSMUSG00000051007 | 1.482695304 | 1.5 | 1.470660762 | 1.464495249 | 1.445123986 | 1.505149978 | 1.444790289 |
| ENSMUSG00000051008 | 0 | 0.5 | 0.396240625 | 0.5 | 0.387761935 | 0.602059991 | 0.224243824 |
| ENSMUSG00000051022 | 0 | 0 | 0 | 0.5 | 0.387761935 | 0.301029996 | 0.224243824 |
| ENSMUSG00000051029 | 1.325351389 | 1.348079356 | 1.26109853 | 1.160964047 | 1 | 1 | 0.969165684 |
| ENSMUSG00000051034 | 1.83262764 | 1.714495249 | 1.634789703 | 1.732684334 | 1.683842353 | 1.84509804 | 1.543390198 |
| ENSMUSG00000051036 | 0 | 0 | 0 | 0 | 0 | 0 | 0.224243824 |
| ENSMUSG00000051037 | 1.036286563 | 1.061981878 | 1.02186571 | 1.25 | 0.955822902 | 0.954242509 | 0.935079929 |
| ENSMUSG00000051038 | 0 | 0 | 0 | 0 | 0 | 0 | 0 |
| ENSMUSG00000051041 | 0 | 0 | 0 | 0 | 0 | 0 | 0 |
| ENSMUSG00000051043 | 1.663535556 | 1.618933358 | 1.552363341 | 0.830482024 | 1.039257769 | 0.84509804 | 0.935079929 |
| ENSMUSG00000051046 | 0 | 0 | 0 | 0 | 0 | 0 | 0 |
| ENSMUSG00000051048 | 2.092368209 | 1.970660762 | 1.959800947 | 1.356566189 | 1.414412788 | 1.505149978 | 1.335191751 |
| ENSMUSG00000051050 | 0 | 0 | 0 | 0 | 0 | 0 | 0 |
| ENSMUSG00000051051 | 0 | 0 | 0 | 0 | 0 | 0 | 0 |
| ENSMUSG00000051052 | 0 | 0 | 0 | 0 | 0 | 0 | 0 |
| ENSMUSG00000051054 | 0.289064826 | 0.5 | 0 | 0.25 | 0.686820904 | 0.602059991 | 0.520678036 |
| ENSMUSG00000051056 | 0 | 0.25 | 0 | 0 | 0.244650542 | 0 | 0.224243824 |
| ENSMUSG00000051062 | 0 | 0 | 0 | 0 | 0 | 0 | 0 |
| ENSMUSG00000051065 | 1.811507563 | 1.77186571 | 1.75835575 | 1.598079356 | 1.533172725 | 1.707570176 | 1.493062805 |
| ENSMUSG00000051067 | 0 | 0 | 0.396240625 | 0.396240625 | 0.489301084 | 0.77815125 | 0.829800754 |

| | | | | | | | |
|---|---|---|---|---|---|---|---|
| ENSMUSG00000051074 | 0 | 0.580482024 | 0.396240625 | 0.25 | 0.244650542 | 0 | 0.448487648 |
| ENSMUSG00000051076 | 0 | 0 | 0 | 0 | 0 | 0 | 0 |
| ENSMUSG00000051079 | 0 | 0 | 0 | 0 | 0 | 0 | 0 |
| ENSMUSG00000051081 | 0 | 0 | 0 | 0 | 0 | 0 | 0 |
| ENSMUSG00000051095 | 0 | 0 | 0 | 0 | 0 | 0 | 0 |
| ENSMUSG00000051098 | 1.069666964 | 1.098079356 | 1.26109853 | 1.557204673 | 1.445123986 | 1.763427994 | 1.369798885 |
| ENSMUSG00000051107 | 0 | 0.5 | 0.5 | 0.25 | 0 | 0 | 0.224243824 |
| ENSMUSG00000051111 | 0.289064826 | 0.396240625 | 0.646240625 | 0.580482024 | 0 | 0.301029996 | 0.224243824 |
| ENSMUSG00000051113 | 0.289064826 | 0 | 0.25 | 0 | 0.244650542 | 0 | 0.355418052 |
| ENSMUSG00000051116 | 0.289064826 | 0 | 0 | 0 | 0 | 0 | 0 |
| ENSMUSG00000051118 | 0 | 0 | 0 | 0 | 0 | 0 | 0 |
| ENSMUSG00000051124 | 0 | 0.25 | 0 | 0.5 | 0.489301084 | 0 | 0.448487648 |
| ENSMUSG00000051133 | 0 | 0 | 0 | 0 | 0 | 0 | 0 |
| ENSMUSG00000051136 | 0 | 0 | 0 | 0 | 0 | 0 | 0 |
| ENSMUSG00000051146 | 1 | 1.098079356 | 0.92510993 | 0.79248125 | 0.686820904 | 0.84509804 | 0.916588299 |
| ENSMUSG00000051147 | 1.663535556 | 1.602347734 | 1.580482024 | 1.339388001 | 1.163285805 | 1.342422681 | 1.078019652 |
| ENSMUSG00000051149 | 0 | 0.580482024 | 0.79248125 | 0.830482024 | 0.632412477 | 0.477121255 | 0.579661877 |
| ENSMUSG00000051153 | 0 | 0 | 0 | 0 | 0 | 0 | 0 |
| ENSMUSG00000051154 | 2.23973002 | 2.086073977 | 2.078220739 | 1.931980114 | 1.934425071 | 2.120573931 | 1.848480858 |
| ENSMUSG00000051156 | 0 | 0 | 0 | 0 | 0 | 0 | 0 |
| ENSMUSG00000051159 | 0.289064826 | 0.5 | 0.701838731 | 0.701838731 | 0.489301084 | 0.602059991 | 0.853775828 |
| ENSMUSG00000051160 | 0 | 0 | 0 | 0 | 0 | 0 | 0 |
| ENSMUSG00000051166 | 1.993049299 | 1.78743678 | 1.747171172 | 2.061981878 | 1.993840051 | 2.250420002 | 1.813563596 |
| ENSMUSG00000051169 | 0.960252568 | 1.17510993 | 1.25 | 1 | 1.121713561 | 1.113943352 | 1.350478789 |
| ENSMUSG00000051172 | 0 | 0 | 0 | 0 | 0 | 0 | 0 |
| ENSMUSG00000051176 | 0 | 0 | 0 | 0 | 0 | 0 | 0 |
| ENSMUSG00000051177 | 1.985980784 | 1.914552871 | 1.864857905 | 1.29248125 | 1.121713561 | 1.361727836 | 1.140832123 |
| ENSMUSG00000051180 | 0 | 0 | 0 | 0 | 0 | 0 | 0 |
| ENSMUSG00000051182 | 0 | 0 | 0 | 0 | 0 | 0 | 0 |
| ENSMUSG00000051184 | 1.069666964 | 1.26109853 | 1.348079356 | 0.830482024 | 1.149965125 | 0.77815125 | 1.185218806 |
| ENSMUSG00000051185 | 1.647796617 | 1.584962501 | 1.54248125 | 1.610735874 | 1.546663136 | 1.672097858 | 1.452143229 |
| ENSMUSG00000051190 | 0 | 0 | 0 | 0 | 0 | 0 | 0 |
| ENSMUSG00000051198 | 0 | 0 | 0 | 0 | 0 | 0 | 0 |
| ENSMUSG00000051200 | 0 | 0 | 0 | 0 | 0 | 0 | 0 |

| | | | | | | | |
|---|---|---|---|---|---|---|---|
| ENSMUSG00000051207 | 0 | 0 | 0 | 0 | 0 | 0 | 0 |
| ENSMUSG00000051209 | 0 | 0 | 0 | 0 | 0 | 0 | 0 |
| ENSMUSG00000051212 | 0 | 0.25 | 0.25 | 0.396240625 | 0.244650542 | 0 | 0.224243824 |
| ENSMUSG00000051219 | 0 | 0 | 0 | 0 | 0 | 0.301029996 | 0 |
| ENSMUSG00000051220 | 2.709994145 | 2.467976309 | 2.439139081 | 2.514836115 | 2.441296923 | 2.828015064 | 2.269400844 |
| ENSMUSG00000051223 | 3.402962068 | 3.092058403 | 3.063166358 | 3.176677961 | 3.066999677 | 3.587374172 | 2.827996137 |
| ENSMUSG00000051224 | 1.325351389 | 1.188721876 | 1.098079356 | 1.445339928 | 1.394615667 | 1.414973348 | 1.302263476 |
| ENSMUSG00000051225 | 0 | 0 | 0.25 | 0.25 | 0.244650542 | 0.301029996 | 0 |
| ENSMUSG00000051228 | 0.289064826 | 0.25 | 0.396240625 | 0.25 | 0.244650542 | 0.301029996 | 0.355418052 |
| ENSMUSG00000051232 | 2.318984359 | 2.146240625 | 2.211372513 | 2.169870025 | 2.116484486 | 2.411619706 | 1.989815042 |
| ENSMUSG00000051234 | 2.496544396 | 2.342671852 | 2.309601185 | 2.344844592 | 2.29040462 | 2.658011397 | 2.157685697 |
| ENSMUSG00000051235 | 2.139333929 | 2.005591953 | 1.992885888 | 2.115881093 | 2.004851707 | 2.281033367 | 1.916588299 |
| ENSMUSG00000051236 | 3.027579933 | 2.763820609 | 2.748234584 | 2.539020769 | 2.456697355 | 2.848804701 | 2.291042162 |
| ENSMUSG00000051237 | 0 | 0 | 0 | 0 | 0 | 0 | 0 |
| ENSMUSG00000051238 | 1.54867984 | 1.470660762 | 1.396240625 | 1.451838731 | 1.489301084 | 1.653212514 | 1.405288179 |
| ENSMUSG00000051242 | 0.578129653 | 0 | 0.580482024 | 0.25 | 0 | 0.301029996 | 0.355418052 |
| ENSMUSG00000051243 | 0 | 0.25 | 0.396240625 | 0.396240625 | 0.733951626 | 0.602059991 | 0.876096088 |
| ENSMUSG00000051246 | 0 | 0.646240625 | 0.396240625 | 0.5 | 0.632412477 | 0 | 0.448487648 |
| ENSMUSG00000051251 | 0 | 0 | 0 | 0 | 0 | 0 | 0.224243824 |
| ENSMUSG00000051255 | 0.289064826 | 0.396240625 | 0 | 0 | 0 | 0.477121255 | 0.224243824 |
| ENSMUSG00000051256 | 2.095120908 | 1.991446071 | 1.92510993 | 2.009729747 | 1.990091809 | 2.176091259 | 1.861046144 |
| ENSMUSG00000051257 | 0 | 0 | 0 | 0 | 0 | 0 | 0 |
| ENSMUSG00000051258 | 0 | 0 | 0 | 0 | 0 | 0 | 0 |
| ENSMUSG00000051262 | 0 | 0.396240625 | 0 | 0 | 0.387761935 | 0.301029996 | 0 |
| ENSMUSG00000051276 | 0 | 0 | 0 | 0 | 0 | 0 | 0 |
| ENSMUSG00000051278 | 2.328498818 | 2.143411797 | 2.122963274 | 2.30357978 | 2.224547964 | 2.472756449 | 2.040177507 |
| ENSMUSG00000051279 | 0 | 0 | 0 | 0.25 | 0.244650542 | 0.301029996 | 0.224243824 |
| ENSMUSG00000051285 | 2.375872521 | 2.225970461 | 2.257666784 | 2.240724001 | 2.179075613 | 2.439332694 | 2.021963573 |
| ENSMUSG00000051295 | 0 | 0.25 | 0 | 0.396240625 | 0 | 0.301029996 | 0.224243824 |
| ENSMUSG00000051297 | 0 | 0 | 0 | 0 | 0 | 0.301029996 | 0 |
| ENSMUSG00000051306 | 2.282114126 | 2.129909063 | 2.087182039 | 1.994319981 | 1.98503142 | 2.209515015 | 1.926805425 |
| ENSMUSG00000051313 | 0 | 0 | 0 | 0 | 0 | 0 | 0 |
| ENSMUSG00000051314 | 0 | 0 | 0 | 0 | 0 | 0 | 0 |
| ENSMUSG00000051316 | 1.482695304 | 1.330482024 | 1.42510993 | 1.566696635 | 1.504538967 | 1.602059991 | 1.459332757 |

| | | | | | | | |
|---|---|---|---|---|---|---|---|
| ENSMUSG00000051319 | 0.960252568 | 0.976722649 | 1.17510993 | 0.951838731 | 0.931471446 | 0.903089987 | 1.041356071 |
| ENSMUSG00000051323 | 2.583267694 | 2.413211243 | 2.405054956 | 2.640560606 | 2.583824821 | 2.933993164 | 2.393225472 |
| ENSMUSG00000051329 | 2.635965117 | 2.412762923 | 2.385757955 | 2.618679987 | 2.556894464 | 2.888740961 | 2.36247044 |
| ENSMUSG00000051331 | 2.305951036 | 2.186208459 | 2.180274797 | 1.953445298 | 1.944572352 | 2.045322979 | 1.873141588 |
| ENSMUSG00000051335 | 2.075459174 | 1.882845365 | 1.937048212 | 1.880890489 | 1.850106709 | 2 | 1.771720588 |
| ENSMUSG00000051339 | 2.788220579 | 2.571061437 | 2.577369088 | 2.429847205 | 2.413656993 | 2.696356389 | 2.3158899 |
| ENSMUSG00000051340 | 0 | 0 | 0 | 0 | 0 | 0 | 0 |
| ENSMUSG00000051341 | 1.671187741 | 1.53743678 | 1.51109853 | 1.505591953 | 1.445123986 | 1.707570176 | 1.405288179 |
| ENSMUSG00000051343 | 2.323768721 | 2.149978211 | 2.210587586 | 2.04248125 | 2.140250123 | 2.336459734 | 2.06186744 |
| ENSMUSG00000051344 | 1.982401085 | 1.894357207 | 1.845926073 | 1.890560606 | 1.895599581 | 2.079181246 | 1.781059827 |
| ENSMUSG00000051346 | 1.753493018 | 1.732684334 | 1.78743678 | 1.880890489 | 1.828942237 | 2.025305865 | 1.723650273 |
| ENSMUSG00000051351 | 2.069666964 | 1.912762923 | 1.940387808 | 1.907339155 | 1.828942237 | 1.959041392 | 1.826389311 |
| ENSMUSG00000051354 | 0 | 0 | 0 | 0 | 0 | 0 | 0 |
| ENSMUSG00000051355 | 2.00683675 | 1.938721876 | 1.926839783 | 2.079353153 | 1.990091809 | 2.274157849 | 1.851668228 |
| ENSMUSG00000051359 | 1.342375483 | 1.282320754 | 1.02186571 | 1.098079356 | 1.074582839 | 1 | 0.74492186 |
| ENSMUSG00000051361 | 0 | 0 | 0 | 0 | 0 | 0 | 0 |
| ENSMUSG00000051362 | 0 | 0 | 0 | 0 | 0 | 0 | 0 |
| ENSMUSG00000051367 | 2.355805236 | 2.222185812 | 2.192046081 | 1.96293726 | 1.862942891 | 2.049218023 | 1.861046144 |
| ENSMUSG00000051373 | 0 | 0 | 0 | 0 | 0 | 0.301029996 | 0 |
| ENSMUSG00000051375 | 2.003432385 | 1.921625132 | 1.995748394 | 1.364857905 | 1.343584837 | 1.301029996 | 1.350478789 |
| ENSMUSG00000051378 | 2.45469604 | 2.30357978 | 2.3184489 | 2.251406137 | 2.365497953 | 2.58546073 | 2.260262214 |
| ENSMUSG00000051379 | 0.867194479 | 0.92510993 | 1.02186571 | 0.92510993 | 0.812711509 | 0.954242509 | 0.803905701 |
| ENSMUSG00000051390 | 1.83262764 | 1.928561379 | 1.995748394 | 1.698603967 | 1.832885921 | 1.954242509 | 1.904245613 |
| ENSMUSG00000051391 | 3.10716689 | 2.892701109 | 2.898231039 | 2.745213402 | 2.755133868 | 3.125481266 | 2.663837308 |
| ENSMUSG00000051392 | 0 | 0 | 0 | 0 | 0 | 0 | 0 |
| ENSMUSG00000051396 | 2.676666078 | 2.466819685 | 2.444929839 | 2.476346751 | 2.434943542 | 2.786751422 | 2.282502732 |
| ENSMUSG00000051397 | 0 | 0.25 | 0.25 | 0.25 | 0.244650542 | 0.301029996 | 0.224243824 |
| ENSMUSG00000051401 | 0 | 0 | 0 | 0 | 0 | 0 | 0 |
| ENSMUSG00000051403 | 1.924654746 | 1.926839783 | 1.984159485 | 1.688721876 | 1.964031554 | 1.875061263 | 2.007270316 |
| ENSMUSG00000051412 | 2.40817504 | 2.268535366 | 2.225970461 | 2.33272922 | 2.193483986 | 2.530199698 | 2.071655208 |
| ENSMUSG00000051413 | 2.139333929 | 1.981203126 | 1.985628626 | 2.14151351 | 2.075570129 | 2.359835482 | 1.954885099 |
| ENSMUSG00000051414 | 0 | 0 | 0 | 0 | 0 | 0 | 0 |
| ENSMUSG00000051424 | 0 | 0 | 0 | 0 | 0 | 0 | 0 |
| ENSMUSG00000051427 | 1.993049299 | 1.858657057 | 1.903677461 | 1.837182039 | 1.929985312 | 2.037426498 | 1.838725884 |

| | | | | | | | |
|---|---|---|---|---|---|---|---|
| ENSMUSG00000051431 | 0 | 0.25 | 0 | 0 | 0 | 0 | 0 |
| ENSMUSG00000051435 | 0 | 0 | 0 | 0 | 0.244650542 | 0.301029996 | 0 |
| ENSMUSG00000051437 | 0 | 0 | 0 | 0 | 0 | 0 | 0 |
| ENSMUSG00000051439 | 1.289064826 | 1.396240625 | 1.470660762 | 1.372963274 | 1.351342427 | 1.462397998 | 1.417653332 |
| ENSMUSG00000051444 | 1.964026642 | 1.80479213 | 1.78743678 | 1.794977273 | 1.726078673 | 1.875061263 | 1.582103514 |
| ENSMUSG00000051451 | 2.390989024 | 2.230460234 | 2.161864607 | 2.170748646 | 2.135290471 | 2.361727836 | 1.955652633 |
| ENSMUSG00000051452 | 0 | 0 | 0 | 0 | 0.244650542 | 0 | 0 |
| ENSMUSG00000051455 | 0.578129653 | 0.25 | 0 | 0.5 | 0.568060967 | 0 | 0.224243824 |
| ENSMUSG00000051456 | 0 | 0 | 0 | 0 | 0 | 0 | 0.224243824 |
| ENSMUSG00000051457 | 0.811507563 | 1 | 1.160964047 | 1.080482024 | 1.121713561 | 0.698970004 | 1 |
| ENSMUSG00000051469 | 2.469490009 | 2.26109853 | 2.252107156 | 2.391037372 | 2.316753004 | 2.62324929 | 2.152303998 |
| ENSMUSG00000051481 | 0 | 0 | 0 | 0 | 0 | 0 | 0 |
| ENSMUSG00000051483 | 1.948720984 | 1.852347734 | 1.841580554 | 1.860735874 | 1.800029087 | 2.045322979 | 1.851668228 |
| ENSMUSG00000051486 | 0 | 0 | 0 | 0 | 0 | 0 | 0 |
| ENSMUSG00000051490 | 0 | 0 | 0 | 0 | 0 | 0 | 0.224243824 |
| ENSMUSG00000051493 | 0 | 0 | 0 | 0 | 0 | 0 | 0 |
| ENSMUSG00000051495 | 2.144240249 | 2.029735268 | 2.03743678 | 1.708222504 | 1.611014646 | 1.851258349 | 1.687112305 |
| ENSMUSG00000051497 | 0 | 0 | 0 | 0 | 0 | 0 | 0 |
| ENSMUSG00000051498 | 1.156259305 | 1.17510993 | 0.951838731 | 0.396240625 | 0.568060967 | 0.602059991 | 0.448487648 |
| ENSMUSG00000051499 | 0.45815691 | 0.646240625 | 0.646240625 | 0.580482024 | 0.568060967 | 0.602059991 | 0.672731473 |
| ENSMUSG00000051502 | 1.227926429 | 1.114857905 | 1.188721876 | 1.214495249 | 1.188507684 | 1.176091259 | 1.20919388 |
| ENSMUSG00000051503 | 0.45815691 | 0.25 | 0.25 | 0 | 0.244650542 | 0 | 0 |
| ENSMUSG00000051504 | 0 | 0 | 0.25 | 0 | 0 | 0.301029996 | 0 |
| ENSMUSG00000051506 | 0 | 0 | 0 | 0.25 | 0 | 0 | 0.224243824 |
| ENSMUSG00000051509 | 0 | 0 | 0 | 0 | 0 | 0 | 0 |
| ENSMUSG00000051510 | 2.659301689 | 2.471805154 | 2.490000483 | 2.257666784 | 2.331380351 | 2.517195898 | 2.199599638 |
| ENSMUSG00000051515 | 0 | 0 | 0 | 0 | 0 | 0 | 0 |
| ENSMUSG00000051517 | 1.837744666 | 1.701838731 | 1.799918086 | 1.527131114 | 1.680838453 | 1.740362689 | 1.657681529 |
| ENSMUSG00000051518 | 2.165631214 | 2.041226732 | 2.063166358 | 1.997171172 | 2.074582839 | 2.220108088 | 1.972093446 |
| ENSMUSG00000051527 | 0.747221736 | 0.951838731 | 0.646240625 | 0.701838731 | 0.686820904 | 0.477121255 | 0.74492186 |
| ENSMUSG00000051528 | 0 | 0 | 0 | 0 | 0 | 0 | 0 |
| ENSMUSG00000051537 | 0.289064826 | 0 | 0.5 | 0.396240625 | 0.632412477 | 0.698970004 | 0.448487648 |
| ENSMUSG00000051548 | 0 | 0 | 0 | 0 | 0 | 0 | 0 |
| ENSMUSG00000051550 | 1.269664473 | 1.27186571 | 1.282320754 | 1.26109853 | 1.254881871 | 1.278753601 | 1.360283038 |

| | | | | | | | |
|---|---|---|---|---|---|---|---|
| ENSMUSG00000051557 | 1.389637216 | 1.410964047 | 1.561981878 | 1.226722649 | 1.407936347 | 1.176091259 | 1.405288179 |
| ENSMUSG00000051579 | 2.720536789 | 2.495034894 | 2.465271726 | 2.469129237 | 2.368520226 | 2.773786445 | 2.246207397 |
| ENSMUSG00000051582 | 0 | 0 | 0 | 0 | 0 | 0 | 0 |
| ENSMUSG00000051586 | 2.267855219 | 2.143411797 | 2.168989258 | 1.966046536 | 2.048957937 | 2.222716471 | 1.977870661 |
| ENSMUSG00000051590 | 0 | 0.25 | 0.25 | 0 | 0.244650542 | 0 | 0.355418052 |
| ENSMUSG00000051591 | 0 | 0 | 0 | 0 | 0 | 0 | 0 |
| ENSMUSG00000051592 | 0 | 0.25 | 0 | 0 | 0 | 0 | 0 |
| ENSMUSG00000051593 | 0 | 0 | 0 | 0 | 0 | 0 | 0 |
| ENSMUSG00000051596 | 0 | 0 | 0 | 0.25 | 0 | 0 | 0.355418052 |
| ENSMUSG00000051599 | 0 | 0 | 0 | 0 | 0 | 0 | 0 |
| ENSMUSG00000051606 | 0 | 0 | 0 | 0 | 0 | 0 | 0 |
| ENSMUSG00000051611 | 0 | 0 | 0 | 0 | 0 | 0 | 0 |
| ENSMUSG00000051615 | 2.105953527 | 1.995748394 | 2.032320754 | 2.027131114 | 1.970729215 | 2.252853031 | 1.897893072 |
| ENSMUSG00000051616 | 0 | 0 | 0 | 0 | 0 | 0 | 0 |
| ENSMUSG00000051617 | 0 | 0 | 0 | 0 | 0 | 0 | 0 |
| ENSMUSG00000051618 | 0 | 0 | 0 | 0 | 0 | 0 | 0 |
| ENSMUSG00000051627 | 0 | 0 | 0.396240625 | 0.25 | 0 | 0.301029996 | 0.74492186 |
| ENSMUSG00000051638 | 0 | 0 | 0 | 0 | 0 | 0 | 0 |
| ENSMUSG00000051639 | 0 | 0 | 0 | 0 | 0 | 0 | 0 |
| ENSMUSG00000051648 | 0 | 0.25 | 0.5 | 0.5 | 0.244650542 | 0.301029996 | 0.224243824 |
| ENSMUSG00000051650 | 0.578129653 | 0.25 | 0.25 | 0.25 | 0.244650542 | 0.301029996 | 0 |
| ENSMUSG00000051652 | 0.45815691 | 0.5 | 0.5 | 1.201838731 | 1.163285805 | 1.255272505 | 1.252393349 |
| ENSMUSG00000051663 | 0 | 0 | 0 | 0 | 0 | 0 | 0 |
| ENSMUSG00000051669 | 1.578129653 | 1.580482024 | 1.494319981 | 0.830482024 | 0.846351821 | 0.77815125 | 0.803905701 |
| ENSMUSG00000051671 | 1.470606563 | 1.476722649 | 1.618933358 | 1.602347734 | 1.489301084 | 1.612783857 | 1.540636858 |
| ENSMUSG00000051674 | 2.283862692 | 2.174241882 | 2.092671852 | 2.099151195 | 2.057362051 | 2.267171728 | 1.895131899 |
| ENSMUSG00000051675 | 2.380049842 | 2.219895812 | 2.226722649 | 2.105516192 | 2.116484486 | 2.269512944 | 1.978585612 |
| ENSMUSG00000051678 | 0.289064826 | 0 | 0 | 0 | 0.244650542 | 0 | 0 |
| ENSMUSG00000051680 | 0 | 0 | 0 | 0 | 0 | 0 | 0 |
| ENSMUSG00000051682 | 0 | 0 | 0 | 0 | 0 | 0 | 0 |
| ENSMUSG00000051687 | 0 | 0 | 0 | 0 | 0 | 0 | 0 |
| ENSMUSG00000051695 | 2.891461985 | 2.639365925 | 2.697383411 | 2.579070383 | 2.582889218 | 2.919078092 | 2.501682732 |
| ENSMUSG00000051703 | 0.289064826 | 0.864857905 | 0.646240625 | 0.25 | 0.387761935 | 0.477121255 | 0.853775828 |
| ENSMUSG00000051705 | 1.876566388 | 1.799918086 | 1.726722649 | 1.779735268 | 1.754126039 | 1.973127854 | 1.605556929 |

| | | | | | | | |
|---|---|---|---|---|---|---|---|
| ENSMUSG00000051706 | 0 | 0 | 0 | 0 | 0 | 0 | 0 |
| ENSMUSG00000051716 | 0.747221736 | 0.396240625 | 0.396240625 | 0.79248125 | 0.877063019 | 0.301029996 | 0.710836105 |
| ENSMUSG00000051721 | 1.527823874 | 1.372963274 | 1.470660762 | 1.356566189 | 1.351342427 | 1.431363764 | 1.324583736 |
| ENSMUSG00000051723 | 0.289064826 | 0.5 | 0.580482024 | 0.5 | 0.387761935 | 0.301029996 | 0.579661877 |
| ENSMUSG00000051726 | 2.381433035 | 2.315523711 | 2.309003548 | 1.372963274 | 1.445123986 | 1.477121255 | 1.413583902 |
| ENSMUSG00000051727 | 0.45815691 | 0.25 | 0.396240625 | 0.396240625 | 0.489301084 | 0.477121255 | 0.448487648 |
| ENSMUSG00000051728 | 0 | 0.25 | 0.396240625 | 0.25 | 0 | 0 | 0.224243824 |
| ENSMUSG00000051729 | 0 | 0 | 0 | 0 | 0 | 0 | 0 |
| ENSMUSG00000051730 | 1.181541736 | 1.431980114 | 1.339388001 | 1.356566189 | 1.343584837 | 1.568201724 | 1.329931221 |
| ENSMUSG00000051732 | 0 | 0 | 0 | 0 | 0 | 0 | 0 |
| ENSMUSG00000051735 | 0.671187741 | 0.5 | 0.396240625 | 0.25 | 0 | 0.301029996 | 0.355418052 |
| ENSMUSG00000051736 | 1.418409478 | 1.27186571 | 1.214495249 | 1.451838731 | 1.319233381 | 1.556302501 | 1.284450788 |
| ENSMUSG00000051747 | 0.578129653 | 0.5 | 0.646240625 | 0.701838731 | 0.686820904 | 0.602059991 | 0.448487648 |
| ENSMUSG00000051748 | 0 | 0 | 0 | 0 | 0 | 0 | 0 |
| ENSMUSG00000051758 | 0 | 0 | 0 | 0 | 0 | 0.301029996 | 0 |
| ENSMUSG00000051768 | 2.163307908 | 2.041226732 | 2.098079356 | 1.942046081 | 2.09661982 | 2.152288344 | 2.052911423 |
| ENSMUSG00000051769 | 0 | 0 | 0 | 0 | 0 | 0 | 0 |
| ENSMUSG00000051777 | 0 | 0 | 0 | 0 | 0 | 0 | 0 |
| ENSMUSG00000051786 | 2.576402808 | 2.357613138 | 2.395770692 | 2.284247778 | 2.363323232 | 2.602059991 | 2.201756416 |
| ENSMUSG00000051788 | 0.289064826 | 0 | 0 | 0 | 0 | 0 | 0 |
| ENSMUSG00000051790 | 2.241665209 | 2.108657057 | 2.149047439 | 2.121960008 | 2.187791022 | 2.418301291 | 2.105204898 |
| ENSMUSG00000051793 | 0 | 0 | 0 | 0 | 0 | 0 | 0 |
| ENSMUSG00000051802 | 0 | 0 | 0 | 0 | 0 | 0 | 0 |
| ENSMUSG00000051804 | 0 | 0 | 0 | 0 | 0 | 0 | 0 |
| ENSMUSG00000051805 | 0 | 0.25 | 0 | 0 | 0 | 0 | 0 |
| ENSMUSG00000051811 | 0 | 0 | 0 | 0.396240625 | 0 | 0.477121255 | 0.355418052 |
| ENSMUSG00000051817 | 1.837744666 | 1.79248125 | 1.848079356 | 1.668106335 | 1.72341484 | 1.832508913 | 1.722075986 |
| ENSMUSG00000051827 | 0 | 0 | 0 | 0 | 0 | 0 | 0 |
| ENSMUSG00000051832 | 1.205378646 | 1.130890489 | 1.201838731 | 0.75 | 0.77552387 | 0.477121255 | 0.829800754 |
| ENSMUSG00000051839 | 0 | 0 | 0 | 0 | 0 | 0 | 0 |
| ENSMUSG00000051844 | 0 | 0.396240625 | 0.396240625 | 0.25 | 0.489301084 | 0.477121255 | 0 |
| ENSMUSG00000051851 | 0.867194479 | 0.580482024 | 0.646240625 | 0.5 | 0.568060967 | 0.698970004 | 0.629532004 |
| ENSMUSG00000051853 | 3.175414803 | 2.908362344 | 2.882723495 | 2.913435195 | 2.881613441 | 3.32756326 | 2.679511242 |
| ENSMUSG00000051855 | 1.156259305 | 1.080482024 | 1.098079356 | 1.438721876 | 1.546663136 | 1.568201724 | 1.313616061 |

| | | | | | | | |
|---|---|---|---|---|---|---|---|
| ENSMUSG00000051860 | 0 | 0 | 0 | 0 | 0 | 0 | 0 |
| ENSMUSG00000051864 | 2.219867581 | 2.075945187 | 2.091580554 | 2.039967834 | 2.135290471 | 2.303196057 | 2.033617522 |
| ENSMUSG00000051877 | 0 | 0 | 0 | 0 | 0 | 0 | 0 |
| ENSMUSG00000051879 | 0 | 0 | 0 | 0 | 0 | 0 | 0 |
| ENSMUSG00000051885 | 0 | 0 | 0 | 0 | 0 | 0 | 0 |
| ENSMUSG00000051890 | 1.269664473 | 1.282320754 | 1.388647213 | 1.339388001 | 1.293076518 | 1.301029996 | 1.272006351 |
| ENSMUSG00000051896 | 0 | 0 | 0 | 0 | 0 | 0 | 0 |
| ENSMUSG00000051900 | 0 | 0 | 0 | 0 | 0.244650542 | 0 | 0 |
| ENSMUSG00000051906 | 0 | 0 | 0 | 0 | 0 | 0 | 0 |
| ENSMUSG00000051910 | 0.289064826 | 0.25 | 0.25 | 0 | 0.244650542 | 0 | 0 |
| ENSMUSG00000051917 | 0 | 0 | 0 | 0 | 0 | 0 | 0 |
| ENSMUSG00000051920 | 1.727821383 | 1.566696635 | 1.614857905 | 0.5 | 0.489301084 | 0.602059991 | 0.224243824 |
| ENSMUSG00000051934 | 2.578129653 | 2.387686696 | 2.352877747 | 2.509386738 | 2.488383918 | 2.816903839 | 2.343157052 |
| ENSMUSG00000051936 | 0 | 0 | 0 | 0 | 0 | 0 | 0 |
| ENSMUSG00000051940 | 0 | 0 | 0 | 0 | 0 | 0 | 0 |
| ENSMUSG00000051950 | 2.355805236 | 2.183677405 | 2.131869252 | 2.269203899 | 2.278401864 | 2.522444234 | 2.098866035 |
| ENSMUSG00000051951 | 0 | 0 | 0 | 0 | 0.244650542 | 0 | 0 |
| ENSMUSG00000051952 | 0 | 0 | 0 | 0 | 0 | 0 | 0 |
| ENSMUSG00000051956 | 0 | 0 | 0 | 0 | 0 | 0 | 0 |
| ENSMUSG00000051965 | 0 | 0 | 0 | 0 | 0 | 0 | 0 |
| ENSMUSG00000051969 | 0 | 0 | 0 | 0 | 0 | 0 | 0 |
| ENSMUSG00000051977 | 1.596667477 | 1.527131114 | 1.458222504 | 1.598079356 | 1.528558853 | 1.643452676 | 1.473249546 |
| ENSMUSG00000051978 | 1.714367558 | 1.766522298 | 1.784887838 | 1.692046081 | 1.55104774 | 1.86332286 | 1.559435575 |
| ENSMUSG00000051980 | 0 | 0 | 0 | 0 | 0 | 0 | 0 |
| ENSMUSG00000051984 | 1 | 0.92510993 | 0.896240625 | 0.5 | 0.489301084 | 0.477121255 | 0.672731473 |
| ENSMUSG00000051985 | 0 | 0 | 0 | 0 | 0 | 0 | 0 |
| ENSMUSG00000051986 | 0 | 0 | 0 | 0 | 0 | 0 | 0 |
| ENSMUSG00000051989 | 1.842799666 | 1.839388001 | 1.779735268 | 1.870953944 | 1.840643511 | 2.096910013 | 1.750751125 |
| ENSMUSG00000051998 | 0.45815691 | 0.75 | 0.580482024 | 0.5 | 0 | 0.477121255 | 0.355418052 |
| ENSMUSG00000052005 | 1.693336259 | 1.482684334 | 1.494319981 | 1.364857905 | 1.163285805 | 1.531478917 | 1.360283038 |
| ENSMUSG00000052012 | 0 | 0 | 0 | 0 | 0 | 0 | 0 |
| ENSMUSG00000052013 | 0 | 0 | 0 | 0 | 0 | 0 | 0 |
| ENSMUSG00000052014 | 0 | 0 | 0 | 0 | 0 | 0 | 0 |
| ENSMUSG00000052025 | 0.289064826 | 0 | 0 | 0 | 0 | 0 | 0 |

| | | | | | | | |
|---|---|---|---|---|---|---|---|
| ENSMUSG00000052026 | 0.91631382 | 0.951838731 | 1.201838731 | 0.396240625 | 0.244650542 | 0 | 0.448487648 |
| ENSMUSG00000052031 | 1.596667477 | 1.552363341 | 1.634789703 | 1.62694866 | 1.665423072 | 1.826074803 | 1.564611933 |
| ENSMUSG00000052033 | 2.042557845 | 1.930274797 | 1.87694866 | 2.12892496 | 2.039257769 | 2.348304863 | 1.874129423 |
| ENSMUSG00000052040 | 1.876566388 | 1.794977273 | 1.848079356 | 1.438721876 | 1.504538967 | 1.579783597 | 1.425641774 |
| ENSMUSG00000052056 | 1.99653913 | 1.799918086 | 1.80479213 | 1.825945187 | 1.877063019 | 2.029383778 | 1.773071385 |
| ENSMUSG00000052058 | 0 | 0 | 0 | 0 | 0 | 0 | 0 |
| ENSMUSG00000052062 | 1.862428343 | 1.705044741 | 1.723704441 | 1.830482024 | 1.763796674 | 2.025305865 | 1.641897157 |
| ENSMUSG00000052075 | 0 | 0 | 0 | 0 | 0 | 0 | 0 |
| ENSMUSG00000052085 | 1.470606563 | 1.339388001 | 1.396240625 | 1.226722649 | 1.327538614 | 1.255272505 | 1.176815335 |
| ENSMUSG00000052087 | 0.289064826 | 0 | 0 | 0.25 | 0 | 0 | 0.224243824 |
| ENSMUSG00000052099 | 0 | 0 | 0 | 0 | 0 | 0 | 0 |
| ENSMUSG00000052102 | 2.177057474 | 2.036164561 | 2.05357978 | 1.834962501 | 1.704182902 | 1.968482949 | 1.667169465 |
| ENSMUSG00000052105 | 1.568542279 | 1.789967834 | 1.744319981 | 1.505591953 | 1.625423019 | 1.62324929 | 1.67819947 |
| ENSMUSG00000052117 | 0.289064826 | 0.25 | 0.25 | 0.25 | 0.244650542 | 0 | 0.224243824 |
| ENSMUSG00000052125 | 1.765759561 | 1.77186571 | 1.732684334 | 1.930274797 | 1.871833944 | 2.045322979 | 1.725216937 |
| ENSMUSG00000052131 | 0.289064826 | 0 | 0.25 | 0 | 0 | 0 | 0 |
| ENSMUSG00000052133 | 0 | 0 | 0.25 | 0.25 | 0 | 0 | 0.224243824 |
| ENSMUSG00000052135 | 1.129344651 | 1.26109853 | 1.146240625 | 0.701838731 | 0.489301084 | 0.903089987 | 0.952571511 |
| ENSMUSG00000052137 | 2.023453784 | 1.898114259 | 1.839388001 | 1.882845365 | 1.828942237 | 2.012837225 | 1.670888075 |
| ENSMUSG00000052139 | 2.185978293 | 2.05357978 | 2.02053726 | 1.987091808 | 1.945998439 | 2.161368002 | 1.909593098 |
| ENSMUSG00000052142 | 0 | 0 | 0 | 0 | 0 | 0.301029996 | 0 |
| ENSMUSG00000052143 | 0.289064826 | 0 | 0.5 | 0.701838731 | 0.686820904 | 0.602059991 | 0.520678036 |
| ENSMUSG00000052144 | 2.934301833 | 2.687257846 | 2.676408097 | 2.659835177 | 2.580303447 | 2.985875357 | 2.387828426 |
| ENSMUSG00000052146 | 2.666334442 | 2.584684096 | 2.757839149 | 2.415889526 | 2.503953633 | 2.755874856 | 2.568901575 |
| ENSMUSG00000052151 | 2.45815691 | 2.282320754 | 2.348079356 | 2.298689214 | 2.368090054 | 2.602059991 | 2.243699513 |
| ENSMUSG00000052154 | 0.289064826 | 0.646240625 | 0 | 0.25 | 0 | 0 | 0.355418052 |
| ENSMUSG00000052155 | 2.131864492 | 1.961372513 | 1.938721876 | 2.001406137 | 1.916320788 | 2.133538908 | 1.770364127 |
| ENSMUSG00000052157 | 0.289064826 | 0 | 0.25 | 0 | 0.387761935 | 0.301029996 | 0.224243824 |
| ENSMUSG00000052160 | 0 | 0.25 | 0.396240625 | 0.25 | 0.244650542 | 0 | 0 |
| ENSMUSG00000052180 | 0.289064826 | 0.25 | 0 | 0 | 0 | 0 | 0 |
| ENSMUSG00000052182 | 0 | 0 | 0 | 0 | 0 | 0 | 0 |
| ENSMUSG00000052187 | 0 | 0 | 0 | 0 | 0 | 0 | 0 |
| ENSMUSG00000052188 | 0 | 0 | 0 | 0 | 0 | 0 | 0.224243824 |
| ENSMUSG00000052192 | 0 | 0 | 0 | 0 | 0 | 0 | 0 |

| | | | | | | | |
|---|---|---|---|---|---|---|---|
| ENSMUSG00000052212 | 0 | 0 | 0 | 0 | 0 | 0 | 0 |
| ENSMUSG00000052214 | 2.373064211 | 2.148114259 | 2.188721876 | 2.266522298 | 2.229681962 | 2.531478917 | 2.15146804 |
| ENSMUSG00000052217 | 0 | 0 | 0 | 0 | 0 | 0 | 0 |
| ENSMUSG00000052221 | 0 | 0 | 0 | 0.396240625 | 0.387761935 | 0 | 0.224243824 |
| ENSMUSG00000052229 | 0 | 0.396240625 | 0.5 | 0 | 0 | 0 | 0 |
| ENSMUSG00000052234 | 0 | 0 | 0 | 0 | 0 | 0 | 0 |
| ENSMUSG00000052236 | 0 | 0 | 0 | 0 | 0 | 0 | 0 |
| ENSMUSG00000052241 | 0 | 0.25 | 0.25 | 0 | 0 | 0 | 0.224243824 |
| ENSMUSG00000052248 | 0 | 0.25 | 0.5 | 0.25 | 0.244650542 | 0.477121255 | 0.355418052 |
| ENSMUSG00000052253 | 2.23778581 | 2.063166358 | 2.100219859 | 2.171625132 | 2.110284345 | 2.367355921 | 1.989124507 |
| ENSMUSG00000052270 | 0 | 0 | 0 | 0 | 0 | 0.301029996 | 0 |
| ENSMUSG00000052271 | 0.45815691 | 0 | 0.25 | 0 | 0.244650542 | 0 | 0.355418052 |
| ENSMUSG00000052273 | 0 | 0 | 0 | 0 | 0 | 0 | 0 |
| ENSMUSG00000052276 | 0 | 0 | 0 | 0 | 0 | 0 | 0 |
| ENSMUSG00000052281 | 0 | 0.25 | 0 | 0 | 0 | 0 | 0 |
| ENSMUSG00000052291 | 0.811507563 | 0.896240625 | 0.92510993 | 1.160964047 | 1 | 0.903089987 | 0.916588299 |
| ENSMUSG00000052293 | 0.671187741 | 0.5 | 0.25 | 0.830482024 | 0.733951626 | 0.602059991 | 0.448487648 |
| ENSMUSG00000052295 | 0 | 0.25 | 0.5 | 0.396240625 | 0 | 0 | 0 |
| ENSMUSG00000052296 | 2.475043119 | 2.341033664 | 2.353406982 | 2.209800947 | 2.326087113 | 2.519827994 | 2.269110043 |
| ENSMUSG00000052298 | 2.481610694 | 2.3184489 | 2.321928095 | 2.438304187 | 2.390196125 | 2.753583059 | 2.2551389 |
| ENSMUSG00000052299 | 2.796978839 | 2.569612365 | 2.541226732 | 2.645770692 | 2.55563165 | 2.965201701 | 2.366155413 |
| ENSMUSG00000052301 | 0 | 0 | 0 | 0 | 0 | 0 | 0 |
| ENSMUSG00000052302 | 0.45815691 | 0.25 | 0 | 1.238549078 | 1.244650542 | 1.278753601 | 1.176815335 |
| ENSMUSG00000052303 | 0 | 0 | 0 | 0 | 0 | 0 | 0 |
| ENSMUSG00000052305 | 0 | 0 | 0 | 0 | 0 | 0 | 0 |
| ENSMUSG00000052310 | 2.604828841 | 2.385274154 | 2.426408097 | 2.493244947 | 2.449252175 | 2.780317312 | 2.349118526 |
| ENSMUSG00000052316 | 1.721148722 | 1.649978211 | 1.580482024 | 0.830482024 | 0.686820904 | 0.698970004 | 0.74492186 |
| ENSMUSG00000052331 | 2.156259305 | 1.994319981 | 1.856566189 | 1.921625132 | 1.86473911 | 2.093421685 | 1.71730663 |
| ENSMUSG00000052334 | 0 | 0 | 0 | 0 | 0.244650542 | 0 | 0 |
| ENSMUSG00000052336 | 0 | 0 | 0 | 0 | 0 | 0 | 0 |
| ENSMUSG00000052337 | 3.062176765 | 2.795132702 | 2.79106961 | 2.830200137 | 2.733798931 | 3.178976947 | 2.543888297 |
| ENSMUSG00000052353 | 1.978790393 | 1.890560606 | 1.77186571 | 1.744319981 | 1.754126039 | 1.806179974 | 1.641897157 |
| ENSMUSG00000052363 | 0 | 0 | 0 | 0 | 0 | 0 | 0 |
| ENSMUSG00000052364 | 1.129344651 | 1.080482024 | 1.146240625 | 1.226722649 | 1.188507684 | 1.301029996 | 1.100339912 |

| | | | | | | | |
|---|---|---|---|---|---|---|---|
| ENSMUSG00000052368 | 0 | 0 | 0 | 0 | 0 | 0 | 0 |
| ENSMUSG00000052369 | 2.467247894 | 2.234894804 | 2.269871196 | 2.159156155 | 2.091002363 | 2.330413773 | 1.997988498 |
| ENSMUSG00000052371 | 0.747221736 | 0.79248125 | 1.04248125 | 0 | 0 | 0 | 0 |
| ENSMUSG00000052372 | 0 | 0 | 0 | 0.25 | 0 | 0 | 0 |
| ENSMUSG00000052373 | 1.100572389 | 1.26109853 | 1.114857905 | 1.214495249 | 1.343584837 | 1.146128036 | 1.224243824 |
| ENSMUSG00000052374 | 0 | 0.5 | 0.5 | 0.25 | 0.489301084 | 0.301029996 | 0.629532004 |
| ENSMUSG00000052376 | 0 | 0 | 0 | 0 | 0 | 0 | 0 |
| ENSMUSG00000052382 | 0 | 0 | 0 | 0 | 0 | 0 | 0 |
| ENSMUSG00000052384 | 0.289064826 | 0.25 | 0.25 | 0.580482024 | 0.632412477 | 0.77815125 | 0.74492186 |
| ENSMUSG00000052387 | 0.671187741 | 0.701838731 | 0.580482024 | 0.396240625 | 0.568060967 | 0.77815125 | 0.74492186 |
| ENSMUSG00000052392 | 0 | 0 | 0 | 0 | 0.244650542 | 0 | 0 |
| ENSMUSG00000052395 | 1.89911417 | 1.62694866 | 1.695339928 | 2.038704527 | 2.003645018 | 2.161368002 | 1.854824501 |
| ENSMUSG00000052396 | 1.505869732 | 1.42510993 | 1.438721876 | 0.79248125 | 0.77552387 | 0.698970004 | 0.710836105 |
| ENSMUSG00000052397 | 2.778209214 | 2.598883534 | 2.65136988 | 2.334962501 | 2.405621873 | 2.668385917 | 2.330172201 |
| ENSMUSG00000052403 | 0 | 0 | 0 | 0.25 | 0 | 0 | 0 |
| ENSMUSG00000052406 | 2.382811656 | 2.207430684 | 2.209012589 | 2.224461364 | 2.218681522 | 2.45484486 | 2.084261301 |
| ENSMUSG00000052407 | 1.249317394 | 1.146240625 | 0.864857905 | 1.098079356 | 0.978602168 | 0.954242509 | 1.041356071 |
| ENSMUSG00000052414 | 0 | 0 | 0 | 0 | 0 | 0 | 0 |
| ENSMUSG00000052415 | 0 | 0 | 0.580482024 | 1.25 | 1.327538614 | 1.431363764 | 1.360283038 |
| ENSMUSG00000052417 | 0 | 0 | 0 | 0 | 0 | 0 | 0 |
| ENSMUSG00000052419 | 2.473938397 | 2.28360658 | 2.293731421 | 2.087182039 | 2.03596422 | 2.357934847 | 1.877074955 |
| ENSMUSG00000052423 | 1.932832114 | 1.896240625 | 1.930274797 | 1.864857905 | 1.875328588 | 2.017033339 | 1.82753046 |
| ENSMUSG00000052426 | 0.289064826 | 0 | 0 | 0 | 0.244650542 | 0 | 0 |
| ENSMUSG00000052428 | 2.487005755 | 2.252806814 | 2.243603647 | 2.269203899 | 2.194189193 | 2.484299839 | 2.098866035 |
| ENSMUSG00000052429 | 2.904270129 | 2.666999634 | 2.660964047 | 2.629663289 | 2.660005804 | 3.044931546 | 2.513787281 |
| ENSMUSG00000052430 | 0.671187741 | 0.75 | 1.02186571 | 0.976722649 | 0.846351821 | 0.77815125 | 0.775756176 |
| ENSMUSG00000052435 | 0 | 0 | 0 | 0 | 0 | 0 | 0 |
| ENSMUSG00000052446 | 2.042557845 | 1.942046081 | 1.953445298 | 2.036164561 | 1.926994141 | 2.23299611 | 1.822941512 |
| ENSMUSG00000052456 | 2.691370667 | 2.519537702 | 2.552363341 | 2.492166698 | 2.56462465 | 2.86569606 | 2.41893749 |
| ENSMUSG00000052459 | 3.053448496 | 2.782642641 | 2.796992648 | 2.774836953 | 2.700005857 | 3.118595365 | 2.509940767 |
| ENSMUSG00000052468 | 0 | 0 | 0 | 0 | 0 | 0 | 0 |
| ENSMUSG00000052469 | 0 | 0 | 0 | 0 | 0 | 0 | 0 |
| ENSMUSG00000052471 | 0 | 0 | 0 | 0 | 0 | 0 | 0 |
| ENSMUSG00000052477 | 0.289064826 | 0 | 0 | 0.25 | 0 | 0 | 0 |

| | | | | | | | |
|---|---|---|---|---|---|---|---|
| ENSMUSG00000052479 | 0 | 0.25 | 0 | 0.25 | 0 | 0 | 0.224243824 |
| ENSMUSG00000052482 | 0 | 0.25 | 0.25 | 0 | 0.387761935 | 0.301029996 | 0 |
| ENSMUSG00000052485 | 1.596667477 | 1.649978211 | 1.638647213 | 1.282320754 | 1.283908311 | 1.414973348 | 1.245582246 |
| ENSMUSG00000052488 | 2.346900291 | 2.282320754 | 2.343216265 | 2.059601185 | 2.257246745 | 2.392696953 | 2.271717885 |
| ENSMUSG00000052496 | 0 | 0 | 0 | 0.75 | 0.489301084 | 0.698970004 | 0.579661877 |
| ENSMUSG00000052504 | 0.671187741 | 0.830482024 | 0.646240625 | 0 | 0.489301084 | 0.698970004 | 0.579661877 |
| ENSMUSG00000052508 | 0 | 0 | 0 | 0 | 0.244650542 | 0 | 0.224243824 |
| ENSMUSG00000052512 | 2.339331437 | 2.186208459 | 2.231203126 | 1.828220739 | 1.870073561 | 2 | 1.871156825 |
| ENSMUSG00000052516 | 1.663535556 | 1.410964047 | 1.516522298 | 1.678561379 | 1.686820904 | 1.785329835 | 1.546120303 |
| ENSMUSG00000052520 | 0 | 0 | 0 | 0 | 0 | 0 | 0 |
| ENSMUSG00000052525 | 1.036286563 | 0.92510993 | 1.061981878 | 1.160964047 | 1.188507684 | 1.079181246 | 1.185218806 |
| ENSMUSG00000052533 | 2.889424695 | 2.647880587 | 2.656198864 | 2.531676118 | 2.561901018 | 2.856124444 | 2.445802857 |
| ENSMUSG00000052534 | 2.750084015 | 2.564641508 | 2.556903986 | 2.28743678 | 2.223252711 | 2.501059262 | 2.068965954 |
| ENSMUSG00000052537 | 0 | 0 | 0 | 0 | 0 | 0 | 0 |
| ENSMUSG00000052539 | 2.780336935 | 2.575374049 | 2.54652856 | 2.635516136 | 2.565117605 | 2.971275849 | 2.375293074 |
| ENSMUSG00000052544 | 0.578129653 | 0.580482024 | 0.580482024 | 1.080482024 | 1.057362051 | 1.079181246 | 1 |
| ENSMUSG00000052549 | 0.289064826 | 0.25 | 0 | 0.5 | 0.632412477 | 0.602059991 | 0.224243824 |
| ENSMUSG00000052551 | 0.289064826 | 0.396240625 | 0 | 0 | 0.244650542 | 0 | 0.224243824 |
| ENSMUSG00000052554 | 0 | 0 | 0 | 0 | 0 | 0 | 0 |
| ENSMUSG00000052557 | 2.40033094 | 2.226722649 | 2.252806814 | 2.192046081 | 2.230318492 | 2.409933123 | 2.096890362 |
| ENSMUSG00000052560 | 3.054824387 | 2.767194569 | 2.725593779 | 2.599685923 | 2.517458689 | 2.870988814 | 2.343850547 |
| ENSMUSG00000052562 | 0 | 0 | 0 | 0 | 0 | 0 | 0 |
| ENSMUSG00000052563 | 0.867194479 | 1 | 0.92510993 | 1.201838731 | 1.136121934 | 1.278753601 | 1.20919388 |
| ENSMUSG00000052565 | 0.289064826 | 0 | 0 | 0 | 0.244650542 | 0.301029996 | 0.224243824 |
| ENSMUSG00000052566 | 1.740854706 | 1.701838731 | 1.720660762 | 1.53743678 | 1.580303447 | 1.707570176 | 1.591689859 |
| ENSMUSG00000052572 | 0 | 0 | 0 | 0.396240625 | 0.387761935 | 0.477121255 | 0 |
| ENSMUSG00000052581 | 0 | 0.25 | 0.25 | 0 | 0 | 0 | 0 |
| ENSMUSG00000052584 | 1.37447073 | 1.146240625 | 1.302363341 | 1.356566189 | 1.456697355 | 1.431363764 | 1.302263476 |
| ENSMUSG00000052593 | 2.578990401 | 2.388647213 | 2.385757955 | 2.459406983 | 2.388981128 | 2.759667845 | 2.235411964 |
| ENSMUSG00000052595 | 0 | 0 | 0.25 | 0 | 0 | 0 | 0 |
| ENSMUSG00000052605 | 0.45815691 | 0.580482024 | 0.864857905 | 0.646240625 | 0.812711509 | 0.77815125 | 0.896975297 |
| ENSMUSG00000052609 | 2.402962068 | 2.237821179 | 2.279735268 | 1.951838731 | 2.06759338 | 2.225309282 | 2.016928211 |
| ENSMUSG00000052613 | 0.289064826 | 0 | 0 | 0.25 | 0 | 0 | 0 |
| ENSMUSG00000052616 | 0.45815691 | 0.25 | 0.25 | 0.25 | 0.489301084 | 0 | 0.224243824 |

| | | | | | | | |
|---|---|---|---|---|---|---|---|
| ENSMUSG00000052625 | 0 | 0 | 0 | 0 | 0 | 0 | 0 |
| ENSMUSG00000052629 | 0.289064826 | 0.25 | 0.396240625 | 0 | 0 | 0 | 0 |
| ENSMUSG00000052631 | 0.671187741 | 0.580482024 | 0.580482024 | 0 | 0 | 0 | 0 |
| ENSMUSG00000052632 | 1.842799666 | 1.755591953 | 1.75835575 | 1.557204673 | 1.618292349 | 1.681241237 | 1.529383012 |
| ENSMUSG00000052642 | 0 | 0 | 0 | 0 | 0 | 0 | 0 |
| ENSMUSG00000052656 | 2.211649928 | 2.001406137 | 2.043731421 | 2.08272922 | 2.008447216 | 2.267171728 | 1.866140981 |
| ENSMUSG00000052658 | 0.289064826 | 0.396240625 | 0.646240625 | 0.396240625 | 0.568060967 | 0.477121255 | 0.520678036 |
| ENSMUSG00000052673 | 0 | 0.25 | 0 | 0 | 0 | 0 | 0 |
| ENSMUSG00000052675 | 1.249317394 | 1.238549078 | 1.188721876 | 1.080482024 | 1.176121988 | 1.113943352 | 1.110947927 |
| ENSMUSG00000052676 | 1.734388958 | 1.678561379 | 1.622963274 | 1.67510993 | 1.639266209 | 1.832508913 | 1.529383012 |
| ENSMUSG00000052681 | 3.008375049 | 2.741986676 | 2.743961992 | 2.832168732 | 2.702851618 | 3.156549151 | 2.531587776 |
| ENSMUSG00000052684 | 2.054824387 | 1.942046081 | 2.095926073 | 1.720660762 | 1.766173492 | 1.880813592 | 1.819456572 |
| ENSMUSG00000052688 | 1.432083896 | 1.527131114 | 1.505591953 | 1.372963274 | 1.293076518 | 1.447158031 | 1.224243824 |
| ENSMUSG00000052698 | 2.401648579 | 2.222185812 | 2.195339928 | 1.982684334 | 2.024758322 | 2.209515015 | 1.97354745 |
| ENSMUSG00000052707 | 2.703665763 | 2.502806814 | 2.475593779 | 2.476346751 | 2.409470941 | 2.761927838 | 2.229218532 |
| ENSMUSG00000052712 | 2.43085913 | 2.208222504 | 2.228222334 | 2.247880462 | 2.201854879 | 2.487138375 | 2.092902448 |
| ENSMUSG00000052713 | 1.89911417 | 1.789967834 | 1.747171172 | 1.79745614 | 1.728722551 | 2.049218023 | 1.657681529 |
| ENSMUSG00000052724 | 0 | 0.396240625 | 0 | 0 | 0 | 0 | 0.448487648 |
| ENSMUSG00000052726 | 1.418409478 | 1.372963274 | 1.380890489 | 1.04248125 | 1.057362051 | 1.230448921 | 0.984950056 |
| ENSMUSG00000052727 | 3.219312402 | 2.923589472 | 2.909948223 | 2.966529948 | 2.909702667 | 3.329804522 | 2.699114618 |
| ENSMUSG00000052730 | 0 | 0 | 0 | 0 | 0 | 0 | 0 |
| ENSMUSG00000052736 | 0 | 0 | 0 | 0 | 0 | 0 | 0 |
| ENSMUSG00000052738 | 2.495495257 | 2.33272922 | 2.343216265 | 2.397646762 | 2.353143227 | 2.671172843 | 2.196701384 |
| ENSMUSG00000052748 | 2.042557845 | 1.888647213 | 1.910964047 | 1.96759118 | 1.903713872 | 2.201397124 | 1.764880683 |
| ENSMUSG00000052749 | 0 | 0 | 0 | 0 | 0 | 0 | 0 |
| ENSMUSG00000052751 | 1.765759561 | 1.630890489 | 1.723704441 | 1.458222504 | 1.595992969 | 1.579783597 | 1.551512352 |
| ENSMUSG00000052752 | 2.813962146 | 2.615625568 | 2.618933358 | 2.597810897 | 2.630166292 | 2.937517892 | 2.481351231 |
| ENSMUSG00000052759 | 0 | 0 | 0 | 0 | 0 | 0 | 0 |
| ENSMUSG00000052760 | 0 | 0.5 | 0 | 0.25 | 0 | 0.477121255 | 0 |
| ENSMUSG00000052763 | 1.964026642 | 1.909156155 | 1.931980114 | 1.825945187 | 1.83872004 | 2.025305865 | 1.829800754 |
| ENSMUSG00000052767 | 0 | 0 | 0 | 0 | 0 | 0 | 0 |
| ENSMUSG00000052769 | 0.578129653 | 0.25 | 0.5 | 0.25 | 0.244650542 | 0.301029996 | 0.355418052 |
| ENSMUSG00000052776 | 0 | 0.25 | 0 | 0.646240625 | 0.686820904 | 0.698970004 | 0.448487648 |
| ENSMUSG00000052779 | 0 | 0 | 0 | 0 | 0 | 0 | 0 |

| | | | | | | | |
|---|---|---|---|---|---|---|---|
| ENSMUSG00000052783 | 1.358731791 | 1.388647213 | 1.396240625 | 1.470660762 | 1.42077253 | 1.568201724 | 1.307989563 |
| ENSMUSG00000052785 | 0 | 0 | 0 | 0 | 0 | 0 | 0 |
| ENSMUSG00000052790 | 0 | 0 | 0 | 0 | 0 | 0 | 0 |
| ENSMUSG00000052794 | 1.721148722 | 1.664552871 | 1.642463902 | 1.62694866 | 1.652586889 | 1.69019608 | 1.535058868 |
| ENSMUSG00000052798 | 2.749513129 | 2.545598588 | 2.544977273 | 2.594302633 | 2.556389881 | 2.945960704 | 2.414141848 |
| ENSMUSG00000052812 | 1.971473847 | 1.830482024 | 1.837182039 | 1.956637122 | 1.906908067 | 2.123851641 | 1.782372264 |
| ENSMUSG00000052818 | 0 | 0 | 0 | 0 | 0 | 0 | 0 |
| ENSMUSG00000052819 | 0.289064826 | 0.25 | 0 | 0 | 0 | 0 | 0.448487648 |
| ENSMUSG00000052821 | 0.289064826 | 0 | 0 | 0 | 0 | 0 | 0.224243824 |
| ENSMUSG00000052825 | 0 | 0.25 | 0 | 0.5 | 0.244650542 | 0 | 0.355418052 |
| ENSMUSG00000052833 | 2.900707419 | 2.649280357 | 2.631869252 | 2.719321033 | 2.628523678 | 3.061829307 | 2.474639684 |
| ENSMUSG00000052837 | 1.53838222 | 1.646240625 | 1.979715809 | 1.238549078 | 1.621875691 | 1.322219295 | 1.925124822 |
| ENSMUSG00000052848 | 0 | 0 | 0 | 0 | 0 | 0 | 0 |
| ENSMUSG00000052850 | 0 | 0 | 0 | 0.25 | 0 | 0 | 0 |
| ENSMUSG00000052852 | 1.759671389 | 1.671625132 | 1.557204673 | 2.089388001 | 1.995080671 | 2.338456494 | 1.932619724 |
| ENSMUSG00000052854 | 1.269664473 | 1.29248125 | 1.330482024 | 1.602347734 | 1.595992969 | 1.653212514 | 1.437266334 |
| ENSMUSG00000052861 | 1.342375483 | 1.080482024 | 1.302363341 | 0.5 | 0.568060967 | 0.477121255 | 0.355418052 |
| ENSMUSG00000052865 | 0 | 0 | 0 | 0 | 0 | 0 | 0 |
| ENSMUSG00000052889 | 0 | 0 | 0 | 0 | 0 | 0 | 0 |
| ENSMUSG00000052906 | 1.948720984 | 1.821350555 | 1.769203899 | 2.04248125 | 1.964031554 | 2.193124598 | 1.881925353 |
| ENSMUSG00000052909 | 0 | 0 | 0 | 0 | 0 | 0 | 0 |
| ENSMUSG00000052911 | 2.696591798 | 2.585518667 | 2.668548067 | 2.310793496 | 2.56854915 | 2.658011397 | 2.525850149 |
| ENSMUSG00000052912 | 0 | 0 | 0 | 0.25 | 0 | 0 | 0 |
| ENSMUSG00000052914 | 2.10861828 | 1.807204673 | 1.837182039 | 1.584962501 | 1.47337555 | 1.531478917 | 1.38802993 |
| ENSMUSG00000052915 | 2.675985209 | 2.421625132 | 2.42510993 | 2.454645544 | 2.386129765 | 2.725094521 | 2.237664194 |
| ENSMUSG00000052917 | 2.217828299 | 2.067865757 | 2.038704527 | 2.038704527 | 1.986303335 | 2.245512668 | 1.824094866 |
| ENSMUSG00000052920 | 1.527823874 | 1.396240625 | 1.396240625 | 1.27186571 | 1.176121988 | 1.462397998 | 1.100339912 |
| ENSMUSG00000052921 | 1.249317394 | 1.282320754 | 1.302363341 | 1.130890489 | 1.039257769 | 1.079181246 | 1.066254157 |
| ENSMUSG00000052922 | 0 | 0 | 0 | 0 | 0 | 0 | 0 |
| ENSMUSG00000052926 | 2.273259602 | 2.088286706 | 2.088286706 | 2.228222334 | 2.273371525 | 2.482873584 | 2.144273809 |
| ENSMUSG00000052928 | 2.167941649 | 2.05479213 | 2.065523711 | 1.903677461 | 1.987570683 | 2.222716471 | 1.825244123 |
| ENSMUSG00000052934 | 1.734388958 | 1.75 | 1.814346961 | 1.774508021 | 1.923977404 | 1.995635195 | 1.877074955 |
| ENSMUSG00000052942 | 2.17479713 | 2.025821952 | 2.001406137 | 2.281030328 | 2.244650542 | 2.519827994 | 2.080113611 |
| ENSMUSG00000052949 | 2.325351389 | 2.17510993 | 2.178561379 | 1.856566189 | 1.91476933 | 2.041392685 | 1.848480858 |

| | | | | | | | |
|---|---|---|---|---|---|---|---|
| ENSMUSG00000052951 | 0 | 0 | 0 | 0.646240625 | 0.77552387 | 0.602059991 | 0.969165684 |
| ENSMUSG00000052955 | 0 | 0 | 0 | 0 | 0 | 0 | 0 |
| ENSMUSG00000052957 | 2.312518611 | 2.138647213 | 2.157339155 | 1.890560606 | 1.826953746 | 2.068185862 | 1.806347338 |
| ENSMUSG00000052962 | 2.121692466 | 1.961372513 | 2.017865591 | 2.139605178 | 2.084333628 | 2.394451681 | 1.959463225 |
| ENSMUSG00000052974 | 0 | 0 | 0 | 0 | 0 | 0 | 0 |
| ENSMUSG00000052981 | 0 | 0 | 0 | 0 | 0 | 0 | 0.224243824 |
| ENSMUSG00000052997 | 2.875723046 | 2.609438018 | 2.586905342 | 2.667221246 | 2.58335733 | 2.973589623 | 2.411904268 |
| ENSMUSG00000053004 | 0.91631382 | 0.5 | 0.830482024 | 0.976722649 | 0.955822902 | 0.954242509 | 0.803905701 |
| ENSMUSG00000053007 | 1.445324132 | 1.348079356 | 1.321350555 | 1.5 | 1.478764298 | 1.62324929 | 1.360283038 |
| ENSMUSG00000053012 | 2.404271433 | 2.267865591 | 2.247171172 | 2.247171172 | 2.139428339 | 2.423245874 | 1.997988498 |
| ENSMUSG00000053024 | 0 | 0 | 0.25 | 0.25 | 0 | 0 | 0.448487648 |
| ENSMUSG00000053025 | 0 | 0 | 0 | 0 | 0 | 0 | 0.224243824 |
| ENSMUSG00000053030 | 0.289064826 | 0 | 0.25 | 0 | 0.387761935 | 0 | 0.224243824 |
| ENSMUSG00000053038 | 1.325351389 | 1.29248125 | 1.188721876 | 1.201838731 | 1.149965125 | 1.204119983 | 1.100339912 |
| ENSMUSG00000053040 | 0.578129653 | 0.896240625 | 0.79248125 | 1.146240625 | 1.074582839 | 1.041392685 | 1.150210039 |
| ENSMUSG00000053044 | 0 | 0.25 | 0 | 0 | 0 | 0 | 0 |
| ENSMUSG00000053046 | 0.289064826 | 0 | 0 | 0 | 0.244650542 | 0 | 0 |
| ENSMUSG00000053049 | 0 | 0 | 0.25 | 0 | 0 | 0 | 0 |
| ENSMUSG00000053054 | 0 | 0 | 0 | 0 | 0 | 0 | 0 |
| ENSMUSG00000053062 | 2.215778996 | 2.069031101 | 2.080482024 | 1.866901388 | 1.824953989 | 2.075546961 | 1.710836105 |
| ENSMUSG00000053063 | 0 | 0.25 | 0.396240625 | 0 | 0 | 0 | 0.224243824 |
| ENSMUSG00000053070 | 1.568542279 | 1.589388001 | 1.660964047 | 1.602347734 | 1.467903253 | 1.633468456 | 1.38802993 |
| ENSMUSG00000053080 | 2.364524 | 2.186208459 | 2.216046536 | 2.05357978 | 2.101233692 | 2.209515015 | 1.995286892 |
| ENSMUSG00000053081 | 0 | 0 | 0 | 0 | 0 | 0 | 0 |
| ENSMUSG00000053091 | 1.734388958 | 1.67510993 | 1.622963274 | 1.71759118 | 1.671670246 | 1.826074803 | 1.574722613 |
| ENSMUSG00000053093 | 0 | 0.25 | 0 | 0 | 0.244650542 | 0 | 0.224243824 |
| ENSMUSG00000053094 | 2.736751754 | 2.505591953 | 2.52219706 | 2.444108258 | 2.398585969 | 2.703291378 | 2.266481009 |
| ENSMUSG00000053101 | 0 | 0.25 | 0 | 0.25 | 0.244650542 | 0 | 0.224243824 |
| ENSMUSG00000053110 | 2.946233352 | 2.677701608 | 2.660061234 | 2.699415381 | 2.656030391 | 3.041392685 | 2.485997583 |
| ENSMUSG00000053111 | 0.289064826 | 0.5 | 0.25 | 0.25 | 0.244650542 | 0.301029996 | 0.448487648 |
| ENSMUSG00000053113 | 1.993049299 | 1.951838731 | 1.942046081 | 1.747171172 | 1.800029087 | 1.949390007 | 1.788855757 |
| ENSMUSG00000053117 | 0.45815691 | 0.396240625 | 0.25 | 0.396240625 | 0.244650542 | 0 | 0.448487648 |
| ENSMUSG00000053119 | 2.759671389 | 2.501406137 | 2.526804269 | 2.53456795 | 2.455355315 | 2.821513528 | 2.288312058 |
| ENSMUSG00000053121 | 0 | 0 | 0 | 0 | 0 | 0 | 0 |

| | | | | | | | |
|---|---|---|---|---|---|---|---|
| ENSMUSG00000053128 | 1.971473847 | 1.860735874 | 1.935366747 | 1.870953944 | 1.895599581 | 2.127104798 | 1.882886767 |
| ENSMUSG00000053129 | 0 | 0 | 0 | 0 | 0 | 0 | 0 |
| ENSMUSG00000053134 | 2.111266113 | 2.013820609 | 1.987091808 | 2.015173983 | 2.025895058 | 2.176091259 | 1.853775828 |
| ENSMUSG00000053137 | 2.10861828 | 1.940387808 | 1.994319981 | 1.898114259 | 1.917865457 | 2.117271296 | 1.848480858 |
| ENSMUSG00000053141 | 0 | 0.25 | 0.5 | 0.25 | 0 | 0 | 0.224243824 |
| ENSMUSG00000053146 | 0 | 0 | 0 | 0 | 0 | 0 | 0 |
| ENSMUSG00000053153 | 0.289064826 | 0 | 0 | 0 | 0 | 0 | 0 |
| ENSMUSG00000053158 | 1.647796617 | 1.54248125 | 1.698603967 | 1.17510993 | 1.283908311 | 1.322219295 | 1.417653332 |
| ENSMUSG00000053161 | 0 | 0 | 0.25 | 0 | 0 | 0 | 0 |
| ENSMUSG00000053164 | 0.45815691 | 0.25 | 0.396240625 | 0.646240625 | 0.568060967 | 0 | 0 |
| ENSMUSG00000053166 | 0 | 0 | 0.25 | 0 | 0 | 0 | 0.448487648 |
| ENSMUSG00000053168 | 0 | 0 | 0 | 0 | 0 | 0 | 0 |
| ENSMUSG00000053173 | 0 | 0 | 0 | 0 | 0 | 0.301029996 | 0 |
| ENSMUSG00000053175 | 1.482695304 | 1.388647213 | 1.571350555 | 1.114857905 | 1.106691885 | 1.301029996 | 1.383567577 |
| ENSMUSG00000053178 | 0.45815691 | 0.701838731 | 0.580482024 | 0.580482024 | 0.568060967 | 0.301029996 | 0.579661877 |
| ENSMUSG00000053181 | 0.671187741 | 0 | 0.396240625 | 0.5 | 0.387761935 | 0.477121255 | 0.224243824 |
| ENSMUSG00000053182 | 0.45815691 | 0 | 0.5 | 0.5 | 0.244650542 | 0 | 0.224243824 |
| ENSMUSG00000053184 | 0 | 0 | 0 | 0 | 0 | 0 | 0 |
| ENSMUSG00000053185 | 0 | 0 | 0 | 0 | 0 | 0 | 0 |
| ENSMUSG00000053192 | 2.144240249 | 1.976722649 | 1.982684334 | 2.039967834 | 1.948833444 | 2.201397124 | 1.838725884 |
| ENSMUSG00000053194 | 0 | 0 | 0 | 0 | 0 | 0 | 0 |
| ENSMUSG00000053198 | 1.920505136 | 1.828220739 | 1.886723615 | 1.653677461 | 1.782377602 | 1.826074803 | 1.769001954 |
| ENSMUSG00000053199 | 0.671187741 | 0.79248125 | 0.830482024 | 0.580482024 | 0.489301084 | 0.602059991 | 0.74492186 |
| ENSMUSG00000053205 | 1.205378646 | 1.146240625 | 1.372963274 | 1.610735874 | 1.427019704 | 1.707570176 | 1.401059159 |
| ENSMUSG00000053208 | 0 | 0 | 0 | 0 | 0 | 0 | 0 |
| ENSMUSG00000053211 | 0 | 0 | 0 | 0 | 0 | 0 | 0 |
| ENSMUSG00000053214 | 0.289064826 | 0.5 | 0.25 | 0 | 0 | 0 | 0 |
| ENSMUSG00000053216 | 0 | 0.25 | 0.25 | 0.5 | 0.686820904 | 0.477121255 | 0.629532004 |
| ENSMUSG00000053217 | 0 | 0 | 0 | 0 | 0 | 0 | 0 |
| ENSMUSG00000053219 | 1.432083896 | 1.356566189 | 1.438721876 | 1.403677461 | 1.293076518 | 1.447158031 | 1.272006351 |
| ENSMUSG00000053226 | 1.227926429 | 1.388647213 | 1.380890489 | 1.226722649 | 1.254881871 | 1.447158031 | 1.379042812 |
| ENSMUSG00000053228 | 0 | 0 | 0 | 0 | 0 | 0 | 0 |
| ENSMUSG00000053251 | 0.289064826 | 0.25 | 0.5 | 0.396240625 | 0.244650542 | 0.301029996 | 0.520678036 |
| ENSMUSG00000053253 | 2.384185734 | 2.173371739 | 2.19697564 | 2.437467357 | 2.385720541 | 2.719331287 | 2.230532302 |

| | | | | | | | |
|---|---|---|---|---|---|---|---|
| ENSMUSG00000053263 | 0.747221736 | 0.951838731 | 0.79248125 | 0.701838731 | 0.489301084 | 0.84509804 | 0.629532004 |
| ENSMUSG00000053268 | 0 | 0 | 0 | 0 | 0 | 0 | 0 |
| ENSMUSG00000053279 | 0.578129653 | 0.580482024 | 0.646240625 | 0.580482024 | 0.387761935 | 0.301029996 | 0.224243824 |
| ENSMUSG00000053286 | 2.23973002 | 2.023189285 | 2.04248125 | 2.169870025 | 2.117361396 | 2.385606274 | 1.991191705 |
| ENSMUSG00000053287 | 0 | 0 | 0 | 0 | 0 | 0 | 0 |
| ENSMUSG00000053289 | 2.348397716 | 2.168989258 | 2.171625132 | 2.349151195 | 2.225838481 | 2.563481085 | 2.059095225 |
| ENSMUSG00000053291 | 0 | 0.25 | 0 | 0.25 | 0 | 0 | 0 |
| ENSMUSG00000053293 | 2.612833548 | 2.490362424 | 2.527457664 | 2.234159485 | 2.382429551 | 2.586587305 | 2.310815043 |
| ENSMUSG00000053297 | 0 | 0.25 | 0.25 | 0.25 | 0.244650542 | 0 | 0 |
| ENSMUSG00000053303 | 0 | 0 | 0 | 0 | 0 | 0 | 0 |
| ENSMUSG00000053310 | 0 | 0 | 0.396240625 | 0 | 0 | 0 | 0.448487648 |
| ENSMUSG00000053317 | 2.054824387 | 1.994319981 | 2.077084758 | 1.945339928 | 1.97205363 | 2.127104798 | 1.980721036 |
| ENSMUSG00000053318 | 0 | 0 | 0 | 0.5 | 0.632412477 | 0 | 0.520678036 |
| ENSMUSG00000053329 | 1.907802659 | 1.75835575 | 1.87694866 | 1.789967834 | 1.94314048 | 2.029383778 | 1.981429723 |
| ENSMUSG00000053332 | 2.721148722 | 2.451435968 | 2.453044326 | 2.475216702 | 2.44234479 | 2.761175813 | 2.278571072 |
| ENSMUSG00000053333 | 1.789260561 | 1.671625132 | 1.630890489 | 1.83272922 | 1.770879644 | 2 | 1.774416565 |
| ENSMUSG00000053334 | 1.671187741 | 1.701838731 | 1.698603967 | 1.321350555 | 1.499532413 | 1.491361694 | 1.437266334 |
| ENSMUSG00000053338 | 0.45815691 | 0 | 0.396240625 | 0 | 0.244650542 | 0 | 0.224243824 |
| ENSMUSG00000053347 | 1.87190676 | 1.732684334 | 1.726722649 | 1.862802778 | 1.718026092 | 1.939519253 | 1.616675393 |
| ENSMUSG00000053353 | 0 | 0 | 0 | 0 | 0 | 0 | 0 |
| ENSMUSG00000053358 | 0.578129653 | 0.701838731 | 0.701838731 | 0.701838731 | 0.812711509 | 0.301029996 | 0.775756176 |
| ENSMUSG00000053360 | 0 | 0 | 0 | 0 | 0 | 0 | 0 |
| ENSMUSG00000053367 | 0 | 0 | 0 | 0 | 0 | 0 | 0 |
| ENSMUSG00000053368 | 0 | 0 | 0 | 0 | 0 | 0 | 0.224243824 |
| ENSMUSG00000053375 | 0 | 0 | 0 | 0 | 0 | 0 | 0 |
| ENSMUSG00000053388 | 0 | 0 | 0 | 0.396240625 | 0.244650542 | 0 | 0 |
| ENSMUSG00000053389 | 0 | 0 | 0 | 0 | 0 | 0 | 0 |
| ENSMUSG00000053390 | 1.83262764 | 1.657339155 | 1.630890489 | 1.850219859 | 1.758994476 | 1.986771734 | 1.641897157 |
| ENSMUSG00000053391 | 0 | 0 | 0 | 0 | 0 | 0 | 0 |
| ENSMUSG00000053395 | 0.671187741 | 0.646240625 | 0.92510993 | 0.25 | 0.489301084 | 0.301029996 | 0.448487648 |
| ENSMUSG00000053398 | 2.761340634 | 2.629417347 | 2.628185116 | 2.452642909 | 2.599585429 | 2.80140371 | 2.498691813 |
| ENSMUSG00000053399 | 1.069666964 | 0.976722649 | 0.951838731 | 0 | 0 | 0 | 0 |
| ENSMUSG00000053411 | 1.862428343 | 1.76109853 | 1.821350555 | 1.766522298 | 1.861137485 | 2.012837225 | 1.834294096 |
| ENSMUSG00000053414 | 0 | 0.5 | 0.25 | 1.339388001 | 1.293076518 | 1.414973348 | 1.307989563 |

| | | | | | | | |
|---|---|---|---|---|---|---|---|
| ENSMUSG00000053420 | 0 | 0 | 0 | 0 | 0 | 0 | 0 |
| ENSMUSG00000053436 | 2.854509768 | 2.625459296 | 2.609697963 | 2.544666213 | 2.513204446 | 2.848189117 | 2.332093605 |
| ENSMUSG00000053441 | 0 | 0 | 0 | 0 | 0 | 0 | 0 |
| ENSMUSG00000053442 | 0 | 0 | 0.25 | 0 | 0 | 0.301029996 | 0.224243824 |
| ENSMUSG00000053453 | 2.010213549 | 1.864857905 | 1.886723615 | 2.167221246 | 2.052132976 | 2.318063335 | 1.929310074 |
| ENSMUSG00000053460 | 1.971473847 | 1.905512955 | 1.83272922 | 1.850219859 | 1.848234258 | 2.100370545 | 1.759302692 |
| ENSMUSG00000053461 | 0.45815691 | 0.646240625 | 0.396240625 | 0 | 0.387761935 | 0.301029996 | 0.224243824 |
| ENSMUSG00000053465 | 0 | 0 | 0 | 0 | 0 | 0 | 0 |
| ENSMUSG00000053469 | 1.771760131 | 1.642463902 | 1.653677461 | 1.451838731 | 1.373641807 | 1.681241237 | 1.290497981 |
| ENSMUSG00000053470 | 2.620687497 | 2.391989019 | 2.406427211 | 2.203445298 | 2.13194489 | 2.419955748 | 2.035419841 |
| ENSMUSG00000053472 | 0 | 0 | 0 | 0 | 0 | 0 | 0 |
| ENSMUSG00000053475 | 1.53838222 | 1.445339928 | 1.470660762 | 1.282320754 | 1.163285805 | 1.278753601 | 0.984950056 |
| ENSMUSG00000053477 | 2.563218389 | 2.38040011 | 2.385274154 | 2.341033664 | 2.283908311 | 2.582063363 | 2.143416814 |
| ENSMUSG00000053483 | 2.267855219 | 2.101285366 | 2.158248799 | 2.125956434 | 2.168627946 | 2.350248018 | 2.10131878 |
| ENSMUSG00000053490 | 0 | 0 | 0 | 0 | 0 | 0 | 0 |
| ENSMUSG00000053499 | 0 | 0 | 0 | 0 | 0 | 0 | 0 |
| ENSMUSG00000053508 | 0 | 0 | 0 | 0 | 0 | 0 | 0 |
| ENSMUSG00000053510 | 2.99653913 | 2.701838731 | 2.707232457 | 2.734894804 | 2.660945767 | 3.037426498 | 2.501257147 |
| ENSMUSG00000053519 | 0 | 0.25 | 0 | 0 | 0 | 0 | 0 |
| ENSMUSG00000053522 | 0 | 0 | 0 | 0 | 0 | 0 | 0 |
| ENSMUSG00000053528 | 0 | 0 | 0 | 0 | 0 | 0 | 0 |
| ENSMUSG00000053536 | 2.721759759 | 2.449009902 | 2.444519282 | 2.459406983 | 2.395811454 | 2.774516966 | 2.259663665 |
| ENSMUSG00000053541 | 0 | 0 | 0 | 0 | 0.244650542 | 0 | 0 |
| ENSMUSG00000053545 | 0 | 0 | 0 | 0 | 0 | 0 | 0 |
| ENSMUSG00000053550 | 0 | 0 | 0 | 0 | 0.244650542 | 0 | 0.224243824 |
| ENSMUSG00000053552 | 0.671187741 | 0.580482024 | 0.864857905 | 0.976722649 | 1.020174412 | 1 | 1.100339912 |
| ENSMUSG00000053553 | 1.862428343 | 1.843759858 | 1.912762923 | 1.668106335 | 1.749189509 | 1.875061263 | 1.800208341 |
| ENSMUSG00000053559 | 0.45815691 | 0.396240625 | 0.5 | 1.302363341 | 1.407936347 | 1.531478917 | 1.272006351 |
| ENSMUSG00000053560 | 1.516991255 | 1.589388001 | 1.714495249 | 1.25 | 1.519146132 | 1.431363764 | 1.720494 |
| ENSMUSG00000053565 | 2.511972754 | 2.358657057 | 2.399978211 | 2.300530956 | 2.338940611 | 2.607455023 | 2.22088434 |
| ENSMUSG00000053574 | 0.91631382 | 0.75 | 0.646240625 | 0.830482024 | 0.733951626 | 0.77815125 | 0.829800754 |
| ENSMUSG00000053580 | 2.46499366 | 2.257666784 | 2.259729747 | 1.979715809 | 1.919403394 | 2.127104798 | 1.843097781 |
| ENSMUSG00000053581 | 2.433305075 | 2.259729747 | 2.234159485 | 2.31611065 | 2.264824954 | 2.5774918 | 2.130728307 |
| ENSMUSG00000053588 | 0 | 0 | 0 | 0 | 0 | 0 | 0 |

| | | | | | | | |
|---|---|---|---|---|---|---|---|
| ENSMUSG00000053593 | 0 | 0 | 0 | 0 | 0 | 0 | 0 |
| ENSMUSG00000053600 | 1.777675583 | 1.67510993 | 1.711372513 | 1.71759118 | 1.66856048 | 1.806179974 | 1.540636858 |
| ENSMUSG00000053603 | 0.289064826 | 0 | 0.25 | 0.25 | 0 | 0 | 0 |
| ENSMUSG00000053604 | 1.789260561 | 1.755591953 | 1.777131114 | 1.75835575 | 1.855665188 | 2.004321374 | 1.743447983 |
| ENSMUSG00000053613 | 0.578129653 | 0 | 0 | 0.25 | 0 | 0.477121255 | 0 |
| ENSMUSG00000053615 | 0.289064826 | 0 | 0 | 0.25 | 0.632412477 | 0.477121255 | 0.579661877 |
| ENSMUSG00000053617 | 3.397683106 | 3.123276223 | 3.137746804 | 2.893648382 | 2.917139396 | 3.29578694 | 2.752322863 |
| ENSMUSG00000053624 | 0 | 0 | 0 | 0 | 0 | 0 | 0 |
| ENSMUSG00000053626 | 2.042557845 | 1.997171172 | 1.92510993 | 0.92510993 | 0.931471446 | 0.903089987 | 0.853775828 |
| ENSMUSG00000053640 | 0 | 0 | 0 | 0 | 0 | 0 | 0 |
| ENSMUSG00000053641 | 2.396352858 | 2.140560606 | 2.187048212 | 2.310197833 | 2.174635855 | 2.511883361 | 2.03780742 |
| ENSMUSG00000053644 | 2.111266113 | 1.990000483 | 2.034887838 | 2.192872367 | 2.203230928 | 2.440909082 | 2.104237736 |
| ENSMUSG00000053646 | 2.342375483 | 2.206637122 | 2.247171172 | 2.098079356 | 2.230318492 | 2.324282455 | 2.128938453 |
| ENSMUSG00000053647 | 0.578129653 | 0.830482024 | 0.75 | 0.701838731 | 0.877063019 | 0.84509804 | 0.876096088 |
| ENSMUSG00000053654 | 0 | 0 | 0 | 0 | 0 | 0 | 0 |
| ENSMUSG00000053656 | 0 | 0 | 0 | 0 | 0 | 0 | 0 |
| ENSMUSG00000053675 | 0.45815691 | 0.646240625 | 0.5 | 0 | 0 | 0.301029996 | 0.629532004 |
| ENSMUSG00000053678 | 0 | 0 | 0 | 0 | 0 | 0 | 0 |
| ENSMUSG00000053684 | 1.53838222 | 1.54248125 | 1.557204673 | 1.403677461 | 1.49445382 | 1.51851394 | 1.444790289 |
| ENSMUSG00000053687 | 0.289064826 | 0.396240625 | 0.396240625 | 0.396240625 | 0 | 0.301029996 | 0.224243824 |
| ENSMUSG00000053693 | 0.747221736 | 0.864857905 | 0.25 | 0.864857905 | 1.039257769 | 0.698970004 | 1.014380832 |
| ENSMUSG00000053695 | 0 | 0 | 0 | 0 | 0 | 0 | 0 |
| ENSMUSG00000053702 | 1.45815691 | 1.431980114 | 1.516522298 | 1.470660762 | 1.387761935 | 1.544068044 | 1.355418052 |
| ENSMUSG00000053706 | 0 | 0 | 0 | 0 | 0 | 0 | 0 |
| ENSMUSG00000053714 | 0.289064826 | 0 | 0.25 | 0 | 0.244650542 | 0.301029996 | 0 |
| ENSMUSG00000053716 | 1.876566388 | 1.850219859 | 1.953445298 | 1.657339155 | 1.836786029 | 1.963787827 | 1.805128823 |
| ENSMUSG00000053719 | 0 | 0 | 0 | 0 | 0 | 0 | 0 |
| ENSMUSG00000053720 | 0 | 0 | 0 | 0 | 0 | 0 | 0 |
| ENSMUSG00000053729 | 0 | 0 | 0 | 0 | 0 | 0 | 0 |
| ENSMUSG00000053730 | 1.707474304 | 1.668106335 | 1.678561379 | 1.388647213 | 1.504538967 | 1.380211242 | 1.48984372 |
| ENSMUSG00000053740 | 0 | 0 | 0.25 | 0.25 | 0.244650542 | 0 | 0 |
| ENSMUSG00000053742 | 0 | 0 | 0 | 0 | 0 | 0 | 0 |
| ENSMUSG00000053746 | 1.568542279 | 1.42510993 | 1.52186571 | 1.146240625 | 1.136121934 | 1.041392685 | 1.20139795 |
| ENSMUSG00000053747 | 0 | 0 | 0 | 0 | 0 | 0 | 0 |

| | | | | | | | |
|---|---|---|---|---|---|---|---|
| ENSMUSG00000053749 | 0.45815691 | 0 | 0.25 | 0.25 | 0.244650542 | 0 | 0 |
| ENSMUSG00000053754 | 2.710621733 | 2.486726569 | 2.458222504 | 2.740000483 | 2.694073682 | 3.054995862 | 2.515285987 |
| ENSMUSG00000053765 | 0 | 0 | 0 | 0 | 0 | 0 | 0 |
| ENSMUSG00000053768 | 2.525875121 | 2.329353153 | 2.368933358 | 2.36434522 | 2.288252441 | 2.640481437 | 2.170957925 |
| ENSMUSG00000053769 | 0.91631382 | 0.896240625 | 0.896240625 | 1.114857905 | 0.846351821 | 0.903089987 | 0.984950056 |
| ENSMUSG00000053773 | 0.289064826 | 0 | 0 | 0 | 0.244650542 | 0.602059991 | 0.224243824 |
| ENSMUSG00000053774 | 2.746646916 | 2.511780978 | 2.459800947 | 2.572504712 | 2.473055988 | 2.814247596 | 2.305919057 |
| ENSMUSG00000053783 | 0 | 0 | 0 | 0.396240625 | 0 | 0 | 0 |
| ENSMUSG00000053790 | 0 | 0 | 0 | 0 | 0 | 0 | 0 |
| ENSMUSG00000053797 | 0 | 0 | 0 | 0 | 0.387761935 | 0.301029996 | 0.355418052 |
| ENSMUSG00000053799 | 2.069666964 | 1.94697564 | 1.905512955 | 2.177701608 | 2.062514787 | 2.376576957 | 1.937521567 |
| ENSMUSG00000053801 | 2.183765914 | 2.105516192 | 2.174241882 | 2.03360658 | 2.209357611 | 2.28780173 | 2.111421942 |
| ENSMUSG00000053815 | 0 | 0 | 0 | 0 | 0 | 0 | 0 |
| ENSMUSG00000053819 | 2.730819472 | 2.523849256 | 2.507321807 | 2.367410454 | 2.32560195 | 2.649334859 | 2.142557543 |
| ENSMUSG00000053820 | 0 | 0 | 0 | 0 | 0 | 0 | 0 |
| ENSMUSG00000053821 | 0 | 0 | 0.25 | 0 | 0 | 0 | 0.355418052 |
| ENSMUSG00000053825 | 0.289064826 | 0.25 | 0 | 0.701838731 | 0.632412477 | 0.698970004 | 0.448487648 |
| ENSMUSG00000053830 | 0.289064826 | 0.25 | 0 | 0 | 0.244650542 | 0.301029996 | 0.224243824 |
| ENSMUSG00000053835 | 1.129344651 | 1.130890489 | 1.098079356 | 1.214495249 | 1.091002363 | 1.230448921 | 1.066254157 |
| ENSMUSG00000053838 | 2.69788693 | 2.480088214 | 2.468361028 | 2.51685859 | 2.48469123 | 2.860936621 | 2.344312053 |
| ENSMUSG00000053841 | 2.755187126 | 2.586351312 | 2.605778978 | 2.524178789 | 2.576508184 | 2.853698212 | 2.471230965 |
| ENSMUSG00000053846 | 1.129344651 | 1.226722649 | 1.146240625 | 1.201838731 | 1.234113756 | 1.041392685 | 1.23151414 |
| ENSMUSG00000053852 | 0 | 0 | 0 | 0 | 0 | 0 | 0 |
| ENSMUSG00000053856 | 0 | 0 | 0 | 0 | 0 | 0 | 0 |
| ENSMUSG00000053861 | 0 | 0 | 0 | 0 | 0 | 0 | 0 |
| ENSMUSG00000053862 | 0 | 0 | 0 | 0 | 0 | 0.301029996 | 0 |
| ENSMUSG00000053863 | 0 | 0 | 0 | 0 | 0 | 0 | 0 |
| ENSMUSG00000053868 | 0 | 0 | 0 | 0 | 0 | 0 | 0 |
| ENSMUSG00000053870 | 1.985980784 | 1.845926073 | 1.78743678 | 1.914552871 | 1.861137485 | 2.056904851 | 1.709198048 |
| ENSMUSG00000053873 | 0 | 0 | 0 | 0 | 0 | 0 | 0 |
| ENSMUSG00000053877 | 1.129344651 | 1.27186571 | 1.403677461 | 1.188721876 | 1.351342427 | 1 | 1.396774124 |
| ENSMUSG00000053886 | 0.811507563 | 0.701838731 | 0.5 | 0 | 0.244650542 | 0.301029996 | 0 |
| ENSMUSG00000053889 | 0.671187741 | 0.396240625 | 0.580482024 | 0.580482024 | 0.686820904 | 0 | 0.579661877 |
| ENSMUSG00000053896 | 0 | 0 | 0 | 0 | 0 | 0 | 0 |

| | | | | | | | |
|---|---|---|---|---|---|---|---|
| ENSMUSG00000053897 | 1.765759561 | 1.685366747 | 1.732684334 | 1.618933358 | 1.456697355 | 1.62324929 | 1.319146374 |
| ENSMUSG00000053898 | 2.097855556 | 1.979715809 | 2.001406137 | 1.850219859 | 1.868304353 | 2.045322979 | 1.803905701 |
| ENSMUSG00000053907 | 3.135049698 | 2.864473459 | 2.841717147 | 2.863189007 | 2.790143872 | 3.202215776 | 2.57876942 |
| ENSMUSG00000053909 | 0 | 0 | 0 | 0 | 0 | 0 | 0 |
| ENSMUSG00000053914 | 0.45815691 | 0.25 | 0.5 | 0 | 0.568060967 | 0.301029996 | 0.520678036 |
| ENSMUSG00000053916 | 0.289064826 | 0 | 0.25 | 0 | 0 | 0.301029996 | 0 |
| ENSMUSG00000053925 | 1.036286563 | 0.92510993 | 0.79248125 | 0.896240625 | 0.905314583 | 0.698970004 | 0.74492186 |
| ENSMUSG00000053929 | 2.545896521 | 2.399047439 | 2.406427211 | 2.365881093 | 2.446505421 | 2.678518379 | 2.329207203 |
| ENSMUSG00000053930 | 0.45815691 | 0.25 | 0 | 0.25 | 0 | 0 | 0 |
| ENSMUSG00000053931 | 3.421613914 | 3.105516192 | 3.06508288 | 2.998765117 | 2.894438858 | 3.409764104 | 2.731826698 |
| ENSMUSG00000053950 | 2.477243823 | 2.311388177 | 2.291226732 | 2.168106335 | 2.153144932 | 2.394451681 | 2.015657029 |
| ENSMUSG00000053957 | 0 | 0 | 0 | 0 | 0 | 0.301029996 | 0 |
| ENSMUSG00000053961 | 0 | 0 | 0 | 0 | 0 | 0 | 0 |
| ENSMUSG00000053963 | 0 | 0 | 0 | 0.396240625 | 0.489301084 | 0 | 0.355418052 |
| ENSMUSG00000053964 | 0.289064826 | 0.25 | 0.701838731 | 0.580482024 | 0.387761935 | 0 | 0.520678036 |
| ENSMUSG00000053965 | 2.048736215 | 1.918106335 | 1.841580554 | 1.614857905 | 1.572189156 | 1.681241237 | 1.496250175 |
| ENSMUSG00000053977 | 0 | 0.25 | 0 | 0.5 | 0.489301084 | 0.301029996 | 0 |
| ENSMUSG00000053979 | 0 | 0 | 0 | 0 | 0 | 0 | 0 |
| ENSMUSG00000053980 | 0 | 0 | 0 | 0 | 0 | 0 | 0 |
| ENSMUSG00000053985 | 0.811507563 | 0.864857905 | 0.92510993 | 0.92510993 | 0.632412477 | 0.77815125 | 0.876096088 |
| ENSMUSG00000054000 | 1.470606563 | 1.29248125 | 1.29248125 | 1.311981878 | 1.136121934 | 1.041392685 | 1.066254157 |
| ENSMUSG00000054003 | 0 | 0 | 0 | 0 | 0 | 0 | 0 |
| ENSMUSG00000054005 | 0 | 0 | 0 | 0 | 0 | 0 | 0.224243824 |
| ENSMUSG00000054006 | 0 | 0 | 0 | 0 | 0 | 0 | 0 |
| ENSMUSG00000054008 | 2.692026894 | 2.509043403 | 2.525493918 | 2.420748646 | 2.441646558 | 2.767155866 | 2.371499364 |
| ENSMUSG00000054013 | 0 | 0.5 | 0.396240625 | 0 | 0 | 0 | 0 |
| ENSMUSG00000054021 | 1.605636262 | 1.431980114 | 1.532320754 | 1.488549078 | 1.533172725 | 1.579783597 | 1.413583902 |
| ENSMUSG00000054027 | 2.545896521 | 2.3843046 | 2.370953944 | 2.323080408 | 2.315755952 | 2.593286067 | 2.14725554 |
| ENSMUSG00000054031 | 0 | 0 | 0 | 0 | 0 | 0 | 0 |
| ENSMUSG00000054033 | 0 | 0 | 0 | 0 | 0 | 0 | 0 |
| ENSMUSG00000054034 | 0 | 0 | 0 | 0 | 0 | 0 | 0 |
| ENSMUSG00000054036 | 0 | 0 | 0 | 0 | 0 | 0 | 0 |
| ENSMUSG00000054046 | 0 | 0 | 0 | 0 | 0 | 0 | 0 |
| ENSMUSG00000054051 | 2.160971586 | 1.982684334 | 2.01109853 | 2.117918804 | 2.093822267 | 2.292256071 | 1.951019875 |

| | | | | | | | |
|---|---|---|---|---|---|---|---|
| ENSMUSG00000054052 | 0 | 0.25 | 0.25 | 0 | 0 | 0 | 0 |
| ENSMUSG00000054054 | 0 | 0 | 0 | 0 | 0 | 0 | 0 |
| ENSMUSG00000054057 | 1.663535556 | 1.561981878 | 1.646240625 | 1.114857905 | 1.136121934 | 1.204119983 | 1.185218806 |
| ENSMUSG00000054061 | 0 | 0 | 0 | 0 | 0 | 0 | 0 |
| ENSMUSG00000054065 | 0.671187741 | 0.580482024 | 0.79248125 | 0.75 | 0.686820904 | 0.602059991 | 0.74492186 |
| ENSMUSG00000054069 | 0 | 0 | 0 | 0 | 0 | 0 | 0 |
| ENSMUSG00000054072 | 0 | 0 | 0 | 0.92510993 | 0.632412477 | 0.698970004 | 0.672731473 |
| ENSMUSG00000054074 | 1.686083339 | 1.618933358 | 1.698603967 | 1.372963274 | 1.234113756 | 1.431363764 | 1.131174228 |
| ENSMUSG00000054079 | 2.519973621 | 2.337734795 | 2.313757142 | 2.402294685 | 2.345425551 | 2.707570176 | 2.179128924 |
| ENSMUSG00000054083 | 0 | 0 | 0.25 | 0 | 0 | 0 | 0 |
| ENSMUSG00000054087 | 0 | 0 | 0 | 0 | 0 | 0 | 0 |
| ENSMUSG00000054091 | 2.273259602 | 2.164552871 | 2.211372513 | 2.185366747 | 2.207327177 | 2.403120521 | 2.105687397 |
| ENSMUSG00000054099 | 2.249317394 | 2.05479213 | 2.028435542 | 2.04248125 | 1.954436039 | 2.209515015 | 1.825244123 |
| ENSMUSG00000054102 | 0 | 0 | 0 | 0 | 0 | 0 | 0 |
| ENSMUSG00000054106 | 0 | 0 | 0 | 0 | 0 | 0 | 0 |
| ENSMUSG00000054115 | 2.688735382 | 2.456637122 | 2.445339928 | 2.498234584 | 2.443389555 | 2.800717078 | 2.30331215 |
| ENSMUSG00000054116 | 0 | 0 | 0 | 0 | 0 | 0 | 0 |
| ENSMUSG00000054117 | 0 | 0 | 0 | 0 | 0 | 0 | 0 |
| ENSMUSG00000054128 | 0 | 0 | 0 | 0.25 | 0.387761935 | 0 | 0 |
| ENSMUSG00000054134 | 0 | 0 | 0 | 0 | 0 | 0 | 0 |
| ENSMUSG00000054135 | 0.578129653 | 0.701838731 | 0.75 | 0.580482024 | 0 | 0.77815125 | 0.448487648 |
| ENSMUSG00000054136 | 0.45815691 | 0.580482024 | 0.79248125 | 0 | 0 | 0 | 0.520678036 |
| ENSMUSG00000054141 | 0 | 0 | 0 | 0 | 0 | 0 | 0 |
| ENSMUSG00000054142 | 0 | 0 | 0 | 0 | 0 | 0 | 0 |
| ENSMUSG00000054146 | 0 | 0 | 0 | 0 | 0 | 0 | 0 |
| ENSMUSG00000054150 | 2.156259305 | 2.069031101 | 2.135757955 | 1.951838731 | 2.001219193 | 2.164352856 | 1.96172822 |
| ENSMUSG00000054156 | 0 | 0 | 0 | 0 | 0 | 0 | 0 |
| ENSMUSG00000054160 | 0 | 0 | 0 | 0 | 0 | 0 | 0 |
| ENSMUSG00000054161 | 0.45815691 | 0.5 | 0.580482024 | 0.25 | 0.387761935 | 0.301029996 | 0.775756176 |
| ENSMUSG00000054162 | 0 | 0 | 0 | 0 | 0 | 0 | 0 |
| ENSMUSG00000054169 | 0 | 0 | 0 | 0 | 0 | 0 | 0 |
| ENSMUSG00000054178 | 1.960252568 | 1.852347734 | 1.83272922 | 1.868933358 | 1.749189509 | 1.838849091 | 1.647906904 |
| ENSMUSG00000054181 | 0.289064826 | 0 | 0.396240625 | 0.396240625 | 0.489301084 | 0.301029996 | 0.448487648 |
| ENSMUSG00000054191 | 0 | 0 | 0.396240625 | 0 | 0 | 0 | 0.355418052 |

| | | | | | | | |
|---|---|---|---|---|---|---|---|
| ENSMUSG00000054196 | 0 | 0.25 | 0 | 2.225216702 | 2.19135987 | 2.511883361 | 2.06842541 |
| ENSMUSG00000054199 | 2.297593393 | 2.136723615 | 2.142463902 | 2.161864607 | 2.19135987 | 2.383815366 | 2.01501956 |
| ENSMUSG00000054200 | 0 | 0 | 0 | 0 | 0 | 0 | 0 |
| ENSMUSG00000054203 | 0.289064826 | 0.646240625 | 0.396240625 | 0.5 | 0.632412477 | 0 | 0.355418052 |
| ENSMUSG00000054204 | 0 | 0.25 | 0 | 0 | 0 | 0 | 0 |
| ENSMUSG00000054206 | 0.747221736 | 0.830482024 | 0.79248125 | 0.580482024 | 0.489301084 | 0.602059991 | 0.74492186 |
| ENSMUSG00000054215 | 0.867194479 | 0.951838731 | 0.75 | 0 | 0.244650542 | 0.477121255 | 0.224243824 |
| ENSMUSG00000054226 | 2.251208701 | 2.061981878 | 2.072504712 | 2.170748646 | 2.063536368 | 2.428134794 | 1.948678378 |
| ENSMUSG00000054236 | 0 | 0 | 0 | 0 | 0 | 0 | 0 |
| ENSMUSG00000054237 | 1.940852214 | 1.75835575 | 1.744319981 | 1.83272922 | 1.763796674 | 1.903089987 | 1.625302983 |
| ENSMUSG00000054247 | 0 | 0 | 0.25 | 0.25 | 0.244650542 | 0.301029996 | 0 |
| ENSMUSG00000054252 | 1.129344651 | 1.26109853 | 1.282320754 | 1.602347734 | 1.632412477 | 1.799340549 | 1.68886578 |
| ENSMUSG00000054256 | 1.389637216 | 1.330482024 | 1.380890489 | 1.339388001 | 1.49445382 | 1.380211242 | 1.396774124 |
| ENSMUSG00000054258 | 0.289064826 | 0.25 | 0.5 | 0 | 0 | 0 | 0 |
| ENSMUSG00000054263 | 2.247417471 | 2.046218836 | 2.05479213 | 2.200224975 | 2.135290471 | 2.418301291 | 2.043115906 |
| ENSMUSG00000054266 | 0 | 0.396240625 | 0.25 | 0.25 | 0 | 0 | 0 |
| ENSMUSG00000054272 | 0 | 0 | 0 | 0 | 0 | 0 | 0 |
| ENSMUSG00000054277 | 2.536482297 | 2.368426437 | 2.367410454 | 2.151832578 | 2.175379703 | 2.451786436 | 2.043700396 |
| ENSMUSG00000054280 | 2.559630991 | 2.370953944 | 2.351816941 | 2.382357639 | 2.369807605 | 2.650307523 | 2.191935631 |
| ENSMUSG00000054293 | 1 | 0.976722649 | 1.080482024 | 1.476722649 | 1.343584837 | 1.62324929 | 1.193409508 |
| ENSMUSG00000054302 | 2.124258828 | 2.004202072 | 2.001406137 | 2.03360658 | 1.94314048 | 2.178976947 | 1.825244123 |
| ENSMUSG00000054304 | 1 | 0.976722649 | 0.75 | 0.646240625 | 0.387761935 | 0.477121255 | 0.520678036 |
| ENSMUSG00000054309 | 2.392336464 | 2.218361028 | 2.234159485 | 2.280383379 | 2.222603297 | 2.537819095 | 2.089878917 |
| ENSMUSG00000054310 | 0 | 0 | 0 | 0 | 0 | 0 | 0 |
| ENSMUSG00000054312 | 2.100572389 | 1.997171172 | 1.961372513 | 1.984159485 | 2.022473809 | 2.292256071 | 1.894206246 |
| ENSMUSG00000054320 | 0 | 0 | 0.25 | 0 | 0 | 0 | 0 |
| ENSMUSG00000054321 | 1.655740331 | 1.54248125 | 1.653677461 | 1.75 | 1.621875691 | 1.72427587 | 1.532233387 |
| ENSMUSG00000054325 | 0 | 0 | 0 | 0 | 0 | 0 | 0 |
| ENSMUSG00000054342 | 0.45815691 | 0.646240625 | 0.5 | 0 | 0.489301084 | 0 | 0.224243824 |
| ENSMUSG00000054351 | 0 | 0 | 0 | 0 | 0 | 0 | 0 |
| ENSMUSG00000054360 | 0 | 0 | 0 | 0 | 0 | 0 | 0 |
| ENSMUSG00000054362 | 0 | 0 | 0 | 0 | 0 | 0 | 0 |
| ENSMUSG00000054364 | 2.477243823 | 2.254897682 | 2.315523711 | 2.246460484 | 2.257835496 | 2.499687083 | 2.178744473 |
| ENSMUSG00000054379 | 0 | 0 | 0 | 0 | 0 | 0 | 0.224243824 |

| | | | | | | | |
|---|---|---|---|---|---|---|---|
| ENSMUSG00000054381 | 1.269664473 | 1.445339928 | 1.388647213 | 1.098079356 | 1.27449559 | 1.230448921 | 1.302263476 |
| ENSMUSG00000054383 | 0.867194479 | 0.951838731 | 1.04248125 | 0.92510993 | 0.686820904 | 1.041392685 | 0.896975297 |
| ENSMUSG00000054385 | 0.45815691 | 0.396240625 | 0.580482024 | 0 | 0.489301084 | 0.301029996 | 0 |
| ENSMUSG00000054387 | 2.623788124 | 2.383818844 | 2.348615674 | 2.530059469 | 2.460030299 | 2.804820679 | 2.267652107 |
| ENSMUSG00000054391 | 0 | 0 | 0 | 0 | 0 | 0 | 0 |
| ENSMUSG00000054400 | 1.54867984 | 1.464495249 | 1.451838731 | 1.5 | 1.439191811 | 1.73239376 | 1.392431569 |
| ENSMUSG00000054404 | 0.289064826 | 0 | 0.25 | 0.580482024 | 0.568060967 | 0.477121255 | 0.672731473 |
| ENSMUSG00000054405 | 2.834495649 | 2.570482507 | 2.58993739 | 2.584405475 | 2.544319634 | 2.944482672 | 2.395004664 |
| ENSMUSG00000054406 | 0 | 0 | 0 | 0 | 0 | 0 | 0 |
| ENSMUSG00000054408 | 3.058380605 | 2.811981878 | 2.762462137 | 2.854990044 | 2.764077132 | 3.190891717 | 2.553692546 |
| ENSMUSG00000054409 | 1.445324132 | 1.458222504 | 1.42510993 | 1.27186571 | 1.074582839 | 0.602059991 | 1.078019652 |
| ENSMUSG00000054412 | 0.45815691 | 0 | 0.25 | 0.25 | 0 | 0 | 0 |
| ENSMUSG00000054414 | 2.494443473 | 2.281676118 | 2.282320754 | 2.451032755 | 2.386538516 | 2.711807229 | 2.252699564 |
| ENSMUSG00000054417 | 0 | 0 | 0 | 0 | 0 | 0 | 0 |
| ENSMUSG00000054418 | 0 | 0 | 0 | 0.5 | 0.905314583 | 0.602059991 | 0.629532004 |
| ENSMUSG00000054422 | 0 | 0 | 0 | 0 | 0 | 0 | 0 |
| ENSMUSG00000054423 | 0 | 0 | 0 | 0 | 0 | 0 | 0 |
| ENSMUSG00000054426 | 1.100572389 | 0.701838731 | 0.976722649 | 1 | 0.77552387 | 0.77815125 | 0.896975297 |
| ENSMUSG00000054428 | 2.434522689 | 2.262462137 | 2.309003548 | 2.132845365 | 2.154724156 | 2.408239965 | 2.148525059 |
| ENSMUSG00000054431 | 0 | 0 | 0 | 0 | 0 | 0 | 0 |
| ENSMUSG00000054434 | 1.721148722 | 1.614857905 | 1.516522298 | 1.527131114 | 1.563883923 | 1.643452676 | 1.562034107 |
| ENSMUSG00000054435 | 0 | 0 | 0 | 0 | 0 | 0.477121255 | 0 |
| ENSMUSG00000054446 | 0 | 0 | 0 | 0 | 0 | 0 | 0 |
| ENSMUSG00000054450 | 0 | 0 | 0 | 0 | 0 | 0 | 0 |
| ENSMUSG00000054452 | 2.711874084 | 2.55205859 | 2.621708755 | 2.480831871 | 2.524713379 | 2.894316063 | 2.519606791 |
| ENSMUSG00000054453 | 1.971473847 | 1.845926073 | 1.830482024 | 0.896240625 | 0.77552387 | 0.602059991 | 0.829800754 |
| ENSMUSG00000054455 | 2.422183552 | 2.26584877 | 2.263820609 | 2.338286706 | 2.313753359 | 2.617000341 | 2.231187192 |
| ENSMUSG00000054457 | 0 | 0 | 0.25 | 0 | 0 | 0 | 0 |
| ENSMUSG00000054459 | 0.578129653 | 0.701838731 | 0.830482024 | 0.5 | 0.489301084 | 0.301029996 | 0.520678036 |
| ENSMUSG00000054469 | 2.438154344 | 2.228222334 | 2.212155735 | 2.289336734 | 2.230953877 | 2.565847819 | 2.075383 |
| ENSMUSG00000054474 | 1 | 1.098079356 | 1.130890489 | 1.130890489 | 1.319233381 | 1.342422681 | 1.265599895 |
| ENSMUSG00000054477 | 0.289064826 | 0 | 0 | 0.396240625 | 0.244650542 | 0 | 0 |
| ENSMUSG00000054484 | 2.100572389 | 1.921625132 | 1.921625132 | 2.194519282 | 2.094757251 | 2.414973348 | 1.921737195 |
| ENSMUSG00000054488 | 0 | 0 | 0 | 0 | 0 | 0 | 0 |

| | | | | | | | |
|---|---|---|---|---|---|---|---|
| ENSMUSG00000054493 | 0 | 0 | 0 | 0 | 0 | 0 | 0 |
| ENSMUSG00000054497 | 0 | 0 | 0 | 0 | 0 | 0 | 0 |
| ENSMUSG00000054498 | 0 | 0 | 0 | 0 | 0 | 0 | 0 |
| ENSMUSG00000054499 | 0.811507563 | 1.02186571 | 1.04248125 | 0.864857905 | 1.091002363 | 1.041392685 | 1.100339912 |
| ENSMUSG00000054509 | 1.89911417 | 1.766522298 | 1.76109853 | 2.132845365 | 2.020174412 | 2.328379603 | 1.939139147 |
| ENSMUSG00000054510 | 0.289064826 | 0 | 0 | 0 | 0 | 0.301029996 | 0 |
| ENSMUSG00000054514 | 0.289064826 | 0.25 | 0.25 | 0.701838731 | 0.244650542 | 0.301029996 | 0.448487648 |
| ENSMUSG00000054517 | 1.45815691 | 1.418106335 | 1.396240625 | 1.146240625 | 1.163285805 | 1.278753601 | 1.121219121 |
| ENSMUSG00000054519 | 1.827447047 | 1.681980114 | 1.692046081 | 1.928561379 | 1.836786029 | 2.10720997 | 1.749303626 |
| ENSMUSG00000054520 | 1.940852214 | 1.892463902 | 1.950224975 | 1.714495249 | 1.770879644 | 1.892094603 | 1.704233519 |
| ENSMUSG00000054523 | 0 | 0 | 0 | 0 | 0 | 0 | 0 |
| ENSMUSG00000054526 | 0 | 0 | 0 | 0 | 0 | 0 | 0 |
| ENSMUSG00000054537 | 0 | 0 | 0 | 0 | 0 | 0 | 0 |
| ENSMUSG00000054545 | 0 | 0.25 | 0 | 0 | 0 | 0 | 0.224243824 |
| ENSMUSG00000054555 | 2.302628051 | 2.167221246 | 2.113831805 | 1.622963274 | 1.607318736 | 1.612783857 | 1.405288179 |
| ENSMUSG00000054556 | 0.747221736 | 0.25 | 0.580482024 | 0.396240625 | 0.244650542 | 0.477121255 | 0 |
| ENSMUSG00000054568 | 0.289064826 | 0.701838731 | 0.580482024 | 0.25 | 0.387761935 | 0.477121255 | 0.448487648 |
| ENSMUSG00000054580 | 2.233869984 | 2.094844592 | 2.109697963 | 1.02186571 | 1.091002363 | 1.146128036 | 0.896975297 |
| ENSMUSG00000054582 | 0.289064826 | 0.5 | 0.25 | 0.646240625 | 0.489301084 | 0 | 0.448487648 |
| ENSMUSG00000054589 | 0.747221736 | 0.75 | 0.5 | 0.5 | 0.568060967 | 0 | 0.629532004 |
| ENSMUSG00000054594 | 0 | 0.396240625 | 0.25 | 0.25 | 0.244650542 | 0 | 0.224243824 |
| ENSMUSG00000054598 | 0 | 0.396240625 | 0.5 | 0.25 | 0.244650542 | 0 | 0.224243824 |
| ENSMUSG00000054604 | 2.88283767 | 2.631869252 | 2.623213655 | 2.68473417 | 2.595767222 | 2.968482949 | 2.388834696 |
| ENSMUSG00000054611 | 3.061095669 | 2.800684011 | 2.785526764 | 2.87246199 | 2.819844621 | 3.243038049 | 2.622387534 |
| ENSMUSG00000054612 | 1.205378646 | 1.302363341 | 1.114857905 | 0.75 | 0.955822902 | 0.77815125 | 1.041356071 |
| ENSMUSG00000054618 | 0 | 0 | 0.25 | 0 | 0 | 0 | 0.224243824 |
| ENSMUSG00000054619 | 1.671187741 | 1.561981878 | 1.575945187 | 1.42510993 | 1.484072009 | 1.643452676 | 1.379042812 |
| ENSMUSG00000054622 | 0.289064826 | 0 | 0.25 | 0 | 0.244650542 | 0 | 0 |
| ENSMUSG00000054626 | 0 | 0 | 0 | 0 | 0 | 0 | 0 |
| ENSMUSG00000054630 | 0 | 0 | 0 | 0 | 0 | 0 | 0 |
| ENSMUSG00000054640 | 1.482695304 | 1.380890489 | 1.356566189 | 0 | 0 | 0 | 0.224243824 |
| ENSMUSG00000054641 | 0 | 0 | 0 | 0 | 0 | 0 | 0 |
| ENSMUSG00000054648 | 1.928763473 | 1.858657057 | 1.841580554 | 1.94697564 | 1.890639929 | 2.100370545 | 1.750751125 |
| ENSMUSG00000054659 | 0 | 0 | 0 | 0.25 | 0.244650542 | 0 | 0 |

| | | | | | | | |
|---|---|---|---|---|---|---|---|
| ENSMUSG00000054662 | 0 | 0 | 0.25 | 0 | 0 | 0 | 0 |
| ENSMUSG00000054666 | 0 | 0 | 0 | 0 | 0 | 0 | 0 |
| ENSMUSG00000054667 | 0 | 0.396240625 | 0.25 | 0.396240625 | 0.244650542 | 0.301029996 | 0.520678036 |
| ENSMUSG00000054672 | 0 | 0 | 0 | 0 | 0 | 0 | 0 |
| ENSMUSG00000054675 | 2.647796617 | 2.499647607 | 2.514497931 | 1.53743678 | 1.701347897 | 1.681241237 | 1.712465909 |
| ENSMUSG00000054676 | 1.505869732 | 1.488549078 | 1.622963274 | 1.5 | 1.618292349 | 1.643452676 | 1.493062805 |
| ENSMUSG00000054679 | 0 | 0 | 0 | 0.951838731 | 0.931471446 | 0.903089987 | 0.579661877 |
| ENSMUSG00000054690 | 0 | 0 | 0 | 0 | 0 | 0 | 0 |
| ENSMUSG00000054693 | 2.949075151 | 2.71585299 | 2.701838731 | 2.773684376 | 2.682431937 | 3.131297797 | 2.48525267 |
| ENSMUSG00000054702 | 1.700465191 | 1.598079356 | 1.552363341 | 1.903677461 | 1.758994476 | 1.973127854 | 1.653807046 |
| ENSMUSG00000054708 | 1.404271433 | 1.29248125 | 1.451838731 | 1.356566189 | 1.439191811 | 1.361727836 | 1.340368108 |
| ENSMUSG00000054715 | 0.867194479 | 0.701838731 | 0.75 | 1.561981878 | 1.514343934 | 1.633468456 | 1.444790289 |
| ENSMUSG00000054716 | 0.747221736 | 0.79248125 | 0.951838731 | 0.864857905 | 0.905314583 | 0.77815125 | 1.078019652 |
| ENSMUSG00000054717 | 2.845984872 | 2.613574823 | 2.63015467 | 2.688930538 | 2.648915262 | 3.025715384 | 2.509249051 |
| ENSMUSG00000054720 | 2.724194991 | 2.486360959 | 2.466819685 | 2.31728167 | 2.278401864 | 2.57634135 | 2.100339912 |
| ENSMUSG00000054723 | 1.482695304 | 1.302363341 | 1.464495249 | 1.54248125 | 1.467903253 | 1.491361694 | 1.433437704 |
| ENSMUSG00000054727 | 0 | 0 | 0 | 0 | 0 | 0 | 0 |
| ENSMUSG00000054728 | 0.91631382 | 0.396240625 | 0.5 | 0.830482024 | 0.733951626 | 0.477121255 | 0.74492186 |
| ENSMUSG00000054733 | 1.527823874 | 1.488549078 | 1.557204673 | 1.593759858 | 1.542223377 | 1.716003344 | 1.499406449 |
| ENSMUSG00000054737 | 0 | 0 | 0 | 1.802363341 | 1.66856048 | 1.880813592 | 1.517723536 |
| ENSMUSG00000054745 | 0 | 0 | 0 | 0 | 0 | 0 | 0 |
| ENSMUSG00000054746 | 0 | 0 | 0 | 0 | 0 | 0 | 0 |
| ENSMUSG00000054752 | 2.016886209 | 1.890560606 | 1.87892496 | 2.006976499 | 1.961316494 | 2.235528447 | 1.771720588 |
| ENSMUSG00000054753 | 0 | 0 | 0.25 | 0.396240625 | 0.244650542 | 0.602059991 | 0.448487648 |
| ENSMUSG00000054757 | 0 | 0 | 0.25 | 0.25 | 0.244650542 | 0.301029996 | 0 |
| ENSMUSG00000054758 | 0 | 0 | 0 | 0 | 0 | 0 | 0 |
| ENSMUSG00000054759 | 0 | 0 | 0 | 0 | 0 | 0 | 0 |
| ENSMUSG00000054763 | 0 | 0.580482024 | 0.396240625 | 0.5 | 0.387761935 | 0.301029996 | 0.355418052 |
| ENSMUSG00000054764 | 0.289064826 | 0 | 0 | 0 | 0 | 0 | 0 |
| ENSMUSG00000054766 | 3.122393951 | 2.848079356 | 2.832449085 | 2.821783797 | 2.763094553 | 3.166430114 | 2.561445361 |
| ENSMUSG00000054770 | 1.890240806 | 1.668106335 | 1.720660762 | 1.774508021 | 1.768534412 | 1.913813852 | 1.655750087 |
| ENSMUSG00000054779 | 0.289064826 | 0.396240625 | 0 | 0 | 0.244650542 | 0 | 0 |
| ENSMUSG00000054792 | 2.042557845 | 1.935366747 | 1.966046536 | 2.001406137 | 2.047893212 | 2.217483944 | 1.993245779 |
| ENSMUSG00000054793 | 1.700465191 | 1.705044741 | 1.782320754 | 1.476722649 | 1.665423072 | 1.698970004 | 1.690609803 |

| | | | | | | | |
|---|---|---|---|---|---|---|---|
| ENSMUSG00000054808 | 3.390129289 | 3.139904021 | 3.160625757 | 2.814935816 | 2.903902568 | 3.261738547 | 2.75968608 |
| ENSMUSG00000054814 | 2.375872521 | 2.17510993 | 2.187886015 | 2.170748646 | 2.091002363 | 2.434568904 | 1.949460761 |
| ENSMUSG00000054822 | 0 | 0.396240625 | 0.25 | 0 | 0 | 0 | 0.224243824 |
| ENSMUSG00000054823 | 2.823638505 | 2.591580554 | 2.594031347 | 2.62019755 | 2.562149491 | 2.940018155 | 2.391037533 |
| ENSMUSG00000054827 | 0 | 0 | 0 | 0 | 0 | 0 | 0 |
| ENSMUSG00000054836 | 1.920505136 | 1.837182039 | 1.811981878 | 1.940387808 | 1.92093466 | 2.133538908 | 1.904245613 |
| ENSMUSG00000054843 | 2.545896521 | 2.385757955 | 2.397178659 | 2.451032755 | 2.375332618 | 2.697229343 | 2.235734673 |
| ENSMUSG00000054850 | 0.960252568 | 0.951838731 | 1.02186571 | 0.701838731 | 0.568060967 | 0.698970004 | 0.803905701 |
| ENSMUSG00000054855 | 2.235832493 | 2.122963274 | 2.105516192 | 2.328220739 | 2.353143227 | 2.660865478 | 2.239900853 |
| ENSMUSG00000054863 | 1.389637216 | 1.410964047 | 1.527131114 | 1.160964047 | 1.091002363 | 0.954242509 | 1.121219121 |
| ENSMUSG00000054871 | 0.811507563 | 0.976722649 | 1.080482024 | 0.396240625 | 0.77552387 | 0.77815125 | 0.829800754 |
| ENSMUSG00000054874 | 2.587501561 | 2.483053688 | 2.591033664 | 2.149047439 | 2.39421617 | 2.439332694 | 2.397361799 |
| ENSMUSG00000054885 | 0 | 0 | 0 | 0 | 0 | 0 | 0 |
| ENSMUSG00000054889 | 1.894701088 | 1.886723615 | 1.825945187 | 2.29745614 | 2.304598364 | 2.549003262 | 2.126687118 |
| ENSMUSG00000054890 | 0 | 0 | 0 | 0 | 0 | 0 | 0 |
| ENSMUSG00000054892 | 0.578129653 | 0 | 0.25 | 0.25 | 0.489301084 | 0.477121255 | 0.224243824 |
| ENSMUSG00000054893 | 1.156259305 | 1.26109853 | 1.321350555 | 1.52186571 | 1.394615667 | 1.556302501 | 1.324583736 |
| ENSMUSG00000054894 | 1.671187741 | 1.53743678 | 1.566696635 | 1.53743678 | 1.47337555 | 1.633468456 | 1.21680636 |
| ENSMUSG00000054901 | 0 | 0.5 | 0.25 | 0 | 0.244650542 | 0.301029996 | 0.355418052 |
| ENSMUSG00000054905 | 0 | 0 | 0 | 0 | 0 | 0 | 0 |
| ENSMUSG00000054909 | 0 | 0 | 0 | 0 | 0 | 0 | 0 |
| ENSMUSG00000054910 | 0.289064826 | 0.396240625 | 0.5 | 0.580482024 | 0.244650542 | 0.698970004 | 0.355418052 |
| ENSMUSG00000054920 | 2.37447073 | 2.197790722 | 2.240000483 | 2.158248799 | 2.136951444 | 2.403120521 | 2.02383181 |
| ENSMUSG00000054931 | 0 | 0 | 0 | 0 | 0 | 0.301029996 | 0 |
| ENSMUSG00000054932 | 0.289064826 | 0.396240625 | 0.25 | 0.25 | 0 | 0.477121255 | 0.74492186 |
| ENSMUSG00000054934 | 0.747221736 | 0.646240625 | 0.701838731 | 1 | 0.955822902 | 1.204119983 | 1.078019652 |
| ENSMUSG00000054938 | 0 | 0.25 | 0 | 0 | 0 | 0 | 0 |
| ENSMUSG00000054939 | 1.342375483 | 1.201838731 | 1.372963274 | 1.130890489 | 1.254881871 | 1.342422681 | 1.176815335 |
| ENSMUSG00000054940 | 0 | 0 | 0 | 0 | 0 | 0 | 0 |
| ENSMUSG00000054942 | 2.292497211 | 2.146240625 | 2.142463902 | 2.214495249 | 2.12771782 | 2.371067862 | 1.983546516 |
| ENSMUSG00000054944 | 0 | 0.25 | 0 | 0 | 0 | 0 | 0 |
| ENSMUSG00000054945 | 1.205378646 | 1.061981878 | 0.896240625 | 1.061981878 | 0.846351821 | 0.954242509 | 0.935079929 |
| ENSMUSG00000054951 | 1.53838222 | 1.476722649 | 1.532320754 | 1.451838731 | 1.514343934 | 1.633468456 | 1.374453863 |
| ENSMUSG00000054957 | 0 | 0 | 0 | 0 | 0 | 0 | 0 |

| | | | | | | | |
|---|---|---|---|---|---|---|---|
| ENSMUSG00000054958 | 0 | 0 | 0 | 0 | 0 | 0 | 0 |
| ENSMUSG00000054966 | 0 | 0 | 0.25 | 0 | 0 | 0 | 0 |
| ENSMUSG00000054967 | 1.404271433 | 1.552363341 | 1.5 | 1.282320754 | 1.439191811 | 1.447158031 | 1.429563222 |
| ENSMUSG00000054976 | 0 | 0 | 0 | 0 | 0 | 0 | 0 |
| ENSMUSG00000054978 | 0 | 0 | 0.25 | 0 | 0.244650542 | 0 | 0.224243824 |
| ENSMUSG00000054986 | 0 | 0 | 0 | 0 | 0 | 0 | 0 |
| ENSMUSG00000054988 | 0 | 0 | 0 | 0 | 0 | 0 | 0 |
| ENSMUSG00000054994 | 0 | 0 | 0 | 0 | 0 | 0 | 0 |
| ENSMUSG00000054999 | 1.358731791 | 1.339388001 | 1.505591953 | 1.302363341 | 1.439191811 | 1.255272505 | 1.441050184 |
| ENSMUSG00000055003 | 0 | 0.5 | 0.396240625 | 0.396240625 | 0.489301084 | 0.301029996 | 0.520678036 |
| ENSMUSG00000055010 | 0 | 0 | 0 | 0 | 0 | 0 | 0 |
| ENSMUSG00000055013 | 2.377269616 | 2.248588359 | 2.257666784 | 2.229715809 | 2.208682096 | 2.396199347 | 2.114720744 |
| ENSMUSG00000055015 | 0 | 0 | 0 | 0 | 0 | 0 | 0 |
| ENSMUSG00000055022 | 0.578129653 | 0.646240625 | 0.5 | 0.701838731 | 0.489301084 | 0.477121255 | 0.355418052 |
| ENSMUSG00000055024 | 2.95713908 | 2.752107156 | 2.731017546 | 2.738003291 | 2.728877458 | 3.095866453 | 2.561209563 |
| ENSMUSG00000055026 | 0 | 0.25 | 0 | 0 | 0 | 0 | 0 |
| ENSMUSG00000055027 | 0.45815691 | 0.5 | 0 | 0.5 | 0.489301084 | 0.602059991 | 0.579661877 |
| ENSMUSG00000055030 | 0.45815691 | 0 | 0 | 0 | 0 | 0 | 0 |
| ENSMUSG00000055033 | 0 | 0 | 0 | 0 | 0 | 0 | 0 |
| ENSMUSG00000055041 | 1.894701088 | 1.850219859 | 1.880890489 | 1.657339155 | 1.78687392 | 1.826074803 | 1.783679399 |
| ENSMUSG00000055044 | 2.731416512 | 2.523189285 | 2.547147211 | 2.432405186 | 2.47337555 | 2.794488047 | 2.354078442 |
| ENSMUSG00000055045 | 0 | 0 | 0 | 0 | 0 | 0 | 0 |
| ENSMUSG00000055048 | 0.289064826 | 0.396240625 | 0.25 | 0 | 0.387761935 | 0.301029996 | 0.224243824 |
| ENSMUSG00000055053 | 2.751223449 | 2.61817271 | 2.652986735 | 2.39151351 | 2.519146132 | 2.752048448 | 2.482406255 |
| ENSMUSG00000055061 | 0 | 0 | 0 | 0.25 | 0 | 0 | 0 |
| ENSMUSG00000055065 | 3.379167233 | 3.056301859 | 3.057655233 | 3.164664448 | 3.111229201 | 3.571242851 | 2.884801065 |
| ENSMUSG00000055067 | 1.783508299 | 1.678561379 | 1.71759118 | 1.769203899 | 1.802174723 | 1.924279286 | 1.692344475 |
| ENSMUSG00000055069 | 0 | 0 | 0 | 0 | 0.632412477 | 0.477121255 | 0.355418052 |
| ENSMUSG00000055072 | 0 | 0.396240625 | 0.5 | 0.25 | 0.387761935 | 0.301029996 | 0.224243824 |
| ENSMUSG00000055078 | 0 | 0 | 0 | 0 | 0 | 0 | 0 |
| ENSMUSG00000055088 | 0 | 0 | 0 | 0 | 0 | 0 | 0 |
| ENSMUSG00000055093 | 0.747221736 | 0.5 | 0 | 0.701838731 | 0.686820904 | 0.477121255 | 0.579661877 |
| ENSMUSG00000055095 | 0 | 0 | 0 | 0 | 0 | 0 | 0 |
| ENSMUSG00000055099 | 0 | 0 | 0 | 0 | 0 | 0 | 0 |

| | | | | | | | |
|---|---|---|---|---|---|---|---|
| ENSMUSG00000055102 | 0.289064826 | 0 | 0 | 0.79248125 | 0.812711509 | 0.954242509 | 0.803905701 |
| ENSMUSG00000055108 | 0 | 0 | 0 | 0 | 0 | 0 | 0.224243824 |
| ENSMUSG00000055109 | 0 | 0 | 0 | 0 | 0 | 0 | 0 |
| ENSMUSG00000055110 | 0 | 0 | 0 | 0 | 0 | 0 | 0 |
| ENSMUSG00000055114 | 0 | 0 | 0 | 0 | 0.387761935 | 0 | 0 |
| ENSMUSG00000055116 | 1.806056082 | 1.701838731 | 1.67510993 | 1.630890489 | 1.655839964 | 1.826074803 | 1.582103514 |
| ENSMUSG00000055125 | 0 | 0 | 0 | 0 | 0 | 0 | 0 |
| ENSMUSG00000055128 | 1.881174527 | 1.714495249 | 1.854463129 | 1.961372513 | 1.90849439 | 2.164352856 | 1.845261772 |
| ENSMUSG00000055134 | 0.671187741 | 0.896240625 | 0.701838731 | 0.830482024 | 0.686820904 | 0.77815125 | 0.672731473 |
| ENSMUSG00000055137 | 0.289064826 | 0.396240625 | 0.396240625 | 0.75 | 0.387761935 | 0.84509804 | 0.710836105 |
| ENSMUSG00000055138 | 0.289064826 | 0 | 0 | 0 | 0 | 0 | 0 |
| ENSMUSG00000055148 | 1.249317394 | 1.02186571 | 1.372963274 | 1.396240625 | 1.484072009 | 1.544068044 | 1.574722613 |
| ENSMUSG00000055150 | 0.289064826 | 0.830482024 | 0.396240625 | 0.79248125 | 0.812711509 | 0.77815125 | 0.710836105 |
| ENSMUSG00000055159 | 0 | 0 | 0 | 0.25 | 0 | 0 | 0 |
| ENSMUSG00000055170 | 0 | 0 | 0 | 0 | 0 | 0 | 0 |
| ENSMUSG00000055172 | 1.671187741 | 1.584962501 | 1.67510993 | 1.356566189 | 1.462344774 | 1.505149978 | 1.335191751 |
| ENSMUSG00000055177 | 0.289064826 | 0 | 0 | 0 | 0.244650542 | 0 | 0 |
| ENSMUSG00000055184 | 1.227926429 | 1.311981878 | 1.388647213 | 1.26109853 | 1.27449559 | 1.322219295 | 1.319146374 |
| ENSMUSG00000055188 | 0 | 0.5 | 0.5 | 0.646240625 | 0.568060967 | 0 | 0.355418052 |
| ENSMUSG00000055193 | 0 | 0 | 0 | 0 | 0 | 0 | 0 |
| ENSMUSG00000055194 | 0 | 0 | 0.25 | 0 | 0 | 0.301029996 | 0 |
| ENSMUSG00000055197 | 0 | 0 | 0 | 0 | 0 | 0 | 0 |
| ENSMUSG00000055198 | 0 | 0 | 0 | 0 | 0 | 0 | 0 |
| ENSMUSG00000055200 | 1.358731791 | 1.380890489 | 1.431980114 | 1.339388001 | 1.343584837 | 1.505149978 | 1.401059159 |
| ENSMUSG00000055202 | 0.578129653 | 0.580482024 | 0.580482024 | 0.25 | 0.387761935 | 0.477121255 | 0.448487648 |
| ENSMUSG00000055204 | 3.296671435 | 2.969800079 | 2.960783978 | 2.903217124 | 2.806924204 | 3.254548077 | 2.618853957 |
| ENSMUSG00000055210 | 0 | 0.396240625 | 0 | 0.5 | 0.686820904 | 0 | 0 |
| ENSMUSG00000055214 | 0.45815691 | 0 | 0 | 0.646240625 | 0.733951626 | 0.602059991 | 0.448487648 |
| ENSMUSG00000055216 | 0.578129653 | 0.25 | 0.646240625 | 0.646240625 | 0.568060967 | 0.301029996 | 0.672731473 |
| ENSMUSG00000055228 | 1.558729299 | 1.339388001 | 1.418106335 | 1.584962501 | 1.456697355 | 1.77815125 | 1.379042812 |
| ENSMUSG00000055235 | 0 | 0 | 0 | 0 | 0.244650542 | 0 | 0 |
| ENSMUSG00000055239 | 2.637462542 | 2.451032755 | 2.439972046 | 2.399047439 | 2.346342321 | 2.608526034 | 2.224243824 |
| ENSMUSG00000055240 | 2.278594844 | 2.113831805 | 2.132845365 | 2.247171172 | 2.169384551 | 2.409933123 | 2.021338422 |
| ENSMUSG00000055254 | 0 | 0 | 0 | 0 | 0 | 0 | 0 |

| | | | | | | | |
|---|---|---|---|---|---|---|---|
| ENSMUSG00000055271 | 0.45815691 | 0 | 0.25 | 0.396240625 | 0.244650542 | 0.301029996 | 0.355418052 |
| ENSMUSG00000055296 | 2.912079952 | 2.672717752 | 2.653447366 | 2.760927716 | 2.682961504 | 3.063708559 | 2.463347644 |
| ENSMUSG00000055298 | 0 | 0 | 0 | 0 | 0 | 0 | 0 |
| ENSMUSG00000055301 | 1.227926429 | 1.061981878 | 1.02186571 | 1.080482024 | 1.074582839 | 1.041392685 | 1.100339912 |
| ENSMUSG00000055302 | 3.236721499 | 2.942769185 | 2.931980114 | 3.065597131 | 2.974610452 | 3.465977368 | 2.782491313 |
| ENSMUSG00000055305 | 1.227926429 | 1.321350555 | 1.188721876 | 1.25 | 1.283908311 | 1.255272505 | 1.140832123 |
| ENSMUSG00000055312 | 0 | 0 | 0 | 0 | 0 | 0 | 0 |
| ENSMUSG00000055313 | 0.811507563 | 0.951838731 | 0.976722649 | 1.114857905 | 1.335652905 | 1.041392685 | 1.140832123 |
| ENSMUSG00000055319 | 2.499676068 | 2.302972074 | 2.25835575 | 2.334962501 | 2.339874358 | 2.621176282 | 2.177588371 |
| ENSMUSG00000055320 | 3.045939207 | 2.798843052 | 2.774508021 | 2.752806814 | 2.68401826 | 3.06483222 | 2.504646253 |
| ENSMUSG00000055322 | 3.033693607 | 2.80479213 | 2.815670536 | 2.504202072 | 2.541697362 | 2.859138297 | 2.41875435 |
| ENSMUSG00000055323 | 0.45815691 | 0.580482024 | 0.396240625 | 0.580482024 | 0.387761935 | 0.301029996 | 0.520678036 |
| ENSMUSG00000055333 | 0.671187741 | 0.646240625 | 0.5 | 0.701838731 | 0.568060967 | 0.903089987 | 0.775756176 |
| ENSMUSG00000055334 | 1.857607105 | 1.657339155 | 1.705044741 | 1.921625132 | 1.892300902 | 2.164352856 | 1.760706232 |
| ENSMUSG00000055341 | 0.578129653 | 0.5 | 0.396240625 | 0.5 | 0.632412477 | 0.477121255 | 0.579661877 |
| ENSMUSG00000055357 | 0 | 0.25 | 0 | 0 | 0 | 0 | 0 |
| ENSMUSG00000055360 | 0 | 0 | 0.5 | 0.25 | 0 | 0.301029996 | 0.224243824 |
| ENSMUSG00000055368 | 0 | 0 | 0.25 | 0.396240625 | 0.387761935 | 0.301029996 | 0 |
| ENSMUSG00000055370 | 0 | 0.5 | 0 | 0.396240625 | 0.489301084 | 0 | 0.355418052 |
| ENSMUSG00000055371 | 2.578129653 | 2.407794264 | 2.357090043 | 2.380890489 | 2.311739339 | 2.596597096 | 2.152721168 |
| ENSMUSG00000055373 | 0.578129653 | 0.580482024 | 0.75 | 0 | 0 | 0 | 0 |
| ENSMUSG00000055385 | 0 | 0 | 0 | 0 | 0 | 0 | 0 |
| ENSMUSG00000055401 | 2.121692466 | 1.990000483 | 2.006976499 | 1.868933358 | 1.919403394 | 2.064457989 | 1.861046144 |
| ENSMUSG00000055403 | 0 | 0.5 | 0.396240625 | 0 | 0.244650542 | 0 | 0 |
| ENSMUSG00000055407 | 2.201144779 | 2.056000419 | 2.067865757 | 2.065523711 | 2.090057366 | 2.278753601 | 1.991191705 |
| ENSMUSG00000055408 | 0 | 0 | 0 | 0 | 0 | 0 | 0 |
| ENSMUSG00000055409 | 0 | 0 | 0 | 0 | 0 | 0 | 0 |
| ENSMUSG00000055413 | 0.289064826 | 0.25 | 0.5 | 0.976722649 | 1.223252711 | 1.176091259 | 1.350478789 |
| ENSMUSG00000055415 | 0 | 0 | 0 | 0 | 0 | 0 | 0 |
| ENSMUSG00000055416 | 0 | 0.25 | 0 | 0 | 0 | 0 | 0.224243824 |
| ENSMUSG00000055421 | 1.358731791 | 1.27186571 | 1.080482024 | 1.431980114 | 1.343584837 | 1.322219295 | 1.272006351 |
| ENSMUSG00000055424 | 0.289064826 | 0 | 0 | 0.25 | 0.244650542 | 0 | 0.355418052 |
| ENSMUSG00000055430 | 0 | 0.5 | 0.25 | 0.646240625 | 0.686820904 | 0.698970004 | 0.579661877 |
| ENSMUSG00000055435 | 1.734388958 | 1.720660762 | 1.54745614 | 1.732684334 | 1.625423019 | 1.681241237 | 1.556816002 |

| | | | | | | | |
|---|---|---|---|---|---|---|---|
| ENSMUSG00000055436 | 2.984682628 | 2.701637405 | 2.668106335 | 2.771202097 | 2.709458383 | 3.089905111 | 2.506886092 |
| ENSMUSG00000055446 | 0 | 0 | 0 | 0 | 0 | 0 | 0 |
| ENSMUSG00000055447 | 2.645603628 | 2.418106335 | 2.381380202 | 2.396709947 | 2.293608479 | 2.630427875 | 2.13295179 |
| ENSMUSG00000055452 | 0 | 0 | 0 | 0 | 0 | 0 | 0 |
| ENSMUSG00000055456 | 0 | 0 | 0 | 0 | 0 | 0 | 0 |
| ENSMUSG00000055471 | 0 | 0 | 0 | 0 | 0 | 0 | 0 |
| ENSMUSG00000055480 | 1.249317394 | 1.17510993 | 1.282320754 | 1.458222504 | 1.335652905 | 1.579783597 | 1.265599895 |
| ENSMUSG00000055485 | 2.163307908 | 2.057204673 | 2.102347734 | 2.015173983 | 2.150762767 | 2.29666519 | 2.124874702 |
| ENSMUSG00000055489 | 0 | 0 | 0.25 | 0 | 0 | 0 | 0 |
| ENSMUSG00000055491 | 2.895909295 | 2.696158711 | 2.689555804 | 2.53360658 | 2.603804521 | 2.8876173 | 2.450818701 |
| ENSMUSG00000055493 | 1.069666964 | 1.02186571 | 0.951838731 | 0.976722649 | 0.931471446 | 1.146128036 | 1.066254157 |
| ENSMUSG00000055494 | 0.45815691 | 0.580482024 | 0.580482024 | 0.79248125 | 0.387761935 | 0.77815125 | 0.629532004 |
| ENSMUSG00000055497 | 0 | 0.5 | 0.25 | 0.25 | 0.244650542 | 0.301029996 | 0 |
| ENSMUSG00000055515 | 0 | 0 | 0 | 0 | 0 | 0 | 0 |
| ENSMUSG00000055523 | 0 | 0 | 0 | 0 | 0 | 0 | 0 |
| ENSMUSG00000055531 | 2.935400733 | 2.675759597 | 2.664552871 | 2.733791262 | 2.700173892 | 3.064083436 | 2.493208369 |
| ENSMUSG00000055538 | 2.732012699 | 2.504897682 | 2.494319981 | 2.641751343 | 2.621875691 | 3.006466042 | 2.463347644 |
| ENSMUSG00000055540 | 0 | 0 | 0 | 0 | 0 | 0 | 0 |
| ENSMUSG00000055541 | 0 | 0 | 0 | 0.25 | 0 | 0 | 0 |
| ENSMUSG00000055546 | 0 | 0 | 0 | 0 | 0 | 0 | 0 |
| ENSMUSG00000055547 | 0 | 0 | 0.25 | 0 | 0 | 0 | 0 |
| ENSMUSG00000055553 | 1.975148165 | 1.77186571 | 1.774508021 | 1.752806814 | 1.770879644 | 2.089905111 | 1.734460864 |
| ENSMUSG00000055555 | 0 | 0 | 0 | 0 | 0 | 0 | 0 |
| ENSMUSG00000055560 | 1.342375483 | 1.160964047 | 1.02186571 | 1.17510993 | 1.264824954 | 1.255272505 | 1.066254157 |
| ENSMUSG00000055561 | 0 | 0 | 0 | 0 | 0 | 0 | 0 |
| ENSMUSG00000055567 | 0 | 0.5 | 0.25 | 0.25 | 0 | 0 | 0.224243824 |
| ENSMUSG00000055571 | 0 | 0 | 0 | 0 | 0 | 0 | 0 |
| ENSMUSG00000055594 | 0 | 0 | 0 | 0 | 0 | 0 | 0 |
| ENSMUSG00000055602 | 0.289064826 | 0 | 0 | 0 | 0 | 0 | 0 |
| ENSMUSG00000055609 | 0 | 0 | 0 | 0 | 0 | 0 | 0 |
| ENSMUSG00000055610 | 0 | 0 | 0 | 0 | 0 | 0 | 0 |
| ENSMUSG00000055612 | 2.330063669 | 2.212155735 | 2.182829758 | 2.211372513 | 2.192777367 | 2.409933123 | 2.060206963 |
| ENSMUSG00000055621 | 0 | 0 | 0 | 0 | 0 | 0 | 0 |
| ENSMUSG00000055629 | 1.181541736 | 1.396240625 | 1.388647213 | 0.976722649 | 1.176121988 | 1.301029996 | 1.185218806 |

| | | | | | | | |
|---|---|---|---|---|---|---|---|
| ENSMUSG00000055632 | 1.156259305 | 1.238549078 | 1.321350555 | 0.646240625 | 0.632412477 | 0 | 0.829800754 |
| ENSMUSG00000055633 | 0.867194479 | 0.951838731 | 1.098079356 | 0.79248125 | 0.846351821 | 0.698970004 | 1.089372236 |
| ENSMUSG00000055639 | 0.289064826 | 0.25 | 0.25 | 0.5 | 0.568060967 | 0 | 0.224243824 |
| ENSMUSG00000055643 | 0 | 0 | 0 | 0 | 0 | 0 | 0 |
| ENSMUSG00000055652 | 1.881174527 | 1.79745614 | 1.821350555 | 1.614857905 | 1.677808768 | 1.86923172 | 1.700880998 |
| ENSMUSG00000055653 | 0 | 0 | 0 | 0.5 | 0.387761935 | 0 | 0 |
| ENSMUSG00000055660 | 1.789260561 | 1.720660762 | 1.71759118 | 1.777131114 | 1.72073075 | 1.973127854 | 1.56716938 |
| ENSMUSG00000055670 | 2.753493018 | 2.521202097 | 2.530383379 | 2.43325383 | 2.42262144 | 2.737192643 | 2.273444839 |
| ENSMUSG00000055675 | 0.671187741 | 0.5 | 0.25 | 0.701838731 | 0.489301084 | 0.301029996 | 0.520678036 |
| ENSMUSG00000055676 | 0.960252568 | 0.830482024 | 0.580482024 | 0.701838731 | 0.978602168 | 0.903089987 | 0.829800754 |
| ENSMUSG00000055679 | 0 | 0 | 0 | 0 | 0 | 0 | 0 |
| ENSMUSG00000055681 | 2.429630757 | 2.323655187 | 2.356566189 | 2.160964047 | 2.28608706 | 2.434568904 | 2.240537069 |
| ENSMUSG00000055691 | 0.289064826 | 0.5 | 0 | 0.396240625 | 0.244650542 | 0 | 0.224243824 |
| ENSMUSG00000055692 | 0.289064826 | 0.25 | 0.396240625 | 0 | 0.489301084 | 0 | 0.672731473 |
| ENSMUSG00000055694 | 0 | 0 | 0 | 0 | 0 | 0 | 0 |
| ENSMUSG00000055700 | 0 | 0 | 0 | 0 | 0 | 0 | 0 |
| ENSMUSG00000055704 | 0.867194479 | 0.864857905 | 0.75 | 0.79248125 | 0.733951626 | 0.477121255 | 0.629532004 |
| ENSMUSG00000055707 | 2.205378646 | 2.083847589 | 2.144357207 | 1.87892496 | 1.945998439 | 2.08278537 | 1.978585612 |
| ENSMUSG00000055717 | 0 | 0.25 | 0 | 1.17510993 | 1.106691885 | 1.041392685 | 1.066254157 |
| ENSMUSG00000055720 | 2.213719572 | 2.046218836 | 2.099151195 | 2.106566189 | 2.119108707 | 2.311753861 | 2.036615833 |
| ENSMUSG00000055723 | 2.262379879 | 2.133818844 | 2.104463129 | 2.275821952 | 2.1962964 | 2.5289167 | 2.02383181 |
| ENSMUSG00000055725 | 1.249317394 | 1.238549078 | 1.29248125 | 1.372963274 | 1.212046813 | 1.301029996 | 1.168187745 |
| ENSMUSG00000055730 | 0 | 0 | 0 | 0 | 0 | 0 | 0 |
| ENSMUSG00000055732 | 0 | 0 | 0 | 0 | 0 | 0 | 0 |
| ENSMUSG00000055733 | 0.91631382 | 0.951838731 | 0.701838731 | 1.29248125 | 1.244650542 | 1.255272505 | 1.054044578 |
| ENSMUSG00000055737 | 2.554191361 | 2.338286706 | 2.377443751 | 2.112802778 | 2.022473809 | 2.260071388 | 1.857950279 |
| ENSMUSG00000055745 | 0.45815691 | 0.5 | 0.396240625 | 1.488549078 | 1.55104774 | 1.740362689 | 1.511732338 |
| ENSMUSG00000055746 | 0 | 0 | 0 | 0 | 0 | 0 | 0 |
| ENSMUSG00000055748 | 0.289064826 | 0.5 | 0.25 | 0 | 0 | 0.602059991 | 0.355418052 |
| ENSMUSG00000055760 | 2.036286563 | 1.919870025 | 1.928561379 | 2.019203899 | 1.941702774 | 2.117271296 | 1.793950594 |
| ENSMUSG00000055761 | 0.289064826 | 0 | 0.396240625 | 0.25 | 0 | 0 | 0.224243824 |
| ENSMUSG00000055762 | 3.077025005 | 2.823511578 | 2.837320307 | 2.825802487 | 2.86251892 | 3.267875419 | 2.668780475 |
| ENSMUSG00000055763 | 0 | 0.25 | 0.25 | 0 | 0 | 0 | 0.224243824 |
| ENSMUSG00000055771 | 0.289064826 | 0 | 0.396240625 | 0 | 0 | 0 | 0 |

| | | | | | | | |
|---|---|---|---|---|---|---|---|
| ENSMUSG00000055775 | 0 | 0.25 | 0.5 | 0.5 | 0.632412477 | 0.301029996 | 0.448487648 |
| ENSMUSG00000055780 | 0 | 0 | 0 | 0 | 0.244650542 | 0 | 0.355418052 |
| ENSMUSG00000055782 | 1.205378646 | 1.080482024 | 1.061981878 | 0.896240625 | 0.877063019 | 1.113943352 | 0.853775828 |
| ENSMUSG00000055789 | 0 | 0 | 0 | 0 | 0 | 0 | 0 |
| ENSMUSG00000055795 | 0.45815691 | 0.646240625 | 0.25 | 0.396240625 | 0.244650542 | 0 | 0.224243824 |
| ENSMUSG00000055799 | 1.89911417 | 1.825945187 | 1.848079356 | 1.575945187 | 1.701347897 | 1.84509804 | 1.74047993 |
| ENSMUSG00000055805 | 0.91631382 | 1.080482024 | 1.26109853 | 1.729715809 | 1.741652793 | 1.924279286 | 1.709198048 |
| ENSMUSG00000055809 | 0.289064826 | 0.396240625 | 0 | 0.864857905 | 0.955822902 | 0.954242509 | 1.100339912 |
| ENSMUSG00000055811 | 0 | 0 | 0 | 0 | 0 | 0 | 0 |
| ENSMUSG00000055817 | 2.199011616 | 2.070192693 | 2.090485943 | 1.938721876 | 1.979897422 | 2.133538908 | 1.982842456 |
| ENSMUSG00000055818 | 0.747221736 | 0.25 | 0.830482024 | 0.646240625 | 0 | 0.301029996 | 0.710836105 |
| ENSMUSG00000055820 | 0 | 0 | 0 | 0 | 0 | 0 | 0 |
| ENSMUSG00000055826 | 0 | 0 | 0 | 0 | 0 | 0 | 0 |
| ENSMUSG00000055827 | 0 | 0 | 0 | 0 | 0 | 0 | 0 |
| ENSMUSG00000055833 | 0.45815691 | 0.5 | 0.92510993 | 0.864857905 | 0.632412477 | 0.698970004 | 0.448487648 |
| ENSMUSG00000055835 | 1.794934559 | 1.692046081 | 1.618933358 | 1.71759118 | 1.701347897 | 1.826074803 | 1.616675393 |
| ENSMUSG00000055838 | 0 | 0 | 0 | 0 | 0 | 0 | 0 |
| ENSMUSG00000055839 | 2.537433341 | 2.477098155 | 2.58412664 | 2.098079356 | 2.330902415 | 2.356025857 | 2.414327616 |
| ENSMUSG00000055850 | 2.402962068 | 2.178561379 | 2.25 | 2.256976499 | 2.253693465 | 2.574031268 | 2.178359564 |
| ENSMUSG00000055862 | 1 | 1 | 1.114857905 | 1.02186571 | 1.057362051 | 0.903089987 | 0.984950056 |
| ENSMUSG00000055865 | 0 | 0 | 0 | 0 | 0 | 0 | 0 |
| ENSMUSG00000055866 | 1.721148722 | 1.610735874 | 1.701838731 | 1.445339928 | 1.546663136 | 1.643452676 | 1.562034107 |
| ENSMUSG00000055872 | 0.289064826 | 0 | 0 | 0 | 0 | 0.477121255 | 0 |
| ENSMUSG00000055874 | 0 | 0 | 0 | 0 | 0 | 0 | 0 |
| ENSMUSG00000055882 | 0 | 0 | 0 | 0.5 | 0.244650542 | 0 | 0.520678036 |
| ENSMUSG00000055884 | 2.343889214 | 2.188721876 | 2.169870025 | 2.23927551 | 2.168627946 | 2.439332694 | 1.997315209 |
| ENSMUSG00000055891 | 0 | 0 | 0.25 | 0 | 0.244650542 | 0 | 0.224243824 |
| ENSMUSG00000055895 | 0 | 0 | 0 | 0 | 0 | 0 | 0 |
| ENSMUSG00000055897 | 1.700465191 | 1.566696635 | 1.638647213 | 1.505591953 | 1.559656853 | 1.72427587 | 1.444790289 |
| ENSMUSG00000055900 | 0.960252568 | 1.080482024 | 0.864857905 | 1.711372513 | 1.559656853 | 1.792391689 | 1.48984372 |
| ENSMUSG00000055912 | 1.842799666 | 1.799918086 | 1.77186571 | 1.418106335 | 1.559656853 | 1.505149978 | 1.473249546 |
| ENSMUSG00000055917 | 2.072573125 | 1.94697564 | 1.923371739 | 1.914552871 | 1.855665188 | 2.113943352 | 1.71890424 |
| ENSMUSG00000055923 | 1.867194479 | 1.726722649 | 1.76109853 | 1.735628626 | 1.683842353 | 1.806179974 | 1.651852264 |
| ENSMUSG00000055926 | 1.289064826 | 1.098079356 | 1.17510993 | 1.671625132 | 1.677808768 | 1.707570176 | 1.559435575 |

| | | | | | | | |
|---|---|---|---|---|---|---|---|
| ENSMUSG00000055932 | 2.634462296 | 2.414106216 | 2.402294685 | 2.442459461 | 2.435299524 | 2.756636108 | 2.259363975 |
| ENSMUSG00000055933 | 0 | 0 | 0 | 0 | 0 | 0 | 0 |
| ENSMUSG00000055937 | 0 | 0 | 0 | 0.25 | 0 | 0 | 0 |
| ENSMUSG00000055942 | 0 | 0 | 0 | 0 | 0 | 0 | 0 |
| ENSMUSG00000055943 | 2.29078455 | 2.149047439 | 2.160061234 | 2.383332433 | 2.307676489 | 2.631443769 | 2.16539364 |
| ENSMUSG00000055944 | 0.289064826 | 0.25 | 0 | 0.396240625 | 0 | 0 | 0.224243824 |
| ENSMUSG00000055945 | 0.91631382 | 0.830482024 | 1.080482024 | 0.5 | 1.020174412 | 0.77815125 | 0.629532004 |
| ENSMUSG00000055958 | 0 | 0.5 | 0.5 | 0 | 0.387761935 | 0 | 0.355418052 |
| ENSMUSG00000055960 | 0.289064826 | 0 | 0 | 0.25 | 0.387761935 | 0 | 0.224243824 |
| ENSMUSG00000055961 | 0 | 0 | 0 | 0 | 0 | 0 | 0 |
| ENSMUSG00000055963 | 1.325351389 | 1.26109853 | 1.214495249 | 1.160964047 | 1.163285805 | 1.301029996 | 1.100339912 |
| ENSMUSG00000055968 | 0 | 0 | 0 | 0 | 0 | 0 | 0 |
| ENSMUSG00000055971 | 0 | 0 | 0 | 0 | 0 | 0 | 0 |
| ENSMUSG00000055972 | 0 | 0 | 0 | 0.25 | 0 | 0 | 0 |
| ENSMUSG00000055976 | 0 | 0 | 0 | 0 | 0 | 0 | 0 |
| ENSMUSG00000055978 | 0.289064826 | 0.25 | 0 | 0 | 0 | 0 | 0 |
| ENSMUSG00000055980 | 2.029919532 | 1.916333979 | 1.896240625 | 2.038704527 | 2.126012715 | 2.238046103 | 2.026921761 |
| ENSMUSG00000055991 | 1.924654746 | 1.735628626 | 1.779735268 | 1.789967834 | 1.756568651 | 1.963787827 | 1.71890424 |
| ENSMUSG00000055994 | 0.671187741 | 0.864857905 | 0.701838731 | 1.476722649 | 1.563883923 | 1.579783597 | 1.448487648 |
| ENSMUSG00000056004 | 1.527823874 | 1.380890489 | 1.527131114 | 0.976722649 | 1.106691885 | 0.903089987 | 0.896975297 |
| ENSMUSG00000056008 | 0 | 0 | 0 | 0 | 0 | 0 | 0 |
| ENSMUSG00000056014 | 1.714367558 | 1.614857905 | 1.62694866 | 1.78743678 | 1.733951626 | 1.86332286 | 1.651852264 |
| ENSMUSG00000056018 | 1.432083896 | 1.464495249 | 1.445339928 | 1.464495249 | 1.254881871 | 1.431363764 | 1.140832123 |
| ENSMUSG00000056019 | 0.91631382 | 0.79248125 | 0.830482024 | 1.26109853 | 1.223252711 | 1.230448921 | 1.21680636 |
| ENSMUSG00000056023 | 0 | 0 | 0 | 0 | 0 | 0 | 0 |
| ENSMUSG00000056025 | 0.671187741 | 1 | 0.830482024 | 0.396240625 | 0.244650542 | 0.602059991 | 0.224243824 |
| ENSMUSG00000056031 | 1.129344651 | 1.160964047 | 1.04248125 | 0.896240625 | 0.877063019 | 0.698970004 | 0.896975297 |
| ENSMUSG00000056032 | 0.45815691 | 0 | 0.25 | 0.25 | 0.244650542 | 0 | 0 |
| ENSMUSG00000056035 | 0 | 0 | 0 | 0 | 0 | 0 | 0 |
| ENSMUSG00000056043 | 0.45815691 | 0.75 | 0.396240625 | 0.580482024 | 0.568060967 | 0.602059991 | 0.629532004 |
| ENSMUSG00000056050 | 2.551444705 | 2.371457327 | 2.377443751 | 2.53199858 | 2.469521578 | 2.77524626 | 2.308247447 |
| ENSMUSG00000056054 | 0 | 0.25 | 0 | 0 | 0 | 0.301029996 | 0 |
| ENSMUSG00000056055 | 1.036286563 | 0.864857905 | 0.79248125 | 0.976722649 | 0.931471446 | 1.041392685 | 0.853775828 |
| ENSMUSG00000056061 | 0 | 0 | 0 | 0 | 0 | 0 | 0 |

| | | | | | | | |
|---|---|---|---|---|---|---|---|
| ENSMUSG00000056069 | 0.578129653 | 0.75 | 0.701838731 | 1.201838731 | 1.039257769 | 1.146128036 | 1.100339912 |
| ENSMUSG00000056071 | 0 | 0 | 0 | 0 | 0 | 0 | 0 |
| ENSMUSG00000056073 | 0.45815691 | 0.701838731 | 0.701838731 | 0.396240625 | 0.686820904 | 0 | 0 |
| ENSMUSG00000056076 | 3.30504739 | 3.004636985 | 3.025083455 | 3.005851961 | 2.983830121 | 3.417471693 | 2.807065219 |
| ENSMUSG00000056078 | 0 | 0.25 | 0 | 0 | 0.244650542 | 0 | 0 |
| ENSMUSG00000056089 | 0 | 0 | 0 | 0 | 0 | 0 | 0 |
| ENSMUSG00000056091 | 2.639698646 | 2.470660762 | 2.466433318 | 2.179419106 | 2.201854879 | 2.457881897 | 2.114720744 |
| ENSMUSG00000056106 | 0 | 0 | 0.396240625 | 0.396240625 | 0.244650542 | 0 | 0 |
| ENSMUSG00000056115 | 0 | 0 | 0 | 0 | 0 | 0 | 0 |
| ENSMUSG00000056116 | 1.936861443 | 1.777131114 | 1.837182039 | 2.063166358 | 2.082404901 | 2.326335861 | 1.964723754 |
| ENSMUSG00000056121 | 3.051791411 | 2.796373731 | 2.79075516 | 2.534247778 | 2.465625059 | 2.863917377 | 2.323847637 |
| ENSMUSG00000056124 | 2.258689303 | 2.151832578 | 2.069031101 | 2.193696765 | 2.112954895 | 2.356025857 | 1.964723754 |
| ENSMUSG00000056128 | 0.289064826 | 0 | 0.25 | 0 | 0 | 0 | 0 |
| ENSMUSG00000056130 | 0.45815691 | 0.580482024 | 0.646240625 | 0.646240625 | 0.489301084 | 0.477121255 | 0.710836105 |
| ENSMUSG00000056131 | 2.042557845 | 1.955044741 | 1.937048212 | 2.012462137 | 1.94314048 | 2.193124598 | 1.842010333 |
| ENSMUSG00000056133 | 0 | 0 | 0 | 0 | 0 | 0 | 0 |
| ENSMUSG00000056144 | 0.45815691 | 0.396240625 | 0.25 | 0.25 | 0.244650542 | 0.301029996 | 0 |
| ENSMUSG00000056145 | 0 | 0 | 0 | 0.25 | 0 | 0 | 0.224243824 |
| ENSMUSG00000056148 | 0.289064826 | 0 | 0 | 0 | 0 | 0 | 0.224243824 |
| ENSMUSG00000056153 | 2.440557995 | 2.284887838 | 2.241446071 | 2.209800947 | 2.192777367 | 2.436162647 | 2.021963573 |
| ENSMUSG00000056155 | 0 | 0.25 | 0 | 0 | 0.244650542 | 0.301029996 | 0.224243824 |
| ENSMUSG00000056158 | 0 | 0 | 0.25 | 0 | 0 | 0 | 0.224243824 |
| ENSMUSG00000056162 | 0 | 0 | 0.25 | 0.25 | 0.244650542 | 0 | 0.355418052 |
| ENSMUSG00000056167 | 2.442947872 | 2.27908599 | 2.259043403 | 2.310197833 | 2.25665701 | 2.571708832 | 2.136039214 |
| ENSMUSG00000056174 | 2.103271637 | 1.985628626 | 1.972185812 | 1.874961472 | 1.947418788 | 2.049218023 | 1.943148064 |
| ENSMUSG00000056184 | 0 | 0 | 0 | 0 | 0 | 0 | 0 |
| ENSMUSG00000056185 | 1.527823874 | 1.42510993 | 1.516522298 | 1.146240625 | 1.254881871 | 1.176091259 | 1.066254157 |
| ENSMUSG00000056197 | 0 | 0 | 0 | 0 | 0 | 0 | 0 |
| ENSMUSG00000056201 | 3.793768556 | 3.453395201 | 3.45920984 | 3.308255107 | 3.259767994 | 3.804003055 | 3.056834735 |
| ENSMUSG00000056203 | 0 | 0 | 0 | 0 | 0 | 0 | 0 |
| ENSMUSG00000056204 | 2.144240249 | 2.112802778 | 2.101285366 | 2.132845365 | 2.136951444 | 2.392696953 | 2.076440241 |
| ENSMUSG00000056209 | 2.330063669 | 2.202642909 | 2.247880462 | 2.269871196 | 2.288252441 | 2.586587305 | 2.173704585 |
| ENSMUSG00000056211 | 2.651426165 | 2.455044741 | 2.405054956 | 2.414106216 | 2.336595379 | 2.675778342 | 2.171745067 |
| ENSMUSG00000056214 | 1.956444026 | 1.825945187 | 1.87892496 | 1.732684334 | 1.766173492 | 1.763427994 | 1.68886578 |

| | | | | | | | |
|---|---|---|---|---|---|---|---|
| ENSMUSG00000056215 | 0.747221736 | 0.79248125 | 0.75 | 0.75 | 0.686820904 | 1 | 0.672731473 |
| ENSMUSG00000056216 | 2.427163103 | 2.259043403 | 2.238549078 | 2.198603967 | 2.150762767 | 2.392696953 | 1.960974982 |
| ENSMUSG00000056219 | 0 | 0 | 0 | 0 | 0 | 0 | 0 |
| ENSMUSG00000056220 | 2.500714755 | 2.30357978 | 2.299918086 | 2.192046081 | 2.097547432 | 2.338456494 | 1.932619724 |
| ENSMUSG00000056222 | 0 | 0 | 0 | 0 | 0 | 0 | 0.224243824 |
| ENSMUSG00000056223 | 0 | 0 | 0 | 0 | 0 | 0 | 0 |
| ENSMUSG00000056228 | 1.924654746 | 1.769203899 | 1.882845365 | 1.688721876 | 1.733951626 | 1.908485019 | 1.699191618 |
| ENSMUSG00000056234 | 1.894701088 | 1.660964047 | 1.695339928 | 1.964495249 | 1.92093466 | 2.130333768 | 1.798966438 |
| ENSMUSG00000056257 | 0 | 0 | 0.25 | 0.25 | 0 | 0 | 0 |
| ENSMUSG00000056258 | 0.45815691 | 0.5 | 0.75 | 0.580482024 | 0 | 0.477121255 | 0.448487648 |
| ENSMUSG00000056260 | 2.170239354 | 1.984159485 | 1.956637122 | 2.117918804 | 2.015530186 | 2.28780173 | 1.909593098 |
| ENSMUSG00000056267 | 1.721148722 | 1.726722649 | 1.708222504 | 1.834962501 | 1.756568651 | 1.86332286 | 1.61448206 |
| ENSMUSG00000056268 | 2.181541736 | 2.025821952 | 2.02186571 | 2.121960008 | 2.039257769 | 2.283301229 | 1.864112664 |
| ENSMUSG00000056270 | 0 | 0.25 | 0 | 0 | 0 | 0 | 0 |
| ENSMUSG00000056271 | 0 | 0 | 0.25 | 0 | 0.387761935 | 0 | 0.448487648 |
| ENSMUSG00000056281 | 0 | 0 | 0 | 0 | 0 | 0 | 0 |
| ENSMUSG00000056288 | 0 | 0 | 0 | 0 | 0 | 0 | 0 |
| ENSMUSG00000056290 | 0 | 0 | 0 | 0 | 0 | 0 | 0 |
| ENSMUSG00000056293 | 0 | 0 | 0 | 0 | 0 | 0 | 0 |
| ENSMUSG00000056296 | 0.289064826 | 0.25 | 0 | 0 | 0.244650542 | 0 | 0.448487648 |
| ENSMUSG00000056300 | 0 | 0.25 | 0.580482024 | 0.396240625 | 0.568060967 | 0.301029996 | 0.224243824 |
| ENSMUSG00000056305 | 2.6088504 | 2.394828984 | 2.372963274 | 2.429419106 | 2.347256716 | 2.703291378 | 2.190083792 |
| ENSMUSG00000056306 | 0 | 0 | 0 | 0.25 | 0 | 0 | 0 |
| ENSMUSG00000056310 | 2.278594844 | 2.145300145 | 2.182829758 | 2.036164561 | 2.10031573 | 2.276461804 | 2.01501956 |
| ENSMUSG00000056313 | 1.558729299 | 1.356566189 | 1.42510993 | 1 | 0.812711509 | 0.954242509 | 0.952571511 |
| ENSMUSG00000056316 | 0 | 0.5 | 0 | 0 | 0 | 0 | 0 |
| ENSMUSG00000056328 | 0 | 0 | 0 | 0 | 0 | 0.301029996 | 0 |
| ENSMUSG00000056329 | 0 | 0 | 0 | 0 | 0 | 0 | 0 |
| ENSMUSG00000056332 | 0.289064826 | 0 | 0 | 0 | 0 | 0 | 0 |
| ENSMUSG00000056342 | 3.124956012 | 2.871457327 | 2.810793496 | 3.01610149 | 2.916866738 | 3.370698093 | 2.67860997 |
| ENSMUSG00000056350 | 0 | 0 | 0 | 0 | 0 | 0 | 0 |
| ENSMUSG00000056359 | 0 | 0 | 0 | 0 | 0 | 0 | 0 |
| ENSMUSG00000056366 | 0 | 0 | 0.25 | 0 | 0 | 0 | 0 |
| ENSMUSG00000056367 | 1.671187741 | 1.598079356 | 1.561981878 | 1.339388001 | 1.27449559 | 1.431363764 | 1.20919388 |

| | | | | | | | |
|---|---|---|---|---|---|---|---|
| ENSMUSG00000056370 | 0.289064826 | 0 | 0.25 | 0 | 0 | 0 | 0 |
| ENSMUSG00000056379 | 0 | 0 | 0 | 0 | 0 | 0 | 0 |
| ENSMUSG00000056380 | 0 | 0.396240625 | 0 | 0.25 | 0.244650542 | 0.301029996 | 0.224243824 |
| ENSMUSG00000056383 | 1.686083339 | 1.598079356 | 1.575945187 | 1.602347734 | 1.433158226 | 1.62324929 | 1.459332757 |
| ENSMUSG00000056394 | 2.697239867 | 2.463717096 | 2.49539182 | 2.444108258 | 2.462015062 | 2.783903579 | 2.343850547 |
| ENSMUSG00000056399 | 0 | 0 | 0 | 0 | 0 | 0 | 0 |
| ENSMUSG00000056411 | 0.578129653 | 0.75 | 0.701838731 | 0.896240625 | 0.686820904 | 0.698970004 | 0.803905701 |
| ENSMUSG00000056412 | 0 | 0 | 0 | 0 | 0 | 0 | 0 |
| ENSMUSG00000056413 | 0.867194479 | 0.701838731 | 0.79248125 | 0.75 | 0.877063019 | 0 | 0.629532004 |
| ENSMUSG00000056418 | 0.45815691 | 0.25 | 0.896240625 | 0.646240625 | 0.568060967 | 0.301029996 | 0.579661877 |
| ENSMUSG00000056423 | 0 | 0 | 0 | 0 | 0 | 0 | 0 |
| ENSMUSG00000056426 | 0 | 0 | 0 | 0 | 0 | 0 | 0 |
| ENSMUSG00000056427 | 0 | 0 | 0 | 0.646240625 | 0.905314583 | 0.903089987 | 0.952571511 |
| ENSMUSG00000056429 | 2.773920925 | 2.526477127 | 2.511780978 | 2.693078643 | 2.607537217 | 3.021189299 | 2.416917233 |
| ENSMUSG00000056436 | 0 | 0 | 0.25 | 0 | 0 | 0 | 0 |
| ENSMUSG00000056445 | 1.227926429 | 1.311981878 | 1.372963274 | 1.146240625 | 1 | 0.84509804 | 1.028149525 |
| ENSMUSG00000056457 | 0.289064826 | 0.396240625 | 0 | 0.25 | 0 | 0 | 0.224243824 |
| ENSMUSG00000056458 | 1.129344651 | 1.282320754 | 1.146240625 | 1.321350555 | 1.283908311 | 1.51851394 | 1.307989563 |
| ENSMUSG00000056459 | 1.885732304 | 1.811981878 | 1.809601185 | 1.646240625 | 1.642643806 | 1.812913357 | 1.511732338 |
| ENSMUSG00000056468 | 1.418409478 | 1.26109853 | 1.188721876 | 0.580482024 | 0.632412477 | 0.903089987 | 0.579661877 |
| ENSMUSG00000056476 | 0.91631382 | 1.04248125 | 0.896240625 | 1.664552871 | 1.677808768 | 1.819543936 | 1.483307832 |
| ENSMUSG00000056481 | 2.322180025 | 2.218361028 | 2.232684334 | 1.649978211 | 1.728722551 | 1.799340549 | 1.777090262 |
| ENSMUSG00000056486 | 0.747221736 | 0.976722649 | 0.951838731 | 1.302363341 | 1.188507684 | 1.322219295 | 1.014380832 |
| ENSMUSG00000056487 | 0 | 0 | 0 | 0 | 0 | 0 | 0 |
| ENSMUSG00000056492 | 0.45815691 | 0.580482024 | 0.25 | 0.976722649 | 0.877063019 | 0.698970004 | 0.803905701 |
| ENSMUSG00000056493 | 2.388281011 | 2.198603967 | 2.316696635 | 2.043731421 | 2.120847411 | 2.252853031 | 2.040177507 |
| ENSMUSG00000056494 | 0 | 0 | 0 | 0 | 0 | 0 | 0 |
| ENSMUSG00000056498 | 0 | 0.25 | 0.25 | 0.701838731 | 0.387761935 | 0.477121255 | 0.710836105 |
| ENSMUSG00000056501 | 1.156259305 | 1.214495249 | 1.282320754 | 1.130890489 | 1.057362051 | 1.113943352 | 1.121219121 |
| ENSMUSG00000056508 | 0.45815691 | 0.75 | 0.580482024 | 0.646240625 | 0.877063019 | 0.602059991 | 0.803905701 |
| ENSMUSG00000056509 | 0 | 0 | 0 | 0 | 0 | 0 | 0 |
| ENSMUSG00000056515 | 3.065135408 | 2.769370839 | 2.789494612 | 2.592126615 | 2.545102531 | 2.931457871 | 2.389035574 |
| ENSMUSG00000056529 | 0.289064826 | 0 | 0.25 | 0.25 | 0.244650542 | 0.477121255 | 0.520678036 |
| ENSMUSG00000056531 | 1.967766868 | 1.864857905 | 1.899978211 | 1.811981878 | 1.782377602 | 1.954242509 | 1.692344475 |

| | | | | | | | |
|---|---|---|---|---|---|---|---|
| ENSMUSG00000056536 | 2.699822119 | 2.469129237 | 2.471424093 | 2.571350555 | 2.499532413 | 2.881954971 | 2.306955966 |
| ENSMUSG00000056537 | 3.015076955 | 2.754202072 | 2.736178015 | 2.728969845 | 2.638666793 | 3.030194785 | 2.442585457 |
| ENSMUSG00000056544 | 0 | 0 | 0 | 0 | 0 | 0 | 0 |
| ENSMUSG00000056553 | 0 | 0 | 0 | 0 | 0 | 0 | 0 |
| ENSMUSG00000056564 | 0 | 0 | 0.25 | 0 | 0 | 0 | 0 |
| ENSMUSG00000056569 | 0 | 0 | 0.25 | 0.25 | 0.387761935 | 0 | 0 |
| ENSMUSG00000056579 | 2.754058486 | 2.536483036 | 2.526149688 | 2.558704358 | 2.472415995 | 2.82672252 | 2.269691383 |
| ENSMUSG00000056586 | 0 | 0 | 0 | 0 | 0 | 0 | 0 |
| ENSMUSG00000056592 | 1.389637216 | 1.464495249 | 1.364857905 | 1.470660762 | 1.47337555 | 1.579783597 | 1.374453863 |
| ENSMUSG00000056596 | 0.867194479 | 1 | 1.146240625 | 0.896240625 | 1.020174412 | 1.113943352 | 1.100339912 |
| ENSMUSG00000056598 | 1.069666964 | 1.02186571 | 0.864857905 | 1.214495249 | 1.293076518 | 1.230448921 | 1.159323753 |
| ENSMUSG00000056600 | 0.289064826 | 0 | 0.25 | 0 | 0 | 0 | 0 |
| ENSMUSG00000056602 | 0.45815691 | 0.5 | 0.5 | 1.17510993 | 1.302012594 | 1.361727836 | 1.185218806 |
| ENSMUSG00000056605 | 0 | 0 | 0 | 0 | 0 | 0 | 0 |
| ENSMUSG00000056606 | 0.289064826 | 0 | 0 | 0 | 0 | 0 | 0.224243824 |
| ENSMUSG00000056608 | 2.579849376 | 2.405512955 | 2.392938161 | 2.540597832 | 2.44581538 | 2.802089258 | 2.266481009 |
| ENSMUSG00000056612 | 2.646335907 | 2.547764804 | 2.615625568 | 2.327084758 | 2.441995847 | 2.6599162 | 2.391635713 |
| ENSMUSG00000056617 | 0 | 0 | 0.25 | 0 | 0.244650542 | 0.301029996 | 0.355418052 |
| ENSMUSG00000056629 | 2.326928073 | 2.189555804 | 2.24929487 | 2.140560606 | 2.218023635 | 2.44870632 | 2.097385413 |
| ENSMUSG00000056632 | 0 | 0 | 0 | 0 | 0 | 0 | 0 |
| ENSMUSG00000056643 | 0 | 0 | 0 | 0.396240625 | 0.244650542 | 0 | 0.224243824 |
| ENSMUSG00000056648 | 1.948720984 | 1.837182039 | 1.864857905 | 1.671625132 | 1.665423072 | 1.857332496 | 1.651852264 |
| ENSMUSG00000056656 | 0.289064826 | 0.580482024 | 0.75 | 0.646240625 | 0.387761935 | 0.301029996 | 0.448487648 |
| ENSMUSG00000056665 | 1.605636262 | 1.566696635 | 1.698603967 | 1.372963274 | 1.652586889 | 1.51851394 | 1.707551656 |
| ENSMUSG00000056666 | 1.91631382 | 1.811981878 | 1.839388001 | 1.860735874 | 1.94314048 | 2.139879086 | 1.756477211 |
| ENSMUSG00000056671 | 1.181541736 | 1.282320754 | 1.226722649 | 1.364857905 | 1.373641807 | 1.380211242 | 1.20139795 |
| ENSMUSG00000056673 | 1.100572389 | 0.75 | 0.92510993 | 0.951838731 | 0.686820904 | 0.84509804 | 0.672731473 |
| ENSMUSG00000056679 | 1.568542279 | 1.566696635 | 1.622963274 | 1.71759118 | 1.773209395 | 1.86332286 | 1.625302983 |
| ENSMUSG00000056682 | 0 | 0.25 | 0 | 0 | 0 | 0 | 0.355418052 |
| ENSMUSG00000056687 | 0.91631382 | 1.146240625 | 0.976722649 | 0.864857905 | 1.039257769 | 1.255272505 | 0.916588299 |
| ENSMUSG00000056692 | 2.785084983 | 2.619439567 | 2.609957721 | 2.573080408 | 2.594409696 | 2.90579588 | 2.445971311 |
| ENSMUSG00000056696 | 0 | 0 | 0 | 0 | 0 | 0 | 0 |
| ENSMUSG00000056698 | 1.707474304 | 1.527131114 | 1.653677461 | 1.649978211 | 1.689774529 | 1.819543936 | 1.577201671 |
| ENSMUSG00000056699 | 0 | 0 | 0 | 0 | 0 | 0 | 0 |

| | | | | | | | |
|---|---|---|---|---|---|---|---|
| ENSMUSG00000056706 | 0 | 0 | 0 | 0 | 0 | 0 | 0 |
| ENSMUSG00000056708 | 2.396352858 | 2.248588359 | 2.278435542 | 2.094844592 | 2.084333628 | 2.281033367 | 2.014380832 |
| ENSMUSG00000056716 | 0 | 0 | 0 | 0 | 0 | 0 | 0 |
| ENSMUSG00000056718 | 0.747221736 | 0.92510993 | 1.146240625 | 0.830482024 | 0.978602168 | 0.903089987 | 0.710836105 |
| ENSMUSG00000056724 | 1.623015126 | 1.5 | 1.642463902 | 1.445339928 | 1.692703642 | 1.69019608 | 1.71890424 |
| ENSMUSG00000056728 | 0 | 0 | 0 | 0 | 0 | 0 | 0 |
| ENSMUSG00000056735 | 1.100572389 | 1.04248125 | 1.114857905 | 1.25 | 1.283908311 | 1.176091259 | 1.252393349 |
| ENSMUSG00000056737 | 2.460448303 | 2.32991803 | 2.416777886 | 1.945339928 | 2.053185007 | 2.252853031 | 2.052911423 |
| ENSMUSG00000056738 | 0 | 0 | 0 | 0 | 0 | 0 | 0 |
| ENSMUSG00000056749 | 1.960252568 | 1.828220739 | 1.862802778 | 1.87892496 | 1.861137485 | 2.037426498 | 1.755055163 |
| ENSMUSG00000056752 | 0 | 0.25 | 0 | 0 | 0 | 0 | 0 |
| ENSMUSG00000056753 | 0.578129653 | 0.79248125 | 0.701838731 | 0.864857905 | 0.905314583 | 0.477121255 | 0.969165684 |
| ENSMUSG00000056755 | 0 | 0 | 0 | 0 | 0 | 0 | 0.355418052 |
| ENSMUSG00000056758 | 3.059468753 | 2.817719824 | 2.827084758 | 2.333288838 | 2.232853197 | 2.553883027 | 2.061314894 |
| ENSMUSG00000056763 | 2.22189694 | 2.070192693 | 2.105516192 | 2.145300145 | 2.122577592 | 2.356025857 | 1.984249047 |
| ENSMUSG00000056770 | 2.556920045 | 2.362802778 | 2.351285366 | 2.312574604 | 2.229044281 | 2.551449998 | 2.122594809 |
| ENSMUSG00000056771 | 0 | 0 | 0.5 | 0.5 | 0.387761935 | 0.698970004 | 0.579661877 |
| ENSMUSG00000056772 | 0.289064826 | 0 | 0 | 0 | 0 | 0 | 0 |
| ENSMUSG00000056777 | 0 | 0 | 0 | 0 | 0 | 0 | 0 |
| ENSMUSG00000056782 | 0 | 0 | 0 | 0 | 0 | 0 | 0 |
| ENSMUSG00000056797 | 0 | 0.25 | 0 | 0 | 0 | 0 | 0 |
| ENSMUSG00000056812 | 0 | 0 | 0 | 0 | 0.244650542 | 0 | 0 |
| ENSMUSG00000056815 | 0 | 0 | 0 | 0 | 0 | 0 | 0 |
| ENSMUSG00000056820 | 2.271465912 | 2.106566189 | 2.111770807 | 2.385757955 | 2.337535342 | 2.668385917 | 2.193041668 |
| ENSMUSG00000056821 | 0 | 0.25 | 0.25 | 0 | 0.244650542 | 0 | 0.224243824 |
| ENSMUSG00000056822 | 0 | 0 | 0 | 0 | 0 | 0 | 0 |
| ENSMUSG00000056824 | 0 | 0 | 0 | 0 | 0 | 0 | 0.224243824 |
| ENSMUSG00000056829 | 0 | 0 | 0 | 0 | 0 | 0 | 0 |
| ENSMUSG00000056832 | 2 | 1.834962501 | 1.890560606 | 1.870953944 | 1.826953746 | 2.041392685 | 1.672731473 |
| ENSMUSG00000056836 | 0 | 0 | 0 | 0 | 0 | 0 | 0 |
| ENSMUSG00000056849 | 0 | 0.396240625 | 0.25 | 0.25 | 0.244650542 | 0 | 0 |
| ENSMUSG00000056851 | 3.26900747 | 2.958518988 | 2.986086509 | 3.016942614 | 2.954517771 | 3.408748606 | 2.753105878 |
| ENSMUSG00000056853 | 0 | 0 | 0 | 0 | 0 | 0 | 0 |
| ENSMUSG00000056854 | 0 | 0 | 0 | 0 | 0 | 0 | 0 |

| | | | | | | | |
|---|---|---|---|---|---|---|---|
| ENSMUSG00000056856 | 0.289064826 | 0.396240625 | 0.580482024 | 0 | 0.244650542 | 0.301029996 | 0.520678036 |
| ENSMUSG00000056858 | 0 | 0 | 0 | 0 | 0 | 0 | 0 |
| ENSMUSG00000056863 | 0 | 0 | 0 | 0 | 0 | 0 | 0 |
| ENSMUSG00000056870 | 2.386920383 | 2.192046081 | 2.113831805 | 2.333847589 | 2.288791717 | 2.588831726 | 2.110473216 |
| ENSMUSG00000056877 | 0.289064826 | 0.25 | 0 | 0 | 0.387761935 | 0.301029996 | 0 |
| ENSMUSG00000056880 | 0 | 0.25 | 0.25 | 0.25 | 0 | 0 | 0 |
| ENSMUSG00000056883 | 0 | 0 | 0 | 0 | 0 | 0 | 0 |
| ENSMUSG00000056885 | 0 | 0 | 0 | 0 | 0 | 0 | 0 |
| ENSMUSG00000056888 | 0.960252568 | 1 | 1.02186571 | 0.25 | 0.568060967 | 0.602059991 | 0.448487648 |
| ENSMUSG00000056895 | 0 | 0 | 0.25 | 0 | 0.489301084 | 0.301029996 | 0.355418052 |
| ENSMUSG00000056899 | 0.960252568 | 1 | 1.04248125 | 0.79248125 | 0.733951626 | 0.903089987 | 0.672731473 |
| ENSMUSG00000056900 | 1.885732304 | 1.726722649 | 1.646240625 | 1.732684334 | 1.726078673 | 1.86332286 | 1.584526862 |
| ENSMUSG00000056901 | 0 | 0 | 0 | 0 | 0 | 0 | 0 |
| ENSMUSG00000056904 | 1.129344651 | 1.201838731 | 1.17510993 | 0.951838731 | 1.212046813 | 1.146128036 | 1.21680636 |
| ENSMUSG00000056910 | 0 | 0 | 0 | 0 | 0 | 0 | 0 |
| ENSMUSG00000056912 | 0 | 0 | 0 | 0 | 0 | 0 | 0 |
| ENSMUSG00000056917 | 2.254965775 | 2.163659007 | 2.264497931 | 2.174241882 | 2.261929482 | 2.482873584 | 2.275448037 |
| ENSMUSG00000056919 | 2.375872521 | 2.224461364 | 2.207430684 | 2.31728167 | 2.230318492 | 2.532754379 | 2.082712142 |
| ENSMUSG00000056921 | 0 | 0 | 0 | 0 | 0 | 0 | 0 |
| ENSMUSG00000056926 | 0 | 0 | 0 | 0 | 0 | 0 | 0 |
| ENSMUSG00000056938 | 1.678702042 | 1.614857905 | 1.720660762 | 1.610735874 | 1.795698282 | 1.832508913 | 1.770364127 |
| ENSMUSG00000056941 | 1.993049299 | 1.898114259 | 1.866901388 | 2.013820609 | 2.025895058 | 2.264817823 | 1.966211184 |
| ENSMUSG00000056943 | 0 | 0 | 0 | 0 | 0 | 0 | 0 |
| ENSMUSG00000056946 | 0 | 0 | 0 | 0 | 0 | 0 | 0 |
| ENSMUSG00000056947 | 0.671187741 | 0.864857905 | 0.79248125 | 0 | 0 | 0.602059991 | 0.224243824 |
| ENSMUSG00000056952 | 2.475043119 | 2.326515422 | 2.277131114 | 2.302363341 | 2.266550905 | 2.589949601 | 2.124874702 |
| ENSMUSG00000056959 | 0 | 0 | 0 | 0 | 0 | 0 | 0 |
| ENSMUSG00000056961 | 0 | 0 | 0 | 0 | 0 | 0 | 0 |
| ENSMUSG00000056962 | 2.412042447 | 2.307805295 | 2.281030328 | 2.308404919 | 2.30971376 | 2.545307116 | 2.212329757 |
| ENSMUSG00000056966 | 0.289064826 | 0.701838731 | 0.396240625 | 0.5 | 0.632412477 | 0.602059991 | 0.629532004 |
| ENSMUSG00000056972 | 0 | 0 | 0 | 0 | 0 | 0 | 0 |
| ENSMUSG00000056973 | 0 | 0 | 0 | 0 | 0 | 0 | 0 |
| ENSMUSG00000056978 | 0 | 0 | 0 | 0 | 0 | 0 | 0 |
| ENSMUSG00000056987 | 0 | 0 | 0 | 0.25 | 0 | 0 | 0 |

| | | | | | | | |
|---|---|---|---|---|---|---|---|
| ENSMUSG00000056995 | 0 | 0 | 0 | 0 | 0 | 0 | 0 |
| ENSMUSG00000056999 | 3.084255469 | 2.800377836 | 2.77401406 | 2.859438018 | 2.765894696 | 3.199480915 | 2.566125594 |
| ENSMUSG00000057000 | 0 | 0.25 | 0 | 0 | 0.244650542 | 0 | 0 |
| ENSMUSG00000057003 | 0 | 0 | 0 | 0 | 0 | 0 | 0 |
| ENSMUSG00000057021 | 0 | 0 | 0 | 0 | 0 | 0 | 0 |
| ENSMUSG00000057036 | 0 | 0 | 0 | 0 | 0 | 0 | 0 |
| ENSMUSG00000057037 | 0 | 0 | 0.25 | 0 | 0 | 0 | 0 |
| ENSMUSG00000057047 | 0 | 0 | 0 | 0 | 0.244650542 | 0 | 0 |
| ENSMUSG00000057050 | 0 | 0 | 0 | 0 | 0 | 0 | 0 |
| ENSMUSG00000057054 | 0.671187741 | 0.976722649 | 0.896240625 | 0.79248125 | 0.812711509 | 0.77815125 | 0.803905701 |
| ENSMUSG00000057058 | 0.811507563 | 0.896240625 | 1 | 1.061981878 | 1.039257769 | 0.954242509 | 0.984950056 |
| ENSMUSG00000057060 | 0 | 0.25 | 0 | 0.396240625 | 0 | 0 | 0.224243824 |
| ENSMUSG00000057067 | 0 | 0 | 0 | 0 | 0 | 0 | 0 |
| ENSMUSG00000057068 | 0 | 0 | 0 | 0 | 0 | 0 | 0 |
| ENSMUSG00000057069 | 1.671187741 | 1.54248125 | 1.575945187 | 2.060793496 | 1.931471446 | 2.264817823 | 1.845261772 |
| ENSMUSG00000057072 | 0 | 0 | 0 | 0 | 0 | 0.301029996 | 0.520678036 |
| ENSMUSG00000057074 | 0.45815691 | 0.5 | 0.25 | 0.864857905 | 0.686820904 | 0.477121255 | 0.520678036 |
| ENSMUSG00000057092 | 0 | 0 | 0 | 0 | 0 | 0 | 0 |
| ENSMUSG00000057093 | 0.91631382 | 1 | 1.02186571 | 1.02186571 | 1 | 1 | 0.876096088 |
| ENSMUSG00000057098 | 2.299278375 | 2.135757955 | 2.095926073 | 1.630890489 | 1.509475496 | 1.73239376 | 1.505628092 |
| ENSMUSG00000057101 | 2.23778581 | 2.071350555 | 2.065523711 | 1.988549078 | 2.016696984 | 2.181843588 | 1.863093717 |
| ENSMUSG00000057103 | 0.811507563 | 0.951838731 | 0.701838731 | 0.830482024 | 0.846351821 | 0.301029996 | 0.803905701 |
| ENSMUSG00000057110 | 2.463861956 | 2.256976499 | 2.296218836 | 2.309003548 | 2.269408865 | 2.575187845 | 2.156863548 |
| ENSMUSG00000057113 | 3.600471251 | 3.270620432 | 3.271534056 | 3.437729075 | 3.306532872 | 3.890588668 | 3.074780752 |
| ENSMUSG00000057116 | 0 | 0.25 | 0.396240625 | 0 | 0 | 0 | 0.224243824 |
| ENSMUSG00000057123 | 0 | 0.25 | 0 | 0 | 0 | 0 | 0.224243824 |
| ENSMUSG00000057130 | 2.325351389 | 2.149047439 | 2.168106335 | 2.084962501 | 2.046825265 | 2.309630167 | 1.972821265 |
| ENSMUSG00000057132 | 0.578129653 | 0.396240625 | 0.396240625 | 0.75 | 0.632412477 | 0.698970004 | 0.520678036 |
| ENSMUSG00000057133 | 2.446507203 | 2.274508021 | 2.204245906 | 2.284247778 | 2.235369829 | 2.536558443 | 2.075383 |
| ENSMUSG00000057134 | 2.358731791 | 2.171625132 | 2.172499493 | 2.146240625 | 2.085294053 | 2.31386722 | 1.958704689 |
| ENSMUSG00000057135 | 0 | 0 | 0 | 0 | 0 | 0 | 0 |
| ENSMUSG00000057137 | 1.129344651 | 1.130890489 | 1.098079356 | 0.976722649 | 0.846351821 | 1 | 0.829800754 |
| ENSMUSG00000057143 | 1.307602651 | 1.302363341 | 1.201838731 | 1.505591953 | 1.47337555 | 1.612783857 | 1.355418052 |
| ENSMUSG00000057147 | 2.069666964 | 1.882845365 | 1.894357207 | 2.019203899 | 1.947418788 | 2.217483944 | 1.839824409 |

| | | | | | | | |
|---|---|---|---|---|---|---|---|
| ENSMUSG00000057149 | 0 | 0 | 0 | 0 | 0 | 0 | 0 |
| ENSMUSG00000057156 | 1.432083896 | 1.348079356 | 1.302363341 | 1.598079356 | 1.533172725 | 1.414973348 | 1.417653332 |
| ENSMUSG00000057157 | 0 | 0 | 0 | 0 | 0 | 0 | 0 |
| ENSMUSG00000057160 | 0 | 0 | 0 | 0 | 0 | 0 | 0 |
| ENSMUSG00000057161 | 0 | 0 | 0 | 0 | 0 | 0 | 0 |
| ENSMUSG00000057163 | 0 | 0 | 0 | 0 | 0 | 0 | 0 |
| ENSMUSG00000057170 | 0 | 0 | 0 | 0 | 0 | 0 | 0 |
| ENSMUSG00000057173 | 0 | 0 | 0 | 0 | 0 | 0.301029996 | 0 |
| ENSMUSG00000057174 | 0 | 0 | 0 | 0 | 0 | 0 | 0 |
| ENSMUSG00000057176 | 1.036286563 | 0.701838731 | 0.896240625 | 0.75 | 0.931471446 | 0.602059991 | 0.935079929 |
| ENSMUSG00000057177 | 2.217828299 | 2.057204673 | 2.070192693 | 2.067865757 | 2.069604532 | 2.303196057 | 1.944737824 |
| ENSMUSG00000057179 | 0 | 0 | 0 | 0 | 0 | 0 | 0 |
| ENSMUSG00000057181 | 2.22189694 | 2.099151195 | 2.072504712 | 2.273189285 | 2.189936658 | 2.544068044 | 2.05629905 |
| ENSMUSG00000057182 | 0.578129653 | 0.25 | 0.580482024 | 0.830482024 | 0.846351821 | 0.903089987 | 0.710836105 |
| ENSMUSG00000057191 | 0.289064826 | 0 | 0.25 | 0.5 | 0.244650542 | 0.477121255 | 0.520678036 |
| ENSMUSG00000057193 | 2.828393772 | 2.562278363 | 2.570192693 | 2.609697963 | 2.588465994 | 2.923244019 | 2.395201749 |
| ENSMUSG00000057195 | 0 | 0 | 0 | 0 | 0 | 0 | 0 |
| ENSMUSG00000057203 | 0 | 0 | 0 | 0 | 0 | 0 | 0 |
| ENSMUSG00000057207 | 0 | 0 | 0 | 0 | 0 | 0 | 0 |
| ENSMUSG00000057215 | 0.45815691 | 0.5 | 0.396240625 | 0.396240625 | 0.244650542 | 0.602059991 | 0.224243824 |
| ENSMUSG00000057216 | 0 | 0 | 0 | 0 | 0 | 0 | 0 |
| ENSMUSG00000057219 | 1.568542279 | 1.488549078 | 1.671625132 | 1.51109853 | 1.576269619 | 1.716003344 | 1.612273755 |
| ENSMUSG00000057223 | 0 | 0 | 0.25 | 0 | 0.244650542 | 0 | 0 |
| ENSMUSG00000057228 | 0 | 0.25 | 0.5 | 0.396240625 | 0.387761935 | 0.301029996 | 0 |
| ENSMUSG00000057229 | 1.45815691 | 1.464495249 | 1.494319981 | 1.431980114 | 1.546663136 | 1.653212514 | 1.455757965 |
| ENSMUSG00000057230 | 2.862428343 | 2.606303976 | 2.601019284 | 2.684311836 | 2.624799605 | 2.985875357 | 2.421490612 |
| ENSMUSG00000057234 | 1.389637216 | 1.339388001 | 1.364857905 | 1.451838731 | 1.433158226 | 1.602059991 | 1.417653332 |
| ENSMUSG00000057236 | 3.239553648 | 2.927809192 | 2.897061538 | 2.958024712 | 2.869346137 | 3.322839273 | 2.661077362 |
| ENSMUSG00000057240 | 0 | 0 | 0 | 0 | 0 | 0 | 0 |
| ENSMUSG00000057244 | 0 | 0 | 0 | 0 | 0 | 0.301029996 | 0 |
| ENSMUSG00000057246 | 0 | 0 | 0 | 0 | 0 | 0 | 0.224243824 |
| ENSMUSG00000057262 | 0 | 0 | 0 | 0 | 0 | 0.301029996 | 0 |
| ENSMUSG00000057265 | 0.91631382 | 1.04248125 | 1.114857905 | 1.02186571 | 0.931471446 | 0.903089987 | 0.775756176 |
| ENSMUSG00000057270 | 0 | 0 | 0 | 0 | 0 | 0 | 0 |

| | | | | | | | |
|---|---|---|---|---|---|---|---|
| ENSMUSG00000057278 | 2.280358197 | 2.110735874 | 2.150906586 | 2.183677405 | 2.103062484 | 2.442479769 | 2.013740841 |
| ENSMUSG00000057280 | 0 | 0 | 0 | 0 | 0 | 0 | 0 |
| ENSMUSG00000057286 | 1.614416215 | 1.476722649 | 1.458222504 | 1.988549078 | 2.003645018 | 2.204119983 | 1.865128412 |
| ENSMUSG00000057290 | 0 | 0 | 0.396240625 | 0.25 | 0.244650542 | 0.477121255 | 0 |
| ENSMUSG00000057315 | 1.505869732 | 1.53743678 | 1.527131114 | 0.79248125 | 0.733951626 | 0.903089987 | 0.952571511 |
| ENSMUSG00000057321 | 0 | 0 | 0 | 0 | 0.244650542 | 0 | 0 |
| ENSMUSG00000057322 | 2.752359777 | 2.542794199 | 2.58272922 | 2.540597832 | 2.520266629 | 2.895422546 | 2.37466387 |
| ENSMUSG00000057329 | 2.051791411 | 1.96759118 | 1.928561379 | 1.819031101 | 1.806427258 | 1.939519253 | 1.651852264 |
| ENSMUSG00000057335 | 2.998430437 | 2.751055117 | 2.720851747 | 2.724650347 | 2.635250738 | 3.048441804 | 2.424384006 |
| ENSMUSG00000057337 | 2.020182925 | 1.903677461 | 1.874961472 | 1.418106335 | 1.310728002 | 1.342422681 | 1.350478789 |
| ENSMUSG00000057342 | 1.596667477 | 1.692046081 | 1.755591953 | 1.364857905 | 1.683842353 | 1.556302501 | 1.729871916 |
| ENSMUSG00000057346 | 0.671187741 | 0.580482024 | 0.864857905 | 1.431980114 | 1.366364104 | 1.556302501 | 1.405288179 |
| ENSMUSG00000057349 | 0 | 0 | 0 | 0 | 0 | 0 | 0 |
| ENSMUSG00000057351 | 0 | 0 | 0 | 0 | 0 | 0 | 0 |
| ENSMUSG00000057359 | 1.587501561 | 1.464495249 | 1.311981878 | 1.438721876 | 1.47337555 | 1.602059991 | 1.355418052 |
| ENSMUSG00000057363 | 2.396352858 | 2.213717096 | 2.215271726 | 2.47934302 | 2.430282804 | 2.771587481 | 2.254226303 |
| ENSMUSG00000057367 | 2.285603957 | 2.056000419 | 2 | 2.242166698 | 2.182004726 | 2.434568904 | 2.022587518 |
| ENSMUSG00000057370 | 0.289064826 | 0.580482024 | 0.646240625 | 0.396240625 | 0.489301084 | 0.602059991 | 0.224243824 |
| ENSMUSG00000057375 | 2.119110214 | 1.874961472 | 1.912762923 | 2.103406982 | 2.013184954 | 2.305351369 | 1.865128412 |
| ENSMUSG00000057378 | 1.289064826 | 1.201838731 | 1.321350555 | 0 | 0.244650542 | 0 | 0.224243824 |
| ENSMUSG00000057381 | 0 | 0 | 0 | 0 | 0 | 0 | 0 |
| ENSMUSG00000057388 | 2.442947872 | 2.273189285 | 2.308404919 | 2.3464656 | 2.314756076 | 2.562292864 | 2.182948513 |
| ENSMUSG00000057396 | 1.129344651 | 1.160964047 | 1.160964047 | 1.311981878 | 1.254881871 | 1.462397998 | 1.245582246 |
| ENSMUSG00000057400 | 0 | 0 | 0.25 | 0 | 0 | 0 | 0 |
| ENSMUSG00000057402 | 0 | 0 | 0 | 0 | 0.244650542 | 0 | 0 |
| ENSMUSG00000057406 | 3.251551655 | 2.928883266 | 2.909156155 | 2.952341552 | 2.859964341 | 3.288696261 | 2.64979592 |
| ENSMUSG00000057409 | 2.081172036 | 1.982684334 | 1.890560606 | 1.928561379 | 1.822942838 | 2.071882007 | 1.707551656 |
| ENSMUSG00000057411 | 1.623015126 | 1.584962501 | 1.678561379 | 1.42510993 | 1.65906333 | 1.51851394 | 1.674564426 |
| ENSMUSG00000057417 | 0 | 0 | 0 | 0 | 0 | 0 | 0 |
| ENSMUSG00000057421 | 2.573799059 | 2.370953944 | 2.353935442 | 2.370953944 | 2.311234033 | 2.643452676 | 2.171351735 |
| ENSMUSG00000057424 | 0 | 0 | 0 | 0 | 0 | 0 | 0 |
| ENSMUSG00000057425 | 0 | 0 | 0 | 0 | 0 | 0 | 0 |
| ENSMUSG00000057439 | 0 | 0 | 0 | 0 | 0 | 0 | 0 |
| ENSMUSG00000057440 | 1.771760131 | 1.634789703 | 1.649978211 | 1.811981878 | 1.808534465 | 2.041392685 | 1.635773653 |

| | | | | | | | |
|---|---|---|---|---|---|---|---|
| ENSMUSG00000057443 | 0 | 0 | 0 | 0 | 0 | 0 | 0 |
| ENSMUSG00000057444 | 0 | 0 | 0 | 0 | 0 | 0 | 0 |
| ENSMUSG00000057446 | 0.289064826 | 0 | 0 | 0 | 0.244650542 | 0 | 0 |
| ENSMUSG00000057447 | 0 | 0 | 0 | 0 | 0 | 0 | 0 |
| ENSMUSG00000057454 | 0 | 0 | 0 | 0 | 0 | 0 | 0 |
| ENSMUSG00000057455 | 0 | 0 | 0 | 0 | 0 | 0 | 0 |
| ENSMUSG00000057457 | 1.289064826 | 1.226722649 | 1.321350555 | 1.214495249 | 1.163285805 | 1.204119983 | 1.078019652 |
| ENSMUSG00000057461 | 0 | 0 | 0 | 0 | 0 | 0 | 0 |
| ENSMUSG00000057464 | 0 | 0 | 0 | 0 | 0 | 0 | 0 |
| ENSMUSG00000057465 | 0.45815691 | 0 | 0 | 0 | 0 | 0 | 0 |
| ENSMUSG00000057469 | 2.273259602 | 2.092671852 | 2.111770807 | 1.921625132 | 1.87878897 | 2.178976947 | 1.798966438 |
| ENSMUSG00000057497 | 2.00683675 | 1.834962501 | 1.852347734 | 2.029735268 | 1.945998439 | 2.170261715 | 1.808770687 |
| ENSMUSG00000057499 | 0 | 0.396240625 | 0 | 0.25 | 0.387761935 | 0 | 0.520678036 |
| ENSMUSG00000057503 | 0 | 0 | 0 | 0 | 0 | 0 | 0 |
| ENSMUSG00000057506 | 2.183765914 | 2.086073977 | 2.048689214 | 2.118933358 | 2.091002363 | 2.334453751 | 1.969165684 |
| ENSMUSG00000057513 | 0 | 0 | 0 | 0 | 0 | 0 | 0 |
| ENSMUSG00000057522 | 2.897114011 | 2.68155454 | 2.689555804 | 2.57508814 | 2.521383579 | 2.920645001 | 2.36247044 |
| ENSMUSG00000057530 | 2.669114668 | 2.455443495 | 2.498234584 | 2.413659007 | 2.435655147 | 2.71432976 | 2.31789776 |
| ENSMUSG00000057531 | 2.405576699 | 2.250703754 | 2.269871196 | 2.229715809 | 2.252501045 | 2.51851394 | 2.142987464 |
| ENSMUSG00000057534 | 0 | 0 | 0 | 0 | 0 | 0 | 0 |
| ENSMUSG00000057540 | 0 | 0 | 0 | 0 | 0 | 0 | 0 |
| ENSMUSG00000057541 | 2.575536697 | 2.31728167 | 2.343759858 | 2.390560606 | 2.349532392 | 2.636487896 | 2.153137801 |
| ENSMUSG00000057551 | 2.116511873 | 1.896240625 | 1.866901388 | 1.969129237 | 1.83872004 | 2.139879086 | 1.735976162 |
| ENSMUSG00000057554 | 2.422183552 | 2.228222334 | 2.200224975 | 2.43325383 | 2.356272688 | 2.674861141 | 2.178359564 |
| ENSMUSG00000057561 | 2.789779581 | 2.577084758 | 2.558404919 | 2.514159429 | 2.446849937 | 2.800717078 | 2.260561074 |
| ENSMUSG00000057564 | 0 | 0 | 0 | 0 | 0 | 0 | 0 |
| ENSMUSG00000057572 | 1.964026642 | 1.884789703 | 1.80479213 | 1.78743678 | 1.706995316 | 1.949390007 | 1.605556929 |
| ENSMUSG00000057577 | 0.960252568 | 0.75 | 0.92510993 | 1.02186571 | 0.978602168 | 0.602059991 | 0.853775828 |
| ENSMUSG00000057580 | 0.578129653 | 0.25 | 0.25 | 0.75 | 0.846351821 | 0.602059991 | 0.672731473 |
| ENSMUSG00000057592 | 0 | 0 | 0 | 0 | 0 | 0 | 0 |
| ENSMUSG00000057594 | 1.777675583 | 1.705044741 | 1.688721876 | 1.850219859 | 1.806427258 | 2.017033339 | 1.71730663 |
| ENSMUSG00000057596 | 0 | 0 | 0 | 0 | 0 | 0 | 0 |
| ENSMUSG00000057604 | 1.912079952 | 1.794977273 | 1.916333979 | 0.896240625 | 1.244650542 | 0.903089987 | 1.23151414 |
| ENSMUSG00000057605 | 0.45815691 | 0.580482024 | 0 | 0.646240625 | 0.568060967 | 0 | 0.448487648 |

| | | | | | | | |
|---|---|---|---|---|---|---|---|
| ENSMUSG00000057606 | 0.289064826 | 0.25 | 0 | 0.25 | 0.387761935 | 0.301029996 | 0.224243824 |
| ENSMUSG00000057609 | 0 | 0 | 0 | 0 | 0 | 0 | 0 |
| ENSMUSG00000057612 | 0 | 0 | 0 | 0 | 0 | 0 | 0 |
| ENSMUSG00000057614 | 2.605636262 | 2.399047439 | 2.354463129 | 1.854463129 | 1.777823267 | 2.08278537 | 1.663407615 |
| ENSMUSG00000057615 | 0 | 0 | 0 | 0 | 0 | 0 | 0 |
| ENSMUSG00000057626 | 0 | 0 | 0 | 0 | 0 | 0 | 0 |
| ENSMUSG00000057637 | 2.489144387 | 2.307805295 | 2.280383379 | 2.326515422 | 2.321205208 | 2.661812686 | 2.164188739 |
| ENSMUSG00000057649 | 2.373064211 | 2.167221246 | 2.180274797 | 2.410061234 | 2.39421617 | 2.72427587 | 2.23151414 |
| ENSMUSG00000057650 | 0 | 0 | 0 | 0 | 0 | 0 | 0 |
| ENSMUSG00000057654 | 0 | 0 | 0 | 0 | 0 | 0 | 0 |
| ENSMUSG00000057657 | 0 | 0.396240625 | 0 | 0 | 0 | 0 | 0.448487648 |
| ENSMUSG00000057666 | 3.395261308 | 3.134243916 | 3.148114259 | 2.859047749 | 2.9199446 | 3.312177356 | 2.803905701 |
| ENSMUSG00000057667 | 1.45815691 | 1.26109853 | 1.302363341 | 1.396240625 | 1.27449559 | 1.505149978 | 1.259064007 |
| ENSMUSG00000057672 | 2.057835465 | 2.004202072 | 2.086073977 | 1.789967834 | 1.917865457 | 1.949390007 | 1.945529784 |
| ENSMUSG00000057674 | 0 | 0 | 0 | 0 | 0 | 0 | 0 |
| ENSMUSG00000057685 | 0 | 0 | 0.396240625 | 0.25 | 0 | 0 | 0 |
| ENSMUSG00000057691 | 2.020182925 | 1.910964047 | 1.919870025 | 1.866901388 | 1.882215755 | 2.08278537 | 1.885753983 |
| ENSMUSG00000057696 | 0 | 0 | 0 | 0 | 0 | 0 | 0 |
| ENSMUSG00000057697 | 0 | 0 | 0 | 0 | 0 | 0 | 0 |
| ENSMUSG00000057699 | 0 | 0 | 0 | 0 | 0 | 0 | 0 |
| ENSMUSG00000057706 | 1.678702042 | 1.602347734 | 1.606566189 | 1.130890489 | 1.234113756 | 1.146128036 | 1.168187745 |
| ENSMUSG00000057710 | 0 | 0 | 0 | 0 | 0 | 0 | 0 |
| ENSMUSG00000057715 | 0.45815691 | 0.25 | 0.396240625 | 0 | 0.387761935 | 0 | 0.224243824 |
| ENSMUSG00000057716 | 0.578129653 | 0 | 0 | 0.25 | 0 | 0.301029996 | 0 |
| ENSMUSG00000057719 | 0.671187741 | 0.396240625 | 0.5 | 1.438721876 | 1.401338844 | 1.633468456 | 1.360283038 |
| ENSMUSG00000057722 | 0 | 0.5 | 0.5 | 0 | 0.489301084 | 0.301029996 | 0.224243824 |
| ENSMUSG00000057723 | 0 | 0 | 0 | 0 | 0 | 0 | 0 |
| ENSMUSG00000057726 | 0.671187741 | 0.396240625 | 0.25 | 0 | 0.387761935 | 0.477121255 | 0 |
| ENSMUSG00000057729 | 0.289064826 | 0.5 | 0.25 | 0 | 0 | 0 | 0.224243824 |
| ENSMUSG00000057735 | 0 | 0 | 0 | 0 | 0 | 0 | 0 |
| ENSMUSG00000057738 | 3.43152763 | 3.128616876 | 3.112222657 | 3.176947624 | 3.188044128 | 3.635885685 | 2.980107736 |
| ENSMUSG00000057751 | 0 | 0 | 0 | 1.561981878 | 1.880506521 | 1.672097858 | 1.975716277 |
| ENSMUSG00000057754 | 0 | 0 | 0 | 0 | 0 | 0 | 0 |
| ENSMUSG00000057761 | 0 | 0 | 0 | 0 | 0 | 0 | 0 |

| | | | | | | | |
|---|---|---|---|---|---|---|---|
| ENSMUSG00000057762 | 0 | 0 | 0 | 0 | 0 | 0 | 0 |
| ENSMUSG00000057766 | 1.249317394 | 1.214495249 | 1.380890489 | 1.649978211 | 1.555378544 | 1.672097858 | 1.505628092 |
| ENSMUSG00000057770 | 0 | 0 | 0 | 0 | 0 | 0 | 0 |
| ENSMUSG00000057777 | 0 | 0 | 0 | 0 | 0 | 0 | 0 |
| ENSMUSG00000057778 | 1.771760131 | 1.681980114 | 1.698603967 | 1.678561379 | 1.652586889 | 1.72427587 | 1.559435575 |
| ENSMUSG00000057788 | 2.092368209 | 1.935366747 | 1.926839783 | 1.735628626 | 1.940259189 | 2.029383778 | 1.927642465 |
| ENSMUSG00000057789 | 2.482695304 | 2.330482024 | 2.333288838 | 2.300530956 | 2.422990063 | 2.63748973 | 2.310046902 |
| ENSMUSG00000057799 | 0 | 0 | 0 | 0 | 0 | 0 | 0 |
| ENSMUSG00000057801 | 0 | 0 | 0 | 0 | 0 | 0 | 0 |
| ENSMUSG00000057802 | 0.578129653 | 0.25 | 0 | 0.5 | 0.733951626 | 0.477121255 | 0.355418052 |
| ENSMUSG00000057805 | 0 | 0 | 0 | 0 | 0 | 0 | 0 |
| ENSMUSG00000057816 | 0.289064826 | 0 | 0 | 0 | 0.244650542 | 0 | 0 |
| ENSMUSG00000057817 | 0 | 0 | 0 | 0 | 0 | 0 | 0 |
| ENSMUSG00000057818 | 0.289064826 | 0 | 0.25 | 0 | 0 | 0.301029996 | 0 |
| ENSMUSG00000057829 | 0 | 0 | 0 | 0 | 0 | 0 | 0 |
| ENSMUSG00000057832 | 0 | 0 | 0 | 0 | 0 | 0 | 0 |
| ENSMUSG00000057835 | 1.494443473 | 1.488549078 | 1.438721876 | 1.482684334 | 1.523883869 | 1.73239376 | 1.340368108 |
| ENSMUSG00000057836 | 0 | 0 | 0 | 0 | 0 | 0 | 0.224243824 |
| ENSMUSG00000057841 | 3.166052246 | 2.87007133 | 2.897880587 | 2.957430684 | 2.854039408 | 3.342817315 | 2.670467652 |
| ENSMUSG00000057842 | 1.205378646 | 1.238549078 | 1.321350555 | 1.527131114 | 1.394615667 | 1.633468456 | 1.296434211 |
| ENSMUSG00000057855 | 0 | 0 | 0 | 0 | 0 | 0 | 0 |
| ENSMUSG00000057858 | 2.334723298 | 2.158248799 | 2.175975893 | 2.174241882 | 2.095689764 | 2.372912003 | 1.962479708 |
| ENSMUSG00000057863 | 2.010213549 | 2.066696635 | 2.24929487 | 1.598079356 | 1.910073614 | 1.812913357 | 2.144701457 |
| ENSMUSG00000057880 | 0.671187741 | 1.02186571 | 0.92510993 | 1.898114259 | 1.806427258 | 2.133538908 | 1.74492186 |
| ENSMUSG00000057886 | 0 | 0 | 0 | 0.25 | 0 | 0 | 0 |
| ENSMUSG00000057890 | 0 | 0 | 0 | 0 | 0 | 0 | 0 |
| ENSMUSG00000057894 | 1.978790393 | 1.80479213 | 1.802363341 | 1.75835575 | 1.706995316 | 1.886490725 | 1.591689859 |
| ENSMUSG00000057895 | 0 | 0 | 0 | 1.226722649 | 1.163285805 | 1.146128036 | 1 |
| ENSMUSG00000057897 | 0.289064826 | 0.5 | 0.396240625 | 0.701838731 | 0.489301084 | 0.477121255 | 0.672731473 |
| ENSMUSG00000057899 | 0 | 0.25 | 0.25 | 0 | 0 | 0.301029996 | 0.355418052 |
| ENSMUSG00000057903 | 0 | 0 | 0 | 0 | 0 | 0 | 0 |
| ENSMUSG00000057913 | 0 | 0 | 0.25 | 0 | 0 | 0 | 0 |
| ENSMUSG00000057914 | 1.578129653 | 1.5 | 1.431980114 | 1.464495249 | 1.414412788 | 1.431363764 | 1.360283038 |
| ENSMUSG00000057924 | 1.358731791 | 1.356566189 | 1.238549078 | 1.438721876 | 1.433158226 | 1.51851394 | 1.238624656 |

| | | | | | | | |
|---|---|---|---|---|---|---|---|
| ENSMUSG00000057933 | 0 | 0 | 0 | 0 | 0 | 0 | 0 |
| ENSMUSG00000057945 | 0 | 0 | 0 | 0 | 0 | 0 | 0 |
| ENSMUSG00000057948 | 1.249317394 | 1.27186571 | 1.372963274 | 1.080482024 | 1.234113756 | 0.954242509 | 1.159323753 |
| ENSMUSG00000057963 | 1.527823874 | 1.476722649 | 1.561981878 | 1.372963274 | 1.427019704 | 1.556302501 | 1.476637173 |
| ENSMUSG00000057967 | 0.578129653 | 0.830482024 | 0.701838731 | 0.864857905 | 1.074582839 | 1 | 1.168187745 |
| ENSMUSG00000057969 | 1.470606563 | 1.451838731 | 1.584962501 | 1.380890489 | 1.49445382 | 1.505149978 | 1.548827561 |
| ENSMUSG00000057977 | 0 | 0 | 0 | 0 | 0 | 0 | 0 |
| ENSMUSG00000057981 | 0 | 0 | 0 | 0 | 0 | 0 | 0 |
| ENSMUSG00000057982 | 1.587501561 | 1.532320754 | 1.516522298 | 1.752806814 | 1.611014646 | 1.740362689 | 1.502532226 |
| ENSMUSG00000057990 | 0 | 0.5 | 0 | 0.396240625 | 0 | 0 | 0.224243824 |
| ENSMUSG00000057997 | 0 | 0 | 0 | 0 | 0 | 0 | 0 |
| ENSMUSG00000058005 | 0 | 0 | 0 | 0 | 0 | 0 | 0 |
| ENSMUSG00000058006 | 2.936861443 | 2.680701883 | 2.661864607 | 2.671187155 | 2.635654347 | 2.967079734 | 2.42257867 |
| ENSMUSG00000058013 | 3.658720998 | 3.304527277 | 3.262589712 | 3.302781955 | 3.226141448 | 3.754195388 | 2.984631604 |
| ENSMUSG00000058014 | 0 | 0 | 0 | 0 | 0 | 0 | 0 |
| ENSMUSG00000058017 | 0 | 0 | 0 | 0 | 0 | 0 | 0 |
| ENSMUSG00000058019 | 0 | 0 | 0.25 | 0 | 0.244650542 | 0 | 0.224243824 |
| ENSMUSG00000058022 | 0 | 0 | 0.5 | 0.25 | 0.632412477 | 0.301029996 | 0.224243824 |
| ENSMUSG00000058028 | 0 | 0 | 0 | 0.25 | 0 | 0 | 0 |
| ENSMUSG00000058030 | 0 | 0 | 0 | 0 | 0 | 0 | 0 |
| ENSMUSG00000058046 | 0.578129653 | 0.580482024 | 0.75 | 0.75 | 0.812711509 | 0.602059991 | 0.579661877 |
| ENSMUSG00000058050 | 0.289064826 | 0 | 0.25 | 0.396240625 | 0.387761935 | 0.477121255 | 0.355418052 |
| ENSMUSG00000058052 | 0 | 0 | 0 | 0 | 0 | 0 | 0 |
| ENSMUSG00000058056 | 2.821721112 | 2.592399337 | 2.571928095 | 2.273189285 | 2.328502976 | 2.550228353 | 2.174874624 |
| ENSMUSG00000058057 | 0.289064826 | 0.5 | 0.5 | 0.5 | 0.387761935 | 0.698970004 | 0.224243824 |
| ENSMUSG00000058063 | 0 | 0 | 0 | 0 | 0 | 0 | 0 |
| ENSMUSG00000058064 | 1.129344651 | 1.02186571 | 1.114857905 | 0.92510993 | 1.176121988 | 0.954242509 | 1.028149525 |
| ENSMUSG00000058070 | 2.623015126 | 2.427270953 | 2.42813175 | 1.985628626 | 1.990091809 | 2.23299611 | 1.89141329 |
| ENSMUSG00000058071 | 0 | 0 | 0 | 0 | 0 | 0 | 0 |
| ENSMUSG00000058073 | 0 | 0 | 0 | 0 | 0 | 0 | 0 |
| ENSMUSG00000058076 | 2.539328946 | 2.381869252 | 2.358135475 | 2.503156135 | 2.432082686 | 2.769377326 | 2.280542874 |
| ENSMUSG00000058078 | 0 | 0 | 0 | 0 | 0 | 0 | 0 |
| ENSMUSG00000058084 | 0 | 0 | 0 | 0 | 0 | 0 | 0 |
| ENSMUSG00000058093 | 1.822201287 | 1.741446071 | 1.67510993 | 1.888647213 | 1.824953989 | 2.017033339 | 1.68886578 |

| | | | | | | | |
|---|---|---|---|---|---|---|---|
| ENSMUSG00000058099 | 0 | 0 | 0 | 0.25 | 0 | 0 | 0 |
| ENSMUSG00000058101 | 0 | 0 | 0.25 | 0 | 0 | 0 | 0.224243824 |
| ENSMUSG00000058114 | 0 | 0 | 0 | 0 | 0 | 0 | 0 |
| ENSMUSG00000058119 | 0 | 0 | 0 | 0 | 0 | 0 | 0 |
| ENSMUSG00000058124 | 0 | 0 | 0 | 0 | 0 | 0 | 0 |
| ENSMUSG00000058126 | 2.516991255 | 2.361253711 | 2.33272922 | 2.211372513 | 2.232221225 | 2.541579244 | 2.087847416 |
| ENSMUSG00000058132 | 0 | 0 | 0 | 0 | 0 | 0 | 0 |
| ENSMUSG00000058135 | 2.457006475 | 2.25 | 2.29248125 | 2.05479213 | 1.991345651 | 2.217483944 | 1.94074868 |
| ENSMUSG00000058145 | 0 | 0 | 0 | 0.25 | 0.387761935 | 0.301029996 | 0.579661877 |
| ENSMUSG00000058147 | 0 | 0 | 0 | 0 | 0 | 0 | 0 |
| ENSMUSG00000058152 | 0 | 0 | 0 | 0.5 | 0.387761935 | 0.477121255 | 0.355418052 |
| ENSMUSG00000058153 | 0 | 0 | 0 | 0 | 0 | 0 | 0 |
| ENSMUSG00000058159 | 0.91631382 | 0.830482024 | 0.25 | 0.396240625 | 0.387761935 | 0 | 0.448487648 |
| ENSMUSG00000058163 | 0.289064826 | 0.580482024 | 0.5 | 0.580482024 | 0.733951626 | 0.602059991 | 0 |
| ENSMUSG00000058172 | 0 | 0 | 0 | 0 | 0 | 0 | 0 |
| ENSMUSG00000058173 | 1.036286563 | 1.160964047 | 1.42510993 | 1.302363341 | 1.310728002 | 1.204119983 | 1.329931221 |
| ENSMUSG00000058174 | 0.289064826 | 0 | 0 | 0 | 0 | 0 | 0.224243824 |
| ENSMUSG00000058176 | 0 | 0 | 0 | 0 | 0 | 0 | 0 |
| ENSMUSG00000058183 | 0.578129653 | 0.580482024 | 0.396240625 | 0.646240625 | 0.686820904 | 0.477121255 | 0.579661877 |
| ENSMUSG00000058186 | 0.289064826 | 0 | 0 | 0.25 | 0 | 0 | 0 |
| ENSMUSG00000058188 | 0 | 0 | 0 | 0 | 0 | 0 | 0 |
| ENSMUSG00000058192 | 1.771760131 | 1.62694866 | 1.602347734 | 1.720660762 | 1.611014646 | 1.770852012 | 1.462868481 |
| ENSMUSG00000058194 | 0 | 0 | 0 | 0 | 0 | 0 | 0 |
| ENSMUSG00000058200 | 0 | 0 | 0 | 0 | 0 | 0 | 0 |
| ENSMUSG00000058207 | 0 | 0.25 | 0.396240625 | 0 | 0 | 0 | 0.224243824 |
| ENSMUSG00000058216 | 0.289064826 | 0 | 0 | 0 | 0 | 0 | 0 |
| ENSMUSG00000058230 | 2.896712826 | 2.666111821 | 2.633332433 | 2.574229052 | 2.499532413 | 2.868056362 | 2.333050044 |
| ENSMUSG00000058239 | 2.116511873 | 2.01109853 | 2.058404919 | 1.80479213 | 1.868304353 | 2.045322979 | 1.838725884 |
| ENSMUSG00000058240 | 2.245508853 | 2.004202072 | 2.066696635 | 2.122963274 | 2.084333628 | 2.372912003 | 1.927642465 |
| ENSMUSG00000058244 | 0 | 0 | 0 | 0 | 0 | 0 | 0 |
| ENSMUSG00000058246 | 0 | 0 | 0 | 0 | 0 | 0 | 0 |
| ENSMUSG00000058248 | 0 | 0.396240625 | 0.396240625 | 0 | 0.244650542 | 0 | 0.355418052 |
| ENSMUSG00000058250 | 0 | 0 | 0 | 0 | 0 | 0 | 0 |
| ENSMUSG00000058251 | 0 | 0 | 0 | 0 | 0 | 0 | 0 |

| | | | | | | | |
|---|---|---|---|---|---|---|---|
| ENSMUSG00000058252 | 0 | 0 | 0 | 0 | 0 | 0 | 0.224243824 |
| ENSMUSG00000058254 | 1.842799666 | 1.711372513 | 1.720660762 | 1.51109853 | 1.401338844 | 1.568201724 | 1.452143229 |
| ENSMUSG00000058258 | 2.572927518 | 2.405512955 | 2.407794264 | 2.660964047 | 2.598242538 | 3.002597981 | 2.395988891 |
| ENSMUSG00000058260 | 0.960252568 | 0.79248125 | 0.701838731 | 0.5 | 0.244650542 | 0.301029996 | 0.224243824 |
| ENSMUSG00000058267 | 2.410757293 | 2.197790722 | 2.260414788 | 2.319612365 | 2.267696867 | 2.606381365 | 2.16539364 |
| ENSMUSG00000058270 | 0 | 0 | 0 | 0 | 0 | 0 | 0 |
| ENSMUSG00000058275 | 0 | 0 | 0 | 0 | 0 | 0 | 0 |
| ENSMUSG00000058281 | 0.289064826 | 0 | 0 | 0 | 0.387761935 | 0 | 0 |
| ENSMUSG00000058287 | 0 | 0 | 0.25 | 0 | 0 | 0.602059991 | 0 |
| ENSMUSG00000058290 | 2.789260561 | 2.585240691 | 2.625459296 | 2.526149688 | 2.659629116 | 2.899820502 | 2.512692936 |
| ENSMUSG00000058291 | 2.544030601 | 2.322504712 | 2.31728167 | 2.362802778 | 2.297309926 | 2.629409599 | 2.113781657 |
| ENSMUSG00000058295 | 0 | 0 | 0 | 0 | 0 | 0 | 0 |
| ENSMUSG00000058297 | 1.445324132 | 1.226722649 | 1.238549078 | 1.575945187 | 1.704182902 | 1.897627091 | 1.596378571 |
| ENSMUSG00000058298 | 2.026699189 | 1.814346961 | 1.811981878 | 1.942046081 | 1.890639929 | 2.08278537 | 1.720494 |
| ENSMUSG00000058301 | 2.56765967 | 2.434523065 | 2.463717096 | 2.298073204 | 2.444777781 | 2.608526034 | 2.338730052 |
| ENSMUSG00000058317 | 1.789260561 | 1.794977273 | 1.811981878 | 1.75 | 1.698489937 | 1.924279286 | 1.676387053 |
| ENSMUSG00000058318 | 2.262379879 | 2.08272922 | 2.044977273 | 2.122963274 | 2.113840608 | 2.374748346 | 1.94074868 |
| ENSMUSG00000058325 | 2.553277818 | 2.350753006 | 2.334405475 | 2.277131114 | 2.228405446 | 2.517195898 | 2.088864761 |
| ENSMUSG00000058328 | 0 | 0 | 0 | 0 | 0 | 0 | 0 |
| ENSMUSG00000058331 | 1.036286563 | 1.02186571 | 1.098079356 | 1.282320754 | 1.136121934 | 1.278753601 | 1.159323753 |
| ENSMUSG00000058349 | 0 | 0 | 0 | 0 | 0 | 0 | 0 |
| ENSMUSG00000058351 | 1.181541736 | 1.26109853 | 1.146240625 | 1.146240625 | 1.27449559 | 1.414973348 | 1.307989563 |
| ENSMUSG00000058354 | 0 | 0 | 0 | 0 | 0 | 0 | 0 |
| ENSMUSG00000058355 | 3.197062863 | 2.932935823 | 2.914441259 | 3.083358729 | 2.97197586 | 3.47070443 | 2.741427268 |
| ENSMUSG00000058357 | 0 | 0 | 0 | 0 | 0.387761935 | 0 | 0.224243824 |
| ENSMUSG00000058360 | 1.069666964 | 1.04248125 | 1 | 1.061981878 | 0.905314583 | 0.903089987 | 0.876096088 |
| ENSMUSG00000058368 | 0 | 0 | 0 | 0 | 0 | 0 | 0 |
| ENSMUSG00000058385 | 0 | 0 | 0 | 0 | 0 | 0 | 0 |
| ENSMUSG00000058388 | 2.481610694 | 2.294354884 | 2.253505118 | 2.253505118 | 2.205967036 | 2.507855872 | 2.089878917 |
| ENSMUSG00000058392 | 2.420929319 | 2.266522298 | 2.250703754 | 2.224461364 | 2.277846464 | 2.501059262 | 2.134278581 |
| ENSMUSG00000058396 | 0 | 0.5 | 0.701838731 | 0.646240625 | 0.686820904 | 0.477121255 | 0.829800754 |
| ENSMUSG00000058398 | 0 | 0 | 0 | 0.25 | 0 | 0 | 0 |
| ENSMUSG00000058399 | 0 | 0 | 0 | 0 | 0 | 0 | 0 |
| ENSMUSG00000058400 | 0 | 0.396240625 | 0.25 | 0 | 0 | 0 | 0.224243824 |

| | | | | | | | |
|---|---|---|---|---|---|---|---|
| ENSMUSG00000058402 | 1.389637216 | 1.226722649 | 1.321350555 | 1.380890489 | 1.27449559 | 1.301029996 | 1.168187745 |
| ENSMUSG00000058407 | 2.66563648 | 2.428990498 | 2.410964047 | 2.588837774 | 2.508030729 | 2.87909588 | 2.323847637 |
| ENSMUSG00000058420 | 1.129344651 | 1.26109853 | 1.188721876 | 0.92510993 | 1.039257769 | 1.113943352 | 1.041356071 |
| ENSMUSG00000058427 | 0.45815691 | 0 | 0.25 | 0.75 | 0.387761935 | 0.84509804 | 0.629532004 |
| ENSMUSG00000058430 | 0 | 0 | 0 | 0 | 0 | 0 | 0 |
| ENSMUSG00000058435 | 0 | 0 | 0 | 0 | 0 | 0 | 0 |
| ENSMUSG00000058440 | 2.309247751 | 2.169870025 | 2.219129237 | 2.168106335 | 2.185632262 | 2.456366033 | 2.085803078 |
| ENSMUSG00000058441 | 0 | 0.25 | 0 | 0 | 0 | 0.301029996 | 0.579661877 |
| ENSMUSG00000058443 | 0.671187741 | 0.580482024 | 0.580482024 | 0.830482024 | 0.812711509 | 0.903089987 | 0.829800754 |
| ENSMUSG00000058444 | 2.17479713 | 1.978222334 | 1.992885888 | 2.016522298 | 1.958580386 | 2.285557309 | 1.87511425 |
| ENSMUSG00000058446 | 2.258689303 | 2.097004321 | 2.058404919 | 2.109697963 | 2.028157605 | 2.307496038 | 1.838725884 |
| ENSMUSG00000058447 | 0 | 0 | 0 | 0 | 0 | 0 | 0 |
| ENSMUSG00000058454 | 1.794934559 | 1.571350555 | 1.695339928 | 1.909156155 | 1.910073614 | 2.093421685 | 1.801445495 |
| ENSMUSG00000058463 | 0 | 0 | 0 | 0 | 0 | 0 | 0 |
| ENSMUSG00000058470 | 0 | 0 | 0 | 0 | 0 | 0 | 0 |
| ENSMUSG00000058486 | 1.794934559 | 1.698603967 | 1.823655187 | 1.62694866 | 1.718026092 | 1.77815125 | 1.73898563 |
| ENSMUSG00000058488 | 0 | 0.25 | 0 | 0.25 | 0.387761935 | 0 | 0.355418052 |
| ENSMUSG00000058491 | 0 | 0 | 0 | 0 | 0 | 0 | 0 |
| ENSMUSG00000058492 | 0.289064826 | 0 | 0 | 0 | 0 | 0 | 0 |
| ENSMUSG00000058498 | 0 | 0.646240625 | 0.646240625 | 0.396240625 | 0.632412477 | 0 | 0.224243824 |
| ENSMUSG00000058499 | 0 | 0 | 0 | 0 | 0 | 0 | 0 |
| ENSMUSG00000058503 | 2.302628051 | 2.123963757 | 2.115881093 | 2.226722649 | 2.179810173 | 2.519827994 | 2.007270316 |
| ENSMUSG00000058508 | 0 | 0 | 0 | 0 | 0 | 0 | 0 |
| ENSMUSG00000058513 | 0 | 0 | 0 | 0 | 0 | 0 | 0 |
| ENSMUSG00000058515 | 0 | 0 | 0 | 0 | 0 | 0 | 0 |
| ENSMUSG00000058521 | 0 | 0 | 0 | 0 | 0 | 0 | 0 |
| ENSMUSG00000058523 | 0 | 0 | 0 | 0 | 0 | 0 | 0 |
| ENSMUSG00000058537 | 0 | 0 | 0 | 0 | 0 | 0 | 0 |
| ENSMUSG00000058542 | 0 | 0.25 | 0.25 | 0 | 0 | 0 | 0 |
| ENSMUSG00000058546 | 0.578129653 | 0.75 | 0.701838731 | 0.896240625 | 0.489301084 | 0.954242509 | 0.775756176 |
| ENSMUSG00000058550 | 0 | 0 | 0 | 0 | 0 | 0 | 0 |
| ENSMUSG00000058558 | 3.264212992 | 2.939659909 | 2.941217898 | 2.869945066 | 2.802803318 | 3.224791956 | 2.599746361 |
| ENSMUSG00000058567 | 0 | 0.396240625 | 0 | 0.25 | 0.244650542 | 0 | 0 |
| ENSMUSG00000058568 | 0 | 0 | 0 | 0 | 0 | 0 | 0 |

| | | | | | | | |
|---|---|---|---|---|---|---|---|
| ENSMUSG00000058569 | 2.736751754 | 2.539967834 | 2.54248125 | 2.627443751 | 2.69373167 | 3.045322979 | 2.541390096 |
| ENSMUSG00000058571 | 2.111266113 | 1.955044741 | 1.975216702 | 1.819031101 | 1.791313678 | 2.012837225 | 1.661510159 |
| ENSMUSG00000058579 | 0 | 0 | 0 | 0 | 0 | 0 | 0 |
| ENSMUSG00000058581 | 0 | 0 | 0.25 | 0 | 0 | 0 | 0 |
| ENSMUSG00000058586 | 1.771760131 | 1.51109853 | 1.642463902 | 1.738549078 | 1.642643806 | 1.919078092 | 1.643912836 |
| ENSMUSG00000058587 | 3.015379043 | 2.779735268 | 2.739819376 | 2.804337976 | 2.709948941 | 3.155336037 | 2.522809941 |
| ENSMUSG00000058588 | 0 | 0 | 0 | 0 | 0 | 0 | 0 |
| ENSMUSG00000058589 | 0.45815691 | 0.701838731 | 0.25 | 0.396240625 | 0.387761935 | 0.602059991 | 0.224243824 |
| ENSMUSG00000058590 | 0 | 0 | 0 | 0 | 0 | 0 | 0 |
| ENSMUSG00000058594 | 2.543094501 | 2.311981878 | 2.326515422 | 2.307805295 | 2.313753359 | 2.57054294 | 2.166993225 |
| ENSMUSG00000058600 | 2.99368599 | 2.729715809 | 2.724083101 | 2.742706225 | 2.668192818 | 3.077004327 | 2.448487648 |
| ENSMUSG00000058603 | 0.289064826 | 0.646240625 | 0.396240625 | 0.396240625 | 0.568060967 | 0.477121255 | 0.74492186 |
| ENSMUSG00000058607 | 0 | 0.25 | 0 | 0.396240625 | 0.244650542 | 0 | 0.224243824 |
| ENSMUSG00000058613 | 0 | 0 | 0 | 0 | 0 | 0 | 0 |
| ENSMUSG00000058618 | 0 | 0 | 0 | 0 | 0 | 0 | 0 |
| ENSMUSG00000058620 | 0 | 0.25 | 0 | 0.25 | 0.244650542 | 0 | 0.355418052 |
| ENSMUSG00000058622 | 0 | 0 | 0 | 0.396240625 | 0.387761935 | 0.301029996 | 0.520678036 |
| ENSMUSG00000058624 | 0.811507563 | 0.25 | 0.396240625 | 0.580482024 | 0.489301084 | 0.301029996 | 0.710836105 |
| ENSMUSG00000058625 | 0.45815691 | 0.396240625 | 0.5 | 0.25 | 0.632412477 | 0 | 0.355418052 |
| ENSMUSG00000058626 | 0 | 0 | 0 | 0 | 0.244650542 | 0 | 0.224243824 |
| ENSMUSG00000058628 | 0 | 0 | 0 | 0 | 0 | 0 | 0 |
| ENSMUSG00000058631 | 0 | 0 | 0 | 0 | 0 | 0 | 0 |
| ENSMUSG00000058638 | 1.99653913 | 1.886723615 | 1.834962501 | 1.94697564 | 1.870073561 | 2.10720997 | 1.723650273 |
| ENSMUSG00000058643 | 0 | 0 | 0.25 | 0 | 0 | 0 | 0 |
| ENSMUSG00000058655 | 3.553277818 | 3.210783978 | 3.204695468 | 3.179044101 | 3.127667787 | 3.611510887 | 2.913069858 |
| ENSMUSG00000058656 | 0.747221736 | 1.098079356 | 0.830482024 | 0.864857905 | 0.686820904 | 0.301029996 | 0.629532004 |
| ENSMUSG00000058659 | 0 | 0 | 0 | 0 | 0 | 0 | 0 |
| ENSMUSG00000058662 | 0 | 0 | 0 | 0 | 0 | 0 | 0 |
| ENSMUSG00000058665 | 1.45815691 | 1.356566189 | 1.580482024 | 1.130890489 | 1.310728002 | 1.204119983 | 1.307989563 |
| ENSMUSG00000058669 | 0 | 0 | 0 | 0 | 0 | 0 | 0.224243824 |
| ENSMUSG00000058670 | 0 | 0 | 0.25 | 0 | 0 | 0.301029996 | 0 |
| ENSMUSG00000058672 | 2.292497211 | 2.097004321 | 2.094844592 | 2.139605178 | 2.182004726 | 2.413299764 | 2.089372236 |
| ENSMUSG00000058685 | 0 | 0 | 0 | 0 | 0 | 0 | 0 |
| ENSMUSG00000058690 | 2.37447073 | 2.201838731 | 2.187048212 | 2.439139081 | 2.334234459 | 2.709269961 | 2.172530299 |

| | | | | | | | |
|---|---|---|---|---|---|---|---|
| ENSMUSG00000058692 | 0 | 0 | 0 | 0 | 0 | 0 | 0 |
| ENSMUSG00000058700 | 0.747221736 | 0 | 0.580482024 | 0.5 | 0.489301084 | 0.301029996 | 0.355418052 |
| ENSMUSG00000058704 | 2.541215974 | 2.364857905 | 2.345926073 | 2.276477127 | 2.204601633 | 2.536558443 | 2.082712142 |
| ENSMUSG00000058706 | 2.160971586 | 2.038704527 | 1.976722649 | 2.05479213 | 1.990091809 | 2.328379603 | 1.906037947 |
| ENSMUSG00000058709 | 2.00683675 | 1.958222504 | 2.102347734 | 1.688721876 | 1.851969278 | 1.924279286 | 1.984950056 |
| ENSMUSG00000058715 | 0.45815691 | 0.5 | 0.25 | 0.5 | 0.632412477 | 0.477121255 | 0.355418052 |
| ENSMUSG00000058717 | 0 | 0 | 0 | 0 | 0 | 0 | 0 |
| ENSMUSG00000058725 | 0 | 0 | 0 | 0 | 0 | 0 | 0 |
| ENSMUSG00000058728 | 0 | 0 | 0 | 0 | 0 | 0 | 0 |
| ENSMUSG00000058729 | 2.3855553 | 2.209012589 | 2.183677405 | 2.338837774 | 2.260180728 | 2.612783857 | 2.107130591 |
| ENSMUSG00000058731 | 0 | 0 | 0 | 0 | 0 | 0 | 0 |
| ENSMUSG00000058740 | 0 | 0.396240625 | 0 | 0.396240625 | 0.568060967 | 0.477121255 | 0.520678036 |
| ENSMUSG00000058741 | 0 | 0 | 0 | 0 | 0 | 0.301029996 | 0 |
| ENSMUSG00000058743 | 0.91631382 | 0.5 | 0.646240625 | 0.646240625 | 0.387761935 | 0.77815125 | 0.629532004 |
| ENSMUSG00000058745 | 0 | 0 | 0 | 0 | 0 | 0 | 0 |
| ENSMUSG00000058748 | 1.747221736 | 1.794977273 | 1.589388001 | 1.606566189 | 1.445123986 | 1.653212514 | 1.413583902 |
| ENSMUSG00000058755 | 0 | 0 | 0 | 0 | 0 | 0 | 0 |
| ENSMUSG00000058756 | 2.469490009 | 2.333847589 | 2.365881093 | 2.156427211 | 2.187791022 | 2.414973348 | 2.161765391 |
| ENSMUSG00000058761 | 2.10861828 | 1.914552871 | 1.894357207 | 1.854463129 | 1.836786029 | 2.113943352 | 1.702561601 |
| ENSMUSG00000058773 | 0 | 0.396240625 | 0.25 | 0.396240625 | 0.489301084 | 0 | 0.520678036 |
| ENSMUSG00000058793 | 2.523917219 | 2.331607372 | 2.327084758 | 2.403677461 | 2.392613642 | 2.688419822 | 2.228889254 |
| ENSMUSG00000058794 | 0.578129653 | 0.79248125 | 0.396240625 | 0.580482024 | 0.77552387 | 0.477121255 | 0.579661877 |
| ENSMUSG00000058799 | 3.369333216 | 3.054413708 | 3.03743678 | 3.123401327 | 3.028290245 | 3.514680544 | 2.777392698 |
| ENSMUSG00000058802 | 0 | 0 | 0 | 0 | 0 | 0 | 0 |
| ENSMUSG00000058806 | 0 | 0 | 0 | 0 | 0.244650542 | 0 | 0 |
| ENSMUSG00000058807 | 0 | 0 | 0 | 0 | 0 | 0 | 0 |
| ENSMUSG00000058809 | 0.289064826 | 0.646240625 | 0.701838731 | 0.976722649 | 0.812711509 | 0.954242509 | 0.448487648 |
| ENSMUSG00000058812 | 0 | 0 | 0 | 0 | 0 | 0 | 0.224243824 |
| ENSMUSG00000058816 | 0 | 0 | 0 | 0 | 0 | 0 | 0 |
| ENSMUSG00000058818 | 0.45815691 | 0.580482024 | 0.25 | 0.25 | 0.568060967 | 0.301029996 | 0.355418052 |
| ENSMUSG00000058820 | 0 | 0 | 0 | 0 | 0 | 0 | 0 |
| ENSMUSG00000058831 | 0 | 0.580482024 | 0.75 | 0.580482024 | 0.632412477 | 0.477121255 | 0.448487648 |
| ENSMUSG00000058833 | 1.671187741 | 1.720660762 | 1.898114259 | 1.29248125 | 1.528558853 | 1.591064607 | 1.803905701 |
| ENSMUSG00000058835 | 2.733202521 | 2.484527332 | 2.481944491 | 2.470660762 | 2.415543455 | 2.731588765 | 2.235411964 |

| | | | | | | | |
|---|---|---|---|---|---|---|---|
| ENSMUSG00000058838 | 0 | 0 | 0 | 0 | 0 | 0 | 0 |
| ENSMUSG00000058856 | 0 | 0 | 0 | 0 | 0 | 0 | 0 |
| ENSMUSG00000058869 | 0 | 0.25 | 0 | 0 | 0 | 0 | 0.224243824 |
| ENSMUSG00000058873 | 0.289064826 | 0 | 0 | 0 | 0 | 0 | 0 |
| ENSMUSG00000058881 | 2.377269616 | 2.219895812 | 2.268535366 | 2.01109853 | 2.059432181 | 2.243038049 | 2.026921761 |
| ENSMUSG00000058883 | 0 | 0 | 0 | 0 | 0 | 0 | 0 |
| ENSMUSG00000058884 | 0 | 0 | 0 | 0 | 0 | 0 | 0 |
| ENSMUSG00000058886 | 1.83262764 | 1.79745614 | 1.732684334 | 1.678561379 | 1.744182955 | 1.763427994 | 1.694069894 |
| ENSMUSG00000058897 | 0 | 0.396240625 | 0.5 | 0.396240625 | 0.244650542 | 0.698970004 | 0.448487648 |
| ENSMUSG00000058900 | 0.811507563 | 1.114857905 | 0.580482024 | 1.25 | 1.212046813 | 1.301029996 | 1.20919388 |
| ENSMUSG00000058904 | 0 | 0 | 0 | 0 | 0 | 0 | 0 |
| ENSMUSG00000058905 | 0 | 0.25 | 0 | 0 | 0 | 0 | 0 |
| ENSMUSG00000058908 | 0 | 0 | 0 | 0 | 0 | 0 | 0 |
| ENSMUSG00000058914 | 0.289064826 | 0.396240625 | 0 | 0 | 0.387761935 | 0.301029996 | 0.224243824 |
| ENSMUSG00000058921 | 0 | 0 | 0 | 0.396240625 | 0 | 0 | 0 |
| ENSMUSG00000058922 | 0 | 0 | 0 | 0 | 0 | 0 | 0 |
| ENSMUSG00000058923 | 0 | 0 | 0 | 0 | 0 | 0 | 0 |
| ENSMUSG00000058925 | 0 | 0 | 0 | 0 | 0 | 0 | 0 |
| ENSMUSG00000058927 | 0 | 0 | 0 | 0 | 0 | 0 | 0 |
| ENSMUSG00000058932 | 0 | 0 | 0 | 0 | 0 | 0 | 0 |
| ENSMUSG00000058933 | 0 | 0 | 0.25 | 0 | 0 | 0 | 0 |
| ENSMUSG00000058934 | 0 | 0 | 0 | 0 | 0.244650542 | 0 | 0 |
| ENSMUSG00000058935 | 0 | 0 | 0 | 0 | 0 | 0 | 0 |
| ENSMUSG00000058952 | 0.289064826 | 0.396240625 | 0 | 0 | 0 | 0 | 0 |
| ENSMUSG00000058953 | 0 | 0 | 0 | 0 | 0 | 0 | 0 |
| ENSMUSG00000058966 | 0 | 0 | 0.25 | 0 | 0.686820904 | 0.477121255 | 0.672731473 |
| ENSMUSG00000058975 | 0 | 0.25 | 0 | 0.25 | 0 | 0.301029996 | 0 |
| ENSMUSG00000058976 | 0 | 0.396240625 | 0 | 0 | 0.244650542 | 0.301029996 | 0 |
| ENSMUSG00000058979 | 1.837744666 | 1.747171172 | 1.784887838 | 1.763820609 | 1.85749873 | 1.968482949 | 1.723650273 |
| ENSMUSG00000058981 | 0 | 0 | 0 | 0 | 0 | 0 | 0 |
| ENSMUSG00000058997 | 2.251208701 | 2.074802005 | 2.04745614 | 2.271202097 | 2.242813819 | 2.507855872 | 2.098866035 |
| ENSMUSG00000059003 | 0.289064826 | 0.25 | 0 | 0.25 | 0.489301084 | 0 | 0 |
| ENSMUSG00000059005 | 3.299889412 | 2.989547546 | 2.988003291 | 3.196158711 | 3.082746031 | 3.597585502 | 2.853775828 |
| ENSMUSG00000059013 | 0.960252568 | 1.214495249 | 1.282320754 | 1 | 1.106691885 | 0.903089987 | 1.319146374 |

| | | | | | | | |
|---|---|---|---|---|---|---|---|
| ENSMUSG00000059014 | 0 | 0 | 0 | 0 | 0 | 0 | 0 |
| ENSMUSG00000059022 | 0.578129653 | 0.701838731 | 0.92510993 | 0.79248125 | 1 | 0.84509804 | 1.20919388 |
| ENSMUSG00000059023 | 0 | 0 | 0 | 0 | 0 | 0 | 0 |
| ENSMUSG00000059027 | 0.811507563 | 0.79248125 | 0.701838731 | 0.396240625 | 0.877063019 | 0.84509804 | 0.803905701 |
| ENSMUSG00000059030 | 0 | 0 | 0 | 0 | 0 | 0 | 0 |
| ENSMUSG00000059031 | 0 | 0 | 0 | 0 | 0 | 0 | 0 |
| ENSMUSG00000059033 | 0.289064826 | 0 | 0.25 | 0 | 0 | 0 | 0.224243824 |
| ENSMUSG00000059040 | 1.777675583 | 1.866901388 | 1.970660762 | 1.380890489 | 1.78687392 | 1.491361694 | 1.869159809 |
| ENSMUSG00000059041 | 0 | 0 | 0 | 0 | 0 | 0 | 0 |
| ENSMUSG00000059042 | 0 | 0 | 0 | 0 | 0 | 0 | 0 |
| ENSMUSG00000059043 | 0 | 0 | 0 | 0 | 0 | 0 | 0 |
| ENSMUSG00000059049 | 0.747221736 | 0.5 | 0.25 | 0.646240625 | 0.244650542 | 0.477121255 | 0.448487648 |
| ENSMUSG00000059058 | 0.91631382 | 0.830482024 | 0.75 | 1.226722649 | 1.106691885 | 1.204119983 | 1.089372236 |
| ENSMUSG00000059060 | 1.181541736 | 1.226722649 | 1.282320754 | 1.25 | 1.136121934 | 1.204119983 | 1.159323753 |
| ENSMUSG00000059064 | 0 | 0.396240625 | 0 | 0 | 0 | 0 | 0.224243824 |
| ENSMUSG00000059069 | 0 | 0 | 0 | 0 | 0 | 0 | 0 |
| ENSMUSG00000059070 | 2.820759099 | 2.638407323 | 2.718745337 | 2.528435542 | 2.640660901 | 2.934498451 | 2.584307307 |
| ENSMUSG00000059077 | 0 | 0 | 0 | 0 | 0 | 0 | 0 |
| ENSMUSG00000059087 | 0 | 0 | 0 | 0 | 0 | 0 | 0 |
| ENSMUSG00000059089 | 0 | 0 | 0 | 0 | 0.244650542 | 0 | 0 |
| ENSMUSG00000059101 | 0 | 0 | 0 | 0 | 0 | 0 | 0 |
| ENSMUSG00000059105 | 0 | 0 | 0 | 0 | 0 | 0 | 0 |
| ENSMUSG00000059106 | 0 | 0 | 0 | 0 | 0 | 0 | 0 |
| ENSMUSG00000059108 | 0 | 0 | 0 | 0 | 0 | 0 | 0 |
| ENSMUSG00000059112 | 0 | 0 | 0 | 0 | 0 | 0 | 0 |
| ENSMUSG00000059113 | 0 | 0 | 0 | 0 | 0 | 0 | 0 |
| ENSMUSG00000059114 | 0 | 0 | 0 | 0 | 0 | 0 | 0 |
| ENSMUSG00000059119 | 2.679378494 | 2.480088214 | 2.483791262 | 2.472185812 | 2.451298303 | 2.767155866 | 2.285558718 |
| ENSMUSG00000059128 | 0 | 0 | 0 | 0 | 0 | 0 | 0 |
| ENSMUSG00000059134 | 0 | 0 | 0 | 0 | 0 | 0 | 0 |
| ENSMUSG00000059136 | 0 | 0 | 0 | 0 | 0 | 0 | 0 |
| ENSMUSG00000059142 | 2.051791411 | 1.870953944 | 1.828220739 | 1.890560606 | 1.830919587 | 2.004321374 | 1.676387053 |
| ENSMUSG00000059146 | 1.069666964 | 1.130890489 | 1.061981878 | 0 | 0 | 0 | 0.224243824 |
| ENSMUSG00000059149 | 1.907802659 | 1.841580554 | 1.821350555 | 1.845926073 | 1.797870327 | 2.004321374 | 1.749303626 |

| | | | | | | | |
|---|---|---|---|---|---|---|---|
| ENSMUSG00000059150 | 0.289064826 | 0 | 0 | 0.25 | 0.387761935 | 0 | 0 |
| ENSMUSG00000059159 | 0 | 0 | 0 | 0 | 0 | 0 | 0 |
| ENSMUSG00000059169 | 0 | 0 | 0 | 0 | 0 | 0 | 0 |
| ENSMUSG00000059173 | 0.289064826 | 0.396240625 | 0.830482024 | 0.75 | 0.632412477 | 0.477121255 | 0.629532004 |
| ENSMUSG00000059179 | 0 | 0 | 0 | 0.25 | 0 | 0 | 0 |
| ENSMUSG00000059182 | 2.528794847 | 2.269203899 | 2.269871196 | 2.19697564 | 2.158641579 | 2.439332694 | 2.001334086 |
| ENSMUSG00000059183 | 1.881174527 | 1.75835575 | 1.741446071 | 1.819031101 | 1.810629166 | 2.041392685 | 1.74196736 |
| ENSMUSG00000059187 | 0 | 0 | 0 | 0 | 0.244650542 | 0 | 0.448487648 |
| ENSMUSG00000059189 | 0 | 0 | 0 | 0 | 0 | 0 | 0 |
| ENSMUSG00000059195 | 0.578129653 | 0.646240625 | 0.646240625 | 0 | 0.387761935 | 0 | 0.579661877 |
| ENSMUSG00000059201 | 0 | 0 | 0 | 0 | 0 | 0 | 0 |
| ENSMUSG00000059203 | 0 | 0 | 0 | 0 | 0 | 0 | 0 |
| ENSMUSG00000059205 | 0 | 0 | 0 | 0 | 0 | 0 | 0 |
| ENSMUSG00000059206 | 0 | 0 | 0 | 0 | 0 | 0 | 0 |
| ENSMUSG00000059208 | 3.203651802 | 2.951939351 | 2.966916212 | 2.947587124 | 2.869761991 | 3.312600439 | 2.746122785 |
| ENSMUSG00000059213 | 0.671187741 | 0.701838731 | 0.701838731 | 0.396240625 | 0.568060967 | 0.477121255 | 0.520678036 |
| ENSMUSG00000059218 | 0 | 0 | 0 | 0 | 0 | 0 | 0 |
| ENSMUSG00000059227 | 0 | 0 | 0 | 0 | 0 | 0 | 0 |
| ENSMUSG00000059229 | 0.578129653 | 0.646240625 | 0.701838731 | 0.580482024 | 0.812711509 | 0.602059991 | 0.520678036 |
| ENSMUSG00000059230 | 0 | 0 | 0 | 0 | 0 | 0 | 0 |
| ENSMUSG00000059237 | 0 | 0 | 0 | 0 | 0 | 0 | 0 |
| ENSMUSG00000059244 | 0 | 0 | 0 | 0 | 0 | 0 | 0 |
| ENSMUSG00000059246 | 0 | 0 | 0 | 0 | 0 | 0 | 0 |
| ENSMUSG00000059248 | 2.997485856 | 2.759558284 | 2.787912707 | 2.564641508 | 2.617443897 | 2.903089987 | 2.496394312 |
| ENSMUSG00000059251 | 0 | 0 | 0 | 0 | 0 | 0 | 0 |
| ENSMUSG00000059252 | 0.289064826 | 0.25 | 0.25 | 0 | 0.244650542 | 0 | 0.448487648 |
| ENSMUSG00000059256 | 0 | 0 | 0 | 0 | 0 | 0 | 0 |
| ENSMUSG00000059263 | 2.939767604 | 2.683889006 | 2.636723615 | 2.722185812 | 2.627905721 | 3.019946682 | 2.437439291 |
| ENSMUSG00000059273 | 2.476144923 | 2.296218836 | 2.347004321 | 2.118933358 | 2.227765453 | 2.42975228 | 2.161359729 |
| ENSMUSG00000059277 | 0.91631382 | 0.701838731 | 0.864857905 | 0.896240625 | 0.812711509 | 0.698970004 | 0.710836105 |
| ENSMUSG00000059278 | 1.418409478 | 1.451838731 | 1.494319981 | 1.339388001 | 1.49445382 | 1.612783857 | 1.607811402 |
| ENSMUSG00000059279 | 0 | 0 | 0 | 0 | 0 | 0 | 0 |
| ENSMUSG00000059280 | 0 | 0 | 0 | 0 | 0 | 0 | 0 |
| ENSMUSG00000059288 | 2.097855556 | 1.951838731 | 1.901832578 | 1.901832578 | 1.871833944 | 2.021189299 | 1.750751125 |

| | | | | | | | |
|---|---|---|---|---|---|---|---|
| ENSMUSG00000059291 | 2.970129664 | 2.743244947 | 2.761610487 | 2.756457917 | 2.760272018 | 3.170261715 | 2.609542797 |
| ENSMUSG00000059301 | 0.289064826 | 0.25 | 0 | 0 | 0 | 0 | 0 |
| ENSMUSG00000059303 | 0 | 0 | 0 | 0 | 0 | 0 | 0 |
| ENSMUSG00000059305 | 0.289064826 | 0.396240625 | 0.396240625 | 0.25 | 0 | 0.477121255 | 0.520678036 |
| ENSMUSG00000059316 | 2.23973002 | 2.152756199 | 2.199415381 | 2.159156155 | 2.285543632 | 2.51054501 | 2.195242462 |
| ENSMUSG00000059319 | 0 | 0 | 0 | 0 | 0 | 0 | 0 |
| ENSMUSG00000059323 | 2.635965117 | 2.467976309 | 2.518200634 | 2.469129237 | 2.558403901 | 2.862131379 | 2.442585457 |
| ENSMUSG00000059325 | 0.960252568 | 0.864857905 | 0.92510993 | 0.864857905 | 0.686820904 | 0.301029996 | 0.876096088 |
| ENSMUSG00000059326 | 0 | 0 | 0 | 0.396240625 | 0.812711509 | 0.602059991 | 0.829800754 |
| ENSMUSG00000059327 | 1.558729299 | 1.403677461 | 1.380890489 | 1.476722649 | 1.484072009 | 1.477121255 | 1.335191751 |
| ENSMUSG00000059334 | 0 | 0 | 0 | 0 | 0 | 0 | 0 |
| ENSMUSG00000059336 | 0 | 0 | 0 | 0.25 | 0.244650542 | 0 | 0 |
| ENSMUSG00000059343 | 0 | 0 | 0.5 | 0.396240625 | 0 | 0 | 0.355418052 |
| ENSMUSG00000059352 | 0 | 0 | 0.25 | 0 | 0 | 0 | 0 |
| ENSMUSG00000059355 | 2.181541736 | 2.006976499 | 2.048689214 | 1.926839783 | 1.982473755 | 2.127104798 | 1.895131899 |
| ENSMUSG00000059361 | 0 | 0 | 0 | 0 | 0 | 0 | 0 |
| ENSMUSG00000059363 | 1.907802659 | 1.834962501 | 1.839388001 | 1.959800947 | 1.903713872 | 2.161368002 | 1.792684387 |
| ENSMUSG00000059366 | 0 | 0 | 0 | 0 | 0 | 0 | 0 |
| ENSMUSG00000059371 | 0 | 0 | 0 | 0 | 0 | 0 | 0 |
| ENSMUSG00000059375 | 0 | 0 | 0 | 0 | 0 | 0 | 0 |
| ENSMUSG00000059379 | 0 | 0 | 0 | 0 | 0 | 0 | 0 |
| ENSMUSG00000059382 | 0 | 0 | 0 | 0 | 0 | 0 | 0 |
| ENSMUSG00000059383 | 0 | 0 | 0 | 0 | 0 | 0 | 0 |
| ENSMUSG00000059395 | 0.45815691 | 0.79248125 | 0.646240625 | 0.701838731 | 0.244650542 | 0.77815125 | 0.74492186 |
| ENSMUSG00000059397 | 0 | 0 | 0 | 0 | 0 | 0 | 0 |
| ENSMUSG00000059401 | 1.936861443 | 1.779735268 | 1.769203899 | 1.54248125 | 1.614672255 | 1.77815125 | 1.523605797 |
| ENSMUSG00000059406 | 0.578129653 | 0.25 | 0.5 | 0.25 | 0.244650542 | 0.698970004 | 0.629532004 |
| ENSMUSG00000059408 | 0 | 0 | 0 | 0.25 | 0 | 0 | 0 |
| ENSMUSG00000059409 | 2.905450967 | 2.666556001 | 2.663882681 | 2.759901129 | 2.720413629 | 3.113943352 | 2.565194949 |
| ENSMUSG00000059410 | 0 | 0 | 0 | 0 | 0 | 0 | 0 |
| ENSMUSG00000059411 | 0 | 0 | 0 | 0 | 0 | 0 | 0 |
| ENSMUSG00000059412 | 0 | 0 | 0 | 0 | 0 | 0 | 0 |
| ENSMUSG00000059422 | 1.069666964 | 1.17510993 | 0.864857905 | 1.418106335 | 1.335652905 | 1.361727836 | 1.259064007 |
| ENSMUSG00000059423 | 1.881174527 | 1.752806814 | 1.738549078 | 1.926839783 | 1.870073561 | 2.100370545 | 1.683576582 |

| | | | | | | | |
|---|---|---|---|---|---|---|---|
| ENSMUSG00000059429 | 0 | 0 | 0 | 0 | 0 | 0 | 0 |
| ENSMUSG00000059430 | 3.120285948 | 2.826800202 | 2.788704527 | 0.396240625 | 0.632412477 | 0 | 0.579661877 |
| ENSMUSG00000059434 | 0 | 0 | 0.396240625 | 0.396240625 | 0.387761935 | 0.301029996 | 0 |
| ENSMUSG00000059436 | 2.502784398 | 2.336073977 | 2.356041572 | 2.03360658 | 2.072599934 | 2.33243846 | 1.986347533 |
| ENSMUSG00000059439 | 2.054824387 | 1.828220739 | 1.860735874 | 1.726722649 | 1.758994476 | 1.892094603 | 1.681794123 |
| ENSMUSG00000059447 | 2.503815381 | 2.311388177 | 2.313166358 | 2.410964047 | 2.321205208 | 2.691965103 | 2.13295179 |
| ENSMUSG00000059448 | 0 | 0 | 0 | 0 | 0 | 0 | 0 |
| ENSMUSG00000059455 | 0 | 0 | 0 | 0 | 0 | 0 | 0.355418052 |
| ENSMUSG00000059456 | 0.45815691 | 0.701838731 | 0.396240625 | 1.098079356 | 1.136121934 | 1.041392685 | 1.089372236 |
| ENSMUSG00000059459 | 0 | 0 | 0 | 0 | 0 | 0 | 0 |
| ENSMUSG00000059460 | 0 | 0 | 0 | 0 | 0 | 0 | 0 |
| ENSMUSG00000059461 | 0 | 0.5 | 0.701838731 | 0.580482024 | 0.568060967 | 0.477121255 | 0.74492186 |
| ENSMUSG00000059463 | 0 | 0 | 0 | 0 | 0 | 0 | 0 |
| ENSMUSG00000059473 | 0 | 0 | 0 | 0 | 0 | 0 | 0 |
| ENSMUSG00000059474 | 2.389637216 | 2.201838731 | 2.205044741 | 2.238549078 | 2.139428339 | 2.44870632 | 2.014380832 |
| ENSMUSG00000059475 | 2.089597219 | 1.975216702 | 1.899978211 | 2.078220739 | 2.029283463 | 2.274157849 | 1.863093717 |
| ENSMUSG00000059479 | 0 | 0 | 0.396240625 | 0 | 0.686820904 | 0.477121255 | 0.629532004 |
| ENSMUSG00000059481 | 0 | 0 | 0 | 0 | 0 | 0 | 0 |
| ENSMUSG00000059482 | 1.978790393 | 1.909156155 | 1.874961472 | 1.912762923 | 1.848234258 | 2.068185862 | 1.73898563 |
| ENSMUSG00000059483 | 0.289064826 | 0 | 0 | 0 | 0 | 0.301029996 | 0.224243824 |
| ENSMUSG00000059486 | 2.544030601 | 2.324229052 | 2.307805295 | 2.27186571 | 2.190648982 | 2.462397998 | 2.036615833 |
| ENSMUSG00000059488 | 0 | 0 | 0 | 0 | 0 | 0 | 0 |
| ENSMUSG00000059493 | 1.389637216 | 1.348079356 | 1.302363341 | 1.098079356 | 1.136121934 | 1 | 1.089372236 |
| ENSMUSG00000059495 | 2.867625075 | 2.616901388 | 2.605253214 | 2.729529463 | 2.665423072 | 3.038222638 | 2.4836078 |
| ENSMUSG00000059498 | 0 | 0 | 0.25 | 0.25 | 0 | 0.301029996 | 0.224243824 |
| ENSMUSG00000059503 | 0 | 0 | 0 | 0 | 0 | 0 | 0 |
| ENSMUSG00000059504 | 0 | 0 | 0 | 0 | 0 | 0 | 0 |
| ENSMUSG00000059511 | 0 | 0 | 0 | 0 | 0.244650542 | 0 | 0 |
| ENSMUSG00000059518 | 1.789260561 | 1.782320754 | 1.852347734 | 1.77186571 | 1.826953746 | 1.84509804 | 1.86815666 |
| ENSMUSG00000059534 | 2.170239354 | 2.048689214 | 2.165444524 | 1.933677405 | 2.055279708 | 2.193124598 | 2.042530357 |
| ENSMUSG00000059540 | 0.960252568 | 0.951838731 | 0.896240625 | 0.830482024 | 1.039257769 | 1.079181246 | 1.054044578 |
| ENSMUSG00000059547 | 0 | 0 | 0 | 0 | 0 | 0 | 0 |
| ENSMUSG00000059552 | 2.774997138 | 2.564052172 | 2.586628433 | 2.68176739 | 2.715300549 | 3.070407322 | 2.580884999 |
| ENSMUSG00000059554 | 1.45815691 | 1.27186571 | 1.364857905 | 1.339388001 | 1.358933174 | 1.477121255 | 1.383567577 |

| | | | | | | | |
|---|---|---|---|---|---|---|---|
| ENSMUSG00000059555 | 1.54867984 | 1.5 | 1.552363341 | 1.476722649 | 1.519146132 | 1.544068044 | 1.473249546 |
| ENSMUSG00000059562 | 0 | 0.25 | 0.5 | 0.25 | 0.632412477 | 0 | 0.224243824 |
| ENSMUSG00000059565 | 0 | 0 | 0 | 0 | 0 | 0 | 0.224243824 |
| ENSMUSG00000059571 | 0 | 0.25 | 0 | 0 | 0 | 0 | 0 |
| ENSMUSG00000059585 | 0 | 0 | 0 | 0.25 | 0.244650542 | 0 | 0.224243824 |
| ENSMUSG00000059586 | 2.536482297 | 2.338286706 | 2.337734795 | 2.383818844 | 2.310728002 | 2.600972896 | 2.128938453 |
| ENSMUSG00000059588 | 1.655740331 | 1.593759858 | 1.470660762 | 1.130890489 | 1.121713561 | 1.146128036 | 1.041356071 |
| ENSMUSG00000059595 | 0 | 0 | 0 | 0 | 0 | 0 | 0 |
| ENSMUSG00000059602 | 0.578129653 | 1.04248125 | 0.976722649 | 0.646240625 | 0.931471446 | 0.903089987 | 0.853775828 |
| ENSMUSG00000059606 | 0 | 0 | 0 | 0 | 0 | 0 | 0 |
| ENSMUSG00000059610 | 0 | 0 | 0 | 0 | 0 | 0 | 0 |
| ENSMUSG00000059623 | 0 | 0 | 0 | 0 | 0 | 0 | 0 |
| ENSMUSG00000059625 | 0 | 0 | 0 | 0 | 0 | 0 | 0.224243824 |
| ENSMUSG00000059626 | 0 | 0 | 0 | 0 | 0 | 0 | 0 |
| ENSMUSG00000059631 | 0 | 0 | 0 | 0 | 0.244650542 | 0 | 0 |
| ENSMUSG00000059632 | 0 | 0 | 0 | 0 | 0 | 0 | 0 |
| ENSMUSG00000059639 | 0 | 0 | 0 | 0 | 0 | 0 | 0 |
| ENSMUSG00000059645 | 0 | 0.25 | 0 | 0 | 0 | 0 | 0 |
| ENSMUSG00000059647 | 0 | 0 | 0 | 0 | 0 | 0 | 0 |
| ENSMUSG00000059648 | 0 | 0 | 0 | 0 | 0 | 0 | 0 |
| ENSMUSG00000059654 | 0 | 0 | 0 | 0 | 0 | 0 | 0 |
| ENSMUSG00000059657 | 0 | 0 | 0 | 0 | 0 | 0 | 0 |
| ENSMUSG00000059658 | 0 | 0 | 0.25 | 0 | 0 | 0.301029996 | 0.224243824 |
| ENSMUSG00000059659 | 0.578129653 | 0.79248125 | 0.79248125 | 0.646240625 | 0.568060967 | 0.698970004 | 0.629532004 |
| ENSMUSG00000059663 | 0 | 0 | 0 | 0 | 0 | 0 | 0 |
| ENSMUSG00000059668 | 0 | 0 | 0 | 0 | 0 | 0 | 0 |
| ENSMUSG00000059669 | 1.811507563 | 1.692046081 | 1.763820609 | 1.738549078 | 1.72341484 | 1.913813852 | 1.598697688 |
| ENSMUSG00000059674 | 1.678702042 | 1.614857905 | 1.714495249 | 1.777131114 | 1.983754904 | 1.949390007 | 1.926805425 |
| ENSMUSG00000059687 | 0 | 0 | 0 | 0 | 0 | 0 | 0 |
| ENSMUSG00000059689 | 2.205378646 | 2.029735268 | 2.019203899 | 1.921625132 | 1.935892666 | 2.181843588 | 1.774416565 |
| ENSMUSG00000059690 | 0 | 0 | 0 | 0 | 0 | 0 | 0 |
| ENSMUSG00000059695 | 0 | 0 | 0 | 0 | 0 | 0 | 0 |
| ENSMUSG00000059706 | 0 | 0 | 0 | 0 | 0 | 0 | 0 |
| ENSMUSG00000059708 | 1.822201287 | 1.71759118 | 1.789967834 | 1.668106335 | 1.559656853 | 1.838849091 | 1.459332757 |

| | | | | | | | |
|---|---|---|---|---|---|---|---|
| ENSMUSG00000059713 | 2.451206209 | 2.229715809 | 2.237091808 | 2.307805295 | 2.223252711 | 2.583198774 | 2.123052075 |
| ENSMUSG00000059714 | 2.048736215 | 1.966046536 | 1.990000483 | 1.79248125 | 1.804307395 | 1.934498451 | 1.829800754 |
| ENSMUSG00000059729 | 0 | 0 | 0 | 0 | 0 | 0 | 0 |
| ENSMUSG00000059731 | 0 | 0 | 0 | 0 | 0 | 0 | 0 |
| ENSMUSG00000059734 | 2.402962068 | 2.259729747 | 2.299304173 | 2.065523711 | 2.254881871 | 2.445604203 | 2.179128924 |
| ENSMUSG00000059741 | 0 | 0.396240625 | 0 | 0 | 0 | 0 | 0 |
| ENSMUSG00000059742 | 1.129344651 | 0.951838731 | 1.061981878 | 1.130890489 | 1.039257769 | 1.113943352 | 0.803905701 |
| ENSMUSG00000059743 | 1.404271433 | 1.470660762 | 1.356566189 | 1.657339155 | 1.649303553 | 1.72427587 | 1.653807046 |
| ENSMUSG00000059751 | 0.671187741 | 0.396240625 | 0.830482024 | 0.864857905 | 0.568060967 | 0.301029996 | 0.579661877 |
| ENSMUSG00000059762 | 0 | 0 | 0 | 0 | 0 | 0 | 0 |
| ENSMUSG00000059763 | 0 | 0 | 0 | 0 | 0 | 0 | 0 |
| ENSMUSG00000059768 | 0 | 0 | 0 | 0 | 0 | 0 | 0 |
| ENSMUSG00000059772 | 1.269664473 | 1.26109853 | 1.372963274 | 1.418106335 | 1.310728002 | 1.301029996 | 1.369798885 |
| ENSMUSG00000059775 | 0 | 0.396240625 | 0 | 0 | 0.244650542 | 0 | 0.224243824 |
| ENSMUSG00000059776 | 0.289064826 | 0 | 0 | 0.25 | 0.387761935 | 0.477121255 | 0.448487648 |
| ENSMUSG00000059791 | 1.912079952 | 1.782320754 | 1.837182039 | 1.807204673 | 1.834841362 | 1.963787827 | 1.812372009 |
| ENSMUSG00000059796 | 3.38543098 | 3.057279805 | 3.043028733 | 3.211470509 | 3.127667787 | 3.645422269 | 2.901620727 |
| ENSMUSG00000059810 | 1.837744666 | 1.837182039 | 1.816696635 | 1.561981878 | 1.618292349 | 1.698970004 | 1.676387053 |
| ENSMUSG00000059811 | 2.595015862 | 2.386723615 | 2.404137211 | 2.443284802 | 2.386129765 | 2.720159303 | 2.213713783 |
| ENSMUSG00000059816 | 0 | 0 | 0.25 | 0 | 0 | 0 | 0.224243824 |
| ENSMUSG00000059820 | 2.196867486 | 2.004202072 | 2.002806814 | 2.14151351 | 2.104881849 | 2.330413773 | 1.94631981 |
| ENSMUSG00000059821 | 0 | 0 | 0 | 0 | 0 | 0 | 0 |
| ENSMUSG00000059824 | 1.129344651 | 1.146240625 | 1.160964047 | 0.701838731 | 0.568060967 | 0.477121255 | 0.710836105 |
| ENSMUSG00000059827 | 0 | 0 | 0 | 0 | 0 | 0 | 0 |
| ENSMUSG00000059830 | 0 | 0.25 | 0 | 0 | 0 | 0 | 0 |
| ENSMUSG00000059832 | 0 | 0 | 0 | 0 | 0 | 0 | 0 |
| ENSMUSG00000059834 | 1.932832114 | 1.823655187 | 1.738549078 | 1.83272922 | 1.698489937 | 1.991226076 | 1.649885598 |
| ENSMUSG00000059835 | 0 | 0 | 0 | 0 | 0 | 0 | 0.224243824 |
| ENSMUSG00000059839 | 1.54867984 | 1.445339928 | 1.418106335 | 1.532320754 | 1.445123986 | 1.531478917 | 1.278288402 |
| ENSMUSG00000059842 | 1.069666964 | 1.02186571 | 1 | 1.04248125 | 1.223252711 | 0.903089987 | 1.245582246 |
| ENSMUSG00000059845 | 0 | 0 | 0 | 0 | 0 | 0 | 0 |
| ENSMUSG00000059851 | 1.789260561 | 1.580482024 | 1.638647213 | 1.527131114 | 1.701347897 | 1.633468456 | 1.635773653 |
| ENSMUSG00000059852 | 0 | 0 | 0 | 0 | 0 | 0 | 0 |
| ENSMUSG00000059854 | 0 | 0 | 0 | 0.25 | 0 | 0.301029996 | 0 |

| | | | | | | | |
|---|---|---|---|---|---|---|---|
| ENSMUSG00000059857 | 0 | 0.396240625 | 0.25 | 0 | 0 | 0.301029996 | 0 |
| ENSMUSG00000059862 | 0 | 0 | 0 | 0 | 0 | 0 | 0 |
| ENSMUSG00000059864 | 0 | 0 | 0.396240625 | 0 | 0 | 0 | 0 |
| ENSMUSG00000059866 | 1.862428343 | 1.681980114 | 1.77186571 | 1.779735268 | 1.782377602 | 1.897627091 | 1.690609803 |
| ENSMUSG00000059867 | 0 | 0 | 0 | 0 | 0 | 0 | 0 |
| ENSMUSG00000059873 | 0 | 0 | 0 | 0 | 0 | 0 | 0 |
| ENSMUSG00000059874 | 0 | 0 | 0 | 0 | 0 | 0 | 0 |
| ENSMUSG00000059877 | 0 | 0 | 0 | 0 | 0 | 0 | 0 |
| ENSMUSG00000059878 | 2.355805236 | 2.19697564 | 2.105516192 | 2.172499493 | 2.101233692 | 2.411619706 | 1.97354745 |
| ENSMUSG00000059883 | 1.940852214 | 1.744319981 | 1.695339928 | 1.823655187 | 1.800029087 | 1.892094603 | 1.771720588 |
| ENSMUSG00000059887 | 0 | 0 | 0 | 0 | 0 | 0 | 0 |
| ENSMUSG00000059890 | 2.666334442 | 2.419870025 | 2.418106335 | 2.597810897 | 2.527190274 | 2.893761762 | 2.334241631 |
| ENSMUSG00000059891 | 0 | 0 | 0 | 0.25 | 0 | 0 | 0 |
| ENSMUSG00000059895 | 2.020182925 | 2.013820609 | 2.121960008 | 1.438721876 | 1.568060967 | 1.361727836 | 1.610050272 |
| ENSMUSG00000059897 | 1.794934559 | 1.76109853 | 1.729715809 | 1.864857905 | 1.777823267 | 1.963787827 | 1.647906904 |
| ENSMUSG00000059898 | 0 | 0 | 0.25 | 0 | 0 | 0 | 0 |
| ENSMUSG00000059900 | 0 | 0 | 0.396240625 | 0.79248125 | 0.733951626 | 0.602059991 | 0.629532004 |
| ENSMUSG00000059901 | 1.794934559 | 1.701838731 | 1.735628626 | 1.380890489 | 1.614672255 | 1.633468456 | 1.523605797 |
| ENSMUSG00000059908 | 0 | 0 | 0 | 0 | 0 | 0 | 0 |
| ENSMUSG00000059910 | 0 | 0 | 0 | 0 | 0 | 0 | 0 |
| ENSMUSG00000059912 | 0.289064826 | 0 | 0.25 | 0.396240625 | 0.387761935 | 0 | 0.355418052 |
| ENSMUSG00000059920 | 2.249317394 | 2.12892496 | 2.015173983 | 2.136723615 | 2.066583491 | 2.334453751 | 1.950241257 |
| ENSMUSG00000059921 | 2.229917041 | 2.031030328 | 2.049918086 | 1.777131114 | 1.810629166 | 1.973127854 | 1.627424404 |
| ENSMUSG00000059923 | 2.680728114 | 2.465659339 | 2.45584181 | 2.472945926 | 2.461685042 | 2.789580712 | 2.324338556 |
| ENSMUSG00000059934 | 0 | 0 | 0 | 0 | 0 | 0 | 0 |
| ENSMUSG00000059939 | 2.342375483 | 2.106566189 | 2.135757955 | 2.169870025 | 2.176121988 | 2.383815366 | 2.03780742 |
| ENSMUSG00000059956 | 0 | 0 | 0 | 0 | 0 | 0 | 0 |
| ENSMUSG00000059964 | 0 | 0 | 0 | 0 | 0 | 0 | 0 |
| ENSMUSG00000059970 | 1.647796617 | 1.708222504 | 1.685366747 | 1.348079356 | 1.489301084 | 1.447158031 | 1.526507301 |
| ENSMUSG00000059972 | 0 | 0 | 0 | 0 | 0 | 0 | 0 |
| ENSMUSG00000059974 | 0 | 0 | 0 | 0.25 | 0 | 0 | 0 |
| ENSMUSG00000059975 | 1.765759561 | 1.653677461 | 1.610735874 | 1.766522298 | 1.671670246 | 1.812913357 | 1.56716938 |
| ENSMUSG00000059981 | 2.371652933 | 2.209800947 | 2.285526764 | 2.167221246 | 2.247084732 | 2.408239965 | 2.199960101 |
| ENSMUSG00000059991 | 0.578129653 | 0 | 0 | 0 | 0 | 0 | 0.224243824 |

| | | | | | | | |
|---|---|---|---|---|---|---|---|
| ENSMUSG00000059994 | 0 | 0 | 0 | 0 | 0 | 0 | 0 |
| ENSMUSG00000059995 | 2.660717759 | 2.460587586 | 2.487821179 | 2.26584877 | 2.308696595 | 2.545307116 | 2.20779034 |
| ENSMUSG00000059996 | 0 | 0 | 0 | 0 | 0 | 0 | 0 |
| ENSMUSG00000060002 | 2.305951036 | 2.145300145 | 2.155512955 | 2.343216265 | 2.254881871 | 2.600972896 | 2.074322292 |
| ENSMUSG00000060012 | 1.83262764 | 1.701838731 | 1.668106335 | 1.937048212 | 1.941702774 | 2.164352856 | 1.853775828 |
| ENSMUSG00000060017 | 0 | 0 | 0 | 0 | 0 | 0 | 0 |
| ENSMUSG00000060019 | 0.671187741 | 0.646240625 | 1 | 0.646240625 | 0.568060967 | 0.602059991 | 0.74492186 |
| ENSMUSG00000060024 | 0 | 0 | 0 | 0 | 0 | 0 | 0 |
| ENSMUSG00000060029 | 0.289064826 | 0.25 | 0.396240625 | 0.396240625 | 0.489301084 | 0 | 0.629532004 |
| ENSMUSG00000060030 | 0 | 0 | 0 | 0 | 0 | 0 | 0 |
| ENSMUSG00000060032 | 1.516991255 | 1.403677461 | 1.557204673 | 1.339388001 | 1.343584837 | 1.414973348 | 1.224243824 |
| ENSMUSG00000060034 | 0.671187741 | 0.580482024 | 0.701838731 | 0.580482024 | 0 | 0.477121255 | 0.520678036 |
| ENSMUSG00000060036 | 3.510777687 | 3.182617534 | 3.177109325 | 3.088079839 | 3.057667245 | 3.512817759 | 2.839575072 |
| ENSMUSG00000060038 | 2.20747957 | 2.073655187 | 2.048689214 | 2.044977273 | 2.052132976 | 2.271841607 | 1.918313719 |
| ENSMUSG00000060044 | 1.527823874 | 1.445339928 | 1.488549078 | 1.160964047 | 0.877063019 | 0.954242509 | 0.829800754 |
| ENSMUSG00000060049 | 0 | 0 | 0 | 0 | 0 | 0 | 0 |
| ENSMUSG00000060057 | 0 | 0 | 0 | 0 | 0 | 0 | 0 |
| ENSMUSG00000060063 | 0.289064826 | 0.25 | 0.25 | 0.25 | 0.244650542 | 0.301029996 | 0.355418052 |
| ENSMUSG00000060068 | 0.578129653 | 0 | 0.396240625 | 0.25 | 0.244650542 | 0.301029996 | 0.448487648 |
| ENSMUSG00000060070 | 0 | 0 | 0 | 0 | 0 | 0 | 0 |
| ENSMUSG00000060073 | 2.960941308 | 2.678776002 | 2.632601584 | 2.68473417 | 2.623966669 | 3.010299957 | 2.42167221 |
| ENSMUSG00000060081 | 0 | 0 | 0 | 0 | 0 | 0 | 0 |
| ENSMUSG00000060084 | 0 | 0 | 0 | 0 | 0 | 0 | 0 |
| ENSMUSG00000060090 | 2.502784398 | 2.331045137 | 2.330482024 | 2.317865757 | 2.26997769 | 2.579783597 | 2.073790632 |
| ENSMUSG00000060093 | 0 | 0 | 0 | 0 | 0 | 0 | 0 |
| ENSMUSG00000060096 | 0 | 0.25 | 0 | 0 | 0.489301084 | 0 | 0 |
| ENSMUSG00000060098 | 2.441754646 | 2.269203899 | 2.313166358 | 2.23927551 | 2.31325093 | 2.541579244 | 2.15146804 |
| ENSMUSG00000060105 | 0 | 0 | 0 | 0 | 0 | 0 | 0 |
| ENSMUSG00000060112 | 0 | 0 | 0 | 0 | 0 | 0 | 0 |
| ENSMUSG00000060114 | 0 | 0 | 0 | 0 | 0 | 0 | 0 |
| ENSMUSG00000060121 | 2.131864492 | 2.041226732 | 1.994319981 | 1.948603967 | 1.871833944 | 2.136720567 | 1.766260282 |
| ENSMUSG00000060126 | 3.348261808 | 3.053655672 | 3.05175358 | 3.270994467 | 3.16078916 | 3.712060142 | 2.939542282 |
| ENSMUSG00000060128 | 0.45815691 | 0 | 0 | 0 | 0 | 0 | 0.224243824 |
| ENSMUSG00000060131 | 0.91631382 | 0.79248125 | 0.896240625 | 0 | 0 | 0.477121255 | 0.224243824 |

| | | | | | | | |
|---|---|---|---|---|---|---|---|
| ENSMUSG00000060143 | 0.578129653 | 0.396240625 | 0.580482024 | 0.396240625 | 0.733951626 | 0.602059991 | 0.803905701 |
| ENSMUSG00000060147 | 3.074935921 | 2.7844079 | 2.800377836 | 2.889724745 | 2.782643683 | 3.263872677 | 2.606481113 |
| ENSMUSG00000060149 | 1.727821383 | 1.735628626 | 1.618933358 | 1.681980114 | 1.677808768 | 1.77815125 | 1.511732338 |
| ENSMUSG00000060152 | 2.033115198 | 1.845926073 | 1.882845365 | 2.08272922 | 2.0510778 | 2.301029996 | 1.914853627 |
| ENSMUSG00000060161 | 0 | 0 | 0 | 0 | 0 | 0 | 0 |
| ENSMUSG00000060166 | 2.349889783 | 2.251406137 | 2.345926073 | 1.951838731 | 2.16097133 | 2.292256071 | 2.15438449 |
| ENSMUSG00000060170 | 0 | 0 | 0 | 0 | 0 | 0 | 0 |
| ENSMUSG00000060176 | 0.811507563 | 0.92510993 | 0.830482024 | 1.26109853 | 1.212046813 | 1.278753601 | 1 |
| ENSMUSG00000060177 | 0 | 0 | 0 | 0 | 0 | 0 | 0 |
| ENSMUSG00000060180 | 0 | 0 | 0 | 0 | 0 | 0 | 0 |
| ENSMUSG00000060181 | 2.375872521 | 2.109697963 | 2.165444524 | 2.195339928 | 2.168627946 | 2.394451681 | 2.007270316 |
| ENSMUSG00000060183 | 0 | 0.396240625 | 0 | 1.080482024 | 1.200473444 | 1.322219295 | 1.238624656 |
| ENSMUSG00000060187 | 0 | 0 | 0 | 0 | 0 | 0 | 0 |
| ENSMUSG00000060188 | 0.289064826 | 0 | 0 | 0 | 0.489301084 | 0 | 0.224243824 |
| ENSMUSG00000060198 | 0 | 0.25 | 0.396240625 | 0 | 0.244650542 | 0 | 0 |
| ENSMUSG00000060201 | 0 | 0 | 0 | 0 | 0 | 0 | 0 |
| ENSMUSG00000060204 | 0 | 0 | 0 | 0 | 0 | 0 | 0.448487648 |
| ENSMUSG00000060205 | 0 | 0 | 0 | 0 | 0 | 0 | 0.224243824 |
| ENSMUSG00000060206 | 2.402962068 | 2.244319981 | 2.205044741 | 2.083847589 | 2.046825265 | 2.324282455 | 1.939944915 |
| ENSMUSG00000060208 | 0 | 0 | 0 | 0 | 0 | 0 | 0 |
| ENSMUSG00000060212 | 0 | 0 | 0.25 | 0 | 0 | 0 | 0 |
| ENSMUSG00000060216 | 2.177057474 | 2.016522298 | 2.025821952 | 2.064346961 | 2.158641579 | 2.31386722 | 2.044283833 |
| ENSMUSG00000060224 | 1.205378646 | 1.302363341 | 1.282320754 | 0.5 | 0.733951626 | 0.903089987 | 0.579661877 |
| ENSMUSG00000060227 | 2.50175086 | 2.337182039 | 2.325374049 | 2.544043287 | 2.462674177 | 2.839478047 | 2.283895396 |
| ENSMUSG00000060240 | 0 | 0 | 0 | 0 | 0 | 0 | 0 |
| ENSMUSG00000060244 | 0.960252568 | 0.92510993 | 0.976722649 | 0.830482024 | 0.846351821 | 0.77815125 | 1 |
| ENSMUSG00000060245 | 0 | 0 | 0 | 0 | 0 | 0 | 0 |
| ENSMUSG00000060254 | 0 | 0 | 0 | 0 | 0 | 0 | 0 |
| ENSMUSG00000060256 | 0 | 0 | 0 | 0 | 0 | 0 | 0 |
| ENSMUSG00000060257 | 0 | 0 | 0 | 0 | 0 | 0 | 0 |
| ENSMUSG00000060260 | 1.623015126 | 1.649978211 | 1.67510993 | 1.130890489 | 1.302012594 | 1.176091259 | 1.345462945 |
| ENSMUSG00000060261 | 3.060283005 | 2.778109877 | 2.786005217 | 2.756630861 | 2.729032298 | 3.104487111 | 2.573135083 |
| ENSMUSG00000060268 | 0.289064826 | 0 | 0.25 | 0 | 0 | 0.301029996 | 0.355418052 |
| ENSMUSG00000060275 | 0.45815691 | 0.5 | 0.580482024 | 0.864857905 | 1.057362051 | 1.204119983 | 1.20919388 |

| | | | | | | | |
|---|---|---|---|---|---|---|---|
| ENSMUSG00000060279 | 2.468370459 | 2.363317659 | 2.418106335 | 2.130890489 | 2.27729019 | 2.414973348 | 2.27027168 |
| ENSMUSG00000060284 | 0.45815691 | 0.25 | 0 | 0 | 0.387761935 | 0 | 0.520678036 |
| ENSMUSG00000060288 | 2.190368148 | 2.001406137 | 1.964495249 | 2.168106335 | 2.103062484 | 2.403120521 | 1.927642465 |
| ENSMUSG00000060301 | 2.364524 | 2.154596376 | 2.12892496 | 2.343759858 | 2.245869735 | 2.588831726 | 2.080113611 |
| ENSMUSG00000060303 | 0 | 0 | 0 | 0 | 0 | 0 | 0 |
| ENSMUSG00000060311 | 0 | 0 | 0 | 0 | 0 | 0 | 0 |
| ENSMUSG00000060314 | 0.671187741 | 0.396240625 | 0.5 | 1.146240625 | 1.188507684 | 1.431363764 | 1.131174228 |
| ENSMUSG00000060317 | 0 | 0 | 0 | 0 | 0 | 0 | 0 |
| ENSMUSG00000060332 | 0.289064826 | 0.396240625 | 0 | 0.25 | 0.244650542 | 0 | 0 |
| ENSMUSG00000060335 | 0 | 0 | 0 | 0 | 0 | 0 | 0 |
| ENSMUSG00000060336 | 1.404271433 | 1.410964047 | 1.214495249 | 1.451838731 | 1.462344774 | 1.491361694 | 1.259064007 |
| ENSMUSG00000060371 | 0 | 0 | 0.25 | 0 | 0 | 0 | 0 |
| ENSMUSG00000060373 | 3.040288275 | 2.793419284 | 2.789020769 | 3.025411863 | 2.913486518 | 3.391816924 | 2.69749337 |
| ENSMUSG00000060375 | 0 | 0 | 0 | 0 | 0 | 0 | 0 |
| ENSMUSG00000060376 | 1.671187741 | 1.769203899 | 1.850219859 | 1.410964047 | 1.652586889 | 1.556302501 | 1.707551656 |
| ENSMUSG00000060377 | 0 | 0 | 0.25 | 0 | 0 | 0 | 0 |
| ENSMUSG00000060380 | 1.069666964 | 1.02186571 | 1.214495249 | 1.188721876 | 1.020174412 | 1.176091259 | 1.121219121 |
| ENSMUSG00000060397 | 1.269664473 | 1.238549078 | 1.238549078 | 1.080482024 | 1.136121934 | 1.278753601 | 1.224243824 |
| ENSMUSG00000060402 | 0 | 0 | 0 | 0 | 0 | 0 | 0.355418052 |
| ENSMUSG00000060404 | 0 | 0 | 0 | 0 | 0 | 0 | 0 |
| ENSMUSG00000060407 | 0 | 0 | 0 | 0 | 0 | 0 | 0 |
| ENSMUSG00000060411 | 0 | 0 | 0 | 0 | 0 | 0 | 0 |
| ENSMUSG00000060412 | 0 | 0 | 0 | 0 | 0 | 0 | 0 |
| ENSMUSG00000060416 | 0 | 0 | 0 | 0 | 0 | 0 | 0 |
| ENSMUSG00000060417 | 0 | 0 | 0 | 0 | 0 | 0 | 0 |
| ENSMUSG00000060419 | 0 | 0 | 0 | 0 | 0 | 0 | 0 |
| ENSMUSG00000060424 | 0 | 0 | 0 | 0 | 0 | 0 | 0 |
| ENSMUSG00000060427 | 2.160971586 | 2.006976499 | 1.959800947 | 2.091580554 | 2.009637626 | 2.274157849 | 1.850609257 |
| ENSMUSG00000060429 | 0 | 0.396240625 | 0.396240625 | 0.25 | 0.244650542 | 0.477121255 | 0.448487648 |
| ENSMUSG00000060438 | 0.289064826 | 0.580482024 | 0.75 | 0.580482024 | 0.568060967 | 0 | 0.579661877 |
| ENSMUSG00000060441 | 0 | 0 | 0 | 0.396240625 | 0 | 0 | 0.520678036 |
| ENSMUSG00000060445 | 0 | 0 | 0 | 0.25 | 0.568060967 | 0 | 0 |
| ENSMUSG00000060450 | 2.655024396 | 2.439555804 | 2.412762923 | 2.449820405 | 2.407551654 | 2.711807229 | 2.23605706 |
| ENSMUSG00000060459 | 0 | 0 | 0 | 0 | 0 | 0 | 0 |

| | | | | | | | |
|---|---|---|---|---|---|---|---|
| ENSMUSG00000060461 | 0 | 0.25 | 0 | 0 | 0 | 0 | 0.224243824 |
| ENSMUSG00000060467 | 0.289064826 | 0.5 | 0.25 | 0 | 0 | 0 | 0.224243824 |
| ENSMUSG00000060469 | 0 | 0 | 0 | 0 | 0 | 0 | 0 |
| ENSMUSG00000060470 | 0.289064826 | 0.396240625 | 0.396240625 | 0 | 0.244650542 | 0 | 0 |
| ENSMUSG00000060475 | 2.706208651 | 2.46293726 | 2.451838731 | 2.508011432 | 2.418540977 | 2.784617293 | 2.242438242 |
| ENSMUSG00000060477 | 2.328498818 | 2.190387808 | 2.137686696 | 1.769203899 | 1.72073075 | 1.86923172 | 1.645916034 |
| ENSMUSG00000060480 | 0 | 0 | 0 | 0 | 0 | 0 | 0 |
| ENSMUSG00000060487 | 0 | 0 | 0 | 0 | 0.244650542 | 0.301029996 | 0 |
| ENSMUSG00000060490 | 0 | 0 | 0 | 0 | 0 | 0 | 0 |
| ENSMUSG00000060491 | 0 | 0 | 0 | 0 | 0 | 0 | 0 |
| ENSMUSG00000060499 | 0 | 0 | 0 | 0 | 0 | 0 | 0 |
| ENSMUSG00000060503 | 0 | 0 | 0 | 0 | 0 | 0 | 0 |
| ENSMUSG00000060508 | 0 | 0 | 0 | 0 | 0 | 0 | 0 |
| ENSMUSG00000060509 | 0.45815691 | 0.396240625 | 0 | 0 | 0 | 0.477121255 | 0.520678036 |
| ENSMUSG00000060510 | 2.631440309 | 2.388647213 | 2.345385738 | 2.453845824 | 2.372793355 | 2.699837726 | 2.204609933 |
| ENSMUSG00000060512 | 0.289064826 | 0.25 | 0.25 | 0.25 | 0 | 0.301029996 | 0.448487648 |
| ENSMUSG00000060519 | 2.029919532 | 1.982684334 | 2.009729747 | 2.14151351 | 2.105788027 | 2.376576957 | 2.065163042 |
| ENSMUSG00000060523 | 0 | 0 | 0 | 0 | 0 | 0 | 0 |
| ENSMUSG00000060530 | 0 | 0 | 0 | 0 | 0.244650542 | 0 | 0 |
| ENSMUSG00000060534 | 0 | 0 | 0 | 0 | 0.387761935 | 0 | 0 |
| ENSMUSG00000060538 | 1.342375483 | 1.364857905 | 1.418106335 | 1.130890489 | 1.366364104 | 1.447158031 | 1.20919388 |
| ENSMUSG00000060548 | 1.432083896 | 1.27186571 | 1.02186571 | 1.668106335 | 1.628935049 | 1.832508913 | 1.633706455 |
| ENSMUSG00000060550 | 0.747221736 | 0.864857905 | 0.830482024 | 1.552363341 | 1.65906333 | 1.770852012 | 1.674564426 |
| ENSMUSG00000060556 | 0 | 0 | 0 | 0 | 0 | 0 | 0 |
| ENSMUSG00000060559 | 0 | 0.396240625 | 0 | 0 | 0 | 0.301029996 | 0 |
| ENSMUSG00000060560 | 0 | 0 | 0 | 0 | 0 | 0 | 0 |
| ENSMUSG00000060565 | 0 | 0 | 0 | 0 | 0 | 0 | 0 |
| ENSMUSG00000060566 | 0.45815691 | 0.396240625 | 0.25 | 0.580482024 | 0.387761935 | 0.477121255 | 0.520678036 |
| ENSMUSG00000060568 | 0.289064826 | 0 | 0 | 0 | 0 | 0 | 0.224243824 |
| ENSMUSG00000060572 | 1.655740331 | 1.630890489 | 1.744319981 | 1.458222504 | 1.739104362 | 1.591064607 | 1.830929954 |
| ENSMUSG00000060579 | 0 | 0 | 0 | 0.25 | 0 | 0 | 0 |
| ENSMUSG00000060583 | 0 | 0 | 0 | 0 | 0 | 0 | 0 |
| ENSMUSG00000060586 | 0 | 0 | 0 | 0.25 | 0.244650542 | 0 | 0 |
| ENSMUSG00000060591 | 2.410757293 | 2.21293726 | 2.228969845 | 2.138647213 | 2.139428339 | 2.469822016 | 2.051774285 |

| | | | | | | | |
|---|---|---|---|---|---|---|---|
| ENSMUSG00000060594 | 1.944805158 | 1.848079356 | 1.843759858 | 1.777131114 | 1.706995316 | 1.913813852 | 1.700880998 |
| ENSMUSG00000060600 | 2.900309677 | 2.687886015 | 2.694724619 | 2.03360658 | 2.072599934 | 2.322219295 | 1.976436 |
| ENSMUSG00000060601 | 1.727821383 | 1.62694866 | 1.841580554 | 1.396240625 | 1.66856048 | 1.579783597 | 1.793950594 |
| ENSMUSG00000060613 | 0 | 0 | 0 | 0 | 0 | 0 | 0 |
| ENSMUSG00000060615 | 0 | 0.25 | 0 | 0 | 0 | 0 | 0 |
| ENSMUSG00000060621 | 0 | 0.396240625 | 0.396240625 | 0.25 | 0.387761935 | 0.301029996 | 0 |
| ENSMUSG00000060630 | 0 | 0 | 0 | 0 | 0 | 0 | 0 |
| ENSMUSG00000060636 | 2.253091469 | 2.063166358 | 2.101285366 | 2.358135475 | 2.268268456 | 2.631443769 | 2.126234965 |
| ENSMUSG00000060639 | 0.811507563 | 0.580482024 | 0.75 | 0.951838731 | 1.020174412 | 1.079181246 | 1.014380832 |
| ENSMUSG00000060647 | 0 | 0 | 0 | 0 | 0 | 0 | 0 |
| ENSMUSG00000060657 | 2.761895567 | 2.536801231 | 2.486360959 | 2.447790722 | 2.406781007 | 2.691081492 | 2.242438242 |
| ENSMUSG00000060663 | 0 | 0 | 0 | 0 | 0 | 0 | 0 |
| ENSMUSG00000060671 | 2.706841958 | 2.502107156 | 2.525493918 | 2.504897682 | 2.48376199 | 2.779596491 | 2.352058568 |
| ENSMUSG00000060673 | 0 | 0 | 0 | 0 | 0 | 0 | 0 |
| ENSMUSG00000060675 | 1.842799666 | 1.811981878 | 1.799918086 | 2 | 2.053185007 | 2.294466226 | 1.94631981 |
| ENSMUSG00000060678 | 0 | 0 | 0 | 0 | 0 | 0 | 0 |
| ENSMUSG00000060679 | 2.192545868 | 2.064346961 | 2.034887838 | 2.048689214 | 2.09661982 | 2.338456494 | 2.015657029 |
| ENSMUSG00000060680 | 0 | 0 | 0 | 0.25 | 0.244650542 | 0 | 0 |
| ENSMUSG00000060681 | 2.360187403 | 2.193696765 | 2.183677405 | 2.306000419 | 2.216704172 | 2.525044807 | 2.06186744 |
| ENSMUSG00000060685 | 0 | 0 | 0 | 0 | 0 | 0 | 0 |
| ENSMUSG00000060688 | 0 | 0 | 0 | 0 | 0 | 0 | 0 |
| ENSMUSG00000060691 | 0 | 0 | 0 | 0 | 0 | 0 | 0 |
| ENSMUSG00000060699 | 0 | 0 | 0 | 0 | 0 | 0 | 0 |
| ENSMUSG00000060701 | 0 | 0 | 0 | 0 | 0 | 0 | 0 |
| ENSMUSG00000060703 | 1.91631382 | 1.807204673 | 1.858657057 | 1.701838731 | 1.568060967 | 1.77815125 | 1.523605797 |
| ENSMUSG00000060708 | 1.470606563 | 1.470660762 | 1.54745614 | 1.54745614 | 1.53772706 | 1.755874856 | 1.537859885 |
| ENSMUSG00000060715 | 0 | 0 | 0 | 0 | 0.244650542 | 0.301029996 | 0 |
| ENSMUSG00000060716 | 1.631440309 | 1.5 | 1.476722649 | 1.53743678 | 1.456697355 | 1.531478917 | 1.433437704 |
| ENSMUSG00000060724 | 0 | 0 | 0 | 0 | 0 | 0 | 0 |
| ENSMUSG00000060726 | 0 | 0 | 0 | 0 | 0 | 0 | 0 |
| ENSMUSG00000060727 | 0.289064826 | 0.25 | 0.25 | 0.396240625 | 0.244650542 | 0 | 0.224243824 |
| ENSMUSG00000060730 | 0.578129653 | 0.75 | 0.5 | 0.79248125 | 0.632412477 | 0.84509804 | 0.935079929 |
| ENSMUSG00000060733 | 2.578990401 | 2.401832578 | 2.38040011 | 2.371457327 | 2.327055443 | 2.67669361 | 2.151886289 |
| ENSMUSG00000060735 | 0.289064826 | 0 | 0 | 0 | 0.244650542 | 0 | 0 |

| | | | | | | | |
|---|---|---|---|---|---|---|---|
| ENSMUSG00000060738 | 0 | 0 | 0 | 0 | 0 | 0 | 0 |
| ENSMUSG00000060739 | 2.750654121 | 2.509729747 | 2.51685859 | 2.716239979 | 2.634239692 | 3.060320029 | 2.447985944 |
| ENSMUSG00000060742 | 0 | 0 | 0 | 0 | 0 | 0 | 0 |
| ENSMUSG00000060743 | 1.907802659 | 1.866901388 | 1.928561379 | 1.892463902 | 1.890639929 | 2.029383778 | 1.87511425 |
| ENSMUSG00000060747 | 0 | 0 | 0 | 0 | 0 | 0 | 0 |
| ENSMUSG00000060756 | 0 | 0 | 0 | 0 | 0.387761935 | 0 | 0 |
| ENSMUSG00000060759 | 0 | 0 | 0 | 0 | 0 | 0 | 0 |
| ENSMUSG00000060765 | 0 | 0 | 0 | 0 | 0 | 0 | 0 |
| ENSMUSG00000060771 | 1.527823874 | 1.348079356 | 1.52186571 | 1.51109853 | 1.533172725 | 1.716003344 | 1.433437704 |
| ENSMUSG00000060777 | 0 | 0 | 0 | 0.580482024 | 0 | 0.301029996 | 0.448487648 |
| ENSMUSG00000060780 | 0 | 0 | 0 | 0 | 0 | 0 | 0 |
| ENSMUSG00000060787 | 0 | 0 | 0 | 0 | 0 | 0 | 0 |
| ENSMUSG00000060791 | 0 | 0 | 0 | 0.5 | 0 | 0.301029996 | 0 |
| ENSMUSG00000060794 | 0 | 0 | 0 | 0 | 0 | 0 | 0 |
| ENSMUSG00000060795 | 0 | 0 | 0 | 0 | 0 | 0 | 0 |
| ENSMUSG00000060798 | 1.912079952 | 1.856566189 | 1.777131114 | 1.809601185 | 1.791313678 | 1.968482949 | 1.669034113 |
| ENSMUSG00000060802 | 3.055921842 | 2.795908845 | 2.804640809 | 2.826372948 | 2.724357314 | 3.187520721 | 2.564261618 |
| ENSMUSG00000060803 | 2.172524468 | 2.02186571 | 2.080482024 | 1.994319981 | 2.00243419 | 2.149219113 | 1.855869786 |
| ENSMUSG00000060807 | 0 | 0 | 0 | 0 | 0 | 0 | 0 |
| ENSMUSG00000060808 | 0 | 0.25 | 0.25 | 0 | 0 | 0.301029996 | 0 |
| ENSMUSG00000060811 | 0 | 0 | 0 | 0 | 0 | 0 | 0 |
| ENSMUSG00000060816 | 0 | 0 | 0 | 0 | 0 | 0 | 0 |
| ENSMUSG00000060827 | 0 | 0 | 0 | 0 | 0 | 0 | 0 |
| ENSMUSG00000060843 | 0 | 0 | 0 | 0.5 | 0.244650542 | 0.301029996 | 0.224243824 |
| ENSMUSG00000060860 | 2.674620126 | 2.480831871 | 2.523189285 | 2.503853763 | 2.519146132 | 2.881954971 | 2.399507523 |
| ENSMUSG00000060862 | 1.989530018 | 1.910964047 | 1.930274797 | 1.75835575 | 1.842556557 | 2.068185862 | 1.811176017 |
| ENSMUSG00000060863 | 0 | 0 | 0 | 0 | 0 | 0 | 0 |
| ENSMUSG00000060878 | 0.289064826 | 0 | 0.25 | 0.396240625 | 0 | 0 | 0 |
| ENSMUSG00000060879 | 0.289064826 | 0 | 0 | 0 | 0.244650542 | 0 | 0 |
| ENSMUSG00000060882 | 0 | 0.25 | 0 | 0.25 | 0.244650542 | 0 | 0 |
| ENSMUSG00000060888 | 0 | 0 | 0 | 0 | 0 | 0 | 0 |
| ENSMUSG00000060890 | 0.289064826 | 0 | 0 | 0 | 0.244650542 | 0 | 0 |
| ENSMUSG00000060904 | 3.085022778 | 2.822072336 | 2.810495787 | 2.870701989 | 2.775795166 | 3.254064453 | 2.601104574 |
| ENSMUSG00000060913 | 0 | 0 | 0 | 0 | 0 | 0 | 0 |

| | | | | | | | |
|---|---|---|---|---|---|---|---|
| ENSMUSG00000060918 | 0 | 0 | 0 | 0 | 0 | 0 | 0 |
| ENSMUSG00000060923 | 1.494443473 | 1.410964047 | 1.396240625 | 1.311981878 | 1.283908311 | 1.380211242 | 1.307989563 |
| ENSMUSG00000060924 | 0 | 0 | 0 | 0.25 | 0 | 0 | 0.224243824 |
| ENSMUSG00000060935 | 2.794934559 | 2.526804269 | 2.507321807 | 2.481944491 | 2.407166541 | 2.757396029 | 2.22088434 |
| ENSMUSG00000060938 | 2.668421347 | 2.418548067 | 2.438304187 | 2.563166358 | 2.531009002 | 2.942008053 | 2.364643151 |
| ENSMUSG00000060950 | 1.605636262 | 1.735628626 | 1.735628626 | 1.410964047 | 1.523883869 | 1.491361694 | 1.586932193 |
| ENSMUSG00000060956 | 0 | 0 | 0 | 0 | 0 | 0 | 0 |
| ENSMUSG00000060961 | 2.23778581 | 2.077084758 | 2.025821952 | 1.988549078 | 1.888971102 | 2.230448921 | 1.788855757 |
| ENSMUSG00000060962 | 0 | 0 | 0 | 0 | 0 | 0 | 0 |
| ENSMUSG00000060967 | 0 | 0 | 0 | 0 | 0 | 0 | 0 |
| ENSMUSG00000060969 | 0 | 0 | 0 | 0.701838731 | 0.905314583 | 0.698970004 | 1.100339912 |
| ENSMUSG00000060974 | 0 | 0 | 0 | 0 | 0 | 0 | 0 |
| ENSMUSG00000060981 | 0.289064826 | 0 | 0 | 0.396240625 | 0 | 0 | 0 |
| ENSMUSG00000060985 | 0 | 0.25 | 0 | 0.5 | 0.568060967 | 0.698970004 | 0.629532004 |
| ENSMUSG00000060988 | 1.894701088 | 1.779735268 | 1.816696635 | 1.695339928 | 1.499532413 | 1.740362689 | 1.448487648 |
| ENSMUSG00000060989 | 0.578129653 | 0.75 | 0.864857905 | 1.098079356 | 0.877063019 | 0.903089987 | 0.579661877 |
| ENSMUSG00000060992 | 2.996854944 | 2.716046536 | 2.721233412 | 2.828928915 | 2.764217277 | 3.196176185 | 2.579550454 |
| ENSMUSG00000061000 | 0 | 0 | 0 | 0 | 0 | 0 | 0 |
| ENSMUSG00000061013 | 1.249317394 | 1.26109853 | 1.388647213 | 1.388647213 | 1.366364104 | 1.342422681 | 1.284450788 |
| ENSMUSG00000061022 | 0 | 0 | 0 | 0 | 0 | 0 | 0 |
| ENSMUSG00000061024 | 2.304292853 | 2.182829758 | 2.195339928 | 2.173371739 | 2.149165677 | 2.357934847 | 2.078544414 |
| ENSMUSG00000061028 | 1.707474304 | 1.729715809 | 1.819031101 | 1.630890489 | 1.72341484 | 1.792391689 | 1.705896842 |
| ENSMUSG00000061032 | 2.81882839 | 2.588837774 | 2.607090043 | 2.664775991 | 2.689081778 | 3.019946682 | 2.550660514 |
| ENSMUSG00000061039 | 0.289064826 | 0 | 0.25 | 0.25 | 0.489301084 | 0.301029996 | 0.224243824 |
| ENSMUSG00000061046 | 1.753493018 | 1.701838731 | 1.816696635 | 1.575945187 | 1.78687392 | 1.77815125 | 1.725216937 |
| ENSMUSG00000061048 | 0.960252568 | 1.04248125 | 0.976722649 | 1.282320754 | 1.366364104 | 1.531478917 | 1.302263476 |
| ENSMUSG00000061062 | 1.936861443 | 1.905512955 | 1.856566189 | 2.02186571 | 1.995080671 | 2.190331698 | 1.864112664 |
| ENSMUSG00000061065 | 0 | 0 | 0 | 0 | 0 | 0 | 0 |
| ENSMUSG00000061068 | 0 | 0 | 0 | 0 | 0 | 0 | 0 |
| ENSMUSG00000061075 | 0 | 0 | 0 | 0 | 0 | 0 | 0 |
| ENSMUSG00000061079 | 2.060824957 | 1.930274797 | 1.938721876 | 2.023189285 | 1.978602168 | 2.204119983 | 1.849546807 |
| ENSMUSG00000061080 | 0 | 0 | 0 | 0.396240625 | 0 | 0 | 0 |
| ENSMUSG00000061082 | 0.578129653 | 0.25 | 0 | 0.830482024 | 0.568060967 | 0.602059991 | 0.579661877 |
| ENSMUSG00000061086 | 0.91631382 | 0.864857905 | 0.92510993 | 0.646240625 | 0.955822902 | 0.698970004 | 0.710836105 |

| | | | | | | | |
|---|---|---|---|---|---|---|---|
| ENSMUSG00000061099 | 0.45815691 | 0.580482024 | 0.25 | 0.396240625 | 0.387761935 | 0.301029996 | 0.520678036 |
| ENSMUSG00000061100 | 0 | 0 | 0 | 0 | 0 | 0 | 0 |
| ENSMUSG00000061104 | 0 | 0.25 | 0.396240625 | 0 | 0.489301084 | 0.698970004 | 0.224243824 |
| ENSMUSG00000061111 | 2.318984359 | 2.228222334 | 2.184523065 | 1.921625132 | 2.153935427 | 2.307496038 | 2.088356488 |
| ENSMUSG00000061115 | 0 | 0 | 0 | 0 | 0 | 0 | 0 |
| ENSMUSG00000061118 | 1.470606563 | 1.388647213 | 1.330482024 | 1.464495249 | 1.542223377 | 1.579783597 | 1.413583902 |
| ENSMUSG00000061119 | 1.700465191 | 1.51109853 | 1.42510993 | 1.602347734 | 1.588235379 | 1.698970004 | 1.425641774 |
| ENSMUSG00000061126 | 0 | 0 | 0 | 0 | 0 | 0 | 0.355418052 |
| ENSMUSG00000061130 | 2.592526109 | 2.356566189 | 2.370953944 | 2.451032755 | 2.342197974 | 2.701567985 | 2.18859465 |
| ENSMUSG00000061132 | 0 | 0.25 | 0.5 | 0.646240625 | 0.733951626 | 0.301029996 | 0.520678036 |
| ENSMUSG00000061136 | 3.034848022 | 2.78199858 | 2.763142013 | 2.859438018 | 2.756138828 | 3.13893394 | 2.528731682 |
| ENSMUSG00000061143 | 1.747221736 | 1.610735874 | 1.685366747 | 1.356566189 | 1.351342427 | 1.51851394 | 1.369798885 |
| ENSMUSG00000061144 | 0 | 0 | 0 | 0 | 0 | 0 | 0 |
| ENSMUSG00000061150 | 0 | 0 | 0 | 0 | 0 | 0 | 0 |
| ENSMUSG00000061165 | 0 | 0 | 0 | 0 | 0 | 0 | 0 |
| ENSMUSG00000061167 | 0 | 0 | 0 | 0 | 0 | 0 | 0 |
| ENSMUSG00000061171 | 0.289064826 | 0 | 0 | 0 | 0 | 0 | 0 |
| ENSMUSG00000061175 | 2.429630757 | 2.228222334 | 2.252806814 | 2.01109853 | 1.993840051 | 2.334453751 | 1.900630878 |
| ENSMUSG00000061184 | 0.289064826 | 0 | 0 | 0 | 0 | 0 | 0 |
| ENSMUSG00000061186 | 1.753493018 | 1.77186571 | 1.794977273 | 1.552363341 | 1.584291694 | 1.568201724 | 1.493062805 |
| ENSMUSG00000061195 | 0 | 0 | 0 | 0 | 0 | 0 | 0 |
| ENSMUSG00000061207 | 1.759671389 | 1.685366747 | 1.779735268 | 1.614857905 | 1.830919587 | 1.86332286 | 1.833176598 |
| ENSMUSG00000061208 | 0 | 0 | 0 | 0 | 0 | 0 | 0 |
| ENSMUSG00000061210 | 0 | 0 | 0 | 0 | 0 | 0 | 0 |
| ENSMUSG00000061232 | 2.247417471 | 2.089388001 | 2.072504712 | 2.605516192 | 2.60666248 | 2.975891136 | 2.44865471 |
| ENSMUSG00000061242 | 0 | 0.25 | 0 | 0 | 0.244650542 | 0.477121255 | 0.224243824 |
| ENSMUSG00000061244 | 2.590857977 | 2.411414608 | 2.391989019 | 2.592126615 | 2.471453824 | 2.839478047 | 2.281384267 |
| ENSMUSG00000061246 | 0 | 0 | 0 | 0 | 0 | 0 | 0 |
| ENSMUSG00000061259 | 0 | 0 | 0.396240625 | 0.25 | 0 | 0 | 0.224243824 |
| ENSMUSG00000061272 | 0 | 0 | 0 | 0 | 0 | 0 | 0 |
| ENSMUSG00000061273 | 2.76355595 | 2.530383379 | 2.492166698 | 2.618426437 | 2.524160589 | 2.921686475 | 2.321629237 |
| ENSMUSG00000061286 | 1.45815691 | 1.5 | 1.638647213 | 1.226722649 | 1.427019704 | 1.380211242 | 1.508694612 |
| ENSMUSG00000061288 | 2.388281011 | 2.216046536 | 2.265173983 | 2.160964047 | 2.144330586 | 2.42975228 | 2.075912052 |
| ENSMUSG00000061292 | 0 | 0 | 0 | 0 | 0 | 0 | 0 |

| | | | | | | | |
|---|---|---|---|---|---|---|---|
| ENSMUSG00000061295 | 0 | 0 | 0 | 0 | 0 | 0 | 0 |
| ENSMUSG00000061296 | 0 | 0 | 0 | 0 | 0 | 0 | 0 |
| ENSMUSG00000061298 | 0.811507563 | 0.951838731 | 0.701838731 | 0.896240625 | 0.77552387 | 0.77815125 | 0.710836105 |
| ENSMUSG00000061305 | 0 | 0 | 0 | 0 | 0 | 0 | 0 |
| ENSMUSG00000061306 | 2.778742163 | 2.631135429 | 2.658475852 | 2.54961126 | 2.665237647 | 2.922725458 | 2.548337014 |
| ENSMUSG00000061311 | 0 | 0 | 0 | 0 | 0.244650542 | 0 | 0 |
| ENSMUSG00000061313 | 2.450036412 | 2.204245906 | 2.248588359 | 2.401832578 | 2.317250476 | 2.673020907 | 2.172137922 |
| ENSMUSG00000061315 | 3.176031563 | 2.908702763 | 2.87246199 | 2.975593779 | 2.884515151 | 3.385248682 | 2.725358985 |
| ENSMUSG00000061322 | 0 | 0 | 0 | 0 | 0.387761935 | 0 | 0.224243824 |
| ENSMUSG00000061330 | 0.45815691 | 0.5 | 0.75 | 0.580482024 | 0.489301084 | 0.602059991 | 0.224243824 |
| ENSMUSG00000061331 | 1.623015126 | 1.488549078 | 1.431980114 | 1.403677461 | 1.319233381 | 1.278753601 | 1.252393349 |
| ENSMUSG00000061336 | 0 | 0 | 0 | 0 | 0 | 0 | 0 |
| ENSMUSG00000061342 | 0 | 0 | 0 | 0 | 0 | 0 | 0 |
| ENSMUSG00000061353 | 3.77436153 | 3.424784656 | 3.368077516 | 2.877814624 | 2.846573806 | 3.248708736 | 2.645461849 |
| ENSMUSG00000061356 | 0 | 0 | 0 | 0.25 | 0.387761935 | 0 | 0 |
| ENSMUSG00000061360 | 2.70430294 | 2.502107156 | 2.450224975 | 2.484159485 | 2.490824419 | 2.84509804 | 2.312345877 |
| ENSMUSG00000061361 | 0 | 0 | 0 | 0 | 0 | 0 | 0 |
| ENSMUSG00000061367 | 0 | 0 | 0 | 0 | 0 | 0 | 0 |
| ENSMUSG00000061371 | 1.714367558 | 1.580482024 | 1.527131114 | 1.580482024 | 1.572189156 | 1.755874856 | 1.448487648 |
| ENSMUSG00000061374 | 1.827447047 | 1.755591953 | 1.809601185 | 1.732684334 | 1.614672255 | 1.826074803 | 1.771720588 |
| ENSMUSG00000061376 | 0 | 0 | 0 | 0 | 0 | 0 | 0 |
| ENSMUSG00000061387 | 0 | 0 | 0 | 0 | 0 | 0 | 0 |
| ENSMUSG00000061388 | 0 | 0 | 0 | 0 | 0 | 0 | 0 |
| ENSMUSG00000061390 | 0.289064826 | 0.25 | 0 | 0 | 0.244650542 | 0 | 0.224243824 |
| ENSMUSG00000061392 | 0 | 0 | 0 | 0 | 0 | 0 | 0 |
| ENSMUSG00000061393 | 0.578129653 | 0.5 | 0.830482024 | 0.25 | 0.568060967 | 0.301029996 | 0.74492186 |
| ENSMUSG00000061397 | 0 | 0 | 0 | 0.25 | 0 | 0.301029996 | 0.355418052 |
| ENSMUSG00000061410 | 2.701107274 | 2.525165585 | 2.55175358 | 2.432405186 | 2.41779398 | 2.706717782 | 2.359623923 |
| ENSMUSG00000061411 | 1.686083339 | 1.695339928 | 1.598079356 | 1.77186571 | 1.789100779 | 1.86923172 | 1.74196736 |
| ENSMUSG00000061414 | 0 | 0 | 0.5 | 0 | 0.387761935 | 0.301029996 | 0.224243824 |
| ENSMUSG00000061436 | 2.544030601 | 2.383332433 | 2.407339155 | 2.149978211 | 2.095689764 | 2.352182518 | 2.032410373 |
| ENSMUSG00000061451 | 1.289064826 | 1.27186571 | 1.438721876 | 1.061981878 | 1.176121988 | 1.113943352 | 1.313616061 |
| ENSMUSG00000061455 | 2.266038081 | 2.057204673 | 2.024508021 | 2.144357207 | 2.10031573 | 2.352182518 | 1.943943921 |
| ENSMUSG00000061457 | 0 | 0 | 0 | 0 | 0 | 0 | 0 |

| | | | | | | | |
|---|---|---|---|---|---|---|---|
| ENSMUSG00000061458 | 2.425923778 | 2.294354884 | 2.298689214 | 2.280383379 | 2.214715681 | 2.551449998 | 2.108089156 |
| ENSMUSG00000061461 | 1.964026642 | 1.872963274 | 1.892463902 | 1.83272922 | 1.846351821 | 2.113943352 | 1.792684387 |
| ENSMUSG00000061462 | 0 | 0 | 0 | 0.701838731 | 0.568060967 | 0 | 0.672731473 |
| ENSMUSG00000061469 | 0 | 0 | 0 | 0 | 0 | 0 | 0 |
| ENSMUSG00000061474 | 1.857607105 | 1.735628626 | 1.701838731 | 1.912762923 | 1.940259189 | 2.127104798 | 1.828667599 |
| ENSMUSG00000061477 | 3.074674049 | 2.866264042 | 2.836351312 | 2.758527787 | 2.753981828 | 3.169968174 | 2.611466982 |
| ENSMUSG00000061479 | 2.129344651 | 2.081607372 | 2.148114259 | 1.874961472 | 1.970729215 | 2.158362492 | 2.057979654 |
| ENSMUSG00000061482 | 0.289064826 | 0 | 0.25 | 0 | 0 | 0 | 0.224243824 |
| ENSMUSG00000061486 | 0 | 0.25 | 0 | 0 | 0 | 0 | 0 |
| ENSMUSG00000061488 | 0.289064826 | 0 | 0.396240625 | 0 | 0 | 0 | 0 |
| ENSMUSG00000061489 | 0 | 0 | 0 | 0 | 0 | 0 | 0 |
| ENSMUSG00000061490 | 0 | 0 | 0 | 0 | 0 | 0 | 0 |
| ENSMUSG00000061501 | 0 | 0 | 0 | 0 | 0 | 0 | 0 |
| ENSMUSG00000061510 | 0.45815691 | 0.864857905 | 0.79248125 | 1.080482024 | 0.978602168 | 1.146128036 | 0.876096088 |
| ENSMUSG00000061517 | 0.289064826 | 0 | 0 | 0.75 | 0.812711509 | 0.301029996 | 0.629532004 |
| ENSMUSG00000061518 | 1.890240806 | 1.777131114 | 1.823655187 | 1.886723615 | 1.916320788 | 2.079181246 | 1.921737195 |
| ENSMUSG00000061520 | 0 | 0 | 0 | 0 | 0 | 0 | 0 |
| ENSMUSG00000061524 | 0.578129653 | 0.5 | 0.396240625 | 1.476722649 | 1.53772706 | 1.612783857 | 1.48984372 |
| ENSMUSG00000061525 | 0 | 0.25 | 0 | 0 | 0 | 0 | 0 |
| ENSMUSG00000061527 | 0.289064826 | 0 | 0 | 0 | 0 | 0 | 0.224243824 |
| ENSMUSG00000061531 | 0.289064826 | 0.396240625 | 0 | 0 | 0 | 0 | 0.224243824 |
| ENSMUSG00000061533 | 2.097855556 | 1.975216702 | 2.005591953 | 2.064346961 | 1.958580386 | 2.220108088 | 1.855869786 |
| ENSMUSG00000061535 | 0 | 0.25 | 0 | 0 | 0 | 0 | 0 |
| ENSMUSG00000061536 | 2.136858952 | 2.002806814 | 1.918106335 | 1.958222504 | 1.940259189 | 2.161368002 | 1.838725884 |
| ENSMUSG00000061540 | 0 | 0 | 0 | 0.25 | 0 | 0 | 0.224243824 |
| ENSMUSG00000061544 | 1.753493018 | 1.638647213 | 1.681980114 | 1.814346961 | 1.741652793 | 1.995635195 | 1.649885598 |
| ENSMUSG00000061549 | 0 | 0 | 0 | 0 | 0 | 0 | 0 |
| ENSMUSG00000061559 | 2.34539747 | 2.194519282 | 2.197790722 | 2.33272922 | 2.284999367 | 2.679427897 | 2.139531992 |
| ENSMUSG00000061561 | 0 | 0 | 0 | 0 | 0 | 0 | 0 |
| ENSMUSG00000061576 | 0.289064826 | 0 | 0.25 | 0 | 0 | 0.301029996 | 0.224243824 |
| ENSMUSG00000061577 | 0.45815691 | 0.580482024 | 0.396240625 | 0.5 | 0.489301084 | 0.477121255 | 0.448487648 |
| ENSMUSG00000061578 | 0 | 0.25 | 0 | 0.25 | 0.244650542 | 0.301029996 | 0.224243824 |
| ENSMUSG00000061584 | 0 | 0 | 0 | 0 | 0 | 0 | 0 |
| ENSMUSG00000061589 | 2.528794847 | 2.413659007 | 2.460587586 | 2.251406137 | 2.389791591 | 2.5774918 | 2.365075947 |

| | | | | | | | |
|---|---|---|---|---|---|---|---|
| ENSMUSG00000061601 | 0.289064826 | 0 | 0 | 0.25 | 0 | 0.301029996 | 0 |
| ENSMUSG00000061602 | 0 | 0 | 0 | 0 | 0 | 0 | 0 |
| ENSMUSG00000061603 | 1.289064826 | 1.188721876 | 1.339388001 | 0.92510993 | 0.978602168 | 0.954242509 | 0.935079929 |
| ENSMUSG00000061607 | 2.91246663 | 2.642938161 | 2.649978211 | 2.640799068 | 2.604466107 | 2.99343623 | 2.445634316 |
| ENSMUSG00000061613 | 2.674620126 | 2.428561379 | 2.449820405 | 2.539336734 | 2.477187895 | 2.840106094 | 2.336139088 |
| ENSMUSG00000061614 | 0 | 0 | 0 | 0 | 0 | 0 | 0 |
| ENSMUSG00000061615 | 0 | 0 | 0 | 0 | 0 | 0 | 0 |
| ENSMUSG00000061616 | 0 | 0 | 0 | 0 | 0 | 0 | 0 |
| ENSMUSG00000061619 | 0 | 0 | 0 | 0 | 0 | 0 | 0 |
| ENSMUSG00000061626 | 0 | 0 | 0 | 0 | 0 | 0 | 0 |
| ENSMUSG00000061633 | 0 | 0.25 | 0 | 0 | 0.387761935 | 0 | 0 |
| ENSMUSG00000061637 | 0 | 0 | 0 | 0 | 0 | 0 | 0 |
| ENSMUSG00000061650 | 1.727821383 | 1.782320754 | 1.802363341 | 1.862802778 | 1.937354184 | 2.10720997 | 1.870159858 |
| ENSMUSG00000061653 | 0 | 0 | 0 | 0 | 0 | 0 | 0 |
| ENSMUSG00000061654 | 1.289064826 | 1.26109853 | 1.080482024 | 1.098079356 | 1 | 1.230448921 | 0.969165684 |
| ENSMUSG00000061665 | 2.909363119 | 2.67619206 | 2.699415381 | 2.805698726 | 2.723886392 | 3.134495856 | 2.502107758 |
| ENSMUSG00000061666 | 2.253091469 | 2.125956434 | 2.119945066 | 2.291226732 | 2.232221225 | 2.572871602 | 2.054609671 |
| ENSMUSG00000061669 | 0.747221736 | 0.5 | 0.701838731 | 0.830482024 | 0.686820904 | 0.698970004 | 0.876096088 |
| ENSMUSG00000061684 | 0.289064826 | 0.646240625 | 0.396240625 | 0.646240625 | 0.244650542 | 0.698970004 | 0.355418052 |
| ENSMUSG00000061689 | 2.179305632 | 2.089388001 | 2.119945066 | 1.919870025 | 2.066583491 | 2.190331698 | 2.037212175 |
| ENSMUSG00000061701 | 0 | 0 | 0 | 0 | 0 | 0 | 0 |
| ENSMUSG00000061702 | 0.45815691 | 0.396240625 | 0.580482024 | 0.396240625 | 0.489301084 | 0.477121255 | 0.355418052 |
| ENSMUSG00000061707 | 0 | 0 | 0 | 0 | 0 | 0 | 0 |
| ENSMUSG00000061718 | 0 | 0.25 | 0 | 0.830482024 | 0.733951626 | 0.84509804 | 0.710836105 |
| ENSMUSG00000061723 | 0.289064826 | 0.580482024 | 0.580482024 | 0 | 0.244650542 | 0.301029996 | 0 |
| ENSMUSG00000061724 | 0 | 0 | 0 | 0 | 0 | 0 | 0 |
| ENSMUSG00000061728 | 0 | 0 | 0 | 0 | 0 | 0 | 0 |
| ENSMUSG00000061731 | 2.987597818 | 2.739094039 | 2.714300868 | 2.459800947 | 2.398190943 | 2.770115295 | 2.236700872 |
| ENSMUSG00000061740 | 0 | 0.25 | 0 | 0.646240625 | 0.489301084 | 0.698970004 | 0.710836105 |
| ENSMUSG00000061742 | 0 | 0 | 0 | 0 | 0 | 0 | 0 |
| ENSMUSG00000061751 | 2.393679564 | 2.277131114 | 2.24929487 | 1.729715809 | 1.782377602 | 1.892094603 | 1.749303626 |
| ENSMUSG00000061755 | 2.911305519 | 2.665667094 | 2.654825739 | 2.647880587 | 2.570495157 | 2.94792362 | 2.386211856 |
| ENSMUSG00000061758 | 1.978790393 | 1.809601185 | 1.816696635 | 1.678561379 | 1.695608647 | 1.913813852 | 1.607811402 |
| ENSMUSG00000061759 | 2.361637952 | 2.145300145 | 2.159156155 | 2.181980114 | 2.111176775 | 2.414973348 | 2.017561934 |

| | | | | | | | |
|---|---|---|---|---|---|---|---|
| ENSMUSG00000061762 | 0 | 0 | 0 | 0 | 0 | 0 | 0 |
| ENSMUSG00000061769 | 0 | 0.25 | 0.580482024 | 0 | 0 | 0 | 0 |
| ENSMUSG00000061777 | 0 | 0 | 0 | 0 | 0 | 0 | 0 |
| ENSMUSG00000061778 | 2.258689303 | 2.078220739 | 2.089388001 | 2.331607372 | 2.263090523 | 2.59439255 | 2.076967568 |
| ENSMUSG00000061780 | 0 | 0 | 0 | 0 | 0 | 0 | 0 |
| ENSMUSG00000061787 | 2.80754993 | 2.589388001 | 2.605778978 | 2.583568321 | 2.590076093 | 2.959518377 | 2.393027179 |
| ENSMUSG00000061798 | 0 | 0 | 0 | 0 | 0 | 0 | 0 |
| ENSMUSG00000061802 | 0 | 0 | 0 | 0 | 0 | 0 | 0 |
| ENSMUSG00000061808 | 0 | 0 | 0 | 0 | 0 | 0 | 0 |
| ENSMUSG00000061809 | 0 | 0 | 0 | 0 | 0 | 0 | 0 |
| ENSMUSG00000061815 | 1.156259305 | 1.114857905 | 1.25 | 0.701838731 | 0.978602168 | 0.84509804 | 0.803905701 |
| ENSMUSG00000061816 | 0 | 0 | 0 | 0 | 0 | 0 | 0 |
| ENSMUSG00000061825 | 0 | 0 | 0 | 0 | 0.244650542 | 0 | 0.224243824 |
| ENSMUSG00000061829 | 0 | 0 | 0 | 0 | 0 | 0 | 0 |
| ENSMUSG00000061833 | 0 | 0.25 | 0 | 0 | 0 | 0 | 0 |
| ENSMUSG00000061838 | 2.481610694 | 2.30479213 | 2.299304173 | 2.313757142 | 2.291475807 | 2.633468456 | 2.129834618 |
| ENSMUSG00000061843 | 0 | 0 | 0 | 0 | 0 | 0 | 0 |
| ENSMUSG00000061845 | 0 | 0 | 0 | 0 | 0 | 0 | 0 |
| ENSMUSG00000061847 | 0 | 0 | 0 | 0 | 0 | 0 | 0 |
| ENSMUSG00000061848 | 0.45815691 | 0 | 0.5 | 0.396240625 | 0.244650542 | 0.301029996 | 0.629532004 |
| ENSMUSG00000061852 | 0 | 0 | 0 | 0 | 0 | 0 | 0 |
| ENSMUSG00000061859 | 2.196867486 | 1.995748394 | 2.057204673 | 1.909156155 | 1.916320788 | 2.176091259 | 1.808770687 |
| ENSMUSG00000061863 | 0.289064826 | 0 | 0 | 0 | 0 | 0 | 0 |
| ENSMUSG00000061875 | 0 | 0 | 0 | 0 | 0 | 0 | 0 |
| ENSMUSG00000061877 | 0 | 0 | 0 | 0 | 0 | 0 | 0 |
| ENSMUSG00000061878 | 0.747221736 | 0.830482024 | 1.080482024 | 1.214495249 | 1.319233381 | 1.380211242 | 1.313616061 |
| ENSMUSG00000061882 | 1.227926429 | 1.061981878 | 1.061981878 | 1.098079356 | 1.106691885 | 0.84509804 | 0.952571511 |
| ENSMUSG00000061887 | 2.616779011 | 2.447790722 | 2.461372513 | 2.234894804 | 2.302531265 | 2.56937391 | 2.199238773 |
| ENSMUSG00000061894 | 1.655740331 | 1.458222504 | 1.53743678 | 1.494319981 | 1.572189156 | 1.662757832 | 1.502532226 |
| ENSMUSG00000061897 | 0.671187741 | 0.79248125 | 0.79248125 | 0.396240625 | 0.733951626 | 0.602059991 | 0.629532004 |
| ENSMUSG00000061898 | 1.940852214 | 1.870953944 | 1.864857905 | 1.880890489 | 1.830919587 | 2.025305865 | 1.723650273 |
| ENSMUSG00000061904 | 3.289847398 | 2.967302063 | 2.979063175 | 3.083498471 | 3.027360714 | 3.520090328 | 2.804462239 |
| ENSMUSG00000061906 | 0 | 0 | 0 | 0 | 0 | 0 | 0 |
| ENSMUSG00000061911 | 0 | 0 | 0 | 0.25 | 0 | 0 | 0 |

| | | | | | | | |
|---|---|---|---|---|---|---|---|
| ENSMUSG00000061923 | 0 | 0.25 | 0 | 0 | 0 | 0.301029996 | 0 |
| ENSMUSG00000061928 | 0.289064826 | 0 | 0 | 0 | 0 | 0 | 0.224243824 |
| ENSMUSG00000061937 | 0 | 0 | 0 | 0 | 0 | 0 | 0.224243824 |
| ENSMUSG00000061947 | 0 | 0 | 0 | 0.25 | 0 | 0 | 0 |
| ENSMUSG00000061950 | 2.750654121 | 2.562574604 | 2.531353368 | 2.510756821 | 2.464316601 | 2.808885867 | 2.307473178 |
| ENSMUSG00000061952 | 0 | 0 | 0 | 0 | 0 | 0 | 0 |
| ENSMUSG00000061958 | 0 | 0 | 0 | 0 | 0 | 0 | 0 |
| ENSMUSG00000061959 | 0 | 0 | 0 | 0 | 0 | 0 | 0 |
| ENSMUSG00000061961 | 0 | 0 | 0 | 0 | 0 | 0 | 0 |
| ENSMUSG00000061969 | 0 | 0 | 0 | 0 | 0 | 0 | 0 |
| ENSMUSG00000061972 | 0.578129653 | 0.5 | 0.646240625 | 0.5 | 0.387761935 | 0.301029996 | 0.224243824 |
| ENSMUSG00000061974 | 0 | 0 | 0 | 0 | 0 | 0 | 0 |
| ENSMUSG00000061975 | 0 | 0 | 0 | 0 | 0 | 0 | 0 |
| ENSMUSG00000061977 | 0 | 0 | 0 | 0 | 0 | 0 | 0 |
| ENSMUSG00000061979 | 2.136858952 | 2.036164561 | 2.069031101 | 1.809601185 | 2.016696984 | 2.029383778 | 1.939944915 |
| ENSMUSG00000061981 | 2.701748369 | 2.533927322 | 2.596195938 | 2.402294685 | 2.477187895 | 2.770115295 | 2.409839468 |
| ENSMUSG00000061983 | 3.119345626 | 2.909948223 | 2.961470509 | 2.754202072 | 2.845907432 | 3.186673867 | 2.743179282 |
| ENSMUSG00000061984 | 0 | 0 | 0 | 0 | 0 | 0 | 0 |
| ENSMUSG00000061988 | 0 | 0.25 | 0 | 0.5 | 0.244650542 | 0.301029996 | 0 |
| ENSMUSG00000061991 | 0 | 0 | 0 | 0 | 0 | 0 | 0 |
| ENSMUSG00000061992 | 0.45815691 | 0.25 | 0.5 | 0.5 | 0.489301084 | 0 | 0.224243824 |
| ENSMUSG00000062006 | 2.945876762 | 2.734343455 | 2.757321807 | 2.738003291 | 2.802426296 | 3.169380495 | 2.600478415 |
| ENSMUSG00000062007 | 0 | 0 | 0 | 0 | 0 | 0 | 0 |
| ENSMUSG00000062012 | 0.45815691 | 0.580482024 | 0.896240625 | 1.505591953 | 1.523883869 | 1.579783597 | 1.520678036 |
| ENSMUSG00000062014 | 2.747795766 | 2.49929487 | 2.460980263 | 2.845385738 | 2.732881367 | 3.148602655 | 2.529252851 |
| ENSMUSG00000062017 | 0 | 0 | 0 | 0 | 0 | 0 | 0 |
| ENSMUSG00000062028 | 0 | 0 | 0 | 0 | 0 | 0 | 0 |
| ENSMUSG00000062031 | 1.948720984 | 1.854463129 | 1.862802778 | 1.714495249 | 1.834841362 | 1.973127854 | 1.71730663 |
| ENSMUSG00000062036 | 0 | 0 | 0 | 0 | 0 | 0 | 0 |
| ENSMUSG00000062037 | 0 | 0 | 0 | 0 | 0 | 0 | 0 |
| ENSMUSG00000062038 | 0 | 0 | 0 | 0 | 0 | 0 | 0 |
| ENSMUSG00000062040 | 1.686083339 | 1.67510993 | 1.688721876 | 1.701838731 | 1.665423072 | 1.799340549 | 1.523605797 |
| ENSMUSG00000062042 | 0 | 0 | 0 | 0 | 0 | 0 | 0 |
| ENSMUSG00000062044 | 0.867194479 | 1.17510993 | 1.226722649 | 0.951838731 | 1.121713561 | 0.698970004 | 1.350478789 |

| | | | | | | | |
|---|---|---|---|---|---|---|---|
| ENSMUSG00000062046 | 0 | 0 | 0 | 0 | 0 | 0 | 0 |
| ENSMUSG00000062054 | 1.389637216 | 1.505591953 | 1.470660762 | 1.339388001 | 1.467903253 | 1.62324929 | 1.502532226 |
| ENSMUSG00000062061 | 0 | 0 | 0 | 0 | 0 | 0 | 0 |
| ENSMUSG00000062064 | 0 | 0 | 0 | 0 | 0 | 0 | 0.224243824 |
| ENSMUSG00000062070 | 3.044251953 | 2.789652422 | 2.74929487 | 2.77236262 | 2.685422363 | 3.135132651 | 2.493353867 |
| ENSMUSG00000062073 | 0 | 0.25 | 0.396240625 | 0 | 0 | 0 | 0 |
| ENSMUSG00000062074 | 0.747221736 | 0.646240625 | 0.646240625 | 0.864857905 | 0.931471446 | 0.602059991 | 0.803905701 |
| ENSMUSG00000062075 | 2.201144779 | 2.177701608 | 2.233422664 | 2.146240625 | 2.301493158 | 2.416640507 | 2.26501115 |
| ENSMUSG00000062077 | 0.671187741 | 0.79248125 | 1.098079356 | 0 | 0 | 0 | 0.224243824 |
| ENSMUSG00000062078 | 3.175003123 | 2.899280357 | 2.874587652 | 2.849819481 | 2.745958094 | 3.18582536 | 2.544261369 |
| ENSMUSG00000062081 | 0 | 0 | 0 | 0 | 0 | 0 | 0 |
| ENSMUSG00000062082 | 0.289064826 | 0 | 0.25 | 0 | 0 | 0 | 0 |
| ENSMUSG00000062083 | 0.289064826 | 0.25 | 0.25 | 0 | 0.387761935 | 0 | 0.629532004 |
| ENSMUSG00000062093 | 0.289064826 | 0.396240625 | 0.580482024 | 0.75 | 0.568060967 | 0.301029996 | 0.448487648 |
| ENSMUSG00000062098 | 1.89911417 | 1.763820609 | 1.79248125 | 1.78743678 | 1.754126039 | 1.959041392 | 1.659601507 |
| ENSMUSG00000062101 | 1.432083896 | 1.403677461 | 1.388647213 | 1.388647213 | 1.394615667 | 1.477121255 | 1.278288402 |
| ENSMUSG00000062103 | 0 | 0 | 0 | 0 | 0 | 0 | 0 |
| ENSMUSG00000062105 | 0 | 0 | 0 | 0 | 0 | 0 | 0 |
| ENSMUSG00000062110 | 1.663535556 | 1.580482024 | 1.593759858 | 1.664552871 | 1.642643806 | 1.880813592 | 1.63986884 |
| ENSMUSG00000062115 | 2.346900291 | 2.233422664 | 2.355516192 | 1.905512955 | 2.128567294 | 2.139879086 | 2.185218806 |
| ENSMUSG00000062116 | 1.505869732 | 1.364857905 | 1.476722649 | 1.516522298 | 1.456697355 | 1.633468456 | 1.441050184 |
| ENSMUSG00000062121 | 0 | 0 | 0 | 0 | 0 | 0 | 0 |
| ENSMUSG00000062124 | 0 | 0 | 0 | 0 | 0 | 0 | 0 |
| ENSMUSG00000062127 | 2.312518611 | 2.211372513 | 2.231203126 | 2.172499493 | 2.171644683 | 2.419955748 | 2.073790632 |
| ENSMUSG00000062128 | 0 | 0 | 0 | 0 | 0 | 0 | 0 |
| ENSMUSG00000062132 | 0 | 0 | 0 | 0 | 0 | 0 | 0 |
| ENSMUSG00000062142 | 0 | 0 | 0 | 0 | 0 | 0 | 0 |
| ENSMUSG00000062148 | 0 | 0 | 0 | 0 | 0 | 0 | 0 |
| ENSMUSG00000062151 | 0.45815691 | 0.646240625 | 0.896240625 | 0.580482024 | 0.244650542 | 0 | 0.355418052 |
| ENSMUSG00000062152 | 0.811507563 | 0.701838731 | 0.951838731 | 0.976722649 | 0.978602168 | 1.041392685 | 0.969165684 |
| ENSMUSG00000062154 | 0 | 0 | 0 | 0 | 0 | 0 | 0 |
| ENSMUSG00000062157 | 0.747221736 | 0.701838731 | 0.75 | 1.201838731 | 1.039257769 | 0.954242509 | 1 |
| ENSMUSG00000062162 | 0 | 0 | 0 | 0 | 0 | 0 | 0 |
| ENSMUSG00000062165 | 0 | 0 | 0 | 0 | 0 | 0 | 0 |

| | | | | | | | |
|---|---|---|---|---|---|---|---|
| ENSMUSG00000062168 | 0.289064826 | 0.25 | 0.25 | 0 | 0.244650542 | 0.301029996 | 0.355418052 |
| ENSMUSG00000062169 | 2.542156295 | 2.345385738 | 2.350753006 | 2.460980263 | 2.386129765 | 2.749736316 | 2.244014063 |
| ENSMUSG00000062170 | 0 | 0 | 0 | 0 | 0 | 0 | 0 |
| ENSMUSG00000062175 | 0.289064826 | 0.701838731 | 0.646240625 | 0.896240625 | 0.632412477 | 0.77815125 | 0.710836105 |
| ENSMUSG00000062181 | 0 | 0 | 0 | 0 | 0 | 0 | 0 |
| ENSMUSG00000062184 | 2.264212992 | 2.184523065 | 2.168989258 | 2.238549078 | 2.182733222 | 2.489958479 | 2.028761664 |
| ENSMUSG00000062186 | 0 | 0 | 0 | 0 | 0 | 0 | 0 |
| ENSMUSG00000062190 | 2.196867486 | 2.060793496 | 2.046218836 | 2.187048212 | 2.09661982 | 2.411619706 | 1.98213686 |
| ENSMUSG00000062198 | 1.789260561 | 1.657339155 | 1.695339928 | 1.744319981 | 1.704182902 | 1.86332286 | 1.479989695 |
| ENSMUSG00000062199 | 0 | 0 | 0 | 0 | 0 | 0 | 0 |
| ENSMUSG00000062200 | 0 | 0 | 0 | 0 | 0 | 0 | 0 |
| ENSMUSG00000062201 | 0 | 0 | 0 | 0 | 0 | 0 | 0 |
| ENSMUSG00000062202 | 2.23973002 | 2.112802778 | 2.057204673 | 2.151832578 | 2.129414728 | 2.414973348 | 2.027536225 |
| ENSMUSG00000062203 | 3.001251411 | 2.725593779 | 2.699009902 | 2.795288064 | 2.710112309 | 3.125806458 | 2.537225416 |
| ENSMUSG00000062204 | 0 | 0 | 0 | 0 | 0 | 0 | 0 |
| ENSMUSG00000062209 | 0.289064826 | 0 | 0.25 | 0 | 0.387761935 | 0 | 0 |
| ENSMUSG00000062210 | 2.163307908 | 2.015173983 | 2.02053726 | 2.134789703 | 2.030405741 | 2.318063335 | 1.930140666 |
| ENSMUSG00000062224 | 0 | 0 | 0 | 0 | 0 | 0 | 0 |
| ENSMUSG00000062226 | 0 | 0 | 0 | 0 | 0 | 0 | 0 |
| ENSMUSG00000062232 | 2.302628051 | 2.090485943 | 2.107613138 | 2.247880462 | 2.194892994 | 2.489958479 | 2.071119143 |
| ENSMUSG00000062234 | 2.91898589 | 2.671843921 | 2.688721876 | 2.567573832 | 2.571464152 | 2.910090546 | 2.401639109 |
| ENSMUSG00000062235 | 0 | 0 | 0 | 0 | 0 | 0 | 0 |
| ENSMUSG00000062245 | 0 | 0 | 0 | 0 | 0 | 0 | 0 |
| ENSMUSG00000062248 | 2.444137694 | 2.26584877 | 2.313757142 | 2.400442697 | 2.339407793 | 2.69019608 | 2.249005722 |
| ENSMUSG00000062252 | 0 | 0 | 0 | 0 | 0 | 0 | 0 |
| ENSMUSG00000062257 | 0.578129653 | 0 | 0.396240625 | 0.396240625 | 0.489301084 | 0.477121255 | 0.355418052 |
| ENSMUSG00000062270 | 3.123094257 | 2.845656007 | 2.79106961 | 2.956637122 | 2.850765209 | 3.342422681 | 2.65820629 |
| ENSMUSG00000062272 | 0 | 0 | 0 | 0 | 0 | 0 | 0 |
| ENSMUSG00000062275 | 0 | 0 | 0 | 0 | 0 | 0 | 0 |
| ENSMUSG00000062278 | 0 | 0 | 0 | 0 | 0 | 0 | 0 |
| ENSMUSG00000062279 | 0.289064826 | 0 | 0 | 0 | 0 | 0 | 0 |
| ENSMUSG00000062280 | 0 | 0 | 0 | 0 | 0 | 0 | 0 |
| ENSMUSG00000062282 | 0 | 0 | 0 | 0 | 0 | 0 | 0 |
| ENSMUSG00000062284 | 0.289064826 | 0 | 0 | 0.396240625 | 0.733951626 | 0 | 0.355418052 |

| | | | | | | | |
|---|---|---|---|---|---|---|---|
| ENSMUSG00000062296 | 0 | 0.25 | 0 | 0.25 | 0.387761935 | 0.301029996 | 0.520678036 |
| ENSMUSG00000062300 | 1.639698646 | 1.67510993 | 1.76109853 | 1.738549078 | 1.842556557 | 1.892094603 | 1.900630878 |
| ENSMUSG00000062309 | 1.156259305 | 1.17510993 | 1.282320754 | 0.896240625 | 0.812711509 | 0.84509804 | 0.984950056 |
| ENSMUSG00000062310 | 0.289064826 | 0.396240625 | 0 | 0 | 0.244650542 | 0.301029996 | 0 |
| ENSMUSG00000062312 | 2.490209605 | 2.319031101 | 2.371457327 | 2.293106877 | 2.378272187 | 2.582063363 | 2.296968506 |
| ENSMUSG00000062314 | 0 | 0 | 0 | 0 | 0 | 0 | 0 |
| ENSMUSG00000062319 | 0 | 0 | 0 | 0 | 0 | 0 | 0 |
| ENSMUSG00000062327 | 0 | 0 | 0.25 | 0 | 0 | 0 | 0 |
| ENSMUSG00000062328 | 2.976144659 | 2.692459461 | 2.697790722 | 2.788546302 | 2.661883234 | 3.127752516 | 2.518935458 |
| ENSMUSG00000062329 | 0 | 0 | 0 | 0 | 0 | 0 | 0 |
| ENSMUSG00000062342 | 0.578129653 | 0.396240625 | 0.580482024 | 0 | 0 | 0 | 0 |
| ENSMUSG00000062345 | 1.069666964 | 1.114857905 | 1.214495249 | 0.396240625 | 0.244650542 | 0 | 0.224243824 |
| ENSMUSG00000062352 | 2.134369199 | 2.025821952 | 2.073655187 | 1.943696765 | 1.906908067 | 2.117271296 | 1.821784031 |
| ENSMUSG00000062353 | 2.425923778 | 2.21293726 | 2.247880462 | 2.421625132 | 2.321696448 | 2.7355989 | 2.218171406 |
| ENSMUSG00000062365 | 0.671187741 | 0.75 | 0.79248125 | 0.830482024 | 0.489301084 | 0.477121255 | 0.775756176 |
| ENSMUSG00000062369 | 0 | 0 | 0 | 0 | 0 | 0 | 0 |
| ENSMUSG00000062372 | 0 | 0 | 0 | 0 | 0 | 0 | 0 |
| ENSMUSG00000062373 | 2.774459379 | 2.551142786 | 2.52053726 | 2.606041572 | 2.514628232 | 2.874481818 | 2.31209124 |
| ENSMUSG00000062376 | 1.647796617 | 1.618933358 | 1.657339155 | 1.75835575 | 1.715300549 | 1.908485019 | 1.681794123 |
| ENSMUSG00000062380 | 2.029919532 | 1.94697564 | 1.951838731 | 1.71759118 | 1.855665188 | 1.968482949 | 1.89141329 |
| ENSMUSG00000062382 | 0.289064826 | 0 | 0.25 | 0 | 0.244650542 | 0 | 0.355418052 |
| ENSMUSG00000062391 | 0 | 0 | 0 | 0 | 0 | 0 | 0 |
| ENSMUSG00000062393 | 0.289064826 | 0.396240625 | 0.5 | 0.951838731 | 0.846351821 | 0.903089987 | 0.876096088 |
| ENSMUSG00000062397 | 3.099092675 | 2.806903986 | 2.796373731 | 2.859438018 | 2.732268329 | 3.193958978 | 2.527425075 |
| ENSMUSG00000062400 | 0 | 0 | 0 | 0 | 0 | 0 | 0 |
| ENSMUSG00000062410 | 0.45815691 | 0 | 0 | 0 | 0 | 0 | 0.224243824 |
| ENSMUSG00000062421 | 2.429630757 | 2.273849256 | 2.265173983 | 2.520869832 | 2.484072009 | 2.831229694 | 2.349118526 |
| ENSMUSG00000062426 | 0 | 0 | 0 | 0 | 0 | 0 | 0 |
| ENSMUSG00000062432 | 0.289064826 | 0.25 | 0.396240625 | 0 | 0.244650542 | 0 | 0.224243824 |
| ENSMUSG00000062433 | 0 | 0 | 0 | 0 | 0 | 0 | 0 |
| ENSMUSG00000062434 | 0 | 0 | 0 | 0 | 0 | 0 | 0 |
| ENSMUSG00000062438 | 0 | 0 | 0 | 0.25 | 0.244650542 | 0 | 0.224243824 |
| ENSMUSG00000062444 | 0 | 0 | 0.25 | 0.396240625 | 0.244650542 | 0.301029996 | 0.355418052 |
| ENSMUSG00000062456 | 0.578129653 | 0 | 0.25 | 0.580482024 | 0.244650542 | 0 | 0.448487648 |

| | | | | | | | |
|---|---|---|---|---|---|---|---|
| ENSMUSG00000062458 | 0 | 0 | 0 | 0 | 0 | 0 | 0 |
| ENSMUSG00000062461 | 0 | 0 | 0 | 0 | 0 | 0 | 0 |
| ENSMUSG00000062464 | 0 | 0 | 0 | 0 | 0 | 0.301029996 | 0 |
| ENSMUSG00000062470 | 0.960252568 | 1.160964047 | 1.098079356 | 1.201838731 | 1.106691885 | 1.079181246 | 1.028149525 |
| ENSMUSG00000062472 | 0 | 0.25 | 0 | 0 | 0 | 0.301029996 | 0.224243824 |
| ENSMUSG00000062477 | 0 | 0.25 | 0 | 0.25 | 0 | 0 | 0 |
| ENSMUSG00000062478 | 0 | 0 | 0 | 0 | 0 | 0 | 0 |
| ENSMUSG00000062480 | 0.671187741 | 0.580482024 | 0.580482024 | 0.580482024 | 0.812711509 | 0.84509804 | 0.829800754 |
| ENSMUSG00000062483 | 0 | 0 | 0 | 0 | 0 | 0 | 0 |
| ENSMUSG00000062488 | 0.578129653 | 0.396240625 | 0.646240625 | 0.396240625 | 0.244650542 | 0.301029996 | 0.448487648 |
| ENSMUSG00000062496 | 0.289064826 | 0 | 0 | 0 | 0 | 0 | 0 |
| ENSMUSG00000062497 | 0 | 0 | 0 | 0 | 0 | 0 | 0 |
| ENSMUSG00000062510 | 2.223916471 | 2.034887838 | 2.031030328 | 2.210587586 | 2.180543208 | 2.44870632 | 2.072190387 |
| ENSMUSG00000062511 | 0.45815691 | 0.25 | 0 | 0 | 0 | 0 | 0.355418052 |
| ENSMUSG00000062515 | 0.578129653 | 0.976722649 | 0.701838731 | 0.25 | 0.489301084 | 0 | 0.224243824 |
| ENSMUSG00000062518 | 0.289064826 | 0 | 0.25 | 0 | 0.244650542 | 0 | 0 |
| ENSMUSG00000062519 | 1.693336259 | 1.692046081 | 1.723704441 | 1.79745614 | 1.806427258 | 2.012837225 | 1.661510159 |
| ENSMUSG00000062524 | 0 | 0 | 0 | 0 | 0 | 0 | 0 |
| ENSMUSG00000062526 | 1.614416215 | 1.364857905 | 1.458222504 | 1.25 | 1.319233381 | 1.431363764 | 1.259064007 |
| ENSMUSG00000062527 | 0 | 0 | 0 | 0 | 0 | 0 | 0 |
| ENSMUSG00000062528 | 0 | 0 | 0 | 0 | 0 | 0 | 0 |
| ENSMUSG00000062542 | 0 | 0 | 0 | 0.396240625 | 0.632412477 | 0.477121255 | 0.355418052 |
| ENSMUSG00000062543 | 0 | 0 | 0 | 0 | 0 | 0 | 0 |
| ENSMUSG00000062545 | 0 | 0 | 0 | 0 | 0 | 0 | 0 |
| ENSMUSG00000062546 | 0 | 0 | 0 | 0 | 0 | 0 | 0 |
| ENSMUSG00000062551 | 0 | 0 | 0 | 0 | 0 | 0 | 0 |
| ENSMUSG00000062553 | 0 | 0 | 0 | 0 | 0 | 0 | 0 |
| ENSMUSG00000062554 | 0 | 0 | 0 | 0 | 0 | 0 | 0 |
| ENSMUSG00000062556 | 0 | 0 | 0 | 0 | 0 | 0 | 0 |
| ENSMUSG00000062561 | 0 | 0 | 0 | 0 | 0 | 0 | 0 |
| ENSMUSG00000062563 | 0 | 0 | 0.25 | 0 | 0 | 0.301029996 | 0.520678036 |
| ENSMUSG00000062564 | 0 | 0 | 0 | 0 | 0 | 0 | 0 |
| ENSMUSG00000062580 | 2.615205302 | 2.440387808 | 2.45584181 | 2.55388325 | 2.479078736 | 2.84260924 | 2.315385981 |
| ENSMUSG00000062582 | 0 | 0 | 0 | 0 | 0 | 0 | 0 |

| | | | | | | | |
|---|---|---|---|---|---|---|---|
| ENSMUSG00000062585 | 0.289064826 | 0.25 | 0.580482024 | 0.646240625 | 0.489301084 | 0.477121255 | 0.710836105 |
| ENSMUSG00000062588 | 0 | 0 | 0 | 0.25 | 0.244650542 | 0.301029996 | 0.224243824 |
| ENSMUSG00000062590 | 2.060824957 | 1.87892496 | 1.951838731 | 1.886723615 | 1.941702774 | 2.10720997 | 1.857950279 |
| ENSMUSG00000062591 | 0.811507563 | 0.896240625 | 0.864857905 | 0.830482024 | 0.955822902 | 1 | 1.014380832 |
| ENSMUSG00000062593 | 0 | 0 | 0 | 0 | 0 | 0 | 0 |
| ENSMUSG00000062598 | 0 | 0 | 0 | 0 | 0 | 0 | 0 |
| ENSMUSG00000062604 | 2.675985209 | 2.483791262 | 2.495034894 | 2.412314044 | 2.35849115 | 2.679427897 | 2.176428123 |
| ENSMUSG00000062608 | 0 | 0 | 0 | 0 | 0 | 0 | 0 |
| ENSMUSG00000062609 | 0 | 0.25 | 0 | 0 | 0.387761935 | 0 | 0.224243824 |
| ENSMUSG00000062611 | 0 | 0 | 0 | 0 | 0.244650542 | 0 | 0 |
| ENSMUSG00000062619 | 1.358731791 | 1.25 | 1.410964047 | 1.114857905 | 1.136121934 | 1.113943352 | 1.401059159 |
| ENSMUSG00000062621 | 0 | 0 | 0 | 0 | 0 | 0 | 0 |
| ENSMUSG00000062624 | 0 | 0 | 0 | 0 | 0 | 0 | 0 |
| ENSMUSG00000062627 | 2.289064826 | 2.154596376 | 2.116901388 | 2.254897682 | 2.181274723 | 2.356025857 | 1.990504106 |
| ENSMUSG00000062629 | 0 | 0 | 0 | 0 | 0 | 0 | 0 |
| ENSMUSG00000062638 | 0 | 0 | 0 | 0 | 0 | 0 | 0 |
| ENSMUSG00000062646 | 1.578129653 | 1.580482024 | 1.53743678 | 1.688721876 | 1.572189156 | 1.698970004 | 1.537859885 |
| ENSMUSG00000062647 | 3.19881715 | 2.916333979 | 2.916445007 | 2.877938163 | 2.905784002 | 3.312811826 | 2.696485654 |
| ENSMUSG00000062649 | 0 | 0 | 0 | 0 | 0 | 0 | 0 |
| ENSMUSG00000062651 | 0 | 0 | 0 | 0 | 0 | 0 | 0 |
| ENSMUSG00000062661 | 2.705574381 | 2.491806565 | 2.471805154 | 2.360217292 | 2.355381384 | 2.645422269 | 2.229218532 |
| ENSMUSG00000062683 | 2.167941649 | 2.145300145 | 2.160061234 | 2.006976499 | 2.114724103 | 2.303196057 | 2.087337541 |
| ENSMUSG00000062687 | 0.289064826 | 0.646240625 | 0.5 | 0.701838731 | 0.568060967 | 0.698970004 | 0.355418052 |
| ENSMUSG00000062691 | 1.727821383 | 1.610735874 | 1.598079356 | 1.649978211 | 1.709785497 | 1.826074803 | 1.532233387 |
| ENSMUSG00000062694 | 0 | 0 | 0 | 0 | 0 | 0 | 0 |
| ENSMUSG00000062704 | 0.45815691 | 0.25 | 0.5 | 0.830482024 | 0.77552387 | 0.477121255 | 0.710836105 |
| ENSMUSG00000062705 | 0 | 0 | 0 | 0 | 0 | 0 | 0 |
| ENSMUSG00000062712 | 0 | 0 | 0 | 0 | 0 | 0 | 0 |
| ENSMUSG00000062713 | 0 | 0 | 0 | 0.396240625 | 0 | 0.477121255 | 0.710836105 |
| ENSMUSG00000062727 | 0 | 0 | 0 | 0 | 0.244650542 | 0 | 0 |
| ENSMUSG00000062729 | 1.978790393 | 1.83272922 | 1.870953944 | 1.856566189 | 1.98503142 | 2.021189299 | 1.886704102 |
| ENSMUSG00000062732 | 0 | 0 | 0 | 0 | 0 | 0 | 0 |
| ENSMUSG00000062737 | 0 | 0 | 0 | 0 | 0 | 0 | 0 |
| ENSMUSG00000062742 | 0.811507563 | 0.830482024 | 0.5 | 0.580482024 | 0.387761935 | 0.477121255 | 0.579661877 |

| | | | | | | | |
|---|---|---|---|---|---|---|---|
| ENSMUSG00000062743 | 1.389637216 | 1.451838731 | 1.54248125 | 1.25 | 1.264824954 | 1.255272505 | 1.176815335 |
| ENSMUSG00000062751 | 0 | 0 | 0 | 0 | 0 | 0 | 0 |
| ENSMUSG00000062753 | 1.605636262 | 1.561981878 | 1.668106335 | 0.92510993 | 1.254881871 | 1.447158031 | 1.21680636 |
| ENSMUSG00000062757 | 0 | 0 | 0 | 0 | 0 | 0 | 0 |
| ENSMUSG00000062758 | 0 | 0 | 0 | 0 | 0 | 0.301029996 | 0 |
| ENSMUSG00000062760 | 0 | 0 | 0.25 | 0.396240625 | 0.568060967 | 0.602059991 | 0.579661877 |
| ENSMUSG00000062761 | 2.529763563 | 2.26109853 | 2.257666784 | 2.284247778 | 2.242813819 | 2.499687083 | 2.126687118 |
| ENSMUSG00000062762 | 2.632197861 | 2.401832578 | 2.37694866 | 2.52908599 | 2.44788147 | 2.818225894 | 2.291856721 |
| ENSMUSG00000062773 | 0 | 0 | 0 | 0 | 0 | 0 | 0 |
| ENSMUSG00000062778 | 0 | 0 | 0 | 0 | 0 | 0 | 0 |
| ENSMUSG00000062782 | 0 | 0 | 0 | 0 | 0 | 0 | 0 |
| ENSMUSG00000062783 | 0 | 0.396240625 | 0 | 0 | 0 | 0 | 0 |
| ENSMUSG00000062785 | 0 | 0.25 | 0.25 | 0.5 | 0.955822902 | 1 | 0.969165684 |
| ENSMUSG00000062791 | 0 | 0 | 0 | 0 | 0 | 0 | 0 |
| ENSMUSG00000062793 | 0 | 0 | 0 | 0 | 0 | 0 | 0 |
| ENSMUSG00000062794 | 1.759671389 | 1.723704441 | 1.732684334 | 1.816696635 | 1.784632921 | 2.004321374 | 1.603286636 |
| ENSMUSG00000062797 | 2.013563224 | 1.890560606 | 1.933677405 | 1.87694866 | 1.861137485 | 2.173186268 | 1.815933683 |
| ENSMUSG00000062814 | 0 | 0 | 0 | 0 | 0 | 0 | 0 |
| ENSMUSG00000062818 | 0 | 0 | 0 | 0 | 0 | 0 | 0 |
| ENSMUSG00000062822 | 2.042557845 | 1.884789703 | 1.896240625 | 1.848079356 | 1.810629166 | 2.071882007 | 1.729871916 |
| ENSMUSG00000062825 | 4.247842172 | 3.79985099 | 3.791648658 | 3.490949803 | 3.454484209 | 4.013553431 | 3.178796925 |
| ENSMUSG00000062826 | 0.289064826 | 0 | 0 | 0.25 | 0 | 0 | 0 |
| ENSMUSG00000062833 | 0 | 0 | 0 | 0 | 0 | 0 | 0 |
| ENSMUSG00000062844 | 0 | 0 | 0 | 0 | 0 | 0 | 0 |
| ENSMUSG00000062846 | 0 | 0 | 0 | 0 | 0 | 0.301029996 | 0 |
| ENSMUSG00000062859 | 0 | 0 | 0 | 0 | 0.244650542 | 0.301029996 | 0.520678036 |
| ENSMUSG00000062861 | 1.470606563 | 1.464495249 | 1.431980114 | 1.458222504 | 1.499532413 | 1.447158031 | 1.335191751 |
| ENSMUSG00000062866 | 1.269664473 | 1.42510993 | 1.188721876 | 1.807204673 | 1.793512788 | 1.991226076 | 1.67819947 |
| ENSMUSG00000062867 | 2.720536789 | 2.47104263 | 2.516522298 | 2.573367912 | 2.553601703 | 2.92788341 | 2.388432563 |
| ENSMUSG00000062868 | 0 | 0 | 0 | 0 | 0 | 0 | 0 |
| ENSMUSG00000062873 | 0 | 0 | 0 | 0 | 0 | 0 | 0 |
| ENSMUSG00000062874 | 0.578129653 | 0.5 | 0.5 | 0.25 | 0.244650542 | 0.301029996 | 0.355418052 |
| ENSMUSG00000062878 | 0 | 0 | 0 | 0 | 0 | 0 | 0 |
| ENSMUSG00000062892 | 0 | 0 | 0 | 0 | 0 | 0 | 0 |

| | | | | | | | |
|---|---|---|---|---|---|---|---|
| ENSMUSG00000062896 | 0.289064826 | 0.5 | 0.396240625 | 0.25 | 0.244650542 | 0.301029996 | 0 |
| ENSMUSG00000062901 | 2.35433422 | 2.132845365 | 2.123963757 | 1.852347734 | 1.749189509 | 1.973127854 | 1.627424404 |
| ENSMUSG00000062905 | 0 | 0.25 | 0 | 0.580482024 | 0.489301084 | 0.477121255 | 0.448487648 |
| ENSMUSG00000062906 | 1.885732304 | 1.816696635 | 1.856566189 | 1.678561379 | 1.887294348 | 1.908485019 | 1.793950594 |
| ENSMUSG00000062908 | 2.536482297 | 2.284887838 | 2.319612365 | 2.363317659 | 2.306653428 | 2.627365857 | 2.15438449 |
| ENSMUSG00000062914 | 0 | 0 | 0 | 0 | 0 | 0 | 0 |
| ENSMUSG00000062929 | 2.987920473 | 2.722566069 | 2.714300868 | 2.696771581 | 2.617231464 | 3.017867719 | 2.416917233 |
| ENSMUSG00000062931 | 1.639698646 | 1.671625132 | 1.744319981 | 1.819031101 | 1.72341484 | 1.875061263 | 1.625302983 |
| ENSMUSG00000062933 | 0 | 0 | 0 | 0 | 0 | 0 | 0 |
| ENSMUSG00000062937 | 3.044533636 | 2.78456795 | 2.756457917 | 2.834684096 | 2.732728207 | 3.207365037 | 2.53823997 |
| ENSMUSG00000062939 | 0 | 0.25 | 0 | 0 | 0.244650542 | 0 | 0 |
| ENSMUSG00000062944 | 1.205378646 | 1.282320754 | 1.25 | 1.330482024 | 1.27449559 | 1.380211242 | 1.278288402 |
| ENSMUSG00000062949 | 2.29078455 | 2.153677461 | 2.125956434 | 2.256284891 | 2.148364414 | 2.487138375 | 2 |
| ENSMUSG00000062952 | 0 | 0 | 0 | 0 | 0 | 0 | 0 |
| ENSMUSG00000062960 | 1.100572389 | 1.02186571 | 1.17510993 | 0.830482024 | 0.686820904 | 0.602059991 | 0.672731473 |
| ENSMUSG00000062961 | 0 | 0 | 0 | 0.25 | 0 | 0 | 0.224243824 |
| ENSMUSG00000062962 | 0 | 0 | 0 | 0 | 0 | 0 | 0 |
| ENSMUSG00000062963 | 2.440557995 | 2.214495249 | 2.272528105 | 2.420748646 | 2.394615667 | 2.7084209 | 2.257861346 |
| ENSMUSG00000062980 | 1.558729299 | 1.445339928 | 1.52186571 | 0.951838731 | 0.812711509 | 0.77815125 | 0.853775828 |
| ENSMUSG00000062981 | 2.297593393 | 2.223704441 | 2.192046081 | 2.31611065 | 2.280614762 | 2.610660163 | 2.104721678 |
| ENSMUSG00000062987 | 0 | 0 | 0 | 0 | 0 | 0 | 0 |
| ENSMUSG00000062991 | 2.229917041 | 2.08272922 | 2.059601185 | 1.741446071 | 1.855665188 | 1.913813852 | 1.709198048 |
| ENSMUSG00000062995 | 2.042557845 | 1.894357207 | 1.938721876 | 1.816696635 | 1.828942237 | 2.064457989 | 1.707551656 |
| ENSMUSG00000062997 | 2.50175086 | 2.314346961 | 2.385274154 | 2.290597832 | 2.313753359 | 2.624282096 | 2.215091912 |
| ENSMUSG00000063001 | 0 | 0 | 0.25 | 0 | 0 | 0 | 0.224243824 |
| ENSMUSG00000063011 | 2.241665209 | 2.019203899 | 2.111770807 | 2.263820609 | 2.371516839 | 2.571708832 | 2.211635518 |
| ENSMUSG00000063015 | 3.105223811 | 2.806452485 | 2.77219706 | 2.74770327 | 2.647942663 | 3.062957834 | 2.471542337 |
| ENSMUSG00000063018 | 1.227926429 | 1.214495249 | 1.25 | 0.864857905 | 0.905314583 | 1 | 0.984950056 |
| ENSMUSG00000063019 | 1.862428343 | 1.862802778 | 1.843759858 | 1.854463129 | 1.903713872 | 2.037426498 | 1.87805087 |
| ENSMUSG00000063020 | 0 | 0 | 0 | 0 | 0 | 0 | 0 |
| ENSMUSG00000063021 | 0 | 0 | 0 | 0 | 0 | 0 | 0 |
| ENSMUSG00000063047 | 1.181541736 | 1.160964047 | 1.238549078 | 1.282320754 | 1.27449559 | 1.322219295 | 1.168187745 |
| ENSMUSG00000063049 | 1.932832114 | 1.752806814 | 1.76109853 | 1.799918086 | 1.715300549 | 1.995635195 | 1.687112305 |
| ENSMUSG00000063052 | 2.528794847 | 2.280383379 | 2.311981878 | 2.444929839 | 2.371090306 | 2.700703717 | 2.218171406 |

| | | | | | | | |
|---|---|---|---|---|---|---|---|
| ENSMUSG00000063058 | 0 | 0 | 0 | 0 | 0 | 0 | 0 |
| ENSMUSG00000063060 | 0.747221736 | 0.580482024 | 0.5 | 0.25 | 0.905314583 | 0.602059991 | 0.629532004 |
| ENSMUSG00000063063 | 0.289064826 | 0.396240625 | 0.25 | 0 | 0.244650542 | 0.301029996 | 0 |
| ENSMUSG00000063065 | 2.707474304 | 2.491085217 | 2.539020769 | 2.411864607 | 2.44581538 | 2.720985744 | 2.338260508 |
| ENSMUSG00000063077 | 3.057562627 | 2.797147211 | 2.775165585 | 2.879786197 | 2.795056894 | 3.256477206 | 2.597012708 |
| ENSMUSG00000063087 | 0.289064826 | 0.75 | 0.92510993 | 0.646240625 | 0.244650542 | 0.903089987 | 0.710836105 |
| ENSMUSG00000063089 | 0 | 0 | 0 | 0 | 0 | 0 | 0 |
| ENSMUSG00000063106 | 0 | 0 | 0 | 0 | 0 | 0 | 0 |
| ENSMUSG00000063108 | 2.158622101 | 2.023189285 | 1.985628626 | 2.243603647 | 2.157079823 | 2.432969291 | 1.991191705 |
| ENSMUSG00000063109 | 0 | 0 | 0 | 0.396240625 | 0 | 0 | 0 |
| ENSMUSG00000063116 | 0 | 0 | 0 | 0 | 0 | 0 | 0 |
| ENSMUSG00000063120 | 0 | 0 | 0 | 0 | 0 | 0 | 0 |
| ENSMUSG00000063129 | 0 | 0 | 0.25 | 0 | 0 | 0 | 0.224243824 |
| ENSMUSG00000063130 | 0 | 0 | 0 | 0 | 0 | 0 | 0 |
| ENSMUSG00000063133 | 0 | 0 | 0 | 0 | 0 | 0 | 0 |
| ENSMUSG00000063137 | 0 | 0 | 0 | 0 | 0 | 0 | 0 |
| ENSMUSG00000063142 | 0 | 0 | 0 | 0.396240625 | 0 | 0.301029996 | 0.224243824 |
| ENSMUSG00000063145 | 1.678702042 | 1.51109853 | 1.552363341 | 1.705044741 | 1.683842353 | 1.832508913 | 1.540636858 |
| ENSMUSG00000063146 | 2.323768721 | 2.26584877 | 2.373963757 | 1.896240625 | 2.106691885 | 2.120573931 | 2.126687118 |
| ENSMUSG00000063157 | 0 | 0 | 0.25 | 0.25 | 0 | 0 | 0 |
| ENSMUSG00000063160 | 1.783508299 | 1.769203899 | 1.828220739 | 1.410964047 | 1.568060967 | 1.505149978 | 1.625302983 |
| ENSMUSG00000063163 | 0 | 0 | 0 | 0 | 0 | 0 | 0 |
| ENSMUSG00000063166 | 0 | 0 | 0 | 0.25 | 0.244650542 | 0 | 0.224243824 |
| ENSMUSG00000063171 | 0 | 0 | 0.25 | 1.146240625 | 1.489301084 | 1.322219295 | 1.753626837 |
| ENSMUSG00000063172 | 1.806056082 | 1.752806814 | 1.695339928 | 1.752806814 | 1.731346772 | 1.934498451 | 1.616675393 |
| ENSMUSG00000063173 | 0 | 0 | 0 | 0 | 0 | 0 | 0 |
| ENSMUSG00000063176 | 0 | 0 | 0 | 0 | 0 | 0 | 0 |
| ENSMUSG00000063177 | 0 | 0 | 0 | 0 | 0 | 0 | 0 |
| ENSMUSG00000063179 | 1.671187741 | 1.54248125 | 1.552363341 | 1.668106335 | 1.533172725 | 1.681241237 | 1.496250175 |
| ENSMUSG00000063180 | 0 | 0.25 | 0.25 | 0 | 0 | 0.301029996 | 0 |
| ENSMUSG00000063188 | 0 | 0 | 0 | 0 | 0 | 0 | 0 |
| ENSMUSG00000063193 | 1.269664473 | 1.098079356 | 1.098079356 | 0.580482024 | 0.489301084 | 0.477121255 | 0.672731473 |
| ENSMUSG00000063200 | 2.380049842 | 2.251406137 | 2.272528105 | 2.481944491 | 2.386538516 | 2.751279104 | 2.225910574 |
| ENSMUSG00000063206 | 0 | 0 | 0 | 0 | 0 | 0 | 0 |

| | | | | | | | |
|---|---|---|---|---|---|---|---|
| ENSMUSG00000063216 | 0 | 0 | 0 | 0 | 0 | 0 | 0 |
| ENSMUSG00000063221 | 0 | 0 | 0 | 0 | 0 | 0 | 0 |
| ENSMUSG00000063225 | 0 | 0 | 0 | 0 | 0 | 0 | 0 |
| ENSMUSG00000063229 | 3.420586599 | 3.124587652 | 3.136060002 | 3.012632226 | 3.005630313 | 3.453165393 | 2.815826328 |
| ENSMUSG00000063230 | 0 | 0 | 0 | 0 | 0 | 0 | 0 |
| ENSMUSG00000063232 | 0.289064826 | 0.25 | 0 | 0 | 0 | 0 | 0 |
| ENSMUSG00000063234 | 0 | 0 | 0 | 0.25 | 0.244650542 | 0 | 0 |
| ENSMUSG00000063235 | 1.765759561 | 1.660964047 | 1.642463902 | 1.995748394 | 1.951645858 | 2.264817823 | 1.817112247 |
| ENSMUSG00000063236 | 1.975148165 | 1.935366747 | 1.943696765 | 1.76109853 | 1.897237431 | 2 | 1.913982791 |
| ENSMUSG00000063239 | 0 | 0 | 0 | 0 | 0.244650542 | 0 | 0.448487648 |
| ENSMUSG00000063240 | 0 | 0 | 0 | 0 | 0 | 0 | 0 |
| ENSMUSG00000063245 | 0.289064826 | 0.396240625 | 0.396240625 | 1.04248125 | 0.632412477 | 0.84509804 | 0.710836105 |
| ENSMUSG00000063251 | 0 | 0 | 0 | 0 | 0 | 0 | 0 |
| ENSMUSG00000063252 | 0 | 0 | 0 | 0 | 0 | 0 | 0 |
| ENSMUSG00000063253 | 2.711248379 | 2.53199858 | 2.524508021 | 2.613574823 | 2.558403901 | 2.936010796 | 2.326294816 |
| ENSMUSG00000063254 | 0.45815691 | 0.396240625 | 0.864857905 | 0.646240625 | 0.489301084 | 0.602059991 | 0.579661877 |
| ENSMUSG00000063260 | 0 | 0 | 0 | 0 | 0 | 0 | 0 |
| ENSMUSG00000063268 | 1.960252568 | 1.837182039 | 1.923371739 | 1.910964047 | 2.048957937 | 2.209515015 | 1.993927582 |
| ENSMUSG00000063273 | 3.014774648 | 2.749118372 | 2.749647607 | 2.908816164 | 2.813808973 | 3.230704314 | 2.560973592 |
| ENSMUSG00000063275 | 2.097855556 | 1.966046536 | 1.938721876 | 1.729715809 | 1.689774529 | 1.880813592 | 1.540636858 |
| ENSMUSG00000063277 | 0 | 0 | 0 | 0 | 0 | 0 | 0 |
| ENSMUSG00000063281 | 2.144240249 | 1.992885888 | 1.979715809 | 2.05479213 | 1.996316946 | 2.230448921 | 1.848480858 |
| ENSMUSG00000063286 | 0 | 0.25 | 0.396240625 | 0 | 0 | 0 | 0 |
| ENSMUSG00000063296 | 0.747221736 | 0.396240625 | 0.864857905 | 1.630890489 | 1.611014646 | 1.643452676 | 1.429563222 |
| ENSMUSG00000063297 | 0 | 0 | 0 | 0 | 0.387761935 | 0 | 0 |
| ENSMUSG00000063305 | 0 | 0 | 0 | 0 | 0 | 0 | 0 |
| ENSMUSG00000063314 | 0.289064826 | 0.25 | 0.25 | 0.396240625 | 0 | 0 | 0.448487648 |
| ENSMUSG00000063316 | 2.742600832 | 2.566989271 | 2.58993739 | 2.497880462 | 2.57000966 | 2.856124444 | 2.434835182 |
| ENSMUSG00000063317 | 2.40033094 | 2.242885888 | 2.243603647 | 2.149978211 | 2.113840608 | 2.369215857 | 1.963977467 |
| ENSMUSG00000063320 | 0.91631382 | 1.188721876 | 1.146240625 | 1 | 1.200473444 | 1.361727836 | 1.238624656 |
| ENSMUSG00000063328 | 0 | 0 | 0 | 0 | 0 | 0 | 0 |
| ENSMUSG00000063334 | 2.472830741 | 2.321928095 | 2.31728167 | 2.402294685 | 2.319727371 | 2.675778342 | 2.166993225 |
| ENSMUSG00000063350 | 0 | 0 | 0 | 0 | 0 | 0 | 0 |
| ENSMUSG00000063354 | 0.578129653 | 0.396240625 | 0 | 0.580482024 | 0.568060967 | 0.301029996 | 0.629532004 |

| | | | | | | | |
|---|---|---|---|---|---|---|---|
| ENSMUSG00000063358 | 3.120285948 | 2.826942508 | 2.80175358 | 2.937153044 | 2.830919587 | 3.287353773 | 2.628288264 |
| ENSMUSG00000063362 | 2.201144779 | 1.994319981 | 1.951838731 | 2.078220739 | 2.004851707 | 2.320146286 | 1.854824501 |
| ENSMUSG00000063364 | 0.671187741 | 0.701838731 | 0.646240625 | 0.864857905 | 0.632412477 | 1.113943352 | 0.520678036 |
| ENSMUSG00000063374 | 0 | 0 | 0 | 0 | 0 | 0 | 0 |
| ENSMUSG00000063376 | 0 | 0 | 0 | 0 | 0 | 0 | 0 |
| ENSMUSG00000063380 | 0 | 0 | 0 | 0 | 0 | 0 | 0 |
| ENSMUSG00000063382 | 2.533616084 | 2.43325383 | 2.530706999 | 1.909156155 | 2.179075613 | 2.127104798 | 2.242438242 |
| ENSMUSG00000063383 | 1.249317394 | 1.201838731 | 1.282320754 | 1.410964047 | 1.343584837 | 1.414973348 | 1.278288402 |
| ENSMUSG00000063386 | 0 | 0 | 0 | 0 | 0 | 0 | 0 |
| ENSMUSG00000063388 | 0 | 0 | 0 | 0.25 | 0 | 0.301029996 | 0 |
| ENSMUSG00000063394 | 0 | 0 | 0 | 0 | 0 | 0 | 0 |
| ENSMUSG00000063406 | 2.869771412 | 2.670748646 | 2.709012589 | 2.769036882 | 2.676014376 | 3.091666958 | 2.459817183 |
| ENSMUSG00000063409 | 0 | 0 | 0.25 | 0 | 0 | 0 | 0 |
| ENSMUSG00000063410 | 2.395018353 | 2.21759118 | 2.209800947 | 2.407794264 | 2.366796383 | 2.672097858 | 2.211982824 |
| ENSMUSG00000063412 | 0 | 0 | 0 | 0 | 0 | 0 | 0 |
| ENSMUSG00000063415 | 0.960252568 | 0.976722649 | 0.864857905 | 0.5 | 0.568060967 | 0.477121255 | 0.520678036 |
| ENSMUSG00000063428 | 0 | 0 | 0 | 0 | 0 | 0 | 0 |
| ENSMUSG00000063430 | 0.45815691 | 0 | 0.25 | 0.896240625 | 1.074582839 | 1 | 1.014380832 |
| ENSMUSG00000063434 | 0 | 0 | 0.25 | 0 | 0 | 0 | 0.355418052 |
| ENSMUSG00000063439 | 1.289064826 | 1.26109853 | 1.42510993 | 1.160964047 | 1.407936347 | 1.278753601 | 1.508694612 |
| ENSMUSG00000063442 | 0.289064826 | 0.701838731 | 0.79248125 | 0.396240625 | 0.686820904 | 0.301029996 | 0.520678036 |
| ENSMUSG00000063445 | 2.35433422 | 2.189555804 | 2.159156155 | 2.134789703 | 2.188507684 | 2.371067862 | 2.088356488 |
| ENSMUSG00000063446 | 0 | 0 | 0 | 0 | 0 | 0 | 0 |
| ENSMUSG00000063447 | 0 | 0 | 0 | 0 | 0 | 0 | 0.224243824 |
| ENSMUSG00000063450 | 3.031955969 | 2.790597832 | 2.787754135 | 2.525165585 | 2.489606278 | 2.804820679 | 2.342462067 |
| ENSMUSG00000063455 | 2.160971586 | 2.031030328 | 2.017865591 | 1.987091808 | 1.94314048 | 2.178976947 | 1.830929954 |
| ENSMUSG00000063457 | 3.079362782 | 2.859307975 | 2.916111821 | 2.709997768 | 2.884913522 | 3.168497484 | 2.760196558 |
| ENSMUSG00000063458 | 0.289064826 | 0.25 | 0 | 0 | 0 | 0.301029996 | 0 |
| ENSMUSG00000063478 | 0 | 0 | 0 | 0 | 0 | 0 | 0 |
| ENSMUSG00000063480 | 2.956096064 | 2.708024712 | 2.664329612 | 2.821061437 | 2.742547864 | 3.196728723 | 2.561798737 |
| ENSMUSG00000063488 | 0.960252568 | 0.92510993 | 0.830482024 | 0.864857905 | 0.686820904 | 0.954242509 | 0.829800754 |
| ENSMUSG00000063506 | 1.45815691 | 1.438721876 | 1.606566189 | 1.482684334 | 1.632412477 | 1.681241237 | 1.720494 |
| ENSMUSG00000063507 | 0 | 0 | 0 | 0 | 0 | 0 | 0 |
| ENSMUSG00000063511 | 2.667031238 | 2.481944491 | 2.552363341 | 2.465659339 | 2.461354714 | 2.767897616 | 2.466839994 |

| | | | | | | | |
|---|---|---|---|---|---|---|---|
| ENSMUSG00000063514 | 0.960252568 | 1.02186571 | 1 | 0.864857905 | 0.812711509 | 0.77815125 | 0.775756176 |
| ENSMUSG00000063522 | 0.960252568 | 0.864857905 | 0.830482024 | 1.04248125 | 0.905314583 | 0.84509804 | 0.952571511 |
| ENSMUSG00000063524 | 3.317962399 | 3.04575375 | 3.074229052 | 2.915778327 | 2.993913149 | 3.349083169 | 2.829286175 |
| ENSMUSG00000063529 | 0 | 0 | 0 | 0 | 0 | 0 | 0 |
| ENSMUSG00000063531 | 2.381433035 | 2.276477127 | 2.289967834 | 2.245034894 | 2.176862714 | 2.460897843 | 2.044283833 |
| ENSMUSG00000063535 | 0 | 0.396240625 | 0.25 | 0.580482024 | 0.489301084 | 0.477121255 | 0.520678036 |
| ENSMUSG00000063543 | 0 | 0 | 0 | 0 | 0 | 0 | 0 |
| ENSMUSG00000063548 | 0 | 0 | 0 | 0.25 | 0 | 0 | 0 |
| ENSMUSG00000063549 | 0 | 0 | 0 | 0 | 0 | 0 | 0 |
| ENSMUSG00000063550 | 2.502784398 | 2.310793496 | 2.291226732 | 2.323655187 | 2.2705456 | 2.559906625 | 2.106650241 |
| ENSMUSG00000063556 | 0 | 0 | 0.25 | 0 | 0 | 0 | 0 |
| ENSMUSG00000063558 | 1.99653913 | 1.845926073 | 1.821350555 | 1.516522298 | 1.445123986 | 1.568201724 | 1.425641774 |
| ENSMUSG00000063564 | 0 | 0.396240625 | 0.396240625 | 1.364857905 | 1.387761935 | 1.544068044 | 1.46982607 |
| ENSMUSG00000063568 | 1.156259305 | 1.29248125 | 1.201838731 | 1.364857905 | 1.42077253 | 1.431363764 | 1.374453863 |
| ENSMUSG00000063576 | 2.499676068 | 2.376452888 | 2.389126514 | 2.267194569 | 2.317747248 | 2.571708832 | 2.279699287 |
| ENSMUSG00000063582 | 0 | 0 | 0 | 0 | 0 | 0 | 0 |
| ENSMUSG00000063583 | 0 | 0 | 0 | 0 | 0 | 0 | 0 |
| ENSMUSG00000063586 | 0.289064826 | 0.79248125 | 0.25 | 0.701838731 | 0.387761935 | 0.954242509 | 0.355418052 |
| ENSMUSG00000063590 | 0 | 0 | 0 | 0 | 0 | 0 | 0 |
| ENSMUSG00000063594 | 0.289064826 | 0.79248125 | 0.396240625 | 0.396240625 | 0 | 0 | 0.448487648 |
| ENSMUSG00000063600 | 0 | 0 | 0 | 0 | 0 | 0 | 0 |
| ENSMUSG00000063605 | 2.054824387 | 2.063166358 | 2.106566189 | 1.76109853 | 1.94314048 | 1.924279286 | 1.988432495 |
| ENSMUSG00000063611 | 0.289064826 | 0.580482024 | 0.25 | 0.25 | 0.387761935 | 0.301029996 | 0.355418052 |
| ENSMUSG00000063615 | 0 | 0 | 0 | 0 | 0 | 0 | 0 |
| ENSMUSG00000063623 | 0.289064826 | 0.396240625 | 0.5 | 0.5 | 0.489301084 | 0.477121255 | 0.224243824 |
| ENSMUSG00000063626 | 0 | 0 | 0 | 0 | 0 | 0 | 0 |
| ENSMUSG00000063628 | 0 | 0 | 0 | 0 | 0 | 0.301029996 | 0 |
| ENSMUSG00000063632 | 2.711874084 | 2.483053688 | 2.461764337 | 2.204245906 | 2.16634835 | 2.445604203 | 2.009225098 |
| ENSMUSG00000063646 | 0 | 0.25 | 0.25 | 0.701838731 | 0 | 0.602059991 | 0 |
| ENSMUSG00000063651 | 0 | 0 | 0 | 0 | 0.244650542 | 0.301029996 | 0 |
| ENSMUSG00000063652 | 1.558729299 | 1.396240625 | 1.494319981 | 1.618933358 | 1.625423019 | 1.84509804 | 1.517723536 |
| ENSMUSG00000063656 | 0 | 0 | 0 | 0 | 0 | 0 | 0 |
| ENSMUSG00000063659 | 2.233869984 | 2.090485943 | 2.102347734 | 1.868933358 | 1.84445929 | 1.995635195 | 1.787569411 |
| ENSMUSG00000063660 | 0 | 0 | 0 | 0 | 0.244650542 | 0 | 0 |

| | | | | | | | |
|---|---|---|---|---|---|---|---|
| ENSMUSG00000063661 | 0 | 0 | 0 | 0 | 0 | 0 | 0 |
| ENSMUSG00000063663 | 2.229917041 | 2.03743678 | 2.00835575 | 2.255591953 | 2.229044281 | 2.498310554 | 2.016293244 |
| ENSMUSG00000063672 | 0 | 0 | 0 | 0 | 0 | 0 | 0 |
| ENSMUSG00000063681 | 0 | 0 | 0.25 | 0.580482024 | 0.244650542 | 0.301029996 | 0.355418052 |
| ENSMUSG00000063683 | 0 | 0.396240625 | 0.25 | 0 | 0.387761935 | 0.301029996 | 0.224243824 |
| ENSMUSG00000063684 | 0 | 0.396240625 | 0 | 0 | 0 | 0 | 0.224243824 |
| ENSMUSG00000063687 | 0.289064826 | 0.580482024 | 0.580482024 | 0.25 | 0.244650542 | 0.477121255 | 0.579661877 |
| ENSMUSG00000063689 | 0 | 0 | 0 | 0 | 0.244650542 | 0 | 0 |
| ENSMUSG00000063694 | 2.241665209 | 2.084962501 | 2.115881093 | 2.189555804 | 2.107593434 | 2.399673721 | 1.991877847 |
| ENSMUSG00000063696 | 2.531694272 | 2.42510993 | 2.488549078 | 1.978222334 | 2.226481975 | 2.309630167 | 2.170168862 |
| ENSMUSG00000063698 | 1.734388958 | 1.610735874 | 1.557204673 | 1.763820609 | 1.665423072 | 1.86332286 | 1.546120303 |
| ENSMUSG00000063704 | 0.289064826 | 0 | 0.580482024 | 0.580482024 | 0.812711509 | 0.477121255 | 0.896975297 |
| ENSMUSG00000063713 | 0 | 0 | 0 | 0 | 0 | 0 | 0 |
| ENSMUSG00000063714 | 0.91631382 | 0.580482024 | 0.79248125 | 0.92510993 | 0.978602168 | 0.84509804 | 0.853775828 |
| ENSMUSG00000063715 | 0 | 0 | 0 | 0 | 0 | 0 | 0 |
| ENSMUSG00000063727 | 1.842799666 | 1.681980114 | 1.685366747 | 0.396240625 | 0.489301084 | 0 | 0.448487648 |
| ENSMUSG00000063728 | 0 | 0 | 0 | 0 | 0 | 0 | 0 |
| ENSMUSG00000063730 | 1.036286563 | 0.976722649 | 0.5 | 1.02186571 | 0.931471446 | 0.698970004 | 0.853775828 |
| ENSMUSG00000063732 | 0 | 0 | 0 | 0 | 0 | 0 | 0 |
| ENSMUSG00000063739 | 0 | 0.396240625 | 0.25 | 0.701838731 | 0.489301084 | 0 | 0 |
| ENSMUSG00000063751 | 0 | 0 | 0 | 0.580482024 | 0.387761935 | 0 | 0 |
| ENSMUSG00000063754 | 0 | 0 | 0.396240625 | 0.5 | 0.244650542 | 0.477121255 | 0.629532004 |
| ENSMUSG00000063757 | 0 | 0.701838731 | 0.25 | 0 | 0.489301084 | 0.301029996 | 0.520678036 |
| ENSMUSG00000063760 | 2.309247751 | 2.149047439 | 2.171625132 | 2.099151195 | 2.006054284 | 2.311753861 | 1.857950279 |
| ENSMUSG00000063762 | 0 | 0 | 0 | 0 | 0 | 0 | 0 |
| ENSMUSG00000063764 | 0 | 0 | 0 | 0 | 0 | 0 | 0 |
| ENSMUSG00000063765 | 0.811507563 | 0.951838731 | 0.951838731 | 0.701838731 | 1.074582839 | 1.041392685 | 1.238624656 |
| ENSMUSG00000063767 | 1.269664473 | 1.330482024 | 1.201838731 | 1.380890489 | 1.254881871 | 1.531478917 | 1.23151414 |
| ENSMUSG00000063777 | 0 | 0 | 0 | 0 | 0 | 0 | 0 |
| ENSMUSG00000063779 | 0 | 0 | 0 | 0 | 0 | 0 | 0 |
| ENSMUSG00000063780 | 0 | 0 | 0 | 0 | 0 | 0 | 0 |
| ENSMUSG00000063785 | 2.370236862 | 2.161864607 | 2.138647213 | 2.130890489 | 2.063536368 | 2.318063335 | 1.902443294 |
| ENSMUSG00000063787 | 2.179305632 | 2.004202072 | 2.099151195 | 2.27908599 | 2.21338377 | 2.588831726 | 2.059095225 |
| ENSMUSG00000063796 | 0 | 0 | 0 | 0 | 0 | 0 | 0.224243824 |

| | | | | | | | |
|---|---|---|---|---|---|---|---|
| ENSMUSG00000063800 | 2.508932407 | 2.313166358 | 2.286164561 | 2.460587586 | 2.378690133 | 2.73479983 | 2.205673547 |
| ENSMUSG00000063801 | 2.304292853 | 2.173371739 | 2.137686696 | 2.304186465 | 2.229044281 | 2.542825427 | 2.117056606 |
| ENSMUSG00000063802 | 1.765759561 | 1.845926073 | 1.96293726 | 1.464495249 | 1.777823267 | 1.672097858 | 1.894206246 |
| ENSMUSG00000063804 | 0.811507563 | 0 | 0.701838731 | 0 | 0.489301084 | 0.477121255 | 0.448487648 |
| ENSMUSG00000063808 | 1.707474304 | 1.685366747 | 1.642463902 | 1.726722649 | 1.686820904 | 1.897627091 | 1.643912836 |
| ENSMUSG00000063810 | 2.331622671 | 2.195339928 | 2.14151351 | 2.144357207 | 2.097547432 | 2.281033367 | 1.938331368 |
| ENSMUSG00000063816 | 0.671187741 | 0.396240625 | 0.580482024 | 0.75 | 0.632412477 | 0.602059991 | 0.579661877 |
| ENSMUSG00000063820 | 0 | 0.5 | 0.5 | 0.580482024 | 0.244650542 | 0.301029996 | 0 |
| ENSMUSG00000063821 | 0 | 0 | 0 | 0 | 0 | 0 | 0 |
| ENSMUSG00000063823 | 0 | 0 | 0 | 0 | 0 | 0 | 0 |
| ENSMUSG00000063827 | 0 | 0 | 0 | 0 | 0 | 0 | 0 |
| ENSMUSG00000063830 | 0.671187741 | 0.25 | 0.396240625 | 0.25 | 0.632412477 | 0.77815125 | 0.710836105 |
| ENSMUSG00000063838 | 0.867194479 | 0.976722649 | 1.02186571 | 0.75 | 0.877063019 | 0.84509804 | 1.054044578 |
| ENSMUSG00000063842 | 0 | 0 | 0 | 0 | 0 | 0 | 0 |
| ENSMUSG00000063844 | 0 | 0 | 0 | 0 | 0 | 0 | 0 |
| ENSMUSG00000063846 | 0 | 0 | 0 | 0 | 0 | 0 | 0 |
| ENSMUSG00000063849 | 1.765759561 | 1.732684334 | 1.76109853 | 1.678561379 | 1.689774529 | 1.73239376 | 1.548827561 |
| ENSMUSG00000063851 | 0 | 0 | 0.25 | 0.396240625 | 0.387761935 | 0 | 0.629532004 |
| ENSMUSG00000063856 | 2.988886945 | 2.80479213 | 2.872838018 | 2.55175358 | 2.692188503 | 2.937517892 | 2.629245411 |
| ENSMUSG00000063867 | 0 | 0 | 0 | 0 | 0 | 0 | 0 |
| ENSMUSG00000063870 | 3.32043997 | 2.997614641 | 3.024672524 | 3.006284891 | 2.980808865 | 3.413299764 | 2.792164961 |
| ENSMUSG00000063873 | 2.294202867 | 2.060793496 | 2.041226732 | 1.622963274 | 1.592135487 | 1.73239376 | 1.559435575 |
| ENSMUSG00000063875 | 0 | 0.580482024 | 0.25 | 0.580482024 | 0.387761935 | 0.301029996 | 0.629532004 |
| ENSMUSG00000063881 | 0 | 0 | 0 | 0 | 0 | 0 | 0 |
| ENSMUSG00000063882 | 2.653588827 | 2.414552871 | 2.415889526 | 2.574802005 | 2.480646735 | 2.88536122 | 2.307731473 |
| ENSMUSG00000063884 | 2.634462296 | 2.416777886 | 2.405054956 | 2.484527332 | 2.389791591 | 2.762678564 | 2.242122155 |
| ENSMUSG00000063885 | 0.811507563 | 0.75 | 0.951838731 | 0.25 | 0.568060967 | 0.477121255 | 0.448487648 |
| ENSMUSG00000063887 | 0.289064826 | 0.25 | 0.396240625 | 0.25 | 0 | 0 | 0.224243824 |
| ENSMUSG00000063888 | 2.887787667 | 2.648581151 | 2.632113528 | 2.749471282 | 2.721680409 | 3.133858125 | 2.544882201 |
| ENSMUSG00000063889 | 1.516991255 | 1.488549078 | 1.348079356 | 1.348079356 | 1.327538614 | 1.491361694 | 1.324583736 |
| ENSMUSG00000063894 | 2.275045611 | 2.12892496 | 2.061981878 | 2.247880462 | 2.174635855 | 2.485721426 | 2.002662694 |
| ENSMUSG00000063895 | 2.461589295 | 2.25835575 | 2.242885888 | 2.476722649 | 2.393816219 | 2.774516966 | 2.264421331 |
| ENSMUSG00000063897 | 1.516991255 | 1.54745614 | 1.735628626 | 1.418106335 | 1.528558853 | 1.447158031 | 1.676387053 |
| ENSMUSG00000063900 | 0 | 0 | 0 | 0 | 0 | 0 | 0 |

| | | | | | | | |
|---|---|---|---|---|---|---|---|
| ENSMUSG00000063902 | 0 | 0.25 | 0.5 | 0.396240625 | 0 | 0.602059991 | 0.448487648 |
| ENSMUSG00000063903 | 0 | 0 | 0.25 | 0 | 0 | 0 | 0.224243824 |
| ENSMUSG00000063904 | 2.791849225 | 2.587182039 | 2.615369862 | 2.539652422 | 2.60819185 | 2.902002891 | 2.488369842 |
| ENSMUSG00000063905 | 0 | 0 | 0 | 0 | 0 | 0 | 0 |
| ENSMUSG00000063911 | 0 | 0 | 0 | 0 | 0 | 0 | 0.224243824 |
| ENSMUSG00000063919 | 1.771760131 | 1.735628626 | 1.777131114 | 0.580482024 | 0.905314583 | 1.041392685 | 0.710836105 |
| ENSMUSG00000063929 | 0 | 0 | 0 | 0 | 0 | 0 | 0 |
| ENSMUSG00000063931 | 2.241665209 | 2.100219859 | 2.12892496 | 1.821350555 | 1.88560959 | 1.977723605 | 1.852723744 |
| ENSMUSG00000063932 | 0 | 0 | 0 | 0 | 0 | 0 | 0 |
| ENSMUSG00000063935 | 0.45815691 | 0.25 | 0.396240625 | 0.25 | 0 | 0.301029996 | 0.355418052 |
| ENSMUSG00000063952 | 2.442947872 | 2.323655187 | 2.32991803 | 2.213717096 | 2.276175004 | 2.482873584 | 2.225910574 |
| ENSMUSG00000063953 | 1 | 0.896240625 | 0.75 | 1.061981878 | 1 | 0.602059991 | 0.952571511 |
| ENSMUSG00000063954 | 0 | 0 | 0 | 0 | 0 | 0 | 0 |
| ENSMUSG00000063971 | 0.578129653 | 0.5 | 0.896240625 | 0.25 | 0.244650542 | 0 | 0 |
| ENSMUSG00000063972 | 1.800532393 | 1.732684334 | 1.634789703 | 1.571350555 | 1.514343934 | 1.653212514 | 1.417653332 |
| ENSMUSG00000063975 | 1 | 0.864857905 | 0.896240625 | 1.339388001 | 1.091002363 | 1.204119983 | 1.238624656 |
| ENSMUSG00000063994 | 0 | 0 | 0 | 0 | 0 | 0 | 0 |
| ENSMUSG00000063998 | 0 | 0 | 0 | 0 | 0 | 0 | 0 |
| ENSMUSG00000064006 | 0 | 0 | 0 | 0 | 0 | 0 | 0 |
| ENSMUSG00000064010 | 0 | 0 | 0 | 0 | 0 | 0 | 0 |
| ENSMUSG00000064016 | 0.289064826 | 0.396240625 | 0.5 | 0.5 | 0 | 0.301029996 | 0.448487648 |
| ENSMUSG00000064023 | 0 | 0 | 0 | 0 | 0.244650542 | 0.602059991 | 0.448487648 |
| ENSMUSG00000064030 | 1.940852214 | 1.903677461 | 1.966046536 | 1.811981878 | 1.958580386 | 2.120573931 | 1.914853627 |
| ENSMUSG00000064032 | 1.205378646 | 1.080482024 | 1.061981878 | 1.098079356 | 0.931471446 | 1.079181246 | 1.041356071 |
| ENSMUSG00000064036 | 0 | 0 | 0 | 0.25 | 0 | 0 | 0 |
| ENSMUSG00000064037 | 2.450036412 | 2.272528105 | 2.219895812 | 2.347542239 | 2.273371525 | 2.599883072 | 2.135599953 |
| ENSMUSG00000064039 | 0 | 0 | 0 | 0 | 0 | 0 | 0 |
| ENSMUSG00000064043 | 2.139333929 | 1.981203126 | 2.072504712 | 1.825945187 | 1.88560959 | 1.995635195 | 1.828667599 |
| ENSMUSG00000064044 | 0 | 0 | 0 | 0 | 0 | 0 | 0 |
| ENSMUSG00000064052 | 0 | 0 | 0 | 0 | 0 | 0 | 0 |
| ENSMUSG00000064057 | 0 | 0 | 0 | 0.25 | 0 | 0 | 0 |
| ENSMUSG00000064061 | 2.750084015 | 2.518535366 | 2.522528105 | 2.650210529 | 2.555378544 | 2.907411361 | 2.349572583 |
| ENSMUSG00000064063 | 0 | 0 | 0 | 0 | 0 | 0 | 0 |
| ENSMUSG00000064065 | 0 | 0 | 0 | 0 | 0 | 0 | 0 |

| | | | | | | | |
|---|---|---|---|---|---|---|---|
| ENSMUSG00000064068 | 2.023453784 | 1.894357207 | 2.005591953 | 1.868933358 | 1.897237431 | 2.029383778 | 1.901538355 |
| ENSMUSG00000064080 | 3.111026095 | 2.918879011 | 2.965465585 | 2.583847589 | 2.711579226 | 2.98721923 | 2.570739083 |
| ENSMUSG00000064081 | 0.289064826 | 0 | 0 | 0 | 0 | 0 | 0 |
| ENSMUSG00000064084 | 0 | 0 | 0 | 0 | 0 | 0 | 0 |
| ENSMUSG00000064090 | 1.940852214 | 1.794977273 | 1.777131114 | 2.02186571 | 1.934425071 | 2.209515015 | 1.788855757 |
| ENSMUSG00000064105 | 1.587501561 | 1.561981878 | 1.54745614 | 1.470660762 | 1.478764298 | 1.681241237 | 1.455757965 |
| ENSMUSG00000064109 | 0 | 0 | 0 | 0 | 0 | 0 | 0 |
| ENSMUSG00000064110 | 0 | 0 | 0 | 0 | 0 | 0 | 0 |
| ENSMUSG00000064115 | 0 | 0 | 0.25 | 0.75 | 0.733951626 | 0.477121255 | 0.355418052 |
| ENSMUSG00000064120 | 1.993049299 | 1.898114259 | 1.912762923 | 1.660964047 | 1.773209395 | 1.73239376 | 1.753626837 |
| ENSMUSG00000064121 | 0 | 0.25 | 0.25 | 0 | 0 | 0 | 0 |
| ENSMUSG00000064125 | 0.45815691 | 0.580482024 | 0.396240625 | 0.580482024 | 0.568060967 | 0.77815125 | 0.448487648 |
| ENSMUSG00000064127 | 2.899513054 | 2.627196291 | 2.599151195 | 2.71816872 | 2.6330226 | 3.026941628 | 2.425821057 |
| ENSMUSG00000064128 | 1.967766868 | 1.874961472 | 1.811981878 | 2.242166698 | 2.182004726 | 2.489958479 | 2.019455688 |
| ENSMUSG00000064129 | 0 | 0 | 0 | 0 | 0 | 0 | 0 |
| ENSMUSG00000064137 | 0 | 0.5 | 0 | 0.5 | 0.244650542 | 0 | 0 |
| ENSMUSG00000064138 | 2.215778996 | 2.046218836 | 2.086073977 | 2.231944491 | 2.085294053 | 2.357934847 | 1.976436 |
| ENSMUSG00000064140 | 0 | 0 | 0 | 0 | 0 | 0 | 0.224243824 |
| ENSMUSG00000064141 | 1.269664473 | 1.27186571 | 1.372963274 | 1.188721876 | 1.264824954 | 1.146128036 | 1.185218806 |
| ENSMUSG00000064145 | 2.489144387 | 2.345926073 | 2.315523711 | 2.403677461 | 2.357161747 | 2.658011397 | 2.239582276 |
| ENSMUSG00000064147 | 0 | 0 | 0 | 0 | 0 | 0 | 0 |
| ENSMUSG00000064156 | 0 | 0 | 0 | 0 | 0 | 0 | 0 |
| ENSMUSG00000064158 | 0 | 0 | 0 | 0 | 0 | 0.477121255 | 0 |
| ENSMUSG00000064165 | 0 | 0 | 0 | 0 | 0 | 0 | 0 |
| ENSMUSG00000064168 | 0 | 0 | 0 | 0 | 0 | 0 | 0 |
| ENSMUSG00000064177 | 0.747221736 | 0.5 | 0.830482024 | 0.580482024 | 0.733951626 | 0.477121255 | 0.629532004 |
| ENSMUSG00000064179 | 0 | 0 | 0 | 0 | 0.733951626 | 0.301029996 | 0.224243824 |
| ENSMUSG00000064181 | 2.295901576 | 2.14151351 | 2.074802005 | 2.207430684 | 2.223252711 | 2.46834733 | 2.064616101 |
| ENSMUSG00000064193 | 0 | 0.25 | 0 | 0 | 0 | 0 | 0 |
| ENSMUSG00000064194 | 0.289064826 | 0.25 | 0 | 0 | 0 | 0.477121255 | 0.224243824 |
| ENSMUSG00000064201 | 0 | 0 | 0 | 0 | 0 | 0 | 0 |
| ENSMUSG00000064202 | 0.578129653 | 0.25 | 0.646240625 | 0.25 | 0 | 0.301029996 | 0.520678036 |
| ENSMUSG00000064208 | 0 | 0.25 | 0 | 0 | 0 | 0 | 0 |
| ENSMUSG00000064210 | 3.143796597 | 2.874836908 | 2.87521047 | 2.758183632 | 2.691672611 | 3.122215878 | 2.517858418 |

| | | | | | | | |
|---|---|---|---|---|---|---|---|
| ENSMUSG00000064213 | 0 | 0 | 0 | 0 | 0 | 0 | 0 |
| ENSMUSG00000064215 | 1.129344651 | 0.896240625 | 0.951838731 | 1.660964047 | 1.744182955 | 1.886490725 | 1.669034113 |
| ENSMUSG00000064220 | 0 | 0 | 0 | 0 | 0 | 0 | 0 |
| ENSMUSG00000064223 | 0 | 0 | 0 | 0 | 0 | 0 | 0 |
| ENSMUSG00000064224 | 0 | 0 | 0 | 0 | 0 | 0 | 0 |
| ENSMUSG00000064225 | 0 | 0.396240625 | 0.25 | 0.830482024 | 0.812711509 | 1 | 0.775756176 |
| ENSMUSG00000064231 | 0.45815691 | 0.396240625 | 0.396240625 | 0.25 | 0.387761935 | 0 | 0.355418052 |
| ENSMUSG00000064232 | 0 | 0 | 0 | 0 | 0 | 0 | 0 |
| ENSMUSG00000064246 | 0 | 0 | 0 | 0 | 0 | 0 | 0.224243824 |
| ENSMUSG00000064247 | 1.181541736 | 0.701838731 | 0.79248125 | 0.75 | 0.686820904 | 0.477121255 | 0.803905701 |
| ENSMUSG00000064252 | 0 | 0 | 0 | 0 | 0 | 0 | 0 |
| ENSMUSG00000064254 | 1.753493018 | 1.752806814 | 1.809601185 | 1.705044741 | 1.866526233 | 1.977723605 | 1.800208341 |
| ENSMUSG00000064259 | 0 | 0 | 0 | 0 | 0 | 0 | 0 |
| ENSMUSG00000064262 | 0 | 0.25 | 0 | 0.396240625 | 0 | 0 | 0.224243824 |
| ENSMUSG00000064263 | 0 | 0 | 0 | 0 | 0 | 0 | 0 |
| ENSMUSG00000064264 | 1.181541736 | 1.348079356 | 1.431980114 | 1.226722649 | 1.234113756 | 1.113943352 | 1.392431569 |
| ENSMUSG00000064267 | 1.181541736 | 1.098079356 | 1.27186571 | 0.25 | 0.568060967 | 0.301029996 | 0.355418052 |
| ENSMUSG00000064272 | 0 | 0 | 0 | 0 | 0 | 0 | 0 |
| ENSMUSG00000064280 | 0.289064826 | 0.25 | 0.25 | 0.5 | 0.568060967 | 0 | 0.355418052 |
| ENSMUSG00000064281 | 0.289064826 | 0.25 | 0.396240625 | 0 | 0.244650542 | 0 | 0.520678036 |
| ENSMUSG00000064284 | 1.639698646 | 1.610735874 | 1.657339155 | 1.451838731 | 1.607318736 | 1.591064607 | 1.618853957 |
| ENSMUSG00000064288 | 0 | 0 | 0 | 0 | 0 | 0.301029996 | 0 |
| ENSMUSG00000064289 | 2.445324132 | 2.245034894 | 2.24929487 | 2.271202097 | 2.204601633 | 2.568201724 | 2.078019652 |
| ENSMUSG00000064293 | 0 | 0 | 0 | 0 | 0 | 0 | 0 |
| ENSMUSG00000064294 | 0 | 0 | 0 | 0 | 0 | 0 | 0 |
| ENSMUSG00000064299 | 0 | 0 | 0 | 0 | 0 | 0 | 0 |
| ENSMUSG00000064302 | 2.789779581 | 2.569902645 | 2.521534056 | 2.524836953 | 2.50483127 | 2.802773725 | 2.333765523 |
| ENSMUSG00000064307 | 1.205378646 | 1.130890489 | 1.238549078 | 0.92510993 | 0.955822902 | 1.230448921 | 1.041356071 |
| ENSMUSG00000064308 | 0 | 0 | 0 | 0.25 | 0 | 0 | 0 |
| ENSMUSG00000064310 | 0 | 0 | 0 | 0 | 0 | 0 | 0 |
| ENSMUSG00000064317 | 0 | 0 | 0 | 0 | 0 | 0 | 0.224243824 |
| ENSMUSG00000064325 | 0 | 0 | 0 | 0.79248125 | 0.955822902 | 0.903089987 | 0.775756176 |
| ENSMUSG00000064326 | 1.928763473 | 1.973704441 | 2.103406982 | 1.458222504 | 1.733951626 | 1.77815125 | 1.830929954 |
| ENSMUSG00000064329 | 0.867194479 | 0.5 | 0.646240625 | 0.580482024 | 0.632412477 | 0.698970004 | 0.355418052 |

| | | | | | | | |
|---|---|---|---|---|---|---|---|
| ENSMUSG00000064330 | 0.289064826 | 0.396240625 | 0.25 | 0.25 | 0.489301084 | 0.301029996 | 0.448487648 |
| ENSMUSG00000064333 | 0 | 0 | 0 | 0 | 0 | 0 | 0 |
| ENSMUSG00000064336 | 0.289064826 | 0 | 0 | 0 | 0 | 0 | 0 |
| ENSMUSG00000064337 | 3.152146313 | 2.89234524 | 2.914106216 | 2.688095162 | 2.521104673 | 2.918554531 | 2.382749586 |
| ENSMUSG00000064338 | 0.289064826 | 0 | 0.25 | 0.25 | 0 | 0.301029996 | 0 |
| ENSMUSG00000064339 | 3.376254011 | 3.049764706 | 3.128308528 | 3.246638287 | 3.110441996 | 3.594060901 | 2.859830095 |
| ENSMUSG00000064340 | 1.181541736 | 1.17510993 | 1.25 | 1.516522298 | 1.572189156 | 1.612783857 | 1.329931221 |
| ENSMUSG00000064341 | 4.14367552 | 3.741006232 | 3.733791262 | 3.78080807 | 3.627128713 | 4.315193376 | 3.360522384 |
| ENSMUSG00000064342 | 0.45815691 | 0.646240625 | 0.646240625 | 0.830482024 | 0.568060967 | 0.77815125 | 0.579661877 |
| ENSMUSG00000064343 | 0.578129653 | 0.864857905 | 0.864857905 | 0.5 | 0.632412477 | 0.602059991 | 0.710836105 |
| ENSMUSG00000064344 | 0.671187741 | 0.580482024 | 0.830482024 | 0.79248125 | 0.489301084 | 0.698970004 | 0.829800754 |
| ENSMUSG00000064345 | 3.91376465 | 3.530929319 | 3.53801178 | 3.54724378 | 3.414701341 | 4.033182414 | 3.153497189 |
| ENSMUSG00000064346 | 0.671187741 | 0.25 | 0.75 | 0.864857905 | 0.733951626 | 0.602059991 | 0.74492186 |
| ENSMUSG00000064347 | 0.960252568 | 0.864857905 | 0.79248125 | 1.04248125 | 0.877063019 | 1.041392685 | 1.028149525 |
| ENSMUSG00000064348 | 1.418409478 | 1.302363341 | 1.372963274 | 1.42510993 | 1.373641807 | 1.414973348 | 1.369798885 |
| ENSMUSG00000064349 | 1.721148722 | 1.527131114 | 1.671625132 | 1.688721876 | 1.683842353 | 1.819543936 | 1.486592281 |
| ENSMUSG00000064350 | 1.671187741 | 1.470660762 | 1.630890489 | 1.614857905 | 1.414412788 | 1.544068044 | 1.345462945 |
| ENSMUSG00000064351 | 4.127607319 | 3.702529929 | 3.711482756 | 3.81960329 | 3.676595914 | 4.352105303 | 3.405210043 |
| ENSMUSG00000064352 | 2.482695304 | 2.345385738 | 2.352877747 | 2.320192693 | 2.331857641 | 2.605305046 | 2.063519437 |
| ENSMUSG00000064353 | 0 | 0 | 0 | 0 | 0 | 0 | 0 |
| ENSMUSG00000064354 | 0.747221736 | 0.830482024 | 0.830482024 | 0.701838731 | 0.686820904 | 0.954242509 | 0 |
| ENSMUSG00000064355 | 0.289064826 | 0.25 | 0.396240625 | 0.396240625 | 0.568060967 | 0.602059991 | 0.224243824 |
| ENSMUSG00000064356 | 2.617563644 | 2.442459461 | 2.435366747 | 2.621457327 | 2.421513257 | 2.858537198 | 2.296701469 |
| ENSMUSG00000064357 | 0.867194479 | 1.080482024 | 1.160964047 | 0.92510993 | 0.733951626 | 0.602059991 | 0.896975297 |
| ENSMUSG00000064358 | 0.671187741 | 0.5 | 0.701838731 | 0.75 | 0.568060967 | 0.301029996 | 0.520678036 |
| ENSMUSG00000064359 | 1.358731791 | 1.17510993 | 1.080482024 | 1.188721876 | 1.335652905 | 1.342422681 | 1.140832123 |
| ENSMUSG00000064360 | 1.227926429 | 1.226722649 | 1.27186571 | 1.160964047 | 1.074582839 | 1.079181246 | 0.803905701 |
| ENSMUSG00000064361 | 0 | 0 | 0.396240625 | 0.25 | 0.244650542 | 0.301029996 | 0.579661877 |
| ENSMUSG00000064363 | 3.747117282 | 3.380062587 | 3.374400597 | 3.375987483 | 3.217092762 | 3.781180721 | 2.983546516 |
| ENSMUSG00000064364 | 0.578129653 | 0.701838731 | 0.396240625 | 0.701838731 | 0.686820904 | 0.477121255 | 0.629532004 |
| ENSMUSG00000064365 | 0 | 0.580482024 | 0.25 | 0.580482024 | 0.77552387 | 0.477121255 | 0.672731473 |
| ENSMUSG00000064366 | 0.45815691 | 0.396240625 | 0 | 0.580482024 | 0.568060967 | 0.602059991 | 0 |
| ENSMUSG00000064367 | 3.951545939 | 3.545870078 | 3.551963302 | 3.548593031 | 3.400508305 | 3.994361152 | 3.148102439 |
| ENSMUSG00000064368 | 3.427612855 | 3.100619793 | 3.121268641 | 3.08993739 | 2.971275164 | 3.453624074 | 2.759366621 |

| | | | | | | | |
|---|---|---|---|---|---|---|---|
| ENSMUSG00000064369 | 1.993049299 | 1.864857905 | 1.834962501 | 1.845926073 | 1.770879644 | 1.924279286 | 1.653807046 |
| ENSMUSG00000064370 | 4.163153844 | 3.756803721 | 3.764011234 | 3.788209844 | 3.645274317 | 4.314014771 | 3.368741247 |
| ENSMUSG00000064371 | 0.45815691 | 0.580482024 | 0.25 | 0.5 | 0.489301084 | 0.698970004 | 0.579661877 |
| ENSMUSG00000064372 | 1.205378646 | 1.061981878 | 1.130890489 | 1.130890489 | 1.057362051 | 1.255272505 | 0.984950056 |
| ENSMUSG00000064373 | 2.066740409 | 1.933677405 | 1.950224975 | 1.76109853 | 1.741652793 | 1.963787827 | 1.661510159 |
| ENSMUSG00000064377 | 0 | 0 | 0 | 0 | 0 | 0 | 0 |
| ENSMUSG00000064380 | 0.289064826 | 0 | 0 | 0.25 | 0 | 0.477121255 | 0 |
| ENSMUSG00000064382 | 0 | 0 | 0 | 0 | 0 | 0 | 0 |
| ENSMUSG00000064386 | 0 | 0 | 0 | 0 | 0 | 0 | 0 |
| ENSMUSG00000064387 | 0 | 0 | 0 | 0 | 0 | 0 | 0 |
| ENSMUSG00000064389 | 0 | 0 | 0 | 0.396240625 | 0 | 0 | 0 |
| ENSMUSG00000064390 | 0 | 0.25 | 0.25 | 0.5 | 0.686820904 | 0 | 0.355418052 |
| ENSMUSG00000064391 | 0 | 0 | 0 | 0 | 0 | 0 | 0 |
| ENSMUSG00000064392 | 0 | 0 | 0 | 0 | 0 | 0 | 0 |
| ENSMUSG00000064393 | 0 | 0 | 0 | 0 | 0 | 0 | 0 |
| ENSMUSG00000064394 | 0 | 0 | 0 | 0 | 0 | 0 | 0 |
| ENSMUSG00000064395 | 0 | 0 | 0 | 0 | 0 | 0 | 0 |
| ENSMUSG00000064397 | 0 | 0 | 0 | 0 | 0 | 0 | 0 |
| ENSMUSG00000064398 | 0 | 0.25 | 0.25 | 0 | 0 | 0.301029996 | 0 |
| ENSMUSG00000064399 | 0 | 0 | 0 | 0 | 0 | 0 | 0 |
| ENSMUSG00000064400 | 0 | 0 | 0 | 0 | 0 | 0 | 0 |
| ENSMUSG00000064403 | 0 | 0 | 0 | 0 | 0 | 0 | 0 |
| ENSMUSG00000064405 | 0 | 0 | 0 | 0 | 0 | 0 | 0 |
| ENSMUSG00000064407 | 0 | 0 | 0 | 0 | 0 | 0 | 0 |
| ENSMUSG00000064408 | 0 | 0 | 0 | 0 | 0 | 0 | 0 |
| ENSMUSG00000064409 | 0 | 0 | 0 | 0.25 | 0 | 0 | 0.224243824 |
| ENSMUSG00000064410 | 0 | 0 | 0 | 0 | 0 | 0 | 0 |
| ENSMUSG00000064412 | 0 | 0 | 0 | 0 | 0 | 0 | 0 |
| ENSMUSG00000064413 | 0 | 0 | 0 | 0 | 0 | 0 | 0 |
| ENSMUSG00000064414 | 0 | 0 | 0 | 0 | 0 | 0 | 0 |
| ENSMUSG00000064416 | 0 | 0 | 0 | 0 | 0 | 0 | 0 |
| ENSMUSG00000064417 | 0 | 0.25 | 0 | 0 | 0 | 0 | 0 |
| ENSMUSG00000064419 | 0 | 0 | 0 | 0 | 0 | 0 | 0 |
| ENSMUSG00000064422 | 0 | 0 | 0 | 0.25 | 0 | 0 | 0 |

| | | | | | | | |
|---|---|---|---|---|---|---|---|
| ENSMUSG00000064424 | 0 | 0 | 0 | 0 | 0 | 0 | 0 |
| ENSMUSG00000064427 | 0.289064826 | 0.396240625 | 0.25 | 0 | 0.244650542 | 0.301029996 | 0 |
| ENSMUSG00000064428 | 0 | 0 | 0 | 0 | 0 | 0 | 0 |
| ENSMUSG00000064429 | 0 | 0 | 0 | 0 | 0 | 0 | 0 |
| ENSMUSG00000064432 | 0 | 0 | 0 | 0 | 0 | 0 | 0 |
| ENSMUSG00000064433 | 0 | 0 | 0 | 0 | 0 | 0 | 0 |
| ENSMUSG00000064435 | 0.289064826 | 0 | 0 | 0 | 0.244650542 | 0 | 0.355418052 |
| ENSMUSG00000064436 | 0 | 0 | 0 | 0 | 0 | 0 | 0 |
| ENSMUSG00000064437 | 0 | 0 | 0 | 0 | 0.244650542 | 0.477121255 | 0.448487648 |
| ENSMUSG00000064440 | 0 | 0 | 0 | 0 | 0 | 0 | 0 |
| ENSMUSG00000064441 | 0 | 0.396240625 | 0.25 | 0 | 0.244650542 | 0 | 0.355418052 |
| ENSMUSG00000064442 | 0 | 0 | 0 | 0 | 0 | 0 | 0 |
| ENSMUSG00000064443 | 0 | 0 | 0 | 0 | 0 | 0 | 0 |
| ENSMUSG00000064444 | 0 | 0 | 0 | 0 | 0 | 0 | 0 |
| ENSMUSG00000064446 | 0 | 0 | 0 | 0 | 0 | 0 | 0 |
| ENSMUSG00000064450 | 0 | 0 | 0 | 0 | 0 | 0 | 0 |
| ENSMUSG00000064451 | 0 | 0.25 | 0 | 0.396240625 | 0 | 0 | 0 |
| ENSMUSG00000064452 | 0 | 0 | 0 | 0 | 0 | 0 | 0 |
| ENSMUSG00000064453 | 0 | 0 | 0 | 0 | 0 | 0 | 0 |
| ENSMUSG00000064455 | 0 | 0 | 0 | 0 | 0 | 0 | 0 |
| ENSMUSG00000064460 | 0 | 0 | 0 | 0 | 0 | 0 | 0 |
| ENSMUSG00000064463 | 0 | 0 | 0 | 0 | 0 | 0 | 0 |
| ENSMUSG00000064464 | 0 | 0 | 0 | 0 | 0 | 0 | 0 |
| ENSMUSG00000064467 | 0 | 0 | 0 | 0 | 0 | 0 | 0 |
| ENSMUSG00000064468 | 0 | 0 | 0 | 0 | 0 | 0 | 0 |
| ENSMUSG00000064471 | 0 | 0 | 0 | 0 | 0 | 0 | 0 |
| ENSMUSG00000064472 | 0 | 0 | 0 | 0 | 0 | 0 | 0 |
| ENSMUSG00000064476 | 0 | 0 | 0.25 | 0 | 0 | 0 | 0 |
| ENSMUSG00000064477 | 0 | 0 | 0 | 0 | 0 | 0 | 0 |
| ENSMUSG00000064481 | 0 | 0 | 0 | 0 | 0 | 0 | 0 |
| ENSMUSG00000064485 | 0 | 0 | 0 | 0 | 0 | 0 | 0 |
| ENSMUSG00000064486 | 0 | 0 | 0 | 0 | 0 | 0 | 0 |
| ENSMUSG00000064487 | 0 | 0 | 0 | 0 | 0 | 0 | 0 |
| ENSMUSG00000064489 | 0 | 0 | 0 | 0 | 0 | 0 | 0 |

| | | | | | | | |
|---|---|---|---|---|---|---|---|
| ENSMUSG00000064490 | 0 | 0 | 0 | 0 | 0 | 0 | 0 |
| ENSMUSG00000064491 | 0 | 0 | 0 | 0 | 0 | 0 | 0 |
| ENSMUSG00000064492 | 0 | 0 | 0 | 0 | 0 | 0 | 0 |
| ENSMUSG00000064493 | 0.960252568 | 0.25 | 0.75 | 0.79248125 | 0.686820904 | 0.477121255 | 0.355418052 |
| ENSMUSG00000064494 | 0.289064826 | 0 | 0.25 | 0 | 0 | 0 | 0 |
| ENSMUSG00000064495 | 0 | 0 | 0 | 0 | 0 | 0.301029996 | 0.224243824 |
| ENSMUSG00000064496 | 0 | 0 | 0 | 0 | 0 | 0 | 0 |
| ENSMUSG00000064497 | 0 | 0 | 0 | 0 | 0 | 0 | 0 |
| ENSMUSG00000064500 | 0 | 0 | 0 | 0 | 0 | 0 | 0 |
| ENSMUSG00000064507 | 0 | 0 | 0 | 0 | 0 | 0 | 0 |
| ENSMUSG00000064508 | 0 | 0 | 0 | 0 | 0 | 0 | 0 |
| ENSMUSG00000064510 | 0 | 0 | 0 | 0 | 0 | 0 | 0 |
| ENSMUSG00000064511 | 0 | 0 | 0 | 0 | 0 | 0 | 0 |
| ENSMUSG00000064512 | 0 | 0 | 0 | 0 | 0 | 0 | 0 |
| ENSMUSG00000064513 | 0 | 0 | 0 | 0 | 0 | 0 | 0 |
| ENSMUSG00000064514 | 0.811507563 | 0 | 0.396240625 | 0.580482024 | 0.568060967 | 0.602059991 | 0.355418052 |
| ENSMUSG00000064515 | 0 | 0 | 0 | 0 | 0 | 0 | 0 |
| ENSMUSG00000064517 | 0 | 0 | 0 | 0 | 0 | 0 | 0 |
| ENSMUSG00000064519 | 0 | 0 | 0 | 0 | 0 | 0 | 0 |
| ENSMUSG00000064522 | 0 | 0 | 0 | 0 | 0 | 0 | 0 |
| ENSMUSG00000064524 | 0 | 0 | 0 | 0 | 0 | 0 | 0 |
| ENSMUSG00000064525 | 0.289064826 | 0.25 | 0 | 0.396240625 | 0.568060967 | 0 | 0.355418052 |
| ENSMUSG00000064526 | 0 | 0 | 0 | 0 | 0 | 0 | 0 |
| ENSMUSG00000064527 | 0 | 0 | 0 | 0 | 0 | 0 | 0 |
| ENSMUSG00000064529 | 0 | 0 | 0 | 0 | 0 | 0 | 0 |
| ENSMUSG00000064530 | 0 | 0.25 | 0 | 0 | 0 | 0 | 0 |
| ENSMUSG00000064532 | 0 | 0 | 0 | 0 | 0 | 0 | 0 |
| ENSMUSG00000064536 | 0 | 0.25 | 0 | 0 | 0 | 0 | 0 |
| ENSMUSG00000064540 | 0 | 0 | 0 | 0 | 0 | 0 | 0 |
| ENSMUSG00000064541 | 0 | 0 | 0 | 0 | 0 | 0 | 0 |
| ENSMUSG00000064542 | 0 | 0 | 0 | 0 | 0.387761935 | 0 | 0 |
| ENSMUSG00000064543 | 0 | 0 | 0 | 0 | 0 | 0 | 0 |
| ENSMUSG00000064545 | 0 | 0 | 0 | 0 | 0 | 0 | 0 |
| ENSMUSG00000064547 | 0 | 0 | 0 | 0 | 0 | 0 | 0 |

| | | | | | | | |
|---|---|---|---|---|---|---|---|
| ENSMUSG00000064548 | 0 | 0 | 0 | 0 | 0 | 0 | 0 |
| ENSMUSG00000064549 | 0 | 0 | 0 | 0 | 0 | 0 | 0 |
| ENSMUSG00000064550 | 0 | 0 | 0 | 0 | 0 | 0 | 0 |
| ENSMUSG00000064551 | 0 | 0 | 0 | 0 | 0 | 0 | 0 |
| ENSMUSG00000064553 | 0 | 0 | 0 | 0 | 0 | 0 | 0 |
| ENSMUSG00000064554 | 0 | 0 | 0 | 0 | 0 | 0 | 0 |
| ENSMUSG00000064555 | 0 | 0 | 0 | 0 | 0 | 0 | 0 |
| ENSMUSG00000064556 | 0 | 0 | 0 | 0 | 0 | 0 | 0 |
| ENSMUSG00000064558 | 0 | 0 | 0 | 0 | 0 | 0 | 0.224243824 |
| ENSMUSG00000064563 | 0 | 0 | 0 | 0 | 0 | 0 | 0 |
| ENSMUSG00000064565 | 0 | 0 | 0 | 0 | 0 | 0 | 0 |
| ENSMUSG00000064566 | 0.289064826 | 0.396240625 | 0.25 | 0.5 | 0 | 0.477121255 | 0.355418052 |
| ENSMUSG00000064567 | 0 | 0 | 0 | 0 | 0 | 0 | 0 |
| ENSMUSG00000064569 | 0 | 0 | 0 | 0 | 0 | 0 | 0 |
| ENSMUSG00000064570 | 0 | 0 | 0 | 0 | 0 | 0 | 0 |
| ENSMUSG00000064572 | 0 | 0 | 0 | 0 | 0 | 0 | 0 |
| ENSMUSG00000064573 | 0 | 0 | 0 | 0 | 0 | 0 | 0 |
| ENSMUSG00000064574 | 0 | 0 | 0 | 0 | 0 | 0 | 0 |
| ENSMUSG00000064575 | 0 | 0 | 0 | 0 | 0 | 0 | 0 |
| ENSMUSG00000064577 | 0 | 0 | 0 | 0 | 0 | 0 | 0 |
| ENSMUSG00000064579 | 0 | 0 | 0 | 0 | 0 | 0 | 0 |
| ENSMUSG00000064580 | 0.289064826 | 0 | 0 | 0 | 0.244650542 | 0 | 0 |
| ENSMUSG00000064581 | 0 | 0 | 0 | 0 | 0 | 0 | 0 |
| ENSMUSG00000064582 | 0 | 0 | 0 | 0 | 0 | 0 | 0 |
| ENSMUSG00000064584 | 0 | 0 | 0 | 0 | 0 | 0 | 0 |
| ENSMUSG00000064585 | 0 | 0 | 0 | 0 | 0 | 0 | 0 |
| ENSMUSG00000064586 | 0 | 0 | 0 | 0 | 0 | 0 | 0 |
| ENSMUSG00000064587 | 0 | 0.25 | 0 | 0 | 0 | 0 | 0 |
| ENSMUSG00000064588 | 0 | 0 | 0 | 0 | 0 | 0 | 0 |
| ENSMUSG00000064590 | 0 | 0.25 | 0 | 0.25 | 0 | 0 | 0 |
| ENSMUSG00000064591 | 0 | 0 | 0 | 0 | 0 | 0 | 0 |
| ENSMUSG00000064595 | 0.45815691 | 0.25 | 0 | 0 | 0.244650542 | 0 | 0 |
| ENSMUSG00000064596 | 0 | 0 | 0 | 0 | 0 | 0 | 0 |
| ENSMUSG00000064597 | 0 | 0 | 0 | 0 | 0 | 0 | 0 |

| | | | | | | | |
|---|---|---|---|---|---|---|---|
| ENSMUSG00000064600 | 0 | 0 | 0 | 0 | 0 | 0 | 0 |
| ENSMUSG00000064601 | 0 | 0 | 0 | 0 | 0 | 0 | 0 |
| ENSMUSG00000064602 | 0.289064826 | 0.396240625 | 0.701838731 | 0.646240625 | 0.686820904 | 0.477121255 | 0.629532004 |
| ENSMUSG00000064604 | 0 | 0 | 0 | 0 | 0 | 0 | 0 |
| ENSMUSG00000064605 | 0 | 0 | 0 | 0 | 0 | 0 | 0 |
| ENSMUSG00000064606 | 0 | 0 | 0 | 0 | 0 | 0 | 0 |
| ENSMUSG00000064609 | 0 | 0 | 0 | 0 | 0 | 0 | 0 |
| ENSMUSG00000064612 | 0.45815691 | 0 | 0.25 | 0 | 0 | 0.301029996 | 0.355418052 |
| ENSMUSG00000064613 | 0 | 0 | 0 | 0 | 0 | 0 | 0 |
| ENSMUSG00000064616 | 0 | 0 | 0 | 0 | 0 | 0 | 0 |
| ENSMUSG00000064617 | 0 | 0 | 0 | 0 | 0 | 0 | 0 |
| ENSMUSG00000064618 | 0 | 0 | 0 | 0 | 0 | 0 | 0 |
| ENSMUSG00000064619 | 0 | 0 | 0 | 0 | 0 | 0 | 0 |
| ENSMUSG00000064620 | 0 | 0 | 0 | 0 | 0 | 0 | 0 |
| ENSMUSG00000064621 | 0 | 0 | 0 | 0 | 0 | 0 | 0 |
| ENSMUSG00000064622 | 0 | 0 | 0 | 0 | 0 | 0 | 0.224243824 |
| ENSMUSG00000064624 | 0 | 0 | 0 | 0 | 0 | 0 | 0 |
| ENSMUSG00000064626 | 0 | 0 | 0 | 0 | 0 | 0 | 0 |
| ENSMUSG00000064630 | 0 | 0 | 0 | 0 | 0 | 0 | 0 |
| ENSMUSG00000064631 | 0 | 0 | 0 | 0 | 0 | 0 | 0 |
| ENSMUSG00000064632 | 0 | 0 | 0 | 0 | 0 | 0 | 0 |
| ENSMUSG00000064634 | 0.91631382 | 0.701838731 | 0.701838731 | 1.080482024 | 1 | 0.954242509 | 0.952571511 |
| ENSMUSG00000064637 | 0.289064826 | 0.25 | 0 | 0 | 0 | 0 | 0.224243824 |
| ENSMUSG00000064642 | 0 | 0 | 0 | 0 | 0 | 0 | 0 |
| ENSMUSG00000064644 | 0 | 0 | 0 | 0 | 0 | 0 | 0 |
| ENSMUSG00000064646 | 0 | 0 | 0 | 0 | 0 | 0 | 0 |
| ENSMUSG00000064647 | 0.671187741 | 0.580482024 | 0.701838731 | 0.396240625 | 0.387761935 | 0.301029996 | 0.74492186 |
| ENSMUSG00000064648 | 0 | 0.25 | 0.25 | 0.25 | 0 | 0 | 0.224243824 |
| ENSMUSG00000064652 | 0 | 0 | 0.25 | 0.25 | 0 | 0 | 0 |
| ENSMUSG00000064653 | 0 | 0 | 0 | 0 | 0 | 0 | 0 |
| ENSMUSG00000064655 | 0 | 0 | 0 | 0 | 0 | 0 | 0 |
| ENSMUSG00000064658 | 0 | 0.5 | 0 | 0.396240625 | 0.489301084 | 0 | 0 |
| ENSMUSG00000064659 | 0 | 0 | 0 | 0 | 0 | 0 | 0 |
| ENSMUSG00000064663 | 0 | 0 | 0 | 0 | 0 | 0 | 0 |

| | | | | | | | |
|---|---|---|---|---|---|---|---|
| ENSMUSG00000064664 | 0 | 0 | 0 | 0 | 0 | 0 | 0 |
| ENSMUSG00000064665 | 0 | 0 | 0 | 0 | 0 | 0 | 0 |
| ENSMUSG00000064666 | 0 | 0 | 0 | 0 | 0 | 0 | 0 |
| ENSMUSG00000064669 | 0 | 0 | 0 | 0 | 0 | 0 | 0 |
| ENSMUSG00000064670 | 0 | 0 | 0 | 0 | 0 | 0 | 0 |
| ENSMUSG00000064671 | 0 | 0 | 0 | 0 | 0 | 0 | 0 |
| ENSMUSG00000064672 | 0.671187741 | 0.5 | 0.580482024 | 0.646240625 | 0.686820904 | 0.84509804 | 0.803905701 |
| ENSMUSG00000064673 | 0 | 0 | 0 | 0 | 0 | 0 | 0 |
| ENSMUSG00000064675 | 0 | 0 | 0 | 0 | 0 | 0 | 0 |
| ENSMUSG00000064677 | 0 | 0 | 0 | 0 | 0 | 0 | 0 |
| ENSMUSG00000064679 | 0.289064826 | 0 | 0 | 0 | 0 | 0 | 0 |
| ENSMUSG00000064682 | 0 | 0 | 0 | 0 | 0 | 0 | 0 |
| ENSMUSG00000064685 | 0 | 0 | 0 | 0 | 0 | 0 | 0 |
| ENSMUSG00000064686 | 0 | 0 | 0 | 0 | 0 | 0 | 0 |
| ENSMUSG00000064687 | 0 | 0 | 0 | 0 | 0 | 0 | 0 |
| ENSMUSG00000064688 | 0 | 0 | 0 | 0 | 0 | 0 | 0 |
| ENSMUSG00000064689 | 0 | 0 | 0 | 0 | 0 | 0 | 0 |
| ENSMUSG00000064691 | 0 | 0 | 0 | 0 | 0 | 0 | 0 |
| ENSMUSG00000064692 | 0 | 0 | 0 | 0 | 0 | 0 | 0.224243824 |
| ENSMUSG00000064693 | 0 | 0 | 0 | 0 | 0 | 0 | 0 |
| ENSMUSG00000064694 | 0 | 0 | 0 | 0 | 0 | 0 | 0 |
| ENSMUSG00000064696 | 0 | 0 | 0 | 0 | 0 | 0 | 0 |
| ENSMUSG00000064697 | 0 | 0 | 0 | 0 | 0 | 0 | 0 |
| ENSMUSG00000064698 | 0 | 0 | 0 | 0 | 0 | 0 | 0 |
| ENSMUSG00000064702 | 0 | 0 | 0 | 0 | 0 | 0 | 0 |
| ENSMUSG00000064704 | 0 | 0 | 0 | 0 | 0 | 0 | 0 |
| ENSMUSG00000064706 | 0 | 0 | 0 | 0 | 0 | 0 | 0 |
| ENSMUSG00000064707 | 0 | 0 | 0 | 0 | 0 | 0 | 0 |
| ENSMUSG00000064709 | 0 | 0 | 0 | 0 | 0 | 0 | 0 |
| ENSMUSG00000064714 | 0 | 0 | 0 | 0 | 0 | 0 | 0 |
| ENSMUSG00000064715 | 0 | 0 | 0 | 0 | 0 | 0 | 0 |
| ENSMUSG00000064717 | 0 | 0 | 0 | 0 | 0 | 0 | 0 |
| ENSMUSG00000064718 | 0 | 0 | 0 | 0 | 0 | 0 | 0 |
| ENSMUSG00000064719 | 0 | 0 | 0 | 0 | 0 | 0 | 0 |

| | | | | | | | |
|---|---|---|---|---|---|---|---|
| ENSMUSG00000064720 | 0 | 0 | 0 | 0 | 0 | 0 | 0 |
| ENSMUSG00000064721 | 1 | 0.92510993 | 0.830482024 | 0.896240625 | 0.955822902 | 0.698970004 | 0.896975297 |
| ENSMUSG00000064722 | 0 | 0 | 0 | 0 | 0 | 0 | 0 |
| ENSMUSG00000064724 | 0 | 0 | 0 | 0 | 0 | 0 | 0 |
| ENSMUSG00000064725 | 0 | 0 | 0 | 0 | 0 | 0 | 0 |
| ENSMUSG00000064726 | 0 | 0 | 0 | 0 | 0 | 0 | 0 |
| ENSMUSG00000064727 | 0 | 0 | 0 | 0 | 0 | 0 | 0 |
| ENSMUSG00000064728 | 0 | 0 | 0 | 0 | 0 | 0 | 0 |
| ENSMUSG00000064731 | 0 | 0 | 0 | 0 | 0 | 0 | 0 |
| ENSMUSG00000064732 | 0 | 0 | 0 | 0 | 0 | 0 | 0 |
| ENSMUSG00000064734 | 0 | 0 | 0 | 0 | 0 | 0 | 0 |
| ENSMUSG00000064735 | 0 | 0 | 0 | 0 | 0 | 0 | 0 |
| ENSMUSG00000064737 | 0 | 0 | 0 | 0 | 0 | 0 | 0 |
| ENSMUSG00000064738 | 0 | 0 | 0 | 0 | 0 | 0 | 0 |
| ENSMUSG00000064739 | 0 | 0 | 0 | 0 | 0 | 0 | 0 |
| ENSMUSG00000064740 | 0 | 0 | 0 | 0 | 0 | 0 | 0 |
| ENSMUSG00000064741 | 0 | 0.5 | 0 | 0 | 0 | 0.301029996 | 0.355418052 |
| ENSMUSG00000064742 | 0 | 0 | 0 | 0 | 0 | 0 | 0 |
| ENSMUSG00000064743 | 0 | 0 | 0 | 0 | 0 | 0 | 0 |
| ENSMUSG00000064745 | 0 | 0 | 0 | 0 | 0 | 0 | 0 |
| ENSMUSG00000064746 | 0 | 0 | 0 | 0 | 0 | 0 | 0 |
| ENSMUSG00000064748 | 0 | 0 | 0 | 0 | 0 | 0 | 0 |
| ENSMUSG00000064751 | 0 | 0 | 0 | 0 | 0 | 0 | 0 |
| ENSMUSG00000064755 | 0 | 0 | 0 | 0 | 0 | 0 | 0 |
| ENSMUSG00000064758 | 0 | 0 | 0 | 0 | 0 | 0 | 0 |
| ENSMUSG00000064759 | 0 | 0 | 0 | 0 | 0 | 0 | 0 |
| ENSMUSG00000064761 | 0 | 0 | 0 | 0 | 0 | 0 | 0 |
| ENSMUSG00000064762 | 0 | 0 | 0 | 0 | 0 | 0 | 0 |
| ENSMUSG00000064763 | 0 | 0 | 0 | 0 | 0 | 0 | 0 |
| ENSMUSG00000064764 | 0 | 0 | 0 | 0 | 0 | 0 | 0 |
| ENSMUSG00000064765 | 0 | 0 | 0 | 0.25 | 0 | 0 | 0 |
| ENSMUSG00000064766 | 0 | 0 | 0 | 0 | 0 | 0 | 0 |
| ENSMUSG00000064767 | 0.289064826 | 0.25 | 0 | 0.5 | 0.387761935 | 0 | 0.224243824 |
| ENSMUSG00000064768 | 0 | 0 | 0 | 0 | 0 | 0 | 0 |

| | | | | | | | |
|---|---|---|---|---|---|---|---|
| ENSMUSG00000064769 | 0 | 0 | 0 | 0 | 0 | 0 | 0 |
| ENSMUSG00000064770 | 0 | 0 | 0.25 | 0 | 0 | 0 | 0 |
| ENSMUSG00000064772 | 0 | 0 | 0 | 0 | 0 | 0 | 0 |
| ENSMUSG00000064775 | 0 | 0 | 0 | 0 | 0 | 0 | 0 |
| ENSMUSG00000064776 | 0 | 0 | 0 | 0 | 0 | 0 | 0 |
| ENSMUSG00000064777 | 0 | 0 | 0 | 0 | 0 | 0 | 0 |
| ENSMUSG00000064778 | 0 | 0 | 0.396240625 | 0.25 | 0 | 0 | 0.355418052 |
| ENSMUSG00000064779 | 0 | 0 | 0 | 0 | 0 | 0 | 0 |
| ENSMUSG00000064780 | 0 | 0 | 0 | 0 | 0 | 0 | 0 |
| ENSMUSG00000064781 | 0 | 0 | 0 | 0 | 0 | 0 | 0 |
| ENSMUSG00000064782 | 0 | 0 | 0 | 0 | 0 | 0 | 0 |
| ENSMUSG00000064783 | 0 | 0 | 0 | 0 | 0 | 0 | 0 |
| ENSMUSG00000064788 | 0 | 0 | 0 | 0 | 0 | 0 | 0 |
| ENSMUSG00000064791 | 0 | 0 | 0 | 0 | 0 | 0 | 0 |
| ENSMUSG00000064792 | 0 | 0 | 0 | 0 | 0 | 0 | 0 |
| ENSMUSG00000064793 | 0 | 0 | 0 | 0 | 0 | 0 | 0 |
| ENSMUSG00000064795 | 0 | 0 | 0 | 0 | 0 | 0 | 0 |
| ENSMUSG00000064796 | 0 | 0 | 0 | 0.25 | 0 | 0 | 0 |
| ENSMUSG00000064797 | 1.036286563 | 1.188721876 | 0.976722649 | 1.238549078 | 1.039257769 | 1.146128036 | 1.014380832 |
| ENSMUSG00000064798 | 0 | 0 | 0 | 0 | 0 | 0 | 0 |
| ENSMUSG00000064800 | 0 | 0 | 0 | 0 | 0 | 0 | 0 |
| ENSMUSG00000064801 | 0 | 0 | 0 | 0 | 0 | 0 | 0 |
| ENSMUSG00000064802 | 0 | 0.25 | 0.25 | 0.25 | 0 | 0 | 0.224243824 |
| ENSMUSG00000064805 | 0 | 0 | 0 | 0 | 0 | 0 | 0 |
| ENSMUSG00000064806 | 0 | 0 | 0 | 0 | 0 | 0 | 0 |
| ENSMUSG00000064811 | 0.289064826 | 0 | 0 | 0 | 0.244650542 | 0 | 0 |
| ENSMUSG00000064814 | 0 | 0 | 0 | 0 | 0 | 0 | 0 |
| ENSMUSG00000064816 | 0 | 0 | 0 | 0 | 0 | 0 | 0 |
| ENSMUSG00000064818 | 0 | 0 | 0 | 0 | 0 | 0 | 0 |
| ENSMUSG00000064822 | 0 | 0 | 0 | 0 | 0 | 0 | 0 |
| ENSMUSG00000064823 | 0 | 0 | 0 | 0 | 0 | 0 | 0 |
| ENSMUSG00000064824 | 0 | 0 | 0 | 0 | 0 | 0 | 0 |
| ENSMUSG00000064829 | 0 | 0 | 0 | 0 | 0 | 0 | 0 |
| ENSMUSG00000064830 | 0 | 0 | 0 | 0 | 0 | 0 | 0 |

| | | | | | | | |
|---|---|---|---|---|---|---|---|
| ENSMUSG00000064831 | 0 | 0 | 0 | 0 | 0 | 0 | 0 |
| ENSMUSG00000064833 | 0 | 0 | 0 | 0 | 0.244650542 | 0 | 0 |
| ENSMUSG00000064834 | 0 | 0 | 0 | 0 | 0 | 0 | 0 |
| ENSMUSG00000064836 | 0 | 0 | 0 | 0 | 0 | 0 | 0 |
| ENSMUSG00000064837 | 0 | 0 | 0 | 0 | 0 | 0 | 0 |
| ENSMUSG00000064840 | 0 | 0 | 0 | 0 | 0 | 0 | 0 |
| ENSMUSG00000064841 | 0 | 0 | 0 | 0 | 0 | 0 | 0 |
| ENSMUSG00000064842 | 0 | 0 | 0 | 0 | 0 | 0 | 0 |
| ENSMUSG00000064843 | 0 | 0 | 0 | 0 | 0 | 0 | 0 |
| ENSMUSG00000064844 | 0.671187741 | 0 | 0.646240625 | 0.396240625 | 0.387761935 | 0.301029996 | 0.448487648 |
| ENSMUSG00000064845 | 0.289064826 | 0 | 0.396240625 | 0.5 | 0.387761935 | 0 | 0.355418052 |
| ENSMUSG00000064850 | 0 | 0 | 0 | 0 | 0 | 0 | 0 |
| ENSMUSG00000064851 | 0 | 0 | 0 | 0 | 0 | 0 | 0 |
| ENSMUSG00000064853 | 0.45815691 | 0 | 0.25 | 0.25 | 0.568060967 | 0 | 0.520678036 |
| ENSMUSG00000064854 | 0 | 0 | 0 | 0 | 0 | 0 | 0 |
| ENSMUSG00000064855 | 0 | 0 | 0 | 0 | 0 | 0 | 0 |
| ENSMUSG00000064856 | 0 | 0 | 0 | 0 | 0 | 0 | 0 |
| ENSMUSG00000064857 | 0 | 0 | 0 | 0 | 0 | 0 | 0 |
| ENSMUSG00000064858 | 0 | 0 | 0 | 0 | 0 | 0 | 0 |
| ENSMUSG00000064859 | 0 | 0 | 0 | 0 | 0 | 0 | 0 |
| ENSMUSG00000064860 | 0 | 0 | 0 | 0 | 0 | 0 | 0 |
| ENSMUSG00000064862 | 0 | 0 | 0 | 0 | 0 | 0 | 0 |
| ENSMUSG00000064867 | 0 | 0 | 0 | 0 | 0 | 0 | 0 |
| ENSMUSG00000064871 | 0 | 0 | 0 | 0 | 0 | 0 | 0 |
| ENSMUSG00000064872 | 0 | 0 | 0 | 0 | 0 | 0 | 0 |
| ENSMUSG00000064876 | 0 | 0 | 0 | 0 | 0 | 0 | 0 |
| ENSMUSG00000064877 | 0 | 0 | 0 | 0 | 0 | 0 | 0 |
| ENSMUSG00000064880 | 0 | 0 | 0 | 0 | 0 | 0 | 0 |
| ENSMUSG00000064882 | 0 | 0 | 0 | 0 | 0 | 0 | 0 |
| ENSMUSG00000064883 | 0 | 0 | 0 | 0 | 0 | 0 | 0 |
| ENSMUSG00000064884 | 0 | 0 | 0 | 0 | 0 | 0 | 0 |
| ENSMUSG00000064887 | 0 | 0 | 0 | 0 | 0 | 0 | 0 |
| ENSMUSG00000064889 | 0 | 0 | 0 | 0 | 0 | 0 | 0 |
| ENSMUSG00000064890 | 0 | 0 | 0 | 0 | 0 | 0 | 0 |

| | | | | | | | |
|---|---|---|---|---|---|---|---|
| ENSMUSG00000064897 | 0 | 0 | 0 | 0 | 0 | 0 | 0 |
| ENSMUSG00000064899 | 0 | 0 | 0 | 0 | 0 | 0 | 0 |
| ENSMUSG00000064900 | 0 | 0 | 0 | 0 | 0 | 0 | 0 |
| ENSMUSG00000064901 | 0.289064826 | 0 | 0.25 | 0.25 | 0.244650542 | 0 | 0.355418052 |
| ENSMUSG00000064904 | 0 | 0 | 0 | 0 | 0 | 0 | 0 |
| ENSMUSG00000064905 | 0 | 0 | 0 | 0 | 0 | 0 | 0 |
| ENSMUSG00000064906 | 0.289064826 | 0 | 0 | 0 | 0 | 0 | 0 |
| ENSMUSG00000064907 | 0 | 0 | 0 | 0 | 0 | 0 | 0 |
| ENSMUSG00000064909 | 0 | 0 | 0 | 0 | 0 | 0 | 0 |
| ENSMUSG00000064915 | 0 | 0 | 0 | 0 | 0 | 0 | 0 |
| ENSMUSG00000064916 | 0 | 0 | 0.25 | 0 | 0 | 0 | 0.224243824 |
| ENSMUSG00000064917 | 0 | 0 | 0 | 0 | 0.244650542 | 0.301029996 | 0.224243824 |
| ENSMUSG00000064918 | 0 | 0.701838731 | 0.580482024 | 0.396240625 | 0.632412477 | 0.301029996 | 0.448487648 |
| ENSMUSG00000064919 | 0.289064826 | 0 | 0.396240625 | 0.396240625 | 0 | 0 | 0.579661877 |
| ENSMUSG00000064921 | 0 | 0 | 0 | 0 | 0 | 0 | 0 |
| ENSMUSG00000064923 | 0 | 0 | 0 | 0 | 0 | 0 | 0 |
| ENSMUSG00000064925 | 0.289064826 | 0.396240625 | 0.25 | 0.25 | 0.244650542 | 0.477121255 | 0.520678036 |
| ENSMUSG00000064926 | 0 | 0 | 0 | 0 | 0 | 0 | 0 |
| ENSMUSG00000064927 | 0 | 0 | 0 | 0 | 0 | 0.301029996 | 0 |
| ENSMUSG00000064928 | 0 | 0 | 0 | 0 | 0 | 0 | 0 |
| ENSMUSG00000064930 | 0 | 0 | 0.25 | 0 | 0.387761935 | 0 | 0.224243824 |
| ENSMUSG00000064932 | 0 | 0 | 0 | 0 | 0 | 0 | 0 |
| ENSMUSG00000064936 | 0 | 0 | 0 | 0 | 0 | 0.477121255 | 0 |
| ENSMUSG00000064938 | 0 | 0 | 0 | 0 | 0 | 0 | 0 |
| ENSMUSG00000064939 | 0 | 0 | 0 | 0 | 0 | 0 | 0 |
| ENSMUSG00000064940 | 0 | 0 | 0 | 0 | 0 | 0 | 0 |
| ENSMUSG00000064941 | 0 | 0 | 0 | 0 | 0 | 0 | 0 |
| ENSMUSG00000064943 | 0 | 0 | 0 | 0 | 0 | 0 | 0 |
| ENSMUSG00000064945 | 0 | 0 | 0 | 0 | 0 | 0 | 0 |
| ENSMUSG00000064948 | 0 | 0 | 0 | 0 | 0 | 0 | 0 |
| ENSMUSG00000064949 | 0 | 0 | 0 | 0 | 0 | 0 | 0 |
| ENSMUSG00000064952 | 0 | 0 | 0 | 0 | 0 | 0 | 0 |
| ENSMUSG00000064954 | 0 | 0 | 0 | 0 | 0 | 0 | 0 |
| ENSMUSG00000064956 | 0 | 0 | 0 | 0 | 0 | 0 | 0 |

| | | | | | | | |
|---|---|---|---|---|---|---|---|
| ENSMUSG00000064958 | 0 | 0 | 0 | 0 | 0 | 0 | 0 |
| ENSMUSG00000064959 | 0 | 0 | 0 | 0 | 0 | 0 | 0 |
| ENSMUSG00000064960 | 0 | 0 | 0 | 0.25 | 0 | 0 | 0 |
| ENSMUSG00000064961 | 0 | 0 | 0 | 0 | 0 | 0 | 0 |
| ENSMUSG00000064964 | 0 | 0 | 0 | 0 | 0 | 0 | 0 |
| ENSMUSG00000064966 | 0 | 0 | 0 | 0 | 0 | 0 | 0 |
| ENSMUSG00000064967 | 0 | 0 | 0 | 0 | 0 | 0 | 0 |
| ENSMUSG00000064968 | 0 | 0 | 0 | 0 | 0 | 0 | 0 |
| ENSMUSG00000064970 | 0 | 0 | 0 | 0 | 0 | 0 | 0 |
| ENSMUSG00000064971 | 0 | 0 | 0 | 0 | 0 | 0 | 0 |
| ENSMUSG00000064972 | 0 | 0 | 0 | 0 | 0 | 0 | 0 |
| ENSMUSG00000064977 | 0 | 0 | 0 | 0 | 0 | 0 | 0 |
| ENSMUSG00000064981 | 0 | 0 | 0 | 0 | 0 | 0 | 0 |
| ENSMUSG00000064982 | 0 | 0 | 0 | 0 | 0 | 0 | 0 |
| ENSMUSG00000064983 | 0 | 0 | 0 | 0 | 0 | 0 | 0 |
| ENSMUSG00000064984 | 0 | 0.25 | 0.396240625 | 0.396240625 | 0 | 0 | 0 |
| ENSMUSG00000064987 | 0 | 0 | 0 | 0 | 0 | 0 | 0 |
| ENSMUSG00000064992 | 0 | 0 | 0 | 0 | 0 | 0 | 0 |
| ENSMUSG00000064994 | 0 | 0.25 | 0 | 0.25 | 0.244650542 | 0.301029996 | 0 |
| ENSMUSG00000064995 | 0 | 0 | 0 | 0 | 0 | 0 | 0 |
| ENSMUSG00000064999 | 0 | 0 | 0 | 0 | 0 | 0 | 0 |
| ENSMUSG00000065004 | 0 | 0 | 0 | 0 | 0 | 0 | 0 |
| ENSMUSG00000065005 | 0 | 0 | 0 | 0 | 0 | 0 | 0 |
| ENSMUSG00000065006 | 0 | 0 | 0 | 0 | 0 | 0 | 0 |
| ENSMUSG00000065007 | 0 | 0 | 0 | 0 | 0 | 0 | 0 |
| ENSMUSG00000065011 | 0 | 0 | 0 | 0 | 0 | 0 | 0 |
| ENSMUSG00000065012 | 0 | 0 | 0 | 0 | 0 | 0 | 0 |
| ENSMUSG00000065013 | 0 | 0 | 0 | 0 | 0 | 0 | 0 |
| ENSMUSG00000065014 | 0 | 0 | 0 | 0 | 0 | 0 | 0 |
| ENSMUSG00000065016 | 0 | 0 | 0 | 0 | 0 | 0 | 0 |
| ENSMUSG00000065017 | 0 | 0 | 0 | 0 | 0 | 0 | 0 |
| ENSMUSG00000065018 | 0 | 0 | 0 | 0 | 0 | 0 | 0 |
| ENSMUSG00000065022 | 0 | 0 | 0 | 0 | 0 | 0 | 0 |
| ENSMUSG00000065027 | 0 | 0 | 0 | 0 | 0 | 0 | 0 |

| | | | | | | | |
|---|---|---|---|---|---|---|---|
| ENSMUSG00000065028 | 0 | 0 | 0 | 0 | 0 | 0 | 0 |
| ENSMUSG00000065029 | 0 | 0 | 0 | 0 | 0 | 0 | 0 |
| ENSMUSG00000065030 | 0 | 0 | 0 | 0 | 0 | 0 | 0 |
| ENSMUSG00000065031 | 0 | 0 | 0 | 0 | 0 | 0 | 0 |
| ENSMUSG00000065034 | 0 | 0 | 0 | 0 | 0 | 0 | 0 |
| ENSMUSG00000065036 | 0.289064826 | 0.25 | 0 | 0 | 0.244650542 | 0 | 0.224243824 |
| ENSMUSG00000065037 | 0 | 0.701838731 | 0.396240625 | 0.701838731 | 0.568060967 | 0.301029996 | 1.100339912 |
| ENSMUSG00000065040 | 0 | 0 | 0 | 0 | 0 | 0 | 0 |
| ENSMUSG00000065041 | 0.671187741 | 0.646240625 | 0.75 | 0.79248125 | 0.955822902 | 0.903089987 | 0.710836105 |
| ENSMUSG00000065042 | 0 | 0 | 0 | 0 | 0 | 0 | 0 |
| ENSMUSG00000065044 | 0 | 0 | 0.25 | 0.25 | 0 | 0.301029996 | 0 |
| ENSMUSG00000065045 | 0 | 0 | 0 | 0 | 0 | 0 | 0 |
| ENSMUSG00000065047 | 0 | 0 | 0.25 | 0 | 0 | 0 | 0 |
| ENSMUSG00000065048 | 0 | 0 | 0 | 0 | 0 | 0 | 0 |
| ENSMUSG00000065049 | 0.289064826 | 0 | 0 | 0 | 0 | 0 | 0 |
| ENSMUSG00000065050 | 0 | 0 | 0 | 0 | 0 | 0 | 0 |
| ENSMUSG00000065053 | 0 | 0 | 0 | 0 | 0 | 0 | 0 |
| ENSMUSG00000065055 | 0 | 0 | 0 | 0 | 0 | 0 | 0 |
| ENSMUSG00000065057 | 0 | 0 | 0 | 0 | 0 | 0 | 0 |
| ENSMUSG00000065061 | 0 | 0 | 0 | 0 | 0 | 0 | 0 |
| ENSMUSG00000065068 | 0 | 0 | 0 | 0 | 0 | 0 | 0 |
| ENSMUSG00000065070 | 0 | 0 | 0 | 0 | 0 | 0 | 0 |
| ENSMUSG00000065072 | 0 | 0 | 0 | 0 | 0 | 0 | 0 |
| ENSMUSG00000065074 | 0 | 0 | 0 | 0 | 0 | 0 | 0 |
| ENSMUSG00000065078 | 0 | 0 | 0 | 0 | 0 | 0 | 0 |
| ENSMUSG00000065079 | 0 | 0 | 0 | 0 | 0 | 0 | 0 |
| ENSMUSG00000065080 | 0 | 0 | 0 | 0 | 0 | 0 | 0 |
| ENSMUSG00000065083 | 0 | 0 | 0 | 0 | 0 | 0 | 0 |
| ENSMUSG00000065084 | 0 | 0 | 0 | 0 | 0 | 0 | 0 |
| ENSMUSG00000065086 | 0 | 0 | 0 | 0 | 0 | 0 | 0 |
| ENSMUSG00000065087 | 1.587501561 | 1.445339928 | 1.516522298 | 1.516522298 | 1.509475496 | 1.568201724 | 1.345462945 |
| ENSMUSG00000065088 | 0 | 0 | 0 | 0 | 0 | 0 | 0 |
| ENSMUSG00000065089 | 0 | 0.25 | 0.25 | 0 | 0 | 0.301029996 | 0 |
| ENSMUSG00000065090 | 0 | 0 | 0 | 0 | 0 | 0 | 0 |

| | | | | | | | |
|---|---|---|---|---|---|---|---|
| ENSMUSG00000065091 | 0 | 0 | 0 | 0 | 0 | 0 | 0 |
| ENSMUSG00000065094 | 0 | 0 | 0 | 0 | 0 | 0 | 0 |
| ENSMUSG00000065095 | 0 | 0 | 0 | 0 | 0 | 0 | 0 |
| ENSMUSG00000065096 | 0 | 0 | 0 | 0 | 0 | 0 | 0 |
| ENSMUSG00000065097 | 0 | 0 | 0 | 0 | 0 | 0 | 0 |
| ENSMUSG00000065100 | 0 | 0 | 0 | 0.25 | 0 | 0 | 0 |
| ENSMUSG00000065102 | 0 | 0 | 0 | 0 | 0 | 0 | 0 |
| ENSMUSG00000065104 | 0 | 0 | 0 | 0 | 0 | 0 | 0 |
| ENSMUSG00000065105 | 0 | 0 | 0 | 0 | 0 | 0 | 0 |
| ENSMUSG00000065107 | 0 | 0 | 0 | 0 | 0 | 0 | 0 |
| ENSMUSG00000065110 | 0 | 0 | 0 | 0 | 0 | 0 | 0 |
| ENSMUSG00000065111 | 0 | 0 | 0 | 0 | 0 | 0 | 0 |
| ENSMUSG00000065115 | 0 | 0 | 0 | 0 | 0 | 0 | 0 |
| ENSMUSG00000065118 | 0 | 0 | 0 | 0.25 | 0 | 0 | 0 |
| ENSMUSG00000065122 | 0 | 0 | 0 | 0 | 0 | 0 | 0 |
| ENSMUSG00000065124 | 0 | 0 | 0 | 0 | 0 | 0 | 0 |
| ENSMUSG00000065126 | 0.289064826 | 0.830482024 | 0.79248125 | 0.580482024 | 0.244650542 | 0.301029996 | 0.629532004 |
| ENSMUSG00000065127 | 0 | 0 | 0 | 0 | 0 | 0 | 0 |
| ENSMUSG00000065130 | 0 | 0 | 0 | 0 | 0 | 0 | 0 |
| ENSMUSG00000065132 | 0 | 0 | 0 | 0 | 0 | 0 | 0 |
| ENSMUSG00000065138 | 0 | 0 | 0 | 0 | 0 | 0 | 0 |
| ENSMUSG00000065141 | 0 | 0 | 0 | 0 | 0 | 0 | 0 |
| ENSMUSG00000065143 | 0 | 0 | 0 | 0 | 0 | 0 | 0 |
| ENSMUSG00000065145 | 0 | 0 | 0 | 0 | 0 | 0 | 0 |
| ENSMUSG00000065147 | 0.747221736 | 0.92510993 | 0.864857905 | 1.098079356 | 0.931471446 | 1.113943352 | 0.952571511 |
| ENSMUSG00000065151 | 0 | 0 | 0 | 0 | 0 | 0 | 0 |
| ENSMUSG00000065155 | 0 | 0 | 0 | 0 | 0 | 0 | 0 |
| ENSMUSG00000065158 | 0 | 0 | 0 | 0 | 0 | 0 | 0 |
| ENSMUSG00000065159 | 0 | 0 | 0 | 0 | 0 | 0 | 0 |
| ENSMUSG00000065160 | 0 | 0 | 0 | 0 | 0 | 0 | 0 |
| ENSMUSG00000065162 | 0 | 0.396240625 | 0.396240625 | 0.396240625 | 0.387761935 | 0.477121255 | 0.355418052 |
| ENSMUSG00000065164 | 0 | 0 | 0 | 0 | 0 | 0 | 0 |
| ENSMUSG00000065165 | 0 | 0 | 0 | 0 | 0 | 0 | 0 |
| ENSMUSG00000065173 | 0 | 0 | 0 | 0 | 0 | 0 | 0 |

| | | | | | | | |
|---|---|---|---|---|---|---|---|
| ENSMUSG00000065176 | 0 | 0 | 0 | 0 | 0 | 0 | 0 |
| ENSMUSG00000065178 | 0 | 0 | 0 | 0 | 0 | 0 | 0 |
| ENSMUSG00000065181 | 0 | 0 | 0 | 0 | 0 | 0 | 0 |
| ENSMUSG00000065182 | 0 | 0 | 0 | 0.25 | 0 | 0 | 0 |
| ENSMUSG00000065185 | 0 | 0 | 0 | 0 | 0 | 0 | 0 |
| ENSMUSG00000065187 | 0 | 0 | 0 | 0 | 0 | 0 | 0 |
| ENSMUSG00000065188 | 0 | 0 | 0 | 0 | 0 | 0 | 0 |
| ENSMUSG00000065191 | 0.671187741 | 0.396240625 | 0.580482024 | 0.396240625 | 0.387761935 | 0.301029996 | 0 |
| ENSMUSG00000065192 | 0 | 0 | 0 | 0 | 0 | 0 | 0 |
| ENSMUSG00000065193 | 0 | 0 | 0 | 0.25 | 0 | 0 | 0 |
| ENSMUSG00000065195 | 0 | 0 | 0 | 0 | 0 | 0 | 0 |
| ENSMUSG00000065196 | 0 | 0 | 0 | 0 | 0 | 0 | 0 |
| ENSMUSG00000065198 | 0 | 0 | 0 | 0 | 0 | 0 | 0 |
| ENSMUSG00000065199 | 0 | 0 | 0 | 0 | 0 | 0 | 0 |
| ENSMUSG00000065200 | 0 | 0 | 0 | 0 | 0 | 0 | 0 |
| ENSMUSG00000065201 | 0 | 0 | 0 | 0 | 0 | 0 | 0 |
| ENSMUSG00000065202 | 0 | 0 | 0 | 0 | 0 | 0 | 0 |
| ENSMUSG00000065203 | 0 | 0 | 0 | 0 | 0 | 0 | 0 |
| ENSMUSG00000065204 | 0 | 0 | 0 | 0 | 0 | 0.301029996 | 0 |
| ENSMUSG00000065206 | 0 | 0 | 0 | 0 | 0 | 0 | 0 |
| ENSMUSG00000065208 | 0.578129653 | 0.79248125 | 0.580482024 | 0.701838731 | 0.77552387 | 0.698970004 | 0.629532004 |
| ENSMUSG00000065211 | 0 | 0 | 0 | 0 | 0 | 0 | 0 |
| ENSMUSG00000065212 | 0 | 0 | 0 | 0 | 0 | 0 | 0 |
| ENSMUSG00000065215 | 0 | 0 | 0 | 0 | 0 | 0 | 0 |
| ENSMUSG00000065219 | 0 | 0 | 0 | 0 | 0 | 0 | 0 |
| ENSMUSG00000065220 | 0 | 0 | 0 | 0 | 0 | 0 | 0 |
| ENSMUSG00000065221 | 0 | 0 | 0 | 0 | 0 | 0 | 0 |
| ENSMUSG00000065223 | 0 | 0 | 0 | 0 | 0 | 0 | 0 |
| ENSMUSG00000065226 | 0.45815691 | 0.580482024 | 0.75 | 0.976722649 | 0.686820904 | 0.602059991 | 0.775756176 |
| ENSMUSG00000065227 | 0 | 0 | 0 | 0 | 0 | 0 | 0 |
| ENSMUSG00000065228 | 0 | 0 | 0 | 0 | 0 | 0 | 0 |
| ENSMUSG00000065231 | 0 | 0 | 0 | 0 | 0 | 0 | 0 |
| ENSMUSG00000065232 | 0 | 0 | 0 | 0 | 0 | 0 | 0 |
| ENSMUSG00000065233 | 0 | 0 | 0 | 0 | 0 | 0 | 0 |

| | | | | | | | |
|---|---|---|---|---|---|---|---|
| ENSMUSG00000065235 | 0 | 0 | 0 | 0 | 0 | 0 | 0 |
| ENSMUSG00000065236 | 0.289064826 | 0.396240625 | 0 | 0.396240625 | 0.244650542 | 0 | 0 |
| ENSMUSG00000065237 | 0 | 0 | 0 | 0 | 0 | 0 | 0 |
| ENSMUSG00000065238 | 0 | 0 | 0 | 0 | 0 | 0 | 0 |
| ENSMUSG00000065239 | 0 | 0 | 0 | 0 | 0 | 0 | 0 |
| ENSMUSG00000065242 | 0 | 0 | 0 | 0 | 0 | 0 | 0 |
| ENSMUSG00000065244 | 0 | 0 | 0 | 0 | 0 | 0 | 0 |
| ENSMUSG00000065245 | 0 | 0 | 0 | 0 | 0 | 0 | 0 |
| ENSMUSG00000065246 | 0 | 0 | 0 | 0 | 0 | 0 | 0 |
| ENSMUSG00000065248 | 0 | 0 | 0 | 0 | 0 | 0 | 0 |
| ENSMUSG00000065251 | 0 | 0 | 0 | 0 | 0 | 0 | 0 |
| ENSMUSG00000065253 | 0 | 0 | 0 | 0 | 0 | 0 | 0 |
| ENSMUSG00000065254 | 0 | 0 | 0 | 0 | 0 | 0 | 0.224243824 |
| ENSMUSG00000065255 | 0 | 0 | 0 | 0 | 0 | 0 | 0 |
| ENSMUSG00000065256 | 0 | 0 | 0 | 0 | 0 | 0 | 0 |
| ENSMUSG00000065258 | 0.45815691 | 0.25 | 0.25 | 0 | 0 | 0.301029996 | 0.224243824 |
| ENSMUSG00000065259 | 0 | 0 | 0.25 | 0.25 | 0 | 0 | 0.355418052 |
| ENSMUSG00000065260 | 0 | 0 | 0 | 0 | 0 | 0 | 0 |
| ENSMUSG00000065261 | 0 | 0 | 0 | 0 | 0 | 0 | 0 |
| ENSMUSG00000065262 | 0 | 0 | 0 | 0 | 0 | 0 | 0 |
| ENSMUSG00000065265 | 1.069666964 | 1.02186571 | 1.098079356 | 1.061981878 | 1.163285805 | 1.176091259 | 0.984950056 |
| ENSMUSG00000065266 | 0 | 0 | 0 | 0 | 0 | 0 | 0 |
| ENSMUSG00000065269 | 0 | 0 | 0 | 0 | 0 | 0 | 0 |
| ENSMUSG00000065270 | 0 | 0 | 0 | 0 | 0 | 0 | 0 |
| ENSMUSG00000065272 | 0 | 0 | 0 | 0 | 0 | 0 | 0 |
| ENSMUSG00000065273 | 0 | 0.25 | 0.25 | 0 | 0 | 0 | 0.355418052 |
| ENSMUSG00000065274 | 0 | 0 | 0 | 0 | 0 | 0 | 0.224243824 |
| ENSMUSG00000065275 | 0 | 0 | 0 | 0 | 0 | 0 | 0 |
| ENSMUSG00000065277 | 0 | 0 | 0 | 0 | 0 | 0 | 0 |
| ENSMUSG00000065279 | 0 | 0 | 0 | 0 | 0 | 0 | 0 |
| ENSMUSG00000065280 | 1.325351389 | 1.17510993 | 1.188721876 | 1.188721876 | 1.212046813 | 1.204119983 | 0.984950056 |
| ENSMUSG00000065281 | 1.181541736 | 1 | 1.080482024 | 1.130890489 | 1.223252711 | 1 | 1.054044578 |
| ENSMUSG00000065282 | 0.91631382 | 1.080482024 | 0.92510993 | 1.04248125 | 0.931471446 | 0.84509804 | 0.952571511 |
| ENSMUSG00000065283 | 0 | 0 | 0 | 0 | 0 | 0 | 0 |

| | | | | | | | |
|---|---|---|---|---|---|---|---|
| ENSMUSG00000065287 | 0 | 0 | 0 | 0 | 0 | 0 | 0 |
| ENSMUSG00000065289 | 0 | 0 | 0 | 0 | 0 | 0 | 0 |
| ENSMUSG00000065291 | 0 | 0 | 0 | 0 | 0 | 0 | 0 |
| ENSMUSG00000065293 | 0 | 0 | 0 | 0 | 0 | 0 | 0 |
| ENSMUSG00000065295 | 0 | 0 | 0 | 0 | 0 | 0 | 0 |
| ENSMUSG00000065296 | 0 | 0 | 0 | 0 | 0 | 0 | 0 |
| ENSMUSG00000065297 | 0 | 0 | 0 | 0 | 0 | 0 | 0 |
| ENSMUSG00000065298 | 0 | 0 | 0 | 0 | 0 | 0 | 0 |
| ENSMUSG00000065299 | 0 | 0 | 0 | 0 | 0 | 0 | 0 |
| ENSMUSG00000065301 | 0 | 0 | 0 | 0 | 0 | 0.301029996 | 0 |
| ENSMUSG00000065303 | 0 | 0 | 0 | 0 | 0 | 0 | 0 |
| ENSMUSG00000065304 | 0.289064826 | 0.5 | 0.580482024 | 0.25 | 0.244650542 | 0.477121255 | 0.448487648 |
| ENSMUSG00000065305 | 1.100572389 | 1.02186571 | 0.92510993 | 0.92510993 | 0.955822902 | 1.079181246 | 1.014380832 |
| ENSMUSG00000065307 | 0 | 0 | 0 | 0 | 0 | 0 | 0 |
| ENSMUSG00000065309 | 0 | 0 | 0 | 0 | 0 | 0 | 0 |
| ENSMUSG00000065311 | 0 | 0 | 0 | 0 | 0 | 0 | 0 |
| ENSMUSG00000065312 | 0 | 0 | 0 | 0 | 0 | 0 | 0 |
| ENSMUSG00000065313 | 0 | 0 | 0 | 0 | 0 | 0 | 0 |
| ENSMUSG00000065314 | 0 | 0 | 0 | 0 | 0 | 0 | 0 |
| ENSMUSG00000065315 | 0 | 0 | 0 | 0.25 | 0.244650542 | 0 | 0.355418052 |
| ENSMUSG00000065319 | 0 | 0 | 0 | 0 | 0 | 0 | 0 |
| ENSMUSG00000065320 | 0 | 0 | 0 | 0 | 0 | 0 | 0 |
| ENSMUSG00000065321 | 0 | 0 | 0 | 0 | 0 | 0 | 0 |
| ENSMUSG00000065322 | 0 | 0 | 0 | 0 | 0 | 0 | 0 |
| ENSMUSG00000065327 | 0 | 0 | 0 | 0 | 0 | 0 | 0 |
| ENSMUSG00000065328 | 0 | 0 | 0 | 0 | 0 | 0 | 0 |
| ENSMUSG00000065329 | 0 | 0 | 0 | 0 | 0 | 0 | 0 |
| ENSMUSG00000065330 | 0 | 0 | 0 | 0 | 0 | 0 | 0 |
| ENSMUSG00000065331 | 0 | 0 | 0 | 0 | 0 | 0 | 0 |
| ENSMUSG00000065333 | 0 | 0 | 0 | 0 | 0 | 0 | 0 |
| ENSMUSG00000065336 | 0.45815691 | 0 | 0.580482024 | 0.5 | 0.244650542 | 0.477121255 | 0.224243824 |
| ENSMUSG00000065337 | 0 | 0 | 0 | 0 | 0 | 0 | 0 |
| ENSMUSG00000065340 | 0 | 0 | 0 | 0 | 0 | 0 | 0 |
| ENSMUSG00000065341 | 0 | 0.580482024 | 0 | 0.396240625 | 0.387761935 | 0 | 0 |

| | | | | | | | |
|---|---|---|---|---|---|---|---|
| ENSMUSG00000065343 | 0.289064826 | 0.25 | 0.25 | 0.5 | 0 | 0 | 0 |
| ENSMUSG00000065344 | 0 | 0 | 0 | 0 | 0 | 0 | 0 |
| ENSMUSG00000065346 | 0 | 0 | 0 | 0 | 0 | 0 | 0 |
| ENSMUSG00000065347 | 0 | 0 | 0 | 0 | 0 | 0 | 0 |
| ENSMUSG00000065348 | 0 | 0 | 0 | 0 | 0 | 0 | 0 |
| ENSMUSG00000065353 | 0 | 0 | 0 | 0 | 0 | 0 | 0 |
| ENSMUSG00000065360 | 0 | 0.25 | 0 | 0 | 0 | 0.301029996 | 0 |
| ENSMUSG00000065361 | 0 | 0 | 0 | 0 | 0 | 0 | 0 |
| ENSMUSG00000065362 | 0 | 0 | 0 | 0 | 0 | 0 | 0 |
| ENSMUSG00000065367 | 0 | 0 | 0 | 0 | 0 | 0 | 0 |
| ENSMUSG00000065368 | 0 | 0 | 0 | 0 | 0 | 0 | 0 |
| ENSMUSG00000065371 | 0 | 0 | 0 | 0 | 0 | 0 | 0 |
| ENSMUSG00000065372 | 0 | 0 | 0 | 0 | 0 | 0 | 0 |
| ENSMUSG00000065373 | 0 | 0 | 0 | 0 | 0 | 0 | 0 |
| ENSMUSG00000065374 | 0 | 0 | 0 | 0.25 | 0 | 0 | 0 |
| ENSMUSG00000065378 | 1.418409478 | 1.146240625 | 1.114857905 | 1.188721876 | 1.176121988 | 1 | 1.100339912 |
| ENSMUSG00000065379 | 0 | 0 | 0 | 0 | 0 | 0 | 0 |
| ENSMUSG00000065380 | 0 | 0 | 0 | 0 | 0 | 0 | 0 |
| ENSMUSG00000065381 | 0 | 0 | 0 | 0 | 0 | 0 | 0 |
| ENSMUSG00000065382 | 0 | 0 | 0 | 0 | 0 | 0 | 0 |
| ENSMUSG00000065384 | 0 | 0 | 0 | 0 | 0 | 0 | 0 |
| ENSMUSG00000065385 | 0 | 0 | 0 | 0 | 0 | 0 | 0 |
| ENSMUSG00000065387 | 0.289064826 | 0 | 0 | 0 | 0 | 0 | 0 |
| ENSMUSG00000065388 | 0 | 0 | 0 | 0 | 0 | 0 | 0 |
| ENSMUSG00000065389 | 0 | 0 | 0 | 0 | 0 | 0 | 0 |
| ENSMUSG00000065392 | 0.747221736 | 0.701838731 | 0.79248125 | 0.896240625 | 0.877063019 | 0.698970004 | 0.803905701 |
| ENSMUSG00000065394 | 0 | 0 | 0 | 0 | 0 | 0 | 0 |
| ENSMUSG00000065395 | 0 | 0 | 0 | 0 | 0 | 0 | 0 |
| ENSMUSG00000065396 | 0 | 0 | 0 | 0 | 0 | 0 | 0 |
| ENSMUSG00000065397 | 0 | 0 | 0 | 0 | 0 | 0 | 0 |
| ENSMUSG00000065398 | 0 | 0 | 0 | 0 | 0 | 0 | 0 |
| ENSMUSG00000065399 | 0 | 0 | 0 | 0.25 | 0 | 0 | 0 |
| ENSMUSG00000065400 | 0 | 0 | 0 | 0 | 0 | 0 | 0 |
| ENSMUSG00000065401 | 0 | 0 | 0 | 0 | 0 | 0 | 0 |

| | | | | | | | |
|---|---|---|---|---|---|---|---|
| ENSMUSG00000065402 | 0 | 0 | 0 | 0 | 0 | 0 | 0 |
| ENSMUSG00000065403 | 0 | 0 | 0 | 0 | 0 | 0 | 0 |
| ENSMUSG00000065404 | 0 | 0 | 0 | 0 | 0 | 0 | 0 |
| ENSMUSG00000065405 | 0 | 0 | 0 | 0 | 0 | 0 | 0 |
| ENSMUSG00000065406 | 0 | 0 | 0 | 0 | 0 | 0 | 0 |
| ENSMUSG00000065407 | 0 | 0 | 0 | 0 | 0 | 0 | 0 |
| ENSMUSG00000065408 | 0 | 0 | 0 | 0 | 0 | 0 | 0 |
| ENSMUSG00000065409 | 0 | 0 | 0 | 0 | 0 | 0 | 0 |
| ENSMUSG00000065410 | 0 | 0 | 0 | 0 | 0 | 0 | 0 |
| ENSMUSG00000065411 | 0 | 0 | 0 | 0 | 0 | 0 | 0 |
| ENSMUSG00000065412 | 0 | 0 | 0 | 0 | 0 | 0 | 0 |
| ENSMUSG00000065413 | 0 | 0 | 0 | 0 | 0 | 0 | 0 |
| ENSMUSG00000065414 | 0 | 0 | 0 | 0 | 0 | 0 | 0 |
| ENSMUSG00000065415 | 0 | 0 | 0 | 0 | 0 | 0 | 0 |
| ENSMUSG00000065416 | 0 | 0 | 0 | 0 | 0 | 0 | 0 |
| ENSMUSG00000065417 | 0 | 0 | 0 | 0 | 0 | 0 | 0 |
| ENSMUSG00000065418 | 0 | 0 | 0 | 0 | 0 | 0 | 0 |
| ENSMUSG00000065420 | 0 | 0 | 0 | 0 | 0 | 0 | 0 |
| ENSMUSG00000065421 | 0 | 0 | 0 | 0 | 0 | 0 | 0 |
| ENSMUSG00000065422 | 0 | 0 | 0 | 0 | 0 | 0.301029996 | 0 |
| ENSMUSG00000065423 | 0 | 0 | 0 | 0 | 0 | 0 | 0 |
| ENSMUSG00000065424 | 0 | 0 | 0 | 0 | 0 | 0 | 0 |
| ENSMUSG00000065425 | 0 | 0 | 0 | 0 | 0 | 0 | 0 |
| ENSMUSG00000065426 | 0 | 0 | 0 | 0 | 0 | 0 | 0 |
| ENSMUSG00000065427 | 0 | 0 | 0 | 0 | 0 | 0 | 0 |
| ENSMUSG00000065428 | 0 | 0 | 0 | 0 | 0 | 0 | 0 |
| ENSMUSG00000065429 | 0 | 0 | 0 | 0 | 0 | 0 | 0 |
| ENSMUSG00000065430 | 0 | 0 | 0 | 0 | 0 | 0 | 0 |
| ENSMUSG00000065431 | 0.289064826 | 0.5 | 0.580482024 | 0.580482024 | 0.77552387 | 0.602059991 | 0.448487648 |
| ENSMUSG00000065432 | 0 | 0 | 0 | 0 | 0 | 0 | 0 |
| ENSMUSG00000065433 | 0 | 0 | 0 | 0 | 0 | 0 | 0 |
| ENSMUSG00000065434 | 0.578129653 | 0.75 | 0 | 0.701838731 | 0.632412477 | 0.477121255 | 0.448487648 |
| ENSMUSG00000065435 | 0 | 0 | 0 | 0 | 0 | 0 | 0 |
| ENSMUSG00000065436 | 0 | 0 | 0 | 0 | 0 | 0 | 0 |

| | | | | | | | |
|---|---|---|---|---|---|---|---|
| ENSMUSG00000065437 | 0 | 0 | 0 | 0 | 0 | 0 | 0 |
| ENSMUSG00000065438 | 0 | 0 | 0 | 0 | 0 | 0 | 0 |
| ENSMUSG00000065439 | 0 | 0 | 0 | 0 | 0 | 0 | 0 |
| ENSMUSG00000065440 | 0 | 0 | 0 | 0 | 0 | 0 | 0 |
| ENSMUSG00000065441 | 0 | 0 | 0 | 0 | 0 | 0 | 0 |
| ENSMUSG00000065442 | 0 | 0 | 0 | 0 | 0 | 0 | 0 |
| ENSMUSG00000065443 | 0 | 0 | 0 | 0 | 0 | 0 | 0 |
| ENSMUSG00000065444 | 0 | 0 | 0 | 0 | 0 | 0 | 0 |
| ENSMUSG00000065445 | 0 | 0.25 | 0 | 0 | 0 | 0 | 0 |
| ENSMUSG00000065446 | 0 | 0 | 0 | 0 | 0 | 0 | 0 |
| ENSMUSG00000065447 | 0 | 0 | 0 | 0 | 0 | 0 | 0 |
| ENSMUSG00000065448 | 0 | 0 | 0 | 0 | 0 | 0 | 0 |
| ENSMUSG00000065449 | 0 | 0 | 0 | 0 | 0 | 0 | 0 |
| ENSMUSG00000065450 | 0 | 0 | 0 | 0 | 0 | 0 | 0 |
| ENSMUSG00000065451 | 0 | 0 | 0 | 0 | 0 | 0 | 0 |
| ENSMUSG00000065452 | 0 | 0 | 0 | 0 | 0 | 0 | 0 |
| ENSMUSG00000065453 | 0 | 0 | 0 | 0 | 0 | 0 | 0 |
| ENSMUSG00000065454 | 0 | 0 | 0 | 0 | 0 | 0 | 0 |
| ENSMUSG00000065455 | 0 | 0 | 0 | 0 | 0 | 0 | 0 |
| ENSMUSG00000065456 | 0 | 0 | 0 | 0 | 0 | 0 | 0 |
| ENSMUSG00000065457 | 0 | 0 | 0 | 0 | 0 | 0 | 0 |
| ENSMUSG00000065458 | 0 | 0 | 0 | 0 | 0 | 0 | 0 |
| ENSMUSG00000065460 | 0 | 0 | 0 | 0 | 0.244650542 | 0 | 0 |
| ENSMUSG00000065462 | 0 | 0 | 0 | 0 | 0 | 0 | 0 |
| ENSMUSG00000065463 | 0 | 0 | 0 | 0 | 0 | 0 | 0 |
| ENSMUSG00000065464 | 0 | 0 | 0 | 0 | 0 | 0 | 0 |
| ENSMUSG00000065465 | 0 | 0 | 0 | 0 | 0 | 0 | 0 |
| ENSMUSG00000065466 | 0 | 0 | 0 | 0 | 0 | 0 | 0 |
| ENSMUSG00000065468 | 0 | 0.25 | 0 | 0.25 | 0 | 0 | 0 |
| ENSMUSG00000065469 | 0 | 0 | 0 | 0 | 0 | 0 | 0 |
| ENSMUSG00000065470 | 0 | 0 | 0 | 0 | 0 | 0 | 0 |
| ENSMUSG00000065471 | 0 | 0 | 0 | 0 | 0 | 0 | 0 |
| ENSMUSG00000065472 | 0 | 0 | 0 | 0 | 0 | 0 | 0 |
| ENSMUSG00000065473 | 0 | 0 | 0 | 0 | 0 | 0 | 0 |

| | | | | | | | |
|---|---|---|---|---|---|---|---|
| ENSMUSG00000065474 | 0 | 0 | 0 | 0 | 0 | 0 | 0 |
| ENSMUSG00000065475 | 0.289064826 | 0 | 0 | 0.25 | 0.387761935 | 0.301029996 | 0 |
| ENSMUSG00000065476 | 0 | 0 | 0 | 0 | 0 | 0 | 0 |
| ENSMUSG00000065477 | 0 | 0 | 0 | 0 | 0 | 0 | 0 |
| ENSMUSG00000065479 | 0 | 0 | 0 | 0 | 0 | 0 | 0 |
| ENSMUSG00000065480 | 0 | 0 | 0 | 0 | 0 | 0 | 0 |
| ENSMUSG00000065481 | 0 | 0 | 0 | 0 | 0 | 0 | 0 |
| ENSMUSG00000065482 | 0 | 0 | 0 | 0 | 0 | 0 | 0 |
| ENSMUSG00000065483 | 0 | 0 | 0 | 0 | 0 | 0.301029996 | 0 |
| ENSMUSG00000065484 | 0 | 0 | 0 | 0 | 0.244650542 | 0 | 0 |
| ENSMUSG00000065485 | 0 | 0 | 0 | 0 | 0 | 0 | 0 |
| ENSMUSG00000065486 | 0 | 0 | 0 | 0 | 0 | 0 | 0 |
| ENSMUSG00000065488 | 0 | 0 | 0 | 0 | 0 | 0 | 0 |
| ENSMUSG00000065489 | 0 | 0.25 | 0 | 0 | 0.244650542 | 0 | 0 |
| ENSMUSG00000065490 | 0 | 0 | 0 | 0 | 0 | 0 | 0 |
| ENSMUSG00000065491 | 0 | 0 | 0 | 0 | 0 | 0 | 0 |
| ENSMUSG00000065492 | 0 | 0 | 0 | 0 | 0 | 0 | 0 |
| ENSMUSG00000065493 | 0 | 0 | 0 | 0 | 0 | 0 | 0 |
| ENSMUSG00000065494 | 0 | 0 | 0 | 0 | 0 | 0 | 0 |
| ENSMUSG00000065495 | 0 | 0 | 0 | 0 | 0 | 0 | 0 |
| ENSMUSG00000065497 | 0 | 0 | 0 | 0 | 0 | 0 | 0 |
| ENSMUSG00000065498 | 0 | 0 | 0 | 0 | 0 | 0 | 0 |
| ENSMUSG00000065499 | 0 | 0 | 0 | 0 | 0 | 0 | 0 |
| ENSMUSG00000065500 | 0 | 0 | 0 | 0 | 0 | 0 | 0 |
| ENSMUSG00000065501 | 0 | 0 | 0 | 0 | 0 | 0 | 0 |
| ENSMUSG00000065502 | 0 | 0 | 0 | 0 | 0 | 0 | 0 |
| ENSMUSG00000065503 | 0 | 0.5 | 0.5 | 0.25 | 0.244650542 | 0 | 0.224243824 |
| ENSMUSG00000065505 | 0 | 0 | 0 | 0 | 0 | 0 | 0 |
| ENSMUSG00000065507 | 0 | 0 | 0 | 0 | 0 | 0 | 0 |
| ENSMUSG00000065508 | 0 | 0 | 0 | 0 | 0 | 0 | 0 |
| ENSMUSG00000065509 | 0 | 0.25 | 0.25 | 0 | 0 | 0 | 0 |
| ENSMUSG00000065510 | 0 | 0 | 0 | 0 | 0 | 0 | 0 |
| ENSMUSG00000065511 | 0 | 0 | 0 | 0 | 0 | 0 | 0 |
| ENSMUSG00000065512 | 0 | 0 | 0 | 0 | 0 | 0 | 0 |

| | | | | | | | |
|---|---|---|---|---|---|---|---|
| ENSMUSG00000065513 | 0 | 0 | 0 | 0 | 0 | 0 | 0 |
| ENSMUSG00000065514 | 0 | 0 | 0 | 0 | 0.244650542 | 0 | 0.224243824 |
| ENSMUSG00000065515 | 0 | 0 | 0 | 0 | 0 | 0 | 0 |
| ENSMUSG00000065516 | 0 | 0 | 0 | 0 | 0 | 0 | 0 |
| ENSMUSG00000065518 | 0 | 0 | 0 | 0 | 0.244650542 | 0 | 0 |
| ENSMUSG00000065519 | 0 | 0 | 0 | 0.25 | 0 | 0 | 0 |
| ENSMUSG00000065520 | 0 | 0 | 0 | 0 | 0 | 0 | 0 |
| ENSMUSG00000065521 | 0 | 0 | 0 | 0 | 0 | 0 | 0 |
| ENSMUSG00000065523 | 0 | 0 | 0 | 0 | 0 | 0 | 0 |
| ENSMUSG00000065524 | 0 | 0 | 0 | 0 | 0 | 0 | 0 |
| ENSMUSG00000065526 | 0 | 0 | 0 | 0 | 0 | 0 | 0 |
| ENSMUSG00000065527 | 0 | 0 | 0 | 0 | 0 | 0 | 0 |
| ENSMUSG00000065528 | 0 | 0 | 0 | 0 | 0 | 0 | 0 |
| ENSMUSG00000065529 | 0 | 0 | 0 | 0 | 0 | 0 | 0 |
| ENSMUSG00000065530 | 0 | 0 | 0 | 0 | 0 | 0 | 0 |
| ENSMUSG00000065532 | 0 | 0 | 0 | 0 | 0 | 0 | 0 |
| ENSMUSG00000065533 | 0 | 0 | 0 | 0 | 0 | 0 | 0 |
| ENSMUSG00000065534 | 0 | 0 | 0 | 0 | 0 | 0 | 0 |
| ENSMUSG00000065536 | 0 | 0 | 0 | 0 | 0 | 0 | 0 |
| ENSMUSG00000065537 | 0 | 0 | 0 | 0 | 0 | 0 | 0 |
| ENSMUSG00000065538 | 0 | 0 | 0 | 0 | 0 | 0 | 0 |
| ENSMUSG00000065539 | 0 | 0 | 0 | 0 | 0 | 0 | 0 |
| ENSMUSG00000065540 | 0 | 0 | 0 | 0 | 0 | 0 | 0 |
| ENSMUSG00000065541 | 0 | 0 | 0 | 0 | 0 | 0 | 0 |
| ENSMUSG00000065542 | 0 | 0 | 0 | 0 | 0.244650542 | 0 | 0 |
| ENSMUSG00000065543 | 0 | 0 | 0 | 0 | 0 | 0 | 0 |
| ENSMUSG00000065544 | 0 | 0 | 0 | 0 | 0.387761935 | 0.301029996 | 0 |
| ENSMUSG00000065546 | 0.811507563 | 1.130890489 | 0.75 | 0.646240625 | 0.846351821 | 0.301029996 | 0.896975297 |
| ENSMUSG00000065547 | 0 | 0 | 0 | 0 | 0.244650542 | 0 | 0 |
| ENSMUSG00000065548 | 0 | 0 | 0 | 0 | 0 | 0 | 0 |
| ENSMUSG00000065549 | 0 | 0 | 0 | 0 | 0 | 0 | 0 |
| ENSMUSG00000065551 | 0 | 0 | 0 | 0 | 0 | 0 | 0 |
| ENSMUSG00000065552 | 0 | 0 | 0 | 0 | 0 | 0 | 0 |
| ENSMUSG00000065553 | 0 | 0 | 0 | 0 | 0 | 0 | 0 |

| | | | | | | | |
|---|---|---|---|---|---|---|---|
| ENSMUSG00000065555 | 0 | 0 | 0 | 0 | 0 | 0 | 0 |
| ENSMUSG00000065556 | 0 | 0 | 0 | 0 | 0 | 0 | 0 |
| ENSMUSG00000065557 | 0 | 0 | 0 | 0 | 0 | 0 | 0 |
| ENSMUSG00000065559 | 0 | 0 | 0 | 0 | 0 | 0 | 0 |
| ENSMUSG00000065560 | 0 | 0 | 0 | 0 | 0 | 0 | 0 |
| ENSMUSG00000065561 | 0 | 0 | 0 | 0 | 0 | 0 | 0 |
| ENSMUSG00000065562 | 0 | 0 | 0 | 0 | 0 | 0 | 0 |
| ENSMUSG00000065563 | 0 | 0 | 0 | 0 | 0 | 0 | 0 |
| ENSMUSG00000065564 | 0 | 0 | 0 | 0 | 0 | 0 | 0 |
| ENSMUSG00000065565 | 0 | 0 | 0 | 0 | 0 | 0 | 0 |
| ENSMUSG00000065567 | 0 | 0 | 0 | 0 | 0 | 0 | 0 |
| ENSMUSG00000065568 | 0 | 0 | 0 | 0 | 0 | 0 | 0 |
| ENSMUSG00000065569 | 0 | 0 | 0 | 0 | 0 | 0 | 0 |
| ENSMUSG00000065570 | 0 | 0 | 0 | 0 | 0 | 0 | 0 |
| ENSMUSG00000065571 | 0 | 0 | 0 | 0 | 0 | 0 | 0 |
| ENSMUSG00000065572 | 0 | 0 | 0 | 0 | 0 | 0 | 0 |
| ENSMUSG00000065573 | 0 | 0 | 0 | 0 | 0 | 0 | 0 |
| ENSMUSG00000065574 | 0 | 0 | 0 | 0 | 0 | 0 | 0 |
| ENSMUSG00000065575 | 0 | 0 | 0.25 | 0 | 0 | 0 | 0 |
| ENSMUSG00000065577 | 0 | 0 | 0 | 0 | 0 | 0 | 0 |
| ENSMUSG00000065578 | 0 | 0 | 0 | 0 | 0 | 0 | 0 |
| ENSMUSG00000065579 | 0 | 0 | 0 | 0 | 0 | 0 | 0 |
| ENSMUSG00000065580 | 0 | 0 | 0 | 0 | 0 | 0 | 0 |
| ENSMUSG00000065581 | 0 | 0.25 | 0.25 | 0.25 | 0.244650542 | 0 | 0.224243824 |
| ENSMUSG00000065582 | 0 | 0 | 0 | 0 | 0 | 0 | 0 |
| ENSMUSG00000065583 | 0 | 0 | 0 | 0 | 0 | 0 | 0 |
| ENSMUSG00000065585 | 0 | 0 | 0 | 0 | 0 | 0 | 0 |
| ENSMUSG00000065586 | 0 | 0 | 0 | 0 | 0 | 0 | 0 |
| ENSMUSG00000065587 | 0 | 0 | 0 | 0 | 0 | 0 | 0 |
| ENSMUSG00000065589 | 0 | 0 | 0 | 0 | 0.244650542 | 0 | 0 |
| ENSMUSG00000065590 | 0 | 0 | 0 | 0 | 0 | 0 | 0 |
| ENSMUSG00000065591 | 0.671187741 | 0 | 0.25 | 0.25 | 0.686820904 | 0.301029996 | 0.520678036 |
| ENSMUSG00000065592 | 0 | 0.396240625 | 0 | 0 | 0 | 0 | 0 |
| ENSMUSG00000065593 | 0 | 0 | 0 | 0 | 0 | 0 | 0 |

| | | | | | | | |
|---|---|---|---|---|---|---|---|
| ENSMUSG00000065594 | 0 | 0 | 0 | 0 | 0 | 0 | 0 |
| ENSMUSG00000065596 | 0 | 0 | 0 | 0 | 0 | 0 | 0 |
| ENSMUSG00000065597 | 0 | 0 | 0 | 0 | 0 | 0 | 0 |
| ENSMUSG00000065599 | 0 | 0 | 0 | 0 | 0 | 0 | 0.224243824 |
| ENSMUSG00000065600 | 0 | 0 | 0 | 0 | 0 | 0 | 0 |
| ENSMUSG00000065601 | 0 | 0 | 0 | 0 | 0 | 0 | 0 |
| ENSMUSG00000065602 | 0 | 0 | 0 | 0 | 0 | 0 | 0 |
| ENSMUSG00000065603 | 0 | 0 | 0 | 0 | 0 | 0 | 0 |
| ENSMUSG00000065604 | 0 | 0 | 0 | 0 | 0 | 0 | 0 |
| ENSMUSG00000065606 | 0 | 0.25 | 0 | 0 | 0 | 0 | 0 |
| ENSMUSG00000065607 | 0 | 0 | 0 | 0 | 0 | 0 | 0 |
| ENSMUSG00000065608 | 0 | 0 | 0 | 0 | 0 | 0.301029996 | 0 |
| ENSMUSG00000065609 | 0 | 0 | 0 | 0 | 0 | 0 | 0 |
| ENSMUSG00000065610 | 0 | 0 | 0 | 0 | 0 | 0 | 0 |
| ENSMUSG00000065611 | 0 | 0 | 0 | 0.25 | 0.244650542 | 0 | 0 |
| ENSMUSG00000065612 | 0 | 0 | 0 | 0 | 0.244650542 | 0 | 0 |
| ENSMUSG00000065613 | 0 | 0 | 0 | 0 | 0 | 0 | 0 |
| ENSMUSG00000065616 | 0 | 0 | 0 | 0 | 0 | 0 | 0 |
| ENSMUSG00000065617 | 0 | 0 | 0 | 0 | 0 | 0 | 0 |
| ENSMUSG00000065619 | 0 | 0 | 0 | 0 | 0 | 0 | 0 |
| ENSMUSG00000065620 | 0 | 0 | 0 | 0 | 0 | 0 | 0 |
| ENSMUSG00000065622 | 0 | 0 | 0 | 0 | 0 | 0 | 0 |
| ENSMUSG00000065623 | 0 | 0 | 0 | 0 | 0 | 0 | 0.224243824 |
| ENSMUSG00000065625 | 0 | 0 | 0 | 0 | 0 | 0 | 0 |
| ENSMUSG00000065627 | 0 | 0 | 0 | 0 | 0 | 0 | 0 |
| ENSMUSG00000065628 | 0 | 0 | 0 | 0 | 0 | 0 | 0 |
| ENSMUSG00000065629 | 0 | 0 | 0 | 0 | 0 | 0 | 0 |
| ENSMUSG00000065630 | 0 | 0 | 0 | 0 | 0 | 0 | 0 |
| ENSMUSG00000065634 | 0 | 0 | 0 | 0 | 0 | 0 | 0 |
| ENSMUSG00000065635 | 0 | 0 | 0 | 0 | 0 | 0 | 0 |
| ENSMUSG00000065636 | 0 | 0 | 0 | 0 | 0 | 0 | 0 |
| ENSMUSG00000065637 | 0 | 0.75 | 0.25 | 0.580482024 | 0.244650542 | 0.301029996 | 0.224243824 |
| ENSMUSG00000065640 | 0 | 0 | 0 | 0 | 0 | 0 | 0 |
| ENSMUSG00000065642 | 0.289064826 | 0 | 0.396240625 | 0.25 | 0.244650542 | 0 | 0.224243824 |

| | | | | | | | |
|---|---|---|---|---|---|---|---|
| ENSMUSG00000065643 | 0 | 0 | 0.25 | 0.25 | 0 | 0 | 0.224243824 |
| ENSMUSG00000065644 | 0 | 0 | 0 | 0 | 0 | 0 | 0 |
| ENSMUSG00000065645 | 0 | 0 | 0 | 0 | 0 | 0 | 0 |
| ENSMUSG00000065647 | 0 | 0 | 0 | 0 | 0 | 0 | 0 |
| ENSMUSG00000065648 | 0 | 0 | 0 | 0 | 0 | 0 | 0 |
| ENSMUSG00000065649 | 0 | 0 | 0 | 0 | 0 | 0 | 0 |
| ENSMUSG00000065651 | 0 | 0 | 0 | 0 | 0 | 0 | 0.224243824 |
| ENSMUSG00000065653 | 0 | 0 | 0 | 0 | 0 | 0 | 0 |
| ENSMUSG00000065656 | 0 | 0 | 0 | 0 | 0 | 0 | 0 |
| ENSMUSG00000065657 | 0 | 0 | 0 | 0 | 0 | 0 | 0 |
| ENSMUSG00000065658 | 0 | 0 | 0 | 0 | 0 | 0 | 0 |
| ENSMUSG00000065659 | 0 | 0 | 0 | 0 | 0 | 0 | 0 |
| ENSMUSG00000065660 | 0 | 0 | 0 | 0 | 0 | 0 | 0 |
| ENSMUSG00000065661 | 0 | 0 | 0 | 0 | 0 | 0 | 0 |
| ENSMUSG00000065662 | 0 | 0 | 0.25 | 0 | 0 | 0 | 0 |
| ENSMUSG00000065663 | 0.747221736 | 0.701838731 | 0.79248125 | 0.951838731 | 0.77552387 | 0.602059991 | 0.853775828 |
| ENSMUSG00000065664 | 0 | 0 | 0 | 0 | 0 | 0 | 0 |
| ENSMUSG00000065665 | 0 | 0 | 0 | 0 | 0 | 0 | 0 |
| ENSMUSG00000065667 | 0 | 0 | 0 | 0.25 | 0 | 0 | 0 |
| ENSMUSG00000065669 | 0 | 0.25 | 0.396240625 | 0.396240625 | 0.244650542 | 0 | 0.224243824 |
| ENSMUSG00000065673 | 0 | 0.25 | 0 | 0 | 0 | 0 | 0 |
| ENSMUSG00000065674 | 0 | 0 | 0 | 0 | 0 | 0 | 0 |
| ENSMUSG00000065675 | 0 | 0 | 0 | 0 | 0 | 0 | 0 |
| ENSMUSG00000065676 | 0.289064826 | 0 | 0 | 0 | 0 | 0 | 0.448487648 |
| ENSMUSG00000065678 | 0 | 0 | 0 | 0 | 0 | 0 | 0 |
| ENSMUSG00000065680 | 0 | 0 | 0 | 0 | 0 | 0 | 0 |
| ENSMUSG00000065683 | 0 | 0 | 0 | 0 | 0 | 0 | 0 |
| ENSMUSG00000065684 | 0 | 0 | 0 | 0 | 0 | 0 | 0 |
| ENSMUSG00000065685 | 0 | 0 | 0 | 0 | 0 | 0 | 0 |
| ENSMUSG00000065686 | 0 | 0.25 | 0 | 0 | 0 | 0 | 0 |
| ENSMUSG00000065687 | 0 | 0 | 0 | 0 | 0 | 0 | 0 |
| ENSMUSG00000065689 | 0 | 0 | 0 | 0 | 0 | 0 | 0 |
| ENSMUSG00000065691 | 0 | 0 | 0 | 0 | 0 | 0 | 0 |
| ENSMUSG00000065692 | 0 | 0 | 0 | 0 | 0 | 0 | 0 |

| | | | | | | | |
|---|---|---|---|---|---|---|---|
| ENSMUSG00000065694 | 0 | 0.396240625 | 0 | 0 | 0 | 0 | 0 |
| ENSMUSG00000065698 | 0 | 0 | 0 | 0 | 0 | 0 | 0 |
| ENSMUSG00000065700 | 0 | 0 | 0 | 0 | 0 | 0 | 0 |
| ENSMUSG00000065701 | 0.289064826 | 0 | 0 | 0 | 0.244650542 | 0.301029996 | 0 |
| ENSMUSG00000065702 | 0 | 0 | 0 | 0 | 0 | 0 | 0 |
| ENSMUSG00000065705 | 0 | 0 | 0 | 0 | 0 | 0 | 0 |
| ENSMUSG00000065706 | 0 | 0 | 0 | 0 | 0 | 0 | 0 |
| ENSMUSG00000065708 | 0 | 0 | 0 | 0 | 0 | 0 | 0 |
| ENSMUSG00000065709 | 0 | 0 | 0 | 0 | 0 | 0 | 0 |
| ENSMUSG00000065714 | 0 | 0 | 0 | 0 | 0 | 0 | 0 |
| ENSMUSG00000065715 | 0.578129653 | 0.701838731 | 0.396240625 | 0.701838731 | 0 | 0.77815125 | 0.579661877 |
| ENSMUSG00000065716 | 0 | 0 | 0 | 0 | 0 | 0 | 0 |
| ENSMUSG00000065718 | 0 | 0 | 0 | 0 | 0 | 0 | 0 |
| ENSMUSG00000065719 | 0 | 0 | 0 | 0 | 0 | 0 | 0 |
| ENSMUSG00000065721 | 0 | 0 | 0 | 0 | 0 | 0 | 0 |
| ENSMUSG00000065722 | 0 | 0 | 0 | 0 | 0 | 0 | 0 |
| ENSMUSG00000065725 | 0 | 0 | 0 | 0.25 | 0 | 0 | 0 |
| ENSMUSG00000065726 | 0 | 0 | 0 | 0 | 0 | 0 | 0 |
| ENSMUSG00000065727 | 0 | 0 | 0 | 0 | 0 | 0 | 0 |
| ENSMUSG00000065728 | 0 | 0.25 | 0 | 0 | 0.387761935 | 0 | 0 |
| ENSMUSG00000065729 | 0 | 0 | 0 | 0 | 0 | 0 | 0 |
| ENSMUSG00000065732 | 0 | 0 | 0 | 0 | 0 | 0 | 0 |
| ENSMUSG00000065733 | 0 | 0 | 0 | 0 | 0 | 0 | 0 |
| ENSMUSG00000065734 | 1.069666964 | 0.75 | 1 | 0.976722649 | 0.978602168 | 0.602059991 | 0.853775828 |
| ENSMUSG00000065735 | 0 | 0 | 0 | 0 | 0 | 0 | 0 |
| ENSMUSG00000065737 | 0 | 0 | 0 | 0 | 0 | 0 | 0 |
| ENSMUSG00000065738 | 0.45815691 | 0.396240625 | 0.25 | 0.5 | 0.568060967 | 0.301029996 | 0.355418052 |
| ENSMUSG00000065739 | 0 | 0 | 0 | 0 | 0 | 0 | 0 |
| ENSMUSG00000065742 | 0 | 0 | 0 | 0 | 0 | 0 | 0 |
| ENSMUSG00000065743 | 0 | 0 | 0 | 0 | 0 | 0 | 0 |
| ENSMUSG00000065745 | 0 | 0 | 0 | 0 | 0 | 0 | 0 |
| ENSMUSG00000065746 | 0 | 0 | 0 | 0 | 0 | 0 | 0 |
| ENSMUSG00000065747 | 0 | 0 | 0 | 0 | 0 | 0 | 0 |
| ENSMUSG00000065749 | 0 | 0 | 0 | 0 | 0 | 0 | 0 |

| | | | | | | | |
|---|---|---|---|---|---|---|---|
| ENSMUSG00000065750 | 0 | 0.25 | 0 | 0 | 0 | 0 | 0 |
| ENSMUSG00000065751 | 0 | 0 | 0 | 0 | 0 | 0 | 0 |
| ENSMUSG00000065752 | 0.45815691 | 0.25 | 0.580482024 | 0 | 0 | 0 | 0.355418052 |
| ENSMUSG00000065757 | 0 | 0 | 0 | 0 | 0 | 0 | 0 |
| ENSMUSG00000065758 | 0 | 0 | 0 | 0 | 0 | 0 | 0 |
| ENSMUSG00000065760 | 0 | 0 | 0 | 0 | 0 | 0 | 0 |
| ENSMUSG00000065763 | 0 | 0 | 0 | 0 | 0 | 0 | 0 |
| ENSMUSG00000065764 | 0 | 0 | 0 | 0 | 0 | 0 | 0 |
| ENSMUSG00000065765 | 0.289064826 | 0.25 | 0.25 | 0.25 | 0.387761935 | 0 | 0.355418052 |
| ENSMUSG00000065767 | 0 | 0.25 | 0.396240625 | 0.25 | 0 | 0.477121255 | 0.224243824 |
| ENSMUSG00000065768 | 0 | 0 | 0 | 0 | 0 | 0 | 0 |
| ENSMUSG00000065769 | 0 | 0 | 0 | 0 | 0 | 0 | 0 |
| ENSMUSG00000065770 | 0 | 0 | 0 | 0 | 0 | 0 | 0 |
| ENSMUSG00000065771 | 0.289064826 | 0 | 0 | 0 | 0 | 0 | 0 |
| ENSMUSG00000065773 | 0 | 0 | 0 | 0 | 0 | 0 | 0 |
| ENSMUSG00000065778 | 0 | 0.25 | 0 | 0 | 0 | 0 | 0 |
| ENSMUSG00000065782 | 1.181541736 | 0.92510993 | 0.896240625 | 0.896240625 | 0.905314583 | 0.903089987 | 0.916588299 |
| ENSMUSG00000065787 | 0 | 0 | 0 | 0 | 0 | 0 | 0 |
| ENSMUSG00000065788 | 0 | 0 | 0 | 0 | 0 | 0 | 0 |
| ENSMUSG00000065789 | 0 | 0 | 0 | 0 | 0 | 0 | 0 |
| ENSMUSG00000065791 | 0 | 0 | 0 | 0 | 0 | 0 | 0 |
| ENSMUSG00000065792 | 0 | 0 | 0 | 0 | 0 | 0 | 0.224243824 |
| ENSMUSG00000065794 | 0 | 0 | 0 | 0 | 0 | 0 | 0 |
| ENSMUSG00000065795 | 0 | 0 | 0 | 0 | 0 | 0 | 0 |
| ENSMUSG00000065797 | 0 | 0 | 0 | 0 | 0 | 0 | 0 |
| ENSMUSG00000065798 | 0 | 0 | 0 | 0 | 0 | 0 | 0 |
| ENSMUSG00000065799 | 0 | 0 | 0 | 0 | 0 | 0 | 0 |
| ENSMUSG00000065800 | 0 | 0 | 0 | 0 | 0 | 0 | 0 |
| ENSMUSG00000065802 | 0 | 0 | 0 | 0 | 0 | 0 | 0 |
| ENSMUSG00000065803 | 0 | 0 | 0 | 0 | 0 | 0 | 0 |
| ENSMUSG00000065804 | 0 | 0 | 0 | 0 | 0 | 0 | 0 |
| ENSMUSG00000065806 | 0.289064826 | 0 | 0 | 0 | 0 | 0 | 0 |
| ENSMUSG00000065807 | 0 | 0 | 0 | 0 | 0 | 0 | 0 |
| ENSMUSG00000065808 | 0.289064826 | 0 | 0 | 0 | 0.244650542 | 0.301029996 | 0 |

| | | | | | | | |
|---|---|---|---|---|---|---|---|
| ENSMUSG00000065811 | 0 | 0 | 0 | 0 | 0 | 0 | 0 |
| ENSMUSG00000065812 | 0 | 0 | 0 | 0 | 0 | 0 | 0 |
| ENSMUSG00000065813 | 0 | 0 | 0 | 0 | 0 | 0 | 0 |
| ENSMUSG00000065815 | 0 | 0 | 0 | 0 | 0 | 0 | 0 |
| ENSMUSG00000065817 | 0 | 0 | 0.25 | 0 | 0 | 0 | 0 |
| ENSMUSG00000065818 | 0.289064826 | 0.5 | 0.75 | 0.25 | 0.387761935 | 0.477121255 | 0.448487648 |
| ENSMUSG00000065819 | 0 | 0 | 0 | 0 | 0 | 0 | 0 |
| ENSMUSG00000065820 | 0 | 0 | 0 | 0 | 0 | 0.301029996 | 0.224243824 |
| ENSMUSG00000065822 | 0 | 0 | 0 | 0 | 0 | 0 | 0 |
| ENSMUSG00000065823 | 0 | 0 | 0 | 0 | 0 | 0 | 0 |
| ENSMUSG00000065824 | 0 | 0 | 0 | 0 | 0 | 0 | 0 |
| ENSMUSG00000065825 | 0 | 0 | 0 | 0 | 0 | 0 | 0 |
| ENSMUSG00000065828 | 0 | 0 | 0 | 0 | 0 | 0 | 0 |
| ENSMUSG00000065836 | 0 | 0 | 0 | 0 | 0 | 0 | 0 |
| ENSMUSG00000065837 | 0 | 0 | 0 | 0 | 0 | 0 | 0 |
| ENSMUSG00000065840 | 0 | 0 | 0 | 0 | 0 | 0 | 0 |
| ENSMUSG00000065843 | 0 | 0 | 0 | 0 | 0 | 0 | 0 |
| ENSMUSG00000065845 | 0 | 0 | 0 | 0 | 0 | 0 | 0 |
| ENSMUSG00000065847 | 0 | 0 | 0 | 0 | 0 | 0 | 0 |
| ENSMUSG00000065851 | 0 | 0 | 0 | 0 | 0 | 0 | 0 |
| ENSMUSG00000065852 | 0 | 0 | 0 | 0 | 0 | 0 | 0 |
| ENSMUSG00000065853 | 0 | 0 | 0 | 0 | 0 | 0 | 0 |
| ENSMUSG00000065858 | 0 | 0 | 0 | 0 | 0 | 0 | 0 |
| ENSMUSG00000065859 | 0 | 0 | 0 | 0 | 0 | 0 | 0 |
| ENSMUSG00000065862 | 0 | 0 | 0 | 0 | 0 | 0 | 0 |
| ENSMUSG00000065864 | 0 | 0 | 0 | 0 | 0 | 0 | 0 |
| ENSMUSG00000065865 | 0 | 0 | 0 | 0 | 0 | 0 | 0 |
| ENSMUSG00000065866 | 0 | 0 | 0 | 0 | 0 | 0 | 0 |
| ENSMUSG00000065869 | 0 | 0 | 0 | 0 | 0 | 0 | 0 |
| ENSMUSG00000065870 | 0 | 0 | 0 | 0 | 0 | 0 | 0 |
| ENSMUSG00000065872 | 0 | 0 | 0 | 0 | 0 | 0 | 0 |
| ENSMUSG00000065873 | 0 | 0.5 | 0 | 0 | 0.387761935 | 0 | 0.355418052 |
| ENSMUSG00000065874 | 0 | 0 | 0 | 0 | 0 | 0 | 0 |
| ENSMUSG00000065875 | 0 | 0 | 0 | 0 | 0 | 0 | 0 |

| | | | | | | | |
|---|---|---|---|---|---|---|---|
| ENSMUSG00000065876 | 0 | 0 | 0 | 0 | 0 | 0 | 0 |
| ENSMUSG00000065878 | 0 | 0 | 0 | 0 | 0 | 0 | 0 |
| ENSMUSG00000065881 | 0 | 0 | 0 | 0 | 0 | 0 | 0 |
| ENSMUSG00000065882 | 0 | 0 | 0 | 0 | 0 | 0 | 0 |
| ENSMUSG00000065883 | 0 | 0 | 0 | 0 | 0 | 0 | 0.355418052 |
| ENSMUSG00000065884 | 0 | 0 | 0 | 0 | 0 | 0 | 0 |
| ENSMUSG00000065887 | 0 | 0 | 0 | 0 | 0 | 0 | 0 |
| ENSMUSG00000065889 | 0 | 0 | 0 | 0 | 0 | 0 | 0 |
| ENSMUSG00000065892 | 0 | 0 | 0 | 0 | 0 | 0 | 0 |
| ENSMUSG00000065893 | 0 | 0 | 0 | 0 | 0 | 0 | 0 |
| ENSMUSG00000065894 | 0 | 0 | 0 | 0 | 0 | 0 | 0 |
| ENSMUSG00000065898 | 0 | 0 | 0 | 0 | 0 | 0 | 0 |
| ENSMUSG00000065899 | 0 | 0 | 0 | 0 | 0 | 0 | 0 |
| ENSMUSG00000065903 | 0 | 0 | 0 | 0 | 0 | 0 | 0 |
| ENSMUSG00000065904 | 0.45815691 | 0 | 0.25 | 0 | 0 | 0 | 0 |
| ENSMUSG00000065905 | 0 | 0 | 0 | 0 | 0 | 0 | 0 |
| ENSMUSG00000065906 | 0 | 0 | 0 | 0 | 0 | 0 | 0 |
| ENSMUSG00000065907 | 0 | 0 | 0 | 0 | 0 | 0 | 0 |
| ENSMUSG00000065911 | 0 | 0 | 0 | 0 | 0 | 0 | 0 |
| ENSMUSG00000065913 | 0 | 0 | 0 | 0 | 0 | 0 | 0 |
| ENSMUSG00000065915 | 0 | 0 | 0 | 0 | 0 | 0 | 0 |
| ENSMUSG00000065916 | 0 | 0 | 0 | 0 | 0 | 0 | 0 |
| ENSMUSG00000065919 | 0 | 0 | 0 | 0 | 0 | 0 | 0 |
| ENSMUSG00000065922 | 0 | 0 | 0 | 0 | 0 | 0 | 0 |
| ENSMUSG00000065924 | 0 | 0 | 0 | 0 | 0 | 0 | 0 |
| ENSMUSG00000065926 | 0 | 0 | 0 | 0 | 0 | 0 | 0 |
| ENSMUSG00000065934 | 0 | 0.25 | 0 | 0 | 0 | 0 | 0 |
| ENSMUSG00000065935 | 0 | 0 | 0 | 0 | 0 | 0 | 0 |
| ENSMUSG00000065936 | 0 | 0 | 0 | 0 | 0 | 0 | 0 |
| ENSMUSG00000065937 | 0 | 0 | 0 | 0 | 0 | 0 | 0 |
| ENSMUSG00000065939 | 0.91631382 | 0.896240625 | 0.830482024 | 0.646240625 | 0.733951626 | 1 | 0.803905701 |
| ENSMUSG00000065940 | 0 | 0 | 0 | 0 | 0 | 0 | 0 |
| ENSMUSG00000065943 | 0 | 0 | 0 | 0 | 0 | 0 | 0 |
| ENSMUSG00000065944 | 0 | 0 | 0 | 0 | 0 | 0 | 0 |

| | | | | | | | |
|---|---|---|---|---|---|---|---|
| ENSMUSG00000065945 | 0 | 0 | 0 | 0 | 0 | 0 | 0 |
| ENSMUSG00000065947 | 0 | 0 | 0 | 0 | 0 | 0 | 0 |
| ENSMUSG00000065950 | 0 | 0 | 0.25 | 0 | 0 | 0 | 0.224243824 |
| ENSMUSG00000065952 | 0 | 0.25 | 0.25 | 0 | 0 | 0 | 0.224243824 |
| ENSMUSG00000065954 | 2.740854706 | 2.517530236 | 2.510414788 | 2.405054956 | 2.366364104 | 2.697229343 | 2.225910574 |
| ENSMUSG00000065956 | 0 | 0 | 0 | 0 | 0 | 0 | 0 |
| ENSMUSG00000065968 | 0 | 0.5 | 0 | 0.25 | 0.387761935 | 0 | 0 |
| ENSMUSG00000065979 | 2.092368209 | 1.950224975 | 2.01109853 | 1.994319981 | 1.938809675 | 2.170261715 | 1.87511425 |
| ENSMUSG00000065987 | 0 | 0 | 0 | 0 | 0 | 0 | 0 |
| ENSMUSG00000065990 | 2.213719572 | 2.154596376 | 2.186208459 | 2.165444524 | 2.157861565 | 2.474216264 | 2.108089156 |
| ENSMUSG00000065999 | 0 | 0 | 0 | 0.25 | 0.568060967 | 0.602059991 | 0.355418052 |
| ENSMUSG00000066000 | 1.307602651 | 1.330482024 | 1.388647213 | 1.238549078 | 1.283908311 | 1.230448921 | 1.159323753 |
| ENSMUSG00000066007 | 0 | 0 | 0 | 0 | 0 | 0 | 0.224243824 |
| ENSMUSG00000066009 | 0 | 0 | 0 | 0 | 0 | 0 | 0 |
| ENSMUSG00000066026 | 1.325351389 | 1.364857905 | 1.114857905 | 1 | 1 | 0.954242509 | 1.014380832 |
| ENSMUSG00000066027 | 0 | 0.25 | 0 | 0 | 0 | 0 | 0 |
| ENSMUSG00000066030 | 0 | 0 | 0 | 0 | 0 | 0 | 0 |
| ENSMUSG00000066031 | 0 | 0 | 0 | 0 | 0 | 0 | 0 |
| ENSMUSG00000066036 | 3.230097535 | 2.94059551 | 2.927486345 | 2.832168732 | 2.870799488 | 3.212720154 | 2.699806959 |
| ENSMUSG00000066037 | 2.915547216 | 2.637205477 | 2.616136438 | 2.722945926 | 2.625007532 | 2.998259338 | 2.420763202 |
| ENSMUSG00000066042 | 1.516991255 | 1.54745614 | 1.584962501 | 1.470660762 | 1.555378544 | 1.602059991 | 1.396774124 |
| ENSMUSG00000066043 | 2.527823874 | 2.28296424 | 2.296218836 | 2.394828984 | 2.354487824 | 2.697229343 | 2.21680636 |
| ENSMUSG00000066057 | 1.693336259 | 1.638647213 | 1.593759858 | 1.614857905 | 1.555378544 | 1.556302501 | 1.455757965 |
| ENSMUSG00000066058 | 0.289064826 | 0.25 | 0 | 0 | 0 | 0 | 0.355418052 |
| ENSMUSG00000066060 | 0 | 0 | 0 | 0 | 0 | 0 | 0 |
| ENSMUSG00000066061 | 0 | 0 | 0 | 0 | 0 | 0 | 0 |
| ENSMUSG00000066068 | 0 | 0 | 0 | 0 | 0 | 0 | 0 |
| ENSMUSG00000066071 | 0 | 0 | 0 | 0 | 0 | 0 | 0 |
| ENSMUSG00000066072 | 0 | 0 | 0 | 0 | 0 | 0 | 0 |
| ENSMUSG00000066090 | 0 | 0 | 0 | 0 | 0 | 0 | 0 |
| ENSMUSG00000066097 | 0 | 0 | 0 | 0 | 0 | 0 | 0 |
| ENSMUSG00000066100 | 0 | 0 | 0 | 0 | 0 | 0 | 0 |
| ENSMUSG00000066101 | 0 | 0 | 0 | 0 | 0 | 0 | 0 |
| ENSMUSG00000066107 | 1.036286563 | 1.061981878 | 1.098079356 | 1.146240625 | 1.234113756 | 1 | 1.089372236 |

| | | | | | | | |
|---|---|---|---|---|---|---|---|
| ENSMUSG00000066108 | 0 | 0 | 0 | 0.25 | 0 | 0 | 0 |
| ENSMUSG00000066113 | 1.205378646 | 1.380890489 | 1.25 | 1 | 1.020174412 | 0.84509804 | 1.20139795 |
| ENSMUSG00000066116 | 0.289064826 | 0 | 0 | 0 | 0 | 0 | 0 |
| ENSMUSG00000066122 | 0 | 0 | 0 | 0 | 0 | 0 | 0 |
| ENSMUSG00000066129 | 0 | 0 | 0.396240625 | 0 | 0.387761935 | 0 | 0 |
| ENSMUSG00000066137 | 0 | 0 | 0 | 0 | 0 | 0 | 0.224243824 |
| ENSMUSG00000066141 | 0 | 0 | 0 | 0 | 0 | 0 | 0 |
| ENSMUSG00000066148 | 2.444137694 | 2.254897682 | 2.259043403 | 2.375956434 | 2.315755952 | 2.661812686 | 2.152303998 |
| ENSMUSG00000066149 | 2.203267085 | 2.080482024 | 2.013820609 | 2.081607372 | 2.034859505 | 2.283301229 | 1.953344547 |
| ENSMUSG00000066150 | 2.685417684 | 2.472945926 | 2.447383411 | 2.636964627 | 2.546663136 | 2.929418926 | 2.34615152 |
| ENSMUSG00000066151 | 2.262379879 | 2.161864607 | 2.130890489 | 2.250703754 | 2.199086582 | 2.499687083 | 2.090889903 |
| ENSMUSG00000066152 | 1.789260561 | 1.71759118 | 1.688721876 | 1.816696635 | 1.777823267 | 1.968482949 | 1.661510159 |
| ENSMUSG00000066153 | 0 | 0 | 0 | 0 | 0 | 0 | 0 |
| ENSMUSG00000066154 | 0 | 0 | 0 | 0 | 0 | 0 | 0 |
| ENSMUSG00000066158 | 0 | 0.396240625 | 0.5 | 0.580482024 | 0.387761935 | 0.301029996 | 0.355418052 |
| ENSMUSG00000066170 | 0.45815691 | 0 | 0.25 | 0.25 | 0.568060967 | 0.301029996 | 0.355418052 |
| ENSMUSG00000066175 | 0.289064826 | 0.396240625 | 0.396240625 | 0.580482024 | 0.632412477 | 0.77815125 | 0.579661877 |
| ENSMUSG00000066176 | 0 | 0 | 0 | 0 | 0 | 0 | 0 |
| ENSMUSG00000066178 | 0.45815691 | 0.701838731 | 0.79248125 | 0.701838731 | 0.686820904 | 0.477121255 | 0.672731473 |
| ENSMUSG00000066180 | 0 | 0.396240625 | 0 | 0 | 0 | 0 | 0 |
| ENSMUSG00000066189 | 0 | 0 | 0 | 0 | 0 | 0 | 0 |
| ENSMUSG00000066191 | 1.707474304 | 1.664552871 | 1.744319981 | 1.458222504 | 1.580303447 | 1.602059991 | 1.661510159 |
| ENSMUSG00000066196 | 0 | 0.5 | 0.646240625 | 0.396240625 | 0.632412477 | 0.301029996 | 0.520678036 |
| ENSMUSG00000066197 | 0 | 0 | 0 | 0 | 0 | 0 | 0 |
| ENSMUSG00000066224 | 0 | 0 | 0 | 0.396240625 | 0.568060967 | 0.602059991 | 0.74492186 |
| ENSMUSG00000066232 | 2.896311254 | 2.663659007 | 2.652294685 | 2.722945926 | 2.630984742 | 3.013258665 | 2.407950873 |
| ENSMUSG00000066233 | 1.227926429 | 1.226722649 | 1.330482024 | 1.160964047 | 1.254881871 | 1.146128036 | 1.296434211 |
| ENSMUSG00000066235 | 1.342375483 | 1.589388001 | 1.610735874 | 1.25 | 1.414412788 | 1.477121255 | 1.586932193 |
| ENSMUSG00000066239 | 0 | 0 | 0 | 0 | 0 | 0 | 0 |
| ENSMUSG00000066240 | 0 | 0 | 0 | 0 | 0 | 0 | 0 |
| ENSMUSG00000066241 | 0 | 0 | 0 | 0 | 0 | 0 | 0 |
| ENSMUSG00000066242 | 0 | 0 | 0 | 0 | 0 | 0 | 0 |
| ENSMUSG00000066245 | 0.45815691 | 0.580482024 | 0 | 0.25 | 0 | 0 | 0 |
| ENSMUSG00000066257 | 0 | 0 | 0 | 0 | 0 | 0 | 0 |

| | | | | | | | |
|---|---|---|---|---|---|---|---|
| ENSMUSG00000066258 | 0 | 0 | 0.25 | 0 | 0.387761935 | 0 | 0.520678036 |
| ENSMUSG00000066262 | 0 | 0 | 0 | 0 | 0 | 0 | 0 |
| ENSMUSG00000066263 | 0 | 0 | 0 | 0 | 0 | 0 | 0 |
| ENSMUSG00000066268 | 0 | 0 | 0 | 0 | 0 | 0 | 0 |
| ENSMUSG00000066269 | 0 | 0 | 0 | 0 | 0 | 0 | 0 |
| ENSMUSG00000066270 | 0 | 0 | 0 | 0 | 0 | 0 | 0 |
| ENSMUSG00000066272 | 0 | 0 | 0 | 0 | 0 | 0 | 0 |
| ENSMUSG00000066273 | 0 | 0 | 0 | 0 | 0 | 0 | 0 |
| ENSMUSG00000066278 | 1.83262764 | 1.688721876 | 1.78743678 | 1.634789703 | 1.706995316 | 1.77815125 | 1.762103709 |
| ENSMUSG00000066279 | 0 | 0 | 0 | 0 | 0.244650542 | 0 | 0 |
| ENSMUSG00000066306 | 2.920505136 | 2.710980263 | 2.741806565 | 2.496104615 | 2.588465994 | 2.857332496 | 2.466682066 |
| ENSMUSG00000066315 | 0 | 0 | 0 | 0 | 0 | 0 | 0 |
| ENSMUSG00000066319 | 0.289064826 | 0.646240625 | 0.646240625 | 0.701838731 | 0.489301084 | 0.477121255 | 0.520678036 |
| ENSMUSG00000066324 | 3.115562993 | 2.832028474 | 2.815523711 | 2.888407323 | 2.792479619 | 3.221674997 | 2.559554144 |
| ENSMUSG00000066357 | 2.890240806 | 2.691425122 | 2.74929487 | 2.537754135 | 2.703684258 | 2.946943271 | 2.599851042 |
| ENSMUSG00000066359 | 0 | 0 | 0 | 0 | 0 | 0 | 0 |
| ENSMUSG00000066361 | 0 | 0 | 0 | 0 | 0 | 0 | 0 |
| ENSMUSG00000066362 | 0.747221736 | 0.25 | 0.5 | 0.75 | 0.733951626 | 0.698970004 | 0.775756176 |
| ENSMUSG00000066363 | 1.623015126 | 1.51109853 | 1.445339928 | 1.17510993 | 1.057362051 | 1.146128036 | 0.896975297 |
| ENSMUSG00000066364 | 0 | 0 | 0 | 0 | 0 | 0 | 0 |
| ENSMUSG00000066366 | 0 | 0 | 0 | 0 | 0 | 0 | 0 |
| ENSMUSG00000066368 | 0 | 0 | 0 | 0 | 0 | 0 | 0 |
| ENSMUSG00000066372 | 0 | 0 | 0 | 0 | 0 | 0 | 0 |
| ENSMUSG00000066378 | 0 | 0 | 0 | 0 | 0 | 0 | 0 |
| ENSMUSG00000066382 | 0 | 0.25 | 0.25 | 0 | 0 | 0 | 0.224243824 |
| ENSMUSG00000066383 | 0 | 0 | 0.5 | 0 | 0 | 0 | 0 |
| ENSMUSG00000066389 | 0 | 0.25 | 0.396240625 | 0.25 | 0.387761935 | 0 | 0.448487648 |
| ENSMUSG00000066392 | 0 | 0.396240625 | 0 | 0.25 | 0 | 0 | 0.224243824 |
| ENSMUSG00000066406 | 2.302628051 | 2.150906586 | 2.12694866 | 2.142463902 | 2.116484486 | 2.334453751 | 1.999330888 |
| ENSMUSG00000066407 | 0 | 0.396240625 | 0.5 | 0 | 0 | 0.301029996 | 0.579661877 |
| ENSMUSG00000066415 | 2.349889783 | 2.192046081 | 2.149047439 | 2.243603647 | 2.13194489 | 2.462397998 | 1.997988498 |
| ENSMUSG00000066438 | 0 | 0 | 0 | 0 | 0 | 0 | 0.224243824 |
| ENSMUSG00000066440 | 2.541215974 | 2.351816941 | 2.384789703 | 2.079353153 | 2.171644683 | 2.338456494 | 2.039586611 |
| ENSMUSG00000066441 | 2.669114668 | 2.449009902 | 2.45584181 | 2.281676118 | 2.187072901 | 2.496929648 | 2.041356071 |

| | | | | | | | |
|---|---|---|---|---|---|---|---|
| ENSMUSG00000066442 | 0.289064826 | 0.396240625 | 0.396240625 | 0.5 | 0.489301084 | 0.84509804 | 0.672731473 |
| ENSMUSG00000066443 | 0 | 0 | 0 | 0 | 0 | 0 | 0 |
| ENSMUSG00000066456 | 2.386920383 | 2.196158711 | 2.222185812 | 2.197790722 | 2.138604637 | 2.450249108 | 2.018825668 |
| ENSMUSG00000066457 | 0.289064826 | 0 | 0 | 0 | 0 | 0 | 0 |
| ENSMUSG00000066463 | 0 | 0 | 0 | 0 | 0 | 0 | 0 |
| ENSMUSG00000066475 | 0.289064826 | 0.25 | 0.25 | 0.396240625 | 0.244650542 | 0 | 0.224243824 |
| ENSMUSG00000066477 | 0.289064826 | 0 | 0.396240625 | 0.25 | 0 | 0.301029996 | 0 |
| ENSMUSG00000066478 | 0 | 0 | 0.25 | 0 | 0 | 0 | 0 |
| ENSMUSG00000066487 | 0.811507563 | 0.75 | 1.02186571 | 0.646240625 | 0.733951626 | 0.477121255 | 0.876096088 |
| ENSMUSG00000066489 | 0.45815691 | 0 | 0.25 | 0.25 | 0.244650542 | 0 | 0.355418052 |
| ENSMUSG00000066491 | 0 | 0 | 0 | 0.25 | 0 | 0 | 0 |
| ENSMUSG00000066500 | 0 | 0 | 0 | 0 | 0 | 0 | 0 |
| ENSMUSG00000066510 | 0 | 0 | 0 | 0 | 0 | 0 | 0 |
| ENSMUSG00000066512 | 0 | 0 | 0 | 0 | 0 | 0 | 0 |
| ENSMUSG00000066513 | 0 | 0 | 0 | 0 | 0 | 0 | 0 |
| ENSMUSG00000066515 | 0 | 0 | 0 | 0 | 0 | 0 | 0 |
| ENSMUSG00000066516 | 0 | 0 | 0 | 0 | 0 | 0 | 0 |
| ENSMUSG00000066537 | 0 | 0 | 0 | 0 | 0 | 0 | 0 |
| ENSMUSG00000066538 | 0 | 0.396240625 | 0 | 0 | 0.244650542 | 0 | 0 |
| ENSMUSG00000066543 | 0 | 0 | 0 | 0 | 0.244650542 | 0 | 0.224243824 |
| ENSMUSG00000066551 | 3.064867306 | 2.79745614 | 2.782159703 | 2.847945152 | 2.766591274 | 3.187802639 | 2.590399237 |
| ENSMUSG00000066553 | 0 | 0 | 0 | 0 | 0 | 0 | 0 |
| ENSMUSG00000066554 | 0.747221736 | 0.580482024 | 0.646240625 | 0.75 | 0.387761935 | 0.602059991 | 0.629532004 |
| ENSMUSG00000066568 | 2.673935906 | 2.505938588 | 2.485261902 | 2.474083101 | 2.410235744 | 2.710963119 | 2.260561074 |
| ENSMUSG00000066571 | 2.158622101 | 2.101285366 | 2.086073977 | 2.001406137 | 1.959951091 | 2.113943352 | 1.860017477 |
| ENSMUSG00000066583 | 0 | 0 | 0 | 0 | 0 | 0 | 0 |
| ENSMUSG00000066584 | 0 | 0 | 0 | 0 | 0 | 0 | 0 |
| ENSMUSG00000066586 | 0 | 0 | 0 | 0 | 0 | 0 | 0 |
| ENSMUSG00000066592 | 0.289064826 | 0.580482024 | 0 | 0.396240625 | 0.387761935 | 0 | 0 |
| ENSMUSG00000066595 | 2.060824957 | 1.938721876 | 1.942046081 | 2.121960008 | 2.064555002 | 2.305351369 | 1.906037947 |
| ENSMUSG00000066607 | 1.389637216 | 1.52186571 | 1.630890489 | 1.418106335 | 1.592135487 | 1.431363764 | 1.647906904 |
| ENSMUSG00000066613 | 1.596667477 | 1.488549078 | 1.476722649 | 1.732684334 | 1.642643806 | 1.707570176 | 1.532233387 |
| ENSMUSG00000066620 | 0 | 0 | 0 | 0 | 0 | 0 | 0.224243824 |
| ENSMUSG00000066621 | 2.023453784 | 1.990000483 | 2.093759858 | 1.766522298 | 1.957204337 | 1.963787827 | 1.951796623 |

| | | | | | | | |
|---|---|---|---|---|---|---|---|
| ENSMUSG00000066626 | 0.578129653 | 0 | 0.396240625 | 0.25 | 0.387761935 | 0.301029996 | 0.355418052 |
| ENSMUSG00000066629 | 0 | 0 | 0.5 | 0 | 0 | 0 | 0.224243824 |
| ENSMUSG00000066632 | 0.45815691 | 0.580482024 | 0.5 | 0.5 | 0.568060967 | 0 | 0.520678036 |
| ENSMUSG00000066637 | 1.129344651 | 1.321350555 | 1.201838731 | 1.372963274 | 1.343584837 | 1.431363764 | 1.121219121 |
| ENSMUSG00000066640 | 2.271465912 | 2.170748646 | 2.213717096 | 1.892463902 | 1.974687643 | 2.086359831 | 1.925124822 |
| ENSMUSG00000066643 | 2.377269616 | 2.158248799 | 2.122963274 | 2.194519282 | 2.149165677 | 2.418301291 | 1.977870661 |
| ENSMUSG00000066647 | 1.325351389 | 1.339388001 | 1.321350555 | 1.25 | 1.200473444 | 1.176091259 | 1.150210039 |
| ENSMUSG00000066652 | 0 | 0 | 0 | 0 | 0 | 0 | 0 |
| ENSMUSG00000066671 | 0 | 0 | 0 | 0 | 0 | 0 | 0 |
| ENSMUSG00000066672 | 0 | 0 | 0 | 0 | 0 | 0 | 0 |
| ENSMUSG00000066677 | 0.45815691 | 0.5 | 0.580482024 | 0.5 | 0.568060967 | 0 | 0.448487648 |
| ENSMUSG00000066680 | 0 | 0 | 0 | 0 | 0 | 0 | 0 |
| ENSMUSG00000066682 | 0 | 0 | 0 | 0 | 0.244650542 | 0.301029996 | 0 |
| ENSMUSG00000066684 | 0 | 0 | 0 | 0 | 0 | 0 | 0 |
| ENSMUSG00000066687 | 0 | 0 | 0 | 0 | 0 | 0.477121255 | 0.355418052 |
| ENSMUSG00000066688 | 0 | 0.580482024 | 0 | 0.25 | 0.489301084 | 0 | 0.224243824 |
| ENSMUSG00000066693 | 0 | 0 | 0 | 0 | 0 | 0 | 0 |
| ENSMUSG00000066697 | 0.289064826 | 0.25 | 0 | 0 | 0 | 0 | 0 |
| ENSMUSG00000066704 | 0 | 0 | 0 | 0 | 0 | 0 | 0 |
| ENSMUSG00000066705 | 0 | 0 | 0 | 0.25 | 0.387761935 | 0.301029996 | 0.224243824 |
| ENSMUSG00000066714 | 0 | 0 | 0 | 0.25 | 0 | 0 | 0 |
| ENSMUSG00000066720 | 0.578129653 | 0.5 | 0.75 | 0 | 0 | 0.602059991 | 0.355418052 |
| ENSMUSG00000066721 | 0.671187741 | 0.580482024 | 0.75 | 0.396240625 | 0.387761935 | 0.698970004 | 0.803905701 |
| ENSMUSG00000066723 | 0 | 0 | 0 | 0 | 0 | 0 | 0 |
| ENSMUSG00000066724 | 0.811507563 | 0.75 | 0.896240625 | 0.976722649 | 0.955822902 | 1.079181246 | 0.935079929 |
| ENSMUSG00000066735 | 2.759113486 | 2.503853763 | 2.491446071 | 2.547764804 | 2.472736137 | 2.822821645 | 2.280261923 |
| ENSMUSG00000066747 | 0 | 0 | 0 | 0 | 0 | 0 | 0 |
| ENSMUSG00000066748 | 0 | 0 | 0 | 0 | 0 | 0 | 0 |
| ENSMUSG00000066749 | 0.289064826 | 0 | 0 | 0.25 | 0 | 0 | 0 |
| ENSMUSG00000066750 | 0 | 0 | 0 | 0 | 0 | 0 | 0 |
| ENSMUSG00000066752 | 0 | 0 | 0 | 0 | 0 | 0 | 0 |
| ENSMUSG00000066755 | 0.289064826 | 0.25 | 0.25 | 0.580482024 | 0.244650542 | 0.301029996 | 0.579661877 |
| ENSMUSG00000066756 | 0 | 0 | 0 | 0 | 0 | 0 | 0 |
| ENSMUSG00000066760 | 1.852729478 | 1.816696635 | 1.777131114 | 1.874961472 | 1.848234258 | 2.004321374 | 1.674564426 |

| | | | | | | | |
|---|---|---|---|---|---|---|---|
| ENSMUSG00000066767 | 0.289064826 | 0.25 | 0 | 0 | 0 | 0.301029996 | 0 |
| ENSMUSG00000066771 | 0 | 0 | 0 | 0 | 0 | 0 | 0.224243824 |
| ENSMUSG00000066772 | 0 | 0 | 0 | 0 | 0 | 0 | 0 |
| ENSMUSG00000066797 | 0 | 0 | 0 | 0 | 0 | 0 | 0 |
| ENSMUSG00000066798 | 2.136858952 | 2.024508021 | 2.067865757 | 2.044977273 | 2.024758322 | 2.243038049 | 1.896054911 |
| ENSMUSG00000066800 | 0.811507563 | 1 | 1.04248125 | 1.380890489 | 1.223252711 | 1.255272505 | 1.284450788 |
| ENSMUSG00000066803 | 0 | 0 | 0 | 0 | 0 | 0 | 0 |
| ENSMUSG00000066804 | 0 | 0 | 0 | 0 | 0 | 0 | 0 |
| ENSMUSG00000066807 | 0 | 0 | 0 | 0 | 0 | 0 | 0 |
| ENSMUSG00000066809 | 0 | 0 | 0 | 0 | 0 | 0 | 0 |
| ENSMUSG00000066810 | 0 | 0 | 0 | 0 | 0 | 0 | 0 |
| ENSMUSG00000066820 | 0 | 0 | 0 | 0 | 0 | 0 | 0 |
| ENSMUSG00000066828 | 0 | 0 | 0 | 0 | 0 | 0 | 0 |
| ENSMUSG00000066829 | 1.445324132 | 1.482684334 | 1.339388001 | 1.5 | 1.55104774 | 1.681241237 | 1.417653332 |
| ENSMUSG00000066838 | 1.404271433 | 1.418106335 | 1.25 | 1.321350555 | 1.27449559 | 1.255272505 | 1.21680636 |
| ENSMUSG00000066839 | 1.912079952 | 1.843759858 | 1.909156155 | 1.79745614 | 1.870073561 | 1.991226076 | 1.836517579 |
| ENSMUSG00000066842 | 1.605636262 | 1.557204673 | 1.321350555 | 1.634789703 | 1.632412477 | 1.62324929 | 1.629532004 |
| ENSMUSG00000066850 | 0.671187741 | 0.646240625 | 0.646240625 | 0.580482024 | 0.733951626 | 0.602059991 | 0.710836105 |
| ENSMUSG00000066861 | 0 | 0 | 0 | 0 | 0 | 0 | 0.224243824 |
| ENSMUSG00000066867 | 0 | 0 | 0 | 0 | 0 | 0 | 0 |
| ENSMUSG00000066877 | 2.165631214 | 2.058404919 | 2.104463129 | 1.774508021 | 1.853822071 | 1.913813852 | 1.834294096 |
| ENSMUSG00000066878 | 0 | 0 | 0 | 0 | 0 | 0 | 0 |
| ENSMUSG00000066880 | 2.00683675 | 1.862802778 | 1.811981878 | 1.860735874 | 1.791313678 | 2.056904851 | 1.702561601 |
| ENSMUSG00000066892 | 1.432083896 | 1.5 | 1.505591953 | 1.458222504 | 1.401338844 | 1.544068044 | 1.46982607 |
| ENSMUSG00000066894 | 2.181541736 | 1.987091808 | 1.995748394 | 2.03743678 | 2.030405741 | 2.204119983 | 1.913109605 |
| ENSMUSG00000066896 | 0 | 0 | 0 | 0 | 0 | 0 | 0 |
| ENSMUSG00000066897 | 0 | 0 | 0 | 0 | 0 | 0 | 0 |
| ENSMUSG00000066899 | 0 | 0 | 0 | 0 | 0 | 0 | 0 |
| ENSMUSG00000066900 | 2.750654121 | 2.528435542 | 2.56168515 | 2.400442697 | 2.418167676 | 2.666517981 | 2.320638377 |
| ENSMUSG00000066902 | 0 | 0 | 0 | 0 | 0 | 0 | 0 |
| ENSMUSG00000066905 | 0 | 0 | 0 | 0 | 0 | 0 | 0 |
| ENSMUSG00000066912 | 0 | 0 | 0 | 0 | 0 | 0 | 0 |
| ENSMUSG00000066914 | 0 | 0 | 0 | 0 | 0 | 0 | 0 |
| ENSMUSG00000066936 | 0 | 0 | 0 | 0 | 0 | 0 | 0 |

| | | | | | | | |
|---|---|---|---|---|---|---|---|
| ENSMUSG00000066938 | 0 | 0 | 0 | 0 | 0 | 0 | 0 |
| ENSMUSG00000066944 | 0.578129653 | 0 | 0.79248125 | 0.5 | 0.387761935 | 0.477121255 | 0.448487648 |
| ENSMUSG00000066952 | 1.482695304 | 1.330482024 | 1.403677461 | 1.364857905 | 1.343584837 | 1.255272505 | 1.284450788 |
| ENSMUSG00000066958 | 0 | 0 | 0 | 0 | 0 | 0 | 0 |
| ENSMUSG00000066964 | 0.289064826 | 0.25 | 0 | 0 | 0 | 0 | 0 |
| ENSMUSG00000066975 | 0.747221736 | 0.830482024 | 0.701838731 | 0.25 | 0.387761935 | 0 | 0.448487648 |
| ENSMUSG00000066979 | 2.713122685 | 2.478969845 | 2.466046536 | 2.418106335 | 2.444431236 | 2.776701184 | 2.319395519 |
| ENSMUSG00000066983 | 0 | 0 | 0 | 0 | 0 | 0 | 0.224243824 |
| ENSMUSG00000066990 | 0 | 0 | 0 | 0 | 0 | 0 | 0 |
| ENSMUSG00000067001 | 0 | 0.25 | 0 | 0 | 0 | 0 | 0 |
| ENSMUSG00000067006 | 0 | 0 | 0 | 0 | 0 | 0 | 0 |
| ENSMUSG00000067010 | 0 | 0 | 0 | 0.25 | 0 | 0 | 0 |
| ENSMUSG00000067017 | 0.289064826 | 0.396240625 | 0.396240625 | 0 | 0.489301084 | 0.301029996 | 0.448487648 |
| ENSMUSG00000067028 | 0 | 0 | 0 | 0 | 0 | 0 | 0 |
| ENSMUSG00000067038 | 2.163307908 | 2.061981878 | 2.069031101 | 1.94697564 | 2.017859938 | 2.198657087 | 1.865128412 |
| ENSMUSG00000067047 | 0 | 0 | 0 | 0 | 0 | 0 | 0 |
| ENSMUSG00000067049 | 0 | 0 | 0.25 | 0.396240625 | 0 | 0 | 0.224243824 |
| ENSMUSG00000067058 | 0.289064826 | 0 | 0 | 0 | 0.387761935 | 0 | 0.224243824 |
| ENSMUSG00000067063 | 0 | 0 | 0 | 0 | 0 | 0 | 0 |
| ENSMUSG00000067064 | 0 | 0 | 0 | 0 | 0 | 0 | 0 |
| ENSMUSG00000067071 | 1.721148722 | 1.622963274 | 1.618933358 | 1.62694866 | 1.649303553 | 1.755874856 | 1.674564426 |
| ENSMUSG00000067081 | 0.578129653 | 0.25 | 0.5 | 0.864857905 | 0.733951626 | 0.903089987 | 0.629532004 |
| ENSMUSG00000067085 | 0 | 0 | 0 | 0 | 0 | 0 | 0 |
| ENSMUSG00000067101 | 0 | 0 | 0.25 | 0 | 0 | 0 | 0 |
| ENSMUSG00000067103 | 0 | 0 | 0 | 0 | 0 | 0 | 0 |
| ENSMUSG00000067106 | 0.289064826 | 0.25 | 0 | 0.25 | 0.489301084 | 0 | 0 |
| ENSMUSG00000067121 | 0.289064826 | 0.25 | 0.5 | 0 | 0.244650542 | 0 | 0.224243824 |
| ENSMUSG00000067122 | 0 | 0 | 0 | 0 | 0 | 0 | 0 |
| ENSMUSG00000067144 | 0 | 0 | 0 | 0.25 | 0 | 0 | 0 |
| ENSMUSG00000067147 | 0 | 0 | 0 | 0 | 0 | 0 | 0 |
| ENSMUSG00000067148 | 2.523917219 | 2.329353153 | 2.324802005 | 2.321350555 | 2.344045917 | 2.664641976 | 2.196701384 |
| ENSMUSG00000067149 | 0 | 0.25 | 0 | 0.25 | 0.244650542 | 0 | 0 |
| ENSMUSG00000067150 | 2.953652167 | 2.669870025 | 2.640560606 | 2.707232457 | 2.627905721 | 2.994756945 | 2.442244914 |
| ENSMUSG00000067156 | 0 | 0.25 | 0 | 0 | 0 | 0.301029996 | 0.355418052 |

| | | | | | | | |
|---|---|---|---|---|---|---|---|
| ENSMUSG00000067158 | 0.289064826 | 0.396240625 | 0.396240625 | 0.701838731 | 0.632412477 | 0.301029996 | 0.448487648 |
| ENSMUSG00000067161 | 0.289064826 | 0 | 0 | 0.5 | 0.244650542 | 0 | 0.224243824 |
| ENSMUSG00000067173 | 0 | 0 | 0 | 0 | 0 | 0 | 0 |
| ENSMUSG00000067186 | 0 | 0 | 0 | 0 | 0 | 0 | 0 |
| ENSMUSG00000067189 | 0 | 0 | 0 | 0 | 0 | 0 | 0 |
| ENSMUSG00000067194 | 2.746071301 | 2.52053726 | 2.532320754 | 2.814494265 | 2.715300549 | 3.13225969 | 2.541264678 |
| ENSMUSG00000067199 | 0 | 0 | 0 | 0.25 | 0 | 0 | 0 |
| ENSMUSG00000067201 | 0 | 0 | 0 | 0 | 0 | 0 | 0 |
| ENSMUSG00000067203 | 0 | 0.25 | 0.25 | 0.5 | 0.632412477 | 0.698970004 | 0.876096088 |
| ENSMUSG00000067206 | 0 | 0 | 0 | 0 | 0 | 0 | 0.224243824 |
| ENSMUSG00000067212 | 1.45815691 | 1.552363341 | 1.630890489 | 1.622963274 | 1.689774529 | 1.770852012 | 1.63986884 |
| ENSMUSG00000067215 | 0.671187741 | 0.701838731 | 0.830482024 | 0.580482024 | 0.686820904 | 0.301029996 | 0 |
| ENSMUSG00000067219 | 1.707474304 | 1.657339155 | 1.671625132 | 1.188721876 | 1.200473444 | 1.462397998 | 1.23151414 |
| ENSMUSG00000067220 | 0 | 0 | 0 | 0 | 0 | 0 | 0 |
| ENSMUSG00000067224 | 0.289064826 | 0.25 | 0.25 | 0.25 | 0.387761935 | 0 | 0.224243824 |
| ENSMUSG00000067225 | 0 | 0 | 0 | 0 | 0 | 0 | 0 |
| ENSMUSG00000067229 | 0 | 0 | 0 | 0 | 0 | 0 | 0 |
| ENSMUSG00000067231 | 0 | 0.25 | 0 | 0.25 | 0.244650542 | 0.301029996 | 0.224243824 |
| ENSMUSG00000067235 | 0.671187741 | 0.830482024 | 0.646240625 | 1.348079356 | 1.427019704 | 1.176091259 | 1.661510159 |
| ENSMUSG00000067242 | 0.289064826 | 0.25 | 0.25 | 0.646240625 | 0.387761935 | 0.301029996 | 0.520678036 |
| ENSMUSG00000067261 | 0 | 0 | 0 | 0 | 0 | 0 | 0 |
| ENSMUSG00000067262 | 0 | 0 | 0 | 0 | 0 | 0 | 0 |
| ENSMUSG00000067274 | 3.577267124 | 3.328078937 | 3.363221174 | 3.049994752 | 3.182776028 | 3.549493713 | 3.026670052 |
| ENSMUSG00000067276 | 0.91631382 | 0.864857905 | 0.92510993 | 0.864857905 | 0.244650542 | 0.477121255 | 0.672731473 |
| ENSMUSG00000067279 | 1.156259305 | 0.864857905 | 1.114857905 | 0.701838731 | 0.77552387 | 0.77815125 | 0.672731473 |
| ENSMUSG00000067285 | 0.578129653 | 0.25 | 0.25 | 0.5 | 0.77552387 | 0.301029996 | 0.579661877 |
| ENSMUSG00000067288 | 2.699178053 | 2.504897682 | 2.58272922 | 2.302363341 | 2.443737124 | 2.663700925 | 2.337083659 |
| ENSMUSG00000067292 | 0 | 0.25 | 0.25 | 0.25 | 0.387761935 | 0 | 0 |
| ENSMUSG00000067297 | 0.45815691 | 0.5 | 0.646240625 | 0.646240625 | 0.686820904 | 0.301029996 | 0.448487648 |
| ENSMUSG00000067299 | 0 | 0 | 0 | 0 | 0 | 0 | 0 |
| ENSMUSG00000067321 | 0 | 0 | 0 | 0 | 0 | 0 | 0 |
| ENSMUSG00000067331 | 0 | 0 | 0 | 0 | 0 | 0 | 0 |
| ENSMUSG00000067336 | 2.850493374 | 2.63503201 | 2.603142462 | 2.768869787 | 2.695948846 | 3.093071306 | 2.463028281 |
| ENSMUSG00000067338 | 0 | 0 | 0 | 0 | 0 | 0 | 0 |

| | | | | | | | |
|---|---|---|---|---|---|---|---|
| ENSMUSG00000067341 | 0 | 0 | 0 | 0 | 0 | 0 | 0 |
| ENSMUSG00000067344 | 0.289064826 | 0 | 0 | 0.396240625 | 0.387761935 | 0.602059991 | 0.355418052 |
| ENSMUSG00000067351 | 0 | 0 | 0 | 0 | 0 | 0 | 0 |
| ENSMUSG00000067352 | 0 | 0.396240625 | 0.5 | 0.25 | 0.686820904 | 0.301029996 | 0.224243824 |
| ENSMUSG00000067356 | 0.811507563 | 0.79248125 | 0.79248125 | 0.580482024 | 0.489301084 | 0.301029996 | 0.710836105 |
| ENSMUSG00000067360 | 0 | 0 | 0 | 0 | 0 | 0 | 0 |
| ENSMUSG00000067365 | 2.31414447 | 2.127938163 | 2.140560606 | 2.255591953 | 2.200473444 | 2.536558443 | 2.086826862 |
| ENSMUSG00000067367 | 2.479432974 | 2.336073977 | 2.365369862 | 2.325374049 | 2.300972958 | 2.613841822 | 2.213713783 |
| ENSMUSG00000067369 | 1.816888701 | 1.75835575 | 1.732684334 | 1.978222334 | 1.92245931 | 2.170261715 | 1.809975587 |
| ENSMUSG00000067370 | 1.129344651 | 1.160964047 | 1.02186571 | 0.896240625 | 1.020174412 | 0.698970004 | 0.803905701 |
| ENSMUSG00000067377 | 1.693336259 | 1.618933358 | 1.584962501 | 1.898114259 | 1.83872004 | 2.103803721 | 1.735976162 |
| ENSMUSG00000067389 | 0.747221736 | 0.75 | 0.896240625 | 0.701838731 | 0.244650542 | 0.477121255 | 0.520678036 |
| ENSMUSG00000067399 | 0.578129653 | 0.646240625 | 0.580482024 | 0.5 | 0.632412477 | 0.698970004 | 0.672731473 |
| ENSMUSG00000067424 | 1.516991255 | 1.51109853 | 1.476722649 | 1.494319981 | 1.462344774 | 1.681241237 | 1.290497981 |
| ENSMUSG00000067430 | 1.936861443 | 1.711372513 | 1.732684334 | 1.77186571 | 1.800029087 | 1.913813852 | 1.616675393 |
| ENSMUSG00000067438 | 0 | 0 | 0 | 0 | 0 | 0 | 0 |
| ENSMUSG00000067441 | 0 | 0 | 0 | 0 | 0 | 0 | 0 |
| ENSMUSG00000067455 | 0 | 0.396240625 | 0.5 | 0.580482024 | 0.387761935 | 0.301029996 | 0.224243824 |
| ENSMUSG00000067482 | 0 | 0 | 0 | 0 | 0 | 0 | 0 |
| ENSMUSG00000067513 | 0 | 0 | 0 | 0 | 0 | 0 | 0 |
| ENSMUSG00000067519 | 0 | 0 | 0 | 0 | 0 | 0 | 0 |
| ENSMUSG00000067520 | 0 | 0 | 0 | 0 | 0 | 0 | 0 |
| ENSMUSG00000067522 | 0 | 0 | 0 | 0 | 0 | 0 | 0 |
| ENSMUSG00000067524 | 0 | 0 | 0 | 0 | 0 | 0 | 0 |
| ENSMUSG00000067525 | 0 | 0 | 0 | 0 | 0 | 0 | 0 |
| ENSMUSG00000067526 | 0 | 0 | 0 | 0 | 0 | 0 | 0 |
| ENSMUSG00000067528 | 0 | 0 | 0 | 0 | 0 | 0 | 0 |
| ENSMUSG00000067529 | 0 | 0 | 0 | 0 | 0 | 0 | 0 |
| ENSMUSG00000067541 | 0.289064826 | 0.396240625 | 0.25 | 0.25 | 0 | 0.602059991 | 0.355418052 |
| ENSMUSG00000067543 | 0 | 0 | 0 | 0 | 0 | 0 | 0 |
| ENSMUSG00000067545 | 0 | 0 | 0 | 0 | 0 | 0 | 0 |
| ENSMUSG00000067547 | 0 | 0 | 0 | 0 | 0 | 0 | 0 |
| ENSMUSG00000067555 | 0 | 0 | 0 | 0 | 0 | 0 | 0 |
| ENSMUSG00000067561 | 0 | 0 | 0 | 0 | 0 | 0 | 0 |

| | | | | | | | |
|---|---|---|---|---|---|---|---|
| ENSMUSG00000067562 | 0 | 0 | 0 | 0 | 0 | 0 | 0 |
| ENSMUSG00000067567 | 1.982401085 | 1.784887838 | 1.816696635 | 1.953445298 | 1.870073561 | 2.075546961 | 1.723650273 |
| ENSMUSG00000067571 | 0.45815691 | 0 | 0 | 0 | 0 | 0 | 0 |
| ENSMUSG00000067575 | 0 | 0 | 0 | 0 | 0 | 0 | 0 |
| ENSMUSG00000067577 | 0 | 0 | 0.396240625 | 0 | 0.244650542 | 0 | 0.224243824 |
| ENSMUSG00000067578 | 0 | 0 | 0 | 0 | 0 | 0 | 0 |
| ENSMUSG00000067586 | 1.734388958 | 1.664552871 | 1.685366747 | 0.701838731 | 0.733951626 | 0.954242509 | 0.672731473 |
| ENSMUSG00000067591 | 0 | 0 | 0 | 0 | 0 | 0 | 0 |
| ENSMUSG00000067594 | 0 | 0 | 0 | 0 | 0 | 0 | 0 |
| ENSMUSG00000067596 | 0 | 0 | 0 | 0 | 0 | 0 | 0 |
| ENSMUSG00000067597 | 0 | 0.25 | 0.25 | 0 | 0 | 0 | 0.224243824 |
| ENSMUSG00000067599 | 0 | 0 | 0 | 0 | 0 | 0 | 0 |
| ENSMUSG00000067604 | 0 | 0 | 0 | 0 | 0 | 0 | 0 |
| ENSMUSG00000067608 | 0.578129653 | 0.5 | 0.25 | 0.5 | 0.387761935 | 0.77815125 | 0.448487648 |
| ENSMUSG00000067610 | 0 | 0 | 0.25 | 0 | 0 | 0 | 0 |
| ENSMUSG00000067613 | 0.289064826 | 0.25 | 0 | 0 | 0 | 0 | 0.224243824 |
| ENSMUSG00000067614 | 0 | 0 | 0 | 0 | 0 | 0 | 0 |
| ENSMUSG00000067615 | 0 | 0 | 0.25 | 0 | 0 | 0 | 0 |
| ENSMUSG00000067616 | 0 | 0 | 0 | 0 | 0 | 0 | 0 |
| ENSMUSG00000067627 | 0.289064826 | 0 | 0 | 0 | 0 | 0 | 0 |
| ENSMUSG00000067629 | 2.136858952 | 2.031030328 | 2.056000419 | 2.065523711 | 2.155511127 | 2.28780173 | 2.055736903 |
| ENSMUSG00000067642 | 0 | 0 | 0 | 0 | 0.244650542 | 0 | 0 |
| ENSMUSG00000067649 | 0 | 0 | 0 | 0 | 0 | 0 | 0 |
| ENSMUSG00000067653 | 0.289064826 | 0.5 | 0.5 | 0.646240625 | 0.686820904 | 0.477121255 | 0.355418052 |
| ENSMUSG00000067656 | 0 | 0 | 0 | 0 | 0 | 0 | 0 |
| ENSMUSG00000067679 | 0 | 0 | 0 | 0 | 0 | 0 | 0 |
| ENSMUSG00000067684 | 0 | 0 | 0 | 0 | 0 | 0 | 0 |
| ENSMUSG00000067698 | 0 | 0 | 0 | 0 | 0 | 0 | 0 |
| ENSMUSG00000067700 | 0 | 0 | 0 | 0 | 0 | 0 | 0 |
| ENSMUSG00000067702 | 0 | 0 | 0 | 0 | 0 | 0 | 0 |
| ENSMUSG00000067704 | 0 | 0 | 0 | 0 | 0 | 0 | 0 |
| ENSMUSG00000067713 | 2.258689303 | 2.123963757 | 2.124961472 | 2.149978211 | 2.210031835 | 2.409933123 | 2.11845007 |
| ENSMUSG00000067714 | 0 | 0 | 0 | 0 | 0 | 0 | 0 |
| ENSMUSG00000067716 | 0 | 0 | 0 | 0 | 0 | 0 | 0 |

| | | | | | | | |
|---|---|---|---|---|---|---|---|
| ENSMUSG00000067719 | 0 | 0 | 0 | 0 | 0 | 0 | 0 |
| ENSMUSG00000067722 | 1.837744666 | 1.76109853 | 1.888647213 | 1.79248125 | 1.770879644 | 1.903089987 | 1.737484395 |
| ENSMUSG00000067724 | 0 | 0.396240625 | 0 | 0.580482024 | 0 | 0.477121255 | 0 |
| ENSMUSG00000067736 | 0 | 0.25 | 0 | 0 | 0 | 0 | 0 |
| ENSMUSG00000067745 | 0 | 0.396240625 | 0 | 0.25 | 0 | 0 | 0 |
| ENSMUSG00000067750 | 0 | 0 | 0 | 0 | 0 | 0 | 0 |
| ENSMUSG00000067764 | 0 | 0 | 0 | 0 | 0 | 0 | 0 |
| ENSMUSG00000067767 | 0 | 0 | 0 | 0 | 0 | 0 | 0 |
| ENSMUSG00000067768 | 0 | 0 | 0 | 0 | 0 | 0 | 0 |
| ENSMUSG00000067771 | 0 | 0 | 0 | 0 | 0 | 0 | 0 |
| ENSMUSG00000067773 | 0 | 0 | 0 | 0 | 0 | 0 | 0 |
| ENSMUSG00000067780 | 0.671187741 | 0.79248125 | 0.5 | 0.396240625 | 0.387761935 | 0.477121255 | 0.579661877 |
| ENSMUSG00000067786 | 2.223916471 | 2.056000419 | 2.106566189 | 2.03360658 | 2.033751322 | 2.334453751 | 2.018825668 |
| ENSMUSG00000067787 | 2.203267085 | 2.005591953 | 2.019203899 | 2.006976499 | 1.973373093 | 2.198657087 | 1.861046144 |
| ENSMUSG00000067795 | 0.289064826 | 0.25 | 0.5 | 0.396240625 | 0.387761935 | 0 | 0.355418052 |
| ENSMUSG00000067813 | 0.289064826 | 0.396240625 | 0.25 | 0 | 0 | 0 | 0 |
| ENSMUSG00000067818 | 3.294048096 | 3.022693514 | 3.032723067 | 1.54248125 | 1.588235379 | 1.579783597 | 1.517723536 |
| ENSMUSG00000067825 | 1.693336259 | 1.642463902 | 1.695339928 | 1.575945187 | 1.568060967 | 1.73239376 | 1.564611933 |
| ENSMUSG00000067835 | 0 | 0 | 0 | 0 | 0 | 0 | 0 |
| ENSMUSG00000067847 | 1.342375483 | 1.53743678 | 1.602347734 | 1.302363341 | 1.467903253 | 1.477121255 | 1.694069894 |
| ENSMUSG00000067848 | 0 | 0 | 0 | 0 | 0 | 0 | 0 |
| ENSMUSG00000067851 | 2.824116479 | 2.621708755 | 2.610217292 | 2.708024712 | 2.624799605 | 2.976349979 | 2.388231309 |
| ENSMUSG00000067855 | 0.289064826 | 0.25 | 0.25 | 0 | 0.387761935 | 0 | 0.224243824 |
| ENSMUSG00000067860 | 0 | 0 | 0 | 0.25 | 0 | 0 | 0 |
| ENSMUSG00000067869 | 0 | 0 | 0 | 0 | 0 | 0 | 0 |
| ENSMUSG00000067870 | 0.867194479 | 0.864857905 | 0.79248125 | 0.976722649 | 0.955822902 | 1.079181246 | 1.028149525 |
| ENSMUSG00000067872 | 0 | 0.5 | 0.396240625 | 0.5 | 0 | 0 | 0.224243824 |
| ENSMUSG00000067873 | 2.58665822 | 2.379909063 | 2.37694866 | 2.409156155 | 2.346342321 | 2.665580991 | 2.17331363 |
| ENSMUSG00000067878 | 0.289064826 | 0.25 | 0.646240625 | 0.646240625 | 0.733951626 | 0.477121255 | 0.520678036 |
| ENSMUSG00000067879 | 0 | 0 | 0 | 0 | 0 | 0 | 0 |
| ENSMUSG00000067882 | 0 | 0 | 0 | 0.396240625 | 0 | 0 | 0 |
| ENSMUSG00000067889 | 0.960252568 | 1.02186571 | 1.146240625 | 1.080482024 | 1.149965125 | 1.278753601 | 1.100339912 |
| ENSMUSG00000067909 | 0 | 0 | 0 | 0 | 0 | 0 | 0 |
| ENSMUSG00000067916 | 1 | 1.04248125 | 0.830482024 | 1.410964047 | 1.42077253 | 1.322219295 | 1.20919388 |

| | | | | | | | |
|---|---|---|---|---|---|---|---|
| ENSMUSG00000067919 | 0 | 0 | 0 | 0 | 0 | 0.301029996 | 0.224243824 |
| ENSMUSG00000067924 | 1.389637216 | 1.364857905 | 1.282320754 | 1.27186571 | 1.380772476 | 1.414973348 | 1.335191751 |
| ENSMUSG00000067925 | 1.307602651 | 1.388647213 | 1.410964047 | 1.380890489 | 1.310728002 | 1.414973348 | 1.340368108 |
| ENSMUSG00000067928 | 2.129344651 | 1.984159485 | 1.931980114 | 1.921625132 | 1.902105869 | 2.123851641 | 1.760706232 |
| ENSMUSG00000067929 | 0.45815691 | 0.580482024 | 0.5 | 0.75 | 0.568060967 | 0.301029996 | 0.520678036 |
| ENSMUSG00000067931 | 2.249317394 | 2.089388001 | 2.155512955 | 2.185366747 | 2.173890435 | 2.369215857 | 1.972093446 |
| ENSMUSG00000067941 | 0 | 0 | 0 | 0.5 | 0.244650542 | 0 | 0.224243824 |
| ENSMUSG00000067942 | 2.249317394 | 2.059601185 | 2.049918086 | 2.081607372 | 2.066583491 | 2.23299611 | 1.87511425 |
| ENSMUSG00000067949 | 0 | 0 | 0 | 0 | 0 | 0 | 0 |
| ENSMUSG00000067951 | 0 | 0 | 0 | 0 | 0 | 0 | 0 |
| ENSMUSG00000067971 | 0 | 0 | 0 | 0 | 0 | 0 | 0 |
| ENSMUSG00000067973 | 0 | 0 | 0 | 0 | 0 | 0 | 0 |
| ENSMUSG00000067995 | 1.960252568 | 1.87892496 | 1.868933358 | 2.129909063 | 2.037065488 | 2.33243846 | 1.980721036 |
| ENSMUSG00000067996 | 0 | 0 | 0 | 0 | 0 | 0 | 0 |
| ENSMUSG00000067998 | 0.289064826 | 0 | 0 | 0 | 0 | 0 | 0 |
| ENSMUSG00000068008 | 0 | 0 | 0 | 0 | 0 | 0 | 0 |
| ENSMUSG00000068009 | 0 | 0 | 0 | 0 | 0.244650542 | 0 | 0 |
| ENSMUSG00000068011 | 0.45815691 | 0.5 | 0.25 | 0.5 | 0.489301084 | 0.301029996 | 0.224243824 |
| ENSMUSG00000068014 | 0 | 0 | 0 | 0 | 0 | 0 | 0 |
| ENSMUSG00000068015 | 1.714367558 | 1.630890489 | 1.701838731 | 1.735628626 | 1.662257525 | 1.799340549 | 1.572224412 |
| ENSMUSG00000068036 | 3.111026095 | 2.808404919 | 2.79853531 | 2.772031423 | 2.718345365 | 3.097951071 | 2.525850149 |
| ENSMUSG00000068037 | 0 | 0.25 | 0.5 | 0 | 0.568060967 | 0.477121255 | 0.224243824 |
| ENSMUSG00000068039 | 3.477941618 | 3.135757955 | 3.119376329 | 3.205194325 | 3.099069969 | 3.635785236 | 2.876006953 |
| ENSMUSG00000068040 | 2.455852858 | 2.27186571 | 2.263820609 | 2.508011432 | 2.469844355 | 2.821513528 | 2.341997913 |
| ENSMUSG00000068048 | 0 | 0 | 0 | 0 | 0 | 0 | 0 |
| ENSMUSG00000068062 | 0 | 0 | 0 | 0 | 0 | 0 | 0 |
| ENSMUSG00000068067 | 0 | 0 | 0 | 0 | 0 | 0 | 0 |
| ENSMUSG00000068068 | 0 | 0 | 0 | 0 | 0 | 0 | 0 |
| ENSMUSG00000068071 | 0 | 0 | 0 | 0 | 0 | 0 | 0 |
| ENSMUSG00000068073 | 0 | 0 | 0 | 0 | 0 | 0 | 0 |
| ENSMUSG00000068074 | 0 | 0 | 0 | 0 | 0 | 0 | 0 |
| ENSMUSG00000068075 | 0 | 0 | 0 | 0 | 0 | 0 | 0 |
| ENSMUSG00000068078 | 0 | 0 | 0 | 0 | 0 | 0 | 0 |
| ENSMUSG00000068079 | 0 | 0 | 0 | 0 | 0 | 0.301029996 | 0 |

| | | | | | | | |
|---|---|---|---|---|---|---|---|
| ENSMUSG00000068082 | 0 | 0 | 0 | 0 | 0 | 0 | 0.224243824 |
| ENSMUSG00000068083 | 0 | 0 | 0 | 0 | 0 | 0 | 0 |
| ENSMUSG00000068085 | 0 | 0 | 0 | 0 | 0 | 0 | 0 |
| ENSMUSG00000068086 | 0 | 0 | 0 | 0 | 0 | 0 | 0 |
| ENSMUSG00000068099 | 0 | 0 | 0 | 0 | 0 | 0 | 0 |
| ENSMUSG00000068101 | 1.759671389 | 1.784887838 | 1.898114259 | 1.589388001 | 1.78687392 | 1.785329835 | 1.900630878 |
| ENSMUSG00000068105 | 0.578129653 | 0.5 | 0 | 0.646240625 | 0.568060967 | 0.84509804 | 0.672731473 |
| ENSMUSG00000068113 | 0 | 0 | 0 | 0 | 0 | 0 | 0 |
| ENSMUSG00000068114 | 1.956444026 | 1.905512955 | 1.882845365 | 1.955044741 | 2.014359518 | 2.184691431 | 1.905143021 |
| ENSMUSG00000068115 | 2.023453784 | 1.942046081 | 1.997171172 | 1.991446071 | 2.052132976 | 2.187520721 | 1.988432495 |
| ENSMUSG00000068117 | 0.289064826 | 0 | 0 | 0.25 | 0 | 0 | 0 |
| ENSMUSG00000068120 | 0 | 0 | 0 | 0 | 0 | 0 | 0 |
| ENSMUSG00000068122 | 0 | 0 | 0 | 0 | 0 | 0 | 0 |
| ENSMUSG00000068129 | 0 | 0 | 0 | 0 | 0 | 0 | 0 |
| ENSMUSG00000068130 | 1.527823874 | 1.282320754 | 1.226722649 | 1.505591953 | 1.484072009 | 1.414973348 | 1.150210039 |
| ENSMUSG00000068134 | 1.956444026 | 1.777131114 | 1.701838731 | 1.901832578 | 1.840643511 | 1.913813852 | 1.692344475 |
| ENSMUSG00000068141 | 0 | 0 | 0 | 0 | 0 | 0 | 0 |
| ENSMUSG00000068149 | 0 | 0 | 0 | 0 | 0 | 0 | 0 |
| ENSMUSG00000068151 | 0 | 0 | 0.25 | 0 | 0 | 0 | 0.448487648 |
| ENSMUSG00000068154 | 0 | 0.25 | 0 | 0 | 0 | 0 | 0 |
| ENSMUSG00000068165 | 0 | 0 | 0 | 0 | 0 | 0 | 0 |
| ENSMUSG00000068167 | 0 | 0 | 0 | 0 | 0 | 0 | 0 |
| ENSMUSG00000068173 | 0 | 0 | 0 | 0 | 0 | 0 | 0 |
| ENSMUSG00000068180 | 0 | 0 | 0 | 0 | 0 | 0 | 0 |
| ENSMUSG00000068181 | 0 | 0 | 0 | 0 | 0 | 0 | 0 |
| ENSMUSG00000068182 | 0 | 0 | 0 | 0 | 0 | 0 | 0 |
| ENSMUSG00000068184 | 1.707474304 | 1.852347734 | 1.814346961 | 1.888647213 | 1.895599581 | 2.086359831 | 1.807561282 |
| ENSMUSG00000068196 | 3.151275227 | 2.897646762 | 2.867918804 | 2.826942508 | 2.794928476 | 3.195622944 | 2.612978034 |
| ENSMUSG00000068200 | 0 | 0 | 0 | 0 | 0 | 0 | 0 |
| ENSMUSG00000068205 | 0.91631382 | 0.976722649 | 1.02186571 | 1.080482024 | 0.846351821 | 1.278753601 | 0.775756176 |
| ENSMUSG00000068206 | 2.092368209 | 1.969129237 | 2.019203899 | 1.921625132 | 1.997548906 | 2.089905111 | 1.960974982 |
| ENSMUSG00000068218 | 0 | 0 | 0 | 0 | 0 | 0 | 0 |
| ENSMUSG00000068219 | 0 | 0 | 0 | 0 | 0 | 0 | 0 |
| ENSMUSG00000068220 | 3.98577453 | 3.59570666 | 3.623307503 | 3.499140439 | 3.480573103 | 4.071697945 | 3.259282193 |

| | | | | | | | |
|---|---|---|---|---|---|---|---|
| ENSMUSG00000068226 | 0.289064826 | 0.25 | 0 | 0 | 0 | 0 | 0 |
| ENSMUSG00000068227 | 0 | 0.396240625 | 0.25 | 0.25 | 0.244650542 | 0 | 0.224243824 |
| ENSMUSG00000068231 | 0 | 0 | 0 | 0 | 0 | 0 | 0 |
| ENSMUSG00000068232 | 0 | 0 | 0 | 0 | 0 | 0.301029996 | 0 |
| ENSMUSG00000068234 | 0 | 0 | 0 | 0 | 0.244650542 | 0 | 0 |
| ENSMUSG00000068240 | 0.45815691 | 0.396240625 | 0.5 | 0.646240625 | 0.568060967 | 0.477121255 | 0.224243824 |
| ENSMUSG00000068243 | 0 | 0 | 0 | 0.25 | 0 | 0 | 0 |
| ENSMUSG00000068245 | 0.45815691 | 0.701838731 | 0.396240625 | 0.646240625 | 0.489301084 | 0.301029996 | 0.224243824 |
| ENSMUSG00000068246 | 0.671187741 | 0.646240625 | 0.701838731 | 0.79248125 | 0.733951626 | 1.146128036 | 1.140832123 |
| ENSMUSG00000068250 | 1.964026642 | 1.814346961 | 1.75835575 | 1.726722649 | 1.680838453 | 1.851258349 | 1.540636858 |
| ENSMUSG00000068252 | 0 | 0 | 0.396240625 | 0 | 0.244650542 | 0 | 0 |
| ENSMUSG00000068259 | 0 | 0 | 0.25 | 0 | 0 | 0.301029996 | 0.224243824 |
| ENSMUSG00000068262 | 0 | 0 | 0 | 0 | 0 | 0 | 0.224243824 |
| ENSMUSG00000068263 | 0.578129653 | 0.396240625 | 0 | 0.25 | 0.686820904 | 0.602059991 | 0.672731473 |
| ENSMUSG00000068264 | 1.418409478 | 1.25 | 1.364857905 | 1.114857905 | 1.380772476 | 1.230448921 | 1.365075947 |
| ENSMUSG00000068267 | 2.45469604 | 2.273849256 | 2.339388001 | 2.246460484 | 2.306653428 | 2.5289167 | 2.293479712 |
| ENSMUSG00000068270 | 2.377269616 | 2.161864607 | 2.100219859 | 2.041226732 | 1.898867717 | 2.167317335 | 1.825244123 |
| ENSMUSG00000068284 | 2.547754129 | 2.37246199 | 2.348079356 | 2.385757955 | 2.28771234 | 2.611723308 | 2.115189268 |
| ENSMUSG00000068289 | 0 | 0 | 0 | 0 | 0 | 0 | 0 |
| ENSMUSG00000068290 | 2.054824387 | 1.933677405 | 2.02053726 | 1.935366747 | 2.048957937 | 2.170261715 | 1.993927582 |
| ENSMUSG00000068299 | 0.747221736 | 0.580482024 | 0.896240625 | 0.79248125 | 0.877063019 | 0.77815125 | 0.672731473 |
| ENSMUSG00000068302 | 0 | 0 | 0 | 0 | 0 | 0 | 0 |
| ENSMUSG00000068303 | 0.747221736 | 0.830482024 | 0.79248125 | 0.92510993 | 0.846351821 | 0.602059991 | 0.579661877 |
| ENSMUSG00000068314 | 0 | 0 | 0 | 0 | 0 | 0 | 0 |
| ENSMUSG00000068323 | 0 | 0.25 | 0 | 0 | 0.244650542 | 0.477121255 | 0 |
| ENSMUSG00000068324 | 0.671187741 | 0.5 | 0.25 | 0.5 | 0.489301084 | 0 | 0 |
| ENSMUSG00000068327 | 0 | 0 | 0 | 0 | 0 | 0 | 0 |
| ENSMUSG00000068328 | 2.415874319 | 2.248588359 | 2.30357978 | 2.110735874 | 2.179075613 | 2.399673721 | 2.121678334 |
| ENSMUSG00000068329 | 2.156259305 | 2.016522298 | 2.115881093 | 1.823655187 | 1.974687643 | 2.041392685 | 1.926805425 |
| ENSMUSG00000068335 | 2.351376531 | 2.235628626 | 2.25 | 1.802363341 | 1.925488995 | 2.096910013 | 1.896054911 |
| ENSMUSG00000068341 | 0 | 0 | 0 | 0 | 0 | 0 | 0 |
| ENSMUSG00000068348 | 0 | 0.25 | 0.25 | 0 | 0 | 0 | 0 |
| ENSMUSG00000068349 | 0 | 0 | 0 | 0 | 0 | 0 | 0 |
| ENSMUSG00000068373 | 0.45815691 | 0.5 | 0 | 0 | 0.244650542 | 0 | 0 |

| | | | | | | | |
|---|---|---|---|---|---|---|---|
| ENSMUSG00000068391 | 2.136858952 | 1.981203126 | 2.005591953 | 1.886723615 | 1.928492895 | 2.181843588 | 1.844181586 |
| ENSMUSG00000068392 | 0 | 0 | 0 | 0 | 0 | 0 | 0 |
| ENSMUSG00000068394 | 1.971473847 | 1.870953944 | 1.884789703 | 1.981203126 | 1.959951091 | 2.225309282 | 1.856911705 |
| ENSMUSG00000068396 | 0 | 0 | 0 | 0 | 0 | 0 | 0 |
| ENSMUSG00000068397 | 0 | 0 | 0 | 0.396240625 | 0 | 0 | 0.224243824 |
| ENSMUSG00000068399 | 0 | 0 | 0.25 | 0 | 0.244650542 | 0 | 0 |
| ENSMUSG00000068407 | 0 | 0 | 0 | 0 | 0 | 0 | 0 |
| ENSMUSG00000068417 | 1.036286563 | 1 | 1 | 1.146240625 | 1.039257769 | 1.431363764 | 1.140832123 |
| ENSMUSG00000068426 | 0 | 0 | 0 | 0 | 0 | 0 | 0 |
| ENSMUSG00000068428 | 0 | 0 | 0 | 0 | 0 | 0 | 0 |
| ENSMUSG00000068431 | 0 | 0 | 0 | 0 | 0 | 0 | 0 |
| ENSMUSG00000068437 | 0 | 0 | 0 | 0 | 0 | 0 | 0 |
| ENSMUSG00000068452 | 0.289064826 | 0.580482024 | 0 | 0 | 0 | 0 | 0 |
| ENSMUSG00000068457 | 0 | 0.25 | 0.396240625 | 0.580482024 | 0.387761935 | 0.301029996 | 0.224243824 |
| ENSMUSG00000068463 | 0.289064826 | 0.25 | 0 | 0 | 0 | 0 | 0 |
| ENSMUSG00000068466 | 0.289064826 | 0 | 0.396240625 | 0.25 | 0.387761935 | 0.477121255 | 0 |
| ENSMUSG00000068479 | 1.932832114 | 1.845926073 | 1.856566189 | 1.951838731 | 1.888971102 | 2.086359831 | 1.753626837 |
| ENSMUSG00000068480 | 0 | 0 | 0 | 0 | 0 | 0 | 0.224243824 |
| ENSMUSG00000068487 | 0 | 0 | 0 | 0 | 0.244650542 | 0 | 0 |
| ENSMUSG00000068506 | 0 | 0 | 0 | 0 | 0 | 0 | 0 |
| ENSMUSG00000068522 | 0 | 0.25 | 0.5 | 0.646240625 | 0.387761935 | 0.301029996 | 0.355418052 |
| ENSMUSG00000068523 | 2.521950082 | 2.370953944 | 2.381869252 | 2.369945066 | 2.323654601 | 2.595496222 | 2.245894972 |
| ENSMUSG00000068535 | 0 | 0 | 0 | 0 | 0 | 0 | 0 |
| ENSMUSG00000068536 | 0 | 0 | 0 | 0 | 0 | 0 | 0 |
| ENSMUSG00000068537 | 0 | 0 | 0 | 0 | 0 | 0 | 0 |
| ENSMUSG00000068547 | 0.289064826 | 0.25 | 0 | 0.580482024 | 0.387761935 | 0 | 0 |
| ENSMUSG00000068551 | 1.663535556 | 1.552363341 | 1.552363341 | 1.061981878 | 0.978602168 | 0.477121255 | 0.935079929 |
| ENSMUSG00000068566 | 2.977469624 | 2.708222504 | 2.722185812 | 2.716626553 | 2.662444522 | 3.100715087 | 2.498262273 |
| ENSMUSG00000068574 | 0 | 0 | 0 | 0 | 0 | 0 | 0 |
| ENSMUSG00000068579 | 0 | 0.5 | 0 | 0 | 0.489301084 | 0 | 0.448487648 |
| ENSMUSG00000068580 | 1.631440309 | 1.630890489 | 1.653677461 | 1.705044741 | 1.890639929 | 1.929418926 | 1.788855757 |
| ENSMUSG00000068587 | 0 | 0.396240625 | 0.25 | 0 | 0 | 0 | 0 |
| ENSMUSG00000068600 | 0 | 0 | 0 | 0 | 0 | 0 | 0 |
| ENSMUSG00000068601 | 0.289064826 | 0 | 0.25 | 0 | 0 | 0.477121255 | 0.224243824 |

| | | | | | | | |
|---|---|---|---|---|---|---|---|
| ENSMUSG00000068604 | 0 | 0 | 0.25 | 0.25 | 0.489301084 | 0.301029996 | 0.355418052 |
| ENSMUSG00000068606 | 0 | 0 | 0.25 | 0 | 0.387761935 | 0 | 0.355418052 |
| ENSMUSG00000068614 | 0 | 0.5 | 0.25 | 0 | 0 | 0 | 0 |
| ENSMUSG00000068615 | 0 | 0 | 0 | 0 | 0 | 0 | 0 |
| ENSMUSG00000068617 | 0 | 0 | 0 | 0 | 0 | 0 | 0 |
| ENSMUSG00000068629 | 0 | 0.25 | 0.25 | 0.25 | 0 | 0 | 0 |
| ENSMUSG00000068631 | 0 | 0 | 0.25 | 0 | 0 | 0 | 0 |
| ENSMUSG00000068638 | 0 | 0 | 0.25 | 0.25 | 0 | 0 | 0 |
| ENSMUSG00000068646 | 0 | 0 | 0 | 0 | 0 | 0 | 0 |
| ENSMUSG00000068647 | 0 | 0 | 0 | 0 | 0 | 0 | 0 |
| ENSMUSG00000068663 | 2.069666964 | 1.956637122 | 1.994319981 | 2.05357978 | 2.046825265 | 2.28780173 | 1.936709733 |
| ENSMUSG00000068674 | 0.811507563 | 0.580482024 | 0.79248125 | 0.580482024 | 0.568060967 | 0 | 0.224243824 |
| ENSMUSG00000068680 | 0 | 0 | 0 | 0 | 0 | 0 | 0 |
| ENSMUSG00000068686 | 1.470606563 | 1.571350555 | 1.516522298 | 1.482684334 | 1.358933174 | 1.477121255 | 1.365075947 |
| ENSMUSG00000068689 | 0 | 0.25 | 0.646240625 | 0 | 0.244650542 | 0.301029996 | 0 |
| ENSMUSG00000068696 | 0 | 0.25 | 0 | 0 | 0.244650542 | 0 | 0 |
| ENSMUSG00000068697 | 0 | 0 | 0 | 0.25 | 0 | 0 | 0 |
| ENSMUSG00000068699 | 3.036860592 | 2.825945187 | 2.900674716 | 2.625459296 | 2.802426296 | 3.025305865 | 2.709496494 |
| ENSMUSG00000068706 | 0.578129653 | 0.646240625 | 0.580482024 | 0.701838731 | 0.632412477 | 0 | 0.579661877 |
| ENSMUSG00000068732 | 2.223916471 | 2.059601185 | 2.088286706 | 2.347004321 | 2.253693465 | 2.609594409 | 2.088864761 |
| ENSMUSG00000068735 | 1.631440309 | 1.54248125 | 1.488549078 | 1.664552871 | 1.782377602 | 1.880813592 | 1.757893036 |
| ENSMUSG00000068739 | 3.037147311 | 2.772031423 | 2.747525991 | 2.748411515 | 2.763796674 | 3.155943018 | 2.614382009 |
| ENSMUSG00000068740 | 0.578129653 | 0.646240625 | 0.580482024 | 1.681980114 | 1.78010778 | 1.755874856 | 1.806347338 |
| ENSMUSG00000068742 | 1.432083896 | 1.339388001 | 1.380890489 | 1.664552871 | 1.758994476 | 1.826074803 | 1.699191618 |
| ENSMUSG00000068744 | 1.842799666 | 1.763820609 | 1.843759858 | 1.593759858 | 1.603583716 | 1.633468456 | 1.526507301 |
| ENSMUSG00000068745 | 0 | 0 | 0 | 0 | 0 | 0 | 0 |
| ENSMUSG00000068747 | 2.054824387 | 1.948603967 | 1.909156155 | 2.464106382 | 2.436365321 | 2.771587481 | 2.280542874 |
| ENSMUSG00000068748 | 2.205378646 | 2.004202072 | 1.976722649 | 0.396240625 | 0.489301084 | 0.301029996 | 0.672731473 |
| ENSMUSG00000068749 | 2.658591846 | 2.47934302 | 2.471805154 | 2.648114259 | 2.612740596 | 3.014100322 | 2.392033886 |
| ENSMUSG00000068758 | 0.91631382 | 1.188721876 | 1.25 | 0.25 | 0.812711509 | 0 | 1 |
| ENSMUSG00000068762 | 0.747221736 | 0.5 | 0 | 0.5 | 0.733951626 | 0.84509804 | 0.520678036 |
| ENSMUSG00000068776 | 0 | 0 | 0 | 0 | 0 | 0 | 0 |
| ENSMUSG00000068794 | 0.289064826 | 0.25 | 0.580482024 | 0 | 0 | 0 | 0 |
| ENSMUSG00000068798 | 2.45815691 | 2.26109853 | 2.271202097 | 2.435787849 | 2.351342427 | 2.683947131 | 2.185218806 |

| | | | | | | | |
|---|---|---|---|---|---|---|---|
| ENSMUSG00000068806 | 0 | 0 | 0 | 0 | 0 | 0 | 0 |
| ENSMUSG00000068808 | 0 | 0 | 0 | 0 | 0 | 0 | 0 |
| ENSMUSG00000068809 | 0 | 0 | 0 | 0 | 0 | 0 | 0 |
| ENSMUSG00000068810 | 0 | 0 | 0 | 0 | 0 | 0 | 0 |
| ENSMUSG00000068814 | 0 | 0 | 0 | 0 | 0 | 0 | 0 |
| ENSMUSG00000068815 | 0 | 0 | 0 | 0 | 0 | 0 | 0 |
| ENSMUSG00000068816 | 0 | 0 | 0 | 0 | 0 | 0 | 0 |
| ENSMUSG00000068817 | 0 | 0 | 0 | 0 | 0 | 0 | 0 |
| ENSMUSG00000068818 | 0 | 0 | 0 | 0 | 0 | 0 | 0 |
| ENSMUSG00000068819 | 0 | 0 | 0 | 0 | 0 | 0 | 0 |
| ENSMUSG00000068823 | 3.227381882 | 2.93325383 | 2.904825739 | 3.012632226 | 2.923620792 | 3.384174139 | 2.701340211 |
| ENSMUSG00000068824 | 0 | 0 | 0 | 0 | 0 | 0 | 0 |
| ENSMUSG00000068854 | 1.156259305 | 1.26109853 | 1.26109853 | 1.348079356 | 1.351342427 | 1.477121255 | 1.23151414 |
| ENSMUSG00000068855 | 0 | 0 | 0 | 0.25 | 0 | 0 | 0 |
| ENSMUSG00000068856 | 2.179305632 | 2.067865757 | 2.171625132 | 1.96293726 | 2.053185007 | 2.149219113 | 2.055736903 |
| ENSMUSG00000068859 | 0 | 0 | 0 | 0 | 0 | 0.301029996 | 0.355418052 |
| ENSMUSG00000068860 | 0.289064826 | 0.5 | 0.396240625 | 0.25 | 0.244650542 | 0 | 0.224243824 |
| ENSMUSG00000068874 | 0 | 0 | 0 | 0.5 | 0 | 0 | 0 |
| ENSMUSG00000068876 | 1.686083339 | 1.62694866 | 1.701838731 | 1.948603967 | 2.037065488 | 2.298853076 | 1.977870661 |
| ENSMUSG00000068877 | 0 | 0 | 0.5 | 0.396240625 | 0.733951626 | 0.301029996 | 0.224243824 |
| ENSMUSG00000068879 | 0 | 0 | 0 | 0 | 0 | 0 | 0 |
| ENSMUSG00000068882 | 2.85804771 | 2.649978211 | 2.635999613 | 2.862802778 | 2.757712293 | 3.203848464 | 2.557413225 |
| ENSMUSG00000068885 | 0 | 0 | 0 | 0 | 0 | 0 | 0 |
| ENSMUSG00000068887 | 0 | 0 | 0 | 0 | 0 | 0 | 0 |
| ENSMUSG00000068888 | 0 | 0 | 0 | 0 | 0 | 0 | 0 |
| ENSMUSG00000068889 | 0 | 0 | 0 | 0 | 0 | 0 | 0 |
| ENSMUSG00000068890 | 0 | 0 | 0 | 0 | 0 | 0 | 0 |
| ENSMUSG00000068893 | 0 | 0 | 0 | 0 | 0 | 0 | 0 |
| ENSMUSG00000068917 | 2.103271637 | 2.044977273 | 2.069031101 | 1.955044741 | 2.017859938 | 2.11058971 | 1.906930404 |
| ENSMUSG00000068921 | 2.483777101 | 2.291226732 | 2.289967834 | 2.341033664 | 2.317250476 | 2.57634135 | 2.178359564 |
| ENSMUSG00000068922 | 2.328498818 | 2.188721876 | 2.218361028 | 2.167221246 | 2.195595395 | 2.340444115 | 2.10278155 |
| ENSMUSG00000068923 | 2.075459174 | 1.890560606 | 1.83272922 | 1.858657057 | 1.892300902 | 2.08278537 | 1.793950594 |
| ENSMUSG00000068940 | 0 | 0 | 0 | 0 | 0 | 0 | 0 |
| ENSMUSG00000068947 | 0 | 0 | 0 | 0 | 0 | 0 | 0 |

| | | | | | | | |
|---|---|---|---|---|---|---|---|
| ENSMUSG00000068948 | 0 | 0 | 0 | 0 | 0 | 0 | 0 |
| ENSMUSG00000068950 | 0 | 0 | 0 | 0 | 0 | 0 | 0 |
| ENSMUSG00000068957 | 0 | 0 | 0 | 0 | 0 | 0.301029996 | 0 |
| ENSMUSG00000068959 | 1.100572389 | 1.146240625 | 1.080482024 | 0.92510993 | 1.057362051 | 0.954242509 | 1.041356071 |
| ENSMUSG00000068962 | 0 | 0 | 0 | 0 | 0.244650542 | 0 | 0 |
| ENSMUSG00000068964 | 0 | 0 | 0 | 0 | 0 | 0 | 0 |
| ENSMUSG00000068966 | 1.960252568 | 1.837182039 | 1.766522298 | 1.854463129 | 1.814781639 | 2.037426498 | 1.753626837 |
| ENSMUSG00000068969 | 0.289064826 | 0.396240625 | 0.396240625 | 0.25 | 0.489301084 | 0 | 0.224243824 |
| ENSMUSG00000068999 | 0 | 0 | 0 | 0 | 0 | 0 | 0 |
| ENSMUSG00000069008 | 0 | 0 | 0 | 0 | 0 | 0 | 0 |
| ENSMUSG00000069011 | 0 | 0 | 0 | 0 | 0 | 0 | 0.224243824 |
| ENSMUSG00000069014 | 0 | 0 | 0 | 0 | 0 | 0 | 0 |
| ENSMUSG00000069020 | 2.103271637 | 1.966046536 | 2.044977273 | 2.019203899 | 2.03596422 | 2.204119983 | 1.941550454 |
| ENSMUSG00000069036 | 0 | 0 | 0 | 0 | 0 | 0 | 0 |
| ENSMUSG00000069038 | 0 | 0 | 0 | 0 | 0 | 0 | 0 |
| ENSMUSG00000069041 | 0 | 0 | 0 | 0.25 | 0 | 0 | 0.355418052 |
| ENSMUSG00000069044 | 0 | 0 | 0.25 | 0 | 0 | 0 | 0 |
| ENSMUSG00000069045 | 0 | 0 | 0 | 0.25 | 0.244650542 | 0 | 0 |
| ENSMUSG00000069049 | 0 | 0 | 0 | 0.25 | 0.244650542 | 0 | 0 |
| ENSMUSG00000069053 | 0.45815691 | 0 | 0.5 | 0.5 | 0.244650542 | 0.477121255 | 0.224243824 |
| ENSMUSG00000069072 | 0 | 0.25 | 0 | 0 | 0.244650542 | 0 | 0 |
| ENSMUSG00000069074 | 0 | 0 | 0 | 0 | 0 | 0 | 0 |
| ENSMUSG00000069080 | 0 | 0 | 0 | 0 | 0 | 0 | 0 |
| ENSMUSG00000069083 | 0 | 0 | 0 | 0 | 0 | 0 | 0.355418052 |
| ENSMUSG00000069085 | 0 | 0 | 0 | 0 | 0 | 0 | 0 |
| ENSMUSG00000069089 | 2.623788124 | 2.38040011 | 2.399047439 | 2.560197833 | 2.508030729 | 2.882524538 | 2.314375784 |
| ENSMUSG00000069094 | 2.414600936 | 2.170748646 | 2.163659007 | 2.280383379 | 2.219993628 | 2.521138084 | 2.033014511 |
| ENSMUSG00000069114 | 2.095120908 | 1.868933358 | 1.886723615 | 1.956637122 | 1.895599581 | 2.037426498 | 1.753626837 |
| ENSMUSG00000069117 | 0 | 0 | 0 | 0 | 0 | 0 | 0 |
| ENSMUSG00000069118 | 0 | 0 | 0 | 0 | 0 | 0 | 0 |
| ENSMUSG00000069125 | 0.289064826 | 0 | 0.25 | 0.25 | 0.244650542 | 0 | 0.224243824 |
| ENSMUSG00000069132 | 0 | 0 | 0 | 0 | 0 | 0 | 0 |
| ENSMUSG00000069135 | 2.585813169 | 2.358657057 | 2.352877747 | 2.251406137 | 2.227124297 | 2.530199698 | 2.068965954 |
| ENSMUSG00000069136 | 0.289064826 | 0 | 0.396240625 | 0.25 | 0 | 0 | 0 |

| | | | | | | | |
|---|---|---|---|---|---|---|---|
| ENSMUSG00000069170 | 0.671187741 | 0.5 | 0.580482024 | 1.610735874 | 1.618292349 | 1.716003344 | 1.486592281 |
| ENSMUSG00000069171 | 1.578129653 | 1.53743678 | 1.488549078 | 1.226722649 | 1.335652905 | 1.230448921 | 1.340368108 |
| ENSMUSG00000069184 | 1.269664473 | 1.201838731 | 1.160964047 | 1.431980114 | 1.176121988 | 1.322219295 | 1.185218806 |
| ENSMUSG00000069188 | 0 | 0 | 0 | 0 | 0 | 0 | 0 |
| ENSMUSG00000069189 | 0 | 0 | 0 | 0 | 0 | 0 | 0.355418052 |
| ENSMUSG00000069196 | 0 | 0 | 0 | 0 | 0 | 0 | 0 |
| ENSMUSG00000069206 | 1.205378646 | 1.396240625 | 1.188721876 | 1.410964047 | 1.373641807 | 1.505149978 | 1.150210039 |
| ENSMUSG00000069208 | 1.771760131 | 1.738549078 | 1.726722649 | 1.87694866 | 1.826953746 | 1.977723605 | 1.669034113 |
| ENSMUSG00000069223 | 0.747221736 | 0.580482024 | 0.396240625 | 0.79248125 | 0.77552387 | 0.602059991 | 0.672731473 |
| ENSMUSG00000069227 | 0 | 0 | 0 | 1.589388001 | 1.572189156 | 1.672097858 | 1.511732338 |
| ENSMUSG00000069236 | 0.289064826 | 0.701838731 | 0.580482024 | 0 | 0.387761935 | 0 | 0.520678036 |
| ENSMUSG00000069237 | 2.552362269 | 2.377443751 | 2.36434522 | 2.443696765 | 2.37023569 | 2.704150517 | 2.187473278 |
| ENSMUSG00000069248 | 0 | 0 | 0 | 0 | 0 | 0 | 0 |
| ENSMUSG00000069255 | 1.482695304 | 1.431980114 | 1.403677461 | 1.311981878 | 1.351342427 | 1.361727836 | 1.302263476 |
| ENSMUSG00000069258 | 0 | 0 | 0 | 0 | 0 | 0 | 0 |
| ENSMUSG00000069259 | 0 | 0 | 0 | 0 | 0 | 0 | 0 |
| ENSMUSG00000069265 | 0 | 0 | 0 | 0.25 | 0.244650542 | 0 | 0 |
| ENSMUSG00000069266 | 0 | 0 | 0.25 | 0 | 0 | 0 | 0 |
| ENSMUSG00000069267 | 0 | 0 | 0 | 0 | 0 | 0 | 0 |
| ENSMUSG00000069268 | 0 | 0 | 0 | 0 | 0 | 0 | 0 |
| ENSMUSG00000069270 | 0 | 0 | 0 | 0 | 0 | 0 | 0 |
| ENSMUSG00000069272 | 0 | 0 | 0.25 | 0 | 0 | 0 | 0.224243824 |
| ENSMUSG00000069273 | 0 | 0 | 0 | 0 | 0 | 0 | 0.224243824 |
| ENSMUSG00000069274 | 0 | 0 | 0.25 | 0 | 0 | 0 | 0 |
| ENSMUSG00000069276 | 0 | 0 | 0 | 0 | 0 | 0 | 0 |
| ENSMUSG00000069280 | 0 | 0 | 0 | 0 | 0 | 0 | 0 |
| ENSMUSG00000069289 | 0 | 0 | 0 | 0 | 0 | 0 | 0 |
| ENSMUSG00000069292 | 0 | 0 | 0 | 0 | 0 | 0 | 0 |
| ENSMUSG00000069294 | 0 | 0 | 0 | 0 | 0 | 0 | 0 |
| ENSMUSG00000069295 | 0 | 0 | 0 | 0 | 0 | 0 | 0 |
| ENSMUSG00000069296 | 0 | 0 | 0 | 0 | 0 | 0 | 0 |
| ENSMUSG00000069297 | 0 | 0 | 0 | 0 | 0 | 0 | 0 |
| ENSMUSG00000069299 | 0 | 0 | 0 | 0 | 0 | 0 | 0 |
| ENSMUSG00000069300 | 0 | 0.396240625 | 0 | 0 | 0 | 0 | 0.224243824 |

| | | | | | | | |
|---|---|---|---|---|---|---|---|
| ENSMUSG00000069301 | 0 | 0 | 0 | 0 | 0 | 0 | 0 |
| ENSMUSG00000069302 | 0 | 0 | 0 | 0 | 0 | 0 | 0 |
| ENSMUSG00000069303 | 0 | 0 | 0 | 0 | 0 | 0 | 0 |
| ENSMUSG00000069305 | 0 | 0 | 0 | 0 | 0 | 0 | 0 |
| ENSMUSG00000069306 | 0 | 0 | 0 | 0 | 0 | 0 | 0 |
| ENSMUSG00000069307 | 0 | 0 | 0 | 0 | 0 | 0 | 0 |
| ENSMUSG00000069308 | 0 | 0 | 0 | 0.25 | 0 | 0 | 0 |
| ENSMUSG00000069309 | 0 | 0 | 0 | 0 | 0 | 0 | 0 |
| ENSMUSG00000069310 | 0.289064826 | 0 | 0.25 | 0 | 0.244650542 | 0 | 0.224243824 |
| ENSMUSG00000069315 | 0 | 0 | 0 | 0 | 0 | 0 | 0 |
| ENSMUSG00000069324 | 0 | 0 | 0 | 0 | 0 | 0 | 0 |
| ENSMUSG00000069355 | 0 | 0 | 0 | 0 | 0 | 0 | 0 |
| ENSMUSG00000069372 | 0 | 0 | 0 | 0 | 0 | 0 | 0 |
| ENSMUSG00000069376 | 0.45815691 | 0.580482024 | 0 | 0 | 0.489301084 | 0 | 0.224243824 |
| ENSMUSG00000069378 | 0 | 0 | 0.25 | 0.396240625 | 0.877063019 | 0.301029996 | 0.829800754 |
| ENSMUSG00000069379 | 0 | 0 | 0 | 0 | 0.244650542 | 0 | 0.224243824 |
| ENSMUSG00000069385 | 0 | 0 | 0 | 0 | 0 | 0 | 0 |
| ENSMUSG00000069390 | 0 | 0 | 0 | 0 | 0 | 0 | 0 |
| ENSMUSG00000069421 | 0 | 0 | 0 | 0 | 0 | 0 | 0 |
| ENSMUSG00000069430 | 0 | 0 | 0 | 0 | 0 | 0 | 0 |
| ENSMUSG00000069439 | 0.289064826 | 0 | 0 | 0 | 0.244650542 | 0 | 0.448487648 |
| ENSMUSG00000069441 | 0 | 0 | 0 | 0 | 0 | 0 | 0 |
| ENSMUSG00000069456 | 0 | 0.5 | 0.646240625 | 0.701838731 | 0.632412477 | 0.698970004 | 0.829800754 |
| ENSMUSG00000069475 | 0 | 0 | 0 | 0 | 0 | 0 | 0 |
| ENSMUSG00000069476 | 0 | 0 | 0 | 0 | 0 | 0 | 0 |
| ENSMUSG00000069495 | 2.479432974 | 2.259729747 | 2.267865591 | 2.26584877 | 2.199780694 | 2.514547753 | 2.028761664 |
| ENSMUSG00000069515 | 0 | 0 | 0 | 0.25 | 0 | 0 | 0 |
| ENSMUSG00000069516 | 0 | 0.25 | 0 | 0.396240625 | 0.244650542 | 0.477121255 | 0.355418052 |
| ENSMUSG00000069518 | 0.45815691 | 0.396240625 | 0.396240625 | 0.580482024 | 0 | 0 | 0.224243824 |
| ENSMUSG00000069520 | 2.336265009 | 2.148114259 | 2.178561379 | 2.144357207 | 2.097547432 | 2.389166084 | 1.978585612 |
| ENSMUSG00000069539 | 2.561428547 | 2.341033664 | 2.312574604 | 2.514836115 | 2.454682377 | 2.788168371 | 2.260561074 |
| ENSMUSG00000069540 | 0.671187741 | 0.25 | 0.25 | 0.580482024 | 0 | 0.698970004 | 0.448487648 |
| ENSMUSG00000069554 | 0 | 0 | 0 | 0.25 | 0.244650542 | 0.301029996 | 0 |
| ENSMUSG00000069565 | 2.519973621 | 2.369945066 | 2.433677405 | 2.313166358 | 2.357605438 | 2.584331224 | 2.323109859 |

| | | | | | | | |
|---|---|---|---|---|---|---|---|
| ENSMUSG00000069581 | 0 | 0 | 0 | 0.580482024 | 0.632412477 | 0.477121255 | 0.672731473 |
| ENSMUSG00000069582 | 0 | 0 | 0 | 0 | 0 | 0 | 0 |
| ENSMUSG00000069583 | 0 | 0 | 0 | 0 | 0 | 0 | 0 |
| ENSMUSG00000069584 | 0 | 0 | 0 | 0 | 0 | 0 | 0 |
| ENSMUSG00000069588 | 0 | 0 | 0 | 0.25 | 0.387761935 | 0 | 0.355418052 |
| ENSMUSG00000069601 | 1.847794126 | 1.75 | 1.823655187 | 1.470660762 | 1.427019704 | 1.681241237 | 1.459332757 |
| ENSMUSG00000069607 | 0 | 0 | 0 | 0 | 0 | 0 | 0 |
| ENSMUSG00000069609 | 0 | 0 | 0 | 0 | 0 | 0 | 0 |
| ENSMUSG00000069620 | 0 | 0 | 0 | 0 | 0 | 0 | 0 |
| ENSMUSG00000069622 | 0 | 0 | 0 | 0 | 0.387761935 | 0.301029996 | 0.355418052 |
| ENSMUSG00000069631 | 2.181541736 | 2.058404919 | 1.972185812 | 1.984159485 | 2.015530186 | 2.206825876 | 1.926805425 |
| ENSMUSG00000069633 | 1.494443473 | 1.364857905 | 1.410964047 | 0.976722649 | 1.020174412 | 1.041392685 | 0.984950056 |
| ENSMUSG00000069662 | 3.102292113 | 2.817719824 | 2.791226732 | 2.562574604 | 2.480333692 | 2.854913022 | 2.313616061 |
| ENSMUSG00000069668 | 0 | 0 | 0 | 0 | 0 | 0 | 0 |
| ENSMUSG00000069670 | 0 | 0.5 | 0.5 | 0 | 0.244650542 | 0 | 0 |
| ENSMUSG00000069678 | 1.307602651 | 1.26109853 | 1.348079356 | 1.321350555 | 1.234113756 | 1.342422681 | 1.238624656 |
| ENSMUSG00000069682 | 0.747221736 | 0.896240625 | 0.75 | 0.864857905 | 0.812711509 | 0.477121255 | 0.829800754 |
| ENSMUSG00000069706 | 0 | 0 | 0 | 0 | 0 | 0 | 0 |
| ENSMUSG00000069707 | 0 | 0 | 0 | 0 | 0 | 0 | 0 |
| ENSMUSG00000069708 | 0 | 0 | 0 | 0 | 0 | 0 | 0 |
| ENSMUSG00000069712 | 0 | 0 | 0.25 | 0 | 0.244650542 | 0.301029996 | 0.224243824 |
| ENSMUSG00000069713 | 0 | 0.580482024 | 0.25 | 0.396240625 | 0.387761935 | 0 | 0.579661877 |
| ENSMUSG00000069717 | 0 | 0 | 0 | 0 | 0 | 0 | 0 |
| ENSMUSG00000069718 | 0 | 0 | 0 | 0 | 0 | 0 | 0 |
| ENSMUSG00000069720 | 0 | 0 | 0 | 0 | 0 | 0 | 0 |
| ENSMUSG00000069721 | 0 | 0.25 | 0 | 0 | 0 | 0 | 0 |
| ENSMUSG00000069722 | 0 | 0 | 0 | 0 | 0 | 0 | 0 |
| ENSMUSG00000069727 | 1.156259305 | 1.17510993 | 1.17510993 | 1.114857905 | 1.244650542 | 1.255272505 | 1.100339912 |
| ENSMUSG00000069729 | 2.4661223 | 2.352347734 | 2.341580554 | 2.112802778 | 2.167869715 | 2.374748346 | 2.082194103 |
| ENSMUSG00000069733 | 0 | 0 | 0 | 0 | 0 | 0 | 0 |
| ENSMUSG00000069743 | 1.903481042 | 1.856566189 | 1.87694866 | 1.943696765 | 1.890639929 | 2.004321374 | 1.729871916 |
| ENSMUSG00000069744 | 2.657168526 | 2.468361028 | 2.51685859 | 2.431980114 | 2.482519186 | 2.757396029 | 2.354525595 |
| ENSMUSG00000069755 | 1.587501561 | 1.54745614 | 1.552363341 | 1.527131114 | 1.523883869 | 1.633468456 | 1.365075947 |
| ENSMUSG00000069763 | 0 | 0 | 0 | 0 | 0.244650542 | 0 | 0 |

| | | | | | | | |
|---|---|---|---|---|---|---|---|
| ENSMUSG00000069769 | 2.375872521 | 2.173371739 | 2.167221246 | 2.145300145 | 2.020174412 | 2.365487985 | 1.920029985 |
| ENSMUSG00000069785 | 0.289064826 | 0 | 0 | 0.5 | 0 | 0 | 0 |
| ENSMUSG00000069792 | 0 | 0 | 0 | 0 | 0 | 0 | 0 |
| ENSMUSG00000069793 | 2.584966402 | 2.404137211 | 2.414106216 | 2.407339155 | 2.297835549 | 2.620136055 | 2.136039214 |
| ENSMUSG00000069796 | 0 | 0 | 0 | 0 | 0 | 0 | 0 |
| ENSMUSG00000069804 | 0.289064826 | 0.396240625 | 0.5 | 0.830482024 | 0.686820904 | 0 | 0.448487648 |
| ENSMUSG00000069805 | 0 | 0 | 0 | 0 | 0 | 0 | 0 |
| ENSMUSG00000069806 | 2.020182925 | 1.919870025 | 1.905512955 | 1.67510993 | 1.736537397 | 1.954242509 | 1.767634022 |
| ENSMUSG00000069808 | 2.266038081 | 2.080482024 | 2.072504712 | 2.148114259 | 2.103062484 | 2.40654018 | 1.962479708 |
| ENSMUSG00000069814 | 0 | 0 | 0 | 0.5 | 0.244650542 | 0 | 0.355418052 |
| ENSMUSG00000069816 | 0 | 0 | 0 | 0 | 0 | 0 | 0 |
| ENSMUSG00000069818 | 0 | 0 | 0 | 0 | 0 | 0 | 0 |
| ENSMUSG00000069823 | 0 | 0 | 0 | 0 | 0 | 0 | 0 |
| ENSMUSG00000069824 | 0 | 0 | 0 | 0 | 0 | 0 | 0 |
| ENSMUSG00000069825 | 0.45815691 | 0.25 | 0.25 | 0 | 0.244650542 | 0 | 0.448487648 |
| ENSMUSG00000069830 | 0 | 0 | 0 | 0 | 0 | 0 | 0 |
| ENSMUSG00000069833 | 3.667979542 | 3.295947609 | 3.299994752 | 3.385425411 | 3.329210388 | 3.784759895 | 3.061239474 |
| ENSMUSG00000069835 | 1.418409478 | 1.25 | 1.226722649 | 1.302363341 | 1.223252711 | 1.301029996 | 1.168187745 |
| ENSMUSG00000069844 | 1.924654746 | 1.799918086 | 1.789967834 | 1.979715809 | 1.882215755 | 2.093421685 | 1.766260282 |
| ENSMUSG00000069855 | 0 | 0 | 0 | 0 | 0 | 0 | 0 |
| ENSMUSG00000069862 | 0 | 0.396240625 | 0 | 0 | 0 | 0 | 0.224243824 |
| ENSMUSG00000069867 | 0 | 0 | 0 | 0 | 0 | 0 | 0 |
| ENSMUSG00000069873 | 0 | 0 | 0 | 0 | 0 | 0 | 0 |
| ENSMUSG00000069874 | 0.811507563 | 0.79248125 | 0.896240625 | 1.470660762 | 1.343584837 | 1.477121255 | 1.252393349 |
| ENSMUSG00000069890 | 0 | 0 | 0 | 0 | 0 | 0 | 0 |
| ENSMUSG00000069892 | 1.747221736 | 1.634789703 | 1.54248125 | 1.42510993 | 1.310728002 | 1.662757832 | 1.265599895 |
| ENSMUSG00000069893 | 0.747221736 | 0.830482024 | 0.896240625 | 0.5 | 0.387761935 | 0 | 0.448487648 |
| ENSMUSG00000069895 | 2.731416512 | 2.501055117 | 2.502457154 | 2.562574604 | 2.50570673 | 2.848189117 | 2.365292129 |
| ENSMUSG00000069899 | 0 | 0.646240625 | 0.951838731 | 0.92510993 | 0.846351821 | 0.698970004 | 0.672731473 |
| ENSMUSG00000069910 | 2.196867486 | 2.109697963 | 2.124961472 | 2.311981878 | 2.26251048 | 2.57054294 | 2.121678334 |
| ENSMUSG00000069911 | 0 | 0 | 0 | 0 | 0 | 0 | 0 |
| ENSMUSG00000069913 | 0 | 0 | 0 | 0.25 | 0 | 0 | 0 |
| ENSMUSG00000069917 | 0 | 0 | 0 | 0 | 0 | 0 | 0 |
| ENSMUSG00000069919 | 0 | 0 | 0 | 0 | 0 | 0 | 0 |

| | | | | | | | |
|---|---|---|---|---|---|---|---|
| ENSMUSG00000069920 | 1.269664473 | 1.302363341 | 1.321350555 | 1.339388001 | 1.427019704 | 1.556302501 | 1.441050184 |
| ENSMUSG00000069922 | 0 | 0 | 0 | 0 | 0 | 0 | 0 |
| ENSMUSG00000069925 | 0.289064826 | 0.25 | 0 | 0 | 0 | 0 | 0 |
| ENSMUSG00000069939 | 0.289064826 | 0.79248125 | 0.580482024 | 0 | 0 | 0 | 0 |
| ENSMUSG00000069962 | 0 | 0.5 | 0 | 0.25 | 0 | 0 | 0 |
| ENSMUSG00000069971 | 0 | 0 | 0 | 0 | 0 | 0 | 0 |
| ENSMUSG00000069972 | 0 | 0 | 0 | 0 | 0 | 0 | 0 |
| ENSMUSG00000069986 | 0 | 0 | 0.25 | 0 | 0 | 0 | 0.355418052 |
| ENSMUSG00000069996 | 0 | 0 | 0 | 0 | 0 | 0 | 0.224243824 |
| ENSMUSG00000069998 | 0 | 0 | 0 | 0 | 0 | 0 | 0 |
| ENSMUSG00000070000 | 0 | 0 | 0 | 0.830482024 | 1.136121934 | 0.477121255 | 1.159323753 |
| ENSMUSG00000070002 | 1.956444026 | 1.862802778 | 1.903677461 | 1.852347734 | 1.962676635 | 2.123851641 | 1.925124822 |
| ENSMUSG00000070003 | 1.129344651 | 1.311981878 | 1.527131114 | 1.321350555 | 1.674752852 | 1.505149978 | 1.855869786 |
| ENSMUSG00000070023 | 0 | 0 | 0 | 0 | 0 | 0 | 0 |
| ENSMUSG00000070025 | 0 | 0 | 0 | 0 | 0 | 0 | 0.224243824 |
| ENSMUSG00000070031 | 1.069666964 | 0.92510993 | 0.830482024 | 0.896240625 | 1.121713561 | 1.146128036 | 0.935079929 |
| ENSMUSG00000070034 | 0 | 0 | 0 | 0 | 0.244650542 | 0 | 0 |
| ENSMUSG00000070044 | 2.392336464 | 2.271202097 | 2.294977273 | 2.389605178 | 2.349985771 | 2.665580991 | 2.213368331 |
| ENSMUSG00000070047 | 3.424001185 | 3.135999613 | 3.114601653 | 2.89939676 | 2.893371499 | 3.284881715 | 2.718614352 |
| ENSMUSG00000070056 | 2.312518611 | 2.160061234 | 2.168106335 | 1.998588359 | 1.987570683 | 2.190331698 | 1.905143021 |
| ENSMUSG00000070063 | 0.867194479 | 0.830482024 | 0.75 | 0.896240625 | 0.77552387 | 0.698970004 | 0.969165684 |
| ENSMUSG00000070065 | 0 | 0 | 0 | 0 | 0 | 0 | 0 |
| ENSMUSG00000070071 | 0 | 0 | 0 | 0 | 0 | 0 | 0 |
| ENSMUSG00000070072 | 0 | 0 | 0 | 0 | 0 | 0 | 0 |
| ENSMUSG00000070073 | 0 | 0 | 0 | 0 | 0 | 0 | 0 |
| ENSMUSG00000070074 | 0 | 0 | 0 | 0.25 | 0 | 0 | 0 |
| ENSMUSG00000070075 | 0 | 0 | 0 | 0 | 0 | 0 | 0 |
| ENSMUSG00000070076 | 0 | 0 | 0 | 0 | 0 | 0 | 0 |
| ENSMUSG00000070077 | 0 | 0 | 0.25 | 0 | 0 | 0 | 0 |
| ENSMUSG00000070078 | 0 | 0 | 0 | 0 | 0 | 0 | 0 |
| ENSMUSG00000070080 | 0 | 0 | 0 | 0 | 0 | 0 | 0 |
| ENSMUSG00000070081 | 0 | 0 | 0 | 0 | 0 | 0 | 0 |
| ENSMUSG00000070084 | 0 | 0 | 0 | 0 | 0 | 0 | 0 |
| ENSMUSG00000070099 | 0 | 0 | 0 | 0 | 0 | 0 | 0 |

| | | | | | | | |
|---|---|---|---|---|---|---|---|
| ENSMUSG00000070100 | 0 | 0 | 0 | 0 | 0 | 0 | 0 |
| ENSMUSG00000070101 | 0 | 0 | 0 | 0 | 0 | 0 | 0 |
| ENSMUSG00000070102 | 0 | 0 | 0 | 0 | 0 | 0 | 0 |
| ENSMUSG00000070103 | 0 | 0 | 0 | 0 | 0 | 0 | 0 |
| ENSMUSG00000070104 | 0 | 0 | 0 | 0 | 0 | 0 | 0 |
| ENSMUSG00000070105 | 0 | 0 | 0 | 0 | 0 | 0 | 0 |
| ENSMUSG00000070106 | 0 | 0 | 0 | 0 | 0 | 0 | 0 |
| ENSMUSG00000070107 | 0 | 0 | 0 | 0 | 0 | 0 | 0 |
| ENSMUSG00000070108 | 0 | 0 | 0 | 0 | 0 | 0 | 0 |
| ENSMUSG00000070109 | 0 | 0 | 0 | 0 | 0 | 0 | 0 |
| ENSMUSG00000070110 | 0 | 0 | 0 | 0 | 0 | 0 | 0 |
| ENSMUSG00000070111 | 0 | 0.396240625 | 0 | 0.25 | 0.244650542 | 0 | 0.224243824 |
| ENSMUSG00000070126 | 0 | 0 | 0 | 0 | 0 | 0 | 0 |
| ENSMUSG00000070127 | 0 | 0 | 0 | 0 | 0 | 0 | 0 |
| ENSMUSG00000070128 | 0 | 0 | 0 | 0 | 0 | 0 | 0 |
| ENSMUSG00000070129 | 0 | 0 | 0 | 0 | 0 | 0 | 0 |
| ENSMUSG00000070130 | 0 | 0 | 0 | 0 | 0 | 0 | 0 |
| ENSMUSG00000070133 | 0 | 0 | 0 | 0 | 0 | 0 | 0 |
| ENSMUSG00000070136 | 0 | 0 | 0 | 0 | 0 | 0 | 0 |
| ENSMUSG00000070138 | 0 | 0 | 0.25 | 0 | 0 | 0 | 0 |
| ENSMUSG00000070139 | 0 | 0 | 0 | 0 | 0 | 0 | 0 |
| ENSMUSG00000070140 | 0 | 0 | 0 | 0 | 0 | 0 | 0 |
| ENSMUSG00000070141 | 0 | 0 | 0 | 0 | 0 | 0 | 0 |
| ENSMUSG00000070144 | 0 | 0 | 0 | 0 | 0 | 0 | 0 |
| ENSMUSG00000070150 | 0 | 0 | 0 | 0 | 0 | 0 | 0 |
| ENSMUSG00000070166 | 0 | 0 | 0 | 0 | 0 | 0 | 0 |
| ENSMUSG00000070167 | 0.671187741 | 0.580482024 | 0.580482024 | 0.396240625 | 0.489301084 | 0.301029996 | 0.629532004 |
| ENSMUSG00000070170 | 0.289064826 | 0 | 0 | 0 | 0.244650542 | 0 | 0 |
| ENSMUSG00000070173 | 0 | 0 | 0 | 0 | 0 | 0 | 0 |
| ENSMUSG00000070175 | 0 | 0 | 0 | 0 | 0 | 0 | 0 |
| ENSMUSG00000070178 | 0 | 0 | 0 | 0 | 0 | 0 | 0 |
| ENSMUSG00000070182 | 0 | 0 | 0 | 0 | 0 | 0 | 0 |
| ENSMUSG00000070192 | 0 | 0 | 0 | 0 | 0 | 0 | 0 |
| ENSMUSG00000070203 | 0 | 0 | 0 | 0 | 0 | 0 | 0 |

| | | | | | | | |
|---|---|---|---|---|---|---|---|
| ENSMUSG00000070209 | 0 | 0 | 0 | 0 | 0 | 0 | 0 |
| ENSMUSG00000070214 | 0 | 0 | 0 | 0 | 0 | 0 | 0 |
| ENSMUSG00000070219 | 0 | 0 | 0 | 0 | 0 | 0 | 0 |
| ENSMUSG00000070225 | 0 | 0 | 0 | 0.25 | 0 | 0 | 0 |
| ENSMUSG00000070227 | 0 | 0 | 0 | 0 | 0 | 0 | 0 |
| ENSMUSG00000070235 | 0 | 0 | 0 | 0 | 0 | 0 | 0 |
| ENSMUSG00000070236 | 0 | 0 | 0 | 0 | 0 | 0 | 0 |
| ENSMUSG00000070240 | 0 | 0 | 0 | 0 | 0 | 0 | 0 |
| ENSMUSG00000070249 | 0 | 0 | 0 | 0 | 0 | 0 | 0 |
| ENSMUSG00000070258 | 0 | 0 | 0 | 0 | 0 | 0 | 0 |
| ENSMUSG00000070263 | 0 | 0 | 0 | 0 | 0 | 0 | 0 |
| ENSMUSG00000070271 | 0.289064826 | 0 | 0 | 0 | 0.244650542 | 0 | 0.224243824 |
| ENSMUSG00000070280 | 0 | 0.25 | 0 | 0.5 | 0.632412477 | 0.84509804 | 0.629532004 |
| ENSMUSG00000070282 | 1.100572389 | 1.330482024 | 1.188721876 | 1.226722649 | 1.302012594 | 1.146128036 | 1.150210039 |
| ENSMUSG00000070283 | 1.205378646 | 1.238549078 | 1.372963274 | 1.311981878 | 1.319233381 | 1.414973348 | 1.360283038 |
| ENSMUSG00000070284 | 1.862428343 | 1.794977273 | 1.71759118 | 1.714495249 | 1.78687392 | 1.826074803 | 1.667169465 |
| ENSMUSG00000070287 | 0.45815691 | 0.646240625 | 0.75 | 0.580482024 | 0.733951626 | 0.698970004 | 0.803905701 |
| ENSMUSG00000070291 | 1.036286563 | 0.701838731 | 0.79248125 | 1.445339928 | 1.176121988 | 1.322219295 | 1.224243824 |
| ENSMUSG00000070298 | 0 | 0.25 | 0 | 0 | 0 | 0 | 0.224243824 |
| ENSMUSG00000070304 | 0 | 0.25 | 0.25 | 0 | 0 | 0.477121255 | 0 |
| ENSMUSG00000070305 | 1.558729299 | 1.464495249 | 1.458222504 | 1.784887838 | 1.78687392 | 1.924279286 | 1.62316756 |
| ENSMUSG00000070306 | 0.289064826 | 0.25 | 0 | 0 | 0.244650542 | 0 | 0 |
| ENSMUSG00000070311 | 0 | 0 | 0 | 0 | 0 | 0 | 0 |
| ENSMUSG00000070313 | 0 | 0 | 0 | 0 | 0 | 0.301029996 | 0 |
| ENSMUSG00000070315 | 1.205378646 | 1.348079356 | 1.160964047 | 0.976722649 | 1.121713561 | 0.84509804 | 0.896975297 |
| ENSMUSG00000070319 | 2.556012466 | 2.370449858 | 2.415889526 | 2.418106335 | 2.501012315 | 2.770852012 | 2.410403904 |
| ENSMUSG00000070323 | 0.289064826 | 0 | 0.25 | 0.25 | 0 | 0 | 0 |
| ENSMUSG00000070324 | 0 | 0 | 0 | 0 | 0 | 0 | 0 |
| ENSMUSG00000070327 | 2.768497801 | 2.51685859 | 2.552363341 | 2.814788692 | 2.788054595 | 3.149527014 | 2.597118276 |
| ENSMUSG00000070330 | 0 | 0.25 | 0 | 0.25 | 0.568060967 | 0.301029996 | 0.520678036 |
| ENSMUSG00000070331 | 0.671187741 | 0.580482024 | 0.5 | 0 | 0.489301084 | 0 | 0.224243824 |
| ENSMUSG00000070332 | 0 | 0 | 0 | 0 | 0 | 0 | 0 |
| ENSMUSG00000070334 | 0 | 0 | 0 | 0 | 0 | 0 | 0 |
| ENSMUSG00000070335 | 0 | 0 | 0 | 0 | 0 | 0 | 0 |

| | | | | | | | |
|---|---|---|---|---|---|---|---|
| ENSMUSG00000070336 | 0.747221736 | 1.02186571 | 1.04248125 | 1.214495249 | 1.176121988 | 1.113943352 | 0.969165684 |
| ENSMUSG00000070337 | 0.578129653 | 0.5 | 0.646240625 | 0.75 | 0.733951626 | 0.77815125 | 0.672731473 |
| ENSMUSG00000070342 | 0.578129653 | 0.75 | 0.25 | 0.396240625 | 0.244650542 | 0.301029996 | 0.224243824 |
| ENSMUSG00000070343 | 0.811507563 | 0.5 | 0.646240625 | 0.830482024 | 0.733951626 | 0.903089987 | 0.520678036 |
| ENSMUSG00000070345 | 0.671187741 | 0.25 | 0.580482024 | 0.25 | 0.568060967 | 0 | 0 |
| ENSMUSG00000070348 | 2.397683106 | 2.300530956 | 2.30357978 | 2.532320754 | 2.496553863 | 2.811575006 | 2.411717103 |
| ENSMUSG00000070354 | 0.289064826 | 0 | 0 | 0 | 0 | 0 | 0.224243824 |
| ENSMUSG00000070360 | 0 | 0 | 0 | 0 | 0 | 0 | 0 |
| ENSMUSG00000070366 | 0 | 0 | 0 | 0 | 0 | 0 | 0 |
| ENSMUSG00000070368 | 0 | 0 | 0 | 0 | 0 | 0 | 0 |
| ENSMUSG00000070369 | 0 | 0 | 0 | 0.25 | 0 | 0 | 0 |
| ENSMUSG00000070371 | 0.91631382 | 0.830482024 | 1.061981878 | 0.92510993 | 1 | 0.698970004 | 1.014380832 |
| ENSMUSG00000070372 | 2.524897319 | 2.353406982 | 2.336073977 | 2.56932185 | 2.495055108 | 2.879669206 | 2.314375784 |
| ENSMUSG00000070374 | 0 | 0 | 0 | 0 | 0 | 0 | 0 |
| ENSMUSG00000070375 | 0 | 0 | 0 | 0 | 0 | 0 | 0 |
| ENSMUSG00000070377 | 0 | 0 | 0 | 0 | 0 | 0 | 0 |
| ENSMUSG00000070378 | 0 | 0 | 0 | 0 | 0 | 0 | 0 |
| ENSMUSG00000070379 | 0 | 0 | 0 | 0 | 0 | 0 | 0 |
| ENSMUSG00000070380 | 0 | 0 | 0 | 0 | 0 | 0 | 0 |
| ENSMUSG00000070382 | 0 | 0 | 0 | 0 | 0 | 0 | 0 |
| ENSMUSG00000070383 | 0 | 0 | 0 | 0 | 0 | 0 | 0 |
| ENSMUSG00000070385 | 0.45815691 | 0.25 | 0 | 0 | 0 | 0.301029996 | 0 |
| ENSMUSG00000070388 | 0 | 0 | 0 | 0 | 0 | 0.301029996 | 0 |
| ENSMUSG00000070390 | 0 | 0 | 0 | 0 | 0 | 0 | 0 |
| ENSMUSG00000070392 | 0 | 0 | 0 | 0 | 0 | 0 | 0 |
| ENSMUSG00000070394 | 2.131864492 | 2.025821952 | 2.092671852 | 1.854463129 | 1.830919587 | 2.017033339 | 1.779742043 |
| ENSMUSG00000070402 | 0 | 0 | 0 | 0 | 0 | 0 | 0 |
| ENSMUSG00000070407 | 0 | 0.25 | 0.5 | 0.79248125 | 0.733951626 | 0.477121255 | 0.710836105 |
| ENSMUSG00000070411 | 0 | 0 | 0 | 0 | 0 | 0 | 0 |
| ENSMUSG00000070417 | 0 | 0 | 0 | 0 | 0 | 0 | 0 |
| ENSMUSG00000070419 | 0 | 0 | 0.25 | 0 | 0 | 0 | 0.224243824 |
| ENSMUSG00000070420 | 1.707474304 | 1.668106335 | 1.809601185 | 1.431980114 | 1.645989387 | 1.602059991 | 1.687112305 |
| ENSMUSG00000070421 | 0 | 0 | 0 | 0 | 0 | 0 | 0 |
| ENSMUSG00000070423 | 0 | 0 | 0 | 0 | 0 | 0 | 0 |

| | | | | | | | |
|---|---|---|---|---|---|---|---|
| ENSMUSG00000070424 | 0.289064826 | 0.5 | 0.25 | 0 | 0 | 0 | 0 |
| ENSMUSG00000070425 | 0.289064826 | 0.25 | 0.25 | 0 | 0.489301084 | 0.301029996 | 0 |
| ENSMUSG00000070426 | 2.190368148 | 2.029735268 | 1.997171172 | 2.116901388 | 2.06149024 | 2.374748346 | 1.918313719 |
| ENSMUSG00000070427 | 1.069666964 | 1.282320754 | 1.130890489 | 1.238549078 | 1.27449559 | 1.204119983 | 1.150210039 |
| ENSMUSG00000070436 | 3.360187403 | 3.13058408 | 3.151022465 | 2.785526764 | 2.892690586 | 3.235275877 | 2.837372575 |
| ENSMUSG00000070438 | 0 | 0 | 0 | 0 | 0 | 0 | 0 |
| ENSMUSG00000070443 | 0.45815691 | 0.5 | 0.5 | 0 | 0.244650542 | 0 | 0 |
| ENSMUSG00000070448 | 0 | 0 | 0 | 0 | 0 | 0 | 0 |
| ENSMUSG00000070458 | 0 | 0 | 0 | 0 | 0 | 0 | 0 |
| ENSMUSG00000070459 | 0 | 0 | 0 | 0 | 0 | 0 | 0 |
| ENSMUSG00000070460 | 0 | 0 | 0 | 0 | 0 | 0 | 0 |
| ENSMUSG00000070461 | 1.482695304 | 1.445339928 | 1.470660762 | 1.445339928 | 1.462344774 | 1.491361694 | 1.392431569 |
| ENSMUSG00000070462 | 1.596667477 | 1.458222504 | 1.622963274 | 1.42510993 | 1.433158226 | 1.568201724 | 1.514741806 |
| ENSMUSG00000070464 | 0 | 0 | 0 | 0 | 0 | 0 | 0 |
| ENSMUSG00000070465 | 0 | 0 | 0 | 0 | 0 | 0 | 0 |
| ENSMUSG00000070469 | 1.358731791 | 1.364857905 | 1.188721876 | 1.930274797 | 1.917865457 | 2.113943352 | 1.800208341 |
| ENSMUSG00000070471 | 0 | 0 | 0 | 0.25 | 0.244650542 | 0.301029996 | 0.224243824 |
| ENSMUSG00000070473 | 0 | 0 | 0 | 0 | 0 | 0.301029996 | 0 |
| ENSMUSG00000070476 | 1.289064826 | 1.26109853 | 1.17510993 | 1.356566189 | 1.234113756 | 1.397940009 | 1.21680636 |
| ENSMUSG00000070489 | 0 | 0 | 0 | 0 | 0 | 0 | 0 |
| ENSMUSG00000070490 | 0.747221736 | 0.646240625 | 0.5 | 0.75 | 0.489301084 | 0.602059991 | 0.629532004 |
| ENSMUSG00000070493 | 2.652148299 | 2.466433318 | 2.46293726 | 2.55539678 | 2.477187895 | 2.872156273 | 2.358301651 |
| ENSMUSG00000070495 | 1.156259305 | 0.896240625 | 0.646240625 | 0.79248125 | 0.77552387 | 0.954242509 | 0.916588299 |
| ENSMUSG00000070498 | 0 | 0 | 0 | 0 | 0 | 0 | 0.224243824 |
| ENSMUSG00000070501 | 0 | 0 | 0 | 0 | 0.244650542 | 0 | 0 |
| ENSMUSG00000070504 | 0 | 0 | 0 | 0 | 0 | 0 | 0 |
| ENSMUSG00000070509 | 1.707474304 | 1.711372513 | 1.738549078 | 1 | 1.106691885 | 1.176091259 | 1.110947927 |
| ENSMUSG00000070511 | 0 | 0 | 0 | 0 | 0 | 0 | 0 |
| ENSMUSG00000070520 | 1.578129653 | 1.494319981 | 1.681980114 | 1.527131114 | 1.49445382 | 1.568201724 | 1.543390198 |
| ENSMUSG00000070522 | 0 | 0 | 0 | 0 | 0 | 0 | 0 |
| ENSMUSG00000070524 | 0 | 0 | 0 | 0 | 0 | 0 | 0 |
| ENSMUSG00000070526 | 1.678702042 | 1.77186571 | 1.708222504 | 1.557204673 | 1.489301084 | 1.716003344 | 1.444790289 |
| ENSMUSG00000070527 | 1.069666964 | 1.146240625 | 1.098079356 | 0.976722649 | 1.106691885 | 0.84509804 | 1.041356071 |
| ENSMUSG00000070529 | 0 | 0 | 0 | 0 | 0 | 0 | 0 |

| | | | | | | | |
|---|---|---|---|---|---|---|---|
| ENSMUSG00000070530 | 0 | 0 | 0 | 0 | 0 | 0 | 0 |
| ENSMUSG00000070531 | 0 | 0 | 0 | 0 | 0 | 0 | 0 |
| ENSMUSG00000070532 | 0 | 0 | 0 | 0 | 0 | 0 | 0 |
| ENSMUSG00000070533 | 0 | 0 | 0 | 0 | 0 | 0 | 0 |
| ENSMUSG00000070544 | 2.72721917 | 2.530383379 | 2.51348147 | 2.70444578 | 2.634037134 | 3.017033339 | 2.421308913 |
| ENSMUSG00000070546 | 0 | 0 | 0 | 0 | 0 | 0 | 0 |
| ENSMUSG00000070547 | 0 | 0.25 | 0.5 | 0 | 0.489301084 | 0 | 0.224243824 |
| ENSMUSG00000070550 | 0 | 0 | 0 | 0 | 0 | 0 | 0 |
| ENSMUSG00000070551 | 0 | 0 | 0 | 0 | 0 | 0 | 0 |
| ENSMUSG00000070552 | 0 | 0 | 0 | 0 | 0 | 0 | 0 |
| ENSMUSG00000070563 | 0 | 0 | 0.25 | 0 | 0.387761935 | 0 | 0 |
| ENSMUSG00000070564 | 0 | 0 | 0 | 0 | 0.244650542 | 0 | 0 |
| ENSMUSG00000070565 | 2.560530738 | 2.373463862 | 2.379909063 | 2.512121718 | 2.454682377 | 2.788875116 | 2.260561074 |
| ENSMUSG00000070568 | 0 | 0 | 0 | 0 | 0 | 0 | 0 |
| ENSMUSG00000070570 | 0.289064826 | 0.396240625 | 0.25 | 1.02186571 | 1.106691885 | 1.146128036 | 1.150210039 |
| ENSMUSG00000070572 | 0 | 0 | 0 | 0 | 0 | 0 | 0 |
| ENSMUSG00000070574 | 0 | 0 | 0 | 0 | 0 | 0 | 0 |
| ENSMUSG00000070576 | 1.714367558 | 1.671625132 | 1.714495249 | 1.160964047 | 1.302012594 | 0.954242509 | 1.290497981 |
| ENSMUSG00000070577 | 0 | 0 | 0 | 0 | 0 | 0 | 0 |
| ENSMUSG00000070583 | 1.671187741 | 1.589388001 | 1.552363341 | 1.614857905 | 1.584291694 | 1.698970004 | 1.429563222 |
| ENSMUSG00000070594 | 0 | 0 | 0 | 0 | 0 | 0 | 0 |
| ENSMUSG00000070601 | 0 | 0 | 0 | 0 | 0 | 0 | 0 |
| ENSMUSG00000070604 | 0.671187741 | 1.02186571 | 1.17510993 | 0.701838731 | 1.057362051 | 0.77815125 | 1.110947927 |
| ENSMUSG00000070605 | 1.827447047 | 1.732684334 | 1.692046081 | 1.931980114 | 1.90849439 | 2.053078443 | 1.709198048 |
| ENSMUSG00000070606 | 0 | 0 | 0 | 0 | 0 | 0 | 0 |
| ENSMUSG00000070609 | 0 | 0 | 0 | 0 | 0 | 0 | 0 |
| ENSMUSG00000070610 | 0 | 0 | 0 | 0 | 0 | 0 | 0 |
| ENSMUSG00000070616 | 0 | 0 | 0 | 0 | 0 | 0 | 0 |
| ENSMUSG00000070617 | 0 | 0 | 0 | 0 | 0 | 0 | 0 |
| ENSMUSG00000070618 | 0 | 0.25 | 0 | 0 | 0 | 0 | 0.224243824 |
| ENSMUSG00000070619 | 0 | 0 | 0 | 0 | 0 | 0.301029996 | 0 |
| ENSMUSG00000070621 | 0 | 0 | 0 | 0 | 0 | 0 | 0 |
| ENSMUSG00000070632 | 0.867194479 | 1.061981878 | 0.92510993 | 1.061981878 | 0.955822902 | 1 | 0.829800754 |
| ENSMUSG00000070637 | 0 | 0 | 0 | 0 | 0 | 0 | 0 |

| | | | | | | | |
|---|---|---|---|---|---|---|---|
| ENSMUSG00000070639 | 2.597490839 | 2.410512923 | 2.412762923 | 2.190387808 | 2.09661982 | 2.36361198 | 1.975716277 |
| ENSMUSG00000070641 | 0 | 0 | 0 | 0.25 | 0 | 0 | 0 |
| ENSMUSG00000070643 | 0.747221736 | 0.701838731 | 0.79248125 | 1.080482024 | 1.254881871 | 1.113943352 | 1.193409508 |
| ENSMUSG00000070644 | 0 | 0 | 0.25 | 0.5 | 0.686820904 | 0.698970004 | 0.448487648 |
| ENSMUSG00000070645 | 0 | 0 | 0 | 0 | 0 | 0 | 0 |
| ENSMUSG00000070661 | 0 | 0 | 0.25 | 0 | 0 | 0 | 0 |
| ENSMUSG00000070667 | 0.289064826 | 0 | 0.25 | 0.396240625 | 0 | 0 | 0.355418052 |
| ENSMUSG00000070677 | 0 | 0 | 0 | 0 | 0 | 0 | 0 |
| ENSMUSG00000070683 | 0 | 0 | 0 | 0 | 0 | 0 | 0 |
| ENSMUSG00000070686 | 0 | 0 | 0 | 0 | 0 | 0 | 0 |
| ENSMUSG00000070687 | 0.867194479 | 0.75 | 0.396240625 | 0.396240625 | 0.489301084 | 0.477121255 | 0.520678036 |
| ENSMUSG00000070690 | 0.289064826 | 0.25 | 0.25 | 0.25 | 0.244650542 | 0.301029996 | 0 |
| ENSMUSG00000070691 | 2.158622101 | 2.00835575 | 2.058404919 | 1.956637122 | 1.978602168 | 2.206825876 | 1.905143021 |
| ENSMUSG00000070692 | 0.747221736 | 0.896240625 | 1.238549078 | 0.896240625 | 0.877063019 | 0.84509804 | 0.896975297 |
| ENSMUSG00000070695 | 0 | 0 | 0.396240625 | 0.25 | 0.244650542 | 0.477121255 | 0 |
| ENSMUSG00000070697 | 2.571178952 | 2.413211243 | 2.412314044 | 2.373963757 | 2.297835549 | 2.622214023 | 2.1908258 |
| ENSMUSG00000070699 | 1.605636262 | 1.51109853 | 1.630890489 | 1.464495249 | 1.662257525 | 1.462397998 | 1.637827726 |
| ENSMUSG00000070702 | 0 | 0 | 0 | 0 | 0 | 0 | 0 |
| ENSMUSG00000070704 | 0 | 0 | 0 | 0 | 0 | 0 | 0 |
| ENSMUSG00000070705 | 1.156259305 | 1.17510993 | 1.02186571 | 0.976722649 | 1.039257769 | 1.041392685 | 0.935079929 |
| ENSMUSG00000070708 | 0 | 0 | 0 | 0 | 0 | 0 | 0 |
| ENSMUSG00000070709 | 1.671187741 | 1.646240625 | 1.580482024 | 1.571350555 | 1.533172725 | 1.591064607 | 1.392431569 |
| ENSMUSG00000070713 | 0.289064826 | 0.5 | 0.5 | 0.396240625 | 0.387761935 | 0.301029996 | 0.448487648 |
| ENSMUSG00000070717 | 0 | 0.396240625 | 0 | 0.25 | 0 | 0 | 0 |
| ENSMUSG00000070719 | 0 | 0 | 0 | 0 | 0.387761935 | 0 | 0 |
| ENSMUSG00000070720 | 0.45815691 | 1 | 0.976722649 | 0.580482024 | 1 | 1.079181246 | 1.121219121 |
| ENSMUSG00000070729 | 0 | 0 | 0.25 | 0 | 0 | 0 | 0.224243824 |
| ENSMUSG00000070730 | 1.753493018 | 1.692046081 | 1.705044741 | 1.959800947 | 1.981187939 | 2.252853031 | 1.941550454 |
| ENSMUSG00000070732 | 0.289064826 | 0 | 0 | 0 | 0 | 0 | 0 |
| ENSMUSG00000070733 | 2.81052167 | 2.55479213 | 2.544977273 | 2.466819685 | 2.419286397 | 2.774516966 | 2.244642249 |
| ENSMUSG00000070737 | 1.129344651 | 1.17510993 | 1.114857905 | 1.098079356 | 1.188507684 | 1.278753601 | 1.089372236 |
| ENSMUSG00000070738 | 2.562324428 | 2.344302633 | 2.288071211 | 2.360735874 | 2.341270363 | 2.595496222 | 2.168981648 |
| ENSMUSG00000070777 | 0.289064826 | 0 | 0 | 0 | 0 | 0 | 0.224243824 |
| ENSMUSG00000070778 | 0.45815691 | 0.25 | 0.25 | 0.396240625 | 0.244650542 | 0.477121255 | 0.448487648 |

| | | | | | | | |
|---|---|---|---|---|---|---|---|
| ENSMUSG00000070780 | 1.307602651 | 1.330482024 | 1.25 | 1.380890489 | 1.394615667 | 1.204119983 | 1.302263476 |
| ENSMUSG00000070794 | 0.289064826 | 0 | 0.5 | 0.25 | 0 | 0.301029996 | 0.224243824 |
| ENSMUSG00000070796 | 0 | 0 | 0 | 0 | 0 | 0 | 0 |
| ENSMUSG00000070797 | 0 | 0 | 0 | 0 | 0 | 0 | 0 |
| ENSMUSG00000070798 | 0 | 0 | 0 | 0 | 0 | 0 | 0 |
| ENSMUSG00000070799 | 0 | 0 | 0 | 0 | 0 | 0 | 0 |
| ENSMUSG00000070802 | 0 | 0 | 0 | 1.061981878 | 1.106691885 | 1.146128036 | 1.131174228 |
| ENSMUSG00000070803 | 0 | 0 | 0 | 0 | 0.244650542 | 0.301029996 | 0.224243824 |
| ENSMUSG00000070806 | 0 | 0 | 0 | 0 | 0.244650542 | 0 | 0 |
| ENSMUSG00000070808 | 1.289064826 | 1.431980114 | 1.622963274 | 1.226722649 | 1.358933174 | 1.414973348 | 1.56970677 |
| ENSMUSG00000070810 | 0 | 0 | 0 | 0 | 0 | 0 | 0 |
| ENSMUSG00000070811 | 0 | 0 | 0 | 0 | 0 | 0 | 0 |
| ENSMUSG00000070814 | 1.727821383 | 1.646240625 | 1.671625132 | 1.606566189 | 1.533172725 | 1.643452676 | 1.554175045 |
| ENSMUSG00000070815 | 0 | 0 | 0 | 0 | 0 | 0 | 0 |
| ENSMUSG00000070816 | 0 | 0 | 0 | 0 | 0 | 0 | 0 |
| ENSMUSG00000070817 | 0 | 0 | 0 | 0 | 0 | 0 | 0 |
| ENSMUSG00000070820 | 0 | 0 | 0 | 0 | 0 | 0 | 0 |
| ENSMUSG00000070821 | 0 | 0 | 0 | 0 | 0 | 0 | 0 |
| ENSMUSG00000070822 | 0.671187741 | 0.646240625 | 0.701838731 | 0.396240625 | 0.812711509 | 0.84509804 | 0.853775828 |
| ENSMUSG00000070823 | 0.289064826 | 0 | 0 | 0 | 0 | 0 | 0 |
| ENSMUSG00000070828 | 0 | 0 | 0 | 0 | 0 | 0 | 0 |
| ENSMUSG00000070834 | 0 | 0 | 0.25 | 0 | 0 | 0 | 0 |
| ENSMUSG00000070837 | 0 | 0 | 0 | 0 | 0 | 0 | 0 |
| ENSMUSG00000070841 | 0 | 0 | 0 | 0 | 0 | 0 | 0 |
| ENSMUSG00000070844 | 0 | 0 | 0 | 0 | 0 | 0 | 0 |
| ENSMUSG00000070847 | 0 | 0 | 0 | 0 | 0 | 0 | 0 |
| ENSMUSG00000070852 | 0 | 0 | 0 | 0 | 0 | 0 | 0 |
| ENSMUSG00000070853 | 0 | 0 | 0 | 0 | 0 | 0 | 0 |
| ENSMUSG00000070855 | 0 | 0 | 0 | 0 | 0 | 0 | 0 |
| ENSMUSG00000070856 | 0 | 0 | 0 | 0 | 0 | 0 | 0 |
| ENSMUSG00000070857 | 0 | 0 | 0 | 0 | 0 | 0 | 0 |
| ENSMUSG00000070858 | 0.578129653 | 0.646240625 | 0.830482024 | 0 | 0.568060967 | 0.602059991 | 0.853775828 |
| ENSMUSG00000070866 | 0 | 0 | 0.25 | 0.25 | 0 | 0 | 0.224243824 |
| ENSMUSG00000070867 | 2.365959569 | 2.173371739 | 2.150906586 | 0.976722649 | 1.106691885 | 1.041392685 | 1.041356071 |

| | | | | | | | |
|---|---|---|---|---|---|---|---|
| ENSMUSG00000070868 | 0 | 0 | 0 | 0 | 0 | 0 | 0 |
| ENSMUSG00000070870 | 0 | 0 | 0 | 0 | 0 | 0 | 0 |
| ENSMUSG00000070871 | 2.352857998 | 2.179419106 | 2.164552871 | 2.25835575 | 2.230953877 | 2.5132176 | 2.085803078 |
| ENSMUSG00000070873 | 0 | 0 | 0 | 0 | 0 | 0 | 0 |
| ENSMUSG00000070875 | 0 | 0 | 0 | 0 | 0 | 0 | 0 |
| ENSMUSG00000070877 | 0 | 0.25 | 0 | 0 | 0 | 0 | 0.355418052 |
| ENSMUSG00000070880 | 0 | 0 | 0.25 | 0 | 0.244650542 | 0 | 0 |
| ENSMUSG00000070883 | 0.867194479 | 1.130890489 | 1 | 1.201838731 | 1.106691885 | 1.204119983 | 1.014380832 |
| ENSMUSG00000070890 | 0 | 0.25 | 0 | 0 | 0 | 0 | 0.224243824 |
| ENSMUSG00000070891 | 0 | 0.25 | 0 | 0 | 0.244650542 | 0 | 0.224243824 |
| ENSMUSG00000070896 | 0 | 0 | 0.25 | 0 | 0 | 0.477121255 | 0.448487648 |
| ENSMUSG00000070900 | 0 | 0 | 0 | 0 | 0 | 0 | 0 |
| ENSMUSG00000070902 | 0 | 0 | 0 | 0 | 0 | 0 | 0 |
| ENSMUSG00000070904 | 0 | 0 | 0 | 0 | 0 | 0 | 0 |
| ENSMUSG00000070908 | 0 | 0 | 0 | 0 | 0 | 0 | 0 |
| ENSMUSG00000070923 | 2.543094501 | 2.345385738 | 2.321350555 | 2.365881093 | 2.28445426 | 2.551449998 | 2.112839835 |
| ENSMUSG00000070933 | 0 | 0 | 0 | 0 | 0 | 0 | 0 |
| ENSMUSG00000070934 | 2.512981297 | 2.344302633 | 2.338286706 | 2.369439567 | 2.287171411 | 2.625312451 | 2.157685697 |
| ENSMUSG00000070939 | 2.309247751 | 2.127938163 | 2.175975893 | 2.199415381 | 2.173890435 | 2.437750563 | 2.065163042 |
| ENSMUSG00000070942 | 0 | 0.396240625 | 0 | 0 | 0 | 0 | 0.355418052 |
| ENSMUSG00000070943 | 0 | 0 | 0 | 0 | 0 | 0 | 0 |
| ENSMUSG00000070953 | 1.993049299 | 1.852347734 | 1.907339155 | 1.864857905 | 1.875328588 | 2.117271296 | 1.792684387 |
| ENSMUSG00000070960 | 0 | 0 | 0 | 0 | 0 | 0 | 0 |
| ENSMUSG00000070972 | 1.960252568 | 1.766522298 | 1.777131114 | 1.890560606 | 1.853822071 | 2.1430148 | 1.747849621 |
| ENSMUSG00000070977 | 0 | 0 | 0 | 0 | 0 | 0 | 0 |
| ENSMUSG00000070979 | 0 | 0.396240625 | 0.396240625 | 0 | 0 | 0 | 0.224243824 |
| ENSMUSG00000070980 | 0 | 0 | 0 | 0 | 0 | 0 | 0 |
| ENSMUSG00000070983 | 0 | 0 | 0 | 0 | 0 | 0 | 0 |
| ENSMUSG00000070984 | 0 | 0 | 0 | 0 | 0 | 0 | 0 |
| ENSMUSG00000070985 | 0 | 0.25 | 0.396240625 | 0.25 | 0 | 0.301029996 | 0.224243824 |
| ENSMUSG00000070990 | 0 | 0 | 0 | 0 | 0 | 0 | 0.224243824 |
| ENSMUSG00000070995 | 0.289064826 | 0.580482024 | 0.646240625 | 0.580482024 | 0.489301084 | 0.602059991 | 0 |
| ENSMUSG00000070997 | 0 | 0.25 | 0.5 | 0.580482024 | 0.568060967 | 0 | 0.448487648 |
| ENSMUSG00000070999 | 0.289064826 | 0.5 | 0.396240625 | 0 | 0.387761935 | 0.477121255 | 0.224243824 |

| | | | | | | | |
|---|---|---|---|---|---|---|---|
| ENSMUSG00000071000 | 0 | 0 | 0 | 0 | 0 | 0 | 0 |
| ENSMUSG00000071001 | 0.960252568 | 1 | 0.976722649 | 0.580482024 | 0.812711509 | 0.477121255 | 0.629532004 |
| ENSMUSG00000071005 | 0 | 0 | 0 | 0 | 0 | 0 | 0 |
| ENSMUSG00000071014 | 1.54867984 | 1.688721876 | 1.834962501 | 1.458222504 | 1.625423019 | 1.544068044 | 1.734460864 |
| ENSMUSG00000071015 | 0 | 0 | 0 | 0 | 0 | 0 | 0 |
| ENSMUSG00000071019 | 0 | 0 | 0 | 0 | 0 | 0 | 0 |
| ENSMUSG00000071033 | 0 | 0 | 0 | 0 | 0 | 0 | 0 |
| ENSMUSG00000071035 | 0 | 0 | 0 | 0 | 0 | 0.301029996 | 0 |
| ENSMUSG00000071036 | 0 | 0 | 0 | 0 | 0 | 0.301029996 | 0.224243824 |
| ENSMUSG00000071037 | 1.249317394 | 1.130890489 | 1.098079356 | 1.201838731 | 1.283908311 | 1.146128036 | 1.168187745 |
| ENSMUSG00000071040 | 0 | 0 | 0 | 0 | 0 | 0 | 0 |
| ENSMUSG00000071041 | 0 | 0.25 | 0 | 0.396240625 | 0.244650542 | 0 | 0 |
| ENSMUSG00000071042 | 2.179305632 | 2.016522298 | 2.009729747 | 1.02186571 | 1 | 1.041392685 | 0.935079929 |
| ENSMUSG00000071047 | 0.45815691 | 0.396240625 | 0.25 | 0 | 0.244650542 | 0 | 0 |
| ENSMUSG00000071052 | 0 | 0 | 0 | 0 | 0 | 0 | 0.224243824 |
| ENSMUSG00000071054 | 2.63895461 | 2.457430684 | 2.501406137 | 2.412314044 | 2.430643515 | 2.706717782 | 2.364426535 |
| ENSMUSG00000071064 | 2.392336464 | 2.240000483 | 2.191217898 | 1.896240625 | 1.883916752 | 2.056904851 | 1.808770687 |
| ENSMUSG00000071065 | 0 | 0 | 0 | 0 | 0 | 0 | 0 |
| ENSMUSG00000071068 | 0 | 0 | 0 | 0 | 0 | 0 | 0 |
| ENSMUSG00000071072 | 3.224648428 | 2.94225283 | 2.883940344 | 2.961862226 | 2.883417309 | 3.336659823 | 2.656541617 |
| ENSMUSG00000071073 | 1 | 1.061981878 | 1.02186571 | 0.830482024 | 1.106691885 | 0.903089987 | 1.159323753 |
| ENSMUSG00000071074 | 2.146671932 | 2.036164561 | 2.105516192 | 1.892463902 | 2.014359518 | 2.146128036 | 1.928477344 |
| ENSMUSG00000071076 | 1.671187741 | 1.638647213 | 1.777131114 | 1.664552871 | 1.595992969 | 1.698970004 | 1.786277931 |
| ENSMUSG00000071078 | 1.181541736 | 1.201838731 | 1.146240625 | 1.146240625 | 1.176121988 | 1.255272505 | 1.089372236 |
| ENSMUSG00000071083 | 1.036286563 | 1 | 0.976722649 | 1.061981878 | 1.176121988 | 0.84509804 | 1.041356071 |
| ENSMUSG00000071084 | 0 | 0 | 0 | 0 | 0 | 0 | 0 |
| ENSMUSG00000071089 | 0 | 0.396240625 | 0 | 0 | 0.244650542 | 0 | 0 |
| ENSMUSG00000071102 | 0.671187741 | 0.25 | 0.25 | 0.25 | 0.632412477 | 0.602059991 | 0.448487648 |
| ENSMUSG00000071103 | 1 | 0.896240625 | 1 | 0.75 | 1.020174412 | 0.903089987 | 0.672731473 |
| ENSMUSG00000071104 | 0 | 0.396240625 | 0 | 0.25 | 0 | 0 | 0 |
| ENSMUSG00000071112 | 0.811507563 | 1.04248125 | 0.896240625 | 0.864857905 | 0.77552387 | 0.84509804 | 1 |
| ENSMUSG00000071113 | 0.578129653 | 0.25 | 0.396240625 | 0.701838731 | 0.387761935 | 0.477121255 | 0.355418052 |
| ENSMUSG00000071138 | 0 | 0 | 0 | 0 | 0 | 0 | 0 |
| ENSMUSG00000071141 | 0.289064826 | 0 | 0 | 0 | 0 | 0 | 0 |

| | | | | | | | |
|---|---|---|---|---|---|---|---|
| ENSMUSG00000071147 | 0 | 0 | 0 | 0 | 0 | 0 | 0 |
| ENSMUSG00000071149 | 0 | 0 | 0 | 0 | 0 | 0 | 0 |
| ENSMUSG00000071150 | 0 | 0 | 0 | 0 | 0 | 0 | 0 |
| ENSMUSG00000071151 | 0 | 0 | 0 | 0 | 0 | 0 | 0 |
| ENSMUSG00000071158 | 0 | 0 | 0 | 0 | 0 | 0 | 0 |
| ENSMUSG00000071165 | 0 | 0 | 0 | 0 | 0 | 0 | 0 |
| ENSMUSG00000071169 | 0 | 0 | 0 | 0 | 0 | 0 | 0 |
| ENSMUSG00000071172 | 3.155396777 | 2.877938163 | 2.869818758 | 2.929847205 | 2.897237431 | 3.325515663 | 2.677129724 |
| ENSMUSG00000071176 | 2.404271433 | 2.269871196 | 2.330482024 | 2.00835575 | 2.141887974 | 2.264817823 | 2.078019652 |
| ENSMUSG00000071177 | 0 | 0 | 0 | 0 | 0 | 0 | 0 |
| ENSMUSG00000071178 | 0 | 0 | 0 | 0 | 0 | 0 | 0 |
| ENSMUSG00000071179 | 0 | 0 | 0 | 0 | 0 | 0 | 0 |
| ENSMUSG00000071180 | 2.469490009 | 2.302972074 | 2.289336734 | 2.302972074 | 2.199086582 | 2.525044807 | 2.040177507 |
| ENSMUSG00000071181 | 0.289064826 | 0.580482024 | 0.646240625 | 0.396240625 | 0.387761935 | 0.301029996 | 0.355418052 |
| ENSMUSG00000071185 | 0 | 0 | 0 | 0 | 0 | 0 | 0 |
| ENSMUSG00000071192 | 0.45815691 | 0.580482024 | 0.646240625 | 0.580482024 | 0.489301084 | 0 | 0.629532004 |
| ENSMUSG00000071193 | 0 | 0 | 0 | 0 | 0 | 0 | 0 |
| ENSMUSG00000071195 | 0 | 0 | 0 | 0 | 0 | 0 | 0 |
| ENSMUSG00000071202 | 0.578129653 | 0.5 | 0.75 | 0.396240625 | 0.931471446 | 0.477121255 | 0.672731473 |
| ENSMUSG00000071203 | 0.578129653 | 0.580482024 | 0.396240625 | 0.701838731 | 0 | 0.301029996 | 0.448487648 |
| ENSMUSG00000071204 | 0 | 0 | 0 | 0 | 0 | 0 | 0 |
| ENSMUSG00000071207 | 0 | 0 | 0 | 0 | 0 | 0 | 0 |
| ENSMUSG00000071217 | 0 | 0 | 0 | 0 | 0 | 0 | 0 |
| ENSMUSG00000071226 | 0.91631382 | 0.580482024 | 0.580482024 | 1.160964047 | 1.136121934 | 1.113943352 | 1.100339912 |
| ENSMUSG00000071229 | 0 | 0 | 0 | 0 | 0 | 0 | 0 |
| ENSMUSG00000071230 | 0 | 0.25 | 0 | 0 | 0 | 0.477121255 | 0.448487648 |
| ENSMUSG00000071234 | 0 | 0 | 0 | 0 | 0 | 0 | 0 |
| ENSMUSG00000071235 | 0 | 0 | 0 | 0 | 0 | 0 | 0 |
| ENSMUSG00000071237 | 0 | 0 | 0 | 0 | 0 | 0 | 0 |
| ENSMUSG00000071252 | 1.389637216 | 1.282320754 | 1.114857905 | 1.561981878 | 1.467903253 | 1.602059991 | 1.40946263 |
| ENSMUSG00000071253 | 2.167941649 | 2.02186571 | 2.015173983 | 2.145300145 | 2.066583491 | 2.385606274 | 1.956418351 |
| ENSMUSG00000071256 | 0.91631382 | 1.188721876 | 1.458222504 | 1 | 1.212046813 | 1 | 1.265599895 |
| ENSMUSG00000071262 | 0 | 0 | 0 | 0.25 | 0.244650542 | 0 | 0 |
| ENSMUSG00000071265 | 0 | 0 | 0 | 0 | 0 | 0 | 0 |

| | | | | | | | |
|---|---|---|---|---|---|---|---|
| ENSMUSG00000071266 | 2.020182925 | 1.868933358 | 1.866901388 | 2.00835575 | 1.898867717 | 2.152288344 | 1.779742043 |
| ENSMUSG00000071267 | 2.185978293 | 2.012462137 | 2.036164561 | 1.978222334 | 1.940259189 | 2.170261715 | 1.781059827 |
| ENSMUSG00000071273 | 0 | 0 | 0 | 0 | 0 | 0 | 0 |
| ENSMUSG00000071275 | 0 | 0 | 0 | 0 | 0 | 0 | 0 |
| ENSMUSG00000071276 | 0 | 0 | 0 | 0 | 0 | 0 | 0 |
| ENSMUSG00000071280 | 0 | 0 | 0 | 0 | 0 | 0 | 0 |
| ENSMUSG00000071281 | 1.734388958 | 1.668106335 | 1.711372513 | 1.950224975 | 1.830919587 | 2.103803721 | 1.725216937 |
| ENSMUSG00000071291 | 1.307602651 | 1.26109853 | 1.26109853 | 1.431980114 | 1.366364104 | 1.397940009 | 1.307989563 |
| ENSMUSG00000071294 | 0 | 0 | 0 | 0 | 0 | 0 | 0 |
| ENSMUSG00000071295 | 0 | 0.396240625 | 0 | 0 | 0 | 0 | 0 |
| ENSMUSG00000071302 | 2.121692466 | 2.044977273 | 2.001406137 | 2.071350555 | 1.951645858 | 2.230448921 | 1.839824409 |
| ENSMUSG00000071303 | 0.45815691 | 0.5 | 0.5 | 0.396240625 | 0.489301084 | 0 | 0.448487648 |
| ENSMUSG00000071311 | 0 | 0 | 0 | 0 | 0 | 0 | 0 |
| ENSMUSG00000071317 | 0.960252568 | 1.080482024 | 0.951838731 | 0.92510993 | 0.877063019 | 0.698970004 | 0.916588299 |
| ENSMUSG00000071322 | 0 | 0 | 0 | 0 | 0 | 0 | 0 |
| ENSMUSG00000071324 | 0.747221736 | 1.17510993 | 1.061981878 | 1.302363341 | 1.327538614 | 1.176091259 | 1.23151414 |
| ENSMUSG00000071332 | 0 | 0.396240625 | 0.396240625 | 0.25 | 0 | 0 | 0 |
| ENSMUSG00000071335 | 0.289064826 | 0.5 | 0.580482024 | 0.25 | 0 | 0 | 0.224243824 |
| ENSMUSG00000071337 | 2.705574381 | 2.485628626 | 2.449820405 | 2.492166698 | 2.456697355 | 2.746634199 | 2.259963078 |
| ENSMUSG00000071340 | 0 | 0.25 | 0 | 0 | 0 | 0 | 0 |
| ENSMUSG00000071341 | 0 | 0 | 0 | 0 | 0 | 0 | 0 |
| ENSMUSG00000071342 | 0 | 0 | 0 | 0.25 | 0 | 0 | 0 |
| ENSMUSG00000071343 | 0.671187741 | 0.75 | 0.951838731 | 0.580482024 | 0.686820904 | 0.301029996 | 0.710836105 |
| ENSMUSG00000071347 | 0 | 0.25 | 0 | 0 | 0 | 0 | 0 |
| ENSMUSG00000071350 | 1.837744666 | 1.769203899 | 1.779735268 | 1.987091808 | 1.926994141 | 2.103803721 | 1.756477211 |
| ENSMUSG00000071356 | 0 | 0 | 0 | 0 | 0 | 0 | 0 |
| ENSMUSG00000071359 | 2.368815967 | 2.179419106 | 2.112802778 | 2.215271726 | 2.118236133 | 2.477121255 | 2.01501956 |
| ENSMUSG00000071361 | 0 | 0 | 0 | 0 | 0 | 0 | 0 |
| ENSMUSG00000071362 | 1.227926429 | 1.17510993 | 1.061981878 | 1.311981878 | 1.176121988 | 1.462397998 | 1.121219121 |
| ENSMUSG00000071369 | 1.181541736 | 1.061981878 | 0.79248125 | 1.53743678 | 1.456697355 | 1.579783597 | 1.413583902 |
| ENSMUSG00000071379 | 2.278594844 | 2.086073977 | 2.121960008 | 1.919870025 | 1.970729215 | 2.161368002 | 1.910475814 |
| ENSMUSG00000071392 | 0 | 0 | 0 | 0.25 | 0 | 0 | 0.224243824 |
| ENSMUSG00000071398 | 0 | 0.5 | 0.25 | 0.25 | 0.568060967 | 0.301029996 | 0.579661877 |
| ENSMUSG00000071414 | 0 | 0 | 0 | 0 | 0 | 0.477121255 | 0.224243824 |

| | | | | | | | |
|---|---|---|---|---|---|---|---|
| ENSMUSG00000071415 | 3.187380084 | 2.907111385 | 2.916556001 | 2.94225283 | 2.881211326 | 3.339053736 | 2.675725467 |
| ENSMUSG00000071419 | 0.45815691 | 0.646240625 | 0.976722649 | 0.396240625 | 0.489301084 | 0.84509804 | 0.775756176 |
| ENSMUSG00000071424 | 0 | 0 | 0.25 | 0 | 0.244650542 | 0 | 0 |
| ENSMUSG00000071428 | 0 | 0 | 0 | 0 | 0 | 0 | 0 |
| ENSMUSG00000071434 | 0 | 0 | 0 | 0 | 0 | 0 | 0 |
| ENSMUSG00000071451 | 1.445324132 | 1.5 | 1.575945187 | 1.380890489 | 1.584291694 | 1.633468456 | 1.543390198 |
| ENSMUSG00000071452 | 0 | 0 | 0 | 0 | 0 | 0 | 0 |
| ENSMUSG00000071454 | 1.876566388 | 1.763820609 | 1.854463129 | 1.905512955 | 1.853822071 | 2.10720997 | 1.852723744 |
| ENSMUSG00000071456 | 1.129344651 | 1.282320754 | 1.17510993 | 1.160964047 | 1.212046813 | 1.204119983 | 1.131174228 |
| ENSMUSG00000071466 | 0 | 0 | 0 | 0 | 0 | 0 | 0 |
| ENSMUSG00000071470 | 0 | 0.5 | 0.5 | 0.25 | 0.568060967 | 0.301029996 | 0.448487648 |
| ENSMUSG00000071471 | 0 | 0 | 0 | 0 | 0 | 0 | 0 |
| ENSMUSG00000071475 | 0.289064826 | 0.396240625 | 0.25 | 0.25 | 0.387761935 | 0.477121255 | 0.520678036 |
| ENSMUSG00000071477 | 1.91631382 | 1.87694866 | 1.866901388 | 1.779735268 | 1.802174723 | 1.954242509 | 1.756477211 |
| ENSMUSG00000071478 | 0 | 0 | 0 | 0 | 0 | 0 | 0 |
| ENSMUSG00000071481 | 0 | 0 | 0 | 0 | 0 | 0 | 0 |
| ENSMUSG00000071489 | 0.289064826 | 0 | 0 | 0 | 0 | 0 | 0 |
| ENSMUSG00000071490 | 0 | 0 | 0 | 0 | 0 | 0 | 0 |
| ENSMUSG00000071491 | 0 | 0 | 0 | 0 | 0 | 0 | 0 |
| ENSMUSG00000071493 | 0 | 0 | 0 | 0 | 0 | 0 | 0 |
| ENSMUSG00000071494 | 0.289064826 | 0 | 0 | 0 | 0 | 0 | 0 |
| ENSMUSG00000071497 | 0 | 0 | 0 | 0 | 0 | 0 | 0 |
| ENSMUSG00000071506 | 0 | 0.25 | 0 | 0 | 0.244650542 | 0.301029996 | 0 |
| ENSMUSG00000071510 | 0 | 0 | 0 | 0 | 0 | 0 | 0 |
| ENSMUSG00000071516 | 0 | 0 | 0 | 0 | 0 | 0 | 0 |
| ENSMUSG00000071517 | 0 | 0 | 0 | 0 | 0 | 0 | 0 |
| ENSMUSG00000071519 | 0 | 0 | 0 | 0 | 0 | 0 | 0 |
| ENSMUSG00000071521 | 0 | 0 | 0 | 0 | 0 | 0 | 0 |
| ENSMUSG00000071522 | 0 | 0 | 0 | 0 | 0 | 0 | 0 |
| ENSMUSG00000071528 | 2.587501561 | 2.3184489 | 2.357090043 | 2.466819685 | 2.360255934 | 2.783188691 | 2.244955885 |
| ENSMUSG00000071531 | 0 | 0 | 0.25 | 0 | 0 | 0.477121255 | 0.355418052 |
| ENSMUSG00000071532 | 0 | 0 | 0 | 0 | 0 | 0 | 0 |
| ENSMUSG00000071533 | 2.716227929 | 2.491085217 | 2.495034894 | 2.601816941 | 2.503660602 | 2.886490725 | 2.327511498 |
| ENSMUSG00000071537 | 0.867194479 | 0.951838731 | 1.02186571 | 1.302363341 | 1.234113756 | 1.431363764 | 1.238624656 |

| | | | | | | | |
|---|---|---|---|---|---|---|---|
| ENSMUSG00000071540 | 0 | 0 | 0 | 0 | 0 | 0 | 0 |
| ENSMUSG00000071547 | 2.553277818 | 2.435366747 | 2.478222334 | 2.391037372 | 2.506289166 | 2.730782276 | 2.440194096 |
| ENSMUSG00000071550 | 0.289064826 | 0 | 0 | 0.580482024 | 0.387761935 | 0 | 0 |
| ENSMUSG00000071551 | 0 | 0 | 0 | 0 | 0 | 0 | 0 |
| ENSMUSG00000071552 | 0.578129653 | 0.75 | 0.25 | 1.146240625 | 0.846351821 | 0.84509804 | 0.896975297 |
| ENSMUSG00000071553 | 0 | 0.646240625 | 0.580482024 | 0.5 | 0.489301084 | 0.698970004 | 0.448487648 |
| ENSMUSG00000071561 | 0 | 0 | 0 | 0 | 0 | 0 | 0 |
| ENSMUSG00000071562 | 0 | 0 | 0 | 0 | 0 | 0 | 0 |
| ENSMUSG00000071567 | 0 | 0 | 0 | 0 | 0 | 0 | 0 |
| ENSMUSG00000071568 | 0 | 0 | 0 | 0 | 0 | 0 | 0 |
| ENSMUSG00000071573 | 1.249317394 | 1.302363341 | 1.29248125 | 1.114857905 | 1.136121934 | 1.079181246 | 1.078019652 |
| ENSMUSG00000071586 | 0 | 0 | 0 | 0 | 0 | 0.301029996 | 0 |
| ENSMUSG00000071591 | 0 | 0 | 0 | 0 | 0 | 0 | 0 |
| ENSMUSG00000071604 | 1.418409478 | 1.29248125 | 1.388647213 | 1.146240625 | 1.020174412 | 1.079181246 | 1.078019652 |
| ENSMUSG00000071613 | 0 | 0 | 0 | 0 | 0 | 0 | 0 |
| ENSMUSG00000071629 | 0 | 0 | 0 | 0 | 0 | 0 | 0 |
| ENSMUSG00000071632 | 1.45815691 | 1.330482024 | 1.42510993 | 1.418106335 | 1.456697355 | 1.544068044 | 1.369798885 |
| ENSMUSG00000071633 | 0 | 0 | 0.25 | 0 | 0 | 0 | 0 |
| ENSMUSG00000071636 | 0 | 0 | 0.25 | 0.25 | 0.489301084 | 0.477121255 | 0.355418052 |
| ENSMUSG00000071637 | 1.129344651 | 1.26109853 | 1.51109853 | 0.75 | 0.812711509 | 0.84509804 | 1.193409508 |
| ENSMUSG00000071640 | 0.289064826 | 0 | 0 | 0 | 0 | 0 | 0 |
| ENSMUSG00000071644 | 3.409350774 | 3.097609422 | 3.092399337 | 3.025329789 | 2.964587954 | 3.427972714 | 2.766885432 |
| ENSMUSG00000071645 | 2.177057474 | 2.145300145 | 2.129909063 | 1.978222334 | 2.09661982 | 2.201397124 | 1.952571511 |
| ENSMUSG00000071646 | 3.054274575 | 2.78199858 | 2.795288064 | 2.757666784 | 2.749043266 | 3.11991541 | 2.571311171 |
| ENSMUSG00000071647 | 2.35727108 | 2.219895812 | 2.247880462 | 2.189555804 | 2.305627392 | 2.424881637 | 2.192673409 |
| ENSMUSG00000071648 | 1.269664473 | 1.214495249 | 1.201838731 | 1.282320754 | 1.394615667 | 1.447158031 | 1.441050184 |
| ENSMUSG00000071649 | 2.302628051 | 2.219895812 | 2.306603048 | 1.907339155 | 2.112954895 | 2.212187604 | 2.069505596 |
| ENSMUSG00000071650 | 3.351646281 | 3.040597832 | 3.030706999 | 3.104529035 | 3.055709431 | 3.538573734 | 2.828099529 |
| ENSMUSG00000071652 | 2.612039956 | 2.398114259 | 2.426839783 | 2.227473271 | 2.288791717 | 2.539076099 | 2.192673409 |
| ENSMUSG00000071653 | 0.671187741 | 0.646240625 | 0.396240625 | 0.701838731 | 0.77552387 | 0.301029996 | 0.224243824 |
| ENSMUSG00000071654 | 1.623015126 | 1.580482024 | 1.646240625 | 1.438721876 | 1.546663136 | 1.612783857 | 1.607811402 |
| ENSMUSG00000071655 | 2.63895461 | 2.512462137 | 2.547147211 | 2.345926073 | 2.440245937 | 2.698970004 | 2.354748941 |
| ENSMUSG00000071656 | 1.432083896 | 1.380890489 | 1.311981878 | 1.201838731 | 1.283908311 | 1.113943352 | 1.265599895 |
| ENSMUSG00000071657 | 1.777675583 | 1.62694866 | 1.681980114 | 1.552363341 | 1.692703642 | 1.681241237 | 1.603286636 |

| | | | | | | | |
|---|---|---|---|---|---|---|---|
| ENSMUSG00000071658 | 0.811507563 | 1.061981878 | 0.976722649 | 0.951838731 | 0.905314583 | 0.602059991 | 1 |
| ENSMUSG00000071659 | 3.174591036 | 2.897061538 | 2.888407323 | 2.963327389 | 2.882316041 | 3.330413773 | 2.692894471 |
| ENSMUSG00000071660 | 2.692026894 | 2.47027849 | 2.469129237 | 2.459800947 | 2.355381384 | 2.747411808 | 2.185972044 |
| ENSMUSG00000071661 | 1.404271433 | 1.403677461 | 1.311981878 | 1.17510993 | 1.283908311 | 1.255272505 | 1.20139795 |
| ENSMUSG00000071662 | 2.687411469 | 2.438304187 | 2.455443495 | 2.527457664 | 2.439543536 | 2.806179974 | 2.285835109 |
| ENSMUSG00000071665 | 0 | 0.25 | 0 | 0 | 0 | 0 | 0 |
| ENSMUSG00000071669 | 1.036286563 | 1.201838731 | 1.282320754 | 1.188721876 | 1.244650542 | 1.146128036 | 1.20139795 |
| ENSMUSG00000071671 | 0 | 0.25 | 0.25 | 0.5 | 0.387761935 | 0.477121255 | 0 |
| ENSMUSG00000071679 | 0 | 0 | 0 | 0 | 0 | 0 | 0 |
| ENSMUSG00000071680 | 0 | 0.25 | 0 | 0 | 0 | 0 | 0 |
| ENSMUSG00000071686 | 0 | 0 | 0 | 0 | 0 | 0 | 0 |
| ENSMUSG00000071691 | 0.45815691 | 0.646240625 | 0.25 | 0.5 | 0.931471446 | 0.602059991 | 0.579661877 |
| ENSMUSG00000071708 | 2.37447073 | 2.240000483 | 2.179419106 | 2.416777886 | 2.318738698 | 2.653212514 | 2.158915016 |
| ENSMUSG00000071711 | 1.663535556 | 1.67510993 | 1.834962501 | 1.575945187 | 1.784632921 | 1.812913357 | 1.889537832 |
| ENSMUSG00000071713 | 0 | 0.25 | 0.25 | 0 | 0 | 0 | 0.224243824 |
| ENSMUSG00000071714 | 0.289064826 | 0 | 0.25 | 0.25 | 0.387761935 | 0 | 0.224243824 |
| ENSMUSG00000071715 | 0 | 0 | 0 | 0 | 0.244650542 | 0 | 0.355418052 |
| ENSMUSG00000071716 | 0 | 0 | 0 | 0 | 0 | 0 | 0 |
| ENSMUSG00000071719 | 0 | 0 | 0 | 0.25 | 0 | 0 | 0 |
| ENSMUSG00000071721 | 0 | 0 | 0 | 0 | 0 | 0 | 0 |
| ENSMUSG00000071722 | 1.777675583 | 1.638647213 | 1.552363341 | 1.711372513 | 1.715300549 | 1.86923172 | 1.537859885 |
| ENSMUSG00000071723 | 1.885732304 | 1.711372513 | 1.711372513 | 1.909156155 | 1.816839698 | 2.029383778 | 1.699191618 |
| ENSMUSG00000071724 | 0.747221736 | 0.830482024 | 1.061981878 | 0.896240625 | 1.020174412 | 0.903089987 | 1.054044578 |
| ENSMUSG00000071726 | 0 | 0 | 0 | 0 | 0 | 0 | 0 |
| ENSMUSG00000071738 | 0 | 0 | 0 | 0 | 0 | 0 | 0 |
| ENSMUSG00000071745 | 0 | 0 | 0 | 0 | 0 | 0 | 0 |
| ENSMUSG00000071748 | 2.194712275 | 2.02053726 | 2.059601185 | 2.278435542 | 2.235369829 | 2.496929648 | 2.03601839 |
| ENSMUSG00000071749 | 0 | 0 | 0 | 0 | 0 | 0 | 0 |
| ENSMUSG00000071750 | 0 | 0 | 0 | 0 | 0 | 0 | 0 |
| ENSMUSG00000071753 | 0 | 0 | 0 | 0 | 0 | 0 | 0 |
| ENSMUSG00000071757 | 1.944805158 | 1.87892496 | 1.880890489 | 0.92510993 | 0.931471446 | 0.477121255 | 0.876096088 |
| ENSMUSG00000071763 | 0 | 0 | 0 | 0 | 0 | 0 | 0 |
| ENSMUSG00000071764 | 0 | 0 | 0 | 0 | 0 | 0 | 0 |
| ENSMUSG00000071766 | 0 | 0 | 0 | 0 | 0 | 0 | 0 |

| | | | | | | | |
|---|---|---|---|---|---|---|---|
| ENSMUSG00000071767 | 0 | 0 | 0 | 0 | 0 | 0 | 0 |
| ENSMUSG00000071769 | 0 | 0 | 0 | 0 | 0 | 0 | 0 |
| ENSMUSG00000071770 | 0 | 0 | 0 | 0 | 0 | 0 | 0 |
| ENSMUSG00000071771 | 0 | 0 | 0 | 0 | 0 | 0 | 0 |
| ENSMUSG00000071772 | 0 | 0 | 0 | 0 | 0 | 0 | 0 |
| ENSMUSG00000071773 | 0 | 0 | 0 | 0 | 0 | 0 | 0 |
| ENSMUSG00000071788 | 0 | 0 | 0 | 0 | 0 | 0 | 0 |
| ENSMUSG00000071793 | 1.771760131 | 1.705044741 | 1.664552871 | 1.638647213 | 1.695608647 | 1.72427587 | 1.670888075 |
| ENSMUSG00000071796 | 1.91631382 | 1.828220739 | 1.816696635 | 2.001406137 | 1.916320788 | 2.178976947 | 1.774416565 |
| ENSMUSG00000071815 | 0 | 0 | 0 | 0 | 0 | 0 | 0 |
| ENSMUSG00000071816 | 0 | 0 | 0 | 0 | 0 | 0 | 0 |
| ENSMUSG00000071818 | 0 | 0 | 0 | 0 | 0 | 0 | 0 |
| ENSMUSG00000071847 | 2.17479713 | 2.039967834 | 2.052363341 | 1.606566189 | 1.655839964 | 1.806179974 | 1.603286636 |
| ENSMUSG00000071855 | 1.783508299 | 1.671625132 | 1.708222504 | 1.916333979 | 1.793512788 | 2.021189299 | 1.680001789 |
| ENSMUSG00000071856 | 2.310886388 | 2.153677461 | 2.17510993 | 2.093759858 | 2.097547432 | 2.303196057 | 1.944737824 |
| ENSMUSG00000071858 | 0 | 0 | 0 | 0 | 0 | 0 | 0 |
| ENSMUSG00000071860 | 0 | 0 | 0.25 | 0 | 0 | 0 | 0 |
| ENSMUSG00000071862 | 0 | 0.5 | 0.396240625 | 0.396240625 | 0.244650542 | 0 | 0.520678036 |
| ENSMUSG00000071866 | 3.428510906 | 3.13804719 | 3.121708755 | 3.16107674 | 3.087319309 | 3.611723308 | 2.893658022 |
| ENSMUSG00000071867 | 0 | 0 | 0 | 0 | 0 | 0.301029996 | 0 |
| ENSMUSG00000071890 | 0 | 0 | 0 | 0 | 0 | 0 | 0 |
| ENSMUSG00000071893 | 0.578129653 | 0.25 | 0.25 | 0.25 | 0.489301084 | 0.301029996 | 0.520678036 |
| ENSMUSG00000071909 | 0 | 0 | 0 | 0 | 0 | 0 | 0 |
| ENSMUSG00000071937 | 0 | 0 | 0 | 0 | 0 | 0 | 0 |
| ENSMUSG00000071952 | 0 | 0 | 0 | 0 | 0 | 0 | 0 |
| ENSMUSG00000071984 | 0 | 0 | 0 | 0 | 0.489301084 | 0 | 0 |
| ENSMUSG00000072049 | 0 | 0 | 0 | 0 | 0 | 0 | 0 |
| ENSMUSG00000072066 | 1.932832114 | 1.837182039 | 1.807204673 | 1.874961472 | 1.816839698 | 1.963787827 | 1.61448206 |
| ENSMUSG00000072068 | 0 | 0 | 0 | 0 | 0 | 0 | 0 |
| ENSMUSG00000072082 | 2.626865868 | 2.417664062 | 2.431980114 | 2.345926073 | 2.404458919 | 2.662757832 | 2.286662868 |
| ENSMUSG00000072100 | 0 | 0 | 0 | 0 | 0 | 0 | 0 |
| ENSMUSG00000072109 | 0.45815691 | 0.5 | 0 | 0.25 | 0.244650542 | 0 | 0.579661877 |
| ENSMUSG00000072115 | 0 | 0.396240625 | 0 | 0.79248125 | 0.733951626 | 0.602059991 | 0.629532004 |
| ENSMUSG00000072145 | 0 | 0 | 0 | 0 | 0 | 0 | 0 |

| | | | | | | | |
|---|---|---|---|---|---|---|---|
| ENSMUSG00000072188 | 0 | 0 | 0 | 0 | 0 | 0 | 0 |
| ENSMUSG00000072214 | 2.243591459 | 2.101285366 | 2.08272922 | 1.912762923 | 2.014359518 | 2.190331698 | 1.927642465 |
| ENSMUSG00000072235 | 3.190764944 | 2.918989258 | 2.921625132 | 2.911302021 | 2.925400257 | 3.335658452 | 2.77153595 |
| ENSMUSG00000072244 | 1.269664473 | 1.130890489 | 1.321350555 | 1.146240625 | 1.188507684 | 1.041392685 | 1.20139795 |
| ENSMUSG00000072249 | 0 | 0 | 0 | 0 | 0 | 0 | 0 |
| ENSMUSG00000072255 | 0 | 0.5 | 0.396240625 | 0.396240625 | 0.387761935 | 0.477121255 | 0.355418052 |
| ENSMUSG00000072258 | 1.948720984 | 1.782320754 | 1.681980114 | 1.823655187 | 1.828942237 | 2.056904851 | 1.69749337 |
| ENSMUSG00000072259 | 0 | 0 | 0 | 0 | 0 | 0 | 0 |
| ENSMUSG00000072294 | 0.91631382 | 1.201838731 | 1.146240625 | 0 | 0.244650542 | 0.477121255 | 0.520678036 |
| ENSMUSG00000072295 | 0 | 0 | 0 | 0 | 0 | 0 | 0 |
| ENSMUSG00000072324 | 0.289064826 | 0.25 | 0 | 0 | 0.387761935 | 0.477121255 | 0.355418052 |
| ENSMUSG00000072381 | 0 | 0 | 0 | 0 | 0.244650542 | 0 | 0 |
| ENSMUSG00000072387 | 0 | 0 | 0 | 0 | 0 | 0 | 0 |
| ENSMUSG00000072407 | 0 | 0 | 0 | 0 | 0 | 0 | 0 |
| ENSMUSG00000072419 | 0 | 0 | 0 | 0.25 | 0 | 0.301029996 | 0 |
| ENSMUSG00000072421 | 0 | 0.396240625 | 0.646240625 | 0.25 | 0.244650542 | 0.698970004 | 0.520678036 |
| ENSMUSG00000072423 | 0.289064826 | 0.25 | 0.25 | 0.580482024 | 0.568060967 | 0.301029996 | 0.355418052 |
| ENSMUSG00000072432 | 0 | 0 | 0 | 0 | 0 | 0 | 0 |
| ENSMUSG00000072437 | 1.816888701 | 1.843759858 | 1.744319981 | 1.868933358 | 1.756568651 | 1.929418926 | 1.645916034 |
| ENSMUSG00000072460 | 0 | 0 | 0 | 0.25 | 0 | 0 | 0 |
| ENSMUSG00000072473 | 0 | 0 | 0 | 0 | 0 | 0 | 0 |
| ENSMUSG00000072476 | 0.289064826 | 0.25 | 0.25 | 0.25 | 0 | 0.301029996 | 0 |
| ENSMUSG00000072477 | 0 | 0 | 0 | 0 | 0 | 0 | 0 |
| ENSMUSG00000072479 | 0 | 0 | 0 | 0 | 0 | 0 | 0 |
| ENSMUSG00000072487 | 0 | 0 | 0.25 | 0 | 0 | 0 | 0 |
| ENSMUSG00000072492 | 0 | 0.580482024 | 0.5 | 0.701838731 | 0.632412477 | 0.301029996 | 0.629532004 |
| ENSMUSG00000072494 | 0.578129653 | 0.646240625 | 0.79248125 | 0.951838731 | 0.733951626 | 0.602059991 | 0.520678036 |
| ENSMUSG00000072501 | 2.840509532 | 2.590211771 | 2.560197833 | 2.635274154 | 2.568305143 | 2.937016107 | 2.355195169 |
| ENSMUSG00000072511 | 0 | 0 | 0 | 0 | 0 | 0 | 0 |
| ENSMUSG00000072541 | 0 | 0 | 0 | 0 | 0 | 0 | 0 |
| ENSMUSG00000072553 | 0 | 0 | 0 | 0 | 0.244650542 | 0 | 0 |
| ENSMUSG00000072568 | 2.302628051 | 2.149978211 | 2.124961472 | 2.03743678 | 1.98503142 | 2.255272505 | 1.86715039 |
| ENSMUSG00000072571 | 0.747221736 | 0.5 | 0.5 | 0.646240625 | 0.733951626 | 0.477121255 | 0.629532004 |
| ENSMUSG00000072572 | 0.578129653 | 0.25 | 0.396240625 | 0.396240625 | 0.632412477 | 0.301029996 | 0.448487648 |

| | | | | | | | |
|---|---|---|---|---|---|---|---|
| ENSMUSG00000072573 | 0 | 0.396240625 | 0.25 | 0.5 | 0.632412477 | 0.301029996 | 0.355418052 |
| ENSMUSG00000072574 | 0 | 0 | 0 | 0 | 0 | 0 | 0 |
| ENSMUSG00000072575 | 0 | 0 | 0 | 0 | 0 | 0 | 0 |
| ENSMUSG00000072576 | 0 | 0 | 0.25 | 0 | 0 | 0 | 0 |
| ENSMUSG00000072582 | 2.409468165 | 2.261780978 | 2.240724001 | 2.293106877 | 2.224547964 | 2.536558443 | 2.078544414 |
| ENSMUSG00000072584 | 0 | 0 | 0 | 0 | 0 | 0 | 0 |
| ENSMUSG00000072589 | 0.671187741 | 0.25 | 0.25 | 0.25 | 0.244650542 | 0 | 0 |
| ENSMUSG00000072591 | 0 | 0 | 0 | 0 | 0.244650542 | 0 | 0.224243824 |
| ENSMUSG00000072592 | 0 | 0.25 | 0 | 0 | 0 | 0 | 0 |
| ENSMUSG00000072594 | 0 | 0 | 0 | 0 | 0 | 0 | 0 |
| ENSMUSG00000072595 | 0 | 0 | 0 | 0 | 0 | 0 | 0 |
| ENSMUSG00000072596 | 0 | 0 | 0 | 0 | 0 | 0 | 0 |
| ENSMUSG00000072598 | 0 | 0 | 0 | 0 | 0 | 0 | 0 |
| ENSMUSG00000072599 | 0 | 0 | 0 | 0 | 0 | 0 | 0 |
| ENSMUSG00000072600 | 0 | 0 | 0 | 0 | 0 | 0 | 0 |
| ENSMUSG00000072601 | 0 | 0 | 0 | 0 | 0 | 0 | 0 |
| ENSMUSG00000072605 | 0 | 0 | 0 | 0 | 0 | 0 | 0 |
| ENSMUSG00000072612 | 0 | 0 | 0 | 0 | 0 | 0 | 0.224243824 |
| ENSMUSG00000072613 | 0 | 0 | 0 | 0 | 0 | 0 | 0 |
| ENSMUSG00000072616 | 0 | 0 | 0 | 0 | 0 | 0 | 0 |
| ENSMUSG00000072618 | 0 | 0 | 0 | 0.25 | 0 | 0.477121255 | 0 |
| ENSMUSG00000072620 | 0 | 0.396240625 | 0.5 | 0 | 0.244650542 | 0 | 0.355418052 |
| ENSMUSG00000072621 | 0 | 0.25 | 0.5 | 0.5 | 0.244650542 | 0 | 0.224243824 |
| ENSMUSG00000072623 | 2.262379879 | 2.123963757 | 2.090485943 | 2.049918086 | 1.962676635 | 2.301029996 | 1.865128412 |
| ENSMUSG00000072624 | 0 | 0 | 0 | 0.25 | 0 | 0 | 0 |
| ENSMUSG00000072625 | 0 | 0 | 0 | 0 | 0 | 0 | 0 |
| ENSMUSG00000072640 | 1.827447047 | 1.634789703 | 1.618933358 | 1.755591953 | 1.655839964 | 1.929418926 | 1.579661877 |
| ENSMUSG00000072647 | 1.100572389 | 1.188721876 | 1 | 1.160964047 | 1.121713561 | 1.079181246 | 1.185218806 |
| ENSMUSG00000072653 | 0 | 0.396240625 | 0.396240625 | 0.5 | 0.489301084 | 0.301029996 | 0.520678036 |
| ENSMUSG00000072660 | 0.289064826 | 0.75 | 0.396240625 | 0.580482024 | 0.244650542 | 0.698970004 | 0.579661877 |
| ENSMUSG00000072662 | 0 | 0.396240625 | 0.5 | 0.25 | 0.244650542 | 0 | 0.224243824 |
| ENSMUSG00000072663 | 0.289064826 | 0 | 0.25 | 0 | 0 | 0 | 0 |
| ENSMUSG00000072664 | 0.45815691 | 0.396240625 | 0.396240625 | 0.25 | 0.387761935 | 0.301029996 | 0.448487648 |
| ENSMUSG00000072672 | 0 | 0 | 0 | 0 | 0 | 0 | 0.224243824 |

| | | | | | | | |
|---|---|---|---|---|---|---|---|
| ENSMUSG00000072673 | 0 | 0 | 0.5 | 0.25 | 0.244650542 | 0.301029996 | 0 |
| ENSMUSG00000072674 | 0 | 0 | 0 | 0 | 0 | 0 | 0 |
| ENSMUSG00000072675 | 0 | 0 | 0 | 0 | 0 | 0 | 0 |
| ENSMUSG00000072676 | 0 | 0 | 0 | 0 | 0 | 0 | 0 |
| ENSMUSG00000072677 | 0 | 0 | 0 | 0 | 0 | 0 | 0 |
| ENSMUSG00000072679 | 0 | 0 | 0 | 0.25 | 0 | 0.301029996 | 0 |
| ENSMUSG00000072680 | 0 | 0 | 0 | 0 | 0 | 0 | 0 |
| ENSMUSG00000072683 | 0 | 0 | 0 | 0.25 | 0 | 0 | 0 |
| ENSMUSG00000072686 | 0 | 0 | 0 | 0 | 0 | 0 | 0 |
| ENSMUSG00000072688 | 0.45815691 | 0.396240625 | 0.5 | 0.396240625 | 0.489301084 | 0.301029996 | 0.672731473 |
| ENSMUSG00000072692 | 1.358731791 | 1.42510993 | 1.130890489 | 1.214495249 | 1.302012594 | 1.230448921 | 1.296434211 |
| ENSMUSG00000072693 | 0 | 0 | 0 | 0 | 0 | 0 | 0 |
| ENSMUSG00000072694 | 1.639698646 | 1.622963274 | 1.54248125 | 1.5 | 1.489301084 | 1.681241237 | 1.42167221 |
| ENSMUSG00000072704 | 2.497590902 | 2.353935442 | 2.354990044 | 2.347004321 | 2.257246745 | 2.580924976 | 2.098373247 |
| ENSMUSG00000072707 | 0 | 0 | 0 | 0 | 0 | 0 | 0 |
| ENSMUSG00000072708 | 0 | 0 | 0 | 0 | 0 | 0 | 0 |
| ENSMUSG00000072709 | 0 | 0 | 0 | 0 | 0 | 0 | 0 |
| ENSMUSG00000072714 | 0 | 0 | 0 | 0 | 0 | 0 | 0 |
| ENSMUSG00000072717 | 0 | 0 | 0 | 0 | 0 | 0 | 0 |
| ENSMUSG00000072718 | 0 | 0 | 0 | 0 | 0 | 0 | 0 |
| ENSMUSG00000072720 | 0 | 0 | 0 | 0 | 0 | 0 | 0 |
| ENSMUSG00000072721 | 0 | 0 | 0 | 0 | 0 | 0 | 0 |
| ENSMUSG00000072722 | 0 | 0 | 0 | 0 | 0 | 0 | 0 |
| ENSMUSG00000072723 | 0 | 0 | 0 | 0 | 0 | 0 | 0 |
| ENSMUSG00000072726 | 0 | 0 | 0 | 0 | 0 | 0 | 0 |
| ENSMUSG00000072731 | 0 | 0 | 0 | 0 | 0 | 0 | 0 |
| ENSMUSG00000072738 | 0 | 0 | 0 | 0 | 0 | 0 | 0 |
| ENSMUSG00000072739 | 0 | 0 | 0 | 0 | 0 | 0 | 0 |
| ENSMUSG00000072753 | 0 | 0 | 0 | 0 | 0 | 0 | 0 |
| ENSMUSG00000072754 | 0 | 0.25 | 0 | 0 | 0 | 0 | 0 |
| ENSMUSG00000072761 | 1.269664473 | 1.080482024 | 1.27186571 | 1.311981878 | 1.136121934 | 1.176091259 | 1.121219121 |
| ENSMUSG00000072762 | 1.783508299 | 1.622963274 | 1.705044741 | 1.79745614 | 1.665423072 | 1.763427994 | 1.598697688 |
| ENSMUSG00000072763 | 1.568542279 | 1.630890489 | 1.571350555 | 1.470660762 | 1.387761935 | 1.568201724 | 1.38802993 |
| ENSMUSG00000072768 | 0 | 0 | 0 | 0 | 0 | 0 | 0 |

| | | | | | | | |
|---|---|---|---|---|---|---|---|
| ENSMUSG00000072769 | 0 | 0.396240625 | 0.25 | 0.75 | 0.632412477 | 0.602059991 | 0.579661877 |
| ENSMUSG00000072770 | 1.069666964 | 0.976722649 | 0.976722649 | 0.830482024 | 0.812711509 | 0.477121255 | 0.74492186 |
| ENSMUSG00000072772 | 1.129344651 | 1.201838731 | 1.114857905 | 1.04248125 | 1.091002363 | 0.954242509 | 1.089372236 |
| ENSMUSG00000072774 | 1.37447073 | 1.388647213 | 1.356566189 | 1.201838731 | 1.039257769 | 1.278753601 | 1.140832123 |
| ENSMUSG00000072778 | 0 | 0 | 0 | 0 | 0 | 0 | 0 |
| ENSMUSG00000072780 | 0 | 0 | 0 | 0 | 0 | 0 | 0 |
| ENSMUSG00000072789 | 0.578129653 | 0.830482024 | 0.701838731 | 0.5 | 0.877063019 | 0.477121255 | 0.579661877 |
| ENSMUSG00000072791 | 0 | 0 | 0 | 0 | 0 | 0 | 0 |
| ENSMUSG00000072809 | 0.45815691 | 0.396240625 | 0.396240625 | 0.580482024 | 0.489301084 | 0.477121255 | 0.355418052 |
| ENSMUSG00000072812 | 2.635965117 | 2.445339928 | 2.485261902 | 2.145300145 | 2.150762767 | 2.359835482 | 2.028149525 |
| ENSMUSG00000072813 | 0 | 0 | 0 | 0 | 0 | 0 | 0 |
| ENSMUSG00000072814 | 0 | 0 | 0 | 0 | 0 | 0 | 0 |
| ENSMUSG00000072821 | 0 | 0 | 0 | 0 | 0 | 0 | 0 |
| ENSMUSG00000072822 | 0 | 0 | 0 | 0 | 0 | 0 | 0 |
| ENSMUSG00000072825 | 2.425923778 | 2.342671852 | 2.443696765 | 2.04248125 | 2.261929482 | 2.29666519 | 2.230859913 |
| ENSMUSG00000072834 | 0 | 0 | 0 | 0 | 0 | 0 | 0 |
| ENSMUSG00000072837 | 0 | 0 | 0 | 0 | 0 | 0 | 0 |
| ENSMUSG00000072844 | 0.578129653 | 0.5 | 0.701838731 | 0.864857905 | 0.489301084 | 0.602059991 | 0.579661877 |
| ENSMUSG00000072845 | 0 | 0 | 0 | 0.25 | 0.244650542 | 0.301029996 | 0 |
| ENSMUSG00000072847 | 1.205378646 | 1.238549078 | 1.238549078 | 1.348079356 | 1.358933174 | 1.414973348 | 1.265599895 |
| ENSMUSG00000072849 | 0 | 0 | 0 | 0 | 0 | 0 | 0 |
| ENSMUSG00000072852 | 0 | 0 | 0 | 0.25 | 0.244650542 | 0 | 0.224243824 |
| ENSMUSG00000072872 | 2.551444705 | 2.359177886 | 2.343216265 | 2.248588359 | 2.146756411 | 2.522444234 | 2.02383181 |
| ENSMUSG00000072874 | 0 | 0 | 0 | 0 | 0 | 0 | 0 |
| ENSMUSG00000072875 | 0 | 0.25 | 0 | 0.580482024 | 0.77552387 | 0.301029996 | 0.520678036 |
| ENSMUSG00000072878 | 0 | 0 | 0 | 0 | 0 | 0 | 0 |
| ENSMUSG00000072884 | 0.289064826 | 0.5 | 0.396240625 | 0.25 | 0.244650542 | 0 | 0 |
| ENSMUSG00000072888 | 0 | 0 | 0 | 0 | 0 | 0 | 0 |
| ENSMUSG00000072889 | 2.405576699 | 2.241446071 | 2.200224975 | 2.273189285 | 2.227765453 | 2.51851394 | 2.055736903 |
| ENSMUSG00000072893 | 1.37447073 | 1.380890489 | 1.311981878 | 1.42510993 | 1.427019704 | 1.531478917 | 1.401059159 |
| ENSMUSG00000072898 | 0 | 0 | 0 | 0 | 0 | 0 | 0 |
| ENSMUSG00000072900 | 0 | 0 | 0 | 0 | 0 | 0 | 0 |
| ENSMUSG00000072902 | 0 | 0 | 0 | 0 | 0 | 0 | 0 |
| ENSMUSG00000072905 | 0 | 0 | 0 | 0 | 0 | 0 | 0.224243824 |

| | | | | | | | |
|---|---|---|---|---|---|---|---|
| ENSMUSG00000072915 | 1.759671389 | 1.54248125 | 1.602347734 | 1.779735268 | 1.686820904 | 1.954242509 | 1.661510159 |
| ENSMUSG00000072919 | 0 | 0.79248125 | 0.25 | 0.396240625 | 0.387761935 | 0.477121255 | 0.579661877 |
| ENSMUSG00000072923 | 0 | 0 | 0 | 0 | 0 | 0 | 0 |
| ENSMUSG00000072929 | 0 | 0 | 0 | 0 | 0 | 0 | 0 |
| ENSMUSG00000072930 | 0 | 0 | 0 | 0 | 0 | 0 | 0 |
| ENSMUSG00000072931 | 0 | 0 | 0 | 0 | 0 | 0 | 0 |
| ENSMUSG00000072934 | 0.289064826 | 0.396240625 | 0.25 | 0.396240625 | 0.244650542 | 0 | 0.355418052 |
| ENSMUSG00000072940 | 0 | 0 | 0 | 0 | 0 | 0 | 0 |
| ENSMUSG00000072941 | 1.771760131 | 1.630890489 | 1.646240625 | 1.160964047 | 1.264824954 | 1.255272505 | 1.185218806 |
| ENSMUSG00000072944 | 0.289064826 | 0.396240625 | 0.5 | 0.79248125 | 0.568060967 | 0.698970004 | 0.579661877 |
| ENSMUSG00000072945 | 0 | 0 | 0 | 0 | 0 | 0 | 0 |
| ENSMUSG00000072946 | 2.131864492 | 2.002806814 | 1.994319981 | 2.02186571 | 1.978602168 | 2.193124598 | 1.833176598 |
| ENSMUSG00000072949 | 0.289064826 | 0 | 0.5 | 0.580482024 | 0.733951626 | 0.477121255 | 0.579661877 |
| ENSMUSG00000072950 | 0 | 0 | 0 | 0 | 0 | 0 | 0 |
| ENSMUSG00000072952 | 0 | 0 | 0 | 0 | 0 | 0 | 0 |
| ENSMUSG00000072955 | 0 | 0 | 0 | 0.25 | 0 | 0.301029996 | 0 |
| ENSMUSG00000072962 | 0 | 0 | 0 | 0 | 0 | 0 | 0 |
| ENSMUSG00000072963 | 0.578129653 | 0 | 0.396240625 | 0.25 | 0.387761935 | 0.301029996 | 0.355418052 |
| ENSMUSG00000072964 | 2.283862692 | 2.12892496 | 2.078220739 | 2.428990498 | 2.298884454 | 2.689308859 | 2.154798988 |
| ENSMUSG00000072966 | 0.671187741 | 0.701838731 | 0.79248125 | 1.380890489 | 1.358933174 | 1.477121255 | 1.265599895 |
| ENSMUSG00000072968 | 0 | 0 | 0 | 0 | 0 | 0 | 0 |
| ENSMUSG00000072969 | 1.605636262 | 1.380890489 | 1.52186571 | 1.782320754 | 1.715300549 | 1.897627091 | 1.63986884 |
| ENSMUSG00000072972 | 0.578129653 | 0.25 | 0.79248125 | 0.25 | 0.812711509 | 0.77815125 | 0.672731473 |
| ENSMUSG00000072974 | 0.671187741 | 0.5 | 0.646240625 | 0.580482024 | 0.686820904 | 0.301029996 | 0.355418052 |
| ENSMUSG00000072978 | 0 | 0 | 0 | 0 | 0 | 0 | 0 |
| ENSMUSG00000072980 | 1.700465191 | 1.606566189 | 1.668106335 | 1.923371739 | 1.814781639 | 2.086359831 | 1.674564426 |
| ENSMUSG00000072982 | 0 | 0 | 0 | 0 | 0 | 0 | 0 |
| ENSMUSG00000072983 | 0 | 0 | 0 | 0 | 0 | 0 | 0 |
| ENSMUSG00000072995 | 0 | 0 | 0 | 0 | 0 | 0 | 0 |
| ENSMUSG00000072999 | 0.289064826 | 0 | 0.25 | 0.25 | 0.387761935 | 0.301029996 | 0.355418052 |
| ENSMUSG00000073000 | 0 | 0.25 | 0 | 0.25 | 0 | 0.602059991 | 0 |
| ENSMUSG00000073001 | 0 | 0 | 0 | 0 | 0 | 0 | 0 |
| ENSMUSG00000073002 | 1.655740331 | 1.614857905 | 1.723704441 | 1.584962501 | 1.611014646 | 1.707570176 | 1.56716938 |
| ENSMUSG00000073006 | 0 | 0 | 0 | 0 | 0 | 0 | 0 |

| | | | | | | | |
|---|---|---|---|---|---|---|---|
| ENSMUSG00000073007 | 0 | 0.25 | 0.25 | 0.25 | 0 | 0.301029996 | 0 |
| ENSMUSG00000073008 | 0 | 0 | 0.25 | 0 | 0 | 0 | 0 |
| ENSMUSG00000073010 | 0 | 0 | 0 | 0 | 0 | 0 | 0 |
| ENSMUSG00000073012 | 0 | 0 | 0 | 0 | 0 | 0 | 0 |
| ENSMUSG00000073016 | 1.993049299 | 1.870953944 | 1.901832578 | 1.886723615 | 1.840643511 | 2.049218023 | 1.699191618 |
| ENSMUSG00000073018 | 0.671187741 | 0.5 | 0.396240625 | 0 | 0.489301084 | 0.301029996 | 0 |
| ENSMUSG00000073027 | 0 | 0 | 0 | 0 | 0 | 0 | 0 |
| ENSMUSG00000073028 | 0 | 0 | 0 | 0 | 0 | 0 | 0 |
| ENSMUSG00000073043 | 0 | 0 | 0 | 0 | 0 | 0 | 0 |
| ENSMUSG00000073045 | 0 | 0 | 0 | 0 | 0 | 0 | 0 |
| ENSMUSG00000073051 | 0 | 0 | 0 | 0 | 0 | 0 | 0 |
| ENSMUSG00000073052 | 0 | 0 | 0 | 0 | 0 | 0 | 0 |
| ENSMUSG00000073060 | 0 | 0 | 0 | 0 | 0 | 0 | 0 |
| ENSMUSG00000073062 | 1.91631382 | 1.79745614 | 1.789967834 | 1.850219859 | 1.806427258 | 2.053078443 | 1.653807046 |
| ENSMUSG00000073063 | 0 | 0 | 0 | 0 | 0 | 0 | 0 |
| ENSMUSG00000073067 | 0 | 0 | 0.396240625 | 0.25 | 0 | 0 | 0 |
| ENSMUSG00000073069 | 0 | 0 | 0 | 0 | 0 | 0 | 0 |
| ENSMUSG00000073072 | 0 | 0 | 0 | 0 | 0 | 0 | 0 |
| ENSMUSG00000073073 | 0.45815691 | 0.25 | 0 | 0.396240625 | 0 | 0.301029996 | 0.355418052 |
| ENSMUSG00000073077 | 0.671187741 | 0.864857905 | 0.830482024 | 0 | 0 | 0 | 0 |
| ENSMUSG00000073079 | 0 | 0 | 0 | 0 | 0 | 0 | 0 |
| ENSMUSG00000073085 | 0 | 0 | 0 | 0 | 0 | 0 | 0 |
| ENSMUSG00000073094 | 0 | 0 | 0 | 0 | 0.244650542 | 0 | 0 |
| ENSMUSG00000073096 | 1.100572389 | 1.146240625 | 1.080482024 | 1.061981878 | 1 | 0.84509804 | 1.028149525 |
| ENSMUSG00000073102 | 0 | 0.25 | 0 | 0 | 0 | 0.477121255 | 0.355418052 |
| ENSMUSG00000073103 | 0 | 0 | 0 | 0 | 0 | 0 | 0 |
| ENSMUSG00000073110 | 0 | 0 | 0 | 0 | 0 | 0 | 0 |
| ENSMUSG00000073111 | 0 | 0 | 0 | 0 | 0 | 0 | 0 |
| ENSMUSG00000073116 | 0 | 0 | 0 | 0 | 0 | 0 | 0 |
| ENSMUSG00000073117 | 0 | 0 | 0 | 0 | 0 | 0 | 0 |
| ENSMUSG00000073119 | 0 | 0 | 0 | 0 | 0 | 0 | 0 |
| ENSMUSG00000073125 | 0 | 0 | 0 | 0.396240625 | 0.244650542 | 0 | 0.520678036 |
| ENSMUSG00000073130 | 0.811507563 | 0.92510993 | 0.701838731 | 0.5 | 0.387761935 | 0.84509804 | 0.672731473 |
| ENSMUSG00000073131 | 2.664237041 | 2.453044326 | 2.451032755 | 2.621457327 | 2.548733265 | 2.92992956 | 2.338260508 |

| | | | | | | | |
|---|---|---|---|---|---|---|---|
| ENSMUSG00000073139 | 2.378662046 | 2.21293726 | 2.170748646 | 2.286801231 | 2.239111448 | 2.57634135 | 2.093904044 |
| ENSMUSG00000073141 | 0 | 0 | 0 | 0 | 0 | 0 | 0 |
| ENSMUSG00000073144 | 0 | 0 | 0 | 0.396240625 | 0 | 0 | 0 |
| ENSMUSG00000073145 | 0 | 0 | 0 | 0 | 0 | 0 | 0 |
| ENSMUSG00000073147 | 1.678702042 | 1.638647213 | 1.735628626 | 1.660964047 | 1.599808748 | 1.819543936 | 1.607811402 |
| ENSMUSG00000073152 | 0 | 0 | 0 | 0 | 0 | 0 | 0 |
| ENSMUSG00000073154 | 0.289064826 | 0 | 0.25 | 0.396240625 | 0.387761935 | 0 | 0.355418052 |
| ENSMUSG00000073155 | 2.413323654 | 2.245034894 | 2.256284891 | 1.961372513 | 1.968065382 | 2.146128036 | 1.926805425 |
| ENSMUSG00000073158 | 0 | 0.25 | 0 | 0.646240625 | 0.244650542 | 0.301029996 | 0.579661877 |
| ENSMUSG00000073164 | 1.227926429 | 1.114857905 | 1.17510993 | 1.527131114 | 1.478764298 | 1.681241237 | 1.437266334 |
| ENSMUSG00000073173 | 0 | 0 | 0 | 0 | 0 | 0 | 0 |
| ENSMUSG00000073174 | 0 | 0 | 0 | 0.396240625 | 0.244650542 | 0 | 0.224243824 |
| ENSMUSG00000073176 | 2.045658471 | 1.923371739 | 1.868933358 | 1.78743678 | 1.733951626 | 1.959041392 | 1.616675393 |
| ENSMUSG00000073177 | 0 | 0 | 0 | 0 | 0 | 0 | 0 |
| ENSMUSG00000073179 | 0.671187741 | 0.646240625 | 0.396240625 | 0.396240625 | 0.632412477 | 0.602059991 | 0.672731473 |
| ENSMUSG00000073184 | 0 | 0 | 0 | 0 | 0.244650542 | 0 | 0 |
| ENSMUSG00000073197 | 1.596667477 | 1.532320754 | 1.488549078 | 1.458222504 | 1.484072009 | 1.602059991 | 1.379042812 |
| ENSMUSG00000073198 | 0 | 0 | 0 | 0 | 0 | 0 | 0 |
| ENSMUSG00000073207 | 0.289064826 | 0 | 0.25 | 0.5 | 0.489301084 | 0.602059991 | 0.355418052 |
| ENSMUSG00000073208 | 0 | 0 | 0 | 0 | 0 | 0 | 0 |
| ENSMUSG00000073209 | 0 | 0 | 0 | 0 | 0 | 0 | 0.224243824 |
| ENSMUSG00000073226 | 0 | 0.396240625 | 0.25 | 0.25 | 0.244650542 | 0.301029996 | 0 |
| ENSMUSG00000073233 | 0 | 0 | 0 | 0 | 0 | 0 | 0 |
| ENSMUSG00000073234 | 1.342375483 | 1.372963274 | 1.364857905 | 1.561981878 | 1.484072009 | 1.681241237 | 1.379042812 |
| ENSMUSG00000073236 | 1.307602651 | 1.098079356 | 1.114857905 | 1.380890489 | 1.302012594 | 1.462397998 | 1.272006351 |
| ENSMUSG00000073242 | 0 | 0 | 0 | 0 | 0.244650542 | 0 | 0 |
| ENSMUSG00000073243 | 0 | 0 | 0 | 0 | 0 | 0 | 0 |
| ENSMUSG00000073245 | 0 | 0 | 0 | 0 | 0 | 0 | 0 |
| ENSMUSG00000073247 | 0 | 0 | 0 | 0 | 0 | 0 | 0 |
| ENSMUSG00000073255 | 0 | 0 | 0 | 0 | 0 | 0 | 0 |
| ENSMUSG00000073257 | 0 | 0 | 0 | 0 | 0 | 0 | 0 |
| ENSMUSG00000073267 | 0 | 0 | 0 | 0 | 0 | 0 | 0 |
| ENSMUSG00000073274 | 0 | 0.25 | 0.396240625 | 0.5 | 0.244650542 | 0.301029996 | 0.224243824 |
| ENSMUSG00000073290 | 0 | 0 | 0 | 0 | 0.387761935 | 0 | 0.224243824 |

| | | | | | | | |
|---|---|---|---|---|---|---|---|
| ENSMUSG00000073291 | 0 | 0 | 0 | 0 | 0 | 0 | 0 |
| ENSMUSG00000073293 | 0 | 0 | 0.25 | 0.25 | 0.244650542 | 0.602059991 | 0.629532004 |
| ENSMUSG00000073294 | 0 | 0 | 0 | 0 | 0 | 0 | 0 |
| ENSMUSG00000073295 | 0 | 0 | 0 | 0.25 | 0.568060967 | 0.301029996 | 0 |
| ENSMUSG00000073367 | 0 | 0 | 0 | 0 | 0 | 0 | 0 |
| ENSMUSG00000073371 | 0.289064826 | 0.25 | 0.25 | 0.396240625 | 0 | 0 | 0.224243824 |
| ENSMUSG00000073374 | 1.036286563 | 1.061981878 | 0.976722649 | 1.098079356 | 0.978602168 | 0.77815125 | 0.916588299 |
| ENSMUSG00000073375 | 0 | 0 | 0 | 0 | 0 | 0 | 0 |
| ENSMUSG00000073377 | 0 | 0 | 0 | 0 | 0 | 0 | 0 |
| ENSMUSG00000073380 | 0 | 0 | 0 | 0 | 0.244650542 | 0 | 0 |
| ENSMUSG00000073386 | 0 | 0 | 0 | 0 | 0 | 0 | 0 |
| ENSMUSG00000073388 | 0 | 0 | 0.25 | 0 | 0 | 0 | 0.224243824 |
| ENSMUSG00000073394 | 0.578129653 | 0.25 | 0.79248125 | 0 | 0.387761935 | 0.301029996 | 0 |
| ENSMUSG00000073396 | 0 | 0 | 0 | 0 | 0 | 0 | 0 |
| ENSMUSG00000073399 | 0 | 0 | 0 | 0 | 0 | 0 | 0.224243824 |
| ENSMUSG00000073400 | 0 | 0 | 0 | 0 | 0 | 0 | 0 |
| ENSMUSG00000073402 | 0.289064826 | 0.580482024 | 0.701838731 | 0.701838731 | 0.733951626 | 0.602059991 | 1.131174228 |
| ENSMUSG00000073403 | 0.747221736 | 0.79248125 | 0.79248125 | 1.080482024 | 1.074582839 | 1.431363764 | 1.278288402 |
| ENSMUSG00000073405 | 0.578129653 | 0.396240625 | 0.580482024 | 0.79248125 | 0.877063019 | 0.477121255 | 0.896975297 |
| ENSMUSG00000073406 | 0.289064826 | 0 | 0.25 | 0.79248125 | 0.387761935 | 0.301029996 | 0.710836105 |
| ENSMUSG00000073407 | 0 | 0 | 0 | 0.25 | 0 | 0.301029996 | 0.355418052 |
| ENSMUSG00000073408 | 0 | 0.25 | 0 | 0 | 0 | 0 | 0 |
| ENSMUSG00000073409 | 1.181541736 | 1.311981878 | 1.330482024 | 1.987091808 | 1.991345651 | 2.206825876 | 1.95794437 |
| ENSMUSG00000073411 | 1.54867984 | 1.388647213 | 1.438721876 | 2.58412664 | 2.611878676 | 2.970346876 | 2.480444177 |
| ENSMUSG00000073412 | 0.289064826 | 0.396240625 | 0.25 | 0.5 | 0.568060967 | 0.301029996 | 0.629532004 |
| ENSMUSG00000073413 | 0 | 0 | 0 | 0 | 0 | 0 | 0 |
| ENSMUSG00000073414 | 0.289064826 | 0.5 | 0.396240625 | 0 | 0.387761935 | 0.301029996 | 0.224243824 |
| ENSMUSG00000073415 | 0.289064826 | 0.580482024 | 0.864857905 | 0.396240625 | 0.489301084 | 0.477121255 | 0.672731473 |
| ENSMUSG00000073418 | 0 | 0.5 | 0.25 | 0.5 | 0.686820904 | 0.301029996 | 0.710836105 |
| ENSMUSG00000073420 | 0 | 0 | 0 | 0 | 0 | 0 | 0 |
| ENSMUSG00000073421 | 1.069666964 | 1.02186571 | 1.226722649 | 0.830482024 | 1.106691885 | 0.698970004 | 1.150210039 |
| ENSMUSG00000073422 | 0 | 0.396240625 | 0.5 | 0.5 | 0.489301084 | 0.477121255 | 0.448487648 |
| ENSMUSG00000073423 | 1.639698646 | 1.741446071 | 1.720660762 | 1.451838731 | 1.607318736 | 1.653212514 | 1.625302983 |
| ENSMUSG00000073424 | 0 | 0 | 0 | 0 | 0 | 0 | 0.224243824 |

| | | | | | | | |
|---|---|---|---|---|---|---|---|
| ENSMUSG00000073427 | 1.307602651 | 1.17510993 | 1.356566189 | 1.451838731 | 1.456697355 | 1.491361694 | 1.302263476 |
| ENSMUSG00000073430 | 0 | 0 | 0 | 0 | 0 | 0 | 0 |
| ENSMUSG00000073433 | 0.747221736 | 0.896240625 | 0.92510993 | 0.951838731 | 1.020174412 | 0.77815125 | 1.028149525 |
| ENSMUSG00000073434 | 2.20747957 | 2.027131114 | 2.123963757 | 2.017865591 | 2.155511127 | 2.209515015 | 2.100829716 |
| ENSMUSG00000073435 | 0 | 0 | 0 | 0 | 0 | 0 | 0 |
| ENSMUSG00000073436 | 1.404271433 | 1.29248125 | 1.302363341 | 1.458222504 | 1.373641807 | 1.462397998 | 1.302263476 |
| ENSMUSG00000073437 | 1.53838222 | 1.458222504 | 1.5 | 1.29248125 | 1.445123986 | 1.301029996 | 1.278288402 |
| ENSMUSG00000073448 | 0.289064826 | 0.580482024 | 0.396240625 | 0.646240625 | 0.244650542 | 0.477121255 | 0.448487648 |
| ENSMUSG00000073457 | 0 | 0 | 0 | 0 | 0 | 0 | 0 |
| ENSMUSG00000073458 | 0 | 0 | 0 | 0 | 0 | 0 | 0 |
| ENSMUSG00000073460 | 0.45815691 | 0.701838731 | 0.646240625 | 0.646240625 | 0.733951626 | 0.77815125 | 0.74492186 |
| ENSMUSG00000073468 | 1.960252568 | 1.779735268 | 1.821350555 | 1.789967834 | 1.810629166 | 1.944482672 | 1.680001789 |
| ENSMUSG00000073471 | 1.069666964 | 1.146240625 | 1.188721876 | 1.080482024 | 1.163285805 | 1.176091259 | 1.078019652 |
| ENSMUSG00000073478 | 0.960252568 | 0.830482024 | 0.951838731 | 1.061981878 | 0.955822902 | 0.84509804 | 1.028149525 |
| ENSMUSG00000073481 | 2.203267085 | 2.056000419 | 2.043731421 | 2.28296424 | 2.184909731 | 2.481442629 | 2.076967568 |
| ENSMUSG00000073482 | 0.289064826 | 0 | 0.25 | 0 | 0 | 0 | 0 |
| ENSMUSG00000073485 | 0.867194479 | 0.646240625 | 0.646240625 | 0.92510993 | 0.568060967 | 0.301029996 | 0.629532004 |
| ENSMUSG00000073486 | 0 | 0 | 0 | 0 | 0 | 0 | 0 |
| ENSMUSG00000073489 | 2 | 1.884789703 | 1.945339928 | 1.802363341 | 1.744182955 | 1.924279286 | 1.594042709 |
| ENSMUSG00000073490 | 0.578129653 | 0.864857905 | 0.830482024 | 0.25 | 0.387761935 | 0.301029996 | 0.224243824 |
| ENSMUSG00000073491 | 0.289064826 | 0 | 0 | 0 | 0 | 0 | 0 |
| ENSMUSG00000073492 | 0.289064826 | 0 | 0 | 0 | 0 | 0 | 0 |
| ENSMUSG00000073494 | 0 | 0 | 0 | 0 | 0 | 0 | 0 |
| ENSMUSG00000073497 | 0 | 0 | 0 | 0 | 0 | 0 | 0 |
| ENSMUSG00000073510 | 0 | 0 | 0 | 0 | 0 | 0 | 0 |
| ENSMUSG00000073514 | 0.289064826 | 0.396240625 | 0.580482024 | 0 | 0 | 0 | 0.224243824 |
| ENSMUSG00000073528 | 0 | 0 | 0 | 0 | 0 | 0 | 0 |
| ENSMUSG00000073529 | 0.45815691 | 0 | 0.25 | 0 | 0 | 0 | 0 |
| ENSMUSG00000073530 | 0.45815691 | 0.830482024 | 0.701838731 | 0.75 | 0.632412477 | 0.698970004 | 0.629532004 |
| ENSMUSG00000073532 | 0 | 0 | 0 | 0 | 0 | 0 | 0 |
| ENSMUSG00000073535 | 0 | 0 | 0.396240625 | 0.25 | 0.244650542 | 0.301029996 | 0 |
| ENSMUSG00000073538 | 0.671187741 | 0.580482024 | 0.580482024 | 0 | 0.686820904 | 0.602059991 | 0.520678036 |
| ENSMUSG00000073542 | 2.287337982 | 2.135757955 | 2.123963757 | 2.154596376 | 2.105788027 | 2.394451681 | 1.943148064 |
| ENSMUSG00000073551 | 0 | 0 | 0 | 0 | 0 | 0 | 0 |

| | | | | | | | |
|---|---|---|---|---|---|---|---|
| ENSMUSG00000073554 | 0 | 0 | 0 | 0 | 0 | 0 | 0 |
| ENSMUSG00000073555 | 0 | 0 | 0.396240625 | 0.5 | 0 | 0.301029996 | 0.224243824 |
| ENSMUSG00000073557 | 2.144240249 | 2.023189285 | 1.956637122 | 2.103406982 | 2.041436517 | 2.23299611 | 1.924281235 |
| ENSMUSG00000073559 | 0.45815691 | 0.646240625 | 0.25 | 0 | 0.387761935 | 0.477121255 | 0.355418052 |
| ENSMUSG00000073563 | 2.453536005 | 2.299918086 | 2.281030328 | 2.256976499 | 2.154724156 | 2.444044796 | 2.001334086 |
| ENSMUSG00000073565 | 0 | 0 | 0 | 0.25 | 0 | 0.477121255 | 0 |
| ENSMUSG00000073568 | 0.289064826 | 0 | 0 | 0.25 | 0 | 0.301029996 | 0.355418052 |
| ENSMUSG00000073569 | 0 | 0 | 0.25 | 0 | 0 | 0 | 0 |
| ENSMUSG00000073572 | 0 | 0 | 0 | 0 | 0 | 0 | 0 |
| ENSMUSG00000073573 | 0 | 0 | 0 | 0 | 0 | 0 | 0 |
| ENSMUSG00000073574 | 0 | 0 | 0 | 0 | 0 | 0 | 0 |
| ENSMUSG00000073590 | 1.605636262 | 1.54745614 | 1.614857905 | 0.580482024 | 0.568060967 | 0.602059991 | 0.355418052 |
| ENSMUSG00000073591 | 1.54867984 | 1.470660762 | 1.445339928 | 1.364857905 | 1.445123986 | 1.505149978 | 1.307989563 |
| ENSMUSG00000073594 | 0 | 0 | 0 | 0 | 0 | 0 | 0 |
| ENSMUSG00000073598 | 0 | 0 | 0 | 0.396240625 | 0 | 0 | 0.224243824 |
| ENSMUSG00000073599 | 0.91631382 | 0.75 | 0.5 | 1.356566189 | 1.310728002 | 1.322219295 | 1.245582246 |
| ENSMUSG00000073600 | 0.867194479 | 0.79248125 | 0.864857905 | 0.864857905 | 0.905314583 | 0.77815125 | 0.935079929 |
| ENSMUSG00000073601 | 0 | 0 | 0 | 0 | 0 | 0 | 0 |
| ENSMUSG00000073602 | 0 | 0 | 0 | 0 | 0 | 0 | 0 |
| ENSMUSG00000073607 | 0 | 0 | 0 | 0.5 | 0.489301084 | 0.954242509 | 0.448487648 |
| ENSMUSG00000073608 | 0.289064826 | 0.396240625 | 0.25 | 0 | 0 | 0 | 0.355418052 |
| ENSMUSG00000073609 | 1.794934559 | 1.692046081 | 1.738549078 | 1.593759858 | 1.632412477 | 1.819543936 | 1.511732338 |
| ENSMUSG00000073610 | 0 | 0 | 0 | 0 | 0 | 0 | 0 |
| ENSMUSG00000073616 | 1.982401085 | 1.87892496 | 1.923371739 | 1.880890489 | 1.873585591 | 2.100370545 | 1.763495175 |
| ENSMUSG00000073627 | 0.867194479 | 1.17510993 | 1.146240625 | 1.02186571 | 1.074582839 | 1.041392685 | 1.014380832 |
| ENSMUSG00000073628 | 0 | 0 | 0 | 0 | 0 | 0 | 0 |
| ENSMUSG00000073631 | 0 | 0 | 0 | 0 | 0 | 0 | 0 |
| ENSMUSG00000073633 | 1.578129653 | 1.396240625 | 1.396240625 | 1.403677461 | 1.387761935 | 1.602059991 | 1.319146374 |
| ENSMUSG00000073639 | 2.874454827 | 2.622712719 | 2.619692405 | 2.633575721 | 2.569766661 | 2.960470778 | 2.362688369 |
| ENSMUSG00000073640 | 0 | 0 | 0 | 0.25 | 0.244650542 | 0 | 0 |
| ENSMUSG00000073643 | 2.713122685 | 2.526149688 | 2.507666784 | 2.186208459 | 2.104881849 | 2.416640507 | 2.00464543 |
| ENSMUSG00000073647 | 1.269664473 | 1.26109853 | 1.238549078 | 1.226722649 | 1.254881871 | 1.342422681 | 1.131174228 |
| ENSMUSG00000073650 | 0.867194479 | 0.580482024 | 0.830482024 | 0 | 0.244650542 | 0.602059991 | 0.224243824 |
| ENSMUSG00000073652 | 0 | 0.396240625 | 0.396240625 | 0.396240625 | 0.244650542 | 0.77815125 | 0.448487648 |

| | | | | | | | |
|---|---|---|---|---|---|---|---|
| ENSMUSG00000073656 | 0 | 0.25 | 0 | 0.25 | 0 | 0 | 0.224243824 |
| ENSMUSG00000073658 | 0 | 0 | 0 | 0 | 0 | 0 | 0 |
| ENSMUSG00000073664 | 2.63295404 | 2.441217898 | 2.439555804 | 2.468745337 | 2.391003805 | 2.688419822 | 2.176815335 |
| ENSMUSG00000073676 | 2.842342644 | 2.622211087 | 2.637927065 | 2.67293588 | 2.631801298 | 2.968015714 | 2.436920143 |
| ENSMUSG00000073678 | 2.149089517 | 2.002806814 | 1.955044741 | 2.100219859 | 2.041436517 | 2.29666519 | 1.900630878 |
| ENSMUSG00000073679 | 0 | 0 | 0.396240625 | 0.25 | 0.244650542 | 0 | 0.224243824 |
| ENSMUSG00000073680 | 0.671187741 | 0.75 | 0.864857905 | 0.701838731 | 0.686820904 | 0.477121255 | 0.520678036 |
| ENSMUSG00000073682 | 0 | 0.5 | 0 | 0.396240625 | 0.244650542 | 0 | 0.448487648 |
| ENSMUSG00000073684 | 1.87190676 | 1.860735874 | 1.888647213 | 1.642463902 | 1.662257525 | 1.748188027 | 1.665294008 |
| ENSMUSG00000073686 | 0 | 0 | 0 | 0 | 0 | 0 | 0 |
| ENSMUSG00000073699 | 0 | 0 | 0 | 0 | 0 | 0 | 0 |
| ENSMUSG00000073700 | 2.201144779 | 2.088286706 | 2.136723615 | 1.976722649 | 2.045754077 | 2.193124598 | 1.95411574 |
| ENSMUSG00000073702 | 2.95990777 | 2.689972046 | 2.694724619 | 2.822792675 | 2.714494897 | 3.176380692 | 2.549194983 |
| ENSMUSG00000073705 | 1.811507563 | 1.779735268 | 1.784887838 | 1.678561379 | 1.793512788 | 1.880813592 | 1.819456572 |
| ENSMUSG00000073716 | 0.289064826 | 0.580482024 | 0.580482024 | 0.396240625 | 0.387761935 | 0.477121255 | 0.224243824 |
| ENSMUSG00000073721 | 0 | 0 | 0 | 0 | 0 | 0 | 0 |
| ENSMUSG00000073722 | 0 | 0 | 0 | 0 | 0 | 0 | 0 |
| ENSMUSG00000073723 | 0 | 0 | 0 | 0 | 0 | 0 | 0 |
| ENSMUSG00000073724 | 0 | 0 | 0 | 0 | 0 | 0 | 0 |
| ENSMUSG00000073725 | 2.29078455 | 2.124961472 | 2.041226732 | 2.284247778 | 2.216042585 | 2.546542663 | 2.091394214 |
| ENSMUSG00000073728 | 0.289064826 | 0.25 | 0.5 | 0 | 0.244650542 | 0.301029996 | 0.448487648 |
| ENSMUSG00000073730 | 0 | 0 | 0 | 0 | 0 | 0 | 0 |
| ENSMUSG00000073731 | 0 | 0 | 0 | 0 | 0 | 0 | 0 |
| ENSMUSG00000073733 | 0.578129653 | 0.92510993 | 0.864857905 | 0.75 | 1.057362051 | 0.698970004 | 0.935079929 |
| ENSMUSG00000073735 | 0 | 0 | 0 | 0 | 0 | 0 | 0 |
| ENSMUSG00000073737 | 0 | 0 | 0 | 0 | 0 | 0 | 0 |
| ENSMUSG00000073739 | 0 | 0.5 | 0 | 0 | 0.387761935 | 0.301029996 | 0.224243824 |
| ENSMUSG00000073747 | 0 | 0 | 0 | 0 | 0 | 0 | 0 |
| ENSMUSG00000073752 | 0 | 0 | 0.25 | 0 | 0 | 0 | 0 |
| ENSMUSG00000073753 | 0.289064826 | 0.5 | 0.396240625 | 0 | 0.244650542 | 0.301029996 | 0.355418052 |
| ENSMUSG00000073755 | 1.482695304 | 1.458222504 | 1.464495249 | 1.130890489 | 1.121713561 | 1.113943352 | 1.140832123 |
| ENSMUSG00000073758 | 0 | 0.75 | 0.646240625 | 0.580482024 | 0.387761935 | 0.77815125 | 0.710836105 |
| ENSMUSG00000073759 | 0 | 0 | 0 | 0 | 0.387761935 | 0.477121255 | 0.355418052 |
| ENSMUSG00000073761 | 0.289064826 | 0.5 | 0.25 | 0.25 | 0 | 0.301029996 | 0.448487648 |

| | | | | | | | |
|---|---|---|---|---|---|---|---|
| ENSMUSG00000073764 | 0 | 0 | 0 | 0 | 0 | 0 | 0 |
| ENSMUSG00000073765 | 0 | 0 | 0 | 0 | 0 | 0 | 0 |
| ENSMUSG00000073768 | 0 | 0 | 0 | 0 | 0 | 0 | 0 |
| ENSMUSG00000073769 | 0 | 0 | 0 | 0 | 0 | 0 | 0 |
| ENSMUSG00000073770 | 0 | 0 | 0 | 0 | 0 | 0 | 0 |
| ENSMUSG00000073771 | 1.876566388 | 1.685366747 | 1.79745614 | 1.598079356 | 1.509475496 | 1.755874856 | 1.551512352 |
| ENSMUSG00000073774 | 0 | 0 | 0 | 0 | 0 | 0 | 0 |
| ENSMUSG00000073775 | 1.903481042 | 1.834962501 | 1.782320754 | 1.794977273 | 1.830919587 | 2 | 1.690609803 |
| ENSMUSG00000073778 | 0.578129653 | 0 | 0.396240625 | 0.5 | 0.632412477 | 0.477121255 | 0.520678036 |
| ENSMUSG00000073779 | 0.289064826 | 0 | 0 | 0 | 0 | 0 | 0 |
| ENSMUSG00000073781 | 0 | 0 | 0 | 0 | 0 | 0 | 0 |
| ENSMUSG00000073785 | 0 | 0 | 0 | 0 | 0 | 0 | 0 |
| ENSMUSG00000073786 | 0 | 0 | 0 | 0 | 0 | 0 | 0 |
| ENSMUSG00000073787 | 0.578129653 | 0.646240625 | 0.864857905 | 0.79248125 | 0 | 0.301029996 | 0.775756176 |
| ENSMUSG00000073789 | 0 | 0 | 0 | 0 | 0 | 0 | 0 |
| ENSMUSG00000073791 | 1.800532393 | 1.714495249 | 1.76109853 | 1.744319981 | 1.614672255 | 1.949390007 | 1.520678036 |
| ENSMUSG00000073792 | 2.235832493 | 2.069031101 | 1.995748394 | 2.21759118 | 2.153935427 | 2.428134794 | 1.978585612 |
| ENSMUSG00000073794 | 0 | 0.25 | 0 | 0.396240625 | 0 | 0 | 0.224243824 |
| ENSMUSG00000073795 | 0 | 0.25 | 0 | 0.25 | 0 | 0 | 0 |
| ENSMUSG00000073797 | 0 | 0 | 0 | 0 | 0 | 0 | 0 |
| ENSMUSG00000073802 | 2.40033094 | 2.256284891 | 2.181128463 | 2.166333979 | 2.103973339 | 2.350248018 | 2.041356071 |
| ENSMUSG00000073803 | 0 | 0 | 0 | 0 | 0 | 0 | 0 |
| ENSMUSG00000073804 | 0 | 0 | 0 | 0 | 0 | 0.301029996 | 0 |
| ENSMUSG00000073805 | 0.289064826 | 0 | 0.25 | 0.396240625 | 0.244650542 | 0 | 0.448487648 |
| ENSMUSG00000073807 | 0 | 0 | 0 | 0 | 0 | 0 | 0 |
| ENSMUSG00000073809 | 0 | 0 | 0 | 0 | 0.244650542 | 0 | 0 |
| ENSMUSG00000073811 | 0 | 0 | 0 | 0 | 0 | 0 | 0 |
| ENSMUSG00000073821 | 0.578129653 | 0.646240625 | 0.396240625 | 0.396240625 | 0.77552387 | 0.477121255 | 0.520678036 |
| ENSMUSG00000073830 | 0 | 0 | 0 | 0 | 0 | 0 | 0 |
| ENSMUSG00000073834 | 0 | 0 | 0 | 0 | 0 | 0 | 0 |
| ENSMUSG00000073835 | 0 | 0 | 0 | 0 | 0 | 0 | 0 |
| ENSMUSG00000073838 | 2.003432385 | 1.837182039 | 1.905512955 | 1.830482024 | 1.892300902 | 2.11058971 | 1.865128412 |
| ENSMUSG00000073840 | 0 | 0 | 0 | 0 | 0 | 0 | 0 |
| ENSMUSG00000073842 | 0 | 0 | 0 | 0 | 0 | 0 | 0 |

| | | | | | | | |
|---|---|---|---|---|---|---|---|
| ENSMUSG00000073843 | 0 | 0 | 0.5 | 0 | 0 | 0 | 0.224243824 |
| ENSMUSG00000073844 | 0 | 0 | 0 | 0 | 0 | 0 | 0 |
| ENSMUSG00000073846 | 0 | 0 | 0 | 0.25 | 0 | 0 | 0 |
| ENSMUSG00000073856 | 0.45815691 | 0.25 | 0.5 | 0.5 | 0.568060967 | 0.602059991 | 0.448487648 |
| ENSMUSG00000073860 | 0 | 0 | 0 | 0 | 0 | 0 | 0 |
| ENSMUSG00000073862 | 0 | 0 | 0 | 0 | 0 | 0 | 0 |
| ENSMUSG00000073867 | 0.45815691 | 0.79248125 | 1 | 0.79248125 | 0.877063019 | 0.84509804 | 1 |
| ENSMUSG00000073876 | 0 | 0 | 0 | 0 | 0 | 0 | 0 |
| ENSMUSG00000073877 | 0 | 0 | 0 | 0 | 0 | 0 | 0 |
| ENSMUSG00000073878 | 0 | 0 | 0 | 0 | 0 | 0 | 0 |
| ENSMUSG00000073879 | 0 | 0 | 0 | 0 | 0 | 0 | 0 |
| ENSMUSG00000073888 | 0.578129653 | 0 | 0.396240625 | 0.5 | 0.632412477 | 0.698970004 | 0.579661877 |
| ENSMUSG00000073889 | 1.482695304 | 1.5 | 1.431980114 | 1.388647213 | 1.674752852 | 1.662757832 | 1.577201671 |
| ENSMUSG00000073893 | 0 | 0 | 0 | 0 | 0 | 0 | 0 |
| ENSMUSG00000073894 | 0 | 0 | 0 | 0 | 0 | 0 | 0 |
| ENSMUSG00000073896 | 0 | 0 | 0 | 0 | 0 | 0 | 0 |
| ENSMUSG00000073897 | 0 | 0 | 0 | 0 | 0 | 0 | 0 |
| ENSMUSG00000073898 | 0 | 0 | 0 | 0 | 0 | 0 | 0 |
| ENSMUSG00000073899 | 0 | 0 | 0 | 0 | 0 | 0 | 0 |
| ENSMUSG00000073900 | 0 | 0 | 0 | 0 | 0 | 0 | 0 |
| ENSMUSG00000073901 | 0 | 0 | 0 | 0 | 0 | 0 | 0 |
| ENSMUSG00000073902 | 0 | 0 | 0 | 0 | 0 | 0 | 0 |
| ENSMUSG00000073906 | 0 | 0.25 | 0 | 0.25 | 0.244650542 | 0 | 0 |
| ENSMUSG00000073907 | 0 | 0 | 0 | 0 | 0 | 0 | 0 |
| ENSMUSG00000073908 | 0 | 0 | 0 | 0 | 0 | 0 | 0 |
| ENSMUSG00000073909 | 0 | 0 | 0 | 0 | 0 | 0 | 0 |
| ENSMUSG00000073910 | 1.269664473 | 1.321350555 | 1.29248125 | 1.418106335 | 1.27449559 | 1.447158031 | 1.259064007 |
| ENSMUSG00000073913 | 0 | 0 | 0 | 0 | 0 | 0 | 0 |
| ENSMUSG00000073914 | 0 | 0 | 0 | 0 | 0 | 0 | 0 |
| ENSMUSG00000073915 | 0 | 0 | 0 | 0 | 0 | 0 | 0 |
| ENSMUSG00000073916 | 0 | 0 | 0 | 0 | 0 | 0 | 0 |
| ENSMUSG00000073917 | 0 | 0 | 0 | 0 | 0 | 0 | 0 |
| ENSMUSG00000073919 | 0 | 0 | 0 | 0 | 0 | 0 | 0 |
| ENSMUSG00000073920 | 0 | 0 | 0 | 0 | 0 | 0 | 0 |

| | | | | | | | |
|---|---|---|---|---|---|---|---|
| ENSMUSG00000073922 | 0 | 0 | 0 | 0 | 0 | 0 | 0 |
| ENSMUSG00000073923 | 0 | 0 | 0 | 0 | 0 | 0 | 0 |
| ENSMUSG00000073924 | 0 | 0 | 0 | 0 | 0 | 0 | 0 |
| ENSMUSG00000073925 | 0 | 0 | 0 | 0 | 0 | 0 | 0 |
| ENSMUSG00000073926 | 0 | 0 | 0 | 0 | 0 | 0 | 0 |
| ENSMUSG00000073927 | 0 | 0 | 0 | 0 | 0 | 0 | 0 |
| ENSMUSG00000073928 | 0 | 0 | 0 | 0 | 0 | 0 | 0 |
| ENSMUSG00000073929 | 0 | 0 | 0 | 0 | 0 | 0 | 0 |
| ENSMUSG00000073931 | 0 | 0 | 0 | 0 | 0 | 0 | 0 |
| ENSMUSG00000073932 | 0 | 0 | 0 | 0 | 0 | 0 | 0 |
| ENSMUSG00000073937 | 0 | 0 | 0 | 0 | 0 | 0 | 0 |
| ENSMUSG00000073938 | 0 | 0 | 0 | 0 | 0 | 0 | 0 |
| ENSMUSG00000073940 | 0 | 0 | 0 | 0 | 0 | 0 | 0 |
| ENSMUSG00000073943 | 0 | 0 | 0 | 0 | 0 | 0 | 0 |
| ENSMUSG00000073944 | 0 | 0 | 0 | 0 | 0 | 0 | 0 |
| ENSMUSG00000073945 | 0 | 0 | 0 | 0 | 0 | 0 | 0 |
| ENSMUSG00000073946 | 0 | 0 | 0 | 0 | 0 | 0 | 0 |
| ENSMUSG00000073947 | 0 | 0 | 0 | 0 | 0 | 0 | 0 |
| ENSMUSG00000073948 | 0 | 0 | 0 | 0 | 0 | 0 | 0 |
| ENSMUSG00000073949 | 0 | 0 | 0 | 0 | 0 | 0 | 0 |
| ENSMUSG00000073950 | 0 | 0 | 0 | 0 | 0 | 0 | 0 |
| ENSMUSG00000073951 | 0 | 0 | 0 | 0 | 0 | 0 | 0 |
| ENSMUSG00000073952 | 0 | 0 | 0 | 0 | 0 | 0 | 0 |
| ENSMUSG00000073953 | 0 | 0 | 0 | 0 | 0 | 0 | 0 |
| ENSMUSG00000073954 | 0 | 0 | 0 | 0 | 0 | 0 | 0 |
| ENSMUSG00000073955 | 0 | 0 | 0 | 0 | 0 | 0 | 0 |
| ENSMUSG00000073956 | 0 | 0 | 0 | 0 | 0 | 0 | 0 |
| ENSMUSG00000073959 | 0 | 0 | 0 | 0 | 0 | 0 | 0 |
| ENSMUSG00000073960 | 0 | 0 | 0 | 0 | 0 | 0 | 0 |
| ENSMUSG00000073961 | 0 | 0 | 0 | 0 | 0 | 0 | 0 |
| ENSMUSG00000073962 | 0 | 0 | 0 | 0 | 0 | 0 | 0 |
| ENSMUSG00000073963 | 0 | 0 | 0 | 0 | 0 | 0 | 0 |
| ENSMUSG00000073964 | 0 | 0 | 0 | 0 | 0 | 0 | 0 |
| ENSMUSG00000073965 | 0 | 0 | 0 | 0 | 0 | 0 | 0 |

| | | | | | | | |
|---|---|---|---|---|---|---|---|
| ENSMUSG00000073966 | 0 | 0 | 0 | 0 | 0 | 0 | 0 |
| ENSMUSG00000073967 | 0 | 0 | 0 | 0 | 0 | 0 | 0 |
| ENSMUSG00000073968 | 1.325351389 | 1.431980114 | 1.403677461 | 1.17510993 | 1.264824954 | 1.342422681 | 1.110947927 |
| ENSMUSG00000073969 | 0 | 0 | 0 | 0 | 0 | 0 | 0 |
| ENSMUSG00000073970 | 0 | 0 | 0 | 0 | 0 | 0 | 0 |
| ENSMUSG00000073971 | 0 | 0 | 0 | 0 | 0 | 0 | 0 |
| ENSMUSG00000073972 | 0 | 0 | 0 | 0 | 0 | 0 | 0 |
| ENSMUSG00000073973 | 0 | 0 | 0 | 0 | 0 | 0 | 0 |
| ENSMUSG00000073974 | 0 | 0 | 0 | 0 | 0 | 0 | 0 |
| ENSMUSG00000073975 | 0 | 0 | 0 | 0 | 0 | 0 | 0 |
| ENSMUSG00000073976 | 0 | 0 | 0 | 0 | 0 | 0 | 0 |
| ENSMUSG00000073977 | 0 | 0 | 0 | 0.25 | 0 | 0 | 0 |
| ENSMUSG00000073978 | 0 | 0 | 0 | 0 | 0 | 0 | 0 |
| ENSMUSG00000073979 | 0 | 0 | 0 | 0 | 0 | 0 | 0 |
| ENSMUSG00000073980 | 0 | 0 | 0 | 0 | 0 | 0 | 0.224243824 |
| ENSMUSG00000073982 | 2.051791411 | 2.004202072 | 2.028435542 | 1.890560606 | 1.905314583 | 2.06069784 | 1.911356129 |
| ENSMUSG00000073985 | 1.227926429 | 1.098079356 | 1.146240625 | 1.02186571 | 0.77552387 | 0.477121255 | 0.775756176 |
| ENSMUSG00000073987 | 1.960252568 | 1.819031101 | 1.830482024 | 1.921625132 | 1.816839698 | 2.093421685 | 1.729871916 |
| ENSMUSG00000073988 | 0.45815691 | 0.79248125 | 0.830482024 | 0.976722649 | 0.877063019 | 0.954242509 | 0.916588299 |
| ENSMUSG00000073991 | 0 | 0 | 0 | 0 | 0 | 0 | 0 |
| ENSMUSG00000073995 | 0 | 0 | 0.396240625 | 0.25 | 0 | 0.301029996 | 0.224243824 |
| ENSMUSG00000073997 | 0 | 0 | 0 | 0 | 0 | 0 | 0 |
| ENSMUSG00000073998 | 0 | 0 | 0 | 0 | 0 | 0 | 0 |
| ENSMUSG00000074001 | 0 | 0 | 0 | 0 | 0 | 0 | 0.224243824 |
| ENSMUSG00000074003 | 0 | 0 | 0 | 0 | 0 | 0 | 0 |
| ENSMUSG00000074004 | 0 | 0.25 | 0.25 | 0 | 0 | 0 | 0 |
| ENSMUSG00000074006 | 0.747221736 | 0.5 | 0.580482024 | 0.396240625 | 0.632412477 | 0.301029996 | 0.629532004 |
| ENSMUSG00000074011 | 0 | 0 | 0.25 | 0 | 0 | 0 | 0 |
| ENSMUSG00000074024 | 1.156259305 | 1.130890489 | 1.098079356 | 1.26109853 | 0.931471446 | 1.079181246 | 1.028149525 |
| ENSMUSG00000074026 | 0 | 0 | 0 | 0 | 0 | 0 | 0 |
| ENSMUSG00000074028 | 0.289064826 | 0 | 0.25 | 0.25 | 0.387761935 | 0 | 0.355418052 |
| ENSMUSG00000074029 | 0 | 0 | 0.25 | 0 | 0 | 0 | 0 |
| ENSMUSG00000074030 | 2.10861828 | 1.959800947 | 1.955044741 | 1.969129237 | 1.882215755 | 2.071882007 | 1.855869786 |
| ENSMUSG00000074034 | 0.671187741 | 0.976722649 | 0.79248125 | 0.396240625 | 0.877063019 | 0.301029996 | 0.803905701 |

| | | | | | | | |
|---|---|---|---|---|---|---|---|
| ENSMUSG00000074037 | 0 | 0.25 | 0 | 0 | 0.244650542 | 0 | 0 |
| ENSMUSG00000074039 | 1.129344651 | 1.130890489 | 1.29248125 | 0.830482024 | 1 | 1.146128036 | 0.829800754 |
| ENSMUSG00000074052 | 0 | 0 | 0.25 | 0.396240625 | 0 | 0 | 0.629532004 |
| ENSMUSG00000074059 | 0 | 0 | 0 | 0 | 0 | 0 | 0 |
| ENSMUSG00000074060 | 0 | 0 | 0 | 0 | 0 | 0 | 0 |
| ENSMUSG00000074061 | 0 | 0.646240625 | 0.5 | 0.5 | 0.387761935 | 0.477121255 | 0.629532004 |
| ENSMUSG00000074062 | 0 | 0 | 0 | 0 | 0 | 0 | 0 |
| ENSMUSG00000074063 | 1.358731791 | 1.584962501 | 1.593759858 | 1 | 1.380772476 | 1.342422681 | 1.517723536 |
| ENSMUSG00000074064 | 1.587501561 | 1.356566189 | 1.494319981 | 1.445339928 | 1.546663136 | 1.568201724 | 1.532233387 |
| ENSMUSG00000074065 | 0 | 0.25 | 0 | 0 | 0 | 0 | 0 |
| ENSMUSG00000074067 | 0 | 0 | 0 | 0 | 0 | 0 | 0 |
| ENSMUSG00000074071 | 0.289064826 | 0.25 | 0 | 0.5 | 0 | 0.301029996 | 0.224243824 |
| ENSMUSG00000074072 | 0 | 0 | 0 | 0 | 0 | 0 | 0 |
| ENSMUSG00000074088 | 2.552362269 | 2.336628433 | 2.350219859 | 2.374462959 | 2.373641807 | 2.678518379 | 2.228229691 |
| ENSMUSG00000074092 | 0 | 0 | 0 | 0.25 | 0 | 0 | 0 |
| ENSMUSG00000074093 | 1.069666964 | 1.146240625 | 1.17510993 | 1.201838731 | 1.039257769 | 1.146128036 | 1.121219121 |
| ENSMUSG00000074102 | 2.564110437 | 2.381380202 | 2.400906586 | 2.357613138 | 2.362449578 | 2.622214023 | 2.248075634 |
| ENSMUSG00000074108 | 0 | 0 | 0.25 | 0 | 0.244650542 | 0 | 0 |
| ENSMUSG00000074109 | 0 | 0 | 0 | 0 | 0 | 0 | 0 |
| ENSMUSG00000074111 | 0 | 0 | 0 | 0 | 0 | 0 | 0 |
| ENSMUSG00000074113 | 0 | 0 | 0 | 0 | 0 | 0 | 0 |
| ENSMUSG00000074115 | 0 | 0 | 0 | 0 | 0 | 0 | 0 |
| ENSMUSG00000074121 | 0.867194479 | 0.5 | 0.646240625 | 0.830482024 | 0.77552387 | 1.146128036 | 0.916588299 |
| ENSMUSG00000074123 | 0 | 0 | 0 | 0 | 0 | 0 | 0 |
| ENSMUSG00000074127 | 0.289064826 | 0.701838731 | 0 | 0.396240625 | 0.568060967 | 0.301029996 | 0.224243824 |
| ENSMUSG00000074128 | 0 | 0 | 0 | 0.25 | 0 | 0 | 0 |
| ENSMUSG00000074129 | 3.01386641 | 2.825945187 | 2.87019755 | 2.682617534 | 2.710765025 | 3.077004327 | 2.628192393 |
| ENSMUSG00000074136 | 1.249317394 | 1.201838731 | 1.02186571 | 1.25 | 1.234113756 | 1.255272505 | 1.066254157 |
| ENSMUSG00000074139 | 0 | 0 | 0 | 0 | 0 | 0 | 0.224243824 |
| ENSMUSG00000074141 | 0.578129653 | 0.864857905 | 0.75 | 0.701838731 | 0.77552387 | 0.602059991 | 0.672731473 |
| ENSMUSG00000074146 | 0 | 0 | 0.25 | 0 | 0.489301084 | 0 | 0 |
| ENSMUSG00000074149 | 0 | 0 | 0 | 0 | 0 | 0 | 0 |
| ENSMUSG00000074151 | 1.289064826 | 1.26109853 | 1.26109853 | 1.606566189 | 1.649303553 | 1.681241237 | 1.543390198 |
| ENSMUSG00000074155 | 0 | 0 | 0 | 0 | 0 | 0 | 0 |

| | | | | | | | |
|---|---|---|---|---|---|---|---|
| ENSMUSG00000074156 | 0 | 0 | 0 | 0 | 0 | 0 | 0 |
| ENSMUSG00000074158 | 1.558729299 | 1.52186571 | 1.54248125 | 1.54248125 | 1.47337555 | 1.602059991 | 1.374453863 |
| ENSMUSG00000074164 | 0.289064826 | 0 | 0.25 | 0 | 0.244650542 | 0 | 0 |
| ENSMUSG00000074165 | 1.777675583 | 1.580482024 | 1.67510993 | 1.566696635 | 1.523883869 | 1.716003344 | 1.479989695 |
| ENSMUSG00000074166 | 1.156259305 | 1.321350555 | 1.25 | 1.372963274 | 1.264824954 | 1.51851394 | 1.131174228 |
| ENSMUSG00000074170 | 1.721148722 | 1.735628626 | 1.789967834 | 1.494319981 | 1.55104774 | 1.681241237 | 1.649885598 |
| ENSMUSG00000074171 | 0 | 0 | 0 | 0 | 0 | 0 | 0 |
| ENSMUSG00000074178 | 0 | 0 | 0.25 | 0 | 0 | 0 | 0.224243824 |
| ENSMUSG00000074179 | 0 | 0.25 | 0 | 0 | 0 | 0 | 0 |
| ENSMUSG00000074182 | 2.066740409 | 1.976722649 | 1.912762923 | 2.173371739 | 2.126866297 | 2.365487985 | 1.983546516 |
| ENSMUSG00000074183 | 0.578129653 | 0.701838731 | 0.396240625 | 0 | 0.244650542 | 0.698970004 | 0.520678036 |
| ENSMUSG00000074184 | 0 | 0 | 0 | 0 | 0 | 0 | 0 |
| ENSMUSG00000074194 | 1.036286563 | 0.951838731 | 0.92510993 | 1.311981878 | 1.200473444 | 1.397940009 | 1.252393349 |
| ENSMUSG00000074195 | 0 | 0 | 0 | 0 | 0 | 0 | 0 |
| ENSMUSG00000074196 | 0 | 0 | 0 | 0 | 0 | 0 | 0 |
| ENSMUSG00000074199 | 0 | 0 | 0 | 0.25 | 0 | 0 | 0 |
| ENSMUSG00000074203 | 0.289064826 | 0.75 | 0.25 | 0 | 0.244650542 | 0.301029996 | 0.520678036 |
| ENSMUSG00000074206 | 0 | 0.25 | 0 | 0 | 0 | 0 | 0 |
| ENSMUSG00000074207 | 0 | 0 | 0 | 0 | 0 | 0 | 0 |
| ENSMUSG00000074210 | 0 | 0 | 0 | 0 | 0 | 0 | 0 |
| ENSMUSG00000074211 | 1.100572389 | 1.27186571 | 1.02186571 | 1.226722649 | 1.057362051 | 1.230448921 | 1.041356071 |
| ENSMUSG00000074212 | 2.294202867 | 2.132845365 | 2.065523711 | 2.29248125 | 2.179810173 | 2.498310554 | 2.028761664 |
| ENSMUSG00000074213 | 0 | 0 | 0 | 0 | 0 | 0 | 0.355418052 |
| ENSMUSG00000074215 | 0.960252568 | 0.396240625 | 0.701838731 | 0.896240625 | 0.846351821 | 1 | 0.853775828 |
| ENSMUSG00000074217 | 0 | 0 | 0 | 0 | 0.244650542 | 0.301029996 | 0.224243824 |
| ENSMUSG00000074218 | 0.45815691 | 0.396240625 | 0.580482024 | 0.580482024 | 0 | 0.477121255 | 0 |
| ENSMUSG00000074219 | 0.578129653 | 0.646240625 | 0.580482024 | 0.75 | 0.686820904 | 0.602059991 | 0.775756176 |
| ENSMUSG00000074220 | 1.639698646 | 1.618933358 | 1.62694866 | 1.552363341 | 1.528558853 | 1.681241237 | 1.42167221 |
| ENSMUSG00000074221 | 2.31414447 | 2.154596376 | 2.118933358 | 2.109697963 | 2.072599934 | 2.357934847 | 1.934261938 |
| ENSMUSG00000074224 | 0 | 0 | 0 | 0 | 0 | 0 | 0 |
| ENSMUSG00000074227 | 0.747221736 | 0.896240625 | 0.951838731 | 1.364857905 | 1.414412788 | 1.591064607 | 1.42167221 |
| ENSMUSG00000074228 | 0 | 0 | 0.25 | 0 | 0 | 0 | 0 |
| ENSMUSG00000074232 | 0.289064826 | 0.396240625 | 0.25 | 0.580482024 | 0.489301084 | 0 | 0.448487648 |
| ENSMUSG00000074235 | 0 | 0 | 0 | 0 | 0 | 0 | 0 |

| | | | | | | | |
|---|---|---|---|---|---|---|---|
| ENSMUSG00000074236 | 0 | 0 | 0 | 0 | 0 | 0 | 0.224243824 |
| ENSMUSG00000074237 | 0 | 0 | 0 | 0 | 0 | 0 | 0 |
| ENSMUSG00000074238 | 2.493389029 | 2.296218836 | 2.289336734 | 2.358657057 | 2.298360391 | 2.62838893 | 2.138226615 |
| ENSMUSG00000074240 | 0 | 0 | 0 | 0 | 0 | 0 | 0 |
| ENSMUSG00000074243 | 0.45815691 | 0 | 0.396240625 | 0.25 | 0 | 0.301029996 | 0.355418052 |
| ENSMUSG00000074247 | 2.245508853 | 2.183677405 | 2.214495249 | 2.058404919 | 2.198391101 | 2.382017043 | 2.214058866 |
| ENSMUSG00000074248 | 0 | 0 | 0 | 0 | 0 | 0 | 0 |
| ENSMUSG00000074250 | 0.747221736 | 0.701838731 | 0.580482024 | 0.396240625 | 0.489301084 | 0.477121255 | 0.579661877 |
| ENSMUSG00000074252 | 0 | 0 | 0 | 0 | 0 | 0 | 0 |
| ENSMUSG00000074254 | 0 | 0 | 0 | 0 | 0 | 0 | 0 |
| ENSMUSG00000074256 | 0 | 0.25 | 0.396240625 | 0 | 0 | 0 | 0 |
| ENSMUSG00000074259 | 1.432083896 | 1.364857905 | 1.431980114 | 1.403677461 | 1.394615667 | 1.531478917 | 1.379042812 |
| ENSMUSG00000074261 | 0 | 0 | 0 | 0 | 0 | 0 | 0.355418052 |
| ENSMUSG00000074264 | 0.578129653 | 0.5 | 0.75 | 0.75 | 0.905314583 | 0.77815125 | 0.579661877 |
| ENSMUSG00000074268 | 0 | 0 | 0.25 | 0 | 0 | 0 | 0 |
| ENSMUSG00000074269 | 0.91631382 | 0.79248125 | 0.580482024 | 0.580482024 | 0.568060967 | 0.698970004 | 0.916588299 |
| ENSMUSG00000074272 | 0 | 0.5 | 0.396240625 | 0.25 | 0 | 0.477121255 | 0.355418052 |
| ENSMUSG00000074274 | 0 | 0 | 0 | 0 | 0 | 0 | 0.224243824 |
| ENSMUSG00000074277 | 0.578129653 | 1.04248125 | 1.098079356 | 1.061981878 | 1.121713561 | 0.84509804 | 1.329931221 |
| ENSMUSG00000074280 | 2.057835465 | 1.816696635 | 1.868933358 | 1.614857905 | 1.635855979 | 1.812913357 | 1.546120303 |
| ENSMUSG00000074281 | 0.45815691 | 0.396240625 | 0.25 | 0.25 | 0.568060967 | 0.301029996 | 0.355418052 |
| ENSMUSG00000074282 | 1.307602651 | 1.188721876 | 1.321350555 | 1.27186571 | 1.335652905 | 1.380211242 | 1.20139795 |
| ENSMUSG00000074283 | 1.205378646 | 1.201838731 | 1.02186571 | 1.17510993 | 1.223252711 | 0.903089987 | 1.014380832 |
| ENSMUSG00000074284 | 0.747221736 | 0.396240625 | 0.646240625 | 0.580482024 | 0.568060967 | 0.477121255 | 0.829800754 |
| ENSMUSG00000074289 | 0 | 0 | 0 | 0 | 0 | 0 | 0 |
| ENSMUSG00000074291 | 0 | 0 | 0 | 0 | 0 | 0 | 0 |
| ENSMUSG00000074292 | 0.671187741 | 0.701838731 | 0.646240625 | 0.646240625 | 0.733951626 | 0.77815125 | 0.448487648 |
| ENSMUSG00000074300 | 0 | 0.396240625 | 0 | 0 | 0 | 0 | 0 |
| ENSMUSG00000074302 | 0 | 0 | 0 | 0 | 0 | 0 | 0 |
| ENSMUSG00000074305 | 2.6088504 | 2.420748646 | 2.376452888 | 2.472185812 | 2.427747449 | 2.766412847 | 2.277155247 |
| ENSMUSG00000074311 | 0 | 0 | 0 | 0 | 0 | 0 | 0 |
| ENSMUSG00000074322 | 0 | 0 | 0 | 0 | 0 | 0 | 0 |
| ENSMUSG00000074336 | 0 | 0 | 0 | 0 | 0 | 0 | 0 |
| ENSMUSG00000074340 | 1.100572389 | 1 | 1.04248125 | 1.02186571 | 0.978602168 | 0.698970004 | 0.829800754 |

| | | | | | | | |
|---|---|---|---|---|---|---|---|
| ENSMUSG00000074342 | 0.289064826 | 0 | 0 | 0 | 0.244650542 | 0.301029996 | 0.448487648 |
| ENSMUSG00000074343 | 0 | 0 | 0 | 0 | 0 | 0 | 0 |
| ENSMUSG00000074344 | 0 | 0 | 0 | 0 | 0 | 0 | 0 |
| ENSMUSG00000074345 | 0 | 0 | 0 | 0 | 0 | 0.301029996 | 0 |
| ENSMUSG00000074346 | 0 | 0 | 0 | 0 | 0 | 0 | 0 |
| ENSMUSG00000074354 | 0 | 0.25 | 0 | 0.25 | 0.244650542 | 0.301029996 | 0.224243824 |
| ENSMUSG00000074355 | 0.671187741 | 0.5 | 0.646240625 | 0.25 | 0.489301084 | 0.301029996 | 0.355418052 |
| ENSMUSG00000074357 | 0.289064826 | 0.25 | 0.5 | 0.396240625 | 0.387761935 | 0.301029996 | 0.448487648 |
| ENSMUSG00000074358 | 1.432083896 | 1.403677461 | 1.571350555 | 0.701838731 | 1.310728002 | 1.278753601 | 1.396774124 |
| ENSMUSG00000074359 | 0.45815691 | 0.25 | 0.701838731 | 0.5 | 0 | 0.301029996 | 0.629532004 |
| ENSMUSG00000074361 | 0.289064826 | 0.25 | 0 | 0 | 0 | 0 | 0.355418052 |
| ENSMUSG00000074364 | 2.812982046 | 2.602877747 | 2.65892953 | 2.453445298 | 2.465298399 | 2.730782276 | 2.426179324 |
| ENSMUSG00000074365 | 0 | 0 | 0 | 0 | 0 | 0 | 0 |
| ENSMUSG00000074366 | 0.91631382 | 0.830482024 | 0.830482024 | 0 | 0 | 0 | 0 |
| ENSMUSG00000074369 | 0 | 0 | 0 | 0 | 0 | 0 | 0 |
| ENSMUSG00000074373 | 0 | 0 | 0 | 0 | 0 | 0 | 0 |
| ENSMUSG00000074375 | 0 | 0 | 0 | 0 | 0 | 0 | 0 |
| ENSMUSG00000074377 | 0 | 0 | 0 | 0 | 0 | 0 | 0 |
| ENSMUSG00000074378 | 0 | 0 | 0 | 0 | 0 | 0 | 0 |
| ENSMUSG00000074384 | 1.249317394 | 1.311981878 | 1.339388001 | 1.25 | 1.200473444 | 1.322219295 | 1.159323753 |
| ENSMUSG00000074385 | 0 | 0 | 0 | 0 | 0 | 0 | 0 |
| ENSMUSG00000074388 | 0.45815691 | 0 | 0.25 | 0 | 0.244650542 | 0 | 0.355418052 |
| ENSMUSG00000074397 | 0 | 0 | 0.25 | 0.396240625 | 0.244650542 | 0.301029996 | 0.224243824 |
| ENSMUSG00000074398 | 0.45815691 | 0.25 | 0.5 | 0.25 | 0 | 0.301029996 | 0.355418052 |
| ENSMUSG00000074401 | 0 | 0 | 0 | 0 | 0 | 0 | 0 |
| ENSMUSG00000074402 | 0 | 0 | 0 | 0 | 0 | 0 | 0 |
| ENSMUSG00000074403 | 0 | 0 | 0 | 0 | 0 | 0 | 0 |
| ENSMUSG00000074404 | 0 | 0 | 0 | 0 | 0 | 0 | 0 |
| ENSMUSG00000074405 | 1.516991255 | 1.418106335 | 1.532320754 | 1.25 | 1.254881871 | 0.903089987 | 1.520678036 |
| ENSMUSG00000074406 | 1.432083896 | 1.380890489 | 1.566696635 | 1.02186571 | 1.335652905 | 1.041392685 | 1.462868481 |
| ENSMUSG00000074412 | 1 | 1.02186571 | 0.92510993 | 0.976722649 | 1.106691885 | 1 | 0.969165684 |
| ENSMUSG00000074415 | 2.636714502 | 2.353935442 | 2.32991803 | 2.255591953 | 2.193483986 | 2.383815366 | 1.995286892 |
| ENSMUSG00000074417 | 0 | 0 | 0 | 0 | 0 | 0 | 0 |
| ENSMUSG00000074418 | 0 | 0 | 0 | 0 | 0 | 0 | 0 |

| | | | | | | | |
|---|---|---|---|---|---|---|---|
| ENSMUSG00000074419 | 0 | 0 | 0 | 0 | 0 | 0 | 0 |
| ENSMUSG00000074420 | 0 | 0 | 0 | 0 | 0 | 0 | 0 |
| ENSMUSG00000074424 | 0 | 0 | 0 | 0 | 0 | 0 | 0 |
| ENSMUSG00000074433 | 0 | 0 | 0 | 0 | 0 | 0 | 0 |
| ENSMUSG00000074434 | 0 | 0 | 0 | 0 | 0 | 0 | 0 |
| ENSMUSG00000074435 | 0 | 0 | 0 | 0 | 0 | 0 | 0 |
| ENSMUSG00000074436 | 0 | 0 | 0 | 0 | 0 | 0 | 0 |
| ENSMUSG00000074437 | 0 | 0 | 0 | 0 | 0 | 0 | 0 |
| ENSMUSG00000074439 | 0 | 0 | 0 | 0 | 0 | 0 | 0 |
| ENSMUSG00000074440 | 0 | 0 | 0 | 0 | 0 | 0 | 0 |
| ENSMUSG00000074441 | 0 | 0 | 0 | 0 | 0 | 0 | 0 |
| ENSMUSG00000074442 | 0 | 0 | 0 | 0 | 0 | 0 | 0 |
| ENSMUSG00000074443 | 0 | 0 | 0 | 0 | 0 | 0 | 0 |
| ENSMUSG00000074444 | 0 | 0 | 0 | 0 | 0 | 0 | 0 |
| ENSMUSG00000074445 | 0 | 0 | 0 | 0 | 0 | 0 | 0 |
| ENSMUSG00000074446 | 0 | 0 | 0 | 0 | 0 | 0 | 0 |
| ENSMUSG00000074447 | 0 | 0 | 0 | 0 | 0 | 0 | 0 |
| ENSMUSG00000074448 | 0 | 0.25 | 0 | 0.25 | 0 | 0.301029996 | 0 |
| ENSMUSG00000074449 | 0 | 0 | 0 | 0 | 0 | 0 | 0 |
| ENSMUSG00000074452 | 0.811507563 | 0.396240625 | 0.701838731 | 0.396240625 | 0.568060967 | 0 | 0.224243824 |
| ENSMUSG00000074454 | 0 | 0 | 0 | 0 | 0 | 0 | 0 |
| ENSMUSG00000074457 | 1.693336259 | 1.584962501 | 1.527131114 | 0.896240625 | 0.877063019 | 0.954242509 | 1.054044578 |
| ENSMUSG00000074461 | 0.289064826 | 0.25 | 0.25 | 0.25 | 0.244650542 | 0.477121255 | 0.579661877 |
| ENSMUSG00000074465 | 0 | 0.25 | 0.396240625 | 0.25 | 0 | 0.477121255 | 0 |
| ENSMUSG00000074466 | 0.671187741 | 0.864857905 | 0.92510993 | 0.701838731 | 0.686820904 | 0.477121255 | 0.803905701 |
| ENSMUSG00000074467 | 0 | 0 | 0 | 0 | 0 | 0 | 0 |
| ENSMUSG00000074469 | 0 | 0 | 0 | 0 | 0 | 0 | 0 |
| ENSMUSG00000074472 | 0 | 0.396240625 | 0 | 0.25 | 0.244650542 | 0 | 0.224243824 |
| ENSMUSG00000074473 | 0 | 0 | 0 | 0 | 0 | 0 | 0 |
| ENSMUSG00000074476 | 2.10861828 | 1.998588359 | 2.071350555 | 2.155512955 | 2.190648982 | 2.428134794 | 2.157274883 |
| ENSMUSG00000074479 | 0 | 0 | 0 | 0 | 0 | 0.301029996 | 0 |
| ENSMUSG00000074480 | 2.598312578 | 2.365369862 | 2.331607372 | 2.320772088 | 2.265401209 | 2.568201724 | 2.148947128 |
| ENSMUSG00000074482 | 0.289064826 | 0 | 0 | 0 | 0 | 0 | 0 |
| ENSMUSG00000074483 | 0 | 0.25 | 0.25 | 0 | 0.244650542 | 0 | 0.224243824 |

| | | | | | | | |
|---|---|---|---|---|---|---|---|
| ENSMUSG00000074485 | 0 | 0 | 0.25 | 0 | 0 | 0 | 0 |
| ENSMUSG00000074486 | 0 | 0 | 0 | 0 | 0 | 0 | 0 |
| ENSMUSG00000074489 | 0.45815691 | 0.396240625 | 0.25 | 0.25 | 0.568060967 | 0.301029996 | 0.224243824 |
| ENSMUSG00000074491 | 0 | 0 | 0 | 0 | 0 | 0 | 0 |
| ENSMUSG00000074497 | 0.811507563 | 0.396240625 | 0 | 0.646240625 | 0.387761935 | 0.698970004 | 0.448487648 |
| ENSMUSG00000074500 | 0.867194479 | 1.214495249 | 1.188721876 | 1.098079356 | 0.77552387 | 0.903089987 | 0.876096088 |
| ENSMUSG00000074501 | 0 | 0 | 0 | 0 | 0 | 0 | 0 |
| ENSMUSG00000074502 | 0.45815691 | 0 | 0 | 0 | 0 | 0 | 0.224243824 |
| ENSMUSG00000074505 | 0 | 0 | 0 | 0 | 0 | 0 | 0 |
| ENSMUSG00000074506 | 0 | 0 | 0 | 0 | 0 | 0 | 0 |
| ENSMUSG00000074507 | 0.289064826 | 0.25 | 0 | 0.5 | 0.244650542 | 0 | 0 |
| ENSMUSG00000074508 | 0 | 0 | 0 | 0 | 0 | 0 | 0 |
| ENSMUSG00000074513 | 1.811507563 | 1.692046081 | 1.685366747 | 1.784887838 | 1.689774529 | 1.832508913 | 1.683576582 |
| ENSMUSG00000074516 | 0 | 0 | 0 | 0 | 0.244650542 | 0 | 0.355418052 |
| ENSMUSG00000074517 | 0 | 0 | 0 | 0 | 0 | 0 | 0.224243824 |
| ENSMUSG00000074519 | 1.289064826 | 1.130890489 | 1.114857905 | 1.321350555 | 1.293076518 | 1.431363764 | 1.176815335 |
| ENSMUSG00000074521 | 0 | 0 | 0 | 0 | 0.244650542 | 0 | 0.355418052 |
| ENSMUSG00000074527 | 1.036286563 | 1.130890489 | 0.830482024 | 1.388647213 | 1.223252711 | 1.361727836 | 1.110947927 |
| ENSMUSG00000074529 | 0 | 0 | 0 | 0.25 | 0 | 0.301029996 | 0.448487648 |
| ENSMUSG00000074547 | 0.289064826 | 0 | 0.580482024 | 0 | 0 | 0 | 0.355418052 |
| ENSMUSG00000074553 | 0 | 0 | 0 | 0 | 0 | 0 | 0 |
| ENSMUSG00000074555 | 0 | 0 | 0 | 0.25 | 0 | 0 | 0 |
| ENSMUSG00000074564 | 0 | 0 | 0 | 0 | 0 | 0 | 0 |
| ENSMUSG00000074569 | 0 | 0.396240625 | 0.396240625 | 0.5 | 0 | 0.602059991 | 0.520678036 |
| ENSMUSG00000074570 | 0 | 0.5 | 0.25 | 0.25 | 0 | 0 | 0 |
| ENSMUSG00000074575 | 0 | 0 | 0 | 0.830482024 | 0.812711509 | 0.602059991 | 0.829800754 |
| ENSMUSG00000074576 | 0.960252568 | 0.951838731 | 0.830482024 | 1.098079356 | 1.039257769 | 1.079181246 | 1.089372236 |
| ENSMUSG00000074577 | 0 | 0 | 0.25 | 0.25 | 0.632412477 | 0.477121255 | 0.876096088 |
| ENSMUSG00000074578 | 2.326928073 | 2.125956434 | 2.121960008 | 2.14151351 | 2.136951444 | 2.385606274 | 1.951019875 |
| ENSMUSG00000074579 | 1.227926429 | 1.27186571 | 1.188721876 | 1.214495249 | 1.136121934 | 1.230448921 | 1.054044578 |
| ENSMUSG00000074580 | 0 | 0.25 | 0.5 | 0.580482024 | 0.568060967 | 0.477121255 | 0.520678036 |
| ENSMUSG00000074582 | 2.590021402 | 2.405512955 | 2.415889526 | 2.447383411 | 2.450958105 | 2.739572344 | 2.276587178 |
| ENSMUSG00000074589 | 0 | 0 | 0 | 0 | 0 | 0 | 0 |
| ENSMUSG00000074591 | 0 | 0 | 0 | 0.25 | 0 | 0 | 0.224243824 |

| | | | | | | | |
|---|---|---|---|---|---|---|---|
| ENSMUSG00000074593 | 0 | 0 | 0 | 0 | 0 | 0 | 0 |
| ENSMUSG00000074594 | 0 | 0 | 0 | 0 | 0 | 0 | 0 |
| ENSMUSG00000074595 | 0 | 0 | 0 | 0 | 0 | 0 | 0 |
| ENSMUSG00000074596 | 0 | 0 | 0 | 0 | 0 | 0 | 0 |
| ENSMUSG00000074603 | 0 | 0.396240625 | 0 | 0 | 0.387761935 | 0 | 0 |
| ENSMUSG00000074604 | 0.45815691 | 0.396240625 | 0.396240625 | 0.25 | 0.244650542 | 0.301029996 | 0.224243824 |
| ENSMUSG00000074607 | 0 | 0 | 0 | 0 | 0 | 0 | 0 |
| ENSMUSG00000074619 | 0 | 0 | 0 | 0.25 | 0.244650542 | 0.301029996 | 0.355418052 |
| ENSMUSG00000074620 | 0 | 0 | 0 | 0 | 0 | 0 | 0 |
| ENSMUSG00000074622 | 0 | 0.25 | 0.25 | 0 | 0.489301084 | 0 | 0.224243824 |
| ENSMUSG00000074623 | 0 | 0 | 0 | 0 | 0 | 0 | 0 |
| ENSMUSG00000074625 | 0 | 0.25 | 0 | 0 | 0 | 0 | 0 |
| ENSMUSG00000074627 | 0 | 0 | 0.25 | 0 | 0 | 0 | 0.224243824 |
| ENSMUSG00000074628 | 0 | 0 | 0.25 | 0 | 0.244650542 | 0 | 0.224243824 |
| ENSMUSG00000074629 | 0.578129653 | 0.646240625 | 0.646240625 | 0.896240625 | 1 | 0.602059991 | 0.853775828 |
| ENSMUSG00000074634 | 0.578129653 | 0.580482024 | 0.5 | 1.25 | 1.121713561 | 1.204119983 | 1.159323753 |
| ENSMUSG00000074635 | 0.578129653 | 0.5 | 0.396240625 | 0.580482024 | 0.568060967 | 0.77815125 | 0.672731473 |
| ENSMUSG00000074637 | 0 | 0 | 0 | 0 | 0 | 0 | 0 |
| ENSMUSG00000074639 | 0 | 0 | 0 | 0 | 0 | 0 | 0 |
| ENSMUSG00000074643 | 0.289064826 | 0 | 0 | 0.396240625 | 0.244650542 | 0.301029996 | 0.224243824 |
| ENSMUSG00000074646 | 0.578129653 | 0.580482024 | 0.5 | 0.580482024 | 0.877063019 | 0.84509804 | 0.629532004 |
| ENSMUSG00000074647 | 0.45815691 | 0.25 | 0 | 0.646240625 | 0.489301084 | 0 | 0.629532004 |
| ENSMUSG00000074649 | 1.623015126 | 1.598079356 | 1.688721876 | 1.956637122 | 1.898867717 | 2.187520721 | 1.921737195 |
| ENSMUSG00000074650 | 0 | 0.396240625 | 0 | 0.580482024 | 0 | 0.77815125 | 0.629532004 |
| ENSMUSG00000074651 | 0 | 0 | 0 | 0 | 0 | 0.301029996 | 0.355418052 |
| ENSMUSG00000074652 | 0 | 0 | 0 | 0.25 | 0.387761935 | 0.477121255 | 0.672731473 |
| ENSMUSG00000074653 | 0.289064826 | 0 | 0 | 0.25 | 0.244650542 | 0 | 0 |
| ENSMUSG00000074655 | 0 | 0 | 0 | 0 | 0 | 0 | 0 |
| ENSMUSG00000074656 | 2.879086229 | 2.666556001 | 2.596735061 | 2.824229052 | 2.720254961 | 3.166726056 | 2.527686819 |
| ENSMUSG00000074657 | 1.069666964 | 1.188721876 | 0.864857905 | 1.380890489 | 1.427019704 | 1.342422681 | 1.462868481 |
| ENSMUSG00000074664 | 0 | 0 | 0 | 0 | 0 | 0 | 0 |
| ENSMUSG00000074665 | 0 | 0 | 0 | 0 | 0 | 0 | 0.224243824 |
| ENSMUSG00000074671 | 0.289064826 | 0.25 | 0.580482024 | 1.388647213 | 1.433158226 | 1.531478917 | 1.360283038 |
| ENSMUSG00000074673 | 0 | 0.25 | 0 | 0.396240625 | 0.632412477 | 0.477121255 | 0.520678036 |

| | | | | | | | |
|---|---|---|---|---|---|---|---|
| ENSMUSG00000074676 | 1.069666964 | 1.282320754 | 1.130890489 | 0 | 0 | 0 | 0 |
| ENSMUSG00000074677 | 0 | 0 | 0 | 0 | 0 | 0 | 0 |
| ENSMUSG00000074678 | 0 | 0 | 0 | 0 | 0 | 0 | 0 |
| ENSMUSG00000074679 | 0 | 0 | 0 | 0 | 0 | 0 | 0 |
| ENSMUSG00000074680 | 0 | 0 | 0 | 0 | 0 | 0 | 0 |
| ENSMUSG00000074681 | 0 | 0 | 0 | 0 | 0 | 0 | 0 |
| ENSMUSG00000074682 | 1.482695304 | 1.380890489 | 1.445339928 | 1.5 | 1.49445382 | 1.633468456 | 1.535058868 |
| ENSMUSG00000074695 | 0 | 0 | 0 | 0 | 0 | 0 | 0 |
| ENSMUSG00000074698 | 2.613625632 | 2.369945066 | 2.339388001 | 2.535845803 | 2.462015062 | 2.892094603 | 2.305399353 |
| ENSMUSG00000074704 | 0 | 0 | 0 | 0 | 0 | 0 | 0 |
| ENSMUSG00000074715 | 0.289064826 | 0 | 0.396240625 | 1.27186571 | 1.163285805 | 1.255272505 | 1.110947927 |
| ENSMUSG00000074720 | 0 | 0 | 0 | 0 | 0 | 0 | 0 |
| ENSMUSG00000074731 | 0.91631382 | 0.951838731 | 0.79248125 | 0.580482024 | 0.489301084 | 0.477121255 | 0.579661877 |
| ENSMUSG00000074733 | 2.029919532 | 1.966046536 | 1.868933358 | 2.01109853 | 1.982473755 | 2.093421685 | 1.840919217 |
| ENSMUSG00000074734 | 0 | 0 | 0 | 0 | 0 | 0 | 0 |
| ENSMUSG00000074735 | 0.671187741 | 0.92510993 | 0.92510993 | 1.330482024 | 1.163285805 | 1.255272505 | 1.131174228 |
| ENSMUSG00000074736 | 0.45815691 | 0.25 | 0.5 | 0 | 0 | 0 | 0 |
| ENSMUSG00000074737 | 0 | 0 | 0 | 0 | 0 | 0 | 0 |
| ENSMUSG00000074738 | 0 | 0 | 0 | 0 | 0 | 0 | 0 |
| ENSMUSG00000074739 | 0 | 0 | 0.25 | 0 | 0 | 0 | 0 |
| ENSMUSG00000074743 | 1.924654746 | 1.769203899 | 1.79745614 | 0.92510993 | 1 | 1 | 1.159323753 |
| ENSMUSG00000074746 | 2.938680166 | 2.690179987 | 2.64881437 | 2.62521047 | 2.562149491 | 2.928395852 | 2.363558619 |
| ENSMUSG00000074748 | 2.940852214 | 2.678346629 | 2.654825739 | 2.597004321 | 2.527738343 | 2.921166051 | 2.368517634 |
| ENSMUSG00000074749 | 1.707474304 | 1.589388001 | 1.618933358 | 1.811981878 | 1.749189509 | 2.012837225 | 1.635773653 |
| ENSMUSG00000074753 | 0 | 0 | 0 | 0 | 0 | 0 | 0 |
| ENSMUSG00000074754 | 1.325351389 | 1.25 | 1.372963274 | 1.348079356 | 1.343584837 | 1.397940009 | 1.290497981 |
| ENSMUSG00000074758 | 0 | 0 | 0 | 0 | 0 | 0 | 0 |
| ENSMUSG00000074763 | 0 | 0 | 0 | 0 | 0 | 0 | 0 |
| ENSMUSG00000074764 | 0 | 0 | 0 | 0 | 0 | 0 | 0 |
| ENSMUSG00000074766 | 0 | 0.25 | 0 | 0.396240625 | 0 | 0 | 0.224243824 |
| ENSMUSG00000074768 | 0 | 0 | 0 | 0.25 | 0 | 0 | 0 |
| ENSMUSG00000074771 | 0.289064826 | 0.25 | 0.396240625 | 0 | 0.244650542 | 0.698970004 | 0 |
| ENSMUSG00000074776 | 0 | 0 | 0 | 0 | 0 | 0 | 0 |
| ENSMUSG00000074780 | 0 | 0 | 0.25 | 0.25 | 0 | 0 | 0 |

| | | | | | | | |
|---|---|---|---|---|---|---|---|
| ENSMUSG00000074781 | 2.855840016 | 2.614857905 | 2.565229883 | 2.760927716 | 2.633834461 | 3.114277297 | 2.467785947 |
| ENSMUSG00000074782 | 0.747221736 | 0.701838731 | 0.864857905 | 0.830482024 | 1 | 0.477121255 | 0.829800754 |
| ENSMUSG00000074783 | 0.289064826 | 0 | 0.25 | 0.25 | 0.387761935 | 0 | 0.224243824 |
| ENSMUSG00000074785 | 0 | 0.25 | 0 | 0.580482024 | 0.686820904 | 0.477121255 | 0.579661877 |
| ENSMUSG00000074793 | 0.811507563 | 1.02186571 | 1.080482024 | 1.214495249 | 1.136121934 | 1.301029996 | 1.245582246 |
| ENSMUSG00000074794 | 1.920505136 | 1.807204673 | 1.80479213 | 1.923371739 | 1.791313678 | 2.113943352 | 1.694069894 |
| ENSMUSG00000074796 | 1.129344651 | 1.27186571 | 1.26109853 | 1.226722649 | 1.264824954 | 1.113943352 | 1.335191751 |
| ENSMUSG00000074797 | 2.317377295 | 2.158248799 | 2.151832578 | 2.286801231 | 2.272243869 | 2.525044807 | 2.093403634 |
| ENSMUSG00000074800 | 0.811507563 | 0.75 | 0.75 | 0.75 | 0.978602168 | 0.477121255 | 0.916588299 |
| ENSMUSG00000074802 | 2.507914013 | 2.288071211 | 2.321350555 | 2.371457327 | 2.303049176 | 2.588831726 | 2.102294694 |
| ENSMUSG00000074804 | 0 | 0 | 0 | 0 | 0 | 0 | 0 |
| ENSMUSG00000074805 | 0 | 0.25 | 0 | 0 | 0 | 0 | 0 |
| ENSMUSG00000074807 | 0.811507563 | 0.976722649 | 0.79248125 | 0.79248125 | 0.733951626 | 1.041392685 | 1.014380832 |
| ENSMUSG00000074811 | 1.822201287 | 1.809601185 | 1.777131114 | 1.561981878 | 1.728722551 | 1.763427994 | 1.791413205 |
| ENSMUSG00000074812 | 0 | 0 | 0 | 0 | 0 | 0 | 0 |
| ENSMUSG00000074813 | 1.389637216 | 1.410964047 | 1.42510993 | 1.527131114 | 1.559656853 | 1.72427587 | 1.459332757 |
| ENSMUSG00000074817 | 0.960252568 | 1.080482024 | 0.79248125 | 1.451838731 | 1.387761935 | 1.477121255 | 1.324583736 |
| ENSMUSG00000074818 | 1.069666964 | 1 | 0.830482024 | 0.75 | 0.877063019 | 0.602059991 | 0.984950056 |
| ENSMUSG00000074824 | 0.45815691 | 0.5 | 0.646240625 | 0.646240625 | 0.632412477 | 0.698970004 | 0.629532004 |
| ENSMUSG00000074825 | 1.944805158 | 1.896240625 | 1.839388001 | 1.909156155 | 1.937354184 | 2.103803721 | 1.829800754 |
| ENSMUSG00000074826 | 0.91631382 | 1.02186571 | 1.02186571 | 1.188721876 | 1 | 0.903089987 | 0.935079929 |
| ENSMUSG00000074829 | 1.837744666 | 1.720660762 | 1.642463902 | 1.598079356 | 1.621875691 | 1.748188027 | 1.499406449 |
| ENSMUSG00000074830 | 0 | 0 | 0 | 0 | 0 | 0 | 0 |
| ENSMUSG00000074832 | 0 | 0.5 | 0.646240625 | 0.646240625 | 0.489301084 | 0 | 0.579661877 |
| ENSMUSG00000074837 | 0 | 0 | 0 | 0 | 0 | 0 | 0 |
| ENSMUSG00000074841 | 0 | 0 | 0 | 0 | 0 | 0 | 0 |
| ENSMUSG00000074846 | 0 | 0 | 0 | 0 | 0 | 0 | 0 |
| ENSMUSG00000074847 | 0.289064826 | 0.396240625 | 0.396240625 | 0.25 | 0.387761935 | 0 | 0.579661877 |
| ENSMUSG00000074849 | 0 | 0 | 0 | 0 | 0 | 0 | 0 |
| ENSMUSG00000074852 | 0 | 0 | 0 | 0 | 0 | 0 | 0 |
| ENSMUSG00000074862 | 0.960252568 | 0.92510993 | 0.951838731 | 0.701838731 | 1 | 1 | 0.896975297 |
| ENSMUSG00000074863 | 1.404271433 | 1.339388001 | 1.396240625 | 1.464495249 | 1.456697355 | 1.62324929 | 1.302263476 |
| ENSMUSG00000074865 | 1.249317394 | 1.438721876 | 1.201838731 | 1.410964047 | 1.414412788 | 1.414973348 | 1.21680636 |
| ENSMUSG00000074867 | 1.827447047 | 1.741446071 | 1.779735268 | 1.834962501 | 1.822942838 | 1.982271233 | 1.637827726 |

| | | | | | | | |
|---|---|---|---|---|---|---|---|
| ENSMUSG00000074868 | 0.578129653 | 0 | 0.25 | 0 | 0.244650542 | 0 | 0 |
| ENSMUSG00000074870 | 0 | 0 | 0 | 0 | 0 | 0 | 0 |
| ENSMUSG00000074871 | 0 | 0 | 0 | 0 | 0 | 0 | 0 |
| ENSMUSG00000074872 | 0 | 0 | 0 | 0 | 0 | 0 | 0 |
| ENSMUSG00000074873 | 1.623015126 | 1.470660762 | 1.476722649 | 1.516522298 | 1.462344774 | 1.477121255 | 1.396774124 |
| ENSMUSG00000074874 | 0.747221736 | 0.580482024 | 0.830482024 | 0.5 | 0.489301084 | 0.301029996 | 0.224243824 |
| ENSMUSG00000074876 | 0.45815691 | 0.580482024 | 0.646240625 | 0.5 | 0.568060967 | 0.477121255 | 0.876096088 |
| ENSMUSG00000074877 | 0 | 0 | 0 | 0 | 0 | 0 | 0 |
| ENSMUSG00000074881 | 0.45815691 | 0.25 | 0.25 | 0 | 0.387761935 | 0 | 0.224243824 |
| ENSMUSG00000074882 | 0 | 0 | 0 | 0 | 0 | 0 | 0 |
| ENSMUSG00000074884 | 2.165631214 | 1.995748394 | 2.013820609 | 1.905512955 | 1.92093466 | 2.053078443 | 1.774416565 |
| ENSMUSG00000074885 | 0 | 0 | 0 | 0 | 0 | 0 | 0 |
| ENSMUSG00000074886 | 2.66563648 | 2.490724001 | 2.488185312 | 2.451838731 | 2.555125257 | 2.80140371 | 2.421490612 |
| ENSMUSG00000074887 | 0 | 0 | 0 | 0 | 0.244650542 | 0 | 0 |
| ENSMUSG00000074890 | 1.516991255 | 1.451838731 | 1.464495249 | 1.445339928 | 1.401338844 | 1.544068044 | 1.340368108 |
| ENSMUSG00000074892 | 0 | 0.25 | 0.25 | 0.396240625 | 0.387761935 | 0.301029996 | 0.224243824 |
| ENSMUSG00000074895 | 0 | 0 | 0 | 0.25 | 0 | 0 | 0 |
| ENSMUSG00000074896 | 0.671187741 | 0.830482024 | 0.79248125 | 0.830482024 | 0.686820904 | 0.602059991 | 0.710836105 |
| ENSMUSG00000074899 | 0 | 0 | 0.25 | 0 | 0.387761935 | 0 | 0.224243824 |
| ENSMUSG00000074903 | 0 | 0 | 0 | 0 | 0 | 0 | 0 |
| ENSMUSG00000074909 | 2.17479713 | 2.015173983 | 1.942046081 | 2.107613138 | 2.042520869 | 2.301029996 | 1.835407748 |
| ENSMUSG00000074912 | 0 | 0 | 0 | 0 | 0 | 0.301029996 | 0.355418052 |
| ENSMUSG00000074913 | 0 | 0 | 0 | 0 | 0 | 0 | 0 |
| ENSMUSG00000074916 | 1.862428343 | 1.78743678 | 1.80479213 | 1.821350555 | 1.866526233 | 2.021189299 | 1.781059827 |
| ENSMUSG00000074918 | 1.765759561 | 1.685366747 | 1.732684334 | 1.959800947 | 1.90849439 | 2.025305865 | 1.771720588 |
| ENSMUSG00000074919 | 0 | 0 | 0 | 0 | 0 | 0 | 0 |
| ENSMUSG00000074922 | 1.53838222 | 1.5 | 1.552363341 | 1.396240625 | 1.358933174 | 1.653212514 | 1.369798885 |
| ENSMUSG00000074923 | 0.578129653 | 0.864857905 | 1.04248125 | 0.79248125 | 1 | 1.079181246 | 1.066254157 |
| ENSMUSG00000074925 | 2.00683675 | 1.821350555 | 1.75 | 1.789967834 | 1.797870327 | 1.949390007 | 1.589319772 |
| ENSMUSG00000074928 | 0 | 0 | 0 | 0 | 0 | 0 | 0 |
| ENSMUSG00000074930 | 0 | 0.25 | 0.25 | 0 | 0 | 0 | 0 |
| ENSMUSG00000074934 | 2.754623188 | 2.519871196 | 2.605516192 | 1.982684334 | 1.970729215 | 2.123851641 | 1.737484395 |
| ENSMUSG00000074939 | 0 | 0 | 0 | 0 | 0 | 0 | 0 |
| ENSMUSG00000074943 | 0 | 0 | 0 | 0 | 0 | 0 | 0 |

| | | | | | | | |
|---|---|---|---|---|---|---|---|
| ENSMUSG00000074945 | 0 | 0 | 0 | 0 | 0 | 0 | 0 |
| ENSMUSG00000074946 | 0 | 0 | 0 | 0 | 0 | 0 | 0 |
| ENSMUSG00000074947 | 0 | 0 | 0 | 0 | 0 | 0 | 0 |
| ENSMUSG00000074951 | 0 | 0 | 0 | 0 | 0 | 0 | 0 |
| ENSMUSG00000074952 | 0 | 0 | 0 | 0 | 0 | 0 | 0 |
| ENSMUSG00000074955 | 0 | 0 | 0 | 0 | 0 | 0 | 0 |
| ENSMUSG00000074957 | 0 | 0 | 0 | 0 | 0 | 0 | 0 |
| ENSMUSG00000074965 | 0 | 0 | 0 | 0 | 0 | 0 | 0 |
| ENSMUSG00000074966 | 0 | 0 | 0 | 0 | 0 | 0 | 0 |
| ENSMUSG00000074968 | 0.747221736 | 0.830482024 | 0.951838731 | 0.25 | 0.244650542 | 0.301029996 | 0 |
| ENSMUSG00000074970 | 0 | 0 | 0 | 0 | 0 | 0 | 0 |
| ENSMUSG00000074971 | 0.747221736 | 0.25 | 0.5 | 0 | 0 | 0 | 0 |
| ENSMUSG00000074973 | 0 | 0.396240625 | 0.396240625 | 0.25 | 0 | 0 | 0 |
| ENSMUSG00000074978 | 0.45815691 | 0.396240625 | 0.396240625 | 0 | 0 | 0 | 0.224243824 |
| ENSMUSG00000074981 | 0 | 0.25 | 0 | 0 | 0 | 0 | 0.224243824 |
| ENSMUSG00000074985 | 0 | 0 | 0 | 0 | 0 | 0 | 0 |
| ENSMUSG00000074987 | 0 | 0 | 0 | 0 | 0 | 0 | 0 |
| ENSMUSG00000074991 | 0 | 0 | 0 | 0 | 0 | 0 | 0 |
| ENSMUSG00000074994 | 2.523917219 | 2.313757142 | 2.294354884 | 2.529410775 | 2.422990063 | 2.796574333 | 2.226574877 |
| ENSMUSG00000074995 | 0 | 0 | 0 | 0 | 0 | 0 | 0 |
| ENSMUSG00000074996 | 0 | 0 | 0 | 0 | 0 | 0 | 0 |
| ENSMUSG00000074997 | 0 | 0 | 0 | 0 | 0 | 0 | 0 |
| ENSMUSG00000074999 | 0 | 0 | 0 | 0.25 | 0 | 0.301029996 | 0.224243824 |
| ENSMUSG00000075000 | 1.932832114 | 1.78743678 | 1.769203899 | 1.830482024 | 1.784632921 | 1.929418926 | 1.732938436 |
| ENSMUSG00000075002 | 0 | 0 | 0 | 0 | 0 | 0 | 0 |
| ENSMUSG00000075006 | 0 | 0 | 0 | 0 | 0 | 0 | 0 |
| ENSMUSG00000075010 | 0 | 0 | 0 | 0 | 0 | 0 | 0 |
| ENSMUSG00000075012 | 1.289064826 | 1.160964047 | 1.27186571 | 0.75 | 1.020174412 | 0.698970004 | 1 |
| ENSMUSG00000075014 | 0 | 0 | 0 | 0 | 0.387761935 | 0 | 0 |
| ENSMUSG00000075015 | 0.289064826 | 0 | 0 | 0 | 0 | 0 | 0.224243824 |
| ENSMUSG00000075020 | 0 | 0 | 0 | 0 | 0 | 0 | 0 |
| ENSMUSG00000075023 | 0 | 0 | 0.25 | 0 | 0 | 0 | 0 |
| ENSMUSG00000075024 | 0 | 0 | 0 | 0 | 0 | 0 | 0 |
| ENSMUSG00000075025 | 0 | 0 | 0.396240625 | 0 | 0 | 0 | 0.224243824 |

| | | | | | | | |
|---|---|---|---|---|---|---|---|
| ENSMUSG00000075027 | 0 | 0 | 0.25 | 0.396240625 | 0 | 0.301029996 | 0.224243824 |
| ENSMUSG00000075028 | 1.404271433 | 1.339388001 | 1.476722649 | 1.482684334 | 1.53772706 | 1.568201724 | 1.543390198 |
| ENSMUSG00000075029 | 0 | 0 | 0 | 0 | 0 | 0 | 0 |
| ENSMUSG00000075031 | 0 | 0 | 0 | 0.396240625 | 0 | 0.477121255 | 0.224243824 |
| ENSMUSG00000075033 | 1.816888701 | 1.708222504 | 1.685366747 | 1.692046081 | 1.689774529 | 1.832508913 | 1.543390198 |
| ENSMUSG00000075036 | 0 | 0 | 0 | 0 | 0 | 0 | 0 |
| ENSMUSG00000075040 | 1.822201287 | 1.777131114 | 1.777131114 | 1.77186571 | 1.828942237 | 1.939519253 | 1.812372009 |
| ENSMUSG00000075042 | 0 | 0 | 0 | 0 | 0 | 0 | 0 |
| ENSMUSG00000075044 | 0 | 0 | 0 | 0 | 0 | 0 | 0 |
| ENSMUSG00000075045 | 0 | 0 | 0 | 0 | 0 | 0 | 0 |
| ENSMUSG00000075046 | 0 | 0 | 0 | 0 | 0.244650542 | 0 | 0 |
| ENSMUSG00000075053 | 0.578129653 | 0.396240625 | 0.25 | 0.396240625 | 0.387761935 | 0.301029996 | 0 |
| ENSMUSG00000075054 | 2.045658471 | 2.03360658 | 1.966046536 | 2.066696635 | 1.969399812 | 2.217483944 | 1.870159858 |
| ENSMUSG00000075061 | 0 | 0 | 0 | 0 | 0 | 0 | 0 |
| ENSMUSG00000075062 | 0 | 0 | 0 | 0 | 0 | 0 | 0 |
| ENSMUSG00000075063 | 0 | 0 | 0 | 0 | 0 | 0 | 0 |
| ENSMUSG00000075064 | 0 | 0 | 0 | 0 | 0 | 0 | 0 |
| ENSMUSG00000075065 | 0 | 0 | 0 | 0 | 0 | 0 | 0 |
| ENSMUSG00000075066 | 0 | 0 | 0 | 0 | 0 | 0 | 0 |
| ENSMUSG00000075067 | 0 | 0 | 0 | 0 | 0 | 0 | 0 |
| ENSMUSG00000075068 | 0 | 0 | 0 | 0 | 0 | 0 | 0 |
| ENSMUSG00000075069 | 0 | 0 | 0 | 0 | 0 | 0 | 0 |
| ENSMUSG00000075070 | 0 | 0 | 0 | 0 | 0 | 0 | 0 |
| ENSMUSG00000075072 | 0 | 0 | 0 | 0 | 0 | 0 | 0 |
| ENSMUSG00000075073 | 0 | 0 | 0 | 0 | 0 | 0 | 0 |
| ENSMUSG00000075074 | 0 | 0 | 0 | 0 | 0 | 0 | 0 |
| ENSMUSG00000075075 | 0 | 0 | 0 | 0 | 0 | 0 | 0 |
| ENSMUSG00000075078 | 0 | 0 | 0 | 0 | 0 | 0 | 0 |
| ENSMUSG00000075079 | 0 | 0 | 0 | 0 | 0 | 0 | 0 |
| ENSMUSG00000075081 | 0 | 0 | 0 | 0 | 0 | 0 | 0 |
| ENSMUSG00000075084 | 0 | 0 | 0 | 0 | 0 | 0 | 0 |
| ENSMUSG00000075085 | 0 | 0 | 0 | 0 | 0 | 0 | 0 |
| ENSMUSG00000075086 | 0 | 0 | 0 | 0 | 0 | 0 | 0 |
| ENSMUSG00000075088 | 0 | 0 | 0 | 0 | 0 | 0 | 0 |

| | | | | | | | |
|---|---|---|---|---|---|---|---|
| ENSMUSG00000075089 | 0 | 0 | 0 | 0 | 0 | 0 | 0 |
| ENSMUSG00000075090 | 0 | 0 | 0 | 0 | 0 | 0 | 0 |
| ENSMUSG00000075091 | 0 | 0 | 0 | 0 | 0 | 0 | 0 |
| ENSMUSG00000075092 | 0 | 0 | 0 | 0 | 0 | 0 | 0 |
| ENSMUSG00000075093 | 0 | 0 | 0 | 0 | 0 | 0 | 0 |
| ENSMUSG00000075094 | 0 | 0 | 0 | 0 | 0 | 0 | 0 |
| ENSMUSG00000075095 | 0 | 0 | 0 | 0 | 0 | 0 | 0 |
| ENSMUSG00000075097 | 0 | 0 | 0 | 0 | 0 | 0 | 0 |
| ENSMUSG00000075099 | 0 | 0 | 0 | 0 | 0 | 0 | 0 |
| ENSMUSG00000075100 | 0 | 0 | 0 | 0 | 0 | 0 | 0 |
| ENSMUSG00000075101 | 0 | 0 | 0 | 0 | 0 | 0 | 0 |
| ENSMUSG00000075102 | 0 | 0 | 0 | 0 | 0 | 0 | 0 |
| ENSMUSG00000075104 | 0 | 0 | 0 | 0 | 0 | 0 | 0 |
| ENSMUSG00000075105 | 0 | 0 | 0 | 0 | 0 | 0 | 0 |
| ENSMUSG00000075107 | 0 | 0 | 0 | 0 | 0 | 0 | 0 |
| ENSMUSG00000075110 | 0 | 0 | 0 | 0 | 0 | 0 | 0 |
| ENSMUSG00000075111 | 0 | 0 | 0 | 0 | 0 | 0 | 0 |
| ENSMUSG00000075112 | 0 | 0 | 0 | 0 | 0 | 0 | 0 |
| ENSMUSG00000075113 | 0 | 0 | 0 | 0 | 0 | 0 | 0 |
| ENSMUSG00000075114 | 0 | 0 | 0 | 0 | 0 | 0 | 0 |
| ENSMUSG00000075115 | 0 | 0 | 0 | 0 | 0 | 0 | 0 |
| ENSMUSG00000075116 | 0 | 0 | 0 | 0 | 0 | 0 | 0 |
| ENSMUSG00000075117 | 0 | 0 | 0 | 0 | 0 | 0 | 0 |
| ENSMUSG00000075119 | 0 | 0 | 0 | 0 | 0 | 0 | 0 |
| ENSMUSG00000075120 | 0 | 0 | 0 | 0 | 0 | 0 | 0 |
| ENSMUSG00000075121 | 0 | 0 | 0 | 0 | 0 | 0 | 0 |
| ENSMUSG00000075122 | 0.867194479 | 0.701838731 | 0.830482024 | 0 | 0.244650542 | 0 | 0 |
| ENSMUSG00000075124 | 0 | 0 | 0 | 0 | 0 | 0 | 0 |
| ENSMUSG00000075125 | 0 | 0 | 0 | 0 | 0 | 0 | 0 |
| ENSMUSG00000075126 | 0 | 0 | 0 | 0 | 0 | 0 | 0 |
| ENSMUSG00000075127 | 0 | 0 | 0 | 0 | 0 | 0 | 0 |
| ENSMUSG00000075128 | 0 | 0 | 0 | 0 | 0 | 0 | 0 |
| ENSMUSG00000075132 | 0 | 0 | 0 | 0 | 0 | 0 | 0 |
| ENSMUSG00000075133 | 0 | 0 | 0 | 0 | 0 | 0 | 0 |

| | | | | | | | |
|---|---|---|---|---|---|---|---|
| ENSMUSG00000075136 | 0 | 0 | 0 | 0 | 0 | 0 | 0 |
| ENSMUSG00000075137 | 0 | 0 | 0 | 0 | 0 | 0 | 0 |
| ENSMUSG00000075139 | 0 | 0 | 0 | 0 | 0 | 0 | 0 |
| ENSMUSG00000075140 | 0 | 0 | 0 | 0 | 0 | 0 | 0 |
| ENSMUSG00000075141 | 0 | 0 | 0 | 0 | 0 | 0 | 0 |
| ENSMUSG00000075142 | 0 | 0 | 0 | 0 | 0 | 0 | 0 |
| ENSMUSG00000075143 | 0 | 0 | 0 | 0 | 0 | 0 | 0 |
| ENSMUSG00000075144 | 0 | 0 | 0 | 0 | 0 | 0 | 0 |
| ENSMUSG00000075145 | 0 | 0 | 0 | 0 | 0 | 0 | 0 |
| ENSMUSG00000075146 | 0 | 0 | 0 | 0 | 0 | 0 | 0 |
| ENSMUSG00000075148 | 0 | 0 | 0 | 0 | 0 | 0 | 0 |
| ENSMUSG00000075149 | 0 | 0 | 0 | 0 | 0 | 0 | 0 |
| ENSMUSG00000075150 | 0 | 0 | 0 | 0 | 0 | 0 | 0 |
| ENSMUSG00000075151 | 0 | 0 | 0 | 0 | 0 | 0 | 0 |
| ENSMUSG00000075153 | 0 | 0 | 0 | 0 | 0 | 0 | 0 |
| ENSMUSG00000075154 | 0 | 0 | 0 | 0 | 0 | 0 | 0 |
| ENSMUSG00000075155 | 0 | 0 | 0 | 0 | 0 | 0 | 0 |
| ENSMUSG00000075156 | 0 | 0 | 0 | 0 | 0 | 0 | 0 |
| ENSMUSG00000075158 | 0 | 0 | 0 | 0 | 0 | 0 | 0 |
| ENSMUSG00000075159 | 0 | 0 | 0 | 0 | 0 | 0 | 0 |
| ENSMUSG00000075160 | 0 | 0 | 0 | 0 | 0 | 0 | 0 |
| ENSMUSG00000075161 | 0 | 0 | 0 | 0 | 0 | 0 | 0 |
| ENSMUSG00000075162 | 0 | 0 | 0 | 0 | 0 | 0 | 0 |
| ENSMUSG00000075163 | 0 | 0 | 0 | 0 | 0 | 0 | 0 |
| ENSMUSG00000075164 | 0 | 0 | 0 | 0 | 0 | 0 | 0 |
| ENSMUSG00000075165 | 0 | 0 | 0 | 0 | 0 | 0 | 0 |
| ENSMUSG00000075166 | 0 | 0 | 0 | 0 | 0 | 0 | 0 |
| ENSMUSG00000075167 | 0 | 0 | 0 | 0 | 0 | 0 | 0 |
| ENSMUSG00000075168 | 0 | 0 | 0 | 0 | 0 | 0 | 0 |
| ENSMUSG00000075169 | 0 | 0 | 0 | 0 | 0 | 0 | 0 |
| ENSMUSG00000075170 | 0 | 0 | 0 | 0 | 0 | 0 | 0 |
| ENSMUSG00000075171 | 0 | 0 | 0 | 0 | 0 | 0 | 0 |
| ENSMUSG00000075172 | 0 | 0 | 0 | 0 | 0 | 0 | 0 |
| ENSMUSG00000075173 | 0 | 0 | 0 | 0 | 0 | 0 | 0 |

| | | | | | | | |
|---|---|---|---|---|---|---|---|
| ENSMUSG00000075174 | 0 | 0 | 0 | 0 | 0 | 0 | 0 |
| ENSMUSG00000075175 | 0 | 0 | 0 | 0 | 0 | 0 | 0 |
| ENSMUSG00000075176 | 0 | 0 | 0 | 0 | 0 | 0 | 0 |
| ENSMUSG00000075177 | 0 | 0 | 0 | 0 | 0 | 0 | 0 |
| ENSMUSG00000075178 | 0 | 0 | 0 | 0 | 0 | 0 | 0 |
| ENSMUSG00000075179 | 0 | 0 | 0 | 0 | 0 | 0 | 0 |
| ENSMUSG00000075180 | 0 | 0 | 0 | 0 | 0 | 0 | 0 |
| ENSMUSG00000075181 | 0 | 0 | 0 | 0 | 0 | 0 | 0 |
| ENSMUSG00000075182 | 0 | 0 | 0 | 0 | 0 | 0 | 0 |
| ENSMUSG00000075184 | 0 | 0 | 0 | 0 | 0 | 0 | 0 |
| ENSMUSG00000075185 | 0 | 0 | 0 | 0 | 0 | 0 | 0 |
| ENSMUSG00000075186 | 0 | 0 | 0 | 0 | 0 | 0 | 0 |
| ENSMUSG00000075187 | 0 | 0 | 0 | 0 | 0 | 0 | 0 |
| ENSMUSG00000075188 | 0 | 0 | 0 | 0 | 0 | 0 | 0 |
| ENSMUSG00000075189 | 0 | 0.25 | 0.25 | 0 | 0 | 0 | 0.224243824 |
| ENSMUSG00000075190 | 0 | 0 | 0 | 0 | 0 | 0 | 0 |
| ENSMUSG00000075192 | 0 | 0 | 0 | 0 | 0 | 0 | 0 |
| ENSMUSG00000075193 | 0 | 0 | 0 | 0 | 0 | 0 | 0 |
| ENSMUSG00000075194 | 0 | 0 | 0 | 0 | 0 | 0 | 0 |
| ENSMUSG00000075195 | 0 | 0 | 0 | 0 | 0 | 0 | 0 |
| ENSMUSG00000075196 | 0 | 0 | 0 | 0 | 0 | 0 | 0 |
| ENSMUSG00000075197 | 0 | 0 | 0 | 0 | 0 | 0 | 0 |
| ENSMUSG00000075198 | 0 | 0 | 0 | 0 | 0 | 0 | 0 |
| ENSMUSG00000075199 | 0 | 0 | 0 | 0 | 0 | 0 | 0 |
| ENSMUSG00000075200 | 0 | 0 | 0 | 0 | 0 | 0 | 0 |
| ENSMUSG00000075201 | 0 | 0 | 0 | 0 | 0 | 0 | 0 |
| ENSMUSG00000075202 | 0 | 0 | 0 | 0 | 0 | 0 | 0 |
| ENSMUSG00000075203 | 0 | 0 | 0 | 0 | 0 | 0 | 0 |
| ENSMUSG00000075204 | 0 | 0 | 0 | 0 | 0 | 0 | 0 |
| ENSMUSG00000075205 | 0 | 0 | 0 | 0 | 0 | 0 | 0 |
| ENSMUSG00000075206 | 0 | 0 | 0 | 0 | 0 | 0 | 0 |
| ENSMUSG00000075208 | 0 | 0 | 0 | 0 | 0 | 0 | 0 |
| ENSMUSG00000075209 | 0 | 0 | 0 | 0 | 0 | 0 | 0 |
| ENSMUSG00000075210 | 0 | 0 | 0 | 0 | 0 | 0 | 0 |

| | | | | | | | |
|---|---|---|---|---|---|---|---|
| ENSMUSG00000075211 | 0 | 0 | 0 | 0 | 0 | 0 | 0 |
| ENSMUSG00000075212 | 0 | 0 | 0 | 0 | 0 | 0 | 0 |
| ENSMUSG00000075214 | 0 | 0 | 0 | 0 | 0 | 0 | 0 |
| ENSMUSG00000075215 | 0 | 0 | 0 | 0 | 0 | 0 | 0 |
| ENSMUSG00000075217 | 0 | 0 | 0 | 0 | 0 | 0 | 0 |
| ENSMUSG00000075218 | 0 | 0 | 0 | 0 | 0 | 0 | 0 |
| ENSMUSG00000075219 | 0 | 0 | 0 | 0 | 0 | 0 | 0 |
| ENSMUSG00000075220 | 0 | 0 | 0 | 0 | 0 | 0 | 0 |
| ENSMUSG00000075221 | 0 | 0 | 0 | 0 | 0 | 0 | 0 |
| ENSMUSG00000075222 | 0 | 0 | 0 | 0 | 0 | 0 | 0 |
| ENSMUSG00000075223 | 0 | 0 | 0 | 0 | 0 | 0 | 0 |
| ENSMUSG00000075224 | 0.289064826 | 0 | 0 | 0.25 | 0.244650542 | 0.301029996 | 0.355418052 |
| ENSMUSG00000075225 | 0 | 0.5 | 0.25 | 1.638647213 | 1.563883923 | 1.755874856 | 1.455757965 |
| ENSMUSG00000075227 | 1.227926429 | 1.238549078 | 1.488549078 | 1.080482024 | 1.283908311 | 1.113943352 | 1.462868481 |
| ENSMUSG00000075228 | 0 | 0 | 0 | 0 | 0 | 0 | 0 |
| ENSMUSG00000075229 | 1.940852214 | 1.834962501 | 1.80479213 | 1.892463902 | 1.826953746 | 2.012837225 | 1.714087544 |
| ENSMUSG00000075232 | 2.317377295 | 2.161864607 | 2.119945066 | 2.296838018 | 2.209357611 | 2.534026106 | 2.034219411 |
| ENSMUSG00000075245 | 0.867194479 | 1 | 1.02186571 | 0.896240625 | 1.057362051 | 0.954242509 | 0.896975297 |
| ENSMUSG00000075249 | 0 | 0 | 0 | 0 | 0 | 0 | 0 |
| ENSMUSG00000075254 | 2.235832493 | 2.103406982 | 2.064346961 | 2.236360959 | 2.19206933 | 2.545307116 | 2.095400649 |
| ENSMUSG00000075256 | 0 | 0 | 0 | 0 | 0.387761935 | 0 | 0.448487648 |
| ENSMUSG00000075265 | 0.91631382 | 0.646240625 | 0.646240625 | 0.79248125 | 0.77552387 | 0.602059991 | 0.775756176 |
| ENSMUSG00000075266 | 1.721148722 | 1.705044741 | 1.79248125 | 1.837182039 | 1.804307395 | 1.995635195 | 1.692344475 |
| ENSMUSG00000075267 | 0.289064826 | 0.25 | 0 | 0.5 | 0 | 0 | 0 |
| ENSMUSG00000075268 | 0 | 0 | 0 | 0 | 0 | 0 | 0 |
| ENSMUSG00000075269 | 0 | 0 | 0 | 0 | 0 | 0 | 0 |
| ENSMUSG00000075270 | 0 | 0.396240625 | 0.25 | 0.25 | 0 | 0 | 0.355418052 |
| ENSMUSG00000075271 | 1.342375483 | 1.282320754 | 1.080482024 | 1.410964047 | 1.343584837 | 1.431363764 | 1.23151414 |
| ENSMUSG00000075272 | 0.45815691 | 0.25 | 0.25 | 0.396240625 | 0.632412477 | 0.477121255 | 0.355418052 |
| ENSMUSG00000075273 | 1.129344651 | 1.188721876 | 1.201838731 | 1.311981878 | 1.283908311 | 1.505149978 | 1.20139795 |
| ENSMUSG00000075276 | 0.289064826 | 0.396240625 | 0.25 | 0 | 0.387761935 | 0.301029996 | 0.224243824 |
| ENSMUSG00000075277 | 0.578129653 | 0.75 | 0.701838731 | 0 | 0 | 0 | 0 |
| ENSMUSG00000075279 | 0 | 0.25 | 0.25 | 0 | 0 | 0 | 0.224243824 |
| ENSMUSG00000075284 | 2.183765914 | 2.098079356 | 2.064346961 | 2.227473271 | 2.203230928 | 2.501059262 | 2.081155529 |

| | | | | | | | |
|---|---|---|---|---|---|---|---|
| ENSMUSG00000075286 | 0 | 0 | 0 | 0.25 | 0 | 0 | 0 |
| ENSMUSG00000075288 | 0.289064826 | 0 | 0 | 0 | 0 | 0 | 0 |
| ENSMUSG00000075289 | 1.100572389 | 1.201838731 | 1.146240625 | 1.146240625 | 1.091002363 | 1.255272505 | 1.259064007 |
| ENSMUSG00000075296 | 0 | 0 | 0 | 0 | 0 | 0 | 0 |
| ENSMUSG00000075297 | 1.54867984 | 1.418106335 | 1.410964047 | 1.061981878 | 1.121713561 | 1.278753601 | 1.20919388 |
| ENSMUSG00000075302 | 0 | 0.580482024 | 0 | 0.701838731 | 0.686820904 | 0.477121255 | 0.775756176 |
| ENSMUSG00000075304 | 0 | 0 | 0 | 0.25 | 0 | 0 | 0.448487648 |
| ENSMUSG00000075307 | 0 | 0.396240625 | 0 | 0.396240625 | 0.489301084 | 0.301029996 | 0.224243824 |
| ENSMUSG00000075312 | 0 | 0 | 0 | 0 | 0 | 0 | 0 |
| ENSMUSG00000075314 | 0 | 0 | 0 | 0 | 0 | 0 | 0 |
| ENSMUSG00000075316 | 1.100572389 | 0.92510993 | 1.080482024 | 0 | 0 | 0 | 0 |
| ENSMUSG00000075318 | 1.671187741 | 1.54745614 | 1.53743678 | 1.146240625 | 1.020174412 | 1.079181246 | 1.066254157 |
| ENSMUSG00000075324 | 1.45815691 | 1.458222504 | 1.29248125 | 1.26109853 | 1.244650542 | 1.278753601 | 1.131174228 |
| ENSMUSG00000075325 | 0 | 0 | 0 | 0 | 0 | 0 | 0 |
| ENSMUSG00000075327 | 1.956444026 | 1.880890489 | 1.888647213 | 1.843759858 | 1.784632921 | 2.053078443 | 1.68886578 |
| ENSMUSG00000075330 | 0 | 0 | 0 | 0 | 0 | 0 | 0 |
| ENSMUSG00000075334 | 0.289064826 | 0.396240625 | 0.75 | 0 | 0 | 0.301029996 | 0 |
| ENSMUSG00000075335 | 0 | 0 | 0 | 0 | 0 | 0 | 0 |
| ENSMUSG00000075359 | 0 | 0 | 0 | 0 | 0 | 0 | 0 |
| ENSMUSG00000075370 | 0 | 0 | 0 | 0 | 0.244650542 | 0 | 0 |
| ENSMUSG00000075376 | 2.752359777 | 2.52053726 | 2.480460234 | 2.541226732 | 2.466277473 | 2.784617293 | 2.270850938 |
| ENSMUSG00000075377 | 0 | 0 | 0 | 0 | 0 | 0 | 0 |
| ENSMUSG00000075378 | 0 | 0 | 0 | 0 | 0 | 0 | 0 |
| ENSMUSG00000075379 | 0 | 0 | 0 | 0 | 0 | 0 | 0 |
| ENSMUSG00000075380 | 0 | 0 | 0 | 0 | 0 | 0 | 0 |
| ENSMUSG00000075382 | 0 | 0 | 0 | 0 | 0 | 0 | 0 |
| ENSMUSG00000075383 | 0 | 0 | 0 | 0 | 0 | 0 | 0 |
| ENSMUSG00000075384 | 0 | 0 | 0 | 0 | 0 | 0 | 0 |
| ENSMUSG00000075387 | 0 | 0 | 0 | 0 | 0 | 0 | 0 |
| ENSMUSG00000075389 | 0.960252568 | 1.130890489 | 1.061981878 | 0.864857905 | 1.057362051 | 1 | 1 |
| ENSMUSG00000075390 | 0 | 0 | 0 | 0 | 0 | 0 | 0 |
| ENSMUSG00000075391 | 0.811507563 | 1 | 0.951838731 | 1.388647213 | 1.467903253 | 1.602059991 | 1.401059159 |
| ENSMUSG00000075394 | 0.45815691 | 0.5 | 0.396240625 | 1.438721876 | 1.414412788 | 1.544068044 | 1.392431569 |
| ENSMUSG00000075395 | 0 | 0.396240625 | 0.25 | 0.25 | 0.244650542 | 0.301029996 | 0.448487648 |

| | | | | | | | |
|---|---|---|---|---|---|---|---|
| ENSMUSG00000075402 | 0 | 0 | 0 | 0 | 0 | 0 | 0 |
| ENSMUSG00000075405 | 0 | 0 | 0 | 0 | 0.244650542 | 0 | 0 |
| ENSMUSG00000075408 | 0 | 0.5 | 0.5 | 0.25 | 0.489301084 | 0 | 0.520678036 |
| ENSMUSG00000075410 | 0 | 0.25 | 0.25 | 0.580482024 | 0.244650542 | 0 | 0.448487648 |
| ENSMUSG00000075415 | 2.53552908 | 2.321928095 | 2.309601185 | 2.273849256 | 2.289867804 | 2.579783597 | 2.16858494 |
| ENSMUSG00000075416 | 0 | 0.25 | 0.396240625 | 0 | 0.244650542 | 0.301029996 | 0 |
| ENSMUSG00000075419 | 2.010213549 | 1.837182039 | 1.930274797 | 1.894357207 | 1.929985312 | 2.152288344 | 1.809975587 |
| ENSMUSG00000075420 | 0.578129653 | 0.25 | 0.5 | 0.5 | 0.733951626 | 0 | 0.355418052 |
| ENSMUSG00000075425 | 0 | 0 | 0 | 0 | 0.244650542 | 0 | 0 |
| ENSMUSG00000075427 | 0 | 0 | 0 | 0 | 0 | 0 | 0 |
| ENSMUSG00000075431 | 0 | 0 | 0 | 0 | 0 | 0 | 0 |
| ENSMUSG00000075437 | 0 | 0 | 0 | 0 | 0 | 0 | 0 |
| ENSMUSG00000075463 | 0.45815691 | 0.701838731 | 0.79248125 | 0.79248125 | 0.77552387 | 0.477121255 | 0.579661877 |
| ENSMUSG00000075465 | 0 | 0 | 0 | 0 | 0 | 0 | 0 |
| ENSMUSG00000075467 | 2.063793172 | 1.914552871 | 1.923371739 | 1.930274797 | 1.897237431 | 2.053078443 | 1.815933683 |
| ENSMUSG00000075469 | 0 | 0 | 0.396240625 | 0.25 | 0 | 0 | 0 |
| ENSMUSG00000075470 | 3.022860988 | 2.783124932 | 2.758699741 | 2.801448312 | 2.722312098 | 3.120244796 | 2.515829254 |
| ENSMUSG00000075478 | 0 | 0 | 0 | 0 | 0 | 0 | 0 |
| ENSMUSG00000075480 | 0 | 0 | 0 | 0 | 0 | 0 | 0 |
| ENSMUSG00000075485 | 0 | 0 | 0 | 0 | 0 | 0 | 0 |
| ENSMUSG00000075486 | 1.811507563 | 1.692046081 | 1.695339928 | 1.956637122 | 1.898867717 | 2.100370545 | 1.755055163 |
| ENSMUSG00000075502 | 0.747221736 | 0.951838731 | 0.79248125 | 1.146240625 | 1.223252711 | 1.041392685 | 1.121219121 |
| ENSMUSG00000075510 | 0 | 0 | 0 | 0 | 0 | 0 | 0 |
| ENSMUSG00000075511 | 0.578129653 | 0.864857905 | 0.75 | 0.701838731 | 0.733951626 | 0.84509804 | 0.916588299 |
| ENSMUSG00000075512 | 0 | 0 | 0 | 0 | 0 | 0 | 0 |
| ENSMUSG00000075514 | 0.45815691 | 0.646240625 | 0.701838731 | 0.25 | 0.244650542 | 0.477121255 | 0.355418052 |
| ENSMUSG00000075517 | 0 | 0 | 0 | 0 | 0 | 0 | 0 |
| ENSMUSG00000075520 | 0 | 0 | 0 | 0 | 0 | 0 | 0 |
| ENSMUSG00000075524 | 0.289064826 | 0 | 0 | 0 | 0 | 0 | 0 |
| ENSMUSG00000075525 | 0 | 0 | 0 | 0 | 0 | 0 | 0 |
| ENSMUSG00000075528 | 0.578129653 | 0.580482024 | 0.646240625 | 0.5 | 0.812711509 | 0.84509804 | 0.448487648 |
| ENSMUSG00000075534 | 0 | 0 | 0 | 0 | 0 | 0.301029996 | 0 |
| ENSMUSG00000075538 | 0 | 0 | 0 | 0 | 0 | 0 | 0 |
| ENSMUSG00000075543 | 0 | 0 | 0 | 0 | 0 | 0 | 0 |

| | | | | | | | |
|---|---|---|---|---|---|---|---|
| ENSMUSG00000075549 | 0 | 0 | 0 | 0 | 0 | 0 | 0 |
| ENSMUSG00000075551 | 0 | 0 | 0 | 0 | 0 | 0 | 0 |
| ENSMUSG00000075552 | 0 | 0 | 0 | 0 | 0 | 0 | 0 |
| ENSMUSG00000075553 | 0 | 0 | 0 | 0.396240625 | 0 | 0 | 0.224243824 |
| ENSMUSG00000075555 | 0 | 0 | 0 | 0.25 | 0 | 0 | 0 |
| ENSMUSG00000075558 | 0 | 0.25 | 0 | 0.5 | 0.244650542 | 0.301029996 | 0.224243824 |
| ENSMUSG00000075566 | 0 | 0 | 0 | 0 | 0 | 0 | 0 |
| ENSMUSG00000075567 | 0 | 0 | 0 | 0 | 0 | 0 | 0 |
| ENSMUSG00000075569 | 0.671187741 | 0.951838731 | 0.830482024 | 0.75 | 0.632412477 | 0.301029996 | 0.984950056 |
| ENSMUSG00000075570 | 0 | 0 | 0 | 0.25 | 0 | 0 | 0 |
| ENSMUSG00000075571 | 0 | 0 | 0 | 0 | 0 | 0 | 0 |
| ENSMUSG00000075572 | 0 | 0 | 0 | 0 | 0 | 0 | 0 |
| ENSMUSG00000075573 | 0 | 0 | 0 | 0 | 0 | 0 | 0 |
| ENSMUSG00000075574 | 0 | 0 | 0 | 0 | 0 | 0 | 0 |
| ENSMUSG00000075576 | 0.91631382 | 0.976722649 | 0.75 | 1.098079356 | 0.846351821 | 0.84509804 | 0.829800754 |
| ENSMUSG00000075581 | 0.747221736 | 0.5 | 0.396240625 | 0 | 0 | 0.301029996 | 0.355418052 |
| ENSMUSG00000075585 | 1.700465191 | 1.598079356 | 1.618933358 | 1.146240625 | 1.176121988 | 1.278753601 | 1.259064007 |
| ENSMUSG00000075586 | 0 | 0 | 0 | 0 | 0 | 0 | 0 |
| ENSMUSG00000075587 | 0.671187741 | 0.396240625 | 0.5 | 0.701838731 | 0.244650542 | 0.84509804 | 0.579661877 |
| ENSMUSG00000075588 | 1.740854706 | 1.660964047 | 1.54248125 | 1.720660762 | 1.72073075 | 1.86923172 | 1.579661877 |
| ENSMUSG00000075589 | 0.45815691 | 0.580482024 | 0.830482024 | 0.5 | 0.686820904 | 0.77815125 | 0.74492186 |
| ENSMUSG00000075590 | 1.800532393 | 1.692046081 | 1.755591953 | 1.610735874 | 1.744182955 | 1.770852012 | 1.594042709 |
| ENSMUSG00000075591 | 0 | 0.25 | 0 | 0 | 0 | 0.477121255 | 0 |
| ENSMUSG00000075592 | 1.156259305 | 0.79248125 | 1.098079356 | 1.664552871 | 1.797870327 | 1.892094603 | 1.750751125 |
| ENSMUSG00000075593 | 0.289064826 | 0 | 0 | 0 | 0 | 0 | 0 |
| ENSMUSG00000075595 | 2.451206209 | 2.267194569 | 2.226722649 | 2.296218836 | 2.239731213 | 2.556302501 | 2.116590781 |
| ENSMUSG00000075596 | 0 | 0.396240625 | 0.25 | 0 | 0.387761935 | 0.301029996 | 0.355418052 |
| ENSMUSG00000075598 | 0 | 0 | 0 | 0 | 0 | 0 | 0 |
| ENSMUSG00000075599 | 0 | 0 | 0 | 0 | 0 | 0 | 0 |
| ENSMUSG00000075600 | 1.418409478 | 1.488549078 | 1.53743678 | 1.238549078 | 1.47337555 | 1.531478917 | 1.577201671 |
| ENSMUSG00000075602 | 2.513987407 | 2.257666784 | 2.320772088 | 1.653677461 | 1.621875691 | 1.806179974 | 1.511732338 |
| ENSMUSG00000075604 | 0 | 0 | 0.25 | 0.25 | 0 | 0 | 0 |
| ENSMUSG00000075605 | 0 | 0 | 0 | 0 | 0 | 0 | 0 |
| ENSMUSG00000075609 | 0.289064826 | 0.396240625 | 0 | 0 | 0 | 0 | 0 |

| | | | | | | | |
|---|---|---|---|---|---|---|---|
| ENSMUSG00000075610 | 0.289064826 | 0 | 0 | 0.396240625 | 0.244650542 | 0 | 0 |
| ENSMUSG00000075611 | 0 | 0 | 0 | 0 | 0 | 0 | 0 |
| ENSMUSG00000075700 | 3.057835465 | 2.815670536 | 2.810495787 | 2.955443495 | 2.873791097 | 3.33405144 | 2.643638709 |
| ENSMUSG00000075701 | 2.282114126 | 2.14151351 | 2.160964047 | 2.167221246 | 2.131103516 | 2.416640507 | 2.020712061 |
| ENSMUSG00000075702 | 1.852729478 | 1.843759858 | 2.051142786 | 1.598079356 | 1.744182955 | 1.86923172 | 1.965468324 |
| ENSMUSG00000075703 | 2.829338353 | 2.595115266 | 2.590211771 | 2.62246199 | 2.532633036 | 2.900913068 | 2.344542559 |
| ENSMUSG00000075704 | 1.777675583 | 1.614857905 | 1.657339155 | 1.642463902 | 1.701347897 | 1.86332286 | 1.520678036 |
| ENSMUSG00000075705 | 2.275045611 | 2.074802005 | 2.115881093 | 2.182829758 | 2.114724103 | 2.536558443 | 2.074853081 |
| ENSMUSG00000075706 | 2.975812759 | 2.759215112 | 2.817573832 | 2.638167274 | 2.665423072 | 2.987666265 | 2.563208393 |
| ENSMUSG00000075707 | 0 | 0 | 0 | 0 | 0 | 0 | 0 |
| ENSMUSG00000075719 | 0 | 0 | 0 | 0 | 0 | 0 | 0 |
| ENSMUSG00000075726 | 0 | 0 | 0 | 0 | 0 | 0 | 0 |
| ENSMUSG00000075760 | 0 | 0 | 0 | 0 | 0 | 0 | 0 |
| ENSMUSG00000075770 | 0 | 0 | 0 | 0 | 0 | 0 | 0 |
| ENSMUSG00000075826 | 0 | 0 | 0 | 0 | 0 | 0 | 0 |
| ENSMUSG00000075829 | 0 | 0 | 0 | 0 | 0 | 0.301029996 | 0 |
| ENSMUSG00000075832 | 0 | 0 | 0 | 0 | 0 | 0 | 0 |
| ENSMUSG00000075833 | 0 | 0 | 0 | 0 | 0 | 0 | 0 |
| ENSMUSG00000075837 | 0 | 0 | 0.396240625 | 0 | 0.489301084 | 0 | 0 |
| ENSMUSG00000075864 | 0 | 0 | 0 | 0 | 0 | 0 | 0 |
| ENSMUSG00000075874 | 0 | 0 | 0 | 0 | 0 | 0 | 0 |
| ENSMUSG00000075893 | 0 | 0 | 0 | 0 | 0 | 0 | 0 |
| ENSMUSG00000075903 | 0 | 0 | 0 | 0 | 0 | 0 | 0 |
| ENSMUSG00000075906 | 0 | 0 | 0 | 0 | 0 | 0 | 0 |
| ENSMUSG00000075908 | 0 | 0.25 | 0 | 0 | 0.244650542 | 0 | 0 |
| ENSMUSG00000075918 | 0 | 0 | 0 | 0 | 0 | 0 | 0 |
| ENSMUSG00000075934 | 0 | 0 | 0 | 0 | 0 | 0 | 0 |
| ENSMUSG00000075952 | 0 | 0 | 0 | 0 | 0 | 0 | 0 |
| ENSMUSG00000075958 | 0 | 0 | 0 | 0 | 0 | 0 | 0 |
| ENSMUSG00000075968 | 0 | 0 | 0 | 0 | 0 | 0 | 0 |
| ENSMUSG00000075973 | 0 | 0 | 0 | 0 | 0 | 0 | 0 |
| ENSMUSG00000075983 | 0 | 0 | 0 | 0 | 0 | 0 | 0 |
| ENSMUSG00000076005 | 0 | 0 | 0 | 0 | 0 | 0 | 0 |
| ENSMUSG00000076006 | 0 | 0 | 0 | 0 | 0 | 0 | 0 |

| | | | | | | | |
|---|---|---|---|---|---|---|---|
| ENSMUSG00000076009 | 0 | 0 | 0 | 0 | 0 | 0 | 0 |
| ENSMUSG00000076010 | 0 | 0 | 0 | 0 | 0 | 0 | 0 |
| ENSMUSG00000076011 | 0 | 0 | 0 | 0 | 0 | 0 | 0 |
| ENSMUSG00000076012 | 0 | 0 | 0 | 0 | 0 | 0 | 0 |
| ENSMUSG00000076014 | 0 | 0 | 0 | 0 | 0 | 0 | 0 |
| ENSMUSG00000076028 | 0 | 0 | 0 | 0 | 0 | 0 | 0 |
| ENSMUSG00000076030 | 0 | 0 | 0 | 0 | 0 | 0 | 0 |
| ENSMUSG00000076031 | 0 | 0 | 0 | 0 | 0 | 0 | 0 |
| ENSMUSG00000076033 | 0 | 0 | 0 | 0 | 0 | 0 | 0 |
| ENSMUSG00000076036 | 0 | 0 | 0 | 0 | 0 | 0 | 0 |
| ENSMUSG00000076037 | 0 | 0 | 0 | 0 | 0 | 0 | 0 |
| ENSMUSG00000076038 | 0 | 0 | 0 | 0 | 0 | 0 | 0 |
| ENSMUSG00000076039 | 0 | 0 | 0 | 0 | 0.244650542 | 0 | 0 |
| ENSMUSG00000076041 | 0 | 0 | 0 | 0 | 0 | 0 | 0 |
| ENSMUSG00000076042 | 0 | 0 | 0 | 0 | 0 | 0 | 0 |
| ENSMUSG00000076049 | 0 | 0 | 0 | 0 | 0 | 0 | 0 |
| ENSMUSG00000076050 | 0 | 0 | 0 | 0 | 0 | 0 | 0 |
| ENSMUSG00000076051 | 0 | 0 | 0 | 0 | 0 | 0 | 0 |
| ENSMUSG00000076052 | 0 | 0 | 0 | 0 | 0 | 0 | 0 |
| ENSMUSG00000076060 | 0 | 0 | 0 | 0.25 | 0 | 0 | 0 |
| ENSMUSG00000076062 | 0 | 0 | 0 | 0 | 0 | 0 | 0 |
| ENSMUSG00000076063 | 0 | 0 | 0 | 0 | 0 | 0 | 0 |
| ENSMUSG00000076064 | 0 | 0 | 0 | 0 | 0 | 0 | 0 |
| ENSMUSG00000076065 | 0 | 0 | 0 | 0 | 0 | 0 | 0 |
| ENSMUSG00000076066 | 0 | 0 | 0 | 0 | 0 | 0 | 0 |
| ENSMUSG00000076067 | 0 | 0 | 0 | 0 | 0 | 0 | 0 |
| ENSMUSG00000076068 | 0 | 0 | 0 | 0 | 0 | 0 | 0 |
| ENSMUSG00000076117 | 0 | 0 | 0 | 0 | 0 | 0 | 0 |
| ENSMUSG00000076118 | 0 | 0 | 0 | 0 | 0 | 0 | 0 |
| ENSMUSG00000076119 | 0 | 0 | 0 | 0 | 0 | 0 | 0 |
| ENSMUSG00000076120 | 0 | 0 | 0 | 0 | 0 | 0 | 0 |
| ENSMUSG00000076121 | 0 | 0 | 0 | 0 | 0 | 0 | 0.224243824 |
| ENSMUSG00000076122 | 0 | 0 | 0.396240625 | 0 | 0 | 0 | 0 |
| ENSMUSG00000076123 | 0 | 0 | 0 | 0 | 0 | 0 | 0 |

| | | | | | | | |
|---|---|---|---|---|---|---|---|
| ENSMUSG00000076126 | 0 | 0 | 0 | 0 | 0 | 0 | 0 |
| ENSMUSG00000076127 | 0 | 0 | 0 | 0 | 0 | 0 | 0 |
| ENSMUSG00000076128 | 0 | 0 | 0 | 0 | 0 | 0 | 0 |
| ENSMUSG00000076129 | 0 | 0 | 0 | 0 | 0 | 0 | 0 |
| ENSMUSG00000076135 | 0 | 0 | 0 | 0 | 0 | 0 | 0.224243824 |
| ENSMUSG00000076136 | 0.289064826 | 0.25 | 0.5 | 0.75 | 0.686820904 | 0.477121255 | 0.224243824 |
| ENSMUSG00000076137 | 0 | 0.396240625 | 0.396240625 | 0 | 0 | 0 | 0 |
| ENSMUSG00000076138 | 0 | 0 | 0 | 0 | 0 | 0 | 0 |
| ENSMUSG00000076140 | 0 | 0 | 0 | 0 | 0 | 0 | 0 |
| ENSMUSG00000076141 | 0 | 0 | 0 | 0 | 0 | 0 | 0 |
| ENSMUSG00000076142 | 0 | 0 | 0 | 0.25 | 0 | 0 | 0 |
| ENSMUSG00000076143 | 0 | 0 | 0 | 0 | 0 | 0 | 0 |
| ENSMUSG00000076144 | 0 | 0 | 0 | 0 | 0 | 0 | 0 |
| ENSMUSG00000076145 | 0 | 0 | 0 | 0 | 0 | 0 | 0 |
| ENSMUSG00000076146 | 0 | 0 | 0 | 0 | 0 | 0 | 0 |
| ENSMUSG00000076147 | 0 | 0 | 0 | 0 | 0 | 0 | 0 |
| ENSMUSG00000076163 | 0 | 0 | 0 | 0 | 0 | 0 | 0 |
| ENSMUSG00000076180 | 0 | 0 | 0 | 0 | 0 | 0 | 0 |
| ENSMUSG00000076190 | 0 | 0 | 0 | 0 | 0 | 0 | 0 |
| ENSMUSG00000076199 | 0 | 0 | 0 | 0 | 0 | 0 | 0 |
| ENSMUSG00000076212 | 0 | 0 | 0 | 0 | 0 | 0 | 0 |
| ENSMUSG00000076214 | 0 | 0 | 0 | 0 | 0 | 0 | 0 |
| ENSMUSG00000076216 | 0 | 0 | 0 | 0 | 0 | 0 | 0 |
| ENSMUSG00000076217 | 0 | 0 | 0 | 0 | 0 | 0 | 0 |
| ENSMUSG00000076218 | 0 | 0 | 0 | 0 | 0 | 0 | 0 |
| ENSMUSG00000076219 | 0 | 0 | 0 | 0 | 0 | 0 | 0 |
| ENSMUSG00000076222 | 0 | 0 | 0 | 0 | 0 | 0 | 0 |
| ENSMUSG00000076227 | 0 | 0 | 0 | 0 | 0 | 0 | 0 |
| ENSMUSG00000076228 | 0 | 0 | 0 | 0 | 0 | 0 | 0 |
| ENSMUSG00000076229 | 0 | 0 | 0 | 0 | 0 | 0 | 0 |
| ENSMUSG00000076230 | 0 | 0 | 0 | 0 | 0 | 0 | 0 |
| ENSMUSG00000076231 | 0 | 0 | 0 | 0 | 0 | 0 | 0 |
| ENSMUSG00000076232 | 0 | 0 | 0 | 0 | 0 | 0 | 0 |
| ENSMUSG00000076233 | 0 | 0 | 0 | 0 | 0 | 0 | 0 |

| | | | | | | | |
|---|---|---|---|---|---|---|---|
| ENSMUSG00000076234 | 0 | 0 | 0 | 0 | 0 | 0 | 0 |
| ENSMUSG00000076236 | 0 | 0 | 0 | 0 | 0 | 0 | 0 |
| ENSMUSG00000076237 | 0 | 0 | 0 | 0 | 0 | 0 | 0 |
| ENSMUSG00000076238 | 0 | 0 | 0 | 0 | 0 | 0 | 0 |
| ENSMUSG00000076240 | 0 | 0 | 0 | 0 | 0 | 0 | 0 |
| ENSMUSG00000076241 | 0 | 0 | 0 | 0 | 0 | 0 | 0 |
| ENSMUSG00000076246 | 0 | 0 | 0 | 0 | 0 | 0 | 0 |
| ENSMUSG00000076251 | 0 | 0 | 0 | 0 | 0 | 0 | 0 |
| ENSMUSG00000076252 | 0 | 0 | 0 | 0 | 0 | 0 | 0 |
| ENSMUSG00000076253 | 0 | 0 | 0 | 0 | 0 | 0 | 0 |
| ENSMUSG00000076255 | 0 | 0 | 0 | 0 | 0 | 0 | 0 |
| ENSMUSG00000076256 | 0 | 0 | 0 | 0 | 0 | 0 | 0 |
| ENSMUSG00000076258 | 0 | 0 | 0.25 | 0 | 0 | 0 | 0.579661877 |
| ENSMUSG00000076269 | 0 | 0 | 0 | 0 | 0 | 0 | 0 |
| ENSMUSG00000076270 | 0 | 0 | 0 | 0 | 0 | 0 | 0 |
| ENSMUSG00000076271 | 0 | 0 | 0 | 0.25 | 0.568060967 | 0 | 0.224243824 |
| ENSMUSG00000076272 | 0 | 0 | 0 | 0 | 0 | 0 | 0 |
| ENSMUSG00000076274 | 0 | 0 | 0 | 0 | 0 | 0 | 0 |
| ENSMUSG00000076275 | 0 | 0 | 0 | 0 | 0 | 0 | 0 |
| ENSMUSG00000076281 | 0 | 0 | 0 | 0 | 0 | 0 | 0.224243824 |
| ENSMUSG00000076282 | 0 | 0 | 0 | 0 | 0 | 0 | 0 |
| ENSMUSG00000076284 | 0 | 0 | 0 | 0 | 0 | 0 | 0 |
| ENSMUSG00000076286 | 0 | 0 | 0 | 0 | 0 | 0 | 0 |
| ENSMUSG00000076287 | 0 | 0 | 0 | 0 | 0 | 0 | 0 |
| ENSMUSG00000076288 | 0 | 0 | 0 | 0 | 0 | 0 | 0 |
| ENSMUSG00000076294 | 0 | 0 | 0 | 0 | 0 | 0 | 0 |
| ENSMUSG00000076300 | 0 | 0 | 0 | 0 | 0 | 0 | 0 |
| ENSMUSG00000076303 | 0 | 0 | 0 | 0 | 0 | 0 | 0 |
| ENSMUSG00000076304 | 0 | 0 | 0 | 0 | 0 | 0 | 0 |
| ENSMUSG00000076305 | 0 | 0 | 0 | 0 | 0 | 0 | 0 |
| ENSMUSG00000076309 | 0 | 0 | 0 | 0 | 0 | 0 | 0 |
| ENSMUSG00000076312 | 0 | 0 | 0 | 0 | 0 | 0 | 0 |
| ENSMUSG00000076313 | 0 | 0 | 0 | 0 | 0 | 0 | 0 |
| ENSMUSG00000076315 | 0 | 0 | 0 | 0 | 0.244650542 | 0.301029996 | 0 |

| | | | | | | | |
|---|---|---|---|---|---|---|---|
| ENSMUSG00000076316 | 0 | 0 | 0 | 0 | 0 | 0 | 0 |
| ENSMUSG00000076318 | 0 | 0 | 0 | 0 | 0 | 0 | 0 |
| ENSMUSG00000076320 | 0 | 0 | 0 | 0 | 0 | 0 | 0 |
| ENSMUSG00000076323 | 0 | 0 | 0 | 0 | 0 | 0 | 0 |
| ENSMUSG00000076327 | 0 | 0 | 0 | 0 | 0 | 0 | 0 |
| ENSMUSG00000076332 | 0 | 0.25 | 0 | 0 | 0 | 0 | 0 |
| ENSMUSG00000076333 | 0 | 0 | 0 | 0.25 | 0 | 0 | 0.224243824 |
| ENSMUSG00000076335 | 0 | 0 | 0 | 0 | 0 | 0 | 0 |
| ENSMUSG00000076337 | 0 | 0 | 0 | 0 | 0 | 0 | 0 |
| ENSMUSG00000076338 | 0 | 0 | 0 | 0 | 0 | 0 | 0 |
| ENSMUSG00000076343 | 0 | 0 | 0 | 0 | 0 | 0 | 0 |
| ENSMUSG00000076355 | 0 | 0 | 0 | 0 | 0 | 0 | 0 |
| ENSMUSG00000076357 | 0 | 0 | 0 | 0 | 0 | 0 | 0 |
| ENSMUSG00000076360 | 0 | 0 | 0 | 0 | 0 | 0 | 0 |
| ENSMUSG00000076361 | 0 | 0 | 0 | 0 | 0 | 0 | 0 |
| ENSMUSG00000076364 | 0 | 0 | 0 | 0 | 0 | 0 | 0 |
| ENSMUSG00000076366 | 0 | 0 | 0 | 0 | 0 | 0 | 0 |
| ENSMUSG00000076370 | 0 | 0 | 0 | 0 | 0 | 0 | 0 |
| ENSMUSG00000076371 | 0 | 0 | 0 | 0 | 0 | 0 | 0 |
| ENSMUSG00000076372 | 0 | 0 | 0 | 0 | 0 | 0 | 0 |
| ENSMUSG00000076376 | 0 | 0 | 0 | 0 | 0.244650542 | 0 | 0 |
| ENSMUSG00000076377 | 0 | 0.25 | 0 | 0 | 0 | 0 | 0 |
| ENSMUSG00000076379 | 0 | 0 | 0 | 0 | 0 | 0 | 0 |
| ENSMUSG00000076382 | 0 | 0 | 0 | 0 | 0 | 0 | 0 |
| ENSMUSG00000076385 | 0 | 0 | 0 | 0 | 0 | 0 | 0 |
| ENSMUSG00000076387 | 0 | 0 | 0 | 0 | 0 | 0 | 0 |
| ENSMUSG00000076389 | 0 | 0 | 0 | 0 | 0 | 0 | 0 |
| ENSMUSG00000076392 | 0 | 0 | 0 | 0 | 0 | 0 | 0 |
| ENSMUSG00000076394 | 0 | 0 | 0 | 0 | 0.387761935 | 0 | 0 |
| ENSMUSG00000076395 | 0 | 0 | 0 | 0 | 0 | 0 | 0 |
| ENSMUSG00000076396 | 0 | 0 | 0 | 0 | 0 | 0 | 0 |
| ENSMUSG00000076398 | 0 | 0 | 0 | 0 | 0 | 0 | 0 |
| ENSMUSG00000076399 | 0 | 0 | 0 | 0 | 0 | 0 | 0 |
| ENSMUSG00000076407 | 0 | 0 | 0 | 0 | 0 | 0 | 0 |

| | | | | | | | |
|---|---|---|---|---|---|---|---|
| ENSMUSG00000076424 | 0 | 0 | 0 | 0 | 0 | 0 | 0 |
| ENSMUSG00000076430 | 0 | 0 | 0 | 0 | 0 | 0 | 0.224243824 |
| ENSMUSG00000076431 | 2.026699189 | 1.950224975 | 1.981203126 | 1.598079356 | 1.72073075 | 1.832508913 | 1.669034113 |
| ENSMUSG00000076432 | 3.104004774 | 2.816550228 | 2.774836953 | 2.889365925 | 2.78661101 | 3.205475037 | 2.583647744 |
| ENSMUSG00000076433 | 1.307602651 | 1.380890489 | 1.339388001 | 1.25 | 1.27449559 | 1.342422681 | 1.193409508 |
| ENSMUSG00000076434 | 0.289064826 | 0.25 | 0.25 | 0.396240625 | 0.244650542 | 0.477121255 | 0.520678036 |
| ENSMUSG00000076435 | 2.334723298 | 2.088286706 | 2.144357207 | 1.995748394 | 2.043601899 | 2.250420002 | 1.951019875 |
| ENSMUSG00000076436 | 0.45815691 | 0 | 0 | 0 | 0.244650542 | 0 | 0.355418052 |
| ENSMUSG00000076437 | 2.072573125 | 2.075945187 | 2.208222504 | 2.006976499 | 2.05001946 | 2.260071388 | 2.153553898 |
| ENSMUSG00000076438 | 0 | 0 | 0 | 0 | 0 | 0.301029996 | 0 |
| ENSMUSG00000076439 | 0.45815691 | 0.25 | 0.5 | 0 | 0 | 0 | 0.355418052 |
| ENSMUSG00000076441 | 2.463861956 | 2.267865591 | 2.334405475 | 1.919870025 | 1.953043706 | 2.033423755 | 1.825244123 |
| ENSMUSG00000076444 | 0 | 0.25 | 0 | 0 | 0 | 0 | 0 |
| ENSMUSG00000076445 | 0 | 0 | 0 | 0 | 0 | 0 | 0 |
| ENSMUSG00000076448 | 0 | 0.25 | 0 | 0 | 0.244650542 | 0 | 0 |
| ENSMUSG00000076450 | 0 | 0 | 0 | 0 | 0 | 0 | 0 |
| ENSMUSG00000076451 | 0 | 0 | 0 | 0 | 0 | 0 | 0 |
| ENSMUSG00000076453 | 0 | 0 | 0 | 0 | 0 | 0 | 0 |
| ENSMUSG00000076454 | 0 | 0 | 0 | 0 | 0 | 0 | 0 |
| ENSMUSG00000076455 | 0 | 0 | 0 | 0 | 0 | 0 | 0 |
| ENSMUSG00000076456 | 0 | 0 | 0 | 0 | 0 | 0 | 0 |
| ENSMUSG00000076457 | 0 | 0 | 0 | 0 | 0 | 0 | 0 |
| ENSMUSG00000076458 | 0 | 0 | 0 | 0 | 0 | 0 | 0 |
| ENSMUSG00000076459 | 0 | 0 | 0 | 0 | 0 | 0 | 0 |
| ENSMUSG00000076460 | 0 | 0 | 0 | 0 | 0 | 0 | 0 |
| ENSMUSG00000076461 | 0 | 0 | 0 | 0 | 0 | 0 | 0 |
| ENSMUSG00000076462 | 0 | 0 | 0 | 0 | 0 | 0 | 0 |
| ENSMUSG00000076463 | 0 | 0 | 0 | 0 | 0 | 0 | 0 |
| ENSMUSG00000076465 | 0 | 0 | 0 | 0 | 0 | 0 | 0 |
| ENSMUSG00000076467 | 0 | 0 | 0 | 0 | 0 | 0 | 0 |
| ENSMUSG00000076469 | 0 | 0 | 0 | 0 | 0 | 0 | 0 |
| ENSMUSG00000076470 | 0 | 0 | 0 | 0 | 0 | 0 | 0 |
| ENSMUSG00000076471 | 0 | 0 | 0 | 0 | 0 | 0 | 0 |
| ENSMUSG00000076472 | 0 | 0 | 0 | 0 | 0 | 0 | 0 |

| | | | | | | | |
|---|---|---|---|---|---|---|---|
| ENSMUSG00000076473 | 0 | 0 | 0 | 0 | 0 | 0 | 0 |
| ENSMUSG00000076474 | 0 | 0 | 0 | 0 | 0 | 0 | 0 |
| ENSMUSG00000076475 | 0 | 0 | 0 | 0 | 0 | 0 | 0 |
| ENSMUSG00000076476 | 0 | 0 | 0 | 0 | 0 | 0 | 0 |
| ENSMUSG00000076477 | 0 | 0 | 0 | 0 | 0 | 0 | 0 |
| ENSMUSG00000076478 | 0 | 0 | 0 | 0 | 0 | 0 | 0 |
| ENSMUSG00000076479 | 0 | 0 | 0 | 0 | 0 | 0 | 0 |
| ENSMUSG00000076480 | 0 | 0 | 0 | 0 | 0 | 0 | 0 |
| ENSMUSG00000076481 | 0 | 0 | 0 | 0 | 0 | 0 | 0 |
| ENSMUSG00000076483 | 0 | 0 | 0 | 0 | 0 | 0 | 0 |
| ENSMUSG00000076484 | 0 | 0 | 0 | 0 | 0 | 0 | 0 |
| ENSMUSG00000076485 | 0 | 0 | 0 | 0 | 0 | 0 | 0 |
| ENSMUSG00000076486 | 0 | 0 | 0 | 0 | 0 | 0 | 0 |
| ENSMUSG00000076487 | 0 | 0 | 0 | 0 | 0 | 0 | 0 |
| ENSMUSG00000076488 | 0 | 0 | 0 | 0 | 0 | 0 | 0 |
| ENSMUSG00000076489 | 0 | 0 | 0 | 0 | 0 | 0 | 0 |
| ENSMUSG00000076490 | 0 | 0 | 0 | 0 | 0 | 0 | 0 |
| ENSMUSG00000076492 | 0 | 0 | 0 | 0 | 0 | 0 | 0 |
| ENSMUSG00000076493 | 0 | 0 | 0 | 0 | 0 | 0 | 0 |
| ENSMUSG00000076494 | 0 | 0 | 0 | 0 | 0 | 0 | 0 |
| ENSMUSG00000076495 | 0 | 0 | 0 | 0 | 0 | 0 | 0 |
| ENSMUSG00000076496 | 0 | 0 | 0 | 0 | 0 | 0 | 0 |
| ENSMUSG00000076497 | 0 | 0 | 0 | 0 | 0 | 0 | 0 |
| ENSMUSG00000076498 | 0 | 0 | 0 | 0 | 0 | 0 | 0 |
| ENSMUSG00000076499 | 0 | 0 | 0 | 0 | 0 | 0 | 0 |
| ENSMUSG00000076500 | 0 | 0 | 0 | 0 | 0 | 0 | 0 |
| ENSMUSG00000076501 | 0 | 0 | 0 | 0 | 0 | 0 | 0 |
| ENSMUSG00000076505 | 0 | 0 | 0 | 0 | 0 | 0 | 0 |
| ENSMUSG00000076508 | 0 | 0 | 0 | 0 | 0 | 0 | 0 |
| ENSMUSG00000076512 | 0 | 0 | 0 | 0 | 0 | 0 | 0 |
| ENSMUSG00000076514 | 0 | 0 | 0 | 0 | 0 | 0 | 0 |
| ENSMUSG00000076518 | 0 | 0 | 0 | 0 | 0 | 0 | 0 |
| ENSMUSG00000076522 | 0 | 0 | 0 | 0 | 0 | 0 | 0 |
| ENSMUSG00000076523 | 0 | 0 | 0 | 0 | 0 | 0 | 0 |

| | | | | | | | |
|---|---|---|---|---|---|---|---|
| ENSMUSG00000076525 | 0 | 0 | 0 | 0 | 0 | 0 | 0 |
| ENSMUSG00000076526 | 0 | 0 | 0 | 0 | 0 | 0 | 0 |
| ENSMUSG00000076530 | 0 | 0 | 0 | 0 | 0 | 0 | 0 |
| ENSMUSG00000076531 | 0 | 0 | 0 | 0 | 0 | 0 | 0 |
| ENSMUSG00000076532 | 0 | 0 | 0 | 0 | 0 | 0 | 0 |
| ENSMUSG00000076533 | 0 | 0 | 0 | 0 | 0 | 0 | 0 |
| ENSMUSG00000076534 | 0 | 0 | 0 | 0 | 0 | 0 | 0 |
| ENSMUSG00000076535 | 0 | 0 | 0 | 0 | 0 | 0 | 0 |
| ENSMUSG00000076536 | 0 | 0 | 0 | 0 | 0 | 0 | 0 |
| ENSMUSG00000076538 | 0 | 0 | 0 | 0 | 0 | 0 | 0 |
| ENSMUSG00000076539 | 0 | 0 | 0 | 0 | 0 | 0 | 0 |
| ENSMUSG00000076540 | 0 | 0 | 0 | 0 | 0 | 0 | 0 |
| ENSMUSG00000076541 | 0 | 0 | 0 | 0 | 0 | 0 | 0 |
| ENSMUSG00000076543 | 0 | 0 | 0 | 0 | 0 | 0 | 0 |
| ENSMUSG00000076545 | 0 | 0 | 0 | 0 | 0 | 0 | 0 |
| ENSMUSG00000076547 | 0 | 0 | 0 | 0 | 0 | 0 | 0 |
| ENSMUSG00000076548 | 0 | 0 | 0 | 0 | 0 | 0 | 0 |
| ENSMUSG00000076549 | 0 | 0 | 0 | 0 | 0 | 0 | 0 |
| ENSMUSG00000076550 | 0 | 0 | 0 | 0 | 0 | 0 | 0 |
| ENSMUSG00000076552 | 0 | 0 | 0 | 0 | 0 | 0 | 0 |
| ENSMUSG00000076555 | 0 | 0 | 0 | 0 | 0 | 0 | 0 |
| ENSMUSG00000076556 | 0 | 0 | 0 | 0 | 0 | 0 | 0 |
| ENSMUSG00000076557 | 0 | 0 | 0 | 0 | 0 | 0 | 0 |
| ENSMUSG00000076562 | 0 | 0 | 0 | 0 | 0 | 0 | 0 |
| ENSMUSG00000076563 | 0 | 0 | 0 | 0 | 0 | 0 | 0 |
| ENSMUSG00000076564 | 0 | 0 | 0 | 0 | 0 | 0 | 0 |
| ENSMUSG00000076569 | 0 | 0 | 0 | 0 | 0 | 0 | 0 |
| ENSMUSG00000076570 | 0 | 0 | 0 | 0 | 0 | 0 | 0 |
| ENSMUSG00000076571 | 0 | 0 | 0 | 0 | 0 | 0 | 0 |
| ENSMUSG00000076572 | 0 | 0 | 0 | 0 | 0 | 0 | 0 |
| ENSMUSG00000076573 | 0 | 0 | 0 | 0 | 0 | 0 | 0 |
| ENSMUSG00000076575 | 0 | 0 | 0 | 0 | 0 | 0 | 0 |
| ENSMUSG00000076576 | 0 | 0 | 0 | 0 | 0 | 0 | 0 |
| ENSMUSG00000076577 | 0 | 0 | 0 | 0 | 0 | 0 | 0 |

| | | | | | | | |
|---|---|---|---|---|---|---|---|
| ENSMUSG00000076578 | 0 | 0 | 0 | 0 | 0 | 0 | 0 |
| ENSMUSG00000076580 | 0 | 0 | 0 | 0 | 0 | 0 | 0 |
| ENSMUSG00000076581 | 0 | 0 | 0 | 0 | 0 | 0 | 0 |
| ENSMUSG00000076583 | 0 | 0 | 0 | 0 | 0 | 0 | 0 |
| ENSMUSG00000076586 | 0 | 0 | 0 | 0 | 0 | 0 | 0 |
| ENSMUSG00000076587 | 0 | 0 | 0 | 0 | 0 | 0 | 0 |
| ENSMUSG00000076589 | 0 | 0 | 0 | 0 | 0 | 0 | 0 |
| ENSMUSG00000076591 | 0 | 0 | 0 | 0 | 0 | 0 | 0 |
| ENSMUSG00000076594 | 0 | 0 | 0 | 0 | 0 | 0 | 0 |
| ENSMUSG00000076596 | 0 | 0 | 0 | 0 | 0 | 0 | 0 |
| ENSMUSG00000076598 | 0 | 0 | 0 | 0 | 0 | 0 | 0 |
| ENSMUSG00000076604 | 0 | 0 | 0 | 0 | 0 | 0 | 0 |
| ENSMUSG00000076605 | 0 | 0 | 0 | 0 | 0 | 0 | 0 |
| ENSMUSG00000076606 | 0 | 0 | 0 | 0 | 0 | 0 | 0 |
| ENSMUSG00000076607 | 0 | 0 | 0 | 0 | 0 | 0 | 0 |
| ENSMUSG00000076608 | 0 | 0 | 0 | 0 | 0 | 0 | 0 |
| ENSMUSG00000076609 | 0 | 0 | 0 | 0 | 0 | 0 | 0 |
| ENSMUSG00000076612 | 0 | 0 | 0 | 0 | 0 | 0 | 0 |
| ENSMUSG00000076613 | 0 | 0 | 0 | 0 | 0 | 0 | 0 |
| ENSMUSG00000076614 | 0 | 0 | 0 | 0 | 0 | 0 | 0 |
| ENSMUSG00000076615 | 0 | 0 | 0 | 0 | 0 | 0 | 0 |
| ENSMUSG00000076617 | 0 | 0 | 0 | 0 | 0 | 0 | 0 |
| ENSMUSG00000076618 | 0 | 0 | 0 | 0 | 0 | 0 | 0 |
| ENSMUSG00000076619 | 0 | 0 | 0 | 0 | 0 | 0 | 0 |
| ENSMUSG00000076620 | 0 | 0 | 0 | 0 | 0 | 0 | 0 |
| ENSMUSG00000076621 | 0 | 0 | 0 | 0 | 0 | 0 | 0 |
| ENSMUSG00000076630 | 0 | 0 | 0 | 0 | 0 | 0 | 0 |
| ENSMUSG00000076632 | 0 | 0 | 0 | 0 | 0 | 0 | 0 |
| ENSMUSG00000076633 | 0 | 0 | 0 | 0 | 0 | 0 | 0 |
| ENSMUSG00000076646 | 0 | 0 | 0 | 0 | 0 | 0 | 0 |
| ENSMUSG00000076652 | 0 | 0 | 0 | 0 | 0 | 0 | 0 |
| ENSMUSG00000076653 | 0 | 0 | 0 | 0 | 0 | 0 | 0 |
| ENSMUSG00000076655 | 0 | 0 | 0 | 0 | 0 | 0 | 0 |
| ENSMUSG00000076661 | 0 | 0 | 0 | 0 | 0 | 0 | 0 |

| | | | | | | | |
|---|---|---|---|---|---|---|---|
| ENSMUSG00000076665 | 0 | 0 | 0 | 0 | 0 | 0 | 0 |
| ENSMUSG00000076666 | 0 | 0 | 0 | 0 | 0 | 0 | 0 |
| ENSMUSG00000076668 | 0 | 0 | 0 | 0 | 0 | 0 | 0 |
| ENSMUSG00000076670 | 0 | 0 | 0 | 0 | 0 | 0 | 0 |
| ENSMUSG00000076671 | 0 | 0 | 0 | 0 | 0 | 0 | 0 |
| ENSMUSG00000076672 | 0 | 0 | 0 | 0 | 0 | 0 | 0 |
| ENSMUSG00000076674 | 0 | 0 | 0 | 0 | 0 | 0 | 0 |
| ENSMUSG00000076676 | 0 | 0 | 0 | 0 | 0 | 0 | 0 |
| ENSMUSG00000076677 | 0 | 0 | 0 | 0 | 0 | 0 | 0 |
| ENSMUSG00000076680 | 0 | 0 | 0 | 0 | 0 | 0 | 0 |
| ENSMUSG00000076688 | 0 | 0 | 0 | 0 | 0 | 0 | 0 |
| ENSMUSG00000076695 | 0 | 0 | 0 | 0 | 0 | 0 | 0 |
| ENSMUSG00000076709 | 0 | 0 | 0 | 0 | 0 | 0 | 0 |
| ENSMUSG00000076710 | 0 | 0 | 0 | 0 | 0 | 0 | 0 |
| ENSMUSG00000076731 | 0 | 0 | 0 | 0 | 0 | 0 | 0 |
| ENSMUSG00000076733 | 0 | 0 | 0 | 0 | 0 | 0 | 0 |
| ENSMUSG00000076744 | 0 | 0 | 0 | 0 | 0 | 0 | 0 |
| ENSMUSG00000076745 | 0 | 0 | 0 | 0 | 0 | 0 | 0 |
| ENSMUSG00000076746 | 0 | 0 | 0 | 0 | 0 | 0 | 0 |
| ENSMUSG00000076747 | 0 | 0 | 0 | 0 | 0 | 0 | 0 |
| ENSMUSG00000076749 | 0 | 0 | 0 | 0 | 0 | 0 | 0 |
| ENSMUSG00000076750 | 0 | 0 | 0 | 0 | 0 | 0 | 0 |
| ENSMUSG00000076752 | 0 | 0 | 0 | 0 | 0 | 0 | 0 |
| ENSMUSG00000076754 | 0 | 0 | 0 | 0 | 0 | 0 | 0 |
| ENSMUSG00000076755 | 0 | 0 | 0 | 0 | 0 | 0 | 0 |
| ENSMUSG00000076757 | 0 | 0 | 0 | 0 | 0 | 0 | 0 |
| ENSMUSG00000076758 | 0 | 0 | 0 | 0 | 0 | 0 | 0 |
| ENSMUSG00000076759 | 0 | 0 | 0 | 0 | 0 | 0 | 0 |
| ENSMUSG00000076760 | 0 | 0 | 0 | 0 | 0 | 0 | 0 |
| ENSMUSG00000076761 | 0 | 0 | 0 | 0 | 0 | 0 | 0 |
| ENSMUSG00000076762 | 0 | 0 | 0 | 0 | 0 | 0 | 0 |
| ENSMUSG00000076764 | 0 | 0 | 0 | 0 | 0 | 0 | 0 |
| ENSMUSG00000076770 | 0 | 0 | 0 | 0 | 0 | 0 | 0 |
| ENSMUSG00000076778 | 0 | 0 | 0 | 0 | 0 | 0 | 0 |

| | | | | | | | |
|---|---|---|---|---|---|---|---|
| ENSMUSG00000076795 | 0 | 0 | 0 | 0 | 0 | 0 | 0 |
| ENSMUSG00000076796 | 0 | 0 | 0 | 0 | 0 | 0 | 0 |
| ENSMUSG00000076800 | 0 | 0 | 0 | 0 | 0 | 0 | 0 |
| ENSMUSG00000076802 | 0 | 0 | 0 | 0 | 0 | 0 | 0 |
| ENSMUSG00000076821 | 0 | 0 | 0 | 0 | 0 | 0 | 0 |
| ENSMUSG00000076822 | 0 | 0 | 0 | 0 | 0 | 0 | 0 |
| ENSMUSG00000076823 | 0 | 0 | 0 | 0 | 0 | 0 | 0 |
| ENSMUSG00000076824 | 0 | 0 | 0 | 0 | 0 | 0 | 0 |
| ENSMUSG00000076826 | 0 | 0 | 0 | 0 | 0 | 0 | 0 |
| ENSMUSG00000076831 | 0 | 0 | 0 | 0 | 0 | 0 | 0 |
| ENSMUSG00000076839 | 0 | 0 | 0 | 0 | 0 | 0 | 0 |
| ENSMUSG00000076840 | 0 | 0 | 0 | 0 | 0 | 0 | 0 |
| ENSMUSG00000076846 | 0 | 0 | 0 | 0.25 | 0 | 0 | 0.224243824 |
| ENSMUSG00000076848 | 0 | 0 | 0 | 0 | 0 | 0 | 0 |
| ENSMUSG00000076855 | 0 | 0 | 0 | 0 | 0 | 0 | 0 |
| ENSMUSG00000076858 | 0 | 0 | 0 | 0 | 0 | 0 | 0 |
| ENSMUSG00000076861 | 0 | 0 | 0 | 0 | 0 | 0 | 0 |
| ENSMUSG00000076862 | 0 | 0 | 0 | 0 | 0 | 0 | 0 |
| ENSMUSG00000076863 | 0 | 0 | 0 | 0 | 0 | 0 | 0 |
| ENSMUSG00000076864 | 0 | 0 | 0 | 0 | 0 | 0 | 0 |
| ENSMUSG00000076865 | 0 | 0 | 0 | 0 | 0 | 0 | 0 |
| ENSMUSG00000076866 | 0 | 0 | 0 | 0 | 0 | 0 | 0 |
| ENSMUSG00000076867 | 0 | 0 | 0 | 0 | 0 | 0 | 0 |
| ENSMUSG00000076870 | 0 | 0 | 0 | 0 | 0 | 0 | 0 |
| ENSMUSG00000076871 | 0 | 0 | 0 | 0 | 0 | 0 | 0 |
| ENSMUSG00000076873 | 0 | 0 | 0 | 0 | 0 | 0 | 0 |
| ENSMUSG00000076874 | 0 | 0 | 0 | 0 | 0 | 0 | 0 |
| ENSMUSG00000076875 | 0 | 0 | 0 | 0 | 0 | 0 | 0 |
| ENSMUSG00000076876 | 0 | 0 | 0 | 0 | 0 | 0 | 0 |
| ENSMUSG00000076877 | 0 | 0 | 0 | 0 | 0 | 0 | 0 |
| ENSMUSG00000076878 | 0 | 0 | 0 | 0 | 0 | 0 | 0 |
| ENSMUSG00000076879 | 0 | 0 | 0 | 0 | 0 | 0 | 0 |
| ENSMUSG00000076880 | 0 | 0 | 0 | 0 | 0 | 0 | 0 |
| ENSMUSG00000076881 | 0 | 0 | 0 | 0 | 0 | 0 | 0 |

| | | | | | | | |
|---|---|---|---|---|---|---|---|
| ENSMUSG00000076882 | 0 | 0 | 0 | 0 | 0 | 0 | 0 |
| ENSMUSG00000076883 | 0 | 0 | 0 | 0 | 0 | 0 | 0 |
| ENSMUSG00000076884 | 0 | 0 | 0 | 0 | 0 | 0 | 0 |
| ENSMUSG00000076885 | 0 | 0 | 0 | 0 | 0 | 0 | 0 |
| ENSMUSG00000076886 | 0 | 0 | 0 | 0 | 0 | 0 | 0 |
| ENSMUSG00000076887 | 0 | 0 | 0 | 0 | 0 | 0 | 0 |
| ENSMUSG00000076888 | 0 | 0 | 0 | 0 | 0 | 0 | 0 |
| ENSMUSG00000076889 | 0 | 0 | 0 | 0 | 0 | 0 | 0 |
| ENSMUSG00000076890 | 0 | 0 | 0 | 0 | 0 | 0 | 0 |
| ENSMUSG00000076891 | 0 | 0 | 0 | 0 | 0 | 0 | 0 |
| ENSMUSG00000076892 | 0 | 0 | 0 | 0 | 0 | 0 | 0 |
| ENSMUSG00000076893 | 0 | 0 | 0 | 0 | 0 | 0 | 0 |
| ENSMUSG00000076894 | 0 | 0 | 0 | 0 | 0 | 0 | 0 |
| ENSMUSG00000076895 | 0 | 0 | 0 | 0 | 0 | 0 | 0 |
| ENSMUSG00000076896 | 0 | 0 | 0 | 0 | 0 | 0 | 0 |
| ENSMUSG00000076897 | 0.578129653 | 0 | 0.25 | 0 | 0 | 0 | 0 |
| ENSMUSG00000076898 | 0 | 0 | 0 | 0 | 0 | 0 | 0 |
| ENSMUSG00000076899 | 0 | 0 | 0 | 0 | 0 | 0 | 0 |
| ENSMUSG00000076900 | 0 | 0 | 0 | 0 | 0 | 0 | 0 |
| ENSMUSG00000076901 | 0 | 0 | 0 | 0 | 0 | 0 | 0 |
| ENSMUSG00000076902 | 0 | 0 | 0 | 0 | 0 | 0 | 0 |
| ENSMUSG00000076903 | 0 | 0.25 | 0 | 0 | 0 | 0 | 0 |
| ENSMUSG00000076904 | 0 | 0 | 0 | 0 | 0 | 0 | 0 |
| ENSMUSG00000076905 | 0 | 0 | 0 | 0 | 0 | 0 | 0 |
| ENSMUSG00000076906 | 0 | 0 | 0 | 0 | 0 | 0 | 0 |
| ENSMUSG00000076907 | 0 | 0 | 0 | 0 | 0 | 0 | 0 |
| ENSMUSG00000076908 | 0 | 0 | 0.25 | 0 | 0 | 0 | 0 |
| ENSMUSG00000076909 | 0 | 0.25 | 0 | 0 | 0 | 0 | 0 |
| ENSMUSG00000076910 | 0 | 0 | 0 | 0 | 0 | 0 | 0 |
| ENSMUSG00000076911 | 0 | 0 | 0 | 0 | 0 | 0 | 0 |
| ENSMUSG00000076912 | 0 | 0 | 0 | 0 | 0 | 0 | 0 |
| ENSMUSG00000076913 | 0 | 0 | 0 | 0 | 0 | 0 | 0 |
| ENSMUSG00000076914 | 0 | 0 | 0 | 0 | 0.244650542 | 0 | 0 |
| ENSMUSG00000076915 | 0 | 0 | 0 | 0 | 0 | 0 | 0 |

| | | | | | | | |
|---|---|---|---|---|---|---|---|
| ENSMUSG00000076916 | 0 | 0 | 0 | 0 | 0 | 0 | 0 |
| ENSMUSG00000076917 | 0 | 0 | 0 | 0 | 0 | 0 | 0 |
| ENSMUSG00000076918 | 0 | 0 | 0 | 0 | 0 | 0 | 0 |
| ENSMUSG00000076919 | 0 | 0 | 0 | 0 | 0 | 0 | 0 |
| ENSMUSG00000076920 | 0 | 0 | 0.25 | 0 | 0 | 0 | 0 |
| ENSMUSG00000076921 | 0 | 0 | 0 | 0 | 0 | 0 | 0 |
| ENSMUSG00000076922 | 0.289064826 | 0 | 0 | 0 | 0 | 0 | 0 |
| ENSMUSG00000076923 | 0 | 0 | 0 | 0 | 0 | 0 | 0 |
| ENSMUSG00000076924 | 0 | 0 | 0 | 0 | 0 | 0 | 0 |
| ENSMUSG00000076925 | 0 | 0 | 0 | 0 | 0 | 0 | 0 |
| ENSMUSG00000076926 | 0 | 0 | 0 | 0 | 0 | 0 | 0 |
| ENSMUSG00000076927 | 0 | 0 | 0 | 0 | 0 | 0 | 0 |
| ENSMUSG00000076928 | 1.693336259 | 1.552363341 | 1.464495249 | 1.080482024 | 1.020174412 | 1.176091259 | 0.916588299 |
| ENSMUSG00000076934 | 0 | 0 | 0 | 0 | 0 | 0 | 0 |
| ENSMUSG00000076937 | 0 | 0 | 0 | 0 | 0 | 0 | 0 |
| ENSMUSG00000076939 | 0 | 0 | 0 | 0 | 0 | 0 | 0 |
| ENSMUSG00000076940 | 0 | 0 | 0 | 0 | 0 | 0 | 0 |
| ENSMUSG00000076941 | 0 | 0 | 0 | 0 | 0 | 0 | 0 |
| ENSMUSG00000076944 | 0 | 0 | 0 | 0 | 0 | 0 | 0 |
| ENSMUSG00000076946 | 0 | 0 | 0 | 0 | 0 | 0 | 0 |
| ENSMUSG00000076948 | 0 | 0 | 0 | 0 | 0 | 0 | 0 |
| ENSMUSG00000076949 | 0 | 0 | 0 | 0 | 0 | 0 | 0 |
| ENSMUSG00000076950 | 0 | 0 | 0 | 0 | 0 | 0 | 0 |
| ENSMUSG00000076951 | 0 | 0 | 0 | 0 | 0 | 0 | 0 |
| ENSMUSG00000076952 | 0 | 0 | 0 | 0 | 0 | 0 | 0 |
| ENSMUSG00000076956 | 0 | 0 | 0 | 0 | 0 | 0 | 0 |
| ENSMUSG00000076958 | 0 | 0 | 0 | 0 | 0 | 0 | 0 |
| ENSMUSG00000076959 | 0 | 0 | 0 | 0 | 0 | 0 | 0 |
| ENSMUSG00000076961 | 0 | 0 | 0 | 0 | 0 | 0 | 0 |
| ENSMUSG00000076964 | 0 | 0 | 0 | 0 | 0 | 0 | 0 |
| ENSMUSG00000076966 | 0 | 0 | 0 | 0 | 0 | 0 | 0 |
| ENSMUSG00000076972 | 0 | 0 | 0 | 0 | 0 | 0 | 0 |
| ENSMUSG00000076976 | 0 | 0 | 0 | 0 | 0 | 0 | 0 |
| ENSMUSG00000076977 | 0 | 0 | 0 | 0 | 0 | 0 | 0 |

| | | | | | | | |
|---|---|---|---|---|---|---|---|
| ENSMUSG00000076983 | 0 | 0 | 0 | 0 | 0 | 0 | 0 |
| ENSMUSG00000076987 | 0 | 0 | 0 | 0 | 0 | 0 | 0 |
| ENSMUSG00000076994 | 0 | 0 | 0 | 0 | 0 | 0 | 0 |
| ENSMUSG00000076999 | 0 | 0 | 0 | 0 | 0 | 0 | 0 |
| ENSMUSG00000077001 | 0 | 0 | 0 | 0 | 0 | 0 | 0 |
| ENSMUSG00000077004 | 0 | 0 | 0 | 0 | 0 | 0 | 0 |
| ENSMUSG00000077010 | 0 | 0 | 0 | 0 | 0 | 0 | 0 |
| ENSMUSG00000077015 | 0 | 0 | 0 | 0 | 0 | 0 | 0 |
| ENSMUSG00000077019 | 0 | 0 | 0 | 0 | 0 | 0 | 0 |
| ENSMUSG00000077021 | 0 | 0 | 0 | 0 | 0 | 0 | 0 |
| ENSMUSG00000077022 | 0 | 0 | 0 | 0 | 0 | 0 | 0 |
| ENSMUSG00000077025 | 0 | 0 | 0 | 0 | 0 | 0 | 0 |
| ENSMUSG00000077038 | 0 | 0 | 0 | 0 | 0 | 0 | 0 |
| ENSMUSG00000077039 | 0 | 0 | 0 | 0 | 0 | 0 | 0 |
| ENSMUSG00000077040 | 0 | 0 | 0 | 0 | 0 | 0 | 0 |
| ENSMUSG00000077042 | 0 | 0 | 0 | 0 | 0 | 0 | 0 |
| ENSMUSG00000077049 | 0 | 0 | 0 | 0 | 0 | 0 | 0 |
| ENSMUSG00000077050 | 0 | 0 | 0 | 0 | 0 | 0 | 0 |
| ENSMUSG00000077055 | 0 | 0 | 0 | 0 | 0 | 0 | 0 |
| ENSMUSG00000077058 | 0 | 0 | 0 | 0 | 0 | 0 | 0 |
| ENSMUSG00000077059 | 0 | 0 | 0 | 0 | 0 | 0 | 0 |
| ENSMUSG00000077065 | 0 | 0 | 0 | 0 | 0 | 0 | 0 |
| ENSMUSG00000077081 | 0 | 0 | 0 | 0 | 0 | 0 | 0 |
| ENSMUSG00000077082 | 0 | 0 | 0 | 0 | 0 | 0 | 0 |
| ENSMUSG00000077085 | 0 | 0 | 0 | 0 | 0 | 0 | 0 |
| ENSMUSG00000077086 | 0 | 0 | 0 | 0 | 0 | 0 | 0 |
| ENSMUSG00000077090 | 0 | 0 | 0 | 0 | 0 | 0 | 0 |
| ENSMUSG00000077092 | 0 | 0 | 0 | 0 | 0 | 0 | 0 |
| ENSMUSG00000077095 | 0 | 0 | 0 | 0 | 0 | 0 | 0 |
| ENSMUSG00000077096 | 0 | 0 | 0 | 0 | 0 | 0 | 0 |
| ENSMUSG00000077104 | 0 | 0 | 0 | 0 | 0 | 0 | 0 |
| ENSMUSG00000077107 | 0 | 0 | 0 | 0 | 0 | 0 | 0 |
| ENSMUSG00000077110 | 0 | 0 | 0 | 0 | 0 | 0 | 0 |
| ENSMUSG00000077111 | 0 | 0 | 0 | 0 | 0 | 0 | 0 |

| | | | | | | | |
|---|---|---|---|---|---|---|---|
| ENSMUSG00000077113 | 0 | 0 | 0 | 0 | 0 | 0 | 0 |
| ENSMUSG00000077119 | 0 | 0 | 0 | 0 | 0 | 0 | 0 |
| ENSMUSG00000077125 | 0 | 0 | 0 | 0 | 0 | 0 | 0 |
| ENSMUSG00000077127 | 0 | 0 | 0 | 0 | 0 | 0 | 0 |
| ENSMUSG00000077131 | 0 | 0 | 0 | 0 | 0 | 0 | 0 |
| ENSMUSG00000077138 | 0 | 0 | 0 | 0 | 0 | 0 | 0 |
| ENSMUSG00000077142 | 0 | 0 | 0 | 0 | 0 | 0 | 0 |
| ENSMUSG00000077143 | 0 | 0 | 0 | 0 | 0 | 0 | 0 |
| ENSMUSG00000077144 | 0 | 0 | 0 | 0 | 0 | 0 | 0 |
| ENSMUSG00000077146 | 0 | 0 | 0 | 0 | 0 | 0 | 0 |
| ENSMUSG00000077148 | 0 | 0.25 | 0 | 0 | 0.489301084 | 0 | 0.224243824 |
| ENSMUSG00000077149 | 0 | 0 | 0 | 0 | 0 | 0 | 0 |
| ENSMUSG00000077150 | 0 | 0 | 0 | 0 | 0 | 0 | 0 |
| ENSMUSG00000077152 | 0 | 0 | 0 | 0 | 0 | 0 | 0 |
| ENSMUSG00000077153 | 0 | 0 | 0 | 0 | 0 | 0 | 0 |
| ENSMUSG00000077155 | 0 | 0 | 0 | 0 | 0 | 0 | 0 |
| ENSMUSG00000077156 | 0 | 0 | 0 | 0 | 0 | 0 | 0 |
| ENSMUSG00000077158 | 0 | 0 | 0 | 0 | 0 | 0 | 0 |
| ENSMUSG00000077160 | 0 | 0 | 0 | 0 | 0 | 0 | 0 |
| ENSMUSG00000077161 | 0 | 0 | 0 | 0 | 0 | 0 | 0 |
| ENSMUSG00000077163 | 0 | 0 | 0 | 0 | 0 | 0 | 0 |
| ENSMUSG00000077166 | 0 | 0 | 0 | 0 | 0 | 0 | 0 |
| ENSMUSG00000077167 | 0.45815691 | 0.864857905 | 0.75 | 0.5 | 0.568060967 | 0.698970004 | 0.629532004 |
| ENSMUSG00000077169 | 0 | 0 | 0 | 0 | 0 | 0 | 0 |
| ENSMUSG00000077171 | 0 | 0 | 0 | 0 | 0 | 0 | 0 |
| ENSMUSG00000077172 | 0 | 0 | 0 | 0 | 0 | 0 | 0 |
| ENSMUSG00000077174 | 0 | 0 | 0 | 0 | 0 | 0 | 0 |
| ENSMUSG00000077175 | 0 | 0 | 0 | 0 | 0 | 0 | 0 |
| ENSMUSG00000077176 | 0 | 0 | 0 | 0 | 0 | 0 | 0 |
| ENSMUSG00000077177 | 0 | 0 | 0 | 0 | 0 | 0 | 0 |
| ENSMUSG00000077178 | 0 | 0 | 0 | 0 | 0 | 0 | 0 |
| ENSMUSG00000077180 | 0 | 0 | 0 | 0 | 0 | 0 | 0 |
| ENSMUSG00000077181 | 0 | 0 | 0 | 0 | 0 | 0 | 0 |
| ENSMUSG00000077184 | 0 | 0 | 0 | 0 | 0 | 0 | 0 |

| | | | | | | | |
|---|---|---|---|---|---|---|---|
| ENSMUSG00000077185 | 0 | 0 | 0 | 0 | 0 | 0 | 0 |
| ENSMUSG00000077188 | 0 | 0 | 0 | 0 | 0 | 0 | 0 |
| ENSMUSG00000077190 | 0 | 0 | 0 | 0 | 0 | 0 | 0 |
| ENSMUSG00000077191 | 0 | 0 | 0 | 0 | 0 | 0 | 0 |
| ENSMUSG00000077192 | 0.289064826 | 0.580482024 | 0 | 0.580482024 | 0.489301084 | 0 | 0.355418052 |
| ENSMUSG00000077194 | 0 | 0 | 0 | 0 | 0 | 0 | 0 |
| ENSMUSG00000077196 | 0 | 0 | 0 | 0 | 0 | 0 | 0 |
| ENSMUSG00000077197 | 0 | 0 | 0 | 0 | 0 | 0 | 0 |
| ENSMUSG00000077199 | 0 | 0 | 0 | 0 | 0 | 0 | 0 |
| ENSMUSG00000077200 | 0 | 0 | 0 | 0 | 0 | 0 | 0 |
| ENSMUSG00000077201 | 0 | 0 | 0 | 0 | 0 | 0 | 0 |
| ENSMUSG00000077202 | 0.578129653 | 0 | 0 | 0 | 0 | 0.301029996 | 0.224243824 |
| ENSMUSG00000077204 | 0 | 0 | 0 | 0 | 0 | 0 | 0 |
| ENSMUSG00000077205 | 0 | 0 | 0 | 0 | 0 | 0 | 0 |
| ENSMUSG00000077207 | 0 | 0 | 0 | 0 | 0 | 0 | 0 |
| ENSMUSG00000077208 | 0 | 0 | 0 | 0 | 0 | 0 | 0 |
| ENSMUSG00000077209 | 0 | 0 | 0 | 0 | 0 | 0 | 0 |
| ENSMUSG00000077212 | 0.671187741 | 0.396240625 | 0.75 | 0.701838731 | 0.77552387 | 0.698970004 | 0.74492186 |
| ENSMUSG00000077214 | 0 | 0 | 0 | 0 | 0 | 0 | 0 |
| ENSMUSG00000077215 | 0 | 0 | 0 | 0 | 0 | 0 | 0 |
| ENSMUSG00000077217 | 0 | 0 | 0 | 0 | 0 | 0 | 0 |
| ENSMUSG00000077218 | 0 | 0 | 0 | 0 | 0 | 0 | 0 |
| ENSMUSG00000077220 | 0 | 0 | 0 | 0 | 0 | 0 | 0 |
| ENSMUSG00000077221 | 0 | 0 | 0 | 0 | 0 | 0 | 0 |
| ENSMUSG00000077222 | 0.867194479 | 0.864857905 | 0.701838731 | 0.396240625 | 0.905314583 | 0.301029996 | 0.775756176 |
| ENSMUSG00000077223 | 0 | 0 | 0 | 0 | 0 | 0 | 0 |
| ENSMUSG00000077224 | 0 | 0 | 0 | 0 | 0 | 0 | 0 |
| ENSMUSG00000077227 | 0 | 0 | 0 | 0 | 0 | 0.301029996 | 0 |
| ENSMUSG00000077228 | 0 | 0 | 0 | 0 | 0 | 0 | 0 |
| ENSMUSG00000077230 | 0 | 0 | 0 | 0 | 0 | 0 | 0 |
| ENSMUSG00000077231 | 0 | 0 | 0 | 0 | 0 | 0 | 0 |
| ENSMUSG00000077235 | 0 | 0 | 0 | 0 | 0 | 0 | 0 |
| ENSMUSG00000077236 | 0 | 0 | 0 | 0 | 0 | 0 | 0 |
| ENSMUSG00000077237 | 0 | 0 | 0 | 0 | 0 | 0 | 0 |

| | | | | | | | |
|---|---|---|---|---|---|---|---|
| ENSMUSG00000077239 | 0 | 0 | 0 | 0 | 0.244650542 | 0 | 0 |
| ENSMUSG00000077240 | 0 | 0 | 0 | 0 | 0 | 0 | 0 |
| ENSMUSG00000077241 | 0 | 0 | 0 | 0 | 0 | 0 | 0 |
| ENSMUSG00000077243 | 0 | 0 | 0 | 0 | 0 | 0 | 0 |
| ENSMUSG00000077244 | 0 | 0 | 0 | 0 | 0 | 0 | 0 |
| ENSMUSG00000077246 | 0 | 0 | 0 | 0 | 0 | 0 | 0 |
| ENSMUSG00000077248 | 0 | 0 | 0 | 0 | 0 | 0 | 0 |
| ENSMUSG00000077250 | 0 | 0 | 0 | 0 | 0 | 0 | 0 |
| ENSMUSG00000077251 | 0 | 0 | 0 | 0 | 0 | 0 | 0 |
| ENSMUSG00000077252 | 0 | 0 | 0 | 0 | 0 | 0 | 0 |
| ENSMUSG00000077253 | 0 | 0 | 0 | 0 | 0 | 0 | 0 |
| ENSMUSG00000077254 | 0 | 0 | 0 | 0 | 0 | 0 | 0 |
| ENSMUSG00000077255 | 0 | 0 | 0 | 0 | 0 | 0 | 0 |
| ENSMUSG00000077256 | 0 | 0 | 0 | 0 | 0 | 0 | 0 |
| ENSMUSG00000077257 | 0 | 0 | 0 | 0 | 0 | 0 | 0 |
| ENSMUSG00000077258 | 0 | 0 | 0 | 0 | 0 | 0 | 0 |
| ENSMUSG00000077259 | 0 | 0 | 0 | 0 | 0 | 0 | 0 |
| ENSMUSG00000077260 | 0 | 0 | 0 | 0 | 0 | 0 | 0 |
| ENSMUSG00000077261 | 0 | 0 | 0 | 0 | 0 | 0 | 0 |
| ENSMUSG00000077262 | 0 | 0 | 0 | 0 | 0 | 0 | 0 |
| ENSMUSG00000077263 | 0 | 0 | 0 | 0 | 0 | 0 | 0 |
| ENSMUSG00000077264 | 0 | 0 | 0 | 0 | 0 | 0 | 0 |
| ENSMUSG00000077265 | 0 | 0.580482024 | 0 | 0.25 | 0 | 0 | 0 |
| ENSMUSG00000077268 | 0 | 0 | 0 | 0 | 0 | 0 | 0 |
| ENSMUSG00000077269 | 0 | 0 | 0 | 0 | 0 | 0 | 0 |
| ENSMUSG00000077270 | 0 | 0 | 0 | 0 | 0 | 0 | 0 |
| ENSMUSG00000077271 | 0 | 0 | 0 | 0 | 0 | 0 | 0 |
| ENSMUSG00000077272 | 0 | 0 | 0 | 0 | 0 | 0 | 0 |
| ENSMUSG00000077273 | 0 | 0 | 0 | 0 | 0 | 0 | 0 |
| ENSMUSG00000077274 | 0 | 0 | 0 | 0 | 0 | 0 | 0 |
| ENSMUSG00000077275 | 0 | 0 | 0 | 0 | 0 | 0 | 0 |
| ENSMUSG00000077276 | 0 | 0 | 0 | 0 | 0 | 0 | 0 |
| ENSMUSG00000077277 | 0 | 0 | 0 | 0 | 0 | 0 | 0 |
| ENSMUSG00000077278 | 0 | 0 | 0 | 0 | 0 | 0 | 0 |

| | | | | | | | |
|---|---|---|---|---|---|---|---|
| ENSMUSG00000077279 | 0 | 0 | 0 | 0 | 0 | 0 | 0 |
| ENSMUSG00000077280 | 0 | 0 | 0 | 0 | 0 | 0 | 0 |
| ENSMUSG00000077281 | 0 | 0 | 0 | 0 | 0 | 0 | 0 |
| ENSMUSG00000077282 | 0 | 0 | 0 | 0 | 0 | 0 | 0 |
| ENSMUSG00000077283 | 0 | 0 | 0 | 0 | 0 | 0 | 0 |
| ENSMUSG00000077285 | 0.45815691 | 0.5 | 0.5 | 0.646240625 | 0.387761935 | 0 | 0.579661877 |
| ENSMUSG00000077287 | 0 | 0 | 0 | 0 | 0 | 0 | 0 |
| ENSMUSG00000077288 | 0 | 0 | 0 | 0 | 0 | 0 | 0 |
| ENSMUSG00000077289 | 0 | 0 | 0 | 0 | 0 | 0 | 0 |
| ENSMUSG00000077290 | 0 | 0 | 0 | 0 | 0 | 0 | 0 |
| ENSMUSG00000077291 | 0 | 0 | 0 | 0 | 0 | 0 | 0 |
| ENSMUSG00000077292 | 0 | 0 | 0 | 0 | 0 | 0 | 0 |
| ENSMUSG00000077293 | 0 | 0 | 0 | 0 | 0 | 0 | 0 |
| ENSMUSG00000077294 | 0 | 0 | 0 | 0 | 0 | 0 | 0 |
| ENSMUSG00000077295 | 0 | 0 | 0 | 0 | 0 | 0 | 0 |
| ENSMUSG00000077298 | 0 | 0 | 0 | 0 | 0 | 0 | 0 |
| ENSMUSG00000077299 | 0 | 0 | 0 | 0 | 0 | 0 | 0 |
| ENSMUSG00000077302 | 0 | 0 | 0 | 0 | 0 | 0 | 0 |
| ENSMUSG00000077303 | 0 | 0 | 0 | 0 | 0 | 0 | 0 |
| ENSMUSG00000077304 | 0 | 0 | 0 | 0 | 0 | 0 | 0 |
| ENSMUSG00000077305 | 0 | 0 | 0 | 0 | 0 | 0 | 0 |
| ENSMUSG00000077306 | 0 | 0 | 0 | 0 | 0 | 0 | 0 |
| ENSMUSG00000077309 | 0.289064826 | 0.25 | 0.5 | 0 | 0.244650542 | 0 | 0.224243824 |
| ENSMUSG00000077312 | 0 | 0 | 0 | 0 | 0 | 0 | 0 |
| ENSMUSG00000077314 | 0 | 0 | 0 | 0 | 0 | 0 | 0 |
| ENSMUSG00000077315 | 0 | 0 | 0 | 0 | 0 | 0 | 0 |
| ENSMUSG00000077316 | 0 | 0 | 0 | 0 | 0 | 0 | 0 |
| ENSMUSG00000077317 | 0 | 0 | 0 | 0 | 0 | 0 | 0 |
| ENSMUSG00000077318 | 0 | 0.25 | 0 | 0 | 0 | 0 | 0 |
| ENSMUSG00000077320 | 0 | 0 | 0 | 0 | 0 | 0 | 0 |
| ENSMUSG00000077323 | 0 | 0 | 0 | 0 | 0 | 0 | 0 |
| ENSMUSG00000077325 | 0 | 0 | 0 | 0 | 0 | 0 | 0 |
| ENSMUSG00000077327 | 0 | 0 | 0 | 0 | 0 | 0 | 0 |
| ENSMUSG00000077328 | 0 | 0 | 0 | 0 | 0 | 0 | 0 |

| | | | | | | | |
|---|---|---|---|---|---|---|---|
| ENSMUSG00000077332 | 0 | 0 | 0 | 0 | 0 | 0 | 0 |
| ENSMUSG00000077333 | 0 | 0 | 0 | 0 | 0 | 0 | 0 |
| ENSMUSG00000077334 | 0 | 0 | 0 | 0 | 0 | 0 | 0 |
| ENSMUSG00000077336 | 0 | 0 | 0 | 0 | 0 | 0 | 0 |
| ENSMUSG00000077337 | 0 | 0 | 0 | 0 | 0 | 0 | 0 |
| ENSMUSG00000077338 | 0 | 0 | 0 | 0 | 0 | 0 | 0 |
| ENSMUSG00000077339 | 0 | 0 | 0 | 0 | 0 | 0 | 0 |
| ENSMUSG00000077342 | 0 | 0 | 0 | 0 | 0 | 0 | 0 |
| ENSMUSG00000077344 | 0 | 0 | 0 | 0 | 0 | 0 | 0 |
| ENSMUSG00000077345 | 0.289064826 | 0.25 | 0 | 0.396240625 | 0.244650542 | 0 | 0.520678036 |
| ENSMUSG00000077347 | 0 | 0 | 0 | 0 | 0 | 0 | 0 |
| ENSMUSG00000077348 | 0 | 0 | 0 | 0 | 0 | 0 | 0 |
| ENSMUSG00000077349 | 0 | 0 | 0 | 0 | 0 | 0 | 0 |
| ENSMUSG00000077350 | 0 | 0.25 | 0 | 0 | 0 | 0 | 0 |
| ENSMUSG00000077351 | 0 | 0 | 0 | 0 | 0 | 0 | 0 |
| ENSMUSG00000077354 | 0 | 0 | 0 | 0 | 0 | 0 | 0.224243824 |
| ENSMUSG00000077356 | 0 | 0 | 0 | 0 | 0 | 0 | 0 |
| ENSMUSG00000077357 | 0 | 0 | 0 | 0 | 0 | 0 | 0 |
| ENSMUSG00000077358 | 0 | 0 | 0 | 0 | 0 | 0 | 0 |
| ENSMUSG00000077359 | 0 | 0 | 0 | 0 | 0 | 0 | 0 |
| ENSMUSG00000077360 | 0 | 0 | 0.25 | 0.25 | 0 | 0 | 0 |
| ENSMUSG00000077361 | 0 | 0 | 0 | 0 | 0 | 0 | 0 |
| ENSMUSG00000077366 | 0 | 0 | 0 | 0 | 0 | 0 | 0 |
| ENSMUSG00000077367 | 0 | 0 | 0 | 0.25 | 0 | 0 | 0 |
| ENSMUSG00000077368 | 0 | 0 | 0 | 0 | 0 | 0 | 0 |
| ENSMUSG00000077369 | 0 | 0 | 0 | 0 | 0 | 0 | 0 |
| ENSMUSG00000077376 | 0 | 0 | 0 | 0 | 0 | 0 | 0 |
| ENSMUSG00000077377 | 0 | 0 | 0 | 0 | 0 | 0 | 0 |
| ENSMUSG00000077378 | 0 | 0 | 0 | 0 | 0 | 0 | 0 |
| ENSMUSG00000077379 | 0 | 0 | 0 | 0 | 0 | 0 | 0 |
| ENSMUSG00000077380 | 0 | 0.25 | 0.25 | 0.5 | 0.733951626 | 0.698970004 | 0 |
| ENSMUSG00000077381 | 0 | 0 | 0 | 0 | 0 | 0 | 0 |
| ENSMUSG00000077382 | 0 | 0 | 0.25 | 0 | 0 | 0.301029996 | 0 |
| ENSMUSG00000077383 | 0 | 0 | 0 | 0 | 0 | 0 | 0 |

| | | | | | | | |
|---|---|---|---|---|---|---|---|
| ENSMUSG00000077384 | 0 | 0 | 0 | 0 | 0 | 0 | 0 |
| ENSMUSG00000077385 | 0 | 0 | 0 | 0 | 0 | 0 | 0 |
| ENSMUSG00000077387 | 0 | 0 | 0 | 0 | 0 | 0 | 0 |
| ENSMUSG00000077391 | 0 | 0 | 0.25 | 0 | 0.244650542 | 0.301029996 | 0 |
| ENSMUSG00000077392 | 0 | 0 | 0 | 0 | 0 | 0 | 0 |
| ENSMUSG00000077393 | 0 | 0 | 0 | 0 | 0 | 0 | 0 |
| ENSMUSG00000077394 | 0 | 0 | 0 | 0 | 0 | 0 | 0.224243824 |
| ENSMUSG00000077396 | 0 | 0 | 0 | 0 | 0 | 0 | 0 |
| ENSMUSG00000077397 | 0 | 0 | 0 | 0 | 0 | 0 | 0 |
| ENSMUSG00000077398 | 0 | 0 | 0 | 0 | 0 | 0 | 0 |
| ENSMUSG00000077399 | 0 | 0 | 0 | 0 | 0 | 0 | 0 |
| ENSMUSG00000077400 | 0 | 0 | 0 | 0 | 0 | 0 | 0 |
| ENSMUSG00000077401 | 0 | 0 | 0 | 0 | 0 | 0 | 0 |
| ENSMUSG00000077402 | 0 | 0 | 0 | 0 | 0 | 0 | 0 |
| ENSMUSG00000077403 | 0 | 0 | 0 | 0 | 0 | 0 | 0 |
| ENSMUSG00000077404 | 0 | 0 | 0 | 0 | 0 | 0 | 0 |
| ENSMUSG00000077406 | 0 | 0 | 0 | 0 | 0 | 0 | 0 |
| ENSMUSG00000077407 | 0 | 0 | 0 | 0 | 0 | 0 | 0 |
| ENSMUSG00000077408 | 0 | 0 | 0 | 0 | 0 | 0 | 0 |
| ENSMUSG00000077409 | 0 | 0 | 0 | 0 | 0 | 0 | 0 |
| ENSMUSG00000077410 | 0 | 0 | 0 | 0 | 0 | 0 | 0 |
| ENSMUSG00000077411 | 0 | 0 | 0 | 0 | 0 | 0 | 0 |
| ENSMUSG00000077413 | 0 | 0 | 0 | 0 | 0 | 0 | 0 |
| ENSMUSG00000077415 | 0 | 0 | 0 | 0 | 0 | 0 | 0 |
| ENSMUSG00000077416 | 0 | 0.25 | 0 | 0.25 | 0 | 0 | 0.224243824 |
| ENSMUSG00000077417 | 0 | 0 | 0 | 0 | 0 | 0 | 0 |
| ENSMUSG00000077418 | 0 | 0 | 0 | 0 | 0 | 0 | 0 |
| ENSMUSG00000077419 | 0 | 0 | 0 | 0 | 0 | 0 | 0 |
| ENSMUSG00000077420 | 0 | 0 | 0 | 0 | 0 | 0 | 0 |
| ENSMUSG00000077421 | 0 | 0 | 0 | 0 | 0 | 0 | 0 |
| ENSMUSG00000077422 | 0 | 0 | 0 | 0 | 0 | 0 | 0 |
| ENSMUSG00000077424 | 0 | 0 | 0 | 0 | 0 | 0 | 0 |
| ENSMUSG00000077425 | 0 | 0 | 0 | 0 | 0 | 0 | 0 |
| ENSMUSG00000077426 | 0.289064826 | 0.646240625 | 0.5 | 0.396240625 | 0.387761935 | 0.477121255 | 0.224243824 |

| | | | | | | | |
|---|---|---|---|---|---|---|---|
| ENSMUSG00000077427 | 0 | 0 | 0 | 0 | 0 | 0 | 0 |
| ENSMUSG00000077428 | 0 | 0 | 0 | 0 | 0 | 0 | 0 |
| ENSMUSG00000077430 | 0 | 0 | 0 | 0 | 0 | 0 | 0 |
| ENSMUSG00000077431 | 0 | 0 | 0 | 0 | 0 | 0 | 0 |
| ENSMUSG00000077433 | 0 | 0 | 0 | 0 | 0 | 0 | 0 |
| ENSMUSG00000077435 | 0 | 0 | 0 | 0 | 0 | 0 | 0 |
| ENSMUSG00000077437 | 0 | 0 | 0 | 0 | 0 | 0 | 0 |
| ENSMUSG00000077438 | 0 | 0 | 0 | 0 | 0 | 0 | 0 |
| ENSMUSG00000077439 | 0 | 0 | 0 | 0 | 0 | 0 | 0 |
| ENSMUSG00000077440 | 0.578129653 | 0.864857905 | 0.864857905 | 1.17510993 | 1 | 0.903089987 | 0.969165684 |
| ENSMUSG00000077441 | 0 | 0 | 0 | 0 | 0 | 0 | 0 |
| ENSMUSG00000077442 | 0 | 0 | 0 | 0 | 0 | 0 | 0 |
| ENSMUSG00000077443 | 0 | 0 | 0 | 0 | 0 | 0 | 0 |
| ENSMUSG00000077445 | 0 | 0 | 0 | 0 | 0 | 0 | 0 |
| ENSMUSG00000077447 | 0 | 0 | 0 | 0 | 0 | 0 | 0 |
| ENSMUSG00000077448 | 0 | 0 | 0 | 0 | 0 | 0 | 0 |
| ENSMUSG00000077449 | 0 | 0.25 | 0 | 0 | 0 | 0.301029996 | 0.224243824 |
| ENSMUSG00000077450 | 2.736751754 | 2.559899632 | 2.582449085 | 2.413211243 | 2.438134528 | 2.736396502 | 2.362688369 |
| ENSMUSG00000077451 | 0 | 0 | 0 | 0 | 0 | 0 | 0 |
| ENSMUSG00000077453 | 0 | 0 | 0 | 0 | 0 | 0 | 0 |
| ENSMUSG00000077454 | 0.45815691 | 0 | 0 | 0.25 | 0 | 0 | 0 |
| ENSMUSG00000077455 | 0 | 0 | 0 | 0 | 0 | 0 | 0 |
| ENSMUSG00000077456 | 0 | 0 | 0 | 0 | 0 | 0 | 0 |
| ENSMUSG00000077457 | 0.960252568 | 0.75 | 0.580482024 | 0.830482024 | 0.686820904 | 0.301029996 | 0.74492186 |
| ENSMUSG00000077459 | 0 | 0 | 0 | 0 | 0 | 0 | 0 |
| ENSMUSG00000077460 | 0 | 0 | 0 | 0 | 0 | 0 | 0 |
| ENSMUSG00000077461 | 0 | 0 | 0 | 0 | 0 | 0 | 0 |
| ENSMUSG00000077462 | 0 | 0 | 0 | 0 | 0 | 0 | 0 |
| ENSMUSG00000077463 | 0 | 0 | 0 | 0 | 0 | 0 | 0 |
| ENSMUSG00000077464 | 0 | 0 | 0 | 0 | 0 | 0 | 0 |
| ENSMUSG00000077465 | 0 | 0 | 0 | 0 | 0 | 0 | 0 |
| ENSMUSG00000077467 | 0 | 0 | 0 | 0 | 0 | 0 | 0 |
| ENSMUSG00000077468 | 0 | 0 | 0 | 0 | 0 | 0 | 0 |
| ENSMUSG00000077469 | 0 | 0 | 0 | 0 | 0 | 0 | 0 |

| | | | | | | | |
|---|---|---|---|---|---|---|---|
| ENSMUSG00000077470 | 0 | 0.25 | 0 | 0 | 0.244650542 | 0.301029996 | 0 |
| ENSMUSG00000077471 | 0 | 0 | 0 | 0 | 0 | 0 | 0 |
| ENSMUSG00000077472 | 0 | 0 | 0 | 0 | 0 | 0 | 0 |
| ENSMUSG00000077473 | 0 | 0 | 0 | 0 | 0 | 0 | 0 |
| ENSMUSG00000077474 | 0 | 0 | 0 | 0 | 0 | 0 | 0 |
| ENSMUSG00000077476 | 0 | 0 | 0 | 0 | 0 | 0 | 0 |
| ENSMUSG00000077477 | 0 | 0 | 0 | 0 | 0 | 0 | 0 |
| ENSMUSG00000077480 | 0 | 0 | 0 | 0 | 0 | 0 | 0 |
| ENSMUSG00000077481 | 0 | 0 | 0 | 0 | 0 | 0 | 0 |
| ENSMUSG00000077483 | 0 | 0 | 0 | 0 | 0 | 0 | 0 |
| ENSMUSG00000077484 | 0 | 0 | 0 | 0 | 0 | 0 | 0 |
| ENSMUSG00000077485 | 0 | 0 | 0 | 0 | 0 | 0 | 0 |
| ENSMUSG00000077487 | 0 | 0 | 0 | 0 | 0 | 0 | 0 |
| ENSMUSG00000077488 | 0 | 0 | 0 | 0 | 0 | 0 | 0 |
| ENSMUSG00000077492 | 0 | 0 | 0 | 0 | 0 | 0 | 0 |
| ENSMUSG00000077493 | 0 | 0.5 | 0.5 | 0.25 | 0.489301084 | 0.301029996 | 0.579661877 |
| ENSMUSG00000077494 | 0 | 0 | 0 | 0 | 0 | 0 | 0 |
| ENSMUSG00000077495 | 0 | 0 | 0 | 0 | 0 | 0 | 0 |
| ENSMUSG00000077497 | 0 | 0 | 0 | 0 | 0 | 0 | 0 |
| ENSMUSG00000077499 | 0 | 0 | 0 | 0 | 0 | 0 | 0 |
| ENSMUSG00000077501 | 0 | 0 | 0 | 0 | 0 | 0 | 0 |
| ENSMUSG00000077502 | 0 | 0 | 0 | 0 | 0 | 0 | 0 |
| ENSMUSG00000077503 | 0 | 0 | 0 | 0 | 0 | 0 | 0 |
| ENSMUSG00000077504 | 0 | 0 | 0 | 0 | 0 | 0 | 0 |
| ENSMUSG00000077505 | 0 | 0 | 0 | 0 | 0 | 0 | 0 |
| ENSMUSG00000077506 | 0.289064826 | 0 | 0 | 0 | 0 | 0 | 0.224243824 |
| ENSMUSG00000077507 | 0 | 0 | 0 | 0 | 0 | 0 | 0 |
| ENSMUSG00000077511 | 0 | 0 | 0 | 0 | 0 | 0 | 0 |
| ENSMUSG00000077514 | 0 | 0 | 0 | 0 | 0 | 0 | 0 |
| ENSMUSG00000077515 | 0 | 0 | 0 | 0 | 0 | 0 | 0 |
| ENSMUSG00000077516 | 0 | 0 | 0 | 0 | 0 | 0 | 0 |
| ENSMUSG00000077517 | 0 | 0 | 0 | 0 | 0 | 0 | 0 |
| ENSMUSG00000077518 | 0 | 0 | 0 | 0 | 0 | 0 | 0 |
| ENSMUSG00000077519 | 0 | 0 | 0 | 0 | 0 | 0 | 0 |

| | | | | | | | |
|---|---|---|---|---|---|---|---|
| ENSMUSG00000077521 | 0 | 0 | 0 | 0 | 0 | 0 | 0 |
| ENSMUSG00000077522 | 0 | 0 | 0 | 0 | 0 | 0 | 0 |
| ENSMUSG00000077524 | 0 | 0.25 | 0 | 0 | 0 | 0 | 0 |
| ENSMUSG00000077526 | 0 | 0 | 0 | 0 | 0.244650542 | 0 | 0 |
| ENSMUSG00000077527 | 0 | 0.25 | 0 | 0 | 0 | 0 | 0 |
| ENSMUSG00000077528 | 0 | 0 | 0 | 0 | 0 | 0 | 0 |
| ENSMUSG00000077535 | 0 | 0 | 0 | 0 | 0 | 0 | 0 |
| ENSMUSG00000077537 | 0 | 0 | 0 | 0 | 0 | 0 | 0 |
| ENSMUSG00000077538 | 0 | 0 | 0 | 0 | 0 | 0 | 0 |
| ENSMUSG00000077539 | 0 | 0 | 0 | 0 | 0 | 0 | 0 |
| ENSMUSG00000077541 | 0 | 0 | 0 | 0 | 0 | 0 | 0 |
| ENSMUSG00000077542 | 0 | 0 | 0 | 0 | 0 | 0 | 0 |
| ENSMUSG00000077543 | 0 | 0 | 0 | 0 | 0 | 0 | 0 |
| ENSMUSG00000077545 | 0 | 0 | 0 | 0 | 0 | 0 | 0 |
| ENSMUSG00000077546 | 0 | 0 | 0 | 0 | 0 | 0 | 0 |
| ENSMUSG00000077547 | 0 | 0 | 0 | 0 | 0 | 0 | 0 |
| ENSMUSG00000077549 | 0.578129653 | 0.396240625 | 0.396240625 | 0.25 | 0 | 0.698970004 | 0.448487648 |
| ENSMUSG00000077552 | 0 | 0 | 0 | 0 | 0 | 0 | 0 |
| ENSMUSG00000077553 | 0 | 0 | 0 | 0 | 0 | 0 | 0 |
| ENSMUSG00000077554 | 0 | 0 | 0 | 0 | 0 | 0 | 0 |
| ENSMUSG00000077555 | 0 | 0 | 0 | 0 | 0 | 0 | 0 |
| ENSMUSG00000077557 | 0 | 0 | 0 | 0 | 0 | 0 | 0 |
| ENSMUSG00000077558 | 0 | 0 | 0 | 0 | 0 | 0 | 0 |
| ENSMUSG00000077559 | 0 | 0 | 0 | 0 | 0 | 0 | 0 |
| ENSMUSG00000077561 | 0 | 0 | 0 | 0 | 0 | 0 | 0 |
| ENSMUSG00000077562 | 0 | 0 | 0 | 0 | 0 | 0 | 0 |
| ENSMUSG00000077563 | 1.100572389 | 1.098079356 | 1.061981878 | 0.896240625 | 1 | 0.903089987 | 0.876096088 |
| ENSMUSG00000077564 | 0 | 0.396240625 | 0 | 0.25 | 0.489301084 | 0 | 0.224243824 |
| ENSMUSG00000077565 | 0 | 0 | 0 | 0 | 0 | 0 | 0 |
| ENSMUSG00000077567 | 0 | 0 | 0 | 0 | 0 | 0 | 0 |
| ENSMUSG00000077569 | 0 | 0.25 | 0 | 0 | 0 | 0 | 0 |
| ENSMUSG00000077571 | 0 | 0 | 0 | 0 | 0 | 0 | 0 |
| ENSMUSG00000077572 | 0 | 0 | 0 | 0 | 0 | 0 | 0 |
| ENSMUSG00000077574 | 0 | 0 | 0 | 0 | 0 | 0 | 0 |

| | | | | | | | |
|---|---|---|---|---|---|---|---|
| ENSMUSG00000077575 | 0 | 0.25 | 0 | 0.25 | 0 | 0.301029996 | 0 |
| ENSMUSG00000077576 | 0 | 0 | 0 | 0 | 0 | 0 | 0 |
| ENSMUSG00000077577 | 0 | 0 | 0 | 0 | 0.244650542 | 0 | 0 |
| ENSMUSG00000077578 | 0 | 0 | 0 | 0 | 0 | 0 | 0 |
| ENSMUSG00000077579 | 0 | 0 | 0 | 0 | 0 | 0 | 0 |
| ENSMUSG00000077581 | 0 | 0 | 0 | 0 | 0 | 0 | 0 |
| ENSMUSG00000077582 | 0 | 0 | 0 | 0 | 0 | 0 | 0 |
| ENSMUSG00000077585 | 0 | 0 | 0 | 0 | 0 | 0 | 0 |
| ENSMUSG00000077586 | 0 | 0 | 0 | 0 | 0 | 0 | 0 |
| ENSMUSG00000077587 | 0 | 0 | 0 | 0 | 0 | 0 | 0 |
| ENSMUSG00000077588 | 0 | 0 | 0 | 0 | 0 | 0 | 0 |
| ENSMUSG00000077590 | 0 | 0 | 0 | 0 | 0 | 0 | 0 |
| ENSMUSG00000077591 | 0 | 0 | 0 | 0 | 0 | 0 | 0 |
| ENSMUSG00000077592 | 0 | 0 | 0 | 0 | 0 | 0 | 0 |
| ENSMUSG00000077593 | 0 | 0 | 0 | 0 | 0 | 0 | 0 |
| ENSMUSG00000077594 | 0 | 0 | 0 | 0 | 0 | 0 | 0 |
| ENSMUSG00000077595 | 0 | 0 | 0 | 0 | 0 | 0 | 0 |
| ENSMUSG00000077597 | 0 | 0 | 0 | 0 | 0 | 0 | 0 |
| ENSMUSG00000077598 | 0 | 0 | 0 | 0 | 0 | 0 | 0 |
| ENSMUSG00000077599 | 0 | 0 | 0 | 0 | 0 | 0 | 0 |
| ENSMUSG00000077601 | 0 | 0 | 0 | 0 | 0 | 0 | 0 |
| ENSMUSG00000077602 | 0 | 0 | 0 | 0 | 0 | 0 | 0 |
| ENSMUSG00000077603 | 0 | 0 | 0 | 0 | 0 | 0 | 0 |
| ENSMUSG00000077604 | 0 | 0 | 0 | 0 | 0 | 0 | 0 |
| ENSMUSG00000077607 | 0 | 0 | 0 | 0 | 0.244650542 | 0.301029996 | 0 |
| ENSMUSG00000077608 | 0 | 0 | 0 | 0 | 0 | 0 | 0 |
| ENSMUSG00000077609 | 0 | 0 | 0 | 0 | 0 | 0 | 0 |
| ENSMUSG00000077611 | 0 | 0 | 0 | 0 | 0.244650542 | 0 | 0 |
| ENSMUSG00000077612 | 0 | 0 | 0 | 0 | 0 | 0 | 0 |
| ENSMUSG00000077613 | 0 | 0 | 0 | 0 | 0 | 0 | 0 |
| ENSMUSG00000077614 | 0.289064826 | 0.580482024 | 0.396240625 | 0.25 | 0 | 0.602059991 | 0.355418052 |
| ENSMUSG00000077615 | 0 | 0 | 0 | 0 | 0 | 0 | 0 |
| ENSMUSG00000077616 | 0 | 0 | 0 | 0 | 0 | 0 | 0 |
| ENSMUSG00000077617 | 0 | 0 | 0 | 0 | 0 | 0 | 0 |

| | | | | | | | |
|---|---|---|---|---|---|---|---|
| ENSMUSG00000077618 | 0 | 0 | 0 | 0 | 0 | 0 | 0 |
| ENSMUSG00000077619 | 0 | 0 | 0 | 0 | 0 | 0 | 0 |
| ENSMUSG00000077620 | 0 | 0 | 0 | 0 | 0 | 0 | 0 |
| ENSMUSG00000077621 | 0 | 0 | 0 | 0 | 0 | 0 | 0 |
| ENSMUSG00000077622 | 0 | 0 | 0 | 0 | 0 | 0 | 0 |
| ENSMUSG00000077624 | 0 | 0 | 0 | 0 | 0.244650542 | 0 | 0 |
| ENSMUSG00000077625 | 0 | 0 | 0 | 0 | 0 | 0 | 0 |
| ENSMUSG00000077627 | 0 | 0 | 0.25 | 0 | 0 | 0 | 0 |
| ENSMUSG00000077628 | 0 | 0 | 0 | 0 | 0 | 0 | 0 |
| ENSMUSG00000077629 | 0 | 0 | 0 | 0 | 0 | 0 | 0 |
| ENSMUSG00000077630 | 0 | 0 | 0 | 0 | 0 | 0 | 0 |
| ENSMUSG00000077633 | 0 | 0 | 0 | 0 | 0 | 0 | 0 |
| ENSMUSG00000077635 | 0 | 0 | 0 | 0 | 0 | 0 | 0 |
| ENSMUSG00000077637 | 0 | 0.5 | 0.396240625 | 0.580482024 | 0.244650542 | 0.301029996 | 0.355418052 |
| ENSMUSG00000077639 | 0.289064826 | 0.25 | 0 | 0.396240625 | 0.387761935 | 0.301029996 | 0 |
| ENSMUSG00000077640 | 0 | 0 | 0 | 0 | 0 | 0 | 0 |
| ENSMUSG00000077641 | 0 | 0 | 0 | 0 | 0 | 0 | 0 |
| ENSMUSG00000077642 | 0 | 0 | 0 | 0 | 0 | 0 | 0 |
| ENSMUSG00000077643 | 0 | 0 | 0 | 0 | 0 | 0 | 0 |
| ENSMUSG00000077644 | 0 | 0 | 0 | 0 | 0 | 0 | 0 |
| ENSMUSG00000077645 | 0 | 0 | 0 | 0 | 0 | 0 | 0 |
| ENSMUSG00000077647 | 0 | 0 | 0 | 0 | 0 | 0 | 0 |
| ENSMUSG00000077648 | 0 | 0 | 0 | 0 | 0 | 0 | 0 |
| ENSMUSG00000077649 | 0 | 0 | 0 | 0 | 0 | 0 | 0 |
| ENSMUSG00000077650 | 0 | 0 | 0 | 0 | 0 | 0 | 0 |
| ENSMUSG00000077652 | 0 | 0 | 0 | 0 | 0 | 0 | 0.224243824 |
| ENSMUSG00000077653 | 0 | 0 | 0 | 0 | 0 | 0 | 0 |
| ENSMUSG00000077655 | 0 | 0 | 0 | 0 | 0 | 0 | 0 |
| ENSMUSG00000077659 | 0 | 0 | 0 | 0 | 0 | 0 | 0 |
| ENSMUSG00000077660 | 0 | 0 | 0 | 0 | 0 | 0 | 0 |
| ENSMUSG00000077661 | 0 | 0 | 0 | 0 | 0 | 0 | 0 |
| ENSMUSG00000077663 | 0 | 0 | 0 | 0 | 0 | 0 | 0 |
| ENSMUSG00000077664 | 0 | 0 | 0 | 0 | 0 | 0 | 0 |
| ENSMUSG00000077666 | 0 | 0 | 0 | 0 | 0 | 0 | 0 |

| | | | | | | | |
|---|---|---|---|---|---|---|---|
| ENSMUSG00000077667 | 0 | 0 | 0 | 0 | 0 | 0 | 0 |
| ENSMUSG00000077668 | 0 | 0 | 0 | 0 | 0 | 0 | 0 |
| ENSMUSG00000077669 | 0 | 0 | 0 | 0 | 0 | 0 | 0 |
| ENSMUSG00000077671 | 0 | 0 | 0 | 0 | 0 | 0 | 0 |
| ENSMUSG00000077672 | 0 | 0 | 0 | 0 | 0 | 0 | 0 |
| ENSMUSG00000077673 | 0 | 0 | 0 | 0 | 0 | 0 | 0 |
| ENSMUSG00000077674 | 0 | 0 | 0 | 0 | 0 | 0 | 0 |
| ENSMUSG00000077675 | 0 | 0 | 0 | 0 | 0 | 0 | 0 |
| ENSMUSG00000077677 | 0 | 0 | 0 | 0 | 0 | 0 | 0 |
| ENSMUSG00000077678 | 0 | 0 | 0 | 0 | 0 | 0 | 0 |
| ENSMUSG00000077680 | 0 | 0 | 0 | 0 | 0 | 0 | 0 |
| ENSMUSG00000077681 | 0 | 0 | 0 | 0 | 0 | 0 | 0 |
| ENSMUSG00000077682 | 0 | 0 | 0 | 0 | 0 | 0 | 0 |
| ENSMUSG00000077683 | 0 | 0 | 0 | 0 | 0 | 0 | 0 |
| ENSMUSG00000077685 | 0 | 0 | 0 | 0 | 0 | 0 | 0 |
| ENSMUSG00000077687 | 0 | 0 | 0 | 0 | 0 | 0 | 0 |
| ENSMUSG00000077689 | 0 | 0 | 0 | 0 | 0 | 0 | 0 |
| ENSMUSG00000077691 | 0 | 0 | 0 | 0 | 0 | 0 | 0 |
| ENSMUSG00000077693 | 0 | 0 | 0 | 0 | 0 | 0 | 0 |
| ENSMUSG00000077696 | 0 | 0.25 | 0 | 0 | 0 | 0 | 0 |
| ENSMUSG00000077697 | 0 | 0 | 0 | 0 | 0 | 0 | 0 |
| ENSMUSG00000077698 | 0 | 0 | 0 | 0 | 0 | 0 | 0 |
| ENSMUSG00000077700 | 0 | 0 | 0 | 0 | 0 | 0 | 0 |
| ENSMUSG00000077701 | 0 | 0 | 0 | 0 | 0 | 0 | 0 |
| ENSMUSG00000077702 | 0 | 0 | 0 | 0 | 0 | 0 | 0 |
| ENSMUSG00000077703 | 0 | 0 | 0 | 0 | 0 | 0 | 0 |
| ENSMUSG00000077704 | 0.578129653 | 0 | 0.25 | 0.396240625 | 0.244650542 | 0 | 0 |
| ENSMUSG00000077705 | 0 | 0 | 0 | 0 | 0 | 0 | 0 |
| ENSMUSG00000077706 | 0 | 0 | 0 | 0 | 0 | 0 | 0 |
| ENSMUSG00000077707 | 0 | 0 | 0 | 0 | 0 | 0 | 0 |
| ENSMUSG00000077708 | 0 | 0 | 0 | 0 | 0 | 0 | 0 |
| ENSMUSG00000077709 | 0 | 0 | 0 | 0 | 0 | 0 | 0 |
| ENSMUSG00000077710 | 0 | 0 | 0 | 0 | 0 | 0 | 0 |
| ENSMUSG00000077711 | 0.45815691 | 0.25 | 0 | 0 | 0 | 0.301029996 | 0.224243824 |

| | | | | | | | |
|---|---|---|---|---|---|---|---|
| ENSMUSG00000077713 | 0 | 0 | 0 | 0 | 0 | 0 | 0 |
| ENSMUSG00000077714 | 0.45815691 | 0.396240625 | 0 | 0.25 | 0.244650542 | 0 | 0.224243824 |
| ENSMUSG00000077716 | 0 | 0 | 0 | 0 | 0 | 0 | 0 |
| ENSMUSG00000077717 | 0 | 0 | 0 | 0 | 0 | 0 | 0 |
| ENSMUSG00000077718 | 0 | 0 | 0 | 0 | 0 | 0 | 0 |
| ENSMUSG00000077719 | 0 | 0 | 0 | 0 | 0 | 0 | 0 |
| ENSMUSG00000077726 | 0 | 0 | 0 | 0 | 0 | 0 | 0 |
| ENSMUSG00000077728 | 0 | 0 | 0 | 0 | 0 | 0 | 0 |
| ENSMUSG00000077730 | 0 | 0 | 0 | 0 | 0 | 0 | 0 |
| ENSMUSG00000077733 | 0 | 0.25 | 0 | 0 | 0 | 0 | 0 |
| ENSMUSG00000077734 | 0.671187741 | 0.580482024 | 0.864857905 | 0.646240625 | 0.568060967 | 0.602059991 | 0.775756176 |
| ENSMUSG00000077735 | 0 | 0 | 0 | 0 | 0 | 0 | 0 |
| ENSMUSG00000077736 | 0 | 0 | 0 | 0 | 0 | 0 | 0 |
| ENSMUSG00000077737 | 0.811507563 | 0.75 | 0.646240625 | 0.830482024 | 0.955822902 | 0.903089987 | 0.775756176 |
| ENSMUSG00000077738 | 0 | 0 | 0 | 0 | 0 | 0 | 0 |
| ENSMUSG00000077741 | 0 | 0 | 0 | 0 | 0 | 0 | 0 |
| ENSMUSG00000077743 | 0 | 0 | 0 | 0 | 0 | 0 | 0 |
| ENSMUSG00000077744 | 0 | 0 | 0 | 0 | 0 | 0 | 0 |
| ENSMUSG00000077745 | 0 | 0 | 0 | 0 | 0 | 0 | 0 |
| ENSMUSG00000077749 | 0 | 0 | 0 | 0 | 0 | 0 | 0 |
| ENSMUSG00000077750 | 0 | 0 | 0 | 0 | 0 | 0 | 0 |
| ENSMUSG00000077752 | 0 | 0 | 0 | 0 | 0 | 0 | 0 |
| ENSMUSG00000077755 | 0 | 0 | 0 | 0 | 0 | 0 | 0 |
| ENSMUSG00000077756 | 0 | 0 | 0 | 0 | 0 | 0 | 0 |
| ENSMUSG00000077757 | 0 | 0 | 0 | 0 | 0 | 0 | 0 |
| ENSMUSG00000077758 | 0 | 0 | 0 | 0 | 0 | 0 | 0 |
| ENSMUSG00000077759 | 0 | 0 | 0 | 0 | 0 | 0 | 0 |
| ENSMUSG00000077761 | 0 | 0 | 0 | 0 | 0 | 0 | 0 |
| ENSMUSG00000077767 | 0 | 0 | 0 | 0 | 0 | 0 | 0 |
| ENSMUSG00000077768 | 0 | 0 | 0 | 0 | 0 | 0 | 0 |
| ENSMUSG00000077769 | 0 | 0 | 0 | 0 | 0 | 0 | 0 |
| ENSMUSG00000077771 | 0 | 0 | 0 | 0 | 0 | 0 | 0 |
| ENSMUSG00000077772 | 0 | 0 | 0 | 0 | 0 | 0 | 0 |
| ENSMUSG00000077776 | 0 | 0 | 0 | 0 | 0 | 0 | 0 |

| | | | | | | | |
|---|---|---|---|---|---|---|---|
| ENSMUSG00000077777 | 0 | 0 | 0 | 0 | 0 | 0 | 0 |
| ENSMUSG00000077780 | 0 | 0 | 0 | 0 | 0 | 0 | 0 |
| ENSMUSG00000077782 | 0 | 0 | 0 | 0 | 0 | 0 | 0 |
| ENSMUSG00000077783 | 0 | 0 | 0 | 0 | 0 | 0 | 0 |
| ENSMUSG00000077784 | 0 | 0 | 0 | 0 | 0 | 0 | 0 |
| ENSMUSG00000077785 | 0 | 0 | 0 | 0 | 0 | 0 | 0.224243824 |
| ENSMUSG00000077786 | 0 | 0 | 0 | 0 | 0 | 0 | 0 |
| ENSMUSG00000077787 | 0.45815691 | 0.25 | 0.396240625 | 0.580482024 | 0.244650542 | 0 | 0.355418052 |
| ENSMUSG00000077789 | 0 | 0 | 0 | 0 | 0 | 0 | 0 |
| ENSMUSG00000077791 | 0 | 0 | 0 | 0 | 0 | 0 | 0 |
| ENSMUSG00000077792 | 0 | 0 | 0 | 0 | 0 | 0 | 0 |
| ENSMUSG00000077793 | 0 | 0 | 0 | 0 | 0 | 0 | 0 |
| ENSMUSG00000077794 | 0 | 0 | 0 | 0 | 0 | 0 | 0 |
| ENSMUSG00000077795 | 0 | 0 | 0 | 0 | 0 | 0 | 0.224243824 |
| ENSMUSG00000077796 | 0 | 0 | 0 | 0 | 0 | 0 | 0 |
| ENSMUSG00000077797 | 0.289064826 | 0.396240625 | 0.396240625 | 0.580482024 | 0.686820904 | 0.477121255 | 0.775756176 |
| ENSMUSG00000077799 | 0 | 0 | 0 | 0 | 0 | 0 | 0 |
| ENSMUSG00000077802 | 0 | 0 | 0 | 0 | 0 | 0 | 0 |
| ENSMUSG00000077804 | 0 | 0 | 0 | 0 | 0 | 0 | 0 |
| ENSMUSG00000077815 | 0 | 0 | 0 | 0 | 0 | 0 | 0 |
| ENSMUSG00000077821 | 0 | 0 | 0 | 0 | 0 | 0 | 0 |
| ENSMUSG00000077829 | 0 | 0 | 0 | 0 | 0 | 0 | 0 |
| ENSMUSG00000077830 | 0 | 0 | 0 | 0 | 0 | 0 | 0 |
| ENSMUSG00000077832 | 0 | 0 | 0 | 0 | 0 | 0 | 0 |
| ENSMUSG00000077834 | 0 | 0 | 0 | 0 | 0 | 0 | 0 |
| ENSMUSG00000077836 | 0 | 0 | 0 | 0 | 0 | 0 | 0 |
| ENSMUSG00000077844 | 0 | 0 | 0 | 0 | 0 | 0 | 0 |
| ENSMUSG00000077846 | 0 | 0 | 0 | 0 | 0 | 0 | 0 |
| ENSMUSG00000077847 | 0 | 0 | 0 | 0 | 0 | 0 | 0 |
| ENSMUSG00000077851 | 0 | 0 | 0 | 0 | 0 | 0 | 0 |
| ENSMUSG00000077857 | 0 | 0 | 0 | 0 | 0 | 0 | 0 |
| ENSMUSG00000077858 | 0 | 0.396240625 | 0 | 0 | 0 | 0 | 0 |
| ENSMUSG00000077869 | 0 | 0 | 0 | 0 | 0 | 0 | 0 |
| ENSMUSG00000077872 | 0 | 0 | 0 | 0 | 0 | 0 | 0 |

| | | | | | | | |
|---|---|---|---|---|---|---|---|
| ENSMUSG00000077879 | 0 | 0 | 0 | 0 | 0 | 0 | 0 |
| ENSMUSG00000077881 | 0 | 0 | 0 | 0 | 0 | 0 | 0 |
| ENSMUSG00000077884 | 0 | 0 | 0 | 0 | 0 | 0 | 0 |
| ENSMUSG00000077886 | 0 | 0 | 0 | 0 | 0 | 0 | 0 |
| ENSMUSG00000077890 | 0 | 0 | 0 | 0.25 | 0 | 0 | 0 |
| ENSMUSG00000077892 | 0 | 0 | 0 | 0 | 0 | 0 | 0 |
| ENSMUSG00000077896 | 0 | 0 | 0 | 0 | 0 | 0 | 0 |
| ENSMUSG00000077897 | 0 | 0 | 0 | 0 | 0 | 0 | 0 |
| ENSMUSG00000077902 | 0 | 0 | 0 | 0 | 0 | 0 | 0 |
| ENSMUSG00000077903 | 0 | 0 | 0 | 0 | 0 | 0 | 0 |
| ENSMUSG00000077921 | 0 | 0 | 0 | 0 | 0 | 0 | 0 |
| ENSMUSG00000077925 | 0 | 0 | 0 | 0 | 0 | 0 | 0 |
| ENSMUSG00000077928 | 0 | 0 | 0 | 0 | 0 | 0 | 0 |
| ENSMUSG00000077931 | 0 | 0 | 0 | 0 | 0 | 0 | 0 |
| ENSMUSG00000077938 | 0 | 0 | 0 | 0 | 0 | 0 | 0 |
| ENSMUSG00000077939 | 0 | 0 | 0 | 0 | 0 | 0 | 0 |
| ENSMUSG00000077941 | 0 | 0 | 0 | 0 | 0 | 0 | 0 |
| ENSMUSG00000077948 | 0 | 0 | 0 | 0 | 0 | 0 | 0 |
| ENSMUSG00000077954 | 0 | 0 | 0 | 0 | 0 | 0 | 0 |
| ENSMUSG00000077962 | 0 | 0 | 0 | 0 | 0 | 0 | 0 |
| ENSMUSG00000077965 | 0 | 0 | 0 | 0 | 0 | 0 | 0 |
| ENSMUSG00000077971 | 0 | 0 | 0 | 0 | 0 | 0 | 0 |
| ENSMUSG00000077972 | 0 | 0 | 0 | 0 | 0 | 0 | 0 |
| ENSMUSG00000077980 | 0 | 0 | 0 | 0 | 0 | 0 | 0 |
| ENSMUSG00000077981 | 0 | 0 | 0 | 0 | 0 | 0 | 0 |
| ENSMUSG00000077990 | 0 | 0 | 0 | 0 | 0 | 0 | 0 |
| ENSMUSG00000077991 | 0 | 0 | 0 | 0 | 0 | 0 | 0 |
| ENSMUSG00000077996 | 0 | 0 | 0 | 0 | 0 | 0 | 0 |
| ENSMUSG00000077998 | 0 | 0 | 0 | 0 | 0 | 0 | 0 |
| ENSMUSG00000078002 | 0 | 0 | 0 | 0 | 0 | 0 | 0 |
| ENSMUSG00000078006 | 0 | 0 | 0 | 0 | 0 | 0 | 0 |
| ENSMUSG00000078008 | 0 | 0 | 0 | 0 | 0 | 0 | 0 |
| ENSMUSG00000078017 | 0 | 0 | 0 | 0 | 0 | 0 | 0 |
| ENSMUSG00000078024 | 0 | 0 | 0 | 0 | 0 | 0 | 0 |

| | | | | | | | |
|---|---|---|---|---|---|---|---|
| ENSMUSG00000078025 | 0 | 0 | 0 | 0 | 0 | 0 | 0 |
| ENSMUSG00000078026 | 0 | 0 | 0 | 0 | 0 | 0 | 0 |
| ENSMUSG00000078027 | 0 | 0 | 0 | 0 | 0 | 0 | 0 |
| ENSMUSG00000078031 | 0 | 0 | 0 | 0 | 0 | 0 | 0 |
| ENSMUSG00000078032 | 0 | 0 | 0 | 0 | 0 | 0 | 0 |
| ENSMUSG00000078034 | 0 | 0 | 0 | 0 | 0 | 0 | 0 |
| ENSMUSG00000078035 | 0 | 0 | 0 | 0 | 0 | 0 | 0 |
| ENSMUSG00000078038 | 0 | 0 | 0 | 0 | 0 | 0 | 0 |
| ENSMUSG00000078041 | 0 | 0 | 0 | 0 | 0 | 0 | 0 |
| ENSMUSG00000078042 | 0 | 0 | 0 | 0 | 0 | 0 | 0 |
| ENSMUSG00000078057 | 0 | 0 | 0 | 0 | 0 | 0 | 0 |
| ENSMUSG00000078060 | 0 | 0 | 0 | 0 | 0 | 0 | 0 |
| ENSMUSG00000078061 | 0 | 0 | 0 | 0 | 0 | 0.301029996 | 0 |
| ENSMUSG00000078080 | 0 | 0 | 0 | 0 | 0 | 0 | 0 |
| ENSMUSG00000078087 | 0 | 0 | 0 | 0 | 0 | 0 | 0 |
| ENSMUSG00000078091 | 0.91631382 | 0.646240625 | 0.701838731 | 0.646240625 | 0.387761935 | 0.301029996 | 0.355418052 |
| ENSMUSG00000078097 | 0 | 0 | 0 | 0 | 0 | 0.301029996 | 0 |
| ENSMUSG00000078116 | 0 | 0 | 0 | 0 | 0 | 0 | 0 |
| ENSMUSG00000078117 | 0 | 0 | 0 | 0 | 0 | 0 | 0 |
| ENSMUSG00000078118 | 0 | 0 | 0 | 0 | 0 | 0 | 0 |
| ENSMUSG00000078122 | 0 | 0 | 0 | 0 | 0 | 0 | 0 |
| ENSMUSG00000078123 | 0 | 0 | 0 | 0 | 0 | 0 | 0 |
| ENSMUSG00000078125 | 0.45815691 | 0.396240625 | 0 | 0 | 0.387761935 | 0.301029996 | 0 |
| ENSMUSG00000078126 | 0.289064826 | 0 | 0 | 0 | 0 | 0 | 0 |
| ENSMUSG00000078127 | 0 | 0 | 0 | 0 | 0 | 0 | 0 |
| ENSMUSG00000078128 | 0 | 0 | 0 | 0 | 0 | 0 | 0 |
| ENSMUSG00000078129 | 0.45815691 | 0.396240625 | 0.25 | 0 | 0.387761935 | 0.301029996 | 0 |
| ENSMUSG00000078130 | 0 | 0 | 0 | 0 | 0 | 0 | 0 |
| ENSMUSG00000078131 | 0 | 0 | 0.25 | 0 | 0 | 0 | 0 |
| ENSMUSG00000078132 | 0 | 0 | 0 | 0 | 0 | 0 | 0 |
| ENSMUSG00000078134 | 0.578129653 | 0.5 | 0.396240625 | 0.396240625 | 0.568060967 | 0.301029996 | 0.448487648 |
| ENSMUSG00000078137 | 0 | 0.25 | 0 | 0.25 | 0 | 0 | 0.224243824 |
| ENSMUSG00000078139 | 0.91631382 | 1.214495249 | 1.04248125 | 1.321350555 | 1.264824954 | 1.079181246 | 1.302263476 |
| ENSMUSG00000078141 | 0.289064826 | 0 | 0 | 0 | 0 | 0 | 0 |

| | | | | | | | |
|---|---|---|---|---|---|---|---|
| ENSMUSG00000078143 | 0 | 0 | 0 | 0 | 0 | 0 | 0 |
| ENSMUSG00000078144 | 0.960252568 | 0.976722649 | 0.976722649 | 0.75 | 0.978602168 | 0.698970004 | 0.853775828 |
| ENSMUSG00000078151 | 0 | 0 | 0 | 0 | 0.244650542 | 0 | 0 |
| ENSMUSG00000078153 | 0.289064826 | 0.5 | 0.396240625 | 0.580482024 | 0.489301084 | 0 | 0 |
| ENSMUSG00000078154 | 0.578129653 | 0.580482024 | 0.396240625 | 0.701838731 | 0.632412477 | 0.77815125 | 0.672731473 |
| ENSMUSG00000078157 | 0 | 0.25 | 0.25 | 0.396240625 | 0 | 0 | 0.224243824 |
| ENSMUSG00000078160 | 0.578129653 | 0.5 | 0.396240625 | 0.79248125 | 0.632412477 | 0.477121255 | 0.629532004 |
| ENSMUSG00000078161 | 0 | 0 | 0 | 0.25 | 0.244650542 | 0 | 0.224243824 |
| ENSMUSG00000078162 | 0 | 0 | 0 | 0 | 0 | 0 | 0 |
| ENSMUSG00000078173 | 0 | 0 | 0 | 0 | 0 | 0 | 0 |
| ENSMUSG00000078177 | 0 | 0 | 0 | 0.25 | 0.244650542 | 0 | 0.224243824 |
| ENSMUSG00000078179 | 0 | 0 | 0 | 0 | 0 | 0 | 0 |
| ENSMUSG00000078180 | 0.289064826 | 0.5 | 0 | 0.25 | 0.489301084 | 0.301029996 | 0.520678036 |
| ENSMUSG00000078182 | 0.289064826 | 0 | 0 | 0.25 | 0 | 0 | 0 |
| ENSMUSG00000078183 | 0.811507563 | 0.864857905 | 0.92510993 | 0.79248125 | 1.091002363 | 0.903089987 | 1 |
| ENSMUSG00000078184 | 0 | 0.25 | 0.396240625 | 0.5 | 0.489301084 | 0.301029996 | 0.355418052 |
| ENSMUSG00000078185 | 2.170239354 | 2.077084758 | 2.02186571 | 2.048689214 | 2.020174412 | 2.220108088 | 1.803905701 |
| ENSMUSG00000078190 | 1.655740331 | 1.614857905 | 1.557204673 | 1.54248125 | 1.528558853 | 1.556302501 | 1.379042812 |
| ENSMUSG00000078192 | 0.289064826 | 0.646240625 | 0 | 0.396240625 | 0.489301084 | 0.301029996 | 0.579661877 |
| ENSMUSG00000078193 | 1 | 0.75 | 0.580482024 | 0.79248125 | 0.905314583 | 0.954242509 | 0.74492186 |
| ENSMUSG00000078197 | 0 | 0 | 0 | 0 | 0 | 0 | 0 |
| ENSMUSG00000078198 | 0 | 0 | 0 | 0 | 0 | 0 | 0 |
| ENSMUSG00000078201 | 0.960252568 | 1.02186571 | 1.02186571 | 0.580482024 | 1 | 0.602059991 | 0.829800754 |
| ENSMUSG00000078202 | 0 | 0 | 0.580482024 | 0 | 0 | 0 | 0.579661877 |
| ENSMUSG00000078206 | 0 | 0 | 0 | 0 | 0 | 0 | 0 |
| ENSMUSG00000078208 | 0 | 0 | 0 | 0 | 0 | 0 | 0 |
| ENSMUSG00000078210 | 0 | 0 | 0 | 0 | 0 | 0 | 0 |
| ENSMUSG00000078212 | 0 | 0 | 0 | 0 | 0 | 0 | 0 |
| ENSMUSG00000078213 | 0 | 0 | 0 | 0 | 0 | 0 | 0 |
| ENSMUSG00000078214 | 0 | 0 | 0 | 0 | 0 | 0 | 0 |
| ENSMUSG00000078218 | 0 | 0 | 0 | 0 | 0 | 0 | 0 |
| ENSMUSG00000078234 | 0.671187741 | 0.396240625 | 0.646240625 | 0.580482024 | 0.568060967 | 0.477121255 | 0.224243824 |
| ENSMUSG00000078235 | 0 | 0 | 0 | 0.396240625 | 0.387761935 | 0.602059991 | 0.224243824 |
| ENSMUSG00000078238 | 0 | 0 | 0.25 | 0 | 0 | 0 | 0.224243824 |

| | | | | | | | |
|---|---|---|---|---|---|---|---|
| ENSMUSG00000078240 | 0.811507563 | 0.75 | 0.864857905 | 0.75 | 0.77552387 | 0.477121255 | 0.853775828 |
| ENSMUSG00000078247 | 1.53838222 | 1.356566189 | 1.418106335 | 1.302363341 | 1.200473444 | 1.230448921 | 1.066254157 |
| ENSMUSG00000078249 | 1.445324132 | 1.516522298 | 1.75 | 0.75 | 1.373641807 | 1.079181246 | 1.657681529 |
| ENSMUSG00000078251 | 0 | 0 | 0 | 0 | 0 | 0 | 0 |
| ENSMUSG00000078252 | 0 | 0 | 0 | 0 | 0 | 0 | 0 |
| ENSMUSG00000078253 | 0 | 0 | 0 | 0 | 0 | 0 | 0 |
| ENSMUSG00000078254 | 0 | 0 | 0 | 0 | 0 | 0 | 0 |
| ENSMUSG00000078255 | 0 | 0 | 0 | 0 | 0 | 0 | 0 |
| ENSMUSG00000078256 | 0 | 0 | 0 | 0 | 0 | 0 | 0 |
| ENSMUSG00000078257 | 0 | 0 | 0 | 0 | 0 | 0 | 0 |
| ENSMUSG00000078258 | 0 | 0 | 0 | 0 | 0 | 0 | 0 |
| ENSMUSG00000078259 | 0 | 0 | 0 | 0 | 0 | 0 | 0 |
| ENSMUSG00000078260 | 0 | 0 | 0 | 0 | 0 | 0 | 0 |
| ENSMUSG00000078261 | 0 | 0 | 0 | 0 | 0 | 0 | 0 |
| ENSMUSG00000078262 | 0 | 0 | 0 | 0 | 0 | 0 | 0 |
| ENSMUSG00000078263 | 0 | 0 | 0 | 0 | 0 | 0 | 0 |
| ENSMUSG00000078269 | 0 | 0 | 0 | 0 | 0 | 0 | 0 |
| ENSMUSG00000078270 | 0 | 0 | 0 | 0 | 0 | 0 | 0 |
| ENSMUSG00000078276 | 0 | 0 | 0 | 0 | 0 | 0 | 0 |
| ENSMUSG00000078280 | 0 | 0 | 0 | 0 | 0 | 0 | 0 |
| ENSMUSG00000078289 | 0.289064826 | 0 | 0.5 | 0 | 0.244650542 | 0.602059991 | 0.579661877 |
| ENSMUSG00000078291 | 0.289064826 | 0 | 0.25 | 0.25 | 0 | 0.301029996 | 0.224243824 |
| ENSMUSG00000078300 | 0 | 0 | 0 | 0 | 0 | 0 | 0.224243824 |
| ENSMUSG00000078302 | 2.129344651 | 1.981203126 | 1.994319981 | 1.969129237 | 1.937354184 | 2.161368002 | 1.943943921 |
| ENSMUSG00000078307 | 0 | 0 | 0.25 | 0 | 0.244650542 | 0 | 0 |
| ENSMUSG00000078308 | 0.289064826 | 0.580482024 | 0.396240625 | 0.25 | 0.244650542 | 0.301029996 | 0.224243824 |
| ENSMUSG00000078314 | 0 | 0 | 0.25 | 0 | 0 | 0 | 0 |
| ENSMUSG00000078315 | 0 | 0 | 0 | 0 | 0 | 0 | 0 |
| ENSMUSG00000078317 | 1.389637216 | 1.080482024 | 1.356566189 | 1.114857905 | 1.106691885 | 1.146128036 | 1.245582246 |
| ENSMUSG00000078320 | 0 | 0 | 0 | 0 | 0 | 0 | 0 |
| ENSMUSG00000078324 | 0 | 0 | 0 | 0 | 0 | 0 | 0 |
| ENSMUSG00000078346 | 0 | 0 | 0 | 0 | 0 | 0 | 0 |
| ENSMUSG00000078348 | 1.928763473 | 1.912762923 | 2.063166358 | 1.51109853 | 1.836786029 | 1.73239376 | 2.016293244 |
| ENSMUSG00000078349 | 0 | 0.701838731 | 0.396240625 | 0.79248125 | 0.905314583 | 0.77815125 | 0.629532004 |

| | | | | | | | |
|---|---|---|---|---|---|---|---|
| ENSMUSG00000078350 | 1.37447073 | 1.311981878 | 1.372963274 | 0.79248125 | 1.020174412 | 0.954242509 | 1.110947927 |
| ENSMUSG00000078354 | 0 | 0 | 0 | 0 | 0 | 0 | 0 |
| ENSMUSG00000078355 | 0 | 0 | 0 | 0 | 0 | 0 | 0 |
| ENSMUSG00000078365 | 0 | 0 | 0 | 0 | 0.244650542 | 0 | 0 |
| ENSMUSG00000078370 | 0.45815691 | 0.646240625 | 0.5 | 0.25 | 0.489301084 | 0.301029996 | 0.629532004 |
| ENSMUSG00000078377 | 0.289064826 | 0 | 0.25 | 0.25 | 0.244650542 | 0 | 0.355418052 |
| ENSMUSG00000078380 | 0 | 0.25 | 0.580482024 | 0.25 | 0.244650542 | 0 | 0.224243824 |
| ENSMUSG00000078384 | 0 | 0 | 0 | 0 | 0 | 0 | 0 |
| ENSMUSG00000078420 | 0 | 0 | 0 | 0 | 0 | 0 | 0 |
| ENSMUSG00000078427 | 2.530730035 | 2.331045137 | 2.337734795 | 2.502107156 | 2.422990063 | 2.789580712 | 2.256956407 |
| ENSMUSG00000078429 | 2.937953628 | 2.704645544 | 2.678561379 | 2.520204382 | 2.555125257 | 2.884795364 | 2.42760844 |
| ENSMUSG00000078430 | 0 | 0 | 0 | 0 | 0 | 0 | 0 |
| ENSMUSG00000078435 | 1.53838222 | 1.418106335 | 1.464495249 | 1.52186571 | 1.47337555 | 1.643452676 | 1.413583902 |
| ENSMUSG00000078439 | 0.289064826 | 0.396240625 | 0.25 | 0.25 | 0.489301084 | 0 | 0.448487648 |
| ENSMUSG00000078440 | 1.964026642 | 1.958222504 | 2.117918804 | 1.726722649 | 1.950242452 | 1.897627091 | 2.101807105 |
| ENSMUSG00000078441 | 0 | 0 | 0.25 | 0 | 0 | 0 | 0.224243824 |
| ENSMUSG00000078442 | 0 | 0 | 0 | 0 | 0 | 0 | 0 |
| ENSMUSG00000078444 | 0 | 0.701838731 | 0.79248125 | 0 | 0.489301084 | 0 | 0.629532004 |
| ENSMUSG00000078451 | 0.289064826 | 0 | 0.396240625 | 0.5 | 0.387761935 | 0.301029996 | 0.224243824 |
| ENSMUSG00000078452 | 1.227926429 | 1.29248125 | 1.146240625 | 1.282320754 | 1.106691885 | 1.477121255 | 1.193409508 |
| ENSMUSG00000078453 | 2.641182749 | 2.413659007 | 2.448603967 | 2.331045137 | 2.247690665 | 2.5774918 | 2.089878917 |
| ENSMUSG00000078480 | 0.811507563 | 0.75 | 0.951838731 | 1.17510993 | 1.091002363 | 1.113943352 | 1.028149525 |
| ENSMUSG00000078481 | 0 | 0 | 0 | 0 | 0 | 0 | 0 |
| ENSMUSG00000078484 | 1.783508299 | 1.646240625 | 1.76109853 | 1.552363341 | 1.736537397 | 1.591064607 | 1.687112305 |
| ENSMUSG00000078485 | 0.960252568 | 1.238549078 | 1.348079356 | 0.951838731 | 1.293076518 | 1.322219295 | 1.21680636 |
| ENSMUSG00000078486 | 0.867194479 | 0.92510993 | 1.080482024 | 1 | 1 | 0.84509804 | 0.916588299 |
| ENSMUSG00000078487 | 0 | 0 | 0 | 0 | 0 | 0 | 0 |
| ENSMUSG00000078488 | 0 | 0 | 0 | 0 | 0 | 0 | 0 |
| ENSMUSG00000078489 | 0.578129653 | 0.396240625 | 0.396240625 | 0.580482024 | 0.244650542 | 0.301029996 | 0 |
| ENSMUSG00000078490 | 1.605636262 | 1.575945187 | 1.5 | 1.445339928 | 1.343584837 | 1.602059991 | 1.272006351 |
| ENSMUSG00000078491 | 0 | 0 | 0 | 0 | 0 | 0 | 0 |
| ENSMUSG00000078492 | 0.289064826 | 0 | 0 | 0 | 0 | 0 | 0 |
| ENSMUSG00000078494 | 0 | 0.25 | 0.25 | 0 | 0.244650542 | 0.301029996 | 0.224243824 |
| ENSMUSG00000078495 | 1.587501561 | 1.649978211 | 1.695339928 | 1.575945187 | 1.53772706 | 1.579783597 | 1.396774124 |

| | | | | | | | |
|---|---|---|---|---|---|---|---|
| ENSMUSG00000078496 | 1.036286563 | 1.02186571 | 1.130890489 | 1.396240625 | 1.467903253 | 1.531478917 | 1.245582246 |
| ENSMUSG00000078497 | 0 | 0 | 0 | 0 | 0 | 0 | 0 |
| ENSMUSG00000078498 | 0 | 0 | 0 | 0 | 0 | 0 | 0 |
| ENSMUSG00000078500 | 0 | 0 | 0 | 0 | 0 | 0 | 0 |
| ENSMUSG00000078502 | 0.747221736 | 0.5 | 0.580482024 | 1 | 0.955822902 | 0.698970004 | 0.629532004 |
| ENSMUSG00000078503 | 0.289064826 | 0.5 | 0 | 0 | 0.244650542 | 0.301029996 | 0 |
| ENSMUSG00000078504 | 0 | 0 | 0 | 0 | 0 | 0 | 0 |
| ENSMUSG00000078505 | 0 | 0 | 0 | 0 | 0 | 0 | 0 |
| ENSMUSG00000078506 | 0 | 0 | 0 | 0 | 0 | 0 | 0 |
| ENSMUSG00000078507 | 0 | 0 | 0 | 0 | 0 | 0 | 0 |
| ENSMUSG00000078508 | 0 | 0 | 0 | 0 | 0 | 0 | 0 |
| ENSMUSG00000078509 | 0 | 0 | 0 | 0 | 0 | 0 | 0 |
| ENSMUSG00000078510 | 0 | 0 | 0 | 0 | 0 | 0 | 0 |
| ENSMUSG00000078511 | 0 | 0 | 0 | 0 | 0.387761935 | 0 | 0 |
| ENSMUSG00000078512 | 0 | 0 | 0 | 0 | 0 | 0 | 0 |
| ENSMUSG00000078513 | 0 | 0 | 0 | 0 | 0 | 0 | 0 |
| ENSMUSG00000078515 | 2.568542279 | 2.40136988 | 2.386723615 | 2.433677405 | 2.404070415 | 2.725094521 | 2.222904214 |
| ENSMUSG00000078517 | 2.433305075 | 2.245748394 | 2.246460484 | 2.215271726 | 2.288252441 | 2.419955748 | 2.153969461 |
| ENSMUSG00000078518 | 0 | 0 | 0 | 0 | 0 | 0 | 0 |
| ENSMUSG00000078519 | 0.45815691 | 0.396240625 | 0.396240625 | 0.396240625 | 0 | 0.602059991 | 0.224243824 |
| ENSMUSG00000078520 | 0 | 0 | 0 | 0 | 0 | 0 | 0 |
| ENSMUSG00000078521 | 2.033115198 | 1.83272922 | 1.890560606 | 1.951838731 | 1.873585591 | 2.176091259 | 1.811176017 |
| ENSMUSG00000078528 | 0.289064826 | 0.25 | 0.396240625 | 0 | 0.387761935 | 0 | 0 |
| ENSMUSG00000078531 | 0.289064826 | 0 | 0 | 0 | 0 | 0 | 0.224243824 |
| ENSMUSG00000078532 | 2.010213549 | 1.928561379 | 1.928561379 | 1.634789703 | 1.834841362 | 1.939519253 | 1.822941512 |
| ENSMUSG00000078537 | 0 | 0 | 0 | 0 | 0 | 0 | 0 |
| ENSMUSG00000078546 | 1.623015126 | 1.51109853 | 1.527131114 | 1.618933358 | 1.584291694 | 1.672097858 | 1.38802993 |
| ENSMUSG00000078552 | 0.578129653 | 0.396240625 | 0.25 | 0.646240625 | 0.568060967 | 0.477121255 | 0.520678036 |
| ENSMUSG00000078554 | 0 | 0 | 0 | 0 | 0 | 0 | 0 |
| ENSMUSG00000078566 | 2.119110214 | 2.04248125 | 2.006976499 | 2.012462137 | 1.940259189 | 2.184691431 | 1.803905701 |
| ENSMUSG00000078570 | 0.671187741 | 1 | 1.130890489 | 0.580482024 | 0.632412477 | 0.477121255 | 0.969165684 |
| ENSMUSG00000078572 | 0.811507563 | 1.238549078 | 1.201838731 | 0.896240625 | 1.020174412 | 1 | 1.245582246 |
| ENSMUSG00000078575 | 0 | 0 | 0 | 0 | 0 | 0 | 0 |
| ENSMUSG00000078576 | 0 | 0 | 0 | 0 | 0 | 0 | 0 |

| | | | | | | | |
|---|---|---|---|---|---|---|---|
| ENSMUSG00000078577 | 0 | 0 | 0 | 0 | 0 | 0 | 0 |
| ENSMUSG00000078578 | 3.050683019 | 2.771534056 | 2.741446071 | 2.935366747 | 2.833922504 | 3.311117843 | 2.633423538 |
| ENSMUSG00000078580 | 1.181541736 | 1.114857905 | 1.146240625 | 0.701838731 | 1.121713561 | 0.954242509 | 1.121219121 |
| ENSMUSG00000078584 | 1.928763473 | 1.87892496 | 1.926839783 | 1.951838731 | 1.905314583 | 2.158362492 | 1.820622395 |
| ENSMUSG00000078586 | 0 | 0 | 0 | 0 | 0 | 0 | 0 |
| ENSMUSG00000078588 | 0.289064826 | 0.646240625 | 0.701838731 | 0.396240625 | 0 | 0.301029996 | 0.224243824 |
| ENSMUSG00000078590 | 0 | 0 | 0 | 0 | 0 | 0 | 0 |
| ENSMUSG00000078591 | 0 | 0 | 0 | 0 | 0 | 0 | 0 |
| ENSMUSG00000078592 | 0.45815691 | 0.25 | 0.25 | 0 | 0.387761935 | 0 | 0.224243824 |
| ENSMUSG00000078593 | 0 | 0 | 0 | 0 | 0 | 0 | 0 |
| ENSMUSG00000078597 | 0 | 0 | 0 | 0 | 0 | 0 | 0 |
| ENSMUSG00000078598 | 0 | 0 | 0.25 | 0 | 0 | 0 | 0.355418052 |
| ENSMUSG00000078599 | 0.960252568 | 0.864857905 | 0.896240625 | 0.896240625 | 0.955822902 | 0.84509804 | 0.829800754 |
| ENSMUSG00000078600 | 0 | 0 | 0 | 0 | 0 | 0 | 0 |
| ENSMUSG00000078601 | 0 | 0 | 0 | 0 | 0.244650542 | 0 | 0 |
| ENSMUSG00000078604 | 0 | 0 | 0.25 | 0.25 | 0 | 0 | 0 |
| ENSMUSG00000078605 | 0 | 0 | 0 | 0 | 0 | 0 | 0 |
| ENSMUSG00000078606 | 0.289064826 | 0 | 0.25 | 0 | 0 | 0 | 0 |
| ENSMUSG00000078607 | 0.671187741 | 0.830482024 | 0.5 | 0.396240625 | 0.733951626 | 0.477121255 | 0.775756176 |
| ENSMUSG00000078611 | 0 | 0 | 0 | 0 | 0 | 0 | 0 |
| ENSMUSG00000078612 | 1.036286563 | 1.146240625 | 1.098079356 | 0.830482024 | 0.955822902 | 0.903089987 | 0.672731473 |
| ENSMUSG00000078616 | 0 | 0 | 0 | 0 | 0 | 0 | 0 |
| ENSMUSG00000078618 | 0 | 0 | 0 | 0 | 0 | 0 | 0 |
| ENSMUSG00000078619 | 2.42468076 | 2.314346961 | 2.279735268 | 2.328220739 | 2.379941013 | 2.711807229 | 2.253616472 |
| ENSMUSG00000078620 | 0 | 0 | 0 | 0 | 0 | 0 | 0 |
| ENSMUSG00000078621 | 0 | 0 | 0 | 0 | 0 | 0 | 0 |
| ENSMUSG00000078622 | 2.794421919 | 2.578787391 | 2.571350555 | 2.66942991 | 2.610365232 | 3.004321374 | 2.414513278 |
| ENSMUSG00000078624 | 1.558729299 | 1.494319981 | 1.388647213 | 1.575945187 | 1.387761935 | 1.491361694 | 1.38802993 |
| ENSMUSG00000078625 | 0.45815691 | 0 | 0.25 | 0 | 0.387761935 | 0 | 0.355418052 |
| ENSMUSG00000078626 | 0 | 0 | 0 | 0 | 0 | 0 | 0 |
| ENSMUSG00000078627 | 0 | 0 | 0.25 | 0 | 0 | 0 | 0 |
| ENSMUSG00000078630 | 0.289064826 | 0.25 | 0.580482024 | 0.5 | 0.387761935 | 0 | 0.579661877 |
| ENSMUSG00000078632 | 0 | 0 | 0 | 0 | 0 | 0 | 0 |
| ENSMUSG00000078635 | 0 | 0.25 | 0.25 | 0 | 0 | 0 | 0 |

| | | | | | | | |
|---|---|---|---|---|---|---|---|
| ENSMUSG00000078636 | 0 | 0 | 0 | 0 | 0 | 0 | 0 |
| ENSMUSG00000078639 | 0 | 0 | 0 | 0 | 0 | 0 | 0 |
| ENSMUSG00000078640 | 0.289064826 | 0.25 | 0.25 | 0.396240625 | 0 | 0 | 0.224243824 |
| ENSMUSG00000078648 | 0 | 0 | 0 | 0 | 0.387761935 | 0.301029996 | 0 |
| ENSMUSG00000078650 | 0 | 0 | 0 | 0 | 0 | 0 | 0 |
| ENSMUSG00000078651 | 1.771760131 | 1.630890489 | 1.646240625 | 1.482684334 | 1.595992969 | 1.681241237 | 1.473249546 |
| ENSMUSG00000078652 | 2.885732304 | 2.644121087 | 2.63162481 | 2.686838457 | 2.640859694 | 3.015778756 | 2.468886073 |
| ENSMUSG00000078653 | 0.289064826 | 0.5 | 0.5 | 0.580482024 | 0.632412477 | 0 | 0.520678036 |
| ENSMUSG00000078655 | 0 | 0 | 0 | 0 | 0 | 0 | 0 |
| ENSMUSG00000078656 | 0.45815691 | 0.25 | 0.5 | 0.5 | 0.489301084 | 0.477121255 | 0.672731473 |
| ENSMUSG00000078657 | 0 | 0 | 0 | 0 | 0 | 0 | 0 |
| ENSMUSG00000078664 | 0 | 0 | 0 | 0 | 0 | 0 | 0 |
| ENSMUSG00000078668 | 0 | 0 | 0 | 0 | 0 | 0 | 0 |
| ENSMUSG00000078670 | 0 | 0 | 0 | 0.396240625 | 0.387761935 | 0.301029996 | 0.672731473 |
| ENSMUSG00000078671 | 2.392336464 | 2.21759118 | 2.174241882 | 2.181128463 | 2.116484486 | 2.350248018 | 2 |
| ENSMUSG00000078672 | 0 | 0 | 0 | 0 | 0 | 0 | 0 |
| ENSMUSG00000078673 | 0 | 0 | 0 | 0 | 0 | 0 | 0 |
| ENSMUSG00000078674 | 0 | 0 | 0 | 0 | 0 | 0 | 0 |
| ENSMUSG00000078675 | 0 | 0 | 0 | 0 | 0 | 0 | 0 |
| ENSMUSG00000078676 | 2.70238848 | 2.481573999 | 2.49929487 | 2.540282971 | 2.506870642 | 2.819543936 | 2.347752519 |
| ENSMUSG00000078677 | 0 | 0.25 | 0 | 0.25 | 0 | 0 | 0 |
| ENSMUSG00000078680 | 0 | 0 | 0 | 0 | 0 | 0 | 0 |
| ENSMUSG00000078681 | 1.578129653 | 1.618933358 | 1.649978211 | 1.339388001 | 1.478764298 | 1.681241237 | 1.586932193 |
| ENSMUSG00000078683 | 0 | 0 | 0 | 0 | 0 | 0 | 0 |
| ENSMUSG00000078684 | 0.289064826 | 0 | 0 | 1.238549078 | 1.27449559 | 1.447158031 | 1.20919388 |
| ENSMUSG00000078685 | 0 | 0 | 0 | 0 | 0 | 0 | 0 |
| ENSMUSG00000078686 | 0 | 0 | 0 | 0 | 0 | 0 | 0 |
| ENSMUSG00000078687 | 0 | 0 | 0 | 0 | 0 | 0 | 0 |
| ENSMUSG00000078688 | 0 | 0 | 0 | 0 | 0 | 0 | 0 |
| ENSMUSG00000078689 | 0 | 0 | 0 | 0 | 0 | 0 | 0 |
| ENSMUSG00000078691 | 0 | 0 | 0 | 0 | 0 | 0 | 0 |
| ENSMUSG00000078695 | 1.568542279 | 1.54248125 | 1.532320754 | 1.646240625 | 1.652586889 | 1.806179974 | 1.601000298 |
| ENSMUSG00000078698 | 0 | 0 | 0 | 0 | 0 | 0 | 0 |
| ENSMUSG00000078700 | 1.578129653 | 1.494319981 | 1.516522298 | 1.566696635 | 1.568060967 | 1.633468456 | 1.540636858 |

| | | | | | | | |
|---|---|---|---|---|---|---|---|
| ENSMUSG00000078706 | 1.903481042 | 1.834962501 | 1.843759858 | 1.53743678 | 1.595992969 | 1.591064607 | 1.582103514 |
| ENSMUSG00000078713 | 2.258689303 | 2.103406982 | 2.103406982 | 2.049918086 | 2.06149024 | 2.307496038 | 1.951796623 |
| ENSMUSG00000078716 | 1.389637216 | 1.214495249 | 1.130890489 | 1.214495249 | 1.433158226 | 1.462397998 | 1.324583736 |
| ENSMUSG00000078719 | 0 | 0 | 0 | 0 | 0 | 0 | 0 |
| ENSMUSG00000078721 | 0 | 0 | 0 | 0 | 0 | 0 | 0 |
| ENSMUSG00000078722 | 0 | 0 | 0 | 0 | 0 | 0 | 0 |
| ENSMUSG00000078735 | 0 | 0 | 0 | 0 | 0 | 0 | 0 |
| ENSMUSG00000078737 | 0 | 0 | 0 | 0 | 0 | 0 | 0 |
| ENSMUSG00000078742 | 0 | 0 | 0 | 0 | 0 | 0 | 0.224243824 |
| ENSMUSG00000078746 | 0 | 0 | 0 | 0 | 0 | 0 | 0 |
| ENSMUSG00000078747 | 0 | 0 | 0 | 0 | 0.489301084 | 0 | 0.355418052 |
| ENSMUSG00000078752 | 0 | 0 | 0 | 0 | 0 | 0 | 0 |
| ENSMUSG00000078753 | 0 | 0 | 0 | 0 | 0 | 0 | 0 |
| ENSMUSG00000078754 | 0 | 0 | 0 | 0 | 0 | 0 | 0 |
| ENSMUSG00000078757 | 0 | 0 | 0 | 0 | 0 | 0 | 0 |
| ENSMUSG00000078759 | 0 | 0 | 0 | 0 | 0 | 0 | 0 |
| ENSMUSG00000078762 | 0.960252568 | 1.302363341 | 1.26109853 | 1.146240625 | 1.254881871 | 1.146128036 | 1.319146374 |
| ENSMUSG00000078763 | 0 | 0 | 0 | 0 | 0 | 0 | 0 |
| ENSMUSG00000078765 | 1.505869732 | 1.51109853 | 1.464495249 | 1.557204673 | 1.655839964 | 1.672097858 | 1.62316756 |
| ENSMUSG00000078768 | 1.325351389 | 1.17510993 | 1.214495249 | 1.330482024 | 1.200473444 | 1.431363764 | 1.245582246 |
| ENSMUSG00000078771 | 0.578129653 | 0.580482024 | 0.396240625 | 0.701838731 | 0.568060967 | 0.903089987 | 0.579661877 |
| ENSMUSG00000078772 | 0.45815691 | 0.396240625 | 0.25 | 0 | 0.387761935 | 0 | 0.224243824 |
| ENSMUSG00000078773 | 2.192545868 | 2.05357978 | 2.015173983 | 2.144357207 | 2.122577592 | 2.322219295 | 1.989815042 |
| ENSMUSG00000078776 | 0.91631382 | 0.830482024 | 0.79248125 | 0.951838731 | 1.020174412 | 0.84509804 | 0.853775828 |
| ENSMUSG00000078779 | 1.036286563 | 1.080482024 | 1.061981878 | 1.080482024 | 1.020174412 | 1.204119983 | 0.896975297 |
| ENSMUSG00000078780 | 0 | 0 | 0 | 0 | 0 | 0 | 0 |
| ENSMUSG00000078783 | 0 | 0 | 0 | 0 | 0 | 0 | 0 |
| ENSMUSG00000078784 | 1.978790393 | 1.901832578 | 1.890560606 | 1.955044741 | 1.893954095 | 2.152288344 | 1.769001954 |
| ENSMUSG00000078786 | 1.45815691 | 1.26109853 | 1.188721876 | 1.282320754 | 1.121713561 | 1.342422681 | 1.23151414 |
| ENSMUSG00000078787 | 0.578129653 | 0.25 | 0.396240625 | 0.396240625 | 0.568060967 | 0 | 0.579661877 |
| ENSMUSG00000078789 | 1.827447047 | 1.653677461 | 1.681980114 | 1.566696635 | 1.514343934 | 1.748188027 | 1.596378571 |
| ENSMUSG00000078793 | 0 | 0 | 0 | 0 | 0 | 0 | 0 |
| ENSMUSG00000078794 | 1.494443473 | 1.282320754 | 1.17510993 | 0.92510993 | 1.149965125 | 1 | 0.969165684 |
| ENSMUSG00000078795 | 0 | 0 | 0 | 0 | 0 | 0 | 0 |

| | | | | | | | |
|---|---|---|---|---|---|---|---|
| ENSMUSG00000078796 | 0 | 0 | 0 | 0 | 0.244650542 | 0 | 0 |
| ENSMUSG00000078798 | 0 | 0 | 0 | 0 | 0 | 0 | 0 |
| ENSMUSG00000078799 | 0 | 0 | 0 | 0 | 0 | 0 | 0 |
| ENSMUSG00000078800 | 0 | 0 | 0 | 0 | 0 | 0 | 0 |
| ENSMUSG00000078808 | 1.325351389 | 1.098079356 | 1.080482024 | 1.380890489 | 1 | 1.176091259 | 1 |
| ENSMUSG00000078810 | 0 | 0 | 0 | 0 | 0 | 0 | 0 |
| ENSMUSG00000078812 | 3.482301227 | 3.202291301 | 3.203995907 | 3.153389819 | 3.16634835 | 3.634678752 | 2.982168966 |
| ENSMUSG00000078813 | 1.470606563 | 1.438721876 | 1.445339928 | 1.356566189 | 1.407936347 | 1.477121255 | 1.48984372 |
| ENSMUSG00000078815 | 0 | 0 | 0 | 0 | 0 | 0 | 0.355418052 |
| ENSMUSG00000078816 | 1.100572389 | 1.04248125 | 1 | 0.701838731 | 0.733951626 | 0.477121255 | 0.829800754 |
| ENSMUSG00000078817 | 0.45815691 | 0.25 | 0.396240625 | 0.25 | 0.489301084 | 0 | 0.520678036 |
| ENSMUSG00000078838 | 0.578129653 | 0.396240625 | 0 | 0.396240625 | 0.489301084 | 0.602059991 | 0.520678036 |
| ENSMUSG00000078840 | 0 | 0 | 0 | 0.25 | 0 | 0 | 0 |
| ENSMUSG00000078847 | 0 | 0 | 0.25 | 0 | 0 | 0 | 0.224243824 |
| ENSMUSG00000078850 | 0 | 0 | 0 | 0 | 0 | 0 | 0 |
| ENSMUSG00000078851 | 0.578129653 | 0.646240625 | 0.701838731 | 1.061981878 | 1.020174412 | 1.041392685 | 0.969165684 |
| ENSMUSG00000078853 | 1.129344651 | 1.061981878 | 1.061981878 | 1.488549078 | 1.467903253 | 1.653212514 | 1.437266334 |
| ENSMUSG00000078861 | 1.227926429 | 1.302363341 | 1.201838731 | 1.356566189 | 1.264824954 | 1.544068044 | 1.20139795 |
| ENSMUSG00000078862 | 0.811507563 | 0.830482024 | 1.080482024 | 0.976722649 | 0.931471446 | 1 | 0.775756176 |
| ENSMUSG00000078864 | 0 | 0.25 | 0 | 0.646240625 | 0.812711509 | 0.77815125 | 0.448487648 |
| ENSMUSG00000078865 | 0 | 0 | 0 | 0 | 0 | 0 | 0.224243824 |
| ENSMUSG00000078866 | 1.307602651 | 1.348079356 | 1.02186571 | 1.726722649 | 1.599808748 | 1.763427994 | 1.486592281 |
| ENSMUSG00000078867 | 0.289064826 | 0 | 0.396240625 | 1 | 0.877063019 | 0.77815125 | 0.829800754 |
| ENSMUSG00000078868 | 0 | 0 | 0 | 0.5 | 0.568060967 | 0.301029996 | 0.448487648 |
| ENSMUSG00000078869 | 0 | 0 | 0 | 0.25 | 0.244650542 | 0.301029996 | 0 |
| ENSMUSG00000078870 | 0.578129653 | 0.75 | 0.580482024 | 0.646240625 | 0.77552387 | 0.84509804 | 0.829800754 |
| ENSMUSG00000078872 | 0.45815691 | 0.580482024 | 0.25 | 0.5 | 0.489301084 | 0 | 0.224243824 |
| ENSMUSG00000078875 | 0 | 0.396240625 | 0.396240625 | 0.25 | 0.244650542 | 0.301029996 | 0.355418052 |
| ENSMUSG00000078876 | 0 | 0 | 0 | 0 | 0 | 0 | 0 |
| ENSMUSG00000078877 | 0.867194479 | 1.061981878 | 0.896240625 | 0.830482024 | 0.686820904 | 0.698970004 | 0 |
| ENSMUSG00000078878 | 0.578129653 | 0.701838731 | 0.396240625 | 0.92510993 | 0.632412477 | 0.477121255 | 0.74492186 |
| ENSMUSG00000078879 | 0.811507563 | 0.580482024 | 0.5 | 0.701838731 | 0.686820904 | 0.698970004 | 0.775756176 |
| ENSMUSG00000078880 | 0 | 0 | 0.25 | 0.580482024 | 0.244650542 | 0 | 0.224243824 |
| ENSMUSG00000078881 | 0 | 0 | 0 | 0 | 0 | 0 | 0 |

| | | | | | | | |
|---|---|---|---|---|---|---|---|
| ENSMUSG00000078886 | 0 | 0 | 0 | 0 | 0 | 0 | 0 |
| ENSMUSG00000078887 | 0.747221736 | 0.830482024 | 0.75 | 0.864857905 | 0.978602168 | 0.77815125 | 0.969165684 |
| ENSMUSG00000078889 | 0 | 0 | 0 | 0 | 0 | 0 | 0 |
| ENSMUSG00000078891 | 0 | 0 | 0 | 0 | 0 | 0 | 0 |
| ENSMUSG00000078894 | 0 | 0 | 0 | 0 | 0 | 0 | 0 |
| ENSMUSG00000078895 | 0 | 0 | 0 | 0 | 0 | 0 | 0 |
| ENSMUSG00000078896 | 0 | 0 | 0 | 0 | 0 | 0 | 0 |
| ENSMUSG00000078897 | 0 | 0 | 0 | 0 | 0 | 0 | 0 |
| ENSMUSG00000078898 | 0.289064826 | 0 | 0 | 0.25 | 0.568060967 | 0.477121255 | 0.448487648 |
| ENSMUSG00000078899 | 2.158622101 | 2.016522298 | 2.015173983 | 2.173371739 | 2.112066955 | 2.338456494 | 1.930140666 |
| ENSMUSG00000078901 | 0 | 0 | 0 | 0.25 | 0.244650542 | 0.301029996 | 0 |
| ENSMUSG00000078902 | 0.867194479 | 0.92510993 | 0.976722649 | 0.951838731 | 1 | 0.698970004 | 1.054044578 |
| ENSMUSG00000078903 | 1.827447047 | 1.614857905 | 1.571350555 | 1.866901388 | 1.695608647 | 1.86923172 | 1.548827561 |
| ENSMUSG00000078905 | 0 | 0.396240625 | 0.396240625 | 0.25 | 0 | 0.301029996 | 0.224243824 |
| ENSMUSG00000078906 | 0.289064826 | 0 | 0 | 0.396240625 | 0.387761935 | 0 | 0.224243824 |
| ENSMUSG00000078907 | 0 | 0.396240625 | 0 | 0 | 0.244650542 | 0 | 0 |
| ENSMUSG00000078908 | 2.023453784 | 1.905512955 | 1.874961472 | 1.763820609 | 1.842556557 | 1.959041392 | 1.757893036 |
| ENSMUSG00000078912 | 0 | 0 | 0 | 0 | 0 | 0 | 0 |
| ENSMUSG00000078915 | 0.289064826 | 0 | 0 | 0 | 0 | 0 | 0 |
| ENSMUSG00000078917 | 0.671187741 | 0.5 | 0.75 | 0.646240625 | 0.877063019 | 0.477121255 | 0.710836105 |
| ENSMUSG00000078919 | 1.432083896 | 1.339388001 | 1.348079356 | 1.527131114 | 1.580303447 | 1.662757832 | 1.401059159 |
| ENSMUSG00000078920 | 1.036286563 | 0.79248125 | 0.701838731 | 1.098079356 | 1.074582839 | 1.204119983 | 1.054044578 |
| ENSMUSG00000078921 | 0.289064826 | 0 | 0 | 0 | 0.244650542 | 0 | 0.224243824 |
| ENSMUSG00000078922 | 0.811507563 | 0.75 | 0.396240625 | 0.25 | 0.244650542 | 0 | 0.520678036 |
| ENSMUSG00000078923 | 0.671187741 | 0.75 | 0.646240625 | 0.646240625 | 0.733951626 | 0.301029996 | 0.355418052 |
| ENSMUSG00000078924 | 0 | 0 | 0 | 0 | 0 | 0 | 0 |
| ENSMUSG00000078925 | 0 | 0.25 | 0 | 0 | 0 | 0 | 0 |
| ENSMUSG00000078926 | 0 | 0 | 0 | 0 | 0 | 0 | 0.224243824 |
| ENSMUSG00000078931 | 0.45815691 | 0.864857905 | 0.5 | 0.646240625 | 0.733951626 | 0.698970004 | 0.579661877 |
| ENSMUSG00000078932 | 0 | 0 | 0 | 0 | 0 | 0 | 0 |
| ENSMUSG00000078934 | 0 | 0 | 0 | 0 | 0 | 0 | 0 |
| ENSMUSG00000078935 | 0 | 0 | 0 | 0 | 0 | 0 | 0 |
| ENSMUSG00000078937 | 0 | 0.5 | 0.25 | 0 | 0.568060967 | 0 | 0.579661877 |
| ENSMUSG00000078938 | 0 | 0 | 0 | 0 | 0 | 0 | 0 |

| | | | | | | | |
|---|---|---|---|---|---|---|---|
| ENSMUSG00000078940 | 0 | 0 | 0 | 0 | 0 | 0 | 0 |
| ENSMUSG00000078941 | 2.256831695 | 2.151832578 | 2.122963274 | 2.256284891 | 2.187072901 | 2.514547753 | 2.040767326 |
| ENSMUSG00000078942 | 0 | 0 | 0 | 0.396240625 | 0.632412477 | 0.477121255 | 0.579661877 |
| ENSMUSG00000078945 | 0.671187741 | 0.79248125 | 0.25 | 1.17510993 | 0.978602168 | 1.041392685 | 0.876096088 |
| ENSMUSG00000078949 | 0 | 0 | 0 | 0 | 0 | 0 | 0 |
| ENSMUSG00000078952 | 1.358731791 | 1.214495249 | 1.330482024 | 1.080482024 | 0.955822902 | 0.954242509 | 0.829800754 |
| ENSMUSG00000078953 | 0 | 0 | 0 | 0 | 0 | 0 | 0 |
| ENSMUSG00000078954 | 1.578129653 | 1.622963274 | 1.610735874 | 1.282320754 | 1.293076518 | 1.397940009 | 1.374453863 |
| ENSMUSG00000078955 | 0 | 0 | 0 | 0 | 0 | 0 | 0 |
| ENSMUSG00000078956 | 0 | 0 | 0 | 0 | 0 | 0 | 0 |
| ENSMUSG00000078957 | 0 | 0 | 0 | 0 | 0 | 0 | 0 |
| ENSMUSG00000078958 | 0 | 0 | 0 | 0 | 0 | 0 | 0 |
| ENSMUSG00000078962 | 0 | 0 | 0 | 0 | 0 | 0 | 0 |
| ENSMUSG00000078963 | 0.45815691 | 0.25 | 0.5 | 0.396240625 | 0.387761935 | 0 | 0 |
| ENSMUSG00000078964 | 0 | 0 | 0 | 0 | 0 | 0 | 0 |
| ENSMUSG00000078965 | 0 | 0 | 0 | 0 | 0 | 0 | 0 |
| ENSMUSG00000078967 | 0.45815691 | 0.646240625 | 0.75 | 0.396240625 | 0.686820904 | 0 | 0.448487648 |
| ENSMUSG00000078970 | 2.086807695 | 1.926839783 | 1.888647213 | 2.061981878 | 1.991345651 | 2.240549248 | 1.884801065 |
| ENSMUSG00000078972 | 0 | 0 | 0 | 0 | 0 | 0 | 0 |
| ENSMUSG00000078974 | 2.072573125 | 1.945339928 | 1.976722649 | 2.219895812 | 2.058398634 | 2.387389826 | 1.966952343 |
| ENSMUSG00000078982 | 0 | 0 | 0 | 0 | 0 | 0 | 0 |
| ENSMUSG00000078984 | 0.45815691 | 0.396240625 | 0 | 0.25 | 0.244650542 | 0.301029996 | 0.355418052 |
| ENSMUSG00000078994 | 1.156259305 | 0.896240625 | 1.080482024 | 1.431980114 | 1.414412788 | 1.51851394 | 1.224243824 |
| ENSMUSG00000078995 | 0.811507563 | 0.701838731 | 0.79248125 | 0.701838731 | 0.733951626 | 0.477121255 | 0.803905701 |
| ENSMUSG00000078998 | 0 | 0 | 0 | 0 | 0 | 0 | 0 |
| ENSMUSG00000079000 | 0 | 0 | 0 | 0 | 0 | 0 | 0.224243824 |
| ENSMUSG00000079001 | 0 | 0 | 0 | 0 | 0 | 0 | 0 |
| ENSMUSG00000079003 | 2.139333929 | 2.001406137 | 2.056000419 | 1.841580554 | 1.929985312 | 2.068185862 | 1.876096088 |
| ENSMUSG00000079005 | 0 | 0 | 0 | 0 | 0 | 0 | 0 |
| ENSMUSG00000079006 | 0 | 0 | 0 | 0 | 0 | 0 | 0 |
| ENSMUSG00000079007 | 0 | 0 | 0 | 0 | 0 | 0 | 0 |
| ENSMUSG00000079008 | 0 | 0.580482024 | 0 | 0.396240625 | 0 | 0 | 0.224243824 |
| ENSMUSG00000079009 | 0 | 0 | 0.25 | 0 | 0.244650542 | 0 | 0 |
| ENSMUSG00000079010 | 0 | 0.25 | 0.646240625 | 0.25 | 0.568060967 | 0 | 0.520678036 |

| | | | | | | | |
|---|---|---|---|---|---|---|---|
| ENSMUSG00000079011 | 0.578129653 | 0.25 | 0.396240625 | 0.79248125 | 0.387761935 | 0.698970004 | 0.520678036 |
| ENSMUSG00000079012 | 0 | 0 | 0 | 0 | 0 | 0 | 0 |
| ENSMUSG00000079013 | 0 | 0 | 0 | 0 | 0 | 0 | 0 |
| ENSMUSG00000079014 | 1.956444026 | 1.819031101 | 1.823655187 | 0.951838731 | 0.77552387 | 1.230448921 | 1.078019652 |
| ENSMUSG00000079015 | 0 | 0 | 0 | 0 | 0 | 0 | 0 |
| ENSMUSG00000079016 | 0 | 0.25 | 0 | 0 | 0.244650542 | 0 | 0 |
| ENSMUSG00000079017 | 0 | 0 | 0 | 0 | 0 | 0 | 0 |
| ENSMUSG00000079018 | 1.482695304 | 1.476722649 | 1.488549078 | 1.061981878 | 0.877063019 | 0.903089987 | 0.935079929 |
| ENSMUSG00000079019 | 0 | 0 | 0 | 0 | 0 | 0 | 0 |
| ENSMUSG00000079020 | 2.457006475 | 2.284247778 | 2.307204673 | 1.912762923 | 2.044679629 | 2.227886705 | 1.982842456 |
| ENSMUSG00000079022 | 0 | 0.580482024 | 0.396240625 | 0.646240625 | 0.568060967 | 0.301029996 | 0.448487648 |
| ENSMUSG00000079024 | 0 | 0.25 | 0 | 0.25 | 0 | 0 | 0.355418052 |
| ENSMUSG00000079025 | 0 | 0 | 0 | 0 | 0 | 0 | 0 |
| ENSMUSG00000079029 | 0 | 0.580482024 | 0.396240625 | 0 | 0.387761935 | 0 | 0 |
| ENSMUSG00000079030 | 0 | 0 | 0 | 0 | 0 | 0 | 0 |
| ENSMUSG00000079031 | 0 | 0 | 0 | 0 | 0 | 0 | 0 |
| ENSMUSG00000079033 | 0 | 0.25 | 0 | 0 | 0 | 0 | 0 |
| ENSMUSG00000079034 | 0.45815691 | 0.646240625 | 0.646240625 | 0 | 0 | 0 | 0 |
| ENSMUSG00000079036 | 2.283862692 | 2.097004321 | 2.051142786 | 2.012462137 | 1.945998439 | 2.127104798 | 1.837623616 |
| ENSMUSG00000079037 | 2.463861956 | 2.256976499 | 2.310197833 | 2.324229052 | 2.302012594 | 2.56937391 | 2.159731975 |
| ENSMUSG00000079038 | 0.578129653 | 0.396240625 | 0.5 | 1.160964047 | 1.057362051 | 0.77815125 | 0.916588299 |
| ENSMUSG00000079039 | 0 | 0 | 0 | 0 | 0 | 0 | 0 |
| ENSMUSG00000079042 | 0 | 0 | 0 | 0 | 0 | 0 | 0 |
| ENSMUSG00000079043 | 1.623015126 | 1.494319981 | 1.42510993 | 1.589388001 | 1.555378544 | 1.84509804 | 1.40946263 |
| ENSMUSG00000079045 | 0 | 0 | 0 | 0 | 0.244650542 | 0 | 0 |
| ENSMUSG00000079048 | 0 | 0 | 0 | 0 | 0 | 0 | 0 |
| ENSMUSG00000079049 | 0.671187741 | 0.830482024 | 0.79248125 | 0.646240625 | 0.632412477 | 0.301029996 | 0.448487648 |
| ENSMUSG00000079051 | 0 | 0 | 0.25 | 0 | 0 | 0 | 0.355418052 |
| ENSMUSG00000079053 | 0 | 0.25 | 0 | 0.25 | 0.387761935 | 0 | 0 |
| ENSMUSG00000079055 | 0 | 0 | 0.396240625 | 0.75 | 0.686820904 | 0.602059991 | 0.579661877 |
| ENSMUSG00000079056 | 0.747221736 | 0.79248125 | 0.896240625 | 0.976722649 | 1.212046813 | 1.255272505 | 0.984950056 |
| ENSMUSG00000079057 | 1.470606563 | 1.396240625 | 1.527131114 | 1.705044741 | 1.572189156 | 1.716003344 | 1.517723536 |
| ENSMUSG00000079058 | 0 | 0 | 0 | 0 | 0 | 0 | 0 |
| ENSMUSG00000079061 | 0 | 0 | 0 | 0 | 0 | 0 | 0 |

| | | | | | | | |
|---|---|---|---|---|---|---|---|
| ENSMUSG00000079064 | 0 | 0 | 0 | 0 | 0 | 0 | 0 |
| ENSMUSG00000079065 | 1.948720984 | 1.87892496 | 1.880890489 | 1.83272922 | 1.758994476 | 1.954242509 | 1.635773653 |
| ENSMUSG00000079067 | 0 | 0 | 0 | 0 | 0 | 0 | 0 |
| ENSMUSG00000079069 | 0.289064826 | 0.396240625 | 0.646240625 | 0.25 | 0.244650542 | 0 | 0.672731473 |
| ENSMUSG00000079070 | 0 | 0 | 0.25 | 0 | 0 | 0 | 0 |
| ENSMUSG00000079071 | 0 | 0 | 0 | 0 | 0 | 0 | 0 |
| ENSMUSG00000079076 | 0 | 0.75 | 0.25 | 0.5 | 0 | 0.477121255 | 0.448487648 |
| ENSMUSG00000079083 | 1.948720984 | 1.874961472 | 1.816696635 | 1.839388001 | 1.766173492 | 1.973127854 | 1.631625962 |
| ENSMUSG00000079084 | 1.91631382 | 1.856566189 | 1.834962501 | 1.981203126 | 1.871833944 | 2.089905111 | 1.783679399 |
| ENSMUSG00000079091 | 0 | 0 | 0 | 0 | 0 | 0 | 0 |
| ENSMUSG00000079092 | 0.867194479 | 0.896240625 | 0.976722649 | 0.646240625 | 0.632412477 | 0.602059991 | 0.579661877 |
| ENSMUSG00000079093 | 0 | 0 | 0 | 0 | 0 | 0 | 0 |
| ENSMUSG00000079101 | 0 | 0 | 0 | 0 | 0 | 0 | 0 |
| ENSMUSG00000079103 | 0 | 0 | 0 | 0 | 0 | 0 | 0 |
| ENSMUSG00000079104 | 2.282114126 | 2.117918804 | 2.115881093 | 2.099151195 | 1.997548906 | 2.278753601 | 1.871156825 |
| ENSMUSG00000079105 | 0 | 0 | 0 | 0 | 0 | 0 | 0 |
| ENSMUSG00000079108 | 2.141794304 | 2.058404919 | 2.066696635 | 2.093759858 | 2.024758322 | 2.235528447 | 1.911356129 |
| ENSMUSG00000079109 | 2.287337982 | 2.106566189 | 2.094844592 | 2.184523065 | 2.135290471 | 2.42975228 | 2.005303651 |
| ENSMUSG00000079110 | 0 | 0 | 0 | 0 | 0.244650542 | 0 | 0.355418052 |
| ENSMUSG00000079111 | 2.930987651 | 2.69961795 | 2.707232457 | 2.617918804 | 2.615740848 | 2.971739591 | 2.500831002 |
| ENSMUSG00000079112 | 0 | 0 | 0 | 0 | 0 | 0 | 0 |
| ENSMUSG00000079113 | 0 | 0 | 0 | 0 | 0 | 0 | 0 |
| ENSMUSG00000079114 | 0 | 0 | 0 | 0 | 0 | 0 | 0 |
| ENSMUSG00000079116 | 0 | 0 | 0 | 0 | 0 | 0 | 0 |
| ENSMUSG00000079120 | 0 | 0 | 0 | 0 | 0 | 0 | 0 |
| ENSMUSG00000079137 | 0 | 0 | 0 | 0 | 0.244650542 | 0 | 0.355418052 |
| ENSMUSG00000079138 | 0 | 0 | 0 | 0 | 0.387761935 | 0 | 0.224243824 |
| ENSMUSG00000079139 | 0.289064826 | 0.396240625 | 0.25 | 0.396240625 | 0.387761935 | 0.477121255 | 0.224243824 |
| ENSMUSG00000079140 | 0 | 0 | 0 | 0 | 0 | 0 | 0 |
| ENSMUSG00000079143 | 0 | 0 | 0 | 0 | 0 | 0.301029996 | 0.224243824 |
| ENSMUSG00000079144 | 1.671187741 | 1.593759858 | 1.602347734 | 1.584962501 | 1.514343934 | 1.591064607 | 1.462868481 |
| ENSMUSG00000079146 | 0 | 0 | 0 | 0 | 0 | 0 | 0 |
| ENSMUSG00000079156 | 0 | 0 | 0 | 0 | 0 | 0 | 0 |
| ENSMUSG00000079157 | 0 | 0 | 0 | 0.25 | 0 | 0 | 0 |

| | | | | | | | |
|---|---|---|---|---|---|---|---|
| ENSMUSG00000079162 | 0.671187741 | 0.5 | 0.79248125 | 0.646240625 | 0.489301084 | 0.698970004 | 0.710836105 |
| ENSMUSG00000079163 | 0 | 0 | 0 | 0 | 0 | 0 | 0 |
| ENSMUSG00000079164 | 0.289064826 | 0.25 | 0.701838731 | 0.646240625 | 0.244650542 | 0.698970004 | 0.829800754 |
| ENSMUSG00000079165 | 0 | 0 | 0 | 0 | 0.244650542 | 0 | 0.224243824 |
| ENSMUSG00000079168 | 0 | 0 | 0 | 0 | 0 | 0 | 0 |
| ENSMUSG00000079169 | 0 | 0 | 0 | 0 | 0 | 0 | 0 |
| ENSMUSG00000079170 | 0 | 0 | 0 | 0 | 0 | 0 | 0 |
| ENSMUSG00000079171 | 0 | 0 | 0 | 0 | 0 | 0 | 0 |
| ENSMUSG00000079173 | 0.867194479 | 0.79248125 | 0.896240625 | 0.25 | 0.489301084 | 0.477121255 | 0.520678036 |
| ENSMUSG00000079174 | 0 | 0 | 0 | 0 | 0 | 0 | 0 |
| ENSMUSG00000079175 | 0 | 0 | 0 | 0 | 0 | 0 | 0 |
| ENSMUSG00000079177 | 0 | 0.396240625 | 0 | 0.396240625 | 0 | 0.301029996 | 0 |
| ENSMUSG00000079179 | 1.852729478 | 1.799918086 | 1.732684334 | 1.614857905 | 1.642643806 | 1.86332286 | 1.621017948 |
| ENSMUSG00000079180 | 0 | 0 | 0 | 0 | 0 | 0 | 0 |
| ENSMUSG00000079183 | 0 | 0 | 0 | 0 | 0 | 0 | 0 |
| ENSMUSG00000079184 | 2.211649928 | 2.100219859 | 2.086073977 | 2.365369862 | 2.2705456 | 2.582063363 | 2.092902448 |
| ENSMUSG00000079186 | 0 | 0 | 0 | 0 | 0 | 0 | 0 |
| ENSMUSG00000079190 | 0 | 0 | 0 | 0 | 0 | 0 | 0 |
| ENSMUSG00000079192 | 0 | 0 | 0 | 0 | 0 | 0 | 0 |
| ENSMUSG00000079197 | 2.253091469 | 2.038704527 | 2.004202072 | 2.555698726 | 2.512061267 | 2.875061263 | 2.338495365 |
| ENSMUSG00000079215 | 2.993049299 | 2.745748394 | 2.724272282 | 2.637446167 | 2.572189156 | 2.967079734 | 2.407192337 |
| ENSMUSG00000079222 | 0 | 0 | 0.25 | 0 | 0 | 0 | 0.224243824 |
| ENSMUSG00000079223 | 0.671187741 | 0.646240625 | 0.396240625 | 0.580482024 | 0.244650542 | 0.602059991 | 0.224243824 |
| ENSMUSG00000079224 | 0 | 0 | 0 | 0 | 0 | 0 | 0 |
| ENSMUSG00000079225 | 0.289064826 | 0.25 | 0.396240625 | 0.25 | 0.244650542 | 0 | 0.448487648 |
| ENSMUSG00000079227 | 0.45815691 | 0.396240625 | 0.25 | 0.580482024 | 0 | 0 | 0.579661877 |
| ENSMUSG00000079230 | 0 | 0 | 0 | 0 | 0 | 0 | 0 |
| ENSMUSG00000079235 | 0.671187741 | 0.5 | 0.701838731 | 0.25 | 0 | 0 | 0.224243824 |
| ENSMUSG00000079239 | 0 | 0 | 0 | 0 | 0 | 0 | 0 |
| ENSMUSG00000079242 | 0.747221736 | 0.646240625 | 0.79248125 | 0.976722649 | 0.686820904 | 0.77815125 | 0.803905701 |
| ENSMUSG00000079243 | 0.289064826 | 0 | 0 | 0 | 0 | 0 | 0 |
| ENSMUSG00000079244 | 0 | 0 | 0 | 0 | 0 | 0 | 0 |
| ENSMUSG00000079247 | 0 | 0 | 0 | 0 | 0 | 0 | 0 |
| ENSMUSG00000079258 | 0 | 0 | 0 | 0 | 0.244650542 | 0 | 0 |

| | | | | | | | |
|---|---|---|---|---|---|---|---|
| ENSMUSG00000079259 | 1.693336259 | 1.664552871 | 1.649978211 | 1.42510993 | 1.445123986 | 1.414973348 | 1.20139795 |
| ENSMUSG00000079260 | 1.894701088 | 1.79248125 | 1.843759858 | 1.87892496 | 1.900490506 | 2.164352856 | 1.826389311 |
| ENSMUSG00000079261 | 0.45815691 | 0 | 0 | 0.396240625 | 0 | 0 | 0 |
| ENSMUSG00000079262 | 0.671187741 | 0.646240625 | 0.580482024 | 0.830482024 | 0.733951626 | 0.602059991 | 0.579661877 |
| ENSMUSG00000079263 | 0 | 0.25 | 0 | 0 | 0 | 0 | 0 |
| ENSMUSG00000079264 | 0 | 0 | 0 | 0 | 0 | 0 | 0 |
| ENSMUSG00000079265 | 0 | 0 | 0 | 0 | 0 | 0 | 0 |
| ENSMUSG00000079266 | 0 | 0 | 0 | 0 | 0 | 0 | 0 |
| ENSMUSG00000079267 | 0 | 0 | 0 | 0 | 0 | 0 | 0 |
| ENSMUSG00000079269 | 0 | 0 | 0 | 0 | 0 | 0 | 0 |
| ENSMUSG00000079271 | 0 | 0 | 0 | 0 | 0 | 0 | 0 |
| ENSMUSG00000079272 | 0 | 0 | 0 | 0 | 0 | 0 | 0 |
| ENSMUSG00000079273 | 0 | 0 | 0 | 0 | 0 | 0 | 0 |
| ENSMUSG00000079276 | 0 | 0 | 0 | 0 | 0 | 0 | 0 |
| ENSMUSG00000079277 | 0.578129653 | 0.5 | 0.5 | 0 | 0 | 0 | 0.224243824 |
| ENSMUSG00000079278 | 0 | 0 | 0 | 0 | 0 | 0 | 0 |
| ENSMUSG00000079283 | 1.100572389 | 0.864857905 | 1.29248125 | 0.951838731 | 1.039257769 | 1.041392685 | 1.121219121 |
| ENSMUSG00000079286 | 0 | 0 | 0 | 0.25 | 0 | 0 | 0 |
| ENSMUSG00000079293 | 0 | 0 | 0 | 0 | 0 | 0 | 0 |
| ENSMUSG00000079295 | 0 | 0 | 0 | 0 | 0 | 0 | 0 |
| ENSMUSG00000079297 | 0.289064826 | 0 | 0 | 0 | 0 | 0.301029996 | 0 |
| ENSMUSG00000079298 | 0 | 0 | 0 | 0 | 0 | 0 | 0 |
| ENSMUSG00000079299 | 0 | 0 | 0 | 0 | 0 | 0 | 0 |
| ENSMUSG00000079304 | 0.45815691 | 0.25 | 0.646240625 | 0.5 | 0.387761935 | 0.301029996 | 0.355418052 |
| ENSMUSG00000079311 | 0.811507563 | 0.580482024 | 0.864857905 | 0.830482024 | 0.489301084 | 0.77815125 | 0.710836105 |
| ENSMUSG00000079316 | 2.111266113 | 1.899978211 | 1.872963274 | 2.039967834 | 1.961316494 | 2.257678575 | 1.872150729 |
| ENSMUSG00000079317 | 1.753493018 | 1.67510993 | 1.685366747 | 1.905512955 | 1.905314583 | 2.049218023 | 1.725216937 |
| ENSMUSG00000079323 | 0 | 0.25 | 0 | 0 | 0 | 0 | 0 |
| ENSMUSG00000079324 | 0 | 0 | 0.25 | 0 | 0 | 0.301029996 | 0.224243824 |
| ENSMUSG00000079326 | 0 | 0 | 0 | 0 | 0 | 0 | 0 |
| ENSMUSG00000079330 | 0.45815691 | 0.646240625 | 0.396240625 | 0.580482024 | 0.568060967 | 0.477121255 | 0.710836105 |
| ENSMUSG00000079333 | 0 | 0.25 | 0 | 0 | 0 | 0 | 0 |
| ENSMUSG00000079334 | 1.505869732 | 1.51109853 | 1.642463902 | 1.364857905 | 1.595992969 | 1.568201724 | 1.63986884 |
| ENSMUSG00000079339 | 0.747221736 | 0.5 | 0.5 | 0.701838731 | 0 | 0.602059991 | 0.224243824 |

| | | | | | | | |
|---|---|---|---|---|---|---|---|
| ENSMUSG00000079342 | 0 | 0 | 0 | 0 | 0 | 0 | 0 |
| ENSMUSG00000079343 | 0 | 0.580482024 | 0.5 | 0.25 | 0 | 0 | 0.224243824 |
| ENSMUSG00000079344 | 0 | 0 | 0 | 0 | 0 | 0 | 0.224243824 |
| ENSMUSG00000079346 | 0 | 0 | 0 | 0 | 0 | 0 | 0 |
| ENSMUSG00000079349 | 0 | 0 | 0 | 0 | 0 | 0 | 0 |
| ENSMUSG00000079350 | 0 | 0 | 0 | 0 | 0 | 0 | 0 |
| ENSMUSG00000079353 | 0 | 0 | 0 | 0 | 0 | 0 | 0 |
| ENSMUSG00000079355 | 0 | 0.25 | 0.25 | 0.5 | 0.489301084 | 0.301029996 | 0.224243824 |
| ENSMUSG00000079356 | 0 | 0 | 0 | 0 | 0 | 0 | 0.224243824 |
| ENSMUSG00000079357 | 0 | 0 | 0 | 0.396240625 | 0.387761935 | 0 | 0.355418052 |
| ENSMUSG00000079359 | 0 | 0 | 0.25 | 0 | 0 | 0 | 0 |
| ENSMUSG00000079362 | 0 | 0 | 0 | 0 | 0 | 0 | 0 |
| ENSMUSG00000079363 | 0.289064826 | 0.396240625 | 0.396240625 | 0.830482024 | 0.733951626 | 0.954242509 | 0.876096088 |
| ENSMUSG00000079364 | 0 | 0 | 0 | 0 | 0 | 0 | 0 |
| ENSMUSG00000079371 | 0 | 0 | 0 | 0 | 0 | 0 | 0 |
| ENSMUSG00000079374 | 0 | 0 | 0 | 0 | 0 | 0 | 0 |
| ENSMUSG00000079378 | 0 | 0 | 0 | 0 | 0 | 0 | 0 |
| ENSMUSG00000079379 | 0 | 0 | 0 | 0 | 0 | 0 | 0 |
| ENSMUSG00000079380 | 0 | 0 | 0 | 0 | 0 | 0 | 0 |
| ENSMUSG00000079382 | 0 | 0 | 0 | 0 | 0.244650542 | 0 | 0 |
| ENSMUSG00000079383 | 0 | 0 | 0 | 0 | 0 | 0 | 0 |
| ENSMUSG00000079386 | 0 | 0 | 0 | 0 | 0 | 0 | 0 |
| ENSMUSG00000079387 | 0 | 0 | 0 | 0 | 0 | 0 | 0 |
| ENSMUSG00000079388 | 0 | 0 | 0 | 0 | 0 | 0 | 0 |
| ENSMUSG00000079389 | 0 | 0 | 0 | 0 | 0 | 0 | 0 |
| ENSMUSG00000079391 | 0 | 0 | 0 | 0 | 0 | 0 | 0.224243824 |
| ENSMUSG00000079392 | 0 | 0 | 0 | 0 | 0 | 0 | 0 |
| ENSMUSG00000079395 | 0 | 0 | 0 | 0 | 0 | 0 | 0 |
| ENSMUSG00000079396 | 0.289064826 | 0.25 | 0.396240625 | 0.396240625 | 0 | 0.301029996 | 0 |
| ENSMUSG00000079402 | 0 | 0 | 0 | 0 | 0 | 0 | 0 |
| ENSMUSG00000079407 | 0.671187741 | 0.580482024 | 0.5 | 0.830482024 | 0.931471446 | 0.84509804 | 0.829800754 |
| ENSMUSG00000079409 | 0 | 0 | 0 | 0 | 0 | 0 | 0 |
| ENSMUSG00000079410 | 0.289064826 | 0 | 0 | 0 | 0 | 0 | 0.224243824 |
| ENSMUSG00000079414 | 1.307602651 | 1.321350555 | 1.311981878 | 1.130890489 | 1 | 1.278753601 | 1.159323753 |

| | | | | | | | |
|---|---|---|---|---|---|---|---|
| ENSMUSG00000079415 | 0.811507563 | 0.951838731 | 0.75 | 0.951838731 | 0.877063019 | 0.903089987 | 0.916588299 |
| ENSMUSG00000079416 | 0 | 0 | 0 | 0 | 0 | 0 | 0 |
| ENSMUSG00000079418 | 1.663535556 | 1.630890489 | 1.610735874 | 1.92510993 | 1.826953746 | 2.089905111 | 1.720494 |
| ENSMUSG00000079419 | 0 | 0 | 0 | 0 | 0 | 0 | 0 |
| ENSMUSG00000079421 | 0 | 0.25 | 0 | 0 | 0 | 0 | 0 |
| ENSMUSG00000079423 | 0 | 0 | 0 | 0 | 0 | 0 | 0 |
| ENSMUSG00000079424 | 0 | 0 | 0 | 0 | 0 | 0 | 0 |
| ENSMUSG00000079426 | 2.717463581 | 2.477473271 | 2.512121718 | 2.388647213 | 2.376174996 | 2.656098202 | 2.250241695 |
| ENSMUSG00000079427 | 0.960252568 | 0.864857905 | 1.114857905 | 0.92510993 | 1.163285805 | 1.204119983 | 0.952571511 |
| ENSMUSG00000079428 | 0 | 0 | 0 | 0 | 0 | 0 | 0.224243824 |
| ENSMUSG00000079429 | 1.639698646 | 1.571350555 | 1.589388001 | 1.711372513 | 1.65906333 | 1.62324929 | 1.586932193 |
| ENSMUSG00000079432 | 0 | 0 | 0 | 0 | 0 | 0 | 0 |
| ENSMUSG00000079433 | 0 | 0 | 0 | 0 | 0 | 0 | 0 |
| ENSMUSG00000079434 | 0 | 0 | 0 | 0 | 0.244650542 | 0 | 0 |
| ENSMUSG00000079435 | 2.609650079 | 2.390560606 | 2.397178659 | 2.632601584 | 2.491431917 | 2.924795996 | 2.338260508 |
| ENSMUSG00000079436 | 0 | 0 | 0 | 0 | 0 | 0 | 0.224243824 |
| ENSMUSG00000079438 | 0 | 0 | 0 | 0 | 0 | 0 | 0 |
| ENSMUSG00000079439 | 0 | 0 | 0 | 0 | 0 | 0 | 0 |
| ENSMUSG00000079440 | 0.867194479 | 0.976722649 | 1 | 0 | 0 | 0 | 0 |
| ENSMUSG00000079442 | 1.956444026 | 1.938721876 | 1.888647213 | 1.953445298 | 1.982473755 | 2.178976947 | 1.965468324 |
| ENSMUSG00000079444 | 0 | 0 | 0 | 0 | 0 | 0 | 0 |
| ENSMUSG00000079445 | 0 | 0 | 0 | 0.25 | 0.632412477 | 0.301029996 | 0.520678036 |
| ENSMUSG00000079450 | 0.671187741 | 0 | 0.396240625 | 0.701838731 | 0.568060967 | 0.477121255 | 0.710836105 |
| ENSMUSG00000079451 | 0 | 0.396240625 | 0.25 | 0 | 0.489301084 | 0.301029996 | 0.224243824 |
| ENSMUSG00000079455 | 0 | 0 | 0 | 0 | 0 | 0 | 0 |
| ENSMUSG00000079457 | 0.747221736 | 0.580482024 | 0.75 | 0.79248125 | 0.877063019 | 0.77815125 | 0.448487648 |
| ENSMUSG00000079460 | 0 | 0 | 0 | 0 | 0 | 0 | 0 |
| ENSMUSG00000079462 | 0 | 0 | 0 | 0 | 0 | 0 | 0 |
| ENSMUSG00000079465 | 0.45815691 | 0.396240625 | 0.5 | 0.25 | 0.387761935 | 0 | 0.355418052 |
| ENSMUSG00000079466 | 0 | 0 | 0 | 0 | 0 | 0 | 0 |
| ENSMUSG00000079467 | 0.45815691 | 0.396240625 | 0.396240625 | 0 | 0.387761935 | 0.301029996 | 0.629532004 |
| ENSMUSG00000079469 | 1.721148722 | 1.714495249 | 1.638647213 | 1.779735268 | 1.78010778 | 1.954242509 | 1.699191618 |
| ENSMUSG00000079470 | 1.952600383 | 1.80479213 | 1.823655187 | 1.880890489 | 1.78687392 | 2.06069784 | 1.663407615 |
| ENSMUSG00000079471 | 0.289064826 | 0 | 0.25 | 0 | 0 | 0 | 0 |

| | | | | | | | |
|---|---|---|---|---|---|---|---|
| ENSMUSG00000079476 | 0 | 0 | 0 | 0 | 0 | 0 | 0 |
| ENSMUSG00000079477 | 3.082459571 | 2.833149014 | 2.842126615 | 2.788546302 | 2.719460551 | 3.149527014 | 2.581106889 |
| ENSMUSG00000079478 | 1.587501561 | 1.638647213 | 1.660964047 | 1.470660762 | 1.572189156 | 1.51851394 | 1.769001954 |
| ENSMUSG00000079479 | 0 | 0 | 0 | 0 | 0 | 0 | 0 |
| ENSMUSG00000079480 | 1.54867984 | 1.610735874 | 1.602347734 | 1.692046081 | 1.559656853 | 1.819543936 | 1.473249546 |
| ENSMUSG00000079481 | 1.655740331 | 1.53743678 | 1.470660762 | 1.348079356 | 1.200473444 | 1.361727836 | 1.23151414 |
| ENSMUSG00000079484 | 0.45815691 | 0.646240625 | 0.75 | 0.396240625 | 0.387761935 | 0.477121255 | 0.224243824 |
| ENSMUSG00000079487 | 2.718080037 | 2.473325383 | 2.499647607 | 2.490362424 | 2.487464361 | 2.770852012 | 2.326782038 |
| ENSMUSG00000079489 | 0.45815691 | 0.701838731 | 0.580482024 | 1.17510993 | 0.955822902 | 0.477121255 | 0.803905701 |
| ENSMUSG00000079491 | 1.307602651 | 1.146240625 | 1.188721876 | 1.488549078 | 1.588235379 | 1.62324929 | 1.405288179 |
| ENSMUSG00000079492 | 0.45815691 | 0.25 | 0.5 | 0.864857905 | 0.77552387 | 0.602059991 | 0.672731473 |
| ENSMUSG00000079494 | 0 | 0 | 0 | 0 | 0 | 0 | 0 |
| ENSMUSG00000079495 | 0.289064826 | 0 | 0.396240625 | 0 | 0 | 0 | 0 |
| ENSMUSG00000079497 | 0 | 0 | 0 | 0 | 0 | 0 | 0 |
| ENSMUSG00000079499 | 0.747221736 | 1.114857905 | 1 | 1.410964047 | 1.414412788 | 1.505149978 | 1.379042812 |
| ENSMUSG00000079501 | 0 | 0 | 0 | 0 | 0 | 0 | 0 |
| ENSMUSG00000079502 | 0 | 0.580482024 | 0 | 0.25 | 0.568060967 | 0.301029996 | 0.355418052 |
| ENSMUSG00000079505 | 0.671187741 | 0.25 | 0.701838731 | 1.02186571 | 0.978602168 | 1.146128036 | 1.014380832 |
| ENSMUSG00000079507 | 0 | 0.396240625 | 0.5 | 0.5 | 0.77552387 | 0.84509804 | 0.74492186 |
| ENSMUSG00000079508 | 1.631440309 | 1.618933358 | 1.566696635 | 1.747171172 | 1.662257525 | 1.755874856 | 1.535058868 |
| ENSMUSG00000079509 | 2.65645504 | 2.473704441 | 2.423807073 | 2.576230417 | 2.501307552 | 2.830588669 | 2.268527657 |
| ENSMUSG00000079511 | 0.671187741 | 0.830482024 | 0.79248125 | 0.75 | 0.846351821 | 0.903089987 | 0.775756176 |
| ENSMUSG00000079513 | 0 | 0 | 0 | 0 | 0 | 0 | 0 |
| ENSMUSG00000079516 | 0 | 0 | 0 | 0 | 0 | 0 | 0 |
| ENSMUSG00000079519 | 0 | 0 | 0 | 0 | 0 | 0 | 0 |
| ENSMUSG00000079521 | 0 | 0 | 0 | 0 | 0 | 0 | 0 |
| ENSMUSG00000079522 | 0 | 0 | 0 | 0 | 0 | 0 | 0 |
| ENSMUSG00000079523 | 2.328498818 | 2.183677405 | 2.221424093 | 2.289336734 | 2.115605392 | 2.527629901 | 2.064068234 |
| ENSMUSG00000079524 | 0 | 0 | 0 | 0 | 0 | 0 | 0 |
| ENSMUSG00000079525 | 0 | 0 | 0 | 0 | 0 | 0 | 0 |
| ENSMUSG00000079528 | 0 | 0 | 0 | 0 | 0 | 0 | 0 |
| ENSMUSG00000079531 | 0 | 0 | 0 | 0 | 0 | 0 | 0 |
| ENSMUSG00000079532 | 0 | 0 | 0 | 0 | 0 | 0 | 0 |
| ENSMUSG00000079534 | 0 | 0 | 0 | 0 | 0 | 0 | 0 |

| | | | | | | | |
|---|---|---|---|---|---|---|---|
| ENSMUSG00000079536 | 0 | 0 | 0 | 0 | 0 | 0 | 0 |
| ENSMUSG00000079537 | 0 | 0 | 0 | 0 | 0 | 0 | 0 |
| ENSMUSG00000079539 | 0 | 0 | 0 | 0 | 0 | 0 | 0 |
| ENSMUSG00000079542 | 0 | 0 | 0 | 0 | 0 | 0 | 0 |
| ENSMUSG00000079543 | 0 | 0 | 0 | 0 | 0 | 0 | 0 |
| ENSMUSG00000079544 | 0 | 0 | 0 | 0 | 0 | 0 | 0 |
| ENSMUSG00000079546 | 0 | 0 | 0 | 0 | 0 | 0 | 0 |
| ENSMUSG00000079547 | 0.45815691 | 0.25 | 0 | 0.25 | 0 | 0.477121255 | 0.448487648 |
| ENSMUSG00000079550 | 0 | 0 | 0.25 | 0.5 | 0 | 0 | 0 |
| ENSMUSG00000079553 | 2.20747957 | 2.051142786 | 2.077084758 | 1.83272922 | 2.010824034 | 2.100370545 | 1.988432495 |
| ENSMUSG00000079554 | 0 | 0.25 | 0 | 0 | 0 | 0.477121255 | 0 |
| ENSMUSG00000079555 | 2.530730035 | 2.370953944 | 2.333847589 | 2.423807073 | 2.328502976 | 2.679427897 | 2.139097452 |
| ENSMUSG00000079557 | 2.149089517 | 2.024508021 | 2.029735268 | 2.016522298 | 1.986303335 | 2.225309282 | 1.952571511 |
| ENSMUSG00000079559 | 0.578129653 | 0.396240625 | 0.701838731 | 0 | 0.244650542 | 0.77815125 | 0.579661877 |
| ENSMUSG00000079560 | 1.307602651 | 1.282320754 | 1.356566189 | 0.830482024 | 1.020174412 | 1.079181246 | 1.028149525 |
| ENSMUSG00000079562 | 2.732608034 | 2.581045137 | 2.586351312 | 2.348079356 | 2.480333692 | 2.673020907 | 2.39794841 |
| ENSMUSG00000079563 | 0 | 0 | 0 | 0.396240625 | 0.244650542 | 0 | 0.448487648 |
| ENSMUSG00000079564 | 0.289064826 | 0 | 0 | 0 | 0.244650542 | 0 | 0.224243824 |
| ENSMUSG00000079566 | 0 | 0 | 0 | 0 | 0 | 0 | 0 |
| ENSMUSG00000079575 | 1.470606563 | 1.27186571 | 1.26109853 | 1.27186571 | 1.254881871 | 1.230448921 | 1.150210039 |
| ENSMUSG00000079577 | 0 | 0 | 0 | 0 | 0 | 0 | 0 |
| ENSMUSG00000079578 | 0 | 0 | 0 | 0 | 0 | 0 | 0 |
| ENSMUSG00000079579 | 0 | 0 | 0 | 0 | 0 | 0 | 0 |
| ENSMUSG00000079580 | 0 | 0 | 0 | 0.580482024 | 0.387761935 | 0 | 0.448487648 |
| ENSMUSG00000079583 | 0 | 0 | 0 | 0 | 0 | 0 | 0 |
| ENSMUSG00000079584 | 0 | 0 | 0 | 0 | 0 | 0 | 0 |
| ENSMUSG00000079588 | 0 | 0 | 0 | 0 | 0 | 0 | 0 |
| ENSMUSG00000079592 | 0 | 0 | 0 | 0 | 0 | 0 | 0 |
| ENSMUSG00000079593 | 0 | 0 | 0 | 0 | 0 | 0 | 0.355418052 |
| ENSMUSG00000079594 | 0 | 0 | 0 | 0 | 0 | 0 | 0 |
| ENSMUSG00000079595 | 0 | 0 | 0 | 0 | 0 | 0 | 0 |
| ENSMUSG00000079597 | 0 | 0 | 0 | 0 | 0 | 0 | 0 |
| ENSMUSG00000079598 | 0.289064826 | 0.701838731 | 0.701838731 | 0.864857905 | 1.074582839 | 0.84509804 | 1.168187745 |
| ENSMUSG00000079600 | 0.578129653 | 0.396240625 | 0.396240625 | 0.396240625 | 0.387761935 | 0 | 0.520678036 |

| | | | | | | | |
|---|---|---|---|---|---|---|---|
| ENSMUSG00000079602 | 0 | 0 | 0 | 0.25 | 0 | 0 | 0 |
| ENSMUSG00000079603 | 0 | 0 | 0 | 0 | 0 | 0 | 0 |
| ENSMUSG00000079604 | 0 | 0 | 0 | 0 | 0 | 0 | 0 |
| ENSMUSG00000079605 | 0 | 0 | 0 | 0 | 0 | 0 | 0 |
| ENSMUSG00000079606 | 0 | 0 | 0 | 0 | 0 | 0 | 0 |
| ENSMUSG00000079608 | 1.181541736 | 1.188721876 | 1.188721876 | 1.114857905 | 1.057362051 | 0.903089987 | 1.121219121 |
| ENSMUSG00000079610 | 1.418409478 | 1.188721876 | 1.418106335 | 1.42510993 | 1.414412788 | 1.301029996 | 1.340368108 |
| ENSMUSG00000079614 | 2.813962146 | 2.579070383 | 2.558105232 | 2.709406983 | 2.637063356 | 3.054995862 | 2.450154398 |
| ENSMUSG00000079619 | 0 | 0 | 0 | 0 | 0 | 0 | 0 |
| ENSMUSG00000079620 | 0 | 0 | 0 | 0 | 0.244650542 | 0 | 0 |
| ENSMUSG00000079625 | 0 | 0 | 0.5 | 0 | 0 | 0 | 0.224243824 |
| ENSMUSG00000079626 | 0 | 0 | 0 | 0 | 0 | 0 | 0 |
| ENSMUSG00000079627 | 0 | 0 | 0 | 0 | 0 | 0 | 0 |
| ENSMUSG00000079628 | 0.289064826 | 0 | 0 | 0 | 0 | 0 | 0 |
| ENSMUSG00000079629 | 0 | 0 | 0 | 0 | 0 | 0 | 0 |
| ENSMUSG00000079630 | 0 | 0 | 0 | 0 | 0 | 0 | 0 |
| ENSMUSG00000079633 | 0 | 0 | 0 | 0 | 0 | 0 | 0 |
| ENSMUSG00000079635 | 0 | 0 | 0 | 0 | 0 | 0 | 0 |
| ENSMUSG00000079636 | 0 | 0 | 0 | 0 | 0 | 0 | 0 |
| ENSMUSG00000079637 | 0 | 0 | 0 | 0 | 0 | 0 | 0 |
| ENSMUSG00000079638 | 0 | 0 | 0 | 0 | 0 | 0 | 0 |
| ENSMUSG00000079639 | 0.289064826 | 0 | 0 | 0 | 0 | 0.301029996 | 0 |
| ENSMUSG00000079641 | 2.710621733 | 2.453845824 | 2.458617762 | 2.54216803 | 2.393014957 | 2.780317312 | 2.218851775 |
| ENSMUSG00000079642 | 0 | 0 | 0 | 0 | 0 | 0 | 0 |
| ENSMUSG00000079644 | 0 | 0 | 0 | 0 | 0.244650542 | 0.477121255 | 0 |
| ENSMUSG00000079645 | 0.867194479 | 0.75 | 0.79248125 | 0.701838731 | 0.877063019 | 0 | 0.224243824 |
| ENSMUSG00000079649 | 0 | 0 | 0 | 0 | 0 | 0 | 0 |
| ENSMUSG00000079652 | 0.289064826 | 0.25 | 0.25 | 0.25 | 0 | 0.301029996 | 0.448487648 |
| ENSMUSG00000079654 | 0 | 0.25 | 0.5 | 0.396240625 | 0.568060967 | 0.301029996 | 0.803905701 |
| ENSMUSG00000079655 | 0 | 0 | 0 | 0 | 0 | 0 | 0 |
| ENSMUSG00000079657 | 0 | 0 | 0 | 0.25 | 0.387761935 | 0 | 0.355418052 |
| ENSMUSG00000079658 | 2.729622822 | 2.545598588 | 2.535207443 | 2.614857905 | 2.505123331 | 2.918554531 | 2.323355972 |
| ENSMUSG00000079659 | 1.505869732 | 1.348079356 | 1.438721876 | 1.372963274 | 1.427019704 | 1.431363764 | 1.401059159 |
| ENSMUSG00000079666 | 0 | 0.25 | 0 | 0.25 | 0.387761935 | 0 | 0 |

| | | | | | | | |
|---|---|---|---|---|---|---|---|
| ENSMUSG00000079669 | 0.45815691 | 0.701838731 | 0.75 | 0.580482024 | 0.568060967 | 0.84509804 | 0.710836105 |
| ENSMUSG00000079671 | 0 | 0 | 0 | 0 | 0.244650542 | 0 | 0 |
| ENSMUSG00000079677 | 1.727821383 | 1.598079356 | 1.708222504 | 1.67510993 | 1.731346772 | 1.934498451 | 1.67819947 |
| ENSMUSG00000079679 | 0.289064826 | 0 | 0 | 0 | 0.244650542 | 0 | 0 |
| ENSMUSG00000079681 | 0.45815691 | 0.864857905 | 0.79248125 | 0.580482024 | 0.733951626 | 0.602059991 | 0.916588299 |
| ENSMUSG00000079685 | 0.747221736 | 0.5 | 0.5 | 2.201032755 | 2.102149272 | 2.376576957 | 1.969165684 |
| ENSMUSG00000079693 | 0 | 0 | 0 | 0 | 0 | 0 | 0 |
| ENSMUSG00000079694 | 0.289064826 | 0.396240625 | 0.25 | 0.25 | 0.387761935 | 0.301029996 | 0 |
| ENSMUSG00000079697 | 0 | 0 | 0 | 0 | 0 | 0 | 0.355418052 |
| ENSMUSG00000079698 | 0 | 0 | 0 | 0 | 0 | 0 | 0 |
| ENSMUSG00000079699 | 0 | 0 | 0 | 0 | 0 | 0 | 0 |
| ENSMUSG00000079700 | 0 | 0 | 0 | 0 | 0 | 0 | 0 |
| ENSMUSG00000079701 | 0 | 0 | 0 | 0 | 0 | 0 | 0 |
| ENSMUSG00000079702 | 0 | 0 | 0 | 0 | 0 | 0 | 0 |
| ENSMUSG00000079703 | 0 | 0 | 0 | 0 | 0 | 0 | 0 |
| ENSMUSG00000079704 | 0 | 0 | 0 | 0 | 0 | 0 | 0 |
| ENSMUSG00000079705 | 0 | 0 | 0.25 | 0 | 0 | 0 | 0 |
| ENSMUSG00000079707 | 0 | 0.25 | 0 | 0.25 | 0 | 0.301029996 | 0 |
| ENSMUSG00000079710 | 0 | 0 | 0 | 0 | 0 | 0 | 0 |
| ENSMUSG00000079711 | 0 | 0 | 0 | 0 | 0 | 0 | 0.224243824 |
| ENSMUSG00000079722 | 0.45815691 | 0.25 | 0 | 0 | 0 | 0.477121255 | 0.224243824 |
| ENSMUSG00000079737 | 1.269664473 | 1.17510993 | 1.356566189 | 1.27186571 | 1.234113756 | 1.204119983 | 1.131174228 |
| ENSMUSG00000079740 | 0 | 0 | 0 | 0 | 0 | 0 | 0 |
| ENSMUSG00000079741 | 0 | 0 | 0 | 0 | 0 | 0 | 0 |
| ENSMUSG00000079764 | 0 | 0 | 0 | 0 | 0 | 0 | 0 |
| ENSMUSG00000079773 | 0 | 0 | 0 | 0 | 0 | 0 | 0 |
| ENSMUSG00000079774 | 0 | 0 | 0 | 0 | 0 | 0 | 0 |
| ENSMUSG00000079794 | 0 | 0 | 0 | 0 | 0 | 0 | 0 |
| ENSMUSG00000079800 | 0 | 0 | 0 | 0 | 0 | 0 | 0 |
| ENSMUSG00000079806 | 0 | 0 | 0 | 0 | 0 | 0 | 0 |
| ENSMUSG00000079808 | 0 | 0.25 | 0 | 0.25 | 0 | 0 | 0 |
| ENSMUSG00000079834 | 2.203267085 | 2.001406137 | 2.017865591 | 1.907339155 | 1.828942237 | 2.053078443 | 1.687112305 |
| ENSMUSG00000079842 | 0 | 0 | 0 | 0 | 0 | 0 | 0 |
| ENSMUSG00000079845 | 0 | 0 | 0 | 0 | 0 | 0 | 0 |

| | | | | | | | |
|---|---|---|---|---|---|---|---|
| ENSMUSG00000079852 | 1.568542279 | 1.52186571 | 1.488549078 | 0 | 0 | 0 | 0 |
| ENSMUSG00000079853 | 0 | 0 | 0 | 0 | 0 | 0 | 0 |
| ENSMUSG00000079871 | 0 | 0 | 0 | 0.25 | 0 | 0 | 0.224243824 |
| ENSMUSG00000079875 | 0 | 0 | 0 | 0 | 0 | 0 | 0 |
| ENSMUSG00000079884 | 0 | 0 | 0 | 0 | 0 | 0 | 0 |
| ENSMUSG00000079889 | 0.289064826 | 0.396240625 | 0 | 0.396240625 | 0 | 0 | 0.224243824 |
| ENSMUSG00000079890 | 0 | 0 | 0.25 | 0.25 | 0 | 0 | 0 |
| ENSMUSG00000079892 | 0 | 0 | 0 | 0 | 0 | 0 | 0 |
| ENSMUSG00000079893 | 0 | 0 | 0 | 0 | 0 | 0 | 0 |
| ENSMUSG00000079906 | 0 | 0.396240625 | 0.25 | 0.25 | 0.387761935 | 0.477121255 | 0.355418052 |
| ENSMUSG00000079915 | 0.45815691 | 0.396240625 | 0 | 0.25 | 0.489301084 | 0.477121255 | 0 |
| ENSMUSG00000079941 | 0 | 0 | 0 | 0 | 0 | 0 | 0 |
| ENSMUSG00000079942 | 0 | 0 | 0 | 0.25 | 0 | 0 | 0.224243824 |
| ENSMUSG00000079962 | 0 | 0.25 | 0 | 0.25 | 0.387761935 | 0 | 0 |
| ENSMUSG00000079963 | 0.289064826 | 0.25 | 0.5 | 0.25 | 0.568060967 | 0.301029996 | 0 |
| ENSMUSG00000079965 | 0 | 0.396240625 | 0 | 0 | 0.244650542 | 0 | 0.224243824 |
| ENSMUSG00000079966 | 0 | 0 | 0 | 0 | 0 | 0 | 0 |
| ENSMUSG00000079988 | 0.289064826 | 0.646240625 | 0.5 | 0.75 | 0.632412477 | 0.903089987 | 0.629532004 |
| ENSMUSG00000079993 | 0 | 0 | 0 | 0 | 0 | 0 | 0 |
| ENSMUSG00000079998 | 0 | 0 | 0 | 0 | 0.244650542 | 0 | 0 |
| ENSMUSG00000080002 | 0 | 0.396240625 | 0.396240625 | 0 | 0 | 0.301029996 | 0.355418052 |
| ENSMUSG00000080006 | 0 | 0 | 0 | 0 | 0 | 0 | 0 |
| ENSMUSG00000080021 | 0 | 0.396240625 | 0.25 | 0.25 | 0.387761935 | 0.301029996 | 0.224243824 |
| ENSMUSG00000080046 | 0 | 0 | 0 | 0 | 0 | 0 | 0 |
| ENSMUSG00000080050 | 0 | 0 | 0 | 0 | 0 | 0 | 0 |
| ENSMUSG00000080051 | 0 | 0 | 0 | 0 | 0 | 0 | 0 |
| ENSMUSG00000080054 | 0 | 0 | 0.25 | 0 | 0 | 0 | 0 |
| ENSMUSG00000080057 | 0 | 0 | 0 | 0.25 | 0 | 0 | 0 |
| ENSMUSG00000080058 | 0 | 0 | 0 | 0 | 0 | 0 | 0 |
| ENSMUSG00000080059 | 0.578129653 | 0.701838731 | 0.701838731 | 0.864857905 | 0.846351821 | 1.041392685 | 0.853775828 |
| ENSMUSG00000080065 | 0 | 0 | 0 | 0 | 0 | 0 | 0 |
| ENSMUSG00000080069 | 0 | 0 | 0 | 0 | 0 | 0 | 0 |
| ENSMUSG00000080071 | 0 | 0 | 0 | 0 | 0 | 0 | 0 |
| ENSMUSG00000080072 | 0 | 0 | 0 | 0 | 0 | 0 | 0 |

| | | | | | | | |
|---|---|---|---|---|---|---|---|
| ENSMUSG00000080073 | 0 | 0 | 0 | 0 | 0 | 0 | 0 |
| ENSMUSG00000080076 | 0 | 0 | 0 | 0 | 0 | 0 | 0 |
| ENSMUSG00000080078 | 0 | 0 | 0 | 0.25 | 0.244650542 | 0 | 0 |
| ENSMUSG00000080115 | 0.671187741 | 0.75 | 0.75 | 0.830482024 | 0.632412477 | 0.301029996 | 0.829800754 |
| ENSMUSG00000080142 | 0.289064826 | 0.25 | 0 | 0.25 | 0 | 0 | 0.355418052 |
| ENSMUSG00000080152 | 0.289064826 | 0 | 0 | 0 | 0 | 0 | 0 |
| ENSMUSG00000080155 | 0 | 0 | 0 | 0 | 0 | 0 | 0 |
| ENSMUSG00000080158 | 0 | 0.25 | 0 | 0 | 0 | 0 | 0 |
| ENSMUSG00000080175 | 0 | 0 | 0 | 0 | 0 | 0 | 0 |
| ENSMUSG00000080180 | 0 | 0.396240625 | 0.396240625 | 0.580482024 | 0.387761935 | 0.698970004 | 0.448487648 |
| ENSMUSG00000080186 | 0.289064826 | 0 | 0.25 | 0.25 | 0 | 0 | 0 |
| ENSMUSG00000080193 | 0 | 0 | 0 | 0 | 0 | 0 | 0 |
| ENSMUSG00000080198 | 0.578129653 | 0.646240625 | 0.396240625 | 0 | 0.244650542 | 0 | 0.355418052 |
| ENSMUSG00000080205 | 0 | 0 | 0 | 0 | 0.244650542 | 0 | 0.224243824 |
| ENSMUSG00000080222 | 0 | 0 | 0 | 0 | 0 | 0 | 0 |
| ENSMUSG00000080223 | 0 | 0 | 0 | 0 | 0 | 0 | 0 |
| ENSMUSG00000080227 | 0 | 0 | 0 | 0 | 0 | 0 | 0 |
| ENSMUSG00000080237 | 0 | 0.25 | 0 | 0 | 0 | 0 | 0 |
| ENSMUSG00000080241 | 0 | 0 | 0 | 0 | 0 | 0 | 0 |
| ENSMUSG00000080242 | 1.404271433 | 1.593759858 | 1.807204673 | 1.188721876 | 1.456697355 | 1.361727836 | 1.659601507 |
| ENSMUSG00000080250 | 0 | 0 | 0 | 0 | 0 | 0 | 0 |
| ENSMUSG00000080254 | 0 | 0 | 0 | 0 | 0 | 0 | 0.224243824 |
| ENSMUSG00000080266 | 0 | 0 | 0 | 0 | 0 | 0 | 0 |
| ENSMUSG00000080268 | 2.156259305 | 2.051142786 | 2.125956434 | 2.088286706 | 2.164820399 | 2.404833717 | 2.098866035 |
| ENSMUSG00000080283 | 0 | 0 | 0 | 0 | 0 | 0 | 0 |
| ENSMUSG00000080298 | 0 | 0 | 0 | 0 | 0 | 0 | 0 |
| ENSMUSG00000080299 | 0 | 0 | 0 | 0 | 0 | 0 | 0 |
| ENSMUSG00000080300 | 0 | 0 | 0 | 0 | 0 | 0 | 0 |
| ENSMUSG00000080303 | 0.747221736 | 0.830482024 | 0.830482024 | 0.5 | 0.931471446 | 0.77815125 | 0.935079929 |
| ENSMUSG00000080316 | 2.477243823 | 2.234159485 | 2.25 | 2.198603967 | 2.153935427 | 2.344392274 | 1.950241257 |
| ENSMUSG00000080317 | 0 | 0 | 0 | 0 | 0 | 0 | 0 |
| ENSMUSG00000080321 | 0 | 0 | 0 | 0 | 0 | 0 | 0 |
| ENSMUSG00000080330 | 0 | 0 | 0 | 0 | 0 | 0 | 0 |
| ENSMUSG00000080331 | 0 | 0 | 0 | 0 | 0 | 0 | 0 |

| | | | | | | | |
|---|---|---|---|---|---|---|---|
| ENSMUSG00000080334 | 0 | 0 | 0 | 0 | 0 | 0 | 0 |
| ENSMUSG00000080336 | 0 | 0 | 0 | 0 | 0 | 0 | 0 |
| ENSMUSG00000080342 | 0 | 0 | 0 | 0 | 0 | 0 | 0 |
| ENSMUSG00000080345 | 0 | 0 | 0 | 0 | 0 | 0 | 0 |
| ENSMUSG00000080346 | 0 | 0 | 0 | 0 | 0 | 0.301029996 | 0 |
| ENSMUSG00000080347 | 0 | 0 | 0 | 0 | 0 | 0 | 0 |
| ENSMUSG00000080348 | 0 | 0 | 0 | 0 | 0 | 0 | 0 |
| ENSMUSG00000080349 | 0 | 0 | 0 | 0 | 0 | 0 | 0 |
| ENSMUSG00000080350 | 0 | 0 | 0 | 0 | 0 | 0 | 0 |
| ENSMUSG00000080352 | 0 | 0 | 0 | 0 | 0 | 0 | 0 |
| ENSMUSG00000080356 | 0 | 0 | 0 | 0 | 0 | 0 | 0 |
| ENSMUSG00000080358 | 0 | 0 | 0 | 0 | 0 | 0 | 0 |
| ENSMUSG00000080359 | 0 | 0 | 0 | 0 | 0 | 0 | 0 |
| ENSMUSG00000080360 | 0 | 0 | 0 | 0 | 0 | 0 | 0 |
| ENSMUSG00000080363 | 0 | 0 | 0 | 0 | 0 | 0 | 0 |
| ENSMUSG00000080364 | 0 | 0.25 | 0 | 0 | 0 | 0 | 0 |
| ENSMUSG00000080365 | 0 | 0 | 0 | 0.396240625 | 0.244650542 | 0 | 0.224243824 |
| ENSMUSG00000080366 | 0 | 0 | 0 | 0 | 0 | 0 | 0 |
| ENSMUSG00000080367 | 0 | 0 | 0 | 0 | 0 | 0 | 0 |
| ENSMUSG00000080368 | 0 | 0 | 0 | 0 | 0 | 0 | 0 |
| ENSMUSG00000080370 | 0.289064826 | 0.25 | 0.25 | 0 | 0.244650542 | 0 | 0.355418052 |
| ENSMUSG00000080374 | 0 | 0 | 0 | 0 | 0 | 0 | 0 |
| ENSMUSG00000080378 | 0 | 0 | 0 | 0 | 0 | 0 | 0 |
| ENSMUSG00000080380 | 0 | 0 | 0 | 0 | 0 | 0 | 0 |
| ENSMUSG00000080383 | 0 | 0 | 0 | 0 | 0 | 0 | 0 |
| ENSMUSG00000080384 | 0 | 0 | 0 | 0 | 0 | 0 | 0 |
| ENSMUSG00000080391 | 0 | 0 | 0 | 0 | 0 | 0 | 0 |
| ENSMUSG00000080394 | 0 | 0 | 0 | 0 | 0 | 0 | 0 |
| ENSMUSG00000080396 | 0.289064826 | 0 | 0 | 0 | 0 | 0 | 0.224243824 |
| ENSMUSG00000080401 | 0 | 0 | 0 | 0 | 0.244650542 | 0 | 0 |
| ENSMUSG00000080402 | 0 | 0 | 0 | 0 | 0 | 0 | 0 |
| ENSMUSG00000080405 | 0 | 0 | 0 | 0 | 0 | 0 | 0 |
| ENSMUSG00000080406 | 0 | 0 | 0 | 0 | 0 | 0 | 0 |
| ENSMUSG00000080408 | 0 | 0 | 0 | 0 | 0 | 0 | 0 |

| | | | | | | | |
|---|---|---|---|---|---|---|---|
| ENSMUSG00000080409 | 0 | 0 | 0 | 0 | 0 | 0 | 0 |
| ENSMUSG00000080410 | 0 | 0 | 0 | 0 | 0 | 0 | 0 |
| ENSMUSG00000080411 | 0 | 0 | 0 | 0 | 0 | 0 | 0 |
| ENSMUSG00000080412 | 0 | 0 | 0 | 0 | 0 | 0 | 0 |
| ENSMUSG00000080413 | 0 | 0 | 0 | 0 | 0 | 0 | 0 |
| ENSMUSG00000080415 | 0 | 0 | 0 | 0 | 0 | 0 | 0 |
| ENSMUSG00000080419 | 0 | 0 | 0 | 0 | 0 | 0 | 0 |
| ENSMUSG00000080424 | 0 | 0 | 0 | 0 | 0 | 0 | 0 |
| ENSMUSG00000080425 | 0 | 0 | 0 | 0 | 0 | 0 | 0 |
| ENSMUSG00000080427 | 0 | 0 | 0 | 0 | 0 | 0 | 0 |
| ENSMUSG00000080428 | 0 | 0 | 0 | 0 | 0 | 0 | 0 |
| ENSMUSG00000080430 | 0 | 0 | 0 | 0 | 0 | 0 | 0 |
| ENSMUSG00000080432 | 0 | 0 | 0 | 0 | 0 | 0 | 0 |
| ENSMUSG00000080434 | 0 | 0 | 0 | 0 | 0 | 0 | 0 |
| ENSMUSG00000080437 | 0 | 0 | 0 | 0 | 0 | 0 | 0 |
| ENSMUSG00000080440 | 0 | 0 | 0 | 0 | 0 | 0 | 0 |
| ENSMUSG00000080441 | 0 | 0 | 0 | 0 | 0 | 0 | 0 |
| ENSMUSG00000080443 | 0 | 0 | 0 | 0 | 0 | 0 | 0 |
| ENSMUSG00000080451 | 0 | 0 | 0 | 0 | 0 | 0 | 0 |
| ENSMUSG00000080452 | 0 | 0 | 0 | 0 | 0 | 0 | 0 |
| ENSMUSG00000080454 | 0 | 0 | 0.25 | 0.396240625 | 0 | 0.301029996 | 0.224243824 |
| ENSMUSG00000080455 | 0 | 0 | 0 | 0 | 0 | 0 | 0 |
| ENSMUSG00000080459 | 0 | 0 | 0 | 0 | 0 | 0 | 0 |
| ENSMUSG00000080461 | 0.289064826 | 0 | 0 | 0 | 0 | 0 | 0.224243824 |
| ENSMUSG00000080463 | 0 | 0 | 0 | 0 | 0 | 0 | 0 |
| ENSMUSG00000080465 | 0.289064826 | 0.5 | 0.396240625 | 0.25 | 0.244650542 | 0.301029996 | 0.448487648 |
| ENSMUSG00000080468 | 0 | 0 | 0 | 0 | 0 | 0 | 0 |
| ENSMUSG00000080469 | 0.289064826 | 0.396240625 | 0.25 | 0.25 | 0 | 0 | 0.355418052 |
| ENSMUSG00000080473 | 0 | 0 | 0 | 0 | 0 | 0 | 0 |
| ENSMUSG00000080474 | 0 | 0 | 0 | 0 | 0 | 0 | 0 |
| ENSMUSG00000080476 | 0 | 0 | 0 | 0 | 0 | 0 | 0 |
| ENSMUSG00000080478 | 0 | 0 | 0 | 0 | 0 | 0 | 0 |
| ENSMUSG00000080480 | 0 | 0 | 0 | 0 | 0 | 0 | 0 |
| ENSMUSG00000080485 | 0 | 0 | 0 | 0 | 0.244650542 | 0.301029996 | 0 |

| | | | | | | | |
|---|---|---|---|---|---|---|---|
| ENSMUSG00000080486 | 1.181541736 | 0.951838731 | 1.130890489 | 1.02186571 | 0.955822902 | 1.079181246 | 0.952571511 |
| ENSMUSG00000080487 | 0 | 0 | 0 | 0 | 0 | 0 | 0 |
| ENSMUSG00000080495 | 0 | 0 | 0 | 0 | 0 | 0 | 0 |
| ENSMUSG00000080498 | 0 | 0 | 0 | 0 | 0 | 0 | 0 |
| ENSMUSG00000080499 | 0 | 0 | 0 | 0 | 0 | 0 | 0 |
| ENSMUSG00000080500 | 0 | 0 | 0 | 0 | 0 | 0 | 0 |
| ENSMUSG00000080504 | 0 | 0 | 0 | 0 | 0 | 0 | 0 |
| ENSMUSG00000080506 | 0 | 0 | 0 | 0 | 0 | 0 | 0 |
| ENSMUSG00000080507 | 0.289064826 | 0 | 0 | 0 | 0 | 0 | 0 |
| ENSMUSG00000080508 | 0 | 0 | 0 | 0 | 0 | 0 | 0 |
| ENSMUSG00000080511 | 0 | 0 | 0 | 0 | 0 | 0 | 0 |
| ENSMUSG00000080515 | 0 | 0 | 0 | 0 | 0 | 0 | 0 |
| ENSMUSG00000080516 | 0 | 0 | 0 | 0 | 0 | 0 | 0 |
| ENSMUSG00000080517 | 0 | 0 | 0 | 0 | 0 | 0 | 0 |
| ENSMUSG00000080518 | 0 | 0 | 0 | 0 | 0 | 0 | 0 |
| ENSMUSG00000080519 | 0 | 0 | 0 | 0 | 0 | 0 | 0 |
| ENSMUSG00000080522 | 0 | 0 | 0 | 0 | 0 | 0 | 0 |
| ENSMUSG00000080524 | 0 | 0 | 0 | 0 | 0 | 0 | 0 |
| ENSMUSG00000080526 | 0 | 0 | 0 | 0 | 0 | 0 | 0 |
| ENSMUSG00000080527 | 0 | 0 | 0 | 0 | 0 | 0 | 0 |
| ENSMUSG00000080529 | 0 | 0 | 0 | 0 | 0 | 0 | 0 |
| ENSMUSG00000080530 | 0 | 0 | 0 | 0 | 0 | 0 | 0 |
| ENSMUSG00000080533 | 0.289064826 | 0.25 | 0.396240625 | 0.396240625 | 0.387761935 | 0 | 0.224243824 |
| ENSMUSG00000080536 | 0 | 0 | 0 | 0 | 0 | 0 | 0 |
| ENSMUSG00000080538 | 0 | 0 | 0 | 0 | 0 | 0 | 0 |
| ENSMUSG00000080539 | 0 | 0 | 0 | 0 | 0 | 0 | 0 |
| ENSMUSG00000080540 | 0 | 0 | 0 | 0 | 0 | 0 | 0 |
| ENSMUSG00000080542 | 0 | 0 | 0 | 0 | 0 | 0 | 0 |
| ENSMUSG00000080543 | 0 | 0 | 0 | 0 | 0 | 0 | 0 |
| ENSMUSG00000080544 | 0 | 0 | 0 | 0 | 0 | 0 | 0 |
| ENSMUSG00000080545 | 0 | 0 | 0 | 0 | 0 | 0 | 0 |
| ENSMUSG00000080549 | 0 | 0 | 0 | 0 | 0 | 0 | 0 |
| ENSMUSG00000080555 | 0 | 0 | 0 | 0 | 0 | 0 | 0 |
| ENSMUSG00000080556 | 0 | 0 | 0 | 0 | 0 | 0 | 0 |

| | | | | | | | |
|---|---|---|---|---|---|---|---|
| ENSMUSG00000080558 | 0 | 0 | 0 | 0 | 0 | 0 | 0 |
| ENSMUSG00000080561 | 0 | 0 | 0 | 0 | 0 | 0 | 0 |
| ENSMUSG00000080562 | 0 | 0 | 0 | 0 | 0 | 0 | 0 |
| ENSMUSG00000080566 | 0 | 0 | 0 | 0 | 0 | 0 | 0 |
| ENSMUSG00000080573 | 0 | 0.25 | 0.25 | 0 | 0.387761935 | 0 | 0 |
| ENSMUSG00000080574 | 0 | 0.25 | 0.25 | 0.25 | 0.244650542 | 0.301029996 | 0.224243824 |
| ENSMUSG00000080575 | 0 | 0 | 0 | 0 | 0 | 0 | 0 |
| ENSMUSG00000080577 | 0 | 0 | 0 | 0 | 0 | 0 | 0 |
| ENSMUSG00000080580 | 0 | 0 | 0 | 0 | 0 | 0 | 0 |
| ENSMUSG00000080582 | 0 | 0 | 0 | 0 | 0 | 0 | 0 |
| ENSMUSG00000080586 | 0 | 0 | 0 | 0 | 0 | 0 | 0 |
| ENSMUSG00000080590 | 0 | 0 | 0 | 0 | 0 | 0 | 0 |
| ENSMUSG00000080592 | 0 | 0 | 0 | 0 | 0 | 0 | 0 |
| ENSMUSG00000080594 | 0 | 0.25 | 0 | 0 | 0 | 0 | 0 |
| ENSMUSG00000080595 | 0 | 0 | 0 | 0 | 0 | 0 | 0 |
| ENSMUSG00000080596 | 0 | 0 | 0 | 0 | 0 | 0 | 0 |
| ENSMUSG00000080597 | 0 | 0 | 0 | 0 | 0 | 0 | 0 |
| ENSMUSG00000080600 | 0 | 0 | 0 | 0 | 0 | 0 | 0 |
| ENSMUSG00000080601 | 0 | 0 | 0 | 0 | 0 | 0 | 0 |
| ENSMUSG00000080607 | 0 | 0 | 0 | 0 | 0 | 0 | 0 |
| ENSMUSG00000080609 | 0 | 0 | 0 | 0 | 0 | 0 | 0 |
| ENSMUSG00000080610 | 0 | 0 | 0.25 | 0 | 0 | 0.301029996 | 0.224243824 |
| ENSMUSG00000080613 | 0 | 0 | 0 | 0.25 | 0 | 0 | 0 |
| ENSMUSG00000080615 | 0 | 0 | 0 | 0 | 0 | 0 | 0 |
| ENSMUSG00000080616 | 0 | 0 | 0 | 0 | 0 | 0 | 0 |
| ENSMUSG00000080622 | 0 | 0 | 0 | 0 | 0 | 0 | 0 |
| ENSMUSG00000080624 | 0 | 0 | 0 | 0 | 0 | 0 | 0 |
| ENSMUSG00000080625 | 0 | 0 | 0 | 0 | 0 | 0 | 0 |
| ENSMUSG00000080626 | 0 | 0 | 0 | 0 | 0 | 0 | 0 |
| ENSMUSG00000080627 | 0 | 0 | 0 | 0 | 0 | 0 | 0 |
| ENSMUSG00000080631 | 0 | 0 | 0 | 0 | 0 | 0 | 0 |
| ENSMUSG00000080635 | 0 | 0 | 0 | 0 | 0 | 0 | 0 |
| ENSMUSG00000080636 | 0 | 0 | 0 | 0 | 0 | 0 | 0 |
| ENSMUSG00000080645 | 0 | 0 | 0 | 0 | 0.244650542 | 0 | 0.224243824 |

| | | | | | | | |
|---|---|---|---|---|---|---|---|
| ENSMUSG00000080649 | 0 | 0 | 0 | 0 | 0 | 0 | 0 |
| ENSMUSG00000080653 | 0 | 0 | 0 | 0 | 0 | 0 | 0 |
| ENSMUSG00000080655 | 0 | 0 | 0 | 0 | 0 | 0 | 0 |
| ENSMUSG00000080657 | 0 | 0 | 0 | 0 | 0 | 0 | 0 |
| ENSMUSG00000080659 | 0 | 0 | 0 | 0 | 0 | 0 | 0 |
| ENSMUSG00000080662 | 0 | 0 | 0 | 0 | 0 | 0 | 0 |
| ENSMUSG00000080663 | 0 | 0 | 0 | 0 | 0 | 0 | 0 |
| ENSMUSG00000080665 | 0 | 0 | 0 | 0 | 0 | 0 | 0 |
| ENSMUSG00000080666 | 0 | 0 | 0 | 0 | 0 | 0 | 0 |
| ENSMUSG00000080668 | 0 | 0 | 0 | 0 | 0 | 0 | 0 |
| ENSMUSG00000080669 | 0 | 0 | 0 | 0 | 0 | 0 | 0 |
| ENSMUSG00000080671 | 0 | 0 | 0 | 0 | 0 | 0 | 0 |
| ENSMUSG00000080673 | 0 | 0 | 0 | 0 | 0 | 0 | 0 |
| ENSMUSG00000080676 | 0 | 0 | 0 | 0 | 0 | 0 | 0 |
| ENSMUSG00000080678 | 0 | 0 | 0 | 0 | 0 | 0 | 0 |
| ENSMUSG00000080680 | 0 | 0 | 0 | 0 | 0 | 0 | 0 |
| ENSMUSG00000080681 | 0 | 0 | 0 | 0 | 0 | 0 | 0 |
| ENSMUSG00000080683 | 0 | 0 | 0 | 0 | 0 | 0 | 0 |
| ENSMUSG00000080684 | 0 | 0 | 0 | 0 | 0 | 0 | 0 |
| ENSMUSG00000080686 | 0 | 0 | 0 | 0 | 0 | 0 | 0 |
| ENSMUSG00000080687 | 0 | 0 | 0 | 0 | 0 | 0 | 0 |
| ENSMUSG00000080689 | 0 | 0 | 0 | 0 | 0 | 0 | 0 |
| ENSMUSG00000080694 | 0 | 0 | 0 | 0 | 0 | 0 | 0 |
| ENSMUSG00000080695 | 0 | 0 | 0 | 0 | 0 | 0 | 0 |
| ENSMUSG00000080696 | 0 | 0 | 0 | 0 | 0 | 0 | 0 |
| ENSMUSG00000080697 | 0 | 0 | 0 | 0 | 0 | 0 | 0 |
| ENSMUSG00000080698 | 0 | 0 | 0 | 0 | 0 | 0 | 0 |
| ENSMUSG00000080700 | 0 | 0 | 0 | 0 | 0 | 0.301029996 | 0.224243824 |
| ENSMUSG00000080701 | 0 | 0.25 | 0 | 0 | 0 | 0 | 0.224243824 |
| ENSMUSG00000080703 | 0 | 0.25 | 0 | 0 | 0.244650542 | 0 | 0.224243824 |
| ENSMUSG00000080704 | 0 | 0 | 0 | 0 | 0 | 0 | 0 |
| ENSMUSG00000080705 | 0 | 0 | 0 | 0 | 0 | 0 | 0 |
| ENSMUSG00000080706 | 0 | 0 | 0 | 0 | 0 | 0 | 0 |
| ENSMUSG00000080707 | 0 | 0 | 0 | 0 | 0 | 0 | 0 |

| | | | | | | | |
|---|---|---|---|---|---|---|---|
| ENSMUSG00000080708 | 0 | 0 | 0 | 0 | 0 | 0 | 0 |
| ENSMUSG00000080709 | 0.578129653 | 0.701838731 | 0.396240625 | 1 | 0.489301084 | 0.602059991 | 0.74492186 |
| ENSMUSG00000080710 | 0 | 0 | 0 | 0 | 0 | 0 | 0 |
| ENSMUSG00000080711 | 0 | 0 | 0 | 0 | 0 | 0 | 0 |
| ENSMUSG00000080712 | 0 | 0 | 0 | 0 | 0 | 0 | 0 |
| ENSMUSG00000080713 | 0 | 0 | 0 | 0 | 0 | 0 | 0 |
| ENSMUSG00000080714 | 0 | 0 | 0 | 0 | 0 | 0 | 0 |
| ENSMUSG00000080715 | 0.289064826 | 0 | 0 | 0 | 0 | 0 | 0 |
| ENSMUSG00000080716 | 0 | 0 | 0 | 0 | 0 | 0 | 0 |
| ENSMUSG00000080717 | 1.036286563 | 1.02186571 | 0.976722649 | 1 | 1.091002363 | 1.113943352 | 1.014380832 |
| ENSMUSG00000080718 | 0.289064826 | 0 | 0 | 0.25 | 0.387761935 | 0 | 0 |
| ENSMUSG00000080719 | 0.45815691 | 0 | 0.396240625 | 0 | 0.244650542 | 0 | 0 |
| ENSMUSG00000080720 | 0 | 0 | 0 | 0 | 0 | 0 | 0 |
| ENSMUSG00000080722 | 0 | 0 | 0 | 0 | 0 | 0 | 0 |
| ENSMUSG00000080723 | 0 | 0 | 0 | 0 | 0 | 0 | 0 |
| ENSMUSG00000080725 | 0 | 0 | 0 | 0 | 0 | 0 | 0 |
| ENSMUSG00000080726 | 0 | 0 | 0 | 0 | 0 | 0 | 0 |
| ENSMUSG00000080727 | 0.747221736 | 0.396240625 | 0.701838731 | 0.830482024 | 0.387761935 | 0.698970004 | 0.803905701 |
| ENSMUSG00000080728 | 0 | 0 | 0 | 0 | 0 | 0 | 0 |
| ENSMUSG00000080729 | 0 | 0 | 0 | 0 | 0 | 0 | 0 |
| ENSMUSG00000080730 | 0.45815691 | 0.580482024 | 0.25 | 0.25 | 0 | 0 | 0 |
| ENSMUSG00000080731 | 0.578129653 | 0.646240625 | 0.701838731 | 0.646240625 | 0.568060967 | 0.477121255 | 0.520678036 |
| ENSMUSG00000080732 | 0 | 0 | 0 | 0 | 0 | 0 | 0 |
| ENSMUSG00000080733 | 0 | 0 | 0 | 0 | 0 | 0 | 0 |
| ENSMUSG00000080734 | 0 | 0 | 0 | 0 | 0 | 0 | 0 |
| ENSMUSG00000080735 | 0 | 0 | 0 | 0 | 0 | 0 | 0 |
| ENSMUSG00000080736 | 0 | 0 | 0 | 0 | 0 | 0 | 0 |
| ENSMUSG00000080737 | 0.747221736 | 0.79248125 | 0.701838731 | 0.976722649 | 0.877063019 | 0.84509804 | 0.672731473 |
| ENSMUSG00000080738 | 0 | 0 | 0 | 0 | 0 | 0 | 0 |
| ENSMUSG00000080741 | 0 | 0 | 0 | 0 | 0 | 0 | 0 |
| ENSMUSG00000080742 | 0 | 0 | 0 | 0 | 0 | 0 | 0 |
| ENSMUSG00000080743 | 0.91631382 | 0.896240625 | 0.951838731 | 0.5 | 0.632412477 | 0 | 0.853775828 |
| ENSMUSG00000080744 | 0 | 0 | 0 | 0 | 0 | 0 | 0 |
| ENSMUSG00000080746 | 0 | 0.646240625 | 0.92510993 | 0.75 | 0.568060967 | 0.477121255 | 0.710836105 |

| | | | | | | | |
|---|---|---|---|---|---|---|---|
| ENSMUSG00000080747 | 0.289064826 | 0.396240625 | 0.5 | 0.5 | 0.568060967 | 0.301029996 | 0.448487648 |
| ENSMUSG00000080748 | 0.289064826 | 0 | 0 | 0 | 0 | 0.301029996 | 0 |
| ENSMUSG00000080749 | 0 | 0 | 0 | 0 | 0 | 0 | 0 |
| ENSMUSG00000080750 | 0 | 0 | 0 | 0 | 0 | 0 | 0 |
| ENSMUSG00000080751 | 0 | 0 | 0 | 0 | 0 | 0 | 0 |
| ENSMUSG00000080752 | 0 | 0 | 0 | 0 | 0 | 0 | 0 |
| ENSMUSG00000080753 | 0 | 0 | 0 | 0 | 0 | 0 | 0 |
| ENSMUSG00000080754 | 0 | 0 | 0 | 0 | 0 | 0 | 0 |
| ENSMUSG00000080755 | 0 | 0 | 0 | 0 | 0 | 0 | 0 |
| ENSMUSG00000080756 | 0 | 0 | 0 | 0 | 0 | 0 | 0 |
| ENSMUSG00000080757 | 0 | 0 | 0 | 0 | 0 | 0 | 0 |
| ENSMUSG00000080758 | 0 | 0 | 0 | 0 | 0 | 0 | 0 |
| ENSMUSG00000080759 | 0 | 0 | 0 | 0 | 0 | 0 | 0.224243824 |
| ENSMUSG00000080760 | 0 | 0.25 | 0 | 0 | 0 | 0 | 0 |
| ENSMUSG00000080762 | 0 | 0 | 0 | 0.25 | 0.244650542 | 0 | 0 |
| ENSMUSG00000080763 | 0 | 0.25 | 0 | 0 | 0 | 0 | 0 |
| ENSMUSG00000080764 | 0 | 0.396240625 | 0 | 0.25 | 0 | 0 | 0 |
| ENSMUSG00000080765 | 0 | 0 | 0 | 0 | 0 | 0 | 0 |
| ENSMUSG00000080766 | 0 | 0 | 0 | 0 | 0 | 0 | 0 |
| ENSMUSG00000080767 | 0 | 0 | 0 | 0 | 0 | 0 | 0 |
| ENSMUSG00000080768 | 0.45815691 | 0.75 | 0.5 | 0.646240625 | 0.77552387 | 0.602059991 | 0.672731473 |
| ENSMUSG00000080770 | 0 | 0 | 0 | 0 | 0 | 0 | 0 |
| ENSMUSG00000080771 | 0 | 0 | 0 | 0 | 0 | 0 | 0 |
| ENSMUSG00000080772 | 0 | 0 | 0 | 0 | 0 | 0.301029996 | 0.224243824 |
| ENSMUSG00000080773 | 0 | 0 | 0 | 0 | 0 | 0 | 0 |
| ENSMUSG00000080774 | 0.289064826 | 0.25 | 0.5 | 0.396240625 | 0.244650542 | 0.301029996 | 0 |
| ENSMUSG00000080775 | 1.036286563 | 1.080482024 | 0.580482024 | 0.976722649 | 1 | 0.903089987 | 0.853775828 |
| ENSMUSG00000080776 | 0 | 0 | 0 | 0.25 | 0 | 0 | 0.224243824 |
| ENSMUSG00000080777 | 0 | 0.25 | 0 | 0 | 0 | 0 | 0 |
| ENSMUSG00000080778 | 0 | 0 | 0 | 0 | 0 | 0 | 0 |
| ENSMUSG00000080779 | 0.289064826 | 0.25 | 0 | 0 | 0 | 0 | 0.224243824 |
| ENSMUSG00000080780 | 0 | 0 | 0 | 0 | 0 | 0 | 0 |
| ENSMUSG00000080781 | 0 | 0 | 0 | 0 | 0 | 0 | 0 |
| ENSMUSG00000080782 | 0 | 0 | 0 | 0 | 0 | 0 | 0 |

| | | | | | | | |
|---|---|---|---|---|---|---|---|
| ENSMUSG00000080783 | 0.45815691 | 0 | 0.25 | 0.25 | 0.387761935 | 0 | 0.579661877 |
| ENSMUSG00000080785 | 0 | 0 | 0 | 0 | 0 | 0 | 0 |
| ENSMUSG00000080788 | 0 | 0 | 0 | 0 | 0 | 0 | 0 |
| ENSMUSG00000080789 | 0 | 0 | 0 | 0 | 0 | 0 | 0 |
| ENSMUSG00000080790 | 0.45815691 | 0.701838731 | 0.5 | 0.396240625 | 0.387761935 | 0 | 0.355418052 |
| ENSMUSG00000080791 | 0 | 0 | 0 | 0 | 0 | 0 | 0.224243824 |
| ENSMUSG00000080792 | 0 | 0 | 0 | 0 | 0 | 0 | 0 |
| ENSMUSG00000080793 | 0 | 0 | 0 | 0 | 0 | 0.301029996 | 0 |
| ENSMUSG00000080794 | 0 | 0 | 0 | 0 | 0 | 0 | 0 |
| ENSMUSG00000080795 | 0 | 0 | 0 | 0 | 0 | 0 | 0.224243824 |
| ENSMUSG00000080796 | 0 | 0 | 0 | 0 | 0 | 0.301029996 | 0 |
| ENSMUSG00000080797 | 0.289064826 | 0.25 | 0 | 0 | 0 | 0 | 0.579661877 |
| ENSMUSG00000080798 | 0 | 0 | 0.25 | 0 | 0 | 0 | 0 |
| ENSMUSG00000080799 | 0 | 0 | 0 | 0 | 0 | 0 | 0 |
| ENSMUSG00000080800 | 0 | 0 | 0 | 0 | 0 | 0 | 0 |
| ENSMUSG00000080801 | 0.289064826 | 0 | 0 | 0.25 | 0 | 0.477121255 | 0.448487648 |
| ENSMUSG00000080802 | 0 | 0 | 0 | 0 | 0 | 0 | 0 |
| ENSMUSG00000080803 | 0 | 0 | 0 | 0 | 0 | 0 | 0 |
| ENSMUSG00000080805 | 0 | 0 | 0 | 0 | 0 | 0 | 0 |
| ENSMUSG00000080806 | 0 | 0 | 0.25 | 0 | 0 | 0 | 0 |
| ENSMUSG00000080807 | 0 | 0 | 0 | 0 | 0 | 0 | 0 |
| ENSMUSG00000080808 | 0.289064826 | 0.580482024 | 0.396240625 | 0.701838731 | 0 | 0.477121255 | 0.448487648 |
| ENSMUSG00000080809 | 0 | 0 | 0 | 0 | 0 | 0 | 0 |
| ENSMUSG00000080810 | 0 | 0 | 0 | 0 | 0 | 0 | 0 |
| ENSMUSG00000080811 | 0 | 0 | 0 | 0.25 | 0 | 0 | 0 |
| ENSMUSG00000080812 | 0 | 0 | 0 | 0 | 0 | 0 | 0 |
| ENSMUSG00000080813 | 0 | 0 | 0 | 0 | 0 | 0 | 0 |
| ENSMUSG00000080814 | 0.289064826 | 0.396240625 | 0 | 0 | 0 | 0 | 0 |
| ENSMUSG00000080815 | 0 | 0.396240625 | 0 | 0 | 0 | 0 | 0 |
| ENSMUSG00000080816 | 0 | 0 | 0 | 0 | 0 | 0 | 0 |
| ENSMUSG00000080817 | 0 | 0 | 0 | 0 | 0 | 0 | 0 |
| ENSMUSG00000080818 | 0 | 0.25 | 0 | 0 | 0.244650542 | 0 | 0 |
| ENSMUSG00000080819 | 0 | 0 | 0 | 0 | 0 | 0 | 0 |
| ENSMUSG00000080820 | 0 | 0 | 0 | 0 | 0 | 0 | 0 |

| | | | | | | | |
|---|---|---|---|---|---|---|---|
| ENSMUSG00000080822 | 0 | 0 | 0 | 0 | 0 | 0 | 0 |
| ENSMUSG00000080823 | 0 | 0 | 0 | 0 | 0 | 0 | 0.224243824 |
| ENSMUSG00000080824 | 0 | 0 | 0 | 0 | 0 | 0 | 0 |
| ENSMUSG00000080825 | 0 | 0 | 0.25 | 0.25 | 0 | 0 | 0.224243824 |
| ENSMUSG00000080826 | 0 | 0 | 0 | 0 | 0 | 0 | 0 |
| ENSMUSG00000080827 | 0 | 0 | 0 | 0 | 0 | 0 | 0 |
| ENSMUSG00000080828 | 0 | 0 | 0 | 0 | 0 | 0 | 0 |
| ENSMUSG00000080829 | 0 | 0.25 | 0 | 0.25 | 0 | 0 | 0 |
| ENSMUSG00000080831 | 0 | 0 | 0 | 0 | 0 | 0 | 0 |
| ENSMUSG00000080832 | 0 | 0.396240625 | 0.25 | 0 | 0.244650542 | 0 | 0.224243824 |
| ENSMUSG00000080833 | 0 | 0 | 0 | 0 | 0 | 0 | 0 |
| ENSMUSG00000080834 | 0 | 0 | 0 | 0 | 0 | 0 | 0 |
| ENSMUSG00000080835 | 0 | 0 | 0 | 0 | 0 | 0 | 0 |
| ENSMUSG00000080836 | 0 | 0.25 | 0.25 | 0 | 0 | 0 | 0 |
| ENSMUSG00000080839 | 0 | 0 | 0.25 | 0 | 0.244650542 | 0 | 0 |
| ENSMUSG00000080840 | 0 | 0 | 0.25 | 0.25 | 0.387761935 | 0 | 0.448487648 |
| ENSMUSG00000080841 | 0 | 0 | 0 | 0 | 0 | 0 | 0 |
| ENSMUSG00000080842 | 0 | 0 | 0 | 0 | 0 | 0 | 0 |
| ENSMUSG00000080843 | 0 | 0 | 0 | 0 | 0 | 0 | 0 |
| ENSMUSG00000080844 | 0 | 0.25 | 0 | 0 | 0 | 0 | 0 |
| ENSMUSG00000080845 | 0.289064826 | 0.396240625 | 0.25 | 0.25 | 0.244650542 | 0 | 0.224243824 |
| ENSMUSG00000080846 | 0.289064826 | 0 | 0 | 0 | 0 | 0 | 0 |
| ENSMUSG00000080847 | 0 | 0 | 0 | 0 | 0 | 0 | 0 |
| ENSMUSG00000080848 | 0.289064826 | 0 | 0.25 | 0 | 0.244650542 | 0.301029996 | 0 |
| ENSMUSG00000080849 | 0 | 0 | 0 | 0.25 | 0 | 0 | 0 |
| ENSMUSG00000080850 | 0.289064826 | 0.396240625 | 0 | 0.5 | 0.568060967 | 0.477121255 | 0.355418052 |
| ENSMUSG00000080851 | 0 | 0.25 | 0 | 0 | 0 | 0 | 0 |
| ENSMUSG00000080852 | 0 | 0 | 0 | 0 | 0 | 0 | 0 |
| ENSMUSG00000080855 | 0 | 0 | 0 | 0 | 0 | 0 | 0 |
| ENSMUSG00000080857 | 0 | 0 | 0 | 0 | 0 | 0 | 0 |
| ENSMUSG00000080859 | 0 | 0 | 0 | 0 | 0 | 0 | 0 |
| ENSMUSG00000080860 | 0 | 0.396240625 | 0.25 | 0.25 | 0.387761935 | 0.477121255 | 0.520678036 |
| ENSMUSG00000080861 | 0 | 0 | 0 | 0 | 0 | 0 | 0 |
| ENSMUSG00000080862 | 0 | 0 | 0 | 0 | 0 | 0 | 0 |

| | | | | | | | |
|---|---|---|---|---|---|---|---|
| ENSMUSG00000080863 | 0 | 0 | 0 | 0 | 0 | 0 | 0 |
| ENSMUSG00000080865 | 0 | 0 | 0 | 0 | 0 | 0 | 0 |
| ENSMUSG00000080866 | 1.036286563 | 1.160964047 | 0.951838731 | 0.92510993 | 1.020174412 | 1.113943352 | 1.041356071 |
| ENSMUSG00000080868 | 0 | 0 | 0 | 0 | 0 | 0 | 0 |
| ENSMUSG00000080869 | 0 | 0 | 0 | 0 | 0 | 0 | 0 |
| ENSMUSG00000080870 | 0.289064826 | 0.25 | 0.646240625 | 0.25 | 0.244650542 | 0 | 0.224243824 |
| ENSMUSG00000080871 | 0 | 0 | 0 | 0 | 0 | 0 | 0 |
| ENSMUSG00000080872 | 0 | 0 | 0 | 0 | 0 | 0 | 0 |
| ENSMUSG00000080873 | 0 | 0.25 | 0 | 0.25 | 0 | 0 | 0 |
| ENSMUSG00000080874 | 0 | 0 | 0 | 0 | 0 | 0 | 0 |
| ENSMUSG00000080875 | 0.289064826 | 0 | 0 | 0 | 0 | 0 | 0 |
| ENSMUSG00000080876 | 0 | 0 | 0 | 0 | 0 | 0 | 0 |
| ENSMUSG00000080877 | 0 | 0.25 | 0.25 | 0.5 | 0.489301084 | 0.301029996 | 0.355418052 |
| ENSMUSG00000080878 | 0 | 0 | 0 | 0 | 0 | 0 | 0 |
| ENSMUSG00000080880 | 0 | 0 | 0 | 0 | 0 | 0 | 0 |
| ENSMUSG00000080884 | 0 | 0 | 0 | 0 | 0 | 0 | 0 |
| ENSMUSG00000080885 | 0.45815691 | 0.580482024 | 0.396240625 | 0 | 0.568060967 | 0 | 0.355418052 |
| ENSMUSG00000080886 | 0 | 0 | 0 | 0 | 0 | 0 | 0 |
| ENSMUSG00000080887 | 0 | 0 | 0 | 0 | 0 | 0 | 0 |
| ENSMUSG00000080888 | 0 | 0.25 | 0.396240625 | 0.580482024 | 0.686820904 | 0.477121255 | 0.520678036 |
| ENSMUSG00000080889 | 0 | 0 | 0 | 0 | 0 | 0 | 0 |
| ENSMUSG00000080890 | 0 | 0 | 0 | 0 | 0 | 0 | 0 |
| ENSMUSG00000080891 | 0 | 0 | 0 | 0 | 0 | 0 | 0 |
| ENSMUSG00000080892 | 0 | 0 | 0 | 0 | 0 | 0 | 0 |
| ENSMUSG00000080893 | 1.54867984 | 1.464495249 | 1.527131114 | 1.339388001 | 1.327538614 | 1.531478917 | 1.284450788 |
| ENSMUSG00000080894 | 0 | 0 | 0 | 0 | 0 | 0 | 0 |
| ENSMUSG00000080895 | 1.156259305 | 1.04248125 | 1.201838731 | 0.25 | 0 | 0.301029996 | 0.224243824 |
| ENSMUSG00000080896 | 0 | 0 | 0 | 0 | 0 | 0 | 0 |
| ENSMUSG00000080897 | 0 | 0 | 0 | 0 | 0 | 0 | 0 |
| ENSMUSG00000080898 | 0 | 0 | 0 | 0 | 0 | 0 | 0 |
| ENSMUSG00000080899 | 0 | 0 | 0 | 0 | 0 | 0 | 0 |
| ENSMUSG00000080902 | 0.289064826 | 0.25 | 0.5 | 0.396240625 | 0 | 0.301029996 | 0.355418052 |
| ENSMUSG00000080903 | 0 | 0 | 0 | 0 | 0 | 0 | 0 |
| ENSMUSG00000080904 | 0 | 0 | 0 | 0 | 0 | 0 | 0 |

| | | | | | | | |
|---|---|---|---|---|---|---|---|
| ENSMUSG00000080906 | 0 | 0 | 0 | 0 | 0 | 0 | 0 |
| ENSMUSG00000080907 | 0 | 0 | 0 | 0 | 0 | 0 | 0 |
| ENSMUSG00000080910 | 0 | 0 | 0 | 0 | 0 | 0 | 0 |
| ENSMUSG00000080911 | 0 | 0 | 0 | 0 | 0 | 0 | 0 |
| ENSMUSG00000080912 | 0 | 0.25 | 0 | 0 | 0 | 0 | 0 |
| ENSMUSG00000080913 | 0 | 0 | 0 | 0 | 0 | 0 | 0 |
| ENSMUSG00000080914 | 0.289064826 | 0.5 | 0 | 0 | 0 | 0 | 0 |
| ENSMUSG00000080915 | 0 | 0 | 0 | 0 | 0 | 0 | 0 |
| ENSMUSG00000080916 | 0 | 0 | 0 | 0 | 0 | 0 | 0 |
| ENSMUSG00000080917 | 0 | 0 | 0 | 0 | 0 | 0 | 0 |
| ENSMUSG00000080918 | 0 | 0 | 0 | 0 | 0 | 0 | 0 |
| ENSMUSG00000080919 | 0 | 0 | 0 | 0 | 0 | 0 | 0 |
| ENSMUSG00000080921 | 0 | 0 | 0.25 | 0 | 0.244650542 | 0 | 0.355418052 |
| ENSMUSG00000080922 | 0 | 0 | 0 | 0 | 0 | 0 | 0 |
| ENSMUSG00000080923 | 0 | 0 | 0 | 0 | 0 | 0 | 0 |
| ENSMUSG00000080924 | 0 | 0 | 0 | 0 | 0 | 0 | 0 |
| ENSMUSG00000080925 | 0 | 0 | 0 | 0 | 0 | 0 | 0 |
| ENSMUSG00000080927 | 0 | 0 | 0 | 0 | 0 | 0 | 0 |
| ENSMUSG00000080928 | 0.289064826 | 0 | 0.25 | 0 | 0 | 0.477121255 | 0 |
| ENSMUSG00000080929 | 0 | 0 | 0 | 0.25 | 0 | 0 | 0 |
| ENSMUSG00000080930 | 0 | 0.25 | 0 | 0 | 0 | 0 | 0 |
| ENSMUSG00000080932 | 0.289064826 | 0.396240625 | 0.25 | 0.396240625 | 0.387761935 | 0.301029996 | 0.579661877 |
| ENSMUSG00000080933 | 0 | 0 | 0 | 0 | 0 | 0 | 0 |
| ENSMUSG00000080934 | 0 | 0 | 0 | 0 | 0 | 0 | 0 |
| ENSMUSG00000080935 | 0 | 0 | 0 | 0 | 0 | 0 | 0 |
| ENSMUSG00000080936 | 0 | 0 | 0 | 0 | 0 | 0 | 0 |
| ENSMUSG00000080937 | 0 | 0 | 0 | 0 | 0 | 0 | 0 |
| ENSMUSG00000080938 | 0.289064826 | 0.25 | 0 | 0.25 | 0 | 0.301029996 | 0 |
| ENSMUSG00000080939 | 0 | 0 | 0 | 0 | 0 | 0 | 0 |
| ENSMUSG00000080941 | 0 | 0 | 0 | 0 | 0 | 0 | 0 |
| ENSMUSG00000080942 | 0 | 0 | 0 | 0 | 0 | 0 | 0 |
| ENSMUSG00000080943 | 0 | 0 | 0 | 0 | 0 | 0 | 0 |
| ENSMUSG00000080944 | 0 | 0.5 | 0.396240625 | 0 | 0.387761935 | 0 | 0.579661877 |
| ENSMUSG00000080946 | 0 | 0 | 0 | 0 | 0 | 0 | 0 |

| | | | | | | | |
|---|---|---|---|---|---|---|---|
| ENSMUSG00000080947 | 0 | 0 | 0 | 0 | 0 | 0 | 0 |
| ENSMUSG00000080948 | 0 | 0 | 0 | 0 | 0 | 0 | 0 |
| ENSMUSG00000080949 | 0 | 0 | 0 | 0 | 0 | 0 | 0 |
| ENSMUSG00000080950 | 0.671187741 | 0.5 | 0.646240625 | 0.396240625 | 0.489301084 | 0 | 0 |
| ENSMUSG00000080951 | 0 | 0.396240625 | 0.701838731 | 0.25 | 0.489301084 | 0.477121255 | 0.355418052 |
| ENSMUSG00000080952 | 0 | 0 | 0 | 0 | 0 | 0 | 0 |
| ENSMUSG00000080953 | 0 | 0 | 0 | 0 | 0 | 0 | 0 |
| ENSMUSG00000080954 | 0 | 0 | 0 | 0 | 0 | 0 | 0 |
| ENSMUSG00000080955 | 0 | 0 | 0.396240625 | 0 | 0 | 0 | 0.448487648 |
| ENSMUSG00000080956 | 0 | 0 | 0 | 0.25 | 0 | 0 | 0 |
| ENSMUSG00000080957 | 0.289064826 | 0 | 0 | 0.25 | 0 | 0.301029996 | 0 |
| ENSMUSG00000080958 | 0 | 0 | 0 | 0 | 0 | 0 | 0 |
| ENSMUSG00000080959 | 0 | 0 | 0 | 0 | 0 | 0 | 0 |
| ENSMUSG00000080960 | 0 | 0 | 0 | 0 | 0 | 0 | 0 |
| ENSMUSG00000080961 | 0 | 0 | 0 | 0 | 0 | 0 | 0 |
| ENSMUSG00000080962 | 0 | 0 | 0.25 | 0 | 0 | 0 | 0 |
| ENSMUSG00000080964 | 0 | 0 | 0 | 0 | 0 | 0 | 0 |
| ENSMUSG00000080965 | 0 | 0.25 | 0 | 0 | 0 | 0 | 0 |
| ENSMUSG00000080966 | 0 | 0 | 0 | 0 | 0.244650542 | 0 | 0 |
| ENSMUSG00000080967 | 0 | 0 | 0 | 0 | 0 | 0 | 0 |
| ENSMUSG00000080968 | 0 | 0 | 0.396240625 | 0 | 0 | 0 | 0 |
| ENSMUSG00000080969 | 0 | 0.25 | 0.25 | 0 | 0 | 0.301029996 | 0 |
| ENSMUSG00000080970 | 0 | 0 | 0 | 0 | 0 | 0 | 0 |
| ENSMUSG00000080971 | 0 | 0 | 0.25 | 0.25 | 0 | 0 | 0 |
| ENSMUSG00000080972 | 0.45815691 | 0.646240625 | 0.79248125 | 0.580482024 | 0.387761935 | 0.477121255 | 0.579661877 |
| ENSMUSG00000080974 | 0 | 0 | 0.396240625 | 0 | 0.244650542 | 0 | 0.355418052 |
| ENSMUSG00000080976 | 0 | 0 | 0 | 0.25 | 0 | 0 | 0 |
| ENSMUSG00000080977 | 0 | 0 | 0 | 0 | 0 | 0 | 0 |
| ENSMUSG00000080978 | 0 | 0 | 0 | 0 | 0 | 0 | 0 |
| ENSMUSG00000080979 | 0 | 0 | 0 | 0 | 0 | 0 | 0 |
| ENSMUSG00000080980 | 0 | 0 | 0 | 0 | 0 | 0 | 0 |
| ENSMUSG00000080981 | 0 | 0 | 0 | 0 | 0 | 0 | 0 |
| ENSMUSG00000080982 | 0 | 0 | 0 | 0 | 0 | 0 | 0 |
| ENSMUSG00000080983 | 0 | 0 | 0 | 0 | 0 | 0 | 0 |

| | | | | | | | |
|---|---|---|---|---|---|---|---|
| ENSMUSG00000080984 | 0 | 0 | 0 | 0 | 0 | 0 | 0 |
| ENSMUSG00000080985 | 0 | 0 | 0.25 | 0 | 0.244650542 | 0 | 0.355418052 |
| ENSMUSG00000080986 | 0 | 0 | 0 | 0 | 0 | 0 | 0 |
| ENSMUSG00000080987 | 0 | 0 | 0 | 0 | 0 | 0 | 0 |
| ENSMUSG00000080989 | 0 | 0 | 0 | 0 | 0 | 0 | 0 |
| ENSMUSG00000080990 | 0 | 0 | 0 | 0 | 0 | 0 | 0 |
| ENSMUSG00000080992 | 0.289064826 | 0 | 0.25 | 0 | 0 | 0 | 0.224243824 |
| ENSMUSG00000080993 | 0 | 0 | 0 | 0 | 0 | 0 | 0 |
| ENSMUSG00000080994 | 0 | 0.25 | 0 | 0 | 0 | 0 | 0 |
| ENSMUSG00000080995 | 0 | 0 | 0.25 | 0 | 0 | 0 | 0 |
| ENSMUSG00000080996 | 0 | 0 | 0 | 0 | 0 | 0 | 0 |
| ENSMUSG00000080997 | 0.45815691 | 0.25 | 0.25 | 0.25 | 0 | 0 | 0 |
| ENSMUSG00000080998 | 0 | 0 | 0 | 0 | 0.244650542 | 0 | 0.224243824 |
| ENSMUSG00000080999 | 0 | 0 | 0 | 0 | 0 | 0 | 0 |
| ENSMUSG00000081000 | 0 | 0 | 0 | 0 | 0 | 0 | 0 |
| ENSMUSG00000081001 | 0 | 0 | 0 | 0 | 0 | 0 | 0 |
| ENSMUSG00000081002 | 0 | 0 | 0 | 0 | 0 | 0 | 0 |
| ENSMUSG00000081003 | 0 | 0 | 0 | 0 | 0 | 0 | 0 |
| ENSMUSG00000081004 | 0.289064826 | 0 | 0.396240625 | 0.25 | 0 | 0 | 0 |
| ENSMUSG00000081005 | 0 | 0 | 0 | 0 | 0 | 0 | 0 |
| ENSMUSG00000081006 | 0 | 0 | 0 | 0 | 0 | 0 | 0 |
| ENSMUSG00000081007 | 0.289064826 | 0 | 0 | 0 | 0 | 0 | 0 |
| ENSMUSG00000081008 | 0 | 0 | 0 | 0 | 0 | 0 | 0 |
| ENSMUSG00000081009 | 0 | 0 | 0 | 0 | 0 | 0 | 0 |
| ENSMUSG00000081010 | 0 | 0 | 0.5 | 0 | 0.244650542 | 0 | 0 |
| ENSMUSG00000081011 | 0.578129653 | 0.5 | 0.5 | 0.396240625 | 0.489301084 | 0.602059991 | 0.520678036 |
| ENSMUSG00000081012 | 0 | 0 | 0 | 0 | 0 | 0 | 0 |
| ENSMUSG00000081013 | 0.289064826 | 0 | 0 | 0 | 0 | 0 | 0 |
| ENSMUSG00000081014 | 0 | 0 | 0 | 0 | 0 | 0 | 0 |
| ENSMUSG00000081016 | 0 | 0.25 | 0 | 0 | 0 | 0 | 0 |
| ENSMUSG00000081017 | 0 | 0 | 0 | 0 | 0 | 0 | 0 |
| ENSMUSG00000081018 | 0 | 0 | 0 | 0 | 0 | 0 | 0 |
| ENSMUSG00000081020 | 0 | 0 | 0 | 0 | 0 | 0 | 0 |
| ENSMUSG00000081021 | 0 | 0 | 0 | 0 | 0 | 0 | 0 |

| | | | | | | | |
|---|---|---|---|---|---|---|---|
| ENSMUSG00000081022 | 0 | 0 | 0 | 0 | 0 | 0 | 0 |
| ENSMUSG00000081023 | 0.289064826 | 0.25 | 0 | 0 | 0.387761935 | 0 | 0.224243824 |
| ENSMUSG00000081024 | 0 | 0 | 0 | 0 | 0 | 0 | 0.224243824 |
| ENSMUSG00000081025 | 0 | 0 | 0 | 0 | 0 | 0 | 0 |
| ENSMUSG00000081026 | 0 | 0.396240625 | 0.25 | 0.25 | 0 | 0 | 0 |
| ENSMUSG00000081027 | 0 | 0 | 0 | 0 | 0 | 0 | 0 |
| ENSMUSG00000081029 | 0 | 0 | 0 | 0 | 0 | 0 | 0.224243824 |
| ENSMUSG00000081030 | 0.289064826 | 0.396240625 | 0 | 0.25 | 0.686820904 | 0.301029996 | 0.355418052 |
| ENSMUSG00000081033 | 0 | 0 | 0 | 0 | 0 | 0 | 0 |
| ENSMUSG00000081034 | 0 | 0 | 0 | 0.25 | 0 | 0 | 0 |
| ENSMUSG00000081035 | 0 | 0 | 0 | 0 | 0 | 0 | 0 |
| ENSMUSG00000081036 | 0 | 0 | 0 | 0 | 0 | 0 | 0 |
| ENSMUSG00000081037 | 0 | 0 | 0 | 0 | 0 | 0 | 0 |
| ENSMUSG00000081038 | 0 | 0 | 0 | 0 | 0 | 0 | 0 |
| ENSMUSG00000081041 | 0 | 0 | 0 | 0 | 0 | 0 | 0.224243824 |
| ENSMUSG00000081042 | 0 | 0.396240625 | 0.25 | 0 | 0.387761935 | 0 | 0.355418052 |
| ENSMUSG00000081043 | 0 | 0.396240625 | 0.25 | 0.5 | 0.568060967 | 0.477121255 | 0.224243824 |
| ENSMUSG00000081044 | 0 | 0.25 | 0 | 0 | 0 | 0 | 0 |
| ENSMUSG00000081045 | 0 | 0 | 0 | 0 | 0 | 0 | 0 |
| ENSMUSG00000081046 | 0 | 0 | 0 | 0 | 0 | 0.477121255 | 0 |
| ENSMUSG00000081047 | 0 | 0 | 0 | 0 | 0 | 0 | 0 |
| ENSMUSG00000081048 | 0.671187741 | 0.25 | 0.396240625 | 0 | 0.568060967 | 0.301029996 | 0.448487648 |
| ENSMUSG00000081049 | 0 | 0 | 0.396240625 | 0.25 | 0.244650542 | 0 | 0 |
| ENSMUSG00000081050 | 0 | 0 | 0 | 0 | 0 | 0 | 0 |
| ENSMUSG00000081051 | 0.45815691 | 1 | 0.75 | 0.396240625 | 0.686820904 | 0.698970004 | 0.710836105 |
| ENSMUSG00000081052 | 0 | 0 | 0 | 0 | 0 | 0 | 0 |
| ENSMUSG00000081053 | 0 | 0 | 0 | 0 | 0 | 0 | 0 |
| ENSMUSG00000081054 | 0 | 0 | 0 | 0 | 0 | 0 | 0 |
| ENSMUSG00000081055 | 0 | 0 | 0 | 0 | 0 | 0 | 0.224243824 |
| ENSMUSG00000081056 | 0 | 0 | 0 | 0 | 0 | 0 | 0 |
| ENSMUSG00000081057 | 0 | 0 | 0 | 0 | 0 | 0 | 0 |
| ENSMUSG00000081058 | 0 | 0 | 0 | 0.25 | 0 | 0 | 0.448487648 |
| ENSMUSG00000081059 | 0 | 0.25 | 0.25 | 0.25 | 0 | 0.301029996 | 0.355418052 |
| ENSMUSG00000081060 | 0 | 0 | 0 | 0 | 0 | 0 | 0 |

| | | | | | | | |
|---|---|---|---|---|---|---|---|
| ENSMUSG00000081061 | 0 | 0 | 0 | 0 | 0 | 0 | 0 |
| ENSMUSG00000081062 | 0 | 0 | 0 | 0 | 0 | 0 | 0 |
| ENSMUSG00000081063 | 0 | 0 | 0 | 0 | 0 | 0 | 0 |
| ENSMUSG00000081064 | 0.289064826 | 0 | 0 | 0 | 0 | 0 | 0.224243824 |
| ENSMUSG00000081065 | 0 | 0 | 0 | 0 | 0 | 0 | 0 |
| ENSMUSG00000081066 | 0 | 0 | 0 | 0 | 0 | 0 | 0 |
| ENSMUSG00000081067 | 1.181541736 | 1.160964047 | 1.188721876 | 1.080482024 | 1.039257769 | 1.397940009 | 1.140832123 |
| ENSMUSG00000081068 | 0 | 0 | 0.25 | 0 | 0.244650542 | 0 | 0 |
| ENSMUSG00000081069 | 0 | 0 | 0 | 0 | 0 | 0 | 0 |
| ENSMUSG00000081070 | 0 | 0 | 0 | 0 | 0 | 0 | 0 |
| ENSMUSG00000081071 | 0 | 0 | 0 | 0 | 0 | 0 | 0 |
| ENSMUSG00000081072 | 0 | 0 | 0 | 0 | 0 | 0 | 0 |
| ENSMUSG00000081073 | 0 | 0 | 0 | 0 | 0 | 0 | 0 |
| ENSMUSG00000081076 | 0.867194479 | 1 | 0.79248125 | 0.5 | 0.812711509 | 0.602059991 | 0.775756176 |
| ENSMUSG00000081077 | 0 | 0 | 0 | 0.25 | 0 | 0 | 0 |
| ENSMUSG00000081078 | 0 | 0 | 0 | 0 | 0 | 0 | 0.224243824 |
| ENSMUSG00000081079 | 0 | 0 | 0 | 0 | 0 | 0 | 0 |
| ENSMUSG00000081080 | 0 | 0 | 0 | 0 | 0 | 0 | 0 |
| ENSMUSG00000081081 | 0 | 0 | 0 | 0 | 0 | 0 | 0 |
| ENSMUSG00000081082 | 0 | 0 | 0 | 0 | 0 | 0 | 0 |
| ENSMUSG00000081083 | 0 | 0 | 0 | 0 | 0 | 0 | 0 |
| ENSMUSG00000081084 | 0.289064826 | 0 | 0.396240625 | 0 | 0.244650542 | 0 | 0.355418052 |
| ENSMUSG00000081085 | 0 | 0 | 0 | 0 | 0 | 0 | 0 |
| ENSMUSG00000081086 | 0 | 0 | 0 | 0 | 0 | 0 | 0.224243824 |
| ENSMUSG00000081087 | 0 | 0 | 0.25 | 0.25 | 0.244650542 | 0.301029996 | 0 |
| ENSMUSG00000081088 | 0 | 0 | 0 | 0 | 0 | 0 | 0 |
| ENSMUSG00000081089 | 0 | 0 | 0 | 0 | 0 | 0 | 0 |
| ENSMUSG00000081092 | 0 | 0 | 0 | 0 | 0 | 0 | 0 |
| ENSMUSG00000081093 | 0.289064826 | 0 | 0.396240625 | 0.25 | 0.244650542 | 0.301029996 | 0.224243824 |
| ENSMUSG00000081094 | 1.036286563 | 1.226722649 | 1.114857905 | 0.864857905 | 1.149965125 | 0.954242509 | 1.21680636 |
| ENSMUSG00000081095 | 0 | 0 | 0 | 0 | 0 | 0 | 0 |
| ENSMUSG00000081096 | 0 | 0 | 0 | 0.25 | 0 | 0 | 0 |
| ENSMUSG00000081097 | 0 | 0 | 0 | 0 | 0 | 0 | 0 |
| ENSMUSG00000081098 | 0 | 0 | 0 | 0 | 0 | 0 | 0 |

| | | | | | | | |
|---|---|---|---|---|---|---|---|
| ENSMUSG00000081099 | 0 | 0 | 0 | 0 | 0 | 0 | 0.224243824 |
| ENSMUSG00000081100 | 0 | 0 | 0 | 0 | 0 | 0 | 0.224243824 |
| ENSMUSG00000081101 | 0 | 0.25 | 0 | 0 | 0 | 0 | 0 |
| ENSMUSG00000081103 | 0.747221736 | 0.580482024 | 0.25 | 0.5 | 0.244650542 | 0 | 0.355418052 |
| ENSMUSG00000081104 | 0 | 0 | 0.396240625 | 0.396240625 | 0.244650542 | 0 | 0.355418052 |
| ENSMUSG00000081106 | 0 | 0 | 0 | 0 | 0 | 0 | 0 |
| ENSMUSG00000081107 | 0 | 0 | 0 | 0 | 0 | 0 | 0 |
| ENSMUSG00000081108 | 0 | 0 | 0 | 0 | 0 | 0 | 0 |
| ENSMUSG00000081110 | 0 | 0 | 0 | 0 | 0 | 0 | 0 |
| ENSMUSG00000081111 | 0 | 0.5 | 0.396240625 | 0.396240625 | 0.244650542 | 0.477121255 | 0.579661877 |
| ENSMUSG00000081112 | 0 | 0 | 0 | 0 | 0.244650542 | 0 | 0.224243824 |
| ENSMUSG00000081113 | 0.45815691 | 0.25 | 0.396240625 | 0.580482024 | 0.568060967 | 0 | 0.224243824 |
| ENSMUSG00000081114 | 0 | 0 | 0 | 0 | 0 | 0 | 0 |
| ENSMUSG00000081115 | 0 | 0 | 0 | 0 | 0 | 0 | 0 |
| ENSMUSG00000081116 | 0 | 0 | 0 | 0 | 0 | 0 | 0 |
| ENSMUSG00000081117 | 0 | 0 | 0 | 0 | 0 | 0 | 0 |
| ENSMUSG00000081118 | 0 | 0 | 0.25 | 0.396240625 | 0.489301084 | 0.301029996 | 0.355418052 |
| ENSMUSG00000081119 | 0 | 0 | 0 | 0 | 0 | 0 | 0 |
| ENSMUSG00000081120 | 0.289064826 | 0 | 0 | 0 | 0 | 0 | 0 |
| ENSMUSG00000081121 | 0 | 0.25 | 0 | 0.25 | 0 | 0 | 0 |
| ENSMUSG00000081122 | 0 | 0 | 0 | 0 | 0 | 0 | 0 |
| ENSMUSG00000081123 | 0 | 0 | 0.25 | 0 | 0 | 0 | 0 |
| ENSMUSG00000081124 | 0 | 0 | 0 | 0 | 0 | 0 | 0 |
| ENSMUSG00000081125 | 0 | 0 | 0 | 0 | 0 | 0 | 0 |
| ENSMUSG00000081126 | 0.811507563 | 1 | 0.396240625 | 0.5 | 0.931471446 | 0.954242509 | 1.100339912 |
| ENSMUSG00000081127 | 0 | 0 | 0 | 0 | 0 | 0.301029996 | 0 |
| ENSMUSG00000081128 | 0 | 0 | 0 | 0 | 0 | 0 | 0 |
| ENSMUSG00000081129 | 0 | 0 | 0 | 0 | 0 | 0 | 0 |
| ENSMUSG00000081130 | 0 | 0 | 0 | 0 | 0 | 0 | 0 |
| ENSMUSG00000081131 | 0 | 0 | 0 | 0 | 0 | 0 | 0 |
| ENSMUSG00000081132 | 0 | 0 | 0 | 0 | 0 | 0 | 0 |
| ENSMUSG00000081133 | 0 | 0 | 0 | 0 | 0 | 0 | 0 |
| ENSMUSG00000081134 | 0 | 0 | 0 | 0 | 0 | 0 | 0 |
| ENSMUSG00000081135 | 0.289064826 | 0.25 | 0 | 0.25 | 0 | 0 | 0 |

| | | | | | | | |
|---|---|---|---|---|---|---|---|
| ENSMUSG00000081136 | 0 | 0.25 | 0 | 0 | 0 | 0 | 0 |
| ENSMUSG00000081137 | 1.342375483 | 1.282320754 | 1.282320754 | 1.311981878 | 1.212046813 | 1.278753601 | 1.193409508 |
| ENSMUSG00000081138 | 0 | 0 | 0.25 | 0.25 | 0.244650542 | 0 | 0 |
| ENSMUSG00000081139 | 0.289064826 | 0 | 0 | 0.396240625 | 0.387761935 | 0 | 0.224243824 |
| ENSMUSG00000081141 | 0 | 0 | 0 | 0 | 0 | 0 | 0 |
| ENSMUSG00000081142 | 0.45815691 | 0.830482024 | 0.396240625 | 0.5 | 0.489301084 | 0.477121255 | 0.224243824 |
| ENSMUSG00000081143 | 0.289064826 | 0.25 | 0 | 0.25 | 0 | 0 | 0 |
| ENSMUSG00000081144 | 0 | 0 | 0 | 0 | 0 | 0 | 0 |
| ENSMUSG00000081145 | 0 | 0 | 0 | 0 | 0 | 0 | 0 |
| ENSMUSG00000081146 | 0 | 0 | 0 | 0 | 0 | 0 | 0 |
| ENSMUSG00000081147 | 0 | 0.25 | 0.25 | 0 | 0 | 0.301029996 | 0 |
| ENSMUSG00000081148 | 0 | 0 | 0 | 0 | 0 | 0 | 0 |
| ENSMUSG00000081150 | 0 | 0 | 0 | 0 | 0 | 0 | 0 |
| ENSMUSG00000081151 | 0 | 0 | 0.25 | 0.25 | 0 | 0 | 0 |
| ENSMUSG00000081152 | 0 | 0 | 0.25 | 0 | 0.244650542 | 0.301029996 | 0 |
| ENSMUSG00000081153 | 0 | 0 | 0 | 0 | 0 | 0.301029996 | 0.224243824 |
| ENSMUSG00000081154 | 0 | 0 | 0 | 0 | 0 | 0 | 0 |
| ENSMUSG00000081155 | 0 | 0 | 0 | 0 | 0 | 0 | 0 |
| ENSMUSG00000081156 | 0.289064826 | 0 | 0.25 | 0 | 0 | 0 | 0 |
| ENSMUSG00000081157 | 0 | 0 | 0.25 | 0 | 0 | 0 | 0 |
| ENSMUSG00000081158 | 0 | 0 | 0 | 0 | 0 | 0 | 0 |
| ENSMUSG00000081159 | 0 | 0 | 0 | 0 | 0 | 0 | 0 |
| ENSMUSG00000081160 | 0 | 0 | 0 | 0 | 0 | 0 | 0 |
| ENSMUSG00000081161 | 0 | 0 | 0 | 0.25 | 0 | 0.301029996 | 0.355418052 |
| ENSMUSG00000081162 | 0 | 0 | 0 | 0 | 0 | 0 | 0 |
| ENSMUSG00000081163 | 0 | 0 | 0 | 0 | 0 | 0 | 0 |
| ENSMUSG00000081164 | 0 | 0.25 | 0 | 0 | 0 | 0 | 0 |
| ENSMUSG00000081165 | 0 | 0 | 0 | 0 | 0 | 0 | 0 |
| ENSMUSG00000081166 | 0 | 0 | 0 | 0 | 0 | 0 | 0 |
| ENSMUSG00000081167 | 0 | 0.25 | 0 | 0 | 0.244650542 | 0 | 0 |
| ENSMUSG00000081168 | 0 | 0 | 0 | 0 | 0 | 0 | 0 |
| ENSMUSG00000081169 | 0 | 0 | 0 | 0.396240625 | 0 | 0 | 0 |
| ENSMUSG00000081170 | 0 | 0.396240625 | 0.396240625 | 0.25 | 0 | 0 | 0.448487648 |
| ENSMUSG00000081171 | 0 | 0 | 0 | 0 | 0 | 0 | 0 |

| | | | | | | | |
|---|---|---|---|---|---|---|---|
| ENSMUSG00000081172 | 0 | 0.25 | 0 | 0 | 0.244650542 | 0 | 0 |
| ENSMUSG00000081173 | 0 | 0 | 0 | 0 | 0 | 0 | 0 |
| ENSMUSG00000081174 | 0 | 0 | 0 | 0 | 0 | 0 | 0 |
| ENSMUSG00000081175 | 0 | 0 | 0 | 0 | 0 | 0 | 0 |
| ENSMUSG00000081176 | 0 | 0.25 | 0 | 0 | 0 | 0 | 0 |
| ENSMUSG00000081177 | 0 | 0 | 0 | 0 | 0 | 0 | 0 |
| ENSMUSG00000081179 | 0 | 0.25 | 0 | 0.25 | 0 | 0 | 0 |
| ENSMUSG00000081180 | 0 | 0 | 0 | 0 | 0 | 0 | 0 |
| ENSMUSG00000081181 | 0 | 0 | 0 | 0 | 0 | 0 | 0 |
| ENSMUSG00000081182 | 0.289064826 | 0 | 0 | 0 | 0.244650542 | 0 | 0 |
| ENSMUSG00000081183 | 0 | 0 | 0 | 0 | 0 | 0 | 0 |
| ENSMUSG00000081184 | 0 | 0 | 0 | 0 | 0 | 0 | 0 |
| ENSMUSG00000081185 | 0 | 0 | 0 | 0 | 0 | 0 | 0 |
| ENSMUSG00000081186 | 0 | 0 | 0 | 0 | 0 | 0 | 0 |
| ENSMUSG00000081187 | 0.289064826 | 0.5 | 0.396240625 | 0.5 | 0.686820904 | 0.698970004 | 0.224243824 |
| ENSMUSG00000081188 | 0 | 0 | 0 | 0 | 0 | 0 | 0 |
| ENSMUSG00000081189 | 0.578129653 | 0.79248125 | 0.580482024 | 0.701838731 | 0.632412477 | 0.77815125 | 0.520678036 |
| ENSMUSG00000081191 | 0 | 0.25 | 0.25 | 0 | 0 | 0 | 0 |
| ENSMUSG00000081194 | 0 | 0 | 0 | 0 | 0.244650542 | 0.301029996 | 0 |
| ENSMUSG00000081195 | 0 | 0 | 0 | 0 | 0 | 0 | 0 |
| ENSMUSG00000081196 | 0 | 0.25 | 0 | 0 | 0 | 0 | 0 |
| ENSMUSG00000081197 | 0 | 0 | 0 | 0 | 0 | 0 | 0 |
| ENSMUSG00000081199 | 0 | 0 | 0 | 0 | 0 | 0 | 0 |
| ENSMUSG00000081200 | 0 | 0 | 0 | 0 | 0 | 0 | 0 |
| ENSMUSG00000081201 | 0 | 0 | 0 | 0 | 0 | 0 | 0 |
| ENSMUSG00000081202 | 0 | 0 | 0 | 0 | 0 | 0 | 0 |
| ENSMUSG00000081205 | 0 | 0 | 0 | 0 | 0 | 0 | 0 |
| ENSMUSG00000081206 | 0 | 0.396240625 | 0 | 0 | 0 | 0 | 0 |
| ENSMUSG00000081207 | 0 | 0 | 0 | 0 | 0 | 0 | 0 |
| ENSMUSG00000081208 | 0.747221736 | 0.701838731 | 0.864857905 | 0.830482024 | 0.905314583 | 0.602059991 | 0.916588299 |
| ENSMUSG00000081209 | 0 | 0 | 0 | 0 | 0 | 0 | 0 |
| ENSMUSG00000081210 | 0.45815691 | 0.25 | 0.580482024 | 0.396240625 | 0.244650542 | 0.301029996 | 0.448487648 |
| ENSMUSG00000081211 | 0 | 0 | 0 | 0 | 0 | 0 | 0 |
| ENSMUSG00000081212 | 0 | 0 | 0 | 0 | 0 | 0 | 0 |

| | | | | | | | |
|---|---|---|---|---|---|---|---|
| ENSMUSG00000081214 | 0 | 0 | 0 | 0 | 0 | 0 | 0 |
| ENSMUSG00000081215 | 0 | 0 | 0 | 0 | 0 | 0 | 0 |
| ENSMUSG00000081216 | 0 | 0 | 0 | 0 | 0 | 0 | 0 |
| ENSMUSG00000081217 | 0.289064826 | 0.396240625 | 0.25 | 0.5 | 0.244650542 | 0 | 0 |
| ENSMUSG00000081218 | 0 | 0 | 0 | 0 | 0 | 0 | 0 |
| ENSMUSG00000081219 | 0.45815691 | 0.25 | 0.396240625 | 0.396240625 | 0.632412477 | 0.602059991 | 0.520678036 |
| ENSMUSG00000081220 | 0 | 0 | 0 | 0 | 0 | 0 | 0 |
| ENSMUSG00000081221 | 0 | 0 | 0.396240625 | 0.25 | 0 | 0 | 0 |
| ENSMUSG00000081222 | 0.289064826 | 0.25 | 0 | 0 | 0.387761935 | 0.477121255 | 0.355418052 |
| ENSMUSG00000081223 | 0.45815691 | 0.25 | 0.396240625 | 0.25 | 0.244650542 | 0 | 0.224243824 |
| ENSMUSG00000081224 | 0 | 0 | 0 | 0 | 0 | 0 | 0 |
| ENSMUSG00000081225 | 0 | 0 | 0 | 0 | 0 | 0 | 0 |
| ENSMUSG00000081226 | 0 | 0 | 0.25 | 0 | 0 | 0 | 0 |
| ENSMUSG00000081227 | 0 | 0 | 0 | 0.25 | 0.244650542 | 0 | 0.224243824 |
| ENSMUSG00000081228 | 0 | 0.25 | 0 | 0 | 0.387761935 | 0 | 0 |
| ENSMUSG00000081229 | 0 | 0 | 0 | 0 | 0 | 0 | 0 |
| ENSMUSG00000081230 | 0 | 0 | 0 | 0 | 0 | 0 | 0 |
| ENSMUSG00000081231 | 0 | 0 | 0 | 0 | 0 | 0 | 0 |
| ENSMUSG00000081232 | 0 | 0.396240625 | 0.396240625 | 0.25 | 0 | 0 | 0 |
| ENSMUSG00000081233 | 0 | 0 | 0 | 0 | 0 | 0 | 0 |
| ENSMUSG00000081234 | 0 | 0 | 0 | 0 | 0 | 0 | 0 |
| ENSMUSG00000081235 | 0 | 0 | 0 | 0 | 0 | 0 | 0.224243824 |
| ENSMUSG00000081236 | 0 | 0 | 0 | 0 | 0 | 0 | 0 |
| ENSMUSG00000081237 | 0 | 0 | 0 | 0 | 0 | 0 | 0 |
| ENSMUSG00000081238 | 0 | 0 | 0.25 | 0 | 0.387761935 | 0 | 0.224243824 |
| ENSMUSG00000081239 | 0.671187741 | 0.701838731 | 0.25 | 0.25 | 0.387761935 | 0.477121255 | 0.710836105 |
| ENSMUSG00000081240 | 0 | 0 | 0 | 0 | 0 | 0 | 0 |
| ENSMUSG00000081242 | 0 | 0 | 0 | 0 | 0.244650542 | 0 | 0 |
| ENSMUSG00000081243 | 0 | 0 | 0.25 | 0.25 | 0 | 0 | 0 |
| ENSMUSG00000081244 | 0.91631382 | 0.646240625 | 0.79248125 | 0.5 | 0.568060967 | 0.77815125 | 0.629532004 |
| ENSMUSG00000081245 | 0 | 0 | 0 | 0 | 0 | 0 | 0 |
| ENSMUSG00000081248 | 0 | 0 | 0 | 0 | 0 | 0 | 0 |
| ENSMUSG00000081249 | 0.578129653 | 0.75 | 0.396240625 | 0.5 | 0.733951626 | 0.301029996 | 0.520678036 |
| ENSMUSG00000081250 | 0.289064826 | 0 | 0 | 0 | 0 | 0.301029996 | 0 |

| | | | | | | | |
|---|---|---|---|---|---|---|---|
| ENSMUSG00000081251 | 1 | 0.79248125 | 0.896240625 | 0 | 0 | 0.301029996 | 0.710836105 |
| ENSMUSG00000081252 | 0 | 0 | 0 | 0 | 0 | 0 | 0 |
| ENSMUSG00000081253 | 0 | 0 | 0 | 0 | 0 | 0 | 0 |
| ENSMUSG00000081254 | 0 | 0 | 0 | 0 | 0 | 0 | 0 |
| ENSMUSG00000081255 | 0.289064826 | 0.5 | 0.396240625 | 0.25 | 0.244650542 | 0.477121255 | 0.224243824 |
| ENSMUSG00000081256 | 0 | 0 | 0 | 0.25 | 0.244650542 | 0 | 0 |
| ENSMUSG00000081257 | 0 | 0 | 0 | 0 | 0 | 0 | 0 |
| ENSMUSG00000081258 | 0 | 0 | 0 | 0 | 0 | 0 | 0 |
| ENSMUSG00000081259 | 0 | 0 | 0 | 0 | 0 | 0 | 0 |
| ENSMUSG00000081260 | 0 | 0 | 0 | 0 | 0 | 0 | 0 |
| ENSMUSG00000081261 | 0 | 0 | 0 | 0 | 0 | 0 | 0 |
| ENSMUSG00000081262 | 0 | 0 | 0.25 | 0.25 | 0 | 0 | 0 |
| ENSMUSG00000081263 | 0 | 0 | 0 | 0 | 0 | 0 | 0 |
| ENSMUSG00000081264 | 0 | 0 | 0 | 0 | 0 | 0 | 0 |
| ENSMUSG00000081265 | 0 | 0.580482024 | 0.25 | 0.25 | 0 | 0 | 0.224243824 |
| ENSMUSG00000081268 | 0 | 0 | 0 | 0 | 0 | 0 | 0 |
| ENSMUSG00000081269 | 0 | 0 | 0 | 0 | 0 | 0.301029996 | 0.224243824 |
| ENSMUSG00000081270 | 0 | 0 | 0 | 0 | 0 | 0 | 0 |
| ENSMUSG00000081271 | 0 | 0 | 0 | 0 | 0 | 0 | 0 |
| ENSMUSG00000081272 | 0 | 0.25 | 0.25 | 0.25 | 0 | 0 | 0 |
| ENSMUSG00000081273 | 0 | 0 | 0 | 0 | 0 | 0 | 0 |
| ENSMUSG00000081274 | 0 | 0.396240625 | 0.396240625 | 0.25 | 0.244650542 | 0.477121255 | 0.224243824 |
| ENSMUSG00000081275 | 0 | 0 | 0 | 0 | 0 | 0 | 0 |
| ENSMUSG00000081276 | 0 | 0 | 0.396240625 | 0.5 | 0.244650542 | 0 | 0.448487648 |
| ENSMUSG00000081277 | 0.289064826 | 0 | 0 | 0.5 | 0.244650542 | 0.477121255 | 0 |
| ENSMUSG00000081278 | 0 | 0 | 0 | 0 | 0 | 0 | 0 |
| ENSMUSG00000081279 | 0 | 0 | 0 | 0 | 0 | 0 | 0 |
| ENSMUSG00000081280 | 0 | 0 | 0.25 | 0 | 0 | 0 | 0 |
| ENSMUSG00000081281 | 0 | 0.25 | 0 | 0.5 | 0.387761935 | 0.301029996 | 0.224243824 |
| ENSMUSG00000081282 | 0 | 0 | 0 | 0 | 0 | 0 | 0 |
| ENSMUSG00000081283 | 0 | 0 | 0 | 0 | 0 | 0 | 0 |
| ENSMUSG00000081284 | 0 | 0 | 0 | 0 | 0 | 0 | 0 |
| ENSMUSG00000081285 | 0 | 0.396240625 | 0 | 0.25 | 0.387761935 | 0.301029996 | 0 |
| ENSMUSG00000081286 | 0 | 0 | 0 | 0 | 0 | 0 | 0 |

| | | | | | | | |
|---|---|---|---|---|---|---|---|
| ENSMUSG00000081287 | 0 | 0 | 0 | 0 | 0 | 0 | 0 |
| ENSMUSG00000081288 | 0 | 0 | 0 | 0 | 0 | 0 | 0 |
| ENSMUSG00000081289 | 0 | 0.396240625 | 0 | 0.396240625 | 0.244650542 | 0 | 0 |
| ENSMUSG00000081290 | 0 | 0 | 0 | 0 | 0 | 0 | 0 |
| ENSMUSG00000081292 | 0 | 0 | 0 | 0 | 0 | 0 | 0 |
| ENSMUSG00000081293 | 0 | 0 | 0 | 0 | 0 | 0 | 0 |
| ENSMUSG00000081295 | 0 | 0.25 | 0 | 0 | 0 | 0 | 0 |
| ENSMUSG00000081296 | 0 | 0 | 0 | 0 | 0 | 0 | 0 |
| ENSMUSG00000081297 | 0 | 0 | 0 | 0 | 0 | 0 | 0 |
| ENSMUSG00000081298 | 0 | 0 | 0 | 0 | 0 | 0 | 0 |
| ENSMUSG00000081299 | 0 | 0 | 0 | 0 | 0 | 0 | 0 |
| ENSMUSG00000081300 | 0 | 0 | 0 | 0 | 0 | 0 | 0 |
| ENSMUSG00000081301 | 0 | 0 | 0 | 0.5 | 0.244650542 | 0 | 0.224243824 |
| ENSMUSG00000081302 | 0.289064826 | 0.646240625 | 0.396240625 | 0.5 | 0.387761935 | 0.301029996 | 0.629532004 |
| ENSMUSG00000081303 | 0 | 0 | 0 | 0 | 0 | 0 | 0 |
| ENSMUSG00000081304 | 0 | 0 | 0 | 0 | 0 | 0 | 0 |
| ENSMUSG00000081305 | 0.578129653 | 0.396240625 | 0.5 | 0.580482024 | 0.244650542 | 0.301029996 | 0.629532004 |
| ENSMUSG00000081306 | 0 | 0 | 0 | 0 | 0 | 0 | 0 |
| ENSMUSG00000081307 | 0 | 0 | 0 | 0 | 0 | 0 | 0 |
| ENSMUSG00000081308 | 0 | 0.25 | 0 | 0.25 | 0 | 0.301029996 | 0.355418052 |
| ENSMUSG00000081309 | 0 | 0 | 0 | 0 | 0 | 0 | 0 |
| ENSMUSG00000081310 | 0 | 0 | 0 | 0 | 0 | 0 | 0 |
| ENSMUSG00000081312 | 0 | 0 | 0 | 0 | 0 | 0 | 0 |
| ENSMUSG00000081313 | 0 | 0 | 0 | 0 | 0 | 0 | 0 |
| ENSMUSG00000081314 | 0 | 0.396240625 | 0.396240625 | 0.25 | 0 | 0 | 0 |
| ENSMUSG00000081315 | 0 | 0 | 0 | 0 | 0 | 0 | 0 |
| ENSMUSG00000081316 | 0 | 0.25 | 0 | 0 | 0 | 0 | 0 |
| ENSMUSG00000081317 | 0.289064826 | 0.396240625 | 0 | 0 | 0 | 0 | 0 |
| ENSMUSG00000081318 | 0 | 0 | 0 | 0 | 0 | 0 | 0 |
| ENSMUSG00000081319 | 0 | 0.25 | 0.25 | 0 | 0 | 0 | 0.355418052 |
| ENSMUSG00000081320 | 0 | 0 | 0 | 0 | 0 | 0 | 0 |
| ENSMUSG00000081321 | 0 | 0.25 | 0 | 0.580482024 | 0.489301084 | 0 | 0.224243824 |
| ENSMUSG00000081323 | 0 | 0 | 0 | 0 | 0 | 0 | 0 |
| ENSMUSG00000081324 | 0 | 0 | 0 | 0 | 0 | 0 | 0 |

| | | | | | | | |
|---|---|---|---|---|---|---|---|
| ENSMUSG00000081325 | 0 | 0 | 0 | 0 | 0 | 0 | 0 |
| ENSMUSG00000081326 | 0 | 0 | 0 | 0 | 0 | 0 | 0 |
| ENSMUSG00000081327 | 0 | 0 | 0 | 0 | 0.244650542 | 0 | 0 |
| ENSMUSG00000081328 | 0 | 0 | 0 | 0 | 0 | 0 | 0 |
| ENSMUSG00000081329 | 0.289064826 | 0.5 | 0.25 | 0.864857905 | 0.244650542 | 0.698970004 | 0.448487648 |
| ENSMUSG00000081330 | 0 | 0 | 0 | 0 | 0 | 0 | 0 |
| ENSMUSG00000081331 | 0 | 0 | 0 | 0 | 0 | 0 | 0 |
| ENSMUSG00000081332 | 0 | 0 | 0 | 0 | 0 | 0.301029996 | 0 |
| ENSMUSG00000081333 | 0 | 0 | 0 | 0 | 0 | 0 | 0 |
| ENSMUSG00000081334 | 0 | 0 | 0 | 0 | 0 | 0 | 0 |
| ENSMUSG00000081335 | 0 | 0 | 0 | 0 | 0 | 0 | 0 |
| ENSMUSG00000081336 | 0 | 0 | 0 | 0 | 0 | 0 | 0 |
| ENSMUSG00000081339 | 0.289064826 | 0 | 0 | 0.25 | 0 | 0.301029996 | 0 |
| ENSMUSG00000081340 | 0 | 0 | 0 | 0.25 | 0 | 0 | 0.448487648 |
| ENSMUSG00000081341 | 0 | 0 | 0 | 0 | 0 | 0 | 0.224243824 |
| ENSMUSG00000081342 | 0 | 0 | 0 | 0 | 0 | 0 | 0 |
| ENSMUSG00000081343 | 0 | 0 | 0 | 0 | 0.244650542 | 0 | 0 |
| ENSMUSG00000081344 | 0 | 0 | 0 | 0 | 0 | 0 | 0 |
| ENSMUSG00000081345 | 0 | 0 | 0 | 0 | 0 | 0 | 0 |
| ENSMUSG00000081346 | 0 | 0 | 0 | 0 | 0 | 0 | 0 |
| ENSMUSG00000081348 | 0 | 0 | 0 | 0 | 0 | 0 | 0 |
| ENSMUSG00000081349 | 0 | 0 | 0 | 0 | 0 | 0 | 0 |
| ENSMUSG00000081350 | 0.45815691 | 0.701838731 | 1.061981878 | 0.396240625 | 0.568060967 | 0.477121255 | 0.579661877 |
| ENSMUSG00000081352 | 0 | 0 | 0 | 0 | 0 | 0 | 0 |
| ENSMUSG00000081353 | 0 | 0 | 0 | 0 | 0 | 0 | 0 |
| ENSMUSG00000081354 | 0 | 0 | 0 | 0 | 0 | 0 | 0 |
| ENSMUSG00000081355 | 0.867194479 | 1.080482024 | 1.098079356 | 0.92510993 | 0.77552387 | 0.903089987 | 0.775756176 |
| ENSMUSG00000081357 | 0 | 0 | 0 | 0 | 0 | 0 | 0 |
| ENSMUSG00000081358 | 0 | 0.396240625 | 0 | 0 | 0 | 0 | 0 |
| ENSMUSG00000081359 | 0 | 0.25 | 0 | 0 | 0 | 0 | 0.224243824 |
| ENSMUSG00000081360 | 0 | 0 | 0 | 0.396240625 | 0.387761935 | 0 | 0 |
| ENSMUSG00000081361 | 0 | 0 | 0 | 0 | 0 | 0 | 0 |
| ENSMUSG00000081362 | 0 | 0 | 0 | 0 | 0 | 0 | 0 |
| ENSMUSG00000081363 | 0 | 0 | 0 | 0 | 0 | 0 | 0 |

| | | | | | | | |
|---|---|---|---|---|---|---|---|
| ENSMUSG00000081364 | 0 | 0 | 0 | 0 | 0 | 0 | 0 |
| ENSMUSG00000081365 | 0 | 0 | 0 | 0 | 0 | 0 | 0 |
| ENSMUSG00000081367 | 0 | 0 | 0 | 0 | 0 | 0 | 0 |
| ENSMUSG00000081369 | 0 | 0 | 0 | 0 | 0 | 0 | 0 |
| ENSMUSG00000081370 | 0 | 0 | 0 | 0 | 0 | 0 | 0 |
| ENSMUSG00000081372 | 0 | 0 | 0 | 0 | 0 | 0 | 0 |
| ENSMUSG00000081373 | 0 | 0.396240625 | 0.5 | 0.396240625 | 0.387761935 | 0.301029996 | 0.629532004 |
| ENSMUSG00000081374 | 0 | 0 | 0 | 0 | 0 | 0 | 0 |
| ENSMUSG00000081375 | 0 | 0 | 0 | 0 | 0 | 0 | 0 |
| ENSMUSG00000081376 | 0 | 0 | 0 | 0.396240625 | 0.244650542 | 0.301029996 | 0 |
| ENSMUSG00000081377 | 0 | 0 | 0 | 0 | 0 | 0 | 0 |
| ENSMUSG00000081378 | 0.289064826 | 0 | 0 | 0 | 0 | 0 | 0.224243824 |
| ENSMUSG00000081379 | 0 | 0 | 0 | 0 | 0 | 0 | 0 |
| ENSMUSG00000081381 | 0 | 0 | 0 | 0 | 0.244650542 | 0 | 0 |
| ENSMUSG00000081382 | 0.289064826 | 0 | 0 | 0 | 0.387761935 | 0 | 0.224243824 |
| ENSMUSG00000081383 | 0 | 0 | 0 | 0 | 0.244650542 | 0 | 0 |
| ENSMUSG00000081384 | 0 | 0 | 0 | 0 | 0 | 0 | 0 |
| ENSMUSG00000081386 | 0 | 0 | 0 | 0 | 0 | 0 | 0 |
| ENSMUSG00000081388 | 0 | 0 | 0 | 0 | 0 | 0 | 0 |
| ENSMUSG00000081389 | 0 | 0 | 0 | 0 | 0 | 0 | 0 |
| ENSMUSG00000081390 | 0 | 0 | 0 | 0 | 0 | 0 | 0 |
| ENSMUSG00000081391 | 0.289064826 | 0 | 0 | 0 | 0 | 0 | 0 |
| ENSMUSG00000081392 | 0 | 0.25 | 0 | 0 | 0 | 0 | 0 |
| ENSMUSG00000081394 | 0 | 0.396240625 | 0 | 0.396240625 | 0.489301084 | 0 | 0.224243824 |
| ENSMUSG00000081395 | 0 | 0 | 0 | 0 | 0 | 0 | 0 |
| ENSMUSG00000081396 | 0 | 0 | 0 | 0 | 0 | 0 | 0 |
| ENSMUSG00000081397 | 0 | 0 | 0 | 0 | 0 | 0 | 0 |
| ENSMUSG00000081398 | 0 | 0 | 0 | 0 | 0 | 0 | 0 |
| ENSMUSG00000081399 | 0 | 0 | 0 | 0 | 0 | 0 | 0 |
| ENSMUSG00000081400 | 0.671187741 | 0.5 | 0.830482024 | 0.75 | 0.846351821 | 0.602059991 | 0.803905701 |
| ENSMUSG00000081402 | 0 | 0 | 0.25 | 0 | 0.387761935 | 0.301029996 | 0 |
| ENSMUSG00000081404 | 0.578129653 | 0.75 | 0.396240625 | 0.580482024 | 0.77552387 | 0.602059991 | 0.448487648 |
| ENSMUSG00000081405 | 0 | 0 | 0 | 0 | 0 | 0 | 0 |
| ENSMUSG00000081406 | 0 | 0 | 0 | 0 | 0 | 0 | 0 |

| | | | | | | | |
|---|---|---|---|---|---|---|---|
| ENSMUSG00000081407 | 0 | 0 | 0 | 0 | 0 | 0 | 0 |
| ENSMUSG00000081408 | 0 | 0 | 0 | 0 | 0 | 0 | 0 |
| ENSMUSG00000081409 | 0 | 0 | 0 | 0 | 0 | 0 | 0 |
| ENSMUSG00000081410 | 0 | 0 | 0 | 0 | 0 | 0 | 0 |
| ENSMUSG00000081412 | 0 | 0 | 0 | 0 | 0.244650542 | 0 | 0 |
| ENSMUSG00000081413 | 0 | 0 | 0 | 0 | 0 | 0 | 0 |
| ENSMUSG00000081414 | 0 | 0 | 0 | 0 | 0 | 0 | 0 |
| ENSMUSG00000081415 | 0 | 0 | 0 | 0 | 0 | 0 | 0 |
| ENSMUSG00000081416 | 0 | 0 | 0 | 0.25 | 0.244650542 | 0 | 0 |
| ENSMUSG00000081417 | 0 | 0.5 | 0.25 | 0 | 0 | 0 | 0 |
| ENSMUSG00000081418 | 0 | 0 | 0.396240625 | 0 | 0.244650542 | 0 | 0.448487648 |
| ENSMUSG00000081420 | 0 | 0 | 0 | 0 | 0 | 0 | 0 |
| ENSMUSG00000081421 | 0 | 0.396240625 | 0 | 0 | 0 | 0 | 0 |
| ENSMUSG00000081422 | 0 | 0 | 0 | 0 | 0 | 0 | 0 |
| ENSMUSG00000081423 | 0 | 0 | 0.25 | 0 | 0 | 0.301029996 | 0 |
| ENSMUSG00000081424 | 0 | 0 | 0 | 0 | 0 | 0 | 0 |
| ENSMUSG00000081426 | 0 | 0 | 0 | 0 | 0 | 0 | 0 |
| ENSMUSG00000081427 | 0 | 0 | 0 | 0 | 0 | 0 | 0 |
| ENSMUSG00000081428 | 0 | 0.25 | 0 | 0 | 0.244650542 | 0 | 0.224243824 |
| ENSMUSG00000081429 | 0 | 0 | 0 | 0 | 0.244650542 | 0.301029996 | 0 |
| ENSMUSG00000081431 | 0.289064826 | 0 | 0 | 0 | 0 | 0 | 0 |
| ENSMUSG00000081432 | 0.289064826 | 0.25 | 0 | 0 | 0 | 0 | 0 |
| ENSMUSG00000081433 | 0 | 0 | 0 | 0 | 0 | 0 | 0 |
| ENSMUSG00000081434 | 2.116511873 | 1.995748394 | 2.034887838 | 1.779735268 | 1.862942891 | 1.954242509 | 1.809975587 |
| ENSMUSG00000081435 | 0 | 0 | 0 | 0 | 0 | 0 | 0 |
| ENSMUSG00000081436 | 0.289064826 | 0 | 0 | 0 | 0 | 0 | 0 |
| ENSMUSG00000081437 | 0 | 0 | 0 | 0 | 0 | 0 | 0 |
| ENSMUSG00000081438 | 0 | 0 | 0 | 0 | 0 | 0 | 0 |
| ENSMUSG00000081439 | 0 | 0 | 0 | 0 | 0 | 0 | 0 |
| ENSMUSG00000081440 | 0 | 0 | 0 | 0 | 0 | 0 | 0 |
| ENSMUSG00000081441 | 0 | 0 | 0 | 0.25 | 0 | 0 | 0 |
| ENSMUSG00000081442 | 0 | 0 | 0 | 0 | 0 | 0.301029996 | 0 |
| ENSMUSG00000081443 | 0 | 0 | 0 | 0 | 0.244650542 | 0 | 0 |
| ENSMUSG00000081444 | 0 | 0 | 0 | 0 | 0 | 0 | 0.224243824 |

| | | | | | | | |
|---|---|---|---|---|---|---|---|
| ENSMUSG00000081445 | 0 | 0 | 0 | 0 | 0 | 0 | 0.224243824 |
| ENSMUSG00000081447 | 0 | 0 | 0.396240625 | 0 | 0.244650542 | 0 | 0.224243824 |
| ENSMUSG00000081449 | 0 | 0 | 0 | 0 | 0 | 0 | 0 |
| ENSMUSG00000081450 | 0 | 0 | 0 | 0 | 0 | 0 | 0 |
| ENSMUSG00000081451 | 0 | 0 | 0 | 0 | 0 | 0 | 0.224243824 |
| ENSMUSG00000081452 | 0 | 0 | 0 | 0 | 0 | 0 | 0 |
| ENSMUSG00000081453 | 0.960252568 | 1.061981878 | 0.864857905 | 0.976722649 | 1.039257769 | 0.84509804 | 0.710836105 |
| ENSMUSG00000081455 | 0.289064826 | 0.25 | 0.396240625 | 0 | 0.387761935 | 0.301029996 | 0.355418052 |
| ENSMUSG00000081456 | 0.289064826 | 0 | 0.396240625 | 0.25 | 0.387761935 | 0 | 0.355418052 |
| ENSMUSG00000081457 | 0 | 0 | 0 | 0 | 0 | 0 | 0 |
| ENSMUSG00000081458 | 0 | 0 | 0 | 0 | 0 | 0 | 0 |
| ENSMUSG00000081459 | 0 | 0.25 | 0.25 | 0 | 0 | 0.301029996 | 0.224243824 |
| ENSMUSG00000081460 | 0 | 0 | 0 | 0 | 0 | 0 | 0 |
| ENSMUSG00000081461 | 0.578129653 | 0 | 0.25 | 0 | 0.387761935 | 0 | 0 |
| ENSMUSG00000081463 | 0 | 0 | 0 | 0 | 0 | 0 | 0 |
| ENSMUSG00000081464 | 0 | 0 | 0 | 0 | 0 | 0 | 0 |
| ENSMUSG00000081465 | 0 | 0 | 0 | 0 | 0 | 0 | 0 |
| ENSMUSG00000081466 | 0 | 0 | 0.25 | 0 | 0 | 0 | 0 |
| ENSMUSG00000081467 | 2.103271637 | 1.918106335 | 1.918106335 | 1.630890489 | 1.695608647 | 1.86332286 | 1.659601507 |
| ENSMUSG00000081469 | 0 | 0 | 0 | 0 | 0 | 0 | 0 |
| ENSMUSG00000081470 | 0 | 0 | 0 | 0 | 0 | 0 | 0 |
| ENSMUSG00000081471 | 1.100572389 | 1.114857905 | 0.976722649 | 1.188721876 | 1.106691885 | 1.041392685 | 1 |
| ENSMUSG00000081473 | 0 | 0 | 0 | 0.25 | 0.244650542 | 0.301029996 | 0 |
| ENSMUSG00000081474 | 0 | 0 | 0 | 0 | 0 | 0 | 0 |
| ENSMUSG00000081475 | 0 | 0 | 0 | 0 | 0 | 0 | 0 |
| ENSMUSG00000081476 | 1.181541736 | 1.201838731 | 1.302363341 | 1.348079356 | 1.387761935 | 1.477121255 | 1.345462945 |
| ENSMUSG00000081477 | 0 | 0 | 0 | 0 | 0 | 0 | 0 |
| ENSMUSG00000081478 | 0 | 0 | 0 | 0.25 | 0 | 0 | 0 |
| ENSMUSG00000081480 | 0.289064826 | 0 | 0 | 0 | 0 | 0 | 0 |
| ENSMUSG00000081481 | 0 | 0 | 0 | 0 | 0 | 0 | 0 |
| ENSMUSG00000081483 | 0.289064826 | 0 | 0.25 | 0.580482024 | 0.244650542 | 0.698970004 | 0.520678036 |
| ENSMUSG00000081484 | 0 | 0 | 0 | 0 | 0 | 0 | 0 |
| ENSMUSG00000081485 | 0 | 0 | 0 | 0 | 0 | 0 | 0 |
| ENSMUSG00000081486 | 0 | 0 | 0 | 0 | 0 | 0 | 0 |

| | | | | | | | |
|---|---|---|---|---|---|---|---|
| ENSMUSG00000081487 | 0.289064826 | 0 | 0 | 0.25 | 0.244650542 | 0 | 0.224243824 |
| ENSMUSG00000081488 | 0 | 0 | 0 | 0 | 0 | 0 | 0 |
| ENSMUSG00000081490 | 0 | 0.25 | 0 | 0.25 | 0 | 0 | 0 |
| ENSMUSG00000081491 | 0 | 0 | 0.396240625 | 0 | 0 | 0 | 0.224243824 |
| ENSMUSG00000081492 | 0.289064826 | 0.25 | 0 | 0.396240625 | 0 | 0 | 0 |
| ENSMUSG00000081494 | 0.289064826 | 0 | 0.396240625 | 0.25 | 0.387761935 | 0 | 0 |
| ENSMUSG00000081495 | 0 | 0 | 0 | 0 | 0 | 0 | 0 |
| ENSMUSG00000081496 | 0 | 0 | 0 | 0 | 0 | 0 | 0 |
| ENSMUSG00000081497 | 0 | 0.25 | 0 | 0 | 0 | 0 | 0 |
| ENSMUSG00000081498 | 0 | 0 | 0 | 0 | 0 | 0 | 0 |
| ENSMUSG00000081499 | 0.289064826 | 0 | 0 | 0.25 | 0 | 0 | 0 |
| ENSMUSG00000081500 | 0 | 0 | 0 | 0 | 0 | 0 | 0 |
| ENSMUSG00000081501 | 0 | 0 | 0 | 0 | 0 | 0 | 0 |
| ENSMUSG00000081502 | 0 | 0 | 0 | 0 | 0 | 0 | 0 |
| ENSMUSG00000081504 | 0 | 0 | 0 | 0 | 0 | 0 | 0 |
| ENSMUSG00000081505 | 0 | 0 | 0 | 0 | 0 | 0 | 0 |
| ENSMUSG00000081506 | 0 | 0 | 0 | 0 | 0 | 0 | 0 |
| ENSMUSG00000081507 | 0 | 0 | 0 | 0 | 0 | 0 | 0 |
| ENSMUSG00000081508 | 0 | 0 | 0 | 0 | 0 | 0 | 0 |
| ENSMUSG00000081510 | 0 | 0 | 0 | 0 | 0 | 0 | 0 |
| ENSMUSG00000081511 | 0 | 0 | 0 | 0 | 0 | 0.301029996 | 0 |
| ENSMUSG00000081512 | 0.289064826 | 0 | 0.396240625 | 0 | 0.244650542 | 0.301029996 | 0 |
| ENSMUSG00000081513 | 0 | 0 | 0 | 0 | 0 | 0 | 0 |
| ENSMUSG00000081514 | 0 | 0 | 0 | 0 | 0 | 0 | 0 |
| ENSMUSG00000081515 | 0 | 0 | 0 | 0 | 0 | 0 | 0 |
| ENSMUSG00000081516 | 0.289064826 | 0 | 0.25 | 0.25 | 0 | 0 | 0 |
| ENSMUSG00000081517 | 0 | 0 | 0.396240625 | 0 | 0 | 0 | 0 |
| ENSMUSG00000081519 | 0 | 0 | 0 | 0 | 0 | 0 | 0 |
| ENSMUSG00000081520 | 0.289064826 | 0 | 0.396240625 | 0 | 0 | 0 | 0 |
| ENSMUSG00000081522 | 0 | 0 | 0 | 0 | 0 | 0 | 0 |
| ENSMUSG00000081523 | 0 | 0 | 0 | 0 | 0 | 0 | 0 |
| ENSMUSG00000081524 | 0 | 0.25 | 0 | 0 | 0 | 0 | 0 |
| ENSMUSG00000081525 | 0 | 0 | 0 | 0 | 0 | 0 | 0 |
| ENSMUSG00000081526 | 0 | 0 | 0 | 0 | 0 | 0 | 0 |

| | | | | | | | |
|---|---|---|---|---|---|---|---|
| ENSMUSG00000081528 | 0 | 0 | 0 | 0 | 0 | 0 | 0 |
| ENSMUSG00000081529 | 0 | 0 | 0 | 0 | 0 | 0 | 0 |
| ENSMUSG00000081530 | 0 | 0 | 0 | 0 | 0 | 0 | 0 |
| ENSMUSG00000081531 | 0 | 0 | 0 | 0 | 0 | 0 | 0 |
| ENSMUSG00000081532 | 0 | 0 | 0 | 0 | 0 | 0 | 0 |
| ENSMUSG00000081533 | 0 | 0 | 0 | 0 | 0 | 0 | 0 |
| ENSMUSG00000081534 | 2.649252229 | 2.420748646 | 2.418989258 | 2.390560606 | 2.376595432 | 2.769377326 | 2.249624299 |
| ENSMUSG00000081536 | 0 | 0 | 0 | 0 | 0 | 0 | 0 |
| ENSMUSG00000081537 | 0.289064826 | 0 | 0 | 0 | 0.244650542 | 0.301029996 | 0 |
| ENSMUSG00000081538 | 0 | 0 | 0 | 0 | 0 | 0 | 0 |
| ENSMUSG00000081539 | 0 | 0.25 | 0 | 0 | 0 | 0 | 0 |
| ENSMUSG00000081540 | 0 | 0 | 0 | 0 | 0 | 0 | 0 |
| ENSMUSG00000081541 | 0 | 0 | 0 | 0 | 0 | 0 | 0 |
| ENSMUSG00000081542 | 0 | 0 | 0 | 0 | 0 | 0 | 0 |
| ENSMUSG00000081543 | 0.289064826 | 0.25 | 0 | 0 | 0 | 0.477121255 | 0.355418052 |
| ENSMUSG00000081544 | 0 | 0 | 0 | 0 | 0 | 0 | 0 |
| ENSMUSG00000081546 | 0 | 0 | 0 | 0 | 0 | 0 | 0 |
| ENSMUSG00000081547 | 0 | 0 | 0 | 0 | 0 | 0 | 0 |
| ENSMUSG00000081548 | 0 | 0 | 0 | 0 | 0 | 0 | 0 |
| ENSMUSG00000081550 | 0.45815691 | 0.5 | 0.396240625 | 0.396240625 | 0 | 0.301029996 | 0.224243824 |
| ENSMUSG00000081551 | 0.671187741 | 0.396240625 | 0.701838731 | 0.646240625 | 0.489301084 | 0.602059991 | 0.775756176 |
| ENSMUSG00000081552 | 0.45815691 | 0.396240625 | 0.25 | 0 | 0 | 0.477121255 | 0.355418052 |
| ENSMUSG00000081553 | 0.45815691 | 0 | 0 | 0 | 0 | 0.301029996 | 0 |
| ENSMUSG00000081555 | 0 | 0 | 0 | 0 | 0 | 0 | 0 |
| ENSMUSG00000081556 | 0 | 0 | 0 | 0 | 0 | 0 | 0.224243824 |
| ENSMUSG00000081557 | 0.578129653 | 0.25 | 0.5 | 0 | 0.489301084 | 0 | 0.629532004 |
| ENSMUSG00000081559 | 0 | 0 | 0 | 0 | 0 | 0 | 0 |
| ENSMUSG00000081560 | 0 | 0 | 0 | 0 | 0 | 0 | 0 |
| ENSMUSG00000081561 | 0 | 0 | 0 | 0 | 0 | 0 | 0 |
| ENSMUSG00000081562 | 0 | 0 | 0 | 0.25 | 0 | 0 | 0 |
| ENSMUSG00000081563 | 0 | 0 | 0 | 0 | 0 | 0 | 0 |
| ENSMUSG00000081564 | 0.578129653 | 0.580482024 | 0.580482024 | 0.396240625 | 0.387761935 | 0 | 0.520678036 |
| ENSMUSG00000081565 | 0 | 0 | 0.396240625 | 0.25 | 0.489301084 | 0 | 0 |
| ENSMUSG00000081566 | 0 | 0 | 0 | 0 | 0 | 0 | 0 |

| | | | | | | | |
|---|---|---|---|---|---|---|---|
| ENSMUSG00000081567 | 0 | 0 | 0 | 0 | 0 | 0 | 0 |
| ENSMUSG00000081568 | 0 | 0.25 | 0.396240625 | 0 | 0.387761935 | 0 | 0.448487648 |
| ENSMUSG00000081569 | 0 | 0 | 0 | 0 | 0 | 0 | 0 |
| ENSMUSG00000081570 | 0 | 0 | 0 | 0.25 | 0 | 0 | 0 |
| ENSMUSG00000081571 | 0 | 0 | 0 | 0 | 0 | 0 | 0 |
| ENSMUSG00000081572 | 0 | 0 | 0 | 0 | 0.244650542 | 0.477121255 | 0 |
| ENSMUSG00000081573 | 0 | 0 | 0 | 0 | 0 | 0 | 0 |
| ENSMUSG00000081574 | 0.289064826 | 0 | 0 | 0 | 0 | 0.602059991 | 0.224243824 |
| ENSMUSG00000081575 | 0.289064826 | 0 | 0.25 | 0.25 | 0.387761935 | 0.301029996 | 0 |
| ENSMUSG00000081576 | 0 | 0 | 0 | 0 | 0 | 0 | 0 |
| ENSMUSG00000081577 | 0 | 0 | 0 | 0 | 0 | 0 | 0 |
| ENSMUSG00000081578 | 0 | 0 | 0 | 0.25 | 0 | 0 | 0 |
| ENSMUSG00000081579 | 0 | 0 | 0 | 0 | 0 | 0 | 0 |
| ENSMUSG00000081580 | 0 | 0 | 0 | 0 | 0 | 0 | 0 |
| ENSMUSG00000081583 | 0 | 0 | 0 | 0 | 0 | 0 | 0 |
| ENSMUSG00000081584 | 0 | 0 | 0 | 0 | 0 | 0 | 0 |
| ENSMUSG00000081585 | 0 | 0 | 0 | 0 | 0 | 0 | 0 |
| ENSMUSG00000081586 | 0 | 0 | 0 | 0 | 0 | 0 | 0 |
| ENSMUSG00000081587 | 0 | 0 | 0.25 | 0.25 | 0.244650542 | 0 | 0.520678036 |
| ENSMUSG00000081588 | 0 | 0 | 0 | 0 | 0 | 0 | 0 |
| ENSMUSG00000081589 | 0 | 0 | 0 | 0 | 0 | 0 | 0 |
| ENSMUSG00000081590 | 0 | 0 | 0 | 0 | 0 | 0 | 0 |
| ENSMUSG00000081591 | 0 | 0 | 0 | 0 | 0 | 0 | 0 |
| ENSMUSG00000081592 | 0 | 0 | 0 | 0 | 0 | 0 | 0 |
| ENSMUSG00000081593 | 0 | 0.25 | 0.5 | 0.25 | 0.244650542 | 0 | 0 |
| ENSMUSG00000081594 | 0.289064826 | 0.25 | 0.25 | 0.25 | 0 | 0 | 0 |
| ENSMUSG00000081595 | 0 | 0 | 0 | 0 | 0 | 0 | 0 |
| ENSMUSG00000081596 | 0 | 0.25 | 0 | 0 | 0 | 0 | 0 |
| ENSMUSG00000081597 | 0.289064826 | 0 | 0.25 | 0 | 0 | 0 | 0 |
| ENSMUSG00000081598 | 0 | 0 | 0 | 0 | 0 | 0 | 0 |
| ENSMUSG00000081599 | 0 | 0 | 0 | 0 | 0.244650542 | 0 | 0 |
| ENSMUSG00000081600 | 0.960252568 | 0.75 | 0.830482024 | 0.5 | 0.686820904 | 0.477121255 | 0.629532004 |
| ENSMUSG00000081601 | 0 | 0 | 0 | 0 | 0 | 0 | 0 |
| ENSMUSG00000081602 | 0 | 0 | 0 | 0 | 0 | 0 | 0 |

| | | | | | | | |
|---|---|---|---|---|---|---|---|
| ENSMUSG00000081603 | 1.734388958 | 1.356566189 | 1.685366747 | 0.896240625 | 0.905314583 | 0.84509804 | 0.853775828 |
| ENSMUSG00000081604 | 0 | 0 | 0 | 0 | 0 | 0 | 0 |
| ENSMUSG00000081605 | 0 | 0.5 | 0.25 | 0.25 | 0.387761935 | 0 | 0.448487648 |
| ENSMUSG00000081606 | 0 | 0 | 0 | 0 | 0 | 0 | 0 |
| ENSMUSG00000081607 | 0 | 0.5 | 0.25 | 0 | 0 | 0.301029996 | 0.224243824 |
| ENSMUSG00000081608 | 0.811507563 | 0.830482024 | 0.79248125 | 0.896240625 | 0.978602168 | 0.698970004 | 1.014380832 |
| ENSMUSG00000081609 | 0 | 0 | 0 | 0 | 0 | 0 | 0 |
| ENSMUSG00000081610 | 0 | 0 | 0 | 0 | 0 | 0 | 0 |
| ENSMUSG00000081611 | 0 | 0 | 0 | 0 | 0.244650542 | 0.301029996 | 0 |
| ENSMUSG00000081612 | 0 | 0 | 0 | 0 | 0 | 0 | 0 |
| ENSMUSG00000081613 | 0.289064826 | 0.25 | 0.701838731 | 0.25 | 0.387761935 | 0.301029996 | 0.579661877 |
| ENSMUSG00000081614 | 0 | 0 | 0 | 0 | 0.244650542 | 0 | 0 |
| ENSMUSG00000081615 | 0 | 0 | 0 | 0 | 0 | 0 | 0 |
| ENSMUSG00000081616 | 0 | 0 | 0 | 0 | 0 | 0 | 0 |
| ENSMUSG00000081617 | 0.289064826 | 0 | 0 | 0 | 0 | 0 | 0 |
| ENSMUSG00000081618 | 0 | 0 | 0 | 0 | 0 | 0 | 0 |
| ENSMUSG00000081619 | 0 | 0.5 | 0.396240625 | 0 | 0.244650542 | 0.301029996 | 0.355418052 |
| ENSMUSG00000081620 | 0 | 0 | 0 | 0 | 0 | 0 | 0.224243824 |
| ENSMUSG00000081621 | 0 | 0 | 0 | 0 | 0 | 0 | 0 |
| ENSMUSG00000081622 | 0 | 0.25 | 0 | 0 | 0.244650542 | 0 | 0 |
| ENSMUSG00000081623 | 0 | 0 | 0 | 0 | 0.244650542 | 0 | 0 |
| ENSMUSG00000081624 | 0 | 0 | 0 | 0 | 0 | 0 | 0 |
| ENSMUSG00000081625 | 0 | 0 | 0 | 0 | 0 | 0 | 0.224243824 |
| ENSMUSG00000081626 | 0 | 0 | 0 | 0 | 0 | 0 | 0 |
| ENSMUSG00000081627 | 0 | 0 | 0 | 0 | 0 | 0 | 0 |
| ENSMUSG00000081628 | 0 | 0 | 0 | 0 | 0 | 0 | 0 |
| ENSMUSG00000081629 | 0 | 0.25 | 0 | 0 | 0 | 0 | 0 |
| ENSMUSG00000081630 | 0 | 0 | 0 | 0 | 0 | 0 | 0 |
| ENSMUSG00000081631 | 0 | 0 | 0 | 0 | 0 | 0 | 0 |
| ENSMUSG00000081632 | 0 | 0 | 0 | 0 | 0 | 0 | 0 |
| ENSMUSG00000081633 | 0 | 0.25 | 0 | 0 | 0 | 0.301029996 | 0.224243824 |
| ENSMUSG00000081635 | 0 | 0 | 0 | 0 | 0 | 0 | 0 |
| ENSMUSG00000081636 | 0 | 0 | 0 | 0 | 0 | 0 | 0 |
| ENSMUSG00000081638 | 0 | 0 | 0 | 0 | 0 | 0 | 0 |

| | | | | | | | |
|---|---|---|---|---|---|---|---|
| ENSMUSG00000081639 | 0 | 0 | 0 | 0 | 0 | 0 | 0 |
| ENSMUSG00000081640 | 1.940852214 | 1.923371739 | 1.942046081 | 1.606566189 | 1.770879644 | 1.84509804 | 1.731408809 |
| ENSMUSG00000081641 | 0 | 0 | 0 | 0 | 0 | 0 | 0.224243824 |
| ENSMUSG00000081642 | 0 | 0.25 | 0.25 | 0.25 | 0.568060967 | 0 | 0.224243824 |
| ENSMUSG00000081643 | 0 | 0 | 0 | 0 | 0 | 0 | 0.224243824 |
| ENSMUSG00000081646 | 0 | 0 | 0 | 0 | 0 | 0 | 0 |
| ENSMUSG00000081647 | 0 | 0 | 0 | 0 | 0 | 0 | 0 |
| ENSMUSG00000081648 | 0.45815691 | 0.396240625 | 0.25 | 0.25 | 0.387761935 | 0.301029996 | 0.448487648 |
| ENSMUSG00000081649 | 0 | 0 | 0 | 0 | 0 | 0 | 0 |
| ENSMUSG00000081650 | 0.289064826 | 0.646240625 | 0 | 0.396240625 | 0.632412477 | 0.477121255 | 0.355418052 |
| ENSMUSG00000081651 | 0 | 0 | 0 | 0 | 0 | 0 | 0.224243824 |
| ENSMUSG00000081652 | 0 | 0 | 0 | 0 | 0 | 0 | 0 |
| ENSMUSG00000081653 | 0 | 0 | 0.25 | 0.25 | 0 | 0 | 0 |
| ENSMUSG00000081654 | 0 | 0 | 0 | 0 | 0 | 0 | 0 |
| ENSMUSG00000081656 | 0 | 0 | 0 | 0 | 0 | 0 | 0 |
| ENSMUSG00000081657 | 0.289064826 | 0 | 0.25 | 0.580482024 | 0.244650542 | 0 | 0.448487648 |
| ENSMUSG00000081658 | 0 | 0 | 0 | 0 | 0 | 0 | 0 |
| ENSMUSG00000081659 | 0.289064826 | 0 | 0 | 0.25 | 0 | 0 | 0.224243824 |
| ENSMUSG00000081660 | 0 | 0 | 0 | 0 | 0 | 0 | 0 |
| ENSMUSG00000081661 | 0 | 0 | 0 | 0.25 | 0 | 0 | 0 |
| ENSMUSG00000081662 | 0.45815691 | 0.396240625 | 0 | 0.25 | 0 | 0 | 0.224243824 |
| ENSMUSG00000081663 | 0 | 0 | 0 | 0 | 0.244650542 | 0 | 0 |
| ENSMUSG00000081664 | 0 | 0 | 0 | 0 | 0 | 0 | 0.224243824 |
| ENSMUSG00000081665 | 0 | 0 | 0 | 0 | 0 | 0 | 0 |
| ENSMUSG00000081666 | 0 | 0 | 0 | 0 | 0 | 0 | 0 |
| ENSMUSG00000081667 | 0 | 0 | 0 | 0 | 0 | 0 | 0 |
| ENSMUSG00000081668 | 0 | 0 | 0 | 0 | 0 | 0 | 0 |
| ENSMUSG00000081669 | 0.45815691 | 0.396240625 | 0 | 0.25 | 0.244650542 | 0 | 0.355418052 |
| ENSMUSG00000081670 | 0 | 0.580482024 | 0 | 0.25 | 0.244650542 | 0.301029996 | 0.448487648 |
| ENSMUSG00000081671 | 0 | 0 | 0 | 0 | 0 | 0 | 0 |
| ENSMUSG00000081672 | 0 | 0 | 0 | 0 | 0 | 0 | 0 |
| ENSMUSG00000081673 | 0 | 0 | 0 | 0 | 0 | 0 | 0 |
| ENSMUSG00000081674 | 0 | 0.25 | 0 | 0 | 0.244650542 | 0 | 0 |
| ENSMUSG00000081675 | 0.289064826 | 0.25 | 0.25 | 0.25 | 0.244650542 | 0 | 0.355418052 |

| | | | | | | | |
|---|---|---|---|---|---|---|---|
| ENSMUSG00000081676 | 0 | 0.396240625 | 0 | 0 | 0 | 0 | 0 |
| ENSMUSG00000081677 | 0 | 0 | 0 | 0 | 0 | 0 | 0 |
| ENSMUSG00000081678 | 0 | 0 | 0 | 0 | 0 | 0 | 0 |
| ENSMUSG00000081679 | 0.45815691 | 0 | 0.25 | 0.25 | 0.489301084 | 0.477121255 | 0.355418052 |
| ENSMUSG00000081680 | 0.289064826 | 0.25 | 0 | 0.25 | 0 | 0 | 0 |
| ENSMUSG00000081681 | 0 | 0 | 0 | 0 | 0 | 0 | 0 |
| ENSMUSG00000081683 | 0 | 0 | 0 | 0.25 | 0.387761935 | 0.301029996 | 0.448487648 |
| ENSMUSG00000081684 | 0 | 0.25 | 0 | 0 | 0 | 0 | 0 |
| ENSMUSG00000081685 | 0 | 0 | 0 | 0 | 0 | 0 | 0 |
| ENSMUSG00000081686 | 0 | 0 | 0 | 0 | 0 | 0 | 0 |
| ENSMUSG00000081687 | 0 | 0 | 0 | 0 | 0 | 0 | 0 |
| ENSMUSG00000081688 | 0 | 0 | 0 | 0 | 0 | 0 | 0 |
| ENSMUSG00000081689 | 0 | 0 | 0 | 0 | 0 | 0 | 0 |
| ENSMUSG00000081690 | 0 | 0 | 0 | 0 | 0 | 0.301029996 | 0 |
| ENSMUSG00000081691 | 0 | 0 | 0 | 0.25 | 0 | 0 | 0 |
| ENSMUSG00000081692 | 0 | 0 | 0 | 0 | 0 | 0.301029996 | 0 |
| ENSMUSG00000081693 | 0 | 0 | 0 | 0 | 0 | 0 | 0 |
| ENSMUSG00000081694 | 0 | 0 | 0 | 0 | 0 | 0 | 0 |
| ENSMUSG00000081695 | 0.671187741 | 0.5 | 0.701838731 | 0.5 | 0.812711509 | 0.477121255 | 0.629532004 |
| ENSMUSG00000081696 | 0 | 0.25 | 0.25 | 0 | 0 | 0 | 0 |
| ENSMUSG00000081697 | 0 | 0 | 0 | 0 | 0 | 0 | 0 |
| ENSMUSG00000081698 | 0 | 0 | 0 | 0 | 0 | 0 | 0 |
| ENSMUSG00000081699 | 0 | 0 | 0 | 0 | 0 | 0 | 0 |
| ENSMUSG00000081700 | 0 | 0 | 0 | 0 | 0 | 0 | 0 |
| ENSMUSG00000081701 | 0 | 0 | 0 | 0 | 0 | 0 | 0 |
| ENSMUSG00000081703 | 0.289064826 | 0 | 0 | 0 | 0 | 0 | 0.224243824 |
| ENSMUSG00000081704 | 0 | 0 | 0 | 0 | 0 | 0 | 0 |
| ENSMUSG00000081705 | 0 | 0 | 0 | 0 | 0 | 0.301029996 | 0 |
| ENSMUSG00000081706 | 0.289064826 | 0.25 | 0 | 0.396240625 | 0 | 0 | 0 |
| ENSMUSG00000081707 | 0 | 0 | 0.25 | 0.5 | 0 | 0 | 0.224243824 |
| ENSMUSG00000081709 | 0 | 0 | 0 | 0 | 0 | 0 | 0 |
| ENSMUSG00000081711 | 0 | 0 | 0 | 0 | 0 | 0 | 0 |
| ENSMUSG00000081712 | 0.289064826 | 0.396240625 | 0 | 0.396240625 | 0 | 0.301029996 | 0.224243824 |
| ENSMUSG00000081713 | 0 | 0 | 0 | 0 | 0 | 0 | 0 |

| | | | | | | | |
|---|---|---|---|---|---|---|---|
| ENSMUSG00000081714 | 0 | 0 | 0 | 0 | 0 | 0 | 0 |
| ENSMUSG00000081715 | 0 | 0.25 | 0 | 0 | 0 | 0 | 0 |
| ENSMUSG00000081716 | 0 | 0 | 0 | 0 | 0 | 0 | 0 |
| ENSMUSG00000081717 | 0 | 0 | 0 | 0 | 0 | 0 | 0 |
| ENSMUSG00000081718 | 0 | 0 | 0.5 | 0.25 | 0 | 0 | 0 |
| ENSMUSG00000081719 | 0.289064826 | 0 | 0 | 0 | 0 | 0 | 0 |
| ENSMUSG00000081720 | 0 | 0 | 0 | 0 | 0.244650542 | 0 | 0.224243824 |
| ENSMUSG00000081721 | 0.289064826 | 0.25 | 0.5 | 0.25 | 0 | 0 | 0.579661877 |
| ENSMUSG00000081722 | 0 | 0 | 0 | 0 | 0 | 0 | 0 |
| ENSMUSG00000081723 | 0 | 0 | 0 | 0 | 0 | 0 | 0 |
| ENSMUSG00000081724 | 0 | 0 | 0 | 0 | 0 | 0 | 0 |
| ENSMUSG00000081728 | 0 | 0 | 0 | 0 | 0 | 0 | 0 |
| ENSMUSG00000081729 | 0 | 0.396240625 | 0.580482024 | 0.396240625 | 0.244650542 | 0 | 0.224243824 |
| ENSMUSG00000081730 | 0.45815691 | 0.25 | 0 | 0.25 | 0.244650542 | 0 | 0 |
| ENSMUSG00000081731 | 0 | 0 | 0 | 0 | 0.244650542 | 0 | 0.224243824 |
| ENSMUSG00000081732 | 0.289064826 | 0 | 0 | 0.25 | 0.244650542 | 0 | 0.355418052 |
| ENSMUSG00000081733 | 0 | 0.25 | 0.396240625 | 0.396240625 | 0 | 0.301029996 | 0.224243824 |
| ENSMUSG00000081734 | 0 | 0 | 0 | 0 | 0 | 0 | 0 |
| ENSMUSG00000081735 | 0 | 0 | 0 | 0.25 | 0 | 0 | 0 |
| ENSMUSG00000081736 | 0 | 0 | 0 | 0 | 0 | 0 | 0 |
| ENSMUSG00000081737 | 0 | 0 | 0 | 0 | 0 | 0 | 0 |
| ENSMUSG00000081738 | 0.289064826 | 0.396240625 | 0 | 0 | 0.244650542 | 0 | 0.355418052 |
| ENSMUSG00000081739 | 0.91631382 | 1.114857905 | 1.04248125 | 1 | 0.846351821 | 1.230448921 | 1.089372236 |
| ENSMUSG00000081740 | 0 | 0 | 0.25 | 0 | 0 | 0 | 0 |
| ENSMUSG00000081741 | 0 | 0 | 0 | 0 | 0 | 0 | 0 |
| ENSMUSG00000081742 | 0 | 0 | 0 | 0 | 0 | 0 | 0 |
| ENSMUSG00000081743 | 0 | 0 | 0 | 0 | 0 | 0 | 0 |
| ENSMUSG00000081744 | 0 | 0 | 0 | 0 | 0 | 0 | 0 |
| ENSMUSG00000081745 | 0.45815691 | 0.25 | 0 | 0.396240625 | 0.244650542 | 0 | 0.224243824 |
| ENSMUSG00000081746 | 0 | 0 | 0 | 0 | 0 | 0 | 0 |
| ENSMUSG00000081747 | 0 | 0.25 | 0 | 0 | 0 | 0 | 0 |
| ENSMUSG00000081748 | 0 | 0 | 0 | 0.25 | 0 | 0 | 0.224243824 |
| ENSMUSG00000081749 | 0 | 0 | 0 | 0 | 0 | 0 | 0 |
| ENSMUSG00000081750 | 0 | 0 | 0 | 0 | 0 | 0 | 0 |

| | | | | | | | |
|---|---|---|---|---|---|---|---|
| ENSMUSG00000081751 | 0 | 0 | 0 | 0 | 0 | 0 | 0 |
| ENSMUSG00000081752 | 0 | 0 | 0 | 0 | 0 | 0 | 0 |
| ENSMUSG00000081755 | 0 | 0 | 0 | 0 | 0 | 0 | 0 |
| ENSMUSG00000081756 | 0 | 0 | 0 | 0 | 0 | 0 | 0 |
| ENSMUSG00000081757 | 0 | 0 | 0 | 0 | 0 | 0 | 0 |
| ENSMUSG00000081758 | 0 | 0 | 0 | 0 | 0 | 0 | 0 |
| ENSMUSG00000081761 | 0 | 0 | 0 | 0 | 0 | 0 | 0 |
| ENSMUSG00000081762 | 0 | 0 | 0 | 0 | 0 | 0 | 0 |
| ENSMUSG00000081763 | 0 | 0.5 | 0.646240625 | 0.396240625 | 0.568060967 | 0.477121255 | 0.224243824 |
| ENSMUSG00000081765 | 0 | 0 | 0 | 0 | 0 | 0 | 0 |
| ENSMUSG00000081766 | 0 | 0 | 0 | 0 | 0 | 0 | 0 |
| ENSMUSG00000081767 | 0 | 0 | 0 | 0 | 0 | 0 | 0 |
| ENSMUSG00000081768 | 0 | 0 | 0 | 0 | 0 | 0 | 0 |
| ENSMUSG00000081769 | 0 | 0 | 0.25 | 0.5 | 0.489301084 | 0.301029996 | 0.355418052 |
| ENSMUSG00000081770 | 0 | 0 | 0 | 0 | 0 | 0 | 0 |
| ENSMUSG00000081771 | 0 | 0.25 | 0 | 0 | 0 | 0 | 0.224243824 |
| ENSMUSG00000081772 | 0 | 0 | 0 | 0 | 0 | 0 | 0 |
| ENSMUSG00000081773 | 0.578129653 | 0.830482024 | 1.04248125 | 0.646240625 | 0.955822902 | 0.602059991 | 0.710836105 |
| ENSMUSG00000081774 | 0 | 0 | 0 | 0 | 0 | 0 | 0 |
| ENSMUSG00000081775 | 0 | 0 | 0 | 0.25 | 0 | 0 | 0 |
| ENSMUSG00000081777 | 0 | 0 | 0 | 0 | 0 | 0 | 0 |
| ENSMUSG00000081778 | 0 | 0 | 0.25 | 0 | 0 | 0 | 0 |
| ENSMUSG00000081779 | 0 | 0 | 0 | 0 | 0 | 0 | 0 |
| ENSMUSG00000081781 | 0 | 0 | 0 | 0 | 0 | 0 | 0.224243824 |
| ENSMUSG00000081782 | 0 | 0 | 0 | 0 | 0 | 0.301029996 | 0 |
| ENSMUSG00000081783 | 0.671187741 | 0.396240625 | 0.25 | 0.5 | 0 | 0.477121255 | 0.224243824 |
| ENSMUSG00000081784 | 0 | 0 | 0 | 0 | 0 | 0 | 0 |
| ENSMUSG00000081786 | 0 | 0 | 0 | 0 | 0 | 0 | 0 |
| ENSMUSG00000081787 | 0 | 0 | 0 | 0 | 0.387761935 | 0 | 0 |
| ENSMUSG00000081788 | 1.516991255 | 1.388647213 | 1.321350555 | 1.438721876 | 1.358933174 | 1.556302501 | 1.307989563 |
| ENSMUSG00000081789 | 0 | 0 | 0 | 0 | 0 | 0 | 0 |
| ENSMUSG00000081791 | 0 | 0 | 0 | 0 | 0 | 0 | 0 |
| ENSMUSG00000081792 | 0.45815691 | 0.396240625 | 0.25 | 0.25 | 0.387761935 | 0.301029996 | 0 |
| ENSMUSG00000081793 | 0 | 0 | 0 | 0 | 0 | 0 | 0 |

| | | | | | | | |
|---|---|---|---|---|---|---|---|
| ENSMUSG00000081796 | 0 | 0 | 0.25 | 0 | 0 | 0 | 0 |
| ENSMUSG00000081797 | 0.289064826 | 0 | 0 | 0 | 0 | 0 | 0.355418052 |
| ENSMUSG00000081798 | 0 | 0 | 0.396240625 | 0.25 | 0 | 0 | 0.355418052 |
| ENSMUSG00000081799 | 0 | 0 | 0 | 0 | 0 | 0 | 0 |
| ENSMUSG00000081800 | 0 | 0 | 0 | 0 | 0 | 0 | 0 |
| ENSMUSG00000081801 | 0 | 0 | 0 | 0 | 0 | 0.301029996 | 0 |
| ENSMUSG00000081803 | 0 | 0.25 | 0 | 0 | 0 | 0.301029996 | 0 |
| ENSMUSG00000081804 | 0 | 0 | 0 | 0 | 0 | 0 | 0 |
| ENSMUSG00000081805 | 0 | 0 | 0 | 0.25 | 0.387761935 | 0 | 0.224243824 |
| ENSMUSG00000081806 | 0 | 0 | 0 | 0 | 0 | 0 | 0 |
| ENSMUSG00000081807 | 0.289064826 | 0 | 0.396240625 | 0 | 0.489301084 | 0.301029996 | 0.224243824 |
| ENSMUSG00000081809 | 0.289064826 | 0.25 | 0.25 | 0 | 0.489301084 | 0.477121255 | 0.520678036 |
| ENSMUSG00000081810 | 0 | 0 | 0 | 0.25 | 0 | 0 | 0.355418052 |
| ENSMUSG00000081811 | 0.289064826 | 0 | 0 | 0 | 0 | 0 | 0 |
| ENSMUSG00000081812 | 0 | 0 | 0 | 0 | 0 | 0 | 0 |
| ENSMUSG00000081813 | 0 | 0 | 0 | 0 | 0 | 0 | 0 |
| ENSMUSG00000081814 | 0 | 0 | 0 | 0 | 0 | 0 | 0 |
| ENSMUSG00000081816 | 0 | 0 | 0 | 0 | 0 | 0 | 0 |
| ENSMUSG00000081817 | 0.45815691 | 0.25 | 0.396240625 | 0.25 | 0.244650542 | 0 | 0 |
| ENSMUSG00000081819 | 0 | 0 | 0 | 0.25 | 0.244650542 | 0 | 0 |
| ENSMUSG00000081820 | 0 | 0 | 0 | 0 | 0 | 0 | 0.224243824 |
| ENSMUSG00000081821 | 0 | 0 | 0 | 0 | 0 | 0 | 0 |
| ENSMUSG00000081822 | 0 | 0 | 0 | 0 | 0 | 0 | 0 |
| ENSMUSG00000081823 | 0 | 0 | 0 | 0 | 0 | 0 | 0 |
| ENSMUSG00000081824 | 0 | 0 | 0 | 0 | 0 | 0 | 0 |
| ENSMUSG00000081827 | 0 | 0 | 0 | 0 | 0 | 0 | 0 |
| ENSMUSG00000081828 | 0 | 0 | 0 | 0 | 0 | 0 | 0 |
| ENSMUSG00000081829 | 0 | 0 | 0 | 0 | 0 | 0 | 0 |
| ENSMUSG00000081830 | 0 | 0 | 0 | 0 | 0 | 0 | 0 |
| ENSMUSG00000081831 | 0 | 0 | 0 | 0.25 | 0 | 0 | 0 |
| ENSMUSG00000081832 | 0 | 0 | 0 | 0 | 0 | 0 | 0 |
| ENSMUSG00000081833 | 0 | 0 | 0 | 0 | 0 | 0 | 0 |
| ENSMUSG00000081834 | 0.289064826 | 0 | 0 | 0 | 0 | 0 | 0 |
| ENSMUSG00000081835 | 0 | 0 | 0 | 0 | 0 | 0 | 0 |

| | | | | | | | |
|---|---|---|---|---|---|---|---|
| ENSMUSG00000081836 | 0 | 0 | 0 | 0 | 0 | 0 | 0 |
| ENSMUSG00000081837 | 0 | 0 | 0 | 0 | 0 | 0 | 0 |
| ENSMUSG00000081838 | 0 | 0 | 0.25 | 0 | 0 | 0 | 0 |
| ENSMUSG00000081839 | 0 | 0 | 0 | 0 | 0 | 0 | 0 |
| ENSMUSG00000081840 | 0.289064826 | 0 | 0 | 0 | 0 | 0 | 0 |
| ENSMUSG00000081841 | 0 | 0 | 0 | 0 | 0 | 0 | 0 |
| ENSMUSG00000081842 | 0 | 0.25 | 0 | 0 | 0 | 0 | 0 |
| ENSMUSG00000081843 | 0 | 0 | 0 | 0 | 0 | 0 | 0 |
| ENSMUSG00000081844 | 0 | 0 | 0 | 0 | 0 | 0 | 0 |
| ENSMUSG00000081845 | 0 | 0 | 0 | 0 | 0 | 0 | 0.224243824 |
| ENSMUSG00000081847 | 0 | 0 | 0 | 0 | 0 | 0 | 0 |
| ENSMUSG00000081848 | 0 | 0 | 0 | 0 | 0 | 0 | 0 |
| ENSMUSG00000081849 | 0 | 0 | 0 | 0 | 0 | 0 | 0 |
| ENSMUSG00000081850 | 0 | 0 | 0.25 | 0 | 0 | 0 | 0.224243824 |
| ENSMUSG00000081851 | 0 | 0.25 | 0.396240625 | 0.396240625 | 0.387761935 | 0 | 0.448487648 |
| ENSMUSG00000081852 | 0 | 0 | 0 | 0 | 0 | 0 | 0 |
| ENSMUSG00000081853 | 0 | 0 | 0 | 0 | 0 | 0 | 0 |
| ENSMUSG00000081854 | 0 | 0 | 0 | 0 | 0 | 0 | 0 |
| ENSMUSG00000081855 | 0 | 0 | 0 | 0.396240625 | 0.387761935 | 0.477121255 | 0 |
| ENSMUSG00000081856 | 0 | 0 | 0 | 0 | 0 | 0 | 0 |
| ENSMUSG00000081857 | 0 | 0 | 0 | 0 | 0 | 0 | 0 |
| ENSMUSG00000081858 | 0 | 0 | 0 | 0 | 0 | 0 | 0 |
| ENSMUSG00000081859 | 0 | 0 | 0 | 0 | 0 | 0 | 0 |
| ENSMUSG00000081860 | 0 | 0.25 | 0 | 0 | 0 | 0 | 0 |
| ENSMUSG00000081861 | 0.289064826 | 0.25 | 0.396240625 | 0.580482024 | 0.244650542 | 0.301029996 | 0 |
| ENSMUSG00000081862 | 0 | 0 | 0 | 0 | 0 | 0 | 0 |
| ENSMUSG00000081863 | 0 | 0 | 0 | 0 | 0 | 0 | 0 |
| ENSMUSG00000081864 | 0 | 0 | 0 | 0 | 0 | 0 | 0 |
| ENSMUSG00000081865 | 0.289064826 | 0.5 | 0.580482024 | 0.396240625 | 0.568060967 | 0.602059991 | 0.355418052 |
| ENSMUSG00000081866 | 0 | 0 | 0 | 0 | 0 | 0 | 0 |
| ENSMUSG00000081868 | 0 | 0 | 0 | 0 | 0 | 0 | 0 |
| ENSMUSG00000081869 | 0 | 0 | 0 | 0 | 0 | 0 | 0 |
| ENSMUSG00000081870 | 0 | 0 | 0 | 0 | 0 | 0 | 0 |
| ENSMUSG00000081871 | 0 | 0.25 | 0.25 | 0.25 | 0 | 0 | 0 |

| | | | | | | | |
|---|---|---|---|---|---|---|---|
| ENSMUSG00000081872 | 0 | 0 | 0 | 0 | 0 | 0 | 0 |
| ENSMUSG00000081873 | 0 | 0.25 | 0 | 0 | 0.244650542 | 0 | 0 |
| ENSMUSG00000081874 | 0 | 0 | 0 | 0 | 0 | 0 | 0 |
| ENSMUSG00000081875 | 0.289064826 | 0 | 0 | 0 | 0 | 0 | 0 |
| ENSMUSG00000081876 | 0 | 0 | 0 | 0 | 0 | 0 | 0 |
| ENSMUSG00000081877 | 0 | 0 | 0 | 0 | 0 | 0 | 0.224243824 |
| ENSMUSG00000081878 | 0 | 0 | 0 | 0 | 0 | 0 | 0 |
| ENSMUSG00000081879 | 0 | 0 | 0 | 0 | 0 | 0 | 0 |
| ENSMUSG00000081880 | 0 | 0 | 0 | 0 | 0 | 0 | 0 |
| ENSMUSG00000081881 | 0 | 0 | 0 | 0 | 0 | 0 | 0 |
| ENSMUSG00000081882 | 0 | 0.25 | 0 | 0 | 0 | 0 | 0 |
| ENSMUSG00000081883 | 0 | 0 | 0 | 0 | 0 | 0.301029996 | 0.448487648 |
| ENSMUSG00000081884 | 0 | 0 | 0 | 0 | 0 | 0 | 0 |
| ENSMUSG00000081885 | 0.289064826 | 0.396240625 | 0.396240625 | 0.79248125 | 0.632412477 | 0.477121255 | 0.629532004 |
| ENSMUSG00000081886 | 0 | 0 | 0 | 0 | 0 | 0 | 0 |
| ENSMUSG00000081887 | 0.671187741 | 0.5 | 0.75 | 0.5 | 0.387761935 | 0.698970004 | 0.579661877 |
| ENSMUSG00000081888 | 0.671187741 | 0.830482024 | 0.646240625 | 0.396240625 | 0.387761935 | 0.602059991 | 0.853775828 |
| ENSMUSG00000081889 | 0 | 0 | 0 | 0 | 0 | 0 | 0 |
| ENSMUSG00000081890 | 0 | 0 | 0 | 0 | 0 | 0 | 0 |
| ENSMUSG00000081891 | 0 | 0 | 0 | 0 | 0 | 0 | 0 |
| ENSMUSG00000081892 | 0 | 0.396240625 | 0.25 | 0.25 | 0.244650542 | 0 | 0.355418052 |
| ENSMUSG00000081893 | 0 | 0 | 0 | 0 | 0 | 0 | 0 |
| ENSMUSG00000081894 | 0 | 0 | 0 | 0 | 0 | 0 | 0 |
| ENSMUSG00000081895 | 0.578129653 | 0.701838731 | 0.646240625 | 0.75 | 0.686820904 | 0.602059991 | 0.672731473 |
| ENSMUSG00000081896 | 0 | 0.396240625 | 0.25 | 0 | 0.244650542 | 0.477121255 | 0.224243824 |
| ENSMUSG00000081897 | 0 | 0 | 0 | 0 | 0 | 0 | 0 |
| ENSMUSG00000081898 | 0.289064826 | 0 | 0 | 0 | 0 | 0.301029996 | 0.355418052 |
| ENSMUSG00000081900 | 0 | 0 | 0 | 0 | 0 | 0 | 0 |
| ENSMUSG00000081901 | 0 | 0 | 0 | 0 | 0 | 0 | 0 |
| ENSMUSG00000081903 | 0 | 0 | 0 | 0 | 0 | 0 | 0 |
| ENSMUSG00000081904 | 0 | 0 | 0 | 0.25 | 0 | 0 | 0 |
| ENSMUSG00000081905 | 0 | 0 | 0 | 0 | 0 | 0 | 0 |
| ENSMUSG00000081906 | 0 | 0 | 0 | 0 | 0 | 0 | 0 |
| ENSMUSG00000081907 | 0 | 0 | 0 | 0 | 0 | 0 | 0 |

| | | | | | | | |
|---|---|---|---|---|---|---|---|
| ENSMUSG00000081908 | 0 | 0 | 0 | 0 | 0 | 0 | 0 |
| ENSMUSG00000081909 | 0 | 0 | 0 | 0 | 0 | 0 | 0 |
| ENSMUSG00000081910 | 0 | 0 | 0 | 0 | 0 | 0 | 0 |
| ENSMUSG00000081911 | 0 | 0 | 0 | 0 | 0.244650542 | 0 | 0 |
| ENSMUSG00000081912 | 0 | 0 | 0 | 0 | 0 | 0 | 0 |
| ENSMUSG00000081913 | 0 | 0 | 0 | 0 | 0 | 0 | 0 |
| ENSMUSG00000081914 | 0 | 0 | 0 | 0 | 0 | 0 | 0 |
| ENSMUSG00000081916 | 0 | 0 | 0 | 0 | 0 | 0 | 0 |
| ENSMUSG00000081917 | 0 | 0 | 0 | 0 | 0 | 0 | 0 |
| ENSMUSG00000081918 | 0 | 0 | 0 | 0 | 0 | 0 | 0 |
| ENSMUSG00000081919 | 0 | 0 | 0 | 0 | 0 | 0 | 0 |
| ENSMUSG00000081920 | 0 | 0 | 0 | 0 | 0 | 0 | 0 |
| ENSMUSG00000081921 | 0 | 0 | 0 | 0 | 0 | 0 | 0 |
| ENSMUSG00000081923 | 0 | 0 | 0 | 0 | 0 | 0 | 0 |
| ENSMUSG00000081924 | 0 | 0 | 0 | 0.25 | 0 | 0 | 0.224243824 |
| ENSMUSG00000081925 | 0 | 0 | 0 | 0 | 0 | 0 | 0 |
| ENSMUSG00000081926 | 0 | 0.396240625 | 0 | 0 | 0 | 0 | 0.224243824 |
| ENSMUSG00000081928 | 0 | 0 | 0 | 0 | 0 | 0 | 0 |
| ENSMUSG00000081929 | 0.289064826 | 0.5 | 0 | 0.396240625 | 0 | 0.301029996 | 0.224243824 |
| ENSMUSG00000081931 | 0 | 0 | 0 | 0 | 0 | 0 | 0 |
| ENSMUSG00000081932 | 0 | 0.25 | 0.25 | 0 | 0.244650542 | 0 | 0.355418052 |
| ENSMUSG00000081933 | 0 | 0 | 0.25 | 0 | 0 | 0 | 0 |
| ENSMUSG00000081934 | 0 | 0 | 0 | 0 | 0 | 0 | 0 |
| ENSMUSG00000081935 | 0 | 0 | 0 | 0 | 0 | 0 | 0 |
| ENSMUSG00000081936 | 0 | 0 | 0 | 0 | 0 | 0 | 0 |
| ENSMUSG00000081937 | 0 | 0 | 0 | 0 | 0 | 0 | 0 |
| ENSMUSG00000081938 | 0 | 0 | 0 | 0 | 0 | 0 | 0 |
| ENSMUSG00000081939 | 0 | 0 | 0 | 0 | 0 | 0 | 0 |
| ENSMUSG00000081940 | 0 | 0 | 0 | 0 | 0 | 0 | 0 |
| ENSMUSG00000081941 | 0 | 0.396240625 | 0 | 0 | 0.387761935 | 0.477121255 | 0.224243824 |
| ENSMUSG00000081942 | 0 | 0 | 0 | 0 | 0 | 0 | 0 |
| ENSMUSG00000081943 | 0 | 0.25 | 0.25 | 0 | 0 | 0 | 0 |
| ENSMUSG00000081944 | 0.578129653 | 0.864857905 | 0.701838731 | 0.830482024 | 0.955822902 | 0.602059991 | 0.952571511 |
| ENSMUSG00000081945 | 0 | 0 | 0 | 0 | 0 | 0 | 0 |

| | | | | | | | |
|---|---|---|---|---|---|---|---|
| ENSMUSG00000081946 | 0.45815691 | 0.396240625 | 0.396240625 | 0.396240625 | 0.632412477 | 0.477121255 | 0.672731473 |
| ENSMUSG00000081947 | 0 | 0 | 0 | 0 | 0 | 0 | 0 |
| ENSMUSG00000081948 | 0 | 0 | 0 | 0 | 0 | 0 | 0 |
| ENSMUSG00000081949 | 0 | 0 | 0 | 0 | 0 | 0 | 0 |
| ENSMUSG00000081950 | 0.289064826 | 0 | 0 | 0 | 0 | 0 | 0 |
| ENSMUSG00000081951 | 0 | 0 | 0 | 0 | 0 | 0 | 0 |
| ENSMUSG00000081952 | 0 | 0 | 0 | 0 | 0 | 0 | 0 |
| ENSMUSG00000081953 | 0 | 0.25 | 0.396240625 | 0.25 | 0 | 0 | 0.224243824 |
| ENSMUSG00000081954 | 0 | 0 | 0 | 0.25 | 0 | 0 | 0 |
| ENSMUSG00000081955 | 0 | 0 | 0 | 0 | 0 | 0 | 0 |
| ENSMUSG00000081957 | 0 | 0 | 0 | 0 | 0 | 0 | 0 |
| ENSMUSG00000081958 | 0 | 0 | 0 | 0 | 0 | 0 | 0 |
| ENSMUSG00000081959 | 0 | 0 | 0 | 0 | 0 | 0 | 0 |
| ENSMUSG00000081960 | 0 | 0 | 0 | 0 | 0 | 0 | 0 |
| ENSMUSG00000081961 | 0 | 0 | 0 | 0 | 0 | 0 | 0 |
| ENSMUSG00000081962 | 0 | 0 | 0 | 0 | 0 | 0 | 0 |
| ENSMUSG00000081964 | 0 | 0 | 0 | 0 | 0 | 0 | 0 |
| ENSMUSG00000081965 | 0 | 0.25 | 0.5 | 0.25 | 0.387761935 | 0 | 0.224243824 |
| ENSMUSG00000081966 | 0 | 0 | 0 | 0 | 0 | 0 | 0 |
| ENSMUSG00000081967 | 0 | 0 | 0.25 | 0 | 0 | 0 | 0 |
| ENSMUSG00000081968 | 0.289064826 | 0 | 0 | 0.25 | 0 | 0.301029996 | 0 |
| ENSMUSG00000081969 | 0 | 0 | 0 | 0 | 0 | 0 | 0 |
| ENSMUSG00000081970 | 0 | 0 | 0 | 0 | 0 | 0 | 0 |
| ENSMUSG00000081971 | 0.289064826 | 0.580482024 | 0.396240625 | 0.25 | 0.387761935 | 0.477121255 | 0.355418052 |
| ENSMUSG00000081972 | 0 | 0 | 0 | 0 | 0 | 0 | 0 |
| ENSMUSG00000081973 | 0 | 0.396240625 | 0.396240625 | 0 | 0 | 0 | 0.224243824 |
| ENSMUSG00000081974 | 1.100572389 | 1.17510993 | 1.061981878 | 1.114857905 | 1.091002363 | 0.903089987 | 1.100339912 |
| ENSMUSG00000081975 | 0.289064826 | 0.396240625 | 0.25 | 0.5 | 0 | 0.477121255 | 0.355418052 |
| ENSMUSG00000081976 | 0 | 0 | 0 | 0 | 0 | 0 | 0 |
| ENSMUSG00000081977 | 0 | 0 | 0 | 0 | 0 | 0 | 0 |
| ENSMUSG00000081978 | 0 | 0 | 0 | 0 | 0 | 0 | 0 |
| ENSMUSG00000081979 | 0 | 0 | 0.25 | 0 | 0 | 0.301029996 | 0.224243824 |
| ENSMUSG00000081980 | 0 | 0 | 0 | 0 | 0 | 0 | 0 |
| ENSMUSG00000081982 | 0 | 0 | 0 | 0 | 0 | 0 | 0 |

| | | | | | | | |
|---|---|---|---|---|---|---|---|
| ENSMUSG00000081984 | 0 | 0 | 0 | 0.25 | 0.244650542 | 0.477121255 | 0.448487648 |
| ENSMUSG00000081985 | 0 | 0 | 0 | 0 | 0 | 0 | 0 |
| ENSMUSG00000081986 | 0 | 0 | 0 | 0 | 0 | 0 | 0 |
| ENSMUSG00000081987 | 0 | 0 | 0 | 0 | 0 | 0 | 0 |
| ENSMUSG00000081988 | 0 | 0 | 0 | 0 | 0 | 0 | 0 |
| ENSMUSG00000081991 | 0 | 0 | 0.396240625 | 0 | 0 | 0 | 0 |
| ENSMUSG00000081992 | 0 | 0.580482024 | 0.5 | 0.5 | 0.244650542 | 0.477121255 | 0.224243824 |
| ENSMUSG00000081993 | 0 | 0 | 0 | 0 | 0 | 0 | 0 |
| ENSMUSG00000081994 | 0 | 0 | 0 | 0.25 | 0 | 0 | 0 |
| ENSMUSG00000081995 | 0 | 0 | 0 | 0 | 0 | 0 | 0 |
| ENSMUSG00000081996 | 0.45815691 | 0.396240625 | 0.701838731 | 0.701838731 | 0.244650542 | 0.301029996 | 0.355418052 |
| ENSMUSG00000081998 | 0.45815691 | 0.701838731 | 0.75 | 0.580482024 | 0.489301084 | 0.477121255 | 0.355418052 |
| ENSMUSG00000081999 | 0 | 0 | 0 | 0.25 | 0 | 0.477121255 | 0 |
| ENSMUSG00000082000 | 0 | 0 | 0 | 0 | 0 | 0 | 0 |
| ENSMUSG00000082001 | 0 | 0 | 0 | 0 | 0 | 0 | 0 |
| ENSMUSG00000082002 | 0 | 0 | 0 | 0 | 0.244650542 | 0 | 0 |
| ENSMUSG00000082005 | 0 | 0 | 0 | 0 | 0 | 0 | 0 |
| ENSMUSG00000082006 | 0.289064826 | 0 | 0 | 0 | 0 | 0 | 0 |
| ENSMUSG00000082007 | 0 | 0 | 0 | 0 | 0 | 0 | 0 |
| ENSMUSG00000082008 | 0 | 0 | 0 | 0.25 | 0 | 0.301029996 | 0 |
| ENSMUSG00000082009 | 0 | 0 | 0 | 0 | 0 | 0 | 0 |
| ENSMUSG00000082010 | 0 | 0 | 0 | 0 | 0 | 0 | 0 |
| ENSMUSG00000082011 | 0 | 0 | 0 | 0 | 0 | 0 | 0 |
| ENSMUSG00000082012 | 0.289064826 | 0.25 | 0.25 | 0.25 | 0 | 0.301029996 | 0.224243824 |
| ENSMUSG00000082013 | 0.289064826 | 0.5 | 0.79248125 | 0.25 | 0.387761935 | 0.477121255 | 0.448487648 |
| ENSMUSG00000082014 | 0 | 0 | 0 | 0 | 0 | 0 | 0 |
| ENSMUSG00000082015 | 0 | 0 | 0 | 0 | 0 | 0 | 0 |
| ENSMUSG00000082016 | 0 | 0 | 0 | 0 | 0 | 0 | 0 |
| ENSMUSG00000082017 | 0 | 0 | 0 | 0 | 0 | 0 | 0 |
| ENSMUSG00000082018 | 0.289064826 | 0 | 0.25 | 0 | 0 | 0 | 0.224243824 |
| ENSMUSG00000082019 | 0 | 0 | 0 | 0 | 0 | 0 | 0 |
| ENSMUSG00000082020 | 0 | 0 | 0 | 0 | 0 | 0 | 0 |
| ENSMUSG00000082021 | 0 | 0 | 0 | 0 | 0 | 0 | 0 |
| ENSMUSG00000082022 | 0 | 0 | 0 | 0 | 0 | 0 | 0 |

| | | | | | | | |
|---|---|---|---|---|---|---|---|
| ENSMUSG00000082023 | 0 | 0 | 0 | 0 | 0 | 0 | 0 |
| ENSMUSG00000082024 | 0 | 0 | 0 | 0 | 0 | 0 | 0 |
| ENSMUSG00000082025 | 0 | 0 | 0 | 0 | 0 | 0 | 0 |
| ENSMUSG00000082026 | 0 | 0.25 | 0 | 0 | 0.489301084 | 0.301029996 | 0.355418052 |
| ENSMUSG00000082027 | 0 | 0 | 0 | 0 | 0 | 0 | 0 |
| ENSMUSG00000082028 | 0 | 0 | 0 | 0 | 0 | 0 | 0 |
| ENSMUSG00000082029 | 0 | 0 | 0 | 0 | 0 | 0 | 0 |
| ENSMUSG00000082030 | 0 | 0 | 0 | 0 | 0 | 0 | 0 |
| ENSMUSG00000082031 | 0 | 0 | 0.25 | 0 | 0 | 0.301029996 | 0.224243824 |
| ENSMUSG00000082032 | 0 | 0 | 0.25 | 0 | 0 | 0 | 0.355418052 |
| ENSMUSG00000082033 | 0 | 0 | 0 | 0 | 0 | 0 | 0 |
| ENSMUSG00000082035 | 0 | 0 | 0 | 0 | 0 | 0 | 0 |
| ENSMUSG00000082036 | 0 | 0 | 0 | 0 | 0 | 0 | 0 |
| ENSMUSG00000082037 | 0 | 0 | 0 | 0 | 0 | 0 | 0 |
| ENSMUSG00000082038 | 0 | 0 | 0 | 0 | 0 | 0 | 0 |
| ENSMUSG00000082039 | 0.289064826 | 0 | 0 | 0 | 0 | 0 | 0 |
| ENSMUSG00000082040 | 0 | 0 | 0 | 0 | 0 | 0 | 0 |
| ENSMUSG00000082042 | 0 | 0 | 0 | 0 | 0 | 0 | 0 |
| ENSMUSG00000082043 | 0 | 0 | 0 | 0.25 | 0 | 0 | 0.224243824 |
| ENSMUSG00000082044 | 0 | 0 | 0.25 | 0.25 | 0.244650542 | 0 | 0 |
| ENSMUSG00000082045 | 0 | 0 | 0 | 0 | 0 | 0 | 0 |
| ENSMUSG00000082046 | 0 | 0 | 0 | 0 | 0 | 0 | 0 |
| ENSMUSG00000082047 | 0 | 0 | 0 | 0 | 0 | 0 | 0 |
| ENSMUSG00000082048 | 0 | 0 | 0 | 0 | 0 | 0 | 0 |
| ENSMUSG00000082049 | 0.45815691 | 0.25 | 0 | 0 | 0.489301084 | 0.301029996 | 0.448487648 |
| ENSMUSG00000082051 | 0 | 0 | 0 | 0 | 0 | 0.301029996 | 0.224243824 |
| ENSMUSG00000082052 | 0.289064826 | 0 | 0.396240625 | 0 | 0.244650542 | 0.477121255 | 0.355418052 |
| ENSMUSG00000082053 | 0 | 0 | 0 | 0 | 0 | 0 | 0 |
| ENSMUSG00000082054 | 0 | 0 | 0 | 0 | 0 | 0 | 0 |
| ENSMUSG00000082056 | 0 | 0 | 0 | 0 | 0 | 0 | 0 |
| ENSMUSG00000082057 | 0.45815691 | 0.580482024 | 0 | 0.396240625 | 0.244650542 | 0.602059991 | 0.224243824 |
| ENSMUSG00000082058 | 0 | 0 | 0 | 0 | 0 | 0 | 0 |
| ENSMUSG00000082059 | 0 | 0 | 0 | 0 | 0 | 0 | 0 |
| ENSMUSG00000082060 | 0 | 0 | 0 | 0 | 0 | 0 | 0 |

| | | | | | | | |
|---|---|---|---|---|---|---|---|
| ENSMUSG00000082061 | 0 | 0 | 0 | 0.25 | 0.387761935 | 0.301029996 | 0.355418052 |
| ENSMUSG00000082062 | 0.289064826 | 0 | 0 | 0 | 0 | 0 | 0 |
| ENSMUSG00000082063 | 0 | 0 | 0 | 0 | 0.387761935 | 0 | 0.224243824 |
| ENSMUSG00000082064 | 1.069666964 | 0.92510993 | 1.061981878 | 0.701838731 | 0.568060967 | 0.301029996 | 0.876096088 |
| ENSMUSG00000082065 | 0 | 0 | 0 | 0 | 0 | 0 | 0 |
| ENSMUSG00000082066 | 0.289064826 | 0 | 0 | 0.396240625 | 0 | 0.301029996 | 0 |
| ENSMUSG00000082068 | 0 | 0 | 0 | 0 | 0 | 0 | 0 |
| ENSMUSG00000082069 | 0 | 0 | 0 | 0 | 0 | 0 | 0 |
| ENSMUSG00000082070 | 0.289064826 | 0 | 0.396240625 | 0.396240625 | 0 | 0 | 0 |
| ENSMUSG00000082071 | 0 | 0 | 0 | 0 | 0 | 0 | 0 |
| ENSMUSG00000082072 | 0.289064826 | 0.396240625 | 0 | 0.25 | 0.244650542 | 0 | 0.448487648 |
| ENSMUSG00000082073 | 0 | 0 | 0 | 0 | 0 | 0 | 0 |
| ENSMUSG00000082074 | 0 | 0 | 0 | 0 | 0 | 0 | 0 |
| ENSMUSG00000082076 | 0 | 0.25 | 0 | 0.396240625 | 0.244650542 | 0 | 0 |
| ENSMUSG00000082077 | 0 | 0 | 0 | 0 | 0 | 0 | 0 |
| ENSMUSG00000082078 | 0.289064826 | 0 | 0 | 0.25 | 0 | 0.301029996 | 0 |
| ENSMUSG00000082079 | 0 | 0 | 0 | 0 | 0 | 0 | 0 |
| ENSMUSG00000082080 | 0 | 0 | 0 | 0 | 0 | 0 | 0 |
| ENSMUSG00000082081 | 0 | 0 | 0 | 0 | 0 | 0 | 0 |
| ENSMUSG00000082082 | 0.289064826 | 0.646240625 | 0.396240625 | 0.396240625 | 0.387761935 | 0.602059991 | 0.448487648 |
| ENSMUSG00000082083 | 0 | 0 | 0 | 0 | 0 | 0 | 0 |
| ENSMUSG00000082084 | 0 | 0 | 0 | 0 | 0 | 0 | 0 |
| ENSMUSG00000082085 | 0.289064826 | 0 | 0 | 0 | 0 | 0 | 0 |
| ENSMUSG00000082087 | 0 | 0 | 0 | 0 | 0 | 0 | 0 |
| ENSMUSG00000082088 | 0 | 0.25 | 0 | 0 | 0.489301084 | 0 | 0 |
| ENSMUSG00000082089 | 0 | 0 | 0 | 0 | 0 | 0 | 0 |
| ENSMUSG00000082090 | 0.289064826 | 0.25 | 0.25 | 0.25 | 0 | 0 | 0 |
| ENSMUSG00000082091 | 0 | 0 | 0 | 0 | 0 | 0 | 0.224243824 |
| ENSMUSG00000082092 | 0 | 0 | 0 | 0 | 0 | 0 | 0 |
| ENSMUSG00000082093 | 0 | 0 | 0 | 0 | 0 | 0 | 0 |
| ENSMUSG00000082095 | 0 | 0 | 0.25 | 0.396240625 | 0 | 0.301029996 | 0.355418052 |
| ENSMUSG00000082096 | 0 | 0 | 0 | 0 | 0 | 0 | 0 |
| ENSMUSG00000082097 | 0 | 0 | 0 | 0 | 0 | 0 | 0 |
| ENSMUSG00000082098 | 0 | 0 | 0 | 0 | 0 | 0 | 0 |

| | | | | | | | |
|---|---|---|---|---|---|---|---|
| ENSMUSG00000082099 | 0 | 0 | 0 | 0 | 0 | 0 | 0 |
| ENSMUSG00000082100 | 0.289064826 | 0 | 0 | 0 | 0.244650542 | 0 | 0 |
| ENSMUSG00000082101 | 0 | 0 | 0 | 0.25 | 0 | 0 | 0 |
| ENSMUSG00000082103 | 0 | 0 | 0 | 0 | 0 | 0 | 0 |
| ENSMUSG00000082104 | 0 | 0 | 0 | 0 | 0 | 0 | 0 |
| ENSMUSG00000082106 | 0 | 0 | 0 | 0 | 0 | 0 | 0 |
| ENSMUSG00000082107 | 0.45815691 | 0.580482024 | 0.5 | 0.646240625 | 0.489301084 | 0 | 0.629532004 |
| ENSMUSG00000082108 | 0 | 0.25 | 0.25 | 0 | 0 | 0 | 0 |
| ENSMUSG00000082109 | 0.45815691 | 0.701838731 | 0.580482024 | 0 | 0.686820904 | 0.477121255 | 0.579661877 |
| ENSMUSG00000082110 | 0 | 0 | 0.25 | 0 | 0 | 0 | 0 |
| ENSMUSG00000082111 | 0 | 0 | 0 | 0 | 0 | 0 | 0 |
| ENSMUSG00000082112 | 0 | 0 | 0 | 0 | 0 | 0 | 0 |
| ENSMUSG00000082113 | 0 | 0 | 0 | 0 | 0 | 0 | 0 |
| ENSMUSG00000082114 | 0 | 0 | 0 | 0 | 0 | 0 | 0 |
| ENSMUSG00000082115 | 0 | 0 | 0 | 0 | 0 | 0 | 0 |
| ENSMUSG00000082116 | 0 | 0 | 0 | 0 | 0 | 0 | 0 |
| ENSMUSG00000082117 | 0.45815691 | 0.25 | 0.25 | 0.5 | 0.244650542 | 0.477121255 | 0.355418052 |
| ENSMUSG00000082118 | 0 | 0.25 | 0 | 0.25 | 0.568060967 | 0 | 0.224243824 |
| ENSMUSG00000082119 | 0 | 0 | 0 | 0 | 0 | 0 | 0 |
| ENSMUSG00000082120 | 0.747221736 | 0.646240625 | 0.75 | 0.5 | 0.387761935 | 0.602059991 | 0.629532004 |
| ENSMUSG00000082121 | 1.432083896 | 1.410964047 | 1.516522298 | 0.896240625 | 1.163285805 | 1.146128036 | 1.140832123 |
| ENSMUSG00000082122 | 0 | 0 | 0 | 0 | 0 | 0 | 0 |
| ENSMUSG00000082123 | 0 | 0 | 0 | 0 | 0 | 0 | 0 |
| ENSMUSG00000082125 | 0 | 0 | 0 | 0 | 0 | 0 | 0 |
| ENSMUSG00000082126 | 0 | 0 | 0 | 0 | 0 | 0 | 0 |
| ENSMUSG00000082127 | 0 | 0 | 0 | 0 | 0 | 0 | 0 |
| ENSMUSG00000082128 | 0.45815691 | 0.25 | 0 | 0.396240625 | 0.489301084 | 0.301029996 | 0.355418052 |
| ENSMUSG00000082129 | 0 | 0 | 0 | 0 | 0 | 0 | 0 |
| ENSMUSG00000082130 | 0.289064826 | 0 | 0 | 0 | 0 | 0 | 0 |
| ENSMUSG00000082132 | 0 | 0 | 0 | 0 | 0 | 0 | 0 |
| ENSMUSG00000082134 | 0 | 0 | 0 | 0 | 0 | 0 | 0 |
| ENSMUSG00000082135 | 0 | 0 | 0 | 0 | 0 | 0 | 0 |
| ENSMUSG00000082136 | 0.289064826 | 0 | 0.5 | 0 | 0 | 0 | 0 |
| ENSMUSG00000082138 | 0 | 0 | 0 | 0 | 0 | 0 | 0 |

| | | | | | | | |
|---|---|---|---|---|---|---|---|
| ENSMUSG00000082139 | 0 | 0 | 0 | 0 | 0 | 0 | 0 |
| ENSMUSG00000082140 | 0 | 0.396240625 | 0.5 | 0.396240625 | 0.244650542 | 0.301029996 | 0.448487648 |
| ENSMUSG00000082141 | 0 | 0 | 0 | 0 | 0 | 0 | 0 |
| ENSMUSG00000082142 | 0 | 0 | 0 | 0 | 0 | 0 | 0 |
| ENSMUSG00000082143 | 0 | 0 | 0 | 0 | 0 | 0 | 0 |
| ENSMUSG00000082144 | 0.289064826 | 0.646240625 | 0.580482024 | 0.75 | 0.77552387 | 0.301029996 | 0.355418052 |
| ENSMUSG00000082145 | 0.45815691 | 0.25 | 0.5 | 0.580482024 | 0.568060967 | 0.301029996 | 0.224243824 |
| ENSMUSG00000082146 | 0 | 0 | 0 | 0 | 0 | 0 | 0 |
| ENSMUSG00000082147 | 0 | 0 | 0 | 0 | 0 | 0 | 0 |
| ENSMUSG00000082148 | 0 | 0 | 0 | 0 | 0 | 0 | 0.224243824 |
| ENSMUSG00000082149 | 0.45815691 | 0.25 | 0 | 0.396240625 | 0.244650542 | 0 | 0 |
| ENSMUSG00000082151 | 0 | 0 | 0 | 0 | 0 | 0.301029996 | 0.224243824 |
| ENSMUSG00000082152 | 0 | 0 | 0 | 0 | 0 | 0 | 0 |
| ENSMUSG00000082154 | 0 | 0 | 0 | 0 | 0 | 0 | 0 |
| ENSMUSG00000082155 | 0 | 0 | 0 | 0 | 0 | 0 | 0 |
| ENSMUSG00000082156 | 0 | 0 | 0 | 0 | 0 | 0 | 0 |
| ENSMUSG00000082157 | 0 | 0 | 0 | 0 | 0 | 0 | 0 |
| ENSMUSG00000082158 | 0 | 0 | 0 | 0 | 0 | 0 | 0 |
| ENSMUSG00000082159 | 0 | 0 | 0 | 0 | 0 | 0 | 0 |
| ENSMUSG00000082160 | 0 | 0 | 0 | 0 | 0 | 0 | 0 |
| ENSMUSG00000082161 | 0 | 0 | 0 | 0 | 0 | 0 | 0.224243824 |
| ENSMUSG00000082162 | 0 | 0 | 0 | 0 | 0 | 0 | 0 |
| ENSMUSG00000082163 | 1.587501561 | 1.418106335 | 1.311981878 | 1.451838731 | 1.509475496 | 1.633468456 | 1.429563222 |
| ENSMUSG00000082164 | 0 | 0 | 0 | 0 | 0 | 0 | 0 |
| ENSMUSG00000082165 | 0 | 0 | 0 | 0.396240625 | 0.244650542 | 0 | 0.355418052 |
| ENSMUSG00000082167 | 0 | 0 | 0 | 0 | 0 | 0 | 0 |
| ENSMUSG00000082168 | 0 | 0 | 0 | 0 | 0 | 0 | 0 |
| ENSMUSG00000082169 | 0 | 0 | 0 | 0 | 0 | 0 | 0 |
| ENSMUSG00000082170 | 0 | 0 | 0 | 0 | 0 | 0 | 0 |
| ENSMUSG00000082171 | 0 | 0 | 0.25 | 0.25 | 0 | 0 | 0 |
| ENSMUSG00000082172 | 0 | 0 | 0 | 0 | 0 | 0 | 0 |
| ENSMUSG00000082173 | 0 | 0 | 0 | 0 | 0 | 0 | 0 |
| ENSMUSG00000082174 | 0 | 0 | 0 | 0 | 0 | 0 | 0 |
| ENSMUSG00000082175 | 0 | 0 | 0 | 0 | 0 | 0 | 0 |

| | | | | | | | |
|---|---|---|---|---|---|---|---|
| ENSMUSG00000082176 | 0 | 0 | 0 | 0 | 0 | 0 | 0 |
| ENSMUSG00000082177 | 0 | 0.25 | 0 | 0 | 0 | 0 | 0 |
| ENSMUSG00000082178 | 0.289064826 | 0 | 0.5 | 0 | 0 | 0 | 0.355418052 |
| ENSMUSG00000082179 | 0.747221736 | 0.79248125 | 0.5 | 0.5 | 0.733951626 | 0.77815125 | 0.448487648 |
| ENSMUSG00000082180 | 0.45815691 | 0 | 0 | 0 | 0.244650542 | 0.301029996 | 0.224243824 |
| ENSMUSG00000082181 | 0 | 0 | 0 | 0 | 0 | 0 | 0 |
| ENSMUSG00000082183 | 0 | 0.25 | 0 | 0 | 0.244650542 | 0.301029996 | 0 |
| ENSMUSG00000082185 | 0.289064826 | 0 | 0 | 0 | 0 | 0 | 0 |
| ENSMUSG00000082186 | 0 | 0.5 | 0.25 | 0.25 | 0.387761935 | 0.301029996 | 0.355418052 |
| ENSMUSG00000082187 | 0 | 0 | 0 | 0 | 0 | 0 | 0.224243824 |
| ENSMUSG00000082188 | 0 | 0 | 0 | 0 | 0 | 0 | 0 |
| ENSMUSG00000082189 | 0.289064826 | 0.580482024 | 0.396240625 | 0.25 | 0.244650542 | 0 | 0 |
| ENSMUSG00000082190 | 0 | 0 | 0.580482024 | 0.5 | 0.568060967 | 0.602059991 | 0.448487648 |
| ENSMUSG00000082191 | 0 | 0 | 0 | 0 | 0 | 0 | 0 |
| ENSMUSG00000082192 | 0 | 0 | 0 | 0 | 0 | 0 | 0 |
| ENSMUSG00000082193 | 0.289064826 | 0.25 | 0 | 0.396240625 | 0.686820904 | 0.301029996 | 0.448487648 |
| ENSMUSG00000082194 | 0.811507563 | 0.701838731 | 0.79248125 | 0.701838731 | 0.632412477 | 0.698970004 | 0.672731473 |
| ENSMUSG00000082195 | 0 | 0 | 0 | 0 | 0 | 0 | 0 |
| ENSMUSG00000082196 | 0 | 0 | 0 | 0.580482024 | 0.387761935 | 0.301029996 | 0.355418052 |
| ENSMUSG00000082197 | 0 | 0 | 0 | 0 | 0 | 0 | 0 |
| ENSMUSG00000082198 | 0 | 0 | 0 | 0 | 0.489301084 | 0 | 0.224243824 |
| ENSMUSG00000082199 | 0 | 0 | 0 | 0 | 0 | 0 | 0 |
| ENSMUSG00000082200 | 0 | 0 | 0 | 0 | 0 | 0 | 0 |
| ENSMUSG00000082202 | 0 | 0 | 0 | 0 | 0 | 0 | 0.224243824 |
| ENSMUSG00000082203 | 0 | 0 | 0 | 0 | 0 | 0 | 0 |
| ENSMUSG00000082204 | 0 | 0.25 | 0 | 0.396240625 | 0.244650542 | 0 | 0.224243824 |
| ENSMUSG00000082205 | 0 | 0 | 0 | 0 | 0 | 0 | 0 |
| ENSMUSG00000082207 | 0 | 0 | 0 | 0 | 0 | 0 | 0 |
| ENSMUSG00000082208 | 0.45815691 | 0.5 | 0.25 | 0.580482024 | 0 | 0.602059991 | 0.355418052 |
| ENSMUSG00000082210 | 0 | 0 | 0 | 0 | 0 | 0 | 0 |
| ENSMUSG00000082211 | 0 | 0 | 0 | 0 | 0 | 0 | 0 |
| ENSMUSG00000082212 | 0 | 0 | 0 | 0 | 0 | 0 | 0 |
| ENSMUSG00000082213 | 0 | 0 | 0 | 0 | 0 | 0 | 0.224243824 |
| ENSMUSG00000082214 | 0 | 0 | 0 | 0 | 0 | 0 | 0 |

| | | | | | | | |
|---|---|---|---|---|---|---|---|
| ENSMUSG00000082215 | 0 | 0 | 0 | 0 | 0 | 0 | 0 |
| ENSMUSG00000082216 | 0 | 0 | 0 | 0 | 0 | 0 | 0 |
| ENSMUSG00000082217 | 0 | 0 | 0 | 0 | 0 | 0 | 0 |
| ENSMUSG00000082218 | 0 | 0.396240625 | 0.25 | 0.25 | 0 | 0 | 0.224243824 |
| ENSMUSG00000082220 | 0 | 0.25 | 0.580482024 | 0 | 0.387761935 | 0 | 0.355418052 |
| ENSMUSG00000082222 | 0 | 0 | 0 | 0 | 0 | 0 | 0 |
| ENSMUSG00000082223 | 0 | 0 | 0 | 0 | 0 | 0 | 0 |
| ENSMUSG00000082225 | 0 | 0 | 0 | 0 | 0 | 0 | 0 |
| ENSMUSG00000082226 | 0 | 0 | 0 | 0 | 0 | 0 | 0.224243824 |
| ENSMUSG00000082227 | 0 | 0 | 0 | 0 | 0 | 0 | 0 |
| ENSMUSG00000082228 | 0 | 0 | 0 | 0 | 0 | 0 | 0 |
| ENSMUSG00000082229 | 0.811507563 | 1.061981878 | 0.701838731 | 1.403677461 | 1.200473444 | 1.278753601 | 1.176815335 |
| ENSMUSG00000082233 | 0.45815691 | 0.396240625 | 0 | 0.396240625 | 0 | 0.301029996 | 0 |
| ENSMUSG00000082234 | 0 | 0 | 0 | 0 | 0 | 0 | 0 |
| ENSMUSG00000082235 | 0.289064826 | 0 | 0 | 0 | 0 | 0 | 0 |
| ENSMUSG00000082236 | 0 | 0 | 0 | 0 | 0 | 0 | 0 |
| ENSMUSG00000082237 | 0 | 0 | 0 | 0 | 0 | 0 | 0 |
| ENSMUSG00000082238 | 0 | 0 | 0 | 0 | 0 | 0 | 0 |
| ENSMUSG00000082239 | 0 | 0 | 0 | 0 | 0 | 0 | 0 |
| ENSMUSG00000082240 | 0 | 0 | 0 | 0 | 0 | 0 | 0 |
| ENSMUSG00000082241 | 0 | 0 | 0 | 0 | 0 | 0 | 0 |
| ENSMUSG00000082242 | 0 | 0 | 0 | 0 | 0 | 0 | 0 |
| ENSMUSG00000082243 | 0 | 0 | 0 | 0.25 | 0 | 0 | 0 |
| ENSMUSG00000082244 | 0.289064826 | 0 | 0.580482024 | 0.396240625 | 0.387761935 | 0 | 0 |
| ENSMUSG00000082245 | 0 | 0 | 0 | 0 | 0 | 0 | 0 |
| ENSMUSG00000082246 | 0.578129653 | 0.580482024 | 0.25 | 0.701838731 | 0.568060967 | 0.301029996 | 0.829800754 |
| ENSMUSG00000082247 | 0 | 0 | 0 | 0 | 0 | 0 | 0.224243824 |
| ENSMUSG00000082248 | 0 | 0 | 0 | 0 | 0 | 0 | 0 |
| ENSMUSG00000082249 | 0 | 0 | 0 | 0 | 0 | 0 | 0 |
| ENSMUSG00000082250 | 0 | 0.396240625 | 0 | 0 | 0 | 0.301029996 | 0.224243824 |
| ENSMUSG00000082251 | 0 | 0 | 0 | 0 | 0 | 0 | 0 |
| ENSMUSG00000082252 | 0 | 0 | 0 | 0 | 0 | 0 | 0 |
| ENSMUSG00000082253 | 0.289064826 | 0 | 0.25 | 0 | 0.244650542 | 0 | 0 |
| ENSMUSG00000082254 | 0 | 0 | 0 | 0 | 0 | 0 | 0.224243824 |

| | | | | | | | |
|---|---|---|---|---|---|---|---|
| ENSMUSG00000082255 | 0 | 0 | 0 | 0 | 0 | 0 | 0 |
| ENSMUSG00000082256 | 0 | 0 | 0 | 0 | 0 | 0 | 0 |
| ENSMUSG00000082258 | 0 | 0 | 0 | 0.25 | 0 | 0 | 0 |
| ENSMUSG00000082259 | 0 | 0 | 0 | 0 | 0 | 0 | 0 |
| ENSMUSG00000082260 | 0 | 0 | 0 | 0 | 0.244650542 | 0 | 0.224243824 |
| ENSMUSG00000082261 | 0 | 0 | 0 | 0 | 0 | 0 | 0 |
| ENSMUSG00000082262 | 0 | 0 | 0.396240625 | 0.396240625 | 0.387761935 | 0.477121255 | 0.355418052 |
| ENSMUSG00000082263 | 0 | 0 | 0 | 0 | 0 | 0 | 0 |
| ENSMUSG00000082264 | 0 | 0 | 0.25 | 0 | 0 | 0.301029996 | 0 |
| ENSMUSG00000082265 | 0.289064826 | 0.25 | 0 | 0.580482024 | 0.489301084 | 0.301029996 | 0.224243824 |
| ENSMUSG00000082266 | 0 | 0 | 0 | 0 | 0 | 0 | 0 |
| ENSMUSG00000082267 | 0 | 0 | 0 | 0 | 0 | 0 | 0 |
| ENSMUSG00000082268 | 0 | 0 | 0 | 0 | 0 | 0 | 0 |
| ENSMUSG00000082269 | 0 | 0 | 0 | 0 | 0 | 0 | 0 |
| ENSMUSG00000082270 | 0 | 0 | 0 | 0 | 0 | 0 | 0 |
| ENSMUSG00000082271 | 0 | 0 | 0 | 0 | 0 | 0 | 0 |
| ENSMUSG00000082272 | 0.289064826 | 0 | 0.25 | 0.25 | 0 | 0.477121255 | 0.224243824 |
| ENSMUSG00000082274 | 0 | 0 | 0.25 | 0 | 0 | 0 | 0 |
| ENSMUSG00000082275 | 0 | 0 | 0 | 0 | 0 | 0 | 0 |
| ENSMUSG00000082276 | 0 | 0 | 0 | 0 | 0.244650542 | 0 | 0 |
| ENSMUSG00000082277 | 0.289064826 | 0.5 | 0.396240625 | 0.25 | 0 | 0.301029996 | 0.355418052 |
| ENSMUSG00000082278 | 0 | 0 | 0 | 0 | 0 | 0 | 0 |
| ENSMUSG00000082279 | 0 | 0.25 | 0.396240625 | 0.25 | 0.244650542 | 0 | 0.355418052 |
| ENSMUSG00000082280 | 0 | 0 | 0.25 | 0 | 0 | 0 | 0 |
| ENSMUSG00000082281 | 0 | 0 | 0 | 0 | 0 | 0 | 0 |
| ENSMUSG00000082283 | 0 | 0 | 0 | 0 | 0 | 0 | 0.224243824 |
| ENSMUSG00000082284 | 0 | 0 | 0.25 | 0 | 0 | 0 | 0 |
| ENSMUSG00000082285 | 0 | 0 | 0 | 0 | 0 | 0 | 0.224243824 |
| ENSMUSG00000082286 | 1.578129653 | 1.575945187 | 1.561981878 | 1.330482024 | 1.572189156 | 1.51851394 | 1.455757965 |
| ENSMUSG00000082287 | 0 | 0 | 0 | 0 | 0 | 0 | 0 |
| ENSMUSG00000082288 | 0 | 0 | 0 | 0 | 0 | 0 | 0 |
| ENSMUSG00000082289 | 0 | 0.25 | 0 | 0 | 0.244650542 | 0 | 0.224243824 |
| ENSMUSG00000082291 | 0 | 0 | 0 | 0 | 0 | 0 | 0 |
| ENSMUSG00000082292 | 0.747221736 | 0.646240625 | 0.25 | 0.75 | 0.812711509 | 0.84509804 | 0.775756176 |

| | | | | | | | |
|---|---|---|---|---|---|---|---|
| ENSMUSG00000082294 | 0 | 0 | 0 | 0 | 0 | 0 | 0 |
| ENSMUSG00000082295 | 0 | 0 | 0 | 0 | 0 | 0 | 0 |
| ENSMUSG00000082296 | 0.45815691 | 0 | 0.396240625 | 0 | 0 | 0 | 0.520678036 |
| ENSMUSG00000082297 | 0 | 0 | 0 | 0 | 0 | 0 | 0 |
| ENSMUSG00000082301 | 0 | 0 | 0 | 0 | 0 | 0 | 0 |
| ENSMUSG00000082302 | 0 | 0 | 0 | 0 | 0 | 0 | 0 |
| ENSMUSG00000082303 | 0 | 0 | 0 | 0 | 0 | 0 | 0 |
| ENSMUSG00000082304 | 0.289064826 | 0 | 0 | 0 | 0.387761935 | 0 | 0 |
| ENSMUSG00000082305 | 0.578129653 | 0.396240625 | 0.5 | 0.396240625 | 0.244650542 | 0.477121255 | 0.520678036 |
| ENSMUSG00000082306 | 0 | 0 | 0 | 0 | 0 | 0 | 0 |
| ENSMUSG00000082307 | 0 | 0 | 0 | 0.25 | 0 | 0 | 0.520678036 |
| ENSMUSG00000082308 | 0 | 0 | 0 | 0 | 0 | 0 | 0 |
| ENSMUSG00000082309 | 0 | 0 | 0 | 0.25 | 0 | 0.301029996 | 0 |
| ENSMUSG00000082310 | 0 | 0 | 0 | 0 | 0 | 0 | 0 |
| ENSMUSG00000082311 | 0 | 0 | 0 | 0 | 0 | 0 | 0 |
| ENSMUSG00000082312 | 0 | 0 | 0.25 | 0 | 0 | 0 | 0 |
| ENSMUSG00000082313 | 0.289064826 | 0 | 0 | 0 | 0.244650542 | 0.301029996 | 0.355418052 |
| ENSMUSG00000082314 | 0 | 0 | 0 | 0 | 0 | 0 | 0 |
| ENSMUSG00000082315 | 0 | 0.396240625 | 0.396240625 | 0.396240625 | 0.244650542 | 0.301029996 | 0.224243824 |
| ENSMUSG00000082316 | 0 | 0 | 0 | 0 | 0 | 0 | 0 |
| ENSMUSG00000082317 | 0.289064826 | 0.396240625 | 0 | 0.25 | 0 | 0 | 0.224243824 |
| ENSMUSG00000082318 | 0 | 0 | 0 | 0 | 0 | 0 | 0.224243824 |
| ENSMUSG00000082319 | 0.747221736 | 0.701838731 | 0 | 0.396240625 | 0.812711509 | 0.698970004 | 0.579661877 |
| ENSMUSG00000082320 | 0 | 0 | 0 | 0 | 0 | 0 | 0 |
| ENSMUSG00000082321 | 0 | 0 | 0 | 0 | 0 | 0 | 0 |
| ENSMUSG00000082322 | 0.671187741 | 0.701838731 | 0.25 | 0.25 | 0.568060967 | 0 | 0.224243824 |
| ENSMUSG00000082323 | 0 | 0 | 0 | 0 | 0 | 0 | 0 |
| ENSMUSG00000082324 | 0 | 0 | 0.25 | 0.5 | 0.387761935 | 0 | 0.355418052 |
| ENSMUSG00000082325 | 0 | 0.396240625 | 0.396240625 | 0.396240625 | 0 | 0 | 0.448487648 |
| ENSMUSG00000082326 | 0 | 0 | 0 | 0 | 0 | 0 | 0 |
| ENSMUSG00000082327 | 0 | 0 | 0 | 0 | 0 | 0 | 0 |
| ENSMUSG00000082328 | 0 | 0 | 0 | 0 | 0 | 0 | 0.224243824 |
| ENSMUSG00000082329 | 0 | 0 | 0 | 0 | 0 | 0 | 0 |
| ENSMUSG00000082330 | 0 | 0.25 | 0 | 0 | 0 | 0 | 0 |

| | | | | | | | |
|---|---|---|---|---|---|---|---|
| ENSMUSG00000082332 | 0 | 0 | 0 | 0 | 0 | 0 | 0 |
| ENSMUSG00000082333 | 0 | 0 | 0 | 0 | 0 | 0 | 0 |
| ENSMUSG00000082334 | 0 | 0 | 0 | 0 | 0 | 0 | 0 |
| ENSMUSG00000082336 | 0 | 0 | 0 | 0 | 0 | 0 | 0 |
| ENSMUSG00000082337 | 0 | 0 | 0 | 0 | 0 | 0 | 0 |
| ENSMUSG00000082338 | 0 | 0 | 0 | 0 | 0 | 0 | 0 |
| ENSMUSG00000082339 | 0 | 0 | 0 | 0 | 0 | 0 | 0 |
| ENSMUSG00000082340 | 0 | 0 | 0 | 0 | 0 | 0 | 0 |
| ENSMUSG00000082341 | 0 | 0 | 0 | 0 | 0 | 0 | 0 |
| ENSMUSG00000082342 | 0 | 0 | 0.25 | 0 | 0.244650542 | 0 | 0 |
| ENSMUSG00000082343 | 0 | 0 | 0 | 0 | 0 | 0 | 0 |
| ENSMUSG00000082344 | 0 | 0 | 0 | 0 | 0 | 0 | 0 |
| ENSMUSG00000082345 | 0.45815691 | 0.646240625 | 0.396240625 | 0.75 | 0.387761935 | 0 | 0.224243824 |
| ENSMUSG00000082346 | 0 | 0 | 0 | 0 | 0 | 0 | 0 |
| ENSMUSG00000082347 | 0 | 0 | 0 | 0 | 0 | 0 | 0 |
| ENSMUSG00000082348 | 0 | 0 | 0.25 | 0 | 0.244650542 | 0 | 0 |
| ENSMUSG00000082350 | 0 | 0 | 0 | 0 | 0 | 0 | 0 |
| ENSMUSG00000082351 | 0 | 0 | 0 | 0 | 0 | 0 | 0 |
| ENSMUSG00000082352 | 0 | 0 | 0 | 0 | 0 | 0 | 0 |
| ENSMUSG00000082353 | 0 | 0 | 0 | 0 | 0 | 0 | 0 |
| ENSMUSG00000082354 | 0 | 0 | 0 | 0 | 0 | 0 | 0 |
| ENSMUSG00000082356 | 0 | 0 | 0 | 0 | 0 | 0 | 0 |
| ENSMUSG00000082357 | 0 | 0.25 | 0 | 0 | 0.387761935 | 0 | 0.355418052 |
| ENSMUSG00000082358 | 0 | 0 | 0 | 0 | 0 | 0 | 0 |
| ENSMUSG00000082359 | 0 | 0 | 0 | 0 | 0 | 0 | 0 |
| ENSMUSG00000082360 | 0 | 0 | 0.25 | 0 | 0 | 0 | 0.224243824 |
| ENSMUSG00000082361 | 1.227926429 | 1.29248125 | 1.160964047 | 0.864857905 | 1 | 0.954242509 | 0.984950056 |
| ENSMUSG00000082362 | 0.45815691 | 0.396240625 | 0.580482024 | 0 | 0.244650542 | 0 | 0.355418052 |
| ENSMUSG00000082363 | 0 | 0 | 0 | 0 | 0 | 0 | 0 |
| ENSMUSG00000082364 | 0 | 0 | 0 | 0 | 0 | 0 | 0 |
| ENSMUSG00000082365 | 0 | 0 | 0 | 0 | 0 | 0 | 0 |
| ENSMUSG00000082367 | 0 | 0 | 0 | 0 | 0 | 0 | 0 |
| ENSMUSG00000082368 | 0 | 0 | 0 | 0 | 0 | 0 | 0 |
| ENSMUSG00000082369 | 0 | 0 | 0 | 0 | 0 | 0 | 0 |

| | | | | | | | |
|---|---|---|---|---|---|---|---|
| ENSMUSG00000082370 | 0 | 0 | 0 | 0 | 0 | 0 | 0 |
| ENSMUSG00000082371 | 0 | 0 | 0 | 0 | 0 | 0 | 0 |
| ENSMUSG00000082372 | 0 | 0 | 0 | 0 | 0 | 0 | 0 |
| ENSMUSG00000082373 | 0 | 0 | 0 | 0 | 0 | 0 | 0 |
| ENSMUSG00000082374 | 0 | 0 | 0 | 0.25 | 0 | 0 | 0 |
| ENSMUSG00000082375 | 0.289064826 | 0.25 | 0 | 0.25 | 0 | 0 | 0 |
| ENSMUSG00000082376 | 0 | 0.25 | 0.25 | 0 | 0.244650542 | 0 | 0.224243824 |
| ENSMUSG00000082377 | 0 | 0 | 0 | 0.25 | 0 | 0.301029996 | 0 |
| ENSMUSG00000082378 | 0 | 0 | 0 | 0 | 0 | 0 | 0 |
| ENSMUSG00000082379 | 0 | 0 | 0 | 0 | 0 | 0 | 0 |
| ENSMUSG00000082380 | 0 | 0 | 0 | 0 | 0 | 0 | 0 |
| ENSMUSG00000082381 | 0 | 0 | 0 | 0 | 0 | 0 | 0 |
| ENSMUSG00000082382 | 0 | 0 | 0.25 | 0 | 0 | 0 | 0 |
| ENSMUSG00000082383 | 0 | 0.5 | 0.25 | 0.25 | 0 | 0.602059991 | 0.224243824 |
| ENSMUSG00000082388 | 0 | 0 | 0 | 0 | 0 | 0 | 0 |
| ENSMUSG00000082389 | 0 | 0 | 0 | 0 | 0 | 0 | 0 |
| ENSMUSG00000082390 | 0 | 0.25 | 0.25 | 0.396240625 | 0 | 0 | 0.224243824 |
| ENSMUSG00000082392 | 0 | 0 | 0 | 0 | 0 | 0 | 0 |
| ENSMUSG00000082393 | 0 | 0 | 0 | 0 | 0 | 0 | 0 |
| ENSMUSG00000082394 | 0 | 0 | 0 | 0 | 0.244650542 | 0 | 0 |
| ENSMUSG00000082395 | 0.289064826 | 0.25 | 0.396240625 | 0 | 0.387761935 | 0 | 0 |
| ENSMUSG00000082396 | 0.45815691 | 0.25 | 0.396240625 | 0.5 | 0.632412477 | 0.301029996 | 0.355418052 |
| ENSMUSG00000082397 | 0 | 0 | 0 | 0 | 0 | 0 | 0 |
| ENSMUSG00000082398 | 0 | 0 | 0 | 0 | 0 | 0 | 0 |
| ENSMUSG00000082399 | 0 | 0 | 0 | 0 | 0 | 0 | 0 |
| ENSMUSG00000082400 | 0 | 0 | 0 | 0 | 0.244650542 | 0 | 0 |
| ENSMUSG00000082402 | 0 | 0 | 0.25 | 0.396240625 | 0 | 0.301029996 | 0.355418052 |
| ENSMUSG00000082403 | 0 | 0 | 0 | 0 | 0 | 0 | 0 |
| ENSMUSG00000082404 | 0.45815691 | 0 | 0 | 0.25 | 0 | 0 | 0 |
| ENSMUSG00000082405 | 0 | 0 | 0 | 0 | 0 | 0 | 0 |
| ENSMUSG00000082406 | 0 | 0.25 | 0 | 0.25 | 0.244650542 | 0 | 0 |
| ENSMUSG00000082407 | 0 | 0 | 0 | 0 | 0 | 0 | 0 |
| ENSMUSG00000082408 | 0 | 0 | 0 | 0 | 0 | 0 | 0 |
| ENSMUSG00000082410 | 0 | 0 | 0 | 0 | 0 | 0 | 0 |

| | | | | | | | |
|---|---|---|---|---|---|---|---|
| ENSMUSG00000082411 | 0 | 0 | 0 | 0 | 0 | 0 | 0 |
| ENSMUSG00000082412 | 0 | 0 | 0 | 0 | 0 | 0 | 0 |
| ENSMUSG00000082413 | 0 | 0 | 0 | 0 | 0 | 0 | 0 |
| ENSMUSG00000082414 | 0 | 0 | 0 | 0 | 0 | 0 | 0 |
| ENSMUSG00000082415 | 0 | 0 | 0 | 0 | 0 | 0 | 0 |
| ENSMUSG00000082416 | 0 | 0 | 0 | 0 | 0 | 0 | 0 |
| ENSMUSG00000082417 | 0 | 0 | 0 | 0 | 0 | 0 | 0 |
| ENSMUSG00000082418 | 0 | 0 | 0 | 0 | 0 | 0 | 0.224243824 |
| ENSMUSG00000082419 | 0 | 0 | 0 | 0 | 0 | 0 | 0 |
| ENSMUSG00000082420 | 0.578129653 | 0.701838731 | 0.580482024 | 0.5 | 0.568060967 | 0.301029996 | 0.355418052 |
| ENSMUSG00000082421 | 0 | 0 | 0 | 0 | 0 | 0 | 0 |
| ENSMUSG00000082422 | 0 | 0 | 0 | 0 | 0 | 0 | 0 |
| ENSMUSG00000082423 | 0.289064826 | 0 | 0 | 0 | 0 | 0.301029996 | 0 |
| ENSMUSG00000082424 | 0 | 0 | 0 | 0 | 0 | 0 | 0 |
| ENSMUSG00000082426 | 0 | 0 | 0 | 0 | 0 | 0 | 0 |
| ENSMUSG00000082427 | 0 | 0 | 0 | 0 | 0 | 0 | 0 |
| ENSMUSG00000082428 | 0 | 0 | 0 | 0 | 0 | 0 | 0 |
| ENSMUSG00000082429 | 0 | 0 | 0 | 0 | 0 | 0 | 0 |
| ENSMUSG00000082430 | 0 | 0 | 0 | 0 | 0 | 0 | 0 |
| ENSMUSG00000082431 | 0 | 0.25 | 0.396240625 | 0.25 | 0.244650542 | 0.301029996 | 0 |
| ENSMUSG00000082432 | 0 | 0 | 0 | 0 | 0.244650542 | 0 | 0 |
| ENSMUSG00000082433 | 0 | 0 | 0 | 0 | 0 | 0 | 0 |
| ENSMUSG00000082435 | 0 | 0 | 0 | 0 | 0 | 0 | 0 |
| ENSMUSG00000082436 | 0.578129653 | 0.5 | 0.646240625 | 0.5 | 0.244650542 | 0 | 0.629532004 |
| ENSMUSG00000082437 | 0 | 0 | 0 | 0 | 0 | 0 | 0 |
| ENSMUSG00000082440 | 0 | 0 | 0 | 0 | 0 | 0 | 0 |
| ENSMUSG00000082442 | 0 | 0 | 0 | 0 | 0 | 0 | 0 |
| ENSMUSG00000082443 | 0 | 0 | 0 | 0 | 0 | 0 | 0 |
| ENSMUSG00000082444 | 0 | 0 | 0 | 0 | 0 | 0 | 0 |
| ENSMUSG00000082446 | 0 | 0 | 0 | 0 | 0 | 0 | 0 |
| ENSMUSG00000082447 | 0 | 0 | 0 | 0 | 0 | 0 | 0 |
| ENSMUSG00000082448 | 0 | 0 | 0 | 0 | 0 | 0 | 0.224243824 |
| ENSMUSG00000082449 | 0 | 0 | 0 | 0 | 0 | 0 | 0 |
| ENSMUSG00000082450 | 0 | 0 | 0 | 0 | 0 | 0 | 0 |

| | | | | | | | |
|---|---|---|---|---|---|---|---|
| ENSMUSG00000082451 | 0 | 0 | 0 | 0 | 0 | 0 | 0 |
| ENSMUSG00000082453 | 0 | 0 | 0 | 0 | 0 | 0 | 0 |
| ENSMUSG00000082454 | 0.867194479 | 0.75 | 0.5 | 0.896240625 | 0.877063019 | 0.602059991 | 0.672731473 |
| ENSMUSG00000082455 | 0 | 0 | 0 | 0 | 0 | 0 | 0 |
| ENSMUSG00000082456 | 0 | 0 | 0 | 0 | 0 | 0 | 0 |
| ENSMUSG00000082457 | 0 | 0.25 | 0 | 0 | 0 | 0 | 0 |
| ENSMUSG00000082458 | 0.289064826 | 0 | 0.25 | 0.25 | 0 | 0 | 0 |
| ENSMUSG00000082459 | 0 | 0 | 0 | 0.25 | 0 | 0 | 0 |
| ENSMUSG00000082460 | 0 | 0 | 0 | 0 | 0 | 0 | 0 |
| ENSMUSG00000082461 | 0 | 0.25 | 0.396240625 | 0.25 | 0.244650542 | 0 | 0.224243824 |
| ENSMUSG00000082462 | 0 | 0 | 0 | 0 | 0 | 0 | 0 |
| ENSMUSG00000082463 | 0 | 0 | 0 | 0 | 0 | 0 | 0 |
| ENSMUSG00000082464 | 0 | 0 | 0 | 0 | 0 | 0 | 0 |
| ENSMUSG00000082465 | 0.289064826 | 0.25 | 0.25 | 0 | 0.244650542 | 0 | 0 |
| ENSMUSG00000082467 | 0 | 0 | 0 | 0.25 | 0 | 0 | 0 |
| ENSMUSG00000082468 | 0 | 0 | 0 | 0 | 0 | 0 | 0 |
| ENSMUSG00000082472 | 0 | 0 | 0 | 0 | 0 | 0.301029996 | 0 |
| ENSMUSG00000082473 | 0 | 0 | 0 | 0 | 0 | 0 | 0 |
| ENSMUSG00000082474 | 0 | 0.396240625 | 0 | 0.25 | 0.387761935 | 0.477121255 | 0.520678036 |
| ENSMUSG00000082475 | 0 | 0 | 0 | 0 | 0.244650542 | 0.301029996 | 0.355418052 |
| ENSMUSG00000082476 | 0 | 0.396240625 | 0.396240625 | 0 | 0 | 0 | 0.224243824 |
| ENSMUSG00000082478 | 0 | 0 | 0 | 0 | 0 | 0 | 0 |
| ENSMUSG00000082479 | 0 | 0 | 0 | 0 | 0 | 0 | 0 |
| ENSMUSG00000082480 | 0.45815691 | 0.25 | 0.25 | 0.396240625 | 0.244650542 | 0 | 0.355418052 |
| ENSMUSG00000082481 | 0.289064826 | 0.25 | 0.396240625 | 0 | 0 | 0 | 0.355418052 |
| ENSMUSG00000082482 | 0 | 0 | 0 | 0 | 0 | 0 | 0 |
| ENSMUSG00000082483 | 0 | 0.646240625 | 0.25 | 0.5 | 0.387761935 | 0 | 0.520678036 |
| ENSMUSG00000082484 | 0 | 0.25 | 0.25 | 0 | 0.244650542 | 0 | 0.224243824 |
| ENSMUSG00000082485 | 0 | 0 | 0 | 0 | 0 | 0 | 0 |
| ENSMUSG00000082486 | 0 | 0 | 0 | 0 | 0 | 0 | 0 |
| ENSMUSG00000082487 | 0 | 0 | 0 | 0 | 0 | 0 | 0 |
| ENSMUSG00000082488 | 0 | 0 | 0 | 0 | 0 | 0 | 0 |
| ENSMUSG00000082489 | 0 | 0 | 0 | 0 | 0 | 0 | 0 |
| ENSMUSG00000082490 | 0 | 0 | 0 | 0 | 0 | 0 | 0 |

| | | | | | | | |
|---|---|---|---|---|---|---|---|
| ENSMUSG00000082491 | 0.289064826 | 0.646240625 | 0.830482024 | 0.896240625 | 0.846351821 | 0.301029996 | 0.579661877 |
| ENSMUSG00000082492 | 0 | 0 | 0 | 0 | 0 | 0 | 0 |
| ENSMUSG00000082494 | 0 | 0 | 0 | 0 | 0.244650542 | 0 | 0.355418052 |
| ENSMUSG00000082495 | 0 | 0 | 0 | 0 | 0 | 0 | 0 |
| ENSMUSG00000082496 | 0 | 0 | 0 | 0 | 0 | 0 | 0 |
| ENSMUSG00000082497 | 0 | 0 | 0 | 0 | 0 | 0 | 0 |
| ENSMUSG00000082498 | 0 | 0 | 0.25 | 0 | 0 | 0 | 0.355418052 |
| ENSMUSG00000082499 | 0.289064826 | 0 | 0.396240625 | 0.396240625 | 0 | 0 | 0.355418052 |
| ENSMUSG00000082500 | 0 | 0 | 0 | 0 | 0 | 0 | 0 |
| ENSMUSG00000082501 | 0 | 0 | 0 | 0 | 0 | 0 | 0 |
| ENSMUSG00000082502 | 0 | 0 | 0 | 0 | 0 | 0 | 0 |
| ENSMUSG00000082503 | 0 | 0 | 0 | 0 | 0 | 0 | 0 |
| ENSMUSG00000082504 | 0 | 0 | 0 | 0 | 0 | 0 | 0 |
| ENSMUSG00000082505 | 0 | 0 | 0 | 0 | 0 | 0 | 0 |
| ENSMUSG00000082507 | 0.289064826 | 0.5 | 0 | 0.5 | 0.244650542 | 0.602059991 | 0.355418052 |
| ENSMUSG00000082508 | 0 | 0 | 0 | 0 | 0 | 0 | 0 |
| ENSMUSG00000082509 | 0 | 0 | 0 | 0 | 0 | 0 | 0 |
| ENSMUSG00000082510 | 0.45815691 | 0.646240625 | 0.5 | 0.25 | 0.244650542 | 0 | 0.629532004 |
| ENSMUSG00000082511 | 0 | 0 | 0.25 | 0 | 0 | 0 | 0 |
| ENSMUSG00000082512 | 0 | 0 | 0.25 | 0.25 | 0 | 0 | 0.224243824 |
| ENSMUSG00000082513 | 0 | 0 | 0 | 0 | 0 | 0 | 0 |
| ENSMUSG00000082514 | 0 | 0 | 0 | 0 | 0 | 0 | 0 |
| ENSMUSG00000082515 | 0.578129653 | 0.646240625 | 0.25 | 0.25 | 0.489301084 | 0 | 0.224243824 |
| ENSMUSG00000082516 | 0 | 0 | 0 | 0 | 0 | 0 | 0 |
| ENSMUSG00000082517 | 0 | 0 | 0 | 0 | 0 | 0 | 0 |
| ENSMUSG00000082519 | 0 | 0 | 0 | 0 | 0 | 0 | 0 |
| ENSMUSG00000082520 | 0 | 0 | 0 | 0 | 0 | 0 | 0 |
| ENSMUSG00000082523 | 0 | 0 | 0 | 0 | 0 | 0 | 0 |
| ENSMUSG00000082524 | 0 | 0 | 0 | 0.25 | 0 | 0 | 0 |
| ENSMUSG00000082526 | 0 | 0 | 0 | 0 | 0 | 0 | 0 |
| ENSMUSG00000082527 | 0 | 0 | 0 | 0 | 0 | 0 | 0 |
| ENSMUSG00000082528 | 0 | 0 | 0 | 0 | 0 | 0 | 0 |
| ENSMUSG00000082529 | 0 | 0 | 0 | 0 | 0 | 0 | 0 |
| ENSMUSG00000082530 | 0 | 0 | 0 | 0 | 0 | 0 | 0.224243824 |

| | | | | | | | |
|---|---|---|---|---|---|---|---|
| ENSMUSG00000082531 | 0 | 0 | 0 | 0 | 0 | 0 | 0 |
| ENSMUSG00000082532 | 0 | 0.25 | 0 | 0.5 | 0.387761935 | 0 | 0.224243824 |
| ENSMUSG00000082533 | 0 | 0 | 0 | 0 | 0 | 0 | 0 |
| ENSMUSG00000082534 | 0 | 0 | 0 | 0 | 0 | 0 | 0 |
| ENSMUSG00000082535 | 0.671187741 | 0.75 | 0.701838731 | 0.5 | 0.686820904 | 0.477121255 | 0.448487648 |
| ENSMUSG00000082536 | 1.614416215 | 1.630890489 | 1.584962501 | 1.692046081 | 1.546663136 | 1.748188027 | 1.502532226 |
| ENSMUSG00000082537 | 0 | 0 | 0 | 0 | 0 | 0 | 0 |
| ENSMUSG00000082538 | 0 | 0 | 0 | 0 | 0 | 0 | 0 |
| ENSMUSG00000082539 | 0 | 0 | 0 | 0 | 0 | 0 | 0 |
| ENSMUSG00000082540 | 0 | 0 | 0 | 0 | 0 | 0 | 0 |
| ENSMUSG00000082541 | 0 | 0 | 0 | 0 | 0 | 0 | 0 |
| ENSMUSG00000082542 | 0 | 0.25 | 0 | 0 | 0 | 0 | 0.224243824 |
| ENSMUSG00000082543 | 0.45815691 | 0.396240625 | 0.25 | 0.25 | 0.489301084 | 0 | 0.224243824 |
| ENSMUSG00000082544 | 0 | 0.25 | 0.396240625 | 0 | 0.244650542 | 0.477121255 | 0 |
| ENSMUSG00000082545 | 0 | 0 | 0 | 0 | 0 | 0 | 0.224243824 |
| ENSMUSG00000082546 | 0 | 0 | 0 | 0 | 0 | 0 | 0 |
| ENSMUSG00000082547 | 0 | 0 | 0 | 0 | 0 | 0 | 0 |
| ENSMUSG00000082548 | 0 | 0 | 0 | 0 | 0 | 0 | 0 |
| ENSMUSG00000082549 | 0 | 0.5 | 0.580482024 | 0.25 | 0.244650542 | 0 | 0.224243824 |
| ENSMUSG00000082550 | 0 | 0 | 0 | 0 | 0 | 0 | 0.224243824 |
| ENSMUSG00000082551 | 0 | 0 | 0 | 0 | 0 | 0 | 0 |
| ENSMUSG00000082552 | 0 | 0 | 0 | 0 | 0 | 0 | 0 |
| ENSMUSG00000082553 | 0 | 0 | 0 | 0 | 0 | 0 | 0 |
| ENSMUSG00000082554 | 0 | 0 | 0 | 0 | 0 | 0 | 0 |
| ENSMUSG00000082555 | 0 | 0 | 0 | 0 | 0 | 0 | 0 |
| ENSMUSG00000082556 | 0 | 0 | 0 | 0 | 0 | 0 | 0 |
| ENSMUSG00000082558 | 0 | 0 | 0 | 0 | 0 | 0 | 0 |
| ENSMUSG00000082560 | 0 | 0 | 0 | 0 | 0 | 0 | 0 |
| ENSMUSG00000082561 | 0.289064826 | 0 | 0 | 0 | 0 | 0 | 0.224243824 |
| ENSMUSG00000082562 | 0 | 0 | 0 | 0 | 0 | 0 | 0 |
| ENSMUSG00000082563 | 0 | 0 | 0 | 0 | 0 | 0 | 0 |
| ENSMUSG00000082564 | 0 | 0 | 0 | 0 | 0 | 0 | 0 |
| ENSMUSG00000082565 | 0 | 0 | 0 | 0 | 0 | 0 | 0 |
| ENSMUSG00000082566 | 0 | 0 | 0 | 0 | 0 | 0 | 0 |

| | | | | | | | |
|---|---|---|---|---|---|---|---|
| ENSMUSG00000082567 | 0 | 0 | 0 | 0 | 0 | 0 | 0 |
| ENSMUSG00000082569 | 0 | 0 | 0 | 0 | 0 | 0 | 0 |
| ENSMUSG00000082570 | 0 | 0 | 0 | 0 | 0 | 0 | 0 |
| ENSMUSG00000082571 | 0 | 0 | 0 | 0 | 0 | 0 | 0 |
| ENSMUSG00000082573 | 0 | 0 | 0 | 0 | 0 | 0 | 0 |
| ENSMUSG00000082574 | 0 | 0 | 0 | 0 | 0 | 0 | 0 |
| ENSMUSG00000082575 | 0.960252568 | 0.976722649 | 0.92510993 | 0.646240625 | 0.846351821 | 0.954242509 | 0.984950056 |
| ENSMUSG00000082576 | 0 | 0 | 0.25 | 0 | 0 | 0 | 0.224243824 |
| ENSMUSG00000082577 | 0 | 0 | 0 | 0 | 0 | 0 | 0 |
| ENSMUSG00000082578 | 0 | 0 | 0 | 0 | 0 | 0 | 0 |
| ENSMUSG00000082579 | 0.289064826 | 0.5 | 0.396240625 | 0.396240625 | 0.244650542 | 0.301029996 | 0.224243824 |
| ENSMUSG00000082580 | 0.289064826 | 0 | 0.25 | 0 | 0 | 0 | 0.224243824 |
| ENSMUSG00000082581 | 0 | 0 | 0 | 0 | 0 | 0 | 0 |
| ENSMUSG00000082583 | 0 | 0 | 0 | 0 | 0 | 0 | 0 |
| ENSMUSG00000082584 | 0 | 0 | 0 | 0 | 0 | 0 | 0 |
| ENSMUSG00000082585 | 0 | 0 | 0 | 0 | 0 | 0 | 0 |
| ENSMUSG00000082586 | 0 | 0 | 0 | 0 | 0 | 0 | 0 |
| ENSMUSG00000082587 | 0.289064826 | 0 | 0.5 | 0.25 | 0.244650542 | 0 | 0.520678036 |
| ENSMUSG00000082588 | 1.249317394 | 1.214495249 | 1.29248125 | 1.26109853 | 1.27449559 | 1.079181246 | 1.176815335 |
| ENSMUSG00000082589 | 0 | 0 | 0 | 0 | 0 | 0 | 0 |
| ENSMUSG00000082590 | 0 | 0 | 0 | 0 | 0 | 0 | 0 |
| ENSMUSG00000082591 | 0 | 0 | 0 | 0 | 0 | 0 | 0 |
| ENSMUSG00000082592 | 0 | 0 | 0 | 0 | 0 | 0 | 0 |
| ENSMUSG00000082593 | 0.289064826 | 0 | 0.5 | 0.864857905 | 0.489301084 | 0.602059991 | 0.520678036 |
| ENSMUSG00000082594 | 0 | 0 | 0 | 0 | 0 | 0 | 0 |
| ENSMUSG00000082595 | 0 | 0 | 0 | 0 | 0 | 0 | 0 |
| ENSMUSG00000082596 | 0 | 0.396240625 | 0.25 | 0 | 0.387761935 | 0 | 0.224243824 |
| ENSMUSG00000082597 | 0 | 0 | 0 | 0 | 0 | 0 | 0 |
| ENSMUSG00000082598 | 0 | 0 | 0 | 0 | 0 | 0 | 0 |
| ENSMUSG00000082599 | 0 | 0 | 0 | 0 | 0.489301084 | 0 | 0.355418052 |
| ENSMUSG00000082600 | 0.289064826 | 0 | 0 | 0 | 0.244650542 | 0 | 0 |
| ENSMUSG00000082601 | 0 | 0 | 0 | 0 | 0 | 0 | 0 |
| ENSMUSG00000082602 | 0 | 0 | 0 | 0 | 0 | 0 | 0 |
| ENSMUSG00000082604 | 0 | 0 | 0.25 | 0 | 0 | 0 | 0 |

| | | | | | | | |
|---|---|---|---|---|---|---|---|
| ENSMUSG00000082605 | 0 | 0.25 | 0 | 0 | 0.244650542 | 0.477121255 | 0.355418052 |
| ENSMUSG00000082606 | 0 | 0 | 0 | 0 | 0 | 0 | 0 |
| ENSMUSG00000082607 | 0 | 0 | 0 | 0 | 0 | 0 | 0 |
| ENSMUSG00000082608 | 0.289064826 | 0 | 0.25 | 0 | 0 | 0.301029996 | 0 |
| ENSMUSG00000082609 | 0 | 0.25 | 0 | 0 | 0 | 0.301029996 | 0 |
| ENSMUSG00000082610 | 0 | 0.25 | 0 | 0 | 0 | 0 | 0 |
| ENSMUSG00000082611 | 0 | 0 | 0 | 0 | 0 | 0 | 0 |
| ENSMUSG00000082612 | 0 | 0 | 0 | 0 | 0 | 0 | 0 |
| ENSMUSG00000082613 | 0 | 0 | 0 | 0 | 0 | 0 | 0 |
| ENSMUSG00000082614 | 0 | 0 | 0 | 0 | 0 | 0 | 0 |
| ENSMUSG00000082615 | 0 | 0 | 0 | 0 | 0 | 0 | 0 |
| ENSMUSG00000082616 | 0 | 0 | 0 | 0 | 0 | 0 | 0 |
| ENSMUSG00000082617 | 0 | 0 | 0 | 0 | 0 | 0 | 0 |
| ENSMUSG00000082618 | 0 | 0.25 | 0 | 0 | 0 | 0 | 0 |
| ENSMUSG00000082619 | 0 | 0 | 0 | 0 | 0 | 0 | 0 |
| ENSMUSG00000082620 | 0 | 0 | 0 | 0 | 0 | 0 | 0 |
| ENSMUSG00000082621 | 0 | 0.25 | 0 | 0 | 0 | 0 | 0.355418052 |
| ENSMUSG00000082622 | 0.578129653 | 0.5 | 0.701838731 | 0.5 | 0.489301084 | 0.698970004 | 0.579661877 |
| ENSMUSG00000082623 | 0 | 0 | 0 | 0 | 0 | 0 | 0 |
| ENSMUSG00000082624 | 0 | 0 | 0 | 0 | 0 | 0 | 0 |
| ENSMUSG00000082625 | 0 | 0.396240625 | 0.25 | 0 | 0 | 0.301029996 | 0 |
| ENSMUSG00000082626 | 0 | 0 | 0 | 0 | 0.244650542 | 0 | 0 |
| ENSMUSG00000082628 | 0 | 0 | 0 | 0 | 0 | 0 | 0 |
| ENSMUSG00000082629 | 0 | 0 | 0 | 0 | 0 | 0 | 0 |
| ENSMUSG00000082630 | 0.289064826 | 0.396240625 | 0.25 | 0.396240625 | 0 | 0.301029996 | 0.520678036 |
| ENSMUSG00000082632 | 0 | 0 | 0 | 0 | 0 | 0 | 0 |
| ENSMUSG00000082633 | 0 | 0 | 0.25 | 0.25 | 0.244650542 | 0 | 0.224243824 |
| ENSMUSG00000082634 | 0 | 0 | 0 | 0.25 | 0 | 0 | 0 |
| ENSMUSG00000082635 | 0 | 0 | 0 | 0 | 0 | 0 | 0 |
| ENSMUSG00000082636 | 0.289064826 | 0 | 0 | 0 | 0 | 0 | 0 |
| ENSMUSG00000082637 | 0 | 0 | 0 | 0 | 0 | 0 | 0 |
| ENSMUSG00000082638 | 0 | 0 | 0 | 0 | 0 | 0 | 0 |
| ENSMUSG00000082639 | 0 | 0 | 0 | 0 | 0 | 0 | 0 |
| ENSMUSG00000082641 | 0.289064826 | 0.5 | 0.646240625 | 0.5 | 0.489301084 | 0.301029996 | 0.520678036 |

| | | | | | | | |
|---|---|---|---|---|---|---|---|
| ENSMUSG00000082644 | 0 | 0 | 0 | 0 | 0 | 0 | 0 |
| ENSMUSG00000082645 | 0 | 0 | 0 | 0 | 0 | 0 | 0 |
| ENSMUSG00000082646 | 0.578129653 | 0 | 0.25 | 0.25 | 0.489301084 | 0.301029996 | 0 |
| ENSMUSG00000082647 | 0 | 0 | 0 | 0 | 0 | 0 | 0 |
| ENSMUSG00000082648 | 0 | 0 | 0 | 0 | 0 | 0 | 0 |
| ENSMUSG00000082649 | 0 | 0 | 0 | 0 | 0 | 0 | 0 |
| ENSMUSG00000082650 | 0 | 0 | 0 | 0 | 0 | 0 | 0 |
| ENSMUSG00000082651 | 0 | 0 | 0.25 | 0 | 0.244650542 | 0 | 0.355418052 |
| ENSMUSG00000082652 | 0 | 0 | 0 | 0 | 0 | 0 | 0 |
| ENSMUSG00000082653 | 0 | 0 | 0 | 0 | 0 | 0 | 0 |
| ENSMUSG00000082655 | 0 | 0 | 0 | 0 | 0 | 0 | 0 |
| ENSMUSG00000082656 | 0 | 0 | 0 | 0 | 0 | 0 | 0 |
| ENSMUSG00000082657 | 0 | 0 | 0 | 0 | 0 | 0 | 0 |
| ENSMUSG00000082658 | 0 | 0 | 0 | 0 | 0 | 0 | 0 |
| ENSMUSG00000082660 | 0 | 0 | 0 | 0 | 0 | 0 | 0 |
| ENSMUSG00000082661 | 0 | 0 | 0 | 0 | 0 | 0 | 0 |
| ENSMUSG00000082662 | 0 | 0 | 0 | 0 | 0 | 0 | 0 |
| ENSMUSG00000082663 | 0 | 0 | 0 | 0 | 0 | 0 | 0 |
| ENSMUSG00000082664 | 0 | 0 | 0 | 0 | 0 | 0 | 0 |
| ENSMUSG00000082665 | 0 | 0.5 | 0.396240625 | 0.5 | 0.387761935 | 0.477121255 | 0.448487648 |
| ENSMUSG00000082666 | 0 | 0 | 0 | 0 | 0 | 0 | 0 |
| ENSMUSG00000082667 | 0 | 0 | 0 | 0 | 0 | 0 | 0 |
| ENSMUSG00000082669 | 0 | 0 | 0 | 0 | 0 | 0 | 0 |
| ENSMUSG00000082670 | 0 | 0 | 0 | 0 | 0 | 0 | 0 |
| ENSMUSG00000082671 | 0.289064826 | 0 | 0 | 0 | 0 | 0 | 0 |
| ENSMUSG00000082672 | 0 | 0 | 0 | 0 | 0 | 0 | 0 |
| ENSMUSG00000082673 | 0 | 0 | 0 | 0.25 | 0 | 0 | 0 |
| ENSMUSG00000082674 | 0 | 0 | 0 | 0 | 0 | 0 | 0 |
| ENSMUSG00000082675 | 0.289064826 | 0 | 0.396240625 | 0 | 0 | 0 | 0 |
| ENSMUSG00000082676 | 0 | 0.25 | 0 | 0 | 0 | 0 | 0 |
| ENSMUSG00000082677 | 0 | 0 | 0 | 0 | 0 | 0 | 0 |
| ENSMUSG00000082678 | 0 | 0.25 | 0 | 0 | 0 | 0 | 0 |
| ENSMUSG00000082679 | 0 | 0 | 0 | 0 | 0 | 0 | 0 |
| ENSMUSG00000082680 | 0 | 0 | 0 | 0 | 0 | 0 | 0 |

| | | | | | | | |
|---|---|---|---|---|---|---|---|
| ENSMUSG00000082681 | 0 | 0 | 0 | 0 | 0 | 0 | 0 |
| ENSMUSG00000082682 | 0 | 0 | 0 | 0 | 0 | 0 | 0 |
| ENSMUSG00000082683 | 0 | 0 | 0 | 0 | 0 | 0 | 0.224243824 |
| ENSMUSG00000082684 | 0 | 0 | 0 | 0 | 0.244650542 | 0 | 0 |
| ENSMUSG00000082686 | 0 | 0 | 0 | 0 | 0.244650542 | 0 | 0 |
| ENSMUSG00000082687 | 0 | 0 | 0 | 0 | 0 | 0 | 0 |
| ENSMUSG00000082688 | 0 | 0 | 0 | 0 | 0 | 0 | 0.224243824 |
| ENSMUSG00000082689 | 0 | 0 | 0 | 0 | 0 | 0 | 0 |
| ENSMUSG00000082690 | 0 | 0 | 0 | 0 | 0 | 0 | 0 |
| ENSMUSG00000082691 | 0.289064826 | 0 | 0 | 0 | 0 | 0 | 0.224243824 |
| ENSMUSG00000082692 | 0 | 0 | 0 | 0 | 0 | 0 | 0 |
| ENSMUSG00000082693 | 0 | 0.25 | 0.25 | 0.5 | 0.244650542 | 0 | 0.224243824 |
| ENSMUSG00000082696 | 0 | 0 | 0 | 0 | 0 | 0 | 0 |
| ENSMUSG00000082698 | 0 | 0 | 0 | 0 | 0 | 0 | 0 |
| ENSMUSG00000082699 | 0 | 0.25 | 0 | 0 | 0.244650542 | 0 | 0 |
| ENSMUSG00000082700 | 0 | 0 | 0 | 0 | 0 | 0 | 0 |
| ENSMUSG00000082701 | 0 | 0 | 0 | 0 | 0 | 0 | 0 |
| ENSMUSG00000082703 | 0 | 0 | 0 | 0 | 0 | 0 | 0.224243824 |
| ENSMUSG00000082704 | 0 | 0 | 0 | 0 | 0 | 0 | 0 |
| ENSMUSG00000082705 | 0 | 0 | 0 | 0 | 0.244650542 | 0 | 0 |
| ENSMUSG00000082706 | 0 | 0 | 0 | 0 | 0 | 0 | 0 |
| ENSMUSG00000082707 | 0 | 0 | 0 | 0 | 0 | 0 | 0 |
| ENSMUSG00000082708 | 0 | 0 | 0 | 0 | 0 | 0 | 0 |
| ENSMUSG00000082709 | 0 | 0 | 0 | 0 | 0 | 0 | 0 |
| ENSMUSG00000082710 | 0 | 0 | 0 | 0 | 0 | 0 | 0 |
| ENSMUSG00000082711 | 0 | 0 | 0.25 | 0 | 0 | 0 | 0 |
| ENSMUSG00000082712 | 0 | 0 | 0 | 0 | 0 | 0 | 0 |
| ENSMUSG00000082713 | 0 | 0 | 0 | 0 | 0 | 0 | 0 |
| ENSMUSG00000082714 | 0 | 0 | 0 | 0 | 0 | 0 | 0 |
| ENSMUSG00000082715 | 0 | 0 | 0 | 0 | 0 | 0 | 0 |
| ENSMUSG00000082716 | 0 | 0 | 0 | 0 | 0 | 0 | 0 |
| ENSMUSG00000082717 | 0 | 0 | 0 | 0 | 0 | 0 | 0 |
| ENSMUSG00000082718 | 0 | 0 | 0 | 0 | 0 | 0 | 0 |
| ENSMUSG00000082720 | 0 | 0 | 0 | 0 | 0 | 0 | 0 |

| | | | | | | | |
|---|---|---|---|---|---|---|---|
| ENSMUSG00000082721 | 0 | 0 | 0 | 0 | 0 | 0 | 0 |
| ENSMUSG00000082722 | 0 | 0 | 0 | 0 | 0 | 0 | 0 |
| ENSMUSG00000082723 | 0 | 0 | 0 | 0 | 0 | 0 | 0 |
| ENSMUSG00000082724 | 0 | 0 | 0 | 0 | 0 | 0 | 0 |
| ENSMUSG00000082725 | 0 | 0 | 0 | 0 | 0 | 0 | 0 |
| ENSMUSG00000082726 | 0 | 0 | 0 | 0 | 0 | 0 | 0 |
| ENSMUSG00000082727 | 0 | 0.25 | 0 | 0 | 0 | 0 | 0.355418052 |
| ENSMUSG00000082728 | 0 | 0 | 0 | 0 | 0 | 0 | 0 |
| ENSMUSG00000082729 | 0 | 0 | 0 | 0 | 0 | 0 | 0 |
| ENSMUSG00000082730 | 0.289064826 | 0 | 0 | 0 | 0 | 0 | 0 |
| ENSMUSG00000082732 | 0 | 0 | 0 | 0 | 0 | 0 | 0 |
| ENSMUSG00000082733 | 0 | 0 | 0 | 0 | 0 | 0 | 0 |
| ENSMUSG00000082734 | 0 | 0 | 0 | 0 | 0 | 0 | 0 |
| ENSMUSG00000082735 | 0 | 0 | 0 | 0 | 0 | 0 | 0 |
| ENSMUSG00000082736 | 0 | 0 | 0 | 0 | 0 | 0 | 0 |
| ENSMUSG00000082737 | 0 | 0.25 | 0 | 0 | 0 | 0 | 0 |
| ENSMUSG00000082738 | 0 | 0 | 0 | 0.25 | 0 | 0 | 0 |
| ENSMUSG00000082739 | 0 | 0 | 0 | 0 | 0 | 0 | 0 |
| ENSMUSG00000082740 | 0 | 0 | 0 | 0 | 0 | 0 | 0 |
| ENSMUSG00000082741 | 0 | 0 | 0 | 0.396240625 | 0.387761935 | 0 | 0 |
| ENSMUSG00000082743 | 0 | 0.25 | 0.25 | 0.25 | 0 | 0 | 0.355418052 |
| ENSMUSG00000082744 | 0 | 0 | 0 | 0 | 0 | 0 | 0 |
| ENSMUSG00000082745 | 0 | 0 | 0 | 0 | 0 | 0 | 0 |
| ENSMUSG00000082746 | 0 | 0 | 0 | 0 | 0 | 0 | 0.224243824 |
| ENSMUSG00000082749 | 0.289064826 | 0 | 0 | 0 | 0 | 0 | 0 |
| ENSMUSG00000082750 | 0 | 0 | 0 | 0 | 0 | 0 | 0 |
| ENSMUSG00000082751 | 0 | 0 | 0 | 0 | 0 | 0 | 0 |
| ENSMUSG00000082752 | 0 | 0 | 0 | 0 | 0 | 0 | 0 |
| ENSMUSG00000082754 | 0 | 0 | 0 | 0 | 0 | 0 | 0 |
| ENSMUSG00000082755 | 0 | 0 | 0.25 | 0 | 0 | 0 | 0.224243824 |
| ENSMUSG00000082757 | 0 | 0 | 0 | 0 | 0 | 0 | 0 |
| ENSMUSG00000082758 | 0 | 0 | 0 | 0 | 0 | 0 | 0.224243824 |
| ENSMUSG00000082759 | 0 | 0 | 0 | 0 | 0 | 0 | 0 |
| ENSMUSG00000082760 | 0 | 0 | 0 | 0 | 0 | 0 | 0 |

| | | | | | | | |
|---|---|---|---|---|---|---|---|
| ENSMUSG00000082762 | 0 | 0 | 0.5 | 0 | 0.244650542 | 0 | 0 |
| ENSMUSG00000082763 | 0 | 0 | 0 | 0 | 0 | 0 | 0 |
| ENSMUSG00000082765 | 0.747221736 | 0 | 0.396240625 | 0.75 | 0.244650542 | 0.602059991 | 0.579661877 |
| ENSMUSG00000082766 | 0 | 0 | 0 | 0 | 0 | 0 | 0 |
| ENSMUSG00000082767 | 0 | 0 | 0 | 0 | 0 | 0 | 0 |
| ENSMUSG00000082768 | 0 | 0 | 0 | 0 | 0 | 0.301029996 | 0 |
| ENSMUSG00000082769 | 0.289064826 | 0 | 0.25 | 0.396240625 | 0 | 0.301029996 | 0 |
| ENSMUSG00000082770 | 0 | 0 | 0 | 0 | 0 | 0 | 0 |
| ENSMUSG00000082771 | 0 | 0 | 0 | 0 | 0 | 0 | 0 |
| ENSMUSG00000082772 | 0 | 0 | 0 | 0 | 0 | 0 | 0 |
| ENSMUSG00000082773 | 0 | 0 | 0 | 0 | 0 | 0 | 0 |
| ENSMUSG00000082774 | 0 | 0 | 0 | 0 | 0 | 0 | 0 |
| ENSMUSG00000082775 | 0 | 0.25 | 0.5 | 0.396240625 | 0.244650542 | 0 | 0.224243824 |
| ENSMUSG00000082776 | 0 | 0 | 0 | 0 | 0.387761935 | 0 | 0.224243824 |
| ENSMUSG00000082777 | 0 | 0 | 0 | 0 | 0 | 0 | 0 |
| ENSMUSG00000082778 | 0.45815691 | 0.25 | 0.25 | 0 | 0.568060967 | 0.301029996 | 0.520678036 |
| ENSMUSG00000082779 | 0 | 0 | 0 | 0 | 0 | 0 | 0 |
| ENSMUSG00000082780 | 0 | 0 | 0 | 0 | 0 | 0 | 0 |
| ENSMUSG00000082781 | 0 | 0 | 0 | 0 | 0 | 0 | 0 |
| ENSMUSG00000082782 | 0 | 0 | 0 | 0 | 0 | 0 | 0 |
| ENSMUSG00000082783 | 0 | 0 | 0 | 0 | 0 | 0 | 0 |
| ENSMUSG00000082784 | 0 | 0 | 0 | 0 | 0 | 0 | 0 |
| ENSMUSG00000082785 | 0 | 0 | 0 | 0 | 0 | 0 | 0 |
| ENSMUSG00000082786 | 0 | 0 | 0 | 0 | 0 | 0 | 0 |
| ENSMUSG00000082788 | 0 | 0 | 0 | 0 | 0 | 0 | 0 |
| ENSMUSG00000082789 | 0 | 0 | 0 | 0 | 0 | 0 | 0 |
| ENSMUSG00000082790 | 0 | 0 | 0 | 0 | 0.244650542 | 0 | 0.224243824 |
| ENSMUSG00000082791 | 1.596667477 | 1.403677461 | 1.52186571 | 1.505591953 | 1.462344774 | 1.602059991 | 1.405288179 |
| ENSMUSG00000082792 | 1.100572389 | 1.17510993 | 1.114857905 | 0.92510993 | 1.163285805 | 1.146128036 | 0.984950056 |
| ENSMUSG00000082793 | 0 | 0 | 0 | 0 | 0 | 0 | 0 |
| ENSMUSG00000082794 | 0.289064826 | 0.25 | 0.5 | 0 | 0.244650542 | 0 | 0 |
| ENSMUSG00000082795 | 0 | 0 | 0 | 0 | 0 | 0 | 0 |
| ENSMUSG00000082796 | 0 | 0 | 0 | 0 | 0 | 0 | 0 |
| ENSMUSG00000082797 | 0 | 0 | 0 | 0 | 0 | 0 | 0 |

| | | | | | | | |
|---|---|---|---|---|---|---|---|
| ENSMUSG00000082798 | 0 | 0 | 0 | 0 | 0 | 0 | 0 |
| ENSMUSG00000082799 | 0 | 0 | 0 | 0 | 0 | 0 | 0 |
| ENSMUSG00000082800 | 0 | 0 | 0 | 0 | 0 | 0 | 0 |
| ENSMUSG00000082802 | 0 | 0 | 0 | 0 | 0 | 0 | 0 |
| ENSMUSG00000082804 | 0.289064826 | 0 | 0 | 0 | 0 | 0 | 0 |
| ENSMUSG00000082806 | 0 | 0 | 0 | 0 | 0 | 0 | 0.224243824 |
| ENSMUSG00000082807 | 0 | 0 | 0 | 0 | 0 | 0 | 0 |
| ENSMUSG00000082809 | 1.036286563 | 1.061981878 | 1.04248125 | 0.896240625 | 1 | 0.954242509 | 1.066254157 |
| ENSMUSG00000082810 | 0 | 0 | 0.25 | 0 | 0 | 0 | 0 |
| ENSMUSG00000082812 | 0.289064826 | 0 | 0 | 0 | 0.244650542 | 0 | 0 |
| ENSMUSG00000082814 | 0 | 0 | 0 | 0 | 0 | 0 | 0 |
| ENSMUSG00000082815 | 0 | 0.646240625 | 0.580482024 | 0 | 0.568060967 | 0.301029996 | 0 |
| ENSMUSG00000082816 | 1.205378646 | 1.26109853 | 1.02186571 | 0.896240625 | 1 | 0.954242509 | 0.896975297 |
| ENSMUSG00000082817 | 0 | 0 | 0 | 0 | 0 | 0 | 0 |
| ENSMUSG00000082818 | 0 | 0 | 0 | 0 | 0 | 0 | 0 |
| ENSMUSG00000082819 | 0 | 0 | 0 | 0 | 0 | 0 | 0 |
| ENSMUSG00000082820 | 0 | 0 | 0 | 0 | 0 | 0 | 0 |
| ENSMUSG00000082821 | 0 | 0 | 0 | 0 | 0 | 0 | 0 |
| ENSMUSG00000082822 | 0 | 0 | 0 | 0 | 0 | 0 | 0 |
| ENSMUSG00000082823 | 0 | 0 | 0 | 0 | 0 | 0 | 0 |
| ENSMUSG00000082824 | 0 | 0 | 0 | 0 | 0 | 0 | 0 |
| ENSMUSG00000082825 | 0 | 0 | 0 | 0 | 0 | 0 | 0 |
| ENSMUSG00000082826 | 0 | 0 | 0 | 0 | 0.244650542 | 0 | 0 |
| ENSMUSG00000082827 | 0 | 0 | 0 | 0 | 0 | 0 | 0 |
| ENSMUSG00000082828 | 0 | 0 | 0 | 0 | 0 | 0 | 0 |
| ENSMUSG00000082829 | 0 | 0 | 0 | 0.396240625 | 0 | 0 | 0 |
| ENSMUSG00000082831 | 0.45815691 | 0.25 | 0.580482024 | 0.79248125 | 0.244650542 | 0.903089987 | 0.355418052 |
| ENSMUSG00000082832 | 0 | 0 | 0 | 0 | 0 | 0 | 0 |
| ENSMUSG00000082833 | 0 | 0 | 0 | 0 | 0 | 0 | 0 |
| ENSMUSG00000082835 | 0 | 0 | 0 | 0 | 0 | 0 | 0 |
| ENSMUSG00000082836 | 0.289064826 | 0 | 0 | 0.25 | 0 | 0 | 0.224243824 |
| ENSMUSG00000082837 | 0 | 0 | 0 | 0 | 0 | 0 | 0 |
| ENSMUSG00000082838 | 0 | 0 | 0 | 0 | 0 | 0 | 0 |
| ENSMUSG00000082839 | 0 | 0 | 0 | 0 | 0 | 0 | 0 |

| | | | | | | | |
|---|---|---|---|---|---|---|---|
| ENSMUSG00000082840 | 0 | 0 | 0 | 0 | 0 | 0 | 0 |
| ENSMUSG00000082841 | 0 | 0 | 0 | 0 | 0 | 0 | 0 |
| ENSMUSG00000082842 | 0 | 0 | 0 | 0 | 0 | 0 | 0 |
| ENSMUSG00000082843 | 0 | 0 | 0 | 0 | 0 | 0 | 0 |
| ENSMUSG00000082844 | 0 | 0 | 0 | 0 | 0 | 0 | 0 |
| ENSMUSG00000082845 | 0 | 0 | 0 | 0.25 | 0 | 0 | 0 |
| ENSMUSG00000082846 | 0 | 0 | 0 | 0 | 0 | 0 | 0 |
| ENSMUSG00000082847 | 0.289064826 | 0 | 0 | 0 | 0 | 0 | 0 |
| ENSMUSG00000082848 | 0 | 0 | 0 | 0.25 | 0 | 0 | 0.224243824 |
| ENSMUSG00000082849 | 0 | 0.25 | 0 | 0 | 0 | 0 | 0 |
| ENSMUSG00000082850 | 0 | 0 | 0 | 0 | 0 | 0 | 0 |
| ENSMUSG00000082851 | 0 | 0 | 0.25 | 0 | 0 | 0.301029996 | 0 |
| ENSMUSG00000082852 | 0.45815691 | 0 | 0.25 | 0 | 0 | 0 | 0 |
| ENSMUSG00000082853 | 0 | 0 | 0 | 0 | 0 | 0 | 0 |
| ENSMUSG00000082855 | 0.289064826 | 0.396240625 | 0 | 0 | 0.244650542 | 0 | 0 |
| ENSMUSG00000082856 | 0 | 0 | 0 | 0 | 0 | 0 | 0 |
| ENSMUSG00000082857 | 0 | 0 | 0 | 0 | 0 | 0 | 0 |
| ENSMUSG00000082858 | 0 | 0 | 0 | 0 | 0 | 0 | 0 |
| ENSMUSG00000082859 | 0 | 0 | 0 | 0 | 0 | 0 | 0 |
| ENSMUSG00000082860 | 1.289064826 | 1.130890489 | 1.27186571 | 1.080482024 | 1.136121934 | 1.113943352 | 1.014380832 |
| ENSMUSG00000082861 | 0 | 0 | 0.25 | 0 | 0.244650542 | 0 | 0 |
| ENSMUSG00000082862 | 0 | 0 | 0 | 0 | 0 | 0 | 0 |
| ENSMUSG00000082863 | 0 | 0 | 0 | 0 | 0 | 0 | 0 |
| ENSMUSG00000082864 | 0 | 0 | 0 | 0 | 0 | 0 | 0 |
| ENSMUSG00000082865 | 0 | 0.25 | 0 | 0 | 0.244650542 | 0.301029996 | 0.224243824 |
| ENSMUSG00000082867 | 0.289064826 | 0.25 | 0 | 0 | 0 | 0.301029996 | 0.448487648 |
| ENSMUSG00000082868 | 0 | 0 | 0 | 0 | 0 | 0 | 0 |
| ENSMUSG00000082869 | 0 | 0 | 0 | 0 | 0 | 0 | 0 |
| ENSMUSG00000082870 | 0 | 0 | 0.25 | 0 | 0 | 0 | 0.448487648 |
| ENSMUSG00000082871 | 0 | 0 | 0 | 0 | 0 | 0 | 0 |
| ENSMUSG00000082872 | 0 | 0.396240625 | 0.5 | 0 | 0 | 0.301029996 | 0 |
| ENSMUSG00000082873 | 0 | 0 | 0.25 | 0 | 0 | 0 | 0 |
| ENSMUSG00000082874 | 0 | 0 | 0.396240625 | 0 | 0.244650542 | 0.477121255 | 0.355418052 |
| ENSMUSG00000082875 | 0 | 0 | 0 | 0 | 0 | 0 | 0 |

| | | | | | | | |
|---|---|---|---|---|---|---|---|
| ENSMUSG00000082876 | 0.45815691 | 0.396240625 | 0 | 0.25 | 0 | 0.301029996 | 0.224243824 |
| ENSMUSG00000082877 | 0 | 0 | 0 | 0 | 0 | 0 | 0 |
| ENSMUSG00000082878 | 0.45815691 | 0.396240625 | 0.5 | 0.580482024 | 0.244650542 | 0.77815125 | 0.448487648 |
| ENSMUSG00000082879 | 0.289064826 | 0.396240625 | 0 | 0.396240625 | 0.387761935 | 0 | 0.224243824 |
| ENSMUSG00000082882 | 0 | 0 | 0 | 0 | 0 | 0 | 0 |
| ENSMUSG00000082883 | 0 | 0 | 0 | 0 | 0 | 0 | 0 |
| ENSMUSG00000082884 | 0 | 0.396240625 | 0.5 | 0 | 0 | 0.477121255 | 0.448487648 |
| ENSMUSG00000082885 | 0 | 0 | 0 | 0 | 0 | 0 | 0 |
| ENSMUSG00000082886 | 0 | 0 | 0 | 0 | 0 | 0 | 0 |
| ENSMUSG00000082887 | 0 | 0 | 0 | 0 | 0 | 0 | 0 |
| ENSMUSG00000082888 | 0 | 0 | 0 | 0 | 0 | 0 | 0 |
| ENSMUSG00000082889 | 0 | 0 | 0 | 0 | 0 | 0 | 0 |
| ENSMUSG00000082890 | 0 | 0 | 0 | 0 | 0 | 0 | 0 |
| ENSMUSG00000082891 | 0 | 0 | 0 | 0 | 0 | 0 | 0 |
| ENSMUSG00000082892 | 0 | 0 | 0 | 0 | 0 | 0 | 0 |
| ENSMUSG00000082893 | 0 | 0 | 0.25 | 0 | 0 | 0.301029996 | 0 |
| ENSMUSG00000082894 | 1.069666964 | 1.098079356 | 0.864857905 | 1.02186571 | 0.846351821 | 0.954242509 | 0.896975297 |
| ENSMUSG00000082895 | 0.867194479 | 0.75 | 0.5 | 0.79248125 | 0.568060967 | 0.903089987 | 0.876096088 |
| ENSMUSG00000082896 | 0.578129653 | 0.5 | 0.580482024 | 0.396240625 | 0.387761935 | 0.301029996 | 0.224243824 |
| ENSMUSG00000082897 | 0 | 0 | 0 | 0 | 0 | 0 | 0 |
| ENSMUSG00000082898 | 0 | 0 | 0 | 0 | 0 | 0 | 0 |
| ENSMUSG00000082899 | 0 | 0 | 0 | 0 | 0 | 0 | 0 |
| ENSMUSG00000082900 | 0 | 0.580482024 | 0.646240625 | 0 | 0.387761935 | 0.301029996 | 0.224243824 |
| ENSMUSG00000082901 | 0 | 0 | 0 | 0.25 | 0 | 0 | 0.224243824 |
| ENSMUSG00000082902 | 0 | 0 | 0 | 0 | 0 | 0 | 0 |
| ENSMUSG00000082903 | 0 | 0 | 0 | 0 | 0 | 0 | 0 |
| ENSMUSG00000082905 | 0 | 0 | 0 | 0 | 0 | 0 | 0 |
| ENSMUSG00000082906 | 0 | 0 | 0 | 0 | 0 | 0 | 0 |
| ENSMUSG00000082907 | 0 | 0 | 0 | 0 | 0 | 0 | 0 |
| ENSMUSG00000082908 | 0 | 0 | 0 | 0 | 0 | 0 | 0 |
| ENSMUSG00000082910 | 0 | 0 | 0 | 0 | 0 | 0 | 0 |
| ENSMUSG00000082911 | 0 | 0 | 0 | 0 | 0 | 0 | 0 |
| ENSMUSG00000082912 | 0.289064826 | 0 | 0 | 0 | 0 | 0 | 0 |
| ENSMUSG00000082913 | 0 | 0 | 0 | 0 | 0 | 0 | 0 |

| | | | | | | | |
|---|---|---|---|---|---|---|---|
| ENSMUSG00000082915 | 0 | 0 | 0 | 0 | 0 | 0 | 0 |
| ENSMUSG00000082917 | 0 | 0 | 0 | 0 | 0 | 0 | 0 |
| ENSMUSG00000082918 | 0 | 0 | 0 | 0 | 0 | 0 | 0 |
| ENSMUSG00000082919 | 0 | 0 | 0 | 0 | 0 | 0 | 0 |
| ENSMUSG00000082920 | 0.747221736 | 0.79248125 | 0.25 | 0.396240625 | 0.632412477 | 0.698970004 | 0.448487648 |
| ENSMUSG00000082922 | 0.45815691 | 0.396240625 | 0.830482024 | 0.396240625 | 0.387761935 | 0.602059991 | 0.672731473 |
| ENSMUSG00000082924 | 0 | 0 | 0 | 0 | 0 | 0 | 0.224243824 |
| ENSMUSG00000082925 | 0 | 0 | 0 | 0 | 0 | 0.301029996 | 0 |
| ENSMUSG00000082926 | 0.289064826 | 0 | 0 | 0 | 0 | 0 | 0 |
| ENSMUSG00000082927 | 0.578129653 | 0.25 | 0.25 | 0.25 | 0.387761935 | 0.477121255 | 0.448487648 |
| ENSMUSG00000082928 | 0 | 0 | 0 | 0 | 0 | 0 | 0 |
| ENSMUSG00000082929 | 0 | 0 | 0 | 0.25 | 0 | 0 | 0 |
| ENSMUSG00000082930 | 0 | 0 | 0 | 0 | 0 | 0 | 0 |
| ENSMUSG00000082931 | 0 | 0 | 0 | 0 | 0.244650542 | 0.301029996 | 0 |
| ENSMUSG00000082932 | 0 | 0 | 0 | 0 | 0 | 0 | 0 |
| ENSMUSG00000082933 | 0 | 0 | 0 | 0 | 0 | 0 | 0 |
| ENSMUSG00000082934 | 0 | 0 | 0 | 0 | 0 | 0 | 0 |
| ENSMUSG00000082935 | 1.639698646 | 1.348079356 | 1.339388001 | 1.711372513 | 1.683842353 | 1.84509804 | 1.618853957 |
| ENSMUSG00000082936 | 0.671187741 | 1.02186571 | 0.830482024 | 0.25 | 0.489301084 | 0.77815125 | 0.355418052 |
| ENSMUSG00000082937 | 0 | 0 | 0 | 0 | 0 | 0 | 0 |
| ENSMUSG00000082938 | 0 | 0 | 0.25 | 0 | 0.244650542 | 0 | 0 |
| ENSMUSG00000082939 | 0 | 0 | 0 | 0 | 0 | 0 | 0 |
| ENSMUSG00000082940 | 0 | 0 | 0 | 0 | 0 | 0 | 0 |
| ENSMUSG00000082941 | 0.289064826 | 0.396240625 | 0.396240625 | 0 | 0 | 0 | 0.355418052 |
| ENSMUSG00000082943 | 0 | 0 | 0 | 0 | 0 | 0 | 0 |
| ENSMUSG00000082945 | 0 | 0 | 0 | 0 | 0 | 0 | 0 |
| ENSMUSG00000082947 | 0 | 0 | 0.25 | 0 | 0 | 0 | 0 |
| ENSMUSG00000082949 | 0 | 0 | 0 | 0 | 0 | 0 | 0 |
| ENSMUSG00000082950 | 0 | 0 | 0.25 | 0.25 | 0 | 0 | 0 |
| ENSMUSG00000082951 | 0 | 0 | 0 | 0.25 | 0 | 0 | 0 |
| ENSMUSG00000082952 | 0 | 0 | 0 | 0 | 0 | 0 | 0 |
| ENSMUSG00000082953 | 0 | 0 | 0 | 0 | 0 | 0 | 0 |
| ENSMUSG00000082954 | 0 | 0 | 0 | 0 | 0 | 0 | 0 |
| ENSMUSG00000082955 | 0 | 0 | 0 | 0 | 0 | 0 | 0 |

| | | | | | | | |
|---|---|---|---|---|---|---|---|
| ENSMUSG00000082956 | 0 | 0 | 0 | 0 | 0 | 0 | 0 |
| ENSMUSG00000082957 | 0 | 0 | 0 | 0 | 0 | 0 | 0 |
| ENSMUSG00000082958 | 0 | 0 | 0 | 0 | 0 | 0 | 0.224243824 |
| ENSMUSG00000082960 | 0 | 0 | 0 | 0 | 0.244650542 | 0 | 0 |
| ENSMUSG00000082961 | 0 | 0.5 | 0.5 | 0.396240625 | 0.387761935 | 0.301029996 | 0.448487648 |
| ENSMUSG00000082962 | 0 | 0 | 0 | 0 | 0 | 0 | 0.224243824 |
| ENSMUSG00000082963 | 0 | 0 | 0 | 0 | 0 | 0 | 0 |
| ENSMUSG00000082964 | 0 | 0 | 0 | 0 | 0.244650542 | 0 | 0 |
| ENSMUSG00000082965 | 0 | 0 | 0 | 0 | 0.387761935 | 0 | 0 |
| ENSMUSG00000082966 | 0 | 0 | 0 | 0 | 0 | 0 | 0 |
| ENSMUSG00000082967 | 0 | 0.25 | 0 | 0 | 0 | 0 | 0 |
| ENSMUSG00000082968 | 0 | 0 | 0 | 0 | 0 | 0 | 0 |
| ENSMUSG00000082969 | 0.289064826 | 0.5 | 0.5 | 0.5 | 0.632412477 | 0.301029996 | 0.355418052 |
| ENSMUSG00000082970 | 0 | 0 | 0 | 0 | 0 | 0 | 0 |
| ENSMUSG00000082971 | 0 | 0 | 0.25 | 0 | 0 | 0 | 0 |
| ENSMUSG00000082973 | 0 | 0 | 0 | 0 | 0 | 0 | 0 |
| ENSMUSG00000082974 | 0 | 0 | 0 | 0 | 0 | 0 | 0 |
| ENSMUSG00000082975 | 0 | 0 | 0 | 0 | 0 | 0 | 0 |
| ENSMUSG00000082976 | 0 | 0 | 0 | 0 | 0 | 0 | 0 |
| ENSMUSG00000082977 | 0 | 0 | 0 | 0 | 0 | 0 | 0 |
| ENSMUSG00000082978 | 0.45815691 | 0.580482024 | 0.5 | 0.25 | 0.733951626 | 0 | 0.355418052 |
| ENSMUSG00000082980 | 0 | 0 | 0 | 0 | 0 | 0 | 0 |
| ENSMUSG00000082981 | 0 | 0 | 0.25 | 0.25 | 0 | 0.301029996 | 0 |
| ENSMUSG00000082982 | 0 | 0 | 0 | 0 | 0 | 0 | 0 |
| ENSMUSG00000082983 | 0 | 0 | 0 | 0 | 0 | 0 | 0 |
| ENSMUSG00000082985 | 0 | 0.25 | 0 | 0.25 | 0.568060967 | 0 | 0.224243824 |
| ENSMUSG00000082986 | 0 | 0 | 0.25 | 0.396240625 | 0 | 0 | 0 |
| ENSMUSG00000082987 | 0 | 0 | 0 | 0 | 0 | 0 | 0.224243824 |
| ENSMUSG00000082988 | 0.45815691 | 0 | 0.5 | 0 | 0 | 0 | 0.224243824 |
| ENSMUSG00000082989 | 0 | 0 | 0 | 0 | 0 | 0 | 0 |
| ENSMUSG00000082990 | 0 | 0 | 0.25 | 0.25 | 0 | 0 | 0.224243824 |
| ENSMUSG00000082991 | 0 | 0 | 0 | 0 | 0 | 0 | 0 |
| ENSMUSG00000082993 | 0 | 0 | 0 | 0 | 0 | 0 | 0 |
| ENSMUSG00000082994 | 0 | 0 | 0 | 0 | 0 | 0 | 0 |

| | | | | | | | |
|---|---|---|---|---|---|---|---|
| ENSMUSG00000082995 | 0 | 0 | 0 | 0 | 0 | 0 | 0 |
| ENSMUSG00000082996 | 0.289064826 | 0 | 0 | 0 | 0 | 0 | 0 |
| ENSMUSG00000082997 | 0 | 0.25 | 0 | 0 | 0 | 0 | 0 |
| ENSMUSG00000082998 | 0 | 0 | 0 | 0 | 0 | 0.301029996 | 0 |
| ENSMUSG00000083000 | 0 | 0 | 0 | 0 | 0 | 0 | 0 |
| ENSMUSG00000083001 | 0 | 0 | 0 | 0 | 0 | 0 | 0 |
| ENSMUSG00000083004 | 0 | 0 | 0 | 0.25 | 0.244650542 | 0 | 0.355418052 |
| ENSMUSG00000083005 | 0 | 0 | 0 | 0 | 0 | 0 | 0 |
| ENSMUSG00000083006 | 0.45815691 | 0 | 0.25 | 0 | 0 | 0 | 0 |
| ENSMUSG00000083007 | 0.45815691 | 0.396240625 | 0.646240625 | 0.25 | 0.244650542 | 0.301029996 | 0.520678036 |
| ENSMUSG00000083008 | 0 | 0 | 0 | 0 | 0 | 0 | 0 |
| ENSMUSG00000083009 | 0 | 0 | 0 | 0 | 0 | 0 | 0 |
| ENSMUSG00000083010 | 0 | 0 | 0 | 0 | 0 | 0 | 0 |
| ENSMUSG00000083011 | 0.747221736 | 0.92510993 | 0.580482024 | 0.75 | 0.568060967 | 0.77815125 | 0.775756176 |
| ENSMUSG00000083012 | 2.201144779 | 2.059601185 | 2.049918086 | 1.975216702 | 1.934425071 | 2.201397124 | 1.87902385 |
| ENSMUSG00000083013 | 0 | 0 | 0 | 0 | 0 | 0 | 0 |
| ENSMUSG00000083014 | 0 | 0 | 0 | 0 | 0 | 0 | 0 |
| ENSMUSG00000083015 | 0 | 0 | 0 | 0 | 0.244650542 | 0 | 0 |
| ENSMUSG00000083016 | 0.289064826 | 0 | 0.25 | 0 | 0.387761935 | 0 | 0.355418052 |
| ENSMUSG00000083017 | 0 | 0 | 0 | 0 | 0 | 0 | 0 |
| ENSMUSG00000083018 | 0 | 0 | 0 | 0 | 0 | 0 | 0 |
| ENSMUSG00000083019 | 0 | 0.396240625 | 0.976722649 | 0.580482024 | 0.387761935 | 0.301029996 | 1.054044578 |
| ENSMUSG00000083020 | 0 | 0 | 0 | 0 | 0 | 0 | 0 |
| ENSMUSG00000083022 | 0.289064826 | 0.396240625 | 0.396240625 | 0.396240625 | 0.387761935 | 0.301029996 | 0.448487648 |
| ENSMUSG00000083023 | 0 | 0 | 0 | 0 | 0 | 0 | 0 |
| ENSMUSG00000083024 | 0 | 0 | 0 | 0 | 0 | 0 | 0 |
| ENSMUSG00000083025 | 0 | 0 | 0.25 | 0 | 0 | 0 | 0 |
| ENSMUSG00000083026 | 0 | 0 | 0 | 0 | 0 | 0 | 0 |
| ENSMUSG00000083027 | 1 | 0.896240625 | 0.976722649 | 1.04248125 | 0.931471446 | 1.041392685 | 0.775756176 |
| ENSMUSG00000083029 | 0 | 0 | 0 | 0 | 0 | 0 | 0 |
| ENSMUSG00000083030 | 0 | 0 | 0 | 0 | 0 | 0 | 0 |
| ENSMUSG00000083031 | 0 | 0 | 0 | 0 | 0 | 0 | 0 |
| ENSMUSG00000083032 | 0 | 0 | 0 | 0 | 0 | 0 | 0 |
| ENSMUSG00000083033 | 0.45815691 | 0.646240625 | 0.5 | 0.5 | 0.686820904 | 0.477121255 | 0.629532004 |

| | | | | | | | |
|---|---|---|---|---|---|---|---|
| ENSMUSG00000083034 | 0 | 0 | 0 | 0 | 0 | 0 | 0 |
| ENSMUSG00000083036 | 0 | 0 | 0 | 0 | 0 | 0 | 0 |
| ENSMUSG00000083037 | 0 | 0 | 0 | 0 | 0 | 0 | 0 |
| ENSMUSG00000083038 | 0 | 0 | 0 | 0 | 0 | 0 | 0 |
| ENSMUSG00000083039 | 0 | 0 | 0 | 0 | 0 | 0 | 0 |
| ENSMUSG00000083041 | 0 | 0 | 0 | 0 | 0 | 0 | 0 |
| ENSMUSG00000083042 | 0 | 0 | 0 | 0 | 0 | 0 | 0 |
| ENSMUSG00000083043 | 0.45815691 | 0.396240625 | 0.5 | 0 | 0.244650542 | 0 | 0.355418052 |
| ENSMUSG00000083044 | 0.867194479 | 0.396240625 | 0.701838731 | 0 | 0.387761935 | 0 | 0 |
| ENSMUSG00000083045 | 0 | 0 | 0 | 0 | 0 | 0 | 0 |
| ENSMUSG00000083046 | 0 | 0 | 0 | 0 | 0 | 0 | 0 |
| ENSMUSG00000083047 | 0 | 0 | 0 | 0 | 0 | 0 | 0 |
| ENSMUSG00000083049 | 0 | 0 | 0 | 0 | 0 | 0 | 0 |
| ENSMUSG00000083050 | 0 | 0 | 0 | 0 | 0 | 0 | 0 |
| ENSMUSG00000083052 | 0 | 0 | 0.25 | 0.25 | 0.244650542 | 0 | 0.224243824 |
| ENSMUSG00000083053 | 0 | 0 | 0 | 0 | 0 | 0 | 0 |
| ENSMUSG00000083054 | 0 | 0 | 0.25 | 0 | 0 | 0 | 0.224243824 |
| ENSMUSG00000083055 | 0 | 0 | 0 | 0 | 0 | 0 | 0 |
| ENSMUSG00000083057 | 0 | 0.25 | 0 | 0 | 0 | 0 | 0 |
| ENSMUSG00000083058 | 0 | 0 | 0 | 0 | 0 | 0 | 0 |
| ENSMUSG00000083059 | 0 | 0 | 0 | 0 | 0 | 0 | 0 |
| ENSMUSG00000083060 | 0 | 0 | 0 | 0 | 0 | 0 | 0 |
| ENSMUSG00000083061 | 0 | 0 | 0 | 0 | 0 | 0 | 0 |
| ENSMUSG00000083062 | 0 | 0 | 0 | 0 | 0 | 0 | 0 |
| ENSMUSG00000083063 | 0 | 0 | 0 | 0 | 0 | 0 | 0 |
| ENSMUSG00000083064 | 0.747221736 | 0.646240625 | 0.79248125 | 0.864857905 | 0.686820904 | 1 | 0.829800754 |
| ENSMUSG00000083065 | 0 | 0 | 0 | 0 | 0 | 0 | 0 |
| ENSMUSG00000083066 | 0 | 0 | 0 | 0 | 0 | 0 | 0 |
| ENSMUSG00000083067 | 0 | 0 | 0 | 0 | 0 | 0 | 0 |
| ENSMUSG00000083068 | 0 | 0 | 0 | 0 | 0 | 0.301029996 | 0.224243824 |
| ENSMUSG00000083069 | 0 | 0 | 0 | 0 | 0 | 0 | 0 |
| ENSMUSG00000083070 | 0 | 0 | 0 | 0 | 0 | 0 | 0 |
| ENSMUSG00000083072 | 0.578129653 | 0 | 0.396240625 | 0 | 0.387761935 | 0 | 0.448487648 |
| ENSMUSG00000083073 | 0 | 0 | 0 | 0 | 0 | 0 | 0 |

| | | | | | | | |
|---|---|---|---|---|---|---|---|
| ENSMUSG00000083074 | 0 | 0 | 0 | 0 | 0 | 0 | 0 |
| ENSMUSG00000083075 | 0 | 0 | 0 | 0 | 0 | 0 | 0 |
| ENSMUSG00000083076 | 0.289064826 | 0.25 | 0 | 0.25 | 0 | 0 | 0 |
| ENSMUSG00000083077 | 0 | 0 | 0 | 0 | 0 | 0 | 0 |
| ENSMUSG00000083078 | 0 | 0 | 0 | 0 | 0 | 0 | 0 |
| ENSMUSG00000083079 | 0 | 0 | 0 | 0 | 0 | 0 | 0 |
| ENSMUSG00000083080 | 0 | 0 | 0 | 0 | 0 | 0 | 0 |
| ENSMUSG00000083081 | 0 | 0 | 0 | 0 | 0 | 0 | 0 |
| ENSMUSG00000083083 | 0.45815691 | 0.25 | 0.25 | 0.25 | 0.568060967 | 0 | 0.355418052 |
| ENSMUSG00000083084 | 0.289064826 | 0.25 | 0 | 0 | 0 | 0.301029996 | 0 |
| ENSMUSG00000083085 | 0 | 0 | 0 | 0 | 0 | 0 | 0 |
| ENSMUSG00000083086 | 0.45815691 | 0.701838731 | 0.5 | 0.396240625 | 0.632412477 | 0.77815125 | 0.74492186 |
| ENSMUSG00000083087 | 0.867194479 | 0.830482024 | 0.976722649 | 0.580482024 | 0.846351821 | 0.301029996 | 0.775756176 |
| ENSMUSG00000083088 | 0 | 0 | 0 | 0 | 0 | 0 | 0 |
| ENSMUSG00000083089 | 0 | 0 | 0 | 0.25 | 0.244650542 | 0 | 0 |
| ENSMUSG00000083090 | 0 | 0.25 | 0 | 0 | 0 | 0 | 0 |
| ENSMUSG00000083091 | 0 | 0 | 0 | 0 | 0 | 0 | 0 |
| ENSMUSG00000083093 | 0 | 0 | 0 | 0 | 0 | 0 | 0 |
| ENSMUSG00000083094 | 0.45815691 | 0 | 0.5 | 0.396240625 | 0.244650542 | 0 | 0 |
| ENSMUSG00000083096 | 0 | 0 | 0 | 0 | 0 | 0 | 0 |
| ENSMUSG00000083097 | 0 | 0.396240625 | 0.5 | 0 | 0.568060967 | 0.602059991 | 0.224243824 |
| ENSMUSG00000083098 | 0 | 0 | 0 | 0 | 0 | 0 | 0 |
| ENSMUSG00000083099 | 0 | 0 | 0 | 0 | 0 | 0 | 0 |
| ENSMUSG00000083100 | 0 | 0 | 0 | 0 | 0 | 0 | 0 |
| ENSMUSG00000083101 | 0 | 0 | 0.25 | 0 | 0 | 0 | 0 |
| ENSMUSG00000083103 | 0 | 0 | 0 | 0 | 0 | 0 | 0 |
| ENSMUSG00000083104 | 0.671187741 | 0.580482024 | 0.646240625 | 0.396240625 | 0.77552387 | 0.301029996 | 0.74492186 |
| ENSMUSG00000083105 | 0 | 0 | 0 | 0 | 0 | 0 | 0 |
| ENSMUSG00000083106 | 0 | 0 | 0 | 0 | 0 | 0 | 0 |
| ENSMUSG00000083107 | 0.578129653 | 0.25 | 0.580482024 | 0.79248125 | 0.955822902 | 1.041392685 | 0.803905701 |
| ENSMUSG00000083109 | 0 | 0 | 0 | 0 | 0 | 0 | 0 |
| ENSMUSG00000083110 | 0 | 0 | 0 | 0 | 0 | 0 | 0 |
| ENSMUSG00000083111 | 0 | 0.396240625 | 0.25 | 0.701838731 | 0.686820904 | 0.602059991 | 0.803905701 |
| ENSMUSG00000083112 | 0 | 0 | 0 | 0 | 0 | 0 | 0 |

| | | | | | | | |
|---|---|---|---|---|---|---|---|
| ENSMUSG00000083113 | 0.289064826 | 0.25 | 0 | 0 | 0 | 0.301029996 | 0 |
| ENSMUSG00000083114 | 0 | 0 | 0 | 0 | 0 | 0 | 0 |
| ENSMUSG00000083115 | 0 | 0 | 0 | 0 | 0 | 0 | 0 |
| ENSMUSG00000083116 | 1.249317394 | 0.92510993 | 1.02186571 | 1.214495249 | 1.149965125 | 1.230448921 | 1.028149525 |
| ENSMUSG00000083117 | 0 | 0 | 0 | 0 | 0 | 0 | 0 |
| ENSMUSG00000083118 | 0 | 0 | 0 | 0 | 0 | 0 | 0 |
| ENSMUSG00000083119 | 0 | 0 | 0 | 0 | 0 | 0 | 0 |
| ENSMUSG00000083120 | 0 | 0 | 0 | 0 | 0 | 0 | 0 |
| ENSMUSG00000083121 | 0.289064826 | 0.25 | 0.25 | 0 | 0.244650542 | 0 | 0 |
| ENSMUSG00000083122 | 0 | 0 | 0 | 0 | 0 | 0 | 0 |
| ENSMUSG00000083123 | 0 | 0 | 0 | 0 | 0 | 0 | 0 |
| ENSMUSG00000083124 | 0 | 0 | 0 | 0 | 0 | 0 | 0 |
| ENSMUSG00000083126 | 0 | 0 | 0 | 0 | 0 | 0 | 0 |
| ENSMUSG00000083127 | 0 | 0 | 0 | 0 | 0 | 0 | 0 |
| ENSMUSG00000083128 | 0 | 0 | 0 | 0 | 0 | 0 | 0 |
| ENSMUSG00000083129 | 0 | 0 | 0 | 0 | 0 | 0 | 0 |
| ENSMUSG00000083130 | 0 | 0 | 0 | 0 | 0 | 0 | 0 |
| ENSMUSG00000083135 | 0 | 0 | 0 | 0 | 0 | 0 | 0 |
| ENSMUSG00000083136 | 0 | 0 | 0 | 0 | 0 | 0 | 0 |
| ENSMUSG00000083137 | 0 | 0 | 0 | 0 | 0 | 0 | 0 |
| ENSMUSG00000083138 | 0 | 0 | 0 | 0 | 0 | 0 | 0 |
| ENSMUSG00000083139 | 0 | 0 | 0 | 0 | 0 | 0 | 0 |
| ENSMUSG00000083140 | 0 | 0 | 0 | 0 | 0 | 0 | 0 |
| ENSMUSG00000083141 | 0 | 0 | 0 | 0 | 0 | 0 | 0 |
| ENSMUSG00000083142 | 0 | 0 | 0 | 0 | 0 | 0 | 0 |
| ENSMUSG00000083143 | 0 | 0 | 0 | 0 | 0 | 0 | 0 |
| ENSMUSG00000083144 | 0 | 0 | 0 | 0 | 0 | 0 | 0 |
| ENSMUSG00000083145 | 0 | 0.25 | 0.25 | 0 | 0 | 0 | 0 |
| ENSMUSG00000083147 | 0 | 0 | 0 | 0 | 0 | 0 | 0 |
| ENSMUSG00000083149 | 0 | 0 | 0 | 0 | 0 | 0.477121255 | 0.224243824 |
| ENSMUSG00000083150 | 0 | 0 | 0 | 0 | 0 | 0 | 0 |
| ENSMUSG00000083151 | 0 | 0 | 0 | 0 | 0 | 0 | 0 |
| ENSMUSG00000083152 | 0.578129653 | 0.25 | 0.580482024 | 0.646240625 | 0.632412477 | 0 | 0.355418052 |
| ENSMUSG00000083153 | 0 | 0 | 0 | 0 | 0 | 0 | 0 |

| | | | | | | | |
|---|---|---|---|---|---|---|---|
| ENSMUSG00000083154 | 0 | 0 | 0 | 0 | 0 | 0 | 0 |
| ENSMUSG00000083155 | 0 | 0 | 0 | 0 | 0 | 0 | 0 |
| ENSMUSG00000083156 | 0 | 0 | 0 | 0.25 | 0 | 0 | 0 |
| ENSMUSG00000083157 | 0 | 0 | 0 | 0 | 0 | 0 | 0 |
| ENSMUSG00000083158 | 0 | 0 | 0 | 0 | 0 | 0 | 0 |
| ENSMUSG00000083159 | 0 | 0 | 0 | 0.25 | 0 | 0.301029996 | 0 |
| ENSMUSG00000083160 | 0 | 0 | 0 | 0 | 0 | 0 | 0 |
| ENSMUSG00000083161 | 0 | 0 | 0 | 0 | 0 | 0 | 0 |
| ENSMUSG00000083162 | 0 | 0 | 0 | 0 | 0 | 0 | 0 |
| ENSMUSG00000083163 | 0 | 0 | 0 | 0 | 0 | 0 | 0 |
| ENSMUSG00000083165 | 0 | 0 | 0 | 0 | 0 | 0 | 0 |
| ENSMUSG00000083166 | 0.289064826 | 0.396240625 | 0.396240625 | 0.646240625 | 0.632412477 | 0.301029996 | 0.579661877 |
| ENSMUSG00000083167 | 0 | 0 | 0 | 0 | 0 | 0 | 0 |
| ENSMUSG00000083168 | 0 | 0 | 0 | 0 | 0 | 0 | 0 |
| ENSMUSG00000083169 | 0 | 0 | 0 | 0 | 0 | 0.301029996 | 0 |
| ENSMUSG00000083170 | 0 | 0 | 0 | 0 | 0 | 0 | 0 |
| ENSMUSG00000083171 | 0 | 0 | 0 | 0 | 0.244650542 | 0 | 0 |
| ENSMUSG00000083172 | 0 | 0 | 0 | 0 | 0 | 0 | 0 |
| ENSMUSG00000083173 | 0 | 0 | 0 | 0 | 0 | 0 | 0 |
| ENSMUSG00000083174 | 0 | 0 | 0 | 0 | 0 | 0 | 0 |
| ENSMUSG00000083175 | 0 | 0 | 0 | 0 | 0 | 0 | 0 |
| ENSMUSG00000083176 | 0 | 0 | 0 | 0 | 0.244650542 | 0 | 0 |
| ENSMUSG00000083177 | 0 | 0 | 0 | 0 | 0 | 0 | 0.224243824 |
| ENSMUSG00000083178 | 0 | 0 | 0 | 0 | 0 | 0 | 0 |
| ENSMUSG00000083179 | 0 | 0.25 | 0.25 | 0 | 0.244650542 | 0 | 0 |
| ENSMUSG00000083180 | 0 | 0 | 0 | 0.25 | 0 | 0 | 0 |
| ENSMUSG00000083181 | 0 | 0 | 0 | 0 | 0 | 0 | 0 |
| ENSMUSG00000083182 | 0 | 0 | 0 | 0 | 0 | 0 | 0 |
| ENSMUSG00000083183 | 0 | 0 | 0.25 | 0 | 0.387761935 | 0 | 0.224243824 |
| ENSMUSG00000083185 | 0 | 0 | 0 | 0 | 0 | 0 | 0 |
| ENSMUSG00000083186 | 0 | 0 | 0 | 0 | 0 | 0 | 0.224243824 |
| ENSMUSG00000083188 | 0 | 0 | 0 | 0 | 0 | 0 | 0 |
| ENSMUSG00000083190 | 0 | 0 | 0 | 0 | 0 | 0 | 0 |
| ENSMUSG00000083191 | 0 | 0 | 0 | 0 | 0 | 0 | 0 |

| | | | | | | | |
|---|---|---|---|---|---|---|---|
| ENSMUSG00000083192 | 0 | 0 | 0 | 0 | 0.244650542 | 0 | 0 |
| ENSMUSG00000083193 | 0.289064826 | 0 | 0 | 0 | 0 | 0 | 0 |
| ENSMUSG00000083195 | 0 | 0 | 0 | 0 | 0 | 0 | 0 |
| ENSMUSG00000083196 | 0 | 0 | 0 | 0 | 0 | 0 | 0 |
| ENSMUSG00000083197 | 0 | 0.25 | 0 | 0 | 0 | 0 | 0 |
| ENSMUSG00000083199 | 0 | 0 | 0 | 0 | 0.244650542 | 0 | 0 |
| ENSMUSG00000083200 | 0.747221736 | 0.79248125 | 0.864857905 | 0.830482024 | 1.057362051 | 1.041392685 | 0.853775828 |
| ENSMUSG00000083201 | 0.289064826 | 0 | 0.5 | 0 | 0 | 0.301029996 | 0.224243824 |
| ENSMUSG00000083202 | 0 | 0 | 0 | 0 | 0.244650542 | 0 | 0 |
| ENSMUSG00000083203 | 0 | 0 | 0 | 0 | 0 | 0 | 0 |
| ENSMUSG00000083204 | 0 | 0 | 0 | 0 | 0 | 0 | 0.224243824 |
| ENSMUSG00000083205 | 0 | 0 | 0 | 0 | 0 | 0 | 0 |
| ENSMUSG00000083206 | 0 | 0 | 0 | 0 | 0 | 0 | 0 |
| ENSMUSG00000083207 | 0 | 0.25 | 0.396240625 | 0 | 0.387761935 | 0.301029996 | 0.448487648 |
| ENSMUSG00000083208 | 0 | 0 | 0 | 0 | 0 | 0 | 0 |
| ENSMUSG00000083209 | 0 | 0.25 | 0.25 | 0 | 0 | 0 | 0 |
| ENSMUSG00000083210 | 0 | 0 | 0 | 0.25 | 0 | 0 | 0 |
| ENSMUSG00000083211 | 0 | 0 | 0 | 0 | 0 | 0 | 0 |
| ENSMUSG00000083212 | 0 | 0 | 0 | 0 | 0 | 0 | 0 |
| ENSMUSG00000083213 | 0 | 0 | 0 | 0 | 0 | 0 | 0 |
| ENSMUSG00000083214 | 0 | 0.25 | 0 | 0 | 0 | 0 | 0 |
| ENSMUSG00000083215 | 0 | 0 | 0 | 0 | 0 | 0 | 0 |
| ENSMUSG00000083216 | 0 | 0 | 0 | 0 | 0 | 0 | 0 |
| ENSMUSG00000083218 | 0.671187741 | 0.580482024 | 0.75 | 0.896240625 | 0.846351821 | 1 | 0.896975297 |
| ENSMUSG00000083219 | 0.747221736 | 0.5 | 0 | 0.396240625 | 0 | 0.477121255 | 0 |
| ENSMUSG00000083220 | 0.578129653 | 0.396240625 | 0.25 | 0.396240625 | 0.387761935 | 0 | 0 |
| ENSMUSG00000083221 | 0 | 0 | 0 | 0 | 0 | 0 | 0 |
| ENSMUSG00000083222 | 0 | 0 | 0 | 0 | 0 | 0 | 0 |
| ENSMUSG00000083223 | 0 | 0.5 | 0.701838731 | 0.646240625 | 0.489301084 | 0 | 0.520678036 |
| ENSMUSG00000083224 | 0 | 0 | 0 | 0 | 0 | 0 | 0 |
| ENSMUSG00000083225 | 0 | 0 | 0 | 0 | 0 | 0 | 0 |
| ENSMUSG00000083226 | 0 | 0 | 0 | 0 | 0 | 0 | 0.224243824 |
| ENSMUSG00000083227 | 0 | 0 | 0 | 0 | 0 | 0 | 0 |
| ENSMUSG00000083228 | 0 | 0 | 0 | 0 | 0 | 0 | 0 |

| | | | | | | | |
|---|---|---|---|---|---|---|---|
| ENSMUSG00000083229 | 0 | 0 | 0 | 0 | 0 | 0 | 0 |
| ENSMUSG00000083230 | 0 | 0 | 0 | 0 | 0 | 0 | 0 |
| ENSMUSG00000083231 | 0 | 0 | 0 | 0 | 0 | 0 | 0 |
| ENSMUSG00000083232 | 0 | 0 | 0 | 0 | 0 | 0 | 0 |
| ENSMUSG00000083233 | 0 | 0 | 0 | 0 | 0 | 0 | 0 |
| ENSMUSG00000083234 | 0 | 0 | 0 | 0 | 0 | 0 | 0 |
| ENSMUSG00000083235 | 0 | 0 | 0 | 0 | 0 | 0 | 0 |
| ENSMUSG00000083236 | 0.289064826 | 0.75 | 0.25 | 0 | 0 | 0.301029996 | 0.224243824 |
| ENSMUSG00000083237 | 0 | 0 | 0 | 0 | 0 | 0 | 0 |
| ENSMUSG00000083238 | 0 | 0 | 0 | 0 | 0 | 0 | 0 |
| ENSMUSG00000083239 | 0 | 0 | 0 | 0 | 0 | 0 | 0 |
| ENSMUSG00000083240 | 0 | 0.25 | 0 | 0 | 0 | 0 | 0 |
| ENSMUSG00000083241 | 0 | 0.25 | 0 | 0 | 0.244650542 | 0 | 0 |
| ENSMUSG00000083242 | 0 | 0 | 0 | 0 | 0 | 0 | 0 |
| ENSMUSG00000083243 | 0 | 0 | 0 | 0 | 0 | 0 | 0 |
| ENSMUSG00000083244 | 0 | 0 | 0 | 0 | 0 | 0 | 0 |
| ENSMUSG00000083245 | 0.45815691 | 0.580482024 | 0.580482024 | 0.25 | 0.568060967 | 0 | 0.579661877 |
| ENSMUSG00000083246 | 0 | 0.5 | 0.5 | 0.25 | 0.632412477 | 0.301029996 | 0.448487648 |
| ENSMUSG00000083248 | 0 | 0 | 0 | 0 | 0 | 0 | 0 |
| ENSMUSG00000083249 | 0 | 0 | 0 | 0 | 0 | 0 | 0 |
| ENSMUSG00000083250 | 0.45815691 | 0 | 0.25 | 0.396240625 | 0.244650542 | 0 | 0 |
| ENSMUSG00000083251 | 0 | 0 | 0 | 0 | 0 | 0 | 0 |
| ENSMUSG00000083252 | 0 | 0 | 0 | 0 | 0 | 0 | 0 |
| ENSMUSG00000083253 | 0 | 0 | 0 | 0.646240625 | 0 | 0 | 0 |
| ENSMUSG00000083254 | 0 | 0 | 0 | 0 | 0 | 0 | 0 |
| ENSMUSG00000083255 | 0 | 0 | 0 | 0 | 0 | 0 | 0 |
| ENSMUSG00000083256 | 0 | 0 | 0 | 0 | 0 | 0 | 0 |
| ENSMUSG00000083257 | 0 | 0 | 0 | 0 | 0 | 0 | 0 |
| ENSMUSG00000083258 | 0 | 0.396240625 | 0.25 | 0.25 | 0 | 0 | 0.224243824 |
| ENSMUSG00000083260 | 0 | 0 | 0 | 0 | 0 | 0 | 0 |
| ENSMUSG00000083261 | 0.289064826 | 0 | 0 | 0 | 0.244650542 | 0.301029996 | 0 |
| ENSMUSG00000083262 | 0 | 0 | 0 | 0 | 0 | 0 | 0 |
| ENSMUSG00000083263 | 0 | 0 | 0 | 0 | 0 | 0 | 0 |
| ENSMUSG00000083264 | 0 | 0 | 0 | 0 | 0 | 0 | 0 |

| | | | | | | | |
|---|---|---|---|---|---|---|---|
| ENSMUSG00000083266 | 0 | 0 | 0 | 0 | 0 | 0 | 0 |
| ENSMUSG00000083267 | 0 | 0 | 0 | 0 | 0 | 0 | 0 |
| ENSMUSG00000083268 | 0.45815691 | 0.580482024 | 0 | 0.5 | 0.489301084 | 0.477121255 | 0.224243824 |
| ENSMUSG00000083269 | 0 | 0 | 0 | 0 | 0 | 0 | 0 |
| ENSMUSG00000083270 | 0 | 0.396240625 | 0.25 | 0.580482024 | 0.387761935 | 0 | 0.224243824 |
| ENSMUSG00000083271 | 0 | 0 | 0 | 0 | 0 | 0 | 0 |
| ENSMUSG00000083272 | 0 | 0 | 0 | 0 | 0 | 0 | 0 |
| ENSMUSG00000083274 | 0 | 0 | 0 | 0 | 0 | 0 | 0 |
| ENSMUSG00000083275 | 0 | 0 | 0 | 0 | 0 | 0.301029996 | 0 |
| ENSMUSG00000083276 | 0 | 0 | 0 | 0 | 0 | 0 | 0 |
| ENSMUSG00000083277 | 0.289064826 | 0 | 0 | 0 | 0 | 0 | 0 |
| ENSMUSG00000083278 | 0 | 0 | 0 | 0 | 0 | 0 | 0 |
| ENSMUSG00000083279 | 0 | 0 | 0 | 0 | 0 | 0 | 0 |
| ENSMUSG00000083280 | 0 | 0 | 0 | 0 | 0 | 0 | 0.224243824 |
| ENSMUSG00000083281 | 0 | 0 | 0 | 0 | 0 | 0.301029996 | 0 |
| ENSMUSG00000083282 | 1.432083896 | 1.339388001 | 1.410964047 | 1.62694866 | 1.686820904 | 1.698970004 | 1.635773653 |
| ENSMUSG00000083283 | 0 | 0 | 0 | 0 | 0 | 0 | 0.224243824 |
| ENSMUSG00000083284 | 0.289064826 | 0.396240625 | 0.25 | 0.25 | 0.244650542 | 0.301029996 | 0.355418052 |
| ENSMUSG00000083285 | 0 | 0 | 0 | 0 | 0 | 0 | 0 |
| ENSMUSG00000083287 | 0 | 0 | 0 | 0 | 0 | 0 | 0 |
| ENSMUSG00000083288 | 0.289064826 | 0.25 | 0 | 0.25 | 0.244650542 | 0.301029996 | 0.355418052 |
| ENSMUSG00000083289 | 0 | 0 | 0 | 0 | 0 | 0 | 0 |
| ENSMUSG00000083290 | 0 | 0 | 0 | 0 | 0 | 0 | 0 |
| ENSMUSG00000083291 | 0 | 0 | 0 | 0 | 0 | 0 | 0 |
| ENSMUSG00000083292 | 0 | 0 | 0 | 0 | 0 | 0 | 0 |
| ENSMUSG00000083293 | 0 | 0 | 0 | 0 | 0 | 0 | 0 |
| ENSMUSG00000083294 | 0 | 0 | 0 | 0 | 0 | 0 | 0 |
| ENSMUSG00000083296 | 0 | 0 | 0 | 0 | 0 | 0 | 0 |
| ENSMUSG00000083297 | 0 | 0 | 0 | 0 | 0 | 0 | 0 |
| ENSMUSG00000083298 | 0 | 0 | 0 | 0 | 0 | 0 | 0 |
| ENSMUSG00000083299 | 0 | 0 | 0 | 0 | 0 | 0 | 0 |
| ENSMUSG00000083300 | 0 | 0.396240625 | 0 | 0 | 0 | 0 | 0 |
| ENSMUSG00000083303 | 0 | 0 | 0 | 0 | 0 | 0 | 0 |
| ENSMUSG00000083304 | 0 | 0 | 0 | 0 | 0 | 0 | 0 |

| | | | | | | | |
|---|---|---|---|---|---|---|---|
| ENSMUSG00000083305 | 0 | 0 | 0.25 | 0 | 0 | 0 | 0 |
| ENSMUSG00000083306 | 0.289064826 | 0 | 0 | 0 | 0 | 0 | 0 |
| ENSMUSG00000083307 | 1.470606563 | 1.380890489 | 1.348079356 | 1.146240625 | 0.846351821 | 1.079181246 | 1.041356071 |
| ENSMUSG00000083309 | 0 | 0 | 0 | 0 | 0 | 0 | 0 |
| ENSMUSG00000083310 | 0 | 0 | 0 | 0 | 0 | 0 | 0 |
| ENSMUSG00000083311 | 0 | 0.75 | 0.396240625 | 0.396240625 | 0.632412477 | 0 | 0.355418052 |
| ENSMUSG00000083312 | 0 | 0 | 0 | 0 | 0 | 0 | 0 |
| ENSMUSG00000083314 | 0 | 0 | 0 | 0 | 0.387761935 | 0.301029996 | 0 |
| ENSMUSG00000083315 | 0 | 0 | 0 | 0 | 0 | 0 | 0 |
| ENSMUSG00000083316 | 0 | 0 | 0 | 0 | 0 | 0.301029996 | 0 |
| ENSMUSG00000083317 | 0 | 0 | 0 | 0 | 0 | 0 | 0 |
| ENSMUSG00000083318 | 0 | 0 | 0 | 0 | 0 | 0 | 0 |
| ENSMUSG00000083319 | 0.289064826 | 0.701838731 | 0.396240625 | 0.580482024 | 0.733951626 | 0.77815125 | 0.710836105 |
| ENSMUSG00000083320 | 0 | 0 | 0 | 0 | 0.244650542 | 0.477121255 | 0 |
| ENSMUSG00000083321 | 0 | 0 | 0 | 0 | 0 | 0 | 0 |
| ENSMUSG00000083324 | 0 | 0 | 0 | 0 | 0 | 0 | 0 |
| ENSMUSG00000083325 | 0 | 0 | 0 | 0 | 0.387761935 | 0 | 0 |
| ENSMUSG00000083326 | 0 | 0 | 0 | 0 | 0 | 0 | 0 |
| ENSMUSG00000083327 | 0.289064826 | 0.396240625 | 0.25 | 0.396240625 | 0.244650542 | 0 | 0.448487648 |
| ENSMUSG00000083328 | 0 | 0 | 0.25 | 0 | 0.244650542 | 0 | 0 |
| ENSMUSG00000083329 | 0 | 0 | 0 | 0 | 0 | 0 | 0 |
| ENSMUSG00000083330 | 0 | 0 | 0 | 0 | 0 | 0 | 0 |
| ENSMUSG00000083331 | 0 | 0 | 0 | 0 | 0 | 0 | 0 |
| ENSMUSG00000083332 | 0 | 0.25 | 0 | 0.25 | 0 | 0.301029996 | 0 |
| ENSMUSG00000083333 | 0 | 0 | 0 | 0 | 0 | 0 | 0 |
| ENSMUSG00000083334 | 0 | 0 | 0 | 0 | 0 | 0 | 0 |
| ENSMUSG00000083335 | 0 | 0.25 | 0 | 0 | 0 | 0 | 0.224243824 |
| ENSMUSG00000083336 | 0 | 0 | 0 | 0 | 0 | 0 | 0 |
| ENSMUSG00000083337 | 0.747221736 | 0.646240625 | 0.646240625 | 0.25 | 0.733951626 | 0.301029996 | 0.579661877 |
| ENSMUSG00000083338 | 0 | 0 | 0 | 0 | 0 | 0 | 0 |
| ENSMUSG00000083339 | 0.289064826 | 0.25 | 0.25 | 0.5 | 0.387761935 | 0 | 0.224243824 |
| ENSMUSG00000083340 | 0 | 0.25 | 0.25 | 0 | 0 | 0 | 0 |
| ENSMUSG00000083341 | 0 | 0 | 0 | 0 | 0 | 0 | 0 |
| ENSMUSG00000083342 | 0 | 0 | 0 | 0 | 0 | 0 | 0 |

| | | | | | | | |
|---|---|---|---|---|---|---|---|
| ENSMUSG00000083343 | 0 | 0 | 0 | 0 | 0 | 0 | 0 |
| ENSMUSG00000083344 | 0.289064826 | 0 | 0.25 | 0 | 0.244650542 | 0.477121255 | 0.224243824 |
| ENSMUSG00000083345 | 0 | 0 | 0 | 0 | 0 | 0 | 0 |
| ENSMUSG00000083346 | 0 | 0 | 0 | 0 | 0 | 0 | 0 |
| ENSMUSG00000083347 | 0 | 0 | 0 | 0 | 0 | 0 | 0 |
| ENSMUSG00000083348 | 0 | 0 | 0 | 0 | 0 | 0 | 0 |
| ENSMUSG00000083350 | 0 | 0 | 0 | 0 | 0 | 0 | 0 |
| ENSMUSG00000083351 | 0 | 0 | 0 | 0 | 0 | 0 | 0 |
| ENSMUSG00000083352 | 0 | 0 | 0 | 0 | 0 | 0 | 0 |
| ENSMUSG00000083353 | 0 | 0 | 0 | 0 | 0.387761935 | 0 | 0.355418052 |
| ENSMUSG00000083355 | 0 | 0 | 0 | 0 | 0 | 0 | 0 |
| ENSMUSG00000083356 | 0 | 0 | 0 | 0 | 0 | 0 | 0 |
| ENSMUSG00000083358 | 0 | 0 | 0 | 0 | 0 | 0 | 0 |
| ENSMUSG00000083359 | 0 | 0.396240625 | 0 | 0 | 0 | 0 | 0.224243824 |
| ENSMUSG00000083361 | 0 | 0 | 0 | 0 | 0 | 0 | 0 |
| ENSMUSG00000083362 | 0.45815691 | 0.25 | 0 | 0 | 0.387761935 | 0 | 0.448487648 |
| ENSMUSG00000083363 | 0 | 0 | 0 | 0 | 0 | 0 | 0 |
| ENSMUSG00000083364 | 0 | 0 | 0 | 0 | 0 | 0 | 0 |
| ENSMUSG00000083365 | 0 | 0 | 0 | 0 | 0 | 0 | 0 |
| ENSMUSG00000083366 | 0 | 0 | 0 | 0 | 0 | 0 | 0 |
| ENSMUSG00000083367 | 0.289064826 | 0.646240625 | 0.5 | 0 | 0.387761935 | 0 | 0.355418052 |
| ENSMUSG00000083368 | 0 | 0 | 0 | 0 | 0 | 0 | 0 |
| ENSMUSG00000083369 | 0 | 0 | 0 | 0 | 0 | 0 | 0 |
| ENSMUSG00000083371 | 0 | 0 | 0 | 0 | 0 | 0 | 0 |
| ENSMUSG00000083372 | 0 | 0 | 0 | 0 | 0 | 0 | 0 |
| ENSMUSG00000083373 | 0 | 0 | 0 | 0 | 0 | 0 | 0 |
| ENSMUSG00000083374 | 0 | 0.396240625 | 0 | 0.396240625 | 0.244650542 | 0 | 0.520678036 |
| ENSMUSG00000083375 | 0.289064826 | 0 | 0 | 0.25 | 0 | 0 | 0 |
| ENSMUSG00000083376 | 0 | 0 | 0 | 0 | 0 | 0 | 0 |
| ENSMUSG00000083377 | 0 | 0.25 | 0 | 0 | 0 | 0 | 0.355418052 |
| ENSMUSG00000083378 | 0 | 0.25 | 0.25 | 0.25 | 0.244650542 | 0.301029996 | 0.224243824 |
| ENSMUSG00000083379 | 0 | 0 | 0 | 0 | 0 | 0 | 0.224243824 |
| ENSMUSG00000083380 | 0 | 0 | 0 | 0 | 0 | 0 | 0 |
| ENSMUSG00000083381 | 0 | 0 | 0 | 0 | 0 | 0 | 0 |

| | | | | | | | |
|---|---|---|---|---|---|---|---|
| ENSMUSG00000083382 | 0 | 0.25 | 0 | 0.25 | 0.387761935 | 0 | 0.224243824 |
| ENSMUSG00000083383 | 0 | 0 | 0 | 0 | 0 | 0 | 0 |
| ENSMUSG00000083384 | 0 | 0 | 0 | 0 | 0 | 0 | 0 |
| ENSMUSG00000083385 | 0 | 0 | 0.25 | 0 | 0 | 0 | 0.224243824 |
| ENSMUSG00000083386 | 0 | 0 | 0 | 0 | 0 | 0 | 0 |
| ENSMUSG00000083387 | 0 | 0 | 0 | 0.25 | 0 | 0 | 0 |
| ENSMUSG00000083388 | 0 | 0 | 0 | 0 | 0.244650542 | 0 | 0 |
| ENSMUSG00000083389 | 0 | 0 | 0 | 0 | 0 | 0 | 0 |
| ENSMUSG00000083390 | 0.289064826 | 0 | 0 | 0 | 0 | 0 | 0 |
| ENSMUSG00000083391 | 1.516991255 | 1.482684334 | 1.488549078 | 1.26109853 | 1.358933174 | 1.230448921 | 1.284450788 |
| ENSMUSG00000083392 | 0 | 0 | 0 | 0.5 | 0 | 0 | 0 |
| ENSMUSG00000083393 | 0 | 0 | 0 | 0 | 0 | 0 | 0 |
| ENSMUSG00000083394 | 0 | 0 | 0 | 0.25 | 0 | 0 | 0.224243824 |
| ENSMUSG00000083395 | 0 | 0 | 0 | 0.25 | 0 | 0 | 0 |
| ENSMUSG00000083396 | 0.289064826 | 0.25 | 0.25 | 0.25 | 0.244650542 | 0 | 0.579661877 |
| ENSMUSG00000083397 | 0.578129653 | 0.25 | 0.25 | 0 | 0.244650542 | 0.301029996 | 0.448487648 |
| ENSMUSG00000083398 | 0 | 0 | 0 | 0 | 0 | 0 | 0 |
| ENSMUSG00000083399 | 0 | 0 | 0 | 0 | 0 | 0 | 0 |
| ENSMUSG00000083401 | 0 | 0 | 0 | 0 | 0 | 0 | 0 |
| ENSMUSG00000083402 | 0 | 0 | 0 | 0 | 0 | 0 | 0 |
| ENSMUSG00000083403 | 0 | 0 | 0 | 0 | 0 | 0 | 0 |
| ENSMUSG00000083404 | 0 | 0 | 0 | 0 | 0 | 0 | 0 |
| ENSMUSG00000083405 | 0 | 0 | 0 | 0 | 0 | 0 | 0 |
| ENSMUSG00000083406 | 0 | 0 | 0 | 0 | 0 | 0 | 0 |
| ENSMUSG00000083407 | 0 | 0 | 0 | 0 | 0 | 0 | 0 |
| ENSMUSG00000083409 | 0 | 0 | 0 | 0 | 0 | 0 | 0 |
| ENSMUSG00000083410 | 0.289064826 | 0 | 0 | 0 | 0 | 0.301029996 | 0 |
| ENSMUSG00000083411 | 0 | 0 | 0 | 0 | 0 | 0 | 0 |
| ENSMUSG00000083412 | 0 | 0 | 0 | 0 | 0 | 0 | 0 |
| ENSMUSG00000083413 | 0 | 0 | 0 | 0 | 0 | 0 | 0 |
| ENSMUSG00000083414 | 0.289064826 | 0.25 | 0.396240625 | 0.5 | 0.387761935 | 0.301029996 | 0.355418052 |
| ENSMUSG00000083416 | 0 | 0 | 0 | 0 | 0 | 0 | 0 |
| ENSMUSG00000083417 | 0 | 0 | 0 | 0 | 0 | 0 | 0 |
| ENSMUSG00000083418 | 0 | 0 | 0 | 0 | 0 | 0 | 0 |

| | | | | | | | |
|---|---|---|---|---|---|---|---|
| ENSMUSG00000083419 | 0 | 0 | 0 | 0 | 0 | 0 | 0 |
| ENSMUSG00000083420 | 0 | 0 | 0 | 0 | 0 | 0 | 0 |
| ENSMUSG00000083421 | 0 | 0 | 0 | 0 | 0 | 0 | 0 |
| ENSMUSG00000083422 | 0.578129653 | 0.25 | 0 | 0.396240625 | 0.387761935 | 0.301029996 | 0.224243824 |
| ENSMUSG00000083423 | 0 | 0 | 0 | 0 | 0 | 0 | 0 |
| ENSMUSG00000083424 | 0.578129653 | 0.396240625 | 0.396240625 | 0.580482024 | 0.568060967 | 0.301029996 | 0 |
| ENSMUSG00000083425 | 0 | 0 | 0 | 0 | 0.244650542 | 0 | 0.224243824 |
| ENSMUSG00000083427 | 0 | 0 | 0 | 0 | 0 | 0 | 0 |
| ENSMUSG00000083428 | 0 | 0 | 0 | 0 | 0 | 0 | 0 |
| ENSMUSG00000083429 | 0 | 0 | 0 | 0 | 0 | 0 | 0.355418052 |
| ENSMUSG00000083430 | 0 | 0 | 0 | 0 | 0 | 0 | 0 |
| ENSMUSG00000083431 | 0 | 0 | 0.25 | 0 | 0.387761935 | 0.301029996 | 0.224243824 |
| ENSMUSG00000083432 | 0 | 0 | 0 | 0 | 0 | 0 | 0 |
| ENSMUSG00000083434 | 0 | 0.396240625 | 0.396240625 | 0.646240625 | 0.387761935 | 0.477121255 | 0.672731473 |
| ENSMUSG00000083435 | 0 | 0 | 0.25 | 0 | 0 | 0 | 0.224243824 |
| ENSMUSG00000083436 | 0 | 0 | 0 | 0 | 0 | 0 | 0 |
| ENSMUSG00000083437 | 0 | 0 | 0 | 0 | 0 | 0 | 0 |
| ENSMUSG00000083438 | 0 | 0 | 0 | 0 | 0 | 0 | 0 |
| ENSMUSG00000083439 | 0 | 0 | 0 | 0 | 0 | 0 | 0 |
| ENSMUSG00000083440 | 0.289064826 | 0 | 0 | 0 | 0 | 0 | 0.448487648 |
| ENSMUSG00000083441 | 0 | 0 | 0 | 0 | 0 | 0 | 0 |
| ENSMUSG00000083443 | 0.289064826 | 0.396240625 | 0.396240625 | 0.646240625 | 0 | 0.301029996 | 0.448487648 |
| ENSMUSG00000083444 | 0 | 0 | 0 | 0 | 0 | 0 | 0 |
| ENSMUSG00000083445 | 0 | 0 | 0 | 0 | 0 | 0 | 0 |
| ENSMUSG00000083446 | 0 | 0 | 0 | 0 | 0 | 0 | 0 |
| ENSMUSG00000083447 | 0 | 0 | 0 | 0 | 0 | 0 | 0 |
| ENSMUSG00000083448 | 0 | 0 | 0 | 0 | 0 | 0 | 0 |
| ENSMUSG00000083449 | 0 | 0 | 0 | 0 | 0 | 0 | 0 |
| ENSMUSG00000083450 | 0 | 0 | 0 | 0 | 0 | 0 | 0 |
| ENSMUSG00000083451 | 0 | 0 | 0 | 0 | 0 | 0 | 0 |
| ENSMUSG00000083452 | 0 | 0 | 0 | 0 | 0 | 0 | 0 |
| ENSMUSG00000083453 | 0 | 0 | 0 | 0 | 0 | 0 | 0 |
| ENSMUSG00000083454 | 0.289064826 | 0.25 | 0 | 0 | 0 | 0 | 0 |
| ENSMUSG00000083455 | 0 | 0 | 0 | 0 | 0 | 0 | 0 |

| | | | | | | | |
|---|---|---|---|---|---|---|---|
| ENSMUSG00000083456 | 0 | 0 | 0 | 0 | 0 | 0 | 0 |
| ENSMUSG00000083457 | 0 | 0 | 0 | 0 | 0 | 0 | 0 |
| ENSMUSG00000083458 | 0 | 0.25 | 0 | 0 | 0 | 0 | 0 |
| ENSMUSG00000083460 | 0.578129653 | 0.92510993 | 0.701838731 | 0 | 0.77552387 | 0.477121255 | 0.224243824 |
| ENSMUSG00000083461 | 0 | 0 | 0 | 0 | 0 | 0 | 0 |
| ENSMUSG00000083462 | 0 | 0 | 0 | 0 | 0 | 0 | 0 |
| ENSMUSG00000083463 | 0 | 0 | 0 | 0 | 0 | 0 | 0 |
| ENSMUSG00000083465 | 0.671187741 | 0.396240625 | 0.580482024 | 0.396240625 | 0.244650542 | 0 | 0 |
| ENSMUSG00000083466 | 0 | 0 | 0 | 0 | 0 | 0 | 0 |
| ENSMUSG00000083467 | 0 | 0 | 0 | 0 | 0 | 0 | 0 |
| ENSMUSG00000083468 | 0.289064826 | 0.25 | 0.25 | 0.580482024 | 0.387761935 | 0.301029996 | 0 |
| ENSMUSG00000083469 | 0 | 0 | 0 | 0 | 0 | 0 | 0 |
| ENSMUSG00000083470 | 0 | 0 | 0.25 | 0 | 0 | 0 | 0.355418052 |
| ENSMUSG00000083471 | 0 | 0.25 | 0 | 0 | 0 | 0 | 0.224243824 |
| ENSMUSG00000083472 | 0 | 0 | 0 | 0 | 0 | 0 | 0 |
| ENSMUSG00000083473 | 0 | 0 | 0 | 0 | 0 | 0 | 0 |
| ENSMUSG00000083474 | 0.671187741 | 0 | 0.580482024 | 0.5 | 0 | 0.301029996 | 0.355418052 |
| ENSMUSG00000083475 | 0 | 0 | 0 | 0 | 0 | 0 | 0 |
| ENSMUSG00000083477 | 0 | 0 | 0 | 0.25 | 0.489301084 | 0 | 0 |
| ENSMUSG00000083478 | 0 | 0 | 0 | 0 | 0 | 0 | 0 |
| ENSMUSG00000083481 | 0 | 0 | 0 | 0 | 0 | 0 | 0 |
| ENSMUSG00000083482 | 0 | 0 | 0 | 0 | 0 | 0 | 0 |
| ENSMUSG00000083483 | 0 | 0 | 0 | 0 | 0 | 0 | 0 |
| ENSMUSG00000083484 | 0 | 0 | 0 | 0 | 0 | 0 | 0 |
| ENSMUSG00000083485 | 0 | 0 | 0 | 0 | 0 | 0 | 0 |
| ENSMUSG00000083486 | 0 | 0 | 0 | 0 | 0 | 0 | 0 |
| ENSMUSG00000083487 | 0 | 0 | 0 | 0 | 0 | 0 | 0 |
| ENSMUSG00000083488 | 0 | 0 | 0 | 0 | 0 | 0 | 0 |
| ENSMUSG00000083489 | 0 | 0 | 0 | 0 | 0 | 0 | 0 |
| ENSMUSG00000083490 | 0 | 0 | 0 | 0 | 0 | 0 | 0 |
| ENSMUSG00000083491 | 0 | 0 | 0 | 0 | 0 | 0 | 0 |
| ENSMUSG00000083492 | 0 | 0 | 0 | 0 | 0 | 0 | 0 |
| ENSMUSG00000083493 | 0 | 0 | 0 | 0 | 0 | 0 | 0 |
| ENSMUSG00000083494 | 0 | 0 | 0 | 0 | 0 | 0 | 0 |

| | | | | | | | |
|---|---|---|---|---|---|---|---|
| ENSMUSG00000083495 | 0 | 0 | 0 | 0 | 0 | 0 | 0 |
| ENSMUSG00000083496 | 0 | 0.580482024 | 0 | 0.25 | 0.489301084 | 0 | 0.224243824 |
| ENSMUSG00000083498 | 0 | 0 | 0 | 0 | 0 | 0 | 0 |
| ENSMUSG00000083499 | 0 | 0 | 0 | 0 | 0 | 0 | 0 |
| ENSMUSG00000083500 | 0.45815691 | 0.580482024 | 0.396240625 | 0.5 | 0.568060967 | 0.301029996 | 0.355418052 |
| ENSMUSG00000083501 | 0 | 0 | 0 | 0 | 0 | 0 | 0 |
| ENSMUSG00000083502 | 0 | 0 | 0 | 0 | 0 | 0 | 0 |
| ENSMUSG00000083503 | 0.289064826 | 0 | 0 | 0 | 0 | 0 | 0.224243824 |
| ENSMUSG00000083505 | 0 | 0.25 | 0.396240625 | 0 | 0.489301084 | 0 | 0.355418052 |
| ENSMUSG00000083507 | 0 | 0.25 | 0 | 0 | 0 | 0.301029996 | 0 |
| ENSMUSG00000083508 | 0.289064826 | 0 | 0 | 0 | 0 | 0 | 0 |
| ENSMUSG00000083509 | 0 | 0 | 0 | 0 | 0 | 0 | 0 |
| ENSMUSG00000083510 | 0 | 0 | 0 | 0 | 0 | 0 | 0 |
| ENSMUSG00000083512 | 0.289064826 | 0.25 | 0.25 | 0.396240625 | 0.387761935 | 0.301029996 | 0 |
| ENSMUSG00000083513 | 0 | 0 | 0 | 0 | 0 | 0 | 0 |
| ENSMUSG00000083514 | 0 | 0 | 0 | 0 | 0 | 0 | 0 |
| ENSMUSG00000083515 | 0 | 0 | 0 | 0 | 0 | 0 | 0 |
| ENSMUSG00000083516 | 0 | 0 | 0 | 0 | 0 | 0 | 0 |
| ENSMUSG00000083517 | 0 | 0 | 0 | 0 | 0 | 0 | 0 |
| ENSMUSG00000083518 | 0 | 0 | 0 | 0 | 0 | 0 | 0 |
| ENSMUSG00000083519 | 0 | 0 | 0 | 0 | 0 | 0 | 0 |
| ENSMUSG00000083520 | 0 | 0 | 0 | 0 | 0 | 0 | 0 |
| ENSMUSG00000083522 | 0 | 0.25 | 0 | 0 | 0 | 0 | 0.224243824 |
| ENSMUSG00000083524 | 0 | 0 | 0 | 0 | 0 | 0 | 0 |
| ENSMUSG00000083527 | 0 | 0 | 0 | 0 | 0 | 0 | 0 |
| ENSMUSG00000083528 | 0 | 0 | 0.25 | 0 | 0.244650542 | 0 | 0.224243824 |
| ENSMUSG00000083530 | 0 | 0 | 0 | 0 | 0 | 0 | 0 |
| ENSMUSG00000083533 | 0 | 0 | 0 | 0 | 0 | 0 | 0 |
| ENSMUSG00000083534 | 0 | 0 | 0 | 0.396240625 | 0 | 0 | 0 |
| ENSMUSG00000083535 | 0 | 0 | 0 | 0 | 0 | 0 | 0 |
| ENSMUSG00000083536 | 0 | 0 | 0 | 0 | 0 | 0 | 0 |
| ENSMUSG00000083537 | 0 | 0 | 0 | 0 | 0 | 0 | 0 |
| ENSMUSG00000083538 | 0 | 0 | 0.25 | 0 | 0.244650542 | 0 | 0.224243824 |
| ENSMUSG00000083539 | 0 | 0.25 | 0.396240625 | 0 | 0.244650542 | 0 | 0.672731473 |

| | | | | | | | |
|---|---|---|---|---|---|---|---|
| ENSMUSG00000083541 | 0 | 0 | 0 | 0 | 0 | 0 | 0 |
| ENSMUSG00000083542 | 0 | 0 | 0 | 0 | 0 | 0 | 0 |
| ENSMUSG00000083543 | 0 | 0 | 0 | 0 | 0 | 0 | 0 |
| ENSMUSG00000083544 | 0 | 0 | 0 | 0 | 0 | 0 | 0 |
| ENSMUSG00000083545 | 0 | 0 | 0 | 0 | 0 | 0 | 0 |
| ENSMUSG00000083546 | 0 | 0 | 0 | 0 | 0 | 0 | 0 |
| ENSMUSG00000083547 | 0 | 0 | 0 | 0.5 | 0.244650542 | 0.301029996 | 0 |
| ENSMUSG00000083548 | 0 | 0 | 0 | 0.25 | 0 | 0 | 0.224243824 |
| ENSMUSG00000083549 | 0 | 0 | 0 | 0 | 0 | 0 | 0 |
| ENSMUSG00000083550 | 0 | 0 | 0 | 0 | 0 | 0 | 0 |
| ENSMUSG00000083551 | 0.289064826 | 0 | 0 | 0 | 0 | 0 | 0 |
| ENSMUSG00000083553 | 0.578129653 | 0.580482024 | 0.701838731 | 0.25 | 0.387761935 | 0.602059991 | 0.355418052 |
| ENSMUSG00000083554 | 0 | 0 | 0 | 0 | 0 | 0 | 0.224243824 |
| ENSMUSG00000083555 | 0 | 0 | 0 | 0 | 0 | 0 | 0 |
| ENSMUSG00000083556 | 0 | 0 | 0 | 0.25 | 0 | 0 | 0 |
| ENSMUSG00000083557 | 0 | 0 | 0 | 0 | 0 | 0 | 0 |
| ENSMUSG00000083558 | 0 | 0 | 0 | 0 | 0 | 0 | 0 |
| ENSMUSG00000083560 | 0 | 0 | 0 | 0 | 0 | 0 | 0 |
| ENSMUSG00000083561 | 0 | 0 | 0 | 0 | 0 | 0 | 0.224243824 |
| ENSMUSG00000083562 | 0 | 0 | 0 | 0 | 0 | 0 | 0 |
| ENSMUSG00000083563 | 1.404271433 | 1.438721876 | 1.42510993 | 1.380890489 | 1.401338844 | 1.505149978 | 1.245582246 |
| ENSMUSG00000083564 | 0 | 0 | 0 | 0 | 0 | 0 | 0 |
| ENSMUSG00000083565 | 0 | 0 | 0 | 0 | 0 | 0 | 0 |
| ENSMUSG00000083566 | 0 | 0 | 0 | 0 | 0 | 0 | 0 |
| ENSMUSG00000083567 | 0 | 0 | 0 | 0.25 | 0 | 0 | 0 |
| ENSMUSG00000083568 | 0 | 0 | 0 | 0 | 0 | 0 | 0 |
| ENSMUSG00000083569 | 0.289064826 | 0 | 0.25 | 0.396240625 | 0 | 0 | 0 |
| ENSMUSG00000083570 | 0 | 0 | 0 | 0 | 0 | 0 | 0 |
| ENSMUSG00000083572 | 0 | 0 | 0 | 0 | 0 | 0 | 0 |
| ENSMUSG00000083573 | 0 | 0 | 0 | 0 | 0 | 0 | 0 |
| ENSMUSG00000083574 | 0 | 0 | 0 | 0 | 0 | 0 | 0 |
| ENSMUSG00000083576 | 0 | 0 | 0 | 0 | 0 | 0 | 0 |
| ENSMUSG00000083577 | 0 | 0 | 0 | 0 | 0 | 0 | 0 |
| ENSMUSG00000083579 | 0 | 0 | 0.396240625 | 0 | 0 | 0 | 0 |

| | | | | | | | |
|---|---|---|---|---|---|---|---|
| ENSMUSG00000083580 | 0.289064826 | 0.580482024 | 0.5 | 0.25 | 0.244650542 | 0.301029996 | 0.448487648 |
| ENSMUSG00000083581 | 0 | 0 | 0 | 0 | 0 | 0 | 0 |
| ENSMUSG00000083582 | 0 | 0 | 0 | 0 | 0 | 0 | 0 |
| ENSMUSG00000083584 | 0 | 0 | 0 | 0 | 0 | 0 | 0 |
| ENSMUSG00000083585 | 0 | 0 | 0 | 0 | 0.244650542 | 0 | 0 |
| ENSMUSG00000083586 | 0 | 0.5 | 0.5 | 0 | 0 | 0 | 0 |
| ENSMUSG00000083587 | 0 | 0 | 0 | 0 | 0.244650542 | 0 | 0 |
| ENSMUSG00000083588 | 0.578129653 | 0.25 | 0.646240625 | 0.5 | 0.387761935 | 0.301029996 | 0.224243824 |
| ENSMUSG00000083590 | 0 | 0 | 0 | 0 | 0 | 0 | 0 |
| ENSMUSG00000083592 | 0 | 0.25 | 0 | 0 | 0 | 0 | 0 |
| ENSMUSG00000083594 | 1.036286563 | 1.080482024 | 1.146240625 | 1.26109853 | 1.136121934 | 1.079181246 | 0.984950056 |
| ENSMUSG00000083595 | 0 | 0 | 0 | 0 | 0 | 0 | 0 |
| ENSMUSG00000083596 | 0 | 0.25 | 0 | 0 | 0.387761935 | 0 | 0 |
| ENSMUSG00000083597 | 0 | 0 | 0 | 0 | 0 | 0 | 0 |
| ENSMUSG00000083598 | 0 | 0 | 0 | 0 | 0 | 0 | 0 |
| ENSMUSG00000083599 | 0 | 0 | 0 | 0 | 0 | 0 | 0 |
| ENSMUSG00000083601 | 0 | 0 | 0 | 0 | 0 | 0 | 0 |
| ENSMUSG00000083602 | 0 | 0 | 0 | 0 | 0 | 0 | 0 |
| ENSMUSG00000083603 | 0 | 0 | 0.25 | 0 | 0 | 0 | 0 |
| ENSMUSG00000083605 | 0 | 0 | 0 | 0 | 0.244650542 | 0 | 0 |
| ENSMUSG00000083606 | 0 | 0 | 0 | 0 | 0 | 0 | 0.224243824 |
| ENSMUSG00000083607 | 0 | 0 | 0 | 0 | 0 | 0 | 0 |
| ENSMUSG00000083608 | 0.671187741 | 0.25 | 0.25 | 0 | 0.568060967 | 0.602059991 | 0.224243824 |
| ENSMUSG00000083609 | 0 | 0 | 0 | 0 | 0 | 0 | 0 |
| ENSMUSG00000083610 | 0.45815691 | 0.396240625 | 0.396240625 | 0.580482024 | 0.489301084 | 0.477121255 | 0.710836105 |
| ENSMUSG00000083611 | 0 | 0 | 0 | 0 | 0 | 0 | 0.224243824 |
| ENSMUSG00000083613 | 0 | 0 | 0 | 0 | 0 | 0 | 0 |
| ENSMUSG00000083614 | 0 | 0 | 0 | 0 | 0 | 0 | 0 |
| ENSMUSG00000083615 | 0 | 0 | 0 | 0 | 0 | 0 | 0 |
| ENSMUSG00000083616 | 0 | 0 | 0 | 0 | 0 | 0 | 0 |
| ENSMUSG00000083617 | 0 | 0 | 0 | 0 | 0 | 0 | 0 |
| ENSMUSG00000083618 | 0 | 0.25 | 0.25 | 0 | 0 | 0 | 0 |
| ENSMUSG00000083619 | 0 | 0 | 0 | 0 | 0 | 0 | 0 |
| ENSMUSG00000083620 | 0 | 0 | 0 | 0 | 0 | 0 | 0 |

| | | | | | | | |
|---|---|---|---|---|---|---|---|
| ENSMUSG00000083621 | 0 | 0.25 | 0 | 0 | 0.244650542 | 0.301029996 | 0 |
| ENSMUSG00000083622 | 0 | 0 | 0 | 0 | 0 | 0 | 0 |
| ENSMUSG00000083623 | 1.069666964 | 0.896240625 | 0.951838731 | 0.864857905 | 1 | 1.301029996 | 1 |
| ENSMUSG00000083624 | 0 | 0 | 0 | 0 | 0 | 0 | 0 |
| ENSMUSG00000083625 | 0 | 0 | 0 | 0 | 0 | 0 | 0 |
| ENSMUSG00000083626 | 0 | 0 | 0 | 0 | 0 | 0 | 0 |
| ENSMUSG00000083627 | 0 | 0 | 0 | 0 | 0 | 0 | 0 |
| ENSMUSG00000083628 | 0 | 0 | 0 | 0 | 0 | 0 | 0 |
| ENSMUSG00000083629 | 0 | 0 | 0 | 0 | 0 | 0 | 0 |
| ENSMUSG00000083630 | 0 | 0 | 0 | 0.25 | 0 | 0 | 0.448487648 |
| ENSMUSG00000083631 | 0 | 0 | 0 | 0 | 0 | 0 | 0 |
| ENSMUSG00000083632 | 0 | 0 | 0 | 0 | 0 | 0 | 0 |
| ENSMUSG00000083633 | 0.45815691 | 0.396240625 | 0 | 0 | 0.387761935 | 0 | 0.224243824 |
| ENSMUSG00000083635 | 0 | 0 | 0 | 0 | 0 | 0 | 0 |
| ENSMUSG00000083636 | 0 | 0 | 0 | 0 | 0 | 0 | 0 |
| ENSMUSG00000083637 | 0 | 0 | 0 | 0 | 0 | 0 | 0 |
| ENSMUSG00000083640 | 0.289064826 | 0 | 0 | 0.25 | 0.244650542 | 0.477121255 | 0.224243824 |
| ENSMUSG00000083641 | 0 | 0 | 0 | 0 | 0 | 0 | 0 |
| ENSMUSG00000083642 | 0 | 0 | 0 | 0 | 0 | 0 | 0 |
| ENSMUSG00000083643 | 0 | 0 | 0 | 0 | 0 | 0 | 0 |
| ENSMUSG00000083644 | 0 | 0 | 0 | 0 | 0 | 0 | 0 |
| ENSMUSG00000083645 | 0 | 0 | 0 | 0 | 0 | 0 | 0 |
| ENSMUSG00000083646 | 0 | 0 | 0 | 0.25 | 0 | 0 | 0.224243824 |
| ENSMUSG00000083648 | 0 | 0 | 0 | 0 | 0 | 0 | 0 |
| ENSMUSG00000083649 | 0 | 0.396240625 | 0.396240625 | 0 | 0 | 0.477121255 | 0 |
| ENSMUSG00000083650 | 0.289064826 | 0 | 0.396240625 | 0 | 0 | 0.301029996 | 0.224243824 |
| ENSMUSG00000083651 | 0 | 0.25 | 0.25 | 0 | 0.568060967 | 0 | 0.520678036 |
| ENSMUSG00000083653 | 0 | 0 | 0 | 0 | 0 | 0 | 0 |
| ENSMUSG00000083654 | 0 | 0 | 0 | 0 | 0 | 0 | 0 |
| ENSMUSG00000083655 | 0 | 0 | 0 | 0 | 0 | 0 | 0 |
| ENSMUSG00000083657 | 0 | 0 | 0 | 0 | 0 | 0 | 0 |
| ENSMUSG00000083658 | 0.45815691 | 0 | 0.25 | 0.25 | 0 | 0 | 0 |
| ENSMUSG00000083659 | 0.289064826 | 0.580482024 | 0.5 | 0.646240625 | 0.568060967 | 0.77815125 | 0.448487648 |
| ENSMUSG00000083660 | 0 | 0 | 0 | 0 | 0 | 0 | 0 |

| | | | | | | | |
|---|---|---|---|---|---|---|---|
| ENSMUSG00000083661 | 0 | 0.25 | 0 | 0 | 0 | 0 | 0 |
| ENSMUSG00000083662 | 0 | 0 | 0 | 0 | 0 | 0 | 0 |
| ENSMUSG00000083663 | 0 | 0 | 0 | 0 | 0 | 0 | 0 |
| ENSMUSG00000083664 | 0 | 0 | 0 | 0 | 0 | 0 | 0 |
| ENSMUSG00000083665 | 0 | 0 | 0 | 0 | 0 | 0 | 0 |
| ENSMUSG00000083666 | 0 | 0 | 0 | 0 | 0 | 0 | 0 |
| ENSMUSG00000083667 | 0 | 0 | 0 | 0 | 0 | 0 | 0 |
| ENSMUSG00000083668 | 0 | 0.25 | 0.25 | 0 | 0 | 0 | 0 |
| ENSMUSG00000083669 | 0 | 0 | 0.25 | 0 | 0 | 0 | 0 |
| ENSMUSG00000083670 | 0 | 0 | 0 | 0 | 0 | 0 | 0 |
| ENSMUSG00000083671 | 0 | 0 | 0 | 0 | 0 | 0 | 0 |
| ENSMUSG00000083672 | 0 | 0 | 0 | 0 | 0 | 0 | 0 |
| ENSMUSG00000083673 | 0 | 0 | 0 | 0 | 0 | 0 | 0 |
| ENSMUSG00000083674 | 0 | 0 | 0.25 | 0.701838731 | 0.387761935 | 0.84509804 | 0.672731473 |
| ENSMUSG00000083676 | 0 | 0 | 0.25 | 0 | 0.244650542 | 0 | 0 |
| ENSMUSG00000083677 | 0 | 0 | 0.25 | 0 | 0 | 0 | 0 |
| ENSMUSG00000083678 | 0 | 0 | 0 | 0.396240625 | 0.387761935 | 0 | 0 |
| ENSMUSG00000083679 | 0 | 0 | 0 | 0 | 0 | 0 | 0.224243824 |
| ENSMUSG00000083680 | 0 | 0 | 0 | 0 | 0 | 0 | 0 |
| ENSMUSG00000083681 | 0 | 0 | 0.25 | 0.25 | 0 | 0 | 0.224243824 |
| ENSMUSG00000083682 | 0 | 0 | 0 | 0 | 0 | 0 | 0 |
| ENSMUSG00000083683 | 0 | 0 | 0 | 0 | 0 | 0 | 0 |
| ENSMUSG00000083684 | 0 | 0 | 0 | 0 | 0 | 0 | 0 |
| ENSMUSG00000083685 | 0 | 0 | 0 | 0 | 0 | 0 | 0 |
| ENSMUSG00000083686 | 0 | 0 | 0.25 | 0 | 0 | 0 | 0 |
| ENSMUSG00000083687 | 0 | 0 | 0 | 0 | 0 | 0 | 0 |
| ENSMUSG00000083688 | 0 | 0 | 0 | 0 | 0.244650542 | 0 | 0 |
| ENSMUSG00000083689 | 0 | 0.25 | 0 | 0 | 0 | 0 | 0.224243824 |
| ENSMUSG00000083691 | 0 | 0 | 0 | 0 | 0 | 0 | 0 |
| ENSMUSG00000083692 | 0.45815691 | 0.25 | 0 | 0 | 0.244650542 | 0.301029996 | 0.224243824 |
| ENSMUSG00000083693 | 0.289064826 | 0 | 0 | 0.580482024 | 0 | 0 | 0.224243824 |
| ENSMUSG00000083695 | 0 | 0 | 0 | 0 | 0 | 0 | 0 |
| ENSMUSG00000083697 | 0 | 0 | 0 | 0 | 0 | 0 | 0 |
| ENSMUSG00000083698 | 0 | 0 | 0 | 0 | 0 | 0 | 0 |

| | | | | | | | |
|---|---|---|---|---|---|---|---|
| ENSMUSG00000083699 | 0 | 0 | 0 | 0 | 0 | 0 | 0 |
| ENSMUSG00000083700 | 0.289064826 | 0.396240625 | 0.396240625 | 0.25 | 0.387761935 | 0.477121255 | 0.448487648 |
| ENSMUSG00000083701 | 0 | 0 | 0 | 0 | 0 | 0 | 0 |
| ENSMUSG00000083702 | 0 | 0 | 0 | 0 | 0 | 0 | 0 |
| ENSMUSG00000083704 | 0 | 0 | 0 | 0 | 0 | 0 | 0 |
| ENSMUSG00000083705 | 1.227926429 | 1.321350555 | 1.160964047 | 1.26109853 | 1.351342427 | 1.431363764 | 1.265599895 |
| ENSMUSG00000083706 | 0 | 0 | 0 | 0 | 0 | 0 | 0 |
| ENSMUSG00000083707 | 0 | 0 | 0 | 0 | 0 | 0.301029996 | 0 |
| ENSMUSG00000083708 | 0.289064826 | 0.396240625 | 0.75 | 0 | 0 | 0.477121255 | 0.520678036 |
| ENSMUSG00000083709 | 0 | 0 | 0 | 0 | 0 | 0 | 0 |
| ENSMUSG00000083710 | 0 | 0 | 0.25 | 0 | 0 | 0 | 0 |
| ENSMUSG00000083711 | 0 | 0 | 0 | 0 | 0 | 0 | 0 |
| ENSMUSG00000083713 | 0.578129653 | 0 | 0.5 | 0.5 | 0.632412477 | 0.301029996 | 0.520678036 |
| ENSMUSG00000083714 | 0 | 0 | 0 | 0 | 0 | 0 | 0 |
| ENSMUSG00000083715 | 0 | 0 | 0 | 0 | 0 | 0 | 0 |
| ENSMUSG00000083716 | 0.45815691 | 0.25 | 0.580482024 | 0.396240625 | 0.489301084 | 0.301029996 | 0.224243824 |
| ENSMUSG00000083717 | 0 | 0 | 0 | 0 | 0 | 0 | 0 |
| ENSMUSG00000083718 | 0.578129653 | 0.646240625 | 0.5 | 0.864857905 | 0.686820904 | 0.602059991 | 0.520678036 |
| ENSMUSG00000083719 | 0 | 0 | 0 | 0 | 0 | 0 | 0 |
| ENSMUSG00000083720 | 0 | 0 | 0 | 0 | 0 | 0 | 0 |
| ENSMUSG00000083721 | 0 | 0 | 0 | 0 | 0 | 0 | 0 |
| ENSMUSG00000083722 | 0 | 0 | 0 | 0 | 0 | 0 | 0 |
| ENSMUSG00000083723 | 0 | 0 | 0 | 0 | 0 | 0 | 0 |
| ENSMUSG00000083725 | 0 | 0 | 0 | 0 | 0.244650542 | 0.301029996 | 0 |
| ENSMUSG00000083728 | 0 | 0 | 0 | 0 | 0 | 0 | 0 |
| ENSMUSG00000083729 | 0 | 0 | 0 | 0 | 0 | 0 | 0 |
| ENSMUSG00000083731 | 0.289064826 | 0 | 0 | 0 | 0 | 0.301029996 | 0.224243824 |
| ENSMUSG00000083732 | 0 | 0 | 0 | 0 | 0.489301084 | 0.477121255 | 0 |
| ENSMUSG00000083733 | 0 | 0 | 0 | 0 | 0 | 0 | 0 |
| ENSMUSG00000083734 | 0 | 0 | 0 | 0.25 | 0.387761935 | 0.477121255 | 0.224243824 |
| ENSMUSG00000083735 | 0 | 0 | 0 | 0 | 0 | 0 | 0 |
| ENSMUSG00000083736 | 0.289064826 | 0 | 0.25 | 0.396240625 | 0 | 0 | 0.355418052 |
| ENSMUSG00000083737 | 0 | 0 | 0 | 0 | 0 | 0 | 0 |
| ENSMUSG00000083738 | 0 | 0 | 0 | 0 | 0 | 0 | 0 |

| | | | | | | | |
|---|---|---|---|---|---|---|---|
| ENSMUSG00000083740 | 0 | 0 | 0 | 0 | 0 | 0 | 0 |
| ENSMUSG00000083741 | 0 | 0 | 0 | 0 | 0 | 0 | 0 |
| ENSMUSG00000083742 | 0 | 0 | 0 | 0 | 0 | 0 | 0 |
| ENSMUSG00000083743 | 0 | 0 | 0.25 | 0 | 0.244650542 | 0.301029996 | 0.355418052 |
| ENSMUSG00000083744 | 0 | 0 | 0 | 0 | 0 | 0 | 0 |
| ENSMUSG00000083745 | 0.289064826 | 0 | 0 | 0 | 0 | 0 | 0 |
| ENSMUSG00000083747 | 0 | 0 | 0 | 0 | 0 | 0 | 0 |
| ENSMUSG00000083748 | 0.45815691 | 0.25 | 0.5 | 0 | 0.244650542 | 0 | 0 |
| ENSMUSG00000083749 | 0 | 0 | 0 | 0 | 0 | 0 | 0 |
| ENSMUSG00000083750 | 0.747221736 | 0.701838731 | 0.75 | 0.864857905 | 0.77552387 | 0.477121255 | 0.520678036 |
| ENSMUSG00000083751 | 0 | 0 | 0 | 0 | 0 | 0 | 0 |
| ENSMUSG00000083752 | 0 | 0 | 0 | 0 | 0 | 0 | 0 |
| ENSMUSG00000083753 | 0 | 0 | 0 | 0 | 0 | 0 | 0 |
| ENSMUSG00000083754 | 0.289064826 | 0.580482024 | 0.396240625 | 0.25 | 0.244650542 | 0 | 0.355418052 |
| ENSMUSG00000083755 | 0 | 0 | 0 | 0 | 0 | 0 | 0 |
| ENSMUSG00000083757 | 0 | 0.396240625 | 0.25 | 0.396240625 | 0 | 0.301029996 | 0 |
| ENSMUSG00000083760 | 0 | 0 | 0 | 0 | 0 | 0 | 0 |
| ENSMUSG00000083761 | 0 | 0.5 | 0 | 0 | 0 | 0 | 0.448487648 |
| ENSMUSG00000083763 | 0.289064826 | 0.25 | 0.25 | 0 | 0 | 0 | 0 |
| ENSMUSG00000083764 | 0.289064826 | 0.396240625 | 0.5 | 0.5 | 0.489301084 | 0.301029996 | 0.520678036 |
| ENSMUSG00000083766 | 0 | 0 | 0 | 0 | 0 | 0 | 0 |
| ENSMUSG00000083767 | 0 | 0.25 | 0.25 | 0 | 0.244650542 | 0 | 0 |
| ENSMUSG00000083768 | 0 | 0 | 0 | 0 | 0 | 0 | 0 |
| ENSMUSG00000083769 | 0 | 0 | 0 | 0 | 0 | 0 | 0 |
| ENSMUSG00000083771 | 0 | 0 | 0 | 0 | 0 | 0 | 0 |
| ENSMUSG00000083772 | 0 | 0 | 0.25 | 0.25 | 0 | 0 | 0.448487648 |
| ENSMUSG00000083773 | 2.583267694 | 2.425543171 | 2.439139081 | 2.360735874 | 2.463003274 | 2.737987326 | 2.348891258 |
| ENSMUSG00000083774 | 0 | 0 | 0 | 0 | 0 | 0 | 0 |
| ENSMUSG00000083775 | 0 | 0 | 0 | 0 | 0 | 0 | 0 |
| ENSMUSG00000083776 | 0 | 0 | 0 | 0 | 0 | 0 | 0 |
| ENSMUSG00000083777 | 0.811507563 | 0.701838731 | 0.79248125 | 0.701838731 | 0.632412477 | 0.903089987 | 0.710836105 |
| ENSMUSG00000083778 | 0 | 0 | 0 | 0 | 0 | 0 | 0 |
| ENSMUSG00000083779 | 0 | 0 | 0.25 | 0.25 | 0 | 0.301029996 | 0 |
| ENSMUSG00000083780 | 0.671187741 | 0.701838731 | 0.864857905 | 0.5 | 0.387761935 | 0.602059991 | 0.672731473 |

| | | | | | | | |
|---|---|---|---|---|---|---|---|
| ENSMUSG00000083781 | 0 | 0 | 0 | 0 | 0 | 0 | 0 |
| ENSMUSG00000083782 | 0 | 0 | 0 | 0 | 0 | 0 | 0 |
| ENSMUSG00000083783 | 0 | 0 | 0.25 | 0.25 | 0.244650542 | 0 | 0 |
| ENSMUSG00000083784 | 0 | 0 | 0 | 0 | 0 | 0 | 0 |
| ENSMUSG00000083785 | 0 | 0 | 0.25 | 0 | 0 | 0 | 0 |
| ENSMUSG00000083786 | 0 | 0 | 0 | 0 | 0 | 0 | 0 |
| ENSMUSG00000083787 | 0 | 0 | 0 | 0 | 0 | 0 | 0 |
| ENSMUSG00000083789 | 0 | 0 | 0 | 0 | 0 | 0 | 0 |
| ENSMUSG00000083791 | 0 | 0 | 0 | 0 | 0 | 0 | 0 |
| ENSMUSG00000083792 | 0 | 0 | 0 | 0 | 0 | 0 | 0 |
| ENSMUSG00000083793 | 0.289064826 | 0.25 | 0.25 | 0 | 0 | 0.301029996 | 0 |
| ENSMUSG00000083794 | 0 | 0 | 0 | 0 | 0.244650542 | 0 | 0.224243824 |
| ENSMUSG00000083795 | 0 | 0 | 0 | 0 | 0 | 0 | 0 |
| ENSMUSG00000083796 | 0.45815691 | 0.25 | 0 | 0.396240625 | 0 | 0 | 0.224243824 |
| ENSMUSG00000083798 | 1.156259305 | 1.188721876 | 1.061981878 | 0.830482024 | 0.846351821 | 1.041392685 | 0.74492186 |
| ENSMUSG00000083801 | 0 | 0 | 0 | 0 | 0 | 0 | 0 |
| ENSMUSG00000083803 | 0 | 0.25 | 0 | 0 | 0 | 0 | 0 |
| ENSMUSG00000083804 | 0 | 0 | 0 | 0.25 | 0 | 0 | 0 |
| ENSMUSG00000083805 | 0.289064826 | 0.396240625 | 0.25 | 0 | 0 | 0 | 0 |
| ENSMUSG00000083806 | 0 | 0 | 0 | 0 | 0 | 0 | 0 |
| ENSMUSG00000083807 | 0 | 0 | 0 | 0 | 0 | 0 | 0 |
| ENSMUSG00000083808 | 0 | 0 | 0 | 0 | 0 | 0 | 0 |
| ENSMUSG00000083809 | 0 | 0 | 0 | 0 | 0 | 0 | 0 |
| ENSMUSG00000083810 | 0 | 0 | 0 | 0 | 0 | 0 | 0 |
| ENSMUSG00000083811 | 0.671187741 | 0.396240625 | 0.580482024 | 0.646240625 | 0.387761935 | 0.301029996 | 0.520678036 |
| ENSMUSG00000083812 | 0.289064826 | 0.25 | 0 | 0.25 | 0 | 0 | 0 |
| ENSMUSG00000083813 | 0 | 0 | 0 | 0 | 0 | 0 | 0 |
| ENSMUSG00000083814 | 0 | 0 | 0 | 0 | 0 | 0 | 0 |
| ENSMUSG00000083815 | 0 | 0 | 0 | 0 | 0.244650542 | 0 | 0 |
| ENSMUSG00000083816 | 0.747221736 | 0.701838731 | 0.864857905 | 0.830482024 | 0.568060967 | 0.477121255 | 0.629532004 |
| ENSMUSG00000083817 | 0 | 0 | 0 | 0.396240625 | 0.387761935 | 0 | 0 |
| ENSMUSG00000083818 | 0 | 0 | 0 | 0 | 0 | 0 | 0 |
| ENSMUSG00000083819 | 0.747221736 | 0.5 | 0.830482024 | 0.976722649 | 0.846351821 | 0.602059991 | 0.803905701 |
| ENSMUSG00000083820 | 0 | 0 | 0 | 0 | 0 | 0 | 0.224243824 |

| | | | | | | | |
|---|---|---|---|---|---|---|---|
| ENSMUSG00000083821 | 0 | 0 | 0 | 0 | 0 | 0 | 0 |
| ENSMUSG00000083822 | 0.578129653 | 0.25 | 0 | 0.396240625 | 0.244650542 | 0.477121255 | 0.355418052 |
| ENSMUSG00000083823 | 0.289064826 | 0 | 0.25 | 0 | 0.244650542 | 0 | 0 |
| ENSMUSG00000083825 | 0 | 0 | 0 | 0 | 0 | 0 | 0 |
| ENSMUSG00000083826 | 0.289064826 | 0 | 0.580482024 | 0.5 | 0.387761935 | 0.301029996 | 0.224243824 |
| ENSMUSG00000083827 | 0 | 0 | 0 | 0 | 0 | 0 | 0 |
| ENSMUSG00000083828 | 0 | 0 | 0 | 0 | 0 | 0 | 0 |
| ENSMUSG00000083829 | 0.45815691 | 0 | 0.25 | 0 | 0.244650542 | 0.301029996 | 0 |
| ENSMUSG00000083830 | 0 | 0.25 | 0.25 | 0 | 0.244650542 | 0 | 0.355418052 |
| ENSMUSG00000083831 | 0 | 0 | 0 | 0 | 0 | 0 | 0 |
| ENSMUSG00000083832 | 0 | 0 | 0.25 | 0.396240625 | 0.244650542 | 0 | 0 |
| ENSMUSG00000083833 | 0 | 0 | 0 | 0 | 0 | 0 | 0 |
| ENSMUSG00000083834 | 0.289064826 | 0 | 0.25 | 0 | 0 | 0 | 0.224243824 |
| ENSMUSG00000083835 | 0 | 0 | 0 | 0 | 0 | 0 | 0 |
| ENSMUSG00000083836 | 0.289064826 | 0 | 0.25 | 0 | 0.387761935 | 0 | 0.224243824 |
| ENSMUSG00000083837 | 0 | 0 | 0 | 0 | 0 | 0 | 0.224243824 |
| ENSMUSG00000083838 | 0 | 0 | 0 | 0.25 | 0 | 0 | 0.224243824 |
| ENSMUSG00000083839 | 0 | 0 | 0 | 0 | 0 | 0 | 0 |
| ENSMUSG00000083841 | 0 | 0 | 0 | 0.5 | 0.387761935 | 0 | 0.355418052 |
| ENSMUSG00000083842 | 0 | 0 | 0 | 0 | 0 | 0 | 0 |
| ENSMUSG00000083843 | 0 | 0 | 0 | 0 | 0 | 0 | 0 |
| ENSMUSG00000083844 | 1.91631382 | 1.782320754 | 1.752806814 | 1.698603967 | 1.728722551 | 1.897627091 | 1.649885598 |
| ENSMUSG00000083845 | 0 | 0 | 0 | 0 | 0 | 0 | 0.355418052 |
| ENSMUSG00000083846 | 0 | 0 | 0 | 0 | 0 | 0 | 0 |
| ENSMUSG00000083847 | 0 | 0 | 0 | 0 | 0 | 0 | 0 |
| ENSMUSG00000083848 | 0 | 0 | 0 | 0 | 0 | 0 | 0 |
| ENSMUSG00000083849 | 0.671187741 | 0.580482024 | 0.79248125 | 0.646240625 | 0.387761935 | 0.477121255 | 0.448487648 |
| ENSMUSG00000083851 | 0 | 0 | 0 | 0 | 0 | 0 | 0 |
| ENSMUSG00000083852 | 0 | 0.25 | 0.25 | 0 | 0 | 0.301029996 | 0 |
| ENSMUSG00000083853 | 0.45815691 | 0.580482024 | 0.75 | 0.580482024 | 0.489301084 | 0.477121255 | 0.629532004 |
| ENSMUSG00000083854 | 0 | 0 | 0 | 0 | 0 | 0 | 0 |
| ENSMUSG00000083855 | 0 | 0 | 0 | 0 | 0 | 0 | 0 |
| ENSMUSG00000083856 | 0 | 0.580482024 | 0.396240625 | 0.396240625 | 0.244650542 | 0.477121255 | 0.224243824 |
| ENSMUSG00000083857 | 0 | 0 | 0 | 0 | 0 | 0 | 0 |

| | | | | | | | |
|---|---|---|---|---|---|---|---|
| ENSMUSG00000083858 | 0 | 0 | 0 | 0 | 0 | 0 | 0 |
| ENSMUSG00000083859 | 0 | 0.25 | 0 | 0 | 0.244650542 | 0.301029996 | 0.224243824 |
| ENSMUSG00000083860 | 0 | 0 | 0 | 0 | 0.244650542 | 0 | 0 |
| ENSMUSG00000083861 | 0 | 0 | 0 | 0 | 0 | 0 | 0 |
| ENSMUSG00000083863 | 0.45815691 | 0.864857905 | 1.114857905 | 0.92510993 | 1.163285805 | 1.113943352 | 0.853775828 |
| ENSMUSG00000083864 | 0 | 0 | 0 | 0 | 0 | 0 | 0.224243824 |
| ENSMUSG00000083866 | 0 | 0 | 0 | 0 | 0 | 0 | 0 |
| ENSMUSG00000083867 | 0 | 0 | 0 | 0 | 0 | 0 | 0 |
| ENSMUSG00000083868 | 0 | 0 | 0 | 0 | 0 | 0 | 0 |
| ENSMUSG00000083869 | 0 | 0 | 0 | 0 | 0 | 0 | 0 |
| ENSMUSG00000083870 | 0 | 0 | 0 | 0 | 0 | 0 | 0 |
| ENSMUSG00000083871 | 0 | 0.25 | 0 | 0 | 0 | 0 | 0 |
| ENSMUSG00000083872 | 0 | 0 | 0 | 0 | 0 | 0 | 0 |
| ENSMUSG00000083873 | 0 | 0 | 0 | 0 | 0 | 0 | 0 |
| ENSMUSG00000083874 | 0 | 0 | 0 | 0 | 0 | 0 | 0 |
| ENSMUSG00000083875 | 0 | 0 | 0 | 0 | 0 | 0 | 0 |
| ENSMUSG00000083876 | 0 | 0 | 0 | 0 | 0 | 0 | 0 |
| ENSMUSG00000083877 | 0 | 0.25 | 0 | 0 | 0 | 0 | 0 |
| ENSMUSG00000083878 | 0 | 0.25 | 0 | 0 | 0 | 0 | 0.224243824 |
| ENSMUSG00000083879 | 0 | 0.25 | 0 | 0 | 0 | 0 | 0 |
| ENSMUSG00000083880 | 1.036286563 | 0.951838731 | 1.188721876 | 1.26109853 | 1.234113756 | 1.113943352 | 1.066254157 |
| ENSMUSG00000083882 | 0 | 0 | 0 | 0 | 0 | 0 | 0 |
| ENSMUSG00000083884 | 0 | 0 | 0 | 0.25 | 0 | 0 | 0 |
| ENSMUSG00000083885 | 0 | 0 | 0.25 | 0 | 0 | 0 | 0.224243824 |
| ENSMUSG00000083886 | 0 | 0 | 0 | 0 | 0 | 0 | 0 |
| ENSMUSG00000083887 | 0 | 0 | 0 | 0 | 0 | 0 | 0 |
| ENSMUSG00000083888 | 0 | 0 | 0 | 0 | 0 | 0 | 0 |
| ENSMUSG00000083889 | 0 | 0 | 0 | 0 | 0 | 0 | 0 |
| ENSMUSG00000083890 | 0.289064826 | 0.79248125 | 0.830482024 | 1.25 | 1.254881871 | 1.447158031 | 1.193409508 |
| ENSMUSG00000083892 | 0 | 0.396240625 | 0.25 | 0.396240625 | 0.387761935 | 0.301029996 | 0 |
| ENSMUSG00000083893 | 0 | 0 | 0 | 0 | 0 | 0 | 0 |
| ENSMUSG00000083894 | 0 | 0 | 0 | 0 | 0 | 0 | 0 |
| ENSMUSG00000083895 | 0 | 0 | 0 | 0 | 0 | 0 | 0 |
| ENSMUSG00000083896 | 0 | 0 | 0 | 0 | 0 | 0 | 0 |

| | | | | | | | |
|---|---|---|---|---|---|---|---|
| ENSMUSG00000083897 | 0 | 0 | 0 | 0 | 0 | 0 | 0.224243824 |
| ENSMUSG00000083898 | 0 | 0 | 0 | 0 | 0 | 0 | 0 |
| ENSMUSG00000083899 | 0.578129653 | 0.5 | 0.580482024 | 0.646240625 | 0.733951626 | 0.477121255 | 0.520678036 |
| ENSMUSG00000083900 | 0 | 0.396240625 | 0.25 | 0 | 0.244650542 | 0 | 0 |
| ENSMUSG00000083901 | 0 | 0 | 0 | 0 | 0 | 0 | 0 |
| ENSMUSG00000083902 | 0.867194479 | 1.17510993 | 1.04248125 | 1.188721876 | 1.074582839 | 1.204119983 | 1.150210039 |
| ENSMUSG00000083903 | 0 | 0 | 0 | 0 | 0 | 0.301029996 | 0 |
| ENSMUSG00000083904 | 0 | 0 | 0 | 0 | 0 | 0 | 0 |
| ENSMUSG00000083905 | 0 | 0 | 0.25 | 0 | 0 | 0 | 0 |
| ENSMUSG00000083906 | 0.45815691 | 0 | 0.5 | 0.5 | 0.244650542 | 0.477121255 | 0.355418052 |
| ENSMUSG00000083907 | 0.91631382 | 0.830482024 | 1.04248125 | 0.830482024 | 1.039257769 | 0.77815125 | 0.896975297 |
| ENSMUSG00000083908 | 0 | 0 | 0 | 0 | 0 | 0 | 0 |
| ENSMUSG00000083909 | 0 | 0 | 0.396240625 | 0 | 0 | 0 | 0 |
| ENSMUSG00000083910 | 0 | 0 | 0.25 | 0.25 | 0 | 0 | 0.579661877 |
| ENSMUSG00000083911 | 0.289064826 | 0 | 0.396240625 | 0 | 0 | 0 | 0.224243824 |
| ENSMUSG00000083912 | 0 | 0.25 | 0 | 0.25 | 0 | 0 | 0 |
| ENSMUSG00000083913 | 0 | 0.5 | 0 | 0 | 0 | 0 | 0 |
| ENSMUSG00000083914 | 0 | 0 | 0.396240625 | 0.25 | 0 | 0 | 0 |
| ENSMUSG00000083915 | 0.289064826 | 0.396240625 | 0 | 0 | 0 | 0 | 0.224243824 |
| ENSMUSG00000083916 | 0 | 0 | 0 | 0 | 0 | 0 | 0 |
| ENSMUSG00000083917 | 0 | 0 | 0.25 | 0.25 | 0.244650542 | 0 | 0 |
| ENSMUSG00000083918 | 0 | 0 | 0 | 0 | 0 | 0 | 0 |
| ENSMUSG00000083919 | 0 | 0 | 0 | 0 | 0 | 0 | 0 |
| ENSMUSG00000083920 | 0 | 0 | 0 | 0 | 0 | 0 | 0 |
| ENSMUSG00000083921 | 0 | 0.5 | 0.25 | 0.25 | 0 | 0 | 0.224243824 |
| ENSMUSG00000083922 | 0.45815691 | 0 | 0 | 0 | 0.244650542 | 0.301029996 | 0.355418052 |
| ENSMUSG00000083923 | 0 | 0 | 0 | 0 | 0 | 0 | 0 |
| ENSMUSG00000083924 | 0 | 0 | 0 | 0 | 0 | 0 | 0 |
| ENSMUSG00000083925 | 0 | 0 | 0 | 0 | 0 | 0 | 0 |
| ENSMUSG00000083926 | 0 | 0 | 0 | 0 | 0 | 0 | 0 |
| ENSMUSG00000083927 | 0.45815691 | 0 | 0 | 0 | 0 | 0 | 0 |
| ENSMUSG00000083928 | 0 | 0.580482024 | 0.25 | 0.25 | 0.244650542 | 0 | 0 |
| ENSMUSG00000083929 | 0 | 0 | 0 | 0 | 0 | 0 | 0 |
| ENSMUSG00000083930 | 0 | 0 | 0 | 0 | 0 | 0 | 0 |

| | | | | | | | |
|---|---|---|---|---|---|---|---|
| ENSMUSG00000083931 | 0 | 0 | 0 | 0 | 0 | 0 | 0 |
| ENSMUSG00000083932 | 0 | 0 | 0 | 0 | 0 | 0 | 0 |
| ENSMUSG00000083933 | 0.289064826 | 0.396240625 | 0.396240625 | 0 | 0.244650542 | 0 | 0.355418052 |
| ENSMUSG00000083934 | 0 | 0 | 0 | 0 | 0 | 0 | 0 |
| ENSMUSG00000083935 | 0 | 0.646240625 | 0 | 0 | 0.632412477 | 0.301029996 | 0 |
| ENSMUSG00000083936 | 0 | 0 | 0 | 0 | 0 | 0 | 0 |
| ENSMUSG00000083937 | 0 | 0.25 | 0 | 0 | 0 | 0 | 0.224243824 |
| ENSMUSG00000083938 | 0 | 0 | 0 | 0 | 0 | 0 | 0 |
| ENSMUSG00000083939 | 0 | 0 | 0 | 0 | 0 | 0 | 0 |
| ENSMUSG00000083940 | 0 | 0 | 0 | 0 | 0 | 0 | 0.224243824 |
| ENSMUSG00000083941 | 0 | 0 | 0 | 0 | 0 | 0 | 0 |
| ENSMUSG00000083942 | 0 | 0 | 0 | 0 | 0 | 0 | 0 |
| ENSMUSG00000083943 | 0 | 0 | 0 | 0.5 | 0.244650542 | 0 | 0.224243824 |
| ENSMUSG00000083944 | 0 | 0 | 0 | 0 | 0 | 0 | 0 |
| ENSMUSG00000083945 | 0 | 0 | 0 | 0 | 0 | 0 | 0 |
| ENSMUSG00000083946 | 0 | 0 | 0 | 0 | 0 | 0 | 0 |
| ENSMUSG00000083947 | 0 | 0 | 0 | 0 | 0 | 0 | 0 |
| ENSMUSG00000083948 | 0 | 0 | 0 | 0 | 0 | 0 | 0 |
| ENSMUSG00000083949 | 0 | 0 | 0 | 0 | 0 | 0 | 0 |
| ENSMUSG00000083950 | 0 | 0 | 0 | 0 | 0 | 0 | 0 |
| ENSMUSG00000083951 | 0 | 0 | 0.25 | 0 | 0.244650542 | 0 | 0 |
| ENSMUSG00000083953 | 0.289064826 | 0.25 | 0 | 0 | 0 | 0.301029996 | 0.224243824 |
| ENSMUSG00000083954 | 0 | 0 | 0 | 0 | 0 | 0 | 0 |
| ENSMUSG00000083955 | 0 | 0 | 0 | 0 | 0 | 0 | 0 |
| ENSMUSG00000083956 | 0 | 0 | 0 | 0 | 0 | 0 | 0 |
| ENSMUSG00000083957 | 0 | 0 | 0 | 0 | 0 | 0 | 0 |
| ENSMUSG00000083958 | 0 | 0 | 0 | 0 | 0 | 0 | 0 |
| ENSMUSG00000083960 | 0 | 0 | 0 | 0 | 0 | 0 | 0 |
| ENSMUSG00000083961 | 0 | 0 | 0 | 0 | 0 | 0 | 0 |
| ENSMUSG00000083962 | 0.289064826 | 0 | 0 | 0 | 0.244650542 | 0 | 0 |
| ENSMUSG00000083963 | 0 | 0 | 0 | 0 | 0 | 0 | 0 |
| ENSMUSG00000083964 | 0.747221736 | 0.701838731 | 0.701838731 | 0.646240625 | 0.77552387 | 0.77815125 | 0.775756176 |
| ENSMUSG00000083966 | 0 | 0 | 0 | 0 | 0 | 0 | 0 |
| ENSMUSG00000083967 | 0 | 0 | 0 | 0 | 0 | 0 | 0 |

| | | | | | | | |
|---|---|---|---|---|---|---|---|
| ENSMUSG00000083969 | 0 | 0 | 0 | 0 | 0 | 0 | 0 |
| ENSMUSG00000083970 | 0 | 0 | 0 | 0 | 0 | 0 | 0 |
| ENSMUSG00000083971 | 0.289064826 | 0 | 0 | 0.25 | 0 | 0 | 0 |
| ENSMUSG00000083972 | 0 | 0 | 0 | 0 | 0 | 0 | 0 |
| ENSMUSG00000083973 | 0 | 0 | 0 | 0 | 0 | 0 | 0 |
| ENSMUSG00000083974 | 0 | 0 | 0 | 0 | 0 | 0 | 0 |
| ENSMUSG00000083975 | 0 | 0 | 0 | 0 | 0 | 0 | 0 |
| ENSMUSG00000083976 | 0 | 0 | 0 | 0 | 0 | 0 | 0 |
| ENSMUSG00000083977 | 0 | 0 | 0 | 0 | 0 | 0 | 0 |
| ENSMUSG00000083978 | 0 | 0.25 | 0 | 0 | 0 | 0 | 0.224243824 |
| ENSMUSG00000083979 | 0 | 0 | 0 | 0 | 0 | 0 | 0 |
| ENSMUSG00000083981 | 0 | 0 | 0 | 0 | 0 | 0 | 0 |
| ENSMUSG00000083982 | 0.289064826 | 0.25 | 0.396240625 | 0.25 | 0.387761935 | 0 | 0.224243824 |
| ENSMUSG00000083983 | 0 | 0 | 0 | 0 | 0 | 0 | 0 |
| ENSMUSG00000083985 | 0 | 0 | 0.25 | 0 | 0 | 0 | 0 |
| ENSMUSG00000083986 | 0 | 0 | 0 | 0 | 0 | 0 | 0 |
| ENSMUSG00000083987 | 0 | 0 | 0 | 0 | 0 | 0 | 0 |
| ENSMUSG00000083988 | 0 | 0 | 0.396240625 | 0 | 0 | 0 | 0 |
| ENSMUSG00000083989 | 0 | 0 | 0 | 0 | 0.244650542 | 0.301029996 | 0 |
| ENSMUSG00000083990 | 0 | 0 | 0 | 0 | 0 | 0 | 0 |
| ENSMUSG00000083991 | 0 | 0 | 0 | 0 | 0 | 0 | 0 |
| ENSMUSG00000083992 | 0.578129653 | 0.701838731 | 0.79248125 | 0.79248125 | 0.77552387 | 0.903089987 | 0.672731473 |
| ENSMUSG00000083993 | 0.45815691 | 0.646240625 | 0.5 | 0.830482024 | 0.568060967 | 0.477121255 | 0.224243824 |
| ENSMUSG00000083994 | 0 | 0 | 0 | 0.25 | 0 | 0.477121255 | 0 |
| ENSMUSG00000083995 | 0 | 0 | 0 | 0 | 0 | 0 | 0 |
| ENSMUSG00000083996 | 0 | 0 | 0 | 0 | 0 | 0 | 0 |
| ENSMUSG00000083997 | 0 | 0 | 0 | 0 | 0 | 0 | 0.448487648 |
| ENSMUSG00000083998 | 0 | 0 | 0.25 | 0.25 | 0.244650542 | 0 | 0.355418052 |
| ENSMUSG00000083999 | 0 | 0 | 0 | 0 | 0 | 0 | 0 |
| ENSMUSG00000084000 | 0 | 0.25 | 0 | 0 | 0 | 0 | 0 |
| ENSMUSG00000084001 | 0 | 0 | 0 | 0 | 0 | 0 | 0 |
| ENSMUSG00000084004 | 0 | 0 | 0 | 0.396240625 | 0 | 0 | 0 |
| ENSMUSG00000084006 | 0 | 0 | 0 | 0 | 0 | 0 | 0 |
| ENSMUSG00000084007 | 0.289064826 | 0 | 0.25 | 0.5 | 0.387761935 | 0 | 0.710836105 |

| | | | | | | | |
|---|---|---|---|---|---|---|---|
| ENSMUSG00000084009 | 0 | 0 | 0 | 0 | 0 | 0 | 0 |
| ENSMUSG00000084010 | 0 | 0 | 0 | 0 | 0 | 0 | 0 |
| ENSMUSG00000084011 | 0 | 0 | 0 | 0.25 | 0.489301084 | 0.477121255 | 0.224243824 |
| ENSMUSG00000084012 | 0 | 0 | 0 | 0 | 0 | 0 | 0 |
| ENSMUSG00000084013 | 0 | 0 | 0 | 0 | 0 | 0 | 0 |
| ENSMUSG00000084014 | 0 | 0 | 0 | 0 | 0 | 0 | 0 |
| ENSMUSG00000084015 | 0 | 0 | 0 | 0 | 0 | 0.301029996 | 0 |
| ENSMUSG00000084016 | 0 | 0 | 0 | 0 | 0 | 0 | 0 |
| ENSMUSG00000084017 | 0.45815691 | 0 | 0.646240625 | 0.580482024 | 0.568060967 | 0.301029996 | 0.355418052 |
| ENSMUSG00000084018 | 0 | 0 | 0 | 0 | 0 | 0 | 0 |
| ENSMUSG00000084019 | 0 | 0 | 0 | 0 | 0 | 0 | 0 |
| ENSMUSG00000084020 | 0 | 0 | 0 | 0 | 0 | 0 | 0 |
| ENSMUSG00000084021 | 0 | 0 | 0 | 0 | 0 | 0.301029996 | 0 |
| ENSMUSG00000084022 | 0.289064826 | 0 | 0 | 0.25 | 0.489301084 | 0.602059991 | 0.448487648 |
| ENSMUSG00000084024 | 0 | 0 | 0 | 0 | 0 | 0 | 0 |
| ENSMUSG00000084025 | 0.45815691 | 0 | 0.25 | 0.25 | 0.244650542 | 0 | 0.224243824 |
| ENSMUSG00000084026 | 0 | 0 | 0 | 0 | 0 | 0 | 0 |
| ENSMUSG00000084027 | 0 | 0 | 0 | 0 | 0 | 0 | 0 |
| ENSMUSG00000084028 | 0 | 0 | 0 | 0 | 0 | 0 | 0 |
| ENSMUSG00000084029 | 0 | 0 | 0 | 0 | 0 | 0 | 0 |
| ENSMUSG00000084030 | 0 | 0 | 0 | 0 | 0 | 0 | 0 |
| ENSMUSG00000084031 | 1.129344651 | 1.098079356 | 0.896240625 | 0.864857905 | 0.812711509 | 0.77815125 | 0.829800754 |
| ENSMUSG00000084032 | 0 | 0 | 0 | 0 | 0 | 0 | 0 |
| ENSMUSG00000084033 | 0.45815691 | 0.396240625 | 0.5 | 0.5 | 0.489301084 | 0.477121255 | 0.710836105 |
| ENSMUSG00000084034 | 0 | 0 | 0 | 0 | 0 | 0 | 0 |
| ENSMUSG00000084038 | 0 | 0 | 0 | 0 | 0 | 0 | 0 |
| ENSMUSG00000084039 | 0 | 0 | 0 | 0 | 0 | 0 | 0 |
| ENSMUSG00000084041 | 0 | 0 | 0 | 0 | 0 | 0 | 0 |
| ENSMUSG00000084043 | 0 | 0 | 0 | 0 | 0 | 0 | 0 |
| ENSMUSG00000084044 | 0 | 0 | 0 | 0 | 0 | 0 | 0 |
| ENSMUSG00000084045 | 0 | 0 | 0 | 0 | 0 | 0 | 0 |
| ENSMUSG00000084046 | 0 | 0 | 0 | 0 | 0 | 0 | 0 |
| ENSMUSG00000084047 | 0 | 0 | 0 | 0 | 0.244650542 | 0 | 0.224243824 |
| ENSMUSG00000084048 | 0 | 0 | 0 | 0 | 0 | 0 | 0 |

| | | | | | | | |
|---|---|---|---|---|---|---|---|
| ENSMUSG00000084050 | 0 | 0 | 0 | 0 | 0 | 0 | 0 |
| ENSMUSG00000084052 | 0 | 0 | 0 | 0 | 0 | 0 | 0 |
| ENSMUSG00000084053 | 0 | 0 | 0 | 0 | 0 | 0 | 0 |
| ENSMUSG00000084054 | 0 | 0 | 0 | 0 | 0 | 0 | 0 |
| ENSMUSG00000084055 | 0 | 0 | 0 | 0 | 0 | 0 | 0 |
| ENSMUSG00000084056 | 0 | 0 | 0 | 0 | 0 | 0 | 0 |
| ENSMUSG00000084057 | 0 | 0 | 0 | 0 | 0 | 0 | 0 |
| ENSMUSG00000084058 | 0 | 0 | 0 | 0 | 0 | 0 | 0 |
| ENSMUSG00000084059 | 0 | 0 | 0 | 0 | 0 | 0 | 0 |
| ENSMUSG00000084060 | 0 | 0 | 0 | 0 | 0 | 0 | 0 |
| ENSMUSG00000084061 | 0 | 0 | 0 | 0 | 0 | 0 | 0 |
| ENSMUSG00000084063 | 0 | 0 | 0 | 0 | 0 | 0 | 0.224243824 |
| ENSMUSG00000084064 | 0 | 0 | 0 | 0 | 0 | 0 | 0 |
| ENSMUSG00000084065 | 0 | 0 | 0 | 0 | 0 | 0 | 0 |
| ENSMUSG00000084066 | 0 | 0 | 0 | 0 | 0 | 0 | 0 |
| ENSMUSG00000084067 | 0 | 0 | 0.396240625 | 0.25 | 0 | 0 | 0.355418052 |
| ENSMUSG00000084068 | 0 | 0 | 0 | 0 | 0 | 0 | 0 |
| ENSMUSG00000084069 | 0 | 0 | 0 | 0 | 0 | 0 | 0.224243824 |
| ENSMUSG00000084070 | 0 | 0 | 0 | 0 | 0 | 0 | 0 |
| ENSMUSG00000084071 | 0 | 0 | 0 | 0 | 0 | 0 | 0 |
| ENSMUSG00000084072 | 0 | 0 | 0 | 0 | 0 | 0 | 0 |
| ENSMUSG00000084073 | 0 | 0 | 0 | 0 | 0 | 0 | 0 |
| ENSMUSG00000084074 | 0 | 0 | 0 | 0 | 0 | 0 | 0 |
| ENSMUSG00000084076 | 0 | 0 | 0 | 0 | 0 | 0 | 0 |
| ENSMUSG00000084077 | 0 | 0 | 0 | 0 | 0 | 0 | 0 |
| ENSMUSG00000084078 | 0 | 0 | 0 | 0 | 0 | 0 | 0 |
| ENSMUSG00000084079 | 0.289064826 | 0.396240625 | 0 | 0 | 0.387761935 | 0 | 0.355418052 |
| ENSMUSG00000084080 | 0 | 0 | 0 | 0 | 0 | 0 | 0 |
| ENSMUSG00000084081 | 0.960252568 | 0.75 | 0.701838731 | 0.830482024 | 0.812711509 | 0.602059991 | 0.775756176 |
| ENSMUSG00000084082 | 0 | 0 | 0 | 0 | 0 | 0 | 0 |
| ENSMUSG00000084083 | 0 | 0.25 | 0 | 0.25 | 0 | 0 | 0 |
| ENSMUSG00000084085 | 0.289064826 | 0 | 0 | 0 | 0 | 0 | 0 |
| ENSMUSG00000084086 | 0 | 0 | 0.25 | 0 | 0 | 0 | 0 |
| ENSMUSG00000084087 | 0.289064826 | 0.396240625 | 0 | 0.25 | 0 | 0.301029996 | 0 |

| | | | | | | | |
|---|---|---|---|---|---|---|---|
| ENSMUSG00000084088 | 0.747221736 | 0.75 | 0.646240625 | 0.701838731 | 0.568060967 | 0.301029996 | 0.224243824 |
| ENSMUSG00000084089 | 0 | 0 | 0 | 0 | 0 | 0 | 0 |
| ENSMUSG00000084090 | 0 | 0 | 0 | 0 | 0 | 0 | 0 |
| ENSMUSG00000084092 | 0 | 0 | 0 | 0 | 0 | 0 | 0 |
| ENSMUSG00000084093 | 0 | 0 | 0 | 0 | 0 | 0 | 0 |
| ENSMUSG00000084094 | 0 | 0 | 0 | 0 | 0.244650542 | 0 | 0 |
| ENSMUSG00000084095 | 0 | 0 | 0 | 0 | 0 | 0 | 0 |
| ENSMUSG00000084096 | 0 | 0 | 0 | 0 | 0.244650542 | 0 | 0 |
| ENSMUSG00000084097 | 0 | 0 | 0 | 0 | 0 | 0 | 0 |
| ENSMUSG00000084098 | 0 | 0 | 0 | 0 | 0.244650542 | 0.301029996 | 0 |
| ENSMUSG00000084099 | 0 | 0 | 0.25 | 0 | 0 | 0 | 0 |
| ENSMUSG00000084101 | 0 | 0 | 0 | 0 | 0 | 0 | 0 |
| ENSMUSG00000084102 | 0 | 0 | 0 | 0 | 0 | 0 | 0 |
| ENSMUSG00000084103 | 0 | 0 | 0 | 0 | 0 | 0 | 0 |
| ENSMUSG00000084104 | 0 | 0 | 0 | 0.25 | 0 | 0 | 0.355418052 |
| ENSMUSG00000084105 | 0 | 0.25 | 0.25 | 0.396240625 | 0 | 0 | 0.224243824 |
| ENSMUSG00000084106 | 0.671187741 | 0.830482024 | 0.92510993 | 0.701838731 | 0.733951626 | 0.301029996 | 0.775756176 |
| ENSMUSG00000084107 | 0 | 0 | 0 | 0 | 0 | 0 | 0 |
| ENSMUSG00000084108 | 0 | 0 | 0 | 0 | 0 | 0 | 0 |
| ENSMUSG00000084110 | 0 | 0 | 0 | 0 | 0 | 0 | 0 |
| ENSMUSG00000084111 | 0 | 0.25 | 0 | 0 | 0 | 0 | 0.355418052 |
| ENSMUSG00000084112 | 0 | 0 | 0 | 0 | 0 | 0 | 0 |
| ENSMUSG00000084113 | 0 | 0.25 | 0.396240625 | 0.5 | 0.244650542 | 0.477121255 | 0.520678036 |
| ENSMUSG00000084114 | 0 | 0 | 0 | 0 | 0 | 0 | 0 |
| ENSMUSG00000084115 | 0.289064826 | 0 | 0 | 0 | 0 | 0 | 0 |
| ENSMUSG00000084116 | 0 | 0 | 0 | 0 | 0 | 0 | 0 |
| ENSMUSG00000084117 | 0 | 0 | 0 | 0 | 0 | 0 | 0 |
| ENSMUSG00000084118 | 0 | 0 | 0 | 0 | 0 | 0 | 0 |
| ENSMUSG00000084119 | 0 | 0 | 0.25 | 0 | 0 | 0 | 0 |
| ENSMUSG00000084120 | 0 | 0 | 0 | 0 | 0 | 0.301029996 | 0.224243824 |
| ENSMUSG00000084123 | 0 | 0 | 0 | 0 | 0 | 0 | 0 |
| ENSMUSG00000084124 | 0 | 0 | 0 | 0 | 0.244650542 | 0 | 0.224243824 |
| ENSMUSG00000084125 | 0 | 0 | 0.25 | 0.396240625 | 0.244650542 | 0 | 0 |
| ENSMUSG00000084126 | 0 | 0 | 0 | 0 | 0 | 0 | 0 |

| | | | | | | | |
|---|---|---|---|---|---|---|---|
| ENSMUSG00000084127 | 0 | 0 | 0 | 0 | 0 | 0 | 0 |
| ENSMUSG00000084128 | 1.129344651 | 1.17510993 | 1.201838731 | 1.302363341 | 1.456697355 | 1.544068044 | 1.392431569 |
| ENSMUSG00000084129 | 0 | 0 | 0 | 0 | 0 | 0 | 0 |
| ENSMUSG00000084130 | 0 | 0 | 0 | 0 | 0 | 0 | 0 |
| ENSMUSG00000084131 | 0.578129653 | 0.79248125 | 0.25 | 0.25 | 0.387761935 | 0 | 0.355418052 |
| ENSMUSG00000084132 | 0 | 0 | 0 | 0.25 | 0 | 0 | 0.224243824 |
| ENSMUSG00000084133 | 0.289064826 | 0 | 0 | 0 | 0.244650542 | 0 | 0.224243824 |
| ENSMUSG00000084134 | 0 | 0 | 0 | 0 | 0 | 0 | 0 |
| ENSMUSG00000084135 | 0 | 0 | 0 | 0 | 0 | 0 | 0 |
| ENSMUSG00000084136 | 0 | 0 | 0 | 0 | 0 | 0 | 0 |
| ENSMUSG00000084137 | 0.289064826 | 0 | 0 | 0 | 0.244650542 | 0 | 0.520678036 |
| ENSMUSG00000084138 | 0 | 0 | 0 | 0 | 0 | 0 | 0 |
| ENSMUSG00000084139 | 0 | 0 | 0.5 | 0 | 0.244650542 | 0.301029996 | 0.224243824 |
| ENSMUSG00000084140 | 0.289064826 | 0 | 0.396240625 | 0.25 | 0 | 0.301029996 | 0.224243824 |
| ENSMUSG00000084142 | 0 | 0 | 0 | 0 | 0 | 0.477121255 | 0 |
| ENSMUSG00000084143 | 0 | 0 | 0 | 0 | 0 | 0 | 0 |
| ENSMUSG00000084144 | 0.289064826 | 0 | 0 | 0 | 0.244650542 | 0 | 0.224243824 |
| ENSMUSG00000084145 | 0 | 0 | 0.25 | 0.396240625 | 0.387761935 | 0 | 0 |
| ENSMUSG00000084146 | 0.45815691 | 0.25 | 0 | 0 | 0.244650542 | 0 | 0.224243824 |
| ENSMUSG00000084147 | 0 | 0 | 0 | 0 | 0 | 0 | 0 |
| ENSMUSG00000084148 | 0.45815691 | 0.25 | 0 | 0.25 | 0.387761935 | 0 | 0.224243824 |
| ENSMUSG00000084149 | 0 | 0.396240625 | 0.396240625 | 0 | 0.387761935 | 0.301029996 | 0.224243824 |
| ENSMUSG00000084150 | 0 | 0 | 0 | 0 | 0 | 0 | 0 |
| ENSMUSG00000084151 | 0 | 0 | 0 | 0 | 0 | 0.301029996 | 0 |
| ENSMUSG00000084152 | 0 | 0 | 0 | 0 | 0 | 0 | 0 |
| ENSMUSG00000084153 | 0 | 0.25 | 0 | 0 | 0 | 0 | 0 |
| ENSMUSG00000084154 | 0 | 0 | 0 | 0 | 0 | 0 | 0 |
| ENSMUSG00000084155 | 0 | 0 | 0 | 0 | 0 | 0 | 0 |
| ENSMUSG00000084157 | 0 | 0 | 0.25 | 0 | 0 | 0 | 0 |
| ENSMUSG00000084158 | 0 | 0 | 0 | 0 | 0 | 0 | 0 |
| ENSMUSG00000084159 | 1.686083339 | 1.681980114 | 1.52186571 | 1.735628626 | 1.686820904 | 1.919078092 | 1.663407615 |
| ENSMUSG00000084160 | 0 | 0 | 0 | 0 | 0 | 0 | 0 |
| ENSMUSG00000084161 | 0 | 0 | 0 | 0 | 0 | 0 | 0 |
| ENSMUSG00000084162 | 0 | 0 | 0 | 0 | 0 | 0 | 0 |

| | | | | | | | |
|---|---|---|---|---|---|---|---|
| ENSMUSG00000084163 | 0 | 0 | 0.25 | 0.25 | 0.244650542 | 0 | 0.224243824 |
| ENSMUSG00000084164 | 0 | 0.25 | 0.646240625 | 0.580482024 | 0.489301084 | 0.301029996 | 0.355418052 |
| ENSMUSG00000084166 | 0.45815691 | 0 | 0.25 | 0 | 0 | 0 | 0.224243824 |
| ENSMUSG00000084167 | 0 | 0 | 0 | 0 | 0 | 0 | 0 |
| ENSMUSG00000084168 | 0 | 0 | 0 | 0 | 0 | 0 | 0 |
| ENSMUSG00000084169 | 0 | 0 | 0 | 0 | 0 | 0 | 0 |
| ENSMUSG00000084170 | 0 | 0 | 0 | 0 | 0 | 0 | 0 |
| ENSMUSG00000084171 | 0 | 0 | 0 | 0 | 0 | 0 | 0 |
| ENSMUSG00000084172 | 0 | 0 | 0 | 0 | 0 | 0 | 0 |
| ENSMUSG00000084173 | 0 | 0 | 0 | 0 | 0 | 0 | 0 |
| ENSMUSG00000084174 | 0 | 0 | 0 | 0 | 0 | 0 | 0 |
| ENSMUSG00000084175 | 0 | 0 | 0 | 0 | 0 | 0 | 0 |
| ENSMUSG00000084177 | 0 | 0.25 | 0 | 0.25 | 0.387761935 | 0 | 0 |
| ENSMUSG00000084178 | 0 | 0.25 | 0 | 0.701838731 | 0.387761935 | 0.301029996 | 0.224243824 |
| ENSMUSG00000084179 | 0 | 0 | 0 | 0 | 0 | 0 | 0 |
| ENSMUSG00000084180 | 0 | 0 | 0 | 0 | 0 | 0 | 0 |
| ENSMUSG00000084181 | 0 | 0 | 0 | 0 | 0 | 0 | 0 |
| ENSMUSG00000084182 | 0 | 0 | 0 | 0 | 0 | 0 | 0 |
| ENSMUSG00000084183 | 0.578129653 | 0.396240625 | 0.5 | 0 | 0.489301084 | 0.301029996 | 0.520678036 |
| ENSMUSG00000084184 | 0 | 0 | 0 | 0 | 0 | 0 | 0.224243824 |
| ENSMUSG00000084185 | 0.289064826 | 0 | 0 | 0 | 0 | 0 | 0 |
| ENSMUSG00000084186 | 0 | 0 | 0 | 0 | 0 | 0 | 0 |
| ENSMUSG00000084188 | 0 | 0 | 0 | 0 | 0 | 0 | 0 |
| ENSMUSG00000084189 | 0 | 0 | 0 | 0 | 0 | 0 | 0 |
| ENSMUSG00000084190 | 0.289064826 | 0.25 | 0.25 | 0 | 0.489301084 | 0.477121255 | 0.224243824 |
| ENSMUSG00000084191 | 0 | 0 | 0 | 0 | 0 | 0 | 0 |
| ENSMUSG00000084192 | 0 | 0 | 0 | 0 | 0 | 0 | 0 |
| ENSMUSG00000084193 | 0 | 0 | 0 | 0 | 0 | 0 | 0 |
| ENSMUSG00000084195 | 0 | 0 | 0 | 0 | 0 | 0 | 0 |
| ENSMUSG00000084196 | 0 | 0 | 0 | 0 | 0 | 0 | 0 |
| ENSMUSG00000084197 | 0 | 0 | 0 | 0 | 0 | 0 | 0 |
| ENSMUSG00000084198 | 0 | 0 | 0 | 0 | 0 | 0 | 0 |
| ENSMUSG00000084199 | 0.289064826 | 0 | 0 | 0 | 0 | 0 | 0 |
| ENSMUSG00000084200 | 0 | 0 | 0 | 0 | 0 | 0 | 0 |

| | | | | | | | |
|---|---|---|---|---|---|---|---|
| ENSMUSG00000084201 | 0 | 0 | 0 | 0 | 0 | 0 | 0 |
| ENSMUSG00000084202 | 0 | 0 | 0 | 0 | 0 | 0 | 0 |
| ENSMUSG00000084203 | 0 | 0.25 | 0.25 | 0 | 0 | 0.301029996 | 0.520678036 |
| ENSMUSG00000084204 | 0 | 0 | 0 | 0 | 0 | 0 | 0 |
| ENSMUSG00000084205 | 0 | 0 | 0 | 0 | 0 | 0 | 0 |
| ENSMUSG00000084206 | 0 | 0 | 0 | 0 | 0 | 0 | 0 |
| ENSMUSG00000084207 | 0 | 0.25 | 0 | 0 | 0 | 0 | 0 |
| ENSMUSG00000084208 | 0 | 0 | 0.25 | 0 | 0 | 0 | 0.224243824 |
| ENSMUSG00000084209 | 0 | 0 | 0 | 0 | 0 | 0 | 0 |
| ENSMUSG00000084210 | 0 | 0 | 0 | 0 | 0 | 0 | 0 |
| ENSMUSG00000084211 | 0 | 0 | 0.396240625 | 0 | 0 | 0 | 0 |
| ENSMUSG00000084212 | 0 | 0 | 0 | 0 | 0 | 0 | 0 |
| ENSMUSG00000084213 | 0 | 0 | 0 | 0.25 | 0 | 0 | 0 |
| ENSMUSG00000084214 | 0 | 0 | 0 | 0 | 0 | 0 | 0 |
| ENSMUSG00000084215 | 0 | 0.25 | 0 | 0 | 0 | 0 | 0.355418052 |
| ENSMUSG00000084216 | 0.289064826 | 0 | 0.25 | 0.396240625 | 0 | 0 | 0.520678036 |
| ENSMUSG00000084217 | 0 | 0 | 0 | 0 | 0.244650542 | 0 | 0.448487648 |
| ENSMUSG00000084218 | 0 | 0.25 | 0 | 0 | 0 | 0 | 0 |
| ENSMUSG00000084219 | 0 | 0 | 0 | 0 | 0 | 0 | 0 |
| ENSMUSG00000084220 | 0 | 0 | 0 | 0 | 0 | 0.301029996 | 0 |
| ENSMUSG00000084221 | 0 | 0.25 | 0 | 0.25 | 0 | 0 | 0.355418052 |
| ENSMUSG00000084222 | 0 | 0 | 0 | 0 | 0 | 0 | 0 |
| ENSMUSG00000084223 | 0 | 0 | 0.5 | 0 | 0.489301084 | 0 | 0.355418052 |
| ENSMUSG00000084224 | 0 | 0 | 0 | 0 | 0 | 0 | 0 |
| ENSMUSG00000084226 | 0 | 0 | 0 | 0.25 | 0 | 0 | 0 |
| ENSMUSG00000084228 | 0 | 0 | 0 | 0 | 0 | 0 | 0 |
| ENSMUSG00000084230 | 0 | 0 | 0 | 0 | 0 | 0 | 0 |
| ENSMUSG00000084231 | 0.289064826 | 0.25 | 0 | 0.25 | 0 | 0.301029996 | 0.355418052 |
| ENSMUSG00000084234 | 0 | 0 | 0 | 0 | 0 | 0 | 0 |
| ENSMUSG00000084235 | 0 | 0 | 0.25 | 0.25 | 0 | 0.301029996 | 0 |
| ENSMUSG00000084236 | 0 | 0 | 0 | 0 | 0 | 0 | 0 |
| ENSMUSG00000084237 | 0 | 0 | 0 | 0 | 0 | 0 | 0 |
| ENSMUSG00000084238 | 0 | 0 | 0 | 0 | 0 | 0 | 0 |
| ENSMUSG00000084239 | 0 | 0 | 0 | 0 | 0 | 0 | 0 |

| | | | | | | | |
|---|---|---|---|---|---|---|---|
| ENSMUSG00000084240 | 0 | 0 | 0 | 0 | 0 | 0 | 0 |
| ENSMUSG00000084241 | 0 | 0 | 0 | 0 | 0 | 0 | 0 |
| ENSMUSG00000084243 | 0 | 0 | 0 | 0.25 | 0 | 0 | 0 |
| ENSMUSG00000084244 | 0 | 0 | 0 | 0 | 0 | 0 | 0 |
| ENSMUSG00000084245 | 0 | 0 | 0 | 0 | 0 | 0 | 0 |
| ENSMUSG00000084246 | 0 | 0 | 0 | 0 | 0 | 0 | 0 |
| ENSMUSG00000084247 | 0 | 0 | 0 | 0 | 0 | 0 | 0 |
| ENSMUSG00000084248 | 0 | 0 | 0 | 0 | 0 | 0 | 0 |
| ENSMUSG00000084249 | 0.289064826 | 0.396240625 | 0 | 0 | 0.489301084 | 0.301029996 | 0.224243824 |
| ENSMUSG00000084250 | 0 | 0 | 0 | 0 | 0 | 0 | 0 |
| ENSMUSG00000084251 | 0 | 0 | 0 | 0 | 0 | 0 | 0 |
| ENSMUSG00000084252 | 0 | 0 | 0 | 0 | 0.244650542 | 0 | 0.224243824 |
| ENSMUSG00000084253 | 0 | 0 | 0 | 0 | 0 | 0 | 0 |
| ENSMUSG00000084254 | 0 | 0 | 0 | 0 | 0 | 0 | 0 |
| ENSMUSG00000084256 | 0 | 0 | 0 | 0 | 0 | 0 | 0 |
| ENSMUSG00000084257 | 0 | 0 | 0 | 0 | 0 | 0 | 0 |
| ENSMUSG00000084259 | 0 | 0 | 0 | 0 | 0 | 0 | 0 |
| ENSMUSG00000084260 | 0 | 0 | 0 | 0 | 0 | 0 | 0 |
| ENSMUSG00000084261 | 0 | 0 | 0 | 0 | 0 | 0 | 0 |
| ENSMUSG00000084262 | 0 | 0 | 0 | 0 | 0 | 0 | 0 |
| ENSMUSG00000084264 | 0 | 0 | 0 | 0 | 0 | 0 | 0 |
| ENSMUSG00000084265 | 0 | 0 | 0 | 0 | 0 | 0 | 0 |
| ENSMUSG00000084267 | 0 | 0 | 0 | 0 | 0 | 0 | 0 |
| ENSMUSG00000084268 | 0 | 0 | 0 | 0 | 0 | 0 | 0 |
| ENSMUSG00000084269 | 0 | 0 | 0 | 0 | 0 | 0 | 0 |
| ENSMUSG00000084270 | 0 | 0 | 0 | 0 | 0 | 0 | 0 |
| ENSMUSG00000084271 | 0.289064826 | 0 | 0.25 | 0 | 0 | 0 | 0 |
| ENSMUSG00000084272 | 0 | 0 | 0 | 0 | 0 | 0 | 0 |
| ENSMUSG00000084273 | 0 | 0 | 0 | 0.25 | 0 | 0 | 0 |
| ENSMUSG00000084274 | 0.91631382 | 1.146240625 | 1 | 1.080482024 | 0.931471446 | 0.84509804 | 1.041356071 |
| ENSMUSG00000084275 | 0 | 0.396240625 | 0 | 0 | 0.686820904 | 0 | 0.224243824 |
| ENSMUSG00000084276 | 0.45815691 | 0 | 0.25 | 0.580482024 | 0.489301084 | 0 | 0 |
| ENSMUSG00000084277 | 0 | 0 | 0 | 0 | 0 | 0 | 0 |
| ENSMUSG00000084278 | 0 | 0 | 0 | 0 | 0 | 0 | 0 |

| | | | | | | | |
|---|---|---|---|---|---|---|---|
| ENSMUSG00000084279 | 0 | 0 | 0 | 0 | 0 | 0 | 0 |
| ENSMUSG00000084280 | 0 | 0 | 0 | 0 | 0 | 0 | 0 |
| ENSMUSG00000084281 | 0 | 0 | 0 | 0 | 0 | 0 | 0 |
| ENSMUSG00000084282 | 0 | 0 | 0 | 0 | 0 | 0 | 0 |
| ENSMUSG00000084283 | 0 | 0.396240625 | 0.25 | 0 | 0.244650542 | 0 | 0.224243824 |
| ENSMUSG00000084284 | 0.289064826 | 0.396240625 | 0.580482024 | 0.25 | 0.489301084 | 0.477121255 | 0.74492186 |
| ENSMUSG00000084285 | 0 | 0 | 0 | 0 | 0 | 0 | 0 |
| ENSMUSG00000084286 | 0 | 0 | 0 | 0 | 0 | 0 | 0 |
| ENSMUSG00000084287 | 0 | 0 | 0 | 0 | 0 | 0 | 0 |
| ENSMUSG00000084288 | 0 | 0 | 0.25 | 0 | 0.244650542 | 0.301029996 | 0.224243824 |
| ENSMUSG00000084289 | 0.289064826 | 0 | 0.396240625 | 0 | 0.244650542 | 0.301029996 | 0 |
| ENSMUSG00000084291 | 0 | 0 | 0 | 0 | 0 | 0 | 0 |
| ENSMUSG00000084292 | 0 | 0 | 0.25 | 0 | 0 | 0 | 0 |
| ENSMUSG00000084293 | 0 | 0 | 0 | 0 | 0 | 0 | 0 |
| ENSMUSG00000084294 | 0 | 0 | 0 | 0 | 0 | 0 | 0 |
| ENSMUSG00000084295 | 0 | 0 | 0 | 0 | 0 | 0 | 0 |
| ENSMUSG00000084296 | 0 | 0 | 0 | 0 | 0 | 0 | 0 |
| ENSMUSG00000084297 | 0 | 0 | 0 | 0 | 0 | 0 | 0 |
| ENSMUSG00000084298 | 0 | 0 | 0 | 0 | 0 | 0 | 0 |
| ENSMUSG00000084299 | 0 | 0 | 0 | 0 | 0 | 0 | 0.224243824 |
| ENSMUSG00000084300 | 0 | 0 | 0 | 0 | 0 | 0 | 0 |
| ENSMUSG00000084301 | 0 | 0 | 0 | 0 | 0 | 0 | 0 |
| ENSMUSG00000084302 | 0.578129653 | 0.25 | 0.396240625 | 0 | 0.244650542 | 0.301029996 | 0.355418052 |
| ENSMUSG00000084303 | 0 | 0.25 | 0.25 | 0.25 | 0.244650542 | 0.301029996 | 0.224243824 |
| ENSMUSG00000084304 | 0 | 0 | 0 | 0 | 0 | 0 | 0.224243824 |
| ENSMUSG00000084305 | 0 | 0 | 0 | 0 | 0 | 0 | 0 |
| ENSMUSG00000084306 | 0 | 0 | 0 | 0.25 | 0 | 0 | 0.224243824 |
| ENSMUSG00000084307 | 0 | 0 | 0 | 0.25 | 0 | 0 | 0 |
| ENSMUSG00000084308 | 0 | 0 | 0 | 0 | 0 | 0 | 0 |
| ENSMUSG00000084309 | 0 | 0 | 0 | 0 | 0 | 0 | 0 |
| ENSMUSG00000084310 | 0 | 0 | 0.25 | 0 | 0 | 0 | 0 |
| ENSMUSG00000084311 | 0 | 0 | 0 | 0 | 0 | 0 | 0 |
| ENSMUSG00000084312 | 0 | 0 | 0 | 0 | 0 | 0 | 0 |
| ENSMUSG00000084313 | 0.289064826 | 0 | 0 | 0 | 0 | 0 | 0 |

| | | | | | | | |
|---|---|---|---|---|---|---|---|
| ENSMUSG00000084314 | 0.45815691 | 0 | 0.5 | 0.580482024 | 0.244650542 | 0.698970004 | 0.224243824 |
| ENSMUSG00000084315 | 0.811507563 | 0.864857905 | 0.79248125 | 0.701838731 | 0.733951626 | 0.903089987 | 0.829800754 |
| ENSMUSG00000084316 | 0 | 0 | 0 | 0 | 0 | 0 | 0 |
| ENSMUSG00000084318 | 0 | 0 | 0 | 0 | 0 | 0 | 0 |
| ENSMUSG00000084319 | 0 | 0.25 | 0.25 | 0.25 | 0 | 0 | 0 |
| ENSMUSG00000084320 | 0 | 0 | 0 | 0 | 0 | 0 | 0 |
| ENSMUSG00000084323 | 0 | 0 | 0 | 0 | 0 | 0 | 0 |
| ENSMUSG00000084324 | 0 | 0 | 0 | 0 | 0 | 0 | 0 |
| ENSMUSG00000084325 | 0 | 0.396240625 | 0 | 0 | 0.244650542 | 0 | 0.224243824 |
| ENSMUSG00000084326 | 0 | 0 | 0 | 0.25 | 0 | 0 | 0 |
| ENSMUSG00000084327 | 0 | 0 | 0 | 0 | 0 | 0 | 0 |
| ENSMUSG00000084328 | 0.867194479 | 0.646240625 | 0.896240625 | 0.580482024 | 0.632412477 | 0.903089987 | 0.520678036 |
| ENSMUSG00000084329 | 1.307602651 | 1.27186571 | 1.238549078 | 1.445339928 | 1.343584837 | 1.544068044 | 1.329931221 |
| ENSMUSG00000084330 | 0 | 0 | 0 | 0 | 0 | 0 | 0 |
| ENSMUSG00000084331 | 0 | 0 | 0 | 0.25 | 0 | 0 | 0 |
| ENSMUSG00000084332 | 0 | 0.396240625 | 0.580482024 | 0 | 0.244650542 | 0 | 0.520678036 |
| ENSMUSG00000084333 | 0 | 0.25 | 0.25 | 0.25 | 0 | 0.301029996 | 0 |
| ENSMUSG00000084334 | 0 | 0 | 0 | 0 | 0 | 0 | 0 |
| ENSMUSG00000084335 | 0 | 0 | 0 | 0 | 0 | 0 | 0 |
| ENSMUSG00000084336 | 0 | 0 | 0 | 0 | 0 | 0 | 0 |
| ENSMUSG00000084337 | 0 | 0 | 0 | 0 | 0 | 0 | 0 |
| ENSMUSG00000084338 | 0 | 0 | 0 | 0 | 0 | 0 | 0 |
| ENSMUSG00000084339 | 0 | 0.25 | 0 | 0 | 0 | 0 | 0 |
| ENSMUSG00000084340 | 0 | 0 | 0 | 0.25 | 0 | 0 | 0 |
| ENSMUSG00000084341 | 0 | 0 | 0 | 0 | 0 | 0 | 0 |
| ENSMUSG00000084342 | 0 | 0 | 0.396240625 | 0.25 | 0 | 0 | 0 |
| ENSMUSG00000084343 | 0 | 0.25 | 0 | 0 | 0 | 0 | 0 |
| ENSMUSG00000084344 | 0 | 0 | 0 | 0 | 0 | 0 | 0 |
| ENSMUSG00000084345 | 0 | 0.25 | 0 | 0.25 | 0.244650542 | 0 | 0.520678036 |
| ENSMUSG00000084346 | 0 | 0 | 0 | 0 | 0 | 0 | 0 |
| ENSMUSG00000084347 | 0.578129653 | 0.580482024 | 0.830482024 | 0.580482024 | 0.846351821 | 0.602059991 | 0.672731473 |
| ENSMUSG00000084348 | 0.289064826 | 0 | 0 | 0 | 0 | 0 | 0 |
| ENSMUSG00000084349 | 1.069666964 | 1.114857905 | 1.146240625 | 1.02186571 | 0.978602168 | 1.113943352 | 1.23151414 |
| ENSMUSG00000084350 | 1.100572389 | 1 | 0.92510993 | 1.26109853 | 1.149965125 | 1.176091259 | 1 |

| | | | | | | | |
|---|---|---|---|---|---|---|---|
| ENSMUSG00000084351 | 0 | 0 | 0 | 0 | 0 | 0 | 0 |
| ENSMUSG00000084352 | 0 | 0 | 0 | 0 | 0 | 0 | 0 |
| ENSMUSG00000084353 | 0 | 0 | 0 | 0 | 0 | 0 | 0 |
| ENSMUSG00000084355 | 0 | 0 | 0 | 0 | 0 | 0 | 0 |
| ENSMUSG00000084356 | 0 | 0 | 0 | 0 | 0 | 0 | 0 |
| ENSMUSG00000084357 | 0 | 0 | 0 | 0 | 0 | 0 | 0 |
| ENSMUSG00000084358 | 0 | 0 | 0 | 0 | 0 | 0 | 0 |
| ENSMUSG00000084359 | 0 | 0 | 0 | 0 | 0 | 0 | 0 |
| ENSMUSG00000084360 | 0 | 0 | 0 | 0 | 0 | 0 | 0 |
| ENSMUSG00000084361 | 0 | 0 | 0 | 0 | 0 | 0 | 0 |
| ENSMUSG00000084362 | 0 | 0 | 0 | 0 | 0 | 0 | 0 |
| ENSMUSG00000084363 | 0 | 0 | 0 | 0 | 0 | 0 | 0 |
| ENSMUSG00000084366 | 0 | 0 | 0 | 0 | 0 | 0 | 0 |
| ENSMUSG00000084367 | 0 | 0 | 0 | 0 | 0 | 0 | 0 |
| ENSMUSG00000084368 | 0 | 0 | 0 | 0 | 0 | 0 | 0 |
| ENSMUSG00000084369 | 0 | 0 | 0 | 0 | 0 | 0 | 0 |
| ENSMUSG00000084370 | 0 | 0 | 0 | 0 | 0 | 0 | 0 |
| ENSMUSG00000084371 | 0 | 0 | 0 | 0 | 0 | 0 | 0 |
| ENSMUSG00000084372 | 0.578129653 | 0.79248125 | 0.5 | 0.79248125 | 0.632412477 | 0.602059991 | 0.579661877 |
| ENSMUSG00000084373 | 0.578129653 | 0.701838731 | 0 | 0.25 | 0.387761935 | 0 | 0.448487648 |
| ENSMUSG00000084374 | 0 | 0 | 0 | 0 | 0.244650542 | 0 | 0 |
| ENSMUSG00000084375 | 0 | 0 | 0 | 0.25 | 0 | 0 | 0 |
| ENSMUSG00000084376 | 0 | 0 | 0 | 0 | 0 | 0 | 0 |
| ENSMUSG00000084377 | 0 | 0 | 0 | 0 | 0 | 0.301029996 | 0 |
| ENSMUSG00000084378 | 0.289064826 | 0.396240625 | 0.580482024 | 0.396240625 | 0.244650542 | 0.301029996 | 0.355418052 |
| ENSMUSG00000084379 | 0 | 0 | 0 | 0 | 0 | 0 | 0 |
| ENSMUSG00000084380 | 0 | 0 | 0 | 0 | 0 | 0 | 0 |
| ENSMUSG00000084381 | 0.45815691 | 0.25 | 0.5 | 0.25 | 0.244650542 | 0 | 0.448487648 |
| ENSMUSG00000084382 | 0 | 0 | 0 | 0 | 0 | 0 | 0 |
| ENSMUSG00000084383 | 0 | 0 | 0 | 0 | 0 | 0 | 0 |
| ENSMUSG00000084384 | 0 | 0.25 | 0 | 0 | 0 | 0 | 0.224243824 |
| ENSMUSG00000084385 | 0 | 0 | 0 | 0 | 0 | 0 | 0 |
| ENSMUSG00000084386 | 0 | 0 | 0 | 0 | 0 | 0 | 0 |
| ENSMUSG00000084387 | 0 | 0 | 0 | 0 | 0 | 0 | 0 |

| | | | | | | | |
|---|---|---|---|---|---|---|---|
| ENSMUSG00000084390 | 0 | 0 | 0 | 0 | 0 | 0 | 0 |
| ENSMUSG00000084391 | 0 | 0 | 0 | 0 | 0 | 0 | 0 |
| ENSMUSG00000084392 | 0 | 0 | 0 | 0 | 0.244650542 | 0 | 0 |
| ENSMUSG00000084393 | 0 | 0 | 0 | 0 | 0 | 0 | 0 |
| ENSMUSG00000084394 | 0 | 0 | 0 | 0 | 0 | 0 | 0 |
| ENSMUSG00000084395 | 0 | 0 | 0 | 0 | 0 | 0 | 0 |
| ENSMUSG00000084396 | 0 | 0 | 0 | 0 | 0 | 0 | 0 |
| ENSMUSG00000084397 | 0 | 0 | 0 | 0 | 0 | 0 | 0 |
| ENSMUSG00000084399 | 0 | 0 | 0 | 0 | 0 | 0 | 0 |
| ENSMUSG00000084400 | 0 | 0 | 0 | 0 | 0 | 0 | 0 |
| ENSMUSG00000084401 | 0 | 0.396240625 | 0 | 0 | 0.489301084 | 0 | 0 |
| ENSMUSG00000084403 | 0.578129653 | 0.5 | 0.646240625 | 0.646240625 | 0.489301084 | 0.602059991 | 0 |
| ENSMUSG00000084404 | 0 | 0 | 0.25 | 0 | 0 | 0.301029996 | 0 |
| ENSMUSG00000084405 | 0 | 0 | 0 | 0 | 0 | 0 | 0 |
| ENSMUSG00000084407 | 0 | 0 | 0 | 0 | 0 | 0 | 0 |
| ENSMUSG00000084408 | 0.811507563 | 0.701838731 | 0.701838731 | 0.580482024 | 0.877063019 | 0.602059991 | 0.448487648 |
| ENSMUSG00000084411 | 0 | 0 | 0 | 0 | 0 | 0 | 0 |
| ENSMUSG00000084412 | 0 | 0 | 0 | 0 | 0 | 0 | 0 |
| ENSMUSG00000084413 | 0.578129653 | 0.25 | 0.5 | 0.25 | 0.568060967 | 0 | 0.355418052 |
| ENSMUSG00000084414 | 0 | 0 | 0 | 0 | 0.244650542 | 0 | 0 |
| ENSMUSG00000084415 | 0 | 0 | 0 | 0 | 0 | 0 | 0 |
| ENSMUSG00000084416 | 0.747221736 | 0.896240625 | 0.701838731 | 0.92510993 | 0.877063019 | 0.954242509 | 0.803905701 |
| ENSMUSG00000084417 | 0 | 0 | 0 | 0 | 0 | 0 | 0 |
| ENSMUSG00000084418 | 0 | 0 | 0 | 0 | 0 | 0 | 0 |
| ENSMUSG00000084419 | 0 | 0 | 0 | 0 | 0.244650542 | 0 | 0.224243824 |
| ENSMUSG00000084421 | 0 | 0 | 0.25 | 0.25 | 0 | 0 | 0 |
| ENSMUSG00000084422 | 0.289064826 | 0.396240625 | 0.5 | 0 | 0.489301084 | 0 | 0.224243824 |
| ENSMUSG00000084423 | 0 | 0 | 0 | 0 | 0 | 0 | 0 |
| ENSMUSG00000084425 | 0 | 0 | 0 | 0 | 0 | 0 | 0 |
| ENSMUSG00000084426 | 0 | 0 | 0 | 0 | 0 | 0 | 0 |
| ENSMUSG00000084428 | 0 | 0 | 0 | 0 | 0 | 0 | 0 |
| ENSMUSG00000084431 | 0 | 0 | 0 | 0 | 0 | 0 | 0 |
| ENSMUSG00000084432 | 0 | 0 | 0 | 0 | 0 | 0 | 0 |
| ENSMUSG00000084433 | 0.289064826 | 0.396240625 | 0.5 | 0.580482024 | 0.489301084 | 0.477121255 | 0.448487648 |

| | | | | | | | |
|---|---|---|---|---|---|---|---|
| ENSMUSG00000084435 | 0 | 0 | 0 | 0 | 0 | 0 | 0 |
| ENSMUSG00000084437 | 0 | 0 | 0 | 0 | 0 | 0 | 0 |
| ENSMUSG00000084439 | 0 | 0 | 0 | 0 | 0 | 0 | 0 |
| ENSMUSG00000084440 | 0 | 0.396240625 | 0 | 0.25 | 0 | 0.301029996 | 0.224243824 |
| ENSMUSG00000084441 | 0 | 0 | 0 | 0 | 0 | 0 | 0 |
| ENSMUSG00000084442 | 0.289064826 | 0 | 0 | 0 | 0 | 0 | 0 |
| ENSMUSG00000084443 | 0 | 0 | 0 | 0 | 0 | 0 | 0 |
| ENSMUSG00000084444 | 0 | 0 | 0 | 0 | 0 | 0 | 0 |
| ENSMUSG00000084445 | 0 | 0 | 0 | 0 | 0 | 0 | 0 |
| ENSMUSG00000084446 | 0 | 0 | 0 | 0 | 0 | 0 | 0 |
| ENSMUSG00000084447 | 0 | 0 | 0 | 0 | 0 | 0 | 0 |
| ENSMUSG00000084448 | 0 | 0 | 0 | 0 | 0 | 0 | 0 |
| ENSMUSG00000084449 | 0 | 0 | 0 | 0 | 0 | 0 | 0 |
| ENSMUSG00000084450 | 0 | 0 | 0 | 0 | 0 | 0 | 0 |
| ENSMUSG00000084451 | 0 | 0 | 0 | 0 | 0 | 0 | 0 |
| ENSMUSG00000084452 | 0 | 0 | 0 | 0 | 0 | 0 | 0 |
| ENSMUSG00000084453 | 0 | 0 | 0 | 0 | 0 | 0 | 0 |
| ENSMUSG00000084454 | 0 | 0 | 0 | 0 | 0 | 0 | 0 |
| ENSMUSG00000084456 | 0 | 0 | 0 | 0 | 0 | 0 | 0 |
| ENSMUSG00000084457 | 0 | 0.25 | 0 | 0 | 0 | 0 | 0 |
| ENSMUSG00000084458 | 0 | 0 | 0 | 0.25 | 0 | 0 | 0 |
| ENSMUSG00000084459 | 0 | 0 | 0 | 0 | 0 | 0 | 0 |
| ENSMUSG00000084460 | 0 | 0 | 0 | 0 | 0 | 0 | 0 |
| ENSMUSG00000084461 | 0 | 0 | 0 | 0 | 0 | 0 | 0 |
| ENSMUSG00000084462 | 0 | 0 | 0.25 | 0 | 0 | 0 | 0 |
| ENSMUSG00000084463 | 0 | 0 | 0 | 0 | 0 | 0 | 0 |
| ENSMUSG00000084465 | 0 | 0 | 0 | 0 | 0 | 0 | 0 |
| ENSMUSG00000084466 | 0 | 0 | 0 | 0 | 0 | 0 | 0 |
| ENSMUSG00000084467 | 0 | 0 | 0 | 0 | 0 | 0 | 0 |
| ENSMUSG00000084468 | 0 | 0 | 0 | 0 | 0 | 0 | 0 |
| ENSMUSG00000084469 | 0 | 0 | 0 | 0 | 0 | 0 | 0 |
| ENSMUSG00000084472 | 0 | 0 | 0 | 0 | 0 | 0 | 0 |
| ENSMUSG00000084473 | 0 | 0 | 0 | 0 | 0 | 0 | 0 |
| ENSMUSG00000084474 | 0 | 0 | 0 | 0 | 0 | 0 | 0 |

| | | | | | | | |
|---|---|---|---|---|---|---|---|
| ENSMUSG00000084475 | 0 | 0 | 0 | 0 | 0 | 0 | 0 |
| ENSMUSG00000084476 | 0 | 0 | 0 | 0 | 0 | 0 | 0 |
| ENSMUSG00000084477 | 0 | 0 | 0 | 0.25 | 0 | 0 | 0 |
| ENSMUSG00000084478 | 0 | 0 | 0.25 | 0 | 0 | 0 | 0 |
| ENSMUSG00000084479 | 0 | 0 | 0 | 0 | 0 | 0 | 0 |
| ENSMUSG00000084480 | 0 | 0 | 0 | 0 | 0 | 0 | 0 |
| ENSMUSG00000084484 | 0 | 0 | 0 | 0 | 0 | 0 | 0 |
| ENSMUSG00000084485 | 0 | 0 | 0 | 0 | 0 | 0 | 0 |
| ENSMUSG00000084487 | 0 | 0 | 0 | 0 | 0 | 0 | 0 |
| ENSMUSG00000084492 | 0 | 0 | 0 | 0 | 0 | 0 | 0 |
| ENSMUSG00000084493 | 0 | 0 | 0 | 0 | 0 | 0 | 0 |
| ENSMUSG00000084496 | 0.289064826 | 0 | 0.396240625 | 0 | 0 | 0 | 0 |
| ENSMUSG00000084497 | 0.289064826 | 0 | 0.396240625 | 0 | 0 | 0 | 0 |
| ENSMUSG00000084498 | 0 | 0 | 0 | 0 | 0 | 0 | 0 |
| ENSMUSG00000084500 | 0 | 0 | 0 | 0 | 0 | 0 | 0 |
| ENSMUSG00000084501 | 0 | 0 | 0 | 0 | 0 | 0 | 0 |
| ENSMUSG00000084502 | 0 | 0 | 0 | 0 | 0 | 0 | 0 |
| ENSMUSG00000084503 | 0 | 0 | 0 | 0 | 0 | 0 | 0 |
| ENSMUSG00000084504 | 0 | 0 | 0 | 0 | 0 | 0 | 0 |
| ENSMUSG00000084505 | 0 | 0 | 0 | 0 | 0 | 0 | 0 |
| ENSMUSG00000084506 | 0 | 0 | 0 | 0 | 0 | 0 | 0 |
| ENSMUSG00000084507 | 0.45815691 | 0.25 | 0 | 0 | 0 | 0 | 0 |
| ENSMUSG00000084508 | 0 | 0 | 0 | 0 | 0 | 0 | 0 |
| ENSMUSG00000084511 | 0 | 0 | 0 | 0 | 0 | 0 | 0 |
| ENSMUSG00000084512 | 0 | 0 | 0 | 0 | 0 | 0 | 0 |
| ENSMUSG00000084513 | 0 | 0 | 0 | 0 | 0 | 0 | 0 |
| ENSMUSG00000084514 | 0 | 0 | 0 | 0 | 0 | 0 | 0 |
| ENSMUSG00000084515 | 0 | 0 | 0 | 0 | 0 | 0 | 0 |
| ENSMUSG00000084516 | 0 | 0 | 0 | 0 | 0 | 0 | 0 |
| ENSMUSG00000084517 | 0 | 0 | 0 | 0 | 0 | 0 | 0 |
| ENSMUSG00000084518 | 0 | 0 | 0 | 0 | 0 | 0 | 0 |
| ENSMUSG00000084519 | 0.578129653 | 0.646240625 | 0.5 | 0.79248125 | 0.846351821 | 0 | 0.629532004 |
| ENSMUSG00000084520 | 0 | 0 | 0 | 0 | 0 | 0 | 0 |
| ENSMUSG00000084521 | 0 | 0 | 0 | 0 | 0 | 0 | 0 |

| | | | | | | | |
|---|---|---|---|---|---|---|---|
| ENSMUSG00000084522 | 0 | 0 | 0 | 0.25 | 0 | 0 | 0 |
| ENSMUSG00000084523 | 0 | 0 | 0 | 0 | 0 | 0 | 0 |
| ENSMUSG00000084524 | 0 | 0 | 0 | 0 | 0 | 0 | 0 |
| ENSMUSG00000084526 | 0 | 0 | 0.25 | 0.25 | 0 | 0 | 0 |
| ENSMUSG00000084527 | 0 | 0 | 0 | 0 | 0 | 0 | 0 |
| ENSMUSG00000084530 | 0 | 0 | 0 | 0 | 0 | 0 | 0 |
| ENSMUSG00000084531 | 0 | 0 | 0 | 0 | 0 | 0 | 0 |
| ENSMUSG00000084532 | 0 | 0 | 0 | 0 | 0 | 0 | 0 |
| ENSMUSG00000084535 | 0 | 0 | 0 | 0 | 0 | 0 | 0 |
| ENSMUSG00000084536 | 0 | 0 | 0 | 0 | 0 | 0 | 0 |
| ENSMUSG00000084537 | 0 | 0 | 0 | 0 | 0 | 0 | 0 |
| ENSMUSG00000084539 | 0 | 0 | 0 | 0 | 0 | 0 | 0 |
| ENSMUSG00000084540 | 0 | 0 | 0 | 0 | 0 | 0 | 0 |
| ENSMUSG00000084541 | 0 | 0 | 0 | 0 | 0 | 0 | 0 |
| ENSMUSG00000084543 | 0 | 0 | 0 | 0 | 0 | 0 | 0.224243824 |
| ENSMUSG00000084544 | 0.289064826 | 0.25 | 0.396240625 | 0 | 0.387761935 | 0 | 0.224243824 |
| ENSMUSG00000084545 | 0 | 0 | 0.25 | 0 | 0 | 0 | 0 |
| ENSMUSG00000084546 | 0 | 0 | 0 | 0 | 0 | 0 | 0 |
| ENSMUSG00000084547 | 0 | 0 | 0 | 0 | 0 | 0 | 0 |
| ENSMUSG00000084548 | 0 | 0 | 0 | 0 | 0 | 0 | 0 |
| ENSMUSG00000084549 | 0 | 0 | 0 | 0 | 0 | 0 | 0 |
| ENSMUSG00000084550 | 0 | 0 | 0 | 0 | 0 | 0 | 0 |
| ENSMUSG00000084551 | 0 | 0 | 0 | 0 | 0 | 0 | 0 |
| ENSMUSG00000084552 | 0 | 0 | 0 | 0 | 0 | 0 | 0 |
| ENSMUSG00000084555 | 0 | 0 | 0 | 0 | 0 | 0 | 0 |
| ENSMUSG00000084557 | 0 | 0 | 0 | 0 | 0 | 0 | 0 |
| ENSMUSG00000084558 | 0 | 0 | 0 | 0 | 0 | 0 | 0 |
| ENSMUSG00000084559 | 0 | 0 | 0 | 0 | 0 | 0 | 0 |
| ENSMUSG00000084560 | 0 | 0 | 0 | 0 | 0 | 0 | 0 |
| ENSMUSG00000084562 | 0 | 0.25 | 0.25 | 0.396240625 | 0 | 0 | 0.355418052 |
| ENSMUSG00000084564 | 0 | 0 | 0 | 0 | 0 | 0 | 0 |
| ENSMUSG00000084565 | 0 | 0 | 0 | 0 | 0 | 0 | 0 |
| ENSMUSG00000084566 | 0 | 0 | 0 | 0 | 0 | 0 | 0 |
| ENSMUSG00000084567 | 0 | 0 | 0 | 0 | 0 | 0 | 0 |

| | | | | | | | |
|---|---|---|---|---|---|---|---|
| ENSMUSG00000084568 | 0 | 0 | 0 | 0 | 0 | 0 | 0 |
| ENSMUSG00000084569 | 0 | 0 | 0 | 0 | 0 | 0 | 0 |
| ENSMUSG00000084571 | 0 | 0 | 0.25 | 0.25 | 0 | 0 | 0 |
| ENSMUSG00000084572 | 0 | 0 | 0 | 0 | 0 | 0 | 0 |
| ENSMUSG00000084573 | 0 | 0 | 0 | 0 | 0 | 0 | 0 |
| ENSMUSG00000084574 | 0 | 0 | 0 | 0 | 0 | 0 | 0 |
| ENSMUSG00000084575 | 0 | 0 | 0 | 0 | 0 | 0 | 0 |
| ENSMUSG00000084576 | 0 | 0 | 0 | 0 | 0 | 0 | 0 |
| ENSMUSG00000084579 | 0 | 0 | 0 | 0 | 0 | 0 | 0 |
| ENSMUSG00000084581 | 0 | 0 | 0 | 0 | 0 | 0 | 0 |
| ENSMUSG00000084582 | 0 | 0 | 0 | 0 | 0 | 0 | 0 |
| ENSMUSG00000084583 | 0 | 0 | 0 | 0 | 0 | 0 | 0 |
| ENSMUSG00000084584 | 0 | 0 | 0.25 | 0.25 | 0 | 0 | 0 |
| ENSMUSG00000084587 | 0 | 0 | 0 | 0 | 0 | 0 | 0 |
| ENSMUSG00000084588 | 0 | 0 | 0 | 0 | 0 | 0 | 0 |
| ENSMUSG00000084589 | 0 | 0 | 0 | 0 | 0 | 0 | 0 |
| ENSMUSG00000084590 | 0 | 0 | 0 | 0 | 0 | 0 | 0 |
| ENSMUSG00000084591 | 0 | 0 | 0 | 0 | 0 | 0 | 0 |
| ENSMUSG00000084592 | 0.671187741 | 0.75 | 0.701838731 | 0.646240625 | 0.244650542 | 0.477121255 | 0.629532004 |
| ENSMUSG00000084593 | 0 | 0 | 0 | 0 | 0 | 0 | 0 |
| ENSMUSG00000084595 | 0 | 0 | 0 | 0 | 0.387761935 | 0 | 0 |
| ENSMUSG00000084596 | 0 | 0 | 0 | 0 | 0 | 0 | 0 |
| ENSMUSG00000084597 | 0 | 0 | 0 | 0 | 0 | 0 | 0 |
| ENSMUSG00000084598 | 0 | 0 | 0 | 0 | 0 | 0 | 0 |
| ENSMUSG00000084599 | 0 | 0 | 0 | 0 | 0 | 0 | 0 |
| ENSMUSG00000084600 | 0 | 0 | 0 | 0 | 0 | 0 | 0 |
| ENSMUSG00000084601 | 0 | 0 | 0 | 0 | 0 | 0 | 0.224243824 |
| ENSMUSG00000084602 | 0 | 0 | 0 | 0 | 0 | 0 | 0 |
| ENSMUSG00000084603 | 0.45815691 | 0 | 0.25 | 0 | 0.568060967 | 0.477121255 | 0.448487648 |
| ENSMUSG00000084604 | 0 | 0 | 0 | 0 | 0 | 0 | 0 |
| ENSMUSG00000084605 | 0 | 0 | 0 | 0 | 0 | 0 | 0 |
| ENSMUSG00000084606 | 0 | 0 | 0 | 0 | 0 | 0 | 0 |
| ENSMUSG00000084607 | 0.289064826 | 0.25 | 0.25 | 0.646240625 | 0.568060967 | 0.301029996 | 0.224243824 |
| ENSMUSG00000084608 | 0 | 0 | 0 | 0 | 0 | 0 | 0 |

| | | | | | | | |
|---|---|---|---|---|---|---|---|
| ENSMUSG00000084609 | 0 | 0 | 0 | 0 | 0 | 0 | 0 |
| ENSMUSG00000084610 | 0 | 0 | 0 | 0 | 0 | 0 | 0 |
| ENSMUSG00000084611 | 0 | 0 | 0 | 0 | 0 | 0 | 0 |
| ENSMUSG00000084612 | 0 | 0 | 0 | 0 | 0 | 0 | 0 |
| ENSMUSG00000084613 | 0 | 0 | 0 | 0 | 0 | 0 | 0 |
| ENSMUSG00000084616 | 0.671187741 | 0.25 | 0.5 | 0.396240625 | 0 | 0.477121255 | 0.355418052 |
| ENSMUSG00000084617 | 0 | 0 | 0 | 0 | 0 | 0 | 0 |
| ENSMUSG00000084619 | 0 | 0 | 0 | 0 | 0 | 0 | 0 |
| ENSMUSG00000084620 | 0 | 0 | 0 | 0 | 0 | 0 | 0 |
| ENSMUSG00000084621 | 0 | 0 | 0 | 0 | 0 | 0 | 0 |
| ENSMUSG00000084622 | 0 | 0 | 0 | 0 | 0 | 0 | 0 |
| ENSMUSG00000084624 | 0 | 0 | 0 | 0.25 | 0 | 0 | 0 |
| ENSMUSG00000084625 | 0 | 0 | 0 | 0 | 0 | 0 | 0 |
| ENSMUSG00000084627 | 0 | 0 | 0 | 0 | 0 | 0 | 0 |
| ENSMUSG00000084628 | 0 | 0 | 0 | 0 | 0.244650542 | 0 | 0 |
| ENSMUSG00000084629 | 0 | 0 | 0 | 0 | 0 | 0 | 0 |
| ENSMUSG00000084630 | 0 | 0 | 0 | 0 | 0 | 0 | 0 |
| ENSMUSG00000084631 | 0 | 0 | 0 | 0 | 0 | 0 | 0 |
| ENSMUSG00000084632 | 0 | 0 | 0 | 0 | 0.244650542 | 0 | 0 |
| ENSMUSG00000084633 | 0 | 0 | 0 | 0 | 0 | 0 | 0 |
| ENSMUSG00000084635 | 0 | 0 | 0 | 0 | 0 | 0 | 0 |
| ENSMUSG00000084636 | 0 | 0 | 0 | 0 | 0 | 0 | 0 |
| ENSMUSG00000084637 | 0 | 0 | 0 | 0 | 0 | 0 | 0 |
| ENSMUSG00000084638 | 0 | 0 | 0 | 0 | 0 | 0 | 0 |
| ENSMUSG00000084639 | 0 | 0 | 0 | 0 | 0 | 0 | 0 |
| ENSMUSG00000084640 | 0 | 0 | 0 | 0 | 0 | 0 | 0 |
| ENSMUSG00000084641 | 0 | 0 | 0 | 0 | 0 | 0 | 0 |
| ENSMUSG00000084642 | 0 | 0 | 0 | 0 | 0 | 0 | 0 |
| ENSMUSG00000084643 | 0 | 0 | 0 | 0 | 0 | 0 | 0 |
| ENSMUSG00000084644 | 0.289064826 | 0.25 | 0 | 0 | 0.244650542 | 0.301029996 | 0.224243824 |
| ENSMUSG00000084645 | 0 | 0 | 0 | 0 | 0 | 0 | 0 |
| ENSMUSG00000084646 | 0 | 0 | 0 | 0 | 0 | 0 | 0 |
| ENSMUSG00000084647 | 0 | 0 | 0 | 0 | 0 | 0 | 0 |
| ENSMUSG00000084648 | 0 | 0 | 0.25 | 0 | 0 | 0 | 0.355418052 |

| | | | | | | | |
|---|---|---|---|---|---|---|---|
| ENSMUSG00000084649 | 0 | 0 | 0 | 0 | 0 | 0 | 0 |
| ENSMUSG00000084650 | 0 | 0 | 0 | 0 | 0 | 0 | 0 |
| ENSMUSG00000084651 | 0 | 0 | 0 | 0 | 0 | 0 | 0 |
| ENSMUSG00000084652 | 0 | 0 | 0 | 0 | 0 | 0 | 0 |
| ENSMUSG00000084653 | 0 | 0 | 0 | 0 | 0 | 0 | 0 |
| ENSMUSG00000084654 | 0 | 0 | 0 | 0 | 0 | 0 | 0 |
| ENSMUSG00000084658 | 0 | 0 | 0 | 0 | 0 | 0 | 0 |
| ENSMUSG00000084659 | 0.289064826 | 0 | 0 | 0 | 0 | 0 | 0.224243824 |
| ENSMUSG00000084660 | 0 | 0 | 0 | 0 | 0 | 0 | 0 |
| ENSMUSG00000084661 | 0 | 0 | 0 | 0 | 0 | 0 | 0 |
| ENSMUSG00000084663 | 0 | 0 | 0 | 0 | 0 | 0 | 0 |
| ENSMUSG00000084664 | 0 | 0 | 0 | 0 | 0 | 0 | 0 |
| ENSMUSG00000084665 | 0 | 0 | 0 | 0 | 0 | 0 | 0 |
| ENSMUSG00000084666 | 0 | 0 | 0 | 0 | 0 | 0 | 0 |
| ENSMUSG00000084667 | 0 | 0 | 0 | 0 | 0 | 0 | 0 |
| ENSMUSG00000084668 | 0 | 0 | 0 | 0 | 0 | 0 | 0 |
| ENSMUSG00000084669 | 0 | 0 | 0 | 0 | 0 | 0 | 0 |
| ENSMUSG00000084671 | 0 | 0 | 0 | 0 | 0 | 0 | 0 |
| ENSMUSG00000084673 | 0.289064826 | 0 | 0 | 0 | 0 | 0 | 0 |
| ENSMUSG00000084674 | 0 | 0 | 0.25 | 0 | 0 | 0 | 0 |
| ENSMUSG00000084675 | 0 | 0 | 0 | 0 | 0 | 0 | 0 |
| ENSMUSG00000084678 | 0.289064826 | 0 | 0 | 0 | 0 | 0 | 0 |
| ENSMUSG00000084682 | 0 | 0 | 0 | 0 | 0 | 0 | 0 |
| ENSMUSG00000084683 | 0 | 0 | 0 | 0 | 0 | 0 | 0 |
| ENSMUSG00000084684 | 0 | 0 | 0 | 0 | 0 | 0 | 0 |
| ENSMUSG00000084686 | 0 | 0.25 | 0 | 0 | 0.244650542 | 0 | 0.224243824 |
| ENSMUSG00000084688 | 0 | 0 | 0 | 0 | 0 | 0 | 0 |
| ENSMUSG00000084689 | 0 | 0 | 0 | 0 | 0 | 0 | 0 |
| ENSMUSG00000084690 | 0 | 0 | 0 | 0 | 0 | 0 | 0 |
| ENSMUSG00000084691 | 0 | 0 | 0 | 0 | 0 | 0 | 0 |
| ENSMUSG00000084693 | 0 | 0.25 | 0 | 0 | 0 | 0 | 0 |
| ENSMUSG00000084694 | 0.289064826 | 0 | 0 | 0 | 0 | 0 | 0 |
| ENSMUSG00000084696 | 0 | 0 | 0 | 0 | 0 | 0 | 0 |
| ENSMUSG00000084700 | 0 | 0 | 0 | 0 | 0 | 0 | 0 |

| | | | | | | | |
|---|---|---|---|---|---|---|---|
| ENSMUSG00000084701 | 0 | 0 | 0 | 0 | 0 | 0 | 0 |
| ENSMUSG00000084702 | 0 | 0 | 0 | 0 | 0 | 0 | 0 |
| ENSMUSG00000084703 | 0 | 0 | 0 | 0 | 0 | 0 | 0 |
| ENSMUSG00000084704 | 0 | 0 | 0 | 0 | 0 | 0 | 0 |
| ENSMUSG00000084705 | 0 | 0 | 0 | 0 | 0 | 0 | 0 |
| ENSMUSG00000084706 | 0.289064826 | 0 | 0 | 0 | 0.244650542 | 0 | 0 |
| ENSMUSG00000084708 | 0 | 0 | 0 | 0 | 0 | 0 | 0 |
| ENSMUSG00000084710 | 0 | 0 | 0 | 0 | 0 | 0 | 0 |
| ENSMUSG00000084711 | 0 | 0 | 0 | 0 | 0 | 0.301029996 | 0 |
| ENSMUSG00000084713 | 0 | 0 | 0 | 0 | 0 | 0 | 0 |
| ENSMUSG00000084714 | 0 | 0 | 0 | 0 | 0 | 0 | 0 |
| ENSMUSG00000084717 | 0 | 0 | 0 | 0 | 0 | 0 | 0 |
| ENSMUSG00000084718 | 0 | 0 | 0 | 0 | 0 | 0 | 0 |
| ENSMUSG00000084719 | 0 | 0 | 0 | 0 | 0 | 0 | 0 |
| ENSMUSG00000084722 | 0 | 0 | 0 | 0 | 0 | 0 | 0 |
| ENSMUSG00000084725 | 0 | 0 | 0 | 0 | 0 | 0 | 0 |
| ENSMUSG00000084728 | 0 | 0 | 0 | 0 | 0 | 0 | 0 |
| ENSMUSG00000084729 | 0 | 0 | 0.396240625 | 0.25 | 0 | 0 | 0.224243824 |
| ENSMUSG00000084731 | 0 | 0 | 0 | 0 | 0 | 0 | 0 |
| ENSMUSG00000084732 | 0 | 0 | 0 | 0 | 0 | 0 | 0 |
| ENSMUSG00000084733 | 0 | 0 | 0 | 0 | 0 | 0 | 0 |
| ENSMUSG00000084734 | 0 | 0 | 0 | 0 | 0 | 0 | 0 |
| ENSMUSG00000084737 | 0 | 0 | 0 | 0 | 0 | 0 | 0 |
| ENSMUSG00000084738 | 0 | 0 | 0 | 0 | 0.387761935 | 0 | 0 |
| ENSMUSG00000084739 | 0 | 0 | 0 | 0 | 0.244650542 | 0 | 0 |
| ENSMUSG00000084740 | 0 | 0 | 0 | 0 | 0 | 0 | 0 |
| ENSMUSG00000084741 | 0 | 0 | 0 | 0 | 0 | 0 | 0 |
| ENSMUSG00000084742 | 0 | 0 | 0 | 0 | 0 | 0 | 0 |
| ENSMUSG00000084743 | 0 | 0 | 0 | 0 | 0 | 0 | 0 |
| ENSMUSG00000084744 | 0 | 0.25 | 0.25 | 0 | 0.244650542 | 0 | 0 |
| ENSMUSG00000084745 | 0 | 0 | 0 | 0 | 0 | 0 | 0 |
| ENSMUSG00000084746 | 0 | 0 | 0 | 0 | 0 | 0 | 0 |
| ENSMUSG00000084747 | 0 | 0 | 0 | 0 | 0 | 0 | 0 |
| ENSMUSG00000084748 | 0 | 0 | 0 | 0 | 0 | 0 | 0 |

| | | | | | | | |
|---|---|---|---|---|---|---|---|
| ENSMUSG00000084749 | 0 | 0 | 0 | 0 | 0 | 0 | 0 |
| ENSMUSG00000084750 | 0 | 0 | 0 | 0 | 0 | 0.301029996 | 0.224243824 |
| ENSMUSG00000084751 | 0 | 0 | 0 | 0 | 0 | 0 | 0 |
| ENSMUSG00000084752 | 0 | 0 | 0 | 0 | 0 | 0 | 0 |
| ENSMUSG00000084753 | 0 | 0 | 0 | 0 | 0 | 0 | 0 |
| ENSMUSG00000084754 | 0 | 0 | 0.25 | 0 | 0 | 0 | 0 |
| ENSMUSG00000084755 | 0.289064826 | 0.25 | 0.25 | 0 | 0.489301084 | 0 | 0.355418052 |
| ENSMUSG00000084756 | 0 | 0 | 0 | 0 | 0 | 0 | 0 |
| ENSMUSG00000084757 | 0 | 0 | 0 | 0 | 0 | 0 | 0 |
| ENSMUSG00000084758 | 0.578129653 | 0 | 0.396240625 | 0.25 | 0 | 0.301029996 | 0 |
| ENSMUSG00000084760 | 0 | 0 | 0 | 0 | 0 | 0 | 0 |
| ENSMUSG00000084761 | 0 | 0 | 0 | 0 | 0 | 0 | 0 |
| ENSMUSG00000084762 | 0 | 0 | 0 | 0 | 0 | 0 | 0 |
| ENSMUSG00000084764 | 0 | 0 | 0 | 0 | 0 | 0 | 0 |
| ENSMUSG00000084766 | 0 | 0 | 0 | 0 | 0 | 0 | 0.224243824 |
| ENSMUSG00000084768 | 0 | 0 | 0 | 0 | 0 | 0 | 0 |
| ENSMUSG00000084769 | 0 | 0 | 0 | 0 | 0 | 0 | 0 |
| ENSMUSG00000084771 | 0.45815691 | 0.25 | 0 | 0 | 0.387761935 | 0.301029996 | 0.448487648 |
| ENSMUSG00000084772 | 0 | 0.396240625 | 0.5 | 0.25 | 0.387761935 | 0.477121255 | 0 |
| ENSMUSG00000084773 | 0 | 0 | 0 | 0 | 0 | 0 | 0 |
| ENSMUSG00000084774 | 0 | 0.25 | 0 | 0 | 0 | 0 | 0 |
| ENSMUSG00000084775 | 0 | 0 | 0 | 0 | 0 | 0 | 0 |
| ENSMUSG00000084776 | 0 | 0 | 0.25 | 0 | 0.244650542 | 0 | 0.224243824 |
| ENSMUSG00000084780 | 0 | 0.25 | 0 | 0.396240625 | 0 | 0 | 0 |
| ENSMUSG00000084781 | 0 | 0 | 0 | 0.5 | 0.568060967 | 0.301029996 | 0.629532004 |
| ENSMUSG00000084782 | 0 | 0 | 0 | 0 | 0 | 0 | 0 |
| ENSMUSG00000084783 | 0 | 0 | 0.25 | 0 | 0.244650542 | 0 | 0.355418052 |
| ENSMUSG00000084785 | 0 | 0 | 0 | 0.25 | 0 | 0 | 0 |
| ENSMUSG00000084786 | 2.412042447 | 2.181980114 | 2.259043403 | 2.391989019 | 2.30971376 | 2.661812686 | 2.165794278 |
| ENSMUSG00000084787 | 0 | 0 | 0 | 0 | 0 | 0 | 0 |
| ENSMUSG00000084788 | 0 | 0 | 0 | 0 | 0 | 0 | 0 |
| ENSMUSG00000084789 | 0 | 0 | 0 | 0 | 0 | 0.301029996 | 0 |
| ENSMUSG00000084790 | 0.811507563 | 0.5 | 0.864857905 | 0.75 | 0.632412477 | 0 | 0.579661877 |
| ENSMUSG00000084791 | 0 | 0 | 0 | 0 | 0.244650542 | 0.477121255 | 0 |

| | | | | | | | |
|---|---|---|---|---|---|---|---|
| ENSMUSG00000084792 | 0.45815691 | 0.701838731 | 0.646240625 | 0.830482024 | 0.77552387 | 0.602059991 | 0.710836105 |
| ENSMUSG00000084793 | 0 | 0 | 0 | 0 | 0 | 0 | 0 |
| ENSMUSG00000084794 | 0 | 0 | 0 | 0 | 0 | 0 | 0 |
| ENSMUSG00000084795 | 0 | 0 | 0 | 0 | 0 | 0 | 0 |
| ENSMUSG00000084796 | 0 | 0 | 0 | 0 | 0 | 0 | 0 |
| ENSMUSG00000084797 | 0.289064826 | 0.5 | 0.396240625 | 0.5 | 0.387761935 | 0.301029996 | 0.448487648 |
| ENSMUSG00000084798 | 0.45815691 | 0.396240625 | 0.580482024 | 0.396240625 | 0.244650542 | 0 | 0.448487648 |
| ENSMUSG00000084799 | 0.960252568 | 1.02186571 | 1 | 1.188721876 | 1.136121934 | 1.113943352 | 1.078019652 |
| ENSMUSG00000084800 | 0 | 0.25 | 0.25 | 0.396240625 | 0.244650542 | 0 | 0.224243824 |
| ENSMUSG00000084802 | 0 | 0 | 0 | 0 | 0 | 0 | 0 |
| ENSMUSG00000084803 | 0 | 0.25 | 0 | 0.396240625 | 0.244650542 | 0 | 0.224243824 |
| ENSMUSG00000084804 | 0 | 0 | 0 | 0 | 0 | 0 | 0.224243824 |
| ENSMUSG00000084805 | 0 | 0 | 0 | 0 | 0 | 0 | 0 |
| ENSMUSG00000084806 | 0.578129653 | 0.701838731 | 0.701838731 | 0.25 | 0 | 0 | 0.224243824 |
| ENSMUSG00000084807 | 0.289064826 | 0.25 | 0.25 | 0.396240625 | 0.489301084 | 0.301029996 | 0.224243824 |
| ENSMUSG00000084808 | 1.227926429 | 1.061981878 | 1.04248125 | 1.321350555 | 1.244650542 | 1.255272505 | 1.041356071 |
| ENSMUSG00000084809 | 0.45815691 | 0 | 0 | 0.25 | 0 | 0 | 0.224243824 |
| ENSMUSG00000084811 | 0 | 0 | 0 | 0 | 0 | 0 | 0 |
| ENSMUSG00000084815 | 0 | 0 | 0 | 0 | 0 | 0 | 0 |
| ENSMUSG00000084816 | 0 | 0 | 0 | 0 | 0 | 0 | 0 |
| ENSMUSG00000084817 | 0.45815691 | 0.396240625 | 0.580482024 | 0.580482024 | 0.387761935 | 0.477121255 | 0.355418052 |
| ENSMUSG00000084818 | 0 | 0 | 0 | 0 | 0 | 0 | 0 |
| ENSMUSG00000084819 | 0 | 0 | 0 | 0 | 0 | 0 | 0 |
| ENSMUSG00000084821 | 0 | 0 | 0 | 0 | 0 | 0 | 0.355418052 |
| ENSMUSG00000084822 | 0 | 0 | 0 | 0.25 | 0.244650542 | 0 | 0 |
| ENSMUSG00000084823 | 0 | 0 | 0 | 0 | 0 | 0 | 0 |
| ENSMUSG00000084824 | 1.100572389 | 1.080482024 | 1.098079356 | 1.201838731 | 1.074582839 | 1.447158031 | 1.140832123 |
| ENSMUSG00000084825 | 0 | 0 | 0 | 0 | 0 | 0 | 0 |
| ENSMUSG00000084826 | 0 | 0 | 0 | 0 | 0 | 0 | 0.224243824 |
| ENSMUSG00000084828 | 0.289064826 | 0 | 0 | 0 | 0 | 0.301029996 | 0.224243824 |
| ENSMUSG00000084829 | 0 | 0 | 0 | 0 | 0 | 0 | 0 |
| ENSMUSG00000084830 | 0 | 0 | 0 | 0 | 0 | 0 | 0 |
| ENSMUSG00000084831 | 0 | 0 | 0 | 0 | 0 | 0 | 0 |
| ENSMUSG00000084833 | 0 | 0 | 0 | 0 | 0 | 0 | 0 |

| | | | | | | | |
|---|---|---|---|---|---|---|---|
| ENSMUSG00000084834 | 1.269664473 | 1.146240625 | 0.79248125 | 1.098079356 | 1.121713561 | 1.113943352 | 0.916588299 |
| ENSMUSG00000084835 | 0.289064826 | 0.25 | 0.75 | 0.396240625 | 0.568060967 | 0.602059991 | 0.579661877 |
| ENSMUSG00000084836 | 0 | 0 | 0 | 0 | 0 | 0 | 0 |
| ENSMUSG00000084837 | 0 | 0 | 0 | 0 | 0 | 0 | 0 |
| ENSMUSG00000084838 | 0 | 0 | 0 | 0 | 0 | 0 | 0 |
| ENSMUSG00000084839 | 0 | 0 | 0 | 0 | 0 | 0 | 0 |
| ENSMUSG00000084840 | 0 | 0 | 0 | 0 | 0 | 0 | 0 |
| ENSMUSG00000084842 | 0 | 0 | 0 | 0.396240625 | 0.387761935 | 0 | 0.448487648 |
| ENSMUSG00000084843 | 0.289064826 | 0.5 | 0.5 | 0.396240625 | 0 | 0.301029996 | 0.355418052 |
| ENSMUSG00000084844 | 0.747221736 | 0.79248125 | 0.951838731 | 1 | 0.905314583 | 0.954242509 | 0.853775828 |
| ENSMUSG00000084845 | 0.578129653 | 0.75 | 0.701838731 | 0.5 | 0.733951626 | 0.698970004 | 0.916588299 |
| ENSMUSG00000084846 | 1.069666964 | 1.114857905 | 0.976722649 | 1.146240625 | 1 | 1.113943352 | 1.041356071 |
| ENSMUSG00000084847 | 0 | 0 | 0.25 | 0 | 0 | 0 | 0 |
| ENSMUSG00000084848 | 0.289064826 | 0 | 0.25 | 0 | 0.244650542 | 0 | 0 |
| ENSMUSG00000084849 | 0 | 0 | 0 | 0 | 0 | 0 | 0 |
| ENSMUSG00000084850 | 0 | 0 | 0 | 0 | 0.244650542 | 0 | 0 |
| ENSMUSG00000084851 | 0 | 0 | 0 | 0 | 0 | 0 | 0 |
| ENSMUSG00000084852 | 0 | 0 | 0 | 0 | 0 | 0 | 0 |
| ENSMUSG00000084853 | 0 | 0 | 0 | 0 | 0 | 0 | 0 |
| ENSMUSG00000084854 | 0 | 0 | 0 | 0 | 0 | 0 | 0 |
| ENSMUSG00000084855 | 0 | 0 | 0 | 0 | 0 | 0 | 0 |
| ENSMUSG00000084856 | 0 | 0 | 0 | 0 | 0 | 0 | 0 |
| ENSMUSG00000084858 | 0 | 0 | 0 | 0.25 | 0 | 0.301029996 | 0 |
| ENSMUSG00000084859 | 0 | 0 | 0 | 0 | 0 | 0 | 0 |
| ENSMUSG00000084860 | 0.289064826 | 0.864857905 | 0.396240625 | 0.25 | 0 | 0 | 0 |
| ENSMUSG00000084861 | 0 | 0 | 0 | 0.5 | 0.244650542 | 0.301029996 | 0.448487648 |
| ENSMUSG00000084862 | 0.289064826 | 0 | 0.580482024 | 0.79248125 | 0.387761935 | 0.301029996 | 0.448487648 |
| ENSMUSG00000084863 | 0 | 0 | 0 | 0 | 0 | 0 | 0 |
| ENSMUSG00000084864 | 0.45815691 | 0.25 | 0.25 | 0.25 | 0.387761935 | 0 | 0 |
| ENSMUSG00000084865 | 0 | 0 | 0 | 0 | 0 | 0 | 0 |
| ENSMUSG00000084866 | 0.671187741 | 0.396240625 | 0.864857905 | 0.75 | 0.733951626 | 0.477121255 | 0.710836105 |
| ENSMUSG00000084869 | 0 | 0 | 0 | 0 | 0 | 0 | 0 |
| ENSMUSG00000084870 | 0 | 0 | 0 | 0.25 | 0.387761935 | 0 | 0 |
| ENSMUSG00000084871 | 0 | 0 | 0 | 0 | 0 | 0 | 0 |

| | | | | | | | |
|---|---|---|---|---|---|---|---|
| ENSMUSG00000084873 | 0 | 0 | 0 | 0 | 0 | 0 | 0 |
| ENSMUSG00000084874 | 0 | 0 | 0 | 0 | 0 | 0 | 0 |
| ENSMUSG00000084875 | 0 | 0 | 0 | 0 | 0 | 0 | 0 |
| ENSMUSG00000084876 | 0 | 0 | 0 | 0 | 0 | 0.301029996 | 0 |
| ENSMUSG00000084878 | 0.289064826 | 0 | 0.25 | 0 | 0.244650542 | 0 | 0 |
| ENSMUSG00000084880 | 0.671187741 | 0.5 | 0.646240625 | 0.25 | 0.244650542 | 0.301029996 | 0.74492186 |
| ENSMUSG00000084881 | 0.289064826 | 0.396240625 | 0.396240625 | 0.580482024 | 0.244650542 | 0.602059991 | 0.579661877 |
| ENSMUSG00000084882 | 0 | 0 | 0 | 0 | 0 | 0 | 0 |
| ENSMUSG00000084883 | 0.747221736 | 0.92510993 | 0.864857905 | 0.580482024 | 0.77552387 | 0.477121255 | 0.935079929 |
| ENSMUSG00000084884 | 0 | 0 | 0 | 0 | 0 | 0 | 0 |
| ENSMUSG00000084885 | 0.747221736 | 0.75 | 0.79248125 | 1.114857905 | 1.163285805 | 1.079181246 | 0.952571511 |
| ENSMUSG00000084887 | 0 | 0 | 0 | 0 | 0 | 0 | 0 |
| ENSMUSG00000084889 | 0 | 0 | 0 | 0 | 0 | 0 | 0 |
| ENSMUSG00000084890 | 0 | 0 | 0.25 | 0 | 0.244650542 | 0 | 0 |
| ENSMUSG00000084891 | 0 | 0 | 0 | 0 | 0 | 0 | 0 |
| ENSMUSG00000084892 | 0 | 0.396240625 | 0 | 0.396240625 | 0 | 0.301029996 | 0 |
| ENSMUSG00000084893 | 0 | 0.25 | 0.396240625 | 0 | 0.244650542 | 0 | 0.355418052 |
| ENSMUSG00000084894 | 0 | 0.25 | 0 | 0 | 0 | 0 | 0 |
| ENSMUSG00000084895 | 0 | 0 | 0 | 0 | 0 | 0 | 0 |
| ENSMUSG00000084896 | 0 | 0 | 0 | 0 | 0 | 0 | 0 |
| ENSMUSG00000084897 | 0.671187741 | 0.75 | 0.92510993 | 0.830482024 | 0.244650542 | 0.954242509 | 0.629532004 |
| ENSMUSG00000084898 | 0 | 0 | 0 | 0 | 0 | 0 | 0 |
| ENSMUSG00000084899 | 0.867194479 | 0.5 | 0.75 | 0.580482024 | 0.632412477 | 0.477121255 | 0.829800754 |
| ENSMUSG00000084901 | 0 | 0 | 0 | 0 | 0 | 0 | 0 |
| ENSMUSG00000084902 | 0 | 0 | 0 | 0 | 0 | 0 | 0 |
| ENSMUSG00000084904 | 0.671187741 | 0.92510993 | 0.896240625 | 0.701838731 | 0.733951626 | 0.77815125 | 0.829800754 |
| ENSMUSG00000084905 | 0 | 0 | 0 | 0 | 0 | 0 | 0 |
| ENSMUSG00000084906 | 0 | 0 | 0 | 0 | 0 | 0 | 0 |
| ENSMUSG00000084907 | 0 | 0 | 0 | 0 | 0 | 0 | 0 |
| ENSMUSG00000084908 | 0 | 0 | 0.25 | 0.396240625 | 0 | 0 | 0 |
| ENSMUSG00000084909 | 0 | 0 | 0 | 0 | 0 | 0 | 0 |
| ENSMUSG00000084910 | 0.867194479 | 1.098079356 | 0.580482024 | 0.75 | 0.812711509 | 0.77815125 | 0.775756176 |
| ENSMUSG00000084911 | 1.822201287 | 1.618933358 | 1.602347734 | 1.714495249 | 1.645989387 | 1.740362689 | 1.473249546 |
| ENSMUSG00000084913 | 0 | 0 | 0 | 0 | 0 | 0 | 0.224243824 |

| | | | | | | | |
|---|---|---|---|---|---|---|---|
| ENSMUSG00000084915 | 0.289064826 | 0.25 | 0.25 | 0.701838731 | 0.489301084 | 0.477121255 | 0.224243824 |
| ENSMUSG00000084916 | 0 | 0 | 0.25 | 0 | 0 | 0 | 0 |
| ENSMUSG00000084917 | 1.227926429 | 1.04248125 | 1.098079356 | 1.114857905 | 1 | 1.041392685 | 1.078019652 |
| ENSMUSG00000084918 | 0 | 0 | 0 | 0 | 0 | 0 | 0 |
| ENSMUSG00000084919 | 0 | 0 | 0 | 0 | 0 | 0 | 0 |
| ENSMUSG00000084920 | 0 | 0 | 0 | 0 | 0 | 0 | 0 |
| ENSMUSG00000084921 | 0 | 0 | 0 | 0 | 0 | 0 | 0 |
| ENSMUSG00000084923 | 0 | 0 | 0 | 0 | 0 | 0 | 0 |
| ENSMUSG00000084924 | 0 | 0 | 0 | 0 | 0 | 0 | 0 |
| ENSMUSG00000084925 | 0.289064826 | 0.25 | 0.25 | 0.75 | 0.489301084 | 0.301029996 | 0.520678036 |
| ENSMUSG00000084927 | 0 | 0 | 0 | 0 | 0 | 0 | 0 |
| ENSMUSG00000084929 | 0 | 0 | 0 | 0 | 0 | 0 | 0 |
| ENSMUSG00000084930 | 0.45815691 | 0.396240625 | 0.396240625 | 0.5 | 0.568060967 | 0.602059991 | 0.355418052 |
| ENSMUSG00000084932 | 0 | 0 | 0 | 0 | 0 | 0 | 0 |
| ENSMUSG00000084933 | 0 | 0.396240625 | 0 | 0.25 | 0 | 0.301029996 | 0 |
| ENSMUSG00000084934 | 0 | 0 | 0 | 0 | 0 | 0 | 0 |
| ENSMUSG00000084935 | 0 | 0 | 0 | 0 | 0 | 0 | 0 |
| ENSMUSG00000084936 | 0 | 0 | 0 | 0 | 0 | 0 | 0 |
| ENSMUSG00000084937 | 0 | 0 | 0 | 0 | 0 | 0 | 0 |
| ENSMUSG00000084938 | 0.578129653 | 0.701838731 | 0.396240625 | 0.5 | 0.244650542 | 0.301029996 | 0.355418052 |
| ENSMUSG00000084939 | 0 | 0 | 0.396240625 | 0.5 | 0 | 0.301029996 | 0.224243824 |
| ENSMUSG00000084940 | 0 | 0 | 0 | 0 | 0 | 0 | 0.224243824 |
| ENSMUSG00000084941 | 0.811507563 | 0.79248125 | 0.864857905 | 0.75 | 0.632412477 | 0.698970004 | 0.629532004 |
| ENSMUSG00000084942 | 0 | 0 | 0.25 | 0 | 0 | 0 | 0 |
| ENSMUSG00000084944 | 0 | 0 | 0 | 0 | 0 | 0 | 0 |
| ENSMUSG00000084945 | 0 | 0.25 | 0 | 0 | 0 | 0 | 0 |
| ENSMUSG00000084946 | 0.747221736 | 0.896240625 | 0.75 | 1.098079356 | 1 | 0.84509804 | 0.803905701 |
| ENSMUSG00000084947 | 0 | 0 | 0 | 0 | 0 | 0 | 0 |
| ENSMUSG00000084948 | 0 | 0 | 0 | 0 | 0.244650542 | 0 | 0 |
| ENSMUSG00000084949 | 0.45815691 | 0.646240625 | 0.5 | 0.701838731 | 0.568060967 | 0.602059991 | 0.520678036 |
| ENSMUSG00000084950 | 1.100572389 | 0.976722649 | 1.114857905 | 1.098079356 | 1.136121934 | 0.954242509 | 0.984950056 |
| ENSMUSG00000084951 | 0 | 0 | 0 | 0 | 0 | 0 | 0.224243824 |
| ENSMUSG00000084954 | 0.289064826 | 0 | 0 | 0 | 0 | 0 | 0 |
| ENSMUSG00000084955 | 0 | 0 | 0 | 0 | 0 | 0 | 0 |

| | | | | | | | |
|---|---|---|---|---|---|---|---|
| ENSMUSG00000084956 | 0 | 0 | 0 | 0 | 0 | 0 | 0 |
| ENSMUSG00000084957 | 2.213719572 | 2.070192693 | 2.049918086 | 2.052363341 | 1.953043706 | 2.290034611 | 1.817112247 |
| ENSMUSG00000084959 | 0 | 0 | 0 | 0 | 0 | 0 | 0 |
| ENSMUSG00000084960 | 0.289064826 | 0.25 | 0 | 0 | 0 | 0 | 0.224243824 |
| ENSMUSG00000084962 | 0 | 0 | 0 | 0 | 0 | 0 | 0 |
| ENSMUSG00000084963 | 0 | 0 | 0 | 0 | 0 | 0 | 0 |
| ENSMUSG00000084964 | 0 | 0 | 0 | 0 | 0.244650542 | 0.301029996 | 0 |
| ENSMUSG00000084965 | 0 | 0 | 0 | 0 | 0 | 0 | 0 |
| ENSMUSG00000084966 | 0 | 0 | 0 | 0 | 0 | 0 | 0 |
| ENSMUSG00000084967 | 0 | 0 | 0 | 0 | 0.244650542 | 0 | 0 |
| ENSMUSG00000084968 | 1.069666964 | 0.580482024 | 0.830482024 | 0.864857905 | 0.489301084 | 0.698970004 | 0.876096088 |
| ENSMUSG00000084969 | 0 | 0 | 0 | 0 | 0.244650542 | 0 | 0 |
| ENSMUSG00000084970 | 0 | 0.25 | 0.25 | 0.396240625 | 0.489301084 | 0.301029996 | 0.355418052 |
| ENSMUSG00000084971 | 0 | 0 | 0 | 0 | 0 | 0 | 0 |
| ENSMUSG00000084973 | 0 | 0 | 0 | 0 | 0 | 0 | 0 |
| ENSMUSG00000084974 | 0.867194479 | 0.396240625 | 0.396240625 | 0.5 | 0.387761935 | 0.477121255 | 0.448487648 |
| ENSMUSG00000084978 | 0 | 0 | 0 | 0 | 0.244650542 | 0.301029996 | 0 |
| ENSMUSG00000084979 | 0.289064826 | 0.396240625 | 0.5 | 0.75 | 0.733951626 | 0.602059991 | 0.355418052 |
| ENSMUSG00000084980 | 0.578129653 | 0.5 | 0.5 | 0 | 0.387761935 | 0.301029996 | 0.224243824 |
| ENSMUSG00000084981 | 0 | 0 | 0.25 | 0 | 0 | 0 | 0 |
| ENSMUSG00000084982 | 0 | 0 | 0 | 0 | 0 | 0 | 0 |
| ENSMUSG00000084983 | 0 | 0 | 0 | 0 | 0 | 0 | 0 |
| ENSMUSG00000084984 | 0.867194479 | 0.864857905 | 0.646240625 | 0.79248125 | 0.686820904 | 0.477121255 | 0.672731473 |
| ENSMUSG00000084985 | 0.45815691 | 0 | 0 | 0.25 | 0 | 0.301029996 | 0.224243824 |
| ENSMUSG00000084986 | 0 | 0 | 0 | 0 | 0 | 0 | 0 |
| ENSMUSG00000084987 | 0 | 0 | 0 | 0 | 0 | 0 | 0 |
| ENSMUSG00000084989 | 0.45815691 | 0.79248125 | 0.580482024 | 0 | 0.387761935 | 0.301029996 | 0.448487648 |
| ENSMUSG00000084990 | 0 | 0 | 0 | 0 | 0 | 0 | 0 |
| ENSMUSG00000084991 | 0 | 0 | 0 | 0 | 0 | 0 | 0 |
| ENSMUSG00000084992 | 0 | 0 | 0 | 0 | 0 | 0 | 0 |
| ENSMUSG00000084993 | 0 | 0 | 0 | 0.25 | 0.244650542 | 0 | 0 |
| ENSMUSG00000084994 | 0 | 0 | 0 | 0 | 0 | 0 | 0 |
| ENSMUSG00000084995 | 0 | 0 | 0 | 0 | 0 | 0 | 0 |
| ENSMUSG00000084996 | 0.289064826 | 0.580482024 | 0.75 | 0.396240625 | 0.489301084 | 0.301029996 | 0.579661877 |

| | | | | | | | |
|---|---|---|---|---|---|---|---|
| ENSMUSG00000084997 | 0 | 0 | 0 | 0 | 0 | 0 | 0 |
| ENSMUSG00000084998 | 1.100572389 | 0.976722649 | 1 | 0.79248125 | 1.020174412 | 1 | 0.984950056 |
| ENSMUSG00000084999 | 0 | 0 | 0 | 0 | 0 | 0.301029996 | 0.448487648 |
| ENSMUSG00000085000 | 0 | 0 | 0 | 0 | 0 | 0 | 0 |
| ENSMUSG00000085001 | 0 | 0 | 0 | 0 | 0 | 0 | 0 |
| ENSMUSG00000085002 | 0 | 0 | 0 | 0 | 0 | 0 | 0 |
| ENSMUSG00000085003 | 0 | 0 | 0 | 0 | 0 | 0 | 0 |
| ENSMUSG00000085004 | 0 | 0 | 0 | 0 | 0 | 0 | 0 |
| ENSMUSG00000085006 | 0.289064826 | 0 | 0 | 0.396240625 | 0.489301084 | 0.301029996 | 0.448487648 |
| ENSMUSG00000085007 | 0 | 0 | 0 | 0 | 0 | 0 | 0.224243824 |
| ENSMUSG00000085008 | 0.289064826 | 0 | 0 | 0.580482024 | 0 | 0.301029996 | 0.224243824 |
| ENSMUSG00000085009 | 0 | 0 | 0.25 | 0.396240625 | 0.387761935 | 0.301029996 | 0.224243824 |
| ENSMUSG00000085010 | 0 | 0 | 0 | 0 | 0 | 0 | 0 |
| ENSMUSG00000085012 | 0 | 0.25 | 0.25 | 0.25 | 0 | 0 | 0 |
| ENSMUSG00000085014 | 0 | 0 | 0 | 0 | 0 | 0 | 0 |
| ENSMUSG00000085015 | 0 | 0 | 0 | 0 | 0 | 0 | 0 |
| ENSMUSG00000085016 | 0.45815691 | 0.396240625 | 0.25 | 0 | 0.568060967 | 0 | 0.355418052 |
| ENSMUSG00000085017 | 0 | 0 | 0 | 0 | 0 | 0 | 0 |
| ENSMUSG00000085019 | 0.289064826 | 0 | 0 | 0 | 0 | 0.301029996 | 0 |
| ENSMUSG00000085020 | 0 | 0 | 0 | 0 | 0 | 0 | 0 |
| ENSMUSG00000085021 | 0 | 0 | 0 | 0.25 | 0 | 0 | 0 |
| ENSMUSG00000085022 | 0.578129653 | 0 | 0.5 | 0.25 | 0 | 0.301029996 | 0.629532004 |
| ENSMUSG00000085023 | 0 | 0 | 0 | 0 | 0 | 0 | 0 |
| ENSMUSG00000085024 | 0.289064826 | 0.646240625 | 0.25 | 0.580482024 | 0.632412477 | 0.602059991 | 0.579661877 |
| ENSMUSG00000085025 | 0 | 0 | 0 | 0 | 0 | 0 | 0 |
| ENSMUSG00000085026 | 0 | 0 | 0 | 0 | 0 | 0 | 0.224243824 |
| ENSMUSG00000085027 | 0 | 0 | 0 | 0 | 0 | 0 | 0 |
| ENSMUSG00000085028 | 1.37447073 | 1.25 | 1.348079356 | 1.410964047 | 1.478764298 | 1.431363764 | 1.340368108 |
| ENSMUSG00000085029 | 0.289064826 | 0 | 0 | 0 | 0 | 0 | 0 |
| ENSMUSG00000085030 | 1.069666964 | 1.02186571 | 1.061981878 | 1.098079356 | 1.121713561 | 1.146128036 | 1.110947927 |
| ENSMUSG00000085032 | 0 | 0 | 0 | 0 | 0 | 0 | 0.224243824 |
| ENSMUSG00000085033 | 0 | 0 | 0 | 0 | 0 | 0.301029996 | 0 |
| ENSMUSG00000085034 | 0 | 0 | 0 | 0 | 0 | 0 | 0 |
| ENSMUSG00000085035 | 0 | 0.25 | 0 | 0 | 0.244650542 | 0 | 0.448487648 |

| | | | | | | | |
|---|---|---|---|---|---|---|---|
| ENSMUSG00000085036 | 0 | 0 | 0 | 0 | 0 | 0 | 0 |
| ENSMUSG00000085037 | 1.249317394 | 1.114857905 | 1.080482024 | 1.188721876 | 1.163285805 | 1.079181246 | 1.159323753 |
| ENSMUSG00000085039 | 0.289064826 | 0.396240625 | 0.580482024 | 0.580482024 | 0.387761935 | 0.477121255 | 0.224243824 |
| ENSMUSG00000085041 | 0 | 0 | 0 | 0 | 0 | 0 | 0 |
| ENSMUSG00000085042 | 0.45815691 | 0 | 0.396240625 | 0 | 0 | 0 | 0.520678036 |
| ENSMUSG00000085043 | 0.289064826 | 0.396240625 | 0 | 0.580482024 | 0.489301084 | 0 | 0 |
| ENSMUSG00000085044 | 0 | 0 | 0 | 0 | 0 | 0 | 0 |
| ENSMUSG00000085045 | 0 | 0 | 0 | 0 | 0 | 0 | 0 |
| ENSMUSG00000085046 | 0 | 0 | 0 | 0 | 0 | 0 | 0 |
| ENSMUSG00000085047 | 0 | 0 | 0 | 0 | 0 | 0 | 0 |
| ENSMUSG00000085048 | 0 | 0 | 0 | 0 | 0 | 0 | 0 |
| ENSMUSG00000085049 | 0 | 0 | 0 | 0 | 0 | 0 | 0 |
| ENSMUSG00000085051 | 0.289064826 | 0 | 0 | 0.25 | 0.387761935 | 0 | 0.224243824 |
| ENSMUSG00000085053 | 0 | 0 | 0 | 0 | 0 | 0 | 0 |
| ENSMUSG00000085054 | 0.578129653 | 0.5 | 0.896240625 | 0.5 | 0.733951626 | 0.77815125 | 0.74492186 |
| ENSMUSG00000085055 | 0.45815691 | 0.5 | 0.25 | 0 | 0 | 0 | 0 |
| ENSMUSG00000085057 | 1.181541736 | 1.098079356 | 1.238549078 | 1.02186571 | 0.978602168 | 1.041392685 | 1.014380832 |
| ENSMUSG00000085058 | 0.45815691 | 0 | 0 | 0 | 0 | 0 | 0 |
| ENSMUSG00000085059 | 0 | 0 | 0 | 0 | 0 | 0 | 0 |
| ENSMUSG00000085060 | 0 | 0 | 0 | 0 | 0 | 0 | 0 |
| ENSMUSG00000085063 | 0 | 0 | 0 | 0.25 | 0 | 0 | 0 |
| ENSMUSG00000085066 | 0.960252568 | 1.02186571 | 0.830482024 | 0.646240625 | 0.387761935 | 0.301029996 | 0.520678036 |
| ENSMUSG00000085067 | 0 | 0 | 0.25 | 0 | 0 | 0 | 0 |
| ENSMUSG00000085068 | 0.289064826 | 0.396240625 | 0.396240625 | 0.25 | 0 | 0 | 0.355418052 |
| ENSMUSG00000085069 | 0 | 0 | 0 | 0 | 0 | 0 | 0 |
| ENSMUSG00000085070 | 0 | 0 | 0 | 0 | 0 | 0 | 0 |
| ENSMUSG00000085071 | 0.289064826 | 0 | 0.25 | 0 | 0 | 0 | 0 |
| ENSMUSG00000085072 | 0.811507563 | 0.646240625 | 0.5 | 0.864857905 | 0.877063019 | 0.698970004 | 0.803905701 |
| ENSMUSG00000085073 | 0 | 0 | 0 | 0 | 0 | 0 | 0 |
| ENSMUSG00000085074 | 0 | 0 | 0 | 0 | 0 | 0 | 0 |
| ENSMUSG00000085075 | 0 | 0 | 0 | 0 | 0 | 0 | 0 |
| ENSMUSG00000085077 | 0 | 0 | 0 | 0 | 0 | 0 | 0 |
| ENSMUSG00000085078 | 0 | 0 | 0.25 | 0 | 0 | 0 | 0 |
| ENSMUSG00000085079 | 0 | 0 | 0 | 0 | 0 | 0 | 0 |

| | | | | | | | |
|---|---|---|---|---|---|---|---|
| ENSMUSG00000085080 | 0 | 0 | 0 | 0 | 0 | 0 | 0 |
| ENSMUSG00000085081 | 0 | 0 | 0 | 0.25 | 0.387761935 | 0 | 0 |
| ENSMUSG00000085082 | 0 | 0 | 0 | 0 | 0 | 0 | 0 |
| ENSMUSG00000085083 | 0 | 0.25 | 0.396240625 | 0.25 | 0.244650542 | 0 | 0.224243824 |
| ENSMUSG00000085084 | 0.671187741 | 0.79248125 | 0.396240625 | 0.25 | 0 | 0.301029996 | 0.355418052 |
| ENSMUSG00000085085 | 1 | 0.79248125 | 0.79248125 | 0 | 0.244650542 | 0.301029996 | 0.448487648 |
| ENSMUSG00000085087 | 0 | 0 | 0 | 0 | 0 | 0 | 0 |
| ENSMUSG00000085088 | 0.578129653 | 0.701838731 | 0.701838731 | 0.5 | 0.387761935 | 0.602059991 | 0.448487648 |
| ENSMUSG00000085089 | 0 | 0 | 0.396240625 | 0 | 0 | 0 | 0 |
| ENSMUSG00000085090 | 0 | 0 | 0 | 0 | 0 | 0 | 0 |
| ENSMUSG00000085091 | 0.578129653 | 0.396240625 | 0.646240625 | 0.25 | 0.387761935 | 0.301029996 | 0.224243824 |
| ENSMUSG00000085093 | 0 | 0.25 | 0 | 0.25 | 0 | 0.301029996 | 0 |
| ENSMUSG00000085094 | 0 | 0 | 0 | 0 | 0 | 0 | 0 |
| ENSMUSG00000085095 | 0 | 0 | 0 | 0 | 0 | 0 | 0 |
| ENSMUSG00000085097 | 0 | 0 | 0 | 0 | 0 | 0 | 0 |
| ENSMUSG00000085099 | 0 | 0.396240625 | 0.646240625 | 0.5 | 0.387761935 | 0.477121255 | 0.448487648 |
| ENSMUSG00000085100 | 0 | 0 | 0 | 0 | 0 | 0 | 0 |
| ENSMUSG00000085101 | 0 | 0 | 0 | 0 | 0 | 0 | 0 |
| ENSMUSG00000085102 | 0 | 0 | 0 | 0 | 0 | 0 | 0 |
| ENSMUSG00000085103 | 0 | 0 | 0 | 0 | 0 | 0 | 0 |
| ENSMUSG00000085104 | 0 | 0 | 0 | 0 | 0 | 0 | 0 |
| ENSMUSG00000085105 | 0.289064826 | 0.25 | 0.25 | 0.5 | 0.387761935 | 0.698970004 | 0.448487648 |
| ENSMUSG00000085106 | 0 | 0 | 0 | 0 | 0 | 0 | 0 |
| ENSMUSG00000085107 | 0 | 0 | 0 | 0 | 0 | 0 | 0 |
| ENSMUSG00000085108 | 0 | 0 | 0 | 0 | 0 | 0 | 0 |
| ENSMUSG00000085109 | 0 | 0.25 | 0 | 0 | 0 | 0 | 0 |
| ENSMUSG00000085110 | 0 | 0 | 0 | 0 | 0 | 0 | 0 |
| ENSMUSG00000085111 | 0 | 0.25 | 0.25 | 0.25 | 0.244650542 | 0.301029996 | 0.355418052 |
| ENSMUSG00000085112 | 0.45815691 | 0.396240625 | 0 | 0 | 0.387761935 | 0.301029996 | 0.224243824 |
| ENSMUSG00000085113 | 0 | 0.25 | 0.25 | 0 | 0 | 0 | 0 |
| ENSMUSG00000085114 | 0 | 0 | 0.25 | 0 | 0 | 0 | 0 |
| ENSMUSG00000085115 | 0 | 0 | 0 | 0 | 0.244650542 | 0 | 0 |
| ENSMUSG00000085116 | 0 | 0 | 0 | 0 | 0 | 0 | 0 |
| ENSMUSG00000085118 | 0.578129653 | 0.646240625 | 0.646240625 | 0.646240625 | 0.568060967 | 0.602059991 | 0.520678036 |

| | | | | | | | |
|---|---|---|---|---|---|---|---|
| ENSMUSG00000085120 | 0 | 0 | 0 | 0 | 0 | 0 | 0 |
| ENSMUSG00000085121 | 0 | 0 | 0 | 0 | 0 | 0 | 0 |
| ENSMUSG00000085122 | 0 | 0 | 0 | 0 | 0 | 0 | 0 |
| ENSMUSG00000085123 | 0 | 0 | 0 | 0 | 0 | 0 | 0 |
| ENSMUSG00000085124 | 0.289064826 | 0.25 | 0.25 | 0 | 0 | 0.301029996 | 0 |
| ENSMUSG00000085125 | 0.289064826 | 0 | 0.396240625 | 0.25 | 0 | 0 | 0 |
| ENSMUSG00000085126 | 0 | 0 | 0.396240625 | 0 | 0 | 0 | 0 |
| ENSMUSG00000085127 | 0.289064826 | 0.25 | 0.396240625 | 0 | 0 | 0.301029996 | 0.355418052 |
| ENSMUSG00000085129 | 0 | 0 | 0 | 0 | 0 | 0 | 0 |
| ENSMUSG00000085130 | 0 | 0 | 0 | 0 | 0 | 0 | 0 |
| ENSMUSG00000085132 | 0 | 0 | 0 | 0 | 0 | 0 | 0 |
| ENSMUSG00000085133 | 1.759671389 | 1.688721876 | 1.610735874 | 1.990000483 | 1.983754904 | 2.113943352 | 1.755055163 |
| ENSMUSG00000085134 | 0.289064826 | 0.25 | 0 | 0 | 0 | 0.301029996 | 0 |
| ENSMUSG00000085135 | 0.578129653 | 0.396240625 | 0 | 0.25 | 0.244650542 | 0 | 0.355418052 |
| ENSMUSG00000085139 | 0 | 0 | 0 | 0 | 0 | 0 | 0 |
| ENSMUSG00000085141 | 0 | 0 | 0 | 0 | 0 | 0 | 0 |
| ENSMUSG00000085142 | 0 | 0 | 0 | 0 | 0 | 0 | 0 |
| ENSMUSG00000085143 | 0 | 0.701838731 | 0.5 | 0.396240625 | 0.632412477 | 0 | 0.710836105 |
| ENSMUSG00000085145 | 0 | 0.25 | 0 | 0.25 | 0 | 0 | 0.448487648 |
| ENSMUSG00000085146 | 0 | 0 | 0 | 0 | 0 | 0 | 0 |
| ENSMUSG00000085147 | 1 | 0.896240625 | 0.951838731 | 1.061981878 | 0.955822902 | 1.079181246 | 0.803905701 |
| ENSMUSG00000085148 | 2.054824387 | 1.935366747 | 1.916333979 | 1.976722649 | 1.983754904 | 2.173186268 | 1.849546807 |
| ENSMUSG00000085150 | 0 | 0 | 0 | 0 | 0 | 0 | 0 |
| ENSMUSG00000085151 | 0 | 0 | 0 | 0.25 | 0 | 0.477121255 | 0 |
| ENSMUSG00000085152 | 0 | 0 | 0 | 0 | 0.244650542 | 0 | 0 |
| ENSMUSG00000085154 | 0.960252568 | 0.92510993 | 1.061981878 | 0.75 | 0.955822902 | 0.77815125 | 0.935079929 |
| ENSMUSG00000085155 | 0 | 0 | 0 | 0 | 0 | 0 | 0 |
| ENSMUSG00000085156 | 1.623015126 | 1.557204673 | 1.660964047 | 1.418106335 | 1.343584837 | 1.380211242 | 1.405288179 |
| ENSMUSG00000085157 | 0 | 0 | 0 | 0 | 0 | 0 | 0 |
| ENSMUSG00000085158 | 0 | 0 | 0 | 0 | 0 | 0 | 0 |
| ENSMUSG00000085160 | 0 | 0 | 0 | 0 | 0 | 0 | 0 |
| ENSMUSG00000085161 | 0 | 0 | 0.25 | 0 | 0 | 0 | 0 |
| ENSMUSG00000085162 | 0.289064826 | 0.25 | 0.580482024 | 0.5 | 0.489301084 | 0.698970004 | 0.224243824 |
| ENSMUSG00000085165 | 0 | 0 | 0.5 | 0.25 | 0 | 0.477121255 | 0.448487648 |

| | | | | | | | |
|---|---|---|---|---|---|---|---|
| ENSMUSG00000085166 | 0 | 0 | 0 | 0 | 0 | 0 | 0 |
| ENSMUSG00000085167 | 0 | 0 | 0 | 0 | 0 | 0 | 0 |
| ENSMUSG00000085168 | 0 | 0 | 0 | 0 | 0 | 0 | 0 |
| ENSMUSG00000085169 | 0.578129653 | 0.5 | 0.75 | 0.830482024 | 0.686820904 | 0.954242509 | 0.775756176 |
| ENSMUSG00000085170 | 0 | 0.25 | 0 | 0.396240625 | 0 | 0 | 0 |
| ENSMUSG00000085171 | 0 | 0 | 0 | 0 | 0 | 0 | 0 |
| ENSMUSG00000085172 | 0.578129653 | 0.5 | 0.25 | 0.396240625 | 0.632412477 | 0.84509804 | 0.579661877 |
| ENSMUSG00000085173 | 0 | 0 | 0 | 0 | 0 | 0 | 0 |
| ENSMUSG00000085174 | 0.671187741 | 0.701838731 | 0.396240625 | 0.580482024 | 0.387761935 | 0.301029996 | 0.520678036 |
| ENSMUSG00000085175 | 0.811507563 | 0.92510993 | 0.864857905 | 0.75 | 0.686820904 | 0.77815125 | 0.520678036 |
| ENSMUSG00000085176 | 0 | 0 | 0 | 0 | 0 | 0 | 0 |
| ENSMUSG00000085177 | 0 | 0 | 0 | 0 | 0 | 0 | 0 |
| ENSMUSG00000085178 | 0.45815691 | 0.5 | 0.646240625 | 0.5 | 0.686820904 | 0.301029996 | 0.224243824 |
| ENSMUSG00000085179 | 0 | 0 | 0 | 0 | 0 | 0 | 0 |
| ENSMUSG00000085180 | 0.91631382 | 0.92510993 | 0.951838731 | 0.864857905 | 0.568060967 | 0.77815125 | 0.579661877 |
| ENSMUSG00000085181 | 0 | 0 | 0 | 0 | 0 | 0 | 0 |
| ENSMUSG00000085182 | 0 | 0 | 0 | 0 | 0 | 0 | 0 |
| ENSMUSG00000085183 | 0.289064826 | 0.5 | 0.396240625 | 0 | 0 | 0 | 0 |
| ENSMUSG00000085184 | 0 | 0.25 | 0 | 0.25 | 0.244650542 | 0 | 0.448487648 |
| ENSMUSG00000085185 | 0.289064826 | 0.25 | 0.580482024 | 0.646240625 | 0.244650542 | 0.477121255 | 0.448487648 |
| ENSMUSG00000085186 | 0 | 0 | 0.5 | 0 | 0 | 0.301029996 | 0 |
| ENSMUSG00000085187 | 0 | 0 | 0 | 0 | 0 | 0 | 0 |
| ENSMUSG00000085188 | 0 | 0 | 0 | 0 | 0 | 0 | 0 |
| ENSMUSG00000085189 | 0 | 0 | 0 | 0 | 0 | 0 | 0 |
| ENSMUSG00000085192 | 0 | 0 | 0 | 0 | 0 | 0 | 0.224243824 |
| ENSMUSG00000085193 | 0 | 0 | 0 | 0 | 0 | 0 | 0 |
| ENSMUSG00000085194 | 0 | 0 | 0 | 0 | 0 | 0 | 0 |
| ENSMUSG00000085196 | 0.671187741 | 0.701838731 | 0.701838731 | 0.701838731 | 0.812711509 | 0.602059991 | 0.448487648 |
| ENSMUSG00000085197 | 0.45815691 | 0.396240625 | 0 | 0 | 0 | 0 | 0 |
| ENSMUSG00000085198 | 0 | 0 | 0 | 0 | 0 | 0 | 0 |
| ENSMUSG00000085199 | 0 | 0 | 0 | 0 | 0 | 0 | 0 |
| ENSMUSG00000085201 | 0.45815691 | 0.25 | 0 | 0 | 0 | 0 | 0 |
| ENSMUSG00000085202 | 0 | 0 | 0 | 0 | 0 | 0 | 0 |
| ENSMUSG00000085203 | 0 | 0 | 0 | 0 | 0 | 0 | 0 |

| | | | | | | | |
|---|---|---|---|---|---|---|---|
| ENSMUSG00000085204 | 0 | 0 | 0 | 0 | 0 | 0 | 0 |
| ENSMUSG00000085205 | 0 | 0 | 0.25 | 0 | 0 | 0 | 0 |
| ENSMUSG00000085206 | 0 | 0 | 0 | 0 | 0 | 0.301029996 | 0 |
| ENSMUSG00000085207 | 0.45815691 | 0 | 0.396240625 | 0.580482024 | 0.244650542 | 0.602059991 | 0.579661877 |
| ENSMUSG00000085208 | 2.958871665 | 2.698400827 | 2.689972046 | 2.741986676 | 2.652009692 | 3.085290578 | 2.45362687 |
| ENSMUSG00000085209 | 0 | 0 | 0 | 0 | 0 | 0 | 0 |
| ENSMUSG00000085211 | 0 | 0.25 | 0 | 0.25 | 0 | 0 | 0.355418052 |
| ENSMUSG00000085213 | 0.578129653 | 0.79248125 | 0.79248125 | 0.646240625 | 0.568060967 | 0.477121255 | 0.579661877 |
| ENSMUSG00000085214 | 0 | 0 | 0.5 | 0.580482024 | 0 | 0.602059991 | 0 |
| ENSMUSG00000085217 | 0 | 0 | 0 | 0 | 0 | 0 | 0 |
| ENSMUSG00000085218 | 0.747221736 | 0.75 | 0.646240625 | 0.396240625 | 0.387761935 | 0 | 0.448487648 |
| ENSMUSG00000085221 | 0 | 0 | 0 | 0 | 0 | 0 | 0 |
| ENSMUSG00000085222 | 0 | 0 | 0 | 0 | 0 | 0 | 0 |
| ENSMUSG00000085223 | 0 | 0 | 0 | 0 | 0 | 0 | 0 |
| ENSMUSG00000085224 | 0 | 0 | 0 | 0 | 0 | 0.301029996 | 0.355418052 |
| ENSMUSG00000085225 | 0 | 0 | 0 | 0 | 0 | 0 | 0 |
| ENSMUSG00000085226 | 0 | 0 | 0 | 0 | 0 | 0 | 0 |
| ENSMUSG00000085227 | 0.578129653 | 0.580482024 | 0.580482024 | 0.830482024 | 0.812711509 | 0.77815125 | 0.829800754 |
| ENSMUSG00000085228 | 0.289064826 | 0.396240625 | 0.396240625 | 0 | 0 | 0 | 0.224243824 |
| ENSMUSG00000085229 | 0 | 0 | 0.25 | 0 | 0 | 0 | 0.224243824 |
| ENSMUSG00000085230 | 0 | 0 | 0 | 0 | 0 | 0 | 0 |
| ENSMUSG00000085232 | 0 | 0 | 0 | 0 | 0 | 0 | 0 |
| ENSMUSG00000085234 | 0.671187741 | 0.701838731 | 0.75 | 0.701838731 | 0.733951626 | 0.77815125 | 0.876096088 |
| ENSMUSG00000085235 | 0 | 0 | 0 | 0 | 0 | 0 | 0 |
| ENSMUSG00000085236 | 0.289064826 | 0 | 0 | 0.25 | 0 | 0 | 0 |
| ENSMUSG00000085237 | 0.289064826 | 0 | 0.396240625 | 0 | 0 | 0 | 0 |
| ENSMUSG00000085238 | 0 | 0 | 0 | 0 | 0 | 0 | 0 |
| ENSMUSG00000085239 | 0 | 0 | 0 | 0 | 0 | 0 | 0 |
| ENSMUSG00000085240 | 0 | 0 | 0 | 0 | 0 | 0 | 0 |
| ENSMUSG00000085241 | 0 | 0 | 0 | 0.25 | 0 | 0 | 0 |
| ENSMUSG00000085242 | 0 | 0.25 | 0.25 | 0.25 | 0.387761935 | 0.602059991 | 0 |
| ENSMUSG00000085244 | 0 | 0 | 0 | 0 | 0 | 0 | 0 |
| ENSMUSG00000085245 | 0 | 0 | 0 | 0 | 0 | 0 | 0 |
| ENSMUSG00000085246 | 0.45815691 | 0.396240625 | 0.396240625 | 0 | 0.244650542 | 0 | 0.224243824 |

| | | | | | | | |
|---|---|---|---|---|---|---|---|
| ENSMUSG00000085247 | 0 | 0 | 0 | 0 | 0 | 0 | 0 |
| ENSMUSG00000085248 | 0 | 0 | 0 | 0 | 0 | 0 | 0 |
| ENSMUSG00000085249 | 0 | 0 | 0 | 0 | 0 | 0 | 0 |
| ENSMUSG00000085250 | 0.289064826 | 0 | 0.396240625 | 0 | 0.244650542 | 0 | 0.224243824 |
| ENSMUSG00000085251 | 0 | 0 | 0 | 0 | 0 | 0 | 0 |
| ENSMUSG00000085252 | 0 | 0 | 0 | 0 | 0 | 0 | 0 |
| ENSMUSG00000085253 | 0 | 0 | 0 | 0 | 0 | 0 | 0 |
| ENSMUSG00000085254 | 0 | 0 | 0 | 0 | 0 | 0 | 0 |
| ENSMUSG00000085255 | 0.578129653 | 0 | 0.25 | 0.25 | 0.568060967 | 0.477121255 | 0.224243824 |
| ENSMUSG00000085256 | 0 | 0 | 0.396240625 | 0 | 0.489301084 | 0 | 0.448487648 |
| ENSMUSG00000085257 | 0 | 0 | 0 | 0 | 0 | 0 | 0 |
| ENSMUSG00000085260 | 0.671187741 | 0.396240625 | 0.396240625 | 0.5 | 0.686820904 | 0.477121255 | 0.775756176 |
| ENSMUSG00000085261 | 0 | 0.25 | 0.396240625 | 0 | 0 | 0 | 0.224243824 |
| ENSMUSG00000085262 | 0 | 0.25 | 0 | 0 | 0 | 0 | 0 |
| ENSMUSG00000085263 | 0 | 0 | 0 | 0 | 0 | 0 | 0 |
| ENSMUSG00000085264 | 0 | 0 | 0 | 0 | 0.244650542 | 0 | 0 |
| ENSMUSG00000085265 | 0 | 0 | 0 | 0 | 0 | 0 | 0 |
| ENSMUSG00000085267 | 0 | 0 | 0 | 0 | 0 | 0 | 0 |
| ENSMUSG00000085269 | 0.811507563 | 1.188721876 | 1.061981878 | 0.864857905 | 0.812711509 | 0.77815125 | 1.028149525 |
| ENSMUSG00000085271 | 0 | 0 | 0 | 0 | 0 | 0 | 0 |
| ENSMUSG00000085272 | 0 | 0 | 0 | 0.5 | 0 | 0.301029996 | 0.355418052 |
| ENSMUSG00000085274 | 0 | 0 | 0 | 0 | 0.387761935 | 0.301029996 | 0.224243824 |
| ENSMUSG00000085275 | 0.578129653 | 0 | 0 | 0 | 0 | 0 | 0 |
| ENSMUSG00000085276 | 0.45815691 | 0 | 0.25 | 0 | 0.244650542 | 0.301029996 | 0.224243824 |
| ENSMUSG00000085277 | 0 | 0 | 0 | 0 | 0 | 0 | 0 |
| ENSMUSG00000085278 | 0 | 0 | 0 | 0 | 0 | 0 | 0 |
| ENSMUSG00000085279 | 1.470606563 | 1.356566189 | 1.431980114 | 1.388647213 | 1.462344774 | 1.531478917 | 1.329931221 |
| ENSMUSG00000085280 | 0 | 0 | 0 | 0 | 0 | 0 | 0 |
| ENSMUSG00000085281 | 0 | 0 | 0 | 0 | 0 | 0 | 0 |
| ENSMUSG00000085282 | 0 | 0.5 | 0.396240625 | 0.396240625 | 0.244650542 | 0 | 0.520678036 |
| ENSMUSG00000085284 | 0 | 0 | 0 | 0 | 0 | 0 | 0 |
| ENSMUSG00000085285 | 0 | 0 | 0 | 0 | 0 | 0 | 0 |
| ENSMUSG00000085286 | 0.45815691 | 0.580482024 | 0.646240625 | 0.580482024 | 0.568060967 | 0 | 0.629532004 |
| ENSMUSG00000085287 | 1 | 1.080482024 | 0.976722649 | 1 | 0.905314583 | 0.954242509 | 0.775756176 |

| | | | | | | | |
|---|---|---|---|---|---|---|---|
| ENSMUSG00000085288 | 0 | 0 | 0 | 0 | 0 | 0 | 0 |
| ENSMUSG00000085289 | 0.671187741 | 0.830482024 | 1.130890489 | 0.951838731 | 0.812711509 | 0.477121255 | 0.672731473 |
| ENSMUSG00000085290 | 0 | 0 | 0 | 0 | 0 | 0 | 0 |
| ENSMUSG00000085292 | 0 | 0 | 0 | 0 | 0 | 0 | 0 |
| ENSMUSG00000085293 | 0 | 0 | 0 | 0 | 0 | 0 | 0 |
| ENSMUSG00000085295 | 0.671187741 | 0.646240625 | 0.25 | 0.5 | 0.244650542 | 0.477121255 | 0 |
| ENSMUSG00000085297 | 0 | 0 | 0 | 0 | 0 | 0 | 0 |
| ENSMUSG00000085298 | 0 | 0 | 0.25 | 0 | 0 | 0 | 0.224243824 |
| ENSMUSG00000085300 | 0 | 0 | 0 | 0 | 0 | 0 | 0 |
| ENSMUSG00000085301 | 0 | 0 | 0 | 0 | 0 | 0 | 0 |
| ENSMUSG00000085302 | 0 | 0 | 0 | 0 | 0 | 0 | 0 |
| ENSMUSG00000085303 | 0 | 0 | 0 | 0 | 0 | 0 | 0 |
| ENSMUSG00000085304 | 0 | 0 | 0 | 0 | 0 | 0 | 0 |
| ENSMUSG00000085305 | 0 | 0 | 0 | 0 | 0 | 0 | 0 |
| ENSMUSG00000085307 | 0.289064826 | 0.951838731 | 0.701838731 | 0.830482024 | 0.489301084 | 0.602059991 | 0.710836105 |
| ENSMUSG00000085310 | 0.671187741 | 0 | 0 | 0.5 | 0 | 0.477121255 | 0 |
| ENSMUSG00000085311 | 0.671187741 | 0.25 | 0.646240625 | 0.5 | 0.733951626 | 0.301029996 | 0.520678036 |
| ENSMUSG00000085312 | 0 | 0.25 | 0.396240625 | 0.25 | 0 | 0 | 0 |
| ENSMUSG00000085314 | 0 | 0 | 0 | 0 | 0 | 0 | 0 |
| ENSMUSG00000085315 | 0.578129653 | 0.5 | 0 | 0.25 | 0.244650542 | 0 | 0.355418052 |
| ENSMUSG00000085316 | 0 | 0 | 0 | 0 | 0 | 0.301029996 | 0 |
| ENSMUSG00000085317 | 0 | 0 | 0 | 0 | 0 | 0 | 0 |
| ENSMUSG00000085318 | 0 | 0 | 0 | 0 | 0 | 0 | 0 |
| ENSMUSG00000085319 | 0 | 0 | 0 | 0 | 0 | 0 | 0 |
| ENSMUSG00000085320 | 0 | 0 | 0 | 0 | 0 | 0 | 0 |
| ENSMUSG00000085321 | 0 | 0 | 0.25 | 0.25 | 0 | 0 | 0 |
| ENSMUSG00000085322 | 0 | 0 | 0 | 0 | 0 | 0 | 0 |
| ENSMUSG00000085323 | 0 | 0 | 0 | 0 | 0 | 0 | 0 |
| ENSMUSG00000085324 | 0 | 0 | 0 | 0 | 0 | 0 | 0 |
| ENSMUSG00000085325 | 0 | 0 | 0 | 0 | 0 | 0 | 0 |
| ENSMUSG00000085326 | 0 | 0 | 0 | 0 | 0 | 0 | 0 |
| ENSMUSG00000085327 | 0.289064826 | 0 | 0.396240625 | 0 | 0 | 0 | 0.224243824 |
| ENSMUSG00000085328 | 1.568542279 | 1.557204673 | 1.470660762 | 1.438721876 | 1.244650542 | 1.462397998 | 1.284450788 |
| ENSMUSG00000085329 | 0 | 0 | 0 | 0 | 0 | 0 | 0 |

| | | | | | | | |
|---|---|---|---|---|---|---|---|
| ENSMUSG00000085330 | 0.289064826 | 0.396240625 | 0.25 | 0 | 0 | 0 | 0 |
| ENSMUSG00000085331 | 0 | 0 | 0 | 0 | 0 | 0 | 0 |
| ENSMUSG00000085333 | 0 | 0 | 0 | 0 | 0 | 0 | 0 |
| ENSMUSG00000085334 | 2.095120908 | 1.958222504 | 1.872963274 | 1.912762923 | 1.850106709 | 2.012837225 | 1.680001789 |
| ENSMUSG00000085335 | 0.289064826 | 0.25 | 0.5 | 0.396240625 | 0.387761935 | 0 | 0.355418052 |
| ENSMUSG00000085336 | 0 | 0 | 0 | 0 | 0 | 0 | 0 |
| ENSMUSG00000085337 | 0.671187741 | 0.75 | 0.646240625 | 0.75 | 0.489301084 | 0.301029996 | 0.672731473 |
| ENSMUSG00000085338 | 0 | 0 | 0 | 0 | 0 | 0 | 0 |
| ENSMUSG00000085340 | 0.289064826 | 0 | 0.25 | 0.25 | 0.632412477 | 0 | 0.224243824 |
| ENSMUSG00000085342 | 0 | 0 | 0 | 0 | 0.244650542 | 0 | 0.355418052 |
| ENSMUSG00000085345 | 0 | 0 | 0 | 0 | 0 | 0 | 0 |
| ENSMUSG00000085347 | 0.671187741 | 0.646240625 | 0.646240625 | 0.701838731 | 0.387761935 | 1.041392685 | 0.829800754 |
| ENSMUSG00000085348 | 0 | 0 | 0 | 0 | 0 | 0 | 0 |
| ENSMUSG00000085350 | 0.289064826 | 0 | 0 | 0 | 0 | 0 | 0.224243824 |
| ENSMUSG00000085351 | 0 | 0 | 0 | 0 | 0.387761935 | 0 | 0.224243824 |
| ENSMUSG00000085352 | 0 | 0 | 0.25 | 0 | 0.244650542 | 0 | 0 |
| ENSMUSG00000085353 | 0 | 0 | 0 | 0 | 0 | 0 | 0 |
| ENSMUSG00000085354 | 0.289064826 | 0 | 0.25 | 0 | 0.244650542 | 0 | 0 |
| ENSMUSG00000085355 | 0.578129653 | 0.646240625 | 0.5 | 0.25 | 0 | 0.477121255 | 0.448487648 |
| ENSMUSG00000085357 | 0 | 0 | 0 | 0 | 0 | 0 | 0 |
| ENSMUSG00000085358 | 0 | 0 | 0 | 0 | 0 | 0 | 0 |
| ENSMUSG00000085359 | 0 | 0 | 0.25 | 0 | 0 | 0 | 0 |
| ENSMUSG00000085360 | 0 | 0 | 0 | 0 | 0 | 0 | 0 |
| ENSMUSG00000085361 | 0 | 0 | 0.5 | 0 | 0.387761935 | 0 | 0.355418052 |
| ENSMUSG00000085363 | 0.45815691 | 0.25 | 0.25 | 0.25 | 0.489301084 | 0 | 0 |
| ENSMUSG00000085366 | 0 | 0 | 0 | 0 | 0 | 0 | 0 |
| ENSMUSG00000085368 | 0 | 0 | 0 | 0 | 0 | 0 | 0 |
| ENSMUSG00000085369 | 0 | 0 | 0 | 0 | 0 | 0 | 0 |
| ENSMUSG00000085371 | 0 | 0 | 0 | 0 | 0 | 0 | 0 |
| ENSMUSG00000085372 | 0 | 0 | 0 | 0 | 0 | 0 | 0 |
| ENSMUSG00000085375 | 0 | 0 | 0 | 0 | 0 | 0 | 0 |
| ENSMUSG00000085376 | 0.91631382 | 0.830482024 | 0.864857905 | 0.646240625 | 0.877063019 | 0.301029996 | 0.935079929 |
| ENSMUSG00000085377 | 0 | 0 | 0 | 0 | 0 | 0 | 0 |
| ENSMUSG00000085378 | 0 | 0 | 0 | 0 | 0 | 0 | 0 |

| | | | | | | | |
|---|---|---|---|---|---|---|---|
| ENSMUSG00000085379 | 0.289064826 | 0.79248125 | 0.701838731 | 0.896240625 | 0.686820904 | 0.698970004 | 0.775756176 |
| ENSMUSG00000085380 | 0 | 0 | 0 | 0 | 0 | 0 | 0 |
| ENSMUSG00000085382 | 0 | 0 | 0 | 0 | 0 | 0 | 0 |
| ENSMUSG00000085383 | 0 | 0 | 0 | 0 | 0 | 0 | 0 |
| ENSMUSG00000085385 | 2.22592627 | 1.994319981 | 1.998588359 | 2.142463902 | 2.146756411 | 2.315970345 | 2.003985867 |
| ENSMUSG00000085386 | 0 | 0 | 0 | 0 | 0 | 0 | 0 |
| ENSMUSG00000085387 | 0 | 0 | 0 | 0 | 0 | 0 | 0 |
| ENSMUSG00000085388 | 0 | 0 | 0 | 0 | 0 | 0 | 0 |
| ENSMUSG00000085389 | 0 | 0.25 | 0 | 0.396240625 | 0.244650542 | 0.477121255 | 0.355418052 |
| ENSMUSG00000085390 | 0 | 0 | 0 | 0 | 0 | 0 | 0 |
| ENSMUSG00000085391 | 0 | 0 | 0 | 0 | 0 | 0 | 0 |
| ENSMUSG00000085393 | 0 | 0 | 0 | 0 | 0 | 0 | 0 |
| ENSMUSG00000085394 | 0 | 0 | 0 | 0 | 0 | 0 | 0 |
| ENSMUSG00000085395 | 1 | 1.226722649 | 1.080482024 | 0.830482024 | 1.039257769 | 1.113943352 | 1.110947927 |
| ENSMUSG00000085396 | 2.170239354 | 1.94697564 | 2.052363341 | 1.926839783 | 1.998776581 | 2.120573931 | 1.805128823 |
| ENSMUSG00000085397 | 0.45815691 | 0 | 0 | 0.25 | 0 | 0.301029996 | 0.224243824 |
| ENSMUSG00000085398 | 0 | 0 | 0 | 0 | 0 | 0 | 0 |
| ENSMUSG00000085399 | 0.867194479 | 1.02186571 | 1 | 1.330482024 | 1.234113756 | 1.176091259 | 1.150210039 |
| ENSMUSG00000085400 | 0 | 0 | 0 | 0 | 0 | 0 | 0 |
| ENSMUSG00000085401 | 0 | 0.25 | 0 | 0 | 0 | 0 | 0.224243824 |
| ENSMUSG00000085402 | 0 | 0 | 0 | 0.5 | 0 | 0 | 0 |
| ENSMUSG00000085403 | 0 | 0.580482024 | 0.646240625 | 0.5 | 0.568060967 | 0 | 0.579661877 |
| ENSMUSG00000085404 | 0 | 0 | 0 | 0.25 | 0 | 0 | 0 |
| ENSMUSG00000085405 | 0 | 0 | 0 | 0 | 0 | 0 | 0 |
| ENSMUSG00000085407 | 0 | 0.5 | 0 | 0.25 | 0.244650542 | 0 | 0 |
| ENSMUSG00000085408 | 0.45815691 | 0.646240625 | 0.25 | 0.396240625 | 0.568060967 | 0.301029996 | 0.710836105 |
| ENSMUSG00000085409 | 0 | 0 | 0 | 0 | 0 | 0 | 0 |
| ENSMUSG00000085411 | 0.45815691 | 0 | 0 | 0 | 0 | 0 | 0 |
| ENSMUSG00000085412 | 0 | 0.580482024 | 0.580482024 | 0 | 0.244650542 | 0 | 0.224243824 |
| ENSMUSG00000085413 | 0 | 0 | 0 | 0 | 0 | 0 | 0 |
| ENSMUSG00000085414 | 0 | 0 | 0.25 | 0 | 0.244650542 | 0 | 0 |
| ENSMUSG00000085415 | 0 | 0.580482024 | 0.396240625 | 0.646240625 | 0.489301084 | 0 | 0.448487648 |
| ENSMUSG00000085416 | 0 | 0 | 0 | 0 | 0 | 0 | 0 |
| ENSMUSG00000085417 | 0 | 0 | 0 | 0 | 0 | 0 | 0 |

| | | | | | | | |
|---|---|---|---|---|---|---|---|
| ENSMUSG00000085418 | 0 | 0 | 0 | 0 | 0 | 0 | 0 |
| ENSMUSG00000085419 | 0 | 0 | 0 | 0 | 0.244650542 | 0.301029996 | 0 |
| ENSMUSG00000085420 | 0 | 0 | 0 | 0 | 0 | 0 | 0 |
| ENSMUSG00000085421 | 0.578129653 | 0 | 0.25 | 0 | 0 | 0 | 0 |
| ENSMUSG00000085423 | 0 | 0 | 0 | 0.25 | 0.244650542 | 0 | 0 |
| ENSMUSG00000085424 | 0 | 0 | 0 | 0 | 0 | 0 | 0 |
| ENSMUSG00000085425 | 0 | 0 | 0 | 0 | 0 | 0 | 0 |
| ENSMUSG00000085426 | 0 | 0 | 0 | 0 | 0 | 0 | 0 |
| ENSMUSG00000085427 | 0 | 0 | 0 | 0 | 0 | 0 | 0 |
| ENSMUSG00000085428 | 0 | 0 | 0 | 0 | 0 | 0 | 0 |
| ENSMUSG00000085429 | 0 | 0.25 | 0 | 0.25 | 0 | 0.301029996 | 0 |
| ENSMUSG00000085430 | 0 | 0 | 0 | 0 | 0 | 0 | 0 |
| ENSMUSG00000085431 | 0 | 0 | 0 | 0 | 0 | 0 | 0 |
| ENSMUSG00000085433 | 1.227926429 | 1.04248125 | 0.864857905 | 1.02186571 | 1.121713561 | 0.903089987 | 1.176815335 |
| ENSMUSG00000085434 | 0.289064826 | 0 | 0.25 | 0 | 0 | 0 | 0.224243824 |
| ENSMUSG00000085435 | 0 | 0 | 0 | 0 | 0 | 0 | 0 |
| ENSMUSG00000085436 | 1.036286563 | 1.02186571 | 1.080482024 | 1.438721876 | 1.366364104 | 1.361727836 | 1.379042812 |
| ENSMUSG00000085437 | 0 | 0.25 | 0 | 0 | 0 | 0 | 0 |
| ENSMUSG00000085438 | 2.692026894 | 2.524836953 | 2.521202097 | 2.82991803 | 2.737747706 | 3.152899596 | 2.523870648 |
| ENSMUSG00000085439 | 0 | 0 | 0 | 0 | 0 | 0 | 0 |
| ENSMUSG00000085440 | 0.578129653 | 0.5 | 0.396240625 | 0 | 0 | 0 | 0.224243824 |
| ENSMUSG00000085441 | 0 | 0 | 0 | 0 | 0 | 0 | 0 |
| ENSMUSG00000085442 | 0.289064826 | 0.580482024 | 0.5 | 0.5 | 0.632412477 | 0.301029996 | 0.579661877 |
| ENSMUSG00000085443 | 0 | 0.25 | 0 | 0 | 0 | 0 | 0 |
| ENSMUSG00000085444 | 0.289064826 | 0.580482024 | 0.646240625 | 0.5 | 0.387761935 | 0 | 0.224243824 |
| ENSMUSG00000085445 | 0 | 0 | 0 | 0 | 0 | 0 | 0 |
| ENSMUSG00000085447 | 0 | 0 | 0 | 0.25 | 0 | 0 | 0 |
| ENSMUSG00000085448 | 0 | 0 | 0 | 0 | 0 | 0 | 0.224243824 |
| ENSMUSG00000085449 | 0 | 0 | 0 | 0 | 0 | 0 | 0 |
| ENSMUSG00000085450 | 0 | 0 | 0 | 0 | 0 | 0 | 0 |
| ENSMUSG00000085451 | 0 | 0.25 | 0 | 0 | 0.244650542 | 0 | 0 |
| ENSMUSG00000085452 | 0 | 0 | 0 | 0.25 | 0 | 0 | 0 |
| ENSMUSG00000085453 | 0.45815691 | 0.75 | 0.580482024 | 0.5 | 0.489301084 | 0.602059991 | 0.579661877 |
| ENSMUSG00000085454 | 0 | 0 | 0.25 | 0 | 0 | 0 | 0 |

| | | | | | | | |
|---|---|---|---|---|---|---|---|
| ENSMUSG00000085456 | 0 | 0 | 0 | 0 | 0 | 0 | 0 |
| ENSMUSG00000085457 | 1.494443473 | 1.431980114 | 1.464495249 | 1.445339928 | 1.234113756 | 1.301029996 | 1.110947927 |
| ENSMUSG00000085459 | 0.289064826 | 0 | 0.25 | 0 | 0 | 0.301029996 | 0.224243824 |
| ENSMUSG00000085463 | 0 | 0 | 0 | 0 | 0 | 0 | 0 |
| ENSMUSG00000085464 | 0 | 0 | 0 | 0 | 0 | 0 | 0 |
| ENSMUSG00000085465 | 0.811507563 | 1 | 0.701838731 | 0.396240625 | 0.812711509 | 0.698970004 | 0.935079929 |
| ENSMUSG00000085466 | 0 | 0 | 0 | 0 | 0 | 0 | 0 |
| ENSMUSG00000085467 | 0 | 0 | 0 | 0 | 0 | 0 | 0 |
| ENSMUSG00000085468 | 0 | 0 | 0 | 0 | 0 | 0 | 0 |
| ENSMUSG00000085469 | 0 | 0 | 0 | 0 | 0 | 0 | 0 |
| ENSMUSG00000085470 | 0 | 0 | 0 | 0 | 0 | 0 | 0 |
| ENSMUSG00000085471 | 0 | 0 | 0.25 | 0 | 0 | 0 | 0 |
| ENSMUSG00000085472 | 0 | 0 | 0.25 | 0 | 0 | 0 | 0 |
| ENSMUSG00000085473 | 0.289064826 | 0 | 0 | 0.25 | 0 | 0 | 0 |
| ENSMUSG00000085476 | 0 | 0 | 0 | 0 | 0 | 0 | 0 |
| ENSMUSG00000085477 | 0 | 0 | 0 | 0 | 0 | 0 | 0 |
| ENSMUSG00000085478 | 0.289064826 | 0.25 | 0.5 | 0.25 | 0 | 0.477121255 | 0.355418052 |
| ENSMUSG00000085479 | 0 | 0 | 0 | 0 | 0 | 0.301029996 | 0 |
| ENSMUSG00000085481 | 0 | 0 | 0 | 0 | 0 | 0 | 0 |
| ENSMUSG00000085482 | 0.45815691 | 0.5 | 0.646240625 | 0.5 | 0.489301084 | 0.301029996 | 0.224243824 |
| ENSMUSG00000085483 | 0.289064826 | 0 | 0 | 0 | 0 | 0 | 0 |
| ENSMUSG00000085484 | 0 | 0 | 0 | 0 | 0 | 0 | 0 |
| ENSMUSG00000085485 | 0 | 0 | 0 | 0 | 0 | 0 | 0 |
| ENSMUSG00000085486 | 0.289064826 | 0 | 0.25 | 0 | 0 | 0 | 0 |
| ENSMUSG00000085487 | 0 | 0.25 | 0.25 | 0.25 | 0 | 0.301029996 | 0 |
| ENSMUSG00000085488 | 0 | 0 | 0 | 0 | 0 | 0 | 0.224243824 |
| ENSMUSG00000085489 | 0 | 0 | 0 | 0 | 0 | 0 | 0 |
| ENSMUSG00000085491 | 0 | 0 | 0 | 0 | 0 | 0 | 0 |
| ENSMUSG00000085492 | 0.747221736 | 1.188721876 | 1.226722649 | 1.130890489 | 0.905314583 | 0.698970004 | 1.131174228 |
| ENSMUSG00000085495 | 0 | 0 | 0.25 | 0.396240625 | 0.244650542 | 0 | 0.355418052 |
| ENSMUSG00000085496 | 0 | 0 | 0 | 0 | 0 | 0 | 0 |
| ENSMUSG00000085497 | 0 | 0 | 0 | 0 | 0 | 0 | 0 |
| ENSMUSG00000085498 | 0 | 0.25 | 0.25 | 0 | 0 | 0.477121255 | 0.224243824 |
| ENSMUSG00000085499 | 0 | 0 | 0 | 0 | 0 | 0 | 0 |

| | | | | | | | |
|---|---|---|---|---|---|---|---|
| ENSMUSG00000085500 | 0 | 0 | 0 | 0 | 0 | 0 | 0 |
| ENSMUSG00000085501 | 0.289064826 | 0.5 | 0.580482024 | 0.25 | 0.733951626 | 0.477121255 | 0.629532004 |
| ENSMUSG00000085502 | 0 | 0 | 0 | 0 | 0 | 0 | 0 |
| ENSMUSG00000085504 | 0 | 0 | 0 | 0 | 0 | 0 | 0 |
| ENSMUSG00000085505 | 0 | 0 | 0 | 0 | 0 | 0 | 0 |
| ENSMUSG00000085507 | 0 | 0 | 0 | 0 | 0 | 0 | 0 |
| ENSMUSG00000085508 | 0 | 0.25 | 0 | 0 | 0 | 0 | 0 |
| ENSMUSG00000085509 | 0 | 0 | 0 | 0 | 0 | 0 | 0 |
| ENSMUSG00000085510 | 0 | 0 | 0 | 0 | 0 | 0 | 0 |
| ENSMUSG00000085511 | 0 | 0 | 0 | 0 | 0.387761935 | 0 | 0.355418052 |
| ENSMUSG00000085512 | 0 | 0 | 0 | 0 | 0 | 0 | 0 |
| ENSMUSG00000085513 | 0 | 0 | 0 | 0 | 0 | 0 | 0 |
| ENSMUSG00000085514 | 0 | 0 | 0 | 0 | 0 | 0 | 0 |
| ENSMUSG00000085515 | 0 | 0 | 0 | 0 | 0 | 0 | 0 |
| ENSMUSG00000085516 | 0 | 0 | 0 | 0 | 0 | 0 | 0 |
| ENSMUSG00000085517 | 0.289064826 | 0.396240625 | 0 | 0.580482024 | 0.568060967 | 0.301029996 | 0.448487648 |
| ENSMUSG00000085519 | 0 | 0 | 0 | 0 | 0 | 0 | 0 |
| ENSMUSG00000085521 | 0 | 0.5 | 0 | 0.25 | 0.387761935 | 0 | 0.520678036 |
| ENSMUSG00000085522 | 0.289064826 | 0.396240625 | 0.396240625 | 0 | 0 | 0.301029996 | 0 |
| ENSMUSG00000085523 | 0 | 0 | 0.25 | 0 | 0 | 0 | 0 |
| ENSMUSG00000085524 | 0 | 0 | 0 | 0 | 0 | 0 | 0 |
| ENSMUSG00000085525 | 0.578129653 | 0.25 | 0.25 | 0.396240625 | 0.568060967 | 0.301029996 | 0.448487648 |
| ENSMUSG00000085526 | 0 | 0 | 0 | 0 | 0 | 0 | 0 |
| ENSMUSG00000085527 | 0.578129653 | 0.701838731 | 0.830482024 | 0.830482024 | 0.931471446 | 0.77815125 | 0.775756176 |
| ENSMUSG00000085528 | 0 | 0 | 0 | 0 | 0 | 0 | 0 |
| ENSMUSG00000085530 | 0 | 0 | 0 | 0 | 0 | 0 | 0 |
| ENSMUSG00000085531 | 0 | 0 | 0 | 0 | 0 | 0 | 0 |
| ENSMUSG00000085532 | 0.289064826 | 0 | 0.396240625 | 0 | 0 | 0 | 0.355418052 |
| ENSMUSG00000085533 | 0 | 0 | 0 | 0 | 0 | 0 | 0 |
| ENSMUSG00000085534 | 0 | 0 | 0 | 0 | 0 | 0 | 0 |
| ENSMUSG00000085536 | 0 | 0 | 0 | 0 | 0 | 0 | 0 |
| ENSMUSG00000085538 | 0.289064826 | 0 | 0 | 0 | 0 | 0.301029996 | 0.224243824 |
| ENSMUSG00000085539 | 0.289064826 | 0.25 | 0.25 | 0 | 0 | 0 | 0 |
| ENSMUSG00000085540 | 0 | 0 | 0 | 0 | 0 | 0 | 0 |

| | | | | | | | |
|---|---|---|---|---|---|---|---|
| ENSMUSG00000085541 | 0 | 0 | 0 | 0 | 0 | 0 | 0 |
| ENSMUSG00000085542 | 0 | 0 | 0 | 0 | 0 | 0 | 0 |
| ENSMUSG00000085543 | 0 | 0 | 0 | 0 | 0 | 0 | 0 |
| ENSMUSG00000085545 | 0 | 0 | 0 | 0 | 0 | 0 | 0 |
| ENSMUSG00000085546 | 0.45815691 | 0.580482024 | 0.396240625 | 0.701838731 | 0.489301084 | 0 | 0.355418052 |
| ENSMUSG00000085547 | 0 | 0 | 0 | 0 | 0 | 0 | 0 |
| ENSMUSG00000085548 | 0 | 0 | 0 | 0.25 | 0 | 0 | 0 |
| ENSMUSG00000085549 | 0 | 0 | 0 | 0 | 0 | 0 | 0 |
| ENSMUSG00000085550 | 0 | 0 | 0 | 0 | 0 | 0 | 0 |
| ENSMUSG00000085551 | 0 | 0 | 0 | 0 | 0 | 0 | 0 |
| ENSMUSG00000085552 | 0 | 0 | 0 | 0 | 0 | 0 | 0 |
| ENSMUSG00000085553 | 0 | 0 | 0 | 0 | 0 | 0 | 0 |
| ENSMUSG00000085555 | 0 | 0 | 0 | 0.5 | 0.686820904 | 0.301029996 | 0.74492186 |
| ENSMUSG00000085556 | 0 | 0 | 0 | 0 | 0 | 0 | 0 |
| ENSMUSG00000085558 | 0.289064826 | 0 | 0.396240625 | 0.396240625 | 0.244650542 | 0 | 0 |
| ENSMUSG00000085559 | 0 | 0 | 0 | 0 | 0 | 0 | 0 |
| ENSMUSG00000085560 | 0 | 0.25 | 0.25 | 0 | 0 | 0 | 0 |
| ENSMUSG00000085561 | 0 | 0 | 0 | 0 | 0 | 0 | 0 |
| ENSMUSG00000085562 | 0 | 0 | 0.25 | 0 | 0 | 0 | 0 |
| ENSMUSG00000085563 | 0.747221736 | 0.396240625 | 0.580482024 | 0.701838731 | 0.489301084 | 0.602059991 | 0.579661877 |
| ENSMUSG00000085564 | 0.289064826 | 0.701838731 | 0.25 | 0 | 0 | 0 | 0 |
| ENSMUSG00000085565 | 0 | 0 | 0.25 | 0 | 0.244650542 | 0 | 0 |
| ENSMUSG00000085566 | 1.069666964 | 0.79248125 | 0.92510993 | 0.79248125 | 0.568060967 | 0.84509804 | 0.74492186 |
| ENSMUSG00000085568 | 0.289064826 | 0.25 | 0.25 | 0.396240625 | 0.244650542 | 0 | 0.224243824 |
| ENSMUSG00000085569 | 1.432083896 | 1.330482024 | 1.311981878 | 0.396240625 | 0.568060967 | 0.301029996 | 0.629532004 |
| ENSMUSG00000085572 | 0.289064826 | 0.25 | 0.396240625 | 0.25 | 0 | 0 | 0.355418052 |
| ENSMUSG00000085573 | 0.45815691 | 0.5 | 0.25 | 0.396240625 | 0.733951626 | 0.301029996 | 0.355418052 |
| ENSMUSG00000085574 | 0 | 0 | 0 | 0 | 0 | 0 | 0 |
| ENSMUSG00000085575 | 0 | 0 | 0 | 0 | 0 | 0 | 0 |
| ENSMUSG00000085576 | 0 | 0 | 0 | 0 | 0 | 0 | 0.224243824 |
| ENSMUSG00000085577 | 0 | 0 | 0 | 0 | 0 | 0 | 0 |
| ENSMUSG00000085578 | 0.289064826 | 0 | 0 | 0 | 0.244650542 | 0 | 0 |
| ENSMUSG00000085579 | 0 | 0 | 0 | 0 | 0 | 0 | 0 |
| ENSMUSG00000085580 | 0 | 0 | 0 | 0 | 0 | 0 | 0 |

| | | | | | | | |
|---|---|---|---|---|---|---|---|
| ENSMUSG00000085581 | 0.289064826 | 0.25 | 0.25 | 0.396240625 | 0.244650542 | 0.301029996 | 0.355418052 |
| ENSMUSG00000085582 | 0 | 0 | 0.25 | 0.25 | 0.244650542 | 0 | 0 |
| ENSMUSG00000085583 | 0.578129653 | 0.25 | 0.396240625 | 0.25 | 0.632412477 | 0.602059991 | 0.829800754 |
| ENSMUSG00000085584 | 0.45815691 | 0 | 0 | 0.25 | 0.244650542 | 0 | 0 |
| ENSMUSG00000085585 | 0 | 0 | 0 | 0 | 0 | 0 | 0 |
| ENSMUSG00000085586 | 1.205378646 | 1.098079356 | 1.17510993 | 1.282320754 | 1.212046813 | 1.414973348 | 1.23151414 |
| ENSMUSG00000085587 | 0 | 0 | 0 | 0 | 0 | 0 | 0 |
| ENSMUSG00000085588 | 0 | 0 | 0 | 0 | 0 | 0 | 0 |
| ENSMUSG00000085589 | 0.671187741 | 0.396240625 | 0.5 | 0 | 0 | 0.301029996 | 0 |
| ENSMUSG00000085591 | 0 | 0 | 0.25 | 0 | 0 | 0 | 0 |
| ENSMUSG00000085592 | 0 | 0 | 0 | 0 | 0 | 0 | 0 |
| ENSMUSG00000085593 | 0 | 0 | 0 | 0 | 0 | 0 | 0 |
| ENSMUSG00000085594 | 0 | 0 | 0 | 0 | 0 | 0 | 0 |
| ENSMUSG00000085595 | 0 | 0 | 0 | 0 | 0 | 0 | 0 |
| ENSMUSG00000085596 | 0.289064826 | 0.580482024 | 0 | 0.396240625 | 0 | 0 | 0 |
| ENSMUSG00000085598 | 0 | 0 | 0 | 0 | 0 | 0 | 0 |
| ENSMUSG00000085599 | 0 | 0 | 0 | 0 | 0 | 0 | 0 |
| ENSMUSG00000085600 | 0 | 0 | 0 | 0 | 0 | 0 | 0 |
| ENSMUSG00000085601 | 0 | 0 | 0 | 0 | 0 | 0 | 0 |
| ENSMUSG00000085603 | 0.45815691 | 0.396240625 | 0.25 | 0.580482024 | 0.489301084 | 0.698970004 | 0.448487648 |
| ENSMUSG00000085604 | 0 | 0.25 | 0 | 0 | 0 | 0 | 0 |
| ENSMUSG00000085606 | 0.747221736 | 0.25 | 0.580482024 | 0.580482024 | 0.812711509 | 0.602059991 | 0.355418052 |
| ENSMUSG00000085608 | 0 | 0 | 0 | 0 | 0 | 0 | 0 |
| ENSMUSG00000085609 | 0 | 0 | 0 | 0 | 0 | 0 | 0.224243824 |
| ENSMUSG00000085610 | 0.45815691 | 0.25 | 0.396240625 | 0.25 | 0 | 0 | 0.355418052 |
| ENSMUSG00000085611 | 0 | 0 | 0.25 | 0 | 0 | 0 | 0 |
| ENSMUSG00000085612 | 0.289064826 | 0 | 0.25 | 0 | 0.489301084 | 0.602059991 | 0.520678036 |
| ENSMUSG00000085614 | 0.671187741 | 0.396240625 | 0.75 | 0.75 | 0.733951626 | 0.602059991 | 0.355418052 |
| ENSMUSG00000085615 | 0.747221736 | 0.92510993 | 0.646240625 | 0.92510993 | 0.812711509 | 0.77815125 | 0.876096088 |
| ENSMUSG00000085616 | 0 | 0 | 0 | 0 | 0 | 0 | 0 |
| ENSMUSG00000085618 | 0 | 0 | 0 | 0 | 0 | 0 | 0 |
| ENSMUSG00000085619 | 0 | 0 | 0 | 0 | 0 | 0 | 0 |
| ENSMUSG00000085620 | 0 | 0.25 | 0 | 0.25 | 0 | 0 | 0 |
| ENSMUSG00000085622 | 1.289064826 | 1.160964047 | 1.061981878 | 1.214495249 | 1.212046813 | 1.176091259 | 1.110947927 |

| | | | | | | | |
|---|---|---|---|---|---|---|---|
| ENSMUSG00000085623 | 0.578129653 | 0.396240625 | 0.396240625 | 0 | 0 | 0.301029996 | 0.224243824 |
| ENSMUSG00000085624 | 0 | 0 | 0 | 0 | 0 | 0 | 0 |
| ENSMUSG00000085625 | 0 | 0 | 0 | 0 | 0 | 0 | 0 |
| ENSMUSG00000085626 | 0 | 0 | 0 | 0 | 0 | 0 | 0 |
| ENSMUSG00000085627 | 0.289064826 | 0 | 0 | 0 | 0 | 0.301029996 | 0 |
| ENSMUSG00000085628 | 0.289064826 | 0.396240625 | 0.396240625 | 0.25 | 0.244650542 | 0 | 0.224243824 |
| ENSMUSG00000085629 | 0 | 0 | 0 | 0 | 0 | 0 | 0 |
| ENSMUSG00000085630 | 0 | 0 | 0 | 0 | 0 | 0 | 0 |
| ENSMUSG00000085631 | 0 | 0 | 0 | 0 | 0 | 0 | 0 |
| ENSMUSG00000085632 | 0 | 0 | 0 | 0 | 0 | 0 | 0 |
| ENSMUSG00000085633 | 0 | 0.25 | 0.25 | 0 | 0.244650542 | 0 | 0 |
| ENSMUSG00000085634 | 0 | 0 | 0 | 0 | 0 | 0 | 0 |
| ENSMUSG00000085635 | 0.578129653 | 0.701838731 | 0.580482024 | 0.5 | 0.387761935 | 0.477121255 | 0.629532004 |
| ENSMUSG00000085636 | 0.578129653 | 0.25 | 0.25 | 0.580482024 | 0.387761935 | 0.301029996 | 0.224243824 |
| ENSMUSG00000085637 | 0.45815691 | 0.25 | 0.580482024 | 0 | 0.489301084 | 0.477121255 | 0.448487648 |
| ENSMUSG00000085638 | 0 | 0 | 0 | 0 | 0 | 0 | 0 |
| ENSMUSG00000085639 | 0.45815691 | 0.580482024 | 0.25 | 0 | 0 | 0.301029996 | 0 |
| ENSMUSG00000085640 | 0 | 0 | 0 | 0 | 0 | 0 | 0 |
| ENSMUSG00000085641 | 0 | 0 | 0 | 0 | 0 | 0 | 0 |
| ENSMUSG00000085642 | 1.181541736 | 1.27186571 | 1.080482024 | 1.02186571 | 1.136121934 | 1.204119983 | 1.066254157 |
| ENSMUSG00000085643 | 0 | 0 | 0 | 0 | 0 | 0 | 0 |
| ENSMUSG00000085645 | 0.578129653 | 0.864857905 | 0.951838731 | 0.580482024 | 0.632412477 | 0.301029996 | 0.710836105 |
| ENSMUSG00000085646 | 0 | 0 | 0 | 0 | 0 | 0 | 0 |
| ENSMUSG00000085648 | 0 | 0 | 0 | 0 | 0 | 0 | 0 |
| ENSMUSG00000085649 | 0 | 0 | 0 | 0 | 0 | 0 | 0 |
| ENSMUSG00000085651 | 0 | 0 | 0 | 0 | 0 | 0 | 0 |
| ENSMUSG00000085652 | 0 | 0 | 0 | 0 | 0 | 0 | 0 |
| ENSMUSG00000085653 | 0 | 0 | 0 | 0 | 0 | 0 | 0 |
| ENSMUSG00000085654 | 0 | 0 | 0 | 0 | 0 | 0 | 0 |
| ENSMUSG00000085655 | 0.671187741 | 0.646240625 | 0.646240625 | 0.701838731 | 0.568060967 | 0.477121255 | 0.629532004 |
| ENSMUSG00000085656 | 0 | 0 | 0 | 0 | 0 | 0 | 0 |
| ENSMUSG00000085657 | 0 | 0 | 0 | 0 | 0 | 0 | 0 |
| ENSMUSG00000085658 | 0.45815691 | 0.25 | 0 | 0 | 0 | 0 | 0.224243824 |
| ENSMUSG00000085659 | 0 | 0 | 0 | 0 | 0 | 0 | 0 |

| | | | | | | | |
|---|---|---|---|---|---|---|---|
| ENSMUSG00000085660 | 0 | 0 | 0 | 0 | 0 | 0 | 0 |
| ENSMUSG00000085662 | 0 | 0 | 0 | 0 | 0 | 0 | 0 |
| ENSMUSG00000085663 | 0 | 0.25 | 0.25 | 0 | 0 | 0 | 0 |
| ENSMUSG00000085664 | 0.289064826 | 0.5 | 0.25 | 0.396240625 | 0.244650542 | 0 | 0.224243824 |
| ENSMUSG00000085665 | 0.811507563 | 0.864857905 | 0.701838731 | 0.864857905 | 0.686820904 | 0.77815125 | 0.710836105 |
| ENSMUSG00000085666 | 1.727821383 | 1.589388001 | 1.62694866 | 1.494319981 | 1.433158226 | 1.591064607 | 1.517723536 |
| ENSMUSG00000085667 | 0.289064826 | 0.25 | 0.646240625 | 0.701838731 | 0.77552387 | 0.84509804 | 0.579661877 |
| ENSMUSG00000085668 | 0 | 0 | 0 | 0 | 0 | 0 | 0 |
| ENSMUSG00000085669 | 0.578129653 | 0.25 | 0.5 | 0.646240625 | 0.387761935 | 0 | 0.224243824 |
| ENSMUSG00000085672 | 0 | 0 | 0 | 0 | 0 | 0 | 0 |
| ENSMUSG00000085673 | 0.289064826 | 0 | 0 | 0 | 0 | 0 | 0 |
| ENSMUSG00000085675 | 0.289064826 | 0.75 | 0.580482024 | 0 | 0 | 0 | 0 |
| ENSMUSG00000085676 | 0 | 0 | 0 | 0 | 0 | 0 | 0 |
| ENSMUSG00000085677 | 0 | 0 | 0 | 0 | 0 | 0 | 0 |
| ENSMUSG00000085678 | 0.289064826 | 0 | 0.5 | 0.396240625 | 0.568060967 | 0 | 0.448487648 |
| ENSMUSG00000085679 | 0 | 0 | 0 | 0 | 0 | 0 | 0 |
| ENSMUSG00000085680 | 0 | 0 | 0 | 0 | 0 | 0 | 0 |
| ENSMUSG00000085681 | 0 | 0 | 0 | 0 | 0 | 0.301029996 | 0 |
| ENSMUSG00000085682 | 0 | 0 | 0 | 0 | 0 | 0 | 0.224243824 |
| ENSMUSG00000085683 | 0 | 0 | 0 | 0 | 0 | 0 | 0 |
| ENSMUSG00000085684 | 0 | 0 | 0 | 0 | 0 | 0 | 0 |
| ENSMUSG00000085686 | 0 | 0 | 0 | 0 | 0 | 0 | 0 |
| ENSMUSG00000085687 | 1.404271433 | 1.311981878 | 1.330482024 | 1.356566189 | 1.244650542 | 1.255272505 | 1.150210039 |
| ENSMUSG00000085688 | 0 | 0 | 0 | 0 | 0 | 0 | 0 |
| ENSMUSG00000085689 | 0.45815691 | 0 | 0 | 0.25 | 0 | 0 | 0 |
| ENSMUSG00000085690 | 0 | 0 | 0 | 0 | 0 | 0 | 0 |
| ENSMUSG00000085691 | 0 | 0 | 0 | 0 | 0 | 0 | 0 |
| ENSMUSG00000085693 | 0.289064826 | 0.25 | 0 | 0.25 | 0.387761935 | 0 | 0.224243824 |
| ENSMUSG00000085694 | 0 | 0 | 0 | 0 | 0 | 0 | 0 |
| ENSMUSG00000085695 | 0 | 0 | 0 | 0 | 0 | 0 | 0 |
| ENSMUSG00000085696 | 1.036286563 | 0.896240625 | 0.951838731 | 0.79248125 | 0.632412477 | 0.602059991 | 0.710836105 |
| ENSMUSG00000085698 | 0.289064826 | 0.25 | 0 | 0.25 | 0.244650542 | 0 | 0 |
| ENSMUSG00000085699 | 0 | 0 | 0 | 0 | 0 | 0 | 0 |
| ENSMUSG00000085700 | 0 | 0 | 0 | 0 | 0 | 0 | 0 |

| | | | | | | | |
|---|---|---|---|---|---|---|---|
| ENSMUSG00000085701 | 0 | 0 | 0 | 0 | 0 | 0 | 0 |
| ENSMUSG00000085702 | 0.45815691 | 0.396240625 | 0.25 | 0.5 | 0.244650542 | 0 | 0 |
| ENSMUSG00000085704 | 0.811507563 | 1.02186571 | 0.92510993 | 1.330482024 | 1.039257769 | 1.113943352 | 0.984950056 |
| ENSMUSG00000085705 | 0 | 0 | 0 | 0.25 | 0 | 0 | 0 |
| ENSMUSG00000085706 | 0 | 0 | 0 | 0 | 0 | 0 | 0 |
| ENSMUSG00000085707 | 0 | 0.25 | 0 | 0 | 0.244650542 | 0 | 0 |
| ENSMUSG00000085708 | 0 | 0 | 0 | 0 | 0 | 0 | 0 |
| ENSMUSG00000085709 | 0 | 0 | 0 | 0 | 0 | 0 | 0 |
| ENSMUSG00000085710 | 0 | 0 | 0 | 0 | 0 | 0 | 0 |
| ENSMUSG00000085711 | 0 | 0 | 0 | 0 | 0 | 0 | 0.224243824 |
| ENSMUSG00000085712 | 0 | 0 | 0 | 0 | 0 | 0 | 0.355418052 |
| ENSMUSG00000085713 | 0 | 0 | 0 | 0 | 0 | 0 | 0 |
| ENSMUSG00000085714 | 0 | 0 | 0 | 0 | 0 | 0 | 0 |
| ENSMUSG00000085715 | 0 | 0.25 | 0 | 0.25 | 0.244650542 | 0 | 0 |
| ENSMUSG00000085716 | 0 | 0 | 0 | 0 | 0 | 0 | 0 |
| ENSMUSG00000085717 | 0 | 0 | 0 | 0 | 0 | 0 | 0 |
| ENSMUSG00000085718 | 0.289064826 | 0.5 | 0.396240625 | 0.396240625 | 0.489301084 | 0 | 0.448487648 |
| ENSMUSG00000085719 | 0 | 0 | 0 | 0 | 0 | 0 | 0 |
| ENSMUSG00000085720 | 0 | 0 | 0 | 0 | 0 | 0 | 0 |
| ENSMUSG00000085721 | 0 | 0 | 0 | 0 | 0 | 0 | 0 |
| ENSMUSG00000085722 | 0 | 0 | 0 | 0 | 0 | 0 | 0 |
| ENSMUSG00000085723 | 0.289064826 | 0 | 0 | 0.25 | 0 | 0 | 0.224243824 |
| ENSMUSG00000085724 | 0 | 0 | 0 | 0 | 0 | 0 | 0 |
| ENSMUSG00000085725 | 0 | 0 | 0 | 0 | 0 | 0 | 0 |
| ENSMUSG00000085728 | 0 | 0 | 0 | 0 | 0 | 0 | 0 |
| ENSMUSG00000085730 | 0 | 0.25 | 0 | 0 | 0 | 0 | 0 |
| ENSMUSG00000085731 | 0 | 0.25 | 0 | 0 | 0 | 0 | 0 |
| ENSMUSG00000085732 | 0 | 0 | 0 | 0 | 0.244650542 | 0 | 0 |
| ENSMUSG00000085733 | 0 | 0 | 0 | 0 | 0 | 0 | 0 |
| ENSMUSG00000085734 | 0 | 0 | 0 | 0 | 0 | 0 | 0 |
| ENSMUSG00000085735 | 0.45815691 | 0.396240625 | 0.25 | 0 | 0.686820904 | 0.477121255 | 0 |
| ENSMUSG00000085737 | 0.289064826 | 0 | 0 | 0 | 0 | 0 | 0.224243824 |
| ENSMUSG00000085738 | 0.289064826 | 0.396240625 | 0.396240625 | 0.646240625 | 0.489301084 | 0.477121255 | 0.448487648 |
| ENSMUSG00000085739 | 0 | 0 | 0 | 0 | 0 | 0 | 0 |

| | | | | | | | |
|---|---|---|---|---|---|---|---|
| ENSMUSG00000085740 | 0 | 0 | 0 | 0 | 0 | 0 | 0 |
| ENSMUSG00000085741 | 0.45815691 | 0.976722649 | 0.79248125 | 0.864857905 | 0.978602168 | 0.301029996 | 0.896975297 |
| ENSMUSG00000085742 | 0 | 0 | 0 | 0 | 0 | 0 | 0 |
| ENSMUSG00000085743 | 0.289064826 | 0.646240625 | 0.580482024 | 0.25 | 0 | 0.301029996 | 0.224243824 |
| ENSMUSG00000085744 | 0 | 0.396240625 | 0.396240625 | 0.5 | 0.387761935 | 0 | 0.355418052 |
| ENSMUSG00000085745 | 0 | 0 | 0 | 0 | 0 | 0 | 0 |
| ENSMUSG00000085747 | 0 | 0 | 0.25 | 0 | 0.244650542 | 0 | 0 |
| ENSMUSG00000085748 | 0.289064826 | 0.396240625 | 0.396240625 | 0 | 0.632412477 | 0 | 0.224243824 |
| ENSMUSG00000085749 | 0.578129653 | 0.25 | 0.25 | 0 | 0.244650542 | 0.301029996 | 0.629532004 |
| ENSMUSG00000085750 | 0 | 0 | 0 | 0.396240625 | 0 | 0 | 0 |
| ENSMUSG00000085751 | 0 | 0 | 0 | 0 | 0 | 0 | 0 |
| ENSMUSG00000085753 | 0 | 0 | 0 | 0 | 0 | 0 | 0 |
| ENSMUSG00000085754 | 0 | 0 | 0 | 0 | 0 | 0 | 0 |
| ENSMUSG00000085755 | 0 | 0 | 0.25 | 0 | 0 | 0 | 0.224243824 |
| ENSMUSG00000085756 | 0 | 0 | 0 | 0 | 0 | 0 | 0 |
| ENSMUSG00000085757 | 0 | 0 | 0 | 0 | 0 | 0 | 0 |
| ENSMUSG00000085759 | 0 | 0 | 0 | 0 | 0 | 0 | 0 |
| ENSMUSG00000085760 | 0 | 0 | 0.25 | 0.5 | 0.387761935 | 0 | 0 |
| ENSMUSG00000085761 | 0 | 0 | 0 | 0 | 0 | 0 | 0 |
| ENSMUSG00000085762 | 0 | 0 | 0 | 0.396240625 | 0 | 0 | 0 |
| ENSMUSG00000085763 | 0.289064826 | 0 | 0.25 | 0 | 0 | 0.301029996 | 0 |
| ENSMUSG00000085765 | 0 | 0 | 0 | 0.25 | 0.244650542 | 0.301029996 | 0 |
| ENSMUSG00000085766 | 1 | 1.04248125 | 1.061981878 | 0.79248125 | 1.074582839 | 1 | 0.952571511 |
| ENSMUSG00000085767 | 1.100572389 | 0.79248125 | 1 | 0.92510993 | 1.057362051 | 0.77815125 | 0.969165684 |
| ENSMUSG00000085768 | 0.289064826 | 0 | 0 | 0 | 0 | 0 | 0 |
| ENSMUSG00000085770 | 0 | 0 | 0 | 0 | 0 | 0 | 0 |
| ENSMUSG00000085771 | 0 | 0 | 0 | 0 | 0 | 0 | 0 |
| ENSMUSG00000085772 | 0 | 0 | 0 | 0 | 0 | 0 | 0 |
| ENSMUSG00000085773 | 0 | 0 | 0 | 0 | 0.244650542 | 0 | 0 |
| ENSMUSG00000085774 | 0.671187741 | 0.396240625 | 0.396240625 | 0.580482024 | 0.632412477 | 0 | 0.355418052 |
| ENSMUSG00000085775 | 0 | 0 | 0 | 0 | 0 | 0 | 0 |
| ENSMUSG00000085776 | 0 | 0 | 0.396240625 | 0.5 | 0 | 0.301029996 | 0 |
| ENSMUSG00000085779 | 0.289064826 | 0.25 | 0 | 0 | 0 | 0 | 0.224243824 |
| ENSMUSG00000085781 | 0.45815691 | 0 | 0.580482024 | 0 | 0 | 0 | 0.224243824 |

| | | | | | | | |
|---|---|---|---|---|---|---|---|
| ENSMUSG00000085782 | 0.578129653 | 0.580482024 | 0.5 | 0.580482024 | 0.489301084 | 0 | 0.355418052 |
| ENSMUSG00000085783 | 0 | 0 | 0 | 0 | 0 | 0 | 0 |
| ENSMUSG00000085784 | 0.671187741 | 0.701838731 | 0.646240625 | 0.864857905 | 0.686820904 | 0.698970004 | 0.710836105 |
| ENSMUSG00000085785 | 0 | 0 | 0 | 0 | 0 | 0 | 0 |
| ENSMUSG00000085786 | 0.45815691 | 0.396240625 | 0 | 0 | 0 | 0 | 0 |
| ENSMUSG00000085787 | 0 | 0 | 0 | 0.25 | 0 | 0 | 0 |
| ENSMUSG00000085788 | 0 | 0.25 | 0 | 0.25 | 0.244650542 | 0 | 0 |
| ENSMUSG00000085790 | 0 | 0.25 | 0 | 0 | 0 | 0 | 0 |
| ENSMUSG00000085791 | 0 | 0 | 0 | 0 | 0 | 0 | 0 |
| ENSMUSG00000085792 | 0 | 0 | 0 | 0 | 0 | 0 | 0 |
| ENSMUSG00000085793 | 1.903481042 | 1.862802778 | 1.723704441 | 1.732684334 | 1.712553795 | 1.897627091 | 1.582103514 |
| ENSMUSG00000085794 | 0 | 0.396240625 | 0.25 | 0.830482024 | 0.905314583 | 0.698970004 | 0.984950056 |
| ENSMUSG00000085795 | 2.121692466 | 2.03360658 | 2.070192693 | 1.747171172 | 1.800029087 | 1.963787827 | 1.762103709 |
| ENSMUSG00000085796 | 0 | 0 | 0 | 0 | 0 | 0 | 0 |
| ENSMUSG00000085797 | 0 | 0 | 0 | 0 | 0 | 0 | 0 |
| ENSMUSG00000085798 | 0 | 0 | 0 | 0 | 0 | 0 | 0 |
| ENSMUSG00000085799 | 0 | 0.25 | 0 | 0 | 0 | 0 | 0 |
| ENSMUSG00000085800 | 0 | 0 | 0.25 | 0.25 | 0 | 0 | 0 |
| ENSMUSG00000085801 | 0 | 0 | 0 | 0 | 0 | 0 | 0 |
| ENSMUSG00000085802 | 0.45815691 | 0 | 0.580482024 | 0.25 | 0.489301084 | 0 | 0.448487648 |
| ENSMUSG00000085803 | 0 | 0.25 | 0.396240625 | 0 | 0 | 0 | 0 |
| ENSMUSG00000085804 | 0 | 0 | 0 | 0 | 0 | 0 | 0 |
| ENSMUSG00000085805 | 0 | 0 | 0 | 0 | 0 | 0 | 0 |
| ENSMUSG00000085806 | 0 | 0 | 0.5 | 0 | 0 | 0 | 0 |
| ENSMUSG00000085807 | 0.289064826 | 0.25 | 0.396240625 | 0.580482024 | 0 | 0 | 0.224243824 |
| ENSMUSG00000085808 | 0 | 0 | 0 | 0 | 0 | 0 | 0.224243824 |
| ENSMUSG00000085810 | 0 | 0 | 0 | 0 | 0.244650542 | 0 | 0 |
| ENSMUSG00000085811 | 1.794934559 | 1.741446071 | 1.638647213 | 1.584962501 | 1.618292349 | 1.755874856 | 1.473249546 |
| ENSMUSG00000085812 | 0 | 0 | 0.25 | 0.25 | 0 | 0 | 0 |
| ENSMUSG00000085813 | 0 | 0 | 0 | 0 | 0 | 0 | 0 |
| ENSMUSG00000085815 | 0 | 0 | 0 | 0 | 0 | 0 | 0.224243824 |
| ENSMUSG00000085816 | 0 | 0 | 0 | 0 | 0 | 0 | 0 |
| ENSMUSG00000085817 | 0 | 0.25 | 0 | 0.25 | 0.244650542 | 0 | 0 |
| ENSMUSG00000085818 | 0 | 0.25 | 0 | 0 | 0.244650542 | 0 | 0 |

| | | | | | | | |
|---|---|---|---|---|---|---|---|
| ENSMUSG00000085819 | 0.747221736 | 0 | 0.25 | 0.5 | 0.387761935 | 0 | 0.224243824 |
| ENSMUSG00000085820 | 0 | 0 | 0 | 0 | 0 | 0 | 0 |
| ENSMUSG00000085822 | 0.289064826 | 0 | 0.396240625 | 0.25 | 0.568060967 | 0 | 0.355418052 |
| ENSMUSG00000085823 | 0 | 0 | 0 | 0 | 0 | 0 | 0 |
| ENSMUSG00000085824 | 0.289064826 | 0.5 | 0 | 0 | 0.244650542 | 0 | 0.224243824 |
| ENSMUSG00000085826 | 0.45815691 | 0.580482024 | 0.25 | 0.396240625 | 0.568060967 | 0.602059991 | 0.520678036 |
| ENSMUSG00000085827 | 0 | 0 | 0 | 0 | 0 | 0 | 0 |
| ENSMUSG00000085828 | 0.45815691 | 0 | 0.25 | 0.5 | 0.489301084 | 0.301029996 | 0.672731473 |
| ENSMUSG00000085829 | 0.671187741 | 0.580482024 | 0.646240625 | 0.701838731 | 0.77552387 | 0.477121255 | 0.355418052 |
| ENSMUSG00000085830 | 0 | 0 | 0 | 0.25 | 0.244650542 | 0 | 0 |
| ENSMUSG00000085831 | 0 | 0 | 0 | 0 | 0 | 0 | 0 |
| ENSMUSG00000085833 | 0 | 0 | 0 | 0 | 0.244650542 | 0 | 0 |
| ENSMUSG00000085834 | 0 | 0 | 0 | 0 | 0 | 0 | 0 |
| ENSMUSG00000085835 | 0 | 0 | 0 | 0 | 0 | 0 | 0 |
| ENSMUSG00000085836 | 0 | 0 | 0 | 0 | 0 | 0 | 0 |
| ENSMUSG00000085837 | 0.289064826 | 0 | 0 | 0.25 | 0.387761935 | 0.301029996 | 0 |
| ENSMUSG00000085838 | 0 | 0 | 0 | 0 | 0 | 0 | 0 |
| ENSMUSG00000085839 | 0 | 0.25 | 0 | 0 | 0 | 0 | 0 |
| ENSMUSG00000085840 | 1.036286563 | 1.02186571 | 0.75 | 0.75 | 0.877063019 | 0.903089987 | 0.74492186 |
| ENSMUSG00000085841 | 0 | 0 | 0 | 0 | 0 | 0 | 0 |
| ENSMUSG00000085842 | 0 | 0 | 0.25 | 0.5 | 0.489301084 | 0 | 0 |
| ENSMUSG00000085843 | 0.45815691 | 0.25 | 0 | 0.25 | 0 | 0.301029996 | 0.355418052 |
| ENSMUSG00000085844 | 0 | 0 | 0 | 0 | 0 | 0 | 0 |
| ENSMUSG00000085845 | 0 | 0 | 0 | 0 | 0 | 0 | 0 |
| ENSMUSG00000085846 | 0 | 0 | 0 | 0 | 0 | 0 | 0 |
| ENSMUSG00000085847 | 0 | 0.25 | 0 | 0 | 0 | 0 | 0 |
| ENSMUSG00000085851 | 0.289064826 | 0.396240625 | 0 | 0.701838731 | 0.244650542 | 0.301029996 | 0.355418052 |
| ENSMUSG00000085852 | 0 | 0.396240625 | 0.396240625 | 0.646240625 | 0.686820904 | 0.301029996 | 0.672731473 |
| ENSMUSG00000085853 | 0 | 0 | 0 | 0 | 0 | 0 | 0 |
| ENSMUSG00000085856 | 0 | 0 | 0 | 0 | 0 | 0 | 0 |
| ENSMUSG00000085857 | 0 | 0 | 0 | 0 | 0 | 0 | 0 |
| ENSMUSG00000085858 | 0 | 0 | 0 | 0 | 0 | 0 | 0 |
| ENSMUSG00000085860 | 0 | 0 | 0 | 0 | 0 | 0 | 0 |
| ENSMUSG00000085861 | 0.289064826 | 0.396240625 | 0.396240625 | 0.25 | 0.686820904 | 0 | 0 |

| | | | | | | | |
|---|---|---|---|---|---|---|---|
| ENSMUSG00000085862 | 0.811507563 | 0.396240625 | 0.92510993 | 0.580482024 | 0.568060967 | 0.477121255 | 0.355418052 |
| ENSMUSG00000085863 | 0 | 0 | 0 | 0 | 0 | 0 | 0 |
| ENSMUSG00000085864 | 0 | 0 | 0 | 0.25 | 0 | 0 | 0 |
| ENSMUSG00000085865 | 0 | 0 | 0 | 0 | 0 | 0 | 0 |
| ENSMUSG00000085866 | 0 | 0 | 0 | 0 | 0 | 0 | 0 |
| ENSMUSG00000085867 | 0 | 0 | 0 | 0 | 0 | 0 | 0 |
| ENSMUSG00000085868 | 0 | 0 | 0.25 | 0 | 0.244650542 | 0.602059991 | 0.355418052 |
| ENSMUSG00000085869 | 0 | 0 | 0 | 0 | 0 | 0 | 0 |
| ENSMUSG00000085870 | 0.289064826 | 0 | 0 | 0 | 0 | 0 | 0 |
| ENSMUSG00000085871 | 0 | 0 | 0 | 0 | 0 | 0 | 0 |
| ENSMUSG00000085872 | 0 | 0 | 0 | 0 | 0 | 0 | 0 |
| ENSMUSG00000085873 | 0.671187741 | 0.646240625 | 0.896240625 | 0.580482024 | 0.568060967 | 0.477121255 | 0.520678036 |
| ENSMUSG00000085875 | 0.45815691 | 0 | 0 | 0 | 0 | 0 | 0 |
| ENSMUSG00000085876 | 0 | 0 | 0 | 0 | 0 | 0 | 0 |
| ENSMUSG00000085877 | 0 | 0 | 0 | 0 | 0 | 0 | 0 |
| ENSMUSG00000085878 | 0 | 0 | 0.25 | 0 | 0 | 0 | 0 |
| ENSMUSG00000085880 | 0 | 0 | 0 | 0 | 0 | 0 | 0 |
| ENSMUSG00000085881 | 0.671187741 | 0.25 | 0 | 0 | 0.387761935 | 0 | 0.355418052 |
| ENSMUSG00000085882 | 0.289064826 | 0.25 | 0.25 | 0.5 | 0.489301084 | 0.602059991 | 0.579661877 |
| ENSMUSG00000085883 | 0 | 0 | 0 | 0 | 0 | 0 | 0 |
| ENSMUSG00000085884 | 0 | 0 | 0.25 | 0.25 | 0.387761935 | 0 | 0.355418052 |
| ENSMUSG00000085885 | 0 | 0 | 0 | 0 | 0 | 0 | 0 |
| ENSMUSG00000085886 | 0 | 0 | 0 | 0 | 0.244650542 | 0 | 0 |
| ENSMUSG00000085887 | 0 | 0.396240625 | 0 | 0 | 0 | 0 | 0 |
| ENSMUSG00000085888 | 0 | 0 | 0 | 0.396240625 | 0 | 0 | 0 |
| ENSMUSG00000085889 | 0 | 0 | 0 | 0 | 0.244650542 | 0 | 0 |
| ENSMUSG00000085890 | 0.289064826 | 0.396240625 | 0.396240625 | 0 | 0.568060967 | 0 | 0.829800754 |
| ENSMUSG00000085891 | 0 | 0 | 0.396240625 | 0.5 | 0.244650542 | 0.477121255 | 0.224243824 |
| ENSMUSG00000085892 | 0 | 0 | 0 | 0 | 0 | 0 | 0 |
| ENSMUSG00000085893 | 0.578129653 | 0.75 | 0.580482024 | 0.5 | 0.846351821 | 0.301029996 | 0.672731473 |
| ENSMUSG00000085894 | 0.747221736 | 0.79248125 | 0.79248125 | 1.02186571 | 0.955822902 | 0.698970004 | 0.935079929 |
| ENSMUSG00000085895 | 0 | 0 | 0 | 0 | 0 | 0 | 0 |
| ENSMUSG00000085896 | 0 | 0 | 0 | 0 | 0 | 0 | 0 |
| ENSMUSG00000085897 | 0 | 0.25 | 0.396240625 | 0.25 | 0.244650542 | 0.301029996 | 0.224243824 |

| | | | | | | | |
|---|---|---|---|---|---|---|---|
| ENSMUSG00000085898 | 0.747221736 | 0.646240625 | 0.701838731 | 0.701838731 | 0.686820904 | 0.698970004 | 0.520678036 |
| ENSMUSG00000085899 | 0 | 0 | 0 | 0 | 0 | 0 | 0 |
| ENSMUSG00000085900 | 0 | 0 | 0 | 0.396240625 | 0.387761935 | 0 | 0.224243824 |
| ENSMUSG00000085902 | 0.289064826 | 0 | 0.25 | 0.25 | 0 | 0 | 0 |
| ENSMUSG00000085903 | 0 | 0 | 0 | 0 | 0 | 0 | 0 |
| ENSMUSG00000085905 | 0 | 0 | 0 | 0 | 0 | 0 | 0 |
| ENSMUSG00000085908 | 0 | 0 | 0 | 0 | 0 | 0 | 0 |
| ENSMUSG00000085909 | 0 | 0 | 0 | 0 | 0 | 0 | 0 |
| ENSMUSG00000085910 | 0 | 0 | 0.25 | 0 | 0 | 0 | 0.224243824 |
| ENSMUSG00000085911 | 0 | 0 | 0 | 0 | 0 | 0 | 0 |
| ENSMUSG00000085912 | 1.036286563 | 0.864857905 | 1.04248125 | 0.25 | 0.387761935 | 0.698970004 | 0.579661877 |
| ENSMUSG00000085913 | 0.747221736 | 0.830482024 | 0.396240625 | 0.896240625 | 0.978602168 | 0.84509804 | 0.803905701 |
| ENSMUSG00000085914 | 0 | 0.396240625 | 0 | 0 | 0 | 0 | 0 |
| ENSMUSG00000085915 | 0 | 0 | 0 | 0 | 0 | 0 | 0 |
| ENSMUSG00000085917 | 0 | 0 | 0 | 0 | 0 | 0 | 0 |
| ENSMUSG00000085918 | 0 | 0 | 0 | 0 | 0 | 0 | 0 |
| ENSMUSG00000085919 | 0 | 0 | 0 | 0 | 0 | 0 | 0 |
| ENSMUSG00000085920 | 0 | 0 | 0 | 0 | 0 | 0 | 0 |
| ENSMUSG00000085921 | 0 | 0 | 0 | 0 | 0 | 0 | 0 |
| ENSMUSG00000085922 | 0 | 0 | 0 | 0 | 0 | 0 | 0 |
| ENSMUSG00000085923 | 0 | 0 | 0 | 0 | 0 | 0 | 0 |
| ENSMUSG00000085924 | 0 | 0 | 0 | 0 | 0.244650542 | 0 | 0 |
| ENSMUSG00000085925 | 0 | 0.580482024 | 0.580482024 | 0.25 | 0 | 0 | 0 |
| ENSMUSG00000085926 | 0 | 0 | 0 | 0 | 0 | 0 | 0 |
| ENSMUSG00000085927 | 0 | 0 | 0 | 0 | 0 | 0 | 0 |
| ENSMUSG00000085928 | 0 | 0 | 0 | 0 | 0 | 0 | 0 |
| ENSMUSG00000085929 | 0.811507563 | 0.75 | 0.75 | 0.75 | 0.846351821 | 0.698970004 | 0.775756176 |
| ENSMUSG00000085931 | 0 | 0 | 0 | 0.25 | 0 | 0.301029996 | 0 |
| ENSMUSG00000085932 | 0 | 0 | 0 | 0 | 0 | 0 | 0 |
| ENSMUSG00000085933 | 0 | 0 | 0 | 0 | 0 | 0 | 0 |
| ENSMUSG00000085934 | 0 | 0 | 0 | 0 | 0 | 0 | 0 |
| ENSMUSG00000085936 | 0.747221736 | 0.701838731 | 0.79248125 | 1.04248125 | 0.905314583 | 0.602059991 | 0.775756176 |
| ENSMUSG00000085937 | 0 | 0 | 0 | 0 | 0.244650542 | 0 | 0 |
| ENSMUSG00000085938 | 0 | 0 | 0.25 | 0 | 0 | 0 | 0 |

| | | | | | | | |
|---|---|---|---|---|---|---|---|
| ENSMUSG00000085939 | 0.811507563 | 0.864857905 | 1.061981878 | 0.976722649 | 0.905314583 | 0.477121255 | 1 |
| ENSMUSG00000085940 | 0 | 0 | 0 | 0 | 0 | 0 | 0 |
| ENSMUSG00000085941 | 0.45815691 | 0.396240625 | 0.396240625 | 0 | 0.244650542 | 0 | 0 |
| ENSMUSG00000085944 | 0 | 0 | 0 | 0 | 0 | 0.301029996 | 0 |
| ENSMUSG00000085945 | 0.289064826 | 0 | 0.396240625 | 0 | 0.244650542 | 0 | 0.224243824 |
| ENSMUSG00000085946 | 0 | 0 | 0 | 0 | 0 | 0 | 0 |
| ENSMUSG00000085949 | 0 | 0 | 0 | 0 | 0 | 0.301029996 | 0.224243824 |
| ENSMUSG00000085950 | 0.289064826 | 0.701838731 | 0.25 | 0.5 | 0.244650542 | 0.301029996 | 0.355418052 |
| ENSMUSG00000085951 | 0 | 0 | 0 | 0 | 0 | 0 | 0 |
| ENSMUSG00000085952 | 0 | 0 | 0 | 0.25 | 0 | 0 | 0 |
| ENSMUSG00000085953 | 0 | 0 | 0 | 0 | 0 | 0 | 0 |
| ENSMUSG00000085955 | 0 | 0 | 0.5 | 0.5 | 0.387761935 | 0 | 0 |
| ENSMUSG00000085956 | 0 | 0.830482024 | 0.5 | 0.25 | 0.733951626 | 0.477121255 | 0.355418052 |
| ENSMUSG00000085957 | 0.811507563 | 0 | 0.580482024 | 0.25 | 0.244650542 | 0 | 0.579661877 |
| ENSMUSG00000085958 | 0.289064826 | 0.25 | 0 | 0 | 0 | 0 | 0.224243824 |
| ENSMUSG00000085961 | 0 | 0.25 | 0 | 0.25 | 0 | 0 | 0 |
| ENSMUSG00000085962 | 0.671187741 | 0.580482024 | 0.396240625 | 0.830482024 | 0.387761935 | 0.477121255 | 0.579661877 |
| ENSMUSG00000085965 | 0 | 0 | 0 | 0 | 0 | 0 | 0 |
| ENSMUSG00000085966 | 0 | 0 | 0 | 0 | 0 | 0 | 0 |
| ENSMUSG00000085967 | 0 | 0 | 0 | 0 | 0 | 0 | 0 |
| ENSMUSG00000085968 | 0 | 0 | 0 | 0 | 0 | 0 | 0 |
| ENSMUSG00000085969 | 0 | 0 | 0 | 0 | 0 | 0 | 0 |
| ENSMUSG00000085970 | 0 | 0 | 0 | 0 | 0 | 0 | 0 |
| ENSMUSG00000085971 | 0.960252568 | 0.75 | 0.646240625 | 0.580482024 | 0.632412477 | 0.477121255 | 0.448487648 |
| ENSMUSG00000085972 | 0.45815691 | 0 | 0 | 0 | 0 | 0 | 0 |
| ENSMUSG00000085973 | 0 | 0 | 0 | 0 | 0 | 0 | 0 |
| ENSMUSG00000085975 | 0 | 0 | 0 | 0 | 0 | 0 | 0 |
| ENSMUSG00000085976 | 0 | 0.396240625 | 0.5 | 0.580482024 | 0.387761935 | 0.698970004 | 0.629532004 |
| ENSMUSG00000085977 | 0 | 0 | 0 | 0 | 0 | 0 | 0 |
| ENSMUSG00000085978 | 0 | 0 | 0 | 0 | 0 | 0 | 0 |
| ENSMUSG00000085979 | 0 | 0 | 0 | 0 | 0 | 0 | 0 |
| ENSMUSG00000085980 | 0 | 0 | 0 | 0 | 0 | 0 | 0 |
| ENSMUSG00000085981 | 0 | 0 | 0 | 0 | 0 | 0 | 0 |
| ENSMUSG00000085982 | 0.289064826 | 0.646240625 | 0.396240625 | 0.25 | 0 | 0.301029996 | 0.355418052 |

| | | | | | | | |
|---|---|---|---|---|---|---|---|
| ENSMUSG00000085984 | 0.45815691 | 0.25 | 0.25 | 0.580482024 | 0.489301084 | 0.698970004 | 0.579661877 |
| ENSMUSG00000085985 | 0 | 0 | 0 | 0 | 0 | 0 | 0 |
| ENSMUSG00000085987 | 0 | 0 | 0 | 0.25 | 0 | 0 | 0 |
| ENSMUSG00000085989 | 0 | 0 | 0 | 0 | 0 | 0 | 0 |
| ENSMUSG00000085990 | 0 | 0 | 0 | 0 | 0 | 0 | 0 |
| ENSMUSG00000085991 | 0 | 0 | 0 | 0 | 0 | 0 | 0 |
| ENSMUSG00000085992 | 0 | 0 | 0.25 | 0 | 0 | 0 | 0.355418052 |
| ENSMUSG00000085993 | 0 | 0 | 0 | 0 | 0 | 0 | 0 |
| ENSMUSG00000085995 | 0.289064826 | 0.25 | 0.396240625 | 0.896240625 | 0.905314583 | 1 | 0.672731473 |
| ENSMUSG00000085996 | 0 | 0 | 0.25 | 0 | 0 | 0 | 0 |
| ENSMUSG00000085998 | 0.289064826 | 0 | 0 | 0 | 0.244650542 | 0 | 0.224243824 |
| ENSMUSG00000085999 | 0 | 0 | 0 | 0 | 0 | 0 | 0 |
| ENSMUSG00000086000 | 0 | 0 | 0 | 0 | 0 | 0 | 0 |
| ENSMUSG00000086001 | 0 | 0 | 0 | 0 | 0.244650542 | 0.301029996 | 0 |
| ENSMUSG00000086003 | 0.289064826 | 0.396240625 | 0.5 | 0 | 0 | 0.301029996 | 0.355418052 |
| ENSMUSG00000086004 | 0 | 0 | 0 | 0.25 | 0 | 0 | 0 |
| ENSMUSG00000086006 | 0.289064826 | 0 | 0.25 | 0 | 0 | 0 | 0 |
| ENSMUSG00000086007 | 0 | 0 | 0 | 0 | 0 | 0 | 0 |
| ENSMUSG00000086008 | 0 | 0.25 | 0.25 | 0 | 0 | 0 | 0 |
| ENSMUSG00000086009 | 0 | 0 | 0 | 0 | 0 | 0 | 0 |
| ENSMUSG00000086010 | 0 | 0 | 0 | 0 | 0 | 0 | 0 |
| ENSMUSG00000086011 | 0 | 0 | 0 | 0 | 0 | 0 | 0 |
| ENSMUSG00000086012 | 0 | 0 | 0 | 0 | 0 | 0 | 0 |
| ENSMUSG00000086013 | 0.91631382 | 1.114857905 | 1.02186571 | 0.92510993 | 0.812711509 | 0.77815125 | 0.829800754 |
| ENSMUSG00000086014 | 0 | 0 | 0 | 0 | 0 | 0 | 0 |
| ENSMUSG00000086015 | 0.747221736 | 0.701838731 | 0.701838731 | 0.646240625 | 0.489301084 | 0.602059991 | 0.803905701 |
| ENSMUSG00000086016 | 0.45815691 | 0 | 0.396240625 | 0 | 0.244650542 | 0 | 0.224243824 |
| ENSMUSG00000086017 | 0 | 0 | 0 | 0 | 0 | 0 | 0 |
| ENSMUSG00000086018 | 0 | 0 | 0 | 0 | 0 | 0 | 0 |
| ENSMUSG00000086019 | 0 | 0 | 0 | 0 | 0 | 0 | 0 |
| ENSMUSG00000086020 | 0 | 0 | 0 | 0 | 0 | 0 | 0 |
| ENSMUSG00000086021 | 0 | 0 | 0 | 0 | 0 | 0 | 0 |
| ENSMUSG00000086022 | 0.45815691 | 0 | 0.396240625 | 0.5 | 0.489301084 | 0 | 0.224243824 |
| ENSMUSG00000086025 | 0.289064826 | 0 | 0.25 | 0 | 0 | 0 | 0.224243824 |

| | | | | | | | |
|---|---|---|---|---|---|---|---|
| ENSMUSG00000086026 | 0 | 0 | 0 | 0 | 0 | 0 | 0 |
| ENSMUSG00000086027 | 0 | 0 | 0 | 0 | 0 | 0 | 0 |
| ENSMUSG00000086028 | 0 | 0 | 0 | 0 | 0 | 0 | 0 |
| ENSMUSG00000086029 | 0 | 0 | 0 | 0 | 0 | 0 | 0.224243824 |
| ENSMUSG00000086030 | 0 | 0 | 0 | 0 | 0 | 0 | 0 |
| ENSMUSG00000086031 | 0 | 0 | 0 | 0 | 0 | 0 | 0 |
| ENSMUSG00000086032 | 0 | 0 | 0 | 0 | 0 | 0 | 0 |
| ENSMUSG00000086034 | 0 | 0 | 0 | 0 | 0 | 0 | 0 |
| ENSMUSG00000086035 | 0.811507563 | 1 | 1.130890489 | 0.976722649 | 0.931471446 | 1 | 0.952571511 |
| ENSMUSG00000086036 | 0 | 0 | 0 | 0 | 0 | 0 | 0 |
| ENSMUSG00000086037 | 0 | 0 | 0 | 0 | 0 | 0 | 0 |
| ENSMUSG00000086039 | 0 | 0 | 0 | 0.25 | 0 | 0 | 0 |
| ENSMUSG00000086040 | 0.289064826 | 0 | 0 | 0.5 | 0.244650542 | 0.77815125 | 0.224243824 |
| ENSMUSG00000086041 | 0.289064826 | 0.5 | 0.25 | 0 | 0 | 0 | 0 |
| ENSMUSG00000086042 | 0 | 0 | 0 | 0 | 0 | 0 | 0 |
| ENSMUSG00000086043 | 0.45815691 | 0.5 | 0.580482024 | 0.25 | 0.244650542 | 0.301029996 | 0.579661877 |
| ENSMUSG00000086044 | 0 | 0 | 0 | 0 | 0 | 0 | 0 |
| ENSMUSG00000086046 | 0 | 0 | 0 | 0 | 0 | 0 | 0 |
| ENSMUSG00000086047 | 0.45815691 | 0.396240625 | 0.25 | 0.25 | 0.244650542 | 0.301029996 | 0.224243824 |
| ENSMUSG00000086048 | 0 | 0 | 0 | 0 | 0 | 0 | 0 |
| ENSMUSG00000086049 | 0 | 0 | 0 | 0 | 0 | 0.301029996 | 0.448487648 |
| ENSMUSG00000086050 | 0 | 0 | 0.25 | 0.25 | 0.387761935 | 0.301029996 | 0.224243824 |
| ENSMUSG00000086052 | 0 | 0 | 0 | 0 | 0 | 0 | 0 |
| ENSMUSG00000086053 | 0 | 0 | 0.25 | 0.25 | 0 | 0 | 0 |
| ENSMUSG00000086054 | 0 | 0 | 0.25 | 0 | 0 | 0 | 0 |
| ENSMUSG00000086055 | 0 | 0 | 0 | 0 | 0 | 0 | 0 |
| ENSMUSG00000086056 | 0 | 0 | 0 | 0 | 0 | 0 | 0 |
| ENSMUSG00000086057 | 0 | 0 | 0 | 0 | 0 | 0 | 0 |
| ENSMUSG00000086058 | 0 | 0 | 0 | 0 | 0 | 0 | 0 |
| ENSMUSG00000086060 | 0 | 0 | 0 | 0 | 0 | 0 | 0 |
| ENSMUSG00000086061 | 0 | 0 | 0 | 0.25 | 0.244650542 | 0 | 0 |
| ENSMUSG00000086063 | 0 | 0 | 0 | 0 | 0 | 0 | 0 |
| ENSMUSG00000086064 | 0 | 0 | 0 | 0 | 0 | 0 | 0 |
| ENSMUSG00000086065 | 0 | 0 | 0 | 0 | 0 | 0 | 0 |

| | | | | | | | |
|---|---|---|---|---|---|---|---|
| ENSMUSG00000086066 | 0 | 0 | 0 | 0 | 0 | 0 | 0 |
| ENSMUSG00000086067 | 0.960252568 | 0.976722649 | 1.02186571 | 0.5 | 0.568060967 | 0.602059991 | 0.448487648 |
| ENSMUSG00000086068 | 0 | 0 | 0 | 0 | 0 | 0 | 0 |
| ENSMUSG00000086069 | 0 | 0 | 0 | 0 | 0 | 0 | 0 |
| ENSMUSG00000086070 | 0 | 0 | 0.396240625 | 0 | 0 | 0 | 0 |
| ENSMUSG00000086071 | 0 | 0 | 0 | 0 | 0 | 0 | 0 |
| ENSMUSG00000086072 | 0 | 0 | 0 | 0 | 0 | 0 | 0 |
| ENSMUSG00000086074 | 0 | 0 | 0 | 0 | 0 | 0 | 0 |
| ENSMUSG00000086075 | 1.129344651 | 1.114857905 | 0.92510993 | 1.02186571 | 0.978602168 | 0.903089987 | 1.078019652 |
| ENSMUSG00000086076 | 0 | 0 | 0 | 0 | 0.489301084 | 0 | 0.224243824 |
| ENSMUSG00000086077 | 0 | 0 | 0 | 0 | 0 | 0 | 0 |
| ENSMUSG00000086078 | 0.289064826 | 0.25 | 0 | 0 | 0 | 0 | 0 |
| ENSMUSG00000086080 | 0 | 0 | 0 | 0 | 0 | 0 | 0 |
| ENSMUSG00000086081 | 0 | 0 | 0 | 0 | 0 | 0 | 0 |
| ENSMUSG00000086083 | 0 | 0 | 0.25 | 0 | 0.387761935 | 0 | 0 |
| ENSMUSG00000086087 | 0 | 0 | 0 | 0 | 0 | 0 | 0 |
| ENSMUSG00000086089 | 0 | 0 | 0 | 0 | 0 | 0 | 0 |
| ENSMUSG00000086090 | 0 | 0 | 0 | 0 | 0 | 0 | 0 |
| ENSMUSG00000086092 | 0 | 0 | 0 | 0 | 0 | 0 | 0 |
| ENSMUSG00000086093 | 0 | 0 | 0 | 0 | 0 | 0 | 0.224243824 |
| ENSMUSG00000086095 | 0.289064826 | 0 | 0.5 | 0 | 0 | 0.477121255 | 0 |
| ENSMUSG00000086096 | 0 | 0 | 0 | 0 | 0 | 0 | 0.224243824 |
| ENSMUSG00000086097 | 0 | 0 | 0 | 0 | 0 | 0 | 0 |
| ENSMUSG00000086098 | 0 | 0 | 0.25 | 0.25 | 0.387761935 | 0 | 0.355418052 |
| ENSMUSG00000086103 | 0.960252568 | 0.646240625 | 0.646240625 | 0.79248125 | 0.733951626 | 0.84509804 | 0.896975297 |
| ENSMUSG00000086104 | 0 | 0 | 0 | 0 | 0 | 0 | 0 |
| ENSMUSG00000086105 | 0.578129653 | 0.864857905 | 0.79248125 | 0.5 | 0.387761935 | 0.602059991 | 0.672731473 |
| ENSMUSG00000086107 | 0 | 0 | 0 | 0 | 0 | 0 | 0 |
| ENSMUSG00000086108 | 0.45815691 | 0.25 | 0 | 0.701838731 | 0.387761935 | 0 | 0.520678036 |
| ENSMUSG00000086109 | 0 | 0 | 0 | 0 | 0 | 0 | 0 |
| ENSMUSG00000086111 | 0 | 0 | 0 | 0 | 0 | 0 | 0 |
| ENSMUSG00000086112 | 0 | 0 | 0 | 0 | 0 | 0 | 0 |
| ENSMUSG00000086114 | 0 | 0 | 0 | 0 | 0 | 0 | 0 |
| ENSMUSG00000086115 | 0 | 0 | 0 | 0 | 0 | 0 | 0 |

| | | | | | | | |
|---|---|---|---|---|---|---|---|
| ENSMUSG00000086117 | 0 | 0 | 0 | 0 | 0 | 0 | 0 |
| ENSMUSG00000086118 | 0 | 0 | 0 | 0 | 0 | 0 | 0 |
| ENSMUSG00000086119 | 0.671187741 | 0.701838731 | 0.701838731 | 0.79248125 | 0.846351821 | 0.954242509 | 0.876096088 |
| ENSMUSG00000086121 | 0 | 0 | 0 | 0 | 0 | 0.301029996 | 0 |
| ENSMUSG00000086122 | 0 | 0 | 0 | 0 | 0 | 0 | 0 |
| ENSMUSG00000086123 | 0 | 0 | 0 | 0 | 0.387761935 | 0 | 0 |
| ENSMUSG00000086124 | 0 | 0 | 0 | 0 | 0 | 0 | 0 |
| ENSMUSG00000086125 | 0 | 0.25 | 0 | 0 | 0 | 0 | 0 |
| ENSMUSG00000086126 | 0 | 0 | 0 | 0 | 0 | 0 | 0 |
| ENSMUSG00000086127 | 0.811507563 | 0.5 | 0.5 | 0.701838731 | 0.632412477 | 0.301029996 | 0.448487648 |
| ENSMUSG00000086128 | 0 | 0 | 0 | 0 | 0 | 0 | 0 |
| ENSMUSG00000086129 | 0 | 0 | 0 | 0 | 0.244650542 | 0 | 0 |
| ENSMUSG00000086130 | 1.527823874 | 1.410964047 | 1.372963274 | 1.348079356 | 1.433158226 | 1.397940009 | 1.224243824 |
| ENSMUSG00000086132 | 0 | 0 | 0 | 0 | 0 | 0 | 0 |
| ENSMUSG00000086133 | 0 | 0 | 0 | 0 | 0 | 0 | 0 |
| ENSMUSG00000086134 | 0.289064826 | 0 | 0 | 0.25 | 0.387761935 | 0 | 0 |
| ENSMUSG00000086135 | 0 | 0 | 0 | 0 | 0 | 0 | 0 |
| ENSMUSG00000086136 | 0 | 0 | 0 | 0 | 0 | 0 | 0 |
| ENSMUSG00000086137 | 0 | 0 | 0 | 0 | 0 | 0 | 0 |
| ENSMUSG00000086138 | 0 | 0 | 0 | 0 | 0 | 0 | 0 |
| ENSMUSG00000086139 | 0 | 0 | 0 | 0 | 0 | 0 | 0 |
| ENSMUSG00000086140 | 0 | 0 | 0 | 0 | 0 | 0 | 0 |
| ENSMUSG00000086141 | 0 | 0 | 0 | 0 | 0 | 0 | 0 |
| ENSMUSG00000086142 | 0 | 0 | 0 | 0 | 0 | 0 | 0 |
| ENSMUSG00000086144 | 0 | 0 | 0 | 0 | 0 | 0 | 0 |
| ENSMUSG00000086145 | 0 | 0.25 | 0 | 0 | 0 | 0 | 0 |
| ENSMUSG00000086146 | 0 | 0 | 0 | 0 | 0.244650542 | 0 | 0 |
| ENSMUSG00000086147 | 0 | 0 | 0 | 0 | 0 | 0 | 0.224243824 |
| ENSMUSG00000086148 | 0 | 0 | 0 | 0 | 0 | 0 | 0 |
| ENSMUSG00000086149 | 0.289064826 | 0.25 | 0 | 0.25 | 0 | 0 | 0 |
| ENSMUSG00000086150 | 0.671187741 | 0.580482024 | 0.75 | 0.25 | 0.568060967 | 0.602059991 | 0.355418052 |
| ENSMUSG00000086151 | 0 | 0 | 0.25 | 0 | 0 | 0 | 0 |
| ENSMUSG00000086152 | 0 | 0 | 0 | 0.25 | 0 | 0 | 0.224243824 |
| ENSMUSG00000086153 | 0 | 0 | 0 | 0 | 0 | 0 | 0 |

| | | | | | | | |
|---|---|---|---|---|---|---|---|
| ENSMUSG00000086154 | 0 | 0 | 0 | 0 | 0.244650542 | 0 | 0 |
| ENSMUSG00000086155 | 0.289064826 | 0 | 0 | 0.25 | 0.244650542 | 0 | 0 |
| ENSMUSG00000086156 | 0 | 0 | 0 | 0 | 0 | 0 | 0 |
| ENSMUSG00000086157 | 0 | 0 | 0 | 0 | 0 | 0 | 0 |
| ENSMUSG00000086158 | 1.156259305 | 1.04248125 | 1.04248125 | 1.146240625 | 1 | 1.079181246 | 0.969165684 |
| ENSMUSG00000086159 | 0 | 0.25 | 0 | 0.396240625 | 0.244650542 | 0 | 0 |
| ENSMUSG00000086160 | 0 | 0 | 0 | 0 | 0 | 0 | 0 |
| ENSMUSG00000086161 | 0 | 0.25 | 0 | 0 | 0.244650542 | 0 | 0 |
| ENSMUSG00000086162 | 0 | 0 | 0 | 0 | 0 | 0 | 0 |
| ENSMUSG00000086163 | 0 | 0 | 0 | 0 | 0 | 0 | 0 |
| ENSMUSG00000086164 | 0.289064826 | 0 | 0 | 0 | 0.387761935 | 0 | 0.224243824 |
| ENSMUSG00000086165 | 0 | 0.5 | 0.396240625 | 0.25 | 0 | 0 | 0 |
| ENSMUSG00000086166 | 0.289064826 | 0.25 | 0 | 0.25 | 0.244650542 | 0.477121255 | 0 |
| ENSMUSG00000086167 | 0 | 0 | 0 | 0 | 0 | 0 | 0 |
| ENSMUSG00000086168 | 0 | 0 | 0 | 0 | 0 | 0 | 0 |
| ENSMUSG00000086170 | 0 | 0 | 0 | 0 | 0 | 0 | 0 |
| ENSMUSG00000086171 | 0 | 0.25 | 0 | 0 | 0 | 0 | 0 |
| ENSMUSG00000086172 | 0 | 0 | 0 | 0.580482024 | 0.244650542 | 0 | 0.224243824 |
| ENSMUSG00000086173 | 0 | 0 | 0 | 0 | 0 | 0.301029996 | 0 |
| ENSMUSG00000086174 | 0 | 0 | 0 | 0 | 0 | 0 | 0 |
| ENSMUSG00000086175 | 0 | 0 | 0 | 0 | 0 | 0 | 0 |
| ENSMUSG00000086176 | 0 | 0 | 0 | 0 | 0 | 0 | 0 |
| ENSMUSG00000086177 | 0 | 0 | 0 | 0 | 0 | 0 | 0 |
| ENSMUSG00000086179 | 0 | 0 | 0 | 0 | 0 | 0 | 0 |
| ENSMUSG00000086181 | 0.578129653 | 0.79248125 | 0.75 | 0 | 0 | 0 | 0.520678036 |
| ENSMUSG00000086184 | 0 | 0 | 0 | 0 | 0.244650542 | 0 | 0 |
| ENSMUSG00000086185 | 0 | 0 | 0 | 0 | 0 | 0 | 0 |
| ENSMUSG00000086186 | 0.45815691 | 0 | 0.25 | 0 | 0 | 0 | 0 |
| ENSMUSG00000086187 | 0 | 0 | 0 | 0.25 | 0 | 0 | 0 |
| ENSMUSG00000086188 | 0.289064826 | 0.25 | 0.5 | 0.25 | 0.489301084 | 0.301029996 | 0.355418052 |
| ENSMUSG00000086189 | 0.289064826 | 0.25 | 0.396240625 | 0.396240625 | 0.568060967 | 0.477121255 | 0.224243824 |
| ENSMUSG00000086190 | 0 | 0 | 0 | 0 | 0 | 0 | 0 |
| ENSMUSG00000086191 | 0.289064826 | 0 | 0.396240625 | 0 | 0.489301084 | 0.477121255 | 0.224243824 |
| ENSMUSG00000086192 | 0 | 0.25 | 0.25 | 0 | 0.244650542 | 0 | 0.224243824 |

| | | | | | | | |
|---|---|---|---|---|---|---|---|
| ENSMUSG00000086193 | 0 | 0 | 0.396240625 | 0 | 0 | 0 | 0 |
| ENSMUSG00000086194 | 0 | 0 | 0 | 0 | 0 | 0 | 0 |
| ENSMUSG00000086195 | 0 | 0 | 0 | 0 | 0 | 0 | 0 |
| ENSMUSG00000086196 | 0.289064826 | 0 | 0 | 0 | 0 | 0 | 0 |
| ENSMUSG00000086197 | 0 | 0 | 0 | 0 | 0 | 0 | 0 |
| ENSMUSG00000086199 | 0 | 0 | 0.25 | 0.396240625 | 0.244650542 | 0.301029996 | 0 |
| ENSMUSG00000086201 | 0 | 0 | 0 | 0 | 0 | 0 | 0 |
| ENSMUSG00000086202 | 0 | 0 | 0 | 0 | 0 | 0 | 0 |
| ENSMUSG00000086203 | 0 | 0.25 | 0 | 0 | 0 | 0 | 0.224243824 |
| ENSMUSG00000086204 | 0 | 0 | 0 | 0 | 0 | 0 | 0 |
| ENSMUSG00000086205 | 0 | 0 | 0 | 0 | 0 | 0 | 0 |
| ENSMUSG00000086206 | 0 | 0 | 0.25 | 0 | 0 | 0 | 0 |
| ENSMUSG00000086207 | 0 | 0 | 0 | 0 | 0 | 0 | 0 |
| ENSMUSG00000086209 | 0 | 0 | 0 | 0 | 0 | 0 | 0 |
| ENSMUSG00000086211 | 0 | 0 | 0 | 0 | 0 | 0 | 0 |
| ENSMUSG00000086212 | 0.747221736 | 0.5 | 0.396240625 | 0.646240625 | 0.632412477 | 0.301029996 | 0.224243824 |
| ENSMUSG00000086213 | 0.578129653 | 0.396240625 | 0.396240625 | 0 | 0 | 0 | 0.448487648 |
| ENSMUSG00000086214 | 0 | 0 | 0 | 0 | 0 | 0 | 0 |
| ENSMUSG00000086216 | 0.289064826 | 0 | 0 | 0.25 | 0 | 0 | 0 |
| ENSMUSG00000086219 | 0.811507563 | 0.646240625 | 0.396240625 | 0 | 0 | 0 | 0 |
| ENSMUSG00000086220 | 0 | 0 | 0 | 0 | 0 | 0 | 0 |
| ENSMUSG00000086221 | 0 | 0 | 0 | 0 | 0 | 0 | 0 |
| ENSMUSG00000086222 | 0 | 0 | 0 | 0 | 0 | 0 | 0 |
| ENSMUSG00000086224 | 0 | 0.25 | 0.25 | 0 | 0.244650542 | 0 | 0.355418052 |
| ENSMUSG00000086225 | 0 | 0 | 0 | 0 | 0 | 0 | 0 |
| ENSMUSG00000086226 | 0 | 0 | 0 | 0 | 0 | 0 | 0 |
| ENSMUSG00000086227 | 0 | 0 | 0 | 0 | 0 | 0 | 0 |
| ENSMUSG00000086228 | 0.289064826 | 0.25 | 0.25 | 0.580482024 | 0 | 0 | 0.629532004 |
| ENSMUSG00000086229 | 0.45815691 | 0 | 0 | 0 | 0 | 0 | 0.224243824 |
| ENSMUSG00000086231 | 0 | 0 | 0 | 0 | 0 | 0 | 0 |
| ENSMUSG00000086232 | 0 | 0 | 0 | 0 | 0 | 0 | 0 |
| ENSMUSG00000086233 | 0.578129653 | 0 | 0.580482024 | 0 | 0 | 0 | 0 |
| ENSMUSG00000086234 | 0 | 0 | 0 | 0 | 0 | 0 | 0 |
| ENSMUSG00000086235 | 0 | 0 | 0 | 0 | 0 | 0 | 0 |

| | | | | | | | |
|---|---|---|---|---|---|---|---|
| ENSMUSG00000086236 | 0 | 0 | 0 | 0.25 | 0 | 0 | 0 |
| ENSMUSG00000086237 | 0.289064826 | 0.396240625 | 0.5 | 0.396240625 | 0.387761935 | 0.477121255 | 0.355418052 |
| ENSMUSG00000086238 | 0 | 0 | 0 | 0.25 | 0 | 0 | 0 |
| ENSMUSG00000086239 | 0 | 0 | 0 | 0 | 0 | 0 | 0 |
| ENSMUSG00000086240 | 0 | 0 | 0.25 | 0 | 0 | 0 | 0 |
| ENSMUSG00000086241 | 0 | 0 | 0 | 0 | 0.77552387 | 0.301029996 | 0.224243824 |
| ENSMUSG00000086243 | 0 | 0 | 0 | 0 | 0 | 0 | 0 |
| ENSMUSG00000086244 | 0 | 0 | 0 | 0 | 0 | 0 | 0 |
| ENSMUSG00000086245 | 0.671187741 | 0.646240625 | 0.79248125 | 0.701838731 | 0.846351821 | 0.477121255 | 0.710836105 |
| ENSMUSG00000086246 | 0 | 0 | 0 | 0 | 0 | 0 | 0 |
| ENSMUSG00000086247 | 0.747221736 | 0.92510993 | 0.701838731 | 0.92510993 | 0.846351821 | 0.301029996 | 0.829800754 |
| ENSMUSG00000086248 | 0 | 0 | 0 | 0 | 0 | 0 | 0 |
| ENSMUSG00000086249 | 0 | 0 | 0 | 0 | 0 | 0 | 0 |
| ENSMUSG00000086250 | 0 | 0 | 0 | 0 | 0 | 0 | 0 |
| ENSMUSG00000086251 | 0 | 0 | 0 | 0 | 0 | 0 | 0.224243824 |
| ENSMUSG00000086253 | 0 | 0 | 0 | 0 | 0 | 0 | 0 |
| ENSMUSG00000086254 | 0 | 0.25 | 0 | 0 | 0 | 0 | 0 |
| ENSMUSG00000086255 | 0 | 0 | 0 | 0 | 0 | 0 | 0 |
| ENSMUSG00000086256 | 0 | 0.25 | 0 | 0 | 0 | 0 | 0 |
| ENSMUSG00000086257 | 0 | 0 | 0 | 0 | 0 | 0 | 0 |
| ENSMUSG00000086258 | 0 | 0 | 0 | 0 | 0 | 0 | 0.224243824 |
| ENSMUSG00000086259 | 0 | 0 | 0 | 0 | 0 | 0 | 0 |
| ENSMUSG00000086260 | 0 | 0 | 0 | 0 | 0 | 0 | 0 |
| ENSMUSG00000086261 | 0 | 0 | 0 | 0 | 0 | 0.301029996 | 0 |
| ENSMUSG00000086262 | 0 | 0 | 0 | 0 | 0 | 0 | 0 |
| ENSMUSG00000086263 | 0 | 0 | 0 | 0 | 0 | 0 | 0 |
| ENSMUSG00000086264 | 0.289064826 | 0.580482024 | 0.25 | 0.25 | 0.387761935 | 0.301029996 | 0 |
| ENSMUSG00000086265 | 0 | 0 | 0 | 0 | 0 | 0 | 0 |
| ENSMUSG00000086266 | 0 | 0 | 0 | 0 | 0 | 0.301029996 | 0 |
| ENSMUSG00000086267 | 0.45815691 | 0.25 | 0.396240625 | 0 | 0.387761935 | 0 | 0.448487648 |
| ENSMUSG00000086268 | 0 | 0 | 0 | 0 | 0 | 0 | 0 |
| ENSMUSG00000086269 | 0.578129653 | 0.25 | 0 | 0.25 | 0.244650542 | 0.301029996 | 0 |
| ENSMUSG00000086272 | 0.45815691 | 0.396240625 | 0.396240625 | 0 | 0 | 0 | 0.224243824 |
| ENSMUSG00000086273 | 0.289064826 | 0.5 | 0.25 | 0 | 0 | 0 | 0.224243824 |

| | | | | | | | |
|---|---|---|---|---|---|---|---|
| ENSMUSG00000086275 | 0 | 0 | 0.25 | 0 | 0.244650542 | 0 | 0 |
| ENSMUSG00000086277 | 0 | 0 | 0 | 0 | 0 | 0 | 0 |
| ENSMUSG00000086278 | 0 | 0 | 0 | 0 | 0 | 0 | 0 |
| ENSMUSG00000086279 | 0 | 0 | 0 | 0 | 0 | 0 | 0 |
| ENSMUSG00000086282 | 0 | 0 | 0 | 0 | 0 | 0 | 0 |
| ENSMUSG00000086283 | 0.747221736 | 0.75 | 0.75 | 0.396240625 | 0.244650542 | 0.477121255 | 0.355418052 |
| ENSMUSG00000086284 | 0 | 0 | 0 | 0 | 0 | 0 | 0 |
| ENSMUSG00000086286 | 0.289064826 | 0 | 0 | 0 | 0 | 0 | 0 |
| ENSMUSG00000086287 | 0 | 0.25 | 0 | 0 | 0 | 0 | 0 |
| ENSMUSG00000086288 | 0.91631382 | 0.830482024 | 0.701838731 | 0.896240625 | 1.020174412 | 0.602059991 | 0.916588299 |
| ENSMUSG00000086289 | 0 | 0 | 0 | 0 | 0 | 0 | 0 |
| ENSMUSG00000086290 | 1.876566388 | 1.695339928 | 1.744319981 | 1.660964047 | 1.680838453 | 1.716003344 | 1.574722613 |
| ENSMUSG00000086291 | 0 | 0.25 | 0 | 0.25 | 0.489301084 | 0.301029996 | 0.520678036 |
| ENSMUSG00000086292 | 0.289064826 | 0.25 | 0 | 0.396240625 | 0.244650542 | 0.301029996 | 0 |
| ENSMUSG00000086293 | 0 | 0 | 0 | 0 | 0 | 0 | 0 |
| ENSMUSG00000086296 | 0.45815691 | 0.396240625 | 0.396240625 | 0.5 | 0.733951626 | 1 | 0.672731473 |
| ENSMUSG00000086297 | 0 | 0 | 0 | 0 | 0 | 0 | 0 |
| ENSMUSG00000086298 | 0 | 0 | 0 | 0 | 0 | 0 | 0 |
| ENSMUSG00000086302 | 0 | 0 | 0 | 0 | 0 | 0 | 0 |
| ENSMUSG00000086306 | 0 | 0 | 0 | 0 | 0 | 0 | 0 |
| ENSMUSG00000086308 | 0.289064826 | 0.396240625 | 0 | 0.25 | 0.244650542 | 0 | 0.224243824 |
| ENSMUSG00000086310 | 0 | 0 | 0 | 0 | 0 | 0 | 0 |
| ENSMUSG00000086311 | 0 | 0 | 0 | 0.25 | 0 | 0 | 0.224243824 |
| ENSMUSG00000086312 | 0 | 0 | 0 | 0 | 0 | 0 | 0 |
| ENSMUSG00000086313 | 0 | 0 | 0 | 0 | 0 | 0 | 0 |
| ENSMUSG00000086315 | 0 | 0 | 0 | 0 | 0 | 0 | 0 |
| ENSMUSG00000086316 | 1.920505136 | 1.884789703 | 1.874961472 | 1.970660762 | 1.937354184 | 2.178976947 | 1.829800754 |
| ENSMUSG00000086319 | 0 | 0 | 0 | 0 | 0 | 0 | 0 |
| ENSMUSG00000086320 | 1.418409478 | 1.356566189 | 1.311981878 | 1.04248125 | 1.149965125 | 1.204119983 | 1.066254157 |
| ENSMUSG00000086321 | 0 | 0 | 0 | 0 | 0 | 0 | 0 |
| ENSMUSG00000086322 | 0 | 0 | 0 | 0 | 0 | 0 | 0.224243824 |
| ENSMUSG00000086324 | 1.404271433 | 1.418106335 | 1.418106335 | 1.356566189 | 1.27449559 | 1.342422681 | 1.252393349 |
| ENSMUSG00000086326 | 0 | 0 | 0 | 0 | 0 | 0 | 0 |
| ENSMUSG00000086327 | 0 | 0 | 0 | 0 | 0 | 0 | 0 |

| | | | | | | | |
|---|---|---|---|---|---|---|---|
| ENSMUSG00000086328 | 0.747221736 | 0 | 0.580482024 | 0.5 | 0.733951626 | 0 | 0.448487648 |
| ENSMUSG00000086329 | 0 | 0 | 0 | 0 | 0 | 0 | 0 |
| ENSMUSG00000086330 | 0.747221736 | 0.75 | 1 | 1 | 0.77552387 | 0.477121255 | 0.952571511 |
| ENSMUSG00000086331 | 1.036286563 | 0.92510993 | 1.02186571 | 0.864857905 | 0.931471446 | 0.84509804 | 0.672731473 |
| ENSMUSG00000086332 | 0 | 0 | 0 | 0 | 0 | 0 | 0 |
| ENSMUSG00000086333 | 0 | 0 | 0 | 0 | 0 | 0 | 0 |
| ENSMUSG00000086335 | 0 | 0 | 0 | 0 | 0.244650542 | 0 | 0 |
| ENSMUSG00000086336 | 0 | 0 | 0 | 0 | 0 | 0 | 0 |
| ENSMUSG00000086337 | 0 | 0 | 0 | 0 | 0 | 0 | 0 |
| ENSMUSG00000086338 | 0 | 0 | 0 | 0 | 0 | 0 | 0 |
| ENSMUSG00000086339 | 0 | 0 | 0 | 0 | 0 | 0 | 0 |
| ENSMUSG00000086340 | 0 | 0.396240625 | 0 | 0.25 | 0.244650542 | 0 | 0 |
| ENSMUSG00000086341 | 0 | 0 | 0.25 | 0.396240625 | 0.387761935 | 0 | 0.448487648 |
| ENSMUSG00000086342 | 0.671187741 | 0.646240625 | 0.580482024 | 0.396240625 | 0.812711509 | 0.77815125 | 0.629532004 |
| ENSMUSG00000086344 | 0 | 0 | 0 | 0.25 | 0 | 0.301029996 | 0.224243824 |
| ENSMUSG00000086346 | 0 | 0 | 0 | 0 | 0 | 0 | 0 |
| ENSMUSG00000086347 | 0 | 0.25 | 0.25 | 0 | 0 | 0 | 0.224243824 |
| ENSMUSG00000086348 | 0 | 0 | 0 | 0 | 0 | 0 | 0 |
| ENSMUSG00000086349 | 0.45815691 | 0 | 0 | 0 | 0.244650542 | 0 | 0.224243824 |
| ENSMUSG00000086350 | 0.289064826 | 0.396240625 | 0.79248125 | 0.5 | 0.632412477 | 0.477121255 | 0.629532004 |
| ENSMUSG00000086352 | 0 | 0 | 0.25 | 0 | 0 | 0 | 0 |
| ENSMUSG00000086353 | 0 | 0 | 0.25 | 0 | 0 | 0 | 0 |
| ENSMUSG00000086354 | 0 | 0 | 0 | 0 | 0 | 0 | 0 |
| ENSMUSG00000086355 | 0 | 0 | 0 | 0 | 0 | 0 | 0 |
| ENSMUSG00000086356 | 0 | 0 | 0 | 0 | 0 | 0 | 0 |
| ENSMUSG00000086358 | 0.91631382 | 0.396240625 | 0.92510993 | 0.79248125 | 0.489301084 | 0.477121255 | 0.629532004 |
| ENSMUSG00000086359 | 0.867194479 | 0.75 | 0.396240625 | 0.580482024 | 0.568060967 | 0.477121255 | 0.672731473 |
| ENSMUSG00000086360 | 0 | 0 | 0 | 0.25 | 0 | 0 | 0 |
| ENSMUSG00000086361 | 0 | 0 | 0 | 0 | 0 | 0 | 0 |
| ENSMUSG00000086363 | 0.289064826 | 0.25 | 0.25 | 0.25 | 0.244650542 | 0 | 0 |
| ENSMUSG00000086364 | 0 | 0 | 0 | 0 | 0 | 0 | 0 |
| ENSMUSG00000086365 | 0 | 0 | 0 | 0 | 0 | 0 | 0 |
| ENSMUSG00000086366 | 0 | 0 | 0 | 0 | 0 | 0 | 0 |
| ENSMUSG00000086367 | 0 | 0 | 0 | 0 | 0 | 0 | 0 |

| | | | | | | | |
|---|---|---|---|---|---|---|---|
| ENSMUSG00000086368 | 0.45815691 | 0.396240625 | 0.396240625 | 0.396240625 | 0.244650542 | 0.477121255 | 0.355418052 |
| ENSMUSG00000086369 | 0 | 0 | 0 | 0 | 0 | 0 | 0 |
| ENSMUSG00000086370 | 1.881174527 | 1.692046081 | 1.777131114 | 1.688721876 | 1.680838453 | 1.819543936 | 1.523605797 |
| ENSMUSG00000086371 | 0 | 0 | 0 | 0 | 0 | 0 | 0 |
| ENSMUSG00000086372 | 0 | 0 | 0 | 0 | 0 | 0 | 0 |
| ENSMUSG00000086373 | 0 | 0 | 0 | 0 | 0 | 0 | 0 |
| ENSMUSG00000086375 | 0 | 0 | 0 | 0 | 0.244650542 | 0 | 0 |
| ENSMUSG00000086376 | 0 | 0 | 0 | 0 | 0 | 0 | 0 |
| ENSMUSG00000086377 | 0 | 0.396240625 | 0.25 | 0 | 0 | 0.301029996 | 0 |
| ENSMUSG00000086378 | 0 | 0 | 0 | 0 | 0 | 0 | 0 |
| ENSMUSG00000086379 | 0 | 0 | 0 | 0 | 0 | 0 | 0 |
| ENSMUSG00000086380 | 0 | 0 | 0 | 0 | 0 | 0 | 0 |
| ENSMUSG00000086381 | 0 | 0 | 0 | 0 | 0 | 0 | 0 |
| ENSMUSG00000086382 | 0.289064826 | 0.5 | 0.396240625 | 0.75 | 0.632412477 | 0.698970004 | 0.775756176 |
| ENSMUSG00000086384 | 0 | 0 | 0 | 0 | 0 | 0 | 0 |
| ENSMUSG00000086385 | 0 | 0 | 0 | 0 | 0 | 0 | 0 |
| ENSMUSG00000086386 | 0 | 0 | 0 | 0 | 0 | 0 | 0 |
| ENSMUSG00000086387 | 0 | 0 | 0 | 0 | 0 | 0.301029996 | 0 |
| ENSMUSG00000086389 | 0.671187741 | 0.79248125 | 0.75 | 0.864857905 | 0.905314583 | 0.77815125 | 0.969165684 |
| ENSMUSG00000086390 | 0.811507563 | 0.396240625 | 0.701838731 | 0.976722649 | 0.632412477 | 0.301029996 | 0.829800754 |
| ENSMUSG00000086391 | 0.289064826 | 0 | 0 | 0 | 0.387761935 | 0.301029996 | 0 |
| ENSMUSG00000086392 | 0.747221736 | 0.580482024 | 0.396240625 | 0.580482024 | 0.77552387 | 0.698970004 | 0.448487648 |
| ENSMUSG00000086394 | 0 | 0 | 0 | 0 | 0 | 0 | 0 |
| ENSMUSG00000086395 | 0 | 0.25 | 0.25 | 0.5 | 0 | 0.301029996 | 0.448487648 |
| ENSMUSG00000086396 | 0 | 0 | 0 | 0 | 0 | 0 | 0 |
| ENSMUSG00000086399 | 0.578129653 | 0.25 | 0.701838731 | 0.864857905 | 0.812711509 | 0.602059991 | 0.74492186 |
| ENSMUSG00000086401 | 0.747221736 | 0.92510993 | 0.864857905 | 0.79248125 | 0.489301084 | 0.954242509 | 0.74492186 |
| ENSMUSG00000086402 | 0 | 0 | 0 | 0 | 0 | 0 | 0 |
| ENSMUSG00000086404 | 0.289064826 | 0.25 | 0 | 0.580482024 | 0.387761935 | 0.301029996 | 0.520678036 |
| ENSMUSG00000086405 | 0 | 0 | 0 | 0 | 0 | 0 | 0 |
| ENSMUSG00000086407 | 0 | 0 | 0 | 0 | 0 | 0 | 0.224243824 |
| ENSMUSG00000086408 | 0 | 0 | 0 | 0 | 0 | 0 | 0 |
| ENSMUSG00000086409 | 0 | 0 | 0 | 0 | 0 | 0 | 0 |
| ENSMUSG00000086410 | 0 | 0 | 0 | 0 | 0 | 0 | 0 |

| | | | | | | | |
|---|---|---|---|---|---|---|---|
| ENSMUSG00000086411 | 0 | 0 | 0 | 0 | 0 | 0 | 0 |
| ENSMUSG00000086413 | 0 | 0 | 0 | 0.25 | 0.489301084 | 0.301029996 | 0.224243824 |
| ENSMUSG00000086414 | 1.156259305 | 1.114857905 | 1.080482024 | 0.701838731 | 0.632412477 | 0.602059991 | 0.896975297 |
| ENSMUSG00000086415 | 0 | 0 | 0 | 0 | 0 | 0 | 0 |
| ENSMUSG00000086416 | 1.069666964 | 1.061981878 | 1 | 0.646240625 | 0.632412477 | 0.477121255 | 0.710836105 |
| ENSMUSG00000086417 | 0 | 0 | 0 | 0 | 0.244650542 | 0 | 0.224243824 |
| ENSMUSG00000086418 | 0 | 0 | 0 | 0 | 0 | 0 | 0 |
| ENSMUSG00000086420 | 0 | 0 | 0 | 0 | 0 | 0 | 0 |
| ENSMUSG00000086421 | 0.45815691 | 0.830482024 | 0.75 | 0.896240625 | 0.686820904 | 0.698970004 | 0.876096088 |
| ENSMUSG00000086422 | 0 | 0 | 0 | 0 | 0 | 0 | 0 |
| ENSMUSG00000086423 | 0 | 0 | 0 | 0 | 0 | 0 | 0 |
| ENSMUSG00000086424 | 0 | 0 | 0 | 0 | 0 | 0.477121255 | 0.224243824 |
| ENSMUSG00000086425 | 0 | 0 | 0 | 0 | 0 | 0 | 0 |
| ENSMUSG00000086426 | 0 | 0 | 0 | 0 | 0 | 0 | 0 |
| ENSMUSG00000086427 | 0.289064826 | 0.25 | 0.580482024 | 0 | 0.387761935 | 0.477121255 | 0.448487648 |
| ENSMUSG00000086428 | 0 | 0 | 0 | 0 | 0 | 0 | 0 |
| ENSMUSG00000086429 | 2.016886209 | 1.982684334 | 1.914552871 | 1.916333979 | 1.86473911 | 2.033423755 | 1.637827726 |
| ENSMUSG00000086430 | 0.289064826 | 0 | 0.396240625 | 0.580482024 | 0 | 0.301029996 | 0.224243824 |
| ENSMUSG00000086431 | 0.289064826 | 0 | 0 | 0 | 0 | 0 | 0 |
| ENSMUSG00000086432 | 0 | 0 | 0 | 0 | 0 | 0 | 0 |
| ENSMUSG00000086433 | 0 | 0 | 0 | 0 | 0 | 0 | 0 |
| ENSMUSG00000086434 | 0 | 0 | 0 | 0 | 0 | 0 | 0 |
| ENSMUSG00000086436 | 0 | 0 | 0 | 0 | 0 | 0 | 0.224243824 |
| ENSMUSG00000086437 | 0 | 0 | 0 | 0 | 0 | 0 | 0 |
| ENSMUSG00000086438 | 0 | 0 | 0 | 0 | 0 | 0.301029996 | 0 |
| ENSMUSG00000086439 | 0 | 0 | 0 | 0 | 0 | 0 | 0 |
| ENSMUSG00000086441 | 0 | 0 | 0 | 0 | 0 | 0 | 0 |
| ENSMUSG00000086442 | 0 | 0 | 0 | 0.25 | 0.387761935 | 0.301029996 | 0 |
| ENSMUSG00000086443 | 0 | 0.25 | 0.396240625 | 0.396240625 | 0 | 0 | 0.224243824 |
| ENSMUSG00000086444 | 0.289064826 | 0.396240625 | 0.5 | 0.25 | 0 | 0.602059991 | 0 |
| ENSMUSG00000086445 | 0 | 0 | 0 | 0 | 0 | 0 | 0 |
| ENSMUSG00000086446 | 0.289064826 | 0.25 | 0.25 | 0.25 | 0.244650542 | 0 | 0.355418052 |
| ENSMUSG00000086447 | 0 | 0 | 0 | 0 | 0 | 0 | 0 |
| ENSMUSG00000086448 | 0 | 0.25 | 0.5 | 0.646240625 | 0.877063019 | 0.84509804 | 0.710836105 |

| | | | | | | | |
|---|---|---|---|---|---|---|---|
| ENSMUSG00000086449 | 0 | 0 | 0 | 0 | 0 | 0 | 0 |
| ENSMUSG00000086450 | 1.069666964 | 1.04248125 | 0.79248125 | 0.79248125 | 0.846351821 | 0.77815125 | 0.876096088 |
| ENSMUSG00000086451 | 0 | 0 | 0 | 0 | 0.387761935 | 0 | 0.224243824 |
| ENSMUSG00000086452 | 0 | 0 | 0 | 0 | 0 | 0 | 0 |
| ENSMUSG00000086453 | 0.45815691 | 0.5 | 0.25 | 0.5 | 0.387761935 | 0.698970004 | 0.579661877 |
| ENSMUSG00000086454 | 0 | 0.396240625 | 0.646240625 | 0.580482024 | 0.489301084 | 0.602059991 | 0.355418052 |
| ENSMUSG00000086455 | 0.578129653 | 0.75 | 0.396240625 | 0.75 | 0.387761935 | 0.77815125 | 0 |
| ENSMUSG00000086458 | 0 | 0 | 0 | 0 | 0 | 0 | 0 |
| ENSMUSG00000086459 | 0 | 0 | 0 | 0 | 0.244650542 | 0 | 0 |
| ENSMUSG00000086460 | 0 | 0 | 0 | 0 | 0 | 0 | 0 |
| ENSMUSG00000086462 | 0 | 0 | 0 | 0 | 0 | 0 | 0 |
| ENSMUSG00000086464 | 0.747221736 | 0.580482024 | 0.396240625 | 0.396240625 | 0.733951626 | 0.477121255 | 0.448487648 |
| ENSMUSG00000086465 | 0 | 0 | 0 | 0 | 0 | 0 | 0 |
| ENSMUSG00000086467 | 0 | 0 | 0 | 0.25 | 0 | 0 | 0 |
| ENSMUSG00000086468 | 0 | 0.79248125 | 0.646240625 | 0.79248125 | 0.387761935 | 0.301029996 | 0.355418052 |
| ENSMUSG00000086470 | 0 | 0 | 0 | 0 | 0 | 0 | 0 |
| ENSMUSG00000086472 | 0.578129653 | 0.5 | 0.5 | 0.580482024 | 0.812711509 | 0 | 0.448487648 |
| ENSMUSG00000086473 | 0.45815691 | 0.5 | 0.25 | 0.5 | 0.244650542 | 0 | 0.448487648 |
| ENSMUSG00000086474 | 0 | 0.396240625 | 0.5 | 0.75 | 0.387761935 | 0.477121255 | 0.355418052 |
| ENSMUSG00000086475 | 0.289064826 | 0.396240625 | 0.5 | 0.5 | 0.244650542 | 0.602059991 | 0.448487648 |
| ENSMUSG00000086477 | 0 | 0 | 0.25 | 0.25 | 0.387761935 | 0 | 0.224243824 |
| ENSMUSG00000086478 | 0 | 0 | 0 | 0 | 0 | 0 | 0 |
| ENSMUSG00000086479 | 0 | 0 | 0 | 0 | 0 | 0 | 0 |
| ENSMUSG00000086480 | 0.747221736 | 1.080482024 | 0.951838731 | 0.830482024 | 1.149965125 | 1 | 1.054044578 |
| ENSMUSG00000086481 | 0 | 0 | 0 | 0 | 0 | 0 | 0 |
| ENSMUSG00000086482 | 0 | 0 | 0 | 0 | 0 | 0 | 0 |
| ENSMUSG00000086483 | 0 | 0 | 0 | 0 | 0 | 0 | 0 |
| ENSMUSG00000086484 | 0 | 0 | 0 | 0 | 0.244650542 | 0 | 0 |
| ENSMUSG00000086485 | 0 | 0 | 0 | 0 | 0 | 0 | 0 |
| ENSMUSG00000086486 | 0 | 0.396240625 | 0 | 0.5 | 0.489301084 | 0.301029996 | 0.224243824 |
| ENSMUSG00000086487 | 0 | 0 | 0 | 0 | 0 | 0 | 0 |
| ENSMUSG00000086489 | 0 | 0 | 0 | 0 | 0 | 0 | 0 |
| ENSMUSG00000086490 | 0 | 0 | 0 | 0 | 0 | 0 | 0 |
| ENSMUSG00000086491 | 0 | 0 | 0 | 0 | 0 | 0 | 0 |

| | | | | | | | |
|---|---|---|---|---|---|---|---|
| ENSMUSG00000086493 | 0 | 0 | 0.25 | 0 | 0 | 0 | 0 |
| ENSMUSG00000086494 | 0 | 0 | 0 | 0 | 0 | 0 | 0 |
| ENSMUSG00000086495 | 0 | 0 | 0 | 0.25 | 0.387761935 | 0 | 0.224243824 |
| ENSMUSG00000086496 | 0 | 0 | 0 | 0.396240625 | 0.244650542 | 0 | 0.224243824 |
| ENSMUSG00000086497 | 0 | 0 | 0 | 0 | 0 | 0 | 0 |
| ENSMUSG00000086498 | 0 | 0.25 | 0.25 | 0 | 0 | 0 | 0 |
| ENSMUSG00000086499 | 0.45815691 | 0 | 0 | 0 | 0 | 0 | 0 |
| ENSMUSG00000086500 | 0 | 0 | 0 | 0 | 0 | 0 | 0 |
| ENSMUSG00000086501 | 0 | 0 | 0 | 0 | 0 | 0 | 0 |
| ENSMUSG00000086502 | 1.269664473 | 1.302363341 | 1.431980114 | 1.458222504 | 1.335652905 | 1.579783597 | 1.383567577 |
| ENSMUSG00000086503 | 0.811507563 | 0.580482024 | 0.701838731 | 0.75 | 0.812711509 | 0.698970004 | 0.775756176 |
| ENSMUSG00000086504 | 0 | 0 | 0 | 0 | 0 | 0 | 0 |
| ENSMUSG00000086505 | 0 | 0 | 0 | 0 | 0 | 0 | 0 |
| ENSMUSG00000086506 | 0.45815691 | 0.580482024 | 0 | 0 | 0.77552387 | 0 | 0.629532004 |
| ENSMUSG00000086507 | 0 | 0 | 0 | 0 | 0.244650542 | 0 | 0.224243824 |
| ENSMUSG00000086508 | 0 | 0 | 0 | 0 | 0 | 0 | 0 |
| ENSMUSG00000086509 | 0 | 0 | 0 | 0 | 0 | 0 | 0 |
| ENSMUSG00000086512 | 0 | 0 | 0 | 0 | 0 | 0 | 0 |
| ENSMUSG00000086513 | 0 | 0 | 0 | 0 | 0 | 0 | 0 |
| ENSMUSG00000086514 | 0.289064826 | 0 | 0 | 0.25 | 0 | 0 | 0.355418052 |
| ENSMUSG00000086515 | 0.289064826 | 0 | 0 | 0.5 | 0.244650542 | 0 | 0.448487648 |
| ENSMUSG00000086517 | 0 | 0 | 0 | 0 | 0 | 0 | 0 |
| ENSMUSG00000086518 | 0 | 0 | 0 | 0 | 0 | 0 | 0 |
| ENSMUSG00000086519 | 0 | 0 | 0 | 0 | 0 | 0 | 0 |
| ENSMUSG00000086521 | 0 | 0 | 0 | 0 | 0 | 0 | 0 |
| ENSMUSG00000086522 | 0.811507563 | 0.830482024 | 0.976722649 | 0.646240625 | 0.244650542 | 0.477121255 | 0.448487648 |
| ENSMUSG00000086523 | 0 | 0 | 0 | 0 | 0 | 0 | 0 |
| ENSMUSG00000086524 | 0 | 0 | 0 | 0 | 0 | 0 | 0 |
| ENSMUSG00000086525 | 0 | 0 | 0 | 0 | 0 | 0 | 0 |
| ENSMUSG00000086526 | 0.289064826 | 0.5 | 0.580482024 | 0.646240625 | 0.387761935 | 0.477121255 | 0.224243824 |
| ENSMUSG00000086527 | 0 | 0 | 0 | 0 | 0 | 0 | 0 |
| ENSMUSG00000086528 | 1.069666964 | 1.080482024 | 0.92510993 | 1.02186571 | 0.77552387 | 0.602059991 | 0.775756176 |
| ENSMUSG00000086529 | 0 | 0 | 0 | 0 | 0 | 0 | 0.224243824 |
| ENSMUSG00000086530 | 0 | 0 | 0 | 0 | 0 | 0 | 0 |

| | | | | | | | |
|---|---|---|---|---|---|---|---|
| ENSMUSG00000086531 | 0.289064826 | 0 | 0 | 0 | 0 | 0 | 0 |
| ENSMUSG00000086533 | 0.289064826 | 0.580482024 | 0.396240625 | 0.25 | 0.244650542 | 0.477121255 | 0.448487648 |
| ENSMUSG00000086534 | 0 | 0 | 0.25 | 0.5 | 0 | 0 | 0.224243824 |
| ENSMUSG00000086536 | 0.747221736 | 0.864857905 | 0.75 | 1.061981878 | 0.955822902 | 0.84509804 | 0.853775828 |
| ENSMUSG00000086537 | 0 | 0.396240625 | 0.25 | 0.5 | 0.733951626 | 0.602059991 | 0.520678036 |
| ENSMUSG00000086539 | 0.578129653 | 0.396240625 | 0.646240625 | 0.580482024 | 0.632412477 | 0.477121255 | 0.355418052 |
| ENSMUSG00000086540 | 0 | 0 | 0 | 0 | 0 | 0 | 0 |
| ENSMUSG00000086541 | 1 | 0.951838731 | 0.976722649 | 0.580482024 | 0.733951626 | 0 | 0.448487648 |
| ENSMUSG00000086542 | 0 | 0 | 0 | 0 | 0 | 0 | 0 |
| ENSMUSG00000086544 | 0 | 0.25 | 0 | 0.25 | 0.244650542 | 0 | 0 |
| ENSMUSG00000086545 | 0 | 0 | 0 | 0 | 0 | 0 | 0 |
| ENSMUSG00000086546 | 0 | 0 | 0 | 0 | 0 | 0 | 0 |
| ENSMUSG00000086547 | 0 | 0 | 0 | 0 | 0 | 0 | 0 |
| ENSMUSG00000086548 | 0 | 0 | 0 | 0 | 0 | 0 | 0 |
| ENSMUSG00000086549 | 0 | 0 | 0 | 0 | 0 | 0 | 0 |
| ENSMUSG00000086550 | 0 | 0 | 0 | 0 | 0 | 0 | 0 |
| ENSMUSG00000086552 | 0 | 0 | 0 | 0.5 | 0.387761935 | 0.301029996 | 0.224243824 |
| ENSMUSG00000086554 | 0.289064826 | 0.25 | 0.396240625 | 0.5 | 0 | 0 | 0.355418052 |
| ENSMUSG00000086555 | 0 | 0 | 0 | 0 | 0 | 0 | 0 |
| ENSMUSG00000086556 | 0 | 0 | 0 | 0 | 0 | 0 | 0 |
| ENSMUSG00000086557 | 0 | 0 | 0 | 0 | 0 | 0 | 0 |
| ENSMUSG00000086558 | 0 | 0 | 0 | 0 | 0 | 0 | 0 |
| ENSMUSG00000086559 | 0 | 0 | 0 | 0 | 0 | 0 | 0 |
| ENSMUSG00000086560 | 1.289064826 | 1.238549078 | 1.188721876 | 0.79248125 | 1 | 0 | 0.935079929 |
| ENSMUSG00000086561 | 0 | 0 | 0 | 0.25 | 0 | 0 | 0 |
| ENSMUSG00000086562 | 0 | 0 | 0 | 0 | 0 | 0 | 0 |
| ENSMUSG00000086563 | 0 | 0 | 0.396240625 | 0.25 | 0.387761935 | 0.477121255 | 0.629532004 |
| ENSMUSG00000086564 | 0.45815691 | 0 | 0 | 0 | 0 | 0 | 0 |
| ENSMUSG00000086565 | 0 | 0 | 0 | 0 | 0 | 0 | 0 |
| ENSMUSG00000086567 | 0.960252568 | 1.201838731 | 1.214495249 | 1.160964047 | 1.136121934 | 1.301029996 | 1.028149525 |
| ENSMUSG00000086569 | 0 | 0 | 0 | 0 | 0 | 0 | 0 |
| ENSMUSG00000086573 | 0.578129653 | 0.396240625 | 0.396240625 | 0.5 | 0.244650542 | 0.301029996 | 0.224243824 |
| ENSMUSG00000086575 | 0 | 0.25 | 0 | 0 | 0 | 0 | 0 |
| ENSMUSG00000086576 | 0 | 0 | 0 | 0 | 0 | 0 | 0 |

| | | | | | | | |
|---|---|---|---|---|---|---|---|
| ENSMUSG00000086577 | 0 | 0 | 0 | 0 | 0 | 0 | 0 |
| ENSMUSG00000086578 | 0.289064826 | 0.25 | 0 | 0.25 | 0 | 0 | 0 |
| ENSMUSG00000086580 | 0.289064826 | 0.580482024 | 0.25 | 0 | 0.489301084 | 0.301029996 | 0.355418052 |
| ENSMUSG00000086582 | 0 | 0 | 0 | 0 | 0 | 0.301029996 | 0 |
| ENSMUSG00000086583 | 0 | 0.396240625 | 0 | 0 | 0 | 0.301029996 | 0 |
| ENSMUSG00000086584 | 0 | 0.396240625 | 0.580482024 | 0 | 0.244650542 | 0 | 0.355418052 |
| ENSMUSG00000086585 | 0 | 0 | 0 | 0 | 0 | 0 | 0 |
| ENSMUSG00000086586 | 0 | 0 | 0.25 | 0 | 0 | 0.301029996 | 0.224243824 |
| ENSMUSG00000086587 | 0.289064826 | 0.25 | 0 | 0.396240625 | 0 | 0 | 0.224243824 |
| ENSMUSG00000086588 | 0 | 0 | 0 | 0 | 0 | 0 | 0 |
| ENSMUSG00000086589 | 0.289064826 | 0.25 | 0.25 | 0.5 | 0.244650542 | 0.301029996 | 0 |
| ENSMUSG00000086591 | 0 | 0 | 0 | 0 | 0 | 0 | 0 |
| ENSMUSG00000086592 | 0 | 0 | 0 | 0 | 0 | 0 | 0 |
| ENSMUSG00000086593 | 0 | 0 | 0 | 0 | 0 | 0 | 0 |
| ENSMUSG00000086594 | 0 | 0 | 0 | 0 | 0 | 0 | 0 |
| ENSMUSG00000086595 | 0 | 0 | 0 | 0 | 0 | 0 | 0 |
| ENSMUSG00000086596 | 0 | 0 | 0.25 | 0.5 | 0.568060967 | 0.698970004 | 0.74492186 |
| ENSMUSG00000086598 | 0.671187741 | 0.396240625 | 0.25 | 0.580482024 | 0.244650542 | 0.301029996 | 0.224243824 |
| ENSMUSG00000086600 | 0 | 0 | 0 | 0 | 0.387761935 | 0 | 0.224243824 |
| ENSMUSG00000086601 | 0 | 0.25 | 0.25 | 0 | 0 | 0 | 0 |
| ENSMUSG00000086602 | 0 | 0 | 0 | 0 | 0 | 0.301029996 | 0 |
| ENSMUSG00000086603 | 0 | 0 | 0 | 0 | 0 | 0 | 0 |
| ENSMUSG00000086604 | 0 | 0.25 | 0.25 | 0.25 | 0 | 0 | 0.224243824 |
| ENSMUSG00000086605 | 0 | 0 | 0 | 0.25 | 0.568060967 | 0.301029996 | 0.224243824 |
| ENSMUSG00000086606 | 0.289064826 | 0.5 | 0.5 | 0.396240625 | 0.489301084 | 0.698970004 | 0.672731473 |
| ENSMUSG00000086607 | 1.069666964 | 1 | 0.976722649 | 0.92510993 | 0.812711509 | 0.77815125 | 0.916588299 |
| ENSMUSG00000086608 | 0 | 0 | 0 | 0 | 0 | 0 | 0 |
| ENSMUSG00000086610 | 0.289064826 | 0.396240625 | 0 | 0.5 | 0 | 0 | 0.224243824 |
| ENSMUSG00000086612 | 0 | 0 | 0 | 0 | 0 | 0 | 0 |
| ENSMUSG00000086613 | 0 | 0 | 0 | 0 | 0 | 0 | 0 |
| ENSMUSG00000086614 | 0 | 0 | 0 | 0 | 0 | 0 | 0 |
| ENSMUSG00000086617 | 0 | 0 | 0 | 0 | 0 | 0 | 0 |
| ENSMUSG00000086618 | 0 | 0 | 0 | 0 | 0 | 0 | 0 |
| ENSMUSG00000086620 | 0 | 0 | 0 | 0.25 | 0.387761935 | 0 | 0 |

| | | | | | | | |
|---|---|---|---|---|---|---|---|
| ENSMUSG00000086621 | 0 | 0 | 0 | 0 | 0 | 0 | 0 |
| ENSMUSG00000086624 | 0 | 0 | 0 | 0 | 0 | 0 | 0 |
| ENSMUSG00000086625 | 0.289064826 | 0 | 0 | 0.396240625 | 0 | 0 | 0 |
| ENSMUSG00000086628 | 0 | 0 | 0 | 0 | 0 | 0 | 0 |
| ENSMUSG00000086629 | 1.156259305 | 1 | 1.146240625 | 1.146240625 | 1.091002363 | 1.204119983 | 1.100339912 |
| ENSMUSG00000086630 | 0 | 0 | 0 | 0 | 0 | 0 | 0 |
| ENSMUSG00000086631 | 0 | 0 | 0 | 0 | 0 | 0 | 0 |
| ENSMUSG00000086632 | 0 | 0 | 0 | 0 | 0 | 0 | 0 |
| ENSMUSG00000086633 | 0 | 0 | 0 | 0 | 0 | 0 | 0 |
| ENSMUSG00000086636 | 0.578129653 | 0.5 | 0.396240625 | 0.646240625 | 0.387761935 | 0.477121255 | 0.448487648 |
| ENSMUSG00000086637 | 0 | 0 | 0 | 0 | 0 | 0 | 0 |
| ENSMUSG00000086638 | 0.578129653 | 0 | 0.396240625 | 0.396240625 | 0.489301084 | 0.77815125 | 0.355418052 |
| ENSMUSG00000086641 | 0 | 0 | 0 | 0 | 0 | 0 | 0 |
| ENSMUSG00000086643 | 0 | 0 | 0 | 0 | 0 | 0 | 0 |
| ENSMUSG00000086644 | 0 | 0.75 | 0.396240625 | 0.25 | 0 | 0 | 0 |
| ENSMUSG00000086645 | 0 | 0 | 0 | 0 | 0 | 0 | 0 |
| ENSMUSG00000086646 | 0 | 0 | 0 | 0 | 0.244650542 | 0 | 0.224243824 |
| ENSMUSG00000086648 | 0 | 0 | 0 | 0 | 0 | 0 | 0 |
| ENSMUSG00000086649 | 0 | 0.25 | 0 | 0 | 0 | 0 | 0 |
| ENSMUSG00000086650 | 0 | 0 | 0 | 0 | 0 | 0 | 0 |
| ENSMUSG00000086652 | 0 | 0 | 0 | 0 | 0 | 0 | 0 |
| ENSMUSG00000086653 | 0 | 0 | 0 | 0 | 0 | 0 | 0 |
| ENSMUSG00000086654 | 0 | 0 | 0 | 0 | 0.244650542 | 0 | 0 |
| ENSMUSG00000086655 | 0 | 0 | 0 | 0 | 0 | 0 | 0 |
| ENSMUSG00000086656 | 0 | 0.25 | 0.396240625 | 0.25 | 0 | 0 | 0 |
| ENSMUSG00000086657 | 0.578129653 | 0.92510993 | 0.580482024 | 0.580482024 | 0.846351821 | 0.77815125 | 0.803905701 |
| ENSMUSG00000086658 | 0 | 0 | 0 | 0 | 0 | 0 | 0 |
| ENSMUSG00000086659 | 0 | 0 | 0 | 0 | 0 | 0 | 0 |
| ENSMUSG00000086660 | 0 | 0 | 0 | 0 | 0 | 0 | 0 |
| ENSMUSG00000086662 | 0.289064826 | 0.25 | 0.25 | 0 | 0 | 0 | 0.224243824 |
| ENSMUSG00000086665 | 0 | 0 | 0.396240625 | 0.25 | 0 | 0 | 0.224243824 |
| ENSMUSG00000086666 | 0 | 0 | 0 | 0 | 0 | 0 | 0 |
| ENSMUSG00000086667 | 0 | 0 | 0 | 0 | 0 | 0 | 0.224243824 |
| ENSMUSG00000086668 | 0 | 0 | 0 | 0 | 0 | 0 | 0 |

| | | | | | | | |
|---|---|---|---|---|---|---|---|
| ENSMUSG00000086669 | 0 | 0 | 0 | 0 | 0 | 0 | 0 |
| ENSMUSG00000086670 | 0 | 0 | 0 | 0 | 0 | 0 | 0 |
| ENSMUSG00000086671 | 0 | 0 | 0.25 | 0.396240625 | 0.387761935 | 0 | 0.224243824 |
| ENSMUSG00000086674 | 0 | 0.25 | 0 | 0 | 0 | 0 | 0.355418052 |
| ENSMUSG00000086675 | 0 | 0 | 0 | 0 | 0 | 0 | 0 |
| ENSMUSG00000086676 | 0.289064826 | 0 | 0.25 | 0.25 | 0 | 0.301029996 | 0.224243824 |
| ENSMUSG00000086677 | 0 | 0 | 0 | 0.25 | 0 | 0 | 0 |
| ENSMUSG00000086679 | 0 | 0 | 0 | 0 | 0 | 0 | 0 |
| ENSMUSG00000086680 | 0.45815691 | 0 | 0.25 | 0.5 | 0 | 0.301029996 | 0.224243824 |
| ENSMUSG00000086681 | 0.289064826 | 0.396240625 | 0.25 | 0 | 0.568060967 | 0 | 0 |
| ENSMUSG00000086682 | 1.156259305 | 0.830482024 | 0.976722649 | 1.02186571 | 1.020174412 | 1.041392685 | 0.803905701 |
| ENSMUSG00000086683 | 0.289064826 | 0.396240625 | 0 | 0 | 0.489301084 | 0 | 0.448487648 |
| ENSMUSG00000086684 | 0 | 0.5 | 0.396240625 | 0 | 0 | 0 | 0.520678036 |
| ENSMUSG00000086685 | 0 | 0 | 0 | 0 | 0 | 0 | 0 |
| ENSMUSG00000086686 | 0 | 0 | 0 | 0 | 0 | 0 | 0 |
| ENSMUSG00000086687 | 0 | 0 | 0 | 0 | 0 | 0 | 0 |
| ENSMUSG00000086688 | 0.45815691 | 0.5 | 0.396240625 | 0.701838731 | 0.489301084 | 0.602059991 | 0.579661877 |
| ENSMUSG00000086689 | 0 | 0 | 0 | 0 | 0 | 0 | 0.224243824 |
| ENSMUSG00000086690 | 0 | 0 | 0 | 0 | 0 | 0.301029996 | 0 |
| ENSMUSG00000086691 | 0 | 0 | 0 | 0 | 0 | 0 | 0 |
| ENSMUSG00000086693 | 0.578129653 | 0.646240625 | 0.5 | 0.5 | 0.686820904 | 0.698970004 | 0.629532004 |
| ENSMUSG00000086694 | 0 | 0 | 0 | 0 | 0 | 0 | 0 |
| ENSMUSG00000086695 | 0.289064826 | 0.25 | 0.396240625 | 0 | 0.387761935 | 0 | 0.520678036 |
| ENSMUSG00000086696 | 0 | 0 | 0 | 0 | 0 | 0 | 0 |
| ENSMUSG00000086697 | 0.289064826 | 0.25 | 0.5 | 0.5 | 0.244650542 | 0.477121255 | 0.224243824 |
| ENSMUSG00000086698 | 0 | 0 | 0 | 0 | 0 | 0 | 0 |
| ENSMUSG00000086699 | 0 | 0 | 0.396240625 | 0 | 0 | 0.301029996 | 0 |
| ENSMUSG00000086700 | 0 | 0 | 0 | 0 | 0 | 0 | 0 |
| ENSMUSG00000086701 | 0 | 0 | 0 | 0 | 0 | 0 | 0 |
| ENSMUSG00000086702 | 0 | 0 | 0 | 0 | 0 | 0 | 0 |
| ENSMUSG00000086703 | 0 | 0 | 0 | 0 | 0 | 0 | 0 |
| ENSMUSG00000086704 | 0 | 0 | 0 | 0 | 0 | 0 | 0 |
| ENSMUSG00000086706 | 0 | 0.25 | 0 | 0 | 0 | 0 | 0 |
| ENSMUSG00000086707 | 0.289064826 | 0.580482024 | 0 | 0.5 | 0.568060967 | 0 | 0.224243824 |

| | | | | | | | |
|---|---|---|---|---|---|---|---|
| ENSMUSG00000086708 | 0 | 0 | 0 | 0 | 0 | 0 | 0 |
| ENSMUSG00000086709 | 0 | 0 | 0 | 0 | 0 | 0 | 0 |
| ENSMUSG00000086710 | 0 | 0 | 0 | 0 | 0 | 0 | 0 |
| ENSMUSG00000086711 | 0 | 0 | 0 | 0.25 | 0 | 0 | 0.224243824 |
| ENSMUSG00000086712 | 0 | 0 | 0 | 0 | 0 | 0 | 0 |
| ENSMUSG00000086713 | 0 | 0 | 0 | 0 | 0 | 0 | 0 |
| ENSMUSG00000086714 | 0.289064826 | 0.25 | 0.396240625 | 0.396240625 | 0.244650542 | 0.301029996 | 0.579661877 |
| ENSMUSG00000086715 | 0 | 0 | 0 | 0 | 0 | 0 | 0 |
| ENSMUSG00000086716 | 0.289064826 | 0 | 0 | 0 | 0 | 0 | 0 |
| ENSMUSG00000086717 | 0 | 0 | 0 | 0 | 0 | 0 | 0 |
| ENSMUSG00000086718 | 0 | 0.25 | 0 | 0.25 | 0 | 0 | 0 |
| ENSMUSG00000086719 | 0 | 0 | 0 | 0 | 0 | 0 | 0 |
| ENSMUSG00000086720 | 0 | 0 | 0 | 0 | 0 | 0 | 0 |
| ENSMUSG00000086721 | 0.578129653 | 0.5 | 0.25 | 0.25 | 0.244650542 | 0.301029996 | 0.355418052 |
| ENSMUSG00000086722 | 0 | 0 | 0 | 0 | 0 | 0 | 0 |
| ENSMUSG00000086724 | 0 | 0 | 0 | 0 | 0 | 0 | 0 |
| ENSMUSG00000086725 | 0.867194479 | 1.061981878 | 1.061981878 | 1 | 1.020174412 | 1.146128036 | 1.131174228 |
| ENSMUSG00000086726 | 0 | 0 | 0 | 0 | 0 | 0 | 0 |
| ENSMUSG00000086727 | 0 | 0.25 | 0 | 0 | 0 | 0 | 0 |
| ENSMUSG00000086728 | 0.289064826 | 0.5 | 0.5 | 0.701838731 | 0.387761935 | 0.301029996 | 0.579661877 |
| ENSMUSG00000086729 | 0 | 0 | 0 | 0 | 0 | 0 | 0 |
| ENSMUSG00000086730 | 0 | 0 | 0 | 0 | 0 | 0 | 0 |
| ENSMUSG00000086731 | 0 | 0 | 0 | 0 | 0 | 0 | 0 |
| ENSMUSG00000086732 | 0 | 0.25 | 0 | 0 | 0 | 0 | 0 |
| ENSMUSG00000086733 | 0 | 0 | 0 | 0 | 0 | 0 | 0 |
| ENSMUSG00000086735 | 0.867194479 | 0.864857905 | 0.864857905 | 1.02186571 | 0.686820904 | 0.477121255 | 0.672731473 |
| ENSMUSG00000086736 | 0 | 0 | 0 | 0 | 0 | 0 | 0 |
| ENSMUSG00000086737 | 0 | 0 | 0 | 0 | 0 | 0 | 0 |
| ENSMUSG00000086738 | 0 | 0 | 0 | 0 | 0 | 0 | 0 |
| ENSMUSG00000086739 | 0 | 0 | 0 | 0 | 0 | 0 | 0 |
| ENSMUSG00000086740 | 1.036286563 | 0.79248125 | 0.976722649 | 1.061981878 | 0.955822902 | 0 | 0.916588299 |
| ENSMUSG00000086741 | 0 | 0 | 0 | 0.25 | 0 | 0 | 0.355418052 |
| ENSMUSG00000086742 | 0.289064826 | 0.5 | 0.396240625 | 0.396240625 | 0.568060967 | 0.477121255 | 0.355418052 |
| ENSMUSG00000086743 | 0.45815691 | 0.5 | 0.25 | 0.5 | 0 | 0.301029996 | 0.355418052 |

| | | | | | | | |
|---|---|---|---|---|---|---|---|
| ENSMUSG00000086745 | 0 | 0.25 | 0.396240625 | 0 | 0.244650542 | 0 | 0 |
| ENSMUSG00000086746 | 0.45815691 | 0 | 0.396240625 | 0 | 0.489301084 | 0 | 0 |
| ENSMUSG00000086748 | 0 | 0 | 0 | 0 | 0 | 0 | 0 |
| ENSMUSG00000086749 | 0 | 0 | 0 | 0 | 0 | 0 | 0 |
| ENSMUSG00000086750 | 0 | 0.25 | 0 | 0 | 0 | 0 | 0 |
| ENSMUSG00000086752 | 0 | 0 | 0 | 0 | 0 | 0 | 0 |
| ENSMUSG00000086753 | 1.342375483 | 1.201838731 | 1.098079356 | 1.311981878 | 1.200473444 | 1.322219295 | 1.23151414 |
| ENSMUSG00000086754 | 0 | 0 | 0 | 0.396240625 | 0 | 0.301029996 | 0.224243824 |
| ENSMUSG00000086755 | 0 | 0 | 0 | 0 | 0 | 0.301029996 | 0.355418052 |
| ENSMUSG00000086756 | 0 | 0 | 0 | 0 | 0 | 0 | 0 |
| ENSMUSG00000086757 | 0 | 0 | 0 | 0 | 0 | 0 | 0 |
| ENSMUSG00000086758 | 0 | 0 | 0 | 0 | 0 | 0 | 0 |
| ENSMUSG00000086760 | 0 | 0 | 0 | 0 | 0 | 0 | 0 |
| ENSMUSG00000086761 | 0.289064826 | 0 | 0.25 | 0 | 0 | 0 | 0 |
| ENSMUSG00000086762 | 0 | 0 | 0 | 0 | 0 | 0 | 0 |
| ENSMUSG00000086763 | 0 | 0 | 0 | 0 | 0 | 0 | 0 |
| ENSMUSG00000086764 | 0 | 0.25 | 0 | 0 | 0 | 0 | 0 |
| ENSMUSG00000086765 | 0.289064826 | 0 | 0 | 0 | 0.244650542 | 0 | 0 |
| ENSMUSG00000086766 | 0 | 0 | 0 | 0 | 0 | 0 | 0 |
| ENSMUSG00000086767 | 0.45815691 | 0.25 | 0 | 0.25 | 0 | 0 | 0.224243824 |
| ENSMUSG00000086769 | 0 | 0 | 0.25 | 0 | 0 | 0 | 0 |
| ENSMUSG00000086770 | 0 | 0 | 0 | 0 | 0 | 0 | 0 |
| ENSMUSG00000086771 | 0 | 0 | 0 | 0 | 0 | 0 | 0 |
| ENSMUSG00000086772 | 0 | 0 | 0 | 0 | 0 | 0 | 0 |
| ENSMUSG00000086773 | 0 | 0 | 0 | 0 | 0 | 0 | 0 |
| ENSMUSG00000086774 | 0 | 0.25 | 0 | 0.25 | 0 | 0 | 0 |
| ENSMUSG00000086775 | 0.289064826 | 0.25 | 0.396240625 | 0.646240625 | 0.686820904 | 0.301029996 | 0.355418052 |
| ENSMUSG00000086777 | 0 | 0 | 0 | 0 | 0.387761935 | 0 | 0 |
| ENSMUSG00000086779 | 0.811507563 | 0.396240625 | 0.25 | 0.830482024 | 0.812711509 | 0.301029996 | 0.579661877 |
| ENSMUSG00000086781 | 0 | 0 | 0 | 0 | 0 | 0 | 0 |
| ENSMUSG00000086782 | 0 | 0.25 | 0.396240625 | 0.396240625 | 0.244650542 | 0 | 0.355418052 |
| ENSMUSG00000086783 | 0 | 0 | 0 | 0 | 0 | 0 | 0 |
| ENSMUSG00000086784 | 1.568542279 | 1.52186571 | 1.505591953 | 1.188721876 | 1.445123986 | 1.322219295 | 1.444790289 |
| ENSMUSG00000086785 | 0 | 0 | 0 | 0 | 0 | 0 | 0 |

| | | | | | | | |
|---|---|---|---|---|---|---|---|
| ENSMUSG00000086786 | 0 | 0 | 0.5 | 0 | 0 | 0 | 0.224243824 |
| ENSMUSG00000086788 | 0 | 0 | 0 | 0 | 0 | 0 | 0 |
| ENSMUSG00000086789 | 0 | 0 | 0 | 0 | 0 | 0 | 0 |
| ENSMUSG00000086790 | 0 | 0 | 0 | 0 | 0 | 0 | 0 |
| ENSMUSG00000086791 | 0 | 0 | 0 | 0 | 0 | 0 | 0 |
| ENSMUSG00000086792 | 0 | 0 | 0 | 0 | 0 | 0 | 0 |
| ENSMUSG00000086793 | 0 | 0 | 0 | 0 | 0 | 0 | 0 |
| ENSMUSG00000086794 | 0.289064826 | 0.646240625 | 0.5 | 0.646240625 | 0.733951626 | 0.602059991 | 0.74492186 |
| ENSMUSG00000086795 | 0 | 0 | 0 | 0 | 0 | 0 | 0 |
| ENSMUSG00000086796 | 0.289064826 | 0.396240625 | 0.25 | 0.701838731 | 0.632412477 | 0.477121255 | 0.520678036 |
| ENSMUSG00000086798 | 0.747221736 | 0.701838731 | 0.701838731 | 0.646240625 | 0.931471446 | 0.602059991 | 0.710836105 |
| ENSMUSG00000086799 | 0.289064826 | 0 | 0.25 | 0.25 | 0 | 0.301029996 | 0.224243824 |
| ENSMUSG00000086800 | 0 | 0 | 0 | 0 | 0 | 0 | 0 |
| ENSMUSG00000086801 | 0 | 0 | 0 | 0 | 0 | 0 | 0 |
| ENSMUSG00000086804 | 0 | 0 | 0 | 0 | 0 | 0 | 0 |
| ENSMUSG00000086805 | 0 | 0 | 0 | 0 | 0 | 0 | 0.224243824 |
| ENSMUSG00000086806 | 0.45815691 | 0 | 0.396240625 | 0 | 0.387761935 | 0 | 0.224243824 |
| ENSMUSG00000086807 | 0 | 0 | 0 | 0.25 | 0 | 0 | 0 |
| ENSMUSG00000086808 | 0 | 0 | 0 | 0 | 0 | 0 | 0 |
| ENSMUSG00000086810 | 0 | 0 | 0 | 0 | 0 | 0 | 0 |
| ENSMUSG00000086812 | 0 | 0 | 0 | 0 | 0 | 0 | 0 |
| ENSMUSG00000086813 | 0 | 0 | 0 | 0 | 0 | 0 | 0.224243824 |
| ENSMUSG00000086814 | 0 | 0 | 0 | 0 | 0 | 0 | 0 |
| ENSMUSG00000086815 | 0.289064826 | 0.396240625 | 0.5 | 0 | 0 | 0 | 0 |
| ENSMUSG00000086816 | 0 | 0 | 0 | 0 | 0.244650542 | 0 | 0 |
| ENSMUSG00000086817 | 0 | 0 | 0 | 0 | 0 | 0 | 0 |
| ENSMUSG00000086818 | 0 | 0.25 | 0 | 0.25 | 0 | 0 | 0 |
| ENSMUSG00000086819 | 0.289064826 | 0.75 | 0.5 | 0.5 | 0.686820904 | 0.477121255 | 0.448487648 |
| ENSMUSG00000086820 | 0.289064826 | 0 | 0.396240625 | 0.25 | 0 | 0.301029996 | 0.224243824 |
| ENSMUSG00000086821 | 0 | 0 | 0 | 0 | 0 | 0 | 0 |
| ENSMUSG00000086822 | 0 | 0.25 | 0 | 0 | 0 | 0 | 0 |
| ENSMUSG00000086823 | 0 | 0 | 0 | 0 | 0 | 0 | 0 |
| ENSMUSG00000086824 | 0 | 0 | 0 | 0 | 0 | 0 | 0 |
| ENSMUSG00000086825 | 0.578129653 | 0.580482024 | 0.830482024 | 0.75 | 0.877063019 | 0.77815125 | 0.710836105 |

| | | | | | | | |
|---|---|---|---|---|---|---|---|
| ENSMUSG00000086826 | 0.960252568 | 1.04248125 | 1.061981878 | 0.896240625 | 1 | 0.698970004 | 0.969165684 |
| ENSMUSG00000086827 | 0 | 0 | 0 | 0 | 0.244650542 | 0 | 0 |
| ENSMUSG00000086828 | 0 | 0 | 0 | 0 | 0 | 0 | 0 |
| ENSMUSG00000086830 | 0 | 0 | 0 | 0 | 0 | 0 | 0 |
| ENSMUSG00000086831 | 0 | 0 | 0 | 0 | 0 | 0 | 0 |
| ENSMUSG00000086833 | 0 | 0 | 0 | 0 | 0 | 0 | 0 |
| ENSMUSG00000086834 | 0 | 0 | 0 | 0 | 0 | 0 | 0 |
| ENSMUSG00000086835 | 0 | 0 | 0 | 0 | 0 | 0 | 0 |
| ENSMUSG00000086836 | 0 | 0 | 0 | 0 | 0 | 0 | 0.224243824 |
| ENSMUSG00000086837 | 0.289064826 | 0.396240625 | 0.75 | 0.580482024 | 0.733951626 | 0.301029996 | 0.579661877 |
| ENSMUSG00000086838 | 0 | 0 | 0 | 0.25 | 0.244650542 | 0 | 0.224243824 |
| ENSMUSG00000086839 | 0 | 0 | 0 | 0 | 0 | 0 | 0 |
| ENSMUSG00000086840 | 0 | 0 | 0 | 0 | 0 | 0 | 0 |
| ENSMUSG00000086841 | 2.097855556 | 1.988549078 | 2.097004321 | 1.825945187 | 1.887294348 | 1.886490725 | 1.971363985 |
| ENSMUSG00000086842 | 0.289064826 | 0 | 0.25 | 0 | 0 | 0 | 0 |
| ENSMUSG00000086843 | 0 | 0 | 0 | 0 | 0 | 0 | 0 |
| ENSMUSG00000086844 | 0 | 0.25 | 0 | 0 | 0 | 0 | 0 |
| ENSMUSG00000086845 | 0 | 0 | 0 | 0 | 0.244650542 | 0 | 0 |
| ENSMUSG00000086846 | 0 | 0.25 | 0 | 0.25 | 0 | 0 | 0 |
| ENSMUSG00000086847 | 0 | 0 | 0 | 0.25 | 0 | 0 | 0 |
| ENSMUSG00000086848 | 0 | 0 | 0 | 0 | 0 | 0 | 0 |
| ENSMUSG00000086849 | 0 | 0 | 0 | 0 | 0 | 0 | 0 |
| ENSMUSG00000086850 | 0.289064826 | 0 | 0.646240625 | 0.25 | 0.489301084 | 0.477121255 | 0 |
| ENSMUSG00000086851 | 0.45815691 | 0.646240625 | 0.580482024 | 0.701838731 | 0.846351821 | 0.301029996 | 0.520678036 |
| ENSMUSG00000086852 | 0.289064826 | 0 | 0 | 0 | 0 | 0 | 0 |
| ENSMUSG00000086853 | 0 | 0 | 0 | 0 | 0 | 0 | 0 |
| ENSMUSG00000086854 | 0 | 0 | 0 | 0 | 0 | 0 | 0 |
| ENSMUSG00000086855 | 0 | 0 | 0 | 0 | 0 | 0 | 0 |
| ENSMUSG00000086856 | 0 | 0 | 0 | 0 | 0 | 0 | 0 |
| ENSMUSG00000086858 | 0 | 0 | 0 | 0 | 0 | 0 | 0 |
| ENSMUSG00000086859 | 1.53838222 | 1.418106335 | 1.494319981 | 1.364857905 | 1.49445382 | 1.301029996 | 1.473249546 |
| ENSMUSG00000086860 | 0 | 0 | 0 | 0 | 0 | 0 | 0 |
| ENSMUSG00000086861 | 0 | 0 | 0 | 0 | 0 | 0 | 0 |
| ENSMUSG00000086862 | 0 | 0 | 0 | 0 | 0 | 0 | 0 |

| | | | | | | | |
|---|---|---|---|---|---|---|---|
| ENSMUSG00000086863 | 0 | 0 | 0 | 0 | 0 | 0 | 0 |
| ENSMUSG00000086864 | 0 | 0 | 0 | 0 | 0 | 0 | 0 |
| ENSMUSG00000086866 | 0 | 0 | 0 | 0 | 0 | 0 | 0 |
| ENSMUSG00000086867 | 0 | 0 | 0 | 0.25 | 0 | 0 | 0 |
| ENSMUSG00000086868 | 0 | 0 | 0 | 0 | 0 | 0 | 0.355418052 |
| ENSMUSG00000086870 | 0 | 0 | 0 | 0.25 | 0 | 0 | 0 |
| ENSMUSG00000086872 | 0 | 0 | 0 | 0 | 0.244650542 | 0 | 0 |
| ENSMUSG00000086873 | 0.578129653 | 0.701838731 | 0.580482024 | 0.396240625 | 0.632412477 | 0.698970004 | 0.520678036 |
| ENSMUSG00000086874 | 0 | 0 | 0 | 0 | 0 | 0 | 0 |
| ENSMUSG00000086875 | 0 | 0 | 0 | 0 | 0 | 0 | 0 |
| ENSMUSG00000086876 | 0 | 0 | 0 | 0 | 0 | 0 | 0 |
| ENSMUSG00000086877 | 1.069666964 | 1 | 1.17510993 | 0.976722649 | 1 | 1.113943352 | 1.028149525 |
| ENSMUSG00000086879 | 0 | 0 | 0 | 0 | 0 | 0 | 0 |
| ENSMUSG00000086881 | 0 | 0 | 0.25 | 0 | 0 | 0 | 0 |
| ENSMUSG00000086882 | 0 | 0 | 0 | 0 | 0 | 0 | 0 |
| ENSMUSG00000086883 | 0 | 0 | 0 | 0 | 0 | 0 | 0 |
| ENSMUSG00000086884 | 0 | 0 | 0 | 0 | 0 | 0 | 0 |
| ENSMUSG00000086885 | 0 | 0 | 0 | 0 | 0 | 0.301029996 | 0 |
| ENSMUSG00000086886 | 0 | 0 | 0 | 0 | 0 | 0 | 0 |
| ENSMUSG00000086887 | 0 | 0 | 0 | 0 | 0 | 0 | 0 |
| ENSMUSG00000086889 | 0.747221736 | 0.5 | 0.396240625 | 0.701838731 | 0.568060967 | 0.602059991 | 0.355418052 |
| ENSMUSG00000086890 | 0 | 0 | 0 | 0 | 0.244650542 | 0 | 0 |
| ENSMUSG00000086891 | 0.289064826 | 0.25 | 0 | 0 | 0.244650542 | 0 | 0 |
| ENSMUSG00000086892 | 0 | 0 | 0 | 0 | 0 | 0 | 0 |
| ENSMUSG00000086893 | 0 | 0 | 0.25 | 0.25 | 0 | 0.301029996 | 0.224243824 |
| ENSMUSG00000086894 | 0.811507563 | 0.646240625 | 0.75 | 0.646240625 | 0.812711509 | 0.477121255 | 0.710836105 |
| ENSMUSG00000086896 | 0.289064826 | 0.25 | 0 | 0 | 0.244650542 | 0 | 0.355418052 |
| ENSMUSG00000086898 | 0 | 0 | 0 | 0 | 0 | 0 | 0 |
| ENSMUSG00000086899 | 0 | 0 | 0.25 | 0 | 0 | 0 | 0 |
| ENSMUSG00000086900 | 0 | 0.25 | 0 | 0.25 | 0.568060967 | 0.301029996 | 0.355418052 |
| ENSMUSG00000086902 | 0 | 0 | 0 | 0 | 0 | 0 | 0 |
| ENSMUSG00000086903 | 0 | 0 | 0 | 0.25 | 0.244650542 | 0 | 0 |
| ENSMUSG00000086904 | 0 | 0 | 0 | 0 | 0 | 0 | 0 |
| ENSMUSG00000086905 | 0 | 0 | 0 | 0 | 0 | 0 | 0 |

| | | | | | | | |
|---|---|---|---|---|---|---|---|
| ENSMUSG00000086906 | 0 | 0 | 0.5 | 0.25 | 0.244650542 | 0.301029996 | 0 |
| ENSMUSG00000086907 | 0.671187741 | 0.646240625 | 0.864857905 | 0.5 | 0.905314583 | 0.903089987 | 0.853775828 |
| ENSMUSG00000086908 | 0 | 0 | 0 | 0 | 0 | 0 | 0 |
| ENSMUSG00000086909 | 0 | 0 | 0 | 0 | 0 | 0 | 0 |
| ENSMUSG00000086910 | 0 | 0 | 0 | 0 | 0 | 0 | 0 |
| ENSMUSG00000086911 | 0 | 0 | 0 | 0 | 0.244650542 | 0 | 0 |
| ENSMUSG00000086912 | 0 | 0 | 0.25 | 0 | 0 | 0 | 0 |
| ENSMUSG00000086913 | 0 | 0 | 0 | 0.25 | 0 | 0 | 0 |
| ENSMUSG00000086914 | 0 | 0.25 | 0.396240625 | 0.701838731 | 0.489301084 | 0 | 0.520678036 |
| ENSMUSG00000086915 | 1.036286563 | 1 | 0.896240625 | 0.79248125 | 0.632412477 | 0.698970004 | 0.74492186 |
| ENSMUSG00000086916 | 0.747221736 | 0.25 | 0.25 | 0.646240625 | 0.387761935 | 0.301029996 | 0.224243824 |
| ENSMUSG00000086917 | 0 | 0 | 0 | 0 | 0 | 0 | 0.355418052 |
| ENSMUSG00000086918 | 0 | 0.25 | 0 | 0 | 0 | 0 | 0 |
| ENSMUSG00000086919 | 0 | 0.25 | 0.5 | 0 | 0.244650542 | 0.301029996 | 0.224243824 |
| ENSMUSG00000086920 | 0.811507563 | 0.701838731 | 0.396240625 | 0.646240625 | 0.489301084 | 0.477121255 | 0.629532004 |
| ENSMUSG00000086921 | 0 | 0 | 0 | 0 | 0 | 0 | 0 |
| ENSMUSG00000086922 | 0 | 0 | 0 | 0.25 | 0 | 0 | 0.224243824 |
| ENSMUSG00000086923 | 0 | 0 | 0 | 0 | 0 | 0 | 0 |
| ENSMUSG00000086924 | 0.811507563 | 0.79248125 | 0.646240625 | 0.396240625 | 0.846351821 | 0.77815125 | 0.916588299 |
| ENSMUSG00000086925 | 0.289064826 | 0.580482024 | 0.396240625 | 0.396240625 | 0.568060967 | 0.477121255 | 0.579661877 |
| ENSMUSG00000086926 | 0 | 0 | 0 | 0 | 0 | 0 | 0 |
| ENSMUSG00000086927 | 0 | 0 | 0 | 0 | 0 | 0 | 0 |
| ENSMUSG00000086928 | 0 | 0 | 0 | 0 | 0 | 0 | 0 |
| ENSMUSG00000086929 | 0 | 0 | 0 | 0 | 0 | 0 | 0 |
| ENSMUSG00000086930 | 0 | 0.5 | 0 | 0 | 0 | 0.301029996 | 0 |
| ENSMUSG00000086931 | 0 | 0 | 0 | 0 | 0 | 0 | 0 |
| ENSMUSG00000086932 | 0 | 0 | 0 | 0 | 0 | 0.301029996 | 0 |
| ENSMUSG00000086933 | 0 | 0 | 0 | 0.25 | 0 | 0 | 0 |
| ENSMUSG00000086935 | 0 | 0.396240625 | 0.396240625 | 0.396240625 | 0.489301084 | 0.477121255 | 0.448487648 |
| ENSMUSG00000086936 | 0.289064826 | 0 | 0.5 | 0.396240625 | 0.489301084 | 0 | 0 |
| ENSMUSG00000086937 | 0.289064826 | 0.396240625 | 0 | 0.396240625 | 0 | 0 | 0 |
| ENSMUSG00000086938 | 0 | 0 | 0 | 0.396240625 | 0.387761935 | 0 | 0 |
| ENSMUSG00000086939 | 0 | 0 | 0 | 0 | 0 | 0 | 0 |
| ENSMUSG00000086940 | 0.45815691 | 0 | 0 | 0.396240625 | 0.244650542 | 0.301029996 | 0.355418052 |

| | | | | | | | |
|---|---|---|---|---|---|---|---|
| ENSMUSG00000086941 | 0 | 0 | 0 | 0 | 0 | 0 | 0 |
| ENSMUSG00000086942 | 0.289064826 | 0.75 | 0.5 | 0.5 | 0.387761935 | 0.477121255 | 0.355418052 |
| ENSMUSG00000086943 | 0.45815691 | 0 | 0.25 | 0 | 0.244650542 | 0.301029996 | 0.224243824 |
| ENSMUSG00000086944 | 0 | 0 | 0 | 0 | 0 | 0 | 0.224243824 |
| ENSMUSG00000086945 | 0 | 0 | 0 | 0 | 0 | 0 | 0.224243824 |
| ENSMUSG00000086946 | 0 | 0 | 0 | 0 | 0.244650542 | 0 | 0 |
| ENSMUSG00000086947 | 0 | 0 | 0 | 0 | 0 | 0 | 0 |
| ENSMUSG00000086948 | 0 | 0 | 0 | 0 | 0 | 0 | 0 |
| ENSMUSG00000086949 | 0 | 0 | 0 | 0 | 0 | 0 | 0 |
| ENSMUSG00000086951 | 0.289064826 | 0 | 0 | 0 | 0 | 0 | 0 |
| ENSMUSG00000086952 | 0 | 0 | 0 | 0 | 0 | 0 | 0 |
| ENSMUSG00000086953 | 0 | 0 | 0 | 0.25 | 0 | 0 | 0 |
| ENSMUSG00000086954 | 0 | 0 | 0 | 0 | 0 | 0 | 0 |
| ENSMUSG00000086955 | 0 | 0 | 0 | 0 | 0 | 0 | 0 |
| ENSMUSG00000086957 | 0 | 0 | 0 | 0 | 0 | 0 | 0 |
| ENSMUSG00000086959 | 0 | 0 | 0 | 0 | 0 | 0 | 0 |
| ENSMUSG00000086960 | 0 | 0 | 0 | 0 | 0 | 0 | 0 |
| ENSMUSG00000086961 | 0 | 0.396240625 | 0 | 0.25 | 0 | 0 | 0 |
| ENSMUSG00000086962 | 0.578129653 | 0 | 0 | 0 | 0 | 0.301029996 | 0.224243824 |
| ENSMUSG00000086963 | 0 | 0 | 0 | 0 | 0 | 0 | 0 |
| ENSMUSG00000086964 | 0 | 0 | 0 | 0 | 0 | 0 | 0 |
| ENSMUSG00000086965 | 1.156259305 | 1.25 | 1.25 | 1 | 1.223252711 | 0.903089987 | 1.159323753 |
| ENSMUSG00000086968 | 1.753493018 | 1.642463902 | 1.646240625 | 1.723704441 | 1.677808768 | 1.903089987 | 1.502532226 |
| ENSMUSG00000086969 | 0.289064826 | 0.396240625 | 0 | 0.580482024 | 0.387761935 | 0 | 0.355418052 |
| ENSMUSG00000086970 | 0 | 0 | 0 | 0 | 0 | 0 | 0 |
| ENSMUSG00000086971 | 0 | 0 | 0 | 0 | 0 | 0 | 0 |
| ENSMUSG00000086974 | 0 | 0 | 0 | 0 | 0 | 0 | 0 |
| ENSMUSG00000086975 | 0 | 0 | 0 | 0 | 0 | 0 | 0 |
| ENSMUSG00000086976 | 0.289064826 | 0.25 | 0 | 0 | 0.244650542 | 0 | 0 |
| ENSMUSG00000086978 | 0 | 0 | 0 | 0 | 0 | 0 | 0 |
| ENSMUSG00000086980 | 0 | 0 | 0 | 0 | 0 | 0 | 0 |
| ENSMUSG00000086981 | 0 | 0 | 0 | 0 | 0 | 0 | 0 |
| ENSMUSG00000086984 | 0 | 0 | 0 | 0 | 0 | 0 | 0 |
| ENSMUSG00000086986 | 0 | 0 | 0 | 0 | 0 | 0 | 0 |

| | | | | | | | |
|---|---|---|---|---|---|---|---|
| ENSMUSG00000086987 | 0.578129653 | 0.5 | 0 | 0.701838731 | 0 | 0 | 0.224243824 |
| ENSMUSG00000086989 | 0 | 0 | 0 | 0 | 0 | 0 | 0 |
| ENSMUSG00000086991 | 0 | 0 | 0 | 0 | 0 | 0 | 0 |
| ENSMUSG00000086992 | 0 | 0 | 0 | 0 | 0 | 0 | 0 |
| ENSMUSG00000086993 | 0 | 0 | 0.396240625 | 0.25 | 0.244650542 | 0 | 0 |
| ENSMUSG00000086994 | 0 | 0 | 0 | 0 | 0 | 0 | 0 |
| ENSMUSG00000086995 | 0 | 0.25 | 0 | 0 | 0 | 0 | 0.224243824 |
| ENSMUSG00000086996 | 0 | 0 | 0 | 0 | 0 | 0 | 0 |
| ENSMUSG00000086997 | 0.45815691 | 0.5 | 0.25 | 0 | 0.387761935 | 0.301029996 | 0.355418052 |
| ENSMUSG00000086998 | 0 | 0 | 0 | 0 | 0 | 0 | 0 |
| ENSMUSG00000086999 | 0 | 0 | 0 | 0 | 0 | 0.301029996 | 0 |
| ENSMUSG00000087001 | 0 | 0.396240625 | 0.580482024 | 0 | 0.244650542 | 0.301029996 | 0 |
| ENSMUSG00000087002 | 0 | 0 | 0 | 0 | 0 | 0 | 0 |
| ENSMUSG00000087003 | 0 | 0 | 0 | 0 | 0 | 0 | 0 |
| ENSMUSG00000087004 | 0.289064826 | 0 | 0 | 0 | 0 | 0 | 0.224243824 |
| ENSMUSG00000087006 | 0.45815691 | 0.75 | 0.92510993 | 0 | 0 | 0 | 0.355418052 |
| ENSMUSG00000087007 | 0 | 0 | 0 | 0 | 0 | 0 | 0 |
| ENSMUSG00000087008 | 0 | 0 | 0 | 0 | 0.387761935 | 0 | 0 |
| ENSMUSG00000087009 | 0 | 0 | 0 | 0 | 0 | 0 | 0 |
| ENSMUSG00000087010 | 0 | 0 | 0 | 0 | 0 | 0 | 0 |
| ENSMUSG00000087011 | 0 | 0 | 0.25 | 0 | 0 | 0 | 0 |
| ENSMUSG00000087012 | 0 | 0 | 0 | 0 | 0 | 0 | 0 |
| ENSMUSG00000087013 | 0 | 0 | 0.396240625 | 0.25 | 0.244650542 | 0 | 0.224243824 |
| ENSMUSG00000087014 | 0.960252568 | 0.79248125 | 0.646240625 | 0.396240625 | 0.244650542 | 0 | 0.579661877 |
| ENSMUSG00000087015 | 0 | 0 | 0 | 0 | 0 | 0 | 0 |
| ENSMUSG00000087016 | 0 | 0 | 0 | 0 | 0 | 0 | 0 |
| ENSMUSG00000087017 | 0.289064826 | 0.25 | 0.396240625 | 0 | 0 | 0 | 0 |
| ENSMUSG00000087018 | 0.289064826 | 0 | 0 | 0 | 0.489301084 | 0.301029996 | 0 |
| ENSMUSG00000087020 | 0 | 0 | 0.25 | 0 | 0 | 0.301029996 | 0 |
| ENSMUSG00000087021 | 0 | 0 | 0 | 0 | 0 | 0 | 0 |
| ENSMUSG00000087022 | 0.747221736 | 0.701838731 | 0.701838731 | 0.830482024 | 0.568060967 | 0.77815125 | 0.829800754 |
| ENSMUSG00000087024 | 0 | 0 | 0 | 0 | 0 | 0 | 0 |
| ENSMUSG00000087026 | 0.91631382 | 0.896240625 | 0.79248125 | 0.951838731 | 0.931471446 | 0.954242509 | 0.710836105 |
| ENSMUSG00000087027 | 0 | 0 | 0 | 0 | 0 | 0 | 0 |

| | | | | | | | |
|---|---|---|---|---|---|---|---|
| ENSMUSG00000087028 | 0.45815691 | 0.75 | 0.580482024 | 0.79248125 | 0.733951626 | 0.954242509 | 0.775756176 |
| ENSMUSG00000087029 | 0 | 0 | 0 | 0 | 0 | 0 | 0 |
| ENSMUSG00000087030 | 0 | 0 | 0 | 0.25 | 0 | 0 | 0 |
| ENSMUSG00000087031 | 0 | 0 | 0 | 0 | 0 | 0 | 0 |
| ENSMUSG00000087032 | 0 | 0 | 0 | 0 | 0 | 0 | 0 |
| ENSMUSG00000087033 | 0 | 0 | 0 | 0 | 0 | 0 | 0 |
| ENSMUSG00000087034 | 0 | 0 | 0 | 0 | 0 | 0 | 0 |
| ENSMUSG00000087035 | 0 | 0 | 0 | 0.25 | 0.244650542 | 0 | 0.224243824 |
| ENSMUSG00000087036 | 0.289064826 | 0.25 | 0 | 0 | 0 | 0 | 0.224243824 |
| ENSMUSG00000087037 | 0.289064826 | 0 | 0.25 | 0 | 0 | 0 | 0 |
| ENSMUSG00000087038 | 0 | 0 | 0 | 0 | 0 | 0 | 0 |
| ENSMUSG00000087039 | 0 | 0 | 0 | 0 | 0 | 0 | 0 |
| ENSMUSG00000087040 | 0 | 0 | 0 | 0 | 0 | 0 | 0 |
| ENSMUSG00000087041 | 0 | 0 | 0 | 0 | 0 | 0 | 0 |
| ENSMUSG00000087042 | 0 | 0 | 0 | 0.25 | 0.244650542 | 0 | 0 |
| ENSMUSG00000087043 | 0 | 0 | 0 | 0 | 0 | 0 | 0 |
| ENSMUSG00000087044 | 0 | 0 | 0 | 0 | 0 | 0 | 0 |
| ENSMUSG00000087045 | 0 | 0 | 0 | 0 | 0 | 0 | 0 |
| ENSMUSG00000087047 | 0 | 0 | 0 | 0 | 0 | 0 | 0 |
| ENSMUSG00000087049 | 0 | 0 | 0 | 0 | 0 | 0 | 0 |
| ENSMUSG00000087050 | 0.578129653 | 0.75 | 0.25 | 0.92510993 | 0.77552387 | 0.954242509 | 0.775756176 |
| ENSMUSG00000087051 | 0 | 0 | 0 | 0 | 0 | 0 | 0 |
| ENSMUSG00000087052 | 0.45815691 | 0.25 | 0.25 | 0 | 0.244650542 | 0.301029996 | 0.448487648 |
| ENSMUSG00000087053 | 0 | 0 | 0 | 0 | 0 | 0 | 0 |
| ENSMUSG00000087054 | 0 | 0 | 0 | 0 | 0 | 0 | 0 |
| ENSMUSG00000087055 | 0 | 0.396240625 | 0 | 0.396240625 | 0.244650542 | 0.477121255 | 0.224243824 |
| ENSMUSG00000087056 | 0 | 0 | 0 | 0 | 0 | 0.301029996 | 0 |
| ENSMUSG00000087057 | 0 | 0 | 0 | 0 | 0 | 0 | 0 |
| ENSMUSG00000087058 | 0 | 0 | 0 | 0 | 0 | 0 | 0 |
| ENSMUSG00000087059 | 0 | 0 | 0 | 0 | 0 | 0 | 0 |
| ENSMUSG00000087060 | 1.631440309 | 1.571350555 | 1.54745614 | 1.54745614 | 1.55104774 | 1.643452676 | 1.437266334 |
| ENSMUSG00000087061 | 0 | 0 | 0 | 0 | 0 | 0 | 0 |
| ENSMUSG00000087062 | 0 | 0 | 0 | 0 | 0 | 0 | 0 |
| ENSMUSG00000087063 | 0 | 0 | 0 | 0 | 0 | 0 | 0.224243824 |

| | | | | | | | |
|---|---|---|---|---|---|---|---|
| ENSMUSG00000087064 | 0.671187741 | 0.646240625 | 0.830482024 | 0.5 | 0 | 0.602059991 | 0.775756176 |
| ENSMUSG00000087066 | 0 | 0.25 | 0 | 0 | 0.244650542 | 0 | 0.224243824 |
| ENSMUSG00000087067 | 0 | 0 | 0.25 | 0.25 | 0 | 0 | 0 |
| ENSMUSG00000087069 | 0 | 0 | 0 | 0 | 0 | 0 | 0 |
| ENSMUSG00000087070 | 0 | 0.25 | 0 | 0 | 0 | 0.301029996 | 0.224243824 |
| ENSMUSG00000087071 | 0 | 0.25 | 0 | 0 | 0 | 0 | 0 |
| ENSMUSG00000087072 | 0 | 0 | 0 | 0 | 0 | 0 | 0 |
| ENSMUSG00000087074 | 0 | 0 | 0 | 0 | 0 | 0 | 0 |
| ENSMUSG00000087075 | 0 | 0 | 0 | 0 | 0.244650542 | 0 | 0 |
| ENSMUSG00000087076 | 0 | 0 | 0 | 0 | 0 | 0 | 0.224243824 |
| ENSMUSG00000087077 | 0.289064826 | 0.25 | 0.5 | 0 | 0 | 0 | 0 |
| ENSMUSG00000087079 | 0 | 0.25 | 0 | 0 | 0 | 0 | 0 |
| ENSMUSG00000087080 | 0 | 0 | 0 | 0 | 0 | 0 | 0 |
| ENSMUSG00000087082 | 0 | 0 | 0 | 0 | 0 | 0 | 0 |
| ENSMUSG00000087083 | 0 | 0 | 0 | 0 | 0 | 0 | 0 |
| ENSMUSG00000087084 | 0 | 0 | 0 | 0 | 0 | 0 | 0 |
| ENSMUSG00000087085 | 0 | 0 | 0 | 0 | 0 | 0 | 0 |
| ENSMUSG00000087086 | 0 | 0 | 0 | 0 | 0 | 0 | 0 |
| ENSMUSG00000087088 | 0.45815691 | 0.396240625 | 0.5 | 0.5 | 0.632412477 | 0.301029996 | 0.448487648 |
| ENSMUSG00000087089 | 0 | 0 | 0 | 0 | 0 | 0 | 0 |
| ENSMUSG00000087090 | 0 | 0 | 0 | 0 | 0 | 0 | 0 |
| ENSMUSG00000087093 | 0 | 0 | 0 | 0 | 0 | 0.301029996 | 0 |
| ENSMUSG00000087094 | 0 | 0 | 0 | 0.25 | 0 | 0 | 0 |
| ENSMUSG00000087095 | 1.269664473 | 1.302363341 | 1.188721876 | 1.17510993 | 1.106691885 | 1.230448921 | 0.984950056 |
| ENSMUSG00000087097 | 0 | 0 | 0 | 0 | 0 | 0 | 0 |
| ENSMUSG00000087098 | 0 | 0 | 0 | 0 | 0 | 0 | 0 |
| ENSMUSG00000087099 | 0 | 0 | 0 | 0 | 0 | 0 | 0 |
| ENSMUSG00000087100 | 0 | 0 | 0 | 0 | 0 | 0 | 0 |
| ENSMUSG00000087102 | 0 | 0 | 0 | 0 | 0 | 0 | 0 |
| ENSMUSG00000087103 | 0 | 0 | 0 | 0 | 0 | 0 | 0 |
| ENSMUSG00000087104 | 0 | 0 | 0 | 0 | 0 | 0 | 0 |
| ENSMUSG00000087106 | 0 | 0.25 | 0.25 | 0.25 | 0 | 0 | 0 |
| ENSMUSG00000087107 | 0 | 0 | 0 | 0 | 0 | 0 | 0 |
| ENSMUSG00000087108 | 0.289064826 | 0.396240625 | 0.396240625 | 0.5 | 0.244650542 | 0.477121255 | 0.224243824 |

| | | | | | | | |
|---|---|---|---|---|---|---|---|
| ENSMUSG00000087109 | 0 | 0 | 0 | 0 | 0 | 0 | 0 |
| ENSMUSG00000087110 | 0.289064826 | 0 | 0 | 0 | 0 | 0 | 0 |
| ENSMUSG00000087111 | 0 | 0.25 | 0 | 0 | 0.244650542 | 0 | 0 |
| ENSMUSG00000087112 | 0 | 0 | 0 | 0 | 0 | 0 | 0.224243824 |
| ENSMUSG00000087113 | 0.45815691 | 0.25 | 0.580482024 | 0 | 0 | 0 | 0.224243824 |
| ENSMUSG00000087114 | 0 | 0 | 0 | 0 | 0 | 0 | 0 |
| ENSMUSG00000087115 | 0 | 0 | 0 | 0 | 0 | 0 | 0 |
| ENSMUSG00000087116 | 0 | 0 | 0 | 0 | 0 | 0 | 0 |
| ENSMUSG00000087117 | 0 | 0 | 0 | 0 | 0 | 0 | 0 |
| ENSMUSG00000087119 | 0 | 0 | 0.25 | 0 | 0.387761935 | 0 | 0.224243824 |
| ENSMUSG00000087120 | 0 | 0.25 | 0 | 0 | 0.387761935 | 0.301029996 | 0 |
| ENSMUSG00000087121 | 0.289064826 | 0.25 | 0.25 | 0.396240625 | 0.244650542 | 0.477121255 | 0.448487648 |
| ENSMUSG00000087122 | 0 | 0 | 0 | 0 | 0 | 0 | 0 |
| ENSMUSG00000087123 | 0.289064826 | 0 | 0.25 | 0 | 0.244650542 | 0 | 0 |
| ENSMUSG00000087124 | 0 | 0 | 0 | 0 | 0 | 0 | 0 |
| ENSMUSG00000087125 | 0 | 0 | 0 | 0 | 0 | 0 | 0 |
| ENSMUSG00000087127 | 0 | 0 | 0 | 0 | 0 | 0 | 0 |
| ENSMUSG00000087128 | 0.289064826 | 0.5 | 0.396240625 | 0.646240625 | 0.387761935 | 0.477121255 | 0.579661877 |
| ENSMUSG00000087129 | 0 | 0 | 0 | 0 | 0 | 0 | 0.224243824 |
| ENSMUSG00000087131 | 0 | 0 | 0 | 0 | 0 | 0 | 0 |
| ENSMUSG00000087132 | 0.578129653 | 0.580482024 | 0.580482024 | 0.5 | 0.733951626 | 0 | 0.224243824 |
| ENSMUSG00000087133 | 0 | 0 | 0 | 0 | 0 | 0 | 0 |
| ENSMUSG00000087135 | 0.289064826 | 0.396240625 | 0.25 | 0 | 0.244650542 | 0 | 0.224243824 |
| ENSMUSG00000087136 | 0 | 0 | 0 | 0 | 0 | 0 | 0 |
| ENSMUSG00000087137 | 0 | 0 | 0 | 0 | 0 | 0 | 0 |
| ENSMUSG00000087138 | 0.91631382 | 1.114857905 | 0.864857905 | 1.080482024 | 1.163285805 | 1 | 1.041356071 |
| ENSMUSG00000087139 | 0 | 0 | 0 | 0 | 0 | 0 | 0 |
| ENSMUSG00000087140 | 0 | 0 | 0 | 0 | 0 | 0 | 0 |
| ENSMUSG00000087141 | 1.920505136 | 1.802363341 | 1.809601185 | 1.766522298 | 1.704182902 | 1.959041392 | 1.645916034 |
| ENSMUSG00000087142 | 0.811507563 | 1.146240625 | 1 | 0.79248125 | 0.905314583 | 0.698970004 | 0.853775828 |
| ENSMUSG00000087143 | 1.307602651 | 1.114857905 | 1.321350555 | 1.201838731 | 1.136121934 | 1 | 1.089372236 |
| ENSMUSG00000087144 | 0 | 0 | 0 | 0 | 0 | 0 | 0 |
| ENSMUSG00000087145 | 0 | 0 | 0 | 0 | 0 | 0 | 0 |
| ENSMUSG00000087146 | 0 | 0 | 0 | 0 | 0 | 0 | 0 |

| | | | | | | | |
|---|---|---|---|---|---|---|---|
| ENSMUSG00000087147 | 0 | 0 | 0 | 0 | 0 | 0 | 0 |
| ENSMUSG00000087148 | 0.289064826 | 0 | 0 | 0 | 0 | 0 | 0 |
| ENSMUSG00000087149 | 0 | 0.25 | 0 | 0 | 0 | 0 | 0 |
| ENSMUSG00000087150 | 0 | 0 | 0 | 0.25 | 0.686820904 | 0.301029996 | 0 |
| ENSMUSG00000087151 | 0.289064826 | 0.396240625 | 0.5 | 0.646240625 | 0 | 0 | 0 |
| ENSMUSG00000087152 | 0 | 0 | 0 | 0 | 0 | 0 | 0 |
| ENSMUSG00000087153 | 2.100572389 | 1.930274797 | 1.964495249 | 2.041226732 | 1.938809675 | 2.193124598 | 1.844181586 |
| ENSMUSG00000087155 | 0 | 0 | 0 | 0 | 0 | 0.301029996 | 0 |
| ENSMUSG00000087157 | 0 | 0 | 0 | 0 | 0 | 0 | 0 |
| ENSMUSG00000087158 | 0 | 0 | 0 | 0 | 0.244650542 | 0 | 0 |
| ENSMUSG00000087159 | 0.747221736 | 0.701838731 | 0.896240625 | 0.79248125 | 0.77552387 | 0.698970004 | 0.672731473 |
| ENSMUSG00000087160 | 0 | 0 | 0 | 0 | 0 | 0 | 0 |
| ENSMUSG00000087162 | 0 | 0 | 0 | 0 | 0 | 0 | 0 |
| ENSMUSG00000087163 | 0.289064826 | 0.25 | 0 | 0 | 0.244650542 | 0 | 0.224243824 |
| ENSMUSG00000087164 | 0 | 0 | 0 | 0 | 0 | 0 | 0 |
| ENSMUSG00000087165 | 1.156259305 | 1 | 0.92510993 | 1.238549078 | 1.302012594 | 1.255272505 | 1.066254157 |
| ENSMUSG00000087166 | 0 | 0 | 0 | 0 | 0 | 0 | 0 |
| ENSMUSG00000087167 | 0 | 0 | 0 | 0 | 0 | 0 | 0 |
| ENSMUSG00000087168 | 0.811507563 | 0.75 | 0.75 | 0.646240625 | 0.568060967 | 0.477121255 | 0.672731473 |
| ENSMUSG00000087170 | 0 | 0 | 0 | 0 | 0 | 0 | 0 |
| ENSMUSG00000087171 | 0 | 0 | 0 | 0 | 0 | 0 | 0 |
| ENSMUSG00000087172 | 0 | 0.580482024 | 0.75 | 0.25 | 0.387761935 | 0 | 0 |
| ENSMUSG00000087173 | 0 | 0 | 0 | 0 | 0 | 0 | 0 |
| ENSMUSG00000087174 | 1.975148165 | 1.828220739 | 1.779735268 | 1.858657057 | 1.818885826 | 1.986771734 | 1.637827726 |
| ENSMUSG00000087175 | 0 | 0 | 0 | 0 | 0 | 0 | 0 |
| ENSMUSG00000087176 | 0.289064826 | 0 | 0 | 0.25 | 0 | 0.301029996 | 0.355418052 |
| ENSMUSG00000087177 | 1.516991255 | 1.451838731 | 1.470660762 | 1.53743678 | 1.427019704 | 1.612783857 | 1.42167221 |
| ENSMUSG00000087178 | 1.156259305 | 1.146240625 | 0.830482024 | 1.146240625 | 1.121713561 | 1 | 1.066254157 |
| ENSMUSG00000087179 | 1.269664473 | 1.418106335 | 1.396240625 | 1.431980114 | 1.484072009 | 1.278753601 | 1.455757965 |
| ENSMUSG00000087180 | 0.289064826 | 0 | 0 | 0.25 | 0 | 0 | 0 |
| ENSMUSG00000087181 | 0 | 0 | 0 | 0 | 0 | 0 | 0 |
| ENSMUSG00000087184 | 0 | 0 | 0 | 0 | 0 | 0 | 0 |
| ENSMUSG00000087185 | 0 | 0 | 0 | 0 | 0 | 0 | 0 |
| ENSMUSG00000087187 | 0.578129653 | 0.646240625 | 0.646240625 | 0 | 0 | 0 | 0 |

| | | | | | | | |
|---|---|---|---|---|---|---|---|
| ENSMUSG00000087190 | 0 | 0 | 0 | 0 | 0 | 0 | 0 |
| ENSMUSG00000087192 | 0.289064826 | 0.396240625 | 0.25 | 0 | 0 | 0 | 0 |
| ENSMUSG00000087193 | 0.289064826 | 0.396240625 | 0 | 0 | 0 | 0.301029996 | 0 |
| ENSMUSG00000087194 | 0 | 0 | 0.25 | 0.25 | 0 | 0 | 0.224243824 |
| ENSMUSG00000087195 | 0 | 0 | 0 | 0 | 0 | 0 | 0 |
| ENSMUSG00000087196 | 0 | 0 | 0 | 0 | 0 | 0 | 0 |
| ENSMUSG00000087197 | 0 | 0 | 0 | 0 | 0 | 0 | 0 |
| ENSMUSG00000087198 | 0 | 0 | 0.396240625 | 0 | 0.387761935 | 0.477121255 | 0 |
| ENSMUSG00000087199 | 0 | 0 | 0 | 0 | 0 | 0 | 0 |
| ENSMUSG00000087200 | 0.289064826 | 0.5 | 0 | 0.580482024 | 0.568060967 | 0.301029996 | 0.355418052 |
| ENSMUSG00000087201 | 0 | 0 | 0 | 0 | 0.387761935 | 0 | 0 |
| ENSMUSG00000087202 | 0 | 0 | 0 | 0 | 0 | 0 | 0 |
| ENSMUSG00000087203 | 0.747221736 | 0.5 | 0.701838731 | 0.25 | 0.387761935 | 0 | 0.224243824 |
| ENSMUSG00000087204 | 0 | 0.25 | 0.25 | 0.5 | 0.489301084 | 0.301029996 | 0 |
| ENSMUSG00000087205 | 0 | 0 | 0 | 0 | 0 | 0 | 0 |
| ENSMUSG00000087206 | 0 | 0.25 | 0.25 | 0 | 0.244650542 | 0 | 0 |
| ENSMUSG00000087207 | 0 | 0 | 0 | 0 | 0.244650542 | 0 | 0 |
| ENSMUSG00000087208 | 0 | 0 | 0 | 0 | 0 | 0 | 0 |
| ENSMUSG00000087210 | 0 | 0 | 0 | 0 | 0 | 0 | 0 |
| ENSMUSG00000087211 | 0 | 0 | 0 | 0.75 | 0 | 0 | 0.710836105 |
| ENSMUSG00000087213 | 1.53838222 | 1.505591953 | 1.488549078 | 1.52186571 | 1.53772706 | 1.755874856 | 1.502532226 |
| ENSMUSG00000087214 | 0 | 0 | 0 | 0 | 0 | 0 | 0 |
| ENSMUSG00000087215 | 0 | 0 | 0 | 0 | 0 | 0 | 0 |
| ENSMUSG00000087216 | 0 | 0 | 0 | 0 | 0 | 0 | 0 |
| ENSMUSG00000087217 | 0 | 0 | 0 | 0 | 0 | 0 | 0 |
| ENSMUSG00000087219 | 0 | 0 | 0 | 0 | 0 | 0 | 0 |
| ENSMUSG00000087220 | 0 | 0 | 0 | 0 | 0.244650542 | 0 | 0.224243824 |
| ENSMUSG00000087221 | 0.45815691 | 0.580482024 | 0.75 | 0.5 | 0.686820904 | 0.477121255 | 0.520678036 |
| ENSMUSG00000087222 | 0 | 0.25 | 0 | 0 | 0 | 0 | 0 |
| ENSMUSG00000087223 | 0 | 0 | 0 | 0 | 0 | 0 | 0 |
| ENSMUSG00000087224 | 0 | 0 | 0 | 0 | 0 | 0 | 0.224243824 |
| ENSMUSG00000087225 | 0 | 0 | 0 | 0 | 0 | 0 | 0 |
| ENSMUSG00000087226 | 0 | 0.25 | 0 | 0 | 0 | 0 | 0 |
| ENSMUSG00000087228 | 0 | 0 | 0 | 0 | 0 | 0 | 0 |

| | | | | | | | |
|---|---|---|---|---|---|---|---|
| ENSMUSG00000087230 | 0.289064826 | 0.25 | 0.5 | 0.5 | 0.387761935 | 0 | 0.224243824 |
| ENSMUSG00000087231 | 0.867194479 | 0.830482024 | 0.92510993 | 1.17510993 | 0.955822902 | 0.77815125 | 0.969165684 |
| ENSMUSG00000087232 | 0 | 0 | 0 | 0 | 0 | 0 | 0 |
| ENSMUSG00000087233 | 0 | 0 | 0 | 0 | 0 | 0 | 0 |
| ENSMUSG00000087234 | 0 | 0 | 0 | 0 | 0 | 0 | 0 |
| ENSMUSG00000087235 | 0 | 0 | 0 | 0 | 0 | 0 | 0 |
| ENSMUSG00000087236 | 0 | 0 | 0 | 0.25 | 0 | 0 | 0 |
| ENSMUSG00000087238 | 0 | 0 | 0 | 0 | 0 | 0 | 0 |
| ENSMUSG00000087239 | 0 | 0 | 0 | 0.396240625 | 0 | 0 | 0 |
| ENSMUSG00000087240 | 0 | 0 | 0.25 | 0 | 0 | 0.301029996 | 0 |
| ENSMUSG00000087241 | 0 | 0 | 0 | 0 | 0 | 0 | 0 |
| ENSMUSG00000087242 | 0.45815691 | 0.396240625 | 0 | 0.396240625 | 0 | 0 | 0 |
| ENSMUSG00000087244 | 0 | 0 | 0 | 0 | 0 | 0 | 0 |
| ENSMUSG00000087245 | 0 | 0 | 0 | 0 | 0 | 0 | 0 |
| ENSMUSG00000087246 | 0 | 0 | 0.25 | 0.25 | 0 | 0 | 0.224243824 |
| ENSMUSG00000087247 | 1.948720984 | 1.919870025 | 1.882845365 | 0.396240625 | 0.489301084 | 0.84509804 | 0.629532004 |
| ENSMUSG00000087248 | 0 | 0 | 0 | 0 | 0 | 0 | 0 |
| ENSMUSG00000087249 | 0.671187741 | 0.830482024 | 0.951838731 | 0.79248125 | 0.877063019 | 0.698970004 | 0.803905701 |
| ENSMUSG00000087250 | 0 | 0 | 0 | 0 | 0 | 0.301029996 | 0 |
| ENSMUSG00000087252 | 0.289064826 | 0 | 0.396240625 | 0.25 | 0.387761935 | 0 | 0.224243824 |
| ENSMUSG00000087253 | 0 | 0.25 | 0 | 0.25 | 0 | 0 | 0 |
| ENSMUSG00000087254 | 0 | 0 | 0 | 0 | 0 | 0 | 0 |
| ENSMUSG00000087255 | 0 | 0 | 0 | 0 | 0 | 0 | 0 |
| ENSMUSG00000087256 | 0 | 0.25 | 0.396240625 | 0.25 | 0.244650542 | 0.301029996 | 0 |
| ENSMUSG00000087258 | 0 | 0 | 0 | 0 | 0 | 0 | 0.224243824 |
| ENSMUSG00000087259 | 1.404271433 | 1.339388001 | 1.388647213 | 1.080482024 | 1.057362051 | 1.079181246 | 0.969165684 |
| ENSMUSG00000087260 | 2.300956576 | 2.089388001 | 2.109697963 | 2.256284891 | 2.150762767 | 2.507855872 | 2.039586611 |
| ENSMUSG00000087261 | 0 | 0 | 0 | 0 | 0 | 0 | 0 |
| ENSMUSG00000087263 | 0.289064826 | 0 | 0 | 0.25 | 0 | 0 | 0 |
| ENSMUSG00000087264 | 0 | 0.25 | 0.25 | 0.25 | 0.489301084 | 0.602059991 | 0.448487648 |
| ENSMUSG00000087265 | 0.289064826 | 0 | 0 | 0 | 0 | 0 | 0 |
| ENSMUSG00000087266 | 0 | 0 | 0 | 0 | 0 | 0 | 0 |
| ENSMUSG00000087267 | 0.289064826 | 0.25 | 0.580482024 | 0.580482024 | 0.846351821 | 0.477121255 | 0.224243824 |
| ENSMUSG00000087268 | 0 | 0 | 0 | 0 | 0 | 0 | 0 |

| | | | | | | | |
|---|---|---|---|---|---|---|---|
| ENSMUSG00000087269 | 0.91631382 | 0.79248125 | 0.951838731 | 0.646240625 | 0.812711509 | 0.477121255 | 0.710836105 |
| ENSMUSG00000087270 | 0 | 0 | 0 | 0 | 0 | 0 | 0 |
| ENSMUSG00000087272 | 0 | 0.5 | 0.580482024 | 0 | 0 | 0 | 0 |
| ENSMUSG00000087273 | 0.747221736 | 0.79248125 | 0.646240625 | 0.646240625 | 0.489301084 | 0.698970004 | 0.448487648 |
| ENSMUSG00000087276 | 0 | 0 | 0 | 0 | 0 | 0 | 0 |
| ENSMUSG00000087277 | 0 | 0 | 0 | 0 | 0 | 0 | 0 |
| ENSMUSG00000087278 | 0 | 0 | 0 | 0 | 0 | 0 | 0 |
| ENSMUSG00000087279 | 0 | 0 | 0 | 0 | 0.244650542 | 0 | 0 |
| ENSMUSG00000087281 | 0 | 0 | 0 | 0.25 | 0 | 0 | 0 |
| ENSMUSG00000087282 | 0 | 0.396240625 | 0.25 | 0 | 0.244650542 | 0 | 0 |
| ENSMUSG00000087283 | 0 | 0 | 0 | 0 | 0 | 0 | 0 |
| ENSMUSG00000087286 | 0.811507563 | 0.830482024 | 0.701838731 | 0.580482024 | 0.733951626 | 0.602059991 | 0.629532004 |
| ENSMUSG00000087288 | 0 | 0 | 0 | 0 | 0 | 0 | 0 |
| ENSMUSG00000087289 | 0 | 0 | 0 | 0 | 0 | 0 | 0 |
| ENSMUSG00000087290 | 0 | 0 | 0 | 0 | 0 | 0 | 0 |
| ENSMUSG00000087291 | 0 | 0.5 | 0.25 | 0 | 0.244650542 | 0 | 0.224243824 |
| ENSMUSG00000087292 | 0 | 0 | 0 | 0 | 0.244650542 | 0.301029996 | 0 |
| ENSMUSG00000087293 | 0 | 0 | 0 | 0 | 0 | 0 | 0 |
| ENSMUSG00000087294 | 0 | 0 | 0 | 0 | 0 | 0 | 0.224243824 |
| ENSMUSG00000087295 | 0 | 0 | 0 | 0 | 0 | 0 | 0 |
| ENSMUSG00000087296 | 0 | 0 | 0 | 0 | 0 | 0 | 0 |
| ENSMUSG00000087297 | 0 | 0 | 0 | 0 | 0 | 0 | 0 |
| ENSMUSG00000087298 | 0.289064826 | 0 | 0.25 | 0.5 | 0.568060967 | 0.301029996 | 0 |
| ENSMUSG00000087299 | 0 | 0 | 0 | 0 | 0 | 0 | 0 |
| ENSMUSG00000087300 | 0 | 0 | 0 | 0 | 0 | 0 | 0 |
| ENSMUSG00000087301 | 0 | 0 | 0 | 0 | 0 | 0 | 0 |
| ENSMUSG00000087302 | 0 | 0 | 0 | 0 | 0 | 0 | 0 |
| ENSMUSG00000087303 | 0.578129653 | 0.580482024 | 0.5 | 0.396240625 | 0.489301084 | 0.301029996 | 0.355418052 |
| ENSMUSG00000087304 | 0 | 0 | 0 | 0 | 0 | 0 | 0 |
| ENSMUSG00000087305 | 0.289064826 | 0.701838731 | 0.396240625 | 0.75 | 0.387761935 | 0.477121255 | 0 |
| ENSMUSG00000087306 | 0.289064826 | 0 | 0.75 | 0 | 0.244650542 | 0 | 0 |
| ENSMUSG00000087307 | 0.289064826 | 0.396240625 | 0.25 | 0 | 0.387761935 | 0 | 0 |
| ENSMUSG00000087308 | 0 | 0 | 0 | 0.25 | 0 | 0 | 0 |
| ENSMUSG00000087309 | 0 | 0 | 0 | 0 | 0 | 0 | 0 |

| | | | | | | | |
|---|---|---|---|---|---|---|---|
| ENSMUSG00000087312 | 0 | 0.25 | 0 | 0 | 0 | 0 | 0 |
| ENSMUSG00000087314 | 0.91631382 | 0.830482024 | 1 | 0.580482024 | 0.733951626 | 0.84509804 | 0.579661877 |
| ENSMUSG00000087316 | 0 | 0 | 0 | 0 | 0 | 0 | 0 |
| ENSMUSG00000087317 | 0 | 0 | 0 | 0 | 0 | 0 | 0 |
| ENSMUSG00000087318 | 0 | 0 | 0 | 0 | 0 | 0 | 0 |
| ENSMUSG00000087319 | 0.289064826 | 0 | 0 | 0 | 0 | 0 | 0 |
| ENSMUSG00000087321 | 0 | 0 | 0.25 | 0 | 0.244650542 | 0 | 0.224243824 |
| ENSMUSG00000087322 | 0 | 0 | 0 | 0 | 0 | 0 | 0 |
| ENSMUSG00000087323 | 0 | 0 | 0 | 0 | 0 | 0 | 0 |
| ENSMUSG00000087324 | 0 | 0 | 0 | 0 | 0 | 0 | 0 |
| ENSMUSG00000087325 | 0 | 0 | 0 | 0 | 0 | 0 | 0 |
| ENSMUSG00000087326 | 0.289064826 | 0 | 0 | 0 | 0 | 0 | 0 |
| ENSMUSG00000087327 | 0.289064826 | 0.25 | 0 | 0 | 0 | 0.301029996 | 0.355418052 |
| ENSMUSG00000087328 | 0 | 0 | 0 | 0 | 0 | 0 | 0 |
| ENSMUSG00000087329 | 0 | 0 | 0 | 0 | 0 | 0 | 0 |
| ENSMUSG00000087330 | 0 | 0 | 0 | 0 | 0 | 0 | 0 |
| ENSMUSG00000087331 | 0.289064826 | 0.701838731 | 0.5 | 0.25 | 0.632412477 | 0.301029996 | 0.355418052 |
| ENSMUSG00000087332 | 0 | 0 | 0 | 0 | 0 | 0 | 0 |
| ENSMUSG00000087333 | 0 | 0 | 0 | 0 | 0 | 0 | 0 |
| ENSMUSG00000087334 | 0 | 0 | 0 | 0 | 0 | 0 | 0 |
| ENSMUSG00000087335 | 0 | 0 | 0 | 0 | 0 | 0 | 0 |
| ENSMUSG00000087336 | 0 | 0.5 | 0 | 0.5 | 0.244650542 | 0.301029996 | 0.355418052 |
| ENSMUSG00000087337 | 0 | 0 | 0 | 0 | 0 | 0 | 0 |
| ENSMUSG00000087338 | 0 | 0 | 0.5 | 0.396240625 | 0.244650542 | 0 | 0.224243824 |
| ENSMUSG00000087339 | 0 | 0 | 0 | 0 | 0 | 0 | 0 |
| ENSMUSG00000087340 | 1.129344651 | 1.114857905 | 1.061981878 | 0 | 0 | 0 | 0 |
| ENSMUSG00000087341 | 0 | 0.25 | 0.25 | 0 | 0 | 0 | 0 |
| ENSMUSG00000087342 | 0 | 0 | 0 | 0 | 0 | 0 | 0 |
| ENSMUSG00000087343 | 0 | 0 | 0 | 0 | 0 | 0 | 0 |
| ENSMUSG00000087344 | 0 | 0 | 0 | 0 | 0 | 0 | 0 |
| ENSMUSG00000087345 | 0 | 0 | 0 | 0 | 0 | 0 | 0 |
| ENSMUSG00000087346 | 0 | 0 | 0 | 0 | 0 | 0 | 0 |
| ENSMUSG00000087347 | 0.289064826 | 0.396240625 | 0.396240625 | 0.25 | 0.568060967 | 0.477121255 | 0.224243824 |
| ENSMUSG00000087348 | 0.289064826 | 0 | 0 | 0 | 0 | 0 | 0.224243824 |

| | | | | | | | |
|---|---|---|---|---|---|---|---|
| ENSMUSG00000087349 | 0 | 0 | 0 | 0 | 0 | 0.301029996 | 0 |
| ENSMUSG00000087350 | 0 | 0 | 0 | 0 | 0 | 0 | 0 |
| ENSMUSG00000087351 | 0 | 0.25 | 0 | 0 | 0 | 0.301029996 | 0 |
| ENSMUSG00000087352 | 0 | 0 | 0 | 0 | 0 | 0 | 0 |
| ENSMUSG00000087353 | 0 | 0 | 0 | 0 | 0 | 0 | 0 |
| ENSMUSG00000087354 | 0.45815691 | 0.75 | 0.646240625 | 0.75 | 0.77552387 | 0.602059991 | 0.775756176 |
| ENSMUSG00000087355 | 0 | 0.25 | 0.396240625 | 0.396240625 | 0.387761935 | 0 | 0.448487648 |
| ENSMUSG00000087356 | 0.289064826 | 0 | 0.25 | 0.396240625 | 0.244650542 | 0 | 0.355418052 |
| ENSMUSG00000087357 | 0 | 0 | 0 | 0 | 0 | 0 | 0.224243824 |
| ENSMUSG00000087358 | 0 | 0 | 0 | 0 | 0 | 0 | 0.224243824 |
| ENSMUSG00000087360 | 0 | 0 | 0 | 0.396240625 | 0.387761935 | 0.301029996 | 0.355418052 |
| ENSMUSG00000087361 | 0 | 0 | 0.25 | 0 | 0 | 0 | 0 |
| ENSMUSG00000087362 | 0.45815691 | 0.396240625 | 0.396240625 | 0.396240625 | 0.244650542 | 0.301029996 | 0.629532004 |
| ENSMUSG00000087363 | 0 | 0 | 0 | 0 | 0 | 0 | 0 |
| ENSMUSG00000087364 | 0 | 0 | 0 | 0.396240625 | 0 | 0 | 0 |
| ENSMUSG00000087365 | 1.614416215 | 1.51109853 | 1.557204673 | 1.226722649 | 1.121713561 | 1.230448921 | 1.078019652 |
| ENSMUSG00000087366 | 0.811507563 | 0.79248125 | 0.92510993 | 0.580482024 | 0.568060967 | 0.602059991 | 0.579661877 |
| ENSMUSG00000087367 | 0 | 0 | 0 | 0 | 0 | 0 | 0 |
| ENSMUSG00000087368 | 1.249317394 | 1.114857905 | 1.098079356 | 1.339388001 | 1.176121988 | 1.380211242 | 1.176815335 |
| ENSMUSG00000087369 | 0 | 0.25 | 0.25 | 0.25 | 0 | 0 | 0 |
| ENSMUSG00000087370 | 1.837744666 | 1.777131114 | 1.657339155 | 1.991446071 | 1.925488995 | 2.113943352 | 1.796468236 |
| ENSMUSG00000087371 | 1 | 0.79248125 | 0.92510993 | 0.864857905 | 0.568060967 | 1 | 0.710836105 |
| ENSMUSG00000087373 | 0.45815691 | 0.580482024 | 0.646240625 | 0.896240625 | 0.812711509 | 0.698970004 | 0.710836105 |
| ENSMUSG00000087374 | 0 | 0 | 0.25 | 0 | 0.244650542 | 0 | 0.224243824 |
| ENSMUSG00000087376 | 0 | 0 | 0 | 0 | 0 | 0 | 0 |
| ENSMUSG00000087377 | 0 | 0.646240625 | 0.5 | 0.25 | 0.387761935 | 0.477121255 | 0.355418052 |
| ENSMUSG00000087378 | 0 | 0 | 0 | 0 | 0 | 0 | 0 |
| ENSMUSG00000087379 | 0 | 0 | 0 | 0 | 0 | 0 | 0 |
| ENSMUSG00000087380 | 0.960252568 | 0.976722649 | 0.79248125 | 0.92510993 | 0.931471446 | 0.477121255 | 0.876096088 |
| ENSMUSG00000087381 | 0.811507563 | 0.864857905 | 0.646240625 | 0.580482024 | 0.244650542 | 0.477121255 | 0.448487648 |
| ENSMUSG00000087382 | 0 | 0 | 0 | 0 | 0 | 0 | 0 |
| ENSMUSG00000087383 | 0 | 0 | 0 | 0 | 0 | 0 | 0 |
| ENSMUSG00000087384 | 0 | 0 | 0 | 0.396240625 | 0 | 0.477121255 | 0.355418052 |
| ENSMUSG00000087385 | 0.811507563 | 0.976722649 | 0.830482024 | 0.92510993 | 0.632412477 | 0.84509804 | 0.710836105 |

| | | | | | | | |
|---|---|---|---|---|---|---|---|
| ENSMUSG00000087386 | 0 | 0 | 0 | 0 | 0 | 0 | 0 |
| ENSMUSG00000087387 | 0 | 0 | 0 | 0 | 0 | 0 | 0 |
| ENSMUSG00000087388 | 0 | 0 | 0 | 0 | 0 | 0 | 0 |
| ENSMUSG00000087389 | 0 | 0 | 0 | 0.25 | 0.244650542 | 0 | 0 |
| ENSMUSG00000087390 | 0.45815691 | 0.951838731 | 0.75 | 0.5 | 0.686820904 | 0.602059991 | 0.829800754 |
| ENSMUSG00000087393 | 0 | 0 | 0 | 0 | 0 | 0 | 0 |
| ENSMUSG00000087394 | 0 | 0.396240625 | 0 | 0 | 0 | 0.301029996 | 0 |
| ENSMUSG00000087395 | 0 | 0.580482024 | 0 | 0 | 0.387761935 | 0 | 0 |
| ENSMUSG00000087396 | 1.783508299 | 1.634789703 | 1.688721876 | 1.614857905 | 1.519146132 | 1.698970004 | 1.493062805 |
| ENSMUSG00000087397 | 0.289064826 | 0.396240625 | 0.5 | 0 | 0 | 0.301029996 | 0.355418052 |
| ENSMUSG00000087398 | 0 | 0 | 0 | 0 | 0 | 0 | 0 |
| ENSMUSG00000087399 | 0 | 0 | 0 | 0 | 0 | 0 | 0 |
| ENSMUSG00000087400 | 0.289064826 | 0.5 | 0 | 0.396240625 | 0.387761935 | 0 | 0.224243824 |
| ENSMUSG00000087401 | 0 | 0 | 0 | 0 | 0 | 0 | 0 |
| ENSMUSG00000087402 | 0 | 0 | 0 | 0 | 0 | 0 | 0 |
| ENSMUSG00000087403 | 0.811507563 | 0.92510993 | 0.830482024 | 0.92510993 | 0.931471446 | 1.146128036 | 0.829800754 |
| ENSMUSG00000087404 | 0 | 0 | 0 | 0 | 0 | 0 | 0 |
| ENSMUSG00000087405 | 0 | 0 | 0 | 0 | 0 | 0 | 0 |
| ENSMUSG00000087406 | 0 | 0 | 0 | 0 | 0 | 0 | 0 |
| ENSMUSG00000087407 | 0 | 0 | 0 | 0 | 0 | 0 | 0 |
| ENSMUSG00000087408 | 1.156259305 | 1 | 1.146240625 | 1.098079356 | 0.905314583 | 0.954242509 | 1.014380832 |
| ENSMUSG00000087409 | 0 | 0 | 0 | 0 | 0 | 0 | 0 |
| ENSMUSG00000087410 | 0 | 0 | 0 | 0 | 0 | 0 | 0 |
| ENSMUSG00000087411 | 0 | 0 | 0 | 0 | 0 | 0 | 0 |
| ENSMUSG00000087412 | 0.45815691 | 0.896240625 | 0.79248125 | 0.701838731 | 0.812711509 | 0.77815125 | 0.876096088 |
| ENSMUSG00000087413 | 0 | 0 | 0 | 0 | 0 | 0.301029996 | 0.355418052 |
| ENSMUSG00000087415 | 0 | 0.25 | 0 | 0 | 0 | 0 | 0 |
| ENSMUSG00000087416 | 0 | 0 | 0 | 0 | 0 | 0 | 0 |
| ENSMUSG00000087417 | 0 | 0 | 0 | 0 | 0 | 0 | 0 |
| ENSMUSG00000087422 | 0 | 0 | 0 | 0 | 0 | 0 | 0 |
| ENSMUSG00000087424 | 0 | 0.396240625 | 0.396240625 | 0.396240625 | 0.568060967 | 0 | 0.355418052 |
| ENSMUSG00000087425 | 0 | 0 | 0 | 0 | 0 | 0.301029996 | 0 |
| ENSMUSG00000087428 | 0 | 0 | 0 | 0 | 0 | 0 | 0 |
| ENSMUSG00000087429 | 0 | 0 | 0 | 0 | 0 | 0 | 0.224243824 |

| | | | | | | | |
|---|---|---|---|---|---|---|---|
| ENSMUSG00000087430 | 0 | 0 | 0.25 | 0 | 0 | 0 | 0 |
| ENSMUSG00000087431 | 0.289064826 | 0.25 | 0.25 | 0 | 0 | 0 | 0 |
| ENSMUSG00000087432 | 0 | 0 | 0 | 0 | 0 | 0 | 0 |
| ENSMUSG00000087433 | 0 | 0.25 | 0.580482024 | 0.25 | 0.244650542 | 0 | 0 |
| ENSMUSG00000087434 | 0 | 0 | 0 | 0 | 0 | 0 | 0 |
| ENSMUSG00000087435 | 0 | 0 | 0 | 0 | 0.244650542 | 0 | 0 |
| ENSMUSG00000087436 | 0.289064826 | 0.580482024 | 0 | 0 | 0.244650542 | 0 | 0.355418052 |
| ENSMUSG00000087437 | 0 | 0 | 0 | 0 | 0 | 0 | 0 |
| ENSMUSG00000087439 | 0.45815691 | 0.5 | 0.5 | 0.646240625 | 0.733951626 | 0.903089987 | 0.629532004 |
| ENSMUSG00000087440 | 0.811507563 | 0.5 | 0.79248125 | 0.79248125 | 0.77552387 | 0.77815125 | 0.629532004 |
| ENSMUSG00000087441 | 0 | 0 | 0 | 0 | 0 | 0 | 0 |
| ENSMUSG00000087442 | 0.289064826 | 0.5 | 0.396240625 | 0 | 0 | 0 | 0.448487648 |
| ENSMUSG00000087443 | 0 | 0 | 0.396240625 | 0.396240625 | 0 | 0 | 0.224243824 |
| ENSMUSG00000087444 | 0 | 0.25 | 0 | 0.25 | 0 | 0.301029996 | 0.448487648 |
| ENSMUSG00000087445 | 0 | 0.396240625 | 0.580482024 | 0.396240625 | 0.387761935 | 0.301029996 | 0.520678036 |
| ENSMUSG00000087449 | 0 | 0 | 0 | 0 | 0 | 0 | 0 |
| ENSMUSG00000087450 | 0 | 0 | 0 | 0 | 0 | 0 | 0 |
| ENSMUSG00000087451 | 0 | 0 | 0 | 0 | 0.244650542 | 0 | 0.224243824 |
| ENSMUSG00000087452 | 0.289064826 | 0.25 | 0 | 0 | 0 | 0.301029996 | 0 |
| ENSMUSG00000087454 | 0 | 0 | 0 | 0 | 0 | 0 | 0 |
| ENSMUSG00000087455 | 0 | 0 | 0 | 0 | 0 | 0 | 0 |
| ENSMUSG00000087456 | 0 | 0 | 0 | 0 | 0 | 0 | 0 |
| ENSMUSG00000087457 | 0 | 0.396240625 | 0.25 | 0 | 0.489301084 | 0 | 0.355418052 |
| ENSMUSG00000087458 | 0.45815691 | 0 | 0 | 0.396240625 | 0 | 0.301029996 | 0 |
| ENSMUSG00000087459 | 0 | 0 | 0 | 0 | 0 | 0 | 0 |
| ENSMUSG00000087460 | 0 | 0 | 0 | 0 | 0 | 0 | 0 |
| ENSMUSG00000087461 | 0.578129653 | 0.396240625 | 0.580482024 | 0.25 | 0 | 0 | 0.224243824 |
| ENSMUSG00000087462 | 0 | 0 | 0 | 0 | 0 | 0 | 0 |
| ENSMUSG00000087464 | 0 | 0 | 0 | 0 | 0 | 0 | 0 |
| ENSMUSG00000087465 | 0 | 0 | 0 | 0 | 0 | 0 | 0 |
| ENSMUSG00000087466 | 0 | 0.25 | 0.396240625 | 0.25 | 0.244650542 | 0.301029996 | 0 |
| ENSMUSG00000087467 | 0 | 0 | 0 | 0 | 0 | 0 | 0 |
| ENSMUSG00000087468 | 0 | 0 | 0 | 0 | 0 | 0 | 0 |
| ENSMUSG00000087469 | 0 | 0.25 | 0.396240625 | 0.5 | 0.387761935 | 0 | 0 |

| | | | | | | | |
|---|---|---|---|---|---|---|---|
| ENSMUSG00000087470 | 0 | 0 | 0 | 0.396240625 | 0 | 0 | 0.355418052 |
| ENSMUSG00000087473 | 0 | 0 | 0 | 0 | 0 | 0 | 0 |
| ENSMUSG00000087474 | 0 | 0 | 0 | 0 | 0 | 0 | 0 |
| ENSMUSG00000087475 | 0 | 0 | 0 | 0 | 0 | 0 | 0 |
| ENSMUSG00000087476 | 0 | 0.25 | 0.25 | 0 | 0 | 0 | 0 |
| ENSMUSG00000087477 | 0 | 0 | 0 | 0 | 0 | 0 | 0 |
| ENSMUSG00000087478 | 0.289064826 | 0.5 | 0 | 0.396240625 | 0.568060967 | 0.477121255 | 0.710836105 |
| ENSMUSG00000087479 | 0.45815691 | 0.5 | 0.5 | 0.396240625 | 0.686820904 | 0.77815125 | 0.672731473 |
| ENSMUSG00000087480 | 1.069666964 | 1 | 1 | 0.79248125 | 0.77552387 | 0.477121255 | 0.672731473 |
| ENSMUSG00000087481 | 0.747221736 | 0.646240625 | 0.701838731 | 0.896240625 | 0.733951626 | 0.602059991 | 0.579661877 |
| ENSMUSG00000087482 | 0 | 0 | 0.25 | 0.5 | 0 | 0 | 0.224243824 |
| ENSMUSG00000087484 | 1.342375483 | 1.282320754 | 1.29248125 | 1.061981878 | 1.020174412 | 0.84509804 | 0.984950056 |
| ENSMUSG00000087485 | 0 | 0 | 0 | 0 | 0 | 0 | 0 |
| ENSMUSG00000087486 | 0 | 0 | 0 | 0 | 0 | 0 | 0 |
| ENSMUSG00000087487 | 0 | 0 | 0 | 0 | 0 | 0 | 0 |
| ENSMUSG00000087489 | 0.289064826 | 0 | 0 | 0 | 0 | 0 | 0 |
| ENSMUSG00000087491 | 0 | 0 | 0 | 0 | 0 | 0 | 0 |
| ENSMUSG00000087492 | 0 | 0 | 0 | 0 | 0 | 0 | 0 |
| ENSMUSG00000087495 | 0 | 0 | 0 | 0 | 0 | 0 | 0 |
| ENSMUSG00000087496 | 0.578129653 | 0.396240625 | 0.580482024 | 0.396240625 | 0 | 0.477121255 | 0.448487648 |
| ENSMUSG00000087497 | 1 | 1.02186571 | 1.080482024 | 1.061981878 | 1.106691885 | 1.301029996 | 1.014380832 |
| ENSMUSG00000087500 | 0 | 0 | 0 | 0 | 0 | 0 | 0 |
| ENSMUSG00000087501 | 0.45815691 | 0.5 | 0.75 | 0.646240625 | 0.632412477 | 0.602059991 | 0.672731473 |
| ENSMUSG00000087502 | 1.777675583 | 1.695339928 | 1.598079356 | 1.482684334 | 1.427019704 | 1.51851394 | 1.23151414 |
| ENSMUSG00000087504 | 0 | 0 | 0 | 0 | 0 | 0 | 0 |
| ENSMUSG00000087505 | 0 | 0 | 0.25 | 0 | 0 | 0 | 0 |
| ENSMUSG00000087507 | 0.91631382 | 0 | 0.646240625 | 0.701838731 | 0.489301084 | 0.301029996 | 0.520678036 |
| ENSMUSG00000087508 | 0 | 0 | 0 | 0 | 0 | 0 | 0 |
| ENSMUSG00000087510 | 0 | 0 | 0 | 0 | 0 | 0 | 0 |
| ENSMUSG00000087512 | 0 | 0 | 0 | 0 | 0 | 0 | 0 |
| ENSMUSG00000087513 | 0 | 0.25 | 0.646240625 | 0.580482024 | 0.387761935 | 0 | 0.355418052 |
| ENSMUSG00000087514 | 0 | 0 | 0 | 0 | 0 | 0 | 0 |
| ENSMUSG00000087515 | 0 | 0 | 0 | 0 | 0 | 0 | 0 |
| ENSMUSG00000087516 | 0 | 0 | 0 | 0 | 0 | 0 | 0.355418052 |

| | | | | | | | |
|---|---|---|---|---|---|---|---|
| ENSMUSG00000087517 | 0 | 0 | 0 | 0 | 0 | 0 | 0 |
| ENSMUSG00000087518 | 0 | 0 | 0.25 | 0 | 0 | 0 | 0 |
| ENSMUSG00000087519 | 0.45815691 | 0.5 | 0.5 | 0.701838731 | 0.489301084 | 0.301029996 | 0.224243824 |
| ENSMUSG00000087520 | 0 | 0 | 0 | 0 | 0 | 0 | 0 |
| ENSMUSG00000087521 | 0 | 0 | 0 | 0 | 0 | 0 | 0 |
| ENSMUSG00000087522 | 0 | 0 | 0 | 0 | 0 | 0 | 0.224243824 |
| ENSMUSG00000087523 | 0.91631382 | 0.92510993 | 0.92510993 | 0.580482024 | 0.733951626 | 0.77815125 | 0.775756176 |
| ENSMUSG00000087524 | 0 | 0 | 0 | 0 | 0 | 0 | 0 |
| ENSMUSG00000087525 | 0 | 0 | 0 | 0 | 0 | 0 | 0 |
| ENSMUSG00000087526 | 0.289064826 | 0 | 0 | 0 | 0 | 0 | 0 |
| ENSMUSG00000087527 | 0 | 0 | 0 | 0 | 0 | 0 | 0 |
| ENSMUSG00000087528 | 0.811507563 | 0.79248125 | 0.976722649 | 1.160964047 | 1.074582839 | 1 | 0.935079929 |
| ENSMUSG00000087530 | 0 | 0.25 | 0.25 | 0 | 0.489301084 | 0 | 0 |
| ENSMUSG00000087531 | 0 | 0 | 0 | 0 | 0 | 0 | 0 |
| ENSMUSG00000087532 | 0.289064826 | 0.25 | 0.396240625 | 0 | 0.244650542 | 0.477121255 | 0 |
| ENSMUSG00000087534 | 0 | 0 | 0 | 0 | 0 | 0 | 0 |
| ENSMUSG00000087535 | 0 | 0.5 | 0.5 | 0.25 | 0 | 0 | 0.448487648 |
| ENSMUSG00000087536 | 0 | 0 | 0 | 0 | 0 | 0 | 0 |
| ENSMUSG00000087537 | 0 | 0 | 0 | 0 | 0 | 0 | 0 |
| ENSMUSG00000087538 | 0 | 0 | 0 | 0 | 0 | 0 | 0 |
| ENSMUSG00000087539 | 0 | 0 | 0 | 0 | 0 | 0 | 0 |
| ENSMUSG00000087540 | 0 | 0 | 0 | 0 | 0 | 0 | 0 |
| ENSMUSG00000087541 | 0 | 0.25 | 0 | 0 | 0 | 0 | 0 |
| ENSMUSG00000087542 | 0.578129653 | 0.5 | 0.25 | 0.701838731 | 0.387761935 | 0.477121255 | 0.224243824 |
| ENSMUSG00000087543 | 1.587501561 | 1.54248125 | 1.52186571 | 1.445339928 | 1.414412788 | 1.414973348 | 1.360283038 |
| ENSMUSG00000087544 | 0 | 0 | 0 | 0 | 0 | 0 | 0 |
| ENSMUSG00000087545 | 0.289064826 | 0.25 | 0.396240625 | 0 | 0.489301084 | 0 | 0 |
| ENSMUSG00000087547 | 0 | 0 | 0 | 0 | 0.244650542 | 0 | 0 |
| ENSMUSG00000087549 | 0.578129653 | 0.25 | 0.580482024 | 0.5 | 0.244650542 | 0.602059991 | 0.355418052 |
| ENSMUSG00000087550 | 0 | 0.25 | 0 | 0 | 0.489301084 | 0 | 0.224243824 |
| ENSMUSG00000087552 | 0.289064826 | 0 | 0.646240625 | 0.25 | 0.244650542 | 0 | 0 |
| ENSMUSG00000087553 | 0 | 0 | 0 | 0 | 0 | 0 | 0 |
| ENSMUSG00000087554 | 0 | 0 | 0 | 0 | 0 | 0 | 0 |
| ENSMUSG00000087555 | 0 | 0 | 0 | 0 | 0 | 0 | 0 |

| | | | | | | | |
|---|---|---|---|---|---|---|---|
| ENSMUSG00000087556 | 0.578129653 | 0.580482024 | 0.79248125 | 0.701838731 | 0.686820904 | 0.301029996 | 0.710836105 |
| ENSMUSG00000087557 | 0.289064826 | 0 | 0.25 | 0 | 0 | 0 | 0.448487648 |
| ENSMUSG00000087559 | 0 | 0 | 0 | 0 | 0 | 0 | 0 |
| ENSMUSG00000087560 | 0.671187741 | 0.25 | 0.5 | 0.75 | 0.686820904 | 0.301029996 | 0.224243824 |
| ENSMUSG00000087561 | 0 | 0 | 0 | 0 | 0 | 0 | 0 |
| ENSMUSG00000087563 | 0 | 0 | 0 | 0 | 0 | 0 | 0 |
| ENSMUSG00000087564 | 0 | 0 | 0 | 0 | 0 | 0 | 0 |
| ENSMUSG00000087565 | 0 | 0 | 0 | 0 | 0 | 0 | 0 |
| ENSMUSG00000087568 | 0 | 0 | 0 | 0 | 0 | 0 | 0 |
| ENSMUSG00000087569 | 0 | 0 | 0 | 0 | 0 | 0 | 0 |
| ENSMUSG00000087570 | 0.289064826 | 0 | 0 | 0 | 0 | 0 | 0 |
| ENSMUSG00000087571 | 0 | 0 | 0 | 0 | 0 | 0 | 0 |
| ENSMUSG00000087574 | 1.205378646 | 1.201838731 | 1.25 | 0.951838731 | 1.074582839 | 1.146128036 | 0.775756176 |
| ENSMUSG00000087575 | 0.45815691 | 0.580482024 | 0.79248125 | 0.75 | 0.568060967 | 0.602059991 | 0.355418052 |
| ENSMUSG00000087577 | 0 | 0 | 0 | 0 | 0 | 0 | 0 |
| ENSMUSG00000087578 | 0.747221736 | 0.701838731 | 0.5 | 0 | 0.489301084 | 0.477121255 | 0.448487648 |
| ENSMUSG00000087579 | 0 | 0.580482024 | 0 | 0 | 0.387761935 | 0 | 0.224243824 |
| ENSMUSG00000087581 | 0 | 0 | 0 | 0 | 0 | 0 | 0 |
| ENSMUSG00000087582 | 0 | 0 | 0 | 0 | 0 | 0 | 0 |
| ENSMUSG00000087584 | 0 | 0 | 0 | 0 | 0 | 0 | 0 |
| ENSMUSG00000087585 | 0 | 0 | 0 | 0 | 0 | 0 | 0 |
| ENSMUSG00000087586 | 0 | 0 | 0 | 0 | 0 | 0 | 0 |
| ENSMUSG00000087587 | 0 | 0 | 0 | 0 | 0 | 0.301029996 | 0 |
| ENSMUSG00000087588 | 0 | 0 | 0 | 0 | 0 | 0 | 0 |
| ENSMUSG00000087589 | 0.45815691 | 0.5 | 0.396240625 | 0.25 | 0.632412477 | 0.477121255 | 0.520678036 |
| ENSMUSG00000087590 | 0.811507563 | 1.061981878 | 0.92510993 | 1 | 0.846351821 | 1.255272505 | 1.121219121 |
| ENSMUSG00000087591 | 0 | 0 | 0 | 0 | 0 | 0 | 0 |
| ENSMUSG00000087592 | 0 | 0 | 0 | 0 | 0 | 0 | 0 |
| ENSMUSG00000087593 | 0.289064826 | 0.25 | 0.396240625 | 0.396240625 | 0.244650542 | 0 | 0.579661877 |
| ENSMUSG00000087594 | 0.747221736 | 0.976722649 | 0.75 | 0.701838731 | 0.846351821 | 1 | 0.775756176 |
| ENSMUSG00000087595 | 0.578129653 | 0.75 | 0.396240625 | 0.75 | 0.733951626 | 0.77815125 | 0.520678036 |
| ENSMUSG00000087596 | 0 | 0 | 0 | 0.25 | 0.387761935 | 0 | 0.355418052 |
| ENSMUSG00000087597 | 0 | 0 | 0 | 0 | 0.244650542 | 0 | 0.224243824 |
| ENSMUSG00000087598 | 2.054824387 | 1.848079356 | 1.852347734 | 1.956637122 | 1.87878897 | 2.181843588 | 1.757893036 |

| | | | | | | | |
|---|---|---|---|---|---|---|---|
| ENSMUSG00000087600 | 0 | 0.25 | 0 | 0 | 0 | 0 | 0 |
| ENSMUSG00000087601 | 0 | 0 | 0 | 0 | 0 | 0 | 0 |
| ENSMUSG00000087602 | 0 | 0 | 0 | 0 | 0 | 0 | 0 |
| ENSMUSG00000087603 | 0 | 0 | 0 | 0 | 0 | 0 | 0 |
| ENSMUSG00000087604 | 0.289064826 | 0 | 0 | 0 | 0 | 0 | 0 |
| ENSMUSG00000087605 | 0 | 0.25 | 0 | 0 | 0 | 0 | 0.355418052 |
| ENSMUSG00000087606 | 0.289064826 | 0 | 0 | 0.25 | 0.387761935 | 0.301029996 | 0.224243824 |
| ENSMUSG00000087607 | 0 | 0 | 0 | 0 | 0 | 0 | 0 |
| ENSMUSG00000087608 | 0 | 0.25 | 0 | 0 | 0 | 0.301029996 | 0 |
| ENSMUSG00000087609 | 0 | 0 | 0 | 0 | 0 | 0 | 0 |
| ENSMUSG00000087610 | 0.45815691 | 0.5 | 0.396240625 | 0 | 0.568060967 | 0 | 0.355418052 |
| ENSMUSG00000087611 | 0.289064826 | 0.75 | 0.5 | 0.864857905 | 0.568060967 | 0.477121255 | 0.629532004 |
| ENSMUSG00000087612 | 0 | 0 | 0 | 0 | 0 | 0 | 0 |
| ENSMUSG00000087613 | 0 | 0 | 0 | 0 | 0 | 0 | 0 |
| ENSMUSG00000087614 | 0.289064826 | 0 | 0.25 | 0 | 0 | 0 | 0 |
| ENSMUSG00000087615 | 0.289064826 | 0 | 0.25 | 0 | 0 | 0.301029996 | 0.224243824 |
| ENSMUSG00000087616 | 0 | 0 | 0 | 0.396240625 | 0.568060967 | 0 | 0.224243824 |
| ENSMUSG00000087617 | 0 | 0 | 0 | 0 | 0 | 0 | 0 |
| ENSMUSG00000087618 | 0 | 0 | 0 | 0 | 0 | 0 | 0 |
| ENSMUSG00000087620 | 0 | 0 | 0 | 0 | 0 | 0 | 0 |
| ENSMUSG00000087621 | 0 | 0 | 0 | 0 | 0.244650542 | 0 | 0 |
| ENSMUSG00000087622 | 0 | 0 | 0 | 0 | 0.244650542 | 0 | 0 |
| ENSMUSG00000087623 | 0.289064826 | 0.396240625 | 0.396240625 | 0 | 0.387761935 | 0.602059991 | 0.448487648 |
| ENSMUSG00000087624 | 0.289064826 | 0.25 | 0.701838731 | 0.864857905 | 0.733951626 | 0.477121255 | 0.629532004 |
| ENSMUSG00000087625 | 0 | 0 | 0 | 0 | 0 | 0 | 0 |
| ENSMUSG00000087626 | 1.205378646 | 1.311981878 | 1.226722649 | 0.951838731 | 0.955822902 | 0.84509804 | 0.969165684 |
| ENSMUSG00000087627 | 0.671187741 | 0.75 | 0.580482024 | 0.75 | 0.387761935 | 0.698970004 | 0.629532004 |
| ENSMUSG00000087628 | 0 | 0 | 0 | 0 | 0 | 0 | 0 |
| ENSMUSG00000087629 | 0 | 0.25 | 0.580482024 | 0.5 | 0.632412477 | 0.477121255 | 0.355418052 |
| ENSMUSG00000087630 | 0 | 0.25 | 0 | 0 | 0 | 0 | 0 |
| ENSMUSG00000087632 | 0 | 0 | 0 | 0 | 0 | 0 | 0 |
| ENSMUSG00000087633 | 0 | 0 | 0.25 | 0 | 0 | 0 | 0 |
| ENSMUSG00000087635 | 0 | 0 | 0 | 0 | 0 | 0 | 0 |
| ENSMUSG00000087636 | 0 | 0 | 0 | 0 | 0 | 0 | 0 |

| | | | | | | | |
|---|---|---|---|---|---|---|---|
| ENSMUSG00000087637 | 0 | 0 | 0 | 0 | 0 | 0 | 0 |
| ENSMUSG00000087638 | 0 | 0.396240625 | 0 | 0.25 | 0 | 0 | 0.224243824 |
| ENSMUSG00000087639 | 0.289064826 | 0.25 | 0 | 0.701838731 | 0.489301084 | 0 | 0.448487648 |
| ENSMUSG00000087640 | 0 | 0 | 0 | 0 | 0.387761935 | 0 | 0 |
| ENSMUSG00000087641 | 0 | 0 | 0 | 0 | 0 | 0 | 0 |
| ENSMUSG00000087642 | 0 | 0 | 0 | 0 | 0 | 0 | 0 |
| ENSMUSG00000087643 | 0 | 0 | 0 | 0 | 0 | 0 | 0 |
| ENSMUSG00000087644 | 0.45815691 | 0.25 | 0 | 0 | 0 | 0 | 0 |
| ENSMUSG00000087646 | 0 | 0 | 0 | 0 | 0 | 0 | 0 |
| ENSMUSG00000087647 | 0 | 0 | 0 | 0 | 0 | 0 | 0 |
| ENSMUSG00000087648 | 0 | 0 | 0 | 0 | 0 | 0.301029996 | 0.224243824 |
| ENSMUSG00000087649 | 0 | 0 | 0 | 0 | 0 | 0 | 0 |
| ENSMUSG00000087651 | 0.578129653 | 0.646240625 | 0.5 | 1.516522298 | 1.366364104 | 1.653212514 | 1.383567577 |
| ENSMUSG00000087652 | 0 | 0 | 0 | 0 | 0 | 0 | 0 |
| ENSMUSG00000087653 | 0 | 0 | 0 | 0 | 0 | 0 | 0 |
| ENSMUSG00000087654 | 0 | 0 | 0 | 0 | 0 | 0 | 0 |
| ENSMUSG00000087657 | 0 | 0 | 0 | 0 | 0 | 0 | 0 |
| ENSMUSG00000087658 | 0.45815691 | 0.5 | 0.701838731 | 0.646240625 | 0.632412477 | 0 | 0.355418052 |
| ENSMUSG00000087659 | 0.289064826 | 0 | 0 | 0.25 | 0 | 0 | 0 |
| ENSMUSG00000087660 | 0 | 0 | 0.25 | 0.25 | 0.387761935 | 0 | 0.224243824 |
| ENSMUSG00000087662 | 0 | 0 | 0 | 0 | 0 | 0 | 0 |
| ENSMUSG00000087663 | 0 | 0 | 0 | 0 | 0 | 0 | 0 |
| ENSMUSG00000087666 | 0 | 0 | 0 | 0 | 0 | 0 | 0 |
| ENSMUSG00000087667 | 0 | 0.25 | 0 | 0.25 | 0.387761935 | 0 | 0.448487648 |
| ENSMUSG00000087668 | 0 | 0 | 0 | 0 | 0 | 0 | 0 |
| ENSMUSG00000087669 | 0 | 0 | 0 | 0 | 0 | 0 | 0 |
| ENSMUSG00000087670 | 0 | 0 | 0 | 0 | 0 | 0 | 0 |
| ENSMUSG00000087672 | 1.129344651 | 1.130890489 | 1.02186571 | 0.896240625 | 0.877063019 | 0.477121255 | 0.896975297 |
| ENSMUSG00000087674 | 0 | 0 | 0 | 0 | 0 | 0 | 0 |
| ENSMUSG00000087675 | 0 | 0 | 0 | 0 | 0 | 0 | 0 |
| ENSMUSG00000087677 | 0 | 0 | 0 | 0 | 0 | 0 | 0 |
| ENSMUSG00000087678 | 0 | 0.25 | 0 | 0 | 0 | 0 | 0 |
| ENSMUSG00000087679 | 2.23778581 | 2.071350555 | 2.067865757 | 1.919870025 | 1.86473911 | 2.029383778 | 1.790137008 |
| ENSMUSG00000087681 | 0 | 0 | 0 | 0 | 0 | 0 | 0 |

| | | | | | | | |
|---|---|---|---|---|---|---|---|
| ENSMUSG00000087682 | 0 | 0 | 0 | 0 | 0 | 0 | 0 |
| ENSMUSG00000087683 | 0 | 0 | 0 | 0 | 0 | 0 | 0 |
| ENSMUSG00000087684 | 0.289064826 | 0.25 | 0 | 0 | 0.244650542 | 0 | 0 |
| ENSMUSG00000087685 | 0.45815691 | 0 | 0 | 0 | 0 | 0 | 0 |
| ENSMUSG00000087687 | 1.445324132 | 1.445339928 | 1.470660762 | 1.330482024 | 1.254881871 | 1.342422681 | 1.224243824 |
| ENSMUSG00000087688 | 0 | 0 | 0 | 0 | 0 | 0 | 0 |
| ENSMUSG00000087689 | 0 | 0 | 0 | 0 | 0 | 0 | 0 |
| ENSMUSG00000087690 | 0 | 0 | 0 | 0 | 0 | 0 | 0 |
| ENSMUSG00000087691 | 0 | 0 | 0 | 0 | 0 | 0 | 0.224243824 |
| ENSMUSG00000087692 | 0.289064826 | 0.396240625 | 0.396240625 | 0 | 0.387761935 | 0 | 0.224243824 |
| ENSMUSG00000087693 | 0 | 0 | 0 | 0 | 0 | 0 | 0 |
| ENSMUSG00000087694 | 0 | 0 | 0 | 0 | 0 | 0 | 0 |
| ENSMUSG00000087695 | 0 | 0 | 0 | 0 | 0 | 0 | 0 |
| ENSMUSG00000087698 | 0 | 0 | 0 | 0 | 0 | 0 | 0 |
| ENSMUSG00000087700 | 0.747221736 | 0.830482024 | 0.896240625 | 0 | 0.244650542 | 0.301029996 | 0.579661877 |
| ENSMUSG00000087701 | 0 | 0.25 | 0 | 0 | 0 | 0 | 0 |
| ENSMUSG00000087702 | 0 | 0 | 0 | 0 | 0 | 0 | 0 |
| ENSMUSG00000087703 | 0 | 0 | 0.25 | 0 | 0 | 0 | 0.224243824 |
| ENSMUSG00000087704 | 0 | 0 | 0 | 0 | 0 | 0 | 0 |
| ENSMUSG00000087707 | 0 | 0 | 0 | 0 | 0 | 0 | 0 |
| ENSMUSG00000087708 | 0 | 0 | 0 | 0 | 0 | 0 | 0 |
| ENSMUSG00000087710 | 0 | 0 | 0 | 0 | 0 | 0 | 0 |
| ENSMUSG00000087712 | 0 | 0 | 0 | 0 | 0 | 0 | 0 |
| ENSMUSG00000087714 | 0 | 0 | 0 | 0 | 0 | 0 | 0 |
| ENSMUSG00000087715 | 0 | 0 | 0 | 0 | 0 | 0 | 0 |
| ENSMUSG00000087716 | 0 | 0 | 0 | 0 | 0 | 0 | 0 |
| ENSMUSG00000087717 | 0 | 0 | 0 | 0 | 0 | 0 | 0 |
| ENSMUSG00000087718 | 0 | 0 | 0 | 0 | 0 | 0 | 0 |
| ENSMUSG00000087719 | 0 | 0 | 0 | 0 | 0 | 0.301029996 | 0 |
| ENSMUSG00000087720 | 0 | 0 | 0 | 0 | 0 | 0 | 0 |
| ENSMUSG00000087721 | 0.578129653 | 0.5 | 0.701838731 | 0.25 | 0.489301084 | 0.602059991 | 0.355418052 |
| ENSMUSG00000087722 | 0 | 0 | 0 | 0 | 0 | 0.301029996 | 0.224243824 |
| ENSMUSG00000087723 | 0 | 0 | 0 | 0 | 0 | 0 | 0 |
| ENSMUSG00000087724 | 0 | 0 | 0 | 0 | 0 | 0 | 0 |

| | | | | | | | |
|---|---|---|---|---|---|---|---|
| ENSMUSG00000087726 | 0 | 0 | 0 | 0 | 0 | 0 | 0 |
| ENSMUSG00000087728 | 0 | 0 | 0 | 0 | 0 | 0 | 0 |
| ENSMUSG00000087730 | 0 | 0 | 0 | 0 | 0 | 0 | 0 |
| ENSMUSG00000087731 | 0.289064826 | 0 | 0 | 0 | 0 | 0 | 0 |
| ENSMUSG00000087732 | 0 | 0 | 0 | 0 | 0 | 0 | 0 |
| ENSMUSG00000087734 | 0 | 0 | 0 | 0 | 0 | 0 | 0 |
| ENSMUSG00000087736 | 0 | 0 | 0 | 0 | 0 | 0 | 0 |
| ENSMUSG00000087737 | 0 | 0 | 0 | 0 | 0 | 0 | 0 |
| ENSMUSG00000087739 | 0 | 0 | 0.25 | 0 | 0.489301084 | 0 | 0.520678036 |
| ENSMUSG00000087741 | 0 | 0 | 0 | 0 | 0 | 0 | 0 |
| ENSMUSG00000087742 | 0 | 0 | 0 | 0 | 0 | 0 | 0 |
| ENSMUSG00000087743 | 0 | 0 | 0 | 0 | 0 | 0 | 0 |
| ENSMUSG00000087744 | 0 | 0 | 0 | 0 | 0 | 0 | 0 |
| ENSMUSG00000087746 | 0 | 0 | 0 | 0 | 0 | 0 | 0 |
| ENSMUSG00000087747 | 0 | 0 | 0 | 0 | 0 | 0 | 0 |
| ENSMUSG00000087748 | 0 | 0 | 0 | 0 | 0 | 0 | 0 |
| ENSMUSG00000087750 | 0.578129653 | 0 | 0.396240625 | 0.580482024 | 0 | 0 | 0.448487648 |
| ENSMUSG00000087752 | 0 | 0 | 0 | 0 | 0 | 0 | 0 |
| ENSMUSG00000087754 | 0 | 0 | 0 | 0 | 0 | 0 | 0 |
| ENSMUSG00000087756 | 0 | 0 | 0 | 0 | 0 | 0 | 0 |
| ENSMUSG00000087758 | 0 | 0 | 0 | 0 | 0 | 0 | 0 |
| ENSMUSG00000087759 | 0 | 0 | 0 | 0 | 0 | 0 | 0 |
| ENSMUSG00000087760 | 0 | 0 | 0 | 0 | 0 | 0 | 0 |
| ENSMUSG00000087761 | 0 | 0 | 0 | 0 | 0 | 0 | 0 |
| ENSMUSG00000087764 | 0 | 0 | 0 | 0 | 0 | 0 | 0 |
| ENSMUSG00000087765 | 0 | 0 | 0 | 0 | 0 | 0 | 0 |
| ENSMUSG00000087766 | 0 | 0 | 0 | 0 | 0 | 0 | 0 |
| ENSMUSG00000087767 | 0 | 0 | 0 | 0.25 | 0 | 0 | 0 |
| ENSMUSG00000087768 | 0 | 0 | 0 | 0 | 0 | 0 | 0 |
| ENSMUSG00000087769 | 0 | 0 | 0 | 0 | 0 | 0 | 0 |
| ENSMUSG00000087772 | 0 | 0 | 0 | 0 | 0 | 0 | 0 |
| ENSMUSG00000087774 | 0 | 0 | 0 | 0 | 0 | 0 | 0 |
| ENSMUSG00000087775 | 0 | 0 | 0 | 0 | 0 | 0 | 0 |
| ENSMUSG00000087776 | 0 | 0 | 0 | 0 | 0 | 0 | 0 |

| | | | | | | | |
|---|---|---|---|---|---|---|---|
| ENSMUSG00000087777 | 0 | 0 | 0 | 0 | 0 | 0 | 0 |
| ENSMUSG00000087779 | 0 | 0 | 0 | 0 | 0 | 0 | 0 |
| ENSMUSG00000087780 | 0 | 0 | 0 | 0 | 0 | 0 | 0 |
| ENSMUSG00000087781 | 0 | 0 | 0 | 0 | 0 | 0 | 0 |
| ENSMUSG00000087782 | 0 | 0 | 0 | 0 | 0 | 0 | 0 |
| ENSMUSG00000087783 | 0 | 0 | 0 | 0 | 0 | 0 | 0 |
| ENSMUSG00000087784 | 0.289064826 | 0.5 | 0 | 0 | 0.244650542 | 0 | 0 |
| ENSMUSG00000087785 | 0 | 0 | 0.25 | 0.25 | 0 | 0 | 0 |
| ENSMUSG00000087786 | 0 | 0 | 0 | 0 | 0 | 0 | 0 |
| ENSMUSG00000087789 | 0 | 0 | 0 | 0 | 0 | 0 | 0 |
| ENSMUSG00000087790 | 0 | 0.25 | 0 | 0 | 0.387761935 | 0 | 0 |
| ENSMUSG00000087791 | 0 | 0 | 0 | 0 | 0 | 0 | 0 |
| ENSMUSG00000087792 | 0 | 0 | 0 | 0 | 0 | 0 | 0 |
| ENSMUSG00000087793 | 0 | 0 | 0 | 0 | 0 | 0 | 0 |
| ENSMUSG00000087794 | 0 | 0 | 0.396240625 | 0.396240625 | 0 | 0 | 0 |
| ENSMUSG00000087795 | 0 | 0.25 | 0 | 0 | 0 | 0.301029996 | 0 |
| ENSMUSG00000087796 | 0 | 0 | 0 | 0 | 0 | 0 | 0 |
| ENSMUSG00000087798 | 0 | 0 | 0 | 0 | 0 | 0 | 0 |
| ENSMUSG00000087799 | 0 | 0 | 0 | 0 | 0 | 0 | 0 |
| ENSMUSG00000087800 | 0 | 0 | 0 | 0 | 0 | 0 | 0 |
| ENSMUSG00000087801 | 0 | 0 | 0 | 0 | 0 | 0 | 0 |
| ENSMUSG00000087802 | 0 | 0 | 0 | 0 | 0 | 0 | 0 |
| ENSMUSG00000087804 | 0 | 0 | 0 | 0 | 0 | 0 | 0 |
| ENSMUSG00000087805 | 0 | 0 | 0 | 0 | 0 | 0 | 0 |
| ENSMUSG00000087806 | 0 | 0 | 0 | 0 | 0 | 0 | 0 |
| ENSMUSG00000087807 | 0.289064826 | 0 | 0 | 0 | 0 | 0 | 0 |
| ENSMUSG00000087808 | 0.289064826 | 0.25 | 0 | 0 | 0 | 0 | 0 |
| ENSMUSG00000087810 | 0 | 0 | 0 | 0 | 0 | 0 | 0 |
| ENSMUSG00000087812 | 0 | 0 | 0 | 0 | 0 | 0 | 0 |
| ENSMUSG00000087813 | 0 | 0 | 0 | 0 | 0 | 0 | 0 |
| ENSMUSG00000087815 | 0 | 0 | 0 | 0 | 0 | 0 | 0 |
| ENSMUSG00000087816 | 0 | 0 | 0 | 0 | 0 | 0 | 0 |
| ENSMUSG00000087817 | 0 | 0 | 0 | 0 | 0 | 0 | 0 |
| ENSMUSG00000087819 | 0.45815691 | 0.396240625 | 0 | 0.25 | 0.244650542 | 0 | 0.355418052 |

| | | | | | | | |
|---|---|---|---|---|---|---|---|
| ENSMUSG00000087820 | 0 | 0 | 0 | 0 | 0 | 0 | 0 |
| ENSMUSG00000087823 | 0 | 0 | 0 | 0 | 0 | 0 | 0 |
| ENSMUSG00000087825 | 0 | 0 | 0 | 0 | 0 | 0 | 0 |
| ENSMUSG00000087829 | 0 | 0 | 0 | 0 | 0 | 0 | 0 |
| ENSMUSG00000087830 | 0 | 0 | 0 | 0 | 0 | 0 | 0 |
| ENSMUSG00000087831 | 0 | 0 | 0 | 0 | 0 | 0 | 0 |
| ENSMUSG00000087832 | 0 | 0 | 0 | 0 | 0 | 0 | 0 |
| ENSMUSG00000087833 | 0 | 0 | 0.25 | 0 | 0 | 0 | 0 |
| ENSMUSG00000087834 | 0 | 0 | 0 | 0 | 0 | 0 | 0 |
| ENSMUSG00000087835 | 0 | 0 | 0 | 0 | 0 | 0 | 0 |
| ENSMUSG00000087836 | 0 | 0 | 0 | 0 | 0 | 0 | 0 |
| ENSMUSG00000087838 | 0 | 0 | 0 | 0 | 0 | 0 | 0 |
| ENSMUSG00000087839 | 0 | 0 | 0 | 0 | 0 | 0 | 0 |
| ENSMUSG00000087841 | 0 | 0 | 0 | 0 | 0 | 0 | 0 |
| ENSMUSG00000087842 | 0 | 0 | 0 | 0 | 0 | 0 | 0 |
| ENSMUSG00000087843 | 0 | 0 | 0 | 0 | 0 | 0 | 0 |
| ENSMUSG00000087844 | 0 | 0 | 0 | 0 | 0 | 0 | 0 |
| ENSMUSG00000087845 | 0 | 0 | 0 | 0 | 0 | 0 | 0 |
| ENSMUSG00000087846 | 0 | 0 | 0 | 0 | 0 | 0 | 0 |
| ENSMUSG00000087847 | 0 | 0 | 0 | 0 | 0 | 0 | 0 |
| ENSMUSG00000087848 | 0 | 0 | 0 | 0 | 0 | 0 | 0 |
| ENSMUSG00000087850 | 0 | 0 | 0 | 0 | 0 | 0 | 0 |
| ENSMUSG00000087852 | 0 | 0 | 0 | 0 | 0 | 0 | 0 |
| ENSMUSG00000087853 | 0 | 0 | 0 | 0 | 0 | 0 | 0 |
| ENSMUSG00000087854 | 0 | 0 | 0 | 0 | 0 | 0 | 0 |
| ENSMUSG00000087855 | 0 | 0 | 0 | 0 | 0 | 0 | 0 |
| ENSMUSG00000087857 | 0 | 0 | 0 | 0 | 0 | 0 | 0 |
| ENSMUSG00000087858 | 0 | 0 | 0 | 0 | 0 | 0 | 0 |
| ENSMUSG00000087859 | 0 | 0 | 0 | 0 | 0 | 0 | 0 |
| ENSMUSG00000087860 | 0 | 0 | 0 | 0 | 0 | 0 | 0 |
| ENSMUSG00000087861 | 0 | 0 | 0 | 0 | 0 | 0 | 0 |
| ENSMUSG00000087862 | 0 | 0 | 0 | 0 | 0 | 0 | 0 |
| ENSMUSG00000087863 | 0 | 0 | 0 | 0 | 0 | 0 | 0 |
| ENSMUSG00000087867 | 0 | 0 | 0 | 0 | 0 | 0 | 0 |

| | | | | | | | |
|---|---|---|---|---|---|---|---|
| ENSMUSG00000087869 | 0 | 0 | 0 | 0 | 0 | 0 | 0 |
| ENSMUSG00000087870 | 0 | 0 | 0 | 0 | 0 | 0 | 0 |
| ENSMUSG00000087871 | 0 | 0 | 0 | 0 | 0 | 0 | 0 |
| ENSMUSG00000087872 | 0 | 0 | 0 | 0 | 0 | 0 | 0 |
| ENSMUSG00000087873 | 0 | 0 | 0 | 0 | 0 | 0 | 0 |
| ENSMUSG00000087875 | 0 | 0 | 0 | 0 | 0 | 0 | 0 |
| ENSMUSG00000087876 | 0.45815691 | 0.580482024 | 0.25 | 0.25 | 0 | 0 | 0.355418052 |
| ENSMUSG00000087877 | 0 | 0 | 0 | 0 | 0 | 0 | 0 |
| ENSMUSG00000087878 | 0 | 0 | 0 | 0 | 0 | 0 | 0 |
| ENSMUSG00000087879 | 0 | 0 | 0 | 0 | 0 | 0 | 0 |
| ENSMUSG00000087880 | 0 | 0 | 0 | 0 | 0 | 0 | 0 |
| ENSMUSG00000087881 | 0 | 0 | 0 | 0 | 0.244650542 | 0 | 0.224243824 |
| ENSMUSG00000087883 | 0 | 0 | 0 | 0 | 0 | 0 | 0 |
| ENSMUSG00000087884 | 0 | 0 | 0 | 0 | 0 | 0 | 0 |
| ENSMUSG00000087885 | 0 | 0.25 | 0 | 0 | 0 | 0 | 0 |
| ENSMUSG00000087886 | 0 | 0 | 0 | 0 | 0 | 0 | 0 |
| ENSMUSG00000087887 | 0 | 0 | 0 | 0 | 0 | 0 | 0 |
| ENSMUSG00000087890 | 0 | 0 | 0 | 0 | 0 | 0 | 0 |
| ENSMUSG00000087891 | 0 | 0 | 0 | 0 | 0 | 0 | 0 |
| ENSMUSG00000087892 | 0 | 0 | 0 | 0 | 0 | 0 | 0 |
| ENSMUSG00000087893 | 0 | 0 | 0 | 0 | 0 | 0 | 0 |
| ENSMUSG00000087894 | 0 | 0 | 0 | 0 | 0 | 0 | 0 |
| ENSMUSG00000087895 | 0 | 0.25 | 0 | 0 | 0 | 0 | 0 |
| ENSMUSG00000087896 | 0 | 0 | 0 | 0 | 0 | 0 | 0 |
| ENSMUSG00000087897 | 0 | 0 | 0 | 0 | 0 | 0 | 0 |
| ENSMUSG00000087898 | 0 | 0 | 0 | 0 | 0 | 0 | 0 |
| ENSMUSG00000087899 | 0.289064826 | 0 | 0 | 0 | 0 | 0 | 0 |
| ENSMUSG00000087902 | 0 | 0 | 0 | 0 | 0 | 0 | 0 |
| ENSMUSG00000087903 | 0 | 0 | 0 | 0 | 0 | 0 | 0 |
| ENSMUSG00000087904 | 0 | 0 | 0 | 0 | 0 | 0 | 0 |
| ENSMUSG00000087905 | 0 | 0 | 0 | 0 | 0 | 0 | 0 |
| ENSMUSG00000087908 | 0 | 0 | 0 | 0 | 0 | 0 | 0 |
| ENSMUSG00000087909 | 0 | 0 | 0 | 0 | 0 | 0 | 0 |
| ENSMUSG00000087910 | 0 | 0 | 0 | 0 | 0 | 0 | 0 |

| | | | | | | | |
|---|---|---|---|---|---|---|---|
| ENSMUSG00000087911 | 0 | 0 | 0 | 0 | 0 | 0 | 0 |
| ENSMUSG00000087912 | 0 | 0.396240625 | 0 | 0.396240625 | 0.568060967 | 0.301029996 | 0.520678036 |
| ENSMUSG00000087913 | 0 | 0 | 0 | 0 | 0 | 0 | 0 |
| ENSMUSG00000087914 | 0 | 0 | 0 | 0 | 0 | 0 | 0 |
| ENSMUSG00000087915 | 0 | 0 | 0 | 0 | 0 | 0 | 0 |
| ENSMUSG00000087916 | 0 | 0 | 0 | 0 | 0 | 0 | 0 |
| ENSMUSG00000087917 | 0 | 0 | 0 | 0 | 0 | 0 | 0 |
| ENSMUSG00000087919 | 0 | 0 | 0 | 0 | 0 | 0 | 0 |
| ENSMUSG00000087921 | 0 | 0 | 0 | 0 | 0 | 0 | 0 |
| ENSMUSG00000087922 | 0 | 0 | 0 | 0 | 0 | 0 | 0 |
| ENSMUSG00000087923 | 0 | 0 | 0 | 0 | 0 | 0 | 0 |
| ENSMUSG00000087924 | 0 | 0 | 0 | 0 | 0 | 0 | 0 |
| ENSMUSG00000087925 | 0.289064826 | 0 | 0 | 0 | 0 | 0 | 0 |
| ENSMUSG00000087926 | 0 | 0 | 0 | 0 | 0 | 0 | 0 |
| ENSMUSG00000087929 | 0 | 0 | 0 | 0 | 0 | 0 | 0 |
| ENSMUSG00000087930 | 0 | 0 | 0 | 0 | 0 | 0 | 0 |
| ENSMUSG00000087932 | 0 | 0 | 0 | 0 | 0 | 0 | 0 |
| ENSMUSG00000087933 | 0 | 0 | 0 | 0 | 0 | 0 | 0 |
| ENSMUSG00000087935 | 0 | 0 | 0 | 0 | 0 | 0 | 0 |
| ENSMUSG00000087936 | 0 | 0 | 0 | 0 | 0 | 0 | 0 |
| ENSMUSG00000087937 | 0 | 0 | 0 | 0 | 0 | 0 | 0 |
| ENSMUSG00000087938 | 0.289064826 | 0 | 0 | 0.25 | 0 | 0 | 0.355418052 |
| ENSMUSG00000087939 | 0 | 0 | 0 | 0 | 0 | 0 | 0 |
| ENSMUSG00000087940 | 0 | 0 | 0 | 0 | 0 | 0 | 0 |
| ENSMUSG00000087941 | 0 | 0 | 0 | 0 | 0 | 0 | 0 |
| ENSMUSG00000087942 | 0 | 0 | 0 | 0 | 0 | 0 | 0 |
| ENSMUSG00000087943 | 0 | 0 | 0 | 0 | 0 | 0 | 0 |
| ENSMUSG00000087945 | 0 | 0 | 0 | 0 | 0 | 0 | 0 |
| ENSMUSG00000087946 | 0 | 0 | 0 | 0 | 0 | 0 | 0 |
| ENSMUSG00000087947 | 0 | 0 | 0 | 0 | 0 | 0 | 0 |
| ENSMUSG00000087949 | 0 | 0 | 0 | 0 | 0 | 0 | 0 |
| ENSMUSG00000087950 | 0 | 0 | 0 | 0 | 0 | 0 | 0 |
| ENSMUSG00000087951 | 0 | 0 | 0 | 0 | 0 | 0 | 0 |
| ENSMUSG00000087954 | 0 | 0 | 0 | 0 | 0 | 0 | 0 |

| | | | | | | | |
|---|---|---|---|---|---|---|---|
| ENSMUSG00000087956 | 0 | 0 | 0 | 0 | 0 | 0 | 0 |
| ENSMUSG00000087957 | 0 | 0 | 0 | 0 | 0 | 0 | 0 |
| ENSMUSG00000087958 | 0 | 0 | 0 | 0 | 0 | 0 | 0 |
| ENSMUSG00000087959 | 0 | 0 | 0 | 0 | 0 | 0 | 0 |
| ENSMUSG00000087960 | 0 | 0 | 0 | 0 | 0 | 0 | 0 |
| ENSMUSG00000087961 | 0 | 0 | 0 | 0 | 0 | 0 | 0 |
| ENSMUSG00000087963 | 0 | 0 | 0 | 0.25 | 0 | 0 | 0 |
| ENSMUSG00000087964 | 0 | 0 | 0 | 0 | 0 | 0 | 0 |
| ENSMUSG00000087965 | 0 | 0 | 0 | 0 | 0 | 0 | 0 |
| ENSMUSG00000087966 | 0 | 0 | 0.25 | 0 | 0 | 0 | 0 |
| ENSMUSG00000087967 | 0 | 0 | 0 | 0 | 0 | 0 | 0 |
| ENSMUSG00000087968 | 0 | 0 | 0 | 0 | 0 | 0 | 0 |
| ENSMUSG00000087973 | 0 | 0 | 0 | 0 | 0 | 0 | 0 |
| ENSMUSG00000087975 | 0 | 0 | 0 | 0 | 0 | 0 | 0 |
| ENSMUSG00000087979 | 0 | 0 | 0 | 0 | 0 | 0 | 0 |
| ENSMUSG00000087980 | 0 | 0 | 0 | 0 | 0 | 0 | 0 |
| ENSMUSG00000087981 | 0 | 0 | 0 | 0 | 0 | 0 | 0 |
| ENSMUSG00000087982 | 0 | 0 | 0.396240625 | 0.25 | 0 | 0 | 0 |
| ENSMUSG00000087983 | 0 | 0 | 0 | 0 | 0 | 0 | 0 |
| ENSMUSG00000087985 | 0 | 0 | 0 | 0 | 0 | 0 | 0 |
| ENSMUSG00000087987 | 0 | 0 | 0 | 0 | 0 | 0 | 0 |
| ENSMUSG00000087989 | 0 | 0 | 0 | 0 | 0 | 0 | 0 |
| ENSMUSG00000087990 | 0 | 0 | 0 | 0 | 0 | 0 | 0 |
| ENSMUSG00000087991 | 0 | 0 | 0 | 0 | 0 | 0 | 0.224243824 |
| ENSMUSG00000087992 | 0 | 0 | 0 | 0 | 0 | 0 | 0 |
| ENSMUSG00000087993 | 0 | 0 | 0 | 0 | 0 | 0 | 0 |
| ENSMUSG00000087994 | 0 | 0 | 0 | 0 | 0 | 0 | 0 |
| ENSMUSG00000087995 | 0 | 0 | 0 | 0 | 0 | 0 | 0 |
| ENSMUSG00000087997 | 0 | 0 | 0 | 0 | 0 | 0 | 0 |
| ENSMUSG00000087999 | 0 | 0 | 0 | 0 | 0 | 0 | 0 |
| ENSMUSG00000088000 | 0 | 0 | 0 | 0 | 0 | 0 | 0 |
| ENSMUSG00000088001 | 0 | 0 | 0 | 0 | 0 | 0 | 0 |
| ENSMUSG00000088002 | 0 | 0 | 0 | 0 | 0 | 0 | 0 |
| ENSMUSG00000088004 | 0 | 0 | 0 | 0 | 0 | 0 | 0 |

| | | | | | | | |
|---|---|---|---|---|---|---|---|
| ENSMUSG00000088005 | 0 | 0 | 0 | 0 | 0 | 0 | 0 |
| ENSMUSG00000088008 | 0.578129653 | 0.396240625 | 0 | 0 | 0.244650542 | 0.301029996 | 0 |
| ENSMUSG00000088009 | 0 | 0 | 0 | 0 | 0 | 0 | 0 |
| ENSMUSG00000088010 | 0 | 0 | 0 | 0 | 0 | 0 | 0 |
| ENSMUSG00000088014 | 0 | 0 | 0 | 0 | 0 | 0 | 0 |
| ENSMUSG00000088015 | 0 | 0 | 0 | 0 | 0 | 0 | 0 |
| ENSMUSG00000088016 | 0 | 0.25 | 0 | 0 | 0 | 0 | 0 |
| ENSMUSG00000088017 | 0 | 0 | 0 | 0 | 0 | 0 | 0 |
| ENSMUSG00000088018 | 0 | 0 | 0 | 0 | 0 | 0 | 0 |
| ENSMUSG00000088019 | 0 | 0 | 0 | 0 | 0 | 0 | 0 |
| ENSMUSG00000088020 | 0 | 0 | 0 | 0 | 0 | 0 | 0 |
| ENSMUSG00000088021 | 0 | 0 | 0 | 0 | 0 | 0 | 0 |
| ENSMUSG00000088022 | 0 | 0 | 0 | 0 | 0 | 0 | 0 |
| ENSMUSG00000088025 | 0 | 0 | 0 | 0 | 0 | 0 | 0 |
| ENSMUSG00000088026 | 0 | 0 | 0 | 0 | 0 | 0 | 0 |
| ENSMUSG00000088027 | 0 | 0 | 0 | 0 | 0 | 0 | 0 |
| ENSMUSG00000088028 | 0 | 0 | 0 | 0 | 0 | 0 | 0 |
| ENSMUSG00000088030 | 0 | 0 | 0.25 | 0 | 0 | 0 | 0 |
| ENSMUSG00000088031 | 0 | 0 | 0 | 0 | 0 | 0 | 0 |
| ENSMUSG00000088036 | 0 | 0 | 0 | 0 | 0 | 0 | 0 |
| ENSMUSG00000088037 | 0 | 0 | 0 | 0 | 0 | 0 | 0 |
| ENSMUSG00000088039 | 0 | 0 | 0 | 0 | 0 | 0 | 0 |
| ENSMUSG00000088040 | 0 | 0 | 0 | 0 | 0 | 0 | 0 |
| ENSMUSG00000088041 | 0 | 0 | 0 | 0 | 0 | 0 | 0 |
| ENSMUSG00000088043 | 0 | 0 | 0 | 0 | 0 | 0 | 0 |
| ENSMUSG00000088044 | 0 | 0 | 0 | 0 | 0 | 0 | 0 |
| ENSMUSG00000088045 | 0 | 0 | 0 | 0 | 0 | 0 | 0 |
| ENSMUSG00000088046 | 0 | 0 | 0 | 0 | 0 | 0 | 0 |
| ENSMUSG00000088047 | 0 | 0 | 0 | 0 | 0 | 0 | 0 |
| ENSMUSG00000088048 | 0 | 0 | 0 | 0 | 0 | 0 | 0 |
| ENSMUSG00000088049 | 0 | 0 | 0 | 0 | 0 | 0 | 0 |
| ENSMUSG00000088050 | 0 | 0 | 0 | 0 | 0 | 0 | 0 |
| ENSMUSG00000088051 | 0 | 0 | 0 | 0 | 0 | 0 | 0 |
| ENSMUSG00000088052 | 0 | 0 | 0 | 0 | 0 | 0 | 0 |

| | | | | | | | |
|---|---|---|---|---|---|---|---|
| ENSMUSG00000088054 | 0 | 0 | 0 | 0 | 0 | 0 | 0 |
| ENSMUSG00000088055 | 0 | 0 | 0 | 0 | 0 | 0 | 0 |
| ENSMUSG00000088056 | 0 | 0 | 0 | 0 | 0 | 0 | 0 |
| ENSMUSG00000088057 | 0 | 0 | 0 | 0 | 0 | 0 | 0 |
| ENSMUSG00000088058 | 0 | 0 | 0 | 0 | 0 | 0 | 0 |
| ENSMUSG00000088059 | 0 | 0 | 0 | 0 | 0 | 0 | 0 |
| ENSMUSG00000088060 | 0 | 0 | 0 | 0 | 0 | 0 | 0 |
| ENSMUSG00000088062 | 0 | 0 | 0 | 0 | 0 | 0 | 0 |
| ENSMUSG00000088063 | 0 | 0 | 0 | 0 | 0 | 0 | 0 |
| ENSMUSG00000088064 | 0 | 0 | 0 | 0 | 0 | 0 | 0 |
| ENSMUSG00000088066 | 0.289064826 | 0 | 0 | 0 | 0 | 0 | 0 |
| ENSMUSG00000088067 | 0 | 0 | 0 | 0 | 0 | 0 | 0 |
| ENSMUSG00000088068 | 0 | 0 | 0 | 0 | 0 | 0 | 0 |
| ENSMUSG00000088069 | 0.289064826 | 0 | 0 | 0.396240625 | 0.387761935 | 0.301029996 | 0 |
| ENSMUSG00000088070 | 0 | 0 | 0 | 0 | 0 | 0 | 0 |
| ENSMUSG00000088071 | 0 | 0 | 0 | 0 | 0 | 0 | 0 |
| ENSMUSG00000088073 | 0 | 0 | 0 | 0 | 0 | 0 | 0 |
| ENSMUSG00000088075 | 0 | 0 | 0 | 0 | 0.244650542 | 0 | 0 |
| ENSMUSG00000088076 | 0 | 0 | 0 | 0 | 0 | 0 | 0 |
| ENSMUSG00000088077 | 0 | 0 | 0 | 0 | 0 | 0 | 0 |
| ENSMUSG00000088083 | 0 | 0 | 0 | 0 | 0 | 0 | 0 |
| ENSMUSG00000088084 | 0 | 0 | 0 | 0 | 0 | 0 | 0 |
| ENSMUSG00000088085 | 0 | 0 | 0 | 0 | 0 | 0 | 0 |
| ENSMUSG00000088086 | 0 | 0 | 0 | 0 | 0 | 0 | 0 |
| ENSMUSG00000088088 | 0.289064826 | 0 | 0.25 | 0 | 0 | 0 | 0.224243824 |
| ENSMUSG00000088090 | 0 | 0 | 0 | 0 | 0 | 0 | 0 |
| ENSMUSG00000088091 | 0 | 0 | 0 | 0 | 0 | 0 | 0 |
| ENSMUSG00000088093 | 0 | 0 | 0 | 0 | 0 | 0 | 0 |
| ENSMUSG00000088094 | 0 | 0 | 0 | 0 | 0 | 0 | 0 |
| ENSMUSG00000088095 | 0 | 0 | 0 | 0 | 0 | 0 | 0 |
| ENSMUSG00000088096 | 0 | 0 | 0 | 0 | 0 | 0 | 0 |
| ENSMUSG00000088097 | 0 | 0 | 0 | 0 | 0 | 0 | 0 |
| ENSMUSG00000088098 | 0 | 0 | 0 | 0 | 0 | 0 | 0 |
| ENSMUSG00000088099 | 0 | 0 | 0 | 0 | 0 | 0 | 0 |

| | | | | | | | |
|---|---|---|---|---|---|---|---|
| ENSMUSG00000088102 | 0 | 0 | 0 | 0 | 0 | 0 | 0 |
| ENSMUSG00000088103 | 0 | 0 | 0 | 0 | 0 | 0 | 0 |
| ENSMUSG00000088104 | 0 | 0 | 0 | 0 | 0 | 0 | 0 |
| ENSMUSG00000088105 | 0 | 0 | 0 | 0 | 0 | 0 | 0 |
| ENSMUSG00000088106 | 0 | 0 | 0 | 0 | 0 | 0 | 0 |
| ENSMUSG00000088108 | 0 | 0.396240625 | 0.5 | 0.25 | 0.489301084 | 0 | 0.448487648 |
| ENSMUSG00000088111 | 0 | 0 | 0 | 0 | 0 | 0 | 0 |
| ENSMUSG00000088112 | 0 | 0 | 0 | 0 | 0 | 0 | 0 |
| ENSMUSG00000088113 | 0 | 0 | 0.25 | 0 | 0 | 0 | 0 |
| ENSMUSG00000088114 | 0 | 0 | 0 | 0 | 0 | 0 | 0 |
| ENSMUSG00000088115 | 0 | 0 | 0 | 0 | 0 | 0 | 0 |
| ENSMUSG00000088116 | 0 | 0 | 0 | 0 | 0 | 0 | 0 |
| ENSMUSG00000088118 | 0 | 0 | 0 | 0 | 0 | 0 | 0 |
| ENSMUSG00000088119 | 0 | 0.5 | 0.5 | 0.580482024 | 0.244650542 | 0 | 0.355418052 |
| ENSMUSG00000088121 | 0 | 0 | 0 | 0 | 0 | 0 | 0 |
| ENSMUSG00000088122 | 0 | 0 | 0 | 0 | 0 | 0 | 0 |
| ENSMUSG00000088123 | 0 | 0 | 0 | 0 | 0 | 0 | 0 |
| ENSMUSG00000088124 | 0 | 0 | 0 | 0 | 0 | 0 | 0 |
| ENSMUSG00000088125 | 0 | 0 | 0 | 0 | 0 | 0 | 0 |
| ENSMUSG00000088126 | 0 | 0 | 0 | 0 | 0 | 0 | 0 |
| ENSMUSG00000088128 | 0 | 0 | 0 | 0 | 0 | 0 | 0 |
| ENSMUSG00000088132 | 0 | 0 | 0 | 0 | 0 | 0 | 0 |
| ENSMUSG00000088133 | 0 | 0 | 0 | 0 | 0 | 0 | 0 |
| ENSMUSG00000088134 | 0 | 0 | 0.396240625 | 0 | 0 | 0 | 0 |
| ENSMUSG00000088135 | 0 | 0 | 0 | 0 | 0 | 0 | 0 |
| ENSMUSG00000088136 | 0 | 0 | 0 | 0 | 0 | 0 | 0 |
| ENSMUSG00000088138 | 0 | 0 | 0 | 0 | 0 | 0 | 0 |
| ENSMUSG00000088139 | 0 | 0 | 0 | 0 | 0 | 0 | 0 |
| ENSMUSG00000088141 | 0 | 0 | 0 | 0 | 0 | 0 | 0 |
| ENSMUSG00000088143 | 0 | 0 | 0 | 0 | 0 | 0 | 0 |
| ENSMUSG00000088144 | 0 | 0 | 0 | 0 | 0 | 0 | 0 |
| ENSMUSG00000088145 | 0 | 0 | 0 | 0 | 0 | 0 | 0 |
| ENSMUSG00000088146 | 0 | 0 | 0 | 0 | 0 | 0 | 0 |
| ENSMUSG00000088147 | 0 | 0 | 0 | 0 | 0 | 0 | 0 |

| | | | | | | | |
|---|---|---|---|---|---|---|---|
| ENSMUSG00000088148 | 0 | 0 | 0 | 0 | 0 | 0 | 0 |
| ENSMUSG00000088149 | 0 | 0 | 0 | 0 | 0 | 0 | 0 |
| ENSMUSG00000088150 | 0 | 0 | 0 | 0 | 0 | 0 | 0 |
| ENSMUSG00000088151 | 0 | 0 | 0.25 | 0.25 | 0.387761935 | 0 | 0 |
| ENSMUSG00000088152 | 0 | 0 | 0 | 0 | 0 | 0 | 0 |
| ENSMUSG00000088153 | 0.289064826 | 0 | 0.25 | 0 | 0.244650542 | 0.301029996 | 0 |
| ENSMUSG00000088154 | 0 | 0 | 0 | 0 | 0 | 0 | 0 |
| ENSMUSG00000088155 | 0 | 0 | 0 | 0 | 0 | 0 | 0 |
| ENSMUSG00000088156 | 0 | 0 | 0 | 0 | 0 | 0 | 0 |
| ENSMUSG00000088157 | 0 | 0 | 0 | 0 | 0 | 0 | 0 |
| ENSMUSG00000088158 | 0 | 0 | 0 | 0 | 0.244650542 | 0 | 0 |
| ENSMUSG00000088159 | 0 | 0 | 0 | 0 | 0 | 0 | 0 |
| ENSMUSG00000088160 | 0 | 0 | 0 | 0 | 0 | 0 | 0 |
| ENSMUSG00000088161 | 0 | 0 | 0 | 0 | 0 | 0 | 0 |
| ENSMUSG00000088162 | 0 | 0 | 0 | 0 | 0 | 0 | 0 |
| ENSMUSG00000088164 | 0 | 0 | 0 | 0 | 0 | 0 | 0 |
| ENSMUSG00000088165 | 0 | 0 | 0 | 0 | 0 | 0 | 0 |
| ENSMUSG00000088166 | 0 | 0 | 0 | 0 | 0 | 0 | 0 |
| ENSMUSG00000088167 | 0 | 0 | 0 | 0 | 0 | 0 | 0 |
| ENSMUSG00000088168 | 0 | 0 | 0 | 0 | 0 | 0 | 0 |
| ENSMUSG00000088169 | 0 | 0 | 0 | 0 | 0 | 0 | 0.224243824 |
| ENSMUSG00000088170 | 0 | 0 | 0.25 | 0.396240625 | 0.244650542 | 0.301029996 | 0.355418052 |
| ENSMUSG00000088171 | 0 | 0 | 0 | 0 | 0.244650542 | 0 | 0 |
| ENSMUSG00000088172 | 0 | 0 | 0 | 0 | 0 | 0 | 0 |
| ENSMUSG00000088175 | 0 | 0 | 0 | 0 | 0 | 0 | 0 |
| ENSMUSG00000088176 | 0 | 0 | 0.25 | 0 | 0 | 0 | 0 |
| ENSMUSG00000088177 | 0 | 0 | 0 | 0 | 0 | 0 | 0 |
| ENSMUSG00000088179 | 0 | 0 | 0 | 0 | 0 | 0 | 0 |
| ENSMUSG00000088180 | 0 | 0 | 0 | 0 | 0 | 0 | 0 |
| ENSMUSG00000088181 | 0 | 0 | 0 | 0 | 0 | 0 | 0 |
| ENSMUSG00000088182 | 0 | 0 | 0 | 0 | 0 | 0 | 0 |
| ENSMUSG00000088183 | 0 | 0 | 0 | 0.25 | 0 | 0 | 0 |
| ENSMUSG00000088184 | 0 | 0 | 0 | 0 | 0 | 0 | 0 |
| ENSMUSG00000088185 | 0 | 0.25 | 0 | 0 | 0 | 0 | 0 |

| | | | | | | | |
|---|---|---|---|---|---|---|---|
| ENSMUSG00000088187 | 0 | 0 | 0 | 0 | 0 | 0 | 0 |
| ENSMUSG00000088188 | 0 | 0 | 0 | 0.25 | 0 | 0 | 0.224243824 |
| ENSMUSG00000088189 | 0 | 0 | 0 | 0 | 0 | 0 | 0 |
| ENSMUSG00000088191 | 0 | 0 | 0 | 0 | 0 | 0 | 0 |
| ENSMUSG00000088192 | 0 | 0 | 0 | 0 | 0 | 0 | 0 |
| ENSMUSG00000088193 | 0 | 0 | 0 | 0 | 0 | 0 | 0 |
| ENSMUSG00000088194 | 0 | 0 | 0 | 0 | 0 | 0 | 0 |
| ENSMUSG00000088195 | 0 | 0 | 0 | 0 | 0 | 0 | 0 |
| ENSMUSG00000088196 | 0 | 0 | 0 | 0 | 0 | 0 | 0 |
| ENSMUSG00000088197 | 0 | 0 | 0 | 0 | 0 | 0 | 0 |
| ENSMUSG00000088198 | 0 | 0 | 0 | 0 | 0 | 0 | 0 |
| ENSMUSG00000088200 | 0 | 0 | 0.25 | 0 | 0 | 0.301029996 | 0 |
| ENSMUSG00000088201 | 0 | 0 | 0 | 0 | 0 | 0 | 0 |
| ENSMUSG00000088203 | 0 | 0 | 0 | 0 | 0 | 0 | 0 |
| ENSMUSG00000088204 | 0 | 0 | 0 | 0 | 0 | 0 | 0 |
| ENSMUSG00000088205 | 0 | 0 | 0 | 0 | 0 | 0 | 0 |
| ENSMUSG00000088207 | 0 | 0 | 0 | 0 | 0 | 0 | 0 |
| ENSMUSG00000088208 | 0.45815691 | 0.25 | 0.25 | 0 | 0 | 0 | 0.224243824 |
| ENSMUSG00000088209 | 0 | 0 | 0 | 0 | 0 | 0 | 0 |
| ENSMUSG00000088210 | 0 | 0 | 0 | 0 | 0 | 0 | 0 |
| ENSMUSG00000088211 | 0 | 0 | 0 | 0 | 0 | 0 | 0 |
| ENSMUSG00000088212 | 0 | 0 | 0 | 0 | 0 | 0 | 0 |
| ENSMUSG00000088213 | 0 | 0.25 | 0 | 0 | 0 | 0 | 0.224243824 |
| ENSMUSG00000088216 | 0 | 0 | 0 | 0 | 0 | 0 | 0 |
| ENSMUSG00000088217 | 0 | 0 | 0.25 | 0 | 0 | 0 | 0.224243824 |
| ENSMUSG00000088218 | 0 | 0 | 0 | 0 | 0 | 0 | 0 |
| ENSMUSG00000088219 | 0 | 0 | 0 | 0 | 0 | 0 | 0 |
| ENSMUSG00000088220 | 0 | 0 | 0 | 0 | 0 | 0 | 0 |
| ENSMUSG00000088221 | 0 | 0 | 0 | 0 | 0 | 0 | 0 |
| ENSMUSG00000088222 | 0 | 0 | 0 | 0 | 0 | 0 | 0 |
| ENSMUSG00000088223 | 0 | 0 | 0 | 0 | 0 | 0 | 0 |
| ENSMUSG00000088224 | 0 | 0 | 0 | 0 | 0 | 0 | 0 |
| ENSMUSG00000088225 | 0 | 0 | 0 | 0 | 0 | 0 | 0 |
| ENSMUSG00000088226 | 0.671187741 | 0.25 | 0.396240625 | 0.396240625 | 0.387761935 | 0.301029996 | 0 |

| | | | | | | | |
|---|---|---|---|---|---|---|---|
| ENSMUSG00000088227 | 0 | 0 | 0 | 0 | 0 | 0 | 0 |
| ENSMUSG00000088228 | 0 | 0 | 0 | 0 | 0 | 0 | 0 |
| ENSMUSG00000088229 | 0 | 0 | 0 | 0 | 0 | 0 | 0 |
| ENSMUSG00000088230 | 0 | 0 | 0 | 0 | 0 | 0 | 0 |
| ENSMUSG00000088231 | 0 | 0 | 0 | 0 | 0 | 0 | 0 |
| ENSMUSG00000088232 | 0 | 0 | 0 | 0 | 0 | 0 | 0 |
| ENSMUSG00000088233 | 0 | 0 | 0.25 | 0 | 0 | 0 | 0 |
| ENSMUSG00000088234 | 0 | 0 | 0 | 0 | 0 | 0 | 0 |
| ENSMUSG00000088236 | 0 | 0 | 0 | 0 | 0 | 0 | 0 |
| ENSMUSG00000088237 | 0 | 0 | 0 | 0 | 0 | 0 | 0 |
| ENSMUSG00000088239 | 0 | 0 | 0 | 0 | 0 | 0 | 0 |
| ENSMUSG00000088240 | 0 | 0 | 0 | 0 | 0 | 0 | 0 |
| ENSMUSG00000088241 | 0 | 0 | 0 | 0 | 0 | 0 | 0 |
| ENSMUSG00000088242 | 0 | 0 | 0 | 0 | 0 | 0 | 0 |
| ENSMUSG00000088243 | 0 | 0 | 0 | 0 | 0 | 0 | 0 |
| ENSMUSG00000088244 | 0 | 0 | 0 | 0 | 0 | 0 | 0 |
| ENSMUSG00000088245 | 0 | 0 | 0 | 0 | 0 | 0 | 0 |
| ENSMUSG00000088246 | 0 | 0 | 0 | 0 | 0 | 0 | 0 |
| ENSMUSG00000088248 | 0 | 0 | 0 | 0 | 0 | 0 | 0 |
| ENSMUSG00000088249 | 0.289064826 | 0 | 0.25 | 0 | 0 | 0 | 0.224243824 |
| ENSMUSG00000088250 | 0 | 0 | 0 | 0 | 0 | 0 | 0.224243824 |
| ENSMUSG00000088252 | 0 | 0 | 0 | 0 | 0 | 0 | 0 |
| ENSMUSG00000088253 | 0 | 0 | 0 | 0 | 0 | 0 | 0 |
| ENSMUSG00000088254 | 0 | 0 | 0.25 | 0 | 0 | 0 | 0 |
| ENSMUSG00000088255 | 0 | 0 | 0 | 0 | 0 | 0 | 0 |
| ENSMUSG00000088257 | 0 | 0 | 0 | 0 | 0 | 0 | 0 |
| ENSMUSG00000088258 | 0 | 0 | 0 | 0 | 0 | 0 | 0 |
| ENSMUSG00000088259 | 0 | 0 | 0 | 0 | 0 | 0 | 0 |
| ENSMUSG00000088260 | 0 | 0 | 0 | 0 | 0 | 0 | 0 |
| ENSMUSG00000088261 | 0 | 0 | 0 | 0 | 0 | 0 | 0 |
| ENSMUSG00000088264 | 0 | 0 | 0 | 0 | 0 | 0 | 0 |
| ENSMUSG00000088265 | 0 | 0 | 0 | 0 | 0 | 0 | 0 |
| ENSMUSG00000088266 | 0 | 0 | 0 | 0 | 0 | 0 | 0 |
| ENSMUSG00000088268 | 0 | 0.5 | 0.396240625 | 0.5 | 0.489301084 | 0.301029996 | 0.355418052 |

| | | | | | | | |
|---|---|---|---|---|---|---|---|
| ENSMUSG00000088269 | 0 | 0 | 0 | 0 | 0 | 0 | 0 |
| ENSMUSG00000088270 | 0 | 0 | 0 | 0 | 0 | 0 | 0 |
| ENSMUSG00000088273 | 0 | 0 | 0 | 0 | 0 | 0 | 0 |
| ENSMUSG00000088275 | 0 | 0 | 0 | 0 | 0 | 0 | 0 |
| ENSMUSG00000088276 | 0 | 0 | 0 | 0 | 0 | 0 | 0 |
| ENSMUSG00000088278 | 0 | 0 | 0 | 0 | 0 | 0 | 0 |
| ENSMUSG00000088279 | 0 | 0 | 0 | 0 | 0 | 0 | 0 |
| ENSMUSG00000088280 | 0 | 0 | 0 | 0 | 0 | 0 | 0 |
| ENSMUSG00000088281 | 0 | 0 | 0 | 0 | 0 | 0 | 0 |
| ENSMUSG00000088282 | 0 | 0 | 0 | 0 | 0 | 0 | 0 |
| ENSMUSG00000088283 | 0 | 0 | 0 | 0 | 0 | 0 | 0 |
| ENSMUSG00000088284 | 0 | 0 | 0 | 0 | 0 | 0 | 0 |
| ENSMUSG00000088285 | 0 | 0 | 0 | 0 | 0 | 0 | 0 |
| ENSMUSG00000088286 | 0 | 0 | 0 | 0 | 0 | 0 | 0 |
| ENSMUSG00000088288 | 0 | 0 | 0 | 0 | 0 | 0 | 0 |
| ENSMUSG00000088291 | 0 | 0 | 0 | 0 | 0 | 0 | 0 |
| ENSMUSG00000088293 | 0 | 0 | 0 | 0 | 0 | 0 | 0 |
| ENSMUSG00000088294 | 0 | 0 | 0 | 0 | 0 | 0 | 0 |
| ENSMUSG00000088295 | 0 | 0 | 0 | 0 | 0 | 0 | 0 |
| ENSMUSG00000088296 | 0 | 0 | 0 | 0 | 0 | 0 | 0 |
| ENSMUSG00000088297 | 0 | 0 | 0 | 0 | 0 | 0 | 0 |
| ENSMUSG00000088298 | 0 | 0 | 0 | 0 | 0 | 0 | 0.224243824 |
| ENSMUSG00000088299 | 0 | 0 | 0 | 0 | 0 | 0 | 0 |
| ENSMUSG00000088301 | 0 | 0 | 0 | 0 | 0 | 0 | 0 |
| ENSMUSG00000088302 | 0 | 0 | 0 | 0 | 0 | 0 | 0 |
| ENSMUSG00000088303 | 0 | 0 | 0 | 0 | 0 | 0 | 0 |
| ENSMUSG00000088304 | 0 | 0 | 0 | 0 | 0 | 0 | 0 |
| ENSMUSG00000088307 | 0 | 0 | 0 | 0 | 0 | 0 | 0 |
| ENSMUSG00000088308 | 0 | 0 | 0 | 0 | 0.244650542 | 0 | 0.448487648 |
| ENSMUSG00000088310 | 0 | 0 | 0 | 0 | 0 | 0 | 0 |
| ENSMUSG00000088311 | 0 | 0 | 0 | 0 | 0 | 0 | 0 |
| ENSMUSG00000088312 | 0 | 0 | 0 | 0 | 0 | 0 | 0 |
| ENSMUSG00000088313 | 0 | 0 | 0 | 0 | 0 | 0 | 0 |
| ENSMUSG00000088314 | 0 | 0 | 0 | 0 | 0 | 0 | 0 |

| | | | | | | | |
|---|---|---|---|---|---|---|---|
| ENSMUSG00000088317 | 0 | 0 | 0 | 0 | 0 | 0 | 0 |
| ENSMUSG00000088318 | 0 | 0 | 0 | 0 | 0 | 0 | 0 |
| ENSMUSG00000088320 | 0 | 0 | 0 | 0 | 0 | 0 | 0 |
| ENSMUSG00000088322 | 0 | 0 | 0 | 0 | 0 | 0 | 0 |
| ENSMUSG00000088323 | 0 | 0 | 0 | 0 | 0 | 0 | 0 |
| ENSMUSG00000088324 | 0 | 0 | 0 | 0 | 0 | 0 | 0 |
| ENSMUSG00000088325 | 0 | 0 | 0 | 0 | 0 | 0 | 0 |
| ENSMUSG00000088326 | 0 | 0 | 0.396240625 | 0 | 0.387761935 | 0.602059991 | 0.355418052 |
| ENSMUSG00000088327 | 0 | 0 | 0 | 0 | 0 | 0 | 0 |
| ENSMUSG00000088329 | 0 | 0 | 0 | 0 | 0 | 0 | 0 |
| ENSMUSG00000088331 | 0 | 0 | 0 | 0 | 0 | 0 | 0 |
| ENSMUSG00000088332 | 0 | 0 | 0 | 0 | 0 | 0.301029996 | 0 |
| ENSMUSG00000088333 | 0 | 0 | 0 | 0 | 0 | 0 | 0 |
| ENSMUSG00000088334 | 0 | 0 | 0 | 0 | 0 | 0 | 0 |
| ENSMUSG00000088335 | 0 | 0 | 0 | 0 | 0 | 0 | 0 |
| ENSMUSG00000088337 | 0 | 0 | 0 | 0 | 0 | 0 | 0 |
| ENSMUSG00000088338 | 0 | 0 | 0 | 0 | 0 | 0 | 0 |
| ENSMUSG00000088339 | 0 | 0 | 0 | 0 | 0 | 0 | 0 |
| ENSMUSG00000088340 | 0 | 0 | 0 | 0 | 0 | 0 | 0 |
| ENSMUSG00000088342 | 0 | 0 | 0 | 0 | 0 | 0 | 0 |
| ENSMUSG00000088343 | 0 | 0 | 0 | 0 | 0 | 0 | 0 |
| ENSMUSG00000088344 | 0 | 0 | 0 | 0 | 0 | 0 | 0 |
| ENSMUSG00000088346 | 0 | 0 | 0 | 0 | 0 | 0 | 0 |
| ENSMUSG00000088348 | 0 | 0 | 0 | 0 | 0 | 0 | 0 |
| ENSMUSG00000088349 | 0 | 0 | 0 | 0 | 0 | 0 | 0 |
| ENSMUSG00000088350 | 0 | 0 | 0 | 0 | 0 | 0 | 0 |
| ENSMUSG00000088351 | 0 | 0 | 0 | 0 | 0 | 0 | 0 |
| ENSMUSG00000088352 | 0 | 0 | 0 | 0 | 0 | 0 | 0 |
| ENSMUSG00000088353 | 0 | 0 | 0 | 0 | 0 | 0 | 0 |
| ENSMUSG00000088354 | 0 | 0 | 0 | 0 | 0 | 0 | 0 |
| ENSMUSG00000088356 | 0 | 0 | 0 | 0 | 0 | 0 | 0 |
| ENSMUSG00000088357 | 0 | 0 | 0 | 0.25 | 0 | 0 | 0 |
| ENSMUSG00000088358 | 0 | 0 | 0 | 0 | 0 | 0 | 0 |
| ENSMUSG00000088359 | 0 | 0 | 0 | 0 | 0 | 0 | 0 |

| | | | | | | | |
|---|---|---|---|---|---|---|---|
| ENSMUSG00000088360 | 0 | 0 | 0 | 0 | 0 | 0 | 0 |
| ENSMUSG00000088361 | 0 | 0 | 0 | 0 | 0 | 0 | 0 |
| ENSMUSG00000088362 | 0 | 0 | 0 | 0 | 0 | 0 | 0 |
| ENSMUSG00000088363 | 0 | 0 | 0 | 0 | 0 | 0 | 0 |
| ENSMUSG00000088364 | 0 | 0 | 0 | 0 | 0 | 0 | 0 |
| ENSMUSG00000088365 | 0 | 0 | 0 | 0 | 0 | 0 | 0 |
| ENSMUSG00000088366 | 0 | 0 | 0 | 0 | 0 | 0 | 0 |
| ENSMUSG00000088367 | 0 | 0 | 0 | 0 | 0 | 0 | 0 |
| ENSMUSG00000088368 | 0 | 0 | 0 | 0 | 0 | 0 | 0 |
| ENSMUSG00000088369 | 0 | 0 | 0 | 0 | 0 | 0 | 0 |
| ENSMUSG00000088370 | 0 | 0 | 0 | 0 | 0 | 0 | 0 |
| ENSMUSG00000088371 | 0 | 0 | 0 | 0 | 0 | 0 | 0 |
| ENSMUSG00000088372 | 0 | 0 | 0 | 0 | 0 | 0 | 0 |
| ENSMUSG00000088373 | 0 | 0 | 0 | 0 | 0 | 0 | 0 |
| ENSMUSG00000088374 | 0 | 0 | 0 | 0 | 0 | 0 | 0 |
| ENSMUSG00000088377 | 0 | 0 | 0 | 0 | 0 | 0 | 0 |
| ENSMUSG00000088378 | 0 | 0 | 0.25 | 0.25 | 0 | 0 | 0 |
| ENSMUSG00000088379 | 0 | 0 | 0 | 0 | 0 | 0 | 0 |
| ENSMUSG00000088380 | 0 | 0 | 0 | 0 | 0 | 0 | 0 |
| ENSMUSG00000088383 | 0 | 0 | 0 | 0 | 0 | 0 | 0 |
| ENSMUSG00000088384 | 0 | 0 | 0 | 0 | 0 | 0 | 0 |
| ENSMUSG00000088385 | 0 | 0 | 0 | 0.25 | 0 | 0 | 0 |
| ENSMUSG00000088386 | 0 | 0 | 0 | 0 | 0 | 0 | 0 |
| ENSMUSG00000088387 | 0 | 0 | 0 | 0 | 0 | 0 | 0 |
| ENSMUSG00000088389 | 0 | 0 | 0 | 0 | 0 | 0 | 0 |
| ENSMUSG00000088391 | 0 | 0 | 0 | 0 | 0 | 0 | 0 |
| ENSMUSG00000088392 | 0 | 0 | 0 | 0 | 0 | 0 | 0 |
| ENSMUSG00000088393 | 0 | 0 | 0 | 0 | 0 | 0 | 0 |
| ENSMUSG00000088394 | 0 | 0 | 0 | 0 | 0 | 0 | 0 |
| ENSMUSG00000088395 | 0 | 0 | 0 | 0 | 0 | 0 | 0 |
| ENSMUSG00000088397 | 0 | 0 | 0 | 0 | 0 | 0 | 0 |
| ENSMUSG00000088398 | 0 | 0 | 0 | 0 | 0 | 0 | 0 |
| ENSMUSG00000088400 | 0 | 0 | 0 | 0 | 0 | 0 | 0 |
| ENSMUSG00000088402 | 0 | 0 | 0 | 0.25 | 0 | 0 | 0 |

| | | | | | | | |
|---|---|---|---|---|---|---|---|
| ENSMUSG00000088403 | 0 | 0 | 0 | 0 | 0 | 0 | 0 |
| ENSMUSG00000088404 | 0 | 0 | 0 | 0 | 0 | 0 | 0 |
| ENSMUSG00000088405 | 0 | 0 | 0 | 0 | 0 | 0 | 0 |
| ENSMUSG00000088406 | 0 | 0 | 0 | 0 | 0 | 0 | 0 |
| ENSMUSG00000088407 | 0 | 0 | 0 | 0 | 0 | 0 | 0 |
| ENSMUSG00000088408 | 0 | 0 | 0 | 0 | 0 | 0 | 0 |
| ENSMUSG00000088409 | 0 | 0 | 0 | 0 | 0 | 0 | 0 |
| ENSMUSG00000088410 | 0 | 0 | 0 | 0 | 0 | 0 | 0 |
| ENSMUSG00000088412 | 0 | 0 | 0 | 0.396240625 | 0 | 0 | 0.224243824 |
| ENSMUSG00000088414 | 0 | 0 | 0 | 0 | 0.244650542 | 0 | 0 |
| ENSMUSG00000088415 | 0 | 0 | 0 | 0 | 0 | 0 | 0 |
| ENSMUSG00000088416 | 0 | 0 | 0 | 0 | 0 | 0 | 0 |
| ENSMUSG00000088417 | 0 | 0 | 0 | 0 | 0 | 0 | 0 |
| ENSMUSG00000088418 | 0 | 0 | 0 | 0 | 0 | 0 | 0 |
| ENSMUSG00000088419 | 0 | 0 | 0 | 0 | 0 | 0 | 0 |
| ENSMUSG00000088420 | 0 | 0 | 0 | 0 | 0 | 0 | 0 |
| ENSMUSG00000088421 | 0 | 0 | 0 | 0 | 0 | 0 | 0 |
| ENSMUSG00000088422 | 0 | 0 | 0 | 0 | 0 | 0 | 0 |
| ENSMUSG00000088423 | 0 | 0 | 0 | 0 | 0 | 0 | 0 |
| ENSMUSG00000088425 | 0 | 0 | 0 | 0 | 0 | 0 | 0 |
| ENSMUSG00000088426 | 0 | 0 | 0 | 0 | 0 | 0 | 0 |
| ENSMUSG00000088428 | 0 | 0 | 0 | 0 | 0 | 0 | 0 |
| ENSMUSG00000088429 | 0 | 0 | 0 | 0 | 0 | 0 | 0 |
| ENSMUSG00000088432 | 0 | 0 | 0 | 0 | 0 | 0 | 0 |
| ENSMUSG00000088433 | 0 | 0 | 0 | 0 | 0 | 0 | 0 |
| ENSMUSG00000088435 | 0 | 0 | 0 | 0 | 0 | 0 | 0 |
| ENSMUSG00000088436 | 0 | 0 | 0 | 0 | 0 | 0 | 0 |
| ENSMUSG00000088437 | 0 | 0 | 0.25 | 0 | 0.244650542 | 0 | 0 |
| ENSMUSG00000088438 | 0 | 0 | 0 | 0 | 0 | 0 | 0 |
| ENSMUSG00000088439 | 0 | 0 | 0 | 0 | 0 | 0 | 0 |
| ENSMUSG00000088440 | 0 | 0 | 0 | 0 | 0 | 0 | 0 |
| ENSMUSG00000088441 | 0 | 0 | 0 | 0 | 0 | 0 | 0 |
| ENSMUSG00000088442 | 0 | 0 | 0 | 0 | 0 | 0 | 0 |
| ENSMUSG00000088443 | 0 | 0 | 0 | 0 | 0 | 0 | 0 |

| | | | | | | | |
|---|---|---|---|---|---|---|---|
| ENSMUSG00000088444 | 0 | 0 | 0 | 0 | 0 | 0 | 0 |
| ENSMUSG00000088445 | 0 | 0 | 0 | 0 | 0 | 0 | 0 |
| ENSMUSG00000088446 | 0 | 0 | 0 | 0 | 0 | 0 | 0 |
| ENSMUSG00000088447 | 0 | 0 | 0 | 0 | 0 | 0 | 0 |
| ENSMUSG00000088448 | 0 | 0 | 0 | 0 | 0 | 0 | 0 |
| ENSMUSG00000088449 | 0 | 0 | 0 | 0 | 0 | 0 | 0 |
| ENSMUSG00000088450 | 0 | 0 | 0 | 0 | 0 | 0 | 0 |
| ENSMUSG00000088451 | 0 | 0 | 0 | 0 | 0 | 0 | 0 |
| ENSMUSG00000088453 | 0 | 0 | 0 | 0.25 | 0 | 0 | 0 |
| ENSMUSG00000088454 | 0 | 0 | 0 | 0 | 0 | 0 | 0 |
| ENSMUSG00000088456 | 0 | 0 | 0 | 0 | 0 | 0 | 0 |
| ENSMUSG00000088458 | 0 | 0 | 0 | 0 | 0 | 0 | 0 |
| ENSMUSG00000088459 | 0 | 0 | 0 | 0 | 0 | 0 | 0.224243824 |
| ENSMUSG00000088460 | 0 | 0 | 0 | 0 | 0 | 0 | 0 |
| ENSMUSG00000088461 | 0 | 0 | 0 | 0 | 0 | 0 | 0 |
| ENSMUSG00000088463 | 0 | 0 | 0 | 0 | 0 | 0 | 0 |
| ENSMUSG00000088464 | 0 | 0.396240625 | 0 | 0.25 | 0.387761935 | 0 | 0.224243824 |
| ENSMUSG00000088465 | 0 | 0 | 0 | 0 | 0 | 0 | 0 |
| ENSMUSG00000088468 | 0 | 0 | 0 | 0 | 0 | 0 | 0 |
| ENSMUSG00000088469 | 0 | 0 | 0 | 0 | 0 | 0 | 0 |
| ENSMUSG00000088470 | 0 | 0 | 0 | 0 | 0 | 0 | 0 |
| ENSMUSG00000088471 | 0 | 0 | 0 | 0 | 0 | 0 | 0 |
| ENSMUSG00000088472 | 0 | 0 | 0 | 0 | 0 | 0 | 0 |
| ENSMUSG00000088473 | 0 | 0 | 0 | 0.5 | 0.387761935 | 0 | 0 |
| ENSMUSG00000088474 | 0 | 0 | 0 | 0 | 0 | 0 | 0 |
| ENSMUSG00000088475 | 0 | 0 | 0.5 | 0 | 0 | 0 | 0.224243824 |
| ENSMUSG00000088476 | 0 | 0 | 0 | 0 | 0 | 0 | 0 |
| ENSMUSG00000088477 | 0 | 0 | 0 | 0 | 0 | 0 | 0 |
| ENSMUSG00000088478 | 0 | 0 | 0 | 0 | 0 | 0 | 0 |
| ENSMUSG00000088479 | 0 | 0 | 0 | 0 | 0.244650542 | 0 | 0 |
| ENSMUSG00000088480 | 0 | 0 | 0 | 0 | 0 | 0 | 0 |
| ENSMUSG00000088481 | 0 | 0 | 0 | 0 | 0 | 0 | 0 |
| ENSMUSG00000088482 | 0 | 0 | 0 | 0 | 0 | 0 | 0 |
| ENSMUSG00000088483 | 0 | 0 | 0 | 0 | 0 | 0 | 0 |

| | | | | | | | |
|---|---|---|---|---|---|---|---|
| ENSMUSG00000088484 | 0 | 0 | 0 | 0 | 0 | 0 | 0 |
| ENSMUSG00000088485 | 0 | 0 | 0 | 0 | 0 | 0 | 0 |
| ENSMUSG00000088487 | 0 | 0 | 0 | 0 | 0 | 0 | 0 |
| ENSMUSG00000088490 | 0.289064826 | 0 | 0 | 0.396240625 | 0.244650542 | 0 | 0.224243824 |
| ENSMUSG00000088491 | 0 | 0 | 0 | 0 | 0 | 0 | 0 |
| ENSMUSG00000088493 | 0 | 0.25 | 0 | 0 | 0 | 0 | 0.224243824 |
| ENSMUSG00000088494 | 0 | 0.25 | 0 | 0 | 0.244650542 | 0 | 0 |
| ENSMUSG00000088495 | 0 | 0 | 0 | 0 | 0 | 0 | 0 |
| ENSMUSG00000088497 | 0 | 0 | 0 | 0 | 0 | 0 | 0 |
| ENSMUSG00000088498 | 0 | 0 | 0 | 0 | 0 | 0 | 0 |
| ENSMUSG00000088499 | 0 | 0 | 0 | 0 | 0 | 0 | 0 |
| ENSMUSG00000088500 | 0 | 0 | 0 | 0 | 0 | 0 | 0 |
| ENSMUSG00000088501 | 0 | 0 | 0 | 0 | 0 | 0 | 0 |
| ENSMUSG00000088502 | 0 | 0 | 0 | 0 | 0 | 0 | 0 |
| ENSMUSG00000088504 | 0 | 0 | 0 | 0 | 0.244650542 | 0 | 0.355418052 |
| ENSMUSG00000088505 | 0 | 0 | 0 | 0 | 0 | 0 | 0 |
| ENSMUSG00000088506 | 0 | 0 | 0 | 0 | 0 | 0 | 0 |
| ENSMUSG00000088509 | 0 | 0 | 0 | 0 | 0 | 0 | 0 |
| ENSMUSG00000088510 | 0 | 0 | 0 | 0 | 0 | 0 | 0 |
| ENSMUSG00000088511 | 0 | 0 | 0 | 0 | 0 | 0 | 0 |
| ENSMUSG00000088512 | 0 | 0 | 0 | 0 | 0 | 0 | 0 |
| ENSMUSG00000088514 | 0 | 0 | 0 | 0 | 0 | 0 | 0 |
| ENSMUSG00000088515 | 0 | 0 | 0 | 0 | 0 | 0 | 0 |
| ENSMUSG00000088516 | 0 | 0 | 0 | 0 | 0 | 0 | 0 |
| ENSMUSG00000088517 | 0 | 0 | 0 | 0 | 0 | 0 | 0 |
| ENSMUSG00000088518 | 0 | 0 | 0 | 0 | 0 | 0 | 0 |
| ENSMUSG00000088519 | 0 | 0 | 0 | 0 | 0 | 0 | 0 |
| ENSMUSG00000088521 | 0 | 0 | 0 | 0 | 0 | 0 | 0 |
| ENSMUSG00000088522 | 0 | 0 | 0 | 0 | 0 | 0 | 0 |
| ENSMUSG00000088523 | 0 | 0 | 0 | 0 | 0 | 0 | 0 |
| ENSMUSG00000088524 | 0 | 0 | 0 | 0 | 0 | 0 | 0 |
| ENSMUSG00000088525 | 0 | 0 | 0 | 0 | 0 | 0 | 0 |
| ENSMUSG00000088526 | 0 | 0 | 0 | 0 | 0 | 0 | 0 |
| ENSMUSG00000088527 | 0 | 0 | 0 | 0 | 0 | 0 | 0 |

| | | | | | | | |
|---|---|---|---|---|---|---|---|
| ENSMUSG00000088528 | 0 | 0 | 0 | 0 | 0 | 0 | 0 |
| ENSMUSG00000088529 | 0 | 0 | 0 | 0 | 0 | 0 | 0 |
| ENSMUSG00000088530 | 0 | 0 | 0 | 0 | 0 | 0 | 0 |
| ENSMUSG00000088532 | 0 | 0 | 0 | 0 | 0 | 0 | 0 |
| ENSMUSG00000088533 | 0 | 0 | 0 | 0 | 0 | 0 | 0 |
| ENSMUSG00000088535 | 0 | 0 | 0 | 0 | 0 | 0 | 0 |
| ENSMUSG00000088536 | 0 | 0 | 0 | 0 | 0 | 0 | 0 |
| ENSMUSG00000088537 | 0 | 0 | 0 | 0 | 0 | 0 | 0 |
| ENSMUSG00000088538 | 0 | 0 | 0 | 0 | 0 | 0 | 0 |
| ENSMUSG00000088539 | 0 | 0 | 0 | 0 | 0 | 0 | 0 |
| ENSMUSG00000088541 | 0 | 0 | 0 | 0 | 0 | 0 | 0 |
| ENSMUSG00000088542 | 0 | 0 | 0 | 0 | 0 | 0 | 0 |
| ENSMUSG00000088543 | 0 | 0 | 0 | 0 | 0 | 0 | 0 |
| ENSMUSG00000088544 | 0 | 0 | 0 | 0 | 0 | 0 | 0 |
| ENSMUSG00000088545 | 0 | 0 | 0 | 0 | 0 | 0 | 0 |
| ENSMUSG00000088546 | 0 | 0 | 0 | 0 | 0 | 0 | 0 |
| ENSMUSG00000088547 | 0 | 0 | 0 | 0 | 0 | 0 | 0 |
| ENSMUSG00000088548 | 0 | 0 | 0 | 0 | 0 | 0 | 0 |
| ENSMUSG00000088549 | 0 | 0 | 0 | 0 | 0 | 0 | 0 |
| ENSMUSG00000088550 | 0 | 0 | 0 | 0 | 0 | 0 | 0 |
| ENSMUSG00000088551 | 0 | 0 | 0 | 0 | 0 | 0 | 0 |
| ENSMUSG00000088552 | 0.289064826 | 0 | 0 | 0 | 0 | 0 | 0 |
| ENSMUSG00000088553 | 0 | 0 | 0 | 0 | 0 | 0 | 0 |
| ENSMUSG00000088554 | 0 | 0 | 0 | 0 | 0 | 0 | 0 |
| ENSMUSG00000088555 | 0 | 0 | 0 | 0 | 0.244650542 | 0 | 0 |
| ENSMUSG00000088558 | 0 | 0 | 0 | 0 | 0 | 0 | 0 |
| ENSMUSG00000088559 | 0 | 0 | 0 | 0 | 0 | 0 | 0 |
| ENSMUSG00000088562 | 0 | 0 | 0 | 0 | 0 | 0 | 0 |
| ENSMUSG00000088563 | 0 | 0 | 0 | 0 | 0 | 0 | 0 |
| ENSMUSG00000088564 | 0 | 0 | 0 | 0 | 0 | 0 | 0 |
| ENSMUSG00000088565 | 0 | 0 | 0 | 0 | 0 | 0 | 0 |
| ENSMUSG00000088566 | 0 | 0 | 0 | 0 | 0 | 0 | 0 |
| ENSMUSG00000088570 | 0 | 0 | 0 | 0 | 0 | 0 | 0 |
| ENSMUSG00000088571 | 0 | 0 | 0 | 0 | 0 | 0 | 0 |

| | | | | | | | |
|---|---|---|---|---|---|---|---|
| ENSMUSG00000088572 | 0 | 0 | 0 | 0 | 0 | 0 | 0 |
| ENSMUSG00000088573 | 0 | 0 | 0 | 0 | 0 | 0 | 0 |
| ENSMUSG00000088574 | 0 | 0 | 0 | 0 | 0 | 0 | 0 |
| ENSMUSG00000088575 | 0 | 0 | 0 | 0 | 0 | 0 | 0 |
| ENSMUSG00000088576 | 0 | 0 | 0 | 0 | 0 | 0 | 0 |
| ENSMUSG00000088577 | 0 | 0 | 0 | 0 | 0 | 0 | 0 |
| ENSMUSG00000088578 | 0 | 0 | 0 | 0 | 0 | 0 | 0 |
| ENSMUSG00000088579 | 0 | 0 | 0 | 0 | 0 | 0 | 0 |
| ENSMUSG00000088580 | 0 | 0 | 0 | 0 | 0 | 0 | 0 |
| ENSMUSG00000088581 | 0 | 0 | 0 | 0 | 0 | 0 | 0 |
| ENSMUSG00000088582 | 0 | 0 | 0 | 0 | 0 | 0 | 0 |
| ENSMUSG00000088583 | 0 | 0 | 0 | 0 | 0 | 0.301029996 | 0 |
| ENSMUSG00000088585 | 0 | 0 | 0 | 0 | 0 | 0 | 0 |
| ENSMUSG00000088587 | 0 | 0 | 0 | 0 | 0 | 0 | 0 |
| ENSMUSG00000088589 | 0 | 0 | 0 | 0 | 0 | 0 | 0 |
| ENSMUSG00000088591 | 0.747221736 | 0.396240625 | 0.25 | 0.701838731 | 0.568060967 | 0.477121255 | 0.448487648 |
| ENSMUSG00000088594 | 0 | 0 | 0 | 0 | 0 | 0 | 0 |
| ENSMUSG00000088595 | 0 | 0 | 0 | 0 | 0 | 0 | 0.224243824 |
| ENSMUSG00000088596 | 0 | 0 | 0 | 0 | 0 | 0 | 0 |
| ENSMUSG00000088597 | 0 | 0 | 0 | 0 | 0 | 0 | 0 |
| ENSMUSG00000088598 | 0 | 0 | 0 | 0 | 0 | 0 | 0 |
| ENSMUSG00000088600 | 0 | 0 | 0 | 0 | 0 | 0 | 0 |
| ENSMUSG00000088601 | 0 | 0 | 0 | 0 | 0 | 0 | 0 |
| ENSMUSG00000088602 | 0 | 0.396240625 | 0 | 0 | 0 | 0 | 0 |
| ENSMUSG00000088603 | 0 | 0 | 0 | 0 | 0 | 0 | 0 |
| ENSMUSG00000088605 | 0 | 0 | 0 | 0.396240625 | 0 | 0.301029996 | 0.224243824 |
| ENSMUSG00000088606 | 0 | 0 | 0 | 0 | 0 | 0 | 0 |
| ENSMUSG00000088608 | 0 | 0 | 0 | 0 | 0 | 0 | 0 |
| ENSMUSG00000088609 | 0 | 0 | 0 | 0 | 0 | 0 | 0 |
| ENSMUSG00000088611 | 0 | 0 | 0 | 0 | 0 | 0 | 0 |
| ENSMUSG00000088614 | 0 | 0.25 | 0 | 0 | 0 | 0 | 0 |
| ENSMUSG00000088615 | 0 | 0.25 | 0.25 | 0 | 0.244650542 | 0 | 0.355418052 |
| ENSMUSG00000088616 | 0 | 0 | 0 | 0 | 0 | 0 | 0 |
| ENSMUSG00000088617 | 0 | 0 | 0 | 0 | 0 | 0 | 0 |

| | | | | | | | |
|---|---|---|---|---|---|---|---|
| ENSMUSG00000088618 | 0 | 0 | 0 | 0 | 0 | 0 | 0 |
| ENSMUSG00000088620 | 0 | 0 | 0 | 0 | 0 | 0 | 0 |
| ENSMUSG00000088621 | 0 | 0 | 0 | 0 | 0 | 0 | 0 |
| ENSMUSG00000088622 | 0 | 0 | 0 | 0 | 0 | 0 | 0 |
| ENSMUSG00000088623 | 0 | 0 | 0 | 0 | 0 | 0 | 0.224243824 |
| ENSMUSG00000088624 | 0 | 0 | 0 | 0 | 0 | 0 | 0 |
| ENSMUSG00000088625 | 0.289064826 | 0 | 0 | 0.396240625 | 0 | 0 | 0.224243824 |
| ENSMUSG00000088626 | 0 | 0 | 0 | 0 | 0 | 0 | 0.224243824 |
| ENSMUSG00000088627 | 0 | 0 | 0 | 0 | 0 | 0 | 0 |
| ENSMUSG00000088628 | 0 | 0 | 0 | 0 | 0 | 0 | 0 |
| ENSMUSG00000088629 | 0 | 0 | 0 | 0 | 0 | 0 | 0 |
| ENSMUSG00000088630 | 0 | 0 | 0 | 0 | 0 | 0 | 0 |
| ENSMUSG00000088631 | 0 | 0 | 0 | 0 | 0 | 0 | 0 |
| ENSMUSG00000088632 | 0 | 0.5 | 0.25 | 0 | 0.489301084 | 0 | 0.448487648 |
| ENSMUSG00000088633 | 0 | 0 | 0 | 0 | 0 | 0 | 0 |
| ENSMUSG00000088634 | 0 | 0 | 0 | 0 | 0 | 0 | 0 |
| ENSMUSG00000088635 | 0 | 0 | 0 | 0 | 0 | 0 | 0 |
| ENSMUSG00000088637 | 0 | 0 | 0 | 0 | 0 | 0 | 0 |
| ENSMUSG00000088638 | 0 | 0 | 0 | 0 | 0 | 0 | 0 |
| ENSMUSG00000088639 | 0 | 0 | 0 | 0 | 0.387761935 | 0 | 0 |
| ENSMUSG00000088640 | 0 | 0 | 0 | 0 | 0 | 0 | 0 |
| ENSMUSG00000088641 | 0 | 0 | 0 | 0 | 0 | 0 | 0 |
| ENSMUSG00000088643 | 0 | 0 | 0 | 0.25 | 0 | 0 | 0.224243824 |
| ENSMUSG00000088644 | 0 | 0 | 0 | 0 | 0 | 0 | 0 |
| ENSMUSG00000088646 | 0 | 0 | 0 | 0 | 0 | 0 | 0 |
| ENSMUSG00000088647 | 0 | 0 | 0 | 0 | 0 | 0 | 0 |
| ENSMUSG00000088649 | 0 | 0 | 0 | 0 | 0 | 0 | 0 |
| ENSMUSG00000088650 | 0 | 0 | 0 | 0 | 0.244650542 | 0 | 0.224243824 |
| ENSMUSG00000088651 | 0 | 0 | 0 | 0 | 0 | 0 | 0 |
| ENSMUSG00000088653 | 0 | 0 | 0 | 0 | 0 | 0 | 0 |
| ENSMUSG00000088654 | 0 | 0 | 0 | 0 | 0 | 0 | 0 |
| ENSMUSG00000088656 | 0 | 0 | 0 | 0 | 0 | 0 | 0 |
| ENSMUSG00000088657 | 0 | 0 | 0 | 0 | 0 | 0 | 0 |
| ENSMUSG00000088658 | 0 | 0 | 0 | 0 | 0 | 0 | 0 |

| | | | | | | | |
|---|---|---|---|---|---|---|---|
| ENSMUSG00000088659 | 0 | 0 | 0 | 0 | 0 | 0 | 0 |
| ENSMUSG00000088660 | 0 | 0 | 0 | 0 | 0 | 0 | 0 |
| ENSMUSG00000088661 | 0 | 0 | 0 | 0 | 0 | 0 | 0 |
| ENSMUSG00000088663 | 0 | 0 | 0 | 0 | 0 | 0 | 0 |
| ENSMUSG00000088666 | 0 | 0 | 0 | 0 | 0 | 0 | 0 |
| ENSMUSG00000088668 | 0 | 0 | 0 | 0 | 0 | 0 | 0 |
| ENSMUSG00000088669 | 0 | 0 | 0 | 0 | 0 | 0 | 0 |
| ENSMUSG00000088670 | 0 | 0 | 0 | 0 | 0 | 0 | 0 |
| ENSMUSG00000088671 | 0 | 0 | 0 | 0 | 0 | 0 | 0 |
| ENSMUSG00000088672 | 0 | 0 | 0 | 0 | 0 | 0 | 0 |
| ENSMUSG00000088674 | 0 | 0 | 0 | 0 | 0 | 0 | 0 |
| ENSMUSG00000088675 | 0.289064826 | 0.25 | 0 | 0 | 0 | 0 | 0 |
| ENSMUSG00000088677 | 0 | 0 | 0 | 0 | 0 | 0 | 0 |
| ENSMUSG00000088678 | 0 | 0 | 0 | 0 | 0 | 0 | 0 |
| ENSMUSG00000088679 | 0 | 0 | 0 | 0 | 0 | 0 | 0 |
| ENSMUSG00000088680 | 0 | 0 | 0 | 0 | 0 | 0 | 0 |
| ENSMUSG00000088682 | 0 | 0 | 0 | 0 | 0 | 0 | 0 |
| ENSMUSG00000088683 | 0 | 0 | 0 | 0 | 0 | 0 | 0 |
| ENSMUSG00000088684 | 0 | 0 | 0 | 0 | 0 | 0 | 0 |
| ENSMUSG00000088685 | 0 | 0 | 0 | 0 | 0 | 0 | 0 |
| ENSMUSG00000088686 | 0 | 0 | 0 | 0 | 0 | 0 | 0 |
| ENSMUSG00000088687 | 0 | 0 | 0 | 0 | 0 | 0 | 0 |
| ENSMUSG00000088688 | 0 | 0 | 0 | 0 | 0 | 0 | 0 |
| ENSMUSG00000088689 | 0 | 0 | 0.25 | 0 | 0 | 0 | 0 |
| ENSMUSG00000088691 | 0 | 0 | 0 | 0 | 0 | 0 | 0 |
| ENSMUSG00000088692 | 0 | 0 | 0 | 0 | 0 | 0 | 0 |
| ENSMUSG00000088693 | 0 | 0 | 0 | 0 | 0 | 0 | 0 |
| ENSMUSG00000088695 | 0 | 0 | 0.396240625 | 0 | 0 | 0 | 0 |
| ENSMUSG00000088696 | 0 | 0 | 0 | 0 | 0 | 0 | 0 |
| ENSMUSG00000088697 | 0 | 0 | 0 | 0 | 0 | 0 | 0 |
| ENSMUSG00000088699 | 0 | 0 | 0 | 0 | 0 | 0 | 0 |
| ENSMUSG00000088700 | 0 | 0 | 0 | 0 | 0 | 0 | 0 |
| ENSMUSG00000088701 | 0 | 0 | 0 | 0 | 0 | 0 | 0 |
| ENSMUSG00000088702 | 0 | 0 | 0 | 0 | 0 | 0 | 0 |

| | | | | | | | |
|---|---|---|---|---|---|---|---|
| ENSMUSG00000088703 | 0 | 0 | 0 | 0 | 0 | 0 | 0 |
| ENSMUSG00000088704 | 0 | 0 | 0 | 0 | 0 | 0 | 0 |
| ENSMUSG00000088705 | 0 | 0 | 0.25 | 0 | 0.244650542 | 0 | 0.224243824 |
| ENSMUSG00000088706 | 0 | 0 | 0 | 0 | 0 | 0 | 0 |
| ENSMUSG00000088707 | 0 | 0 | 0 | 0 | 0 | 0 | 0 |
| ENSMUSG00000088708 | 0 | 0 | 0 | 0 | 0.244650542 | 0 | 0 |
| ENSMUSG00000088709 | 0 | 0 | 0 | 0 | 0 | 0 | 0 |
| ENSMUSG00000088710 | 0 | 0 | 0 | 0 | 0 | 0 | 0 |
| ENSMUSG00000088711 | 0 | 0 | 0 | 0 | 0 | 0 | 0 |
| ENSMUSG00000088712 | 0.45815691 | 0 | 0 | 0 | 0.244650542 | 0 | 0.355418052 |
| ENSMUSG00000088713 | 0 | 0 | 0 | 0 | 0 | 0 | 0 |
| ENSMUSG00000088714 | 0 | 0 | 0 | 0 | 0 | 0 | 0 |
| ENSMUSG00000088715 | 0 | 0 | 0 | 0 | 0 | 0 | 0 |
| ENSMUSG00000088716 | 0 | 0 | 0 | 0 | 0 | 0 | 0 |
| ENSMUSG00000088717 | 0 | 0 | 0 | 0 | 0 | 0 | 0 |
| ENSMUSG00000088718 | 0 | 0 | 0 | 0 | 0 | 0 | 0 |
| ENSMUSG00000088719 | 0 | 0 | 0 | 0 | 0 | 0 | 0 |
| ENSMUSG00000088721 | 0 | 0 | 0 | 0 | 0 | 0 | 0 |
| ENSMUSG00000088722 | 0 | 0 | 0 | 0 | 0 | 0 | 0 |
| ENSMUSG00000088726 | 0 | 0 | 0 | 0 | 0 | 0 | 0 |
| ENSMUSG00000088730 | 0 | 0 | 0 | 0 | 0 | 0 | 0 |
| ENSMUSG00000088731 | 0 | 0 | 0 | 0 | 0 | 0 | 0 |
| ENSMUSG00000088732 | 0 | 0 | 0 | 0 | 0 | 0 | 0 |
| ENSMUSG00000088733 | 0 | 0 | 0 | 0 | 0 | 0 | 0 |
| ENSMUSG00000088734 | 0 | 0 | 0 | 0 | 0 | 0 | 0 |
| ENSMUSG00000088735 | 0 | 0 | 0 | 0 | 0 | 0 | 0 |
| ENSMUSG00000088736 | 0 | 0 | 0 | 0 | 0 | 0 | 0 |
| ENSMUSG00000088737 | 0 | 0 | 0 | 0 | 0 | 0 | 0 |
| ENSMUSG00000088738 | 0 | 0 | 0 | 0 | 0 | 0 | 0 |
| ENSMUSG00000088739 | 0 | 0 | 0 | 0 | 0 | 0 | 0 |
| ENSMUSG00000088740 | 0 | 0 | 0 | 0 | 0 | 0 | 0 |
| ENSMUSG00000088742 | 0 | 0 | 0 | 0 | 0 | 0 | 0 |
| ENSMUSG00000088743 | 0 | 0 | 0 | 0 | 0 | 0 | 0 |
| ENSMUSG00000088744 | 0 | 0 | 0 | 0 | 0 | 0 | 0 |

| | | | | | | | |
|---|---|---|---|---|---|---|---|
| ENSMUSG00000088746 | 0 | 0 | 0 | 0 | 0 | 0 | 0 |
| ENSMUSG00000088747 | 0 | 0 | 0 | 0 | 0 | 0 | 0 |
| ENSMUSG00000088748 | 0 | 0 | 0 | 0 | 0 | 0 | 0 |
| ENSMUSG00000088749 | 0 | 0 | 0 | 0 | 0 | 0 | 0 |
| ENSMUSG00000088750 | 0 | 0 | 0 | 0 | 0 | 0 | 0 |
| ENSMUSG00000088751 | 0 | 0 | 0 | 0 | 0 | 0 | 0 |
| ENSMUSG00000088752 | 0 | 0 | 0 | 0 | 0 | 0 | 0 |
| ENSMUSG00000088753 | 0 | 0 | 0 | 0 | 0 | 0 | 0 |
| ENSMUSG00000088754 | 0 | 0 | 0 | 0 | 0 | 0 | 0 |
| ENSMUSG00000088756 | 0 | 0 | 0 | 0 | 0 | 0 | 0 |
| ENSMUSG00000088758 | 0 | 0 | 0 | 0 | 0 | 0 | 0 |
| ENSMUSG00000088760 | 0 | 0 | 0 | 0 | 0 | 0 | 0 |
| ENSMUSG00000088761 | 0 | 0 | 0 | 0 | 0 | 0 | 0 |
| ENSMUSG00000088763 | 0 | 0 | 0 | 0 | 0 | 0 | 0.224243824 |
| ENSMUSG00000088764 | 0 | 0 | 0 | 0 | 0 | 0 | 0 |
| ENSMUSG00000088766 | 0 | 0 | 0 | 0 | 0 | 0 | 0 |
| ENSMUSG00000088767 | 0 | 0 | 0 | 0 | 0 | 0 | 0 |
| ENSMUSG00000088768 | 0 | 0 | 0 | 0 | 0 | 0 | 0 |
| ENSMUSG00000088769 | 0.289064826 | 0.75 | 0 | 0.396240625 | 0.387761935 | 0.477121255 | 0 |
| ENSMUSG00000088772 | 0 | 0 | 0 | 0 | 0 | 0 | 0 |
| ENSMUSG00000088773 | 0 | 0 | 0 | 0 | 0 | 0 | 0 |
| ENSMUSG00000088774 | 0 | 0 | 0 | 0 | 0 | 0 | 0 |
| ENSMUSG00000088775 | 0 | 0 | 0 | 0 | 0 | 0 | 0 |
| ENSMUSG00000088776 | 0 | 0 | 0 | 0 | 0 | 0 | 0 |
| ENSMUSG00000088777 | 0 | 0.25 | 0 | 0 | 0.244650542 | 0 | 0.448487648 |
| ENSMUSG00000088779 | 0 | 0 | 0 | 0 | 0 | 0 | 0 |
| ENSMUSG00000088781 | 0 | 0 | 0 | 0 | 0 | 0 | 0 |
| ENSMUSG00000088782 | 0 | 0 | 0 | 0 | 0 | 0 | 0 |
| ENSMUSG00000088783 | 0 | 0 | 0 | 0 | 0 | 0 | 0 |
| ENSMUSG00000088784 | 0 | 0 | 0 | 0 | 0 | 0 | 0 |
| ENSMUSG00000088786 | 0 | 0 | 0 | 0 | 0 | 0 | 0 |
| ENSMUSG00000088787 | 0 | 0 | 0 | 0 | 0 | 0 | 0 |
| ENSMUSG00000088789 | 0 | 0.5 | 0.396240625 | 0 | 0.244650542 | 0 | 0.448487648 |
| ENSMUSG00000088790 | 0 | 0 | 0 | 0 | 0 | 0 | 0 |

| | | | | | | | |
|---|---|---|---|---|---|---|---|
| ENSMUSG00000088791 | 0 | 0 | 0 | 0 | 0 | 0 | 0 |
| ENSMUSG00000088792 | 0 | 0 | 0 | 0 | 0 | 0 | 0 |
| ENSMUSG00000088793 | 0 | 0.25 | 0 | 0 | 0 | 0 | 0 |
| ENSMUSG00000088794 | 0 | 0 | 0 | 0 | 0 | 0 | 0 |
| ENSMUSG00000088796 | 0 | 0 | 0 | 0 | 0 | 0 | 0 |
| ENSMUSG00000088797 | 0 | 0 | 0 | 0 | 0 | 0 | 0 |
| ENSMUSG00000088798 | 0 | 0 | 0 | 0 | 0 | 0 | 0 |
| ENSMUSG00000088799 | 0 | 0 | 0 | 0 | 0 | 0 | 0 |
| ENSMUSG00000088800 | 0 | 0 | 0 | 0 | 0 | 0 | 0 |
| ENSMUSG00000088802 | 0 | 0 | 0 | 0 | 0 | 0 | 0 |
| ENSMUSG00000088803 | 0 | 0 | 0 | 0 | 0 | 0 | 0 |
| ENSMUSG00000088804 | 0 | 0 | 0 | 0 | 0 | 0 | 0 |
| ENSMUSG00000088808 | 0 | 0 | 0 | 0 | 0 | 0 | 0 |
| ENSMUSG00000088809 | 0 | 0 | 0 | 0 | 0 | 0 | 0 |
| ENSMUSG00000088810 | 0 | 0 | 0 | 0 | 0 | 0 | 0 |
| ENSMUSG00000088811 | 0 | 0 | 0 | 0 | 0 | 0 | 0 |
| ENSMUSG00000088812 | 0 | 0 | 0 | 0 | 0 | 0 | 0 |
| ENSMUSG00000088813 | 0 | 0 | 0 | 0 | 0 | 0 | 0 |
| ENSMUSG00000088814 | 0 | 0 | 0 | 0 | 0 | 0 | 0 |
| ENSMUSG00000088816 | 0 | 0 | 0 | 0 | 0 | 0 | 0 |
| ENSMUSG00000088817 | 0 | 0 | 0 | 0 | 0 | 0 | 0 |
| ENSMUSG00000088818 | 0 | 0 | 0 | 0 | 0 | 0 | 0 |
| ENSMUSG00000088819 | 0 | 0 | 0 | 0.25 | 0 | 0 | 0 |
| ENSMUSG00000088820 | 0 | 0 | 0 | 0 | 0 | 0 | 0 |
| ENSMUSG00000088822 | 0 | 0 | 0 | 0 | 0 | 0 | 0 |
| ENSMUSG00000088823 | 0 | 0 | 0 | 0 | 0 | 0 | 0 |
| ENSMUSG00000088824 | 0 | 0 | 0 | 0 | 0 | 0 | 0 |
| ENSMUSG00000088825 | 0 | 0 | 0 | 0 | 0 | 0 | 0 |
| ENSMUSG00000088829 | 0 | 0 | 0 | 0 | 0 | 0 | 0 |
| ENSMUSG00000088830 | 0 | 0 | 0 | 0 | 0 | 0 | 0 |
| ENSMUSG00000088832 | 0 | 0.25 | 0 | 0 | 0.244650542 | 0 | 0.355418052 |
| ENSMUSG00000088833 | 0 | 0 | 0 | 0 | 0 | 0 | 0 |
| ENSMUSG00000088835 | 0.289064826 | 0.25 | 0.396240625 | 0.5 | 0.632412477 | 0.477121255 | 0.448487648 |
| ENSMUSG00000088836 | 0 | 0 | 0 | 0 | 0 | 0 | 0 |

| | | | | | | | |
|---|---|---|---|---|---|---|---|
| ENSMUSG00000088839 | 0 | 0 | 0 | 0 | 0 | 0 | 0 |
| ENSMUSG00000088840 | 0 | 0 | 0 | 0 | 0 | 0 | 0 |
| ENSMUSG00000088842 | 0 | 0 | 0 | 0 | 0 | 0 | 0 |
| ENSMUSG00000088843 | 0 | 0 | 0 | 0 | 0 | 0 | 0 |
| ENSMUSG00000088845 | 0 | 0 | 0 | 0 | 0 | 0 | 0 |
| ENSMUSG00000088846 | 0 | 0 | 0 | 0 | 0 | 0 | 0 |
| ENSMUSG00000088848 | 0 | 0 | 0 | 0 | 0 | 0 | 0 |
| ENSMUSG00000088849 | 0 | 0 | 0 | 0 | 0 | 0 | 0 |
| ENSMUSG00000088850 | 0 | 0 | 0 | 0 | 0 | 0 | 0 |
| ENSMUSG00000088851 | 0 | 0 | 0 | 0 | 0 | 0 | 0 |
| ENSMUSG00000088852 | 0 | 0 | 0 | 0 | 0 | 0 | 0 |
| ENSMUSG00000088853 | 0 | 0 | 0 | 0 | 0 | 0 | 0 |
| ENSMUSG00000088854 | 0 | 0 | 0 | 0 | 0 | 0 | 0 |
| ENSMUSG00000088855 | 0 | 0 | 0 | 0 | 0 | 0 | 0 |
| ENSMUSG00000088856 | 0.289064826 | 0 | 0 | 0.396240625 | 0 | 0 | 0 |
| ENSMUSG00000088857 | 0 | 0 | 0 | 0 | 0 | 0 | 0 |
| ENSMUSG00000088858 | 0 | 0 | 0 | 0 | 0 | 0 | 0 |
| ENSMUSG00000088860 | 0 | 0 | 0 | 0 | 0 | 0 | 0 |
| ENSMUSG00000088861 | 0 | 0 | 0 | 0 | 0 | 0 | 0 |
| ENSMUSG00000088862 | 0 | 0 | 0 | 0 | 0 | 0 | 0 |
| ENSMUSG00000088863 | 0 | 0 | 0 | 0 | 0 | 0 | 0 |
| ENSMUSG00000088864 | 0 | 0 | 0 | 0 | 0 | 0 | 0 |
| ENSMUSG00000088865 | 0 | 0 | 0 | 0 | 0 | 0 | 0 |
| ENSMUSG00000088867 | 0 | 0 | 0 | 0 | 0 | 0 | 0 |
| ENSMUSG00000088868 | 0 | 0 | 0 | 0 | 0 | 0 | 0 |
| ENSMUSG00000088870 | 0 | 0 | 0 | 0 | 0 | 0 | 0 |
| ENSMUSG00000088872 | 0 | 0 | 0 | 0 | 0 | 0 | 0 |
| ENSMUSG00000088873 | 0 | 0 | 0 | 0 | 0 | 0 | 0 |
| ENSMUSG00000088874 | 0 | 0 | 0 | 0 | 0 | 0 | 0 |
| ENSMUSG00000088875 | 0 | 0 | 0 | 0 | 0 | 0 | 0 |
| ENSMUSG00000088876 | 0 | 0 | 0 | 0 | 0 | 0 | 0 |
| ENSMUSG00000088877 | 0 | 0 | 0 | 0 | 0 | 0 | 0 |
| ENSMUSG00000088878 | 0 | 0 | 0 | 0 | 0 | 0 | 0 |
| ENSMUSG00000088879 | 0 | 0 | 0 | 0 | 0 | 0 | 0 |

| | | | | | | | |
|---|---|---|---|---|---|---|---|
| ENSMUSG00000088880 | 0 | 0 | 0 | 0 | 0 | 0 | 0 |
| ENSMUSG00000088883 | 0 | 0 | 0 | 0 | 0 | 0 | 0 |
| ENSMUSG00000088884 | 0 | 0 | 0 | 0 | 0 | 0 | 0 |
| ENSMUSG00000088885 | 0 | 0 | 0 | 0 | 0 | 0 | 0 |
| ENSMUSG00000088888 | 0 | 0 | 0 | 0 | 0 | 0 | 0 |
| ENSMUSG00000088889 | 0 | 0 | 0 | 0 | 0 | 0 | 0 |
| ENSMUSG00000088890 | 0 | 0 | 0 | 0 | 0 | 0 | 0 |
| ENSMUSG00000088891 | 0 | 0.25 | 0 | 0.25 | 0 | 0 | 0.224243824 |
| ENSMUSG00000088892 | 0 | 0 | 0 | 0 | 0 | 0 | 0 |
| ENSMUSG00000088894 | 0 | 0 | 0 | 0 | 0 | 0 | 0 |
| ENSMUSG00000088895 | 0 | 0 | 0 | 0 | 0 | 0 | 0 |
| ENSMUSG00000088896 | 0 | 0 | 0 | 0 | 0 | 0 | 0 |
| ENSMUSG00000088897 | 0 | 0 | 0 | 0 | 0 | 0 | 0 |
| ENSMUSG00000088898 | 0 | 0 | 0 | 0 | 0 | 0 | 0 |
| ENSMUSG00000088900 | 0 | 0 | 0 | 0 | 0 | 0 | 0 |
| ENSMUSG00000088901 | 0 | 0 | 0 | 0.25 | 0 | 0.301029996 | 0 |
| ENSMUSG00000088902 | 0 | 0 | 0 | 0 | 0 | 0 | 0 |
| ENSMUSG00000088903 | 0 | 0 | 0 | 0 | 0 | 0 | 0 |
| ENSMUSG00000088904 | 0 | 0 | 0 | 0 | 0 | 0 | 0 |
| ENSMUSG00000088905 | 0 | 0 | 0 | 0 | 0 | 0 | 0 |
| ENSMUSG00000088906 | 0 | 0 | 0 | 0 | 0 | 0 | 0 |
| ENSMUSG00000088907 | 0 | 0 | 0 | 0 | 0 | 0 | 0 |
| ENSMUSG00000088908 | 0 | 0 | 0 | 0 | 0 | 0 | 0 |
| ENSMUSG00000088909 | 0 | 0 | 0 | 0 | 0 | 0 | 0 |
| ENSMUSG00000088910 | 0 | 0 | 0 | 0 | 0 | 0 | 0 |
| ENSMUSG00000088911 | 0 | 0 | 0 | 0 | 0 | 0 | 0 |
| ENSMUSG00000088913 | 0 | 0 | 0.25 | 0 | 0 | 0 | 0 |
| ENSMUSG00000088914 | 0 | 0 | 0 | 0 | 0 | 0 | 0 |
| ENSMUSG00000088915 | 0 | 0 | 0 | 0 | 0 | 0 | 0 |
| ENSMUSG00000088916 | 0 | 0 | 0 | 0 | 0 | 0 | 0 |
| ENSMUSG00000088918 | 0 | 0 | 0 | 0 | 0 | 0 | 0 |
| ENSMUSG00000088920 | 0 | 0 | 0 | 0 | 0 | 0 | 0 |
| ENSMUSG00000088922 | 0 | 0 | 0 | 0 | 0 | 0 | 0 |
| ENSMUSG00000088923 | 0 | 0 | 0 | 0 | 0 | 0 | 0 |

| | | | | | | | |
|---|---|---|---|---|---|---|---|
| ENSMUSG00000088924 | 0 | 0 | 0 | 0 | 0 | 0 | 0 |
| ENSMUSG00000088925 | 0 | 0 | 0 | 0 | 0 | 0 | 0 |
| ENSMUSG00000088926 | 0 | 0 | 0 | 0 | 0 | 0 | 0 |
| ENSMUSG00000088928 | 0 | 0 | 0 | 0 | 0 | 0 | 0 |
| ENSMUSG00000088929 | 0.671187741 | 0.896240625 | 0.896240625 | 0.92510993 | 1.106691885 | 1.113943352 | 0.952571511 |
| ENSMUSG00000088930 | 0 | 0 | 0 | 0 | 0 | 0 | 0 |
| ENSMUSG00000088932 | 0 | 0 | 0 | 0 | 0 | 0 | 0 |
| ENSMUSG00000088933 | 0 | 0 | 0 | 0 | 0 | 0 | 0 |
| ENSMUSG00000088935 | 0 | 0 | 0 | 0 | 0 | 0 | 0 |
| ENSMUSG00000088937 | 0 | 0 | 0 | 0 | 0 | 0 | 0 |
| ENSMUSG00000088938 | 0 | 0 | 0 | 0 | 0 | 0 | 0 |
| ENSMUSG00000088939 | 0 | 0 | 0 | 0 | 0 | 0 | 0 |
| ENSMUSG00000088940 | 0 | 0 | 0 | 0 | 0 | 0 | 0 |
| ENSMUSG00000088941 | 0 | 0 | 0 | 0 | 0 | 0 | 0 |
| ENSMUSG00000088942 | 0 | 0 | 0 | 0 | 0 | 0 | 0 |
| ENSMUSG00000088943 | 0 | 0 | 0 | 0 | 0 | 0 | 0 |
| ENSMUSG00000088944 | 0 | 0 | 0 | 0 | 0 | 0 | 0 |
| ENSMUSG00000088947 | 0 | 0 | 0 | 0 | 0 | 0 | 0 |
| ENSMUSG00000088948 | 0.289064826 | 0 | 0.25 | 0 | 0.244650542 | 0 | 0 |
| ENSMUSG00000088949 | 0 | 0 | 0 | 0 | 0 | 0 | 0 |
| ENSMUSG00000088950 | 0 | 0 | 0 | 0 | 0 | 0 | 0 |
| ENSMUSG00000088951 | 0 | 0 | 0 | 0 | 0 | 0 | 0.224243824 |
| ENSMUSG00000088953 | 0 | 0 | 0 | 0 | 0 | 0 | 0 |
| ENSMUSG00000088954 | 0 | 0 | 0 | 0 | 0 | 0 | 0 |
| ENSMUSG00000088955 | 0 | 0 | 0 | 0 | 0 | 0 | 0 |
| ENSMUSG00000088956 | 0 | 0 | 0 | 0 | 0 | 0 | 0 |
| ENSMUSG00000088957 | 0 | 0 | 0 | 0 | 0 | 0 | 0 |
| ENSMUSG00000088958 | 0 | 0 | 0.25 | 0 | 0 | 0.301029996 | 0 |
| ENSMUSG00000088959 | 0 | 0 | 0 | 0 | 0 | 0 | 0 |
| ENSMUSG00000088960 | 0 | 0 | 0 | 0 | 0 | 0 | 0 |
| ENSMUSG00000088961 | 0 | 0.5 | 0 | 0.25 | 0 | 0 | 0 |
| ENSMUSG00000088964 | 0 | 0 | 0 | 0 | 0 | 0 | 0 |
| ENSMUSG00000088966 | 0 | 0 | 0 | 0 | 0 | 0 | 0 |
| ENSMUSG00000088969 | 0.289064826 | 0 | 0.25 | 0 | 0 | 0 | 0 |

| | | | | | | | |
|---|---|---|---|---|---|---|---|
| ENSMUSG00000088970 | 0 | 0 | 0 | 0 | 0 | 0 | 0 |
| ENSMUSG00000088971 | 0 | 0 | 0 | 0 | 0 | 0 | 0 |
| ENSMUSG00000088972 | 0 | 0 | 0 | 0 | 0 | 0 | 0 |
| ENSMUSG00000088973 | 0 | 0 | 0 | 0 | 0 | 0 | 0 |
| ENSMUSG00000088975 | 0 | 0 | 0 | 0 | 0 | 0 | 0 |
| ENSMUSG00000088976 | 0 | 0 | 0 | 0 | 0 | 0 | 0 |
| ENSMUSG00000088978 | 0 | 0 | 0 | 0 | 0 | 0 | 0 |
| ENSMUSG00000088979 | 0 | 0 | 0 | 0 | 0 | 0 | 0 |
| ENSMUSG00000088980 | 0 | 0 | 0 | 0 | 0 | 0 | 0 |
| ENSMUSG00000088981 | 0 | 0 | 0 | 0 | 0 | 0 | 0 |
| ENSMUSG00000088982 | 0 | 0 | 0 | 0 | 0.387761935 | 0 | 0.224243824 |
| ENSMUSG00000088983 | 0 | 0 | 0 | 0 | 0 | 0 | 0 |
| ENSMUSG00000088984 | 0.289064826 | 0 | 0.25 | 0.25 | 0.244650542 | 0 | 0.355418052 |
| ENSMUSG00000088985 | 0 | 0 | 0.25 | 0 | 0 | 0 | 0 |
| ENSMUSG00000088986 | 0 | 0 | 0 | 0 | 0 | 0 | 0 |
| ENSMUSG00000088987 | 0 | 0 | 0 | 0 | 0 | 0 | 0 |
| ENSMUSG00000088988 | 0 | 0 | 0 | 0 | 0 | 0 | 0 |
| ENSMUSG00000088989 | 0 | 0 | 0 | 0 | 0 | 0 | 0 |
| ENSMUSG00000088990 | 0.289064826 | 0 | 0 | 0.25 | 0.244650542 | 0 | 0.224243824 |
| ENSMUSG00000088991 | 0 | 0 | 0 | 0 | 0 | 0 | 0 |
| ENSMUSG00000088992 | 0 | 0 | 0 | 0 | 0 | 0 | 0 |
| ENSMUSG00000088993 | 0 | 0 | 0 | 0.580482024 | 0.244650542 | 0 | 0.224243824 |
| ENSMUSG00000088994 | 0 | 0 | 0 | 0 | 0 | 0 | 0 |
| ENSMUSG00000088995 | 0 | 0 | 0 | 0 | 0 | 0 | 0 |
| ENSMUSG00000088996 | 0 | 0 | 0 | 0 | 0 | 0 | 0 |
| ENSMUSG00000088997 | 0 | 0 | 0 | 0 | 0 | 0 | 0 |
| ENSMUSG00000088999 | 0 | 0 | 0 | 0 | 0 | 0 | 0 |
| ENSMUSG00000089000 | 0 | 0 | 0 | 0 | 0 | 0 | 0 |
| ENSMUSG00000089001 | 0 | 0 | 0 | 0 | 0 | 0 | 0 |
| ENSMUSG00000089002 | 0 | 0 | 0 | 0 | 0 | 0 | 0 |
| ENSMUSG00000089003 | 0 | 0 | 0 | 0 | 0 | 0 | 0 |
| ENSMUSG00000089004 | 0 | 0 | 0 | 0 | 0 | 0 | 0 |
| ENSMUSG00000089005 | 0 | 0 | 0 | 0 | 0 | 0 | 0 |
| ENSMUSG00000089007 | 0 | 0 | 0 | 0 | 0 | 0 | 0 |

| | | | | | | | |
|---|---|---|---|---|---|---|---|
| ENSMUSG00000089008 | 0 | 0 | 0 | 0 | 0 | 0 | 0 |
| ENSMUSG00000089009 | 0 | 0 | 0 | 0 | 0 | 0 | 0 |
| ENSMUSG00000089011 | 0 | 0 | 0 | 0 | 0 | 0 | 0 |
| ENSMUSG00000089014 | 0 | 0 | 0 | 0.25 | 0 | 0 | 0 |
| ENSMUSG00000089015 | 0 | 0 | 0 | 0 | 0 | 0 | 0 |
| ENSMUSG00000089016 | 0 | 0 | 0 | 0 | 0 | 0 | 0 |
| ENSMUSG00000089017 | 0 | 0 | 0 | 0 | 0 | 0 | 0 |
| ENSMUSG00000089018 | 0 | 0 | 0 | 0 | 0 | 0 | 0 |
| ENSMUSG00000089019 | 0 | 0 | 0 | 0 | 0 | 0 | 0 |
| ENSMUSG00000089021 | 0 | 0 | 0 | 0 | 0 | 0 | 0 |
| ENSMUSG00000089023 | 0 | 0 | 0 | 0 | 0 | 0 | 0 |
| ENSMUSG00000089024 | 0 | 0 | 0 | 0 | 0 | 0 | 0 |
| ENSMUSG00000089025 | 0 | 0 | 0 | 0 | 0 | 0 | 0 |
| ENSMUSG00000089026 | 0 | 0.25 | 0.25 | 0 | 0 | 0.301029996 | 0 |
| ENSMUSG00000089027 | 0 | 0 | 0 | 0 | 0 | 0 | 0 |
| ENSMUSG00000089028 | 0 | 0 | 0 | 0 | 0 | 0 | 0 |
| ENSMUSG00000089029 | 0 | 0 | 0 | 0 | 0 | 0 | 0 |
| ENSMUSG00000089030 | 0 | 0 | 0 | 0 | 0 | 0 | 0 |
| ENSMUSG00000089033 | 0 | 0 | 0 | 0 | 0 | 0 | 0 |
| ENSMUSG00000089034 | 0 | 0 | 0 | 0 | 0 | 0 | 0 |
| ENSMUSG00000089035 | 0 | 0 | 0 | 0 | 0 | 0 | 0 |
| ENSMUSG00000089036 | 0 | 0 | 0 | 0 | 0 | 0 | 0 |
| ENSMUSG00000089037 | 0 | 0 | 0 | 0 | 0 | 0 | 0 |
| ENSMUSG00000089038 | 0 | 0 | 0 | 0 | 0 | 0 | 0 |
| ENSMUSG00000089039 | 0 | 0 | 0 | 0 | 0 | 0 | 0 |
| ENSMUSG00000089040 | 0 | 0 | 0 | 0 | 0 | 0 | 0 |
| ENSMUSG00000089041 | 0 | 0 | 0 | 0 | 0 | 0 | 0 |
| ENSMUSG00000089043 | 0 | 0 | 0 | 0 | 0 | 0 | 0 |
| ENSMUSG00000089044 | 0 | 0 | 0 | 0 | 0 | 0 | 0 |
| ENSMUSG00000089045 | 0 | 0 | 0 | 0 | 0.244650542 | 0 | 0 |
| ENSMUSG00000089046 | 0 | 0 | 0 | 0 | 0 | 0 | 0 |
| ENSMUSG00000089047 | 0 | 0 | 0 | 0 | 0 | 0 | 0 |
| ENSMUSG00000089048 | 0 | 0 | 0 | 0 | 0 | 0.301029996 | 0 |
| ENSMUSG00000089049 | 0 | 0 | 0 | 0 | 0 | 0 | 0 |

| | | | | | | | |
|---|---|---|---|---|---|---|---|
| ENSMUSG00000089050 | 0 | 0 | 0 | 0 | 0 | 0 | 0 |
| ENSMUSG00000089051 | 0 | 0 | 0 | 0 | 0 | 0 | 0 |
| ENSMUSG00000089052 | 0 | 0 | 0 | 0 | 0 | 0 | 0 |
| ENSMUSG00000089053 | 0 | 0 | 0 | 0 | 0 | 0 | 0 |
| ENSMUSG00000089054 | 0 | 0 | 0 | 0 | 0 | 0 | 0 |
| ENSMUSG00000089055 | 0 | 0 | 0 | 0 | 0 | 0 | 0 |
| ENSMUSG00000089057 | 0 | 0 | 0 | 0 | 0 | 0 | 0 |
| ENSMUSG00000089060 | 0 | 0 | 0 | 0 | 0 | 0 | 0 |
| ENSMUSG00000089061 | 0 | 0 | 0 | 0 | 0 | 0 | 0 |
| ENSMUSG00000089062 | 0 | 0 | 0 | 0 | 0 | 0 | 0 |
| ENSMUSG00000089065 | 0 | 0 | 0 | 0 | 0 | 0 | 0 |
| ENSMUSG00000089066 | 0 | 0 | 0 | 0 | 0 | 0 | 0 |
| ENSMUSG00000089068 | 0 | 0 | 0 | 0 | 0 | 0 | 0 |
| ENSMUSG00000089069 | 0 | 0 | 0 | 0 | 0 | 0 | 0 |
| ENSMUSG00000089070 | 0 | 0 | 0 | 0 | 0 | 0 | 0 |
| ENSMUSG00000089071 | 0 | 0 | 0 | 0 | 0 | 0 | 0 |
| ENSMUSG00000089072 | 0.289064826 | 0.25 | 0.25 | 0.5 | 0.632412477 | 0 | 0.579661877 |
| ENSMUSG00000089073 | 0 | 0 | 0 | 0 | 0 | 0 | 0 |
| ENSMUSG00000089074 | 0 | 0 | 0 | 0 | 0 | 0 | 0 |
| ENSMUSG00000089075 | 0 | 0 | 0 | 0 | 0 | 0 | 0 |
| ENSMUSG00000089077 | 0 | 0 | 0 | 0 | 0 | 0 | 0 |
| ENSMUSG00000089078 | 0 | 0 | 0 | 0 | 0 | 0 | 0 |
| ENSMUSG00000089079 | 0 | 0 | 0 | 0 | 0 | 0 | 0 |
| ENSMUSG00000089080 | 0 | 0 | 0 | 0 | 0 | 0 | 0 |
| ENSMUSG00000089081 | 0 | 0 | 0 | 0 | 0 | 0 | 0 |
| ENSMUSG00000089082 | 0 | 0 | 0 | 0 | 0 | 0 | 0 |
| ENSMUSG00000089083 | 0 | 0 | 0 | 0 | 0 | 0 | 0 |
| ENSMUSG00000089084 | 0 | 0 | 0 | 0 | 0 | 0 | 0 |
| ENSMUSG00000089085 | 0 | 0 | 0 | 0 | 0 | 0 | 0 |
| ENSMUSG00000089087 | 0 | 0 | 0 | 0 | 0 | 0 | 0 |
| ENSMUSG00000089088 | 0 | 0 | 0 | 0 | 0 | 0 | 0 |
| ENSMUSG00000089090 | 0 | 0 | 0 | 0 | 0 | 0 | 0 |
| ENSMUSG00000089091 | 0 | 0 | 0 | 0 | 0 | 0 | 0 |
| ENSMUSG00000089092 | 0 | 0 | 0 | 0 | 0 | 0 | 0 |

| | | | | | | | |
|---|---|---|---|---|---|---|---|
| ENSMUSG00000089093 | 0 | 0 | 0 | 0 | 0 | 0 | 0 |
| ENSMUSG00000089094 | 0 | 0 | 0 | 0 | 0 | 0 | 0 |
| ENSMUSG00000089095 | 0 | 0 | 0 | 0 | 0 | 0 | 0 |
| ENSMUSG00000089096 | 0 | 0 | 0 | 0 | 0 | 0 | 0 |
| ENSMUSG00000089097 | 0 | 0 | 0 | 0 | 0 | 0 | 0 |
| ENSMUSG00000089098 | 0 | 0 | 0 | 0 | 0 | 0 | 0 |
| ENSMUSG00000089102 | 0 | 0 | 0 | 0 | 0 | 0 | 0 |
| ENSMUSG00000089103 | 0 | 0 | 0 | 0 | 0 | 0 | 0 |
| ENSMUSG00000089105 | 0 | 0.396240625 | 0 | 0 | 0 | 0 | 0.224243824 |
| ENSMUSG00000089106 | 0 | 0 | 0 | 0 | 0 | 0 | 0 |
| ENSMUSG00000089107 | 0 | 0 | 0 | 0 | 0 | 0 | 0 |
| ENSMUSG00000089109 | 0 | 0 | 0 | 0 | 0 | 0 | 0 |
| ENSMUSG00000089111 | 0 | 0 | 0 | 0 | 0 | 0 | 0 |
| ENSMUSG00000089113 | 0 | 0.25 | 0 | 0 | 0 | 0 | 0 |
| ENSMUSG00000089114 | 0 | 0 | 0 | 0 | 0 | 0 | 0 |
| ENSMUSG00000089115 | 0 | 0 | 0 | 0 | 0 | 0 | 0 |
| ENSMUSG00000089116 | 0 | 0 | 0 | 0 | 0 | 0 | 0 |
| ENSMUSG00000089117 | 0 | 0 | 0 | 0 | 0 | 0 | 0 |
| ENSMUSG00000089118 | 0 | 0 | 0 | 0 | 0 | 0 | 0 |
| ENSMUSG00000089119 | 0 | 0 | 0 | 0 | 0 | 0 | 0 |
| ENSMUSG00000089120 | 0 | 0 | 0 | 0 | 0 | 0 | 0 |
| ENSMUSG00000089123 | 0 | 0 | 0 | 0 | 0 | 0 | 0 |
| ENSMUSG00000089124 | 0 | 0 | 0 | 0 | 0 | 0 | 0 |
| ENSMUSG00000089125 | 0 | 0.25 | 0 | 0 | 0.244650542 | 0.301029996 | 0.355418052 |
| ENSMUSG00000089127 | 0 | 0 | 0 | 0 | 0 | 0 | 0 |
| ENSMUSG00000089128 | 0 | 0 | 0 | 0 | 0 | 0 | 0 |
| ENSMUSG00000089129 | 0.45815691 | 0.25 | 0.646240625 | 0.580482024 | 0.568060967 | 0.477121255 | 0.448487648 |
| ENSMUSG00000089130 | 0 | 0 | 0 | 0 | 0 | 0 | 0 |
| ENSMUSG00000089131 | 0 | 0 | 0 | 0 | 0 | 0 | 0 |
| ENSMUSG00000089132 | 0 | 0 | 0 | 0 | 0 | 0 | 0 |
| ENSMUSG00000089133 | 0 | 0 | 0 | 0 | 0 | 0 | 0 |
| ENSMUSG00000089134 | 0 | 0 | 0 | 0 | 0 | 0 | 0 |
| ENSMUSG00000089136 | 0 | 0 | 0 | 0 | 0 | 0 | 0 |
| ENSMUSG00000089137 | 0 | 0 | 0 | 0 | 0 | 0 | 0 |

| | | | | | | | |
|---|---|---|---|---|---|---|---|
| ENSMUSG00000089138 | 0 | 0 | 0 | 0 | 0 | 0 | 0 |
| ENSMUSG00000089139 | 0 | 0 | 0 | 0 | 0 | 0 | 0 |
| ENSMUSG00000089141 | 0 | 0 | 0 | 0 | 0 | 0 | 0 |
| ENSMUSG00000089143 | 0 | 0 | 0 | 0 | 0 | 0 | 0 |
| ENSMUSG00000089144 | 0 | 0 | 0 | 0 | 0 | 0 | 0 |
| ENSMUSG00000089145 | 0 | 0 | 0 | 0 | 0 | 0 | 0 |
| ENSMUSG00000089146 | 0 | 0 | 0 | 0 | 0 | 0 | 0 |
| ENSMUSG00000089147 | 0 | 0 | 0 | 0 | 0 | 0 | 0 |
| ENSMUSG00000089148 | 0 | 0 | 0 | 0 | 0 | 0 | 0 |
| ENSMUSG00000089149 | 0 | 0 | 0 | 0 | 0 | 0 | 0 |
| ENSMUSG00000089151 | 0 | 0 | 0 | 0.396240625 | 0 | 0 | 0 |
| ENSMUSG00000089152 | 0 | 0 | 0 | 0 | 0 | 0 | 0 |
| ENSMUSG00000089153 | 0 | 0 | 0 | 0 | 0 | 0 | 0 |
| ENSMUSG00000089155 | 0 | 0 | 0 | 0 | 0 | 0 | 0.224243824 |
| ENSMUSG00000089156 | 0 | 0 | 0 | 0 | 0 | 0 | 0 |
| ENSMUSG00000089157 | 0 | 0 | 0 | 0 | 0 | 0 | 0 |
| ENSMUSG00000089158 | 0 | 0 | 0 | 0 | 0 | 0 | 0 |
| ENSMUSG00000089161 | 0 | 0 | 0 | 0 | 0 | 0 | 0 |
| ENSMUSG00000089162 | 0 | 0 | 0 | 0 | 0 | 0 | 0 |
| ENSMUSG00000089163 | 0 | 0 | 0 | 0 | 0 | 0 | 0 |
| ENSMUSG00000089164 | 0 | 0 | 0 | 0 | 0 | 0 | 0 |
| ENSMUSG00000089168 | 0 | 0 | 0 | 0 | 0 | 0 | 0 |
| ENSMUSG00000089169 | 0 | 0 | 0 | 0 | 0 | 0 | 0 |
| ENSMUSG00000089170 | 0 | 0.25 | 0 | 0 | 0 | 0.301029996 | 0.224243824 |
| ENSMUSG00000089171 | 0 | 0 | 0 | 0 | 0 | 0 | 0 |
| ENSMUSG00000089174 | 0 | 0 | 0 | 0 | 0 | 0 | 0 |
| ENSMUSG00000089175 | 0 | 0 | 0 | 0 | 0 | 0 | 0 |
| ENSMUSG00000089177 | 0 | 0 | 0 | 0 | 0 | 0 | 0 |
| ENSMUSG00000089178 | 0 | 0 | 0 | 0 | 0 | 0 | 0 |
| ENSMUSG00000089179 | 0 | 0 | 0 | 0 | 0 | 0 | 0.224243824 |
| ENSMUSG00000089182 | 0 | 0 | 0 | 0 | 0 | 0 | 0 |
| ENSMUSG00000089183 | 0 | 0 | 0 | 0 | 0 | 0 | 0 |
| ENSMUSG00000089184 | 0 | 0 | 0 | 0 | 0 | 0 | 0 |
| ENSMUSG00000089185 | 0 | 0 | 0 | 0 | 0.244650542 | 0 | 0 |

| | | | | | | | |
|---|---|---|---|---|---|---|---|
| ENSMUSG00000089186 | 0 | 0 | 0 | 0 | 0 | 0 | 0 |
| ENSMUSG00000089187 | 0 | 0 | 0.25 | 0 | 0 | 0 | 0 |
| ENSMUSG00000089188 | 0 | 0 | 0 | 0 | 0 | 0 | 0 |
| ENSMUSG00000089189 | 0 | 0 | 0 | 0 | 0 | 0 | 0 |
| ENSMUSG00000089190 | 0 | 0 | 0 | 0 | 0 | 0 | 0 |
| ENSMUSG00000089191 | 0 | 0 | 0 | 0 | 0 | 0 | 0 |
| ENSMUSG00000089194 | 0 | 0 | 0 | 0 | 0 | 0 | 0 |
| ENSMUSG00000089195 | 0 | 0 | 0 | 0 | 0 | 0 | 0 |
| ENSMUSG00000089197 | 0 | 0 | 0 | 0 | 0 | 0 | 0 |
| ENSMUSG00000089199 | 0 | 0 | 0 | 0 | 0 | 0 | 0 |
| ENSMUSG00000089203 | 0 | 0 | 0 | 0 | 0 | 0 | 0 |
| ENSMUSG00000089204 | 0 | 0 | 0 | 0 | 0 | 0 | 0 |
| ENSMUSG00000089205 | 0 | 0 | 0 | 0 | 0 | 0 | 0 |
| ENSMUSG00000089206 | 0 | 0 | 0 | 0 | 0 | 0 | 0 |
| ENSMUSG00000089207 | 0 | 0 | 0 | 0 | 0 | 0 | 0 |
| ENSMUSG00000089208 | 0 | 0 | 0 | 0 | 0 | 0 | 0 |
| ENSMUSG00000089209 | 0 | 0 | 0 | 0 | 0 | 0 | 0 |
| ENSMUSG00000089210 | 0 | 0 | 0 | 0 | 0 | 0 | 0 |
| ENSMUSG00000089211 | 0 | 0 | 0 | 0 | 0 | 0 | 0 |
| ENSMUSG00000089213 | 0 | 0 | 0 | 0 | 0 | 0 | 0 |
| ENSMUSG00000089214 | 0 | 0 | 0 | 0 | 0 | 0 | 0 |
| ENSMUSG00000089217 | 0 | 0 | 0 | 0 | 0 | 0 | 0 |
| ENSMUSG00000089218 | 0 | 0 | 0 | 0 | 0 | 0 | 0 |
| ENSMUSG00000089219 | 0 | 0 | 0 | 0 | 0 | 0 | 0 |
| ENSMUSG00000089220 | 0 | 0 | 0 | 0 | 0 | 0 | 0 |
| ENSMUSG00000089222 | 0 | 0 | 0 | 0 | 0 | 0 | 0 |
| ENSMUSG00000089223 | 0 | 0 | 0 | 0 | 0 | 0 | 0 |
| ENSMUSG00000089224 | 0.289064826 | 0 | 0 | 0 | 0 | 0 | 0 |
| ENSMUSG00000089226 | 0 | 0 | 0 | 0 | 0 | 0 | 0 |
| ENSMUSG00000089227 | 0 | 0 | 0 | 0 | 0 | 0 | 0 |
| ENSMUSG00000089228 | 0 | 0 | 0 | 0 | 0 | 0 | 0 |
| ENSMUSG00000089229 | 0.578129653 | 0.396240625 | 0 | 0.396240625 | 0.244650542 | 0.602059991 | 0.520678036 |
| ENSMUSG00000089230 | 0 | 0 | 0 | 0 | 0 | 0 | 0 |
| ENSMUSG00000089231 | 0 | 0 | 0 | 0 | 0 | 0 | 0 |

| | | | | | | | |
|---|---|---|---|---|---|---|---|
| ENSMUSG00000089233 | 0 | 0 | 0 | 0 | 0 | 0 | 0 |
| ENSMUSG00000089234 | 0 | 0 | 0 | 0 | 0 | 0 | 0 |
| ENSMUSG00000089235 | 0 | 0 | 0 | 0 | 0.244650542 | 0 | 0.224243824 |
| ENSMUSG00000089237 | 0 | 0 | 0 | 0 | 0 | 0 | 0 |
| ENSMUSG00000089239 | 0 | 0.396240625 | 0 | 0 | 0 | 0 | 0 |
| ENSMUSG00000089241 | 0 | 0 | 0 | 0 | 0 | 0 | 0 |
| ENSMUSG00000089242 | 0 | 0 | 0 | 0 | 0 | 0 | 0 |
| ENSMUSG00000089244 | 0 | 0 | 0 | 0 | 0 | 0 | 0 |
| ENSMUSG00000089245 | 0 | 0 | 0 | 0 | 0 | 0 | 0 |
| ENSMUSG00000089247 | 0 | 0 | 0 | 0 | 0 | 0 | 0 |
| ENSMUSG00000089248 | 0 | 0.25 | 0 | 0.25 | 0 | 0 | 0.224243824 |
| ENSMUSG00000089250 | 0 | 0 | 0.25 | 0 | 0 | 0 | 0 |
| ENSMUSG00000089251 | 0 | 0 | 0 | 0 | 0 | 0 | 0 |
| ENSMUSG00000089252 | 0 | 0 | 0 | 0 | 0 | 0 | 0 |
| ENSMUSG00000089253 | 0 | 0.396240625 | 0 | 0 | 0 | 0 | 0.224243824 |
| ENSMUSG00000089255 | 0 | 0.396240625 | 0.25 | 0 | 0.244650542 | 0 | 0.520678036 |
| ENSMUSG00000089256 | 0 | 0 | 0 | 0 | 0 | 0 | 0 |
| ENSMUSG00000089257 | 0 | 0 | 0 | 0 | 0 | 0 | 0 |
| ENSMUSG00000089258 | 0 | 0 | 0 | 0 | 0 | 0 | 0 |
| ENSMUSG00000089259 | 0 | 0 | 0 | 0 | 0 | 0 | 0 |
| ENSMUSG00000089260 | 0 | 0 | 0 | 0.25 | 0.244650542 | 0 | 0.224243824 |
| ENSMUSG00000089261 | 0 | 0.25 | 0 | 0 | 0 | 0 | 0 |
| ENSMUSG00000089262 | 0 | 0 | 0 | 0 | 0 | 0 | 0 |
| ENSMUSG00000089263 | 0 | 0 | 0 | 0 | 0 | 0 | 0 |
| ENSMUSG00000089264 | 0 | 0 | 0 | 0 | 0 | 0 | 0 |
| ENSMUSG00000089265 | 0 | 0 | 0 | 0 | 0 | 0 | 0 |
| ENSMUSG00000089266 | 0 | 0 | 0 | 0 | 0 | 0 | 0 |
| ENSMUSG00000089267 | 0 | 0 | 0.25 | 0 | 0 | 0 | 0 |
| ENSMUSG00000089268 | 0 | 0 | 0 | 0 | 0 | 0 | 0 |
| ENSMUSG00000089269 | 0 | 0 | 0 | 0 | 0 | 0 | 0 |
| ENSMUSG00000089270 | 0 | 0 | 0 | 0 | 0 | 0 | 0 |
| ENSMUSG00000089271 | 0 | 0 | 0 | 0 | 0 | 0 | 0 |
| ENSMUSG00000089272 | 0 | 0 | 0 | 0 | 0 | 0 | 0 |
| ENSMUSG00000089273 | 0 | 0 | 0 | 0 | 0 | 0 | 0 |

| | | | | | | | |
|---|---|---|---|---|---|---|---|
| ENSMUSG00000089274 | 0 | 0 | 0 | 0 | 0 | 0 | 0 |
| ENSMUSG00000089275 | 0 | 0 | 0 | 0 | 0 | 0 | 0 |
| ENSMUSG00000089276 | 0 | 0 | 0 | 0 | 0 | 0 | 0 |
| ENSMUSG00000089277 | 0 | 0 | 0 | 0 | 0 | 0 | 0 |
| ENSMUSG00000089278 | 0 | 0 | 0 | 0 | 0 | 0 | 0 |
| ENSMUSG00000089279 | 0 | 0 | 0 | 0 | 0 | 0 | 0 |
| ENSMUSG00000089280 | 0 | 0 | 0 | 0 | 0 | 0 | 0 |
| ENSMUSG00000089281 | 0 | 0 | 0.25 | 0.25 | 0 | 0.301029996 | 0 |
| ENSMUSG00000089282 | 0 | 0 | 0 | 0 | 0 | 0 | 0 |
| ENSMUSG00000089284 | 0 | 0 | 0 | 0 | 0 | 0 | 0 |
| ENSMUSG00000089285 | 0 | 0 | 0 | 0 | 0 | 0 | 0 |
| ENSMUSG00000089286 | 0 | 0 | 0 | 0 | 0 | 0 | 0 |
| ENSMUSG00000089287 | 0 | 0 | 0 | 0 | 0 | 0 | 0 |
| ENSMUSG00000089288 | 0 | 0 | 0 | 0 | 0 | 0 | 0 |
| ENSMUSG00000089289 | 0 | 0 | 0 | 0 | 0 | 0 | 0 |
| ENSMUSG00000089290 | 0 | 0 | 0 | 0 | 0 | 0 | 0 |
| ENSMUSG00000089291 | 0 | 0 | 0 | 0 | 0 | 0 | 0 |
| ENSMUSG00000089292 | 0 | 0 | 0 | 0 | 0 | 0 | 0.224243824 |
| ENSMUSG00000089293 | 0 | 0 | 0 | 0 | 0 | 0 | 0 |
| ENSMUSG00000089294 | 0 | 0 | 0 | 0 | 0 | 0 | 0 |
| ENSMUSG00000089296 | 0.289064826 | 0.25 | 0.396240625 | 0.5 | 0.489301084 | 0 | 0.224243824 |
| ENSMUSG00000089297 | 0 | 0 | 0 | 0 | 0 | 0 | 0 |
| ENSMUSG00000089298 | 0 | 0 | 0 | 0.25 | 0 | 0 | 0 |
| ENSMUSG00000089299 | 0 | 0 | 0 | 0 | 0 | 0 | 0 |
| ENSMUSG00000089301 | 0 | 0 | 0 | 0 | 0 | 0 | 0 |
| ENSMUSG00000089302 | 0 | 0 | 0 | 0 | 0 | 0 | 0 |
| ENSMUSG00000089304 | 0 | 0 | 0 | 0 | 0 | 0 | 0 |
| ENSMUSG00000089305 | 0 | 0 | 0 | 0 | 0 | 0 | 0 |
| ENSMUSG00000089306 | 0 | 0 | 0 | 0 | 0 | 0 | 0 |
| ENSMUSG00000089307 | 0 | 0 | 0 | 0 | 0 | 0 | 0 |
| ENSMUSG00000089308 | 0 | 0 | 0 | 0 | 0 | 0 | 0.224243824 |
| ENSMUSG00000089309 | 0 | 0 | 0 | 0 | 0 | 0 | 0 |
| ENSMUSG00000089310 | 0 | 0 | 0 | 0 | 0 | 0 | 0 |
| ENSMUSG00000089311 | 0 | 0 | 0 | 0 | 0 | 0 | 0 |

| | | | | | | | |
|---|---|---|---|---|---|---|---|
| ENSMUSG00000089312 | 0 | 0 | 0 | 0 | 0 | 0 | 0 |
| ENSMUSG00000089313 | 0 | 0.25 | 0 | 0 | 0 | 0 | 0 |
| ENSMUSG00000089314 | 0 | 0 | 0 | 0 | 0 | 0 | 0 |
| ENSMUSG00000089316 | 0 | 0 | 0 | 0 | 0 | 0 | 0 |
| ENSMUSG00000089317 | 0.45815691 | 0.396240625 | 0 | 0 | 0 | 0 | 0 |
| ENSMUSG00000089318 | 0 | 0 | 0 | 0 | 0 | 0 | 0 |
| ENSMUSG00000089320 | 0 | 0 | 0 | 0 | 0 | 0 | 0 |
| ENSMUSG00000089321 | 0 | 0 | 0 | 0 | 0 | 0 | 0 |
| ENSMUSG00000089322 | 0 | 0 | 0 | 0 | 0 | 0 | 0 |
| ENSMUSG00000089323 | 0 | 0 | 0 | 0 | 0 | 0 | 0 |
| ENSMUSG00000089324 | 0 | 0 | 0 | 0 | 0 | 0 | 0 |
| ENSMUSG00000089326 | 0 | 0 | 0 | 0 | 0 | 0 | 0 |
| ENSMUSG00000089327 | 0 | 0 | 0 | 0 | 0 | 0 | 0 |
| ENSMUSG00000089331 | 0 | 0 | 0 | 0 | 0 | 0 | 0 |
| ENSMUSG00000089333 | 0 | 0 | 0 | 0 | 0 | 0 | 0 |
| ENSMUSG00000089334 | 0 | 0 | 0 | 0 | 0 | 0 | 0 |
| ENSMUSG00000089335 | 0 | 0 | 0 | 0 | 0 | 0 | 0 |
| ENSMUSG00000089336 | 0 | 0 | 0 | 0 | 0 | 0 | 0 |
| ENSMUSG00000089337 | 0 | 0 | 0.25 | 0 | 0.244650542 | 0.301029996 | 0 |
| ENSMUSG00000089338 | 0 | 0 | 0 | 0 | 0 | 0 | 0 |
| ENSMUSG00000089341 | 0 | 0 | 0 | 0 | 0 | 0 | 0 |
| ENSMUSG00000089342 | 0 | 0 | 0 | 0 | 0 | 0 | 0 |
| ENSMUSG00000089343 | 0 | 0 | 0.25 | 0 | 0 | 0 | 0 |
| ENSMUSG00000089344 | 0 | 0 | 0 | 0 | 0 | 0 | 0 |
| ENSMUSG00000089345 | 0 | 0 | 0 | 0 | 0 | 0 | 0 |
| ENSMUSG00000089346 | 0 | 0 | 0 | 0 | 0 | 0 | 0 |
| ENSMUSG00000089348 | 0 | 0 | 0 | 0 | 0 | 0 | 0 |
| ENSMUSG00000089349 | 0 | 0 | 0 | 0 | 0 | 0 | 0 |
| ENSMUSG00000089350 | 0 | 0 | 0 | 0 | 0 | 0 | 0 |
| ENSMUSG00000089351 | 0 | 0 | 0 | 0 | 0 | 0 | 0 |
| ENSMUSG00000089352 | 0 | 0 | 0 | 0 | 0 | 0 | 0 |
| ENSMUSG00000089353 | 0 | 0 | 0 | 0 | 0 | 0 | 0 |
| ENSMUSG00000089357 | 0 | 0 | 0 | 0 | 0 | 0 | 0 |
| ENSMUSG00000089358 | 0 | 0 | 0 | 0 | 0 | 0 | 0 |

| | | | | | | | |
|---|---|---|---|---|---|---|---|
| ENSMUSG00000089359 | 0 | 0 | 0 | 0 | 0 | 0 | 0 |
| ENSMUSG00000089363 | 0 | 0 | 0 | 0 | 0 | 0 | 0 |
| ENSMUSG00000089364 | 0 | 0 | 0 | 0 | 0 | 0 | 0 |
| ENSMUSG00000089365 | 0 | 0.25 | 0 | 0 | 0 | 0.301029996 | 0 |
| ENSMUSG00000089366 | 0 | 0 | 0 | 0 | 0 | 0 | 0 |
| ENSMUSG00000089369 | 0 | 0 | 0 | 0 | 0 | 0 | 0 |
| ENSMUSG00000089370 | 0 | 0 | 0 | 0 | 0 | 0 | 0 |
| ENSMUSG00000089371 | 0 | 0 | 0 | 0 | 0 | 0 | 0 |
| ENSMUSG00000089372 | 0 | 0 | 0 | 0 | 0 | 0 | 0 |
| ENSMUSG00000089373 | 0 | 0 | 0 | 0 | 0 | 0 | 0 |
| ENSMUSG00000089375 | 0 | 0 | 0 | 0 | 0 | 0 | 0 |
| ENSMUSG00000089376 | 0 | 0 | 0 | 0 | 0 | 0 | 0 |
| ENSMUSG00000089377 | 0 | 0 | 0 | 0 | 0 | 0 | 0 |
| ENSMUSG00000089380 | 0 | 0 | 0 | 0 | 0 | 0 | 0 |
| ENSMUSG00000089381 | 0 | 0.25 | 0 | 0 | 0 | 0 | 0 |
| ENSMUSG00000089382 | 0 | 0 | 0 | 0 | 0 | 0 | 0 |
| ENSMUSG00000089383 | 0 | 0 | 0 | 0 | 0 | 0 | 0 |
| ENSMUSG00000089384 | 0 | 0 | 0 | 0 | 0 | 0 | 0 |
| ENSMUSG00000089385 | 0 | 0 | 0 | 0 | 0 | 0 | 0 |
| ENSMUSG00000089387 | 0 | 0 | 0 | 0 | 0 | 0 | 0 |
| ENSMUSG00000089389 | 0.289064826 | 0 | 0 | 0 | 0 | 0 | 0 |
| ENSMUSG00000089390 | 0 | 0 | 0 | 0 | 0 | 0 | 0 |
| ENSMUSG00000089391 | 0 | 0 | 0 | 0 | 0 | 0 | 0 |
| ENSMUSG00000089392 | 0 | 0 | 0 | 0 | 0 | 0 | 0 |
| ENSMUSG00000089393 | 0 | 0 | 0 | 0 | 0 | 0 | 0 |
| ENSMUSG00000089394 | 0 | 0 | 0 | 0 | 0 | 0 | 0 |
| ENSMUSG00000089395 | 0 | 0 | 0 | 0 | 0 | 0 | 0 |
| ENSMUSG00000089396 | 0 | 0.5 | 0 | 0.396240625 | 0.387761935 | 0.301029996 | 0.224243824 |
| ENSMUSG00000089397 | 0 | 0 | 0 | 0 | 0 | 0 | 0 |
| ENSMUSG00000089398 | 0 | 0 | 0 | 0 | 0 | 0 | 0 |
| ENSMUSG00000089399 | 0 | 0 | 0 | 0 | 0 | 0 | 0 |
| ENSMUSG00000089401 | 0 | 0 | 0 | 0 | 0 | 0 | 0 |
| ENSMUSG00000089404 | 0 | 0 | 0.25 | 0 | 0 | 0 | 0 |
| ENSMUSG00000089405 | 0 | 0 | 0 | 0 | 0 | 0 | 0 |

| | | | | | | | |
|---|---|---|---|---|---|---|---|
| ENSMUSG00000089406 | 0 | 0 | 0 | 0 | 0 | 0 | 0 |
| ENSMUSG00000089407 | 0 | 0 | 0 | 0 | 0 | 0 | 0 |
| ENSMUSG00000089408 | 0 | 0 | 0 | 0 | 0 | 0 | 0 |
| ENSMUSG00000089409 | 0 | 0 | 0 | 0 | 0 | 0 | 0 |
| ENSMUSG00000089410 | 0 | 0 | 0 | 0 | 0 | 0 | 0 |
| ENSMUSG00000089413 | 0 | 0 | 0 | 0 | 0 | 0 | 0 |
| ENSMUSG00000089414 | 0 | 0 | 0 | 0 | 0 | 0 | 0 |
| ENSMUSG00000089415 | 0 | 0 | 0 | 0 | 0 | 0 | 0 |
| ENSMUSG00000089416 | 0 | 0 | 0 | 0 | 0 | 0 | 0 |
| ENSMUSG00000089417 | 0.45815691 | 0.25 | 0 | 0.646240625 | 0.387761935 | 0.301029996 | 0.224243824 |
| ENSMUSG00000089418 | 0 | 0 | 0 | 0 | 0 | 0 | 0 |
| ENSMUSG00000089420 | 0 | 0 | 0 | 0 | 0 | 0 | 0 |
| ENSMUSG00000089422 | 0 | 0 | 0 | 0 | 0 | 0 | 0 |
| ENSMUSG00000089423 | 0 | 0 | 0 | 0 | 0 | 0 | 0 |
| ENSMUSG00000089424 | 0 | 0 | 0 | 0 | 0 | 0 | 0 |
| ENSMUSG00000089425 | 0 | 0 | 0 | 0 | 0 | 0 | 0 |
| ENSMUSG00000089427 | 0 | 0 | 0 | 0 | 0 | 0 | 0 |
| ENSMUSG00000089429 | 0 | 0 | 0 | 0 | 0 | 0 | 0 |
| ENSMUSG00000089431 | 0.289064826 | 0 | 0 | 0.25 | 0 | 0 | 0 |
| ENSMUSG00000089432 | 0 | 0.25 | 0 | 0 | 0 | 0 | 0 |
| ENSMUSG00000089434 | 0 | 0 | 0 | 0 | 0 | 0 | 0 |
| ENSMUSG00000089436 | 0 | 0 | 0 | 0 | 0 | 0 | 0 |
| ENSMUSG00000089437 | 0.289064826 | 0 | 0 | 0 | 0 | 0.477121255 | 0.355418052 |
| ENSMUSG00000089438 | 0 | 0 | 0 | 0 | 0 | 0 | 0 |
| ENSMUSG00000089439 | 0 | 0 | 0 | 0 | 0 | 0 | 0 |
| ENSMUSG00000089441 | 0 | 0 | 0 | 0 | 0 | 0 | 0 |
| ENSMUSG00000089442 | 0 | 0 | 0 | 0 | 0 | 0 | 0 |
| ENSMUSG00000089443 | 0 | 0 | 0 | 0 | 0 | 0 | 0 |
| ENSMUSG00000089444 | 0 | 0 | 0 | 0 | 0 | 0 | 0 |
| ENSMUSG00000089445 | 0 | 0 | 0 | 0 | 0 | 0 | 0 |
| ENSMUSG00000089446 | 0 | 0 | 0 | 0 | 0 | 0 | 0 |
| ENSMUSG00000089447 | 0 | 0 | 0 | 0 | 0 | 0 | 0 |
| ENSMUSG00000089450 | 0 | 0 | 0 | 0 | 0 | 0 | 0 |
| ENSMUSG00000089451 | 0 | 0 | 0 | 0 | 0 | 0 | 0 |

| | | | | | | | |
|---|---|---|---|---|---|---|---|
| ENSMUSG00000089452 | 0 | 0 | 0 | 0 | 0 | 0 | 0 |
| ENSMUSG00000089453 | 0 | 0 | 0.25 | 0 | 0.244650542 | 0 | 0 |
| ENSMUSG00000089454 | 0 | 0 | 0 | 0 | 0 | 0 | 0 |
| ENSMUSG00000089455 | 0 | 0 | 0 | 0 | 0 | 0 | 0 |
| ENSMUSG00000089458 | 0 | 0 | 0 | 0 | 0 | 0 | 0 |
| ENSMUSG00000089459 | 0 | 0 | 0 | 0 | 0 | 0 | 0 |
| ENSMUSG00000089460 | 0 | 0 | 0 | 0 | 0 | 0 | 0 |
| ENSMUSG00000089461 | 0 | 0 | 0 | 0 | 0 | 0 | 0 |
| ENSMUSG00000089462 | 0 | 0 | 0 | 0 | 0 | 0 | 0 |
| ENSMUSG00000089464 | 0 | 0 | 0 | 0 | 0 | 0 | 0 |
| ENSMUSG00000089465 | 0 | 0 | 0 | 0 | 0 | 0 | 0 |
| ENSMUSG00000089467 | 0 | 0 | 0 | 0 | 0 | 0 | 0 |
| ENSMUSG00000089468 | 0 | 0 | 0 | 0 | 0 | 0 | 0 |
| ENSMUSG00000089469 | 0 | 0 | 0 | 0 | 0 | 0 | 0 |
| ENSMUSG00000089470 | 0 | 0 | 0 | 0 | 0 | 0 | 0 |
| ENSMUSG00000089471 | 0 | 0 | 0 | 0 | 0 | 0 | 0 |
| ENSMUSG00000089472 | 0 | 0 | 0 | 0 | 0 | 0 | 0 |
| ENSMUSG00000089473 | 0 | 0 | 0 | 0 | 0 | 0 | 0 |
| ENSMUSG00000089475 | 0 | 0 | 0 | 0 | 0 | 0 | 0.224243824 |
| ENSMUSG00000089476 | 0.289064826 | 0.5 | 0.396240625 | 0 | 0.387761935 | 0 | 0.224243824 |
| ENSMUSG00000089477 | 0 | 0 | 0 | 0 | 0 | 0 | 0 |
| ENSMUSG00000089479 | 0 | 0 | 0 | 0 | 0 | 0 | 0 |
| ENSMUSG00000089480 | 0 | 0 | 0.25 | 0 | 0 | 0 | 0 |
| ENSMUSG00000089481 | 0 | 0 | 0 | 0 | 0 | 0 | 0 |
| ENSMUSG00000089482 | 0 | 0 | 0 | 0 | 0 | 0 | 0 |
| ENSMUSG00000089484 | 0 | 0 | 0 | 0 | 0 | 0 | 0 |
| ENSMUSG00000089485 | 0 | 0 | 0 | 0 | 0 | 0 | 0 |
| ENSMUSG00000089486 | 0 | 0 | 0 | 0 | 0 | 0 | 0 |
| ENSMUSG00000089487 | 0 | 0 | 0 | 0 | 0 | 0 | 0 |
| ENSMUSG00000089488 | 0 | 0 | 0 | 0 | 0 | 0 | 0 |
| ENSMUSG00000089489 | 0 | 0 | 0 | 0 | 0 | 0 | 0 |
| ENSMUSG00000089490 | 0 | 0 | 0 | 0 | 0 | 0 | 0 |
| ENSMUSG00000089491 | 0 | 0 | 0.25 | 0 | 0 | 0 | 0 |
| ENSMUSG00000089493 | 0 | 0 | 0 | 0 | 0 | 0 | 0.224243824 |

| | | | | | | | |
|---|---|---|---|---|---|---|---|
| ENSMUSG00000089497 | 0 | 0 | 0 | 0 | 0 | 0 | 0 |
| ENSMUSG00000089498 | 0 | 0.25 | 0 | 0 | 0.244650542 | 0 | 0 |
| ENSMUSG00000089499 | 0 | 0 | 0 | 0 | 0 | 0 | 0 |
| ENSMUSG00000089501 | 0 | 0 | 0 | 0 | 0 | 0 | 0 |
| ENSMUSG00000089504 | 0 | 0 | 0 | 0 | 0 | 0 | 0 |
| ENSMUSG00000089506 | 0 | 0 | 0 | 0 | 0 | 0 | 0 |
| ENSMUSG00000089507 | 0 | 0 | 0 | 0 | 0 | 0 | 0 |
| ENSMUSG00000089508 | 0 | 0 | 0 | 0 | 0 | 0 | 0 |
| ENSMUSG00000089509 | 0 | 0 | 0 | 0 | 0 | 0 | 0 |
| ENSMUSG00000089510 | 0 | 0 | 0 | 0 | 0 | 0 | 0 |
| ENSMUSG00000089512 | 0 | 0.25 | 0.25 | 0 | 0 | 0.301029996 | 0 |
| ENSMUSG00000089513 | 0 | 0 | 0 | 0 | 0 | 0 | 0 |
| ENSMUSG00000089514 | 0 | 0 | 0 | 0 | 0 | 0 | 0 |
| ENSMUSG00000089516 | 0 | 0 | 0 | 0 | 0 | 0 | 0 |
| ENSMUSG00000089517 | 0 | 0 | 0 | 0 | 0 | 0 | 0 |
| ENSMUSG00000089520 | 0 | 0 | 0 | 0 | 0 | 0 | 0 |
| ENSMUSG00000089522 | 0 | 0 | 0 | 0 | 0 | 0 | 0 |
| ENSMUSG00000089524 | 0 | 0 | 0 | 0 | 0 | 0 | 0 |
| ENSMUSG00000089525 | 0.289064826 | 0.580482024 | 0.396240625 | 0.396240625 | 0 | 0.301029996 | 0.224243824 |
| ENSMUSG00000089528 | 0 | 0 | 0 | 0 | 0 | 0 | 0 |
| ENSMUSG00000089529 | 0 | 0 | 0 | 0 | 0 | 0 | 0 |
| ENSMUSG00000089530 | 0 | 0 | 0 | 0 | 0 | 0 | 0 |
| ENSMUSG00000089531 | 0 | 0 | 0 | 0 | 0 | 0 | 0 |
| ENSMUSG00000089533 | 0 | 0 | 0 | 0 | 0 | 0 | 0 |
| ENSMUSG00000089534 | 0 | 0 | 0 | 0 | 0 | 0 | 0 |
| ENSMUSG00000089536 | 0 | 0 | 0 | 0 | 0 | 0 | 0 |
| ENSMUSG00000089538 | 0 | 0 | 0 | 0 | 0 | 0 | 0 |
| ENSMUSG00000089539 | 0 | 0 | 0 | 0 | 0 | 0 | 0 |
| ENSMUSG00000089541 | 0 | 0 | 0 | 0 | 0 | 0 | 0 |
| ENSMUSG00000089542 | 0 | 0.5 | 0.75 | 0.701838731 | 0.568060967 | 0.602059991 | 0.448487648 |
| ENSMUSG00000089543 | 0 | 0 | 0 | 0 | 0 | 0 | 0 |
| ENSMUSG00000089544 | 0 | 0 | 0 | 0 | 0 | 0 | 0 |
| ENSMUSG00000089545 | 0 | 0 | 0 | 0 | 0 | 0.301029996 | 0.224243824 |
| ENSMUSG00000089546 | 0 | 0 | 0 | 0 | 0 | 0 | 0 |

| | | | | | | | |
|---|---|---|---|---|---|---|---|
| ENSMUSG00000089548 | 0 | 0 | 0 | 0 | 0 | 0 | 0 |
| ENSMUSG00000089549 | 0 | 0 | 0 | 0 | 0 | 0 | 0 |
| ENSMUSG00000089550 | 0 | 0 | 0 | 0 | 0 | 0 | 0 |
| ENSMUSG00000089551 | 0 | 0 | 0 | 0 | 0 | 0 | 0 |
| ENSMUSG00000089552 | 0 | 0 | 0 | 0 | 0 | 0 | 0 |
| ENSMUSG00000089553 | 0 | 0 | 0 | 0 | 0 | 0 | 0 |
| ENSMUSG00000089555 | 0 | 0 | 0 | 0 | 0 | 0 | 0 |
| ENSMUSG00000089556 | 0 | 0 | 0 | 0 | 0 | 0 | 0 |
| ENSMUSG00000089558 | 0 | 0 | 0 | 0 | 0 | 0 | 0 |
| ENSMUSG00000089560 | 0 | 0 | 0 | 0 | 0 | 0 | 0 |
| ENSMUSG00000089561 | 0 | 0 | 0 | 0 | 0 | 0 | 0 |
| ENSMUSG00000089562 | 0 | 0 | 0 | 0 | 0 | 0 | 0 |
| ENSMUSG00000089563 | 0 | 0 | 0 | 0.25 | 0 | 0 | 0.224243824 |
| ENSMUSG00000089564 | 0.289064826 | 0 | 0 | 0 | 0 | 0 | 0 |
| ENSMUSG00000089570 | 0 | 0 | 0 | 0 | 0 | 0 | 0 |
| ENSMUSG00000089571 | 0.45815691 | 0.396240625 | 0 | 0.25 | 0 | 0 | 0.224243824 |
| ENSMUSG00000089575 | 0 | 0 | 0 | 0 | 0 | 0 | 0 |
| ENSMUSG00000089576 | 0 | 0 | 0 | 0 | 0 | 0 | 0 |
| ENSMUSG00000089578 | 0 | 0 | 0 | 0 | 0 | 0 | 0 |
| ENSMUSG00000089580 | 0 | 0 | 0 | 0 | 0 | 0 | 0 |
| ENSMUSG00000089581 | 0.45815691 | 0.5 | 0.396240625 | 0.25 | 0.387761935 | 0.301029996 | 0.448487648 |
| ENSMUSG00000089583 | 0 | 0 | 0 | 0 | 0 | 0 | 0 |
| ENSMUSG00000089584 | 0 | 0 | 0 | 0 | 0 | 0 | 0 |
| ENSMUSG00000089585 | 0 | 0 | 0 | 0 | 0 | 0 | 0 |
| ENSMUSG00000089586 | 0 | 0 | 0 | 0 | 0 | 0 | 0 |
| ENSMUSG00000089587 | 0 | 0 | 0 | 0 | 0 | 0 | 0 |
| ENSMUSG00000089588 | 0 | 0 | 0 | 0.25 | 0 | 0 | 0 |
| ENSMUSG00000089589 | 0 | 0 | 0 | 0 | 0 | 0 | 0 |
| ENSMUSG00000089590 | 0 | 0 | 0 | 0 | 0 | 0 | 0 |
| ENSMUSG00000089591 | 0 | 0 | 0 | 0 | 0 | 0 | 0 |
| ENSMUSG00000089593 | 0 | 0 | 0 | 0 | 0 | 0 | 0 |
| ENSMUSG00000089596 | 0 | 0 | 0 | 0 | 0 | 0 | 0 |
| ENSMUSG00000089597 | 0 | 0 | 0 | 0 | 0 | 0 | 0 |
| ENSMUSG00000089598 | 0 | 0 | 0 | 0 | 0 | 0 | 0 |

| | | | | | | | |
|---|---|---|---|---|---|---|---|
| ENSMUSG00000089599 | 0 | 0 | 0 | 0 | 0 | 0 | 0 |
| ENSMUSG00000089601 | 0 | 0 | 0 | 0 | 0 | 0 | 0 |
| ENSMUSG00000089603 | 0 | 0 | 0 | 0 | 0 | 0 | 0 |
| ENSMUSG00000089605 | 0 | 0 | 0 | 0 | 0 | 0 | 0 |
| ENSMUSG00000089606 | 0 | 0 | 0 | 0 | 0 | 0 | 0 |
| ENSMUSG00000089607 | 0 | 0 | 0 | 0 | 0 | 0 | 0 |
| ENSMUSG00000089608 | 0 | 0 | 0 | 0 | 0 | 0 | 0 |
| ENSMUSG00000089609 | 0 | 0 | 0 | 0 | 0 | 0 | 0 |
| ENSMUSG00000089610 | 0 | 0 | 0 | 0 | 0 | 0 | 0 |
| ENSMUSG00000089612 | 0 | 0 | 0 | 0 | 0 | 0 | 0 |
| ENSMUSG00000089613 | 0 | 0 | 0 | 0 | 0 | 0 | 0 |
| ENSMUSG00000089614 | 0 | 0 | 0 | 0 | 0 | 0 | 0 |
| ENSMUSG00000089615 | 0 | 0 | 0 | 0 | 0 | 0 | 0 |
| ENSMUSG00000089616 | 0 | 0 | 0 | 0 | 0 | 0 | 0 |
| ENSMUSG00000089617 | 0 | 0 | 0 | 0 | 0 | 0 | 0 |
| ENSMUSG00000089618 | 0 | 0 | 0 | 0 | 0 | 0 | 0 |
| ENSMUSG00000089619 | 0 | 0 | 0 | 0 | 0 | 0 | 0 |
| ENSMUSG00000089621 | 0 | 0 | 0 | 0 | 0 | 0 | 0 |
| ENSMUSG00000089622 | 0 | 0 | 0 | 0 | 0 | 0 | 0 |
| ENSMUSG00000089624 | 0 | 0 | 0 | 0 | 0 | 0 | 0 |
| ENSMUSG00000089625 | 0 | 0 | 0 | 0 | 0 | 0 | 0 |
| ENSMUSG00000089627 | 0 | 0 | 0 | 0 | 0 | 0 | 0 |
| ENSMUSG00000089628 | 0 | 0 | 0 | 0 | 0 | 0 | 0 |
| ENSMUSG00000089630 | 0 | 0 | 0 | 0 | 0.244650542 | 0 | 0 |
| ENSMUSG00000089631 | 0.45815691 | 0.396240625 | 0.25 | 0.25 | 0.387761935 | 0 | 0.355418052 |
| ENSMUSG00000089633 | 0 | 0.25 | 0 | 0 | 0 | 0 | 0 |
| ENSMUSG00000089634 | 0 | 0 | 0 | 0 | 0.244650542 | 0 | 0.224243824 |
| ENSMUSG00000089635 | 0 | 0 | 0.25 | 0 | 0 | 0 | 0 |
| ENSMUSG00000089636 | 0.960252568 | 0.701838731 | 0.701838731 | 0.396240625 | 0.489301084 | 0.301029996 | 0.629532004 |
| ENSMUSG00000089639 | 0.289064826 | 0 | 0 | 0 | 0.244650542 | 0 | 0 |
| ENSMUSG00000089640 | 0 | 0 | 0 | 0 | 0 | 0 | 0.224243824 |
| ENSMUSG00000089642 | 0 | 0 | 0 | 0 | 0 | 0 | 0 |
| ENSMUSG00000089643 | 0 | 0 | 0 | 0 | 0.244650542 | 0 | 0 |
| ENSMUSG00000089645 | 0.671187741 | 0.646240625 | 0.5 | 0.5 | 0.387761935 | 0.301029996 | 0.520678036 |

| | | | | | | | |
|---|---|---|---|---|---|---|---|
| ENSMUSG00000089646 | 0.289064826 | 0.580482024 | 0.580482024 | 0.25 | 0.877063019 | 0.477121255 | 0.672731473 |
| ENSMUSG00000089647 | 0.960252568 | 0.951838731 | 0.646240625 | 1 | 0.877063019 | 0.602059991 | 0.672731473 |
| ENSMUSG00000089648 | 0 | 0.25 | 0 | 0 | 0 | 0 | 0 |
| ENSMUSG00000089651 | 0.289064826 | 0.25 | 0 | 0.580482024 | 0.387761935 | 0 | 0.224243824 |
| ENSMUSG00000089652 | 0 | 0 | 0 | 0 | 0 | 0 | 0 |
| ENSMUSG00000089653 | 0 | 0 | 0 | 0 | 0 | 0 | 0 |
| ENSMUSG00000089654 | 0 | 0 | 0 | 0 | 0 | 0 | 0 |
| ENSMUSG00000089655 | 0 | 0 | 0 | 0 | 0 | 0 | 0 |
| ENSMUSG00000089656 | 0 | 0 | 0 | 0 | 0 | 0 | 0 |
| ENSMUSG00000089657 | 0.45815691 | 0.396240625 | 0 | 0.5 | 0 | 0 | 0.224243824 |
| ENSMUSG00000089659 | 0 | 0 | 0 | 0 | 0 | 0 | 0 |
| ENSMUSG00000089661 | 0 | 0.25 | 0 | 0 | 0 | 0 | 0 |
| ENSMUSG00000089662 | 0 | 0 | 0 | 0 | 0.244650542 | 0 | 0 |
| ENSMUSG00000089663 | 0 | 0 | 0 | 0 | 0 | 0 | 0 |
| ENSMUSG00000089664 | 0.289064826 | 0 | 0 | 0 | 0.489301084 | 0 | 0.224243824 |
| ENSMUSG00000089665 | 0.289064826 | 0.580482024 | 0.396240625 | 0.25 | 0.733951626 | 0.602059991 | 0.629532004 |
| ENSMUSG00000089666 | 0.289064826 | 0.396240625 | 0 | 0.25 | 0 | 0 | 0.224243824 |
| ENSMUSG00000089667 | 0.289064826 | 0.5 | 0 | 0 | 0 | 0 | 0 |
| ENSMUSG00000089669 | 0.289064826 | 0 | 0.701838731 | 0.396240625 | 0.387761935 | 0 | 0.579661877 |
| ENSMUSG00000089670 | 0.289064826 | 0.580482024 | 0.701838731 | 0.5 | 0.244650542 | 0 | 0.224243824 |
| ENSMUSG00000089671 | 0.289064826 | 0.396240625 | 0.396240625 | 0.25 | 0.244650542 | 0 | 0.224243824 |
| ENSMUSG00000089672 | 0 | 0 | 0 | 0 | 0 | 0 | 0 |
| ENSMUSG00000089673 | 0 | 0 | 0 | 0 | 0 | 0 | 0 |
| ENSMUSG00000089675 | 0 | 0 | 0 | 0 | 0 | 0 | 0 |
| ENSMUSG00000089678 | 0 | 0 | 0 | 0 | 0 | 0 | 0 |
| ENSMUSG00000089679 | 0 | 0.396240625 | 0 | 0 | 0 | 0 | 0 |
| ENSMUSG00000089680 | 0 | 0 | 0.25 | 0 | 0 | 0 | 0.224243824 |
| ENSMUSG00000089682 | 2.10861828 | 1.990000483 | 1.96759118 | 2.143411797 | 2.068600388 | 2.303196057 | 1.928477344 |
| ENSMUSG00000089683 | 0 | 0 | 0 | 0 | 0 | 0 | 0 |
| ENSMUSG00000089684 | 0 | 0 | 0 | 0 | 0 | 0 | 0 |
| ENSMUSG00000089685 | 0 | 0 | 0 | 0 | 0 | 0 | 0 |
| ENSMUSG00000089686 | 0 | 0 | 0 | 0 | 0 | 0 | 0 |
| ENSMUSG00000089687 | 0.289064826 | 0.396240625 | 0.25 | 0 | 0.244650542 | 0 | 0.224243824 |
| ENSMUSG00000089689 | 0 | 0 | 0 | 0 | 0 | 0 | 0 |

| | | | | | | | |
|---|---|---|---|---|---|---|---|
| ENSMUSG00000089690 | 0 | 0 | 0 | 0.25 | 0 | 0 | 0 |
| ENSMUSG00000089694 | 0 | 0 | 0 | 0 | 0 | 0 | 0 |
| ENSMUSG00000089695 | 0 | 0 | 0.25 | 0.580482024 | 0.244650542 | 0 | 0.355418052 |
| ENSMUSG00000089696 | 0 | 0 | 0 | 0 | 0 | 0 | 0 |
| ENSMUSG00000089697 | 0 | 0.396240625 | 0 | 0.25 | 0.244650542 | 0 | 0 |
| ENSMUSG00000089698 | 0 | 0 | 0 | 0 | 0 | 0 | 0 |
| ENSMUSG00000089699 | 0 | 0 | 0 | 0 | 0 | 0 | 0 |
| ENSMUSG00000089700 | 0 | 0 | 0 | 0 | 0 | 0 | 0 |
| ENSMUSG00000089701 | 0.289064826 | 0 | 0 | 0 | 0.244650542 | 0 | 0 |
| ENSMUSG00000089702 | 0 | 0 | 0 | 0 | 0 | 0 | 0 |
| ENSMUSG00000089703 | 0 | 0 | 0 | 0 | 0 | 0 | 0 |
| ENSMUSG00000089704 | 0 | 0 | 0 | 0 | 0 | 0 | 0 |
| ENSMUSG00000089705 | 0 | 0 | 0 | 0 | 0 | 0 | 0 |
| ENSMUSG00000089706 | 0.578129653 | 0.5 | 0.5 | 0.5 | 0.877063019 | 0.602059991 | 0.74492186 |
| ENSMUSG00000089707 | 0 | 0 | 0 | 0 | 0.244650542 | 0 | 0 |
| ENSMUSG00000089708 | 0 | 0 | 0 | 0.25 | 0 | 0 | 0 |
| ENSMUSG00000089709 | 0 | 0 | 0 | 0 | 0 | 0 | 0.224243824 |
| ENSMUSG00000089710 | 0 | 0 | 0 | 0 | 0 | 0 | 0 |
| ENSMUSG00000089711 | 0 | 0 | 0.25 | 0.25 | 0.244650542 | 0 | 0 |
| ENSMUSG00000089712 | 0 | 0 | 0 | 0 | 0 | 0 | 0 |
| ENSMUSG00000089713 | 0 | 0 | 0 | 0 | 0 | 0 | 0 |
| ENSMUSG00000089714 | 0 | 0 | 0 | 0 | 0 | 0 | 0 |
| ENSMUSG00000089715 | 3.008682029 | 2.796218836 | 2.848481669 | 2.525493918 | 2.64026298 | 2.860338007 | 2.505768111 |
| ENSMUSG00000089716 | 0 | 0.5 | 0.580482024 | 0.396240625 | 0 | 0.301029996 | 0.672731473 |
| ENSMUSG00000089717 | 0 | 0 | 0 | 0 | 0 | 0 | 0 |
| ENSMUSG00000089718 | 0 | 0 | 0 | 0 | 0 | 0 | 0 |
| ENSMUSG00000089719 | 0.45815691 | 0 | 0.646240625 | 0.580482024 | 0.632412477 | 0.301029996 | 0.74492186 |
| ENSMUSG00000089720 | 0 | 0 | 0 | 0 | 0 | 0 | 0 |
| ENSMUSG00000089722 | 0 | 0 | 0 | 0 | 0 | 0 | 0 |
| ENSMUSG00000089723 | 0 | 0 | 0 | 0 | 0 | 0 | 0 |
| ENSMUSG00000089724 | 0 | 0 | 0 | 0 | 0 | 0 | 0 |
| ENSMUSG00000089726 | 1.418409478 | 1.356566189 | 1.25 | 1.532320754 | 1.533172725 | 1.477121255 | 1.302263476 |
| ENSMUSG00000089727 | 0 | 0 | 0 | 0 | 0 | 0 | 0 |
| ENSMUSG00000089728 | 0 | 0 | 0 | 0 | 0 | 0 | 0 |

| | | | | | | | |
|---|---|---|---|---|---|---|---|
| ENSMUSG00000089729 | 0 | 0 | 0 | 0 | 0 | 0 | 0 |
| ENSMUSG00000089730 | 0.45815691 | 0.396240625 | 0.580482024 | 0.580482024 | 0.568060967 | 0.77815125 | 0.448487648 |
| ENSMUSG00000089731 | 0 | 0 | 0 | 0 | 0 | 0 | 0 |
| ENSMUSG00000089732 | 0 | 0 | 0 | 0 | 0 | 0 | 0 |
| ENSMUSG00000089733 | 0 | 0 | 0 | 0 | 0 | 0 | 0 |
| ENSMUSG00000089735 | 0.45815691 | 0.396240625 | 0.5 | 0.25 | 0.244650542 | 0 | 0 |
| ENSMUSG00000089736 | 1.036286563 | 1.04248125 | 1 | 0.951838731 | 0.955822902 | 1 | 1.028149525 |
| ENSMUSG00000089737 | 0 | 0 | 0 | 0 | 0 | 0 | 0 |
| ENSMUSG00000089739 | 0 | 0 | 0 | 0 | 0 | 0 | 0 |
| ENSMUSG00000089742 | 0 | 0.25 | 0.5 | 0.25 | 0.387761935 | 0 | 0.355418052 |
| ENSMUSG00000089743 | 0 | 0 | 0 | 0 | 0 | 0 | 0 |
| ENSMUSG00000089744 | 0 | 0 | 0 | 0 | 0.244650542 | 0 | 0 |
| ENSMUSG00000089745 | 0 | 0.25 | 0 | 0 | 0 | 0 | 0 |
| ENSMUSG00000089746 | 0 | 0 | 0 | 0.25 | 0 | 0 | 0 |
| ENSMUSG00000089748 | 0 | 0 | 0.25 | 0 | 0 | 0 | 0 |
| ENSMUSG00000089749 | 0 | 0 | 0 | 0 | 0 | 0 | 0 |
| ENSMUSG00000089751 | 0 | 0 | 0 | 0 | 0 | 0 | 0 |
| ENSMUSG00000089753 | 0 | 0 | 0 | 0 | 0 | 0 | 0 |
| ENSMUSG00000089754 | 0 | 0 | 0 | 0 | 0 | 0 | 0 |
| ENSMUSG00000089755 | 0 | 0 | 0 | 0 | 0 | 0 | 0 |
| ENSMUSG00000089756 | 1 | 1.160964047 | 1.04248125 | 1.226722649 | 1.074582839 | 1.204119983 | 1.066254157 |
| ENSMUSG00000089757 | 0 | 0 | 0 | 0 | 0 | 0 | 0 |
| ENSMUSG00000089759 | 0.289064826 | 0.396240625 | 0 | 0.396240625 | 0.568060967 | 0.301029996 | 0.224243824 |
| ENSMUSG00000089760 | 0 | 0 | 0 | 0.25 | 0 | 0 | 0 |
| ENSMUSG00000089761 | 0 | 0 | 0 | 0 | 0 | 0 | 0 |
| ENSMUSG00000089762 | 1.249317394 | 1.380890489 | 1.552363341 | 0.976722649 | 0.931471446 | 0.698970004 | 1.21680636 |
| ENSMUSG00000089763 | 0 | 0 | 0 | 0 | 0 | 0 | 0 |
| ENSMUSG00000089764 | 0.289064826 | 0.646240625 | 0.396240625 | 0.396240625 | 0.686820904 | 0 | 0.520678036 |
| ENSMUSG00000089766 | 0 | 0 | 0.25 | 0 | 0.244650542 | 0 | 0.224243824 |
| ENSMUSG00000089767 | 0 | 0 | 0 | 0 | 0 | 0 | 0 |
| ENSMUSG00000089768 | 0 | 0 | 0.25 | 0.396240625 | 0 | 0.301029996 | 0.520678036 |
| ENSMUSG00000089769 | 0.45815691 | 0.646240625 | 0.976722649 | 0.5 | 0.733951626 | 0 | 0.579661877 |
| ENSMUSG00000089771 | 0 | 0 | 0 | 0 | 0 | 0 | 0 |
| ENSMUSG00000089773 | 0 | 0 | 0 | 0 | 0 | 0 | 0 |

| | | | | | | | |
|---|---|---|---|---|---|---|---|
| ENSMUSG00000089774 | 2.787177998 | 2.596735061 | 2.575659731 | 2.438721876 | 2.371090306 | 2.654176542 | 2.158505762 |
| ENSMUSG00000089775 | 0 | 0 | 0 | 0 | 0 | 0 | 0 |
| ENSMUSG00000089776 | 0 | 0 | 0.396240625 | 0 | 0 | 0 | 0 |
| ENSMUSG00000089779 | 0 | 0 | 0 | 0 | 0 | 0 | 0 |
| ENSMUSG00000089780 | 0 | 0.25 | 0.25 | 0 | 0 | 0 | 0.224243824 |
| ENSMUSG00000089781 | 0 | 0 | 0 | 0 | 0 | 0 | 0 |
| ENSMUSG00000089782 | 0.289064826 | 0.396240625 | 0.580482024 | 0.701838731 | 0.244650542 | 0.301029996 | 0.672731473 |
| ENSMUSG00000089783 | 0 | 0.25 | 0 | 0 | 0 | 0 | 0 |
| ENSMUSG00000089785 | 0 | 0 | 0 | 0 | 0.387761935 | 0.301029996 | 0.355418052 |
| ENSMUSG00000089787 | 0 | 0 | 0 | 0 | 0 | 0 | 0 |
| ENSMUSG00000089788 | 2.081172036 | 1.890560606 | 1.837182039 | 1.942046081 | 1.919403394 | 2.103803721 | 1.680001789 |
| ENSMUSG00000089789 | 0.867194479 | 0.75 | 0.79248125 | 0.79248125 | 0.733951626 | 0.954242509 | 0.803905701 |
| ENSMUSG00000089790 | 0.578129653 | 0.5 | 0.5 | 0.396240625 | 0.387761935 | 0.698970004 | 0 |
| ENSMUSG00000089791 | 0.867194479 | 0.864857905 | 0.92510993 | 0.976722649 | 0.846351821 | 0.698970004 | 0.853775828 |
| ENSMUSG00000089792 | 0 | 0 | 0 | 0 | 0 | 0 | 0 |
| ENSMUSG00000089793 | 0 | 0 | 0 | 0 | 0 | 0 | 0 |
| ENSMUSG00000089794 | 0 | 0 | 0.25 | 0 | 0 | 0 | 0 |
| ENSMUSG00000089795 | 0 | 0 | 0 | 0 | 0 | 0 | 0 |
| ENSMUSG00000089796 | 0.578129653 | 0.25 | 0 | 0.25 | 0 | 0.301029996 | 0.224243824 |
| ENSMUSG00000089797 | 0 | 0 | 0 | 0 | 0 | 0 | 0 |
| ENSMUSG00000089798 | 0 | 0 | 0.25 | 0.396240625 | 0.387761935 | 0 | 0.224243824 |
| ENSMUSG00000089800 | 0.289064826 | 0.396240625 | 0.25 | 0 | 0.244650542 | 0 | 0 |
| ENSMUSG00000089802 | 0 | 0 | 0 | 0 | 0 | 0 | 0 |
| ENSMUSG00000089803 | 0 | 0 | 0 | 0 | 0 | 0 | 0 |
| ENSMUSG00000089804 | 0 | 0 | 0 | 0 | 0 | 0 | 0 |
| ENSMUSG00000089806 | 0 | 0 | 0 | 0 | 0 | 0 | 0 |
| ENSMUSG00000089807 | 0 | 0 | 0.25 | 0 | 0.244650542 | 0 | 0.448487648 |
| ENSMUSG00000089808 | 0.45815691 | 0.5 | 0.646240625 | 0.580482024 | 0.568060967 | 0.698970004 | 0.74492186 |
| ENSMUSG00000089809 | 0 | 0 | 0 | 0.25 | 0.244650542 | 0 | 0 |
| ENSMUSG00000089810 | 1.655740331 | 1.396240625 | 1.516522298 | 1.566696635 | 1.467903253 | 1.633468456 | 1.360283038 |
| ENSMUSG00000089811 | 0 | 0.396240625 | 0 | 0 | 0 | 0 | 0 |
| ENSMUSG00000089812 | 0.747221736 | 0.75 | 0.580482024 | 0.396240625 | 0.244650542 | 0.301029996 | 0.672731473 |
| ENSMUSG00000089814 | 0 | 0 | 0 | 0 | 0 | 0 | 0 |
| ENSMUSG00000089815 | 0 | 0 | 0 | 0 | 0 | 0 | 0 |

| | | | | | | | |
|---|---|---|---|---|---|---|---|
| ENSMUSG00000089816 | 0 | 0 | 0 | 0 | 0 | 0 | 0 |
| ENSMUSG00000089817 | 0.867194479 | 0.864857905 | 0.896240625 | 0.646240625 | 0.686820904 | 0.477121255 | 0.672731473 |
| ENSMUSG00000089818 | 0.578129653 | 0.646240625 | 0.79248125 | 0.580482024 | 0.812711509 | 0.301029996 | 0.710836105 |
| ENSMUSG00000089819 | 0 | 0 | 0 | 0 | 0 | 0 | 0 |
| ENSMUSG00000089820 | 0.45815691 | 0.396240625 | 0.701838731 | 0 | 0.632412477 | 0 | 0 |
| ENSMUSG00000089821 | 0 | 0 | 0 | 0 | 0 | 0 | 0 |
| ENSMUSG00000089822 | 0 | 0 | 0 | 0 | 0 | 0 | 0 |
| ENSMUSG00000089824 | 2.100572389 | 1.973704441 | 1.990000483 | 2.112802778 | 2.075570129 | 2.281033367 | 1.925124822 |
| ENSMUSG00000089826 | 0 | 0 | 0 | 0 | 0 | 0 | 0 |
| ENSMUSG00000089827 | 0.289064826 | 0 | 0 | 0 | 0 | 0 | 0 |
| ENSMUSG00000089828 | 0 | 0 | 0 | 0 | 0 | 0 | 0 |
| ENSMUSG00000089829 | 0 | 0 | 0.25 | 0 | 0.244650542 | 0 | 0 |
| ENSMUSG00000089830 | 0 | 0 | 0 | 0 | 0 | 0 | 0 |
| ENSMUSG00000089831 | 0 | 0 | 0 | 0 | 0 | 0 | 0 |
| ENSMUSG00000089832 | 1.857607105 | 1.816696635 | 1.78743678 | 1.372963274 | 1.671670246 | 1.491361694 | 1.757893036 |
| ENSMUSG00000089833 | 0 | 0 | 0 | 0 | 0 | 0 | 0 |
| ENSMUSG00000089834 | 0 | 0 | 0 | 0 | 0 | 0 | 0 |
| ENSMUSG00000089835 | 0 | 0 | 0 | 0 | 0 | 0 | 0 |
| ENSMUSG00000089836 | 0 | 0 | 0 | 0 | 0 | 0 | 0 |
| ENSMUSG00000089837 | 0 | 0 | 0 | 0 | 0 | 0 | 0 |
| ENSMUSG00000089838 | 0 | 0 | 0 | 0 | 0 | 0 | 0 |
| ENSMUSG00000089839 | 0 | 0 | 0 | 0 | 0.244650542 | 0 | 0 |
| ENSMUSG00000089840 | 0 | 0 | 0 | 0 | 0 | 0 | 0 |
| ENSMUSG00000089842 | 0.289064826 | 0.396240625 | 0.396240625 | 0.396240625 | 0.387761935 | 0.77815125 | 0.355418052 |
| ENSMUSG00000089844 | 1.100572389 | 1.02186571 | 0.976722649 | 1.080482024 | 1.091002363 | 1.079181246 | 0.984950056 |
| ENSMUSG00000089847 | 1.069666964 | 1.29248125 | 1.348079356 | 1.114857905 | 1.188507684 | 1.342422681 | 1.150210039 |
| ENSMUSG00000089849 | 0 | 0.396240625 | 0.25 | 0.25 | 0.489301084 | 0.477121255 | 0.355418052 |
| ENSMUSG00000089851 | 0 | 0.5 | 0.25 | 0.5 | 0.387761935 | 0.477121255 | 0.355418052 |
| ENSMUSG00000089853 | 0.289064826 | 0.396240625 | 0.5 | 0.396240625 | 0 | 0.301029996 | 0 |
| ENSMUSG00000089854 | 0 | 0.5 | 0.5 | 0 | 0 | 0 | 0.224243824 |
| ENSMUSG00000089855 | 1.389637216 | 1.302363341 | 1.372963274 | 1.17510993 | 1.244650542 | 1.255272505 | 1.401059159 |
| ENSMUSG00000089857 | 0.289064826 | 0.75 | 0.75 | 0.79248125 | 1 | 0.954242509 | 0.935079929 |
| ENSMUSG00000089859 | 0 | 0 | 0 | 0 | 0 | 0 | 0 |
| ENSMUSG00000089861 | 0 | 0 | 0 | 0 | 0 | 0 | 0 |

| | | | | | | | |
|---|---|---|---|---|---|---|---|
| ENSMUSG00000089862 | 1.956444026 | 1.848079356 | 1.814346961 | 1.75835575 | 1.66856048 | 1.913813852 | 1.603286636 |
| ENSMUSG00000089865 | 0 | 0 | 0 | 0 | 0 | 0 | 0 |
| ENSMUSG00000089866 | 0 | 0 | 0 | 0 | 0 | 0 | 0 |
| ENSMUSG00000089868 | 0 | 0 | 0 | 0 | 0 | 0 | 0.224243824 |
| ENSMUSG00000089870 | 0.289064826 | 0 | 0 | 0 | 0.244650542 | 0 | 0 |
| ENSMUSG00000089871 | 0 | 0.25 | 0 | 0 | 0 | 0 | 0.224243824 |
| ENSMUSG00000089872 | 2.219867581 | 2.08272922 | 2.12892496 | 2.095926073 | 2.072599934 | 2.322219295 | 1.997315209 |
| ENSMUSG00000089873 | 0 | 0 | 0 | 0 | 0 | 0 | 0 |
| ENSMUSG00000089874 | 0 | 0 | 0 | 0 | 0 | 0 | 0 |
| ENSMUSG00000089875 | 0.91631382 | 0.92510993 | 0.951838731 | 1.146240625 | 0.812711509 | 0.954242509 | 0.916588299 |
| ENSMUSG00000089876 | 0.289064826 | 0 | 0.25 | 0.580482024 | 0.387761935 | 0.602059991 | 0.74492186 |
| ENSMUSG00000089877 | 0 | 0 | 0.25 | 0.25 | 0.244650542 | 0.301029996 | 0 |
| ENSMUSG00000089879 | 0 | 0 | 0 | 0 | 0 | 0 | 0 |
| ENSMUSG00000089881 | 0 | 0 | 0 | 0 | 0 | 0 | 0 |
| ENSMUSG00000089883 | 0.289064826 | 0.5 | 0.5 | 0.580482024 | 0.244650542 | 0 | 0.355418052 |
| ENSMUSG00000089884 | 0 | 0 | 0 | 0 | 0 | 0 | 0 |
| ENSMUSG00000089885 | 0 | 0 | 0 | 0.25 | 0 | 0 | 0 |
| ENSMUSG00000089886 | 0.960252568 | 0.5 | 0.701838731 | 0.25 | 0.387761935 | 0.477121255 | 0.520678036 |
| ENSMUSG00000089887 | 0 | 0 | 0 | 0 | 0 | 0 | 0 |
| ENSMUSG00000089888 | 0 | 0 | 0 | 0 | 0 | 0 | 0 |
| ENSMUSG00000089889 | 0.671187741 | 0.79248125 | 0.701838731 | 0.92510993 | 0.846351821 | 0.602059991 | 0.520678036 |
| ENSMUSG00000089890 | 0 | 0 | 0 | 0 | 0 | 0 | 0 |
| ENSMUSG00000089891 | 0 | 0 | 0 | 0 | 0 | 0 | 0 |
| ENSMUSG00000089892 | 0 | 0 | 0 | 0 | 0 | 0 | 0 |
| ENSMUSG00000089893 | 0 | 0 | 0 | 0 | 0 | 0 | 0 |
| ENSMUSG00000089894 | 0 | 0 | 0 | 0 | 0 | 0 | 0 |
| ENSMUSG00000089898 | 0 | 0 | 0 | 0 | 0 | 0 | 0 |
| ENSMUSG00000089900 | 0 | 0.396240625 | 0.25 | 0.396240625 | 0.568060967 | 0 | 0.224243824 |
| ENSMUSG00000089901 | 2.167941649 | 1.990000483 | 1.933677405 | 1.766522298 | 1.680838453 | 1.763427994 | 1.535058868 |
| ENSMUSG00000089902 | 0.289064826 | 0.25 | 0.25 | 0 | 0.387761935 | 0 | 0 |
| ENSMUSG00000089904 | 0 | 0 | 0 | 0 | 0 | 0 | 0 |
| ENSMUSG00000089905 | 0 | 0 | 0 | 0 | 0 | 0 | 0 |
| ENSMUSG00000089906 | 0 | 0 | 0 | 0 | 0 | 0 | 0 |
| ENSMUSG00000089907 | 0 | 0 | 0 | 0.25 | 0.489301084 | 0 | 0.224243824 |

| | | | | | | | |
|---|---|---|---|---|---|---|---|
| ENSMUSG00000089908 | 0 | 0 | 0 | 0.25 | 0.244650542 | 0 | 0 |
| ENSMUSG00000089909 | 0 | 0 | 0 | 0 | 0 | 0 | 0 |
| ENSMUSG00000089910 | 0 | 0 | 0 | 0 | 0 | 0 | 0 |
| ENSMUSG00000089911 | 2.360187403 | 2.186208459 | 2.129909063 | 2.33993739 | 2.289330171 | 2.57634135 | 2.130728307 |
| ENSMUSG00000089914 | 0 | 0 | 0 | 0 | 0 | 0 | 0 |
| ENSMUSG00000089916 | 0 | 0 | 0 | 0 | 0 | 0 | 0 |
| ENSMUSG00000089917 | 1.678702042 | 1.634789703 | 1.653677461 | 1.566696635 | 1.563883923 | 1.633468456 | 1.517723536 |
| ENSMUSG00000089922 | 1.129344651 | 1.130890489 | 1.114857905 | 1.04248125 | 1.091002363 | 0.954242509 | 0.896975297 |
| ENSMUSG00000089924 | 0 | 0.25 | 0 | 0 | 0 | 0 | 0 |
| ENSMUSG00000089926 | 0 | 0 | 0 | 0 | 0 | 0 | 0 |
| ENSMUSG00000089927 | 0 | 0 | 0 | 0 | 0 | 0 | 0 |
| ENSMUSG00000089929 | 0 | 0 | 0 | 0 | 0 | 0 | 0 |
| ENSMUSG00000089931 | 0 | 0 | 0 | 0 | 0 | 0 | 0 |
| ENSMUSG00000089932 | 0 | 0 | 0 | 0 | 0 | 0 | 0 |
| ENSMUSG00000089933 | 0 | 0 | 0 | 0 | 0 | 0 | 0 |
| ENSMUSG00000089934 | 0 | 0 | 0 | 0 | 0 | 0 | 0 |
| ENSMUSG00000089935 | 0 | 0 | 0 | 0 | 0 | 0 | 0 |
| ENSMUSG00000089936 | 0 | 0 | 0 | 0 | 0 | 0 | 0 |
| ENSMUSG00000089937 | 0 | 0 | 0 | 0 | 0 | 0 | 0 |
| ENSMUSG00000089938 | 0 | 0 | 0.25 | 0 | 0.244650542 | 0 | 0 |
| ENSMUSG00000089940 | 1.800532393 | 1.705044741 | 1.646240625 | 1.723704441 | 1.592135487 | 1.770852012 | 1.479989695 |
| ENSMUSG00000089941 | 0 | 0 | 0 | 0 | 0 | 0 | 0 |
| ENSMUSG00000089942 | 0 | 0 | 0 | 0 | 0 | 0 | 0 |
| ENSMUSG00000089943 | 0 | 0 | 0 | 0 | 0 | 0 | 0 |
| ENSMUSG00000089945 | 0 | 0 | 0 | 0 | 0 | 0 | 0 |
| ENSMUSG00000089947 | 0 | 0 | 0 | 0 | 0 | 0 | 0 |
| ENSMUSG00000089948 | 0.578129653 | 0 | 0.25 | 0.396240625 | 0.387761935 | 0.698970004 | 0.629532004 |
| ENSMUSG00000089949 | 0 | 0 | 0 | 0 | 0 | 0 | 0 |
| ENSMUSG00000089950 | 0 | 0 | 0 | 0 | 0 | 0 | 0 |
| ENSMUSG00000089951 | 0.578129653 | 0.864857905 | 0.896240625 | 0.701838731 | 0.877063019 | 0.77815125 | 0.74492186 |
| ENSMUSG00000089952 | 0.671187741 | 0.396240625 | 0.701838731 | 0.864857905 | 0.686820904 | 0.77815125 | 0.629532004 |
| ENSMUSG00000089953 | 0 | 0 | 0.25 | 0 | 0 | 0 | 0 |
| ENSMUSG00000089956 | 0 | 0 | 0 | 0 | 0 | 0 | 0 |
| ENSMUSG00000089957 | 0 | 0 | 0 | 0 | 0 | 0 | 0 |

| | | | | | | | |
|---|---|---|---|---|---|---|---|
| ENSMUSG00000089958 | 0 | 0.396240625 | 0.5 | 0 | 0 | 0.301029996 | 0.224243824 |
| ENSMUSG00000089959 | 0 | 0 | 0.25 | 0.396240625 | 0.244650542 | 0.477121255 | 0.224243824 |
| ENSMUSG00000089960 | 0 | 0 | 0 | 0 | 0 | 0 | 0 |
| ENSMUSG00000089961 | 0 | 0 | 0 | 0 | 0 | 0 | 0 |
| ENSMUSG00000089963 | 0 | 0 | 0 | 0 | 0 | 0 | 0 |
| ENSMUSG00000089964 | 0 | 0 | 0 | 0 | 0 | 0 | 0 |
| ENSMUSG00000089965 | 0 | 0.25 | 0.396240625 | 0 | 0 | 0 | 0.224243824 |
| ENSMUSG00000089966 | 0 | 0 | 0 | 0 | 0 | 0 | 0 |
| ENSMUSG00000089967 | 0 | 0 | 0 | 0 | 0 | 0 | 0 |
| ENSMUSG00000089968 | 0.289064826 | 0.396240625 | 0.580482024 | 0.396240625 | 0.244650542 | 0.602059991 | 0.579661877 |
| ENSMUSG00000089972 | 0 | 0 | 0 | 0 | 0 | 0 | 0 |
| ENSMUSG00000089973 | 0 | 0 | 0 | 0.25 | 0 | 0 | 0 |
| ENSMUSG00000089975 | 0.289064826 | 0.25 | 0.25 | 0 | 0.244650542 | 0 | 0 |
| ENSMUSG00000089976 | 0 | 0 | 0 | 0 | 0 | 0 | 0 |
| ENSMUSG00000089978 | 0 | 0 | 0 | 0 | 0 | 0 | 0 |
| ENSMUSG00000089979 | 0 | 0.25 | 0.580482024 | 0.25 | 0.244650542 | 0.477121255 | 0.224243824 |
| ENSMUSG00000089982 | 0 | 0 | 0 | 0 | 0 | 0 | 0 |
| ENSMUSG00000089983 | 0 | 0 | 0 | 0 | 0 | 0 | 0 |
| ENSMUSG00000089984 | 0.45815691 | 0.75 | 0.580482024 | 0.580482024 | 0.632412477 | 0.84509804 | 0.853775828 |
| ENSMUSG00000089987 | 0 | 0 | 0 | 0 | 0 | 0 | 0 |
| ENSMUSG00000089988 | 0.811507563 | 0.580482024 | 0.701838731 | 0.701838731 | 0.77552387 | 0.477121255 | 0.710836105 |
| ENSMUSG00000089989 | 0.289064826 | 0.79248125 | 0.896240625 | 1.098079356 | 1.057362051 | 0.77815125 | 0.952571511 |
| ENSMUSG00000089990 | 0 | 0 | 0 | 0 | 0 | 0 | 0 |
| ENSMUSG00000089991 | 0 | 0 | 0 | 0 | 0 | 0 | 0 |
| ENSMUSG00000089992 | 0 | 0 | 0 | 0 | 0.244650542 | 0 | 0.224243824 |
| ENSMUSG00000089993 | 1.227926429 | 1.226722649 | 1.160964047 | 1.226722649 | 1.234113756 | 1.255272505 | 1.121219121 |
| ENSMUSG00000089994 | 0.578129653 | 0.580482024 | 0 | 0.396240625 | 0 | 0.301029996 | 0.448487648 |
| ENSMUSG00000089995 | 0 | 0 | 0 | 0 | 0.244650542 | 0 | 0 |
| ENSMUSG00000089996 | 0 | 0 | 0 | 0 | 0 | 0 | 0 |
| ENSMUSG00000089997 | 0 | 0 | 0 | 0 | 0 | 0 | 0 |
| ENSMUSG00000089998 | 0.867194479 | 0.646240625 | 0.646240625 | 0.646240625 | 0.568060967 | 0.602059991 | 0.803905701 |
| ENSMUSG00000089999 | 0 | 0 | 0 | 0 | 0.244650542 | 0 | 0.224243824 |
| ENSMUSG00000090000 | 2.40033094 | 2.142463902 | 2.127938163 | 2.102347734 | 2.006054284 | 2.352182518 | 1.903345709 |
| ENSMUSG00000090001 | 0.578129653 | 0.580482024 | 0.396240625 | 0.396240625 | 0.489301084 | 0 | 0 |

| | | | | | | | |
|---|---|---|---|---|---|---|---|
| ENSMUSG00000090002 | 0 | 0.25 | 0 | 0.396240625 | 0 | 0 | 0.224243824 |
| ENSMUSG00000090003 | 0 | 0 | 0 | 0 | 0 | 0 | 0 |
| ENSMUSG00000090004 | 0.289064826 | 0.25 | 0 | 0 | 0 | 0.301029996 | 0.224243824 |
| ENSMUSG00000090005 | 0.289064826 | 0.25 | 0.25 | 0 | 0 | 0 | 0 |
| ENSMUSG00000090006 | 0 | 0 | 0 | 0 | 0 | 0 | 0 |
| ENSMUSG00000090007 | 0 | 0.25 | 0.25 | 0 | 0 | 0 | 0 |
| ENSMUSG00000090008 | 0.45815691 | 0.646240625 | 0.580482024 | 0 | 0 | 0 | 0 |
| ENSMUSG00000090009 | 0 | 0 | 0 | 0 | 0 | 0 | 0 |
| ENSMUSG00000090012 | 0 | 0.25 | 0 | 0 | 0 | 0 | 0 |
| ENSMUSG00000090013 | 0 | 0 | 0 | 0 | 0 | 0 | 0 |
| ENSMUSG00000090015 | 1.527823874 | 1.470660762 | 1.580482024 | 1.464495249 | 1.373641807 | 1.556302501 | 1.340368108 |
| ENSMUSG00000090016 | 0 | 0 | 0 | 0 | 0 | 0 | 0 |
| ENSMUSG00000090017 | 0 | 0 | 0 | 0 | 0 | 0 | 0 |
| ENSMUSG00000090018 | 0 | 0 | 0 | 0 | 0 | 0 | 0 |
| ENSMUSG00000090019 | 0.289064826 | 0 | 0.396240625 | 0.396240625 | 0.244650542 | 0 | 0 |
| ENSMUSG00000090021 | 0 | 0 | 0.25 | 0 | 0.387761935 | 0 | 0.224243824 |
| ENSMUSG00000090023 | 0.289064826 | 0.25 | 0.25 | 0 | 0.244650542 | 0.477121255 | 0.224243824 |
| ENSMUSG00000090024 | 0.45815691 | 0.396240625 | 0.5 | 0.25 | 0.244650542 | 0 | 0.448487648 |
| ENSMUSG00000090026 | 0.45815691 | 0 | 0 | 0.25 | 0 | 0 | 0.224243824 |
| ENSMUSG00000090027 | 0 | 0 | 0 | 0 | 0 | 0 | 0 |
| ENSMUSG00000090030 | 0 | 0 | 0.25 | 0 | 0 | 0 | 0.224243824 |
| ENSMUSG00000090031 | 1.342375483 | 1.330482024 | 1.356566189 | 1.226722649 | 1.351342427 | 1.204119983 | 1.100339912 |
| ENSMUSG00000090033 | 0 | 0 | 0 | 0 | 0 | 0 | 0 |
| ENSMUSG00000090034 | 0 | 0 | 0.25 | 0.25 | 0 | 0 | 0.224243824 |
| ENSMUSG00000090035 | 2.185978293 | 2.059601185 | 1.976722649 | 2.12892496 | 2.070605826 | 2.294466226 | 1.938331368 |
| ENSMUSG00000090036 | 0 | 0 | 0 | 0 | 0 | 0 | 0 |
| ENSMUSG00000090038 | 0 | 0 | 0 | 0.25 | 0 | 0 | 0 |
| ENSMUSG00000090039 | 0 | 0 | 0 | 0 | 0.387761935 | 0 | 0.672731473 |
| ENSMUSG00000090040 | 0 | 0 | 0 | 0 | 0 | 0 | 0 |
| ENSMUSG00000090041 | 0 | 0 | 0.25 | 0 | 0.387761935 | 0 | 0.355418052 |
| ENSMUSG00000090042 | 0 | 0 | 0 | 0 | 0 | 0 | 0 |
| ENSMUSG00000090043 | 0 | 0 | 0 | 0 | 0 | 0 | 0.224243824 |
| ENSMUSG00000090044 | 0 | 0 | 0 | 0 | 0 | 0 | 0 |
| ENSMUSG00000090047 | 0 | 0 | 0 | 0 | 0 | 0 | 0 |

| | | | | | | | |
|---|---|---|---|---|---|---|---|
| ENSMUSG00000090048 | 0 | 0.25 | 0.25 | 0 | 0.244650542 | 0 | 0 |
| ENSMUSG00000090050 | 0 | 0 | 0 | 0 | 0 | 0 | 0 |
| ENSMUSG00000090051 | 0 | 0 | 0 | 0 | 0 | 0 | 0 |
| ENSMUSG00000090053 | 1.647796617 | 1.571350555 | 1.52186571 | 1.348079356 | 1.254881871 | 1.278753601 | 1.140832123 |
| ENSMUSG00000090054 | 0 | 0 | 0 | 0 | 0 | 0 | 0 |
| ENSMUSG00000090057 | 0 | 0 | 0 | 0 | 0 | 0 | 0 |
| ENSMUSG00000090059 | 0 | 0 | 0 | 0 | 0 | 0 | 0 |
| ENSMUSG00000090061 | 0 | 0.25 | 0 | 0.25 | 0.244650542 | 0 | 0 |
| ENSMUSG00000090062 | 0 | 0 | 0 | 0 | 0 | 0 | 0 |
| ENSMUSG00000090063 | 0.867194479 | 0.92510993 | 0.976722649 | 0.951838731 | 1.020174412 | 1 | 0.984950056 |
| ENSMUSG00000090066 | 0 | 0 | 0 | 0.701838731 | 0.77552387 | 0.477121255 | 0.629532004 |
| ENSMUSG00000090067 | 0 | 0.25 | 0 | 0 | 0 | 0 | 0.224243824 |
| ENSMUSG00000090069 | 0.578129653 | 0.646240625 | 0.396240625 | 0.646240625 | 0.632412477 | 0.301029996 | 0.520678036 |
| ENSMUSG00000090070 | 0.578129653 | 0.25 | 0.5 | 0.396240625 | 0.489301084 | 0.301029996 | 0.355418052 |
| ENSMUSG00000090071 | 0 | 0 | 0 | 0.396240625 | 0.387761935 | 0.301029996 | 0.629532004 |
| ENSMUSG00000090073 | 0 | 0 | 0 | 0 | 0 | 0 | 0 |
| ENSMUSG00000090074 | 0 | 0 | 0 | 0 | 0 | 0 | 0 |
| ENSMUSG00000090075 | 0 | 0 | 0 | 0 | 0 | 0 | 0 |
| ENSMUSG00000090077 | 0.289064826 | 0.396240625 | 0.396240625 | 0 | 0.77552387 | 0.477121255 | 0.579661877 |
| ENSMUSG00000090078 | 0 | 0.25 | 0.396240625 | 0 | 0.244650542 | 0 | 0.224243824 |
| ENSMUSG00000090079 | 0 | 0 | 0.25 | 0 | 0 | 0 | 0 |
| ENSMUSG00000090080 | 0 | 0 | 0 | 0 | 0 | 0 | 0 |
| ENSMUSG00000090081 | 0 | 0 | 0 | 0 | 0 | 0 | 0 |
| ENSMUSG00000090083 | 1.587501561 | 1.575945187 | 1.488549078 | 1.431980114 | 1.499532413 | 1.591064607 | 1.455757965 |
| ENSMUSG00000090084 | 0 | 0 | 0 | 0.25 | 0 | 0 | 0 |
| ENSMUSG00000090086 | 1.036286563 | 1.098079356 | 1.080482024 | 1.02186571 | 0.978602168 | 0.698970004 | 0.896975297 |
| ENSMUSG00000090087 | 0.289064826 | 0 | 0.25 | 0 | 0 | 0 | 0 |
| ENSMUSG00000090088 | 0 | 0 | 0 | 0 | 0 | 0 | 0 |
| ENSMUSG00000090089 | 0 | 0 | 0 | 0 | 0 | 0 | 0 |
| ENSMUSG00000090090 | 0 | 0 | 0 | 0 | 0 | 0 | 0 |
| ENSMUSG00000090093 | 0.91631382 | 0.951838731 | 0.864857905 | 0.864857905 | 0.877063019 | 0.77815125 | 0.896975297 |
| ENSMUSG00000090095 | 0.289064826 | 0 | 0 | 0 | 0 | 0 | 0 |
| ENSMUSG00000090097 | 0 | 0 | 0 | 0 | 0 | 0 | 0 |
| ENSMUSG00000090098 | 0 | 0 | 0 | 0 | 0 | 0 | 0 |

| | | | | | | | |
|---|---|---|---|---|---|---|---|
| ENSMUSG00000090099 | 0 | 0 | 0 | 0 | 0 | 0 | 0 |
| ENSMUSG00000090100 | 2.35727108 | 2.183677405 | 2.173371739 | 2.281676118 | 2.164820399 | 2.463892989 | 2.041943747 |
| ENSMUSG00000090101 | 0 | 0.25 | 0.396240625 | 0.396240625 | 0 | 0 | 0.224243824 |
| ENSMUSG00000090102 | 0 | 0 | 0 | 0 | 0 | 0 | 0 |
| ENSMUSG00000090103 | 0 | 0 | 0 | 0 | 0 | 0 | 0 |
| ENSMUSG00000090104 | 1.289064826 | 1.282320754 | 1.321350555 | 1.188721876 | 1.149965125 | 1.255272505 | 1.014380832 |
| ENSMUSG00000090105 | 0 | 0 | 0 | 0 | 0 | 0 | 0.224243824 |
| ENSMUSG00000090106 | 0 | 0 | 0 | 0 | 0 | 0 | 0 |
| ENSMUSG00000090107 | 0 | 0 | 0 | 0 | 0 | 0 | 0 |
| ENSMUSG00000090108 | 0 | 0.646240625 | 0.25 | 0.25 | 0 | 0.602059991 | 0 |
| ENSMUSG00000090109 | 0 | 0 | 0 | 0 | 0 | 0 | 0 |
| ENSMUSG00000090110 | 1.389637216 | 1.282320754 | 1.29248125 | 1.380890489 | 1.42077253 | 1.322219295 | 1.238624656 |
| ENSMUSG00000090111 | 0 | 0 | 0 | 0 | 0 | 0 | 0 |
| ENSMUSG00000090112 | 2.576402808 | 2.393411797 | 2.354463129 | 2.491446071 | 2.422990063 | 2.760422483 | 2.21302251 |
| ENSMUSG00000090113 | 0 | 0.25 | 0.25 | 0 | 0 | 0 | 0.224243824 |
| ENSMUSG00000090114 | 0 | 0 | 0 | 0 | 0 | 0 | 0 |
| ENSMUSG00000090115 | 2.086807695 | 1.916333979 | 1.950224975 | 2.051142786 | 2.022473809 | 2.225309282 | 1.863093717 |
| ENSMUSG00000090116 | 0 | 0 | 0 | 0 | 0 | 0 | 0 |
| ENSMUSG00000090117 | 0 | 0 | 0 | 0 | 0 | 0 | 0 |
| ENSMUSG00000090118 | 0 | 0 | 0 | 0 | 0 | 0 | 0 |
| ENSMUSG00000090119 | 0 | 0.25 | 0 | 0.25 | 0 | 0 | 0 |
| ENSMUSG00000090120 | 0.671187741 | 0.580482024 | 0.25 | 0.396240625 | 0.387761935 | 0.698970004 | 0.520678036 |
| ENSMUSG00000090121 | 0 | 0 | 0 | 0 | 0 | 0 | 0 |
| ENSMUSG00000090122 | 0 | 0 | 0 | 0 | 0 | 0 | 0 |
| ENSMUSG00000090124 | 0.289064826 | 0 | 0 | 0 | 0 | 0 | 0 |
| ENSMUSG00000090125 | 0.289064826 | 0.25 | 0.25 | 1.27186571 | 1.293076518 | 1.477121255 | 1.429563222 |
| ENSMUSG00000090126 | 0 | 0.25 | 0 | 0 | 0 | 0 | 0.224243824 |
| ENSMUSG00000090128 | 0 | 0 | 0 | 0 | 0 | 0 | 0 |
| ENSMUSG00000090129 | 0 | 0 | 0 | 0.396240625 | 0 | 0 | 0 |
| ENSMUSG00000090131 | 0 | 0 | 0 | 0 | 0 | 0 | 0 |
| ENSMUSG00000090132 | 0 | 0 | 0 | 0 | 0 | 0 | 0 |
| ENSMUSG00000090135 | 0 | 0 | 0 | 0 | 0 | 0 | 0 |
| ENSMUSG00000090136 | 0.289064826 | 0.25 | 0.580482024 | 0.396240625 | 0.244650542 | 0 | 0.448487648 |
| ENSMUSG00000090137 | 1.069666964 | 0.951838731 | 1 | 1.061981878 | 1 | 1 | 0.829800754 |

| | | | | | | | |
|---|---|---|---|---|---|---|---|
| ENSMUSG00000090138 | 0 | 0 | 0 | 0 | 0 | 0 | 0 |
| ENSMUSG00000090141 | 0 | 0 | 0 | 0 | 0 | 0 | 0 |
| ENSMUSG00000090142 | 0 | 0 | 0 | 0 | 0 | 0 | 0.355418052 |
| ENSMUSG00000090145 | 0 | 0 | 0 | 0 | 0 | 0 | 0 |
| ENSMUSG00000090146 | 0 | 0 | 0 | 0 | 0 | 0 | 0 |
| ENSMUSG00000090147 | 0 | 0 | 0 | 0 | 0 | 0 | 0 |
| ENSMUSG00000090150 | 1.975148165 | 1.763820609 | 1.766522298 | 1.830482024 | 1.795698282 | 2.103803721 | 1.705896842 |
| ENSMUSG00000090151 | 0 | 0 | 0 | 0 | 0 | 0 | 0 |
| ENSMUSG00000090152 | 0 | 0 | 0 | 0 | 0 | 0 | 0 |
| ENSMUSG00000090157 | 0 | 0 | 0 | 0 | 0 | 0 | 0 |
| ENSMUSG00000090159 | 0 | 0 | 0 | 0 | 0 | 0 | 0 |
| ENSMUSG00000090160 | 0 | 0 | 0 | 0.25 | 0 | 0 | 0 |
| ENSMUSG00000090162 | 0 | 0 | 0.25 | 0 | 0 | 0 | 0 |
| ENSMUSG00000090163 | 0 | 0 | 0 | 0 | 0 | 0 | 0 |
| ENSMUSG00000090164 | 0 | 0 | 0 | 0 | 0 | 0 | 0 |
| ENSMUSG00000090165 | 0 | 0 | 0 | 0 | 0 | 0 | 0 |
| ENSMUSG00000090166 | 0 | 0 | 0 | 0 | 0 | 0 | 0 |
| ENSMUSG00000090168 | 0 | 0 | 0 | 0 | 0 | 0 | 0 |
| ENSMUSG00000090171 | 0 | 0 | 0 | 0 | 0 | 0 | 0 |
| ENSMUSG00000090173 | 0.289064826 | 0.25 | 0.580482024 | 0.5 | 0.489301084 | 0.301029996 | 0.448487648 |
| ENSMUSG00000090174 | 0 | 0 | 0 | 0 | 0 | 0 | 0 |
| ENSMUSG00000090175 | 0 | 0 | 0 | 0 | 0 | 0 | 0 |
| ENSMUSG00000090176 | 0 | 0 | 0 | 0 | 0.244650542 | 0 | 0.224243824 |
| ENSMUSG00000090179 | 0 | 0 | 0 | 0 | 0 | 0 | 0 |
| ENSMUSG00000090180 | 0 | 0 | 0 | 0 | 0 | 0 | 0 |
| ENSMUSG00000090185 | 0 | 0 | 0 | 0 | 0 | 0 | 0 |
| ENSMUSG00000090186 | 0 | 0 | 0 | 0 | 0 | 0 | 0 |
| ENSMUSG00000090188 | 0.45815691 | 0.646240625 | 0.646240625 | 0.396240625 | 0.733951626 | 0.698970004 | 0.629532004 |
| ENSMUSG00000090191 | 0 | 0 | 0.396240625 | 0.396240625 | 0.244650542 | 0.477121255 | 0 |
| ENSMUSG00000090192 | 0 | 0 | 0.25 | 0 | 0 | 0 | 0 |
| ENSMUSG00000090193 | 0 | 0 | 0 | 0 | 0 | 0 | 0 |
| ENSMUSG00000090194 | 0.578129653 | 0.396240625 | 0.5 | 0.5 | 0 | 0.602059991 | 0.355418052 |
| ENSMUSG00000090197 | 0 | 0 | 0 | 0.25 | 0 | 0 | 0 |
| ENSMUSG00000090199 | 0 | 0 | 0 | 0 | 0 | 0 | 0 |

| | | | | | | | |
|---|---|---|---|---|---|---|---|
| ENSMUSG00000090200 | 0 | 0 | 0.25 | 0 | 0 | 0 | 0.224243824 |
| ENSMUSG00000090202 | 0 | 0 | 0 | 0 | 0 | 0 | 0 |
| ENSMUSG00000090203 | 0 | 0 | 0 | 0 | 0 | 0 | 0 |
| ENSMUSG00000090206 | 0.45815691 | 0.701838731 | 0.5 | 0.75 | 0.77552387 | 0.77815125 | 0.803905701 |
| ENSMUSG00000090207 | 0.289064826 | 0 | 0 | 0 | 0 | 0 | 0 |
| ENSMUSG00000090208 | 0 | 0 | 0 | 0 | 0 | 0 | 0 |
| ENSMUSG00000090210 | 1 | 1 | 0.830482024 | 1.114857905 | 1.074582839 | 1.079181246 | 0.935079929 |
| ENSMUSG00000090211 | 0.289064826 | 0.25 | 0.396240625 | 0.25 | 0.244650542 | 0.477121255 | 0 |
| ENSMUSG00000090212 | 0 | 0 | 0 | 0 | 0 | 0 | 0 |
| ENSMUSG00000090213 | 2.113897241 | 1.997171172 | 2.073655187 | 2.034887838 | 2.103973339 | 2.320146286 | 2.02383181 |
| ENSMUSG00000090214 | 0 | 0 | 0 | 0 | 0 | 0 | 0 |
| ENSMUSG00000090215 | 0 | 0.25 | 0.396240625 | 0 | 0 | 0 | 0.355418052 |
| ENSMUSG00000090216 | 0 | 0 | 0 | 0 | 0 | 0 | 0 |
| ENSMUSG00000090217 | 0.45815691 | 0.25 | 0.25 | 0 | 0 | 0.477121255 | 0 |
| ENSMUSG00000090219 | 0 | 0 | 0 | 0 | 0 | 0 | 0 |
| ENSMUSG00000090220 | 0 | 0 | 0 | 0.25 | 0 | 0 | 0.224243824 |
| ENSMUSG00000090222 | 0 | 0 | 0 | 0 | 0 | 0.301029996 | 0 |
| ENSMUSG00000090223 | 0 | 0 | 0 | 0 | 0 | 0 | 0 |
| ENSMUSG00000090224 | 0 | 0 | 0 | 0.25 | 0 | 0 | 0.224243824 |
| ENSMUSG00000090225 | 0 | 0 | 0 | 0 | 0 | 0 | 0 |
| ENSMUSG00000090226 | 0 | 0 | 0 | 0 | 0 | 0 | 0 |
| ENSMUSG00000090227 | 0 | 0 | 0 | 0 | 0 | 0 | 0 |
| ENSMUSG00000090228 | 0 | 0 | 0 | 0 | 0 | 0 | 0 |
| ENSMUSG00000090230 | 0 | 0 | 0.5 | 0 | 0.244650542 | 0 | 0 |
| ENSMUSG00000090231 | 0 | 0.25 | 0 | 0.396240625 | 0.387761935 | 0.301029996 | 0.355418052 |
| ENSMUSG00000090232 | 0 | 0 | 0 | 0 | 0 | 0 | 0 |
| ENSMUSG00000090235 | 0.91631382 | 0.830482024 | 0.646240625 | 0.79248125 | 0.846351821 | 0.84509804 | 0.448487648 |
| ENSMUSG00000090236 | 0 | 0 | 0.25 | 0.396240625 | 0 | 0.301029996 | 0.224243824 |
| ENSMUSG00000090237 | 0 | 0 | 0 | 0 | 0 | 0 | 0 |
| ENSMUSG00000090242 | 0 | 0.25 | 0 | 0 | 0 | 0 | 0 |
| ENSMUSG00000090243 | 1.714367558 | 1.606566189 | 1.62694866 | 1.53743678 | 1.580303447 | 1.653212514 | 1.486592281 |
| ENSMUSG00000090244 | 0 | 0.25 | 0 | 0 | 0 | 0 | 0.224243824 |
| ENSMUSG00000090246 | 0 | 0 | 0 | 0 | 0 | 0 | 0 |
| ENSMUSG00000090247 | 0.45815691 | 0.5 | 0.580482024 | 0.646240625 | 0.489301084 | 0.602059991 | 0.629532004 |

| | | | | | | | |
|---|---|---|---|---|---|---|---|
| ENSMUSG00000090248 | 0.578129653 | 0.25 | 0.701838731 | 0 | 0.568060967 | 0.477121255 | 0.224243824 |
| ENSMUSG00000090249 | 0 | 0 | 0 | 0 | 0 | 0 | 0 |
| ENSMUSG00000090252 | 0 | 0 | 0 | 0 | 0 | 0 | 0 |
| ENSMUSG00000090254 | 0 | 0 | 0 | 0 | 0 | 0 | 0 |
| ENSMUSG00000090255 | 0 | 0 | 0 | 0 | 0 | 0 | 0 |
| ENSMUSG00000090256 | 0.289064826 | 0 | 0.5 | 0.396240625 | 0 | 0 | 0 |
| ENSMUSG00000090257 | 1.036286563 | 0.79248125 | 0.896240625 | 1.02186571 | 0.686820904 | 0.84509804 | 0.876096088 |
| ENSMUSG00000090258 | 1.671187741 | 1.602347734 | 1.589388001 | 1.622963274 | 1.528558853 | 1.763427994 | 1.462868481 |
| ENSMUSG00000090259 | 0 | 0 | 0 | 0 | 0 | 0 | 0 |
| ENSMUSG00000090260 | 0.289064826 | 0.25 | 0 | 0 | 0 | 0 | 0.224243824 |
| ENSMUSG00000090263 | 0 | 0 | 0.25 | 0 | 0 | 0 | 0 |
| ENSMUSG00000090264 | 0.45815691 | 0.396240625 | 0.580482024 | 0 | 0.244650542 | 0.301029996 | 0.672731473 |
| ENSMUSG00000090265 | 0 | 0 | 0 | 0 | 0 | 0 | 0 |
| ENSMUSG00000090266 | 1.827447047 | 1.488549078 | 1.575945187 | 1.660964047 | 1.519146132 | 1.672097858 | 1.401059159 |
| ENSMUSG00000090268 | 0 | 0 | 0 | 0 | 0 | 0 | 0 |
| ENSMUSG00000090270 | 0 | 0 | 0 | 0 | 0.387761935 | 0 | 0.224243824 |
| ENSMUSG00000090272 | 0.747221736 | 0.896240625 | 0.864857905 | 0.75 | 0.489301084 | 0.301029996 | 0.579661877 |
| ENSMUSG00000090273 | 0.747221736 | 0.646240625 | 0.92510993 | 0.396240625 | 0.632412477 | 0.477121255 | 1.131174228 |
| ENSMUSG00000090277 | 0 | 0 | 0 | 0 | 0 | 0 | 0 |
| ENSMUSG00000090284 | 0 | 0.25 | 0 | 0 | 0 | 0 | 0.224243824 |
| ENSMUSG00000090286 | 0.671187741 | 0.75 | 0.701838731 | 0.646240625 | 0.568060967 | 0.477121255 | 0.355418052 |
| ENSMUSG00000090290 | 1.989530018 | 1.896240625 | 1.890560606 | 1.671625132 | 1.692703642 | 1.851258349 | 1.610050272 |
| ENSMUSG00000090291 | 0 | 0.25 | 0 | 0.396240625 | 0.244650542 | 0.301029996 | 0.355418052 |
| ENSMUSG00000090292 | 0 | 0 | 0 | 0 | 0 | 0 | 0 |
| ENSMUSG00000090293 | 0 | 0.396240625 | 0 | 0 | 0 | 0 | 0 |
| ENSMUSG00000090298 | 0 | 0 | 0 | 0 | 0 | 0 | 0 |
| ENSMUSG00000090300 | 0 | 0 | 0 | 0 | 0 | 0 | 0 |
| ENSMUSG00000090302 | 0 | 0 | 0 | 0 | 0 | 0 | 0 |
| ENSMUSG00000090303 | 0.289064826 | 0 | 0 | 0 | 0 | 0 | 0 |
| ENSMUSG00000090304 | 0 | 0 | 0 | 0 | 0 | 0 | 0 |
| ENSMUSG00000090305 | 0.289064826 | 0 | 0.5 | 0.396240625 | 0.387761935 | 0 | 0.355418052 |
| ENSMUSG00000090306 | 0 | 0 | 0 | 0 | 0 | 0 | 0 |
| ENSMUSG00000090307 | 0.289064826 | 0 | 0 | 0.25 | 0 | 0 | 0.224243824 |
| ENSMUSG00000090308 | 0 | 0 | 0 | 0 | 0 | 0 | 0 |

| | | | | | | | |
|---|---|---|---|---|---|---|---|
| ENSMUSG00000090314 | 0 | 0 | 0 | 0 | 0 | 0 | 0 |
| ENSMUSG00000090315 | 0 | 0 | 0 | 0 | 0 | 0 | 0 |
| ENSMUSG00000090317 | 0 | 0 | 0 | 0.25 | 0 | 0 | 0 |
| ENSMUSG00000090318 | 0 | 0 | 0 | 0 | 0 | 0 | 0 |
| ENSMUSG00000090319 | 1.069666964 | 0.864857905 | 1.02186571 | 1.02186571 | 0.955822902 | 1.113943352 | 0.916588299 |
| ENSMUSG00000090322 | 0 | 0 | 0.25 | 0 | 0 | 0 | 0 |
| ENSMUSG00000090323 | 0 | 0 | 0 | 0 | 0 | 0 | 0 |
| ENSMUSG00000090326 | 0 | 0 | 0 | 0 | 0 | 0 | 0 |
| ENSMUSG00000090327 | 0.747221736 | 0.580482024 | 0.25 | 0.396240625 | 0.244650542 | 0 | 0.448487648 |
| ENSMUSG00000090328 | 0 | 0 | 0 | 0 | 0 | 0 | 0 |
| ENSMUSG00000090329 | 0 | 0 | 0 | 0 | 0.244650542 | 0 | 0 |
| ENSMUSG00000090330 | 1.289064826 | 1.146240625 | 1.372963274 | 1.098079356 | 1.106691885 | 1.462397998 | 1.238624656 |
| ENSMUSG00000090331 | 0 | 0 | 0 | 0 | 0 | 0 | 0 |
| ENSMUSG00000090332 | 0 | 0 | 0 | 0 | 0 | 0 | 0 |
| ENSMUSG00000090333 | 0 | 0 | 0 | 0 | 0 | 0 | 0 |
| ENSMUSG00000090334 | 0.45815691 | 0 | 0 | 0.25 | 0.244650542 | 0 | 0 |
| ENSMUSG00000090336 | 0 | 0.25 | 0 | 0 | 0 | 0 | 0 |
| ENSMUSG00000090338 | 0 | 0 | 0 | 0 | 0 | 0 | 0 |
| ENSMUSG00000090341 | 0 | 0 | 0 | 0 | 0 | 0 | 0 |
| ENSMUSG00000090342 | 0 | 0 | 0 | 0 | 0 | 0 | 0 |
| ENSMUSG00000090346 | 0 | 0 | 0 | 0 | 0 | 0 | 0 |
| ENSMUSG00000090350 | 0 | 0 | 0 | 0 | 0 | 0 | 0 |
| ENSMUSG00000090351 | 0 | 0 | 0 | 0 | 0 | 0 | 0 |
| ENSMUSG00000090352 | 0 | 0 | 0 | 0 | 0 | 0 | 0 |
| ENSMUSG00000090353 | 0.811507563 | 0.864857905 | 0.92510993 | 0.951838731 | 0.877063019 | 1 | 0.853775828 |
| ENSMUSG00000090355 | 0 | 0 | 0 | 0 | 0 | 0 | 0 |
| ENSMUSG00000090356 | 0 | 0 | 0 | 0 | 0 | 0 | 0 |
| ENSMUSG00000090357 | 0 | 0 | 0 | 0 | 0 | 0 | 0.224243824 |
| ENSMUSG00000090358 | 0.578129653 | 0.25 | 0.5 | 0.5 | 0.632412477 | 0.602059991 | 0.579661877 |
| ENSMUSG00000090362 | 0 | 0 | 0 | 0 | 0 | 0 | 0 |
| ENSMUSG00000090363 | 0 | 0.25 | 0.25 | 0 | 0 | 0 | 0.224243824 |
| ENSMUSG00000090364 | 0 | 0 | 0 | 0 | 0 | 0 | 0 |
| ENSMUSG00000090365 | 0 | 0 | 0 | 0 | 0 | 0 | 0 |
| ENSMUSG00000090369 | 0.671187741 | 0.580482024 | 0.646240625 | 0.92510993 | 0.733951626 | 0.477121255 | 0.74492186 |

| | | | | | | | |
|---|---|---|---|---|---|---|---|
| ENSMUSG00000090370 | 0 | 0 | 0 | 0 | 0 | 0 | 0 |
| ENSMUSG00000090374 | 0 | 0 | 0 | 0 | 0 | 0 | 0 |
| ENSMUSG00000090379 | 0 | 0 | 0 | 0 | 0 | 0 | 0 |
| ENSMUSG00000090381 | 0 | 0 | 0 | 0 | 0 | 0 | 0 |
| ENSMUSG00000090383 | 0 | 0 | 0 | 0 | 0 | 0 | 0 |
| ENSMUSG00000090385 | 0 | 0 | 0 | 0 | 0 | 0 | 0 |
| ENSMUSG00000090386 | 0.289064826 | 0.951838731 | 0.701838731 | 1.575945187 | 1.528558853 | 1.643452676 | 1.383567577 |
| ENSMUSG00000090387 | 0 | 0 | 0 | 0.5 | 0.489301084 | 0 | 0.224243824 |
| ENSMUSG00000090389 | 0 | 0.580482024 | 0 | 0.25 | 0.489301084 | 0.301029996 | 0.448487648 |
| ENSMUSG00000090394 | 1.325351389 | 1.238549078 | 1.226722649 | 1.080482024 | 0.77552387 | 1.041392685 | 0.829800754 |
| ENSMUSG00000090395 | 0 | 0 | 0 | 0 | 0 | 0 | 0 |
| ENSMUSG00000090396 | 0 | 0 | 0 | 0 | 0 | 0 | 0 |
| ENSMUSG00000090397 | 0.578129653 | 0.5 | 0.396240625 | 0.580482024 | 0.568060967 | 0.698970004 | 0.629532004 |
| ENSMUSG00000090399 | 1.036286563 | 1.04248125 | 1.214495249 | 1.098079356 | 1 | 1 | 1.140832123 |
| ENSMUSG00000090404 | 0 | 0 | 0 | 0 | 0 | 0 | 0 |
| ENSMUSG00000090406 | 0.811507563 | 0.25 | 0.25 | 0 | 0 | 0 | 0 |
| ENSMUSG00000090408 | 0 | 0 | 0 | 0 | 0.244650542 | 0 | 0 |
| ENSMUSG00000090411 | 0 | 0 | 0 | 0 | 0 | 0 | 0 |
| ENSMUSG00000090417 | 0 | 0 | 0 | 0 | 0 | 0 | 0 |
| ENSMUSG00000090427 | 0 | 0 | 0 | 0 | 0 | 0 | 0 |
| ENSMUSG00000090429 | 0 | 0 | 0 | 0 | 0 | 0 | 0 |
| ENSMUSG00000090436 | 0 | 0 | 0 | 0 | 0 | 0 | 0 |
| ENSMUSG00000090439 | 0 | 0 | 0 | 0 | 0 | 0 | 0 |
| ENSMUSG00000090440 | 0 | 0.25 | 0 | 0.25 | 0 | 0 | 0.355418052 |
| ENSMUSG00000090441 | 0 | 0 | 0 | 0 | 0 | 0 | 0 |
| ENSMUSG00000090448 | 0 | 0 | 0 | 0 | 0 | 0 | 0 |
| ENSMUSG00000090449 | 0 | 0 | 0 | 0 | 0 | 0 | 0 |
| ENSMUSG00000090451 | 0 | 0 | 0.396240625 | 0.646240625 | 0.387761935 | 0.602059991 | 0 |
| ENSMUSG00000090455 | 0 | 0 | 0 | 0 | 0 | 0 | 0 |
| ENSMUSG00000090457 | 0 | 0 | 0 | 0 | 0 | 0 | 0 |
| ENSMUSG00000090458 | 0 | 0 | 0 | 0 | 0 | 0 | 0 |
| ENSMUSG00000090459 | 0 | 0 | 0.5 | 0.25 | 0.489301084 | 0.477121255 | 0.448487648 |
| ENSMUSG00000090460 | 0 | 0 | 0 | 0 | 0 | 0 | 0.224243824 |
| ENSMUSG00000090461 | 0 | 0 | 0 | 0 | 0 | 0 | 0 |

| | | | | | | | |
|---|---|---|---|---|---|---|---|
| ENSMUSG00000090462 | 0 | 0 | 0 | 0 | 0 | 0 | 0 |
| ENSMUSG00000090464 | 0 | 0 | 0 | 0 | 0 | 0 | 0 |
| ENSMUSG00000090465 | 0 | 0 | 0 | 0 | 0 | 0 | 0 |
| ENSMUSG00000090466 | 0 | 0 | 0 | 0 | 0 | 0 | 0 |
| ENSMUSG00000090467 | 0 | 0 | 0 | 0 | 0 | 0 | 0 |
| ENSMUSG00000090468 | 0 | 0 | 0 | 0 | 0 | 0 | 0 |
| ENSMUSG00000090470 | 0 | 0 | 0 | 0 | 0 | 0 | 0 |
| ENSMUSG00000090471 | 0 | 0 | 0 | 0 | 0 | 0 | 0 |
| ENSMUSG00000090472 | 0 | 0 | 0 | 0 | 0 | 0 | 0 |
| ENSMUSG00000090474 | 0 | 0 | 0 | 0 | 0 | 0 | 0 |
| ENSMUSG00000090475 | 0 | 0 | 0 | 0 | 0.244650542 | 0 | 0 |
| ENSMUSG00000090476 | 0 | 0 | 0 | 0 | 0 | 0 | 0 |
| ENSMUSG00000090478 | 0 | 0 | 0 | 0 | 0 | 0 | 0 |
| ENSMUSG00000090481 | 0 | 0 | 0 | 0 | 0 | 0 | 0 |
| ENSMUSG00000090482 | 0 | 0 | 0 | 0 | 0 | 0 | 0 |
| ENSMUSG00000090485 | 0 | 0 | 0 | 0 | 0 | 0 | 0 |
| ENSMUSG00000090486 | 1.069666964 | 1.130890489 | 1.098079356 | 0.864857905 | 0.568060967 | 0.602059991 | 0.775756176 |
| ENSMUSG00000090487 | 0 | 0 | 0 | 0 | 0 | 0 | 0 |
| ENSMUSG00000090489 | 0.671187741 | 0.396240625 | 0.396240625 | 0.396240625 | 0.632412477 | 0 | 0.355418052 |
| ENSMUSG00000090494 | 0 | 0 | 0 | 0 | 0 | 0 | 0 |
| ENSMUSG00000090497 | 0 | 0 | 0 | 0 | 0 | 0 | 0 |
| ENSMUSG00000090500 | 0.45815691 | 0.396240625 | 0.701838731 | 0.396240625 | 0.387761935 | 0.77815125 | 0.448487648 |
| ENSMUSG00000090505 | 0 | 0 | 0 | 0 | 0 | 0 | 0 |
| ENSMUSG00000090509 | 0 | 0 | 0 | 0.25 | 0.244650542 | 0.477121255 | 0.224243824 |
| ENSMUSG00000090512 | 0 | 0 | 0 | 0 | 0 | 0 | 0 |
| ENSMUSG00000090515 | 0 | 0 | 0 | 0 | 0 | 0 | 0 |
| ENSMUSG00000090516 | 0 | 0 | 0 | 0.396240625 | 0 | 0 | 0 |
| ENSMUSG00000090519 | 0 | 0 | 0 | 0 | 0 | 0 | 0 |
| ENSMUSG00000090523 | 2.629921064 | 2.423807073 | 2.392938161 | 2.218361028 | 2.164820399 | 2.428134794 | 2.052911423 |
| ENSMUSG00000090527 | 0 | 0 | 0 | 0 | 0 | 0 | 0 |
| ENSMUSG00000090533 | 0 | 0 | 0 | 0 | 0 | 0 | 0 |
| ENSMUSG00000090534 | 0 | 0 | 0 | 0.25 | 0 | 0 | 0 |
| ENSMUSG00000090535 | 0 | 0 | 0 | 0 | 0 | 0 | 0 |
| ENSMUSG00000090538 | 0 | 0 | 0 | 0 | 0 | 0 | 0.224243824 |

| | | | | | | | |
|---|---|---|---|---|---|---|---|
| ENSMUSG00000090539 | 0 | 0 | 0 | 0 | 0 | 0 | 0 |
| ENSMUSG00000090544 | 0.289064826 | 0.830482024 | 0.75 | 0.5 | 0.387761935 | 0.301029996 | 0.520678036 |
| ENSMUSG00000090546 | 0 | 0 | 0 | 0 | 0 | 0 | 0 |
| ENSMUSG00000090547 | 0 | 0 | 0 | 0 | 0 | 0 | 0 |
| ENSMUSG00000090549 | 0 | 0 | 0 | 0 | 0 | 0 | 0 |
| ENSMUSG00000090550 | 0 | 0 | 0 | 0 | 0 | 0 | 0 |
| ENSMUSG00000090551 | 0.578129653 | 0.75 | 0.5 | 0.25 | 0 | 0.301029996 | 0.355418052 |
| ENSMUSG00000090552 | 0 | 0 | 0 | 0 | 0 | 0 | 0 |
| ENSMUSG00000090553 | 2.583267694 | 2.363317659 | 2.382357639 | 2.474461364 | 2.399768403 | 2.755112266 | 2.232493008 |
| ENSMUSG00000090555 | 0 | 0 | 0 | 0 | 0 | 0 | 0 |
| ENSMUSG00000090556 | 0 | 0 | 0 | 0 | 0 | 0 | 0 |
| ENSMUSG00000090558 | 0 | 0 | 0 | 0 | 0 | 0 | 0 |
| ENSMUSG00000090559 | 0 | 0 | 0.25 | 0.25 | 0.387761935 | 0 | 0.355418052 |
| ENSMUSG00000090561 | 0 | 0 | 0 | 0 | 0 | 0 | 0 |
| ENSMUSG00000090564 | 0.747221736 | 0.864857905 | 0.92510993 | 0.646240625 | 0.632412477 | 0.477121255 | 0.520678036 |
| ENSMUSG00000090565 | 0 | 0 | 0.25 | 0 | 0.387761935 | 0 | 0 |
| ENSMUSG00000090570 | 0 | 0 | 0 | 0 | 0 | 0 | 0 |
| ENSMUSG00000090572 | 0 | 0 | 0 | 0 | 0 | 0 | 0 |
| ENSMUSG00000090574 | 0 | 0 | 0 | 0 | 0 | 0 | 0 |
| ENSMUSG00000090576 | 0.747221736 | 0.75 | 0.830482024 | 0.75 | 0.686820904 | 0.477121255 | 0.74492186 |
| ENSMUSG00000090577 | 0 | 0 | 0 | 0 | 0 | 0 | 0 |
| ENSMUSG00000090578 | 0 | 0 | 0 | 0 | 0 | 0 | 0 |
| ENSMUSG00000090581 | 0 | 0 | 0 | 0 | 0 | 0 | 0 |
| ENSMUSG00000090582 | 0.578129653 | 0.951838731 | 0.951838731 | 0.92510993 | 0.846351821 | 0.954242509 | 0.355418052 |
| ENSMUSG00000090584 | 0 | 0 | 0 | 0 | 0 | 0 | 0 |
| ENSMUSG00000090585 | 0 | 0 | 0 | 0 | 0 | 0 | 0 |
| ENSMUSG00000090588 | 0 | 0 | 0.25 | 0 | 0 | 0 | 0.224243824 |
| ENSMUSG00000090589 | 0 | 0 | 0 | 0 | 0 | 0 | 0 |
| ENSMUSG00000090592 | 0 | 0 | 0 | 0.25 | 0 | 0 | 0 |
| ENSMUSG00000090594 | 0 | 0 | 0 | 0 | 0 | 0 | 0 |
| ENSMUSG00000090598 | 0 | 0 | 0 | 0 | 0 | 0 | 0 |
| ENSMUSG00000090602 | 0 | 0.25 | 0 | 0.5 | 0.387761935 | 0.301029996 | 0.355418052 |
| ENSMUSG00000090603 | 0 | 0 | 0 | 0 | 0 | 0 | 0 |
| ENSMUSG00000090607 | 0 | 0 | 0 | 0 | 0 | 0 | 0 |

| | | | | | | | |
|---|---|---|---|---|---|---|---|
| ENSMUSG00000090608 | 0 | 0.5 | 0 | 0 | 0.244650542 | 0 | 0.224243824 |
| ENSMUSG00000090610 | 0 | 0.25 | 0.580482024 | 0 | 0.244650542 | 0.477121255 | 0.224243824 |
| ENSMUSG00000090618 | 0 | 0 | 0 | 0 | 0 | 0 | 0 |
| ENSMUSG00000090619 | 0 | 0 | 0 | 0 | 0 | 0 | 0 |
| ENSMUSG00000090622 | 1.100572389 | 1.061981878 | 1 | 0.864857905 | 0.77552387 | 1 | 0.672731473 |
| ENSMUSG00000090623 | 0 | 0 | 0 | 0 | 0 | 0 | 0 |
| ENSMUSG00000090624 | 0 | 0 | 0 | 0.396240625 | 0 | 0.477121255 | 0 |
| ENSMUSG00000090625 | 0 | 0 | 0 | 0 | 0 | 0 | 0 |
| ENSMUSG00000090626 | 1.759671389 | 1.622963274 | 1.685366747 | 1.802363341 | 1.731346772 | 1.908485019 | 1.61448206 |
| ENSMUSG00000090627 | 0 | 0 | 0 | 0 | 0 | 0 | 0 |
| ENSMUSG00000090628 | 0.867194479 | 0 | 0.5 | 0.701838731 | 0.489301084 | 0 | 0.224243824 |
| ENSMUSG00000090629 | 0 | 0 | 0 | 0 | 0 | 0 | 0 |
| ENSMUSG00000090631 | 0 | 0 | 0 | 0.25 | 0.489301084 | 0.477121255 | 0.448487648 |
| ENSMUSG00000090634 | 0 | 0 | 0 | 0 | 0 | 0 | 0 |
| ENSMUSG00000090635 | 0 | 0 | 0 | 0 | 0 | 0 | 0 |
| ENSMUSG00000090636 | 0 | 0 | 0 | 0 | 0 | 0 | 0 |
| ENSMUSG00000090637 | 0 | 0 | 0 | 0 | 0 | 0 | 0 |
| ENSMUSG00000090638 | 1.069666964 | 1.25 | 1.25 | 0 | 0 | 0 | 0 |
| ENSMUSG00000090639 | 0 | 0 | 0 | 0 | 0 | 0 | 0 |
| ENSMUSG00000090641 | 0.811507563 | 1 | 1.02186571 | 1.04248125 | 0.955822902 | 1.204119983 | 0.853775828 |
| ENSMUSG00000090642 | 0 | 0 | 0 | 0 | 0 | 0 | 0 |
| ENSMUSG00000090643 | 0 | 0 | 0 | 0 | 0 | 0 | 0 |
| ENSMUSG00000090655 | 0 | 0 | 0 | 0 | 0 | 0 | 0 |
| ENSMUSG00000090657 | 0 | 0 | 0 | 0 | 0 | 0 | 0 |
| ENSMUSG00000090658 | 0 | 0 | 0 | 0 | 0 | 0 | 0 |
| ENSMUSG00000090659 | 1.069666964 | 1.080482024 | 0.951838731 | 0.951838731 | 1.057362051 | 1.176091259 | 0.952571511 |
| ENSMUSG00000090660 | 0 | 0 | 0 | 0 | 0 | 0 | 0 |
| ENSMUSG00000090662 | 0 | 0 | 0 | 0 | 0 | 0 | 0 |
| ENSMUSG00000090665 | 0 | 0 | 0 | 0 | 0 | 0 | 0 |
| ENSMUSG00000090666 | 0 | 0 | 0 | 0.25 | 0 | 0.477121255 | 0.355418052 |
| ENSMUSG00000090667 | 0 | 0 | 0 | 0 | 0 | 0 | 0 |
| ENSMUSG00000090671 | 0 | 0 | 0 | 0 | 0 | 0 | 0 |
| ENSMUSG00000090673 | 2.045658471 | 1.864857905 | 1.92510993 | 1.969129237 | 1.905314583 | 2.133538908 | 1.782372264 |
| ENSMUSG00000090674 | 0.289064826 | 0 | 0 | 0.25 | 0.244650542 | 0 | 0.579661877 |

| | | | | | | | |
|---|---|---|---|---|---|---|---|
| ENSMUSG00000090675 | 0 | 0 | 0 | 0 | 0 | 0 | 0 |
| ENSMUSG00000090681 | 0 | 0 | 0 | 0 | 0 | 0 | 0 |
| ENSMUSG00000090682 | 0 | 0 | 0 | 0 | 0 | 0 | 0 |
| ENSMUSG00000090684 | 0 | 0 | 0 | 0 | 0 | 0 | 0 |
| ENSMUSG00000090685 | 0 | 0 | 0 | 0 | 0 | 0 | 0 |
| ENSMUSG00000090688 | 0 | 0 | 0 | 0 | 0 | 0 | 0 |
| ENSMUSG00000090690 | 0 | 0 | 0 | 0 | 0 | 0 | 0 |
| ENSMUSG00000090691 | 0 | 0 | 0 | 0 | 0.244650542 | 0 | 0.224243824 |
| ENSMUSG00000090692 | 0 | 0 | 0 | 0 | 0 | 0 | 0 |
| ENSMUSG00000090693 | 0.45815691 | 0.25 | 0.5 | 0.25 | 0.387761935 | 0 | 0.355418052 |
| ENSMUSG00000090698 | 0.91631382 | 0.75 | 0.75 | 0.25 | 0.387761935 | 0.301029996 | 0.448487648 |
| ENSMUSG00000090699 | 0 | 0 | 0 | 0 | 0 | 0 | 0 |
| ENSMUSG00000090700 | 0 | 0 | 0 | 0 | 0 | 0 | 0 |
| ENSMUSG00000090701 | 0 | 0 | 0 | 0 | 0 | 0 | 0 |
| ENSMUSG00000090704 | 0 | 0 | 0 | 0 | 0 | 0 | 0 |
| ENSMUSG00000090706 | 0 | 0 | 0 | 0 | 0 | 0 | 0 |
| ENSMUSG00000090707 | 0 | 0 | 0 | 0 | 0 | 0.301029996 | 0 |
| ENSMUSG00000090708 | 0 | 0 | 0 | 0 | 0.244650542 | 0 | 0.224243824 |
| ENSMUSG00000090709 | 0.578129653 | 0 | 0.25 | 0 | 0 | 0 | 0 |
| ENSMUSG00000090710 | 0 | 0 | 0 | 0 | 0 | 0 | 0 |
| ENSMUSG00000090713 | 0 | 0.25 | 0 | 0 | 0 | 0 | 0 |
| ENSMUSG00000090714 | 0 | 0 | 0.25 | 0 | 0 | 0 | 0 |
| ENSMUSG00000090715 | 0 | 0 | 0 | 0 | 0 | 0 | 0 |
| ENSMUSG00000090716 | 0 | 0 | 0 | 0 | 0 | 0 | 0 |
| ENSMUSG00000090718 | 0 | 0 | 0 | 0 | 0 | 0 | 0 |
| ENSMUSG00000090722 | 0 | 0 | 0 | 0 | 0 | 0 | 0 |
| ENSMUSG00000090724 | 0 | 0 | 0 | 0 | 0 | 0 | 0 |
| ENSMUSG00000090726 | 0 | 0 | 0 | 0 | 0 | 0 | 0 |
| ENSMUSG00000090727 | 0 | 0 | 0 | 0 | 0 | 0 | 0 |
| ENSMUSG00000090728 | 0 | 0 | 0 | 0 | 0 | 0 | 0 |
| ENSMUSG00000090731 | 0 | 0 | 0 | 0 | 0 | 0 | 0 |
| ENSMUSG00000090732 | 0 | 0 | 0 | 0 | 0 | 0 | 0 |
| ENSMUSG00000090733 | 2.669114668 | 2.429419106 | 2.446567407 | 2.59727338 | 2.44788147 | 2.890979597 | 2.324828731 |
| ENSMUSG00000090736 | 0 | 0 | 0 | 0 | 0 | 0 | 0 |

| | | | | | | | |
|---|---|---|---|---|---|---|---|
| ENSMUSG00000090737 | 0 | 0 | 0 | 0 | 0 | 0 | 0.224243824 |
| ENSMUSG00000090738 | 0 | 0 | 0 | 0 | 0 | 0 | 0 |
| ENSMUSG00000090740 | 0 | 0 | 0 | 0 | 0 | 0 | 0 |
| ENSMUSG00000090742 | 0 | 0 | 0 | 0 | 0 | 0 | 0 |
| ENSMUSG00000090743 | 0 | 0 | 0 | 0 | 0 | 0 | 0 |
| ENSMUSG00000090744 | 0.578129653 | 0.896240625 | 0.580482024 | 0.580482024 | 0.77552387 | 0.84509804 | 0.74492186 |
| ENSMUSG00000090747 | 0 | 0 | 0 | 0 | 0 | 0 | 0 |
| ENSMUSG00000090750 | 0 | 0 | 0 | 0 | 0 | 0 | 0 |
| ENSMUSG00000090751 | 0 | 0 | 0 | 0 | 0 | 0 | 0 |
| ENSMUSG00000090753 | 0 | 0 | 0 | 0.25 | 0 | 0 | 0 |
| ENSMUSG00000090758 | 0 | 0 | 0 | 0 | 0 | 0 | 0 |
| ENSMUSG00000090761 | 1.100572389 | 1.114857905 | 1.061981878 | 1.04248125 | 1.244650542 | 1.204119983 | 1.23151414 |
| ENSMUSG00000090762 | 0 | 0 | 0 | 0 | 0 | 0 | 0 |
| ENSMUSG00000090764 | 0 | 0 | 0 | 0 | 0 | 0 | 0 |
| ENSMUSG00000090765 | 0 | 0 | 0 | 0 | 0 | 0 | 0 |
| ENSMUSG00000090772 | 0 | 0 | 0 | 0 | 0 | 0 | 0 |
| ENSMUSG00000090773 | 0 | 0 | 0 | 0 | 0 | 0 | 0 |
| ENSMUSG00000090774 | 0 | 0 | 0 | 0 | 0 | 0 | 0 |
| ENSMUSG00000090777 | 0.45815691 | 0 | 0 | 0.25 | 0 | 0 | 0.224243824 |
| ENSMUSG00000090778 | 0.289064826 | 0.396240625 | 0 | 0.25 | 0 | 0 | 0 |
| ENSMUSG00000090779 | 0 | 0 | 0 | 0 | 0 | 0 | 0.224243824 |
| ENSMUSG00000090781 | 0 | 0 | 0 | 0 | 0 | 0 | 0 |
| ENSMUSG00000090782 | 0 | 0 | 0 | 0 | 0 | 0 | 0 |
| ENSMUSG00000090785 | 0 | 0 | 0 | 0 | 0.244650542 | 0 | 0 |
| ENSMUSG00000090788 | 0 | 0 | 0 | 0 | 0 | 0 | 0 |
| ENSMUSG00000090790 | 0 | 0 | 0.25 | 0 | 0 | 0 | 0 |
| ENSMUSG00000090793 | 0.289064826 | 0.25 | 0 | 0 | 0 | 0 | 0 |
| ENSMUSG00000090794 | 0 | 0 | 0 | 0 | 0 | 0 | 0 |
| ENSMUSG00000090795 | 0 | 0 | 0 | 0 | 0 | 0 | 0 |
| ENSMUSG00000090796 | 0 | 0 | 0 | 0 | 0 | 0 | 0 |
| ENSMUSG00000090798 | 0 | 0 | 0 | 0 | 0 | 0 | 0 |
| ENSMUSG00000090799 | 0.289064826 | 0.396240625 | 0.25 | 0.396240625 | 0.686820904 | 0 | 0.355418052 |
| ENSMUSG00000090801 | 0 | 0 | 0 | 0 | 0 | 0 | 0 |
| ENSMUSG00000090802 | 0 | 0 | 0 | 0 | 0 | 0 | 0 |

| | | | | | | | |
|---|---|---|---|---|---|---|---|
| ENSMUSG00000090805 | 0 | 0 | 0 | 0 | 0 | 0 | 0 |
| ENSMUSG00000090806 | 0 | 0 | 0 | 0 | 0 | 0 | 0 |
| ENSMUSG00000090808 | 0 | 0 | 0 | 0 | 0 | 0 | 0 |
| ENSMUSG00000090812 | 0.578129653 | 0.646240625 | 0.580482024 | 0.580482024 | 0.568060967 | 0.84509804 | 0.520678036 |
| ENSMUSG00000090816 | 0 | 0 | 0 | 0 | 0 | 0 | 0 |
| ENSMUSG00000090817 | 0 | 0 | 0 | 0 | 0 | 0 | 0 |
| ENSMUSG00000090818 | 0 | 0 | 0 | 0 | 0 | 0 | 0 |
| ENSMUSG00000090821 | 0 | 0 | 0 | 0 | 0 | 0 | 0 |
| ENSMUSG00000090824 | 0 | 0 | 0 | 0 | 0 | 0 | 0 |
| ENSMUSG00000090827 | 0 | 0 | 0 | 0 | 0 | 0 | 0 |
| ENSMUSG00000090839 | 0 | 0 | 0 | 0 | 0 | 0 | 0 |
| ENSMUSG00000090840 | 0.45815691 | 0.5 | 0.5 | 0.396240625 | 0.387761935 | 0 | 0.579661877 |
| ENSMUSG00000090841 | 3.391111698 | 3.095520898 | 3.084196369 | 2.854990044 | 2.838946875 | 3.284881715 | 2.671307952 |
| ENSMUSG00000090843 | 0 | 0 | 0 | 0 | 0 | 0.301029996 | 0.224243824 |
| ENSMUSG00000090849 | 0 | 0 | 0 | 0 | 0 | 0 | 0 |
| ENSMUSG00000090853 | 0 | 0 | 0 | 0 | 0 | 0 | 0 |
| ENSMUSG00000090854 | 0 | 0 | 0 | 0 | 0 | 0 | 0 |
| ENSMUSG00000090855 | 0 | 0 | 0 | 0 | 0 | 0 | 0 |
| ENSMUSG00000090860 | 0 | 0 | 0 | 0 | 0 | 0 | 0 |
| ENSMUSG00000090861 | 0 | 0 | 0 | 0 | 0.244650542 | 0 | 0.224243824 |
| ENSMUSG00000090862 | 2.546826359 | 2.351285366 | 2.42813175 | 2.381869252 | 2.337065673 | 2.655138435 | 2.275733199 |
| ENSMUSG00000090863 | 0.45815691 | 0.25 | 0.701838731 | 0.5 | 0.244650542 | 0.698970004 | 0.224243824 |
| ENSMUSG00000090864 | 0 | 0 | 0 | 0 | 0 | 0 | 0 |
| ENSMUSG00000090865 | 0 | 0 | 0 | 0 | 0 | 0 | 0 |
| ENSMUSG00000090868 | 0 | 0 | 0 | 0 | 0 | 0 | 0 |
| ENSMUSG00000090872 | 0 | 0 | 0 | 0 | 0 | 0 | 0 |
| ENSMUSG00000090873 | 0.289064826 | 0.580482024 | 0.580482024 | 0.396240625 | 0.568060967 | 0.301029996 | 0.448487648 |
| ENSMUSG00000090874 | 0 | 0 | 0 | 0 | 0 | 0 | 0 |
| ENSMUSG00000090877 | 1.714367558 | 1.634789703 | 1.720660762 | 1.685366747 | 1.689774529 | 1.643452676 | 1.594042709 |
| ENSMUSG00000090881 | 0 | 0 | 0.25 | 0 | 0 | 0.301029996 | 0.224243824 |
| ENSMUSG00000090882 | 0 | 0 | 0 | 0 | 0 | 0 | 0 |
| ENSMUSG00000090889 | 0 | 0 | 0 | 0 | 0 | 0 | 0 |
| ENSMUSG00000090891 | 0.45815691 | 0.5 | 0.580482024 | 0.580482024 | 0.387761935 | 0.301029996 | 0.672731473 |
| ENSMUSG00000090892 | 0 | 0 | 0 | 0 | 0 | 0 | 0 |

| | | | | | | | |
|---|---|---|---|---|---|---|---|
| ENSMUSG00000090894 | 0 | 0 | 0 | 0 | 0 | 0 | 0 |
| ENSMUSG00000090897 | 0 | 0 | 0 | 0 | 0 | 0 | 0 |
| ENSMUSG00000090900 | 0 | 0 | 0 | 0 | 0 | 0 | 0 |
| ENSMUSG00000090907 | 0 | 0 | 0 | 0 | 0 | 0 | 0.224243824 |
| ENSMUSG00000090919 | 0.671187741 | 0.830482024 | 0.701838731 | 0.75 | 0.846351821 | 0.77815125 | 0.74492186 |
| ENSMUSG00000090925 | 0.289064826 | 0.396240625 | 0.580482024 | 0.580482024 | 0.387761935 | 0.602059991 | 0.579661877 |
| ENSMUSG00000090927 | 0 | 0 | 0 | 0 | 0 | 0 | 0 |
| ENSMUSG00000090935 | 2.231898196 | 2.046218836 | 2.00835575 | 2.084962501 | 2.022473809 | 2.298853076 | 1.846338363 |
| ENSMUSG00000090936 | 0.289064826 | 0.580482024 | 0.701838731 | 0.25 | 0.244650542 | 0.301029996 | 0.520678036 |
| ENSMUSG00000090939 | 0 | 0.25 | 0.25 | 0 | 0 | 0 | 0.224243824 |
| ENSMUSG00000090941 | 0 | 0.25 | 0 | 0 | 0 | 0 | 0 |
| ENSMUSG00000090942 | 0.867194479 | 0.701838731 | 0.396240625 | 0.5 | 0.244650542 | 0.477121255 | 0.355418052 |
| ENSMUSG00000090946 | 1.800532393 | 1.720660762 | 1.660964047 | 1.470660762 | 1.433158226 | 1.602059991 | 1.340368108 |
| ENSMUSG00000090947 | 0 | 0 | 0 | 0 | 0 | 0 | 0 |
| ENSMUSG00000090949 | 0 | 0 | 0 | 0 | 0 | 0 | 0 |
| ENSMUSG00000090951 | 0 | 0 | 0 | 0 | 0 | 0 | 0 |
| ENSMUSG00000090952 | 0.747221736 | 0.976722649 | 1.080482024 | 0.830482024 | 0.812711509 | 0.602059991 | 0.853775828 |
| ENSMUSG00000090955 | 0.289064826 | 0.25 | 0.396240625 | 0 | 0.568060967 | 0.477121255 | 0.448487648 |
| ENSMUSG00000090957 | 0 | 0 | 0 | 0 | 0 | 0.301029996 | 0 |
| ENSMUSG00000090958 | 2.550525119 | 2.387205477 | 2.392463902 | 1.348079356 | 1.49445382 | 1.51851394 | 1.562034107 |
| ENSMUSG00000090961 | 0 | 0 | 0 | 0 | 0 | 0 | 0 |
| ENSMUSG00000090963 | 0.747221736 | 0.896240625 | 1.114857905 | 0.92510993 | 0.955822902 | 1.176091259 | 0.829800754 |
| ENSMUSG00000090964 | 0.289064826 | 0.25 | 0 | 0 | 0 | 0 | 0 |
| ENSMUSG00000090965 | 0.45815691 | 0.5 | 0.580482024 | 0.646240625 | 0.812711509 | 0.301029996 | 0.520678036 |
| ENSMUSG00000090966 | 0 | 0 | 0 | 0 | 0 | 0 | 0 |
| ENSMUSG00000090967 | 0 | 0 | 0 | 0 | 0 | 0 | 0 |
| ENSMUSG00000090968 | 0 | 0 | 0 | 0 | 0 | 0 | 0 |
| ENSMUSG00000090975 | 0 | 0 | 0 | 0 | 0 | 0 | 0 |
| ENSMUSG00000090981 | 0 | 0 | 0 | 0 | 0 | 0 | 0 |
| ENSMUSG00000090987 | 0 | 0 | 0 | 0 | 0 | 0 | 0 |
| ENSMUSG00000090989 | 0 | 0 | 0 | 0 | 0 | 0 | 0 |
| ENSMUSG00000090990 | 0 | 0 | 0 | 0 | 0 | 0 | 0 |
| ENSMUSG00000090991 | 0 | 0 | 0 | 0 | 0 | 0 | 0 |
| ENSMUSG00000090994 | 0.289064826 | 0 | 0.25 | 0.396240625 | 0 | 0 | 0 |

| | | | | | | | |
|---|---|---|---|---|---|---|---|
| ENSMUSG00000090996 | 0 | 0 | 0 | 0 | 0 | 0 | 0 |
| ENSMUSG00000091002 | 0 | 0 | 0 | 0 | 0 | 0 | 0 |
| ENSMUSG00000091003 | 0 | 0 | 0 | 0 | 0 | 0 | 0 |
| ENSMUSG00000091004 | 0 | 0 | 0 | 0 | 0 | 0 | 0 |
| ENSMUSG00000091005 | 0 | 0 | 0 | 0 | 0 | 0 | 0 |
| ENSMUSG00000091006 | 0 | 0 | 0 | 0 | 0 | 0 | 0 |
| ENSMUSG00000091007 | 0 | 0.25 | 0 | 0 | 0 | 0 | 0 |
| ENSMUSG00000091008 | 0 | 0 | 0 | 0 | 0 | 0 | 0 |
| ENSMUSG00000091010 | 0 | 0 | 0 | 0 | 0 | 0 | 0 |
| ENSMUSG00000091011 | 0 | 0 | 0 | 0 | 0 | 0 | 0 |
| ENSMUSG00000091012 | 0 | 0 | 0 | 0 | 0 | 0 | 0 |
| ENSMUSG00000091013 | 0 | 0.25 | 0 | 0 | 0 | 0 | 0 |
| ENSMUSG00000091014 | 0 | 0 | 0 | 0.396240625 | 0 | 0 | 0 |
| ENSMUSG00000091016 | 0 | 0 | 0 | 0 | 0 | 0 | 0 |
| ENSMUSG00000091017 | 0 | 0 | 0 | 0 | 0 | 0 | 0 |
| ENSMUSG00000091019 | 0.289064826 | 0.396240625 | 0.396240625 | 0.25 | 0.489301084 | 0 | 0.224243824 |
| ENSMUSG00000091020 | 0.671187741 | 0.830482024 | 0.830482024 | 1 | 1.020174412 | 1.113943352 | 0.876096088 |
| ENSMUSG00000091021 | 1.289064826 | 1.061981878 | 1.130890489 | 1.080482024 | 0.955822902 | 0.77815125 | 0.952571511 |
| ENSMUSG00000091022 | 0 | 0 | 0 | 0 | 0 | 0 | 0 |
| ENSMUSG00000091027 | 0 | 0 | 0 | 0 | 0 | 0 | 0 |
| ENSMUSG00000091028 | 0 | 0 | 0 | 0 | 0 | 0 | 0 |
| ENSMUSG00000091031 | 0 | 0 | 0 | 0 | 0 | 0 | 0 |
| ENSMUSG00000091034 | 0 | 0 | 0 | 0 | 0 | 0 | 0 |
| ENSMUSG00000091037 | 0 | 0 | 0 | 0 | 0 | 0 | 0 |
| ENSMUSG00000091039 | 0 | 0 | 0 | 0 | 0 | 0 | 0 |
| ENSMUSG00000091040 | 0 | 0 | 0 | 0 | 0 | 0 | 0.224243824 |
| ENSMUSG00000091041 | 0 | 0 | 0 | 0 | 0 | 0 | 0 |
| ENSMUSG00000091043 | 0 | 0 | 0 | 0 | 0.387761935 | 0.301029996 | 0.224243824 |
| ENSMUSG00000091045 | 0.45815691 | 0.25 | 0.25 | 0.5 | 0.387761935 | 0 | 0 |
| ENSMUSG00000091048 | 0 | 0 | 0 | 0 | 0 | 0 | 0 |
| ENSMUSG00000091049 | 0 | 0 | 0 | 0 | 0 | 0 | 0 |
| ENSMUSG00000091050 | 0.747221736 | 0.830482024 | 0.92510993 | 0.830482024 | 0.489301084 | 0.602059991 | 0.710836105 |
| ENSMUSG00000091053 | 0 | 0 | 0 | 0 | 0 | 0 | 0 |
| ENSMUSG00000091055 | 0 | 0 | 0 | 0 | 0 | 0 | 0 |

| | | | | | | | |
|---|---|---|---|---|---|---|---|
| ENSMUSG00000091058 | 0 | 0 | 0 | 0.25 | 0 | 0 | 0 |
| ENSMUSG00000091059 | 0 | 0 | 0 | 0 | 0 | 0 | 0 |
| ENSMUSG00000091060 | 0 | 0 | 0 | 0.25 | 0 | 0 | 0 |
| ENSMUSG00000091061 | 0.289064826 | 0 | 0 | 0 | 0 | 0 | 0 |
| ENSMUSG00000091071 | 0 | 0 | 0 | 0 | 0 | 0 | 0 |
| ENSMUSG00000091076 | 0 | 0 | 0 | 0 | 0 | 0 | 0 |
| ENSMUSG00000091077 | 0 | 0 | 0 | 0 | 0 | 0 | 0 |
| ENSMUSG00000091078 | 0 | 0 | 0 | 0 | 0 | 0 | 0 |
| ENSMUSG00000091079 | 0 | 0 | 0 | 0 | 0 | 0 | 0 |
| ENSMUSG00000091080 | 0 | 0 | 0 | 0 | 0 | 0 | 0 |
| ENSMUSG00000091083 | 0 | 0 | 0 | 0.25 | 0.387761935 | 0 | 0 |
| ENSMUSG00000091086 | 0 | 0.396240625 | 0.580482024 | 0 | 0.387761935 | 0.301029996 | 0 |
| ENSMUSG00000091087 | 0 | 0 | 0 | 0 | 0 | 0 | 0 |
| ENSMUSG00000091089 | 0.289064826 | 0.396240625 | 0.396240625 | 0 | 0.489301084 | 0 | 0.355418052 |
| ENSMUSG00000091091 | 0 | 0 | 0 | 0 | 0 | 0 | 0 |
| ENSMUSG00000091095 | 0 | 0 | 0 | 0 | 0 | 0 | 0 |
| ENSMUSG00000091096 | 0 | 0 | 0 | 0 | 0 | 0 | 0 |
| ENSMUSG00000091099 | 0 | 0 | 0 | 0 | 0 | 0 | 0 |
| ENSMUSG00000091101 | 0 | 0 | 0 | 0 | 0 | 0 | 0 |
| ENSMUSG00000091102 | 0.289064826 | 0.396240625 | 0.396240625 | 0.396240625 | 0.489301084 | 0.477121255 | 0.520678036 |
| ENSMUSG00000091105 | 1.578129653 | 1.445339928 | 1.42510993 | 1.364857905 | 1.523883869 | 1.612783857 | 1.42167221 |
| ENSMUSG00000091109 | 0.289064826 | 0 | 0.25 | 0 | 0.244650542 | 0 | 0.355418052 |
| ENSMUSG00000091110 | 0 | 0 | 0 | 0 | 0 | 0 | 0 |
| ENSMUSG00000091114 | 0 | 0 | 0 | 0 | 0 | 0 | 0 |
| ENSMUSG00000091116 | 0 | 0 | 0 | 0 | 0 | 0 | 0 |
| ENSMUSG00000091117 | 0 | 0 | 0 | 0 | 0 | 0 | 0.224243824 |
| ENSMUSG00000091119 | 0 | 0.25 | 0.25 | 0.25 | 0.244650542 | 0 | 0.448487648 |
| ENSMUSG00000091122 | 0 | 0 | 0 | 0 | 0 | 0 | 0 |
| ENSMUSG00000091123 | 0 | 0 | 0 | 0 | 0 | 0 | 0 |
| ENSMUSG00000091129 | 0 | 0 | 0 | 0 | 0 | 0 | 0 |
| ENSMUSG00000091131 | 0 | 0 | 0 | 0 | 0 | 0 | 0 |
| ENSMUSG00000091132 | 0 | 0 | 0 | 0 | 0 | 0 | 0 |
| ENSMUSG00000091133 | 0.289064826 | 0.580482024 | 0.5 | 0.580482024 | 0.632412477 | 0.602059991 | 0.629532004 |
| ENSMUSG00000091135 | 0 | 0 | 0 | 0 | 0 | 0 | 0 |

| | | | | | | | |
|---|---|---|---|---|---|---|---|
| ENSMUSG00000091140 | 0 | 0 | 0 | 0 | 0 | 0 | 0 |
| ENSMUSG00000091142 | 0 | 0 | 0 | 0 | 0 | 0 | 0 |
| ENSMUSG00000091144 | 1.156259305 | 1.02186571 | 1.130890489 | 1.311981878 | 1.136121934 | 1.342422681 | 1.20919388 |
| ENSMUSG00000091146 | 0 | 0 | 0 | 0 | 0 | 0 | 0 |
| ENSMUSG00000091147 | 0 | 0 | 0 | 0.25 | 0 | 0 | 0 |
| ENSMUSG00000091148 | 0 | 0 | 0 | 0 | 0 | 0 | 0 |
| ENSMUSG00000091149 | 0 | 0 | 0 | 0 | 0 | 0 | 0 |
| ENSMUSG00000091151 | 0 | 0 | 0 | 0 | 0 | 0 | 0 |
| ENSMUSG00000091154 | 0.289064826 | 0.25 | 0 | 0.396240625 | 0 | 0 | 0 |
| ENSMUSG00000091155 | 0.289064826 | 0.25 | 0.25 | 0.79248125 | 0.77552387 | 0.477121255 | 0.74492186 |
| ENSMUSG00000091157 | 0 | 0 | 0 | 0 | 0 | 0 | 0 |
| ENSMUSG00000091159 | 0 | 0 | 0 | 0 | 0 | 0 | 0 |
| ENSMUSG00000091160 | 0.289064826 | 0.5 | 0.396240625 | 0 | 0.244650542 | 0 | 0 |
| ENSMUSG00000091163 | 0 | 0 | 0 | 0 | 0 | 0 | 0 |
| ENSMUSG00000091165 | 0 | 0 | 0 | 0 | 0 | 0 | 0 |
| ENSMUSG00000091172 | 0 | 0 | 0 | 0 | 0 | 0 | 0 |
| ENSMUSG00000091174 | 0 | 0 | 0 | 0 | 0 | 0 | 0 |
| ENSMUSG00000091175 | 0 | 0 | 0 | 0 | 0 | 0 | 0 |
| ENSMUSG00000091177 | 0 | 0 | 0 | 0 | 0 | 0 | 0 |
| ENSMUSG00000091183 | 1.693336259 | 1.51109853 | 1.571350555 | 1.614857905 | 1.542223377 | 1.602059991 | 1.392431569 |
| ENSMUSG00000091184 | 0.289064826 | 0 | 0 | 0.5 | 0.244650542 | 0.301029996 | 0 |
| ENSMUSG00000091185 | 0 | 0 | 0 | 0 | 0 | 0 | 0 |
| ENSMUSG00000091189 | 0 | 0 | 0 | 0 | 0 | 0 | 0 |
| ENSMUSG00000091191 | 0 | 0.396240625 | 0.5 | 0.580482024 | 0.632412477 | 0.602059991 | 0.672731473 |
| ENSMUSG00000091192 | 0 | 0 | 0 | 0 | 0 | 0 | 0 |
| ENSMUSG00000091193 | 0 | 0 | 0 | 0 | 0 | 0 | 0 |
| ENSMUSG00000091194 | 0 | 0.580482024 | 0.396240625 | 0.396240625 | 0.387761935 | 0 | 0.224243824 |
| ENSMUSG00000091195 | 0 | 0 | 0 | 0 | 0 | 0 | 0 |
| ENSMUSG00000091198 | 0 | 0 | 0 | 0 | 0.244650542 | 0 | 0 |
| ENSMUSG00000091199 | 0 | 0 | 0 | 0 | 0 | 0 | 0 |
| ENSMUSG00000091204 | 0 | 0 | 0 | 0 | 0.244650542 | 0 | 0 |
| ENSMUSG00000091205 | 0 | 0 | 0 | 0 | 0 | 0 | 0 |
| ENSMUSG00000091206 | 0 | 0 | 0 | 0 | 0 | 0 | 0 |
| ENSMUSG00000091212 | 0 | 0 | 0 | 0 | 0 | 0 | 0 |

| | | | | | | | |
|---|---|---|---|---|---|---|---|
| ENSMUSG00000091215 | 0 | 0 | 0 | 0 | 0 | 0 | 0.355418052 |
| ENSMUSG00000091216 | 0 | 0 | 0 | 0 | 0 | 0 | 0 |
| ENSMUSG00000091221 | 0 | 0 | 0 | 0 | 0 | 0 | 0 |
| ENSMUSG00000091223 | 0 | 0 | 0.25 | 0.25 | 0.244650542 | 0.477121255 | 0.224243824 |
| ENSMUSG00000091227 | 0 | 0 | 0.396240625 | 0 | 0 | 0 | 0.355418052 |
| ENSMUSG00000091228 | 0 | 0 | 0 | 0 | 0 | 0 | 0 |
| ENSMUSG00000091230 | 0.578129653 | 0.25 | 0.5 | 0.25 | 0.489301084 | 0 | 0.224243824 |
| ENSMUSG00000091233 | 0 | 0 | 0.25 | 0 | 0 | 0 | 0 |
| ENSMUSG00000091237 | 0 | 0 | 0 | 0 | 0 | 0 | 0 |
| ENSMUSG00000091238 | 0.289064826 | 0.25 | 0.25 | 0 | 0 | 0 | 0.355418052 |
| ENSMUSG00000091239 | 0 | 0 | 0 | 0 | 0 | 0 | 0 |
| ENSMUSG00000091243 | 2.553277818 | 2.325945187 | 2.333847589 | 2.229715809 | 2.173143437 | 2.451786436 | 2.057420422 |
| ENSMUSG00000091248 | 0 | 0 | 0 | 0 | 0 | 0 | 0 |
| ENSMUSG00000091255 | 0 | 0 | 0 | 0 | 0 | 0 | 0 |
| ENSMUSG00000091259 | 0 | 0 | 0 | 0 | 0 | 0 | 0 |
| ENSMUSG00000091260 | 0 | 0.25 | 0 | 0 | 0 | 0 | 0 |
| ENSMUSG00000091264 | 2.131864492 | 2.009729747 | 1.955044741 | 2.046218836 | 2.001219193 | 2.31386722 | 1.870159858 |
| ENSMUSG00000091268 | 0 | 0 | 0 | 0 | 0 | 0 | 0 |
| ENSMUSG00000091269 | 0 | 0 | 0 | 0 | 0 | 0 | 0 |
| ENSMUSG00000091271 | 0 | 0 | 0 | 0 | 0 | 0 | 0 |
| ENSMUSG00000091272 | 0 | 0 | 0 | 0 | 0 | 0 | 0 |
| ENSMUSG00000091273 | 0 | 0 | 0 | 0 | 0 | 0 | 0 |
| ENSMUSG00000091275 | 0 | 0 | 0 | 0 | 0 | 0 | 0 |
| ENSMUSG00000091277 | 0 | 0.396240625 | 0 | 0.396240625 | 0.568060967 | 0.602059991 | 0.775756176 |
| ENSMUSG00000091280 | 0 | 0 | 0 | 0 | 0 | 0 | 0 |
| ENSMUSG00000091281 | 0 | 0 | 0 | 0 | 0 | 0 | 0 |
| ENSMUSG00000091283 | 0 | 0 | 0 | 0 | 0 | 0 | 0 |
| ENSMUSG00000091285 | 0.867194479 | 0.79248125 | 0.75 | 0.646240625 | 0.632412477 | 0.477121255 | 0.710836105 |
| ENSMUSG00000091286 | 0 | 0 | 0 | 0 | 0 | 0 | 0 |
| ENSMUSG00000091288 | 0 | 0 | 0 | 0 | 0 | 0 | 0 |
| ENSMUSG00000091289 | 0 | 0 | 0 | 0 | 0 | 0 | 0 |
| ENSMUSG00000091290 | 0 | 0 | 0 | 0 | 0 | 0 | 0 |
| ENSMUSG00000091292 | 0 | 0 | 0 | 0 | 0 | 0 | 0 |
| ENSMUSG00000091293 | 0 | 0.580482024 | 0.396240625 | 0.25 | 0.387761935 | 0 | 0 |

| | | | | | | | |
|---|---|---|---|---|---|---|---|
| ENSMUSG00000091296 | 0 | 0 | 0 | 0 | 0 | 0 | 0 |
| ENSMUSG00000091297 | 0 | 0 | 0 | 0 | 0 | 0 | 0 |
| ENSMUSG00000091304 | 0 | 0 | 0 | 0 | 0 | 0 | 0 |
| ENSMUSG00000091305 | 0 | 0 | 0 | 0 | 0 | 0 | 0 |
| ENSMUSG00000091306 | 0 | 0 | 0 | 0 | 0 | 0 | 0 |
| ENSMUSG00000091307 | 0 | 0 | 0 | 0 | 0 | 0 | 0 |
| ENSMUSG00000091311 | 0 | 0 | 0 | 0 | 0 | 0 | 0 |
| ENSMUSG00000091312 | 0 | 0 | 0 | 0 | 0 | 0 | 0 |
| ENSMUSG00000091313 | 0.289064826 | 0 | 0 | 0.25 | 0.244650542 | 0.301029996 | 0.355418052 |
| ENSMUSG00000091315 | 0 | 0 | 0 | 0 | 0 | 0 | 0 |
| ENSMUSG00000091318 | 0.578129653 | 0.25 | 0.25 | 0.396240625 | 0 | 0 | 0.224243824 |
| ENSMUSG00000091325 | 0 | 0 | 0 | 0 | 0 | 0 | 0 |
| ENSMUSG00000091326 | 0 | 0 | 0 | 0 | 0 | 0 | 0 |
| ENSMUSG00000091329 | 0.289064826 | 0.5 | 0 | 0 | 0.244650542 | 0 | 0.355418052 |
| ENSMUSG00000091336 | 0 | 0 | 0.25 | 0 | 0 | 0 | 0 |
| ENSMUSG00000091337 | 2.638209243 | 2.414106216 | 2.416777886 | 2.402294685 | 2.307676489 | 2.613841822 | 2.162979334 |
| ENSMUSG00000091345 | 0.578129653 | 0.25 | 0 | 1.431980114 | 1.462344774 | 1.431363764 | 1.374453863 |
| ENSMUSG00000091347 | 0 | 0 | 0 | 0 | 0 | 0 | 0 |
| ENSMUSG00000091350 | 0 | 0 | 0 | 0 | 0 | 0 | 0 |
| ENSMUSG00000091351 | 0 | 0 | 0 | 0 | 0 | 0.301029996 | 0 |
| ENSMUSG00000091355 | 0 | 0 | 0 | 0 | 0 | 0 | 0 |
| ENSMUSG00000091358 | 0 | 0.25 | 0 | 0 | 0 | 0 | 0 |
| ENSMUSG00000091361 | 0 | 0 | 0.25 | 0 | 0 | 0.301029996 | 0.224243824 |
| ENSMUSG00000091363 | 0 | 0.25 | 0.580482024 | 0 | 0 | 0.477121255 | 0 |
| ENSMUSG00000091366 | 0 | 0 | 0 | 0 | 0.387761935 | 0 | 0 |
| ENSMUSG00000091367 | 0 | 0 | 0 | 0 | 0 | 0 | 0 |
| ENSMUSG00000091368 | 0 | 0 | 0 | 0 | 0 | 0 | 0 |
| ENSMUSG00000091370 | 0 | 0 | 0 | 0 | 0.244650542 | 0 | 0 |
| ENSMUSG00000091373 | 0.811507563 | 1.098079356 | 1.130890489 | 1.201838731 | 1.373641807 | 1.477121255 | 1.345462945 |
| ENSMUSG00000091375 | 0 | 0 | 0 | 0 | 0 | 0 | 0 |
| ENSMUSG00000091376 | 0 | 0 | 0 | 0 | 0 | 0 | 0 |
| ENSMUSG00000091378 | 0.289064826 | 0 | 0 | 0 | 0 | 0.301029996 | 0 |
| ENSMUSG00000091381 | 0 | 0 | 0 | 0 | 0 | 0 | 0 |
| ENSMUSG00000091382 | 0 | 0 | 0 | 0 | 0 | 0 | 0 |

| | | | | | | | |
|---|---|---|---|---|---|---|---|
| ENSMUSG00000091383 | 0 | 0 | 0 | 0 | 0 | 0 | 0 |
| ENSMUSG00000091386 | 0 | 0 | 0 | 0 | 0 | 0 | 0 |
| ENSMUSG00000091387 | 1.816888701 | 1.705044741 | 1.726722649 | 1.990000483 | 1.877063019 | 2.11058971 | 1.67819947 |
| ENSMUSG00000091388 | 0 | 0 | 0 | 0 | 0 | 0 | 0 |
| ENSMUSG00000091390 | 0.289064826 | 0.25 | 0.25 | 0 | 0 | 0.301029996 | 0.224243824 |
| ENSMUSG00000091393 | 0 | 0.580482024 | 0 | 0.5 | 0.733951626 | 0.602059991 | 0.710836105 |
| ENSMUSG00000091396 | 0 | 0 | 0 | 0 | 0 | 0 | 0 |
| ENSMUSG00000091400 | 0 | 0 | 0 | 0 | 0 | 0 | 0 |
| ENSMUSG00000091401 | 0.45815691 | 0.396240625 | 0.25 | 0.25 | 0 | 0.301029996 | 0 |
| ENSMUSG00000091402 | 0 | 0 | 0 | 0 | 0 | 0 | 0 |
| ENSMUSG00000091404 | 0 | 0 | 0 | 0 | 0 | 0 | 0 |
| ENSMUSG00000091405 | 0.289064826 | 0.25 | 0.25 | 0 | 0.244650542 | 0 | 0 |
| ENSMUSG00000091407 | 0 | 0 | 0 | 0 | 0 | 0 | 0 |
| ENSMUSG00000091408 | 0.289064826 | 0.25 | 0.396240625 | 0.5 | 0 | 0.301029996 | 0.520678036 |
| ENSMUSG00000091410 | 0.671187741 | 0.75 | 0.951838731 | 0.25 | 0.632412477 | 0.477121255 | 0.710836105 |
| ENSMUSG00000091411 | 0 | 0 | 0 | 0 | 0 | 0 | 0 |
| ENSMUSG00000091412 | 0 | 0 | 0 | 0 | 0 | 0 | 0 |
| ENSMUSG00000091415 | 0.289064826 | 0.25 | 0.25 | 0.5 | 0.244650542 | 0.477121255 | 0.355418052 |
| ENSMUSG00000091416 | 0 | 0 | 0 | 0 | 0 | 0 | 0 |
| ENSMUSG00000091418 | 0 | 0.25 | 0 | 0 | 0 | 0 | 0 |
| ENSMUSG00000091419 | 0 | 0 | 0.396240625 | 0 | 0 | 0 | 0 |
| ENSMUSG00000091421 | 0 | 0 | 0 | 0 | 0 | 0 | 0 |
| ENSMUSG00000091422 | 0 | 0 | 0 | 0 | 0 | 0 | 0 |
| ENSMUSG00000091423 | 0.45815691 | 0.580482024 | 0.5 | 0.580482024 | 0.244650542 | 0 | 0.224243824 |
| ENSMUSG00000091425 | 0 | 0 | 0 | 0 | 0 | 0 | 0 |
| ENSMUSG00000091426 | 0.45815691 | 0 | 0.396240625 | 0 | 0 | 0.301029996 | 0.355418052 |
| ENSMUSG00000091429 | 0 | 0 | 0 | 0 | 0 | 0 | 0 |
| ENSMUSG00000091430 | 0 | 0 | 0 | 0 | 0 | 0 | 0 |
| ENSMUSG00000091435 | 0 | 0 | 0 | 0 | 0 | 0 | 0 |
| ENSMUSG00000091438 | 0.289064826 | 0 | 0 | 0 | 0 | 0 | 0 |
| ENSMUSG00000091441 | 0 | 0 | 0 | 0 | 0 | 0 | 0 |
| ENSMUSG00000091442 | 0 | 0 | 0 | 0.25 | 0 | 0 | 0 |
| ENSMUSG00000091443 | 0.45815691 | 0.5 | 0.5 | 0.396240625 | 0 | 0 | 0.224243824 |
| ENSMUSG00000091449 | 0.289064826 | 0 | 0.5 | 0.25 | 0.244650542 | 0 | 0 |

| | | | | | | | |
|---|---|---|---|---|---|---|---|
| ENSMUSG00000091450 | 0 | 0 | 0 | 0 | 0 | 0 | 0 |
| ENSMUSG00000091453 | 0 | 0 | 0 | 0 | 0 | 0 | 0 |
| ENSMUSG00000091455 | 0 | 0.396240625 | 0 | 0.5 | 0.387761935 | 0 | 0.579661877 |
| ENSMUSG00000091457 | 0 | 0 | 0 | 0 | 0 | 0 | 0 |
| ENSMUSG00000091459 | 0 | 0 | 0 | 0 | 0 | 0 | 0 |
| ENSMUSG00000091460 | 0 | 0.25 | 0.580482024 | 0 | 0 | 0 | 0 |
| ENSMUSG00000091462 | 0 | 0.25 | 0 | 0.396240625 | 0.489301084 | 0.301029996 | 0.520678036 |
| ENSMUSG00000091466 | 0 | 0 | 0 | 0 | 0 | 0 | 0 |
| ENSMUSG00000091468 | 0 | 0 | 0 | 0 | 0 | 0 | 0 |
| ENSMUSG00000091470 | 0 | 0 | 0 | 0 | 0 | 0 | 0 |
| ENSMUSG00000091471 | 0 | 0 | 0 | 0 | 0 | 0 | 0 |
| ENSMUSG00000091472 | 0 | 0 | 0 | 0.25 | 0 | 0 | 0.224243824 |
| ENSMUSG00000091474 | 1.777675583 | 1.726722649 | 1.741446071 | 1.752806814 | 1.635855979 | 1.799340549 | 1.508694612 |
| ENSMUSG00000091475 | 0.289064826 | 0.5 | 0.396240625 | 1.061981878 | 0.978602168 | 1 | 1.028149525 |
| ENSMUSG00000091476 | 0 | 0 | 0.25 | 0.25 | 0 | 0 | 0 |
| ENSMUSG00000091477 | 0 | 0 | 0 | 0 | 0 | 0 | 0 |
| ENSMUSG00000091478 | 1.45815691 | 1.418106335 | 1.403677461 | 1.226722649 | 1.489301084 | 1.505149978 | 1.401059159 |
| ENSMUSG00000091479 | 0 | 0 | 0 | 0 | 0 | 0 | 0 |
| ENSMUSG00000091481 | 0 | 0 | 0 | 0 | 0 | 0 | 0 |
| ENSMUSG00000091483 | 0 | 0 | 0 | 0 | 0 | 0 | 0 |
| ENSMUSG00000091486 | 0 | 0 | 0 | 0 | 0 | 0 | 0 |
| ENSMUSG00000091487 | 0 | 0 | 0 | 0 | 0 | 0 | 0 |
| ENSMUSG00000091490 | 0 | 0 | 0 | 0 | 0 | 0 | 0 |
| ENSMUSG00000091491 | 0 | 0 | 0 | 0 | 0 | 0 | 0 |
| ENSMUSG00000091492 | 0 | 0 | 0 | 0 | 0 | 0 | 0 |
| ENSMUSG00000091494 | 0 | 0 | 0 | 0 | 0 | 0 | 0 |
| ENSMUSG00000091496 | 0 | 0 | 0 | 0 | 0 | 0 | 0 |
| ENSMUSG00000091497 | 0 | 0 | 0 | 0 | 0 | 0 | 0 |
| ENSMUSG00000091498 | 0 | 0 | 0 | 0 | 0 | 0 | 0 |
| ENSMUSG00000091504 | 0 | 0 | 0.25 | 0 | 0 | 0 | 0.224243824 |
| ENSMUSG00000091507 | 0 | 0.5 | 0.5 | 0.25 | 0 | 0 | 0.448487648 |
| ENSMUSG00000091509 | 1.100572389 | 1.201838731 | 1.26109853 | 1.445339928 | 1.407936347 | 1.568201724 | 1.355418052 |
| ENSMUSG00000091510 | 0.289064826 | 0.25 | 0.25 | 0 | 0 | 0 | 0.448487648 |
| ENSMUSG00000091511 | 0 | 0 | 0 | 0 | 0 | 0 | 0 |

| | | | | | | | |
|---|---|---|---|---|---|---|---|
| ENSMUSG00000091512 | 2.063793172 | 1.981203126 | 2.004202072 | 2.017865591 | 2 | 2.245512668 | 1.895131899 |
| ENSMUSG00000091514 | 0.747221736 | 0.701838731 | 0.976722649 | 0.864857905 | 0.846351821 | 0.698970004 | 0.829800754 |
| ENSMUSG00000091515 | 0.91631382 | 0.896240625 | 1.02186571 | 0.79248125 | 0.955822902 | 0.954242509 | 1.041356071 |
| ENSMUSG00000091519 | 0 | 0 | 0 | 0 | 0 | 0 | 0 |
| ENSMUSG00000091523 | 0 | 0 | 0 | 0 | 0.244650542 | 0 | 0.224243824 |
| ENSMUSG00000091526 | 0 | 0 | 0 | 0 | 0 | 0 | 0 |
| ENSMUSG00000091528 | 0 | 0 | 0 | 0 | 0 | 0 | 0 |
| ENSMUSG00000091530 | 0.578129653 | 0.396240625 | 0.396240625 | 0.580482024 | 0.244650542 | 0.477121255 | 0.579661877 |
| ENSMUSG00000091531 | 0 | 0.25 | 0 | 0 | 0 | 0 | 0 |
| ENSMUSG00000091534 | 0.747221736 | 0.951838731 | 0.79248125 | 0.646240625 | 0.568060967 | 0.477121255 | 0.579661877 |
| ENSMUSG00000091537 | 2.558729299 | 2.390083209 | 2.392463902 | 2.556000419 | 2.522218979 | 2.909020854 | 2.353854633 |
| ENSMUSG00000091539 | 0 | 0 | 0 | 0 | 0 | 0 | 0 |
| ENSMUSG00000091541 | 0 | 0 | 0 | 0 | 0 | 0 | 0 |
| ENSMUSG00000091542 | 0.578129653 | 0.830482024 | 0.79248125 | 0.75 | 0.686820904 | 0.698970004 | 0.672731473 |
| ENSMUSG00000091548 | 0 | 0 | 0 | 0 | 0 | 0 | 0 |
| ENSMUSG00000091549 | 2.020182925 | 1.899978211 | 1.914552871 | 1.880890489 | 1.746695109 | 1.982271233 | 1.667169465 |
| ENSMUSG00000091550 | 0 | 0 | 0 | 0 | 0 | 0 | 0 |
| ENSMUSG00000091552 | 0 | 0 | 0 | 0 | 0 | 0 | 0 |
| ENSMUSG00000091553 | 1.578129653 | 1.54745614 | 1.494319981 | 0.864857905 | 0.77552387 | 0.77815125 | 0.803905701 |
| ENSMUSG00000091556 | 0 | 0 | 0 | 0.25 | 0 | 0 | 0 |
| ENSMUSG00000091557 | 0 | 0.25 | 0 | 0 | 0 | 0 | 0 |
| ENSMUSG00000091558 | 0 | 0.25 | 0 | 0 | 0.489301084 | 0 | 0.355418052 |
| ENSMUSG00000091561 | 0.289064826 | 0.25 | 0.25 | 0 | 0.244650542 | 0.301029996 | 0.224243824 |
| ENSMUSG00000091562 | 1.614416215 | 1.532320754 | 1.494319981 | 1.692046081 | 1.639266209 | 1.851258349 | 1.499406449 |
| ENSMUSG00000091563 | 0 | 0 | 0 | 0 | 0 | 0 | 0 |
| ENSMUSG00000091568 | 0 | 0 | 0 | 0 | 0 | 0 | 0 |
| ENSMUSG00000091569 | 0 | 0 | 0 | 0 | 0 | 0 | 0 |
| ENSMUSG00000091570 | 0 | 0 | 0 | 0 | 0 | 0 | 0 |
| ENSMUSG00000091572 | 0 | 0 | 0 | 0 | 0 | 0 | 0 |
| ENSMUSG00000091573 | 0 | 0 | 0 | 0 | 0 | 0 | 0 |
| ENSMUSG00000091575 | 0.289064826 | 0 | 0 | 0.25 | 0 | 0.301029996 | 0.355418052 |
| ENSMUSG00000091577 | 0.91631382 | 0.701838731 | 0.646240625 | 0.896240625 | 0.686820904 | 0.698970004 | 0.448487648 |
| ENSMUSG00000091580 | 0.45815691 | 0.646240625 | 0.646240625 | 0.5 | 0.244650542 | 0.602059991 | 0.520678036 |
| ENSMUSG00000091583 | 0 | 0 | 0 | 0.25 | 0 | 0 | 0 |

| | | | | | | | |
|---|---|---|---|---|---|---|---|
| ENSMUSG00000091584 | 0 | 0 | 0 | 0 | 0 | 0 | 0 |
| ENSMUSG00000091585 | 0 | 0 | 0 | 0 | 0 | 0 | 0 |
| ENSMUSG00000091586 | 1.470606563 | 1.26109853 | 1.488549078 | 1.098079356 | 1.136121934 | 1.079181246 | 1.224243824 |
| ENSMUSG00000091587 | 0.289064826 | 0.396240625 | 0 | 0.25 | 0.244650542 | 0 | 0 |
| ENSMUSG00000091594 | 0 | 0.25 | 0 | 0 | 0 | 0 | 0 |
| ENSMUSG00000091595 | 0 | 0 | 0 | 0 | 0 | 0 | 0 |
| ENSMUSG00000091601 | 0 | 0 | 0 | 0 | 0 | 0 | 0 |
| ENSMUSG00000091604 | 0 | 0.25 | 0 | 0 | 0 | 0.301029996 | 0 |
| ENSMUSG00000091605 | 0 | 0 | 0 | 0 | 0 | 0 | 0 |
| ENSMUSG00000091606 | 0 | 0 | 0 | 0 | 0 | 0 | 0 |
| ENSMUSG00000091613 | 0 | 0.25 | 0 | 0 | 0.244650542 | 0 | 0.224243824 |
| ENSMUSG00000091614 | 0 | 0 | 0 | 0 | 0 | 0 | 0.224243824 |
| ENSMUSG00000091615 | 0 | 0 | 0 | 0 | 0 | 0 | 0 |
| ENSMUSG00000091617 | 0 | 0 | 0 | 0 | 0 | 0 | 0 |
| ENSMUSG00000091618 | 1 | 1.04248125 | 1.04248125 | 0.5 | 0.387761935 | 0.477121255 | 0.355418052 |
| ENSMUSG00000091619 | 0 | 0 | 0 | 0 | 0 | 0 | 0 |
| ENSMUSG00000091620 | 0 | 0 | 0 | 0 | 0 | 0 | 0 |
| ENSMUSG00000091623 | 0.747221736 | 0.646240625 | 0.701838731 | 0 | 0.387761935 | 0.301029996 | 0.520678036 |
| ENSMUSG00000091624 | 0 | 0 | 0 | 0 | 0 | 0 | 0 |
| ENSMUSG00000091625 | 2.066740409 | 1.940387808 | 1.938721876 | 1.991446071 | 1.950242452 | 2.176091259 | 1.812372009 |
| ENSMUSG00000091627 | 0 | 0 | 0 | 0 | 0 | 0 | 0 |
| ENSMUSG00000091631 | 0 | 0 | 0 | 0 | 0 | 0 | 0 |
| ENSMUSG00000091635 | 0 | 0 | 0.25 | 0 | 0 | 0 | 0 |
| ENSMUSG00000091636 | 0 | 0 | 0 | 0 | 0 | 0 | 0 |
| ENSMUSG00000091638 | 0 | 0 | 0 | 0 | 0 | 0 | 0 |
| ENSMUSG00000091639 | 0 | 0 | 0 | 0 | 0.489301084 | 0 | 0.224243824 |
| ENSMUSG00000091648 | 0.747221736 | 0.396240625 | 0 | 0 | 0 | 0 | 0 |
| ENSMUSG00000091649 | 0 | 0 | 0 | 0 | 0.244650542 | 0 | 0 |
| ENSMUSG00000091650 | 0 | 0 | 0 | 0 | 0 | 0 | 0 |
| ENSMUSG00000091651 | 0 | 0 | 0 | 0 | 0 | 0 | 0 |
| ENSMUSG00000091652 | 0 | 0 | 0 | 0 | 0 | 0 | 0 |
| ENSMUSG00000091655 | 0 | 0 | 0 | 0 | 0 | 0 | 0 |
| ENSMUSG00000091656 | 0 | 0 | 0 | 0 | 0 | 0 | 0 |
| ENSMUSG00000091657 | 0 | 0 | 0 | 0 | 0 | 0 | 0 |

| | | | | | | | |
|---|---|---|---|---|---|---|---|
| ENSMUSG00000091659 | 0 | 0 | 0 | 0 | 0 | 0 | 0 |
| ENSMUSG00000091661 | 0 | 0 | 0 | 0 | 0 | 0 | 0 |
| ENSMUSG00000091662 | 0 | 0 | 0 | 0 | 0 | 0 | 0 |
| ENSMUSG00000091665 | 0 | 0 | 0 | 0 | 0 | 0 | 0 |
| ENSMUSG00000091668 | 0 | 0 | 0 | 0 | 0 | 0 | 0 |
| ENSMUSG00000091669 | 0 | 0 | 0 | 0 | 0 | 0 | 0 |
| ENSMUSG00000091670 | 0 | 0 | 0 | 0 | 0 | 0 | 0 |
| ENSMUSG00000091671 | 0 | 0 | 0 | 0 | 0 | 0 | 0 |
| ENSMUSG00000091672 | 0 | 0 | 0 | 0 | 0 | 0 | 0 |
| ENSMUSG00000091676 | 0 | 0 | 0 | 0 | 0 | 0 | 0 |
| ENSMUSG00000091679 | 0 | 0 | 0 | 0 | 0 | 0 | 0 |
| ENSMUSG00000091680 | 0 | 0 | 0 | 0 | 0 | 0 | 0 |
| ENSMUSG00000091682 | 0 | 0 | 0 | 0 | 0 | 0 | 0 |
| ENSMUSG00000091685 | 0 | 0 | 0 | 0 | 0 | 0 | 0 |
| ENSMUSG00000091686 | 0 | 0 | 0 | 0 | 0.244650542 | 0 | 0 |
| ENSMUSG00000091692 | 0 | 0 | 0 | 0 | 0 | 0 | 0 |
| ENSMUSG00000091694 | 0 | 0.580482024 | 0.396240625 | 0 | 0 | 0 | 0 |
| ENSMUSG00000091695 | 0 | 0 | 0 | 0 | 0 | 0 | 0 |
| ENSMUSG00000091697 | 0 | 0.25 | 0 | 0 | 0 | 0 | 0 |
| ENSMUSG00000091698 | 0 | 0 | 0 | 0 | 0 | 0 | 0 |
| ENSMUSG00000091700 | 0 | 0 | 0 | 0 | 0 | 0 | 0 |
| ENSMUSG00000091703 | 0 | 0 | 0 | 0 | 0 | 0 | 0 |
| ENSMUSG00000091705 | 0.747221736 | 0.92510993 | 1 | 1.557204673 | 1.621875691 | 1.662757832 | 1.674564426 |
| ENSMUSG00000091707 | 0 | 0 | 0 | 0 | 0 | 0 | 0 |
| ENSMUSG00000091709 | 0 | 0 | 0 | 0 | 0 | 0 | 0 |
| ENSMUSG00000091712 | 0.289064826 | 0 | 0.396240625 | 0 | 0.568060967 | 0 | 0.224243824 |
| ENSMUSG00000091713 | 0 | 0 | 0 | 0 | 0 | 0 | 0 |
| ENSMUSG00000091718 | 0 | 0 | 0 | 0 | 0 | 0 | 0 |
| ENSMUSG00000091721 | 0 | 0 | 0 | 0 | 0 | 0 | 0 |
| ENSMUSG00000091722 | 0 | 0 | 0 | 0 | 0 | 0 | 0 |
| ENSMUSG00000091723 | 0 | 0 | 0 | 0 | 0 | 0 | 0 |
| ENSMUSG00000091725 | 0 | 0 | 0 | 0 | 0 | 0.301029996 | 0 |
| ENSMUSG00000091730 | 0.45815691 | 0 | 0.396240625 | 0 | 0 | 0 | 0 |
| ENSMUSG00000091731 | 0 | 0 | 0 | 0 | 0 | 0 | 0 |

| | | | | | | | |
|---|---|---|---|---|---|---|---|
| ENSMUSG00000091732 | 0.811507563 | 0.701838731 | 0.701838731 | 0.5 | 0.77552387 | 0.602059991 | 0.672731473 |
| ENSMUSG00000091733 | 0 | 0 | 0 | 0 | 0 | 0 | 0 |
| ENSMUSG00000091734 | 0 | 0 | 0 | 0 | 0 | 0 | 0 |
| ENSMUSG00000091735 | 0.45815691 | 0 | 0.580482024 | 0.5 | 0.489301084 | 0.477121255 | 0.520678036 |
| ENSMUSG00000091736 | 0 | 0 | 0 | 0 | 0 | 0 | 0 |
| ENSMUSG00000091740 | 0 | 0 | 0 | 0 | 0 | 0 | 0 |
| ENSMUSG00000091741 | 0 | 0 | 0 | 0 | 0 | 0 | 0 |
| ENSMUSG00000091742 | 0.289064826 | 0 | 0.25 | 0.25 | 0.387761935 | 0 | 0.224243824 |
| ENSMUSG00000091743 | 0 | 0 | 0 | 0 | 0 | 0 | 0.224243824 |
| ENSMUSG00000091744 | 0 | 0.396240625 | 0 | 0.396240625 | 0.877063019 | 0.84509804 | 0.579661877 |
| ENSMUSG00000091745 | 0 | 0 | 0 | 0 | 0 | 0 | 0 |
| ENSMUSG00000091749 | 0 | 0 | 0 | 0 | 0 | 0 | 0 |
| ENSMUSG00000091752 | 0 | 0 | 0 | 0 | 0 | 0 | 0 |
| ENSMUSG00000091754 | 0.867194479 | 0.830482024 | 0.951838731 | 0.92510993 | 0.905314583 | 0.301029996 | 0.876096088 |
| ENSMUSG00000091756 | 0 | 0 | 0 | 0 | 0 | 0 | 0 |
| ENSMUSG00000091757 | 0 | 0 | 0 | 0 | 0 | 0 | 0 |
| ENSMUSG00000091762 | 0 | 0 | 0 | 0 | 0 | 0 | 0 |
| ENSMUSG00000091764 | 1.404271433 | 1.380890489 | 1.348079356 | 1.418106335 | 1.358933174 | 1.414973348 | 1.224243824 |
| ENSMUSG00000091768 | 0 | 0 | 0 | 0 | 0 | 0 | 0 |
| ENSMUSG00000091771 | 0 | 0 | 0 | 0 | 0 | 0 | 0 |
| ENSMUSG00000091775 | 0 | 0 | 0 | 0 | 0 | 0 | 0 |
| ENSMUSG00000091776 | 0 | 0 | 0 | 0 | 0 | 0 | 0 |
| ENSMUSG00000091779 | 0 | 0 | 0 | 0 | 0 | 0 | 0 |
| ENSMUSG00000091780 | 0 | 0 | 0 | 0 | 0 | 0 | 0 |
| ENSMUSG00000091784 | 0.747221736 | 0.896240625 | 0.896240625 | 0.830482024 | 0.846351821 | 0.77815125 | 0.710836105 |
| ENSMUSG00000091785 | 0 | 0.25 | 0 | 0 | 0 | 0 | 0.224243824 |
| ENSMUSG00000091786 | 0.867194479 | 0.92510993 | 0.976722649 | 0.830482024 | 1 | 0.84509804 | 0.935079929 |
| ENSMUSG00000091788 | 0 | 0 | 0 | 0 | 0 | 0 | 0 |
| ENSMUSG00000091792 | 0 | 0 | 0 | 0 | 0 | 0 | 0 |
| ENSMUSG00000091794 | 0 | 0 | 0 | 0.25 | 0 | 0 | 0 |
| ENSMUSG00000091800 | 0 | 0 | 0 | 0 | 0 | 0 | 0 |
| ENSMUSG00000091803 | 0.289064826 | 0.396240625 | 0.646240625 | 0.5 | 0.387761935 | 0.301029996 | 0.579661877 |
| ENSMUSG00000091805 | 0 | 0 | 0 | 0 | 0 | 0 | 0 |
| ENSMUSG00000091806 | 0 | 0 | 0 | 0 | 0 | 0 | 0 |

| | | | | | | | |
|---|---|---|---|---|---|---|---|
| ENSMUSG00000091808 | 0 | 0 | 0 | 0 | 0 | 0 | 0 |
| ENSMUSG00000091809 | 0 | 0 | 0 | 0 | 0 | 0 | 0 |
| ENSMUSG00000091811 | 1.307602651 | 1.302363341 | 1.372963274 | 1.311981878 | 1.244650542 | 1.380211242 | 1.245582246 |
| ENSMUSG00000091812 | 0 | 0 | 0 | 0 | 0 | 0 | 0 |
| ENSMUSG00000091813 | 0 | 0 | 0 | 0 | 0 | 0 | 0 |
| ENSMUSG00000091814 | 0 | 0 | 0.25 | 0 | 0 | 0 | 0 |
| ENSMUSG00000091815 | 0 | 0 | 0 | 0 | 0 | 0 | 0 |
| ENSMUSG00000091816 | 0 | 0 | 0 | 0 | 0 | 0 | 0 |
| ENSMUSG00000091817 | 0 | 0 | 0 | 0 | 0 | 0 | 0 |
| ENSMUSG00000091818 | 0 | 0.580482024 | 0.396240625 | 0.25 | 0.568060967 | 0 | 0.224243824 |
| ENSMUSG00000091821 | 0 | 0 | 0 | 0 | 0 | 0 | 0 |
| ENSMUSG00000091825 | 0.289064826 | 0.396240625 | 0.79248125 | 0.5 | 0.489301084 | 0.477121255 | 0.520678036 |
| ENSMUSG00000091827 | 0 | 0 | 0 | 0 | 0 | 0 | 0 |
| ENSMUSG00000091831 | 0.578129653 | 0 | 0.580482024 | 0 | 0.77552387 | 0.477121255 | 0.520678036 |
| ENSMUSG00000091832 | 0 | 0 | 0 | 0 | 0 | 0 | 0 |
| ENSMUSG00000091834 | 0 | 0 | 0 | 0 | 0 | 0 | 0 |
| ENSMUSG00000091842 | 0 | 0 | 0 | 0 | 0 | 0 | 0 |
| ENSMUSG00000091844 | 0.671187741 | 0.646240625 | 0.75 | 0.75 | 0.812711509 | 1.079181246 | 0.579661877 |
| ENSMUSG00000091845 | 0.747221736 | 1.080482024 | 1.130890489 | 0.396240625 | 0.955822902 | 0 | 1.089372236 |
| ENSMUSG00000091847 | 0 | 0 | 0 | 0 | 0 | 0 | 0 |
| ENSMUSG00000091849 | 0 | 0 | 0 | 0 | 0 | 0 | 0 |
| ENSMUSG00000091859 | 0 | 0 | 0 | 0 | 0 | 0 | 0 |
| ENSMUSG00000091863 | 0 | 0 | 0 | 0 | 0 | 0 | 0 |
| ENSMUSG00000091864 | 0 | 0.580482024 | 0.5 | 0.396240625 | 0.244650542 | 0.301029996 | 0.448487648 |
| ENSMUSG00000091866 | 0.289064826 | 0.25 | 0.25 | 0.646240625 | 0.387761935 | 0.698970004 | 0.448487648 |
| ENSMUSG00000091867 | 0 | 0 | 0 | 0 | 0 | 0 | 0 |
| ENSMUSG00000091873 | 0 | 0 | 0 | 0 | 0 | 0 | 0 |
| ENSMUSG00000091874 | 0 | 0 | 0 | 0 | 0 | 0 | 0 |
| ENSMUSG00000091876 | 0 | 0 | 0 | 0 | 0 | 0 | 0 |
| ENSMUSG00000091879 | 0 | 0 | 0 | 0 | 0 | 0 | 0 |
| ENSMUSG00000091881 | 0.289064826 | 0.396240625 | 0.396240625 | 0.396240625 | 0.489301084 | 0.477121255 | 0.355418052 |
| ENSMUSG00000091882 | 0 | 0 | 0 | 0 | 0 | 0 | 0 |
| ENSMUSG00000091887 | 0 | 0 | 0 | 0 | 0.244650542 | 0 | 0 |
| ENSMUSG00000091888 | 0 | 0 | 0 | 0 | 0 | 0 | 0 |

| | | | | | | | |
|---|---|---|---|---|---|---|---|
| ENSMUSG00000091890 | 0 | 0 | 0 | 0 | 0 | 0 | 0 |
| ENSMUSG00000091894 | 0 | 0 | 0 | 0 | 0 | 0 | 0 |
| ENSMUSG00000091896 | 2.942293987 | 2.667442722 | 2.645535495 | 2.697179583 | 2.599361968 | 3.02325246 | 2.405288179 |
| ENSMUSG00000091897 | 0 | 0 | 0 | 0 | 0 | 0 | 0 |
| ENSMUSG00000091898 | 0.747221736 | 0.701838731 | 0.646240625 | 0.580482024 | 0.905314583 | 0.77815125 | 0.969165684 |
| ENSMUSG00000091900 | 1.069666964 | 0.951838731 | 0.896240625 | 1.188721876 | 1.200473444 | 1.176091259 | 1.185218806 |
| ENSMUSG00000091902 | 0 | 0 | 0 | 0 | 0 | 0 | 0 |
| ENSMUSG00000091903 | 0 | 0 | 0 | 0 | 0 | 0 | 0 |
| ENSMUSG00000091905 | 0 | 0.396240625 | 0 | 0 | 0 | 0 | 0 |
| ENSMUSG00000091906 | 0 | 0.25 | 0 | 0 | 0 | 0 | 0 |
| ENSMUSG00000091908 | 0 | 0 | 0 | 0 | 0 | 0 | 0 |
| ENSMUSG00000091911 | 0 | 0 | 0 | 0 | 0 | 0 | 0 |
| ENSMUSG00000091922 | 0 | 0 | 0 | 0 | 0 | 0 | 0 |
| ENSMUSG00000091923 | 0 | 0 | 0 | 0 | 0 | 0 | 0 |
| ENSMUSG00000091924 | 0 | 0 | 0 | 0 | 0 | 0 | 0 |
| ENSMUSG00000091926 | 0 | 0 | 0 | 0 | 0 | 0 | 0 |
| ENSMUSG00000091929 | 0 | 0 | 0 | 0 | 0 | 0 | 0 |
| ENSMUSG00000091930 | 0.289064826 | 0 | 0 | 0 | 0 | 0 | 0 |
| ENSMUSG00000091931 | 1.587501561 | 1.575945187 | 1.614857905 | 1.494319981 | 1.394615667 | 1.62324929 | 1.379042812 |
| ENSMUSG00000091933 | 0 | 0 | 0 | 0 | 0 | 0 | 0 |
| ENSMUSG00000091936 | 0 | 0 | 0 | 0 | 0 | 0 | 0 |
| ENSMUSG00000091937 | 0 | 0 | 0 | 0 | 0 | 0 | 0 |
| ENSMUSG00000091938 | 0 | 0 | 0 | 0 | 0.244650542 | 0 | 0 |
| ENSMUSG00000091941 | 0 | 0.5 | 0 | 0.5 | 0 | 0.301029996 | 0.224243824 |
| ENSMUSG00000091942 | 0 | 0 | 0 | 0 | 0 | 0.301029996 | 0.224243824 |
| ENSMUSG00000091945 | 0 | 0 | 0 | 0 | 0 | 0 | 0 |
| ENSMUSG00000091947 | 0 | 0.396240625 | 0 | 0.5 | 0.244650542 | 0 | 0.448487648 |
| ENSMUSG00000091950 | 0 | 0 | 0 | 0.25 | 0 | 0 | 0 |
| ENSMUSG00000091952 | 0 | 0 | 0.5 | 0.5 | 0.387761935 | 0 | 0.355418052 |
| ENSMUSG00000091955 | 0.578129653 | 0.25 | 0 | 0.25 | 0.387761935 | 0.301029996 | 0.224243824 |
| ENSMUSG00000091956 | 0 | 0 | 0 | 0 | 0 | 0 | 0 |
| ENSMUSG00000091957 | 1.205378646 | 1.26109853 | 1.238549078 | 1.282320754 | 1.327538614 | 1.322219295 | 1.278288402 |
| ENSMUSG00000091960 | 0 | 0 | 0 | 0 | 0 | 0 | 0 |
| ENSMUSG00000091961 | 0 | 0 | 0 | 0 | 0 | 0 | 0 |

| | | | | | | | |
|---|---|---|---|---|---|---|---|
| ENSMUSG00000091962 | 0 | 0 | 0 | 0 | 0 | 0 | 0 |
| ENSMUSG00000091967 | 0 | 0 | 0 | 0 | 0 | 0 | 0 |
| ENSMUSG00000091968 | 0 | 0.25 | 0.5 | 0.396240625 | 0.686820904 | 0.301029996 | 0.448487648 |
| ENSMUSG00000091970 | 0 | 0 | 0 | 0 | 0 | 0 | 0 |
| ENSMUSG00000091971 | 1.156259305 | 1.146240625 | 1.226722649 | 0.896240625 | 1 | 1.041392685 | 0.952571511 |
| ENSMUSG00000091972 | 0 | 0 | 0 | 0 | 0 | 0 | 0 |
| ENSMUSG00000091980 | 0 | 0 | 0 | 0 | 0 | 0 | 0 |
| ENSMUSG00000091983 | 0 | 0 | 0 | 0 | 0 | 0 | 0 |
| ENSMUSG00000091985 | 1.325351389 | 1.080482024 | 1.188721876 | 1.356566189 | 1.343584837 | 1.447158031 | 1.20139795 |
| ENSMUSG00000091987 | 0 | 0 | 0 | 0 | 0 | 0 | 0 |
| ENSMUSG00000091989 | 0 | 0 | 0 | 0 | 0 | 0 | 0 |
| ENSMUSG00000091990 | 0 | 0 | 0 | 0 | 0 | 0 | 0 |
| ENSMUSG00000091993 | 0.671187741 | 0.79248125 | 0.896240625 | 0.580482024 | 0.686820904 | 0.477121255 | 0.579661877 |
| ENSMUSG00000091994 | 1.470606563 | 1.388647213 | 1.445339928 | 1.396240625 | 1.42077253 | 1.397940009 | 1.38802993 |
| ENSMUSG00000091995 | 0 | 0 | 0 | 0 | 0 | 0 | 0 |
| ENSMUSG00000091996 | 0 | 0 | 0 | 0 | 0 | 0 | 0 |
| ENSMUSG00000091997 | 0 | 0 | 0 | 0.5 | 0.387761935 | 0 | 0.355418052 |
| ENSMUSG00000091998 | 0 | 0 | 0 | 0 | 0 | 0 | 0 |
| ENSMUSG00000092004 | 0 | 0 | 0 | 0 | 0 | 0 | 0 |
| ENSMUSG00000092006 | 0 | 0 | 0 | 0 | 0 | 0 | 0 |
| ENSMUSG00000092008 | 0 | 0 | 0 | 0 | 0 | 0 | 0 |
| ENSMUSG00000092009 | 0 | 0 | 0.25 | 0.25 | 0 | 0.301029996 | 0.224243824 |
| ENSMUSG00000092014 | 0 | 0 | 0 | 0 | 0 | 0 | 0 |
| ENSMUSG00000092019 | 0.289064826 | 0 | 0 | 0 | 0 | 0 | 0 |
| ENSMUSG00000092021 | 0 | 0 | 0 | 0 | 0 | 0 | 0.224243824 |
| ENSMUSG00000092027 | 0 | 0 | 0 | 0 | 0 | 0 | 0 |
| ENSMUSG00000092029 | 0 | 0 | 0 | 0.25 | 0 | 0 | 0 |
| ENSMUSG00000092031 | 0 | 0 | 0 | 0 | 0 | 0 | 0 |
| ENSMUSG00000092032 | 0 | 0 | 0 | 0 | 0 | 0 | 0 |
| ENSMUSG00000092035 | 3.995359894 | 3.612786676 | 3.55133377 | 2.974272282 | 2.888084401 | 3.343408594 | 2.705670528 |
| ENSMUSG00000092036 | 0 | 0 | 0 | 0.25 | 0 | 0 | 0.224243824 |
| ENSMUSG00000092039 | 0 | 0 | 0 | 0 | 0 | 0 | 0 |
| ENSMUSG00000092040 | 0 | 0 | 0 | 0 | 0 | 0 | 0 |
| ENSMUSG00000092042 | 0 | 0 | 0 | 0 | 0 | 0 | 0 |

| | | | | | | | |
|---|---|---|---|---|---|---|---|
| ENSMUSG00000092043 | 0.289064826 | 0 | 0 | 0 | 0 | 0 | 0 |
| ENSMUSG00000092045 | 0 | 0 | 0 | 0 | 0 | 0 | 0 |
| ENSMUSG00000092047 | 0 | 0 | 0 | 0 | 0 | 0 | 0 |
| ENSMUSG00000092048 | 0 | 0 | 0 | 0 | 0 | 0 | 0 |
| ENSMUSG00000092049 | 0 | 0 | 0 | 0 | 0 | 0 | 0 |
| ENSMUSG00000092051 | 0 | 0 | 0 | 0 | 0 | 0 | 0 |
| ENSMUSG00000092054 | 0 | 0 | 0 | 0 | 0 | 0 | 0 |
| ENSMUSG00000092055 | 0 | 0 | 0 | 0 | 0 | 0 | 0 |
| ENSMUSG00000092056 | 0 | 0 | 0.396240625 | 0 | 0 | 0 | 0.224243824 |
| ENSMUSG00000092060 | 2.315764015 | 2.21759118 | 2.168106335 | 2.409156155 | 2.31625483 | 2.618048097 | 2.194143936 |
| ENSMUSG00000092062 | 0 | 0.5 | 0.25 | 0.396240625 | 0.244650542 | 0.301029996 | 0.355418052 |
| ENSMUSG00000092066 | 0 | 0 | 0 | 0 | 0 | 0 | 0 |
| ENSMUSG00000092071 | 0.45815691 | 0.5 | 0.580482024 | 0.580482024 | 0 | 0 | 0.355418052 |
| ENSMUSG00000092072 | 0.578129653 | 0.5 | 0.25 | 0.5 | 0.244650542 | 0.477121255 | 0.448487648 |
| ENSMUSG00000092073 | 0 | 0 | 0 | 0 | 0 | 0 | 0 |
| ENSMUSG00000092074 | 0 | 0.25 | 0 | 0 | 0 | 0 | 0.224243824 |
| ENSMUSG00000092075 | 0 | 0 | 0 | 0 | 0 | 0 | 0 |
| ENSMUSG00000092077 | 0 | 0.25 | 0.5 | 0 | 0 | 0 | 0 |
| ENSMUSG00000092080 | 0 | 0 | 0 | 0 | 0 | 0 | 0.224243824 |
| ENSMUSG00000092083 | 0 | 0.25 | 0 | 0.25 | 0 | 0 | 0.224243824 |
| ENSMUSG00000092085 | 0 | 0 | 0 | 0 | 0 | 0.301029996 | 0 |
| ENSMUSG00000092086 | 0.671187741 | 0.580482024 | 0.830482024 | 0.580482024 | 0.733951626 | 0.698970004 | 0.672731473 |
| ENSMUSG00000092090 | 0 | 0.25 | 0 | 0 | 0 | 0 | 0 |
| ENSMUSG00000092092 | 0 | 0 | 0 | 0 | 0 | 0 | 0 |
| ENSMUSG00000092093 | 0 | 0 | 0 | 0 | 0 | 0 | 0 |
| ENSMUSG00000092094 | 0 | 0 | 0 | 0 | 0 | 0 | 0 |
| ENSMUSG00000092097 | 0 | 0.25 | 0.25 | 0.396240625 | 0.244650542 | 0 | 0.224243824 |
| ENSMUSG00000092103 | 0 | 0 | 0 | 0.25 | 0 | 0 | 0 |
| ENSMUSG00000092107 | 0 | 0 | 0 | 0 | 0 | 0 | 0 |
| ENSMUSG00000092109 | 0 | 0 | 0 | 0 | 0 | 0 | 0 |
| ENSMUSG00000092111 | 0 | 0 | 0 | 0 | 0 | 0 | 0 |
| ENSMUSG00000092116 | 0 | 0.25 | 0 | 0 | 0 | 0.301029996 | 0 |
| ENSMUSG00000092118 | 1.249317394 | 1.380890489 | 1.214495249 | 1.17510993 | 1.176121988 | 1.113943352 | 1.028149525 |
| ENSMUSG00000092120 | 0 | 0 | 0 | 0 | 0 | 0 | 0 |

| | | | | | | | |
|---|---|---|---|---|---|---|---|
| ENSMUSG00000092123 | 0 | 0 | 0 | 0 | 0 | 0 | 0 |
| ENSMUSG00000092124 | 0.578129653 | 0.580482024 | 0.646240625 | 0.5 | 0.244650542 | 0.477121255 | 0.520678036 |
| ENSMUSG00000092125 | 0 | 0 | 0 | 0 | 0 | 0 | 0 |
| ENSMUSG00000092130 | 0.671187741 | 0.701838731 | 0.92510993 | 0 | 0 | 0.301029996 | 0.355418052 |
| ENSMUSG00000092131 | 0 | 0.25 | 0.25 | 0.25 | 0.244650542 | 0.301029996 | 0.224243824 |
| ENSMUSG00000092134 | 0 | 0.396240625 | 0.25 | 0.25 | 0.244650542 | 0.301029996 | 0 |
| ENSMUSG00000092137 | 0 | 0 | 0 | 0 | 0 | 0 | 0 |
| ENSMUSG00000092142 | 0 | 0 | 0 | 0 | 0 | 0.301029996 | 0 |
| ENSMUSG00000092144 | 0 | 0 | 0 | 0 | 0 | 0 | 0 |
| ENSMUSG00000092147 | 0 | 0 | 0 | 0 | 0 | 0 | 0 |
| ENSMUSG00000092148 | 0 | 0 | 0 | 0 | 0 | 0 | 0 |
| ENSMUSG00000092152 | 0 | 0 | 0 | 0 | 0 | 0 | 0 |
| ENSMUSG00000092157 | 0 | 0 | 0 | 0 | 0 | 0 | 0 |
| ENSMUSG00000092162 | 0 | 0 | 0 | 0 | 0 | 0 | 0 |
| ENSMUSG00000092164 | 0 | 0 | 0 | 0 | 0 | 0 | 0 |
| ENSMUSG00000092165 | 0 | 0 | 0 | 0 | 0 | 0 | 0 |
| ENSMUSG00000092166 | 0 | 0 | 0 | 0 | 0 | 0 | 0 |
| ENSMUSG00000092167 | 0.289064826 | 0.25 | 0.25 | 0 | 0 | 0 | 0 |
| ENSMUSG00000092168 | 0 | 0 | 0 | 0 | 0 | 0 | 0 |
| ENSMUSG00000092171 | 0.289064826 | 0.396240625 | 0.396240625 | 0.25 | 0.489301084 | 0 | 0 |
| ENSMUSG00000092172 | 0 | 0 | 0 | 0 | 0 | 0 | 0 |
| ENSMUSG00000092173 | 0 | 0 | 0 | 0 | 0 | 0 | 0 |
| ENSMUSG00000092174 | 0 | 0 | 0 | 0 | 0 | 0 | 0 |
| ENSMUSG00000092175 | 0 | 0 | 0 | 0 | 0 | 0 | 0 |
| ENSMUSG00000092176 | 0 | 0 | 0 | 0 | 0 | 0 | 0 |
| ENSMUSG00000092178 | 1.129344651 | 1.080482024 | 1 | 1.061981878 | 0.978602168 | 1.113943352 | 1.014380832 |
| ENSMUSG00000092179 | 0 | 0 | 0 | 0 | 0 | 0 | 0 |
| ENSMUSG00000092181 | 0 | 0 | 0 | 0 | 0 | 0 | 0 |
| ENSMUSG00000092182 | 0 | 0 | 0 | 0 | 0 | 0 | 0 |
| ENSMUSG00000092183 | 0.578129653 | 0.396240625 | 0.701838731 | 0.75 | 0.877063019 | 0.84509804 | 0.579661877 |
| ENSMUSG00000092184 | 0 | 0 | 0 | 0 | 0 | 0 | 0 |
| ENSMUSG00000092185 | 0 | 0 | 0 | 0 | 0 | 0 | 0 |
| ENSMUSG00000092186 | 0 | 0 | 0.5 | 0.5 | 0.387761935 | 0.301029996 | 0.224243824 |
| ENSMUSG00000092187 | 0.671187741 | 0.79248125 | 0.580482024 | 0.580482024 | 0.905314583 | 0.477121255 | 0.579661877 |

| | | | | | | | |
|---|---|---|---|---|---|---|---|
| ENSMUSG00000092188 | 0 | 0 | 0 | 0.25 | 0 | 0 | 0 |
| ENSMUSG00000092189 | 0 | 0 | 0 | 0 | 0 | 0 | 0 |
| ENSMUSG00000092190 | 0 | 0 | 0 | 0.25 | 0.489301084 | 0 | 0.224243824 |
| ENSMUSG00000092191 | 0 | 0 | 0 | 0 | 0 | 0 | 0 |
| ENSMUSG00000092192 | 0 | 0 | 0 | 0 | 0 | 0 | 0 |
| ENSMUSG00000092193 | 0 | 0 | 0 | 0 | 0 | 0 | 0 |
| ENSMUSG00000092195 | 0 | 0 | 0 | 0 | 0 | 0 | 0 |
| ENSMUSG00000092196 | 0 | 0 | 0 | 0 | 0 | 0 | 0 |
| ENSMUSG00000092197 | 0 | 0 | 0 | 0 | 0 | 0 | 0 |
| ENSMUSG00000092199 | 0 | 0 | 0 | 0 | 0 | 0 | 0 |
| ENSMUSG00000092200 | 0 | 0 | 0 | 0 | 0 | 0 | 0 |
| ENSMUSG00000092201 | 0 | 0 | 0 | 0 | 0 | 0 | 0 |
| ENSMUSG00000092202 | 0.289064826 | 0.396240625 | 0.5 | 0.25 | 0.244650542 | 0.602059991 | 0.74492186 |
| ENSMUSG00000092203 | 2.617563644 | 2.408702763 | 2.490724001 | 2.192046081 | 2.266550905 | 2.42975228 | 2.197791274 |
| ENSMUSG00000092204 | 0 | 0 | 0 | 0 | 0 | 0 | 0 |
| ENSMUSG00000092208 | 0 | 0 | 0 | 0 | 0 | 0 | 0 |
| ENSMUSG00000092210 | 0 | 0 | 0 | 0 | 0 | 0 | 0 |
| ENSMUSG00000092211 | 0 | 0 | 0 | 0 | 0 | 0 | 0 |
| ENSMUSG00000092212 | 0 | 0.25 | 0 | 0.25 | 0.387761935 | 0.698970004 | 0.224243824 |
| ENSMUSG00000092213 | 0 | 0 | 0 | 0 | 0 | 0 | 0 |
| ENSMUSG00000092214 | 0 | 0 | 0 | 0 | 0 | 0 | 0 |
| ENSMUSG00000092216 | 0 | 0 | 0 | 0 | 0 | 0 | 0 |
| ENSMUSG00000092217 | 0.289064826 | 0 | 0 | 0 | 0 | 0 | 0 |
| ENSMUSG00000092218 | 0 | 0 | 0 | 0 | 0 | 0 | 0 |
| ENSMUSG00000092219 | 0 | 0 | 0 | 0 | 0 | 0 | 0 |
| ENSMUSG00000092220 | 0 | 0 | 0 | 0 | 0 | 0 | 0 |
| ENSMUSG00000092221 | 0 | 0 | 0 | 0 | 0 | 0 | 0 |
| ENSMUSG00000092222 | 0 | 0 | 0 | 0 | 0 | 0 | 0 |
| ENSMUSG00000092223 | 0 | 0 | 0 | 0 | 0 | 0 | 0.355418052 |
| ENSMUSG00000092224 | 0 | 0 | 0 | 0 | 0 | 0 | 0 |
| ENSMUSG00000092225 | 0.289064826 | 0 | 0.25 | 0.25 | 0 | 0.301029996 | 0 |
| ENSMUSG00000092227 | 0 | 0 | 0 | 0 | 0 | 0 | 0 |
| ENSMUSG00000092228 | 0.289064826 | 0 | 0.396240625 | 0 | 0 | 0 | 0 |
| ENSMUSG00000092229 | 0 | 0.25 | 0.25 | 0 | 0 | 0 | 0 |

| | | | | | | | |
|---|---|---|---|---|---|---|---|
| ENSMUSG00000092230 | 0 | 0 | 0 | 0.25 | 0 | 0 | 0 |
| ENSMUSG00000092231 | 0 | 0 | 0 | 0 | 0 | 0 | 0 |
| ENSMUSG00000092232 | 0 | 0 | 0 | 0 | 0 | 0 | 0 |
| ENSMUSG00000092233 | 0 | 0 | 0 | 0 | 0.244650542 | 0 | 0.224243824 |
| ENSMUSG00000092237 | 0 | 0 | 0 | 0 | 0 | 0 | 0 |
| ENSMUSG00000092238 | 0 | 0 | 0 | 0 | 0 | 0 | 0 |
| ENSMUSG00000092239 | 0.45815691 | 0 | 0.25 | 0.25 | 0.387761935 | 0.301029996 | 0 |
| ENSMUSG00000092240 | 0 | 0 | 0 | 0 | 0 | 0 | 0.224243824 |
| ENSMUSG00000092241 | 0.289064826 | 0.25 | 0.5 | 0 | 0 | 0 | 0.520678036 |
| ENSMUSG00000092242 | 0 | 0 | 0 | 0 | 0 | 0 | 0 |
| ENSMUSG00000092243 | 0 | 0 | 0 | 0 | 0 | 0 | 0 |
| ENSMUSG00000092244 | 0 | 0 | 0 | 0 | 0 | 0 | 0 |
| ENSMUSG00000092245 | 0 | 0 | 0 | 0 | 0 | 0 | 0 |
| ENSMUSG00000092246 | 0.45815691 | 0.25 | 0 | 0 | 0 | 0 | 0 |
| ENSMUSG00000092247 | 0 | 0.25 | 0 | 0 | 0 | 0 | 0 |
| ENSMUSG00000092248 | 0 | 0.25 | 0 | 0 | 0.244650542 | 0 | 0 |
| ENSMUSG00000092250 | 0 | 0.25 | 0 | 0 | 0 | 0 | 0.355418052 |
| ENSMUSG00000092251 | 0 | 0 | 0 | 0 | 0 | 0 | 0 |
| ENSMUSG00000092252 | 0 | 0 | 0 | 0 | 0 | 0 | 0 |
| ENSMUSG00000092253 | 0 | 0 | 0 | 0 | 0 | 0 | 0 |
| ENSMUSG00000092255 | 0 | 0 | 0 | 0 | 0 | 0 | 0 |
| ENSMUSG00000092256 | 0 | 0 | 0 | 0 | 0 | 0 | 0 |
| ENSMUSG00000092257 | 0 | 0.396240625 | 0.25 | 0.25 | 0.244650542 | 0 | 0 |
| ENSMUSG00000092258 | 0 | 0 | 0 | 0 | 0 | 0 | 0 |
| ENSMUSG00000092259 | 0 | 0 | 0 | 0 | 0 | 0 | 0 |
| ENSMUSG00000092260 | 1.289064826 | 1.330482024 | 1.238549078 | 1.364857905 | 1.188507684 | 1.491361694 | 1.324583736 |
| ENSMUSG00000092261 | 0 | 0.25 | 0.25 | 0.396240625 | 0.244650542 | 0 | 0.224243824 |
| ENSMUSG00000092263 | 0 | 0 | 0 | 0 | 0 | 0 | 0 |
| ENSMUSG00000092264 | 0 | 0 | 0 | 0 | 0 | 0 | 0 |
| ENSMUSG00000092265 | 0 | 0 | 0 | 0.25 | 0 | 0.301029996 | 0 |
| ENSMUSG00000092266 | 0 | 0 | 0 | 0 | 0 | 0 | 0 |
| ENSMUSG00000092267 | 0 | 0 | 0 | 0 | 0 | 0 | 0 |
| ENSMUSG00000092268 | 0 | 0 | 0.25 | 0 | 0 | 0 | 0.224243824 |
| ENSMUSG00000092269 | 0 | 0 | 0 | 0 | 0 | 0 | 0 |

| | | | | | | | |
|---|---|---|---|---|---|---|---|
| ENSMUSG00000092270 | 0 | 0 | 0 | 0 | 0 | 0 | 0 |
| ENSMUSG00000092271 | 0 | 0 | 0 | 0 | 0 | 0 | 0 |
| ENSMUSG00000092272 | 0 | 0 | 0 | 0 | 0 | 0 | 0 |
| ENSMUSG00000092274 | 2.963342983 | 2.693078643 | 2.703244868 | 2.378431899 | 2.41479008 | 2.633468456 | 2.275733199 |
| ENSMUSG00000092275 | 0 | 0 | 0.25 | 0 | 0 | 0 | 0 |
| ENSMUSG00000092276 | 0 | 0 | 0 | 0 | 0 | 0 | 0 |
| ENSMUSG00000092277 | 0 | 0 | 0 | 0 | 0 | 0 | 0 |
| ENSMUSG00000092278 | 0 | 0.25 | 0.25 | 0.25 | 0.387761935 | 0 | 0.448487648 |
| ENSMUSG00000092281 | 0 | 0 | 0.5 | 0.396240625 | 0.489301084 | 0.301029996 | 0.224243824 |
| ENSMUSG00000092283 | 0 | 0 | 0 | 0 | 0 | 0 | 0 |
| ENSMUSG00000092284 | 0 | 0.25 | 0.396240625 | 0 | 0 | 0 | 0 |
| ENSMUSG00000092285 | 0 | 0 | 0 | 0 | 0 | 0 | 0 |
| ENSMUSG00000092286 | 0.671187741 | 0.646240625 | 0.75 | 0.830482024 | 0.905314583 | 0 | 0.629532004 |
| ENSMUSG00000092287 | 0 | 0 | 0 | 0 | 0 | 0 | 0 |
| ENSMUSG00000092289 | 0 | 0 | 0 | 0 | 0 | 0.301029996 | 0 |
| ENSMUSG00000092290 | 0 | 0 | 0.25 | 0 | 0 | 0 | 0 |
| ENSMUSG00000092292 | 0 | 0 | 0 | 0 | 0 | 0 | 0 |
| ENSMUSG00000092293 | 0 | 0 | 0 | 0 | 0 | 0 | 0 |
| ENSMUSG00000092294 | 0 | 0 | 0 | 0 | 0 | 0 | 0 |
| ENSMUSG00000092295 | 0 | 0 | 0 | 0 | 0 | 0 | 0 |
| ENSMUSG00000092296 | 0 | 0 | 0 | 0 | 0 | 0 | 0 |
| ENSMUSG00000092297 | 0 | 0 | 0 | 0 | 0 | 0 | 0 |
| ENSMUSG00000092298 | 0 | 0 | 0 | 0 | 0 | 0 | 0 |
| ENSMUSG00000092300 | 0.45815691 | 0.79248125 | 0.864857905 | 0.646240625 | 0.686820904 | 0.301029996 | 0.710836105 |
| ENSMUSG00000092301 | 0.289064826 | 0 | 0.25 | 0 | 0 | 0 | 0.224243824 |
| ENSMUSG00000092302 | 0 | 0 | 0 | 0 | 0 | 0 | 0 |
| ENSMUSG00000092303 | 0 | 0 | 0 | 0 | 0 | 0 | 0 |
| ENSMUSG00000092304 | 0 | 0 | 0 | 0 | 0 | 0 | 0 |
| ENSMUSG00000092305 | 0 | 0 | 0 | 0 | 0 | 0 | 0 |
| ENSMUSG00000092306 | 0 | 0 | 0 | 0 | 0 | 0 | 0 |
| ENSMUSG00000092307 | 0 | 0 | 0 | 0 | 0 | 0 | 0 |
| ENSMUSG00000092308 | 0 | 0 | 0 | 0 | 0 | 0 | 0 |
| ENSMUSG00000092309 | 0 | 0 | 0 | 0 | 0 | 0 | 0 |
| ENSMUSG00000092310 | 0 | 0 | 0 | 0 | 0 | 0 | 0 |

| | | | | | | | |
|---|---|---|---|---|---|---|---|
| ENSMUSG00000092311 | 0 | 0 | 0 | 0 | 0 | 0 | 0 |
| ENSMUSG00000092312 | 0 | 0 | 0 | 0 | 0.244650542 | 0 | 0 |
| ENSMUSG00000092313 | 0 | 0 | 0 | 0 | 0 | 0 | 0 |
| ENSMUSG00000092314 | 0 | 0 | 0 | 0 | 0 | 0 | 0 |
| ENSMUSG00000092316 | 0 | 0 | 0 | 0 | 0 | 0 | 0 |
| ENSMUSG00000092317 | 0 | 0 | 0 | 0 | 0 | 0 | 0.224243824 |
| ENSMUSG00000092318 | 0 | 0 | 0 | 0 | 0 | 0 | 0 |
| ENSMUSG00000092322 | 0 | 0 | 0 | 0 | 0 | 0 | 0 |
| ENSMUSG00000092323 | 0 | 0 | 0 | 0 | 0 | 0 | 0 |
| ENSMUSG00000092325 | 0.578129653 | 0.25 | 0.5 | 0.580482024 | 0.568060967 | 0.602059991 | 0.355418052 |
| ENSMUSG00000092326 | 0 | 0 | 0 | 0 | 0 | 0 | 0 |
| ENSMUSG00000092328 | 0 | 0 | 0 | 0 | 0 | 0 | 0 |
| ENSMUSG00000092329 | 0 | 0 | 0 | 0 | 0 | 0 | 0 |
| ENSMUSG00000092330 | 0 | 0 | 0 | 0 | 0 | 0 | 0 |
| ENSMUSG00000092331 | 0 | 0 | 0 | 0 | 0 | 0 | 0 |
| ENSMUSG00000092332 | 0 | 0 | 0 | 0 | 0 | 0 | 0 |
| ENSMUSG00000092335 | 0 | 0 | 0 | 0.580482024 | 0.489301084 | 0.477121255 | 0.448487648 |
| ENSMUSG00000092336 | 0 | 0 | 0 | 0 | 0 | 0 | 0 |
| ENSMUSG00000092338 | 0 | 0 | 0 | 0 | 0 | 0 | 0 |
| ENSMUSG00000092339 | 0 | 0 | 0 | 0 | 0 | 0 | 0 |
| ENSMUSG00000092340 | 0 | 0 | 0 | 0 | 0 | 0 | 0 |
| ENSMUSG00000092341 | 3.502220968 | 3.127691042 | 3.104924222 | 3.071711626 | 3.058398634 | 3.36078269 | 2.77398916 |
| ENSMUSG00000092342 | 0 | 0 | 0 | 0 | 0 | 0 | 0 |
| ENSMUSG00000092343 | 0 | 0 | 0 | 0 | 0 | 0 | 0 |
| ENSMUSG00000092344 | 0.289064826 | 0.396240625 | 0.580482024 | 0.5 | 0.387761935 | 0 | 0.520678036 |
| ENSMUSG00000092345 | 0 | 0 | 0 | 0 | 0 | 0 | 0 |
| ENSMUSG00000092346 | 0 | 0 | 0 | 0 | 0 | 0 | 0 |
| ENSMUSG00000092349 | 0.45815691 | 0.646240625 | 0.5 | 0.701838731 | 0.77552387 | 0.477121255 | 0.448487648 |
| ENSMUSG00000092350 | 0 | 0 | 0 | 0 | 0 | 0 | 0 |
| ENSMUSG00000092351 | 0 | 0 | 0 | 0 | 0 | 0 | 0 |
| ENSMUSG00000092352 | 0 | 0 | 0 | 0 | 0 | 0 | 0 |
| ENSMUSG00000092353 | 0 | 0.25 | 0 | 0.25 | 0.244650542 | 0 | 0.224243824 |
| ENSMUSG00000092354 | 0.671187741 | 0.701838731 | 0.830482024 | 0.646240625 | 0.489301084 | 0.602059991 | 0.853775828 |
| ENSMUSG00000092356 | 0 | 0 | 0 | 0 | 0 | 0 | 0 |

| | | | | | | | |
|---|---|---|---|---|---|---|---|
| ENSMUSG00000092357 | 0 | 0 | 0 | 0 | 0 | 0 | 0 |
| ENSMUSG00000092358 | 0.289064826 | 0 | 0 | 0.25 | 0.244650542 | 0 | 0.448487648 |
| ENSMUSG00000092359 | 0 | 0 | 0 | 0 | 0 | 0 | 0 |
| ENSMUSG00000092360 | 0 | 0 | 0 | 0 | 0 | 0 | 0 |
| ENSMUSG00000092362 | 0 | 0 | 0 | 0 | 0 | 0 | 0 |
| ENSMUSG00000092363 | 0 | 0 | 0 | 0 | 0 | 0 | 0 |
| ENSMUSG00000092364 | 0 | 0 | 0 | 0 | 0 | 0.477121255 | 0 |
| ENSMUSG00000092365 | 0 | 0 | 0 | 0 | 0 | 0 | 0 |
| ENSMUSG00000092366 | 0 | 0 | 0 | 0.25 | 0.489301084 | 0 | 0 |
| ENSMUSG00000092367 | 0 | 0 | 0 | 0 | 0 | 0 | 0 |
| ENSMUSG00000092368 | 0 | 0 | 0 | 0 | 0 | 0 | 0 |
| ENSMUSG00000092369 | 0 | 0 | 0 | 0 | 0 | 0 | 0 |
| ENSMUSG00000092370 | 0 | 0 | 0 | 0 | 0 | 0 | 0 |
| ENSMUSG00000092371 | 0 | 0 | 0 | 0 | 0.244650542 | 0 | 0 |
| ENSMUSG00000092372 | 0 | 0 | 0 | 0 | 0 | 0 | 0 |
| ENSMUSG00000092373 | 0.45815691 | 0 | 0 | 0 | 0.244650542 | 0.301029996 | 0.224243824 |
| ENSMUSG00000092374 | 0 | 0 | 0 | 0 | 0 | 0 | 0 |
| ENSMUSG00000092375 | 0 | 0.25 | 0.25 | 0 | 0 | 0 | 0 |
| ENSMUSG00000092376 | 0 | 0 | 0 | 0 | 0 | 0.301029996 | 0 |
| ENSMUSG00000092377 | 0 | 0 | 0 | 0 | 0 | 0 | 0 |
| ENSMUSG00000092381 | 0.289064826 | 0 | 0 | 0 | 0 | 0 | 0.224243824 |
| ENSMUSG00000092382 | 0 | 0 | 0 | 0 | 0 | 0 | 0 |
| ENSMUSG00000092383 | 0 | 0 | 0 | 0 | 0 | 0 | 0 |
| ENSMUSG00000092384 | 0.289064826 | 0.701838731 | 0.580482024 | 0.25 | 0.489301084 | 0 | 0.672731473 |
| ENSMUSG00000092385 | 0 | 0 | 0 | 0 | 0 | 0 | 0.224243824 |
| ENSMUSG00000092386 | 0 | 0 | 0 | 0 | 0 | 0 | 0 |
| ENSMUSG00000092387 | 0 | 0 | 0 | 0 | 0 | 0 | 0 |
| ENSMUSG00000092388 | 0 | 0 | 0 | 0 | 0 | 0 | 0 |
| ENSMUSG00000092389 | 0 | 0 | 0 | 0 | 0 | 0 | 0 |
| ENSMUSG00000092390 | 0.45815691 | 0.25 | 0.396240625 | 0.830482024 | 0.846351821 | 0.602059991 | 0.672731473 |
| ENSMUSG00000092392 | 0.747221736 | 0.5 | 0.79248125 | 0.396240625 | 0.489301084 | 0 | 0.672731473 |
| ENSMUSG00000092393 | 0 | 0 | 0 | 0 | 0 | 0 | 0 |
| ENSMUSG00000092394 | 0 | 0 | 0.25 | 0.25 | 0 | 0 | 0 |
| ENSMUSG00000092395 | 0 | 0 | 0 | 0 | 0 | 0 | 0 |

| | | | | | | | |
|---|---|---|---|---|---|---|---|
| ENSMUSG00000092396 | 0 | 0 | 0 | 0 | 0 | 0 | 0 |
| ENSMUSG00000092397 | 0 | 0 | 0 | 0 | 0 | 0 | 0 |
| ENSMUSG00000092398 | 0 | 0 | 0 | 0 | 0 | 0 | 0 |
| ENSMUSG00000092399 | 0 | 0.396240625 | 0 | 0 | 0 | 0 | 0 |
| ENSMUSG00000092401 | 0 | 0 | 0 | 0 | 0 | 0 | 0 |
| ENSMUSG00000092402 | 0 | 0 | 0 | 0 | 0 | 0 | 0 |
| ENSMUSG00000092403 | 0 | 0 | 0 | 0 | 0 | 0 | 0 |
| ENSMUSG00000092404 | 0 | 0.25 | 0 | 0 | 0 | 0 | 0 |
| ENSMUSG00000092405 | 0 | 0 | 0 | 0 | 0 | 0 | 0 |
| ENSMUSG00000092406 | 0 | 0 | 0 | 0 | 0 | 0 | 0 |
| ENSMUSG00000092408 | 0 | 0 | 0 | 0 | 0 | 0 | 0 |
| ENSMUSG00000092409 | 0 | 0 | 0 | 0 | 0 | 0 | 0 |
| ENSMUSG00000092410 | 0 | 0 | 0 | 0 | 0 | 0 | 0 |
| ENSMUSG00000092411 | 0 | 0 | 0 | 0 | 0 | 0 | 0 |
| ENSMUSG00000092412 | 0.671187741 | 0.701838731 | 0.25 | 0.646240625 | 0.686820904 | 0.301029996 | 0.629532004 |
| ENSMUSG00000092413 | 0 | 0 | 0 | 0 | 0 | 0 | 0 |
| ENSMUSG00000092414 | 0 | 0 | 0 | 0 | 0 | 0 | 0 |
| ENSMUSG00000092415 | 0 | 0 | 0 | 0 | 0 | 0 | 0 |
| ENSMUSG00000092416 | 1.605636262 | 1.593759858 | 1.552363341 | 1.54745614 | 1.456697355 | 1.556302501 | 1.448487648 |
| ENSMUSG00000092417 | 1.734388958 | 1.711372513 | 1.708222504 | 1.657339155 | 1.674752852 | 1.72427587 | 1.647906904 |
| ENSMUSG00000092418 | 0 | 0 | 0 | 0 | 0.244650542 | 0 | 0 |
| ENSMUSG00000092419 | 0 | 0 | 0 | 0 | 0 | 0 | 0 |
| ENSMUSG00000092422 | 0 | 0 | 0 | 0 | 0 | 0 | 0 |
| ENSMUSG00000092425 | 0 | 0 | 0 | 0 | 0 | 0 | 0 |
| ENSMUSG00000092426 | 0 | 0 | 0 | 0 | 0 | 0 | 0 |
| ENSMUSG00000092427 | 0 | 0 | 0 | 0 | 0 | 0 | 0 |
| ENSMUSG00000092428 | 0.289064826 | 0 | 0 | 0 | 0.244650542 | 0 | 0.224243824 |
| ENSMUSG00000092429 | 0 | 0 | 0 | 0 | 0 | 0 | 0 |
| ENSMUSG00000092431 | 0 | 0 | 0 | 0 | 0 | 0 | 0 |
| ENSMUSG00000092433 | 0 | 0 | 0 | 0 | 0 | 0 | 0 |
| ENSMUSG00000092435 | 0 | 0 | 0 | 0 | 0 | 0 | 0 |
| ENSMUSG00000092437 | 0 | 0 | 0 | 0 | 0 | 0 | 0.224243824 |
| ENSMUSG00000092438 | 0 | 0.25 | 0.25 | 0 | 0.244650542 | 0 | 0 |
| ENSMUSG00000092439 | 0 | 0 | 0 | 0 | 0 | 0 | 0 |

| | | | | | | | |
|---|---|---|---|---|---|---|---|
| ENSMUSG00000092440 | 0 | 0 | 0 | 0 | 0 | 0 | 0 |
| ENSMUSG00000092441 | 0 | 0 | 0 | 0 | 0 | 0 | 0 |
| ENSMUSG00000092443 | 0 | 0 | 0.25 | 0 | 0.387761935 | 0 | 0 |
| ENSMUSG00000092444 | 0.747221736 | 0.951838731 | 0.5 | 0.75 | 0.931471446 | 0.903089987 | 0.969165684 |
| ENSMUSG00000092447 | 0 | 0 | 0 | 0 | 0 | 0 | 0 |
| ENSMUSG00000092448 | 0 | 0.25 | 0.25 | 0 | 0 | 0 | 0 |
| ENSMUSG00000092450 | 0 | 0 | 0 | 0 | 0 | 0 | 0 |
| ENSMUSG00000092451 | 0 | 0 | 0 | 0 | 0 | 0 | 0 |
| ENSMUSG00000092452 | 0 | 0 | 0 | 0 | 0 | 0 | 0 |
| ENSMUSG00000092454 | 0 | 0 | 0 | 0 | 0 | 0 | 0 |
| ENSMUSG00000092455 | 0 | 0 | 0 | 0 | 0 | 0 | 0 |
| ENSMUSG00000092456 | 0 | 0 | 0 | 0 | 0 | 0 | 0 |
| ENSMUSG00000092457 | 0 | 0 | 0 | 0.580482024 | 0.387761935 | 0.698970004 | 0.448487648 |
| ENSMUSG00000092458 | 0 | 0 | 0 | 0 | 0 | 0 | 0 |
| ENSMUSG00000092460 | 0 | 0 | 0 | 0 | 0 | 0 | 0 |
| ENSMUSG00000092461 | 0 | 0 | 0 | 0 | 0 | 0 | 0 |
| ENSMUSG00000092462 | 0 | 0 | 0 | 0 | 0 | 0 | 0 |
| ENSMUSG00000092463 | 0 | 0 | 0 | 0 | 0 | 0 | 0 |
| ENSMUSG00000092464 | 0.671187741 | 0.75 | 0.830482024 | 0.75 | 0.733951626 | 0.77815125 | 0.775756176 |
| ENSMUSG00000092465 | 0.960252568 | 0.580482024 | 0.25 | 0 | 0 | 0.301029996 | 0 |
| ENSMUSG00000092466 | 0 | 0 | 0 | 0 | 0 | 0 | 0 |
| ENSMUSG00000092467 | 0 | 0 | 0 | 0 | 0 | 0 | 0 |
| ENSMUSG00000092468 | 0 | 0 | 0 | 0 | 0 | 0 | 0 |
| ENSMUSG00000092469 | 0 | 0 | 0 | 0 | 0 | 0 | 0 |
| ENSMUSG00000092470 | 0.289064826 | 0.25 | 0.25 | 0.25 | 0.244650542 | 0 | 0.224243824 |
| ENSMUSG00000092471 | 0 | 0 | 0 | 0 | 0 | 0 | 0 |
| ENSMUSG00000092472 | 0 | 0 | 0 | 0 | 0 | 0 | 0 |
| ENSMUSG00000092473 | 0 | 0 | 0 | 0 | 0 | 0 | 0 |
| ENSMUSG00000092474 | 0 | 0 | 0 | 0 | 0 | 0.477121255 | 0.448487648 |
| ENSMUSG00000092476 | 0 | 0 | 0 | 0 | 0 | 0 | 0 |
| ENSMUSG00000092477 | 0 | 0 | 0 | 0 | 0 | 0 | 0 |
| ENSMUSG00000092480 | 0 | 0 | 0 | 0 | 0 | 0 | 0 |
| ENSMUSG00000092482 | 0.289064826 | 0 | 0 | 0 | 0 | 0 | 0 |
| ENSMUSG00000092483 | 0 | 0 | 0 | 0 | 0 | 0 | 0 |

| | | | | | | | |
|---|---|---|---|---|---|---|---|
| ENSMUSG00000092485 | 0 | 0 | 0 | 0 | 0 | 0 | 0 |
| ENSMUSG00000092486 | 1.639698646 | 1.561981878 | 1.614857905 | 1.852347734 | 1.822942838 | 2.049218023 | 1.707551656 |
| ENSMUSG00000092487 | 0 | 0 | 0 | 0 | 0 | 0 | 0 |
| ENSMUSG00000092488 | 0 | 0 | 0 | 0 | 0 | 0 | 0 |
| ENSMUSG00000092489 | 0 | 0 | 0 | 0 | 0 | 0 | 0 |
| ENSMUSG00000092490 | 0.45815691 | 0.396240625 | 0 | 0.396240625 | 0.568060967 | 0.301029996 | 0.355418052 |
| ENSMUSG00000092492 | 0 | 0.25 | 0 | 0 | 0.244650542 | 0 | 0 |
| ENSMUSG00000092493 | 0 | 0 | 0 | 0 | 0 | 0 | 0 |
| ENSMUSG00000092494 | 0 | 0 | 0 | 0 | 0 | 0 | 0 |
| ENSMUSG00000092495 | 0.578129653 | 0 | 0.5 | 0.25 | 0.387761935 | 0 | 0.224243824 |
| ENSMUSG00000092496 | 0 | 0 | 0 | 0 | 0 | 0 | 0 |
| ENSMUSG00000092499 | 0 | 0 | 0 | 0 | 0 | 0 | 0 |
| ENSMUSG00000092500 | 0 | 0.5 | 0.5 | 0.580482024 | 0.632412477 | 0.602059991 | 0.448487648 |
| ENSMUSG00000092502 | 0 | 0 | 0 | 0 | 0 | 0 | 0 |
| ENSMUSG00000092503 | 0 | 0 | 0 | 0 | 0 | 0 | 0 |
| ENSMUSG00000092504 | 0 | 0 | 0 | 0 | 0 | 0 | 0 |
| ENSMUSG00000092505 | 0 | 0 | 0 | 0 | 0 | 0 | 0 |
| ENSMUSG00000092506 | 0 | 0 | 0 | 0 | 0 | 0 | 0 |
| ENSMUSG00000092507 | 0 | 0 | 0 | 0 | 0 | 0 | 0 |
| ENSMUSG00000092508 | 0 | 0 | 0 | 0 | 0 | 0 | 0 |
| ENSMUSG00000092509 | 0 | 0 | 0.25 | 0.5 | 0.77552387 | 0.477121255 | 0.579661877 |
| ENSMUSG00000092510 | 0 | 0 | 0 | 0 | 0 | 0 | 0 |
| ENSMUSG00000092511 | 0 | 0 | 0 | 0 | 0 | 0 | 0 |
| ENSMUSG00000092512 | 0 | 0 | 0 | 0 | 0 | 0 | 0 |
| ENSMUSG00000092513 | 0 | 0 | 0 | 0 | 0 | 0 | 0 |
| ENSMUSG00000092514 | 0 | 0 | 0 | 0 | 0 | 0 | 0 |
| ENSMUSG00000092515 | 0 | 0 | 0 | 0.25 | 0.387761935 | 0 | 0.520678036 |
| ENSMUSG00000092516 | 0 | 0 | 0 | 0 | 0 | 0 | 0 |
| ENSMUSG00000092517 | 0 | 0 | 0 | 0 | 0 | 0 | 0 |
| ENSMUSG00000092518 | 0 | 0 | 0 | 0 | 0.244650542 | 0 | 0 |
| ENSMUSG00000092519 | 0 | 0 | 0 | 0 | 0 | 0 | 0 |
| ENSMUSG00000092520 | 0 | 0 | 0 | 0 | 0 | 0 | 0 |
| ENSMUSG00000092521 | 0 | 0 | 0 | 0 | 0.244650542 | 0 | 0 |
| ENSMUSG00000092522 | 0 | 0 | 0 | 0 | 0 | 0 | 0 |

| | | | | | | | |
|---|---|---|---|---|---|---|---|
| ENSMUSG00000092523 | 0 | 0 | 0 | 0 | 0 | 0 | 0 |
| ENSMUSG00000092524 | 0.578129653 | 0.25 | 0.25 | 0.396240625 | 0 | 0.301029996 | 0 |
| ENSMUSG00000092525 | 0 | 0 | 0 | 0 | 0 | 0 | 0 |
| ENSMUSG00000092526 | 0 | 0.396240625 | 0.25 | 0 | 0 | 0 | 0.355418052 |
| ENSMUSG00000092528 | 0 | 0 | 0 | 0 | 0 | 0 | 0 |
| ENSMUSG00000092529 | 0 | 0 | 0 | 0 | 0 | 0 | 0 |
| ENSMUSG00000092530 | 0 | 0 | 0 | 0 | 0 | 0 | 0 |
| ENSMUSG00000092532 | 0 | 0 | 0 | 0 | 0 | 0 | 0 |
| ENSMUSG00000092533 | 0 | 0 | 0 | 0 | 0 | 0 | 0 |
| ENSMUSG00000092534 | 0 | 0 | 0 | 0 | 0 | 0 | 0 |
| ENSMUSG00000092535 | 0 | 0 | 0 | 0 | 0 | 0 | 0 |
| ENSMUSG00000092536 | 0 | 0 | 0 | 0 | 0 | 0 | 0.224243824 |
| ENSMUSG00000092538 | 0 | 0 | 0 | 0 | 0 | 0 | 0 |
| ENSMUSG00000092539 | 0 | 0.25 | 0.396240625 | 0 | 0.244650542 | 0 | 0.224243824 |
| ENSMUSG00000092540 | 0 | 0 | 0 | 0 | 0 | 0.477121255 | 0.224243824 |
| ENSMUSG00000092541 | 0 | 0 | 0 | 0 | 0 | 0 | 0 |
| ENSMUSG00000092542 | 0 | 0 | 0 | 0 | 0 | 0 | 0 |
| ENSMUSG00000092543 | 0 | 0.25 | 0.25 | 0 | 0.244650542 | 0 | 0 |
| ENSMUSG00000092544 | 0 | 0 | 0 | 0 | 0 | 0 | 0 |
| ENSMUSG00000092545 | 0 | 0 | 0.25 | 0 | 0 | 0 | 0 |
| ENSMUSG00000092547 | 0 | 0 | 0 | 0 | 0 | 0 | 0 |
| ENSMUSG00000092548 | 0 | 0 | 0 | 0 | 0 | 0 | 0 |
| ENSMUSG00000092549 | 0 | 0 | 0 | 0 | 0 | 0 | 0 |
| ENSMUSG00000092550 | 0 | 0 | 0 | 0.25 | 0.387761935 | 0.477121255 | 0 |
| ENSMUSG00000092551 | 0 | 0 | 0 | 0 | 0 | 0 | 0 |
| ENSMUSG00000092552 | 0 | 0 | 0 | 0 | 0 | 0 | 0 |
| ENSMUSG00000092553 | 0 | 0 | 0 | 0.25 | 0 | 0 | 0 |
| ENSMUSG00000092554 | 0 | 0 | 0 | 0 | 0 | 0 | 0 |
| ENSMUSG00000092555 | 0 | 0 | 0 | 0 | 0 | 0 | 0 |
| ENSMUSG00000092556 | 0 | 0.5 | 0.396240625 | 0.396240625 | 0 | 0.301029996 | 0 |
| ENSMUSG00000092557 | 0.289064826 | 0 | 0 | 0 | 0.244650542 | 0 | 0 |
| ENSMUSG00000092558 | 2.209569962 | 2.088286706 | 2.074802005 | 2.120953944 | 2.09661982 | 2.269512944 | 2.005960535 |
| ENSMUSG00000092559 | 0 | 0 | 0 | 0.25 | 0 | 0 | 0 |
| ENSMUSG00000092560 | 0 | 0 | 0 | 0 | 0 | 0 | 0 |

| | | | | | | | |
|---|---|---|---|---|---|---|---|
| ENSMUSG00000092561 | 0 | 0 | 0 | 0 | 0 | 0 | 0 |
| ENSMUSG00000092562 | 0 | 0 | 0 | 0 | 0 | 0 | 0 |
| ENSMUSG00000092563 | 0 | 0 | 0 | 0 | 0.244650542 | 0 | 0 |
| ENSMUSG00000092564 | 0 | 0.646240625 | 0.75 | 0.396240625 | 0.387761935 | 0 | 0.710836105 |
| ENSMUSG00000092565 | 0 | 0 | 0 | 0 | 0 | 0 | 0 |
| ENSMUSG00000092566 | 0 | 0 | 0 | 0 | 0 | 0 | 0 |
| ENSMUSG00000092567 | 0 | 0 | 0 | 0 | 0 | 0 | 0 |
| ENSMUSG00000092568 | 0 | 0 | 0 | 0 | 0 | 0 | 0 |
| ENSMUSG00000092569 | 1.227926429 | 1.226722649 | 1.146240625 | 0.830482024 | 0.489301084 | 0.477121255 | 0.803905701 |
| ENSMUSG00000092570 | 0 | 0 | 0 | 0 | 0 | 0 | 0 |
| ENSMUSG00000092571 | 0 | 0.25 | 0 | 0 | 0 | 0 | 0.224243824 |
| ENSMUSG00000092572 | 0 | 0 | 0 | 0 | 0 | 0 | 0.224243824 |
| ENSMUSG00000092573 | 0 | 0 | 0 | 0 | 0 | 0 | 0 |
| ENSMUSG00000092574 | 0 | 0 | 0 | 0 | 0 | 0 | 0 |
| ENSMUSG00000092575 | 0 | 0 | 0 | 0 | 0 | 0 | 0 |
| ENSMUSG00000092576 | 0 | 0 | 0 | 0 | 0 | 0 | 0 |
| ENSMUSG00000092577 | 0.289064826 | 0.25 | 0.25 | 0.25 | 0.244650542 | 0.301029996 | 0.448487648 |
| ENSMUSG00000092578 | 0 | 0 | 0 | 0 | 0 | 0 | 0 |
| ENSMUSG00000092579 | 0 | 0 | 0 | 0 | 0 | 0 | 0 |
| ENSMUSG00000092580 | 0 | 0 | 0 | 0 | 0 | 0 | 0 |
| ENSMUSG00000092581 | 0 | 0 | 0 | 0 | 0 | 0 | 0 |
| ENSMUSG00000092583 | 0 | 0 | 0 | 0 | 0 | 0 | 0 |
| ENSMUSG00000092584 | 0 | 0 | 0 | 0 | 0 | 0 | 0 |
| ENSMUSG00000092586 | 0.289064826 | 0.646240625 | 0.646240625 | 0.396240625 | 0.489301084 | 0.301029996 | 0.448487648 |
| ENSMUSG00000092587 | 0 | 0 | 0 | 0 | 0 | 0 | 0 |
| ENSMUSG00000092588 | 0 | 0 | 0 | 0 | 0 | 0 | 0 |
| ENSMUSG00000092591 | 0 | 0 | 0 | 0 | 0 | 0 | 0 |
| ENSMUSG00000092592 | 0.578129653 | 0.580482024 | 0.701838731 | 0.25 | 0.387761935 | 0.477121255 | 0.520678036 |
| ENSMUSG00000092593 | 0.578129653 | 0 | 0.396240625 | 0 | 0.244650542 | 0.301029996 | 0 |
| ENSMUSG00000092595 | 0 | 0.396240625 | 0.25 | 0.396240625 | 0.244650542 | 0 | 0.224243824 |
| ENSMUSG00000092596 | 0 | 0 | 0 | 0 | 0 | 0 | 0 |
| ENSMUSG00000092597 | 0 | 0 | 0 | 0 | 0 | 0 | 0 |
| ENSMUSG00000092599 | 0 | 0 | 0 | 0 | 0 | 0 | 0 |
| ENSMUSG00000092600 | 0.289064826 | 0.25 | 0.25 | 0.25 | 0.244650542 | 0 | 0.224243824 |

| | | | | | | | |
|---|---|---|---|---|---|---|---|
| ENSMUSG00000092601 | 0 | 0 | 0 | 0 | 0 | 0 | 0 |
| ENSMUSG00000092602 | 0 | 0 | 0 | 0 | 0 | 0 | 0 |
| ENSMUSG00000092603 | 0 | 0 | 0 | 0 | 0 | 0 | 0 |
| ENSMUSG00000092604 | 0 | 0 | 0 | 0 | 0 | 0 | 0 |
| ENSMUSG00000092606 | 0 | 0 | 0 | 0 | 0 | 0 | 0 |
| ENSMUSG00000092607 | 1.54867984 | 1.571350555 | 1.505591953 | 1.752806814 | 1.758994476 | 1.995635195 | 1.714087544 |
| ENSMUSG00000092608 | 0 | 0 | 0 | 0 | 0 | 0 | 0 |
| ENSMUSG00000092609 | 0 | 0 | 0 | 0 | 0 | 0 | 0 |
| ENSMUSG00000092610 | 0 | 0 | 0 | 0 | 0 | 0 | 0 |
| ENSMUSG00000092611 | 0 | 0 | 0 | 0 | 0 | 0 | 0 |
| ENSMUSG00000092612 | 0 | 0 | 0 | 0 | 0 | 0 | 0 |
| ENSMUSG00000092614 | 0 | 0 | 0 | 0 | 0 | 0 | 0 |
| ENSMUSG00000092615 | 0 | 0 | 0 | 0 | 0 | 0 | 0 |
| ENSMUSG00000092616 | 0 | 0 | 0 | 0 | 0 | 0 | 0 |
| ENSMUSG00000092617 | 0 | 0 | 0.25 | 0.25 | 0 | 0 | 0.224243824 |
| ENSMUSG00000092618 | 0 | 0 | 0 | 0 | 0 | 0 | 0 |
| ENSMUSG00000092619 | 0 | 0 | 0.25 | 0 | 0 | 0 | 0 |
| ENSMUSG00000092620 | 0 | 0 | 0 | 0 | 0 | 0 | 0 |
| ENSMUSG00000092621 | 0 | 0 | 0 | 0 | 0 | 0 | 0 |
| ENSMUSG00000092622 | 0 | 0 | 0 | 0 | 0 | 0 | 0 |
| ENSMUSG00000092624 | 0 | 0 | 0 | 0 | 0 | 0.301029996 | 0 |
| ENSMUSG00000092625 | 0.289064826 | 0.396240625 | 0.396240625 | 0.396240625 | 0.387761935 | 0.301029996 | 0.224243824 |
| ENSMUSG00000092626 | 0.45815691 | 0.25 | 0.25 | 0.396240625 | 0 | 0.301029996 | 0.355418052 |
| ENSMUSG00000092627 | 0 | 0 | 0.5 | 0.701838731 | 0.905314583 | 0.77815125 | 0.853775828 |
| ENSMUSG00000092628 | 0 | 0 | 0 | 0 | 0 | 0 | 0 |
| ENSMUSG00000092629 | 0 | 0 | 0 | 0 | 0 | 0 | 0 |
| ENSMUSG00000092631 | 0 | 0 | 0 | 0 | 0 | 0 | 0 |
| ENSMUSG00000092633 | 0 | 0 | 0 | 0 | 0 | 0 | 0 |
| ENSMUSG00000092636 | 0 | 0 | 0 | 0 | 0 | 0 | 0 |
| ENSMUSG00000092638 | 0 | 0 | 0 | 0 | 0 | 0 | 0 |
| ENSMUSG00000092639 | 0 | 0 | 0 | 0 | 0 | 0 | 0 |
| ENSMUSG00000092641 | 0 | 0.25 | 0.25 | 0.396240625 | 0 | 0 | 0.520678036 |
| ENSMUSG00000092643 | 0 | 0 | 0 | 0 | 0 | 0 | 0 |
| ENSMUSG00000092644 | 0.45815691 | 0 | 0 | 0.396240625 | 0.244650542 | 0 | 0.448487648 |

| | | | | | | | |
|---|---|---|---|---|---|---|---|
| ENSMUSG00000092647 | 0 | 0 | 0 | 0 | 0 | 0 | 0 |
| ENSMUSG00000092648 | 0 | 0 | 0.25 | 0 | 0 | 0 | 0 |
| ENSMUSG00000092650 | 0 | 0 | 0 | 0 | 0 | 0 | 0 |
| ENSMUSG00000092652 | 0 | 0 | 0 | 0 | 0 | 0 | 0 |
| ENSMUSG00000092653 | 0 | 0 | 0 | 0 | 0 | 0 | 0 |
| ENSMUSG00000092655 | 0 | 0 | 0 | 0 | 0 | 0 | 0 |
| ENSMUSG00000092656 | 0 | 0 | 0 | 0 | 0 | 0 | 0 |
| ENSMUSG00000092657 | 0 | 0 | 0.25 | 0 | 0 | 0 | 0 |
| ENSMUSG00000092659 | 0 | 0 | 0.25 | 0 | 0 | 0 | 0 |
| ENSMUSG00000092660 | 0.45815691 | 0 | 0 | 0 | 0.489301084 | 0 | 0.355418052 |
| ENSMUSG00000092661 | 0 | 0 | 0 | 0 | 0 | 0 | 0 |
| ENSMUSG00000092664 | 0 | 0 | 0 | 0 | 0 | 0 | 0 |
| ENSMUSG00000092666 | 0 | 0 | 0 | 0 | 0 | 0 | 0 |
| ENSMUSG00000092668 | 0 | 0 | 0 | 0 | 0 | 0 | 0 |
| ENSMUSG00000092671 | 0 | 0 | 0 | 0 | 0 | 0 | 0 |
| ENSMUSG00000092673 | 0 | 0.25 | 0 | 0 | 0 | 0 | 0 |
| ENSMUSG00000092674 | 0 | 0 | 0 | 0 | 0 | 0 | 0 |
| ENSMUSG00000092675 | 0 | 0 | 0 | 0 | 0 | 0 | 0 |
| ENSMUSG00000092677 | 0 | 0 | 0 | 0 | 0 | 0 | 0 |
| ENSMUSG00000092679 | 0 | 0 | 0 | 0 | 0 | 0 | 0 |
| ENSMUSG00000092680 | 0 | 0 | 0 | 0 | 0 | 0 | 0 |
| ENSMUSG00000092683 | 0 | 0 | 0 | 0 | 0 | 0 | 0 |
| ENSMUSG00000092685 | 0 | 0 | 0 | 0 | 0 | 0 | 0 |
| ENSMUSG00000092686 | 0 | 0 | 0 | 0 | 0 | 0 | 0 |
| ENSMUSG00000092687 | 0 | 0 | 0 | 0 | 0 | 0 | 0 |
| ENSMUSG00000092693 | 0 | 0 | 0 | 0 | 0 | 0 | 0 |
| ENSMUSG00000092695 | 0 | 0 | 0 | 0 | 0 | 0 | 0 |
| ENSMUSG00000092696 | 0 | 0 | 0 | 0 | 0 | 0 | 0 |
| ENSMUSG00000092697 | 0 | 0 | 0 | 0 | 0 | 0 | 0 |
| ENSMUSG00000092698 | 0 | 0 | 0 | 0 | 0 | 0 | 0 |
| ENSMUSG00000092701 | 0 | 0 | 0 | 0 | 0 | 0 | 0 |
| ENSMUSG00000092702 | 0 | 0 | 0 | 0 | 0 | 0 | 0 |
| ENSMUSG00000092704 | 0 | 0 | 0 | 0 | 0 | 0 | 0 |
| ENSMUSG00000092705 | 0 | 0 | 0 | 0 | 0 | 0 | 0 |

| | | | | | | | |
|---|---|---|---|---|---|---|---|
| ENSMUSG00000092706 | 0 | 0 | 0 | 0 | 0 | 0 | 0 |
| ENSMUSG00000092710 | 0 | 0 | 0 | 0 | 0 | 0 | 0 |
| ENSMUSG00000092713 | 0.289064826 | 0 | 0.25 | 0 | 0 | 0 | 0 |
| ENSMUSG00000092714 | 0 | 0 | 0 | 0 | 0 | 0 | 0 |
| ENSMUSG00000092715 | 0 | 0 | 0 | 0 | 0 | 0 | 0 |
| ENSMUSG00000092718 | 0 | 0 | 0 | 0 | 0 | 0 | 0 |
| ENSMUSG00000092719 | 0 | 0 | 0 | 0 | 0.244650542 | 0 | 0 |
| ENSMUSG00000092720 | 0 | 0 | 0 | 0 | 0 | 0 | 0 |
| ENSMUSG00000092723 | 0 | 0 | 0 | 0 | 0 | 0 | 0 |
| ENSMUSG00000092724 | 0 | 0 | 0 | 0 | 0 | 0 | 0 |
| ENSMUSG00000092725 | 0 | 0 | 0 | 0 | 0 | 0 | 0 |
| ENSMUSG00000092727 | 0 | 0 | 0 | 0 | 0.387761935 | 0 | 0 |
| ENSMUSG00000092730 | 0 | 0 | 0 | 0 | 0 | 0 | 0 |
| ENSMUSG00000092731 | 0 | 0 | 0 | 0 | 0 | 0 | 0 |
| ENSMUSG00000092732 | 0 | 0 | 0 | 0 | 0 | 0 | 0 |
| ENSMUSG00000092734 | 0 | 0 | 0 | 0 | 0 | 0 | 0.224243824 |
| ENSMUSG00000092737 | 0 | 0 | 0 | 0 | 0 | 0 | 0 |
| ENSMUSG00000092738 | 0 | 0 | 0 | 0 | 0 | 0 | 0 |
| ENSMUSG00000092741 | 0 | 0 | 0 | 0 | 0.244650542 | 0 | 0 |
| ENSMUSG00000092743 | 0 | 0 | 0 | 0 | 0 | 0 | 0 |
| ENSMUSG00000092745 | 0 | 0 | 0 | 0 | 0 | 0 | 0 |
| ENSMUSG00000092746 | 0 | 0 | 0 | 0 | 0 | 0 | 0 |
| ENSMUSG00000092747 | 0 | 0 | 0 | 0 | 0 | 0 | 0 |
| ENSMUSG00000092748 | 0 | 0 | 0 | 0 | 0 | 0 | 0 |
| ENSMUSG00000092751 | 0 | 0 | 0 | 0 | 0 | 0 | 0 |
| ENSMUSG00000092752 | 0 | 0 | 0 | 0 | 0 | 0 | 0 |
| ENSMUSG00000092753 | 0 | 0 | 0 | 0 | 0 | 0 | 0 |
| ENSMUSG00000092758 | 0 | 0 | 0 | 0 | 0 | 0 | 0 |
| ENSMUSG00000092759 | 0 | 0 | 0 | 0 | 0 | 0 | 0 |
| ENSMUSG00000092760 | 0 | 0 | 0 | 0 | 0 | 0 | 0.448487648 |
| ENSMUSG00000092761 | 0 | 0 | 0 | 0 | 0 | 0 | 0 |
| ENSMUSG00000092762 | 0 | 0 | 0 | 0 | 0 | 0 | 0 |
| ENSMUSG00000092763 | 0 | 0 | 0 | 0 | 0 | 0 | 0 |
| ENSMUSG00000092764 | 0 | 0 | 0 | 0 | 0 | 0 | 0 |

| | | | | | | | |
|---|---|---|---|---|---|---|---|
| ENSMUSG00000092765 | 0 | 0 | 0 | 0 | 0 | 0 | 0 |
| ENSMUSG00000092767 | 0 | 0 | 0 | 0 | 0 | 0 | 0 |
| ENSMUSG00000092770 | 0 | 0 | 0 | 0 | 0 | 0 | 0 |
| ENSMUSG00000092774 | 0 | 0 | 0 | 0 | 0 | 0 | 0 |
| ENSMUSG00000092776 | 0 | 0 | 0 | 0 | 0 | 0 | 0 |
| ENSMUSG00000092777 | 0 | 0 | 0 | 0 | 0 | 0 | 0 |
| ENSMUSG00000092781 | 0 | 0 | 0.5 | 0.25 | 0 | 0 | 0.355418052 |
| ENSMUSG00000092784 | 0 | 0 | 0 | 0 | 0 | 0 | 0 |
| ENSMUSG00000092787 | 0 | 0 | 0 | 0 | 0 | 0 | 0 |
| ENSMUSG00000092790 | 0 | 0 | 0 | 0 | 0 | 0 | 0 |
| ENSMUSG00000092792 | 0 | 0 | 0 | 0 | 0 | 0 | 0 |
| ENSMUSG00000092793 | 0 | 0 | 0 | 0 | 0 | 0 | 0 |
| ENSMUSG00000092794 | 0 | 0 | 0 | 0 | 0 | 0 | 0 |
| ENSMUSG00000092795 | 0.289064826 | 0 | 0 | 0.396240625 | 0 | 0 | 0.224243824 |
| ENSMUSG00000092796 | 0 | 0 | 0 | 0 | 0 | 0 | 0 |
| ENSMUSG00000092799 | 0 | 0 | 0 | 0 | 0 | 0 | 0 |
| ENSMUSG00000092802 | 0 | 0 | 0.25 | 0.5 | 0 | 0 | 0 |
| ENSMUSG00000092805 | 0 | 0.25 | 0.25 | 0.25 | 0 | 0 | 0 |
| ENSMUSG00000092806 | 0 | 0 | 0 | 0 | 0 | 0 | 0 |
| ENSMUSG00000092807 | 0 | 0 | 0 | 0 | 0 | 0 | 0 |
| ENSMUSG00000092811 | 0 | 0 | 0 | 0 | 0 | 0 | 0 |
| ENSMUSG00000092812 | 0 | 0 | 0 | 0 | 0 | 0 | 0 |
| ENSMUSG00000092813 | 0 | 0 | 0 | 0 | 0 | 0 | 0 |
| ENSMUSG00000092814 | 0 | 0 | 0 | 0 | 0 | 0 | 0 |
| ENSMUSG00000092815 | 0 | 0 | 0 | 0 | 0 | 0 | 0 |
| ENSMUSG00000092816 | 0 | 0 | 0 | 0 | 0 | 0.301029996 | 0 |
| ENSMUSG00000092818 | 0 | 0 | 0 | 0 | 0 | 0 | 0 |
| ENSMUSG00000092819 | 0 | 0 | 0.396240625 | 0.25 | 0 | 0 | 0 |
| ENSMUSG00000092821 | 0 | 0 | 0 | 0 | 0 | 0 | 0 |
| ENSMUSG00000092822 | 0 | 0 | 0 | 0 | 0 | 0 | 0 |
| ENSMUSG00000092825 | 0 | 0 | 0 | 0 | 0 | 0 | 0 |
| ENSMUSG00000092826 | 0 | 0 | 0 | 0 | 0 | 0 | 0 |
| ENSMUSG00000092827 | 0 | 0 | 0 | 0 | 0 | 0 | 0.224243824 |
| ENSMUSG00000092829 | 0 | 0 | 0 | 0 | 0 | 0 | 0 |

| | | | | | | | |
|---|---|---|---|---|---|---|---|
| ENSMUSG00000092830 | 0 | 0 | 0 | 0 | 0 | 0 | 0 |
| ENSMUSG00000092832 | 0 | 0 | 0 | 0 | 0 | 0 | 0 |
| ENSMUSG00000092833 | 0 | 0 | 0 | 0 | 0 | 0 | 0 |
| ENSMUSG00000092834 | 0 | 0 | 0 | 0 | 0 | 0 | 0 |
| ENSMUSG00000092835 | 0 | 0 | 0 | 0 | 0 | 0 | 0 |
| ENSMUSG00000092837 | 0 | 0 | 0 | 0 | 0.244650542 | 0 | 0 |
| ENSMUSG00000092838 | 0 | 0 | 0 | 0 | 0 | 0 | 0 |
| ENSMUSG00000092840 | 0 | 0 | 0 | 0 | 0 | 0 | 0 |
| ENSMUSG00000092841 | 0 | 0 | 0 | 0 | 0 | 0 | 0 |
| ENSMUSG00000092842 | 0 | 0 | 0 | 0 | 0 | 0 | 0 |
| ENSMUSG00000092843 | 0 | 0 | 0 | 0 | 0 | 0 | 0 |
| ENSMUSG00000092845 | 0 | 0 | 0 | 0 | 0 | 0 | 0 |
| ENSMUSG00000092847 | 0 | 0 | 0 | 0 | 0 | 0 | 0 |
| ENSMUSG00000092848 | 0 | 0 | 0 | 0 | 0 | 0 | 0 |
| ENSMUSG00000092849 | 0 | 0 | 0.25 | 0 | 0 | 0 | 0 |
| ENSMUSG00000092850 | 0 | 0 | 0 | 0 | 0 | 0 | 0 |
| ENSMUSG00000092851 | 0 | 0 | 0 | 0 | 0 | 0 | 0 |
| ENSMUSG00000092852 | 0 | 0 | 0 | 0 | 0 | 0 | 0 |
| ENSMUSG00000092853 | 0 | 0 | 0 | 0 | 0 | 0 | 0 |
| ENSMUSG00000092855 | 0 | 0 | 0 | 0 | 0 | 0 | 0 |
| ENSMUSG00000092856 | 0 | 0 | 0 | 0 | 0 | 0 | 0 |
| ENSMUSG00000092862 | 0 | 0 | 0 | 0 | 0 | 0 | 0 |
| ENSMUSG00000092866 | 0 | 0 | 0 | 0 | 0 | 0 | 0 |
| ENSMUSG00000092867 | 0.671187741 | 0.396240625 | 0.25 | 0 | 0 | 0 | 0 |
| ENSMUSG00000092868 | 0 | 0 | 0 | 0 | 0 | 0 | 0 |
| ENSMUSG00000092870 | 0 | 0 | 0 | 0 | 0 | 0 | 0 |
| ENSMUSG00000092871 | 0 | 0 | 0 | 0 | 0 | 0 | 0 |
| ENSMUSG00000092875 | 0 | 0 | 0 | 0 | 0 | 0 | 0 |
| ENSMUSG00000092876 | 0 | 0 | 0 | 0 | 0 | 0 | 0 |
| ENSMUSG00000092878 | 0 | 0 | 0 | 0 | 0 | 0 | 0 |
| ENSMUSG00000092879 | 0 | 0 | 0 | 0 | 0 | 0 | 0 |
| ENSMUSG00000092880 | 0 | 0 | 0 | 0 | 0 | 0 | 0 |
| ENSMUSG00000092885 | 0 | 0 | 0 | 0 | 0 | 0 | 0 |
| ENSMUSG00000092887 | 0 | 0 | 0.5 | 0 | 0.244650542 | 0 | 0.224243824 |

| | | | | | | | |
|---|---|---|---|---|---|---|---|
| ENSMUSG00000092888 | 0 | 0 | 0 | 0 | 0 | 0 | 0 |
| ENSMUSG00000092889 | 0.45815691 | 0.25 | 0 | 0.25 | 0.244650542 | 0 | 0.224243824 |
| ENSMUSG00000092891 | 0 | 0 | 0 | 0 | 0 | 0 | 0 |
| ENSMUSG00000092894 | 0 | 0 | 0 | 0 | 0 | 0 | 0 |
| ENSMUSG00000092896 | 0 | 0 | 0 | 0 | 0 | 0 | 0 |
| ENSMUSG00000092897 | 0 | 0 | 0 | 0 | 0 | 0 | 0 |
| ENSMUSG00000092898 | 0 | 0 | 0 | 0 | 0 | 0 | 0 |
| ENSMUSG00000092901 | 0 | 0 | 0 | 0 | 0 | 0 | 0 |
| ENSMUSG00000092903 | 0 | 0 | 0 | 0.25 | 0.244650542 | 0.301029996 | 0 |
| ENSMUSG00000092905 | 0 | 0 | 0 | 0.396240625 | 0.387761935 | 0.301029996 | 0.224243824 |
| ENSMUSG00000092906 | 0 | 0 | 0 | 0 | 0 | 0 | 0 |
| ENSMUSG00000092907 | 0 | 0 | 0 | 0 | 0 | 0 | 0 |
| ENSMUSG00000092908 | 0 | 0 | 0 | 0 | 0 | 0 | 0 |
| ENSMUSG00000092909 | 0.289064826 | 0.25 | 0.25 | 0 | 0 | 0 | 0.224243824 |
| ENSMUSG00000092912 | 0 | 0 | 0 | 0 | 0 | 0 | 0 |
| ENSMUSG00000092913 | 0 | 0 | 0 | 0 | 0 | 0 | 0 |
| ENSMUSG00000092914 | 0 | 0 | 0 | 0 | 0 | 0 | 0 |
| ENSMUSG00000092915 | 0 | 0 | 0 | 0 | 0 | 0 | 0 |
| ENSMUSG00000092916 | 0 | 0 | 0 | 0 | 0 | 0 | 0 |
| ENSMUSG00000092917 | 0 | 0 | 0 | 0 | 0 | 0 | 0 |
| ENSMUSG00000092918 | 0 | 0 | 0 | 0 | 0 | 0 | 0.355418052 |
| ENSMUSG00000092919 | 0 | 0 | 0 | 0 | 0 | 0 | 0 |
| ENSMUSG00000092920 | 0 | 0 | 0 | 0 | 0 | 0 | 0 |
| ENSMUSG00000092921 | 0 | 0 | 0 | 0 | 0 | 0 | 0 |
| ENSMUSG00000092922 | 0 | 0 | 0 | 0 | 0 | 0 | 0 |
| ENSMUSG00000092923 | 0 | 0 | 0 | 0 | 0 | 0 | 0 |
| ENSMUSG00000092928 | 0 | 0 | 0 | 0 | 0 | 0 | 0 |
| ENSMUSG00000092931 | 0 | 0 | 0 | 0 | 0 | 0 | 0 |
| ENSMUSG00000092932 | 0 | 0.396240625 | 0.5 | 0.25 | 0.244650542 | 0 | 0 |
| ENSMUSG00000092934 | 0 | 0 | 0 | 0 | 0 | 0 | 0 |
| ENSMUSG00000092935 | 0 | 0 | 0 | 0 | 0 | 0 | 0 |
| ENSMUSG00000092939 | 0 | 0 | 0 | 0 | 0 | 0 | 0 |
| ENSMUSG00000092948 | 0 | 0 | 0 | 0 | 0 | 0 | 0 |
| ENSMUSG00000092949 | 0 | 0 | 0 | 0 | 0 | 0 | 0 |

| | | | | | | | |
|---|---|---|---|---|---|---|---|
| ENSMUSG00000092952 | 0 | 0 | 0 | 0 | 0 | 0 | 0 |
| ENSMUSG00000092953 | 0 | 0 | 0 | 0 | 0 | 0 | 0 |
| ENSMUSG00000092956 | 0 | 0 | 0 | 0 | 0 | 0 | 0 |
| ENSMUSG00000092957 | 0 | 0 | 0 | 0 | 0 | 0 | 0 |
| ENSMUSG00000092958 | 0 | 0 | 0 | 0 | 0 | 0 | 0 |
| ENSMUSG00000092961 | 0 | 0 | 0 | 0 | 0 | 0 | 0 |
| ENSMUSG00000092963 | 0 | 0 | 0 | 0 | 0 | 0 | 0 |
| ENSMUSG00000092965 | 0 | 0 | 0 | 0 | 0 | 0 | 0 |
| ENSMUSG00000092966 | 0 | 0 | 0 | 0 | 0 | 0 | 0 |
| ENSMUSG00000092972 | 0 | 0 | 0 | 0 | 0 | 0 | 0 |
| ENSMUSG00000092973 | 0 | 0 | 0 | 0 | 0 | 0 | 0 |
| ENSMUSG00000092974 | 0 | 0 | 0 | 0 | 0 | 0 | 0 |
| ENSMUSG00000092978 | 0 | 0 | 0 | 0 | 0 | 0 | 0 |
| ENSMUSG00000092979 | 0 | 0 | 0 | 0 | 0 | 0 | 0 |
| ENSMUSG00000092981 | 0 | 0.75 | 0.830482024 | 0.25 | 0.489301084 | 0 | 0.710836105 |
| ENSMUSG00000092985 | 0 | 0 | 0 | 0 | 0 | 0 | 0 |
| ENSMUSG00000092986 | 0 | 0.25 | 0 | 0.25 | 0.244650542 | 0 | 0 |
| ENSMUSG00000092988 | 0 | 0 | 0 | 0 | 0 | 0 | 0 |
| ENSMUSG00000092990 | 0 | 0.25 | 0 | 0 | 0 | 0 | 0 |
| ENSMUSG00000092994 | 0 | 0 | 0 | 0 | 0 | 0 | 0 |
| ENSMUSG00000092995 | 0 | 0 | 0 | 0 | 0 | 0 | 0 |
| ENSMUSG00000092997 | 0 | 0 | 0 | 0 | 0 | 0 | 0 |
| ENSMUSG00000092998 | 0 | 0 | 0 | 0 | 0 | 0 | 0 |
| ENSMUSG00000092999 | 0 | 0 | 0 | 0 | 0 | 0 | 0 |
| ENSMUSG00000093000 | 0 | 0 | 0 | 0 | 0 | 0 | 0 |
| ENSMUSG00000093002 | 0 | 0 | 0 | 0 | 0 | 0 | 0 |
| ENSMUSG00000093003 | 0 | 0 | 0 | 0 | 0 | 0 | 0 |
| ENSMUSG00000093004 | 0 | 0 | 0 | 0 | 0 | 0 | 0 |
| ENSMUSG00000093005 | 0 | 0 | 0 | 0 | 0 | 0 | 0 |
| ENSMUSG00000093006 | 0 | 0 | 0 | 0 | 0 | 0 | 0 |
| ENSMUSG00000093007 | 0 | 0 | 0 | 0 | 0 | 0 | 0 |
| ENSMUSG00000093009 | 0 | 0 | 0 | 0 | 0 | 0 | 0 |
| ENSMUSG00000093010 | 0 | 0 | 0 | 0 | 0 | 0 | 0 |
| ENSMUSG00000093011 | 0 | 0 | 0 | 0 | 0 | 0 | 0 |

| | | | | | | | |
|---|---|---|---|---|---|---|---|
| ENSMUSG00000093013 | 0 | 0 | 0 | 0 | 0 | 0 | 0 |
| ENSMUSG00000093014 | 0 | 0 | 0 | 0 | 0 | 0 | 0 |
| ENSMUSG00000093015 | 0 | 0 | 0 | 0 | 0 | 0 | 0 |
| ENSMUSG00000093017 | 0 | 0 | 0 | 0 | 0 | 0 | 0 |
| ENSMUSG00000093019 | 0 | 0 | 0 | 0 | 0 | 0 | 0 |
| ENSMUSG00000093021 | 0 | 0 | 0 | 0 | 0 | 0 | 0 |
| ENSMUSG00000093023 | 0.671187741 | 0.25 | 0.25 | 0.25 | 0.244650542 | 0.301029996 | 0 |
| ENSMUSG00000093024 | 0 | 0 | 0 | 0 | 0 | 0 | 0 |
| ENSMUSG00000093025 | 0 | 0 | 0 | 0 | 0 | 0 | 0 |
| ENSMUSG00000093026 | 0 | 0 | 0 | 0 | 0 | 0 | 0 |
| ENSMUSG00000093029 | 0 | 0 | 0 | 0 | 0 | 0 | 0 |
| ENSMUSG00000093030 | 0 | 0.25 | 0 | 0 | 0 | 0 | 0 |
| ENSMUSG00000093031 | 0 | 0 | 0 | 0 | 0 | 0 | 0 |
| ENSMUSG00000093032 | 0 | 0 | 0 | 0.25 | 0 | 0 | 0 |
| ENSMUSG00000093033 | 0 | 0 | 0 | 0 | 0 | 0 | 0 |
| ENSMUSG00000093034 | 0 | 0 | 0 | 0 | 0 | 0 | 0 |
| ENSMUSG00000093036 | 0 | 0 | 0 | 0 | 0 | 0 | 0 |
| ENSMUSG00000093037 | 0 | 0 | 0 | 0 | 0 | 0 | 0 |
| ENSMUSG00000093039 | 0 | 0 | 0 | 0 | 0 | 0 | 0 |
| ENSMUSG00000093040 | 0 | 0 | 0 | 0 | 0 | 0 | 0 |
| ENSMUSG00000093041 | 0 | 0 | 0 | 0 | 0 | 0 | 0 |
| ENSMUSG00000093042 | 0 | 0 | 0 | 0 | 0 | 0 | 0 |
| ENSMUSG00000093044 | 0.289064826 | 0 | 0.25 | 0.396240625 | 0 | 0 | 0.224243824 |
| ENSMUSG00000093046 | 0 | 0 | 0 | 0 | 0 | 0 | 0 |
| ENSMUSG00000093051 | 0 | 0 | 0 | 0 | 0 | 0 | 0 |
| ENSMUSG00000093052 | 0 | 0 | 0 | 0 | 0 | 0 | 0 |
| ENSMUSG00000093054 | 0 | 0 | 0 | 0 | 0 | 0 | 0 |
| ENSMUSG00000093056 | 0 | 0 | 0 | 0 | 0 | 0 | 0 |
| ENSMUSG00000093058 | 0 | 0 | 0 | 0 | 0 | 0 | 0 |
| ENSMUSG00000093059 | 0.289064826 | 0.396240625 | 0.396240625 | 0.580482024 | 0.387761935 | 0.301029996 | 0 |
| ENSMUSG00000093060 | 0 | 0 | 0 | 0 | 0 | 0 | 0 |
| ENSMUSG00000093061 | 0 | 0 | 0 | 0 | 0 | 0 | 0 |
| ENSMUSG00000093063 | 0 | 0 | 0 | 0 | 0 | 0 | 0 |
| ENSMUSG00000093064 | 0 | 0 | 0 | 0 | 0 | 0 | 0 |

| | | | | | | | |
|---|---|---|---|---|---|---|---|
| ENSMUSG00000093065 | 0 | 0 | 0 | 0 | 0 | 0 | 0 |
| ENSMUSG00000093067 | 0 | 0 | 0 | 0 | 0 | 0 | 0 |
| ENSMUSG00000093070 | 0 | 0 | 0 | 0 | 0 | 0 | 0 |
| ENSMUSG00000093071 | 0 | 0 | 0 | 0 | 0 | 0 | 0 |
| ENSMUSG00000093072 | 0 | 0 | 0 | 0 | 0 | 0 | 0 |
| ENSMUSG00000093073 | 0 | 0 | 0 | 0 | 0 | 0 | 0 |
| ENSMUSG00000093074 | 0 | 0 | 0 | 0 | 0 | 0 | 0 |
| ENSMUSG00000093075 | 0 | 0 | 0 | 0 | 0.387761935 | 0 | 0.224243824 |
| ENSMUSG00000093076 | 0 | 0 | 0 | 0 | 0 | 0 | 0 |
| ENSMUSG00000093079 | 0 | 0 | 0 | 0 | 0 | 0 | 0 |
| ENSMUSG00000093080 | 0.45815691 | 0 | 0.396240625 | 0.25 | 0 | 0 | 0.224243824 |
| ENSMUSG00000093081 | 0 | 0 | 0 | 0 | 0 | 0 | 0 |
| ENSMUSG00000093082 | 0 | 0 | 0 | 0 | 0 | 0 | 0 |
| ENSMUSG00000093083 | 0 | 0 | 0 | 0 | 0 | 0 | 0 |
| ENSMUSG00000093085 | 0 | 0.25 | 0 | 0.25 | 0.244650542 | 0 | 0 |
| ENSMUSG00000093087 | 0 | 0 | 0 | 0 | 0 | 0 | 0 |
| ENSMUSG00000093089 | 0 | 0 | 0 | 0 | 0 | 0 | 0 |
| ENSMUSG00000093090 | 0 | 0 | 0 | 0 | 0 | 0 | 0 |
| ENSMUSG00000093091 | 0 | 0 | 0 | 0 | 0 | 0 | 0 |
| ENSMUSG00000093092 | 0 | 0 | 0 | 0 | 0 | 0 | 0 |
| ENSMUSG00000093093 | 0.289064826 | 0.25 | 0 | 0.25 | 0 | 0 | 0.224243824 |
| ENSMUSG00000093096 | 0 | 0 | 0 | 0 | 0 | 0 | 0 |
| ENSMUSG00000093097 | 0 | 0 | 0 | 0 | 0 | 0 | 0 |
| ENSMUSG00000093100 | 0 | 0 | 0 | 0 | 0 | 0 | 0 |
| ENSMUSG00000093101 | 0 | 0 | 0 | 0 | 0 | 0 | 0 |
| ENSMUSG00000093102 | 0 | 0 | 0 | 0 | 0 | 0 | 0 |
| ENSMUSG00000093105 | 0.289064826 | 0 | 0 | 0 | 0.244650542 | 0 | 0.224243824 |
| ENSMUSG00000093106 | 0 | 0 | 0 | 0 | 0 | 0 | 0 |
| ENSMUSG00000093107 | 0 | 0 | 0 | 0 | 0 | 0 | 0 |
| ENSMUSG00000093108 | 0 | 0 | 0 | 0 | 0 | 0 | 0 |
| ENSMUSG00000093112 | 0 | 0 | 0 | 0 | 0 | 0 | 0 |
| ENSMUSG00000093113 | 0 | 0 | 0 | 0 | 0 | 0 | 0 |
| ENSMUSG00000093114 | 0 | 0 | 0 | 0 | 0 | 0 | 0 |
| ENSMUSG00000093115 | 0 | 0 | 0 | 0 | 0 | 0 | 0 |

| | | | | | | | |
|---|---|---|---|---|---|---|---|
| ENSMUSG00000093116 | 0 | 0 | 0 | 0 | 0 | 0 | 0 |
| ENSMUSG00000093117 | 0 | 0 | 0 | 0 | 0 | 0 | 0 |
| ENSMUSG00000093118 | 0 | 0 | 0 | 0 | 0 | 0 | 0 |
| ENSMUSG00000093119 | 0.289064826 | 0 | 0.25 | 0.580482024 | 0.244650542 | 0 | 0 |
| ENSMUSG00000093120 | 0 | 0 | 0 | 0 | 0 | 0 | 0.224243824 |
| ENSMUSG00000093122 | 0 | 0 | 0 | 0 | 0 | 0 | 0 |
| ENSMUSG00000093124 | 0.289064826 | 0.25 | 0 | 0 | 0 | 0 | 0.224243824 |
| ENSMUSG00000093125 | 0 | 0 | 0 | 0 | 0 | 0 | 0 |
| ENSMUSG00000093128 | 0.289064826 | 0 | 0 | 0 | 0 | 0 | 0 |
| ENSMUSG00000093129 | 0 | 0 | 0 | 0 | 0 | 0 | 0 |
| ENSMUSG00000093132 | 0 | 0 | 0 | 0 | 0 | 0 | 0 |
| ENSMUSG00000093133 | 0 | 0 | 0 | 0 | 0 | 0 | 0 |
| ENSMUSG00000093134 | 0 | 0 | 0.25 | 0.25 | 0 | 0 | 0 |
| ENSMUSG00000093137 | 0 | 0 | 0 | 0 | 0 | 0 | 0 |
| ENSMUSG00000093138 | 0 | 0 | 0 | 0 | 0 | 0 | 0 |
| ENSMUSG00000093139 | 0 | 0 | 0 | 0 | 0 | 0 | 0 |
| ENSMUSG00000093140 | 0 | 0 | 0 | 0 | 0 | 0 | 0 |
| ENSMUSG00000093142 | 0 | 0 | 0 | 0 | 0 | 0 | 0 |
| ENSMUSG00000093147 | 0 | 0 | 0 | 0 | 0 | 0.301029996 | 0.224243824 |
| ENSMUSG00000093148 | 0 | 0 | 0 | 0 | 0 | 0 | 0 |
| ENSMUSG00000093149 | 0 | 0 | 0 | 0 | 0 | 0 | 0 |
| ENSMUSG00000093152 | 0 | 0 | 0 | 0 | 0 | 0 | 0 |
| ENSMUSG00000093153 | 0 | 0 | 0 | 0 | 0 | 0 | 0 |
| ENSMUSG00000093155 | 0 | 0 | 0 | 0.396240625 | 0 | 0 | 0.224243824 |
| ENSMUSG00000093156 | 0 | 0 | 0 | 0 | 0 | 0 | 0.224243824 |
| ENSMUSG00000093157 | 0 | 0 | 0 | 0 | 0 | 0 | 0 |
| ENSMUSG00000093158 | 0 | 0 | 0 | 0 | 0 | 0 | 0 |
| ENSMUSG00000093159 | 0 | 0 | 0 | 0 | 0 | 0 | 0 |
| ENSMUSG00000093160 | 0 | 0 | 0 | 0 | 0 | 0 | 0 |
| ENSMUSG00000093161 | 0 | 0 | 0 | 0 | 0 | 0 | 0.224243824 |
| ENSMUSG00000093162 | 0 | 0 | 0 | 0 | 0 | 0 | 0 |
| ENSMUSG00000093163 | 0 | 0 | 0 | 0.25 | 0 | 0 | 0 |
| ENSMUSG00000093164 | 0 | 0 | 0 | 0 | 0.244650542 | 0.301029996 | 0 |
| ENSMUSG00000093167 | 0 | 0 | 0 | 0 | 0 | 0 | 0 |

| | | | | | | | |
|---|---|---|---|---|---|---|---|
| ENSMUSG00000093168 | 0 | 0 | 0 | 0 | 0.244650542 | 0.301029996 | 0 |
| ENSMUSG00000093169 | 0 | 0 | 0 | 0 | 0 | 0 | 0 |
| ENSMUSG00000093170 | 0 | 0 | 0 | 0 | 0 | 0 | 0 |
| ENSMUSG00000093172 | 0 | 0 | 0 | 0 | 0 | 0 | 0 |
| ENSMUSG00000093173 | 0 | 0 | 0 | 0 | 0 | 0 | 0 |
| ENSMUSG00000093176 | 0 | 0 | 0.25 | 0.25 | 0 | 0 | 0 |
| ENSMUSG00000093178 | 0.289064826 | 0.396240625 | 0.646240625 | 0.5 | 0.568060967 | 0.301029996 | 0.355418052 |
| ENSMUSG00000093180 | 0 | 0 | 0.25 | 0 | 0 | 0 | 0 |
| ENSMUSG00000093183 | 0 | 0 | 0 | 0 | 0 | 0 | 0 |
| ENSMUSG00000093184 | 0 | 0 | 0 | 0 | 0 | 0 | 0 |
| ENSMUSG00000093185 | 0 | 0 | 0 | 0 | 0 | 0 | 0 |
| ENSMUSG00000093187 | 0 | 0 | 0 | 0 | 0 | 0 | 0 |
| ENSMUSG00000093189 | 0 | 0 | 0 | 0 | 0 | 0 | 0 |
| ENSMUSG00000093190 | 0.45815691 | 0.25 | 0.25 | 0.75 | 0.489301084 | 0.477121255 | 0.224243824 |
| ENSMUSG00000093191 | 0 | 0 | 0 | 0 | 0 | 0 | 0 |
| ENSMUSG00000093192 | 0 | 0 | 0 | 0 | 0 | 0 | 0 |
| ENSMUSG00000093202 | 0 | 0 | 0 | 0 | 0 | 0 | 0 |
| ENSMUSG00000093203 | 0 | 0 | 0.25 | 0 | 0.387761935 | 0 | 0 |
| ENSMUSG00000093206 | 0 | 0 | 0 | 0 | 0 | 0 | 0 |
| ENSMUSG00000093207 | 0 | 0 | 0 | 0 | 0 | 0 | 0 |
| ENSMUSG00000093208 | 0 | 0 | 0 | 0 | 0 | 0 | 0 |
| ENSMUSG00000093211 | 0 | 0 | 0 | 0 | 0 | 0 | 0 |
| ENSMUSG00000093212 | 0 | 0 | 0 | 0 | 0 | 0 | 0 |
| ENSMUSG00000093214 | 0 | 0 | 0 | 0 | 0 | 0 | 0 |
| ENSMUSG00000093215 | 0 | 0 | 0 | 0 | 0 | 0 | 0 |
| ENSMUSG00000093216 | 0 | 0 | 0 | 0 | 0 | 0 | 0 |
| ENSMUSG00000093217 | 0 | 0 | 0 | 0 | 0 | 0 | 0 |
| ENSMUSG00000093218 | 0 | 0 | 0 | 0 | 0 | 0 | 0 |
| ENSMUSG00000093219 | 0 | 0 | 0 | 0 | 0 | 0 | 0 |
| ENSMUSG00000093221 | 0 | 0 | 0 | 0 | 0 | 0 | 0 |
| ENSMUSG00000093223 | 0 | 0 | 0 | 0 | 0 | 0 | 0 |
| ENSMUSG00000093224 | 0 | 0 | 0 | 0 | 0 | 0 | 0 |
| ENSMUSG00000093226 | 0 | 0 | 0 | 0 | 0 | 0 | 0 |
| ENSMUSG00000093227 | 0 | 0 | 0 | 0 | 0 | 0 | 0 |

| | | | | | | | |
|---|---|---|---|---|---|---|---|
| ENSMUSG00000093228 | 0 | 0 | 0 | 0 | 0 | 0 | 0 |
| ENSMUSG00000093230 | 0 | 0 | 0 | 0 | 0 | 0 | 0 |
| ENSMUSG00000093231 | 0.289064826 | 0.5 | 0 | 0.25 | 0.244650542 | 0.301029996 | 0.355418052 |
| ENSMUSG00000093232 | 0 | 0 | 0 | 0 | 0 | 0 | 0 |
| ENSMUSG00000093234 | 0 | 0 | 0 | 0 | 0 | 0 | 0 |
| ENSMUSG00000093236 | 0 | 0 | 0 | 0 | 0 | 0 | 0 |
| ENSMUSG00000093238 | 0 | 0 | 0 | 0 | 0 | 0 | 0 |
| ENSMUSG00000093239 | 0 | 0 | 0 | 0 | 0 | 0 | 0 |
| ENSMUSG00000093240 | 0 | 0 | 0 | 0 | 0 | 0 | 0 |
| ENSMUSG00000093241 | 0 | 0 | 0 | 0 | 0 | 0 | 0 |
| ENSMUSG00000093242 | 0 | 0 | 0 | 0 | 0 | 0 | 0 |
| ENSMUSG00000093243 | 0 | 0 | 0 | 0 | 0 | 0 | 0 |
| ENSMUSG00000093245 | 0 | 0 | 0 | 0 | 0 | 0 | 0 |
| ENSMUSG00000093246 | 0 | 0 | 0 | 0 | 0 | 0 | 0 |
| ENSMUSG00000093249 | 0 | 0 | 0.25 | 0.25 | 0 | 0.301029996 | 0 |
| ENSMUSG00000093251 | 0 | 0 | 0 | 0 | 0 | 0 | 0 |
| ENSMUSG00000093254 | 0 | 0 | 0 | 0 | 0 | 0 | 0 |
| ENSMUSG00000093256 | 0 | 0 | 0 | 0 | 0 | 0 | 0 |
| ENSMUSG00000093258 | 0 | 0 | 0 | 0 | 0 | 0 | 0 |
| ENSMUSG00000093260 | 0 | 0 | 0 | 0.25 | 0 | 0 | 0 |
| ENSMUSG00000093261 | 0 | 0 | 0 | 0 | 0 | 0 | 0 |
| ENSMUSG00000093262 | 0 | 0 | 0 | 0 | 0 | 0 | 0 |
| ENSMUSG00000093264 | 0.45815691 | 0 | 0.25 | 0 | 0.244650542 | 0 | 0 |
| ENSMUSG00000093266 | 0 | 0 | 0 | 0 | 0 | 0 | 0 |
| ENSMUSG00000093269 | 0 | 0 | 0 | 0 | 0 | 0 | 0 |
| ENSMUSG00000093270 | 0 | 0 | 0.25 | 0 | 0 | 0 | 0 |
| ENSMUSG00000093271 | 0 | 0 | 0 | 0 | 0 | 0 | 0 |
| ENSMUSG00000093273 | 0 | 0 | 0 | 0 | 0 | 0 | 0 |
| ENSMUSG00000093274 | 0 | 0 | 0 | 0 | 0 | 0 | 0 |
| ENSMUSG00000093276 | 0 | 0 | 0 | 0 | 0.244650542 | 0 | 0 |
| ENSMUSG00000093277 | 0 | 0 | 0 | 0 | 0 | 0 | 0 |
| ENSMUSG00000093278 | 0 | 0 | 0 | 0 | 0 | 0 | 0 |
| ENSMUSG00000093279 | 0 | 0 | 0 | 0 | 0 | 0 | 0 |
| ENSMUSG00000093280 | 0 | 0 | 0 | 0 | 0 | 0 | 0 |

| | | | | | | | |
|---|---|---|---|---|---|---|---|
| ENSMUSG00000093281 | 0 | 0 | 0 | 0 | 0 | 0 | 0 |
| ENSMUSG00000093282 | 0 | 0 | 0.25 | 0 | 0 | 0 | 0 |
| ENSMUSG00000093283 | 0 | 0 | 0 | 0 | 0 | 0 | 0 |
| ENSMUSG00000093286 | 0 | 0 | 0 | 0 | 0 | 0 | 0 |
| ENSMUSG00000093289 | 0 | 0 | 0 | 0 | 0 | 0 | 0 |
| ENSMUSG00000093290 | 0 | 0 | 0 | 0 | 0 | 0 | 0 |
| ENSMUSG00000093291 | 0 | 0 | 0 | 0 | 0 | 0 | 0 |
| ENSMUSG00000093292 | 0 | 0 | 0 | 0 | 0 | 0 | 0 |
| ENSMUSG00000093293 | 0 | 0 | 0 | 0 | 0 | 0 | 0 |
| ENSMUSG00000093294 | 0 | 0 | 0 | 0 | 0 | 0 | 0 |
| ENSMUSG00000093296 | 0 | 0 | 0 | 0 | 0 | 0 | 0 |
| ENSMUSG00000093298 | 0 | 0 | 0 | 0 | 0 | 0 | 0 |
| ENSMUSG00000093300 | 0 | 0 | 0 | 0 | 0 | 0 | 0 |
| ENSMUSG00000093301 | 0 | 0 | 0 | 0 | 0 | 0 | 0 |
| ENSMUSG00000093302 | 0 | 0 | 0 | 0 | 0 | 0 | 0 |
| ENSMUSG00000093303 | 0 | 0 | 0 | 0 | 0 | 0 | 0 |
| ENSMUSG00000093306 | 0 | 0 | 0 | 0 | 0 | 0 | 0 |
| ENSMUSG00000093311 | 0 | 0 | 0 | 0 | 0 | 0 | 0 |
| ENSMUSG00000093314 | 0 | 0 | 0.25 | 0.25 | 0 | 0 | 0 |
| ENSMUSG00000093315 | 0 | 0 | 0 | 0 | 0 | 0 | 0 |
| ENSMUSG00000093317 | 0 | 0 | 0 | 0 | 0 | 0 | 0 |
| ENSMUSG00000093319 | 0 | 0.25 | 0 | 0.396240625 | 0 | 0 | 0.355418052 |
| ENSMUSG00000093322 | 0 | 0 | 0 | 0 | 0 | 0 | 0 |
| ENSMUSG00000093323 | 0 | 0 | 0 | 0 | 0 | 0 | 0 |
| ENSMUSG00000093325 | 0 | 0 | 0 | 0 | 0 | 0 | 0 |
| ENSMUSG00000093327 | 0 | 0 | 0 | 0 | 0 | 0 | 0 |
| ENSMUSG00000093329 | 0 | 0 | 0 | 0 | 0 | 0 | 0 |
| ENSMUSG00000093331 | 0 | 0 | 0 | 0 | 0 | 0 | 0 |
| ENSMUSG00000093334 | 0 | 0 | 0 | 0 | 0 | 0 | 0 |
| ENSMUSG00000093335 | 0 | 0 | 0 | 0 | 0 | 0 | 0 |
| ENSMUSG00000093336 | 0 | 0 | 0 | 0 | 0 | 0 | 0 |
| ENSMUSG00000093338 | 0 | 0 | 0 | 0 | 0 | 0 | 0 |
| ENSMUSG00000093340 | 0 | 0 | 0 | 0 | 0 | 0 | 0 |
| ENSMUSG00000093341 | 0 | 0 | 0 | 0 | 0 | 0 | 0 |

| | | | | | | | |
|---|---|---|---|---|---|---|---|
| ENSMUSG00000093342 | 0 | 0 | 0 | 0 | 0 | 0 | 0 |
| ENSMUSG00000093344 | 0 | 0 | 0 | 0 | 0 | 0 | 0 |
| ENSMUSG00000093345 | 0 | 0 | 0 | 0 | 0 | 0 | 0.224243824 |
| ENSMUSG00000093346 | 0 | 0 | 0 | 0 | 0 | 0 | 0 |
| ENSMUSG00000093348 | 0 | 0 | 0 | 0 | 0 | 0 | 0 |
| ENSMUSG00000093351 | 0 | 0 | 0.25 | 0 | 0 | 0 | 0 |
| ENSMUSG00000093352 | 0 | 0 | 0 | 0 | 0 | 0 | 0 |
| ENSMUSG00000093354 | 0 | 0 | 0 | 0 | 0 | 0 | 0 |
| ENSMUSG00000093355 | 0 | 0 | 0 | 0 | 0 | 0 | 0 |
| ENSMUSG00000093356 | 0 | 0 | 0 | 0 | 0 | 0 | 0 |
| ENSMUSG00000093358 | 0 | 0 | 0 | 0 | 0 | 0 | 0 |
| ENSMUSG00000093359 | 0 | 0 | 0 | 0 | 0 | 0 | 0 |
| ENSMUSG00000093362 | 0 | 0 | 0.25 | 0 | 0 | 0 | 0 |
| ENSMUSG00000093363 | 0 | 0.396240625 | 0 | 0 | 0 | 0.301029996 | 0 |
| ENSMUSG00000093365 | 0 | 0 | 0 | 0 | 0 | 0 | 0 |
| ENSMUSG00000093368 | 0 | 0 | 0 | 0 | 0 | 0 | 0 |
| ENSMUSG00000093373 | 0 | 0 | 0 | 0 | 0 | 0 | 0 |
| ENSMUSG00000093376 | 0 | 0 | 0 | 0 | 0 | 0 | 0 |
| ENSMUSG00000093378 | 0.578129653 | 0.25 | 0.5 | 0.25 | 0.489301084 | 0.301029996 | 0.355418052 |
| ENSMUSG00000093379 | 0 | 0 | 0 | 0 | 0 | 0 | 0 |
| ENSMUSG00000093380 | 0 | 0 | 0 | 0 | 0 | 0 | 0 |
| ENSMUSG00000093381 | 0 | 0 | 0 | 0 | 0 | 0 | 0 |
| ENSMUSG00000093382 | 0 | 0 | 0 | 0 | 0 | 0 | 0 |
| ENSMUSG00000093383 | 0 | 0 | 0 | 0 | 0 | 0 | 0 |
| ENSMUSG00000093384 | 0 | 0 | 0 | 0.25 | 0 | 0 | 0 |
| ENSMUSG00000093385 | 1.289064826 | 1.098079356 | 1.061981878 | 1.25 | 1.188507684 | 1.301029996 | 1.014380832 |
| ENSMUSG00000093386 | 0 | 0 | 0 | 0 | 0 | 0 | 0 |
| ENSMUSG00000093387 | 0 | 0 | 0 | 0 | 0 | 0 | 0 |
| ENSMUSG00000093388 | 0 | 0.25 | 0 | 0 | 0 | 0 | 0 |
| ENSMUSG00000093390 | 0 | 0 | 0 | 0 | 0.244650542 | 0 | 0.224243824 |
| ENSMUSG00000093392 | 0 | 0 | 0 | 0 | 0 | 0 | 0 |
| ENSMUSG00000093394 | 0.289064826 | 0 | 0.25 | 0 | 0.489301084 | 0.301029996 | 0.224243824 |
| ENSMUSG00000093395 | 0 | 0 | 0 | 0 | 0 | 0 | 0 |
| ENSMUSG00000093397 | 0 | 0 | 0 | 0 | 0 | 0 | 0 |

| | | | | | | | |
|---|---|---|---|---|---|---|---|
| ENSMUSG00000093398 | 0 | 0 | 0 | 0 | 0 | 0 | 0 |
| ENSMUSG00000093400 | 0 | 0 | 0 | 0 | 0 | 0 | 0 |
| ENSMUSG00000093401 | 0 | 0 | 0 | 0 | 0 | 0 | 0 |
| ENSMUSG00000093402 | 0 | 0 | 0 | 0 | 0 | 0 | 0 |
| ENSMUSG00000093403 | 0 | 0 | 0 | 0 | 0 | 0 | 0 |
| ENSMUSG00000093404 | 0 | 0 | 0.25 | 0.25 | 0 | 0 | 0 |
| ENSMUSG00000093405 | 0 | 0 | 0 | 0 | 0 | 0 | 0 |
| ENSMUSG00000093406 | 0 | 0 | 0 | 0 | 0 | 0 | 0 |
| ENSMUSG00000093408 | 0 | 0 | 0 | 0 | 0 | 0 | 0 |
| ENSMUSG00000093409 | 0 | 0.25 | 0 | 0 | 0 | 0.301029996 | 0 |
| ENSMUSG00000093410 | 0 | 0 | 0 | 0 | 0 | 0 | 0 |
| ENSMUSG00000093411 | 0 | 0 | 0 | 0 | 0 | 0 | 0 |
| ENSMUSG00000093413 | 0 | 0.25 | 0 | 0 | 0 | 0 | 0 |
| ENSMUSG00000093414 | 0 | 0 | 0 | 0 | 0 | 0 | 0 |
| ENSMUSG00000093416 | 0.578129653 | 0.25 | 0.5 | 0.646240625 | 0.733951626 | 0 | 0.520678036 |
| ENSMUSG00000093418 | 0.289064826 | 0 | 0 | 0 | 0 | 0 | 0 |
| ENSMUSG00000093419 | 0 | 0 | 0 | 0 | 0 | 0 | 0 |
| ENSMUSG00000093420 | 0 | 0 | 0 | 0 | 0 | 0 | 0 |
| ENSMUSG00000093421 | 0 | 0 | 0 | 0 | 0 | 0 | 0 |
| ENSMUSG00000093424 | 0.289064826 | 1.04248125 | 0.830482024 | 0.646240625 | 0.77552387 | 0.602059991 | 0.803905701 |
| ENSMUSG00000093426 | 0 | 0 | 0 | 0 | 0 | 0 | 0 |
| ENSMUSG00000093427 | 0 | 0 | 0 | 0 | 0 | 0 | 0 |
| ENSMUSG00000093428 | 0 | 0 | 0 | 0 | 0 | 0 | 0 |
| ENSMUSG00000093429 | 0 | 0.25 | 0 | 0 | 0 | 0 | 0 |
| ENSMUSG00000093433 | 0 | 0 | 0 | 0 | 0 | 0 | 0 |
| ENSMUSG00000093434 | 0 | 0 | 0 | 0 | 0 | 0 | 0 |
| ENSMUSG00000093436 | 0 | 0 | 0 | 0 | 0 | 0 | 0 |
| ENSMUSG00000093437 | 0 | 0.25 | 0 | 0.396240625 | 0.568060967 | 0.301029996 | 0 |
| ENSMUSG00000093441 | 0 | 0 | 0 | 0 | 0 | 0 | 0 |
| ENSMUSG00000093443 | 0 | 0 | 0 | 0 | 0 | 0 | 0 |
| ENSMUSG00000093444 | 0 | 0 | 0 | 0 | 0 | 0 | 0 |
| ENSMUSG00000093445 | 0 | 0.25 | 0.25 | 0 | 0.568060967 | 0 | 0 |
| ENSMUSG00000093446 | 0 | 0 | 0 | 0 | 0 | 0 | 0 |
| ENSMUSG00000093447 | 0.289064826 | 0.25 | 0.25 | 0.25 | 0.387761935 | 0.477121255 | 0.355418052 |

| | | | | | | | |
|---|---|---|---|---|---|---|---|
| ENSMUSG00000093449 | 0 | 0 | 0 | 0 | 0 | 0 | 0 |
| ENSMUSG00000093451 | 0 | 0 | 0 | 0 | 0 | 0 | 0 |
| ENSMUSG00000093452 | 0 | 0 | 0 | 0 | 0 | 0 | 0 |
| ENSMUSG00000093453 | 0 | 0 | 0 | 0 | 0 | 0 | 0 |
| ENSMUSG00000093454 | 0 | 0 | 0 | 0 | 0 | 0 | 0 |
| ENSMUSG00000093456 | 0 | 0 | 0 | 0 | 0 | 0 | 0 |
| ENSMUSG00000093458 | 0 | 0 | 0 | 0.396240625 | 0 | 0 | 0.224243824 |
| ENSMUSG00000093460 | 0.289064826 | 0 | 0 | 0.5 | 0.77552387 | 0.84509804 | 0.520678036 |
| ENSMUSG00000093461 | 0 | 0 | 0 | 0 | 0 | 0 | 0 |
| ENSMUSG00000093465 | 0 | 0.25 | 0 | 0 | 0.244650542 | 0 | 0 |
| ENSMUSG00000093467 | 0 | 0 | 0 | 0 | 0 | 0 | 0 |
| ENSMUSG00000093469 | 0 | 0 | 0 | 0 | 0 | 0 | 0 |
| ENSMUSG00000093472 | 0 | 0 | 0 | 0 | 0 | 0 | 0 |
| ENSMUSG00000093473 | 0 | 0 | 0 | 0 | 0 | 0 | 0 |
| ENSMUSG00000093474 | 0 | 0 | 0 | 0 | 0 | 0 | 0 |
| ENSMUSG00000093476 | 0 | 0 | 0 | 0 | 0.244650542 | 0.301029996 | 0.224243824 |
| ENSMUSG00000093479 | 0 | 0 | 0 | 0 | 0 | 0 | 0 |
| ENSMUSG00000093480 | 0 | 0 | 0 | 0 | 0 | 0 | 0 |
| ENSMUSG00000093482 | 0 | 0 | 0 | 0 | 0 | 0 | 0 |
| ENSMUSG00000093483 | 1.069666964 | 1.061981878 | 0.951838731 | 1.04248125 | 1.020174412 | 1 | 1.014380832 |
| ENSMUSG00000093484 | 0 | 0 | 0 | 0.25 | 0 | 0 | 0 |
| ENSMUSG00000093485 | 0 | 0 | 0 | 0 | 0 | 0 | 0.224243824 |
| ENSMUSG00000093487 | 0 | 0 | 0 | 0 | 0 | 0 | 0 |
| ENSMUSG00000093489 | 0 | 0 | 0 | 0 | 0 | 0 | 0 |
| ENSMUSG00000093490 | 0 | 0 | 0 | 0 | 0 | 0 | 0 |
| ENSMUSG00000093492 | 0 | 0 | 0 | 0 | 0 | 0 | 0 |
| ENSMUSG00000093493 | 0 | 0 | 0 | 0 | 0 | 0 | 0 |
| ENSMUSG00000093495 | 0.289064826 | 0 | 0.396240625 | 0 | 0.244650542 | 0.301029996 | 0 |
| ENSMUSG00000093497 | 0 | 0 | 0 | 0 | 0 | 0 | 0 |
| ENSMUSG00000093498 | 0 | 0 | 0 | 0 | 0 | 0 | 0 |
| ENSMUSG00000093499 | 0 | 0 | 0 | 0 | 0 | 0 | 0 |
| ENSMUSG00000093501 | 0 | 0 | 0 | 0 | 0 | 0 | 0 |
| ENSMUSG00000093502 | 0 | 0 | 0 | 0 | 0 | 0 | 0 |
| ENSMUSG00000093503 | 0 | 0 | 0 | 0 | 0 | 0 | 0 |

| | | | | | | | |
|---|---|---|---|---|---|---|---|
| ENSMUSG00000093504 | 0 | 0 | 0 | 0 | 0 | 0 | 0 |
| ENSMUSG00000093505 | 0 | 0 | 0 | 0 | 0 | 0 | 0 |
| ENSMUSG00000093507 | 0 | 0.5 | 0 | 0 | 0 | 0 | 0 |
| ENSMUSG00000093508 | 0 | 0 | 0 | 0 | 0 | 0 | 0 |
| ENSMUSG00000093509 | 0 | 0 | 0 | 0 | 0 | 0 | 0 |
| ENSMUSG00000093510 | 0 | 0 | 0 | 0 | 0 | 0 | 0 |
| ENSMUSG00000093511 | 0 | 0 | 0 | 0 | 0 | 0 | 0 |
| ENSMUSG00000093513 | 0 | 0 | 0 | 0 | 0 | 0 | 0 |
| ENSMUSG00000093514 | 0.45815691 | 0.396240625 | 0 | 0.25 | 0 | 0 | 0 |
| ENSMUSG00000093515 | 0.289064826 | 0 | 0 | 0 | 0 | 0 | 0 |
| ENSMUSG00000093516 | 0 | 0 | 0 | 0 | 0 | 0 | 0 |
| ENSMUSG00000093518 | 0 | 0 | 0 | 0 | 0 | 0 | 0 |
| ENSMUSG00000093519 | 0 | 0 | 0 | 0 | 0 | 0 | 0 |
| ENSMUSG00000093520 | 0 | 0 | 0 | 0 | 0 | 0 | 0 |
| ENSMUSG00000093522 | 0 | 0 | 0 | 0 | 0 | 0 | 0 |
| ENSMUSG00000093523 | 0 | 0 | 0 | 0 | 0 | 0 | 0 |
| ENSMUSG00000093525 | 0 | 0 | 0 | 0 | 0 | 0 | 0 |
| ENSMUSG00000093528 | 0 | 0 | 0 | 0 | 0 | 0 | 0 |
| ENSMUSG00000093529 | 0 | 0 | 0 | 0 | 0 | 0 | 0 |
| ENSMUSG00000093530 | 0 | 0 | 0 | 0 | 0 | 0 | 0 |
| ENSMUSG00000093531 | 0 | 0 | 0 | 0 | 0 | 0 | 0 |
| ENSMUSG00000093533 | 0 | 0 | 0 | 0 | 0 | 0 | 0 |
| ENSMUSG00000093534 | 0 | 0 | 0 | 0 | 0 | 0 | 0 |
| ENSMUSG00000093535 | 0 | 0.396240625 | 0.25 | 0.580482024 | 0.489301084 | 0.301029996 | 0.355418052 |
| ENSMUSG00000093536 | 0 | 0 | 0.25 | 0.396240625 | 0.489301084 | 0 | 0.448487648 |
| ENSMUSG00000093537 | 0 | 0 | 0 | 0 | 0 | 0 | 0 |
| ENSMUSG00000093538 | 0 | 0.25 | 0 | 0.396240625 | 0.387761935 | 0.301029996 | 0.224243824 |
| ENSMUSG00000093539 | 0 | 0 | 0 | 0 | 0 | 0 | 0 |
| ENSMUSG00000093540 | 0 | 0 | 0 | 0 | 0 | 0 | 0 |
| ENSMUSG00000093541 | 0.671187741 | 0.396240625 | 0.5 | 0.25 | 0.568060967 | 0 | 0.520678036 |
| ENSMUSG00000093543 | 0 | 0 | 0 | 0 | 0 | 0 | 0.224243824 |
| ENSMUSG00000093544 | 0 | 0 | 0 | 0 | 0 | 0 | 0 |
| ENSMUSG00000093545 | 0 | 0.25 | 0 | 0 | 0 | 0.301029996 | 0 |
| ENSMUSG00000093546 | 0 | 0 | 0 | 0 | 0 | 0 | 0 |

| | | | | | | | |
|---|---|---|---|---|---|---|---|
| ENSMUSG00000093547 | 0 | 0 | 0 | 0 | 0 | 0 | 0 |
| ENSMUSG00000093548 | 0 | 0.5 | 0.396240625 | 0 | 0 | 0 | 0 |
| ENSMUSG00000093549 | 0 | 0 | 0 | 0 | 0 | 0 | 0 |
| ENSMUSG00000093550 | 0 | 0 | 0 | 0 | 0 | 0 | 0 |
| ENSMUSG00000093551 | 0 | 0 | 0 | 0 | 0 | 0 | 0 |
| ENSMUSG00000093552 | 0.289064826 | 0 | 0.396240625 | 0.25 | 0 | 0.477121255 | 0.224243824 |
| ENSMUSG00000093553 | 0.578129653 | 0.896240625 | 0.75 | 0.396240625 | 0.387761935 | 0.301029996 | 0.448487648 |
| ENSMUSG00000093554 | 0 | 0 | 0 | 0 | 0 | 0 | 0 |
| ENSMUSG00000093556 | 0 | 0 | 0 | 0 | 0 | 0 | 0 |
| ENSMUSG00000093557 | 0 | 0 | 0 | 0 | 0 | 0 | 0 |
| ENSMUSG00000093559 | 0.289064826 | 0.396240625 | 0 | 0 | 0 | 0 | 0 |
| ENSMUSG00000093560 | 0 | 0 | 0.25 | 0.25 | 0 | 0 | 0 |
| ENSMUSG00000093561 | 0 | 0 | 0 | 0.25 | 0.387761935 | 0.301029996 | 0 |
| ENSMUSG00000093562 | 0 | 0 | 0 | 0 | 0 | 0 | 0 |
| ENSMUSG00000093563 | 0 | 0 | 0 | 0 | 0 | 0 | 0 |
| ENSMUSG00000093564 | 0 | 0 | 0 | 0 | 0 | 0 | 0 |
| ENSMUSG00000093565 | 1.249317394 | 1.201838731 | 1.364857905 | 1.282320754 | 1.200473444 | 1.380211242 | 1.150210039 |
| ENSMUSG00000093567 | 0 | 0 | 0 | 0 | 0 | 0 | 0 |
| ENSMUSG00000093568 | 0 | 0 | 0 | 0 | 0 | 0 | 0 |
| ENSMUSG00000093570 | 0 | 0 | 0 | 0 | 0 | 0 | 0 |
| ENSMUSG00000093571 | 0 | 0 | 0 | 0 | 0 | 0 | 0 |
| ENSMUSG00000093572 | 0 | 0 | 0 | 0 | 0 | 0 | 0.224243824 |
| ENSMUSG00000093573 | 0 | 0 | 0 | 0.25 | 0 | 0 | 0 |
| ENSMUSG00000093574 | 0 | 0 | 0 | 0 | 0 | 0 | 0 |
| ENSMUSG00000093575 | 0.289064826 | 0 | 0 | 0 | 0 | 0 | 0 |
| ENSMUSG00000093577 | 0.578129653 | 0.580482024 | 0.701838731 | 0.646240625 | 0 | 0 | 0.448487648 |
| ENSMUSG00000093579 | 0.578129653 | 0.701838731 | 0.646240625 | 0.646240625 | 0.568060967 | 0.477121255 | 0.74492186 |
| ENSMUSG00000093580 | 0 | 0 | 0 | 0 | 0 | 0 | 0 |
| ENSMUSG00000093581 | 0 | 0 | 0 | 0 | 0 | 0 | 0 |
| ENSMUSG00000093582 | 0 | 0 | 0 | 0 | 0 | 0 | 0 |
| ENSMUSG00000093584 | 0 | 0 | 0 | 0 | 0 | 0 | 0 |
| ENSMUSG00000093587 | 0.671187741 | 0.25 | 0.646240625 | 0.976722649 | 0.846351821 | 0.602059991 | 0.74492186 |
| ENSMUSG00000093588 | 0 | 0 | 0 | 0 | 0 | 0 | 0 |
| ENSMUSG00000093589 | 0.45815691 | 0 | 0.5 | 0.25 | 0.632412477 | 0.477121255 | 0.579661877 |

| | | | | | | | |
|---|---|---|---|---|---|---|---|
| ENSMUSG00000093590 | 0 | 0 | 0 | 0 | 0 | 0 | 0 |
| ENSMUSG00000093592 | 0 | 0 | 0 | 0 | 0 | 0 | 0 |
| ENSMUSG00000093593 | 0 | 0 | 0.25 | 0 | 0 | 0 | 0 |
| ENSMUSG00000093594 | 0.811507563 | 0.830482024 | 1.098079356 | 0.896240625 | 0.877063019 | 0.954242509 | 0.710836105 |
| ENSMUSG00000093595 | 0 | 0 | 0 | 0 | 0 | 0 | 0 |
| ENSMUSG00000093598 | 0 | 0 | 0 | 0 | 0 | 0 | 0 |
| ENSMUSG00000093599 | 0 | 0 | 0 | 0 | 0 | 0 | 0 |
| ENSMUSG00000093601 | 0 | 0 | 0 | 0 | 0 | 0 | 0 |
| ENSMUSG00000093602 | 0 | 0 | 0 | 0 | 0 | 0 | 0 |
| ENSMUSG00000093606 | 0.811507563 | 0.79248125 | 1.04248125 | 1.04248125 | 0.877063019 | 1.079181246 | 0.74492186 |
| ENSMUSG00000093607 | 0 | 0 | 0 | 0 | 0 | 0 | 0 |
| ENSMUSG00000093610 | 0 | 0 | 0 | 0 | 0 | 0 | 0 |
| ENSMUSG00000093614 | 0 | 0 | 0 | 0 | 0 | 0 | 0.224243824 |
| ENSMUSG00000093615 | 0 | 0 | 0 | 0 | 0 | 0 | 0 |
| ENSMUSG00000093616 | 0 | 0 | 0 | 0 | 0 | 0 | 0 |
| ENSMUSG00000093617 | 0 | 0 | 0 | 0 | 0 | 0 | 0 |
| ENSMUSG00000093618 | 0 | 0 | 0 | 0 | 0 | 0 | 0 |
| ENSMUSG00000093619 | 0 | 0 | 0 | 0 | 0 | 0 | 0 |
| ENSMUSG00000093621 | 0 | 0 | 0 | 0 | 0 | 0 | 0 |
| ENSMUSG00000093622 | 0 | 0.25 | 0 | 0 | 0 | 0 | 0 |
| ENSMUSG00000093623 | 0 | 0 | 0 | 0 | 0 | 0 | 0 |
| ENSMUSG00000093624 | 0 | 0 | 0 | 0 | 0 | 0 | 0 |
| ENSMUSG00000093625 | 0 | 0 | 0 | 0 | 0 | 0 | 0 |
| ENSMUSG00000093626 | 0 | 0 | 0 | 0 | 0 | 0 | 0 |
| ENSMUSG00000093627 | 0 | 0 | 0 | 0 | 0 | 0 | 0 |
| ENSMUSG00000093628 | 0 | 0 | 0 | 0 | 0 | 0 | 0 |
| ENSMUSG00000093629 | 0.578129653 | 0.5 | 0.830482024 | 0.580482024 | 0.387761935 | 0 | 0.579661877 |
| ENSMUSG00000093632 | 0 | 0 | 0 | 0 | 0 | 0 | 0 |
| ENSMUSG00000093634 | 0 | 0 | 0 | 0.25 | 0 | 0 | 0 |
| ENSMUSG00000093635 | 0 | 0 | 0 | 0 | 0 | 0 | 0 |
| ENSMUSG00000093637 | 0.45815691 | 0.5 | 0.396240625 | 0.396240625 | 0.387761935 | 0.301029996 | 0.355418052 |
| ENSMUSG00000093639 | 0 | 0 | 0 | 0 | 0 | 0 | 0 |
| ENSMUSG00000093642 | 0 | 0 | 0 | 0 | 0 | 0 | 0 |
| ENSMUSG00000093644 | 0 | 0 | 0 | 0 | 0 | 0 | 0 |

| | | | | | | | |
|---|---|---|---|---|---|---|---|
| ENSMUSG00000093645 | 0 | 0 | 0 | 0 | 0 | 0 | 0 |
| ENSMUSG00000093648 | 0 | 0 | 0 | 0 | 0 | 0 | 0 |
| ENSMUSG00000093649 | 0 | 0.25 | 0 | 0 | 0 | 0 | 0 |
| ENSMUSG00000093650 | 0.747221736 | 0.75 | 0.646240625 | 0.701838731 | 0.877063019 | 0.77815125 | 0.579661877 |
| ENSMUSG00000093651 | 0 | 0 | 0.25 | 0 | 0 | 0 | 0 |
| ENSMUSG00000093652 | 0 | 0 | 0 | 0 | 0 | 0 | 0 |
| ENSMUSG00000093656 | 0.45815691 | 0.25 | 0 | 0 | 0 | 0 | 0.224243824 |
| ENSMUSG00000093658 | 0 | 0 | 0 | 0 | 0 | 0 | 0 |
| ENSMUSG00000093660 | 0 | 0 | 0 | 0 | 0 | 0 | 0 |
| ENSMUSG00000093661 | 1.45815691 | 1.29248125 | 1.201838731 | 1.445339928 | 1.394615667 | 1.62324929 | 1.392431569 |
| ENSMUSG00000093662 | 0 | 0 | 0 | 0 | 0 | 0 | 0 |
| ENSMUSG00000093663 | 0 | 0 | 0 | 0 | 0 | 0 | 0 |
| ENSMUSG00000093664 | 0 | 0 | 0 | 0 | 0 | 0 | 0 |
| ENSMUSG00000093665 | 0 | 0 | 0 | 0 | 0 | 0 | 0 |
| ENSMUSG00000093666 | 0 | 0 | 0 | 0 | 0 | 0 | 0 |
| ENSMUSG00000093668 | 0.289064826 | 0.396240625 | 0.25 | 0.25 | 0.244650542 | 0 | 0 |
| ENSMUSG00000093669 | 0 | 0.25 | 0 | 0 | 0 | 0 | 0 |
| ENSMUSG00000093671 | 0 | 0 | 0 | 0 | 0 | 0 | 0 |
| ENSMUSG00000093672 | 0 | 0.5 | 0.79248125 | 0.396240625 | 0.244650542 | 0.698970004 | 0 |
| ENSMUSG00000093673 | 0.289064826 | 0.25 | 0.580482024 | 0.580482024 | 0.489301084 | 0.301029996 | 0.520678036 |
| ENSMUSG00000093674 | 2.85804771 | 2.666333979 | 2.747525991 | 2.698806992 | 2.696288717 | 3.053846427 | 2.591475112 |
| ENSMUSG00000093675 | 0 | 0 | 0 | 0 | 0 | 0 | 0 |
| ENSMUSG00000093677 | 1 | 1 | 1.114857905 | 1.098079356 | 0.877063019 | 1.041392685 | 0.984950056 |
| ENSMUSG00000093679 | 0 | 0 | 0 | 0 | 0 | 0 | 0 |
| ENSMUSG00000093681 | 0 | 0 | 0 | 0 | 0 | 0 | 0 |
| ENSMUSG00000093682 | 0 | 0.25 | 0 | 0 | 0.387761935 | 0.301029996 | 0.520678036 |
| ENSMUSG00000093683 | 0 | 0 | 0 | 0 | 0 | 0 | 0 |
| ENSMUSG00000093684 | 0 | 0 | 0 | 0 | 0 | 0 | 0 |
| ENSMUSG00000093685 | 0 | 0 | 0 | 0 | 0 | 0 | 0 |
| ENSMUSG00000093686 | 0 | 0 | 0 | 0 | 0 | 0 | 0 |
| ENSMUSG00000093687 | 0.578129653 | 0 | 0 | 0.25 | 0 | 0 | 0.224243824 |
| ENSMUSG00000093688 | 0 | 0 | 0 | 0 | 0 | 0 | 0 |
| ENSMUSG00000093689 | 0 | 0 | 0 | 0 | 0 | 0 | 0 |
| ENSMUSG00000093690 | 0 | 0 | 0 | 0 | 0 | 0 | 0 |

| | | | | | | | |
|---|---|---|---|---|---|---|---|
| ENSMUSG00000093692 | 0 | 0 | 0 | 0 | 0 | 0 | 0 |
| ENSMUSG00000093695 | 0 | 0.396240625 | 0.396240625 | 0.25 | 0.244650542 | 0 | 0 |
| ENSMUSG00000093696 | 0 | 0 | 0 | 0 | 0 | 0 | 0 |
| ENSMUSG00000093697 | 0 | 0 | 0 | 0 | 0 | 0 | 0 |
| ENSMUSG00000093699 | 0 | 0 | 0 | 0 | 0 | 0 | 0 |
| ENSMUSG00000093701 | 0 | 0 | 0 | 0 | 0 | 0 | 0 |
| ENSMUSG00000093702 | 0 | 0 | 0 | 0 | 0 | 0 | 0 |
| ENSMUSG00000093703 | 0 | 0 | 0 | 0 | 0 | 0 | 0 |
| ENSMUSG00000093705 | 0 | 0 | 0 | 0 | 0 | 0 | 0 |
| ENSMUSG00000093707 | 0 | 0 | 0 | 0 | 0 | 0 | 0 |
| ENSMUSG00000093708 | 0 | 0 | 0 | 0 | 0 | 0 | 0 |
| ENSMUSG00000093710 | 0 | 0 | 0 | 0 | 0 | 0 | 0 |
| ENSMUSG00000093711 | 0 | 0 | 0 | 0 | 0 | 0 | 0 |
| ENSMUSG00000093714 | 0 | 0 | 0 | 0 | 0 | 0 | 0 |
| ENSMUSG00000093716 | 0 | 0 | 0.396240625 | 0 | 0.387761935 | 0 | 0 |
| ENSMUSG00000093719 | 0 | 0 | 0 | 0 | 0 | 0 | 0 |
| ENSMUSG00000093720 | 0 | 0 | 0 | 0 | 0 | 0 | 0.224243824 |
| ENSMUSG00000093721 | 0 | 0 | 0 | 0 | 0 | 0 | 0 |
| ENSMUSG00000093722 | 0 | 0 | 0 | 0 | 0 | 0 | 0 |
| ENSMUSG00000093724 | 0 | 0 | 0 | 0 | 0 | 0 | 0 |
| ENSMUSG00000093725 | 0 | 0 | 0 | 0 | 0 | 0 | 0 |
| ENSMUSG00000093726 | 1.205378646 | 1.388647213 | 1.17510993 | 1.54745614 | 1.519146132 | 1.591064607 | 1.360283038 |
| ENSMUSG00000093727 | 0 | 0 | 0 | 0 | 0 | 0 | 0 |
| ENSMUSG00000093728 | 0 | 0 | 0 | 0 | 0 | 0 | 0 |
| ENSMUSG00000093729 | 0 | 0 | 0 | 0 | 0 | 0 | 0 |
| ENSMUSG00000093730 | 0 | 0 | 0 | 0 | 0 | 0 | 0 |
| ENSMUSG00000093731 | 0 | 0 | 0 | 0 | 0 | 0 | 0 |
| ENSMUSG00000093732 | 0 | 0 | 0 | 0 | 0 | 0 | 0 |
| ENSMUSG00000093734 | 0.289064826 | 0.5 | 0.396240625 | 0.580482024 | 0.244650542 | 0.477121255 | 0.448487648 |
| ENSMUSG00000093736 | 0 | 0 | 0 | 0 | 0 | 0 | 0 |
| ENSMUSG00000093738 | 0 | 0 | 0 | 0 | 0 | 0 | 0 |
| ENSMUSG00000093739 | 0.289064826 | 0 | 0.5 | 0.396240625 | 0.632412477 | 0 | 0.520678036 |
| ENSMUSG00000093740 | 0 | 0 | 0.25 | 0 | 0 | 0 | 0 |
| ENSMUSG00000093741 | 0 | 0 | 0 | 0 | 0 | 0 | 0 |

| | | | | | | | |
|---|---|---|---|---|---|---|---|
| ENSMUSG00000093743 | 0 | 0 | 0 | 0 | 0 | 0 | 0 |
| ENSMUSG00000093744 | 0 | 0 | 0 | 0 | 0 | 0 | 0 |
| ENSMUSG00000093745 | 0 | 0 | 0 | 0 | 0 | 0 | 0 |
| ENSMUSG00000093746 | 0 | 0 | 0 | 0 | 0 | 0 | 0 |
| ENSMUSG00000093747 | 0 | 0 | 0 | 0 | 0 | 0.477121255 | 0 |
| ENSMUSG00000093748 | 0 | 0 | 0 | 0 | 0 | 0 | 0 |
| ENSMUSG00000093749 | 0 | 0 | 0.396240625 | 0 | 0.244650542 | 0 | 0.448487648 |
| ENSMUSG00000093751 | 0.289064826 | 0.25 | 0 | 0.646240625 | 0 | 0 | 0.224243824 |
| ENSMUSG00000093752 | 0 | 0 | 0 | 0 | 0 | 0 | 0 |
| ENSMUSG00000093753 | 0 | 0 | 0 | 0 | 0 | 0 | 0 |
| ENSMUSG00000093755 | 0 | 0 | 0 | 0 | 0 | 0 | 0 |
| ENSMUSG00000093756 | 0 | 0 | 0 | 0 | 0 | 0 | 0 |
| ENSMUSG00000093757 | 0 | 0 | 0 | 0 | 0 | 0 | 0 |
| ENSMUSG00000093758 | 0 | 0 | 0 | 0 | 0 | 0.301029996 | 0 |
| ENSMUSG00000093759 | 0 | 0 | 0 | 0 | 0 | 0 | 0 |
| ENSMUSG00000093760 | 1.389637216 | 1.372963274 | 1.282320754 | 1.282320754 | 1.091002363 | 1.176091259 | 1.110947927 |
| ENSMUSG00000093761 | 0 | 0 | 0 | 0 | 0 | 0 | 0 |
| ENSMUSG00000093762 | 0 | 0 | 0 | 0 | 0 | 0 | 0 |
| ENSMUSG00000093763 | 0 | 0 | 0 | 0 | 0 | 0 | 0 |
| ENSMUSG00000093764 | 0 | 0 | 0 | 0 | 0 | 0 | 0 |
| ENSMUSG00000093765 | 0 | 0 | 0 | 0 | 0 | 0 | 0 |
| ENSMUSG00000093766 | 0 | 0 | 0 | 0 | 0 | 0 | 0 |
| ENSMUSG00000093767 | 0 | 0 | 0 | 0 | 0 | 0 | 0 |
| ENSMUSG00000093768 | 0 | 0 | 0 | 0 | 0 | 0 | 0 |
| ENSMUSG00000093769 | 0 | 0 | 0 | 0 | 0 | 0 | 0 |
| ENSMUSG00000093770 | 0 | 0 | 0 | 0 | 0 | 0 | 0 |
| ENSMUSG00000093771 | 0 | 0 | 0 | 0 | 0 | 0 | 0 |
| ENSMUSG00000093772 | 0 | 0 | 0 | 0.25 | 0 | 0 | 0 |
| ENSMUSG00000093773 | 0 | 0 | 0 | 0 | 0 | 0 | 0 |
| ENSMUSG00000093774 | 0.289064826 | 0 | 0.25 | 0.25 | 0 | 0 | 0.224243824 |
| ENSMUSG00000093775 | 0 | 0 | 0 | 0 | 0 | 0 | 0 |
| ENSMUSG00000093776 | 0 | 0 | 0 | 0 | 0 | 0 | 0 |
| ENSMUSG00000093779 | 0 | 0 | 0 | 0 | 0 | 0 | 0 |
| ENSMUSG00000093780 | 0.45815691 | 0 | 0.396240625 | 0 | 0 | 0 | 0 |

| | | | | | | | |
|---|---|---|---|---|---|---|---|
| ENSMUSG00000093781 | 0 | 0 | 0 | 0 | 0 | 0 | 0 |
| ENSMUSG00000093782 | 0 | 0 | 0 | 0 | 0 | 0 | 0 |
| ENSMUSG00000093784 | 0 | 0 | 0 | 0 | 0 | 0 | 0 |
| ENSMUSG00000093785 | 0 | 0 | 0 | 0 | 0 | 0 | 0 |
| ENSMUSG00000093787 | 0 | 0 | 0 | 0 | 0 | 0 | 0 |
| ENSMUSG00000093788 | 0 | 0 | 0 | 0 | 0 | 0 | 0 |
| ENSMUSG00000093789 | 0 | 0 | 0 | 0 | 0 | 0 | 0 |
| ENSMUSG00000093791 | 0 | 0 | 0 | 0 | 0 | 0 | 0 |
| ENSMUSG00000093793 | 0 | 0 | 0 | 0 | 0 | 0 | 0 |
| ENSMUSG00000093795 | 0 | 0 | 0 | 0 | 0 | 0 | 0 |
| ENSMUSG00000093796 | 0 | 0 | 0 | 0 | 0 | 0 | 0 |
| ENSMUSG00000093798 | 0 | 0 | 0 | 0 | 0 | 0 | 0 |
| ENSMUSG00000093800 | 0.289064826 | 0 | 0.396240625 | 0.25 | 0.244650542 | 0.301029996 | 0.224243824 |
| ENSMUSG00000093801 | 0 | 0 | 0 | 0 | 0 | 0 | 0 |
| ENSMUSG00000093802 | 0 | 0 | 0 | 0 | 0 | 0 | 0 |
| ENSMUSG00000093803 | 0.91631382 | 1.26109853 | 1.403677461 | 0.79248125 | 1.254881871 | 0.903089987 | 1.383567577 |
| ENSMUSG00000093804 | 0 | 0 | 0 | 0 | 0 | 0 | 0 |
| ENSMUSG00000093805 | 0 | 0.5 | 0.396240625 | 0 | 0.387761935 | 0 | 0 |
| ENSMUSG00000093806 | 0 | 0 | 0 | 0 | 0 | 0 | 0 |
| ENSMUSG00000093807 | 0 | 0 | 0 | 0 | 0 | 0 | 0 |
| ENSMUSG00000093808 | 0 | 0 | 0 | 0 | 0 | 0 | 0 |
| ENSMUSG00000093812 | 0 | 0 | 0 | 0 | 0 | 0 | 0 |
| ENSMUSG00000093813 | 0 | 0 | 0 | 0 | 0 | 0 | 0 |
| ENSMUSG00000093814 | 0 | 0 | 0 | 0 | 0 | 0 | 0 |
| ENSMUSG00000093815 | 0 | 0 | 0 | 0 | 0 | 0 | 0 |
| ENSMUSG00000093816 | 0 | 0 | 0 | 0 | 0 | 0 | 0 |
| ENSMUSG00000093817 | 0 | 0 | 0 | 0 | 0 | 0 | 0 |
| ENSMUSG00000093818 | 0 | 0 | 0 | 0 | 0 | 0 | 0 |
| ENSMUSG00000093820 | 0 | 0 | 0 | 0 | 0 | 0 | 0 |
| ENSMUSG00000093822 | 0 | 0 | 0 | 0 | 0 | 0 | 0 |
| ENSMUSG00000093823 | 0 | 0 | 0 | 0 | 0 | 0 | 0 |
| ENSMUSG00000093824 | 0 | 0 | 0 | 0 | 0 | 0 | 0 |
| ENSMUSG00000093825 | 0 | 0 | 0 | 0 | 0 | 0 | 0 |
| ENSMUSG00000093826 | 0 | 0 | 0 | 0 | 0 | 0 | 0 |

| | | | | | | | |
|---|---|---|---|---|---|---|---|
| ENSMUSG00000093827 | 0 | 0 | 0 | 0 | 0 | 0 | 0 |
| ENSMUSG00000093828 | 0 | 0 | 0 | 0 | 0 | 0 | 0 |
| ENSMUSG00000093829 | 0 | 0 | 0 | 0 | 0 | 0 | 0 |
| ENSMUSG00000093830 | 0 | 0 | 0 | 0 | 0 | 0 | 0 |
| ENSMUSG00000093831 | 0 | 0 | 0 | 0 | 0 | 0 | 0 |
| ENSMUSG00000093833 | 0 | 0 | 0 | 0 | 0 | 0 | 0 |
| ENSMUSG00000093834 | 0 | 0 | 0 | 0 | 0 | 0 | 0 |
| ENSMUSG00000093835 | 0 | 0 | 0 | 0 | 0 | 0 | 0 |
| ENSMUSG00000093838 | 0 | 0 | 0 | 0 | 0 | 0 | 0 |
| ENSMUSG00000093839 | 0 | 0 | 0 | 0 | 0 | 0 | 0 |
| ENSMUSG00000093843 | 0 | 0.25 | 0 | 0 | 0 | 0 | 0 |
| ENSMUSG00000093844 | 0 | 0 | 0 | 0 | 0 | 0 | 0 |
| ENSMUSG00000093846 | 1.100572389 | 1.061981878 | 0.976722649 | 0.896240625 | 0.733951626 | 0.477121255 | 0.579661877 |
| ENSMUSG00000093847 | 0 | 0.25 | 0 | 0 | 0 | 0 | 0 |
| ENSMUSG00000093848 | 0 | 0 | 0 | 0 | 0 | 0 | 0 |
| ENSMUSG00000093849 | 0 | 0.396240625 | 0.25 | 0 | 0 | 0 | 0 |
| ENSMUSG00000093851 | 0 | 0 | 0 | 0 | 0 | 0 | 0 |
| ENSMUSG00000093852 | 0 | 0 | 0 | 0 | 0 | 0 | 0 |
| ENSMUSG00000093853 | 0 | 0 | 0 | 0 | 0 | 0 | 0 |
| ENSMUSG00000093854 | 0 | 0 | 0 | 0 | 0 | 0 | 0 |
| ENSMUSG00000093856 | 0 | 0 | 0 | 0 | 0 | 0 | 0 |
| ENSMUSG00000093857 | 0 | 0 | 0 | 0 | 0 | 0 | 0 |
| ENSMUSG00000093858 | 0 | 0.25 | 0.396240625 | 0.5 | 0 | 0.301029996 | 0.355418052 |
| ENSMUSG00000093859 | 0 | 0 | 0 | 0.396240625 | 0 | 0 | 0.355418052 |
| ENSMUSG00000093860 | 0 | 0 | 0 | 0 | 0 | 0 | 0 |
| ENSMUSG00000093861 | 0 | 0 | 0 | 0 | 0 | 0 | 0 |
| ENSMUSG00000093862 | 0.289064826 | 0.396240625 | 0.5 | 0 | 0.244650542 | 0.477121255 | 0.224243824 |
| ENSMUSG00000093863 | 0 | 0 | 0 | 0 | 0 | 0 | 0 |
| ENSMUSG00000093864 | 0 | 0 | 0 | 0 | 0 | 0 | 0 |
| ENSMUSG00000093865 | 0.289064826 | 0 | 0 | 0 | 0 | 0 | 0.224243824 |
| ENSMUSG00000093866 | 0 | 0 | 0 | 0 | 0 | 0 | 0 |
| ENSMUSG00000093867 | 0.45815691 | 0.396240625 | 0.25 | 0.396240625 | 0 | 0.477121255 | 0.224243824 |
| ENSMUSG00000093868 | 0 | 0 | 0 | 0 | 0 | 0 | 0 |
| ENSMUSG00000093869 | 0 | 0 | 0 | 0 | 0 | 0 | 0 |

| | | | | | | | |
|---|---|---|---|---|---|---|---|
| ENSMUSG00000093870 | 0 | 0 | 0 | 0 | 0 | 0 | 0 |
| ENSMUSG00000093871 | 0 | 0 | 0 | 0 | 0 | 0 | 0 |
| ENSMUSG00000093873 | 0 | 0 | 0 | 0 | 0 | 0 | 0 |
| ENSMUSG00000093874 | 0 | 0 | 0 | 0 | 0 | 0 | 0 |
| ENSMUSG00000093875 | 0 | 0 | 0 | 0 | 0 | 0 | 0 |
| ENSMUSG00000093876 | 0 | 0 | 0 | 0 | 0 | 0 | 0 |
| ENSMUSG00000093877 | 0 | 0 | 0 | 0 | 0 | 0 | 0 |
| ENSMUSG00000093878 | 0 | 0 | 0 | 0 | 0 | 0 | 0 |
| ENSMUSG00000093879 | 0 | 0 | 0 | 0 | 0 | 0 | 0 |
| ENSMUSG00000093880 | 0.578129653 | 0.5 | 0.25 | 0.864857905 | 0.955822902 | 0.77815125 | 0.853775828 |
| ENSMUSG00000093883 | 0 | 0 | 0 | 0 | 0 | 0 | 0 |
| ENSMUSG00000093884 | 0 | 0 | 0 | 0 | 0 | 0 | 0 |
| ENSMUSG00000093885 | 0 | 0 | 0.25 | 0 | 0 | 0 | 0.224243824 |
| ENSMUSG00000093886 | 0 | 0 | 0 | 0 | 0 | 0 | 0.224243824 |
| ENSMUSG00000093887 | 0 | 0 | 0 | 0 | 0 | 0 | 0 |
| ENSMUSG00000093889 | 0 | 0 | 0 | 0 | 0 | 0 | 0 |
| ENSMUSG00000093890 | 0 | 0 | 0 | 0 | 0 | 0 | 0 |
| ENSMUSG00000093891 | 0 | 0 | 0 | 0 | 0 | 0 | 0 |
| ENSMUSG00000093892 | 0 | 0 | 0 | 0 | 0 | 0 | 0 |
| ENSMUSG00000093893 | 0 | 0 | 0 | 0 | 0 | 0 | 0 |
| ENSMUSG00000093894 | 0 | 0 | 0 | 0 | 0 | 0 | 0 |
| ENSMUSG00000093895 | 0 | 0 | 0 | 0 | 0 | 0 | 0 |
| ENSMUSG00000093896 | 0 | 0 | 0 | 0 | 0 | 0 | 0 |
| ENSMUSG00000093897 | 0 | 0 | 0 | 0.25 | 0 | 0 | 0 |
| ENSMUSG00000093898 | 0 | 0 | 0 | 0 | 0 | 0 | 0 |
| ENSMUSG00000093899 | 0 | 0 | 0 | 0 | 0 | 0 | 0 |
| ENSMUSG00000093900 | 0 | 0 | 0 | 0 | 0 | 0 | 0 |
| ENSMUSG00000093901 | 0 | 0 | 0 | 0 | 0 | 0 | 0 |
| ENSMUSG00000093902 | 0 | 0 | 0 | 0 | 0 | 0 | 0 |
| ENSMUSG00000093903 | 0 | 0 | 0 | 0 | 0 | 0 | 0 |
| ENSMUSG00000093904 | 3.120755314 | 2.83076369 | 2.813609537 | 2.914775991 | 2.805431261 | 3.282395505 | 2.592118924 |
| ENSMUSG00000093905 | 0 | 0 | 0 | 0 | 0 | 0 | 0 |
| ENSMUSG00000093906 | 0 | 0 | 0 | 0 | 0 | 0 | 0 |
| ENSMUSG00000093907 | 0 | 0 | 0 | 0 | 0 | 0 | 0 |

| | | | | | | | |
|---|---|---|---|---|---|---|---|
| ENSMUSG00000093908 | 0.45815691 | 0.830482024 | 0.79248125 | 0.830482024 | 0.77552387 | 0.77815125 | 0.579661877 |
| ENSMUSG00000093909 | 0 | 0 | 0 | 0 | 0 | 0 | 0 |
| ENSMUSG00000093910 | 0 | 0.25 | 0 | 0 | 0 | 0 | 0.355418052 |
| ENSMUSG00000093912 | 0 | 0 | 0 | 0 | 0.244650542 | 0.301029996 | 0 |
| ENSMUSG00000093913 | 0 | 0 | 0 | 0 | 0 | 0 | 0 |
| ENSMUSG00000093914 | 0 | 0 | 0 | 0 | 0 | 0 | 0 |
| ENSMUSG00000093915 | 0 | 0 | 0 | 0 | 0 | 0 | 0 |
| ENSMUSG00000093916 | 0 | 0 | 0 | 0 | 0 | 0 | 0 |
| ENSMUSG00000093917 | 0 | 0 | 0 | 0 | 0 | 0 | 0 |
| ENSMUSG00000093918 | 0 | 0 | 0 | 0 | 0 | 0 | 0 |
| ENSMUSG00000093919 | 0 | 0 | 0 | 0 | 0 | 0 | 0 |
| ENSMUSG00000093920 | 0 | 0 | 0 | 0 | 0 | 0 | 0 |
| ENSMUSG00000093921 | 0.578129653 | 0.25 | 0 | 0 | 0.244650542 | 0 | 0.224243824 |
| ENSMUSG00000093922 | 0 | 0 | 0 | 0 | 0 | 0 | 0 |
| ENSMUSG00000093923 | 0 | 0 | 0 | 0 | 0 | 0 | 0 |
| ENSMUSG00000093924 | 0 | 0 | 0 | 0 | 0 | 0 | 0 |
| ENSMUSG00000093926 | 0 | 0 | 0 | 0 | 0 | 0 | 0 |
| ENSMUSG00000093927 | 0 | 0 | 0 | 0 | 0 | 0 | 0 |
| ENSMUSG00000093928 | 0 | 0 | 0 | 0 | 0 | 0 | 0 |
| ENSMUSG00000093929 | 0 | 0 | 0 | 0 | 0 | 0 | 0 |
| ENSMUSG00000093930 | 2.81052167 | 2.566696635 | 2.567865757 | 2.736178015 | 2.686820904 | 3.075546961 | 2.510768879 |
| ENSMUSG00000093931 | 0 | 0 | 0 | 0 | 0 | 0 | 0 |
| ENSMUSG00000093932 | 0 | 0 | 0 | 0 | 0 | 0 | 0 |
| ENSMUSG00000093933 | 0.289064826 | 0 | 0.25 | 0.580482024 | 0.244650542 | 0.301029996 | 0.448487648 |
| ENSMUSG00000093934 | 0 | 0 | 0 | 0 | 0 | 0 | 0 |
| ENSMUSG00000093935 | 0 | 0 | 0 | 0 | 0 | 0 | 0 |
| ENSMUSG00000093937 | 0 | 0 | 0 | 0 | 0 | 0 | 0 |
| ENSMUSG00000093938 | 0 | 0 | 0 | 0 | 0 | 0 | 0 |
| ENSMUSG00000093939 | 0 | 0 | 0 | 0 | 0 | 0 | 0 |
| ENSMUSG00000093940 | 0 | 0 | 0 | 0 | 0 | 0 | 0 |
| ENSMUSG00000093941 | 0 | 0 | 0 | 0 | 0 | 0 | 0 |
| ENSMUSG00000093942 | 0 | 0 | 0 | 0 | 0 | 0 | 0 |
| ENSMUSG00000093943 | 0 | 0 | 0 | 0 | 0 | 0 | 0 |
| ENSMUSG00000093944 | 0 | 0 | 0 | 0 | 0 | 0 | 0 |

| | | | | | | | |
|---|---|---|---|---|---|---|---|
| ENSMUSG00000093945 | 0 | 0 | 0 | 0 | 0 | 0 | 0 |
| ENSMUSG00000093946 | 0 | 0 | 0 | 0 | 0 | 0 | 0 |
| ENSMUSG00000093947 | 0 | 0 | 0 | 0 | 0 | 0 | 0 |
| ENSMUSG00000093948 | 0 | 0 | 0 | 0 | 0 | 0 | 0 |
| ENSMUSG00000093949 | 0 | 0 | 0 | 0 | 0 | 0 | 0 |
| ENSMUSG00000093950 | 0 | 0 | 0 | 0 | 0 | 0 | 0 |
| ENSMUSG00000093952 | 0 | 0 | 0 | 0 | 0 | 0 | 0 |
| ENSMUSG00000093953 | 0 | 0 | 0 | 0 | 0 | 0 | 0 |
| ENSMUSG00000093954 | 0 | 0 | 0 | 0 | 0 | 0 | 0 |
| ENSMUSG00000093955 | 0 | 0 | 0 | 0 | 0 | 0 | 0 |
| ENSMUSG00000093956 | 0 | 0 | 0 | 0 | 0 | 0 | 0 |
| ENSMUSG00000093957 | 0 | 0 | 0 | 0 | 0 | 0 | 0 |
| ENSMUSG00000093958 | 0 | 0 | 0 | 0 | 0 | 0 | 0 |
| ENSMUSG00000093959 | 0 | 0 | 0 | 0 | 0 | 0 | 0 |
| ENSMUSG00000093960 | 0 | 0 | 0 | 0 | 0 | 0 | 0 |
| ENSMUSG00000093961 | 0 | 0 | 0 | 0 | 0 | 0 | 0 |
| ENSMUSG00000093962 | 0 | 0 | 0 | 0 | 0 | 0 | 0 |
| ENSMUSG00000093963 | 0 | 0 | 0 | 0 | 0 | 0 | 0 |
| ENSMUSG00000093964 | 0 | 0 | 0 | 0 | 0 | 0 | 0 |
| ENSMUSG00000093965 | 0 | 0 | 0 | 0 | 0 | 0 | 0 |
| ENSMUSG00000093966 | 0 | 0 | 0 | 0 | 0 | 0 | 0 |
| ENSMUSG00000093967 | 0.45815691 | 0.5 | 0 | 0.396240625 | 0 | 0.301029996 | 0.448487648 |
| ENSMUSG00000093968 | 0 | 0 | 0 | 0 | 0 | 0 | 0 |
| ENSMUSG00000093969 | 0 | 0 | 0 | 0 | 0 | 0 | 0 |
| ENSMUSG00000093970 | 0 | 0 | 0 | 0 | 0 | 0 | 0 |
| ENSMUSG00000093971 | 0 | 0 | 0 | 0 | 0 | 0 | 0 |
| ENSMUSG00000093973 | 0 | 0 | 0 | 0 | 0 | 0 | 0 |
| ENSMUSG00000093974 | 0 | 0 | 0 | 0 | 0 | 0 | 0 |
| ENSMUSG00000093975 | 0 | 0 | 0.25 | 0 | 0 | 0 | 0 |
| ENSMUSG00000093976 | 0 | 0 | 0 | 0 | 0 | 0 | 0 |
| ENSMUSG00000093977 | 0 | 0 | 0 | 0 | 0 | 0 | 0 |
| ENSMUSG00000093978 | 0 | 0 | 0 | 0 | 0 | 0 | 0 |
| ENSMUSG00000093979 | 0.578129653 | 0.580482024 | 0.79248125 | 0.75 | 0.387761935 | 0.477121255 | 0.448487648 |
| ENSMUSG00000093980 | 0 | 0 | 0 | 0 | 0 | 0 | 0 |

| | | | | | | | |
|---|---|---|---|---|---|---|---|
| ENSMUSG00000093981 | 0 | 0 | 0 | 0 | 0 | 0 | 0 |
| ENSMUSG00000093983 | 0 | 0 | 0 | 0 | 0 | 0 | 0 |
| ENSMUSG00000093984 | 0 | 0 | 0 | 0 | 0 | 0 | 0 |
| ENSMUSG00000093985 | 0 | 0 | 0 | 0 | 0 | 0 | 0 |
| ENSMUSG00000093986 | 0 | 0 | 0 | 0 | 0 | 0 | 0 |
| ENSMUSG00000093987 | 0 | 0 | 0 | 0 | 0 | 0 | 0 |
| ENSMUSG00000093988 | 0 | 0 | 0 | 0 | 0 | 0 | 0 |
| ENSMUSG00000093989 | 0.289064826 | 0.5 | 0.646240625 | 0 | 0.568060967 | 0.698970004 | 0.520678036 |
| ENSMUSG00000093990 | 0 | 0 | 0 | 0 | 0 | 0 | 0 |
| ENSMUSG00000093992 | 0 | 0 | 0 | 0 | 0 | 0 | 0 |
| ENSMUSG00000093993 | 0 | 0 | 0 | 0 | 0 | 0 | 0 |
| ENSMUSG00000093994 | 0 | 0 | 0 | 0 | 0 | 0 | 0 |
| ENSMUSG00000093995 | 0 | 0 | 0 | 0 | 0 | 0 | 0 |
| ENSMUSG00000093996 | 0 | 0 | 0 | 0 | 0 | 0 | 0 |
| ENSMUSG00000093997 | 0 | 0 | 0 | 0 | 0 | 0 | 0 |
| ENSMUSG00000093998 | 0 | 0 | 0 | 0 | 0 | 0 | 0 |
| ENSMUSG00000093999 | 0 | 0 | 0 | 0 | 0 | 0 | 0 |
| ENSMUSG00000094001 | 0 | 0 | 0 | 0 | 0 | 0 | 0 |
| ENSMUSG00000094002 | 0 | 0 | 0 | 0 | 0 | 0 | 0 |
| ENSMUSG00000094003 | 0 | 0 | 0 | 0 | 0 | 0 | 0 |
| ENSMUSG00000094004 | 0 | 0 | 0 | 0 | 0 | 0 | 0 |
| ENSMUSG00000094005 | 0 | 0 | 0 | 0 | 0 | 0 | 0 |
| ENSMUSG00000094006 | 0 | 0 | 0 | 0 | 0 | 0 | 0 |
| ENSMUSG00000094007 | 0 | 0 | 0 | 0 | 0 | 0 | 0 |
| ENSMUSG00000094008 | 0 | 0 | 0 | 0 | 0 | 0 | 0 |
| ENSMUSG00000094009 | 0 | 0 | 0 | 0 | 0 | 0 | 0 |
| ENSMUSG00000094010 | 0 | 0 | 0 | 0 | 0 | 0 | 0 |
| ENSMUSG00000094011 | 0 | 0 | 0 | 0 | 0 | 0 | 0 |
| ENSMUSG00000094012 | 0 | 0 | 0 | 0 | 0 | 0 | 0 |
| ENSMUSG00000094013 | 0 | 0 | 0 | 0 | 0 | 0 | 0 |
| ENSMUSG00000094015 | 0 | 0 | 0 | 0 | 0 | 0 | 0 |
| ENSMUSG00000094016 | 0 | 0 | 0 | 0 | 0 | 0 | 0 |
| ENSMUSG00000094017 | 0 | 0 | 0.396240625 | 0.25 | 0 | 0.301029996 | 0 |
| ENSMUSG00000094018 | 0 | 0 | 0 | 0.25 | 0 | 0 | 0 |

| | | | | | | | |
|---|---|---|---|---|---|---|---|
| ENSMUSG00000094020 | 0 | 0 | 0 | 0 | 0 | 0 | 0 |
| ENSMUSG00000094021 | 0 | 0 | 0 | 0 | 0 | 0 | 0 |
| ENSMUSG00000094023 | 0 | 0 | 0 | 0 | 0 | 0 | 0 |
| ENSMUSG00000094024 | 0 | 0 | 0 | 0 | 0 | 0 | 0 |
| ENSMUSG00000094025 | 0 | 0 | 0 | 0 | 0 | 0 | 0 |
| ENSMUSG00000094026 | 0 | 0 | 0 | 0 | 0 | 0 | 0 |
| ENSMUSG00000094027 | 0 | 0.25 | 0 | 0 | 0 | 0 | 0 |
| ENSMUSG00000094028 | 0 | 0 | 0 | 0 | 0 | 0 | 0 |
| ENSMUSG00000094029 | 0 | 0 | 0 | 0 | 0 | 0 | 0 |
| ENSMUSG00000094030 | 0.45815691 | 0.580482024 | 0.75 | 0.75 | 0.632412477 | 0.477121255 | 0.520678036 |
| ENSMUSG00000094031 | 0 | 0 | 0 | 0 | 0 | 0 | 0 |
| ENSMUSG00000094032 | 0 | 0 | 0 | 0 | 0 | 0 | 0 |
| ENSMUSG00000094033 | 0 | 0.25 | 0 | 0 | 0 | 0 | 0 |
| ENSMUSG00000094034 | 0 | 0 | 0 | 0 | 0 | 0 | 0 |
| ENSMUSG00000094035 | 0 | 0 | 0 | 0 | 0 | 0 | 0.224243824 |
| ENSMUSG00000094036 | 0 | 0 | 0 | 0 | 0 | 0 | 0 |
| ENSMUSG00000094037 | 0 | 0 | 0 | 0 | 0 | 0 | 0 |
| ENSMUSG00000094038 | 0 | 0 | 0 | 0 | 0 | 0 | 0 |
| ENSMUSG00000094039 | 0 | 0 | 0 | 0 | 0 | 0 | 0 |
| ENSMUSG00000094041 | 0 | 0 | 0 | 0 | 0 | 0 | 0 |
| ENSMUSG00000094043 | 0 | 0 | 0 | 0 | 0 | 0 | 0 |
| ENSMUSG00000094044 | 0 | 0 | 0 | 0 | 0 | 0 | 0 |
| ENSMUSG00000094048 | 0 | 0 | 0 | 0 | 0 | 0 | 0 |
| ENSMUSG00000094049 | 0 | 0 | 0 | 0 | 0 | 0 | 0 |
| ENSMUSG00000094050 | 0 | 0 | 0 | 0 | 0 | 0 | 0 |
| ENSMUSG00000094051 | 0 | 0 | 0 | 0 | 0 | 0 | 0 |
| ENSMUSG00000094052 | 0 | 0 | 0 | 0 | 0 | 0 | 0 |
| ENSMUSG00000094053 | 0 | 0 | 0 | 0 | 0 | 0 | 0 |
| ENSMUSG00000094054 | 0 | 0 | 0 | 0 | 0 | 0 | 0 |
| ENSMUSG00000094057 | 0 | 0 | 0 | 0 | 0 | 0 | 0 |
| ENSMUSG00000094059 | 0 | 0 | 0 | 0 | 0 | 0 | 0 |
| ENSMUSG00000094060 | 0 | 0 | 0 | 0 | 0 | 0 | 0 |
| ENSMUSG00000094063 | 0 | 0 | 0 | 0 | 0 | 0 | 0 |
| ENSMUSG00000094064 | 0 | 0 | 0 | 0 | 0 | 0 | 0 |

| | | | | | | | |
|---|---|---|---|---|---|---|---|
| ENSMUSG00000094065 | 0 | 0 | 0 | 0 | 0 | 0 | 0 |
| ENSMUSG00000094066 | 0 | 0 | 0 | 0 | 0 | 0 | 0 |
| ENSMUSG00000094067 | 0 | 0 | 0 | 0 | 0 | 0 | 0 |
| ENSMUSG00000094068 | 0 | 0 | 0 | 0 | 0 | 0 | 0 |
| ENSMUSG00000094069 | 0 | 0 | 0 | 0 | 0 | 0 | 0 |
| ENSMUSG00000094071 | 0 | 0 | 0 | 0 | 0 | 0 | 0 |
| ENSMUSG00000094072 | 0 | 0 | 0 | 0 | 0 | 0 | 0 |
| ENSMUSG00000094074 | 0 | 0 | 0 | 0 | 0 | 0 | 0 |
| ENSMUSG00000094075 | 0 | 0 | 0 | 0 | 0 | 0 | 0 |
| ENSMUSG00000094076 | 0 | 0 | 0 | 0 | 0 | 0 | 0 |
| ENSMUSG00000094077 | 0 | 0 | 0 | 0 | 0 | 0 | 0 |
| ENSMUSG00000094078 | 0 | 0 | 0 | 0 | 0 | 0 | 0 |
| ENSMUSG00000094079 | 0 | 0 | 0 | 0 | 0 | 0 | 0 |
| ENSMUSG00000094080 | 0 | 0 | 0 | 0 | 0 | 0 | 0 |
| ENSMUSG00000094081 | 0 | 0 | 0 | 0 | 0 | 0 | 0 |
| ENSMUSG00000094082 | 0 | 0 | 0 | 0 | 0 | 0 | 0 |
| ENSMUSG00000094083 | 0 | 0 | 0.25 | 0 | 0 | 0 | 0 |
| ENSMUSG00000094084 | 0 | 0 | 0 | 0 | 0 | 0 | 0 |
| ENSMUSG00000094085 | 0 | 0 | 0 | 0 | 0 | 0 | 0 |
| ENSMUSG00000094086 | 0 | 0 | 0 | 0.396240625 | 0.489301084 | 0 | 0.448487648 |
| ENSMUSG00000094087 | 0 | 0 | 0 | 0 | 0 | 0 | 0 |
| ENSMUSG00000094088 | 0 | 0 | 0 | 0 | 0 | 0 | 0 |
| ENSMUSG00000094090 | 1.069666964 | 0.864857905 | 1 | 0.896240625 | 0.931471446 | 0.954242509 | 0.984950056 |
| ENSMUSG00000094091 | 0 | 0 | 0 | 0 | 0.244650542 | 0 | 0.224243824 |
| ENSMUSG00000094092 | 0.578129653 | 0 | 0 | 0.25 | 0.244650542 | 0 | 0.224243824 |
| ENSMUSG00000094093 | 0 | 0 | 0 | 0 | 0 | 0 | 0 |
| ENSMUSG00000094094 | 0 | 0 | 0 | 0 | 0 | 0 | 0 |
| ENSMUSG00000094095 | 0 | 0 | 0 | 0 | 0 | 0 | 0 |
| ENSMUSG00000094096 | 0 | 0 | 0 | 0 | 0 | 0 | 0 |
| ENSMUSG00000094097 | 0 | 0 | 0 | 0 | 0 | 0 | 0 |
| ENSMUSG00000094098 | 0 | 0 | 0 | 0 | 0 | 0 | 0 |
| ENSMUSG00000094099 | 0 | 0 | 0 | 0 | 0 | 0 | 0 |
| ENSMUSG00000094101 | 0 | 0 | 0 | 0 | 0 | 0 | 0 |
| ENSMUSG00000094102 | 0 | 0 | 0 | 0 | 0 | 0 | 0 |

| | | | | | | | |
|---|---|---|---|---|---|---|---|
| ENSMUSG00000094103 | 0 | 0 | 0 | 0 | 0 | 0 | 0 |
| ENSMUSG00000094104 | 0 | 0 | 0 | 0 | 0 | 0 | 0 |
| ENSMUSG00000094105 | 0 | 0 | 0 | 0 | 0 | 0 | 0 |
| ENSMUSG00000094107 | 0 | 0 | 0 | 0 | 0 | 0 | 0 |
| ENSMUSG00000094108 | 0 | 0 | 0 | 0 | 0 | 0 | 0 |
| ENSMUSG00000094109 | 0 | 0 | 0 | 0 | 0 | 0 | 0 |
| ENSMUSG00000094111 | 0 | 0 | 0 | 0 | 0 | 0 | 0 |
| ENSMUSG00000094112 | 0 | 0 | 0 | 0 | 0 | 0 | 0 |
| ENSMUSG00000094113 | 0 | 0 | 0 | 0 | 0 | 0 | 0 |
| ENSMUSG00000094114 | 1.289064826 | 1.311981878 | 1.25 | 1.950224975 | 1.916320788 | 2.298853076 | 1.826389311 |
| ENSMUSG00000094115 | 0 | 0 | 0 | 0 | 0 | 0 | 0 |
| ENSMUSG00000094116 | 0 | 0 | 0 | 0 | 0 | 0 | 0 |
| ENSMUSG00000094117 | 0 | 0 | 0 | 0 | 0 | 0 | 0 |
| ENSMUSG00000094118 | 0 | 0 | 0 | 0 | 0 | 0 | 0 |
| ENSMUSG00000094119 | 0 | 0 | 0 | 0 | 0 | 0 | 0 |
| ENSMUSG00000094120 | 0 | 0 | 0 | 0 | 0 | 0 | 0 |
| ENSMUSG00000094121 | 0 | 0 | 0 | 0 | 0 | 0 | 0 |
| ENSMUSG00000094122 | 0 | 0 | 0 | 0 | 0 | 0 | 0.224243824 |
| ENSMUSG00000094123 | 0 | 0 | 0 | 0 | 0 | 0 | 0 |
| ENSMUSG00000094124 | 0 | 0 | 0 | 0 | 0 | 0 | 0 |
| ENSMUSG00000094125 | 0 | 0 | 0 | 0 | 0 | 0 | 0 |
| ENSMUSG00000094127 | 0 | 0 | 0 | 0 | 0 | 0 | 0 |
| ENSMUSG00000094128 | 0 | 0 | 0 | 0 | 0 | 0 | 0 |
| ENSMUSG00000094129 | 0 | 0 | 0 | 0 | 0 | 0 | 0 |
| ENSMUSG00000094131 | 0 | 0 | 0 | 0 | 0 | 0 | 0 |
| ENSMUSG00000094132 | 0 | 0 | 0 | 0 | 0 | 0 | 0 |
| ENSMUSG00000094133 | 0 | 0 | 0 | 0 | 0 | 0 | 0 |
| ENSMUSG00000094134 | 0 | 0 | 0 | 0 | 0 | 0 | 0 |
| ENSMUSG00000094135 | 0 | 0 | 0 | 0 | 0 | 0 | 0 |
| ENSMUSG00000094136 | 0 | 0 | 0 | 0 | 0 | 0 | 0 |
| ENSMUSG00000094137 | 0 | 0 | 0 | 0 | 0 | 0 | 0 |
| ENSMUSG00000094139 | 0 | 0 | 0 | 0 | 0 | 0 | 0 |
| ENSMUSG00000094140 | 0 | 0 | 0 | 0 | 0 | 0 | 0 |
| ENSMUSG00000094144 | 0 | 0 | 0 | 0 | 0 | 0 | 0 |

| | | | | | | | |
|---|---|---|---|---|---|---|---|
| ENSMUSG00000094145 | 0 | 0 | 0 | 0 | 0 | 0 | 0 |
| ENSMUSG00000094146 | 0 | 0 | 0 | 0 | 0 | 0 | 0 |
| ENSMUSG00000094147 | 0 | 0 | 0 | 0 | 0 | 0 | 0 |
| ENSMUSG00000094148 | 0 | 0 | 0 | 0 | 0 | 0 | 0 |
| ENSMUSG00000094149 | 0 | 0 | 0 | 0 | 0 | 0 | 0 |
| ENSMUSG00000094150 | 0 | 0 | 0 | 0 | 0 | 0 | 0 |
| ENSMUSG00000094151 | 0 | 0 | 0 | 0 | 0 | 0 | 0 |
| ENSMUSG00000094152 | 0 | 0 | 0 | 0 | 0 | 0 | 0 |
| ENSMUSG00000094153 | 0 | 0.25 | 0 | 0 | 0 | 0 | 0 |
| ENSMUSG00000094154 | 0 | 0 | 0 | 0 | 0 | 0 | 0 |
| ENSMUSG00000094155 | 0 | 0 | 0 | 0 | 0 | 0 | 0 |
| ENSMUSG00000094156 | 0 | 0 | 0 | 0 | 0 | 0 | 0 |
| ENSMUSG00000094157 | 0 | 0 | 0 | 0 | 0 | 0 | 0 |
| ENSMUSG00000094158 | 0 | 0.396240625 | 0.396240625 | 0.25 | 0 | 0.301029996 | 0 |
| ENSMUSG00000094159 | 0 | 0 | 0 | 0 | 0 | 0 | 0 |
| ENSMUSG00000094160 | 0 | 0 | 0 | 0 | 0 | 0 | 0 |
| ENSMUSG00000094161 | 0 | 0 | 0 | 0 | 0 | 0 | 0 |
| ENSMUSG00000094162 | 0 | 0 | 0 | 0 | 0 | 0 | 0 |
| ENSMUSG00000094163 | 0 | 0 | 0 | 0 | 0 | 0 | 0 |
| ENSMUSG00000094164 | 0 | 0 | 0 | 0 | 0 | 0 | 0 |
| ENSMUSG00000094165 | 0 | 0 | 0 | 0 | 0 | 0 | 0 |
| ENSMUSG00000094167 | 0 | 0 | 0 | 0 | 0 | 0 | 0 |
| ENSMUSG00000094168 | 0 | 0 | 0 | 0 | 0 | 0 | 0 |
| ENSMUSG00000094170 | 0 | 0 | 0 | 0 | 0 | 0 | 0 |
| ENSMUSG00000094172 | 0 | 0 | 0 | 0 | 0 | 0 | 0 |
| ENSMUSG00000094173 | 0 | 0 | 0 | 0 | 0 | 0 | 0 |
| ENSMUSG00000094174 | 0 | 0 | 0 | 0 | 0 | 0 | 0 |
| ENSMUSG00000094175 | 0 | 0 | 0 | 0 | 0 | 0 | 0 |
| ENSMUSG00000094176 | 0 | 0 | 0 | 0 | 0 | 0 | 0 |
| ENSMUSG00000094177 | 0 | 0 | 0 | 0 | 0 | 0 | 0 |
| ENSMUSG00000094178 | 0 | 0 | 0 | 0 | 0 | 0 | 0 |
| ENSMUSG00000094179 | 0 | 0 | 0 | 0 | 0 | 0 | 0 |
| ENSMUSG00000094180 | 0 | 0 | 0 | 0 | 0 | 0 | 0 |
| ENSMUSG00000094181 | 0 | 0 | 0 | 0 | 0 | 0 | 0 |

| | | | | | | | |
|---|---|---|---|---|---|---|---|
| ENSMUSG00000094182 | 0 | 0 | 0 | 0 | 0 | 0 | 0 |
| ENSMUSG00000094183 | 0 | 0 | 0 | 0 | 0 | 0 | 0 |
| ENSMUSG00000094184 | 0 | 0 | 0 | 0 | 0 | 0 | 0 |
| ENSMUSG00000094185 | 0.289064826 | 0 | 0 | 0.25 | 0 | 0 | 0 |
| ENSMUSG00000094186 | 0 | 0 | 0 | 0 | 0 | 0 | 0 |
| ENSMUSG00000094187 | 0 | 0 | 0 | 0 | 0 | 0 | 0 |
| ENSMUSG00000094188 | 0 | 0 | 0 | 0 | 0 | 0 | 0 |
| ENSMUSG00000094189 | 0 | 0 | 0 | 0 | 0 | 0 | 0 |
| ENSMUSG00000094190 | 0 | 0 | 0 | 0 | 0 | 0 | 0 |
| ENSMUSG00000094191 | 0.289064826 | 0 | 0.580482024 | 0.396240625 | 0 | 0.301029996 | 0.355418052 |
| ENSMUSG00000094192 | 0 | 0 | 0 | 0 | 0 | 0 | 0 |
| ENSMUSG00000094194 | 0 | 0 | 0 | 0 | 0 | 0 | 0 |
| ENSMUSG00000094195 | 0 | 0 | 0 | 0 | 0 | 0 | 0 |
| ENSMUSG00000094196 | 0 | 0 | 0 | 0 | 0 | 0 | 0 |
| ENSMUSG00000094197 | 0 | 0 | 0 | 0 | 0 | 0 | 0 |
| ENSMUSG00000094198 | 0 | 0 | 0 | 0 | 0 | 0 | 0 |
| ENSMUSG00000094199 | 0 | 0 | 0 | 0 | 0 | 0 | 0 |
| ENSMUSG00000094200 | 0 | 0 | 0 | 0 | 0 | 0 | 0 |
| ENSMUSG00000094201 | 0 | 0 | 0 | 0 | 0 | 0 | 0 |
| ENSMUSG00000094202 | 0 | 0 | 0 | 0 | 0 | 0 | 0 |
| ENSMUSG00000094203 | 0 | 0 | 0 | 0 | 0 | 0 | 0 |
| ENSMUSG00000094204 | 0 | 0 | 0 | 0 | 0 | 0 | 0 |
| ENSMUSG00000094205 | 0 | 0 | 0 | 0 | 0 | 0 | 0 |
| ENSMUSG00000094206 | 0 | 0 | 0 | 0 | 0 | 0 | 0 |
| ENSMUSG00000094208 | 0 | 0 | 0 | 0 | 0 | 0 | 0 |
| ENSMUSG00000094210 | 0 | 0 | 0 | 0 | 0 | 0 | 0 |
| ENSMUSG00000094211 | 0 | 0 | 0 | 0 | 0 | 0 | 0 |
| ENSMUSG00000094212 | 0 | 0 | 0 | 0.25 | 0 | 0 | 0 |
| ENSMUSG00000094213 | 0 | 0 | 0 | 0 | 0 | 0 | 0 |
| ENSMUSG00000094214 | 0 | 0 | 0 | 0 | 0 | 0 | 0 |
| ENSMUSG00000094215 | 0 | 0 | 0 | 0 | 0 | 0 | 0 |
| ENSMUSG00000094217 | 0 | 0 | 0 | 0 | 0 | 0 | 0 |
| ENSMUSG00000094218 | 0 | 0 | 0 | 0 | 0 | 0 | 0 |
| ENSMUSG00000094219 | 0 | 0.25 | 0 | 0 | 0 | 0 | 0 |

| | | | | | | | |
|---|---|---|---|---|---|---|---|
| ENSMUSG00000094220 | 0 | 0 | 0 | 0 | 0 | 0 | 0 |
| ENSMUSG00000094221 | 0 | 0 | 0 | 0 | 0 | 0 | 0 |
| ENSMUSG00000094222 | 0 | 0 | 0 | 0 | 0 | 0 | 0 |
| ENSMUSG00000094226 | 0 | 0 | 0 | 0 | 0 | 0 | 0 |
| ENSMUSG00000094227 | 0 | 0 | 0 | 0 | 0 | 0 | 0 |
| ENSMUSG00000094228 | 0 | 0 | 0 | 0 | 0 | 0 | 0 |
| ENSMUSG00000094229 | 0 | 0 | 0 | 0 | 0 | 0 | 0 |
| ENSMUSG00000094230 | 0 | 0 | 0 | 0 | 0 | 0 | 0 |
| ENSMUSG00000094231 | 0 | 0 | 0 | 0 | 0 | 0 | 0 |
| ENSMUSG00000094232 | 0 | 0 | 0 | 0 | 0 | 0 | 0 |
| ENSMUSG00000094233 | 0 | 0 | 0 | 0 | 0 | 0 | 0 |
| ENSMUSG00000094235 | 0 | 0 | 0 | 0 | 0 | 0 | 0 |
| ENSMUSG00000094236 | 0 | 0 | 0 | 0 | 0 | 0 | 0 |
| ENSMUSG00000094237 | 0 | 0 | 0 | 0 | 0 | 0 | 0 |
| ENSMUSG00000094238 | 0 | 0 | 0 | 0 | 0 | 0 | 0 |
| ENSMUSG00000094239 | 0 | 0 | 0 | 0 | 0 | 0 | 0 |
| ENSMUSG00000094240 | 0 | 0 | 0 | 0 | 0 | 0 | 0 |
| ENSMUSG00000094241 | 0 | 0 | 0 | 0 | 0 | 0 | 0 |
| ENSMUSG00000094242 | 0 | 0 | 0 | 0 | 0 | 0 | 0 |
| ENSMUSG00000094243 | 0 | 0 | 0 | 0 | 0 | 0 | 0 |
| ENSMUSG00000094244 | 0 | 0 | 0 | 0 | 0 | 0 | 0 |
| ENSMUSG00000094245 | 0.289064826 | 0 | 0 | 0.5 | 0.244650542 | 0 | 0 |
| ENSMUSG00000094247 | 0 | 0 | 0 | 0 | 0 | 0 | 0 |
| ENSMUSG00000094248 | 0 | 0 | 0 | 0 | 0 | 0 | 0 |
| ENSMUSG00000094249 | 0 | 0 | 0 | 0 | 0 | 0 | 0 |
| ENSMUSG00000094250 | 0 | 0.25 | 0 | 0 | 0 | 0 | 0 |
| ENSMUSG00000094251 | 0 | 0.396240625 | 0 | 0 | 0 | 0 | 0 |
| ENSMUSG00000094252 | 0 | 0 | 0 | 0 | 0 | 0 | 0 |
| ENSMUSG00000094253 | 0 | 0 | 0 | 0 | 0 | 0 | 0 |
| ENSMUSG00000094254 | 0 | 0 | 0 | 0 | 0 | 0 | 0 |
| ENSMUSG00000094255 | 0 | 0 | 0 | 0 | 0 | 0 | 0 |
| ENSMUSG00000094256 | 0 | 0.25 | 0 | 0 | 0 | 0 | 0 |
| ENSMUSG00000094257 | 0 | 0 | 0 | 0.25 | 0 | 0 | 0 |
| ENSMUSG00000094258 | 0 | 0 | 0 | 0 | 0 | 0 | 0 |

| | | | | | | | |
|---|---|---|---|---|---|---|---|
| ENSMUSG00000094259 | 0 | 0 | 0 | 0 | 0 | 0 | 0 |
| ENSMUSG00000094261 | 0 | 0 | 0 | 0 | 0 | 0 | 0 |
| ENSMUSG00000094262 | 0 | 0 | 0 | 0 | 0 | 0 | 0 |
| ENSMUSG00000094263 | 0 | 0 | 0 | 0 | 0 | 0 | 0 |
| ENSMUSG00000094264 | 0 | 0 | 0 | 0 | 0 | 0 | 0 |
| ENSMUSG00000094265 | 0 | 0 | 0 | 0 | 0 | 0 | 0 |
| ENSMUSG00000094266 | 0 | 0 | 0 | 0 | 0 | 0 | 0 |
| ENSMUSG00000094267 | 0 | 0 | 0 | 0 | 0 | 0 | 0 |
| ENSMUSG00000094268 | 0 | 0 | 0 | 0 | 0 | 0 | 0 |
| ENSMUSG00000094269 | 0 | 0 | 0 | 0 | 0 | 0 | 0 |
| ENSMUSG00000094270 | 0 | 0 | 0 | 0 | 0 | 0 | 0 |
| ENSMUSG00000094271 | 0 | 0 | 0 | 0 | 0 | 0 | 0 |
| ENSMUSG00000094273 | 0 | 0 | 0 | 0 | 0 | 0 | 0 |
| ENSMUSG00000094274 | 0 | 0 | 0 | 0 | 0 | 0 | 0 |
| ENSMUSG00000094275 | 0 | 0 | 0 | 0 | 0 | 0 | 0 |
| ENSMUSG00000094276 | 0 | 0 | 0 | 0 | 0 | 0 | 0 |
| ENSMUSG00000094277 | 0 | 0 | 0 | 0 | 0 | 0 | 0 |
| ENSMUSG00000094278 | 0 | 0 | 0 | 0 | 0 | 0 | 0 |
| ENSMUSG00000094280 | 0 | 0 | 0 | 0 | 0 | 0 | 0 |
| ENSMUSG00000094281 | 0 | 0.25 | 0.396240625 | 0.396240625 | 0.812711509 | 0.477121255 | 0.448487648 |
| ENSMUSG00000094282 | 0 | 0 | 0 | 0 | 0.387761935 | 0 | 0 |
| ENSMUSG00000094283 | 0 | 0 | 0 | 0 | 0 | 0 | 0 |
| ENSMUSG00000094284 | 0 | 0 | 0 | 0 | 0 | 0 | 0 |
| ENSMUSG00000094285 | 0 | 0 | 0 | 0 | 0 | 0 | 0 |
| ENSMUSG00000094286 | 0.289064826 | 0.396240625 | 0.396240625 | 0.580482024 | 0.244650542 | 0.602059991 | 0.672731473 |
| ENSMUSG00000094287 | 0 | 0 | 0 | 0 | 0 | 0 | 0 |
| ENSMUSG00000094288 | 0 | 0 | 0 | 0 | 0 | 0 | 0 |
| ENSMUSG00000094289 | 0 | 0 | 0 | 0 | 0 | 0 | 0 |
| ENSMUSG00000094290 | 0 | 0 | 0 | 0 | 0 | 0 | 0 |
| ENSMUSG00000094291 | 0 | 0 | 0 | 0 | 0 | 0 | 0 |
| ENSMUSG00000094292 | 0 | 0 | 0 | 0 | 0 | 0 | 0 |
| ENSMUSG00000094293 | 0 | 0 | 0 | 0 | 0 | 0 | 0 |
| ENSMUSG00000094294 | 0 | 0 | 0 | 0 | 0 | 0 | 0 |
| ENSMUSG00000094295 | 0 | 0 | 0 | 0 | 0 | 0 | 0 |

| | | | | | | | |
|---|---|---|---|---|---|---|---|
| ENSMUSG00000094296 | 0.45815691 | 0.396240625 | 0.646240625 | 0.646240625 | 0.877063019 | 0.77815125 | 0.629532004 |
| ENSMUSG00000094297 | 0 | 0 | 0 | 0 | 0 | 0 | 0 |
| ENSMUSG00000094298 | 0 | 0 | 0 | 0 | 0 | 0 | 0 |
| ENSMUSG00000094301 | 0 | 0 | 0 | 0 | 0 | 0 | 0 |
| ENSMUSG00000094302 | 0 | 0 | 0 | 0 | 0 | 0 | 0 |
| ENSMUSG00000094303 | 0 | 0 | 0 | 0 | 0 | 0 | 0 |
| ENSMUSG00000094304 | 0 | 0 | 0 | 0 | 0 | 0 | 0 |
| ENSMUSG00000094305 | 0 | 0 | 0 | 0 | 0 | 0 | 0 |
| ENSMUSG00000094306 | 0 | 0 | 0 | 0 | 0 | 0 | 0 |
| ENSMUSG00000094307 | 0 | 0 | 0 | 0 | 0 | 0 | 0 |
| ENSMUSG00000094308 | 0 | 0 | 0 | 0 | 0 | 0 | 0 |
| ENSMUSG00000094309 | 0 | 0 | 0 | 0 | 0 | 0 | 0 |
| ENSMUSG00000094310 | 0 | 0 | 0 | 0 | 0 | 0 | 0 |
| ENSMUSG00000094311 | 0 | 0 | 0 | 0 | 0 | 0 | 0 |
| ENSMUSG00000094312 | 0 | 0 | 0 | 0 | 0 | 0 | 0 |
| ENSMUSG00000094313 | 0 | 0 | 0 | 0 | 0 | 0 | 0 |
| ENSMUSG00000094314 | 0 | 0 | 0 | 0 | 0 | 0 | 0 |
| ENSMUSG00000094315 | 0 | 0 | 0 | 0 | 0 | 0 | 0 |
| ENSMUSG00000094316 | 0 | 0 | 0 | 0 | 0 | 0 | 0 |
| ENSMUSG00000094317 | 0 | 0 | 0 | 0 | 0 | 0 | 0 |
| ENSMUSG00000094318 | 0 | 0 | 0 | 0 | 0 | 0 | 0 |
| ENSMUSG00000094319 | 0 | 0 | 0 | 0 | 0 | 0 | 0 |
| ENSMUSG00000094320 | 0 | 0 | 0 | 0 | 0 | 0 | 0 |
| ENSMUSG00000094321 | 0 | 0 | 0 | 0 | 0 | 0 | 0 |
| ENSMUSG00000094322 | 0 | 0 | 0 | 0 | 0 | 0 | 0 |
| ENSMUSG00000094323 | 0 | 0 | 0 | 0 | 0 | 0 | 0 |
| ENSMUSG00000094325 | 0 | 0 | 0 | 0 | 0 | 0 | 0 |
| ENSMUSG00000094326 | 0 | 0 | 0 | 0 | 0 | 0 | 0 |
| ENSMUSG00000094327 | 0 | 0 | 0 | 0 | 0 | 0 | 0 |
| ENSMUSG00000094328 | 0 | 0 | 0 | 0 | 0 | 0 | 0 |
| ENSMUSG00000094330 | 0 | 0 | 0 | 0 | 0 | 0 | 0 |
| ENSMUSG00000094334 | 0.289064826 | 0 | 0.25 | 0 | 0 | 0 | 0.224243824 |
| ENSMUSG00000094335 | 0 | 0 | 0 | 0 | 0 | 0 | 0 |
| ENSMUSG00000094336 | 0 | 0 | 0 | 0 | 0 | 0 | 0 |

| | | | | | | | |
|---|---|---|---|---|---|---|---|
| ENSMUSG00000094337 | 0 | 0 | 0 | 0 | 0 | 0 | 0 |
| ENSMUSG00000094338 | 0 | 0 | 0 | 0 | 0 | 0 | 0 |
| ENSMUSG00000094340 | 0 | 0 | 0 | 0 | 0 | 0 | 0 |
| ENSMUSG00000094342 | 0 | 0 | 0 | 0 | 0 | 0 | 0 |
| ENSMUSG00000094344 | 0 | 0 | 0 | 0 | 0.387761935 | 0 | 0.224243824 |
| ENSMUSG00000094345 | 0 | 0 | 0 | 0 | 0 | 0 | 0 |
| ENSMUSG00000094346 | 0 | 0 | 0 | 0 | 0 | 0 | 0 |
| ENSMUSG00000094347 | 0 | 0 | 0 | 0 | 0 | 0 | 0 |
| ENSMUSG00000094349 | 0 | 0 | 0 | 0 | 0 | 0 | 0 |
| ENSMUSG00000094350 | 0 | 0 | 0 | 0 | 0 | 0 | 0 |
| ENSMUSG00000094351 | 0 | 0 | 0 | 0 | 0 | 0 | 0 |
| ENSMUSG00000094352 | 0 | 0 | 0 | 0 | 0 | 0 | 0 |
| ENSMUSG00000094353 | 0 | 0 | 0 | 0 | 0 | 0 | 0 |
| ENSMUSG00000094354 | 0 | 0 | 0 | 0 | 0 | 0 | 0 |
| ENSMUSG00000094356 | 0 | 0 | 0 | 0 | 0 | 0 | 0 |
| ENSMUSG00000094357 | 0 | 0 | 0 | 0 | 0 | 0 | 0 |
| ENSMUSG00000094358 | 0 | 0 | 0 | 0 | 0 | 0 | 0 |
| ENSMUSG00000094359 | 0 | 0 | 0 | 0 | 0 | 0 | 0 |
| ENSMUSG00000094360 | 0 | 0 | 0 | 0 | 0 | 0 | 0 |
| ENSMUSG00000094361 | 0 | 0 | 0 | 0 | 0 | 0 | 0 |
| ENSMUSG00000094362 | 0 | 0 | 0 | 0 | 0 | 0 | 0 |
| ENSMUSG00000094363 | 0 | 0 | 0 | 0 | 0 | 0 | 0 |
| ENSMUSG00000094364 | 0 | 0 | 0 | 0 | 0 | 0 | 0 |
| ENSMUSG00000094365 | 0 | 0 | 0 | 0 | 0 | 0 | 0 |
| ENSMUSG00000094366 | 0 | 0 | 0 | 0 | 0 | 0 | 0 |
| ENSMUSG00000094367 | 0 | 0 | 0 | 0 | 0 | 0 | 0 |
| ENSMUSG00000094369 | 0 | 0 | 0 | 0 | 0 | 0 | 0 |
| ENSMUSG00000094370 | 0 | 0 | 0 | 0 | 0 | 0 | 0 |
| ENSMUSG00000094371 | 0 | 0 | 0 | 0 | 0 | 0 | 0 |
| ENSMUSG00000094372 | 0 | 0 | 0 | 0 | 0 | 0 | 0 |
| ENSMUSG00000094374 | 0 | 0 | 0 | 0 | 0 | 0 | 0 |
| ENSMUSG00000094377 | 0 | 0 | 0 | 0 | 0 | 0 | 0 |
| ENSMUSG00000094378 | 0 | 0 | 0 | 0 | 0 | 0 | 0 |
| ENSMUSG00000094379 | 0 | 0 | 0 | 0 | 0 | 0 | 0 |

| | | | | | | | |
|---|---|---|---|---|---|---|---|
| ENSMUSG00000094380 | 0 | 0 | 0 | 0 | 0 | 0 | 0 |
| ENSMUSG00000094381 | 0 | 0 | 0 | 0 | 0 | 0 | 0 |
| ENSMUSG00000094382 | 0 | 0 | 0 | 0 | 0 | 0 | 0.355418052 |
| ENSMUSG00000094383 | 0 | 0 | 0 | 0.396240625 | 0 | 0 | 0 |
| ENSMUSG00000094384 | 0 | 0 | 0 | 0 | 0 | 0 | 0 |
| ENSMUSG00000094385 | 0 | 0 | 0 | 0 | 0 | 0 | 0 |
| ENSMUSG00000094386 | 0 | 0 | 0 | 0 | 0 | 0 | 0 |
| ENSMUSG00000094388 | 0 | 0 | 0 | 0 | 0 | 0 | 0 |
| ENSMUSG00000094389 | 0 | 0 | 0 | 0 | 0 | 0 | 0 |
| ENSMUSG00000094390 | 0 | 0 | 0 | 0 | 0 | 0 | 0 |
| ENSMUSG00000094391 | 0 | 0 | 0 | 0 | 0 | 0 | 0 |
| ENSMUSG00000094392 | 0 | 0 | 0 | 0 | 0 | 0 | 0 |
| ENSMUSG00000094394 | 0 | 0 | 0 | 0 | 0 | 0 | 0 |
| ENSMUSG00000094396 | 0 | 0 | 0 | 0 | 0 | 0 | 0 |
| ENSMUSG00000094399 | 0 | 0 | 0 | 0 | 0 | 0 | 0 |
| ENSMUSG00000094400 | 0 | 0 | 0 | 0 | 0 | 0 | 0 |
| ENSMUSG00000094401 | 0 | 0 | 0 | 0 | 0 | 0 | 0 |
| ENSMUSG00000094402 | 0 | 0 | 0 | 0 | 0 | 0 | 0 |
| ENSMUSG00000094403 | 0 | 0 | 0 | 0 | 0 | 0 | 0 |
| ENSMUSG00000094404 | 0 | 0 | 0 | 0 | 0 | 0 | 0 |
| ENSMUSG00000094405 | 0 | 0 | 0 | 0 | 0 | 0 | 0 |
| ENSMUSG00000094406 | 0 | 0 | 0 | 0 | 0 | 0 | 0 |
| ENSMUSG00000094407 | 0 | 0 | 0 | 0 | 0 | 0 | 0 |
| ENSMUSG00000094408 | 0 | 0 | 0 | 0 | 0 | 0 | 0 |
| ENSMUSG00000094409 | 0 | 0 | 0 | 0 | 0 | 0 | 0 |
| ENSMUSG00000094410 | 1.655740331 | 1.664552871 | 1.630890489 | 1.642463902 | 1.695608647 | 1.763427994 | 1.548827561 |
| ENSMUSG00000094411 | 0.671187741 | 1.061981878 | 0.701838731 | 0.646240625 | 0.931471446 | 0.698970004 | 0.803905701 |
| ENSMUSG00000094412 | 0 | 0 | 0 | 0 | 0 | 0 | 0 |
| ENSMUSG00000094413 | 0 | 0 | 0 | 0 | 0 | 0 | 0 |
| ENSMUSG00000094415 | 0 | 0 | 0 | 0 | 0 | 0 | 0 |
| ENSMUSG00000094416 | 0 | 0 | 0 | 0 | 0 | 0 | 0 |
| ENSMUSG00000094417 | 0 | 0 | 0 | 0 | 0 | 0 | 0 |
| ENSMUSG00000094419 | 0 | 0 | 0 | 0 | 0 | 0 | 0 |
| ENSMUSG00000094420 | 0 | 0 | 0 | 0 | 0 | 0 | 0 |

| | | | | | | | |
|---|---|---|---|---|---|---|---|
| ENSMUSG00000094421 | 0 | 0 | 0 | 0 | 0 | 0 | 0 |
| ENSMUSG00000094422 | 0 | 0 | 0 | 0 | 0 | 0 | 0 |
| ENSMUSG00000094423 | 0 | 0 | 0 | 0 | 0 | 0 | 0 |
| ENSMUSG00000094424 | 0 | 0 | 0 | 0 | 0 | 0 | 0 |
| ENSMUSG00000094426 | 0 | 0 | 0 | 0 | 0 | 0 | 0 |
| ENSMUSG00000094427 | 0 | 0 | 0 | 0 | 0 | 0 | 0 |
| ENSMUSG00000094428 | 0 | 0 | 0 | 0 | 0 | 0 | 0 |
| ENSMUSG00000094429 | 0 | 0 | 0 | 0 | 0 | 0 | 0 |
| ENSMUSG00000094430 | 0 | 0 | 0 | 0 | 0 | 0 | 0 |
| ENSMUSG00000094431 | 0 | 0 | 0 | 0 | 0 | 0 | 0 |
| ENSMUSG00000094432 | 0 | 0 | 0 | 0 | 0 | 0 | 0 |
| ENSMUSG00000094433 | 0 | 0 | 0 | 0 | 0 | 0 | 0 |
| ENSMUSG00000094435 | 0 | 0 | 0 | 0 | 0 | 0 | 0.224243824 |
| ENSMUSG00000094436 | 0 | 0 | 0 | 0 | 0 | 0 | 0 |
| ENSMUSG00000094437 | 0 | 0 | 0 | 0 | 0 | 0 | 0 |
| ENSMUSG00000094438 | 0 | 0 | 0 | 0 | 0 | 0 | 0 |
| ENSMUSG00000094439 | 0 | 0 | 0 | 0 | 0 | 0 | 0 |
| ENSMUSG00000094440 | 0 | 0 | 0 | 0 | 0 | 0 | 0 |
| ENSMUSG00000094441 | 1.45815691 | 1.403677461 | 1.27186571 | 1.527131114 | 1.49445382 | 1.531478917 | 1.272006351 |
| ENSMUSG00000094442 | 0 | 0 | 0 | 0 | 0 | 0 | 0 |
| ENSMUSG00000094443 | 0 | 0 | 0 | 0 | 0.244650542 | 0 | 0.224243824 |
| ENSMUSG00000094444 | 0 | 0 | 0 | 0 | 0 | 0 | 0 |
| ENSMUSG00000094445 | 0 | 0 | 0 | 0 | 0 | 0 | 0 |
| ENSMUSG00000094446 | 0 | 0 | 0 | 0 | 0 | 0 | 0 |
| ENSMUSG00000094447 | 0 | 0 | 0 | 0 | 0 | 0 | 0 |
| ENSMUSG00000094449 | 0 | 0 | 0 | 0 | 0 | 0 | 0 |
| ENSMUSG00000094452 | 0 | 0 | 0 | 0 | 0 | 0 | 0 |
| ENSMUSG00000094454 | 0 | 0 | 0 | 0 | 0 | 0 | 0 |
| ENSMUSG00000094455 | 0 | 0 | 0 | 0 | 0 | 0 | 0 |
| ENSMUSG00000094456 | 0 | 0 | 0 | 0 | 0 | 0 | 0 |
| ENSMUSG00000094457 | 0 | 0 | 0 | 0 | 0 | 0 | 0 |
| ENSMUSG00000094458 | 0 | 0 | 0 | 0 | 0 | 0 | 0 |
| ENSMUSG00000094460 | 0 | 0.25 | 0 | 0 | 0 | 0 | 0 |
| ENSMUSG00000094461 | 0 | 0 | 0 | 0 | 0 | 0 | 0 |

| | | | | | | | |
|---|---|---|---|---|---|---|---|
| ENSMUSG00000094462 | 0 | 0 | 0 | 0 | 0 | 0 | 0 |
| ENSMUSG00000094463 | 0 | 0 | 0 | 0 | 0 | 0 | 0 |
| ENSMUSG00000094464 | 0 | 0 | 0 | 0 | 0 | 0 | 0 |
| ENSMUSG00000094465 | 0 | 0 | 0 | 0 | 0 | 0 | 0 |
| ENSMUSG00000094466 | 0 | 0 | 0 | 0 | 0 | 0 | 0 |
| ENSMUSG00000094467 | 0 | 0 | 0 | 0.25 | 0 | 0 | 0 |
| ENSMUSG00000094468 | 0 | 0 | 0 | 0 | 0 | 0 | 0 |
| ENSMUSG00000094470 | 0 | 0 | 0 | 0 | 0 | 0 | 0 |
| ENSMUSG00000094471 | 0 | 0 | 0 | 0 | 0 | 0 | 0 |
| ENSMUSG00000094472 | 1.605636262 | 1.557204673 | 1.532320754 | 1.54248125 | 1.450958105 | 1.491361694 | 1.307989563 |
| ENSMUSG00000094473 | 0 | 0 | 0 | 0 | 0 | 0 | 0 |
| ENSMUSG00000094474 | 0 | 0 | 0 | 0 | 0 | 0 | 0 |
| ENSMUSG00000094475 | 0 | 0 | 0 | 0 | 0 | 0 | 0 |
| ENSMUSG00000094476 | 0 | 0 | 0 | 0 | 0 | 0 | 0 |
| ENSMUSG00000094477 | 0 | 0 | 0 | 0 | 0 | 0 | 0 |
| ENSMUSG00000094478 | 0 | 0 | 0 | 0 | 0 | 0 | 0 |
| ENSMUSG00000094481 | 0 | 0 | 0 | 0 | 0 | 0 | 0 |
| ENSMUSG00000094482 | 0 | 0 | 0 | 0 | 0 | 0 | 0 |
| ENSMUSG00000094483 | 3.010519179 | 2.741446071 | 2.713911792 | 2.826800202 | 2.732881367 | 3.127428778 | 2.501540933 |
| ENSMUSG00000094484 | 0 | 0 | 0 | 0 | 0 | 0 | 0 |
| ENSMUSG00000094485 | 0 | 0 | 0 | 0 | 0 | 0 | 0 |
| ENSMUSG00000094486 | 0 | 0 | 0.25 | 0 | 0 | 0 | 0.355418052 |
| ENSMUSG00000094487 | 0 | 0 | 0 | 0 | 0 | 0 | 0 |
| ENSMUSG00000094488 | 0 | 0 | 0 | 0 | 0 | 0 | 0 |
| ENSMUSG00000094489 | 0 | 0 | 0 | 0 | 0 | 0 | 0 |
| ENSMUSG00000094491 | 0 | 0 | 0 | 0 | 0 | 0 | 0 |
| ENSMUSG00000094493 | 0 | 0 | 0 | 0 | 0 | 0 | 0 |
| ENSMUSG00000094494 | 0 | 0 | 0 | 0 | 0 | 0 | 0 |
| ENSMUSG00000094496 | 0 | 0 | 0 | 0 | 0 | 0 | 0 |
| ENSMUSG00000094497 | 0 | 0 | 0 | 0 | 0 | 0 | 0 |
| ENSMUSG00000094498 | 0 | 0 | 0 | 0 | 0 | 0 | 0 |
| ENSMUSG00000094499 | 0 | 0 | 0 | 0 | 0 | 0 | 0 |
| ENSMUSG00000094500 | 0 | 0.25 | 0.25 | 0 | 0.387761935 | 0 | 0 |
| ENSMUSG00000094501 | 0 | 0 | 0 | 0 | 0 | 0 | 0 |

| | | | | | | | |
|---|---|---|---|---|---|---|---|
| ENSMUSG00000094502 | 0 | 0 | 0 | 0 | 0 | 0 | 0 |
| ENSMUSG00000094504 | 0 | 0 | 0 | 0 | 0 | 0 | 0 |
| ENSMUSG00000094505 | 0 | 0 | 0 | 0 | 0 | 0 | 0 |
| ENSMUSG00000094506 | 0 | 0 | 0 | 0 | 0 | 0 | 0 |
| ENSMUSG00000094507 | 0 | 0 | 0 | 0 | 0 | 0 | 0 |
| ENSMUSG00000094508 | 0 | 0 | 0 | 0 | 0 | 0 | 0 |
| ENSMUSG00000094509 | 0 | 0 | 0 | 0 | 0 | 0 | 0 |
| ENSMUSG00000094510 | 0 | 0 | 0 | 0 | 0 | 0 | 0 |
| ENSMUSG00000094511 | 0 | 0 | 0 | 0 | 0 | 0 | 0 |
| ENSMUSG00000094513 | 0 | 0 | 0 | 0 | 0 | 0 | 0 |
| ENSMUSG00000094514 | 0 | 0 | 0 | 0 | 0 | 0 | 0 |
| ENSMUSG00000094515 | 0 | 0 | 0 | 0 | 0 | 0 | 0 |
| ENSMUSG00000094516 | 0 | 0 | 0 | 0 | 0 | 0 | 0 |
| ENSMUSG00000094517 | 0 | 0 | 0 | 0 | 0 | 0 | 0 |
| ENSMUSG00000094518 | 0.45815691 | 0.580482024 | 0.580482024 | 0 | 0.244650542 | 0.77815125 | 0.520678036 |
| ENSMUSG00000094519 | 0 | 0 | 0 | 0 | 0 | 0 | 0 |
| ENSMUSG00000094520 | 0 | 0 | 0 | 0 | 0 | 0 | 0 |
| ENSMUSG00000094521 | 0 | 0 | 0 | 0 | 0 | 0 | 0 |
| ENSMUSG00000094522 | 0 | 0 | 0 | 0 | 0 | 0 | 0 |
| ENSMUSG00000094523 | 0 | 0 | 0 | 0 | 0 | 0 | 0 |
| ENSMUSG00000094524 | 0 | 0 | 0 | 0 | 0 | 0 | 0 |
| ENSMUSG00000094525 | 0 | 0 | 0 | 0 | 0 | 0 | 0 |
| ENSMUSG00000094526 | 0 | 0 | 0 | 0 | 0 | 0 | 0 |
| ENSMUSG00000094527 | 0 | 0 | 0 | 0 | 0 | 0 | 0 |
| ENSMUSG00000094528 | 0 | 0 | 0 | 0 | 0 | 0 | 0 |
| ENSMUSG00000094529 | 0 | 0 | 0 | 0 | 0 | 0 | 0 |
| ENSMUSG00000094530 | 0 | 0.5 | 0.25 | 0.396240625 | 0 | 0.477121255 | 0.224243824 |
| ENSMUSG00000094531 | 0 | 0 | 0 | 0 | 0 | 0 | 0 |
| ENSMUSG00000094532 | 0 | 0 | 0 | 0 | 0 | 0 | 0 |
| ENSMUSG00000094533 | 0 | 0 | 0 | 0 | 0 | 0 | 0 |
| ENSMUSG00000094534 | 0 | 0 | 0 | 0 | 0 | 0 | 0 |
| ENSMUSG00000094535 | 0 | 0 | 0 | 0 | 0 | 0 | 0 |
| ENSMUSG00000094537 | 0 | 0 | 0 | 0 | 0 | 0 | 0 |
| ENSMUSG00000094538 | 0 | 0 | 0 | 0 | 0 | 0 | 0 |

| | | | | | | | |
|---|---|---|---|---|---|---|---|
| ENSMUSG00000094539 | 0 | 0 | 0 | 0 | 0 | 0 | 0 |
| ENSMUSG00000094540 | 0 | 0 | 0 | 0 | 0 | 0 | 0 |
| ENSMUSG00000094541 | 0 | 0 | 0 | 0 | 0 | 0 | 0 |
| ENSMUSG00000094542 | 0 | 0 | 0 | 0 | 0 | 0 | 0 |
| ENSMUSG00000094543 | 0 | 0 | 0 | 0 | 0 | 0 | 0 |
| ENSMUSG00000094544 | 0 | 0 | 0 | 0 | 0 | 0 | 0 |
| ENSMUSG00000094545 | 0 | 0 | 0 | 0 | 0 | 0 | 0 |
| ENSMUSG00000094546 | 0 | 0 | 0 | 0 | 0 | 0 | 0 |
| ENSMUSG00000094548 | 0 | 0 | 0 | 0 | 0 | 0 | 0 |
| ENSMUSG00000094550 | 0 | 0 | 0 | 0 | 0 | 0 | 0 |
| ENSMUSG00000094551 | 0.811507563 | 0.646240625 | 0.830482024 | 0.701838731 | 0.812711509 | 0.602059991 | 0.803905701 |
| ENSMUSG00000094552 | 0 | 0 | 0 | 0 | 0 | 0 | 0 |
| ENSMUSG00000094553 | 0 | 0 | 0 | 0 | 0 | 0 | 0 |
| ENSMUSG00000094554 | 0 | 0 | 0 | 0 | 0 | 0 | 0 |
| ENSMUSG00000094555 | 0 | 0 | 0 | 0 | 0 | 0 | 0 |
| ENSMUSG00000094556 | 0 | 0 | 0 | 0 | 0 | 0 | 0 |
| ENSMUSG00000094557 | 0 | 0.25 | 0 | 0.25 | 0.244650542 | 0 | 0 |
| ENSMUSG00000094558 | 0 | 0 | 0 | 0 | 0 | 0 | 0 |
| ENSMUSG00000094559 | 0 | 0 | 0 | 0 | 0 | 0 | 0 |
| ENSMUSG00000094560 | 0 | 0 | 0 | 0 | 0 | 0 | 0 |
| ENSMUSG00000094561 | 0 | 0 | 0 | 0 | 0 | 0 | 0 |
| ENSMUSG00000094562 | 0 | 0 | 0 | 0 | 0 | 0 | 0 |
| ENSMUSG00000094563 | 0 | 0 | 0 | 0 | 0 | 0 | 0 |
| ENSMUSG00000094564 | 0 | 0 | 0 | 0 | 0 | 0 | 0 |
| ENSMUSG00000094565 | 0 | 0 | 0 | 0 | 0 | 0 | 0 |
| ENSMUSG00000094566 | 0 | 0 | 0 | 0 | 0 | 0 | 0 |
| ENSMUSG00000094567 | 0 | 0 | 0 | 0 | 0 | 0 | 0 |
| ENSMUSG00000094568 | 0 | 0.25 | 0 | 0 | 0 | 0.301029996 | 0 |
| ENSMUSG00000094569 | 0 | 0 | 0 | 0 | 0 | 0 | 0 |
| ENSMUSG00000094570 | 0 | 0 | 0 | 0 | 0 | 0 | 0 |
| ENSMUSG00000094571 | 0 | 0 | 0 | 0 | 0 | 0 | 0 |
| ENSMUSG00000094572 | 0 | 0 | 0 | 0 | 0 | 0 | 0 |
| ENSMUSG00000094573 | 0 | 0 | 0 | 0 | 0 | 0 | 0 |
| ENSMUSG00000094574 | 0 | 0 | 0 | 0 | 0 | 0 | 0 |

| | | | | | | | |
|---|---|---|---|---|---|---|---|
| ENSMUSG00000094575 | 0 | 0 | 0 | 0 | 0 | 0 | 0 |
| ENSMUSG00000094576 | 0 | 0 | 0 | 0 | 0 | 0 | 0 |
| ENSMUSG00000094577 | 0 | 0 | 0 | 0 | 0 | 0 | 0 |
| ENSMUSG00000094578 | 0 | 0 | 0 | 0 | 0 | 0 | 0 |
| ENSMUSG00000094579 | 0 | 0 | 0 | 0 | 0 | 0 | 0 |
| ENSMUSG00000094581 | 0 | 0 | 0 | 0 | 0 | 0 | 0 |
| ENSMUSG00000094582 | 0 | 0 | 0 | 0 | 0 | 0 | 0 |
| ENSMUSG00000094584 | 0 | 0 | 0 | 0 | 0 | 0 | 0 |
| ENSMUSG00000094585 | 0 | 0 | 0 | 0 | 0 | 0 | 0 |
| ENSMUSG00000094586 | 0 | 0 | 0 | 0 | 0 | 0 | 0 |
| ENSMUSG00000094587 | 0 | 0 | 0 | 0 | 0 | 0 | 0 |
| ENSMUSG00000094588 | 0 | 0 | 0 | 0 | 0 | 0 | 0 |
| ENSMUSG00000094589 | 0 | 0 | 0 | 0 | 0 | 0 | 0 |
| ENSMUSG00000094590 | 0 | 0 | 0 | 0.25 | 0 | 0 | 0 |
| ENSMUSG00000094592 | 0 | 0 | 0 | 0 | 0 | 0 | 0 |
| ENSMUSG00000094593 | 0 | 0 | 0 | 0 | 0 | 0 | 0 |
| ENSMUSG00000094594 | 0 | 0 | 0 | 0 | 0 | 0.301029996 | 0 |
| ENSMUSG00000094595 | 1.404271433 | 1.25 | 1.330482024 | 1.42510993 | 1.407936347 | 1.462397998 | 1.245582246 |
| ENSMUSG00000094596 | 0 | 0 | 0 | 0 | 0 | 0 | 0 |
| ENSMUSG00000094597 | 0 | 0 | 0 | 0 | 0 | 0.301029996 | 0 |
| ENSMUSG00000094598 | 0 | 0 | 0 | 0 | 0 | 0 | 0 |
| ENSMUSG00000094599 | 0 | 0 | 0 | 0 | 0 | 0 | 0 |
| ENSMUSG00000094600 | 0 | 0 | 0 | 0 | 0 | 0 | 0 |
| ENSMUSG00000094601 | 0 | 0 | 0 | 0 | 0 | 0 | 0 |
| ENSMUSG00000094602 | 0 | 0 | 0 | 0 | 0 | 0 | 0 |
| ENSMUSG00000094603 | 0.289064826 | 0 | 0 | 0 | 0.244650542 | 0 | 0 |
| ENSMUSG00000094604 | 0 | 0 | 0 | 0 | 0 | 0 | 0 |
| ENSMUSG00000094605 | 0 | 0 | 0 | 0 | 0 | 0 | 0 |
| ENSMUSG00000094606 | 0 | 0 | 0 | 0 | 0 | 0 | 0 |
| ENSMUSG00000094607 | 0 | 0 | 0 | 0 | 0 | 0 | 0 |
| ENSMUSG00000094608 | 0 | 0 | 0 | 0 | 0 | 0 | 0 |
| ENSMUSG00000094609 | 0 | 0 | 0 | 0 | 0 | 0 | 0 |
| ENSMUSG00000094610 | 0 | 0 | 0 | 0 | 0 | 0 | 0 |
| ENSMUSG00000094612 | 0 | 0 | 0 | 0 | 0 | 0 | 0 |

| | | | | | | | |
|---|---|---|---|---|---|---|---|
| ENSMUSG00000094613 | 0 | 0 | 0 | 0 | 0 | 0 | 0 |
| ENSMUSG00000094614 | 1.837744666 | 1.774508021 | 1.593759858 | 1.886723615 | 1.808534465 | 2.056904851 | 1.695786161 |
| ENSMUSG00000094615 | 0.289064826 | 0.646240625 | 0 | 0 | 0.244650542 | 0 | 0.448487648 |
| ENSMUSG00000094616 | 0 | 0 | 0 | 0 | 0 | 0 | 0 |
| ENSMUSG00000094618 | 0 | 0.5 | 0 | 0 | 0.387761935 | 0 | 0.224243824 |
| ENSMUSG00000094619 | 0 | 0 | 0 | 0 | 0 | 0 | 0 |
| ENSMUSG00000094620 | 0 | 0 | 0 | 0 | 0 | 0 | 0 |
| ENSMUSG00000094621 | 0 | 0 | 0 | 0 | 0 | 0 | 0 |
| ENSMUSG00000094622 | 0 | 0 | 0 | 0 | 0 | 0 | 0 |
| ENSMUSG00000094623 | 0 | 0 | 0 | 0 | 0 | 0 | 0 |
| ENSMUSG00000094624 | 0 | 0 | 0 | 0 | 0 | 0 | 0 |
| ENSMUSG00000094625 | 0 | 0 | 0 | 0 | 0 | 0 | 0 |
| ENSMUSG00000094626 | 0.289064826 | 0 | 0 | 0 | 0 | 0 | 0 |
| ENSMUSG00000094627 | 2.377269616 | 2.194519282 | 2.159156155 | 2.410964047 | 2.335180723 | 2.680335513 | 2.179128924 |
| ENSMUSG00000094628 | 0 | 0 | 0 | 0 | 0 | 0 | 0 |
| ENSMUSG00000094630 | 0 | 0 | 0 | 0 | 0 | 0 | 0 |
| ENSMUSG00000094631 | 0 | 0 | 0 | 0 | 0 | 0 | 0 |
| ENSMUSG00000094632 | 0 | 0 | 0 | 0 | 0 | 0 | 0 |
| ENSMUSG00000094634 | 0.671187741 | 0.396240625 | 0.396240625 | 0.79248125 | 0.244650542 | 0.698970004 | 0.448487648 |
| ENSMUSG00000094635 | 0.289064826 | 0 | 0 | 0 | 0 | 0 | 0 |
| ENSMUSG00000094636 | 0 | 0 | 0 | 0 | 0 | 0 | 0 |
| ENSMUSG00000094637 | 0 | 0 | 0 | 0 | 0 | 0 | 0 |
| ENSMUSG00000094638 | 0 | 0 | 0 | 0 | 0 | 0 | 0 |
| ENSMUSG00000094639 | 0 | 0 | 0 | 0 | 0 | 0 | 0 |
| ENSMUSG00000094641 | 0 | 0 | 0 | 0 | 0 | 0 | 0 |
| ENSMUSG00000094642 | 0 | 0 | 0 | 0 | 0 | 0 | 0 |
| ENSMUSG00000094643 | 0 | 0 | 0 | 0 | 0 | 0 | 0 |
| ENSMUSG00000094647 | 0 | 0 | 0 | 0 | 0 | 0 | 0 |
| ENSMUSG00000094648 | 0 | 0 | 0 | 0 | 0 | 0 | 0 |
| ENSMUSG00000094649 | 0.45815691 | 0.580482024 | 0.79248125 | 0.25 | 0.489301084 | 0.301029996 | 0.629532004 |
| ENSMUSG00000094650 | 0.578129653 | 0 | 0 | 0 | 0 | 0 | 0 |
| ENSMUSG00000094651 | 0 | 0 | 0 | 0 | 0 | 0 | 0 |
| ENSMUSG00000094652 | 0 | 0 | 0 | 0 | 0 | 0 | 0 |
| ENSMUSG00000094654 | 0 | 0 | 0 | 0 | 0 | 0 | 0 |

| | | | | | | | |
|---|---|---|---|---|---|---|---|
| ENSMUSG00000094655 | 0 | 0 | 0 | 0.25 | 0 | 0 | 0 |
| ENSMUSG00000094656 | 0 | 0 | 0 | 0 | 0 | 0 | 0 |
| ENSMUSG00000094657 | 0 | 0 | 0 | 0 | 0 | 0 | 0 |
| ENSMUSG00000094658 | 0 | 0 | 0 | 0 | 0 | 0 | 0 |
| ENSMUSG00000094659 | 0 | 0 | 0 | 0 | 0 | 0 | 0 |
| ENSMUSG00000094660 | 0 | 0 | 0 | 0 | 0 | 0 | 0 |
| ENSMUSG00000094661 | 0 | 0 | 0 | 0 | 0 | 0 | 0 |
| ENSMUSG00000094662 | 0 | 0 | 0 | 0 | 0 | 0 | 0 |
| ENSMUSG00000094663 | 0 | 0 | 0 | 0 | 0 | 0 | 0 |
| ENSMUSG00000094664 | 0 | 0 | 0 | 0 | 0 | 0 | 0 |
| ENSMUSG00000094665 | 0 | 0 | 0 | 0 | 0 | 0 | 0 |
| ENSMUSG00000094667 | 0 | 0 | 0 | 0 | 0 | 0 | 0 |
| ENSMUSG00000094668 | 0 | 0 | 0 | 0 | 0 | 0 | 0 |
| ENSMUSG00000094669 | 0 | 0 | 0 | 0 | 0 | 0 | 0 |
| ENSMUSG00000094670 | 0 | 0 | 0 | 0 | 0 | 0 | 0 |
| ENSMUSG00000094672 | 0 | 0 | 0 | 0 | 0 | 0 | 0 |
| ENSMUSG00000094673 | 0 | 0 | 0 | 0 | 0 | 0 | 0 |
| ENSMUSG00000094675 | 0 | 0 | 0 | 0 | 0 | 0 | 0 |
| ENSMUSG00000094676 | 0 | 0 | 0 | 0 | 0 | 0 | 0 |
| ENSMUSG00000094677 | 0.289064826 | 0 | 0 | 0 | 0 | 0 | 0 |
| ENSMUSG00000094678 | 0 | 0 | 0 | 0 | 0 | 0 | 0 |
| ENSMUSG00000094679 | 0 | 0 | 0 | 0 | 0 | 0 | 0 |
| ENSMUSG00000094680 | 0 | 0 | 0 | 0 | 0 | 0 | 0 |
| ENSMUSG00000094681 | 0 | 0 | 0 | 0 | 0 | 0 | 0 |
| ENSMUSG00000094682 | 0 | 0 | 0 | 0 | 0 | 0 | 0 |
| ENSMUSG00000094683 | 0 | 0 | 0 | 0 | 0.244650542 | 0.301029996 | 0 |
| ENSMUSG00000094684 | 0 | 0 | 0 | 0 | 0 | 0 | 0 |
| ENSMUSG00000094685 | 1.587501561 | 1.51109853 | 1.552363341 | 1.410964047 | 1.433158226 | 1.414973348 | 1.265599895 |
| ENSMUSG00000094686 | 0 | 0 | 0 | 0 | 0 | 0 | 0 |
| ENSMUSG00000094687 | 0 | 0 | 0 | 0 | 0 | 0 | 0 |
| ENSMUSG00000094688 | 0 | 0 | 0 | 0 | 0 | 0 | 0 |
| ENSMUSG00000094689 | 0 | 0 | 0 | 0 | 0 | 0 | 0 |
| ENSMUSG00000094690 | 0.578129653 | 0 | 0 | 0 | 0 | 0 | 0 |
| ENSMUSG00000094691 | 0 | 0 | 0 | 0 | 0 | 0 | 0 |

| | | | | | | | |
|---|---|---|---|---|---|---|---|
| ENSMUSG00000094692 | 0 | 0 | 0 | 0 | 0 | 0 | 0 |
| ENSMUSG00000094693 | 0 | 0 | 0 | 0 | 0 | 0 | 0 |
| ENSMUSG00000094694 | 0 | 0 | 0 | 0 | 0 | 0 | 0 |
| ENSMUSG00000094695 | 0 | 0 | 0 | 0 | 0 | 0 | 0 |
| ENSMUSG00000094696 | 0 | 0 | 0 | 0 | 0 | 0 | 0 |
| ENSMUSG00000094697 | 0 | 0 | 0 | 0 | 0 | 0 | 0 |
| ENSMUSG00000094698 | 0 | 0 | 0 | 0 | 0 | 0 | 0 |
| ENSMUSG00000094699 | 0 | 0 | 0 | 0 | 0 | 0 | 0 |
| ENSMUSG00000094700 | 0 | 0 | 0 | 0 | 0 | 0 | 0 |
| ENSMUSG00000094701 | 0 | 0 | 0 | 0 | 0 | 0 | 0 |
| ENSMUSG00000094702 | 0 | 0 | 0 | 0 | 0 | 0 | 0 |
| ENSMUSG00000094703 | 0 | 0 | 0 | 0 | 0 | 0 | 0 |
| ENSMUSG00000094704 | 0 | 0 | 0 | 0 | 0 | 0 | 0 |
| ENSMUSG00000094706 | 0 | 0 | 0 | 0 | 0 | 0 | 0 |
| ENSMUSG00000094707 | 0 | 0.25 | 0 | 0 | 0 | 0 | 0 |
| ENSMUSG00000094708 | 0 | 0 | 0 | 0 | 0 | 0 | 0 |
| ENSMUSG00000094709 | 0 | 0 | 0 | 0 | 0 | 0 | 0 |
| ENSMUSG00000094710 | 0 | 0 | 0 | 0 | 0 | 0 | 0 |
| ENSMUSG00000094712 | 0 | 0 | 0 | 0 | 0 | 0 | 0 |
| ENSMUSG00000094713 | 0 | 0 | 0 | 0 | 0 | 0 | 0 |
| ENSMUSG00000094714 | 0 | 0 | 0 | 0 | 0 | 0 | 0 |
| ENSMUSG00000094715 | 0 | 0 | 0 | 0 | 0 | 0 | 0 |
| ENSMUSG00000094717 | 0 | 0 | 0 | 0 | 0 | 0 | 0 |
| ENSMUSG00000094718 | 0 | 0 | 0 | 0 | 0 | 0 | 0 |
| ENSMUSG00000094719 | 0 | 0 | 0.25 | 0.25 | 0.387761935 | 0 | 0 |
| ENSMUSG00000094720 | 0 | 0 | 0 | 0 | 0 | 0 | 0 |
| ENSMUSG00000094721 | 0 | 0 | 0 | 0 | 0 | 0 | 0 |
| ENSMUSG00000094722 | 0 | 0 | 0 | 0 | 0 | 0 | 0 |
| ENSMUSG00000094723 | 0 | 0 | 0 | 0 | 0 | 0 | 0 |
| ENSMUSG00000094724 | 0.867194479 | 0.701838731 | 0.5 | 0.646240625 | 0.387761935 | 0.602059991 | 0.579661877 |
| ENSMUSG00000094725 | 0 | 0 | 0 | 0 | 0 | 0 | 0 |
| ENSMUSG00000094726 | 0 | 0 | 0 | 0 | 0 | 0 | 0 |
| ENSMUSG00000094727 | 0 | 0 | 0 | 0 | 0 | 0 | 0 |
| ENSMUSG00000094728 | 0 | 0 | 0 | 0 | 0 | 0 | 0 |

| | | | | | | | |
|---|---|---|---|---|---|---|---|
| ENSMUSG00000094729 | 0 | 0 | 0 | 0 | 0 | 0 | 0 |
| ENSMUSG00000094730 | 0 | 0 | 0 | 0 | 0 | 0 | 0 |
| ENSMUSG00000094732 | 0.289064826 | 0 | 0 | 0 | 0 | 0 | 0 |
| ENSMUSG00000094733 | 0 | 0 | 0 | 0 | 0 | 0 | 0 |
| ENSMUSG00000094734 | 0 | 0 | 0 | 0 | 0 | 0 | 0 |
| ENSMUSG00000094735 | 0 | 0 | 0 | 0 | 0 | 0 | 0 |
| ENSMUSG00000094736 | 0 | 0 | 0 | 0 | 0 | 0 | 0 |
| ENSMUSG00000094737 | 0 | 0 | 0 | 0 | 0 | 0 | 0 |
| ENSMUSG00000094738 | 0 | 0 | 0 | 0 | 0 | 0 | 0 |
| ENSMUSG00000094739 | 0 | 0 | 0 | 0 | 0 | 0 | 0 |
| ENSMUSG00000094740 | 0 | 0 | 0 | 0 | 0 | 0 | 0 |
| ENSMUSG00000094741 | 0 | 0 | 0 | 0 | 0 | 0 | 0 |
| ENSMUSG00000094743 | 0 | 0 | 0 | 0 | 0 | 0 | 0 |
| ENSMUSG00000094745 | 0 | 0 | 0 | 0 | 0 | 0 | 0 |
| ENSMUSG00000094746 | 0 | 0 | 0 | 0 | 0 | 0 | 0 |
| ENSMUSG00000094747 | 0 | 0 | 0 | 0 | 0 | 0 | 0 |
| ENSMUSG00000094748 | 0 | 0 | 0 | 0 | 0 | 0 | 0 |
| ENSMUSG00000094750 | 0 | 0 | 0 | 0 | 0 | 0 | 0 |
| ENSMUSG00000094751 | 0 | 0 | 0 | 0 | 0 | 0 | 0 |
| ENSMUSG00000094752 | 0 | 0 | 0 | 0 | 0 | 0 | 0 |
| ENSMUSG00000094753 | 0 | 0 | 0 | 0 | 0 | 0 | 0 |
| ENSMUSG00000094754 | 0 | 0 | 0 | 0 | 0 | 0 | 0 |
| ENSMUSG00000094755 | 0 | 0 | 0 | 0 | 0 | 0 | 0 |
| ENSMUSG00000094757 | 0 | 0 | 0 | 0 | 0 | 0 | 0 |
| ENSMUSG00000094758 | 0 | 0 | 0 | 0 | 0 | 0 | 0 |
| ENSMUSG00000094759 | 0 | 0 | 0 | 0 | 0 | 0 | 0 |
| ENSMUSG00000094760 | 0 | 0 | 0 | 0 | 0 | 0 | 0 |
| ENSMUSG00000094762 | 0 | 0 | 0 | 0 | 0 | 0 | 0 |
| ENSMUSG00000094763 | 0 | 0 | 0 | 0 | 0 | 0 | 0 |
| ENSMUSG00000094764 | 0 | 0 | 0 | 0 | 0 | 0 | 0 |
| ENSMUSG00000094765 | 0 | 0 | 0 | 0 | 0 | 0 | 0 |
| ENSMUSG00000094766 | 0 | 0 | 0 | 0 | 0 | 0 | 0 |
| ENSMUSG00000094767 | 0 | 0 | 0 | 0 | 0 | 0 | 0 |
| ENSMUSG00000094768 | 0 | 0 | 0 | 0 | 0 | 0 | 0 |

| | | | | | | | |
|---|---|---|---|---|---|---|---|
| ENSMUSG00000094771 | 0 | 0 | 0 | 0 | 0 | 0 | 0 |
| ENSMUSG00000094772 | 0.289064826 | 0.396240625 | 0.25 | 0 | 0 | 0.301029996 | 0.224243824 |
| ENSMUSG00000094773 | 0 | 0 | 0 | 0 | 0 | 0 | 0 |
| ENSMUSG00000094775 | 0 | 0 | 0 | 0 | 0 | 0 | 0 |
| ENSMUSG00000094777 | 0 | 0 | 0 | 0 | 0 | 0 | 0 |
| ENSMUSG00000094778 | 0 | 0 | 0 | 0 | 0 | 0 | 0 |
| ENSMUSG00000094779 | 0 | 0 | 0 | 0 | 0 | 0 | 0 |
| ENSMUSG00000094780 | 0 | 0 | 0 | 0 | 0 | 0 | 0 |
| ENSMUSG00000094782 | 0 | 0 | 0 | 0 | 0 | 0 | 0 |
| ENSMUSG00000094784 | 0 | 0 | 0 | 0 | 0 | 0 | 0 |
| ENSMUSG00000094785 | 0 | 0 | 0 | 0 | 0 | 0 | 0 |
| ENSMUSG00000094786 | 0.867194479 | 0.75 | 0.580482024 | 1.238549078 | 1.264824954 | 1.278753601 | 1.110947927 |
| ENSMUSG00000094787 | 0 | 0 | 0 | 0 | 0 | 0 | 0 |
| ENSMUSG00000094789 | 0 | 0 | 0 | 0 | 0 | 0 | 0 |
| ENSMUSG00000094790 | 0 | 0.5 | 0.396240625 | 0.5 | 0.244650542 | 0 | 0.448487648 |
| ENSMUSG00000094791 | 0 | 0 | 0 | 0 | 0 | 0 | 0 |
| ENSMUSG00000094792 | 0 | 0 | 0 | 0 | 0 | 0 | 0 |
| ENSMUSG00000094793 | 0 | 0 | 0 | 0 | 0 | 0 | 0 |
| ENSMUSG00000094794 | 0 | 0 | 0 | 0 | 0 | 0 | 0 |
| ENSMUSG00000094796 | 0 | 0 | 0 | 0 | 0 | 0 | 0 |
| ENSMUSG00000094797 | 0 | 0 | 0 | 0 | 0 | 0 | 0 |
| ENSMUSG00000094798 | 0 | 0 | 0 | 0 | 0 | 0 | 0 |
| ENSMUSG00000094799 | 0 | 0 | 0 | 0 | 0 | 0 | 0 |
| ENSMUSG00000094800 | 0.289064826 | 0 | 0.25 | 0 | 0 | 0 | 0 |
| ENSMUSG00000094801 | 0 | 0 | 0 | 0 | 0 | 0 | 0 |
| ENSMUSG00000094802 | 0 | 0 | 0 | 0 | 0 | 0 | 0 |
| ENSMUSG00000094803 | 0 | 0 | 0 | 0 | 0 | 0 | 0 |
| ENSMUSG00000094804 | 0 | 0 | 0 | 0 | 0 | 0 | 0 |
| ENSMUSG00000094805 | 0 | 0 | 0 | 0 | 0 | 0 | 0 |
| ENSMUSG00000094806 | 0 | 0 | 0 | 0 | 0 | 0 | 0 |
| ENSMUSG00000094807 | 0 | 0 | 0 | 0 | 0 | 0 | 0 |
| ENSMUSG00000094808 | 0 | 0 | 0.25 | 0 | 0 | 0 | 0 |
| ENSMUSG00000094810 | 0 | 0 | 0 | 0 | 0 | 0 | 0 |
| ENSMUSG00000094811 | 0 | 0 | 0 | 0 | 0 | 0 | 0 |

| | | | | | | | |
|---|---|---|---|---|---|---|---|
| ENSMUSG00000094812 | 0 | 0 | 0 | 0 | 0 | 0 | 0 |
| ENSMUSG00000094813 | 0 | 0 | 0 | 0 | 0 | 0 | 0 |
| ENSMUSG00000094814 | 0 | 0 | 0 | 0 | 0 | 0 | 0 |
| ENSMUSG00000094815 | 0 | 0 | 0 | 0 | 0 | 0 | 0 |
| ENSMUSG00000094817 | 0 | 0 | 0 | 0 | 0 | 0 | 0 |
| ENSMUSG00000094818 | 0 | 0 | 0 | 0 | 0 | 0 | 0 |
| ENSMUSG00000094819 | 0 | 0 | 0 | 0 | 0 | 0 | 0 |
| ENSMUSG00000094820 | 0 | 0 | 0 | 0 | 0 | 0.301029996 | 0 |
| ENSMUSG00000094821 | 0 | 0 | 0 | 0 | 0 | 0 | 0 |
| ENSMUSG00000094822 | 0 | 0 | 0 | 0 | 0 | 0 | 0 |
| ENSMUSG00000094823 | 0 | 0 | 0 | 0 | 0 | 0 | 0 |
| ENSMUSG00000094824 | 0 | 0 | 0 | 0 | 0 | 0 | 0 |
| ENSMUSG00000094825 | 0 | 0 | 0 | 0 | 0 | 0 | 0 |
| ENSMUSG00000094826 | 0 | 0 | 0 | 0 | 0 | 0 | 0 |
| ENSMUSG00000094827 | 0 | 0 | 0 | 0 | 0 | 0 | 0 |
| ENSMUSG00000094828 | 0 | 0 | 0 | 0 | 0 | 0 | 0 |
| ENSMUSG00000094829 | 0 | 0 | 0 | 0 | 0 | 0 | 0 |
| ENSMUSG00000094830 | 0 | 0 | 0 | 0 | 0 | 0 | 0 |
| ENSMUSG00000094831 | 0 | 0 | 0 | 0 | 0 | 0 | 0 |
| ENSMUSG00000094832 | 0.289064826 | 0 | 0.25 | 0.25 | 0 | 0 | 0 |
| ENSMUSG00000094833 | 0 | 0 | 0 | 0 | 0 | 0 | 0 |
| ENSMUSG00000094834 | 0 | 0.25 | 0 | 0.25 | 0 | 0.477121255 | 0 |
| ENSMUSG00000094836 | 0 | 0 | 0 | 0 | 0 | 0 | 0 |
| ENSMUSG00000094837 | 0 | 0 | 0 | 0 | 0 | 0 | 0 |
| ENSMUSG00000094838 | 0 | 0 | 0 | 0 | 0 | 0 | 0 |
| ENSMUSG00000094840 | 0 | 0 | 0 | 0 | 0 | 0 | 0.224243824 |
| ENSMUSG00000094841 | 0 | 0 | 0 | 0 | 0 | 0 | 0 |
| ENSMUSG00000094842 | 0 | 0 | 0 | 0 | 0.244650542 | 0 | 0 |
| ENSMUSG00000094843 | 0 | 0 | 0 | 0 | 0 | 0 | 0 |
| ENSMUSG00000094844 | 0 | 0 | 0 | 0 | 0 | 0 | 0 |
| ENSMUSG00000094845 | 0 | 0.701838731 | 0 | 0.646240625 | 0.244650542 | 0.602059991 | 0.448487648 |
| ENSMUSG00000094846 | 0 | 0 | 0 | 0 | 0 | 0 | 0 |
| ENSMUSG00000094847 | 0 | 0 | 0 | 0 | 0 | 0 | 0 |
| ENSMUSG00000094848 | 0 | 0 | 0 | 0 | 0 | 0 | 0 |

| | | | | | | | |
|---|---|---|---|---|---|---|---|
| ENSMUSG00000094849 | 0 | 0 | 0 | 0.25 | 0 | 0 | 0 |
| ENSMUSG00000094850 | 0 | 0 | 0 | 0 | 0 | 0 | 0 |
| ENSMUSG00000094852 | 0 | 0 | 0.25 | 0 | 0 | 0 | 0 |
| ENSMUSG00000094854 | 0 | 0 | 0 | 0 | 0 | 0 | 0 |
| ENSMUSG00000094855 | 0 | 0 | 0 | 0 | 0 | 0 | 0 |
| ENSMUSG00000094856 | 0 | 0.25 | 0 | 0.396240625 | 0.489301084 | 0 | 0.224243824 |
| ENSMUSG00000094858 | 0 | 0 | 0 | 0 | 0 | 0 | 0 |
| ENSMUSG00000094860 | 0 | 0 | 0 | 0 | 0 | 0 | 0 |
| ENSMUSG00000094861 | 0 | 0 | 0 | 0 | 0 | 0 | 0 |
| ENSMUSG00000094862 | 0 | 0 | 0 | 0 | 0 | 0 | 0 |
| ENSMUSG00000094864 | 0 | 0 | 0 | 0 | 0 | 0 | 0 |
| ENSMUSG00000094865 | 0 | 0 | 0 | 0.25 | 0 | 0 | 0 |
| ENSMUSG00000094866 | 0 | 0 | 0 | 0 | 0 | 0 | 0 |
| ENSMUSG00000094867 | 0 | 0 | 0 | 0 | 0 | 0 | 0 |
| ENSMUSG00000094868 | 0 | 0 | 0 | 0 | 0 | 0 | 0 |
| ENSMUSG00000094869 | 0 | 0 | 0 | 0 | 0 | 0 | 0 |
| ENSMUSG00000094870 | 2.481610694 | 2.339388001 | 2.302363341 | 2.441217898 | 2.32316608 | 2.702430536 | 2.156863548 |
| ENSMUSG00000094872 | 0 | 0 | 0 | 0 | 0 | 0 | 0 |
| ENSMUSG00000094873 | 0 | 0 | 0 | 0 | 0 | 0 | 0 |
| ENSMUSG00000094874 | 0 | 0 | 0 | 0 | 0 | 0 | 0 |
| ENSMUSG00000094875 | 0 | 0 | 0 | 0 | 0 | 0 | 0 |
| ENSMUSG00000094876 | 0 | 0 | 0 | 0 | 0 | 0 | 0 |
| ENSMUSG00000094877 | 0 | 0 | 0 | 0 | 0 | 0 | 0 |
| ENSMUSG00000094878 | 0 | 0 | 0 | 0 | 0 | 0 | 0 |
| ENSMUSG00000094879 | 0 | 0 | 0 | 0 | 0 | 0 | 0 |
| ENSMUSG00000094880 | 0 | 0 | 0 | 0 | 0 | 0 | 0 |
| ENSMUSG00000094881 | 0 | 0 | 0 | 0 | 0 | 0 | 0 |
| ENSMUSG00000094883 | 0 | 0 | 0 | 0 | 0 | 0 | 0 |
| ENSMUSG00000094884 | 0 | 0 | 0 | 0 | 0 | 0 | 0 |
| ENSMUSG00000094885 | 0 | 0 | 0 | 0 | 0 | 0 | 0 |
| ENSMUSG00000094886 | 0 | 0 | 0 | 0 | 0 | 0 | 0 |
| ENSMUSG00000094887 | 0 | 0 | 0 | 0 | 0 | 0 | 0 |
| ENSMUSG00000094888 | 0 | 0 | 0 | 0 | 0 | 0 | 0 |
| ENSMUSG00000094889 | 0 | 0 | 0 | 0 | 0 | 0 | 0 |

| | | | | | | | |
|---|---|---|---|---|---|---|---|
| ENSMUSG00000094890 | 0 | 0 | 0 | 0 | 0 | 0 | 0 |
| ENSMUSG00000094891 | 0.289064826 | 0 | 0 | 0 | 0 | 0 | 0 |
| ENSMUSG00000094892 | 0 | 0 | 0 | 0 | 0 | 0 | 0 |
| ENSMUSG00000094893 | 0 | 0.25 | 0.25 | 0.396240625 | 0 | 0.301029996 | 0.224243824 |
| ENSMUSG00000094894 | 0 | 0 | 0 | 0 | 0 | 0 | 0 |
| ENSMUSG00000094895 | 0 | 0 | 0 | 0 | 0 | 0 | 0 |
| ENSMUSG00000094896 | 0 | 0 | 0 | 0 | 0 | 0 | 0 |
| ENSMUSG00000094897 | 0 | 0 | 0 | 0 | 0 | 0 | 0 |
| ENSMUSG00000094898 | 0 | 0 | 0 | 0 | 0 | 0 | 0 |
| ENSMUSG00000094901 | 0 | 0 | 0 | 0 | 0 | 0 | 0 |
| ENSMUSG00000094902 | 0 | 0 | 0 | 0 | 0 | 0 | 0 |
| ENSMUSG00000094903 | 0 | 0 | 0 | 0 | 0 | 0 | 0 |
| ENSMUSG00000094904 | 0 | 0 | 0 | 0 | 0 | 0 | 0 |
| ENSMUSG00000094905 | 0 | 0 | 0 | 0 | 0 | 0 | 0 |
| ENSMUSG00000094906 | 0 | 0 | 0 | 0 | 0 | 0 | 0 |
| ENSMUSG00000094910 | 0 | 0 | 0 | 0.396240625 | 0 | 0.301029996 | 0.355418052 |
| ENSMUSG00000094911 | 0 | 0 | 0 | 0 | 0 | 0 | 0 |
| ENSMUSG00000094912 | 0 | 0 | 0 | 0 | 0 | 0 | 0 |
| ENSMUSG00000094913 | 0 | 0 | 0 | 0 | 0 | 0 | 0 |
| ENSMUSG00000094914 | 0 | 0 | 0 | 0 | 0 | 0 | 0 |
| ENSMUSG00000094915 | 0 | 0 | 0 | 0 | 0 | 0 | 0 |
| ENSMUSG00000094916 | 0 | 0 | 0 | 0 | 0 | 0 | 0 |
| ENSMUSG00000094917 | 0 | 0 | 0 | 0 | 0 | 0 | 0 |
| ENSMUSG00000094918 | 0 | 0 | 0 | 0 | 0 | 0 | 0 |
| ENSMUSG00000094919 | 0 | 0 | 0 | 0 | 0 | 0 | 0 |
| ENSMUSG00000094920 | 0 | 0 | 0 | 0 | 0 | 0 | 0 |
| ENSMUSG00000094921 | 0 | 0 | 0 | 0 | 0 | 0 | 0 |
| ENSMUSG00000094922 | 0 | 0 | 0 | 0 | 0 | 0 | 0 |
| ENSMUSG00000094923 | 0 | 0 | 0 | 0 | 0 | 0 | 0 |
| ENSMUSG00000094925 | 0 | 0 | 0 | 0 | 0 | 0 | 0 |
| ENSMUSG00000094926 | 0 | 0 | 0 | 0 | 0 | 0 | 0 |
| ENSMUSG00000094927 | 0 | 0 | 0 | 0 | 0 | 0 | 0 |
| ENSMUSG00000094928 | 0 | 0 | 0 | 0 | 0 | 0 | 0 |
| ENSMUSG00000094929 | 0 | 0 | 0 | 0 | 0 | 0 | 0 |

| | | | | | | | |
|---|---|---|---|---|---|---|---|
| ENSMUSG00000094930 | 0 | 0 | 0 | 0 | 0 | 0 | 0 |
| ENSMUSG00000094931 | 0 | 0 | 0 | 0 | 0 | 0 | 0 |
| ENSMUSG00000094932 | 0 | 0 | 0 | 0 | 0 | 0 | 0 |
| ENSMUSG00000094933 | 0 | 0 | 0 | 0 | 0 | 0 | 0 |
| ENSMUSG00000094934 | 0 | 0 | 0 | 0 | 0 | 0 | 0 |
| ENSMUSG00000094935 | 0.45815691 | 0.396240625 | 0.396240625 | 0.25 | 0 | 0 | 0.355418052 |
| ENSMUSG00000094936 | 1.647796617 | 1.552363341 | 1.580482024 | 1.566696635 | 1.632412477 | 1.643452676 | 1.452143229 |
| ENSMUSG00000094937 | 0 | 0 | 0 | 0 | 0 | 0 | 0 |
| ENSMUSG00000094938 | 0 | 0 | 0 | 0 | 0 | 0 | 0 |
| ENSMUSG00000094939 | 0 | 0 | 0 | 0 | 0 | 0 | 0 |
| ENSMUSG00000094940 | 0 | 0 | 0 | 0 | 0 | 0 | 0 |
| ENSMUSG00000094941 | 0 | 0 | 0 | 0 | 0 | 0 | 0 |
| ENSMUSG00000094942 | 1.494443473 | 1.356566189 | 1.282320754 | 1.527131114 | 1.394615667 | 1.477121255 | 1.296434211 |
| ENSMUSG00000094943 | 0 | 0 | 0 | 0 | 0 | 0 | 0 |
| ENSMUSG00000094945 | 0 | 0 | 0 | 0 | 0 | 0 | 0 |
| ENSMUSG00000094946 | 0 | 0 | 0 | 0 | 0 | 0 | 0 |
| ENSMUSG00000094947 | 0 | 0 | 0 | 0 | 0 | 0 | 0 |
| ENSMUSG00000094948 | 0 | 0 | 0 | 0 | 0 | 0 | 0 |
| ENSMUSG00000094949 | 0 | 0 | 0 | 0 | 0 | 0 | 0 |
| ENSMUSG00000094950 | 0 | 0 | 0 | 0 | 0 | 0 | 0 |
| ENSMUSG00000094951 | 0 | 0 | 0 | 0 | 0 | 0 | 0 |
| ENSMUSG00000094952 | 0 | 0 | 0 | 0 | 0 | 0 | 0 |
| ENSMUSG00000094954 | 0 | 0 | 0 | 0 | 0 | 0 | 0 |
| ENSMUSG00000094955 | 0 | 0 | 0 | 0 | 0 | 0 | 0 |
| ENSMUSG00000094956 | 0 | 0 | 0 | 0 | 0 | 0 | 0 |
| ENSMUSG00000094957 | 0 | 0 | 0 | 0 | 0 | 0 | 0 |
| ENSMUSG00000094958 | 0 | 0.25 | 0 | 0 | 0.244650542 | 0.301029996 | 0 |
| ENSMUSG00000094959 | 0 | 0 | 0 | 0 | 0 | 0 | 0 |
| ENSMUSG00000094960 | 0 | 0 | 0 | 0 | 0 | 0 | 0 |
| ENSMUSG00000094962 | 0 | 0 | 0 | 0 | 0.244650542 | 0 | 0 |
| ENSMUSG00000094963 | 0 | 0 | 0 | 0 | 0 | 0 | 0 |
| ENSMUSG00000094964 | 0 | 0 | 0 | 0 | 0 | 0 | 0 |
| ENSMUSG00000094965 | 0 | 0 | 0 | 0 | 0 | 0 | 0 |
| ENSMUSG00000094966 | 0 | 0 | 0 | 0 | 0 | 0 | 0 |

| | | | | | | | |
|---|---|---|---|---|---|---|---|
| ENSMUSG00000094967 | 0 | 0 | 0 | 0 | 0 | 0 | 0 |
| ENSMUSG00000094968 | 0 | 0 | 0 | 0 | 0 | 0 | 0 |
| ENSMUSG00000094969 | 0.289064826 | 0 | 0 | 0.25 | 0 | 0 | 0.224243824 |
| ENSMUSG00000094970 | 0 | 0 | 0 | 0 | 0 | 0 | 0 |
| ENSMUSG00000094971 | 0 | 0 | 0 | 0 | 0 | 0 | 0 |
| ENSMUSG00000094972 | 0 | 0 | 0 | 0 | 0 | 0 | 0 |
| ENSMUSG00000094973 | 0 | 0 | 0 | 0 | 0 | 0.301029996 | 0.520678036 |
| ENSMUSG00000094974 | 0 | 0 | 0 | 0 | 0 | 0 | 0 |
| ENSMUSG00000094975 | 0 | 0 | 0 | 0 | 0 | 0 | 0 |
| ENSMUSG00000094976 | 0 | 0 | 0 | 0 | 0 | 0 | 0 |
| ENSMUSG00000094977 | 0 | 0 | 0 | 0 | 0 | 0 | 0 |
| ENSMUSG00000094978 | 0 | 0 | 0 | 0 | 0 | 0 | 0 |
| ENSMUSG00000094979 | 0 | 0 | 0 | 0 | 0 | 0 | 0 |
| ENSMUSG00000094980 | 0 | 0 | 0 | 0 | 0 | 0 | 0 |
| ENSMUSG00000094981 | 0 | 0 | 0 | 0 | 0 | 0 | 0 |
| ENSMUSG00000094982 | 0 | 0 | 0 | 0 | 0 | 0 | 0 |
| ENSMUSG00000094983 | 0 | 0 | 0 | 0 | 0 | 0 | 0 |
| ENSMUSG00000094985 | 0 | 0 | 0 | 0 | 0 | 0 | 0 |
| ENSMUSG00000094986 | 0 | 0 | 0 | 0 | 0 | 0 | 0 |
| ENSMUSG00000094987 | 0 | 0 | 0 | 0 | 0 | 0 | 0 |
| ENSMUSG00000094989 | 0 | 0 | 0.25 | 0 | 0 | 0 | 0 |
| ENSMUSG00000094991 | 0 | 0 | 0 | 0 | 0 | 0 | 0 |
| ENSMUSG00000094992 | 0 | 0 | 0 | 0 | 0 | 0 | 0 |
| ENSMUSG00000094993 | 0 | 0 | 0 | 0 | 0 | 0 | 0 |
| ENSMUSG00000094994 | 0 | 0 | 0 | 0 | 0 | 0 | 0 |
| ENSMUSG00000094995 | 0.289064826 | 0 | 0 | 0 | 0 | 0 | 0 |
| ENSMUSG00000094997 | 0 | 0 | 0 | 0 | 0 | 0 | 0 |
| ENSMUSG00000094998 | 0 | 0 | 0 | 0 | 0 | 0 | 0 |
| ENSMUSG00000094999 | 0 | 0 | 0 | 0 | 0 | 0 | 0 |
| ENSMUSG00000095000 | 0 | 0 | 0 | 0 | 0 | 0 | 0 |
| ENSMUSG00000095002 | 0 | 0 | 0 | 0 | 0 | 0 | 0 |
| ENSMUSG00000095003 | 0 | 0 | 0 | 0 | 0 | 0 | 0 |
| ENSMUSG00000095005 | 0 | 0 | 0 | 0 | 0 | 0 | 0 |
| ENSMUSG00000095006 | 0 | 0 | 0 | 0 | 0 | 0 | 0 |

| | | | | | | | |
|---|---|---|---|---|---|---|---|
| ENSMUSG00000095007 | 0 | 0 | 0 | 0 | 0 | 0 | 0 |
| ENSMUSG00000095008 | 0 | 0 | 0 | 0 | 0 | 0 | 0 |
| ENSMUSG00000095009 | 0 | 0 | 0 | 0 | 0 | 0 | 0 |
| ENSMUSG00000095010 | 0 | 0 | 0 | 0 | 0 | 0 | 0.224243824 |
| ENSMUSG00000095011 | 0 | 0 | 0 | 0 | 0 | 0 | 0 |
| ENSMUSG00000095015 | 0 | 0 | 0 | 0 | 0 | 0 | 0 |
| ENSMUSG00000095016 | 0 | 0 | 0 | 0 | 0 | 0 | 0 |
| ENSMUSG00000095017 | 0 | 0 | 0 | 0 | 0 | 0 | 0 |
| ENSMUSG00000095019 | 0 | 0 | 0 | 0 | 0 | 0 | 0 |
| ENSMUSG00000095020 | 0 | 0 | 0 | 0 | 0 | 0 | 0 |
| ENSMUSG00000095021 | 0 | 0 | 0 | 0 | 0 | 0 | 0 |
| ENSMUSG00000095022 | 0 | 0 | 0 | 0 | 0 | 0 | 0 |
| ENSMUSG00000095023 | 0 | 0 | 0 | 0 | 0 | 0 | 0 |
| ENSMUSG00000095024 | 0 | 0 | 0 | 0 | 0 | 0 | 0 |
| ENSMUSG00000095025 | 0 | 0 | 0 | 0 | 0 | 0 | 0 |
| ENSMUSG00000095026 | 0 | 0 | 0 | 0.25 | 0 | 0 | 0 |
| ENSMUSG00000095027 | 0 | 0 | 0 | 0 | 0 | 0 | 0 |
| ENSMUSG00000095028 | 0 | 0 | 0 | 0 | 0 | 0 | 0 |
| ENSMUSG00000095029 | 0 | 0 | 0 | 0 | 0 | 0 | 0 |
| ENSMUSG00000095030 | 0 | 0 | 0 | 0 | 0 | 0 | 0 |
| ENSMUSG00000095031 | 0 | 0 | 0 | 0 | 0 | 0 | 0 |
| ENSMUSG00000095032 | 0 | 0 | 0 | 0 | 0 | 0 | 0 |
| ENSMUSG00000095033 | 0 | 0 | 0 | 0 | 0 | 0 | 0 |
| ENSMUSG00000095034 | 0 | 0 | 0 | 0 | 0 | 0 | 0 |
| ENSMUSG00000095035 | 0 | 0 | 0 | 0 | 0 | 0 | 0 |
| ENSMUSG00000095037 | 0 | 0 | 0 | 0 | 0 | 0 | 0 |
| ENSMUSG00000095040 | 0 | 0 | 0 | 0 | 0 | 0 | 0 |
| ENSMUSG00000095041 | 2.898315257 | 2.629171237 | 2.638167274 | 2.583568321 | 2.609931625 | 2.933487288 | 2.420763202 |
| ENSMUSG00000095042 | 0 | 0 | 0 | 0 | 0 | 0 | 0 |
| ENSMUSG00000095044 | 0 | 0 | 0 | 0 | 0 | 0 | 0 |
| ENSMUSG00000095045 | 0 | 0 | 0 | 0 | 0 | 0 | 0 |
| ENSMUSG00000095047 | 0 | 0 | 0.25 | 0 | 0 | 0.301029996 | 0 |
| ENSMUSG00000095048 | 0 | 0 | 0 | 0 | 0 | 0 | 0 |
| ENSMUSG00000095049 | 0 | 0 | 0 | 0 | 0 | 0 | 0 |

| | | | | | | | |
|---|---|---|---|---|---|---|---|
| ENSMUSG00000095051 | 0 | 0 | 0 | 0 | 0 | 0 | 0 |
| ENSMUSG00000095052 | 0 | 0 | 0 | 0 | 0 | 0 | 0 |
| ENSMUSG00000095053 | 0 | 0 | 0 | 0 | 0 | 0 | 0 |
| ENSMUSG00000095054 | 0 | 0 | 0.25 | 0 | 0 | 0 | 0 |
| ENSMUSG00000095056 | 0 | 0 | 0 | 0 | 0 | 0 | 0 |
| ENSMUSG00000095057 | 0 | 0.25 | 0 | 0.25 | 0 | 0 | 0 |
| ENSMUSG00000095059 | 0 | 0 | 0 | 0 | 0 | 0 | 0 |
| ENSMUSG00000095060 | 0 | 0 | 0 | 0 | 0 | 0 | 0 |
| ENSMUSG00000095061 | 0 | 0 | 0 | 0 | 0 | 0 | 0 |
| ENSMUSG00000095063 | 0 | 0 | 0 | 0 | 0 | 0 | 0 |
| ENSMUSG00000095064 | 0 | 0 | 0 | 0 | 0 | 0 | 0 |
| ENSMUSG00000095065 | 0 | 0 | 0 | 0 | 0 | 0 | 0 |
| ENSMUSG00000095066 | 0 | 0 | 0 | 0 | 0 | 0 | 0 |
| ENSMUSG00000095068 | 0 | 0 | 0.25 | 0 | 0 | 0 | 0 |
| ENSMUSG00000095069 | 0 | 0 | 0 | 0 | 0 | 0 | 0 |
| ENSMUSG00000095070 | 0 | 0 | 0 | 0 | 0 | 0 | 0 |
| ENSMUSG00000095071 | 0 | 0 | 0 | 0 | 0 | 0 | 0 |
| ENSMUSG00000095072 | 0 | 0 | 0 | 0 | 0 | 0 | 0 |
| ENSMUSG00000095073 | 0 | 0 | 0 | 0 | 0 | 0 | 0 |
| ENSMUSG00000095074 | 0 | 0 | 0 | 0 | 0 | 0 | 0 |
| ENSMUSG00000095075 | 0 | 0 | 0 | 0 | 0 | 0 | 0 |
| ENSMUSG00000095076 | 0 | 0 | 0 | 0 | 0 | 0 | 0 |
| ENSMUSG00000095078 | 0 | 0 | 0 | 0 | 0 | 0 | 0 |
| ENSMUSG00000095079 | 0 | 0 | 0 | 0 | 0 | 0 | 0 |
| ENSMUSG00000095081 | 0 | 0 | 0 | 0 | 0 | 0 | 0 |
| ENSMUSG00000095082 | 0 | 0 | 0 | 0 | 0 | 0 | 0 |
| ENSMUSG00000095085 | 0 | 0 | 0 | 0 | 0 | 0 | 0 |
| ENSMUSG00000095086 | 0 | 0 | 0 | 0 | 0 | 0 | 0 |
| ENSMUSG00000095087 | 0 | 0 | 0.396240625 | 0.396240625 | 0.387761935 | 0 | 0.355418052 |
| ENSMUSG00000095088 | 0 | 0 | 0 | 0 | 0 | 0 | 0 |
| ENSMUSG00000095089 | 0 | 0 | 0 | 0 | 0 | 0 | 0 |
| ENSMUSG00000095091 | 0 | 0 | 0 | 0 | 0 | 0 | 0 |
| ENSMUSG00000095092 | 0 | 0 | 0 | 0 | 0 | 0 | 0 |
| ENSMUSG00000095093 | 0 | 0 | 0 | 0 | 0 | 0 | 0 |

| | | | | | | | |
|---|---|---|---|---|---|---|---|
| ENSMUSG00000095094 | 0 | 0 | 0 | 0 | 0 | 0 | 0 |
| ENSMUSG00000095095 | 0 | 0 | 0 | 0 | 0 | 0 | 0 |
| ENSMUSG00000095096 | 0 | 0 | 0 | 0 | 0 | 0 | 0 |
| ENSMUSG00000095097 | 0 | 0 | 0 | 0 | 0 | 0 | 0 |
| ENSMUSG00000095098 | 0.671187741 | 1.04248125 | 1.364857905 | 0.701838731 | 1.163285805 | 0.477121255 | 1.313616061 |
| ENSMUSG00000095100 | 0 | 0 | 0 | 0 | 0 | 0 | 0 |
| ENSMUSG00000095101 | 0 | 0 | 0 | 0 | 0 | 0 | 0 |
| ENSMUSG00000095102 | 0 | 0 | 0 | 0 | 0 | 0 | 0 |
| ENSMUSG00000095103 | 0 | 0 | 0 | 0 | 0 | 0 | 0 |
| ENSMUSG00000095104 | 0 | 0 | 0 | 0 | 0 | 0 | 0 |
| ENSMUSG00000095105 | 0.45815691 | 0.580482024 | 0.580482024 | 1.04248125 | 1.074582839 | 1.113943352 | 1.100339912 |
| ENSMUSG00000095107 | 0 | 0 | 0 | 0 | 0 | 0 | 0 |
| ENSMUSG00000095108 | 0 | 0 | 0 | 0 | 0 | 0 | 0 |
| ENSMUSG00000095110 | 0 | 0 | 0 | 0 | 0 | 0 | 0 |
| ENSMUSG00000095111 | 0 | 0 | 0 | 0 | 0 | 0 | 0 |
| ENSMUSG00000095112 | 0 | 0 | 0 | 0 | 0 | 0 | 0 |
| ENSMUSG00000095113 | 0 | 0 | 0 | 0 | 0 | 0 | 0 |
| ENSMUSG00000095115 | 2.654307229 | 2.491085217 | 2.480088214 | 2.415889526 | 2.357605438 | 2.649334859 | 2.164590871 |
| ENSMUSG00000095116 | 0 | 0 | 0 | 0 | 0 | 0 | 0 |
| ENSMUSG00000095117 | 0 | 0 | 0 | 0 | 0 | 0 | 0 |
| ENSMUSG00000095118 | 0 | 0 | 0 | 0 | 0 | 0 | 0 |
| ENSMUSG00000095119 | 0 | 0 | 0 | 0 | 0 | 0 | 0 |
| ENSMUSG00000095121 | 0 | 0 | 0 | 0 | 0 | 0 | 0 |
| ENSMUSG00000095122 | 0 | 0 | 0 | 0 | 0 | 0 | 0 |
| ENSMUSG00000095123 | 1.777675583 | 1.688721876 | 1.606566189 | 1.78743678 | 1.72073075 | 1.86332286 | 1.621017948 |
| ENSMUSG00000095125 | 0 | 0 | 0 | 0 | 0 | 0 | 0 |
| ENSMUSG00000095126 | 0 | 0 | 0 | 0 | 0 | 0 | 0 |
| ENSMUSG00000095127 | 0 | 0 | 0 | 0 | 0 | 0 | 0 |
| ENSMUSG00000095128 | 0 | 0 | 0 | 0 | 0 | 0 | 0 |
| ENSMUSG00000095129 | 0 | 0 | 0 | 0 | 0 | 0 | 0 |
| ENSMUSG00000095130 | 0 | 0 | 0 | 0 | 0 | 0 | 0 |
| ENSMUSG00000095131 | 0 | 0 | 0 | 0 | 0 | 0 | 0 |
| ENSMUSG00000095132 | 0 | 0 | 0 | 0 | 0 | 0 | 0 |
| ENSMUSG00000095133 | 0 | 0 | 0 | 0 | 0 | 0 | 0 |

| | | | | | | | |
|---|---|---|---|---|---|---|---|
| ENSMUSG00000095134 | 0 | 0 | 0 | 0 | 0 | 0 | 0 |
| ENSMUSG00000095135 | 0 | 0 | 0 | 0 | 0 | 0 | 0 |
| ENSMUSG00000095136 | 0 | 0 | 0 | 0 | 0 | 0 | 0 |
| ENSMUSG00000095137 | 0 | 0 | 0 | 0 | 0 | 0 | 0 |
| ENSMUSG00000095138 | 0 | 0 | 0 | 0 | 0 | 0 | 0 |
| ENSMUSG00000095139 | 0 | 0.25 | 0.396240625 | 0.25 | 0 | 0.301029996 | 0.224243824 |
| ENSMUSG00000095140 | 0 | 0 | 0 | 0 | 0 | 0 | 0 |
| ENSMUSG00000095141 | 0 | 0 | 0 | 0 | 0 | 0 | 0 |
| ENSMUSG00000095142 | 0 | 0 | 0 | 0 | 0 | 0 | 0 |
| ENSMUSG00000095143 | 0 | 0 | 0 | 0 | 0 | 0 | 0 |
| ENSMUSG00000095146 | 0 | 0 | 0 | 0 | 0 | 0 | 0 |
| ENSMUSG00000095147 | 0 | 0 | 0 | 0 | 0 | 0 | 0 |
| ENSMUSG00000095148 | 0 | 0 | 0 | 0 | 0 | 0 | 0 |
| ENSMUSG00000095149 | 0 | 0 | 0 | 0 | 0 | 0 | 0 |
| ENSMUSG00000095150 | 0.289064826 | 0 | 0.25 | 0.25 | 0.387761935 | 0 | 0.448487648 |
| ENSMUSG00000095151 | 0 | 0 | 0 | 0 | 0 | 0 | 0 |
| ENSMUSG00000095152 | 0 | 0 | 0 | 0 | 0 | 0 | 0 |
| ENSMUSG00000095153 | 0 | 0 | 0 | 0 | 0 | 0 | 0 |
| ENSMUSG00000095154 | 0 | 0 | 0 | 0 | 0 | 0 | 0 |
| ENSMUSG00000095155 | 0 | 0 | 0 | 0 | 0 | 0 | 0 |
| ENSMUSG00000095156 | 0 | 0 | 0 | 0 | 0 | 0 | 0 |
| ENSMUSG00000095159 | 0 | 0.25 | 0.25 | 0.25 | 0.387761935 | 0 | 0.355418052 |
| ENSMUSG00000095160 | 0 | 0 | 0 | 0 | 0 | 0 | 0 |
| ENSMUSG00000095161 | 0 | 0 | 0 | 0 | 0 | 0 | 0 |
| ENSMUSG00000095163 | 0 | 0 | 0 | 0 | 0 | 0 | 0 |
| ENSMUSG00000095165 | 0 | 0 | 0 | 0 | 0 | 0 | 0 |
| ENSMUSG00000095168 | 0 | 0.5 | 0.701838731 | 0 | 0.387761935 | 0.301029996 | 0 |
| ENSMUSG00000095169 | 0 | 0 | 0 | 0 | 0 | 0 | 0 |
| ENSMUSG00000095170 | 0 | 0 | 0 | 0 | 0 | 0 | 0 |
| ENSMUSG00000095171 | 0 | 0 | 0 | 0 | 0 | 0 | 0 |
| ENSMUSG00000095172 | 0 | 0 | 0 | 0 | 0 | 0 | 0 |
| ENSMUSG00000095174 | 0 | 0 | 0 | 0 | 0 | 0 | 0 |
| ENSMUSG00000095175 | 0 | 0 | 0 | 0 | 0 | 0 | 0 |
| ENSMUSG00000095176 | 0 | 0 | 0 | 0 | 0 | 0 | 0 |

| | | | | | | | |
|---|---|---|---|---|---|---|---|
| ENSMUSG00000095178 | 0.289064826 | 0.396240625 | 0 | 0 | 0.244650542 | 0 | 0 |
| ENSMUSG00000095179 | 0 | 0 | 0 | 0 | 0 | 0 | 0 |
| ENSMUSG00000095180 | 0 | 0.25 | 0.25 | 0 | 0.244650542 | 0.477121255 | 0.448487648 |
| ENSMUSG00000095182 | 0 | 0 | 0 | 0 | 0 | 0 | 0 |
| ENSMUSG00000095183 | 0 | 0 | 0 | 0 | 0 | 0 | 0 |
| ENSMUSG00000095184 | 0 | 0 | 0.25 | 0 | 0 | 0 | 0 |
| ENSMUSG00000095185 | 0 | 0 | 0 | 0 | 0 | 0 | 0 |
| ENSMUSG00000095186 | 0 | 0 | 0 | 0 | 0 | 0 | 0 |
| ENSMUSG00000095187 | 0 | 0 | 0 | 0 | 0 | 0 | 0 |
| ENSMUSG00000095188 | 0 | 0 | 0 | 0 | 0 | 0 | 0 |
| ENSMUSG00000095189 | 0 | 0 | 0 | 0 | 0 | 0 | 0 |
| ENSMUSG00000095190 | 0 | 0 | 0 | 0 | 0 | 0 | 0 |
| ENSMUSG00000095191 | 0 | 0 | 0 | 0 | 0 | 0 | 0 |
| ENSMUSG00000095192 | 0 | 0 | 0 | 0 | 0 | 0 | 0 |
| ENSMUSG00000095193 | 1.734388958 | 1.606566189 | 1.52186571 | 1.584962501 | 1.559656853 | 1.73239376 | 1.345462945 |
| ENSMUSG00000095194 | 0 | 0 | 0 | 0 | 0 | 0 | 0 |
| ENSMUSG00000095195 | 0 | 0 | 0 | 0 | 0.244650542 | 0 | 0 |
| ENSMUSG00000095196 | 0 | 0 | 0 | 0 | 0 | 0 | 0 |
| ENSMUSG00000095197 | 0 | 0 | 0 | 0 | 0 | 0 | 0 |
| ENSMUSG00000095198 | 0 | 0 | 0 | 0 | 0 | 0 | 0 |
| ENSMUSG00000095199 | 0 | 0 | 0 | 0 | 0 | 0 | 0 |
| ENSMUSG00000095200 | 0 | 0 | 0 | 0 | 0 | 0 | 0 |
| ENSMUSG00000095201 | 0 | 0 | 0 | 0 | 0 | 0 | 0 |
| ENSMUSG00000095202 | 0 | 0 | 0.25 | 0 | 0 | 0 | 0 |
| ENSMUSG00000095203 | 0 | 0 | 0 | 0 | 0 | 0 | 0 |
| ENSMUSG00000095204 | 0 | 0 | 0 | 0 | 0 | 0 | 0 |
| ENSMUSG00000095205 | 0.671187741 | 0.646240625 | 0.396240625 | 0.396240625 | 0.489301084 | 0.301029996 | 0.520678036 |
| ENSMUSG00000095206 | 0 | 0 | 0 | 0 | 0 | 0 | 0 |
| ENSMUSG00000095207 | 0 | 0 | 0 | 0 | 0 | 0 | 0 |
| ENSMUSG00000095209 | 0 | 0 | 0 | 0 | 0 | 0 | 0 |
| ENSMUSG00000095210 | 0 | 0 | 0 | 0 | 0 | 0 | 0 |
| ENSMUSG00000095212 | 0 | 0 | 0 | 0 | 0 | 0 | 0 |
| ENSMUSG00000095213 | 0 | 0 | 0 | 0 | 0 | 0 | 0 |
| ENSMUSG00000095214 | 0 | 0 | 0 | 0 | 0 | 0 | 0 |

| | | | | | | | |
|---|---|---|---|---|---|---|---|
| ENSMUSG00000095217 | 0 | 0 | 0 | 0.25 | 0 | 0 | 0 |
| ENSMUSG00000095218 | 0 | 0 | 0 | 0 | 0 | 0 | 0 |
| ENSMUSG00000095220 | 0 | 0 | 0 | 0 | 0 | 0 | 0 |
| ENSMUSG00000095221 | 0 | 0 | 0 | 0 | 0 | 0 | 0 |
| ENSMUSG00000095222 | 0 | 0 | 0 | 0 | 0 | 0 | 0 |
| ENSMUSG00000095224 | 0 | 0 | 0 | 0 | 0 | 0 | 0 |
| ENSMUSG00000095226 | 0 | 0 | 0 | 0 | 0 | 0 | 0 |
| ENSMUSG00000095228 | 0 | 0 | 0 | 0 | 0 | 0 | 0 |
| ENSMUSG00000095229 | 0 | 0 | 0 | 0 | 0 | 0 | 0 |
| ENSMUSG00000095231 | 0 | 0 | 0 | 0 | 0 | 0 | 0 |
| ENSMUSG00000095232 | 0 | 0 | 0 | 0 | 0 | 0 | 0 |
| ENSMUSG00000095233 | 0 | 0 | 0 | 0 | 0 | 0 | 0 |
| ENSMUSG00000095234 | 0 | 0 | 0 | 0 | 0 | 0 | 0 |
| ENSMUSG00000095236 | 0 | 0 | 0 | 0 | 0 | 0 | 0 |
| ENSMUSG00000095238 | 0 | 0 | 0 | 0 | 0 | 0 | 0 |
| ENSMUSG00000095239 | 0 | 0 | 0 | 0 | 0 | 0 | 0 |
| ENSMUSG00000095240 | 0 | 0 | 0 | 0 | 0 | 0 | 0 |
| ENSMUSG00000095241 | 0 | 0 | 0 | 0 | 0 | 0 | 0 |
| ENSMUSG00000095242 | 0 | 0 | 0 | 0 | 0 | 0 | 0 |
| ENSMUSG00000095243 | 0 | 0 | 0 | 0 | 0 | 0 | 0 |
| ENSMUSG00000095245 | 0 | 0 | 0 | 0 | 0 | 0 | 0 |
| ENSMUSG00000095246 | 0 | 0 | 0 | 0 | 0 | 0 | 0 |
| ENSMUSG00000095247 | 0 | 0 | 0 | 0 | 0 | 0 | 0 |
| ENSMUSG00000095248 | 0 | 0 | 0 | 0 | 0 | 0 | 0 |
| ENSMUSG00000095249 | 0 | 0 | 0 | 0 | 0 | 0 | 0 |
| ENSMUSG00000095250 | 0 | 0 | 0 | 0 | 0 | 0 | 0 |
| ENSMUSG00000095251 | 0 | 0 | 0 | 0 | 0 | 0 | 0 |
| ENSMUSG00000095253 | 1.894701088 | 1.726722649 | 1.735628626 | 1.774508021 | 1.739104362 | 2 | 1.586932193 |
| ENSMUSG00000095254 | 0 | 0 | 0 | 0 | 0 | 0 | 0 |
| ENSMUSG00000095255 | 0.45815691 | 0 | 0 | 0 | 0 | 0 | 0 |
| ENSMUSG00000095256 | 0 | 0 | 0 | 0 | 0 | 0 | 0 |
| ENSMUSG00000095257 | 0 | 0 | 0 | 0 | 0 | 0 | 0 |
| ENSMUSG00000095258 | 0 | 0 | 0 | 0 | 0 | 0 | 0 |
| ENSMUSG00000095259 | 0 | 0 | 0 | 0 | 0 | 0 | 0 |

| | | | | | | | |
|---|---|---|---|---|---|---|---|
| ENSMUSG00000095260 | 0 | 0 | 0 | 0 | 0 | 0 | 0 |
| ENSMUSG00000095261 | 0 | 0 | 0 | 0 | 0 | 0 | 0 |
| ENSMUSG00000095262 | 0 | 0 | 0 | 0 | 0.244650542 | 0 | 0 |
| ENSMUSG00000095263 | 0 | 0 | 0 | 0 | 0 | 0 | 0 |
| ENSMUSG00000095264 | 0 | 0 | 0 | 0 | 0 | 0 | 0 |
| ENSMUSG00000095265 | 0 | 0 | 0 | 0 | 0 | 0 | 0 |
| ENSMUSG00000095266 | 0 | 0 | 0.25 | 0 | 0 | 0 | 0 |
| ENSMUSG00000095267 | 0 | 0 | 0 | 0 | 0 | 0 | 0 |
| ENSMUSG00000095269 | 0 | 0 | 0 | 0 | 0 | 0 | 0 |
| ENSMUSG00000095270 | 0 | 0 | 0 | 0 | 0 | 0 | 0 |
| ENSMUSG00000095272 | 0 | 0 | 0 | 0 | 0 | 0 | 0 |
| ENSMUSG00000095273 | 0 | 0 | 0 | 0 | 0 | 0 | 0 |
| ENSMUSG00000095274 | 0 | 0 | 0 | 0 | 0 | 0 | 0 |
| ENSMUSG00000095275 | 0 | 0 | 0 | 0 | 0 | 0 | 0 |
| ENSMUSG00000095276 | 0 | 0 | 0 | 0 | 0 | 0 | 0 |
| ENSMUSG00000095277 | 0 | 0 | 0 | 0 | 0 | 0 | 0 |
| ENSMUSG00000095278 | 0 | 0 | 0.25 | 0 | 0 | 0 | 0 |
| ENSMUSG00000095279 | 0 | 0 | 0 | 0 | 0 | 0 | 0 |
| ENSMUSG00000095280 | 0 | 0 | 0 | 0 | 0 | 0 | 0 |
| ENSMUSG00000095281 | 0 | 0 | 0 | 0 | 0 | 0 | 0 |
| ENSMUSG00000095282 | 0 | 0 | 0 | 0 | 0 | 0 | 0 |
| ENSMUSG00000095283 | 0 | 0 | 0 | 0 | 0 | 0 | 0 |
| ENSMUSG00000095284 | 0 | 0 | 0 | 0 | 0 | 0 | 0 |
| ENSMUSG00000095285 | 0 | 0 | 0 | 0 | 0 | 0 | 0 |
| ENSMUSG00000095286 | 0 | 0 | 0 | 0 | 0 | 0 | 0 |
| ENSMUSG00000095287 | 0 | 0 | 0 | 0 | 0 | 0 | 0 |
| ENSMUSG00000095288 | 0 | 0 | 0 | 0 | 0 | 0 | 0 |
| ENSMUSG00000095289 | 0 | 0 | 0 | 0 | 0 | 0 | 0 |
| ENSMUSG00000095290 | 0 | 0 | 0 | 0 | 0 | 0 | 0 |
| ENSMUSG00000095291 | 0 | 0 | 0 | 0 | 0 | 0 | 0 |
| ENSMUSG00000095292 | 0 | 0 | 0 | 0 | 0 | 0 | 0 |
| ENSMUSG00000095293 | 0 | 0 | 0 | 0 | 0 | 0 | 0 |
| ENSMUSG00000095294 | 0 | 0 | 0 | 0 | 0 | 0 | 0 |
| ENSMUSG00000095295 | 0 | 0 | 0 | 0 | 0 | 0 | 0 |

| | | | | | | | |
|---|---|---|---|---|---|---|---|
| ENSMUSG00000095296 | 0 | 0 | 0 | 0 | 0 | 0 | 0 |
| ENSMUSG00000095297 | 0 | 0 | 0 | 0 | 0 | 0 | 0 |
| ENSMUSG00000095298 | 0 | 0 | 0 | 0 | 0 | 0 | 0 |
| ENSMUSG00000095299 | 0 | 0 | 0 | 0 | 0 | 0 | 0 |
| ENSMUSG00000095300 | 0 | 0 | 0 | 0 | 0 | 0 | 0 |
| ENSMUSG00000095301 | 0 | 0 | 0 | 0 | 0 | 0 | 0 |
| ENSMUSG00000095302 | 0 | 0 | 0 | 0 | 0 | 0 | 0 |
| ENSMUSG00000095304 | 0 | 0 | 0 | 0 | 0 | 0 | 0 |
| ENSMUSG00000095305 | 0 | 0 | 0 | 0 | 0 | 0 | 0 |
| ENSMUSG00000095306 | 0 | 0 | 0 | 0 | 0.244650542 | 0 | 0 |
| ENSMUSG00000095307 | 0.289064826 | 0 | 0 | 0 | 0 | 0 | 0 |
| ENSMUSG00000095309 | 0 | 0 | 0 | 0 | 0 | 0 | 0 |
| ENSMUSG00000095310 | 0 | 0 | 0 | 0 | 0 | 0 | 0 |
| ENSMUSG00000095311 | 0 | 0 | 0 | 0 | 0 | 0.301029996 | 0 |
| ENSMUSG00000095312 | 0 | 0 | 0 | 0 | 0 | 0 | 0 |
| ENSMUSG00000095314 | 0 | 0 | 0 | 0 | 0 | 0 | 0 |
| ENSMUSG00000095315 | 0 | 0 | 0.396240625 | 0.646240625 | 0.733951626 | 0.477121255 | 0.74492186 |
| ENSMUSG00000095316 | 0 | 0 | 0 | 0 | 0 | 0 | 0 |
| ENSMUSG00000095318 | 0 | 0 | 0 | 0 | 0 | 0 | 0 |
| ENSMUSG00000095319 | 0 | 0 | 0 | 0 | 0 | 0 | 0 |
| ENSMUSG00000095320 | 0 | 0 | 0 | 0 | 0 | 0 | 0 |
| ENSMUSG00000095321 | 0 | 0 | 0 | 0 | 0 | 0 | 0 |
| ENSMUSG00000095322 | 0 | 0 | 0 | 0 | 0 | 0 | 0 |
| ENSMUSG00000095323 | 0 | 0 | 0 | 0 | 0 | 0 | 0 |
| ENSMUSG00000095324 | 0 | 0 | 0 | 0 | 0 | 0 | 0 |
| ENSMUSG00000095325 | 1.587501561 | 1.552363341 | 1.516522298 | 1.774508021 | 1.726078673 | 1.968482949 | 1.610050272 |
| ENSMUSG00000095327 | 0 | 0 | 0 | 0 | 0 | 0 | 0 |
| ENSMUSG00000095328 | 0 | 0 | 0 | 0 | 0 | 0 | 0 |
| ENSMUSG00000095329 | 0 | 0 | 0 | 0 | 0.244650542 | 0 | 0 |
| ENSMUSG00000095330 | 0 | 0 | 0 | 0 | 0 | 0 | 0 |
| ENSMUSG00000095331 | 0 | 0 | 0 | 0 | 0 | 0 | 0 |
| ENSMUSG00000095332 | 0 | 0.580482024 | 0.580482024 | 0.701838731 | 0.632412477 | 0.602059991 | 0.520678036 |
| ENSMUSG00000095333 | 0 | 0 | 0 | 0 | 0 | 0 | 0 |
| ENSMUSG00000095334 | 0 | 0 | 0 | 0 | 0 | 0 | 0 |

| | | | | | | | |
|---|---|---|---|---|---|---|---|
| ENSMUSG00000095335 | 0 | 0 | 0 | 0 | 0 | 0 | 0 |
| ENSMUSG00000095336 | 0 | 0 | 0 | 0 | 0 | 0 | 0 |
| ENSMUSG00000095338 | 0 | 0 | 0 | 0 | 0 | 0 | 0 |
| ENSMUSG00000095339 | 0 | 0 | 0 | 0 | 0 | 0 | 0 |
| ENSMUSG00000095340 | 0 | 0 | 0 | 0 | 0 | 0 | 0 |
| ENSMUSG00000095341 | 0 | 0 | 0 | 0 | 0 | 0 | 0 |
| ENSMUSG00000095342 | 0 | 0 | 0 | 0 | 0 | 0 | 0 |
| ENSMUSG00000095344 | 0 | 0 | 0.396240625 | 0 | 0.387761935 | 0 | 0 |
| ENSMUSG00000095345 | 0 | 0 | 0 | 0 | 0 | 0 | 0 |
| ENSMUSG00000095346 | 0 | 0 | 0 | 0 | 0 | 0 | 0 |
| ENSMUSG00000095347 | 0 | 0 | 0 | 0 | 0 | 0 | 0 |
| ENSMUSG00000095348 | 0 | 0 | 0 | 0 | 0 | 0 | 0 |
| ENSMUSG00000095349 | 0 | 0 | 0 | 0 | 0 | 0 | 0 |
| ENSMUSG00000095350 | 0 | 0 | 0 | 0 | 0 | 0 | 0 |
| ENSMUSG00000095351 | 0 | 0 | 0 | 0 | 0 | 0 | 0 |
| ENSMUSG00000095353 | 0 | 0 | 0 | 0 | 0 | 0 | 0 |
| ENSMUSG00000095356 | 0 | 0 | 0 | 0 | 0 | 0 | 0 |
| ENSMUSG00000095357 | 0 | 0 | 0 | 0 | 0 | 0 | 0 |
| ENSMUSG00000095358 | 0 | 0 | 0 | 0 | 0 | 0 | 0 |
| ENSMUSG00000095360 | 0 | 0 | 0 | 0 | 0 | 0 | 0 |
| ENSMUSG00000095361 | 0 | 0 | 0 | 0 | 0 | 0 | 0 |
| ENSMUSG00000095362 | 1.100572389 | 0.896240625 | 0.75 | 1.04248125 | 1.039257769 | 1.176091259 | 1.054044578 |
| ENSMUSG00000095363 | 0 | 0 | 0 | 0 | 0 | 0 | 0 |
| ENSMUSG00000095365 | 0 | 0 | 0 | 0 | 0 | 0 | 0 |
| ENSMUSG00000095366 | 0 | 0 | 0 | 0 | 0 | 0 | 0 |
| ENSMUSG00000095367 | 0 | 0 | 0 | 0 | 0 | 0 | 0 |
| ENSMUSG00000095368 | 0 | 0 | 0 | 0 | 0 | 0 | 0 |
| ENSMUSG00000095369 | 0 | 0 | 0 | 0 | 0 | 0 | 0 |
| ENSMUSG00000095370 | 0 | 0 | 0 | 0 | 0 | 0 | 0 |
| ENSMUSG00000095371 | 0 | 0 | 0 | 0 | 0 | 0 | 0 |
| ENSMUSG00000095372 | 0 | 0 | 0 | 0 | 0 | 0 | 0 |
| ENSMUSG00000095373 | 0 | 0 | 0 | 0 | 0 | 0 | 0 |
| ENSMUSG00000095374 | 0 | 0 | 0 | 0 | 0 | 0 | 0 |
| ENSMUSG00000095376 | 0 | 0 | 0 | 0 | 0 | 0 | 0 |

| | | | | | | | |
|---|---|---|---|---|---|---|---|
| ENSMUSG00000095377 | 0 | 0 | 0 | 0 | 0.244650542 | 0 | 0 |
| ENSMUSG00000095379 | 0 | 0 | 0 | 0 | 0 | 0 | 0 |
| ENSMUSG00000095380 | 0 | 0 | 0 | 0 | 0 | 0 | 0 |
| ENSMUSG00000095382 | 0 | 0 | 0 | 0 | 0 | 0 | 0 |
| ENSMUSG00000095383 | 0 | 0 | 0 | 0 | 0 | 0 | 0 |
| ENSMUSG00000095384 | 0 | 0 | 0 | 0 | 0 | 0 | 0 |
| ENSMUSG00000095385 | 0 | 0 | 0.25 | 0 | 0 | 0 | 0 |
| ENSMUSG00000095386 | 0 | 0 | 0 | 0 | 0 | 0 | 0 |
| ENSMUSG00000095387 | 0 | 0 | 0 | 0 | 0 | 0 | 0 |
| ENSMUSG00000095388 | 0 | 0 | 0 | 0 | 0 | 0 | 0 |
| ENSMUSG00000095389 | 0 | 0 | 0 | 0 | 0 | 0 | 0 |
| ENSMUSG00000095390 | 0 | 0 | 0 | 0 | 0 | 0 | 0 |
| ENSMUSG00000095392 | 0 | 0 | 0 | 0 | 0 | 0 | 0 |
| ENSMUSG00000095393 | 0 | 0 | 0 | 0 | 0 | 0 | 0 |
| ENSMUSG00000095394 | 0 | 0 | 0 | 0 | 0 | 0 | 0 |
| ENSMUSG00000095395 | 0 | 0 | 0 | 0 | 0 | 0 | 0 |
| ENSMUSG00000095396 | 0 | 0 | 0 | 0 | 0 | 0.301029996 | 0 |
| ENSMUSG00000095398 | 0 | 0 | 0 | 0 | 0 | 0 | 0 |
| ENSMUSG00000095399 | 0 | 0 | 0 | 0 | 0 | 0 | 0 |
| ENSMUSG00000095401 | 0 | 0 | 0 | 0 | 0 | 0 | 0 |
| ENSMUSG00000095402 | 0 | 0 | 0 | 0 | 0 | 0 | 0 |
| ENSMUSG00000095403 | 2.163307908 | 2.087182039 | 2.075945187 | 2.240724001 | 2.176121988 | 2.411619706 | 2.018825668 |
| ENSMUSG00000095406 | 0 | 0 | 0.25 | 0 | 0 | 0 | 0 |
| ENSMUSG00000095407 | 0 | 0 | 0 | 0 | 0 | 0 | 0 |
| ENSMUSG00000095408 | 0 | 0 | 0 | 0 | 0 | 0 | 0 |
| ENSMUSG00000095409 | 0 | 0 | 0 | 0 | 0 | 0 | 0 |
| ENSMUSG00000095410 | 0 | 0 | 0 | 0 | 0 | 0 | 0 |
| ENSMUSG00000095411 | 0 | 0 | 0 | 0 | 0 | 0 | 0 |
| ENSMUSG00000095412 | 0 | 0 | 0 | 0 | 0 | 0 | 0 |
| ENSMUSG00000095413 | 0 | 0 | 0 | 0 | 0 | 0 | 0 |
| ENSMUSG00000095414 | 0 | 0 | 0 | 0 | 0 | 0 | 0 |
| ENSMUSG00000095416 | 0 | 0 | 0 | 0 | 0 | 0 | 0 |
| ENSMUSG00000095418 | 0 | 0 | 0 | 0 | 0 | 0 | 0 |
| ENSMUSG00000095419 | 0 | 0 | 0 | 0 | 0 | 0 | 0.224243824 |

| | | | | | | | |
|---|---|---|---|---|---|---|---|
| ENSMUSG00000095420 | 0 | 0 | 0 | 0 | 0 | 0 | 0 |
| ENSMUSG00000095422 | 0 | 0 | 0 | 0 | 0 | 0 | 0 |
| ENSMUSG00000095423 | 0 | 0 | 0 | 0 | 0 | 0 | 0 |
| ENSMUSG00000095424 | 0 | 0 | 0 | 0 | 0 | 0 | 0 |
| ENSMUSG00000095425 | 0 | 0 | 0 | 0 | 0 | 0 | 0 |
| ENSMUSG00000095426 | 0 | 0 | 0 | 0 | 0 | 0 | 0 |
| ENSMUSG00000095427 | 0 | 0.25 | 0 | 0.25 | 0.244650542 | 0.301029996 | 0 |
| ENSMUSG00000095428 | 0 | 0 | 0 | 0 | 0 | 0 | 0 |
| ENSMUSG00000095429 | 0 | 0 | 0 | 0 | 0 | 0 | 0 |
| ENSMUSG00000095430 | 0 | 0 | 0 | 0 | 0 | 0 | 0 |
| ENSMUSG00000095431 | 0 | 0 | 0 | 0 | 0 | 0 | 0 |
| ENSMUSG00000095432 | 1.857607105 | 1.845926073 | 1.809601185 | 2.072504712 | 1.992595054 | 2.269512944 | 1.815933683 |
| ENSMUSG00000095433 | 0 | 0 | 0 | 0 | 0 | 0 | 0 |
| ENSMUSG00000095434 | 0 | 0 | 0 | 0 | 0 | 0 | 0 |
| ENSMUSG00000095435 | 0 | 0 | 0 | 0 | 0 | 0 | 0 |
| ENSMUSG00000095436 | 0 | 0 | 0 | 0 | 0 | 0 | 0 |
| ENSMUSG00000095437 | 0 | 0 | 0 | 0 | 0 | 0 | 0 |
| ENSMUSG00000095438 | 0.289064826 | 0.25 | 0 | 0.25 | 0 | 0.301029996 | 0.224243824 |
| ENSMUSG00000095440 | 0.671187741 | 1.080482024 | 0.951838731 | 1.321350555 | 1.223252711 | 1.322219295 | 1.296434211 |
| ENSMUSG00000095442 | 0 | 0 | 0 | 0 | 0 | 0 | 0 |
| ENSMUSG00000095444 | 0 | 0 | 0 | 0 | 0 | 0 | 0 |
| ENSMUSG00000095445 | 0 | 0 | 0 | 0 | 0 | 0 | 0 |
| ENSMUSG00000095446 | 0 | 0 | 0 | 0 | 0 | 0 | 0 |
| ENSMUSG00000095447 | 0 | 0 | 0 | 0 | 0 | 0 | 0 |
| ENSMUSG00000095448 | 0 | 0 | 0 | 0 | 0 | 0 | 0 |
| ENSMUSG00000095449 | 0 | 0 | 0 | 0 | 0 | 0 | 0 |
| ENSMUSG00000095450 | 0 | 0 | 0 | 0 | 0 | 0 | 0 |
| ENSMUSG00000095452 | 0 | 0 | 0 | 0 | 0 | 0 | 0 |
| ENSMUSG00000095453 | 0 | 0 | 0 | 0 | 0 | 0 | 0 |
| ENSMUSG00000095455 | 0 | 0 | 0 | 0 | 0 | 0 | 0 |
| ENSMUSG00000095456 | 0 | 0 | 0 | 0 | 0 | 0 | 0 |
| ENSMUSG00000095457 | 0 | 0 | 0 | 0 | 0 | 0 | 0 |
| ENSMUSG00000095458 | 0 | 0 | 0 | 0 | 0 | 0 | 0 |
| ENSMUSG00000095460 | 0 | 0 | 0 | 0 | 0 | 0 | 0 |

| | | | | | | | |
|---|---|---|---|---|---|---|---|
| ENSMUSG00000095462 | 0 | 0 | 0 | 0 | 0 | 0 | 0 |
| ENSMUSG00000095463 | 0 | 0 | 0 | 0 | 0 | 0 | 0 |
| ENSMUSG00000095464 | 0 | 0 | 0 | 0 | 0 | 0 | 0 |
| ENSMUSG00000095465 | 0 | 0 | 0 | 0 | 0 | 0 | 0 |
| ENSMUSG00000095466 | 0 | 0 | 0 | 0 | 0 | 0 | 0 |
| ENSMUSG00000095467 | 0 | 0 | 0 | 0 | 0 | 0 | 0 |
| ENSMUSG00000095469 | 0 | 0 | 0 | 0 | 0 | 0 | 0 |
| ENSMUSG00000095470 | 0 | 0 | 0 | 0 | 0 | 0 | 0 |
| ENSMUSG00000095472 | 0 | 0 | 0 | 0 | 0 | 0 | 0 |
| ENSMUSG00000095474 | 0 | 0 | 0 | 0 | 0 | 0 | 0 |
| ENSMUSG00000095475 | 0 | 0 | 0 | 0 | 0 | 0 | 0 |
| ENSMUSG00000095476 | 0 | 0 | 0 | 0 | 0 | 0 | 0 |
| ENSMUSG00000095477 | 0 | 0 | 0.25 | 0 | 0 | 0 | 0.224243824 |
| ENSMUSG00000095478 | 0 | 0 | 0 | 0 | 0 | 0 | 0.224243824 |
| ENSMUSG00000095479 | 0 | 0 | 0 | 0 | 0 | 0 | 0 |
| ENSMUSG00000095482 | 0 | 0 | 0 | 0 | 0 | 0 | 0 |
| ENSMUSG00000095483 | 0 | 0 | 0 | 0 | 0 | 0 | 0 |
| ENSMUSG00000095484 | 0 | 0 | 0 | 0 | 0 | 0 | 0 |
| ENSMUSG00000095486 | 0 | 0 | 0 | 0 | 0 | 0 | 0 |
| ENSMUSG00000095487 | 0 | 0 | 0 | 0 | 0 | 0 | 0 |
| ENSMUSG00000095489 | 0 | 0 | 0 | 0 | 0 | 0 | 0 |
| ENSMUSG00000095490 | 0 | 0 | 0 | 0 | 0 | 0 | 0 |
| ENSMUSG00000095491 | 0 | 0 | 0 | 0 | 0 | 0 | 0 |
| ENSMUSG00000095492 | 0 | 0 | 0 | 0 | 0 | 0 | 0 |
| ENSMUSG00000095493 | 0.289064826 | 0 | 0.396240625 | 0.25 | 0 | 0 | 0.224243824 |
| ENSMUSG00000095494 | 0 | 0 | 0 | 0 | 0 | 0 | 0 |
| ENSMUSG00000095495 | 0 | 0 | 0 | 0 | 0 | 0 | 0 |
| ENSMUSG00000095497 | 0 | 0 | 0 | 0 | 0 | 0 | 0 |
| ENSMUSG00000095498 | 0 | 0 | 0 | 0 | 0 | 0 | 0 |
| ENSMUSG00000095499 | 0 | 0 | 0 | 0 | 0 | 0 | 0 |
| ENSMUSG00000095500 | 0 | 0 | 0 | 0 | 0 | 0 | 0 |
| ENSMUSG00000095501 | 0 | 0 | 0 | 0 | 0 | 0 | 0 |
| ENSMUSG00000095502 | 0 | 0 | 0 | 0 | 0 | 0 | 0 |
| ENSMUSG00000095503 | 0 | 0 | 0 | 0 | 0 | 0 | 0 |

| | | | | | | | |
|---|---|---|---|---|---|---|---|
| ENSMUSG00000095504 | 0 | 0 | 0 | 0 | 0 | 0 | 0 |
| ENSMUSG00000095505 | 0 | 0 | 0 | 0 | 0 | 0 | 0 |
| ENSMUSG00000095506 | 0 | 0 | 0 | 0 | 0 | 0 | 0 |
| ENSMUSG00000095507 | 0 | 0 | 0 | 0 | 0 | 0 | 0 |
| ENSMUSG00000095508 | 0 | 0 | 0 | 0 | 0 | 0 | 0 |
| ENSMUSG00000095509 | 0 | 0 | 0 | 0 | 0 | 0 | 0 |
| ENSMUSG00000095510 | 0 | 0 | 0 | 0 | 0 | 0 | 0 |
| ENSMUSG00000095511 | 0 | 0 | 0 | 0 | 0 | 0 | 0 |
| ENSMUSG00000095512 | 0 | 0 | 0 | 0 | 0 | 0 | 0 |
| ENSMUSG00000095513 | 0 | 0 | 0 | 0 | 0 | 0 | 0 |
| ENSMUSG00000095514 | 0.289064826 | 0.25 | 0 | 0 | 0 | 0 | 0 |
| ENSMUSG00000095516 | 0 | 0 | 0 | 0 | 0 | 0 | 0 |
| ENSMUSG00000095517 | 0 | 0 | 0 | 0 | 0 | 0 | 0 |
| ENSMUSG00000095518 | 0 | 0 | 0 | 0 | 0 | 0 | 0 |
| ENSMUSG00000095519 | 0 | 0 | 0 | 0 | 0 | 0 | 0 |
| ENSMUSG00000095520 | 0 | 0 | 0 | 0 | 0 | 0 | 0 |
| ENSMUSG00000095521 | 0 | 0 | 0 | 0 | 0 | 0 | 0 |
| ENSMUSG00000095522 | 0 | 0 | 0 | 0 | 0 | 0 | 0 |
| ENSMUSG00000095523 | 0 | 0 | 0 | 0 | 0 | 0 | 0 |
| ENSMUSG00000095524 | 0.45815691 | 0.396240625 | 0.5 | 0.25 | 0.387761935 | 0 | 0.629532004 |
| ENSMUSG00000095525 | 0 | 0 | 0 | 0 | 0 | 0 | 0 |
| ENSMUSG00000095526 | 0 | 0 | 0 | 0 | 0 | 0 | 0 |
| ENSMUSG00000095527 | 0 | 0 | 0 | 0 | 0 | 0 | 0 |
| ENSMUSG00000095528 | 0 | 0 | 0 | 0 | 0 | 0 | 0 |
| ENSMUSG00000095529 | 0 | 0 | 0 | 0 | 0 | 0 | 0 |
| ENSMUSG00000095530 | 0 | 0.25 | 0 | 0.25 | 0 | 0 | 0 |
| ENSMUSG00000095531 | 0 | 0 | 0 | 0 | 0 | 0 | 0 |
| ENSMUSG00000095532 | 0 | 0 | 0 | 0 | 0 | 0 | 0 |
| ENSMUSG00000095533 | 0 | 0 | 0 | 0 | 0 | 0 | 0 |
| ENSMUSG00000095534 | 0 | 0 | 0 | 0 | 0 | 0 | 0 |
| ENSMUSG00000095536 | 0 | 0 | 0 | 0 | 0 | 0 | 0 |
| ENSMUSG00000095537 | 0 | 0 | 0 | 0 | 0 | 0 | 0 |
| ENSMUSG00000095538 | 0 | 0 | 0 | 0 | 0 | 0 | 0 |
| ENSMUSG00000095539 | 0 | 0 | 0 | 0 | 0 | 0 | 0 |

| | | | | | | | |
|---|---|---|---|---|---|---|---|
| ENSMUSG00000095540 | 0 | 0 | 0 | 0 | 0 | 0 | 0 |
| ENSMUSG00000095541 | 0 | 0 | 0 | 0 | 0 | 0 | 0 |
| ENSMUSG00000095542 | 0 | 0 | 0 | 0 | 0 | 0 | 0 |
| ENSMUSG00000095543 | 0 | 0 | 0 | 0 | 0 | 0 | 0 |
| ENSMUSG00000095544 | 0 | 0 | 0 | 0 | 0 | 0 | 0 |
| ENSMUSG00000095545 | 0 | 0 | 0 | 0 | 0 | 0 | 0 |
| ENSMUSG00000095546 | 0 | 0 | 0 | 0 | 0 | 0 | 0 |
| ENSMUSG00000095547 | 0 | 0 | 0 | 0 | 0 | 0 | 0 |
| ENSMUSG00000095549 | 0 | 0 | 0 | 0 | 0 | 0 | 0 |
| ENSMUSG00000095550 | 0 | 0 | 0 | 0 | 0 | 0 | 0 |
| ENSMUSG00000095551 | 0 | 0 | 0 | 0 | 0 | 0 | 0 |
| ENSMUSG00000095552 | 0 | 0 | 0 | 0 | 0 | 0 | 0 |
| ENSMUSG00000095553 | 0 | 0 | 0 | 0 | 0 | 0 | 0 |
| ENSMUSG00000095554 | 0 | 0 | 0 | 0 | 0 | 0 | 0 |
| ENSMUSG00000095555 | 0 | 0 | 0 | 0 | 0 | 0 | 0 |
| ENSMUSG00000095557 | 0 | 0 | 0 | 0 | 0 | 0 | 0 |
| ENSMUSG00000095558 | 0 | 0 | 0 | 0 | 0 | 0 | 0 |
| ENSMUSG00000095559 | 0 | 0 | 0 | 0 | 0 | 0 | 0 |
| ENSMUSG00000095560 | 0 | 0 | 0 | 0 | 0 | 0 | 0 |
| ENSMUSG00000095561 | 0 | 0 | 0 | 0 | 0 | 0 | 0 |
| ENSMUSG00000095562 | 0 | 0 | 0 | 0 | 0 | 0 | 0.224243824 |
| ENSMUSG00000095563 | 0 | 0 | 0 | 0 | 0 | 0 | 0 |
| ENSMUSG00000095564 | 0 | 0 | 0 | 0 | 0 | 0 | 0 |
| ENSMUSG00000095565 | 0 | 0 | 0 | 0 | 0 | 0 | 0 |
| ENSMUSG00000095567 | 2.569423023 | 2.413211243 | 2.342126615 | 2.293106877 | 2.32316608 | 2.588831726 | 2.17331363 |
| ENSMUSG00000095569 | 0 | 0 | 0 | 0 | 0 | 0 | 0 |
| ENSMUSG00000095570 | 0 | 0 | 0 | 0 | 0 | 0 | 0 |
| ENSMUSG00000095571 | 0 | 0 | 0 | 0 | 0 | 0 | 0 |
| ENSMUSG00000095572 | 0 | 0 | 0 | 0 | 0 | 0 | 0 |
| ENSMUSG00000095573 | 0 | 0 | 0 | 0 | 0 | 0 | 0 |
| ENSMUSG00000095574 | 0 | 0 | 0 | 0 | 0 | 0 | 0 |
| ENSMUSG00000095576 | 0 | 0 | 0 | 0 | 0 | 0 | 0 |
| ENSMUSG00000095577 | 0 | 0 | 0 | 0 | 0 | 0 | 0 |
| ENSMUSG00000095578 | 0 | 0 | 0 | 0 | 0 | 0 | 0 |

| | | | | | | | |
|---|---|---|---|---|---|---|---|
| ENSMUSG00000095580 | 0 | 0 | 0 | 0 | 0 | 0 | 0 |
| ENSMUSG00000095581 | 0 | 0 | 0 | 0 | 0 | 0 | 0 |
| ENSMUSG00000095583 | 0 | 0 | 0 | 0 | 0 | 0 | 0 |
| ENSMUSG00000095584 | 0 | 0 | 0 | 0 | 0 | 0 | 0 |
| ENSMUSG00000095585 | 0 | 0 | 0 | 0 | 0 | 0 | 0 |
| ENSMUSG00000095586 | 0 | 0 | 0 | 0 | 0 | 0 | 0 |
| ENSMUSG00000095587 | 0 | 0 | 0 | 0 | 0 | 0 | 0 |
| ENSMUSG00000095588 | 0 | 0 | 0 | 0 | 0 | 0 | 0.224243824 |
| ENSMUSG00000095589 | 0 | 0 | 0 | 0 | 0 | 0 | 0 |
| ENSMUSG00000095590 | 0 | 0 | 0 | 0 | 0 | 0 | 0 |
| ENSMUSG00000095591 | 0 | 0 | 0 | 0 | 0 | 0 | 0 |
| ENSMUSG00000095592 | 0 | 0 | 0 | 0 | 0 | 0 | 0 |
| ENSMUSG00000095593 | 0 | 0 | 0 | 0 | 0 | 0 | 0 |
| ENSMUSG00000095594 | 0 | 0 | 0 | 0 | 0 | 0 | 0 |
| ENSMUSG00000095595 | 2.205378646 | 2.049918086 | 2.015173983 | 2.411864607 | 2.29993025 | 2.673020907 | 2.135160095 |
| ENSMUSG00000095596 | 0 | 0 | 0 | 0 | 0 | 0 | 0 |
| ENSMUSG00000095597 | 1.940852214 | 1.901832578 | 1.973704441 | 1.146240625 | 1.47337555 | 1.447158031 | 1.452143229 |
| ENSMUSG00000095599 | 0 | 0 | 0 | 0 | 0 | 0 | 0 |
| ENSMUSG00000095600 | 0 | 0.25 | 0 | 0.396240625 | 0 | 0 | 0.224243824 |
| ENSMUSG00000095601 | 0 | 0 | 0 | 0 | 0 | 0 | 0 |
| ENSMUSG00000095602 | 0 | 0 | 0 | 0 | 0 | 0 | 0 |
| ENSMUSG00000095603 | 0 | 0 | 0 | 0 | 0 | 0 | 0 |
| ENSMUSG00000095604 | 0 | 0 | 0 | 0 | 0 | 0 | 0 |
| ENSMUSG00000095605 | 0 | 0 | 0 | 0 | 0 | 0 | 0 |
| ENSMUSG00000095606 | 0 | 0 | 0 | 0 | 0 | 0 | 0 |
| ENSMUSG00000095607 | 0 | 0 | 0 | 0 | 0 | 0 | 0 |
| ENSMUSG00000095608 | 0 | 0 | 0 | 0 | 0 | 0 | 0 |
| ENSMUSG00000095609 | 0 | 0 | 0 | 0 | 0 | 0 | 0 |
| ENSMUSG00000095610 | 0 | 0 | 0 | 0 | 0 | 0 | 0 |
| ENSMUSG00000095612 | 0 | 0 | 0 | 0 | 0 | 0 | 0 |
| ENSMUSG00000095614 | 0 | 0 | 0 | 0 | 0.244650542 | 0 | 0.224243824 |
| ENSMUSG00000095615 | 0 | 0 | 0 | 0 | 0 | 0 | 0 |
| ENSMUSG00000095616 | 0 | 0 | 0 | 0 | 0 | 0 | 0 |
| ENSMUSG00000095618 | 0 | 0 | 0 | 0 | 0 | 0 | 0 |

| | | | | | | | |
|---|---|---|---|---|---|---|---|
| ENSMUSG00000095619 | 0 | 0 | 0 | 0 | 0 | 0 | 0 |
| ENSMUSG00000095620 | 0 | 0 | 0 | 0.25 | 0 | 0 | 0 |
| ENSMUSG00000095621 | 0 | 0 | 0 | 0 | 0 | 0 | 0 |
| ENSMUSG00000095623 | 0 | 0 | 0 | 0 | 0 | 0 | 0 |
| ENSMUSG00000095624 | 0 | 0 | 0 | 0 | 0 | 0 | 0 |
| ENSMUSG00000095625 | 0 | 0 | 0 | 0 | 0 | 0 | 0 |
| ENSMUSG00000095626 | 0 | 0 | 0 | 0 | 0 | 0 | 0 |
| ENSMUSG00000095627 | 0 | 0 | 0 | 0 | 0 | 0 | 0 |
| ENSMUSG00000095628 | 0 | 0 | 0 | 0 | 0 | 0 | 0 |
| ENSMUSG00000095629 | 0 | 0 | 0 | 0 | 0 | 0 | 0 |
| ENSMUSG00000095630 | 0 | 0 | 0 | 0 | 0 | 0 | 0 |
| ENSMUSG00000095631 | 0 | 0 | 0 | 0 | 0 | 0 | 0 |
| ENSMUSG00000095632 | 0 | 0 | 0 | 0 | 0 | 0 | 0 |
| ENSMUSG00000095633 | 0 | 0 | 0 | 0 | 0 | 0 | 0 |
| ENSMUSG00000095634 | 0 | 0 | 0 | 0 | 0 | 0 | 0 |
| ENSMUSG00000095635 | 0 | 0.25 | 0 | 0 | 0 | 0 | 0 |
| ENSMUSG00000095636 | 0 | 0 | 0 | 0 | 0 | 0 | 0 |
| ENSMUSG00000095637 | 0 | 0 | 0 | 0 | 0 | 0 | 0 |
| ENSMUSG00000095638 | 0 | 0 | 0 | 0 | 0 | 0 | 0 |
| ENSMUSG00000095639 | 0 | 0 | 0 | 0 | 0 | 0 | 0 |
| ENSMUSG00000095640 | 0 | 0 | 0 | 0 | 0 | 0 | 0 |
| ENSMUSG00000095641 | 0 | 0 | 0 | 0 | 0 | 0 | 0 |
| ENSMUSG00000095642 | 0 | 0 | 0 | 0 | 0 | 0 | 0 |
| ENSMUSG00000095643 | 0 | 0 | 0 | 0 | 0 | 0 | 0 |
| ENSMUSG00000095645 | 0 | 0 | 0 | 0 | 0 | 0 | 0 |
| ENSMUSG00000095646 | 0 | 0 | 0 | 0 | 0 | 0 | 0 |
| ENSMUSG00000095647 | 0 | 0 | 0 | 0 | 0 | 0 | 0 |
| ENSMUSG00000095648 | 0 | 0 | 0 | 0 | 0 | 0 | 0 |
| ENSMUSG00000095649 | 0 | 0 | 0 | 0 | 0 | 0 | 0 |
| ENSMUSG00000095650 | 0 | 0 | 0 | 0 | 0 | 0 | 0 |
| ENSMUSG00000095651 | 0.671187741 | 0.75 | 0.646240625 | 0.701838731 | 0.877063019 | 0.698970004 | 0.896975297 |
| ENSMUSG00000095652 | 0 | 0 | 0 | 0 | 0 | 0 | 0 |
| ENSMUSG00000095653 | 0 | 0.25 | 0 | 0 | 0 | 0 | 0 |
| ENSMUSG00000095654 | 0 | 0 | 0 | 0 | 0 | 0 | 0 |

| | | | | | | | |
|---|---|---|---|---|---|---|---|
| ENSMUSG00000095655 | 0 | 0 | 0 | 0 | 0 | 0 | 0 |
| ENSMUSG00000095656 | 0 | 0 | 0 | 0 | 0 | 0 | 0 |
| ENSMUSG00000095657 | 0 | 0 | 0 | 0 | 0 | 0 | 0 |
| ENSMUSG00000095658 | 0 | 0 | 0 | 0 | 0 | 0 | 0 |
| ENSMUSG00000095659 | 0 | 0 | 0 | 0 | 0 | 0 | 0 |
| ENSMUSG00000095660 | 0 | 0 | 0 | 0 | 0 | 0 | 0 |
| ENSMUSG00000095661 | 0 | 0 | 0 | 0.5 | 0 | 0 | 0 |
| ENSMUSG00000095662 | 0 | 0 | 0 | 0 | 0 | 0 | 0 |
| ENSMUSG00000095663 | 0 | 0 | 0 | 0 | 0 | 0 | 0 |
| ENSMUSG00000095664 | 0 | 0 | 0 | 0 | 0 | 0 | 0 |
| ENSMUSG00000095666 | 0 | 0 | 0 | 0 | 0 | 0 | 0 |
| ENSMUSG00000095667 | 0 | 0 | 0 | 0 | 0 | 0 | 0 |
| ENSMUSG00000095668 | 0 | 0 | 0 | 0 | 0 | 0 | 0 |
| ENSMUSG00000095670 | 0 | 0 | 0 | 0 | 0 | 0 | 0 |
| ENSMUSG00000095671 | 0 | 0 | 0 | 0 | 0 | 0 | 0 |
| ENSMUSG00000095672 | 0 | 0 | 0 | 0 | 0 | 0 | 0 |
| ENSMUSG00000095673 | 0 | 0 | 0 | 0 | 0 | 0 | 0 |
| ENSMUSG00000095674 | 0 | 0 | 0 | 0 | 0 | 0 | 0 |
| ENSMUSG00000095675 | 0 | 0 | 0 | 0 | 0 | 0 | 0 |
| ENSMUSG00000095676 | 0 | 0 | 0 | 0 | 0 | 0 | 0 |
| ENSMUSG00000095677 | 1.800532393 | 1.593759858 | 1.575945187 | 1.566696635 | 1.625423019 | 1.681241237 | 1.554175045 |
| ENSMUSG00000095678 | 0 | 0 | 0 | 0 | 0 | 0 | 0 |
| ENSMUSG00000095680 | 0 | 0 | 0 | 0 | 0 | 0 | 0 |
| ENSMUSG00000095681 | 0 | 0 | 0 | 0 | 0 | 0 | 0 |
| ENSMUSG00000095682 | 0 | 0 | 0 | 0 | 0 | 0 | 0 |
| ENSMUSG00000095683 | 0 | 0 | 0 | 0 | 0 | 0 | 0 |
| ENSMUSG00000095684 | 0 | 0 | 0 | 0 | 0 | 0 | 0 |
| ENSMUSG00000095685 | 0 | 0 | 0 | 0 | 0 | 0 | 0 |
| ENSMUSG00000095686 | 0 | 0 | 0 | 0 | 0 | 0 | 0 |
| ENSMUSG00000095687 | 1.069666964 | 1.114857905 | 1.282320754 | 0.864857905 | 1.039257769 | 1.230448921 | 1.078019652 |
| ENSMUSG00000095689 | 0 | 0 | 0 | 0 | 0 | 0 | 0 |
| ENSMUSG00000095690 | 0 | 0 | 0 | 0 | 0 | 0 | 0 |
| ENSMUSG00000095691 | 0 | 0 | 0 | 0 | 0 | 0 | 0 |
| ENSMUSG00000095692 | 0 | 0 | 0 | 0 | 0 | 0 | 0 |

| | | | | | | | |
|---|---|---|---|---|---|---|---|
| ENSMUSG00000095693 | 0 | 0 | 0 | 0 | 0 | 0 | 0 |
| ENSMUSG00000095696 | 0 | 0 | 0 | 0 | 0 | 0 | 0 |
| ENSMUSG00000095697 | 0 | 0 | 0 | 0 | 0 | 0 | 0 |
| ENSMUSG00000095698 | 0 | 0 | 0 | 0 | 0 | 0 | 0 |
| ENSMUSG00000095699 | 0 | 0 | 0 | 0 | 0 | 0 | 0 |
| ENSMUSG00000095700 | 0 | 0 | 0 | 0 | 0 | 0 | 0 |
| ENSMUSG00000095701 | 0 | 0 | 0 | 0 | 0 | 0 | 0 |
| ENSMUSG00000095702 | 0 | 0 | 0 | 0 | 0 | 0 | 0 |
| ENSMUSG00000095703 | 0 | 0 | 0 | 0 | 0 | 0 | 0 |
| ENSMUSG00000095704 | 0 | 0 | 0 | 0 | 0 | 0 | 0 |
| ENSMUSG00000095705 | 0 | 0 | 0 | 0 | 0 | 0 | 0 |
| ENSMUSG00000095706 | 0 | 0 | 0 | 0 | 0 | 0 | 0 |
| ENSMUSG00000095709 | 0 | 0 | 0 | 0 | 0 | 0 | 0 |
| ENSMUSG00000095710 | 0 | 0 | 0 | 0 | 0 | 0 | 0 |
| ENSMUSG00000095711 | 0 | 0 | 0 | 0 | 0 | 0 | 0 |
| ENSMUSG00000095712 | 0 | 0 | 0 | 0 | 0 | 0 | 0 |
| ENSMUSG00000095713 | 0 | 0 | 0 | 0 | 0 | 0 | 0 |
| ENSMUSG00000095714 | 0 | 0 | 0 | 0 | 0 | 0 | 0 |
| ENSMUSG00000095716 | 0 | 0 | 0 | 0 | 0 | 0 | 0 |
| ENSMUSG00000095717 | 0 | 0 | 0 | 0 | 0 | 0 | 0 |
| ENSMUSG00000095718 | 0 | 0 | 0 | 0 | 0 | 0 | 0 |
| ENSMUSG00000095720 | 0 | 0 | 0 | 0 | 0 | 0 | 0 |
| ENSMUSG00000095721 | 0 | 0 | 0 | 0 | 0 | 0 | 0 |
| ENSMUSG00000095722 | 0 | 0 | 0 | 0 | 0 | 0 | 0 |
| ENSMUSG00000095723 | 0 | 0 | 0 | 0 | 0 | 0 | 0 |
| ENSMUSG00000095724 | 0 | 0.25 | 0.25 | 0.5 | 0.244650542 | 0.301029996 | 0 |
| ENSMUSG00000095725 | 0 | 0 | 0 | 0 | 0 | 0 | 0 |
| ENSMUSG00000095727 | 0 | 0 | 0 | 0 | 0 | 0 | 0 |
| ENSMUSG00000095728 | 0 | 0 | 0 | 0 | 0 | 0 | 0 |
| ENSMUSG00000095729 | 0 | 0 | 0 | 0 | 0 | 0 | 0 |
| ENSMUSG00000095730 | 0.289064826 | 0.25 | 0 | 0 | 0 | 0 | 0 |
| ENSMUSG00000095731 | 0 | 0 | 0 | 0 | 0 | 0 | 0 |
| ENSMUSG00000095732 | 0 | 0 | 0 | 0 | 0 | 0 | 0 |
| ENSMUSG00000095734 | 0 | 0 | 0 | 0 | 0 | 0 | 0 |

| | | | | | | | |
|---|---|---|---|---|---|---|---|
| ENSMUSG00000095736 | 0 | 0 | 0 | 0 | 0 | 0 | 0 |
| ENSMUSG00000095737 | 0 | 0 | 0 | 0 | 0 | 0 | 0 |
| ENSMUSG00000095738 | 0 | 0 | 0 | 0 | 0 | 0 | 0 |
| ENSMUSG00000095739 | 0 | 0 | 0 | 0 | 0 | 0 | 0 |
| ENSMUSG00000095740 | 0 | 0 | 0 | 0 | 0.244650542 | 0 | 0 |
| ENSMUSG00000095741 | 0 | 0 | 0 | 0 | 0 | 0 | 0 |
| ENSMUSG00000095742 | 0 | 0.5 | 0.646240625 | 0.25 | 0.244650542 | 0 | 0.896975297 |
| ENSMUSG00000095743 | 0 | 0 | 0 | 0 | 0 | 0 | 0 |
| ENSMUSG00000095744 | 0 | 0 | 0 | 0 | 0 | 0 | 0 |
| ENSMUSG00000095745 | 0 | 0 | 0 | 0 | 0 | 0 | 0 |
| ENSMUSG00000095747 | 0 | 0 | 0 | 0 | 0 | 0 | 0 |
| ENSMUSG00000095749 | 0 | 0 | 0 | 0 | 0 | 0 | 0 |
| ENSMUSG00000095752 | 0 | 0 | 0 | 0 | 0 | 0 | 0 |
| ENSMUSG00000095753 | 0 | 0 | 0 | 0 | 0 | 0 | 0 |
| ENSMUSG00000095754 | 0 | 0 | 0 | 0 | 0 | 0 | 0 |
| ENSMUSG00000095755 | 0 | 0 | 0 | 0 | 0 | 0 | 0 |
| ENSMUSG00000095756 | 0 | 0 | 0 | 0 | 0 | 0 | 0 |
| ENSMUSG00000095757 | 0 | 0 | 0 | 0 | 0 | 0 | 0 |
| ENSMUSG00000095758 | 0 | 0 | 0 | 0 | 0 | 0 | 0 |
| ENSMUSG00000095759 | 0 | 0 | 0 | 0 | 0 | 0 | 0 |
| ENSMUSG00000095761 | 0 | 0 | 0 | 0 | 0 | 0 | 0 |
| ENSMUSG00000095762 | 0 | 0 | 0.25 | 0.25 | 0.244650542 | 0 | 0 |
| ENSMUSG00000095763 | 0 | 0 | 0 | 0 | 0 | 0 | 0 |
| ENSMUSG00000095764 | 0 | 0 | 0 | 0 | 0 | 0 | 0 |
| ENSMUSG00000095765 | 0 | 0 | 0 | 0 | 0 | 0 | 0 |
| ENSMUSG00000095766 | 0 | 0 | 0 | 0 | 0 | 0 | 0 |
| ENSMUSG00000095768 | 0 | 0 | 0 | 0 | 0 | 0 | 0 |
| ENSMUSG00000095769 | 0 | 0 | 0 | 0 | 0 | 0 | 0 |
| ENSMUSG00000095770 | 0 | 0 | 0 | 0 | 0 | 0 | 0 |
| ENSMUSG00000095771 | 0 | 0 | 0 | 0 | 0 | 0 | 0 |
| ENSMUSG00000095772 | 0 | 0 | 0 | 0 | 0 | 0 | 0 |
| ENSMUSG00000095773 | 0 | 0 | 0 | 0 | 0 | 0 | 0 |
| ENSMUSG00000095774 | 0 | 0 | 0 | 0 | 0 | 0 | 0 |
| ENSMUSG00000095776 | 0 | 0 | 0 | 0 | 0 | 0 | 0 |

| | | | | | | | |
|---|---|---|---|---|---|---|---|
| ENSMUSG00000095778 | 0 | 0 | 0 | 0 | 0 | 0 | 0 |
| ENSMUSG00000095779 | 0 | 0 | 0 | 0 | 0 | 0 | 0 |
| ENSMUSG00000095780 | 0 | 0 | 0.25 | 0 | 0 | 0 | 0 |
| ENSMUSG00000095782 | 0 | 0 | 0 | 0 | 0 | 0 | 0 |
| ENSMUSG00000095785 | 0 | 0 | 0 | 0 | 0 | 0 | 0 |
| ENSMUSG00000095786 | 0 | 0 | 0 | 0 | 0 | 0 | 0 |
| ENSMUSG00000095787 | 0 | 0 | 0 | 0 | 0 | 0 | 0 |
| ENSMUSG00000095788 | 0 | 0 | 0.25 | 0 | 0 | 0.477121255 | 0.520678036 |
| ENSMUSG00000095789 | 0 | 0.580482024 | 0.5 | 0.25 | 0.489301084 | 0.84509804 | 0.520678036 |
| ENSMUSG00000095791 | 0 | 0 | 0 | 0 | 0 | 0 | 0 |
| ENSMUSG00000095792 | 0 | 0 | 0 | 0 | 0 | 0 | 0 |
| ENSMUSG00000095793 | 0 | 0 | 0 | 0 | 0 | 0 | 0 |
| ENSMUSG00000095794 | 0 | 0 | 0 | 0 | 0 | 0 | 0 |
| ENSMUSG00000095796 | 0 | 0 | 0 | 0 | 0 | 0 | 0 |
| ENSMUSG00000095797 | 0 | 0 | 0 | 0 | 0 | 0 | 0 |
| ENSMUSG00000095798 | 0 | 0 | 0 | 0 | 0 | 0 | 0 |
| ENSMUSG00000095799 | 0 | 0 | 0 | 0 | 0 | 0 | 0 |
| ENSMUSG00000095802 | 0 | 0 | 0 | 0 | 0 | 0 | 0 |
| ENSMUSG00000095803 | 0 | 0 | 0 | 0 | 0 | 0 | 0 |
| ENSMUSG00000095804 | 0 | 0 | 0 | 0 | 0 | 0 | 0 |
| ENSMUSG00000095805 | 0 | 0 | 0 | 0 | 0 | 0 | 0 |
| ENSMUSG00000095806 | 0 | 0 | 0 | 0 | 0 | 0 | 0 |
| ENSMUSG00000095807 | 0 | 0 | 0 | 0 | 0 | 0 | 0 |
| ENSMUSG00000095809 | 0 | 0 | 0 | 0 | 0 | 0 | 0 |
| ENSMUSG00000095812 | 0 | 0.25 | 0 | 0 | 0.244650542 | 0.301029996 | 0.224243824 |
| ENSMUSG00000095813 | 0 | 0 | 0 | 0 | 0 | 0 | 0 |
| ENSMUSG00000095814 | 0 | 0 | 0 | 0 | 0 | 0 | 0 |
| ENSMUSG00000095816 | 0 | 0 | 0 | 0 | 0 | 0 | 0 |
| ENSMUSG00000095817 | 0 | 0 | 0 | 0 | 0 | 0 | 0 |
| ENSMUSG00000095818 | 0 | 0 | 0 | 0 | 0 | 0 | 0 |
| ENSMUSG00000095820 | 0 | 0 | 0 | 0 | 0 | 0 | 0 |
| ENSMUSG00000095821 | 0 | 0 | 0 | 0 | 0 | 0 | 0 |
| ENSMUSG00000095822 | 0 | 0 | 0 | 0 | 0 | 0 | 0 |
| ENSMUSG00000095824 | 0 | 0 | 0 | 0 | 0 | 0 | 0 |

| | | | | | | | |
|---|---|---|---|---|---|---|---|
| ENSMUSG00000095825 | 0 | 0 | 0 | 0 | 0 | 0 | 0 |
| ENSMUSG00000095826 | 0 | 0.25 | 0.5 | 0 | 0 | 0 | 0 |
| ENSMUSG00000095827 | 0 | 0 | 0 | 0 | 0 | 0 | 0 |
| ENSMUSG00000095828 | 0 | 0 | 0 | 0 | 0 | 0 | 0 |
| ENSMUSG00000095829 | 0 | 0 | 0 | 0 | 0 | 0 | 0 |
| ENSMUSG00000095831 | 0 | 0 | 0 | 0 | 0 | 0 | 0 |
| ENSMUSG00000095834 | 0 | 0 | 0 | 0 | 0 | 0 | 0 |
| ENSMUSG00000095836 | 0.45815691 | 0.580482024 | 0.580482024 | 0.580482024 | 0.877063019 | 0.301029996 | 0.520678036 |
| ENSMUSG00000095837 | 0 | 0 | 0 | 0 | 0 | 0 | 0 |
| ENSMUSG00000095839 | 0 | 0 | 0 | 0 | 0 | 0 | 0 |
| ENSMUSG00000095841 | 0 | 0 | 0 | 0 | 0 | 0 | 0 |
| ENSMUSG00000095842 | 0 | 0 | 0 | 0 | 0 | 0 | 0 |
| ENSMUSG00000095843 | 0 | 0 | 0 | 0 | 0 | 0 | 0 |
| ENSMUSG00000095845 | 0 | 0.25 | 0.25 | 0.25 | 0.244650542 | 0 | 0.355418052 |
| ENSMUSG00000095846 | 0 | 0 | 0 | 0 | 0 | 0 | 0 |
| ENSMUSG00000095847 | 0 | 0.396240625 | 0 | 0.25 | 0 | 0.301029996 | 0.224243824 |
| ENSMUSG00000095849 | 0 | 0 | 0 | 0 | 0 | 0 | 0 |
| ENSMUSG00000095850 | 0 | 0 | 0 | 0 | 0 | 0 | 0 |
| ENSMUSG00000095851 | 0 | 0 | 0 | 0 | 0 | 0 | 0 |
| ENSMUSG00000095852 | 0 | 0 | 0 | 0 | 0 | 0 | 0 |
| ENSMUSG00000095853 | 0 | 0 | 0 | 0 | 0 | 0 | 0 |
| ENSMUSG00000095854 | 0 | 0 | 0 | 0 | 0 | 0 | 0 |
| ENSMUSG00000095856 | 0 | 0 | 0 | 0 | 0 | 0 | 0 |
| ENSMUSG00000095857 | 0 | 0 | 0 | 0 | 0 | 0 | 0 |
| ENSMUSG00000095858 | 0 | 0 | 0 | 0 | 0 | 0 | 0 |
| ENSMUSG00000095859 | 0 | 0 | 0 | 0 | 0 | 0 | 0 |
| ENSMUSG00000095860 | 0 | 0 | 0 | 0 | 0 | 0 | 0 |
| ENSMUSG00000095862 | 0 | 0 | 0 | 0 | 0 | 0 | 0 |
| ENSMUSG00000095863 | 0 | 0 | 0 | 0 | 0 | 0 | 0 |
| ENSMUSG00000095864 | 0 | 0 | 0 | 0 | 0 | 0 | 0 |
| ENSMUSG00000095865 | 0 | 0 | 0 | 0 | 0 | 0 | 0 |
| ENSMUSG00000095866 | 0 | 0 | 0 | 0 | 0 | 0 | 0 |
| ENSMUSG00000095867 | 0 | 0 | 0 | 0 | 0 | 0 | 0 |
| ENSMUSG00000095868 | 0.45815691 | 0.25 | 0 | 0.25 | 0 | 0 | 0 |

| | | | | | | | |
|---|---|---|---|---|---|---|---|
| ENSMUSG00000095869 | 0 | 0 | 0 | 0 | 0 | 0 | 0 |
| ENSMUSG00000095870 | 0 | 0 | 0 | 0 | 0 | 0 | 0 |
| ENSMUSG00000095871 | 0 | 0 | 0 | 0 | 0 | 0 | 0 |
| ENSMUSG00000095872 | 0 | 0 | 0 | 0 | 0 | 0 | 0 |
| ENSMUSG00000095873 | 0 | 0 | 0 | 0 | 0 | 0 | 0 |
| ENSMUSG00000095874 | 0 | 0 | 0 | 0 | 0 | 0 | 0 |
| ENSMUSG00000095876 | 0 | 0 | 0 | 0 | 0 | 0 | 0 |
| ENSMUSG00000095878 | 0 | 0 | 0.25 | 0 | 0 | 0 | 0 |
| ENSMUSG00000095879 | 0 | 0 | 0 | 0 | 0 | 0 | 0 |
| ENSMUSG00000095880 | 0 | 0 | 0 | 0 | 0 | 0 | 0 |
| ENSMUSG00000095883 | 0 | 0 | 0 | 0 | 0 | 0 | 0 |
| ENSMUSG00000095884 | 0 | 0 | 0 | 0 | 0 | 0 | 0 |
| ENSMUSG00000095885 | 0 | 0 | 0 | 0 | 0 | 0 | 0 |
| ENSMUSG00000095886 | 0 | 0 | 0 | 0 | 0 | 0 | 0 |
| ENSMUSG00000095887 | 0 | 0 | 0 | 0 | 0 | 0 | 0 |
| ENSMUSG00000095889 | 0 | 0 | 0 | 0 | 0 | 0 | 0 |
| ENSMUSG00000095890 | 0 | 0 | 0 | 0 | 0 | 0 | 0 |
| ENSMUSG00000095891 | 0 | 0 | 0.25 | 0 | 0 | 0 | 0 |
| ENSMUSG00000095892 | 0 | 0 | 0.25 | 0 | 0 | 0 | 0 |
| ENSMUSG00000095893 | 0 | 0 | 0 | 0 | 0 | 0 | 0 |
| ENSMUSG00000095894 | 0 | 0 | 0 | 0 | 0 | 0 | 0 |
| ENSMUSG00000095895 | 0 | 0 | 0 | 0 | 0.244650542 | 0 | 0 |
| ENSMUSG00000095896 | 0 | 0 | 0 | 0 | 0 | 0 | 0 |
| ENSMUSG00000095897 | 0 | 0 | 0 | 0 | 0 | 0 | 0 |
| ENSMUSG00000095898 | 0 | 0 | 0 | 0 | 0 | 0 | 0 |
| ENSMUSG00000095899 | 0 | 0 | 0 | 0 | 0 | 0 | 0 |
| ENSMUSG00000095900 | 0 | 0 | 0 | 0 | 0 | 0 | 0 |
| ENSMUSG00000095901 | 0 | 0 | 0 | 0 | 0 | 0 | 0 |
| ENSMUSG00000095902 | 0 | 0 | 0 | 0 | 0 | 0 | 0 |
| ENSMUSG00000095903 | 0 | 0 | 0 | 0 | 0 | 0 | 0 |
| ENSMUSG00000095905 | 0 | 0 | 0 | 0 | 0 | 0 | 0 |
| ENSMUSG00000095906 | 0 | 0 | 0 | 0 | 0 | 0 | 0 |
| ENSMUSG00000095907 | 0 | 0 | 0 | 0 | 0 | 0 | 0 |
| ENSMUSG00000095909 | 0 | 0 | 0 | 0.396240625 | 0.387761935 | 0.301029996 | 0.224243824 |

| | | | | | | | |
|---|---|---|---|---|---|---|---|
| ENSMUSG00000095910 | 0 | 0 | 0 | 0 | 0 | 0 | 0 |
| ENSMUSG00000095911 | 0 | 0 | 0 | 0 | 0 | 0 | 0 |
| ENSMUSG00000095912 | 0 | 0 | 0 | 0 | 0 | 0 | 0 |
| ENSMUSG00000095913 | 0 | 0 | 0.396240625 | 0.25 | 0.387761935 | 0.698970004 | 0.355418052 |
| ENSMUSG00000095914 | 0 | 0 | 0 | 0 | 0 | 0 | 0 |
| ENSMUSG00000095915 | 0 | 0 | 0 | 0 | 0 | 0 | 0 |
| ENSMUSG00000095916 | 0 | 0 | 0 | 0 | 0 | 0 | 0 |
| ENSMUSG00000095917 | 0 | 0 | 0 | 0 | 0 | 0 | 0 |
| ENSMUSG00000095918 | 0 | 0 | 0 | 0 | 0 | 0 | 0 |
| ENSMUSG00000095920 | 0 | 0 | 0 | 0.25 | 0 | 0 | 0 |
| ENSMUSG00000095922 | 0 | 0 | 0 | 0 | 0 | 0 | 0 |
| ENSMUSG00000095923 | 0 | 0 | 0 | 0 | 0 | 0 | 0 |
| ENSMUSG00000095924 | 0 | 0 | 0 | 0 | 0 | 0 | 0 |
| ENSMUSG00000095925 | 0 | 0 | 0 | 0 | 0 | 0 | 0 |
| ENSMUSG00000095926 | 0 | 0 | 0 | 0 | 0 | 0 | 0 |
| ENSMUSG00000095927 | 0 | 0 | 0 | 0 | 0 | 0 | 0 |
| ENSMUSG00000095928 | 0 | 0 | 0 | 0 | 0 | 0 | 0 |
| ENSMUSG00000095929 | 0 | 0 | 0 | 0 | 0 | 0 | 0 |
| ENSMUSG00000095930 | 1.325351389 | 1.396240625 | 1.372963274 | 1.561981878 | 1.433158226 | 1.73239376 | 1.448487648 |
| ENSMUSG00000095931 | 0 | 0 | 0 | 0 | 0 | 0 | 0 |
| ENSMUSG00000095932 | 0 | 0 | 0 | 0 | 0 | 0 | 0 |
| ENSMUSG00000095933 | 0 | 0 | 0 | 0 | 0 | 0 | 0 |
| ENSMUSG00000095934 | 0 | 0 | 0 | 0 | 0 | 0 | 0 |
| ENSMUSG00000095935 | 0 | 0 | 0 | 0 | 0 | 0 | 0 |
| ENSMUSG00000095936 | 0 | 0 | 0 | 0 | 0 | 0 | 0 |
| ENSMUSG00000095937 | 0 | 0 | 0 | 0 | 0 | 0 | 0 |
| ENSMUSG00000095938 | 0 | 0 | 0 | 0 | 0 | 0 | 0 |
| ENSMUSG00000095939 | 0 | 0 | 0 | 0 | 0 | 0 | 0 |
| ENSMUSG00000095940 | 0 | 0.25 | 0 | 0 | 0 | 0 | 0.224243824 |
| ENSMUSG00000095941 | 0 | 0 | 0 | 0 | 0 | 0 | 0 |
| ENSMUSG00000095942 | 0 | 0 | 0 | 0 | 0 | 0 | 0 |
| ENSMUSG00000095943 | 0 | 0.396240625 | 0 | 0 | 0.244650542 | 0 | 0 |
| ENSMUSG00000095944 | 0 | 0 | 0 | 0 | 0 | 0 | 0 |
| ENSMUSG00000095945 | 0 | 0 | 0 | 0 | 0 | 0 | 0 |

| | | | | | | | |
|---|---|---|---|---|---|---|---|
| ENSMUSG00000095946 | 0 | 0 | 0 | 0 | 0 | 0 | 0 |
| ENSMUSG00000095947 | 0 | 0 | 0 | 0 | 0 | 0 | 0 |
| ENSMUSG00000095948 | 0 | 0 | 0 | 0 | 0 | 0 | 0 |
| ENSMUSG00000095950 | 0 | 0 | 0 | 0 | 0 | 0 | 0 |
| ENSMUSG00000095951 | 0 | 0 | 0 | 0 | 0 | 0 | 0 |
| ENSMUSG00000095952 | 0 | 0 | 0 | 0 | 0 | 0 | 0 |
| ENSMUSG00000095953 | 0 | 0 | 0 | 0 | 0 | 0 | 0 |
| ENSMUSG00000095954 | 0 | 0 | 0 | 0 | 0 | 0 | 0 |
| ENSMUSG00000095955 | 0 | 0 | 0 | 0 | 0 | 0 | 0 |
| ENSMUSG00000095956 | 0 | 0 | 0 | 0 | 0 | 0 | 0 |
| ENSMUSG00000095957 | 0 | 0 | 0 | 0 | 0 | 0 | 0 |
| ENSMUSG00000095958 | 0 | 0 | 0 | 0 | 0 | 0 | 0 |
| ENSMUSG00000095959 | 0 | 0 | 0 | 0 | 0 | 0 | 0 |
| ENSMUSG00000095960 | 0 | 0 | 0 | 0 | 0 | 0 | 0 |
| ENSMUSG00000095961 | 0 | 0 | 0 | 0 | 0 | 0 | 0 |
| ENSMUSG00000095962 | 0 | 0 | 0 | 0 | 0 | 0 | 0 |
| ENSMUSG00000095963 | 0 | 0 | 0 | 0 | 0 | 0 | 0 |
| ENSMUSG00000095964 | 0 | 0 | 0 | 0 | 0 | 0 | 0 |
| ENSMUSG00000095965 | 0 | 0 | 0 | 0 | 0 | 0 | 0 |
| ENSMUSG00000095966 | 0 | 0 | 0 | 0 | 0 | 0 | 0 |
| ENSMUSG00000095968 | 0 | 0 | 0 | 0 | 0 | 0 | 0 |
| ENSMUSG00000095969 | 0 | 0 | 0 | 0 | 0 | 0 | 0 |
| ENSMUSG00000095970 | 0 | 0 | 0 | 0 | 0 | 0 | 0 |
| ENSMUSG00000095971 | 0 | 0 | 0 | 0 | 0 | 0 | 0 |
| ENSMUSG00000095972 | 0 | 0 | 0 | 0 | 0 | 0 | 0 |
| ENSMUSG00000095973 | 0 | 0 | 0 | 0 | 0 | 0 | 0 |
| ENSMUSG00000095974 | 0 | 0 | 0 | 0 | 0 | 0 | 0 |
| ENSMUSG00000095975 | 0 | 0 | 0 | 0 | 0 | 0 | 0 |
| ENSMUSG00000095976 | 0 | 0 | 0 | 0 | 0 | 0 | 0 |
| ENSMUSG00000095977 | 0 | 0 | 0 | 0 | 0 | 0 | 0 |
| ENSMUSG00000095978 | 0 | 0 | 0 | 0 | 0 | 0 | 0 |
| ENSMUSG00000095979 | 0 | 0 | 0 | 0 | 0 | 0 | 0 |
| ENSMUSG00000095980 | 0 | 0 | 0 | 0 | 0 | 0 | 0 |
| ENSMUSG00000095981 | 0 | 0 | 0 | 0 | 0 | 0 | 0 |

| | | | | | | | |
|---|---|---|---|---|---|---|---|
| ENSMUSG00000095982 | 0 | 0 | 0 | 0 | 0 | 0 | 0 |
| ENSMUSG00000095984 | 0 | 0 | 0 | 0 | 0 | 0 | 0 |
| ENSMUSG00000095985 | 0 | 0 | 0 | 0 | 0 | 0 | 0 |
| ENSMUSG00000095986 | 0 | 0 | 0 | 0 | 0 | 0 | 0 |
| ENSMUSG00000095987 | 0 | 0 | 0 | 0 | 0 | 0 | 0 |
| ENSMUSG00000095988 | 0 | 0 | 0 | 0 | 0 | 0 | 0 |
| ENSMUSG00000095989 | 0 | 0 | 0 | 0 | 0 | 0 | 0 |
| ENSMUSG00000095990 | 1 | 1.188721876 | 1.201838731 | 1.188721876 | 1.302012594 | 1.041392685 | 1.014380832 |
| ENSMUSG00000095991 | 0 | 0 | 0 | 0 | 0 | 0 | 0 |
| ENSMUSG00000095992 | 0 | 0 | 0 | 0 | 0 | 0 | 0 |
| ENSMUSG00000095993 | 0 | 0 | 0.25 | 0.25 | 0.244650542 | 0 | 0.224243824 |
| ENSMUSG00000095994 | 0 | 0 | 0 | 0 | 0 | 0 | 0 |
| ENSMUSG00000095996 | 0 | 0 | 0 | 0 | 0 | 0 | 0 |
| ENSMUSG00000095997 | 0 | 0 | 0 | 0 | 0 | 0 | 0 |
| ENSMUSG00000095998 | 0 | 0 | 0 | 0 | 0 | 0 | 0 |
| ENSMUSG00000096000 | 0 | 0 | 0 | 0 | 0 | 0 | 0 |
| ENSMUSG00000096001 | 1 | 1.160964047 | 1 | 0.976722649 | 1.106691885 | 1.204119983 | 1.041356071 |
| ENSMUSG00000096002 | 0 | 0 | 0 | 0 | 0 | 0 | 0 |
| ENSMUSG00000096003 | 0 | 0 | 0 | 0 | 0 | 0 | 0 |
| ENSMUSG00000096005 | 0 | 0 | 0 | 0 | 0 | 0 | 0 |
| ENSMUSG00000096006 | 0.747221736 | 0.75 | 0.830482024 | 0.75 | 0.686820904 | 0.77815125 | 0.520678036 |
| ENSMUSG00000096008 | 0 | 0 | 0 | 0 | 0 | 0 | 0 |
| ENSMUSG00000096009 | 0 | 0 | 0 | 0 | 0 | 0 | 0 |
| ENSMUSG00000096010 | 0.289064826 | 0.5 | 0.25 | 0.25 | 0.387761935 | 0.301029996 | 0.448487648 |
| ENSMUSG00000096011 | 0 | 0 | 0 | 0 | 0 | 0 | 0 |
| ENSMUSG00000096014 | 0 | 0 | 0 | 0 | 0 | 0 | 0 |
| ENSMUSG00000096015 | 0 | 0 | 0 | 0 | 0 | 0 | 0 |
| ENSMUSG00000096016 | 0 | 0 | 0 | 0 | 0 | 0 | 0 |
| ENSMUSG00000096017 | 0 | 0 | 0 | 0 | 0 | 0 | 0 |
| ENSMUSG00000096018 | 0 | 0 | 0 | 0 | 0 | 0 | 0 |
| ENSMUSG00000096019 | 0 | 0 | 0 | 0 | 0 | 0 | 0 |
| ENSMUSG00000096020 | 0 | 0 | 0 | 0 | 0 | 0 | 0 |
| ENSMUSG00000096021 | 0 | 0 | 0 | 0 | 0 | 0 | 0 |
| ENSMUSG00000096022 | 0 | 0 | 0 | 0 | 0 | 0 | 0 |

| | | | | | | | |
|---|---|---|---|---|---|---|---|
| ENSMUSG00000096023 | 0 | 0 | 0 | 0 | 0 | 0 | 0 |
| ENSMUSG00000096024 | 0.289064826 | 0 | 0 | 0 | 0 | 0.301029996 | 0 |
| ENSMUSG00000096025 | 0 | 0.25 | 0.25 | 0 | 0 | 0 | 0 |
| ENSMUSG00000096027 | 0 | 0 | 0 | 0 | 0 | 0 | 0 |
| ENSMUSG00000096028 | 0 | 0 | 0 | 0 | 0 | 0 | 0 |
| ENSMUSG00000096029 | 0 | 0 | 0 | 0 | 0 | 0 | 0 |
| ENSMUSG00000096030 | 0 | 0 | 0 | 0 | 0 | 0 | 0 |
| ENSMUSG00000096031 | 0 | 0 | 0 | 0 | 0 | 0 | 0 |
| ENSMUSG00000096033 | 0 | 0 | 0 | 0.25 | 0 | 0 | 0 |
| ENSMUSG00000096035 | 0.289064826 | 0.25 | 0.580482024 | 0.5 | 0.244650542 | 0.477121255 | 0.448487648 |
| ENSMUSG00000096036 | 0 | 0 | 0 | 0 | 0 | 0 | 0 |
| ENSMUSG00000096037 | 0 | 0 | 0 | 0 | 0 | 0 | 0 |
| ENSMUSG00000096038 | 0 | 0 | 0 | 0 | 0 | 0 | 0 |
| ENSMUSG00000096039 | 0 | 0.25 | 0 | 0.5 | 0.387761935 | 0 | 0.448487648 |
| ENSMUSG00000096041 | 0 | 0 | 0 | 0 | 0 | 0 | 0 |
| ENSMUSG00000096042 | 0 | 0 | 0 | 0 | 0 | 0 | 0 |
| ENSMUSG00000096043 | 0 | 0 | 0 | 0 | 0 | 0 | 0 |
| ENSMUSG00000096044 | 0 | 0 | 0 | 0 | 0.387761935 | 0 | 0.224243824 |
| ENSMUSG00000096045 | 0 | 0 | 0 | 0 | 0 | 0 | 0 |
| ENSMUSG00000096046 | 0 | 0 | 0 | 0 | 0 | 0 | 0 |
| ENSMUSG00000096047 | 0 | 0 | 0 | 0 | 0 | 0 | 0 |
| ENSMUSG00000096048 | 0 | 0 | 0 | 0 | 0 | 0 | 0 |
| ENSMUSG00000096049 | 0 | 0 | 0 | 0 | 0 | 0 | 0 |
| ENSMUSG00000096051 | 0 | 0 | 0 | 0 | 0 | 0 | 0 |
| ENSMUSG00000096052 | 0 | 0.25 | 0.25 | 0.396240625 | 0 | 0 | 0.224243824 |
| ENSMUSG00000096053 | 0 | 0 | 0 | 0 | 0 | 0 | 0 |
| ENSMUSG00000096054 | 2.016886209 | 1.868933358 | 1.866901388 | 1.747171172 | 1.733951626 | 1.851258349 | 1.670888075 |
| ENSMUSG00000096055 | 0 | 0 | 0 | 0 | 0 | 0 | 0 |
| ENSMUSG00000096056 | 0.578129653 | 0.75 | 0.79248125 | 0.701838731 | 0.931471446 | 0.903089987 | 0.74492186 |
| ENSMUSG00000096057 | 0 | 0 | 0 | 0 | 0 | 0 | 0 |
| ENSMUSG00000096061 | 0 | 0 | 0 | 0 | 0 | 0 | 0 |
| ENSMUSG00000096065 | 0 | 0 | 0 | 0 | 0 | 0 | 0 |
| ENSMUSG00000096066 | 0 | 0 | 0 | 0 | 0 | 0 | 0 |
| ENSMUSG00000096067 | 0 | 0 | 0 | 0 | 0 | 0 | 0 |

| | | | | | | | |
|---|---|---|---|---|---|---|---|
| ENSMUSG00000096068 | 0 | 0 | 0 | 0 | 0 | 0 | 0 |
| ENSMUSG00000096069 | 0 | 0 | 0 | 0 | 0 | 0 | 0 |
| ENSMUSG00000096070 | 0 | 0 | 0 | 0 | 0 | 0 | 0 |
| ENSMUSG00000096071 | 0 | 0 | 0 | 0 | 0 | 0 | 0 |
| ENSMUSG00000096072 | 0 | 0 | 0 | 0 | 0 | 0 | 0 |
| ENSMUSG00000096073 | 0 | 0 | 0 | 0 | 0 | 0 | 0 |
| ENSMUSG00000096074 | 0 | 0 | 0 | 0 | 0 | 0 | 0 |
| ENSMUSG00000096075 | 0 | 0 | 0 | 0 | 0 | 0 | 0 |
| ENSMUSG00000096076 | 0 | 0 | 0 | 0 | 0 | 0 | 0 |
| ENSMUSG00000096078 | 0 | 0 | 0 | 0 | 0 | 0 | 0 |
| ENSMUSG00000096079 | 0 | 0 | 0 | 0 | 0 | 0 | 0 |
| ENSMUSG00000096080 | 0 | 0 | 0 | 0 | 0 | 0 | 0 |
| ENSMUSG00000096081 | 0 | 0 | 0 | 0 | 0 | 0 | 0 |
| ENSMUSG00000096082 | 0 | 0 | 0 | 0 | 0 | 0 | 0 |
| ENSMUSG00000096083 | 0 | 0 | 0 | 0 | 0 | 0 | 0 |
| ENSMUSG00000096084 | 0 | 0 | 0 | 0 | 0 | 0 | 0 |
| ENSMUSG00000096085 | 0 | 0 | 0 | 0 | 0 | 0 | 0 |
| ENSMUSG00000096086 | 0.289064826 | 0 | 0.5 | 0.396240625 | 0.244650542 | 0 | 0.448487648 |
| ENSMUSG00000096087 | 0 | 0 | 0 | 0 | 0 | 0 | 0 |
| ENSMUSG00000096088 | 0 | 0 | 0 | 0 | 0 | 0 | 0 |
| ENSMUSG00000096090 | 0 | 0 | 0 | 0 | 0 | 0 | 0 |
| ENSMUSG00000096091 | 0 | 0 | 0 | 0 | 0 | 0 | 0 |
| ENSMUSG00000096092 | 0 | 0 | 0 | 0 | 0 | 0 | 0 |
| ENSMUSG00000096094 | 0.289064826 | 0 | 0 | 0 | 0 | 0.301029996 | 0.224243824 |
| ENSMUSG00000096095 | 0 | 0 | 0 | 0 | 0 | 0 | 0 |
| ENSMUSG00000096096 | 0 | 0 | 0 | 0.25 | 0 | 0 | 0 |
| ENSMUSG00000096097 | 0 | 0 | 0 | 0 | 0 | 0 | 0 |
| ENSMUSG00000096098 | 0 | 0 | 0 | 0 | 0 | 0 | 0 |
| ENSMUSG00000096099 | 0 | 0 | 0 | 0 | 0 | 0 | 0 |
| ENSMUSG00000096100 | 0 | 0 | 0 | 0 | 0 | 0 | 0 |
| ENSMUSG00000096102 | 0 | 0 | 0 | 0 | 0 | 0.301029996 | 0 |
| ENSMUSG00000096105 | 0 | 0 | 0 | 0 | 0 | 0 | 0 |
| ENSMUSG00000096106 | 0 | 0 | 0 | 0 | 0 | 0 | 0 |
| ENSMUSG00000096107 | 0 | 0 | 0 | 0 | 0 | 0 | 0 |

| | | | | | | | |
|---|---|---|---|---|---|---|---|
| ENSMUSG00000096108 | 0 | 0 | 0 | 0 | 0 | 0 | 0 |
| ENSMUSG00000096109 | 0 | 0 | 0 | 0 | 0 | 0 | 0 |
| ENSMUSG00000096110 | 0 | 0 | 0 | 0 | 0 | 0 | 0 |
| ENSMUSG00000096111 | 0.289064826 | 0.25 | 0 | 0.580482024 | 0 | 0.301029996 | 0.224243824 |
| ENSMUSG00000096112 | 0 | 0 | 0 | 0 | 0 | 0 | 0 |
| ENSMUSG00000096113 | 0 | 0 | 0 | 0 | 0 | 0 | 0 |
| ENSMUSG00000096114 | 0 | 0 | 0 | 0 | 0 | 0 | 0 |
| ENSMUSG00000096117 | 0 | 0 | 0 | 0 | 0 | 0 | 0 |
| ENSMUSG00000096119 | 0 | 0 | 0 | 0 | 0 | 0 | 0 |
| ENSMUSG00000096120 | 0 | 0 | 0 | 0 | 0 | 0 | 0 |
| ENSMUSG00000096122 | 0 | 0 | 0 | 0 | 0 | 0 | 0 |
| ENSMUSG00000096124 | 0 | 0 | 0 | 0 | 0 | 0 | 0 |
| ENSMUSG00000096126 | 0 | 0 | 0 | 0 | 0 | 0 | 0 |
| ENSMUSG00000096127 | 0 | 0 | 0 | 0 | 0 | 0 | 0 |
| ENSMUSG00000096129 | 0 | 0 | 0 | 0 | 0 | 0 | 0 |
| ENSMUSG00000096130 | 0 | 0 | 0 | 0 | 0 | 0 | 0 |
| ENSMUSG00000096131 | 0 | 0 | 0 | 0 | 0 | 0 | 0 |
| ENSMUSG00000096132 | 0 | 0 | 0 | 0 | 0 | 0 | 0 |
| ENSMUSG00000096133 | 0 | 0 | 0 | 0 | 0 | 0 | 0 |
| ENSMUSG00000096134 | 0 | 0 | 0 | 0 | 0 | 0 | 0 |
| ENSMUSG00000096135 | 0 | 0 | 0 | 0 | 0 | 0 | 0 |
| ENSMUSG00000096136 | 0 | 0 | 0 | 0 | 0 | 0 | 0 |
| ENSMUSG00000096137 | 0 | 0 | 0 | 0 | 0 | 0 | 0 |
| ENSMUSG00000096138 | 0 | 0 | 0 | 0 | 0 | 0 | 0 |
| ENSMUSG00000096139 | 0 | 0 | 0 | 0 | 0 | 0 | 0 |
| ENSMUSG00000096140 | 0 | 0.25 | 0 | 0 | 0.244650542 | 0.477121255 | 0 |
| ENSMUSG00000096141 | 0 | 0 | 0.25 | 0 | 0 | 0 | 0 |
| ENSMUSG00000096142 | 0 | 0 | 0 | 0 | 0 | 0 | 0 |
| ENSMUSG00000096144 | 0 | 0.25 | 0 | 0 | 0 | 0 | 0 |
| ENSMUSG00000096145 | 1.37447073 | 1.29248125 | 1.330482024 | 1.410964047 | 1.302012594 | 1.531478917 | 1.324583736 |
| ENSMUSG00000096146 | 0 | 0 | 0 | 0 | 0 | 0 | 0.224243824 |
| ENSMUSG00000096147 | 0 | 0 | 0 | 0 | 0 | 0 | 0 |
| ENSMUSG00000096148 | 0 | 0 | 0 | 0 | 0 | 0 | 0 |
| ENSMUSG00000096149 | 0 | 0 | 0 | 0 | 0 | 0 | 0 |

| | | | | | | | |
|---|---|---|---|---|---|---|---|
| ENSMUSG00000096150 | 0 | 0 | 0 | 0 | 0 | 0 | 0 |
| ENSMUSG00000096151 | 0 | 0 | 0 | 0 | 0 | 0 | 0 |
| ENSMUSG00000096152 | 0 | 0 | 0 | 0 | 0 | 0 | 0 |
| ENSMUSG00000096153 | 0 | 0 | 0 | 0 | 0 | 0 | 0 |
| ENSMUSG00000096154 | 0 | 0 | 0 | 0 | 0 | 0 | 0 |
| ENSMUSG00000096156 | 0 | 0 | 0 | 0 | 0 | 0 | 0 |
| ENSMUSG00000096157 | 0.289064826 | 0 | 0 | 0 | 0 | 0 | 0.224243824 |
| ENSMUSG00000096158 | 0 | 0 | 0 | 0 | 0 | 0 | 0 |
| ENSMUSG00000096159 | 0 | 0 | 0 | 0 | 0 | 0 | 0 |
| ENSMUSG00000096160 | 0 | 0 | 0 | 0 | 0 | 0 | 0.224243824 |
| ENSMUSG00000096161 | 0 | 0 | 0 | 0 | 0 | 0 | 0 |
| ENSMUSG00000096163 | 0 | 0 | 0 | 0 | 0 | 0 | 0 |
| ENSMUSG00000096164 | 0 | 0 | 0 | 0 | 0 | 0 | 0 |
| ENSMUSG00000096166 | 0 | 0 | 0 | 0 | 0 | 0 | 0 |
| ENSMUSG00000096167 | 0 | 0 | 0 | 0 | 0 | 0 | 0 |
| ENSMUSG00000096168 | 0 | 0 | 0 | 0 | 0 | 0 | 0 |
| ENSMUSG00000096169 | 0 | 0 | 0.25 | 0 | 0 | 0 | 0.224243824 |
| ENSMUSG00000096170 | 0 | 0 | 0 | 0 | 0 | 0 | 0 |
| ENSMUSG00000096171 | 0 | 0 | 0 | 0.25 | 0 | 0 | 0.355418052 |
| ENSMUSG00000096172 | 0 | 0 | 0 | 0 | 0 | 0 | 0 |
| ENSMUSG00000096173 | 0 | 0 | 0 | 0 | 0 | 0 | 0 |
| ENSMUSG00000096174 | 0 | 0 | 0 | 0 | 0 | 0 | 0 |
| ENSMUSG00000096175 | 0 | 0 | 0 | 0 | 0 | 0 | 0 |
| ENSMUSG00000096176 | 0 | 0 | 0 | 0 | 0 | 0 | 0 |
| ENSMUSG00000096177 | 0.289064826 | 0 | 0.396240625 | 0.25 | 0 | 0 | 0.355418052 |
| ENSMUSG00000096178 | 0 | 0 | 0 | 0 | 0 | 0 | 0 |
| ENSMUSG00000096179 | 0 | 0 | 0 | 0 | 0 | 0 | 0 |
| ENSMUSG00000096180 | 0 | 0 | 0 | 0 | 0 | 0 | 0 |
| ENSMUSG00000096181 | 0 | 0 | 0 | 0 | 0 | 0 | 0 |
| ENSMUSG00000096182 | 0 | 0 | 0 | 0 | 0 | 0 | 0 |
| ENSMUSG00000096183 | 0 | 0 | 0 | 0 | 0 | 0 | 0 |
| ENSMUSG00000096184 | 0 | 0 | 0 | 0 | 0 | 0 | 0 |
| ENSMUSG00000096185 | 0 | 0 | 0 | 0 | 0 | 0 | 0 |
| ENSMUSG00000096186 | 0 | 0 | 0 | 0 | 0 | 0 | 0 |

| | | | | | | | |
|---|---|---|---|---|---|---|---|
| ENSMUSG00000096188 | 3.002187511 | 2.738185312 | 2.736178015 | 2.70083098 | 2.658874533 | 3.030599722 | 2.472784836 |
| ENSMUSG00000096189 | 0 | 0 | 0 | 0 | 0 | 0 | 0 |
| ENSMUSG00000096190 | 0 | 0 | 0 | 0 | 0 | 0 | 0 |
| ENSMUSG00000096191 | 0 | 0 | 0 | 0 | 0 | 0 | 0 |
| ENSMUSG00000096192 | 0 | 0 | 0 | 0 | 0.244650542 | 0 | 0 |
| ENSMUSG00000096193 | 0 | 0 | 0 | 0 | 0 | 0 | 0 |
| ENSMUSG00000096194 | 0 | 0 | 0 | 0 | 0 | 0 | 0 |
| ENSMUSG00000096195 | 0 | 0.396240625 | 0.25 | 0 | 0 | 0 | 0 |
| ENSMUSG00000096196 | 0 | 0 | 0 | 0 | 0 | 0 | 0 |
| ENSMUSG00000096197 | 0 | 0 | 0 | 0 | 0 | 0 | 0 |
| ENSMUSG00000096198 | 0 | 0 | 0 | 0 | 0 | 0 | 0 |
| ENSMUSG00000096199 | 1.516991255 | 1.458222504 | 1.575945187 | 1.566696635 | 1.489301084 | 1.602059991 | 1.594042709 |
| ENSMUSG00000096200 | 0 | 0 | 0 | 0 | 0 | 0 | 0 |
| ENSMUSG00000096201 | 0 | 0 | 0 | 0 | 0 | 0 | 0 |
| ENSMUSG00000096202 | 0 | 0 | 0 | 0 | 0 | 0 | 0 |
| ENSMUSG00000096205 | 0 | 0 | 0 | 0 | 0 | 0 | 0 |
| ENSMUSG00000096206 | 0 | 0 | 0 | 0 | 0 | 0 | 0 |
| ENSMUSG00000096208 | 0 | 0 | 0 | 0 | 0 | 0 | 0 |
| ENSMUSG00000096209 | 0 | 0 | 0 | 0 | 0 | 0 | 0 |
| ENSMUSG00000096210 | 3.130720997 | 2.838837774 | 2.877320042 | 2.920309603 | 2.875226296 | 3.31302311 | 2.734805874 |
| ENSMUSG00000096211 | 0 | 0 | 0 | 0 | 0 | 0 | 0 |
| ENSMUSG00000096212 | 0 | 0 | 0 | 0 | 0 | 0 | 0 |
| ENSMUSG00000096213 | 0 | 0 | 0 | 0 | 0 | 0 | 0 |
| ENSMUSG00000096214 | 0 | 0 | 0 | 0 | 0 | 0 | 0 |
| ENSMUSG00000096215 | 0 | 0 | 0 | 0 | 0 | 0 | 0 |
| ENSMUSG00000096216 | 0 | 0 | 0 | 0 | 0 | 0 | 0 |
| ENSMUSG00000096217 | 0 | 0 | 0 | 0 | 0 | 0 | 0 |
| ENSMUSG00000096218 | 0 | 0 | 0 | 0 | 0 | 0 | 0 |
| ENSMUSG00000096219 | 0 | 0 | 0 | 0 | 0 | 0 | 0 |
| ENSMUSG00000096220 | 0 | 0 | 0 | 0 | 0 | 0 | 0 |
| ENSMUSG00000096221 | 1.205378646 | 1.098079356 | 1.146240625 | 1.130890489 | 1 | 1.041392685 | 1.168187745 |
| ENSMUSG00000096222 | 0 | 0 | 0 | 0 | 0 | 0 | 0 |
| ENSMUSG00000096223 | 0 | 0 | 0 | 0 | 0 | 0 | 0 |
| ENSMUSG00000096225 | 0 | 0 | 0 | 0 | 0 | 0 | 0 |

| | | | | | | | |
|---|---|---|---|---|---|---|---|
| ENSMUSG00000096228 | 0 | 0 | 0 | 0 | 0 | 0 | 0 |
| ENSMUSG00000096229 | 0 | 0 | 0 | 0 | 0 | 0 | 0 |
| ENSMUSG00000096230 | 0 | 0 | 0 | 0 | 0 | 0 | 0 |
| ENSMUSG00000096231 | 0 | 0 | 0 | 0 | 0 | 0 | 0 |
| ENSMUSG00000096232 | 0 | 0 | 0 | 0 | 0 | 0 | 0 |
| ENSMUSG00000096233 | 0 | 0 | 0 | 0 | 0 | 0 | 0 |
| ENSMUSG00000096235 | 0 | 0 | 0 | 0 | 0 | 0 | 0 |
| ENSMUSG00000096236 | 0 | 0 | 0 | 0 | 0 | 0 | 0 |
| ENSMUSG00000096237 | 0 | 0 | 0 | 0 | 0 | 0 | 0 |
| ENSMUSG00000096238 | 0 | 0 | 0 | 0 | 0 | 0 | 0 |
| ENSMUSG00000096239 | 0 | 0 | 0 | 0 | 0 | 0 | 0 |
| ENSMUSG00000096240 | 0 | 0 | 0 | 0 | 0 | 0 | 0 |
| ENSMUSG00000096241 | 0 | 0 | 0 | 0 | 0 | 0 | 0 |
| ENSMUSG00000096243 | 0 | 0 | 0 | 0 | 0 | 0 | 0.224243824 |
| ENSMUSG00000096244 | 0 | 0 | 0 | 0 | 0 | 0 | 0 |
| ENSMUSG00000096245 | 0 | 0 | 0 | 0 | 0 | 0 | 0 |
| ENSMUSG00000096247 | 0 | 0 | 0 | 0 | 0 | 0 | 0 |
| ENSMUSG00000096248 | 0 | 0 | 0 | 0 | 0 | 0 | 0 |
| ENSMUSG00000096249 | 0 | 0 | 0 | 0 | 0 | 0 | 0 |
| ENSMUSG00000096250 | 0 | 0 | 0 | 0 | 0 | 0 | 0 |
| ENSMUSG00000096252 | 0.289064826 | 0 | 0 | 0.25 | 0 | 0 | 0 |
| ENSMUSG00000096254 | 0 | 0 | 0 | 0 | 0 | 0 | 0 |
| ENSMUSG00000096255 | 0.578129653 | 0.79248125 | 0.580482024 | 0.92510993 | 0.632412477 | 0.602059991 | 0.520678036 |
| ENSMUSG00000096256 | 0 | 0 | 0 | 0 | 0 | 0 | 0 |
| ENSMUSG00000096257 | 0 | 0 | 0 | 0.25 | 0 | 0 | 0 |
| ENSMUSG00000096258 | 0 | 0 | 0 | 0 | 0 | 0 | 0 |
| ENSMUSG00000096259 | 0 | 0 | 0 | 0 | 0 | 0 | 0 |
| ENSMUSG00000096263 | 0 | 0 | 0 | 0 | 0 | 0 | 0 |
| ENSMUSG00000096264 | 0 | 0 | 0 | 0 | 0 | 0 | 0 |
| ENSMUSG00000096265 | 0 | 0 | 0 | 0 | 0 | 0 | 0 |
| ENSMUSG00000096268 | 0 | 0 | 0 | 0 | 0 | 0 | 0 |
| ENSMUSG00000096269 | 0 | 0 | 0 | 0 | 0 | 0 | 0.224243824 |
| ENSMUSG00000096270 | 0 | 0 | 0 | 0 | 0 | 0 | 0 |
| ENSMUSG00000096271 | 0 | 0 | 0 | 0 | 0 | 0 | 0 |

| | | | | | | | |
|---|---|---|---|---|---|---|---|
| ENSMUSG00000096272 | 0 | 0 | 0 | 0 | 0 | 0 | 0 |
| ENSMUSG00000096273 | 0 | 0 | 0 | 0 | 0 | 0 | 0 |
| ENSMUSG00000096274 | 0 | 0 | 0 | 0 | 0 | 0 | 0 |
| ENSMUSG00000096275 | 0 | 0 | 0 | 0 | 0 | 0 | 0 |
| ENSMUSG00000096276 | 0 | 0 | 0 | 0 | 0 | 0 | 0 |
| ENSMUSG00000096277 | 0 | 0 | 0 | 0 | 0 | 0 | 0 |
| ENSMUSG00000096278 | 0 | 0 | 0 | 0 | 0 | 0 | 0 |
| ENSMUSG00000096279 | 0 | 0 | 0 | 0 | 0 | 0 | 0 |
| ENSMUSG00000096280 | 0 | 0 | 0 | 0 | 0 | 0 | 0 |
| ENSMUSG00000096281 | 0 | 0 | 0 | 0 | 0 | 0 | 0 |
| ENSMUSG00000096282 | 0 | 0 | 0 | 0 | 0 | 0 | 0 |
| ENSMUSG00000096283 | 0 | 0 | 0 | 0 | 0 | 0 | 0 |
| ENSMUSG00000096284 | 0 | 0.25 | 0 | 0 | 0.244650542 | 0 | 0 |
| ENSMUSG00000096285 | 0 | 0 | 0 | 0 | 0 | 0 | 0 |
| ENSMUSG00000096286 | 0 | 0 | 0 | 0 | 0 | 0 | 0 |
| ENSMUSG00000096287 | 0 | 0 | 0 | 0 | 0 | 0 | 0 |
| ENSMUSG00000096288 | 0 | 0 | 0 | 0 | 0 | 0 | 0 |
| ENSMUSG00000096289 | 0 | 0 | 0 | 0 | 0 | 0 | 0 |
| ENSMUSG00000096290 | 0 | 0 | 0 | 0 | 0 | 0 | 0 |
| ENSMUSG00000096291 | 0 | 0 | 0 | 0 | 0 | 0 | 0 |
| ENSMUSG00000096292 | 0 | 0 | 0 | 0 | 0 | 0 | 0 |
| ENSMUSG00000096293 | 0 | 0 | 0 | 0 | 0 | 0 | 0 |
| ENSMUSG00000096294 | 0 | 0 | 0 | 0 | 0 | 0 | 0 |
| ENSMUSG00000096295 | 0 | 0 | 0 | 0 | 0 | 0 | 0 |
| ENSMUSG00000096296 | 0.289064826 | 0 | 0.25 | 0.396240625 | 0.244650542 | 0.301029996 | 0.224243824 |
| ENSMUSG00000096297 | 0 | 0 | 0 | 0 | 0 | 0 | 0 |
| ENSMUSG00000096298 | 0 | 0 | 0 | 0 | 0 | 0 | 0 |
| ENSMUSG00000096299 | 0.289064826 | 0.25 | 0.396240625 | 0.646240625 | 0.387761935 | 0.477121255 | 0.579661877 |
| ENSMUSG00000096300 | 0 | 0 | 0 | 0 | 0 | 0 | 0 |
| ENSMUSG00000096301 | 0 | 0 | 0 | 0 | 0 | 0 | 0 |
| ENSMUSG00000096302 | 0 | 0 | 0 | 0 | 0 | 0 | 0 |
| ENSMUSG00000096303 | 0 | 0 | 0 | 0 | 0 | 0 | 0 |
| ENSMUSG00000096304 | 0 | 0 | 0 | 0 | 0 | 0 | 0 |
| ENSMUSG00000096305 | 0 | 0 | 0 | 0 | 0 | 0 | 0 |

| | | | | | | | |
|---|---|---|---|---|---|---|---|
| ENSMUSG00000096306 | 0 | 0 | 0 | 0 | 0 | 0 | 0 |
| ENSMUSG00000096307 | 0 | 0 | 0 | 0 | 0 | 0 | 0 |
| ENSMUSG00000096308 | 0 | 0 | 0 | 0 | 0 | 0 | 0 |
| ENSMUSG00000096309 | 0 | 0 | 0 | 0 | 0 | 0 | 0 |
| ENSMUSG00000096310 | 0 | 0 | 0 | 0 | 0 | 0 | 0 |
| ENSMUSG00000096311 | 0 | 0 | 0 | 0 | 0 | 0 | 0 |
| ENSMUSG00000096313 | 0 | 0 | 0 | 0 | 0 | 0 | 0 |
| ENSMUSG00000096314 | 0 | 0 | 0 | 0 | 0 | 0 | 0 |
| ENSMUSG00000096316 | 0 | 0 | 0 | 0 | 0 | 0 | 0 |
| ENSMUSG00000096318 | 0 | 0 | 0 | 0 | 0 | 0 | 0 |
| ENSMUSG00000096320 | 0 | 0 | 0 | 0 | 0 | 0 | 0 |
| ENSMUSG00000096322 | 0 | 0 | 0 | 0 | 0 | 0 | 0 |
| ENSMUSG00000096323 | 0 | 0 | 0 | 0 | 0 | 0 | 0 |
| ENSMUSG00000096324 | 0 | 0 | 0 | 0 | 0 | 0 | 0 |
| ENSMUSG00000096326 | 0 | 0 | 0 | 0 | 0 | 0 | 0 |
| ENSMUSG00000096327 | 0 | 0 | 0 | 0 | 0 | 0 | 0 |
| ENSMUSG00000096328 | 0 | 0 | 0 | 0 | 0 | 0 | 0 |
| ENSMUSG00000096329 | 0 | 0 | 0 | 0 | 0 | 0 | 0 |
| ENSMUSG00000096330 | 0 | 0 | 0 | 0.25 | 0.244650542 | 0 | 0.224243824 |
| ENSMUSG00000096331 | 0 | 0 | 0 | 0 | 0 | 0 | 0 |
| ENSMUSG00000096332 | 0 | 0 | 0 | 0 | 0 | 0 | 0 |
| ENSMUSG00000096333 | 0 | 0 | 0 | 0 | 0 | 0 | 0 |
| ENSMUSG00000096335 | 0 | 0 | 0 | 0 | 0 | 0 | 0 |
| ENSMUSG00000096336 | 0 | 0 | 0 | 0 | 0 | 0 | 0 |
| ENSMUSG00000096337 | 0 | 0 | 0 | 0 | 0 | 0 | 0 |
| ENSMUSG00000096338 | 0 | 0 | 0 | 0 | 0 | 0 | 0 |
| ENSMUSG00000096339 | 0 | 0 | 0 | 0 | 0 | 0 | 0 |
| ENSMUSG00000096340 | 0 | 0 | 0 | 0 | 0 | 0 | 0 |
| ENSMUSG00000096341 | 0 | 0 | 0 | 0 | 0 | 0 | 0 |
| ENSMUSG00000096342 | 0 | 0 | 0 | 0 | 0 | 0 | 0 |
| ENSMUSG00000096343 | 0 | 0 | 0 | 0 | 0 | 0 | 0 |
| ENSMUSG00000096344 | 0 | 0 | 0 | 0 | 0 | 0 | 0 |
| ENSMUSG00000096345 | 0 | 0 | 0 | 0 | 0 | 0 | 0 |
| ENSMUSG00000096346 | 0 | 0 | 0 | 0 | 0 | 0 | 0 |

| | | | | | | | |
|---|---|---|---|---|---|---|---|
| ENSMUSG00000096347 | 0 | 0 | 0 | 0 | 0 | 0 | 0 |
| ENSMUSG00000096348 | 0 | 0 | 0 | 0 | 0 | 0 | 0 |
| ENSMUSG00000096349 | 0 | 0 | 0 | 0 | 0 | 0 | 0 |
| ENSMUSG00000096350 | 0 | 0 | 0 | 0 | 0 | 0 | 0 |
| ENSMUSG00000096351 | 0.578129653 | 0.25 | 0.25 | 0 | 0.244650542 | 0 | 0.224243824 |
| ENSMUSG00000096352 | 0 | 0 | 0 | 0 | 0 | 0 | 0 |
| ENSMUSG00000096353 | 0 | 0 | 0 | 0 | 0 | 0 | 0 |
| ENSMUSG00000096354 | 0 | 0 | 0 | 0 | 0 | 0 | 0 |
| ENSMUSG00000096355 | 0 | 0 | 0 | 0 | 0 | 0 | 0 |
| ENSMUSG00000096356 | 0 | 0 | 0 | 0 | 0 | 0 | 0 |
| ENSMUSG00000096357 | 0 | 0 | 0 | 0 | 0 | 0 | 0 |
| ENSMUSG00000096358 | 0 | 0 | 0 | 0 | 0 | 0 | 0 |
| ENSMUSG00000096359 | 0 | 0 | 0 | 0 | 0 | 0 | 0 |
| ENSMUSG00000096360 | 0 | 0 | 0 | 0 | 0 | 0 | 0 |
| ENSMUSG00000096361 | 0 | 0 | 0 | 0 | 0 | 0 | 0 |
| ENSMUSG00000096362 | 0 | 0 | 0 | 0 | 0 | 0 | 0 |
| ENSMUSG00000096363 | 0 | 0 | 0 | 0 | 0 | 0 | 0 |
| ENSMUSG00000096365 | 0 | 0 | 0 | 0 | 0 | 0 | 0 |
| ENSMUSG00000096366 | 0 | 0 | 0 | 0 | 0 | 0 | 0 |
| ENSMUSG00000096367 | 0 | 0 | 0 | 0 | 0 | 0 | 0 |
| ENSMUSG00000096368 | 0 | 0 | 0 | 0 | 0 | 0 | 0 |
| ENSMUSG00000096370 | 1.205378646 | 1.02186571 | 1.098079356 | 1.201838731 | 1.176121988 | 1.342422681 | 1.20139795 |
| ENSMUSG00000096371 | 0 | 0 | 0 | 0 | 0 | 0 | 0 |
| ENSMUSG00000096372 | 0 | 0 | 0 | 0 | 0 | 0 | 0 |
| ENSMUSG00000096373 | 0 | 0 | 0 | 0 | 0 | 0 | 0 |
| ENSMUSG00000096374 | 0 | 0 | 0 | 0 | 0 | 0 | 0 |
| ENSMUSG00000096375 | 0 | 0 | 0 | 0 | 0 | 0 | 0 |
| ENSMUSG00000096376 | 0 | 0 | 0 | 0 | 0 | 0 | 0 |
| ENSMUSG00000096377 | 0 | 0 | 0 | 0 | 0 | 0 | 0 |
| ENSMUSG00000096379 | 0 | 0 | 0 | 0 | 0 | 0 | 0 |
| ENSMUSG00000096380 | 0 | 0 | 0 | 0 | 0 | 0 | 0 |
| ENSMUSG00000096381 | 0 | 0 | 0 | 0 | 0 | 0 | 0 |
| ENSMUSG00000096383 | 0 | 0 | 0 | 0 | 0 | 0 | 0 |
| ENSMUSG00000096384 | 0 | 0 | 0 | 0 | 0 | 0 | 0 |

| | | | | | | | |
|---|---|---|---|---|---|---|---|
| ENSMUSG00000096385 | 0 | 0 | 0 | 0 | 0 | 0 | 0 |
| ENSMUSG00000096386 | 0 | 0 | 0 | 0 | 0 | 0 | 0 |
| ENSMUSG00000096387 | 0 | 0 | 0 | 0 | 0 | 0 | 0 |
| ENSMUSG00000096388 | 0 | 0 | 0 | 0 | 0 | 0 | 0 |
| ENSMUSG00000096390 | 0 | 0 | 0 | 0 | 0 | 0 | 0 |
| ENSMUSG00000096391 | 0 | 0 | 0 | 0 | 0 | 0 | 0 |
| ENSMUSG00000096392 | 0 | 0 | 0 | 0 | 0 | 0 | 0 |
| ENSMUSG00000096393 | 0 | 0 | 0 | 0 | 0 | 0 | 0 |
| ENSMUSG00000096396 | 0 | 0 | 0 | 0 | 0 | 0 | 0 |
| ENSMUSG00000096397 | 0 | 0 | 0 | 0 | 0 | 0 | 0 |
| ENSMUSG00000096398 | 0 | 0 | 0 | 0 | 0 | 0 | 0 |
| ENSMUSG00000096399 | 0 | 0 | 0 | 0 | 0 | 0 | 0 |
| ENSMUSG00000096400 | 0 | 0 | 0 | 0.25 | 0.244650542 | 0 | 0.355418052 |
| ENSMUSG00000096401 | 1.069666964 | 1.061981878 | 1.02186571 | 1.27186571 | 1.244650542 | 1.204119983 | 1.140832123 |
| ENSMUSG00000096402 | 0 | 0 | 0 | 0 | 0 | 0 | 0 |
| ENSMUSG00000096403 | 1.847794126 | 1.841580554 | 1.850219859 | 1.698603967 | 1.816839698 | 1.963787827 | 1.743447983 |
| ENSMUSG00000096405 | 0 | 0 | 0 | 0 | 0 | 0 | 0 |
| ENSMUSG00000096406 | 0 | 0 | 0 | 0 | 0 | 0 | 0 |
| ENSMUSG00000096407 | 0 | 0 | 0 | 0 | 0 | 0 | 0 |
| ENSMUSG00000096408 | 0 | 0 | 0 | 0 | 0 | 0 | 0 |
| ENSMUSG00000096409 | 0 | 0 | 0 | 0 | 0 | 0 | 0 |
| ENSMUSG00000096410 | 0 | 0 | 0 | 0 | 0 | 0 | 0 |
| ENSMUSG00000096411 | 0 | 0 | 0 | 0 | 0 | 0 | 0 |
| ENSMUSG00000096412 | 0 | 0 | 0 | 0 | 0 | 0 | 0 |
| ENSMUSG00000096413 | 0 | 0 | 0 | 0 | 0 | 0 | 0 |
| ENSMUSG00000096414 | 0 | 0 | 0 | 0 | 0 | 0 | 0 |
| ENSMUSG00000096415 | 0 | 0 | 0 | 0 | 0 | 0 | 0 |
| ENSMUSG00000096417 | 0 | 0 | 0 | 0 | 0 | 0 | 0 |
| ENSMUSG00000096420 | 0 | 0 | 0 | 0 | 0 | 0 | 0 |
| ENSMUSG00000096421 | 0 | 0 | 0 | 0 | 0 | 0 | 0 |
| ENSMUSG00000096422 | 0 | 0 | 0 | 0 | 0 | 0 | 0 |
| ENSMUSG00000096423 | 0 | 0 | 0 | 0 | 0 | 0 | 0 |
| ENSMUSG00000096424 | 0 | 0 | 0 | 0 | 0 | 0 | 0 |
| ENSMUSG00000096425 | 0 | 0 | 0 | 0 | 0 | 0 | 0 |

| | | | | | | | |
|---|---|---|---|---|---|---|---|
| ENSMUSG00000096426 | 0 | 0 | 0 | 0 | 0 | 0 | 0 |
| ENSMUSG00000096427 | 0 | 0 | 0 | 0 | 0 | 0 | 0 |
| ENSMUSG00000096429 | 0 | 0 | 0 | 0 | 0 | 0 | 0 |
| ENSMUSG00000096430 | 0 | 0 | 0 | 0 | 0 | 0 | 0 |
| ENSMUSG00000096431 | 0 | 0 | 0 | 0 | 0 | 0 | 0 |
| ENSMUSG00000096432 | 0 | 0.25 | 0 | 0 | 0 | 0 | 0 |
| ENSMUSG00000096433 | 1.985980784 | 1.990000483 | 1.959800947 | 1.998588359 | 1.94314048 | 2.120573931 | 1.74196736 |
| ENSMUSG00000096434 | 0 | 0 | 0 | 0 | 0 | 0 | 0 |
| ENSMUSG00000096435 | 0 | 0 | 0 | 0 | 0 | 0 | 0 |
| ENSMUSG00000096436 | 0 | 0 | 0 | 0 | 0 | 0 | 0 |
| ENSMUSG00000096437 | 0 | 0 | 0 | 0.25 | 0 | 0 | 0 |
| ENSMUSG00000096438 | 0 | 0 | 0 | 0 | 0 | 0 | 0 |
| ENSMUSG00000096439 | 0 | 0 | 0 | 0 | 0 | 0 | 0 |
| ENSMUSG00000096440 | 0.289064826 | 0 | 0.25 | 0.396240625 | 0 | 0 | 0 |
| ENSMUSG00000096441 | 0 | 0 | 0 | 0 | 0 | 0 | 0 |
| ENSMUSG00000096442 | 0 | 0 | 0 | 0 | 0 | 0 | 0 |
| ENSMUSG00000096444 | 0 | 0 | 0 | 0 | 0 | 0 | 0 |
| ENSMUSG00000096445 | 0 | 0 | 0 | 0 | 0 | 0 | 0 |
| ENSMUSG00000096446 | 0 | 0 | 0 | 0 | 0 | 0 | 0 |
| ENSMUSG00000096447 | 0 | 0 | 0 | 0 | 0 | 0 | 0 |
| ENSMUSG00000096448 | 0 | 0 | 0.396240625 | 0 | 0 | 0 | 0 |
| ENSMUSG00000096449 | 0 | 0.5 | 0.580482024 | 0.25 | 0.632412477 | 0 | 0.224243824 |
| ENSMUSG00000096450 | 0 | 0 | 0 | 0 | 0 | 0 | 0 |
| ENSMUSG00000096451 | 0 | 0 | 0 | 0 | 0 | 0 | 0 |
| ENSMUSG00000096452 | 0 | 0 | 0 | 0 | 0 | 0 | 0 |
| ENSMUSG00000096453 | 0 | 0 | 0 | 0 | 0 | 0 | 0 |
| ENSMUSG00000096454 | 0 | 0 | 0 | 0 | 0 | 0 | 0 |
| ENSMUSG00000096455 | 0 | 0 | 0 | 0 | 0 | 0 | 0 |
| ENSMUSG00000096457 | 0 | 0 | 0 | 0 | 0 | 0 | 0 |
| ENSMUSG00000096458 | 0 | 0 | 0 | 0 | 0 | 0 | 0 |
| ENSMUSG00000096459 | 0 | 0 | 0 | 0 | 0 | 0 | 0 |
| ENSMUSG00000096461 | 0 | 0 | 0 | 0 | 0 | 0 | 0 |
| ENSMUSG00000096462 | 0 | 0 | 0 | 0 | 0 | 0 | 0 |
| ENSMUSG00000096463 | 0.811507563 | 0.5 | 0 | 0.646240625 | 0.489301084 | 0.77815125 | 0.355418052 |

| | | | | | | | |
|---|---|---|---|---|---|---|---|
| ENSMUSG00000096464 | 0 | 0 | 0 | 0 | 0 | 0 | 0 |
| ENSMUSG00000096465 | 0 | 0 | 0 | 0 | 0 | 0 | 0 |
| ENSMUSG00000096467 | 0 | 0 | 0 | 0 | 0 | 0 | 0 |
| ENSMUSG00000096468 | 0 | 0 | 0 | 0 | 0 | 0 | 0 |
| ENSMUSG00000096470 | 0 | 0 | 0 | 0 | 0 | 0 | 0 |
| ENSMUSG00000096472 | 1.912079952 | 1.870953944 | 1.901832578 | 1.864857905 | 1.929985312 | 2.1430148 | 1.923435442 |
| ENSMUSG00000096473 | 0 | 0 | 0 | 0 | 0 | 0 | 0 |
| ENSMUSG00000096474 | 0 | 0 | 0.25 | 0 | 0.387761935 | 0 | 0 |
| ENSMUSG00000096475 | 0 | 0 | 0 | 0 | 0 | 0 | 0 |
| ENSMUSG00000096476 | 0 | 0 | 0 | 0 | 0 | 0 | 0 |
| ENSMUSG00000096477 | 0 | 0 | 0 | 0 | 0 | 0 | 0 |
| ENSMUSG00000096478 | 0 | 0 | 0 | 0 | 0 | 0 | 0 |
| ENSMUSG00000096479 | 0 | 0 | 0 | 0 | 0 | 0 | 0 |
| ENSMUSG00000096480 | 0 | 0 | 0 | 0 | 0 | 0 | 0 |
| ENSMUSG00000096481 | 0 | 0 | 0 | 0 | 0 | 0 | 0 |
| ENSMUSG00000096482 | 0 | 0 | 0 | 0 | 0 | 0 | 0 |
| ENSMUSG00000096483 | 0 | 0 | 0 | 0 | 0 | 0 | 0 |
| ENSMUSG00000096484 | 0 | 0 | 0 | 0 | 0 | 0 | 0 |
| ENSMUSG00000096485 | 0 | 0 | 0 | 0 | 0 | 0 | 0 |
| ENSMUSG00000096486 | 0 | 0 | 0 | 0 | 0 | 0 | 0 |
| ENSMUSG00000096487 | 0 | 0 | 0 | 0 | 0 | 0 | 0 |
| ENSMUSG00000096488 | 0 | 0 | 0 | 0 | 0 | 0 | 0 |
| ENSMUSG00000096490 | 0 | 0 | 0 | 0 | 0 | 0 | 0 |
| ENSMUSG00000096492 | 0 | 0 | 0 | 0 | 0 | 0 | 0 |
| ENSMUSG00000096496 | 0 | 0 | 0 | 0 | 0 | 0 | 0 |
| ENSMUSG00000096497 | 0 | 0 | 0 | 0 | 0 | 0 | 0 |
| ENSMUSG00000096498 | 0 | 0 | 0 | 0 | 0 | 0 | 0 |
| ENSMUSG00000096499 | 0 | 0 | 0 | 0 | 0 | 0 | 0 |
| ENSMUSG00000096500 | 0 | 0 | 0 | 0 | 0 | 0 | 0 |
| ENSMUSG00000096502 | 0 | 0 | 0 | 0 | 0 | 0 | 0 |
| ENSMUSG00000096504 | 0 | 0 | 0 | 0 | 0 | 0 | 0 |
| ENSMUSG00000096505 | 0 | 0 | 0 | 0 | 0 | 0 | 0 |
| ENSMUSG00000096506 | 0 | 0 | 0 | 0 | 0 | 0 | 0 |
| ENSMUSG00000096507 | 0 | 0 | 0 | 0 | 0 | 0 | 0 |

| | | | | | | | |
|---|---|---|---|---|---|---|---|
| ENSMUSG00000096508 | 0 | 0 | 0 | 0 | 0 | 0 | 0 |
| ENSMUSG00000096510 | 0 | 0 | 0 | 0 | 0 | 0 | 0 |
| ENSMUSG00000096512 | 0 | 0 | 0 | 0 | 0 | 0 | 0 |
| ENSMUSG00000096513 | 0 | 0 | 0 | 0 | 0 | 0 | 0 |
| ENSMUSG00000096514 | 0 | 0 | 0 | 0 | 0 | 0 | 0 |
| ENSMUSG00000096515 | 0 | 0 | 0 | 0 | 0 | 0 | 0 |
| ENSMUSG00000096516 | 0 | 0 | 0 | 0 | 0 | 0 | 0 |
| ENSMUSG00000096517 | 0 | 0 | 0 | 0 | 0 | 0 | 0 |
| ENSMUSG00000096518 | 0 | 0 | 0 | 0 | 0 | 0 | 0 |
| ENSMUSG00000096519 | 0 | 0 | 0 | 0 | 0 | 0 | 0 |
| ENSMUSG00000096520 | 0 | 0 | 0 | 0 | 0 | 0 | 0 |
| ENSMUSG00000096521 | 0 | 0 | 0 | 0 | 0 | 0 | 0 |
| ENSMUSG00000096522 | 0 | 0 | 0 | 0 | 0 | 0 | 0 |
| ENSMUSG00000096523 | 0 | 0 | 0 | 0 | 0 | 0 | 0 |
| ENSMUSG00000096524 | 0 | 0 | 0 | 0 | 0 | 0 | 0 |
| ENSMUSG00000096525 | 0 | 0 | 0 | 0 | 0 | 0 | 0 |
| ENSMUSG00000096527 | 0 | 0 | 0 | 0 | 0 | 0 | 0 |
| ENSMUSG00000096528 | 0.289064826 | 0.5 | 0.580482024 | 0.5 | 0.632412477 | 0.602059991 | 0.520678036 |
| ENSMUSG00000096529 | 0 | 0 | 0 | 0 | 0 | 0 | 0 |
| ENSMUSG00000096530 | 0 | 0 | 0 | 0 | 0 | 0 | 0 |
| ENSMUSG00000096531 | 0 | 0 | 0 | 0 | 0 | 0 | 0 |
| ENSMUSG00000096532 | 0 | 0 | 0 | 0 | 0 | 0 | 0 |
| ENSMUSG00000096533 | 0 | 0 | 0 | 0 | 0 | 0 | 0 |
| ENSMUSG00000096534 | 0 | 0 | 0 | 0 | 0 | 0 | 0 |
| ENSMUSG00000096535 | 0 | 0 | 0 | 0 | 0 | 0 | 0 |
| ENSMUSG00000096536 | 0 | 0 | 0 | 0 | 0 | 0 | 0 |
| ENSMUSG00000096537 | 0 | 0 | 0 | 0 | 0 | 0 | 0 |
| ENSMUSG00000096538 | 0 | 0 | 0 | 0 | 0 | 0 | 0 |
| ENSMUSG00000096539 | 0 | 0 | 0 | 0 | 0 | 0 | 0 |
| ENSMUSG00000096541 | 0 | 0 | 0 | 0 | 0 | 0 | 0 |
| ENSMUSG00000096544 | 0 | 0 | 0 | 0 | 0 | 0 | 0 |
| ENSMUSG00000096546 | 0 | 0 | 0 | 0 | 0 | 0 | 0 |
| ENSMUSG00000096547 | 0 | 0 | 0 | 0 | 0.244650542 | 0 | 0 |
| ENSMUSG00000096549 | 0 | 0 | 0 | 0 | 0 | 0 | 0 |

| | | | | | | | |
|---|---|---|---|---|---|---|---|
| ENSMUSG00000096550 | 0 | 0 | 0 | 0 | 0 | 0 | 0 |
| ENSMUSG00000096551 | 0 | 0 | 0 | 0 | 0 | 0 | 0 |
| ENSMUSG00000096553 | 0 | 0 | 0 | 0 | 0 | 0 | 0 |
| ENSMUSG00000096554 | 0 | 0 | 0 | 0 | 0 | 0 | 0 |
| ENSMUSG00000096555 | 0 | 0 | 0 | 0 | 0 | 0 | 0 |
| ENSMUSG00000096557 | 0 | 0 | 0 | 0 | 0 | 0 | 0 |
| ENSMUSG00000096558 | 0 | 0 | 0 | 0 | 0 | 0 | 0 |
| ENSMUSG00000096559 | 0 | 0 | 0 | 0 | 0 | 0 | 0 |
| ENSMUSG00000096560 | 0 | 0 | 0 | 0 | 0 | 0 | 0 |
| ENSMUSG00000096561 | 0 | 0 | 0 | 0 | 0 | 0 | 0 |
| ENSMUSG00000096563 | 0 | 0 | 0 | 0 | 0 | 0 | 0 |
| ENSMUSG00000096564 | 0 | 0 | 0 | 0 | 0 | 0 | 0 |
| ENSMUSG00000096565 | 0 | 0 | 0 | 0 | 0 | 0 | 0 |
| ENSMUSG00000096566 | 0 | 0 | 0 | 0 | 0 | 0 | 0 |
| ENSMUSG00000096567 | 0 | 0 | 0 | 0 | 0 | 0 | 0 |
| ENSMUSG00000096568 | 0 | 0 | 0 | 0 | 0 | 0 | 0 |
| ENSMUSG00000096569 | 0 | 0 | 0 | 0 | 0 | 0 | 0 |
| ENSMUSG00000096573 | 0 | 0 | 0 | 0 | 0 | 0 | 0 |
| ENSMUSG00000096574 | 0 | 0 | 0 | 0 | 0 | 0 | 0 |
| ENSMUSG00000096576 | 0 | 0 | 0 | 0 | 0 | 0 | 0 |
| ENSMUSG00000096577 | 0 | 0 | 0 | 0 | 0 | 0 | 0 |
| ENSMUSG00000096578 | 0 | 0 | 0 | 0 | 0 | 0 | 0 |
| ENSMUSG00000096579 | 0 | 0 | 0 | 0 | 0 | 0 | 0 |
| ENSMUSG00000096580 | 0 | 0 | 0 | 0 | 0 | 0 | 0 |
| ENSMUSG00000096582 | 0 | 0 | 0 | 0 | 0 | 0 | 0 |
| ENSMUSG00000096583 | 0 | 0 | 0 | 0 | 0 | 0 | 0 |
| ENSMUSG00000096584 | 0 | 0 | 0 | 0 | 0 | 0 | 0 |
| ENSMUSG00000096585 | 0 | 0 | 0 | 0 | 0 | 0 | 0 |
| ENSMUSG00000096586 | 0 | 0 | 0 | 0 | 0 | 0 | 0 |
| ENSMUSG00000096587 | 0 | 0 | 0 | 0 | 0 | 0 | 0 |
| ENSMUSG00000096589 | 0 | 0 | 0 | 0 | 0 | 0 | 0 |
| ENSMUSG00000096591 | 0 | 0 | 0 | 0.25 | 0 | 0 | 0 |
| ENSMUSG00000096592 | 0 | 0 | 0 | 0 | 0 | 0 | 0 |
| ENSMUSG00000096593 | 0 | 0 | 0 | 0 | 0 | 0 | 0 |

| | | | | | | | |
|---|---|---|---|---|---|---|---|
| ENSMUSG00000096594 | 0 | 0 | 0 | 0 | 0 | 0 | 0 |
| ENSMUSG00000096595 | 0 | 0 | 0 | 0 | 0 | 0 | 0 |
| ENSMUSG00000096596 | 0 | 0 | 0 | 0 | 0 | 0 | 0 |
| ENSMUSG00000096597 | 0 | 0 | 0 | 0 | 0 | 0 | 0 |
| ENSMUSG00000096598 | 0 | 0 | 0 | 0 | 0 | 0 | 0 |
| ENSMUSG00000096599 | 0 | 0 | 0 | 0 | 0 | 0 | 0 |
| ENSMUSG00000096600 | 0 | 0 | 0 | 0 | 0 | 0 | 0 |
| ENSMUSG00000096601 | 0 | 0 | 0 | 0 | 0 | 0 | 0 |
| ENSMUSG00000096604 | 0 | 0 | 0 | 0 | 0 | 0 | 0 |
| ENSMUSG00000096605 | 0 | 0 | 0 | 0 | 0 | 0 | 0 |
| ENSMUSG00000096606 | 0 | 0 | 0 | 0.25 | 0.387761935 | 0.301029996 | 0.355418052 |
| ENSMUSG00000096607 | 0 | 0 | 0 | 0.25 | 0 | 0 | 0.224243824 |
| ENSMUSG00000096612 | 0 | 0 | 0 | 0 | 0 | 0 | 0 |
| ENSMUSG00000096613 | 0 | 0 | 0 | 0 | 0 | 0 | 0 |
| ENSMUSG00000096614 | 0 | 0 | 0 | 0 | 0 | 0 | 0 |
| ENSMUSG00000096615 | 0 | 0 | 0 | 0 | 0 | 0 | 0 |
| ENSMUSG00000096617 | 0 | 0 | 0 | 0 | 0 | 0 | 0 |
| ENSMUSG00000096618 | 0 | 0 | 0 | 0 | 0 | 0 | 0 |
| ENSMUSG00000096619 | 0 | 0.25 | 0 | 0 | 0 | 0 | 0 |
| ENSMUSG00000096620 | 0 | 0 | 0 | 0 | 0 | 0 | 0 |
| ENSMUSG00000096621 | 0 | 0 | 0 | 0 | 0 | 0 | 0 |
| ENSMUSG00000096623 | 0.289064826 | 0 | 0 | 0.25 | 0.568060967 | 0.301029996 | 0.224243824 |
| ENSMUSG00000096624 | 0 | 0 | 0 | 0 | 0 | 0 | 0 |
| ENSMUSG00000096626 | 0 | 0 | 0 | 0 | 0 | 0 | 0 |
| ENSMUSG00000096627 | 0 | 0 | 0 | 0 | 0 | 0 | 0 |
| ENSMUSG00000096628 | 0 | 0 | 0 | 0 | 0 | 0 | 0 |
| ENSMUSG00000096629 | 0 | 0 | 0 | 0 | 0.244650542 | 0 | 0 |
| ENSMUSG00000096630 | 0 | 0 | 0 | 0 | 0 | 0 | 0 |
| ENSMUSG00000096631 | 0 | 0 | 0 | 0 | 0 | 0 | 0 |
| ENSMUSG00000096632 | 0 | 0 | 0 | 0 | 0 | 0 | 0 |
| ENSMUSG00000096633 | 0 | 0 | 0 | 0 | 0 | 0 | 0 |
| ENSMUSG00000096634 | 0 | 0 | 0 | 0 | 0 | 0 | 0 |
| ENSMUSG00000096635 | 0 | 0.25 | 0.25 | 0 | 0.244650542 | 0 | 0.224243824 |
| ENSMUSG00000096638 | 0 | 0 | 0 | 0 | 0 | 0 | 0 |

| | | | | | | | |
|---|---|---|---|---|---|---|---|
| ENSMUSG00000096639 | 0 | 0 | 0 | 0 | 0 | 0 | 0 |
| ENSMUSG00000096640 | 0 | 0 | 0 | 0 | 0 | 0 | 0 |
| ENSMUSG00000096641 | 0 | 0.396240625 | 0 | 0 | 0 | 0 | 0 |
| ENSMUSG00000096642 | 0 | 0 | 0 | 0 | 0 | 0 | 0 |
| ENSMUSG00000096644 | 0 | 0 | 0 | 0 | 0 | 0 | 0 |
| ENSMUSG00000096645 | 0 | 0 | 0 | 0 | 0 | 0 | 0 |
| ENSMUSG00000096646 | 0 | 0 | 0 | 0 | 0 | 0 | 0 |
| ENSMUSG00000096647 | 0 | 0 | 0 | 0 | 0 | 0 | 0.224243824 |
| ENSMUSG00000096648 | 0 | 0 | 0 | 0 | 0 | 0 | 0 |
| ENSMUSG00000096649 | 0 | 0 | 0 | 0 | 0 | 0 | 0 |
| ENSMUSG00000096650 | 0 | 0 | 0 | 0 | 0 | 0 | 0 |
| ENSMUSG00000096651 | 0 | 0 | 0 | 0 | 0 | 0 | 0 |
| ENSMUSG00000096652 | 0 | 0 | 0 | 0 | 0 | 0 | 0 |
| ENSMUSG00000096653 | 0 | 0 | 0 | 0 | 0 | 0 | 0 |
| ENSMUSG00000096654 | 0 | 0 | 0 | 0 | 0 | 0 | 0 |
| ENSMUSG00000096655 | 0.289064826 | 0.396240625 | 0.396240625 | 0.5 | 0.686820904 | 0.301029996 | 0 |
| ENSMUSG00000096656 | 0 | 0 | 0 | 0 | 0 | 0 | 0 |
| ENSMUSG00000096658 | 0 | 0 | 0 | 0 | 0 | 0 | 0 |
| ENSMUSG00000096659 | 0 | 0 | 0 | 0 | 0 | 0 | 0 |
| ENSMUSG00000096660 | 0 | 0 | 0 | 0 | 0 | 0 | 0 |
| ENSMUSG00000096661 | 0 | 0 | 0 | 0 | 0 | 0 | 0 |
| ENSMUSG00000096663 | 0 | 0 | 0 | 0 | 0 | 0 | 0 |
| ENSMUSG00000096664 | 0 | 0 | 0 | 0 | 0 | 0 | 0 |
| ENSMUSG00000096666 | 0 | 0 | 0 | 0 | 0 | 0 | 0 |
| ENSMUSG00000096667 | 0 | 0 | 0 | 0 | 0 | 0 | 0 |
| ENSMUSG00000096668 | 0 | 0 | 0 | 0 | 0 | 0 | 0 |
| ENSMUSG00000096670 | 0 | 0 | 0 | 0 | 0 | 0 | 0 |
| ENSMUSG00000096672 | 0 | 0 | 0 | 0 | 0 | 0 | 0 |
| ENSMUSG00000096673 | 0 | 0 | 0 | 0 | 0 | 0 | 0 |
| ENSMUSG00000096674 | 0 | 0 | 0 | 0 | 0 | 0 | 0 |
| ENSMUSG00000096675 | 0 | 0 | 0 | 0 | 0 | 0 | 0 |
| ENSMUSG00000096676 | 0 | 0 | 0 | 0 | 0 | 0 | 0 |
| ENSMUSG00000096677 | 0 | 0 | 0 | 0 | 0 | 0 | 0 |
| ENSMUSG00000096678 | 0 | 0 | 0 | 0 | 0 | 0 | 0 |

| | | | | | | | |
|---|---|---|---|---|---|---|---|
| ENSMUSG00000096679 | 0 | 0 | 0 | 0 | 0 | 0 | 0 |
| ENSMUSG00000096680 | 0 | 0 | 0 | 0 | 0 | 0 | 0 |
| ENSMUSG00000096681 | 0 | 0 | 0 | 0 | 0 | 0 | 0 |
| ENSMUSG00000096682 | 0 | 0 | 0 | 0 | 0 | 0 | 0 |
| ENSMUSG00000096683 | 0 | 0 | 0 | 0 | 0 | 0 | 0 |
| ENSMUSG00000096684 | 0 | 0 | 0 | 0 | 0 | 0 | 0 |
| ENSMUSG00000096685 | 0 | 0 | 0 | 0 | 0 | 0 | 0 |
| ENSMUSG00000096686 | 0 | 0 | 0 | 0 | 0 | 0 | 0 |
| ENSMUSG00000096687 | 1.806056082 | 1.642463902 | 1.657339155 | 1.664552871 | 1.542223377 | 1.698970004 | 1.496250175 |
| ENSMUSG00000096688 | 0 | 0 | 0 | 0 | 0 | 0 | 0 |
| ENSMUSG00000096689 | 0 | 0 | 0 | 0 | 0 | 0 | 0 |
| ENSMUSG00000096690 | 0 | 0 | 0 | 0 | 0 | 0 | 0 |
| ENSMUSG00000096691 | 0 | 0 | 0 | 0 | 0 | 0 | 0 |
| ENSMUSG00000096692 | 0 | 0 | 0 | 0 | 0 | 0 | 0 |
| ENSMUSG00000096694 | 0 | 0 | 0 | 0 | 0 | 0 | 0 |
| ENSMUSG00000096695 | 0 | 0 | 0 | 0 | 0 | 0 | 0 |
| ENSMUSG00000096696 | 1.129344651 | 1.188721876 | 1.02186571 | 1.29248125 | 1.149965125 | 1.204119983 | 1.054044578 |
| ENSMUSG00000096697 | 0 | 0 | 0 | 0 | 0 | 0 | 0 |
| ENSMUSG00000096698 | 0 | 0 | 0 | 0 | 0 | 0 | 0 |
| ENSMUSG00000096699 | 0.45815691 | 0.25 | 0.396240625 | 0 | 0.568060967 | 0.602059991 | 0.629532004 |
| ENSMUSG00000096700 | 0 | 0 | 0 | 0 | 0 | 0 | 0 |
| ENSMUSG00000096701 | 0 | 0 | 0 | 0.646240625 | 0 | 0.301029996 | 0.355418052 |
| ENSMUSG00000096703 | 0 | 0 | 0 | 0 | 0 | 0 | 0 |
| ENSMUSG00000096704 | 0 | 0 | 0 | 0 | 0 | 0 | 0 |
| ENSMUSG00000096705 | 0 | 0 | 0 | 0 | 0 | 0 | 0 |
| ENSMUSG00000096706 | 0 | 0 | 0 | 0 | 0 | 0 | 0 |
| ENSMUSG00000096707 | 0 | 0 | 0 | 0 | 0 | 0 | 0 |
| ENSMUSG00000096708 | 0 | 0 | 0 | 0 | 0 | 0 | 0 |
| ENSMUSG00000096709 | 0 | 0 | 0 | 0 | 0 | 0 | 0 |
| ENSMUSG00000096710 | 0 | 0 | 0 | 0 | 0 | 0 | 0 |
| ENSMUSG00000096711 | 0 | 0 | 0 | 0 | 0 | 0 | 0 |
| ENSMUSG00000096712 | 0 | 0.25 | 0 | 0 | 0 | 0 | 0 |
| ENSMUSG00000096713 | 0 | 0 | 0 | 0 | 0 | 0 | 0 |
| ENSMUSG00000096714 | 0 | 0 | 0 | 0 | 0 | 0 | 0 |

| | | | | | | | |
|---|---|---|---|---|---|---|---|
| ENSMUSG00000096715 | 0 | 0 | 0 | 0 | 0 | 0 | 0 |
| ENSMUSG00000096716 | 0 | 0 | 0 | 0 | 0 | 0 | 0 |
| ENSMUSG00000096717 | 0 | 0 | 0 | 0 | 0 | 0 | 0 |
| ENSMUSG00000096718 | 0.289064826 | 0.25 | 0 | 0 | 0 | 0.477121255 | 0 |
| ENSMUSG00000096719 | 0 | 0 | 0 | 0 | 0 | 0 | 0 |
| ENSMUSG00000096721 | 0 | 0 | 0 | 0 | 0 | 0 | 0 |
| ENSMUSG00000096722 | 0 | 0.396240625 | 0 | 0.25 | 0.244650542 | 0.301029996 | 0.355418052 |
| ENSMUSG00000096724 | 0 | 0 | 0 | 0 | 0 | 0 | 0 |
| ENSMUSG00000096726 | 0.747221736 | 0.701838731 | 0.701838731 | 0.79248125 | 0.632412477 | 0.77815125 | 0.672731473 |
| ENSMUSG00000096727 | 0.671187741 | 0 | 0.25 | 1.51109853 | 1.484072009 | 1.51851394 | 1.508694612 |
| ENSMUSG00000096728 | 0 | 0 | 0 | 0 | 0 | 0 | 0 |
| ENSMUSG00000096729 | 0 | 0 | 0 | 0 | 0 | 0 | 0 |
| ENSMUSG00000096730 | 0 | 0 | 0 | 0 | 0 | 0 | 0 |
| ENSMUSG00000096731 | 0 | 0 | 0 | 0 | 0 | 0 | 0 |
| ENSMUSG00000096732 | 0 | 0 | 0 | 0 | 0 | 0 | 0 |
| ENSMUSG00000096733 | 0 | 0 | 0 | 0 | 0 | 0 | 0 |
| ENSMUSG00000096734 | 0 | 0 | 0 | 0 | 0 | 0 | 0 |
| ENSMUSG00000096735 | 0 | 0 | 0 | 0 | 0 | 0 | 0 |
| ENSMUSG00000096736 | 0 | 0 | 0 | 0 | 0 | 0 | 0 |
| ENSMUSG00000096737 | 0 | 0 | 0 | 0 | 0 | 0 | 0 |
| ENSMUSG00000096738 | 0 | 0 | 0 | 0 | 0 | 0 | 0 |
| ENSMUSG00000096740 | 1.759671389 | 1.62694866 | 1.561981878 | 1.668106335 | 1.655839964 | 1.86332286 | 1.548827561 |
| ENSMUSG00000096741 | 0 | 0 | 0 | 0 | 0 | 0 | 0 |
| ENSMUSG00000096742 | 0 | 0 | 0 | 0 | 0 | 0 | 0 |
| ENSMUSG00000096743 | 0 | 0 | 0 | 0 | 0 | 0 | 0 |
| ENSMUSG00000096744 | 0 | 0 | 0 | 0 | 0 | 0 | 0 |
| ENSMUSG00000096745 | 0 | 0 | 0 | 0 | 0 | 0 | 0 |
| ENSMUSG00000096746 | 0 | 0 | 0 | 0 | 0 | 0 | 0 |
| ENSMUSG00000096747 | 0 | 0 | 0 | 0 | 0 | 0 | 0 |
| ENSMUSG00000096748 | 0 | 0 | 0 | 0 | 0 | 0 | 0 |
| ENSMUSG00000096749 | 0 | 0 | 0 | 0 | 0 | 0 | 0 |
| ENSMUSG00000096750 | 0 | 0 | 0 | 0 | 0 | 0 | 0 |
| ENSMUSG00000096751 | 0 | 0 | 0 | 0.25 | 0 | 0.301029996 | 0.224243824 |
| ENSMUSG00000096753 | 0 | 0 | 0 | 0 | 0 | 0 | 0 |

| | | | | | | | |
|---|---|---|---|---|---|---|---|
| ENSMUSG00000096755 | 0.45815691 | 0.396240625 | 0.396240625 | 0 | 0.244650542 | 0.301029996 | 0.355418052 |
| ENSMUSG00000096756 | 0 | 0 | 0 | 0 | 0 | 0 | 0 |
| ENSMUSG00000096757 | 0 | 0 | 0 | 0 | 0 | 0 | 0 |
| ENSMUSG00000096758 | 0 | 0 | 0 | 0 | 0 | 0 | 0 |
| ENSMUSG00000096759 | 0 | 0 | 0 | 0 | 0 | 0 | 0 |
| ENSMUSG00000096760 | 0 | 0 | 0 | 0 | 0 | 0 | 0 |
| ENSMUSG00000096761 | 0 | 0 | 0 | 0 | 0 | 0 | 0 |
| ENSMUSG00000096763 | 0 | 0 | 0 | 0 | 0 | 0 | 0 |
| ENSMUSG00000096764 | 0 | 0 | 0 | 0 | 0 | 0 | 0.224243824 |
| ENSMUSG00000096765 | 0 | 0 | 0 | 0 | 0 | 0 | 0 |
| ENSMUSG00000096766 | 0 | 0 | 0 | 0 | 0 | 0 | 0 |
| ENSMUSG00000096767 | 0 | 0 | 0 | 0 | 0 | 0 | 0 |
| ENSMUSG00000096768 | 0.91631382 | 1.02186571 | 1.17510993 | 1.114857905 | 1.293076518 | 1.176091259 | 1.40946263 |
| ENSMUSG00000096769 | 0 | 0 | 0 | 0 | 0 | 0 | 0 |
| ENSMUSG00000096770 | 0 | 0 | 0 | 0 | 0 | 0 | 0 |
| ENSMUSG00000096771 | 0 | 0 | 0 | 0 | 0 | 0 | 0 |
| ENSMUSG00000096772 | 0 | 0 | 0 | 0 | 0 | 0 | 0 |
| ENSMUSG00000096773 | 0 | 0 | 0 | 0 | 0 | 0 | 0 |
| ENSMUSG00000096774 | 0 | 0 | 0 | 0 | 0 | 0 | 0 |
| ENSMUSG00000096775 | 0 | 0 | 0 | 0 | 0.244650542 | 0 | 0 |
| ENSMUSG00000096776 | 0 | 0 | 0 | 0 | 0 | 0 | 0 |
| ENSMUSG00000096777 | 0 | 0 | 0 | 0 | 0 | 0 | 0 |
| ENSMUSG00000096779 | 0 | 0 | 0 | 0 | 0 | 0 | 0 |
| ENSMUSG00000096780 | 1.783508299 | 1.685366747 | 1.726722649 | 1.649978211 | 1.751666404 | 1.819543936 | 1.657681529 |
| ENSMUSG00000096782 | 0 | 0 | 0 | 0 | 0 | 0 | 0 |
| ENSMUSG00000096784 | 0 | 0 | 0 | 0 | 0 | 0 | 0 |
| ENSMUSG00000096785 | 0 | 0 | 0 | 0 | 0 | 0 | 0 |
| ENSMUSG00000096786 | 0 | 0 | 0 | 0 | 0 | 0 | 0 |
| ENSMUSG00000096788 | 0 | 0 | 0 | 0 | 0 | 0 | 0 |
| ENSMUSG00000096789 | 0 | 0 | 0 | 0 | 0 | 0 | 0 |
| ENSMUSG00000096790 | 0 | 0 | 0 | 0 | 0 | 0 | 0 |
| ENSMUSG00000096791 | 0 | 0 | 0 | 0 | 0 | 0 | 0 |
| ENSMUSG00000096792 | 0 | 0 | 0 | 0 | 0 | 0 | 0 |
| ENSMUSG00000096793 | 0 | 0 | 0 | 0 | 0 | 0 | 0 |

| | | | | | | | |
|---|---|---|---|---|---|---|---|
| ENSMUSG00000096794 | 0 | 0 | 0 | 0 | 0 | 0 | 0 |
| ENSMUSG00000096795 | 1.747221736 | 1.685366747 | 1.593759858 | 1.921625132 | 1.870073561 | 2.120573931 | 1.726776051 |
| ENSMUSG00000096796 | 0 | 0 | 0 | 0 | 0 | 0 | 0 |
| ENSMUSG00000096797 | 0 | 0 | 0 | 0 | 0 | 0 | 0 |
| ENSMUSG00000096798 | 0 | 0 | 0 | 0 | 0 | 0 | 0 |
| ENSMUSG00000096799 | 0 | 0 | 0 | 0 | 0 | 0 | 0 |
| ENSMUSG00000096800 | 0 | 0 | 0 | 0 | 0 | 0 | 0 |
| ENSMUSG00000096801 | 0 | 0 | 0 | 0 | 0 | 0 | 0 |
| ENSMUSG00000096802 | 0 | 0 | 0 | 0 | 0 | 0 | 0 |
| ENSMUSG00000096803 | 0.289064826 | 0 | 0 | 0 | 0 | 0 | 0 |
| ENSMUSG00000096804 | 0 | 0 | 0 | 0 | 0 | 0 | 0 |
| ENSMUSG00000096805 | 0 | 0 | 0 | 0 | 0 | 0 | 0 |
| ENSMUSG00000096806 | 0 | 0 | 0 | 0 | 0 | 0 | 0 |
| ENSMUSG00000096807 | 0 | 0.396240625 | 0 | 0 | 0 | 0 | 0 |
| ENSMUSG00000096808 | 0.578129653 | 0.580482024 | 0.75 | 0.5 | 0.387761935 | 0.477121255 | 0.355418052 |
| ENSMUSG00000096809 | 0 | 0 | 0 | 0 | 0 | 0 | 0 |
| ENSMUSG00000096810 | 0 | 0.25 | 0 | 0 | 0.244650542 | 0 | 0 |
| ENSMUSG00000096811 | 0 | 0 | 0 | 0 | 0 | 0 | 0 |
| ENSMUSG00000096812 | 0 | 0 | 0 | 0 | 0 | 0 | 0 |
| ENSMUSG00000096813 | 0 | 0 | 0 | 0 | 0 | 0 | 0 |
| ENSMUSG00000096814 | 0 | 0 | 0 | 0 | 0 | 0 | 0 |
| ENSMUSG00000096815 | 0 | 0 | 0 | 0 | 0 | 0 | 0 |
| ENSMUSG00000096817 | 0 | 0 | 0 | 0 | 0 | 0 | 0 |
| ENSMUSG00000096818 | 0 | 0 | 0 | 0 | 0 | 0 | 0 |
| ENSMUSG00000096819 | 0 | 0 | 0 | 0 | 0 | 0 | 0 |
| ENSMUSG00000096820 | 0 | 0 | 0 | 0 | 0 | 0 | 0 |
| ENSMUSG00000096821 | 0 | 0 | 0 | 0 | 0 | 0 | 0 |
| ENSMUSG00000096822 | 0 | 0 | 0 | 0 | 0 | 0 | 0 |
| ENSMUSG00000096823 | 0 | 0 | 0 | 0 | 0 | 0 | 0 |
| ENSMUSG00000096824 | 0 | 0 | 0 | 0 | 0 | 0 | 0 |
| ENSMUSG00000096825 | 0 | 0 | 0 | 0 | 0 | 0 | 0 |
| ENSMUSG00000096826 | 0 | 0 | 0 | 0 | 0 | 0 | 0 |
| ENSMUSG00000096827 | 0 | 0 | 0 | 0 | 0 | 0 | 0 |
| ENSMUSG00000096828 | 0 | 0 | 0 | 0 | 0 | 0 | 0 |

| | | | | | | | |
|---|---|---|---|---|---|---|---|
| ENSMUSG00000096829 | 0 | 0 | 0 | 0 | 0 | 0 | 0 |
| ENSMUSG00000096830 | 0 | 0 | 0 | 0 | 0 | 0 | 0 |
| ENSMUSG00000096831 | 0 | 0 | 0 | 0 | 0 | 0 | 0 |
| ENSMUSG00000096832 | 0 | 0 | 0 | 0 | 0 | 0 | 0 |
| ENSMUSG00000096833 | 0 | 0 | 0 | 0 | 0 | 0 | 0 |
| ENSMUSG00000096834 | 0 | 0 | 0 | 0 | 0 | 0 | 0 |
| ENSMUSG00000096835 | 0 | 0 | 0 | 0 | 0 | 0 | 0 |
| ENSMUSG00000096838 | 0 | 0 | 0 | 0 | 0 | 0 | 0 |
| ENSMUSG00000096839 | 0 | 0 | 0 | 0 | 0 | 0 | 0 |
| ENSMUSG00000096840 | 0 | 0 | 0 | 0 | 0 | 0 | 0 |
| ENSMUSG00000096841 | 0.45815691 | 0 | 0.396240625 | 0.580482024 | 0 | 0.301029996 | 0.224243824 |
| ENSMUSG00000096842 | 0 | 0 | 0 | 0 | 0 | 0 | 0 |
| ENSMUSG00000096844 | 0 | 0 | 0 | 0 | 0 | 0 | 0 |
| ENSMUSG00000096845 | 0 | 0 | 0 | 0 | 0 | 0 | 0 |
| ENSMUSG00000096846 | 0 | 0 | 0 | 0 | 0 | 0 | 0 |
| ENSMUSG00000096847 | 0 | 0.5 | 0.396240625 | 0.864857905 | 1 | 1.113943352 | 1.110947927 |
| ENSMUSG00000096848 | 0 | 0 | 0 | 0 | 0 | 0 | 0 |
| ENSMUSG00000096849 | 0 | 0 | 0 | 0 | 0 | 0 | 0 |
| ENSMUSG00000096850 | 0 | 0 | 0 | 0 | 0 | 0 | 0 |
| ENSMUSG00000096851 | 0 | 0 | 0 | 0 | 0 | 0 | 0 |
| ENSMUSG00000096852 | 0 | 0 | 0 | 0 | 0 | 0 | 0 |
| ENSMUSG00000096853 | 0 | 0 | 0 | 0 | 0 | 0 | 0 |
| ENSMUSG00000096854 | 0.289064826 | 0.92510993 | 0.701838731 | 0.864857905 | 0.387761935 | 0 | 0.672731473 |
| ENSMUSG00000096855 | 0 | 0 | 0 | 0 | 0 | 0 | 0 |
| ENSMUSG00000096856 | 0 | 0.25 | 0 | 0 | 0.244650542 | 0.602059991 | 0.224243824 |
| ENSMUSG00000096857 | 0 | 0 | 0 | 0 | 0 | 0 | 0 |
| ENSMUSG00000096858 | 0 | 0 | 0 | 0 | 0 | 0 | 0 |
| ENSMUSG00000096859 | 0 | 0 | 0 | 0 | 0 | 0 | 0 |
| ENSMUSG00000096861 | 0 | 0 | 0 | 0 | 0 | 0 | 0 |
| ENSMUSG00000096862 | 0 | 0 | 0 | 0 | 0 | 0 | 0 |
| ENSMUSG00000096863 | 0 | 0 | 0 | 0 | 0 | 0 | 0 |
| ENSMUSG00000096864 | 0 | 0 | 0 | 0 | 0 | 0 | 0 |
| ENSMUSG00000096865 | 0 | 0 | 0 | 0 | 0 | 0 | 0 |
| ENSMUSG00000096866 | 0 | 0 | 0 | 0 | 0 | 0 | 0 |

| | | | | | | | |
|---|---|---|---|---|---|---|---|
| ENSMUSG00000096867 | 0 | 0.25 | 0 | 0 | 0 | 0 | 0 |
| ENSMUSG00000096868 | 0 | 0 | 0 | 0 | 0 | 0 | 0 |
| ENSMUSG00000096869 | 0 | 0 | 0 | 0 | 0 | 0 | 0 |
| ENSMUSG00000096870 | 0 | 0 | 0 | 0 | 0 | 0 | 0 |
| ENSMUSG00000096871 | 0 | 0 | 0 | 0 | 0 | 0 | 0 |
| ENSMUSG00000096872 | 0 | 0 | 0 | 0 | 0 | 0 | 0 |
| ENSMUSG00000096873 | 0 | 0 | 0 | 0 | 0 | 0 | 0 |
| ENSMUSG00000096874 | 0 | 0 | 0 | 0 | 0 | 0 | 0 |
| ENSMUSG00000096875 | 0 | 0 | 0 | 0 | 0 | 0 | 0 |
| ENSMUSG00000096876 | 0 | 0 | 0 | 0 | 0 | 0 | 0 |
| ENSMUSG00000096878 | 0 | 0 | 0 | 0 | 0 | 0 | 0 |
| ENSMUSG00000096879 | 0.289064826 | 0 | 0 | 0 | 0 | 0 | 0 |
| ENSMUSG00000096880 | 0 | 0 | 0 | 0 | 0 | 0 | 0 |
| ENSMUSG00000096881 | 0 | 0 | 0 | 0 | 0 | 0 | 0 |
| ENSMUSG00000096882 | 0 | 0 | 0 | 0 | 0 | 0 | 0 |
| ENSMUSG00000096883 | 0 | 0 | 0.25 | 0 | 0.489301084 | 0 | 0.448487648 |
| ENSMUSG00000096884 | 0 | 0 | 0 | 0 | 0 | 0 | 0 |
| ENSMUSG00000096885 | 0 | 0 | 0 | 0 | 0 | 0 | 0 |
| ENSMUSG00000096887 | 0 | 0 | 0 | 0.25 | 0 | 0 | 0 |
| ENSMUSG00000096889 | 0 | 0 | 0 | 0 | 0 | 0 | 0 |
| ENSMUSG00000096890 | 0 | 0 | 0 | 0 | 0 | 0 | 0 |
| ENSMUSG00000096891 | 0 | 0 | 0 | 0 | 0 | 0 | 0 |
| ENSMUSG00000096893 | 0 | 0 | 0 | 0 | 0 | 0 | 0 |
| ENSMUSG00000096894 | 0 | 0 | 0 | 0 | 0 | 0 | 0 |
| ENSMUSG00000096895 | 0 | 0 | 0 | 0 | 0 | 0 | 0 |
| ENSMUSG00000096896 | 0 | 0 | 0 | 0 | 0 | 0 | 0 |
| ENSMUSG00000096897 | 0 | 0 | 0 | 0 | 0 | 0 | 0 |
| ENSMUSG00000096898 | 0 | 0 | 0 | 0 | 0 | 0 | 0 |
| ENSMUSG00000096899 | 0 | 0 | 0 | 0 | 0 | 0 | 0 |
| ENSMUSG00000096900 | 0 | 0 | 0 | 0 | 0 | 0 | 0 |
| ENSMUSG00000096901 | 0 | 0 | 0 | 0 | 0 | 0 | 0 |
| ENSMUSG00000096902 | 0 | 0 | 0 | 0 | 0 | 0 | 0 |
| ENSMUSG00000096903 | 0 | 0 | 0 | 0 | 0 | 0 | 0 |
| ENSMUSG00000096904 | 0 | 0 | 0 | 0 | 0 | 0 | 0 |

| | | | | | | | |
|---|---|---|---|---|---|---|---|
| ENSMUSG00000096905 | 0 | 0 | 0 | 0 | 0 | 0 | 0 |
| ENSMUSG00000096906 | 0 | 0 | 0 | 0 | 0 | 0 | 0 |
| ENSMUSG00000096907 | 0 | 0 | 0 | 0 | 0 | 0 | 0 |
| ENSMUSG00000096908 | 0 | 0 | 0 | 0 | 0 | 0 | 0 |
| ENSMUSG00000096910 | 1.578129653 | 1.431980114 | 1.561981878 | 1.488549078 | 1.433158226 | 1.579783597 | 1.329931221 |
| ENSMUSG00000096911 | 0 | 0 | 0 | 0 | 0 | 0 | 0 |
| ENSMUSG00000096912 | 0 | 0 | 0 | 0 | 0 | 0 | 0 |
| ENSMUSG00000096913 | 0 | 0 | 0 | 0 | 0 | 0 | 0 |
| ENSMUSG00000096914 | 0 | 0.396240625 | 0.25 | 0 | 0 | 0 | 0.355418052 |
| ENSMUSG00000096915 | 0 | 0 | 0 | 0 | 0 | 0 | 0 |
| ENSMUSG00000096916 | 1.876566388 | 1.723704441 | 1.75 | 1.880890489 | 1.816839698 | 2.064457989 | 1.625302983 |
| ENSMUSG00000096917 | 1.181541736 | 1.188721876 | 1.114857905 | 1.146240625 | 1.149965125 | 0.903089987 | 0.829800754 |
| ENSMUSG00000096918 | 0 | 0 | 0 | 0 | 0 | 0 | 0 |
| ENSMUSG00000096919 | 0 | 0.25 | 0 | 0 | 0 | 0 | 0 |
| ENSMUSG00000096920 | 0 | 0 | 0.25 | 0 | 0.387761935 | 0 | 0.224243824 |
| ENSMUSG00000096921 | 0 | 0 | 0 | 0 | 0.244650542 | 0 | 0 |
| ENSMUSG00000096922 | 0 | 0 | 0 | 0 | 0 | 0 | 0 |
| ENSMUSG00000096923 | 0.45815691 | 0.25 | 0.25 | 0.25 | 0.244650542 | 0 | 0.355418052 |
| ENSMUSG00000096924 | 0.867194479 | 0.646240625 | 0.396240625 | 0.580482024 | 0.568060967 | 0 | 0.579661877 |
| ENSMUSG00000096926 | 0 | 0.646240625 | 0.580482024 | 0.646240625 | 0.686820904 | 0.301029996 | 0.710836105 |
| ENSMUSG00000096927 | 0 | 0 | 0 | 0 | 0 | 0 | 0 |
| ENSMUSG00000096929 | 0 | 0.25 | 0.396240625 | 0.25 | 0 | 0 | 0.355418052 |
| ENSMUSG00000096930 | 0 | 0 | 0 | 0 | 0 | 0 | 0 |
| ENSMUSG00000096931 | 0 | 0 | 0 | 0 | 0 | 0 | 0 |
| ENSMUSG00000096932 | 0.289064826 | 0.396240625 | 0.396240625 | 0.5 | 0 | 0 | 0.448487648 |
| ENSMUSG00000096934 | 0 | 0 | 0 | 0 | 0 | 0 | 0 |
| ENSMUSG00000096935 | 0 | 0.396240625 | 0.25 | 0.646240625 | 0.632412477 | 0.477121255 | 0.672731473 |
| ENSMUSG00000096936 | 0 | 0 | 0 | 0 | 0 | 0 | 0 |
| ENSMUSG00000096937 | 0 | 0 | 0 | 0 | 0.244650542 | 0 | 0 |
| ENSMUSG00000096938 | 0.671187741 | 0.5 | 0.25 | 0.25 | 0.387761935 | 0 | 0.224243824 |
| ENSMUSG00000096939 | 0 | 0 | 0 | 0 | 0 | 0 | 0 |
| ENSMUSG00000096940 | 0.45815691 | 0.25 | 0 | 0 | 0 | 0 | 0.224243824 |
| ENSMUSG00000096941 | 0 | 0 | 0 | 0 | 0 | 0 | 0 |
| ENSMUSG00000096942 | 0 | 0 | 0 | 0 | 0 | 0 | 0 |

| | | | | | | | |
|---|---|---|---|---|---|---|---|
| ENSMUSG00000096943 | 0 | 0 | 0 | 0 | 0 | 0 | 0 |
| ENSMUSG00000096944 | 0 | 0.580482024 | 0 | 0.25 | 0 | 0 | 0.355418052 |
| ENSMUSG00000096945 | 0.289064826 | 0 | 0.25 | 0.396240625 | 0.244650542 | 0.301029996 | 0.355418052 |
| ENSMUSG00000096946 | 0.578129653 | 0.75 | 0.646240625 | 0.396240625 | 0.686820904 | 0.301029996 | 0.224243824 |
| ENSMUSG00000096947 | 0 | 0 | 0 | 0 | 0 | 0 | 0 |
| ENSMUSG00000096948 | 0.289064826 | 0.580482024 | 0.396240625 | 0.25 | 0.244650542 | 0.477121255 | 0.448487648 |
| ENSMUSG00000096949 | 0 | 0 | 0 | 0 | 0 | 0 | 0 |
| ENSMUSG00000096950 | 0.45815691 | 0.701838731 | 0.75 | 0.5 | 0.244650542 | 0 | 0.448487648 |
| ENSMUSG00000096953 | 0 | 0 | 0 | 0 | 0 | 0 | 0 |
| ENSMUSG00000096954 | 1.558729299 | 1.321350555 | 1.476722649 | 1.26109853 | 1.121713561 | 1.113943352 | 1.066254157 |
| ENSMUSG00000096955 | 0 | 0 | 0 | 0 | 0 | 0 | 0 |
| ENSMUSG00000096956 | 1.605636262 | 1.482684334 | 1.566696635 | 0.830482024 | 0.686820904 | 1.041392685 | 0.896975297 |
| ENSMUSG00000096957 | 0.671187741 | 0.396240625 | 0.25 | 0.396240625 | 0.489301084 | 0.301029996 | 0.355418052 |
| ENSMUSG00000096958 | 0 | 0 | 0.25 | 0 | 0.244650542 | 0 | 0 |
| ENSMUSG00000096959 | 0.289064826 | 0.79248125 | 0.396240625 | 0.646240625 | 0.632412477 | 0.84509804 | 0.803905701 |
| ENSMUSG00000096960 | 0.289064826 | 0.25 | 0.646240625 | 0 | 0.244650542 | 0 | 0.224243824 |
| ENSMUSG00000096962 | 0 | 0.25 | 0 | 0 | 0 | 0 | 0.224243824 |
| ENSMUSG00000096963 | 0 | 0 | 0 | 0 | 0 | 0 | 0 |
| ENSMUSG00000096964 | 0 | 0 | 0 | 0.25 | 0.244650542 | 0 | 0.224243824 |
| ENSMUSG00000096965 | 1.100572389 | 1.188721876 | 1.17510993 | 0.75 | 0.568060967 | 0.477121255 | 0.710836105 |
| ENSMUSG00000096966 | 0 | 0 | 0 | 0 | 0 | 0 | 0 |
| ENSMUSG00000096967 | 0.960252568 | 1.02186571 | 0.976722649 | 1.04248125 | 0.931471446 | 1.041392685 | 0.935079929 |
| ENSMUSG00000096968 | 0.671187741 | 0.25 | 0.396240625 | 0.5 | 0 | 0.301029996 | 0.579661877 |
| ENSMUSG00000096969 | 0 | 0 | 0 | 0 | 0 | 0 | 0 |
| ENSMUSG00000096971 | 0.960252568 | 1.130890489 | 1.25 | 0.646240625 | 0.905314583 | 0.903089987 | 0.829800754 |
| ENSMUSG00000096972 | 0 | 0 | 0 | 0 | 0 | 0 | 0 |
| ENSMUSG00000096973 | 0.289064826 | 0.25 | 0 | 0.25 | 0 | 0 | 0 |
| ENSMUSG00000096974 | 0 | 0.25 | 0 | 0 | 0 | 0.477121255 | 0 |
| ENSMUSG00000096975 | 0.578129653 | 0.79248125 | 0.830482024 | 0.75 | 0.877063019 | 0.84509804 | 0.775756176 |
| ENSMUSG00000096976 | 0 | 0.396240625 | 0.25 | 0 | 0.387761935 | 0 | 0.224243824 |
| ENSMUSG00000096977 | 0.747221736 | 0.5 | 0.5 | 0.701838731 | 0.77552387 | 0.477121255 | 0.829800754 |
| ENSMUSG00000096978 | 0 | 0.25 | 0 | 0.396240625 | 0 | 0.301029996 | 0 |
| ENSMUSG00000096979 | 1.181541736 | 1.130890489 | 1.302363341 | 0.79248125 | 0.733951626 | 0.602059991 | 0.448487648 |
| ENSMUSG00000096980 | 0.289064826 | 0.25 | 0 | 0 | 0.244650542 | 0 | 0 |

| | | | | | | | |
|---|---|---|---|---|---|---|---|
| ENSMUSG00000096981 | 0.289064826 | 0.5 | 0.25 | 0.580482024 | 0.387761935 | 0.477121255 | 0.579661877 |
| ENSMUSG00000096982 | 0 | 0 | 0 | 0 | 0 | 0 | 0 |
| ENSMUSG00000096983 | 0.671187741 | 0.830482024 | 0.830482024 | 0.79248125 | 0.812711509 | 1 | 0.74492186 |
| ENSMUSG00000096984 | 0 | 0 | 0.396240625 | 0 | 0.244650542 | 0 | 0.448487648 |
| ENSMUSG00000096986 | 0.289064826 | 0 | 0.25 | 0.5 | 0.244650542 | 0 | 0 |
| ENSMUSG00000096987 | 0.867194479 | 1.04248125 | 0.79248125 | 0.701838731 | 0.846351821 | 0.477121255 | 0.803905701 |
| ENSMUSG00000096988 | 0.289064826 | 0.396240625 | 0.396240625 | 0.396240625 | 0.244650542 | 0 | 0.520678036 |
| ENSMUSG00000096990 | 0.289064826 | 0.25 | 0.396240625 | 0 | 0.244650542 | 0.301029996 | 0 |
| ENSMUSG00000096991 | 1.129344651 | 1.02186571 | 1 | 0.951838731 | 0.905314583 | 0.84509804 | 0.803905701 |
| ENSMUSG00000096992 | 0.671187741 | 0.864857905 | 0.580482024 | 0.701838731 | 0.686820904 | 0.301029996 | 0.672731473 |
| ENSMUSG00000096993 | 0 | 0 | 0 | 0 | 0 | 0.301029996 | 0 |
| ENSMUSG00000096995 | 1.129344651 | 0.951838731 | 1.130890489 | 1.098079356 | 0.877063019 | 1 | 1.054044578 |
| ENSMUSG00000096996 | 1.432083896 | 1.226722649 | 1.494319981 | 1.17510993 | 1.293076518 | 1.342422681 | 1.20919388 |
| ENSMUSG00000096997 | 0 | 0 | 0 | 0 | 0 | 0 | 0 |
| ENSMUSG00000096999 | 0 | 0 | 0 | 0 | 0 | 0 | 0 |
| ENSMUSG00000097000 | 1.568542279 | 1.451838731 | 1.476722649 | 1.418106335 | 1.401338844 | 1.51851394 | 1.245582246 |
| ENSMUSG00000097001 | 0 | 0 | 0 | 0 | 0 | 0 | 0 |
| ENSMUSG00000097002 | 0.45815691 | 0.646240625 | 0.5 | 0 | 0 | 0 | 0.224243824 |
| ENSMUSG00000097003 | 0.289064826 | 0 | 0 | 0 | 0.244650542 | 0.301029996 | 0 |
| ENSMUSG00000097004 | 0.289064826 | 0.5 | 0.646240625 | 0.396240625 | 0.632412477 | 0.602059991 | 0.520678036 |
| ENSMUSG00000097005 | 0 | 0 | 0 | 0 | 0 | 0 | 0 |
| ENSMUSG00000097006 | 2.23973002 | 2.091580554 | 2.04248125 | 1.981203126 | 2.023617914 | 2.195899652 | 1.917452159 |
| ENSMUSG00000097007 | 0.45815691 | 0 | 0.25 | 0.25 | 0 | 0 | 0.355418052 |
| ENSMUSG00000097008 | 0.289064826 | 0 | 0.5 | 0.25 | 0.387761935 | 0 | 0.224243824 |
| ENSMUSG00000097009 | 0.289064826 | 0.25 | 0 | 0 | 0 | 0 | 0 |
| ENSMUSG00000097010 | 0 | 0 | 0 | 0 | 0 | 0.301029996 | 0 |
| ENSMUSG00000097011 | 0 | 0 | 0.25 | 0 | 0 | 0 | 0 |
| ENSMUSG00000097012 | 0 | 0 | 0 | 0 | 0 | 0 | 0 |
| ENSMUSG00000097013 | 0.671187741 | 0.5 | 0.5 | 0.5 | 0.733951626 | 0 | 0.224243824 |
| ENSMUSG00000097014 | 0 | 0.25 | 0.396240625 | 0.396240625 | 0 | 0 | 0.355418052 |
| ENSMUSG00000097015 | 0.45815691 | 0.5 | 0.5 | 0.580482024 | 0.489301084 | 0.698970004 | 0.520678036 |
| ENSMUSG00000097016 | 0.289064826 | 0.396240625 | 0 | 0.25 | 0.244650542 | 0 | 0 |
| ENSMUSG00000097017 | 0.45815691 | 0.25 | 0.580482024 | 0.701838731 | 0.733951626 | 0.602059991 | 0.520678036 |
| ENSMUSG00000097018 | 0 | 0 | 0 | 0 | 0 | 0 | 0 |

| | | | | | | | |
|---|---|---|---|---|---|---|---|
| ENSMUSG00000097019 | 0.289064826 | 0 | 0 | 0 | 0 | 0 | 0 |
| ENSMUSG00000097020 | 0.578129653 | 0.646240625 | 0.5 | 0.79248125 | 0.632412477 | 0.698970004 | 0.355418052 |
| ENSMUSG00000097021 | 0.289064826 | 0.396240625 | 0 | 0 | 0 | 0.602059991 | 0.224243824 |
| ENSMUSG00000097022 | 0.578129653 | 0.5 | 0.25 | 0.25 | 0.489301084 | 0 | 0 |
| ENSMUSG00000097023 | 0 | 0 | 0 | 0 | 0 | 0 | 0 |
| ENSMUSG00000097024 | 0 | 0 | 0 | 0 | 0 | 0 | 0 |
| ENSMUSG00000097025 | 0 | 0 | 0 | 0 | 0 | 0 | 0.224243824 |
| ENSMUSG00000097026 | 1.389637216 | 1.27186571 | 1.17510993 | 1.04248125 | 1.074582839 | 1 | 0.935079929 |
| ENSMUSG00000097027 | 0.867194479 | 0.646240625 | 0.864857905 | 0.951838731 | 0.846351821 | 0.477121255 | 0.629532004 |
| ENSMUSG00000097028 | 1.693336259 | 1.602347734 | 1.589388001 | 1 | 0.931471446 | 1 | 1.014380832 |
| ENSMUSG00000097029 | 0 | 0 | 0 | 0 | 0 | 0 | 0 |
| ENSMUSG00000097030 | 0 | 0 | 0 | 0 | 0 | 0 | 0 |
| ENSMUSG00000097031 | 0 | 0 | 0 | 0 | 0 | 0 | 0 |
| ENSMUSG00000097032 | 0.578129653 | 0.396240625 | 0.5 | 0.75 | 0.686820904 | 0.84509804 | 0 |
| ENSMUSG00000097033 | 1.389637216 | 1.311981878 | 1.348079356 | 1.214495249 | 1.254881871 | 1.255272505 | 1.238624656 |
| ENSMUSG00000097035 | 0 | 0 | 0.25 | 0 | 0 | 0 | 0.448487648 |
| ENSMUSG00000097036 | 0 | 0 | 0 | 0 | 0 | 0.301029996 | 0 |
| ENSMUSG00000097037 | 0 | 0 | 0.25 | 0 | 0 | 0 | 0.224243824 |
| ENSMUSG00000097038 | 0 | 0 | 0 | 0 | 0 | 0 | 0 |
| ENSMUSG00000097039 | 1.982401085 | 1.896240625 | 1.860735874 | 1.845926073 | 1.848234258 | 2.041392685 | 1.687112305 |
| ENSMUSG00000097040 | 0.578129653 | 0.580482024 | 0.396240625 | 0.5 | 0.489301084 | 0.477121255 | 0.224243824 |
| ENSMUSG00000097041 | 0 | 0 | 0 | 0 | 0 | 0 | 0 |
| ENSMUSG00000097042 | 1.249317394 | 1.114857905 | 1.114857905 | 1.061981878 | 1.106691885 | 1.230448921 | 1.089372236 |
| ENSMUSG00000097043 | 0 | 0 | 0 | 0 | 0 | 0 | 0 |
| ENSMUSG00000097044 | 0 | 0 | 0 | 0 | 0 | 0 | 0 |
| ENSMUSG00000097045 | 0 | 0 | 0 | 0.5 | 0 | 0 | 0 |
| ENSMUSG00000097047 | 0.45815691 | 0.580482024 | 0.5 | 0 | 0 | 0.301029996 | 0.448487648 |
| ENSMUSG00000097048 | 1.342375483 | 1.25 | 1.282320754 | 1.403677461 | 1.484072009 | 1.505149978 | 1.374453863 |
| ENSMUSG00000097049 | 0 | 0 | 0 | 0 | 0 | 0 | 0 |
| ENSMUSG00000097050 | 0 | 0 | 0 | 0 | 0.244650542 | 0 | 0.224243824 |
| ENSMUSG00000097051 | 0.578129653 | 0.701838731 | 0.646240625 | 0.646240625 | 0.686820904 | 0.698970004 | 0.448487648 |
| ENSMUSG00000097052 | 0.289064826 | 0.396240625 | 0 | 0 | 0 | 0 | 0 |
| ENSMUSG00000097053 | 0 | 0 | 0 | 0 | 0 | 0 | 0 |
| ENSMUSG00000097054 | 0 | 0 | 0 | 0 | 0 | 0 | 0 |

| | | | | | | | |
|---|---|---|---|---|---|---|---|
| ENSMUSG00000097055 | 0.45815691 | 0.396240625 | 0.396240625 | 0.5 | 0.244650542 | 0 | 0.520678036 |
| ENSMUSG00000097056 | 1.249317394 | 1.130890489 | 1.02186571 | 1.061981878 | 0.877063019 | 0.84509804 | 0.803905701 |
| ENSMUSG00000097057 | 0.91631382 | 1.098079356 | 1.061981878 | 1.061981878 | 1.212046813 | 1.113943352 | 1.041356071 |
| ENSMUSG00000097058 | 0.45815691 | 0.5 | 0 | 0 | 0 | 0.301029996 | 0.520678036 |
| ENSMUSG00000097059 | 1.036286563 | 0.79248125 | 0.79248125 | 1.061981878 | 1.057362051 | 1.176091259 | 0.916588299 |
| ENSMUSG00000097060 | 0 | 0 | 0 | 0 | 0 | 0 | 0 |
| ENSMUSG00000097061 | 1.45815691 | 1.321350555 | 1.282320754 | 1.238549078 | 1.188507684 | 1.255272505 | 1.078019652 |
| ENSMUSG00000097062 | 1.516991255 | 1.188721876 | 1.464495249 | 1.282320754 | 1.200473444 | 1.278753601 | 1.265599895 |
| ENSMUSG00000097063 | 0 | 0 | 0 | 0 | 0 | 0 | 0 |
| ENSMUSG00000097065 | 0.289064826 | 0.5 | 0.25 | 0.646240625 | 0 | 0 | 0.224243824 |
| ENSMUSG00000097066 | 0 | 0 | 0 | 0 | 0 | 0 | 0 |
| ENSMUSG00000097067 | 0 | 0 | 0 | 0.25 | 0 | 0 | 0 |
| ENSMUSG00000097068 | 0 | 0 | 0 | 0 | 0 | 0 | 0 |
| ENSMUSG00000097069 | 0 | 0.25 | 0 | 0 | 0.244650542 | 0 | 0 |
| ENSMUSG00000097071 | 0 | 0 | 0 | 0.396240625 | 0.244650542 | 0 | 0 |
| ENSMUSG00000097072 | 0 | 0 | 0.25 | 0.646240625 | 0.489301084 | 0.698970004 | 0.355418052 |
| ENSMUSG00000097073 | 1.289064826 | 0.951838731 | 1.114857905 | 1.02186571 | 1.091002363 | 0.903089987 | 0.853775828 |
| ENSMUSG00000097074 | 0 | 0 | 0 | 0 | 0 | 0 | 0 |
| ENSMUSG00000097075 | 0 | 0 | 0 | 0 | 0.244650542 | 0.477121255 | 0 |
| ENSMUSG00000097076 | 0 | 0 | 0 | 0 | 0 | 0 | 0 |
| ENSMUSG00000097077 | 0.289064826 | 0.25 | 0.396240625 | 0.25 | 0.489301084 | 0.301029996 | 0.224243824 |
| ENSMUSG00000097078 | 0 | 0 | 0 | 0 | 0 | 0 | 0 |
| ENSMUSG00000097079 | 0.289064826 | 0.25 | 0 | 0.25 | 0 | 0 | 0.355418052 |
| ENSMUSG00000097080 | 1.445324132 | 1.311981878 | 1.396240625 | 0.951838731 | 1.074582839 | 1.230448921 | 1.014380832 |
| ENSMUSG00000097081 | 1.036286563 | 0.951838731 | 1.080482024 | 0.25 | 0.846351821 | 0.477121255 | 0.803905701 |
| ENSMUSG00000097082 | 0 | 0 | 0 | 0 | 0 | 0 | 0 |
| ENSMUSG00000097083 | 0.289064826 | 0.5 | 0 | 0 | 0.244650542 | 0.477121255 | 0.448487648 |
| ENSMUSG00000097084 | 1.358731791 | 1.061981878 | 1.160964047 | 0.75 | 0.568060967 | 0 | 0.829800754 |
| ENSMUSG00000097085 | 0.578129653 | 0.25 | 0.25 | 0.396240625 | 0.387761935 | 0 | 0.579661877 |
| ENSMUSG00000097086 | 0 | 0 | 0 | 0 | 0 | 0 | 0 |
| ENSMUSG00000097087 | 0 | 0 | 0 | 0 | 0 | 0 | 0 |
| ENSMUSG00000097088 | 0.289064826 | 0.25 | 0 | 0 | 0.244650542 | 0 | 0 |
| ENSMUSG00000097089 | 0 | 0 | 0 | 0 | 0 | 0 | 0 |
| ENSMUSG00000097090 | 1.156259305 | 1.061981878 | 1.098079356 | 1.130890489 | 1.163285805 | 0.477121255 | 1.110947927 |

| | | | | | | | |
|---|---|---|---|---|---|---|---|
| ENSMUSG00000097091 | 0 | 0 | 0.25 | 0.25 | 0.244650542 | 0.301029996 | 0 |
| ENSMUSG00000097092 | 0 | 0 | 0 | 0 | 0 | 0 | 0 |
| ENSMUSG00000097093 | 0.45815691 | 0.5 | 0.580482024 | 0 | 0.387761935 | 0.301029996 | 0.74492186 |
| ENSMUSG00000097094 | 0 | 0 | 0 | 0 | 0 | 0 | 0 |
| ENSMUSG00000097095 | 0 | 0.25 | 0.396240625 | 0 | 0.387761935 | 0 | 0.355418052 |
| ENSMUSG00000097097 | 0 | 0 | 0 | 0 | 0 | 0 | 0 |
| ENSMUSG00000097098 | 0 | 0 | 0.25 | 0 | 0.244650542 | 0 | 0 |
| ENSMUSG00000097099 | 1.227926429 | 1.061981878 | 1.061981878 | 1.061981878 | 1.039257769 | 0.954242509 | 1.121219121 |
| ENSMUSG00000097100 | 0 | 0.396240625 | 0.25 | 0 | 0 | 0 | 0 |
| ENSMUSG00000097101 | 1.036286563 | 0.75 | 0.92510993 | 0.896240625 | 0.846351821 | 0.602059991 | 0.853775828 |
| ENSMUSG00000097102 | 0 | 0 | 0 | 0 | 0 | 0.301029996 | 0 |
| ENSMUSG00000097103 | 0.289064826 | 0.396240625 | 0.396240625 | 0.580482024 | 0.568060967 | 0.477121255 | 0.355418052 |
| ENSMUSG00000097104 | 0.45815691 | 0 | 0.25 | 0 | 0 | 0 | 0 |
| ENSMUSG00000097106 | 0 | 0 | 0 | 0 | 0 | 0 | 0 |
| ENSMUSG00000097107 | 0 | 0 | 0 | 0 | 0 | 0 | 0 |
| ENSMUSG00000097108 | 0.289064826 | 0 | 0 | 0.25 | 0 | 0.477121255 | 0.224243824 |
| ENSMUSG00000097109 | 0 | 0 | 0 | 0 | 0 | 0 | 0 |
| ENSMUSG00000097110 | 0 | 0 | 0 | 0 | 0 | 0 | 0 |
| ENSMUSG00000097111 | 0 | 0 | 0 | 0 | 0.244650542 | 0.301029996 | 0 |
| ENSMUSG00000097112 | 0 | 0 | 0 | 0 | 0 | 0 | 0 |
| ENSMUSG00000097113 | 1.227926429 | 1.098079356 | 1.17510993 | 1.311981878 | 1.302012594 | 1.176091259 | 1.159323753 |
| ENSMUSG00000097115 | 0 | 0 | 0 | 0 | 0 | 0.301029996 | 0.355418052 |
| ENSMUSG00000097116 | 0 | 0 | 0 | 0 | 0 | 0 | 0 |
| ENSMUSG00000097117 | 0.578129653 | 0.25 | 0.5 | 0 | 0.632412477 | 0 | 0.579661877 |
| ENSMUSG00000097118 | 0 | 0 | 0 | 0 | 0 | 0 | 0 |
| ENSMUSG00000097119 | 2.310886388 | 2.114857905 | 2.123963757 | 2.19697564 | 2.09661982 | 2.357934847 | 1.952571511 |
| ENSMUSG00000097120 | 0.289064826 | 0 | 0 | 0 | 0 | 0 | 0 |
| ENSMUSG00000097121 | 0.578129653 | 0.830482024 | 0.5 | 0.396240625 | 0.387761935 | 0.477121255 | 0.520678036 |
| ENSMUSG00000097122 | 0.671187741 | 0.646240625 | 0.396240625 | 0.396240625 | 0.244650542 | 0.301029996 | 0.520678036 |
| ENSMUSG00000097123 | 0.45815691 | 0.580482024 | 0.396240625 | 0.25 | 0.632412477 | 0.477121255 | 0.520678036 |
| ENSMUSG00000097124 | 0 | 0 | 0 | 0 | 0 | 0 | 0 |
| ENSMUSG00000097125 | 0 | 0 | 0 | 0 | 0 | 0 | 0 |
| ENSMUSG00000097127 | 0 | 0 | 0 | 0 | 0 | 0 | 0 |
| ENSMUSG00000097128 | 0.45815691 | 0.25 | 0.5 | 0.25 | 0.244650542 | 0.477121255 | 0.579661877 |

| | | | | | | | |
|---|---|---|---|---|---|---|---|
| ENSMUSG00000097129 | 0.747221736 | 0.864857905 | 0.896240625 | 0.646240625 | 0.733951626 | 0.477121255 | 0.629532004 |
| ENSMUSG00000097130 | 0.45815691 | 0 | 0 | 0 | 0.244650542 | 0 | 0.355418052 |
| ENSMUSG00000097131 | 1 | 0.896240625 | 0.976722649 | 1.061981878 | 1.020174412 | 1.079181246 | 1 |
| ENSMUSG00000097133 | 0.811507563 | 1.04248125 | 1.061981878 | 1.114857905 | 0.877063019 | 0.903089987 | 0.876096088 |
| ENSMUSG00000097134 | 0.45815691 | 0.5 | 0.580482024 | 0.396240625 | 0 | 0.602059991 | 0.579661877 |
| ENSMUSG00000097135 | 0 | 0 | 0 | 0 | 0 | 0 | 0 |
| ENSMUSG00000097136 | 0.811507563 | 0.580482024 | 0.580482024 | 0.646240625 | 0.733951626 | 0.602059991 | 0.448487648 |
| ENSMUSG00000097137 | 0 | 0 | 0 | 0 | 0 | 0 | 0 |
| ENSMUSG00000097138 | 0.289064826 | 0.646240625 | 0.396240625 | 0.396240625 | 0.568060967 | 0 | 0.579661877 |
| ENSMUSG00000097139 | 0 | 0.396240625 | 0 | 0.25 | 0.244650542 | 0.602059991 | 0.224243824 |
| ENSMUSG00000097140 | 0.960252568 | 1.061981878 | 0.976722649 | 1.061981878 | 0.955822902 | 1 | 0.969165684 |
| ENSMUSG00000097141 | 1.482695304 | 1.451838731 | 1.431980114 | 1.464495249 | 1.358933174 | 1.447158031 | 1.307989563 |
| ENSMUSG00000097142 | 0 | 0 | 0 | 0 | 0 | 0 | 0 |
| ENSMUSG00000097145 | 1.558729299 | 1.494319981 | 1.476722649 | 1.356566189 | 1.234113756 | 1.230448921 | 1.131174228 |
| ENSMUSG00000097146 | 0 | 0 | 0.25 | 0 | 0 | 0 | 0.224243824 |
| ENSMUSG00000097147 | 0 | 0 | 0.25 | 0 | 0 | 0 | 0.224243824 |
| ENSMUSG00000097148 | 1.181541736 | 1.188721876 | 0.976722649 | 1 | 0.931471446 | 1.146128036 | 1.028149525 |
| ENSMUSG00000097149 | 0 | 0 | 0 | 0 | 0 | 0 | 0 |
| ENSMUSG00000097150 | 0.747221736 | 0.5 | 0.25 | 0.396240625 | 0.77552387 | 0.301029996 | 0.520678036 |
| ENSMUSG00000097151 | 1.227926429 | 0.951838731 | 1.02186571 | 1.160964047 | 1.149965125 | 1.204119983 | 1.041356071 |
| ENSMUSG00000097152 | 0 | 0.396240625 | 0.396240625 | 0.396240625 | 0 | 0 | 0.355418052 |
| ENSMUSG00000097154 | 0.747221736 | 0.25 | 0.396240625 | 0 | 0 | 0 | 0.224243824 |
| ENSMUSG00000097155 | 0 | 0 | 0.25 | 0 | 0.387761935 | 0 | 0 |
| ENSMUSG00000097156 | 0.289064826 | 0 | 0.396240625 | 0.25 | 0 | 0.301029996 | 0 |
| ENSMUSG00000097157 | 0 | 0 | 0 | 0 | 0 | 0 | 0 |
| ENSMUSG00000097158 | 0.671187741 | 0.896240625 | 0.75 | 1 | 0.733951626 | 0.698970004 | 0.853775828 |
| ENSMUSG00000097159 | 0.289064826 | 0.5 | 0.396240625 | 0.25 | 0.489301084 | 0.77815125 | 0.224243824 |
| ENSMUSG00000097160 | 0 | 0 | 0 | 0 | 0 | 0 | 0 |
| ENSMUSG00000097162 | 0.289064826 | 0 | 0.396240625 | 0.25 | 0.244650542 | 0 | 0.448487648 |
| ENSMUSG00000097163 | 1.505869732 | 1.464495249 | 1.445339928 | 1.403677461 | 1.244650542 | 1.51851394 | 1.278288402 |
| ENSMUSG00000097164 | 1.527823874 | 1.527131114 | 1.557204673 | 1.75 | 1.72073075 | 1.977723605 | 1.603286636 |
| ENSMUSG00000097165 | 1.036286563 | 0.864857905 | 0.646240625 | 0 | 0.489301084 | 0.477121255 | 0.629532004 |
| ENSMUSG00000097166 | 0 | 0.5 | 0.25 | 0.25 | 0.387761935 | 0.602059991 | 0.224243824 |
| ENSMUSG00000097167 | 0.578129653 | 0.646240625 | 0.5 | 0.75 | 0.931471446 | 0.602059991 | 0.775756176 |

| | | | | | | | |
|---|---|---|---|---|---|---|---|
| ENSMUSG00000097168 | 0.289064826 | 0.580482024 | 0.25 | 0.580482024 | 0.387761935 | 0.301029996 | 0.579661877 |
| ENSMUSG00000097169 | 0 | 0 | 0 | 0 | 0 | 0 | 0 |
| ENSMUSG00000097170 | 0.45815691 | 0.5 | 0.75 | 0.646240625 | 0.387761935 | 0.602059991 | 0.672731473 |
| ENSMUSG00000097171 | 0.289064826 | 0 | 0 | 0.5 | 0 | 0.477121255 | 0.224243824 |
| ENSMUSG00000097172 | 0 | 0 | 0 | 0 | 0.387761935 | 0 | 0 |
| ENSMUSG00000097173 | 0 | 0 | 0 | 0.25 | 0 | 0 | 0 |
| ENSMUSG00000097174 | 0.671187741 | 0.830482024 | 0.25 | 0.5 | 0.489301084 | 0.477121255 | 0.74492186 |
| ENSMUSG00000097175 | 0 | 0 | 0 | 0 | 0 | 0 | 0 |
| ENSMUSG00000097176 | 0.289064826 | 0.646240625 | 0.25 | 0.396240625 | 0.244650542 | 0 | 0.355418052 |
| ENSMUSG00000097177 | 1.578129653 | 1.431980114 | 1.42510993 | 1.445339928 | 1.450958105 | 1.477121255 | 1.296434211 |
| ENSMUSG00000097178 | 0 | 0 | 0.25 | 0.25 | 0 | 0.301029996 | 0 |
| ENSMUSG00000097180 | 1.794934559 | 1.634789703 | 1.668106335 | 1.747171172 | 1.78010778 | 1.934498451 | 1.594042709 |
| ENSMUSG00000097181 | 0 | 0 | 0 | 0 | 0 | 0 | 0 |
| ENSMUSG00000097182 | 0 | 0 | 0 | 0.25 | 0 | 0 | 0 |
| ENSMUSG00000097183 | 1.727821383 | 1.688721876 | 1.671625132 | 1.62694866 | 1.639266209 | 1.832508913 | 1.586932193 |
| ENSMUSG00000097184 | 0.289064826 | 0.25 | 0.5 | 0.5 | 0.387761935 | 0.301029996 | 0.710836105 |
| ENSMUSG00000097185 | 0 | 0.25 | 0.396240625 | 0 | 0 | 0 | 0.224243824 |
| ENSMUSG00000097186 | 0 | 0.25 | 0 | 0.25 | 0.244650542 | 0 | 0 |
| ENSMUSG00000097187 | 1.342375483 | 1.146240625 | 1.226722649 | 0.951838731 | 1.136121934 | 0.77815125 | 1.121219121 |
| ENSMUSG00000097189 | 0.45815691 | 0.580482024 | 0.5 | 0.5 | 0.686820904 | 0 | 0.448487648 |
| ENSMUSG00000097190 | 0 | 0 | 0 | 0 | 0 | 0 | 0 |
| ENSMUSG00000097191 | 0.45815691 | 0 | 0.5 | 0.25 | 0.244650542 | 0.301029996 | 0 |
| ENSMUSG00000097193 | 0.578129653 | 0.75 | 0.646240625 | 0.580482024 | 0.387761935 | 0.477121255 | 0.355418052 |
| ENSMUSG00000097194 | 0.45815691 | 0.396240625 | 0 | 0.25 | 0.489301084 | 0.301029996 | 0.520678036 |
| ENSMUSG00000097195 | 2.346900291 | 2.117918804 | 2.122963274 | 2.123963757 | 2.113840608 | 2.250420002 | 1.962479708 |
| ENSMUSG00000097196 | 0.289064826 | 0 | 0 | 0.25 | 0.387761935 | 0 | 0 |
| ENSMUSG00000097197 | 0.289064826 | 0.5 | 0.646240625 | 0.396240625 | 0.632412477 | 0 | 0.448487648 |
| ENSMUSG00000097199 | 0 | 0 | 0.25 | 0 | 0 | 0 | 0 |
| ENSMUSG00000097201 | 0 | 0 | 0 | 0 | 0 | 0 | 0 |
| ENSMUSG00000097202 | 0 | 0 | 0 | 0 | 0 | 0 | 0 |
| ENSMUSG00000097203 | 0.578129653 | 0.580482024 | 0.5 | 0.5 | 0.632412477 | 0.301029996 | 0.355418052 |
| ENSMUSG00000097204 | 1 | 1 | 1.098079356 | 0.896240625 | 1.020174412 | 1.079181246 | 1 |
| ENSMUSG00000097205 | 0 | 0.25 | 0 | 0.25 | 0.244650542 | 0 | 0.355418052 |
| ENSMUSG00000097206 | 0 | 0 | 0 | 0 | 0 | 0 | 0 |

| | | | | | | | |
|---|---|---|---|---|---|---|---|
| ENSMUSG00000097207 | 1.269664473 | 1.302363341 | 1.26109853 | 1.114857905 | 1.254881871 | 1.176091259 | 1.078019652 |
| ENSMUSG00000097208 | 0.45815691 | 0.580482024 | 0.396240625 | 0.25 | 0.387761935 | 0 | 0 |
| ENSMUSG00000097209 | 0 | 0 | 0 | 0 | 0.244650542 | 0.301029996 | 0.355418052 |
| ENSMUSG00000097211 | 0 | 0 | 0 | 0 | 0 | 0 | 0.224243824 |
| ENSMUSG00000097213 | 0 | 0 | 0 | 0 | 0 | 0.301029996 | 0 |
| ENSMUSG00000097214 | 0 | 0 | 0.25 | 0 | 0 | 0 | 0 |
| ENSMUSG00000097215 | 0 | 0 | 0 | 0 | 0 | 0 | 0 |
| ENSMUSG00000097216 | 0 | 0.646240625 | 0.646240625 | 0.25 | 0.489301084 | 0.301029996 | 0.520678036 |
| ENSMUSG00000097217 | 0.91631382 | 0.92510993 | 0.701838731 | 1.04248125 | 0.812711509 | 0.954242509 | 0.896975297 |
| ENSMUSG00000097218 | 0 | 0 | 0 | 0 | 0 | 0 | 0 |
| ENSMUSG00000097219 | 1.129344651 | 0.951838731 | 0.864857905 | 1 | 0.955822902 | 0.84509804 | 0.969165684 |
| ENSMUSG00000097220 | 0.578129653 | 0.396240625 | 0.25 | 0.396240625 | 0.489301084 | 0.602059991 | 0.520678036 |
| ENSMUSG00000097221 | 0 | 0 | 0 | 0 | 0 | 0 | 0 |
| ENSMUSG00000097222 | 0 | 0.25 | 0.25 | 0.580482024 | 0.387761935 | 0.301029996 | 0.448487648 |
| ENSMUSG00000097223 | 0 | 0.25 | 0.25 | 0.396240625 | 0.244650542 | 0 | 0.224243824 |
| ENSMUSG00000097224 | 0 | 0 | 0 | 0 | 0 | 0 | 0 |
| ENSMUSG00000097226 | 0 | 0.25 | 0.396240625 | 0.580482024 | 0.568060967 | 0.301029996 | 0.224243824 |
| ENSMUSG00000097228 | 0 | 0 | 0 | 0 | 0 | 0 | 0 |
| ENSMUSG00000097229 | 0 | 0 | 0 | 0 | 0 | 0 | 0 |
| ENSMUSG00000097230 | 1.45815691 | 1.302363341 | 1.17510993 | 1.685366747 | 1.559656853 | 1.672097858 | 1.40946263 |
| ENSMUSG00000097231 | 0.91631382 | 1.02186571 | 1.098079356 | 1.188721876 | 1.212046813 | 1.278753601 | 1.110947927 |
| ENSMUSG00000097232 | 0.45815691 | 0 | 0.580482024 | 0 | 0.244650542 | 0.477121255 | 0 |
| ENSMUSG00000097233 | 0.671187741 | 0.396240625 | 0.75 | 0.646240625 | 0.632412477 | 0.903089987 | 0.74492186 |
| ENSMUSG00000097234 | 1.307602651 | 1.114857905 | 0.92510993 | 1.080482024 | 1.074582839 | 1.041392685 | 0.916588299 |
| ENSMUSG00000097235 | 0 | 0 | 0 | 0 | 0 | 0 | 0 |
| ENSMUSG00000097236 | 0 | 0.396240625 | 0.5 | 0 | 0 | 0 | 0.224243824 |
| ENSMUSG00000097237 | 0.289064826 | 0 | 0 | 0 | 0 | 0 | 0.224243824 |
| ENSMUSG00000097239 | 0.289064826 | 0.580482024 | 0.75 | 0.5 | 0.387761935 | 0.602059991 | 0.629532004 |
| ENSMUSG00000097240 | 0 | 0 | 0 | 0.25 | 0 | 0 | 0 |
| ENSMUSG00000097241 | 0 | 0 | 0 | 0 | 0 | 0 | 0 |
| ENSMUSG00000097242 | 0.671187741 | 0.646240625 | 0.701838731 | 0.646240625 | 0.812711509 | 0.903089987 | 0.829800754 |
| ENSMUSG00000097243 | 0.289064826 | 0.25 | 0.580482024 | 0.75 | 0.489301084 | 0.477121255 | 0.448487648 |
| ENSMUSG00000097244 | 0 | 0 | 0 | 0.25 | 0.244650542 | 0.301029996 | 0 |
| ENSMUSG00000097245 | 0.867194479 | 0.951838731 | 0.79248125 | 1 | 0.686820904 | 0.84509804 | 0.853775828 |

| | | | | | | | |
|---|---|---|---|---|---|---|---|
| ENSMUSG00000097246 | 0.867194479 | 0.396240625 | 0.580482024 | 0.951838731 | 0.846351821 | 0.698970004 | 0.672731473 |
| ENSMUSG00000097247 | 0 | 0.25 | 0 | 0 | 0 | 0.301029996 | 0.224243824 |
| ENSMUSG00000097248 | 0.578129653 | 0.5 | 0.5 | 0.864857905 | 0.905314583 | 0.954242509 | 0.829800754 |
| ENSMUSG00000097249 | 0 | 0 | 0 | 0 | 0 | 0 | 0 |
| ENSMUSG00000097250 | 0.45815691 | 0.396240625 | 0.5 | 0 | 0 | 0 | 0.355418052 |
| ENSMUSG00000097251 | 0.671187741 | 0.580482024 | 0.25 | 0.79248125 | 0.846351821 | 0.903089987 | 0.803905701 |
| ENSMUSG00000097252 | 1 | 0.92510993 | 1.114857905 | 1.098079356 | 1.074582839 | 1.146128036 | 1 |
| ENSMUSG00000097253 | 0.960252568 | 0.830482024 | 0.79248125 | 0.5 | 0.489301084 | 0.698970004 | 0.579661877 |
| ENSMUSG00000097254 | 0 | 0 | 0.25 | 0.25 | 0.244650542 | 0 | 0.355418052 |
| ENSMUSG00000097255 | 0 | 0.25 | 0 | 0 | 0 | 0 | 0 |
| ENSMUSG00000097256 | 0 | 0 | 0 | 0 | 0 | 0 | 0 |
| ENSMUSG00000097257 | 0.289064826 | 0.580482024 | 0 | 0 | 0.244650542 | 0.477121255 | 0.355418052 |
| ENSMUSG00000097258 | 0.960252568 | 0.864857905 | 0.79248125 | 1.114857905 | 1.020174412 | 0.84509804 | 0.935079929 |
| ENSMUSG00000097259 | 0.45815691 | 0.25 | 0.5 | 0.25 | 0.387761935 | 0 | 0.579661877 |
| ENSMUSG00000097260 | 0 | 0 | 0.25 | 0 | 0 | 0.301029996 | 0 |
| ENSMUSG00000097261 | 0 | 0.25 | 0 | 0.396240625 | 0.387761935 | 0 | 0.355418052 |
| ENSMUSG00000097262 | 0 | 0 | 0 | 0 | 0 | 0 | 0.448487648 |
| ENSMUSG00000097263 | 0.91631382 | 0.896240625 | 1 | 1.29248125 | 1.149965125 | 1.176091259 | 1.131174228 |
| ENSMUSG00000097264 | 0 | 0 | 0 | 0 | 0 | 0 | 0 |
| ENSMUSG00000097265 | 0.45815691 | 0.79248125 | 0.396240625 | 0.5 | 0.489301084 | 0.477121255 | 0.520678036 |
| ENSMUSG00000097266 | 0 | 0.25 | 0 | 0 | 0 | 0 | 0.224243824 |
| ENSMUSG00000097267 | 1.614416215 | 1.438721876 | 1.54745614 | 1.42510993 | 1.343584837 | 1.380211242 | 1.176815335 |
| ENSMUSG00000097268 | 0 | 0.25 | 0 | 0.5 | 0.489301084 | 0.301029996 | 0 |
| ENSMUSG00000097271 | 0.578129653 | 0.701838731 | 0.701838731 | 0.580482024 | 0.387761935 | 0.301029996 | 0.224243824 |
| ENSMUSG00000097272 | 0 | 0 | 0 | 0 | 0 | 0 | 0 |
| ENSMUSG00000097274 | 0 | 0.25 | 0.396240625 | 0 | 0.244650542 | 0 | 0 |
| ENSMUSG00000097275 | 0.960252568 | 0.75 | 0.896240625 | 0.951838731 | 0.877063019 | 0.301029996 | 0.916588299 |
| ENSMUSG00000097276 | 0 | 0.396240625 | 0 | 0 | 0.244650542 | 0 | 0.224243824 |
| ENSMUSG00000097277 | 0.91631382 | 1.04248125 | 1.130890489 | 1 | 1 | 1.176091259 | 1 |
| ENSMUSG00000097278 | 0 | 0 | 0 | 0 | 0.244650542 | 0 | 0 |
| ENSMUSG00000097279 | 0 | 0.396240625 | 0.396240625 | 0.25 | 0.489301084 | 0.301029996 | 0.355418052 |
| ENSMUSG00000097280 | 0 | 0 | 0.25 | 0.25 | 0 | 0 | 0 |
| ENSMUSG00000097281 | 0 | 0.25 | 0 | 0 | 0 | 0 | 0.224243824 |
| ENSMUSG00000097282 | 0.671187741 | 0.646240625 | 0.646240625 | 0.396240625 | 0.244650542 | 0.477121255 | 0.224243824 |

| | | | | | | | |
|---|---|---|---|---|---|---|---|
| ENSMUSG00000097283 | 0 | 0.25 | 0 | 0.25 | 0.387761935 | 0 | 0.355418052 |
| ENSMUSG00000097284 | 0.747221736 | 0.864857905 | 0.701838731 | 1.061981878 | 0.905314583 | 0.698970004 | 0.74492186 |
| ENSMUSG00000097285 | 0 | 0 | 0.396240625 | 0.25 | 0.686820904 | 0 | 0 |
| ENSMUSG00000097286 | 0 | 0 | 0 | 0 | 0 | 0 | 0 |
| ENSMUSG00000097287 | 1.494443473 | 1.396240625 | 1.214495249 | 1.226722649 | 1.039257769 | 1.041392685 | 1.140832123 |
| ENSMUSG00000097288 | 0 | 0.396240625 | 0.25 | 0 | 0 | 0 | 0 |
| ENSMUSG00000097289 | 1.505869732 | 1.403677461 | 1.464495249 | 1.52186571 | 1.489301084 | 1.579783597 | 1.433437704 |
| ENSMUSG00000097290 | 1.249317394 | 1.25 | 1.226722649 | 1.330482024 | 1.293076518 | 1.380211242 | 1.185218806 |
| ENSMUSG00000097291 | 0 | 0 | 0 | 0 | 0 | 0 | 0 |
| ENSMUSG00000097292 | 0.671187741 | 0.75 | 0.25 | 0.701838731 | 0.489301084 | 0.301029996 | 0.520678036 |
| ENSMUSG00000097293 | 0 | 0 | 0 | 0 | 0 | 0 | 0 |
| ENSMUSG00000097294 | 0 | 0 | 0.396240625 | 0.396240625 | 0.387761935 | 0.301029996 | 0.448487648 |
| ENSMUSG00000097295 | 0.289064826 | 0 | 0 | 0 | 0 | 0 | 0 |
| ENSMUSG00000097296 | 1.156259305 | 1.214495249 | 1.302363341 | 1.146240625 | 1.057362051 | 1.176091259 | 1.100339912 |
| ENSMUSG00000097298 | 0.289064826 | 0 | 0 | 0.396240625 | 0.244650542 | 0.301029996 | 0 |
| ENSMUSG00000097299 | 0 | 0 | 0 | 0 | 0 | 0 | 0 |
| ENSMUSG00000097300 | 0 | 0 | 0 | 0 | 0 | 0 | 0.224243824 |
| ENSMUSG00000097301 | 0 | 0 | 0 | 0 | 0 | 0 | 0 |
| ENSMUSG00000097302 | 0 | 0 | 0 | 0 | 0 | 0 | 0 |
| ENSMUSG00000097303 | 0 | 0.580482024 | 0.396240625 | 0.580482024 | 0.489301084 | 0.477121255 | 0.448487648 |
| ENSMUSG00000097304 | 0 | 0 | 0 | 0 | 0 | 0 | 0 |
| ENSMUSG00000097305 | 0 | 0.396240625 | 0 | 0 | 0 | 0.301029996 | 0 |
| ENSMUSG00000097307 | 0 | 0 | 0.396240625 | 0.25 | 0 | 0 | 0 |
| ENSMUSG00000097308 | 0.45815691 | 0.646240625 | 0.646240625 | 0.5 | 0.812711509 | 0 | 0.829800754 |
| ENSMUSG00000097309 | 0 | 0.396240625 | 0.701838731 | 0.5 | 0.632412477 | 0.84509804 | 0.520678036 |
| ENSMUSG00000097310 | 0 | 0 | 0 | 0 | 0.244650542 | 0 | 0 |
| ENSMUSG00000097311 | 0 | 0 | 0 | 0 | 0 | 0 | 0 |
| ENSMUSG00000097312 | 0 | 0 | 0 | 0 | 0 | 0 | 0 |
| ENSMUSG00000097313 | 0 | 0.5 | 0.5 | 0.396240625 | 0.244650542 | 0.477121255 | 0 |
| ENSMUSG00000097314 | 0 | 0 | 0 | 0 | 0 | 0 | 0 |
| ENSMUSG00000097315 | 0 | 0 | 0 | 0 | 0 | 0 | 0 |
| ENSMUSG00000097316 | 0.91631382 | 1.02186571 | 0.864857905 | 1.146240625 | 1.163285805 | 1.204119983 | 0.984950056 |
| ENSMUSG00000097317 | 0.45815691 | 0.701838731 | 0.646240625 | 0.5 | 0.244650542 | 0.301029996 | 0.629532004 |
| ENSMUSG00000097318 | 0.578129653 | 0.701838731 | 0.701838731 | 0.396240625 | 0.568060967 | 0.477121255 | 0.629532004 |

| | | | | | | | |
|---|---|---|---|---|---|---|---|
| ENSMUSG00000097320 | 0.671187741 | 0.5 | 0.580482024 | 0.580482024 | 0.387761935 | 0.698970004 | 0.629532004 |
| ENSMUSG00000097321 | 0.289064826 | 0.25 | 0 | 0.5 | 0.686820904 | 0.698970004 | 0.579661877 |
| ENSMUSG00000097322 | 0 | 0.396240625 | 0 | 0.396240625 | 0 | 0.301029996 | 0 |
| ENSMUSG00000097323 | 0 | 0.5 | 0.5 | 0.396240625 | 0.244650542 | 0 | 0.224243824 |
| ENSMUSG00000097324 | 0.811507563 | 1 | 0.951838731 | 0.5 | 0.244650542 | 0.602059991 | 0.224243824 |
| ENSMUSG00000097325 | 0 | 0 | 0.25 | 0.25 | 0.387761935 | 0.477121255 | 0 |
| ENSMUSG00000097326 | 0 | 0 | 0 | 0 | 0 | 0 | 0 |
| ENSMUSG00000097327 | 1.181541736 | 1.146240625 | 1.080482024 | 1.146240625 | 1.200473444 | 1.146128036 | 1.054044578 |
| ENSMUSG00000097328 | 1.205378646 | 1.29248125 | 1.451838731 | 1.451838731 | 1.439191811 | 1.681241237 | 1.486592281 |
| ENSMUSG00000097329 | 0 | 0 | 0.396240625 | 0 | 0.489301084 | 0.301029996 | 0.224243824 |
| ENSMUSG00000097330 | 0.671187741 | 0.79248125 | 0.79248125 | 0.580482024 | 0.489301084 | 0.301029996 | 0.224243824 |
| ENSMUSG00000097331 | 1.069666964 | 1.02186571 | 1.214495249 | 1.130890489 | 1.074582839 | 1.041392685 | 0.876096088 |
| ENSMUSG00000097332 | 0 | 0 | 0 | 0 | 0 | 0 | 0 |
| ENSMUSG00000097333 | 1.816888701 | 1.701838731 | 1.695339928 | 1.852347734 | 1.804307395 | 2.045322979 | 1.647906904 |
| ENSMUSG00000097334 | 0 | 0 | 0 | 0 | 0 | 0 | 0.448487648 |
| ENSMUSG00000097335 | 0.747221736 | 0.580482024 | 0.79248125 | 0.580482024 | 0.632412477 | 0.602059991 | 0.710836105 |
| ENSMUSG00000097336 | 0 | 0.25 | 0 | 1.02186571 | 0.846351821 | 1.041392685 | 0.803905701 |
| ENSMUSG00000097337 | 1.227926429 | 1.160964047 | 1.02186571 | 1.080482024 | 0.931471446 | 1 | 0.952571511 |
| ENSMUSG00000097338 | 0 | 0 | 0 | 0 | 0 | 0 | 0 |
| ENSMUSG00000097339 | 0 | 0.25 | 0 | 0.25 | 0 | 0 | 0 |
| ENSMUSG00000097340 | 1.568542279 | 1.532320754 | 1.505591953 | 1.54248125 | 1.504538967 | 1.568201724 | 1.413583902 |
| ENSMUSG00000097342 | 0 | 0 | 0 | 0 | 0 | 0 | 0 |
| ENSMUSG00000097343 | 0.811507563 | 0.92510993 | 1.098079356 | 0.830482024 | 1 | 0.77815125 | 0.876096088 |
| ENSMUSG00000097344 | 0.671187741 | 0.580482024 | 0.580482024 | 0.25 | 0.244650542 | 0.301029996 | 0 |
| ENSMUSG00000097345 | 0 | 0.5 | 0.396240625 | 0.5 | 0.244650542 | 0 | 0.520678036 |
| ENSMUSG00000097346 | 0 | 0 | 0 | 0 | 0 | 0 | 0 |
| ENSMUSG00000097347 | 1.505869732 | 1.330482024 | 1.356566189 | 1.388647213 | 1.264824954 | 1.431363764 | 1.20139795 |
| ENSMUSG00000097348 | 0.289064826 | 0.25 | 0 | 0 | 0 | 0 | 0.224243824 |
| ENSMUSG00000097349 | 0 | 0 | 0 | 0 | 0 | 0 | 0 |
| ENSMUSG00000097350 | 1.156259305 | 1.04248125 | 1.061981878 | 0.976722649 | 0.955822902 | 0.954242509 | 1 |
| ENSMUSG00000097352 | 1.432083896 | 1.356566189 | 1.451838731 | 1.160964047 | 1.039257769 | 1 | 0.803905701 |
| ENSMUSG00000097353 | 1.181541736 | 1.061981878 | 1 | 1.061981878 | 1.121713561 | 1.301029996 | 0.935079929 |
| ENSMUSG00000097354 | 0.811507563 | 0.79248125 | 0.75 | 0.580482024 | 0.387761935 | 0.477121255 | 0 |
| ENSMUSG00000097355 | 0 | 0 | 0 | 0 | 0 | 0 | 0 |

| | | | | | | | |
|---|---|---|---|---|---|---|---|
| ENSMUSG00000097356 | 0.289064826 | 0.25 | 0.396240625 | 0.25 | 0.568060967 | 0 | 0.224243824 |
| ENSMUSG00000097357 | 0.747221736 | 0.5 | 0.5 | 0.646240625 | 0.489301084 | 0.477121255 | 0.448487648 |
| ENSMUSG00000097358 | 0.289064826 | 0.5 | 0.580482024 | 1.04248125 | 1.057362051 | 0.698970004 | 0.579661877 |
| ENSMUSG00000097360 | 0 | 0 | 0 | 0.25 | 0.244650542 | 0.301029996 | 0.224243824 |
| ENSMUSG00000097361 | 0 | 0 | 0 | 0 | 0 | 0 | 0 |
| ENSMUSG00000097362 | 0 | 0 | 0 | 0 | 0 | 0 | 0 |
| ENSMUSG00000097363 | 0 | 0 | 0 | 0 | 0 | 0 | 0 |
| ENSMUSG00000097364 | 0.289064826 | 0 | 0.25 | 0.25 | 0 | 0 | 0 |
| ENSMUSG00000097365 | 0 | 0 | 0 | 0 | 0.244650542 | 0 | 0.224243824 |
| ENSMUSG00000097366 | 0 | 0.396240625 | 0.25 | 0 | 0 | 0.301029996 | 0.355418052 |
| ENSMUSG00000097367 | 0 | 0 | 0 | 0 | 0 | 0 | 0 |
| ENSMUSG00000097368 | 0 | 0.396240625 | 0.396240625 | 0.25 | 0 | 0.301029996 | 0.224243824 |
| ENSMUSG00000097369 | 0 | 0 | 0 | 0.5 | 0.387761935 | 0 | 0 |
| ENSMUSG00000097370 | 0 | 0 | 0 | 0.25 | 0 | 0 | 0 |
| ENSMUSG00000097371 | 0 | 0 | 0 | 0 | 0 | 0 | 0 |
| ENSMUSG00000097372 | 0 | 0.25 | 0.25 | 0.396240625 | 0.244650542 | 0.301029996 | 0.448487648 |
| ENSMUSG00000097373 | 0.289064826 | 0.25 | 0 | 0 | 0.244650542 | 0.301029996 | 0.355418052 |
| ENSMUSG00000097374 | 0 | 0 | 0 | 0 | 0 | 0 | 0 |
| ENSMUSG00000097375 | 1.342375483 | 1.302363341 | 1.214495249 | 1.431980114 | 1.439191811 | 1.544068044 | 1.278288402 |
| ENSMUSG00000097376 | 0.289064826 | 0.396240625 | 0 | 0 | 0.244650542 | 0.301029996 | 0.448487648 |
| ENSMUSG00000097377 | 0.578129653 | 0.25 | 0.25 | 0.396240625 | 0.489301084 | 0.301029996 | 0.448487648 |
| ENSMUSG00000097378 | 0.289064826 | 0.646240625 | 0.701838731 | 0.580482024 | 0.812711509 | 0.602059991 | 0.672731473 |
| ENSMUSG00000097379 | 1.325351389 | 1.160964047 | 1.04248125 | 1.160964047 | 1.212046813 | 1.041392685 | 1.159323753 |
| ENSMUSG00000097380 | 0.91631382 | 1.146240625 | 0.92510993 | 1.114857905 | 0.877063019 | 0.84509804 | 0.896975297 |
| ENSMUSG00000097381 | 0 | 0.25 | 0.25 | 0.396240625 | 0 | 0.301029996 | 0.224243824 |
| ENSMUSG00000097382 | 0 | 0 | 0 | 0 | 0 | 0 | 0 |
| ENSMUSG00000097383 | 0.960252568 | 0.92510993 | 0.701838731 | 0.951838731 | 0.812711509 | 0.77815125 | 0.520678036 |
| ENSMUSG00000097384 | 0.289064826 | 0.25 | 0 | 0.25 | 0 | 0 | 0.224243824 |
| ENSMUSG00000097385 | 0.289064826 | 0 | 0 | 0 | 0 | 0 | 0 |
| ENSMUSG00000097386 | 0.289064826 | 0 | 0.25 | 0 | 0 | 0.301029996 | 0.355418052 |
| ENSMUSG00000097387 | 0 | 0 | 0.25 | 0 | 0 | 0 | 0 |
| ENSMUSG00000097388 | 0.289064826 | 0.5 | 0.5 | 0.396240625 | 0.632412477 | 0.301029996 | 0.448487648 |
| ENSMUSG00000097390 | 1 | 0.951838731 | 1.04248125 | 0.864857905 | 1 | 0.84509804 | 0.935079929 |
| ENSMUSG00000097391 | 1.516991255 | 1.410964047 | 1.403677461 | 0.25 | 0.489301084 | 0 | 0.224243824 |

| | | | | | | | |
|---|---|---|---|---|---|---|---|
| ENSMUSG00000097392 | 1.928763473 | 1.856566189 | 1.845926073 | 1.752806814 | 1.706995316 | 1.812913357 | 1.594042709 |
| ENSMUSG00000097393 | 0.867194479 | 0.5 | 0.5 | 0.5 | 0.387761935 | 0.301029996 | 0.448487648 |
| ENSMUSG00000097396 | 0 | 0 | 0 | 0 | 0 | 0 | 0 |
| ENSMUSG00000097397 | 1 | 1 | 1.04248125 | 0.701838731 | 1 | 0.301029996 | 0.775756176 |
| ENSMUSG00000097398 | 0 | 0.25 | 0 | 0.5 | 0.244650542 | 0.477121255 | 0 |
| ENSMUSG00000097399 | 0.578129653 | 0.75 | 0.580482024 | 0.830482024 | 1 | 1.041392685 | 0.896975297 |
| ENSMUSG00000097401 | 0.578129653 | 0.25 | 0.396240625 | 0 | 0.387761935 | 0 | 0 |
| ENSMUSG00000097402 | 0.578129653 | 0.580482024 | 0.701838731 | 0.396240625 | 0.632412477 | 0.477121255 | 0.355418052 |
| ENSMUSG00000097403 | 0.671187741 | 0.580482024 | 0.864857905 | 0.701838731 | 0.733951626 | 0.477121255 | 0.448487648 |
| ENSMUSG00000097404 | 1.069666964 | 0.646240625 | 0.75 | 0.701838731 | 0.686820904 | 0.77815125 | 0.710836105 |
| ENSMUSG00000097405 | 0.811507563 | 0.79248125 | 0.92510993 | 0.701838731 | 1.106691885 | 1.041392685 | 1.078019652 |
| ENSMUSG00000097406 | 0 | 0 | 0 | 0 | 0 | 0 | 0 |
| ENSMUSG00000097407 | 0 | 0 | 0 | 0 | 0 | 0 | 0.224243824 |
| ENSMUSG00000097408 | 0.289064826 | 0.5 | 0.396240625 | 0.25 | 0.387761935 | 0 | 0.448487648 |
| ENSMUSG00000097409 | 0 | 0 | 0 | 0 | 0 | 0 | 0 |
| ENSMUSG00000097410 | 0 | 0.396240625 | 0.5 | 0.75 | 0.77552387 | 0.698970004 | 0.629532004 |
| ENSMUSG00000097411 | 0.45815691 | 0.25 | 0 | 0 | 0.387761935 | 0 | 0 |
| ENSMUSG00000097412 | 0.960252568 | 1.02186571 | 1.02186571 | 0.976722649 | 1.057362051 | 0.954242509 | 1.054044578 |
| ENSMUSG00000097413 | 0 | 0 | 0 | 0.25 | 0 | 0.477121255 | 0.224243824 |
| ENSMUSG00000097414 | 0.747221736 | 0.79248125 | 0.92510993 | 1.02186571 | 1.106691885 | 0.77815125 | 0.935079929 |
| ENSMUSG00000097415 | 1.558729299 | 1.372963274 | 1.282320754 | 1.364857905 | 1.509475496 | 1.568201724 | 1.245582246 |
| ENSMUSG00000097416 | 0.289064826 | 0.396240625 | 0 | 0 | 0.632412477 | 0.477121255 | 0.355418052 |
| ENSMUSG00000097417 | 0 | 0 | 0 | 0 | 0 | 0 | 0 |
| ENSMUSG00000097418 | 1.156259305 | 1.080482024 | 0.976722649 | 1.098079356 | 1.136121934 | 0.954242509 | 0.969165684 |
| ENSMUSG00000097419 | 0.289064826 | 0.25 | 0 | 0 | 0 | 0 | 0 |
| ENSMUSG00000097420 | 0 | 0 | 0 | 0 | 0 | 0 | 0 |
| ENSMUSG00000097421 | 0 | 0 | 0 | 0 | 0 | 0 | 0 |
| ENSMUSG00000097422 | 0 | 0 | 0.396240625 | 0.5 | 0 | 0.301029996 | 0.224243824 |
| ENSMUSG00000097423 | 0.289064826 | 0.25 | 0.396240625 | 0 | 0 | 0.301029996 | 0.224243824 |
| ENSMUSG00000097424 | 0.289064826 | 0 | 0 | 0 | 0 | 0 | 0 |
| ENSMUSG00000097425 | 0 | 0 | 0 | 0 | 0 | 0 | 0 |
| ENSMUSG00000097426 | 0 | 0.25 | 0.25 | 0 | 0 | 0 | 0 |
| ENSMUSG00000097427 | 0 | 0 | 0 | 0 | 0 | 0 | 0 |
| ENSMUSG00000097428 | 0 | 0.25 | 0 | 0 | 0.489301084 | 0.301029996 | 0.224243824 |

| | | | | | | | |
|---|---|---|---|---|---|---|---|
| ENSMUSG00000097429 | 0.289064826 | 0 | 0.5 | 0.25 | 0.244650542 | 0 | 0 |
| ENSMUSG00000097430 | 0 | 0 | 0.25 | 0 | 0 | 0 | 0 |
| ENSMUSG00000097431 | 1.036286563 | 0.896240625 | 1.061981878 | 1.04248125 | 0.955822902 | 0.903089987 | 0.896975297 |
| ENSMUSG00000097433 | 0 | 0.396240625 | 0 | 0.25 | 0.387761935 | 0.477121255 | 0.355418052 |
| ENSMUSG00000097434 | 0.289064826 | 0.396240625 | 0 | 0.646240625 | 0.489301084 | 0.301029996 | 0.579661877 |
| ENSMUSG00000097435 | 0 | 0 | 0 | 0 | 0 | 0 | 0 |
| ENSMUSG00000097437 | 0 | 0 | 0 | 0 | 0 | 0 | 0 |
| ENSMUSG00000097438 | 0 | 0 | 0 | 0 | 0 | 0 | 0 |
| ENSMUSG00000097439 | 0.91631382 | 0.951838731 | 0.864857905 | 0.92510993 | 1 | 0.77815125 | 0.520678036 |
| ENSMUSG00000097440 | 0.289064826 | 0 | 0 | 0.25 | 0.244650542 | 0 | 0.224243824 |
| ENSMUSG00000097441 | 0 | 0 | 0 | 0 | 0 | 0 | 0 |
| ENSMUSG00000097442 | 0 | 0 | 0 | 0.25 | 0 | 0 | 0 |
| ENSMUSG00000097443 | 0.747221736 | 0.701838731 | 0.580482024 | 0.580482024 | 0.568060967 | 0.84509804 | 0.520678036 |
| ENSMUSG00000097444 | 0 | 0 | 0 | 0 | 0 | 0.301029996 | 0 |
| ENSMUSG00000097445 | 0.747221736 | 0.951838731 | 0.75 | 0.79248125 | 0.846351821 | 0.954242509 | 0.672731473 |
| ENSMUSG00000097446 | 0.45815691 | 0.25 | 0 | 0.5 | 0 | 0.301029996 | 0.355418052 |
| ENSMUSG00000097447 | 0 | 0 | 0 | 0 | 0.244650542 | 0 | 0 |
| ENSMUSG00000097448 | 0.671187741 | 0.25 | 0.79248125 | 0.701838731 | 0.846351821 | 0.903089987 | 0.853775828 |
| ENSMUSG00000097450 | 0 | 0 | 0 | 0 | 0 | 0 | 0 |
| ENSMUSG00000097451 | 3.065671095 | 2.778435542 | 2.759215112 | 1.282320754 | 1.319233381 | 1.361727836 | 1.272006351 |
| ENSMUSG00000097452 | 0.45815691 | 0.75 | 0.646240625 | 0.646240625 | 0.733951626 | 0.301029996 | 0.579661877 |
| ENSMUSG00000097453 | 0 | 0 | 0 | 0 | 0 | 0 | 0 |
| ENSMUSG00000097454 | 0 | 0 | 0 | 0 | 0 | 0 | 0 |
| ENSMUSG00000097455 | 0.289064826 | 0.396240625 | 0.25 | 0.580482024 | 0.387761935 | 0.301029996 | 0.224243824 |
| ENSMUSG00000097456 | 0 | 0 | 0 | 0.25 | 0 | 0.477121255 | 0 |
| ENSMUSG00000097457 | 0 | 0.25 | 0.25 | 0.25 | 0.387761935 | 0.301029996 | 0.448487648 |
| ENSMUSG00000097458 | 0 | 0 | 0 | 0 | 0 | 0 | 0 |
| ENSMUSG00000097459 | 0 | 0 | 0 | 0 | 0 | 0 | 0 |
| ENSMUSG00000097460 | 0 | 0.79248125 | 0.580482024 | 0.25 | 0.244650542 | 0.602059991 | 0.672731473 |
| ENSMUSG00000097461 | 0 | 0 | 0 | 0 | 0 | 0 | 0 |
| ENSMUSG00000097462 | 0.289064826 | 0.25 | 0.25 | 0.580482024 | 0.387761935 | 0.602059991 | 0.224243824 |
| ENSMUSG00000097463 | 0 | 0 | 0 | 0 | 0 | 0 | 0 |
| ENSMUSG00000097464 | 0.289064826 | 0 | 0 | 0.5 | 0.244650542 | 0 | 0 |
| ENSMUSG00000097465 | 0 | 0 | 0 | 0 | 0 | 0 | 0 |

| | | | | | | | |
|---|---|---|---|---|---|---|---|
| ENSMUSG00000097466 | 0 | 0 | 0 | 0 | 0 | 0 | 0 |
| ENSMUSG00000097467 | 1.100572389 | 0.951838731 | 1 | 1.380890489 | 1.358933174 | 1.397940009 | 1.265599895 |
| ENSMUSG00000097468 | 0 | 0 | 0 | 0 | 0 | 0 | 0 |
| ENSMUSG00000097469 | 0 | 0.646240625 | 0.580482024 | 0.75 | 0.77552387 | 0.77815125 | 0.579661877 |
| ENSMUSG00000097470 | 1 | 0.864857905 | 0.896240625 | 1 | 1.020174412 | 1.041392685 | 0.853775828 |
| ENSMUSG00000097471 | 0.289064826 | 0 | 0 | 0.25 | 0.244650542 | 0 | 0.355418052 |
| ENSMUSG00000097472 | 1.100572389 | 1.061981878 | 1.114857905 | 0.830482024 | 0.978602168 | 0.602059991 | 0.853775828 |
| ENSMUSG00000097473 | 0 | 0 | 0 | 0 | 0 | 0 | 0 |
| ENSMUSG00000097474 | 0 | 0 | 0.25 | 0.5 | 0 | 0 | 0 |
| ENSMUSG00000097476 | 0.289064826 | 0 | 0 | 0.396240625 | 0 | 0 | 0.224243824 |
| ENSMUSG00000097479 | 0.45815691 | 0.79248125 | 0.646240625 | 0.701838731 | 0.686820904 | 0 | 0.803905701 |
| ENSMUSG00000097480 | 0 | 0 | 0 | 0 | 0 | 0 | 0 |
| ENSMUSG00000097481 | 0 | 0 | 0 | 0.25 | 0 | 0 | 0 |
| ENSMUSG00000097482 | 0 | 0 | 0 | 0 | 0 | 0 | 0 |
| ENSMUSG00000097483 | 0 | 0 | 0 | 0 | 0 | 0 | 0 |
| ENSMUSG00000097484 | 0 | 0 | 0 | 0 | 0 | 0 | 0 |
| ENSMUSG00000097486 | 0 | 0 | 0 | 0 | 0 | 0 | 0 |
| ENSMUSG00000097487 | 1.307602651 | 1.445339928 | 1.418106335 | 1.146240625 | 1.234113756 | 1.397940009 | 1.078019652 |
| ENSMUSG00000097488 | 1.289064826 | 1.238549078 | 1 | 1.130890489 | 1.244650542 | 1.322219295 | 1.121219121 |
| ENSMUSG00000097489 | 0.289064826 | 0 | 0.25 | 0 | 0 | 0 | 0.224243824 |
| ENSMUSG00000097491 | 0.747221736 | 0 | 0.5 | 0.25 | 0 | 0.477121255 | 0.355418052 |
| ENSMUSG00000097493 | 1.418409478 | 1.26109853 | 1.302363341 | 0.79248125 | 0.733951626 | 1.041392685 | 0.853775828 |
| ENSMUSG00000097494 | 0.45815691 | 0 | 0.25 | 0.396240625 | 0.387761935 | 0 | 0.224243824 |
| ENSMUSG00000097495 | 0.867194479 | 1 | 0.896240625 | 0.580482024 | 1.057362051 | 0.602059991 | 1 |
| ENSMUSG00000097496 | 0 | 0 | 0 | 0 | 0 | 0 | 0 |
| ENSMUSG00000097497 | 0 | 0 | 0 | 0.25 | 0 | 0 | 0.224243824 |
| ENSMUSG00000097500 | 0 | 0 | 0 | 0 | 0 | 0 | 0 |
| ENSMUSG00000097501 | 0 | 0 | 0.25 | 0 | 0 | 0 | 0 |
| ENSMUSG00000097502 | 0.289064826 | 0.25 | 0.580482024 | 0.396240625 | 0.244650542 | 0.301029996 | 0 |
| ENSMUSG00000097503 | 1.249317394 | 1.302363341 | 1.311981878 | 1.114857905 | 1.163285805 | 0.954242509 | 0.952571511 |
| ENSMUSG00000097504 | 0.45815691 | 0.396240625 | 0.25 | 0.646240625 | 0.244650542 | 0 | 0.448487648 |
| ENSMUSG00000097505 | 0 | 0 | 0 | 0 | 0 | 0 | 0 |
| ENSMUSG00000097507 | 0.960252568 | 0.830482024 | 0.646240625 | 0.75 | 0.632412477 | 0.77815125 | 0.775756176 |
| ENSMUSG00000097509 | 0 | 0 | 0.25 | 0 | 0 | 0 | 0 |

| | | | | | | | |
|---|---|---|---|---|---|---|---|
| ENSMUSG00000097511 | 0 | 0 | 0 | 0 | 0 | 0 | 0 |
| ENSMUSG00000097513 | 0 | 0.25 | 0 | 0 | 0.387761935 | 0 | 0 |
| ENSMUSG00000097514 | 0 | 0 | 0 | 0 | 0 | 0 | 0 |
| ENSMUSG00000097515 | 0.747221736 | 0.830482024 | 0.25 | 0.646240625 | 0.387761935 | 0.301029996 | 0.355418052 |
| ENSMUSG00000097516 | 0 | 0 | 0 | 0 | 0 | 0 | 0 |
| ENSMUSG00000097517 | 0 | 0 | 0 | 0 | 0 | 0 | 0 |
| ENSMUSG00000097518 | 1.289064826 | 1.214495249 | 1.27186571 | 1.226722649 | 1.478764298 | 1.322219295 | 1.383567577 |
| ENSMUSG00000097519 | 1.342375483 | 1.238549078 | 1.130890489 | 1.302363341 | 1.264824954 | 1.505149978 | 1.131174228 |
| ENSMUSG00000097520 | 0.578129653 | 0 | 0.25 | 0 | 0 | 0.301029996 | 0.520678036 |
| ENSMUSG00000097522 | 0 | 0 | 0 | 0 | 0 | 0 | 0 |
| ENSMUSG00000097523 | 0.578129653 | 0.580482024 | 0.580482024 | 0.646240625 | 0.387761935 | 0.602059991 | 0.355418052 |
| ENSMUSG00000097524 | 0 | 0 | 0 | 0 | 0 | 0 | 0 |
| ENSMUSG00000097525 | 0 | 0.25 | 0 | 0 | 0.244650542 | 0 | 0 |
| ENSMUSG00000097526 | 0.289064826 | 0.25 | 0.646240625 | 0.580482024 | 0.632412477 | 0.301029996 | 0.629532004 |
| ENSMUSG00000097527 | 0 | 0.396240625 | 0.396240625 | 0.5 | 0.387761935 | 0 | 0.448487648 |
| ENSMUSG00000097529 | 0.289064826 | 0 | 0.25 | 0 | 0 | 0.301029996 | 0 |
| ENSMUSG00000097530 | 0 | 0 | 0.25 | 0.25 | 0 | 0 | 0 |
| ENSMUSG00000097532 | 0.289064826 | 0.5 | 0.396240625 | 0.580482024 | 0.568060967 | 0.602059991 | 0.520678036 |
| ENSMUSG00000097533 | 0.289064826 | 0 | 0.25 | 0.25 | 0 | 0.477121255 | 0.224243824 |
| ENSMUSG00000097534 | 0.45815691 | 0.580482024 | 0.701838731 | 0.75 | 0.686820904 | 0.602059991 | 0.672731473 |
| ENSMUSG00000097535 | 0 | 0 | 0.25 | 0 | 0 | 0 | 0.224243824 |
| ENSMUSG00000097536 | 1.342375483 | 1.146240625 | 1.29248125 | 1.27186571 | 1.254881871 | 1.322219295 | 1.131174228 |
| ENSMUSG00000097537 | 1.269664473 | 1.080482024 | 1.201838731 | 1.201838731 | 1.176121988 | 1.361727836 | 1.089372236 |
| ENSMUSG00000097538 | 0 | 0 | 0 | 0 | 0 | 0 | 0 |
| ENSMUSG00000097539 | 0 | 0 | 0 | 0 | 0 | 0 | 0 |
| ENSMUSG00000097540 | 0 | 0 | 0 | 0 | 0.244650542 | 0 | 0.224243824 |
| ENSMUSG00000097541 | 0 | 0 | 0 | 0.25 | 0 | 0 | 0.224243824 |
| ENSMUSG00000097542 | 0 | 0 | 0 | 0 | 0 | 0 | 0 |
| ENSMUSG00000097543 | 0 | 0.25 | 0 | 0.25 | 0 | 0.301029996 | 0 |
| ENSMUSG00000097544 | 0 | 0 | 0 | 0 | 0 | 0 | 0 |
| ENSMUSG00000097545 | 0 | 0 | 0 | 0 | 0 | 0 | 0 |
| ENSMUSG00000097546 | 0 | 0 | 0.396240625 | 0.25 | 0 | 0.477121255 | 0 |
| ENSMUSG00000097547 | 0 | 0 | 0 | 0 | 0 | 0 | 0 |
| ENSMUSG00000097548 | 0.45815691 | 0.396240625 | 0.25 | 0.580482024 | 0.387761935 | 0.477121255 | 0 |

| | | | | | | | |
|---|---|---|---|---|---|---|---|
| ENSMUSG00000097550 | 0 | 0 | 0 | 0 | 0 | 0 | 0 |
| ENSMUSG00000097551 | 0 | 0 | 0 | 0 | 0 | 0 | 0 |
| ENSMUSG00000097552 | 0 | 0.396240625 | 0 | 0 | 0 | 0 | 0 |
| ENSMUSG00000097553 | 0 | 0 | 0 | 0 | 0 | 0 | 0 |
| ENSMUSG00000097554 | 2.423434025 | 2.223704441 | 2.171625132 | 2.234159485 | 2.28445426 | 2.541579244 | 2.11845007 |
| ENSMUSG00000097557 | 0.289064826 | 0.25 | 0.5 | 0.25 | 0.244650542 | 0.301029996 | 0.448487648 |
| ENSMUSG00000097558 | 1.156259305 | 1.214495249 | 1.356566189 | 0.830482024 | 0.978602168 | 1.041392685 | 1 |
| ENSMUSG00000097559 | 0.960252568 | 1.114857905 | 1.061981878 | 0.646240625 | 0.846351821 | 0.903089987 | 0.672731473 |
| ENSMUSG00000097560 | 0.578129653 | 0.396240625 | 0.396240625 | 0.396240625 | 0.244650542 | 0.301029996 | 0 |
| ENSMUSG00000097561 | 0.671187741 | 0.5 | 0 | 0.25 | 0.489301084 | 0.602059991 | 0.224243824 |
| ENSMUSG00000097562 | 0 | 0 | 0 | 0 | 0 | 0 | 0 |
| ENSMUSG00000097563 | 0.289064826 | 0.25 | 0 | 0.25 | 0 | 0 | 0 |
| ENSMUSG00000097564 | 0.671187741 | 0.75 | 0.646240625 | 0.5 | 0.568060967 | 0.301029996 | 0.355418052 |
| ENSMUSG00000097565 | 0 | 0 | 0 | 0 | 0 | 0 | 0 |
| ENSMUSG00000097566 | 0 | 0 | 0 | 0 | 0 | 0 | 0 |
| ENSMUSG00000097567 | 0.671187741 | 0.580482024 | 0.79248125 | 0.580482024 | 0.632412477 | 0.301029996 | 0.520678036 |
| ENSMUSG00000097569 | 0.289064826 | 0.25 | 0 | 0.25 | 0 | 0 | 0 |
| ENSMUSG00000097571 | 1.036286563 | 0.951838731 | 0.92510993 | 0.976722649 | 0.632412477 | 1 | 0.829800754 |
| ENSMUSG00000097572 | 0.578129653 | 0.701838731 | 0.580482024 | 0.5 | 0.387761935 | 0.698970004 | 0.448487648 |
| ENSMUSG00000097573 | 0.747221736 | 0.5 | 0.396240625 | 0.580482024 | 0.489301084 | 0.477121255 | 0.520678036 |
| ENSMUSG00000097574 | 0.289064826 | 0.396240625 | 0.25 | 0.25 | 0.387761935 | 0 | 0 |
| ENSMUSG00000097575 | 0 | 0 | 0 | 0 | 0.244650542 | 0 | 0 |
| ENSMUSG00000097576 | 0.289064826 | 0.5 | 0.580482024 | 0.25 | 0.387761935 | 0.301029996 | 0.224243824 |
| ENSMUSG00000097577 | 0.811507563 | 0.79248125 | 0.75 | 1.061981878 | 1.020174412 | 1.041392685 | 0.952571511 |
| ENSMUSG00000097578 | 0.45815691 | 0.396240625 | 0.646240625 | 0.646240625 | 0.686820904 | 0 | 0.629532004 |
| ENSMUSG00000097579 | 0.578129653 | 0.701838731 | 0.580482024 | 0.5 | 0.686820904 | 0.477121255 | 0.672731473 |
| ENSMUSG00000097580 | 0 | 0.25 | 0 | 0 | 0 | 0 | 0 |
| ENSMUSG00000097581 | 0.45815691 | 0.25 | 0.646240625 | 0 | 0 | 0 | 0 |
| ENSMUSG00000097582 | 0 | 0 | 0 | 0 | 0 | 0 | 0 |
| ENSMUSG00000097583 | 0.578129653 | 0.5 | 0.25 | 0.701838731 | 0.387761935 | 0.602059991 | 0.629532004 |
| ENSMUSG00000097584 | 0 | 0.396240625 | 0 | 0 | 0.244650542 | 0 | 0.224243824 |
| ENSMUSG00000097585 | 0 | 0 | 0.25 | 0 | 0.244650542 | 0.301029996 | 0.355418052 |
| ENSMUSG00000097586 | 0 | 0 | 0 | 0.25 | 0 | 0 | 0 |
| ENSMUSG00000097587 | 0.289064826 | 0 | 0 | 0 | 0 | 0 | 0 |

| | | | | | | | |
|---|---|---|---|---|---|---|---|
| ENSMUSG00000097589 | 1.765759561 | 1.622963274 | 1.681980114 | 1.940387808 | 1.851969278 | 2.100370545 | 1.712465909 |
| ENSMUSG00000097590 | 0 | 0 | 0 | 0 | 0 | 0 | 0 |
| ENSMUSG00000097591 | 0.45815691 | 0.25 | 0.580482024 | 0.580482024 | 0.568060967 | 0.698970004 | 0.224243824 |
| ENSMUSG00000097592 | 0 | 0 | 0 | 0.25 | 0.489301084 | 0.301029996 | 0.579661877 |
| ENSMUSG00000097593 | 0 | 0 | 0 | 0 | 0 | 0 | 0 |
| ENSMUSG00000097594 | 0 | 0 | 0 | 0 | 0 | 0 | 0 |
| ENSMUSG00000097595 | 0 | 0 | 0 | 0 | 0 | 0.301029996 | 0.355418052 |
| ENSMUSG00000097596 | 0 | 0 | 0 | 0 | 0 | 0 | 0 |
| ENSMUSG00000097597 | 0 | 0 | 0 | 0 | 0.387761935 | 0 | 0 |
| ENSMUSG00000097598 | 0 | 0 | 0 | 0 | 0 | 0 | 0 |
| ENSMUSG00000097599 | 0 | 0 | 0 | 0 | 0 | 0 | 0 |
| ENSMUSG00000097601 | 0.289064826 | 0.5 | 0 | 0 | 0.244650542 | 0 | 0 |
| ENSMUSG00000097602 | 0 | 0 | 0 | 0.25 | 0 | 0 | 0 |
| ENSMUSG00000097603 | 0.289064826 | 0.25 | 0.396240625 | 0 | 0.244650542 | 0 | 0.355418052 |
| ENSMUSG00000097604 | 0 | 0 | 0 | 0.25 | 0 | 0 | 0 |
| ENSMUSG00000097605 | 0 | 0 | 0 | 0 | 0 | 0 | 0 |
| ENSMUSG00000097608 | 0 | 0 | 0 | 0 | 0 | 0 | 0 |
| ENSMUSG00000097609 | 0.747221736 | 1.098079356 | 1.146240625 | 1.238549078 | 1.149965125 | 1.176091259 | 1.168187745 |
| ENSMUSG00000097610 | 0.578129653 | 0.75 | 0.5 | 0.646240625 | 0.568060967 | 0.602059991 | 0.448487648 |
| ENSMUSG00000097612 | 0 | 0 | 0 | 0 | 0 | 0 | 0.224243824 |
| ENSMUSG00000097613 | 0.289064826 | 0.5 | 0.396240625 | 0 | 0.489301084 | 0.301029996 | 0.355418052 |
| ENSMUSG00000097614 | 0 | 0 | 0 | 0 | 0 | 0 | 0 |
| ENSMUSG00000097615 | 0 | 0 | 0 | 0 | 0.244650542 | 0 | 0 |
| ENSMUSG00000097616 | 1.129344651 | 1.061981878 | 1.114857905 | 0.830482024 | 0.568060967 | 0.954242509 | 0.710836105 |
| ENSMUSG00000097617 | 0.960252568 | 0.830482024 | 0.896240625 | 0.5 | 0.244650542 | 0.602059991 | 0.520678036 |
| ENSMUSG00000097618 | 0 | 0.25 | 0.396240625 | 0 | 0.244650542 | 0 | 0 |
| ENSMUSG00000097619 | 0.671187741 | 0.580482024 | 0.5 | 0.5 | 0.568060967 | 0.477121255 | 0.448487648 |
| ENSMUSG00000097620 | 0.811507563 | 0.830482024 | 0.396240625 | 0.75 | 0.846351821 | 0.77815125 | 0.952571511 |
| ENSMUSG00000097621 | 0.578129653 | 0.25 | 0.580482024 | 0.396240625 | 0.387761935 | 0.301029996 | 0.579661877 |
| ENSMUSG00000097622 | 0 | 0 | 0 | 0 | 0 | 0 | 0 |
| ENSMUSG00000097623 | 0 | 0 | 0 | 0 | 0 | 0 | 0 |
| ENSMUSG00000097624 | 0 | 0 | 0 | 0 | 0 | 0 | 0 |
| ENSMUSG00000097625 | 0 | 0 | 0 | 0 | 0.244650542 | 0 | 0 |
| ENSMUSG00000097626 | 0.45815691 | 0.701838731 | 1.080482024 | 0 | 0.244650542 | 0 | 0.448487648 |

| | | | | | | | |
|---|---|---|---|---|---|---|---|
| ENSMUSG00000097627 | 0 | 0 | 0 | 0 | 0 | 0 | 0 |
| ENSMUSG00000097628 | 0.671187741 | 0.396240625 | 0.5 | 0 | 0.568060967 | 0.301029996 | 0.579661877 |
| ENSMUSG00000097630 | 0 | 0 | 0 | 0 | 0 | 0 | 0 |
| ENSMUSG00000097631 | 0 | 0 | 0 | 0 | 0 | 0 | 0 |
| ENSMUSG00000097632 | 0 | 0 | 0 | 0 | 0.244650542 | 0 | 0 |
| ENSMUSG00000097633 | 0.289064826 | 0.79248125 | 0.580482024 | 0.701838731 | 0.812711509 | 0.77815125 | 0.775756176 |
| ENSMUSG00000097634 | 0 | 0 | 0 | 0 | 0 | 0 | 0 |
| ENSMUSG00000097635 | 0.45815691 | 0.580482024 | 0.646240625 | 0.25 | 0.244650542 | 0 | 0 |
| ENSMUSG00000097636 | 0.578129653 | 0.646240625 | 0.396240625 | 1.214495249 | 1.234113756 | 1.255272505 | 1.238624656 |
| ENSMUSG00000097637 | 0 | 0.5 | 0.5 | 0.25 | 0.387761935 | 0.301029996 | 0.629532004 |
| ENSMUSG00000097638 | 0 | 0 | 0 | 0 | 0.244650542 | 0 | 0.224243824 |
| ENSMUSG00000097639 | 0 | 0 | 0.5 | 0 | 0.244650542 | 0 | 0 |
| ENSMUSG00000097640 | 0.578129653 | 0.25 | 0.25 | 0 | 0 | 0 | 0.224243824 |
| ENSMUSG00000097641 | 0 | 0 | 0 | 0 | 0 | 0 | 0 |
| ENSMUSG00000097642 | 0 | 0 | 0 | 0 | 0 | 0 | 0 |
| ENSMUSG00000097643 | 1.036286563 | 0.75 | 0.646240625 | 0.646240625 | 0.812711509 | 0.602059991 | 0.355418052 |
| ENSMUSG00000097644 | 0 | 0 | 0 | 0 | 0 | 0 | 0 |
| ENSMUSG00000097645 | 0 | 0 | 0.396240625 | 0.25 | 0 | 0.301029996 | 0.224243824 |
| ENSMUSG00000097646 | 0 | 0 | 0 | 0 | 0 | 0 | 0 |
| ENSMUSG00000097648 | 0 | 0 | 0 | 0 | 0 | 0 | 0 |
| ENSMUSG00000097649 | 1.069666964 | 1.130890489 | 1.188721876 | 1.160964047 | 1.106691885 | 0.954242509 | 1.041356071 |
| ENSMUSG00000097650 | 0 | 0.25 | 0 | 0.25 | 0.244650542 | 0 | 0.224243824 |
| ENSMUSG00000097651 | 0.289064826 | 0.5 | 0.25 | 0.396240625 | 0 | 0 | 0.355418052 |
| ENSMUSG00000097652 | 0 | 0 | 0 | 0 | 0 | 0 | 0 |
| ENSMUSG00000097653 | 0 | 0 | 0 | 0 | 0 | 0 | 0 |
| ENSMUSG00000097654 | 0 | 0 | 0 | 0 | 0 | 0 | 0 |
| ENSMUSG00000097655 | 0 | 0 | 0 | 0 | 0 | 0 | 0 |
| ENSMUSG00000097656 | 0 | 0 | 0 | 0 | 0 | 0 | 0 |
| ENSMUSG00000097657 | 1.249317394 | 1.282320754 | 1.17510993 | 1.080482024 | 1.200473444 | 1.342422681 | 1.265599895 |
| ENSMUSG00000097658 | 0.671187741 | 0.701838731 | 0.646240625 | 0.580482024 | 0.387761935 | 0.698970004 | 0.74492186 |
| ENSMUSG00000097659 | 0 | 0 | 0 | 0 | 0 | 0 | 0 |
| ENSMUSG00000097660 | 0.289064826 | 0.646240625 | 0.75 | 0.396240625 | 0.733951626 | 0.301029996 | 0.775756176 |
| ENSMUSG00000097661 | 0.45815691 | 0 | 0.25 | 0 | 0 | 0 | 0 |
| ENSMUSG00000097662 | 0.45815691 | 0.396240625 | 0.25 | 0 | 0.244650542 | 0 | 0.448487648 |

| | | | | | | | |
|---|---|---|---|---|---|---|---|
| ENSMUSG00000097664 | 0.578129653 | 0.701838731 | 0.646240625 | 0.830482024 | 0.846351821 | 0.602059991 | 0.829800754 |
| ENSMUSG00000097665 | 0 | 0.25 | 0.25 | 0.646240625 | 0.489301084 | 0 | 0.448487648 |
| ENSMUSG00000097666 | 0 | 0 | 0 | 0.25 | 0 | 0 | 0 |
| ENSMUSG00000097667 | 0.45815691 | 0.864857905 | 0.79248125 | 0.5 | 0.632412477 | 0.477121255 | 0.355418052 |
| ENSMUSG00000097668 | 0 | 0 | 0.25 | 0 | 0 | 0 | 0 |
| ENSMUSG00000097669 | 0 | 0 | 0 | 0 | 0.244650542 | 0 | 0 |
| ENSMUSG00000097670 | 0 | 0 | 0 | 0 | 0 | 0 | 0 |
| ENSMUSG00000097671 | 0 | 0 | 0 | 0 | 0 | 0 | 0 |
| ENSMUSG00000097672 | 0.289064826 | 0 | 0.396240625 | 0.25 | 0 | 0 | 0 |
| ENSMUSG00000097673 | 0.289064826 | 0.25 | 0 | 0 | 0.632412477 | 0.477121255 | 0.448487648 |
| ENSMUSG00000097674 | 0.671187741 | 0.646240625 | 0.396240625 | 0.75 | 0.387761935 | 0.77815125 | 0.672731473 |
| ENSMUSG00000097675 | 0.289064826 | 0 | 0 | 0 | 0 | 0 | 0 |
| ENSMUSG00000097676 | 0 | 0.25 | 0 | 0 | 0 | 0 | 0 |
| ENSMUSG00000097677 | 0 | 0 | 0 | 0 | 0 | 0 | 0.224243824 |
| ENSMUSG00000097679 | 0 | 0 | 0 | 0 | 0 | 0 | 0 |
| ENSMUSG00000097680 | 0.811507563 | 0.92510993 | 0.701838731 | 0.830482024 | 1.039257769 | 0.84509804 | 0.710836105 |
| ENSMUSG00000097681 | 0 | 0 | 0 | 0 | 0 | 0 | 0.224243824 |
| ENSMUSG00000097682 | 0 | 0 | 0 | 0 | 0 | 0 | 0 |
| ENSMUSG00000097683 | 0 | 0.25 | 0 | 0 | 0 | 0 | 0 |
| ENSMUSG00000097684 | 0.578129653 | 0.580482024 | 0.830482024 | 0.75 | 0.77552387 | 0.698970004 | 0.710836105 |
| ENSMUSG00000097685 | 0 | 0 | 0 | 0 | 0.244650542 | 0 | 0.224243824 |
| ENSMUSG00000097686 | 0 | 0.580482024 | 0 | 0.396240625 | 0.244650542 | 0 | 0 |
| ENSMUSG00000097689 | 0.289064826 | 0 | 0 | 0 | 0 | 0 | 0 |
| ENSMUSG00000097690 | 0 | 0 | 0 | 0 | 0 | 0 | 0.224243824 |
| ENSMUSG00000097691 | 0 | 0 | 0 | 0 | 0 | 0 | 0 |
| ENSMUSG00000097692 | 0.578129653 | 0.396240625 | 0.5 | 0.396240625 | 0 | 0 | 0.224243824 |
| ENSMUSG00000097693 | 0 | 0 | 0 | 0 | 0 | 0 | 0 |
| ENSMUSG00000097694 | 0 | 0 | 0 | 0 | 0.686820904 | 0.301029996 | 0.224243824 |
| ENSMUSG00000097695 | 1.671187741 | 1.671625132 | 1.482684334 | 1.54248125 | 1.584291694 | 1.643452676 | 1.401059159 |
| ENSMUSG00000097697 | 1.036286563 | 1.04248125 | 1 | 0 | 0 | 0 | 0 |
| ENSMUSG00000097698 | 0 | 0 | 0 | 0 | 0 | 0 | 0 |
| ENSMUSG00000097699 | 0 | 0 | 0.25 | 0.396240625 | 0.244650542 | 0.301029996 | 0.448487648 |
| ENSMUSG00000097700 | 0 | 0 | 0 | 0 | 0 | 0 | 0 |
| ENSMUSG00000097701 | 0.289064826 | 0 | 0 | 0.25 | 0 | 0 | 0 |

| | | | | | | | |
|---|---|---|---|---|---|---|---|
| ENSMUSG00000097702 | 0 | 0.25 | 0 | 0 | 0.244650542 | 0 | 0 |
| ENSMUSG00000097703 | 0.811507563 | 0.580482024 | 0.646240625 | 0.75 | 0.686820904 | 0.84509804 | 0.853775828 |
| ENSMUSG00000097704 | 0 | 0 | 0 | 0 | 0 | 0 | 0 |
| ENSMUSG00000097705 | 0.289064826 | 0.396240625 | 0 | 0.25 | 0.244650542 | 0.301029996 | 0 |
| ENSMUSG00000097706 | 0 | 0.25 | 0 | 0 | 0 | 0.301029996 | 0 |
| ENSMUSG00000097707 | 0 | 0 | 0 | 0 | 0.244650542 | 0 | 0 |
| ENSMUSG00000097709 | 2.160971586 | 2.057204673 | 2.032320754 | 2.03360658 | 1.993840051 | 2.225309282 | 1.769001954 |
| ENSMUSG00000097710 | 0 | 0 | 0 | 0 | 0 | 0 | 0 |
| ENSMUSG00000097711 | 0.45815691 | 0.580482024 | 0.5 | 0.396240625 | 0.244650542 | 0.301029996 | 0.355418052 |
| ENSMUSG00000097712 | 0.289064826 | 0 | 0 | 0 | 0 | 0 | 0 |
| ENSMUSG00000097713 | 0 | 0.396240625 | 0 | 0 | 0.244650542 | 0 | 0 |
| ENSMUSG00000097714 | 0 | 0 | 0 | 0 | 0 | 0 | 0 |
| ENSMUSG00000097715 | 1.747221736 | 1.552363341 | 1.54745614 | 1.431980114 | 1.293076518 | 1.447158031 | 1.272006351 |
| ENSMUSG00000097716 | 0 | 0 | 0 | 0 | 0 | 0 | 0 |
| ENSMUSG00000097717 | 0.45815691 | 0 | 0 | 0.25 | 0.244650542 | 0 | 0 |
| ENSMUSG00000097718 | 0.289064826 | 0.5 | 0.5 | 0.79248125 | 0.632412477 | 0.602059991 | 0.579661877 |
| ENSMUSG00000097719 | 0 | 0 | 0 | 0 | 0.244650542 | 0 | 0.224243824 |
| ENSMUSG00000097720 | 0 | 0 | 0.25 | 0.5 | 0.244650542 | 0.477121255 | 0.355418052 |
| ENSMUSG00000097722 | 0.747221736 | 0.25 | 0.580482024 | 0.646240625 | 0.568060967 | 0.477121255 | 0.520678036 |
| ENSMUSG00000097723 | 1.734388958 | 1.51109853 | 1.356566189 | 1.584962501 | 1.394615667 | 1.653212514 | 1.417653332 |
| ENSMUSG00000097724 | 0 | 0.25 | 0.396240625 | 0.25 | 0.387761935 | 0.477121255 | 0.448487648 |
| ENSMUSG00000097725 | 0 | 0 | 0 | 0 | 0 | 0 | 0 |
| ENSMUSG00000097726 | 0.289064826 | 0 | 0 | 0 | 0 | 0.301029996 | 0 |
| ENSMUSG00000097727 | 0.671187741 | 0.646240625 | 0.646240625 | 0 | 0 | 0 | 0.355418052 |
| ENSMUSG00000097728 | 0.289064826 | 0 | 0 | 0.25 | 0.489301084 | 0.84509804 | 0.579661877 |
| ENSMUSG00000097729 | 1.45815691 | 1.339388001 | 1.321350555 | 0.976722649 | 0.931471446 | 1.113943352 | 1.014380832 |
| ENSMUSG00000097730 | 0 | 0 | 0 | 0 | 0 | 0 | 0 |
| ENSMUSG00000097731 | 0 | 0.5 | 0.396240625 | 0 | 0.387761935 | 0.477121255 | 0.448487648 |
| ENSMUSG00000097732 | 0.45815691 | 0 | 0 | 0.25 | 0 | 0 | 0.224243824 |
| ENSMUSG00000097734 | 0.289064826 | 0.830482024 | 0.646240625 | 0.864857905 | 0.489301084 | 0.698970004 | 0.74492186 |
| ENSMUSG00000097735 | 0 | 0.396240625 | 0 | 0 | 0 | 0 | 0.224243824 |
| ENSMUSG00000097736 | 0 | 0 | 0 | 0.25 | 0 | 0 | 0.224243824 |
| ENSMUSG00000097737 | 1 | 0.951838731 | 0.92510993 | 0.951838731 | 0.812711509 | 1.113943352 | 0.876096088 |
| ENSMUSG00000097738 | 0.578129653 | 0.25 | 0.396240625 | 0.25 | 0.387761935 | 0.301029996 | 0 |

| | | | | | | | |
|---|---|---|---|---|---|---|---|
| ENSMUSG00000097739 | 0.811507563 | 0.396240625 | 0.830482024 | 0.951838731 | 0.733951626 | 0.301029996 | 0.629532004 |
| ENSMUSG00000097740 | 0 | 0 | 0 | 0 | 0 | 0 | 0 |
| ENSMUSG00000097742 | 0 | 0 | 0 | 0 | 0 | 0 | 0 |
| ENSMUSG00000097743 | 1.227926429 | 1.26109853 | 1.17510993 | 1.494319981 | 1.414412788 | 1.204119983 | 1.350478789 |
| ENSMUSG00000097744 | 0 | 0.580482024 | 0.25 | 0.580482024 | 0 | 0 | 0.355418052 |
| ENSMUSG00000097745 | 0 | 0 | 0 | 0 | 0 | 0 | 0.224243824 |
| ENSMUSG00000097746 | 0.747221736 | 0 | 0.396240625 | 0.396240625 | 0.244650542 | 0 | 0.224243824 |
| ENSMUSG00000097747 | 0.578129653 | 0.580482024 | 0.5 | 0.396240625 | 0.387761935 | 0.301029996 | 0.355418052 |
| ENSMUSG00000097748 | 0.289064826 | 0 | 0 | 0.25 | 0.244650542 | 0.301029996 | 0.355418052 |
| ENSMUSG00000097749 | 0.578129653 | 0.580482024 | 0.646240625 | 0.646240625 | 0.489301084 | 0.602059991 | 0.775756176 |
| ENSMUSG00000097750 | 1 | 0.976722649 | 0.896240625 | 0.951838731 | 0.812711509 | 0.954242509 | 0.876096088 |
| ENSMUSG00000097751 | 1.069666964 | 1.04248125 | 0.976722649 | 1.02186571 | 1.163285805 | 1 | 1.054044578 |
| ENSMUSG00000097752 | 0 | 0 | 0 | 0 | 0 | 0 | 0 |
| ENSMUSG00000097753 | 0 | 0 | 0 | 0 | 0 | 0 | 0 |
| ENSMUSG00000097754 | 0 | 0 | 0 | 0 | 0 | 0 | 0 |
| ENSMUSG00000097755 | 1.069666964 | 1.17510993 | 1.061981878 | 1.25 | 1.188507684 | 1.431363764 | 1.140832123 |
| ENSMUSG00000097756 | 0.289064826 | 0.25 | 0 | 0 | 0 | 0 | 0 |
| ENSMUSG00000097757 | 0.671187741 | 0.701838731 | 0.75 | 0.701838731 | 0.489301084 | 0.602059991 | 0.672731473 |
| ENSMUSG00000097759 | 0.45815691 | 0.396240625 | 0.5 | 0 | 0.489301084 | 0 | 0.448487648 |
| ENSMUSG00000097760 | 0.671187741 | 0.5 | 0.79248125 | 0.79248125 | 0.632412477 | 0.698970004 | 0.672731473 |
| ENSMUSG00000097761 | 0 | 0 | 0.25 | 0.25 | 0.489301084 | 0.301029996 | 0.520678036 |
| ENSMUSG00000097762 | 0.289064826 | 0.396240625 | 0.646240625 | 0.396240625 | 0.387761935 | 0 | 0.448487648 |
| ENSMUSG00000097763 | 0 | 0 | 0 | 0 | 0 | 0 | 0 |
| ENSMUSG00000097764 | 0 | 0 | 0 | 0 | 0 | 0 | 0 |
| ENSMUSG00000097766 | 0.289064826 | 0 | 0 | 0 | 0 | 0 | 0 |
| ENSMUSG00000097767 | 0 | 0 | 0 | 0.25 | 0 | 0 | 0.224243824 |
| ENSMUSG00000097768 | 0.289064826 | 0.25 | 0 | 0.25 | 0.244650542 | 0.602059991 | 0.355418052 |
| ENSMUSG00000097769 | 2.075459174 | 1.87892496 | 1.910964047 | 1.958222504 | 1.919403394 | 2.120573931 | 1.784981274 |
| ENSMUSG00000097770 | 0 | 0 | 0 | 0 | 0 | 0 | 0 |
| ENSMUSG00000097772 | 2.105953527 | 1.901832578 | 1.880890489 | 1.789967834 | 1.797870327 | 1.838849091 | 1.621017948 |
| ENSMUSG00000097773 | 0.289064826 | 0 | 0.5 | 0.396240625 | 0.244650542 | 0.301029996 | 0.448487648 |
| ENSMUSG00000097774 | 0 | 0 | 0 | 0 | 0 | 0 | 0 |
| ENSMUSG00000097775 | 0 | 0 | 0 | 0 | 0 | 0.301029996 | 0 |
| ENSMUSG00000097776 | 0 | 0 | 0 | 0 | 0 | 0 | 0 |

| | | | | | | | |
|---|---|---|---|---|---|---|---|
| ENSMUSG00000097777 | 0.578129653 | 0.646240625 | 0.646240625 | 0.976722649 | 0.812711509 | 0.698970004 | 0.74492186 |
| ENSMUSG00000097779 | 0.289064826 | 0 | 0 | 0 | 0 | 0 | 0 |
| ENSMUSG00000097780 | 0 | 0 | 0 | 0 | 0 | 0 | 0 |
| ENSMUSG00000097781 | 0.867194479 | 0.79248125 | 0.92510993 | 0.75 | 0.568060967 | 0 | 0.448487648 |
| ENSMUSG00000097783 | 0 | 0 | 0 | 0 | 0 | 0 | 0 |
| ENSMUSG00000097784 | 0.671187741 | 0.5 | 0.580482024 | 0.396240625 | 0.632412477 | 0.301029996 | 0.579661877 |
| ENSMUSG00000097785 | 0.960252568 | 0.92510993 | 0.951838731 | 1.02186571 | 0.733951626 | 0.602059991 | 0.853775828 |
| ENSMUSG00000097786 | 0 | 0 | 0 | 0 | 0.244650542 | 0 | 0 |
| ENSMUSG00000097787 | 0.867194479 | 0.396240625 | 0.864857905 | 0.701838731 | 0.632412477 | 0.602059991 | 0.775756176 |
| ENSMUSG00000097788 | 0.671187741 | 0.580482024 | 0.864857905 | 0.25 | 0.244650542 | 0.301029996 | 0.672731473 |
| ENSMUSG00000097789 | 2.235832493 | 2.114857905 | 2.103406982 | 1.532320754 | 1.499532413 | 1.62324929 | 1.396774124 |
| ENSMUSG00000097790 | 0 | 0 | 0 | 0 | 0 | 0 | 0 |
| ENSMUSG00000097791 | 0.289064826 | 0.75 | 0.580482024 | 0.25 | 0.77552387 | 0.77815125 | 0.224243824 |
| ENSMUSG00000097792 | 1.325351389 | 1.146240625 | 1.04248125 | 1.282320754 | 1.244650542 | 1.414973348 | 1.078019652 |
| ENSMUSG00000097793 | 0.671187741 | 0.701838731 | 0.896240625 | 0.896240625 | 0.931471446 | 1 | 1.054044578 |
| ENSMUSG00000097794 | 0 | 0 | 0 | 0 | 0 | 0 | 0 |
| ENSMUSG00000097795 | 0 | 0.25 | 0 | 0 | 0 | 0.301029996 | 0 |
| ENSMUSG00000097796 | 1.342375483 | 1.226722649 | 1.160964047 | 1.160964047 | 1.121713561 | 1.041392685 | 1.150210039 |
| ENSMUSG00000097797 | 0.289064826 | 0.646240625 | 0.75 | 0.701838731 | 0.812711509 | 0.301029996 | 0.672731473 |
| ENSMUSG00000097798 | 0 | 0 | 0 | 0 | 0 | 0 | 0 |
| ENSMUSG00000097799 | 0.289064826 | 0.25 | 0 | 0 | 0 | 0 | 0 |
| ENSMUSG00000097800 | 0 | 0 | 0 | 0.25 | 0 | 0 | 0 |
| ENSMUSG00000097801 | 0 | 0 | 0 | 0 | 0 | 0 | 0 |
| ENSMUSG00000097803 | 0 | 0 | 0.25 | 0 | 0.244650542 | 0 | 0 |
| ENSMUSG00000097804 | 0.45815691 | 0.646240625 | 0.646240625 | 0.79248125 | 0.686820904 | 0.903089987 | 0.74492186 |
| ENSMUSG00000097805 | 0 | 0 | 0 | 0 | 0 | 0.301029996 | 0 |
| ENSMUSG00000097806 | 0.747221736 | 0.580482024 | 0.580482024 | 0.79248125 | 0.77552387 | 0.602059991 | 0.672731473 |
| ENSMUSG00000097807 | 0 | 0 | 0 | 0 | 0 | 0 | 0 |
| ENSMUSG00000097809 | 0 | 0 | 0 | 0 | 0 | 0 | 0 |
| ENSMUSG00000097810 | 0 | 0.25 | 0.25 | 0 | 0 | 0 | 0 |
| ENSMUSG00000097811 | 0 | 0 | 0.25 | 0 | 0 | 0.301029996 | 0 |
| ENSMUSG00000097812 | 0.45815691 | 0 | 0 | 0 | 0.244650542 | 0 | 0.224243824 |
| ENSMUSG00000097814 | 0.45815691 | 0.25 | 0.396240625 | 0.396240625 | 0.387761935 | 0.477121255 | 0 |
| ENSMUSG00000097815 | 2.768497801 | 2.58272922 | 2.560793496 | 2.407339155 | 2.341734473 | 2.7084209 | 2.19633727 |

| | | | | | | | |
|---|---|---|---|---|---|---|---|
| ENSMUSG00000097816 | 1.227926429 | 1.080482024 | 1.02186571 | 0.976722649 | 0.846351821 | 0.954242509 | 0.74492186 |
| ENSMUSG00000097817 | 0 | 0 | 0 | 0 | 0 | 0 | 0 |
| ENSMUSG00000097818 | 0 | 0 | 0 | 0 | 0 | 0 | 0 |
| ENSMUSG00000097819 | 0.578129653 | 0.646240625 | 0.5 | 0.864857905 | 0.632412477 | 0.301029996 | 0.710836105 |
| ENSMUSG00000097820 | 1 | 0.864857905 | 1.080482024 | 0.646240625 | 0.489301084 | 0.477121255 | 0.803905701 |
| ENSMUSG00000097821 | 0 | 0 | 0 | 0 | 0 | 0 | 0 |
| ENSMUSG00000097822 | 0 | 0 | 0 | 0 | 0 | 0 | 0 |
| ENSMUSG00000097823 | 0 | 0 | 0 | 0 | 0 | 0 | 0 |
| ENSMUSG00000097824 | 0 | 0 | 0 | 0 | 0 | 0 | 0 |
| ENSMUSG00000097825 | 0 | 0 | 0 | 0 | 0 | 0.301029996 | 0.355418052 |
| ENSMUSG00000097827 | 0 | 0 | 0.25 | 0.396240625 | 0 | 0 | 0 |
| ENSMUSG00000097828 | 0.578129653 | 0.396240625 | 0.5 | 0.830482024 | 0.905314583 | 0.477121255 | 0.629532004 |
| ENSMUSG00000097831 | 0 | 0.25 | 0.5 | 0.75 | 0 | 0.698970004 | 0.224243824 |
| ENSMUSG00000097832 | 0 | 0.396240625 | 0.646240625 | 0 | 0.387761935 | 0.301029996 | 0.355418052 |
| ENSMUSG00000097833 | 0.289064826 | 0.25 | 0.25 | 0.396240625 | 0 | 0 | 0.224243824 |
| ENSMUSG00000097834 | 0 | 0 | 0 | 0 | 0 | 0 | 0 |
| ENSMUSG00000097835 | 0.45815691 | 0.396240625 | 0.580482024 | 0.25 | 0.489301084 | 0 | 0.448487648 |
| ENSMUSG00000097836 | 0 | 0 | 0 | 0 | 0 | 0 | 0 |
| ENSMUSG00000097837 | 0 | 0 | 0 | 0 | 0 | 0 | 0 |
| ENSMUSG00000097838 | 1 | 0.896240625 | 0.79248125 | 1.02186571 | 1 | 1.176091259 | 0.74492186 |
| ENSMUSG00000097839 | 0.671187741 | 0.79248125 | 0.75 | 0.864857905 | 0.568060967 | 0.954242509 | 0.775756176 |
| ENSMUSG00000097840 | 0 | 0 | 0 | 0.25 | 0.387761935 | 0 | 0 |
| ENSMUSG00000097841 | 0 | 0 | 0 | 0 | 0 | 0 | 0 |
| ENSMUSG00000097842 | 0.867194479 | 0.646240625 | 0.75 | 0.75 | 0.77552387 | 0 | 0.629532004 |
| ENSMUSG00000097843 | 0.289064826 | 0 | 0 | 0 | 0 | 0 | 0 |
| ENSMUSG00000097845 | 0 | 0 | 0.25 | 0 | 0 | 0 | 0.355418052 |
| ENSMUSG00000097847 | 0.45815691 | 0.701838731 | 0.396240625 | 0.5 | 0.244650542 | 0.301029996 | 0.355418052 |
| ENSMUSG00000097848 | 0 | 0 | 0 | 0 | 0 | 0 | 0 |
| ENSMUSG00000097849 | 0 | 0 | 0 | 0 | 0 | 0 | 0 |
| ENSMUSG00000097850 | 0 | 0.25 | 0 | 0 | 0 | 0 | 0 |
| ENSMUSG00000097851 | 0 | 0 | 0 | 0 | 0 | 0 | 0 |
| ENSMUSG00000097852 | 0.45815691 | 0.25 | 0.25 | 0.5 | 0 | 0.602059991 | 0.224243824 |
| ENSMUSG00000097853 | 0 | 0 | 0 | 0 | 0 | 0 | 0 |
| ENSMUSG00000097854 | 0 | 0 | 0 | 0 | 0 | 0 | 0 |

| | | | | | | | |
|---|---|---|---|---|---|---|---|
| ENSMUSG00000097855 | 1.036286563 | 1.160964047 | 1.238549078 | 0.396240625 | 0.77552387 | 0.698970004 | 0.672731473 |
| ENSMUSG00000097857 | 0 | 0 | 0.25 | 0.396240625 | 0 | 0 | 0.355418052 |
| ENSMUSG00000097858 | 0.747221736 | 0.701838731 | 0.701838731 | 0.896240625 | 0.846351821 | 1 | 0.672731473 |
| ENSMUSG00000097859 | 0.289064826 | 0.396240625 | 0 | 0.25 | 0.632412477 | 0 | 0.520678036 |
| ENSMUSG00000097860 | 0 | 0 | 0 | 0.396240625 | 0 | 0 | 0.355418052 |
| ENSMUSG00000097861 | 0 | 0 | 0 | 0 | 0 | 0 | 0 |
| ENSMUSG00000097862 | 0 | 0 | 0 | 0 | 0 | 0 | 0 |
| ENSMUSG00000097863 | 0 | 0.25 | 0.25 | 0.25 | 0 | 0.301029996 | 0 |
| ENSMUSG00000097864 | 0 | 0 | 0 | 0.25 | 0 | 0 | 0.355418052 |
| ENSMUSG00000097867 | 0.91631382 | 1.061981878 | 1 | 1.061981878 | 1.176121988 | 1.322219295 | 1.121219121 |
| ENSMUSG00000097868 | 0.671187741 | 1.080482024 | 0.951838731 | 1.02186571 | 0.905314583 | 0.954242509 | 0.969165684 |
| ENSMUSG00000097869 | 0.289064826 | 0.25 | 0.25 | 0.396240625 | 0.387761935 | 0.301029996 | 0.224243824 |
| ENSMUSG00000097870 | 0 | 0 | 0 | 0 | 0 | 0 | 0 |
| ENSMUSG00000097871 | 0 | 0 | 0 | 0 | 0 | 0 | 0 |
| ENSMUSG00000097872 | 0 | 0 | 0 | 0 | 0 | 0 | 0 |
| ENSMUSG00000097873 | 0.867194479 | 0.951838731 | 0.75 | 0.580482024 | 0.846351821 | 0.84509804 | 0.672731473 |
| ENSMUSG00000097874 | 0.45815691 | 0 | 0.25 | 0.25 | 0 | 0.301029996 | 0.355418052 |
| ENSMUSG00000097875 | 0.671187741 | 0.92510993 | 0.92510993 | 0.830482024 | 0.489301084 | 0.698970004 | 0.710836105 |
| ENSMUSG00000097876 | 0.671187741 | 0.92510993 | 0.92510993 | 1.146240625 | 1.091002363 | 1.079181246 | 0.935079929 |
| ENSMUSG00000097877 | 0.45815691 | 0.5 | 0 | 0 | 0 | 0.477121255 | 0.355418052 |
| ENSMUSG00000097878 | 0 | 0 | 0 | 0 | 0 | 0 | 0 |
| ENSMUSG00000097879 | 1.505869732 | 1.451838731 | 1.52186571 | 1.634789703 | 1.635855979 | 1.672097858 | 1.520678036 |
| ENSMUSG00000097881 | 0 | 0.25 | 0.25 | 0 | 0 | 0 | 0 |
| ENSMUSG00000097882 | 0.867194479 | 0.646240625 | 0.701838731 | 0.75 | 0.686820904 | 0.698970004 | 0.710836105 |
| ENSMUSG00000097883 | 0 | 0.25 | 0 | 0 | 0 | 0.301029996 | 0.224243824 |
| ENSMUSG00000097884 | 0 | 0 | 0.25 | 0 | 0 | 0 | 0 |
| ENSMUSG00000097885 | 0.671187741 | 0.864857905 | 0.75 | 0.646240625 | 0.568060967 | 0.84509804 | 0.876096088 |
| ENSMUSG00000097886 | 0 | 0 | 0 | 0 | 0 | 0 | 0 |
| ENSMUSG00000097887 | 0.45815691 | 0.701838731 | 0.75 | 0.5 | 0.686820904 | 0.698970004 | 0.710836105 |
| ENSMUSG00000097888 | 0.578129653 | 0.646240625 | 0.646240625 | 0.25 | 0.568060967 | 0.301029996 | 0.224243824 |
| ENSMUSG00000097889 | 0.45815691 | 0.580482024 | 0.79248125 | 0.896240625 | 0.905314583 | 0.903089987 | 0.969165684 |
| ENSMUSG00000097890 | 0.671187741 | 0.5 | 0.580482024 | 0.5 | 0.387761935 | 0.301029996 | 0.520678036 |
| ENSMUSG00000097891 | 0.747221736 | 0.646240625 | 0.830482024 | 0.75 | 0.77552387 | 0.698970004 | 0.803905701 |
| ENSMUSG00000097892 | 0 | 0 | 0.25 | 0 | 0.244650542 | 0 | 0 |

| | | | | | | | |
|---|---|---|---|---|---|---|---|
| ENSMUSG00000097893 | 0 | 0.25 | 0.25 | 0.5 | 0.686820904 | 0.77815125 | 0.579661877 |
| ENSMUSG00000097895 | 0 | 0 | 0 | 0 | 0 | 0 | 0 |
| ENSMUSG00000097896 | 0 | 0 | 0 | 0 | 0 | 0 | 0 |
| ENSMUSG00000097897 | 0 | 0 | 0 | 0 | 0 | 0 | 0 |
| ENSMUSG00000097898 | 0 | 0 | 0 | 0 | 0 | 0 | 0 |
| ENSMUSG00000097899 | 0 | 0.5 | 0 | 0 | 0.244650542 | 0 | 0 |
| ENSMUSG00000097900 | 0.45815691 | 0.396240625 | 0.580482024 | 0.5 | 0 | 0 | 0 |
| ENSMUSG00000097901 | 0.45815691 | 0.5 | 0.25 | 0.396240625 | 0.387761935 | 0.301029996 | 0.355418052 |
| ENSMUSG00000097902 | 0.289064826 | 0.396240625 | 0.25 | 0.25 | 0.489301084 | 0 | 0.355418052 |
| ENSMUSG00000097903 | 0.289064826 | 0 | 0 | 0 | 0 | 0 | 0 |
| ENSMUSG00000097904 | 0.289064826 | 0.580482024 | 0.75 | 0.75 | 0.632412477 | 0.602059991 | 0.224243824 |
| ENSMUSG00000097906 | 2.063793172 | 1.982684334 | 1.975216702 | 1.819031101 | 1.848234258 | 1.973127854 | 1.680001789 |
| ENSMUSG00000097907 | 0 | 0 | 0 | 0 | 0 | 0 | 0 |
| ENSMUSG00000097908 | 2.361637952 | 2.192872367 | 2.186208459 | 1.918106335 | 1.910073614 | 2.068185862 | 1.74196736 |
| ENSMUSG00000097910 | 0 | 0 | 0 | 0 | 0 | 0 | 0 |
| ENSMUSG00000097911 | 1.129344651 | 1.098079356 | 1.17510993 | 1.26109853 | 1.188507684 | 1 | 1.110947927 |
| ENSMUSG00000097912 | 0.289064826 | 0.396240625 | 0.396240625 | 0.396240625 | 0.387761935 | 0.477121255 | 0.355418052 |
| ENSMUSG00000097913 | 0 | 0 | 0 | 0 | 0 | 0 | 0 |
| ENSMUSG00000097914 | 0.289064826 | 0 | 0 | 0.580482024 | 0.387761935 | 0 | 0.448487648 |
| ENSMUSG00000097915 | 0.289064826 | 0.646240625 | 0.646240625 | 0.646240625 | 0.489301084 | 0.954242509 | 0.853775828 |
| ENSMUSG00000097917 | 0 | 0 | 0 | 0 | 0 | 0 | 0 |
| ENSMUSG00000097918 | 0 | 0 | 0 | 0 | 0 | 0 | 0 |
| ENSMUSG00000097919 | 0 | 0 | 0 | 0 | 0 | 0 | 0 |
| ENSMUSG00000097920 | 0 | 0.25 | 0.396240625 | 0.25 | 0.244650542 | 0.301029996 | 0 |
| ENSMUSG00000097921 | 0 | 0.396240625 | 0.25 | 0 | 0.244650542 | 0 | 0.224243824 |
| ENSMUSG00000097922 | 0 | 0 | 0 | 0 | 0 | 0 | 0 |
| ENSMUSG00000097923 | 0 | 0 | 0 | 0 | 0 | 0 | 0 |
| ENSMUSG00000097924 | 1.389637216 | 1.348079356 | 1.410964047 | 1.29248125 | 0.905314583 | 1.322219295 | 1.078019652 |
| ENSMUSG00000097925 | 0.289064826 | 0.79248125 | 0.396240625 | 0.5 | 0 | 0 | 0 |
| ENSMUSG00000097926 | 0 | 0 | 0.396240625 | 0.396240625 | 0 | 0 | 0 |
| ENSMUSG00000097927 | 0 | 0 | 0 | 0 | 0 | 0 | 0 |
| ENSMUSG00000097928 | 0.811507563 | 0.92510993 | 1 | 0.5 | 0.489301084 | 0.602059991 | 0.896975297 |
| ENSMUSG00000097929 | 0 | 0 | 0 | 0 | 0.244650542 | 0.301029996 | 0 |
| ENSMUSG00000097930 | 0.289064826 | 0.5 | 0.25 | 0.25 | 0.244650542 | 0 | 0 |

| | | | | | | | |
|---|---|---|---|---|---|---|---|
| ENSMUSG00000097931 | 0 | 0 | 0 | 0 | 0 | 0 | 0 |
| ENSMUSG00000097932 | 0 | 0 | 0 | 0 | 0 | 0 | 0 |
| ENSMUSG00000097933 | 0.289064826 | 0 | 0 | 0 | 0 | 0 | 0 |
| ENSMUSG00000097934 | 0.289064826 | 0 | 0.5 | 0 | 0 | 0 | 0.224243824 |
| ENSMUSG00000097935 | 0.45815691 | 0 | 0 | 0 | 0 | 0 | 0 |
| ENSMUSG00000097936 | 0.289064826 | 0.25 | 0.5 | 0.25 | 0 | 0.301029996 | 0 |
| ENSMUSG00000097937 | 0 | 0 | 0 | 0 | 0 | 0 | 0.224243824 |
| ENSMUSG00000097938 | 0 | 0 | 0 | 0 | 0 | 0 | 0 |
| ENSMUSG00000097939 | 0 | 0 | 0 | 0 | 0 | 0 | 0 |
| ENSMUSG00000097940 | 0 | 0 | 0 | 0 | 0 | 0 | 0 |
| ENSMUSG00000097941 | 0.578129653 | 0.701838731 | 0.646240625 | 0.5 | 0.568060967 | 0 | 0.520678036 |
| ENSMUSG00000097942 | 0 | 0 | 0 | 0 | 0 | 0 | 0 |
| ENSMUSG00000097943 | 0 | 0.25 | 0 | 0 | 0 | 0 | 0.224243824 |
| ENSMUSG00000097944 | 1.249317394 | 1.080482024 | 1.146240625 | 1.17510993 | 1.188507684 | 0.954242509 | 1.159323753 |
| ENSMUSG00000097945 | 0 | 0.396240625 | 0.5 | 0.25 | 0.244650542 | 0.301029996 | 0.224243824 |
| ENSMUSG00000097946 | 0 | 0.396240625 | 0.5 | 0.25 | 0.244650542 | 0.301029996 | 0.224243824 |
| ENSMUSG00000097947 | 0 | 0 | 0 | 0 | 0 | 0 | 0 |
| ENSMUSG00000097948 | 0 | 0 | 0 | 0 | 0 | 0 | 0 |
| ENSMUSG00000097949 | 0 | 0 | 0 | 0 | 0 | 0 | 0 |
| ENSMUSG00000097950 | 0 | 0 | 0 | 0 | 0 | 0 | 0 |
| ENSMUSG00000097951 | 0 | 0 | 0.25 | 0.396240625 | 0 | 0 | 0.224243824 |
| ENSMUSG00000097952 | 0 | 0 | 0 | 0 | 0 | 0 | 0 |
| ENSMUSG00000097953 | 0 | 0.5 | 0.25 | 0 | 0 | 0 | 0 |
| ENSMUSG00000097954 | 0.747221736 | 0.951838731 | 0.864857905 | 0.5 | 0.733951626 | 0.698970004 | 0.520678036 |
| ENSMUSG00000097955 | 0 | 0 | 0 | 0 | 0 | 0 | 0 |
| ENSMUSG00000097957 | 0 | 0 | 0 | 0 | 0 | 0 | 0 |
| ENSMUSG00000097958 | 0.578129653 | 0.75 | 0.646240625 | 0 | 0.489301084 | 0.301029996 | 0.710836105 |
| ENSMUSG00000097959 | 0 | 0 | 0 | 0 | 0 | 0 | 0 |
| ENSMUSG00000097960 | 1.777675583 | 1.642463902 | 1.67510993 | 1.282320754 | 1.319233381 | 1.447158031 | 1.296434211 |
| ENSMUSG00000097961 | 0 | 0 | 0 | 0 | 0 | 0 | 0 |
| ENSMUSG00000097962 | 0 | 0 | 0 | 0 | 0 | 0 | 0 |
| ENSMUSG00000097963 | 0 | 0 | 0 | 0 | 0 | 0 | 0 |
| ENSMUSG00000097964 | 0 | 0 | 0 | 0 | 0 | 0 | 0 |
| ENSMUSG00000097965 | 0 | 0 | 0.25 | 0.25 | 0.244650542 | 0 | 0.224243824 |

| | | | | | | | |
|---|---|---|---|---|---|---|---|
| ENSMUSG00000097966 | 0 | 0 | 0 | 0 | 0 | 0 | 0.224243824 |
| ENSMUSG00000097967 | 0 | 0.25 | 0.396240625 | 0.25 | 0.244650542 | 0 | 0.355418052 |
| ENSMUSG00000097968 | 0 | 0 | 0 | 0 | 0 | 0 | 0 |
| ENSMUSG00000097969 | 0 | 0 | 0 | 0 | 0 | 0 | 0 |
| ENSMUSG00000097970 | 0.578129653 | 0.5 | 0 | 0 | 0.244650542 | 0 | 0.355418052 |
| ENSMUSG00000097971 | 1.578129653 | 1.747171172 | 1.923371739 | 1.348079356 | 1.53772706 | 1.255272505 | 1.835407748 |
| ENSMUSG00000097972 | 0 | 0 | 0 | 0 | 0 | 0 | 0 |
| ENSMUSG00000097973 | 0.811507563 | 0.976722649 | 1.080482024 | 0.896240625 | 0.77552387 | 0.84509804 | 0.896975297 |
| ENSMUSG00000097974 | 0 | 0.25 | 0 | 0 | 0 | 0 | 0 |
| ENSMUSG00000097975 | 0 | 0.396240625 | 0.25 | 0 | 0 | 0 | 0 |
| ENSMUSG00000097976 | 0 | 0 | 0 | 0 | 0 | 0 | 0 |
| ENSMUSG00000097977 | 0.578129653 | 0.79248125 | 0.864857905 | 0.580482024 | 0.387761935 | 0 | 0.935079929 |
| ENSMUSG00000097979 | 0.578129653 | 0.396240625 | 0.580482024 | 0.25 | 0.244650542 | 0 | 0.224243824 |
| ENSMUSG00000097980 | 0 | 0 | 0 | 0 | 0 | 0 | 0 |
| ENSMUSG00000097981 | 0.289064826 | 0.25 | 0 | 0.25 | 0.244650542 | 0 | 0.224243824 |
| ENSMUSG00000097982 | 0 | 0 | 0 | 0 | 0 | 0 | 0 |
| ENSMUSG00000097983 | 0 | 0.396240625 | 0 | 0 | 0 | 0 | 0 |
| ENSMUSG00000097984 | 0 | 0 | 0 | 0 | 0 | 0 | 0 |
| ENSMUSG00000097985 | 0.289064826 | 0 | 0.396240625 | 0.25 | 0 | 0.301029996 | 0.448487648 |
| ENSMUSG00000097986 | 0 | 0 | 0 | 0 | 0 | 0 | 0 |
| ENSMUSG00000097987 | 0 | 0 | 0.25 | 0 | 0 | 0 | 0 |
| ENSMUSG00000097988 | 0.289064826 | 0 | 0 | 0 | 0 | 0 | 0 |
| ENSMUSG00000097989 | 1.036286563 | 1.098079356 | 0.830482024 | 0.646240625 | 0.568060967 | 0.84509804 | 0.916588299 |
| ENSMUSG00000097990 | 0.578129653 | 0.701838731 | 0.646240625 | 0.701838731 | 0.686820904 | 0.602059991 | 0.74492186 |
| ENSMUSG00000097991 | 0 | 0.25 | 0 | 0 | 0 | 0 | 0 |
| ENSMUSG00000097992 | 0 | 0 | 0 | 0.396240625 | 0.244650542 | 0.301029996 | 0.224243824 |
| ENSMUSG00000097993 | 0.671187741 | 0.5 | 0 | 0 | 0 | 0 | 0.520678036 |
| ENSMUSG00000097994 | 1.307602651 | 1.330482024 | 1.226722649 | 1.311981878 | 1.439191811 | 1.278753601 | 1.272006351 |
| ENSMUSG00000097995 | 0.289064826 | 0.396240625 | 0 | 0.396240625 | 0.489301084 | 0.698970004 | 0.355418052 |
| ENSMUSG00000097996 | 0 | 0 | 0.25 | 0 | 0.244650542 | 0 | 0.224243824 |
| ENSMUSG00000097997 | 0 | 0 | 0 | 0 | 0 | 0 | 0 |
| ENSMUSG00000097998 | 0 | 0 | 0 | 0 | 0 | 0 | 0 |
| ENSMUSG00000097999 | 0.289064826 | 0.25 | 0.25 | 0 | 0.244650542 | 0 | 0 |
| ENSMUSG00000098000 | 0 | 0 | 0.25 | 0 | 0.244650542 | 0 | 0 |

| | | | | | | | |
|---|---|---|---|---|---|---|---|
| ENSMUSG00000098001 | 0.45815691 | 0.396240625 | 0.25 | 0.25 | 0 | 0.301029996 | 0.224243824 |
| ENSMUSG00000098002 | 0 | 0 | 0 | 0.25 | 0 | 0.301029996 | 0 |
| ENSMUSG00000098003 | 0 | 0 | 0 | 0 | 0.244650542 | 0 | 0 |
| ENSMUSG00000098004 | 0 | 0 | 0 | 0 | 0 | 0 | 0 |
| ENSMUSG00000098005 | 0 | 0.396240625 | 0 | 0 | 0 | 0 | 0 |
| ENSMUSG00000098006 | 0.867194479 | 0.976722649 | 0.79248125 | 0.701838731 | 0.77552387 | 0.301029996 | 0.448487648 |
| ENSMUSG00000098007 | 0 | 0 | 0 | 0 | 0 | 0 | 0 |
| ENSMUSG00000098008 | 0 | 0 | 0 | 0 | 0 | 0 | 0 |
| ENSMUSG00000098009 | 0 | 0 | 0 | 0 | 0 | 0 | 0.224243824 |
| ENSMUSG00000098010 | 0 | 0.25 | 0 | 0 | 0 | 0 | 0 |
| ENSMUSG00000098011 | 0 | 0 | 0 | 0 | 0 | 0 | 0 |
| ENSMUSG00000098012 | 0 | 0 | 0 | 0 | 0 | 0 | 0 |
| ENSMUSG00000098013 | 0 | 0 | 0 | 0 | 0 | 0 | 0 |
| ENSMUSG00000098014 | 0 | 0 | 0 | 0.25 | 0 | 0 | 0 |
| ENSMUSG00000098015 | 0 | 0 | 0 | 0 | 0 | 0 | 0 |
| ENSMUSG00000098016 | 0 | 0.25 | 0 | 0 | 0 | 0 | 0 |
| ENSMUSG00000098017 | 0 | 0 | 0 | 0 | 0 | 0 | 0 |
| ENSMUSG00000098019 | 0.289064826 | 0.25 | 0.5 | 0 | 0.244650542 | 0.301029996 | 0.355418052 |
| ENSMUSG00000098021 | 1.069666964 | 1.25 | 1.396240625 | 0.951838731 | 1.039257769 | 0.698970004 | 1.20139795 |
| ENSMUSG00000098022 | 0.289064826 | 0 | 0.580482024 | 0.396240625 | 0.244650542 | 0.602059991 | 0.355418052 |
| ENSMUSG00000098023 | 0 | 0 | 0 | 0 | 0 | 0 | 0 |
| ENSMUSG00000098024 | 0.289064826 | 0.396240625 | 0.396240625 | 0.79248125 | 0.77552387 | 0 | 0.579661877 |
| ENSMUSG00000098025 | 0 | 0 | 0 | 0 | 0 | 0 | 0 |
| ENSMUSG00000098026 | 0 | 0 | 0 | 0 | 0 | 0 | 0 |
| ENSMUSG00000098027 | 0 | 0 | 0 | 0 | 0 | 0 | 0 |
| ENSMUSG00000098028 | 0.671187741 | 0.25 | 0.25 | 0 | 0 | 0 | 0.355418052 |
| ENSMUSG00000098029 | 0 | 0.25 | 0 | 0 | 0 | 0 | 0 |
| ENSMUSG00000098030 | 0 | 0 | 0 | 0 | 0 | 0 | 0 |
| ENSMUSG00000098031 | 0.289064826 | 0.396240625 | 0.580482024 | 0.25 | 0.244650542 | 0 | 0 |
| ENSMUSG00000098033 | 0 | 0 | 0.25 | 0 | 0 | 0 | 0 |
| ENSMUSG00000098035 | 0.289064826 | 0.25 | 0.646240625 | 0.396240625 | 0.244650542 | 0.301029996 | 0.672731473 |
| ENSMUSG00000098036 | 0 | 0 | 0 | 0 | 0 | 0 | 0.224243824 |
| ENSMUSG00000098037 | 0 | 0 | 0 | 0 | 0 | 0 | 0 |
| ENSMUSG00000098038 | 0 | 0 | 0 | 0 | 0 | 0 | 0 |

| | | | | | | | |
|---|---|---|---|---|---|---|---|
| ENSMUSG00000098039 | 0 | 0 | 0.25 | 0 | 0 | 0 | 0 |
| ENSMUSG00000098040 | 0 | 0 | 0 | 0 | 0 | 0 | 0 |
| ENSMUSG00000098041 | 0.867194479 | 0.79248125 | 0.75 | 0.951838731 | 0.877063019 | 0.602059991 | 0.829800754 |
| ENSMUSG00000098042 | 0 | 0 | 0 | 0 | 0 | 0.301029996 | 0 |
| ENSMUSG00000098043 | 0 | 0 | 0.25 | 0 | 0 | 0 | 0 |
| ENSMUSG00000098044 | 0 | 0 | 0 | 0 | 0 | 0 | 0 |
| ENSMUSG00000098045 | 0 | 0 | 0 | 0 | 0 | 0 | 0 |
| ENSMUSG00000098046 | 0 | 0 | 0.25 | 0.25 | 0.244650542 | 0 | 0 |
| ENSMUSG00000098047 | 0 | 0 | 0 | 0 | 0 | 0 | 0 |
| ENSMUSG00000098048 | 0.289064826 | 0.25 | 0 | 0.25 | 0 | 0 | 0.224243824 |
| ENSMUSG00000098050 | 0.289064826 | 0.5 | 0.25 | 0 | 0 | 0 | 0.224243824 |
| ENSMUSG00000098051 | 0 | 0 | 0 | 0 | 0 | 0 | 0 |
| ENSMUSG00000098052 | 0.289064826 | 0.396240625 | 0 | 0 | 0 | 0 | 0 |
| ENSMUSG00000098053 | 0 | 0 | 0 | 0 | 0 | 0 | 0 |
| ENSMUSG00000098054 | 0 | 0 | 0 | 0 | 0 | 0 | 0 |
| ENSMUSG00000098055 | 0.747221736 | 0.896240625 | 0.701838731 | 0 | 0 | 0.301029996 | 0.355418052 |
| ENSMUSG00000098056 | 0.289064826 | 0 | 0 | 0 | 0.244650542 | 0 | 0 |
| ENSMUSG00000098057 | 0.289064826 | 0.25 | 0.25 | 0.25 | 0.244650542 | 0 | 0.224243824 |
| ENSMUSG00000098058 | 0 | 0 | 0 | 0 | 0 | 0 | 0 |
| ENSMUSG00000098059 | 0 | 0 | 0 | 0 | 0 | 0 | 0 |
| ENSMUSG00000098060 | 0 | 0 | 0 | 0 | 0 | 0 | 0 |
| ENSMUSG00000098061 | 1.036286563 | 0.951838731 | 0.951838731 | 0 | 0 | 0 | 0 |
| ENSMUSG00000098062 | 0.960252568 | 0.646240625 | 0.864857905 | 0.75 | 0.733951626 | 0.84509804 | 0.629532004 |
| ENSMUSG00000098064 | 0 | 0 | 0 | 0 | 0 | 0 | 0 |
| ENSMUSG00000098065 | 0 | 0 | 0 | 0 | 0 | 0 | 0 |
| ENSMUSG00000098066 | 0.289064826 | 0.396240625 | 0.5 | 0.5 | 0.244650542 | 0.301029996 | 0.629532004 |
| ENSMUSG00000098067 | 0 | 0.396240625 | 0 | 0 | 0 | 0 | 0 |
| ENSMUSG00000098068 | 0 | 0 | 0 | 0 | 0 | 0 | 0 |
| ENSMUSG00000098069 | 0 | 0 | 0 | 0 | 0 | 0 | 0 |
| ENSMUSG00000098070 | 0 | 0 | 0 | 0 | 0 | 0 | 0 |
| ENSMUSG00000098071 | 0 | 0 | 0 | 0 | 0 | 0 | 0 |
| ENSMUSG00000098072 | 0 | 0 | 0 | 0 | 0 | 0 | 0 |
| ENSMUSG00000098073 | 0.45815691 | 0 | 0.396240625 | 0.25 | 0.387761935 | 0 | 0 |
| ENSMUSG00000098074 | 0 | 0 | 0 | 0 | 0 | 0 | 0 |

| | | | | | | | |
|---|---|---|---|---|---|---|---|
| ENSMUSG00000098075 | 0 | 0 | 0 | 0 | 0 | 0 | 0 |
| ENSMUSG00000098076 | 0.289064826 | 0 | 0 | 0 | 0.489301084 | 0 | 0.448487648 |
| ENSMUSG00000098077 | 0 | 0 | 0 | 0 | 0.244650542 | 0 | 0.224243824 |
| ENSMUSG00000098078 | 0 | 0 | 0 | 0 | 0 | 0 | 0 |
| ENSMUSG00000098079 | 0 | 0.25 | 0 | 0 | 0 | 0 | 0 |
| ENSMUSG00000098080 | 0.289064826 | 0 | 0 | 0 | 0 | 0 | 0 |
| ENSMUSG00000098081 | 0 | 0 | 0 | 0 | 0 | 0 | 0 |
| ENSMUSG00000098082 | 0.45815691 | 0.396240625 | 0.701838731 | 0.25 | 0.632412477 | 0.477121255 | 0.448487648 |
| ENSMUSG00000098083 | 0 | 0 | 0 | 0 | 0 | 0 | 0.224243824 |
| ENSMUSG00000098084 | 0 | 0 | 0 | 0 | 0 | 0 | 0 |
| ENSMUSG00000098085 | 0 | 0 | 0 | 0 | 0 | 0 | 0 |
| ENSMUSG00000098086 | 0 | 0 | 0 | 0 | 0 | 0 | 0 |
| ENSMUSG00000098087 | 0 | 0 | 0 | 0 | 0 | 0 | 0 |
| ENSMUSG00000098088 | 0 | 0 | 0 | 0 | 0 | 0 | 0 |
| ENSMUSG00000098089 | 0 | 0 | 0 | 0 | 0 | 0 | 0 |
| ENSMUSG00000098090 | 1.967766868 | 1.882845365 | 1.821350555 | 1.899978211 | 1.848234258 | 2.033423755 | 1.737484395 |
| ENSMUSG00000098091 | 0.578129653 | 0.580482024 | 0.701838731 | 0.25 | 0.489301084 | 0.301029996 | 0.629532004 |
| ENSMUSG00000098092 | 0 | 0 | 0.25 | 0 | 0.387761935 | 0 | 0.355418052 |
| ENSMUSG00000098093 | 0 | 0 | 0 | 0 | 0 | 0 | 0.224243824 |
| ENSMUSG00000098094 | 0 | 0 | 0 | 0 | 0 | 0 | 0 |
| ENSMUSG00000098095 | 0 | 0 | 0 | 0 | 0 | 0 | 0 |
| ENSMUSG00000098096 | 0 | 0 | 0 | 0 | 0 | 0 | 0 |
| ENSMUSG00000098097 | 0 | 0 | 0 | 0 | 0 | 0 | 0 |
| ENSMUSG00000098098 | 1.389637216 | 1.348079356 | 1.364857905 | 0.580482024 | 0.244650542 | 0.698970004 | 0.629532004 |
| ENSMUSG00000098099 | 0 | 0.396240625 | 0 | 0 | 0 | 0 | 0.224243824 |
| ENSMUSG00000098100 | 0 | 0 | 0 | 0.25 | 0 | 0 | 0 |
| ENSMUSG00000098101 | 0 | 0 | 0.396240625 | 0.646240625 | 0.244650542 | 0.477121255 | 0.355418052 |
| ENSMUSG00000098102 | 0 | 0 | 0 | 0.25 | 0 | 0 | 0 |
| ENSMUSG00000098103 | 0 | 0 | 0 | 0 | 0 | 0 | 0 |
| ENSMUSG00000098104 | 0 | 0 | 0 | 0 | 0 | 0 | 0 |
| ENSMUSG00000098105 | 0 | 0.25 | 0.396240625 | 0 | 0 | 0 | 0.224243824 |
| ENSMUSG00000098106 | 0 | 0.5 | 0 | 0.25 | 0.244650542 | 0.301029996 | 0.355418052 |
| ENSMUSG00000098107 | 0 | 0 | 0 | 0 | 0 | 0 | 0 |
| ENSMUSG00000098108 | 0 | 0 | 0 | 0 | 0 | 0 | 0 |

| | | | | | | | |
|---|---|---|---|---|---|---|---|
| ENSMUSG00000098109 | 0 | 0 | 0.25 | 0 | 0 | 0 | 0 |
| ENSMUSG00000098110 | 0 | 0 | 0 | 0 | 0 | 0 | 0.224243824 |
| ENSMUSG00000098111 | 0.45815691 | 0.701838731 | 0.75 | 0 | 0.387761935 | 0.477121255 | 0.579661877 |
| ENSMUSG00000098112 | 0.289064826 | 0 | 0.25 | 0.25 | 0.387761935 | 0 | 0.224243824 |
| ENSMUSG00000098113 | 0.747221736 | 0.75 | 0.701838731 | 0.5 | 0.632412477 | 0.477121255 | 0.520678036 |
| ENSMUSG00000098114 | 0 | 0 | 0 | 0 | 0 | 0 | 0 |
| ENSMUSG00000098115 | 0 | 0 | 0 | 0 | 0 | 0 | 0 |
| ENSMUSG00000098116 | 0.289064826 | 0.25 | 0 | 0.25 | 0 | 0 | 0 |
| ENSMUSG00000098118 | 0 | 0 | 0 | 0 | 0 | 0 | 0 |
| ENSMUSG00000098119 | 0 | 0 | 0 | 0 | 0 | 0 | 0 |
| ENSMUSG00000098120 | 0.289064826 | 0.25 | 0.5 | 0.25 | 0.244650542 | 0 | 0.224243824 |
| ENSMUSG00000098121 | 0 | 0 | 0 | 0 | 0 | 0 | 0 |
| ENSMUSG00000098122 | 0 | 0 | 0 | 0 | 0 | 0 | 0 |
| ENSMUSG00000098123 | 0 | 0.5 | 0.25 | 0.25 | 0.244650542 | 0.301029996 | 0.355418052 |
| ENSMUSG00000098124 | 0 | 0 | 0 | 0 | 0 | 0 | 0 |
| ENSMUSG00000098125 | 0 | 0.396240625 | 0.25 | 0 | 0 | 0 | 0 |
| ENSMUSG00000098126 | 0 | 0 | 0 | 0 | 0 | 0 | 0 |
| ENSMUSG00000098127 | 0 | 0 | 0 | 0 | 0 | 0.301029996 | 0 |
| ENSMUSG00000098128 | 0 | 0 | 0 | 0 | 0 | 0 | 0 |
| ENSMUSG00000098129 | 0 | 0 | 0 | 0 | 0 | 0 | 0 |
| ENSMUSG00000098130 | 0 | 0 | 0 | 0 | 0 | 0 | 0 |
| ENSMUSG00000098131 | 0 | 0 | 0 | 0 | 0 | 0 | 0 |
| ENSMUSG00000098132 | 0.289064826 | 0 | 0 | 0 | 0 | 0 | 0.224243824 |
| ENSMUSG00000098134 | 0 | 0 | 0 | 0 | 0 | 0 | 0 |
| ENSMUSG00000098135 | 0 | 0 | 0.25 | 0 | 0 | 0 | 0 |
| ENSMUSG00000098136 | 0 | 0 | 0.25 | 0 | 0 | 0 | 0 |
| ENSMUSG00000098137 | 0 | 0 | 0 | 0 | 0 | 0 | 0 |
| ENSMUSG00000098138 | 0 | 0.25 | 0.25 | 0 | 0 | 0 | 0.224243824 |
| ENSMUSG00000098139 | 0 | 0 | 0 | 0 | 0 | 0 | 0 |
| ENSMUSG00000098140 | 0 | 0 | 0 | 0 | 0 | 0 | 0 |
| ENSMUSG00000098141 | 0.289064826 | 0.646240625 | 0.396240625 | 0.25 | 0.387761935 | 0.301029996 | 0 |
| ENSMUSG00000098142 | 0.289064826 | 0.5 | 0.580482024 | 0 | 0.387761935 | 0.301029996 | 0.224243824 |
| ENSMUSG00000098143 | 0 | 0 | 0 | 0 | 0 | 0 | 0 |
| ENSMUSG00000098144 | 0 | 0 | 0 | 0 | 0 | 0 | 0 |

| | | | | | | | |
|---|---|---|---|---|---|---|---|
| ENSMUSG00000098145 | 0 | 0 | 0 | 0 | 0 | 0 | 0 |
| ENSMUSG00000098146 | 0.289064826 | 0.25 | 0.646240625 | 0.396240625 | 0 | 0.602059991 | 0 |
| ENSMUSG00000098147 | 0 | 0 | 0 | 0 | 0 | 0 | 0 |
| ENSMUSG00000098149 | 0.578129653 | 0.396240625 | 0.580482024 | 0.25 | 0.568060967 | 0.301029996 | 0.520678036 |
| ENSMUSG00000098150 | 0.289064826 | 0.25 | 0 | 0 | 0 | 0 | 0 |
| ENSMUSG00000098151 | 0 | 0 | 0 | 0 | 0 | 0 | 0 |
| ENSMUSG00000098152 | 0 | 0 | 0 | 0 | 0 | 0 | 0 |
| ENSMUSG00000098153 | 0 | 0 | 0 | 0 | 0 | 0 | 0 |
| ENSMUSG00000098154 | 0.578129653 | 0.25 | 0.25 | 0 | 0.244650542 | 0 | 0.224243824 |
| ENSMUSG00000098155 | 0 | 0 | 0 | 0 | 0 | 0 | 0 |
| ENSMUSG00000098156 | 0.45815691 | 0.25 | 0 | 0.396240625 | 0 | 0 | 0 |
| ENSMUSG00000098157 | 0 | 0 | 0 | 0 | 0 | 0 | 0 |
| ENSMUSG00000098158 | 0 | 0 | 0 | 0.25 | 0 | 0 | 0 |
| ENSMUSG00000098159 | 0 | 0 | 0 | 0 | 0 | 0 | 0 |
| ENSMUSG00000098160 | 0 | 0 | 0 | 0 | 0 | 0 | 0 |
| ENSMUSG00000098161 | 0 | 0 | 0 | 0 | 0 | 0 | 0 |
| ENSMUSG00000098162 | 0 | 0.25 | 0 | 0.396240625 | 0 | 0 | 0 |
| ENSMUSG00000098163 | 0 | 0 | 0 | 0 | 0 | 0 | 0 |
| ENSMUSG00000098164 | 0.45815691 | 0.396240625 | 0 | 0.701838731 | 0.632412477 | 0.698970004 | 0.355418052 |
| ENSMUSG00000098165 | 0 | 0 | 0 | 0 | 0 | 0 | 0 |
| ENSMUSG00000098166 | 0 | 0 | 0 | 0 | 0 | 0 | 0 |
| ENSMUSG00000098167 | 0 | 0 | 0 | 0 | 0 | 0 | 0 |
| ENSMUSG00000098169 | 0 | 0 | 0 | 0 | 0 | 0 | 0.224243824 |
| ENSMUSG00000098170 | 0 | 0 | 0 | 0 | 0 | 0 | 0 |
| ENSMUSG00000098171 | 0.289064826 | 0 | 0 | 0 | 0 | 0 | 0 |
| ENSMUSG00000098172 | 0 | 0 | 0 | 0 | 0 | 0 | 0 |
| ENSMUSG00000098173 | 0 | 0 | 0 | 0 | 0 | 0 | 0 |
| ENSMUSG00000098174 | 0 | 0 | 0 | 0 | 0 | 0 | 0 |
| ENSMUSG00000098175 | 0 | 0 | 0 | 0 | 0 | 0.301029996 | 0 |
| ENSMUSG00000098176 | 0.45815691 | 1.080482024 | 0.896240625 | 1.02186571 | 1.020174412 | 0.903089987 | 1.028149525 |
| ENSMUSG00000098177 | 0 | 0 | 0 | 0 | 0 | 0 | 0 |
| ENSMUSG00000098178 | 0.671187741 | 1.146240625 | 1.470660762 | 0.701838731 | 0.931471446 | 0.477121255 | 1.350478789 |
| ENSMUSG00000098179 | 0 | 0 | 0 | 0 | 0 | 0 | 0 |
| ENSMUSG00000098180 | 0.867194479 | 1.080482024 | 1.146240625 | 0.396240625 | 0.877063019 | 0.301029996 | 0.74492186 |

| | | | | | | | |
|---|---|---|---|---|---|---|---|
| ENSMUSG00000098181 | 0 | 0.25 | 0 | 0 | 0 | 0.301029996 | 0 |
| ENSMUSG00000098183 | 1.404271433 | 1.311981878 | 1.348079356 | 1.410964047 | 1.254881871 | 1.505149978 | 1.313616061 |
| ENSMUSG00000098184 | 0 | 0.5 | 0 | 0 | 0 | 0 | 0.224243824 |
| ENSMUSG00000098185 | 0 | 0 | 0 | 0 | 0 | 0 | 0 |
| ENSMUSG00000098186 | 0 | 0 | 0 | 0 | 0 | 0 | 0 |
| ENSMUSG00000098187 | 0.289064826 | 0.25 | 0.25 | 0 | 0 | 0 | 0 |
| ENSMUSG00000098188 | 0 | 0 | 0 | 0 | 0 | 0 | 0 |
| ENSMUSG00000098189 | 0 | 0 | 0 | 0 | 0 | 0 | 0 |
| ENSMUSG00000098190 | 0 | 0 | 0 | 0 | 0.244650542 | 0 | 0.224243824 |
| ENSMUSG00000098191 | 0 | 0 | 0 | 0 | 0 | 0 | 0 |
| ENSMUSG00000098192 | 0.289064826 | 0.396240625 | 0 | 0 | 0 | 0.301029996 | 0 |
| ENSMUSG00000098193 | 0 | 0 | 0 | 0 | 0 | 0 | 0 |
| ENSMUSG00000098194 | 0 | 0 | 0.396240625 | 0.25 | 0 | 0.301029996 | 0.355418052 |
| ENSMUSG00000098195 | 0 | 0.25 | 0 | 0 | 0.244650542 | 0 | 0 |
| ENSMUSG00000098196 | 0 | 0 | 0 | 0.25 | 0.244650542 | 0 | 0 |
| ENSMUSG00000098197 | 0 | 0 | 0 | 0 | 0 | 0 | 0 |
| ENSMUSG00000098198 | 0 | 0 | 0 | 0.25 | 0 | 0 | 0 |
| ENSMUSG00000098199 | 0.45815691 | 0.646240625 | 0.75 | 0.25 | 0.387761935 | 0.477121255 | 0.355418052 |
| ENSMUSG00000098200 | 0 | 0 | 0 | 0 | 0.244650542 | 0 | 0 |
| ENSMUSG00000098201 | 0 | 0 | 0 | 0.25 | 0 | 0.301029996 | 0 |
| ENSMUSG00000098202 | 0 | 0.25 | 0 | 0 | 0 | 0 | 0 |
| ENSMUSG00000098203 | 0.289064826 | 0.25 | 0 | 0 | 0 | 0 | 0 |
| ENSMUSG00000098204 | 0.289064826 | 0 | 0 | 0 | 0 | 0 | 0 |
| ENSMUSG00000098205 | 0 | 0 | 0 | 0 | 0 | 0 | 0 |
| ENSMUSG00000098206 | 0.45815691 | 0.396240625 | 0.701838731 | 0.864857905 | 0.905314583 | 0.602059991 | 0.579661877 |
| ENSMUSG00000098207 | 0.289064826 | 0 | 0 | 0 | 0 | 0 | 0 |
| ENSMUSG00000098208 | 0.747221736 | 0.75 | 0.830482024 | 0.396240625 | 0.568060967 | 0.301029996 | 0.629532004 |
| ENSMUSG00000098209 | 0 | 0 | 0 | 0 | 0 | 0 | 0 |
| ENSMUSG00000098210 | 0.45815691 | 0.5 | 0.396240625 | 0 | 0.244650542 | 0.602059991 | 0.448487648 |
| ENSMUSG00000098211 | 0 | 0 | 0 | 0 | 0 | 0 | 0 |
| ENSMUSG00000098212 | 0 | 0 | 0 | 0 | 0 | 0 | 0 |
| ENSMUSG00000098213 | 0 | 0 | 0 | 0 | 0 | 0 | 0 |
| ENSMUSG00000098214 | 0 | 0 | 0 | 0 | 0 | 0 | 0 |
| ENSMUSG00000098215 | 0 | 0.25 | 0 | 0 | 0 | 0 | 0 |

| | | | | | | | |
|---|---|---|---|---|---|---|---|
| ENSMUSG00000098216 | 0 | 0 | 0 | 0 | 0 | 0 | 0 |
| ENSMUSG00000098217 | 0 | 0 | 0 | 0 | 0 | 0 | 0 |
| ENSMUSG00000098218 | 0 | 0 | 0 | 0 | 0 | 0 | 0 |
| ENSMUSG00000098219 | 0 | 0.25 | 0 | 0 | 0.244650542 | 0 | 0 |
| ENSMUSG00000098221 | 0.289064826 | 0.25 | 0 | 0.396240625 | 0.244650542 | 0.477121255 | 0.579661877 |
| ENSMUSG00000098222 | 0.289064826 | 0 | 0.646240625 | 0 | 0.244650542 | 0 | 0.224243824 |
| ENSMUSG00000098223 | 0 | 0 | 0 | 0 | 0 | 0 | 0 |
| ENSMUSG00000098224 | 0.671187741 | 0.701838731 | 0.701838731 | 0.25 | 0.489301084 | 0.77815125 | 0.710836105 |
| ENSMUSG00000098225 | 0 | 0 | 0 | 0 | 0 | 0 | 0 |
| ENSMUSG00000098226 | 0 | 0 | 0 | 0 | 0 | 0 | 0 |
| ENSMUSG00000098227 | 0 | 0 | 0 | 0 | 0 | 0.301029996 | 0 |
| ENSMUSG00000098228 | 0 | 0 | 0 | 0 | 0 | 0 | 0 |
| ENSMUSG00000098229 | 0 | 0 | 0 | 0 | 0 | 0 | 0 |
| ENSMUSG00000098230 | 0 | 0 | 0 | 0 | 0 | 0 | 0 |
| ENSMUSG00000098231 | 0 | 0 | 0.25 | 0 | 0 | 0 | 0 |
| ENSMUSG00000098232 | 0 | 0.25 | 0.25 | 0 | 0.244650542 | 0 | 0.224243824 |
| ENSMUSG00000098233 | 0 | 0 | 0 | 0 | 0 | 0 | 0 |
| ENSMUSG00000098234 | 1.920505136 | 1.735628626 | 1.78743678 | 1.769203899 | 1.741652793 | 1.919078092 | 1.655750087 |
| ENSMUSG00000098235 | 0.289064826 | 0.396240625 | 0.25 | 0.5 | 0.387761935 | 0.301029996 | 0.448487648 |
| ENSMUSG00000098236 | 0 | 0.25 | 0 | 0 | 0 | 0 | 0 |
| ENSMUSG00000098237 | 0 | 0 | 0 | 0 | 0 | 0 | 0 |
| ENSMUSG00000098238 | 0.91631382 | 1.061981878 | 1.04248125 | 0.75 | 0.846351821 | 0.84509804 | 0.74492186 |
| ENSMUSG00000098240 | 0.289064826 | 0 | 0 | 0 | 0 | 0 | 0 |
| ENSMUSG00000098241 | 0 | 0 | 0 | 0 | 0 | 0 | 0 |
| ENSMUSG00000098242 | 0 | 0 | 0 | 0 | 0 | 0 | 0 |
| ENSMUSG00000098243 | 0 | 0 | 0 | 0.25 | 0.244650542 | 0 | 0.224243824 |
| ENSMUSG00000098244 | 0 | 0 | 0 | 0 | 0 | 0 | 0 |
| ENSMUSG00000098245 | 0 | 0 | 0 | 0 | 0 | 0 | 0 |
| ENSMUSG00000098246 | 0 | 0.25 | 0 | 0 | 0.244650542 | 0 | 0 |
| ENSMUSG00000098247 | 0 | 0 | 0 | 0 | 0 | 0 | 0 |
| ENSMUSG00000098248 | 0 | 0 | 0 | 0 | 0 | 0 | 0 |
| ENSMUSG00000098249 | 0 | 0 | 0 | 0 | 0 | 0 | 0 |
| ENSMUSG00000098251 | 0 | 0 | 0 | 0 | 0 | 0 | 0 |
| ENSMUSG00000098252 | 0 | 0 | 0 | 0 | 0 | 0 | 0 |

| | | | | | | | |
|---|---|---|---|---|---|---|---|
| ENSMUSG00000098253 | 0 | 0 | 0 | 0 | 0 | 0 | 0 |
| ENSMUSG00000098254 | 0 | 0 | 0 | 0 | 0 | 0 | 0 |
| ENSMUSG00000098255 | 0 | 0 | 0 | 0 | 0 | 0 | 0 |
| ENSMUSG00000098257 | 0 | 0 | 0 | 0 | 0 | 0 | 0 |
| ENSMUSG00000098258 | 0 | 0 | 0 | 0 | 0 | 0 | 0 |
| ENSMUSG00000098259 | 0.289064826 | 0 | 0.580482024 | 0.396240625 | 0.387761935 | 0 | 0.355418052 |
| ENSMUSG00000098260 | 0 | 0 | 0 | 0 | 0 | 0 | 0 |
| ENSMUSG00000098261 | 0 | 0 | 0 | 0 | 0 | 0 | 0 |
| ENSMUSG00000098262 | 0 | 0 | 0 | 0 | 0 | 0 | 0 |
| ENSMUSG00000098263 | 0 | 0 | 0 | 0 | 0 | 0 | 0 |
| ENSMUSG00000098264 | 0 | 0 | 0 | 0 | 0 | 0 | 0 |
| ENSMUSG00000098265 | 0 | 0 | 0 | 0 | 0 | 0 | 0 |
| ENSMUSG00000098266 | 0 | 0 | 0 | 0 | 0 | 0 | 0 |
| ENSMUSG00000098267 | 0 | 0 | 0.25 | 0 | 0 | 0 | 0.355418052 |
| ENSMUSG00000098268 | 0 | 0 | 0 | 0 | 0 | 0 | 0 |
| ENSMUSG00000098269 | 0 | 0 | 0 | 0 | 0 | 0 | 0 |
| ENSMUSG00000098270 | 0 | 0 | 0 | 0 | 0 | 0 | 0 |
| ENSMUSG00000098271 | 0.289064826 | 0 | 0 | 0 | 0 | 0 | 0 |
| ENSMUSG00000098272 | 0 | 0 | 0 | 0 | 0 | 0 | 0 |
| ENSMUSG00000098273 | 0 | 0 | 0 | 0 | 0 | 0 | 0 |
| ENSMUSG00000098274 | 2.249317394 | 2.116901388 | 2.118933358 | 2.095926073 | 2.136121934 | 2.356025857 | 1.89141329 |
| ENSMUSG00000098275 | 0 | 0 | 0.25 | 0 | 0.244650542 | 0 | 0 |
| ENSMUSG00000098276 | 0 | 0 | 0 | 0 | 0 | 0 | 0 |
| ENSMUSG00000098278 | 0 | 0 | 0 | 0 | 0 | 0 | 0 |
| ENSMUSG00000098279 | 0 | 0 | 0 | 0 | 0 | 0 | 0 |
| ENSMUSG00000098280 | 0 | 0 | 0 | 0 | 0 | 0 | 0 |
| ENSMUSG00000098281 | 0 | 0 | 0 | 0 | 0 | 0 | 0 |
| ENSMUSG00000098282 | 0 | 0 | 0 | 0 | 0 | 0 | 0 |
| ENSMUSG00000098284 | 0 | 0 | 0 | 0 | 0 | 0 | 0 |
| ENSMUSG00000098285 | 0 | 0 | 0 | 0 | 0 | 0 | 0 |
| ENSMUSG00000098286 | 0.45815691 | 0.5 | 0.580482024 | 0.580482024 | 0.733951626 | 0.602059991 | 0.520678036 |
| ENSMUSG00000098287 | 0 | 0 | 0 | 0 | 0 | 0 | 0 |
| ENSMUSG00000098288 | 0 | 0 | 0 | 0 | 0 | 0 | 0 |
| ENSMUSG00000098289 | 0 | 0 | 0 | 0 | 0 | 0 | 0 |

| | | | | | | | |
|---|---|---|---|---|---|---|---|
| ENSMUSG00000098290 | 0 | 0 | 0 | 0 | 0 | 0 | 0 |
| ENSMUSG00000098291 | 0 | 0 | 0 | 0 | 0.489301084 | 0 | 0 |
| ENSMUSG00000098292 | 0 | 0 | 0 | 0.396240625 | 0 | 0 | 0 |
| ENSMUSG00000098293 | 0 | 0.25 | 0.396240625 | 0 | 0 | 0 | 0.224243824 |
| ENSMUSG00000098294 | 0 | 0 | 0 | 0 | 0 | 0 | 0 |
| ENSMUSG00000098295 | 0 | 0 | 0 | 0 | 0 | 0 | 0 |
| ENSMUSG00000098296 | 0 | 0 | 0 | 0 | 0 | 0 | 0 |
| ENSMUSG00000098297 | 0 | 0 | 0 | 0 | 0 | 0 | 0 |
| ENSMUSG00000098298 | 0 | 0 | 0 | 0 | 0 | 0 | 0 |
| ENSMUSG00000098299 | 0 | 0 | 0 | 0 | 0 | 0 | 0 |
| ENSMUSG00000098300 | 0 | 0 | 0 | 0 | 0 | 0 | 0 |
| ENSMUSG00000098302 | 0 | 0 | 0 | 0 | 0 | 0 | 0 |
| ENSMUSG00000098303 | 0 | 0 | 0 | 0 | 0 | 0 | 0 |
| ENSMUSG00000098304 | 0 | 0 | 0.396240625 | 0 | 0 | 0 | 0 |
| ENSMUSG00000098305 | 0 | 0.25 | 0 | 0 | 0 | 0 | 0 |
| ENSMUSG00000098306 | 0 | 0 | 0 | 0 | 0 | 0 | 0 |
| ENSMUSG00000098307 | 0 | 0 | 0 | 0 | 0 | 0 | 0 |
| ENSMUSG00000098308 | 0 | 0 | 0 | 0 | 0 | 0 | 0 |
| ENSMUSG00000098309 | 0 | 0 | 0 | 0 | 0 | 0 | 0 |
| ENSMUSG00000098310 | 0 | 0 | 0 | 0 | 0 | 0 | 0 |
| ENSMUSG00000098311 | 0 | 0 | 0 | 0 | 0 | 0 | 0 |
| ENSMUSG00000098312 | 0 | 0 | 0 | 0 | 0 | 0 | 0 |
| ENSMUSG00000098313 | 0 | 0 | 0 | 0 | 0 | 0 | 0 |
| ENSMUSG00000098314 | 0 | 0 | 0 | 0 | 0 | 0 | 0 |
| ENSMUSG00000098315 | 0 | 0 | 0 | 0 | 0 | 0 | 0 |
| ENSMUSG00000098316 | 0 | 0 | 0 | 0 | 0 | 0 | 0 |
| ENSMUSG00000098317 | 0 | 0 | 0 | 0 | 0 | 0 | 0 |
| ENSMUSG00000098318 | 1.568542279 | 1.505591953 | 1.614857905 | 1.25 | 1.414412788 | 1.322219295 | 1.324583736 |
| ENSMUSG00000098319 | 0 | 0 | 0 | 0 | 0 | 0 | 0 |
| ENSMUSG00000098320 | 0 | 0 | 0 | 0 | 0 | 0 | 0 |
| ENSMUSG00000098322 | 0 | 0 | 0 | 0 | 0 | 0 | 0 |
| ENSMUSG00000098323 | 0 | 0 | 0 | 0 | 0 | 0 | 0 |
| ENSMUSG00000098325 | 0 | 0 | 0 | 0 | 0 | 0 | 0 |
| ENSMUSG00000098326 | 0 | 0 | 0 | 0 | 0 | 0 | 0 |

| | | | | | | | |
|---|---|---|---|---|---|---|---|
| ENSMUSG00000098327 | 0 | 0 | 0 | 0 | 0 | 0 | 0 |
| ENSMUSG00000098328 | 0 | 0 | 0 | 0 | 0 | 0 | 0.224243824 |
| ENSMUSG00000098329 | 0 | 0 | 0 | 0 | 0 | 0 | 0 |
| ENSMUSG00000098330 | 0 | 0 | 0 | 0 | 0 | 0 | 0 |
| ENSMUSG00000098331 | 0 | 0 | 0 | 0 | 0 | 0 | 0.224243824 |
| ENSMUSG00000098332 | 1 | 1.188721876 | 1.226722649 | 0.896240625 | 1.091002363 | 1.278753601 | 1.21680636 |
| ENSMUSG00000098333 | 0 | 0 | 0 | 0 | 0 | 0 | 0 |
| ENSMUSG00000098334 | 0 | 0 | 0 | 0 | 0 | 0 | 0 |
| ENSMUSG00000098335 | 0 | 0 | 0 | 0 | 0 | 0 | 0 |
| ENSMUSG00000098336 | 0 | 0 | 0 | 0 | 0 | 0 | 0 |
| ENSMUSG00000098338 | 0 | 0 | 0 | 0 | 0 | 0 | 0 |
| ENSMUSG00000098339 | 0 | 0 | 0 | 0 | 0 | 0 | 0 |
| ENSMUSG00000098340 | 0 | 0 | 0 | 0 | 0.244650542 | 0 | 0 |
| ENSMUSG00000098341 | 0 | 0 | 0 | 0 | 0 | 0 | 0.224243824 |
| ENSMUSG00000098342 | 0 | 0 | 0 | 0 | 0 | 0 | 0 |
| ENSMUSG00000098343 | 0.578129653 | 0.580482024 | 0.25 | 0.396240625 | 0.489301084 | 0.301029996 | 0.355418052 |
| ENSMUSG00000098344 | 0.289064826 | 0 | 0.25 | 0.396240625 | 0 | 0 | 0 |
| ENSMUSG00000098345 | 0 | 0 | 0 | 0 | 0 | 0 | 0 |
| ENSMUSG00000098346 | 0 | 0 | 0.25 | 0 | 0 | 0 | 0 |
| ENSMUSG00000098347 | 0 | 0 | 0 | 0 | 0 | 0 | 0 |
| ENSMUSG00000098348 | 0 | 0 | 0 | 0 | 0 | 0 | 0 |
| ENSMUSG00000098349 | 0 | 0 | 0 | 0 | 0 | 0 | 0 |
| ENSMUSG00000098350 | 0 | 0 | 0 | 0 | 0 | 0 | 0 |
| ENSMUSG00000098351 | 0 | 0 | 0 | 0 | 0 | 0 | 0 |
| ENSMUSG00000098353 | 0 | 0 | 0 | 0 | 0 | 0 | 0 |
| ENSMUSG00000098354 | 0.289064826 | 0.25 | 0 | 0 | 0.387761935 | 0 | 0 |
| ENSMUSG00000098355 | 0 | 0 | 0 | 0 | 0 | 0 | 0 |
| ENSMUSG00000098356 | 0 | 0 | 0 | 0 | 0 | 0 | 0 |
| ENSMUSG00000098357 | 0 | 0 | 0 | 0 | 0 | 0 | 0 |
| ENSMUSG00000098358 | 0 | 0 | 0 | 0 | 0 | 0 | 0 |
| ENSMUSG00000098359 | 0 | 0 | 0 | 0 | 0 | 0 | 0 |
| ENSMUSG00000098360 | 0 | 0 | 0 | 0 | 0 | 0 | 0 |
| ENSMUSG00000098361 | 0 | 0 | 0 | 0 | 0 | 0 | 0 |
| ENSMUSG00000098362 | 0 | 0 | 0 | 0 | 0 | 0 | 0 |

| | | | | | | | |
|---|---|---|---|---|---|---|---|
| ENSMUSG00000098363 | 0 | 0.396240625 | 0.396240625 | 0.25 | 0 | 0.477121255 | 0 |
| ENSMUSG00000098364 | 0 | 0 | 0 | 0 | 0 | 0 | 0 |
| ENSMUSG00000098365 | 0 | 0 | 0 | 0.25 | 0 | 0 | 0 |
| ENSMUSG00000098366 | 0 | 0 | 0 | 0 | 0 | 0 | 0 |
| ENSMUSG00000098367 | 0 | 0 | 0 | 0 | 0 | 0 | 0 |
| ENSMUSG00000098368 | 0 | 0 | 0 | 0 | 0 | 0 | 0 |
| ENSMUSG00000098369 | 0 | 0 | 0 | 0 | 0 | 0 | 0 |
| ENSMUSG00000098370 | 0 | 0 | 0 | 0 | 0 | 0 | 0 |
| ENSMUSG00000098371 | 0 | 0 | 0 | 0 | 0 | 0 | 0 |
| ENSMUSG00000098373 | 0 | 0 | 0 | 0 | 0 | 0 | 0 |
| ENSMUSG00000098374 | 1.269664473 | 1.188721876 | 1.302363341 | 1.482684334 | 1.433158226 | 1.51851394 | 1.259064007 |
| ENSMUSG00000098376 | 0 | 0 | 0 | 0 | 0.244650542 | 0 | 0.224243824 |
| ENSMUSG00000098378 | 0 | 0 | 0 | 0 | 0 | 0 | 0 |
| ENSMUSG00000098379 | 0 | 0 | 0 | 0 | 0 | 0 | 0 |
| ENSMUSG00000098380 | 0 | 0 | 0 | 0 | 0 | 0 | 0 |
| ENSMUSG00000098381 | 0 | 0 | 0 | 0 | 0 | 0 | 0 |
| ENSMUSG00000098382 | 0 | 0 | 0 | 0 | 0 | 0 | 0 |
| ENSMUSG00000098383 | 0 | 0 | 0 | 0 | 0 | 0 | 0 |
| ENSMUSG00000098385 | 0 | 0 | 0 | 0 | 0.244650542 | 0 | 0 |
| ENSMUSG00000098386 | 0 | 0 | 0 | 0 | 0 | 0 | 0 |
| ENSMUSG00000098387 | 0 | 0 | 0 | 0 | 0 | 0 | 0 |
| ENSMUSG00000098388 | 0 | 0 | 0 | 0 | 0 | 0 | 0 |
| ENSMUSG00000098389 | 0 | 0 | 0 | 0 | 0 | 0 | 0 |
| ENSMUSG00000098390 | 0 | 0 | 0 | 0 | 0 | 0 | 0 |
| ENSMUSG00000098391 | 0 | 0 | 0 | 0 | 0 | 0 | 0 |
| ENSMUSG00000098392 | 0 | 0 | 0 | 0 | 0 | 0 | 0 |
| ENSMUSG00000098393 | 0 | 0 | 0 | 0 | 0 | 0 | 0 |
| ENSMUSG00000098394 | 0 | 0 | 0 | 0 | 0 | 0 | 0 |
| ENSMUSG00000098395 | 0 | 0 | 0 | 0 | 0 | 0 | 0 |
| ENSMUSG00000098396 | 0 | 0 | 0 | 0 | 0 | 0 | 0 |
| ENSMUSG00000098397 | 0 | 0 | 0 | 0 | 0.489301084 | 0 | 0.224243824 |
| ENSMUSG00000098399 | 0 | 0 | 0 | 0 | 0 | 0 | 0 |
| ENSMUSG00000098400 | 0 | 0 | 0 | 0 | 0 | 0 | 0 |
| ENSMUSG00000098401 | 0 | 0 | 0 | 0 | 0 | 0 | 0 |

| | | | | | | | |
|---|---|---|---|---|---|---|---|
| ENSMUSG00000098402 | 0 | 0 | 0 | 0 | 0 | 0 | 0 |
| ENSMUSG00000098403 | 0 | 0.25 | 0 | 0.25 | 0.387761935 | 0.301029996 | 0 |
| ENSMUSG00000098404 | 0 | 0 | 0 | 0 | 0 | 0 | 0 |
| ENSMUSG00000098405 | 0 | 0 | 0 | 0 | 0 | 0 | 0 |
| ENSMUSG00000098406 | 0 | 0 | 0 | 0 | 0 | 0 | 0 |
| ENSMUSG00000098407 | 0 | 0 | 0 | 0 | 0 | 0 | 0 |
| ENSMUSG00000098408 | 0 | 0 | 0 | 0 | 0 | 0 | 0 |
| ENSMUSG00000098409 | 0 | 0 | 0 | 0 | 0 | 0 | 0 |
| ENSMUSG00000098410 | 0 | 0 | 0 | 0 | 0 | 0 | 0 |
| ENSMUSG00000098412 | 0 | 0.25 | 0 | 0.5 | 0.387761935 | 0 | 0.224243824 |
| ENSMUSG00000098413 | 0 | 0 | 0 | 0 | 0 | 0 | 0 |
| ENSMUSG00000098414 | 0 | 0.25 | 0 | 0 | 0.244650542 | 0 | 0 |
| ENSMUSG00000098415 | 0 | 0 | 0 | 0 | 0.244650542 | 0 | 0 |
| ENSMUSG00000098416 | 0 | 0 | 0 | 0 | 0 | 0 | 0 |
| ENSMUSG00000098417 | 0 | 0 | 0 | 0 | 0 | 0 | 0 |
| ENSMUSG00000098418 | 0 | 0 | 0 | 0 | 0 | 0 | 0 |
| ENSMUSG00000098419 | 0 | 0.25 | 0.5 | 0 | 0.244650542 | 0 | 0 |
| ENSMUSG00000098420 | 0.289064826 | 0 | 0 | 0 | 0 | 0 | 0 |
| ENSMUSG00000098421 | 0 | 0 | 0 | 0 | 0 | 0 | 0 |
| ENSMUSG00000098422 | 0 | 0 | 0 | 0 | 0 | 0 | 0 |
| ENSMUSG00000098423 | 0 | 0 | 0 | 0 | 0 | 0 | 0 |
| ENSMUSG00000098424 | 0 | 0 | 0 | 0 | 0 | 0 | 0 |
| ENSMUSG00000098425 | 0 | 0 | 0 | 0 | 0 | 0 | 0 |
| ENSMUSG00000098426 | 0 | 0 | 0 | 0 | 0 | 0 | 0 |
| ENSMUSG00000098428 | 0 | 0 | 0 | 0 | 0 | 0 | 0 |
| ENSMUSG00000098429 | 0 | 0 | 0 | 0 | 0 | 0 | 0 |
| ENSMUSG00000098430 | 0 | 0 | 0 | 0 | 0 | 0 | 0 |
| ENSMUSG00000098431 | 0 | 0 | 0 | 0 | 0 | 0 | 0 |
| ENSMUSG00000098432 | 0 | 0 | 0 | 0 | 0 | 0 | 0 |
| ENSMUSG00000098433 | 0 | 0 | 0 | 0 | 0 | 0 | 0 |
| ENSMUSG00000098434 | 0.289064826 | 0.25 | 0 | 0.25 | 0.244650542 | 0 | 0.224243824 |
| ENSMUSG00000098435 | 0 | 0 | 0 | 0 | 0 | 0 | 0 |
| ENSMUSG00000098436 | 0 | 0 | 0 | 0 | 0 | 0 | 0 |
| ENSMUSG00000098437 | 0 | 0 | 0 | 0 | 0 | 0 | 0 |

| | | | | | | | |
|---|---|---|---|---|---|---|---|
| ENSMUSG00000098438 | 0 | 0 | 0 | 0 | 0 | 0 | 0 |
| ENSMUSG00000098439 | 0 | 0 | 0 | 0 | 0 | 0 | 0 |
| ENSMUSG00000098441 | 0 | 0 | 0 | 0 | 0 | 0 | 0 |
| ENSMUSG00000098442 | 0 | 0 | 0 | 0 | 0 | 0 | 0 |
| ENSMUSG00000098443 | 0 | 0 | 0 | 0 | 0 | 0 | 0 |
| ENSMUSG00000098444 | 0 | 0 | 0 | 0 | 0 | 0 | 0 |
| ENSMUSG00000098445 | 0.45815691 | 0 | 0.25 | 0.25 | 0.244650542 | 0.301029996 | 0.579661877 |
| ENSMUSG00000098446 | 0 | 0 | 0 | 0 | 0 | 0 | 0 |
| ENSMUSG00000098447 | 0 | 0 | 0 | 0.25 | 0.244650542 | 0 | 0.448487648 |
| ENSMUSG00000098448 | 0 | 0 | 0 | 0 | 0 | 0 | 0 |
| ENSMUSG00000098449 | 0 | 0 | 0.25 | 0 | 0 | 0 | 0.224243824 |
| ENSMUSG00000098451 | 0 | 0 | 0 | 0 | 0 | 0 | 0 |
| ENSMUSG00000098452 | 0 | 0 | 0 | 0 | 0 | 0 | 0 |
| ENSMUSG00000098453 | 0 | 0 | 0 | 0 | 0 | 0 | 0 |
| ENSMUSG00000098454 | 0 | 0 | 0 | 0 | 0 | 0 | 0 |
| ENSMUSG00000098455 | 0 | 0 | 0 | 0 | 0 | 0 | 0 |
| ENSMUSG00000098456 | 0 | 0 | 0.5 | 0 | 0.387761935 | 0 | 0 |
| ENSMUSG00000098457 | 0 | 0 | 0 | 0 | 0 | 0 | 0 |
| ENSMUSG00000098458 | 0 | 0 | 0 | 0 | 0 | 0 | 0 |
| ENSMUSG00000098459 | 0 | 0 | 0 | 0 | 0 | 0 | 0 |
| ENSMUSG00000098460 | 0 | 0 | 0 | 0 | 0 | 0 | 0 |
| ENSMUSG00000098462 | 0 | 0 | 0 | 0 | 0 | 0 | 0 |
| ENSMUSG00000098463 | 0 | 0 | 0 | 0 | 0 | 0 | 0 |
| ENSMUSG00000098464 | 0 | 0 | 0 | 0 | 0 | 0 | 0 |
| ENSMUSG00000098465 | 0 | 0 | 0.25 | 0 | 0 | 0 | 0 |
| ENSMUSG00000098466 | 0 | 0 | 0 | 0 | 0 | 0 | 0 |
| ENSMUSG00000098467 | 0 | 0 | 0 | 0 | 0 | 0 | 0 |
| ENSMUSG00000098468 | 0 | 0 | 0 | 0 | 0 | 0 | 0 |
| ENSMUSG00000098469 | 0 | 0 | 0 | 0 | 0 | 0 | 0 |
| ENSMUSG00000098470 | 0.578129653 | 0.79248125 | 0.5 | 0.580482024 | 0.812711509 | 0.903089987 | 0.579661877 |
| ENSMUSG00000098471 | 0 | 0 | 0 | 0 | 0 | 0 | 0 |
| ENSMUSG00000098472 | 0 | 0 | 0 | 0 | 0 | 0 | 0 |
| ENSMUSG00000098473 | 0 | 0 | 0 | 0 | 0 | 0 | 0 |
| ENSMUSG00000098474 | 0 | 0 | 0 | 0 | 0 | 0 | 0 |

| | | | | | | | |
|---|---|---|---|---|---|---|---|
| ENSMUSG00000098475 | 0 | 0 | 0 | 0 | 0 | 0 | 0 |
| ENSMUSG00000098476 | 0 | 0 | 0 | 0 | 0 | 0 | 0 |
| ENSMUSG00000098477 | 0 | 0 | 0 | 0 | 0 | 0 | 0 |
| ENSMUSG00000098478 | 0 | 0 | 0 | 0 | 0 | 0 | 0 |
| ENSMUSG00000098479 | 0 | 0 | 0 | 0 | 0 | 0 | 0 |
| ENSMUSG00000098480 | 0 | 0 | 0 | 0 | 0 | 0 | 0 |
| ENSMUSG00000098481 | 0 | 0 | 0 | 0 | 0 | 0 | 0 |
| ENSMUSG00000098482 | 0 | 0 | 0 | 0 | 0 | 0 | 0 |
| ENSMUSG00000098483 | 0.45815691 | 0 | 0.25 | 0 | 0 | 0 | 0 |
| ENSMUSG00000098484 | 0 | 0 | 0 | 0 | 0 | 0 | 0 |
| ENSMUSG00000098485 | 0 | 0 | 0 | 0 | 0 | 0 | 0 |
| ENSMUSG00000098486 | 0 | 0 | 0 | 0 | 0 | 0 | 0 |
| ENSMUSG00000098487 | 0 | 0 | 0 | 0 | 0 | 0 | 0 |
| ENSMUSG00000098488 | 0.45815691 | 0.396240625 | 0.5 | 0.79248125 | 0.905314583 | 0.698970004 | 0.710836105 |
| ENSMUSG00000098489 | 0 | 0 | 0 | 0 | 0 | 0 | 0 |
| ENSMUSG00000098490 | 0 | 0.25 | 0.25 | 0.25 | 0.244650542 | 0.301029996 | 0.224243824 |
| ENSMUSG00000098491 | 0 | 0 | 0 | 0 | 0 | 0 | 0 |
| ENSMUSG00000098492 | 0 | 0 | 0.25 | 0 | 0 | 0 | 0 |
| ENSMUSG00000098493 | 0 | 0 | 0 | 0 | 0 | 0 | 0 |
| ENSMUSG00000098494 | 0 | 0 | 0 | 0 | 0.244650542 | 0 | 0.224243824 |
| ENSMUSG00000098496 | 0.867194479 | 0.75 | 0.79248125 | 0.79248125 | 0.568060967 | 0.77815125 | 0.710836105 |
| ENSMUSG00000098497 | 0 | 0 | 0 | 0 | 0 | 0 | 0 |
| ENSMUSG00000098498 | 0 | 0 | 0 | 0 | 0 | 0 | 0 |
| ENSMUSG00000098499 | 0 | 0 | 0 | 0 | 0 | 0 | 0 |
| ENSMUSG00000098500 | 0 | 0 | 0 | 0 | 0 | 0 | 0 |
| ENSMUSG00000098501 | 0 | 0 | 0 | 0 | 0 | 0 | 0 |
| ENSMUSG00000098502 | 0 | 0 | 0 | 0 | 0 | 0 | 0 |
| ENSMUSG00000098503 | 0 | 0 | 0 | 0 | 0 | 0 | 0 |
| ENSMUSG00000098504 | 0 | 0 | 0 | 0 | 0 | 0 | 0 |
| ENSMUSG00000098505 | 0 | 0 | 0 | 0 | 0 | 0 | 0 |
| ENSMUSG00000098506 | 0.45815691 | 0.580482024 | 0.396240625 | 0 | 0.733951626 | 0 | 0.629532004 |
| ENSMUSG00000098507 | 0 | 0 | 0 | 0 | 0 | 0 | 0 |
| ENSMUSG00000098508 | 0 | 0 | 0 | 0 | 0 | 0 | 0 |
| ENSMUSG00000098509 | 0 | 0 | 0 | 0 | 0 | 0 | 0 |

| | | | | | | | |
|---|---|---|---|---|---|---|---|
| ENSMUSG00000098510 | 0 | 0 | 0 | 0.25 | 0.244650542 | 0 | 0.224243824 |
| ENSMUSG00000098511 | 0 | 0 | 0 | 0 | 0 | 0 | 0 |
| ENSMUSG00000098513 | 0 | 0 | 0 | 0 | 0 | 0 | 0 |
| ENSMUSG00000098514 | 0 | 0 | 0 | 0 | 0 | 0 | 0 |
| ENSMUSG00000098515 | 0.578129653 | 0.5 | 0 | 0.75 | 0.489301084 | 0.301029996 | 0.629532004 |
| ENSMUSG00000098516 | 0 | 0 | 0.25 | 0 | 0 | 0 | 0 |
| ENSMUSG00000098518 | 0 | 0 | 0 | 0 | 0 | 0 | 0 |
| ENSMUSG00000098519 | 0 | 0 | 0 | 0 | 0 | 0 | 0 |
| ENSMUSG00000098520 | 0.289064826 | 0 | 0 | 0 | 0.244650542 | 0 | 0.224243824 |
| ENSMUSG00000098521 | 0 | 0 | 0 | 0 | 0 | 0 | 0 |
| ENSMUSG00000098522 | 0 | 0 | 0 | 0 | 0 | 0 | 0 |
| ENSMUSG00000098524 | 0 | 0 | 0 | 0 | 0 | 0 | 0 |
| ENSMUSG00000098525 | 0 | 0 | 0 | 0 | 0 | 0 | 0 |
| ENSMUSG00000098526 | 0 | 0 | 0 | 0 | 0 | 0 | 0 |
| ENSMUSG00000098527 | 0 | 0 | 0 | 0 | 0 | 0 | 0 |
| ENSMUSG00000098528 | 0 | 0 | 0 | 0 | 0 | 0 | 0 |
| ENSMUSG00000098529 | 0 | 0 | 0 | 0 | 0 | 0 | 0 |
| ENSMUSG00000098530 | 0 | 0 | 0 | 0 | 0 | 0 | 0 |
| ENSMUSG00000098531 | 0 | 0 | 0 | 0 | 0 | 0 | 0 |
| ENSMUSG00000098532 | 0 | 0 | 0 | 0.25 | 0 | 0 | 0 |
| ENSMUSG00000098533 | 0 | 0 | 0 | 0 | 0 | 0 | 0 |
| ENSMUSG00000098534 | 0 | 0 | 0 | 0.25 | 0.387761935 | 0 | 0.224243824 |
| ENSMUSG00000098535 | 0 | 0 | 0 | 0 | 0 | 0 | 0 |
| ENSMUSG00000098536 | 0 | 0 | 0 | 0 | 0 | 0 | 0 |
| ENSMUSG00000098537 | 0 | 0 | 0 | 0 | 0 | 0 | 0 |
| ENSMUSG00000098538 | 0 | 0 | 0 | 0 | 0 | 0 | 0 |
| ENSMUSG00000098539 | 0 | 0 | 0 | 0 | 0 | 0 | 0 |
| ENSMUSG00000098540 | 0 | 0 | 0 | 0 | 0 | 0 | 0 |
| ENSMUSG00000098541 | 0 | 0 | 0 | 0 | 0 | 0 | 0 |
| ENSMUSG00000098542 | 0 | 0 | 0 | 0 | 0 | 0 | 0 |
| ENSMUSG00000098543 | 0 | 0 | 0 | 0 | 0 | 0 | 0 |
| ENSMUSG00000098544 | 0 | 0 | 0 | 0 | 0 | 0 | 0 |
| ENSMUSG00000098547 | 0.289064826 | 0 | 0 | 0 | 0.244650542 | 0 | 0.224243824 |
| ENSMUSG00000098548 | 0 | 0 | 0 | 0 | 0 | 0 | 0 |

| | | | | | | | |
|---|---|---|---|---|---|---|---|
| ENSMUSG00000098549 | 0.45815691 | 0 | 0 | 0 | 0.244650542 | 0 | 0.448487648 |
| ENSMUSG00000098550 | 0 | 0 | 0 | 0 | 0 | 0 | 0 |
| ENSMUSG00000098552 | 0 | 0 | 0 | 0 | 0 | 0 | 0 |
| ENSMUSG00000098553 | 0 | 0 | 0 | 0 | 0 | 0 | 0 |
| ENSMUSG00000098555 | 0 | 0 | 0 | 0 | 0 | 0 | 0 |
| ENSMUSG00000098557 | 0.289064826 | 0 | 0.5 | 0.92510993 | 0.931471446 | 1.113943352 | 0.952571511 |
| ENSMUSG00000098558 | 0 | 0 | 0 | 0 | 0 | 0 | 0 |
| ENSMUSG00000098559 | 0.289064826 | 0.396240625 | 0.396240625 | 0.580482024 | 0.686820904 | 0.477121255 | 0.803905701 |
| ENSMUSG00000098560 | 0 | 0 | 0 | 0 | 0 | 0 | 0 |
| ENSMUSG00000098561 | 0 | 0 | 0 | 0 | 0 | 0 | 0 |
| ENSMUSG00000098562 | 0 | 0 | 0 | 0 | 0 | 0 | 0 |
| ENSMUSG00000098563 | 0 | 0 | 0 | 0 | 0 | 0 | 0 |
| ENSMUSG00000098564 | 0 | 0 | 0 | 0 | 0 | 0 | 0 |
| ENSMUSG00000098565 | 0 | 0 | 0 | 0 | 0 | 0 | 0 |
| ENSMUSG00000098566 | 0 | 0 | 0 | 0 | 0 | 0 | 0 |
| ENSMUSG00000098567 | 0 | 0 | 0 | 0 | 0 | 0 | 0 |
| ENSMUSG00000098568 | 0 | 0 | 0 | 0 | 0 | 0 | 0 |
| ENSMUSG00000098569 | 0 | 0 | 0 | 0 | 0 | 0 | 0 |
| ENSMUSG00000098570 | 0 | 0 | 0 | 0 | 0 | 0 | 0 |
| ENSMUSG00000098572 | 0 | 0 | 0 | 0 | 0 | 0 | 0 |
| ENSMUSG00000098573 | 0 | 0 | 0 | 0 | 0 | 0 | 0 |
| ENSMUSG00000098574 | 0 | 0 | 0 | 0 | 0 | 0 | 0 |
| ENSMUSG00000098575 | 0 | 0 | 0 | 0 | 0 | 0 | 0 |
| ENSMUSG00000098577 | 0 | 0 | 0 | 0 | 0 | 0 | 0 |
| ENSMUSG00000098578 | 0 | 0 | 0 | 0 | 0 | 0 | 0 |
| ENSMUSG00000098579 | 0 | 0 | 0 | 0 | 0 | 0 | 0 |
| ENSMUSG00000098580 | 0 | 0 | 0 | 0 | 0 | 0 | 0 |
| ENSMUSG00000098582 | 0 | 0 | 0 | 0 | 0 | 0 | 0 |
| ENSMUSG00000098583 | 0 | 0 | 0 | 0 | 0 | 0 | 0 |
| ENSMUSG00000098585 | 0 | 0 | 0 | 0 | 0 | 0 | 0 |
| ENSMUSG00000098586 | 0 | 0 | 0 | 0 | 0 | 0 | 0 |
| ENSMUSG00000098588 | 0 | 0 | 0 | 0 | 0 | 0 | 0 |
| ENSMUSG00000098589 | 0 | 0 | 0 | 0 | 0.244650542 | 0.301029996 | 0.224243824 |
| ENSMUSG00000098590 | 0 | 0 | 0 | 0 | 0 | 0 | 0 |

| | | | | | | | |
|---|---|---|---|---|---|---|---|
| ENSMUSG00000098591 | 0 | 0 | 0 | 0 | 0 | 0 | 0 |
| ENSMUSG00000098592 | 0 | 0 | 0 | 0 | 0 | 0 | 0 |
| ENSMUSG00000098593 | 0 | 0 | 0 | 0 | 0 | 0 | 0 |
| ENSMUSG00000098594 | 0 | 0 | 0 | 0 | 0 | 0 | 0 |
| ENSMUSG00000098595 | 0 | 0 | 0 | 0 | 0 | 0 | 0 |
| ENSMUSG00000098596 | 0 | 0 | 0 | 0 | 0 | 0 | 0 |
| ENSMUSG00000098597 | 0 | 0 | 0 | 0 | 0 | 0 | 0 |
| ENSMUSG00000098598 | 0 | 0 | 0 | 0 | 0 | 0 | 0 |
| ENSMUSG00000098599 | 0 | 0 | 0 | 0 | 0 | 0 | 0 |
| ENSMUSG00000098600 | 0 | 0.25 | 0 | 0 | 0.244650542 | 0 | 0.224243824 |
| ENSMUSG00000098601 | 0 | 0 | 0 | 0 | 0 | 0 | 0 |
| ENSMUSG00000098602 | 0 | 0 | 0 | 0 | 0 | 0 | 0 |
| ENSMUSG00000098603 | 0 | 0 | 0 | 0 | 0 | 0 | 0 |
| ENSMUSG00000098604 | 0 | 0 | 0 | 0 | 0 | 0 | 0 |
| ENSMUSG00000098606 | 0 | 0 | 0 | 0 | 0 | 0 | 0 |
| ENSMUSG00000098607 | 0 | 0 | 0 | 0 | 0 | 0 | 0 |
| ENSMUSG00000098608 | 0 | 0 | 0 | 0 | 0 | 0 | 0 |
| ENSMUSG00000098609 | 0 | 0.396240625 | 0 | 0.396240625 | 0 | 0.602059991 | 0.355418052 |
| ENSMUSG00000098611 | 0 | 0 | 0 | 0 | 0 | 0 | 0 |
| ENSMUSG00000098612 | 0 | 0 | 0 | 0 | 0 | 0 | 0 |
| ENSMUSG00000098613 | 0 | 0 | 0 | 0 | 0 | 0 | 0 |
| ENSMUSG00000098614 | 0 | 0 | 0 | 0.25 | 0 | 0 | 0 |
| ENSMUSG00000098617 | 0 | 0 | 0 | 0 | 0 | 0 | 0 |
| ENSMUSG00000098618 | 0 | 0 | 0 | 0 | 0 | 0 | 0 |
| ENSMUSG00000098619 | 0 | 0 | 0 | 0 | 0 | 0 | 0 |
| ENSMUSG00000098620 | 0.867194479 | 0.75 | 0.646240625 | 0.864857905 | 0.77552387 | 0.301029996 | 0.579661877 |
| ENSMUSG00000098621 | 0 | 0 | 0 | 0 | 0 | 0 | 0 |
| ENSMUSG00000098622 | 0 | 0 | 0 | 0 | 0 | 0 | 0 |
| ENSMUSG00000098623 | 0 | 0 | 0 | 0 | 0 | 0 | 0 |
| ENSMUSG00000098624 | 0 | 0 | 0 | 0 | 0 | 0 | 0 |
| ENSMUSG00000098625 | 0 | 0 | 0 | 0 | 0 | 0 | 0 |
| ENSMUSG00000098626 | 0 | 0 | 0 | 0 | 0 | 0 | 0 |
| ENSMUSG00000098627 | 0 | 0 | 0 | 0 | 0 | 0 | 0 |
| ENSMUSG00000098628 | 0 | 0 | 0 | 0 | 0 | 0 | 0 |

| | | | | | | | |
|---|---|---|---|---|---|---|---|
| ENSMUSG00000098629 | 0 | 0 | 0 | 0 | 0 | 0 | 0 |
| ENSMUSG00000098630 | 0 | 0 | 0 | 0 | 0 | 0 | 0 |
| ENSMUSG00000098631 | 0 | 0 | 0 | 0 | 0 | 0 | 0 |
| ENSMUSG00000098632 | 0 | 0 | 0 | 0 | 0 | 0 | 0 |
| ENSMUSG00000098634 | 0 | 0 | 0 | 0 | 0 | 0 | 0 |
| ENSMUSG00000098635 | 0 | 0 | 0 | 0 | 0 | 0 | 0 |
| ENSMUSG00000098636 | 0 | 0 | 0 | 0 | 0 | 0 | 0 |
| ENSMUSG00000098638 | 0 | 0 | 0 | 0 | 0 | 0 | 0 |
| ENSMUSG00000098640 | 0 | 0.25 | 0 | 0 | 0 | 0 | 0 |
| ENSMUSG00000098641 | 0 | 0 | 0 | 0 | 0 | 0 | 0 |
| ENSMUSG00000098642 | 0 | 0 | 0 | 0 | 0 | 0 | 0 |
| ENSMUSG00000098644 | 0 | 0 | 0 | 0 | 0 | 0 | 0 |
| ENSMUSG00000098645 | 0 | 0 | 0 | 0 | 0 | 0 | 0 |
| ENSMUSG00000098646 | 0 | 0 | 0 | 0 | 0 | 0 | 0 |
| ENSMUSG00000098647 | 0 | 0 | 0 | 0 | 0 | 0 | 0 |
| ENSMUSG00000098648 | 0 | 0 | 0 | 0 | 0 | 0 | 0 |
| ENSMUSG00000098649 | 0 | 0 | 0.25 | 0 | 0 | 0 | 0 |
| ENSMUSG00000098650 | 0 | 0 | 0 | 0 | 0 | 0 | 0 |
| ENSMUSG00000098651 | 0 | 0 | 0 | 0 | 0 | 0 | 0 |
| ENSMUSG00000098652 | 0 | 0 | 0 | 0 | 0 | 0 | 0 |
| ENSMUSG00000098653 | 0 | 0 | 0 | 0 | 0 | 0 | 0 |
| ENSMUSG00000098654 | 0 | 0 | 0 | 0 | 0 | 0 | 0 |
| ENSMUSG00000098655 | 0 | 0 | 0 | 0 | 0 | 0 | 0 |
| ENSMUSG00000098656 | 0.289064826 | 0 | 0 | 0 | 0 | 0 | 0 |
| ENSMUSG00000098657 | 0 | 0 | 0 | 0 | 0 | 0 | 0 |
| ENSMUSG00000098658 | 0 | 0 | 0 | 0 | 0 | 0 | 0 |
| ENSMUSG00000098659 | 0 | 0 | 0 | 0 | 0 | 0 | 0 |
| ENSMUSG00000098660 | 0 | 0 | 0 | 0 | 0 | 0 | 0 |
| ENSMUSG00000098661 | 0.289064826 | 0.25 | 0 | 0 | 0 | 0 | 0 |
| ENSMUSG00000098662 | 0 | 0 | 0 | 0 | 0 | 0 | 0 |
| ENSMUSG00000098663 | 0 | 0 | 0 | 0 | 0.244650542 | 0 | 0 |
| ENSMUSG00000098664 | 0 | 0 | 0 | 0 | 0 | 0 | 0 |
| ENSMUSG00000098665 | 0 | 0 | 0 | 0 | 0 | 0 | 0 |
| ENSMUSG00000098666 | 0 | 0.25 | 0.25 | 0 | 0 | 0 | 0.224243824 |

| | | | | | | | |
|---|---|---|---|---|---|---|---|
| ENSMUSG00000098667 | 0 | 0 | 0 | 0 | 0 | 0 | 0 |
| ENSMUSG00000098668 | 0 | 0 | 0 | 0 | 0 | 0 | 0 |
| ENSMUSG00000098669 | 0.289064826 | 0.25 | 0 | 0 | 0.387761935 | 0.301029996 | 0.355418052 |
| ENSMUSG00000098670 | 0 | 0.25 | 0.396240625 | 0 | 0 | 0 | 0 |
| ENSMUSG00000098671 | 0 | 0 | 0 | 0 | 0 | 0 | 0 |
| ENSMUSG00000098672 | 0 | 0 | 0 | 0 | 0 | 0 | 0 |
| ENSMUSG00000098674 | 0 | 0 | 0 | 0 | 0 | 0 | 0 |
| ENSMUSG00000098675 | 0 | 0 | 0.396240625 | 0.25 | 0 | 0 | 0 |
| ENSMUSG00000098676 | 0 | 0 | 0 | 0 | 0 | 0 | 0 |
| ENSMUSG00000098677 | 0 | 0 | 0 | 0 | 0 | 0 | 0 |
| ENSMUSG00000098678 | 0 | 0.25 | 0.701838731 | 0.5 | 0.387761935 | 0 | 0.629532004 |
| ENSMUSG00000098679 | 0 | 0 | 0 | 0 | 0.387761935 | 0.698970004 | 0.355418052 |
| ENSMUSG00000098680 | 0 | 0 | 0 | 0 | 0 | 0 | 0 |
| ENSMUSG00000098682 | 0 | 0 | 0 | 0 | 0 | 0 | 0 |
| ENSMUSG00000098683 | 0 | 0 | 0 | 0 | 0 | 0.301029996 | 0 |
| ENSMUSG00000098684 | 0 | 0 | 0 | 0 | 0 | 0 | 0 |
| ENSMUSG00000098685 | 0 | 0.25 | 0.5 | 0 | 0.387761935 | 0.477121255 | 0.355418052 |
| ENSMUSG00000098686 | 0 | 0 | 0 | 0 | 0 | 0 | 0 |
| ENSMUSG00000098687 | 0 | 0 | 0 | 0 | 0 | 0 | 0 |
| ENSMUSG00000098688 | 0 | 0 | 0 | 0 | 0 | 0 | 0 |
| ENSMUSG00000098689 | 0 | 0 | 0 | 0 | 0 | 0 | 0 |
| ENSMUSG00000098690 | 0.289064826 | 0.25 | 0.580482024 | 0.396240625 | 0.387761935 | 0.477121255 | 0 |
| ENSMUSG00000098691 | 0 | 0 | 0 | 0 | 0 | 0 | 0 |
| ENSMUSG00000098692 | 0.289064826 | 0 | 0 | 0.25 | 0 | 0.301029996 | 0 |
| ENSMUSG00000098693 | 0 | 0 | 0 | 0 | 0 | 0 | 0 |
| ENSMUSG00000098694 | 0 | 0 | 0 | 0 | 0 | 0 | 0 |
| ENSMUSG00000098695 | 0 | 0 | 0 | 0 | 0 | 0 | 0 |
| ENSMUSG00000098696 | 0 | 0 | 0 | 0 | 0 | 0 | 0 |
| ENSMUSG00000098697 | 0 | 0 | 0 | 0 | 0 | 0 | 0 |
| ENSMUSG00000098698 | 0.289064826 | 0 | 0.25 | 0.396240625 | 0.244650542 | 0 | 0.224243824 |
| ENSMUSG00000098700 | 0 | 0 | 0 | 0 | 0 | 0 | 0 |
| ENSMUSG00000098701 | 0 | 0 | 0 | 0 | 0 | 0 | 0 |
| ENSMUSG00000098702 | 1.156259305 | 1.214495249 | 1.061981878 | 0.92510993 | 0.877063019 | 0.954242509 | 0.952571511 |
| ENSMUSG00000098703 | 0 | 0 | 0 | 0 | 0 | 0 | 0 |

| | | | | | | | |
|---|---|---|---|---|---|---|---|
| ENSMUSG00000098704 | 0 | 0 | 0 | 0 | 0 | 0 | 0 |
| ENSMUSG00000098705 | 0 | 0 | 0 | 0 | 0 | 0 | 0 |
| ENSMUSG00000098706 | 0 | 0 | 0 | 0 | 0 | 0 | 0 |
| ENSMUSG00000098707 | 0 | 0 | 0 | 0 | 0 | 0 | 0 |
| ENSMUSG00000098708 | 0 | 0 | 0 | 0 | 0 | 0 | 0 |
| ENSMUSG00000098709 | 0 | 0 | 0 | 0 | 0 | 0 | 0 |
| ENSMUSG00000098710 | 0 | 0 | 0.25 | 0 | 0 | 0 | 0 |
| ENSMUSG00000098711 | 0 | 0 | 0 | 0 | 0 | 0 | 0 |
| ENSMUSG00000098712 | 0 | 0 | 0 | 0 | 0 | 0 | 0 |
| ENSMUSG00000098713 | 0.747221736 | 0.25 | 0.396240625 | 0.25 | 0.387761935 | 0.301029996 | 0.355418052 |
| ENSMUSG00000098714 | 0 | 0 | 0 | 0.25 | 0.244650542 | 0 | 0.355418052 |
| ENSMUSG00000098715 | 0 | 0 | 0 | 0 | 0 | 0 | 0 |
| ENSMUSG00000098716 | 0 | 0 | 0 | 0 | 0 | 0 | 0 |
| ENSMUSG00000098717 | 0 | 0 | 0 | 0 | 0 | 0 | 0 |
| ENSMUSG00000098718 | 0 | 0 | 0 | 0 | 0 | 0 | 0 |
| ENSMUSG00000098719 | 0 | 0 | 0 | 0 | 0 | 0 | 0 |
| ENSMUSG00000098720 | 0 | 0 | 0 | 0 | 0 | 0 | 0 |
| ENSMUSG00000098721 | 0 | 0 | 0 | 0 | 0 | 0 | 0 |
| ENSMUSG00000098722 | 0 | 0 | 0 | 0 | 0 | 0 | 0 |
| ENSMUSG00000098723 | 0 | 0 | 0 | 0 | 0 | 0 | 0 |
| ENSMUSG00000098725 | 0 | 0 | 0 | 0 | 0 | 0 | 0 |
| ENSMUSG00000098726 | 0 | 0 | 0 | 0 | 0 | 0 | 0 |
| ENSMUSG00000098727 | 0 | 0 | 0 | 0 | 0 | 0 | 0 |
| ENSMUSG00000098728 | 0 | 0 | 0 | 0 | 0 | 0 | 0 |
| ENSMUSG00000098729 | 0 | 0 | 0 | 0 | 0 | 0 | 0 |
| ENSMUSG00000098730 | 0 | 0 | 0 | 0 | 0 | 0 | 0 |
| ENSMUSG00000098731 | 0.289064826 | 0.580482024 | 0.701838731 | 0 | 0.244650542 | 0 | 0.672731473 |
| ENSMUSG00000098732 | 0 | 0 | 0 | 0 | 0 | 0 | 0 |
| ENSMUSG00000098733 | 0 | 0 | 0 | 0 | 0 | 0 | 0 |
| ENSMUSG00000098734 | 0 | 0 | 0 | 0 | 0 | 0 | 0 |
| ENSMUSG00000098735 | 0 | 0 | 0 | 0 | 0 | 0 | 0 |
| ENSMUSG00000098736 | 0 | 0 | 0 | 0 | 0 | 0 | 0 |
| ENSMUSG00000098737 | 0 | 0 | 0 | 0 | 0 | 0 | 0 |
| ENSMUSG00000098739 | 0 | 0 | 0 | 0 | 0 | 0 | 0 |

| | | | | | | | |
|---|---|---|---|---|---|---|---|
| ENSMUSG00000098740 | 0 | 0 | 0 | 0 | 0 | 0 | 0 |
| ENSMUSG00000098741 | 0 | 0 | 0 | 0 | 0 | 0 | 0 |
| ENSMUSG00000098742 | 0 | 0 | 0 | 0 | 0 | 0 | 0 |
| ENSMUSG00000098743 | 0 | 0 | 0 | 0 | 0 | 0 | 0 |
| ENSMUSG00000098744 | 0 | 0 | 0 | 0 | 0 | 0 | 0 |
| ENSMUSG00000098745 | 0 | 0 | 0 | 0 | 0 | 0 | 0 |
| ENSMUSG00000098747 | 0 | 0 | 0 | 0 | 0 | 0 | 0 |
| ENSMUSG00000098748 | 0 | 0 | 0 | 0 | 0 | 0 | 0 |
| ENSMUSG00000098749 | 0.45815691 | 0.896240625 | 0.646240625 | 0.896240625 | 0.244650542 | 0.477121255 | 0.710836105 |
| ENSMUSG00000098750 | 0 | 0 | 0 | 0 | 0 | 0.301029996 | 0 |
| ENSMUSG00000098751 | 0 | 0 | 0 | 0 | 0 | 0 | 0 |
| ENSMUSG00000098752 | 0 | 0 | 0 | 0 | 0 | 0 | 0 |
| ENSMUSG00000098754 | 0 | 0 | 0 | 0 | 0 | 0 | 0 |
| ENSMUSG00000098755 | 0 | 0 | 0.25 | 0.25 | 0.489301084 | 0.301029996 | 0 |
| ENSMUSG00000098756 | 0 | 0 | 0 | 0 | 0 | 0 | 0 |
| ENSMUSG00000098757 | 0 | 0 | 0 | 0 | 0 | 0 | 0 |
| ENSMUSG00000098758 | 0 | 0 | 0 | 0 | 0 | 0 | 0 |
| ENSMUSG00000098759 | 0 | 0.25 | 0 | 0.25 | 0 | 0 | 0 |
| ENSMUSG00000098760 | 0 | 0 | 0 | 0 | 0 | 0.301029996 | 0 |
| ENSMUSG00000098761 | 0 | 0 | 0 | 0 | 0.387761935 | 0 | 0 |
| ENSMUSG00000098762 | 0 | 0 | 0 | 0 | 0 | 0 | 0 |
| ENSMUSG00000098763 | 0 | 0 | 0 | 0 | 0 | 0 | 0 |
| ENSMUSG00000098764 | 0 | 0 | 0 | 0 | 0 | 0 | 0 |
| ENSMUSG00000098765 | 0 | 0 | 0 | 0 | 0 | 0 | 0 |
| ENSMUSG00000098766 | 0 | 0.396240625 | 0.5 | 0.580482024 | 0.489301084 | 0.477121255 | 0.224243824 |
| ENSMUSG00000098767 | 0 | 0 | 0 | 0 | 0 | 0 | 0 |
| ENSMUSG00000098768 | 0 | 0 | 0 | 0 | 0 | 0 | 0 |
| ENSMUSG00000098769 | 0 | 0 | 0 | 0 | 0 | 0 | 0.224243824 |
| ENSMUSG00000098770 | 0 | 0 | 0 | 0 | 0 | 0 | 0 |
| ENSMUSG00000098771 | 0 | 0 | 0 | 0 | 0 | 0 | 0 |
| ENSMUSG00000098772 | 0 | 0 | 0 | 0 | 0 | 0 | 0 |
| ENSMUSG00000098773 | 0 | 0 | 0 | 0 | 0 | 0 | 0 |
| ENSMUSG00000098774 | 0 | 0 | 0 | 0 | 0 | 0 | 0 |
| ENSMUSG00000098775 | 0 | 0 | 0 | 0 | 0 | 0 | 0 |

| | | | | | | | |
|---|---|---|---|---|---|---|---|
| ENSMUSG00000098776 | 0 | 0 | 0 | 0 | 0 | 0 | 0 |
| ENSMUSG00000098777 | 0 | 0 | 0 | 0 | 0 | 0 | 0 |
| ENSMUSG00000098778 | 0 | 0 | 0 | 0 | 0 | 0 | 0 |
| ENSMUSG00000098779 | 0 | 0 | 0.396240625 | 0 | 0 | 0 | 0 |
| ENSMUSG00000098780 | 0 | 0 | 0 | 0 | 0 | 0 | 0 |
| ENSMUSG00000098781 | 0 | 0 | 0 | 0 | 0 | 0 | 0 |
| ENSMUSG00000098782 | 0 | 0 | 0 | 0 | 0 | 0 | 0 |
| ENSMUSG00000098783 | 0 | 0 | 0 | 0 | 0 | 0 | 0 |
| ENSMUSG00000098784 | 0 | 0 | 0 | 0 | 0 | 0 | 0 |
| ENSMUSG00000098785 | 0 | 0 | 0 | 0 | 0 | 0 | 0 |
| ENSMUSG00000098786 | 0.289064826 | 0 | 0 | 0.25 | 0 | 0 | 0 |
| ENSMUSG00000098787 | 0 | 0 | 0 | 0 | 0 | 0 | 0 |
| ENSMUSG00000098788 | 0 | 0 | 0 | 0 | 0 | 0 | 0 |
| ENSMUSG00000098789 | 0 | 0 | 0 | 0.580482024 | 0 | 0 | 0.448487648 |
| ENSMUSG00000098790 | 0.289064826 | 0.396240625 | 0.25 | 0.396240625 | 0.387761935 | 0 | 0 |
| ENSMUSG00000098791 | 0 | 0 | 0 | 0 | 0 | 0 | 0 |
| ENSMUSG00000098792 | 0 | 0 | 0 | 0 | 0 | 0 | 0 |
| ENSMUSG00000098793 | 0 | 0 | 0 | 0 | 0 | 0 | 0 |
| ENSMUSG00000098794 | 0 | 0 | 0 | 0 | 0 | 0 | 0 |
| ENSMUSG00000098795 | 0 | 0 | 0 | 0 | 0 | 0 | 0 |
| ENSMUSG00000098796 | 0 | 0 | 0 | 0 | 0 | 0 | 0 |
| ENSMUSG00000098797 | 0 | 0 | 0 | 0 | 0 | 0 | 0 |
| ENSMUSG00000098798 | 0 | 0 | 0 | 0 | 0 | 0 | 0 |
| ENSMUSG00000098799 | 0 | 0 | 0 | 0 | 0 | 0 | 0 |
| ENSMUSG00000098800 | 0 | 0 | 0 | 0 | 0 | 0 | 0 |
| ENSMUSG00000098801 | 0 | 0 | 0 | 0 | 0 | 0 | 0 |
| ENSMUSG00000098802 | 0 | 0 | 0 | 0 | 0 | 0 | 0 |
| ENSMUSG00000098803 | 0 | 0 | 0 | 0 | 0.244650542 | 0 | 0 |
| ENSMUSG00000098805 | 0 | 0 | 0 | 0 | 0 | 0 | 0 |
| ENSMUSG00000098807 | 0 | 0 | 0 | 0 | 0 | 0 | 0 |
| ENSMUSG00000098808 | 0 | 0 | 0 | 0 | 0 | 0 | 0 |
| ENSMUSG00000098809 | 0 | 0 | 0 | 0 | 0 | 0 | 0 |
| ENSMUSG00000098811 | 0 | 0 | 0 | 0.396240625 | 0.244650542 | 0 | 0.224243824 |
| ENSMUSG00000098812 | 0 | 0.25 | 0 | 0 | 0 | 0 | 0 |

| | | | | | | | |
|---|---|---|---|---|---|---|---|
| ENSMUSG00000098813 | 0.289064826 | 0.646240625 | 0.646240625 | 0.25 | 0.632412477 | 0.602059991 | 0.448487648 |
| ENSMUSG00000098814 | 0 | 0 | 0 | 0 | 0 | 0 | 0 |
| ENSMUSG00000098815 | 0 | 0 | 0 | 0 | 0 | 0 | 0 |
| ENSMUSG00000098816 | 0 | 0 | 0 | 0 | 0 | 0 | 0 |
| ENSMUSG00000098817 | 0 | 0 | 0 | 0 | 0 | 0 | 0 |
| ENSMUSG00000098818 | 0 | 0 | 0 | 0 | 0 | 0 | 0 |
| ENSMUSG00000098819 | 0 | 0 | 0 | 0 | 0 | 0 | 0 |
| ENSMUSG00000098820 | 0 | 0 | 0 | 0.25 | 0 | 0 | 0 |
| ENSMUSG00000098821 | 0 | 0 | 0 | 0 | 0 | 0 | 0 |
| ENSMUSG00000098822 | 0 | 0 | 0 | 0 | 0 | 0 | 0 |
| ENSMUSG00000098823 | 0 | 0 | 0 | 0 | 0 | 0 | 0 |
| ENSMUSG00000098824 | 0 | 0 | 0 | 0 | 0 | 0 | 0 |
| ENSMUSG00000098825 | 0 | 0 | 0 | 0 | 0 | 0 | 0 |
| ENSMUSG00000098826 | 0 | 0 | 0 | 0 | 0 | 0 | 0 |
| ENSMUSG00000098827 | 0 | 0 | 0 | 0 | 0 | 0 | 0 |
| ENSMUSG00000098828 | 0 | 0 | 0 | 0 | 0 | 0 | 0 |
| ENSMUSG00000098829 | 0 | 0 | 0 | 0 | 0 | 0 | 0 |
| ENSMUSG00000098830 | 0 | 0 | 0 | 0 | 0 | 0 | 0.224243824 |
| ENSMUSG00000098831 | 0 | 0 | 0 | 0 | 0 | 0 | 0 |
| ENSMUSG00000098832 | 0 | 0 | 0 | 0.25 | 0 | 0 | 0.448487648 |
| ENSMUSG00000098833 | 0 | 0 | 0 | 0 | 0 | 0 | 0 |
| ENSMUSG00000098834 | 0 | 0 | 0 | 0 | 0 | 0 | 0 |
| ENSMUSG00000098836 | 0 | 0 | 0 | 0 | 0 | 0 | 0 |
| ENSMUSG00000098837 | 0 | 0 | 0 | 0 | 0 | 0 | 0 |
| ENSMUSG00000098838 | 0 | 0 | 0 | 0 | 0 | 0 | 0 |
| ENSMUSG00000098839 | 0 | 0 | 0 | 0 | 0 | 0 | 0 |
| ENSMUSG00000098840 | 0.45815691 | 0.25 | 0.25 | 0 | 0 | 0.301029996 | 0 |
| ENSMUSG00000098842 | 0 | 0 | 0 | 0 | 0 | 0 | 0 |
| ENSMUSG00000098843 | 0 | 0.396240625 | 0.25 | 0.646240625 | 0.387761935 | 0.301029996 | 0.448487648 |
| ENSMUSG00000098844 | 0 | 0 | 0 | 0 | 0 | 0 | 0 |
| ENSMUSG00000098845 | 0 | 0 | 0 | 0 | 0 | 0 | 0 |
| ENSMUSG00000098846 | 0 | 0 | 0 | 0 | 0 | 0 | 0 |
| ENSMUSG00000098847 | 0 | 0 | 0 | 0 | 0 | 0 | 0 |
| ENSMUSG00000098848 | 0 | 0 | 0 | 0 | 0 | 0 | 0 |

| | | | | | | | |
|---|---|---|---|---|---|---|---|
| ENSMUSG00000098849 | 0 | 0 | 0 | 0 | 0 | 0 | 0 |
| ENSMUSG00000098850 | 0 | 0 | 0 | 0 | 0 | 0 | 0 |
| ENSMUSG00000098851 | 0 | 0 | 0 | 0 | 0 | 0 | 0 |
| ENSMUSG00000098852 | 0 | 0 | 0 | 0 | 0.244650542 | 0 | 0 |
| ENSMUSG00000098853 | 0 | 0 | 0 | 0 | 0 | 0 | 0 |
| ENSMUSG00000098854 | 0 | 0 | 0 | 0 | 0 | 0 | 0 |
| ENSMUSG00000098855 | 0 | 0 | 0 | 0 | 0 | 0 | 0 |
| ENSMUSG00000098856 | 0 | 0 | 0 | 0 | 0 | 0 | 0 |
| ENSMUSG00000098857 | 0 | 0 | 0 | 0 | 0 | 0 | 0 |
| ENSMUSG00000098858 | 0 | 0 | 0 | 0 | 0 | 0 | 0 |
| ENSMUSG00000098859 | 0 | 0 | 0 | 0 | 0 | 0 | 0 |
| ENSMUSG00000098860 | 0 | 0 | 0 | 0 | 0 | 0 | 0 |
| ENSMUSG00000098861 | 0 | 0 | 0 | 0 | 0 | 0 | 0 |
| ENSMUSG00000098862 | 0 | 0 | 0 | 0 | 0 | 0 | 0 |
| ENSMUSG00000098863 | 0 | 0 | 0 | 0 | 0 | 0 | 0 |
| ENSMUSG00000098864 | 0 | 0 | 0 | 0 | 0 | 0 | 0 |
| ENSMUSG00000098865 | 0 | 0 | 0 | 0 | 0 | 0 | 0 |
| ENSMUSG00000098866 | 0 | 0 | 0 | 0 | 0 | 0 | 0 |
| ENSMUSG00000098867 | 0 | 0 | 0 | 0 | 0 | 0 | 0 |
| ENSMUSG00000098868 | 0 | 0 | 0 | 0 | 0 | 0 | 0 |
| ENSMUSG00000098869 | 0 | 0 | 0 | 0 | 0 | 0 | 0 |
| ENSMUSG00000098871 | 0 | 0 | 0 | 0 | 0 | 0 | 0 |
| ENSMUSG00000098873 | 0 | 0 | 0 | 0 | 0 | 0 | 0 |
| ENSMUSG00000098874 | 0 | 0 | 0.5 | 0 | 0 | 0 | 0 |
| ENSMUSG00000098875 | 0 | 0 | 0 | 0 | 0 | 0 | 0 |
| ENSMUSG00000098876 | 0 | 0 | 0 | 0 | 0 | 0 | 0 |
| ENSMUSG00000098877 | 0 | 0 | 0 | 0 | 0 | 0 | 0 |
| ENSMUSG00000098878 | 0 | 0 | 0 | 0 | 0 | 0 | 0 |
| ENSMUSG00000098879 | 0 | 0 | 0 | 0 | 0 | 0 | 0 |
| ENSMUSG00000098880 | 0 | 0 | 0 | 0 | 0 | 0 | 0 |
| ENSMUSG00000098881 | 0 | 0 | 0 | 0 | 0 | 0 | 0 |
| ENSMUSG00000098882 | 0 | 0 | 0 | 0 | 0 | 0 | 0 |
| ENSMUSG00000098883 | 0 | 0 | 0 | 0 | 0 | 0 | 0 |
| ENSMUSG00000098884 | 0 | 0 | 0 | 0 | 0 | 0 | 0 |

| | | | | | | | |
|---|---|---|---|---|---|---|---|
| ENSMUSG00000098885 | 0 | 0 | 0 | 0 | 0 | 0 | 0 |
| ENSMUSG00000098886 | 0 | 0 | 0 | 0 | 0 | 0 | 0 |
| ENSMUSG00000098887 | 0 | 0 | 0 | 0 | 0 | 0 | 0 |
| ENSMUSG00000098888 | 0 | 0 | 0 | 0 | 0 | 0 | 0 |
| ENSMUSG00000098889 | 0.289064826 | 0.396240625 | 0.396240625 | 0.25 | 0.387761935 | 0.301029996 | 0 |
| ENSMUSG00000098890 | 0 | 0 | 0 | 0 | 0 | 0 | 0 |
| ENSMUSG00000098891 | 0 | 0 | 0 | 0 | 0 | 0 | 0 |
| ENSMUSG00000098893 | 0.289064826 | 0 | 0.25 | 0 | 0.244650542 | 0 | 0.224243824 |
| ENSMUSG00000098894 | 0 | 0 | 0 | 0 | 0 | 0 | 0 |
| ENSMUSG00000098895 | 0 | 0 | 0 | 0 | 0 | 0 | 0 |
| ENSMUSG00000098896 | 0 | 0 | 0 | 0 | 0 | 0 | 0 |
| ENSMUSG00000098897 | 0 | 0 | 0 | 0 | 0 | 0 | 0 |
| ENSMUSG00000098898 | 0 | 0.25 | 0 | 0 | 0 | 0 | 0 |
| ENSMUSG00000098899 | 0 | 0 | 0 | 0 | 0 | 0 | 0 |
| ENSMUSG00000098900 | 0 | 0 | 0 | 0 | 0 | 0 | 0 |
| ENSMUSG00000098901 | 0 | 0 | 0 | 0 | 0 | 0 | 0 |
| ENSMUSG00000098903 | 0 | 0 | 0 | 0 | 0 | 0 | 0 |
| ENSMUSG00000098904 | 0 | 0 | 0 | 0 | 0.489301084 | 0 | 0.224243824 |
| ENSMUSG00000098905 | 1.100572389 | 1.17510993 | 1.27186571 | 1.410964047 | 1.283908311 | 1.342422681 | 1.140832123 |
| ENSMUSG00000098906 | 0 | 0 | 0 | 0 | 0 | 0 | 0 |
| ENSMUSG00000098907 | 0 | 0 | 0 | 0 | 0.244650542 | 0 | 0.224243824 |
| ENSMUSG00000098908 | 0 | 0 | 0 | 0 | 0 | 0 | 0 |
| ENSMUSG00000098909 | 0 | 0 | 0 | 0 | 0 | 0 | 0 |
| ENSMUSG00000098910 | 0 | 0 | 0 | 0 | 0 | 0 | 0 |
| ENSMUSG00000098912 | 1.129344651 | 0.951838731 | 0.896240625 | 1.25 | 1.223252711 | 1.041392685 | 1.110947927 |
| ENSMUSG00000098914 | 0 | 0 | 0 | 0 | 0 | 0 | 0 |
| ENSMUSG00000098915 | 0.289064826 | 0.580482024 | 0 | 0.580482024 | 0.387761935 | 0.301029996 | 0.520678036 |
| ENSMUSG00000098916 | 0 | 0 | 0 | 0 | 0 | 0 | 0 |
| ENSMUSG00000098917 | 0 | 0 | 0 | 0 | 0 | 0 | 0 |
| ENSMUSG00000098918 | 0 | 0.396240625 | 0 | 0.25 | 0 | 0 | 0 |
| ENSMUSG00000098919 | 0 | 0 | 0 | 0 | 0 | 0 | 0 |
| ENSMUSG00000098920 | 0 | 0 | 0 | 0 | 0 | 0 | 0.224243824 |
| ENSMUSG00000098922 | 0 | 0 | 0 | 0 | 0 | 0 | 0 |
| ENSMUSG00000098923 | 0.289064826 | 0 | 0.396240625 | 0.25 | 0 | 0.301029996 | 0.224243824 |

| | | | | | | | |
|---|---|---|---|---|---|---|---|
| ENSMUSG00000098924 | 0 | 0 | 0 | 0 | 0 | 0 | 0 |
| ENSMUSG00000098925 | 0 | 0 | 0 | 0 | 0 | 0 | 0 |
| ENSMUSG00000098926 | 0 | 0 | 0 | 0 | 0 | 0 | 0.224243824 |
| ENSMUSG00000098928 | 0 | 0 | 0 | 0 | 0 | 0 | 0 |
| ENSMUSG00000098929 | 0 | 0 | 0 | 0 | 0 | 0 | 0 |
| ENSMUSG00000098931 | 0 | 0 | 0 | 0 | 0 | 0 | 0 |
| ENSMUSG00000098932 | 0 | 0 | 0 | 0 | 0 | 0 | 0 |
| ENSMUSG00000098934 | 0 | 0 | 0 | 0 | 0 | 0 | 0 |
| ENSMUSG00000098935 | 0 | 0 | 0 | 0 | 0 | 0 | 0 |
| ENSMUSG00000098937 | 0 | 0 | 0 | 0 | 0 | 0 | 0 |
| ENSMUSG00000098938 | 0 | 0 | 0 | 0 | 0 | 0 | 0 |
| ENSMUSG00000098939 | 0 | 0 | 0 | 0 | 0 | 0 | 0 |
| ENSMUSG00000098940 | 0 | 0 | 0 | 0 | 0 | 0 | 0 |
| ENSMUSG00000098941 | 0 | 0 | 0 | 0 | 0 | 0 | 0 |
| ENSMUSG00000098942 | 0 | 0 | 0 | 0 | 0 | 0 | 0 |
| ENSMUSG00000098943 | 0 | 0 | 0 | 0 | 0 | 0 | 0 |
| ENSMUSG00000098944 | 0 | 0 | 0 | 0 | 0 | 0 | 0 |
| ENSMUSG00000098945 | 0 | 0 | 0 | 0 | 0 | 0 | 0 |
| ENSMUSG00000098946 | 0 | 0 | 0 | 0 | 0 | 0 | 0 |
| ENSMUSG00000098947 | 0 | 0 | 0 | 0 | 0 | 0 | 0 |
| ENSMUSG00000098948 | 0 | 0 | 0 | 0 | 0 | 0 | 0 |
| ENSMUSG00000098949 | 0 | 0 | 0 | 0 | 0 | 0 | 0 |
| ENSMUSG00000098950 | 0 | 0 | 0 | 0 | 0 | 0 | 0 |
| ENSMUSG00000098951 | 0 | 0.25 | 0 | 0 | 0 | 0 | 0 |
| ENSMUSG00000098952 | 0 | 0 | 0 | 0 | 0 | 0 | 0 |
| ENSMUSG00000098953 | 0 | 0 | 0 | 0 | 0 | 0 | 0 |
| ENSMUSG00000098954 | 0 | 0 | 0 | 0 | 0 | 0 | 0 |
| ENSMUSG00000098955 | 0 | 0 | 0 | 0 | 0 | 0 | 0 |
| ENSMUSG00000098956 | 0 | 0 | 0 | 0 | 0 | 0 | 0 |
| ENSMUSG00000098957 | 0 | 0 | 0 | 0 | 0 | 0 | 0 |
| ENSMUSG00000098958 | 0 | 0 | 0 | 0 | 0 | 0 | 0 |
| ENSMUSG00000098959 | 0 | 0 | 0 | 0 | 0 | 0 | 0 |
| ENSMUSG00000098960 | 0 | 0 | 0 | 0 | 0 | 0 | 0 |
| ENSMUSG00000098961 | 0 | 0 | 0 | 0 | 0 | 0 | 0 |

| | | | | | | | |
|---|---|---|---|---|---|---|---|
| ENSMUSG00000098962 | 0 | 0 | 0 | 0 | 0 | 0 | 0 |
| ENSMUSG00000098963 | 0 | 0 | 0 | 0 | 0 | 0 | 0 |
| ENSMUSG00000098964 | 0 | 0 | 0 | 0 | 0 | 0 | 0 |
| ENSMUSG00000098965 | 0 | 0 | 0 | 0 | 0 | 0 | 0 |
| ENSMUSG00000098966 | 0 | 0 | 0 | 0 | 0 | 0 | 0 |
| ENSMUSG00000098967 | 0 | 0 | 0 | 0 | 0 | 0 | 0 |
| ENSMUSG00000098968 | 0 | 0 | 0 | 0 | 0 | 0 | 0 |
| ENSMUSG00000098969 | 0 | 0 | 0 | 0 | 0 | 0 | 0 |
| ENSMUSG00000098970 | 0 | 0 | 0 | 0 | 0 | 0 | 0 |
| ENSMUSG00000098972 | 0 | 0 | 0 | 0 | 0 | 0 | 0 |
| ENSMUSG00000098973 | 1.714367558 | 1.823655187 | 1.882845365 | 1.527131114 | 1.614672255 | 1.255272505 | 1.759302692 |
| ENSMUSG00000098974 | 0 | 0 | 0 | 0 | 0 | 0 | 0 |
| ENSMUSG00000098975 | 0 | 0 | 0 | 0 | 0 | 0 | 0 |
| ENSMUSG00000098976 | 0 | 0 | 0 | 0.25 | 0.244650542 | 0 | 0 |
| ENSMUSG00000098977 | 0 | 0 | 0 | 0 | 0 | 0 | 0 |
| ENSMUSG00000098978 | 0 | 0 | 0 | 0 | 0 | 0 | 0 |
| ENSMUSG00000098979 | 0 | 0 | 0 | 0.25 | 0 | 0 | 0 |
| ENSMUSG00000098980 | 0 | 0 | 0 | 0.25 | 0 | 0 | 0 |
| ENSMUSG00000098981 | 0 | 0 | 0 | 0 | 0 | 0 | 0 |
| ENSMUSG00000098982 | 0 | 0 | 0 | 0 | 0 | 0 | 0 |
| ENSMUSG00000098983 | 0 | 0 | 0 | 0 | 0 | 0 | 0 |
| ENSMUSG00000098984 | 0 | 0 | 0 | 0 | 0 | 0 | 0 |
| ENSMUSG00000098985 | 0 | 0.396240625 | 0 | 0.25 | 0.244650542 | 0.301029996 | 0.355418052 |
| ENSMUSG00000098986 | 0 | 0 | 0 | 0 | 0 | 0 | 0 |
| ENSMUSG00000098987 | 0 | 0 | 0 | 0 | 0 | 0 | 0 |
| ENSMUSG00000098988 | 0 | 0 | 0 | 0 | 0 | 0 | 0 |
| ENSMUSG00000098989 | 0 | 0.25 | 0 | 0 | 0 | 0 | 0 |
| ENSMUSG00000098990 | 0 | 0 | 0 | 0 | 0 | 0 | 0 |
| ENSMUSG00000098991 | 0 | 0 | 0 | 0 | 0 | 0 | 0 |
| ENSMUSG00000098992 | 0 | 0 | 0 | 0 | 0 | 0 | 0 |
| ENSMUSG00000098993 | 0 | 0 | 0 | 0 | 0 | 0 | 0 |
| ENSMUSG00000098994 | 0 | 0 | 0 | 0 | 0 | 0 | 0 |
| ENSMUSG00000098995 | 0 | 0 | 0 | 0 | 0 | 0 | 0 |
| ENSMUSG00000098996 | 0 | 0 | 0 | 0 | 0 | 0 | 0 |

| | | | | | | | |
|---|---|---|---|---|---|---|---|
| ENSMUSG00000098997 | 0 | 0 | 0 | 0 | 0 | 0 | 0 |
| ENSMUSG00000098998 | 0 | 0 | 0 | 0 | 0 | 0 | 0 |
| ENSMUSG00000098999 | 0 | 0 | 0 | 0 | 0 | 0 | 0 |
| ENSMUSG00000099000 | 0 | 0 | 0 | 0 | 0 | 0 | 0 |
| ENSMUSG00000099001 | 0 | 0 | 0 | 0 | 0 | 0 | 0 |
| ENSMUSG00000099002 | 0 | 0 | 0 | 0 | 0 | 0 | 0 |
| ENSMUSG00000099003 | 0 | 0 | 0 | 0 | 0 | 0 | 0 |
| ENSMUSG00000099004 | 0 | 0 | 0.396240625 | 0 | 0 | 0 | 0 |
| ENSMUSG00000099005 | 0 | 0 | 0 | 0 | 0 | 0 | 0 |
| ENSMUSG00000099006 | 0 | 0 | 0 | 0 | 0 | 0 | 0 |
| ENSMUSG00000099008 | 0.289064826 | 0 | 0 | 0.25 | 0 | 0 | 0 |
| ENSMUSG00000099009 | 0 | 0 | 0 | 0 | 0 | 0 | 0 |
| ENSMUSG00000099010 | 0 | 0 | 0 | 0 | 0 | 0 | 0 |
| ENSMUSG00000099011 | 0 | 0 | 0 | 0 | 0 | 0 | 0 |
| ENSMUSG00000099012 | 0 | 0 | 0 | 0 | 0 | 0 | 0 |
| ENSMUSG00000099013 | 0 | 0 | 0 | 0 | 0 | 0 | 0 |
| ENSMUSG00000099014 | 0 | 0 | 0 | 0 | 0 | 0 | 0 |
| ENSMUSG00000099015 | 0 | 0 | 0 | 0 | 0 | 0 | 0 |
| ENSMUSG00000099016 | 0 | 0 | 0 | 0 | 0 | 0 | 0 |
| ENSMUSG00000099017 | 0 | 0 | 0 | 0 | 0 | 0 | 0 |
| ENSMUSG00000099018 | 0 | 0 | 0 | 0 | 0 | 0 | 0 |
| ENSMUSG00000099019 | 0.289064826 | 0 | 0 | 0 | 0.244650542 | 0 | 0.355418052 |
| ENSMUSG00000099020 | 0 | 0 | 0 | 0 | 0 | 0 | 0 |
| ENSMUSG00000099021 | 0 | 0 | 0 | 0 | 0 | 0 | 0 |
| ENSMUSG00000099022 | 0 | 0 | 0 | 0 | 0 | 0 | 0 |
| ENSMUSG00000099023 | 0 | 0 | 0 | 0 | 0 | 0 | 0 |
| ENSMUSG00000099024 | 0 | 0 | 0 | 0 | 0 | 0 | 0 |
| ENSMUSG00000099025 | 0 | 0 | 0 | 0 | 0 | 0 | 0 |
| ENSMUSG00000099027 | 0 | 0 | 0 | 0 | 0 | 0 | 0 |
| ENSMUSG00000099030 | 0 | 0 | 0 | 0 | 0 | 0 | 0 |
| ENSMUSG00000099031 | 0 | 0 | 0 | 0 | 0 | 0 | 0 |
| ENSMUSG00000099032 | 1.289064826 | 1.26109853 | 1.26109853 | 1.992885888 | 1.916320788 | 2.190331698 | 1.782372264 |
| ENSMUSG00000099033 | 0.289064826 | 0 | 0.25 | 0.5 | 0.244650542 | 0 | 0.355418052 |
| ENSMUSG00000099034 | 1.342375483 | 1.188721876 | 1.27186571 | 1.27186571 | 1.136121934 | 1.230448921 | 1.21680636 |

| | | | | | | | |
|---|---|---|---|---|---|---|---|
| ENSMUSG00000099035 | 0 | 0 | 0 | 0 | 0 | 0 | 0 |
| ENSMUSG00000099036 | 0 | 0 | 0 | 0 | 0 | 0 | 0 |
| ENSMUSG00000099037 | 0 | 0 | 0 | 0 | 0 | 0 | 0 |
| ENSMUSG00000099039 | 0 | 0 | 0 | 0 | 0 | 0 | 0 |
| ENSMUSG00000099041 | 0.289064826 | 0 | 0 | 0 | 0 | 0 | 0 |
| ENSMUSG00000099042 | 0 | 0 | 0 | 0 | 0 | 0 | 0 |
| ENSMUSG00000099043 | 0 | 0 | 0 | 0 | 0 | 0 | 0 |
| ENSMUSG00000099045 | 0 | 0 | 0 | 0 | 0 | 0 | 0 |
| ENSMUSG00000099047 | 0 | 0 | 0 | 0 | 0 | 0 | 0 |
| ENSMUSG00000099049 | 0 | 0 | 0 | 0 | 0 | 0 | 0 |
| ENSMUSG00000099050 | 0 | 0 | 0 | 0 | 0 | 0 | 0 |
| ENSMUSG00000099051 | 0.289064826 | 0 | 0 | 0.25 | 0.568060967 | 0 | 0 |
| ENSMUSG00000099052 | 0 | 0.25 | 0 | 0 | 0 | 0 | 0 |
| ENSMUSG00000099053 | 0 | 0 | 0 | 0 | 0 | 0 | 0 |
| ENSMUSG00000099054 | 0 | 0 | 0 | 0 | 0 | 0 | 0 |
| ENSMUSG00000099055 | 0 | 0 | 0 | 0 | 0 | 0.301029996 | 0 |
| ENSMUSG00000099056 | 0 | 0 | 0 | 0 | 0 | 0 | 0 |
| ENSMUSG00000099057 | 0 | 0 | 0 | 0 | 0 | 0 | 0 |
| ENSMUSG00000099058 | 0 | 0 | 0 | 0 | 0 | 0 | 0.224243824 |
| ENSMUSG00000099059 | 0 | 0 | 0 | 0 | 0 | 0 | 0 |
| ENSMUSG00000099060 | 0.289064826 | 0 | 0 | 0 | 0 | 0 | 0 |
| ENSMUSG00000099061 | 0.578129653 | 0.396240625 | 0.25 | 0.396240625 | 0.489301084 | 0 | 0.355418052 |
| ENSMUSG00000099062 | 0 | 0 | 0 | 0 | 0 | 0 | 0 |
| ENSMUSG00000099063 | 0 | 0 | 0 | 0 | 0 | 0 | 0 |
| ENSMUSG00000099064 | 0 | 0 | 0 | 0 | 0 | 0 | 0 |
| ENSMUSG00000099066 | 0 | 0 | 0 | 0 | 0 | 0 | 0 |
| ENSMUSG00000099068 | 0.289064826 | 0.396240625 | 0.580482024 | 0.5 | 0.244650542 | 0.477121255 | 0.355418052 |
| ENSMUSG00000099069 | 0 | 0 | 0 | 0 | 0 | 0 | 0 |
| ENSMUSG00000099070 | 0 | 0 | 0 | 0 | 0 | 0 | 0 |
| ENSMUSG00000099071 | 0 | 0 | 0 | 0 | 0 | 0 | 0 |
| ENSMUSG00000099072 | 0 | 0 | 0 | 0 | 0 | 0 | 0 |
| ENSMUSG00000099073 | 0 | 0 | 0 | 0 | 0 | 0 | 0 |
| ENSMUSG00000099074 | 0 | 0 | 0 | 0 | 0 | 0 | 0 |
| ENSMUSG00000099075 | 0 | 0 | 0 | 0 | 0 | 0 | 0 |

| | | | | | | | |
|---|---|---|---|---|---|---|---|
| ENSMUSG00000099076 | 0 | 0 | 0 | 0 | 0 | 0 | 0.224243824 |
| ENSMUSG00000099078 | 0 | 0 | 0 | 0 | 0 | 0 | 0 |
| ENSMUSG00000099079 | 0 | 0 | 0 | 0 | 0 | 0 | 0 |
| ENSMUSG00000099080 | 0 | 0 | 0 | 0 | 0 | 0 | 0 |
| ENSMUSG00000099081 | 0 | 0 | 0 | 0 | 0 | 0 | 0 |
| ENSMUSG00000099082 | 0 | 0 | 0 | 0 | 0 | 0 | 0 |
| ENSMUSG00000099083 | 2.839589945 | 2.575659731 | 2.574802005 | 2.600753006 | 2.504246421 | 2.852479994 | 2.330894065 |
| ENSMUSG00000099085 | 0 | 0 | 0 | 0 | 0 | 0 | 0 |
| ENSMUSG00000099086 | 0 | 0 | 0 | 0 | 0 | 0 | 0 |
| ENSMUSG00000099087 | 0 | 0 | 0 | 0 | 0 | 0 | 0 |
| ENSMUSG00000099088 | 0 | 0 | 0 | 0 | 0 | 0 | 0 |
| ENSMUSG00000099089 | 0 | 0 | 0 | 0 | 0 | 0 | 0 |
| ENSMUSG00000099091 | 0 | 0 | 0 | 0 | 0 | 0 | 0 |
| ENSMUSG00000099092 | 0 | 0 | 0 | 0 | 0 | 0 | 0 |
| ENSMUSG00000099093 | 0 | 0 | 0 | 0 | 0 | 0 | 0 |
| ENSMUSG00000099094 | 0.671187741 | 0.701838731 | 0.79248125 | 0.580482024 | 0.568060967 | 0.698970004 | 0.74492186 |
| ENSMUSG00000099095 | 0 | 0 | 0 | 0 | 0 | 0 | 0 |
| ENSMUSG00000099096 | 0 | 0 | 0 | 0 | 0 | 0 | 0 |
| ENSMUSG00000099097 | 0 | 0 | 0 | 0 | 0 | 0 | 0 |
| ENSMUSG00000099098 | 0 | 0.25 | 0 | 0.25 | 0.387761935 | 0 | 0.355418052 |
| ENSMUSG00000099099 | 0 | 0 | 0 | 0 | 0 | 0 | 0 |
| ENSMUSG00000099101 | 0 | 0 | 0 | 0 | 0 | 0 | 0 |
| ENSMUSG00000099102 | 0 | 0 | 0.25 | 0 | 0 | 0 | 0 |
| ENSMUSG00000099103 | 0 | 0 | 0 | 0 | 0 | 0 | 0 |
| ENSMUSG00000099104 | 0 | 0 | 0 | 0 | 0.244650542 | 0 | 0.355418052 |
| ENSMUSG00000099105 | 0 | 0 | 0 | 0 | 0 | 0 | 0 |
| ENSMUSG00000099106 | 0 | 0 | 0 | 0 | 0 | 0 | 0 |
| ENSMUSG00000099107 | 0.289064826 | 0 | 0.25 | 0.25 | 0.244650542 | 0 | 0.224243824 |
| ENSMUSG00000099108 | 0 | 0 | 0 | 0 | 0 | 0 | 0 |
| ENSMUSG00000099109 | 0 | 0 | 0 | 0 | 0 | 0 | 0 |
| ENSMUSG00000099110 | 0 | 0 | 0 | 0 | 0 | 0 | 0 |
| ENSMUSG00000099111 | 0 | 0 | 0 | 0 | 0 | 0 | 0 |
| ENSMUSG00000099112 | 0.45815691 | 0.396240625 | 0.396240625 | 0 | 0.387761935 | 0 | 0.579661877 |
| ENSMUSG00000099114 | 0 | 0 | 0 | 0 | 0 | 0 | 0 |

| | | | | | | | |
|---|---|---|---|---|---|---|---|
| ENSMUSG00000099115 | 0.289064826 | 0 | 0 | 0.25 | 0 | 0 | 0 |
| ENSMUSG00000099117 | 0 | 0 | 0 | 0 | 0 | 0 | 0 |
| ENSMUSG00000099118 | 0 | 0 | 0 | 0 | 0 | 0 | 0 |
| ENSMUSG00000099119 | 0 | 0 | 0 | 0 | 0 | 0 | 0 |
| ENSMUSG00000099120 | 0 | 0 | 0 | 0 | 0 | 0 | 0 |
| ENSMUSG00000099123 | 0 | 0 | 0.396240625 | 0 | 0 | 0 | 0 |
| ENSMUSG00000099124 | 0 | 0 | 0 | 0 | 0 | 0 | 0 |
| ENSMUSG00000099125 | 0 | 0 | 0 | 0 | 0 | 0 | 0 |
| ENSMUSG00000099126 | 0 | 0 | 0 | 0 | 0 | 0 | 0 |
| ENSMUSG00000099127 | 0 | 0 | 0 | 0 | 0 | 0.301029996 | 0 |
| ENSMUSG00000099128 | 0 | 0 | 0 | 0 | 0 | 0 | 0 |
| ENSMUSG00000099129 | 0 | 0 | 0 | 0 | 0 | 0 | 0 |
| ENSMUSG00000099130 | 0.289064826 | 0 | 0 | 0 | 0 | 0 | 0 |
| ENSMUSG00000099131 | 0.289064826 | 0 | 0 | 0.25 | 0.387761935 | 0 | 0 |
| ENSMUSG00000099132 | 0 | 0 | 0 | 0 | 0 | 0 | 0 |
| ENSMUSG00000099133 | 0 | 0 | 0 | 0 | 0.387761935 | 0 | 0.520678036 |
| ENSMUSG00000099134 | 0 | 0 | 0 | 0 | 0 | 0 | 0 |
| ENSMUSG00000099135 | 0 | 0 | 0 | 0 | 0 | 0 | 0 |
| ENSMUSG00000099136 | 0 | 0 | 0 | 0 | 0 | 0 | 0 |
| ENSMUSG00000099137 | 0.747221736 | 0.5 | 0.864857905 | 0.79248125 | 0.686820904 | 1 | 0.629532004 |
| ENSMUSG00000099138 | 0 | 0 | 0 | 0 | 0 | 0 | 0 |
| ENSMUSG00000099139 | 0 | 0 | 0 | 0 | 0 | 0 | 0 |
| ENSMUSG00000099140 | 0 | 0 | 0.396240625 | 0 | 0 | 0 | 0.224243824 |
| ENSMUSG00000099141 | 0 | 0 | 0 | 0 | 0 | 0 | 0 |
| ENSMUSG00000099142 | 0 | 0 | 0 | 0 | 0 | 0 | 0 |
| ENSMUSG00000099143 | 0 | 0 | 0 | 0 | 0 | 0 | 0 |
| ENSMUSG00000099144 | 0 | 0 | 0 | 0 | 0 | 0 | 0 |
| ENSMUSG00000099145 | 0 | 0 | 0 | 0 | 0 | 0 | 0 |
| ENSMUSG00000099146 | 0 | 0 | 0.25 | 0 | 0 | 0.301029996 | 0 |
| ENSMUSG00000099147 | 0 | 0 | 0 | 0 | 0 | 0 | 0 |
| ENSMUSG00000099148 | 0.45815691 | 0.580482024 | 0.75 | 0.396240625 | 0.489301084 | 0.698970004 | 0.672731473 |
| ENSMUSG00000099149 | 0 | 0 | 0 | 0 | 0 | 0 | 0 |
| ENSMUSG00000099150 | 0 | 0 | 0 | 0 | 0 | 0 | 0 |
| ENSMUSG00000099151 | 0 | 0 | 0 | 0 | 0 | 0 | 0 |

| | | | | | | | |
|---|---|---|---|---|---|---|---|
| ENSMUSG00000099152 | 0 | 0 | 0 | 0 | 0 | 0 | 0 |
| ENSMUSG00000099153 | 0 | 0 | 0 | 0 | 0 | 0 | 0 |
| ENSMUSG00000099154 | 0 | 0 | 0 | 0 | 0 | 0 | 0 |
| ENSMUSG00000099155 | 0 | 0 | 0 | 0 | 0 | 0 | 0 |
| ENSMUSG00000099157 | 0 | 0 | 0 | 0 | 0 | 0 | 0 |
| ENSMUSG00000099158 | 0 | 0 | 0 | 0 | 0 | 0 | 0 |
| ENSMUSG00000099159 | 0 | 0 | 0 | 0 | 0 | 0 | 0 |
| ENSMUSG00000099160 | 0 | 0 | 0 | 0 | 0 | 0 | 0 |
| ENSMUSG00000099161 | 0 | 0 | 0 | 0 | 0 | 0 | 0 |
| ENSMUSG00000099162 | 0 | 0 | 0 | 0.25 | 0.244650542 | 0.301029996 | 0 |
| ENSMUSG00000099163 | 0 | 0 | 0 | 0.25 | 0 | 0 | 0 |
| ENSMUSG00000099165 | 0 | 0 | 0 | 0 | 0 | 0 | 0 |
| ENSMUSG00000099166 | 0 | 0 | 0 | 0 | 0 | 0 | 0 |
| ENSMUSG00000099168 | 0 | 0 | 0 | 0 | 0 | 0 | 0 |
| ENSMUSG00000099169 | 0 | 0 | 0 | 0 | 0 | 0 | 0 |
| ENSMUSG00000099170 | 0 | 0 | 0.25 | 0.396240625 | 0.244650542 | 0 | 0 |
| ENSMUSG00000099172 | 0 | 0 | 0 | 0 | 0 | 0 | 0 |
| ENSMUSG00000099173 | 0 | 0 | 0 | 0 | 0 | 0 | 0 |
| ENSMUSG00000099174 | 0 | 0 | 0.25 | 0 | 0 | 0.301029996 | 0 |
| ENSMUSG00000099176 | 0 | 0 | 0 | 0 | 0 | 0 | 0 |
| ENSMUSG00000099178 | 0 | 0 | 0 | 0 | 0.244650542 | 0 | 0 |
| ENSMUSG00000099179 | 0 | 0 | 0 | 0 | 0 | 0 | 0 |
| ENSMUSG00000099180 | 0 | 0 | 0 | 0 | 0 | 0 | 0 |
| ENSMUSG00000099181 | 0 | 0 | 0 | 0 | 0 | 0 | 0 |
| ENSMUSG00000099182 | 0 | 0 | 0 | 0 | 0 | 0 | 0 |
| ENSMUSG00000099183 | 0 | 0 | 0 | 0 | 0 | 0 | 0 |
| ENSMUSG00000099184 | 0 | 0 | 0 | 0 | 0 | 0 | 0 |
| ENSMUSG00000099185 | 0 | 0 | 0 | 0 | 0 | 0 | 0 |
| ENSMUSG00000099186 | 0 | 0 | 0 | 0 | 0 | 0 | 0 |
| ENSMUSG00000099188 | 0 | 0 | 0 | 0 | 0 | 0 | 0.224243824 |
| ENSMUSG00000099189 | 0 | 0 | 0 | 0 | 0 | 0 | 0 |
| ENSMUSG00000099190 | 0 | 0 | 0 | 0 | 0 | 0 | 0 |
| ENSMUSG00000099191 | 0 | 0 | 0 | 0 | 0 | 0 | 0 |
| ENSMUSG00000099192 | 0 | 0 | 0 | 0 | 0 | 0 | 0 |

| | | | | | | | |
|---|---|---|---|---|---|---|---|
| ENSMUSG00000099193 | 0 | 0.396240625 | 0 | 0 | 0 | 0 | 0 |
| ENSMUSG00000099196 | 0 | 0.25 | 0 | 0 | 0 | 0 | 0 |
| ENSMUSG00000099197 | 0 | 0 | 0 | 0 | 0 | 0 | 0 |
| ENSMUSG00000099198 | 0 | 0 | 0 | 0 | 0 | 0 | 0 |
| ENSMUSG00000099199 | 0 | 0 | 0 | 0 | 0 | 0 | 0 |
| ENSMUSG00000099200 | 0 | 0 | 0 | 0 | 0 | 0 | 0 |
| ENSMUSG00000099201 | 0 | 0 | 0 | 0 | 0 | 0 | 0 |
| ENSMUSG00000099202 | 0 | 0 | 0 | 0 | 0 | 0 | 0 |
| ENSMUSG00000099203 | 0 | 0 | 0 | 0 | 0 | 0 | 0 |
| ENSMUSG00000099204 | 0 | 0 | 0 | 0 | 0 | 0 | 0 |
| ENSMUSG00000099205 | 0 | 0 | 0 | 0 | 0 | 0 | 0 |
| ENSMUSG00000099206 | 0 | 0 | 0 | 0.25 | 0 | 0 | 0 |
| ENSMUSG00000099207 | 0 | 0 | 0 | 0 | 0 | 0 | 0 |
| ENSMUSG00000099208 | 0 | 0 | 0 | 0 | 0 | 0 | 0 |
| ENSMUSG00000099209 | 0 | 0 | 0 | 0 | 0 | 0 | 0 |
| ENSMUSG00000099210 | 0.289064826 | 0.25 | 0 | 0 | 0 | 0 | 0 |
| ENSMUSG00000099211 | 0 | 0 | 0 | 0 | 0 | 0 | 0 |
| ENSMUSG00000099212 | 0 | 0 | 0 | 0 | 0 | 0 | 0 |
| ENSMUSG00000099213 | 0 | 0 | 0 | 0 | 0 | 0 | 0 |
| ENSMUSG00000099214 | 0 | 0 | 0 | 0 | 0 | 0 | 0 |
| ENSMUSG00000099215 | 0 | 0 | 0 | 0 | 0 | 0 | 0 |
| ENSMUSG00000099216 | 0 | 0 | 0 | 0 | 0 | 0 | 0 |
| ENSMUSG00000099218 | 0 | 0 | 0 | 0 | 0 | 0 | 0 |
| ENSMUSG00000099219 | 0 | 0 | 0 | 0 | 0 | 0 | 0 |
| ENSMUSG00000099220 | 0 | 0 | 0 | 0 | 0 | 0 | 0 |
| ENSMUSG00000099221 | 0 | 0 | 0 | 0 | 0 | 0 | 0 |
| ENSMUSG00000099222 | 0 | 0 | 0 | 0 | 0 | 0 | 0 |
| ENSMUSG00000099223 | 0 | 0 | 0 | 0 | 0 | 0 | 0 |
| ENSMUSG00000099224 | 0 | 0 | 0 | 0 | 0 | 0 | 0 |
| ENSMUSG00000099225 | 0 | 0 | 0 | 0 | 0 | 0 | 0 |
| ENSMUSG00000099226 | 0 | 0 | 0.25 | 0 | 0 | 0.301029996 | 0 |
| ENSMUSG00000099227 | 0 | 0 | 0 | 0 | 0 | 0 | 0 |
| ENSMUSG00000099228 | 0 | 0 | 0 | 0 | 0 | 0 | 0 |
| ENSMUSG00000099229 | 0 | 0 | 0 | 0 | 0 | 0 | 0 |

| | | | | | | | |
|---|---|---|---|---|---|---|---|
| ENSMUSG00000099230 | 0.289064826 | 0 | 0 | 0 | 0 | 0 | 0 |
| ENSMUSG00000099232 | 0 | 0 | 0 | 0 | 0 | 0 | 0 |
| ENSMUSG00000099233 | 0 | 0 | 0 | 0 | 0 | 0 | 0 |
| ENSMUSG00000099235 | 0 | 0 | 0 | 0 | 0 | 0 | 0 |
| ENSMUSG00000099236 | 0 | 0 | 0 | 0 | 0 | 0 | 0 |
| ENSMUSG00000099237 | 0 | 0 | 0 | 0 | 0 | 0 | 0 |
| ENSMUSG00000099238 | 0.289064826 | 0.396240625 | 0 | 0 | 0.244650542 | 0 | 0.224243824 |
| ENSMUSG00000099239 | 0 | 0 | 0 | 0 | 0 | 0 | 0 |
| ENSMUSG00000099240 | 0 | 0 | 0 | 0 | 0 | 0 | 0 |
| ENSMUSG00000099241 | 0 | 0 | 0 | 0 | 0 | 0 | 0 |
| ENSMUSG00000099242 | 0 | 0 | 0 | 0 | 0.244650542 | 0 | 0 |
| ENSMUSG00000099243 | 0 | 0 | 0 | 0 | 0 | 0 | 0 |
| ENSMUSG00000099244 | 0 | 0 | 0 | 0 | 0 | 0 | 0 |
| ENSMUSG00000099245 | 0 | 0 | 0 | 0 | 0 | 0 | 0 |
| ENSMUSG00000099246 | 0 | 0 | 0 | 0 | 0 | 0.301029996 | 0 |
| ENSMUSG00000099247 | 0 | 0 | 0 | 0 | 0 | 0 | 0 |
| ENSMUSG00000099248 | 0 | 0 | 0 | 0 | 0 | 0 | 0 |
| ENSMUSG00000099250 | 0 | 0 | 0 | 0 | 0 | 0 | 0 |
| ENSMUSG00000099252 | 0 | 0 | 0 | 0 | 0 | 0 | 0 |
| ENSMUSG00000099253 | 0 | 0 | 0 | 0 | 0 | 0 | 0.224243824 |
| ENSMUSG00000099254 | 0 | 0.25 | 0.25 | 0.396240625 | 0.244650542 | 0 | 0.355418052 |
| ENSMUSG00000099255 | 0 | 0 | 0 | 0 | 0 | 0 | 0 |
| ENSMUSG00000099256 | 0 | 0 | 0 | 0 | 0 | 0 | 0 |
| ENSMUSG00000099257 | 0 | 0 | 0 | 0 | 0 | 0 | 0 |
| ENSMUSG00000099258 | 0 | 0 | 0 | 0 | 0 | 0 | 0 |
| ENSMUSG00000099259 | 0 | 0 | 0 | 0 | 0 | 0 | 0 |
| ENSMUSG00000099260 | 0 | 0 | 0 | 0 | 0 | 0 | 0 |
| ENSMUSG00000099261 | 0 | 0 | 0.25 | 0 | 0.244650542 | 0.301029996 | 0 |
| ENSMUSG00000099263 | 0 | 0 | 0 | 0 | 0 | 0 | 0 |
| ENSMUSG00000099264 | 0 | 0 | 0 | 0 | 0 | 0 | 0 |
| ENSMUSG00000099266 | 0 | 0 | 0 | 0 | 0 | 0 | 0 |
| ENSMUSG00000099267 | 0.289064826 | 0.5 | 0.25 | 0.396240625 | 0.244650542 | 0 | 0 |
| ENSMUSG00000099268 | 0 | 0 | 0.25 | 0 | 0 | 0 | 0 |
| ENSMUSG00000099269 | 0 | 0 | 0 | 0 | 0 | 0 | 0 |

| | | | | | | | |
|---|---|---|---|---|---|---|---|
| ENSMUSG00000099270 | 0 | 0 | 0 | 0 | 0 | 0 | 0 |
| ENSMUSG00000099272 | 0 | 0 | 0 | 0 | 0 | 0 | 0 |
| ENSMUSG00000099273 | 0 | 0 | 0 | 0 | 0 | 0 | 0 |
| ENSMUSG00000099274 | 0 | 0 | 0 | 0 | 0 | 0 | 0 |
| ENSMUSG00000099275 | 0 | 0 | 0 | 0 | 0 | 0 | 0 |
| ENSMUSG00000099276 | 0 | 0 | 0 | 0 | 0 | 0 | 0 |
| ENSMUSG00000099278 | 0 | 0 | 0 | 0 | 0 | 0 | 0 |
| ENSMUSG00000099279 | 0 | 0 | 0 | 0 | 0 | 0 | 0 |
| ENSMUSG00000099280 | 0 | 0 | 0 | 0 | 0 | 0 | 0.224243824 |
| ENSMUSG00000099283 | 0 | 0 | 0 | 0 | 0.244650542 | 0 | 0.224243824 |
| ENSMUSG00000099284 | 0.578129653 | 0.396240625 | 0.5 | 0.25 | 0.387761935 | 0 | 0.224243824 |
| ENSMUSG00000099285 | 0 | 0 | 0 | 0 | 0 | 0 | 0 |
| ENSMUSG00000099286 | 0 | 0 | 0 | 0 | 0 | 0 | 0 |
| ENSMUSG00000099287 | 0.289064826 | 0.25 | 0 | 0 | 0 | 0 | 0.224243824 |
| ENSMUSG00000099289 | 0 | 0 | 0 | 0 | 0 | 0 | 0 |
| ENSMUSG00000099290 | 0 | 0 | 0 | 0 | 0 | 0 | 0 |
| ENSMUSG00000099291 | 0 | 0 | 0 | 0 | 0 | 0 | 0 |
| ENSMUSG00000099292 | 0.289064826 | 0 | 0 | 0 | 0 | 0 | 0 |
| ENSMUSG00000099293 | 0 | 0 | 0 | 0 | 0 | 0 | 0 |
| ENSMUSG00000099294 | 0.289064826 | 0 | 0 | 0 | 0 | 0 | 0.224243824 |
| ENSMUSG00000099295 | 0 | 0 | 0 | 0 | 0 | 0 | 0 |
| ENSMUSG00000099298 | 0 | 0 | 0 | 0 | 0 | 0 | 0 |
| ENSMUSG00000099299 | 0 | 0 | 0 | 0 | 0 | 0 | 0 |
| ENSMUSG00000099300 | 0 | 0 | 0 | 0 | 0 | 0 | 0 |
| ENSMUSG00000099301 | 0 | 0 | 0 | 0 | 0 | 0 | 0.224243824 |
| ENSMUSG00000099302 | 0 | 0 | 0 | 0 | 0 | 0 | 0 |
| ENSMUSG00000099303 | 0 | 0 | 0 | 0 | 0.244650542 | 0 | 0 |
| ENSMUSG00000099304 | 0 | 0 | 0 | 0 | 0 | 0 | 0 |
| ENSMUSG00000099307 | 0 | 0 | 0 | 0 | 0 | 0 | 0 |
| ENSMUSG00000099308 | 0 | 0 | 0 | 0 | 0 | 0 | 0 |
| ENSMUSG00000099309 | 0 | 0 | 0 | 0 | 0 | 0 | 0 |
| ENSMUSG00000099310 | 0 | 0 | 0 | 0 | 0.244650542 | 0 | 0 |
| ENSMUSG00000099311 | 0 | 0 | 0 | 0 | 0 | 0 | 0 |
| ENSMUSG00000099312 | 0 | 0 | 0 | 0 | 0 | 0 | 0 |

| | | | | | | | |
|---|---|---|---|---|---|---|---|
| ENSMUSG00000099313 | 0 | 0.25 | 0.25 | 0.396240625 | 0 | 0 | 0.224243824 |
| ENSMUSG00000099314 | 0 | 0 | 0 | 0 | 0 | 0.301029996 | 0 |
| ENSMUSG00000099315 | 0 | 0 | 0 | 0 | 0 | 0 | 0 |
| ENSMUSG00000099316 | 0 | 0 | 0 | 0 | 0 | 0 | 0 |
| ENSMUSG00000099317 | 0 | 0 | 0 | 0 | 0 | 0 | 0 |
| ENSMUSG00000099318 | 0 | 0 | 0 | 0 | 0 | 0 | 0 |
| ENSMUSG00000099319 | 0 | 0 | 0 | 0 | 0 | 0 | 0 |
| ENSMUSG00000099320 | 0 | 0 | 0 | 0 | 0 | 0 | 0 |
| ENSMUSG00000099321 | 0 | 0 | 0 | 0 | 0 | 0 | 0 |
| ENSMUSG00000099322 | 0 | 0 | 0 | 0 | 0 | 0 | 0 |
| ENSMUSG00000099323 | 0 | 0 | 0 | 0 | 0 | 0 | 0 |
| ENSMUSG00000099324 | 0 | 0 | 0 | 0 | 0 | 0 | 0 |
| ENSMUSG00000099325 | 0 | 0 | 0 | 0 | 0 | 0 | 0 |
| ENSMUSG00000099326 | 0 | 0 | 0 | 0 | 0 | 0 | 0 |
| ENSMUSG00000099328 | 0 | 0 | 0 | 0 | 0 | 0 | 0 |
| ENSMUSG00000099329 | 0 | 0 | 0 | 0 | 0 | 0 | 0 |
| ENSMUSG00000099330 | 0 | 0 | 0 | 0 | 0 | 0 | 0 |
| ENSMUSG00000099331 | 0 | 0 | 0 | 0 | 0 | 0 | 0 |
| ENSMUSG00000099332 | 0 | 0 | 0 | 0 | 0 | 0 | 0 |
| ENSMUSG00000099334 | 0 | 0 | 0 | 0 | 0 | 0 | 0 |
| ENSMUSG00000099335 | 0 | 0 | 0 | 0 | 0 | 0 | 0 |
| ENSMUSG00000099336 | 0 | 0 | 0 | 0 | 0 | 0 | 0 |
| ENSMUSG00000099337 | 0 | 0 | 0 | 0 | 0.244650542 | 0 | 0 |
| ENSMUSG00000099338 | 0 | 0.396240625 | 0 | 0 | 0 | 0 | 0.224243824 |
| ENSMUSG00000099339 | 0 | 0 | 0 | 0 | 0 | 0 | 0 |
| ENSMUSG00000099340 | 0 | 0 | 0 | 0 | 0 | 0 | 0 |
| ENSMUSG00000099341 | 0 | 0 | 0 | 0 | 0 | 0 | 0 |
| ENSMUSG00000099342 | 0.671187741 | 0.701838731 | 0.75 | 0.580482024 | 0.733951626 | 0.602059991 | 0.448487648 |
| ENSMUSG00000099343 | 0 | 0 | 0 | 0 | 0 | 0 | 0 |
| ENSMUSG00000099344 | 0 | 0 | 0 | 0 | 0 | 0 | 0 |
| ENSMUSG00000099346 | 0 | 0 | 0 | 0 | 0 | 0 | 0 |
| ENSMUSG00000099347 | 0 | 0 | 0 | 0 | 0 | 0 | 0 |
| ENSMUSG00000099348 | 0 | 0 | 0 | 0 | 0 | 0 | 0 |
| ENSMUSG00000099349 | 0 | 0 | 0 | 0 | 0 | 0 | 0 |

| | | | | | | | |
|---|---|---|---|---|---|---|---|
| ENSMUSG00000099350 | 0 | 0 | 0 | 0 | 0 | 0 | 0 |
| ENSMUSG00000099351 | 0 | 0 | 0 | 0 | 0 | 0 | 0 |
| ENSMUSG00000099352 | 0 | 0 | 0 | 0 | 0 | 0 | 0 |
| ENSMUSG00000099353 | 0 | 0 | 0 | 0 | 0 | 0 | 0 |
| ENSMUSG00000099354 | 0.289064826 | 0.396240625 | 0.396240625 | 0.25 | 0.632412477 | 0.477121255 | 0.520678036 |
| ENSMUSG00000099355 | 0 | 0 | 0 | 0 | 0 | 0 | 0 |
| ENSMUSG00000099356 | 0 | 0 | 0 | 0 | 0 | 0 | 0 |
| ENSMUSG00000099357 | 0 | 0 | 0 | 0 | 0 | 0 | 0 |
| ENSMUSG00000099359 | 0 | 0 | 0 | 0 | 0 | 0 | 0 |
| ENSMUSG00000099360 | 0 | 0 | 0.25 | 0 | 0 | 0 | 0 |
| ENSMUSG00000099361 | 0 | 0 | 0 | 0 | 0 | 0 | 0 |
| ENSMUSG00000099362 | 0 | 0 | 0 | 0 | 0 | 0 | 0 |
| ENSMUSG00000099363 | 0 | 0 | 0 | 0 | 0 | 0 | 0 |
| ENSMUSG00000099364 | 0.289064826 | 0 | 0 | 0 | 0 | 0 | 0 |
| ENSMUSG00000099366 | 0 | 0 | 0 | 0 | 0 | 0 | 0 |
| ENSMUSG00000099367 | 0 | 0 | 0 | 0 | 0 | 0 | 0 |
| ENSMUSG00000099368 | 0 | 0 | 0 | 0 | 0 | 0 | 0 |
| ENSMUSG00000099369 | 0 | 0 | 0 | 0 | 0 | 0 | 0 |
| ENSMUSG00000099370 | 0.45815691 | 0.580482024 | 0 | 0 | 0 | 0.301029996 | 0 |
| ENSMUSG00000099371 | 0 | 0 | 0 | 0 | 0 | 0 | 0 |
| ENSMUSG00000099372 | 0 | 0 | 0 | 0 | 0 | 0 | 0 |
| ENSMUSG00000099373 | 0 | 0 | 0.396240625 | 0 | 0.387761935 | 0 | 0 |
| ENSMUSG00000099374 | 0 | 0 | 0 | 0 | 0 | 0 | 0 |
| ENSMUSG00000099375 | 0.289064826 | 0.396240625 | 0 | 0.396240625 | 0.244650542 | 0.301029996 | 0 |
| ENSMUSG00000099376 | 0 | 0 | 0 | 0 | 0 | 0 | 0 |
| ENSMUSG00000099377 | 0 | 0 | 0 | 0 | 0.244650542 | 0 | 0.224243824 |
| ENSMUSG00000099378 | 0 | 0 | 0 | 0 | 0 | 0 | 0 |
| ENSMUSG00000099379 | 0 | 0 | 0 | 0 | 0 | 0 | 0 |
| ENSMUSG00000099380 | 0 | 0 | 0 | 0 | 0 | 0 | 0 |
| ENSMUSG00000099381 | 0 | 0 | 0 | 0 | 0 | 0 | 0 |
| ENSMUSG00000099382 | 0 | 0 | 0 | 0 | 0 | 0 | 0 |
| ENSMUSG00000099383 | 0 | 0 | 0 | 0 | 0 | 0 | 0 |
| ENSMUSG00000099384 | 0.289064826 | 0 | 0 | 0 | 0 | 0 | 0.224243824 |
| ENSMUSG00000099385 | 0 | 0 | 0 | 0 | 0 | 0 | 0 |

| | | | | | | | |
|---|---|---|---|---|---|---|---|
| ENSMUSG00000099386 | 0 | 0 | 0 | 0 | 0 | 0 | 0 |
| ENSMUSG00000099387 | 0 | 0 | 0 | 0 | 0 | 0 | 0 |
| ENSMUSG00000099388 | 0 | 0 | 0 | 0 | 0 | 0 | 0 |
| ENSMUSG00000099389 | 0 | 0 | 0 | 0 | 0 | 0 | 0 |
| ENSMUSG00000099390 | 0 | 0 | 0 | 0 | 0 | 0 | 0 |
| ENSMUSG00000099391 | 0 | 0 | 0 | 0 | 0.244650542 | 0 | 0 |
| ENSMUSG00000099392 | 0 | 0 | 0 | 0 | 0 | 0 | 0 |
| ENSMUSG00000099393 | 0 | 0 | 0 | 0 | 0 | 0 | 0 |
| ENSMUSG00000099394 | 0 | 0 | 0 | 0 | 0 | 0 | 0 |
| ENSMUSG00000099395 | 0 | 0 | 0 | 0 | 0 | 0 | 0 |
| ENSMUSG00000099396 | 0 | 0 | 0 | 0 | 0 | 0 | 0 |
| ENSMUSG00000099397 | 2.231898196 | 2.077084758 | 2.109697963 | 1.955044741 | 1.948833444 | 2.103803721 | 1.880961074 |
| ENSMUSG00000099398 | 0 | 0 | 0 | 0 | 0 | 0 | 0 |
| ENSMUSG00000099399 | 0 | 0 | 0 | 0 | 0 | 0 | 0 |
| ENSMUSG00000099400 | 0 | 0 | 0 | 0 | 0 | 0 | 0 |
| ENSMUSG00000099401 | 0 | 0 | 0.25 | 0 | 0 | 0 | 0 |
| ENSMUSG00000099403 | 0 | 0 | 0 | 0 | 0 | 0 | 0 |
| ENSMUSG00000099404 | 0 | 0 | 0 | 0 | 0 | 0 | 0 |
| ENSMUSG00000099405 | 0 | 0 | 0 | 0 | 0 | 0 | 0.224243824 |
| ENSMUSG00000099406 | 0.289064826 | 0.951838731 | 0.830482024 | 0.580482024 | 0.632412477 | 0 | 0.710836105 |
| ENSMUSG00000099407 | 0 | 0 | 0 | 0 | 0 | 0 | 0 |
| ENSMUSG00000099408 | 0 | 0 | 0 | 0 | 0 | 0 | 0 |
| ENSMUSG00000099409 | 0 | 0 | 0 | 0 | 0 | 0 | 0 |
| ENSMUSG00000099410 | 0 | 0 | 0 | 0 | 0 | 0 | 0 |
| ENSMUSG00000099411 | 0 | 0 | 0.25 | 0 | 0 | 0 | 0 |
| ENSMUSG00000099412 | 0 | 0 | 0 | 0 | 0 | 0 | 0 |
| ENSMUSG00000099413 | 0.578129653 | 0 | 0 | 0.25 | 0 | 0.301029996 | 0.224243824 |
| ENSMUSG00000099414 | 0 | 0 | 0 | 0 | 0 | 0 | 0 |
| ENSMUSG00000099415 | 0 | 0 | 0 | 0 | 0 | 0 | 0 |
| ENSMUSG00000099416 | 0 | 0 | 0 | 0 | 0 | 0 | 0 |
| ENSMUSG00000099417 | 0.289064826 | 0 | 0 | 0 | 0 | 0.301029996 | 0.355418052 |
| ENSMUSG00000099418 | 0 | 0.25 | 0 | 0 | 0 | 0 | 0 |
| ENSMUSG00000099419 | 0 | 0 | 0 | 0 | 0 | 0 | 0 |
| ENSMUSG00000099420 | 0 | 0 | 0 | 0 | 0 | 0 | 0 |

| | | | | | | | |
|---|---|---|---|---|---|---|---|
| ENSMUSG00000099421 | 0 | 0 | 0 | 0 | 0 | 0 | 0 |
| ENSMUSG00000099422 | 0 | 0 | 0 | 0 | 0 | 0 | 0 |
| ENSMUSG00000099423 | 0 | 0 | 0 | 0 | 0.387761935 | 0.301029996 | 0.224243824 |
| ENSMUSG00000099424 | 0.289064826 | 0.25 | 0 | 0 | 0.387761935 | 0 | 0.224243824 |
| ENSMUSG00000099425 | 0 | 0 | 0 | 0 | 0 | 0 | 0 |
| ENSMUSG00000099426 | 0 | 0 | 0 | 0 | 0 | 0 | 0 |
| ENSMUSG00000099427 | 0 | 0 | 0 | 0 | 0 | 0 | 0 |
| ENSMUSG00000099428 | 0.289064826 | 0 | 0 | 0 | 0 | 0 | 0 |
| ENSMUSG00000099429 | 0 | 0.396240625 | 0.396240625 | 0 | 0 | 0 | 0 |
| ENSMUSG00000099430 | 0 | 0 | 0 | 0 | 0 | 0 | 0 |
| ENSMUSG00000099431 | 0 | 0 | 0 | 0 | 0 | 0 | 0 |
| ENSMUSG00000099432 | 0 | 0 | 0 | 0 | 0 | 0 | 0 |
| ENSMUSG00000099433 | 0 | 0 | 0 | 0 | 0 | 0 | 0 |
| ENSMUSG00000099434 | 0 | 0 | 0 | 0 | 0 | 0 | 0 |
| ENSMUSG00000099435 | 0 | 0 | 0 | 0 | 0 | 0 | 0 |
| ENSMUSG00000099436 | 0 | 0 | 0 | 0 | 0 | 0 | 0 |
| ENSMUSG00000099437 | 0 | 0 | 0 | 0 | 0 | 0 | 0 |
| ENSMUSG00000099438 | 0 | 0 | 0 | 0 | 0 | 0 | 0 |
| ENSMUSG00000099439 | 0 | 0 | 0 | 0 | 0 | 0 | 0 |
| ENSMUSG00000099440 | 0 | 0 | 0 | 0 | 0 | 0 | 0 |
| ENSMUSG00000099441 | 0 | 0 | 0 | 0 | 0 | 0 | 0 |
| ENSMUSG00000099442 | 0 | 0 | 0 | 0 | 0 | 0 | 0 |
| ENSMUSG00000099443 | 0 | 0 | 0 | 0 | 0 | 0 | 0 |
| ENSMUSG00000099444 | 0.45815691 | 0.396240625 | 0.25 | 0.396240625 | 0.387761935 | 0.477121255 | 0.448487648 |
| ENSMUSG00000099445 | 0 | 0 | 0 | 0 | 0 | 0 | 0 |
| ENSMUSG00000099446 | 0 | 0.5 | 0.5 | 0.580482024 | 0 | 0 | 0 |
| ENSMUSG00000099447 | 0 | 0 | 0 | 0 | 0 | 0 | 0 |
| ENSMUSG00000099448 | 0 | 0 | 0 | 0 | 0 | 0 | 0 |
| ENSMUSG00000099449 | 0 | 0 | 0 | 0 | 0 | 0 | 0 |
| ENSMUSG00000099450 | 0 | 0 | 0 | 0 | 0 | 0 | 0 |
| ENSMUSG00000099451 | 0 | 0 | 0 | 0 | 0 | 0 | 0 |
| ENSMUSG00000099452 | 0 | 0 | 0 | 0 | 0 | 0 | 0 |
| ENSMUSG00000099453 | 0 | 0 | 0 | 0 | 0 | 0 | 0 |
| ENSMUSG00000099454 | 0 | 0 | 0 | 0 | 0 | 0 | 0 |

| | | | | | | | |
|---|---|---|---|---|---|---|---|
| ENSMUSG00000099455 | 0 | 0 | 0 | 0 | 0 | 0 | 0 |
| ENSMUSG00000099456 | 0 | 0 | 0 | 0 | 0 | 0 | 0 |
| ENSMUSG00000099457 | 0 | 0 | 0 | 0 | 0 | 0 | 0 |
| ENSMUSG00000099458 | 0 | 0 | 0 | 0 | 0 | 0 | 0 |
| ENSMUSG00000099459 | 0 | 0 | 0 | 0 | 0 | 0 | 0 |
| ENSMUSG00000099460 | 0 | 0 | 0 | 0 | 0 | 0 | 0 |
| ENSMUSG00000099461 | 0 | 0 | 0 | 0 | 0 | 0 | 0 |
| ENSMUSG00000099462 | 0 | 0 | 0 | 0 | 0 | 0 | 0 |
| ENSMUSG00000099463 | 0 | 0 | 0 | 0 | 0 | 0 | 0 |
| ENSMUSG00000099464 | 0 | 0 | 0 | 0 | 0 | 0 | 0 |
| ENSMUSG00000099465 | 0 | 0 | 0.25 | 0 | 0 | 0 | 0 |
| ENSMUSG00000099466 | 0 | 0 | 0 | 0 | 0 | 0 | 0 |
| ENSMUSG00000099467 | 0 | 0 | 0 | 0 | 0 | 0 | 0 |
| ENSMUSG00000099468 | 0 | 0 | 0 | 0 | 0 | 0 | 0 |
| ENSMUSG00000099469 | 0 | 0 | 0 | 0 | 0 | 0 | 0 |
| ENSMUSG00000099470 | 0 | 0.396240625 | 0 | 0 | 0.244650542 | 0.301029996 | 0.224243824 |
| ENSMUSG00000099471 | 0.289064826 | 0 | 0 | 0.25 | 0 | 0 | 0.224243824 |
| ENSMUSG00000099472 | 0 | 0 | 0 | 0 | 0 | 0 | 0 |
| ENSMUSG00000099473 | 0 | 0 | 0 | 0 | 0 | 0 | 0 |
| ENSMUSG00000099474 | 0 | 0 | 0 | 0 | 0 | 0 | 0 |
| ENSMUSG00000099476 | 0 | 0 | 0 | 0 | 0 | 0 | 0 |
| ENSMUSG00000099477 | 0 | 0 | 0 | 0 | 0 | 0 | 0 |
| ENSMUSG00000099478 | 1.205378646 | 1.226722649 | 1.27186571 | 1.330482024 | 1.27449559 | 1.146128036 | 1.140832123 |
| ENSMUSG00000099479 | 0 | 0 | 0 | 0 | 0 | 0 | 0 |
| ENSMUSG00000099480 | 0 | 0 | 0 | 0 | 0 | 0 | 0 |
| ENSMUSG00000099481 | 1.505869732 | 1.311981878 | 1.458222504 | 1.458222504 | 1.414412788 | 1.505149978 | 1.355418052 |
| ENSMUSG00000099482 | 0 | 0 | 0 | 0 | 0 | 0 | 0 |
| ENSMUSG00000099483 | 0 | 0 | 0 | 0 | 0 | 0 | 0 |
| ENSMUSG00000099484 | 0 | 0 | 0 | 0 | 0 | 0 | 0 |
| ENSMUSG00000099485 | 0 | 0 | 0 | 0 | 0 | 0 | 0 |
| ENSMUSG00000099486 | 0 | 0 | 0 | 0 | 0 | 0 | 0 |
| ENSMUSG00000099487 | 0 | 0 | 0 | 0 | 0 | 0 | 0 |
| ENSMUSG00000099488 | 0 | 0 | 0 | 0 | 0 | 0 | 0 |
| ENSMUSG00000099489 | 0 | 0 | 0 | 0 | 0 | 0 | 0 |

| | | | | | | | |
|---|---|---|---|---|---|---|---|
| ENSMUSG00000099490 | 0 | 0 | 0 | 0 | 0 | 0 | 0 |
| ENSMUSG00000099491 | 0 | 0.25 | 0 | 0 | 0 | 0 | 0 |
| ENSMUSG00000099492 | 0.578129653 | 0.580482024 | 0.830482024 | 0.25 | 0.244650542 | 0.602059991 | 0.775756176 |
| ENSMUSG00000099493 | 0 | 0 | 0 | 0 | 0 | 0 | 0 |
| ENSMUSG00000099494 | 0 | 0 | 0 | 0 | 0 | 0 | 0 |
| ENSMUSG00000099495 | 0 | 0 | 0 | 0 | 0 | 0 | 0 |
| ENSMUSG00000099496 | 0 | 0 | 0 | 0 | 0 | 0 | 0 |
| ENSMUSG00000099497 | 0 | 0 | 0 | 0 | 0 | 0 | 0 |
| ENSMUSG00000099498 | 0 | 0 | 0 | 0 | 0 | 0 | 0 |
| ENSMUSG00000099499 | 0 | 0 | 0 | 0 | 0 | 0 | 0 |
| ENSMUSG00000099500 | 0 | 0 | 0 | 0 | 0 | 0 | 0 |
| ENSMUSG00000099501 | 0 | 0 | 0 | 0 | 0 | 0 | 0 |
| ENSMUSG00000099502 | 0 | 0 | 0 | 0 | 0 | 0 | 0 |
| ENSMUSG00000099503 | 0 | 0 | 0.25 | 0 | 0 | 0 | 0 |
| ENSMUSG00000099504 | 0 | 0 | 0 | 0 | 0 | 0 | 0 |
| ENSMUSG00000099505 | 0 | 0 | 0 | 0 | 0 | 0 | 0 |
| ENSMUSG00000099506 | 0 | 0 | 0 | 0 | 0 | 0 | 0 |
| ENSMUSG00000099507 | 0 | 0 | 0 | 0 | 0 | 0 | 0 |
| ENSMUSG00000099508 | 0 | 0 | 0 | 0 | 0 | 0 | 0 |
| ENSMUSG00000099509 | 0 | 0 | 0 | 0 | 0 | 0 | 0 |
| ENSMUSG00000099510 | 0 | 0 | 0 | 0 | 0 | 0 | 0 |
| ENSMUSG00000099511 | 0 | 0 | 0 | 0 | 0 | 0 | 0 |
| ENSMUSG00000099512 | 0 | 0 | 0 | 0 | 0 | 0 | 0 |
| ENSMUSG00000099513 | 0 | 0 | 0 | 0 | 0 | 0 | 0 |
| ENSMUSG00000099514 | 0 | 0 | 0 | 0 | 0 | 0 | 0 |
| ENSMUSG00000099516 | 0 | 0 | 0 | 0.396240625 | 0 | 0 | 0.224243824 |
| ENSMUSG00000099517 | 0 | 0 | 0 | 0 | 0 | 0 | 0.224243824 |
| ENSMUSG00000099518 | 0 | 0 | 0 | 0 | 0 | 0 | 0 |
| ENSMUSG00000099519 | 0 | 0 | 0 | 0 | 0 | 0 | 0 |
| ENSMUSG00000099520 | 0 | 0 | 0 | 0 | 0 | 0 | 0 |
| ENSMUSG00000099521 | 0 | 0 | 0 | 0 | 0 | 0 | 0 |
| ENSMUSG00000099522 | 0 | 0 | 0 | 0 | 0 | 0 | 0 |
| ENSMUSG00000099523 | 0 | 0 | 0 | 0 | 0 | 0 | 0 |
| ENSMUSG00000099524 | 0 | 0 | 0 | 0 | 0 | 0 | 0 |

| | | | | | | | |
|---|---|---|---|---|---|---|---|
| ENSMUSG00000099525 | 0 | 0 | 0 | 0 | 0 | 0 | 0 |
| ENSMUSG00000099526 | 0 | 0 | 0 | 0 | 0 | 0 | 0 |
| ENSMUSG00000099527 | 0 | 0 | 0 | 0 | 0 | 0 | 0 |
| ENSMUSG00000099528 | 0 | 0 | 0 | 0 | 0 | 0 | 0 |
| ENSMUSG00000099529 | 0 | 0 | 0 | 0 | 0 | 0 | 0 |
| ENSMUSG00000099530 | 0 | 0 | 0 | 0 | 0 | 0 | 0 |
| ENSMUSG00000099531 | 0 | 0 | 0 | 0 | 0 | 0 | 0 |
| ENSMUSG00000099532 | 0 | 0 | 0 | 0 | 0 | 0 | 0 |
| ENSMUSG00000099534 | 0 | 0 | 0 | 0 | 0 | 0 | 0 |
| ENSMUSG00000099535 | 0 | 0 | 0 | 0 | 0 | 0 | 0 |
| ENSMUSG00000099536 | 0 | 0 | 0 | 0 | 0 | 0 | 0 |
| ENSMUSG00000099537 | 0 | 0 | 0 | 0 | 0 | 0 | 0 |
| ENSMUSG00000099539 | 0 | 0 | 0 | 0 | 0 | 0 | 0 |
| ENSMUSG00000099541 | 0 | 0 | 0 | 0 | 0 | 0 | 0 |
| ENSMUSG00000099542 | 0 | 0 | 0 | 0 | 0 | 0 | 0 |
| ENSMUSG00000099543 | 0 | 0 | 0 | 0 | 0 | 0 | 0 |
| ENSMUSG00000099544 | 0 | 0 | 0 | 0 | 0 | 0 | 0 |
| ENSMUSG00000099545 | 0 | 0 | 0 | 0 | 0 | 0 | 0 |
| ENSMUSG00000099546 | 0 | 0 | 0 | 0 | 0 | 0 | 0 |
| ENSMUSG00000099547 | 0 | 0 | 0 | 0 | 0 | 0 | 0 |
| ENSMUSG00000099548 | 0 | 0 | 0 | 0 | 0 | 0 | 0 |
| ENSMUSG00000099549 | 0 | 0 | 0 | 0 | 0 | 0 | 0 |
| ENSMUSG00000099550 | 0 | 0 | 0 | 0 | 0 | 0 | 0 |
| ENSMUSG00000099551 | 0 | 0 | 0 | 0 | 0 | 0 | 0 |
| ENSMUSG00000099552 | 0.289064826 | 0.396240625 | 0.25 | 0 | 0.387761935 | 0.602059991 | 0.448487648 |
| ENSMUSG00000099553 | 0 | 0 | 0 | 0 | 0 | 0 | 0 |
| ENSMUSG00000099554 | 0 | 0 | 0 | 0 | 0 | 0 | 0 |
| ENSMUSG00000099555 | 0 | 0 | 0 | 0 | 0 | 0 | 0 |
| ENSMUSG00000099556 | 0 | 0 | 0 | 0 | 0 | 0 | 0 |
| ENSMUSG00000099558 | 0 | 0 | 0 | 0 | 0 | 0 | 0 |
| ENSMUSG00000099559 | 0 | 0 | 0 | 0 | 0 | 0 | 0 |
| ENSMUSG00000099560 | 0 | 0 | 0 | 0 | 0 | 0 | 0 |
| ENSMUSG00000099561 | 0 | 0 | 0 | 0.25 | 0 | 0 | 0.224243824 |
| ENSMUSG00000099562 | 0 | 0 | 0 | 0 | 0.387761935 | 0 | 0 |

| | | | | | | | |
|---|---|---|---|---|---|---|---|
| ENSMUSG00000099563 | 0 | 0 | 0 | 0 | 0 | 0 | 0 |
| ENSMUSG00000099564 | 0 | 0 | 0 | 0 | 0 | 0 | 0 |
| ENSMUSG00000099565 | 0 | 0 | 0 | 0 | 0 | 0 | 0 |
| ENSMUSG00000099566 | 0 | 0 | 0 | 0 | 0 | 0 | 0 |
| ENSMUSG00000099568 | 0 | 0 | 0 | 0 | 0 | 0 | 0 |
| ENSMUSG00000099569 | 0 | 0 | 0 | 0 | 0 | 0 | 0 |
| ENSMUSG00000099570 | 0 | 0 | 0 | 0 | 0 | 0 | 0 |
| ENSMUSG00000099571 | 0 | 0.701838731 | 0.396240625 | 0.25 | 0.244650542 | 0.301029996 | 0.355418052 |
| ENSMUSG00000099572 | 0 | 0 | 0 | 0 | 0 | 0 | 0 |
| ENSMUSG00000099573 | 0 | 0 | 0 | 0 | 0 | 0 | 0 |
| ENSMUSG00000099574 | 0 | 0 | 0 | 0 | 0 | 0 | 0 |
| ENSMUSG00000099575 | 0 | 0 | 0 | 0 | 0 | 0 | 0 |
| ENSMUSG00000099576 | 0 | 0 | 0.25 | 0 | 0 | 0 | 0 |
| ENSMUSG00000099577 | 0 | 0 | 0 | 0 | 0 | 0 | 0 |
| ENSMUSG00000099578 | 0 | 0 | 0 | 0 | 0 | 0 | 0 |
| ENSMUSG00000099579 | 0.45815691 | 0.396240625 | 0.580482024 | 0.5 | 0.387761935 | 0.602059991 | 0.629532004 |
| ENSMUSG00000099581 | 0 | 0 | 0 | 0 | 0 | 0 | 0 |
| ENSMUSG00000099582 | 0 | 0 | 0 | 0 | 0 | 0 | 0 |
| ENSMUSG00000099583 | 0 | 0 | 0.25 | 0 | 0.387761935 | 0.301029996 | 0.224243824 |
| ENSMUSG00000099584 | 0 | 0 | 0.25 | 0 | 0 | 0 | 0 |
| ENSMUSG00000099585 | 0 | 0 | 0 | 0 | 0 | 0 | 0 |
| ENSMUSG00000099586 | 0 | 0 | 0 | 0 | 0 | 0 | 0 |
| ENSMUSG00000099587 | 0 | 0 | 0 | 0 | 0 | 0 | 0 |
| ENSMUSG00000099588 | 0 | 0 | 0 | 0 | 0 | 0 | 0 |
| ENSMUSG00000099589 | 0 | 0 | 0 | 0 | 0 | 0 | 0 |
| ENSMUSG00000099590 | 0 | 0 | 0 | 0 | 0 | 0 | 0 |
| ENSMUSG00000099591 | 0 | 0 | 0 | 0 | 0.244650542 | 0 | 0 |
| ENSMUSG00000099592 | 0 | 0 | 0 | 0 | 0 | 0 | 0 |
| ENSMUSG00000099593 | 0 | 0 | 0 | 0 | 0 | 0 | 0 |
| ENSMUSG00000099594 | 0 | 0 | 0 | 0 | 0 | 0 | 0 |
| ENSMUSG00000099595 | 0 | 0 | 0 | 0 | 0 | 0 | 0 |
| ENSMUSG00000099596 | 0 | 0 | 0 | 0.25 | 0 | 0 | 0.224243824 |
| ENSMUSG00000099597 | 1.777675583 | 1.606566189 | 1.692046081 | 1.470660762 | 1.489301084 | 1.568201724 | 1.345462945 |
| ENSMUSG00000099598 | 0 | 0 | 0 | 0 | 0 | 0 | 0 |

| | | | | | | | |
|---|---|---|---|---|---|---|---|
| ENSMUSG00000099599 | 0 | 0 | 0 | 0 | 0 | 0 | 0 |
| ENSMUSG00000099600 | 0 | 0 | 0 | 0 | 0 | 0 | 0 |
| ENSMUSG00000099601 | 0 | 0.25 | 0 | 0 | 0 | 0 | 0 |
| ENSMUSG00000099603 | 0 | 0 | 0 | 0 | 0 | 0 | 0 |
| ENSMUSG00000099604 | 0 | 0 | 0 | 0 | 0 | 0 | 0 |
| ENSMUSG00000099605 | 0 | 0 | 0 | 0 | 0 | 0 | 0 |
| ENSMUSG00000099606 | 0 | 0 | 0 | 0 | 0 | 0 | 0 |
| ENSMUSG00000099607 | 0 | 0 | 0 | 0 | 0 | 0 | 0 |
| ENSMUSG00000099608 | 1.181541736 | 1.114857905 | 1 | 1.114857905 | 1.106691885 | 1.041392685 | 1.014380832 |
| ENSMUSG00000099609 | 0 | 0 | 0 | 0 | 0 | 0 | 0 |
| ENSMUSG00000099610 | 0 | 0 | 0 | 0 | 0 | 0 | 0 |
| ENSMUSG00000099611 | 0 | 0 | 0 | 0 | 0 | 0 | 0 |
| ENSMUSG00000099612 | 0 | 0 | 0 | 0 | 0 | 0 | 0 |
| ENSMUSG00000099613 | 0 | 0 | 0 | 0 | 0 | 0 | 0 |
| ENSMUSG00000099614 | 0 | 0 | 0 | 0 | 0 | 0 | 0 |
| ENSMUSG00000099615 | 0.91631382 | 0.92510993 | 0.864857905 | 0.864857905 | 0.733951626 | 0.602059991 | 0.710836105 |
| ENSMUSG00000099616 | 0 | 0 | 0 | 0 | 0 | 0 | 0 |
| ENSMUSG00000099617 | 0 | 0 | 0 | 0 | 0 | 0 | 0 |
| ENSMUSG00000099618 | 0 | 0 | 0 | 0 | 0 | 0 | 0 |
| ENSMUSG00000099619 | 0 | 0 | 0 | 0 | 0 | 0 | 0 |
| ENSMUSG00000099620 | 0 | 0 | 0 | 0 | 0 | 0 | 0 |
| ENSMUSG00000099621 | 0 | 0 | 0 | 0 | 0 | 0 | 0 |
| ENSMUSG00000099622 | 0 | 0 | 0 | 0 | 0 | 0 | 0 |
| ENSMUSG00000099623 | 0 | 0 | 0 | 0 | 0 | 0 | 0 |
| ENSMUSG00000099624 | 0 | 0.646240625 | 0.396240625 | 0.25 | 0.489301084 | 0.602059991 | 0.520678036 |
| ENSMUSG00000099625 | 0.289064826 | 0.396240625 | 0 | 0.25 | 0.387761935 | 0.301029996 | 0.224243824 |
| ENSMUSG00000099626 | 0 | 0 | 0 | 0 | 0 | 0 | 0 |
| ENSMUSG00000099627 | 0 | 0 | 0 | 0 | 0 | 0 | 0 |
| ENSMUSG00000099628 | 0 | 0 | 0 | 0 | 0 | 0 | 0 |
| ENSMUSG00000099629 | 0 | 0 | 0 | 0 | 0 | 0 | 0 |
| ENSMUSG00000099630 | 0 | 0 | 0 | 0 | 0 | 0 | 0 |
| ENSMUSG00000099631 | 0 | 0.25 | 0.396240625 | 0.5 | 0 | 0 | 0.355418052 |
| ENSMUSG00000099632 | 0.811507563 | 0.701838731 | 0.864857905 | 0 | 0.568060967 | 0.477121255 | 0.74492186 |
| ENSMUSG00000099633 | 0 | 0 | 0 | 0 | 0 | 0 | 0 |

| | | | | | | | |
|---|---|---|---|---|---|---|---|
| ENSMUSG00000099634 | 0 | 0 | 0 | 0 | 0 | 0 | 0 |
| ENSMUSG00000099635 | 0 | 0 | 0 | 0 | 0.244650542 | 0 | 0.224243824 |
| ENSMUSG00000099636 | 0 | 0 | 0 | 0 | 0 | 0 | 0 |
| ENSMUSG00000099637 | 0 | 0 | 0 | 0 | 0 | 0 | 0 |
| ENSMUSG00000099638 | 0 | 0 | 0 | 0 | 0 | 0 | 0 |
| ENSMUSG00000099639 | 0 | 0 | 0 | 0 | 0 | 0 | 0 |
| ENSMUSG00000099640 | 0 | 0 | 0 | 0 | 0 | 0 | 0 |
| ENSMUSG00000099641 | 0 | 0 | 0 | 0 | 0 | 0 | 0 |
| ENSMUSG00000099642 | 0 | 0 | 0 | 0 | 0 | 0 | 0 |
| ENSMUSG00000099643 | 0 | 0 | 0 | 0 | 0 | 0 | 0 |
| ENSMUSG00000099644 | 0 | 0 | 0 | 0 | 0 | 0 | 0 |
| ENSMUSG00000099645 | 0 | 0 | 0 | 0 | 0 | 0 | 0 |
| ENSMUSG00000099646 | 0 | 0 | 0 | 0 | 0 | 0 | 0 |
| ENSMUSG00000099647 | 0 | 0 | 0 | 0 | 0 | 0 | 0 |
| ENSMUSG00000099648 | 0 | 0 | 0 | 0 | 0 | 0 | 0 |
| ENSMUSG00000099649 | 0 | 0 | 0 | 0 | 0 | 0 | 0 |
| ENSMUSG00000099650 | 0 | 0 | 0 | 0 | 0 | 0 | 0 |
| ENSMUSG00000099651 | 0 | 0 | 0 | 0 | 0 | 0 | 0 |
| ENSMUSG00000099652 | 0 | 0 | 0 | 0 | 0 | 0 | 0 |
| ENSMUSG00000099653 | 0 | 0 | 0 | 0 | 0 | 0 | 0 |
| ENSMUSG00000099654 | 0 | 0 | 0 | 0 | 0 | 0 | 0 |
| ENSMUSG00000099655 | 0.45815691 | 0.646240625 | 0.701838731 | 0.701838731 | 0.387761935 | 0.301029996 | 0.448487648 |
| ENSMUSG00000099656 | 0 | 0 | 0 | 0 | 0 | 0 | 0 |
| ENSMUSG00000099657 | 0 | 0 | 0 | 0 | 0 | 0 | 0 |
| ENSMUSG00000099658 | 0 | 0 | 0 | 0 | 0 | 0 | 0 |
| ENSMUSG00000099659 | 0 | 0 | 0 | 0 | 0 | 0 | 0 |
| ENSMUSG00000099660 | 0 | 0 | 0 | 0 | 0 | 0 | 0 |
| ENSMUSG00000099661 | 0 | 0 | 0 | 0 | 0 | 0 | 0 |
| ENSMUSG00000099662 | 0 | 0 | 0 | 0 | 0 | 0 | 0 |
| ENSMUSG00000099663 | 0 | 0 | 0 | 0 | 0 | 0 | 0 |
| ENSMUSG00000099664 | 0 | 0 | 0 | 0 | 0 | 0 | 0 |
| ENSMUSG00000099665 | 0 | 0 | 0 | 0 | 0 | 0 | 0 |
| ENSMUSG00000099666 | 0 | 0 | 0 | 0 | 0 | 0 | 0 |
| ENSMUSG00000099667 | 0 | 0 | 0 | 0 | 0 | 0 | 0 |

| | | | | | | | |
|---|---|---|---|---|---|---|---|
| ENSMUSG00000099668 | 0 | 0 | 0 | 0 | 0 | 0 | 0 |
| ENSMUSG00000099669 | 0 | 0 | 0 | 0 | 0 | 0 | 0 |
| ENSMUSG00000099670 | 0 | 0 | 0 | 0 | 0 | 0 | 0 |
| ENSMUSG00000099671 | 0 | 0 | 0 | 0 | 0 | 0 | 0 |
| ENSMUSG00000099672 | 0 | 0 | 0 | 0 | 0 | 0 | 0 |
| ENSMUSG00000099673 | 0 | 0 | 0 | 0 | 0 | 0 | 0 |
| ENSMUSG00000099674 | 0 | 0 | 0 | 0 | 0 | 0 | 0 |
| ENSMUSG00000099675 | 0 | 0 | 0 | 0 | 0 | 0 | 0 |
| ENSMUSG00000099676 | 0 | 0 | 0 | 0 | 0 | 0 | 0 |
| ENSMUSG00000099677 | 0 | 0 | 0 | 0 | 0 | 0 | 0 |
| ENSMUSG00000099678 | 0 | 0 | 0 | 0 | 0 | 0 | 0 |
| ENSMUSG00000099679 | 0 | 0 | 0 | 0 | 0 | 0 | 0 |
| ENSMUSG00000099681 | 1.663535556 | 1.580482024 | 1.516522298 | 1.622963274 | 1.478764298 | 1.602059991 | 1.444790289 |
| ENSMUSG00000099682 | 0 | 0 | 0 | 0 | 0 | 0 | 0 |
| ENSMUSG00000099683 | 0 | 0 | 0 | 0 | 0 | 0 | 0 |
| ENSMUSG00000099684 | 0 | 0 | 0 | 0 | 0 | 0 | 0 |
| ENSMUSG00000099685 | 0 | 0 | 0 | 0 | 0 | 0 | 0 |
| ENSMUSG00000099686 | 0 | 0 | 0 | 0 | 0 | 0 | 0 |
| ENSMUSG00000099687 | 0 | 0 | 0 | 0 | 0 | 0 | 0 |
| ENSMUSG00000099688 | 0 | 0 | 0 | 0 | 0 | 0 | 0 |
| ENSMUSG00000099689 | 1.771760131 | 1.649978211 | 1.634789703 | 1.735628626 | 1.652586889 | 1.755874856 | 1.502532226 |
| ENSMUSG00000099690 | 0 | 0 | 0 | 0 | 0 | 0 | 0 |
| ENSMUSG00000099691 | 0 | 0 | 0 | 0 | 0 | 0 | 0 |
| ENSMUSG00000099692 | 0 | 0 | 0 | 0 | 0 | 0 | 0.224243824 |
| ENSMUSG00000099693 | 0 | 0 | 0 | 0 | 0 | 0 | 0 |
| ENSMUSG00000099694 | 0 | 0.25 | 0 | 0 | 0.244650542 | 0 | 0.355418052 |
| ENSMUSG00000099695 | 0 | 0 | 0 | 0 | 0 | 0 | 0 |
| ENSMUSG00000099696 | 0 | 0 | 0 | 0 | 0.244650542 | 0 | 0 |
| ENSMUSG00000099697 | 1.129344651 | 1.201838731 | 0.951838731 | 1.02186571 | 1.057362051 | 1.041392685 | 1.054044578 |
| ENSMUSG00000099698 | 0 | 0 | 0 | 0 | 0 | 0 | 0 |
| ENSMUSG00000099699 | 0 | 0 | 0 | 0 | 0 | 0 | 0 |
| ENSMUSG00000099700 | 0 | 0 | 0 | 0 | 0 | 0 | 0 |
| ENSMUSG00000099701 | 0 | 0 | 0.25 | 0 | 0 | 0 | 0.224243824 |
| ENSMUSG00000099702 | 0 | 0 | 0 | 0 | 0 | 0 | 0 |

| | | | | | | | |
|---|---|---|---|---|---|---|---|
| ENSMUSG00000099703 | 0.289064826 | 0.25 | 0.25 | 0.396240625 | 0 | 0 | 0.224243824 |
| ENSMUSG00000099704 | 0 | 0 | 0 | 0 | 0 | 0 | 0 |
| ENSMUSG00000099706 | 0 | 0 | 0 | 0 | 0 | 0 | 0 |
| ENSMUSG00000099707 | 0.747221736 | 0.646240625 | 0.25 | 0.25 | 0.489301084 | 0.301029996 | 0 |
| ENSMUSG00000099708 | 0.45815691 | 0.75 | 0.92510993 | 0.701838731 | 0.812711509 | 0.301029996 | 0.579661877 |
| ENSMUSG00000099709 | 0 | 0 | 0 | 0 | 0 | 0 | 0 |
| ENSMUSG00000099711 | 0 | 0 | 0 | 0 | 0 | 0 | 0 |
| ENSMUSG00000099712 | 0 | 0 | 0 | 0 | 0 | 0 | 0 |
| ENSMUSG00000099713 | 0 | 0 | 0 | 0 | 0 | 0 | 0 |
| ENSMUSG00000099714 | 0 | 0 | 0 | 0 | 0 | 0 | 0 |
| ENSMUSG00000099715 | 0 | 0 | 0 | 0 | 0 | 0 | 0 |
| ENSMUSG00000099716 | 0 | 0 | 0 | 0 | 0 | 0 | 0 |
| ENSMUSG00000099717 | 0 | 0 | 0 | 0 | 0 | 0 | 0 |
| ENSMUSG00000099718 | 0 | 0 | 0 | 0 | 0 | 0 | 0 |
| ENSMUSG00000099719 | 0 | 0.25 | 0.5 | 0.25 | 0.489301084 | 0.301029996 | 0.448487648 |
| ENSMUSG00000099721 | 0 | 0 | 0 | 0 | 0 | 0 | 0 |
| ENSMUSG00000099722 | 0 | 0 | 0 | 0 | 0 | 0 | 0 |
| ENSMUSG00000099723 | 0 | 0 | 0 | 0 | 0 | 0 | 0 |
| ENSMUSG00000099724 | 0 | 0 | 0 | 0 | 0 | 0 | 0 |
| ENSMUSG00000099725 | 0 | 0 | 0 | 0 | 0 | 0 | 0 |
| ENSMUSG00000099726 | 0 | 0 | 0 | 0 | 0 | 0 | 0 |
| ENSMUSG00000099727 | 0 | 0 | 0 | 0 | 0 | 0 | 0 |
| ENSMUSG00000099728 | 0 | 0 | 0 | 0 | 0 | 0 | 0 |
| ENSMUSG00000099729 | 0 | 0 | 0 | 0 | 0 | 0 | 0 |
| ENSMUSG00000099730 | 0 | 0 | 0 | 0 | 0 | 0 | 0 |
| ENSMUSG00000099731 | 0 | 0 | 0 | 0 | 0 | 0 | 0 |
| ENSMUSG00000099732 | 0 | 0 | 0 | 0 | 0 | 0 | 0 |
| ENSMUSG00000099733 | 0 | 0 | 0 | 0 | 0 | 0 | 0 |
| ENSMUSG00000099734 | 0 | 0 | 0 | 0 | 0 | 0 | 0 |
| ENSMUSG00000099735 | 0 | 0 | 0 | 0 | 0 | 0 | 0 |
| ENSMUSG00000099736 | 0 | 0 | 0 | 0 | 0 | 0 | 0 |
| ENSMUSG00000099737 | 0 | 0 | 0 | 0 | 0 | 0 | 0 |
| ENSMUSG00000099738 | 0 | 0 | 0 | 0 | 0 | 0 | 0 |
| ENSMUSG00000099739 | 0 | 0 | 0 | 0 | 0 | 0 | 0 |

| | | | | | | | |
|---|---|---|---|---|---|---|---|
| ENSMUSG00000099740 | 0 | 0 | 0 | 0 | 0 | 0 | 0 |
| ENSMUSG00000099741 | 0 | 0 | 0 | 0 | 0 | 0 | 0 |
| ENSMUSG00000099742 | 0 | 0 | 0 | 0 | 0 | 0 | 0 |
| ENSMUSG00000099743 | 0 | 0 | 0 | 0 | 0 | 0 | 0 |
| ENSMUSG00000099744 | 0 | 0 | 0 | 0 | 0 | 0 | 0 |
| ENSMUSG00000099745 | 0 | 0 | 0 | 0 | 0 | 0 | 0 |
| ENSMUSG00000099746 | 0.671187741 | 0.25 | 0 | 0.25 | 0.489301084 | 0 | 0.224243824 |
| ENSMUSG00000099747 | 0 | 0 | 0 | 0 | 0 | 0 | 0 |
| ENSMUSG00000099748 | 0 | 0 | 0 | 0 | 0 | 0 | 0 |
| ENSMUSG00000099749 | 0 | 0.25 | 0 | 0 | 0 | 0 | 0 |
| ENSMUSG00000099750 | 0 | 0 | 0 | 0 | 0 | 0 | 0 |
| ENSMUSG00000099751 | 0 | 0 | 0 | 0 | 0 | 0 | 0.224243824 |
| ENSMUSG00000099752 | 0 | 0 | 0 | 0 | 0 | 0 | 0 |
| ENSMUSG00000099753 | 0 | 0 | 0 | 0 | 0 | 0 | 0 |
| ENSMUSG00000099754 | 0 | 0 | 0 | 0 | 0 | 0 | 0 |
| ENSMUSG00000099755 | 0 | 0 | 0 | 0 | 0 | 0 | 0 |
| ENSMUSG00000099756 | 0 | 0 | 0 | 0 | 0 | 0 | 0 |
| ENSMUSG00000099757 | 0 | 0 | 0 | 0 | 0 | 0 | 0 |
| ENSMUSG00000099758 | 0 | 0 | 0 | 0 | 0 | 0 | 0 |
| ENSMUSG00000099759 | 0.747221736 | 0.701838731 | 0.5 | 0.75 | 0.387761935 | 0.602059991 | 0.579661877 |
| ENSMUSG00000099760 | 0.289064826 | 0.5 | 0.25 | 0.25 | 0.387761935 | 0.301029996 | 0.448487648 |
| ENSMUSG00000099761 | 0 | 0 | 0 | 0 | 0 | 0 | 0 |
| ENSMUSG00000099762 | 0 | 0 | 0 | 0 | 0 | 0 | 0 |
| ENSMUSG00000099763 | 0 | 0 | 0 | 0 | 0 | 0 | 0 |
| ENSMUSG00000099764 | 0 | 0.25 | 0 | 0 | 0.244650542 | 0.301029996 | 0.224243824 |
| ENSMUSG00000099765 | 0 | 0 | 0 | 0 | 0 | 0 | 0 |
| ENSMUSG00000099766 | 0 | 0 | 0 | 0 | 0 | 0 | 0 |
| ENSMUSG00000099767 | 0 | 0 | 0 | 0 | 0 | 0 | 0 |
| ENSMUSG00000099768 | 0 | 0 | 0 | 0 | 0 | 0 | 0 |
| ENSMUSG00000099770 | 0 | 0 | 0 | 0 | 0 | 0 | 0 |
| ENSMUSG00000099771 | 0 | 0 | 0 | 0 | 0 | 0 | 0 |
| ENSMUSG00000099772 | 0 | 0 | 0 | 0 | 0 | 0 | 0 |
| ENSMUSG00000099773 | 0.578129653 | 0.396240625 | 0.580482024 | 0.580482024 | 0.244650542 | 0 | 0.224243824 |
| ENSMUSG00000099774 | 0 | 0 | 0 | 0 | 0 | 0 | 0 |

| | | | | | | | |
|---|---|---|---|---|---|---|---|
| ENSMUSG00000099775 | 0 | 0 | 0 | 0 | 0 | 0 | 0 |
| ENSMUSG00000099776 | 0 | 0 | 0 | 0 | 0 | 0 | 0 |
| ENSMUSG00000099777 | 0 | 0 | 0 | 0 | 0 | 0 | 0 |
| ENSMUSG00000099778 | 0 | 0 | 0 | 0 | 0 | 0 | 0 |
| ENSMUSG00000099779 | 0.811507563 | 0.580482024 | 0.75 | 0.646240625 | 0.812711509 | 0.77815125 | 0.853775828 |
| ENSMUSG00000099780 | 0 | 0 | 0 | 0 | 0 | 0 | 0 |
| ENSMUSG00000099781 | 0 | 0 | 0 | 0 | 0 | 0 | 0 |
| ENSMUSG00000099782 | 0 | 0 | 0 | 0 | 0 | 0 | 0 |
| ENSMUSG00000099783 | 0 | 0 | 0 | 0 | 0 | 0 | 0 |
| ENSMUSG00000099784 | 0.289064826 | 0 | 0 | 0.25 | 0 | 0 | 0 |
| ENSMUSG00000099785 | 0 | 0 | 0 | 0 | 0 | 0 | 0 |
| ENSMUSG00000099786 | 0 | 0 | 0 | 0 | 0 | 0 | 0 |
| ENSMUSG00000099787 | 0 | 0 | 0 | 0 | 0 | 0 | 0 |
| ENSMUSG00000099788 | 0 | 0 | 0 | 0 | 0 | 0 | 0 |
| ENSMUSG00000099789 | 0 | 0 | 0 | 0 | 0 | 0 | 0 |
| ENSMUSG00000099790 | 0 | 0 | 0 | 0 | 0 | 0 | 0 |
| ENSMUSG00000099791 | 0 | 0 | 0 | 0 | 0 | 0 | 0 |
| ENSMUSG00000099792 | 0 | 0 | 0 | 0 | 0 | 0 | 0 |
| ENSMUSG00000099793 | 0 | 0 | 0 | 0 | 0 | 0 | 0 |
| ENSMUSG00000099794 | 0 | 0 | 0 | 0 | 0 | 0 | 0 |
| ENSMUSG00000099795 | 0 | 0 | 0 | 0 | 0 | 0 | 0 |
| ENSMUSG00000099796 | 0 | 0 | 0 | 0 | 0 | 0 | 0 |
| ENSMUSG00000099797 | 0 | 0.580482024 | 0.5 | 0 | 0.244650542 | 0 | 0.355418052 |
| ENSMUSG00000099798 | 0 | 0 | 0 | 0 | 0 | 0 | 0 |
| ENSMUSG00000099799 | 0 | 0 | 0 | 0 | 0 | 0 | 0 |
| ENSMUSG00000099800 | 0 | 0 | 0 | 0 | 0 | 0 | 0 |
| ENSMUSG00000099801 | 0 | 0 | 0 | 0 | 0 | 0 | 0 |
| ENSMUSG00000099802 | 0 | 0 | 0 | 0 | 0 | 0 | 0 |
| ENSMUSG00000099803 | 0 | 0 | 0 | 0 | 0 | 0 | 0 |
| ENSMUSG00000099804 | 0 | 0.25 | 0 | 0.25 | 0 | 0 | 0.224243824 |
| ENSMUSG00000099805 | 0 | 0 | 0 | 0 | 0 | 0 | 0 |
| ENSMUSG00000099806 | 0 | 0 | 0 | 0 | 0 | 0 | 0 |
| ENSMUSG00000099807 | 0 | 0 | 0 | 0 | 0 | 0 | 0 |
| ENSMUSG00000099808 | 0 | 0 | 0 | 0 | 0 | 0 | 0 |

| | | | | | | | |
|---|---|---|---|---|---|---|---|
| ENSMUSG00000099809 | 0 | 0 | 0 | 0 | 0 | 0 | 0 |
| ENSMUSG00000099810 | 0 | 0 | 0 | 0 | 0 | 0 | 0 |
| ENSMUSG00000099811 | 0 | 0 | 0 | 0 | 0 | 0 | 0 |
| ENSMUSG00000099812 | 0.289064826 | 0 | 0 | 0.25 | 0 | 0 | 0 |
| ENSMUSG00000099813 | 0 | 0 | 0 | 0 | 0 | 0 | 0 |
| ENSMUSG00000099814 | 0 | 0 | 0 | 0 | 0 | 0 | 0 |
| ENSMUSG00000099815 | 0 | 0 | 0 | 0 | 0 | 0 | 0 |
| ENSMUSG00000099816 | 0 | 0 | 0 | 0 | 0 | 0 | 0 |
| ENSMUSG00000099817 | 0 | 0 | 0 | 0 | 0 | 0 | 0 |
| ENSMUSG00000099818 | 0 | 0 | 0 | 0 | 0 | 0 | 0 |
| ENSMUSG00000099819 | 0 | 0 | 0 | 0 | 0 | 0 | 0 |
| ENSMUSG00000099820 | 0 | 0 | 0 | 0 | 0 | 0 | 0 |
| ENSMUSG00000099821 | 0 | 0 | 0 | 0 | 0 | 0 | 0 |
| ENSMUSG00000099822 | 0 | 0 | 0 | 0 | 0 | 0 | 0 |
| ENSMUSG00000099823 | 0 | 0 | 0 | 0 | 0 | 0 | 0 |
| ENSMUSG00000099824 | 0 | 0 | 0 | 0 | 0 | 0 | 0 |
| ENSMUSG00000099825 | 0 | 0.396240625 | 0 | 0.25 | 0.244650542 | 0.477121255 | 0 |
| ENSMUSG00000099826 | 0 | 0 | 0 | 0 | 0 | 0 | 0 |
| ENSMUSG00000099827 | 0 | 0 | 0 | 0 | 0 | 0 | 0 |
| ENSMUSG00000099828 | 0 | 0 | 0 | 0 | 0 | 0 | 0 |
| ENSMUSG00000099829 | 0 | 0 | 0 | 0 | 0 | 0 | 0 |
| ENSMUSG00000099831 | 0 | 0 | 0 | 0 | 0 | 0 | 0 |
| ENSMUSG00000099832 | 0 | 0 | 0 | 0 | 0 | 0 | 0 |
| ENSMUSG00000099833 | 0 | 0 | 0 | 0 | 0 | 0 | 0 |
| ENSMUSG00000099834 | 0 | 0 | 0 | 0 | 0 | 0 | 0 |
| ENSMUSG00000099835 | 0.747221736 | 0.5 | 0.75 | 0.25 | 0.489301084 | 0.301029996 | 0.355418052 |
| ENSMUSG00000099836 | 0 | 0 | 0 | 0 | 0 | 0 | 0 |
| ENSMUSG00000099838 | 0 | 0 | 0 | 0 | 0 | 0 | 0 |
| ENSMUSG00000099839 | 0 | 0 | 0 | 0 | 0 | 0 | 0 |
| ENSMUSG00000099840 | 0 | 0 | 0 | 0 | 0 | 0 | 0 |
| ENSMUSG00000099841 | 0 | 0 | 0 | 0 | 0 | 0 | 0 |
| ENSMUSG00000099842 | 0.289064826 | 0 | 0 | 0.25 | 0.686820904 | 0.301029996 | 0.224243824 |
| ENSMUSG00000099843 | 0.747221736 | 0.92510993 | 0.92510993 | 0.25 | 0.733951626 | 0.698970004 | 0.74492186 |
| ENSMUSG00000099845 | 0 | 0 | 0 | 0 | 0 | 0 | 0 |

| | | | | | | | |
|---|---|---|---|---|---|---|---|
| ENSMUSG00000099846 | 0 | 0 | 0 | 0 | 0 | 0 | 0 |
| ENSMUSG00000099847 | 0 | 0 | 0 | 0 | 0 | 0 | 0 |
| ENSMUSG00000099848 | 0 | 0 | 0 | 0.25 | 0 | 0.301029996 | 0.355418052 |
| ENSMUSG00000099849 | 0 | 0 | 0 | 0 | 0 | 0 | 0 |
| ENSMUSG00000099850 | 0 | 0.396240625 | 0 | 0.25 | 0 | 0 | 0.224243824 |
| ENSMUSG00000099851 | 0 | 0 | 0 | 0 | 0 | 0 | 0 |
| ENSMUSG00000099852 | 0 | 0 | 0 | 0 | 0 | 0 | 0 |
| ENSMUSG00000099853 | 0 | 0 | 0 | 0 | 0 | 0 | 0 |
| ENSMUSG00000099854 | 0.289064826 | 0 | 0 | 0 | 0 | 0 | 0 |
| ENSMUSG00000099855 | 0 | 0 | 0 | 0 | 0 | 0 | 0 |
| ENSMUSG00000099856 | 0 | 0 | 0 | 0 | 0 | 0 | 0 |
| ENSMUSG00000099858 | 0 | 0 | 0 | 0 | 0 | 0 | 0 |
| ENSMUSG00000099860 | 0 | 0 | 0 | 0 | 0 | 0 | 0 |
| ENSMUSG00000099861 | 0 | 0 | 0 | 0 | 0 | 0 | 0 |
| ENSMUSG00000099862 | 0 | 0 | 0 | 0 | 0 | 0 | 0 |
| ENSMUSG00000099863 | 0 | 0 | 0 | 0 | 0 | 0 | 0 |
| ENSMUSG00000099864 | 0 | 0 | 0 | 0 | 0 | 0 | 0 |
| ENSMUSG00000099865 | 0 | 0 | 0 | 0 | 0 | 0 | 0 |
| ENSMUSG00000099866 | 0 | 0 | 0 | 0 | 0 | 0 | 0 |
| ENSMUSG00000099867 | 0 | 0 | 0 | 0 | 0 | 0 | 0 |
| ENSMUSG00000099868 | 0 | 0 | 0 | 0 | 0 | 0 | 0 |
| ENSMUSG00000099869 | 0 | 0 | 0 | 0 | 0 | 0 | 0 |
| ENSMUSG00000099870 | 0 | 0 | 0 | 0 | 0 | 0 | 0 |
| ENSMUSG00000099871 | 0 | 0 | 0 | 0 | 0 | 0 | 0 |
| ENSMUSG00000099872 | 0 | 0 | 0 | 0 | 0 | 0 | 0.355418052 |
| ENSMUSG00000099873 | 0 | 0 | 0 | 0 | 0 | 0 | 0 |
| ENSMUSG00000099874 | 0 | 0 | 0 | 0 | 0 | 0.301029996 | 0 |
| ENSMUSG00000099875 | 0.45815691 | 0.25 | 0.5 | 0 | 0.244650542 | 0.602059991 | 0.224243824 |
| ENSMUSG00000099876 | 0 | 0 | 0 | 0 | 0 | 0 | 0 |
| ENSMUSG00000099877 | 0 | 0.396240625 | 0 | 0 | 0 | 0.301029996 | 0 |
| ENSMUSG00000099878 | 0 | 0 | 0 | 0 | 0 | 0.477121255 | 0 |
| ENSMUSG00000099879 | 0.671187741 | 0.396240625 | 0.5 | 0.646240625 | 0.846351821 | 0.477121255 | 0.853775828 |
| ENSMUSG00000099880 | 0 | 0 | 0 | 0 | 0 | 0 | 0 |
| ENSMUSG00000099881 | 1.639698646 | 1.52186571 | 1.660964047 | 1.589388001 | 1.523883869 | 1.763427994 | 1.466365979 |

| | | | | | | | |
|---|---|---|---|---|---|---|---|
| ENSMUSG00000099882 | 0 | 0 | 0 | 0 | 0 | 0 | 0 |
| ENSMUSG00000099883 | 0 | 0 | 0 | 0 | 0 | 0 | 0 |
| ENSMUSG00000099884 | 0 | 0 | 0 | 0 | 0 | 0 | 0 |
| ENSMUSG00000099886 | 0 | 0 | 0 | 0 | 0 | 0 | 0 |
| ENSMUSG00000099887 | 0 | 0 | 0 | 0 | 0 | 0 | 0 |
| ENSMUSG00000099888 | 0 | 0 | 0 | 0 | 0 | 0 | 0 |
| ENSMUSG00000099889 | 0 | 0 | 0 | 0 | 0 | 0 | 0 |
| ENSMUSG00000099890 | 0.867194479 | 0.79248125 | 0.976722649 | 0.92510993 | 0.877063019 | 1.041392685 | 0.853775828 |
| ENSMUSG00000099891 | 0 | 0.25 | 0 | 0.396240625 | 0 | 0.301029996 | 0.224243824 |
| ENSMUSG00000099892 | 0 | 0 | 0 | 0 | 0 | 0 | 0 |
| ENSMUSG00000099893 | 0 | 0 | 0 | 0.25 | 0 | 0 | 0 |
| ENSMUSG00000099894 | 0 | 0 | 0 | 0 | 0 | 0 | 0 |
| ENSMUSG00000099895 | 0 | 0 | 0 | 0 | 0 | 0 | 0 |
| ENSMUSG00000099896 | 0 | 0 | 0 | 0 | 0 | 0 | 0 |
| ENSMUSG00000099897 | 0 | 0 | 0 | 0 | 0 | 0 | 0 |
| ENSMUSG00000099898 | 0 | 0 | 0 | 0 | 0 | 0 | 0 |
| ENSMUSG00000099899 | 0 | 0 | 0 | 0 | 0 | 0 | 0 |
| ENSMUSG00000099900 | 0 | 0 | 0 | 0 | 0 | 0 | 0 |
| ENSMUSG00000099901 | 0 | 0 | 0 | 0 | 0 | 0 | 0 |
| ENSMUSG00000099902 | 0 | 0 | 0 | 0 | 0 | 0 | 0 |
| ENSMUSG00000099903 | 0 | 0 | 0 | 0 | 0 | 0 | 0 |
| ENSMUSG00000099904 | 0 | 0 | 0 | 0 | 0 | 0 | 0 |
| ENSMUSG00000099905 | 0 | 0 | 0 | 0 | 0 | 0 | 0 |
| ENSMUSG00000099906 | 0 | 0.25 | 0 | 0 | 0 | 0 | 0 |
| ENSMUSG00000099907 | 0 | 0 | 0 | 0 | 0 | 0 | 0 |
| ENSMUSG00000099908 | 0 | 0 | 0 | 0 | 0 | 0 | 0 |
| ENSMUSG00000099909 | 0 | 0 | 0 | 0 | 0 | 0 | 0 |
| ENSMUSG00000099910 | 0 | 0 | 0 | 0 | 0 | 0 | 0 |
| ENSMUSG00000099911 | 0 | 0 | 0 | 0 | 0 | 0 | 0 |
| ENSMUSG00000099912 | 0 | 0 | 0 | 0 | 0 | 0 | 0 |
| ENSMUSG00000099913 | 0.45815691 | 0.580482024 | 0.5 | 0.396240625 | 0.387761935 | 0.301029996 | 0.448487648 |
| ENSMUSG00000099914 | 0 | 0 | 0 | 0 | 0 | 0 | 0 |
| ENSMUSG00000099915 | 0 | 0 | 0 | 0 | 0 | 0 | 0 |
| ENSMUSG00000099916 | 0 | 0 | 0 | 0 | 0 | 0 | 0 |

| | | | | | | | |
|---|---|---|---|---|---|---|---|
| ENSMUSG00000099917 | 0 | 0 | 0 | 0 | 0 | 0 | 0 |
| ENSMUSG00000099918 | 0 | 0 | 0 | 0 | 0 | 0 | 0 |
| ENSMUSG00000099920 | 0 | 0 | 0 | 0 | 0 | 0 | 0 |
| ENSMUSG00000099921 | 0 | 0 | 0 | 0 | 0 | 0 | 0 |
| ENSMUSG00000099922 | 0 | 0 | 0 | 0 | 0 | 0 | 0 |
| ENSMUSG00000099923 | 0 | 0.25 | 0 | 0.25 | 0.244650542 | 0.301029996 | 0.224243824 |
| ENSMUSG00000099924 | 0 | 0.396240625 | 0.25 | 0.25 | 0 | 0.301029996 | 0 |
| ENSMUSG00000099925 | 0 | 0 | 0 | 0 | 0 | 0 | 0 |
| ENSMUSG00000099926 | 0 | 0 | 0 | 0 | 0 | 0 | 0 |
| ENSMUSG00000099927 | 0 | 0 | 0 | 0 | 0 | 0 | 0 |
| ENSMUSG00000099928 | 0 | 0 | 0.25 | 0.25 | 0.244650542 | 0 | 0.448487648 |
| ENSMUSG00000099929 | 0 | 0 | 0 | 0 | 0 | 0 | 0 |
| ENSMUSG00000099930 | 0 | 0 | 0 | 0 | 0 | 0 | 0.355418052 |
| ENSMUSG00000099931 | 0 | 0 | 0 | 0 | 0 | 0 | 0 |
| ENSMUSG00000099932 | 0 | 0 | 0 | 0 | 0 | 0 | 0 |
| ENSMUSG00000099933 | 0 | 0.25 | 0 | 0 | 0 | 0 | 0 |
| ENSMUSG00000099934 | 0 | 0 | 0 | 0 | 0.244650542 | 0 | 0 |
| ENSMUSG00000099935 | 0 | 0 | 0 | 0 | 0 | 0 | 0 |
| ENSMUSG00000099936 | 0 | 0 | 0.25 | 0.25 | 0.244650542 | 0 | 0.224243824 |
| ENSMUSG00000099937 | 0 | 0 | 0 | 0 | 0 | 0 | 0 |
| ENSMUSG00000099938 | 0 | 0 | 0 | 0 | 0 | 0 | 0 |
| ENSMUSG00000099939 | 0 | 0 | 0 | 0 | 0 | 0 | 0 |
| ENSMUSG00000099940 | 0 | 0 | 0 | 0 | 0 | 0 | 0 |
| ENSMUSG00000099941 | 0 | 0 | 0 | 0 | 0 | 0 | 0 |
| ENSMUSG00000099942 | 0 | 0 | 0 | 0 | 0 | 0 | 0 |
| ENSMUSG00000099943 | 0 | 0 | 0 | 0 | 0 | 0 | 0 |
| ENSMUSG00000099944 | 0 | 0 | 0 | 0 | 0 | 0 | 0 |
| ENSMUSG00000099945 | 0 | 0 | 0 | 0 | 0 | 0 | 0 |
| ENSMUSG00000099946 | 0 | 0 | 0 | 0 | 0 | 0 | 0.224243824 |
| ENSMUSG00000099947 | 0 | 0 | 0.25 | 0 | 0 | 0 | 0 |
| ENSMUSG00000099948 | 0 | 0 | 0 | 0 | 0 | 0 | 0 |
| ENSMUSG00000099949 | 0 | 0 | 0.25 | 0 | 0 | 0.301029996 | 0.224243824 |
| ENSMUSG00000099950 | 0 | 0 | 0 | 0 | 0 | 0 | 0 |
| ENSMUSG00000099951 | 0 | 0 | 0 | 0 | 0 | 0 | 0 |

| | | | | | | | |
|---|---|---|---|---|---|---|---|
| ENSMUSG00000099952 | 0 | 0 | 0 | 0 | 0 | 0 | 0 |
| ENSMUSG00000099953 | 0 | 0 | 0 | 0 | 0 | 0 | 0 |
| ENSMUSG00000099954 | 0 | 0 | 0 | 0 | 0.244650542 | 0 | 0 |
| ENSMUSG00000099955 | 0 | 0 | 0 | 0 | 0 | 0 | 0 |
| ENSMUSG00000099956 | 0 | 0 | 0 | 0 | 0 | 0 | 0 |
| ENSMUSG00000099957 | 0 | 0 | 0 | 0 | 0 | 0 | 0 |
| ENSMUSG00000099958 | 0.289064826 | 0 | 0.396240625 | 0 | 0 | 0 | 0 |
| ENSMUSG00000099959 | 0 | 0 | 0 | 0 | 0 | 0 | 0 |
| ENSMUSG00000099960 | 0 | 0 | 0 | 0 | 0 | 0 | 0 |
| ENSMUSG00000099961 | 0 | 0 | 0 | 0 | 0 | 0 | 0 |
| ENSMUSG00000099962 | 0 | 0 | 0 | 0 | 0 | 0 | 0 |
| ENSMUSG00000099963 | 0 | 0 | 0 | 0 | 0 | 0 | 0 |
| ENSMUSG00000099964 | 0.289064826 | 0.396240625 | 0.25 | 0.25 | 0.387761935 | 0 | 0.355418052 |
| ENSMUSG00000099965 | 0 | 0 | 0 | 0 | 0 | 0 | 0 |
| ENSMUSG00000099966 | 1.558729299 | 1.482684334 | 1.451838731 | 1.720660762 | 1.635855979 | 1.770852012 | 1.546120303 |
| ENSMUSG00000099967 | 0 | 0 | 0 | 0 | 0 | 0 | 0 |
| ENSMUSG00000099968 | 0 | 0 | 0 | 0 | 0 | 0 | 0 |
| ENSMUSG00000099969 | 0.45815691 | 0.580482024 | 0.5 | 0.25 | 0.244650542 | 0 | 0.355418052 |
| ENSMUSG00000099970 | 0 | 0 | 0 | 0 | 0 | 0 | 0 |
| ENSMUSG00000099971 | 0 | 0 | 0 | 0 | 0 | 0 | 0 |
| ENSMUSG00000099972 | 0 | 0 | 0 | 0 | 0 | 0 | 0 |
| ENSMUSG00000099973 | 0 | 0 | 0 | 0 | 0 | 0 | 0 |
| ENSMUSG00000099974 | 0 | 0 | 0 | 0 | 0 | 0 | 0 |
| ENSMUSG00000099975 | 0 | 0 | 0 | 0 | 0 | 0 | 0 |
| ENSMUSG00000099976 | 0 | 0 | 0 | 0 | 0 | 0 | 0 |
| ENSMUSG00000099977 | 0 | 0 | 0 | 0 | 0 | 0 | 0 |
| ENSMUSG00000099978 | 0 | 0 | 0 | 0 | 0 | 0 | 0 |
| ENSMUSG00000099979 | 0 | 0 | 0.25 | 0.5 | 0.244650542 | 0 | 0.224243824 |
| ENSMUSG00000099980 | 0 | 0.5 | 0.25 | 0 | 0.568060967 | 0.602059991 | 0.448487648 |
| ENSMUSG00000099981 | 0 | 0 | 0 | 0 | 0 | 0 | 0 |
| ENSMUSG00000099982 | 0 | 0 | 0 | 0 | 0 | 0 | 0 |
| ENSMUSG00000099983 | 0 | 0 | 0 | 0 | 0 | 0 | 0 |
| ENSMUSG00000099985 | 0 | 0 | 0 | 0 | 0 | 0 | 0 |
| ENSMUSG00000099986 | 0 | 0 | 0 | 0 | 0 | 0 | 0 |

| | | | | | | | |
|---|---|---|---|---|---|---|---|
| ENSMUSG00000099988 | 0 | 0 | 0 | 0 | 0 | 0 | 0 |
| ENSMUSG00000099989 | 0 | 0 | 0 | 0 | 0 | 0 | 0 |
| ENSMUSG00000099990 | 0 | 0 | 0 | 0 | 0 | 0 | 0 |
| ENSMUSG00000099991 | 0 | 0 | 0 | 0 | 0 | 0 | 0 |
| ENSMUSG00000099992 | 0 | 0 | 0 | 0 | 0 | 0 | 0 |
| ENSMUSG00000099993 | 0 | 0 | 0 | 0 | 0 | 0 | 0 |
| ENSMUSG00000099994 | 0 | 0 | 0 | 0 | 0 | 0 | 0 |
| ENSMUSG00000099995 | 0 | 0 | 0 | 0 | 0 | 0 | 0 |
| ENSMUSG00000099996 | 0 | 0 | 0 | 0 | 0 | 0 | 0 |
| ENSMUSG00000099997 | 0 | 0.396240625 | 0.25 | 0.25 | 0.387761935 | 0 | 0.355418052 |
| ENSMUSG00000099998 | 0 | 0 | 0 | 0 | 0.244650542 | 0 | 0 |
| ENSMUSG00000099999 | 0 | 0 | 0 | 0 | 0 | 0 | 0 |
| ENSMUSG00000100000 | 0 | 0 | 0 | 0 | 0 | 0 | 0 |
| ENSMUSG00000100001 | 0 | 0 | 0 | 0 | 0 | 0 | 0 |
| ENSMUSG00000100002 | 0 | 0 | 0 | 0.25 | 0.244650542 | 0 | 0 |
| ENSMUSG00000100003 | 0 | 0 | 0 | 0 | 0 | 0 | 0 |
| ENSMUSG00000100004 | 0 | 0 | 0 | 0 | 0 | 0 | 0 |
| ENSMUSG00000100005 | 1 | 0.646240625 | 0.951838731 | 0.646240625 | 0.387761935 | 0.301029996 | 0.74492186 |
| ENSMUSG00000100006 | 0 | 0 | 0 | 0 | 0 | 0 | 0 |
| ENSMUSG00000100007 | 0 | 0 | 0 | 0 | 0 | 0 | 0.224243824 |
| ENSMUSG00000100008 | 0 | 0 | 0 | 0 | 0 | 0 | 0 |
| ENSMUSG00000100009 | 0.45815691 | 0 | 0.5 | 0.25 | 0.568060967 | 0.602059991 | 0.448487648 |
| ENSMUSG00000100010 | 0 | 0 | 0.25 | 0 | 0.244650542 | 0 | 0 |
| ENSMUSG00000100011 | 0 | 0 | 0 | 0 | 0 | 0 | 0 |
| ENSMUSG00000100012 | 0 | 0 | 0 | 0 | 0 | 0 | 0 |
| ENSMUSG00000100013 | 0 | 0 | 0 | 0 | 0 | 0 | 0 |
| ENSMUSG00000100014 | 0 | 0 | 0 | 0 | 0 | 0 | 0 |
| ENSMUSG00000100015 | 0 | 0 | 0 | 0 | 0 | 0 | 0 |
| ENSMUSG00000100016 | 0 | 0 | 0 | 0 | 0 | 0 | 0 |
| ENSMUSG00000100017 | 0.45815691 | 0.396240625 | 0.580482024 | 0.701838731 | 0.686820904 | 0.698970004 | 0.629532004 |
| ENSMUSG00000100018 | 0 | 0 | 0 | 0 | 0 | 0 | 0 |
| ENSMUSG00000100019 | 0 | 0 | 0 | 0 | 0 | 0 | 0 |
| ENSMUSG00000100020 | 0 | 0 | 0 | 0 | 0 | 0 | 0 |
| ENSMUSG00000100021 | 0 | 0 | 0 | 0 | 0 | 0 | 0 |

| | | | | | | | |
|---|---|---|---|---|---|---|---|
| ENSMUSG00000100022 | 0 | 0 | 0.25 | 0 | 0 | 0 | 0 |
| ENSMUSG00000100023 | 0 | 0 | 0 | 0 | 0 | 0 | 0 |
| ENSMUSG00000100024 | 0 | 0 | 0 | 0 | 0 | 0 | 0 |
| ENSMUSG00000100025 | 0.578129653 | 0.396240625 | 0.75 | 0.79248125 | 0.686820904 | 0.698970004 | 0.74492186 |
| ENSMUSG00000100026 | 0 | 0 | 0 | 0 | 0 | 0 | 0 |
| ENSMUSG00000100027 | 0 | 0 | 0 | 0.25 | 0 | 0 | 0 |
| ENSMUSG00000100028 | 0 | 0 | 0 | 0 | 0 | 0 | 0 |
| ENSMUSG00000100029 | 0 | 0 | 0 | 0 | 0 | 0 | 0 |
| ENSMUSG00000100031 | 0 | 0 | 0 | 0 | 0 | 0 | 0 |
| ENSMUSG00000100032 | 0 | 0 | 0 | 0 | 0 | 0 | 0 |
| ENSMUSG00000100033 | 0.747221736 | 0.5 | 0.830482024 | 0.896240625 | 1.106691885 | 0.84509804 | 0.876096088 |
| ENSMUSG00000100034 | 0 | 0 | 0 | 0 | 0 | 0 | 0 |
| ENSMUSG00000100035 | 0 | 0 | 0 | 0 | 0 | 0 | 0 |
| ENSMUSG00000100036 | 0 | 0 | 0 | 0 | 0 | 0 | 0 |
| ENSMUSG00000100037 | 0 | 0 | 0 | 0 | 0 | 0 | 0 |
| ENSMUSG00000100038 | 0 | 0 | 0 | 0 | 0 | 0 | 0 |
| ENSMUSG00000100039 | 0 | 0 | 0 | 0 | 0 | 0 | 0 |
| ENSMUSG00000100040 | 0 | 0 | 0 | 0 | 0 | 0 | 0 |
| ENSMUSG00000100041 | 0.289064826 | 0 | 0 | 0 | 0.244650542 | 0 | 0.355418052 |
| ENSMUSG00000100042 | 0 | 0 | 0 | 0 | 0 | 0 | 0 |
| ENSMUSG00000100043 | 0 | 0 | 0 | 0 | 0 | 0 | 0 |
| ENSMUSG00000100044 | 0 | 0 | 0 | 0 | 0 | 0 | 0 |
| ENSMUSG00000100045 | 0 | 0 | 0 | 0 | 0 | 0 | 0 |
| ENSMUSG00000100046 | 0 | 0 | 0 | 0 | 0 | 0 | 0 |
| ENSMUSG00000100047 | 0 | 0 | 0 | 0 | 0 | 0 | 0 |
| ENSMUSG00000100048 | 0 | 0 | 0 | 0 | 0 | 0 | 0 |
| ENSMUSG00000100049 | 0 | 0 | 0 | 0 | 0 | 0 | 0 |
| ENSMUSG00000100050 | 0 | 0 | 0 | 0 | 0 | 0 | 0 |
| ENSMUSG00000100051 | 0 | 0 | 0 | 0 | 0 | 0 | 0 |
| ENSMUSG00000100052 | 0 | 0 | 0 | 0 | 0 | 0 | 0 |
| ENSMUSG00000100053 | 0 | 0 | 0 | 0 | 0 | 0 | 0 |
| ENSMUSG00000100054 | 0 | 0 | 0 | 0 | 0 | 0 | 0 |
| ENSMUSG00000100055 | 0 | 0 | 0 | 0 | 0 | 0 | 0 |
| ENSMUSG00000100056 | 0 | 0 | 0 | 0 | 0 | 0 | 0 |

| | | | | | | | |
|---|---|---|---|---|---|---|---|
| ENSMUSG00000100057 | 0 | 0 | 0 | 0 | 0 | 0 | 0 |
| ENSMUSG00000100058 | 0 | 0 | 0 | 0 | 0 | 0 | 0 |
| ENSMUSG00000100059 | 0 | 0 | 0 | 0 | 0 | 0 | 0 |
| ENSMUSG00000100060 | 0 | 0 | 0 | 0 | 0 | 0 | 0 |
| ENSMUSG00000100061 | 0 | 0 | 0 | 0 | 0 | 0 | 0 |
| ENSMUSG00000100062 | 0 | 0 | 0 | 0 | 0 | 0 | 0 |
| ENSMUSG00000100063 | 0 | 0 | 0 | 0 | 0 | 0 | 0 |
| ENSMUSG00000100064 | 0 | 0 | 0 | 0 | 0 | 0 | 0 |
| ENSMUSG00000100065 | 0 | 0 | 0 | 0 | 0 | 0 | 0.355418052 |
| ENSMUSG00000100066 | 0 | 0 | 0 | 0 | 0 | 0 | 0 |
| ENSMUSG00000100067 | 0 | 0 | 0 | 0 | 0 | 0 | 0 |
| ENSMUSG00000100068 | 0 | 0 | 0 | 0 | 0 | 0 | 0 |
| ENSMUSG00000100069 | 0 | 0 | 0 | 0 | 0 | 0 | 0.224243824 |
| ENSMUSG00000100070 | 0 | 0 | 0 | 0 | 0 | 0 | 0 |
| ENSMUSG00000100071 | 0 | 0 | 0 | 0.25 | 0 | 0 | 0 |
| ENSMUSG00000100072 | 0 | 0 | 0 | 0 | 0 | 0 | 0 |
| ENSMUSG00000100073 | 0 | 0 | 0 | 0 | 0 | 0 | 0 |
| ENSMUSG00000100074 | 0 | 0 | 0 | 0 | 0 | 0 | 0 |
| ENSMUSG00000100075 | 0.289064826 | 0.5 | 0.5 | 0.396240625 | 0.244650542 | 0 | 0.520678036 |
| ENSMUSG00000100076 | 0 | 0 | 0 | 0 | 0 | 0 | 0 |
| ENSMUSG00000100077 | 0 | 0 | 0 | 0 | 0 | 0 | 0 |
| ENSMUSG00000100078 | 1.605636262 | 1.330482024 | 1.494319981 | 1.51109853 | 1.401338844 | 1.62324929 | 1.360283038 |
| ENSMUSG00000100079 | 0 | 0 | 0 | 0 | 0 | 0 | 0 |
| ENSMUSG00000100080 | 0 | 0 | 0 | 0 | 0 | 0 | 0 |
| ENSMUSG00000100081 | 0 | 0 | 0 | 0 | 0 | 0 | 0 |
| ENSMUSG00000100082 | 0 | 0 | 0 | 0 | 0 | 0 | 0 |
| ENSMUSG00000100083 | 0 | 0 | 0 | 0 | 0 | 0 | 0 |
| ENSMUSG00000100084 | 0.289064826 | 0.25 | 0.580482024 | 0.396240625 | 0.489301084 | 0.477121255 | 0.448487648 |
| ENSMUSG00000100085 | 0 | 0 | 0 | 0 | 0 | 0 | 0 |
| ENSMUSG00000100086 | 0 | 0 | 0 | 0 | 0 | 0 | 0 |
| ENSMUSG00000100087 | 0 | 0 | 0 | 0 | 0 | 0 | 0 |
| ENSMUSG00000100088 | 0 | 0 | 0 | 0 | 0 | 0 | 0 |
| ENSMUSG00000100089 | 0 | 0 | 0 | 0 | 0 | 0 | 0 |
| ENSMUSG00000100090 | 0.289064826 | 0 | 0 | 0.396240625 | 0 | 0 | 0 |

| | | | | | | | |
|---|---|---|---|---|---|---|---|
| ENSMUSG00000100091 | 0 | 0 | 0 | 0 | 0 | 0 | 0 |
| ENSMUSG00000100092 | 0 | 0 | 0 | 0 | 0 | 0 | 0 |
| ENSMUSG00000100093 | 0 | 0 | 0 | 0 | 0 | 0 | 0 |
| ENSMUSG00000100094 | 0 | 0 | 0 | 0 | 0 | 0.301029996 | 0 |
| ENSMUSG00000100095 | 0 | 0 | 0 | 0 | 0 | 0 | 0 |
| ENSMUSG00000100096 | 0 | 0 | 0 | 0 | 0 | 0 | 0.224243824 |
| ENSMUSG00000100097 | 0 | 0 | 0 | 0 | 0 | 0 | 0 |
| ENSMUSG00000100098 | 0 | 0 | 0 | 0 | 0 | 0 | 0 |
| ENSMUSG00000100099 | 0 | 0 | 0 | 0 | 0 | 0 | 0 |
| ENSMUSG00000100100 | 0 | 0 | 0 | 0 | 0 | 0 | 0 |
| ENSMUSG00000100101 | 0 | 0 | 0 | 0 | 0 | 0 | 0 |
| ENSMUSG00000100103 | 0 | 0 | 0 | 0 | 0 | 0 | 0 |
| ENSMUSG00000100104 | 0.289064826 | 0.5 | 0.25 | 0 | 0.489301084 | 0 | 0.224243824 |
| ENSMUSG00000100105 | 0 | 0 | 0 | 0 | 0 | 0 | 0 |
| ENSMUSG00000100106 | 0 | 0 | 0 | 0 | 0 | 0.301029996 | 0 |
| ENSMUSG00000100107 | 0 | 0 | 0 | 0 | 0 | 0 | 0 |
| ENSMUSG00000100108 | 0 | 0 | 0 | 0.25 | 0.244650542 | 0 | 0 |
| ENSMUSG00000100109 | 0 | 0 | 0 | 0 | 0 | 0 | 0 |
| ENSMUSG00000100110 | 0 | 0 | 0 | 0 | 0 | 0 | 0 |
| ENSMUSG00000100111 | 0 | 0 | 0 | 0 | 0 | 0 | 0 |
| ENSMUSG00000100112 | 0 | 0 | 0 | 0 | 0 | 0 | 0 |
| ENSMUSG00000100113 | 0 | 0 | 0 | 0 | 0 | 0 | 0 |
| ENSMUSG00000100115 | 0 | 0 | 0 | 0 | 0 | 0 | 0 |
| ENSMUSG00000100116 | 0 | 0 | 0 | 0 | 0 | 0 | 0 |
| ENSMUSG00000100117 | 0 | 0 | 0 | 0 | 0 | 0 | 0 |
| ENSMUSG00000100118 | 0 | 0 | 0 | 0 | 0 | 0 | 0 |
| ENSMUSG00000100119 | 0 | 0.25 | 0 | 0 | 0 | 0 | 0 |
| ENSMUSG00000100120 | 0 | 0 | 0 | 0 | 0 | 0 | 0 |
| ENSMUSG00000100121 | 0.289064826 | 0.25 | 0.396240625 | 0.5 | 0 | 0 | 0.224243824 |
| ENSMUSG00000100122 | 0 | 0 | 0 | 0 | 0 | 0 | 0 |
| ENSMUSG00000100123 | 0 | 0 | 0 | 0.25 | 0 | 0 | 0 |
| ENSMUSG00000100124 | 0 | 0 | 0 | 0 | 0 | 0 | 0 |
| ENSMUSG00000100125 | 0 | 0 | 0 | 0 | 0 | 0 | 0 |
| ENSMUSG00000100126 | 0 | 0 | 0 | 0 | 0 | 0 | 0 |

| | | | | | | | |
|---|---|---|---|---|---|---|---|
| ENSMUSG00000100127 | 0 | 0 | 0 | 0 | 0.387761935 | 0 | 0 |
| ENSMUSG00000100128 | 0 | 0 | 0 | 0 | 0 | 0 | 0 |
| ENSMUSG00000100129 | 0 | 0 | 0 | 0 | 0 | 0 | 0 |
| ENSMUSG00000100130 | 0 | 0 | 0 | 0 | 0 | 0 | 0 |
| ENSMUSG00000100131 | 0 | 0 | 0 | 0.396240625 | 0 | 0 | 0 |
| ENSMUSG00000100132 | 0 | 0 | 0 | 0 | 0 | 0 | 0 |
| ENSMUSG00000100133 | 0 | 0 | 0 | 0 | 0 | 0 | 0 |
| ENSMUSG00000100134 | 0 | 0 | 0 | 0 | 0 | 0 | 0 |
| ENSMUSG00000100135 | 0 | 0 | 0 | 0 | 0 | 0 | 0 |
| ENSMUSG00000100136 | 0 | 0 | 0 | 0 | 0 | 0 | 0 |
| ENSMUSG00000100137 | 0 | 0 | 0 | 0 | 0 | 0 | 0 |
| ENSMUSG00000100138 | 0 | 0 | 0 | 0 | 0 | 0 | 0 |
| ENSMUSG00000100140 | 0 | 0 | 0 | 0 | 0 | 0 | 0 |
| ENSMUSG00000100141 | 0 | 0 | 0.396240625 | 0.25 | 0.244650542 | 0.301029996 | 0.224243824 |
| ENSMUSG00000100142 | 0 | 0 | 0 | 0 | 0 | 0 | 0 |
| ENSMUSG00000100143 | 0 | 0 | 0 | 0 | 0 | 0 | 0 |
| ENSMUSG00000100144 | 0 | 0 | 0 | 0 | 0 | 0 | 0 |
| ENSMUSG00000100145 | 0 | 0 | 0 | 0 | 0 | 0 | 0 |
| ENSMUSG00000100146 | 0 | 0 | 0 | 0 | 0 | 0 | 0 |
| ENSMUSG00000100147 | 0 | 0 | 0 | 0 | 0 | 0 | 0 |
| ENSMUSG00000100148 | 0 | 0 | 0 | 0 | 0 | 0 | 0 |
| ENSMUSG00000100149 | 0 | 0 | 0 | 0 | 0 | 0 | 0 |
| ENSMUSG00000100150 | 0 | 0 | 0 | 0 | 0 | 0 | 0 |
| ENSMUSG00000100151 | 0 | 0 | 0 | 0 | 0 | 0 | 0 |
| ENSMUSG00000100152 | 0 | 0 | 0 | 0 | 0 | 0 | 0 |
| ENSMUSG00000100153 | 0 | 0 | 0 | 0 | 0 | 0 | 0 |
| ENSMUSG00000100154 | 0 | 0 | 0 | 0 | 0 | 0 | 0 |
| ENSMUSG00000100155 | 0 | 0 | 0 | 0 | 0 | 0 | 0 |
| ENSMUSG00000100156 | 0 | 0 | 0 | 0 | 0 | 0 | 0 |
| ENSMUSG00000100157 | 0 | 0 | 0.25 | 0 | 0 | 0 | 0 |
| ENSMUSG00000100158 | 0 | 0.580482024 | 0.75 | 0.396240625 | 0.568060967 | 0 | 0.579661877 |
| ENSMUSG00000100159 | 0 | 0 | 0 | 0 | 0 | 0 | 0 |
| ENSMUSG00000100160 | 0 | 0.25 | 0 | 0 | 0 | 0 | 0.224243824 |
| ENSMUSG00000100161 | 0 | 0 | 0 | 0 | 0 | 0 | 0 |

| | | | | | | | |
|---|---|---|---|---|---|---|---|
| ENSMUSG00000100162 | 0 | 0 | 0 | 0 | 0 | 0 | 0 |
| ENSMUSG00000100164 | 1.036286563 | 1.098079356 | 1.201838731 | 1.17510993 | 1.234113756 | 1.278753601 | 1.20919388 |
| ENSMUSG00000100165 | 0 | 0 | 0 | 0 | 0 | 0 | 0 |
| ENSMUSG00000100166 | 0 | 0 | 0 | 0 | 0 | 0 | 0 |
| ENSMUSG00000100167 | 0 | 0 | 0 | 0 | 0 | 0 | 0 |
| ENSMUSG00000100168 | 0 | 0 | 0 | 0 | 0 | 0 | 0 |
| ENSMUSG00000100169 | 0 | 0 | 0 | 0 | 0 | 0 | 0 |
| ENSMUSG00000100170 | 0 | 0 | 0 | 0 | 0 | 0 | 0 |
| ENSMUSG00000100171 | 0 | 0 | 0 | 0 | 0 | 0 | 0 |
| ENSMUSG00000100172 | 0 | 0 | 0 | 0 | 0 | 0 | 0 |
| ENSMUSG00000100174 | 0 | 0 | 0 | 0 | 0 | 0 | 0 |
| ENSMUSG00000100175 | 0 | 0 | 0 | 0 | 0 | 0 | 0 |
| ENSMUSG00000100176 | 0 | 0 | 0 | 0 | 0 | 0 | 0.355418052 |
| ENSMUSG00000100177 | 0 | 0 | 0 | 0 | 0 | 0 | 0 |
| ENSMUSG00000100178 | 0 | 0.25 | 0 | 0 | 0 | 0 | 0 |
| ENSMUSG00000100179 | 0 | 0 | 0 | 0 | 0 | 0 | 0 |
| ENSMUSG00000100180 | 0 | 0 | 0 | 0 | 0 | 0 | 0 |
| ENSMUSG00000100181 | 0 | 0 | 0 | 0 | 0.244650542 | 0 | 0 |
| ENSMUSG00000100182 | 0 | 0 | 0 | 0 | 0 | 0 | 0 |
| ENSMUSG00000100183 | 0.671187741 | 0.580482024 | 0.396240625 | 0.580482024 | 0.489301084 | 0.301029996 | 0.579661877 |
| ENSMUSG00000100184 | 0 | 0 | 0 | 0 | 0 | 0 | 0 |
| ENSMUSG00000100185 | 0 | 0 | 0 | 0 | 0 | 0 | 0 |
| ENSMUSG00000100186 | 0 | 0 | 0 | 0 | 0 | 0 | 0 |
| ENSMUSG00000100187 | 0 | 0 | 0 | 0 | 0 | 0 | 0 |
| ENSMUSG00000100188 | 0 | 0 | 0.25 | 0 | 0 | 0 | 0 |
| ENSMUSG00000100189 | 0 | 0 | 0.25 | 0 | 0 | 0 | 0 |
| ENSMUSG00000100190 | 0 | 0 | 0 | 0 | 0 | 0 | 0 |
| ENSMUSG00000100191 | 0 | 0 | 0 | 0 | 0 | 0 | 0 |
| ENSMUSG00000100192 | 0 | 0 | 0 | 0 | 0 | 0 | 0 |
| ENSMUSG00000100193 | 0 | 0 | 0 | 0 | 0 | 0 | 0 |
| ENSMUSG00000100194 | 0 | 0 | 0 | 0 | 0 | 0 | 0 |
| ENSMUSG00000100195 | 0 | 0 | 0 | 0 | 0 | 0 | 0 |
| ENSMUSG00000100196 | 0 | 0 | 0 | 0 | 0 | 0 | 0 |
| ENSMUSG00000100197 | 0 | 0 | 0 | 0 | 0 | 0 | 0 |

| | | | | | | | |
|---|---|---|---|---|---|---|---|
| ENSMUSG00000100198 | 0 | 0 | 0 | 0 | 0 | 0 | 0 |
| ENSMUSG00000100199 | 1 | 0.864857905 | 0.830482024 | 0.896240625 | 0.955822902 | 1.041392685 | 0.775756176 |
| ENSMUSG00000100200 | 0 | 0 | 0 | 0 | 0.244650542 | 0 | 0 |
| ENSMUSG00000100201 | 0.867194479 | 0.951838731 | 0.701838731 | 1.02186571 | 0.905314583 | 0.698970004 | 0.896975297 |
| ENSMUSG00000100202 | 0 | 0 | 0 | 0 | 0 | 0 | 0 |
| ENSMUSG00000100203 | 0 | 0 | 0 | 0 | 0 | 0 | 0 |
| ENSMUSG00000100204 | 0.289064826 | 0 | 0.25 | 0 | 0.387761935 | 0 | 0.224243824 |
| ENSMUSG00000100205 | 0 | 0 | 0 | 0 | 0 | 0 | 0 |
| ENSMUSG00000100206 | 0 | 0 | 0 | 0 | 0 | 0 | 0 |
| ENSMUSG00000100207 | 0 | 0 | 0 | 0 | 0 | 0 | 0 |
| ENSMUSG00000100208 | 0 | 0 | 0 | 0 | 0 | 0 | 0 |
| ENSMUSG00000100209 | 0 | 0 | 0 | 0 | 0 | 0 | 0 |
| ENSMUSG00000100210 | 0 | 0 | 0 | 0 | 0 | 0 | 0 |
| ENSMUSG00000100211 | 0 | 0 | 0 | 0.25 | 0 | 0 | 0 |
| ENSMUSG00000100212 | 0 | 0 | 0 | 0 | 0 | 0 | 0 |
| ENSMUSG00000100213 | 0.91631382 | 0.580482024 | 0.75 | 0.396240625 | 0.387761935 | 0.477121255 | 0.629532004 |
| ENSMUSG00000100214 | 1.596667477 | 1.51109853 | 1.52186571 | 1.451838731 | 1.414412788 | 1.431363764 | 1.23151414 |
| ENSMUSG00000100215 | 2.323768721 | 2.170748646 | 2.176839783 | 2.361770807 | 2.287171411 | 2.69019608 | 2.143416814 |
| ENSMUSG00000100216 | 0 | 0 | 0 | 0 | 0 | 0 | 0 |
| ENSMUSG00000100217 | 0 | 0 | 0 | 0 | 0 | 0 | 0 |
| ENSMUSG00000100218 | 0 | 0 | 0 | 0 | 0 | 0 | 0 |
| ENSMUSG00000100219 | 0 | 0 | 0 | 0 | 0 | 0 | 0 |
| ENSMUSG00000100220 | 0 | 0 | 0 | 0 | 0 | 0 | 0 |
| ENSMUSG00000100221 | 0 | 0 | 0 | 0 | 0 | 0 | 0 |
| ENSMUSG00000100222 | 0 | 0 | 0 | 0 | 0 | 0 | 0 |
| ENSMUSG00000100223 | 0 | 0 | 0 | 0 | 0 | 0 | 0 |
| ENSMUSG00000100224 | 0 | 0 | 0 | 0 | 0 | 0 | 0 |
| ENSMUSG00000100225 | 0 | 0 | 0 | 0 | 0 | 0 | 0 |
| ENSMUSG00000100226 | 0 | 0 | 0 | 0 | 0 | 0 | 0 |
| ENSMUSG00000100227 | 0 | 0 | 0 | 0 | 0 | 0 | 0 |
| ENSMUSG00000100228 | 0.289064826 | 0 | 0.580482024 | 0.580482024 | 0.244650542 | 0.477121255 | 0.224243824 |
| ENSMUSG00000100229 | 0 | 0 | 0 | 0 | 0 | 0 | 0 |
| ENSMUSG00000100230 | 0.289064826 | 0 | 0 | 0 | 0 | 0 | 0 |
| ENSMUSG00000100231 | 0 | 0 | 0 | 0 | 0 | 0 | 0 |

| | | | | | | | |
|---|---|---|---|---|---|---|---|
| ENSMUSG00000100232 | 0 | 0 | 0 | 0 | 0 | 0 | 0 |
| ENSMUSG00000100233 | 0 | 0 | 0 | 0 | 0 | 0 | 0 |
| ENSMUSG00000100234 | 0 | 0 | 0 | 0 | 0 | 0 | 0 |
| ENSMUSG00000100235 | 0.289064826 | 0.79248125 | 0.396240625 | 0.396240625 | 0.77552387 | 0.477121255 | 0.775756176 |
| ENSMUSG00000100236 | 0 | 0 | 0 | 0 | 0 | 0 | 0 |
| ENSMUSG00000100237 | 0 | 0 | 0 | 0 | 0 | 0 | 0 |
| ENSMUSG00000100238 | 0 | 0 | 0 | 0 | 0 | 0 | 0 |
| ENSMUSG00000100239 | 0 | 0 | 0 | 0 | 0 | 0 | 0 |
| ENSMUSG00000100240 | 0 | 0 | 0 | 0 | 0 | 0 | 0 |
| ENSMUSG00000100241 | 0 | 0 | 0 | 0 | 0 | 0 | 0 |
| ENSMUSG00000100242 | 0 | 0 | 0 | 0 | 0 | 0 | 0 |
| ENSMUSG00000100243 | 0 | 0 | 0 | 0 | 0 | 0 | 0 |
| ENSMUSG00000100244 | 0 | 0 | 0 | 0 | 0 | 0 | 0 |
| ENSMUSG00000100245 | 0 | 0 | 0 | 0 | 0 | 0 | 0 |
| ENSMUSG00000100246 | 0 | 0 | 0 | 0 | 0 | 0 | 0 |
| ENSMUSG00000100247 | 0.578129653 | 0 | 0.5 | 0.25 | 0.244650542 | 0 | 0.224243824 |
| ENSMUSG00000100248 | 0 | 0 | 0 | 0 | 0 | 0 | 0 |
| ENSMUSG00000100249 | 0 | 0 | 0 | 0 | 0 | 0 | 0 |
| ENSMUSG00000100250 | 0 | 0 | 0 | 0 | 0 | 0 | 0 |
| ENSMUSG00000100251 | 0 | 0 | 0 | 0 | 0 | 0 | 0 |
| ENSMUSG00000100252 | 0.289064826 | 0 | 0 | 0.25 | 0 | 0 | 0 |
| ENSMUSG00000100253 | 0 | 0 | 0 | 0 | 0 | 0 | 0 |
| ENSMUSG00000100254 | 0 | 0 | 0 | 0 | 0 | 0 | 0 |
| ENSMUSG00000100255 | 0.671187741 | 0.5 | 0.646240625 | 0 | 0.77552387 | 0 | 0.355418052 |
| ENSMUSG00000100256 | 0 | 0 | 0 | 0 | 0 | 0 | 0 |
| ENSMUSG00000100257 | 0.289064826 | 0 | 0 | 0 | 0 | 0 | 0 |
| ENSMUSG00000100258 | 0 | 0 | 0 | 0 | 0 | 0 | 0 |
| ENSMUSG00000100259 | 0 | 0 | 0.25 | 0 | 0 | 0 | 0 |
| ENSMUSG00000100260 | 0 | 0 | 0 | 0 | 0 | 0 | 0 |
| ENSMUSG00000100261 | 0 | 0 | 0.25 | 0 | 0.244650542 | 0 | 0.224243824 |
| ENSMUSG00000100262 | 0 | 0 | 0 | 0 | 0 | 0 | 0 |
| ENSMUSG00000100263 | 0 | 0 | 0 | 0 | 0 | 0 | 0 |
| ENSMUSG00000100264 | 0 | 0 | 0 | 0 | 0 | 0 | 0 |
| ENSMUSG00000100265 | 0 | 0 | 0 | 0 | 0 | 0 | 0 |

| | | | | | | | |
|---|---|---|---|---|---|---|---|
| ENSMUSG00000100266 | 0 | 0 | 0 | 0 | 0 | 0 | 0 |
| ENSMUSG00000100267 | 0 | 0 | 0 | 0 | 0 | 0 | 0 |
| ENSMUSG00000100268 | 0 | 0 | 0 | 0 | 0 | 0 | 0 |
| ENSMUSG00000100269 | 0 | 0 | 0 | 0 | 0 | 0 | 0 |
| ENSMUSG00000100270 | 0 | 0 | 0 | 0 | 0 | 0 | 0 |
| ENSMUSG00000100271 | 0 | 0 | 0 | 0 | 0 | 0 | 0 |
| ENSMUSG00000100272 | 0 | 0 | 0 | 0 | 0 | 0 | 0 |
| ENSMUSG00000100273 | 0 | 0 | 0 | 0 | 0 | 0 | 0 |
| ENSMUSG00000100274 | 0 | 0 | 0 | 0 | 0 | 0 | 0 |
| ENSMUSG00000100275 | 0 | 0 | 0 | 0 | 0 | 0 | 0 |
| ENSMUSG00000100276 | 0 | 0 | 0 | 0 | 0 | 0 | 0 |
| ENSMUSG00000100277 | 0 | 0 | 0 | 0 | 0 | 0 | 0 |
| ENSMUSG00000100278 | 0 | 0 | 0 | 0 | 0 | 0 | 0 |
| ENSMUSG00000100279 | 0 | 0 | 0 | 0 | 0 | 0 | 0 |
| ENSMUSG00000100280 | 0.960252568 | 0.830482024 | 0.79248125 | 0.896240625 | 0.686820904 | 0.84509804 | 0.710836105 |
| ENSMUSG00000100281 | 0.45815691 | 0.580482024 | 0.396240625 | 0.396240625 | 0.568060967 | 0.84509804 | 0.579661877 |
| ENSMUSG00000100282 | 0 | 0 | 0 | 0 | 0 | 0 | 0 |
| ENSMUSG00000100283 | 0 | 0 | 0 | 0 | 0 | 0 | 0 |
| ENSMUSG00000100284 | 0 | 0 | 0 | 0 | 0 | 0 | 0 |
| ENSMUSG00000100285 | 0 | 0 | 0 | 0 | 0 | 0 | 0 |
| ENSMUSG00000100286 | 0 | 0 | 0 | 0 | 0 | 0 | 0 |
| ENSMUSG00000100287 | 0.289064826 | 0 | 0 | 0.25 | 0.244650542 | 0 | 0.224243824 |
| ENSMUSG00000100288 | 0 | 0 | 0 | 0 | 0 | 0 | 0 |
| ENSMUSG00000100289 | 0 | 0 | 0 | 0 | 0 | 0 | 0 |
| ENSMUSG00000100290 | 0 | 0 | 0 | 0 | 0 | 0 | 0 |
| ENSMUSG00000100291 | 0 | 0 | 0 | 0 | 0 | 0 | 0 |
| ENSMUSG00000100292 | 0 | 0 | 0 | 0 | 0 | 0 | 0 |
| ENSMUSG00000100294 | 0.289064826 | 0 | 0 | 0 | 0 | 0 | 0 |
| ENSMUSG00000100295 | 0 | 0 | 0 | 0 | 0 | 0 | 0 |
| ENSMUSG00000100296 | 0 | 0 | 0 | 0 | 0 | 0 | 0 |
| ENSMUSG00000100297 | 0 | 0 | 0 | 0 | 0 | 0 | 0 |
| ENSMUSG00000100298 | 0 | 0 | 0 | 0 | 0 | 0 | 0 |
| ENSMUSG00000100299 | 0 | 0 | 0 | 0 | 0 | 0 | 0 |
| ENSMUSG00000100300 | 0 | 0 | 0 | 0 | 0 | 0 | 0 |

| | | | | | | | |
|---|---|---|---|---|---|---|---|
| ENSMUSG00000100301 | 0 | 0 | 0 | 0 | 0 | 0 | 0 |
| ENSMUSG00000100302 | 0.289064826 | 0.25 | 0.25 | 0 | 0 | 0 | 0 |
| ENSMUSG00000100303 | 0 | 0 | 0 | 0.25 | 0.244650542 | 0 | 0 |
| ENSMUSG00000100304 | 0 | 0 | 0 | 0 | 0 | 0 | 0 |
| ENSMUSG00000100305 | 0 | 0 | 0 | 0 | 0 | 0 | 0 |
| ENSMUSG00000100306 | 0 | 0 | 0 | 0 | 0 | 0 | 0 |
| ENSMUSG00000100307 | 0 | 0 | 0 | 0 | 0 | 0 | 0 |
| ENSMUSG00000100308 | 0 | 0 | 0 | 0 | 0 | 0 | 0 |
| ENSMUSG00000100309 | 0 | 0 | 0 | 0 | 0 | 0 | 0 |
| ENSMUSG00000100310 | 0 | 0 | 0 | 0 | 0 | 0 | 0 |
| ENSMUSG00000100311 | 0 | 0 | 0 | 0 | 0 | 0 | 0 |
| ENSMUSG00000100312 | 0 | 0 | 0 | 0 | 0 | 0 | 0 |
| ENSMUSG00000100313 | 0 | 0 | 0 | 0 | 0 | 0 | 0 |
| ENSMUSG00000100314 | 0 | 0 | 0 | 0 | 0 | 0 | 0 |
| ENSMUSG00000100315 | 0 | 0 | 0 | 0.25 | 0 | 0 | 0 |
| ENSMUSG00000100316 | 0 | 0 | 0 | 0 | 0 | 0 | 0 |
| ENSMUSG00000100317 | 0 | 0 | 0 | 0 | 0 | 0 | 0 |
| ENSMUSG00000100318 | 0 | 0 | 0 | 0 | 0 | 0 | 0.224243824 |
| ENSMUSG00000100319 | 0 | 0.25 | 0.25 | 0.25 | 0 | 0 | 0 |
| ENSMUSG00000100320 | 0.289064826 | 0.25 | 0 | 0.396240625 | 0 | 0 | 0 |
| ENSMUSG00000100321 | 0 | 0 | 0 | 0 | 0 | 0 | 0 |
| ENSMUSG00000100322 | 0 | 0 | 0 | 0 | 0 | 0 | 0 |
| ENSMUSG00000100323 | 0 | 0 | 0 | 0 | 0 | 0 | 0 |
| ENSMUSG00000100324 | 0 | 0 | 0 | 0 | 0 | 0 | 0 |
| ENSMUSG00000100325 | 0 | 0 | 0 | 0 | 0 | 0 | 0 |
| ENSMUSG00000100326 | 0 | 0 | 0 | 0 | 0 | 0 | 0 |
| ENSMUSG00000100328 | 0 | 0 | 0 | 0 | 0 | 0 | 0 |
| ENSMUSG00000100329 | 0 | 0 | 0 | 0 | 0 | 0 | 0 |
| ENSMUSG00000100330 | 0 | 0 | 0 | 0 | 0 | 0 | 0 |
| ENSMUSG00000100331 | 0 | 0 | 0 | 0 | 0 | 0 | 0 |
| ENSMUSG00000100332 | 0 | 0 | 0 | 0 | 0 | 0 | 0 |
| ENSMUSG00000100333 | 0 | 0 | 0 | 0 | 0 | 0 | 0 |
| ENSMUSG00000100334 | 0 | 0 | 0.25 | 0 | 0.244650542 | 0.301029996 | 0 |
| ENSMUSG00000100335 | 0 | 0.25 | 0 | 0 | 0 | 0 | 0 |

| | | | | | | | |
|---|---|---|---|---|---|---|---|
| ENSMUSG00000100336 | 0 | 0 | 0 | 0 | 0 | 0 | 0 |
| ENSMUSG00000100337 | 0 | 0 | 0 | 0 | 0.244650542 | 0 | 0 |
| ENSMUSG00000100338 | 0 | 0 | 0 | 0 | 0 | 0 | 0 |
| ENSMUSG00000100339 | 0 | 0 | 0 | 0 | 0 | 0 | 0 |
| ENSMUSG00000100340 | 0 | 0 | 0 | 0 | 0 | 0 | 0 |
| ENSMUSG00000100341 | 0 | 0 | 0 | 0 | 0 | 0 | 0 |
| ENSMUSG00000100343 | 0 | 0 | 0 | 0 | 0 | 0 | 0 |
| ENSMUSG00000100344 | 0 | 0 | 0 | 0 | 0 | 0 | 0 |
| ENSMUSG00000100345 | 0 | 0 | 0 | 0.25 | 0 | 0 | 0 |
| ENSMUSG00000100346 | 0 | 0 | 0 | 0 | 0 | 0 | 0 |
| ENSMUSG00000100347 | 0 | 0.25 | 0 | 0 | 0 | 0 | 0.224243824 |
| ENSMUSG00000100348 | 0 | 0 | 0 | 0 | 0 | 0 | 0 |
| ENSMUSG00000100349 | 0 | 0 | 0 | 0 | 0 | 0 | 0 |
| ENSMUSG00000100350 | 0 | 0 | 0 | 0 | 0 | 0 | 0 |
| ENSMUSG00000100351 | 0.578129653 | 0.396240625 | 0.25 | 0.396240625 | 0 | 0.301029996 | 0.355418052 |
| ENSMUSG00000100352 | 0 | 0 | 0 | 0 | 0 | 0 | 0 |
| ENSMUSG00000100353 | 0 | 0 | 0 | 0 | 0 | 0 | 0 |
| ENSMUSG00000100354 | 0 | 0 | 0 | 0 | 0 | 0.477121255 | 0 |
| ENSMUSG00000100355 | 0 | 0 | 0 | 0 | 0 | 0 | 0 |
| ENSMUSG00000100356 | 0 | 0 | 0 | 0 | 0 | 0 | 0 |
| ENSMUSG00000100357 | 0 | 0 | 0 | 0 | 0 | 0 | 0 |
| ENSMUSG00000100358 | 0 | 0 | 0 | 0 | 0 | 0 | 0 |
| ENSMUSG00000100359 | 0 | 0 | 0 | 0 | 0 | 0 | 0 |
| ENSMUSG00000100360 | 0 | 0 | 0 | 0 | 0 | 0 | 0 |
| ENSMUSG00000100361 | 0.289064826 | 0.25 | 0.396240625 | 0.25 | 0.244650542 | 0.477121255 | 0.448487648 |
| ENSMUSG00000100362 | 0 | 0 | 0 | 0 | 0 | 0 | 0 |
| ENSMUSG00000100363 | 0 | 0 | 0 | 0 | 0 | 0 | 0 |
| ENSMUSG00000100364 | 0 | 0 | 0 | 0 | 0 | 0 | 0 |
| ENSMUSG00000100365 | 0 | 0 | 0 | 0 | 0 | 0 | 0 |
| ENSMUSG00000100366 | 0 | 0 | 0 | 0 | 0 | 0 | 0 |
| ENSMUSG00000100367 | 0 | 0 | 0 | 0 | 0 | 0 | 0 |
| ENSMUSG00000100368 | 0 | 0 | 0 | 0 | 0 | 0 | 0 |
| ENSMUSG00000100369 | 0 | 0 | 0 | 0 | 0 | 0 | 0 |
| ENSMUSG00000100370 | 1.862428343 | 1.692046081 | 1.642463902 | 1.714495249 | 1.603583716 | 1.77815125 | 1.476637173 |

| | | | | | | | |
|---|---|---|---|---|---|---|---|
| ENSMUSG00000100371 | 0.289064826 | 0 | 0 | 0 | 0 | 0 | 0 |
| ENSMUSG00000100372 | 0 | 0 | 0 | 0 | 0 | 0 | 0 |
| ENSMUSG00000100373 | 0 | 0 | 0 | 0 | 0 | 0 | 0 |
| ENSMUSG00000100374 | 0 | 0 | 0 | 0 | 0 | 0 | 0 |
| ENSMUSG00000100375 | 0 | 0 | 0 | 0 | 0 | 0 | 0 |
| ENSMUSG00000100376 | 0 | 0 | 0 | 0 | 0 | 0 | 0 |
| ENSMUSG00000100377 | 0 | 0 | 0 | 0 | 0 | 0 | 0 |
| ENSMUSG00000100378 | 0 | 0 | 0 | 0 | 0 | 0 | 0 |
| ENSMUSG00000100379 | 0 | 0 | 0 | 0 | 0 | 0 | 0 |
| ENSMUSG00000100380 | 0 | 0 | 0 | 0 | 0 | 0 | 0 |
| ENSMUSG00000100381 | 0 | 0 | 0 | 0 | 0 | 0 | 0 |
| ENSMUSG00000100382 | 0 | 0 | 0.25 | 0 | 0 | 0 | 0 |
| ENSMUSG00000100383 | 0.45815691 | 0.25 | 0.25 | 0 | 0 | 0.301029996 | 0 |
| ENSMUSG00000100384 | 0 | 0 | 0 | 0 | 0 | 0 | 0 |
| ENSMUSG00000100385 | 0 | 0 | 0 | 0 | 0 | 0 | 0 |
| ENSMUSG00000100386 | 0 | 0 | 0 | 0 | 0 | 0 | 0 |
| ENSMUSG00000100387 | 0 | 0 | 0 | 0 | 0 | 0 | 0 |
| ENSMUSG00000100388 | 0 | 0 | 0 | 0 | 0 | 0 | 0 |
| ENSMUSG00000100389 | 0 | 0 | 0 | 0 | 0 | 0 | 0 |
| ENSMUSG00000100390 | 0 | 0 | 0 | 0 | 0 | 0 | 0 |
| ENSMUSG00000100391 | 0 | 0 | 0 | 0 | 0 | 0 | 0 |
| ENSMUSG00000100392 | 0 | 0 | 0 | 0 | 0 | 0 | 0 |
| ENSMUSG00000100393 | 0 | 0.396240625 | 0 | 0 | 0.244650542 | 0 | 0 |
| ENSMUSG00000100394 | 0.747221736 | 0.396240625 | 0.864857905 | 0.646240625 | 0.77552387 | 0.602059991 | 0.448487648 |
| ENSMUSG00000100395 | 0 | 0 | 0 | 0 | 0 | 0 | 0 |
| ENSMUSG00000100396 | 0 | 0 | 0.396240625 | 0 | 0.244650542 | 0.301029996 | 0.355418052 |
| ENSMUSG00000100397 | 0 | 0 | 0 | 0 | 0 | 0 | 0 |
| ENSMUSG00000100398 | 0 | 0 | 0 | 0 | 0 | 0 | 0 |
| ENSMUSG00000100399 | 0 | 0 | 0 | 0 | 0 | 0 | 0 |
| ENSMUSG00000100400 | 0 | 0 | 0.25 | 0 | 0 | 0 | 0 |
| ENSMUSG00000100401 | 0 | 0 | 0 | 0 | 0 | 0 | 0 |
| ENSMUSG00000100402 | 0 | 0 | 0 | 0 | 0 | 0 | 0 |
| ENSMUSG00000100403 | 0 | 0 | 0 | 0 | 0 | 0 | 0 |
| ENSMUSG00000100404 | 0 | 0 | 0 | 0 | 0 | 0 | 0 |

| | | | | | | | |
|---|---|---|---|---|---|---|---|
| ENSMUSG00000100405 | 0 | 0 | 0 | 0 | 0 | 0 | 0 |
| ENSMUSG00000100406 | 0 | 0 | 0 | 0 | 0 | 0 | 0 |
| ENSMUSG00000100407 | 0 | 0 | 0 | 0 | 0 | 0 | 0 |
| ENSMUSG00000100408 | 0 | 0 | 0 | 0 | 0 | 0 | 0 |
| ENSMUSG00000100409 | 0 | 0 | 0 | 0 | 0 | 0 | 0 |
| ENSMUSG00000100410 | 0 | 0 | 0 | 0 | 0 | 0 | 0 |
| ENSMUSG00000100411 | 0 | 0 | 0 | 0 | 0 | 0 | 0 |
| ENSMUSG00000100412 | 0 | 0 | 0 | 0 | 0 | 0 | 0 |
| ENSMUSG00000100413 | 0 | 0 | 0 | 0 | 0 | 0 | 0 |
| ENSMUSG00000100415 | 0 | 0 | 0 | 0 | 0 | 0 | 0 |
| ENSMUSG00000100416 | 0.289064826 | 0 | 0 | 0 | 0.244650542 | 0 | 0 |
| ENSMUSG00000100417 | 0.289064826 | 0.25 | 0 | 0.25 | 0 | 0 | 0.355418052 |
| ENSMUSG00000100418 | 0 | 0 | 0 | 0 | 0 | 0 | 0 |
| ENSMUSG00000100420 | 0 | 0 | 0 | 0 | 0 | 0 | 0 |
| ENSMUSG00000100421 | 0 | 0 | 0 | 0 | 0 | 0 | 0 |
| ENSMUSG00000100422 | 0 | 0 | 0 | 0 | 0 | 0 | 0 |
| ENSMUSG00000100423 | 0 | 0 | 0 | 0 | 0 | 0 | 0 |
| ENSMUSG00000100424 | 0 | 0 | 0 | 0 | 0 | 0 | 0 |
| ENSMUSG00000100425 | 0 | 0 | 0 | 0 | 0 | 0 | 0 |
| ENSMUSG00000100426 | 0 | 0 | 0 | 0 | 0.244650542 | 0 | 0 |
| ENSMUSG00000100428 | 0 | 0 | 0 | 0 | 0 | 0 | 0 |
| ENSMUSG00000100429 | 0 | 0 | 0 | 0 | 0 | 0 | 0 |
| ENSMUSG00000100430 | 0 | 0 | 0 | 0 | 0 | 0 | 0 |
| ENSMUSG00000100431 | 0 | 0 | 0 | 0 | 0 | 0 | 0 |
| ENSMUSG00000100432 | 0 | 0 | 0 | 0 | 0 | 0 | 0.224243824 |
| ENSMUSG00000100433 | 0 | 0 | 0 | 0 | 0 | 0 | 0 |
| ENSMUSG00000100434 | 0 | 0 | 0 | 0 | 0 | 0 | 0 |
| ENSMUSG00000100435 | 0 | 0 | 0 | 0 | 0 | 0 | 0 |
| ENSMUSG00000100436 | 0 | 0 | 0 | 0 | 0 | 0 | 0 |
| ENSMUSG00000100437 | 0 | 0 | 0 | 0 | 0 | 0 | 0 |
| ENSMUSG00000100438 | 0 | 0 | 0 | 0 | 0 | 0 | 0 |
| ENSMUSG00000100439 | 0 | 0 | 0 | 0 | 0 | 0 | 0 |
| ENSMUSG00000100440 | 0 | 0 | 0 | 0 | 0 | 0 | 0 |
| ENSMUSG00000100441 | 0 | 0 | 0 | 0.25 | 0 | 0 | 0 |

| | | | | | | | |
|---|---|---|---|---|---|---|---|
| ENSMUSG00000100442 | 0 | 0 | 0 | 0 | 0 | 0 | 0 |
| ENSMUSG00000100443 | 0 | 0 | 0 | 0 | 0 | 0 | 0 |
| ENSMUSG00000100444 | 0.289064826 | 0.5 | 0.5 | 0 | 0 | 0 | 0 |
| ENSMUSG00000100445 | 0 | 0 | 0 | 0 | 0 | 0 | 0 |
| ENSMUSG00000100446 | 0 | 0 | 0 | 0 | 0 | 0 | 0 |
| ENSMUSG00000100447 | 0 | 0 | 0 | 0 | 0 | 0 | 0 |
| ENSMUSG00000100448 | 0 | 0 | 0 | 0 | 0 | 0 | 0 |
| ENSMUSG00000100449 | 0 | 0 | 0 | 0 | 0 | 0 | 0 |
| ENSMUSG00000100450 | 0 | 0 | 0 | 0 | 0 | 0 | 0 |
| ENSMUSG00000100451 | 0 | 0 | 0 | 0 | 0 | 0 | 0 |
| ENSMUSG00000100452 | 0 | 0 | 0 | 0 | 0 | 0 | 0 |
| ENSMUSG00000100453 | 0 | 0 | 0 | 0 | 0 | 0 | 0 |
| ENSMUSG00000100454 | 0 | 0 | 0 | 0 | 0 | 0 | 0 |
| ENSMUSG00000100455 | 1.156259305 | 1.226722649 | 1.17510993 | 1.396240625 | 1.200473444 | 1.204119983 | 1.272006351 |
| ENSMUSG00000100456 | 0 | 0 | 0 | 0 | 0 | 0 | 0 |
| ENSMUSG00000100457 | 0 | 0 | 0 | 0 | 0 | 0 | 0 |
| ENSMUSG00000100458 | 0 | 0 | 0 | 0 | 0 | 0 | 0 |
| ENSMUSG00000100459 | 0 | 0 | 0 | 0 | 0 | 0 | 0 |
| ENSMUSG00000100460 | 0 | 0 | 0 | 0 | 0 | 0 | 0 |
| ENSMUSG00000100461 | 0 | 0 | 0 | 0 | 0 | 0 | 0 |
| ENSMUSG00000100462 | 0 | 0 | 0 | 0 | 0 | 0 | 0 |
| ENSMUSG00000100463 | 0 | 0 | 0 | 0 | 0 | 0 | 0 |
| ENSMUSG00000100464 | 0 | 0 | 0 | 0 | 0 | 0 | 0 |
| ENSMUSG00000100465 | 0 | 0 | 0 | 0 | 0.244650542 | 0 | 0 |
| ENSMUSG00000100466 | 0 | 0 | 0 | 0 | 0 | 0 | 0 |
| ENSMUSG00000100467 | 0 | 0 | 0 | 0 | 0 | 0 | 0 |
| ENSMUSG00000100468 | 0 | 0 | 0 | 0 | 0 | 0 | 0 |
| ENSMUSG00000100469 | 0 | 0 | 0 | 0 | 0 | 0 | 0 |
| ENSMUSG00000100470 | 0 | 0 | 0 | 0 | 0 | 0 | 0 |
| ENSMUSG00000100471 | 0 | 0 | 0 | 0 | 0 | 0 | 0 |
| ENSMUSG00000100472 | 0 | 0 | 0 | 0 | 0 | 0 | 0 |
| ENSMUSG00000100473 | 0 | 0 | 0 | 0 | 0 | 0 | 0 |
| ENSMUSG00000100474 | 0 | 0 | 0 | 0 | 0 | 0 | 0 |
| ENSMUSG00000100475 | 0 | 0 | 0 | 0 | 0 | 0 | 0 |

| | | | | | | | |
|---|---|---|---|---|---|---|---|
| ENSMUSG00000100476 | 0 | 0 | 0 | 0 | 0 | 0 | 0 |
| ENSMUSG00000100477 | 0 | 0 | 0 | 0 | 0 | 0 | 0 |
| ENSMUSG00000100478 | 0 | 0 | 0 | 0 | 0 | 0 | 0 |
| ENSMUSG00000100479 | 0 | 0 | 0 | 0 | 0 | 0 | 0 |
| ENSMUSG00000100480 | 0 | 0.396240625 | 0.5 | 0.75 | 0.632412477 | 0.698970004 | 0.710836105 |
| ENSMUSG00000100481 | 0 | 0.25 | 0 | 0 | 0 | 0 | 0 |
| ENSMUSG00000100483 | 0 | 0 | 0.25 | 0.25 | 0.244650542 | 0 | 0 |
| ENSMUSG00000100484 | 0 | 0 | 0 | 0 | 0 | 0 | 0 |
| ENSMUSG00000100485 | 0 | 0 | 0 | 0 | 0 | 0 | 0 |
| ENSMUSG00000100486 | 0 | 0 | 0 | 0 | 0 | 0 | 0 |
| ENSMUSG00000100487 | 0 | 0 | 0 | 0 | 0 | 0 | 0 |
| ENSMUSG00000100488 | 0 | 0 | 0 | 0 | 0 | 0 | 0 |
| ENSMUSG00000100489 | 0 | 0 | 0 | 0 | 0 | 0 | 0 |
| ENSMUSG00000100490 | 0 | 0 | 0.25 | 0 | 0.244650542 | 0.477121255 | 0.448487648 |
| ENSMUSG00000100491 | 0 | 0 | 0 | 0 | 0 | 0 | 0 |
| ENSMUSG00000100492 | 0 | 0 | 0 | 0 | 0 | 0 | 0 |
| ENSMUSG00000100494 | 0 | 0 | 0 | 0 | 0 | 0 | 0 |
| ENSMUSG00000100495 | 0 | 0 | 0 | 0 | 0 | 0 | 0 |
| ENSMUSG00000100496 | 0 | 0 | 0 | 0 | 0 | 0 | 0 |
| ENSMUSG00000100497 | 0 | 0 | 0 | 0 | 0 | 0 | 0 |
| ENSMUSG00000100499 | 0 | 0 | 0 | 0 | 0 | 0 | 0 |
| ENSMUSG00000100500 | 0 | 0 | 0 | 0 | 0 | 0 | 0 |
| ENSMUSG00000100501 | 0 | 0 | 0 | 0 | 0 | 0 | 0 |
| ENSMUSG00000100502 | 0 | 0 | 0 | 0 | 0 | 0 | 0 |
| ENSMUSG00000100503 | 0 | 0 | 0 | 0 | 0 | 0 | 0 |
| ENSMUSG00000100504 | 0 | 0 | 0 | 0 | 0 | 0 | 0 |
| ENSMUSG00000100505 | 0 | 0 | 0 | 0 | 0 | 0 | 0 |
| ENSMUSG00000100506 | 0 | 0 | 0 | 0 | 0 | 0 | 0 |
| ENSMUSG00000100507 | 0 | 0 | 0 | 0 | 0 | 0 | 0 |
| ENSMUSG00000100508 | 0 | 0 | 0 | 0 | 0 | 0 | 0 |
| ENSMUSG00000100509 | 0 | 0 | 0 | 0 | 0 | 0 | 0 |
| ENSMUSG00000100510 | 0.289064826 | 0.25 | 0.5 | 0 | 0 | 0.602059991 | 0 |
| ENSMUSG00000100511 | 0 | 0 | 0 | 0 | 0 | 0 | 0 |
| ENSMUSG00000100512 | 0 | 0 | 0 | 0 | 0 | 0 | 0 |

| | | | | | | | |
|---|---|---|---|---|---|---|---|
| ENSMUSG00000100513 | 0 | 0 | 0 | 0 | 0 | 0 | 0 |
| ENSMUSG00000100514 | 0.45815691 | 0 | 0 | 0 | 0 | 0 | 0 |
| ENSMUSG00000100515 | 0 | 0 | 0 | 0 | 0 | 0 | 0 |
| ENSMUSG00000100516 | 0 | 0 | 0 | 0 | 0 | 0 | 0 |
| ENSMUSG00000100517 | 0 | 0 | 0 | 0 | 0 | 0 | 0 |
| ENSMUSG00000100518 | 0 | 0 | 0 | 0 | 0 | 0 | 0 |
| ENSMUSG00000100519 | 0 | 0 | 0 | 0 | 0 | 0 | 0 |
| ENSMUSG00000100520 | 0 | 0 | 0 | 0 | 0 | 0 | 0 |
| ENSMUSG00000100521 | 0 | 0 | 0 | 0 | 0 | 0 | 0 |
| ENSMUSG00000100522 | 0 | 0 | 0 | 0 | 0 | 0 | 0 |
| ENSMUSG00000100523 | 0 | 0 | 0 | 0 | 0 | 0 | 0 |
| ENSMUSG00000100524 | 0 | 0 | 0 | 0 | 0 | 0 | 0 |
| ENSMUSG00000100525 | 0 | 0 | 0 | 0 | 0 | 0 | 0 |
| ENSMUSG00000100526 | 0 | 0 | 0 | 0 | 0 | 0 | 0 |
| ENSMUSG00000100527 | 0 | 0 | 0 | 0 | 0 | 0 | 0 |
| ENSMUSG00000100528 | 0 | 0 | 0 | 0 | 0 | 0 | 0 |
| ENSMUSG00000100529 | 0 | 0 | 0 | 0 | 0 | 0 | 0 |
| ENSMUSG00000100530 | 0 | 0 | 0 | 0 | 0 | 0 | 0 |
| ENSMUSG00000100531 | 0 | 0 | 0 | 0 | 0 | 0 | 0 |
| ENSMUSG00000100532 | 0 | 0 | 0 | 0 | 0 | 0 | 0 |
| ENSMUSG00000100533 | 0 | 0 | 0 | 0 | 0 | 0 | 0 |
| ENSMUSG00000100534 | 0 | 0.5 | 0.396240625 | 0 | 0.244650542 | 0 | 0 |
| ENSMUSG00000100535 | 0 | 0 | 0 | 0 | 0 | 0 | 0 |
| ENSMUSG00000100536 | 0 | 0 | 0 | 0.25 | 0.387761935 | 0.301029996 | 0.448487648 |
| ENSMUSG00000100537 | 0 | 0 | 0 | 0 | 0 | 0 | 0 |
| ENSMUSG00000100538 | 0 | 0 | 0 | 0 | 0 | 0 | 0 |
| ENSMUSG00000100539 | 0 | 0 | 0 | 0 | 0 | 0 | 0 |
| ENSMUSG00000100540 | 0 | 0 | 0 | 0 | 0 | 0 | 0 |
| ENSMUSG00000100541 | 0 | 0 | 0 | 0 | 0 | 0 | 0 |
| ENSMUSG00000100542 | 0 | 0 | 0 | 0 | 0 | 0 | 0 |
| ENSMUSG00000100543 | 0 | 0 | 0 | 0 | 0 | 0 | 0 |
| ENSMUSG00000100544 | 0 | 0 | 0 | 0 | 0 | 0 | 0 |
| ENSMUSG00000100546 | 0 | 0 | 0 | 0 | 0 | 0 | 0 |
| ENSMUSG00000100547 | 0 | 0 | 0 | 0 | 0 | 0 | 0 |

| | | | | | | | |
|---|---|---|---|---|---|---|---|
| ENSMUSG00000100548 | 0.45815691 | 0.580482024 | 0.701838731 | 0.580482024 | 0.568060967 | 0.477121255 | 0.579661877 |
| ENSMUSG00000100549 | 0 | 0 | 0 | 0 | 0 | 0 | 0 |
| ENSMUSG00000100550 | 0.289064826 | 0.25 | 0.396240625 | 0.396240625 | 0 | 0.301029996 | 0 |
| ENSMUSG00000100551 | 0 | 0 | 0 | 0.25 | 0 | 0 | 0 |
| ENSMUSG00000100552 | 0.289064826 | 0 | 0 | 0 | 0.244650542 | 0.301029996 | 0 |
| ENSMUSG00000100553 | 0 | 0 | 0 | 0 | 0 | 0 | 0 |
| ENSMUSG00000100554 | 0 | 0 | 0 | 0 | 0 | 0 | 0.224243824 |
| ENSMUSG00000100555 | 0.289064826 | 0 | 0 | 0 | 0.244650542 | 0 | 0.224243824 |
| ENSMUSG00000100556 | 0 | 0 | 0 | 0 | 0 | 0 | 0 |
| ENSMUSG00000100557 | 0 | 0 | 0 | 0 | 0 | 0 | 0 |
| ENSMUSG00000100558 | 0 | 0 | 0 | 0 | 0 | 0 | 0 |
| ENSMUSG00000100559 | 0 | 0 | 0 | 0 | 0 | 0 | 0 |
| ENSMUSG00000100560 | 0 | 0 | 0 | 0 | 0 | 0 | 0 |
| ENSMUSG00000100561 | 0.747221736 | 0.92510993 | 1 | 0.646240625 | 0.931471446 | 0.477121255 | 0.916588299 |
| ENSMUSG00000100562 | 0 | 0 | 0 | 0 | 0 | 0 | 0 |
| ENSMUSG00000100563 | 0 | 0 | 0 | 0 | 0 | 0 | 0 |
| ENSMUSG00000100564 | 0 | 0 | 0 | 0 | 0 | 0 | 0 |
| ENSMUSG00000100565 | 0 | 0 | 0 | 0 | 0 | 0 | 0 |
| ENSMUSG00000100566 | 0 | 0 | 0 | 0 | 0 | 0 | 0 |
| ENSMUSG00000100567 | 0 | 0 | 0 | 0 | 0 | 0 | 0 |
| ENSMUSG00000100568 | 0 | 0 | 0 | 0.25 | 0.244650542 | 0 | 0 |
| ENSMUSG00000100569 | 0 | 0 | 0 | 0 | 0 | 0 | 0 |
| ENSMUSG00000100570 | 0 | 0 | 0 | 0 | 0 | 0 | 0 |
| ENSMUSG00000100571 | 0 | 0 | 0 | 0 | 0 | 0 | 0 |
| ENSMUSG00000100572 | 0.671187741 | 0.701838731 | 0.701838731 | 0.701838731 | 0.632412477 | 0.698970004 | 0.448487648 |
| ENSMUSG00000100573 | 0 | 0 | 0 | 0 | 0 | 0 | 0 |
| ENSMUSG00000100574 | 0 | 0 | 0 | 0 | 0 | 0 | 0 |
| ENSMUSG00000100575 | 0 | 0 | 0 | 0 | 0 | 0 | 0 |
| ENSMUSG00000100576 | 0 | 0 | 0 | 0 | 0 | 0 | 0 |
| ENSMUSG00000100577 | 0 | 0 | 0 | 0 | 0 | 0 | 0 |
| ENSMUSG00000100578 | 0 | 0 | 0 | 0 | 0 | 0 | 0 |
| ENSMUSG00000100579 | 0 | 0 | 0 | 0 | 0 | 0 | 0 |
| ENSMUSG00000100580 | 0.747221736 | 0.5 | 0.580482024 | 0.580482024 | 0.244650542 | 0.301029996 | 0.520678036 |
| ENSMUSG00000100581 | 0 | 0 | 0 | 0 | 0 | 0 | 0 |

| | | | | | | | |
|---|---|---|---|---|---|---|---|
| ENSMUSG00000100582 | 0 | 0 | 0 | 0 | 0 | 0 | 0 |
| ENSMUSG00000100583 | 0 | 0 | 0.25 | 0 | 0 | 0 | 0.224243824 |
| ENSMUSG00000100584 | 0 | 0 | 0 | 0 | 0 | 0 | 0 |
| ENSMUSG00000100585 | 0 | 0 | 0 | 0 | 0 | 0 | 0 |
| ENSMUSG00000100586 | 0 | 0 | 0 | 0 | 0 | 0 | 0 |
| ENSMUSG00000100587 | 0 | 0 | 0 | 0 | 0 | 0 | 0 |
| ENSMUSG00000100588 | 0 | 0 | 0 | 0 | 0 | 0 | 0 |
| ENSMUSG00000100589 | 0 | 0 | 0 | 0 | 0 | 0 | 0 |
| ENSMUSG00000100590 | 0 | 0 | 0 | 0 | 0 | 0 | 0 |
| ENSMUSG00000100591 | 0 | 0 | 0 | 0 | 0 | 0 | 0 |
| ENSMUSG00000100592 | 0 | 0 | 0 | 0 | 0 | 0 | 0 |
| ENSMUSG00000100593 | 0 | 0 | 0 | 0.25 | 0 | 0 | 0 |
| ENSMUSG00000100594 | 0.811507563 | 1 | 0.830482024 | 0.864857905 | 0.733951626 | 0.84509804 | 0.579661877 |
| ENSMUSG00000100595 | 0 | 0 | 0 | 0.25 | 0 | 0 | 0.224243824 |
| ENSMUSG00000100596 | 0 | 0 | 0 | 0 | 0 | 0 | 0.224243824 |
| ENSMUSG00000100597 | 0 | 0 | 0 | 0 | 0 | 0 | 0 |
| ENSMUSG00000100598 | 0 | 0 | 0 | 0 | 0 | 0 | 0 |
| ENSMUSG00000100599 | 0.867194479 | 0.92510993 | 0.864857905 | 0.580482024 | 0.77552387 | 1 | 0.829800754 |
| ENSMUSG00000100600 | 0.811507563 | 0.580482024 | 0.701838731 | 0.92510993 | 0.686820904 | 0.477121255 | 0.775756176 |
| ENSMUSG00000100601 | 0 | 0 | 0 | 0 | 0 | 0 | 0 |
| ENSMUSG00000100602 | 0.289064826 | 0 | 0 | 0.25 | 0.244650542 | 0 | 0 |
| ENSMUSG00000100603 | 0 | 0 | 0 | 0 | 0 | 0 | 0 |
| ENSMUSG00000100604 | 0 | 0 | 0 | 0 | 0 | 0 | 0 |
| ENSMUSG00000100605 | 0 | 0 | 0 | 0.25 | 0 | 0 | 0 |
| ENSMUSG00000100606 | 0 | 0 | 0 | 0 | 0 | 0 | 0 |
| ENSMUSG00000100607 | 0 | 0 | 0 | 0 | 0 | 0 | 0 |
| ENSMUSG00000100608 | 0 | 0 | 0 | 0 | 0 | 0 | 0 |
| ENSMUSG00000100609 | 0 | 0 | 0 | 0 | 0 | 0 | 0 |
| ENSMUSG00000100610 | 0 | 0 | 0 | 0 | 0 | 0 | 0 |
| ENSMUSG00000100611 | 0 | 0 | 0 | 0 | 0 | 0 | 0 |
| ENSMUSG00000100612 | 0 | 0 | 0 | 0 | 0 | 0 | 0 |
| ENSMUSG00000100613 | 0 | 0 | 0 | 0 | 0 | 0 | 0 |
| ENSMUSG00000100614 | 0 | 0 | 0 | 0 | 0 | 0 | 0 |
| ENSMUSG00000100615 | 0 | 0.580482024 | 0.25 | 0.396240625 | 0.686820904 | 0.602059991 | 0.579661877 |

| | | | | | | | |
|---|---|---|---|---|---|---|---|
| ENSMUSG00000100616 | 0 | 0 | 0 | 0 | 0 | 0 | 0 |
| ENSMUSG00000100617 | 0 | 0 | 0 | 0 | 0 | 0 | 0 |
| ENSMUSG00000100618 | 0 | 0 | 0 | 0 | 0 | 0 | 0 |
| ENSMUSG00000100619 | 0.289064826 | 0.25 | 0.396240625 | 0.25 | 0.489301084 | 0.301029996 | 0.224243824 |
| ENSMUSG00000100620 | 0 | 0.25 | 0.5 | 0.5 | 0 | 0 | 0.355418052 |
| ENSMUSG00000100621 | 0 | 0 | 0 | 0 | 0 | 0 | 0 |
| ENSMUSG00000100622 | 1.181541736 | 1.311981878 | 1.061981878 | 1.476722649 | 1.504538967 | 1.643452676 | 1.459332757 |
| ENSMUSG00000100623 | 0 | 0 | 0 | 0 | 0 | 0 | 0 |
| ENSMUSG00000100624 | 0 | 0 | 0 | 0 | 0 | 0 | 0 |
| ENSMUSG00000100625 | 0 | 0 | 0 | 0 | 0 | 0 | 0 |
| ENSMUSG00000100626 | 0 | 0 | 0 | 0 | 0 | 0 | 0 |
| ENSMUSG00000100627 | 0 | 0.5 | 0.25 | 0.25 | 0.244650542 | 0.301029996 | 0.520678036 |
| ENSMUSG00000100628 | 0 | 0 | 0 | 0 | 0 | 0 | 0 |
| ENSMUSG00000100629 | 1.181541736 | 1.061981878 | 1.04248125 | 0.976722649 | 1.188507684 | 0.954242509 | 1.066254157 |
| ENSMUSG00000100630 | 0 | 0 | 0 | 0 | 0 | 0 | 0 |
| ENSMUSG00000100631 | 0 | 0 | 0 | 0 | 0 | 0 | 0 |
| ENSMUSG00000100632 | 0 | 0.25 | 0.396240625 | 0.5 | 0.387761935 | 0.477121255 | 0.355418052 |
| ENSMUSG00000100634 | 0 | 0 | 0 | 0 | 0 | 0 | 0 |
| ENSMUSG00000100635 | 0 | 0.5 | 0.25 | 0.25 | 0 | 0.602059991 | 0.224243824 |
| ENSMUSG00000100636 | 0 | 0 | 0.25 | 0 | 0 | 0 | 0 |
| ENSMUSG00000100637 | 0 | 0 | 0 | 0 | 0 | 0 | 0 |
| ENSMUSG00000100639 | 0 | 0 | 0 | 0 | 0 | 0 | 0 |
| ENSMUSG00000100640 | 0 | 0 | 0 | 0 | 0 | 0 | 0 |
| ENSMUSG00000100641 | 0 | 0 | 0 | 0 | 0 | 0 | 0 |
| ENSMUSG00000100642 | 0.578129653 | 0.5 | 0.646240625 | 0 | 0 | 0 | 0 |
| ENSMUSG00000100643 | 0 | 0 | 0 | 0 | 0 | 0 | 0 |
| ENSMUSG00000100644 | 0 | 0 | 0 | 0 | 0 | 0 | 0 |
| ENSMUSG00000100645 | 0 | 0 | 0 | 0 | 0 | 0 | 0 |
| ENSMUSG00000100646 | 0 | 0 | 0 | 0 | 0 | 0 | 0 |
| ENSMUSG00000100647 | 0 | 0 | 0 | 0 | 0 | 0 | 0 |
| ENSMUSG00000100648 | 0.747221736 | 0 | 0.25 | 0 | 0.244650542 | 0.602059991 | 0.448487648 |
| ENSMUSG00000100649 | 0 | 0 | 0 | 0 | 0 | 0 | 0 |
| ENSMUSG00000100650 | 0 | 0 | 0 | 0 | 0 | 0 | 0 |
| ENSMUSG00000100651 | 0 | 0 | 0 | 0 | 0 | 0 | 0 |

| | | | | | | | |
|---|---|---|---|---|---|---|---|
| ENSMUSG00000100652 | 0.747221736 | 0.580482024 | 0.646240625 | 0.701838731 | 0.686820904 | 0.602059991 | 0.520678036 |
| ENSMUSG00000100653 | 0 | 0 | 0 | 0 | 0 | 0 | 0 |
| ENSMUSG00000100654 | 0 | 0 | 0 | 0 | 0 | 0 | 0 |
| ENSMUSG00000100655 | 0 | 0 | 0 | 0 | 0 | 0 | 0 |
| ENSMUSG00000100656 | 0 | 0 | 0 | 0 | 0 | 0 | 0 |
| ENSMUSG00000100657 | 0 | 0 | 0 | 0 | 0 | 0 | 0 |
| ENSMUSG00000100658 | 0.578129653 | 0.864857905 | 0.951838731 | 0.646240625 | 0.733951626 | 0.301029996 | 0.355418052 |
| ENSMUSG00000100659 | 0 | 0 | 0 | 0 | 0 | 0 | 0 |
| ENSMUSG00000100660 | 0 | 0 | 0 | 0 | 0 | 0 | 0 |
| ENSMUSG00000100661 | 0 | 0 | 0 | 0 | 0 | 0 | 0 |
| ENSMUSG00000100662 | 0 | 0.25 | 0 | 0 | 0 | 0 | 0.224243824 |
| ENSMUSG00000100663 | 0 | 0 | 0 | 0 | 0 | 0 | 0 |
| ENSMUSG00000100664 | 0 | 0 | 0 | 0 | 0 | 0 | 0 |
| ENSMUSG00000100665 | 0 | 0 | 0 | 0 | 0 | 0 | 0 |
| ENSMUSG00000100666 | 0 | 0 | 0 | 0 | 0 | 0 | 0 |
| ENSMUSG00000100667 | 0 | 0 | 0 | 0 | 0 | 0 | 0 |
| ENSMUSG00000100668 | 0 | 0 | 0 | 0 | 0 | 0 | 0 |
| ENSMUSG00000100669 | 0 | 0 | 0 | 0 | 0 | 0 | 0 |
| ENSMUSG00000100670 | 0 | 0 | 0 | 0 | 0 | 0 | 0 |
| ENSMUSG00000100671 | 0 | 0 | 0 | 0 | 0 | 0 | 0 |
| ENSMUSG00000100672 | 0.289064826 | 0.396240625 | 0.396240625 | 0.646240625 | 0.387761935 | 0.477121255 | 0.520678036 |
| ENSMUSG00000100673 | 0 | 0 | 0 | 0 | 0 | 0 | 0 |
| ENSMUSG00000100674 | 0 | 0 | 0 | 0 | 0 | 0 | 0 |
| ENSMUSG00000100675 | 0 | 0 | 0 | 0 | 0 | 0 | 0 |
| ENSMUSG00000100676 | 0 | 0 | 0 | 0 | 0 | 0 | 0 |
| ENSMUSG00000100677 | 0 | 0 | 0 | 0 | 0 | 0 | 0 |
| ENSMUSG00000100678 | 0 | 0 | 0 | 0 | 0 | 0 | 0 |
| ENSMUSG00000100679 | 0 | 0 | 0 | 0 | 0 | 0 | 0 |
| ENSMUSG00000100680 | 0.578129653 | 0.75 | 0.79248125 | 0.646240625 | 0.632412477 | 0.301029996 | 0.520678036 |
| ENSMUSG00000100681 | 0 | 0 | 0 | 0 | 0 | 0 | 0 |
| ENSMUSG00000100682 | 0 | 0 | 0 | 0 | 0 | 0 | 0 |
| ENSMUSG00000100683 | 0 | 0 | 0 | 0 | 0 | 0 | 0 |
| ENSMUSG00000100684 | 0 | 0 | 0 | 0 | 0 | 0 | 0 |
| ENSMUSG00000100685 | 0 | 0 | 0 | 0 | 0.244650542 | 0 | 0 |

| | | | | | | | |
|---|---|---|---|---|---|---|---|
| ENSMUSG00000100687 | 0 | 0 | 0 | 0 | 0 | 0 | 0 |
| ENSMUSG00000100689 | 0 | 0 | 0 | 0 | 0 | 0 | 0 |
| ENSMUSG00000100690 | 0 | 0 | 0 | 0 | 0 | 0 | 0 |
| ENSMUSG00000100691 | 1.289064826 | 1.330482024 | 1.418106335 | 0.951838731 | 1.254881871 | 1.176091259 | 1.302263476 |
| ENSMUSG00000100692 | 0 | 0 | 0 | 0 | 0 | 0 | 0 |
| ENSMUSG00000100693 | 0 | 0 | 0 | 0 | 0 | 0 | 0 |
| ENSMUSG00000100694 | 0 | 0 | 0 | 0 | 0 | 0 | 0 |
| ENSMUSG00000100695 | 0 | 0 | 0 | 0 | 0 | 0 | 0 |
| ENSMUSG00000100696 | 0 | 0 | 0 | 0.25 | 0.244650542 | 0.301029996 | 0 |
| ENSMUSG00000100697 | 0 | 0 | 0 | 0 | 0 | 0 | 0 |
| ENSMUSG00000100698 | 0 | 0 | 0 | 0 | 0 | 0 | 0 |
| ENSMUSG00000100699 | 0 | 0 | 0 | 0 | 0 | 0 | 0 |
| ENSMUSG00000100700 | 0 | 0 | 0 | 0 | 0 | 0 | 0 |
| ENSMUSG00000100701 | 1.181541736 | 1.080482024 | 1.214495249 | 1.356566189 | 1.327538614 | 1.431363764 | 1.238624656 |
| ENSMUSG00000100702 | 0.747221736 | 0.864857905 | 0.701838731 | 0.25 | 0.812711509 | 0.602059991 | 0.448487648 |
| ENSMUSG00000100703 | 0 | 0 | 0 | 0 | 0 | 0 | 0 |
| ENSMUSG00000100704 | 0 | 0 | 0 | 0 | 0 | 0 | 0 |
| ENSMUSG00000100705 | 0 | 0 | 0 | 0 | 0 | 0 | 0 |
| ENSMUSG00000100706 | 0 | 0 | 0 | 0.79248125 | 0.387761935 | 0.477121255 | 0.629532004 |
| ENSMUSG00000100707 | 0 | 0 | 0 | 0 | 0 | 0 | 0 |
| ENSMUSG00000100708 | 0 | 0 | 0 | 0 | 0 | 0 | 0 |
| ENSMUSG00000100709 | 0 | 0 | 0 | 0 | 0 | 0 | 0 |
| ENSMUSG00000100710 | 0 | 0 | 0 | 0 | 0 | 0 | 0 |
| ENSMUSG00000100711 | 0 | 0 | 0 | 0 | 0 | 0 | 0 |
| ENSMUSG00000100712 | 0 | 0 | 0 | 0 | 0 | 0 | 0 |
| ENSMUSG00000100713 | 0 | 0 | 0 | 0 | 0 | 0 | 0 |
| ENSMUSG00000100714 | 0 | 0.396240625 | 0.396240625 | 0 | 0 | 0 | 0 |
| ENSMUSG00000100715 | 0 | 0 | 0 | 0 | 0 | 0 | 0 |
| ENSMUSG00000100716 | 0 | 0 | 0 | 0 | 0 | 0 | 0 |
| ENSMUSG00000100717 | 0 | 0 | 0 | 0 | 0 | 0 | 0 |
| ENSMUSG00000100718 | 0 | 0 | 0 | 0 | 0 | 0 | 0 |
| ENSMUSG00000100719 | 0 | 0 | 0 | 0 | 0 | 0 | 0 |
| ENSMUSG00000100720 | 0 | 0.25 | 0 | 0.396240625 | 0 | 0 | 0 |
| ENSMUSG00000100721 | 0 | 0 | 0 | 0 | 0 | 0 | 0 |

| | | | | | | | |
|---|---|---|---|---|---|---|---|
| ENSMUSG00000100722 | 0 | 0 | 0 | 0 | 0 | 0 | 0 |
| ENSMUSG00000100723 | 0 | 0 | 0 | 0 | 0 | 0 | 0 |
| ENSMUSG00000100724 | 0 | 0 | 0 | 0 | 0 | 0 | 0 |
| ENSMUSG00000100725 | 0 | 0 | 0 | 0 | 0 | 0 | 0 |
| ENSMUSG00000100726 | 0 | 0 | 0 | 0 | 0 | 0 | 0 |
| ENSMUSG00000100727 | 0 | 0 | 0 | 0 | 0 | 0 | 0 |
| ENSMUSG00000100728 | 0 | 0 | 0 | 0 | 0 | 0 | 0 |
| ENSMUSG00000100729 | 0 | 0 | 0 | 0 | 0 | 0 | 0 |
| ENSMUSG00000100730 | 0 | 0 | 0 | 0 | 0 | 0 | 0 |
| ENSMUSG00000100731 | 0 | 0 | 0 | 0 | 0 | 0 | 0 |
| ENSMUSG00000100732 | 0 | 0 | 0 | 0 | 0 | 0 | 0 |
| ENSMUSG00000100733 | 0.289064826 | 0 | 0 | 0 | 0 | 0 | 0 |
| ENSMUSG00000100734 | 0 | 0 | 0 | 0 | 0 | 0 | 0 |
| ENSMUSG00000100736 | 0.747221736 | 0.25 | 0.646240625 | 0.396240625 | 0 | 0.602059991 | 0.448487648 |
| ENSMUSG00000100737 | 0 | 0 | 0 | 0 | 0 | 0 | 0 |
| ENSMUSG00000100738 | 0 | 0 | 0 | 0 | 0 | 0 | 0 |
| ENSMUSG00000100739 | 0 | 0 | 0 | 0 | 0 | 0 | 0 |
| ENSMUSG00000100740 | 0 | 0 | 0 | 0 | 0 | 0 | 0 |
| ENSMUSG00000100741 | 0 | 0 | 0 | 0 | 0 | 0 | 0 |
| ENSMUSG00000100742 | 0 | 0 | 0 | 0 | 0 | 0 | 0 |
| ENSMUSG00000100743 | 0 | 0 | 0 | 0 | 0 | 0 | 0 |
| ENSMUSG00000100744 | 0 | 0 | 0.25 | 0 | 0.244650542 | 0 | 0.224243824 |
| ENSMUSG00000100745 | 0 | 0 | 0 | 0 | 0 | 0 | 0 |
| ENSMUSG00000100746 | 0 | 0 | 0 | 0 | 0 | 0 | 0 |
| ENSMUSG00000100747 | 0.289064826 | 0.646240625 | 0.25 | 0 | 0 | 0 | 0.355418052 |
| ENSMUSG00000100748 | 0 | 0 | 0 | 0 | 0 | 0 | 0 |
| ENSMUSG00000100749 | 0 | 0 | 0 | 0 | 0 | 0 | 0 |
| ENSMUSG00000100750 | 1.470606563 | 1.418106335 | 1.364857905 | 1.27186571 | 1.27449559 | 1.204119983 | 1.168187745 |
| ENSMUSG00000100751 | 0 | 0 | 0 | 0 | 0 | 0 | 0 |
| ENSMUSG00000100752 | 0 | 0 | 0 | 0 | 0 | 0 | 0 |
| ENSMUSG00000100753 | 0 | 0 | 0 | 0 | 0 | 0 | 0 |
| ENSMUSG00000100754 | 0 | 0 | 0 | 0 | 0 | 0 | 0 |
| ENSMUSG00000100755 | 0 | 0 | 0 | 0 | 0 | 0 | 0 |
| ENSMUSG00000100756 | 0 | 0 | 0 | 0 | 0 | 0 | 0 |

| | | | | | | | |
|---|---|---|---|---|---|---|---|
| ENSMUSG00000100757 | 0 | 0 | 0 | 0 | 0 | 0 | 0 |
| ENSMUSG00000100758 | 0 | 0 | 0 | 0 | 0 | 0 | 0 |
| ENSMUSG00000100759 | 0 | 0 | 0 | 0 | 0 | 0 | 0 |
| ENSMUSG00000100760 | 0 | 0 | 0 | 0 | 0 | 0 | 0 |
| ENSMUSG00000100761 | 0 | 0 | 0 | 0 | 0 | 0 | 0 |
| ENSMUSG00000100762 | 0 | 0 | 0 | 0 | 0 | 0 | 0 |
| ENSMUSG00000100763 | 0 | 0 | 0.25 | 0.25 | 0.387761935 | 0.301029996 | 0 |
| ENSMUSG00000100764 | 0 | 0 | 0 | 0.5 | 0.489301084 | 0.477121255 | 0.448487648 |
| ENSMUSG00000100765 | 0 | 0 | 0 | 0 | 0 | 0 | 0 |
| ENSMUSG00000100767 | 0 | 0 | 0.396240625 | 0.25 | 0 | 0 | 0 |
| ENSMUSG00000100768 | 1 | 1.061981878 | 1.114857905 | 0.92510993 | 1.091002363 | 1.041392685 | 0.803905701 |
| ENSMUSG00000100769 | 0.578129653 | 0.25 | 0.25 | 0.396240625 | 0.568060967 | 0 | 0.224243824 |
| ENSMUSG00000100770 | 0 | 0 | 0 | 0 | 0 | 0 | 0 |
| ENSMUSG00000100771 | 0 | 0 | 0 | 0 | 0 | 0 | 0 |
| ENSMUSG00000100772 | 0 | 0 | 0 | 0 | 0 | 0 | 0 |
| ENSMUSG00000100773 | 0 | 0 | 0 | 0 | 0 | 0 | 0 |
| ENSMUSG00000100774 | 0 | 0 | 0.25 | 0 | 0.244650542 | 0 | 0 |
| ENSMUSG00000100775 | 0.289064826 | 0.396240625 | 0.5 | 0.396240625 | 0.387761935 | 0.301029996 | 0.520678036 |
| ENSMUSG00000100776 | 0 | 0 | 0 | 0 | 0 | 0 | 0 |
| ENSMUSG00000100777 | 0 | 0 | 0 | 0 | 0 | 0 | 0 |
| ENSMUSG00000100778 | 0 | 0 | 0 | 0 | 0 | 0 | 0 |
| ENSMUSG00000100779 | 0.289064826 | 0 | 0.25 | 0.5 | 0 | 0 | 0 |
| ENSMUSG00000100780 | 0 | 0 | 0 | 0 | 0 | 0 | 0 |
| ENSMUSG00000100781 | 0 | 0 | 0 | 0 | 0 | 0.301029996 | 0 |
| ENSMUSG00000100782 | 0.289064826 | 0 | 0.25 | 0 | 0 | 0 | 0 |
| ENSMUSG00000100783 | 0.289064826 | 0 | 0.25 | 0.25 | 0.244650542 | 0 | 0 |
| ENSMUSG00000100784 | 0 | 0 | 0 | 0 | 0 | 0 | 0 |
| ENSMUSG00000100785 | 0 | 0 | 0 | 0 | 0 | 0 | 0 |
| ENSMUSG00000100786 | 0 | 0 | 0 | 0 | 0 | 0 | 0 |
| ENSMUSG00000100787 | 0 | 0 | 0 | 0 | 0 | 0 | 0 |
| ENSMUSG00000100788 | 0 | 0 | 0 | 0 | 0 | 0 | 0 |
| ENSMUSG00000100789 | 0 | 0 | 0 | 0.25 | 0.244650542 | 0 | 0 |
| ENSMUSG00000100790 | 0 | 0 | 0 | 0 | 0 | 0 | 0 |
| ENSMUSG00000100791 | 0 | 0 | 0 | 0 | 0 | 0 | 0 |

| | | | | | | | |
|---|---|---|---|---|---|---|---|
| ENSMUSG00000100792 | 0 | 0 | 0 | 0 | 0 | 0 | 0 |
| ENSMUSG00000100793 | 0 | 0 | 0 | 0 | 0 | 0 | 0 |
| ENSMUSG00000100794 | 1.129344651 | 1.04248125 | 0.896240625 | 0.864857905 | 0.931471446 | 1.146128036 | 0.803905701 |
| ENSMUSG00000100795 | 0 | 0 | 0 | 0 | 0 | 0 | 0 |
| ENSMUSG00000100796 | 0 | 0 | 0 | 0 | 0 | 0 | 0 |
| ENSMUSG00000100797 | 0 | 0 | 0 | 0 | 0 | 0 | 0 |
| ENSMUSG00000100798 | 0 | 0.5 | 0 | 0.580482024 | 0.568060967 | 0.602059991 | 0.448487648 |
| ENSMUSG00000100799 | 0 | 0 | 0 | 0 | 0 | 0 | 0 |
| ENSMUSG00000100800 | 0 | 0 | 0 | 0 | 0 | 0 | 0 |
| ENSMUSG00000100801 | 0 | 0 | 0 | 0 | 0 | 0 | 0 |
| ENSMUSG00000100802 | 0 | 0 | 0 | 0 | 0 | 0 | 0 |
| ENSMUSG00000100803 | 0 | 0 | 0 | 0 | 0 | 0 | 0 |
| ENSMUSG00000100804 | 0 | 0 | 0 | 0 | 0 | 0 | 0 |
| ENSMUSG00000100805 | 0 | 0 | 0 | 0 | 0 | 0 | 0 |
| ENSMUSG00000100806 | 0 | 0 | 0 | 0 | 0 | 0 | 0 |
| ENSMUSG00000100807 | 0 | 0 | 0 | 0 | 0 | 0 | 0 |
| ENSMUSG00000100808 | 0 | 0 | 0 | 0 | 0 | 0 | 0 |
| ENSMUSG00000100809 | 0 | 0 | 0 | 0 | 0 | 0 | 0 |
| ENSMUSG00000100810 | 0 | 0 | 0 | 0 | 0 | 0 | 0 |
| ENSMUSG00000100811 | 0.289064826 | 0 | 0 | 0 | 0 | 0 | 0.224243824 |
| ENSMUSG00000100812 | 0 | 0 | 0 | 0 | 0 | 0 | 0 |
| ENSMUSG00000100813 | 0.867194479 | 0.75 | 0.646240625 | 0.701838731 | 0.77552387 | 0.77815125 | 0.520678036 |
| ENSMUSG00000100814 | 0 | 0 | 0 | 0 | 0 | 0 | 0 |
| ENSMUSG00000100815 | 0 | 0 | 0 | 0 | 0 | 0 | 0 |
| ENSMUSG00000100816 | 0.960252568 | 0.646240625 | 0.896240625 | 0.396240625 | 0.489301084 | 0.77815125 | 0.579661877 |
| ENSMUSG00000100817 | 0 | 0 | 0 | 0 | 0 | 0 | 0 |
| ENSMUSG00000100818 | 0 | 0 | 0 | 0 | 0 | 0 | 0 |
| ENSMUSG00000100819 | 0.289064826 | 0 | 0 | 0 | 0 | 0 | 0 |
| ENSMUSG00000100820 | 0 | 0 | 0 | 0 | 0 | 0 | 0 |
| ENSMUSG00000100821 | 0 | 0 | 0 | 0 | 0 | 0 | 0 |
| ENSMUSG00000100822 | 0 | 0 | 0 | 0 | 0 | 0 | 0 |
| ENSMUSG00000100823 | 0 | 0 | 0 | 0 | 0 | 0 | 0 |
| ENSMUSG00000100824 | 0 | 0 | 0 | 0 | 0 | 0 | 0 |
| ENSMUSG00000100825 | 0 | 0 | 0 | 0 | 0 | 0 | 0.355418052 |

| | | | | | | | |
|---|---|---|---|---|---|---|---|
| ENSMUSG00000100826 | 1.714367558 | 1.580482024 | 1.54745614 | 1.580482024 | 1.625423019 | 1.755874856 | 1.459332757 |
| ENSMUSG00000100827 | 0 | 0 | 0 | 0 | 0 | 0 | 0 |
| ENSMUSG00000100828 | 0 | 0 | 0 | 0 | 0 | 0 | 0 |
| ENSMUSG00000100830 | 0 | 0.396240625 | 0.25 | 0.580482024 | 0.244650542 | 0.301029996 | 0.448487648 |
| ENSMUSG00000100831 | 0 | 0 | 0 | 0 | 0 | 0 | 0 |
| ENSMUSG00000100832 | 0 | 0 | 0 | 0 | 0 | 0 | 0 |
| ENSMUSG00000100833 | 0.289064826 | 0.396240625 | 0 | 0.580482024 | 0.387761935 | 0.301029996 | 0.224243824 |
| ENSMUSG00000100834 | 0 | 0 | 0 | 0 | 0 | 0 | 0 |
| ENSMUSG00000100835 | 0 | 0 | 0 | 0 | 0 | 0 | 0 |
| ENSMUSG00000100836 | 0 | 0 | 0 | 0 | 0 | 0 | 0 |
| ENSMUSG00000100837 | 0.671187741 | 0.701838731 | 0.5 | 0.701838731 | 0.489301084 | 0.698970004 | 0.672731473 |
| ENSMUSG00000100838 | 0 | 0 | 0 | 0 | 0 | 0 | 0 |
| ENSMUSG00000100839 | 0 | 0 | 0 | 0 | 0.387761935 | 0.301029996 | 0.224243824 |
| ENSMUSG00000100841 | 0 | 0 | 0 | 0 | 0 | 0 | 0 |
| ENSMUSG00000100842 | 0 | 0 | 0 | 0 | 0 | 0 | 0 |
| ENSMUSG00000100843 | 0 | 0 | 0 | 0 | 0 | 0 | 0 |
| ENSMUSG00000100844 | 0 | 0 | 0 | 0 | 0 | 0 | 0 |
| ENSMUSG00000100845 | 0 | 0 | 0 | 0 | 0 | 0 | 0 |
| ENSMUSG00000100846 | 0 | 0 | 0 | 0 | 0 | 0 | 0 |
| ENSMUSG00000100847 | 0 | 0 | 0 | 0 | 0 | 0 | 0 |
| ENSMUSG00000100848 | 0 | 0 | 0 | 0 | 0 | 0 | 0 |
| ENSMUSG00000100849 | 0 | 0 | 0 | 0 | 0 | 0 | 0 |
| ENSMUSG00000100850 | 0 | 0 | 0 | 0 | 0 | 0 | 0 |
| ENSMUSG00000100851 | 0 | 0 | 0 | 0 | 0 | 0 | 0 |
| ENSMUSG00000100852 | 0 | 0 | 0 | 0 | 0 | 0 | 0 |
| ENSMUSG00000100853 | 0 | 0 | 0 | 0 | 0 | 0 | 0 |
| ENSMUSG00000100854 | 0 | 0 | 0 | 0 | 0 | 0 | 0 |
| ENSMUSG00000100855 | 0 | 0 | 0.25 | 0 | 0 | 0 | 0 |
| ENSMUSG00000100856 | 0 | 0 | 0 | 0 | 0 | 0 | 0 |
| ENSMUSG00000100857 | 0 | 0 | 0 | 0 | 0 | 0 | 0 |
| ENSMUSG00000100858 | 0 | 0 | 0 | 0 | 0 | 0 | 0 |
| ENSMUSG00000100859 | 0 | 0 | 0 | 0 | 0 | 0 | 0 |
| ENSMUSG00000100860 | 0 | 0 | 0 | 0 | 0 | 0 | 0 |
| ENSMUSG00000100861 | 0 | 0 | 0 | 0 | 0 | 0 | 0 |

| | | | | | | | |
|---|---|---|---|---|---|---|---|
| ENSMUSG00000100862 | 0 | 0.25 | 0.396240625 | 0 | 0.387761935 | 0.477121255 | 0 |
| ENSMUSG00000100863 | 0.289064826 | 0 | 0 | 0 | 0 | 0 | 0 |
| ENSMUSG00000100864 | 0 | 0 | 0 | 0 | 0 | 0 | 0 |
| ENSMUSG00000100865 | 0.289064826 | 0 | 0.396240625 | 0 | 0.387761935 | 0.477121255 | 0.355418052 |
| ENSMUSG00000100866 | 0 | 0 | 0 | 0 | 0 | 0 | 0 |
| ENSMUSG00000100867 | 0 | 0 | 0 | 0 | 0 | 0 | 0 |
| ENSMUSG00000100868 | 0 | 0.25 | 0 | 0 | 0 | 0 | 0 |
| ENSMUSG00000100869 | 0 | 0 | 0 | 0 | 0 | 0 | 0 |
| ENSMUSG00000100870 | 0 | 0 | 0 | 0 | 0 | 0 | 0 |
| ENSMUSG00000100871 | 0 | 0 | 0 | 0 | 0 | 0 | 0 |
| ENSMUSG00000100872 | 0.578129653 | 0.25 | 0 | 0 | 0.244650542 | 0.301029996 | 0 |
| ENSMUSG00000100873 | 0 | 0 | 0 | 0 | 0 | 0 | 0 |
| ENSMUSG00000100874 | 0 | 0 | 0 | 0 | 0 | 0 | 0 |
| ENSMUSG00000100875 | 0 | 0 | 0 | 0 | 0 | 0 | 0 |
| ENSMUSG00000100876 | 0.960252568 | 0.864857905 | 0.864857905 | 0.79248125 | 0.846351821 | 0.84509804 | 0.579661877 |
| ENSMUSG00000100877 | 0 | 0 | 0 | 0 | 0 | 0 | 0 |
| ENSMUSG00000100878 | 0 | 0 | 0 | 0 | 0 | 0 | 0 |
| ENSMUSG00000100879 | 0 | 0 | 0 | 0 | 0 | 0 | 0 |
| ENSMUSG00000100880 | 0 | 0 | 0 | 0 | 0.244650542 | 0 | 0 |
| ENSMUSG00000100881 | 0 | 0 | 0 | 0 | 0 | 0 | 0 |
| ENSMUSG00000100882 | 0 | 0 | 0 | 0 | 0 | 0 | 0 |
| ENSMUSG00000100883 | 0 | 0 | 0 | 0 | 0 | 0 | 0 |
| ENSMUSG00000100884 | 0 | 0 | 0 | 0 | 0 | 0 | 0 |
| ENSMUSG00000100885 | 0 | 0 | 0 | 0 | 0 | 0 | 0 |
| ENSMUSG00000100886 | 0 | 0 | 0 | 0 | 0 | 0 | 0 |
| ENSMUSG00000100887 | 0 | 0 | 0 | 0 | 0 | 0 | 0 |
| ENSMUSG00000100888 | 0 | 0 | 0 | 0 | 0 | 0 | 0 |
| ENSMUSG00000100889 | 0 | 0 | 0 | 0 | 0 | 0 | 0 |
| ENSMUSG00000100890 | 0 | 0 | 0 | 0 | 0 | 0 | 0 |
| ENSMUSG00000100891 | 0 | 0 | 0 | 0 | 0.244650542 | 0 | 0 |
| ENSMUSG00000100892 | 0 | 0 | 0 | 0 | 0 | 0 | 0 |
| ENSMUSG00000100893 | 0 | 0 | 0 | 0 | 0 | 0 | 0 |
| ENSMUSG00000100894 | 0 | 0 | 0 | 0 | 0.244650542 | 0 | 0.355418052 |
| ENSMUSG00000100895 | 0 | 0 | 0 | 0 | 0 | 0 | 0 |

| | | | | | | | |
|---|---|---|---|---|---|---|---|
| ENSMUSG00000100896 | 0 | 0 | 0 | 0 | 0 | 0 | 0 |
| ENSMUSG00000100897 | 0 | 0 | 0 | 0 | 0 | 0 | 0 |
| ENSMUSG00000100898 | 0 | 0 | 0 | 0 | 0 | 0 | 0 |
| ENSMUSG00000100899 | 0 | 0 | 0 | 0 | 0 | 0 | 0 |
| ENSMUSG00000100900 | 0 | 0 | 0 | 0 | 0 | 0 | 0 |
| ENSMUSG00000100901 | 0 | 0 | 0 | 0 | 0 | 0 | 0 |
| ENSMUSG00000100902 | 0 | 0 | 0 | 0 | 0 | 0 | 0 |
| ENSMUSG00000100903 | 0 | 0 | 0 | 0 | 0 | 0 | 0 |
| ENSMUSG00000100904 | 0 | 0 | 0 | 0 | 0 | 0 | 0 |
| ENSMUSG00000100905 | 0 | 0 | 0 | 0 | 0 | 0 | 0 |
| ENSMUSG00000100906 | 0 | 0 | 0 | 0 | 0 | 0 | 0 |
| ENSMUSG00000100907 | 0 | 0 | 0 | 0 | 0 | 0 | 0 |
| ENSMUSG00000100908 | 0 | 0 | 0 | 0 | 0 | 0 | 0 |
| ENSMUSG00000100909 | 0 | 0 | 0 | 0 | 0 | 0 | 0 |
| ENSMUSG00000100910 | 0 | 0 | 0 | 0 | 0 | 0 | 0 |
| ENSMUSG00000100911 | 0 | 0 | 0 | 0 | 0 | 0 | 0 |
| ENSMUSG00000100912 | 0 | 0 | 0 | 0 | 0 | 0 | 0 |
| ENSMUSG00000100913 | 0 | 0 | 0 | 0 | 0.244650542 | 0 | 0 |
| ENSMUSG00000100914 | 0 | 0 | 0 | 0 | 0 | 0 | 0 |
| ENSMUSG00000100915 | 0 | 0 | 0 | 0 | 0 | 0 | 0 |
| ENSMUSG00000100916 | 0 | 0 | 0 | 0 | 0 | 0 | 0 |
| ENSMUSG00000100917 | 0 | 0 | 0 | 0 | 0 | 0 | 0 |
| ENSMUSG00000100918 | 0 | 0 | 0 | 0 | 0 | 0 | 0 |
| ENSMUSG00000100919 | 0 | 0 | 0 | 0 | 0 | 0 | 0 |
| ENSMUSG00000100920 | 0 | 0 | 0 | 0 | 0 | 0 | 0 |
| ENSMUSG00000100922 | 0.960252568 | 1.238549078 | 1.27186571 | 0.701838731 | 1.039257769 | 0.84509804 | 1.110947927 |
| ENSMUSG00000100923 | 0 | 0 | 0 | 0 | 0 | 0 | 0 |
| ENSMUSG00000100924 | 0 | 0 | 0 | 0 | 0 | 0 | 0 |
| ENSMUSG00000100925 | 0 | 0 | 0 | 0 | 0 | 0 | 0 |
| ENSMUSG00000100926 | 0 | 0 | 0 | 0 | 0 | 0 | 0 |
| ENSMUSG00000100927 | 0 | 0 | 0 | 0 | 0 | 0 | 0 |
| ENSMUSG00000100928 | 0 | 0 | 0 | 0 | 0 | 0 | 0 |
| ENSMUSG00000100929 | 0 | 0 | 0 | 0 | 0 | 0 | 0 |
| ENSMUSG00000100930 | 0 | 0 | 0 | 0 | 0 | 0 | 0 |

| | | | | | | | |
|---|---|---|---|---|---|---|---|
| ENSMUSG00000100931 | 0.960252568 | 0.75 | 0.580482024 | 0.75 | 0.568060967 | 0.602059991 | 0.629532004 |
| ENSMUSG00000100932 | 0 | 0 | 0 | 0 | 0 | 0 | 0 |
| ENSMUSG00000100933 | 0 | 0 | 0 | 0 | 0 | 0 | 0 |
| ENSMUSG00000100935 | 0 | 0 | 0 | 0 | 0 | 0 | 0 |
| ENSMUSG00000100936 | 0 | 0 | 0 | 0 | 0 | 0 | 0 |
| ENSMUSG00000100937 | 0.578129653 | 0.92510993 | 0.701838731 | 0.25 | 0.568060967 | 0.301029996 | 0.355418052 |
| ENSMUSG00000100938 | 0 | 0 | 0 | 0 | 0 | 0 | 0 |
| ENSMUSG00000100939 | 0 | 0 | 0 | 0 | 0 | 0 | 0 |
| ENSMUSG00000100940 | 0 | 0 | 0 | 0 | 0 | 0 | 0 |
| ENSMUSG00000100941 | 0 | 0 | 0 | 0 | 0 | 0 | 0 |
| ENSMUSG00000100942 | 0 | 0 | 0 | 0 | 0 | 0 | 0 |
| ENSMUSG00000100943 | 0 | 0 | 0 | 0 | 0 | 0 | 0 |
| ENSMUSG00000100944 | 0 | 0 | 0 | 0 | 0 | 0 | 0 |
| ENSMUSG00000100945 | 0 | 0 | 0 | 0 | 0 | 0 | 0 |
| ENSMUSG00000100946 | 0 | 0 | 0 | 0 | 0 | 0 | 0 |
| ENSMUSG00000100947 | 0 | 0 | 0 | 0 | 0 | 0 | 0 |
| ENSMUSG00000100948 | 0 | 0 | 0 | 0 | 0 | 0 | 0 |
| ENSMUSG00000100949 | 0 | 0 | 0 | 0 | 0 | 0 | 0 |
| ENSMUSG00000100950 | 0 | 0 | 0 | 0 | 0 | 0 | 0 |
| ENSMUSG00000100951 | 0 | 0 | 0 | 0 | 0 | 0 | 0 |
| ENSMUSG00000100952 | 0 | 0 | 0 | 0 | 0 | 0 | 0 |
| ENSMUSG00000100953 | 0 | 0 | 0 | 0 | 0 | 0 | 0 |
| ENSMUSG00000100954 | 0.289064826 | 0.580482024 | 0.646240625 | 0.79248125 | 0.489301084 | 0.477121255 | 0.448487648 |
| ENSMUSG00000100955 | 0 | 0 | 0 | 0 | 0 | 0 | 0 |
| ENSMUSG00000100956 | 0 | 0 | 0 | 0 | 0 | 0 | 0 |
| ENSMUSG00000100957 | 0 | 0 | 0 | 0 | 0 | 0 | 0 |
| ENSMUSG00000100958 | 0 | 0 | 0 | 0 | 0 | 0 | 0 |
| ENSMUSG00000100959 | 0.289064826 | 0.25 | 0 | 0 | 0 | 0 | 0.224243824 |
| ENSMUSG00000100960 | 0 | 0 | 0 | 0 | 0 | 0 | 0 |
| ENSMUSG00000100961 | 0 | 0 | 0 | 0 | 0 | 0 | 0 |
| ENSMUSG00000100962 | 0 | 0 | 0 | 0 | 0 | 0 | 0 |
| ENSMUSG00000100963 | 0 | 0.25 | 0 | 0 | 0 | 0 | 0 |
| ENSMUSG00000100964 | 0 | 0 | 0 | 0 | 0 | 0 | 0 |
| ENSMUSG00000100965 | 0 | 0 | 0 | 0 | 0 | 0 | 0 |

| | | | | | | | |
|---|---|---|---|---|---|---|---|
| ENSMUSG00000100966 | 0 | 0 | 0 | 0 | 0 | 0 | 0 |
| ENSMUSG00000100967 | 1.289064826 | 1.201838731 | 1.214495249 | 1.410964047 | 1.366364104 | 1.301029996 | 1.20139795 |
| ENSMUSG00000100968 | 0 | 0 | 0 | 0 | 0 | 0 | 0 |
| ENSMUSG00000100969 | 0 | 0.25 | 0 | 0 | 0.244650542 | 0 | 0 |
| ENSMUSG00000100970 | 0 | 0 | 0 | 0 | 0 | 0 | 0 |
| ENSMUSG00000100971 | 0 | 0 | 0 | 0 | 0 | 0 | 0 |
| ENSMUSG00000100972 | 0 | 0 | 0 | 0 | 0 | 0 | 0 |
| ENSMUSG00000100973 | 0 | 0 | 0 | 0 | 0 | 0 | 0 |
| ENSMUSG00000100974 | 0 | 0 | 0 | 0 | 0 | 0 | 0 |
| ENSMUSG00000100975 | 1.156259305 | 1.04248125 | 1.04248125 | 0.976722649 | 0.905314583 | 1.176091259 | 0.952571511 |
| ENSMUSG00000100976 | 0 | 0 | 0 | 0 | 0 | 0 | 0 |
| ENSMUSG00000100977 | 0 | 0 | 0 | 0 | 0 | 0 | 0 |
| ENSMUSG00000100978 | 0 | 0 | 0 | 0 | 0 | 0 | 0 |
| ENSMUSG00000100979 | 0 | 0 | 0 | 0 | 0 | 0 | 0 |
| ENSMUSG00000100980 | 1.036286563 | 1.098079356 | 1.114857905 | 0.864857905 | 1.121713561 | 0.954242509 | 0.935079929 |
| ENSMUSG00000100981 | 0 | 0 | 0 | 0 | 0 | 0 | 0 |
| ENSMUSG00000100982 | 0 | 0 | 0 | 0 | 0 | 0 | 0 |
| ENSMUSG00000100983 | 0 | 0 | 0 | 0 | 0 | 0 | 0 |
| ENSMUSG00000100984 | 0 | 0 | 0 | 0 | 0 | 0 | 0 |
| ENSMUSG00000100985 | 0 | 0 | 0 | 0.25 | 0 | 0 | 0 |
| ENSMUSG00000100986 | 0 | 0 | 0 | 0 | 0 | 0 | 0 |
| ENSMUSG00000100987 | 0.45815691 | 0.5 | 0.701838731 | 0 | 0 | 0.301029996 | 0.224243824 |
| ENSMUSG00000100988 | 0 | 0 | 0 | 0 | 0 | 0 | 0 |
| ENSMUSG00000100989 | 0 | 0 | 0 | 0 | 0 | 0 | 0 |
| ENSMUSG00000100990 | 0 | 0 | 0 | 0 | 0 | 0 | 0.224243824 |
| ENSMUSG00000100991 | 0 | 0 | 0 | 0 | 0 | 0 | 0 |
| ENSMUSG00000100992 | 0 | 0 | 0 | 0 | 0 | 0 | 0 |
| ENSMUSG00000100993 | 0 | 0 | 0 | 0 | 0 | 0 | 0 |
| ENSMUSG00000100994 | 0 | 0 | 0 | 0 | 0 | 0 | 0 |
| ENSMUSG00000100995 | 0 | 0 | 0 | 0 | 0 | 0 | 0 |
| ENSMUSG00000100996 | 0 | 0 | 0 | 0 | 0 | 0 | 0 |
| ENSMUSG00000100998 | 0 | 0 | 0 | 0 | 0 | 0 | 0 |
| ENSMUSG00000100999 | 0 | 0 | 0 | 0 | 0 | 0 | 0 |
| ENSMUSG00000101000 | 0 | 0 | 0 | 0 | 0 | 0 | 0 |

| | | | | | | | |
|---|---|---|---|---|---|---|---|
| ENSMUSG00000101001 | 0 | 0 | 0 | 0 | 0 | 0 | 0 |
| ENSMUSG00000101002 | 0 | 0 | 0 | 0 | 0 | 0 | 0 |
| ENSMUSG00000101003 | 0 | 0 | 0 | 0 | 0 | 0 | 0 |
| ENSMUSG00000101004 | 0 | 0 | 0.25 | 0 | 0 | 0.301029996 | 0 |
| ENSMUSG00000101005 | 0 | 0 | 0 | 0 | 0 | 0 | 0 |
| ENSMUSG00000101006 | 0 | 0 | 0 | 0 | 0 | 0 | 0 |
| ENSMUSG00000101007 | 0 | 0.25 | 0.25 | 0 | 0 | 0 | 0 |
| ENSMUSG00000101008 | 0 | 0 | 0 | 0 | 0 | 0 | 0 |
| ENSMUSG00000101009 | 0 | 0.396240625 | 0.25 | 0.396240625 | 0.244650542 | 0 | 0.355418052 |
| ENSMUSG00000101010 | 0 | 0 | 0 | 0 | 0 | 0 | 0 |
| ENSMUSG00000101011 | 0 | 0.396240625 | 0 | 0 | 0.244650542 | 0.301029996 | 0.224243824 |
| ENSMUSG00000101012 | 0 | 0 | 0 | 0 | 0 | 0 | 0 |
| ENSMUSG00000101013 | 0 | 0.25 | 0.25 | 0 | 0.244650542 | 0 | 0.448487648 |
| ENSMUSG00000101014 | 0 | 0 | 0 | 0 | 0 | 0 | 0 |
| ENSMUSG00000101015 | 0 | 0 | 0 | 0 | 0 | 0 | 0 |
| ENSMUSG00000101016 | 0 | 0 | 0 | 0 | 0 | 0 | 0 |
| ENSMUSG00000101017 | 0 | 0 | 0 | 0 | 0 | 0 | 0 |
| ENSMUSG00000101018 | 0 | 0 | 0 | 0 | 0 | 0 | 0 |
| ENSMUSG00000101019 | 0 | 0 | 0 | 0 | 0 | 0 | 0 |
| ENSMUSG00000101020 | 0 | 0 | 0 | 0 | 0 | 0 | 0 |
| ENSMUSG00000101021 | 0 | 0 | 0 | 0 | 0 | 0 | 0 |
| ENSMUSG00000101022 | 0 | 0 | 0 | 0 | 0 | 0 | 0 |
| ENSMUSG00000101023 | 0 | 0 | 0 | 0 | 0 | 0 | 0 |
| ENSMUSG00000101024 | 0 | 0 | 0 | 0 | 0 | 0 | 0 |
| ENSMUSG00000101025 | 0 | 0 | 0 | 0 | 0 | 0 | 0 |
| ENSMUSG00000101026 | 0 | 0 | 0 | 0 | 0 | 0 | 0 |
| ENSMUSG00000101027 | 0 | 0 | 0 | 0 | 0 | 0 | 0 |
| ENSMUSG00000101028 | 0 | 0 | 0 | 0 | 0 | 0 | 0 |
| ENSMUSG00000101029 | 0 | 0 | 0 | 0 | 0 | 0 | 0 |
| ENSMUSG00000101030 | 0 | 0 | 0 | 0 | 0 | 0 | 0 |
| ENSMUSG00000101031 | 0 | 0 | 0 | 0 | 0 | 0 | 0 |
| ENSMUSG00000101032 | 0 | 0 | 0 | 0 | 0 | 0 | 0 |
| ENSMUSG00000101033 | 0 | 0 | 0 | 0 | 0 | 0 | 0 |
| ENSMUSG00000101035 | 0 | 0 | 0 | 0 | 0 | 0 | 0 |

| | | | | | | | |
|---|---|---|---|---|---|---|---|
| ENSMUSG00000101036 | 0 | 0 | 0 | 0 | 0 | 0 | 0.224243824 |
| ENSMUSG00000101037 | 0.289064826 | 0.580482024 | 0.580482024 | 0.580482024 | 0.733951626 | 0.698970004 | 0.448487648 |
| ENSMUSG00000101038 | 0 | 0 | 0 | 0 | 0 | 0 | 0 |
| ENSMUSG00000101039 | 0 | 0 | 0 | 0 | 0 | 0 | 0 |
| ENSMUSG00000101040 | 0 | 0 | 0 | 0 | 0 | 0 | 0 |
| ENSMUSG00000101041 | 0 | 0 | 0 | 0 | 0 | 0 | 0 |
| ENSMUSG00000101042 | 0 | 0 | 0 | 0 | 0 | 0 | 0 |
| ENSMUSG00000101043 | 0 | 0 | 0 | 0 | 0 | 0 | 0 |
| ENSMUSG00000101044 | 0 | 0 | 0 | 0 | 0 | 0 | 0 |
| ENSMUSG00000101045 | 0 | 0 | 0 | 0 | 0 | 0 | 0 |
| ENSMUSG00000101046 | 0 | 0 | 0 | 0 | 0 | 0 | 0 |
| ENSMUSG00000101049 | 0 | 0 | 0 | 0 | 0.244650542 | 0 | 0 |
| ENSMUSG00000101050 | 0 | 0 | 0 | 0 | 0 | 0 | 0 |
| ENSMUSG00000101051 | 0 | 0 | 0 | 0 | 0 | 0 | 0 |
| ENSMUSG00000101052 | 0 | 0 | 0 | 0 | 0 | 0 | 0 |
| ENSMUSG00000101053 | 0 | 0 | 0 | 0 | 0 | 0 | 0 |
| ENSMUSG00000101054 | 0 | 0 | 0 | 0 | 0 | 0 | 0 |
| ENSMUSG00000101055 | 0 | 0 | 0 | 0 | 0 | 0 | 0 |
| ENSMUSG00000101056 | 0 | 0 | 0 | 0 | 0 | 0 | 0 |
| ENSMUSG00000101057 | 0 | 0 | 0 | 0 | 0 | 0 | 0 |
| ENSMUSG00000101058 | 0 | 0 | 0 | 0 | 0 | 0 | 0 |
| ENSMUSG00000101059 | 0 | 0 | 0 | 0 | 0 | 0 | 0.448487648 |
| ENSMUSG00000101060 | 0 | 0 | 0 | 0 | 0 | 0 | 0 |
| ENSMUSG00000101061 | 0.289064826 | 0 | 0.25 | 0.25 | 0.387761935 | 0.477121255 | 0.224243824 |
| ENSMUSG00000101062 | 0 | 0 | 0 | 0 | 0 | 0 | 0 |
| ENSMUSG00000101063 | 0 | 0 | 0 | 0 | 0 | 0 | 0 |
| ENSMUSG00000101064 | 0 | 0 | 0 | 0 | 0 | 0 | 0 |
| ENSMUSG00000101065 | 0 | 0 | 0 | 0 | 0 | 0 | 0 |
| ENSMUSG00000101066 | 0 | 0 | 0 | 0 | 0 | 0 | 0 |
| ENSMUSG00000101067 | 0 | 0 | 0 | 0 | 0 | 0 | 0 |
| ENSMUSG00000101068 | 0 | 0 | 0 | 0 | 0 | 0 | 0 |
| ENSMUSG00000101069 | 0 | 0 | 0 | 0 | 0 | 0 | 0 |
| ENSMUSG00000101070 | 0 | 0 | 0 | 0 | 0 | 0 | 0 |
| ENSMUSG00000101071 | 0 | 0 | 0 | 0 | 0 | 0 | 0 |

| | | | | | | | |
|---|---|---|---|---|---|---|---|
| ENSMUSG00000101072 | 0 | 0 | 0 | 0 | 0 | 0 | 0 |
| ENSMUSG00000101073 | 0 | 0 | 0 | 0 | 0 | 0 | 0 |
| ENSMUSG00000101074 | 0 | 0 | 0 | 0 | 0 | 0 | 0 |
| ENSMUSG00000101075 | 0 | 0 | 0 | 0 | 0 | 0 | 0 |
| ENSMUSG00000101076 | 0 | 0 | 0 | 0 | 0 | 0 | 0 |
| ENSMUSG00000101077 | 0 | 0 | 0.25 | 0 | 0 | 0 | 0 |
| ENSMUSG00000101078 | 0 | 0 | 0 | 0 | 0 | 0 | 0 |
| ENSMUSG00000101079 | 0 | 0 | 0 | 0 | 0 | 0 | 0 |
| ENSMUSG00000101080 | 0 | 0 | 0 | 0 | 0 | 0 | 0 |
| ENSMUSG00000101081 | 0 | 0 | 0 | 0 | 0 | 0 | 0 |
| ENSMUSG00000101082 | 0 | 0 | 0 | 0 | 0 | 0 | 0 |
| ENSMUSG00000101083 | 0 | 0 | 0 | 0 | 0 | 0 | 0 |
| ENSMUSG00000101084 | 0 | 0 | 0 | 0 | 0 | 0 | 0 |
| ENSMUSG00000101085 | 0 | 0 | 0 | 0 | 0 | 0 | 0 |
| ENSMUSG00000101086 | 0 | 0 | 0 | 0 | 0 | 0 | 0 |
| ENSMUSG00000101087 | 0 | 0 | 0 | 0 | 0 | 0 | 0 |
| ENSMUSG00000101088 | 0.289064826 | 0 | 0 | 0 | 0.244650542 | 0 | 0 |
| ENSMUSG00000101089 | 0 | 0 | 0 | 0 | 0 | 0 | 0 |
| ENSMUSG00000101090 | 0 | 0 | 0 | 0 | 0 | 0 | 0 |
| ENSMUSG00000101091 | 0 | 0 | 0 | 0 | 0 | 0 | 0 |
| ENSMUSG00000101092 | 0 | 0 | 0 | 0 | 0 | 0 | 0 |
| ENSMUSG00000101093 | 0 | 0 | 0 | 0 | 0 | 0 | 0.224243824 |
| ENSMUSG00000101094 | 0 | 0 | 0 | 0 | 0 | 0 | 0 |
| ENSMUSG00000101095 | 0 | 0 | 0 | 0 | 0 | 0 | 0 |
| ENSMUSG00000101096 | 0 | 0 | 0 | 0 | 0 | 0 | 0 |
| ENSMUSG00000101097 | 0 | 0 | 0 | 0 | 0.244650542 | 0 | 0.224243824 |
| ENSMUSG00000101098 | 0 | 0 | 0 | 0 | 0 | 0 | 0 |
| ENSMUSG00000101099 | 0 | 0 | 0 | 0 | 0 | 0 | 0 |
| ENSMUSG00000101100 | 0 | 0 | 0 | 0 | 0 | 0 | 0 |
| ENSMUSG00000101101 | 0 | 0 | 0 | 0 | 0 | 0 | 0 |
| ENSMUSG00000101102 | 0 | 0.25 | 0 | 0 | 0 | 0 | 0 |
| ENSMUSG00000101103 | 0 | 0 | 0 | 0 | 0 | 0 | 0 |
| ENSMUSG00000101104 | 0 | 0 | 0 | 0 | 0 | 0 | 0 |
| ENSMUSG00000101105 | 0 | 0 | 0 | 0 | 0 | 0 | 0 |

| | | | | | | | |
|---|---|---|---|---|---|---|---|
| ENSMUSG00000101106 | 0 | 0 | 0 | 0 | 0 | 0 | 0 |
| ENSMUSG00000101107 | 0 | 0 | 0 | 0.25 | 0 | 0 | 0 |
| ENSMUSG00000101108 | 0 | 0 | 0 | 0 | 0 | 0 | 0 |
| ENSMUSG00000101109 | 0 | 0 | 0 | 0 | 0 | 0 | 0 |
| ENSMUSG00000101110 | 0 | 0 | 0 | 0 | 0 | 0 | 0 |
| ENSMUSG00000101111 | 0 | 0.396240625 | 0 | 0.25 | 0 | 0 | 0 |
| ENSMUSG00000101112 | 0 | 0 | 0 | 0 | 0 | 0 | 0 |
| ENSMUSG00000101113 | 0 | 0 | 0 | 0 | 0 | 0 | 0 |
| ENSMUSG00000101114 | 0 | 0 | 0 | 0 | 0 | 0 | 0 |
| ENSMUSG00000101115 | 0 | 0 | 0 | 0 | 0 | 0 | 0 |
| ENSMUSG00000101116 | 0 | 0 | 0 | 0 | 0 | 0 | 0 |
| ENSMUSG00000101117 | 0 | 0 | 0 | 0 | 0 | 0 | 0 |
| ENSMUSG00000101118 | 0 | 0 | 0 | 0 | 0 | 0 | 0 |
| ENSMUSG00000101119 | 0 | 0 | 0 | 0 | 0 | 0 | 0 |
| ENSMUSG00000101120 | 0 | 0 | 0 | 0 | 0 | 0 | 0 |
| ENSMUSG00000101122 | 0.578129653 | 0.951838731 | 0.92510993 | 0.646240625 | 0.686820904 | 0.477121255 | 0.853775828 |
| ENSMUSG00000101123 | 0 | 0 | 0 | 0 | 0 | 0 | 0 |
| ENSMUSG00000101124 | 0 | 0 | 0 | 0 | 0 | 0 | 0 |
| ENSMUSG00000101125 | 0 | 0 | 0 | 0 | 0 | 0 | 0 |
| ENSMUSG00000101126 | 0 | 0 | 0 | 0 | 0 | 0 | 0 |
| ENSMUSG00000101127 | 0 | 0 | 0 | 0 | 0 | 0 | 0 |
| ENSMUSG00000101128 | 0 | 0 | 0 | 0 | 0 | 0 | 0 |
| ENSMUSG00000101129 | 0 | 0 | 0 | 0 | 0.244650542 | 0 | 0 |
| ENSMUSG00000101130 | 0 | 0 | 0 | 0 | 0 | 0 | 0 |
| ENSMUSG00000101131 | 0 | 0 | 0 | 0 | 0 | 0 | 0 |
| ENSMUSG00000101132 | 0 | 0 | 0 | 0 | 0 | 0 | 0 |
| ENSMUSG00000101133 | 0.578129653 | 0.79248125 | 0.701838731 | 0.701838731 | 0.489301084 | 0 | 0.520678036 |
| ENSMUSG00000101134 | 0 | 0 | 0 | 0 | 0 | 0 | 0 |
| ENSMUSG00000101135 | 0.289064826 | 0 | 0 | 0 | 0 | 0 | 0 |
| ENSMUSG00000101136 | 0.289064826 | 0 | 0 | 0 | 0 | 0.301029996 | 0 |
| ENSMUSG00000101137 | 0 | 0 | 0 | 0 | 0 | 0 | 0 |
| ENSMUSG00000101138 | 0 | 0 | 0 | 0 | 0 | 0 | 0 |
| ENSMUSG00000101139 | 0 | 0 | 0 | 0 | 0 | 0 | 0 |
| ENSMUSG00000101140 | 0 | 0 | 0 | 0 | 0 | 0 | 0 |

| | | | | | | | |
|---|---|---|---|---|---|---|---|
| ENSMUSG00000101141 | 0 | 0 | 0 | 0 | 0 | 0 | 0 |
| ENSMUSG00000101142 | 0 | 0 | 0 | 0 | 0 | 0 | 0 |
| ENSMUSG00000101143 | 0 | 0 | 0 | 0 | 0 | 0 | 0 |
| ENSMUSG00000101144 | 0 | 0 | 0.25 | 0 | 0 | 0 | 0 |
| ENSMUSG00000101145 | 0 | 0 | 0 | 0 | 0 | 0 | 0 |
| ENSMUSG00000101146 | 0 | 0 | 0 | 0 | 0 | 0 | 0 |
| ENSMUSG00000101147 | 0 | 0 | 0 | 0 | 0 | 0 | 0 |
| ENSMUSG00000101148 | 0 | 0 | 0 | 0 | 0 | 0 | 0 |
| ENSMUSG00000101149 | 0 | 0 | 0 | 0 | 0 | 0 | 0 |
| ENSMUSG00000101150 | 0 | 0 | 0 | 0 | 0 | 0 | 0 |
| ENSMUSG00000101151 | 0 | 0 | 0 | 0 | 0 | 0 | 0 |
| ENSMUSG00000101152 | 0 | 0 | 0 | 0 | 0 | 0 | 0 |
| ENSMUSG00000101153 | 0 | 0 | 0 | 0 | 0 | 0 | 0 |
| ENSMUSG00000101154 | 0 | 0 | 0 | 0 | 0 | 0 | 0 |
| ENSMUSG00000101155 | 0 | 0 | 0 | 0 | 0 | 0 | 0 |
| ENSMUSG00000101156 | 0 | 0.396240625 | 0 | 0.25 | 0.244650542 | 0 | 0 |
| ENSMUSG00000101157 | 0 | 0 | 0 | 0 | 0 | 0 | 0 |
| ENSMUSG00000101158 | 0 | 0 | 0 | 0 | 0 | 0 | 0 |
| ENSMUSG00000101159 | 0 | 0 | 0 | 0 | 0 | 0 | 0 |
| ENSMUSG00000101160 | 0 | 0 | 0 | 0 | 0 | 0 | 0 |
| ENSMUSG00000101161 | 0 | 0 | 0 | 0 | 0 | 0 | 0 |
| ENSMUSG00000101162 | 0.45815691 | 0 | 0 | 0 | 0 | 0 | 0 |
| ENSMUSG00000101163 | 0 | 0 | 0 | 0 | 0 | 0 | 0 |
| ENSMUSG00000101164 | 0 | 0 | 0 | 0 | 0 | 0 | 0 |
| ENSMUSG00000101166 | 0.289064826 | 0 | 0 | 0 | 0 | 0 | 0 |
| ENSMUSG00000101167 | 0.45815691 | 0.396240625 | 0.25 | 0.396240625 | 0.244650542 | 0 | 0.355418052 |
| ENSMUSG00000101168 | 0 | 0.396240625 | 0 | 0 | 0 | 0 | 0.224243824 |
| ENSMUSG00000101170 | 0 | 0 | 0 | 0 | 0 | 0 | 0 |
| ENSMUSG00000101171 | 0 | 0.25 | 0.25 | 0.396240625 | 0.244650542 | 0 | 0.224243824 |
| ENSMUSG00000101172 | 0 | 0 | 0.25 | 0.25 | 0.632412477 | 0 | 0.224243824 |
| ENSMUSG00000101173 | 0 | 0 | 0 | 0 | 0 | 0 | 0 |
| ENSMUSG00000101174 | 0.578129653 | 0.830482024 | 0.79248125 | 0 | 0 | 0 | 0.224243824 |
| ENSMUSG00000101176 | 0 | 0 | 0 | 0 | 0 | 0 | 0 |
| ENSMUSG00000101177 | 0 | 0 | 0 | 0 | 0 | 0 | 0 |

| | | | | | | | |
|---|---|---|---|---|---|---|---|
| ENSMUSG00000101178 | 0 | 0 | 0 | 0 | 0 | 0 | 0 |
| ENSMUSG00000101179 | 0 | 0 | 0 | 0.396240625 | 0.244650542 | 0 | 0.448487648 |
| ENSMUSG00000101180 | 0 | 0 | 0 | 0 | 0 | 0 | 0 |
| ENSMUSG00000101181 | 0 | 0 | 0 | 0 | 0 | 0 | 0 |
| ENSMUSG00000101182 | 0 | 0 | 0 | 0 | 0 | 0 | 0 |
| ENSMUSG00000101183 | 0 | 0 | 0 | 0 | 0 | 0 | 0 |
| ENSMUSG00000101184 | 0 | 0 | 0 | 0 | 0 | 0 | 0 |
| ENSMUSG00000101185 | 0 | 0 | 0 | 0 | 0 | 0 | 0 |
| ENSMUSG00000101186 | 0 | 0 | 0 | 0 | 0 | 0 | 0 |
| ENSMUSG00000101188 | 0 | 0 | 0.396240625 | 0.25 | 0.489301084 | 0.477121255 | 0.224243824 |
| ENSMUSG00000101189 | 0 | 0 | 0 | 0 | 0 | 0 | 0 |
| ENSMUSG00000101190 | 0 | 0 | 0 | 0 | 0 | 0 | 0 |
| ENSMUSG00000101191 | 0.45815691 | 0.976722649 | 0.864857905 | 0.92510993 | 0.846351821 | 0.84509804 | 0.916588299 |
| ENSMUSG00000101192 | 0 | 0 | 0 | 0 | 0 | 0 | 0 |
| ENSMUSG00000101193 | 0 | 0 | 0 | 0 | 0 | 0 | 0 |
| ENSMUSG00000101194 | 0 | 0 | 0 | 0 | 0 | 0 | 0 |
| ENSMUSG00000101195 | 0 | 0 | 0 | 0 | 0 | 0 | 0 |
| ENSMUSG00000101196 | 0 | 0 | 0 | 0 | 0 | 0 | 0 |
| ENSMUSG00000101197 | 0 | 0 | 0 | 0 | 0 | 0 | 0 |
| ENSMUSG00000101198 | 0 | 0 | 0 | 0 | 0 | 0 | 0 |
| ENSMUSG00000101199 | 0.289064826 | 0.25 | 0 | 0 | 0 | 0 | 0 |
| ENSMUSG00000101200 | 0 | 0 | 0 | 0 | 0 | 0 | 0 |
| ENSMUSG00000101201 | 0 | 0 | 0 | 0 | 0 | 0 | 0 |
| ENSMUSG00000101202 | 0 | 0 | 0 | 0 | 0 | 0 | 0 |
| ENSMUSG00000101203 | 0 | 0 | 0 | 0 | 0 | 0 | 0 |
| ENSMUSG00000101204 | 0 | 0 | 0 | 0 | 0 | 0 | 0 |
| ENSMUSG00000101205 | 0 | 0 | 0 | 0 | 0 | 0 | 0 |
| ENSMUSG00000101206 | 0 | 0 | 0 | 0.25 | 0.244650542 | 0 | 0.355418052 |
| ENSMUSG00000101207 | 0 | 0 | 0 | 0 | 0 | 0 | 0 |
| ENSMUSG00000101208 | 0 | 0 | 0 | 0 | 0 | 0 | 0 |
| ENSMUSG00000101209 | 0 | 0 | 0.25 | 0 | 0 | 0 | 0.224243824 |
| ENSMUSG00000101210 | 0.289064826 | 0.396240625 | 0.25 | 0.646240625 | 0.686820904 | 0.602059991 | 0.224243824 |
| ENSMUSG00000101211 | 0.747221736 | 0.5 | 0.5 | 0.396240625 | 0.387761935 | 0 | 0.224243824 |
| ENSMUSG00000101212 | 0 | 0 | 0 | 0 | 0 | 0 | 0 |

| | | | | | | | |
|---|---|---|---|---|---|---|---|
| ENSMUSG00000101213 | 0 | 0 | 0 | 0 | 0 | 0 | 0 |
| ENSMUSG00000101214 | 0 | 0 | 0 | 0 | 0 | 0 | 0 |
| ENSMUSG00000101215 | 0 | 0 | 0 | 0 | 0 | 0 | 0 |
| ENSMUSG00000101216 | 0 | 0 | 0 | 0 | 0 | 0 | 0 |
| ENSMUSG00000101218 | 0 | 0 | 0 | 0 | 0 | 0 | 0 |
| ENSMUSG00000101219 | 0 | 0.25 | 0 | 0.25 | 0 | 0 | 0.224243824 |
| ENSMUSG00000101220 | 0.45815691 | 0 | 0 | 0 | 0 | 0 | 0.224243824 |
| ENSMUSG00000101221 | 0 | 0 | 0 | 0 | 0 | 0 | 0 |
| ENSMUSG00000101222 | 0 | 0 | 0 | 0 | 0 | 0 | 0 |
| ENSMUSG00000101223 | 0 | 0 | 0 | 0.25 | 0 | 0 | 0 |
| ENSMUSG00000101224 | 0 | 0 | 0.25 | 0 | 0 | 0 | 0 |
| ENSMUSG00000101225 | 0.747221736 | 0.92510993 | 0.830482024 | 0.864857905 | 0.812711509 | 0.477121255 | 0.74492186 |
| ENSMUSG00000101226 | 0 | 0 | 0 | 0 | 0 | 0 | 0 |
| ENSMUSG00000101227 | 0 | 0 | 0 | 0 | 0 | 0 | 0 |
| ENSMUSG00000101228 | 0 | 0 | 0 | 0 | 0 | 0 | 0 |
| ENSMUSG00000101229 | 0 | 0 | 0 | 0 | 0 | 0 | 0 |
| ENSMUSG00000101230 | 0 | 0 | 0 | 0 | 0 | 0 | 0 |
| ENSMUSG00000101231 | 0 | 0 | 0 | 0 | 0 | 0 | 0 |
| ENSMUSG00000101232 | 0 | 0 | 0 | 0 | 0 | 0 | 0 |
| ENSMUSG00000101233 | 0 | 0 | 0 | 0 | 0 | 0 | 0 |
| ENSMUSG00000101234 | 0.747221736 | 0.25 | 0.580482024 | 0.25 | 0 | 0 | 0.224243824 |
| ENSMUSG00000101235 | 0 | 0 | 0 | 0 | 0 | 0 | 0 |
| ENSMUSG00000101236 | 0 | 0 | 0 | 0 | 0 | 0 | 0 |
| ENSMUSG00000101237 | 0 | 0 | 0 | 0 | 0 | 0 | 0 |
| ENSMUSG00000101238 | 0 | 0 | 0 | 0 | 0 | 0 | 0 |
| ENSMUSG00000101239 | 0 | 0 | 0 | 0 | 0 | 0 | 0 |
| ENSMUSG00000101240 | 0 | 0.25 | 0 | 0 | 0 | 0 | 0.224243824 |
| ENSMUSG00000101241 | 0 | 0 | 0 | 0 | 0 | 0 | 0 |
| ENSMUSG00000101242 | 0 | 0 | 0 | 0 | 0.387761935 | 0.301029996 | 0 |
| ENSMUSG00000101243 | 0 | 0 | 0 | 0 | 0 | 0 | 0 |
| ENSMUSG00000101244 | 0 | 0 | 0 | 0 | 0 | 0 | 0 |
| ENSMUSG00000101245 | 0 | 0 | 0 | 0 | 0 | 0 | 0 |
| ENSMUSG00000101246 | 0 | 0 | 0 | 0 | 0 | 0 | 0 |
| ENSMUSG00000101247 | 0 | 0 | 0 | 0 | 0 | 0 | 0 |

| | | | | | | | |
|---|---|---|---|---|---|---|---|
| ENSMUSG00000101248 | 0 | 0 | 0 | 0 | 0 | 0 | 0 |
| ENSMUSG00000101249 | 0 | 0 | 0 | 0.5 | 0.387761935 | 0 | 0.224243824 |
| ENSMUSG00000101250 | 0 | 0 | 0 | 0 | 0 | 0 | 0 |
| ENSMUSG00000101251 | 0 | 0 | 0 | 0 | 0 | 0 | 0 |
| ENSMUSG00000101252 | 0 | 0 | 0 | 0 | 0 | 0 | 0 |
| ENSMUSG00000101253 | 0 | 0 | 0 | 0 | 0 | 0 | 0 |
| ENSMUSG00000101254 | 0 | 0 | 0 | 0 | 0 | 0 | 0 |
| ENSMUSG00000101255 | 0 | 0 | 0 | 0 | 0 | 0 | 0 |
| ENSMUSG00000101256 | 0 | 0 | 0 | 0 | 0 | 0 | 0 |
| ENSMUSG00000101257 | 0.289064826 | 0.25 | 0 | 0.25 | 0.244650542 | 0 | 0 |
| ENSMUSG00000101258 | 0 | 0 | 0 | 0 | 0 | 0 | 0 |
| ENSMUSG00000101259 | 0 | 0 | 0 | 0 | 0 | 0 | 0 |
| ENSMUSG00000101260 | 0 | 0 | 0 | 0 | 0 | 0 | 0 |
| ENSMUSG00000101261 | 0 | 0 | 0 | 0 | 0 | 0 | 0 |
| ENSMUSG00000101262 | 1 | 0.951838731 | 0.92510993 | 1 | 0.905314583 | 1.113943352 | 0.74492186 |
| ENSMUSG00000101263 | 0 | 0 | 0 | 0 | 0 | 0 | 0 |
| ENSMUSG00000101264 | 0 | 0 | 0 | 0 | 0 | 0 | 0 |
| ENSMUSG00000101265 | 0 | 0 | 0 | 0 | 0 | 0 | 0 |
| ENSMUSG00000101266 | 0 | 0 | 0 | 0 | 0 | 0 | 0 |
| ENSMUSG00000101267 | 0 | 0 | 0 | 0 | 0 | 0 | 0 |
| ENSMUSG00000101268 | 0.45815691 | 0.25 | 0 | 0 | 0 | 0 | 0 |
| ENSMUSG00000101270 | 0 | 0 | 0 | 0 | 0 | 0 | 0 |
| ENSMUSG00000101271 | 0 | 0 | 0 | 0 | 0 | 0 | 0 |
| ENSMUSG00000101272 | 0 | 0 | 0 | 0 | 0 | 0 | 0 |
| ENSMUSG00000101273 | 0 | 0 | 0 | 0 | 0 | 0 | 0 |
| ENSMUSG00000101275 | 0 | 0 | 0 | 0 | 0 | 0 | 0 |
| ENSMUSG00000101276 | 0 | 0 | 0 | 0 | 0 | 0 | 0 |
| ENSMUSG00000101277 | 0 | 0 | 0 | 0 | 0 | 0 | 0 |
| ENSMUSG00000101278 | 0 | 0 | 0 | 0 | 0 | 0 | 0 |
| ENSMUSG00000101279 | 0 | 0 | 0 | 0 | 0 | 0 | 0 |
| ENSMUSG00000101280 | 0 | 0 | 0 | 0 | 0 | 0 | 0 |
| ENSMUSG00000101281 | 0 | 0 | 0 | 0.25 | 0 | 0 | 0 |
| ENSMUSG00000101282 | 0 | 0 | 0 | 0 | 0 | 0 | 0 |
| ENSMUSG00000101283 | 0 | 0 | 0 | 0 | 0 | 0 | 0 |

| | | | | | | | |
|---|---|---|---|---|---|---|---|
| ENSMUSG00000101284 | 0 | 0 | 0 | 0 | 0 | 0 | 0 |
| ENSMUSG00000101285 | 0 | 0 | 0 | 0 | 0 | 0 | 0 |
| ENSMUSG00000101286 | 0 | 0 | 0 | 0 | 0 | 0 | 0 |
| ENSMUSG00000101287 | 0 | 0 | 0 | 0 | 0 | 0 | 0 |
| ENSMUSG00000101288 | 0 | 0 | 0 | 0 | 0 | 0 | 0 |
| ENSMUSG00000101289 | 0 | 0 | 0 | 0 | 0 | 0 | 0 |
| ENSMUSG00000101290 | 0 | 0 | 0 | 0 | 0 | 0 | 0 |
| ENSMUSG00000101291 | 0 | 0 | 0 | 0 | 0 | 0 | 0 |
| ENSMUSG00000101292 | 0 | 0 | 0 | 0 | 0 | 0 | 0 |
| ENSMUSG00000101294 | 0 | 0 | 0 | 0 | 0 | 0 | 0 |
| ENSMUSG00000101295 | 0 | 0 | 0 | 0 | 0 | 0 | 0 |
| ENSMUSG00000101296 | 0 | 0 | 0 | 0 | 0 | 0 | 0 |
| ENSMUSG00000101297 | 0 | 0 | 0 | 0 | 0 | 0 | 0 |
| ENSMUSG00000101298 | 0 | 0 | 0 | 0 | 0 | 0 | 0 |
| ENSMUSG00000101299 | 0 | 0 | 0 | 0 | 0 | 0 | 0 |
| ENSMUSG00000101300 | 0 | 0 | 0 | 0 | 0 | 0 | 0 |
| ENSMUSG00000101301 | 0.289064826 | 0 | 0 | 0.25 | 0 | 0 | 0 |
| ENSMUSG00000101302 | 0 | 0 | 0 | 0 | 0 | 0 | 0 |
| ENSMUSG00000101303 | 0 | 0 | 0 | 0.25 | 0 | 0 | 0 |
| ENSMUSG00000101304 | 0 | 0 | 0 | 0 | 0 | 0 | 0 |
| ENSMUSG00000101305 | 0 | 0 | 0 | 0 | 0 | 0 | 0 |
| ENSMUSG00000101306 | 0 | 0 | 0 | 0 | 0 | 0 | 0 |
| ENSMUSG00000101307 | 0 | 0 | 0 | 0 | 0 | 0 | 0 |
| ENSMUSG00000101308 | 0.289064826 | 0.25 | 0.396240625 | 0.75 | 0.489301084 | 0 | 0.448487648 |
| ENSMUSG00000101309 | 0 | 0 | 0 | 0 | 0 | 0 | 0 |
| ENSMUSG00000101310 | 0 | 0 | 0 | 0 | 0 | 0 | 0 |
| ENSMUSG00000101311 | 0 | 0 | 0 | 0 | 0 | 0 | 0 |
| ENSMUSG00000101312 | 0 | 0 | 0 | 0 | 0 | 0 | 0 |
| ENSMUSG00000101313 | 0 | 0 | 0 | 0 | 0 | 0 | 0 |
| ENSMUSG00000101314 | 0 | 0 | 0 | 0 | 0 | 0 | 0 |
| ENSMUSG00000101315 | 0 | 0 | 0 | 0 | 0 | 0 | 0 |
| ENSMUSG00000101316 | 1.036286563 | 1.02186571 | 1.061981878 | 1.238549078 | 1.121713561 | 1.079181246 | 1.054044578 |
| ENSMUSG00000101317 | 0 | 0 | 0 | 0 | 0 | 0 | 0 |
| ENSMUSG00000101318 | 0 | 0 | 0 | 0 | 0 | 0 | 0 |

| | | | | | | | |
|---|---|---|---|---|---|---|---|
| ENSMUSG00000101319 | 0 | 0 | 0 | 0 | 0 | 0 | 0 |
| ENSMUSG00000101320 | 0 | 0 | 0 | 0.396240625 | 0 | 0 | 0.224243824 |
| ENSMUSG00000101321 | 0 | 0 | 0 | 0 | 0 | 0 | 0 |
| ENSMUSG00000101322 | 0 | 0 | 0 | 0 | 0 | 0 | 0 |
| ENSMUSG00000101323 | 0 | 0 | 0 | 0 | 0 | 0 | 0 |
| ENSMUSG00000101324 | 0 | 0 | 0 | 0 | 0 | 0 | 0 |
| ENSMUSG00000101325 | 0 | 0 | 0 | 0 | 0 | 0 | 0 |
| ENSMUSG00000101326 | 0 | 0 | 0 | 0 | 0 | 0 | 0 |
| ENSMUSG00000101327 | 0 | 0 | 0 | 0 | 0 | 0 | 0 |
| ENSMUSG00000101328 | 0 | 0.396240625 | 0 | 0.25 | 0 | 0 | 0 |
| ENSMUSG00000101329 | 0 | 0 | 0 | 0 | 0 | 0 | 0 |
| ENSMUSG00000101330 | 0 | 0 | 0.25 | 0 | 0 | 0 | 0 |
| ENSMUSG00000101331 | 0 | 0 | 0 | 0 | 0 | 0 | 0 |
| ENSMUSG00000101332 | 0 | 0 | 0 | 0 | 0 | 0 | 0 |
| ENSMUSG00000101333 | 0 | 0 | 0 | 0 | 0 | 0 | 0 |
| ENSMUSG00000101334 | 0 | 0 | 0 | 0 | 0 | 0 | 0 |
| ENSMUSG00000101335 | 1.036286563 | 0.976722649 | 1.114857905 | 1.348079356 | 1.163285805 | 1.361727836 | 1.20139795 |
| ENSMUSG00000101336 | 0 | 0 | 0 | 0 | 0 | 0 | 0 |
| ENSMUSG00000101337 | 0 | 0.25 | 0.25 | 0.396240625 | 0.244650542 | 0.301029996 | 0 |
| ENSMUSG00000101338 | 0 | 0 | 0 | 0 | 0 | 0 | 0 |
| ENSMUSG00000101339 | 0 | 0 | 0 | 0 | 0 | 0 | 0 |
| ENSMUSG00000101340 | 0 | 0 | 0 | 0 | 0 | 0 | 0 |
| ENSMUSG00000101341 | 0 | 0 | 0 | 0 | 0 | 0 | 0 |
| ENSMUSG00000101342 | 0 | 0 | 0 | 0 | 0 | 0 | 0 |
| ENSMUSG00000101343 | 0 | 0 | 0 | 0 | 0 | 0 | 0 |
| ENSMUSG00000101344 | 0 | 0 | 0 | 0 | 0 | 0 | 0 |
| ENSMUSG00000101345 | 0 | 0 | 0 | 0 | 0 | 0 | 0 |
| ENSMUSG00000101346 | 0 | 0 | 0 | 0 | 0 | 0 | 0 |
| ENSMUSG00000101347 | 0 | 0 | 0 | 0 | 0 | 0 | 0 |
| ENSMUSG00000101348 | 0 | 0 | 0 | 0 | 0 | 0 | 0 |
| ENSMUSG00000101349 | 0 | 0 | 0 | 0 | 0 | 0 | 0 |
| ENSMUSG00000101350 | 0 | 0 | 0 | 0 | 0 | 0 | 0 |
| ENSMUSG00000101351 | 0 | 0 | 0 | 0 | 0 | 0 | 0 |
| ENSMUSG00000101352 | 0 | 0 | 0 | 0 | 0 | 0 | 0 |

| | | | | | | | |
|---|---|---|---|---|---|---|---|
| ENSMUSG00000101353 | 0 | 0 | 0 | 0 | 0 | 0 | 0 |
| ENSMUSG00000101354 | 0 | 0 | 0 | 0 | 0 | 0 | 0 |
| ENSMUSG00000101355 | 0 | 0 | 0 | 0 | 0 | 0 | 0 |
| ENSMUSG00000101356 | 0 | 0.396240625 | 0.396240625 | 0.25 | 0.686820904 | 0 | 0.355418052 |
| ENSMUSG00000101357 | 0 | 0 | 0 | 0 | 0 | 0 | 0 |
| ENSMUSG00000101358 | 0 | 0 | 0 | 0 | 0 | 0 | 0 |
| ENSMUSG00000101359 | 0 | 0 | 0 | 0 | 0 | 0 | 0 |
| ENSMUSG00000101360 | 0 | 0 | 0 | 0 | 0 | 0 | 0 |
| ENSMUSG00000101361 | 0 | 0 | 0 | 0 | 0 | 0 | 0 |
| ENSMUSG00000101362 | 0 | 0 | 0 | 0 | 0 | 0 | 0 |
| ENSMUSG00000101363 | 0 | 0 | 0 | 0 | 0 | 0 | 0 |
| ENSMUSG00000101364 | 0 | 0 | 0 | 0 | 0 | 0 | 0 |
| ENSMUSG00000101365 | 0.671187741 | 0.25 | 0.580482024 | 0 | 0.244650542 | 0 | 0.224243824 |
| ENSMUSG00000101366 | 0 | 0 | 0 | 0 | 0 | 0 | 0 |
| ENSMUSG00000101367 | 0 | 0 | 0 | 0 | 0 | 0 | 0 |
| ENSMUSG00000101368 | 0 | 0 | 0 | 0 | 0 | 0 | 0 |
| ENSMUSG00000101369 | 0 | 0 | 0 | 0 | 0 | 0 | 0 |
| ENSMUSG00000101370 | 0 | 0 | 0 | 0 | 0 | 0 | 0 |
| ENSMUSG00000101371 | 0 | 0 | 0 | 0 | 0 | 0 | 0 |
| ENSMUSG00000101372 | 0 | 0 | 0 | 0 | 0 | 0 | 0.224243824 |
| ENSMUSG00000101373 | 0 | 0 | 0 | 0 | 0 | 0 | 0 |
| ENSMUSG00000101374 | 0 | 0 | 0 | 0 | 0 | 0 | 0 |
| ENSMUSG00000101375 | 0 | 0 | 0 | 0 | 0 | 0 | 0 |
| ENSMUSG00000101376 | 0 | 0 | 0 | 0 | 0 | 0 | 0 |
| ENSMUSG00000101377 | 0 | 0 | 0 | 0 | 0 | 0 | 0 |
| ENSMUSG00000101378 | 0 | 0 | 0 | 0 | 0 | 0 | 0 |
| ENSMUSG00000101379 | 0 | 0 | 0 | 0 | 0 | 0 | 0 |
| ENSMUSG00000101380 | 0 | 0 | 0 | 0 | 0 | 0 | 0 |
| ENSMUSG00000101381 | 0 | 0 | 0 | 0 | 0 | 0 | 0 |
| ENSMUSG00000101382 | 0 | 0 | 0 | 0 | 0 | 0 | 0 |
| ENSMUSG00000101383 | 0 | 0 | 0 | 0 | 0 | 0 | 0 |
| ENSMUSG00000101385 | 0 | 0 | 0 | 0 | 0 | 0 | 0 |
| ENSMUSG00000101386 | 0.45815691 | 0 | 0.396240625 | 0.25 | 0 | 0.301029996 | 0.224243824 |
| ENSMUSG00000101387 | 0 | 0.25 | 0.396240625 | 0.396240625 | 0 | 0 | 0.448487648 |

| | | | | | | | |
|---|---|---|---|---|---|---|---|
| ENSMUSG00000101388 | 0 | 0 | 0 | 0 | 0 | 0 | 0 |
| ENSMUSG00000101389 | 0 | 0 | 0 | 0 | 0 | 0 | 0 |
| ENSMUSG00000101390 | 0 | 0 | 0 | 0 | 0 | 0 | 0 |
| ENSMUSG00000101391 | 0 | 0 | 0 | 0 | 0 | 0 | 0 |
| ENSMUSG00000101392 | 0 | 0 | 0 | 0 | 0 | 0 | 0 |
| ENSMUSG00000101393 | 0 | 0 | 0 | 0 | 0 | 0 | 0 |
| ENSMUSG00000101394 | 0 | 0 | 0 | 0 | 0 | 0.477121255 | 0 |
| ENSMUSG00000101395 | 0 | 0 | 0 | 0 | 0 | 0 | 0 |
| ENSMUSG00000101396 | 0 | 0 | 0 | 0 | 0 | 0 | 0 |
| ENSMUSG00000101397 | 0 | 0 | 0 | 0 | 0 | 0 | 0 |
| ENSMUSG00000101398 | 0 | 0 | 0 | 0 | 0 | 0 | 0 |
| ENSMUSG00000101399 | 0 | 0 | 0 | 0 | 0 | 0 | 0 |
| ENSMUSG00000101400 | 0.289064826 | 0.396240625 | 0 | 0.396240625 | 0.489301084 | 0.301029996 | 0.355418052 |
| ENSMUSG00000101401 | 0 | 0 | 0 | 0 | 0 | 0 | 0 |
| ENSMUSG00000101402 | 0 | 0 | 0 | 0 | 0 | 0 | 0 |
| ENSMUSG00000101403 | 0 | 0 | 0 | 0 | 0 | 0 | 0 |
| ENSMUSG00000101404 | 0 | 0 | 0 | 0 | 0 | 0 | 0 |
| ENSMUSG00000101405 | 0 | 0 | 0 | 0 | 0 | 0 | 0 |
| ENSMUSG00000101406 | 0 | 0 | 0 | 0 | 0 | 0 | 0 |
| ENSMUSG00000101407 | 0 | 0 | 0 | 0 | 0 | 0 | 0 |
| ENSMUSG00000101408 | 0 | 0 | 0 | 0 | 0 | 0 | 0 |
| ENSMUSG00000101409 | 0 | 0 | 0 | 0 | 0 | 0 | 0 |
| ENSMUSG00000101410 | 0 | 0 | 0 | 0 | 0 | 0 | 0 |
| ENSMUSG00000101411 | 0 | 0 | 0 | 0 | 0 | 0 | 0 |
| ENSMUSG00000101412 | 0 | 0 | 0 | 0 | 0 | 0 | 0 |
| ENSMUSG00000101413 | 0 | 0 | 0 | 0 | 0 | 0 | 0 |
| ENSMUSG00000101414 | 0.45815691 | 0.646240625 | 0.646240625 | 0.580482024 | 0.632412477 | 0.698970004 | 0.448487648 |
| ENSMUSG00000101415 | 0 | 0 | 0 | 0 | 0 | 0 | 0 |
| ENSMUSG00000101416 | 0 | 0 | 0 | 0 | 0 | 0 | 0 |
| ENSMUSG00000101417 | 0 | 0 | 0 | 0 | 0 | 0 | 0 |
| ENSMUSG00000101418 | 0 | 0 | 0 | 0 | 0 | 0 | 0 |
| ENSMUSG00000101419 | 0 | 0 | 0 | 0 | 0 | 0 | 0 |
| ENSMUSG00000101420 | 0 | 0 | 0 | 0 | 0 | 0 | 0 |
| ENSMUSG00000101421 | 0 | 0 | 0 | 0 | 0 | 0 | 0 |

| | | | | | | | |
|---|---|---|---|---|---|---|---|
| ENSMUSG00000101422 | 0 | 0 | 0 | 0 | 0 | 0 | 0 |
| ENSMUSG00000101423 | 0 | 0 | 0 | 0 | 0 | 0 | 0 |
| ENSMUSG00000101424 | 0 | 0 | 0 | 0 | 0 | 0 | 0 |
| ENSMUSG00000101425 | 0 | 0 | 0 | 0 | 0 | 0 | 0 |
| ENSMUSG00000101426 | 0 | 0 | 0 | 0 | 0 | 0 | 0 |
| ENSMUSG00000101427 | 0 | 0 | 0 | 0 | 0 | 0 | 0 |
| ENSMUSG00000101428 | 0 | 0 | 0 | 0 | 0 | 0 | 0 |
| ENSMUSG00000101429 | 0 | 0 | 0 | 0 | 0.244650542 | 0 | 0 |
| ENSMUSG00000101431 | 1.847794126 | 1.87892496 | 1.909156155 | 1.282320754 | 1.607318736 | 1.414973348 | 1.61448206 |
| ENSMUSG00000101433 | 0 | 0 | 0 | 0 | 0 | 0 | 0 |
| ENSMUSG00000101434 | 0 | 0 | 0 | 0 | 0 | 0 | 0 |
| ENSMUSG00000101435 | 0 | 0 | 0 | 0 | 0 | 0 | 0 |
| ENSMUSG00000101436 | 0 | 0 | 0 | 0 | 0 | 0 | 0 |
| ENSMUSG00000101437 | 0 | 0 | 0 | 0 | 0 | 0 | 0 |
| ENSMUSG00000101438 | 0 | 0 | 0.25 | 0 | 0 | 0 | 0.224243824 |
| ENSMUSG00000101439 | 0 | 0 | 0 | 0 | 0 | 0 | 0 |
| ENSMUSG00000101440 | 0 | 0 | 0 | 0 | 0 | 0 | 0 |
| ENSMUSG00000101441 | 0 | 0 | 0 | 0 | 0 | 0 | 0 |
| ENSMUSG00000101442 | 0 | 0 | 0 | 0 | 0 | 0 | 0 |
| ENSMUSG00000101443 | 0 | 0.25 | 0 | 0 | 0 | 0 | 0 |
| ENSMUSG00000101444 | 0 | 0 | 0 | 0 | 0 | 0 | 0 |
| ENSMUSG00000101445 | 0 | 0 | 0 | 0 | 0 | 0 | 0 |
| ENSMUSG00000101446 | 0 | 0 | 0 | 0 | 0 | 0 | 0 |
| ENSMUSG00000101447 | 0 | 0.580482024 | 0.5 | 0 | 0.686820904 | 0.301029996 | 0.520678036 |
| ENSMUSG00000101448 | 0 | 0 | 0 | 0 | 0 | 0 | 0 |
| ENSMUSG00000101449 | 0 | 0.25 | 0 | 0.25 | 0 | 0 | 0 |
| ENSMUSG00000101450 | 0.671187741 | 0.92510993 | 1.02186571 | 1.02186571 | 0.846351821 | 0.698970004 | 0.896975297 |
| ENSMUSG00000101451 | 0.289064826 | 0.25 | 0.5 | 0.5 | 0.244650542 | 0 | 0 |
| ENSMUSG00000101452 | 0.578129653 | 0.701838731 | 0.896240625 | 0.646240625 | 0.733951626 | 0.602059991 | 0.579661877 |
| ENSMUSG00000101453 | 0 | 0 | 0 | 0 | 0 | 0 | 0 |
| ENSMUSG00000101454 | 0 | 0 | 0 | 0 | 0 | 0 | 0 |
| ENSMUSG00000101455 | 0 | 0 | 0 | 0 | 0 | 0 | 0 |
| ENSMUSG00000101456 | 0 | 0 | 0 | 0 | 0 | 0 | 0 |
| ENSMUSG00000101457 | 0 | 0 | 0 | 0 | 0 | 0 | 0 |

| | | | | | | | |
|---|---|---|---|---|---|---|---|
| ENSMUSG00000101458 | 0.671187741 | 0.580482024 | 0.701838731 | 0.92510993 | 0.846351821 | 0.698970004 | 0.803905701 |
| ENSMUSG00000101459 | 0 | 0 | 0 | 0 | 0 | 0 | 0 |
| ENSMUSG00000101462 | 1.036286563 | 1.26109853 | 1.25 | 0.830482024 | 0.812711509 | 0.77815125 | 0.916588299 |
| ENSMUSG00000101463 | 0 | 0 | 0 | 0 | 0 | 0 | 0 |
| ENSMUSG00000101464 | 0 | 0 | 0 | 0 | 0 | 0 | 0 |
| ENSMUSG00000101465 | 0 | 0 | 0 | 0 | 0 | 0 | 0 |
| ENSMUSG00000101466 | 0 | 0 | 0 | 0 | 0 | 0 | 0 |
| ENSMUSG00000101467 | 0 | 0 | 0 | 0 | 0 | 0 | 0 |
| ENSMUSG00000101468 | 0 | 0 | 0 | 0 | 0 | 0 | 0 |
| ENSMUSG00000101469 | 0 | 0 | 0 | 0 | 0 | 0 | 0 |
| ENSMUSG00000101470 | 0 | 0 | 0 | 0 | 0 | 0 | 0 |
| ENSMUSG00000101471 | 0 | 0 | 0 | 0 | 0 | 0 | 0 |
| ENSMUSG00000101472 | 0 | 0 | 0 | 0 | 0 | 0 | 0 |
| ENSMUSG00000101473 | 0 | 0 | 0 | 0 | 0 | 0 | 0 |
| ENSMUSG00000101474 | 0 | 0 | 0 | 0 | 0 | 0 | 0 |
| ENSMUSG00000101475 | 0 | 0 | 0 | 0 | 0 | 0 | 0 |
| ENSMUSG00000101476 | 0 | 0 | 0 | 0 | 0.244650542 | 0 | 0 |
| ENSMUSG00000101477 | 0 | 0 | 0 | 0 | 0 | 0 | 0 |
| ENSMUSG00000101478 | 0 | 0 | 0 | 0 | 0 | 0 | 0 |
| ENSMUSG00000101479 | 0 | 0 | 0 | 0 | 0 | 0 | 0 |
| ENSMUSG00000101480 | 0 | 0 | 0 | 0 | 0 | 0 | 0 |
| ENSMUSG00000101481 | 0 | 0 | 0 | 0 | 0 | 0 | 0 |
| ENSMUSG00000101482 | 0 | 0 | 0 | 0 | 0 | 0 | 0 |
| ENSMUSG00000101483 | 0 | 0 | 0 | 0 | 0.244650542 | 0 | 0 |
| ENSMUSG00000101484 | 0 | 0 | 0 | 0 | 0 | 0 | 0 |
| ENSMUSG00000101485 | 0 | 0 | 0 | 0 | 0 | 0 | 0 |
| ENSMUSG00000101486 | 0 | 0 | 0 | 0 | 0 | 0 | 0 |
| ENSMUSG00000101487 | 0 | 0 | 0 | 0 | 0 | 0 | 0 |
| ENSMUSG00000101488 | 0 | 0 | 0 | 0 | 0 | 0 | 0 |
| ENSMUSG00000101489 | 0 | 0 | 0 | 0 | 0 | 0 | 0 |
| ENSMUSG00000101490 | 0 | 0 | 0 | 0 | 0 | 0 | 0 |
| ENSMUSG00000101491 | 0 | 0 | 0 | 0 | 0 | 0 | 0 |
| ENSMUSG00000101492 | 0 | 0 | 0 | 0 | 0 | 0 | 0 |
| ENSMUSG00000101493 | 0.289064826 | 0.580482024 | 0.5 | 0.646240625 | 0.632412477 | 0.477121255 | 0.579661877 |

| | | | | | | | |
|---|---|---|---|---|---|---|---|
| ENSMUSG00000101494 | 0 | 0 | 0 | 0 | 0 | 0 | 0 |
| ENSMUSG00000101495 | 0 | 0 | 0 | 0 | 0 | 0 | 0 |
| ENSMUSG00000101496 | 0 | 0.396240625 | 0.25 | 0 | 0.489301084 | 0 | 0.355418052 |
| ENSMUSG00000101497 | 0 | 0 | 0 | 0 | 0 | 0 | 0 |
| ENSMUSG00000101498 | 0 | 0 | 0 | 0 | 0 | 0 | 0 |
| ENSMUSG00000101499 | 0 | 0 | 0 | 0 | 0 | 0 | 0 |
| ENSMUSG00000101500 | 0 | 0 | 0 | 0 | 0 | 0 | 0 |
| ENSMUSG00000101501 | 0 | 0 | 0.25 | 0 | 0 | 0.301029996 | 0 |
| ENSMUSG00000101502 | 0 | 0.25 | 0 | 0 | 0 | 0.301029996 | 0 |
| ENSMUSG00000101503 | 0 | 0 | 0 | 0 | 0 | 0 | 0 |
| ENSMUSG00000101504 | 0 | 0 | 0 | 0 | 0 | 0 | 0 |
| ENSMUSG00000101505 | 0 | 0 | 0.25 | 0 | 0 | 0.301029996 | 0 |
| ENSMUSG00000101506 | 0 | 0 | 0 | 0 | 0 | 0 | 0 |
| ENSMUSG00000101507 | 0 | 0 | 0 | 0 | 0 | 0 | 0 |
| ENSMUSG00000101508 | 0 | 0 | 0 | 0 | 0 | 0 | 0 |
| ENSMUSG00000101509 | 0 | 0 | 0 | 0 | 0 | 0 | 0 |
| ENSMUSG00000101510 | 0 | 0 | 0 | 0 | 0 | 0 | 0 |
| ENSMUSG00000101511 | 0 | 0 | 0 | 0 | 0 | 0 | 0 |
| ENSMUSG00000101512 | 0 | 0 | 0 | 0 | 0 | 0 | 0 |
| ENSMUSG00000101513 | 0 | 0 | 0 | 0 | 0 | 0 | 0 |
| ENSMUSG00000101514 | 0 | 0 | 0 | 0 | 0 | 0 | 0 |
| ENSMUSG00000101515 | 0 | 0 | 0 | 0 | 0 | 0 | 0 |
| ENSMUSG00000101516 | 0 | 0 | 0 | 0 | 0 | 0 | 0 |
| ENSMUSG00000101517 | 0 | 0 | 0 | 0.396240625 | 0 | 0 | 0.224243824 |
| ENSMUSG00000101518 | 0 | 0 | 0 | 0 | 0 | 0 | 0 |
| ENSMUSG00000101519 | 0 | 0 | 0 | 0 | 0 | 0.301029996 | 0 |
| ENSMUSG00000101520 | 0 | 0 | 0 | 0 | 0 | 0 | 0 |
| ENSMUSG00000101521 | 0 | 0 | 0 | 0 | 0 | 0 | 0 |
| ENSMUSG00000101522 | 0 | 0 | 0 | 0 | 0 | 0 | 0 |
| ENSMUSG00000101523 | 0 | 0 | 0 | 0 | 0 | 0 | 0 |
| ENSMUSG00000101525 | 0 | 0 | 0 | 0 | 0 | 0 | 0 |
| ENSMUSG00000101526 | 0 | 0 | 0 | 0 | 0 | 0 | 0 |
| ENSMUSG00000101527 | 0 | 0 | 0 | 0 | 0 | 0 | 0 |
| ENSMUSG00000101528 | 0 | 0 | 0 | 0 | 0 | 0 | 0 |

| | | | | | | | |
|---|---|---|---|---|---|---|---|
| ENSMUSG00000101529 | 0 | 0 | 0 | 0 | 0 | 0 | 0 |
| ENSMUSG00000101530 | 0 | 0 | 0 | 0 | 0 | 0 | 0 |
| ENSMUSG00000101531 | 0 | 0 | 0 | 0 | 0 | 0 | 0 |
| ENSMUSG00000101532 | 0 | 0 | 0.25 | 0 | 0 | 0 | 0.224243824 |
| ENSMUSG00000101533 | 0 | 0 | 0 | 0 | 0 | 0 | 0 |
| ENSMUSG00000101534 | 0 | 0 | 0 | 0 | 0 | 0 | 0 |
| ENSMUSG00000101535 | 0 | 0 | 0 | 0 | 0 | 0 | 0 |
| ENSMUSG00000101536 | 0 | 0 | 0 | 0 | 0 | 0 | 0 |
| ENSMUSG00000101537 | 0 | 0 | 0 | 0 | 0 | 0 | 0 |
| ENSMUSG00000101538 | 0 | 0 | 0 | 0 | 0 | 0 | 0 |
| ENSMUSG00000101539 | 0 | 0 | 0 | 0 | 0 | 0 | 0 |
| ENSMUSG00000101540 | 0.289064826 | 0.396240625 | 0.25 | 0.25 | 0.244650542 | 0.301029996 | 0.448487648 |
| ENSMUSG00000101541 | 0 | 0 | 0 | 0 | 0 | 0 | 0 |
| ENSMUSG00000101542 | 0 | 0 | 0 | 0 | 0 | 0 | 0 |
| ENSMUSG00000101543 | 0 | 0 | 0 | 0 | 0 | 0 | 0 |
| ENSMUSG00000101544 | 0 | 0 | 0 | 0 | 0 | 0 | 0 |
| ENSMUSG00000101545 | 0 | 0.25 | 0.396240625 | 0.396240625 | 0 | 0.301029996 | 0 |
| ENSMUSG00000101546 | 0 | 0 | 0 | 0 | 0 | 0 | 0 |
| ENSMUSG00000101547 | 0 | 0 | 0 | 0 | 0 | 0 | 0 |
| ENSMUSG00000101548 | 0 | 0 | 0 | 0 | 0 | 0 | 0 |
| ENSMUSG00000101549 | 0.671187741 | 0.79248125 | 0.864857905 | 0.92510993 | 0.978602168 | 0.698970004 | 0.74492186 |
| ENSMUSG00000101550 | 0 | 0 | 0 | 0 | 0 | 0 | 0 |
| ENSMUSG00000101551 | 0.289064826 | 0 | 0.5 | 0.396240625 | 0.244650542 | 0 | 0.448487648 |
| ENSMUSG00000101552 | 0 | 0 | 0 | 0 | 0 | 0 | 0 |
| ENSMUSG00000101553 | 0 | 0 | 0 | 0 | 0 | 0 | 0 |
| ENSMUSG00000101554 | 0 | 0 | 0 | 0 | 0 | 0 | 0 |
| ENSMUSG00000101555 | 0 | 0 | 0 | 0.25 | 0 | 0 | 0.355418052 |
| ENSMUSG00000101556 | 0 | 0.25 | 0 | 0.25 | 0.244650542 | 0.301029996 | 0.224243824 |
| ENSMUSG00000101557 | 0 | 0 | 0 | 0 | 0 | 0 | 0 |
| ENSMUSG00000101558 | 0 | 0 | 0 | 0 | 0 | 0 | 0 |
| ENSMUSG00000101559 | 0.289064826 | 0 | 0 | 0.25 | 0 | 0 | 0 |
| ENSMUSG00000101560 | 0 | 0 | 0 | 0 | 0 | 0 | 0 |
| ENSMUSG00000101562 | 0 | 0 | 0 | 0 | 0 | 0 | 0 |
| ENSMUSG00000101563 | 0 | 0 | 0 | 0 | 0 | 0 | 0 |

| | | | | | | | |
|---|---|---|---|---|---|---|---|
| ENSMUSG00000101564 | 0 | 0 | 0 | 0 | 0 | 0 | 0 |
| ENSMUSG00000101565 | 0 | 0 | 0 | 0 | 0 | 0 | 0 |
| ENSMUSG00000101566 | 0 | 0 | 0 | 0 | 0 | 0 | 0 |
| ENSMUSG00000101567 | 0 | 0 | 0.25 | 0.25 | 0.244650542 | 0 | 0 |
| ENSMUSG00000101568 | 0 | 0.25 | 0 | 0.396240625 | 0 | 0 | 0 |
| ENSMUSG00000101569 | 0 | 0 | 0 | 0 | 0 | 0 | 0 |
| ENSMUSG00000101570 | 0 | 0 | 0 | 0 | 0 | 0 | 0 |
| ENSMUSG00000101571 | 0.289064826 | 0 | 0.25 | 0.25 | 0 | 0 | 0 |
| ENSMUSG00000101573 | 0 | 0 | 0 | 0 | 0 | 0 | 0 |
| ENSMUSG00000101574 | 0.289064826 | 0.25 | 0 | 0 | 0 | 0 | 0 |
| ENSMUSG00000101575 | 0 | 0 | 0 | 0 | 0 | 0 | 0 |
| ENSMUSG00000101576 | 0 | 0 | 0 | 0 | 0 | 0 | 0 |
| ENSMUSG00000101578 | 0 | 0 | 0 | 0 | 0 | 0 | 0 |
| ENSMUSG00000101579 | 0 | 0 | 0 | 0 | 0 | 0 | 0 |
| ENSMUSG00000101580 | 0 | 0 | 0 | 0 | 0 | 0 | 0 |
| ENSMUSG00000101581 | 0 | 0.396240625 | 0.25 | 0.25 | 0 | 0.301029996 | 0.355418052 |
| ENSMUSG00000101582 | 0 | 0 | 0 | 0 | 0 | 0 | 0 |
| ENSMUSG00000101583 | 0 | 0 | 0 | 0 | 0 | 0 | 0 |
| ENSMUSG00000101584 | 0 | 0 | 0 | 0 | 0 | 0 | 0 |
| ENSMUSG00000101585 | 0 | 0.25 | 0 | 0 | 0.489301084 | 0.698970004 | 0.355418052 |
| ENSMUSG00000101586 | 0.867194479 | 0.896240625 | 0.896240625 | 0.75 | 0.632412477 | 0.77815125 | 0.775756176 |
| ENSMUSG00000101587 | 0.289064826 | 0.25 | 0.25 | 0.25 | 0.244650542 | 0 | 0 |
| ENSMUSG00000101588 | 0 | 0 | 0 | 0.396240625 | 0 | 0 | 0 |
| ENSMUSG00000101589 | 0.578129653 | 0 | 0.396240625 | 0.5 | 0.387761935 | 0.301029996 | 0.224243824 |
| ENSMUSG00000101590 | 0 | 0 | 0 | 0 | 0 | 0 | 0 |
| ENSMUSG00000101591 | 0 | 0 | 0 | 0 | 0 | 0 | 0 |
| ENSMUSG00000101592 | 0 | 0 | 0 | 0 | 0 | 0 | 0 |
| ENSMUSG00000101593 | 0 | 0 | 0 | 0 | 0 | 0 | 0 |
| ENSMUSG00000101594 | 0 | 0 | 0 | 0 | 0 | 0 | 0 |
| ENSMUSG00000101595 | 0 | 0 | 0 | 0 | 0 | 0 | 0 |
| ENSMUSG00000101596 | 0 | 0 | 0 | 0 | 0 | 0 | 0 |
| ENSMUSG00000101597 | 0 | 0 | 0 | 0 | 0 | 0 | 0 |
| ENSMUSG00000101598 | 0 | 0 | 0 | 0 | 0 | 0 | 0 |
| ENSMUSG00000101599 | 1.325351389 | 1.238549078 | 1.339388001 | 1.488549078 | 1.433158226 | 1.643452676 | 1.319146374 |

| | | | | | | | |
|---|---|---|---|---|---|---|---|
| ENSMUSG00000101600 | 0 | 0 | 0 | 0 | 0 | 0 | 0 |
| ENSMUSG00000101601 | 0 | 0 | 0 | 0 | 0 | 0 | 0 |
| ENSMUSG00000101602 | 0 | 0 | 0 | 0 | 0 | 0 | 0 |
| ENSMUSG00000101603 | 0.578129653 | 0.25 | 0 | 0.25 | 0 | 0 | 0.355418052 |
| ENSMUSG00000101605 | 0 | 0 | 0 | 0 | 0 | 0 | 0 |
| ENSMUSG00000101606 | 0 | 0 | 0 | 0 | 0 | 0 | 0 |
| ENSMUSG00000101607 | 0 | 0 | 0 | 0 | 0 | 0 | 0 |
| ENSMUSG00000101608 | 0.289064826 | 0.396240625 | 0.396240625 | 0 | 0.387761935 | 0 | 0.355418052 |
| ENSMUSG00000101609 | 2.165631214 | 1.994319981 | 2.012462137 | 2.084962501 | 1.965381292 | 2.204119983 | 1.788855757 |
| ENSMUSG00000101610 | 0 | 0.25 | 0 | 0.5 | 0 | 0 | 0 |
| ENSMUSG00000101611 | 0 | 0 | 0 | 0 | 0 | 0 | 0 |
| ENSMUSG00000101612 | 0 | 0 | 0 | 0 | 0 | 0 | 0 |
| ENSMUSG00000101613 | 0 | 0 | 0 | 0 | 0 | 0 | 0 |
| ENSMUSG00000101614 | 0 | 0 | 0 | 0 | 0 | 0 | 0 |
| ENSMUSG00000101615 | 0 | 0 | 0 | 0 | 0 | 0 | 0 |
| ENSMUSG00000101616 | 0 | 0 | 0 | 0 | 0 | 0 | 0 |
| ENSMUSG00000101617 | 0 | 0 | 0 | 0 | 0 | 0 | 0 |
| ENSMUSG00000101618 | 0 | 0 | 0 | 0 | 0 | 0 | 0 |
| ENSMUSG00000101619 | 0 | 0 | 0 | 0 | 0 | 0 | 0 |
| ENSMUSG00000101621 | 0 | 0 | 0 | 0 | 0 | 0 | 0 |
| ENSMUSG00000101622 | 0 | 0 | 0 | 0 | 0 | 0 | 0 |
| ENSMUSG00000101623 | 0 | 0 | 0 | 0 | 0 | 0 | 0 |
| ENSMUSG00000101624 | 0 | 0 | 0 | 0 | 0 | 0 | 0 |
| ENSMUSG00000101625 | 0.578129653 | 0 | 0.701838731 | 0.646240625 | 0.489301084 | 0.698970004 | 0.355418052 |
| ENSMUSG00000101626 | 0 | 0 | 0 | 0 | 0 | 0 | 0 |
| ENSMUSG00000101627 | 0 | 0 | 0 | 0 | 0 | 0 | 0 |
| ENSMUSG00000101628 | 0 | 0 | 0 | 0 | 0 | 0 | 0.224243824 |
| ENSMUSG00000101629 | 0 | 0 | 0 | 0 | 0 | 0 | 0 |
| ENSMUSG00000101630 | 0.289064826 | 0 | 0 | 0 | 0 | 0 | 0 |
| ENSMUSG00000101631 | 0 | 0 | 0 | 0 | 0 | 0 | 0 |
| ENSMUSG00000101632 | 0 | 0 | 0 | 0 | 0 | 0 | 0 |
| ENSMUSG00000101633 | 0 | 0 | 0 | 0 | 0 | 0 | 0 |
| ENSMUSG00000101634 | 0 | 0 | 0 | 0 | 0 | 0 | 0 |
| ENSMUSG00000101636 | 0 | 0 | 0 | 0 | 0 | 0 | 0 |

| | | | | | | | |
|---|---|---|---|---|---|---|---|
| ENSMUSG00000101637 | 0 | 0 | 0 | 0 | 0 | 0 | 0 |
| ENSMUSG00000101638 | 0 | 0 | 0 | 0 | 0 | 0 | 0 |
| ENSMUSG00000101639 | 0 | 0.25 | 0 | 0 | 0 | 0 | 0 |
| ENSMUSG00000101640 | 0 | 0 | 0 | 0 | 0 | 0 | 0 |
| ENSMUSG00000101641 | 0 | 0 | 0 | 0 | 0.244650542 | 0.301029996 | 0.224243824 |
| ENSMUSG00000101642 | 0 | 0 | 0 | 0 | 0 | 0 | 0 |
| ENSMUSG00000101643 | 0 | 0 | 0 | 0 | 0 | 0 | 0 |
| ENSMUSG00000101644 | 0 | 0 | 0 | 0 | 0 | 0 | 0 |
| ENSMUSG00000101645 | 0 | 0 | 0 | 0 | 0 | 0 | 0 |
| ENSMUSG00000101646 | 0 | 0 | 0 | 0 | 0 | 0 | 0 |
| ENSMUSG00000101647 | 0 | 0 | 0 | 0 | 0 | 0 | 0 |
| ENSMUSG00000101648 | 0 | 0 | 0 | 0 | 0 | 0 | 0 |
| ENSMUSG00000101649 | 0 | 0 | 0 | 0 | 0 | 0 | 0 |
| ENSMUSG00000101650 | 0 | 0 | 0 | 0 | 0 | 0 | 0 |
| ENSMUSG00000101651 | 0 | 0 | 0 | 0 | 0 | 0 | 0 |
| ENSMUSG00000101652 | 0.289064826 | 0.580482024 | 0.830482024 | 0.25 | 0.632412477 | 0.477121255 | 0.520678036 |
| ENSMUSG00000101653 | 0 | 0 | 0 | 0 | 0 | 0 | 0 |
| ENSMUSG00000101654 | 0 | 0 | 0 | 0 | 0 | 0 | 0 |
| ENSMUSG00000101655 | 0.289064826 | 0 | 0 | 0 | 0 | 0 | 0 |
| ENSMUSG00000101656 | 0 | 0 | 0 | 0 | 0 | 0 | 0 |
| ENSMUSG00000101657 | 0 | 0 | 0 | 0 | 0 | 0 | 0 |
| ENSMUSG00000101658 | 0 | 0 | 0 | 0 | 0 | 0 | 0 |
| ENSMUSG00000101659 | 0 | 0 | 0 | 0 | 0 | 0 | 0 |
| ENSMUSG00000101660 | 0 | 0 | 0 | 0 | 0 | 0 | 0 |
| ENSMUSG00000101661 | 0 | 0 | 0 | 0 | 0 | 0 | 0 |
| ENSMUSG00000101662 | 0 | 0 | 0 | 0 | 0.244650542 | 0 | 0.224243824 |
| ENSMUSG00000101664 | 0 | 0 | 0 | 0 | 0 | 0 | 0 |
| ENSMUSG00000101665 | 0 | 0 | 0 | 0 | 0 | 0 | 0 |
| ENSMUSG00000101666 | 0 | 0 | 0 | 0 | 0 | 0 | 0 |
| ENSMUSG00000101667 | 0 | 0 | 0 | 0 | 0 | 0 | 0 |
| ENSMUSG00000101668 | 0 | 0 | 0 | 0 | 0 | 0 | 0 |
| ENSMUSG00000101669 | 0 | 0 | 0 | 0 | 0 | 0 | 0 |
| ENSMUSG00000101670 | 0 | 0 | 0 | 0 | 0 | 0 | 0 |
| ENSMUSG00000101671 | 0 | 0 | 0 | 0 | 0 | 0 | 0 |

| | | | | | | | |
|---|---|---|---|---|---|---|---|
| ENSMUSG00000101672 | 0 | 0 | 0 | 0 | 0 | 0 | 0 |
| ENSMUSG00000101673 | 0 | 0 | 0 | 0 | 0 | 0 | 0 |
| ENSMUSG00000101674 | 0 | 0 | 0 | 0 | 0 | 0 | 0 |
| ENSMUSG00000101675 | 0 | 0 | 0 | 0 | 0 | 0 | 0 |
| ENSMUSG00000101676 | 0 | 0 | 0 | 0 | 0 | 0 | 0 |
| ENSMUSG00000101677 | 0 | 0 | 0 | 0 | 0 | 0.301029996 | 0 |
| ENSMUSG00000101678 | 0.289064826 | 0.5 | 0.701838731 | 0.580482024 | 0.244650542 | 0.698970004 | 0.629532004 |
| ENSMUSG00000101679 | 0 | 0 | 0 | 0 | 0 | 0 | 0 |
| ENSMUSG00000101680 | 0 | 0 | 0 | 0 | 0 | 0 | 0 |
| ENSMUSG00000101681 | 0 | 0 | 0 | 0 | 0 | 0 | 0 |
| ENSMUSG00000101682 | 0.578129653 | 0.580482024 | 0.646240625 | 0.646240625 | 0.568060967 | 0.602059991 | 0.448487648 |
| ENSMUSG00000101683 | 0 | 0 | 0 | 0 | 0 | 0 | 0 |
| ENSMUSG00000101684 | 0 | 0 | 0 | 0 | 0 | 0 | 0 |
| ENSMUSG00000101685 | 0 | 0 | 0.25 | 0 | 0 | 0 | 0 |
| ENSMUSG00000101686 | 0 | 0 | 0 | 0 | 0 | 0 | 0 |
| ENSMUSG00000101687 | 0 | 0 | 0 | 0 | 0 | 0 | 0 |
| ENSMUSG00000101688 | 0 | 0 | 0 | 0 | 0 | 0 | 0 |
| ENSMUSG00000101689 | 0 | 0 | 0 | 0 | 0 | 0 | 0 |
| ENSMUSG00000101690 | 0 | 0 | 0 | 0 | 0 | 0 | 0 |
| ENSMUSG00000101691 | 0 | 0 | 0 | 0 | 0 | 0 | 0 |
| ENSMUSG00000101692 | 0 | 0 | 0 | 0 | 0 | 0 | 0 |
| ENSMUSG00000101693 | 0 | 0 | 0 | 0 | 0 | 0 | 0 |
| ENSMUSG00000101694 | 0 | 0 | 0 | 0 | 0 | 0 | 0 |
| ENSMUSG00000101695 | 0 | 0 | 0 | 0 | 0 | 0 | 0 |
| ENSMUSG00000101696 | 0 | 0 | 0 | 0 | 0 | 0 | 0 |
| ENSMUSG00000101698 | 1.655740331 | 1.602347734 | 1.678561379 | 1.723704441 | 1.642643806 | 1.832508913 | 1.572224412 |
| ENSMUSG00000101699 | 0 | 0 | 0 | 0 | 0 | 0 | 0 |
| ENSMUSG00000101700 | 0 | 0 | 0 | 0 | 0 | 0 | 0 |
| ENSMUSG00000101701 | 0.289064826 | 0.25 | 0 | 0.25 | 0 | 0.301029996 | 0 |
| ENSMUSG00000101702 | 0 | 0 | 0 | 0 | 0 | 0 | 0 |
| ENSMUSG00000101703 | 0 | 0 | 0 | 0 | 0 | 0 | 0 |
| ENSMUSG00000101704 | 0 | 0 | 0 | 0 | 0 | 0 | 0 |
| ENSMUSG00000101705 | 0 | 0 | 0 | 0 | 0 | 0 | 0 |
| ENSMUSG00000101706 | 0 | 0 | 0 | 0 | 0 | 0 | 0 |

| | | | | | | | |
|---|---|---|---|---|---|---|---|
| ENSMUSG00000101707 | 0 | 0 | 0 | 0 | 0 | 0 | 0 |
| ENSMUSG00000101708 | 0 | 0 | 0 | 0 | 0 | 0 | 0 |
| ENSMUSG00000101709 | 0 | 0 | 0 | 0 | 0 | 0 | 0 |
| ENSMUSG00000101710 | 0 | 0 | 0 | 0 | 0 | 0 | 0 |
| ENSMUSG00000101711 | 0 | 0.25 | 0 | 0.25 | 0.489301084 | 0.602059991 | 0.355418052 |
| ENSMUSG00000101712 | 0 | 0 | 0 | 0 | 0 | 0 | 0 |
| ENSMUSG00000101713 | 0 | 0 | 0 | 0 | 0 | 0 | 0 |
| ENSMUSG00000101714 | 0 | 0 | 0 | 0 | 0 | 0 | 0 |
| ENSMUSG00000101715 | 0 | 0 | 0 | 0 | 0 | 0 | 0 |
| ENSMUSG00000101716 | 0.289064826 | 0.25 | 0 | 0 | 0 | 0 | 0 |
| ENSMUSG00000101717 | 0 | 0 | 0 | 0 | 0 | 0 | 0 |
| ENSMUSG00000101718 | 0 | 0 | 0 | 0 | 0 | 0 | 0 |
| ENSMUSG00000101719 | 0 | 0 | 0 | 0 | 0 | 0 | 0 |
| ENSMUSG00000101720 | 0 | 0 | 0 | 0 | 0 | 0 | 0 |
| ENSMUSG00000101721 | 0 | 0 | 0 | 0 | 0 | 0 | 0 |
| ENSMUSG00000101722 | 0 | 0 | 0 | 0 | 0.244650542 | 0 | 0.224243824 |
| ENSMUSG00000101723 | 0 | 0 | 0 | 0 | 0 | 0 | 0 |
| ENSMUSG00000101724 | 0 | 0 | 0 | 0 | 0 | 0 | 0 |
| ENSMUSG00000101725 | 0 | 0 | 0 | 0 | 0 | 0 | 0 |
| ENSMUSG00000101726 | 0.289064826 | 0 | 0 | 0 | 0 | 0 | 0 |
| ENSMUSG00000101727 | 0 | 0 | 0 | 0 | 0 | 0 | 0 |
| ENSMUSG00000101728 | 0 | 0 | 0 | 0 | 0 | 0 | 0 |
| ENSMUSG00000101729 | 0 | 0.25 | 0 | 0 | 0 | 0 | 0 |
| ENSMUSG00000101730 | 0 | 0.25 | 0.25 | 0 | 0 | 0 | 0 |
| ENSMUSG00000101731 | 0.289064826 | 0 | 0 | 0 | 0 | 0.301029996 | 0.448487648 |
| ENSMUSG00000101732 | 0 | 0 | 0 | 0 | 0 | 0 | 0 |
| ENSMUSG00000101733 | 0 | 0 | 0 | 0 | 0 | 0 | 0 |
| ENSMUSG00000101734 | 0 | 0 | 0 | 0 | 0 | 0 | 0 |
| ENSMUSG00000101735 | 0 | 0 | 0 | 0 | 0 | 0 | 0 |
| ENSMUSG00000101736 | 0.289064826 | 0 | 0.25 | 0 | 0 | 0 | 0.224243824 |
| ENSMUSG00000101737 | 0 | 0 | 0 | 0 | 0 | 0 | 0 |
| ENSMUSG00000101738 | 0 | 0 | 0 | 0 | 0 | 0 | 0 |
| ENSMUSG00000101739 | 0 | 0 | 0 | 0 | 0 | 0 | 0 |
| ENSMUSG00000101740 | 0 | 0 | 0 | 0 | 0 | 0 | 0 |

| | | | | | | | |
|---|---|---|---|---|---|---|---|
| ENSMUSG00000101741 | 0 | 0 | 0 | 0 | 0 | 0 | 0 |
| ENSMUSG00000101742 | 0 | 0 | 0 | 0 | 0 | 0 | 0 |
| ENSMUSG00000101743 | 0 | 0 | 0 | 0 | 0 | 0 | 0 |
| ENSMUSG00000101744 | 0 | 0 | 0 | 0 | 0 | 0 | 0 |
| ENSMUSG00000101745 | 0 | 0 | 0 | 0 | 0 | 0 | 0 |
| ENSMUSG00000101746 | 0 | 0 | 0.25 | 0 | 0 | 0.301029996 | 0 |
| ENSMUSG00000101747 | 0 | 0 | 0 | 0 | 0 | 0 | 0 |
| ENSMUSG00000101748 | 0 | 0 | 0 | 0 | 0 | 0 | 0 |
| ENSMUSG00000101750 | 0 | 0 | 0.25 | 0 | 0 | 0 | 0 |
| ENSMUSG00000101751 | 0 | 0 | 0 | 0 | 0 | 0 | 0.224243824 |
| ENSMUSG00000101752 | 0 | 0 | 0 | 0 | 0 | 0 | 0 |
| ENSMUSG00000101753 | 0 | 0 | 0 | 0 | 0 | 0 | 0 |
| ENSMUSG00000101754 | 0 | 0 | 0 | 0 | 0 | 0 | 0 |
| ENSMUSG00000101755 | 0 | 0 | 0 | 0 | 0 | 0 | 0 |
| ENSMUSG00000101756 | 0 | 0 | 0 | 0 | 0 | 0 | 0 |
| ENSMUSG00000101757 | 0 | 0 | 0 | 0 | 0 | 0 | 0 |
| ENSMUSG00000101758 | 0 | 0 | 0 | 0 | 0 | 0 | 0 |
| ENSMUSG00000101759 | 0 | 0 | 0 | 0 | 0 | 0 | 0 |
| ENSMUSG00000101760 | 0 | 0 | 0 | 0 | 0 | 0 | 0 |
| ENSMUSG00000101761 | 0 | 0 | 0 | 0 | 0 | 0 | 0 |
| ENSMUSG00000101762 | 0 | 0 | 0 | 0 | 0 | 0 | 0 |
| ENSMUSG00000101763 | 0 | 0 | 0 | 0 | 0 | 0 | 0 |
| ENSMUSG00000101764 | 0 | 0 | 0 | 0 | 0 | 0 | 0 |
| ENSMUSG00000101765 | 0 | 0 | 0 | 0 | 0 | 0 | 0 |
| ENSMUSG00000101766 | 0 | 0 | 0 | 0 | 0 | 0 | 0 |
| ENSMUSG00000101767 | 0 | 0 | 0 | 0 | 0 | 0 | 0 |
| ENSMUSG00000101768 | 0 | 0.25 | 0 | 0 | 0 | 0 | 0 |
| ENSMUSG00000101769 | 0 | 0 | 0 | 0 | 0 | 0 | 0 |
| ENSMUSG00000101770 | 0 | 0 | 0 | 0 | 0 | 0 | 0 |
| ENSMUSG00000101771 | 0 | 0 | 0 | 0 | 0 | 0 | 0 |
| ENSMUSG00000101772 | 0 | 0 | 0 | 0 | 0 | 0 | 0 |
| ENSMUSG00000101773 | 0 | 0 | 0 | 0 | 0 | 0 | 0 |
| ENSMUSG00000101774 | 0 | 0 | 0 | 0 | 0 | 0 | 0 |
| ENSMUSG00000101775 | 0 | 0 | 0 | 0 | 0 | 0 | 0 |

| | | | | | | | |
|---|---|---|---|---|---|---|---|
| ENSMUSG00000101776 | 1.470606563 | 1.380890489 | 1.364857905 | 1.380890489 | 1.343584837 | 1.230448921 | 1.278288402 |
| ENSMUSG00000101777 | 0.289064826 | 0 | 0 | 0 | 0 | 0 | 0 |
| ENSMUSG00000101778 | 0 | 0 | 0 | 0 | 0 | 0.301029996 | 0 |
| ENSMUSG00000101779 | 0 | 0 | 0 | 0 | 0 | 0 | 0 |
| ENSMUSG00000101780 | 0 | 0 | 0 | 0 | 0 | 0 | 0 |
| ENSMUSG00000101781 | 0 | 0 | 0 | 0 | 0 | 0 | 0 |
| ENSMUSG00000101782 | 0 | 0 | 0 | 0 | 0 | 0 | 0 |
| ENSMUSG00000101783 | 0 | 0 | 0 | 0 | 0 | 0 | 0 |
| ENSMUSG00000101784 | 0 | 0 | 0 | 0 | 0 | 0 | 0 |
| ENSMUSG00000101785 | 0 | 0 | 0 | 0 | 0 | 0 | 0 |
| ENSMUSG00000101786 | 0 | 0 | 0 | 0 | 0 | 0 | 0 |
| ENSMUSG00000101787 | 0 | 0 | 0 | 0 | 0 | 0 | 0 |
| ENSMUSG00000101788 | 0 | 0 | 0 | 0 | 0 | 0 | 0 |
| ENSMUSG00000101789 | 0 | 0 | 0 | 0 | 0 | 0 | 0 |
| ENSMUSG00000101790 | 0 | 0 | 0 | 0 | 0 | 0 | 0 |
| ENSMUSG00000101791 | 0.578129653 | 0.396240625 | 0.580482024 | 0.5 | 0.387761935 | 0 | 0.629532004 |
| ENSMUSG00000101792 | 0 | 0 | 0 | 0 | 0.244650542 | 0 | 0 |
| ENSMUSG00000101793 | 0 | 0 | 0 | 0 | 0 | 0 | 0 |
| ENSMUSG00000101794 | 0 | 0 | 0 | 0 | 0 | 0 | 0 |
| ENSMUSG00000101795 | 0.45815691 | 0.580482024 | 0.580482024 | 0 | 0.489301084 | 0.602059991 | 0.355418052 |
| ENSMUSG00000101796 | 0 | 0 | 0 | 0 | 0 | 0 | 0 |
| ENSMUSG00000101797 | 0 | 0 | 0 | 0 | 0 | 0 | 0 |
| ENSMUSG00000101798 | 0 | 0.25 | 0 | 0 | 0 | 0 | 0 |
| ENSMUSG00000101799 | 0.289064826 | 0 | 0.25 | 0 | 0 | 0 | 0 |
| ENSMUSG00000101800 | 0 | 0 | 0 | 0 | 0 | 0 | 0 |
| ENSMUSG00000101801 | 0 | 0 | 0 | 0 | 0 | 0 | 0 |
| ENSMUSG00000101802 | 0 | 0 | 0 | 0 | 0 | 0 | 0 |
| ENSMUSG00000101803 | 0 | 0 | 0 | 0 | 0 | 0 | 0 |
| ENSMUSG00000101804 | 0 | 0 | 0 | 0 | 0 | 0 | 0 |
| ENSMUSG00000101805 | 0 | 0 | 0 | 0 | 0 | 0 | 0 |
| ENSMUSG00000101806 | 0 | 0 | 0 | 0 | 0 | 0 | 0 |
| ENSMUSG00000101807 | 0 | 0 | 0 | 0 | 0 | 0 | 0 |
| ENSMUSG00000101808 | 0 | 0 | 0 | 0 | 0 | 0 | 0 |
| ENSMUSG00000101809 | 0 | 0 | 0 | 0 | 0 | 0 | 0 |

| | | | | | | | |
|---|---|---|---|---|---|---|---|
| ENSMUSG00000101810 | 0 | 0 | 0 | 0 | 0 | 0 | 0.224243824 |
| ENSMUSG00000101811 | 0 | 0 | 0 | 0 | 0 | 0 | 0 |
| ENSMUSG00000101812 | 0 | 0 | 0 | 0 | 0 | 0 | 0 |
| ENSMUSG00000101813 | 0 | 0 | 0 | 0 | 0 | 0 | 0 |
| ENSMUSG00000101814 | 0 | 0 | 0 | 0 | 0 | 0 | 0 |
| ENSMUSG00000101815 | 0 | 0 | 0 | 0 | 0 | 0 | 0 |
| ENSMUSG00000101816 | 0 | 0 | 0 | 0 | 0 | 0 | 0 |
| ENSMUSG00000101817 | 0 | 0 | 0 | 0 | 0 | 0 | 0 |
| ENSMUSG00000101818 | 0 | 0 | 0 | 0 | 0 | 0 | 0 |
| ENSMUSG00000101819 | 0 | 0 | 0 | 0 | 0 | 0 | 0 |
| ENSMUSG00000101820 | 0 | 0 | 0 | 0 | 0 | 0 | 0 |
| ENSMUSG00000101821 | 0 | 0 | 0 | 0 | 0 | 0 | 0 |
| ENSMUSG00000101822 | 0 | 0 | 0 | 0 | 0 | 0 | 0 |
| ENSMUSG00000101823 | 0.747221736 | 0.75 | 0.79248125 | 0.75 | 0.686820904 | 0.77815125 | 0.629532004 |
| ENSMUSG00000101824 | 0 | 0 | 0 | 0 | 0 | 0 | 0 |
| ENSMUSG00000101825 | 0 | 0 | 0 | 0 | 0 | 0 | 0 |
| ENSMUSG00000101826 | 0 | 0.25 | 0 | 0 | 0 | 0 | 0 |
| ENSMUSG00000101827 | 0 | 0 | 0 | 0 | 0.244650542 | 0 | 0 |
| ENSMUSG00000101828 | 0 | 0.25 | 0.5 | 0.830482024 | 0.387761935 | 0 | 0.520678036 |
| ENSMUSG00000101829 | 0 | 0 | 0 | 0 | 0 | 0 | 0 |
| ENSMUSG00000101830 | 0 | 0 | 0 | 0 | 0 | 0 | 0 |
| ENSMUSG00000101831 | 0 | 0 | 0 | 0 | 0 | 0 | 0 |
| ENSMUSG00000101832 | 0 | 0 | 0 | 0 | 0 | 0 | 0.224243824 |
| ENSMUSG00000101833 | 0 | 0 | 0 | 0 | 0 | 0 | 0 |
| ENSMUSG00000101834 | 0 | 0 | 0 | 0 | 0 | 0 | 0 |
| ENSMUSG00000101835 | 0 | 0 | 0 | 0 | 0 | 0 | 0 |
| ENSMUSG00000101836 | 0 | 0 | 0 | 0 | 0 | 0 | 0 |
| ENSMUSG00000101837 | 0 | 0 | 0 | 0 | 0 | 0 | 0 |
| ENSMUSG00000101838 | 0 | 0.396240625 | 0 | 0 | 0.387761935 | 0 | 0 |
| ENSMUSG00000101839 | 0 | 0 | 0 | 0 | 0 | 0 | 0 |
| ENSMUSG00000101840 | 0 | 0 | 0 | 0 | 0 | 0 | 0 |
| ENSMUSG00000101841 | 0 | 0 | 0 | 0.396240625 | 0 | 0 | 0 |
| ENSMUSG00000101842 | 0 | 0 | 0 | 0 | 0 | 0 | 0 |
| ENSMUSG00000101843 | 0 | 0 | 0 | 0 | 0 | 0 | 0 |

| | | | | | | | |
|---|---|---|---|---|---|---|---|
| ENSMUSG00000101844 | 0 | 0 | 0 | 0 | 0 | 0 | 0 |
| ENSMUSG00000101845 | 0 | 0.396240625 | 0.25 | 0 | 0.632412477 | 0 | 0.355418052 |
| ENSMUSG00000101846 | 0 | 0 | 0 | 0 | 0 | 0 | 0 |
| ENSMUSG00000101847 | 0 | 0 | 0 | 0 | 0 | 0 | 0 |
| ENSMUSG00000101848 | 0 | 0.25 | 0.25 | 0 | 0 | 0 | 0 |
| ENSMUSG00000101849 | 0.45815691 | 0.701838731 | 0 | 0 | 0 | 0 | 0 |
| ENSMUSG00000101850 | 0 | 0 | 0 | 0 | 0 | 0 | 0 |
| ENSMUSG00000101851 | 0 | 0 | 0 | 0 | 0 | 0 | 0 |
| ENSMUSG00000101852 | 0 | 0 | 0 | 0 | 0 | 0 | 0 |
| ENSMUSG00000101853 | 0 | 0 | 0 | 0 | 0 | 0 | 0 |
| ENSMUSG00000101854 | 0 | 0 | 0 | 0 | 0 | 0 | 0 |
| ENSMUSG00000101855 | 0 | 0 | 0 | 0 | 0 | 0 | 0 |
| ENSMUSG00000101856 | 0.811507563 | 0.701838731 | 0.896240625 | 0.79248125 | 1 | 0.903089987 | 0.803905701 |
| ENSMUSG00000101857 | 0 | 0 | 0 | 0 | 0 | 0 | 0 |
| ENSMUSG00000101858 | 0 | 0 | 0 | 0 | 0 | 0 | 0 |
| ENSMUSG00000101859 | 0.289064826 | 0.25 | 0 | 0 | 0.387761935 | 0 | 0 |
| ENSMUSG00000101860 | 0 | 0 | 0 | 0 | 0 | 0 | 0 |
| ENSMUSG00000101861 | 0 | 0 | 0 | 0 | 0 | 0 | 0 |
| ENSMUSG00000101862 | 0 | 0 | 0 | 0.25 | 0 | 0 | 0 |
| ENSMUSG00000101863 | 0 | 0 | 0 | 0 | 0 | 0 | 0 |
| ENSMUSG00000101864 | 0 | 0 | 0 | 0 | 0 | 0 | 0 |
| ENSMUSG00000101865 | 0 | 0 | 0 | 0 | 0 | 0 | 0 |
| ENSMUSG00000101866 | 0 | 0 | 0 | 0 | 0 | 0 | 0 |
| ENSMUSG00000101867 | 0 | 0 | 0 | 0 | 0 | 0 | 0 |
| ENSMUSG00000101868 | 0 | 0 | 0 | 0 | 0 | 0 | 0 |
| ENSMUSG00000101869 | 0 | 0 | 0 | 0 | 0 | 0 | 0 |
| ENSMUSG00000101870 | 0 | 0 | 0 | 0 | 0 | 0 | 0 |
| ENSMUSG00000101871 | 0 | 0 | 0 | 0 | 0 | 0 | 0 |
| ENSMUSG00000101872 | 1.036286563 | 0.92510993 | 0.976722649 | 1 | 0.846351821 | 0.477121255 | 0.672731473 |
| ENSMUSG00000101873 | 0 | 0 | 0 | 0 | 0 | 0 | 0 |
| ENSMUSG00000101874 | 0 | 0 | 0 | 0 | 0 | 0 | 0 |
| ENSMUSG00000101875 | 0.289064826 | 0.5 | 0 | 0.25 | 0.244650542 | 0 | 0.355418052 |
| ENSMUSG00000101877 | 0 | 0 | 0.25 | 0 | 0 | 0 | 0 |
| ENSMUSG00000101878 | 0 | 0 | 0 | 0 | 0 | 0 | 0 |

| | | | | | | | |
|---|---|---|---|---|---|---|---|
| ENSMUSG00000101879 | 0 | 0 | 0 | 0 | 0 | 0 | 0 |
| ENSMUSG00000101880 | 0 | 0 | 0 | 0 | 0 | 0.301029996 | 0 |
| ENSMUSG00000101881 | 0 | 0 | 0 | 0 | 0 | 0 | 0 |
| ENSMUSG00000101882 | 0 | 0 | 0 | 0 | 0 | 0 | 0 |
| ENSMUSG00000101883 | 0 | 0 | 0 | 0 | 0 | 0 | 0 |
| ENSMUSG00000101884 | 0 | 0 | 0 | 0 | 0 | 0 | 0 |
| ENSMUSG00000101885 | 0 | 0 | 0 | 0 | 0 | 0 | 0 |
| ENSMUSG00000101886 | 0 | 0 | 0 | 0 | 0 | 0 | 0 |
| ENSMUSG00000101887 | 0 | 0 | 0 | 0 | 0 | 0 | 0 |
| ENSMUSG00000101888 | 0 | 0 | 0 | 0 | 0 | 0 | 0 |
| ENSMUSG00000101890 | 0 | 0 | 0 | 0 | 0 | 0 | 0 |
| ENSMUSG00000101891 | 0 | 0 | 0 | 0 | 0 | 0 | 0 |
| ENSMUSG00000101892 | 1.912079952 | 1.912762923 | 1.928561379 | 1.814346961 | 1.814781639 | 1.986771734 | 1.781059827 |
| ENSMUSG00000101893 | 0 | 0 | 0 | 0 | 0 | 0 | 0 |
| ENSMUSG00000101894 | 0 | 0 | 0 | 0 | 0 | 0 | 0 |
| ENSMUSG00000101895 | 0 | 0 | 0 | 0 | 0 | 0.301029996 | 0 |
| ENSMUSG00000101896 | 0.45815691 | 0.396240625 | 0.25 | 0 | 0 | 0.301029996 | 0 |
| ENSMUSG00000101897 | 0 | 0 | 0 | 0 | 0 | 0 | 0 |
| ENSMUSG00000101898 | 0 | 0 | 0 | 0 | 0 | 0 | 0 |
| ENSMUSG00000101899 | 0 | 0 | 0 | 0 | 0 | 0 | 0 |
| ENSMUSG00000101900 | 0 | 0 | 0 | 0 | 0 | 0 | 0 |
| ENSMUSG00000101901 | 0 | 0 | 0 | 0 | 0 | 0 | 0 |
| ENSMUSG00000101902 | 0 | 0 | 0 | 0 | 0 | 0 | 0 |
| ENSMUSG00000101903 | 0 | 0 | 0 | 0 | 0 | 0 | 0 |
| ENSMUSG00000101904 | 0 | 0 | 0 | 0 | 0 | 0 | 0 |
| ENSMUSG00000101905 | 0 | 0 | 0 | 0 | 0 | 0 | 0 |
| ENSMUSG00000101906 | 0.960252568 | 0.92510993 | 0.75 | 1.02186571 | 0.931471446 | 0.602059991 | 0.853775828 |
| ENSMUSG00000101907 | 0 | 0 | 0 | 0 | 0 | 0 | 0 |
| ENSMUSG00000101908 | 0 | 0 | 0 | 0 | 0 | 0 | 0 |
| ENSMUSG00000101909 | 0 | 0 | 0 | 0 | 0 | 0 | 0 |
| ENSMUSG00000101910 | 0 | 0 | 0 | 0 | 0 | 0 | 0 |
| ENSMUSG00000101911 | 0 | 0 | 0 | 0 | 0 | 0 | 0 |
| ENSMUSG00000101912 | 0 | 0 | 0 | 0 | 0 | 0 | 0 |
| ENSMUSG00000101913 | 0 | 0 | 0 | 0 | 0 | 0 | 0 |

| | | | | | | | |
|---|---|---|---|---|---|---|---|
| ENSMUSG00000101914 | 0 | 0 | 0 | 0 | 0 | 0 | 0 |
| ENSMUSG00000101915 | 0 | 0 | 0 | 0 | 0 | 0 | 0 |
| ENSMUSG00000101916 | 0 | 0 | 0 | 0 | 0 | 0 | 0 |
| ENSMUSG00000101917 | 0 | 0 | 0 | 0 | 0 | 0 | 0 |
| ENSMUSG00000101918 | 0 | 0 | 0 | 0 | 0 | 0 | 0 |
| ENSMUSG00000101919 | 0 | 0 | 0 | 0 | 0 | 0 | 0 |
| ENSMUSG00000101920 | 0 | 0 | 0 | 0 | 0 | 0 | 0 |
| ENSMUSG00000101921 | 0 | 0 | 0 | 0 | 0 | 0 | 0 |
| ENSMUSG00000101922 | 0 | 0 | 0 | 0 | 0 | 0 | 0 |
| ENSMUSG00000101923 | 0 | 0 | 0 | 0 | 0 | 0 | 0 |
| ENSMUSG00000101924 | 0 | 0 | 0 | 0 | 0 | 0 | 0 |
| ENSMUSG00000101925 | 0 | 0.5 | 0 | 0 | 0 | 0 | 0 |
| ENSMUSG00000101926 | 0 | 0 | 0 | 0 | 0 | 0 | 0 |
| ENSMUSG00000101927 | 0 | 0 | 0 | 0 | 0 | 0 | 0 |
| ENSMUSG00000101928 | 0 | 0 | 0 | 0 | 0 | 0 | 0 |
| ENSMUSG00000101929 | 0 | 0 | 0 | 0 | 0 | 0 | 0 |
| ENSMUSG00000101930 | 0 | 0 | 0 | 0 | 0 | 0 | 0 |
| ENSMUSG00000101931 | 0 | 0 | 0 | 0 | 0 | 0 | 0 |
| ENSMUSG00000101932 | 0 | 0.25 | 0 | 0 | 0 | 0 | 0 |
| ENSMUSG00000101933 | 0 | 0 | 0 | 0 | 0 | 0 | 0 |
| ENSMUSG00000101934 | 0 | 0 | 0 | 0 | 0 | 0 | 0 |
| ENSMUSG00000101935 | 0 | 0 | 0 | 0 | 0 | 0 | 0 |
| ENSMUSG00000101936 | 0 | 0 | 0 | 0 | 0 | 0 | 0 |
| ENSMUSG00000101937 | 0 | 0 | 0 | 0 | 0 | 0 | 0 |
| ENSMUSG00000101938 | 0 | 0 | 0 | 0 | 0 | 0 | 0 |
| ENSMUSG00000101939 | 1.470606563 | 1.372963274 | 1.445339928 | 1.372963274 | 1.373641807 | 1.342422681 | 1.20139795 |
| ENSMUSG00000101940 | 0 | 0 | 0 | 0 | 0 | 0 | 0 |
| ENSMUSG00000101941 | 0 | 0 | 0 | 0 | 0 | 0 | 0 |
| ENSMUSG00000101942 | 0 | 0 | 0 | 0 | 0 | 0 | 0 |
| ENSMUSG00000101943 | 0 | 0 | 0 | 0 | 0 | 0 | 0 |
| ENSMUSG00000101944 | 0 | 0 | 0 | 0 | 0 | 0 | 0 |
| ENSMUSG00000101945 | 0 | 0 | 0 | 0 | 0 | 0 | 0 |
| ENSMUSG00000101946 | 0 | 0 | 0 | 0 | 0 | 0 | 0 |
| ENSMUSG00000101947 | 0 | 0 | 0 | 0 | 0 | 0 | 0 |

| | | | | | | | |
|---|---|---|---|---|---|---|---|
| ENSMUSG00000101948 | 0 | 0 | 0 | 0 | 0 | 0 | 0 |
| ENSMUSG00000101949 | 0 | 0 | 0 | 0 | 0 | 0 | 0 |
| ENSMUSG00000101950 | 0 | 0 | 0 | 0 | 0 | 0 | 0 |
| ENSMUSG00000101951 | 0 | 0 | 0 | 0 | 0 | 0 | 0 |
| ENSMUSG00000101952 | 0 | 0 | 0 | 0.25 | 0.244650542 | 0 | 0.224243824 |
| ENSMUSG00000101953 | 0.578129653 | 0.396240625 | 0.25 | 0 | 0.489301084 | 0 | 0.224243824 |
| ENSMUSG00000101954 | 0 | 0 | 0 | 0 | 0 | 0 | 0 |
| ENSMUSG00000101955 | 0 | 0 | 0 | 0 | 0 | 0 | 0 |
| ENSMUSG00000101956 | 0 | 0 | 0 | 0 | 0 | 0 | 0 |
| ENSMUSG00000101957 | 0 | 0 | 0 | 0 | 0 | 0 | 0 |
| ENSMUSG00000101958 | 0.289064826 | 0.25 | 0 | 0.396240625 | 0.387761935 | 0.301029996 | 0.355418052 |
| ENSMUSG00000101959 | 0 | 0 | 0 | 0 | 0 | 0 | 0 |
| ENSMUSG00000101960 | 0 | 0 | 0 | 0 | 0 | 0 | 0 |
| ENSMUSG00000101961 | 0 | 0 | 0 | 0 | 0 | 0 | 0 |
| ENSMUSG00000101962 | 0 | 0 | 0 | 0 | 0 | 0 | 0 |
| ENSMUSG00000101963 | 1.249317394 | 1.02186571 | 1.26109853 | 1.114857905 | 1.091002363 | 1.278753601 | 0.969165684 |
| ENSMUSG00000101964 | 0 | 0 | 0 | 0 | 0 | 0 | 0 |
| ENSMUSG00000101965 | 0 | 0.25 | 0 | 0 | 0 | 0 | 0 |
| ENSMUSG00000101966 | 0 | 0.25 | 0.396240625 | 0.25 | 0.244650542 | 0.477121255 | 0.448487648 |
| ENSMUSG00000101967 | 0 | 0 | 0 | 0 | 0 | 0 | 0 |
| ENSMUSG00000101968 | 0 | 0 | 0 | 0 | 0 | 0 | 0 |
| ENSMUSG00000101969 | 0 | 0.25 | 0 | 0 | 0 | 0 | 0 |
| ENSMUSG00000101970 | 1.289064826 | 1.114857905 | 1.311981878 | 1.098079356 | 0.955822902 | 0.954242509 | 1.041356071 |
| ENSMUSG00000101971 | 0 | 0 | 0 | 0 | 0 | 0 | 0 |
| ENSMUSG00000101972 | 0 | 0 | 0 | 0 | 0 | 0 | 0 |
| ENSMUSG00000101973 | 0 | 0.25 | 0.396240625 | 0.25 | 0 | 0 | 0 |
| ENSMUSG00000101974 | 0 | 0 | 0 | 0 | 0 | 0 | 0 |
| ENSMUSG00000101975 | 0 | 0 | 0 | 0 | 0 | 0 | 0 |
| ENSMUSG00000101976 | 0 | 0 | 0 | 0 | 0 | 0 | 0 |
| ENSMUSG00000101977 | 0 | 0 | 0 | 0 | 0 | 0 | 0 |
| ENSMUSG00000101978 | 0 | 0 | 0 | 0 | 0 | 0 | 0 |
| ENSMUSG00000101980 | 0 | 0 | 0 | 0 | 0 | 0 | 0 |
| ENSMUSG00000101981 | 0 | 0 | 0 | 0 | 0 | 0 | 0 |
| ENSMUSG00000101982 | 0 | 0 | 0 | 0 | 0 | 0 | 0 |

| | | | | | | | |
|---|---|---|---|---|---|---|---|
| ENSMUSG00000101983 | 0 | 0 | 0 | 0 | 0 | 0 | 0 |
| ENSMUSG00000101984 | 0 | 0 | 0 | 0 | 0 | 0 | 0 |
| ENSMUSG00000101985 | 0 | 0 | 0 | 0 | 0 | 0 | 0 |
| ENSMUSG00000101986 | 0 | 0 | 0 | 0 | 0 | 0 | 0 |
| ENSMUSG00000101987 | 0 | 0 | 0 | 0 | 0 | 0 | 0 |
| ENSMUSG00000101988 | 0 | 0 | 0 | 0 | 0 | 0 | 0 |
| ENSMUSG00000101989 | 0 | 0 | 0 | 0 | 0 | 0 | 0 |
| ENSMUSG00000101990 | 0 | 0 | 0 | 0 | 0 | 0 | 0 |
| ENSMUSG00000101991 | 0 | 0 | 0 | 0 | 0 | 0 | 0 |
| ENSMUSG00000101992 | 0 | 0 | 0 | 0 | 0 | 0 | 0 |
| ENSMUSG00000101993 | 0 | 0 | 0 | 0 | 0 | 0 | 0 |
| ENSMUSG00000101994 | 0 | 0 | 0 | 0 | 0 | 0 | 0 |
| ENSMUSG00000101995 | 0 | 0.396240625 | 0 | 0.25 | 0.244650542 | 0.301029996 | 0.224243824 |
| ENSMUSG00000101996 | 0 | 0 | 0 | 0 | 0 | 0 | 0 |
| ENSMUSG00000101997 | 0 | 0 | 0 | 0 | 0 | 0 | 0 |
| ENSMUSG00000101998 | 0 | 0 | 0 | 0 | 0 | 0 | 0 |
| ENSMUSG00000102000 | 0 | 0 | 0 | 0 | 0 | 0 | 0 |
| ENSMUSG00000102001 | 0 | 0 | 0 | 0 | 0 | 0 | 0 |
| ENSMUSG00000102002 | 0 | 0 | 0 | 0 | 0 | 0 | 0 |
| ENSMUSG00000102003 | 0 | 0 | 0 | 0 | 0 | 0 | 0 |
| ENSMUSG00000102004 | 0 | 0 | 0 | 0 | 0 | 0 | 0 |
| ENSMUSG00000102005 | 0 | 0 | 0 | 0 | 0 | 0 | 0 |
| ENSMUSG00000102006 | 0 | 0 | 0 | 0 | 0 | 0 | 0 |
| ENSMUSG00000102008 | 0 | 0 | 0 | 0 | 0 | 0 | 0 |
| ENSMUSG00000102009 | 0 | 0.25 | 0.25 | 0.25 | 0 | 0 | 0.224243824 |
| ENSMUSG00000102010 | 0 | 0 | 0 | 0 | 0 | 0 | 0 |
| ENSMUSG00000102011 | 0 | 0 | 0 | 0 | 0 | 0 | 0 |
| ENSMUSG00000102012 | 0 | 0 | 0 | 0.25 | 0 | 0 | 0 |
| ENSMUSG00000102013 | 0 | 0 | 0 | 0 | 0 | 0.301029996 | 0 |
| ENSMUSG00000102014 | 0.45815691 | 0 | 0.396240625 | 0.5 | 0.489301084 | 0.301029996 | 0.355418052 |
| ENSMUSG00000102015 | 0 | 0 | 0 | 0 | 0 | 0 | 0 |
| ENSMUSG00000102016 | 0 | 0 | 0 | 0 | 0 | 0 | 0 |
| ENSMUSG00000102017 | 0 | 0 | 0 | 0 | 0 | 0 | 0 |
| ENSMUSG00000102018 | 0.578129653 | 0.5 | 0.646240625 | 0.951838731 | 1.039257769 | 0.903089987 | 1.054044578 |

| | | | | | | | |
|---|---|---|---|---|---|---|---|
| ENSMUSG00000102019 | 0 | 0 | 0 | 0 | 0 | 0 | 0 |
| ENSMUSG00000102020 | 0 | 0 | 0 | 0 | 0 | 0 | 0 |
| ENSMUSG00000102022 | 0 | 0 | 0 | 0 | 0 | 0 | 0 |
| ENSMUSG00000102023 | 0 | 0 | 0 | 0 | 0 | 0 | 0 |
| ENSMUSG00000102025 | 0 | 0 | 0 | 0 | 0 | 0 | 0 |
| ENSMUSG00000102026 | 0 | 0 | 0 | 0 | 0 | 0 | 0 |
| ENSMUSG00000102027 | 0.45815691 | 0 | 0.25 | 0 | 0.244650542 | 0 | 0 |
| ENSMUSG00000102028 | 0 | 0 | 0 | 0 | 0 | 0 | 0 |
| ENSMUSG00000102029 | 0 | 0 | 0 | 0 | 0 | 0 | 0 |
| ENSMUSG00000102030 | 0.811507563 | 0.646240625 | 0 | 0.701838731 | 0.387761935 | 0.477121255 | 0.448487648 |
| ENSMUSG00000102031 | 0 | 0 | 0 | 0 | 0 | 0 | 0 |
| ENSMUSG00000102032 | 0 | 0 | 0 | 0 | 0 | 0 | 0 |
| ENSMUSG00000102033 | 0 | 0 | 0 | 0 | 0 | 0 | 0 |
| ENSMUSG00000102035 | 0 | 0 | 0 | 0 | 0 | 0 | 0 |
| ENSMUSG00000102036 | 0 | 0 | 0 | 0 | 0 | 0 | 0 |
| ENSMUSG00000102037 | 0 | 0 | 0 | 0 | 0 | 0 | 0 |
| ENSMUSG00000102038 | 0.289064826 | 0.25 | 0.396240625 | 0.396240625 | 0 | 0 | 0.448487648 |
| ENSMUSG00000102039 | 0 | 0 | 0 | 0 | 0 | 0 | 0 |
| ENSMUSG00000102041 | 0 | 0 | 0 | 0 | 0 | 0 | 0 |
| ENSMUSG00000102042 | 0 | 0 | 0 | 0 | 0 | 0 | 0 |
| ENSMUSG00000102043 | 0 | 0 | 0 | 0 | 0 | 0 | 0 |
| ENSMUSG00000102044 | 0 | 0 | 0 | 0 | 0 | 0 | 0 |
| ENSMUSG00000102045 | 0 | 0 | 0 | 0 | 0 | 0 | 0 |
| ENSMUSG00000102046 | 0 | 0 | 0 | 0.25 | 0 | 0 | 0 |
| ENSMUSG00000102047 | 0 | 0 | 0 | 0 | 0 | 0 | 0 |
| ENSMUSG00000102048 | 0.289064826 | 0 | 0 | 0.580482024 | 0 | 0 | 0 |
| ENSMUSG00000102049 | 0 | 0 | 0 | 0 | 0 | 0 | 0 |
| ENSMUSG00000102050 | 0 | 0 | 0 | 0 | 0 | 0 | 0 |
| ENSMUSG00000102051 | 0.289064826 | 0.25 | 0 | 0 | 0 | 0 | 0.224243824 |
| ENSMUSG00000102052 | 0 | 0 | 0 | 0 | 0 | 0 | 0 |
| ENSMUSG00000102053 | 0 | 0 | 0 | 0 | 0 | 0 | 0 |
| ENSMUSG00000102054 | 0 | 0 | 0 | 0 | 0 | 0 | 0 |
| ENSMUSG00000102055 | 0.45815691 | 0.396240625 | 0.396240625 | 0.25 | 0.489301084 | 0.301029996 | 0 |
| ENSMUSG00000102056 | 0 | 0 | 0 | 0 | 0 | 0 | 0 |

| | | | | | | | |
|---|---|---|---|---|---|---|---|
| ENSMUSG00000102057 | 0 | 0 | 0 | 0 | 0 | 0 | 0 |
| ENSMUSG00000102058 | 0 | 0 | 0 | 0 | 0 | 0 | 0 |
| ENSMUSG00000102059 | 1.036286563 | 0.951838731 | 0.79248125 | 0.951838731 | 0.978602168 | 1.041392685 | 0.74492186 |
| ENSMUSG00000102060 | 0 | 0 | 0 | 0 | 0 | 0 | 0 |
| ENSMUSG00000102061 | 0 | 0 | 0 | 0 | 0 | 0 | 0 |
| ENSMUSG00000102062 | 0 | 0 | 0 | 0 | 0 | 0 | 0 |
| ENSMUSG00000102063 | 0 | 0 | 0 | 0 | 0 | 0 | 0 |
| ENSMUSG00000102064 | 0.289064826 | 0 | 0.25 | 0.25 | 0 | 0 | 0.224243824 |
| ENSMUSG00000102065 | 0 | 0 | 0 | 0 | 0 | 0 | 0 |
| ENSMUSG00000102067 | 0 | 0 | 0 | 0 | 0 | 0 | 0 |
| ENSMUSG00000102068 | 0 | 0 | 0 | 0 | 0 | 0 | 0 |
| ENSMUSG00000102069 | 0 | 0 | 0 | 0 | 0 | 0 | 0 |
| ENSMUSG00000102070 | 0 | 0.396240625 | 0.25 | 0 | 0.244650542 | 0.301029996 | 0 |
| ENSMUSG00000102071 | 0 | 0 | 0 | 0 | 0 | 0 | 0 |
| ENSMUSG00000102072 | 0 | 0 | 0 | 0 | 0 | 0 | 0 |
| ENSMUSG00000102073 | 0 | 0 | 0 | 0 | 0 | 0 | 0 |
| ENSMUSG00000102074 | 0 | 0 | 0 | 0 | 0 | 0 | 0 |
| ENSMUSG00000102075 | 0 | 0 | 0 | 0 | 0 | 0 | 0 |
| ENSMUSG00000102076 | 0 | 0 | 0 | 0 | 0 | 0 | 0 |
| ENSMUSG00000102077 | 0 | 0 | 0 | 0 | 0 | 0 | 0 |
| ENSMUSG00000102078 | 0 | 0 | 0 | 0 | 0 | 0 | 0 |
| ENSMUSG00000102079 | 0 | 0 | 0 | 0 | 0 | 0 | 0 |
| ENSMUSG00000102080 | 0 | 0 | 0 | 0 | 0 | 0 | 0 |
| ENSMUSG00000102081 | 0 | 0 | 0 | 0 | 0 | 0 | 0 |
| ENSMUSG00000102082 | 0 | 0 | 0 | 0 | 0 | 0 | 0 |
| ENSMUSG00000102083 | 0 | 0 | 0 | 0.25 | 0 | 0 | 0 |
| ENSMUSG00000102084 | 0 | 0 | 0 | 0 | 0 | 0 | 0 |
| ENSMUSG00000102085 | 0 | 0 | 0 | 0.396240625 | 0.244650542 | 0 | 0.224243824 |
| ENSMUSG00000102087 | 0 | 0 | 0 | 0 | 0 | 0 | 0 |
| ENSMUSG00000102088 | 0 | 0 | 0 | 0 | 0 | 0 | 0 |
| ENSMUSG00000102090 | 0 | 0 | 0.25 | 0 | 0 | 0 | 0 |
| ENSMUSG00000102091 | 0 | 0 | 0 | 0 | 0 | 0 | 0 |
| ENSMUSG00000102092 | 0 | 0 | 0 | 0 | 0 | 0 | 0 |
| ENSMUSG00000102093 | 0 | 0 | 0 | 0 | 0 | 0 | 0 |

| | | | | | | | |
|---|---|---|---|---|---|---|---|
| ENSMUSG00000102095 | 0 | 0 | 0 | 0.25 | 0.244650542 | 0 | 0 |
| ENSMUSG00000102096 | 0 | 0 | 0 | 0 | 0 | 0 | 0 |
| ENSMUSG00000102097 | 0 | 0.5 | 0.25 | 0.5 | 0 | 0 | 0 |
| ENSMUSG00000102098 | 0 | 0 | 0 | 0 | 0 | 0 | 0 |
| ENSMUSG00000102099 | 0 | 0 | 0 | 0 | 0 | 0 | 0 |
| ENSMUSG00000102100 | 0 | 0 | 0 | 0 | 0 | 0 | 0 |
| ENSMUSG00000102101 | 0.289064826 | 0.5 | 0.646240625 | 0.396240625 | 0.244650542 | 0.301029996 | 0.520678036 |
| ENSMUSG00000102102 | 0 | 0 | 0 | 0 | 0 | 0 | 0 |
| ENSMUSG00000102103 | 0 | 0 | 0 | 0 | 0 | 0 | 0 |
| ENSMUSG00000102104 | 0 | 0 | 0 | 0 | 0 | 0 | 0 |
| ENSMUSG00000102105 | 0 | 0 | 0 | 0 | 0 | 0 | 0 |
| ENSMUSG00000102106 | 0 | 0 | 0 | 0 | 0 | 0 | 0 |
| ENSMUSG00000102107 | 0 | 0 | 0 | 0 | 0 | 0 | 0 |
| ENSMUSG00000102108 | 0 | 0 | 0 | 0 | 0 | 0 | 0 |
| ENSMUSG00000102109 | 0 | 0 | 0 | 0 | 0 | 0 | 0 |
| ENSMUSG00000102110 | 0 | 0 | 0 | 0 | 0 | 0 | 0 |
| ENSMUSG00000102111 | 0 | 0 | 0 | 0 | 0 | 0 | 0 |
| ENSMUSG00000102112 | 0 | 0 | 0 | 0.25 | 0.489301084 | 0.301029996 | 0.224243824 |
| ENSMUSG00000102113 | 0 | 0 | 0 | 0 | 0 | 0 | 0 |
| ENSMUSG00000102114 | 0 | 0 | 0 | 0 | 0 | 0 | 0 |
| ENSMUSG00000102115 | 0 | 0 | 0 | 0 | 0 | 0 | 0 |
| ENSMUSG00000102116 | 0 | 0 | 0 | 0 | 0 | 0 | 0 |
| ENSMUSG00000102117 | 0.671187741 | 0.646240625 | 0.864857905 | 0.396240625 | 0.877063019 | 0.903089987 | 0.710836105 |
| ENSMUSG00000102118 | 0 | 0 | 0 | 0 | 0 | 0 | 0 |
| ENSMUSG00000102119 | 0 | 0 | 0 | 0 | 0 | 0 | 0 |
| ENSMUSG00000102120 | 0 | 0 | 0 | 0 | 0 | 0 | 0 |
| ENSMUSG00000102121 | 0 | 0 | 0 | 0 | 0 | 0 | 0 |
| ENSMUSG00000102122 | 0 | 0 | 0 | 0 | 0 | 0 | 0 |
| ENSMUSG00000102123 | 0 | 0.25 | 0 | 0 | 0 | 0 | 0 |
| ENSMUSG00000102124 | 0 | 0 | 0 | 0 | 0 | 0 | 0 |
| ENSMUSG00000102125 | 0 | 0 | 0 | 0 | 0 | 0 | 0 |
| ENSMUSG00000102126 | 0 | 0 | 0 | 0 | 0 | 0 | 0 |
| ENSMUSG00000102127 | 0 | 0 | 0 | 0 | 0 | 0 | 0 |
| ENSMUSG00000102128 | 0 | 0 | 0 | 0 | 0 | 0 | 0 |

| | | | | | | | |
|---|---|---|---|---|---|---|---|
| ENSMUSG00000102129 | 0 | 0 | 0 | 0 | 0 | 0 | 0 |
| ENSMUSG00000102130 | 0 | 0 | 0 | 0 | 0 | 0 | 0 |
| ENSMUSG00000102131 | 0 | 0 | 0 | 0 | 0 | 0 | 0 |
| ENSMUSG00000102132 | 0 | 0 | 0 | 0 | 0 | 0 | 0 |
| ENSMUSG00000102133 | 0 | 0.580482024 | 0.580482024 | 0.396240625 | 0.387761935 | 0 | 0 |
| ENSMUSG00000102134 | 0.289064826 | 0.396240625 | 0 | 0 | 0.244650542 | 0 | 0 |
| ENSMUSG00000102135 | 1 | 0.646240625 | 0.701838731 | 0.830482024 | 0.733951626 | 0.301029996 | 0.916588299 |
| ENSMUSG00000102136 | 0 | 0 | 0 | 0 | 0 | 0 | 0 |
| ENSMUSG00000102137 | 0.45815691 | 0.25 | 0.5 | 0 | 0 | 0.301029996 | 0.224243824 |
| ENSMUSG00000102138 | 0 | 0 | 0 | 0 | 0 | 0 | 0 |
| ENSMUSG00000102139 | 0 | 0 | 0 | 0 | 0 | 0 | 0.224243824 |
| ENSMUSG00000102140 | 0.289064826 | 0.25 | 0 | 0 | 0 | 0 | 0 |
| ENSMUSG00000102141 | 0 | 0 | 0 | 0 | 0 | 0 | 0.224243824 |
| ENSMUSG00000102142 | 0.289064826 | 0 | 0 | 0.25 | 0.244650542 | 0 | 0 |
| ENSMUSG00000102143 | 1 | 1.080482024 | 1.080482024 | 1.061981878 | 0.931471446 | 0.954242509 | 0.672731473 |
| ENSMUSG00000102144 | 0 | 0 | 0 | 0 | 0 | 0 | 0 |
| ENSMUSG00000102145 | 0.45815691 | 0.79248125 | 0.396240625 | 0.580482024 | 0.489301084 | 0.301029996 | 0.520678036 |
| ENSMUSG00000102146 | 0 | 0 | 0 | 0 | 0 | 0 | 0 |
| ENSMUSG00000102147 | 0 | 0 | 0 | 0 | 0 | 0 | 0 |
| ENSMUSG00000102148 | 0 | 0 | 0.25 | 0 | 0 | 0 | 0 |
| ENSMUSG00000102149 | 0 | 0 | 0 | 0 | 0 | 0 | 0 |
| ENSMUSG00000102150 | 0 | 0 | 0 | 0 | 0 | 0 | 0 |
| ENSMUSG00000102151 | 0 | 0 | 0 | 0.25 | 0 | 0 | 0 |
| ENSMUSG00000102152 | 0 | 0 | 0 | 0.646240625 | 0.568060967 | 0.698970004 | 0.448487648 |
| ENSMUSG00000102153 | 0 | 0.25 | 0.25 | 0 | 0 | 0.301029996 | 0.355418052 |
| ENSMUSG00000102154 | 0 | 0 | 0 | 0 | 0 | 0 | 0 |
| ENSMUSG00000102155 | 0 | 0 | 0 | 0 | 0 | 0 | 0 |
| ENSMUSG00000102156 | 0 | 0 | 0 | 0 | 0 | 0 | 0 |
| ENSMUSG00000102157 | 0.289064826 | 0 | 0 | 0 | 0.387761935 | 0 | 0 |
| ENSMUSG00000102158 | 0 | 0.396240625 | 0.25 | 0.25 | 0 | 0 | 0.355418052 |
| ENSMUSG00000102159 | 0 | 0 | 0 | 0 | 0 | 0 | 0 |
| ENSMUSG00000102160 | 0 | 0 | 0 | 0 | 0 | 0 | 0 |
| ENSMUSG00000102162 | 0 | 0.25 | 0 | 0.396240625 | 0.244650542 | 0.301029996 | 0 |
| ENSMUSG00000102163 | 0 | 0 | 0 | 0 | 0 | 0 | 0 |

| | | | | | | | |
|---|---|---|---|---|---|---|---|
| ENSMUSG00000102164 | 0 | 0 | 0 | 0 | 0 | 0 | 0 |
| ENSMUSG00000102166 | 0 | 0 | 0 | 0 | 0 | 0 | 0 |
| ENSMUSG00000102167 | 0 | 0 | 0 | 0 | 0 | 0 | 0 |
| ENSMUSG00000102168 | 0 | 0 | 0 | 0 | 0 | 0 | 0 |
| ENSMUSG00000102169 | 0 | 0 | 0 | 0 | 0 | 0 | 0 |
| ENSMUSG00000102170 | 0 | 0 | 0 | 0 | 0 | 0 | 0 |
| ENSMUSG00000102171 | 1 | 1.02186571 | 0.976722649 | 1.372963274 | 1.310728002 | 1.591064607 | 1.140832123 |
| ENSMUSG00000102172 | 0 | 0 | 0 | 0 | 0.244650542 | 0 | 0 |
| ENSMUSG00000102173 | 0 | 0 | 0 | 0 | 0 | 0 | 0 |
| ENSMUSG00000102174 | 0 | 0 | 0 | 0 | 0 | 0 | 0 |
| ENSMUSG00000102175 | 0 | 0 | 0 | 0 | 0 | 0 | 0 |
| ENSMUSG00000102176 | 0 | 0 | 0 | 0 | 0 | 0 | 0 |
| ENSMUSG00000102177 | 0 | 0 | 0 | 0 | 0 | 0 | 0 |
| ENSMUSG00000102178 | 0 | 0 | 0 | 0 | 0 | 0 | 0 |
| ENSMUSG00000102179 | 0 | 0.396240625 | 0.580482024 | 0.25 | 0.244650542 | 0.602059991 | 0.579661877 |
| ENSMUSG00000102180 | 0 | 0 | 0 | 0 | 0 | 0 | 0 |
| ENSMUSG00000102181 | 0 | 0 | 0 | 0 | 0 | 0 | 0 |
| ENSMUSG00000102182 | 0 | 0 | 0 | 0 | 0 | 0 | 0 |
| ENSMUSG00000102183 | 0.289064826 | 0 | 0.5 | 0.5 | 0.244650542 | 0.602059991 | 0.355418052 |
| ENSMUSG00000102184 | 0 | 0 | 0 | 0 | 0 | 0 | 0 |
| ENSMUSG00000102185 | 0 | 0 | 0 | 0 | 0 | 0 | 0 |
| ENSMUSG00000102186 | 0 | 0 | 0.25 | 0 | 0.244650542 | 0 | 0 |
| ENSMUSG00000102187 | 0 | 0 | 0 | 0 | 0 | 0 | 0 |
| ENSMUSG00000102188 | 0 | 0 | 0 | 0 | 0 | 0 | 0 |
| ENSMUSG00000102189 | 0.289064826 | 0.25 | 0 | 0.5 | 0.632412477 | 0.301029996 | 0.448487648 |
| ENSMUSG00000102190 | 0 | 0 | 0 | 0 | 0 | 0 | 0 |
| ENSMUSG00000102191 | 0 | 0 | 0 | 0 | 0 | 0 | 0 |
| ENSMUSG00000102192 | 0 | 0 | 0 | 0 | 0 | 0 | 0 |
| ENSMUSG00000102193 | 0.45815691 | 0.25 | 0 | 0.396240625 | 0.244650542 | 0 | 0.520678036 |
| ENSMUSG00000102194 | 0 | 0 | 0 | 0 | 0 | 0 | 0 |
| ENSMUSG00000102195 | 0 | 0 | 0 | 0 | 0 | 0 | 0 |
| ENSMUSG00000102196 | 0 | 0 | 0 | 0 | 0.244650542 | 0 | 0 |
| ENSMUSG00000102197 | 0 | 0 | 0 | 0 | 0 | 0.477121255 | 0.224243824 |
| ENSMUSG00000102198 | 0 | 0 | 0 | 0 | 0 | 0 | 0 |

| | | | | | | | |
|---|---|---|---|---|---|---|---|
| ENSMUSG00000102199 | 0 | 0 | 0 | 0 | 0 | 0 | 0 |
| ENSMUSG00000102200 | 0.289064826 | 0 | 0.25 | 0 | 0 | 0 | 0 |
| ENSMUSG00000102201 | 0 | 0 | 0 | 0 | 0 | 0 | 0 |
| ENSMUSG00000102202 | 0.811507563 | 1 | 0.92510993 | 1.302363341 | 1.223252711 | 1.230448921 | 1.131174228 |
| ENSMUSG00000102203 | 0 | 0 | 0 | 0 | 0 | 0 | 0 |
| ENSMUSG00000102204 | 0 | 0 | 0.396240625 | 0.25 | 0.632412477 | 0 | 0.355418052 |
| ENSMUSG00000102205 | 0 | 0 | 0 | 0 | 0 | 0 | 0 |
| ENSMUSG00000102206 | 0 | 0 | 0 | 0 | 0 | 0 | 0 |
| ENSMUSG00000102207 | 0 | 0.25 | 0.25 | 0 | 0.387761935 | 0 | 0 |
| ENSMUSG00000102208 | 0 | 0 | 0 | 0 | 0 | 0 | 0 |
| ENSMUSG00000102211 | 0.289064826 | 0 | 0.25 | 0.25 | 0 | 0 | 0 |
| ENSMUSG00000102212 | 0 | 0 | 0.25 | 0 | 0 | 0 | 0 |
| ENSMUSG00000102213 | 0 | 0 | 0 | 0 | 0 | 0 | 0 |
| ENSMUSG00000102214 | 0 | 0 | 0 | 0 | 0 | 0 | 0 |
| ENSMUSG00000102215 | 0 | 0 | 0 | 0 | 0 | 0 | 0 |
| ENSMUSG00000102216 | 0 | 0 | 0 | 0 | 0 | 0 | 0 |
| ENSMUSG00000102217 | 0 | 0 | 0 | 0 | 0 | 0 | 0 |
| ENSMUSG00000102218 | 0 | 0 | 0 | 0 | 0 | 0 | 0 |
| ENSMUSG00000102219 | 0 | 0 | 0 | 0 | 0 | 0 | 0 |
| ENSMUSG00000102220 | 0.671187741 | 0.896240625 | 0.951838731 | 0.92510993 | 0.812711509 | 0.954242509 | 0.935079929 |
| ENSMUSG00000102221 | 0 | 0 | 0 | 0 | 0 | 0 | 0 |
| ENSMUSG00000102222 | 0 | 0 | 0 | 0 | 0 | 0 | 0 |
| ENSMUSG00000102223 | 0 | 0 | 0 | 0 | 0 | 0 | 0 |
| ENSMUSG00000102224 | 1.100572389 | 1.04248125 | 1.238549078 | 1.445339928 | 1.445123986 | 1.556302501 | 1.396774124 |
| ENSMUSG00000102225 | 0.45815691 | 0.646240625 | 0.5 | 0.580482024 | 0.489301084 | 0.698970004 | 0.629532004 |
| ENSMUSG00000102227 | 0 | 0 | 0 | 0 | 0 | 0 | 0 |
| ENSMUSG00000102229 | 0 | 0 | 0 | 0 | 0 | 0 | 0 |
| ENSMUSG00000102230 | 0 | 0 | 0 | 0 | 0 | 0 | 0 |
| ENSMUSG00000102231 | 0 | 0 | 0 | 0 | 0 | 0 | 0 |
| ENSMUSG00000102232 | 0 | 0 | 0 | 0 | 0 | 0 | 0 |
| ENSMUSG00000102233 | 0 | 0 | 0 | 0 | 0 | 0 | 0 |
| ENSMUSG00000102234 | 0.289064826 | 0 | 0.25 | 0 | 0 | 0 | 0 |
| ENSMUSG00000102235 | 0 | 0.5 | 0.25 | 0 | 0.489301084 | 0.477121255 | 0.224243824 |
| ENSMUSG00000102236 | 0 | 0 | 0 | 0 | 0 | 0 | 0 |

| | | | | | | | |
|---|---|---|---|---|---|---|---|
| ENSMUSG00000102237 | 0 | 0 | 0 | 0 | 0 | 0 | 0 |
| ENSMUSG00000102238 | 0 | 0 | 0 | 0 | 0 | 0 | 0 |
| ENSMUSG00000102239 | 0 | 0 | 0 | 0 | 0 | 0 | 0 |
| ENSMUSG00000102240 | 0 | 0 | 0 | 0 | 0 | 0 | 0 |
| ENSMUSG00000102241 | 0 | 0.5 | 0.396240625 | 0.25 | 0.244650542 | 0 | 0.355418052 |
| ENSMUSG00000102242 | 0 | 0 | 0 | 0 | 0 | 0 | 0 |
| ENSMUSG00000102243 | 0.45815691 | 0.25 | 0.25 | 0.580482024 | 0.632412477 | 0.301029996 | 0.355418052 |
| ENSMUSG00000102244 | 0.289064826 | 0 | 0 | 0 | 0 | 0 | 0 |
| ENSMUSG00000102245 | 0 | 0 | 0 | 0 | 0 | 0 | 0 |
| ENSMUSG00000102246 | 0.45815691 | 0.396240625 | 0.646240625 | 0.5 | 0.568060967 | 1 | 0.710836105 |
| ENSMUSG00000102247 | 0 | 0 | 0 | 0 | 0 | 0 | 0 |
| ENSMUSG00000102248 | 0.289064826 | 0.25 | 0 | 0.25 | 0 | 0 | 0 |
| ENSMUSG00000102249 | 0 | 0 | 0 | 0 | 0 | 0 | 0 |
| ENSMUSG00000102250 | 0 | 0 | 0 | 0 | 0 | 0 | 0 |
| ENSMUSG00000102251 | 0 | 0 | 0 | 0 | 0 | 0 | 0 |
| ENSMUSG00000102252 | 1.837744666 | 1.714495249 | 1.71759118 | 1.685366747 | 1.649303553 | 1.944482672 | 1.610050272 |
| ENSMUSG00000102253 | 0 | 0 | 0 | 0 | 0 | 0 | 0 |
| ENSMUSG00000102254 | 0 | 0 | 0 | 0 | 0 | 0.301029996 | 0 |
| ENSMUSG00000102255 | 0 | 0 | 0 | 0 | 0 | 0 | 0 |
| ENSMUSG00000102256 | 0 | 0 | 0 | 0 | 0 | 0 | 0 |
| ENSMUSG00000102257 | 0 | 0 | 0 | 0 | 0 | 0 | 0 |
| ENSMUSG00000102258 | 0.45815691 | 0.25 | 0.5 | 0.5 | 0.387761935 | 0.301029996 | 0.629532004 |
| ENSMUSG00000102259 | 0 | 0 | 0 | 0 | 0 | 0 | 0.224243824 |
| ENSMUSG00000102260 | 0 | 0 | 0 | 0 | 0 | 0 | 0 |
| ENSMUSG00000102261 | 0 | 0 | 0 | 0 | 0 | 0 | 0 |
| ENSMUSG00000102262 | 0 | 0 | 0 | 0 | 0 | 0 | 0 |
| ENSMUSG00000102263 | 0 | 0 | 0 | 0 | 0 | 0 | 0 |
| ENSMUSG00000102264 | 0 | 0 | 0 | 0 | 0 | 0 | 0 |
| ENSMUSG00000102265 | 0 | 0 | 0 | 0 | 0 | 0 | 0 |
| ENSMUSG00000102266 | 0 | 0 | 0 | 0.396240625 | 0 | 0 | 0 |
| ENSMUSG00000102267 | 0 | 0 | 0 | 0.25 | 0.244650542 | 0.698970004 | 0.224243824 |
| ENSMUSG00000102268 | 0.45815691 | 0.25 | 0.396240625 | 0.25 | 0.387761935 | 0.477121255 | 0 |
| ENSMUSG00000102269 | 0 | 0 | 0.25 | 0 | 0 | 0 | 0 |
| ENSMUSG00000102270 | 0 | 0 | 0 | 0 | 0 | 0 | 0 |

| | | | | | | | |
|---|---|---|---|---|---|---|---|
| ENSMUSG00000102271 | 0 | 0 | 0 | 0 | 0 | 0 | 0 |
| ENSMUSG00000102272 | 0 | 0 | 0 | 0 | 0 | 0 | 0 |
| ENSMUSG00000102273 | 0 | 0 | 0 | 0 | 0 | 0 | 0 |
| ENSMUSG00000102274 | 0 | 0 | 0 | 0 | 0 | 0 | 0 |
| ENSMUSG00000102275 | 0.45815691 | 0.864857905 | 0.75 | 0.830482024 | 0.632412477 | 0.903089987 | 0.579661877 |
| ENSMUSG00000102276 | 0 | 0 | 0 | 0 | 0 | 0 | 0 |
| ENSMUSG00000102277 | 0 | 0 | 0 | 0 | 0 | 0 | 0 |
| ENSMUSG00000102278 | 0.747221736 | 0.580482024 | 0.830482024 | 0.701838731 | 0.77552387 | 0.903089987 | 0.629532004 |
| ENSMUSG00000102279 | 0 | 0 | 0 | 0 | 0 | 0 | 0 |
| ENSMUSG00000102280 | 0 | 0 | 0 | 0 | 0 | 0 | 0 |
| ENSMUSG00000102281 | 0 | 0 | 0 | 0 | 0 | 0 | 0 |
| ENSMUSG00000102282 | 0 | 0 | 0 | 0.25 | 0.244650542 | 0 | 0.224243824 |
| ENSMUSG00000102283 | 0 | 0 | 0 | 0 | 0 | 0 | 0 |
| ENSMUSG00000102284 | 0 | 0 | 0 | 0 | 0 | 0 | 0 |
| ENSMUSG00000102285 | 0 | 0 | 0 | 0 | 0 | 0 | 0 |
| ENSMUSG00000102286 | 0 | 0 | 0 | 0 | 0 | 0 | 0 |
| ENSMUSG00000102287 | 0 | 0 | 0 | 0 | 0 | 0 | 0 |
| ENSMUSG00000102288 | 0 | 0 | 0 | 0 | 0 | 0 | 0 |
| ENSMUSG00000102289 | 0.960252568 | 1.061981878 | 1.130890489 | 0.976722649 | 1.074582839 | 1.079181246 | 0.916588299 |
| ENSMUSG00000102290 | 0 | 0 | 0 | 0 | 0 | 0 | 0 |
| ENSMUSG00000102291 | 0 | 0 | 0 | 0 | 0 | 0 | 0 |
| ENSMUSG00000102293 | 0 | 0.25 | 0 | 0 | 0 | 0 | 0 |
| ENSMUSG00000102294 | 0 | 0 | 0 | 0 | 0 | 0 | 0 |
| ENSMUSG00000102295 | 0 | 0 | 0 | 0 | 0 | 0 | 0 |
| ENSMUSG00000102296 | 0 | 0 | 0 | 0 | 0 | 0 | 0 |
| ENSMUSG00000102297 | 0 | 0 | 0 | 0.25 | 0 | 0 | 0.355418052 |
| ENSMUSG00000102298 | 0.578129653 | 0.646240625 | 0.25 | 0.701838731 | 0.387761935 | 0.477121255 | 0.448487648 |
| ENSMUSG00000102299 | 0.289064826 | 0.25 | 0 | 0 | 0 | 0 | 0 |
| ENSMUSG00000102300 | 0 | 0.25 | 0 | 0 | 0.244650542 | 0 | 0.224243824 |
| ENSMUSG00000102301 | 0 | 0 | 0 | 0 | 0 | 0 | 0 |
| ENSMUSG00000102302 | 1.156259305 | 0.976722649 | 1.282320754 | 1.04248125 | 1.074582839 | 0.903089987 | 1.054044578 |
| ENSMUSG00000102303 | 0 | 0 | 0 | 0 | 0 | 0 | 0 |
| ENSMUSG00000102304 | 0 | 0 | 0 | 0 | 0 | 0 | 0 |
| ENSMUSG00000102305 | 0.671187741 | 0.701838731 | 0.896240625 | 0.79248125 | 0.877063019 | 0.84509804 | 0.876096088 |

| | | | | | | | |
|---|---|---|---|---|---|---|---|
| ENSMUSG00000102306 | 0 | 0 | 0 | 0 | 0 | 0 | 0 |
| ENSMUSG00000102307 | 2.283862692 | 2.059601185 | 2.004202072 | 1.910964047 | 1.842556557 | 2.075546961 | 1.74492186 |
| ENSMUSG00000102308 | 0 | 0 | 0 | 0 | 0 | 0 | 0 |
| ENSMUSG00000102309 | 0 | 0 | 0 | 0 | 0 | 0 | 0 |
| ENSMUSG00000102310 | 0 | 0 | 0 | 0 | 0 | 0 | 0 |
| ENSMUSG00000102311 | 0 | 0 | 0 | 0 | 0 | 0 | 0 |
| ENSMUSG00000102312 | 0 | 0 | 0 | 0 | 0 | 0 | 0 |
| ENSMUSG00000102313 | 0 | 0 | 0 | 0 | 0 | 0 | 0 |
| ENSMUSG00000102314 | 0 | 0 | 0 | 0 | 0 | 0 | 0 |
| ENSMUSG00000102315 | 0 | 0 | 0 | 0 | 0 | 0 | 0 |
| ENSMUSG00000102316 | 0 | 0 | 0 | 0 | 0 | 0 | 0 |
| ENSMUSG00000102317 | 0.289064826 | 0.5 | 0.25 | 0.396240625 | 0.568060967 | 0 | 0.448487648 |
| ENSMUSG00000102318 | 0 | 0 | 0 | 0 | 0 | 0 | 0 |
| ENSMUSG00000102319 | 0.289064826 | 0.5 | 0 | 0.396240625 | 0.568060967 | 0 | 0.448487648 |
| ENSMUSG00000102320 | 0 | 0 | 0 | 0 | 0 | 0 | 0 |
| ENSMUSG00000102321 | 0 | 0 | 0 | 0 | 0 | 0.301029996 | 0 |
| ENSMUSG00000102322 | 0 | 0 | 0 | 0 | 0 | 0 | 0 |
| ENSMUSG00000102323 | 0.747221736 | 0.646240625 | 0.396240625 | 0.864857905 | 0.77552387 | 0.903089987 | 0.803905701 |
| ENSMUSG00000102324 | 0.671187741 | 0.646240625 | 0.701838731 | 0.701838731 | 0.568060967 | 0.698970004 | 0.775756176 |
| ENSMUSG00000102325 | 0.811507563 | 0.396240625 | 0.701838731 | 0.830482024 | 0.632412477 | 0.477121255 | 0.74492186 |
| ENSMUSG00000102326 | 1.036286563 | 1 | 1.02186571 | 1.061981878 | 1.106691885 | 0.954242509 | 0.969165684 |
| ENSMUSG00000102327 | 0 | 0 | 0 | 0 | 0 | 0 | 0 |
| ENSMUSG00000102328 | 0 | 0 | 0 | 0 | 0 | 0 | 0 |
| ENSMUSG00000102329 | 0 | 0 | 0.25 | 0 | 0 | 0 | 0 |
| ENSMUSG00000102330 | 0 | 0.396240625 | 0.580482024 | 0.396240625 | 0.244650542 | 0.301029996 | 0.355418052 |
| ENSMUSG00000102331 | 0.289064826 | 0 | 0 | 0.25 | 0.244650542 | 0.477121255 | 0.355418052 |
| ENSMUSG00000102332 | 0 | 0 | 0 | 0 | 0.244650542 | 0 | 0.224243824 |
| ENSMUSG00000102333 | 0 | 0 | 0 | 0 | 0 | 0 | 0 |
| ENSMUSG00000102334 | 0 | 0 | 0 | 0 | 0 | 0 | 0 |
| ENSMUSG00000102335 | 0 | 0 | 0 | 0 | 0 | 0 | 0 |
| ENSMUSG00000102336 | 2 | 1.958222504 | 1.872963274 | 1.566696635 | 1.47337555 | 1.556302501 | 1.401059159 |
| ENSMUSG00000102337 | 0 | 0 | 0 | 0 | 0 | 0 | 0 |
| ENSMUSG00000102338 | 0 | 0 | 0 | 0 | 0 | 0 | 0 |
| ENSMUSG00000102339 | 0 | 0 | 0 | 0 | 0 | 0 | 0 |

| | | | | | | | |
|---|---|---|---|---|---|---|---|
| ENSMUSG00000102340 | 0.671187741 | 0.976722649 | 0.864857905 | 1.061981878 | 1.121713561 | 0.77815125 | 1.014380832 |
| ENSMUSG00000102341 | 0 | 0 | 0 | 0 | 0 | 0 | 0 |
| ENSMUSG00000102342 | 0 | 0 | 0 | 0 | 0 | 0 | 0 |
| ENSMUSG00000102343 | 0.578129653 | 0.5 | 0.580482024 | 0.75 | 0.568060967 | 0.477121255 | 0.579661877 |
| ENSMUSG00000102344 | 0 | 0 | 0 | 0 | 0 | 0 | 0 |
| ENSMUSG00000102345 | 0 | 0 | 0 | 0 | 0 | 0 | 0 |
| ENSMUSG00000102346 | 0 | 0 | 0 | 0 | 0 | 0 | 0 |
| ENSMUSG00000102347 | 0 | 0 | 0 | 0 | 0 | 0 | 0 |
| ENSMUSG00000102348 | 0 | 0 | 0 | 0 | 0 | 0 | 0 |
| ENSMUSG00000102349 | 0 | 0 | 0 | 0 | 0 | 0 | 0 |
| ENSMUSG00000102350 | 0 | 0 | 0 | 0 | 0 | 0 | 0 |
| ENSMUSG00000102352 | 0 | 0 | 0 | 0 | 0 | 0 | 0 |
| ENSMUSG00000102353 | 1.156259305 | 0.79248125 | 0.976722649 | 0.976722649 | 0.905314583 | 1.113943352 | 0.969165684 |
| ENSMUSG00000102354 | 0 | 0 | 0 | 0 | 0 | 0 | 0 |
| ENSMUSG00000102355 | 0 | 0 | 0 | 0 | 0 | 0 | 0 |
| ENSMUSG00000102356 | 0 | 0 | 0 | 0 | 0 | 0 | 0 |
| ENSMUSG00000102357 | 0 | 0 | 0 | 0 | 0 | 0 | 0 |
| ENSMUSG00000102358 | 0 | 0 | 0 | 0 | 0 | 0 | 0 |
| ENSMUSG00000102359 | 0 | 0 | 0 | 0 | 0 | 0 | 0.224243824 |
| ENSMUSG00000102360 | 0 | 0 | 0 | 0 | 0 | 0 | 0.224243824 |
| ENSMUSG00000102362 | 0 | 0 | 0 | 0 | 0 | 0 | 0.224243824 |
| ENSMUSG00000102363 | 0 | 0 | 0 | 0 | 0 | 0 | 0 |
| ENSMUSG00000102364 | 0 | 0 | 0 | 0 | 0 | 0 | 0 |
| ENSMUSG00000102365 | 0.45815691 | 0.396240625 | 0 | 0.25 | 0 | 0.477121255 | 0.224243824 |
| ENSMUSG00000102366 | 0 | 0 | 0 | 0 | 0 | 0 | 0 |
| ENSMUSG00000102367 | 0 | 0 | 0 | 0 | 0 | 0 | 0 |
| ENSMUSG00000102368 | 0 | 0.25 | 0 | 0 | 0 | 0 | 0 |
| ENSMUSG00000102369 | 0 | 0 | 0 | 0.396240625 | 0 | 0 | 0 |
| ENSMUSG00000102370 | 0.289064826 | 0.5 | 0.5 | 0 | 0.244650542 | 0.477121255 | 0.355418052 |
| ENSMUSG00000102371 | 0 | 0 | 0 | 0 | 0.244650542 | 0 | 0 |
| ENSMUSG00000102372 | 0 | 0 | 0 | 0 | 0.244650542 | 0 | 0 |
| ENSMUSG00000102373 | 0 | 0 | 0 | 0 | 0 | 0 | 0 |
| ENSMUSG00000102374 | 0.671187741 | 0.79248125 | 0.701838731 | 0.830482024 | 0.244650542 | 0.477121255 | 0.520678036 |
| ENSMUSG00000102375 | 0 | 0.25 | 0 | 0 | 0 | 0 | 0.224243824 |

| | | | | | | | |
|---|---|---|---|---|---|---|---|
| ENSMUSG00000102376 | 0 | 0.25 | 0 | 0.25 | 0 | 0.301029996 | 0.224243824 |
| ENSMUSG00000102377 | 0 | 0 | 0 | 0 | 0 | 0 | 0 |
| ENSMUSG00000102378 | 0 | 0 | 0 | 0 | 0 | 0 | 0 |
| ENSMUSG00000102379 | 0 | 0 | 0.25 | 0 | 0 | 0 | 0.224243824 |
| ENSMUSG00000102380 | 0 | 0 | 0 | 0.25 | 0 | 0 | 0.224243824 |
| ENSMUSG00000102381 | 0 | 0 | 0 | 0 | 0 | 0 | 0 |
| ENSMUSG00000102382 | 0 | 0 | 0 | 0 | 0 | 0 | 0 |
| ENSMUSG00000102383 | 0.578129653 | 0 | 0.25 | 0.396240625 | 0.244650542 | 0.301029996 | 0.224243824 |
| ENSMUSG00000102384 | 0.289064826 | 0 | 0.25 | 0.580482024 | 0.244650542 | 0 | 0.355418052 |
| ENSMUSG00000102385 | 0 | 0 | 0 | 0 | 0 | 0 | 0 |
| ENSMUSG00000102386 | 0 | 0 | 0.25 | 0 | 0 | 0 | 0 |
| ENSMUSG00000102387 | 0 | 0 | 0 | 0 | 0 | 0 | 0 |
| ENSMUSG00000102388 | 0 | 0 | 0 | 0 | 0 | 0 | 0 |
| ENSMUSG00000102389 | 0 | 0 | 0 | 0 | 0 | 0 | 0.224243824 |
| ENSMUSG00000102390 | 0 | 0 | 0 | 0 | 0 | 0 | 0 |
| ENSMUSG00000102391 | 0 | 0 | 0 | 0 | 0 | 0 | 0 |
| ENSMUSG00000102392 | 0 | 0 | 0 | 0 | 0 | 0 | 0 |
| ENSMUSG00000102393 | 0 | 0 | 0 | 0 | 0 | 0 | 0 |
| ENSMUSG00000102394 | 0 | 0 | 0 | 0.25 | 0.244650542 | 0 | 0 |
| ENSMUSG00000102395 | 0 | 0 | 0 | 0 | 0 | 0 | 0 |
| ENSMUSG00000102396 | 0 | 0 | 0 | 0 | 0 | 0 | 0 |
| ENSMUSG00000102397 | 0 | 0 | 0 | 0 | 0 | 0 | 0 |
| ENSMUSG00000102398 | 0 | 0 | 0 | 0.25 | 0 | 0.301029996 | 0 |
| ENSMUSG00000102399 | 0 | 0 | 0 | 0 | 0 | 0 | 0 |
| ENSMUSG00000102400 | 0 | 0 | 0 | 0 | 0 | 0 | 0 |
| ENSMUSG00000102401 | 0 | 0.25 | 0 | 0 | 0 | 0 | 0.224243824 |
| ENSMUSG00000102402 | 0 | 0 | 0 | 0 | 0 | 0 | 0 |
| ENSMUSG00000102403 | 0.289064826 | 0.25 | 0 | 0.396240625 | 0.387761935 | 0 | 0.224243824 |
| ENSMUSG00000102404 | 0.867194479 | 0.864857905 | 0.701838731 | 0.75 | 0.632412477 | 0.77815125 | 0.355418052 |
| ENSMUSG00000102405 | 0.289064826 | 0 | 0 | 0 | 0 | 0 | 0 |
| ENSMUSG00000102406 | 0 | 0 | 0 | 0 | 0 | 0 | 0 |
| ENSMUSG00000102407 | 0 | 0 | 0 | 0 | 0 | 0 | 0 |
| ENSMUSG00000102408 | 0 | 0 | 0 | 0 | 0 | 0 | 0 |
| ENSMUSG00000102409 | 0 | 0 | 0 | 0 | 0 | 0 | 0 |

| | | | | | | | |
|---|---|---|---|---|---|---|---|
| ENSMUSG00000102410 | 0 | 0 | 0 | 0 | 0 | 0 | 0 |
| ENSMUSG00000102411 | 1.45815691 | 1.505591953 | 1.388647213 | 1.527131114 | 1.358933174 | 1.51851394 | 1.355418052 |
| ENSMUSG00000102413 | 0.289064826 | 0 | 0 | 0 | 0 | 0 | 0 |
| ENSMUSG00000102414 | 0 | 0.396240625 | 0.25 | 0.25 | 0.489301084 | 0 | 0.355418052 |
| ENSMUSG00000102415 | 0.45815691 | 0.396240625 | 0.396240625 | 0 | 0.489301084 | 0 | 0.355418052 |
| ENSMUSG00000102416 | 0 | 0 | 0 | 0 | 0 | 0 | 0 |
| ENSMUSG00000102417 | 0 | 0 | 0 | 0 | 0 | 0 | 0 |
| ENSMUSG00000102418 | 0.747221736 | 0 | 0.75 | 0.75 | 0.244650542 | 0.301029996 | 0.520678036 |
| ENSMUSG00000102419 | 0.289064826 | 0 | 0 | 0 | 0 | 0 | 0 |
| ENSMUSG00000102420 | 0 | 0 | 0 | 0 | 0 | 0 | 0 |
| ENSMUSG00000102421 | 0 | 0 | 0 | 0 | 0 | 0 | 0 |
| ENSMUSG00000102422 | 0 | 0 | 0 | 0 | 0 | 0 | 0 |
| ENSMUSG00000102423 | 0 | 0 | 0 | 0.396240625 | 0 | 0 | 0 |
| ENSMUSG00000102424 | 0 | 0 | 0 | 0 | 0 | 0 | 0 |
| ENSMUSG00000102425 | 0.578129653 | 0.25 | 0.25 | 0.396240625 | 0.632412477 | 0.301029996 | 0.224243824 |
| ENSMUSG00000102426 | 1.227926429 | 1.214495249 | 1.27186571 | 1.451838731 | 1.439191811 | 1.643452676 | 1.340368108 |
| ENSMUSG00000102427 | 0 | 0 | 0 | 0 | 0 | 0 | 0 |
| ENSMUSG00000102428 | 1.129344651 | 0.75 | 0.864857905 | 1 | 1.057362051 | 1.113943352 | 1.168187745 |
| ENSMUSG00000102429 | 0 | 0 | 0 | 0 | 0 | 0 | 0 |
| ENSMUSG00000102430 | 0 | 0 | 0 | 0 | 0 | 0 | 0 |
| ENSMUSG00000102431 | 0 | 0 | 0 | 0 | 0 | 0 | 0 |
| ENSMUSG00000102432 | 0 | 0 | 0 | 0 | 0 | 0 | 0 |
| ENSMUSG00000102433 | 0.289064826 | 0.646240625 | 0.5 | 0.396240625 | 0.244650542 | 0 | 0 |
| ENSMUSG00000102434 | 0 | 0 | 0 | 0 | 0 | 0 | 0.224243824 |
| ENSMUSG00000102436 | 0.45815691 | 0.79248125 | 0.5 | 0.5 | 0.387761935 | 0.698970004 | 0.579661877 |
| ENSMUSG00000102437 | 0.289064826 | 0.580482024 | 0.396240625 | 0 | 0 | 0 | 0 |
| ENSMUSG00000102438 | 0 | 0 | 0 | 0 | 0 | 0 | 0 |
| ENSMUSG00000102439 | 0 | 0 | 0 | 0 | 0 | 0 | 0 |
| ENSMUSG00000102440 | 1.404271433 | 1.51109853 | 1.388647213 | 0.646240625 | 0.387761935 | 0.602059991 | 0.355418052 |
| ENSMUSG00000102441 | 0 | 0 | 0 | 0 | 0 | 0 | 0 |
| ENSMUSG00000102442 | 0 | 0 | 0 | 0 | 0 | 0 | 0 |
| ENSMUSG00000102443 | 0.578129653 | 0.92510993 | 0.830482024 | 0.701838731 | 0.846351821 | 0.698970004 | 0.579661877 |
| ENSMUSG00000102445 | 0 | 0 | 0 | 0 | 0 | 0 | 0 |
| ENSMUSG00000102446 | 0 | 0 | 0 | 0 | 0 | 0 | 0 |

| | | | | | | | |
|---|---|---|---|---|---|---|---|
| ENSMUSG00000102447 | 0.289064826 | 0.25 | 0 | 0 | 0 | 0 | 0.224243824 |
| ENSMUSG00000102448 | 0.289064826 | 0 | 0.25 | 0 | 0.244650542 | 0 | 0 |
| ENSMUSG00000102449 | 0 | 0 | 0 | 0 | 0 | 0 | 0 |
| ENSMUSG00000102450 | 0 | 0 | 0 | 0 | 0 | 0 | 0 |
| ENSMUSG00000102451 | 0 | 0 | 0 | 0 | 0.387761935 | 0 | 0 |
| ENSMUSG00000102452 | 0.289064826 | 0.396240625 | 0.580482024 | 0 | 0.387761935 | 0 | 0 |
| ENSMUSG00000102453 | 0 | 0 | 0 | 0 | 0 | 0 | 0 |
| ENSMUSG00000102456 | 0 | 0 | 0 | 0 | 0 | 0 | 0 |
| ENSMUSG00000102457 | 0.289064826 | 0 | 0.25 | 0 | 0 | 0 | 0 |
| ENSMUSG00000102458 | 0 | 0 | 0 | 0 | 0 | 0 | 0 |
| ENSMUSG00000102459 | 0 | 0.25 | 0 | 0.25 | 0 | 0 | 0 |
| ENSMUSG00000102460 | 0 | 0 | 0 | 0 | 0 | 0 | 0 |
| ENSMUSG00000102461 | 0.289064826 | 0.25 | 0.25 | 0.25 | 0.244650542 | 0 | 0 |
| ENSMUSG00000102462 | 0 | 0 | 0 | 0 | 0 | 0 | 0 |
| ENSMUSG00000102463 | 0.289064826 | 0 | 0 | 0 | 0 | 0 | 0 |
| ENSMUSG00000102464 | 0 | 0 | 0 | 0 | 0 | 0 | 0 |
| ENSMUSG00000102465 | 0.578129653 | 0.79248125 | 0.396240625 | 0.5 | 0.568060967 | 0.77815125 | 0.448487648 |
| ENSMUSG00000102466 | 0.45815691 | 0.5 | 0.25 | 0.25 | 0.387761935 | 0 | 0.355418052 |
| ENSMUSG00000102467 | 0 | 0 | 0 | 0 | 0 | 0 | 0 |
| ENSMUSG00000102469 | 0 | 0 | 0 | 0 | 0 | 0 | 0 |
| ENSMUSG00000102470 | 0.91631382 | 0.830482024 | 0.75 | 0.5 | 0.387761935 | 0.477121255 | 0.448487648 |
| ENSMUSG00000102471 | 0 | 0 | 0 | 0.25 | 0 | 0 | 0 |
| ENSMUSG00000102473 | 0 | 0 | 0 | 0 | 0 | 0 | 0 |
| ENSMUSG00000102474 | 0.289064826 | 0 | 0.25 | 0.25 | 0 | 0 | 0 |
| ENSMUSG00000102475 | 0.289064826 | 0 | 0 | 0 | 0 | 0 | 0 |
| ENSMUSG00000102476 | 0 | 0 | 0 | 0 | 0 | 0 | 0 |
| ENSMUSG00000102478 | 0.578129653 | 0.396240625 | 0.396240625 | 0.5 | 0 | 0.602059991 | 0.224243824 |
| ENSMUSG00000102479 | 0 | 0.25 | 0 | 0 | 0.489301084 | 0.301029996 | 0 |
| ENSMUSG00000102480 | 0.45815691 | 0.5 | 0.25 | 0.396240625 | 0.387761935 | 0.477121255 | 0.224243824 |
| ENSMUSG00000102481 | 0 | 0 | 0 | 0 | 0.244650542 | 0 | 0 |
| ENSMUSG00000102482 | 0 | 0.25 | 0 | 0.396240625 | 0.244650542 | 0 | 0.224243824 |
| ENSMUSG00000102483 | 0 | 0 | 0 | 0 | 0 | 0 | 0 |
| ENSMUSG00000102484 | 0 | 0 | 0 | 0 | 0 | 0 | 0 |
| ENSMUSG00000102485 | 0 | 0 | 0 | 0 | 0 | 0 | 0 |

| | | | | | | | |
|---|---|---|---|---|---|---|---|
| ENSMUSG00000102486 | 0 | 0 | 0 | 0 | 0 | 0 | 0 |
| ENSMUSG00000102487 | 0 | 0 | 0 | 0 | 0 | 0 | 0 |
| ENSMUSG00000102488 | 0 | 0 | 0 | 0 | 0 | 0 | 0 |
| ENSMUSG00000102489 | 0 | 0 | 0 | 0 | 0 | 0 | 0 |
| ENSMUSG00000102490 | 0 | 0 | 0.25 | 0.25 | 0.387761935 | 0 | 0.355418052 |
| ENSMUSG00000102491 | 0 | 0 | 0 | 0 | 0 | 0 | 0 |
| ENSMUSG00000102492 | 0 | 0 | 0 | 0 | 0 | 0 | 0 |
| ENSMUSG00000102493 | 0.867194479 | 0.580482024 | 0.79248125 | 1.25 | 1.200473444 | 1.146128036 | 1.078019652 |
| ENSMUSG00000102494 | 0.671187741 | 0.646240625 | 0.75 | 0.79248125 | 0.686820904 | 0.477121255 | 0.629532004 |
| ENSMUSG00000102495 | 0 | 0 | 0 | 0 | 0 | 0 | 0 |
| ENSMUSG00000102496 | 1.325351389 | 1.302363341 | 1 | 1.214495249 | 1.074582839 | 0.954242509 | 0.916588299 |
| ENSMUSG00000102497 | 0 | 0 | 0 | 0 | 0 | 0 | 0 |
| ENSMUSG00000102498 | 0 | 0 | 0 | 0 | 0 | 0 | 0 |
| ENSMUSG00000102499 | 0 | 0 | 0 | 0 | 0 | 0 | 0 |
| ENSMUSG00000102500 | 0 | 0 | 0 | 0 | 0 | 0 | 0 |
| ENSMUSG00000102501 | 0.289064826 | 0.25 | 0.5 | 0.396240625 | 0.244650542 | 0 | 0.355418052 |
| ENSMUSG00000102502 | 0 | 0 | 0 | 0 | 0 | 0 | 0 |
| ENSMUSG00000102503 | 0.289064826 | 0.396240625 | 0 | 0 | 0 | 0 | 0 |
| ENSMUSG00000102504 | 0 | 0.25 | 0 | 0.396240625 | 0 | 0 | 0.355418052 |
| ENSMUSG00000102505 | 0 | 0 | 0 | 0 | 0 | 0 | 0 |
| ENSMUSG00000102506 | 0 | 0 | 0 | 0 | 0 | 0 | 0 |
| ENSMUSG00000102507 | 0 | 0 | 0 | 0 | 0 | 0 | 0 |
| ENSMUSG00000102508 | 0 | 0 | 0 | 0 | 0 | 0 | 0 |
| ENSMUSG00000102509 | 0 | 0.396240625 | 0 | 0 | 0.244650542 | 0 | 0.448487648 |
| ENSMUSG00000102510 | 0 | 0 | 0 | 0 | 0 | 0 | 0 |
| ENSMUSG00000102511 | 0 | 0 | 0 | 0 | 0 | 0 | 0 |
| ENSMUSG00000102512 | 0 | 0 | 0 | 0 | 0 | 0 | 0 |
| ENSMUSG00000102513 | 0 | 0 | 0 | 0 | 0 | 0 | 0 |
| ENSMUSG00000102514 | 0 | 0 | 0 | 0.25 | 0.244650542 | 0 | 0 |
| ENSMUSG00000102515 | 0 | 0 | 0 | 0 | 0 | 0 | 0 |
| ENSMUSG00000102516 | 0 | 0.25 | 0 | 0.25 | 0 | 0 | 0 |
| ENSMUSG00000102517 | 0.578129653 | 0.396240625 | 0.25 | 0.580482024 | 0.244650542 | 0.301029996 | 0.448487648 |
| ENSMUSG00000102518 | 0.45815691 | 0 | 0.396240625 | 0.5 | 0.244650542 | 0.301029996 | 0.224243824 |
| ENSMUSG00000102520 | 0 | 0 | 0 | 0 | 0 | 0 | 0 |

| | | | | | | | |
|---|---|---|---|---|---|---|---|
| ENSMUSG00000102521 | 0.289064826 | 0.25 | 0 | 0 | 0 | 0.301029996 | 0 |
| ENSMUSG00000102522 | 0 | 0 | 0 | 0 | 0 | 0 | 0 |
| ENSMUSG00000102523 | 0.289064826 | 0 | 0.25 | 0.396240625 | 0 | 0.301029996 | 0.355418052 |
| ENSMUSG00000102524 | 0 | 0 | 0 | 0 | 0 | 0 | 0 |
| ENSMUSG00000102525 | 0 | 0 | 0 | 0.25 | 0 | 0 | 0 |
| ENSMUSG00000102526 | 0.578129653 | 0.5 | 0.25 | 0.25 | 0.387761935 | 0.477121255 | 0.520678036 |
| ENSMUSG00000102527 | 0 | 0 | 0 | 0 | 0 | 0 | 0 |
| ENSMUSG00000102528 | 0 | 0 | 0 | 0 | 0 | 0 | 0 |
| ENSMUSG00000102529 | 0 | 0 | 0 | 0 | 0 | 0 | 0 |
| ENSMUSG00000102530 | 0 | 0 | 0 | 0 | 0 | 0 | 0 |
| ENSMUSG00000102531 | 0 | 0 | 0 | 0 | 0 | 0 | 0 |
| ENSMUSG00000102532 | 0 | 0 | 0 | 0 | 0 | 0 | 0 |
| ENSMUSG00000102533 | 0.289064826 | 0.5 | 0.25 | 0.25 | 0 | 0 | 0.224243824 |
| ENSMUSG00000102534 | 0 | 0 | 0 | 0 | 0 | 0 | 0 |
| ENSMUSG00000102535 | 0 | 0 | 0 | 0 | 0 | 0 | 0 |
| ENSMUSG00000102536 | 0 | 0 | 0 | 0 | 0 | 0 | 0 |
| ENSMUSG00000102537 | 0 | 0 | 0 | 0 | 0 | 0 | 0 |
| ENSMUSG00000102538 | 0 | 0 | 0 | 0 | 0 | 0 | 0 |
| ENSMUSG00000102539 | 0 | 0 | 0.25 | 0 | 0 | 0 | 0 |
| ENSMUSG00000102540 | 0.289064826 | 0 | 0.25 | 0 | 0.244650542 | 0.301029996 | 0.224243824 |
| ENSMUSG00000102541 | 0 | 0 | 0 | 0 | 0 | 0 | 0 |
| ENSMUSG00000102542 | 0 | 0 | 0 | 0 | 0 | 0 | 0 |
| ENSMUSG00000102543 | 1.928763473 | 1.809601185 | 1.807204673 | 1.27186571 | 1.366364104 | 1.301029996 | 1.252393349 |
| ENSMUSG00000102544 | 0.289064826 | 0 | 0 | 0 | 0.244650542 | 0 | 0 |
| ENSMUSG00000102545 | 0 | 0.396240625 | 0.25 | 0 | 0.244650542 | 0 | 0 |
| ENSMUSG00000102546 | 0.289064826 | 0 | 0.25 | 0.25 | 0 | 0 | 0 |
| ENSMUSG00000102547 | 0 | 0 | 0 | 0 | 0 | 0 | 0 |
| ENSMUSG00000102548 | 0 | 0 | 0 | 0 | 0 | 0 | 0 |
| ENSMUSG00000102549 | 0 | 0 | 0 | 0 | 0 | 0 | 0 |
| ENSMUSG00000102550 | 0.671187741 | 0.951838731 | 0.92510993 | 0.646240625 | 0.77552387 | 0.84509804 | 0.829800754 |
| ENSMUSG00000102551 | 0 | 0 | 0 | 0 | 0 | 0 | 0 |
| ENSMUSG00000102552 | 0 | 0 | 0 | 0 | 0 | 0 | 0 |
| ENSMUSG00000102553 | 1.129344651 | 0.79248125 | 0.830482024 | 0.830482024 | 0.812711509 | 0.602059991 | 0.672731473 |
| ENSMUSG00000102554 | 0 | 0.25 | 0 | 0.25 | 0.387761935 | 0 | 0.224243824 |

| | | | | | | | |
|---|---|---|---|---|---|---|---|
| ENSMUSG00000102555 | 0.811507563 | 1.02186571 | 1.098079356 | 1.160964047 | 1.121713561 | 1.230448921 | 0.984950056 |
| ENSMUSG00000102556 | 0 | 0 | 0 | 0.25 | 0 | 0 | 0 |
| ENSMUSG00000102558 | 0 | 0 | 0 | 0 | 0 | 0 | 0 |
| ENSMUSG00000102559 | 1 | 0.701838731 | 0.896240625 | 0.75 | 0.846351821 | 0.698970004 | 0.775756176 |
| ENSMUSG00000102561 | 0 | 0.25 | 0.25 | 0 | 0.244650542 | 0 | 0 |
| ENSMUSG00000102562 | 0 | 0 | 0 | 1.146240625 | 0.733951626 | 0.954242509 | 0.916588299 |
| ENSMUSG00000102563 | 0 | 0 | 0 | 0 | 0 | 0 | 0 |
| ENSMUSG00000102564 | 0 | 0 | 0 | 0 | 0 | 0 | 0 |
| ENSMUSG00000102565 | 0 | 0 | 0 | 0 | 0.244650542 | 0 | 0 |
| ENSMUSG00000102566 | 0 | 0 | 0 | 0 | 0 | 0 | 0 |
| ENSMUSG00000102567 | 0 | 0 | 0 | 0 | 0 | 0 | 0 |
| ENSMUSG00000102568 | 0.45815691 | 0 | 0.396240625 | 0.25 | 0.387761935 | 0.301029996 | 0 |
| ENSMUSG00000102569 | 0 | 0 | 0 | 0 | 0 | 0 | 0 |
| ENSMUSG00000102570 | 0.747221736 | 0.92510993 | 0.79248125 | 0.75 | 0.846351821 | 0.602059991 | 0.876096088 |
| ENSMUSG00000102571 | 0 | 0 | 0.25 | 0 | 0 | 0 | 0 |
| ENSMUSG00000102572 | 0 | 0 | 0 | 0 | 0 | 0 | 0 |
| ENSMUSG00000102573 | 0.960252568 | 0.79248125 | 0.830482024 | 0.75 | 0.877063019 | 0.477121255 | 0.672731473 |
| ENSMUSG00000102574 | 0 | 0 | 0 | 0 | 0 | 0 | 0 |
| ENSMUSG00000102575 | 0 | 0.25 | 0 | 0 | 0 | 0 | 0 |
| ENSMUSG00000102576 | 0 | 0 | 0 | 0 | 0 | 0 | 0 |
| ENSMUSG00000102577 | 0.289064826 | 0.25 | 0.75 | 0.396240625 | 0.387761935 | 0.77815125 | 0.448487648 |
| ENSMUSG00000102578 | 0 | 0 | 0.396240625 | 0 | 0 | 0 | 0 |
| ENSMUSG00000102579 | 0 | 0 | 0 | 0 | 0 | 0 | 0 |
| ENSMUSG00000102580 | 0 | 0 | 0 | 0 | 0 | 0 | 0 |
| ENSMUSG00000102581 | 0 | 0.25 | 0 | 0 | 0 | 0 | 0 |
| ENSMUSG00000102582 | 0 | 0 | 0 | 0 | 0 | 0 | 0 |
| ENSMUSG00000102583 | 0 | 0 | 0 | 0 | 0 | 0 | 0 |
| ENSMUSG00000102584 | 0 | 0 | 0 | 0 | 0 | 0 | 0 |
| ENSMUSG00000102585 | 0 | 0 | 0 | 0 | 0 | 0 | 0 |
| ENSMUSG00000102586 | 0 | 0 | 0 | 0 | 0 | 0 | 0 |
| ENSMUSG00000102587 | 0 | 0 | 0 | 0 | 0 | 0 | 0 |
| ENSMUSG00000102588 | 0 | 0 | 0 | 0 | 0 | 0 | 0 |
| ENSMUSG00000102589 | 0 | 0 | 0 | 0 | 0 | 0 | 0 |
| ENSMUSG00000102590 | 0 | 0 | 0 | 0 | 0.244650542 | 0 | 0 |

| | | | | | | | |
|---|---|---|---|---|---|---|---|
| ENSMUSG00000102591 | 1.129344651 | 0.951838731 | 1.04248125 | 1.25 | 0.931471446 | 1.146128036 | 1.028149525 |
| ENSMUSG00000102592 | 0 | 0.25 | 0 | 0 | 0 | 0 | 0.224243824 |
| ENSMUSG00000102593 | 0.811507563 | 0.864857905 | 0.5 | 0.92510993 | 0.632412477 | 0.698970004 | 0.629532004 |
| ENSMUSG00000102594 | 0 | 0.25 | 0 | 0 | 0.387761935 | 0 | 0 |
| ENSMUSG00000102595 | 0 | 0 | 0 | 0 | 0 | 0 | 0 |
| ENSMUSG00000102596 | 0 | 0 | 0 | 0 | 0 | 0 | 0 |
| ENSMUSG00000102597 | 0 | 0 | 0 | 0 | 0 | 0 | 0 |
| ENSMUSG00000102598 | 0 | 0 | 0 | 0 | 0 | 0 | 0 |
| ENSMUSG00000102599 | 0.45815691 | 0.646240625 | 0.25 | 0.396240625 | 0.489301084 | 0.698970004 | 0.224243824 |
| ENSMUSG00000102600 | 0 | 0 | 0.396240625 | 0.25 | 0 | 0.301029996 | 0.224243824 |
| ENSMUSG00000102601 | 0 | 0 | 0 | 0 | 0 | 0 | 0 |
| ENSMUSG00000102602 | 2.039433992 | 1.830482024 | 1.814346961 | 1.940387808 | 1.789100779 | 2.056904851 | 1.669034113 |
| ENSMUSG00000102603 | 0 | 0 | 0 | 0 | 0 | 0 | 0 |
| ENSMUSG00000102605 | 0 | 0 | 0 | 0 | 0 | 0 | 0 |
| ENSMUSG00000102606 | 0 | 0 | 0 | 0.25 | 0 | 0 | 0 |
| ENSMUSG00000102607 | 0 | 0 | 0 | 0 | 0 | 0 | 0 |
| ENSMUSG00000102608 | 0 | 0 | 0 | 0 | 0 | 0 | 0 |
| ENSMUSG00000102609 | 0 | 0 | 0 | 0 | 0 | 0 | 0 |
| ENSMUSG00000102610 | 0.747221736 | 0.830482024 | 1.02186571 | 1.130890489 | 0.812711509 | 0.954242509 | 0.969165684 |
| ENSMUSG00000102611 | 0 | 0 | 0 | 0 | 0 | 0 | 0 |
| ENSMUSG00000102612 | 0 | 0 | 0 | 0 | 0 | 0 | 0 |
| ENSMUSG00000102613 | 0 | 0 | 0 | 0 | 0 | 0 | 0 |
| ENSMUSG00000102614 | 0 | 0 | 0 | 0 | 0 | 0 | 0.224243824 |
| ENSMUSG00000102615 | 0 | 0.25 | 0 | 0 | 0 | 0 | 0 |
| ENSMUSG00000102616 | 0 | 0 | 0 | 0 | 0 | 0 | 0 |
| ENSMUSG00000102617 | 0 | 0 | 0 | 0 | 0 | 0 | 0 |
| ENSMUSG00000102618 | 0 | 0 | 0 | 0 | 0 | 0 | 0 |
| ENSMUSG00000102619 | 0 | 0 | 0 | 0.5 | 0 | 0.301029996 | 0 |
| ENSMUSG00000102620 | 0 | 0 | 0 | 0 | 0 | 0 | 0 |
| ENSMUSG00000102621 | 0 | 0.25 | 0.25 | 0.25 | 0 | 0.301029996 | 0.448487648 |
| ENSMUSG00000102622 | 0 | 0 | 0 | 0 | 0 | 0 | 0 |
| ENSMUSG00000102623 | 0 | 0.646240625 | 0.25 | 0 | 0.244650542 | 0.301029996 | 0.224243824 |
| ENSMUSG00000102624 | 0 | 0 | 0 | 0 | 0 | 0 | 0 |
| ENSMUSG00000102625 | 0 | 0.25 | 0 | 0 | 0 | 0 | 0 |

| | | | | | | | |
|---|---|---|---|---|---|---|---|
| ENSMUSG00000102626 | 0 | 0 | 0 | 0 | 0 | 0 | 0 |
| ENSMUSG00000102627 | 0 | 0 | 0 | 0 | 0 | 0 | 0 |
| ENSMUSG00000102628 | 0 | 0 | 0 | 0 | 0 | 0 | 0 |
| ENSMUSG00000102629 | 0 | 0 | 0.25 | 0 | 0 | 0 | 0 |
| ENSMUSG00000102630 | 0 | 0.25 | 0.25 | 0.396240625 | 0 | 0 | 0 |
| ENSMUSG00000102632 | 0.289064826 | 0.25 | 0 | 0.25 | 0.244650542 | 0 | 0.355418052 |
| ENSMUSG00000102633 | 0.747221736 | 0.701838731 | 0.646240625 | 0.701838731 | 0.733951626 | 0.84509804 | 0.672731473 |
| ENSMUSG00000102634 | 0 | 0 | 0 | 0 | 0 | 0 | 0 |
| ENSMUSG00000102635 | 0.289064826 | 0 | 0 | 0 | 0 | 0 | 0 |
| ENSMUSG00000102636 | 0 | 0 | 0 | 0 | 0 | 0 | 0 |
| ENSMUSG00000102637 | 0 | 0.25 | 0 | 0 | 0 | 0 | 0 |
| ENSMUSG00000102639 | 0 | 0 | 0 | 0 | 0 | 0 | 0 |
| ENSMUSG00000102641 | 0 | 0 | 0 | 0 | 0 | 0 | 0 |
| ENSMUSG00000102642 | 0 | 0 | 0 | 0 | 0 | 0 | 0.224243824 |
| ENSMUSG00000102643 | 0 | 0 | 0 | 0 | 0 | 0 | 0 |
| ENSMUSG00000102644 | 1.494443473 | 1.451838731 | 1.527131114 | 1.557204673 | 1.49445382 | 1.591064607 | 1.340368108 |
| ENSMUSG00000102645 | 0 | 0 | 0 | 0 | 0 | 0 | 0 |
| ENSMUSG00000102646 | 0 | 0 | 0 | 0 | 0 | 0 | 0 |
| ENSMUSG00000102647 | 0.45815691 | 0.25 | 0.396240625 | 0.25 | 0.489301084 | 0.477121255 | 0.520678036 |
| ENSMUSG00000102648 | 0 | 0.25 | 0.25 | 0 | 0 | 0 | 0.224243824 |
| ENSMUSG00000102649 | 0 | 0 | 0.25 | 0 | 0.387761935 | 0.477121255 | 0.224243824 |
| ENSMUSG00000102650 | 0 | 0 | 0 | 0 | 0 | 0 | 0 |
| ENSMUSG00000102651 | 0 | 0 | 0 | 0.25 | 0 | 0 | 0 |
| ENSMUSG00000102652 | 0 | 0.25 | 0 | 0.396240625 | 0 | 0 | 0 |
| ENSMUSG00000102653 | 0 | 0 | 0 | 0 | 0 | 0 | 0 |
| ENSMUSG00000102654 | 0 | 0 | 0 | 0 | 0 | 0 | 0 |
| ENSMUSG00000102655 | 0 | 0 | 0 | 0 | 0 | 0 | 0 |
| ENSMUSG00000102656 | 0 | 0 | 0 | 0 | 0 | 0 | 0 |
| ENSMUSG00000102657 | 0.289064826 | 0.701838731 | 0.5 | 0.75 | 0 | 0 | 0.224243824 |
| ENSMUSG00000102658 | 0 | 0 | 0 | 0 | 0 | 0 | 0 |
| ENSMUSG00000102659 | 0.45815691 | 0.79248125 | 0 | 0.701838731 | 0.686820904 | 0.84509804 | 0.710836105 |
| ENSMUSG00000102660 | 0 | 0 | 0 | 0 | 0 | 0 | 0 |
| ENSMUSG00000102661 | 0 | 0 | 0 | 0 | 0 | 0 | 0 |
| ENSMUSG00000102662 | 0 | 0 | 0 | 0.5 | 0.244650542 | 0 | 0.355418052 |

| | | | | | | | |
|---|---|---|---|---|---|---|---|
| ENSMUSG00000102663 | 0 | 0 | 0 | 0 | 0 | 0 | 0.224243824 |
| ENSMUSG00000102664 | 0.289064826 | 0.646240625 | 0.396240625 | 0.25 | 0.489301084 | 0 | 0.355418052 |
| ENSMUSG00000102665 | 0 | 0 | 0.25 | 0.396240625 | 0 | 0.301029996 | 0.355418052 |
| ENSMUSG00000102666 | 0 | 0.396240625 | 0 | 0 | 0 | 0 | 0 |
| ENSMUSG00000102667 | 0 | 0 | 0 | 0 | 0 | 0 | 0 |
| ENSMUSG00000102668 | 0 | 0 | 0 | 0 | 0 | 0 | 0 |
| ENSMUSG00000102669 | 0 | 0 | 0 | 0 | 0 | 0 | 0 |
| ENSMUSG00000102670 | 0 | 0.25 | 0.25 | 0.25 | 0.244650542 | 0 | 0 |
| ENSMUSG00000102672 | 1.069666964 | 1 | 1.04248125 | 1.02186571 | 0.846351821 | 0.903089987 | 1.078019652 |
| ENSMUSG00000102673 | 0.671187741 | 0.5 | 0.646240625 | 0 | 0.244650542 | 0.698970004 | 0.520678036 |
| ENSMUSG00000102674 | 0.45815691 | 0 | 0 | 0 | 0 | 0 | 0.224243824 |
| ENSMUSG00000102676 | 0 | 0.25 | 0 | 0 | 0 | 0 | 0.224243824 |
| ENSMUSG00000102677 | 0 | 0 | 0 | 0 | 0 | 0 | 0 |
| ENSMUSG00000102678 | 0 | 0 | 0 | 0 | 0 | 0 | 0 |
| ENSMUSG00000102679 | 0 | 0 | 0 | 0 | 0 | 0 | 0 |
| ENSMUSG00000102680 | 0 | 0 | 0 | 0.25 | 0 | 0 | 0 |
| ENSMUSG00000102681 | 0 | 0 | 0 | 0 | 0 | 0 | 0 |
| ENSMUSG00000102682 | 0 | 0 | 0 | 0 | 0 | 0 | 0 |
| ENSMUSG00000102683 | 0 | 0 | 0 | 0 | 0 | 0 | 0 |
| ENSMUSG00000102684 | 0 | 0.25 | 0 | 0 | 0 | 0 | 0 |
| ENSMUSG00000102685 | 0.747221736 | 0.5 | 0.75 | 1.02186571 | 0.632412477 | 0.602059991 | 0.710836105 |
| ENSMUSG00000102686 | 0 | 0 | 0 | 0 | 0 | 0 | 0 |
| ENSMUSG00000102687 | 0 | 0 | 0 | 0 | 0 | 0.301029996 | 0 |
| ENSMUSG00000102688 | 0 | 0 | 0 | 0 | 0 | 0 | 0 |
| ENSMUSG00000102689 | 0 | 0 | 0 | 0 | 0 | 0 | 0 |
| ENSMUSG00000102690 | 0.45815691 | 0 | 0.396240625 | 0.25 | 0.244650542 | 0 | 0.355418052 |
| ENSMUSG00000102691 | 0.45815691 | 0.92510993 | 0.646240625 | 1.201838731 | 1.091002363 | 1.361727836 | 1.089372236 |
| ENSMUSG00000102692 | 0.960252568 | 0.830482024 | 0.864857905 | 1.302363341 | 1.200473444 | 1.477121255 | 1.23151414 |
| ENSMUSG00000102693 | 0 | 0 | 0 | 0 | 0 | 0 | 0 |
| ENSMUSG00000102694 | 0.811507563 | 0.580482024 | 0.79248125 | 0.864857905 | 0.733951626 | 0.77815125 | 0.579661877 |
| ENSMUSG00000102696 | 0 | 0 | 0 | 0 | 0 | 0 | 0 |
| ENSMUSG00000102697 | 0 | 0 | 0.25 | 0.25 | 0 | 0 | 0 |
| ENSMUSG00000102698 | 0 | 0 | 0 | 0 | 0 | 0 | 0.224243824 |
| ENSMUSG00000102699 | 0 | 0 | 0 | 0 | 0 | 0 | 0 |

| | | | | | | | |
|---|---|---|---|---|---|---|---|
| ENSMUSG00000102700 | 3.087570326 | 2.818740118 | 2.83272922 | 2.733975421 | 2.651238624 | 3.001300933 | 2.394807458 |
| ENSMUSG00000102701 | 0 | 0 | 0 | 0 | 0.244650542 | 0 | 0.224243824 |
| ENSMUSG00000102702 | 0 | 0.25 | 0.396240625 | 0.25 | 0.244650542 | 0.602059991 | 0.448487648 |
| ENSMUSG00000102703 | 0 | 0 | 0 | 0 | 0 | 0 | 0 |
| ENSMUSG00000102704 | 0.578129653 | 0.79248125 | 0.5 | 0.5 | 0.632412477 | 0 | 0.224243824 |
| ENSMUSG00000102705 | 1.37447073 | 1.311981878 | 1.188721876 | 1.061981878 | 1.212046813 | 1.176091259 | 1.131174228 |
| ENSMUSG00000102706 | 0.671187741 | 0.864857905 | 0.5 | 0 | 0 | 0 | 0 |
| ENSMUSG00000102707 | 0 | 0 | 0 | 0 | 0 | 0 | 0 |
| ENSMUSG00000102709 | 0.289064826 | 0.25 | 0 | 0.25 | 0 | 0 | 0 |
| ENSMUSG00000102710 | 0 | 0 | 0 | 0 | 0 | 0 | 0 |
| ENSMUSG00000102712 | 0.45815691 | 0.25 | 0.580482024 | 0 | 0.387761935 | 0.477121255 | 0.448487648 |
| ENSMUSG00000102713 | 0.45815691 | 0.396240625 | 0.396240625 | 0 | 0.387761935 | 0 | 0.355418052 |
| ENSMUSG00000102714 | 0 | 0.25 | 0.25 | 0 | 0.244650542 | 0 | 0 |
| ENSMUSG00000102715 | 0 | 0.25 | 0 | 0.25 | 0.387761935 | 0 | 0.448487648 |
| ENSMUSG00000102716 | 0 | 0 | 0 | 0 | 0 | 0 | 0 |
| ENSMUSG00000102717 | 0 | 0.25 | 0.5 | 0.580482024 | 0.244650542 | 0 | 0.224243824 |
| ENSMUSG00000102718 | 1.269664473 | 1.098079356 | 1.080482024 | 1.226722649 | 1.136121934 | 1.113943352 | 0.916588299 |
| ENSMUSG00000102719 | 0.578129653 | 0.864857905 | 0.646240625 | 0.646240625 | 0.686820904 | 0.602059991 | 0.775756176 |
| ENSMUSG00000102721 | 0 | 0 | 0 | 0 | 0 | 0 | 0 |
| ENSMUSG00000102722 | 0.289064826 | 0 | 0.396240625 | 0 | 0 | 0 | 0 |
| ENSMUSG00000102723 | 0 | 0 | 0 | 0 | 0 | 0 | 0 |
| ENSMUSG00000102724 | 0 | 0 | 0 | 0 | 0 | 0 | 0 |
| ENSMUSG00000102725 | 0 | 0 | 0 | 0 | 0 | 0 | 0 |
| ENSMUSG00000102726 | 0 | 0 | 0 | 0 | 0 | 0 | 0 |
| ENSMUSG00000102727 | 0 | 0 | 0 | 0 | 0 | 0 | 0 |
| ENSMUSG00000102728 | 0 | 0.25 | 0 | 0.25 | 0 | 0 | 0 |
| ENSMUSG00000102729 | 0 | 0 | 0 | 0 | 0.244650542 | 0 | 0 |
| ENSMUSG00000102730 | 0 | 0 | 0 | 0 | 0 | 0 | 0 |
| ENSMUSG00000102731 | 0 | 0 | 0 | 0 | 0 | 0 | 0 |
| ENSMUSG00000102732 | 0 | 0.25 | 0 | 0 | 0.244650542 | 0 | 0 |
| ENSMUSG00000102733 | 0 | 0 | 0 | 0 | 0 | 0 | 0 |
| ENSMUSG00000102734 | 0 | 0 | 0 | 0 | 0 | 0 | 0 |
| ENSMUSG00000102735 | 0.45815691 | 0 | 0.580482024 | 0.25 | 0.489301084 | 0 | 0 |
| ENSMUSG00000102737 | 0 | 0 | 0 | 0 | 0 | 0 | 0 |

| | | | | | | | |
|---|---|---|---|---|---|---|---|
| ENSMUSG00000102738 | 0 | 0 | 0 | 0 | 0 | 0 | 0 |
| ENSMUSG00000102739 | 0 | 0 | 0 | 0 | 0 | 0 | 0 |
| ENSMUSG00000102740 | 0 | 0 | 0 | 0 | 0 | 0 | 0 |
| ENSMUSG00000102741 | 0 | 0 | 0 | 0 | 0 | 0 | 0 |
| ENSMUSG00000102742 | 0.578129653 | 0.701838731 | 0.5 | 0.646240625 | 0.387761935 | 0.477121255 | 0.355418052 |
| ENSMUSG00000102743 | 0 | 0 | 0 | 0 | 0 | 0 | 0 |
| ENSMUSG00000102744 | 0 | 0 | 0 | 0 | 0 | 0 | 0.224243824 |
| ENSMUSG00000102745 | 0 | 0 | 0 | 0 | 0 | 0 | 0 |
| ENSMUSG00000102746 | 0.578129653 | 0.580482024 | 0.580482024 | 0.864857905 | 0.733951626 | 0.477121255 | 0.853775828 |
| ENSMUSG00000102747 | 0 | 0 | 0 | 0 | 0 | 0 | 0 |
| ENSMUSG00000102748 | 1.404271433 | 1.356566189 | 1.339388001 | 1.226722649 | 1.200473444 | 1.322219295 | 1.110947927 |
| ENSMUSG00000102749 | 0 | 0.25 | 0 | 0 | 0 | 0 | 0 |
| ENSMUSG00000102750 | 0 | 0 | 0 | 0 | 0.244650542 | 0 | 0 |
| ENSMUSG00000102751 | 0 | 0 | 0 | 0 | 0 | 0 | 0 |
| ENSMUSG00000102752 | 1 | 1 | 0.976722649 | 1 | 1.074582839 | 1 | 0.984950056 |
| ENSMUSG00000102753 | 0 | 0.25 | 0 | 0 | 0 | 0 | 0 |
| ENSMUSG00000102754 | 0 | 0 | 0 | 0 | 0 | 0 | 0 |
| ENSMUSG00000102755 | 0 | 0 | 0 | 0 | 0 | 0 | 0 |
| ENSMUSG00000102756 | 0 | 0 | 0 | 0 | 0 | 0 | 0 |
| ENSMUSG00000102757 | 0 | 0 | 0 | 0 | 0 | 0 | 0 |
| ENSMUSG00000102758 | 0.867194479 | 0.79248125 | 0.976722649 | 1.27186571 | 1.106691885 | 1.255272505 | 1 |
| ENSMUSG00000102759 | 0.811507563 | 0.75 | 0.646240625 | 0.396240625 | 0.387761935 | 0.602059991 | 0.672731473 |
| ENSMUSG00000102760 | 0 | 0.25 | 0.25 | 0 | 0.244650542 | 0.301029996 | 0 |
| ENSMUSG00000102761 | 0 | 0 | 0 | 0 | 0 | 0 | 0 |
| ENSMUSG00000102762 | 0 | 0.25 | 0 | 0 | 0.489301084 | 0 | 0 |
| ENSMUSG00000102763 | 0 | 0 | 0 | 0 | 0 | 0 | 0 |
| ENSMUSG00000102764 | 0 | 0 | 0.25 | 0.25 | 0 | 0 | 0 |
| ENSMUSG00000102765 | 0 | 0 | 0 | 0 | 0 | 0 | 0 |
| ENSMUSG00000102766 | 0 | 0 | 0 | 0 | 0 | 0 | 0 |
| ENSMUSG00000102768 | 0 | 0 | 0 | 0 | 0 | 0 | 0 |
| ENSMUSG00000102769 | 0 | 0 | 0 | 0 | 0 | 0 | 0 |
| ENSMUSG00000102770 | 0 | 0 | 0 | 0 | 0.244650542 | 0 | 0 |
| ENSMUSG00000102771 | 0 | 0 | 0 | 0 | 0 | 0 | 0 |
| ENSMUSG00000102772 | 0 | 0 | 0 | 0 | 0 | 0 | 0 |

| | | | | | | | |
|---|---|---|---|---|---|---|---|
| ENSMUSG00000102773 | 0 | 0 | 0 | 0 | 0 | 0 | 0 |
| ENSMUSG00000102774 | 0 | 0 | 0 | 0 | 0 | 0 | 0 |
| ENSMUSG00000102775 | 0 | 0.25 | 0.25 | 0.580482024 | 0.244650542 | 0 | 0.448487648 |
| ENSMUSG00000102776 | 0.747221736 | 0.864857905 | 0.701838731 | 0.79248125 | 0.686820904 | 0.77815125 | 0.803905701 |
| ENSMUSG00000102777 | 0 | 0 | 0 | 0 | 0 | 0 | 0 |
| ENSMUSG00000102778 | 0 | 0 | 0 | 0 | 0 | 0 | 0 |
| ENSMUSG00000102779 | 0 | 0 | 0 | 0.25 | 0 | 0 | 0 |
| ENSMUSG00000102780 | 1.307602651 | 1.146240625 | 1.27186571 | 1.372963274 | 1.27449559 | 1.301029996 | 1.185218806 |
| ENSMUSG00000102782 | 0.811507563 | 0.701838731 | 0.646240625 | 0.75 | 0.568060967 | 0.698970004 | 0.710836105 |
| ENSMUSG00000102783 | 0 | 0 | 0 | 0 | 0 | 0 | 0 |
| ENSMUSG00000102784 | 0 | 0.396240625 | 0 | 0.25 | 0 | 0 | 0 |
| ENSMUSG00000102785 | 0 | 0 | 0 | 0 | 0 | 0 | 0 |
| ENSMUSG00000102786 | 0 | 0 | 0 | 0 | 0 | 0 | 0 |
| ENSMUSG00000102787 | 0 | 0 | 0 | 0 | 0 | 0 | 0 |
| ENSMUSG00000102788 | 0 | 0 | 0 | 0 | 0 | 0 | 0 |
| ENSMUSG00000102789 | 0 | 0 | 0 | 0 | 0 | 0 | 0 |
| ENSMUSG00000102790 | 0 | 0 | 0 | 0 | 0 | 0 | 0 |
| ENSMUSG00000102791 | 0 | 0 | 0 | 0 | 0 | 0 | 0 |
| ENSMUSG00000102792 | 0 | 0 | 0 | 0 | 0 | 0.301029996 | 0 |
| ENSMUSG00000102793 | 0 | 0 | 0 | 0 | 0 | 0 | 0 |
| ENSMUSG00000102794 | 0 | 0 | 0 | 0 | 0 | 0 | 0 |
| ENSMUSG00000102795 | 0 | 0 | 0 | 0 | 0 | 0 | 0 |
| ENSMUSG00000102796 | 0 | 0 | 0 | 0 | 0 | 0 | 0.224243824 |
| ENSMUSG00000102797 | 0 | 0 | 0 | 0 | 0 | 0 | 0 |
| ENSMUSG00000102798 | 0.747221736 | 0.25 | 0.5 | 0.396240625 | 0.632412477 | 0.477121255 | 0.520678036 |
| ENSMUSG00000102799 | 0 | 0 | 0 | 0 | 0 | 0 | 0 |
| ENSMUSG00000102800 | 0.289064826 | 0.396240625 | 0 | 0.25 | 0 | 0 | 0.224243824 |
| ENSMUSG00000102801 | 0 | 0 | 0 | 0 | 0 | 0 | 0 |
| ENSMUSG00000102802 | 0 | 0 | 0 | 0 | 0 | 0 | 0 |
| ENSMUSG00000102803 | 0 | 0 | 0 | 0.396240625 | 0 | 0 | 0.224243824 |
| ENSMUSG00000102804 | 0 | 0 | 0 | 0 | 0 | 0 | 0 |
| ENSMUSG00000102805 | 0 | 0 | 0 | 0 | 0 | 0 | 0 |
| ENSMUSG00000102806 | 0 | 0 | 0 | 0 | 0 | 0 | 0 |
| ENSMUSG00000102807 | 0 | 0 | 0 | 0 | 0 | 0 | 0 |

| | | | | | | | |
|---|---|---|---|---|---|---|---|
| ENSMUSG00000102808 | 0 | 0 | 0 | 0 | 0 | 0 | 0.224243824 |
| ENSMUSG00000102809 | 0.578129653 | 0.5 | 0.646240625 | 0.5 | 0.568060967 | 0 | 0.520678036 |
| ENSMUSG00000102810 | 0 | 0 | 0 | 0 | 0 | 0 | 0 |
| ENSMUSG00000102813 | 1.358731791 | 1.42510993 | 1.330482024 | 1.098079356 | 1.223252711 | 1.230448921 | 1.140832123 |
| ENSMUSG00000102814 | 0 | 0 | 0 | 0 | 0.244650542 | 0 | 0 |
| ENSMUSG00000102815 | 0 | 0 | 0.25 | 0.25 | 0.244650542 | 0.477121255 | 0.355418052 |
| ENSMUSG00000102816 | 0 | 0 | 0 | 0 | 0 | 0 | 0 |
| ENSMUSG00000102817 | 0.289064826 | 0 | 0 | 0 | 0.244650542 | 0.477121255 | 0 |
| ENSMUSG00000102818 | 0 | 0 | 0 | 0 | 0 | 0 | 0 |
| ENSMUSG00000102819 | 0 | 0 | 0 | 0 | 0 | 0 | 0 |
| ENSMUSG00000102820 | 0 | 0 | 0 | 0 | 0 | 0 | 0 |
| ENSMUSG00000102821 | 0 | 0 | 0 | 0 | 0 | 0 | 0 |
| ENSMUSG00000102822 | 0 | 0 | 0 | 0 | 0 | 0 | 0 |
| ENSMUSG00000102824 | 0 | 0 | 0 | 0 | 0 | 0 | 0 |
| ENSMUSG00000102825 | 0 | 0 | 0 | 0 | 0 | 0.301029996 | 0 |
| ENSMUSG00000102826 | 0 | 0 | 0 | 0 | 0 | 0 | 0 |
| ENSMUSG00000102827 | 0 | 0 | 0 | 0 | 0 | 0 | 0.224243824 |
| ENSMUSG00000102828 | 0 | 0 | 0 | 0 | 0 | 0 | 0 |
| ENSMUSG00000102829 | 0 | 0 | 0 | 0 | 0 | 0 | 0 |
| ENSMUSG00000102830 | 0 | 0 | 0.25 | 0.5 | 0.244650542 | 0 | 0.224243824 |
| ENSMUSG00000102831 | 0 | 0 | 0 | 0 | 0 | 0 | 0 |
| ENSMUSG00000102832 | 0 | 0 | 0 | 0 | 0 | 0 | 0 |
| ENSMUSG00000102833 | 0.289064826 | 0.25 | 0 | 0 | 0 | 0 | 0.448487648 |
| ENSMUSG00000102834 | 0 | 0 | 0 | 0 | 0 | 0 | 0 |
| ENSMUSG00000102835 | 0 | 0 | 0 | 0 | 0 | 0 | 0 |
| ENSMUSG00000102836 | 0 | 0 | 0 | 0 | 0 | 0 | 0 |
| ENSMUSG00000102837 | 0 | 0 | 0 | 0 | 0 | 0 | 0.224243824 |
| ENSMUSG00000102838 | 0 | 0 | 0 | 0 | 0 | 0 | 0 |
| ENSMUSG00000102839 | 0 | 0 | 0 | 0 | 0 | 0 | 0 |
| ENSMUSG00000102840 | 0.747221736 | 0.79248125 | 0.951838731 | 0.580482024 | 0.632412477 | 0.477121255 | 0.876096088 |
| ENSMUSG00000102841 | 0 | 0 | 0.25 | 0 | 0.244650542 | 0 | 0.355418052 |
| ENSMUSG00000102842 | 0 | 0 | 0 | 0 | 0 | 0 | 0 |
| ENSMUSG00000102843 | 0 | 0 | 0 | 0 | 0 | 0 | 0 |
| ENSMUSG00000102844 | 0.578129653 | 0.580482024 | 0.580482024 | 0.5 | 0.489301084 | 0.477121255 | 0.579661877 |

| | | | | | | | |
|---|---|---|---|---|---|---|---|
| ENSMUSG00000102845 | 0 | 0 | 0 | 0 | 0 | 0 | 0 |
| ENSMUSG00000102846 | 0 | 0 | 0 | 0 | 0 | 0 | 0 |
| ENSMUSG00000102847 | 0 | 0 | 0 | 0 | 0 | 0 | 0 |
| ENSMUSG00000102848 | 0 | 0 | 0 | 0 | 0 | 0 | 0 |
| ENSMUSG00000102849 | 0 | 0 | 0 | 0 | 0 | 0 | 0 |
| ENSMUSG00000102850 | 0.45815691 | 0.5 | 0 | 0.646240625 | 0.244650542 | 0.698970004 | 0.224243824 |
| ENSMUSG00000102851 | 0 | 0 | 0 | 0 | 0 | 0 | 0 |
| ENSMUSG00000102852 | 0 | 0 | 0 | 0.396240625 | 0 | 0 | 0.224243824 |
| ENSMUSG00000102853 | 0 | 0 | 0 | 0 | 0 | 0 | 0 |
| ENSMUSG00000102854 | 0.747221736 | 0.5 | 0.5 | 0.976722649 | 0.846351821 | 0.301029996 | 0.672731473 |
| ENSMUSG00000102855 | 0 | 0 | 0 | 0 | 0 | 0 | 0 |
| ENSMUSG00000102856 | 0.578129653 | 0.646240625 | 0 | 0.580482024 | 0.387761935 | 0.301029996 | 0 |
| ENSMUSG00000102857 | 0 | 0 | 0 | 0 | 0 | 0 | 0 |
| ENSMUSG00000102858 | 1.505869732 | 1.364857905 | 1.372963274 | 1.42510993 | 1.343584837 | 1.397940009 | 1.131174228 |
| ENSMUSG00000102859 | 0 | 0.25 | 0 | 0 | 0.387761935 | 0 | 0 |
| ENSMUSG00000102860 | 0 | 0 | 0 | 0 | 0 | 0 | 0 |
| ENSMUSG00000102861 | 1 | 0.864857905 | 0.92510993 | 1 | 0.955822902 | 0.903089987 | 1.041356071 |
| ENSMUSG00000102862 | 0 | 0 | 0 | 0 | 0.244650542 | 0 | 0 |
| ENSMUSG00000102863 | 1.069666964 | 0.92510993 | 0.830482024 | 0.79248125 | 0.733951626 | 0.698970004 | 0.520678036 |
| ENSMUSG00000102864 | 0 | 0 | 0 | 0 | 0 | 0 | 0 |
| ENSMUSG00000102865 | 0 | 0 | 0 | 0 | 0 | 0 | 0 |
| ENSMUSG00000102866 | 0 | 0 | 0 | 0 | 0 | 0 | 0 |
| ENSMUSG00000102867 | 0 | 0 | 0 | 0 | 0 | 0 | 0 |
| ENSMUSG00000102868 | 0 | 0.25 | 0 | 0.25 | 0 | 0 | 0 |
| ENSMUSG00000102869 | 1.036286563 | 1.080482024 | 1.114857905 | 1.26109853 | 1.149965125 | 1.278753601 | 1.350478789 |
| ENSMUSG00000102870 | 0 | 0 | 0 | 0 | 0 | 0 | 0 |
| ENSMUSG00000102871 | 0 | 0 | 0 | 0 | 0 | 0 | 0 |
| ENSMUSG00000102872 | 0.289064826 | 0.701838731 | 0.5 | 0.5 | 0.489301084 | 0.602059991 | 0.448487648 |
| ENSMUSG00000102873 | 0 | 0 | 0 | 0 | 0 | 0 | 0 |
| ENSMUSG00000102874 | 0.289064826 | 0 | 0 | 0 | 0 | 0 | 0.224243824 |
| ENSMUSG00000102875 | 0.671187741 | 0.646240625 | 0.396240625 | 0.25 | 0.812711509 | 0.301029996 | 0.710836105 |
| ENSMUSG00000102876 | 0 | 0 | 0 | 0 | 0 | 0 | 0 |
| ENSMUSG00000102877 | 0 | 0 | 0 | 0 | 0 | 0 | 0 |
| ENSMUSG00000102878 | 0 | 0 | 0 | 0 | 0 | 0 | 0 |

| | | | | | | | |
|---|---|---|---|---|---|---|---|
| ENSMUSG00000102879 | 0 | 0 | 0 | 0 | 0 | 0 | 0 |
| ENSMUSG00000102880 | 0 | 0 | 0 | 0 | 0 | 0 | 0 |
| ENSMUSG00000102881 | 0 | 0 | 0.396240625 | 0.25 | 0.244650542 | 0 | 0.224243824 |
| ENSMUSG00000102882 | 0.867194479 | 0.580482024 | 0.896240625 | 0.864857905 | 0.244650542 | 0.903089987 | 0.74492186 |
| ENSMUSG00000102883 | 0 | 0 | 0 | 0 | 0 | 0 | 0 |
| ENSMUSG00000102884 | 0 | 0 | 0 | 0 | 0 | 0 | 0 |
| ENSMUSG00000102885 | 0 | 0 | 0 | 0 | 0 | 0 | 0 |
| ENSMUSG00000102886 | 0 | 0 | 0 | 0 | 0 | 0 | 0 |
| ENSMUSG00000102887 | 0 | 0 | 0 | 0 | 0 | 0 | 0 |
| ENSMUSG00000102888 | 0 | 0 | 0 | 0 | 0 | 0 | 0 |
| ENSMUSG00000102889 | 0 | 0 | 0 | 0 | 0 | 0 | 0 |
| ENSMUSG00000102890 | 0 | 0 | 0.25 | 0 | 0 | 0 | 0 |
| ENSMUSG00000102891 | 0 | 0 | 0 | 0 | 0 | 0 | 0 |
| ENSMUSG00000102892 | 0 | 0.25 | 0.396240625 | 0.580482024 | 0.489301084 | 0 | 0.355418052 |
| ENSMUSG00000102893 | 0.578129653 | 0.75 | 0.864857905 | 0.646240625 | 0.568060967 | 0.77815125 | 0.579661877 |
| ENSMUSG00000102894 | 0 | 0 | 0 | 0 | 0 | 0 | 0 |
| ENSMUSG00000102895 | 0.45815691 | 0 | 0 | 0.25 | 0.387761935 | 0 | 0.520678036 |
| ENSMUSG00000102896 | 0 | 0 | 0 | 0 | 0 | 0 | 0 |
| ENSMUSG00000102897 | 0 | 0 | 0 | 0 | 0 | 0 | 0 |
| ENSMUSG00000102898 | 0 | 0 | 0 | 0 | 0 | 0 | 0 |
| ENSMUSG00000102899 | 0.578129653 | 0.5 | 0.396240625 | 0.580482024 | 0.686820904 | 0.602059991 | 0.448487648 |
| ENSMUSG00000102900 | 0 | 0 | 0 | 0 | 0 | 0 | 0 |
| ENSMUSG00000102901 | 0 | 0 | 0 | 0 | 0 | 0 | 0 |
| ENSMUSG00000102902 | 0 | 0.25 | 0 | 0 | 0 | 0 | 0.355418052 |
| ENSMUSG00000102903 | 0 | 0 | 0 | 0 | 0 | 0 | 0 |
| ENSMUSG00000102904 | 0 | 0 | 0.25 | 0 | 0 | 0.301029996 | 0.224243824 |
| ENSMUSG00000102905 | 0 | 0 | 0 | 0 | 0 | 0 | 0 |
| ENSMUSG00000102906 | 0 | 0 | 0 | 0 | 0 | 0 | 0 |
| ENSMUSG00000102907 | 0 | 0 | 0 | 0 | 0 | 0 | 0 |
| ENSMUSG00000102908 | 0.578129653 | 0.580482024 | 0.396240625 | 0.396240625 | 0 | 0.301029996 | 0.224243824 |
| ENSMUSG00000102909 | 0.289064826 | 0 | 0 | 0 | 0.244650542 | 0 | 0.224243824 |
| ENSMUSG00000102910 | 0 | 0 | 0 | 0 | 0 | 0 | 0 |
| ENSMUSG00000102911 | 0 | 0 | 0 | 0 | 0 | 0 | 0 |
| ENSMUSG00000102912 | 0.867194479 | 0.5 | 0 | 0.25 | 0.733951626 | 0.602059991 | 0 |

| | | | | | | | |
|---|---|---|---|---|---|---|---|
| ENSMUSG00000102913 | 0 | 0 | 0 | 0 | 0 | 0 | 0 |
| ENSMUSG00000102914 | 0.289064826 | 0 | 0 | 0 | 0 | 0 | 0 |
| ENSMUSG00000102915 | 0 | 0 | 0 | 0 | 0 | 0 | 0 |
| ENSMUSG00000102916 | 0 | 0 | 0 | 0 | 0 | 0 | 0 |
| ENSMUSG00000102917 | 1.069666964 | 1.04248125 | 0.830482024 | 1.02186571 | 1.020174412 | 1 | 0.952571511 |
| ENSMUSG00000102918 | 2.227926429 | 2.115881093 | 2.179419106 | 1.910964047 | 1.903713872 | 2.08278537 | 1.905143021 |
| ENSMUSG00000102919 | 0 | 0.580482024 | 0.25 | 0.580482024 | 0.244650542 | 0 | 0 |
| ENSMUSG00000102920 | 0 | 0 | 0 | 0 | 0 | 0 | 0 |
| ENSMUSG00000102921 | 0 | 0 | 0 | 0 | 0 | 0 | 0 |
| ENSMUSG00000102922 | 0 | 0 | 0 | 0 | 0 | 0 | 0 |
| ENSMUSG00000102923 | 0 | 0 | 0 | 0 | 0.244650542 | 0 | 0.224243824 |
| ENSMUSG00000102924 | 0 | 0 | 0 | 0 | 0 | 0 | 0 |
| ENSMUSG00000102925 | 0 | 0 | 0 | 0 | 0 | 0 | 0 |
| ENSMUSG00000102926 | 0 | 0 | 0 | 0 | 0 | 0 | 0 |
| ENSMUSG00000102927 | 0.289064826 | 0.396240625 | 0.5 | 0.580482024 | 0.387761935 | 0.301029996 | 0.448487648 |
| ENSMUSG00000102928 | 0 | 0 | 0 | 0 | 0 | 0 | 0 |
| ENSMUSG00000102929 | 0.867194479 | 0.701838731 | 0.75 | 0.896240625 | 0.877063019 | 0.698970004 | 0.775756176 |
| ENSMUSG00000102930 | 0 | 0 | 0 | 0 | 0.244650542 | 0 | 0.355418052 |
| ENSMUSG00000102931 | 0.671187741 | 0.396240625 | 0.580482024 | 0.5 | 0.244650542 | 0 | 0.448487648 |
| ENSMUSG00000102932 | 0 | 0 | 0 | 0 | 0 | 0 | 0 |
| ENSMUSG00000102933 | 0 | 0 | 0 | 0 | 0 | 0 | 0 |
| ENSMUSG00000102934 | 0 | 0 | 0 | 0 | 0 | 0 | 0 |
| ENSMUSG00000102935 | 0.45815691 | 0.701838731 | 0.580482024 | 0.701838731 | 0.489301084 | 0.301029996 | 0.224243824 |
| ENSMUSG00000102936 | 0 | 0 | 0 | 0 | 0 | 0 | 0 |
| ENSMUSG00000102937 | 0 | 0 | 0 | 0 | 0 | 0 | 0 |
| ENSMUSG00000102938 | 0 | 0 | 0 | 0 | 0 | 0 | 0 |
| ENSMUSG00000102939 | 0.289064826 | 0 | 0 | 0.25 | 0 | 0 | 0.355418052 |
| ENSMUSG00000102940 | 0 | 0.25 | 0 | 0 | 0 | 0 | 0 |
| ENSMUSG00000102941 | 0 | 0 | 0 | 0 | 0 | 0 | 0 |
| ENSMUSG00000102942 | 0 | 0 | 0 | 0 | 0 | 0 | 0 |
| ENSMUSG00000102943 | 0 | 0 | 0 | 0 | 0 | 0 | 0 |
| ENSMUSG00000102945 | 0.289064826 | 0 | 0 | 0 | 0 | 0 | 0 |
| ENSMUSG00000102946 | 0 | 0 | 0 | 0 | 0 | 0 | 0 |
| ENSMUSG00000102948 | 0 | 0 | 0 | 0 | 0 | 0 | 0 |

| | | | | | | | |
|---|---|---|---|---|---|---|---|
| ENSMUSG00000102949 | 0 | 0 | 0 | 0 | 0 | 0 | 0 |
| ENSMUSG00000102950 | 0 | 0.25 | 0 | 0.25 | 0.387761935 | 0 | 0 |
| ENSMUSG00000102951 | 0 | 0.396240625 | 0 | 0 | 0 | 0 | 0.224243824 |
| ENSMUSG00000102952 | 0 | 0 | 0 | 0 | 0 | 0 | 0 |
| ENSMUSG00000102953 | 0 | 0.396240625 | 0.5 | 0 | 0.244650542 | 0 | 0 |
| ENSMUSG00000102955 | 0 | 0 | 0 | 0 | 0 | 0 | 0 |
| ENSMUSG00000102956 | 0.289064826 | 0.25 | 0.25 | 0.25 | 0.244650542 | 0.301029996 | 0.224243824 |
| ENSMUSG00000102957 | 0 | 0 | 0 | 0 | 0 | 0 | 0 |
| ENSMUSG00000102958 | 0 | 0 | 0 | 0 | 0 | 0 | 0 |
| ENSMUSG00000102959 | 0 | 0 | 0 | 0 | 0 | 0 | 0 |
| ENSMUSG00000102960 | 0.289064826 | 0.25 | 0 | 0.396240625 | 0 | 0.301029996 | 0 |
| ENSMUSG00000102961 | 0 | 0 | 0 | 0 | 0 | 0 | 0 |
| ENSMUSG00000102962 | 0 | 0 | 0 | 0 | 0 | 0 | 0 |
| ENSMUSG00000102963 | 0 | 0 | 0 | 0 | 0 | 0.301029996 | 0 |
| ENSMUSG00000102964 | 0.671187741 | 0.79248125 | 0.396240625 | 0.79248125 | 0.686820904 | 0.698970004 | 0.672731473 |
| ENSMUSG00000102965 | 0 | 0 | 0 | 0 | 0 | 0 | 0 |
| ENSMUSG00000102966 | 0 | 0 | 0 | 0 | 0 | 0 | 0 |
| ENSMUSG00000102967 | 0 | 0 | 0.25 | 0 | 0 | 0 | 0 |
| ENSMUSG00000102968 | 0 | 0 | 0 | 0 | 0 | 0 | 0 |
| ENSMUSG00000102969 | 0 | 0 | 0 | 0.25 | 0.244650542 | 0.477121255 | 0 |
| ENSMUSG00000102970 | 0.289064826 | 0.396240625 | 0.25 | 0.25 | 0.244650542 | 0.301029996 | 0.224243824 |
| ENSMUSG00000102971 | 0 | 0 | 0 | 0 | 0 | 0 | 0 |
| ENSMUSG00000102972 | 0 | 0 | 0 | 0 | 0 | 0 | 0 |
| ENSMUSG00000102973 | 0 | 0 | 0 | 0 | 0 | 0 | 0 |
| ENSMUSG00000102975 | 0 | 0 | 0 | 0 | 0 | 0 | 0 |
| ENSMUSG00000102976 | 2.160971586 | 1.972185812 | 1.970660762 | 2.005591953 | 1.978602168 | 2.23299611 | 1.793950594 |
| ENSMUSG00000102977 | 0 | 0 | 0 | 0 | 0 | 0 | 0 |
| ENSMUSG00000102979 | 0 | 0.25 | 0.25 | 0 | 0 | 0 | 0.224243824 |
| ENSMUSG00000102980 | 0 | 0 | 0 | 0 | 0 | 0 | 0 |
| ENSMUSG00000102981 | 0 | 0 | 0 | 0 | 0 | 0 | 0 |
| ENSMUSG00000102982 | 0.91631382 | 0.646240625 | 0.396240625 | 1 | 0.568060967 | 0.84509804 | 0.448487648 |
| ENSMUSG00000102983 | 0.289064826 | 0 | 0 | 0 | 0.244650542 | 0 | 0 |
| ENSMUSG00000102984 | 0 | 0 | 0 | 0 | 0 | 0 | 0 |
| ENSMUSG00000102985 | 0 | 0 | 0.25 | 0 | 0 | 0.301029996 | 0 |

| | | | | | | | |
|---|---|---|---|---|---|---|---|
| ENSMUSG00000102986 | 0 | 0 | 0 | 0 | 0 | 0 | 0 |
| ENSMUSG00000102987 | 0 | 0 | 0 | 0 | 0 | 0 | 0 |
| ENSMUSG00000102988 | 0 | 0 | 0 | 0 | 0 | 0 | 0 |
| ENSMUSG00000102989 | 0 | 0 | 0 | 0 | 0 | 0 | 0 |
| ENSMUSG00000102990 | 0 | 0 | 0 | 0 | 0.244650542 | 0 | 0 |
| ENSMUSG00000102991 | 0 | 0 | 0 | 0 | 0 | 0 | 0 |
| ENSMUSG00000102992 | 0 | 0 | 0 | 0 | 0 | 0 | 0 |
| ENSMUSG00000102993 | 0 | 0 | 0 | 0 | 0 | 0 | 0 |
| ENSMUSG00000102994 | 0.671187741 | 0.5 | 0 | 0.580482024 | 0.387761935 | 0 | 0.448487648 |
| ENSMUSG00000102995 | 0 | 0 | 0 | 0 | 0 | 0 | 0 |
| ENSMUSG00000102997 | 0.578129653 | 0 | 0.396240625 | 0.25 | 0 | 0.301029996 | 0 |
| ENSMUSG00000102998 | 0 | 0 | 0 | 0 | 0 | 0 | 0 |
| ENSMUSG00000102999 | 0.289064826 | 0 | 0 | 0.25 | 0 | 0 | 0 |
| ENSMUSG00000103000 | 0.960252568 | 0.396240625 | 0 | 0.580482024 | 0.686820904 | 0.301029996 | 0.710836105 |
| ENSMUSG00000103001 | 0 | 0 | 0 | 0 | 0 | 0 | 0 |
| ENSMUSG00000103002 | 0 | 0 | 0 | 0 | 0.244650542 | 0.301029996 | 0 |
| ENSMUSG00000103003 | 0 | 0 | 0 | 0 | 0 | 0 | 0 |
| ENSMUSG00000103004 | 0 | 0 | 0 | 0 | 0 | 0 | 0 |
| ENSMUSG00000103005 | 0 | 0 | 0 | 0 | 0 | 0 | 0 |
| ENSMUSG00000103006 | 0 | 0.5 | 0.5 | 0 | 0.568060967 | 0.301029996 | 0.710836105 |
| ENSMUSG00000103007 | 0 | 0 | 0 | 0 | 0 | 0 | 0 |
| ENSMUSG00000103008 | 0 | 0 | 0 | 0 | 0 | 0 | 0 |
| ENSMUSG00000103009 | 1.289064826 | 1.04248125 | 1.130890489 | 1.226722649 | 1.074582839 | 1.113943352 | 1.054044578 |
| ENSMUSG00000103010 | 0.289064826 | 0 | 0 | 0 | 0 | 0 | 0 |
| ENSMUSG00000103011 | 0 | 0 | 0 | 0.396240625 | 0 | 0 | 0.355418052 |
| ENSMUSG00000103012 | 0 | 0 | 0 | 0 | 0 | 0 | 0 |
| ENSMUSG00000103013 | 0.289064826 | 0.25 | 0.396240625 | 0 | 0.489301084 | 0 | 0.224243824 |
| ENSMUSG00000103015 | 0.289064826 | 0 | 0 | 0.25 | 0.244650542 | 0 | 0 |
| ENSMUSG00000103016 | 0 | 0 | 0 | 0 | 0 | 0 | 0 |
| ENSMUSG00000103017 | 0 | 0 | 0 | 0 | 0.244650542 | 0 | 0 |
| ENSMUSG00000103018 | 0 | 0 | 0 | 0 | 0 | 0 | 0 |
| ENSMUSG00000103020 | 0 | 0 | 0 | 0 | 0 | 0 | 0 |
| ENSMUSG00000103021 | 0 | 0 | 0 | 0 | 0.244650542 | 0 | 0 |
| ENSMUSG00000103023 | 0 | 0 | 0 | 0 | 0 | 0 | 0 |

| | | | | | | | |
|---|---|---|---|---|---|---|---|
| ENSMUSG00000103024 | 0 | 0 | 0.396240625 | 0.396240625 | 0 | 0 | 0.224243824 |
| ENSMUSG00000103025 | 0 | 0 | 0 | 0 | 0 | 0 | 0 |
| ENSMUSG00000103026 | 0 | 0 | 0 | 0 | 0 | 0 | 0 |
| ENSMUSG00000103027 | 0 | 0 | 0 | 0 | 0 | 0 | 0 |
| ENSMUSG00000103028 | 0 | 0 | 0 | 0 | 0 | 0 | 0 |
| ENSMUSG00000103029 | 0 | 0 | 0 | 0 | 0 | 0 | 0 |
| ENSMUSG00000103030 | 0.289064826 | 0.5 | 0.396240625 | 0.25 | 0.244650542 | 0 | 0.579661877 |
| ENSMUSG00000103031 | 0 | 0 | 0 | 0 | 0 | 0 | 0 |
| ENSMUSG00000103032 | 1.227926429 | 1.080482024 | 1.080482024 | 0.830482024 | 0.931471446 | 0.602059991 | 0.579661877 |
| ENSMUSG00000103033 | 0 | 0 | 0 | 0 | 0 | 0 | 0 |
| ENSMUSG00000103034 | 0.867194479 | 0.5 | 0.646240625 | 0.951838731 | 0.905314583 | 0.84509804 | 1.041356071 |
| ENSMUSG00000103035 | 0 | 0 | 0 | 0 | 0 | 0 | 0 |
| ENSMUSG00000103037 | 0.45815691 | 0.25 | 0 | 0 | 0 | 0 | 0.520678036 |
| ENSMUSG00000103038 | 0 | 0 | 0 | 0 | 0 | 0 | 0 |
| ENSMUSG00000103039 | 1.036286563 | 1.04248125 | 1.080482024 | 0.864857905 | 0.978602168 | 0.903089987 | 0.896975297 |
| ENSMUSG00000103040 | 0 | 0 | 0 | 0 | 0 | 0 | 0 |
| ENSMUSG00000103041 | 1.445324132 | 1.42510993 | 1.348079356 | 1.464495249 | 1.380772476 | 1.414973348 | 1.259064007 |
| ENSMUSG00000103042 | 0 | 0 | 0 | 0 | 0 | 0 | 0 |
| ENSMUSG00000103043 | 0 | 0 | 0 | 0 | 0 | 0 | 0 |
| ENSMUSG00000103044 | 0 | 0 | 0 | 0 | 0 | 0 | 0 |
| ENSMUSG00000103045 | 0 | 0 | 0 | 0 | 0 | 0 | 0 |
| ENSMUSG00000103046 | 1.205378646 | 1.04248125 | 1.061981878 | 1.130890489 | 1 | 0.903089987 | 1.028149525 |
| ENSMUSG00000103047 | 0.289064826 | 0.701838731 | 0.396240625 | 0.25 | 0 | 0 | 0.224243824 |
| ENSMUSG00000103049 | 0 | 0 | 0 | 0 | 0 | 0 | 0 |
| ENSMUSG00000103050 | 0 | 0 | 0.25 | 0 | 0 | 0 | 0 |
| ENSMUSG00000103051 | 0 | 0 | 0 | 0 | 0 | 0 | 0 |
| ENSMUSG00000103052 | 0 | 0 | 0 | 0 | 0 | 0 | 0 |
| ENSMUSG00000103053 | 1.307602651 | 1.17510993 | 1 | 1.02186571 | 1.176121988 | 1.380211242 | 1.066254157 |
| ENSMUSG00000103054 | 0 | 0 | 0 | 0 | 0 | 0 | 0 |
| ENSMUSG00000103055 | 0 | 0 | 0 | 0 | 0 | 0 | 0 |
| ENSMUSG00000103056 | 0 | 0.25 | 0 | 0 | 0.387761935 | 0.301029996 | 0.224243824 |
| ENSMUSG00000103058 | 0 | 0 | 0 | 0 | 0 | 0 | 0 |
| ENSMUSG00000103059 | 0 | 0 | 0 | 0 | 0 | 0 | 0 |
| ENSMUSG00000103060 | 0 | 0 | 0 | 0 | 0 | 0 | 0 |

| | | | | | | | |
|---|---|---|---|---|---|---|---|
| ENSMUSG00000103061 | 0 | 0 | 0 | 0 | 0 | 0 | 0 |
| ENSMUSG00000103062 | 0.289064826 | 0 | 0.396240625 | 0.580482024 | 0.244650542 | 0.602059991 | 0.448487648 |
| ENSMUSG00000103063 | 0 | 0 | 0 | 0 | 0 | 0 | 0 |
| ENSMUSG00000103064 | 0 | 0 | 0 | 0 | 0 | 0 | 0 |
| ENSMUSG00000103065 | 0 | 0 | 0 | 0 | 0 | 0 | 0 |
| ENSMUSG00000103066 | 0 | 0 | 0 | 0 | 0 | 0 | 0 |
| ENSMUSG00000103067 | 0.578129653 | 0.646240625 | 0.396240625 | 0.701838731 | 0.686820904 | 0.301029996 | 0.520678036 |
| ENSMUSG00000103069 | 0 | 0 | 0 | 0 | 0 | 0 | 0 |
| ENSMUSG00000103070 | 1.156259305 | 1 | 1.17510993 | 1.17510993 | 1.039257769 | 1.079181246 | 0.952571511 |
| ENSMUSG00000103071 | 0.671187741 | 0.79248125 | 0.75 | 0.646240625 | 0.77552387 | 1.041392685 | 0.775756176 |
| ENSMUSG00000103072 | 0 | 0 | 0 | 0 | 0 | 0 | 0 |
| ENSMUSG00000103073 | 0 | 0 | 0 | 0 | 0 | 0 | 0 |
| ENSMUSG00000103074 | 0 | 0 | 0 | 0 | 0 | 0 | 0 |
| ENSMUSG00000103075 | 0 | 0 | 0.25 | 0 | 0 | 0 | 0 |
| ENSMUSG00000103076 | 0.960252568 | 0.79248125 | 0.646240625 | 0.5 | 0.489301084 | 0.698970004 | 0.710836105 |
| ENSMUSG00000103077 | 0 | 0 | 0 | 0 | 0 | 0 | 0 |
| ENSMUSG00000103078 | 0 | 0 | 0 | 0 | 0 | 0 | 0 |
| ENSMUSG00000103079 | 0 | 0 | 0 | 0 | 0 | 0 | 0 |
| ENSMUSG00000103080 | 0.578129653 | 0.396240625 | 0.580482024 | 0.25 | 0.686820904 | 0.477121255 | 0.579661877 |
| ENSMUSG00000103081 | 1.036286563 | 0.864857905 | 0.830482024 | 0.701838731 | 0.568060967 | 0.602059991 | 0.672731473 |
| ENSMUSG00000103082 | 0.45815691 | 0 | 0 | 0.25 | 0.244650542 | 0 | 0 |
| ENSMUSG00000103083 | 0.289064826 | 0 | 0.25 | 0.25 | 0.244650542 | 0 | 0.224243824 |
| ENSMUSG00000103084 | 0 | 0 | 0 | 0 | 0 | 0 | 0 |
| ENSMUSG00000103085 | 0.45815691 | 0.580482024 | 0.396240625 | 0 | 0.686820904 | 0.301029996 | 0.579661877 |
| ENSMUSG00000103086 | 0 | 0 | 0 | 0 | 0 | 0 | 0 |
| ENSMUSG00000103087 | 0 | 0 | 0.396240625 | 0 | 0 | 0 | 0 |
| ENSMUSG00000103088 | 1.249317394 | 1.188721876 | 1.311981878 | 0.830482024 | 0.733951626 | 0.77815125 | 0.579661877 |
| ENSMUSG00000103089 | 0 | 0 | 0.25 | 0 | 0 | 0 | 0.224243824 |
| ENSMUSG00000103090 | 0 | 0 | 0 | 0 | 0 | 0 | 0 |
| ENSMUSG00000103091 | 0 | 0 | 0.25 | 0 | 0 | 0 | 0 |
| ENSMUSG00000103092 | 0 | 0 | 0.25 | 0 | 0 | 0 | 0 |
| ENSMUSG00000103094 | 0.289064826 | 0.580482024 | 0.25 | 0.580482024 | 0.244650542 | 0.301029996 | 0.355418052 |
| ENSMUSG00000103095 | 0 | 0 | 0 | 0 | 0 | 0 | 0 |
| ENSMUSG00000103096 | 0.289064826 | 0 | 0 | 0.25 | 0.244650542 | 0.301029996 | 0 |

| | | | | | | | |
|---|---|---|---|---|---|---|---|
| ENSMUSG00000103097 | 0 | 0 | 0 | 0 | 0 | 0 | 0 |
| ENSMUSG00000103098 | 0.289064826 | 0.5 | 0 | 0.396240625 | 0 | 0 | 0.448487648 |
| ENSMUSG00000103099 | 0 | 0 | 0.25 | 0 | 0 | 0.477121255 | 0.224243824 |
| ENSMUSG00000103100 | 0 | 0.25 | 0.25 | 0 | 0.244650542 | 0.301029996 | 0 |
| ENSMUSG00000103101 | 0.289064826 | 0.25 | 0.396240625 | 0.25 | 0.387761935 | 0.477121255 | 0.355418052 |
| ENSMUSG00000103102 | 0.289064826 | 0.580482024 | 0.25 | 0.396240625 | 0 | 0.301029996 | 0 |
| ENSMUSG00000103103 | 0.45815691 | 0.646240625 | 0.396240625 | 0 | 0 | 0 | 0 |
| ENSMUSG00000103105 | 0 | 0 | 0 | 0 | 0 | 0 | 0 |
| ENSMUSG00000103106 | 0 | 0 | 0 | 0 | 0 | 0 | 0 |
| ENSMUSG00000103107 | 0.289064826 | 0.25 | 0 | 0.396240625 | 0.244650542 | 0.477121255 | 0 |
| ENSMUSG00000103108 | 0 | 0.25 | 0 | 0 | 0 | 0 | 0 |
| ENSMUSG00000103110 | 0 | 0 | 0 | 0 | 0 | 0 | 0 |
| ENSMUSG00000103111 | 0 | 0 | 0 | 0 | 0 | 0 | 0 |
| ENSMUSG00000103112 | 0 | 0 | 0 | 0 | 0 | 0 | 0 |
| ENSMUSG00000103113 | 0 | 0 | 0 | 0 | 0 | 0 | 0 |
| ENSMUSG00000103114 | 0 | 0 | 0.25 | 0.396240625 | 0 | 0 | 0 |
| ENSMUSG00000103115 | 0.289064826 | 0.864857905 | 0.701838731 | 0.701838731 | 0.733951626 | 0.77815125 | 0.579661877 |
| ENSMUSG00000103116 | 0 | 0 | 0 | 0 | 0 | 0 | 0 |
| ENSMUSG00000103118 | 0 | 0 | 0 | 0 | 0 | 0 | 0 |
| ENSMUSG00000103119 | 0 | 0.25 | 0 | 0 | 0.244650542 | 0.477121255 | 0.224243824 |
| ENSMUSG00000103120 | 0 | 0 | 0 | 0 | 0 | 0 | 0 |
| ENSMUSG00000103121 | 0 | 0 | 0 | 0 | 0 | 0 | 0 |
| ENSMUSG00000103122 | 0 | 0 | 0 | 0 | 0.244650542 | 0 | 0 |
| ENSMUSG00000103123 | 0.289064826 | 0.25 | 0.5 | 0.396240625 | 0.387761935 | 0 | 0.224243824 |
| ENSMUSG00000103124 | 0 | 0 | 0 | 0 | 0 | 0 | 0 |
| ENSMUSG00000103125 | 0 | 0 | 0 | 0 | 0 | 0 | 0 |
| ENSMUSG00000103126 | 0 | 0 | 0 | 0 | 0 | 0 | 0 |
| ENSMUSG00000103127 | 0 | 0 | 0 | 0 | 0 | 0 | 0 |
| ENSMUSG00000103129 | 0 | 0 | 0 | 0 | 0 | 0 | 0 |
| ENSMUSG00000103130 | 0 | 0 | 0 | 0 | 0 | 0 | 0 |
| ENSMUSG00000103131 | 0 | 0 | 0 | 0 | 0 | 0 | 0 |
| ENSMUSG00000103132 | 0 | 0 | 0 | 0.25 | 0.244650542 | 0 | 0.224243824 |
| ENSMUSG00000103133 | 0.91631382 | 0.976722649 | 0.701838731 | 1 | 1.057362051 | 0.84509804 | 0.935079929 |
| ENSMUSG00000103135 | 0.45815691 | 0.25 | 0.25 | 0 | 0.568060967 | 0.301029996 | 0.355418052 |

| | | | | | | | |
|---|---|---|---|---|---|---|---|
| ENSMUSG00000103136 | 0 | 0 | 0 | 0 | 0 | 0 | 0 |
| ENSMUSG00000103137 | 0 | 0 | 0 | 0 | 0 | 0 | 0 |
| ENSMUSG00000103138 | 0.578129653 | 0.646240625 | 0.701838731 | 0.5 | 0 | 0 | 0.579661877 |
| ENSMUSG00000103139 | 0.289064826 | 0 | 0 | 0 | 0.244650542 | 0 | 0 |
| ENSMUSG00000103140 | 0 | 0 | 0 | 0 | 0 | 0 | 0 |
| ENSMUSG00000103141 | 0 | 0 | 0.25 | 0 | 0 | 0 | 0 |
| ENSMUSG00000103142 | 0 | 0 | 0 | 0 | 0 | 0 | 0 |
| ENSMUSG00000103143 | 0 | 0.25 | 0 | 0 | 0 | 0 | 0.224243824 |
| ENSMUSG00000103144 | 1.100572389 | 0.976722649 | 0.896240625 | 1.17510993 | 1 | 1.301029996 | 1.089372236 |
| ENSMUSG00000103145 | 0 | 0.25 | 0 | 0.25 | 0 | 0 | 0.224243824 |
| ENSMUSG00000103146 | 0.289064826 | 0.396240625 | 0.5 | 0.396240625 | 0.489301084 | 0.301029996 | 0.355418052 |
| ENSMUSG00000103147 | 0 | 0 | 0 | 0 | 0 | 0 | 0 |
| ENSMUSG00000103148 | 0.578129653 | 0.646240625 | 0.5 | 0.75 | 0.244650542 | 0.301029996 | 0.224243824 |
| ENSMUSG00000103149 | 0.578129653 | 0.830482024 | 0.25 | 0.396240625 | 0.489301084 | 0.698970004 | 0.520678036 |
| ENSMUSG00000103150 | 0 | 0 | 0 | 0 | 0 | 0 | 0 |
| ENSMUSG00000103151 | 0 | 0 | 0 | 0 | 0 | 0 | 0.224243824 |
| ENSMUSG00000103152 | 0 | 0 | 0 | 0 | 0 | 0 | 0 |
| ENSMUSG00000103153 | 0 | 0 | 0 | 0 | 0 | 0 | 0 |
| ENSMUSG00000103154 | 0 | 0 | 0 | 0 | 0 | 0 | 0 |
| ENSMUSG00000103155 | 0 | 0 | 0 | 0 | 0 | 0.301029996 | 0 |
| ENSMUSG00000103156 | 0.289064826 | 0 | 0.25 | 0 | 0 | 0 | 0 |
| ENSMUSG00000103157 | 0 | 0 | 0 | 0 | 0 | 0 | 0 |
| ENSMUSG00000103158 | 0 | 0 | 0 | 0 | 0 | 0 | 0 |
| ENSMUSG00000103159 | 0 | 0 | 0.5 | 0 | 0.387761935 | 0 | 0.224243824 |
| ENSMUSG00000103160 | 0.289064826 | 0.396240625 | 0 | 0.25 | 0.387761935 | 0 | 0.579661877 |
| ENSMUSG00000103161 | 1.036286563 | 1.098079356 | 1.02186571 | 1.114857905 | 1.057362051 | 1.113943352 | 1.089372236 |
| ENSMUSG00000103162 | 0 | 0.25 | 0 | 0.25 | 0.244650542 | 0 | 0 |
| ENSMUSG00000103163 | 0 | 0 | 0 | 0 | 0 | 0 | 0 |
| ENSMUSG00000103164 | 0.45815691 | 0.580482024 | 0.5 | 1.04248125 | 0.955822902 | 1.041392685 | 0.74492186 |
| ENSMUSG00000103165 | 0 | 0 | 0 | 0 | 0 | 0 | 0 |
| ENSMUSG00000103166 | 0 | 0 | 0.25 | 0 | 0 | 0 | 0 |
| ENSMUSG00000103167 | 0 | 0 | 0 | 0 | 0 | 0 | 0 |
| ENSMUSG00000103168 | 0.867194479 | 0.864857905 | 0.75 | 0.580482024 | 0.686820904 | 0.602059991 | 0.672731473 |
| ENSMUSG00000103169 | 0 | 0 | 0 | 0 | 0 | 0 | 0 |

| | | | | | | | |
|---|---|---|---|---|---|---|---|
| ENSMUSG00000103170 | 0.289064826 | 0.25 | 0.396240625 | 0 | 0 | 0.301029996 | 0 |
| ENSMUSG00000103172 | 0 | 0 | 0 | 0 | 0 | 0 | 0 |
| ENSMUSG00000103174 | 0 | 0.396240625 | 0.396240625 | 0.396240625 | 0.489301084 | 0.477121255 | 0.579661877 |
| ENSMUSG00000103175 | 0.671187741 | 0.25 | 0.5 | 0.396240625 | 0.387761935 | 0 | 0.520678036 |
| ENSMUSG00000103176 | 0 | 0 | 0 | 0 | 0 | 0 | 0 |
| ENSMUSG00000103177 | 0 | 0 | 0 | 0 | 0 | 0 | 0 |
| ENSMUSG00000103178 | 0 | 0 | 0 | 0 | 0 | 0 | 0 |
| ENSMUSG00000103179 | 0 | 0 | 0 | 0 | 0 | 0 | 0 |
| ENSMUSG00000103180 | 0 | 0 | 0 | 0 | 0 | 0 | 0 |
| ENSMUSG00000103181 | 0 | 0 | 0.25 | 0 | 0 | 0 | 0 |
| ENSMUSG00000103182 | 0.747221736 | 0.896240625 | 0.951838731 | 1.061981878 | 0.846351821 | 0.903089987 | 0.829800754 |
| ENSMUSG00000103183 | 1 | 0.896240625 | 0.92510993 | 0.79248125 | 0.568060967 | 0.602059991 | 0.520678036 |
| ENSMUSG00000103184 | 0 | 0 | 0 | 0 | 0 | 0 | 0 |
| ENSMUSG00000103185 | 0 | 0 | 0 | 0 | 0 | 0 | 0 |
| ENSMUSG00000103186 | 0.45815691 | 0 | 0.25 | 0.25 | 0.387761935 | 0.698970004 | 0.224243824 |
| ENSMUSG00000103187 | 0 | 0 | 0 | 0 | 0 | 0 | 0 |
| ENSMUSG00000103188 | 0 | 0 | 0 | 0 | 0 | 0 | 0 |
| ENSMUSG00000103189 | 0 | 0 | 0 | 0 | 0 | 0 | 0 |
| ENSMUSG00000103190 | 0 | 0 | 0 | 0 | 0 | 0 | 0 |
| ENSMUSG00000103191 | 0 | 0.396240625 | 0.25 | 0.25 | 0 | 0 | 0 |
| ENSMUSG00000103192 | 0 | 0 | 0 | 0 | 0 | 0 | 0 |
| ENSMUSG00000103193 | 0 | 0 | 0 | 0.25 | 0 | 0 | 0 |
| ENSMUSG00000103194 | 0 | 0 | 0 | 0 | 0 | 0 | 0 |
| ENSMUSG00000103195 | 0.45815691 | 0.25 | 0 | 0 | 0 | 0 | 0 |
| ENSMUSG00000103196 | 0 | 0 | 0 | 0 | 0 | 0 | 0 |
| ENSMUSG00000103197 | 0 | 0 | 0 | 0 | 0 | 0 | 0 |
| ENSMUSG00000103198 | 0 | 0 | 0 | 0 | 0 | 0 | 0 |
| ENSMUSG00000103199 | 0 | 0 | 0.25 | 0.25 | 0 | 0 | 0.448487648 |
| ENSMUSG00000103200 | 0 | 0.396240625 | 0 | 0 | 0 | 0 | 0.355418052 |
| ENSMUSG00000103201 | 0 | 0 | 0 | 0 | 0 | 0 | 0 |
| ENSMUSG00000103202 | 0 | 0 | 0 | 0 | 0 | 0 | 0 |
| ENSMUSG00000103203 | 0 | 0 | 0 | 0 | 0 | 0 | 0 |
| ENSMUSG00000103204 | 0 | 0 | 0 | 0 | 0 | 0 | 0 |
| ENSMUSG00000103206 | 0 | 0 | 0 | 0 | 0 | 0 | 0 |

| | | | | | | | |
|---|---|---|---|---|---|---|---|
| ENSMUSG00000103207 | 0 | 0 | 0.580482024 | 0.580482024 | 0.568060967 | 0.477121255 | 0.520678036 |
| ENSMUSG00000103208 | 0 | 0 | 0 | 0 | 0 | 0 | 0 |
| ENSMUSG00000103209 | 0 | 0 | 0 | 0 | 0 | 0 | 0 |
| ENSMUSG00000103210 | 0 | 0 | 0 | 0 | 0 | 0 | 0 |
| ENSMUSG00000103211 | 0 | 0 | 0 | 0 | 0 | 0 | 0 |
| ENSMUSG00000103213 | 0 | 0 | 0 | 0 | 0 | 0 | 0 |
| ENSMUSG00000103214 | 0 | 0 | 0 | 0 | 0 | 0 | 0 |
| ENSMUSG00000103215 | 0 | 0 | 0 | 0 | 0 | 0 | 0 |
| ENSMUSG00000103216 | 0 | 0 | 0 | 0 | 0 | 0 | 0.224243824 |
| ENSMUSG00000103217 | 0 | 0 | 0 | 0 | 0 | 0 | 0 |
| ENSMUSG00000103218 | 0 | 0 | 0 | 0 | 0 | 0 | 0 |
| ENSMUSG00000103219 | 0 | 0 | 0 | 0 | 0 | 0 | 0 |
| ENSMUSG00000103220 | 0 | 0 | 0 | 0.396240625 | 0 | 0 | 0 |
| ENSMUSG00000103221 | 0 | 0 | 0 | 0 | 0 | 0 | 0 |
| ENSMUSG00000103222 | 0.289064826 | 0.25 | 0.396240625 | 0.646240625 | 0.489301084 | 0.602059991 | 0.224243824 |
| ENSMUSG00000103223 | 0.45815691 | 0 | 0 | 0 | 0 | 0 | 0 |
| ENSMUSG00000103224 | 0 | 0 | 0 | 0 | 0 | 0 | 0 |
| ENSMUSG00000103225 | 0 | 0 | 0 | 0 | 0 | 0 | 0 |
| ENSMUSG00000103226 | 0 | 0 | 0 | 0 | 0 | 0 | 0 |
| ENSMUSG00000103227 | 0.578129653 | 0.396240625 | 0.25 | 0.580482024 | 0.244650542 | 0.477121255 | 0.629532004 |
| ENSMUSG00000103228 | 0 | 0 | 0 | 0 | 0 | 0 | 0 |
| ENSMUSG00000103230 | 0 | 0 | 0.25 | 0 | 0 | 0 | 0 |
| ENSMUSG00000103231 | 0 | 0 | 0 | 0 | 0 | 0 | 0 |
| ENSMUSG00000103232 | 0.289064826 | 0 | 0.25 | 0.25 | 0 | 0 | 0 |
| ENSMUSG00000103233 | 0.289064826 | 0 | 0.396240625 | 0.25 | 0 | 0.301029996 | 0 |
| ENSMUSG00000103234 | 1.358731791 | 1.364857905 | 1.188721876 | 1.380890489 | 1.200473444 | 1.255272505 | 1.193409508 |
| ENSMUSG00000103236 | 0 | 0 | 0 | 0 | 0 | 0 | 0 |
| ENSMUSG00000103237 | 0 | 0 | 0 | 0 | 0 | 0 | 0 |
| ENSMUSG00000103238 | 0.578129653 | 0 | 0.5 | 0.396240625 | 0.244650542 | 0 | 0.224243824 |
| ENSMUSG00000103239 | 0 | 0 | 0 | 0 | 0 | 0 | 0 |
| ENSMUSG00000103240 | 0 | 0 | 0 | 0 | 0 | 0 | 0 |
| ENSMUSG00000103241 | 0.578129653 | 0.396240625 | 0 | 0.25 | 0 | 0.301029996 | 0 |
| ENSMUSG00000103242 | 0 | 0 | 0 | 0 | 0 | 0 | 0 |
| ENSMUSG00000103243 | 0 | 0 | 0 | 0 | 0 | 0 | 0 |

| | | | | | | | |
|---|---|---|---|---|---|---|---|
| ENSMUSG00000103244 | 0 | 0 | 0 | 0 | 0 | 0 | 0 |
| ENSMUSG00000103245 | 0 | 0 | 0 | 0 | 0 | 0 | 0 |
| ENSMUSG00000103246 | 0 | 0 | 0 | 0 | 0 | 0 | 0 |
| ENSMUSG00000103247 | 0 | 0 | 0 | 0 | 0.244650542 | 0.301029996 | 0.448487648 |
| ENSMUSG00000103248 | 0 | 0 | 0 | 0 | 0 | 0 | 0 |
| ENSMUSG00000103249 | 0 | 0 | 0 | 0 | 0 | 0 | 0 |
| ENSMUSG00000103250 | 0 | 0 | 0 | 0 | 0 | 0 | 0 |
| ENSMUSG00000103251 | 0.289064826 | 0.5 | 0 | 0 | 0.244650542 | 0 | 0.355418052 |
| ENSMUSG00000103252 | 0 | 0.396240625 | 0.396240625 | 0.25 | 0.244650542 | 0 | 0.448487648 |
| ENSMUSG00000103253 | 0 | 0 | 0 | 0 | 0 | 0 | 0 |
| ENSMUSG00000103254 | 0 | 0 | 0 | 0 | 0 | 0 | 0 |
| ENSMUSG00000103255 | 0 | 0 | 0 | 0 | 0 | 0 | 0 |
| ENSMUSG00000103256 | 0 | 0 | 0 | 0 | 0 | 0 | 0 |
| ENSMUSG00000103257 | 0 | 0 | 0 | 0 | 0 | 0 | 0.224243824 |
| ENSMUSG00000103258 | 0.289064826 | 0 | 0 | 0.25 | 0.244650542 | 0 | 0.355418052 |
| ENSMUSG00000103259 | 0 | 0 | 0 | 0 | 0 | 0 | 0 |
| ENSMUSG00000103260 | 0.671187741 | 0.580482024 | 0.701838731 | 0.701838731 | 0.846351821 | 0.301029996 | 0.896975297 |
| ENSMUSG00000103261 | 0 | 0 | 0.25 | 0 | 0 | 0.301029996 | 0 |
| ENSMUSG00000103262 | 0 | 0 | 0 | 0 | 0 | 0 | 0 |
| ENSMUSG00000103263 | 0 | 0 | 0.25 | 0 | 0.244650542 | 0 | 0 |
| ENSMUSG00000103264 | 0 | 0 | 0 | 0 | 0 | 0 | 0 |
| ENSMUSG00000103265 | 0 | 0.25 | 0 | 0 | 0 | 0 | 0 |
| ENSMUSG00000103266 | 0 | 0 | 0 | 0 | 0 | 0 | 0 |
| ENSMUSG00000103267 | 0 | 0 | 0 | 0 | 0 | 0 | 0 |
| ENSMUSG00000103269 | 0 | 0 | 0 | 0 | 0 | 0 | 0 |
| ENSMUSG00000103270 | 0 | 0 | 0 | 0.25 | 0 | 0 | 0.355418052 |
| ENSMUSG00000103271 | 0 | 0 | 0 | 0 | 0 | 0 | 0 |
| ENSMUSG00000103272 | 0.811507563 | 0.79248125 | 0.830482024 | 0.75 | 0.846351821 | 0.602059991 | 0.355418052 |
| ENSMUSG00000103273 | 0 | 0 | 0 | 0 | 0 | 0 | 0 |
| ENSMUSG00000103274 | 0 | 0 | 0 | 0 | 0 | 0 | 0 |
| ENSMUSG00000103275 | 0 | 0 | 0 | 0 | 0 | 0 | 0 |
| ENSMUSG00000103276 | 1.227926429 | 1.25 | 1.201838731 | 1.25 | 1.343584837 | 1.361727836 | 1.324583736 |
| ENSMUSG00000103277 | 0.289064826 | 0 | 0.25 | 0.25 | 0.244650542 | 0 | 0 |
| ENSMUSG00000103279 | 0 | 0 | 0 | 0 | 0 | 0 | 0 |

| | | | | | | | |
|---|---|---|---|---|---|---|---|
| ENSMUSG00000103280 | 0 | 0 | 0.25 | 0.5 | 0.489301084 | 0.301029996 | 0.629532004 |
| ENSMUSG00000103281 | 0 | 0 | 0 | 0 | 0 | 0 | 0 |
| ENSMUSG00000103282 | 0 | 0 | 0 | 0 | 0 | 0 | 0 |
| ENSMUSG00000103283 | 0.811507563 | 1 | 0.951838731 | 1.061981878 | 1.020174412 | 1.079181246 | 1.078019652 |
| ENSMUSG00000103284 | 0 | 0 | 0 | 0 | 0 | 0 | 0 |
| ENSMUSG00000103285 | 0.747221736 | 0.5 | 0.701838731 | 0.79248125 | 0.686820904 | 0.77815125 | 0.579661877 |
| ENSMUSG00000103286 | 0 | 0 | 0 | 0 | 0 | 0 | 0 |
| ENSMUSG00000103287 | 0 | 0 | 0 | 0.25 | 0 | 0 | 0 |
| ENSMUSG00000103288 | 0 | 0 | 0 | 0 | 0 | 0 | 0 |
| ENSMUSG00000103289 | 0 | 0 | 0 | 0 | 0 | 0 | 0 |
| ENSMUSG00000103290 | 0 | 0 | 0 | 0 | 0 | 0 | 0 |
| ENSMUSG00000103291 | 0 | 0 | 0 | 0 | 0 | 0.301029996 | 0 |
| ENSMUSG00000103292 | 0.45815691 | 0 | 0.396240625 | 0.646240625 | 0.489301084 | 0.301029996 | 0.520678036 |
| ENSMUSG00000103293 | 0 | 0 | 0 | 0.396240625 | 0 | 0.301029996 | 0.224243824 |
| ENSMUSG00000103294 | 0 | 0.25 | 0 | 0 | 0 | 0 | 0 |
| ENSMUSG00000103295 | 0 | 0 | 0 | 0 | 0 | 0 | 0 |
| ENSMUSG00000103296 | 0 | 0 | 0 | 0 | 0 | 0 | 0 |
| ENSMUSG00000103297 | 0 | 0 | 0 | 0 | 0 | 0 | 0 |
| ENSMUSG00000103298 | 0 | 0 | 0 | 0 | 0 | 0 | 0 |
| ENSMUSG00000103299 | 0 | 0 | 0 | 0 | 0 | 0 | 0 |
| ENSMUSG00000103300 | 0 | 0 | 0 | 0 | 0 | 0 | 0 |
| ENSMUSG00000103301 | 0 | 0.25 | 0 | 0 | 0 | 0 | 0 |
| ENSMUSG00000103302 | 0 | 0 | 0 | 0 | 0 | 0 | 0 |
| ENSMUSG00000103303 | 0 | 0 | 0.5 | 0.396240625 | 0 | 0 | 0 |
| ENSMUSG00000103304 | 0 | 0 | 0 | 0.25 | 0 | 0 | 0 |
| ENSMUSG00000103305 | 0 | 0 | 0 | 0 | 0 | 0 | 0 |
| ENSMUSG00000103306 | 0 | 0 | 0 | 0 | 0 | 0 | 0 |
| ENSMUSG00000103307 | 0 | 0 | 0 | 0 | 0 | 0 | 0 |
| ENSMUSG00000103308 | 0.45815691 | 0.646240625 | 0.79248125 | 0.864857905 | 0.978602168 | 0.84509804 | 0.876096088 |
| ENSMUSG00000103309 | 0.671187741 | 1.04248125 | 0.646240625 | 0.5 | 0.244650542 | 0.301029996 | 0.520678036 |
| ENSMUSG00000103310 | 0 | 0 | 0 | 0 | 0 | 0 | 0 |
| ENSMUSG00000103311 | 0 | 0 | 0 | 0 | 0 | 0 | 0 |
| ENSMUSG00000103312 | 0 | 0 | 0 | 0.25 | 0 | 0 | 0 |
| ENSMUSG00000103313 | 1.686083339 | 1.482684334 | 1.488549078 | 1.488549078 | 1.49445382 | 1.612783857 | 1.365075947 |

| | | | | | | | |
|---|---|---|---|---|---|---|---|
| ENSMUSG00000103314 | 0.289064826 | 0 | 0 | 0 | 0.244650542 | 0.301029996 | 0 |
| ENSMUSG00000103315 | 0 | 0 | 0 | 0 | 0 | 0 | 0 |
| ENSMUSG00000103317 | 0 | 0 | 0.396240625 | 0 | 0 | 0 | 0 |
| ENSMUSG00000103318 | 0 | 0 | 0 | 0 | 0 | 0 | 0 |
| ENSMUSG00000103320 | 0.289064826 | 0.25 | 0 | 0 | 0 | 0 | 0 |
| ENSMUSG00000103321 | 0 | 0 | 0 | 0 | 0 | 0 | 0 |
| ENSMUSG00000103322 | 0.747221736 | 0.396240625 | 0.75 | 0.5 | 0.568060967 | 0.698970004 | 0.520678036 |
| ENSMUSG00000103323 | 0.867194479 | 0.701838731 | 0.646240625 | 0.5 | 0.387761935 | 0.84509804 | 0.629532004 |
| ENSMUSG00000103324 | 0 | 0 | 0 | 0 | 0 | 0.301029996 | 0 |
| ENSMUSG00000103325 | 0 | 0 | 0 | 0 | 0 | 0 | 0 |
| ENSMUSG00000103327 | 0 | 0 | 0 | 0 | 0 | 0 | 0 |
| ENSMUSG00000103328 | 0 | 0 | 0 | 0 | 0 | 0 | 0 |
| ENSMUSG00000103329 | 0 | 0 | 0 | 0 | 0 | 0 | 0 |
| ENSMUSG00000103330 | 0 | 0 | 0 | 0 | 0 | 0 | 0 |
| ENSMUSG00000103331 | 0 | 0 | 0 | 0 | 0.244650542 | 0 | 0.224243824 |
| ENSMUSG00000103332 | 0.671187741 | 0.580482024 | 0.646240625 | 0.5 | 0.387761935 | 0.301029996 | 0.355418052 |
| ENSMUSG00000103333 | 0 | 0 | 0 | 0 | 0 | 0 | 0 |
| ENSMUSG00000103334 | 0 | 0 | 0 | 0 | 0 | 0 | 0 |
| ENSMUSG00000103335 | 0 | 0 | 0 | 0 | 0 | 0 | 0 |
| ENSMUSG00000103336 | 0 | 0 | 0 | 0 | 0 | 0 | 0 |
| ENSMUSG00000103338 | 0 | 0 | 0 | 0 | 0 | 0 | 0 |
| ENSMUSG00000103339 | 0 | 0 | 0 | 0 | 0 | 0 | 0 |
| ENSMUSG00000103340 | 0 | 0 | 0 | 0 | 0 | 0 | 0 |
| ENSMUSG00000103341 | 0 | 0 | 0 | 0 | 0 | 0 | 0 |
| ENSMUSG00000103342 | 0 | 0 | 0 | 0 | 0 | 0 | 0 |
| ENSMUSG00000103343 | 0 | 0.25 | 0 | 0 | 0 | 0 | 0 |
| ENSMUSG00000103345 | 0.289064826 | 0 | 0 | 0 | 0 | 0 | 0 |
| ENSMUSG00000103346 | 0 | 0 | 0 | 0 | 0 | 0 | 0 |
| ENSMUSG00000103347 | 0 | 0 | 0 | 0 | 0 | 0 | 0 |
| ENSMUSG00000103348 | 0 | 0 | 0.25 | 0 | 0 | 0 | 0 |
| ENSMUSG00000103349 | 0 | 0 | 0 | 0 | 0.387761935 | 0 | 0 |
| ENSMUSG00000103350 | 0 | 0 | 0 | 0 | 0 | 0 | 0 |
| ENSMUSG00000103351 | 0 | 0 | 0 | 0 | 0 | 0 | 0.224243824 |
| ENSMUSG00000103352 | 0 | 0 | 0 | 0 | 0 | 0 | 0 |

| | | | | | | | |
|---|---|---|---|---|---|---|---|
| ENSMUSG00000103353 | 0 | 0 | 0 | 0 | 0 | 0 | 0 |
| ENSMUSG00000103354 | 0 | 0 | 0 | 0 | 0 | 0 | 0 |
| ENSMUSG00000103355 | 0.578129653 | 0.701838731 | 0.5 | 0.25 | 0 | 0 | 0.224243824 |
| ENSMUSG00000103356 | 0 | 0 | 0 | 0 | 0 | 0 | 0 |
| ENSMUSG00000103357 | 0 | 0 | 0 | 0 | 0 | 0 | 0 |
| ENSMUSG00000103358 | 0 | 0 | 0 | 0 | 0 | 0 | 0 |
| ENSMUSG00000103360 | 0.289064826 | 0.25 | 0 | 0.25 | 0 | 0 | 0 |
| ENSMUSG00000103361 | 0 | 0 | 0 | 0 | 0 | 0 | 0 |
| ENSMUSG00000103362 | 0 | 0 | 0 | 0 | 0 | 0 | 0 |
| ENSMUSG00000103364 | 0 | 0.25 | 0 | 0.25 | 0.387761935 | 0 | 0.224243824 |
| ENSMUSG00000103365 | 0 | 0 | 0 | 0 | 0 | 0 | 0 |
| ENSMUSG00000103366 | 0 | 0 | 0 | 0.25 | 0 | 0 | 0 |
| ENSMUSG00000103367 | 0.867194479 | 0.75 | 0.896240625 | 0.646240625 | 0.568060967 | 0.602059991 | 0.448487648 |
| ENSMUSG00000103368 | 0 | 0 | 0 | 0 | 0 | 0 | 0 |
| ENSMUSG00000103369 | 0 | 0 | 0 | 0 | 0 | 0 | 0 |
| ENSMUSG00000103370 | 0 | 0 | 0 | 0 | 0 | 0 | 0 |
| ENSMUSG00000103371 | 0 | 0 | 0 | 0 | 0 | 0 | 0 |
| ENSMUSG00000103372 | 0 | 0 | 0 | 0 | 0 | 0 | 0 |
| ENSMUSG00000103373 | 0 | 0.25 | 0.25 | 0.5 | 0.568060967 | 0.477121255 | 0 |
| ENSMUSG00000103374 | 0 | 0 | 0 | 0 | 0 | 0 | 0 |
| ENSMUSG00000103375 | 0 | 0 | 0 | 0 | 0.244650542 | 0 | 0 |
| ENSMUSG00000103376 | 0 | 0.25 | 0.396240625 | 0 | 0.387761935 | 0 | 0.224243824 |
| ENSMUSG00000103377 | 0 | 0 | 0 | 0 | 0 | 0 | 0 |
| ENSMUSG00000103378 | 0 | 0 | 0 | 0 | 0.244650542 | 0 | 0 |
| ENSMUSG00000103379 | 0 | 0 | 0 | 0 | 0 | 0 | 0 |
| ENSMUSG00000103380 | 0 | 0 | 0 | 0 | 0 | 0 | 0 |
| ENSMUSG00000103382 | 0.671187741 | 0.864857905 | 0.701838731 | 0.580482024 | 0.632412477 | 0.301029996 | 0.520678036 |
| ENSMUSG00000103383 | 0 | 0 | 0.25 | 0 | 0 | 0 | 0.224243824 |
| ENSMUSG00000103384 | 0 | 0 | 0 | 0 | 0 | 0 | 0 |
| ENSMUSG00000103385 | 0 | 0 | 0 | 0 | 0 | 0 | 0 |
| ENSMUSG00000103386 | 0 | 0 | 0 | 0 | 0 | 0 | 0.224243824 |
| ENSMUSG00000103387 | 0 | 0 | 0 | 0 | 0 | 0 | 0 |
| ENSMUSG00000103388 | 0.811507563 | 0.92510993 | 0.701838731 | 1.098079356 | 0.846351821 | 0.477121255 | 0.896975297 |
| ENSMUSG00000103389 | 0 | 0 | 0 | 0 | 0 | 0 | 0 |

| | | | | | | | |
|---|---|---|---|---|---|---|---|
| ENSMUSG00000103390 | 0 | 0 | 0 | 0 | 0 | 0 | 0 |
| ENSMUSG00000103391 | 0 | 0.25 | 0.25 | 0.25 | 0 | 0.301029996 | 0.355418052 |
| ENSMUSG00000103392 | 0 | 0 | 0 | 0 | 0 | 0 | 0 |
| ENSMUSG00000103393 | 0 | 0 | 0 | 0 | 0 | 0 | 0 |
| ENSMUSG00000103394 | 0 | 0 | 0 | 0 | 0 | 0 | 0 |
| ENSMUSG00000103395 | 0 | 0 | 0 | 0 | 0 | 0 | 0 |
| ENSMUSG00000103396 | 0 | 0 | 0 | 0 | 0 | 0 | 0 |
| ENSMUSG00000103397 | 0 | 0 | 0 | 0 | 0 | 0 | 0 |
| ENSMUSG00000103398 | 0 | 0.396240625 | 0.25 | 0.396240625 | 0 | 0 | 0 |
| ENSMUSG00000103399 | 0 | 0 | 0 | 0 | 0.387761935 | 0 | 0 |
| ENSMUSG00000103400 | 0 | 0.79248125 | 0.580482024 | 0.580482024 | 0.387761935 | 0.602059991 | 0.74492186 |
| ENSMUSG00000103401 | 0 | 0 | 0 | 0 | 0 | 0 | 0 |
| ENSMUSG00000103402 | 0 | 0 | 0 | 0 | 0 | 0 | 0 |
| ENSMUSG00000103403 | 0.578129653 | 0.646240625 | 0.25 | 0.580482024 | 0.489301084 | 0.477121255 | 0.355418052 |
| ENSMUSG00000103404 | 0.671187741 | 0.5 | 0.701838731 | 0.701838731 | 0.568060967 | 0.301029996 | 0.448487648 |
| ENSMUSG00000103406 | 0 | 0 | 0 | 0 | 0 | 0 | 0 |
| ENSMUSG00000103407 | 0 | 0 | 0 | 0 | 0 | 0 | 0 |
| ENSMUSG00000103408 | 0.289064826 | 0 | 0.396240625 | 0.25 | 0.244650542 | 0.602059991 | 0 |
| ENSMUSG00000103409 | 0.578129653 | 0.646240625 | 0.5 | 0.646240625 | 0.686820904 | 0.698970004 | 0.775756176 |
| ENSMUSG00000103410 | 0 | 0 | 0 | 0 | 0 | 0 | 0 |
| ENSMUSG00000103411 | 0 | 0 | 0 | 0 | 0 | 0 | 0 |
| ENSMUSG00000103412 | 0 | 0 | 0 | 0 | 0 | 0 | 0 |
| ENSMUSG00000103413 | 0 | 0 | 0 | 0 | 0 | 0 | 0 |
| ENSMUSG00000103415 | 0 | 0 | 0 | 0 | 0.244650542 | 0 | 0 |
| ENSMUSG00000103416 | 0 | 0 | 0 | 0 | 0 | 0 | 0 |
| ENSMUSG00000103417 | 0 | 0 | 0 | 0 | 0 | 0 | 0 |
| ENSMUSG00000103419 | 0 | 0 | 0.25 | 0 | 0 | 0 | 0 |
| ENSMUSG00000103420 | 0 | 0 | 0 | 0 | 0 | 0 | 0 |
| ENSMUSG00000103421 | 0 | 0 | 0 | 0 | 0 | 0 | 0 |
| ENSMUSG00000103422 | 0 | 0.25 | 0 | 0.5 | 0.244650542 | 0.301029996 | 0.355418052 |
| ENSMUSG00000103423 | 0.289064826 | 0.701838731 | 0.396240625 | 0.701838731 | 0.812711509 | 0.84509804 | 0.520678036 |
| ENSMUSG00000103424 | 0 | 0 | 0 | 0 | 0.244650542 | 0 | 0 |
| ENSMUSG00000103425 | 0 | 0 | 0 | 0 | 0 | 0 | 0 |
| ENSMUSG00000103426 | 0 | 0 | 0 | 0 | 0 | 0 | 0 |

| | | | | | | | |
|---|---|---|---|---|---|---|---|
| ENSMUSG00000103427 | 0.289064826 | 0.396240625 | 0.25 | 0.580482024 | 0.568060967 | 0.602059991 | 0.224243824 |
| ENSMUSG00000103428 | 0 | 0 | 0 | 0 | 0 | 0 | 0 |
| ENSMUSG00000103430 | 0 | 0 | 0 | 0 | 0 | 0 | 0 |
| ENSMUSG00000103431 | 0 | 0 | 0 | 0 | 0 | 0 | 0 |
| ENSMUSG00000103432 | 0 | 0.25 | 0 | 0.25 | 0 | 0.301029996 | 0 |
| ENSMUSG00000103433 | 0 | 0 | 0 | 0 | 0 | 0 | 0 |
| ENSMUSG00000103434 | 0 | 0 | 0 | 0 | 0.387761935 | 0 | 0 |
| ENSMUSG00000103435 | 0 | 0.396240625 | 0.25 | 0 | 0.244650542 | 0.301029996 | 0 |
| ENSMUSG00000103436 | 0.45815691 | 0.580482024 | 0.25 | 0.396240625 | 0.387761935 | 0.301029996 | 0.355418052 |
| ENSMUSG00000103437 | 0 | 0 | 0 | 0 | 0 | 0 | 0 |
| ENSMUSG00000103438 | 0 | 0 | 0 | 0 | 0 | 0 | 0 |
| ENSMUSG00000103439 | 0 | 0 | 0 | 0 | 0 | 0 | 0.224243824 |
| ENSMUSG00000103440 | 0 | 0 | 0 | 0 | 0 | 0 | 0 |
| ENSMUSG00000103441 | 0 | 0.25 | 0 | 0 | 0 | 0 | 0 |
| ENSMUSG00000103442 | 0 | 0 | 0 | 0 | 0 | 0 | 0 |
| ENSMUSG00000103443 | 0 | 0.25 | 0 | 0.396240625 | 0.489301084 | 0.301029996 | 0.579661877 |
| ENSMUSG00000103444 | 0 | 0 | 0 | 0 | 0 | 0 | 0 |
| ENSMUSG00000103445 | 0.671187741 | 0.79248125 | 0.646240625 | 0.79248125 | 0.846351821 | 1.041392685 | 0.775756176 |
| ENSMUSG00000103446 | 0 | 0 | 0 | 0 | 0 | 0 | 0 |
| ENSMUSG00000103448 | 0 | 0 | 0 | 0 | 0 | 0 | 0 |
| ENSMUSG00000103449 | 0 | 0 | 0 | 0 | 0 | 0 | 0 |
| ENSMUSG00000103451 | 0 | 0 | 0 | 0 | 0 | 0 | 0 |
| ENSMUSG00000103452 | 0 | 0 | 0 | 0 | 0 | 0 | 0 |
| ENSMUSG00000103454 | 0 | 0 | 0 | 0 | 0 | 0 | 0 |
| ENSMUSG00000103456 | 0 | 0 | 0 | 0 | 0 | 0 | 0 |
| ENSMUSG00000103457 | 0 | 0 | 0 | 0 | 0 | 0 | 0 |
| ENSMUSG00000103458 | 0 | 0 | 0 | 0 | 0 | 0 | 0 |
| ENSMUSG00000103459 | 0 | 0.396240625 | 0 | 0.25 | 0 | 0 | 0 |
| ENSMUSG00000103460 | 0 | 0 | 0 | 0 | 0 | 0 | 0 |
| ENSMUSG00000103461 | 0 | 0 | 0 | 0 | 0 | 0 | 0 |
| ENSMUSG00000103462 | 0 | 0 | 0 | 0 | 0 | 0 | 0 |
| ENSMUSG00000103463 | 0 | 0 | 0 | 0 | 0 | 0 | 0 |
| ENSMUSG00000103464 | 0.578129653 | 0.25 | 0.396240625 | 0.396240625 | 0.387761935 | 0 | 0.355418052 |
| ENSMUSG00000103465 | 0 | 0 | 0 | 0 | 0 | 0 | 0 |

| | | | | | | | |
|---|---|---|---|---|---|---|---|
| ENSMUSG00000103466 | 0 | 0 | 0 | 0 | 0 | 0 | 0 |
| ENSMUSG00000103467 | 0.289064826 | 0.580482024 | 0.580482024 | 0.5 | 0.568060967 | 0.602059991 | 0.355418052 |
| ENSMUSG00000103468 | 0 | 0 | 0 | 0 | 0 | 0 | 0 |
| ENSMUSG00000103469 | 0 | 0 | 0 | 0 | 0 | 0 | 0 |
| ENSMUSG00000103470 | 0 | 0 | 0 | 0 | 0 | 0 | 0 |
| ENSMUSG00000103471 | 0 | 0 | 0 | 0 | 0 | 0 | 0 |
| ENSMUSG00000103472 | 1.156259305 | 1.098079356 | 1.061981878 | 0 | 0 | 0.602059991 | 0 |
| ENSMUSG00000103473 | 0.289064826 | 0.25 | 0.5 | 0.580482024 | 0.387761935 | 0.301029996 | 0.520678036 |
| ENSMUSG00000103474 | 0 | 0 | 0 | 0 | 0 | 0 | 0 |
| ENSMUSG00000103475 | 2.571178952 | 2.377443751 | 2.375956434 | 2.395770692 | 2.302012594 | 2.564666064 | 2.106650241 |
| ENSMUSG00000103476 | 0 | 0 | 0 | 0 | 0 | 0 | 0 |
| ENSMUSG00000103477 | 0 | 0 | 0.25 | 0 | 0 | 0.301029996 | 0 |
| ENSMUSG00000103478 | 0 | 0 | 0 | 0 | 0 | 0 | 0 |
| ENSMUSG00000103479 | 0 | 0 | 0 | 0 | 0 | 0 | 0 |
| ENSMUSG00000103481 | 0 | 0 | 0 | 0 | 0 | 0 | 0 |
| ENSMUSG00000103482 | 0 | 0 | 0 | 0 | 0 | 0 | 0 |
| ENSMUSG00000103483 | 0 | 0 | 0 | 0 | 0 | 0 | 0 |
| ENSMUSG00000103484 | 0 | 0 | 0 | 0 | 0 | 0 | 0 |
| ENSMUSG00000103485 | 0 | 0 | 0 | 0 | 0 | 0 | 0.224243824 |
| ENSMUSG00000103486 | 0 | 0 | 0.396240625 | 0.25 | 0.244650542 | 0.301029996 | 0 |
| ENSMUSG00000103487 | 0 | 0 | 0 | 0 | 0 | 0 | 0 |
| ENSMUSG00000103488 | 0 | 0 | 0 | 0 | 0 | 0 | 0 |
| ENSMUSG00000103489 | 0 | 0 | 0 | 0 | 0 | 0 | 0 |
| ENSMUSG00000103491 | 0 | 0 | 0 | 0 | 0 | 0 | 0 |
| ENSMUSG00000103492 | 0.289064826 | 0.25 | 0 | 0 | 0.244650542 | 0 | 0 |
| ENSMUSG00000103493 | 0.91631382 | 0.75 | 0.79248125 | 0.646240625 | 0.686820904 | 0.477121255 | 0.629532004 |
| ENSMUSG00000103494 | 0 | 0.25 | 0 | 0 | 0.387761935 | 0.301029996 | 0.224243824 |
| ENSMUSG00000103495 | 0 | 0 | 0 | 0.25 | 0 | 0.301029996 | 0 |
| ENSMUSG00000103496 | 0 | 0 | 0 | 0 | 0 | 0 | 0 |
| ENSMUSG00000103497 | 0 | 0.25 | 0 | 0 | 0 | 0 | 0 |
| ENSMUSG00000103498 | 0 | 0 | 0 | 0 | 0 | 0 | 0 |
| ENSMUSG00000103499 | 0 | 0 | 0 | 0 | 0 | 0 | 0 |
| ENSMUSG00000103500 | 0 | 0 | 0 | 0 | 0 | 0 | 0 |
| ENSMUSG00000103501 | 0 | 0 | 0 | 0 | 0 | 0 | 0 |

| | | | | | | | |
|---|---|---|---|---|---|---|---|
| ENSMUSG00000103502 | 0 | 0 | 0 | 0 | 0 | 0 | 0 |
| ENSMUSG00000103503 | 0 | 0 | 0 | 0 | 0 | 0 | 0 |
| ENSMUSG00000103504 | 0 | 0 | 0 | 0 | 0 | 0 | 0 |
| ENSMUSG00000103505 | 0.289064826 | 0.5 | 0 | 0.580482024 | 0.733951626 | 0 | 0.448487648 |
| ENSMUSG00000103506 | 0.45815691 | 0.25 | 0.25 | 0.396240625 | 0.489301084 | 0.301029996 | 0 |
| ENSMUSG00000103507 | 1.227926429 | 1.238549078 | 0.896240625 | 1.17510993 | 1.212046813 | 0.954242509 | 1.041356071 |
| ENSMUSG00000103509 | 1.714367558 | 1.584962501 | 1.410964047 | 1.494319981 | 1.445123986 | 1.531478917 | 1.265599895 |
| ENSMUSG00000103510 | 0 | 0 | 0 | 0 | 0 | 0 | 0 |
| ENSMUSG00000103511 | 0 | 0 | 0 | 0 | 0 | 0 | 0 |
| ENSMUSG00000103512 | 0 | 0 | 0 | 0 | 0 | 0 | 0 |
| ENSMUSG00000103513 | 0 | 0 | 0 | 0 | 0 | 0 | 0 |
| ENSMUSG00000103514 | 0 | 0 | 0 | 0 | 0 | 0 | 0 |
| ENSMUSG00000103515 | 1.227926429 | 1.238549078 | 1.17510993 | 1.160964047 | 1.106691885 | 1.255272505 | 1.150210039 |
| ENSMUSG00000103516 | 0 | 0 | 0 | 0 | 0 | 0 | 0 |
| ENSMUSG00000103517 | 0 | 0 | 0 | 0 | 0 | 0 | 0 |
| ENSMUSG00000103518 | 0 | 0 | 0 | 0 | 0 | 0 | 0 |
| ENSMUSG00000103519 | 0 | 0.25 | 0 | 0 | 0 | 0 | 0 |
| ENSMUSG00000103520 | 0.91631382 | 0.701838731 | 0.79248125 | 0.580482024 | 0.877063019 | 0.301029996 | 0.672731473 |
| ENSMUSG00000103521 | 0 | 0 | 0 | 0 | 0 | 0 | 0 |
| ENSMUSG00000103522 | 0 | 0.25 | 0 | 0 | 0 | 0 | 0 |
| ENSMUSG00000103523 | 0 | 0 | 0 | 0 | 0 | 0 | 0 |
| ENSMUSG00000103524 | 0 | 0 | 0 | 0 | 0 | 0 | 0 |
| ENSMUSG00000103525 | 0.289064826 | 0 | 0.25 | 0 | 0 | 0.301029996 | 0 |
| ENSMUSG00000103526 | 0 | 0 | 0 | 0 | 0 | 0 | 0 |
| ENSMUSG00000103527 | 0 | 0 | 0 | 0 | 0 | 0 | 0 |
| ENSMUSG00000103528 | 0 | 0 | 0 | 0 | 0 | 0 | 0 |
| ENSMUSG00000103529 | 0.289064826 | 0.25 | 0.5 | 0 | 0 | 0 | 0.224243824 |
| ENSMUSG00000103530 | 0 | 0 | 0 | 0 | 0 | 0 | 0 |
| ENSMUSG00000103531 | 0 | 0 | 0 | 0 | 0 | 0 | 0 |
| ENSMUSG00000103532 | 0.747221736 | 0.5 | 0.896240625 | 0.646240625 | 0.905314583 | 1.113943352 | 0.448487648 |
| ENSMUSG00000103533 | 0 | 0 | 0 | 0 | 0 | 0 | 0 |
| ENSMUSG00000103534 | 0 | 0 | 0 | 0 | 0 | 0.301029996 | 0 |
| ENSMUSG00000103535 | 0 | 0 | 0 | 0 | 0 | 0 | 0 |
| ENSMUSG00000103536 | 0 | 0 | 0 | 0 | 0 | 0 | 0 |

| | | | | | | | |
|---|---|---|---|---|---|---|---|
| ENSMUSG00000103537 | 0 | 0 | 0 | 0 | 0 | 0 | 0 |
| ENSMUSG00000103538 | 0 | 0 | 0 | 0 | 0 | 0 | 0 |
| ENSMUSG00000103539 | 0.867194479 | 0.580482024 | 0.701838731 | 0.896240625 | 0.686820904 | 0.903089987 | 0.74492186 |
| ENSMUSG00000103540 | 0 | 0 | 0 | 0 | 0 | 0 | 0 |
| ENSMUSG00000103541 | 0 | 0 | 0 | 0 | 0 | 0 | 0 |
| ENSMUSG00000103542 | 0 | 0 | 0 | 0 | 0 | 0 | 0 |
| ENSMUSG00000103543 | 0.45815691 | 0.5 | 0.701838731 | 0.79248125 | 0.846351821 | 0.698970004 | 0.629532004 |
| ENSMUSG00000103544 | 0.289064826 | 0.5 | 0.5 | 0.580482024 | 0.632412477 | 0.77815125 | 0.803905701 |
| ENSMUSG00000103545 | 0 | 0 | 0 | 0 | 0 | 0 | 0 |
| ENSMUSG00000103546 | 0 | 0 | 0.25 | 0 | 0 | 0.301029996 | 0 |
| ENSMUSG00000103547 | 0 | 0 | 0 | 0 | 0 | 0 | 0 |
| ENSMUSG00000103548 | 0 | 0 | 0 | 0 | 0 | 0 | 0 |
| ENSMUSG00000103549 | 0 | 0 | 0 | 0 | 0 | 0 | 0 |
| ENSMUSG00000103550 | 0 | 0 | 0 | 0 | 0 | 0 | 0 |
| ENSMUSG00000103552 | 0 | 0.25 | 0 | 0 | 0 | 0 | 0 |
| ENSMUSG00000103553 | 0.45815691 | 0.580482024 | 0.396240625 | 0.580482024 | 0 | 0 | 0.355418052 |
| ENSMUSG00000103554 | 0 | 0 | 0 | 0 | 0 | 0 | 0 |
| ENSMUSG00000103555 | 0 | 0 | 0 | 0 | 0 | 0 | 0 |
| ENSMUSG00000103556 | 0.578129653 | 0.5 | 0.830482024 | 0.396240625 | 0.77552387 | 0.477121255 | 0.803905701 |
| ENSMUSG00000103557 | 0 | 0 | 0 | 0 | 0 | 0 | 0 |
| ENSMUSG00000103558 | 0 | 0.25 | 0 | 0.25 | 0 | 0 | 0 |
| ENSMUSG00000103559 | 0 | 0 | 0 | 0 | 0 | 0 | 0 |
| ENSMUSG00000103560 | 0.867194479 | 0.951838731 | 0.864857905 | 0.830482024 | 0.931471446 | 0.77815125 | 0.853775828 |
| ENSMUSG00000103561 | 0 | 0 | 0 | 0 | 0 | 0 | 0 |
| ENSMUSG00000103562 | 0 | 0 | 0 | 0 | 0 | 0 | 0 |
| ENSMUSG00000103563 | 0 | 0 | 0 | 0 | 0 | 0 | 0.224243824 |
| ENSMUSG00000103564 | 0 | 0 | 0 | 0 | 0 | 0 | 0 |
| ENSMUSG00000103565 | 0 | 0 | 0 | 0 | 0 | 0 | 0 |
| ENSMUSG00000103566 | 0 | 0 | 0 | 0 | 0 | 0 | 0 |
| ENSMUSG00000103567 | 0.91631382 | 0.864857905 | 0.951838731 | 1.226722649 | 1.293076518 | 1.176091259 | 1.245582246 |
| ENSMUSG00000103568 | 0 | 0 | 0 | 0 | 0 | 0 | 0 |
| ENSMUSG00000103569 | 0.811507563 | 0.5 | 0.580482024 | 0.701838731 | 0.568060967 | 0 | 0.629532004 |
| ENSMUSG00000103570 | 0.289064826 | 0 | 0.25 | 0.396240625 | 0 | 0 | 0 |
| ENSMUSG00000103571 | 0 | 0 | 0 | 0 | 0 | 0 | 0 |

| | | | | | | | |
|---|---|---|---|---|---|---|---|
| ENSMUSG00000103572 | 0.289064826 | 0.25 | 0.396240625 | 0 | 0 | 0 | 0 |
| ENSMUSG00000103573 | 0 | 0 | 0 | 0 | 0 | 0 | 0 |
| ENSMUSG00000103574 | 0.578129653 | 0 | 0.25 | 0.25 | 0.489301084 | 0 | 0.520678036 |
| ENSMUSG00000103575 | 0 | 0 | 0 | 0 | 0 | 0 | 0 |
| ENSMUSG00000103576 | 0.289064826 | 0.701838731 | 0.75 | 0.580482024 | 0.733951626 | 0.477121255 | 0.448487648 |
| ENSMUSG00000103577 | 0 | 0 | 0 | 0 | 0 | 0 | 0 |
| ENSMUSG00000103579 | 0 | 0 | 0 | 0 | 0 | 0 | 0 |
| ENSMUSG00000103580 | 0 | 0 | 0.25 | 0.25 | 0.244650542 | 0 | 0.355418052 |
| ENSMUSG00000103581 | 0.289064826 | 0 | 0 | 0 | 0 | 0 | 0.224243824 |
| ENSMUSG00000103583 | 0 | 0 | 0 | 0 | 0 | 0 | 0 |
| ENSMUSG00000103584 | 0 | 0 | 0 | 0 | 0 | 0 | 0 |
| ENSMUSG00000103585 | 0.671187741 | 0.830482024 | 0.75 | 0.5 | 0.733951626 | 0.77815125 | 0.629532004 |
| ENSMUSG00000103586 | 0 | 0 | 0 | 0 | 0 | 0 | 0 |
| ENSMUSG00000103587 | 0 | 0 | 0 | 0 | 0 | 0 | 0 |
| ENSMUSG00000103588 | 0.671187741 | 0.701838731 | 0.25 | 0 | 0.244650542 | 0.477121255 | 0.355418052 |
| ENSMUSG00000103589 | 0 | 0 | 0 | 0 | 0 | 0 | 0 |
| ENSMUSG00000103590 | 0.289064826 | 0.396240625 | 0.396240625 | 0.25 | 0.387761935 | 0.477121255 | 0.224243824 |
| ENSMUSG00000103591 | 0 | 0.5 | 0 | 0.25 | 0.244650542 | 0.698970004 | 0.579661877 |
| ENSMUSG00000103592 | 0.289064826 | 0 | 0.396240625 | 0 | 0 | 0.602059991 | 0 |
| ENSMUSG00000103593 | 0.578129653 | 0.5 | 0.580482024 | 0 | 0.387761935 | 0.301029996 | 0.355418052 |
| ENSMUSG00000103594 | 0 | 0 | 0 | 0 | 0 | 0 | 0 |
| ENSMUSG00000103595 | 0 | 0 | 0 | 0 | 0 | 0 | 0 |
| ENSMUSG00000103596 | 0 | 0 | 0 | 0.25 | 0 | 0 | 0.224243824 |
| ENSMUSG00000103597 | 0 | 0 | 0 | 0 | 0 | 0 | 0 |
| ENSMUSG00000103598 | 0 | 0 | 0 | 0 | 0 | 0 | 0 |
| ENSMUSG00000103599 | 0 | 0 | 0 | 0.25 | 0 | 0 | 0 |
| ENSMUSG00000103600 | 0 | 0 | 0 | 0 | 0 | 0 | 0 |
| ENSMUSG00000103601 | 0 | 0 | 0 | 0 | 0 | 0 | 0 |
| ENSMUSG00000103602 | 0 | 0 | 0 | 0 | 0 | 0 | 0 |
| ENSMUSG00000103603 | 0 | 0 | 0 | 0 | 0 | 0 | 0 |
| ENSMUSG00000103604 | 0 | 0 | 0 | 0 | 0 | 0 | 0 |
| ENSMUSG00000103605 | 0 | 0 | 0 | 0 | 0 | 0 | 0 |
| ENSMUSG00000103606 | 0 | 0 | 0 | 0 | 0 | 0 | 0 |
| ENSMUSG00000103607 | 0 | 0 | 0 | 0 | 0 | 0 | 0 |

| | | | | | | | |
|---|---|---|---|---|---|---|---|
| ENSMUSG00000103608 | 0 | 0 | 0 | 0 | 0 | 0 | 0 |
| ENSMUSG00000103609 | 0 | 0.79248125 | 0.25 | 0.580482024 | 0.387761935 | 0.477121255 | 0.629532004 |
| ENSMUSG00000103610 | 0 | 0 | 0 | 0 | 0 | 0 | 0 |
| ENSMUSG00000103611 | 0.289064826 | 0 | 0.25 | 0.25 | 0.244650542 | 0 | 0.224243824 |
| ENSMUSG00000103612 | 0 | 0 | 0 | 0 | 0 | 0 | 0 |
| ENSMUSG00000103613 | 0.289064826 | 0.396240625 | 0.396240625 | 0.25 | 0.686820904 | 0.602059991 | 0.355418052 |
| ENSMUSG00000103614 | 0 | 0 | 0.25 | 0.396240625 | 0.489301084 | 0.301029996 | 0.224243824 |
| ENSMUSG00000103615 | 0 | 0 | 0 | 0 | 0 | 0 | 0 |
| ENSMUSG00000103616 | 0 | 0 | 0.25 | 0 | 0 | 0 | 0.224243824 |
| ENSMUSG00000103617 | 0 | 0 | 0 | 0 | 0 | 0 | 0 |
| ENSMUSG00000103618 | 0.289064826 | 0 | 0 | 0 | 0 | 0 | 0 |
| ENSMUSG00000103620 | 0 | 0 | 0 | 0 | 0.244650542 | 0 | 0 |
| ENSMUSG00000103621 | 0 | 0.5 | 0.580482024 | 0.75 | 0.568060967 | 0.698970004 | 0.74492186 |
| ENSMUSG00000103622 | 1.156259305 | 1.02186571 | 1.061981878 | 0.951838731 | 0.978602168 | 1.079181246 | 0.969165684 |
| ENSMUSG00000103624 | 0 | 0 | 0.25 | 0.25 | 0.244650542 | 0 | 0 |
| ENSMUSG00000103625 | 0.671187741 | 0.701838731 | 0.75 | 0.580482024 | 0.632412477 | 0 | 0.520678036 |
| ENSMUSG00000103626 | 0 | 0 | 0 | 0 | 0 | 0 | 0 |
| ENSMUSG00000103628 | 0 | 0 | 0.25 | 0 | 0 | 0 | 0 |
| ENSMUSG00000103629 | 0 | 0 | 0 | 0 | 0 | 0 | 0 |
| ENSMUSG00000103630 | 1.069666964 | 1.098079356 | 0.864857905 | 1.080482024 | 1.091002363 | 1.113943352 | 1.100339912 |
| ENSMUSG00000103631 | 0 | 0 | 0 | 0 | 0 | 0 | 0 |
| ENSMUSG00000103632 | 0.747221736 | 0.5 | 0.5 | 0.646240625 | 0.489301084 | 0.477121255 | 0.448487648 |
| ENSMUSG00000103633 | 0 | 0 | 0 | 0 | 0 | 0 | 0 |
| ENSMUSG00000103634 | 0.45815691 | 0.25 | 0.75 | 0.580482024 | 0.387761935 | 0.301029996 | 0.224243824 |
| ENSMUSG00000103635 | 0 | 0 | 0 | 0 | 0 | 0 | 0 |
| ENSMUSG00000103636 | 0.289064826 | 0.5 | 0.25 | 0 | 0.244650542 | 0.301029996 | 0.224243824 |
| ENSMUSG00000103637 | 0 | 0 | 0 | 0 | 0 | 0 | 0 |
| ENSMUSG00000103638 | 0 | 0 | 0 | 0 | 0 | 0 | 0 |
| ENSMUSG00000103639 | 0 | 0 | 0 | 0 | 0 | 0 | 0 |
| ENSMUSG00000103640 | 0 | 0 | 0 | 0 | 0 | 0 | 0 |
| ENSMUSG00000103641 | 0 | 0 | 0.25 | 0 | 0.244650542 | 0 | 0 |
| ENSMUSG00000103642 | 0 | 0 | 0 | 0 | 0 | 0 | 0 |
| ENSMUSG00000103643 | 0 | 0 | 0 | 0 | 0 | 0 | 0 |
| ENSMUSG00000103644 | 0 | 0 | 0 | 0 | 0 | 0 | 0 |

| | | | | | | | |
|---|---|---|---|---|---|---|---|
| ENSMUSG00000103645 | 0 | 0 | 0 | 0 | 0 | 0 | 0 |
| ENSMUSG00000103646 | 0 | 0.25 | 0.25 | 0 | 0.489301084 | 0 | 0 |
| ENSMUSG00000103647 | 0 | 0 | 0 | 0 | 0 | 0 | 0 |
| ENSMUSG00000103648 | 0 | 0 | 0 | 0 | 0 | 0 | 0 |
| ENSMUSG00000103649 | 1.227926429 | 1.04248125 | 0.976722649 | 0.976722649 | 1.136121934 | 1.204119983 | 1.110947927 |
| ENSMUSG00000103650 | 0 | 0 | 0 | 0 | 0 | 0 | 0 |
| ENSMUSG00000103651 | 0 | 0 | 0 | 0 | 0 | 0 | 0 |
| ENSMUSG00000103652 | 0 | 0 | 0 | 0 | 0 | 0 | 0.224243824 |
| ENSMUSG00000103653 | 0 | 0 | 0 | 0 | 0 | 0 | 0.355418052 |
| ENSMUSG00000103654 | 0 | 0 | 0 | 0 | 0 | 0 | 0 |
| ENSMUSG00000103655 | 0 | 0 | 0 | 0 | 0 | 0 | 0 |
| ENSMUSG00000103656 | 0 | 0 | 0 | 0 | 0 | 0 | 0 |
| ENSMUSG00000103657 | 1.100572389 | 0.864857905 | 0.92510993 | 1.080482024 | 0.77552387 | 0.903089987 | 0.803905701 |
| ENSMUSG00000103658 | 0 | 0 | 0 | 0 | 0 | 0 | 0.224243824 |
| ENSMUSG00000103659 | 0 | 0 | 0 | 0 | 0 | 0 | 0 |
| ENSMUSG00000103660 | 0 | 0 | 0 | 0 | 0 | 0 | 0 |
| ENSMUSG00000103661 | 0 | 0 | 0 | 0 | 0 | 0 | 0 |
| ENSMUSG00000103662 | 0 | 0 | 0 | 0 | 0 | 0 | 0 |
| ENSMUSG00000103663 | 0 | 0 | 0 | 0 | 0 | 0 | 0 |
| ENSMUSG00000103664 | 0 | 0 | 0 | 0 | 0 | 0 | 0 |
| ENSMUSG00000103665 | 0 | 0 | 0 | 0 | 0 | 0 | 0 |
| ENSMUSG00000103666 | 0 | 0 | 0 | 0 | 0 | 0 | 0 |
| ENSMUSG00000103667 | 0 | 0 | 0 | 0 | 0 | 0 | 0 |
| ENSMUSG00000103668 | 0 | 0 | 0.25 | 0 | 0 | 0 | 0.224243824 |
| ENSMUSG00000103669 | 0 | 0.396240625 | 0.25 | 0.25 | 0.244650542 | 0 | 0 |
| ENSMUSG00000103670 | 0.671187741 | 0.646240625 | 0.701838731 | 0.646240625 | 0.686820904 | 0.602059991 | 0.448487648 |
| ENSMUSG00000103672 | 1.069666964 | 1.160964047 | 1.080482024 | 1.160964047 | 1.039257769 | 1.204119983 | 1.066254157 |
| ENSMUSG00000103673 | 0 | 0 | 0 | 0 | 0 | 0 | 0 |
| ENSMUSG00000103674 | 0.289064826 | 0 | 0 | 0 | 0 | 0 | 0 |
| ENSMUSG00000103675 | 0 | 0 | 0 | 0 | 0 | 0 | 0 |
| ENSMUSG00000103676 | 0 | 0 | 0 | 0 | 0 | 0 | 0 |
| ENSMUSG00000103677 | 1.418409478 | 1.410964047 | 1.505591953 | 0.396240625 | 0.244650542 | 0.301029996 | 0.224243824 |
| ENSMUSG00000103678 | 0 | 0 | 0 | 0 | 0 | 0 | 0 |
| ENSMUSG00000103679 | 0 | 0 | 0 | 0 | 0 | 0 | 0 |

| | | | | | | | |
|---|---|---|---|---|---|---|---|
| ENSMUSG00000103680 | 0 | 0.25 | 0 | 0 | 0 | 0.301029996 | 0 |
| ENSMUSG00000103681 | 0 | 0 | 0 | 0 | 0 | 0 | 0 |
| ENSMUSG00000103682 | 0 | 0 | 0 | 0.5 | 0.387761935 | 0.301029996 | 0 |
| ENSMUSG00000103684 | 0 | 0 | 0 | 0 | 0 | 0 | 0 |
| ENSMUSG00000103685 | 0.671187741 | 0.79248125 | 0.580482024 | 0.701838731 | 0.846351821 | 0.77815125 | 0.710836105 |
| ENSMUSG00000103686 | 0 | 0 | 0 | 0 | 0 | 0 | 0 |
| ENSMUSG00000103688 | 0 | 0 | 0 | 0 | 0 | 0 | 0 |
| ENSMUSG00000103689 | 0 | 0 | 0 | 0 | 0 | 0 | 0 |
| ENSMUSG00000103690 | 0 | 0 | 0 | 0 | 0 | 0 | 0 |
| ENSMUSG00000103691 | 0 | 0 | 0 | 0 | 0 | 0 | 0 |
| ENSMUSG00000103692 | 0 | 0 | 0 | 0 | 0 | 0 | 0 |
| ENSMUSG00000103693 | 0.45815691 | 0.396240625 | 0.5 | 0 | 0 | 0.301029996 | 0 |
| ENSMUSG00000103694 | 0 | 0 | 0 | 0.25 | 0.244650542 | 0 | 0 |
| ENSMUSG00000103695 | 0 | 0 | 0 | 0 | 0 | 0 | 0 |
| ENSMUSG00000103696 | 0.45815691 | 0.25 | 0.396240625 | 0.396240625 | 0.387761935 | 0.301029996 | 0.224243824 |
| ENSMUSG00000103697 | 1.181541736 | 0.951838731 | 0.92510993 | 0.951838731 | 0.905314583 | 0.84509804 | 0.896975297 |
| ENSMUSG00000103698 | 0 | 0 | 0 | 0 | 0 | 0 | 0 |
| ENSMUSG00000103699 | 0 | 0 | 0 | 0 | 0 | 0 | 0 |
| ENSMUSG00000103700 | 0 | 0 | 0 | 0 | 0 | 0 | 0 |
| ENSMUSG00000103701 | 0 | 0 | 0 | 0 | 0 | 0 | 0 |
| ENSMUSG00000103702 | 0 | 0 | 0 | 0.25 | 0 | 0 | 0 |
| ENSMUSG00000103703 | 0.578129653 | 0.830482024 | 0.25 | 1 | 0.931471446 | 0.698970004 | 0.74492186 |
| ENSMUSG00000103704 | 0 | 0 | 0 | 0 | 0 | 0 | 0 |
| ENSMUSG00000103705 | 0 | 0 | 0 | 0 | 0 | 0 | 0 |
| ENSMUSG00000103706 | 0.45815691 | 0.580482024 | 0.580482024 | 0.79248125 | 0.686820904 | 0.602059991 | 0.448487648 |
| ENSMUSG00000103707 | 0 | 0 | 0 | 0 | 0 | 0 | 0 |
| ENSMUSG00000103708 | 0 | 0 | 0 | 0 | 0 | 0 | 0 |
| ENSMUSG00000103709 | 0 | 0 | 0 | 0 | 0 | 0 | 0 |
| ENSMUSG00000103710 | 0.289064826 | 0.396240625 | 0.701838731 | 0.75 | 0.489301084 | 0.301029996 | 0.629532004 |
| ENSMUSG00000103711 | 0.578129653 | 0.75 | 0.830482024 | 0.79248125 | 0.955822902 | 0.602059991 | 0.74492186 |
| ENSMUSG00000103712 | 0 | 0 | 0 | 0 | 0 | 0 | 0 |
| ENSMUSG00000103713 | 0 | 0 | 0 | 0 | 0 | 0 | 0 |
| ENSMUSG00000103714 | 0 | 0 | 0 | 0 | 0 | 0 | 0 |
| ENSMUSG00000103715 | 0.867194479 | 1.201838731 | 1.02186571 | 1.098079356 | 0.877063019 | 1.204119983 | 0.984950056 |

| | | | | | | | |
|---|---|---|---|---|---|---|---|
| ENSMUSG00000103716 | 0 | 0 | 0 | 0 | 0 | 0 | 0 |
| ENSMUSG00000103717 | 0.289064826 | 0 | 0 | 0 | 0 | 0 | 0.224243824 |
| ENSMUSG00000103718 | 0 | 0 | 0 | 0 | 0 | 0 | 0 |
| ENSMUSG00000103719 | 0 | 0 | 0 | 0 | 0 | 0 | 0 |
| ENSMUSG00000103720 | 0 | 0 | 0 | 0 | 0 | 0 | 0 |
| ENSMUSG00000103722 | 0.289064826 | 0.25 | 0.396240625 | 0 | 0.489301084 | 0.477121255 | 0.224243824 |
| ENSMUSG00000103723 | 0 | 0 | 0.25 | 0 | 0 | 0 | 0 |
| ENSMUSG00000103724 | 0 | 0 | 0 | 0 | 0 | 0 | 0 |
| ENSMUSG00000103725 | 0 | 0 | 0 | 0 | 0 | 0 | 0 |
| ENSMUSG00000103726 | 0.289064826 | 0.25 | 0.25 | 0.25 | 0 | 0 | 0.224243824 |
| ENSMUSG00000103727 | 0 | 0 | 0 | 0 | 0 | 0 | 0.224243824 |
| ENSMUSG00000103728 | 0.289064826 | 0 | 0 | 0 | 0 | 0.301029996 | 0.224243824 |
| ENSMUSG00000103729 | 0 | 0 | 0 | 0 | 0 | 0 | 0 |
| ENSMUSG00000103730 | 0 | 0 | 0 | 0 | 0 | 0 | 0 |
| ENSMUSG00000103731 | 0.867194479 | 0.864857905 | 0.701838731 | 0.830482024 | 0.931471446 | 0.84509804 | 0.952571511 |
| ENSMUSG00000103732 | 0 | 0 | 0 | 0 | 0 | 0 | 0 |
| ENSMUSG00000103733 | 0 | 0 | 0 | 0 | 0 | 0 | 0 |
| ENSMUSG00000103734 | 0 | 0 | 0 | 0 | 0 | 0.301029996 | 0 |
| ENSMUSG00000103735 | 0 | 0 | 0 | 0 | 0 | 0 | 0.224243824 |
| ENSMUSG00000103736 | 0 | 0 | 0 | 0 | 0 | 0 | 0 |
| ENSMUSG00000103737 | 0 | 0 | 0 | 0 | 0 | 0 | 0 |
| ENSMUSG00000103738 | 0 | 0 | 0 | 0 | 0 | 0 | 0 |
| ENSMUSG00000103739 | 0 | 0.25 | 0 | 0 | 0 | 0.301029996 | 0 |
| ENSMUSG00000103740 | 0 | 0 | 0 | 0 | 0 | 0 | 0 |
| ENSMUSG00000103741 | 0 | 0 | 0 | 0 | 0 | 0 | 0.355418052 |
| ENSMUSG00000103742 | 0 | 0 | 0 | 0 | 0 | 0 | 0 |
| ENSMUSG00000103744 | 0 | 0 | 0 | 0 | 0 | 0 | 0 |
| ENSMUSG00000103745 | 0 | 0 | 0 | 0 | 0 | 0 | 0 |
| ENSMUSG00000103746 | 0 | 0.25 | 0.79248125 | 0 | 0.632412477 | 0.477121255 | 0.448487648 |
| ENSMUSG00000103747 | 0.289064826 | 0 | 0 | 0.25 | 0 | 0 | 0 |
| ENSMUSG00000103748 | 0.578129653 | 0.25 | 0.5 | 0.646240625 | 0.568060967 | 0.301029996 | 0.448487648 |
| ENSMUSG00000103749 | 0.811507563 | 0.5 | 0.5 | 0.701838731 | 0.632412477 | 0.301029996 | 0.520678036 |
| ENSMUSG00000103750 | 0 | 0 | 0 | 0 | 0 | 0 | 0 |
| ENSMUSG00000103751 | 0 | 0 | 0 | 0 | 0 | 0 | 0 |

| | | | | | | | |
|---|---|---|---|---|---|---|---|
| ENSMUSG00000103752 | 0 | 0 | 0 | 0 | 0 | 0 | 0 |
| ENSMUSG00000103753 | 0 | 0 | 0 | 0 | 0.244650542 | 0 | 0 |
| ENSMUSG00000103754 | 0.289064826 | 0 | 0.25 | 0.25 | 0.489301084 | 0.301029996 | 0.448487648 |
| ENSMUSG00000103755 | 0 | 0.25 | 0.396240625 | 0 | 0.489301084 | 0.301029996 | 0.448487648 |
| ENSMUSG00000103756 | 0.867194479 | 0.701838731 | 0.864857905 | 0.79248125 | 0.846351821 | 0.602059991 | 0.579661877 |
| ENSMUSG00000103757 | 0 | 0 | 0 | 0 | 0 | 0 | 0 |
| ENSMUSG00000103758 | 0 | 0 | 0 | 0 | 0 | 0 | 0 |
| ENSMUSG00000103759 | 0 | 0 | 0 | 0 | 0 | 0.477121255 | 0.224243824 |
| ENSMUSG00000103760 | 0.289064826 | 0 | 0 | 0 | 0 | 0 | 0.355418052 |
| ENSMUSG00000103761 | 0 | 0 | 0 | 0 | 0 | 0 | 0 |
| ENSMUSG00000103762 | 0.289064826 | 0.25 | 0 | 0 | 0 | 0 | 0 |
| ENSMUSG00000103763 | 0 | 0.396240625 | 0.25 | 0.25 | 0.244650542 | 0.477121255 | 0.355418052 |
| ENSMUSG00000103764 | 0 | 0 | 0 | 0 | 0 | 0 | 0 |
| ENSMUSG00000103765 | 0 | 0 | 0 | 0 | 0 | 0 | 0 |
| ENSMUSG00000103766 | 0 | 0 | 0 | 0 | 0 | 0 | 0 |
| ENSMUSG00000103767 | 0 | 0 | 0 | 0 | 0 | 0 | 0 |
| ENSMUSG00000103768 | 0 | 0 | 0 | 0 | 0 | 0 | 0 |
| ENSMUSG00000103769 | 0 | 0 | 0 | 0 | 0 | 0 | 0 |
| ENSMUSG00000103770 | 0 | 0 | 0 | 0 | 0 | 0 | 0 |
| ENSMUSG00000103771 | 0 | 0.25 | 0 | 0.396240625 | 0 | 0 | 0.355418052 |
| ENSMUSG00000103772 | 0 | 0 | 0 | 0.25 | 0 | 0 | 0 |
| ENSMUSG00000103773 | 0 | 0 | 0 | 0 | 0 | 0 | 0 |
| ENSMUSG00000103774 | 0 | 0 | 0.396240625 | 0 | 0 | 0.477121255 | 0 |
| ENSMUSG00000103775 | 0 | 0 | 0 | 0 | 0 | 0 | 0 |
| ENSMUSG00000103776 | 0 | 0 | 0 | 0 | 0 | 0 | 0 |
| ENSMUSG00000103777 | 0 | 0 | 0 | 0 | 0 | 0 | 0 |
| ENSMUSG00000103778 | 0.289064826 | 0 | 0 | 0 | 0 | 0 | 0 |
| ENSMUSG00000103779 | 0 | 0 | 0.25 | 0 | 0 | 0 | 0 |
| ENSMUSG00000103780 | 0 | 0 | 0 | 0 | 0 | 0 | 0 |
| ENSMUSG00000103781 | 0 | 0 | 0 | 0 | 0 | 0 | 0 |
| ENSMUSG00000103782 | 0 | 0 | 0 | 0 | 0 | 0 | 0 |
| ENSMUSG00000103783 | 0 | 0 | 0 | 0.396240625 | 0 | 0.301029996 | 0.355418052 |
| ENSMUSG00000103784 | 0 | 0 | 0 | 0 | 0 | 0 | 0 |
| ENSMUSG00000103785 | 0 | 0 | 0 | 0 | 0.244650542 | 0.301029996 | 0 |

| | | | | | | | |
|---|---|---|---|---|---|---|---|
| ENSMUSG00000103786 | 0 | 0 | 0 | 0 | 0.387761935 | 0 | 0.355418052 |
| ENSMUSG00000103787 | 0 | 0 | 0 | 0 | 0 | 0 | 0 |
| ENSMUSG00000103788 | 0 | 0 | 0 | 0 | 0 | 0 | 0 |
| ENSMUSG00000103789 | 0.289064826 | 0 | 0.25 | 0.25 | 0.568060967 | 0.301029996 | 0 |
| ENSMUSG00000103790 | 0 | 0 | 0 | 0 | 0 | 0 | 0 |
| ENSMUSG00000103791 | 0 | 0 | 0 | 0 | 0 | 0 | 0 |
| ENSMUSG00000103792 | 0 | 0 | 0.396240625 | 0 | 0 | 0 | 0.224243824 |
| ENSMUSG00000103793 | 0 | 0.25 | 0.25 | 0.5 | 0.568060967 | 0 | 0.355418052 |
| ENSMUSG00000103794 | 0 | 0 | 0 | 0 | 0 | 0 | 0 |
| ENSMUSG00000103795 | 0 | 0 | 0 | 0 | 0 | 0 | 0 |
| ENSMUSG00000103796 | 0 | 0 | 0 | 0 | 0 | 0 | 0 |
| ENSMUSG00000103797 | 0 | 0 | 0 | 0 | 0 | 0 | 0 |
| ENSMUSG00000103798 | 0 | 0 | 0 | 0 | 0 | 0 | 0 |
| ENSMUSG00000103799 | 0 | 0 | 0 | 0 | 0 | 0 | 0 |
| ENSMUSG00000103800 | 0 | 0 | 0 | 0 | 0 | 0 | 0 |
| ENSMUSG00000103801 | 0 | 0 | 0 | 0 | 0.387761935 | 0 | 0 |
| ENSMUSG00000103802 | 0 | 0 | 0 | 0 | 0 | 0 | 0 |
| ENSMUSG00000103803 | 0 | 0 | 0 | 0 | 0.244650542 | 0 | 0 |
| ENSMUSG00000103804 | 0.45815691 | 0.25 | 0.5 | 0.5 | 0.77552387 | 0.477121255 | 0.629532004 |
| ENSMUSG00000103805 | 0 | 0 | 0 | 0 | 0 | 0 | 0 |
| ENSMUSG00000103807 | 0.289064826 | 0 | 0.25 | 0 | 0 | 0 | 0 |
| ENSMUSG00000103808 | 0 | 0 | 0.25 | 0 | 0.387761935 | 0 | 0.224243824 |
| ENSMUSG00000103809 | 0 | 0 | 0 | 0 | 0 | 0 | 0 |
| ENSMUSG00000103810 | 0.289064826 | 0.75 | 0.646240625 | 0.951838731 | 1.039257769 | 1.113943352 | 0.896975297 |
| ENSMUSG00000103811 | 0 | 0 | 0 | 0 | 0 | 0 | 0 |
| ENSMUSG00000103812 | 0 | 0 | 0 | 0 | 0 | 0 | 0 |
| ENSMUSG00000103813 | 0 | 0 | 0 | 0 | 0 | 0 | 0 |
| ENSMUSG00000103814 | 0 | 0 | 0 | 0 | 0 | 0 | 0 |
| ENSMUSG00000103815 | 0 | 0 | 0 | 0 | 0 | 0 | 0 |
| ENSMUSG00000103816 | 0 | 0 | 0 | 0 | 0 | 0 | 0 |
| ENSMUSG00000103817 | 0 | 0 | 0 | 0 | 0 | 0 | 0 |
| ENSMUSG00000103818 | 0.289064826 | 0 | 0 | 0 | 0 | 0 | 0 |
| ENSMUSG00000103819 | 0 | 0 | 0 | 0 | 0 | 0 | 0 |
| ENSMUSG00000103820 | 0 | 0 | 0 | 0 | 0 | 0 | 0 |

| | | | | | | | |
|---|---|---|---|---|---|---|---|
| ENSMUSG00000103821 | 0.578129653 | 0.864857905 | 0.92510993 | 0.976722649 | 0.77552387 | 0.903089987 | 0.896975297 |
| ENSMUSG00000103822 | 0 | 0.5 | 0 | 0.25 | 0.489301084 | 0 | 0 |
| ENSMUSG00000103823 | 0 | 0.25 | 0.25 | 0 | 0 | 0 | 0 |
| ENSMUSG00000103824 | 0 | 0 | 0 | 0 | 0 | 0 | 0 |
| ENSMUSG00000103825 | 0.289064826 | 0.396240625 | 0.25 | 0 | 0.244650542 | 0.602059991 | 0.355418052 |
| ENSMUSG00000103826 | 0 | 0 | 0 | 0 | 0 | 0 | 0 |
| ENSMUSG00000103827 | 0 | 0 | 0 | 0 | 0 | 0 | 0 |
| ENSMUSG00000103828 | 0 | 0 | 0 | 0.25 | 0 | 0 | 0 |
| ENSMUSG00000103829 | 0 | 0 | 0 | 0 | 0 | 0 | 0 |
| ENSMUSG00000103830 | 0 | 0 | 0 | 0 | 0 | 0 | 0 |
| ENSMUSG00000103831 | 0 | 0 | 0 | 0 | 0 | 0 | 0 |
| ENSMUSG00000103832 | 0 | 0 | 0 | 0 | 0 | 0 | 0 |
| ENSMUSG00000103833 | 0 | 0.396240625 | 0 | 0.25 | 0.387761935 | 0 | 0 |
| ENSMUSG00000103834 | 0 | 0 | 0 | 0 | 0 | 0 | 0 |
| ENSMUSG00000103835 | 0.289064826 | 0.5 | 0.580482024 | 0.5 | 0.632412477 | 0.602059991 | 0.520678036 |
| ENSMUSG00000103836 | 0 | 0 | 0 | 0 | 0 | 0 | 0 |
| ENSMUSG00000103837 | 0 | 0 | 0 | 0 | 0 | 0 | 0 |
| ENSMUSG00000103839 | 1.404271433 | 1.29248125 | 1.372963274 | 1.282320754 | 1.223252711 | 1.397940009 | 1.168187745 |
| ENSMUSG00000103840 | 0 | 0 | 0 | 0 | 0 | 0 | 0 |
| ENSMUSG00000103841 | 0 | 0 | 0 | 0 | 0 | 0 | 0 |
| ENSMUSG00000103842 | 0 | 0 | 0 | 0 | 0 | 0 | 0 |
| ENSMUSG00000103843 | 0.289064826 | 0.25 | 0 | 0 | 0 | 0 | 0 |
| ENSMUSG00000103844 | 0 | 0 | 0 | 0 | 0 | 0 | 0 |
| ENSMUSG00000103845 | 0.45815691 | 0 | 0 | 0.25 | 0.244650542 | 0 | 0 |
| ENSMUSG00000103846 | 0 | 0 | 0 | 0 | 0 | 0 | 0 |
| ENSMUSG00000103847 | 0.45815691 | 0.25 | 0.5 | 0.25 | 0.77552387 | 0.477121255 | 0.355418052 |
| ENSMUSG00000103848 | 0 | 0 | 0 | 0 | 0 | 0 | 0 |
| ENSMUSG00000103849 | 0 | 0 | 0 | 0 | 0 | 0 | 0.224243824 |
| ENSMUSG00000103850 | 0 | 0 | 0 | 0 | 0 | 0 | 0 |
| ENSMUSG00000103851 | 0 | 0 | 0.25 | 0.396240625 | 0.387761935 | 0 | 0.448487648 |
| ENSMUSG00000103852 | 0.289064826 | 0 | 0 | 0 | 0 | 0 | 0 |
| ENSMUSG00000103854 | 0 | 0.25 | 0.25 | 0 | 0 | 0 | 0.448487648 |
| ENSMUSG00000103855 | 0 | 0 | 0 | 0 | 0 | 0 | 0 |
| ENSMUSG00000103856 | 0 | 0 | 0 | 0 | 0 | 0 | 0 |

| | | | | | | | |
|---|---|---|---|---|---|---|---|
| ENSMUSG00000103857 | 0.91631382 | 0.75 | 0.701838731 | 0.92510993 | 0.812711509 | 0.698970004 | 0.710836105 |
| ENSMUSG00000103858 | 0.289064826 | 0.396240625 | 0.396240625 | 0 | 0 | 0 | 0 |
| ENSMUSG00000103859 | 0 | 0 | 0 | 0 | 0 | 0 | 0 |
| ENSMUSG00000103860 | 0 | 0 | 0 | 0 | 0 | 0 | 0 |
| ENSMUSG00000103861 | 0 | 0 | 0 | 0 | 0 | 0 | 0.224243824 |
| ENSMUSG00000103862 | 0 | 0 | 0 | 0 | 0 | 0 | 0 |
| ENSMUSG00000103863 | 0 | 0 | 0 | 0 | 0 | 0 | 0 |
| ENSMUSG00000103864 | 0 | 0.5 | 0 | 0 | 0 | 0 | 0.355418052 |
| ENSMUSG00000103865 | 0.671187741 | 0.79248125 | 0.79248125 | 1 | 1 | 0.84509804 | 1.014380832 |
| ENSMUSG00000103867 | 0 | 0 | 0 | 0.25 | 0.387761935 | 0 | 0.224243824 |
| ENSMUSG00000103869 | 0 | 0 | 0 | 0 | 0.489301084 | 0.301029996 | 0.355418052 |
| ENSMUSG00000103870 | 0 | 0 | 0 | 0 | 0 | 0 | 0 |
| ENSMUSG00000103871 | 0 | 0 | 0 | 0 | 0 | 0 | 0 |
| ENSMUSG00000103872 | 0 | 0 | 0 | 0 | 0 | 0 | 0 |
| ENSMUSG00000103873 | 0 | 0 | 0 | 0 | 0 | 0 | 0 |
| ENSMUSG00000103874 | 0 | 0 | 0 | 0 | 0 | 0 | 0 |
| ENSMUSG00000103875 | 0 | 0 | 0 | 0 | 0 | 0 | 0 |
| ENSMUSG00000103876 | 0 | 0 | 0 | 0 | 0 | 0 | 0 |
| ENSMUSG00000103877 | 0 | 0 | 0.396240625 | 0 | 0 | 0 | 0 |
| ENSMUSG00000103878 | 0 | 0 | 0 | 0 | 0 | 0 | 0 |
| ENSMUSG00000103880 | 0.289064826 | 0.5 | 0.25 | 0 | 0.387761935 | 0 | 0.355418052 |
| ENSMUSG00000103881 | 0.289064826 | 0.25 | 0 | 0.25 | 0 | 0 | 0 |
| ENSMUSG00000103882 | 0 | 0 | 0.396240625 | 0 | 0 | 0 | 0 |
| ENSMUSG00000103883 | 0.91631382 | 0.92510993 | 0.951838731 | 0.864857905 | 0.812711509 | 0.698970004 | 0.803905701 |
| ENSMUSG00000103884 | 0 | 0 | 0 | 0 | 0 | 0 | 0 |
| ENSMUSG00000103885 | 0 | 0 | 0 | 0.25 | 0 | 0 | 0 |
| ENSMUSG00000103886 | 0 | 0 | 0 | 0 | 0 | 0 | 0 |
| ENSMUSG00000103887 | 0 | 0 | 0 | 0 | 0 | 0 | 0 |
| ENSMUSG00000103888 | 0 | 0 | 0 | 0 | 0 | 0 | 0 |
| ENSMUSG00000103889 | 0.289064826 | 0.580482024 | 0.25 | 0 | 0.387761935 | 0.698970004 | 0.355418052 |
| ENSMUSG00000103890 | 0 | 0 | 0 | 0 | 0 | 0 | 0 |
| ENSMUSG00000103891 | 0 | 0.396240625 | 0 | 0.25 | 0.244650542 | 0.301029996 | 0.355418052 |
| ENSMUSG00000103892 | 0 | 0 | 0 | 0 | 0 | 0 | 0 |
| ENSMUSG00000103893 | 0 | 0 | 0 | 0 | 0 | 0 | 0 |

| | | | | | | | |
|---|---|---|---|---|---|---|---|
| ENSMUSG00000103894 | 0 | 0 | 0 | 0 | 0 | 0 | 0 |
| ENSMUSG00000103895 | 0 | 0 | 0 | 0 | 0 | 0 | 0 |
| ENSMUSG00000103896 | 0 | 0 | 0 | 0 | 0 | 0 | 0 |
| ENSMUSG00000103897 | 0.91631382 | 0.896240625 | 0.75 | 0.5 | 0.387761935 | 0.301029996 | 0.224243824 |
| ENSMUSG00000103898 | 0.867194479 | 0.951838731 | 0.75 | 0.5 | 0.632412477 | 0.903089987 | 0.448487648 |
| ENSMUSG00000103899 | 0.289064826 | 0 | 0.25 | 0.25 | 0.244650542 | 0.698970004 | 0.224243824 |
| ENSMUSG00000103900 | 0 | 0.396240625 | 0 | 0 | 0.387761935 | 0 | 0.355418052 |
| ENSMUSG00000103901 | 1.181541736 | 1.130890489 | 1.214495249 | 1.311981878 | 1.234113756 | 1.278753601 | 1 |
| ENSMUSG00000103902 | 0 | 0 | 0 | 0 | 0 | 0 | 0 |
| ENSMUSG00000103903 | 0 | 0 | 0 | 0.25 | 0 | 0 | 0.224243824 |
| ENSMUSG00000103904 | 0 | 0 | 0.5 | 0 | 0.244650542 | 0.301029996 | 0.355418052 |
| ENSMUSG00000103905 | 0 | 0 | 0 | 0 | 0 | 0 | 0 |
| ENSMUSG00000103906 | 0.578129653 | 0.580482024 | 0.580482024 | 0.951838731 | 0.489301084 | 0.477121255 | 0.672731473 |
| ENSMUSG00000103907 | 0 | 0 | 0 | 0 | 0.244650542 | 0 | 0 |
| ENSMUSG00000103908 | 0 | 0.25 | 0 | 0 | 0 | 0 | 0.224243824 |
| ENSMUSG00000103909 | 0 | 0 | 0 | 0 | 0 | 0 | 0 |
| ENSMUSG00000103911 | 0 | 0 | 0 | 0 | 0 | 0 | 0 |
| ENSMUSG00000103912 | 0 | 0 | 0 | 0.25 | 0 | 0 | 0.355418052 |
| ENSMUSG00000103913 | 0 | 0 | 0.25 | 0 | 0 | 0 | 0 |
| ENSMUSG00000103914 | 0 | 0 | 0 | 0 | 0 | 0 | 0 |
| ENSMUSG00000103915 | 0 | 0 | 0 | 0 | 0 | 0 | 0 |
| ENSMUSG00000103916 | 0 | 0 | 0 | 0 | 0 | 0 | 0 |
| ENSMUSG00000103917 | 0 | 0 | 0 | 0 | 0 | 0 | 0 |
| ENSMUSG00000103918 | 0 | 0 | 0 | 0 | 0 | 0 | 0 |
| ENSMUSG00000103919 | 0 | 0 | 0 | 0 | 0 | 0 | 0 |
| ENSMUSG00000103920 | 0.45815691 | 0 | 0.25 | 0 | 0 | 0.301029996 | 0.224243824 |
| ENSMUSG00000103921 | 0 | 0.25 | 0 | 0 | 0 | 0 | 0 |
| ENSMUSG00000103922 | 0 | 0 | 0 | 0 | 0 | 0 | 0 |
| ENSMUSG00000103923 | 0 | 0.25 | 0 | 0.25 | 0 | 0 | 0 |
| ENSMUSG00000103924 | 0 | 0 | 0 | 0 | 0 | 0 | 0 |
| ENSMUSG00000103925 | 0.289064826 | 0 | 0 | 0.25 | 0 | 0.301029996 | 0.355418052 |
| ENSMUSG00000103926 | 0 | 0 | 0.25 | 0.396240625 | 0.387761935 | 0.301029996 | 0.355418052 |
| ENSMUSG00000103927 | 0 | 0.396240625 | 0 | 0 | 0.244650542 | 0.301029996 | 0 |
| ENSMUSG00000103928 | 1.249317394 | 1.214495249 | 1.330482024 | 1.25 | 1.343584837 | 1.431363764 | 1.078019652 |

| | | | | | | | |
|---|---|---|---|---|---|---|---|
| ENSMUSG00000103929 | 0 | 0 | 0 | 0 | 0 | 0 | 0 |
| ENSMUSG00000103930 | 0 | 0 | 0 | 0 | 0 | 0 | 0 |
| ENSMUSG00000103931 | 0 | 0 | 0 | 0 | 0 | 0 | 0 |
| ENSMUSG00000103932 | 0 | 0 | 0 | 0 | 0.244650542 | 0.301029996 | 0.448487648 |
| ENSMUSG00000103933 | 0.747221736 | 0.830482024 | 0.5 | 0.396240625 | 0.686820904 | 0.602059991 | 0.672731473 |
| ENSMUSG00000103934 | 0 | 0 | 0 | 0 | 0 | 0 | 0 |
| ENSMUSG00000103935 | 0 | 0 | 0 | 0 | 0 | 0 | 0 |
| ENSMUSG00000103936 | 0 | 0 | 0 | 0 | 0 | 0 | 0 |
| ENSMUSG00000103937 | 0 | 0 | 0.25 | 0.396240625 | 0 | 0.301029996 | 0 |
| ENSMUSG00000103938 | 0 | 0 | 0 | 0 | 0 | 0 | 0 |
| ENSMUSG00000103939 | 0 | 0 | 0 | 0 | 0 | 0 | 0 |
| ENSMUSG00000103940 | 0 | 0 | 0 | 0 | 0.244650542 | 0.301029996 | 0.224243824 |
| ENSMUSG00000103942 | 0 | 0 | 0 | 0 | 0 | 0 | 0 |
| ENSMUSG00000103943 | 0 | 0 | 0 | 0 | 0 | 0 | 0 |
| ENSMUSG00000103944 | 0 | 0 | 0 | 0 | 0 | 0 | 0 |
| ENSMUSG00000103945 | 1.100572389 | 1.188721876 | 0.92510993 | 1.02186571 | 0.978602168 | 0.84509804 | 0.629532004 |
| ENSMUSG00000103947 | 0 | 0 | 0 | 0 | 0 | 0 | 0 |
| ENSMUSG00000103948 | 0.289064826 | 0.25 | 0 | 0 | 0 | 0 | 0.224243824 |
| ENSMUSG00000103949 | 0.578129653 | 0 | 0.25 | 0.646240625 | 0.568060967 | 0.602059991 | 0.448487648 |
| ENSMUSG00000103950 | 0 | 0 | 0 | 0 | 0 | 0 | 0 |
| ENSMUSG00000103951 | 0 | 0 | 0 | 0 | 0.244650542 | 0 | 0 |
| ENSMUSG00000103952 | 0.578129653 | 0.646240625 | 0 | 0.25 | 0.387761935 | 0 | 0.224243824 |
| ENSMUSG00000103953 | 0 | 0 | 0 | 0 | 0.244650542 | 0.301029996 | 0 |
| ENSMUSG00000103954 | 0 | 0 | 0 | 0 | 0 | 0 | 0 |
| ENSMUSG00000103955 | 0 | 0 | 0 | 0 | 0 | 0 | 0 |
| ENSMUSG00000103956 | 0 | 0 | 0 | 0 | 0 | 0 | 0 |
| ENSMUSG00000103957 | 0 | 0.25 | 0 | 0 | 0 | 0 | 0 |
| ENSMUSG00000103958 | 0 | 0 | 0 | 0 | 0 | 0 | 0 |
| ENSMUSG00000103959 | 0 | 0.25 | 0 | 0.25 | 0 | 0 | 0 |
| ENSMUSG00000103960 | 0 | 0 | 0 | 0 | 0 | 0 | 0 |
| ENSMUSG00000103961 | 0 | 0 | 0 | 0 | 0 | 0 | 0.224243824 |
| ENSMUSG00000103962 | 1 | 0.75 | 0.864857905 | 1.04248125 | 0.955822902 | 0.84509804 | 0.520678036 |
| ENSMUSG00000103963 | 0 | 0 | 0 | 0 | 0 | 0 | 0 |
| ENSMUSG00000103964 | 0 | 0 | 0.396240625 | 0 | 0 | 0 | 0.224243824 |

| | | | | | | | |
|---|---|---|---|---|---|---|---|
| ENSMUSG00000103965 | 0 | 0.25 | 0 | 0 | 0 | 0 | 0 |
| ENSMUSG00000103966 | 0.671187741 | 0.580482024 | 0.580482024 | 0.396240625 | 0.77552387 | 0.477121255 | 0.448487648 |
| ENSMUSG00000103967 | 0.289064826 | 0 | 0 | 0.5 | 0 | 0.477121255 | 0.224243824 |
| ENSMUSG00000103968 | 0 | 0 | 0 | 0 | 0 | 0 | 0 |
| ENSMUSG00000103969 | 0 | 0 | 0 | 0 | 0 | 0 | 0 |
| ENSMUSG00000103970 | 0 | 0.25 | 0.396240625 | 0.396240625 | 0.387761935 | 0 | 0.520678036 |
| ENSMUSG00000103971 | 0 | 0 | 0 | 0 | 0 | 0 | 0 |
| ENSMUSG00000103972 | 0 | 0 | 0 | 0 | 0 | 0 | 0 |
| ENSMUSG00000103973 | 0.45815691 | 0.396240625 | 0.5 | 0.396240625 | 0.77552387 | 0.301029996 | 0.629532004 |
| ENSMUSG00000103975 | 0 | 0 | 0 | 0 | 0 | 0 | 0 |
| ENSMUSG00000103976 | 0.45815691 | 0.701838731 | 0.830482024 | 0.92510993 | 0.686820904 | 0.954242509 | 0.775756176 |
| ENSMUSG00000103977 | 0 | 0 | 0 | 0 | 0 | 0 | 0 |
| ENSMUSG00000103978 | 0 | 0 | 0 | 0 | 0 | 0 | 0 |
| ENSMUSG00000103979 | 0 | 0 | 0 | 0 | 0 | 0 | 0 |
| ENSMUSG00000103980 | 0 | 0 | 0 | 0 | 0 | 0 | 0 |
| ENSMUSG00000103981 | 0 | 0 | 0 | 0 | 0 | 0 | 0 |
| ENSMUSG00000103982 | 0 | 0 | 0 | 0 | 0 | 0 | 0 |
| ENSMUSG00000103983 | 1.269664473 | 1.214495249 | 1.098079356 | 1.364857905 | 1.149965125 | 1.531478917 | 1.259064007 |
| ENSMUSG00000103984 | 1.069666964 | 0.646240625 | 0.75 | 0.5 | 0.812711509 | 0.698970004 | 0.579661877 |
| ENSMUSG00000103985 | 0.289064826 | 0.25 | 0 | 0 | 0 | 0 | 0 |
| ENSMUSG00000103986 | 0 | 0.25 | 0.396240625 | 0 | 0 | 0 | 0.448487648 |
| ENSMUSG00000103987 | 0 | 0 | 0 | 0 | 0 | 0 | 0 |
| ENSMUSG00000103989 | 0 | 0 | 0 | 0 | 0 | 0 | 0 |
| ENSMUSG00000103990 | 0 | 0 | 0 | 0 | 0 | 0 | 0 |
| ENSMUSG00000103991 | 0 | 0 | 0 | 0 | 0 | 0 | 0 |
| ENSMUSG00000103992 | 0.289064826 | 0 | 0.25 | 0 | 0 | 0.301029996 | 0.224243824 |
| ENSMUSG00000103993 | 0 | 0 | 0 | 0 | 0 | 0 | 0 |
| ENSMUSG00000103994 | 0 | 0 | 0 | 0 | 0 | 0 | 0 |
| ENSMUSG00000103995 | 0 | 0 | 0 | 0 | 0 | 0 | 0 |
| ENSMUSG00000103996 | 0 | 0.25 | 0 | 0 | 0 | 0 | 0 |
| ENSMUSG00000103997 | 0 | 0 | 0 | 0 | 0 | 0 | 0 |
| ENSMUSG00000103998 | 0.578129653 | 0.646240625 | 0.396240625 | 0.5 | 0.489301084 | 0.602059991 | 0.579661877 |
| ENSMUSG00000103999 | 0 | 0.5 | 0.25 | 0.396240625 | 0 | 0.301029996 | 0 |
| ENSMUSG00000104000 | 0.289064826 | 0 | 0 | 0 | 0 | 0 | 0 |

| | | | | | | | |
|---|---|---|---|---|---|---|---|
| ENSMUSG00000104001 | 0 | 0 | 0 | 0 | 0 | 0 | 0 |
| ENSMUSG00000104002 | 0.289064826 | 0.701838731 | 0.5 | 0.396240625 | 0.489301084 | 0.301029996 | 0.224243824 |
| ENSMUSG00000104003 | 0.45815691 | 0.701838731 | 0.396240625 | 1.080482024 | 0.846351821 | 0.698970004 | 0.710836105 |
| ENSMUSG00000104004 | 0 | 0 | 0 | 0 | 0 | 0 | 0.224243824 |
| ENSMUSG00000104006 | 0 | 0 | 0 | 0 | 0 | 0 | 0 |
| ENSMUSG00000104007 | 0 | 0 | 0 | 0 | 0 | 0 | 0 |
| ENSMUSG00000104009 | 0 | 0 | 0 | 0 | 0 | 0 | 0 |
| ENSMUSG00000104010 | 0.289064826 | 0 | 0.396240625 | 0.25 | 0.686820904 | 0.301029996 | 0.448487648 |
| ENSMUSG00000104011 | 0 | 0 | 0.396240625 | 0 | 0.244650542 | 0.301029996 | 0 |
| ENSMUSG00000104012 | 0 | 0 | 0 | 0 | 0 | 0 | 0 |
| ENSMUSG00000104013 | 0 | 0 | 0 | 0 | 0 | 0 | 0 |
| ENSMUSG00000104014 | 0 | 0 | 0 | 0 | 0 | 0 | 0 |
| ENSMUSG00000104015 | 0 | 0 | 0.25 | 0.25 | 0 | 0 | 0 |
| ENSMUSG00000104016 | 0.289064826 | 0 | 0.25 | 0.25 | 0 | 0.477121255 | 0 |
| ENSMUSG00000104017 | 0 | 0 | 0 | 0 | 0 | 0 | 0 |
| ENSMUSG00000104018 | 0.671187741 | 0.75 | 0.701838731 | 0 | 0.632412477 | 0.77815125 | 0.710836105 |
| ENSMUSG00000104020 | 0 | 0.25 | 0 | 0 | 0 | 0 | 0 |
| ENSMUSG00000104021 | 0 | 0 | 0 | 0 | 0 | 0 | 0 |
| ENSMUSG00000104022 | 0.45815691 | 0.75 | 0.701838731 | 0.25 | 0.686820904 | 0.698970004 | 0.224243824 |
| ENSMUSG00000104023 | 0 | 0 | 0 | 0 | 0 | 0.301029996 | 0 |
| ENSMUSG00000104024 | 0 | 0 | 0 | 0 | 0 | 0 | 0 |
| ENSMUSG00000104025 | 0 | 0 | 0 | 0 | 0 | 0 | 0 |
| ENSMUSG00000104026 | 0 | 0 | 0 | 0.25 | 0.244650542 | 0 | 0 |
| ENSMUSG00000104027 | 0 | 0 | 0 | 0 | 0.244650542 | 0 | 0 |
| ENSMUSG00000104028 | 0 | 0 | 0 | 0 | 0 | 0 | 0 |
| ENSMUSG00000104029 | 0 | 0 | 0 | 0 | 0 | 0 | 0 |
| ENSMUSG00000104030 | 0 | 0 | 0 | 0 | 0 | 0 | 0.224243824 |
| ENSMUSG00000104031 | 0 | 0 | 0 | 0.25 | 0.387761935 | 0.477121255 | 0 |
| ENSMUSG00000104032 | 0 | 0 | 0 | 0 | 0 | 0 | 0 |
| ENSMUSG00000104033 | 0 | 0.25 | 0 | 0 | 0 | 0 | 0 |
| ENSMUSG00000104034 | 0 | 0 | 0.25 | 0 | 0 | 0 | 0 |
| ENSMUSG00000104035 | 0 | 0 | 0 | 0 | 0 | 0 | 0 |
| ENSMUSG00000104036 | 0.747221736 | 0.701838731 | 0.701838731 | 0.5 | 0.632412477 | 0.301029996 | 0.448487648 |
| ENSMUSG00000104037 | 0.867194479 | 0.701838731 | 0.701838731 | 0.75 | 0.686820904 | 0.84509804 | 0.448487648 |

| | | | | | | | |
|---|---|---|---|---|---|---|---|
| ENSMUSG00000104038 | 0 | 0 | 0 | 0 | 0 | 0 | 0 |
| ENSMUSG00000104039 | 0 | 0 | 0 | 0 | 0 | 0 | 0 |
| ENSMUSG00000104040 | 0 | 0 | 0 | 0 | 0.244650542 | 0 | 0 |
| ENSMUSG00000104041 | 0 | 0 | 0 | 0 | 0 | 0 | 0 |
| ENSMUSG00000104042 | 0 | 0 | 0 | 0 | 0 | 0 | 0 |
| ENSMUSG00000104043 | 0.811507563 | 0.864857905 | 0.864857905 | 0.580482024 | 0.632412477 | 0.477121255 | 0.710836105 |
| ENSMUSG00000104044 | 0 | 0.396240625 | 0 | 0.25 | 0.244650542 | 0 | 0 |
| ENSMUSG00000104045 | 0.747221736 | 0.646240625 | 0.580482024 | 0 | 0 | 0 | 0 |
| ENSMUSG00000104046 | 1.794934559 | 1.606566189 | 1.614857905 | 1.67510993 | 1.72073075 | 1.755874856 | 1.537859885 |
| ENSMUSG00000104048 | 0 | 0 | 0 | 0 | 0 | 0 | 0 |
| ENSMUSG00000104050 | 0 | 0 | 0 | 0 | 0 | 0 | 0 |
| ENSMUSG00000104051 | 0.91631382 | 0.896240625 | 0.864857905 | 0.951838731 | 0.931471446 | 0.84509804 | 0.916588299 |
| ENSMUSG00000104052 | 0 | 0.25 | 0 | 0 | 0.244650542 | 0 | 0.224243824 |
| ENSMUSG00000104053 | 0 | 0 | 0 | 0 | 0 | 0 | 0.224243824 |
| ENSMUSG00000104054 | 0 | 0 | 0 | 0 | 0 | 0 | 0 |
| ENSMUSG00000104055 | 0 | 0 | 0 | 0 | 0 | 0 | 0 |
| ENSMUSG00000104056 | 0 | 0 | 0 | 0 | 0 | 0.477121255 | 0 |
| ENSMUSG00000104057 | 0 | 0 | 0.5 | 0.25 | 0.244650542 | 0 | 0.224243824 |
| ENSMUSG00000104058 | 0 | 0 | 0 | 0 | 0 | 0 | 0 |
| ENSMUSG00000104059 | 0 | 0 | 0 | 0 | 0 | 0 | 0 |
| ENSMUSG00000104060 | 1.389637216 | 1.321350555 | 1.380890489 | 1.348079356 | 1.319233381 | 1.447158031 | 1.20919388 |
| ENSMUSG00000104061 | 0.289064826 | 0.25 | 0.25 | 0.25 | 0.568060967 | 0.477121255 | 0.579661877 |
| ENSMUSG00000104062 | 0 | 0 | 0 | 0 | 0 | 0 | 0 |
| ENSMUSG00000104063 | 0.91631382 | 0.896240625 | 1 | 0.580482024 | 0.77552387 | 0.301029996 | 0.629532004 |
| ENSMUSG00000104064 | 0 | 0 | 0 | 0 | 0 | 0 | 0 |
| ENSMUSG00000104065 | 0 | 0 | 0 | 0 | 0 | 0 | 0 |
| ENSMUSG00000104066 | 0.289064826 | 0 | 0.25 | 0 | 0 | 0 | 0 |
| ENSMUSG00000104067 | 0 | 0 | 0 | 0 | 0 | 0 | 0 |
| ENSMUSG00000104068 | 0 | 0 | 0 | 0 | 0 | 0 | 0 |
| ENSMUSG00000104069 | 0.289064826 | 0.5 | 0.25 | 0.5 | 0.568060967 | 0.301029996 | 0.520678036 |
| ENSMUSG00000104070 | 0 | 0 | 0 | 0 | 0 | 0 | 0 |
| ENSMUSG00000104071 | 0 | 0 | 0 | 0 | 0 | 0 | 0 |
| ENSMUSG00000104072 | 0 | 0 | 0 | 0 | 0 | 0 | 0 |
| ENSMUSG00000104073 | 0 | 0 | 0 | 0 | 0 | 0 | 0 |

| | | | | | | | |
|---|---|---|---|---|---|---|---|
| ENSMUSG00000104074 | 0.289064826 | 0.396240625 | 0.79248125 | 0.396240625 | 0.489301084 | 0 | 0.520678036 |
| ENSMUSG00000104075 | 0 | 0 | 0 | 0 | 0 | 0 | 0 |
| ENSMUSG00000104076 | 0 | 0 | 0 | 0.25 | 0 | 0 | 0 |
| ENSMUSG00000104077 | 0 | 0 | 0 | 0 | 0 | 0 | 0 |
| ENSMUSG00000104078 | 0 | 0 | 0.25 | 0 | 0 | 0 | 0 |
| ENSMUSG00000104079 | 0 | 0 | 0 | 0 | 0 | 0 | 0 |
| ENSMUSG00000104080 | 1 | 0.951838731 | 1.160964047 | 0.92510993 | 1.188507684 | 0.954242509 | 0.710836105 |
| ENSMUSG00000104081 | 0 | 0 | 0 | 0 | 0 | 0 | 0 |
| ENSMUSG00000104082 | 0 | 0 | 0.25 | 0 | 0 | 0.301029996 | 0 |
| ENSMUSG00000104083 | 0.289064826 | 0.396240625 | 0.25 | 0 | 0 | 0.301029996 | 0 |
| ENSMUSG00000104084 | 0 | 0 | 0 | 0 | 0 | 0 | 0 |
| ENSMUSG00000104085 | 0 | 0 | 0 | 0 | 0 | 0 | 0 |
| ENSMUSG00000104086 | 0 | 0 | 0 | 0 | 0 | 0 | 0 |
| ENSMUSG00000104087 | 0 | 0 | 0 | 0 | 0 | 0.301029996 | 0 |
| ENSMUSG00000104088 | 0.45815691 | 0.25 | 0.5 | 0 | 0.244650542 | 0 | 0.224243824 |
| ENSMUSG00000104089 | 0 | 0 | 0 | 0 | 0 | 0 | 0 |
| ENSMUSG00000104090 | 0.671187741 | 0.25 | 0.396240625 | 0.5 | 0.632412477 | 0.477121255 | 0.224243824 |
| ENSMUSG00000104091 | 0.289064826 | 0 | 0.396240625 | 0.396240625 | 0 | 0.477121255 | 0 |
| ENSMUSG00000104092 | 0 | 0 | 0.25 | 0 | 0 | 0.301029996 | 0 |
| ENSMUSG00000104093 | 0.45815691 | 0.580482024 | 0.646240625 | 0.396240625 | 0.244650542 | 0.301029996 | 0.355418052 |
| ENSMUSG00000104094 | 0.747221736 | 0.830482024 | 0.830482024 | 0.580482024 | 0.489301084 | 0 | 0.448487648 |
| ENSMUSG00000104095 | 0.867194479 | 0.864857905 | 1 | 0.951838731 | 0.978602168 | 1 | 0.803905701 |
| ENSMUSG00000104096 | 0 | 0 | 0 | 0 | 0 | 0 | 0 |
| ENSMUSG00000104097 | 0 | 0 | 0.5 | 0 | 0 | 0 | 0.224243824 |
| ENSMUSG00000104098 | 0 | 0 | 0 | 0 | 0 | 0 | 0 |
| ENSMUSG00000104099 | 0 | 0 | 0 | 0 | 0 | 0 | 0 |
| ENSMUSG00000104100 | 0 | 0 | 0.25 | 0.25 | 0 | 0 | 0 |
| ENSMUSG00000104101 | 0 | 0 | 0 | 0 | 0.244650542 | 0 | 0 |
| ENSMUSG00000104102 | 0.289064826 | 0.25 | 0.25 | 0.25 | 0.244650542 | 0 | 0.224243824 |
| ENSMUSG00000104103 | 0 | 0 | 0.25 | 0 | 0 | 0 | 0 |
| ENSMUSG00000104104 | 0 | 0.396240625 | 0 | 0 | 0 | 0 | 0.224243824 |
| ENSMUSG00000104105 | 0 | 0 | 0 | 0 | 0 | 0 | 0 |
| ENSMUSG00000104106 | 0 | 0 | 0 | 0 | 0 | 0 | 0 |
| ENSMUSG00000104107 | 0 | 0 | 0 | 0 | 0 | 0.301029996 | 0 |

| | | | | | | | |
|---|---|---|---|---|---|---|---|
| ENSMUSG00000104108 | 0 | 0.5 | 0 | 0 | 0 | 0 | 0.224243824 |
| ENSMUSG00000104109 | 0.45815691 | 0 | 0 | 0.25 | 0.244650542 | 0 | 0 |
| ENSMUSG00000104110 | 0 | 0 | 0 | 0 | 0 | 0 | 0 |
| ENSMUSG00000104111 | 0 | 0 | 0 | 0 | 0 | 0 | 0 |
| ENSMUSG00000104112 | 0 | 0 | 0 | 0 | 0 | 0 | 0 |
| ENSMUSG00000104113 | 0.289064826 | 0.646240625 | 0.580482024 | 0.25 | 0.568060967 | 0.301029996 | 0.355418052 |
| ENSMUSG00000104114 | 0 | 0.25 | 0.25 | 0 | 0 | 0.301029996 | 0.224243824 |
| ENSMUSG00000104115 | 0 | 0 | 0 | 0 | 0 | 0 | 0 |
| ENSMUSG00000104116 | 0.289064826 | 0.25 | 0.25 | 0 | 0 | 0 | 0 |
| ENSMUSG00000104117 | 0 | 0 | 0 | 0 | 0 | 0.301029996 | 0 |
| ENSMUSG00000104118 | 0 | 0 | 0 | 0 | 0 | 0 | 0 |
| ENSMUSG00000104120 | 0 | 0.25 | 0 | 0.25 | 0.244650542 | 0 | 0 |
| ENSMUSG00000104121 | 0.289064826 | 0 | 0.396240625 | 0.25 | 0.244650542 | 0 | 0 |
| ENSMUSG00000104122 | 0 | 0 | 0.25 | 0.25 | 0 | 0 | 0.224243824 |
| ENSMUSG00000104123 | 0 | 0 | 0 | 0 | 0 | 0 | 0 |
| ENSMUSG00000104125 | 0 | 0.396240625 | 0.396240625 | 0 | 0.244650542 | 0 | 0.355418052 |
| ENSMUSG00000104126 | 0 | 0 | 0 | 0 | 0 | 0.301029996 | 0 |
| ENSMUSG00000104127 | 0 | 0 | 0 | 0 | 0 | 0 | 0 |
| ENSMUSG00000104129 | 0 | 0 | 0 | 0 | 0 | 0 | 0 |
| ENSMUSG00000104130 | 0 | 0 | 0 | 0 | 0 | 0 | 0.224243824 |
| ENSMUSG00000104131 | 0 | 0 | 0 | 0 | 0 | 0 | 0 |
| ENSMUSG00000104132 | 0 | 0 | 0 | 0 | 0 | 0 | 0 |
| ENSMUSG00000104133 | 0 | 0 | 0 | 0 | 0 | 0 | 0 |
| ENSMUSG00000104134 | 0 | 0 | 0 | 0 | 0 | 0 | 0 |
| ENSMUSG00000104135 | 0 | 0 | 0 | 0 | 0 | 0 | 0 |
| ENSMUSG00000104136 | 0.671187741 | 0.92510993 | 0.701838731 | 0.646240625 | 0.489301084 | 0.77815125 | 0.520678036 |
| ENSMUSG00000104137 | 0 | 0 | 0 | 0 | 0 | 0 | 0 |
| ENSMUSG00000104138 | 0 | 0 | 0 | 0.25 | 0 | 0 | 0 |
| ENSMUSG00000104140 | 0 | 0.396240625 | 0 | 0.25 | 0.244650542 | 0.477121255 | 0.224243824 |
| ENSMUSG00000104141 | 0 | 0.25 | 0 | 0.25 | 0 | 0.301029996 | 0 |
| ENSMUSG00000104142 | 0 | 0 | 0 | 0 | 0 | 0 | 0 |
| ENSMUSG00000104144 | 0 | 0 | 0 | 0 | 0 | 0 | 0 |
| ENSMUSG00000104145 | 0 | 0 | 0.25 | 0.396240625 | 0.387761935 | 0 | 0 |
| ENSMUSG00000104146 | 0 | 0 | 0 | 0 | 0 | 0 | 0 |

| | | | | | | | |
|---|---|---|---|---|---|---|---|
| ENSMUSG00000104147 | 0 | 0.25 | 0 | 0 | 0.244650542 | 0.301029996 | 0.224243824 |
| ENSMUSG00000104148 | 0 | 0 | 0 | 0 | 0 | 0 | 0 |
| ENSMUSG00000104149 | 0.91631382 | 0.896240625 | 0.580482024 | 0.864857905 | 0.733951626 | 0.602059991 | 0.803905701 |
| ENSMUSG00000104150 | 0 | 0 | 0 | 0 | 0 | 0 | 0 |
| ENSMUSG00000104152 | 0 | 0 | 0 | 0 | 0 | 0 | 0 |
| ENSMUSG00000104153 | 0 | 0 | 0 | 0 | 0 | 0 | 0 |
| ENSMUSG00000104154 | 0.747221736 | 0.75 | 0.701838731 | 0.701838731 | 0.77552387 | 0.698970004 | 0.803905701 |
| ENSMUSG00000104155 | 0 | 0 | 0 | 0 | 0 | 0 | 0 |
| ENSMUSG00000104156 | 0.45815691 | 0.646240625 | 0.896240625 | 0.75 | 0.632412477 | 0.301029996 | 0.579661877 |
| ENSMUSG00000104157 | 0 | 0 | 0 | 0 | 0 | 0 | 0 |
| ENSMUSG00000104158 | 0 | 0 | 0 | 0 | 0 | 0 | 0 |
| ENSMUSG00000104159 | 0.811507563 | 0.75 | 0.75 | 0.92510993 | 0.877063019 | 0.954242509 | 0.829800754 |
| ENSMUSG00000104161 | 0 | 0 | 0 | 0 | 0 | 0 | 0 |
| ENSMUSG00000104162 | 0 | 0 | 0 | 0 | 0 | 0 | 0 |
| ENSMUSG00000104163 | 0 | 0 | 0 | 0 | 0 | 0 | 0 |
| ENSMUSG00000104164 | 1.516991255 | 1.188721876 | 1.380890489 | 1.282320754 | 1.373641807 | 1.255272505 | 1.176815335 |
| ENSMUSG00000104165 | 0 | 0 | 0 | 0 | 0 | 0 | 0 |
| ENSMUSG00000104166 | 0 | 0.25 | 0.25 | 0 | 0 | 0.301029996 | 0 |
| ENSMUSG00000104167 | 0 | 0 | 0 | 0 | 0 | 0 | 0 |
| ENSMUSG00000104168 | 0.289064826 | 0 | 0.5 | 0.396240625 | 0 | 0.698970004 | 0.520678036 |
| ENSMUSG00000104169 | 0 | 0 | 0 | 0 | 0 | 0 | 0 |
| ENSMUSG00000104170 | 0 | 0.25 | 0 | 0.25 | 0 | 0 | 0 |
| ENSMUSG00000104171 | 0 | 0 | 0 | 0 | 0 | 0 | 0 |
| ENSMUSG00000104172 | 0 | 0 | 0 | 0 | 0 | 0 | 0 |
| ENSMUSG00000104173 | 0 | 0.25 | 0 | 0 | 0.244650542 | 0.301029996 | 0 |
| ENSMUSG00000104174 | 0 | 0 | 0 | 0 | 0 | 0 | 0 |
| ENSMUSG00000104175 | 0 | 0 | 0 | 0 | 0 | 0 | 0 |
| ENSMUSG00000104176 | 0 | 0 | 0 | 0 | 0 | 0 | 0 |
| ENSMUSG00000104177 | 0 | 0 | 0 | 0 | 0 | 0 | 0 |
| ENSMUSG00000104178 | 0 | 0 | 0 | 0 | 0 | 0 | 0 |
| ENSMUSG00000104180 | 0 | 0 | 0 | 0 | 0 | 0 | 0 |
| ENSMUSG00000104181 | 0.45815691 | 0.25 | 0.25 | 0.5 | 0.489301084 | 0.477121255 | 0.224243824 |
| ENSMUSG00000104182 | 0 | 0 | 0 | 0 | 0 | 0 | 0 |
| ENSMUSG00000104183 | 0.671187741 | 0.5 | 0.580482024 | 0.5 | 0.387761935 | 0 | 0.355418052 |

| | | | | | | | |
|---|---|---|---|---|---|---|---|
| ENSMUSG00000104184 | 0.867194479 | 0.830482024 | 0.79248125 | 0.396240625 | 0.568060967 | 0 | 0 |
| ENSMUSG00000104185 | 0 | 0 | 0 | 0 | 0 | 0 | 0 |
| ENSMUSG00000104186 | 0 | 0.396240625 | 0.25 | 0.396240625 | 0 | 0.301029996 | 0 |
| ENSMUSG00000104187 | 0 | 0 | 0 | 0 | 0 | 0.301029996 | 0 |
| ENSMUSG00000104188 | 0 | 0 | 0 | 0 | 0 | 0 | 0 |
| ENSMUSG00000104189 | 0 | 0.25 | 0 | 0 | 0 | 0 | 0.224243824 |
| ENSMUSG00000104190 | 0.289064826 | 0 | 0 | 0 | 0 | 0 | 0.355418052 |
| ENSMUSG00000104191 | 0 | 0 | 0 | 0 | 0 | 0 | 0 |
| ENSMUSG00000104192 | 0 | 0 | 0 | 0 | 0 | 0 | 0 |
| ENSMUSG00000104194 | 0 | 0 | 0 | 0 | 0 | 0 | 0 |
| ENSMUSG00000104195 | 0.578129653 | 0.580482024 | 0.396240625 | 0.580482024 | 0.387761935 | 0.301029996 | 0.710836105 |
| ENSMUSG00000104197 | 0 | 0 | 0 | 0.25 | 0 | 0 | 0 |
| ENSMUSG00000104198 | 0 | 0 | 0 | 0 | 0 | 0 | 0 |
| ENSMUSG00000104199 | 0 | 0 | 0 | 0 | 0 | 0 | 0 |
| ENSMUSG00000104200 | 0 | 0 | 0 | 0 | 0 | 0 | 0 |
| ENSMUSG00000104201 | 0 | 0 | 0 | 0 | 0 | 0 | 0 |
| ENSMUSG00000104202 | 0 | 0 | 0 | 0 | 0 | 0 | 0 |
| ENSMUSG00000104203 | 0 | 0 | 0 | 0 | 0 | 0 | 0 |
| ENSMUSG00000104204 | 0.578129653 | 0.5 | 0.701838731 | 0.5 | 0.387761935 | 0.602059991 | 0.224243824 |
| ENSMUSG00000104205 | 0 | 0 | 0 | 0 | 0 | 0 | 0 |
| ENSMUSG00000104206 | 0 | 0 | 0 | 0 | 0 | 0 | 0 |
| ENSMUSG00000104207 | 0 | 0 | 0 | 0 | 0 | 0 | 0 |
| ENSMUSG00000104208 | 0 | 0 | 0 | 0 | 0 | 0 | 0 |
| ENSMUSG00000104209 | 0 | 0 | 0 | 0 | 0 | 0 | 0 |
| ENSMUSG00000104210 | 0 | 0 | 0 | 0.25 | 0 | 0 | 0 |
| ENSMUSG00000104211 | 1 | 0.92510993 | 0.864857905 | 0.79248125 | 0.77552387 | 0.954242509 | 0.710836105 |
| ENSMUSG00000104212 | 0 | 0 | 0 | 0.25 | 0.387761935 | 0 | 0.448487648 |
| ENSMUSG00000104213 | 0 | 0 | 0 | 0 | 0 | 0 | 0 |
| ENSMUSG00000104214 | 0.289064826 | 0.25 | 0 | 0.580482024 | 0 | 0.301029996 | 0 |
| ENSMUSG00000104215 | 0 | 0 | 0 | 0 | 0 | 0 | 0 |
| ENSMUSG00000104216 | 0 | 0 | 0 | 0 | 0 | 0 | 0 |
| ENSMUSG00000104217 | 0 | 0 | 0 | 0 | 0 | 0 | 0 |
| ENSMUSG00000104218 | 0 | 0 | 0 | 0 | 0 | 0 | 0 |
| ENSMUSG00000104219 | 0 | 0 | 0 | 0 | 0 | 0 | 0 |

| | | | | | | | |
|---|---|---|---|---|---|---|---|
| ENSMUSG00000104220 | 0 | 0 | 0 | 0 | 0 | 0 | 0 |
| ENSMUSG00000104221 | 0 | 0 | 0 | 0 | 0 | 0 | 0 |
| ENSMUSG00000104222 | 2.451206209 | 2.213717096 | 2.175975893 | 2.412314044 | 2.324629618 | 2.678518379 | 2.160546876 |
| ENSMUSG00000104223 | 0 | 0 | 0 | 0 | 0 | 0 | 0 |
| ENSMUSG00000104224 | 0 | 0 | 0 | 0 | 0 | 0 | 0 |
| ENSMUSG00000104225 | 0 | 0 | 0 | 0 | 0 | 0 | 0 |
| ENSMUSG00000104226 | 0 | 0 | 0 | 0 | 0 | 0 | 0 |
| ENSMUSG00000104227 | 0 | 0 | 0 | 0 | 0 | 0 | 0 |
| ENSMUSG00000104228 | 0 | 0 | 0 | 0 | 0 | 0 | 0 |
| ENSMUSG00000104229 | 0 | 0 | 0 | 0 | 0 | 0 | 0 |
| ENSMUSG00000104230 | 0 | 0 | 0 | 0 | 0 | 0 | 0 |
| ENSMUSG00000104231 | 0.578129653 | 0.580482024 | 0.396240625 | 0.75 | 0.568060967 | 0.602059991 | 0.448487648 |
| ENSMUSG00000104232 | 0 | 0 | 0 | 0 | 0 | 0 | 0 |
| ENSMUSG00000104233 | 0 | 0 | 0 | 0 | 0 | 0 | 0 |
| ENSMUSG00000104234 | 0 | 0 | 0 | 0 | 0 | 0 | 0 |
| ENSMUSG00000104235 | 0.45815691 | 0.701838731 | 0.5 | 0.580482024 | 0.244650542 | 0.301029996 | 0.224243824 |
| ENSMUSG00000104236 | 0 | 0 | 0 | 0 | 0 | 0 | 0 |
| ENSMUSG00000104237 | 0.45815691 | 0 | 0 | 0.25 | 0 | 0 | 0 |
| ENSMUSG00000104238 | 0 | 0 | 0 | 0 | 0 | 0 | 0 |
| ENSMUSG00000104239 | 0 | 0 | 0 | 0 | 0 | 0 | 0 |
| ENSMUSG00000104240 | 0 | 0 | 0 | 0 | 0 | 0 | 0 |
| ENSMUSG00000104241 | 0 | 0 | 0.25 | 0.5 | 0.387761935 | 0.301029996 | 0.355418052 |
| ENSMUSG00000104242 | 0 | 0 | 0 | 0 | 0 | 0 | 0 |
| ENSMUSG00000104243 | 0 | 0 | 0.25 | 0 | 0 | 0 | 0 |
| ENSMUSG00000104244 | 0.671187741 | 0.646240625 | 0.896240625 | 0.25 | 0 | 0 | 0.355418052 |
| ENSMUSG00000104245 | 1.269664473 | 1.27186571 | 1.238549078 | 1.146240625 | 1.106691885 | 1.113943352 | 1 |
| ENSMUSG00000104247 | 0 | 0 | 0 | 0 | 0 | 0 | 0 |
| ENSMUSG00000104248 | 0.289064826 | 0.25 | 0 | 0.25 | 0.387761935 | 0 | 0.224243824 |
| ENSMUSG00000104249 | 0 | 0 | 0 | 0 | 0 | 0 | 0 |
| ENSMUSG00000104251 | 0 | 0 | 0 | 0 | 0 | 0 | 0 |
| ENSMUSG00000104252 | 0 | 0 | 0 | 0 | 0 | 0 | 0 |
| ENSMUSG00000104253 | 0.747221736 | 0.830482024 | 0.701838731 | 0.580482024 | 0.568060967 | 0.698970004 | 0.775756176 |
| ENSMUSG00000104254 | 0 | 0 | 0 | 0 | 0 | 0 | 0 |
| ENSMUSG00000104255 | 0 | 0 | 0 | 0 | 0 | 0 | 0 |

| | | | | | | | |
|---|---|---|---|---|---|---|---|
| ENSMUSG00000104256 | 0 | 0 | 0 | 0 | 0 | 0 | 0 |
| ENSMUSG00000104257 | 0 | 0 | 0.25 | 0 | 0 | 0 | 0.355418052 |
| ENSMUSG00000104258 | 0 | 0 | 0 | 0 | 0 | 0 | 0 |
| ENSMUSG00000104259 | 0 | 0 | 0 | 0 | 0 | 0 | 0 |
| ENSMUSG00000104260 | 0 | 0 | 0 | 0 | 0 | 0 | 0 |
| ENSMUSG00000104261 | 0 | 0 | 0 | 0 | 0 | 0 | 0 |
| ENSMUSG00000104262 | 0 | 0 | 0 | 0 | 0 | 0 | 0 |
| ENSMUSG00000104263 | 0.91631382 | 0.701838731 | 0.75 | 0.79248125 | 0.812711509 | 0.903089987 | 0.74492186 |
| ENSMUSG00000104264 | 0 | 0 | 0 | 0 | 0 | 0 | 0 |
| ENSMUSG00000104265 | 0 | 0 | 0 | 0 | 0 | 0 | 0 |
| ENSMUSG00000104266 | 0.289064826 | 0.25 | 0.25 | 0 | 0 | 0 | 0 |
| ENSMUSG00000104267 | 0 | 0 | 0 | 0 | 0 | 0 | 0 |
| ENSMUSG00000104268 | 0 | 0 | 0 | 0 | 0 | 0 | 0 |
| ENSMUSG00000104269 | 0 | 0 | 0 | 0 | 0 | 0 | 0 |
| ENSMUSG00000104270 | 0 | 0 | 0 | 0 | 0 | 0 | 0 |
| ENSMUSG00000104271 | 0 | 0.25 | 0 | 0.580482024 | 0.387761935 | 0.301029996 | 0 |
| ENSMUSG00000104272 | 0 | 0 | 0.25 | 0.396240625 | 0 | 0 | 0.355418052 |
| ENSMUSG00000104273 | 0.867194479 | 0.79248125 | 0.701838731 | 0.79248125 | 0.812711509 | 1.041392685 | 0.629532004 |
| ENSMUSG00000104274 | 0 | 0 | 0 | 0.25 | 0 | 0 | 0.224243824 |
| ENSMUSG00000104275 | 0 | 0.5 | 0.396240625 | 0.25 | 0.489301084 | 0.477121255 | 0.224243824 |
| ENSMUSG00000104276 | 0 | 0 | 0 | 0 | 0 | 0 | 0 |
| ENSMUSG00000104277 | 0 | 0.396240625 | 0.396240625 | 0.5 | 0.489301084 | 0 | 0.224243824 |
| ENSMUSG00000104278 | 0 | 0 | 0 | 0 | 0 | 0 | 0 |
| ENSMUSG00000104279 | 0.45815691 | 0.25 | 0.396240625 | 0 | 0.489301084 | 0.301029996 | 0.448487648 |
| ENSMUSG00000104280 | 0.811507563 | 0.79248125 | 0.830482024 | 0.396240625 | 0.733951626 | 0.301029996 | 0.710836105 |
| ENSMUSG00000104281 | 0 | 0 | 0 | 0 | 0 | 0 | 0 |
| ENSMUSG00000104282 | 0 | 0 | 0 | 0 | 0 | 0 | 0 |
| ENSMUSG00000104283 | 0 | 0 | 0 | 0 | 0 | 0 | 0 |
| ENSMUSG00000104284 | 0 | 0 | 0 | 0 | 0 | 0 | 0 |
| ENSMUSG00000104285 | 0 | 0 | 0 | 0 | 0 | 0 | 0 |
| ENSMUSG00000104286 | 0 | 0 | 0 | 0 | 0 | 0 | 0.224243824 |
| ENSMUSG00000104287 | 0 | 0 | 0 | 0 | 0 | 0 | 0 |
| ENSMUSG00000104288 | 0 | 0 | 0 | 0 | 0 | 0 | 0 |
| ENSMUSG00000104289 | 0 | 0 | 0 | 0 | 0 | 0 | 0 |

| | | | | | | | |
|---|---|---|---|---|---|---|---|
| ENSMUSG00000104290 | 0 | 0 | 0 | 0 | 0 | 0 | 0 |
| ENSMUSG00000104291 | 0.289064826 | 0 | 0 | 0 | 0 | 0 | 0 |
| ENSMUSG00000104292 | 0 | 0 | 0.5 | 0.396240625 | 0.244650542 | 0 | 0.224243824 |
| ENSMUSG00000104293 | 0.811507563 | 0.646240625 | 0.396240625 | 0.25 | 0.568060967 | 0.602059991 | 0.579661877 |
| ENSMUSG00000104294 | 0 | 0 | 0 | 0 | 0 | 0 | 0 |
| ENSMUSG00000104295 | 0 | 0 | 0 | 0 | 0 | 0 | 0 |
| ENSMUSG00000104296 | 0 | 0 | 0 | 0 | 0 | 0 | 0 |
| ENSMUSG00000104297 | 0 | 0 | 0 | 0 | 0.244650542 | 0 | 0 |
| ENSMUSG00000104298 | 0 | 0.25 | 0 | 0 | 0 | 0.301029996 | 0 |
| ENSMUSG00000104299 | 0 | 0 | 0.396240625 | 0.25 | 0.244650542 | 0 | 0.355418052 |
| ENSMUSG00000104301 | 0.289064826 | 0 | 0 | 0 | 0 | 0 | 0 |
| ENSMUSG00000104302 | 0 | 0 | 0 | 0 | 0 | 0 | 0 |
| ENSMUSG00000104303 | 0.960252568 | 0.896240625 | 1.080482024 | 0.951838731 | 0.812711509 | 1 | 0.829800754 |
| ENSMUSG00000104304 | 0 | 0 | 0 | 0 | 0 | 0 | 0 |
| ENSMUSG00000104306 | 0 | 0 | 0 | 0 | 0.244650542 | 0 | 0 |
| ENSMUSG00000104307 | 0 | 0 | 0 | 0 | 0 | 0 | 0 |
| ENSMUSG00000104308 | 0 | 0 | 0 | 0 | 0 | 0 | 0 |
| ENSMUSG00000104309 | 0 | 0.25 | 0.25 | 0 | 0 | 0 | 0.448487648 |
| ENSMUSG00000104310 | 0 | 0 | 0 | 0 | 0 | 0 | 0 |
| ENSMUSG00000104311 | 0 | 0 | 0 | 0 | 0 | 0 | 0 |
| ENSMUSG00000104312 | 0.289064826 | 0 | 0 | 0.25 | 0 | 0 | 0 |
| ENSMUSG00000104313 | 0 | 0 | 0 | 0 | 0 | 0 | 0 |
| ENSMUSG00000104314 | 0 | 0.25 | 0 | 0 | 0 | 0 | 0 |
| ENSMUSG00000104315 | 0.289064826 | 0.580482024 | 0.25 | 0.701838731 | 0.244650542 | 0 | 0.224243824 |
| ENSMUSG00000104316 | 0 | 0 | 0 | 0.25 | 0 | 0 | 0.224243824 |
| ENSMUSG00000104317 | 0 | 0 | 0 | 0 | 0 | 0 | 0 |
| ENSMUSG00000104318 | 0 | 0 | 0 | 0 | 0 | 0 | 0 |
| ENSMUSG00000104319 | 0 | 0 | 0 | 0 | 0 | 0 | 0 |
| ENSMUSG00000104320 | 0 | 0 | 0 | 0 | 0 | 0 | 0 |
| ENSMUSG00000104321 | 0 | 0 | 0 | 0 | 0 | 0 | 0 |
| ENSMUSG00000104323 | 0 | 0 | 0 | 0 | 0 | 0 | 0 |
| ENSMUSG00000104324 | 0.45815691 | 0.646240625 | 0.5 | 0.5 | 0.568060967 | 0.698970004 | 0.520678036 |
| ENSMUSG00000104325 | 0.289064826 | 0.396240625 | 0.396240625 | 0.580482024 | 0.387761935 | 0 | 0.448487648 |
| ENSMUSG00000104326 | 0 | 0 | 0 | 0 | 0 | 0 | 0 |

| | | | | | | | |
|---|---|---|---|---|---|---|---|
| ENSMUSG00000104327 | 0 | 0.25 | 0 | 0 | 0 | 0 | 0 |
| ENSMUSG00000104328 | 0 | 0 | 0 | 0 | 0 | 0 | 0 |
| ENSMUSG00000104329 | 1.227926429 | 1.27186571 | 1.238549078 | 0.864857905 | 0.905314583 | 0.903089987 | 0.896975297 |
| ENSMUSG00000104330 | 0.289064826 | 0 | 0 | 0 | 0 | 0 | 0 |
| ENSMUSG00000104331 | 0.45815691 | 0 | 0.396240625 | 0.5 | 0.686820904 | 0 | 0.448487648 |
| ENSMUSG00000104332 | 0 | 0 | 0 | 0 | 0 | 0 | 0 |
| ENSMUSG00000104333 | 0 | 0 | 0 | 0 | 0 | 0 | 0 |
| ENSMUSG00000104334 | 0 | 0 | 0 | 0 | 0 | 0 | 0 |
| ENSMUSG00000104335 | 0 | 0 | 0 | 0 | 0 | 0 | 0 |
| ENSMUSG00000104336 | 0 | 0 | 0 | 0 | 0 | 0 | 0 |
| ENSMUSG00000104337 | 0 | 0 | 0 | 0 | 0 | 0 | 0 |
| ENSMUSG00000104338 | 0 | 0 | 0 | 0 | 0 | 0 | 0 |
| ENSMUSG00000104339 | 0 | 0 | 0 | 0.25 | 0 | 0 | 0.224243824 |
| ENSMUSG00000104340 | 0 | 0 | 0 | 0.25 | 0 | 0 | 0 |
| ENSMUSG00000104341 | 0 | 0.580482024 | 0.646240625 | 0 | 0.244650542 | 0.602059991 | 0.448487648 |
| ENSMUSG00000104342 | 1.432083896 | 1.380890489 | 1.372963274 | 1.610735874 | 1.621875691 | 1.662757832 | 1.40946263 |
| ENSMUSG00000104343 | 0 | 0 | 0 | 0 | 0 | 0 | 0 |
| ENSMUSG00000104344 | 1.325351389 | 1.29248125 | 1.146240625 | 1.388647213 | 1.283908311 | 1.342422681 | 1.265599895 |
| ENSMUSG00000104345 | 0 | 0 | 0 | 0 | 0 | 0 | 0 |
| ENSMUSG00000104346 | 0.578129653 | 0.75 | 0.701838731 | 0.25 | 0.733951626 | 0.301029996 | 0.710836105 |
| ENSMUSG00000104347 | 0.289064826 | 0 | 0 | 0 | 0 | 0 | 0 |
| ENSMUSG00000104348 | 0 | 0.396240625 | 0.396240625 | 0 | 0.387761935 | 0 | 0 |
| ENSMUSG00000104349 | 0 | 0 | 0 | 0 | 0.244650542 | 0 | 0 |
| ENSMUSG00000104350 | 0 | 0.25 | 0 | 0.25 | 0.387761935 | 0.477121255 | 0.355418052 |
| ENSMUSG00000104351 | 0.289064826 | 0 | 0 | 0.25 | 0 | 0.301029996 | 0.224243824 |
| ENSMUSG00000104352 | 0 | 0 | 0 | 0 | 0 | 0 | 0 |
| ENSMUSG00000104354 | 0 | 0 | 0 | 0 | 0 | 0 | 0 |
| ENSMUSG00000104355 | 0 | 0 | 0 | 0 | 0 | 0 | 0 |
| ENSMUSG00000104356 | 0 | 0 | 0 | 0 | 0 | 0 | 0 |
| ENSMUSG00000104358 | 1 | 1.02186571 | 0.976722649 | 1.061981878 | 1.106691885 | 1.079181246 | 0.935079929 |
| ENSMUSG00000104360 | 0 | 0 | 0 | 0 | 0 | 0 | 0.224243824 |
| ENSMUSG00000104361 | 0.289064826 | 0.396240625 | 0.396240625 | 0.25 | 0.387761935 | 0.301029996 | 0.448487648 |
| ENSMUSG00000104362 | 0 | 0 | 0 | 0 | 0 | 0 | 0 |
| ENSMUSG00000104363 | 0 | 0 | 0 | 0.25 | 0.387761935 | 0 | 0 |

| | | | | | | | |
|---|---|---|---|---|---|---|---|
| ENSMUSG00000104364 | 0 | 0 | 0 | 0 | 0 | 0 | 0 |
| ENSMUSG00000104365 | 0 | 0 | 0 | 0 | 0 | 0 | 0 |
| ENSMUSG00000104366 | 0 | 0 | 0 | 0 | 0 | 0 | 0 |
| ENSMUSG00000104367 | 0 | 0 | 0 | 0 | 0 | 0 | 0.224243824 |
| ENSMUSG00000104368 | 0 | 0 | 0.5 | 0.396240625 | 0.244650542 | 0 | 0.224243824 |
| ENSMUSG00000104369 | 0.578129653 | 0.830482024 | 0.75 | 0.864857905 | 0.632412477 | 0.698970004 | 0.803905701 |
| ENSMUSG00000104370 | 0.289064826 | 0 | 0.25 | 0.25 | 0 | 0 | 0 |
| ENSMUSG00000104371 | 0 | 0 | 0 | 0 | 0 | 0 | 0 |
| ENSMUSG00000104373 | 0 | 0 | 0 | 0 | 0 | 0 | 0 |
| ENSMUSG00000104374 | 0 | 0 | 0 | 0 | 0 | 0 | 0 |
| ENSMUSG00000104375 | 0 | 0 | 0 | 0 | 0 | 0 | 0 |
| ENSMUSG00000104376 | 0 | 0 | 0 | 0 | 0 | 0 | 0 |
| ENSMUSG00000104377 | 1.614416215 | 1.610735874 | 1.494319981 | 1.584962501 | 1.467903253 | 1.531478917 | 1.355418052 |
| ENSMUSG00000104378 | 0 | 0 | 0.25 | 0 | 0 | 0 | 0.355418052 |
| ENSMUSG00000104379 | 0 | 0 | 0 | 0 | 0.244650542 | 0 | 0 |
| ENSMUSG00000104380 | 0 | 0 | 0 | 0 | 0 | 0 | 0 |
| ENSMUSG00000104381 | 0 | 0 | 0 | 0 | 0 | 0 | 0 |
| ENSMUSG00000104382 | 0 | 0 | 0 | 0 | 0 | 0 | 0 |
| ENSMUSG00000104383 | 0.289064826 | 0.701838731 | 0.580482024 | 0 | 0.387761935 | 0 | 0 |
| ENSMUSG00000104384 | 0 | 0.25 | 0.25 | 0.396240625 | 0.244650542 | 0.477121255 | 0 |
| ENSMUSG00000104385 | 0 | 0 | 0 | 0.25 | 0 | 0 | 0 |
| ENSMUSG00000104386 | 0 | 0 | 0 | 0 | 0 | 0 | 0 |
| ENSMUSG00000104388 | 0 | 0.396240625 | 0.25 | 0.5 | 0.489301084 | 0.301029996 | 0.355418052 |
| ENSMUSG00000104389 | 0 | 0 | 0 | 0.5 | 0.387761935 | 0.477121255 | 0.224243824 |
| ENSMUSG00000104390 | 0 | 0 | 0 | 0 | 0 | 0 | 0 |
| ENSMUSG00000104391 | 0.578129653 | 0 | 0.580482024 | 0 | 0.244650542 | 0 | 0.355418052 |
| ENSMUSG00000104392 | 0.45815691 | 0.580482024 | 0.646240625 | 0.396240625 | 0.686820904 | 0.602059991 | 0.520678036 |
| ENSMUSG00000104393 | 0 | 0 | 0 | 0 | 0 | 0 | 0 |
| ENSMUSG00000104394 | 1.156259305 | 1.160964047 | 1.188721876 | 1.418106335 | 1.373641807 | 1.322219295 | 1.290497981 |
| ENSMUSG00000104395 | 0 | 0 | 0 | 0 | 0 | 0 | 0 |
| ENSMUSG00000104396 | 0 | 0 | 0 | 0 | 0 | 0.301029996 | 0.224243824 |
| ENSMUSG00000104397 | 0 | 0 | 0 | 0 | 0 | 0 | 0 |
| ENSMUSG00000104398 | 0 | 0 | 0 | 0 | 0 | 0 | 0 |
| ENSMUSG00000104399 | 1.069666964 | 1.04248125 | 0.701838731 | 0.976722649 | 1 | 1 | 0.984950056 |

| | | | | | | | |
|---|---|---|---|---|---|---|---|
| ENSMUSG00000104400 | 0 | 0.25 | 0 | 0 | 0 | 0 | 0 |
| ENSMUSG00000104401 | 0 | 0 | 0 | 0 | 0 | 0 | 0.224243824 |
| ENSMUSG00000104402 | 0.289064826 | 0 | 0 | 0 | 0.387761935 | 0 | 0.355418052 |
| ENSMUSG00000104403 | 0 | 0 | 0 | 0 | 0 | 0 | 0 |
| ENSMUSG00000104404 | 0 | 0 | 0 | 0 | 0 | 0 | 0 |
| ENSMUSG00000104405 | 0 | 0 | 0 | 0 | 0 | 0 | 0 |
| ENSMUSG00000104406 | 0 | 0.25 | 0.396240625 | 0.396240625 | 0.489301084 | 0.477121255 | 0 |
| ENSMUSG00000104407 | 0 | 0 | 0 | 0 | 0 | 0 | 0 |
| ENSMUSG00000104408 | 0.811507563 | 0.5 | 0.830482024 | 0.830482024 | 0.686820904 | 0.602059991 | 0.896975297 |
| ENSMUSG00000104409 | 0 | 0 | 0 | 0 | 0 | 0 | 0 |
| ENSMUSG00000104410 | 0 | 0 | 0 | 0 | 0 | 0 | 0 |
| ENSMUSG00000104411 | 0 | 0 | 0 | 0 | 0 | 0 | 0 |
| ENSMUSG00000104412 | 0 | 0 | 0 | 0 | 0 | 0 | 0 |
| ENSMUSG00000104413 | 0.671187741 | 0.701838731 | 0.75 | 0.701838731 | 0.568060967 | 0 | 0.629532004 |
| ENSMUSG00000104414 | 0 | 0 | 0 | 0 | 0 | 0 | 0 |
| ENSMUSG00000104415 | 0.45815691 | 0.5 | 0.646240625 | 0.396240625 | 0.489301084 | 0.602059991 | 0.355418052 |
| ENSMUSG00000104416 | 0 | 0.25 | 0.580482024 | 0 | 0.489301084 | 0.301029996 | 0.224243824 |
| ENSMUSG00000104417 | 0 | 0 | 0 | 0 | 0 | 0 | 0 |
| ENSMUSG00000104418 | 0 | 0.396240625 | 0 | 0.396240625 | 0.244650542 | 0 | 0.224243824 |
| ENSMUSG00000104419 | 0 | 0 | 0 | 0 | 0 | 0 | 0 |
| ENSMUSG00000104420 | 0 | 0 | 0.25 | 0.5 | 0 | 0 | 0.224243824 |
| ENSMUSG00000104422 | 0 | 0 | 0 | 0 | 0 | 0 | 0 |
| ENSMUSG00000104423 | 0 | 0 | 0.25 | 0 | 0 | 0 | 0 |
| ENSMUSG00000104424 | 0 | 0 | 0 | 0 | 0 | 0 | 0 |
| ENSMUSG00000104425 | 0 | 0 | 0 | 0 | 0 | 0 | 0 |
| ENSMUSG00000104426 | 0 | 0 | 0 | 0 | 0 | 0 | 0 |
| ENSMUSG00000104427 | 0 | 0 | 0 | 0 | 0 | 0 | 0 |
| ENSMUSG00000104428 | 0 | 0 | 0 | 0 | 0 | 0 | 0 |
| ENSMUSG00000104429 | 0 | 0 | 0 | 0 | 0 | 0 | 0 |
| ENSMUSG00000104430 | 0 | 0 | 0 | 0 | 0 | 0 | 0 |
| ENSMUSG00000104431 | 0 | 0 | 0 | 0 | 0 | 0 | 0 |
| ENSMUSG00000104432 | 0.289064826 | 0.5 | 0 | 0 | 0.632412477 | 0.301029996 | 0.224243824 |
| ENSMUSG00000104433 | 0 | 0.396240625 | 0.5 | 0 | 0.244650542 | 0 | 0.224243824 |
| ENSMUSG00000104434 | 0.747221736 | 0.5 | 0.646240625 | 0.580482024 | 0.905314583 | 0.698970004 | 0.74492186 |

| | | | | | | | |
|---|---|---|---|---|---|---|---|
| ENSMUSG00000104435 | 1.445324132 | 1.418106335 | 1.396240625 | 1.302363341 | 1.439191811 | 1.397940009 | 1.296434211 |
| ENSMUSG00000104436 | 0 | 0.5 | 0 | 0.396240625 | 0.244650542 | 0 | 0.224243824 |
| ENSMUSG00000104437 | 0 | 0 | 0 | 0 | 0 | 0 | 0 |
| ENSMUSG00000104438 | 0 | 0 | 0 | 0 | 0 | 0 | 0 |
| ENSMUSG00000104439 | 0 | 0 | 0 | 0 | 0 | 0 | 0 |
| ENSMUSG00000104440 | 0 | 0 | 0 | 0 | 0 | 0 | 0 |
| ENSMUSG00000104441 | 0 | 0 | 0 | 0 | 0 | 0 | 0 |
| ENSMUSG00000104442 | 0 | 0 | 0 | 0 | 0 | 0 | 0 |
| ENSMUSG00000104443 | 0.867194479 | 0.646240625 | 0.830482024 | 0.75 | 0.489301084 | 0.698970004 | 0.710836105 |
| ENSMUSG00000104444 | 0 | 0 | 0 | 0 | 0 | 0 | 0 |
| ENSMUSG00000104445 | 0 | 0 | 0 | 0 | 0.244650542 | 0 | 0.224243824 |
| ENSMUSG00000104446 | 0 | 0 | 0 | 0 | 0 | 0 | 0 |
| ENSMUSG00000104448 | 0 | 0 | 0 | 0 | 0 | 0 | 0 |
| ENSMUSG00000104449 | 0.289064826 | 0.580482024 | 0.5 | 0.25 | 0.387761935 | 0 | 0.448487648 |
| ENSMUSG00000104450 | 0 | 0 | 0 | 0 | 0 | 0 | 0 |
| ENSMUSG00000104451 | 0 | 0 | 0 | 0 | 0 | 0 | 0 |
| ENSMUSG00000104452 | 0 | 0 | 0 | 0 | 0 | 0 | 0 |
| ENSMUSG00000104453 | 0.578129653 | 0.646240625 | 0.79248125 | 0.646240625 | 0.387761935 | 0.477121255 | 0.448487648 |
| ENSMUSG00000104454 | 0.289064826 | 0 | 0 | 0 | 0 | 0 | 0 |
| ENSMUSG00000104455 | 0 | 0 | 0 | 0 | 0 | 0 | 0 |
| ENSMUSG00000104456 | 0 | 0 | 0 | 0 | 0 | 0 | 0 |
| ENSMUSG00000104457 | 0 | 0 | 0.25 | 0 | 0 | 0 | 0 |
| ENSMUSG00000104458 | 0 | 0 | 0 | 0 | 0 | 0 | 0 |
| ENSMUSG00000104459 | 0.578129653 | 0.396240625 | 0.396240625 | 0.646240625 | 0.244650542 | 0.301029996 | 0.355418052 |
| ENSMUSG00000104460 | 0 | 0 | 0 | 0 | 0 | 0 | 0 |
| ENSMUSG00000104461 | 0 | 0 | 0 | 0 | 0 | 0 | 0 |
| ENSMUSG00000104462 | 0 | 0 | 0 | 0 | 0 | 0 | 0 |
| ENSMUSG00000104464 | 0 | 0 | 0 | 0 | 0 | 0 | 0 |
| ENSMUSG00000104465 | 0 | 0 | 0 | 0 | 0 | 0 | 0 |
| ENSMUSG00000104466 | 0 | 0 | 0 | 0 | 0 | 0 | 0 |
| ENSMUSG00000104467 | 0.747221736 | 0.646240625 | 0.830482024 | 0.864857905 | 0.978602168 | 0.77815125 | 1 |
| ENSMUSG00000104468 | 0 | 0 | 0 | 0 | 0 | 0 | 0 |
| ENSMUSG00000104469 | 0.45815691 | 0.580482024 | 0.580482024 | 0.396240625 | 0.244650542 | 0 | 0.224243824 |
| ENSMUSG00000104470 | 0 | 0.25 | 0.25 | 0 | 0 | 0 | 0.224243824 |

| | | | | | | | |
|---|---|---|---|---|---|---|---|
| ENSMUSG00000104471 | 0 | 0 | 0 | 0 | 0 | 0 | 0 |
| ENSMUSG00000104472 | 0 | 0 | 0 | 0 | 0 | 0 | 0 |
| ENSMUSG00000104473 | 0 | 0 | 0 | 0 | 0 | 0 | 0 |
| ENSMUSG00000104474 | 0 | 0 | 0 | 0 | 0 | 0 | 0 |
| ENSMUSG00000104475 | 1.578129653 | 1.51109853 | 1.464495249 | 1.747171172 | 1.642643806 | 1.826074803 | 1.526507301 |
| ENSMUSG00000104476 | 0.289064826 | 0.25 | 0 | 0.5 | 0.489301084 | 0.698970004 | 0.224243824 |
| ENSMUSG00000104477 | 0 | 0 | 0 | 0 | 0 | 0 | 0 |
| ENSMUSG00000104478 | 0 | 0 | 0 | 0 | 0 | 0 | 0 |
| ENSMUSG00000104479 | 0 | 0.25 | 0.396240625 | 0.25 | 0 | 0 | 0 |
| ENSMUSG00000104480 | 0 | 0 | 0 | 0 | 0 | 0 | 0 |
| ENSMUSG00000104481 | 0 | 0 | 0 | 0 | 0 | 0 | 0 |
| ENSMUSG00000104482 | 0 | 0 | 0 | 0 | 0 | 0 | 0 |
| ENSMUSG00000104484 | 0 | 0 | 0 | 0 | 0 | 0 | 0 |
| ENSMUSG00000104485 | 0.578129653 | 0.396240625 | 0.25 | 0 | 0.244650542 | 0.301029996 | 0.224243824 |
| ENSMUSG00000104486 | 0.289064826 | 0 | 0 | 0 | 0 | 0 | 0.224243824 |
| ENSMUSG00000104487 | 0 | 0 | 0 | 0 | 0 | 0 | 0 |
| ENSMUSG00000104488 | 0 | 0 | 0 | 0 | 0 | 0 | 0 |
| ENSMUSG00000104489 | 0 | 0 | 0 | 0 | 0 | 0 | 0 |
| ENSMUSG00000104490 | 0 | 0 | 0 | 0 | 0 | 0 | 0 |
| ENSMUSG00000104491 | 0 | 0 | 0 | 0 | 0 | 0 | 0 |
| ENSMUSG00000104493 | 0 | 0 | 0.25 | 0.25 | 0 | 0 | 0 |
| ENSMUSG00000104494 | 0.289064826 | 0 | 0 | 0 | 0.244650542 | 0 | 0 |
| ENSMUSG00000104496 | 0 | 0.25 | 0 | 0.25 | 0 | 0 | 0.355418052 |
| ENSMUSG00000104497 | 0 | 0 | 0 | 0 | 0 | 0 | 0 |
| ENSMUSG00000104498 | 0 | 0 | 0 | 0 | 0 | 0 | 0 |
| ENSMUSG00000104499 | 0 | 0 | 0 | 0 | 0 | 0 | 0 |
| ENSMUSG00000104500 | 0 | 0 | 0 | 0 | 0 | 0 | 0 |
| ENSMUSG00000104501 | 0 | 0 | 0 | 0 | 0 | 0 | 0 |
| ENSMUSG00000104502 | 0 | 0 | 0 | 0 | 0 | 0 | 0 |
| ENSMUSG00000104503 | 0.45815691 | 1.130890489 | 0.864857905 | 0.976722649 | 0.632412477 | 0.602059991 | 0.803905701 |
| ENSMUSG00000104504 | 0 | 0 | 0 | 0 | 0 | 0 | 0 |
| ENSMUSG00000104505 | 0 | 0 | 0 | 0 | 0 | 0 | 0 |
| ENSMUSG00000104506 | 0 | 0 | 0 | 0 | 0 | 0 | 0 |
| ENSMUSG00000104507 | 0 | 0 | 0.25 | 0 | 0 | 0 | 0 |

| | | | | | | | |
|---|---|---|---|---|---|---|---|
| ENSMUSG00000104508 | 0 | 0 | 0 | 0 | 0 | 0 | 0.224243824 |
| ENSMUSG00000104509 | 0.289064826 | 0 | 0 | 0 | 0 | 0 | 0 |
| ENSMUSG00000104510 | 0 | 0 | 0 | 0 | 0 | 0 | 0 |
| ENSMUSG00000104511 | 0 | 0 | 0 | 0 | 0 | 0 | 0 |
| ENSMUSG00000104512 | 0 | 0 | 0 | 0 | 0 | 0 | 0 |
| ENSMUSG00000104513 | 0 | 0 | 0 | 0 | 0 | 0 | 0 |
| ENSMUSG00000104514 | 0 | 0 | 0.25 | 0.396240625 | 0.244650542 | 0.602059991 | 0.355418052 |
| ENSMUSG00000104515 | 0.747221736 | 0.396240625 | 0.25 | 0.5 | 0.244650542 | 0.301029996 | 0.224243824 |
| ENSMUSG00000104516 | 0 | 0 | 0 | 0 | 0 | 0 | 0 |
| ENSMUSG00000104517 | 0 | 0 | 0 | 0 | 0 | 0 | 0 |
| ENSMUSG00000104518 | 0 | 0 | 0 | 0 | 0 | 0 | 0 |
| ENSMUSG00000104519 | 0 | 0.580482024 | 0 | 0 | 0 | 0 | 0 |
| ENSMUSG00000104520 | 0 | 0 | 0 | 0 | 0 | 0 | 0 |
| ENSMUSG00000104521 | 0 | 0 | 0 | 0 | 0 | 0 | 0 |
| ENSMUSG00000104522 | 0 | 0 | 0 | 0 | 0 | 0 | 0 |
| ENSMUSG00000104523 | 0 | 0 | 0 | 0 | 0 | 0 | 0 |
| ENSMUSG00000104524 | 0 | 0.25 | 0 | 0 | 0 | 0 | 0.355418052 |
| ENSMUSG00000104525 | 0 | 0 | 0 | 0 | 0.244650542 | 0 | 0.224243824 |
| ENSMUSG00000104526 | 0 | 0 | 0 | 0 | 0 | 0 | 0 |
| ENSMUSG00000104527 | 0 | 0 | 0 | 0 | 0 | 0 | 0 |
| ENSMUSG00000104528 | 0.45815691 | 0.25 | 0.5 | 0 | 0.387761935 | 0.301029996 | 0 |
| ENSMUSG00000104529 | 0 | 0 | 0 | 0 | 0 | 0 | 0 |
| ENSMUSG00000104530 | 0 | 0 | 0 | 0 | 0 | 0 | 0 |
| ENSMUSG00000104531 | 0 | 0 | 0 | 0 | 0 | 0 | 0 |
| ENSMUSG00000104532 | 0 | 0 | 0 | 0 | 0 | 0 | 0 |
| ENSMUSG00000104533 | 0 | 0 | 0 | 0 | 0 | 0 | 0 |
| ENSMUSG00000104534 | 0 | 0 | 0 | 0 | 0 | 0 | 0 |
| ENSMUSG00000104535 | 0.45815691 | 0.701838731 | 0.5 | 0.646240625 | 0.489301084 | 0.477121255 | 0.448487648 |
| ENSMUSG00000104536 | 0 | 0 | 0 | 0 | 0 | 0 | 0 |
| ENSMUSG00000104537 | 0 | 0 | 0 | 0 | 0 | 0 | 0 |
| ENSMUSG00000104538 | 0 | 0 | 0 | 0 | 0 | 0 | 0 |
| ENSMUSG00000104539 | 0 | 0 | 0 | 0 | 0 | 0 | 0 |
| ENSMUSG00000104540 | 0 | 0 | 0 | 0 | 0 | 0 | 0 |
| ENSMUSG00000104541 | 0 | 0 | 0 | 0 | 0 | 0 | 0 |

| | | | | | | | |
|---|---|---|---|---|---|---|---|
| ENSMUSG00000104542 | 0 | 0 | 0 | 0 | 0 | 0 | 0 |
| ENSMUSG00000104543 | 0 | 0 | 0.25 | 0 | 0 | 0 | 0.448487648 |
| ENSMUSG00000104544 | 0 | 0 | 0 | 0 | 0 | 0 | 0 |
| ENSMUSG00000104545 | 0.289064826 | 0 | 0 | 0.396240625 | 0.244650542 | 0.477121255 | 0 |
| ENSMUSG00000104546 | 1 | 0.976722649 | 1 | 1.098079356 | 1.176121988 | 1.146128036 | 1.054044578 |
| ENSMUSG00000104547 | 0 | 0 | 0 | 0 | 0 | 0 | 0 |
| ENSMUSG00000104548 | 0.578129653 | 0.701838731 | 0.75 | 1.061981878 | 1 | 0.954242509 | 0.672731473 |
| ENSMUSG00000104549 | 0 | 0 | 0 | 0 | 0 | 0 | 0 |
| ENSMUSG00000104550 | 0 | 0.396240625 | 0 | 0.25 | 0.387761935 | 0.477121255 | 0.520678036 |
| ENSMUSG00000104551 | 0 | 0 | 0 | 0 | 0 | 0 | 0 |
| ENSMUSG00000104552 | 0 | 0 | 0 | 0 | 0 | 0 | 0 |
| ENSMUSG00000104553 | 0 | 0 | 0 | 0 | 0 | 0 | 0 |
| ENSMUSG00000104554 | 0 | 0 | 0 | 0.396240625 | 0.244650542 | 0.301029996 | 0 |
| ENSMUSG00000104555 | 0 | 0 | 0 | 0 | 0 | 0 | 0.224243824 |
| ENSMUSG00000104556 | 0.289064826 | 0.25 | 0 | 0.25 | 0 | 0 | 0.355418052 |
| ENSMUSG00000104557 | 0 | 0 | 0 | 0 | 0 | 0 | 0 |
| ENSMUSG00000104558 | 0 | 0 | 0 | 0 | 0 | 0 | 0 |
| ENSMUSG00000104559 | 0 | 0 | 0 | 0 | 0 | 0 | 0 |
| ENSMUSG00000104560 | 0 | 0 | 0.25 | 0 | 0 | 0 | 0 |
| ENSMUSG00000104561 | 0 | 0 | 0 | 0 | 0 | 0 | 0 |
| ENSMUSG00000104562 | 0 | 0 | 0 | 0 | 0 | 0 | 0 |
| ENSMUSG00000104563 | 2.363083473 | 2.221424093 | 2.234159485 | 1.985628626 | 1.992595054 | 2.214843848 | 1.889537832 |
| ENSMUSG00000104564 | 0 | 0 | 0 | 0 | 0 | 0.301029996 | 0 |
| ENSMUSG00000104566 | 0 | 0 | 0 | 0 | 0 | 0 | 0 |
| ENSMUSG00000104567 | 0 | 0 | 0 | 0 | 0.244650542 | 0 | 0.224243824 |
| ENSMUSG00000104568 | 0 | 0.25 | 0.25 | 0 | 0 | 0 | 0.224243824 |
| ENSMUSG00000104569 | 0.578129653 | 0.25 | 0.5 | 0.646240625 | 0.632412477 | 0.698970004 | 0.355418052 |
| ENSMUSG00000104570 | 0 | 0 | 0 | 0 | 0 | 0 | 0 |
| ENSMUSG00000104571 | 0 | 0 | 0 | 0 | 0.244650542 | 0 | 0 |
| ENSMUSG00000104572 | 0 | 0 | 0 | 0 | 0 | 0 | 0 |
| ENSMUSG00000104573 | 0 | 0 | 0 | 0 | 0 | 0 | 0 |
| ENSMUSG00000104574 | 0 | 0 | 0 | 0 | 0 | 0 | 0 |
| ENSMUSG00000104575 | 0 | 0 | 0 | 0 | 0 | 0 | 0 |
| ENSMUSG00000104576 | 0 | 0.25 | 0.25 | 0 | 0 | 0.301029996 | 0 |

| | | | | | | | |
|---|---|---|---|---|---|---|---|
| ENSMUSG00000104577 | 0.289064826 | 0.396240625 | 0 | 0.396240625 | 0.244650542 | 0.602059991 | 0 |
| ENSMUSG00000104578 | 0 | 0 | 0 | 0 | 0 | 0 | 0 |
| ENSMUSG00000104579 | 0 | 0 | 0 | 0 | 0 | 0 | 0 |
| ENSMUSG00000104580 | 0 | 0 | 0 | 0 | 0 | 0 | 0 |
| ENSMUSG00000104581 | 0 | 0 | 0.25 | 0 | 0.244650542 | 0 | 0.224243824 |
| ENSMUSG00000104582 | 0 | 0 | 0 | 0 | 0 | 0 | 0 |
| ENSMUSG00000104583 | 0 | 0 | 0 | 0.396240625 | 0.387761935 | 0.301029996 | 0 |
| ENSMUSG00000104584 | 0 | 0 | 0 | 0 | 0 | 0 | 0 |
| ENSMUSG00000104585 | 0 | 0 | 0.25 | 0 | 0 | 0 | 0 |
| ENSMUSG00000104586 | 0 | 0 | 0 | 0 | 0 | 0 | 0 |
| ENSMUSG00000104587 | 0.671187741 | 0.25 | 0.25 | 0.5 | 0.489301084 | 0.301029996 | 0.224243824 |
| ENSMUSG00000104588 | 0 | 0 | 0 | 0 | 0.244650542 | 0 | 0 |
| ENSMUSG00000104589 | 0 | 0 | 0 | 0 | 0 | 0 | 0 |
| ENSMUSG00000104590 | 0 | 0 | 0 | 0 | 0 | 0 | 0 |
| ENSMUSG00000104591 | 0.671187741 | 0 | 0 | 0 | 0 | 0 | 0 |
| ENSMUSG00000104592 | 0.578129653 | 0.75 | 0.396240625 | 0.79248125 | 0.733951626 | 0.602059991 | 0.710836105 |
| ENSMUSG00000104593 | 0 | 0 | 0 | 0 | 0 | 0 | 0 |
| ENSMUSG00000104594 | 0 | 0 | 0 | 0 | 0 | 0 | 0 |
| ENSMUSG00000104595 | 0 | 0 | 0 | 0 | 0 | 0 | 0 |
| ENSMUSG00000104596 | 0 | 0 | 0 | 0 | 0 | 0 | 0 |
| ENSMUSG00000104597 | 0.811507563 | 1.02186571 | 0.701838731 | 0.646240625 | 0.387761935 | 0 | 0.355418052 |
| ENSMUSG00000104598 | 0 | 0 | 0 | 0 | 0 | 0 | 0 |
| ENSMUSG00000104599 | 0 | 0 | 0 | 0 | 0 | 0 | 0 |
| ENSMUSG00000104600 | 0 | 0 | 0 | 0 | 0 | 0 | 0 |
| ENSMUSG00000104601 | 0 | 0 | 0 | 0 | 0 | 0 | 0 |
| ENSMUSG00000104602 | 0 | 0 | 0 | 0 | 0 | 0 | 0 |
| ENSMUSG00000104603 | 0 | 0 | 0 | 0 | 0 | 0 | 0 |
| ENSMUSG00000104604 | 0 | 0 | 0 | 0 | 0.244650542 | 0 | 0 |
| ENSMUSG00000104605 | 0 | 0 | 0 | 0 | 0 | 0 | 0 |
| ENSMUSG00000104606 | 0 | 0 | 0.25 | 0.5 | 0.244650542 | 0.477121255 | 0.224243824 |
| ENSMUSG00000104607 | 0 | 0 | 0 | 0 | 0 | 0 | 0 |
| ENSMUSG00000104608 | 0 | 0 | 0 | 0 | 0 | 0 | 0 |
| ENSMUSG00000104609 | 0 | 0 | 0 | 0 | 0 | 0 | 0 |
| ENSMUSG00000104610 | 0.45815691 | 0.896240625 | 0.79248125 | 0.92510993 | 0.686820904 | 0.77815125 | 0.672731473 |

| | | | | | | | |
|---|---|---|---|---|---|---|---|
| ENSMUSG00000104612 | 0 | 0 | 0 | 0 | 0 | 0 | 0 |
| ENSMUSG00000104613 | 0 | 0 | 0 | 0 | 0 | 0 | 0 |
| ENSMUSG00000104614 | 0 | 0 | 0 | 0 | 0 | 0 | 0 |
| ENSMUSG00000104615 | 0 | 0 | 0 | 0 | 0.244650542 | 0 | 0 |
| ENSMUSG00000104616 | 0.289064826 | 0 | 0 | 0.25 | 0 | 0.301029996 | 0 |
| ENSMUSG00000104617 | 0 | 0 | 0 | 0 | 0 | 0 | 0 |
| ENSMUSG00000104618 | 0 | 0 | 0 | 0 | 0 | 0 | 0 |
| ENSMUSG00000104619 | 0 | 0 | 0 | 0 | 0 | 0 | 0 |
| ENSMUSG00000104620 | 0 | 0 | 0 | 0 | 0 | 0 | 0 |
| ENSMUSG00000104621 | 1.307602651 | 1.146240625 | 1.160964047 | 1 | 0.905314583 | 0.903089987 | 0.952571511 |
| ENSMUSG00000104622 | 0 | 0 | 0.396240625 | 0.25 | 0 | 0.301029996 | 0 |
| ENSMUSG00000104623 | 0.289064826 | 0.396240625 | 0.396240625 | 0 | 0 | 0.301029996 | 0.355418052 |
| ENSMUSG00000104624 | 0 | 0 | 0 | 0 | 0 | 0 | 0 |
| ENSMUSG00000104625 | 0.45815691 | 0.25 | 0.396240625 | 0.25 | 0.686820904 | 0.301029996 | 0.448487648 |
| ENSMUSG00000104626 | 0.45815691 | 0.25 | 0 | 0 | 0.244650542 | 0 | 0 |
| ENSMUSG00000104627 | 0.289064826 | 0 | 0 | 0.25 | 0 | 0 | 0 |
| ENSMUSG00000104628 | 0 | 0 | 0 | 0 | 0 | 0 | 0 |
| ENSMUSG00000104629 | 0 | 0 | 0 | 0 | 0 | 0 | 0 |
| ENSMUSG00000104630 | 0 | 0 | 0 | 0 | 0 | 0 | 0 |
| ENSMUSG00000104631 | 0 | 0 | 0 | 0 | 0 | 0 | 0 |
| ENSMUSG00000104632 | 0.45815691 | 0 | 0 | 0.396240625 | 0.244650542 | 0 | 0 |
| ENSMUSG00000104633 | 0 | 0 | 0 | 0 | 0 | 0 | 0 |
| ENSMUSG00000104634 | 0 | 0 | 0.25 | 0.25 | 0 | 0 | 0 |
| ENSMUSG00000104635 | 0 | 0 | 0 | 0 | 0 | 0 | 0 |
| ENSMUSG00000104636 | 0 | 0 | 0 | 0 | 0 | 0 | 0 |
| ENSMUSG00000104637 | 0 | 0 | 0 | 0 | 0 | 0 | 0 |
| ENSMUSG00000104638 | 0 | 0 | 0 | 0 | 0 | 0 | 0 |
| ENSMUSG00000104639 | 0 | 0 | 0 | 0 | 0 | 0 | 0.224243824 |
| ENSMUSG00000104640 | 0 | 0 | 0 | 0 | 0 | 0 | 0 |
| ENSMUSG00000104641 | 0.289064826 | 0.646240625 | 0.646240625 | 0.580482024 | 0.877063019 | 0.301029996 | 0.672731473 |
| ENSMUSG00000104642 | 0 | 0 | 0 | 0 | 0 | 0 | 0 |
| ENSMUSG00000104643 | 0.289064826 | 0 | 0 | 0 | 0 | 0 | 0.224243824 |
| ENSMUSG00000104644 | 0 | 0 | 0 | 0 | 0 | 0 | 0 |
| ENSMUSG00000104645 | 0 | 0 | 0 | 0 | 0 | 0 | 0 |

| | | | | | | | |
|---|---|---|---|---|---|---|---|
| ENSMUSG00000104646 | 0 | 0 | 0 | 0 | 0 | 0 | 0 |
| ENSMUSG00000104647 | 0 | 0.25 | 0 | 0.25 | 0.244650542 | 0 | 0 |
| ENSMUSG00000104648 | 0 | 0 | 0 | 0 | 0 | 0 | 0 |
| ENSMUSG00000104649 | 0.578129653 | 0.79248125 | 0.646240625 | 0.864857905 | 0.812711509 | 1.041392685 | 0.629532004 |
| ENSMUSG00000104650 | 0 | 0 | 0 | 0 | 0 | 0 | 0 |
| ENSMUSG00000104651 | 0 | 0 | 0 | 0 | 0 | 0 | 0 |
| ENSMUSG00000104652 | 0 | 0 | 0 | 0 | 0 | 0 | 0 |
| ENSMUSG00000104653 | 0 | 0 | 0 | 0 | 0 | 0 | 0 |
| ENSMUSG00000104654 | 0 | 0 | 0 | 0 | 0 | 0 | 0 |
| ENSMUSG00000104655 | 0 | 0 | 0 | 0 | 0 | 0 | 0 |
| ENSMUSG00000104656 | 0 | 0 | 0 | 0 | 0 | 0 | 0 |
| ENSMUSG00000104657 | 0 | 0 | 0 | 0 | 0 | 0 | 0 |
| ENSMUSG00000104658 | 0 | 0 | 0 | 0 | 0 | 0 | 0 |
| ENSMUSG00000104659 | 0 | 0 | 0 | 0 | 0 | 0 | 0 |
| ENSMUSG00000104660 | 0 | 0 | 0 | 0 | 0 | 0 | 0 |
| ENSMUSG00000104661 | 0.811507563 | 0.5 | 0.580482024 | 0.830482024 | 0.632412477 | 0.602059991 | 0.224243824 |
| ENSMUSG00000104662 | 0.289064826 | 0.396240625 | 0.25 | 0.396240625 | 0.489301084 | 0.477121255 | 0 |
| ENSMUSG00000104663 | 0 | 0 | 0 | 0 | 0.244650542 | 0 | 0 |
| ENSMUSG00000104664 | 0 | 0 | 0 | 0 | 0 | 0 | 0 |
| ENSMUSG00000104665 | 0 | 0.396240625 | 0.396240625 | 0.25 | 0.489301084 | 0.301029996 | 0.448487648 |
| ENSMUSG00000104666 | 0 | 0 | 0 | 0 | 0 | 0 | 0 |
| ENSMUSG00000104667 | 0 | 0.5 | 0.396240625 | 0.5 | 0.244650542 | 0.477121255 | 0.448487648 |
| ENSMUSG00000104668 | 0 | 0 | 0 | 0 | 0 | 0 | 0 |
| ENSMUSG00000104669 | 0 | 0.25 | 0.396240625 | 0.25 | 0 | 0 | 0.224243824 |
| ENSMUSG00000104670 | 0 | 0 | 0.5 | 0 | 0.244650542 | 0 | 0 |
| ENSMUSG00000104671 | 0.578129653 | 0.580482024 | 0.580482024 | 0.951838731 | 0.931471446 | 0.77815125 | 0.672731473 |
| ENSMUSG00000104672 | 0 | 0 | 0 | 0 | 0 | 0 | 0 |
| ENSMUSG00000104673 | 0 | 0 | 0 | 0 | 0 | 0 | 0 |
| ENSMUSG00000104674 | 0.671187741 | 0.580482024 | 0.25 | 0.396240625 | 0.244650542 | 0.301029996 | 0.629532004 |
| ENSMUSG00000104675 | 0 | 0 | 0 | 0 | 0 | 0.301029996 | 0 |
| ENSMUSG00000104676 | 0 | 0.25 | 0 | 0.25 | 0.489301084 | 0 | 0 |
| ENSMUSG00000104677 | 0.289064826 | 0 | 0 | 0 | 0.244650542 | 0 | 0 |
| ENSMUSG00000104678 | 0 | 0.396240625 | 0 | 0 | 0 | 0 | 0 |
| ENSMUSG00000104679 | 0 | 0 | 0 | 0 | 0 | 0 | 0 |

| | | | | | | | |
|---|---|---|---|---|---|---|---|
| ENSMUSG00000104680 | 0 | 0 | 0 | 0 | 0 | 0 | 0.224243824 |
| ENSMUSG00000104681 | 0 | 0 | 0 | 0 | 0 | 0 | 0 |
| ENSMUSG00000104682 | 0.960252568 | 0.79248125 | 0.79248125 | 1 | 0.931471446 | 0.903089987 | 0.984950056 |
| ENSMUSG00000104683 | 0 | 0 | 0 | 0 | 0 | 0 | 0 |
| ENSMUSG00000104684 | 0.289064826 | 0 | 0 | 0 | 0 | 0 | 0 |
| ENSMUSG00000104685 | 0.45815691 | 0.396240625 | 0.25 | 0.5 | 0.244650542 | 0.477121255 | 0.448487648 |
| ENSMUSG00000104686 | 0.45815691 | 0.580482024 | 0.646240625 | 0.580482024 | 0.244650542 | 0.477121255 | 0.224243824 |
| ENSMUSG00000104687 | 0.289064826 | 0.25 | 0 | 0 | 0.489301084 | 0 | 0 |
| ENSMUSG00000104688 | 0 | 0 | 0 | 0 | 0 | 0 | 0 |
| ENSMUSG00000104689 | 0 | 0 | 0 | 0 | 0 | 0 | 0 |
| ENSMUSG00000104690 | 0 | 0 | 0 | 0 | 0 | 0 | 0 |
| ENSMUSG00000104691 | 0 | 0 | 0 | 0 | 0 | 0 | 0 |
| ENSMUSG00000104692 | 0 | 0 | 0 | 0 | 0 | 0 | 0 |
| ENSMUSG00000104693 | 0.289064826 | 0 | 0.396240625 | 0.396240625 | 0 | 0.301029996 | 0 |
| ENSMUSG00000104694 | 0 | 0 | 0 | 0 | 0.387761935 | 0 | 0.224243824 |
| ENSMUSG00000104695 | 0.289064826 | 0.25 | 0.396240625 | 0.25 | 0.387761935 | 0.301029996 | 0.355418052 |
| ENSMUSG00000104696 | 0 | 0 | 0 | 0 | 0 | 0 | 0.224243824 |
| ENSMUSG00000104697 | 0 | 0 | 0 | 0 | 0 | 0 | 0 |
| ENSMUSG00000104698 | 0 | 0 | 0 | 0 | 0 | 0 | 0 |
| ENSMUSG00000104699 | 0.45815691 | 0.5 | 0.396240625 | 0.25 | 0.489301084 | 0.477121255 | 0.710836105 |
| ENSMUSG00000104700 | 0 | 0 | 0 | 0 | 0 | 0 | 0 |
| ENSMUSG00000104701 | 0 | 0 | 0 | 0 | 0 | 0 | 0 |
| ENSMUSG00000104702 | 0 | 0 | 0 | 0 | 0 | 0 | 0 |
| ENSMUSG00000104703 | 0 | 0 | 0 | 0 | 0 | 0 | 0 |
| ENSMUSG00000104704 | 0.289064826 | 0.5 | 0.25 | 0.5 | 0.244650542 | 0 | 0.224243824 |
| ENSMUSG00000104705 | 0 | 0 | 0 | 0 | 0 | 0 | 0 |
| ENSMUSG00000104706 | 0.45815691 | 0.701838731 | 0.79248125 | 0.580482024 | 0.244650542 | 0.602059991 | 0.448487648 |
| ENSMUSG00000104707 | 0 | 0 | 0 | 0 | 0 | 0 | 0.224243824 |
| ENSMUSG00000104708 | 1.325351389 | 1.17510993 | 1.188721876 | 1.29248125 | 1.302012594 | 1.531478917 | 1.20139795 |
| ENSMUSG00000104709 | 0 | 0.25 | 0 | 0 | 0 | 0 | 0 |
| ENSMUSG00000104710 | 0 | 0 | 0 | 0 | 0 | 0 | 0 |
| ENSMUSG00000104711 | 0 | 0 | 0 | 0 | 0 | 0 | 0 |
| ENSMUSG00000104712 | 0 | 0 | 0 | 0 | 0 | 0 | 0 |
| ENSMUSG00000104713 | 0 | 0 | 0 | 0 | 0.489301084 | 0 | 0.355418052 |

| | | | | | | | |
|---|---|---|---|---|---|---|---|
| ENSMUSG00000104714 | 0 | 0 | 0 | 0 | 0 | 0 | 0 |
| ENSMUSG00000104715 | 0 | 0 | 0 | 0 | 0 | 0 | 0 |
| ENSMUSG00000104716 | 0 | 0 | 0 | 0 | 0 | 0 | 0 |
| ENSMUSG00000104717 | 0 | 0 | 0 | 0 | 0 | 0 | 0 |
| ENSMUSG00000104718 | 0 | 0 | 0 | 0 | 0 | 0 | 0 |
| ENSMUSG00000104719 | 0.45815691 | 0.5 | 0.5 | 0.25 | 0 | 0.301029996 | 0.355418052 |
| ENSMUSG00000104720 | 0.578129653 | 0.25 | 0.701838731 | 0.75 | 0.686820904 | 0.477121255 | 0.579661877 |
| ENSMUSG00000104721 | 0.45815691 | 0 | 0.25 | 0.396240625 | 0 | 0 | 0.224243824 |
| ENSMUSG00000104722 | 0 | 0 | 0 | 0 | 0 | 0 | 0 |
| ENSMUSG00000104723 | 0 | 0 | 0 | 0 | 0 | 0 | 0 |
| ENSMUSG00000104724 | 0.747221736 | 0.951838731 | 0.976722649 | 0.79248125 | 0.905314583 | 0.903089987 | 0.672731473 |
| ENSMUSG00000104725 | 0.289064826 | 0 | 0 | 0 | 0 | 0 | 0 |
| ENSMUSG00000104726 | 0 | 0 | 0 | 0 | 0 | 0 | 0 |
| ENSMUSG00000104727 | 0 | 0 | 0 | 0 | 0 | 0 | 0 |
| ENSMUSG00000104728 | 0.289064826 | 0.646240625 | 0.79248125 | 0.701838731 | 0.812711509 | 0.698970004 | 0.672731473 |
| ENSMUSG00000104729 | 0 | 0 | 0 | 0 | 0 | 0 | 0 |
| ENSMUSG00000104730 | 0 | 0 | 0 | 0 | 0 | 0 | 0 |
| ENSMUSG00000104731 | 0 | 0 | 0 | 0 | 0 | 0 | 0 |
| ENSMUSG00000104732 | 0 | 0 | 0.25 | 0 | 0 | 0 | 0 |
| ENSMUSG00000104733 | 0 | 0 | 0 | 0 | 0 | 0 | 0 |
| ENSMUSG00000104734 | 0 | 0 | 0 | 0 | 0 | 0 | 0 |
| ENSMUSG00000104735 | 0 | 0 | 0 | 0.25 | 0 | 0 | 0.448487648 |
| ENSMUSG00000104736 | 0 | 0 | 0 | 0 | 0 | 0 | 0 |
| ENSMUSG00000104737 | 0.289064826 | 0.396240625 | 0.25 | 0.25 | 0.244650542 | 0.602059991 | 0.224243824 |
| ENSMUSG00000104738 | 0 | 0 | 0 | 0 | 0 | 0 | 0 |
| ENSMUSG00000104739 | 0 | 0 | 0 | 0 | 0 | 0 | 0 |
| ENSMUSG00000104740 | 0 | 0 | 0 | 0 | 0 | 0 | 0 |
| ENSMUSG00000104741 | 0 | 0 | 0 | 0 | 0 | 0 | 0 |
| ENSMUSG00000104742 | 0 | 0 | 0 | 0 | 0 | 0 | 0 |
| ENSMUSG00000104743 | 0 | 0 | 0 | 0 | 0 | 0 | 0 |
| ENSMUSG00000104744 | 0 | 0 | 0 | 0 | 0 | 0 | 0 |
| ENSMUSG00000104745 | 0 | 0 | 0 | 0 | 0 | 0 | 0.224243824 |
| ENSMUSG00000104746 | 0 | 0 | 0 | 0 | 0 | 0 | 0 |
| ENSMUSG00000104747 | 0 | 0 | 0 | 0 | 0 | 0 | 0 |

| | | | | | | | |
|---|---|---|---|---|---|---|---|
| ENSMUSG00000104748 | 0 | 0 | 0.25 | 0 | 0 | 0 | 0.224243824 |
| ENSMUSG00000104749 | 0 | 0 | 0 | 0 | 0 | 0 | 0 |
| ENSMUSG00000104750 | 0 | 0 | 0 | 0 | 0 | 0 | 0 |
| ENSMUSG00000104751 | 0 | 0 | 0 | 0 | 0 | 0 | 0 |
| ENSMUSG00000104752 | 0 | 0 | 0 | 0 | 0 | 0 | 0 |
| ENSMUSG00000104753 | 0 | 0 | 0 | 0 | 0 | 0 | 0 |
| ENSMUSG00000104754 | 0 | 0 | 0 | 0 | 0 | 0 | 0 |
| ENSMUSG00000104755 | 0 | 0.25 | 0 | 0.25 | 0 | 0 | 0 |
| ENSMUSG00000104756 | 0 | 0.5 | 0.701838731 | 0.580482024 | 0.244650542 | 0 | 0.520678036 |
| ENSMUSG00000104757 | 0 | 0 | 0 | 0 | 0 | 0 | 0 |
| ENSMUSG00000104758 | 0 | 0 | 0 | 0 | 0.244650542 | 0 | 0 |
| ENSMUSG00000104759 | 0 | 0 | 0 | 0 | 0 | 0 | 0 |
| ENSMUSG00000104760 | 0 | 0 | 0 | 0 | 0 | 0 | 0 |
| ENSMUSG00000104761 | 0.91631382 | 1.02186571 | 0.896240625 | 0.79248125 | 1.020174412 | 0.77815125 | 0.829800754 |
| ENSMUSG00000104762 | 0 | 0 | 0 | 0 | 0.244650542 | 0 | 0 |
| ENSMUSG00000104763 | 0 | 0 | 0 | 0 | 0 | 0 | 0 |
| ENSMUSG00000104764 | 0 | 0 | 0 | 0 | 0 | 0 | 0 |
| ENSMUSG00000104765 | 0.289064826 | 0.580482024 | 0.701838731 | 0.92510993 | 0.632412477 | 0.477121255 | 0.74492186 |
| ENSMUSG00000104766 | 0 | 0 | 0.25 | 0 | 0 | 0 | 0 |
| ENSMUSG00000104767 | 0 | 0 | 0 | 0 | 0 | 0 | 0 |
| ENSMUSG00000104768 | 0 | 0.25 | 0 | 0.5 | 0 | 0 | 0 |
| ENSMUSG00000104769 | 0 | 0 | 0 | 0 | 0 | 0 | 0 |
| ENSMUSG00000104770 | 0 | 0 | 0 | 0 | 0 | 0 | 0 |
| ENSMUSG00000104771 | 0 | 0 | 0 | 0 | 0 | 0 | 0 |
| ENSMUSG00000104772 | 0 | 0 | 0 | 0 | 0 | 0 | 0 |
| ENSMUSG00000104773 | 0 | 0 | 0 | 0 | 0 | 0 | 0 |
| ENSMUSG00000104774 | 0 | 0 | 0 | 0 | 0 | 0 | 0 |
| ENSMUSG00000104775 | 0 | 0.25 | 0 | 0.25 | 0 | 0 | 0 |
| ENSMUSG00000104776 | 0 | 0 | 0 | 0 | 0 | 0 | 0 |
| ENSMUSG00000104777 | 0 | 0 | 0 | 0 | 0 | 0 | 0 |
| ENSMUSG00000104778 | 0 | 0 | 0 | 0 | 0 | 0 | 0 |
| ENSMUSG00000104779 | 0 | 0 | 0 | 0 | 0 | 0 | 0 |
| ENSMUSG00000104780 | 0 | 0 | 0 | 0 | 0 | 0 | 0.224243824 |
| ENSMUSG00000104781 | 0 | 0 | 0.25 | 0.25 | 0 | 0.301029996 | 0.224243824 |

| | | | | | | | |
|---|---|---|---|---|---|---|---|
| ENSMUSG00000104782 | 0 | 0 | 0 | 0 | 0 | 0 | 0 |
| ENSMUSG00000104783 | 0 | 0 | 0 | 0 | 0 | 0 | 0 |
| ENSMUSG00000104784 | 0 | 0 | 0 | 0.25 | 0 | 0.301029996 | 0 |
| ENSMUSG00000104785 | 0 | 0 | 0 | 0 | 0 | 0 | 0 |
| ENSMUSG00000104786 | 0.289064826 | 0 | 0 | 0.25 | 0.244650542 | 0.301029996 | 0.224243824 |
| ENSMUSG00000104787 | 0 | 0 | 0 | 0 | 0 | 0 | 0 |
| ENSMUSG00000104788 | 0 | 0.25 | 0 | 0 | 0 | 0.477121255 | 0 |
| ENSMUSG00000104789 | 0 | 0 | 0 | 0 | 0 | 0 | 0 |
| ENSMUSG00000104790 | 0 | 0 | 0 | 0 | 0 | 0 | 0 |
| ENSMUSG00000104791 | 0 | 0 | 0 | 0 | 0 | 0 | 0 |
| ENSMUSG00000104792 | 0 | 0 | 0 | 0 | 0 | 0 | 0 |
| ENSMUSG00000104793 | 0.289064826 | 0.646240625 | 0.646240625 | 0.5 | 0.489301084 | 0.698970004 | 0.355418052 |
| ENSMUSG00000104794 | 0 | 0 | 0 | 0 | 0 | 0 | 0 |
| ENSMUSG00000104795 | 0.289064826 | 0.396240625 | 0.25 | 0.5 | 0.77552387 | 0 | 0.803905701 |
| ENSMUSG00000104796 | 0 | 0 | 0 | 0 | 0 | 0 | 0 |
| ENSMUSG00000104797 | 0 | 0 | 0 | 0 | 0 | 0 | 0 |
| ENSMUSG00000104798 | 0 | 0 | 0 | 0 | 0 | 0 | 0 |
| ENSMUSG00000104799 | 0 | 0 | 0 | 0 | 0 | 0 | 0 |
| ENSMUSG00000104800 | 0 | 0 | 0 | 0 | 0 | 0 | 0 |
| ENSMUSG00000104801 | 0 | 0 | 0 | 0 | 0 | 0 | 0 |
| ENSMUSG00000104802 | 1.325351389 | 1.160964047 | 1.330482024 | 1.29248125 | 1.149965125 | 1.414973348 | 1.21680636 |
| ENSMUSG00000104803 | 0 | 0 | 0 | 0 | 0 | 0 | 0 |
| ENSMUSG00000104804 | 0 | 0 | 0 | 0 | 0 | 0 | 0 |
| ENSMUSG00000104805 | 0 | 0 | 0 | 0 | 0 | 0 | 0 |
| ENSMUSG00000104806 | 0.671187741 | 0.864857905 | 0.75 | 0.830482024 | 0.812711509 | 1 | 0.579661877 |
| ENSMUSG00000104807 | 0 | 0 | 0 | 0 | 0 | 0 | 0 |
| ENSMUSG00000104808 | 0 | 0 | 0 | 0 | 0 | 0 | 0 |
| ENSMUSG00000104809 | 0 | 0 | 0 | 0 | 0 | 0 | 0 |
| ENSMUSG00000104810 | 0 | 0 | 0 | 0 | 0 | 0 | 0 |
| ENSMUSG00000104811 | 0 | 0 | 0 | 0 | 0 | 0 | 0 |
| ENSMUSG00000104812 | 0 | 0 | 0 | 0 | 0 | 0 | 0 |
| ENSMUSG00000104813 | 0 | 0 | 0 | 0 | 0 | 0 | 0 |
| ENSMUSG00000104814 | 1.847794126 | 1.622963274 | 1.646240625 | 1.584962501 | 1.584291694 | 1.491361694 | 1.345462945 |
| ENSMUSG00000104815 | 0 | 0 | 0 | 0 | 0 | 0 | 0 |

| | | | | | | | |
|---|---|---|---|---|---|---|---|
| ENSMUSG00000104816 | 0 | 0.25 | 0 | 0.25 | 0 | 0 | 0.355418052 |
| ENSMUSG00000104817 | 0 | 0 | 0 | 0 | 0 | 0 | 0 |
| ENSMUSG00000104818 | 0 | 0.5 | 0.396240625 | 0.396240625 | 0.489301084 | 0.301029996 | 0 |
| ENSMUSG00000104819 | 0 | 0 | 0 | 0 | 0 | 0 | 0 |
| ENSMUSG00000104820 | 0 | 0 | 0 | 0 | 0 | 0 | 0 |
| ENSMUSG00000104821 | 0.289064826 | 0.396240625 | 0 | 0 | 0.244650542 | 0.301029996 | 0 |
| ENSMUSG00000104822 | 0.578129653 | 0.25 | 0.75 | 0.580482024 | 0.812711509 | 0.602059991 | 0.520678036 |
| ENSMUSG00000104823 | 0 | 0 | 0 | 0 | 0 | 0 | 0 |
| ENSMUSG00000104824 | 0 | 0 | 0 | 0 | 0 | 0 | 0 |
| ENSMUSG00000104825 | 0 | 0 | 0 | 0 | 0 | 0 | 0 |
| ENSMUSG00000104826 | 0 | 0 | 0 | 0 | 0 | 0 | 0 |
| ENSMUSG00000104827 | 0 | 0 | 0 | 0 | 0 | 0 | 0 |
| ENSMUSG00000104828 | 0.45815691 | 0 | 0 | 0 | 0 | 0 | 0 |
| ENSMUSG00000104829 | 0 | 0 | 0 | 0 | 0 | 0 | 0 |
| ENSMUSG00000104830 | 0.960252568 | 0.580482024 | 0.79248125 | 0.646240625 | 0.489301084 | 0.602059991 | 0.629532004 |
| ENSMUSG00000104831 | 0 | 0 | 0 | 0 | 0 | 0 | 0 |
| ENSMUSG00000104832 | 0 | 0 | 0 | 0 | 0 | 0 | 0 |
| ENSMUSG00000104833 | 0 | 0 | 0 | 0 | 0 | 0 | 0 |
| ENSMUSG00000104834 | 0 | 0 | 0 | 0 | 0 | 0 | 0 |
| ENSMUSG00000104835 | 0 | 0 | 0 | 0 | 0 | 0 | 0 |
| ENSMUSG00000104836 | 0 | 0 | 0 | 0 | 0 | 0 | 0 |
| ENSMUSG00000104837 | 0.811507563 | 0.896240625 | 1.061981878 | 1.17510993 | 1.091002363 | 1.146128036 | 0.969165684 |
| ENSMUSG00000104838 | 0 | 0 | 0 | 0 | 0 | 0 | 0 |
| ENSMUSG00000104839 | 0 | 0 | 0 | 0 | 0 | 0 | 0 |
| ENSMUSG00000104840 | 0 | 0 | 0 | 0 | 0 | 0 | 0 |
| ENSMUSG00000104841 | 0 | 0.25 | 0 | 0 | 0 | 0 | 0 |
| ENSMUSG00000104842 | 0 | 0 | 0 | 0 | 0 | 0 | 0 |
| ENSMUSG00000104843 | 0.811507563 | 0.646240625 | 0.5 | 0.646240625 | 0.733951626 | 0.698970004 | 0.629532004 |
| ENSMUSG00000104844 | 0 | 0 | 0 | 0 | 0 | 0 | 0 |
| ENSMUSG00000104845 | 0 | 0 | 0 | 0 | 0 | 0 | 0 |
| ENSMUSG00000104846 | 0 | 0 | 0 | 0 | 0 | 0 | 0 |
| ENSMUSG00000104847 | 0 | 0 | 0 | 0 | 0 | 0 | 0 |
| ENSMUSG00000104848 | 0 | 0 | 0 | 0 | 0 | 0 | 0 |
| ENSMUSG00000104849 | 0 | 0 | 0 | 0 | 0 | 0 | 0 |

| | | | | | | | |
|---|---|---|---|---|---|---|---|
| ENSMUSG00000104850 | 0.45815691 | 0.396240625 | 0.580482024 | 0.25 | 0.244650542 | 0 | 0.224243824 |
| ENSMUSG00000104851 | 0 | 0 | 0 | 0 | 0 | 0 | 0 |
| ENSMUSG00000104852 | 0.289064826 | 0.25 | 0.5 | 0.396240625 | 0.244650542 | 0.477121255 | 0.448487648 |
| ENSMUSG00000104853 | 0 | 0 | 0 | 0 | 0 | 0 | 0 |
| ENSMUSG00000104854 | 0 | 0 | 0 | 0.25 | 0 | 0 | 0 |
| ENSMUSG00000104855 | 0 | 0 | 0 | 0 | 0 | 0 | 0 |
| ENSMUSG00000104856 | 0 | 0 | 0 | 0 | 0 | 0 | 0 |
| ENSMUSG00000104857 | 0 | 0 | 0 | 0 | 0 | 0 | 0 |
| ENSMUSG00000104858 | 0 | 0 | 0 | 0 | 0 | 0 | 0 |
| ENSMUSG00000104859 | 0.289064826 | 0.25 | 0 | 0 | 0.387761935 | 0 | 0 |
| ENSMUSG00000104860 | 0.289064826 | 0 | 0 | 0.396240625 | 0 | 0.301029996 | 0.224243824 |
| ENSMUSG00000104861 | 1.693336259 | 1.685366747 | 1.747171172 | 1.593759858 | 1.595992969 | 1.73239376 | 1.479989695 |
| ENSMUSG00000104862 | 0.289064826 | 0.25 | 0 | 0 | 0 | 0 | 0.355418052 |
| ENSMUSG00000104863 | 0 | 0 | 0 | 0 | 0 | 0 | 0 |
| ENSMUSG00000104864 | 0.578129653 | 0.25 | 0.25 | 0 | 0.489301084 | 0 | 0.224243824 |
| ENSMUSG00000104865 | 0 | 0 | 0 | 0 | 0 | 0 | 0 |
| ENSMUSG00000104866 | 0 | 0 | 0 | 0 | 0 | 0 | 0 |
| ENSMUSG00000104867 | 0 | 0.25 | 0.5 | 0.396240625 | 0.244650542 | 0.301029996 | 0 |
| ENSMUSG00000104868 | 0 | 0 | 0 | 0 | 0 | 0 | 0 |
| ENSMUSG00000104869 | 0.45815691 | 0.396240625 | 0.25 | 0 | 0 | 0 | 0.355418052 |
| ENSMUSG00000104870 | 0 | 0 | 0 | 0 | 0 | 0 | 0 |
| ENSMUSG00000104871 | 0 | 0.25 | 0.25 | 0 | 0 | 0 | 0.224243824 |
| ENSMUSG00000104872 | 0.289064826 | 0.5 | 0.396240625 | 0.396240625 | 0.244650542 | 0.301029996 | 0.224243824 |
| ENSMUSG00000104873 | 0 | 0.396240625 | 0.25 | 0 | 0 | 0 | 0.224243824 |
| ENSMUSG00000104874 | 0 | 0.25 | 0 | 0.25 | 0 | 0 | 0 |
| ENSMUSG00000104875 | 0 | 0 | 0 | 0 | 0 | 0 | 0 |
| ENSMUSG00000104876 | 0 | 0 | 0 | 0 | 0 | 0 | 0 |
| ENSMUSG00000104877 | 0.289064826 | 0.25 | 0 | 0 | 0.244650542 | 0.301029996 | 0.224243824 |
| ENSMUSG00000104878 | 0 | 0 | 0 | 0 | 0 | 0 | 0 |
| ENSMUSG00000104879 | 0 | 0 | 0 | 0 | 0 | 0 | 0 |
| ENSMUSG00000104880 | 0 | 0 | 0 | 0 | 0 | 0 | 0 |
| ENSMUSG00000104881 | 0 | 0 | 0 | 0.25 | 0 | 0.301029996 | 0 |
| ENSMUSG00000104882 | 0 | 0 | 0 | 0.25 | 0 | 0 | 0 |
| ENSMUSG00000104883 | 0 | 0 | 0 | 0 | 0 | 0 | 0 |

| | | | | | | | |
|---|---|---|---|---|---|---|---|
| ENSMUSG00000104884 | 0 | 0 | 0 | 0 | 0 | 0 | 0 |
| ENSMUSG00000104885 | 0 | 0 | 0.396240625 | 0 | 0.387761935 | 0 | 0.224243824 |
| ENSMUSG00000104886 | 0 | 0 | 0 | 0 | 0 | 0 | 0 |
| ENSMUSG00000104887 | 0 | 0 | 0 | 0 | 0 | 0 | 0 |
| ENSMUSG00000104888 | 0 | 0 | 0 | 0.25 | 0 | 0 | 0.355418052 |
| ENSMUSG00000104889 | 0 | 0 | 0 | 0 | 0 | 0 | 0 |
| ENSMUSG00000104890 | 0 | 0 | 0 | 0 | 0 | 0 | 0 |
| ENSMUSG00000104891 | 0.91631382 | 0.646240625 | 0.830482024 | 0.79248125 | 0.568060967 | 0.602059991 | 0.775756176 |
| ENSMUSG00000104892 | 0 | 0.25 | 0 | 0 | 0 | 0 | 0 |
| ENSMUSG00000104893 | 0 | 0.580482024 | 0 | 0 | 0.244650542 | 0.301029996 | 0.224243824 |
| ENSMUSG00000104894 | 0 | 0 | 0 | 0 | 0 | 0 | 0 |
| ENSMUSG00000104895 | 0 | 0 | 0 | 0.25 | 0 | 0 | 0 |
| ENSMUSG00000104896 | 0 | 0 | 0 | 0 | 0 | 0 | 0 |
| ENSMUSG00000104897 | 0.289064826 | 0.5 | 0.580482024 | 0.5 | 0.489301084 | 0.602059991 | 0.355418052 |
| ENSMUSG00000104898 | 0 | 0 | 0 | 0 | 0 | 0 | 0 |
| ENSMUSG00000104899 | 0 | 0 | 0 | 0 | 0 | 0 | 0 |
| ENSMUSG00000104900 | 0 | 0 | 0 | 0 | 0 | 0 | 0 |
| ENSMUSG00000104901 | 0 | 0 | 0 | 0 | 0 | 0 | 0 |
| ENSMUSG00000104902 | 0 | 0 | 0 | 0 | 0 | 0 | 0 |
| ENSMUSG00000104903 | 0 | 0 | 0 | 0 | 0.387761935 | 0 | 0.224243824 |
| ENSMUSG00000104904 | 0.289064826 | 0.25 | 0.580482024 | 0 | 0.387761935 | 0.301029996 | 0 |
| ENSMUSG00000104905 | 0 | 0 | 0 | 0 | 0 | 0 | 0 |
| ENSMUSG00000104906 | 0 | 0 | 0 | 0 | 0 | 0 | 0.224243824 |
| ENSMUSG00000104908 | 0 | 0 | 0 | 0 | 0 | 0 | 0 |
| ENSMUSG00000104909 | 0 | 0 | 0 | 0 | 0 | 0 | 0 |
| ENSMUSG00000104910 | 0 | 0.25 | 0 | 0 | 0.244650542 | 0 | 0.224243824 |
| ENSMUSG00000104911 | 0 | 0 | 0 | 0 | 0 | 0 | 0 |
| ENSMUSG00000104912 | 0.289064826 | 0 | 0.396240625 | 0.25 | 0 | 0 | 0 |
| ENSMUSG00000104913 | 0.811507563 | 0.79248125 | 0.830482024 | 0.5 | 0.686820904 | 0.954242509 | 0.829800754 |
| ENSMUSG00000104914 | 0.289064826 | 0 | 0 | 0 | 0 | 0.301029996 | 0.224243824 |
| ENSMUSG00000104915 | 0 | 0 | 0 | 0 | 0 | 0 | 0 |
| ENSMUSG00000104916 | 0 | 0 | 0.25 | 0 | 0 | 0 | 0 |
| ENSMUSG00000104917 | 0 | 0.580482024 | 0.25 | 0 | 0.244650542 | 0 | 0.448487648 |
| ENSMUSG00000104918 | 0 | 0.25 | 0.25 | 0.25 | 0 | 0 | 0 |

| | | | | | | | |
|---|---|---|---|---|---|---|---|
| ENSMUSG00000104919 | 0 | 0 | 0 | 0 | 0 | 0 | 0 |
| ENSMUSG00000104920 | 0 | 0.25 | 0 | 0 | 0 | 0 | 0.224243824 |
| ENSMUSG00000104921 | 0 | 0 | 0 | 0 | 0 | 0 | 0 |
| ENSMUSG00000104922 | 0 | 0 | 0 | 0 | 0 | 0 | 0 |
| ENSMUSG00000104923 | 0 | 0 | 0 | 0 | 0 | 0.301029996 | 0 |
| ENSMUSG00000104924 | 0.289064826 | 0 | 0 | 0.25 | 0 | 0.301029996 | 0.355418052 |
| ENSMUSG00000104925 | 0.578129653 | 0.580482024 | 0.396240625 | 0.5 | 0.489301084 | 0.477121255 | 0.224243824 |
| ENSMUSG00000104926 | 0 | 0 | 0 | 0 | 0.244650542 | 0 | 0 |
| ENSMUSG00000104927 | 0 | 0 | 0 | 0 | 0 | 0 | 0 |
| ENSMUSG00000104928 | 0 | 0 | 0 | 0 | 0 | 0 | 0 |
| ENSMUSG00000104929 | 0 | 0 | 0 | 0 | 0 | 0 | 0 |
| ENSMUSG00000104930 | 0 | 0 | 0 | 0 | 0 | 0 | 0 |
| ENSMUSG00000104931 | 0 | 0 | 0 | 0 | 0 | 0 | 0 |
| ENSMUSG00000104932 | 0.289064826 | 0.396240625 | 0.25 | 0 | 0 | 0 | 0.224243824 |
| ENSMUSG00000104933 | 0 | 0 | 0 | 0 | 0 | 0 | 0 |
| ENSMUSG00000104934 | 0 | 0 | 0 | 0 | 0 | 0 | 0 |
| ENSMUSG00000104935 | 0 | 0 | 0 | 0 | 0 | 0 | 0 |
| ENSMUSG00000104936 | 0 | 0 | 0 | 0 | 0 | 0 | 0 |
| ENSMUSG00000104937 | 0.289064826 | 0 | 0.396240625 | 0 | 0 | 0 | 0 |
| ENSMUSG00000104938 | 0 | 0 | 0 | 0.25 | 0.244650542 | 0.301029996 | 0.224243824 |
| ENSMUSG00000104939 | 0 | 0 | 0 | 0 | 0 | 0 | 0 |
| ENSMUSG00000104940 | 0.45815691 | 0 | 0 | 0 | 0 | 0 | 0 |
| ENSMUSG00000104941 | 0 | 0 | 0 | 0 | 0 | 0 | 0 |
| ENSMUSG00000104942 | 0 | 0 | 0 | 0 | 0 | 0 | 0 |
| ENSMUSG00000104943 | 0.45815691 | 0.25 | 0 | 0.25 | 0.387761935 | 0.477121255 | 0.448487648 |
| ENSMUSG00000104944 | 0 | 0 | 0 | 0 | 0 | 0 | 0 |
| ENSMUSG00000104945 | 0 | 0 | 0 | 0.25 | 0 | 0 | 0 |
| ENSMUSG00000104946 | 0 | 0 | 0 | 0 | 0 | 0 | 0 |
| ENSMUSG00000104947 | 0 | 0 | 0 | 0.25 | 0 | 0 | 0 |
| ENSMUSG00000104948 | 0 | 0 | 0 | 0 | 0 | 0 | 0 |
| ENSMUSG00000104949 | 0 | 0 | 0 | 0 | 0 | 0 | 0 |
| ENSMUSG00000104950 | 1.129344651 | 1.130890489 | 1.098079356 | 1.17510993 | 1 | 1.204119983 | 0.803905701 |
| ENSMUSG00000104951 | 0.811507563 | 0.25 | 0.701838731 | 0.5 | 0.244650542 | 0.77815125 | 0.520678036 |
| ENSMUSG00000104952 | 0 | 0 | 0 | 0 | 0 | 0 | 0 |

| | | | | | | | |
|---|---|---|---|---|---|---|---|
| ENSMUSG00000104953 | 0 | 0 | 0 | 0 | 0 | 0 | 0 |
| ENSMUSG00000104954 | 0 | 0 | 0 | 0 | 0 | 0 | 0 |
| ENSMUSG00000104955 | 0.289064826 | 0 | 0 | 0 | 0 | 0 | 0.224243824 |
| ENSMUSG00000104956 | 0 | 0 | 0 | 0 | 0 | 0 | 0 |
| ENSMUSG00000104957 | 0 | 0 | 0 | 0 | 0 | 0 | 0 |
| ENSMUSG00000104958 | 0 | 0 | 0 | 0 | 0 | 0 | 0 |
| ENSMUSG00000104959 | 0 | 0 | 0 | 0 | 0 | 0 | 0 |
| ENSMUSG00000104960 | 1.700465191 | 1.75 | 1.921625132 | 1.42510993 | 1.66856048 | 1.591064607 | 1.759302692 |
| ENSMUSG00000104961 | 0 | 0 | 0 | 0 | 0 | 0 | 0 |
| ENSMUSG00000104962 | 0 | 0 | 0 | 0 | 0 | 0 | 0 |
| ENSMUSG00000104963 | 0.289064826 | 0 | 0 | 0.25 | 0.489301084 | 0 | 0.448487648 |
| ENSMUSG00000104964 | 0 | 0 | 0 | 0 | 0 | 0 | 0 |
| ENSMUSG00000104965 | 1.100572389 | 0.580482024 | 0.92510993 | 1.114857905 | 1.212046813 | 1.380211242 | 1.20139795 |
| ENSMUSG00000104966 | 0 | 0 | 0 | 0 | 0 | 0 | 0 |
| ENSMUSG00000104967 | 0.289064826 | 0 | 0 | 0 | 0.244650542 | 0 | 0.224243824 |
| ENSMUSG00000104968 | 0 | 0 | 0 | 0 | 0 | 0 | 0 |
| ENSMUSG00000104969 | 0.578129653 | 0.79248125 | 0 | 0.396240625 | 0.568060967 | 0.477121255 | 0 |
| ENSMUSG00000104970 | 0 | 0 | 0 | 0 | 0 | 0 | 0 |
| ENSMUSG00000104971 | 0 | 0 | 0 | 0.25 | 0 | 0 | 0 |
| ENSMUSG00000104972 | 0 | 0.25 | 0 | 0 | 0 | 0 | 0 |
| ENSMUSG00000104973 | 0.289064826 | 0 | 0 | 0.25 | 0 | 0.301029996 | 0.448487648 |
| ENSMUSG00000104974 | 0 | 0 | 0 | 0 | 0 | 0 | 0 |
| ENSMUSG00000104975 | 0 | 0 | 0 | 0 | 0 | 0 | 0 |
| ENSMUSG00000104976 | 0 | 0 | 0 | 0 | 0 | 0 | 0 |
| ENSMUSG00000104978 | 0 | 0 | 0 | 0 | 0 | 0 | 0 |
| ENSMUSG00000104979 | 0 | 0 | 0 | 0 | 0 | 0 | 0 |
| ENSMUSG00000104980 | 0 | 0 | 0 | 0 | 0 | 0 | 0 |
| ENSMUSG00000104981 | 0 | 0 | 0 | 0 | 0 | 0 | 0 |
| ENSMUSG00000104982 | 0 | 0 | 0 | 0 | 0 | 0 | 0 |
| ENSMUSG00000104983 | 0 | 0 | 0 | 0 | 0 | 0.301029996 | 0 |
| ENSMUSG00000104984 | 0 | 0 | 0 | 0 | 0 | 0 | 0 |
| ENSMUSG00000104985 | 0 | 0 | 0 | 0 | 0 | 0 | 0 |
| ENSMUSG00000104986 | 0.289064826 | 0.25 | 0.396240625 | 0.5 | 0.489301084 | 0.477121255 | 0.448487648 |
| ENSMUSG00000104987 | 0 | 0 | 0 | 0 | 0 | 0 | 0 |

| | | | | | | | |
|---|---|---|---|---|---|---|---|
| ENSMUSG00000104988 | 1.342375483 | 1.311981878 | 1.25 | 1.356566189 | 1.327538614 | 1.301029996 | 1.176815335 |
| ENSMUSG00000104989 | 0 | 0 | 0 | 0 | 0 | 0 | 0 |
| ENSMUSG00000104990 | 0 | 0.25 | 0 | 0 | 0.244650542 | 0 | 0 |
| ENSMUSG00000104991 | 0 | 0 | 0 | 0 | 0 | 0 | 0 |
| ENSMUSG00000104992 | 0 | 0 | 0 | 0 | 0 | 0 | 0 |
| ENSMUSG00000104993 | 0 | 0 | 0 | 0 | 0 | 0 | 0 |
| ENSMUSG00000104994 | 0 | 0 | 0 | 0 | 0 | 0 | 0 |
| ENSMUSG00000104995 | 0 | 0 | 0 | 0 | 0 | 0 | 0 |
| ENSMUSG00000104996 | 0 | 0 | 0 | 0 | 0 | 0 | 0 |
| ENSMUSG00000104997 | 0 | 0 | 0 | 0 | 0 | 0 | 0 |
| ENSMUSG00000104998 | 0 | 0 | 0 | 0 | 0 | 0 | 0 |
| ENSMUSG00000104999 | 0 | 0 | 0.396240625 | 0 | 0.244650542 | 0 | 0 |
| ENSMUSG00000105000 | 0 | 0 | 0 | 0 | 0 | 0 | 0 |
| ENSMUSG00000105001 | 0 | 0 | 0 | 0 | 0 | 0 | 0 |
| ENSMUSG00000105002 | 0 | 0 | 0 | 0 | 0 | 0 | 0 |
| ENSMUSG00000105003 | 0 | 0.25 | 0 | 0 | 0 | 0 | 0 |
| ENSMUSG00000105004 | 0.289064826 | 0.396240625 | 0.25 | 0.396240625 | 0.244650542 | 0 | 0.224243824 |
| ENSMUSG00000105005 | 0 | 0 | 0 | 0 | 0 | 0 | 0 |
| ENSMUSG00000105006 | 0.45815691 | 0.25 | 0.396240625 | 0 | 0.244650542 | 0.301029996 | 0 |
| ENSMUSG00000105007 | 0 | 0 | 0 | 0 | 0 | 0 | 0 |
| ENSMUSG00000105008 | 0.289064826 | 0.25 | 0.25 | 0.25 | 0 | 0 | 0 |
| ENSMUSG00000105009 | 0 | 0 | 0 | 0 | 0 | 0 | 0 |
| ENSMUSG00000105010 | 0 | 0 | 0 | 0 | 0 | 0 | 0 |
| ENSMUSG00000105011 | 0 | 0 | 0.25 | 0.25 | 0 | 0.477121255 | 0.224243824 |
| ENSMUSG00000105012 | 0 | 0 | 0 | 0 | 0 | 0 | 0 |
| ENSMUSG00000105013 | 0 | 0 | 0 | 0 | 0 | 0 | 0 |
| ENSMUSG00000105014 | 0 | 0 | 0 | 0 | 0 | 0 | 0 |
| ENSMUSG00000105015 | 0 | 0 | 0 | 0 | 0.244650542 | 0 | 0 |
| ENSMUSG00000105016 | 0 | 0 | 0 | 0 | 0 | 0 | 0 |
| ENSMUSG00000105017 | 0 | 0 | 0 | 0 | 0 | 0 | 0 |
| ENSMUSG00000105018 | 0 | 0 | 0 | 0 | 0 | 0 | 0 |
| ENSMUSG00000105019 | 0 | 0 | 0 | 0.25 | 0 | 0 | 0 |
| ENSMUSG00000105020 | 0 | 0 | 0 | 0 | 0 | 0 | 0 |
| ENSMUSG00000105021 | 0 | 0 | 0 | 0.646240625 | 0.632412477 | 0.77815125 | 0.355418052 |

| | | | | | | | |
|---|---|---|---|---|---|---|---|
| ENSMUSG00000105022 | 0.578129653 | 0.646240625 | 0.25 | 0.396240625 | 0.387761935 | 0.477121255 | 0.448487648 |
| ENSMUSG00000105023 | 0 | 0 | 0 | 0 | 0 | 0 | 0 |
| ENSMUSG00000105024 | 0 | 0 | 0 | 0 | 0 | 0 | 0 |
| ENSMUSG00000105025 | 0 | 0 | 0 | 0 | 0 | 0 | 0 |
| ENSMUSG00000105026 | 0 | 0 | 0 | 0 | 0 | 0 | 0 |
| ENSMUSG00000105027 | 0 | 0 | 0 | 0 | 0 | 0 | 0 |
| ENSMUSG00000105028 | 0 | 0 | 0 | 0 | 0 | 0 | 0 |
| ENSMUSG00000105029 | 0 | 0 | 0 | 0 | 0 | 0 | 0 |
| ENSMUSG00000105030 | 0 | 0 | 0 | 0 | 0 | 0 | 0 |
| ENSMUSG00000105031 | 0 | 0 | 0 | 0 | 0 | 0.301029996 | 0.224243824 |
| ENSMUSG00000105032 | 0 | 0 | 0 | 0 | 0 | 0 | 0 |
| ENSMUSG00000105033 | 0 | 0.5 | 0.396240625 | 0.25 | 0 | 0.602059991 | 0.355418052 |
| ENSMUSG00000105034 | 0 | 0 | 0 | 0 | 0 | 0 | 0 |
| ENSMUSG00000105035 | 0 | 0 | 0 | 0 | 0 | 0 | 0 |
| ENSMUSG00000105036 | 0 | 0.25 | 0 | 0 | 0 | 0 | 0 |
| ENSMUSG00000105037 | 0 | 0 | 0 | 0 | 0 | 0 | 0 |
| ENSMUSG00000105038 | 0 | 0 | 0 | 0 | 0 | 0 | 0 |
| ENSMUSG00000105039 | 0 | 0 | 0 | 0.25 | 0 | 0 | 0.355418052 |
| ENSMUSG00000105040 | 0 | 0.396240625 | 0 | 0.396240625 | 0 | 0 | 0.224243824 |
| ENSMUSG00000105041 | 0 | 0 | 0 | 0 | 0 | 0 | 0.355418052 |
| ENSMUSG00000105042 | 0.747221736 | 0.646240625 | 0.646240625 | 0.79248125 | 0.812711509 | 0.77815125 | 0.672731473 |
| ENSMUSG00000105043 | 0 | 0 | 0 | 0 | 0 | 0 | 0 |
| ENSMUSG00000105044 | 0.578129653 | 0.5 | 0.75 | 0.75 | 0.77552387 | 0.77815125 | 0.74492186 |
| ENSMUSG00000105045 | 0.289064826 | 0.25 | 0.25 | 0.646240625 | 0.244650542 | 0.602059991 | 0.579661877 |
| ENSMUSG00000105046 | 0 | 0 | 0 | 0 | 0 | 0 | 0 |
| ENSMUSG00000105047 | 0 | 0 | 0 | 0 | 0 | 0 | 0 |
| ENSMUSG00000105048 | 0 | 0 | 0 | 0 | 0 | 0 | 0 |
| ENSMUSG00000105049 | 0 | 0 | 0 | 0 | 0 | 0.301029996 | 0 |
| ENSMUSG00000105050 | 0 | 0 | 0 | 0 | 0 | 0 | 0 |
| ENSMUSG00000105051 | 0 | 0 | 0 | 0 | 0 | 0 | 0.224243824 |
| ENSMUSG00000105052 | 0 | 0 | 0 | 0 | 0 | 0 | 0 |
| ENSMUSG00000105053 | 0 | 0 | 0.25 | 0 | 0 | 0 | 0 |
| ENSMUSG00000105054 | 0 | 0 | 0 | 0 | 0 | 0 | 0 |
| ENSMUSG00000105055 | 0 | 0 | 0 | 0 | 0 | 0 | 0 |

| | | | | | | | |
|---|---|---|---|---|---|---|---|
| ENSMUSG00000105056 | 0 | 0 | 0 | 0 | 0 | 0 | 0 |
| ENSMUSG00000105057 | 0 | 0 | 0 | 0 | 0 | 0 | 0 |
| ENSMUSG00000105058 | 0 | 0 | 0 | 0 | 0 | 0 | 0 |
| ENSMUSG00000105059 | 0 | 0 | 0 | 0 | 0 | 0 | 0 |
| ENSMUSG00000105060 | 0 | 0 | 0 | 0 | 0 | 0 | 0 |
| ENSMUSG00000105061 | 0 | 0 | 0 | 0 | 0 | 0 | 0 |
| ENSMUSG00000105062 | 0.45815691 | 0 | 0 | 0 | 0 | 0.301029996 | 0 |
| ENSMUSG00000105063 | 0.289064826 | 0.396240625 | 0 | 0 | 0.489301084 | 0.301029996 | 0.355418052 |
| ENSMUSG00000105064 | 0 | 0 | 0 | 0 | 0.244650542 | 0 | 0 |
| ENSMUSG00000105065 | 1.342375483 | 1.29248125 | 1.282320754 | 1.226722649 | 1.136121934 | 1.230448921 | 1.131174228 |
| ENSMUSG00000105066 | 0 | 0 | 0 | 0 | 0 | 0 | 0 |
| ENSMUSG00000105067 | 0 | 0 | 0 | 0 | 0 | 0 | 0 |
| ENSMUSG00000105068 | 0 | 0 | 0 | 0 | 0 | 0 | 0 |
| ENSMUSG00000105070 | 0 | 0 | 0 | 0 | 0 | 0 | 0 |
| ENSMUSG00000105071 | 0.45815691 | 0.25 | 0.646240625 | 0.5 | 0.244650542 | 0.301029996 | 0.355418052 |
| ENSMUSG00000105072 | 0 | 0 | 0 | 0 | 0 | 0 | 0 |
| ENSMUSG00000105073 | 0 | 0 | 0 | 0 | 0 | 0 | 0 |
| ENSMUSG00000105074 | 0 | 0 | 0 | 0 | 0 | 0 | 0 |
| ENSMUSG00000105075 | 0 | 0 | 0 | 0 | 0 | 0 | 0 |
| ENSMUSG00000105076 | 0 | 0.396240625 | 0 | 0 | 0.244650542 | 0.301029996 | 0 |
| ENSMUSG00000105077 | 0.45815691 | 0.396240625 | 0.25 | 0 | 0.387761935 | 0.477121255 | 0.629532004 |
| ENSMUSG00000105078 | 0 | 0 | 0 | 0 | 0 | 0 | 0 |
| ENSMUSG00000105079 | 0 | 0 | 0 | 0 | 0 | 0.477121255 | 0.355418052 |
| ENSMUSG00000105080 | 0 | 0 | 0 | 0 | 0 | 0 | 0 |
| ENSMUSG00000105081 | 0 | 0 | 0 | 0 | 0 | 0 | 0 |
| ENSMUSG00000105082 | 0.289064826 | 0 | 0.25 | 0 | 0.387761935 | 0.301029996 | 0.224243824 |
| ENSMUSG00000105083 | 0.45815691 | 0.75 | 0.75 | 0.75 | 0.387761935 | 0.477121255 | 0.520678036 |
| ENSMUSG00000105084 | 0.289064826 | 0 | 0 | 0 | 0.244650542 | 0 | 0 |
| ENSMUSG00000105085 | 0.289064826 | 0 | 0 | 0 | 0 | 0 | 0 |
| ENSMUSG00000105086 | 0 | 0 | 0 | 0 | 0 | 0 | 0 |
| ENSMUSG00000105087 | 0 | 0 | 0 | 0 | 0 | 0 | 0 |
| ENSMUSG00000105088 | 0 | 0 | 0 | 0 | 0 | 0 | 0 |
| ENSMUSG00000105089 | 0 | 0 | 0 | 0 | 0 | 0 | 0 |
| ENSMUSG00000105090 | 0 | 0 | 0 | 0 | 0 | 0 | 0 |

| | | | | | | | |
|---|---|---|---|---|---|---|---|
| ENSMUSG00000105091 | 0 | 0 | 0 | 0 | 0 | 0 | 0 |
| ENSMUSG00000105092 | 0 | 0 | 0 | 0 | 0 | 0 | 0 |
| ENSMUSG00000105093 | 0 | 0 | 0 | 0.5 | 0.244650542 | 0.301029996 | 0.355418052 |
| ENSMUSG00000105094 | 0 | 0 | 0 | 0 | 0 | 0 | 0 |
| ENSMUSG00000105095 | 0.45815691 | 0.830482024 | 0.79248125 | 0.75 | 0.632412477 | 0.477121255 | 0.74492186 |
| ENSMUSG00000105096 | 0 | 0 | 0 | 0 | 0 | 0 | 0 |
| ENSMUSG00000105097 | 0 | 0 | 0 | 0 | 0 | 0 | 0 |
| ENSMUSG00000105098 | 0 | 0 | 0 | 0 | 0 | 0 | 0 |
| ENSMUSG00000105099 | 0 | 0 | 0 | 0 | 0 | 0 | 0 |
| ENSMUSG00000105100 | 0 | 0.25 | 0 | 0 | 0.244650542 | 0 | 0.224243824 |
| ENSMUSG00000105101 | 0 | 0 | 0 | 0 | 0 | 0 | 0 |
| ENSMUSG00000105102 | 0 | 0 | 0 | 0 | 0.244650542 | 0 | 0.224243824 |
| ENSMUSG00000105103 | 0 | 0 | 0 | 0 | 0 | 0 | 0 |
| ENSMUSG00000105104 | 0 | 0.25 | 0.25 | 0 | 0 | 0 | 0 |
| ENSMUSG00000105105 | 0 | 0 | 0 | 0 | 0 | 0 | 0 |
| ENSMUSG00000105106 | 0 | 0 | 0 | 0 | 0 | 0 | 0 |
| ENSMUSG00000105107 | 0 | 0.580482024 | 0.396240625 | 0.580482024 | 0.387761935 | 0.84509804 | 0.629532004 |
| ENSMUSG00000105108 | 0 | 0 | 0 | 0 | 0 | 0 | 0 |
| ENSMUSG00000105109 | 0 | 0 | 0 | 0 | 0 | 0 | 0 |
| ENSMUSG00000105110 | 0 | 0 | 0 | 0 | 0 | 0 | 0 |
| ENSMUSG00000105111 | 0 | 0 | 0 | 0 | 0 | 0 | 0 |
| ENSMUSG00000105112 | 1.249317394 | 1.282320754 | 1.17510993 | 1.061981878 | 0.955822902 | 0.954242509 | 0.896975297 |
| ENSMUSG00000105113 | 0 | 0 | 0 | 0 | 0 | 0 | 0 |
| ENSMUSG00000105114 | 0 | 0 | 0 | 0 | 0 | 0 | 0 |
| ENSMUSG00000105115 | 0 | 0 | 0 | 0 | 0 | 0 | 0 |
| ENSMUSG00000105116 | 0 | 0 | 0 | 0 | 0 | 0 | 0 |
| ENSMUSG00000105117 | 0 | 0 | 0 | 0 | 0 | 0 | 0 |
| ENSMUSG00000105118 | 0 | 0 | 0 | 0 | 0 | 0 | 0 |
| ENSMUSG00000105119 | 0 | 0 | 0.396240625 | 0 | 0.244650542 | 0.477121255 | 0.355418052 |
| ENSMUSG00000105120 | 0 | 0 | 0 | 0 | 0 | 0 | 0 |
| ENSMUSG00000105121 | 0 | 0 | 0 | 0 | 0 | 0 | 0 |
| ENSMUSG00000105122 | 0 | 0 | 0 | 0 | 0 | 0 | 0 |
| ENSMUSG00000105123 | 0 | 0 | 0 | 0 | 0 | 0 | 0 |
| ENSMUSG00000105124 | 0 | 0.396240625 | 0 | 0 | 0 | 0 | 0 |

| | | | | | | | |
|---|---|---|---|---|---|---|---|
| ENSMUSG00000105125 | 0 | 0 | 0.25 | 0 | 0 | 0 | 0 |
| ENSMUSG00000105126 | 0 | 0.25 | 0 | 0 | 0 | 0 | 0 |
| ENSMUSG00000105127 | 0 | 0 | 0 | 0.25 | 0 | 0 | 0.224243824 |
| ENSMUSG00000105128 | 0 | 0 | 0 | 0 | 0 | 0 | 0 |
| ENSMUSG00000105129 | 0 | 0 | 0 | 0 | 0 | 0 | 0 |
| ENSMUSG00000105130 | 0 | 0.580482024 | 0.5 | 0.25 | 0.568060967 | 0.301029996 | 0 |
| ENSMUSG00000105131 | 0 | 0 | 0 | 0 | 0 | 0 | 0 |
| ENSMUSG00000105132 | 0 | 0 | 0 | 0 | 0 | 0 | 0 |
| ENSMUSG00000105133 | 0 | 0 | 0 | 0 | 0 | 0 | 0 |
| ENSMUSG00000105134 | 1.747221736 | 1.714495249 | 1.688721876 | 1.774508021 | 1.671670246 | 1.86332286 | 1.551512352 |
| ENSMUSG00000105135 | 0.578129653 | 0.646240625 | 0.580482024 | 0.79248125 | 1.039257769 | 1.079181246 | 0.935079929 |
| ENSMUSG00000105136 | 0 | 0 | 0 | 0 | 0 | 0 | 0 |
| ENSMUSG00000105137 | 0.578129653 | 0 | 0.396240625 | 0.25 | 0.387761935 | 0.301029996 | 0 |
| ENSMUSG00000105138 | 0 | 0 | 0 | 0 | 0 | 0 | 0 |
| ENSMUSG00000105139 | 1.307602651 | 1.27186571 | 1.146240625 | 1.214495249 | 1.234113756 | 1.176091259 | 1.131174228 |
| ENSMUSG00000105140 | 0.289064826 | 0.25 | 0.75 | 0.646240625 | 0.733951626 | 0.477121255 | 0.896975297 |
| ENSMUSG00000105141 | 0 | 0 | 0 | 0 | 0 | 0 | 0 |
| ENSMUSG00000105142 | 0 | 0 | 0 | 0 | 0 | 0 | 0 |
| ENSMUSG00000105143 | 0 | 0 | 0 | 0 | 0 | 0 | 0 |
| ENSMUSG00000105144 | 0 | 0 | 0 | 0 | 0 | 0 | 0 |
| ENSMUSG00000105145 | 0 | 0 | 0 | 0 | 0 | 0 | 0 |
| ENSMUSG00000105146 | 0 | 0 | 0 | 0 | 0 | 0 | 0 |
| ENSMUSG00000105147 | 0 | 0 | 0 | 0 | 0 | 0 | 0 |
| ENSMUSG00000105148 | 0 | 0 | 0 | 0 | 0 | 0 | 0 |
| ENSMUSG00000105149 | 0 | 0 | 0 | 0 | 0 | 0 | 0 |
| ENSMUSG00000105150 | 0 | 0 | 0 | 0 | 0.244650542 | 0 | 0 |
| ENSMUSG00000105151 | 0 | 0 | 0 | 0 | 0 | 0 | 0 |
| ENSMUSG00000105152 | 0 | 0 | 0 | 0 | 0.244650542 | 0 | 0 |
| ENSMUSG00000105153 | 0 | 0 | 0 | 0 | 0 | 0.477121255 | 0 |
| ENSMUSG00000105154 | 0 | 0.25 | 0 | 0 | 0 | 0 | 0 |
| ENSMUSG00000105155 | 0.289064826 | 0.396240625 | 0 | 0.5 | 0.568060967 | 0.698970004 | 0.355418052 |
| ENSMUSG00000105156 | 0.671187741 | 0.580482024 | 0.701838731 | 0.580482024 | 0.489301084 | 0.477121255 | 0.520678036 |
| ENSMUSG00000105157 | 0 | 0 | 0 | 0 | 0 | 0 | 0 |
| ENSMUSG00000105158 | 0 | 0 | 0 | 0 | 0 | 0 | 0 |

| | | | | | | | |
|---|---|---|---|---|---|---|---|
| ENSMUSG00000105159 | 0 | 0 | 0 | 0 | 0 | 0 | 0 |
| ENSMUSG00000105160 | 0 | 0 | 0 | 0.25 | 0 | 0 | 0.355418052 |
| ENSMUSG00000105161 | 1.249317394 | 1.130890489 | 1.201838731 | 1.25 | 1.254881871 | 1.204119983 | 1.066254157 |
| ENSMUSG00000105162 | 0 | 0.25 | 0.25 | 0.25 | 0.244650542 | 0 | 0 |
| ENSMUSG00000105163 | 0 | 0 | 0.25 | 0 | 0 | 0 | 0.355418052 |
| ENSMUSG00000105164 | 0 | 0 | 0 | 0 | 0 | 0 | 0 |
| ENSMUSG00000105165 | 0 | 0 | 0 | 0 | 0 | 0 | 0 |
| ENSMUSG00000105166 | 0.811507563 | 0.5 | 0.5 | 0.580482024 | 0.568060967 | 0 | 0.672731473 |
| ENSMUSG00000105167 | 0 | 0.396240625 | 0.25 | 0.25 | 0.489301084 | 0 | 0.355418052 |
| ENSMUSG00000105168 | 0 | 0 | 0 | 0 | 0 | 0 | 0 |
| ENSMUSG00000105169 | 0 | 0 | 0 | 0 | 0 | 0 | 0 |
| ENSMUSG00000105170 | 0 | 0 | 0 | 0 | 0 | 0 | 0 |
| ENSMUSG00000105171 | 0.578129653 | 0.5 | 0.396240625 | 0.864857905 | 0.733951626 | 0.77815125 | 0.629532004 |
| ENSMUSG00000105172 | 0.289064826 | 0.396240625 | 0 | 0.25 | 0.387761935 | 0.301029996 | 0.224243824 |
| ENSMUSG00000105173 | 0 | 0 | 0 | 0 | 0 | 0 | 0 |
| ENSMUSG00000105174 | 0 | 0 | 0 | 0 | 0 | 0 | 0 |
| ENSMUSG00000105175 | 0 | 0 | 0 | 0 | 0 | 0.477121255 | 0 |
| ENSMUSG00000105176 | 0.811507563 | 1.330482024 | 1.17510993 | 1.410964047 | 1.27449559 | 1.414973348 | 1.21680636 |
| ENSMUSG00000105177 | 0 | 0 | 0 | 0 | 0 | 0 | 0 |
| ENSMUSG00000105178 | 0 | 0 | 0 | 0 | 0 | 0 | 0 |
| ENSMUSG00000105179 | 0 | 0.25 | 0 | 0 | 0.244650542 | 0.301029996 | 0 |
| ENSMUSG00000105180 | 0 | 0 | 0 | 0 | 0 | 0 | 0 |
| ENSMUSG00000105181 | 0 | 0.25 | 0 | 0.5 | 0 | 0.301029996 | 0.355418052 |
| ENSMUSG00000105182 | 0 | 0 | 0 | 0 | 0 | 0 | 0 |
| ENSMUSG00000105183 | 0.289064826 | 0 | 0.5 | 0 | 0 | 0 | 0 |
| ENSMUSG00000105184 | 0 | 0 | 0 | 0 | 0 | 0 | 0 |
| ENSMUSG00000105185 | 0 | 0 | 0 | 0 | 0 | 0 | 0 |
| ENSMUSG00000105186 | 0.289064826 | 0.75 | 0 | 0.25 | 0 | 0.301029996 | 0.355418052 |
| ENSMUSG00000105187 | 0 | 0 | 0 | 0 | 0 | 0 | 0 |
| ENSMUSG00000105188 | 0 | 0 | 0 | 0 | 0 | 0.301029996 | 0 |
| ENSMUSG00000105189 | 0 | 0 | 0 | 0 | 0 | 0 | 0 |
| ENSMUSG00000105190 | 0 | 0 | 0 | 0 | 0 | 0 | 0 |
| ENSMUSG00000105191 | 0 | 0 | 0 | 0 | 0 | 0 | 0 |
| ENSMUSG00000105192 | 0 | 0 | 0 | 0 | 0 | 0 | 0 |

| | | | | | | | |
|---|---|---|---|---|---|---|---|
| ENSMUSG00000105193 | 0 | 0 | 0 | 0 | 0 | 0 | 0 |
| ENSMUSG00000105194 | 0 | 0 | 0 | 0 | 0 | 0 | 0 |
| ENSMUSG00000105195 | 0 | 0 | 0 | 0 | 0 | 0 | 0 |
| ENSMUSG00000105196 | 0 | 0 | 0 | 0 | 0 | 0 | 0 |
| ENSMUSG00000105197 | 0 | 0 | 0 | 0 | 0 | 0 | 0 |
| ENSMUSG00000105198 | 0.578129653 | 0.5 | 0.580482024 | 0.701838731 | 0.489301084 | 0.698970004 | 0.629532004 |
| ENSMUSG00000105199 | 0.289064826 | 0 | 0.25 | 0.25 | 0.489301084 | 0.477121255 | 0.520678036 |
| ENSMUSG00000105200 | 0 | 0 | 0 | 0 | 0 | 0 | 0 |
| ENSMUSG00000105201 | 0.867194479 | 0.79248125 | 0.5 | 1.098079356 | 0.877063019 | 0.84509804 | 0.775756176 |
| ENSMUSG00000105202 | 0.45815691 | 0 | 0.25 | 0.25 | 0 | 0 | 0.224243824 |
| ENSMUSG00000105203 | 0 | 0 | 0 | 0.25 | 0 | 0.301029996 | 0.224243824 |
| ENSMUSG00000105204 | 0 | 0 | 0 | 0 | 0 | 0 | 0 |
| ENSMUSG00000105205 | 0 | 0 | 0 | 0 | 0 | 0 | 0 |
| ENSMUSG00000105206 | 0.289064826 | 0.25 | 0 | 0.396240625 | 0.387761935 | 0 | 0.224243824 |
| ENSMUSG00000105207 | 2.251208701 | 2.092671852 | 2.077084758 | 2.106566189 | 2.014359518 | 2.222716471 | 1.822941512 |
| ENSMUSG00000105208 | 0 | 0 | 0 | 0 | 0 | 0 | 0 |
| ENSMUSG00000105209 | 0 | 0 | 0 | 0 | 0 | 0 | 0 |
| ENSMUSG00000105210 | 0 | 0 | 0 | 0 | 0 | 0 | 0 |
| ENSMUSG00000105211 | 0 | 0 | 0 | 0 | 0 | 0 | 0 |
| ENSMUSG00000105212 | 0 | 0 | 0 | 0 | 0 | 0 | 0 |
| ENSMUSG00000105213 | 0 | 0 | 0 | 0 | 0 | 0 | 0 |
| ENSMUSG00000105214 | 0 | 0 | 0 | 0 | 0 | 0 | 0 |
| ENSMUSG00000105215 | 0 | 0 | 0 | 0 | 0 | 0 | 0 |
| ENSMUSG00000105216 | 0 | 0 | 0 | 0 | 0 | 0 | 0 |
| ENSMUSG00000105217 | 0 | 0 | 0 | 0 | 0.244650542 | 0.301029996 | 0 |
| ENSMUSG00000105218 | 0 | 0 | 0 | 0 | 0 | 0 | 0 |
| ENSMUSG00000105219 | 0 | 0 | 0 | 0 | 0 | 0 | 0 |
| ENSMUSG00000105220 | 0 | 0 | 0 | 0 | 0 | 0 | 0 |
| ENSMUSG00000105221 | 0 | 0 | 0 | 0.25 | 0.244650542 | 0 | 0 |
| ENSMUSG00000105222 | 0 | 0 | 0 | 0 | 0 | 0 | 0 |
| ENSMUSG00000105223 | 0 | 0 | 0 | 0 | 0 | 0 | 0 |
| ENSMUSG00000105224 | 0 | 0 | 0 | 0 | 0 | 0 | 0 |
| ENSMUSG00000105225 | 0 | 0 | 0 | 0 | 0 | 0 | 0 |
| ENSMUSG00000105226 | 0 | 0 | 0 | 0 | 0.244650542 | 0 | 0 |

| | | | | | | | |
|---|---|---|---|---|---|---|---|
| ENSMUSG00000105227 | 0 | 0 | 0 | 0 | 0 | 0 | 0 |
| ENSMUSG00000105228 | 0.289064826 | 0.25 | 0.580482024 | 0 | 0.489301084 | 0.477121255 | 0.520678036 |
| ENSMUSG00000105229 | 0 | 0 | 0 | 0 | 0 | 0 | 0 |
| ENSMUSG00000105230 | 0 | 0 | 0 | 0 | 0.244650542 | 0 | 0 |
| ENSMUSG00000105231 | 0 | 0 | 0 | 0 | 0 | 0 | 0 |
| ENSMUSG00000105232 | 0 | 0 | 0 | 0 | 0.244650542 | 0 | 0 |
| ENSMUSG00000105233 | 0 | 0.25 | 0.25 | 0 | 0.244650542 | 0 | 0 |
| ENSMUSG00000105234 | 0 | 0 | 0 | 0 | 0 | 0 | 0 |
| ENSMUSG00000105235 | 0 | 0 | 0 | 0 | 0 | 0 | 0 |
| ENSMUSG00000105236 | 0 | 0 | 0 | 0 | 0 | 0 | 0 |
| ENSMUSG00000105237 | 0 | 0 | 0 | 0 | 0 | 0 | 0 |
| ENSMUSG00000105238 | 0 | 0 | 0 | 0 | 0 | 0 | 0 |
| ENSMUSG00000105239 | 0 | 0 | 0 | 0 | 0 | 0 | 0 |
| ENSMUSG00000105240 | 0 | 0 | 0 | 0 | 0 | 0 | 0 |
| ENSMUSG00000105241 | 0 | 0 | 0 | 0 | 0 | 0 | 0 |
| ENSMUSG00000105242 | 0 | 0 | 0 | 0 | 0 | 0 | 0 |
| ENSMUSG00000105243 | 0 | 0 | 0 | 0 | 0 | 0 | 0.224243824 |
| ENSMUSG00000105244 | 0 | 0 | 0 | 0 | 0.244650542 | 0 | 0 |
| ENSMUSG00000105245 | 0.45815691 | 0.25 | 0.396240625 | 0.25 | 0 | 0 | 0.355418052 |
| ENSMUSG00000105246 | 0 | 0 | 0 | 0 | 0 | 0 | 0 |
| ENSMUSG00000105247 | 1.289064826 | 1.130890489 | 1.418106335 | 1.311981878 | 1.302012594 | 1.278753601 | 1.21680636 |
| ENSMUSG00000105248 | 0 | 0 | 0 | 0 | 0 | 0 | 0 |
| ENSMUSG00000105249 | 0 | 0 | 0 | 0 | 0 | 0 | 0 |
| ENSMUSG00000105250 | 0 | 0 | 0 | 0 | 0 | 0 | 0 |
| ENSMUSG00000105251 | 0 | 0 | 0 | 0 | 0 | 0 | 0 |
| ENSMUSG00000105252 | 0 | 0 | 0 | 0 | 0 | 0 | 0 |
| ENSMUSG00000105253 | 0.289064826 | 0.25 | 0 | 0.5 | 0.489301084 | 0.301029996 | 0.355418052 |
| ENSMUSG00000105254 | 0 | 0.25 | 0.5 | 0 | 0 | 0 | 0.224243824 |
| ENSMUSG00000105255 | 0 | 0 | 0 | 0 | 0 | 0 | 0 |
| ENSMUSG00000105256 | 0 | 0 | 0 | 0 | 0 | 0 | 0 |
| ENSMUSG00000105257 | 0 | 0 | 0 | 0 | 0 | 0 | 0 |
| ENSMUSG00000105258 | 0 | 0.25 | 0 | 0 | 0 | 0 | 0 |
| ENSMUSG00000105259 | 0 | 0.25 | 0.25 | 0.5 | 0.387761935 | 0 | 0.224243824 |
| ENSMUSG00000105260 | 0 | 0 | 0 | 0 | 0 | 0 | 0 |

| | | | | | | | |
|---|---|---|---|---|---|---|---|
| ENSMUSG00000105261 | 0 | 0 | 0 | 0 | 0 | 0 | 0.224243824 |
| ENSMUSG00000105262 | 0 | 0 | 0 | 0 | 0 | 0 | 0 |
| ENSMUSG00000105263 | 2.627631768 | 2.424241882 | 2.391989019 | 2.533927322 | 2.501012315 | 2.834420704 | 2.354972132 |
| ENSMUSG00000105264 | 0 | 0 | 0.396240625 | 0.580482024 | 0.387761935 | 0 | 0.224243824 |
| ENSMUSG00000105265 | 0.289064826 | 0.396240625 | 0.396240625 | 0.396240625 | 0 | 0 | 0.355418052 |
| ENSMUSG00000105266 | 0 | 0 | 0 | 0 | 0 | 0 | 0 |
| ENSMUSG00000105267 | 0 | 0 | 0 | 0 | 0 | 0 | 0 |
| ENSMUSG00000105269 | 0 | 0 | 0 | 0 | 0 | 0 | 0 |
| ENSMUSG00000105270 | 1.482695304 | 1.364857905 | 1.464495249 | 1.26109853 | 1.373641807 | 1.361727836 | 1.259064007 |
| ENSMUSG00000105271 | 0 | 0 | 0 | 0 | 0 | 0 | 0 |
| ENSMUSG00000105272 | 0.811507563 | 0.951838731 | 0.92510993 | 1.080482024 | 0.846351821 | 0.84509804 | 0.829800754 |
| ENSMUSG00000105273 | 0 | 0 | 0 | 0 | 0 | 0 | 0 |
| ENSMUSG00000105274 | 0 | 0 | 0 | 0 | 0 | 0 | 0 |
| ENSMUSG00000105275 | 0.289064826 | 0 | 0 | 0.25 | 0.244650542 | 0 | 0 |
| ENSMUSG00000105276 | 0 | 0 | 0 | 0.25 | 0 | 0.301029996 | 0 |
| ENSMUSG00000105277 | 0 | 0 | 0 | 0 | 0 | 0 | 0 |
| ENSMUSG00000105278 | 0.289064826 | 0 | 0.25 | 0 | 0.387761935 | 0 | 0.224243824 |
| ENSMUSG00000105279 | 0 | 0 | 0 | 0 | 0 | 0 | 0 |
| ENSMUSG00000105280 | 0 | 0 | 0 | 0 | 0 | 0 | 0 |
| ENSMUSG00000105281 | 0 | 0 | 0 | 0 | 0 | 0 | 0 |
| ENSMUSG00000105282 | 0 | 0 | 0 | 0 | 0 | 0 | 0 |
| ENSMUSG00000105283 | 0 | 0 | 0 | 0 | 0 | 0 | 0 |
| ENSMUSG00000105284 | 0 | 0 | 0 | 0 | 0 | 0 | 0 |
| ENSMUSG00000105285 | 1.036286563 | 0.976722649 | 0.896240625 | 0.92510993 | 0.846351821 | 0.954242509 | 0.803905701 |
| ENSMUSG00000105286 | 0 | 0.25 | 0 | 0 | 0 | 0 | 0 |
| ENSMUSG00000105287 | 0.960252568 | 0.92510993 | 0.92510993 | 1.02186571 | 1.149965125 | 0.954242509 | 0.952571511 |
| ENSMUSG00000105288 | 0 | 0 | 0 | 0 | 0 | 0 | 0 |
| ENSMUSG00000105289 | 0 | 0 | 0.25 | 0 | 0 | 0 | 0 |
| ENSMUSG00000105290 | 0 | 0 | 0 | 0 | 0 | 0 | 0 |
| ENSMUSG00000105291 | 0 | 0 | 0 | 0 | 0.244650542 | 0 | 0.224243824 |
| ENSMUSG00000105292 | 0 | 0 | 0 | 0 | 0 | 0 | 0 |
| ENSMUSG00000105293 | 0.289064826 | 0.396240625 | 0 | 0 | 0 | 0 | 0 |
| ENSMUSG00000105294 | 0.289064826 | 0 | 0 | 0 | 0 | 0 | 0 |
| ENSMUSG00000105295 | 0 | 0 | 0 | 0 | 0 | 0 | 0 |

| | | | | | | | |
|---|---|---|---|---|---|---|---|
| ENSMUSG00000105296 | 0 | 0 | 0 | 0 | 0 | 0 | 0 |
| ENSMUSG00000105297 | 0 | 0 | 0 | 0 | 0 | 0 | 0 |
| ENSMUSG00000105298 | 0 | 0 | 0 | 0 | 0 | 0 | 0 |
| ENSMUSG00000105299 | 0 | 0 | 0 | 0 | 0 | 0 | 0 |
| ENSMUSG00000105300 | 0 | 0 | 0 | 0 | 0 | 0 | 0 |
| ENSMUSG00000105301 | 0 | 0 | 0 | 0 | 0 | 0 | 0 |
| ENSMUSG00000105302 | 0.289064826 | 0 | 0.25 | 0.25 | 0.244650542 | 0 | 0 |
| ENSMUSG00000105303 | 0.578129653 | 0.5 | 0.25 | 0 | 0 | 0 | 0 |
| ENSMUSG00000105304 | 0.747221736 | 0.25 | 0.396240625 | 0.701838731 | 0.568060967 | 0.698970004 | 0.629532004 |
| ENSMUSG00000105305 | 0 | 0 | 0 | 0 | 0 | 0 | 0 |
| ENSMUSG00000105306 | 0 | 0 | 0 | 0 | 0 | 0 | 0 |
| ENSMUSG00000105307 | 0 | 0 | 0 | 0 | 0 | 0 | 0 |
| ENSMUSG00000105308 | 0 | 0 | 0 | 0 | 0 | 0 | 0 |
| ENSMUSG00000105309 | 0 | 0 | 0 | 0 | 0 | 0 | 0 |
| ENSMUSG00000105310 | 0 | 0 | 0 | 0 | 0 | 0 | 0 |
| ENSMUSG00000105311 | 0 | 0 | 0 | 0 | 0 | 0 | 0 |
| ENSMUSG00000105312 | 0 | 0 | 0 | 0 | 0 | 0 | 0 |
| ENSMUSG00000105313 | 0 | 0 | 0 | 0 | 0 | 0 | 0 |
| ENSMUSG00000105314 | 0 | 0 | 0 | 0 | 0 | 0 | 0 |
| ENSMUSG00000105315 | 0 | 0 | 0 | 0 | 0 | 0 | 0 |
| ENSMUSG00000105316 | 0 | 0 | 0 | 0 | 0 | 0 | 0 |
| ENSMUSG00000105317 | 0 | 0 | 0 | 0 | 0 | 0 | 0 |
| ENSMUSG00000105318 | 0 | 0 | 0 | 0 | 0 | 0 | 0 |
| ENSMUSG00000105319 | 0 | 0 | 0 | 0 | 0 | 0 | 0 |
| ENSMUSG00000105320 | 0 | 0 | 0 | 0 | 0 | 0 | 0 |
| ENSMUSG00000105321 | 0 | 0 | 0 | 0 | 0 | 0 | 0 |
| ENSMUSG00000105322 | 0 | 0 | 0 | 0 | 0 | 0 | 0 |
| ENSMUSG00000105323 | 0 | 0 | 0 | 0 | 0 | 0 | 0 |
| ENSMUSG00000105324 | 0 | 0 | 0 | 0 | 0 | 0 | 0 |
| ENSMUSG00000105325 | 0 | 0 | 0 | 0 | 0 | 0 | 0 |
| ENSMUSG00000105326 | 0 | 0.25 | 0.396240625 | 0.25 | 0 | 0.477121255 | 0.224243824 |
| ENSMUSG00000105327 | 0 | 0 | 0.5 | 0 | 0.387761935 | 0.301029996 | 0.224243824 |
| ENSMUSG00000105328 | 0.45815691 | 0.25 | 0.5 | 0.396240625 | 0 | 0.301029996 | 0 |
| ENSMUSG00000105329 | 0 | 0 | 0 | 0 | 0 | 0 | 0 |

| | | | | | | | |
|---|---|---|---|---|---|---|---|
| ENSMUSG00000105330 | 0 | 0 | 0 | 0 | 0 | 0 | 0 |
| ENSMUSG00000105331 | 0 | 0 | 0 | 0 | 0 | 0 | 0 |
| ENSMUSG00000105332 | 0 | 0 | 0.396240625 | 0.25 | 0 | 0 | 0 |
| ENSMUSG00000105333 | 0 | 0 | 0 | 0 | 0.244650542 | 0 | 0 |
| ENSMUSG00000105334 | 0 | 0 | 0 | 0 | 0 | 0 | 0 |
| ENSMUSG00000105335 | 0 | 0 | 0 | 0 | 0 | 0 | 0 |
| ENSMUSG00000105336 | 0 | 0 | 0.25 | 0 | 0.244650542 | 0.301029996 | 0 |
| ENSMUSG00000105337 | 0 | 0 | 0 | 0 | 0 | 0 | 0 |
| ENSMUSG00000105338 | 0 | 0 | 0 | 0 | 0 | 0 | 0 |
| ENSMUSG00000105339 | 0 | 0 | 0 | 0 | 0 | 0 | 0 |
| ENSMUSG00000105340 | 0.960252568 | 0.976722649 | 1.04248125 | 1.080482024 | 1.091002363 | 0.954242509 | 0.952571511 |
| ENSMUSG00000105341 | 0.45815691 | 0.25 | 0.396240625 | 0 | 0 | 0 | 0.224243824 |
| ENSMUSG00000105342 | 0 | 0 | 0 | 0 | 0 | 0 | 0 |
| ENSMUSG00000105343 | 0 | 0 | 0 | 0 | 0.244650542 | 0 | 0.224243824 |
| ENSMUSG00000105344 | 0 | 0 | 0 | 0 | 0 | 0 | 0 |
| ENSMUSG00000105345 | 1 | 1.130890489 | 1.146240625 | 1.02186571 | 0.931471446 | 0.477121255 | 0.803905701 |
| ENSMUSG00000105346 | 0 | 0 | 0 | 0 | 0 | 0 | 0 |
| ENSMUSG00000105347 | 0 | 0.5 | 0.580482024 | 0.5 | 0.387761935 | 0.301029996 | 0.448487648 |
| ENSMUSG00000105348 | 0 | 0 | 0 | 0 | 0 | 0 | 0 |
| ENSMUSG00000105349 | 1.069666964 | 0.646240625 | 0.75 | 0.896240625 | 0.812711509 | 0.77815125 | 0.710836105 |
| ENSMUSG00000105350 | 0 | 0.25 | 0 | 0 | 0 | 0 | 0 |
| ENSMUSG00000105351 | 0 | 0 | 0 | 0 | 0 | 0 | 0 |
| ENSMUSG00000105352 | 0 | 0.25 | 0 | 0 | 0 | 0 | 0 |
| ENSMUSG00000105353 | 1.100572389 | 1.188721876 | 0.92510993 | 1.311981878 | 1.223252711 | 1.255272505 | 1.121219121 |
| ENSMUSG00000105354 | 0 | 0 | 0 | 0 | 0 | 0 | 0 |
| ENSMUSG00000105355 | 0 | 0 | 0 | 0 | 0 | 0 | 0 |
| ENSMUSG00000105356 | 0.671187741 | 0.75 | 0.5 | 0.646240625 | 0.686820904 | 0.698970004 | 0.579661877 |
| ENSMUSG00000105357 | 0 | 0 | 0 | 0 | 0 | 0 | 0 |
| ENSMUSG00000105358 | 0.289064826 | 0 | 0.580482024 | 0.5 | 0.812711509 | 0.77815125 | 0.74492186 |
| ENSMUSG00000105359 | 0.289064826 | 0.701838731 | 0.864857905 | 0.79248125 | 0.812711509 | 0.698970004 | 0.672731473 |
| ENSMUSG00000105360 | 0 | 0 | 0 | 0 | 0 | 0 | 0 |
| ENSMUSG00000105361 | 0 | 0 | 0 | 0 | 0 | 0 | 0 |
| ENSMUSG00000105362 | 0 | 0 | 0 | 0 | 0 | 0 | 0 |
| ENSMUSG00000105363 | 0 | 0 | 0 | 0 | 0 | 0 | 0 |

| | | | | | | | |
|---|---|---|---|---|---|---|---|
| ENSMUSG00000105364 | 0 | 0 | 0 | 0 | 0.244650542 | 0 | 0 |
| ENSMUSG00000105365 | 0 | 0 | 0 | 0 | 0 | 0 | 0 |
| ENSMUSG00000105366 | 1.404271433 | 1.348079356 | 1.311981878 | 0.701838731 | 0.846351821 | 0.602059991 | 0.579661877 |
| ENSMUSG00000105367 | 0 | 0 | 0 | 0 | 0 | 0 | 0 |
| ENSMUSG00000105368 | 0.45815691 | 0.701838731 | 0.864857905 | 0.396240625 | 0.733951626 | 0.698970004 | 0.74492186 |
| ENSMUSG00000105369 | 0 | 0 | 0 | 0 | 0 | 0 | 0 |
| ENSMUSG00000105370 | 1.482695304 | 1.311981878 | 1.418106335 | 1.42510993 | 1.343584837 | 1.361727836 | 1.23151414 |
| ENSMUSG00000105371 | 0 | 0 | 0 | 0 | 0 | 0 | 0 |
| ENSMUSG00000105372 | 0 | 0 | 0 | 0 | 0 | 0 | 0 |
| ENSMUSG00000105373 | 0 | 0 | 0 | 0 | 0 | 0 | 0 |
| ENSMUSG00000105374 | 0 | 0 | 0 | 0 | 0 | 0 | 0 |
| ENSMUSG00000105375 | 0 | 0 | 0 | 0 | 0 | 0 | 0 |
| ENSMUSG00000105376 | 0.289064826 | 0.79248125 | 0.896240625 | 0.5 | 0.244650542 | 0 | 0.224243824 |
| ENSMUSG00000105377 | 0 | 0 | 0 | 0 | 0 | 0 | 0 |
| ENSMUSG00000105378 | 0.289064826 | 0.396240625 | 0.25 | 0.396240625 | 0 | 0.477121255 | 0.224243824 |
| ENSMUSG00000105379 | 0 | 0 | 0.25 | 0 | 0 | 0 | 0 |
| ENSMUSG00000105380 | 0 | 0 | 0 | 0 | 0 | 0 | 0 |
| ENSMUSG00000105381 | 0 | 0 | 0 | 0 | 0 | 0 | 0 |
| ENSMUSG00000105382 | 1.269664473 | 1.130890489 | 0.976722649 | 1.146240625 | 1.020174412 | 1.113943352 | 1.014380832 |
| ENSMUSG00000105383 | 0.289064826 | 0.646240625 | 0.976722649 | 0.701838731 | 0.568060967 | 0.954242509 | 0.829800754 |
| ENSMUSG00000105384 | 0 | 0 | 0 | 0 | 0 | 0 | 0 |
| ENSMUSG00000105385 | 0 | 0 | 0 | 0 | 0 | 0 | 0 |
| ENSMUSG00000105386 | 0 | 0 | 0 | 0 | 0 | 0 | 0 |
| ENSMUSG00000105387 | 0 | 0.5 | 0.580482024 | 0 | 0.244650542 | 0.477121255 | 0.355418052 |
| ENSMUSG00000105388 | 0.578129653 | 0.5 | 0 | 0.5 | 0.244650542 | 0.602059991 | 0.355418052 |
| ENSMUSG00000105389 | 0.289064826 | 0.25 | 0 | 0 | 0 | 0 | 0 |
| ENSMUSG00000105390 | 0 | 0 | 0 | 0 | 0 | 0 | 0 |
| ENSMUSG00000105391 | 0 | 0.25 | 0 | 0 | 0 | 0 | 0 |
| ENSMUSG00000105392 | 0 | 0 | 0 | 0 | 0 | 0 | 0 |
| ENSMUSG00000105393 | 0 | 0 | 0 | 0 | 0 | 0 | 0 |
| ENSMUSG00000105394 | 0 | 0 | 0 | 0 | 0 | 0 | 0 |
| ENSMUSG00000105395 | 0 | 0.25 | 0.580482024 | 0.5 | 0.568060967 | 0.477121255 | 0.579661877 |
| ENSMUSG00000105396 | 0.289064826 | 0 | 0 | 0 | 0 | 0 | 0 |
| ENSMUSG00000105397 | 0 | 0.396240625 | 0.396240625 | 0.396240625 | 0 | 0 | 0 |

| | | | | | | | |
|---|---|---|---|---|---|---|---|
| ENSMUSG00000105398 | 0 | 0 | 0 | 0 | 0 | 0 | 0 |
| ENSMUSG00000105399 | 0 | 0 | 0 | 0 | 0 | 0 | 0 |
| ENSMUSG00000105400 | 0 | 0 | 0 | 0 | 0 | 0 | 0 |
| ENSMUSG00000105401 | 0.45815691 | 0.25 | 0.25 | 0 | 0 | 0 | 0.520678036 |
| ENSMUSG00000105402 | 1.747221736 | 1.671625132 | 1.681980114 | 1.17510993 | 1.188507684 | 1.079181246 | 1.066254157 |
| ENSMUSG00000105403 | 0.289064826 | 0 | 0 | 0 | 0 | 0 | 0 |
| ENSMUSG00000105404 | 0.289064826 | 0.25 | 0.25 | 0 | 0 | 0 | 0 |
| ENSMUSG00000105405 | 0 | 0 | 0 | 0 | 0 | 0 | 0 |
| ENSMUSG00000105406 | 0.45815691 | 0 | 0 | 0 | 0.244650542 | 0 | 0.520678036 |
| ENSMUSG00000105407 | 0 | 0 | 0 | 0 | 0 | 0 | 0 |
| ENSMUSG00000105408 | 0.671187741 | 0.25 | 0.580482024 | 0.5 | 0.387761935 | 0.477121255 | 0.629532004 |
| ENSMUSG00000105409 | 0 | 0 | 0 | 0 | 0 | 0 | 0 |
| ENSMUSG00000105410 | 0 | 0 | 0 | 0 | 0 | 0 | 0 |
| ENSMUSG00000105411 | 0 | 0 | 0 | 0 | 0 | 0 | 0 |
| ENSMUSG00000105412 | 0 | 0.25 | 0 | 0.25 | 0 | 0 | 0 |
| ENSMUSG00000105413 | 0 | 0 | 0 | 0 | 0 | 0 | 0 |
| ENSMUSG00000105415 | 0 | 0 | 0 | 0 | 0 | 0 | 0 |
| ENSMUSG00000105416 | 0.45815691 | 0.396240625 | 0.701838731 | 0.396240625 | 0.244650542 | 0.477121255 | 0.448487648 |
| ENSMUSG00000105418 | 0 | 0 | 0 | 0 | 0 | 0 | 0 |
| ENSMUSG00000105419 | 0.747221736 | 0.92510993 | 0.896240625 | 0.864857905 | 0.905314583 | 0.84509804 | 1.140832123 |
| ENSMUSG00000105420 | 0 | 0 | 0 | 0 | 0 | 0 | 0 |
| ENSMUSG00000105421 | 0 | 0 | 0 | 0 | 0 | 0 | 0 |
| ENSMUSG00000105422 | 0 | 0 | 0 | 0 | 0 | 0 | 0.224243824 |
| ENSMUSG00000105423 | 0 | 0 | 0 | 0 | 0 | 0 | 0 |
| ENSMUSG00000105424 | 0 | 0 | 0 | 0 | 0 | 0 | 0 |
| ENSMUSG00000105425 | 0.578129653 | 0.580482024 | 0.396240625 | 0.396240625 | 0.387761935 | 0 | 0 |
| ENSMUSG00000105426 | 0 | 0 | 0 | 0 | 0 | 0 | 0 |
| ENSMUSG00000105427 | 0 | 0 | 0 | 0 | 0 | 0 | 0 |
| ENSMUSG00000105428 | 0 | 0 | 0.25 | 0 | 0 | 0 | 0 |
| ENSMUSG00000105429 | 0 | 0.830482024 | 0.79248125 | 0.75 | 0.632412477 | 0.477121255 | 0.672731473 |
| ENSMUSG00000105430 | 0 | 0 | 0 | 0 | 0 | 0.301029996 | 0 |
| ENSMUSG00000105431 | 0 | 0 | 0.25 | 0.25 | 0 | 0 | 0.224243824 |
| ENSMUSG00000105432 | 0 | 0 | 0 | 0 | 0 | 0 | 0 |
| ENSMUSG00000105433 | 0 | 0 | 0 | 0 | 0 | 0 | 0 |

| | | | | | | | |
|---|---|---|---|---|---|---|---|
| ENSMUSG00000105434 | 0.578129653 | 0.25 | 0.580482024 | 0.646240625 | 0.387761935 | 0 | 0.520678036 |
| ENSMUSG00000105435 | 0 | 0 | 0 | 0 | 0 | 0 | 0 |
| ENSMUSG00000105436 | 0 | 0 | 0 | 0 | 0 | 0 | 0 |
| ENSMUSG00000105437 | 0.289064826 | 0.25 | 0 | 0.25 | 0 | 0 | 0 |
| ENSMUSG00000105438 | 0 | 0 | 0 | 0 | 0 | 0 | 0 |
| ENSMUSG00000105439 | 0 | 0 | 0 | 0 | 0 | 0 | 0 |
| ENSMUSG00000105440 | 0.289064826 | 0.580482024 | 0.25 | 0.396240625 | 0.387761935 | 0.301029996 | 0.355418052 |
| ENSMUSG00000105441 | 0.289064826 | 0 | 0.25 | 0.25 | 0.244650542 | 0 | 0 |
| ENSMUSG00000105442 | 0 | 0 | 0 | 0 | 0 | 0 | 0 |
| ENSMUSG00000105443 | 0 | 0 | 0 | 0 | 0 | 0 | 0 |
| ENSMUSG00000105444 | 0 | 0 | 0 | 0 | 0 | 0 | 0 |
| ENSMUSG00000105445 | 0 | 0 | 0 | 0 | 0 | 0 | 0 |
| ENSMUSG00000105446 | 0 | 0 | 0 | 0 | 0 | 0 | 0 |
| ENSMUSG00000105447 | 0 | 0 | 0 | 0 | 0 | 0 | 0.224243824 |
| ENSMUSG00000105448 | 0 | 0 | 0 | 0 | 0 | 0 | 0 |
| ENSMUSG00000105449 | 1.069666964 | 1.160964047 | 1.114857905 | 1.348079356 | 1.264824954 | 1.278753601 | 1.159323753 |
| ENSMUSG00000105450 | 0 | 0 | 0 | 0 | 0 | 0 | 0 |
| ENSMUSG00000105451 | 0 | 0 | 0 | 0 | 0 | 0 | 0 |
| ENSMUSG00000105452 | 0 | 0.580482024 | 0.396240625 | 0.25 | 0.244650542 | 0.301029996 | 0.355418052 |
| ENSMUSG00000105453 | 0.671187741 | 0.646240625 | 0.701838731 | 0.5 | 0.846351821 | 0.77815125 | 0.629532004 |
| ENSMUSG00000105454 | 0.811507563 | 1.098079356 | 0.976722649 | 1.061981878 | 0.931471446 | 1.113943352 | 0.853775828 |
| ENSMUSG00000105455 | 0 | 0 | 0 | 0 | 0 | 0 | 0 |
| ENSMUSG00000105456 | 0.289064826 | 0 | 0.25 | 0 | 0 | 0 | 0 |
| ENSMUSG00000105457 | 0 | 0.75 | 0.580482024 | 0.25 | 0 | 0.477121255 | 0.224243824 |
| ENSMUSG00000105458 | 0 | 0 | 0 | 0 | 0 | 0 | 0 |
| ENSMUSG00000105459 | 0 | 0 | 0 | 0 | 0 | 0 | 0 |
| ENSMUSG00000105460 | 0 | 0.25 | 0 | 0 | 0 | 0 | 0 |
| ENSMUSG00000105461 | 1.227926429 | 1.339388001 | 1.27186571 | 0.896240625 | 0.931471446 | 0.903089987 | 0.710836105 |
| ENSMUSG00000105462 | 0 | 0 | 0 | 0 | 0 | 0 | 0 |
| ENSMUSG00000105463 | 0 | 0 | 0 | 0 | 0 | 0 | 0 |
| ENSMUSG00000105464 | 0 | 0 | 0 | 0 | 0 | 0 | 0 |
| ENSMUSG00000105465 | 0 | 0 | 0 | 0 | 0 | 0 | 0 |
| ENSMUSG00000105466 | 0.289064826 | 0 | 0.5 | 0.25 | 0.568060967 | 0.602059991 | 0.579661877 |
| ENSMUSG00000105467 | 0 | 0 | 0 | 0 | 0 | 0 | 0 |

| | | | | | | | |
|---|---|---|---|---|---|---|---|
| ENSMUSG00000105468 | 0 | 0 | 0 | 0 | 0 | 0 | 0 |
| ENSMUSG00000105469 | 0.289064826 | 0 | 0 | 0 | 0 | 0 | 0 |
| ENSMUSG00000105470 | 0 | 0 | 0 | 0 | 0 | 0 | 0 |
| ENSMUSG00000105471 | 1 | 0.976722649 | 0.864857905 | 0.896240625 | 1.039257769 | 0.903089987 | 0.876096088 |
| ENSMUSG00000105472 | 0 | 0 | 0 | 0.25 | 0.244650542 | 0 | 0 |
| ENSMUSG00000105473 | 0 | 0 | 0 | 0 | 0 | 0 | 0 |
| ENSMUSG00000105474 | 0 | 0 | 0 | 0 | 0 | 0 | 0 |
| ENSMUSG00000105475 | 0 | 0 | 0 | 0 | 0 | 0 | 0 |
| ENSMUSG00000105476 | 0.578129653 | 0.830482024 | 0.976722649 | 0.830482024 | 0.905314583 | 1 | 0.775756176 |
| ENSMUSG00000105477 | 0 | 0 | 0 | 0 | 0 | 0 | 0 |
| ENSMUSG00000105478 | 0.578129653 | 0.25 | 0.396240625 | 0 | 0 | 0 | 0.448487648 |
| ENSMUSG00000105479 | 0 | 0 | 0 | 0 | 0 | 0 | 0 |
| ENSMUSG00000105480 | 0 | 0 | 0 | 0 | 0 | 0 | 0 |
| ENSMUSG00000105481 | 0 | 0 | 0 | 0 | 0 | 0.301029996 | 0 |
| ENSMUSG00000105482 | 0 | 0 | 0.25 | 0 | 0 | 0 | 0 |
| ENSMUSG00000105483 | 0.671187741 | 0.79248125 | 0.580482024 | 1 | 1.091002363 | 1.041392685 | 1.066254157 |
| ENSMUSG00000105484 | 0 | 0 | 0 | 0 | 0 | 0 | 0 |
| ENSMUSG00000105485 | 0 | 0 | 0 | 0 | 0 | 0 | 0 |
| ENSMUSG00000105486 | 0 | 0 | 0 | 0 | 0 | 0 | 0 |
| ENSMUSG00000105487 | 0 | 0 | 0 | 0 | 0 | 0 | 0 |
| ENSMUSG00000105488 | 0 | 0 | 0 | 0 | 0 | 0 | 0 |
| ENSMUSG00000105489 | 0 | 0 | 0 | 0 | 0 | 0 | 0 |
| ENSMUSG00000105490 | 0 | 0 | 0 | 0 | 0 | 0 | 0 |
| ENSMUSG00000105491 | 0 | 0 | 0 | 0 | 0 | 0 | 0 |
| ENSMUSG00000105492 | 0 | 0 | 0 | 0 | 0 | 0 | 0 |
| ENSMUSG00000105493 | 0 | 0 | 0 | 0 | 0 | 0 | 0 |
| ENSMUSG00000105494 | 0 | 0 | 0 | 0 | 0 | 0 | 0 |
| ENSMUSG00000105495 | 0 | 0 | 0 | 0 | 0 | 0 | 0 |
| ENSMUSG00000105496 | 0 | 0 | 0 | 0 | 0 | 0 | 0 |
| ENSMUSG00000105497 | 0 | 0 | 0 | 0 | 0 | 0 | 0 |
| ENSMUSG00000105498 | 0 | 0 | 0 | 0 | 0 | 0 | 0 |
| ENSMUSG00000105499 | 0 | 0 | 0 | 0 | 0 | 0 | 0 |
| ENSMUSG00000105500 | 0 | 0 | 0 | 0 | 0 | 0 | 0 |
| ENSMUSG00000105501 | 0 | 0 | 0.25 | 0.396240625 | 0 | 0.301029996 | 0.224243824 |

| | | | | | | | |
|---|---|---|---|---|---|---|---|
| ENSMUSG00000105502 | 0 | 0 | 0 | 0.25 | 0 | 0 | 0.224243824 |
| ENSMUSG00000105503 | 0 | 0 | 0 | 0 | 0 | 0 | 0 |
| ENSMUSG00000105504 | 0 | 0.580482024 | 0.25 | 0.646240625 | 0.387761935 | 0.698970004 | 0.672731473 |
| ENSMUSG00000105505 | 0 | 0 | 0 | 0 | 0 | 0 | 0 |
| ENSMUSG00000105506 | 0 | 0.25 | 0 | 0.396240625 | 0 | 0 | 0.224243824 |
| ENSMUSG00000105507 | 0 | 0 | 0 | 0 | 0 | 0 | 0 |
| ENSMUSG00000105508 | 0 | 0 | 0 | 0 | 0 | 0 | 0 |
| ENSMUSG00000105509 | 1.036286563 | 1.02186571 | 0.951838731 | 1.114857905 | 1.136121934 | 1.146128036 | 0.969165684 |
| ENSMUSG00000105510 | 0.578129653 | 0.5 | 0 | 0.701838731 | 0.632412477 | 0.698970004 | 0.448487648 |
| ENSMUSG00000105511 | 0 | 0 | 0 | 0 | 0 | 0 | 0 |
| ENSMUSG00000105512 | 0 | 0 | 0 | 0 | 0 | 0 | 0.224243824 |
| ENSMUSG00000105513 | 0 | 0 | 0 | 0 | 0 | 0 | 0 |
| ENSMUSG00000105514 | 0.747221736 | 0.580482024 | 0.646240625 | 0.646240625 | 0.568060967 | 0.477121255 | 0.672731473 |
| ENSMUSG00000105515 | 0 | 0 | 0 | 0 | 0 | 0 | 0 |
| ENSMUSG00000105516 | 0.289064826 | 0.396240625 | 0.580482024 | 0.701838731 | 0.632412477 | 0.301029996 | 0.710836105 |
| ENSMUSG00000105517 | 0 | 0.396240625 | 0 | 0.396240625 | 0.568060967 | 0 | 0.579661877 |
| ENSMUSG00000105518 | 0 | 0 | 0 | 0 | 0 | 0 | 0 |
| ENSMUSG00000105519 | 0 | 0 | 0.25 | 0.25 | 0.244650542 | 0.301029996 | 0 |
| ENSMUSG00000105520 | 0 | 0 | 0 | 0 | 0 | 0 | 0 |
| ENSMUSG00000105521 | 0 | 0 | 0 | 0 | 0 | 0 | 0 |
| ENSMUSG00000105522 | 0 | 0 | 0 | 0 | 0 | 0 | 0 |
| ENSMUSG00000105523 | 0 | 0 | 0 | 0 | 0 | 0 | 0 |
| ENSMUSG00000105524 | 0 | 0 | 0 | 0 | 0 | 0 | 0 |
| ENSMUSG00000105525 | 0 | 0 | 0 | 0 | 0 | 0 | 0 |
| ENSMUSG00000105526 | 0 | 0 | 0 | 0 | 0 | 0.301029996 | 0 |
| ENSMUSG00000105527 | 0 | 0 | 0 | 0 | 0 | 0 | 0 |
| ENSMUSG00000105528 | 0 | 0.25 | 0 | 0 | 0 | 0 | 0 |
| ENSMUSG00000105529 | 0 | 0.5 | 0.25 | 0.5 | 0.568060967 | 0.477121255 | 0.355418052 |
| ENSMUSG00000105530 | 0 | 0 | 0 | 0 | 0 | 0 | 0.355418052 |
| ENSMUSG00000105531 | 0 | 0 | 0 | 0 | 0 | 0 | 0 |
| ENSMUSG00000105532 | 0 | 0 | 0 | 0 | 0 | 0 | 0 |
| ENSMUSG00000105533 | 0 | 0 | 0 | 0 | 0 | 0 | 0 |
| ENSMUSG00000105534 | 0 | 0 | 0.25 | 0.396240625 | 0.244650542 | 0.301029996 | 0.224243824 |
| ENSMUSG00000105535 | 0 | 0 | 0 | 0 | 0 | 0 | 0 |

| | | | | | | | |
|---|---|---|---|---|---|---|---|
| ENSMUSG00000105536 | 0.45815691 | 0.25 | 0.396240625 | 0.25 | 0.387761935 | 0.301029996 | 0 |
| ENSMUSG00000105537 | 0 | 0 | 0 | 0 | 0 | 0 | 0 |
| ENSMUSG00000105538 | 0 | 0 | 0 | 0 | 0 | 0 | 0 |
| ENSMUSG00000105539 | 0 | 0 | 0 | 0 | 0 | 0 | 0 |
| ENSMUSG00000105540 | 0 | 0 | 0 | 0 | 0 | 0 | 0 |
| ENSMUSG00000105541 | 0.289064826 | 0 | 0 | 0.25 | 0.387761935 | 0.301029996 | 0.355418052 |
| ENSMUSG00000105542 | 0 | 0 | 0.25 | 0 | 0.387761935 | 0 | 0 |
| ENSMUSG00000105543 | 0 | 0 | 0 | 0 | 0 | 0 | 0 |
| ENSMUSG00000105544 | 0 | 0 | 0 | 0 | 0 | 0 | 0 |
| ENSMUSG00000105545 | 0 | 0 | 0 | 0 | 0 | 0 | 0 |
| ENSMUSG00000105546 | 0 | 0 | 0 | 0 | 0 | 0 | 0 |
| ENSMUSG00000105547 | 0 | 0 | 0 | 0 | 0 | 0 | 0 |
| ENSMUSG00000105548 | 0 | 0 | 0 | 0 | 0 | 0 | 0 |
| ENSMUSG00000105549 | 1.069666964 | 1.214495249 | 0.92510993 | 1.098079356 | 0.955822902 | 1 | 0.853775828 |
| ENSMUSG00000105550 | 0 | 0 | 0 | 0 | 0 | 0 | 0 |
| ENSMUSG00000105551 | 0 | 0 | 0 | 0 | 0 | 0 | 0 |
| ENSMUSG00000105552 | 1.249317394 | 1 | 1.348079356 | 0.976722649 | 1.176121988 | 0.602059991 | 1.110947927 |
| ENSMUSG00000105553 | 0.289064826 | 0.5 | 0.25 | 0.396240625 | 0.387761935 | 0 | 0.355418052 |
| ENSMUSG00000105554 | 0.289064826 | 0.580482024 | 0.580482024 | 0.5 | 0.387761935 | 0 | 0.579661877 |
| ENSMUSG00000105555 | 0 | 0 | 0 | 0 | 0 | 0 | 0 |
| ENSMUSG00000105556 | 1 | 1.04248125 | 0.92510993 | 0.951838731 | 0.931471446 | 0.903089987 | 0.853775828 |
| ENSMUSG00000105557 | 0 | 0 | 0 | 0 | 0 | 0 | 0 |
| ENSMUSG00000105558 | 1.623015126 | 1.580482024 | 1.705044741 | 1.25 | 1.580303447 | 1.579783597 | 1.42167221 |
| ENSMUSG00000105559 | 0 | 0 | 0 | 0 | 0 | 0 | 0 |
| ENSMUSG00000105560 | 0 | 0 | 0 | 0 | 0 | 0 | 0 |
| ENSMUSG00000105561 | 0.45815691 | 0.580482024 | 0.396240625 | 0.396240625 | 0.244650542 | 0.477121255 | 0.355418052 |
| ENSMUSG00000105562 | 0 | 0 | 0 | 0 | 0 | 0 | 0 |
| ENSMUSG00000105563 | 0 | 0 | 0 | 0 | 0 | 0 | 0 |
| ENSMUSG00000105564 | 0 | 0 | 0 | 0 | 0 | 0 | 0 |
| ENSMUSG00000105565 | 0.45815691 | 0.5 | 0.5 | 0 | 0.387761935 | 0 | 0.710836105 |
| ENSMUSG00000105566 | 0 | 0 | 0 | 0 | 0 | 0 | 0 |
| ENSMUSG00000105567 | 0 | 0 | 0 | 0 | 0 | 0 | 0 |
| ENSMUSG00000105568 | 0 | 0 | 0 | 0 | 0 | 0 | 0 |
| ENSMUSG00000105569 | 0 | 0 | 0 | 0 | 0 | 0 | 0 |

| | | | | | | | |
|---|---|---|---|---|---|---|---|
| ENSMUSG00000105570 | 0 | 0 | 0 | 0.25 | 0.244650542 | 0.301029996 | 0.355418052 |
| ENSMUSG00000105571 | 0 | 0 | 0 | 0 | 0 | 0 | 0 |
| ENSMUSG00000105572 | 1.325351389 | 1.25 | 1.201838731 | 1.282320754 | 1.293076518 | 1.301029996 | 1.089372236 |
| ENSMUSG00000105573 | 0 | 0 | 0 | 0 | 0 | 0 | 0 |
| ENSMUSG00000105574 | 0 | 0 | 0 | 0 | 0 | 0 | 0 |
| ENSMUSG00000105575 | 0 | 0 | 0 | 0 | 0 | 0 | 0 |
| ENSMUSG00000105576 | 0 | 0 | 0 | 0 | 0 | 0 | 0 |
| ENSMUSG00000105577 | 0 | 0 | 0 | 0 | 0 | 0 | 0 |
| ENSMUSG00000105578 | 1.325351389 | 1.238549078 | 1.17510993 | 1.410964047 | 1.136121934 | 1.176091259 | 1.054044578 |
| ENSMUSG00000105579 | 0 | 0.25 | 0 | 0 | 0 | 0 | 0 |
| ENSMUSG00000105580 | 0 | 0 | 0 | 0 | 0 | 0 | 0 |
| ENSMUSG00000105581 | 0 | 0 | 0 | 0 | 0 | 0 | 0 |
| ENSMUSG00000105582 | 0 | 0 | 0 | 0 | 0 | 0 | 0 |
| ENSMUSG00000105583 | 0 | 0 | 0 | 0 | 0 | 0 | 0 |
| ENSMUSG00000105584 | 0 | 0 | 0 | 0 | 0 | 0 | 0 |
| ENSMUSG00000105585 | 0 | 0 | 0 | 0 | 0 | 0 | 0 |
| ENSMUSG00000105586 | 0.289064826 | 0 | 0.25 | 0.5 | 0.244650542 | 0.301029996 | 0.629532004 |
| ENSMUSG00000105588 | 0 | 0 | 0 | 0 | 0 | 0 | 0 |
| ENSMUSG00000105589 | 0 | 0 | 0 | 0 | 0 | 0 | 0 |
| ENSMUSG00000105590 | 0.289064826 | 0 | 0 | 0 | 0 | 0 | 0 |
| ENSMUSG00000105591 | 0.289064826 | 0.5 | 0.396240625 | 0.25 | 0.489301084 | 0.301029996 | 0.355418052 |
| ENSMUSG00000105592 | 0 | 0 | 0 | 0 | 0 | 0 | 0 |
| ENSMUSG00000105593 | 0 | 0 | 0 | 0 | 0 | 0 | 0 |
| ENSMUSG00000105594 | 0 | 0 | 0 | 0.396240625 | 0.632412477 | 0 | 0.355418052 |
| ENSMUSG00000105595 | 0 | 0 | 0 | 0 | 0 | 0 | 0 |
| ENSMUSG00000105596 | 0 | 0 | 0 | 0 | 0 | 0 | 0 |
| ENSMUSG00000105597 | 0.747221736 | 0.646240625 | 0.580482024 | 0.580482024 | 0.846351821 | 0.477121255 | 0.448487648 |
| ENSMUSG00000105598 | 0 | 0 | 0 | 0 | 0 | 0 | 0 |
| ENSMUSG00000105599 | 0 | 0.25 | 0 | 0 | 0 | 0 | 0 |
| ENSMUSG00000105600 | 0.289064826 | 0.396240625 | 0 | 0 | 0.244650542 | 0 | 0.520678036 |
| ENSMUSG00000105601 | 0.45815691 | 0.646240625 | 0.79248125 | 0.646240625 | 0.686820904 | 0.602059991 | 0.629532004 |
| ENSMUSG00000105602 | 0 | 0 | 0 | 0.25 | 0.244650542 | 0 | 0.224243824 |
| ENSMUSG00000105603 | 0 | 0 | 0 | 0 | 0 | 0 | 0 |
| ENSMUSG00000105604 | 0 | 0 | 0 | 0 | 0 | 0 | 0 |

| | | | | | | | |
|---|---|---|---|---|---|---|---|
| ENSMUSG00000105605 | 0 | 0 | 0 | 0 | 0 | 0 | 0.224243824 |
| ENSMUSG00000105606 | 0 | 0 | 0 | 0 | 0 | 0 | 0 |
| ENSMUSG00000105607 | 0.811507563 | 0.79248125 | 0.951838731 | 0.92510993 | 1.039257769 | 0.602059991 | 0.672731473 |
| ENSMUSG00000105608 | 0 | 0 | 0 | 0 | 0 | 0 | 0 |
| ENSMUSG00000105609 | 0 | 0.25 | 0.25 | 0 | 0.244650542 | 0 | 0 |
| ENSMUSG00000105610 | 0 | 0 | 0 | 0 | 0 | 0 | 0 |
| ENSMUSG00000105611 | 0.45815691 | 0.25 | 0.25 | 0.396240625 | 0.387761935 | 0.602059991 | 0.224243824 |
| ENSMUSG00000105612 | 0 | 0 | 0 | 0 | 0 | 0 | 0 |
| ENSMUSG00000105613 | 0.747221736 | 0.75 | 0.896240625 | 0.92510993 | 0.877063019 | 0.84509804 | 0.74492186 |
| ENSMUSG00000105614 | 0 | 0 | 0 | 0 | 0 | 0 | 0 |
| ENSMUSG00000105615 | 0 | 0 | 0 | 0 | 0 | 0 | 0 |
| ENSMUSG00000105616 | 0 | 0 | 0 | 0 | 0 | 0 | 0 |
| ENSMUSG00000105617 | 0 | 0 | 0 | 0 | 0 | 0 | 0 |
| ENSMUSG00000105618 | 0 | 0 | 0 | 0 | 0 | 0 | 0 |
| ENSMUSG00000105619 | 0 | 0 | 0 | 0 | 0 | 0 | 0 |
| ENSMUSG00000105620 | 0 | 0 | 0 | 0 | 0 | 0 | 0 |
| ENSMUSG00000105621 | 0 | 0 | 0 | 0 | 0 | 0 | 0 |
| ENSMUSG00000105622 | 0 | 0.25 | 0.5 | 0 | 0 | 0 | 0 |
| ENSMUSG00000105624 | 0.45815691 | 0.5 | 0.580482024 | 0.25 | 0.244650542 | 0.477121255 | 0.355418052 |
| ENSMUSG00000105625 | 0 | 0 | 0 | 0 | 0 | 0 | 0 |
| ENSMUSG00000105626 | 0 | 0 | 0 | 0 | 0 | 0 | 0 |
| ENSMUSG00000105627 | 0 | 0 | 0 | 0 | 0 | 0 | 0 |
| ENSMUSG00000105628 | 0 | 0 | 0 | 0 | 0 | 0 | 0 |
| ENSMUSG00000105629 | 0 | 0 | 0 | 0 | 0 | 0 | 0 |
| ENSMUSG00000105630 | 0 | 0 | 0 | 0 | 0 | 0 | 0 |
| ENSMUSG00000105632 | 0 | 0.396240625 | 0.25 | 0.396240625 | 0.244650542 | 0 | 0 |
| ENSMUSG00000105633 | 0.289064826 | 0.25 | 0 | 0 | 0 | 0 | 0 |
| ENSMUSG00000105634 | 0 | 0 | 0 | 0 | 0 | 0 | 0 |
| ENSMUSG00000105635 | 0 | 0 | 0 | 0 | 0 | 0 | 0 |
| ENSMUSG00000105636 | 0 | 0.580482024 | 0.396240625 | 0.25 | 0.244650542 | 0 | 0.520678036 |
| ENSMUSG00000105637 | 0 | 0.396240625 | 0 | 0 | 0 | 0 | 0 |
| ENSMUSG00000105638 | 0 | 0 | 0 | 0 | 0 | 0 | 0 |
| ENSMUSG00000105639 | 0 | 0 | 0 | 0 | 0 | 0 | 0 |
| ENSMUSG00000105640 | 0 | 0 | 0 | 0 | 0 | 0 | 0 |

| | | | | | | | |
|---|---|---|---|---|---|---|---|
| ENSMUSG00000105641 | 0 | 0 | 0 | 0.396240625 | 0 | 0 | 0.224243824 |
| ENSMUSG00000105642 | 0 | 0 | 0 | 0 | 0 | 0 | 0 |
| ENSMUSG00000105643 | 0.747221736 | 0.5 | 0.701838731 | 0.830482024 | 0.812711509 | 0.602059991 | 0.579661877 |
| ENSMUSG00000105644 | 0 | 0.580482024 | 0.25 | 0.396240625 | 0 | 0.301029996 | 0.355418052 |
| ENSMUSG00000105645 | 0 | 0 | 0 | 0 | 0 | 0 | 0 |
| ENSMUSG00000105646 | 0 | 0 | 0 | 0 | 0 | 0 | 0 |
| ENSMUSG00000105647 | 0 | 0 | 0 | 0 | 0 | 0 | 0 |
| ENSMUSG00000105648 | 0 | 0 | 0 | 0 | 0 | 0 | 0 |
| ENSMUSG00000105649 | 0 | 0 | 0 | 0 | 0 | 0 | 0 |
| ENSMUSG00000105650 | 0 | 0 | 0 | 0 | 0 | 0.301029996 | 0 |
| ENSMUSG00000105651 | 0 | 0.25 | 0 | 0 | 0 | 0 | 0 |
| ENSMUSG00000105652 | 0.289064826 | 0 | 0.25 | 0 | 0.244650542 | 0 | 0 |
| ENSMUSG00000105653 | 0 | 0.25 | 0 | 0 | 0 | 0 | 0.224243824 |
| ENSMUSG00000105654 | 0 | 0 | 0 | 0 | 0 | 0 | 0 |
| ENSMUSG00000105655 | 1.069666964 | 0.92510993 | 0.830482024 | 1.061981878 | 1.020174412 | 1.041392685 | 0.829800754 |
| ENSMUSG00000105656 | 0 | 0 | 0 | 0 | 0 | 0 | 0 |
| ENSMUSG00000105657 | 0.45815691 | 0.396240625 | 0.830482024 | 0.5 | 0.632412477 | 0 | 0.355418052 |
| ENSMUSG00000105658 | 0 | 0 | 0 | 0.25 | 0 | 0 | 0 |
| ENSMUSG00000105659 | 0 | 0.25 | 0 | 0.396240625 | 0.244650542 | 0.301029996 | 0 |
| ENSMUSG00000105660 | 0 | 0 | 0 | 0 | 0 | 0 | 0 |
| ENSMUSG00000105661 | 0 | 0 | 0 | 0 | 0 | 0 | 0 |
| ENSMUSG00000105662 | 0 | 0 | 0 | 0 | 0 | 0 | 0 |
| ENSMUSG00000105663 | 0.45815691 | 0.580482024 | 0 | 0 | 0.387761935 | 0 | 0 |
| ENSMUSG00000105664 | 0 | 0 | 0 | 0 | 0 | 0 | 0.224243824 |
| ENSMUSG00000105665 | 0 | 0 | 0 | 0 | 0 | 0 | 0 |
| ENSMUSG00000105666 | 0 | 0 | 0 | 0 | 0 | 0 | 0 |
| ENSMUSG00000105667 | 0 | 0 | 0 | 0 | 0 | 0 | 0 |
| ENSMUSG00000105668 | 0 | 0 | 0 | 0 | 0 | 0 | 0 |
| ENSMUSG00000105669 | 0 | 0 | 0 | 0 | 0 | 0 | 0 |
| ENSMUSG00000105670 | 0 | 0 | 0.25 | 0 | 0 | 0.301029996 | 0 |
| ENSMUSG00000105671 | 0 | 0 | 0 | 0 | 0 | 0 | 0 |
| ENSMUSG00000105672 | 0 | 0 | 0 | 0 | 0 | 0 | 0 |
| ENSMUSG00000105673 | 0 | 0 | 0 | 0 | 0 | 0 | 0 |
| ENSMUSG00000105674 | 0 | 0 | 0 | 0 | 0 | 0 | 0 |

| | | | | | | | |
|---|---|---|---|---|---|---|---|
| ENSMUSG00000105675 | 0 | 0 | 0 | 0 | 0 | 0 | 0 |
| ENSMUSG00000105676 | 0 | 0 | 0.25 | 0.25 | 0 | 0.301029996 | 0.224243824 |
| ENSMUSG00000105677 | 1.069666964 | 0.951838731 | 0.701838731 | 0.951838731 | 0.77552387 | 0.477121255 | 0.896975297 |
| ENSMUSG00000105678 | 0 | 0.25 | 0 | 0.25 | 0.244650542 | 0 | 0 |
| ENSMUSG00000105679 | 0.811507563 | 0.701838731 | 0.646240625 | 0.864857905 | 0.489301084 | 0.477121255 | 0.448487648 |
| ENSMUSG00000105680 | 0 | 0 | 0 | 0 | 0 | 0 | 0 |
| ENSMUSG00000105681 | 0.671187741 | 0.396240625 | 0.25 | 0.396240625 | 0 | 0.477121255 | 0.579661877 |
| ENSMUSG00000105682 | 0 | 0 | 0 | 0 | 0 | 0 | 0 |
| ENSMUSG00000105683 | 0 | 0 | 0 | 0 | 0 | 0 | 0 |
| ENSMUSG00000105684 | 0 | 0 | 0 | 0 | 0 | 0 | 0 |
| ENSMUSG00000105685 | 0 | 0 | 0 | 0 | 0 | 0 | 0 |
| ENSMUSG00000105686 | 0 | 0 | 0 | 0 | 0 | 0 | 0 |
| ENSMUSG00000105687 | 0 | 0 | 0 | 0 | 0 | 0 | 0.224243824 |
| ENSMUSG00000105688 | 0 | 0 | 0 | 0 | 0 | 0 | 0 |
| ENSMUSG00000105689 | 0 | 0 | 0 | 0 | 0 | 0.301029996 | 0.224243824 |
| ENSMUSG00000105690 | 0 | 0 | 0 | 0 | 0 | 0 | 0 |
| ENSMUSG00000105691 | 0.45815691 | 0.5 | 0.5 | 0.5 | 0.686820904 | 0 | 0.448487648 |
| ENSMUSG00000105692 | 1.816888701 | 1.52186571 | 1.630890489 | 1.571350555 | 1.555378544 | 1.568201724 | 1.429563222 |
| ENSMUSG00000105693 | 0 | 0 | 0 | 0 | 0 | 0 | 0 |
| ENSMUSG00000105694 | 0 | 0.396240625 | 0 | 0 | 0 | 0 | 0 |
| ENSMUSG00000105695 | 1 | 0.864857905 | 1.061981878 | 0.976722649 | 0.877063019 | 0.954242509 | 0.829800754 |
| ENSMUSG00000105696 | 0 | 0 | 0 | 0 | 0 | 0 | 0 |
| ENSMUSG00000105697 | 0 | 0 | 0 | 0 | 0 | 0 | 0 |
| ENSMUSG00000105698 | 0 | 0 | 0.25 | 0 | 0 | 0 | 0 |
| ENSMUSG00000105699 | 0.578129653 | 0.5 | 0.864857905 | 0.646240625 | 0.686820904 | 0 | 0.448487648 |
| ENSMUSG00000105700 | 0 | 0 | 0.25 | 0 | 0 | 0 | 0.355418052 |
| ENSMUSG00000105701 | 0 | 0 | 0 | 0 | 0 | 0 | 0 |
| ENSMUSG00000105702 | 0 | 0 | 0 | 0 | 0 | 0 | 0 |
| ENSMUSG00000105703 | 2.188178997 | 2.078220739 | 2.091580554 | 2.044977273 | 2.058398634 | 2.23299611 | 1.958704689 |
| ENSMUSG00000105704 | 0 | 0 | 0 | 0 | 0 | 0 | 0 |
| ENSMUSG00000105705 | 0 | 0 | 0 | 0 | 0 | 0 | 0 |
| ENSMUSG00000105706 | 0 | 0 | 0.25 | 0 | 0 | 0 | 0 |
| ENSMUSG00000105707 | 0 | 0 | 0 | 0 | 0 | 0 | 0 |
| ENSMUSG00000105708 | 0.811507563 | 0.75 | 0.580482024 | 0 | 0.489301084 | 0 | 0 |

| | | | | | | | |
|---|---|---|---|---|---|---|---|
| ENSMUSG00000105709 | 0 | 0 | 0 | 0 | 0 | 0 | 0 |
| ENSMUSG00000105710 | 0.91631382 | 0.896240625 | 1 | 0.951838731 | 0.955822902 | 1.041392685 | 0.829800754 |
| ENSMUSG00000105711 | 0 | 0.25 | 0.25 | 0.25 | 0.387761935 | 0 | 0 |
| ENSMUSG00000105712 | 0 | 0 | 0 | 0 | 0 | 0 | 0 |
| ENSMUSG00000105713 | 0 | 0 | 0 | 0 | 0 | 0 | 0 |
| ENSMUSG00000105714 | 0 | 0 | 0 | 0 | 0 | 0 | 0 |
| ENSMUSG00000105715 | 0 | 0 | 0 | 0 | 0 | 0 | 0 |
| ENSMUSG00000105716 | 0 | 0 | 0 | 0 | 0 | 0 | 0 |
| ENSMUSG00000105717 | 0 | 0.396240625 | 0.25 | 0.396240625 | 0.387761935 | 0.477121255 | 0.579661877 |
| ENSMUSG00000105718 | 0 | 0 | 0 | 0 | 0 | 0 | 0 |
| ENSMUSG00000105719 | 0 | 0.396240625 | 0.25 | 0.701838731 | 0.733951626 | 0.602059991 | 0.629532004 |
| ENSMUSG00000105720 | 0 | 0 | 0 | 0 | 0.244650542 | 0 | 0 |
| ENSMUSG00000105721 | 0 | 0 | 0 | 0 | 0 | 0 | 0 |
| ENSMUSG00000105722 | 0.289064826 | 0 | 0 | 0 | 0 | 0 | 0.448487648 |
| ENSMUSG00000105723 | 0 | 0 | 0 | 0 | 0 | 0 | 0 |
| ENSMUSG00000105724 | 0 | 0 | 0 | 0 | 0 | 0 | 0 |
| ENSMUSG00000105725 | 0 | 0 | 0 | 0 | 0 | 0 | 0 |
| ENSMUSG00000105726 | 0 | 0 | 0 | 0 | 0 | 0 | 0 |
| ENSMUSG00000105727 | 0 | 0 | 0 | 0 | 0 | 0 | 0 |
| ENSMUSG00000105728 | 0.289064826 | 0.25 | 0.25 | 0.396240625 | 0.387761935 | 0 | 0.355418052 |
| ENSMUSG00000105729 | 0 | 0 | 0 | 0 | 0 | 0 | 0 |
| ENSMUSG00000105730 | 0 | 0 | 0 | 0 | 0 | 0 | 0 |
| ENSMUSG00000105731 | 0.45815691 | 0.25 | 0.25 | 0.5 | 0.387761935 | 0.477121255 | 0.224243824 |
| ENSMUSG00000105732 | 0 | 0 | 0 | 0 | 0 | 0 | 0 |
| ENSMUSG00000105733 | 0 | 0 | 0 | 0 | 0 | 0 | 0 |
| ENSMUSG00000105734 | 0 | 0 | 0 | 0 | 0 | 0 | 0.224243824 |
| ENSMUSG00000105735 | 0 | 0 | 0 | 0 | 0 | 0 | 0 |
| ENSMUSG00000105736 | 0 | 0 | 0 | 0 | 0 | 0 | 0 |
| ENSMUSG00000105737 | 0 | 0 | 0 | 0 | 0 | 0 | 0 |
| ENSMUSG00000105738 | 0 | 0 | 0 | 0 | 0 | 0 | 0 |
| ENSMUSG00000105739 | 0 | 0 | 0 | 0 | 0 | 0 | 0 |
| ENSMUSG00000105740 | 0.45815691 | 0.25 | 0 | 0.25 | 0.244650542 | 0 | 0.224243824 |
| ENSMUSG00000105741 | 0.867194479 | 0.79248125 | 0.79248125 | 0.92510993 | 0.733951626 | 1 | 0.803905701 |
| ENSMUSG00000105742 | 1.100572389 | 1.061981878 | 1.02186571 | 0.79248125 | 0.77552387 | 0.477121255 | 0.853775828 |

| | | | | | | | |
|---|---|---|---|---|---|---|---|
| ENSMUSG00000105743 | 0 | 0 | 0 | 0 | 0 | 0 | 0 |
| ENSMUSG00000105744 | 0 | 0 | 0 | 0 | 0 | 0 | 0 |
| ENSMUSG00000105745 | 0.45815691 | 0.646240625 | 0.646240625 | 0.580482024 | 0.568060967 | 0.301029996 | 0 |
| ENSMUSG00000105746 | 0.289064826 | 0 | 0 | 0 | 0 | 0 | 0 |
| ENSMUSG00000105747 | 0 | 0 | 0 | 0 | 0 | 0 | 0 |
| ENSMUSG00000105748 | 0 | 0.701838731 | 0.580482024 | 0.79248125 | 0.77552387 | 0.602059991 | 0.74492186 |
| ENSMUSG00000105749 | 0 | 0 | 0 | 0 | 0 | 0 | 0 |
| ENSMUSG00000105750 | 0 | 0.25 | 0.25 | 0 | 0 | 0 | 0 |
| ENSMUSG00000105751 | 0 | 0 | 0 | 0 | 0 | 0 | 0 |
| ENSMUSG00000105752 | 0 | 0 | 0 | 0 | 0 | 0 | 0 |
| ENSMUSG00000105753 | 0.45815691 | 0.5 | 0.396240625 | 0.396240625 | 0.244650542 | 0.698970004 | 0.520678036 |
| ENSMUSG00000105754 | 0 | 0 | 0 | 0 | 0 | 0 | 0 |
| ENSMUSG00000105755 | 0.289064826 | 0 | 0 | 0 | 0 | 0.301029996 | 0 |
| ENSMUSG00000105756 | 0 | 0 | 0 | 0 | 0 | 0 | 0 |
| ENSMUSG00000105757 | 0 | 0 | 0 | 0 | 0 | 0 | 0 |
| ENSMUSG00000105758 | 0 | 0 | 0 | 0 | 0 | 0 | 0 |
| ENSMUSG00000105759 | 0 | 0.25 | 0 | 0 | 0 | 0 | 0 |
| ENSMUSG00000105760 | 0 | 0 | 0 | 0 | 0 | 0 | 0 |
| ENSMUSG00000105761 | 0 | 0.25 | 0.25 | 0 | 0.387761935 | 0 | 0 |
| ENSMUSG00000105762 | 0 | 0.5 | 0.25 | 0 | 0 | 0 | 0 |
| ENSMUSG00000105763 | 0 | 0 | 0 | 0 | 0 | 0 | 0 |
| ENSMUSG00000105764 | 0 | 0 | 0 | 0 | 0 | 0 | 0 |
| ENSMUSG00000105765 | 0 | 0 | 0 | 0 | 0 | 0 | 0 |
| ENSMUSG00000105766 | 0 | 0 | 0 | 0 | 0 | 0.301029996 | 0 |
| ENSMUSG00000105767 | 0 | 0 | 0 | 0 | 0 | 0 | 0 |
| ENSMUSG00000105768 | 0 | 0 | 0 | 0 | 0 | 0 | 0 |
| ENSMUSG00000105769 | 0 | 0 | 0 | 0 | 0 | 0 | 0 |
| ENSMUSG00000105770 | 0 | 0 | 0 | 0 | 0 | 0 | 0 |
| ENSMUSG00000105771 | 0.289064826 | 0.396240625 | 0.580482024 | 0 | 0 | 0 | 0.579661877 |
| ENSMUSG00000105772 | 0 | 0 | 0 | 0.25 | 0 | 0.301029996 | 0.224243824 |
| ENSMUSG00000105773 | 0 | 0 | 0 | 0 | 0 | 0 | 0 |
| ENSMUSG00000105774 | 0 | 0 | 0.25 | 0 | 0 | 0 | 0 |
| ENSMUSG00000105775 | 0 | 0 | 0.25 | 0 | 0 | 0.301029996 | 0.224243824 |
| ENSMUSG00000105776 | 0.45815691 | 0.25 | 0.5 | 0.396240625 | 0.489301084 | 0.301029996 | 0.579661877 |

| | | | | | | | |
|---|---|---|---|---|---|---|---|
| ENSMUSG00000105777 | 0 | 0 | 0 | 0 | 0 | 0 | 0 |
| ENSMUSG00000105778 | 0 | 0 | 0 | 0 | 0 | 0 | 0 |
| ENSMUSG00000105779 | 0 | 0 | 0 | 0 | 0.244650542 | 0 | 0 |
| ENSMUSG00000105780 | 0 | 0 | 0 | 0 | 0 | 0 | 0 |
| ENSMUSG00000105781 | 0 | 0 | 0 | 0 | 0 | 0 | 0 |
| ENSMUSG00000105782 | 0 | 0 | 0.25 | 0 | 0 | 0 | 0 |
| ENSMUSG00000105783 | 0 | 0 | 0 | 0 | 0 | 0 | 0 |
| ENSMUSG00000105784 | 0 | 0 | 0 | 0 | 0 | 0 | 0 |
| ENSMUSG00000105785 | 0 | 0 | 0 | 0 | 0 | 0 | 0 |
| ENSMUSG00000105786 | 0 | 0 | 0 | 0 | 0 | 0 | 0 |
| ENSMUSG00000105787 | 0.289064826 | 0.25 | 0 | 0 | 0 | 0 | 0.224243824 |
| ENSMUSG00000105788 | 0 | 0 | 0 | 0 | 0 | 0 | 0 |
| ENSMUSG00000105789 | 0.289064826 | 0 | 0 | 0.5 | 0.387761935 | 0 | 0.355418052 |
| ENSMUSG00000105790 | 0 | 0 | 0 | 0 | 0 | 0 | 0 |
| ENSMUSG00000105791 | 0 | 0 | 0 | 0 | 0 | 0 | 0 |
| ENSMUSG00000105792 | 0 | 0 | 0 | 0 | 0 | 0 | 0 |
| ENSMUSG00000105793 | 0 | 0.25 | 0 | 0.25 | 0.244650542 | 0 | 0 |
| ENSMUSG00000105794 | 0 | 0 | 0 | 0 | 0 | 0 | 0 |
| ENSMUSG00000105795 | 0 | 0 | 0 | 0 | 0 | 0 | 0 |
| ENSMUSG00000105796 | 0 | 0 | 0 | 0 | 0 | 0 | 0 |
| ENSMUSG00000105797 | 0 | 0 | 0.25 | 0 | 0 | 0 | 0 |
| ENSMUSG00000105798 | 0 | 0 | 0 | 0 | 0 | 0 | 0 |
| ENSMUSG00000105799 | 0 | 0 | 0 | 0 | 0 | 0 | 0 |
| ENSMUSG00000105800 | 0 | 0 | 0 | 0 | 0 | 0 | 0 |
| ENSMUSG00000105801 | 0 | 0 | 0 | 0.25 | 0 | 0 | 0 |
| ENSMUSG00000105802 | 0.289064826 | 0.580482024 | 0.396240625 | 0.25 | 0 | 0 | 0 |
| ENSMUSG00000105803 | 0 | 0 | 0 | 0 | 0 | 0 | 0 |
| ENSMUSG00000105804 | 0.45815691 | 0.646240625 | 0.396240625 | 0.830482024 | 0.846351821 | 0.698970004 | 0.355418052 |
| ENSMUSG00000105805 | 0 | 0 | 0 | 0 | 0 | 0 | 0 |
| ENSMUSG00000105806 | 0 | 0 | 0 | 0 | 0 | 0 | 0 |
| ENSMUSG00000105807 | 0 | 0 | 0 | 0 | 0 | 0 | 0 |
| ENSMUSG00000105808 | 0 | 0 | 0 | 0 | 0 | 0 | 0 |
| ENSMUSG00000105809 | 0 | 0 | 0 | 0 | 0 | 0 | 0 |
| ENSMUSG00000105810 | 0 | 0.25 | 0 | 0 | 0.387761935 | 0.301029996 | 0.224243824 |

| | | | | | | | |
|---|---|---|---|---|---|---|---|
| ENSMUSG00000105811 | 0 | 0 | 0 | 0 | 0 | 0 | 0 |
| ENSMUSG00000105812 | 0 | 0 | 0 | 0 | 0 | 0 | 0 |
| ENSMUSG00000105813 | 0.747221736 | 0.75 | 0.79248125 | 0.646240625 | 0.568060967 | 0.602059991 | 0.448487648 |
| ENSMUSG00000105814 | 1.558729299 | 1.51109853 | 1.339388001 | 1.571350555 | 1.456697355 | 1.643452676 | 1.401059159 |
| ENSMUSG00000105815 | 0 | 0 | 0.25 | 0 | 0 | 0.301029996 | 0 |
| ENSMUSG00000105816 | 0 | 0 | 0 | 0 | 0 | 0 | 0 |
| ENSMUSG00000105817 | 0 | 0 | 0 | 0 | 0 | 0 | 0 |
| ENSMUSG00000105818 | 0 | 0.25 | 0.396240625 | 0.396240625 | 0 | 0.477121255 | 0.355418052 |
| ENSMUSG00000105819 | 0 | 0 | 0 | 0 | 0 | 0 | 0 |
| ENSMUSG00000105820 | 0 | 0 | 0 | 0 | 0 | 0 | 0 |
| ENSMUSG00000105821 | 0 | 0 | 0 | 0 | 0 | 0 | 0 |
| ENSMUSG00000105822 | 0 | 0 | 0 | 0 | 0 | 0 | 0 |
| ENSMUSG00000105823 | 0 | 0 | 0 | 0 | 0 | 0 | 0 |
| ENSMUSG00000105824 | 0.289064826 | 0.396240625 | 0.25 | 0.396240625 | 0.244650542 | 0 | 0.224243824 |
| ENSMUSG00000105825 | 0 | 0 | 0 | 0 | 0 | 0 | 0 |
| ENSMUSG00000105826 | 0.960252568 | 0.646240625 | 0.75 | 0.830482024 | 0.568060967 | 1 | 0.803905701 |
| ENSMUSG00000105827 | 0 | 0 | 0 | 0 | 0 | 0 | 0 |
| ENSMUSG00000105828 | 0 | 0 | 0 | 0 | 0 | 0 | 0 |
| ENSMUSG00000105829 | 0 | 0 | 0 | 0 | 0 | 0 | 0.224243824 |
| ENSMUSG00000105830 | 0 | 0 | 0 | 0 | 0 | 0 | 0 |
| ENSMUSG00000105831 | 0 | 0 | 0 | 0 | 0 | 0 | 0 |
| ENSMUSG00000105832 | 0.960252568 | 1.17510993 | 1.214495249 | 1.098079356 | 1.188507684 | 1.113943352 | 0.969165684 |
| ENSMUSG00000105833 | 0 | 0 | 0.5 | 0.25 | 0.244650542 | 0 | 0.448487648 |
| ENSMUSG00000105834 | 0 | 0 | 0 | 0 | 0 | 0 | 0.355418052 |
| ENSMUSG00000105835 | 0 | 0 | 0 | 0 | 0 | 0 | 0 |
| ENSMUSG00000105836 | 0 | 0 | 0.25 | 0.25 | 0 | 0.301029996 | 0 |
| ENSMUSG00000105837 | 0.289064826 | 0 | 0 | 0.25 | 0.244650542 | 0 | 0.224243824 |
| ENSMUSG00000105838 | 0.289064826 | 0 | 0 | 0 | 0 | 0 | 0 |
| ENSMUSG00000105839 | 0 | 0 | 0 | 0 | 0 | 0 | 0 |
| ENSMUSG00000105840 | 0 | 0 | 0 | 0 | 0 | 0 | 0 |
| ENSMUSG00000105841 | 0 | 0 | 0 | 0 | 0 | 0 | 0 |
| ENSMUSG00000105842 | 0.289064826 | 0.5 | 0 | 0.5 | 0.568060967 | 0.301029996 | 0.448487648 |
| ENSMUSG00000105843 | 0 | 0.396240625 | 0 | 0 | 0 | 0 | 0 |
| ENSMUSG00000105844 | 0 | 0 | 0 | 0 | 0 | 0 | 0 |

| | | | | | | | |
|---|---|---|---|---|---|---|---|
| ENSMUSG00000105845 | 0 | 0 | 0 | 0 | 0 | 0 | 0.224243824 |
| ENSMUSG00000105846 | 0.45815691 | 0.646240625 | 0.646240625 | 0.5 | 0.568060967 | 0.477121255 | 0.448487648 |
| ENSMUSG00000105847 | 0 | 0 | 0 | 0 | 0 | 0 | 0 |
| ENSMUSG00000105848 | 0.289064826 | 0.25 | 0.25 | 0.25 | 0 | 0 | 0 |
| ENSMUSG00000105849 | 0 | 0 | 0 | 0 | 0 | 0 | 0 |
| ENSMUSG00000105850 | 0 | 0 | 0 | 0 | 0 | 0 | 0 |
| ENSMUSG00000105851 | 0.671187741 | 0.864857905 | 0.92510993 | 0.75 | 0.387761935 | 0.602059991 | 0.629532004 |
| ENSMUSG00000105852 | 0.578129653 | 0.580482024 | 0.396240625 | 0.396240625 | 0.387761935 | 0 | 0.74492186 |
| ENSMUSG00000105853 | 0 | 0 | 0 | 0 | 0 | 0 | 0 |
| ENSMUSG00000105854 | 0 | 0.25 | 0.646240625 | 0 | 0.632412477 | 0.301029996 | 0.224243824 |
| ENSMUSG00000105855 | 0.867194479 | 1.04248125 | 0.896240625 | 0.701838731 | 0.877063019 | 0.84509804 | 0.829800754 |
| ENSMUSG00000105856 | 0 | 0 | 0 | 0 | 0 | 0 | 0 |
| ENSMUSG00000105857 | 0 | 0 | 0 | 0 | 0 | 0 | 0 |
| ENSMUSG00000105858 | 0.289064826 | 0.75 | 0.396240625 | 0 | 0.244650542 | 0 | 0.355418052 |
| ENSMUSG00000105859 | 0 | 0 | 0 | 0 | 0 | 0 | 0 |
| ENSMUSG00000105860 | 0 | 0 | 0 | 0.25 | 0.387761935 | 0.301029996 | 0 |
| ENSMUSG00000105861 | 0 | 0 | 0 | 0 | 0 | 0 | 0 |
| ENSMUSG00000105862 | 0 | 0 | 0 | 0 | 0 | 0 | 0 |
| ENSMUSG00000105863 | 0 | 0 | 0 | 0 | 0 | 0 | 0 |
| ENSMUSG00000105864 | 0 | 0.25 | 0 | 0.25 | 0 | 0 | 0.355418052 |
| ENSMUSG00000105865 | 0.45815691 | 0.25 | 0 | 0.25 | 0.387761935 | 0 | 0.224243824 |
| ENSMUSG00000105866 | 0 | 0.5 | 0 | 0.5 | 0.387761935 | 0 | 0 |
| ENSMUSG00000105867 | 0 | 0.396240625 | 0.25 | 0 | 0.244650542 | 0 | 0.448487648 |
| ENSMUSG00000105868 | 0.867194479 | 0.830482024 | 0.396240625 | 0.864857905 | 0.686820904 | 0.477121255 | 0.803905701 |
| ENSMUSG00000105869 | 0 | 0 | 0 | 0 | 0 | 0 | 0 |
| ENSMUSG00000105870 | 0 | 0.25 | 0.396240625 | 0 | 0 | 0 | 0 |
| ENSMUSG00000105871 | 0 | 0 | 0 | 0 | 0 | 0 | 0 |
| ENSMUSG00000105872 | 0 | 0 | 0 | 0 | 0 | 0 | 0 |
| ENSMUSG00000105873 | 0 | 0 | 0 | 0 | 0 | 0 | 0 |
| ENSMUSG00000105874 | 0 | 0 | 0 | 0 | 0 | 0 | 0 |
| ENSMUSG00000105875 | 0 | 0.25 | 0 | 0 | 0 | 0 | 0 |
| ENSMUSG00000105876 | 0 | 0 | 0 | 0 | 0 | 0 | 0 |
| ENSMUSG00000105877 | 0 | 0.25 | 0.25 | 0 | 0.244650542 | 0 | 0.224243824 |
| ENSMUSG00000105878 | 0 | 0 | 0 | 0 | 0 | 0 | 0 |

| | | | | | | | |
|---|---|---|---|---|---|---|---|
| ENSMUSG00000105879 | 0 | 0.25 | 0.25 | 0 | 0 | 0 | 0 |
| ENSMUSG00000105880 | 0 | 0 | 0 | 0 | 0 | 0 | 0 |
| ENSMUSG00000105881 | 0.811507563 | 0.976722649 | 1.080482024 | 0.5 | 0.77552387 | 0.301029996 | 0.74492186 |
| ENSMUSG00000105882 | 0 | 0 | 0 | 0 | 0 | 0 | 0 |
| ENSMUSG00000105883 | 0 | 0 | 0 | 0 | 0 | 0 | 0 |
| ENSMUSG00000105884 | 0 | 0 | 0 | 0 | 0 | 0 | 0 |
| ENSMUSG00000105885 | 0 | 0 | 0 | 0 | 0 | 0 | 0 |
| ENSMUSG00000105886 | 0 | 0 | 0 | 0 | 0 | 0 | 0 |
| ENSMUSG00000105887 | 0 | 0 | 0 | 0 | 0 | 0 | 0 |
| ENSMUSG00000105888 | 0 | 0 | 0 | 0 | 0 | 0 | 0 |
| ENSMUSG00000105889 | 0 | 0 | 0 | 0 | 0 | 0 | 0 |
| ENSMUSG00000105890 | 0 | 0 | 0 | 0 | 0 | 0 | 0 |
| ENSMUSG00000105891 | 0 | 0 | 0.25 | 0 | 0 | 0 | 0 |
| ENSMUSG00000105892 | 0 | 0 | 0 | 0.25 | 0 | 0 | 0 |
| ENSMUSG00000105893 | 0 | 0 | 0.25 | 0.25 | 0.244650542 | 0 | 0 |
| ENSMUSG00000105894 | 0 | 0 | 0 | 0 | 0 | 0 | 0 |
| ENSMUSG00000105895 | 0.289064826 | 0.79248125 | 0.396240625 | 0.896240625 | 0.905314583 | 0.77815125 | 0.579661877 |
| ENSMUSG00000105896 | 0.747221736 | 0.5 | 0.5 | 0.580482024 | 0.568060967 | 0.698970004 | 0.629532004 |
| ENSMUSG00000105897 | 0 | 0 | 0 | 0 | 0 | 0 | 0 |
| ENSMUSG00000105898 | 0 | 0 | 0 | 0 | 0 | 0 | 0 |
| ENSMUSG00000105899 | 0 | 0 | 0 | 0 | 0 | 0 | 0 |
| ENSMUSG00000105900 | 0 | 0 | 0 | 0 | 0 | 0 | 0 |
| ENSMUSG00000105901 | 0 | 0 | 0 | 0 | 0 | 0 | 0 |
| ENSMUSG00000105902 | 0 | 0 | 0 | 0 | 0 | 0 | 0 |
| ENSMUSG00000105903 | 0 | 0 | 0 | 0 | 0 | 0 | 0 |
| ENSMUSG00000105904 | 0 | 0 | 0 | 0 | 0 | 0 | 0 |
| ENSMUSG00000105905 | 0 | 0 | 0 | 0.25 | 0 | 0.301029996 | 0.224243824 |
| ENSMUSG00000105906 | 0 | 0 | 0 | 0 | 0 | 0 | 0 |
| ENSMUSG00000105907 | 0 | 0 | 0 | 0 | 0 | 0 | 0 |
| ENSMUSG00000105908 | 0 | 0 | 0 | 0 | 0 | 0 | 0 |
| ENSMUSG00000105909 | 0 | 0.25 | 0 | 0 | 0 | 0 | 0 |
| ENSMUSG00000105910 | 0 | 0 | 0 | 0 | 0 | 0 | 0 |
| ENSMUSG00000105911 | 0 | 0 | 0 | 0 | 0 | 0 | 0 |
| ENSMUSG00000105912 | 0 | 0 | 0 | 0 | 0 | 0 | 0 |

| | | | | | | | |
|---|---|---|---|---|---|---|---|
| ENSMUSG00000105913 | 0 | 0 | 0 | 0 | 0 | 0 | 0 |
| ENSMUSG00000105914 | 0 | 0 | 0 | 0 | 0 | 0 | 0 |
| ENSMUSG00000105915 | 0 | 0 | 0 | 0 | 0 | 0 | 0 |
| ENSMUSG00000105916 | 0.289064826 | 0 | 0 | 0 | 0 | 0.301029996 | 0 |
| ENSMUSG00000105917 | 0 | 0 | 0.25 | 0 | 0 | 0 | 0 |
| ENSMUSG00000105918 | 0 | 0 | 0 | 0 | 0 | 0 | 0 |
| ENSMUSG00000105919 | 0 | 0 | 0 | 0 | 0 | 0 | 0 |
| ENSMUSG00000105920 | 0 | 0 | 0 | 0 | 0 | 0 | 0 |
| ENSMUSG00000105921 | 0 | 0 | 0 | 0 | 0 | 0 | 0 |
| ENSMUSG00000105922 | 0.289064826 | 0 | 0 | 0.25 | 0.387761935 | 0 | 0.355418052 |
| ENSMUSG00000105923 | 0 | 0 | 0 | 0 | 0 | 0 | 0 |
| ENSMUSG00000105924 | 0 | 0 | 0 | 0 | 0 | 0 | 0 |
| ENSMUSG00000105925 | 0 | 0 | 0 | 0 | 0.244650542 | 0 | 0 |
| ENSMUSG00000105926 | 0.289064826 | 0 | 0 | 0 | 0 | 0 | 0 |
| ENSMUSG00000105927 | 0 | 0 | 0 | 0 | 0 | 0 | 0 |
| ENSMUSG00000105928 | 0 | 0 | 0 | 0 | 0 | 0 | 0 |
| ENSMUSG00000105929 | 0.45815691 | 0.646240625 | 0.864857905 | 0.5 | 0.489301084 | 0.602059991 | 0.355418052 |
| ENSMUSG00000105930 | 0 | 0 | 0 | 0 | 0 | 0 | 0 |
| ENSMUSG00000105931 | 0.747221736 | 0.580482024 | 0.580482024 | 0.396240625 | 0.387761935 | 0.602059991 | 0.74492186 |
| ENSMUSG00000105932 | 0 | 0 | 0 | 0.25 | 0 | 0 | 0 |
| ENSMUSG00000105933 | 0 | 0.25 | 0.5 | 0 | 0 | 0 | 0.224243824 |
| ENSMUSG00000105934 | 0 | 0 | 0 | 0 | 0 | 0 | 0 |
| ENSMUSG00000105935 | 0 | 0.396240625 | 0 | 0.396240625 | 0.632412477 | 0 | 0.579661877 |
| ENSMUSG00000105936 | 0 | 0 | 0 | 0.25 | 0.244650542 | 0.301029996 | 0.224243824 |
| ENSMUSG00000105937 | 0 | 0 | 0 | 0 | 0 | 0 | 0 |
| ENSMUSG00000105938 | 0 | 0 | 0 | 0 | 0 | 0 | 0 |
| ENSMUSG00000105939 | 0 | 0.25 | 0.396240625 | 0.5 | 0.489301084 | 0.477121255 | 0.355418052 |
| ENSMUSG00000105940 | 0.671187741 | 0.75 | 0.75 | 0.701838731 | 0.244650542 | 0.698970004 | 0.224243824 |
| ENSMUSG00000105941 | 0 | 0.25 | 0 | 0 | 0 | 0.477121255 | 0 |
| ENSMUSG00000105942 | 0 | 0 | 0 | 0.396240625 | 0 | 0 | 0.224243824 |
| ENSMUSG00000105943 | 0 | 0 | 0 | 0 | 0 | 0 | 0 |
| ENSMUSG00000105944 | 0.45815691 | 0 | 0.25 | 0 | 0 | 0 | 0.224243824 |
| ENSMUSG00000105945 | 0 | 0.25 | 0.25 | 0.25 | 0.387761935 | 0.477121255 | 0.448487648 |
| ENSMUSG00000105946 | 0 | 0 | 0.25 | 0 | 0 | 0 | 0 |

| | | | | | | | |
|---|---|---|---|---|---|---|---|
| ENSMUSG00000105947 | 0.578129653 | 0 | 0 | 0.25 | 0.244650542 | 0 | 0 |
| ENSMUSG00000105948 | 0 | 0 | 0 | 0.25 | 0 | 0 | 0 |
| ENSMUSG00000105949 | 0 | 0 | 0 | 0 | 0 | 0 | 0 |
| ENSMUSG00000105950 | 1.181541736 | 1.114857905 | 1.17510993 | 1.160964047 | 1.039257769 | 1.146128036 | 1.078019652 |
| ENSMUSG00000105951 | 0 | 0 | 0 | 0 | 0 | 0 | 0 |
| ENSMUSG00000105952 | 0 | 0 | 0 | 0 | 0 | 0 | 0 |
| ENSMUSG00000105953 | 0 | 0 | 0 | 0 | 0 | 0 | 0 |
| ENSMUSG00000105954 | 0 | 0.580482024 | 0.25 | 0 | 0 | 0 | 0 |
| ENSMUSG00000105955 | 0 | 0 | 0 | 0 | 0 | 0 | 0 |
| ENSMUSG00000105956 | 0 | 0 | 0 | 0 | 0 | 0 | 0 |
| ENSMUSG00000105957 | 0.747221736 | 0.864857905 | 0.646240625 | 0.701838731 | 0.846351821 | 0.954242509 | 0.629532004 |
| ENSMUSG00000105958 | 0.289064826 | 0 | 0 | 0 | 0 | 0 | 0 |
| ENSMUSG00000105959 | 0 | 0 | 0 | 0 | 0 | 0 | 0 |
| ENSMUSG00000105960 | 0 | 0 | 0 | 0 | 0 | 0.301029996 | 0 |
| ENSMUSG00000105961 | 0 | 0 | 0 | 0 | 0 | 0.301029996 | 0 |
| ENSMUSG00000105962 | 0.289064826 | 0.396240625 | 0 | 0.5 | 0.632412477 | 0.602059991 | 0.629532004 |
| ENSMUSG00000105963 | 0 | 0 | 0 | 0 | 0 | 0 | 0 |
| ENSMUSG00000105964 | 0 | 0 | 0 | 0 | 0 | 0 | 0 |
| ENSMUSG00000105965 | 0 | 0 | 0 | 0 | 0.244650542 | 0 | 0 |
| ENSMUSG00000105966 | 0 | 0 | 0 | 0 | 0 | 0 | 0 |
| ENSMUSG00000105967 | 0 | 0 | 0 | 0 | 0 | 0 | 0 |
| ENSMUSG00000105968 | 0 | 0 | 0 | 0 | 0 | 0 | 0 |
| ENSMUSG00000105969 | 0 | 0 | 0 | 0 | 0 | 0 | 0 |
| ENSMUSG00000105970 | 0.289064826 | 0.25 | 0.396240625 | 0.5 | 0 | 0.301029996 | 0.224243824 |
| ENSMUSG00000105971 | 0 | 0 | 0 | 0 | 0 | 0 | 0 |
| ENSMUSG00000105972 | 0 | 0 | 0 | 0 | 0 | 0 | 0 |
| ENSMUSG00000105973 | 0.289064826 | 0 | 0 | 0.25 | 0.489301084 | 0 | 0.224243824 |
| ENSMUSG00000105974 | 0.45815691 | 0 | 0.396240625 | 0.396240625 | 0.387761935 | 0.301029996 | 0 |
| ENSMUSG00000105975 | 0 | 0 | 0.25 | 0.25 | 0 | 0 | 0 |
| ENSMUSG00000105976 | 0 | 0.25 | 0.580482024 | 0.25 | 0 | 0.301029996 | 0.448487648 |
| ENSMUSG00000105977 | 0 | 0 | 0 | 0 | 0 | 0 | 0 |
| ENSMUSG00000105978 | 0 | 0 | 0 | 0 | 0 | 0 | 0 |
| ENSMUSG00000105979 | 0 | 0 | 0 | 0 | 0 | 0 | 0 |
| ENSMUSG00000105980 | 0 | 0 | 0.25 | 0.396240625 | 0 | 0 | 0 |

| | | | | | | | |
|---|---|---|---|---|---|---|---|
| ENSMUSG00000105981 | 0.45815691 | 0.92510993 | 0.79248125 | 0.951838731 | 0.846351821 | 0.77815125 | 0.876096088 |
| ENSMUSG00000105982 | 0 | 0 | 0 | 0 | 0 | 0 | 0 |
| ENSMUSG00000105983 | 0 | 0 | 0 | 0 | 0 | 0 | 0 |
| ENSMUSG00000105984 | 0 | 0 | 0 | 0 | 0 | 0 | 0 |
| ENSMUSG00000105985 | 0 | 0 | 0 | 0 | 0 | 0 | 0 |
| ENSMUSG00000105986 | 0.289064826 | 0.25 | 0 | 0 | 0 | 0 | 0 |
| ENSMUSG00000105987 | 2.848245215 | 2.618426437 | 2.624961472 | 2.605778978 | 2.626873383 | 2.957128198 | 2.451150342 |
| ENSMUSG00000105988 | 0 | 0 | 0 | 0 | 0 | 0 | 0 |
| ENSMUSG00000105989 | 0 | 0 | 0 | 0 | 0 | 0 | 0 |
| ENSMUSG00000105990 | 0.289064826 | 0.701838731 | 0.646240625 | 0.864857905 | 0.846351821 | 0.698970004 | 0.710836105 |
| ENSMUSG00000105991 | 0 | 0 | 0 | 0 | 0 | 0 | 0 |
| ENSMUSG00000105992 | 0 | 0.25 | 0 | 0.396240625 | 0.387761935 | 0 | 0.224243824 |
| ENSMUSG00000105993 | 0 | 0 | 0 | 0 | 0 | 0 | 0 |
| ENSMUSG00000105994 | 0 | 0 | 0 | 0 | 0 | 0 | 0 |
| ENSMUSG00000105995 | 0 | 0 | 0 | 0 | 0 | 0 | 0 |
| ENSMUSG00000105996 | 0 | 0 | 0 | 0 | 0 | 0 | 0.224243824 |
| ENSMUSG00000105997 | 0 | 0 | 0 | 0 | 0 | 0 | 0 |
| ENSMUSG00000105998 | 0 | 0 | 0 | 0 | 0 | 0 | 0 |
| ENSMUSG00000105999 | 0 | 0 | 0 | 0 | 0.244650542 | 0 | 0.224243824 |
| ENSMUSG00000106000 | 0.45815691 | 0.396240625 | 0.580482024 | 0.25 | 0.244650542 | 0 | 0 |
| ENSMUSG00000106001 | 0.45815691 | 0.580482024 | 0.396240625 | 0 | 0.244650542 | 0.698970004 | 0.224243824 |
| ENSMUSG00000106002 | 0 | 0 | 0 | 0 | 0 | 0 | 0 |
| ENSMUSG00000106003 | 0 | 0 | 0 | 0 | 0 | 0 | 0 |
| ENSMUSG00000106004 | 0 | 0 | 0 | 0 | 0 | 0 | 0 |
| ENSMUSG00000106005 | 0 | 0 | 0 | 0.25 | 0 | 0 | 0 |
| ENSMUSG00000106006 | 0 | 0 | 0 | 0 | 0 | 0 | 0 |
| ENSMUSG00000106007 | 0 | 0 | 0 | 0 | 0 | 0 | 0 |
| ENSMUSG00000106008 | 0 | 0 | 0 | 0 | 0 | 0 | 0 |
| ENSMUSG00000106009 | 0.289064826 | 0.25 | 0.580482024 | 0 | 0 | 0 | 0.355418052 |
| ENSMUSG00000106010 | 0 | 0 | 0.396240625 | 0 | 0 | 0 | 0 |
| ENSMUSG00000106011 | 0.289064826 | 0 | 0 | 0 | 0 | 0 | 0 |
| ENSMUSG00000106012 | 0 | 0.25 | 0.25 | 0 | 0 | 0 | 0 |
| ENSMUSG00000106013 | 0 | 0 | 0 | 0 | 0 | 0 | 0 |
| ENSMUSG00000106014 | 0 | 0 | 0 | 0 | 0 | 0 | 0 |

| | | | | | | | |
|---|---|---|---|---|---|---|---|
| ENSMUSG00000106015 | 0.289064826 | 0 | 0 | 0 | 0 | 0 | 0 |
| ENSMUSG00000106016 | 0 | 0 | 0 | 0 | 0 | 0 | 0 |
| ENSMUSG00000106017 | 0 | 0 | 0 | 0 | 0 | 0 | 0 |
| ENSMUSG00000106018 | 0 | 0 | 0 | 0 | 0 | 0 | 0 |
| ENSMUSG00000106019 | 1.181541736 | 1.188721876 | 0.830482024 | 1.114857905 | 0.846351821 | 1.146128036 | 0.984950056 |
| ENSMUSG00000106020 | 0 | 0 | 0 | 0 | 0 | 0 | 0 |
| ENSMUSG00000106021 | 0 | 0 | 0 | 0 | 0 | 0 | 0 |
| ENSMUSG00000106022 | 0.91631382 | 0.951838731 | 0.830482024 | 0.79248125 | 0.931471446 | 0.77815125 | 0.74492186 |
| ENSMUSG00000106023 | 0 | 0 | 0 | 0 | 0 | 0 | 0 |
| ENSMUSG00000106024 | 0 | 0 | 0 | 0 | 0 | 0 | 0 |
| ENSMUSG00000106025 | 0.289064826 | 0.580482024 | 0 | 0 | 0.489301084 | 0.301029996 | 0.579661877 |
| ENSMUSG00000106026 | 0 | 0 | 0 | 0 | 0 | 0 | 0 |
| ENSMUSG00000106027 | 0 | 0 | 0 | 0 | 0 | 0 | 0 |
| ENSMUSG00000106028 | 0 | 0 | 0.5 | 0 | 0 | 0 | 0.224243824 |
| ENSMUSG00000106029 | 0 | 0 | 0 | 0 | 0 | 0 | 0 |
| ENSMUSG00000106030 | 0 | 0 | 0 | 0 | 0 | 0 | 0 |
| ENSMUSG00000106031 | 0 | 0 | 0 | 0 | 0 | 0 | 0 |
| ENSMUSG00000106032 | 1 | 1.160964047 | 1 | 0.864857905 | 0.489301084 | 0.477121255 | 0.672731473 |
| ENSMUSG00000106033 | 0 | 0 | 0 | 0 | 0 | 0 | 0 |
| ENSMUSG00000106034 | 0 | 0 | 0 | 0 | 0 | 0 | 0 |
| ENSMUSG00000106035 | 0 | 0.580482024 | 0.701838731 | 0.580482024 | 0 | 0 | 0.355418052 |
| ENSMUSG00000106036 | 0.289064826 | 0.25 | 0.25 | 0 | 0 | 0 | 0 |
| ENSMUSG00000106037 | 0.578129653 | 0.5 | 0.646240625 | 0.580482024 | 0.489301084 | 0 | 0.520678036 |
| ENSMUSG00000106038 | 0 | 0 | 0 | 0 | 0.244650542 | 0.301029996 | 0.224243824 |
| ENSMUSG00000106039 | 0 | 0 | 0 | 0 | 0 | 0 | 0 |
| ENSMUSG00000106040 | 0 | 0 | 0 | 0 | 0 | 0 | 0 |
| ENSMUSG00000106041 | 0 | 0 | 0 | 0 | 0 | 0 | 0 |
| ENSMUSG00000106042 | 0 | 0.25 | 0 | 0 | 0 | 0 | 0.224243824 |
| ENSMUSG00000106043 | 0 | 0 | 0 | 0 | 0 | 0 | 0 |
| ENSMUSG00000106044 | 2.036286563 | 1.961372513 | 1.921625132 | 1.921625132 | 1.883916752 | 2.017033339 | 1.700880998 |
| ENSMUSG00000106045 | 0 | 0 | 0 | 0 | 0 | 0 | 0 |
| ENSMUSG00000106046 | 0 | 0 | 0 | 0 | 0 | 0 | 0 |
| ENSMUSG00000106047 | 0 | 0 | 0 | 0 | 0 | 0 | 0 |
| ENSMUSG00000106048 | 0.289064826 | 0 | 0 | 0.396240625 | 0 | 0 | 0 |

| | | | | | | | |
|---|---|---|---|---|---|---|---|
| ENSMUSG00000106049 | 0 | 0 | 0 | 0 | 0 | 0 | 0 |
| ENSMUSG00000106050 | 0 | 0 | 0 | 0 | 0 | 0 | 0 |
| ENSMUSG00000106051 | 0 | 0 | 0 | 0 | 0 | 0 | 0 |
| ENSMUSG00000106052 | 0 | 0.25 | 0 | 0 | 0 | 0 | 0 |
| ENSMUSG00000106053 | 0.289064826 | 0.25 | 0 | 0.396240625 | 0.489301084 | 0 | 0.355418052 |
| ENSMUSG00000106054 | 0 | 0 | 0 | 0 | 0 | 0 | 0 |
| ENSMUSG00000106056 | 0 | 0 | 0 | 0 | 0 | 0 | 0 |
| ENSMUSG00000106057 | 0 | 0 | 0.25 | 0 | 0 | 0 | 0 |
| ENSMUSG00000106058 | 0.578129653 | 0 | 0 | 0.25 | 0.244650542 | 0.301029996 | 0.520678036 |
| ENSMUSG00000106059 | 0 | 0 | 0 | 0 | 0 | 0 | 0 |
| ENSMUSG00000106060 | 0.671187741 | 0.396240625 | 0.5 | 0.25 | 0.244650542 | 0 | 0.224243824 |
| ENSMUSG00000106061 | 0 | 0.25 | 0.25 | 0 | 0 | 0 | 0 |
| ENSMUSG00000106062 | 0 | 0.25 | 0 | 0 | 0 | 0 | 0 |
| ENSMUSG00000106063 | 0 | 0 | 0 | 0 | 0.244650542 | 0 | 0 |
| ENSMUSG00000106064 | 0 | 0 | 0 | 0.25 | 0 | 0 | 0 |
| ENSMUSG00000106065 | 0 | 0 | 0 | 0 | 0 | 0 | 0 |
| ENSMUSG00000106066 | 0 | 0 | 0 | 0 | 0 | 0 | 0 |
| ENSMUSG00000106067 | 0.578129653 | 0.701838731 | 0.580482024 | 0.5 | 0.568060967 | 0.77815125 | 0.579661877 |
| ENSMUSG00000106068 | 0 | 0 | 0 | 0.25 | 0.244650542 | 0 | 0 |
| ENSMUSG00000106069 | 0 | 0 | 0 | 0 | 0 | 0 | 0 |
| ENSMUSG00000106070 | 0.289064826 | 0 | 0 | 0 | 0 | 0 | 0 |
| ENSMUSG00000106071 | 0 | 0 | 0 | 0 | 0 | 0 | 0 |
| ENSMUSG00000106072 | 0 | 0 | 0 | 0 | 0 | 0 | 0 |
| ENSMUSG00000106073 | 0 | 0 | 0 | 0 | 0 | 0.301029996 | 0 |
| ENSMUSG00000106074 | 0 | 0 | 0 | 0 | 0 | 0 | 0 |
| ENSMUSG00000106075 | 0 | 0 | 0 | 0 | 0 | 0 | 0 |
| ENSMUSG00000106076 | 0 | 0 | 0 | 0 | 0 | 0 | 0 |
| ENSMUSG00000106077 | 0 | 0 | 0 | 0 | 0.244650542 | 0.301029996 | 0.224243824 |
| ENSMUSG00000106078 | 0 | 0 | 0 | 0 | 0 | 0 | 0 |
| ENSMUSG00000106079 | 0 | 0 | 0 | 0 | 0 | 0 | 0 |
| ENSMUSG00000106080 | 0 | 0 | 0 | 0 | 0 | 0 | 0 |
| ENSMUSG00000106081 | 0 | 0 | 0 | 0 | 0 | 0 | 0 |
| ENSMUSG00000106082 | 0 | 0 | 0 | 0 | 0 | 0 | 0 |
| ENSMUSG00000106083 | 0 | 0 | 0 | 0 | 0 | 0 | 0 |

| | | | | | | | |
|---|---|---|---|---|---|---|---|
| ENSMUSG00000106084 | 0 | 0 | 0 | 0 | 0 | 0 | 0 |
| ENSMUSG00000106085 | 0 | 0 | 0 | 0 | 0 | 0 | 0 |
| ENSMUSG00000106086 | 0 | 0 | 0 | 0 | 0 | 0 | 0 |
| ENSMUSG00000106087 | 0.867194479 | 0.896240625 | 0.75 | 0.864857905 | 0.905314583 | 0.77815125 | 0.775756176 |
| ENSMUSG00000106088 | 0 | 0 | 0 | 0 | 0 | 0 | 0 |
| ENSMUSG00000106089 | 0.289064826 | 0.25 | 0.25 | 0.25 | 0.244650542 | 0.301029996 | 0 |
| ENSMUSG00000106090 | 0.45815691 | 0 | 0 | 0 | 0 | 0 | 0 |
| ENSMUSG00000106091 | 0 | 0 | 0 | 0 | 0 | 0 | 0 |
| ENSMUSG00000106092 | 0 | 0 | 0 | 0 | 0.244650542 | 0 | 0.224243824 |
| ENSMUSG00000106093 | 0 | 0 | 0 | 0.25 | 0 | 0 | 0 |
| ENSMUSG00000106094 | 0 | 0.25 | 0 | 0 | 0 | 0.301029996 | 0 |
| ENSMUSG00000106095 | 0 | 0 | 0 | 0 | 0 | 0 | 0 |
| ENSMUSG00000106096 | 0 | 0 | 0 | 0 | 0 | 0 | 0 |
| ENSMUSG00000106097 | 0 | 0 | 0 | 0 | 0 | 0 | 0 |
| ENSMUSG00000106098 | 0 | 0 | 0 | 0 | 0 | 0 | 0 |
| ENSMUSG00000106099 | 1.505869732 | 1.464495249 | 1.482684334 | 1.552363341 | 1.559656853 | 1.556302501 | 1.383567577 |
| ENSMUSG00000106100 | 0 | 0.25 | 0 | 0 | 0.387761935 | 0 | 0.355418052 |
| ENSMUSG00000106101 | 0 | 0 | 0 | 0 | 0 | 0 | 0 |
| ENSMUSG00000106102 | 0 | 0 | 0 | 0 | 0 | 0 | 0 |
| ENSMUSG00000106103 | 0 | 0.580482024 | 0.25 | 0.396240625 | 0.244650542 | 0.301029996 | 0.355418052 |
| ENSMUSG00000106104 | 0 | 0 | 0 | 0 | 0 | 0 | 0 |
| ENSMUSG00000106105 | 0 | 0 | 0 | 0 | 0 | 0 | 0 |
| ENSMUSG00000106106 | 3.363083473 | 3.182564459 | 3.215659339 | 3.108526732 | 2.828358549 | 3.189770956 | 3.051385709 |
| ENSMUSG00000106107 | 1.036286563 | 1.061981878 | 1.061981878 | 1.17510993 | 1.163285805 | 1.079181246 | 0.969165684 |
| ENSMUSG00000106108 | 0.289064826 | 0.396240625 | 0.580482024 | 0.25 | 0.244650542 | 0.301029996 | 0.520678036 |
| ENSMUSG00000106109 | 0 | 0 | 0 | 0 | 0 | 0 | 0 |
| ENSMUSG00000106110 | 0.45815691 | 0.25 | 0.25 | 0 | 0.387761935 | 0.477121255 | 0 |
| ENSMUSG00000106111 | 0 | 0 | 0 | 0 | 0 | 0 | 0 |
| ENSMUSG00000106112 | 0 | 0 | 0 | 0.396240625 | 0 | 0 | 0 |
| ENSMUSG00000106113 | 0 | 0 | 0 | 0 | 0 | 0 | 0 |
| ENSMUSG00000106114 | 0 | 0 | 0 | 0 | 0 | 0 | 0 |
| ENSMUSG00000106115 | 0.45815691 | 0.396240625 | 0.25 | 0.25 | 0 | 0.477121255 | 0.224243824 |
| ENSMUSG00000106116 | 0 | 0 | 0 | 0 | 0 | 0 | 0 |
| ENSMUSG00000106117 | 0 | 0 | 0 | 0 | 0 | 0 | 0.355418052 |

| | | | | | | | |
|---|---|---|---|---|---|---|---|
| ENSMUSG00000106118 | 0 | 0 | 0 | 0 | 0 | 0 | 0 |
| ENSMUSG00000106119 | 0 | 0 | 0 | 0 | 0 | 0 | 0 |
| ENSMUSG00000106120 | 0 | 0 | 0 | 0.396240625 | 0 | 0 | 0 |
| ENSMUSG00000106121 | 0.867194479 | 0.580482024 | 0.79248125 | 0.864857905 | 0.877063019 | 0.602059991 | 0.896975297 |
| ENSMUSG00000106122 | 0 | 0 | 0 | 0 | 0 | 0 | 0 |
| ENSMUSG00000106123 | 0 | 0.396240625 | 0 | 0 | 0 | 0 | 0.224243824 |
| ENSMUSG00000106124 | 0 | 0 | 0 | 0 | 0 | 0 | 0 |
| ENSMUSG00000106125 | 0 | 0.25 | 0 | 0 | 0 | 0 | 0 |
| ENSMUSG00000106126 | 0 | 0 | 0 | 0 | 0 | 0 | 0 |
| ENSMUSG00000106127 | 0 | 0.396240625 | 0.396240625 | 0 | 0 | 0 | 0 |
| ENSMUSG00000106128 | 0 | 0 | 0 | 0 | 0 | 0 | 0 |
| ENSMUSG00000106129 | 0 | 0 | 0 | 0 | 0 | 0 | 0 |
| ENSMUSG00000106130 | 0 | 0 | 0 | 0.25 | 0 | 0.301029996 | 0 |
| ENSMUSG00000106131 | 0 | 0 | 0.25 | 0 | 0 | 0 | 0.224243824 |
| ENSMUSG00000106132 | 0 | 0 | 0 | 0 | 0 | 0 | 0 |
| ENSMUSG00000106133 | 0 | 0 | 0 | 0 | 0 | 0 | 0 |
| ENSMUSG00000106134 | 0.289064826 | 0 | 0 | 0.25 | 0.244650542 | 0 | 0 |
| ENSMUSG00000106135 | 0.811507563 | 0.830482024 | 0.864857905 | 1.02186571 | 0.905314583 | 0.84509804 | 1 |
| ENSMUSG00000106136 | 0 | 0 | 0 | 0 | 0 | 0 | 0 |
| ENSMUSG00000106137 | 0 | 0 | 0 | 0 | 0 | 0 | 0 |
| ENSMUSG00000106138 | 0 | 0 | 0 | 0 | 0 | 0 | 0 |
| ENSMUSG00000106139 | 0 | 0 | 0.396240625 | 0 | 0 | 0 | 0 |
| ENSMUSG00000106140 | 0 | 0.25 | 0 | 0 | 0 | 0 | 0 |
| ENSMUSG00000106141 | 0 | 0 | 0 | 0 | 0 | 0 | 0 |
| ENSMUSG00000106142 | 0 | 0 | 0.25 | 0 | 0 | 0 | 0 |
| ENSMUSG00000106143 | 0.289064826 | 0 | 0 | 0.396240625 | 0 | 0.301029996 | 0 |
| ENSMUSG00000106144 | 0 | 0 | 0 | 0 | 0 | 0 | 0 |
| ENSMUSG00000106145 | 0 | 0 | 0 | 0 | 0 | 0 | 0 |
| ENSMUSG00000106146 | 0 | 0 | 0 | 0 | 0 | 0 | 0 |
| ENSMUSG00000106147 | 0 | 0 | 0 | 0 | 0 | 0 | 0 |
| ENSMUSG00000106148 | 0 | 0 | 0 | 0 | 0 | 0 | 0 |
| ENSMUSG00000106149 | 0.45815691 | 0.25 | 0.25 | 0.396240625 | 0.568060967 | 0 | 0 |
| ENSMUSG00000106150 | 0.671187741 | 0.396240625 | 0.5 | 0.396240625 | 0.489301084 | 0.602059991 | 0.520678036 |
| ENSMUSG00000106151 | 0 | 0 | 0 | 0.25 | 0 | 0 | 0 |

| | | | | | | | |
|---|---|---|---|---|---|---|---|
| ENSMUSG00000106152 | 0 | 0 | 0 | 0 | 0 | 0 | 0 |
| ENSMUSG00000106153 | 0 | 0.25 | 0 | 0.396240625 | 0 | 0 | 0.224243824 |
| ENSMUSG00000106154 | 0 | 0 | 0 | 0 | 0 | 0 | 0 |
| ENSMUSG00000106155 | 0 | 0 | 0 | 0 | 0 | 0 | 0 |
| ENSMUSG00000106156 | 0 | 0 | 0 | 0 | 0 | 0 | 0 |
| ENSMUSG00000106157 | 0 | 0 | 0 | 0 | 0 | 0 | 0 |
| ENSMUSG00000106158 | 0 | 0.25 | 0.25 | 0 | 0 | 0 | 0.224243824 |
| ENSMUSG00000106159 | 0 | 0 | 0 | 0 | 0 | 0 | 0 |
| ENSMUSG00000106160 | 0 | 0 | 0 | 0 | 0 | 0 | 0 |
| ENSMUSG00000106161 | 0 | 0.25 | 0 | 0 | 0 | 0.301029996 | 0 |
| ENSMUSG00000106162 | 0 | 0 | 0 | 0 | 0 | 0 | 0 |
| ENSMUSG00000106163 | 0 | 0 | 0 | 0 | 0 | 0 | 0 |
| ENSMUSG00000106164 | 0.289064826 | 0.5 | 0 | 0 | 0.244650542 | 0 | 0 |
| ENSMUSG00000106165 | 0 | 0 | 0 | 0 | 0 | 0 | 0 |
| ENSMUSG00000106166 | 0 | 0.25 | 0 | 0.396240625 | 0.568060967 | 0 | 0 |
| ENSMUSG00000106167 | 0 | 0 | 0 | 0 | 0 | 0 | 0 |
| ENSMUSG00000106168 | 0 | 0 | 0 | 0 | 0 | 0 | 0 |
| ENSMUSG00000106169 | 0 | 0.25 | 0 | 0.25 | 0.244650542 | 0 | 0 |
| ENSMUSG00000106170 | 0 | 0 | 0 | 0 | 0 | 0 | 0 |
| ENSMUSG00000106172 | 0 | 0 | 0 | 0 | 0 | 0 | 0 |
| ENSMUSG00000106173 | 0.747221736 | 0.864857905 | 0.896240625 | 0.896240625 | 0.812711509 | 0.698970004 | 0.803905701 |
| ENSMUSG00000106174 | 0 | 0 | 0 | 0 | 0 | 0 | 0 |
| ENSMUSG00000106175 | 0 | 0 | 0 | 0 | 0 | 0 | 0 |
| ENSMUSG00000106176 | 0.289064826 | 0 | 0 | 0 | 0 | 0 | 0 |
| ENSMUSG00000106177 | 0 | 0 | 0 | 0 | 0 | 0 | 0 |
| ENSMUSG00000106178 | 0.289064826 | 0 | 0 | 0.25 | 0.387761935 | 0 | 0.448487648 |
| ENSMUSG00000106179 | 0 | 0 | 0 | 0 | 0 | 0 | 0 |
| ENSMUSG00000106180 | 0 | 0 | 0 | 0 | 0 | 0 | 0 |
| ENSMUSG00000106181 | 0 | 0.396240625 | 0.25 | 0 | 0.244650542 | 0 | 0.224243824 |
| ENSMUSG00000106182 | 0.289064826 | 0 | 0 | 0 | 0.244650542 | 0 | 0 |
| ENSMUSG00000106183 | 0 | 0 | 0 | 0 | 0 | 0 | 0 |
| ENSMUSG00000106184 | 0.289064826 | 0.25 | 0.25 | 0.25 | 0.244650542 | 0.301029996 | 0.224243824 |
| ENSMUSG00000106185 | 0 | 0 | 0 | 0 | 0 | 0 | 0 |
| ENSMUSG00000106186 | 0.671187741 | 0.396240625 | 0.25 | 0.5 | 0.244650542 | 0.301029996 | 0 |

| | | | | | | | |
|---|---|---|---|---|---|---|---|
| ENSMUSG00000106187 | 0 | 0 | 0 | 0 | 0 | 0 | 0 |
| ENSMUSG00000106188 | 0 | 0 | 0.396240625 | 0 | 0.489301084 | 0.301029996 | 0.224243824 |
| ENSMUSG00000106189 | 0 | 0 | 0 | 0 | 0 | 0 | 0 |
| ENSMUSG00000106190 | 0.747221736 | 0.75 | 0.701838731 | 0.646240625 | 0.905314583 | 0.954242509 | 0.74492186 |
| ENSMUSG00000106191 | 0 | 0 | 0 | 0 | 0 | 0 | 0 |
| ENSMUSG00000106192 | 0 | 0 | 0 | 0 | 0 | 0 | 0 |
| ENSMUSG00000106193 | 0 | 0 | 0 | 0 | 0 | 0 | 0 |
| ENSMUSG00000106194 | 0 | 0 | 0 | 0 | 0 | 0 | 0 |
| ENSMUSG00000106195 | 0 | 0 | 0 | 0 | 0 | 0 | 0 |
| ENSMUSG00000106196 | 0 | 0 | 0 | 0 | 0 | 0 | 0 |
| ENSMUSG00000106197 | 0.867194479 | 0.5 | 0.396240625 | 0.951838731 | 0.387761935 | 0.698970004 | 0.520678036 |
| ENSMUSG00000106198 | 0 | 0 | 0 | 0 | 0 | 0 | 0 |
| ENSMUSG00000106199 | 0 | 0 | 0 | 0 | 0 | 0 | 0 |
| ENSMUSG00000106200 | 0 | 0 | 0 | 0 | 0 | 0 | 0 |
| ENSMUSG00000106201 | 0 | 0 | 0 | 0 | 0 | 0 | 0 |
| ENSMUSG00000106202 | 0 | 0 | 0 | 0 | 0.244650542 | 0 | 0 |
| ENSMUSG00000106203 | 0 | 0 | 0 | 0 | 0 | 0 | 0 |
| ENSMUSG00000106204 | 0 | 0 | 0 | 0 | 0 | 0 | 0 |
| ENSMUSG00000106205 | 0.747221736 | 0.646240625 | 0.79248125 | 0.25 | 0.568060967 | 0.477121255 | 0.448487648 |
| ENSMUSG00000106206 | 0 | 0 | 0 | 0 | 0 | 0 | 0 |
| ENSMUSG00000106207 | 0 | 0 | 0 | 0 | 0 | 0 | 0 |
| ENSMUSG00000106208 | 0 | 0.25 | 0 | 0.25 | 0.387761935 | 0 | 0.355418052 |
| ENSMUSG00000106209 | 0.45815691 | 0.396240625 | 0.25 | 0.580482024 | 0.244650542 | 0 | 0.355418052 |
| ENSMUSG00000106210 | 0 | 0 | 0 | 0 | 0 | 0 | 0.224243824 |
| ENSMUSG00000106211 | 0 | 0 | 0 | 0 | 0.244650542 | 0.301029996 | 0 |
| ENSMUSG00000106212 | 0 | 0 | 0 | 0 | 0 | 0 | 0 |
| ENSMUSG00000106213 | 0 | 0 | 0 | 0 | 0 | 0 | 0 |
| ENSMUSG00000106214 | 0 | 0 | 0 | 0 | 0 | 0 | 0 |
| ENSMUSG00000106215 | 0 | 0 | 0 | 0 | 0 | 0 | 0 |
| ENSMUSG00000106216 | 0 | 0 | 0 | 0 | 0 | 0 | 0 |
| ENSMUSG00000106217 | 0 | 0 | 0 | 0.25 | 0 | 0 | 0 |
| ENSMUSG00000106218 | 0 | 0 | 0 | 0 | 0 | 0 | 0 |
| ENSMUSG00000106219 | 0 | 0.25 | 0 | 0.396240625 | 0 | 0 | 0 |
| ENSMUSG00000106220 | 0 | 0 | 0 | 0 | 0 | 0 | 0 |

| | | | | | | | |
|---|---|---|---|---|---|---|---|
| ENSMUSG00000106221 | 0 | 0 | 0 | 0 | 0 | 0 | 0 |
| ENSMUSG00000106222 | 0 | 0 | 0 | 0 | 0 | 0 | 0 |
| ENSMUSG00000106223 | 0 | 0 | 0 | 0 | 0 | 0 | 0 |
| ENSMUSG00000106224 | 0.747221736 | 0.701838731 | 0.5 | 0.396240625 | 0.244650542 | 0.301029996 | 0.224243824 |
| ENSMUSG00000106225 | 0 | 0 | 0 | 0 | 0 | 0 | 0 |
| ENSMUSG00000106226 | 0 | 0 | 0 | 0 | 0 | 0 | 0 |
| ENSMUSG00000106227 | 0 | 0 | 0 | 0 | 0 | 0 | 0 |
| ENSMUSG00000106228 | 0 | 0 | 0 | 0 | 0 | 0 | 0 |
| ENSMUSG00000106229 | 0.289064826 | 0.646240625 | 0.5 | 0.646240625 | 0.568060967 | 0.301029996 | 0.520678036 |
| ENSMUSG00000106230 | 0 | 0 | 0 | 0 | 0 | 0 | 0 |
| ENSMUSG00000106231 | 0 | 0 | 0 | 0 | 0 | 0 | 0 |
| ENSMUSG00000106232 | 0 | 0 | 0 | 0 | 0 | 0 | 0 |
| ENSMUSG00000106233 | 0 | 0 | 0 | 0 | 0 | 0 | 0 |
| ENSMUSG00000106234 | 0 | 0 | 0 | 0 | 0 | 0 | 0 |
| ENSMUSG00000106235 | 0 | 0 | 0 | 0 | 0 | 0 | 0 |
| ENSMUSG00000106236 | 0 | 0 | 0 | 0 | 0 | 0 | 0 |
| ENSMUSG00000106237 | 0.578129653 | 0 | 0.25 | 0.396240625 | 0 | 0 | 0.355418052 |
| ENSMUSG00000106238 | 0 | 0 | 0 | 0 | 0 | 0 | 0 |
| ENSMUSG00000106239 | 0 | 0 | 0 | 0.25 | 0 | 0 | 0 |
| ENSMUSG00000106240 | 0 | 0 | 0 | 0 | 0.387761935 | 0 | 0 |
| ENSMUSG00000106241 | 1.069666964 | 1 | 0.896240625 | 0.976722649 | 1.020174412 | 1.079181246 | 0.896975297 |
| ENSMUSG00000106242 | 0 | 0 | 0 | 0 | 0 | 0 | 0 |
| ENSMUSG00000106243 | 0.671187741 | 0.701838731 | 0.580482024 | 0.396240625 | 0.632412477 | 0.301029996 | 0.629532004 |
| ENSMUSG00000106244 | 0 | 0 | 0 | 0 | 0 | 0 | 0 |
| ENSMUSG00000106245 | 0 | 0 | 0 | 0 | 0 | 0 | 0 |
| ENSMUSG00000106246 | 0 | 0 | 0 | 0 | 0 | 0 | 0 |
| ENSMUSG00000106247 | 0 | 0 | 0 | 0 | 0 | 0 | 0 |
| ENSMUSG00000106248 | 0.45815691 | 0.25 | 0.25 | 0 | 0.244650542 | 0 | 0.520678036 |
| ENSMUSG00000106249 | 0 | 0 | 0 | 0 | 0 | 0 | 0 |
| ENSMUSG00000106250 | 0 | 0 | 0 | 0 | 0 | 0 | 0 |
| ENSMUSG00000106251 | 0.747221736 | 0.75 | 0.864857905 | 0.830482024 | 0.812711509 | 0.602059991 | 0.829800754 |
| ENSMUSG00000106252 | 0 | 0 | 0 | 0.25 | 0 | 0 | 0 |
| ENSMUSG00000106253 | 0 | 0 | 0 | 0 | 0 | 0 | 0 |
| ENSMUSG00000106254 | 0.289064826 | 0.25 | 0.580482024 | 0.5 | 0.244650542 | 0.698970004 | 0.355418052 |

| | | | | | | | |
|---|---|---|---|---|---|---|---|
| ENSMUSG00000106255 | 0 | 0 | 0 | 0 | 0 | 0 | 0 |
| ENSMUSG00000106256 | 0 | 0 | 0 | 0 | 0 | 0 | 0 |
| ENSMUSG00000106257 | 0 | 0 | 0 | 0 | 0 | 0.301029996 | 0 |
| ENSMUSG00000106258 | 0 | 0 | 0 | 0 | 0 | 0.301029996 | 0 |
| ENSMUSG00000106259 | 0 | 0 | 0 | 0 | 0 | 0 | 0 |
| ENSMUSG00000106260 | 0 | 0 | 0 | 0 | 0 | 0 | 0 |
| ENSMUSG00000106261 | 0 | 0 | 0 | 0 | 0 | 0 | 0 |
| ENSMUSG00000106262 | 0.960252568 | 1.02186571 | 0.830482024 | 1.114857905 | 1.136121934 | 1.079181246 | 0.916588299 |
| ENSMUSG00000106263 | 0 | 0 | 0 | 0 | 0 | 0 | 0 |
| ENSMUSG00000106264 | 0 | 0 | 0 | 0 | 0 | 0 | 0 |
| ENSMUSG00000106265 | 0 | 0 | 0 | 0.25 | 0 | 0 | 0 |
| ENSMUSG00000106266 | 0 | 0 | 0 | 0 | 0 | 0 | 0 |
| ENSMUSG00000106267 | 0 | 0 | 0 | 0 | 0 | 0 | 0 |
| ENSMUSG00000106268 | 0.289064826 | 0.396240625 | 0.580482024 | 0 | 0.244650542 | 0.301029996 | 0.224243824 |
| ENSMUSG00000106269 | 0.578129653 | 0.830482024 | 0.396240625 | 0.5 | 0.632412477 | 0.602059991 | 0.355418052 |
| ENSMUSG00000106270 | 0 | 0 | 0 | 0 | 0 | 0 | 0 |
| ENSMUSG00000106271 | 0 | 0.25 | 0.396240625 | 0 | 0.387761935 | 0.301029996 | 0 |
| ENSMUSG00000106272 | 0 | 0 | 0 | 0 | 0 | 0 | 0 |
| ENSMUSG00000106273 | 0 | 0 | 0.25 | 0 | 0 | 0 | 0 |
| ENSMUSG00000106274 | 0 | 0 | 0 | 0 | 0 | 0 | 0 |
| ENSMUSG00000106275 | 0 | 0 | 0 | 0 | 0 | 0 | 0 |
| ENSMUSG00000106276 | 0 | 0 | 0 | 0 | 0 | 0 | 0 |
| ENSMUSG00000106278 | 0 | 0 | 0 | 0 | 0 | 0 | 0 |
| ENSMUSG00000106279 | 0 | 0 | 0 | 0 | 0 | 0 | 0 |
| ENSMUSG00000106280 | 0 | 0 | 0 | 0 | 0 | 0 | 0 |
| ENSMUSG00000106281 | 0 | 0 | 0 | 0 | 0 | 0 | 0 |
| ENSMUSG00000106282 | 0 | 0 | 0 | 0 | 0 | 0.301029996 | 0 |
| ENSMUSG00000106283 | 0 | 0.25 | 0 | 0 | 0 | 0 | 0 |
| ENSMUSG00000106284 | 0 | 0 | 0 | 0 | 0 | 0 | 0 |
| ENSMUSG00000106285 | 0 | 0 | 0.25 | 0.25 | 0.244650542 | 0 | 0 |
| ENSMUSG00000106286 | 0 | 0 | 0 | 0 | 0 | 0 | 0 |
| ENSMUSG00000106287 | 0 | 0 | 0 | 0.25 | 0.244650542 | 0 | 0.224243824 |
| ENSMUSG00000106288 | 1.069666964 | 0.79248125 | 0.830482024 | 0.75 | 0.77552387 | 0.477121255 | 0.520678036 |
| ENSMUSG00000106289 | 0 | 0 | 0 | 0 | 0 | 0 | 0 |

| | | | | | | | |
|---|---|---|---|---|---|---|---|
| ENSMUSG00000106290 | 0.747221736 | 1.02186571 | 0.896240625 | 0.75 | 0.905314583 | 1.079181246 | 0.829800754 |
| ENSMUSG00000106291 | 0.671187741 | 0.701838731 | 0.5 | 0.5 | 0.489301084 | 0.698970004 | 0.579661877 |
| ENSMUSG00000106292 | 0 | 0 | 0 | 0 | 0 | 0 | 0 |
| ENSMUSG00000106293 | 0.45815691 | 0 | 0 | 0.25 | 0 | 0 | 0 |
| ENSMUSG00000106294 | 0 | 0 | 0.25 | 0 | 0 | 0 | 0 |
| ENSMUSG00000106295 | 0 | 0 | 0 | 0 | 0 | 0 | 0 |
| ENSMUSG00000106296 | 0.289064826 | 0.25 | 0 | 0 | 0.244650542 | 0 | 0 |
| ENSMUSG00000106297 | 0 | 0 | 0 | 0 | 0 | 0 | 0 |
| ENSMUSG00000106298 | 0 | 0 | 0 | 0 | 0 | 0 | 0 |
| ENSMUSG00000106299 | 0 | 0 | 0 | 0 | 0 | 0 | 0 |
| ENSMUSG00000106300 | 0.289064826 | 0 | 0 | 0.25 | 0 | 0 | 0.224243824 |
| ENSMUSG00000106301 | 0 | 0 | 0.25 | 0 | 0 | 0 | 0 |
| ENSMUSG00000106302 | 0 | 0 | 0 | 0 | 0 | 0 | 0 |
| ENSMUSG00000106303 | 0 | 0 | 0 | 0 | 0 | 0 | 0 |
| ENSMUSG00000106304 | 0 | 0 | 0 | 0 | 0 | 0 | 0 |
| ENSMUSG00000106305 | 0 | 0 | 0 | 0 | 0 | 0 | 0 |
| ENSMUSG00000106306 | 0 | 0 | 0 | 0 | 0 | 0 | 0 |
| ENSMUSG00000106307 | 0 | 0 | 0 | 0 | 0 | 0 | 0 |
| ENSMUSG00000106308 | 0 | 0 | 0 | 0 | 0 | 0 | 0 |
| ENSMUSG00000106309 | 0 | 0.396240625 | 0.25 | 0.396240625 | 0.489301084 | 0.301029996 | 0.355418052 |
| ENSMUSG00000106310 | 0 | 0.25 | 0 | 0.25 | 0.244650542 | 0.301029996 | 0.224243824 |
| ENSMUSG00000106311 | 0 | 0 | 0 | 0 | 0.244650542 | 0 | 0 |
| ENSMUSG00000106312 | 0 | 0 | 0 | 0 | 0 | 0 | 0 |
| ENSMUSG00000106313 | 0.289064826 | 0 | 0.25 | 0 | 0 | 0 | 0 |
| ENSMUSG00000106314 | 0 | 0.25 | 0 | 0 | 0 | 0 | 0 |
| ENSMUSG00000106315 | 0 | 0 | 0 | 0 | 0 | 0 | 0 |
| ENSMUSG00000106316 | 0 | 0 | 0 | 0 | 0 | 0 | 0 |
| ENSMUSG00000106317 | 0.811507563 | 0.79248125 | 0.896240625 | 0.976722649 | 0.905314583 | 1.079181246 | 1 |
| ENSMUSG00000106318 | 0 | 0 | 0 | 0 | 0 | 0 | 0 |
| ENSMUSG00000106319 | 0 | 0 | 0 | 0 | 0 | 0 | 0 |
| ENSMUSG00000106320 | 0 | 0.25 | 0 | 0.396240625 | 0 | 0 | 0.224243824 |
| ENSMUSG00000106321 | 0 | 0.396240625 | 0 | 0 | 0.244650542 | 0 | 0.224243824 |
| ENSMUSG00000106322 | 0 | 0.25 | 0 | 0.580482024 | 0 | 0 | 0 |
| ENSMUSG00000106323 | 0 | 0 | 0 | 0 | 0 | 0 | 0 |

| | | | | | | | |
|---|---|---|---|---|---|---|---|
| ENSMUSG00000106324 | 0 | 0 | 0 | 0 | 0 | 0 | 0 |
| ENSMUSG00000106325 | 0 | 0 | 0 | 0 | 0 | 0 | 0 |
| ENSMUSG00000106326 | 0 | 0 | 0 | 0 | 0 | 0 | 0 |
| ENSMUSG00000106327 | 0 | 0 | 0 | 0 | 0 | 0 | 0 |
| ENSMUSG00000106328 | 0 | 0 | 0 | 0 | 0 | 0 | 0 |
| ENSMUSG00000106329 | 0.45815691 | 0 | 0 | 0.25 | 0.244650542 | 0.301029996 | 0.224243824 |
| ENSMUSG00000106330 | 0 | 0 | 0 | 0.25 | 0 | 0 | 0 |
| ENSMUSG00000106331 | 0 | 0 | 0 | 0 | 0.244650542 | 0 | 0 |
| ENSMUSG00000106332 | 0 | 0.396240625 | 0.25 | 0.5 | 0.244650542 | 0 | 0 |
| ENSMUSG00000106333 | 0 | 0 | 0 | 0 | 0 | 0 | 0 |
| ENSMUSG00000106334 | 1.903481042 | 1.698603967 | 1.67510993 | 2.718745337 | 2.66038209 | 3.024485668 | 2.456573893 |
| ENSMUSG00000106335 | 0 | 0 | 0 | 0 | 0 | 0 | 0 |
| ENSMUSG00000106336 | 0 | 0 | 0 | 0 | 0 | 0 | 0 |
| ENSMUSG00000106337 | 0 | 0.25 | 0 | 0.25 | 0 | 0 | 0 |
| ENSMUSG00000106338 | 0 | 0 | 0 | 0 | 0 | 0 | 0 |
| ENSMUSG00000106339 | 0 | 0.25 | 0 | 0 | 0 | 0 | 0 |
| ENSMUSG00000106340 | 0 | 0 | 0 | 0 | 0 | 0 | 0 |
| ENSMUSG00000106341 | 0 | 0 | 0 | 0 | 0 | 0 | 0 |
| ENSMUSG00000106342 | 0 | 0 | 0 | 0 | 0 | 0 | 0 |
| ENSMUSG00000106343 | 0 | 0 | 0 | 0 | 0 | 0 | 0 |
| ENSMUSG00000106344 | 0 | 0 | 0 | 0 | 0 | 0 | 0 |
| ENSMUSG00000106345 | 0 | 0 | 0 | 0 | 0 | 0 | 0 |
| ENSMUSG00000106346 | 0 | 0 | 0 | 0 | 0 | 0 | 0 |
| ENSMUSG00000106347 | 0 | 0 | 0 | 0 | 0 | 0 | 0 |
| ENSMUSG00000106348 | 0 | 0 | 0 | 0 | 0 | 0 | 0 |
| ENSMUSG00000106349 | 0 | 0 | 0 | 0 | 0 | 0 | 0 |
| ENSMUSG00000106350 | 0 | 0 | 0 | 0 | 0 | 0 | 0 |
| ENSMUSG00000106351 | 0 | 0 | 0 | 0 | 0 | 0 | 0 |
| ENSMUSG00000106352 | 0 | 0 | 0 | 0 | 0 | 0 | 0 |
| ENSMUSG00000106353 | 0 | 0 | 0 | 0.25 | 0 | 0 | 0.355418052 |
| ENSMUSG00000106354 | 0 | 0 | 0 | 0.25 | 0 | 0.301029996 | 0 |
| ENSMUSG00000106355 | 0 | 0 | 0 | 0 | 0 | 0 | 0 |
| ENSMUSG00000106356 | 0 | 0 | 0 | 0 | 0 | 0 | 0 |
| ENSMUSG00000106357 | 0 | 0 | 0 | 0 | 0 | 0 | 0 |

| | | | | | | | |
|---|---|---|---|---|---|---|---|
| ENSMUSG00000106358 | 0 | 0.25 | 0 | 0.25 | 0 | 0 | 0 |
| ENSMUSG00000106359 | 0 | 0 | 0 | 0 | 0 | 0 | 0 |
| ENSMUSG00000106360 | 0 | 0 | 0 | 0 | 0 | 0 | 0 |
| ENSMUSG00000106361 | 0.289064826 | 0.25 | 0 | 0.25 | 0.387761935 | 0 | 0 |
| ENSMUSG00000106362 | 0 | 0 | 0 | 0 | 0 | 0 | 0 |
| ENSMUSG00000106363 | 0 | 0 | 0 | 0 | 0 | 0 | 0 |
| ENSMUSG00000106364 | 0.289064826 | 0 | 0 | 0.396240625 | 0 | 0 | 0.355418052 |
| ENSMUSG00000106365 | 0 | 0 | 0 | 0 | 0.244650542 | 0 | 0 |
| ENSMUSG00000106366 | 0 | 0 | 0 | 0 | 0 | 0 | 0 |
| ENSMUSG00000106367 | 0 | 0 | 0 | 0 | 0 | 0 | 0 |
| ENSMUSG00000106368 | 0 | 0.25 | 0 | 0 | 0 | 0 | 0 |
| ENSMUSG00000106369 | 0 | 0 | 0 | 0 | 0 | 0 | 0 |
| ENSMUSG00000106370 | 0 | 0 | 0 | 0 | 0 | 0 | 0 |
| ENSMUSG00000106371 | 0 | 0 | 0 | 0.25 | 0 | 0 | 0 |
| ENSMUSG00000106372 | 0 | 0 | 0 | 0 | 0 | 0 | 0 |
| ENSMUSG00000106373 | 0 | 0 | 0 | 0 | 0 | 0 | 0 |
| ENSMUSG00000106374 | 0 | 0 | 0 | 0 | 0 | 0 | 0 |
| ENSMUSG00000106375 | 0.578129653 | 0.580482024 | 0.25 | 0.5 | 0.387761935 | 0 | 0.355418052 |
| ENSMUSG00000106376 | 0 | 0 | 0 | 0 | 0 | 0 | 0 |
| ENSMUSG00000106377 | 0 | 0 | 0 | 0 | 0 | 0 | 0 |
| ENSMUSG00000106378 | 0 | 0 | 0 | 0 | 0 | 0 | 0 |
| ENSMUSG00000106379 | 0 | 0 | 0 | 0 | 0 | 0 | 0 |
| ENSMUSG00000106380 | 0.45815691 | 0.25 | 0.396240625 | 0.701838731 | 0 | 0.477121255 | 0.629532004 |
| ENSMUSG00000106381 | 0 | 0 | 0 | 0 | 0 | 0 | 0 |
| ENSMUSG00000106382 | 0 | 0 | 0 | 0 | 0 | 0 | 0 |
| ENSMUSG00000106383 | 0.747221736 | 1.080482024 | 1.130890489 | 1.348079356 | 1.27449559 | 1.301029996 | 1.23151414 |
| ENSMUSG00000106384 | 0 | 0 | 0 | 0 | 0 | 0 | 0 |
| ENSMUSG00000106385 | 0.289064826 | 0.25 | 0.25 | 0 | 0 | 0 | 0.448487648 |
| ENSMUSG00000106386 | 0 | 0.25 | 0 | 0 | 0 | 0 | 0 |
| ENSMUSG00000106387 | 0 | 0 | 0 | 0 | 0 | 0 | 0 |
| ENSMUSG00000106388 | 0 | 0 | 0 | 0 | 0 | 0 | 0 |
| ENSMUSG00000106389 | 0 | 0 | 0 | 0 | 0 | 0 | 0 |
| ENSMUSG00000106390 | 0 | 0 | 0 | 0 | 0 | 0 | 0 |
| ENSMUSG00000106391 | 1 | 0.830482024 | 0.646240625 | 0.5 | 0.846351821 | 0.698970004 | 0.672731473 |

| | | | | | | | |
|---|---|---|---|---|---|---|---|
| ENSMUSG00000106392 | 0 | 0 | 0 | 0 | 0 | 0 | 0 |
| ENSMUSG00000106393 | 0 | 0 | 0 | 0 | 0 | 0 | 0 |
| ENSMUSG00000106394 | 0 | 0 | 0 | 0 | 0 | 0 | 0 |
| ENSMUSG00000106395 | 0 | 0 | 0 | 0 | 0 | 0 | 0 |
| ENSMUSG00000106396 | 0 | 0 | 0 | 0.25 | 0.244650542 | 0 | 0.355418052 |
| ENSMUSG00000106397 | 0 | 0 | 0 | 0 | 0 | 0 | 0 |
| ENSMUSG00000106398 | 0 | 0 | 0 | 0 | 0 | 0 | 0 |
| ENSMUSG00000106399 | 0 | 0 | 0 | 0 | 0 | 0 | 0 |
| ENSMUSG00000106400 | 0 | 0 | 0 | 0 | 0 | 0 | 0 |
| ENSMUSG00000106401 | 0 | 0 | 0 | 0 | 0 | 0 | 0 |
| ENSMUSG00000106402 | 0 | 0 | 0 | 0 | 0 | 0 | 0 |
| ENSMUSG00000106403 | 0 | 0 | 0 | 0 | 0 | 0 | 0 |
| ENSMUSG00000106404 | 0 | 0 | 0 | 0 | 0 | 0 | 0 |
| ENSMUSG00000106405 | 0 | 0 | 0 | 0 | 0 | 0 | 0 |
| ENSMUSG00000106406 | 0 | 0.25 | 0 | 0 | 0 | 0 | 0 |
| ENSMUSG00000106407 | 0 | 0 | 0 | 0 | 0 | 0 | 0 |
| ENSMUSG00000106408 | 0.671187741 | 0.830482024 | 0.701838731 | 0.896240625 | 0.489301084 | 1 | 0.853775828 |
| ENSMUSG00000106409 | 0 | 0 | 0 | 0 | 0 | 0 | 0 |
| ENSMUSG00000106410 | 0 | 0 | 0 | 0 | 0 | 0 | 0 |
| ENSMUSG00000106411 | 1.432083896 | 1.388647213 | 1.27186571 | 1.410964047 | 1.394615667 | 1.301029996 | 1.307989563 |
| ENSMUSG00000106412 | 0 | 0 | 0 | 0 | 0 | 0 | 0 |
| ENSMUSG00000106413 | 0 | 0 | 0 | 0 | 0 | 0 | 0 |
| ENSMUSG00000106414 | 0 | 0 | 0 | 0 | 0 | 0 | 0 |
| ENSMUSG00000106415 | 0 | 0 | 0 | 0 | 0 | 0 | 0 |
| ENSMUSG00000106416 | 0.289064826 | 0.396240625 | 0.396240625 | 0.646240625 | 0.733951626 | 0.77815125 | 0.710836105 |
| ENSMUSG00000106417 | 0 | 0.5 | 0 | 0.25 | 0.244650542 | 0.477121255 | 0.224243824 |
| ENSMUSG00000106418 | 0 | 0 | 0 | 0 | 0 | 0 | 0 |
| ENSMUSG00000106419 | 0.45815691 | 0.25 | 0.5 | 0.396240625 | 0.387761935 | 0.301029996 | 0.355418052 |
| ENSMUSG00000106420 | 0 | 0 | 0 | 0 | 0 | 0 | 0 |
| ENSMUSG00000106421 | 0 | 0 | 0.25 | 0 | 0.244650542 | 0 | 0.520678036 |
| ENSMUSG00000106422 | 0 | 0 | 0 | 0 | 0 | 0 | 0 |
| ENSMUSG00000106423 | 0 | 0 | 0 | 0 | 0 | 0 | 0 |
| ENSMUSG00000106424 | 0 | 0 | 0 | 0 | 0 | 0 | 0 |
| ENSMUSG00000106425 | 0 | 0 | 0 | 0 | 0 | 0 | 0 |

| | | | | | | | |
|---|---|---|---|---|---|---|---|
| ENSMUSG00000106426 | 0.747221736 | 0.92510993 | 0.79248125 | 0.5 | 0.686820904 | 0.477121255 | 0.448487648 |
| ENSMUSG00000106427 | 0.671187741 | 0.25 | 0.646240625 | 0.701838731 | 0.244650542 | 0 | 0.629532004 |
| ENSMUSG00000106428 | 0 | 0 | 0 | 0 | 0 | 0 | 0 |
| ENSMUSG00000106429 | 0 | 0 | 0 | 0 | 0 | 0 | 0 |
| ENSMUSG00000106430 | 0 | 0 | 0 | 0.396240625 | 0 | 0.301029996 | 0 |
| ENSMUSG00000106431 | 0 | 0 | 0 | 0 | 0 | 0 | 0 |
| ENSMUSG00000106432 | 0 | 0.25 | 0 | 0 | 0.489301084 | 0 | 0.448487648 |
| ENSMUSG00000106433 | 0 | 0 | 0 | 0 | 0 | 0 | 0 |
| ENSMUSG00000106434 | 0 | 0 | 0 | 0 | 0 | 0 | 0 |
| ENSMUSG00000106435 | 0 | 0 | 0 | 0 | 0 | 0 | 0 |
| ENSMUSG00000106436 | 0 | 0 | 0 | 0 | 0 | 0 | 0 |
| ENSMUSG00000106437 | 0.289064826 | 0.396240625 | 0 | 0 | 0 | 0.477121255 | 0 |
| ENSMUSG00000106438 | 0 | 0 | 0 | 0.25 | 0.244650542 | 0 | 0 |
| ENSMUSG00000106439 | 0 | 0 | 0 | 0 | 0 | 0 | 0 |
| ENSMUSG00000106440 | 0 | 0 | 0 | 0 | 0 | 0 | 0 |
| ENSMUSG00000106441 | 0 | 0 | 0 | 0 | 0 | 0 | 0 |
| ENSMUSG00000106442 | 0 | 0 | 0 | 0 | 0 | 0 | 0 |
| ENSMUSG00000106443 | 0.45815691 | 0.646240625 | 0.646240625 | 0.701838731 | 0.632412477 | 0.602059991 | 0.629532004 |
| ENSMUSG00000106444 | 0 | 0 | 0 | 0 | 0 | 0 | 0 |
| ENSMUSG00000106445 | 0 | 0 | 0 | 0 | 0 | 0 | 0 |
| ENSMUSG00000106446 | 0.91631382 | 0.75 | 0.646240625 | 0.75 | 0.489301084 | 0 | 0.579661877 |
| ENSMUSG00000106447 | 0 | 0 | 0 | 0 | 0 | 0 | 0 |
| ENSMUSG00000106448 | 0 | 0 | 0 | 0 | 0 | 0 | 0 |
| ENSMUSG00000106450 | 0 | 0 | 0 | 0 | 0 | 0 | 0 |
| ENSMUSG00000106451 | 0 | 0 | 0 | 0 | 0 | 0.301029996 | 0 |
| ENSMUSG00000106452 | 0 | 0 | 0 | 0 | 0 | 0 | 0 |
| ENSMUSG00000106453 | 0 | 0 | 0 | 0 | 0 | 0 | 0 |
| ENSMUSG00000106454 | 0 | 0 | 0 | 0 | 0.244650542 | 0 | 0.224243824 |
| ENSMUSG00000106455 | 0 | 0.396240625 | 0.396240625 | 0.25 | 0.244650542 | 0 | 0.224243824 |
| ENSMUSG00000106456 | 0 | 0 | 0 | 0 | 0 | 0 | 0 |
| ENSMUSG00000106457 | 0.289064826 | 0.396240625 | 0 | 0.25 | 0 | 0.301029996 | 0.355418052 |
| ENSMUSG00000106458 | 0.45815691 | 0.646240625 | 0.580482024 | 0.5 | 0.244650542 | 0.954242509 | 0.629532004 |
| ENSMUSG00000106459 | 0 | 0 | 0 | 0 | 0 | 0 | 0 |
| ENSMUSG00000106460 | 0 | 0 | 0 | 0 | 0 | 0 | 0 |

| | | | | | | | |
|---|---|---|---|---|---|---|---|
| ENSMUSG00000106461 | 0 | 0 | 0 | 0 | 0 | 0 | 0 |
| ENSMUSG00000106462 | 0 | 0 | 0 | 0 | 0 | 0 | 0 |
| ENSMUSG00000106463 | 0.671187741 | 0.396240625 | 0.396240625 | 0.25 | 0.489301084 | 0.477121255 | 0.355418052 |
| ENSMUSG00000106464 | 1.470606563 | 1.464495249 | 1.445339928 | 1.302363341 | 1.176121988 | 1.204119983 | 1.014380832 |
| ENSMUSG00000106465 | 0 | 0 | 0 | 0 | 0 | 0 | 0 |
| ENSMUSG00000106466 | 0 | 0 | 0 | 0 | 0 | 0 | 0 |
| ENSMUSG00000106467 | 0 | 0 | 0 | 0 | 0 | 0 | 0 |
| ENSMUSG00000106468 | 0 | 0 | 0 | 0 | 0 | 0 | 0 |
| ENSMUSG00000106469 | 0 | 0 | 0 | 0 | 0 | 0 | 0 |
| ENSMUSG00000106470 | 0 | 0 | 0 | 0 | 0 | 0 | 0 |
| ENSMUSG00000106471 | 0 | 0 | 0 | 0 | 0 | 0 | 0 |
| ENSMUSG00000106472 | 0 | 0 | 0 | 0 | 0 | 0 | 0 |
| ENSMUSG00000106473 | 0 | 0 | 0 | 0 | 0 | 0 | 0 |
| ENSMUSG00000106474 | 0 | 0 | 0 | 0 | 0 | 0 | 0 |
| ENSMUSG00000106475 | 0 | 0.396240625 | 0 | 0.396240625 | 0.387761935 | 0 | 0.224243824 |
| ENSMUSG00000106476 | 0 | 0.25 | 0 | 0 | 0 | 0 | 0 |
| ENSMUSG00000106477 | 0 | 0 | 0 | 0 | 0 | 0 | 0 |
| ENSMUSG00000106478 | 0.289064826 | 0 | 0.25 | 0.25 | 0 | 0 | 0.224243824 |
| ENSMUSG00000106479 | 0 | 0 | 0 | 0 | 0 | 0 | 0 |
| ENSMUSG00000106480 | 0 | 0 | 0 | 0 | 0 | 0 | 0 |
| ENSMUSG00000106481 | 0 | 0 | 0 | 0 | 0 | 0 | 0 |
| ENSMUSG00000106482 | 0 | 0 | 0 | 0 | 0 | 0 | 0 |
| ENSMUSG00000106483 | 0 | 0 | 0 | 0 | 0 | 0 | 0 |
| ENSMUSG00000106484 | 0.289064826 | 0.5 | 0.25 | 0.701838731 | 0.244650542 | 0.477121255 | 0.224243824 |
| ENSMUSG00000106485 | 0 | 0 | 0 | 0 | 0 | 0 | 0 |
| ENSMUSG00000106486 | 0 | 0 | 0 | 0 | 0 | 0 | 0 |
| ENSMUSG00000106487 | 0 | 0 | 0 | 0 | 0 | 0 | 0 |
| ENSMUSG00000106488 | 0 | 0 | 0 | 0 | 0 | 0 | 0 |
| ENSMUSG00000106489 | 0 | 0 | 0 | 0 | 0 | 0 | 0 |
| ENSMUSG00000106490 | 0.289064826 | 0.25 | 0 | 0 | 0 | 0 | 0 |
| ENSMUSG00000106491 | 0 | 0 | 0.396240625 | 0.396240625 | 0 | 0.301029996 | 0 |
| ENSMUSG00000106492 | 0.289064826 | 0.25 | 0.396240625 | 0 | 0 | 0 | 0 |
| ENSMUSG00000106493 | 0 | 0 | 0 | 0 | 0 | 0 | 0 |
| ENSMUSG00000106494 | 0 | 0 | 0 | 0 | 0 | 0 | 0 |

| | | | | | | | |
|---|---|---|---|---|---|---|---|
| ENSMUSG00000106495 | 0.578129653 | 0.396240625 | 0.75 | 0.396240625 | 0.955822902 | 0.77815125 | 0.448487648 |
| ENSMUSG00000106496 | 0 | 0 | 0 | 0 | 0 | 0 | 0 |
| ENSMUSG00000106497 | 0 | 0.396240625 | 0 | 0 | 0.244650542 | 0 | 0 |
| ENSMUSG00000106498 | 0.747221736 | 0.864857905 | 0.976722649 | 0 | 0 | 0 | 0.224243824 |
| ENSMUSG00000106499 | 0 | 0 | 0 | 0 | 0 | 0.477121255 | 0.448487648 |
| ENSMUSG00000106500 | 0 | 0.25 | 0.25 | 0 | 0 | 0 | 0.224243824 |
| ENSMUSG00000106501 | 0 | 0 | 0 | 0 | 0 | 0 | 0 |
| ENSMUSG00000106502 | 0 | 0 | 0 | 0 | 0 | 0 | 0 |
| ENSMUSG00000106503 | 0 | 0 | 0 | 0 | 0 | 0 | 0 |
| ENSMUSG00000106504 | 0 | 0 | 0 | 0 | 0 | 0 | 0 |
| ENSMUSG00000106505 | 0.289064826 | 0 | 0 | 0 | 0 | 0 | 0 |
| ENSMUSG00000106506 | 0 | 0 | 0 | 0 | 0 | 0 | 0 |
| ENSMUSG00000106507 | 0 | 0 | 0 | 0.25 | 0.489301084 | 0 | 0.520678036 |
| ENSMUSG00000106508 | 0 | 0 | 0 | 0.25 | 0 | 0 | 0 |
| ENSMUSG00000106509 | 0 | 0 | 0 | 0 | 0 | 0 | 0 |
| ENSMUSG00000106510 | 0 | 0 | 0 | 0 | 0 | 0 | 0 |
| ENSMUSG00000106511 | 0 | 0 | 0 | 0 | 0 | 0 | 0 |
| ENSMUSG00000106512 | 0.289064826 | 0 | 0.396240625 | 0 | 0.244650542 | 0 | 0.224243824 |
| ENSMUSG00000106513 | 0 | 0 | 0 | 0 | 0 | 0 | 0 |
| ENSMUSG00000106514 | 0 | 0.25 | 0.396240625 | 0 | 0 | 0 | 0.355418052 |
| ENSMUSG00000106515 | 0 | 0 | 0 | 0 | 0 | 0 | 0 |
| ENSMUSG00000106516 | 0 | 0 | 0 | 0 | 0 | 0 | 0 |
| ENSMUSG00000106517 | 0 | 0 | 0 | 0 | 0 | 0 | 0 |
| ENSMUSG00000106518 | 0.289064826 | 0.25 | 0 | 0 | 0 | 0 | 0 |
| ENSMUSG00000106519 | 0 | 0.25 | 0 | 0 | 0 | 0 | 0 |
| ENSMUSG00000106520 | 0 | 0 | 0 | 0 | 0 | 0 | 0 |
| ENSMUSG00000106521 | 0 | 0 | 0 | 0 | 0 | 0 | 0 |
| ENSMUSG00000106522 | 0 | 0 | 0 | 0 | 0 | 0 | 0 |
| ENSMUSG00000106523 | 0 | 0 | 0 | 0 | 0 | 0 | 0 |
| ENSMUSG00000106524 | 0 | 0 | 0 | 0 | 0 | 0 | 0 |
| ENSMUSG00000106525 | 0 | 0 | 0 | 0 | 0 | 0 | 0 |
| ENSMUSG00000106526 | 0 | 0.25 | 0 | 0 | 0 | 0 | 0 |
| ENSMUSG00000106527 | 0 | 0 | 0 | 0 | 0 | 0 | 0 |
| ENSMUSG00000106528 | 0 | 0.25 | 0 | 0 | 0 | 0 | 0 |

| | | | | | | | |
|---|---|---|---|---|---|---|---|
| ENSMUSG00000106529 | 0 | 0.25 | 0.25 | 0.646240625 | 0.686820904 | 0.903089987 | 0.520678036 |
| ENSMUSG00000106530 | 0 | 0 | 0 | 0 | 0 | 0 | 0 |
| ENSMUSG00000106531 | 0 | 0 | 0 | 0 | 0 | 0 | 0.355418052 |
| ENSMUSG00000106532 | 0 | 0 | 0 | 0 | 0 | 0 | 0 |
| ENSMUSG00000106533 | 0 | 0 | 0 | 0 | 0 | 0 | 0 |
| ENSMUSG00000106534 | 0 | 0 | 0 | 0 | 0.244650542 | 0 | 0.224243824 |
| ENSMUSG00000106535 | 0.867194479 | 0.896240625 | 0.92510993 | 1.061981878 | 0.978602168 | 1.146128036 | 1.014380832 |
| ENSMUSG00000106536 | 0.747221736 | 0.646240625 | 0.580482024 | 0.646240625 | 0.387761935 | 0.301029996 | 0.355418052 |
| ENSMUSG00000106537 | 0 | 0 | 0 | 0 | 0 | 0 | 0 |
| ENSMUSG00000106538 | 0 | 0 | 0.25 | 0 | 0 | 0 | 0 |
| ENSMUSG00000106539 | 0 | 0 | 0 | 0 | 0 | 0 | 0 |
| ENSMUSG00000106540 | 0 | 0 | 0 | 0 | 0 | 0 | 0 |
| ENSMUSG00000106541 | 0 | 0 | 0.25 | 0 | 0 | 0 | 0.224243824 |
| ENSMUSG00000106542 | 0 | 0 | 0 | 0 | 0 | 0 | 0 |
| ENSMUSG00000106543 | 0.45815691 | 0.580482024 | 0.580482024 | 0.396240625 | 0.387761935 | 0 | 0.579661877 |
| ENSMUSG00000106544 | 0 | 0 | 0 | 0 | 0 | 0 | 0 |
| ENSMUSG00000106545 | 0.289064826 | 0 | 0 | 0 | 0 | 0 | 0 |
| ENSMUSG00000106546 | 0 | 0 | 0 | 0 | 0 | 0 | 0 |
| ENSMUSG00000106547 | 0 | 0.5 | 0.396240625 | 0.580482024 | 0.568060967 | 0.477121255 | 0.579661877 |
| ENSMUSG00000106548 | 0 | 0 | 0 | 0 | 0 | 0 | 0 |
| ENSMUSG00000106549 | 1.307602651 | 1.26109853 | 1.114857905 | 1.17510993 | 1.121713561 | 1.278753601 | 1.100339912 |
| ENSMUSG00000106550 | 0 | 0.25 | 0 | 0 | 0 | 0 | 0 |
| ENSMUSG00000106551 | 0 | 0 | 0.25 | 0 | 0 | 0 | 0 |
| ENSMUSG00000106552 | 0 | 0 | 0 | 0 | 0 | 0 | 0 |
| ENSMUSG00000106553 | 0 | 0 | 0 | 0 | 0 | 0 | 0 |
| ENSMUSG00000106554 | 0 | 0 | 0 | 0 | 0 | 0 | 0 |
| ENSMUSG00000106555 | 0 | 0 | 0 | 0.25 | 0 | 0 | 0 |
| ENSMUSG00000106556 | 0 | 0 | 0 | 0 | 0 | 0 | 0 |
| ENSMUSG00000106557 | 0 | 0 | 0 | 0 | 0 | 0 | 0 |
| ENSMUSG00000106558 | 0 | 0 | 0 | 0 | 0 | 0 | 0 |
| ENSMUSG00000106559 | 0 | 0 | 0 | 0 | 0 | 0 | 0 |
| ENSMUSG00000106560 | 0 | 0.580482024 | 0.396240625 | 0.580482024 | 0.489301084 | 0.602059991 | 0.579661877 |
| ENSMUSG00000106561 | 0 | 0 | 0 | 0 | 0 | 0 | 0 |
| ENSMUSG00000106562 | 0.578129653 | 0 | 0.396240625 | 0 | 0 | 0.301029996 | 0.224243824 |

| | | | | | | | |
|---|---|---|---|---|---|---|---|
| ENSMUSG00000106563 | 0 | 0 | 0 | 0 | 0 | 0 | 0 |
| ENSMUSG00000106564 | 0 | 0 | 0 | 0 | 0 | 0 | 0 |
| ENSMUSG00000106565 | 0.289064826 | 0 | 0 | 0.25 | 0 | 0.477121255 | 0.355418052 |
| ENSMUSG00000106566 | 0 | 0 | 0 | 0 | 0 | 0 | 0 |
| ENSMUSG00000106567 | 0 | 0 | 0 | 0 | 0 | 0 | 0 |
| ENSMUSG00000106568 | 0.960252568 | 0.864857905 | 1.04248125 | 0.79248125 | 0.978602168 | 1 | 0.775756176 |
| ENSMUSG00000106569 | 0 | 0 | 0 | 0 | 0 | 0 | 0 |
| ENSMUSG00000106570 | 0 | 0 | 0 | 0 | 0 | 0 | 0 |
| ENSMUSG00000106571 | 0 | 0 | 0 | 0 | 0 | 0 | 0 |
| ENSMUSG00000106572 | 0 | 0 | 0.25 | 0 | 0 | 0 | 0 |
| ENSMUSG00000106573 | 0 | 0 | 0 | 0 | 0 | 0 | 0 |
| ENSMUSG00000106574 | 0 | 0 | 0.25 | 0 | 0 | 0.301029996 | 0 |
| ENSMUSG00000106575 | 0 | 0.25 | 0 | 0 | 0 | 0 | 0 |
| ENSMUSG00000106576 | 0.289064826 | 0.580482024 | 0.396240625 | 0 | 0.489301084 | 0.301029996 | 0.224243824 |
| ENSMUSG00000106577 | 0 | 0.5 | 0 | 0.25 | 0 | 0 | 0 |
| ENSMUSG00000106578 | 0 | 0 | 0 | 0 | 0 | 0 | 0 |
| ENSMUSG00000106579 | 0 | 0 | 0 | 0 | 0 | 0 | 0.224243824 |
| ENSMUSG00000106580 | 0 | 0 | 0.25 | 0 | 0 | 0 | 0 |
| ENSMUSG00000106581 | 0 | 0 | 0 | 0 | 0 | 0 | 0.224243824 |
| ENSMUSG00000106582 | 0 | 0 | 0 | 0 | 0 | 0 | 0 |
| ENSMUSG00000106583 | 0 | 0 | 0 | 0 | 0 | 0 | 0 |
| ENSMUSG00000106584 | 0 | 0 | 0 | 0 | 0 | 0 | 0 |
| ENSMUSG00000106585 | 0 | 0 | 0 | 0 | 0 | 0 | 0 |
| ENSMUSG00000106586 | 0 | 0 | 0 | 0 | 0 | 0 | 0 |
| ENSMUSG00000106587 | 0 | 0 | 0 | 0 | 0 | 0 | 0 |
| ENSMUSG00000106588 | 0.289064826 | 0.25 | 0.25 | 0 | 0 | 0 | 0.224243824 |
| ENSMUSG00000106589 | 0 | 0 | 0 | 0 | 0 | 0 | 0 |
| ENSMUSG00000106590 | 0.960252568 | 0.896240625 | 0.580482024 | 0.396240625 | 0.686820904 | 0 | 0.224243824 |
| ENSMUSG00000106591 | 0 | 0 | 0 | 0 | 0 | 0 | 0 |
| ENSMUSG00000106592 | 0 | 0 | 0 | 0 | 0.244650542 | 0 | 0 |
| ENSMUSG00000106593 | 0 | 0 | 0 | 0 | 0 | 0 | 0 |
| ENSMUSG00000106594 | 0 | 0 | 0 | 0 | 0 | 0 | 0 |
| ENSMUSG00000106595 | 0 | 0 | 0 | 0 | 0 | 0 | 0 |
| ENSMUSG00000106596 | 0 | 0 | 0 | 0 | 0 | 0 | 0.224243824 |

| | | | | | | | |
|---|---|---|---|---|---|---|---|
| ENSMUSG00000106597 | 0 | 0 | 0 | 0 | 0 | 0 | 0 |
| ENSMUSG00000106598 | 0 | 0 | 0 | 0.25 | 0 | 0 | 0.224243824 |
| ENSMUSG00000106599 | 0 | 0 | 0 | 0 | 0 | 0 | 0 |
| ENSMUSG00000106600 | 0 | 0.25 | 0 | 0.25 | 0.387761935 | 0 | 0 |
| ENSMUSG00000106601 | 0 | 0 | 0 | 0 | 0 | 0 | 0 |
| ENSMUSG00000106602 | 0 | 0 | 0 | 0 | 0 | 0 | 0 |
| ENSMUSG00000106603 | 0.578129653 | 0.396240625 | 0.5 | 0 | 0.387761935 | 0 | 0 |
| ENSMUSG00000106604 | 0 | 0 | 0 | 0 | 0 | 0 | 0 |
| ENSMUSG00000106605 | 0 | 0 | 0 | 0 | 0 | 0 | 0 |
| ENSMUSG00000106606 | 0 | 0 | 0 | 0 | 0 | 0 | 0 |
| ENSMUSG00000106607 | 0 | 0.25 | 0 | 0 | 0.489301084 | 0 | 0.224243824 |
| ENSMUSG00000106608 | 0 | 0 | 0 | 0 | 0 | 0 | 0 |
| ENSMUSG00000106609 | 0.289064826 | 0.396240625 | 0.25 | 0.396240625 | 0.387761935 | 0 | 0.224243824 |
| ENSMUSG00000106610 | 0 | 0.25 | 0 | 0 | 0 | 0 | 0 |
| ENSMUSG00000106611 | 0 | 0 | 0 | 0 | 0 | 0 | 0 |
| ENSMUSG00000106612 | 0 | 0.25 | 0.25 | 0 | 0 | 0 | 0 |
| ENSMUSG00000106613 | 0 | 0 | 0 | 0 | 0 | 0 | 0 |
| ENSMUSG00000106614 | 0 | 0 | 0.25 | 0 | 0 | 0 | 0 |
| ENSMUSG00000106615 | 0 | 0 | 0 | 0 | 0 | 0 | 0 |
| ENSMUSG00000106616 | 0 | 0 | 0 | 0 | 0 | 0 | 0 |
| ENSMUSG00000106617 | 0.45815691 | 0.646240625 | 0.396240625 | 0.646240625 | 0.77552387 | 0.301029996 | 0.579661877 |
| ENSMUSG00000106618 | 0 | 0 | 0 | 0 | 0 | 0 | 0 |
| ENSMUSG00000106619 | 0 | 0 | 0 | 0 | 0 | 0.301029996 | 0 |
| ENSMUSG00000106620 | 0 | 0 | 0 | 0 | 0 | 0 | 0 |
| ENSMUSG00000106621 | 0 | 0 | 0 | 0 | 0 | 0 | 0 |
| ENSMUSG00000106622 | 0 | 0 | 0 | 0 | 0.244650542 | 0 | 0 |
| ENSMUSG00000106623 | 0.45815691 | 0.25 | 0.25 | 0.25 | 0.387761935 | 0 | 0.224243824 |
| ENSMUSG00000106624 | 0 | 0 | 0 | 0 | 0 | 0 | 0 |
| ENSMUSG00000106625 | 0 | 0 | 0.396240625 | 0.25 | 0 | 0 | 0 |
| ENSMUSG00000106626 | 0 | 0 | 0 | 0 | 0 | 0 | 0 |
| ENSMUSG00000106627 | 0 | 0 | 0 | 0 | 0 | 0 | 0 |
| ENSMUSG00000106628 | 0.289064826 | 0.25 | 0.25 | 0.25 | 0.568060967 | 0 | 0.224243824 |
| ENSMUSG00000106629 | 0 | 0 | 0 | 0 | 0.244650542 | 0 | 0 |
| ENSMUSG00000106630 | 0 | 0 | 0 | 0 | 0 | 0 | 0 |

| | | | | | | | |
|---|---|---|---|---|---|---|---|
| ENSMUSG00000106631 | 0.811507563 | 0.580482024 | 0.79248125 | 0.396240625 | 0.846351821 | 0.698970004 | 0.74492186 |
| ENSMUSG00000106632 | 0.671187741 | 0.25 | 0.75 | 0.646240625 | 0.77552387 | 0.301029996 | 0.355418052 |
| ENSMUSG00000106633 | 0 | 0 | 0 | 0 | 0 | 0 | 0 |
| ENSMUSG00000106634 | 1.269664473 | 1.17510993 | 1.130890489 | 1.188721876 | 1.254881871 | 1.322219295 | 1.140832123 |
| ENSMUSG00000106635 | 0 | 0 | 0 | 0 | 0 | 0 | 0 |
| ENSMUSG00000106636 | 0.671187741 | 0.830482024 | 0.580482024 | 0.5 | 0.632412477 | 0.301029996 | 0.629532004 |
| ENSMUSG00000106637 | 0.45815691 | 0.396240625 | 0.580482024 | 0.580482024 | 0.686820904 | 0.301029996 | 0.224243824 |
| ENSMUSG00000106639 | 0 | 0 | 0 | 0 | 0 | 0 | 0 |
| ENSMUSG00000106640 | 0 | 0 | 0.25 | 0 | 0 | 0 | 0 |
| ENSMUSG00000106641 | 0 | 0 | 0 | 0.646240625 | 0.244650542 | 0.301029996 | 0.355418052 |
| ENSMUSG00000106642 | 0 | 0 | 0 | 0 | 0 | 0 | 0 |
| ENSMUSG00000106643 | 0 | 0.5 | 0 | 0 | 0 | 0 | 0.224243824 |
| ENSMUSG00000106644 | 0 | 0 | 0 | 0 | 0 | 0 | 0 |
| ENSMUSG00000106645 | 0 | 0 | 0 | 0 | 0 | 0 | 0 |
| ENSMUSG00000106646 | 0.289064826 | 0.396240625 | 0 | 0.25 | 0 | 0 | 0.520678036 |
| ENSMUSG00000106647 | 0.289064826 | 0.396240625 | 0.25 | 0 | 0.244650542 | 0 | 0 |
| ENSMUSG00000106648 | 0.289064826 | 0 | 0.396240625 | 0.5 | 0.244650542 | 0 | 0.355418052 |
| ENSMUSG00000106649 | 0 | 0 | 0 | 0 | 0 | 0 | 0 |
| ENSMUSG00000106650 | 0.289064826 | 0 | 0 | 0.25 | 0 | 0 | 0.355418052 |
| ENSMUSG00000106651 | 0.289064826 | 0.79248125 | 0.646240625 | 0.646240625 | 0.489301084 | 0.698970004 | 0 |
| ENSMUSG00000106652 | 0 | 0 | 0 | 0 | 0 | 0 | 0 |
| ENSMUSG00000106653 | 0 | 0 | 0 | 0 | 0 | 0 | 0 |
| ENSMUSG00000106654 | 0 | 0 | 0 | 0.25 | 0.387761935 | 0.301029996 | 0 |
| ENSMUSG00000106655 | 0 | 0 | 0 | 0 | 0 | 0 | 0 |
| ENSMUSG00000106656 | 0 | 0 | 0 | 0 | 0 | 0 | 0 |
| ENSMUSG00000106657 | 0 | 0 | 0 | 0 | 0.244650542 | 0 | 0 |
| ENSMUSG00000106658 | 0 | 0 | 0 | 0.25 | 0.244650542 | 0 | 0 |
| ENSMUSG00000106659 | 0 | 0 | 0 | 0 | 0 | 0 | 0 |
| ENSMUSG00000106660 | 0 | 0 | 0 | 0 | 0 | 0 | 0 |
| ENSMUSG00000106661 | 0 | 0 | 0 | 0 | 0 | 0 | 0 |
| ENSMUSG00000106662 | 1.129344651 | 1.04248125 | 0.896240625 | 1 | 0.978602168 | 1.146128036 | 1 |
| ENSMUSG00000106663 | 0.45815691 | 0 | 0 | 0 | 0 | 0 | 0 |
| ENSMUSG00000106664 | 0.45815691 | 0.5 | 0.5 | 0.580482024 | 0.489301084 | 0.301029996 | 0.355418052 |
| ENSMUSG00000106665 | 0.747221736 | 0.5 | 0.646240625 | 0.580482024 | 0.632412477 | 0.602059991 | 0.629532004 |

| | | | | | | | |
|---|---|---|---|---|---|---|---|
| ENSMUSG00000106666 | 0 | 0 | 0 | 0 | 0 | 0 | 0 |
| ENSMUSG00000106667 | 0 | 0 | 0 | 0 | 0 | 0 | 0 |
| ENSMUSG00000106668 | 0 | 0 | 0 | 0 | 0 | 0 | 0 |
| ENSMUSG00000106669 | 0 | 0 | 0 | 0 | 0 | 0 | 0 |
| ENSMUSG00000106670 | 0 | 0 | 0 | 0 | 0 | 0 | 0 |
| ENSMUSG00000106671 | 0.91631382 | 0.976722649 | 0.75 | 0.701838731 | 0.955822902 | 0.77815125 | 0.935079929 |
| ENSMUSG00000106673 | 0 | 0 | 0 | 0 | 0 | 0 | 0 |
| ENSMUSG00000106674 | 0 | 0 | 0 | 0.25 | 0.244650542 | 0 | 0 |
| ENSMUSG00000106675 | 0 | 0 | 0 | 0 | 0 | 0 | 0 |
| ENSMUSG00000106676 | 0.289064826 | 0 | 0 | 0 | 0.244650542 | 0 | 0 |
| ENSMUSG00000106677 | 0 | 0 | 0 | 0 | 0 | 0 | 0 |
| ENSMUSG00000106678 | 0.867194479 | 0.896240625 | 0.864857905 | 1 | 1.020174412 | 0.954242509 | 1 |
| ENSMUSG00000106679 | 0 | 0 | 0 | 0 | 0 | 0 | 0 |
| ENSMUSG00000106680 | 0 | 0.396240625 | 0.396240625 | 0 | 0 | 0.301029996 | 0.355418052 |
| ENSMUSG00000106681 | 0 | 0 | 0 | 0 | 0 | 0 | 0 |
| ENSMUSG00000106682 | 1.129344651 | 0.896240625 | 0.701838731 | 0.646240625 | 0.489301084 | 0.301029996 | 0.579661877 |
| ENSMUSG00000106683 | 0 | 0 | 0 | 0.25 | 0.244650542 | 0 | 0 |
| ENSMUSG00000106686 | 0 | 0 | 0 | 0 | 0 | 0 | 0 |
| ENSMUSG00000106688 | 0.671187741 | 0.701838731 | 0.5 | 0.580482024 | 0.244650542 | 0.477121255 | 0.355418052 |
| ENSMUSG00000106689 | 0 | 0 | 0 | 0 | 0 | 0 | 0 |
| ENSMUSG00000106690 | 0 | 0 | 0 | 0 | 0 | 0 | 0 |
| ENSMUSG00000106691 | 0.747221736 | 0.5 | 0.701838731 | 0.396240625 | 0.568060967 | 0.477121255 | 0.224243824 |
| ENSMUSG00000106692 | 0 | 0.25 | 0 | 0 | 0.244650542 | 0 | 0.224243824 |
| ENSMUSG00000106693 | 0 | 0 | 0 | 0 | 0 | 0 | 0 |
| ENSMUSG00000106694 | 0 | 0 | 0.25 | 0 | 0 | 0 | 0 |
| ENSMUSG00000106696 | 0.671187741 | 0.25 | 0.5 | 0.25 | 0.489301084 | 0.301029996 | 0.355418052 |
| ENSMUSG00000106697 | 0 | 0 | 0 | 0 | 0 | 0 | 0 |
| ENSMUSG00000106698 | 0 | 0 | 0.25 | 0 | 0.244650542 | 0.301029996 | 0 |
| ENSMUSG00000106699 | 0 | 0 | 0 | 0 | 0 | 0 | 0 |
| ENSMUSG00000106701 | 0 | 0 | 0 | 0 | 0 | 0 | 0 |
| ENSMUSG00000106702 | 0 | 0 | 0 | 0 | 0 | 0 | 0 |
| ENSMUSG00000106703 | 0 | 0 | 0 | 0 | 0 | 0.301029996 | 0 |
| ENSMUSG00000106704 | 0 | 0 | 0 | 0 | 0 | 0 | 0 |
| ENSMUSG00000106705 | 0 | 0 | 0 | 0 | 0 | 0 | 0 |

| | | | | | | | |
|---|---|---|---|---|---|---|---|
| ENSMUSG00000106706 | 0.289064826 | 0.25 | 0.396240625 | 0.25 | 0.489301084 | 0.602059991 | 0.579661877 |
| ENSMUSG00000106707 | 0.289064826 | 0.25 | 0.25 | 0.25 | 0 | 0 | 0 |
| ENSMUSG00000106708 | 0 | 0 | 0 | 0 | 0 | 0 | 0 |
| ENSMUSG00000106709 | 0 | 0.396240625 | 0 | 0 | 0 | 0 | 0 |
| ENSMUSG00000106710 | 0 | 0.25 | 0.25 | 0.25 | 0 | 0 | 0 |
| ENSMUSG00000106712 | 0 | 0 | 0 | 0 | 0 | 0 | 0 |
| ENSMUSG00000106713 | 0 | 0 | 0 | 0 | 0 | 0 | 0 |
| ENSMUSG00000106714 | 1.358731791 | 1.188721876 | 1.238549078 | 1.25 | 1.244650542 | 1.255272505 | 1.150210039 |
| ENSMUSG00000106715 | 0 | 0 | 0 | 0 | 0 | 0 | 0 |
| ENSMUSG00000106717 | 0.671187741 | 0.5 | 0.646240625 | 0.580482024 | 0.955822902 | 0.698970004 | 0.710836105 |
| ENSMUSG00000106718 | 0 | 0 | 0 | 0 | 0 | 0 | 0 |
| ENSMUSG00000106719 | 0 | 0.25 | 0.25 | 0 | 0.387761935 | 0.301029996 | 0.448487648 |
| ENSMUSG00000106720 | 0.289064826 | 0.25 | 0 | 0 | 0.244650542 | 0 | 0 |
| ENSMUSG00000106721 | 0.578129653 | 0.396240625 | 0.25 | 0 | 0 | 0 | 0 |
| ENSMUSG00000106724 | 0 | 0 | 0 | 0 | 0 | 0 | 0 |
| ENSMUSG00000106725 | 0 | 0 | 0 | 0 | 0.244650542 | 0.477121255 | 0 |
| ENSMUSG00000106726 | 0 | 0 | 0 | 0 | 0 | 0 | 0 |
| ENSMUSG00000106729 | 0 | 0 | 0 | 0 | 0 | 0 | 0 |
| ENSMUSG00000106730 | 0 | 0 | 0 | 0 | 0 | 0 | 0 |
| ENSMUSG00000106732 | 0 | 0.25 | 0.396240625 | 0 | 0 | 0 | 0 |
| ENSMUSG00000106733 | 0 | 0 | 0.25 | 0.396240625 | 0 | 0 | 0 |
| ENSMUSG00000106734 | 1.181541736 | 1.214495249 | 1.25 | 1.282320754 | 1.091002363 | 1.255272505 | 1.168187745 |
| ENSMUSG00000106735 | 0 | 0 | 0 | 0 | 0 | 0 | 0 |
| ENSMUSG00000106736 | 0.45815691 | 0.580482024 | 0 | 0.396240625 | 0.244650542 | 0.301029996 | 0.224243824 |
| ENSMUSG00000106737 | 0 | 0 | 0 | 0 | 0 | 0 | 0 |
| ENSMUSG00000106738 | 0 | 0 | 0 | 0 | 0 | 0 | 0 |
| ENSMUSG00000106739 | 0 | 0.25 | 0.5 | 0 | 0.387761935 | 0.301029996 | 0.224243824 |
| ENSMUSG00000106740 | 0 | 0 | 0 | 0 | 0 | 0 | 0 |
| ENSMUSG00000106741 | 0 | 0 | 0 | 0 | 0 | 0 | 0 |
| ENSMUSG00000106742 | 0 | 0 | 0 | 0 | 0 | 0 | 0 |
| ENSMUSG00000106743 | 0.671187741 | 0.951838731 | 0.830482024 | 0.976722649 | 0.931471446 | 0.903089987 | 1.028149525 |
| ENSMUSG00000106744 | 2.418409478 | 2.202642909 | 2.180274797 | 2.293731421 | 2.214715681 | 2.491361694 | 2.070582188 |
| ENSMUSG00000106745 | 0 | 0 | 0 | 0 | 0 | 0 | 0 |
| ENSMUSG00000106746 | 0 | 0 | 0 | 0 | 0 | 0 | 0 |

| | | | | | | | |
|---|---|---|---|---|---|---|---|
| ENSMUSG00000106747 | 0.578129653 | 0.5 | 0.75 | 0.701838731 | 0.489301084 | 0 | 0.629532004 |
| ENSMUSG00000106748 | 0 | 0 | 0 | 0 | 0 | 0 | 0 |
| ENSMUSG00000106749 | 0 | 0 | 0 | 0 | 0 | 0 | 0 |
| ENSMUSG00000106750 | 0.289064826 | 0 | 0 | 0 | 0 | 0 | 0 |
| ENSMUSG00000106751 | 0.289064826 | 0.25 | 0.25 | 0.5 | 0.568060967 | 0.301029996 | 0.520678036 |
| ENSMUSG00000106752 | 0 | 0 | 0 | 0 | 0 | 0 | 0 |
| ENSMUSG00000106753 | 0 | 0 | 0 | 0 | 0 | 0 | 0 |
| ENSMUSG00000106754 | 0 | 0 | 0 | 0 | 0 | 0 | 0 |
| ENSMUSG00000106755 | 0.289064826 | 0 | 0.25 | 0.5 | 0 | 0.301029996 | 0.224243824 |
| ENSMUSG00000106756 | 0 | 0 | 0 | 0 | 0 | 0 | 0 |
| ENSMUSG00000106757 | 0.671187741 | 0.701838731 | 0.75 | 0.896240625 | 0.568060967 | 0.602059991 | 0.74492186 |
| ENSMUSG00000106758 | 0 | 0 | 0 | 0 | 0 | 0 | 0 |
| ENSMUSG00000106759 | 0.289064826 | 0 | 0 | 0 | 0 | 0 | 0 |
| ENSMUSG00000106760 | 0 | 0 | 0 | 0 | 0 | 0 | 0 |
| ENSMUSG00000106762 | 0 | 0 | 0 | 0 | 0 | 0 | 0 |
| ENSMUSG00000106763 | 0 | 0 | 0 | 0 | 0 | 0 | 0 |
| ENSMUSG00000106764 | 0 | 0 | 0 | 0 | 0 | 0 | 0 |
| ENSMUSG00000106766 | 0 | 0 | 0 | 0 | 0 | 0 | 0 |
| ENSMUSG00000106767 | 1.036286563 | 0.830482024 | 0.75 | 0.646240625 | 0.632412477 | 0.477121255 | 0.672731473 |
| ENSMUSG00000106768 | 0 | 0 | 0 | 0 | 0 | 0 | 0 |
| ENSMUSG00000106769 | 0 | 0 | 0 | 0 | 0 | 0 | 0 |
| ENSMUSG00000106770 | 0 | 0 | 0 | 0 | 0 | 0 | 0 |
| ENSMUSG00000106771 | 0 | 0 | 0 | 0 | 0 | 0 | 0 |
| ENSMUSG00000106772 | 0.289064826 | 0 | 0 | 0 | 0 | 0 | 0 |
| ENSMUSG00000106773 | 0 | 0 | 0 | 0 | 0 | 0 | 0 |
| ENSMUSG00000106775 | 0 | 0.25 | 0 | 0.396240625 | 0 | 0 | 0 |
| ENSMUSG00000106776 | 0 | 0 | 0 | 0 | 0.244650542 | 0 | 0 |
| ENSMUSG00000106777 | 0.671187741 | 0.580482024 | 0.396240625 | 0.701838731 | 0.489301084 | 0.477121255 | 0.520678036 |
| ENSMUSG00000106778 | 0 | 0 | 0 | 0 | 0.244650542 | 0 | 0.224243824 |
| ENSMUSG00000106779 | 0 | 0 | 0 | 0 | 0 | 0 | 0 |
| ENSMUSG00000106780 | 0 | 0 | 0 | 0 | 0 | 0 | 0 |
| ENSMUSG00000106781 | 0 | 0 | 0 | 0 | 0 | 0 | 0 |
| ENSMUSG00000106782 | 0 | 0 | 0 | 0 | 0 | 0 | 0 |
| ENSMUSG00000106783 | 0 | 0 | 0 | 0 | 0 | 0 | 0 |

| | | | | | | | |
|---|---|---|---|---|---|---|---|
| ENSMUSG00000106784 | 0 | 0 | 0 | 0 | 0 | 0 | 0 |
| ENSMUSG00000106785 | 0 | 0 | 0 | 0 | 0 | 0 | 0 |
| ENSMUSG00000106786 | 0 | 0 | 0 | 0 | 0 | 0 | 0 |
| ENSMUSG00000106788 | 0 | 0 | 0 | 0 | 0 | 0 | 0 |
| ENSMUSG00000106790 | 0 | 0 | 0 | 0 | 0 | 0 | 0 |
| ENSMUSG00000106791 | 0 | 0 | 0.25 | 0 | 0.568060967 | 0 | 0.355418052 |
| ENSMUSG00000106793 | 0 | 0.25 | 0 | 0.25 | 0 | 0 | 0 |
| ENSMUSG00000106794 | 0 | 0 | 0 | 0 | 0 | 0 | 0 |
| ENSMUSG00000106795 | 0 | 0 | 0 | 0 | 0 | 0 | 0 |
| ENSMUSG00000106797 | 0 | 0 | 0 | 0 | 0 | 0 | 0 |
| ENSMUSG00000106798 | 0 | 0.25 | 0 | 0 | 0 | 0 | 0 |
| ENSMUSG00000106799 | 0 | 0 | 0 | 0 | 0 | 0 | 0.224243824 |
| ENSMUSG00000106800 | 0 | 0 | 0 | 0 | 0 | 0 | 0 |
| ENSMUSG00000106801 | 0 | 0 | 0 | 0 | 0 | 0 | 0 |
| ENSMUSG00000106802 | 0.289064826 | 0 | 0.25 | 0.396240625 | 0.489301084 | 0 | 0.355418052 |
| ENSMUSG00000106803 | 0 | 0 | 0 | 0 | 0 | 0 | 0 |
| ENSMUSG00000106804 | 0 | 0 | 0 | 0 | 0 | 0 | 0 |
| ENSMUSG00000106805 | 0 | 0 | 0 | 0 | 0 | 0 | 0 |
| ENSMUSG00000106806 | 0.289064826 | 0 | 0 | 0 | 0 | 0 | 0 |
| ENSMUSG00000106807 | 0 | 0 | 0 | 0.25 | 0.244650542 | 0.301029996 | 0 |
| ENSMUSG00000106808 | 0 | 0 | 0 | 0 | 0 | 0 | 0 |
| ENSMUSG00000106810 | 0 | 0 | 0 | 0 | 0 | 0 | 0 |
| ENSMUSG00000106811 | 0 | 0 | 0 | 0 | 0 | 0 | 0 |
| ENSMUSG00000106812 | 0 | 0 | 0 | 0 | 0 | 0 | 0 |
| ENSMUSG00000106813 | 0 | 0 | 0 | 0 | 0 | 0 | 0 |
| ENSMUSG00000106814 | 0 | 0 | 0 | 0 | 0 | 0 | 0 |
| ENSMUSG00000106815 | 0.45815691 | 0.396240625 | 0.5 | 0.25 | 0.244650542 | 0.301029996 | 0 |
| ENSMUSG00000106816 | 0 | 0 | 0 | 0 | 0 | 0 | 0 |
| ENSMUSG00000106817 | 0 | 0 | 0 | 0 | 0 | 0 | 0 |
| ENSMUSG00000106818 | 0.811507563 | 0.580482024 | 0.580482024 | 0.75 | 0.632412477 | 0.301029996 | 0.520678036 |
| ENSMUSG00000106819 | 0.867194479 | 0.951838731 | 0.896240625 | 1.061981878 | 0.877063019 | 0.698970004 | 1 |
| ENSMUSG00000106820 | 0.811507563 | 0.830482024 | 0.864857905 | 0.396240625 | 0.244650542 | 0.477121255 | 0.520678036 |
| ENSMUSG00000106821 | 0 | 0 | 0.25 | 0 | 0 | 0 | 0 |
| ENSMUSG00000106822 | 0 | 0.25 | 0 | 0 | 0 | 0 | 0 |

| | | | | | | | |
|---|---|---|---|---|---|---|---|
| ENSMUSG00000106824 | 0 | 0 | 0 | 0 | 0 | 0 | 0 |
| ENSMUSG00000106825 | 1.069666964 | 0.896240625 | 0.951838731 | 0.646240625 | 0.632412477 | 0.84509804 | 0.775756176 |
| ENSMUSG00000106826 | 0.578129653 | 0.396240625 | 0.396240625 | 0.580482024 | 0.489301084 | 0.602059991 | 0.355418052 |
| ENSMUSG00000106827 | 0 | 0 | 0.25 | 0.396240625 | 0 | 0 | 0 |
| ENSMUSG00000106828 | 0.289064826 | 0.396240625 | 0.25 | 0 | 0 | 0 | 0 |
| ENSMUSG00000106829 | 0 | 0 | 0 | 0 | 0 | 0 | 0 |
| ENSMUSG00000106831 | 0 | 0.25 | 0 | 0 | 0 | 0 | 0 |
| ENSMUSG00000106832 | 0 | 0.25 | 0 | 0.25 | 0 | 0 | 0 |
| ENSMUSG00000106833 | 0 | 0 | 0 | 0 | 0 | 0 | 0 |
| ENSMUSG00000106834 | 0.289064826 | 0 | 0 | 0 | 0 | 0 | 0 |
| ENSMUSG00000106835 | 0 | 0 | 0 | 0 | 0 | 0 | 0 |
| ENSMUSG00000106836 | 0 | 0 | 0 | 0 | 0 | 0 | 0 |
| ENSMUSG00000106837 | 0 | 0 | 0 | 0 | 0 | 0 | 0 |
| ENSMUSG00000106838 | 0 | 0 | 0 | 0 | 0 | 0 | 0 |
| ENSMUSG00000106839 | 0 | 0 | 0 | 0 | 0 | 0 | 0 |
| ENSMUSG00000106840 | 0.671187741 | 0.396240625 | 0.25 | 0.646240625 | 0.387761935 | 0.301029996 | 0 |
| ENSMUSG00000106841 | 0.289064826 | 0.25 | 0.25 | 0.5 | 0 | 0 | 0.224243824 |
| ENSMUSG00000106843 | 0 | 0 | 0 | 0.25 | 0.489301084 | 0 | 0 |
| ENSMUSG00000106844 | 0 | 0 | 0 | 0 | 0 | 0 | 0 |
| ENSMUSG00000106845 | 0 | 0 | 0 | 0 | 0 | 0 | 0 |
| ENSMUSG00000106846 | 0 | 0 | 0 | 0 | 0 | 0 | 0 |
| ENSMUSG00000106847 | 1.971473847 | 1.79248125 | 1.729715809 | 1.653677461 | 1.662257525 | 1.799340549 | 1.612273755 |
| ENSMUSG00000106849 | 0 | 0 | 0 | 0 | 0 | 0 | 0 |
| ENSMUSG00000106851 | 0.289064826 | 0 | 0 | 0 | 0 | 0 | 0 |
| ENSMUSG00000106852 | 0 | 0 | 0 | 0 | 0 | 0 | 0 |
| ENSMUSG00000106853 | 0 | 0 | 0 | 0 | 0 | 0 | 0 |
| ENSMUSG00000106854 | 0 | 0 | 0 | 0 | 0 | 0 | 0 |
| ENSMUSG00000106855 | 0 | 0 | 0.25 | 0.25 | 0 | 0 | 0 |
| ENSMUSG00000106856 | 0 | 0.396240625 | 0 | 0 | 0 | 0 | 0 |
| ENSMUSG00000106857 | 0 | 0 | 0 | 0 | 0 | 0 | 0 |
| ENSMUSG00000106858 | 0.811507563 | 0.79248125 | 0.580482024 | 0.396240625 | 0.77552387 | 0.602059991 | 0.448487648 |
| ENSMUSG00000106859 | 0 | 0 | 0 | 0 | 0 | 0 | 0 |
| ENSMUSG00000106860 | 0 | 0 | 0.396240625 | 0.580482024 | 0.244650542 | 0.301029996 | 0.355418052 |
| ENSMUSG00000106861 | 0 | 0 | 0 | 0 | 0 | 0 | 0 |

| | | | | | | | |
|---|---|---|---|---|---|---|---|
| ENSMUSG00000106862 | 0.45815691 | 0 | 0.396240625 | 0 | 0.244650542 | 0 | 0 |
| ENSMUSG00000106863 | 0 | 0.25 | 0.25 | 0 | 0.387761935 | 0 | 0 |
| ENSMUSG00000106864 | 2.402962068 | 2.237821179 | 2.224461364 | 2.347542239 | 2.279510047 | 2.571708832 | 2.110947927 |
| ENSMUSG00000106865 | 0 | 0 | 0 | 0 | 0 | 0 | 0 |
| ENSMUSG00000106866 | 0.45815691 | 0.25 | 0.25 | 0 | 0.244650542 | 0 | 0 |
| ENSMUSG00000106867 | 0 | 0.25 | 0.5 | 0.396240625 | 0 | 0 | 0.224243824 |
| ENSMUSG00000106868 | 0 | 0 | 0 | 0 | 0 | 0 | 0.355418052 |
| ENSMUSG00000106869 | 0 | 0 | 0.25 | 0.25 | 0 | 0 | 0 |
| ENSMUSG00000106870 | 1.358731791 | 1.356566189 | 1.27186571 | 1.17510993 | 1.091002363 | 1 | 1.089372236 |
| ENSMUSG00000106871 | 0 | 0.25 | 0 | 0 | 0 | 0 | 0 |
| ENSMUSG00000106872 | 0 | 0 | 0 | 0 | 0 | 0 | 0 |
| ENSMUSG00000106873 | 0 | 0 | 0 | 0 | 0 | 0 | 0 |
| ENSMUSG00000106874 | 1.418409478 | 1.356566189 | 1.380890489 | 1.26109853 | 1.212046813 | 1.079181246 | 1.028149525 |
| ENSMUSG00000106875 | 0 | 0.25 | 0.25 | 0 | 0 | 0 | 0 |
| ENSMUSG00000106876 | 0 | 0 | 0 | 0 | 0 | 0 | 0 |
| ENSMUSG00000106878 | 0.45815691 | 0.396240625 | 0.25 | 0.25 | 0.387761935 | 0 | 0 |
| ENSMUSG00000106879 | 0 | 0 | 0 | 0 | 0 | 0 | 0 |
| ENSMUSG00000106880 | 0 | 0 | 0 | 0 | 0 | 0 | 0 |
| ENSMUSG00000106881 | 0 | 0 | 0 | 0 | 0 | 0 | 0 |
| ENSMUSG00000106883 | 0 | 0 | 0 | 0.25 | 0.244650542 | 0 | 0.224243824 |
| ENSMUSG00000106884 | 0 | 0 | 0 | 0 | 0 | 0 | 0 |
| ENSMUSG00000106885 | 0.289064826 | 0 | 0.25 | 0.25 | 0.244650542 | 0.602059991 | 0.224243824 |
| ENSMUSG00000106887 | 0.289064826 | 0.396240625 | 0 | 0.5 | 0.387761935 | 0 | 0 |
| ENSMUSG00000106888 | 0 | 0.25 | 0 | 0 | 0.244650542 | 0.301029996 | 0 |
| ENSMUSG00000106889 | 0 | 0 | 0 | 0 | 0 | 0 | 0 |
| ENSMUSG00000106890 | 0 | 0 | 0 | 0 | 0 | 0 | 0 |
| ENSMUSG00000106891 | 0.289064826 | 0.646240625 | 0.79248125 | 0.79248125 | 0.632412477 | 0.698970004 | 0.775756176 |
| ENSMUSG00000106892 | 0.45815691 | 0.5 | 0.5 | 0.25 | 0 | 0 | 0.224243824 |
| ENSMUSG00000106893 | 0 | 0 | 0 | 0 | 0 | 0 | 0 |
| ENSMUSG00000106894 | 0 | 0.25 | 0 | 0.25 | 0 | 0 | 0 |
| ENSMUSG00000106895 | 0.45815691 | 0.25 | 0.396240625 | 0.5 | 0.244650542 | 0.301029996 | 0.520678036 |
| ENSMUSG00000106896 | 0 | 0 | 0 | 0 | 0 | 0 | 0.224243824 |
| ENSMUSG00000106897 | 0 | 0.396240625 | 0.25 | 0 | 0.387761935 | 0.301029996 | 0.355418052 |
| ENSMUSG00000106898 | 0 | 0 | 0 | 0 | 0 | 0 | 0 |

| | | | | | | | |
|---|---|---|---|---|---|---|---|
| ENSMUSG00000106899 | 0 | 0 | 0 | 0 | 0 | 0 | 0 |
| ENSMUSG00000106900 | 0 | 0 | 0 | 0 | 0 | 0 | 0 |
| ENSMUSG00000106902 | 0 | 0 | 0 | 0 | 0 | 0 | 0 |
| ENSMUSG00000106904 | 0.960252568 | 0.701838731 | 0.646240625 | 0.830482024 | 0.686820904 | 0.903089987 | 0.355418052 |
| ENSMUSG00000106905 | 0 | 0 | 0 | 0 | 0 | 0 | 0 |
| ENSMUSG00000106906 | 0.289064826 | 0 | 0 | 0 | 0 | 0 | 0 |
| ENSMUSG00000106909 | 0 | 0 | 0 | 0 | 0 | 0 | 0 |
| ENSMUSG00000106910 | 0 | 0 | 0.25 | 0 | 0 | 0 | 0 |
| ENSMUSG00000106912 | 0 | 0 | 0 | 0 | 0 | 0 | 0 |
| ENSMUSG00000106913 | 0 | 0 | 0 | 0 | 0 | 0 | 0 |
| ENSMUSG00000106914 | 0 | 0 | 0 | 0 | 0 | 0 | 0 |
| ENSMUSG00000106915 | 0 | 0 | 0 | 0 | 0 | 0 | 0 |
| ENSMUSG00000106916 | 0 | 0 | 0 | 0 | 0 | 0 | 0 |
| ENSMUSG00000106917 | 0 | 0.25 | 0 | 0 | 0 | 0 | 0 |
| ENSMUSG00000106918 | 2.401648579 | 2.196158711 | 2.220660762 | 2.209012589 | 2.136951444 | 2.498310554 | 2.005303651 |
| ENSMUSG00000106920 | 0 | 0.25 | 0.25 | 0 | 0 | 0 | 0.224243824 |
| ENSMUSG00000106924 | 0 | 0.25 | 0 | 0 | 0 | 0 | 0 |
| ENSMUSG00000106925 | 0 | 0 | 0.25 | 0 | 0 | 0 | 0 |
| ENSMUSG00000106926 | 0.578129653 | 0.701838731 | 0.701838731 | 0.701838731 | 0.733951626 | 0.77815125 | 0.629532004 |
| ENSMUSG00000106927 | 0.45815691 | 0.25 | 0.5 | 0 | 0 | 0.698970004 | 0.355418052 |
| ENSMUSG00000106928 | 1.036286563 | 1.160964047 | 1.04248125 | 1.160964047 | 1.200473444 | 1.041392685 | 1.140832123 |
| ENSMUSG00000106930 | 0 | 0.25 | 0.25 | 0 | 0 | 0 | 0.224243824 |
| ENSMUSG00000106931 | 0 | 0 | 0 | 0 | 0 | 0 | 0 |
| ENSMUSG00000106932 | 0 | 0 | 0 | 0 | 0 | 0 | 0 |
| ENSMUSG00000106933 | 0.747221736 | 0.396240625 | 0.580482024 | 0.25 | 0 | 0 | 0.355418052 |
| ENSMUSG00000106934 | 0 | 0 | 0 | 0 | 0 | 0 | 0 |
| ENSMUSG00000106935 | 0 | 0 | 0 | 0 | 0 | 0 | 0 |
| ENSMUSG00000106936 | 0 | 0 | 0 | 0 | 0 | 0 | 0 |
| ENSMUSG00000106937 | 0 | 0 | 0 | 0 | 0 | 0 | 0 |
| ENSMUSG00000106938 | 0 | 0 | 0 | 0 | 0 | 0 | 0 |
| ENSMUSG00000106939 | 0 | 0 | 0 | 0 | 0 | 0 | 0 |
| ENSMUSG00000106940 | 0.289064826 | 0.25 | 0.396240625 | 0.25 | 0.568060967 | 0.477121255 | 0.448487648 |
| ENSMUSG00000106941 | 0 | 0 | 0 | 0 | 0 | 0 | 0 |
| ENSMUSG00000106943 | 0 | 0 | 0 | 0.25 | 0 | 0.301029996 | 0.224243824 |

| | | | | | | | |
|---|---|---|---|---|---|---|---|
| ENSMUSG00000106944 | 0 | 0.25 | 0 | 0 | 0 | 0.477121255 | 0.224243824 |
| ENSMUSG00000106945 | 0 | 0 | 0 | 0 | 0 | 0 | 0 |
| ENSMUSG00000106946 | 0.45815691 | 0.580482024 | 0.25 | 0.25 | 0.387761935 | 0 | 0.224243824 |
| ENSMUSG00000106947 | 0.289064826 | 0 | 0 | 0 | 0 | 0 | 0 |
| ENSMUSG00000106948 | 1.639698646 | 1.557204673 | 1.482684334 | 1.516522298 | 1.484072009 | 1.556302501 | 1.284450788 |
| ENSMUSG00000106950 | 0 | 0 | 0 | 0 | 0 | 0 | 0 |
| ENSMUSG00000106951 | 1.842799666 | 1.726722649 | 1.657339155 | 1.575945187 | 1.559656853 | 1.681241237 | 1.437266334 |
| ENSMUSG00000106952 | 0 | 0 | 0 | 0 | 0 | 0 | 0 |
| ENSMUSG00000106954 | 0 | 0 | 0 | 0 | 0 | 0 | 0 |
| ENSMUSG00000106955 | 0 | 0 | 0 | 0 | 0 | 0 | 0 |
| ENSMUSG00000106956 | 0 | 0 | 0 | 0 | 0 | 0 | 0 |
| ENSMUSG00000106957 | 0.289064826 | 0.396240625 | 0.25 | 0 | 0 | 0 | 0 |
| ENSMUSG00000106958 | 0 | 0 | 0 | 0 | 0 | 0 | 0 |
| ENSMUSG00000106959 | 0.289064826 | 0.5 | 0.580482024 | 0.646240625 | 0.632412477 | 0.698970004 | 0.629532004 |
| ENSMUSG00000106960 | 0 | 0 | 0 | 0 | 0 | 0 | 0 |
| ENSMUSG00000106961 | 1.639698646 | 1.445339928 | 1.505591953 | 1.130890489 | 1.200473444 | 1.301029996 | 1.140832123 |
| ENSMUSG00000106962 | 0.289064826 | 0.701838731 | 0 | 0.79248125 | 0.632412477 | 0.477121255 | 0.520678036 |
| ENSMUSG00000106963 | 0.960252568 | 0.646240625 | 0.701838731 | 0.75 | 0.632412477 | 0.477121255 | 0.710836105 |
| ENSMUSG00000106965 | 0 | 0 | 0.396240625 | 0.5 | 0.387761935 | 0 | 0.224243824 |
| ENSMUSG00000106967 | 0.289064826 | 0 | 0 | 0 | 0 | 0 | 0 |
| ENSMUSG00000106968 | 0.289064826 | 0.25 | 0.396240625 | 0 | 0 | 0 | 0 |
| ENSMUSG00000106969 | 0 | 0 | 0 | 0 | 0 | 0 | 0 |
| ENSMUSG00000106970 | 0 | 0 | 0 | 0 | 0 | 0 | 0 |
| ENSMUSG00000106971 | 0 | 0 | 0 | 0 | 0 | 0 | 0 |
| ENSMUSG00000106973 | 0 | 0 | 0 | 0 | 0 | 0 | 0 |
| ENSMUSG00000106974 | 0 | 0 | 0 | 0 | 0 | 0 | 0 |
| ENSMUSG00000106975 | 0 | 0 | 0 | 0 | 0 | 0 | 0 |
| ENSMUSG00000106976 | 0 | 0 | 0 | 0 | 0 | 0 | 0 |
| ENSMUSG00000106978 | 0.91631382 | 1.188721876 | 1.04248125 | 0.896240625 | 0.77552387 | 1.041392685 | 0.896975297 |
| ENSMUSG00000106979 | 0 | 0 | 0 | 0 | 0 | 0 | 0 |
| ENSMUSG00000106980 | 0 | 0 | 0.25 | 0.25 | 0.244650542 | 0 | 0 |
| ENSMUSG00000106981 | 1.129344651 | 1.238549078 | 1.04248125 | 1.146240625 | 1 | 1.146128036 | 1.131174228 |
| ENSMUSG00000106982 | 0 | 0 | 0 | 0 | 0 | 0 | 0 |
| ENSMUSG00000106985 | 0 | 0 | 0 | 0 | 0 | 0 | 0 |

| | | | | | | | |
|---|---|---|---|---|---|---|---|
| ENSMUSG00000106986 | 0 | 0 | 0 | 0 | 0 | 0 | 0 |
| ENSMUSG00000106987 | 0 | 0 | 0 | 0.25 | 0 | 0 | 0 |
| ENSMUSG00000106988 | 0 | 0.25 | 0 | 0.25 | 0 | 0 | 0.224243824 |
| ENSMUSG00000106989 | 0 | 0 | 0 | 0 | 0 | 0 | 0 |
| ENSMUSG00000106990 | 1.505869732 | 1.476722649 | 1.482684334 | 1.589388001 | 1.607318736 | 1.51851394 | 1.413583902 |
| ENSMUSG00000106991 | 0 | 0 | 0 | 0 | 0 | 0 | 0 |
| ENSMUSG00000106992 | 0 | 0 | 0.25 | 0 | 0 | 0.301029996 | 0.224243824 |
| ENSMUSG00000106993 | 0 | 0 | 0 | 0 | 0 | 0 | 0 |
| ENSMUSG00000106994 | 0 | 0 | 0.25 | 0 | 0 | 0 | 0 |
| ENSMUSG00000106995 | 0.578129653 | 0 | 0.5 | 0.79248125 | 0.489301084 | 0.77815125 | 0.672731473 |
| ENSMUSG00000106996 | 0 | 0 | 0 | 0 | 0 | 0 | 0 |
| ENSMUSG00000106997 | 1.100572389 | 0.976722649 | 1.098079356 | 0.951838731 | 0.955822902 | 0.84509804 | 0.952571511 |
| ENSMUSG00000106998 | 0 | 0 | 0 | 0 | 0 | 0 | 0 |
| ENSMUSG00000106999 | 0 | 0 | 0 | 0 | 0 | 0 | 0 |
| ENSMUSG00000107000 | 1.418409478 | 1.380890489 | 1.410964047 | 1.339388001 | 1.327538614 | 1.301029996 | 1.272006351 |
| ENSMUSG00000107002 | 1.686083339 | 1.660964047 | 1.71759118 | 1.671625132 | 1.607318736 | 1.770852012 | 1.579661877 |
| ENSMUSG00000107004 | 0 | 0 | 0 | 0 | 0 | 0 | 0 |
| ENSMUSG00000107005 | 0.45815691 | 0.5 | 0 | 0.5 | 0.489301084 | 0.301029996 | 0.355418052 |
| ENSMUSG00000107006 | 0 | 0 | 0 | 0 | 0 | 0 | 0.224243824 |
| ENSMUSG00000107008 | 0 | 0 | 0 | 0 | 0 | 0.301029996 | 0.224243824 |
| ENSMUSG00000107009 | 0 | 0.25 | 0 | 0 | 0 | 0 | 0 |
| ENSMUSG00000107010 | 0 | 0 | 0 | 0 | 0 | 0 | 0 |
| ENSMUSG00000107011 | 0 | 0 | 0 | 0 | 0 | 0 | 0 |
| ENSMUSG00000107012 | 0 | 0 | 0.25 | 0 | 0 | 0 | 0 |
| ENSMUSG00000107013 | 0 | 0 | 0 | 0 | 0 | 0 | 0 |
| ENSMUSG00000107014 | 0 | 0 | 0 | 0 | 0 | 0 | 0 |
| ENSMUSG00000107015 | 0 | 0 | 0 | 0 | 0.244650542 | 0 | 0 |
| ENSMUSG00000107016 | 0 | 0 | 0 | 0 | 0 | 0 | 0 |
| ENSMUSG00000107017 | 0 | 0 | 0.396240625 | 0.25 | 0.489301084 | 0 | 0 |
| ENSMUSG00000107018 | 0 | 0 | 0 | 0 | 0 | 0.301029996 | 0 |
| ENSMUSG00000107019 | 1.494443473 | 1.42510993 | 1.330482024 | 1.552363341 | 1.366364104 | 1.544068044 | 1.379042812 |
| ENSMUSG00000107020 | 0 | 0 | 0 | 0 | 0.244650542 | 0 | 0 |
| ENSMUSG00000107021 | 0 | 0 | 0 | 0 | 0.244650542 | 0 | 0 |
| ENSMUSG00000107022 | 0 | 0.25 | 0 | 0 | 0 | 0 | 0 |

| | | | | | | | |
|---|---|---|---|---|---|---|---|
| ENSMUSG00000107023 | 0.867194479 | 0.864857905 | 0.896240625 | 1.02186571 | 1.091002363 | 1 | 0.952571511 |
| ENSMUSG00000107024 | 0 | 0 | 0 | 0 | 0 | 0 | 0 |
| ENSMUSG00000107026 | 0.578129653 | 0.396240625 | 0.25 | 0.5 | 0.387761935 | 0 | 0.448487648 |
| ENSMUSG00000107027 | 0 | 0 | 0 | 0 | 0 | 0 | 0 |
| ENSMUSG00000107028 | 0 | 0 | 0 | 0 | 0 | 0 | 0 |
| ENSMUSG00000107029 | 0.289064826 | 0 | 0.25 | 0.25 | 0 | 0 | 0.224243824 |
| ENSMUSG00000107030 | 0 | 0.25 | 0 | 0 | 0 | 0 | 0 |
| ENSMUSG00000107031 | 0 | 0 | 0 | 0 | 0 | 0 | 0 |
| ENSMUSG00000107032 | 0 | 0 | 0 | 0 | 0 | 0 | 0 |
| ENSMUSG00000107033 | 0 | 0 | 0 | 0 | 0 | 0.301029996 | 0 |
| ENSMUSG00000107034 | 0 | 0 | 0 | 0 | 0 | 0 | 0 |
| ENSMUSG00000107035 | 0 | 0.25 | 0.25 | 0 | 0 | 0 | 0.224243824 |
| ENSMUSG00000107036 | 0 | 0.25 | 0 | 0 | 0 | 0 | 0 |
| ENSMUSG00000107037 | 0 | 0 | 0 | 0 | 0 | 0 | 0 |
| ENSMUSG00000107040 | 0 | 0 | 0 | 0 | 0 | 0 | 0 |
| ENSMUSG00000107041 | 0 | 0 | 0.25 | 0 | 0.387761935 | 0.602059991 | 0 |
| ENSMUSG00000107042 | 0 | 0 | 0 | 0 | 0 | 0 | 0 |
| ENSMUSG00000107043 | 0.289064826 | 0.396240625 | 0 | 0 | 0 | 0 | 0 |
| ENSMUSG00000107044 | 0 | 0 | 0 | 0 | 0 | 0 | 0 |
| ENSMUSG00000107045 | 0 | 0 | 0 | 0.396240625 | 0 | 0.301029996 | 0 |
| ENSMUSG00000107046 | 0 | 0 | 0 | 0 | 0 | 0 | 0 |
| ENSMUSG00000107047 | 0 | 0 | 0 | 0 | 0 | 0 | 0 |
| ENSMUSG00000107048 | 0 | 0 | 0.25 | 0 | 0 | 0.477121255 | 0.224243824 |
| ENSMUSG00000107049 | 0 | 0 | 0.25 | 0 | 0.244650542 | 0 | 0 |
| ENSMUSG00000107050 | 0.289064826 | 0 | 0 | 0.396240625 | 0.244650542 | 0.301029996 | 0.224243824 |
| ENSMUSG00000107053 | 0 | 0 | 0 | 0 | 0 | 0 | 0 |
| ENSMUSG00000107054 | 0 | 0.25 | 0 | 0.25 | 0.387761935 | 0 | 0 |
| ENSMUSG00000107056 | 0 | 0 | 0 | 0 | 0 | 0 | 0 |
| ENSMUSG00000107058 | 0 | 0 | 0 | 0 | 0 | 0 | 0 |
| ENSMUSG00000107059 | 0 | 0 | 0 | 0 | 0 | 0 | 0 |
| ENSMUSG00000107060 | 0 | 0 | 0 | 0 | 0 | 0 | 0 |
| ENSMUSG00000107061 | 0 | 0 | 0 | 0 | 0 | 0 | 0 |
| ENSMUSG00000107062 | 0 | 0 | 0 | 0 | 0 | 0 | 0 |
| ENSMUSG00000107063 | 0 | 0 | 0 | 0 | 0.244650542 | 0 | 0 |

| | | | | | | | |
|---|---|---|---|---|---|---|---|
| ENSMUSG00000107064 | 0 | 0 | 0 | 0 | 0 | 0 | 0 |
| ENSMUSG00000107065 | 0.289064826 | 0 | 0 | 0 | 0 | 0 | 0.224243824 |
| ENSMUSG00000107066 | 0 | 0 | 0 | 0 | 0 | 0 | 0 |
| ENSMUSG00000107067 | 0 | 0 | 0 | 0 | 0 | 0 | 0 |
| ENSMUSG00000107068 | 0 | 0 | 0 | 0 | 0 | 0 | 0 |
| ENSMUSG00000107069 | 0 | 0 | 0 | 0 | 0 | 0 | 0 |
| ENSMUSG00000107070 | 0 | 0.25 | 0 | 0.25 | 0 | 0 | 0.355418052 |
| ENSMUSG00000107071 | 0 | 0 | 0 | 0 | 0 | 0 | 0 |
| ENSMUSG00000107072 | 0 | 0 | 0 | 0 | 0 | 0 | 0 |
| ENSMUSG00000107073 | 0 | 0 | 0 | 0 | 0 | 0 | 0 |
| ENSMUSG00000107074 | 0 | 0 | 0 | 0 | 0 | 0 | 0 |
| ENSMUSG00000107075 | 0 | 0 | 0 | 0 | 0 | 0 | 0 |
| ENSMUSG00000107076 | 0 | 0 | 0 | 0 | 0.244650542 | 0.477121255 | 0.224243824 |
| ENSMUSG00000107078 | 0 | 0 | 0 | 0 | 0 | 0 | 0 |
| ENSMUSG00000107079 | 0 | 0.25 | 0 | 0.396240625 | 0.387761935 | 0.301029996 | 0.224243824 |
| ENSMUSG00000107081 | 0 | 0 | 0 | 0 | 0 | 0 | 0 |
| ENSMUSG00000107082 | 0 | 0 | 0 | 0 | 0 | 0 | 0 |
| ENSMUSG00000107083 | 0.289064826 | 0.5 | 0.396240625 | 0.396240625 | 0.387761935 | 0 | 0.224243824 |
| ENSMUSG00000107084 | 0 | 0 | 0 | 0 | 0 | 0 | 0 |
| ENSMUSG00000107085 | 0 | 0 | 0 | 0 | 0 | 0 | 0 |
| ENSMUSG00000107086 | 0.45815691 | 0.646240625 | 0.396240625 | 0.701838731 | 0.686820904 | 0.477121255 | 0.710836105 |
| ENSMUSG00000107087 | 0 | 0 | 0 | 0 | 0 | 0 | 0 |
| ENSMUSG00000107089 | 0 | 0 | 0 | 0 | 0 | 0 | 0 |
| ENSMUSG00000107090 | 0 | 0 | 0 | 0 | 0 | 0 | 0 |
| ENSMUSG00000107091 | 0 | 0.580482024 | 0 | 0.25 | 0.244650542 | 0 | 0 |
| ENSMUSG00000107092 | 0.578129653 | 0.25 | 0.25 | 0.25 | 0.244650542 | 0.301029996 | 0.355418052 |
| ENSMUSG00000107093 | 0 | 0 | 0 | 0 | 0 | 0 | 0 |
| ENSMUSG00000107094 | 0.91631382 | 0.75 | 0.5 | 0.580482024 | 0.387761935 | 0.602059991 | 0.629532004 |
| ENSMUSG00000107096 | 1.505869732 | 1.42510993 | 1.470660762 | 1.356566189 | 1.327538614 | 1.397940009 | 1.290497981 |
| ENSMUSG00000107097 | 0 | 0 | 0 | 0 | 0 | 0 | 0 |
| ENSMUSG00000107098 | 0 | 0 | 0 | 0 | 0 | 0 | 0 |
| ENSMUSG00000107100 | 0 | 0 | 0 | 0 | 0 | 0 | 0 |
| ENSMUSG00000107102 | 0.747221736 | 0.5 | 0.580482024 | 0.646240625 | 0.77552387 | 0.477121255 | 0.672731473 |
| ENSMUSG00000107103 | 0 | 0.25 | 0.25 | 0 | 0 | 0 | 0 |

| | | | | | | | |
|---|---|---|---|---|---|---|---|
| ENSMUSG00000107105 | 0 | 0 | 0 | 0 | 0 | 0 | 0 |
| ENSMUSG00000107106 | 0 | 0 | 0.25 | 0 | 0 | 0 | 0 |
| ENSMUSG00000107107 | 0.289064826 | 0 | 0 | 0 | 0 | 0 | 0.224243824 |
| ENSMUSG00000107108 | 0 | 0.25 | 0 | 0.396240625 | 0.244650542 | 0 | 0 |
| ENSMUSG00000107109 | 0 | 0 | 0 | 0 | 0 | 0 | 0 |
| ENSMUSG00000107110 | 0 | 0 | 0 | 0 | 0 | 0 | 0 |
| ENSMUSG00000107111 | 0 | 0 | 0 | 0 | 0 | 0 | 0 |
| ENSMUSG00000107112 | 0 | 0 | 0 | 0 | 0 | 0 | 0 |
| ENSMUSG00000107113 | 0 | 0 | 0 | 0 | 0 | 0 | 0 |
| ENSMUSG00000107114 | 0 | 0 | 0 | 0 | 0 | 0 | 0 |
| ENSMUSG00000107115 | 0 | 0.25 | 0 | 0 | 0 | 0 | 0 |
| ENSMUSG00000107116 | 0.578129653 | 0.25 | 0.580482024 | 0.580482024 | 0.489301084 | 0 | 0.448487648 |
| ENSMUSG00000107117 | 0.45815691 | 0.396240625 | 0.25 | 0.646240625 | 0.244650542 | 0.477121255 | 0.520678036 |
| ENSMUSG00000107118 | 0.578129653 | 0.646240625 | 0.75 | 0.396240625 | 0.387761935 | 0.477121255 | 0.355418052 |
| ENSMUSG00000107120 | 1.100572389 | 0.896240625 | 1 | 0.951838731 | 0.931471446 | 0.903089987 | 0.775756176 |
| ENSMUSG00000107121 | 0.91631382 | 0.646240625 | 1.02186571 | 0.25 | 0.568060967 | 0.477121255 | 0.803905701 |
| ENSMUSG00000107122 | 0 | 0 | 0 | 0 | 0 | 0 | 0 |
| ENSMUSG00000107123 | 0 | 0 | 0 | 0 | 0 | 0 | 0 |
| ENSMUSG00000107125 | 0 | 0 | 0 | 0 | 0 | 0 | 0 |
| ENSMUSG00000107126 | 0 | 0 | 0 | 0.25 | 0 | 0 | 0 |
| ENSMUSG00000107127 | 0 | 0 | 0 | 0 | 0 | 0 | 0 |
| ENSMUSG00000107128 | 0 | 0 | 0 | 0 | 0 | 0 | 0 |
| ENSMUSG00000107129 | 0 | 0 | 0 | 0 | 0 | 0 | 0 |
| ENSMUSG00000107130 | 0 | 0 | 0 | 0 | 0 | 0 | 0 |
| ENSMUSG00000107131 | 0 | 0 | 0 | 0 | 0 | 0 | 0 |
| ENSMUSG00000107132 | 0 | 0 | 0 | 0 | 0 | 0 | 0 |
| ENSMUSG00000107133 | 0 | 0.396240625 | 0 | 0 | 0 | 0 | 0 |
| ENSMUSG00000107134 | 0 | 0 | 0 | 0 | 0 | 0 | 0 |
| ENSMUSG00000107135 | 0 | 0 | 0 | 0 | 0 | 0 | 0 |
| ENSMUSG00000107136 | 0.289064826 | 0 | 0 | 0 | 0.244650542 | 0 | 0 |
| ENSMUSG00000107139 | 0 | 0 | 0 | 0 | 0 | 0 | 0 |
| ENSMUSG00000107140 | 0 | 0 | 0 | 0 | 0 | 0 | 0 |
| ENSMUSG00000107141 | 0 | 0 | 0 | 0 | 0 | 0 | 0 |
| ENSMUSG00000107142 | 0 | 0 | 0 | 0 | 0 | 0 | 0 |

| | | | | | | | |
|---|---|---|---|---|---|---|---|
| ENSMUSG00000107143 | 0.578129653 | 0.896240625 | 0.75 | 0.896240625 | 0.812711509 | 0.77815125 | 0.74492186 |
| ENSMUSG00000107144 | 0 | 0 | 0 | 0 | 0 | 0 | 0 |
| ENSMUSG00000107145 | 0 | 0.25 | 0 | 0 | 0 | 0 | 0 |
| ENSMUSG00000107146 | 0 | 0 | 0 | 0 | 0 | 0 | 0 |
| ENSMUSG00000107148 | 0 | 0 | 0 | 0.25 | 0 | 0 | 0 |
| ENSMUSG00000107149 | 0 | 0 | 0.25 | 0.396240625 | 0.387761935 | 0 | 0 |
| ENSMUSG00000107151 | 0 | 0.25 | 0.25 | 0.25 | 0 | 0 | 0 |
| ENSMUSG00000107152 | 0 | 0 | 0 | 0.396240625 | 0.244650542 | 0 | 0 |
| ENSMUSG00000107153 | 0 | 0 | 0 | 0 | 0 | 0 | 0 |
| ENSMUSG00000107154 | 0 | 0 | 0.25 | 0 | 0 | 0 | 0 |
| ENSMUSG00000107155 | 0.289064826 | 0.396240625 | 0.396240625 | 0 | 0.244650542 | 0 | 0 |
| ENSMUSG00000107157 | 0 | 0 | 0 | 0 | 0 | 0 | 0 |
| ENSMUSG00000107158 | 0 | 0 | 0 | 0 | 0 | 0 | 0 |
| ENSMUSG00000107159 | 0 | 0 | 0 | 0 | 0 | 0 | 0 |
| ENSMUSG00000107160 | 0.747221736 | 0.5 | 0.5 | 0.580482024 | 0.686820904 | 0 | 0.355418052 |
| ENSMUSG00000107161 | 0 | 0.25 | 0.396240625 | 0.646240625 | 0.568060967 | 0 | 0 |
| ENSMUSG00000107162 | 0.289064826 | 0 | 0 | 0 | 0 | 0 | 0 |
| ENSMUSG00000107163 | 0 | 0.396240625 | 0 | 0 | 0.244650542 | 0.301029996 | 0 |
| ENSMUSG00000107164 | 0.671187741 | 0.92510993 | 0.646240625 | 0.25 | 0.387761935 | 0.477121255 | 0.629532004 |
| ENSMUSG00000107165 | 0.811507563 | 1.080482024 | 0.79248125 | 0.896240625 | 0.632412477 | 0.477121255 | 0.520678036 |
| ENSMUSG00000107167 | 0 | 0 | 0 | 0 | 0 | 0 | 0 |
| ENSMUSG00000107168 | 0.289064826 | 0.5 | 0.396240625 | 0.5 | 0.387761935 | 0 | 0 |
| ENSMUSG00000107169 | 0 | 0 | 0 | 0 | 0 | 0 | 0 |
| ENSMUSG00000107171 | 0.91631382 | 0.79248125 | 0.79248125 | 0.79248125 | 0.568060967 | 0.77815125 | 0.629532004 |
| ENSMUSG00000107172 | 0 | 0 | 0 | 0 | 0 | 0 | 0 |
| ENSMUSG00000107173 | 0.289064826 | 0.25 | 0.25 | 0.396240625 | 0.489301084 | 0.602059991 | 0.448487648 |
| ENSMUSG00000107175 | 0 | 0 | 0 | 0 | 0 | 0 | 0 |
| ENSMUSG00000107176 | 0.289064826 | 0.25 | 0.25 | 0 | 0 | 0.301029996 | 0.355418052 |
| ENSMUSG00000107177 | 0 | 0 | 0 | 0 | 0 | 0 | 0 |
| ENSMUSG00000107178 | 0.289064826 | 0.580482024 | 0.396240625 | 0.5 | 0.387761935 | 0 | 0.355418052 |
| ENSMUSG00000107179 | 0 | 0 | 0 | 0 | 0 | 0 | 0 |
| ENSMUSG00000107180 | 0 | 0 | 0 | 0 | 0 | 0 | 0 |
| ENSMUSG00000107181 | 0 | 0 | 0 | 0 | 0 | 0 | 0 |
| ENSMUSG00000107182 | 1.432083896 | 1.396240625 | 1.494319981 | 1.653677461 | 1.625423019 | 1.662757832 | 1.444790289 |

| | | | | | | | |
|---|---|---|---|---|---|---|---|
| ENSMUSG00000107183 | 0 | 0 | 0 | 0 | 0 | 0 | 0 |
| ENSMUSG00000107184 | 0.289064826 | 0 | 0 | 0 | 0.387761935 | 0 | 0 |
| ENSMUSG00000107185 | 0 | 0 | 0 | 0 | 0 | 0 | 0 |
| ENSMUSG00000107187 | 0 | 0 | 0 | 0 | 0 | 0 | 0 |
| ENSMUSG00000107189 | 0 | 0 | 0 | 0 | 0 | 0 | 0 |
| ENSMUSG00000107190 | 0 | 0 | 0 | 0 | 0 | 0 | 0 |
| ENSMUSG00000107191 | 0 | 0 | 0 | 0 | 0 | 0 | 0 |
| ENSMUSG00000107192 | 0 | 0 | 0 | 0 | 0 | 0 | 0 |
| ENSMUSG00000107193 | 0 | 0 | 0 | 0 | 0 | 0 | 0 |
| ENSMUSG00000107194 | 0 | 0 | 0 | 0 | 0 | 0 | 0.224243824 |
| ENSMUSG00000107195 | 0.289064826 | 0.25 | 0 | 0.396240625 | 0.489301084 | 0.477121255 | 0.448487648 |
| ENSMUSG00000107197 | 0 | 0 | 0 | 0 | 0.244650542 | 0 | 0 |
| ENSMUSG00000107198 | 0 | 0.25 | 0.25 | 0.25 | 0.244650542 | 0 | 0 |
| ENSMUSG00000107199 | 0 | 0 | 0 | 0 | 0 | 0 | 0 |
| ENSMUSG00000107200 | 1.036286563 | 1.061981878 | 0.951838731 | 0.976722649 | 0.978602168 | 0.954242509 | 0.803905701 |
| ENSMUSG00000107201 | 1.227926429 | 1.188721876 | 1.130890489 | 1.146240625 | 1.121713561 | 1.204119983 | 1.089372236 |
| ENSMUSG00000107202 | 0 | 0 | 0 | 0 | 0 | 0 | 0 |
| ENSMUSG00000107203 | 0 | 0 | 0 | 0.5 | 0 | 0 | 0 |
| ENSMUSG00000107204 | 0 | 0 | 0 | 0 | 0 | 0.301029996 | 0 |
| ENSMUSG00000107205 | 0.578129653 | 0.5 | 0.396240625 | 0.5 | 0.244650542 | 0.477121255 | 0.355418052 |
| ENSMUSG00000107206 | 0 | 0.25 | 0.25 | 0 | 0.244650542 | 0 | 0 |
| ENSMUSG00000107207 | 0 | 0 | 0 | 0 | 0.244650542 | 0 | 0.224243824 |
| ENSMUSG00000107208 | 0 | 0 | 0 | 0 | 0 | 0 | 0 |
| ENSMUSG00000107209 | 0 | 0 | 0 | 0 | 0 | 0 | 0 |
| ENSMUSG00000107211 | 0 | 0.25 | 0 | 0 | 0 | 0.301029996 | 0 |
| ENSMUSG00000107212 | 0 | 0.396240625 | 0 | 0.25 | 0 | 0.301029996 | 0.224243824 |
| ENSMUSG00000107213 | 0 | 0 | 0 | 0 | 0 | 0 | 0 |
| ENSMUSG00000107214 | 0 | 0 | 0 | 0 | 0 | 0 | 0 |
| ENSMUSG00000107215 | 1.985980784 | 1.763820609 | 1.701838731 | 1.593759858 | 1.546663136 | 1.707570176 | 1.42167221 |
| ENSMUSG00000107216 | 0.289064826 | 0.396240625 | 0.580482024 | 0.25 | 0 | 0.602059991 | 0.224243824 |
| ENSMUSG00000107217 | 0 | 0 | 0 | 0 | 0 | 0 | 0 |
| ENSMUSG00000107218 | 0 | 0.25 | 0 | 0 | 0 | 0 | 0 |
| ENSMUSG00000107219 | 0 | 0 | 0 | 0 | 0 | 0 | 0 |
| ENSMUSG00000107220 | 0 | 0 | 0 | 0 | 0 | 0 | 0 |

| | | | | | | | |
|---|---|---|---|---|---|---|---|
| ENSMUSG00000107222 | 0 | 0 | 0 | 0 | 0 | 0 | 0 |
| ENSMUSG00000107223 | 0 | 0 | 0 | 0 | 0 | 0 | 0 |
| ENSMUSG00000107224 | 0 | 0 | 0 | 0 | 0 | 0 | 0 |
| ENSMUSG00000107225 | 0.289064826 | 0 | 0.5 | 0.25 | 0.244650542 | 0.477121255 | 0.355418052 |
| ENSMUSG00000107226 | 0 | 0 | 0 | 0 | 0 | 0 | 0 |
| ENSMUSG00000107227 | 1.036286563 | 0.976722649 | 0.864857905 | 0.896240625 | 0.955822902 | 0.903089987 | 0.710836105 |
| ENSMUSG00000107228 | 0 | 0 | 0 | 0 | 0 | 0 | 0 |
| ENSMUSG00000107229 | 0 | 0 | 0 | 0 | 0 | 0 | 0 |
| ENSMUSG00000107230 | 0.578129653 | 0.701838731 | 0.580482024 | 0.580482024 | 0.686820904 | 0.602059991 | 0.448487648 |
| ENSMUSG00000107231 | 0 | 0 | 0 | 0 | 0 | 0 | 0 |
| ENSMUSG00000107232 | 0 | 0 | 0 | 0 | 0 | 0 | 0 |
| ENSMUSG00000107233 | 1.205378646 | 1.26109853 | 1.130890489 | 1.04248125 | 0.955822902 | 0.77815125 | 0.952571511 |
| ENSMUSG00000107234 | 0 | 0.25 | 0 | 0 | 0 | 0 | 0 |
| ENSMUSG00000107237 | 0 | 0 | 0 | 0 | 0 | 0 | 0 |
| ENSMUSG00000107238 | 0 | 0 | 0 | 0 | 0 | 0 | 0 |
| ENSMUSG00000107240 | 0 | 0 | 0 | 0.25 | 0 | 0 | 0.224243824 |
| ENSMUSG00000107241 | 0 | 0 | 0 | 0 | 0 | 0 | 0 |
| ENSMUSG00000107242 | 0 | 0 | 0 | 0 | 0 | 0 | 0 |
| ENSMUSG00000107243 | 0 | 0 | 0 | 0 | 0 | 0 | 0 |
| ENSMUSG00000107244 | 0.671187741 | 0.396240625 | 0.75 | 0.5 | 0.568060967 | 0.477121255 | 0.579661877 |
| ENSMUSG00000107245 | 0 | 0 | 0 | 0 | 0 | 0 | 0 |
| ENSMUSG00000107246 | 0.289064826 | 0 | 0.396240625 | 0 | 0.244650542 | 0.301029996 | 0.448487648 |
| ENSMUSG00000107247 | 0 | 0 | 0 | 0 | 0 | 0 | 0 |
| ENSMUSG00000107248 | 0 | 0 | 0 | 0 | 0 | 0 | 0 |
| ENSMUSG00000107251 | 0 | 0 | 0 | 0 | 0 | 0 | 0 |
| ENSMUSG00000107252 | 0 | 0.25 | 0 | 0 | 0 | 0 | 0 |
| ENSMUSG00000107253 | 0 | 0 | 0.396240625 | 0 | 0 | 0.301029996 | 0.224243824 |
| ENSMUSG00000107254 | 0.45815691 | 0.25 | 0.25 | 0.5 | 0.568060967 | 0.477121255 | 0.520678036 |
| ENSMUSG00000107255 | 0 | 0 | 0 | 0 | 0 | 0 | 0 |
| ENSMUSG00000107256 | 0.578129653 | 0 | 0.5 | 0.580482024 | 0.489301084 | 0 | 0.448487648 |
| ENSMUSG00000107257 | 0 | 0 | 0 | 0 | 0 | 0 | 0 |
| ENSMUSG00000107258 | 0 | 0 | 0 | 0 | 0 | 0 | 0 |
| ENSMUSG00000107259 | 0 | 0 | 0 | 0.25 | 0 | 0 | 0.224243824 |
| ENSMUSG00000107260 | 0 | 0 | 0 | 0 | 0 | 0 | 0 |

| | | | | | | | |
|---|---|---|---|---|---|---|---|
| ENSMUSG00000107261 | 0 | 0 | 0 | 0 | 0 | 0 | 0 |
| ENSMUSG00000107262 | 0 | 0 | 0 | 0 | 0 | 0 | 0 |
| ENSMUSG00000107263 | 0.45815691 | 0.396240625 | 0.5 | 0.25 | 0 | 0.301029996 | 0.224243824 |
| ENSMUSG00000107264 | 0 | 0 | 0 | 0 | 0 | 0 | 0 |
| ENSMUSG00000107265 | 0 | 0 | 0 | 0 | 0 | 0 | 0 |
| ENSMUSG00000107266 | 0 | 0 | 0 | 0.25 | 0 | 0 | 0.224243824 |
| ENSMUSG00000107270 | 0 | 0 | 0 | 0 | 0 | 0 | 0 |
| ENSMUSG00000107271 | 0.289064826 | 0.396240625 | 0.25 | 0.75 | 0.387761935 | 0.602059991 | 0.448487648 |
| ENSMUSG00000107272 | 0 | 0 | 0 | 0 | 0 | 0 | 0.224243824 |
| ENSMUSG00000107273 | 0 | 0 | 0 | 0 | 0 | 0 | 0 |
| ENSMUSG00000107276 | 0.45815691 | 0.396240625 | 0.580482024 | 0.864857905 | 0.387761935 | 0.477121255 | 0.448487648 |
| ENSMUSG00000107277 | 0.867194479 | 0.75 | 0.75 | 0.580482024 | 0.812711509 | 0.602059991 | 0.629532004 |
| ENSMUSG00000107278 | 1.181541736 | 1.04248125 | 1.114857905 | 1 | 1.149965125 | 1.079181246 | 0.829800754 |
| ENSMUSG00000107279 | 0 | 0 | 0 | 0 | 0 | 0 | 0 |
| ENSMUSG00000107280 | 0 | 0 | 0 | 0 | 0 | 0 | 0 |
| ENSMUSG00000107281 | 0 | 0 | 0 | 0 | 0 | 0 | 0 |
| ENSMUSG00000107282 | 0 | 0 | 0 | 0 | 0 | 0 | 0 |
| ENSMUSG00000107283 | 2.17479713 | 2.016522298 | 2.048689214 | 2.173371739 | 2.106691885 | 2.491361694 | 1.981429723 |
| ENSMUSG00000107284 | 0 | 0 | 0 | 0 | 0 | 0 | 0 |
| ENSMUSG00000107285 | 0 | 0 | 0 | 0 | 0 | 0 | 0 |
| ENSMUSG00000107286 | 0 | 0 | 0 | 0 | 0 | 0 | 0 |
| ENSMUSG00000107287 | 0 | 0 | 0 | 0 | 0 | 0 | 0 |
| ENSMUSG00000107288 | 0.289064826 | 0.25 | 0.25 | 0 | 0 | 0 | 0 |
| ENSMUSG00000107289 | 0 | 0 | 0 | 0 | 0 | 0 | 0 |
| ENSMUSG00000107290 | 0 | 0 | 0 | 0 | 0 | 0 | 0 |
| ENSMUSG00000107291 | 0 | 0 | 0 | 0 | 0 | 0 | 0 |
| ENSMUSG00000107292 | 0 | 0 | 0 | 0 | 0 | 0 | 0 |
| ENSMUSG00000107294 | 0 | 0 | 0 | 0 | 0 | 0 | 0 |
| ENSMUSG00000107295 | 0 | 0 | 0 | 0.25 | 0 | 0 | 0 |
| ENSMUSG00000107296 | 0 | 0 | 0 | 0 | 0 | 0 | 0 |
| ENSMUSG00000107298 | 0 | 0 | 0 | 0 | 0 | 0 | 0 |
| ENSMUSG00000107299 | 0 | 0 | 0 | 0 | 0 | 0 | 0 |
| ENSMUSG00000107300 | 0.289064826 | 0 | 0.396240625 | 0.580482024 | 0.632412477 | 0.301029996 | 0.224243824 |
| ENSMUSG00000107301 | 0 | 0 | 0 | 0 | 0 | 0 | 0 |

| | | | | | | | |
|---|---|---|---|---|---|---|---|
| ENSMUSG00000107303 | 0 | 0 | 0 | 0 | 0 | 0 | 0 |
| ENSMUSG00000107304 | 0 | 0.25 | 0 | 0.396240625 | 0 | 0.301029996 | 0.224243824 |
| ENSMUSG00000107305 | 0 | 0 | 0 | 0 | 0 | 0 | 0 |
| ENSMUSG00000107306 | 1.527823874 | 1.27186571 | 1.388647213 | 1.339388001 | 1.244650542 | 1.301029996 | 1.23151414 |
| ENSMUSG00000107307 | 0 | 0.25 | 0 | 0 | 0 | 0 | 0 |
| ENSMUSG00000107308 | 0 | 0 | 0 | 0 | 0 | 0 | 0 |
| ENSMUSG00000107309 | 0 | 0 | 0 | 0 | 0 | 0 | 0 |
| ENSMUSG00000107310 | 1.289064826 | 1.02186571 | 1.02186571 | 1.061981878 | 1.057362051 | 1.079181246 | 1.078019652 |
| ENSMUSG00000107312 | 0 | 0 | 0.25 | 0 | 0 | 0 | 0 |
| ENSMUSG00000107313 | 0.747221736 | 0.580482024 | 0 | 0.396240625 | 0.387761935 | 0 | 0.448487648 |
| ENSMUSG00000107314 | 0 | 0 | 0 | 0 | 0.244650542 | 0 | 0 |
| ENSMUSG00000107315 | 0 | 0 | 0 | 0 | 0 | 0 | 0 |
| ENSMUSG00000107316 | 0 | 0 | 0 | 0 | 0 | 0 | 0 |
| ENSMUSG00000107317 | 0 | 0 | 0 | 0 | 0 | 0 | 0 |
| ENSMUSG00000107318 | 0 | 0 | 0 | 0 | 0 | 0 | 0 |
| ENSMUSG00000107319 | 0.289064826 | 0.25 | 0.25 | 0.580482024 | 0.387761935 | 0.698970004 | 0.224243824 |
| ENSMUSG00000107320 | 1.100572389 | 1.188721876 | 0.976722649 | 1.214495249 | 1.149965125 | 1.041392685 | 1.028149525 |
| ENSMUSG00000107322 | 0 | 0 | 0 | 0 | 0 | 0 | 0 |
| ENSMUSG00000107323 | 0 | 0 | 0 | 0 | 0 | 0 | 0 |
| ENSMUSG00000107324 | 0 | 0 | 0 | 0 | 0 | 0 | 0 |
| ENSMUSG00000107325 | 0 | 0 | 0 | 0 | 0 | 0 | 0 |
| ENSMUSG00000107326 | 0 | 0 | 0 | 0 | 0 | 0 | 0 |
| ENSMUSG00000107327 | 0 | 0 | 0 | 0 | 0.387761935 | 0 | 0 |
| ENSMUSG00000107329 | 0 | 0 | 0 | 0 | 0 | 0 | 0 |
| ENSMUSG00000107331 | 1.307602651 | 1.339388001 | 1.356566189 | 1.201838731 | 1.121713561 | 1.230448921 | 1.066254157 |
| ENSMUSG00000107332 | 0 | 0 | 0 | 0 | 0 | 0 | 0 |
| ENSMUSG00000107333 | 0.45815691 | 0.25 | 0.396240625 | 0.25 | 0 | 0.301029996 | 0 |
| ENSMUSG00000107334 | 0 | 0 | 0 | 0 | 0 | 0 | 0 |
| ENSMUSG00000107335 | 0 | 0.25 | 0 | 0 | 0 | 0 | 0.520678036 |
| ENSMUSG00000107336 | 0.45815691 | 0.25 | 0.5 | 0.580482024 | 0.244650542 | 0.602059991 | 0.579661877 |
| ENSMUSG00000107337 | 0 | 0.25 | 0.25 | 0 | 0.244650542 | 0.301029996 | 0 |
| ENSMUSG00000107338 | 0.289064826 | 0.25 | 0.25 | 0.25 | 0.244650542 | 0.477121255 | 0.355418052 |
| ENSMUSG00000107340 | 0.289064826 | 0 | 0.25 | 0 | 0 | 0.301029996 | 0 |
| ENSMUSG00000107341 | 0 | 0 | 0 | 0 | 0 | 0 | 0 |

| | | | | | | | |
|---|---|---|---|---|---|---|---|
| ENSMUSG00000107344 | 0 | 0 | 0 | 0 | 0 | 0 | 0 |
| ENSMUSG00000107345 | 0 | 0 | 0 | 0 | 0 | 0 | 0 |
| ENSMUSG00000107346 | 0.289064826 | 0 | 0 | 0 | 0.244650542 | 0 | 0 |
| ENSMUSG00000107347 | 0 | 0 | 0 | 0 | 0 | 0 | 0 |
| ENSMUSG00000107348 | 0 | 0 | 0 | 0 | 0 | 0 | 0 |
| ENSMUSG00000107349 | 0 | 0 | 0.25 | 0 | 0 | 0 | 0 |
| ENSMUSG00000107350 | 0.578129653 | 0.646240625 | 0.864857905 | 0.396240625 | 0.489301084 | 0.301029996 | 0 |
| ENSMUSG00000107351 | 0 | 0 | 0 | 0 | 0 | 0 | 0 |
| ENSMUSG00000107352 | 1.37447073 | 1.214495249 | 1.080482024 | 1.238549078 | 1.121713561 | 1.255272505 | 1.089372236 |
| ENSMUSG00000107353 | 0 | 0 | 0 | 0 | 0 | 0 | 0 |
| ENSMUSG00000107354 | 0 | 0 | 0.25 | 0.25 | 0 | 0.301029996 | 0.224243824 |
| ENSMUSG00000107355 | 1.181541736 | 1.061981878 | 0.976722649 | 0.79248125 | 0.632412477 | 0.84509804 | 0.520678036 |
| ENSMUSG00000107356 | 0 | 0 | 0 | 0 | 0 | 0 | 0 |
| ENSMUSG00000107357 | 0 | 0 | 0 | 0 | 0.244650542 | 0 | 0 |
| ENSMUSG00000107358 | 0 | 0 | 0 | 0 | 0 | 0 | 0.224243824 |
| ENSMUSG00000107359 | 0 | 0 | 0.25 | 0 | 0 | 0 | 0 |
| ENSMUSG00000107360 | 0 | 0 | 0 | 0 | 0 | 0 | 0 |
| ENSMUSG00000107362 | 0 | 0.5 | 0.25 | 0.25 | 0 | 0.477121255 | 0.448487648 |
| ENSMUSG00000107363 | 0 | 0 | 0 | 0 | 0 | 0 | 0 |
| ENSMUSG00000107364 | 0 | 0 | 0 | 0 | 0 | 0 | 0 |
| ENSMUSG00000107365 | 0 | 0 | 0 | 0 | 0 | 0 | 0 |
| ENSMUSG00000107368 | 0 | 0.396240625 | 0.25 | 0 | 0 | 0 | 0 |
| ENSMUSG00000107369 | 0 | 0 | 0 | 0 | 0 | 0 | 0 |
| ENSMUSG00000107370 | 0.45815691 | 0.396240625 | 0.25 | 0.5 | 0.244650542 | 0 | 0.355418052 |
| ENSMUSG00000107371 | 0 | 0 | 0 | 0 | 0 | 0 | 0 |
| ENSMUSG00000107372 | 0 | 0 | 0 | 0 | 0 | 0 | 0 |
| ENSMUSG00000107373 | 0 | 0 | 0 | 0 | 0 | 0 | 0 |
| ENSMUSG00000107374 | 1.205378646 | 1.146240625 | 1.146240625 | 0.75 | 0.846351821 | 0.903089987 | 0.916588299 |
| ENSMUSG00000107375 | 0 | 0 | 0 | 0 | 0 | 0 | 0 |
| ENSMUSG00000107376 | 0 | 0 | 0 | 0 | 0 | 0 | 0 |
| ENSMUSG00000107377 | 0 | 0.5 | 0 | 0.25 | 0.244650542 | 0 | 0.448487648 |
| ENSMUSG00000107378 | 0 | 0.25 | 0 | 0 | 0 | 0 | 0 |
| ENSMUSG00000107379 | 0 | 0 | 0 | 0 | 0 | 0 | 0 |
| ENSMUSG00000107380 | 0 | 0 | 0 | 0 | 0 | 0 | 0 |

| | | | | | | | |
|---|---|---|---|---|---|---|---|
| ENSMUSG00000107383 | 2.213719572 | 2.058404919 | 2.073655187 | 1.990000483 | 2.126012715 | 2.222716471 | 2.001999072 |
| ENSMUSG00000107384 | 0 | 0 | 0 | 0 | 0 | 0 | 0 |
| ENSMUSG00000107385 | 0.671187741 | 0.79248125 | 0.701838731 | 0.75 | 0.686820904 | 0.77815125 | 0.629532004 |
| ENSMUSG00000107386 | 0 | 0 | 0 | 0 | 0 | 0 | 0 |
| ENSMUSG00000107387 | 0 | 0 | 0 | 0 | 0 | 0 | 0 |
| ENSMUSG00000107388 | 0 | 0 | 0.25 | 0.396240625 | 0.244650542 | 0.301029996 | 0 |
| ENSMUSG00000107389 | 0 | 0.396240625 | 0.25 | 0 | 0 | 0 | 0.520678036 |
| ENSMUSG00000107390 | 0.45815691 | 0.396240625 | 0.25 | 0.25 | 0 | 0.301029996 | 0.355418052 |
| ENSMUSG00000107392 | 0 | 0 | 0 | 0 | 0 | 0 | 0 |
| ENSMUSG00000107394 | 0 | 0 | 0 | 0 | 0 | 0 | 0 |
| ENSMUSG00000107395 | 0 | 0 | 0 | 0 | 0 | 0 | 0.224243824 |
| ENSMUSG00000107396 | 0 | 0 | 0 | 0 | 0 | 0 | 0 |
| ENSMUSG00000107397 | 0.289064826 | 0.396240625 | 0.25 | 0 | 0 | 0.301029996 | 0.224243824 |
| ENSMUSG00000107398 | 0 | 0 | 0 | 0 | 0 | 0 | 0 |
| ENSMUSG00000107399 | 0 | 0 | 0 | 0 | 0 | 0 | 0 |
| ENSMUSG00000107400 | 0 | 0 | 0 | 0 | 0 | 0 | 0 |
| ENSMUSG00000107401 | 0 | 0 | 0 | 0 | 0 | 0 | 0 |
| ENSMUSG00000107402 | 0.960252568 | 0.92510993 | 0.864857905 | 0.976722649 | 0.846351821 | 0.602059991 | 0.896975297 |
| ENSMUSG00000107403 | 0 | 0 | 0 | 0 | 0 | 0 | 0 |
| ENSMUSG00000107404 | 0 | 0 | 0 | 0 | 0 | 0 | 0 |
| ENSMUSG00000107405 | 0 | 0 | 0 | 0 | 0 | 0.301029996 | 0 |
| ENSMUSG00000107406 | 0 | 0 | 0.25 | 0.396240625 | 0 | 0 | 0 |
| ENSMUSG00000107407 | 0 | 0 | 0 | 0 | 0 | 0 | 0 |
| ENSMUSG00000107408 | 0 | 0 | 0 | 0 | 0 | 0 | 0 |
| ENSMUSG00000107409 | 0 | 0 | 0 | 0 | 0 | 0 | 0 |
| ENSMUSG00000107410 | 0 | 0 | 0 | 0 | 0 | 0 | 0 |
| ENSMUSG00000107411 | 0 | 0.5 | 0.25 | 0.5 | 0.387761935 | 0.301029996 | 0 |
| ENSMUSG00000107412 | 0 | 0 | 0 | 0 | 0 | 0 | 0 |
| ENSMUSG00000107413 | 0 | 0 | 0 | 0 | 0 | 0 | 0 |
| ENSMUSG00000107414 | 0 | 0 | 0 | 0 | 0 | 0 | 0 |
| ENSMUSG00000107415 | 0 | 0 | 0 | 0 | 0 | 0 | 0 |
| ENSMUSG00000107416 | 0 | 0 | 0 | 0 | 0 | 0 | 0 |
| ENSMUSG00000107417 | 0 | 0 | 0.25 | 0 | 0 | 0 | 0 |
| ENSMUSG00000107418 | 0 | 0 | 0 | 0 | 0 | 0 | 0 |

| | | | | | | | |
|---|---|---|---|---|---|---|---|
| ENSMUSG00000107419 | 0 | 0 | 0 | 0 | 0 | 0 | 0 |
| ENSMUSG00000107420 | 0.289064826 | 0 | 0 | 0 | 0 | 0 | 0 |
| ENSMUSG00000107421 | 0 | 0 | 0 | 0 | 0 | 0 | 0 |
| ENSMUSG00000107422 | 0 | 0 | 0 | 0 | 0 | 0 | 0 |
| ENSMUSG00000107423 | 0 | 0 | 0 | 0 | 0 | 0 | 0 |
| ENSMUSG00000107424 | 0 | 0 | 0 | 0 | 0 | 0 | 0 |
| ENSMUSG00000107425 | 0 | 0 | 0 | 0 | 0 | 0 | 0 |
| ENSMUSG00000107427 | 0 | 0 | 0 | 0 | 0 | 0 | 0 |
| ENSMUSG00000107428 | 0 | 0 | 0.25 | 0 | 0 | 0 | 0 |
| ENSMUSG00000107429 | 1.205378646 | 1.25 | 1.410964047 | 1.410964047 | 1.244650542 | 1.380211242 | 1.193409508 |
| ENSMUSG00000107430 | 0 | 0.25 | 0 | 0.25 | 0 | 0 | 0.224243824 |
| ENSMUSG00000107431 | 0 | 0 | 0 | 0 | 0 | 0 | 0 |
| ENSMUSG00000107432 | 0.867194479 | 0.75 | 0.75 | 1.02186571 | 0.812711509 | 0.84509804 | 0.74492186 |
| ENSMUSG00000107433 | 0 | 0 | 0 | 0 | 0 | 0 | 0 |
| ENSMUSG00000107434 | 0.578129653 | 0.701838731 | 0.646240625 | 0.5 | 0 | 0 | 0.224243824 |
| ENSMUSG00000107435 | 0 | 0 | 0 | 0 | 0 | 0 | 0 |
| ENSMUSG00000107436 | 0.289064826 | 0 | 0.25 | 0 | 0 | 0.301029996 | 0.355418052 |
| ENSMUSG00000107438 | 0 | 0 | 0 | 0 | 0 | 0 | 0 |
| ENSMUSG00000107439 | 0 | 0 | 0 | 0 | 0 | 0 | 0 |
| ENSMUSG00000107440 | 0 | 0 | 0.5 | 0.25 | 0 | 0.477121255 | 0 |
| ENSMUSG00000107441 | 0 | 0 | 0 | 0 | 0 | 0 | 0 |
| ENSMUSG00000107442 | 0 | 0 | 0 | 0 | 0 | 0 | 0 |
| ENSMUSG00000107443 | 0 | 0.5 | 0 | 0 | 0.244650542 | 0.301029996 | 0.224243824 |
| ENSMUSG00000107444 | 0 | 0 | 0 | 0 | 0 | 0 | 0 |
| ENSMUSG00000107445 | 0 | 0 | 0 | 0 | 0 | 0 | 0 |
| ENSMUSG00000107446 | 0 | 0 | 0 | 0 | 0 | 0 | 0 |
| ENSMUSG00000107447 | 0 | 0 | 0 | 0 | 0 | 0 | 0 |
| ENSMUSG00000107448 | 0 | 0 | 0 | 0 | 0 | 0 | 0 |
| ENSMUSG00000107449 | 0 | 0 | 0 | 0 | 0 | 0 | 0 |
| ENSMUSG00000107450 | 0 | 0 | 0 | 0 | 0 | 0 | 0 |
| ENSMUSG00000107451 | 0 | 0 | 0 | 0 | 0 | 0 | 0 |
| ENSMUSG00000107452 | 0 | 0 | 0 | 0 | 0 | 0 | 0 |
| ENSMUSG00000107453 | 0 | 0 | 0 | 0 | 0 | 0 | 0 |
| ENSMUSG00000107454 | 0 | 0 | 0 | 0 | 0 | 0 | 0 |

| | | | | | | | |
|---|---|---|---|---|---|---|---|
| ENSMUSG00000107455 | 0 | 0.25 | 0.25 | 0.25 | 0.244650542 | 0.301029996 | 0 |
| ENSMUSG00000107456 | 0 | 0 | 0 | 0 | 0 | 0 | 0 |
| ENSMUSG00000107457 | 0 | 0 | 0 | 0 | 0 | 0 | 0 |
| ENSMUSG00000107458 | 0.289064826 | 0 | 0 | 0 | 0 | 0 | 0 |
| ENSMUSG00000107459 | 0.45815691 | 0.396240625 | 0.701838731 | 0.25 | 0.489301084 | 0.301029996 | 0.224243824 |
| ENSMUSG00000107460 | 0 | 0.396240625 | 0.5 | 0.396240625 | 0 | 0.301029996 | 0 |
| ENSMUSG00000107461 | 0 | 0 | 0 | 0.396240625 | 0.489301084 | 0.477121255 | 0.224243824 |
| ENSMUSG00000107462 | 0 | 0.396240625 | 0.580482024 | 0 | 0 | 0 | 0.224243824 |
| ENSMUSG00000107463 | 0 | 0 | 0 | 0 | 0 | 0 | 0 |
| ENSMUSG00000107464 | 0 | 0 | 0 | 0 | 0 | 0 | 0 |
| ENSMUSG00000107465 | 0 | 0 | 0 | 0 | 0 | 0 | 0 |
| ENSMUSG00000107466 | 0 | 0 | 0 | 0 | 0 | 0 | 0 |
| ENSMUSG00000107467 | 0 | 0 | 0 | 0 | 0 | 0 | 0 |
| ENSMUSG00000107468 | 0.289064826 | 0 | 0 | 0.25 | 0.244650542 | 0 | 0.355418052 |
| ENSMUSG00000107469 | 0 | 0 | 0 | 0 | 0 | 0 | 0 |
| ENSMUSG00000107470 | 1.932832114 | 1.814346961 | 1.850219859 | 2.02053726 | 2.029283463 | 2.257678575 | 1.95718226 |
| ENSMUSG00000107471 | 0 | 0 | 0 | 0 | 0 | 0 | 0 |
| ENSMUSG00000107472 | 0 | 0 | 0 | 0 | 0 | 0 | 0.224243824 |
| ENSMUSG00000107473 | 0 | 0 | 0 | 0 | 0 | 0 | 0 |
| ENSMUSG00000107474 | 0 | 0 | 0 | 0 | 0 | 0 | 0 |
| ENSMUSG00000107475 | 0 | 0 | 0.396240625 | 0 | 0 | 0 | 0 |
| ENSMUSG00000107476 | 1.99653913 | 1.912762923 | 1.860735874 | 1.87694866 | 1.868304353 | 1.924279286 | 1.802677937 |
| ENSMUSG00000107477 | 0 | 0.25 | 0.25 | 0 | 0.489301084 | 0 | 0.355418052 |
| ENSMUSG00000107478 | 0 | 0.646240625 | 0.396240625 | 0 | 0.568060967 | 0 | 0.224243824 |
| ENSMUSG00000107479 | 0 | 0 | 0 | 0 | 0 | 0 | 0 |
| ENSMUSG00000107480 | 0.289064826 | 0.25 | 0 | 0.25 | 0.244650542 | 0.301029996 | 0 |
| ENSMUSG00000107481 | 0 | 0 | 0 | 0 | 0 | 0 | 0 |
| ENSMUSG00000107482 | 0 | 0 | 0 | 0 | 0 | 0 | 0 |
| ENSMUSG00000107483 | 0 | 0 | 0 | 0.25 | 0 | 0 | 0 |
| ENSMUSG00000107484 | 0 | 0 | 0 | 0 | 0 | 0 | 0 |
| ENSMUSG00000107486 | 0 | 0 | 0 | 0 | 0 | 0 | 0 |
| ENSMUSG00000107487 | 0 | 0 | 0 | 0 | 0 | 0 | 0.224243824 |
| ENSMUSG00000107488 | 0 | 0 | 0 | 0 | 0 | 0 | 0 |
| ENSMUSG00000107489 | 0 | 0 | 0 | 0 | 0 | 0 | 0 |

| | | | | | | | |
|---|---|---|---|---|---|---|---|
| ENSMUSG00000107490 | 0 | 0 | 0 | 0 | 0 | 0 | 0 |
| ENSMUSG00000107491 | 0 | 0.25 | 0 | 0.396240625 | 0.387761935 | 0 | 0 |
| ENSMUSG00000107492 | 0 | 0 | 0 | 0 | 0 | 0 | 0 |
| ENSMUSG00000107493 | 0 | 0 | 0 | 0 | 0 | 0 | 0 |
| ENSMUSG00000107494 | 0 | 0 | 0 | 0 | 0 | 0 | 0 |
| ENSMUSG00000107495 | 2.22592627 | 2.034887838 | 2.009729747 | 2.027131114 | 1.929985312 | 2.155336037 | 1.752192176 |
| ENSMUSG00000107496 | 0 | 0 | 0 | 0 | 0 | 0 | 0 |
| ENSMUSG00000107497 | 0 | 0.25 | 0.25 | 0.25 | 0 | 0.301029996 | 0 |
| ENSMUSG00000107498 | 0 | 0 | 0 | 0 | 0 | 0 | 0 |
| ENSMUSG00000107499 | 0 | 0 | 0 | 0 | 0.387761935 | 0.301029996 | 0.579661877 |
| ENSMUSG00000107500 | 0 | 0 | 0 | 0 | 0 | 0 | 0 |
| ENSMUSG00000107501 | 0 | 0.396240625 | 0 | 0 | 0 | 0 | 0.224243824 |
| ENSMUSG00000107502 | 0.289064826 | 0 | 0 | 0 | 0.244650542 | 0 | 0.520678036 |
| ENSMUSG00000107503 | 0 | 0 | 0 | 0 | 0 | 0 | 0 |
| ENSMUSG00000107504 | 0 | 0 | 0 | 0 | 0 | 0 | 0.355418052 |
| ENSMUSG00000107505 | 0.960252568 | 0.646240625 | 0.396240625 | 0.580482024 | 0.489301084 | 0.301029996 | 0.579661877 |
| ENSMUSG00000107506 | 0 | 0 | 0 | 0 | 0 | 0 | 0 |
| ENSMUSG00000107507 | 0 | 0 | 0 | 0 | 0 | 0 | 0 |
| ENSMUSG00000107508 | 0.289064826 | 0 | 0.25 | 0 | 0 | 0.301029996 | 0 |
| ENSMUSG00000107509 | 1.100572389 | 0.79248125 | 0.701838731 | 0.580482024 | 0.632412477 | 0 | 0.710836105 |
| ENSMUSG00000107510 | 0 | 0 | 0 | 0 | 0 | 0 | 0 |
| ENSMUSG00000107511 | 0.747221736 | 0.396240625 | 0.646240625 | 0.79248125 | 0.632412477 | 0.477121255 | 0.520678036 |
| ENSMUSG00000107512 | 0 | 0 | 0 | 0 | 0 | 0 | 0 |
| ENSMUSG00000107513 | 0 | 0 | 0 | 0 | 0 | 0 | 0 |
| ENSMUSG00000107514 | 0 | 0 | 0 | 0 | 0 | 0 | 0 |
| ENSMUSG00000107515 | 0.289064826 | 0 | 0.25 | 0 | 0 | 0 | 0 |
| ENSMUSG00000107516 | 0 | 0 | 0 | 0 | 0 | 0 | 0 |
| ENSMUSG00000107517 | 0 | 0 | 0 | 0 | 0.387761935 | 0 | 0 |
| ENSMUSG00000107518 | 0 | 0 | 0 | 0 | 0 | 0 | 0 |
| ENSMUSG00000107519 | 0 | 0 | 0 | 0 | 0 | 0 | 0 |
| ENSMUSG00000107520 | 0 | 0 | 0 | 0 | 0 | 0 | 0 |
| ENSMUSG00000107521 | 0 | 0 | 0 | 0 | 0 | 0 | 0.224243824 |
| ENSMUSG00000107522 | 0 | 0 | 0 | 0 | 0 | 0 | 0 |
| ENSMUSG00000107523 | 0 | 0 | 0 | 0 | 0 | 0 | 0 |

| | | | | | | | |
|---|---|---|---|---|---|---|---|
| ENSMUSG00000107524 | 0 | 0 | 0.25 | 0.396240625 | 0.568060967 | 0.301029996 | 0.448487648 |
| ENSMUSG00000107525 | 0 | 0 | 0 | 0 | 0 | 0 | 0 |
| ENSMUSG00000107526 | 0.45815691 | 0.25 | 0.25 | 0 | 0.244650542 | 0 | 0 |
| ENSMUSG00000107527 | 0 | 0 | 0 | 0 | 0 | 0 | 0 |
| ENSMUSG00000107528 | 0 | 0 | 0 | 0 | 0 | 0 | 0 |
| ENSMUSG00000107529 | 0 | 0 | 0 | 0.25 | 0 | 0 | 0.224243824 |
| ENSMUSG00000107530 | 0 | 0 | 0 | 0 | 0 | 0 | 0 |
| ENSMUSG00000107531 | 0.960252568 | 0.92510993 | 0.75 | 0.580482024 | 0.877063019 | 0.77815125 | 0.710836105 |
| ENSMUSG00000107532 | 0 | 0 | 0 | 0 | 0 | 0 | 0 |
| ENSMUSG00000107533 | 0 | 0 | 0 | 0 | 0 | 0 | 0 |
| ENSMUSG00000107534 | 0 | 0 | 0 | 0.396240625 | 0.244650542 | 0 | 0.355418052 |
| ENSMUSG00000107535 | 0 | 0 | 0 | 0 | 0 | 0 | 0 |
| ENSMUSG00000107536 | 0 | 0 | 0 | 0 | 0 | 0 | 0 |
| ENSMUSG00000107538 | 0 | 0 | 0 | 0 | 0 | 0 | 0 |
| ENSMUSG00000107539 | 0 | 0 | 0 | 0 | 0 | 0 | 0 |
| ENSMUSG00000107540 | 0 | 0 | 0 | 0 | 0 | 0 | 0 |
| ENSMUSG00000107541 | 0 | 0 | 0 | 0 | 0 | 0 | 0 |
| ENSMUSG00000107542 | 0 | 0 | 0 | 0 | 0 | 0 | 0 |
| ENSMUSG00000107543 | 0 | 0 | 0 | 0 | 0 | 0 | 0 |
| ENSMUSG00000107544 | 0 | 0 | 0 | 0 | 0 | 0 | 0 |
| ENSMUSG00000107545 | 0 | 0 | 0 | 0 | 0 | 0 | 0 |
| ENSMUSG00000107546 | 0 | 0 | 0 | 0 | 0 | 0.301029996 | 0 |
| ENSMUSG00000107548 | 0 | 0 | 0 | 0 | 0 | 0 | 0 |
| ENSMUSG00000107549 | 0.811507563 | 0.701838731 | 0.701838731 | 0.580482024 | 0.733951626 | 0.301029996 | 0.579661877 |
| ENSMUSG00000107550 | 0 | 0 | 0 | 0 | 0 | 0 | 0 |
| ENSMUSG00000107551 | 0 | 0 | 0 | 0 | 0 | 0.301029996 | 0 |
| ENSMUSG00000107552 | 0 | 0 | 0.5 | 0.25 | 0.568060967 | 0.301029996 | 0.520678036 |
| ENSMUSG00000107553 | 0 | 0 | 0 | 0 | 0 | 0 | 0 |
| ENSMUSG00000107554 | 0 | 0 | 0 | 0 | 0 | 0 | 0 |
| ENSMUSG00000107555 | 0 | 0 | 0 | 0 | 0 | 0 | 0 |
| ENSMUSG00000107556 | 0.45815691 | 0.396240625 | 0 | 0.25 | 0 | 0 | 0 |
| ENSMUSG00000107557 | 0 | 0 | 0 | 0 | 0 | 0 | 0 |
| ENSMUSG00000107558 | 0 | 0 | 0 | 0 | 0 | 0 | 0 |
| ENSMUSG00000107559 | 0 | 0.25 | 0.25 | 0 | 0 | 0 | 0.224243824 |

| | | | | | | | |
|---|---|---|---|---|---|---|---|
| ENSMUSG00000107560 | 0.91631382 | 1 | 0.951838731 | 1 | 0.905314583 | 0.903089987 | 0.803905701 |
| ENSMUSG00000107561 | 0 | 0 | 0 | 0 | 0 | 0 | 0 |
| ENSMUSG00000107562 | 0 | 0 | 0 | 0 | 0 | 0 | 0 |
| ENSMUSG00000107563 | 0 | 0 | 0 | 0 | 0 | 0 | 0 |
| ENSMUSG00000107564 | 0 | 0 | 0 | 0 | 0 | 0 | 0 |
| ENSMUSG00000107565 | 0 | 0 | 0 | 0 | 0 | 0 | 0 |
| ENSMUSG00000107566 | 0 | 0 | 0 | 0 | 0 | 0 | 0 |
| ENSMUSG00000107567 | 0.578129653 | 0.396240625 | 0.396240625 | 0 | 0.387761935 | 0.301029996 | 0.355418052 |
| ENSMUSG00000107568 | 0.289064826 | 0 | 0.396240625 | 0.580482024 | 0 | 0.477121255 | 0 |
| ENSMUSG00000107569 | 0 | 0 | 0 | 0 | 0 | 0 | 0 |
| ENSMUSG00000107570 | 0.867194479 | 0.864857905 | 0.976722649 | 0.75 | 0.733951626 | 0.77815125 | 0.74492186 |
| ENSMUSG00000107571 | 0 | 0 | 0 | 0 | 0 | 0 | 0 |
| ENSMUSG00000107572 | 0 | 0 | 0 | 0 | 0 | 0 | 0 |
| ENSMUSG00000107573 | 0 | 0 | 0 | 0 | 0 | 0 | 0 |
| ENSMUSG00000107574 | 0 | 0 | 0 | 0 | 0 | 0 | 0 |
| ENSMUSG00000107575 | 0 | 0 | 0 | 0 | 0 | 0 | 0 |
| ENSMUSG00000107576 | 0 | 0 | 0.25 | 0.25 | 0.387761935 | 0.477121255 | 0 |
| ENSMUSG00000107577 | 1.129344651 | 1.214495249 | 1.160964047 | 1.146240625 | 1 | 1 | 1.014380832 |
| ENSMUSG00000107578 | 0 | 0 | 0 | 0 | 0 | 0 | 0 |
| ENSMUSG00000107579 | 0 | 0 | 0 | 0 | 0 | 0 | 0 |
| ENSMUSG00000107580 | 0 | 0 | 0 | 0 | 0 | 0 | 0 |
| ENSMUSG00000107581 | 0 | 0 | 0 | 0 | 0 | 0 | 0 |
| ENSMUSG00000107582 | 0 | 0 | 0 | 0 | 0 | 0 | 0 |
| ENSMUSG00000107583 | 0 | 0 | 0 | 0 | 0.244650542 | 0.301029996 | 0 |
| ENSMUSG00000107584 | 0.45815691 | 0.5 | 0.396240625 | 0 | 0.489301084 | 0.301029996 | 0.355418052 |
| ENSMUSG00000107585 | 0 | 0 | 0 | 0 | 0 | 0 | 0 |
| ENSMUSG00000107586 | 0 | 0.25 | 0.25 | 0 | 0 | 0 | 0 |
| ENSMUSG00000107587 | 0 | 0 | 0 | 0 | 0 | 0 | 0 |
| ENSMUSG00000107588 | 0 | 0 | 0 | 0 | 0 | 0 | 0 |
| ENSMUSG00000107590 | 0 | 0 | 0 | 0 | 0 | 0 | 0 |
| ENSMUSG00000107591 | 0 | 0 | 0 | 0.25 | 0 | 0 | 0 |
| ENSMUSG00000107592 | 0 | 0 | 0 | 0 | 0 | 0 | 0 |
| ENSMUSG00000107593 | 0 | 0 | 0.25 | 0 | 0 | 0 | 0 |
| ENSMUSG00000107594 | 0 | 0 | 0 | 0 | 0 | 0 | 0 |

| | | | | | | | |
|---|---|---|---|---|---|---|---|
| ENSMUSG00000107595 | 0 | 0 | 0.25 | 0 | 0 | 0 | 0 |
| ENSMUSG00000107596 | 0 | 0.25 | 0 | 0 | 0 | 0 | 0 |
| ENSMUSG00000107597 | 0 | 0 | 0 | 0 | 0 | 0 | 0 |
| ENSMUSG00000107598 | 0 | 0 | 0 | 0 | 0 | 0 | 0 |
| ENSMUSG00000107599 | 0.45815691 | 0.396240625 | 0.580482024 | 0.5 | 0 | 0 | 0.355418052 |
| ENSMUSG00000107600 | 0 | 0 | 0 | 0 | 0 | 0 | 0 |
| ENSMUSG00000107601 | 0 | 0 | 0 | 0 | 0 | 0 | 0 |
| ENSMUSG00000107602 | 0 | 0 | 0 | 0 | 0 | 0 | 0 |
| ENSMUSG00000107603 | 0 | 0 | 0 | 0.580482024 | 0.244650542 | 0 | 0.224243824 |
| ENSMUSG00000107604 | 0 | 0.25 | 0.25 | 0 | 0 | 0 | 0 |
| ENSMUSG00000107605 | 0 | 0 | 0 | 0 | 0 | 0 | 0 |
| ENSMUSG00000107606 | 0 | 0.25 | 0.396240625 | 0 | 0 | 0.301029996 | 0 |
| ENSMUSG00000107608 | 0.289064826 | 0 | 0.396240625 | 0.25 | 0.489301084 | 0.301029996 | 0.520678036 |
| ENSMUSG00000107609 | 0 | 0 | 0 | 0 | 0 | 0 | 0 |
| ENSMUSG00000107610 | 0.289064826 | 0.580482024 | 0.5 | 0.396240625 | 0.568060967 | 0.477121255 | 0.629532004 |
| ENSMUSG00000107611 | 0 | 0 | 0 | 0 | 0 | 0 | 0 |
| ENSMUSG00000107612 | 0.289064826 | 0 | 0 | 0 | 0 | 0 | 0.224243824 |
| ENSMUSG00000107613 | 0 | 0 | 0 | 0 | 0 | 0 | 0 |
| ENSMUSG00000107614 | 0 | 0 | 0 | 0 | 0 | 0 | 0 |
| ENSMUSG00000107615 | 0 | 0 | 0 | 0 | 0 | 0 | 0 |
| ENSMUSG00000107616 | 0 | 0 | 0 | 0 | 0 | 0 | 0 |
| ENSMUSG00000107618 | 0 | 0 | 0 | 0 | 0 | 0 | 0 |
| ENSMUSG00000107619 | 0 | 0 | 0 | 0 | 0 | 0 | 0 |
| ENSMUSG00000107620 | 0.289064826 | 0.25 | 0.25 | 0.25 | 0.632412477 | 0.477121255 | 0.355418052 |
| ENSMUSG00000107622 | 0 | 0 | 0 | 0 | 0 | 0 | 0 |
| ENSMUSG00000107623 | 0 | 0 | 0 | 0 | 0 | 0 | 0.224243824 |
| ENSMUSG00000107624 | 0 | 0 | 0 | 0 | 0 | 0 | 0 |
| ENSMUSG00000107625 | 0 | 0 | 0 | 0 | 0 | 0 | 0 |
| ENSMUSG00000107626 | 0 | 0 | 0 | 0 | 0 | 0 | 0 |
| ENSMUSG00000107627 | 0 | 0 | 0 | 0 | 0 | 0 | 0 |
| ENSMUSG00000107628 | 0 | 0.5 | 0 | 0 | 0 | 0 | 0.224243824 |
| ENSMUSG00000107629 | 0 | 0 | 0.25 | 0 | 0 | 0 | 0 |
| ENSMUSG00000107630 | 0 | 0 | 0 | 0 | 0 | 0 | 0 |
| ENSMUSG00000107631 | 0 | 0 | 0 | 0 | 0 | 0 | 0 |

| | | | | | | | |
|---|---|---|---|---|---|---|---|
| ENSMUSG00000107632 | 0.867194479 | 0.79248125 | 1 | 0.75 | 0.77552387 | 0.84509804 | 0.710836105 |
| ENSMUSG00000107633 | 1.753493018 | 1.705044741 | 1.584962501 | 1.561981878 | 1.49445382 | 1.612783857 | 1.38802993 |
| ENSMUSG00000107634 | 0 | 0.25 | 0.5 | 0 | 0 | 0 | 0 |
| ENSMUSG00000107635 | 0 | 0 | 0 | 0 | 0.244650542 | 0 | 0 |
| ENSMUSG00000107637 | 0 | 0 | 0 | 0 | 0 | 0 | 0 |
| ENSMUSG00000107638 | 0 | 0 | 0 | 0 | 0 | 0 | 0 |
| ENSMUSG00000107639 | 0 | 0.25 | 0 | 0.396240625 | 0.244650542 | 0.301029996 | 0 |
| ENSMUSG00000107640 | 0 | 0 | 0 | 0 | 0 | 0 | 0 |
| ENSMUSG00000107641 | 0 | 0 | 0 | 0 | 0 | 0 | 0 |
| ENSMUSG00000107642 | 0 | 0 | 0 | 0 | 0 | 0 | 0 |
| ENSMUSG00000107643 | 0 | 0 | 0 | 0 | 0 | 0 | 0 |
| ENSMUSG00000107644 | 0 | 0 | 0 | 0 | 0.244650542 | 0 | 0.224243824 |
| ENSMUSG00000107645 | 0 | 0 | 0 | 0 | 0 | 0 | 0 |
| ENSMUSG00000107646 | 0 | 0 | 0 | 0 | 0 | 0 | 0 |
| ENSMUSG00000107647 | 0 | 0 | 0.396240625 | 0 | 0.244650542 | 0 | 0 |
| ENSMUSG00000107648 | 0 | 0 | 0 | 0 | 0 | 0 | 0 |
| ENSMUSG00000107649 | 0 | 0 | 0 | 0 | 0 | 0 | 0 |
| ENSMUSG00000107650 | 0 | 0 | 0 | 0 | 0 | 0 | 0 |
| ENSMUSG00000107651 | 0 | 0 | 0 | 0 | 0 | 0 | 0 |
| ENSMUSG00000107652 | 0 | 0 | 0 | 0 | 0 | 0 | 0 |
| ENSMUSG00000107653 | 0.671187741 | 0.75 | 0.580482024 | 0.701838731 | 0.632412477 | 0.602059991 | 0.74492186 |
| ENSMUSG00000107655 | 0 | 0.25 | 0.25 | 0 | 0 | 0 | 0 |
| ENSMUSG00000107656 | 0 | 0 | 0 | 0 | 0 | 0 | 0 |
| ENSMUSG00000107657 | 0 | 0 | 0 | 0 | 0 | 0 | 0 |
| ENSMUSG00000107658 | 0.289064826 | 0.5 | 0.580482024 | 0.79248125 | 0.489301084 | 0.602059991 | 0.448487648 |
| ENSMUSG00000107659 | 0 | 0 | 0 | 0 | 0 | 0 | 0 |
| ENSMUSG00000107660 | 0 | 0 | 0 | 0 | 0 | 0 | 0 |
| ENSMUSG00000107661 | 0 | 0 | 0 | 0 | 0 | 0 | 0 |
| ENSMUSG00000107662 | 0 | 0 | 0 | 0 | 0 | 0 | 0 |
| ENSMUSG00000107663 | 0.289064826 | 0 | 0 | 0 | 0 | 0 | 0 |
| ENSMUSG00000107664 | 0 | 0 | 0 | 0 | 0 | 0 | 0 |
| ENSMUSG00000107666 | 0 | 0 | 0 | 0 | 0 | 0 | 0 |
| ENSMUSG00000107667 | 0 | 0 | 0 | 0 | 0 | 0 | 0 |
| ENSMUSG00000107668 | 0.45815691 | 0.5 | 0.25 | 0 | 0.489301084 | 0.301029996 | 0.355418052 |

| | | | | | | | |
|---|---|---|---|---|---|---|---|
| ENSMUSG00000107669 | 0 | 0 | 0 | 0 | 0 | 0 | 0 |
| ENSMUSG00000107670 | 0 | 0 | 0 | 0 | 0 | 0 | 0 |
| ENSMUSG00000107671 | 0 | 0 | 0 | 0 | 0 | 0 | 0 |
| ENSMUSG00000107672 | 0 | 0 | 0 | 0 | 0 | 0 | 0 |
| ENSMUSG00000107673 | 0 | 0 | 0 | 0 | 0 | 0 | 0 |
| ENSMUSG00000107674 | 0 | 0 | 0 | 0 | 0 | 0 | 0 |
| ENSMUSG00000107675 | 0 | 0 | 0.25 | 0 | 0 | 0 | 0 |
| ENSMUSG00000107676 | 0 | 0.25 | 0.25 | 0 | 0 | 0 | 0.224243824 |
| ENSMUSG00000107677 | 0 | 0 | 0 | 0 | 0 | 0 | 0 |
| ENSMUSG00000107678 | 0 | 0 | 0 | 0 | 0 | 0 | 0 |
| ENSMUSG00000107679 | 0 | 0 | 0 | 0 | 0 | 0 | 0 |
| ENSMUSG00000107680 | 0 | 0 | 0 | 0 | 0 | 0 | 0 |
| ENSMUSG00000107681 | 0 | 0 | 0.396240625 | 0 | 0 | 0 | 0 |
| ENSMUSG00000107682 | 0 | 0.25 | 0 | 0.396240625 | 0 | 0 | 0 |
| ENSMUSG00000107683 | 0 | 0 | 0 | 0 | 0 | 0 | 0 |
| ENSMUSG00000107684 | 0 | 0 | 0 | 0 | 0 | 0 | 0 |
| ENSMUSG00000107685 | 0 | 0 | 0.25 | 0 | 0 | 0.301029996 | 0 |
| ENSMUSG00000107686 | 0 | 0 | 0 | 0 | 0 | 0 | 0 |
| ENSMUSG00000107687 | 0 | 0 | 0 | 0 | 0 | 0 | 0 |
| ENSMUSG00000107688 | 0 | 0 | 0.5 | 0.25 | 0 | 0 | 0 |
| ENSMUSG00000107689 | 0 | 0 | 0 | 0 | 0 | 0 | 0 |
| ENSMUSG00000107690 | 0.289064826 | 0.25 | 0.25 | 0 | 0 | 0 | 0 |
| ENSMUSG00000107691 | 0 | 0 | 0 | 0 | 0 | 0 | 0 |
| ENSMUSG00000107692 | 0 | 0 | 0 | 0 | 0 | 0 | 0 |
| ENSMUSG00000107693 | 0 | 0 | 0 | 0 | 0 | 0 | 0 |
| ENSMUSG00000107694 | 0 | 0 | 0 | 0 | 0 | 0 | 0 |
| ENSMUSG00000107695 | 0 | 0 | 0 | 0 | 0 | 0 | 0 |
| ENSMUSG00000107696 | 0 | 0 | 0 | 0 | 0 | 0 | 0 |
| ENSMUSG00000107697 | 0 | 0 | 0 | 0 | 0 | 0 | 0 |
| ENSMUSG00000107698 | 0.45815691 | 0.5 | 0.25 | 0 | 0.632412477 | 0.602059991 | 0 |
| ENSMUSG00000107699 | 0 | 0 | 0 | 0.5 | 0.686820904 | 0.698970004 | 0.672731473 |
| ENSMUSG00000107700 | 0 | 0 | 0 | 0 | 0 | 0 | 0 |
| ENSMUSG00000107701 | 0 | 0 | 0 | 0 | 0 | 0 | 0 |
| ENSMUSG00000107702 | 0 | 0 | 0 | 0 | 0 | 0 | 0 |

| | | | | | | | |
|---|---|---|---|---|---|---|---|
| ENSMUSG00000107703 | 0 | 0 | 0 | 0 | 0 | 0 | 0.355418052 |
| ENSMUSG00000107704 | 0 | 0 | 0 | 0.25 | 0 | 0 | 0 |
| ENSMUSG00000107705 | 0 | 0 | 0 | 0 | 0 | 0 | 0 |
| ENSMUSG00000107706 | 0 | 0 | 0 | 0 | 0 | 0 | 0 |
| ENSMUSG00000107707 | 0 | 0 | 0 | 0 | 0 | 0 | 0 |
| ENSMUSG00000107708 | 0 | 0.396240625 | 0 | 0 | 0 | 0 | 0 |
| ENSMUSG00000107709 | 0.45815691 | 0.75 | 0.396240625 | 0.396240625 | 0.686820904 | 0.301029996 | 0.448487648 |
| ENSMUSG00000107710 | 0 | 0 | 0 | 0 | 0 | 0 | 0 |
| ENSMUSG00000107711 | 0 | 0 | 0 | 0 | 0 | 0 | 0 |
| ENSMUSG00000107712 | 0.45815691 | 0 | 0 | 0.25 | 0 | 0 | 0.355418052 |
| ENSMUSG00000107713 | 0 | 0 | 0 | 0 | 0 | 0 | 0 |
| ENSMUSG00000107714 | 0 | 0 | 0 | 0 | 0 | 0 | 0 |
| ENSMUSG00000107715 | 0 | 0 | 0 | 0 | 0 | 0 | 0 |
| ENSMUSG00000107716 | 0 | 0 | 0 | 0 | 0 | 0 | 0 |
| ENSMUSG00000107718 | 0 | 0 | 0 | 0 | 0 | 0 | 0 |
| ENSMUSG00000107719 | 0 | 0 | 0 | 0 | 0 | 0 | 0 |
| ENSMUSG00000107720 | 0.289064826 | 0 | 0.25 | 0 | 0 | 0 | 0 |
| ENSMUSG00000107721 | 0 | 0 | 0 | 0 | 0 | 0 | 0 |
| ENSMUSG00000107722 | 0 | 0 | 0 | 0 | 0 | 0 | 0.224243824 |
| ENSMUSG00000107723 | 0 | 0 | 0 | 0 | 0 | 0 | 0 |
| ENSMUSG00000107724 | 0 | 0 | 0 | 0 | 0 | 0 | 0 |
| ENSMUSG00000107725 | 0 | 0 | 0 | 0 | 0 | 0 | 0 |
| ENSMUSG00000107726 | 0.671187741 | 0.646240625 | 0.580482024 | 0.396240625 | 0.244650542 | 0 | 0.224243824 |
| ENSMUSG00000107727 | 0 | 0 | 0 | 0 | 0 | 0 | 0 |
| ENSMUSG00000107728 | 0 | 0 | 0 | 0 | 0.244650542 | 0.301029996 | 0.448487648 |
| ENSMUSG00000107729 | 0 | 0 | 0 | 0 | 0 | 0 | 0 |
| ENSMUSG00000107730 | 0 | 0.25 | 0 | 0 | 0 | 0 | 0 |
| ENSMUSG00000107731 | 0 | 0 | 0 | 0 | 0 | 0 | 0 |
| ENSMUSG00000107732 | 0 | 0 | 0 | 0 | 0 | 0 | 0 |
| ENSMUSG00000107733 | 0 | 0 | 0 | 0 | 0 | 0 | 0.224243824 |
| ENSMUSG00000107734 | 0.45815691 | 0 | 0 | 0 | 0.244650542 | 0 | 0 |
| ENSMUSG00000107735 | 0 | 0 | 0 | 0 | 0 | 0 | 0 |
| ENSMUSG00000107736 | 0 | 0 | 0 | 0 | 0 | 0 | 0 |
| ENSMUSG00000107737 | 0 | 0 | 0 | 0 | 0 | 0 | 0 |

| | | | | | | | |
|---|---|---|---|---|---|---|---|
| ENSMUSG00000107738 | 0 | 0 | 0 | 0 | 0 | 0 | 0 |
| ENSMUSG00000107739 | 0 | 0 | 0 | 0 | 0 | 0 | 0 |
| ENSMUSG00000107740 | 0 | 0 | 0 | 0 | 0 | 0 | 0 |
| ENSMUSG00000107741 | 0.91631382 | 0.951838731 | 1 | 1.098079356 | 1.121713561 | 1.146128036 | 1.028149525 |
| ENSMUSG00000107742 | 0.811507563 | 1.061981878 | 1.160964047 | 1.098079356 | 1.057362051 | 1.041392685 | 0.969165684 |
| ENSMUSG00000107743 | 0.960252568 | 1 | 0.951838731 | 1.080482024 | 1 | 1 | 1.131174228 |
| ENSMUSG00000107744 | 0 | 0 | 0 | 0 | 0 | 0 | 0 |
| ENSMUSG00000107745 | 0 | 0.25 | 0 | 0 | 0 | 0 | 0.224243824 |
| ENSMUSG00000107746 | 0 | 0 | 0 | 0 | 0 | 0 | 0 |
| ENSMUSG00000107747 | 0 | 0 | 0.25 | 0 | 0 | 0 | 0 |
| ENSMUSG00000107748 | 0 | 0 | 0 | 0 | 0 | 0 | 0 |
| ENSMUSG00000107749 | 1.205378646 | 1.114857905 | 0.951838731 | 0.951838731 | 1.020174412 | 0.77815125 | 0.710836105 |
| ENSMUSG00000107750 | 0.671187741 | 0.396240625 | 0.396240625 | 0.396240625 | 0.387761935 | 0.301029996 | 0.355418052 |
| ENSMUSG00000107751 | 0.289064826 | 0.25 | 0.396240625 | 0 | 0 | 0.301029996 | 0 |
| ENSMUSG00000107752 | 0 | 0 | 0 | 0 | 0 | 0 | 0 |
| ENSMUSG00000107753 | 0 | 0 | 0 | 0 | 0 | 0 | 0 |
| ENSMUSG00000107754 | 0 | 0 | 0 | 0 | 0 | 0 | 0 |
| ENSMUSG00000107755 | 0 | 0 | 0 | 0 | 0 | 0 | 0 |
| ENSMUSG00000107756 | 0.747221736 | 1.02186571 | 0.896240625 | 0.830482024 | 1.020174412 | 0.77815125 | 0.710836105 |
| ENSMUSG00000107757 | 0 | 0 | 0 | 0 | 0 | 0 | 0 |
| ENSMUSG00000107758 | 0 | 0 | 0 | 0 | 0 | 0 | 0 |
| ENSMUSG00000107759 | 0 | 0 | 0 | 0 | 0 | 0 | 0 |
| ENSMUSG00000107760 | 0.45815691 | 0.396240625 | 0.646240625 | 0 | 0.244650542 | 0.301029996 | 0.224243824 |
| ENSMUSG00000107761 | 0 | 0 | 0 | 0 | 0 | 0 | 0 |
| ENSMUSG00000107762 | 0 | 0 | 0 | 0 | 0 | 0 | 0 |
| ENSMUSG00000107763 | 0.578129653 | 0 | 0 | 0.25 | 0 | 0 | 0 |
| ENSMUSG00000107764 | 0 | 0 | 0 | 0 | 0 | 0 | 0.224243824 |
| ENSMUSG00000107765 | 0 | 0 | 0 | 0 | 0 | 0 | 0 |
| ENSMUSG00000107766 | 0 | 0 | 0 | 0 | 0 | 0 | 0 |
| ENSMUSG00000107767 | 0 | 0 | 0 | 0 | 0 | 0 | 0 |
| ENSMUSG00000107768 | 1.069666964 | 0.951838731 | 0.896240625 | 0.951838731 | 0.77552387 | 0.477121255 | 0.520678036 |
| ENSMUSG00000107769 | 0 | 0 | 0 | 0 | 0 | 0 | 0 |
| ENSMUSG00000107770 | 0 | 0 | 0 | 0 | 0 | 0 | 0 |
| ENSMUSG00000107771 | 0.578129653 | 0.951838731 | 0.79248125 | 0.79248125 | 0.812711509 | 0.698970004 | 0.775756176 |

| | | | | | | | |
|---|---|---|---|---|---|---|---|
| ENSMUSG00000107772 | 0.289064826 | 0.25 | 0 | 0.25 | 0.387761935 | 0 | 0 |
| ENSMUSG00000107773 | 0 | 0 | 0 | 0 | 0 | 0 | 0 |
| ENSMUSG00000107774 | 0 | 0 | 0 | 0 | 0 | 0 | 0 |
| ENSMUSG00000107775 | 0 | 0 | 0 | 0 | 0 | 0 | 0.224243824 |
| ENSMUSG00000107776 | 0 | 0 | 0 | 0 | 0 | 0 | 0 |
| ENSMUSG00000107777 | 0 | 0 | 0 | 0 | 0 | 0 | 0 |
| ENSMUSG00000107778 | 0 | 0 | 0 | 0 | 0 | 0 | 0 |
| ENSMUSG00000107779 | 0 | 0 | 0 | 0.580482024 | 0.244650542 | 0.301029996 | 0.224243824 |
| ENSMUSG00000107780 | 0 | 0 | 0 | 0 | 0 | 0 | 0 |
| ENSMUSG00000107781 | 0 | 0 | 0 | 0 | 0 | 0 | 0 |
| ENSMUSG00000107782 | 0 | 0 | 0 | 0 | 0 | 0 | 0 |
| ENSMUSG00000107783 | 0 | 0 | 0 | 0 | 0 | 0 | 0 |
| ENSMUSG00000107784 | 0 | 0 | 0 | 0 | 0 | 0 | 0 |
| ENSMUSG00000107785 | 0 | 0 | 0.25 | 0 | 0 | 0 | 0 |
| ENSMUSG00000107786 | 0 | 0 | 0 | 0 | 0 | 0 | 0 |
| ENSMUSG00000107787 | 0 | 0 | 0 | 0 | 0 | 0 | 0 |
| ENSMUSG00000107788 | 0.45815691 | 0.75 | 0.79248125 | 0 | 0.387761935 | 0.698970004 | 0.224243824 |
| ENSMUSG00000107789 | 0 | 0 | 0 | 0 | 0 | 0 | 0 |
| ENSMUSG00000107790 | 0 | 0 | 0 | 0 | 0 | 0 | 0 |
| ENSMUSG00000107791 | 0 | 0 | 0 | 0 | 0 | 0 | 0 |
| ENSMUSG00000107792 | 0 | 0 | 0 | 0 | 0 | 0 | 0 |
| ENSMUSG00000107793 | 0 | 0 | 0 | 0 | 0 | 0 | 0 |
| ENSMUSG00000107794 | 0.289064826 | 0 | 0 | 0 | 0 | 0 | 0 |
| ENSMUSG00000107795 | 0 | 0 | 0 | 0 | 0 | 0 | 0 |
| ENSMUSG00000107796 | 0.747221736 | 0.580482024 | 0.580482024 | 0.580482024 | 0.812711509 | 0.77815125 | 0.448487648 |
| ENSMUSG00000107797 | 0 | 0 | 0 | 0 | 0 | 0 | 0 |
| ENSMUSG00000107798 | 0 | 0 | 0 | 0 | 0 | 0 | 0 |
| ENSMUSG00000107799 | 0 | 0 | 0 | 0 | 0 | 0 | 0 |
| ENSMUSG00000107800 | 0 | 0 | 0 | 0 | 0 | 0 | 0 |
| ENSMUSG00000107801 | 0.289064826 | 0 | 0 | 0 | 0 | 0 | 0 |
| ENSMUSG00000107802 | 0.867194479 | 0.580482024 | 0.75 | 0.25 | 0.387761935 | 0.477121255 | 0.672731473 |
| ENSMUSG00000107803 | 0 | 0 | 0 | 0 | 0 | 0 | 0 |
| ENSMUSG00000107804 | 0 | 0.25 | 0 | 0 | 0 | 0 | 0 |
| ENSMUSG00000107805 | 0 | 0 | 0 | 0 | 0 | 0 | 0 |

| | | | | | | | |
|---|---|---|---|---|---|---|---|
| ENSMUSG00000107806 | 0 | 0 | 0 | 0 | 0 | 0 | 0 |
| ENSMUSG00000107808 | 0 | 0 | 0 | 0 | 0 | 0 | 0 |
| ENSMUSG00000107809 | 0.289064826 | 0.25 | 0 | 0 | 0 | 0.301029996 | 0.224243824 |
| ENSMUSG00000107810 | 0.811507563 | 0.646240625 | 0.896240625 | 0.92510993 | 0.632412477 | 0.84509804 | 0.710836105 |
| ENSMUSG00000107811 | 0 | 0 | 0 | 0 | 0 | 0 | 0 |
| ENSMUSG00000107812 | 0 | 0.25 | 0.25 | 0.25 | 0 | 0 | 0.224243824 |
| ENSMUSG00000107813 | 0 | 0 | 0 | 0.25 | 0 | 0.301029996 | 0.224243824 |
| ENSMUSG00000107814 | 0.289064826 | 0.396240625 | 0.396240625 | 0.25 | 0 | 0.301029996 | 0.448487648 |
| ENSMUSG00000107815 | 0 | 0 | 0 | 0 | 0 | 0 | 0 |
| ENSMUSG00000107816 | 0 | 0 | 0 | 0 | 0 | 0 | 0 |
| ENSMUSG00000107817 | 0 | 0 | 0 | 0 | 0 | 0 | 0 |
| ENSMUSG00000107818 | 0 | 0 | 0 | 0 | 0 | 0 | 0 |
| ENSMUSG00000107819 | 0 | 0 | 0 | 0 | 0 | 0 | 0 |
| ENSMUSG00000107821 | 0 | 0 | 0 | 0 | 0 | 0 | 0 |
| ENSMUSG00000107822 | 0 | 0 | 0 | 0 | 0 | 0 | 0 |
| ENSMUSG00000107823 | 0 | 0 | 0 | 0 | 0 | 0 | 0 |
| ENSMUSG00000107824 | 0.747221736 | 0.580482024 | 0.5 | 0 | 0.244650542 | 0 | 0.520678036 |
| ENSMUSG00000107825 | 0 | 0 | 0 | 0 | 0 | 0 | 0 |
| ENSMUSG00000107826 | 0 | 0 | 0 | 0 | 0 | 0 | 0 |
| ENSMUSG00000107828 | 0 | 0 | 0 | 0 | 0 | 0 | 0 |
| ENSMUSG00000107829 | 0 | 0 | 0 | 0 | 0 | 0 | 0 |
| ENSMUSG00000107830 | 0 | 0 | 0 | 0 | 0 | 0 | 0 |
| ENSMUSG00000107831 | 0 | 0 | 0 | 0 | 0 | 0 | 0 |
| ENSMUSG00000107832 | 0 | 0 | 0 | 0 | 0 | 0 | 0 |
| ENSMUSG00000107833 | 0 | 0 | 0.25 | 0 | 0.244650542 | 0 | 0 |
| ENSMUSG00000107834 | 0 | 0 | 0 | 0 | 0 | 0 | 0 |
| ENSMUSG00000107835 | 0 | 0 | 0 | 0 | 0 | 0 | 0 |
| ENSMUSG00000107836 | 0.671187741 | 0.396240625 | 0.646240625 | 0.701838731 | 0.733951626 | 0.698970004 | 0.710836105 |
| ENSMUSG00000107837 | 0 | 0 | 0 | 0 | 0 | 0 | 0 |
| ENSMUSG00000107838 | 0.867194479 | 0.646240625 | 0.75 | 0.75 | 0.489301084 | 0.77815125 | 0.520678036 |
| ENSMUSG00000107839 | 0 | 0 | 0 | 0 | 0 | 0 | 0 |
| ENSMUSG00000107840 | 0 | 0 | 0 | 0 | 0 | 0 | 0 |
| ENSMUSG00000107841 | 0 | 0.25 | 0 | 0 | 0 | 0 | 0 |
| ENSMUSG00000107842 | 0 | 0 | 0 | 0 | 0.244650542 | 0 | 0 |

| | | | | | | | |
|---|---|---|---|---|---|---|---|
| ENSMUSG00000107844 | 0 | 0 | 0 | 0 | 0 | 0 | 0 |
| ENSMUSG00000107845 | 0 | 0 | 0.25 | 0.25 | 0 | 0 | 0 |
| ENSMUSG00000107846 | 0.289064826 | 0.25 | 0 | 0 | 0 | 0.301029996 | 0.355418052 |
| ENSMUSG00000107847 | 0 | 0 | 0 | 0 | 0 | 0 | 0 |
| ENSMUSG00000107848 | 0 | 0 | 0 | 0 | 0 | 0 | 0 |
| ENSMUSG00000107849 | 0 | 0 | 0 | 0 | 0 | 0 | 0 |
| ENSMUSG00000107850 | 0 | 0 | 0 | 0 | 0 | 0 | 0 |
| ENSMUSG00000107851 | 0 | 0 | 0 | 0 | 0 | 0 | 0 |
| ENSMUSG00000107852 | 0 | 0.25 | 0 | 0.25 | 0 | 0 | 0 |
| ENSMUSG00000107853 | 0.289064826 | 0 | 0.25 | 0 | 0.387761935 | 0.477121255 | 0 |
| ENSMUSG00000107854 | 0 | 0 | 0 | 0 | 0 | 0 | 0 |
| ENSMUSG00000107855 | 0 | 0 | 0 | 0 | 0 | 0 | 0 |
| ENSMUSG00000107856 | 0 | 0.25 | 0 | 0 | 0 | 0 | 0 |
| ENSMUSG00000107857 | 0 | 0 | 0 | 0 | 0 | 0 | 0 |
| ENSMUSG00000107858 | 0 | 0 | 0 | 0 | 0 | 0 | 0 |
| ENSMUSG00000107859 | 0 | 0 | 0 | 0 | 0 | 0 | 0 |
| ENSMUSG00000107861 | 0 | 0 | 0 | 0 | 0.244650542 | 0 | 0.224243824 |
| ENSMUSG00000107862 | 0 | 0 | 0 | 0 | 0 | 0 | 0 |
| ENSMUSG00000107863 | 0 | 0 | 0 | 0 | 0 | 0 | 0 |
| ENSMUSG00000107864 | 0 | 0 | 0.25 | 0.25 | 0 | 0 | 0.224243824 |
| ENSMUSG00000107865 | 0.289064826 | 0.25 | 0 | 0 | 0 | 0.301029996 | 0.224243824 |
| ENSMUSG00000107866 | 0 | 0 | 0 | 0 | 0 | 0 | 0 |
| ENSMUSG00000107867 | 0 | 0 | 0 | 0 | 0 | 0 | 0 |
| ENSMUSG00000107868 | 0.91631382 | 1.160964047 | 1.04248125 | 0.951838731 | 0.733951626 | 0.903089987 | 0.74492186 |
| ENSMUSG00000107869 | 0 | 0 | 0 | 0 | 0 | 0 | 0 |
| ENSMUSG00000107870 | 0.578129653 | 0.580482024 | 0 | 0.864857905 | 0.489301084 | 0.954242509 | 0.710836105 |
| ENSMUSG00000107872 | 0 | 0 | 0 | 0 | 0 | 0 | 0 |
| ENSMUSG00000107873 | 0 | 0.25 | 0 | 0 | 0 | 0 | 0 |
| ENSMUSG00000107874 | 0 | 0 | 0 | 0 | 0 | 0 | 0 |
| ENSMUSG00000107875 | 0 | 0 | 0 | 0.25 | 0 | 0 | 0 |
| ENSMUSG00000107876 | 0 | 0 | 0 | 0 | 0 | 0 | 0 |
| ENSMUSG00000107877 | 0.747221736 | 0.79248125 | 0.830482024 | 0.830482024 | 0.931471446 | 0.77815125 | 0.916588299 |
| ENSMUSG00000107878 | 0 | 0 | 0 | 0 | 0.244650542 | 0 | 0 |
| ENSMUSG00000107879 | 0.91631382 | 0.951838731 | 0.896240625 | 1.061981878 | 1.057362051 | 1.176091259 | 1.089372236 |

| | | | | | | | |
|---|---|---|---|---|---|---|---|
| ENSMUSG00000107880 | 0.289064826 | 0 | 0 | 0 | 0 | 0 | 0 |
| ENSMUSG00000107881 | 2.504843822 | 2.288071211 | 2.313757142 | 1.593759858 | 1.478764298 | 1.643452676 | 1.441050184 |
| ENSMUSG00000107882 | 0.289064826 | 0.396240625 | 0 | 0 | 0.244650542 | 0.301029996 | 0 |
| ENSMUSG00000107883 | 0.289064826 | 0.396240625 | 0 | 0 | 0 | 0 | 0.448487648 |
| ENSMUSG00000107884 | 0 | 0.25 | 0 | 0 | 0 | 0 | 0.224243824 |
| ENSMUSG00000107885 | 0 | 0 | 0 | 0 | 0 | 0 | 0 |
| ENSMUSG00000107886 | 0.578129653 | 0.5 | 0.646240625 | 0.580482024 | 0.568060967 | 0.301029996 | 0.448487648 |
| ENSMUSG00000107887 | 0 | 0 | 0 | 0 | 0 | 0 | 0 |
| ENSMUSG00000107888 | 0 | 0 | 0 | 0 | 0 | 0 | 0 |
| ENSMUSG00000107889 | 0 | 0 | 0 | 0 | 0 | 0 | 0 |
| ENSMUSG00000107890 | 0 | 0 | 0 | 0 | 0 | 0 | 0 |
| ENSMUSG00000107891 | 0.289064826 | 0 | 0 | 0 | 0 | 0 | 0.224243824 |
| ENSMUSG00000107892 | 0 | 0 | 0 | 0.396240625 | 0.387761935 | 0.301029996 | 0.224243824 |
| ENSMUSG00000107893 | 0 | 0 | 0 | 0 | 0 | 0 | 0 |
| ENSMUSG00000107894 | 0.747221736 | 0.79248125 | 0.646240625 | 0.646240625 | 0.77552387 | 0 | 0.520678036 |
| ENSMUSG00000107895 | 0 | 0 | 0 | 0 | 0 | 0 | 0 |
| ENSMUSG00000107896 | 0 | 0 | 0 | 0 | 0 | 0 | 0 |
| ENSMUSG00000107898 | 0 | 0 | 0 | 0 | 0 | 0 | 0 |
| ENSMUSG00000107899 | 0 | 0 | 0 | 0 | 0 | 0 | 0 |
| ENSMUSG00000107900 | 0 | 0 | 0 | 0 | 0 | 0 | 0 |
| ENSMUSG00000107901 | 0 | 0 | 0 | 0 | 0 | 0 | 0 |
| ENSMUSG00000107902 | 0.45815691 | 0.396240625 | 0 | 0 | 0.489301084 | 0 | 0.224243824 |
| ENSMUSG00000107903 | 0.289064826 | 0 | 0 | 0 | 0 | 0 | 0 |
| ENSMUSG00000107904 | 0 | 0 | 0 | 0 | 0 | 0 | 0 |
| ENSMUSG00000107905 | 0 | 0 | 0 | 0 | 0 | 0 | 0 |
| ENSMUSG00000107906 | 0 | 0 | 0 | 0 | 0 | 0 | 0.224243824 |
| ENSMUSG00000107907 | 0 | 0 | 0 | 0 | 0 | 0 | 0 |
| ENSMUSG00000107908 | 0 | 0 | 0 | 0 | 0 | 0 | 0 |
| ENSMUSG00000107909 | 0.45815691 | 0.25 | 0.830482024 | 0.580482024 | 0.632412477 | 0.477121255 | 0.520678036 |
| ENSMUSG00000107911 | 0.45815691 | 0.396240625 | 0.25 | 0.396240625 | 0.387761935 | 0.602059991 | 0.224243824 |
| ENSMUSG00000107913 | 0 | 0 | 0 | 0 | 0 | 0 | 0 |
| ENSMUSG00000107914 | 0 | 0 | 0 | 0 | 0 | 0 | 0 |
| ENSMUSG00000107915 | 0.289064826 | 0.396240625 | 0 | 0.25 | 0.489301084 | 0 | 0 |
| ENSMUSG00000107916 | 0 | 0 | 0 | 0 | 0 | 0 | 0 |

| | | | | | | | |
|---|---|---|---|---|---|---|---|
| ENSMUSG00000107917 | 0 | 0 | 0 | 0 | 0 | 0 | 0 |
| ENSMUSG00000107918 | 0 | 0 | 0 | 0 | 0 | 0 | 0 |
| ENSMUSG00000107919 | 0 | 0 | 0 | 0 | 0 | 0 | 0 |
| ENSMUSG00000107920 | 0 | 0 | 0 | 0 | 0 | 0 | 0 |
| ENSMUSG00000107921 | 0 | 0 | 0 | 0 | 0 | 0 | 0 |
| ENSMUSG00000107922 | 0 | 0 | 0.25 | 0 | 0 | 0.301029996 | 0 |
| ENSMUSG00000107923 | 0 | 0 | 0 | 0 | 0 | 0 | 0 |
| ENSMUSG00000107924 | 0 | 0 | 0 | 0 | 0 | 0 | 0 |
| ENSMUSG00000107925 | 0 | 0 | 0 | 0 | 0 | 0 | 0 |
| ENSMUSG00000107926 | 0 | 0 | 0 | 0 | 0 | 0 | 0 |
| ENSMUSG00000107927 | 0 | 0 | 0.25 | 0.25 | 0.244650542 | 0.301029996 | 0 |
| ENSMUSG00000107928 | 0 | 0.25 | 0 | 0.396240625 | 0 | 0.301029996 | 0 |
| ENSMUSG00000107929 | 0 | 0 | 0 | 0 | 0 | 0 | 0 |
| ENSMUSG00000107930 | 0 | 0 | 0 | 0 | 0 | 0 | 0 |
| ENSMUSG00000107931 | 0 | 0 | 0 | 0 | 0 | 0 | 0 |
| ENSMUSG00000107932 | 0.671187741 | 0.396240625 | 0.92510993 | 0.5 | 0.387761935 | 0.477121255 | 0.629532004 |
| ENSMUSG00000107933 | 0 | 0 | 0 | 0 | 0 | 0 | 0 |
| ENSMUSG00000107934 | 0 | 0 | 0 | 0 | 0 | 0 | 0 |
| ENSMUSG00000107935 | 0 | 0 | 0 | 0 | 0 | 0 | 0 |
| ENSMUSG00000107936 | 0 | 0.25 | 0 | 0 | 0 | 0 | 0 |
| ENSMUSG00000107937 | 0.91631382 | 0.79248125 | 0.75 | 0.75 | 0.812711509 | 0.698970004 | 0.74492186 |
| ENSMUSG00000107938 | 0 | 0 | 0 | 0 | 0 | 0 | 0 |
| ENSMUSG00000107940 | 0 | 0 | 0 | 0 | 0 | 0 | 0 |
| ENSMUSG00000107941 | 0 | 0 | 0 | 0 | 0 | 0 | 0 |
| ENSMUSG00000107942 | 0 | 0.25 | 0 | 0 | 0 | 0 | 0 |
| ENSMUSG00000107943 | 0 | 0 | 0 | 0 | 0 | 0 | 0 |
| ENSMUSG00000107944 | 0 | 0.25 | 0.396240625 | 0.396240625 | 0.489301084 | 0 | 0.629532004 |
| ENSMUSG00000107945 | 0 | 0 | 0 | 0 | 0 | 0.301029996 | 0.224243824 |
| ENSMUSG00000107946 | 0 | 0.25 | 0.580482024 | 0 | 0.387761935 | 0 | 0.224243824 |
| ENSMUSG00000107947 | 0 | 0 | 0 | 0 | 0 | 0 | 0 |
| ENSMUSG00000107948 | 0 | 0 | 0 | 0 | 0 | 0 | 0 |
| ENSMUSG00000107949 | 0 | 0 | 0.25 | 0 | 0.244650542 | 0 | 0 |
| ENSMUSG00000107950 | 1.036286563 | 1.080482024 | 0.92510993 | 0.580482024 | 0.686820904 | 0.602059991 | 0.803905701 |
| ENSMUSG00000107951 | 1 | 1.114857905 | 0.951838731 | 1.146240625 | 1.121713561 | 1.380211242 | 0.896975297 |

| | | | | | | | |
|---|---|---|---|---|---|---|---|
| ENSMUSG00000107952 | 0 | 0.25 | 0 | 0 | 0 | 0.301029996 | 0 |
| ENSMUSG00000107953 | 0 | 0 | 0 | 0 | 0 | 0 | 0 |
| ENSMUSG00000107954 | 0 | 0.396240625 | 0 | 0.25 | 0.244650542 | 0 | 0.224243824 |
| ENSMUSG00000107955 | 0 | 0.396240625 | 0.25 | 0.25 | 0.244650542 | 0.301029996 | 0 |
| ENSMUSG00000107956 | 0 | 0 | 0 | 0 | 0 | 0 | 0 |
| ENSMUSG00000107957 | 0 | 0 | 0 | 0 | 0 | 0 | 0 |
| ENSMUSG00000107959 | 0 | 0 | 0 | 0 | 0 | 0 | 0 |
| ENSMUSG00000107960 | 0 | 0 | 0 | 0 | 0 | 0 | 0 |
| ENSMUSG00000107961 | 0.289064826 | 0 | 0 | 0.25 | 0 | 0 | 0 |
| ENSMUSG00000107962 | 1.783508299 | 1.618933358 | 1.505591953 | 1.606566189 | 1.563883923 | 1.633468456 | 1.335191751 |
| ENSMUSG00000107964 | 0 | 0 | 0 | 0 | 0 | 0 | 0 |
| ENSMUSG00000107965 | 0 | 0 | 0 | 0 | 0 | 0 | 0 |
| ENSMUSG00000107966 | 0 | 0 | 0 | 0 | 0 | 0 | 0 |
| ENSMUSG00000107967 | 0 | 0 | 0.25 | 0.25 | 0 | 0.477121255 | 0 |
| ENSMUSG00000107968 | 0 | 0 | 0 | 0 | 0 | 0 | 0 |
| ENSMUSG00000107969 | 0 | 0 | 0 | 0 | 0 | 0 | 0 |
| ENSMUSG00000107970 | 0.45815691 | 0 | 0.25 | 0 | 0.489301084 | 0.301029996 | 0.355418052 |
| ENSMUSG00000107971 | 0.960252568 | 0.896240625 | 1.061981878 | 0.951838731 | 1.020174412 | 0.84509804 | 0.853775828 |
| ENSMUSG00000107972 | 0 | 0 | 0 | 0 | 0 | 0.301029996 | 0 |
| ENSMUSG00000107973 | 0 | 0 | 0 | 0 | 0 | 0 | 0 |
| ENSMUSG00000107975 | 0 | 0 | 0 | 0 | 0 | 0 | 0 |
| ENSMUSG00000107976 | 0 | 0.25 | 0 | 0.25 | 0.489301084 | 0.477121255 | 0.355418052 |
| ENSMUSG00000107977 | 0 | 0.25 | 0.5 | 0.25 | 0.489301084 | 0 | 0.355418052 |
| ENSMUSG00000107978 | 0 | 0 | 0 | 0 | 0 | 0 | 0 |
| ENSMUSG00000107979 | 0 | 0 | 0 | 0 | 0 | 0 | 0 |
| ENSMUSG00000107980 | 0 | 0 | 0 | 0 | 0 | 0 | 0 |
| ENSMUSG00000107981 | 0 | 0 | 0 | 0.25 | 0 | 0 | 0 |
| ENSMUSG00000107982 | 0 | 0 | 0 | 0 | 0 | 0 | 0 |
| ENSMUSG00000107983 | 0 | 0 | 0 | 0 | 0 | 0 | 0 |
| ENSMUSG00000107984 | 0 | 0.25 | 0 | 0.25 | 0 | 0 | 0 |
| ENSMUSG00000107985 | 0 | 0 | 0.396240625 | 0.580482024 | 0.387761935 | 0.698970004 | 0.448487648 |
| ENSMUSG00000107986 | 0 | 0 | 0 | 0 | 0 | 0 | 0 |
| ENSMUSG00000107987 | 0 | 0 | 0 | 0 | 0 | 0 | 0 |
| ENSMUSG00000107988 | 0 | 0 | 0 | 0 | 0 | 0 | 0 |

| | | | | | | | |
|---|---|---|---|---|---|---|---|
| ENSMUSG00000107989 | 0.811507563 | 0 | 0.5 | 0.580482024 | 0.387761935 | 0 | 0.448487648 |
| ENSMUSG00000107990 | 0.289064826 | 0.25 | 0.396240625 | 0.25 | 0.489301084 | 0.301029996 | 0.448487648 |
| ENSMUSG00000107991 | 0.289064826 | 0 | 0 | 0.25 | 0 | 0.301029996 | 0 |
| ENSMUSG00000107992 | 0 | 0 | 0 | 0 | 0 | 0 | 0 |
| ENSMUSG00000107993 | 0 | 0 | 0 | 0 | 0 | 0 | 0 |
| ENSMUSG00000107994 | 0.45815691 | 0.5 | 0.396240625 | 0.25 | 0 | 0 | 0.355418052 |
| ENSMUSG00000107995 | 0 | 0 | 0 | 0 | 0.244650542 | 0 | 0 |
| ENSMUSG00000107996 | 0 | 0 | 0.5 | 0.25 | 0 | 0 | 0.224243824 |
| ENSMUSG00000107997 | 0.45815691 | 0.5 | 0.396240625 | 0 | 0.568060967 | 0.301029996 | 0.224243824 |
| ENSMUSG00000107998 | 0 | 0 | 0 | 0 | 0 | 0 | 0 |
| ENSMUSG00000107999 | 0 | 0 | 0 | 0 | 0 | 0 | 0 |
| ENSMUSG00000108000 | 0 | 0 | 0 | 0 | 0 | 0 | 0 |
| ENSMUSG00000108002 | 0 | 0 | 0 | 0 | 0 | 0 | 0 |
| ENSMUSG00000108003 | 0 | 0 | 0 | 0 | 0 | 0 | 0 |
| ENSMUSG00000108004 | 0.578129653 | 0.580482024 | 0.396240625 | 0.580482024 | 0.387761935 | 0.602059991 | 0.224243824 |
| ENSMUSG00000108005 | 0 | 0 | 0 | 0 | 0 | 0 | 0 |
| ENSMUSG00000108006 | 0 | 0.25 | 0 | 0 | 0 | 0 | 0 |
| ENSMUSG00000108007 | 0 | 0 | 0 | 0 | 0 | 0 | 0 |
| ENSMUSG00000108008 | 0 | 0 | 0.25 | 0.25 | 0 | 0.301029996 | 0 |
| ENSMUSG00000108009 | 0 | 0 | 0 | 0 | 0 | 0 | 0.224243824 |
| ENSMUSG00000108010 | 0.578129653 | 0.646240625 | 0.5 | 0.25 | 0.387761935 | 0 | 0.448487648 |
| ENSMUSG00000108011 | 0 | 0 | 0 | 0 | 0 | 0 | 0 |
| ENSMUSG00000108012 | 0 | 0 | 0 | 0 | 0 | 0 | 0 |
| ENSMUSG00000108013 | 0 | 0 | 0 | 0 | 0 | 0 | 0 |
| ENSMUSG00000108014 | 0 | 0 | 0 | 0 | 0 | 0 | 0 |
| ENSMUSG00000108015 | 0.289064826 | 0 | 0 | 0 | 0 | 0 | 0 |
| ENSMUSG00000108016 | 0 | 0 | 0 | 0 | 0 | 0 | 0 |
| ENSMUSG00000108017 | 0 | 0 | 0 | 0 | 0 | 0 | 0 |
| ENSMUSG00000108018 | 0 | 0 | 0 | 0 | 0 | 0 | 0 |
| ENSMUSG00000108019 | 0 | 0 | 0 | 0 | 0 | 0 | 0 |
| ENSMUSG00000108020 | 0.747221736 | 0.701838731 | 0.701838731 | 0.396240625 | 0.244650542 | 0.301029996 | 0.224243824 |
| ENSMUSG00000108021 | 0 | 0 | 0 | 0 | 0 | 0 | 0 |
| ENSMUSG00000108022 | 0 | 0 | 0 | 0 | 0 | 0 | 0 |
| ENSMUSG00000108023 | 0 | 0.25 | 0 | 0 | 0 | 0 | 0 |

| | | | | | | | |
|---|---|---|---|---|---|---|---|
| ENSMUSG00000108024 | 0.289064826 | 0.25 | 0.25 | 0 | 0.489301084 | 0.301029996 | 0 |
| ENSMUSG00000108025 | 0 | 0 | 0 | 0 | 0 | 0 | 0 |
| ENSMUSG00000108026 | 0.867194479 | 0.75 | 0.864857905 | 1 | 0.846351821 | 0.602059991 | 0.775756176 |
| ENSMUSG00000108027 | 0.289064826 | 0.5 | 0.25 | 0.580482024 | 0.686820904 | 0.301029996 | 0.629532004 |
| ENSMUSG00000108028 | 0 | 0 | 0 | 0 | 0 | 0 | 0 |
| ENSMUSG00000108029 | 0 | 0 | 0 | 0 | 0 | 0 | 0 |
| ENSMUSG00000108030 | 0.289064826 | 0.5 | 0.25 | 0.646240625 | 0.387761935 | 0 | 0.448487648 |
| ENSMUSG00000108032 | 0 | 0 | 0 | 0 | 0 | 0 | 0 |
| ENSMUSG00000108033 | 0.45815691 | 0.25 | 0.396240625 | 0.5 | 0.489301084 | 0 | 0.224243824 |
| ENSMUSG00000108034 | 0.45815691 | 0.5 | 0.646240625 | 0.396240625 | 0.568060967 | 0.301029996 | 0.579661877 |
| ENSMUSG00000108035 | 0 | 0 | 0 | 0 | 0 | 0 | 0 |
| ENSMUSG00000108036 | 0 | 0 | 0 | 0 | 0 | 0 | 0 |
| ENSMUSG00000108037 | 0.45815691 | 0.396240625 | 0.5 | 0.396240625 | 0.632412477 | 0.477121255 | 0.448487648 |
| ENSMUSG00000108038 | 0 | 0 | 0 | 0 | 0 | 0 | 0 |
| ENSMUSG00000108039 | 0 | 0 | 0 | 0 | 0 | 0 | 0 |
| ENSMUSG00000108040 | 0.45815691 | 0 | 0 | 0 | 0 | 0 | 0 |
| ENSMUSG00000108041 | 0 | 0 | 0 | 0 | 0 | 0 | 0 |
| ENSMUSG00000108042 | 0.811507563 | 0.830482024 | 0.92510993 | 0.896240625 | 0.846351821 | 0.954242509 | 0.803905701 |
| ENSMUSG00000108043 | 0 | 0 | 0 | 0 | 0 | 0 | 0.224243824 |
| ENSMUSG00000108044 | 0 | 0 | 0 | 0 | 0 | 0 | 0 |
| ENSMUSG00000108045 | 0 | 0 | 0.25 | 0 | 0 | 0 | 0 |
| ENSMUSG00000108046 | 0.747221736 | 0.396240625 | 0.79248125 | 0.830482024 | 0.686820904 | 0.477121255 | 0.520678036 |
| ENSMUSG00000108047 | 0 | 0 | 0 | 0 | 0 | 0 | 0 |
| ENSMUSG00000108048 | 0 | 0 | 0.396240625 | 0.25 | 0.244650542 | 0 | 0.355418052 |
| ENSMUSG00000108049 | 1.289064826 | 1.26109853 | 1.29248125 | 1.201838731 | 1.106691885 | 1.041392685 | 0.916588299 |
| ENSMUSG00000108050 | 0 | 0 | 0 | 0.25 | 0 | 0 | 0 |
| ENSMUSG00000108052 | 0 | 0 | 0 | 0 | 0 | 0 | 0 |
| ENSMUSG00000108053 | 1.181541736 | 1.04248125 | 1 | 0.976722649 | 0.931471446 | 0.903089987 | 0.775756176 |
| ENSMUSG00000108054 | 0 | 0 | 0 | 0 | 0 | 0 | 0 |
| ENSMUSG00000108055 | 0.45815691 | 0.79248125 | 0.396240625 | 0.701838731 | 0.387761935 | 0 | 0.448487648 |
| ENSMUSG00000108056 | 0.289064826 | 0 | 0.25 | 0.25 | 0.387761935 | 0.301029996 | 0.224243824 |
| ENSMUSG00000108057 | 0 | 0.25 | 0 | 0 | 0 | 0 | 0 |
| ENSMUSG00000108058 | 0 | 0 | 0 | 0 | 0 | 0 | 0 |
| ENSMUSG00000108059 | 0.578129653 | 0.580482024 | 0 | 0.25 | 0 | 0.602059991 | 0.355418052 |

| | | | | | | | |
|---|---|---|---|---|---|---|---|
| ENSMUSG00000108060 | 0 | 0 | 0 | 0.25 | 0 | 0 | 0 |
| ENSMUSG00000108061 | 0 | 0 | 0 | 0 | 0 | 0 | 0 |
| ENSMUSG00000108062 | 0 | 0 | 0 | 0 | 0 | 0 | 0 |
| ENSMUSG00000108063 | 0 | 0 | 0 | 0 | 0 | 0 | 0 |
| ENSMUSG00000108064 | 0.867194479 | 0.5 | 0.5 | 0.951838731 | 0.931471446 | 0.77815125 | 0.579661877 |
| ENSMUSG00000108065 | 0 | 0 | 0 | 0 | 0 | 0 | 0 |
| ENSMUSG00000108066 | 0 | 0 | 0 | 0 | 0 | 0 | 0 |
| ENSMUSG00000108067 | 0 | 0 | 0 | 0 | 0 | 0 | 0 |
| ENSMUSG00000108068 | 0 | 0 | 0 | 0.25 | 0 | 0.477121255 | 0.224243824 |
| ENSMUSG00000108069 | 0 | 0 | 0 | 0 | 0 | 0 | 0 |
| ENSMUSG00000108070 | 0 | 0 | 0 | 0 | 0 | 0 | 0 |
| ENSMUSG00000108071 | 0 | 0 | 0 | 0 | 0 | 0 | 0 |
| ENSMUSG00000108072 | 0.45815691 | 0.5 | 0.646240625 | 0.580482024 | 0.77552387 | 0.77815125 | 0.520678036 |
| ENSMUSG00000108073 | 0 | 0 | 0 | 0 | 0 | 0 | 0 |
| ENSMUSG00000108074 | 0 | 0 | 0 | 0 | 0 | 0 | 0 |
| ENSMUSG00000108075 | 0 | 0 | 0 | 0 | 0 | 0 | 0 |
| ENSMUSG00000108076 | 0 | 0 | 0 | 0 | 0 | 0 | 0 |
| ENSMUSG00000108077 | 0 | 0 | 0 | 0 | 0 | 0 | 0.224243824 |
| ENSMUSG00000108078 | 0 | 0 | 0 | 0 | 0 | 0 | 0 |
| ENSMUSG00000108079 | 0 | 0 | 0.25 | 0 | 0.244650542 | 0 | 0.520678036 |
| ENSMUSG00000108080 | 0 | 0 | 0.25 | 0.25 | 0.387761935 | 0.477121255 | 0.224243824 |
| ENSMUSG00000108081 | 0 | 0 | 0 | 0 | 0 | 0 | 0 |
| ENSMUSG00000108082 | 0 | 0 | 0 | 0 | 0 | 0 | 0 |
| ENSMUSG00000108083 | 0 | 0 | 0 | 0 | 0 | 0 | 0 |
| ENSMUSG00000108084 | 0.45815691 | 0.396240625 | 0 | 0.25 | 0 | 0.301029996 | 0.224243824 |
| ENSMUSG00000108085 | 0 | 0 | 0 | 0 | 0 | 0.301029996 | 0 |
| ENSMUSG00000108086 | 0 | 0.396240625 | 0 | 0 | 0 | 0 | 0 |
| ENSMUSG00000108087 | 0 | 0 | 0 | 0 | 0 | 0 | 0 |
| ENSMUSG00000108088 | 0 | 0 | 0 | 0 | 0 | 0 | 0 |
| ENSMUSG00000108089 | 0.671187741 | 0.396240625 | 0.396240625 | 0 | 0.568060967 | 0.301029996 | 0.224243824 |
| ENSMUSG00000108090 | 0 | 0 | 0 | 0 | 0 | 0 | 0 |
| ENSMUSG00000108091 | 0 | 0 | 0 | 0 | 0 | 0 | 0 |
| ENSMUSG00000108092 | 1.036286563 | 0.951838731 | 0.701838731 | 0.92510993 | 1 | 0.477121255 | 0.803905701 |
| ENSMUSG00000108093 | 0 | 0 | 0 | 0 | 0 | 0 | 0 |

| | | | | | | | |
|---|---|---|---|---|---|---|---|
| ENSMUSG00000108094 | 0 | 0 | 0 | 0 | 0 | 0 | 0 |
| ENSMUSG00000108095 | 0 | 0 | 0 | 0 | 0 | 0 | 0 |
| ENSMUSG00000108096 | 0 | 0 | 0 | 0 | 0 | 0 | 0 |
| ENSMUSG00000108097 | 0 | 0 | 0 | 0 | 0 | 0 | 0 |
| ENSMUSG00000108098 | 0 | 0 | 0 | 0 | 0 | 0 | 0 |
| ENSMUSG00000108099 | 0 | 0 | 0 | 0 | 0 | 0 | 0 |
| ENSMUSG00000108100 | 0 | 0 | 0.580482024 | 0.396240625 | 0 | 0.301029996 | 0.224243824 |
| ENSMUSG00000108101 | 0 | 0.396240625 | 0.396240625 | 0.396240625 | 0.568060967 | 0.301029996 | 0.355418052 |
| ENSMUSG00000108102 | 0 | 0.25 | 0 | 0 | 0.244650542 | 0.301029996 | 0 |
| ENSMUSG00000108103 | 0 | 0.25 | 0.396240625 | 0 | 0 | 0 | 0 |
| ENSMUSG00000108104 | 0 | 0 | 0 | 0 | 0 | 0 | 0 |
| ENSMUSG00000108105 | 0 | 0.396240625 | 0.25 | 0 | 0 | 0 | 0.448487648 |
| ENSMUSG00000108106 | 0 | 0 | 0 | 0 | 0 | 0 | 0 |
| ENSMUSG00000108107 | 0 | 0 | 0 | 0 | 0.244650542 | 0 | 0.224243824 |
| ENSMUSG00000108108 | 0.45815691 | 0.396240625 | 0 | 0 | 0.244650542 | 0 | 0.224243824 |
| ENSMUSG00000108109 | 0 | 0 | 0.25 | 0.396240625 | 0 | 0 | 0 |
| ENSMUSG00000108110 | 0 | 0 | 0 | 0 | 0 | 0 | 0 |
| ENSMUSG00000108111 | 0 | 0 | 0 | 0 | 0 | 0 | 0 |
| ENSMUSG00000108112 | 0 | 0 | 0 | 0 | 0 | 0 | 0 |
| ENSMUSG00000108113 | 0 | 0 | 0 | 0 | 0 | 0 | 0 |
| ENSMUSG00000108114 | 0 | 0 | 0 | 0 | 0 | 0 | 0 |
| ENSMUSG00000108115 | 0.289064826 | 0.25 | 0 | 0.396240625 | 0.489301084 | 0 | 0 |
| ENSMUSG00000108117 | 0 | 0 | 0 | 0 | 0 | 0 | 0 |
| ENSMUSG00000108118 | 0 | 0 | 0 | 0 | 0 | 0 | 0 |
| ENSMUSG00000108119 | 0 | 0 | 0 | 0 | 0.244650542 | 0 | 0 |
| ENSMUSG00000108120 | 0.289064826 | 0.25 | 0.25 | 0.25 | 0 | 0.301029996 | 0.224243824 |
| ENSMUSG00000108121 | 0 | 0 | 0 | 0 | 0 | 0 | 0 |
| ENSMUSG00000108122 | 0 | 0.5 | 0 | 0.396240625 | 0 | 0.477121255 | 0.520678036 |
| ENSMUSG00000108123 | 0 | 0 | 0 | 0 | 0 | 0 | 0 |
| ENSMUSG00000108124 | 0 | 0 | 0 | 0.5 | 0 | 0 | 0 |
| ENSMUSG00000108125 | 0.289064826 | 0.25 | 0.25 | 0.25 | 0.387761935 | 0 | 0.355418052 |
| ENSMUSG00000108126 | 0.45815691 | 0.25 | 0.396240625 | 0 | 0 | 0 | 0 |
| ENSMUSG00000108127 | 0 | 0 | 0 | 0 | 0 | 0 | 0 |
| ENSMUSG00000108128 | 0 | 0 | 0 | 0 | 0 | 0 | 0 |

| | | | | | | | |
|---|---|---|---|---|---|---|---|
| ENSMUSG00000108129 | 0 | 0.396240625 | 0.396240625 | 0 | 0.489301084 | 0 | 0 |
| ENSMUSG00000108130 | 0 | 0 | 0 | 0 | 0 | 0 | 0 |
| ENSMUSG00000108131 | 0 | 0 | 0 | 0 | 0 | 0 | 0 |
| ENSMUSG00000108132 | 0.578129653 | 0.5 | 0.5 | 0 | 0 | 0 | 0.355418052 |
| ENSMUSG00000108133 | 0 | 0 | 0 | 0 | 0 | 0 | 0 |
| ENSMUSG00000108134 | 0 | 0 | 0 | 0 | 0 | 0 | 0 |
| ENSMUSG00000108135 | 0 | 0 | 0 | 0 | 0 | 0 | 0 |
| ENSMUSG00000108136 | 0 | 0 | 0 | 0 | 0 | 0 | 0 |
| ENSMUSG00000108137 | 0.867194479 | 0.580482024 | 0.896240625 | 0.864857905 | 0.77552387 | 0.477121255 | 0.672731473 |
| ENSMUSG00000108138 | 0 | 0 | 0 | 0 | 0 | 0 | 0 |
| ENSMUSG00000108139 | 0 | 0 | 0 | 0 | 0 | 0 | 0 |
| ENSMUSG00000108140 | 0 | 0 | 0 | 0 | 0 | 0 | 0 |
| ENSMUSG00000108141 | 0 | 0 | 0.25 | 0 | 0 | 0 | 0 |
| ENSMUSG00000108142 | 0 | 0 | 0.396240625 | 0.396240625 | 0.244650542 | 0.301029996 | 0.224243824 |
| ENSMUSG00000108143 | 0.289064826 | 0 | 0 | 0.396240625 | 0.387761935 | 0 | 0.448487648 |
| ENSMUSG00000108144 | 0 | 0 | 0 | 0 | 0 | 0 | 0 |
| ENSMUSG00000108145 | 0 | 0 | 0 | 0 | 0 | 0 | 0 |
| ENSMUSG00000108146 | 0 | 0 | 0 | 0 | 0 | 0 | 0 |
| ENSMUSG00000108147 | 0 | 0 | 0 | 0 | 0 | 0 | 0 |
| ENSMUSG00000108148 | 0 | 0 | 0 | 0 | 0 | 0 | 0 |
| ENSMUSG00000108149 | 0 | 0 | 0 | 0 | 0 | 0 | 0 |
| ENSMUSG00000108150 | 0 | 0 | 0 | 0 | 0 | 0 | 0 |
| ENSMUSG00000108151 | 0 | 0 | 0 | 0 | 0 | 0 | 0 |
| ENSMUSG00000108152 | 0 | 0 | 0 | 0 | 0 | 0 | 0.224243824 |
| ENSMUSG00000108153 | 0 | 0 | 0 | 0 | 0 | 0 | 0 |
| ENSMUSG00000108154 | 0.45815691 | 0.646240625 | 0.646240625 | 0.25 | 0.244650542 | 0.301029996 | 0.520678036 |
| ENSMUSG00000108155 | 0 | 0.25 | 0.396240625 | 0.25 | 0.387761935 | 0.301029996 | 0.448487648 |
| ENSMUSG00000108156 | 0 | 0 | 0 | 0 | 0 | 0 | 0 |
| ENSMUSG00000108157 | 0 | 0 | 0 | 0 | 0 | 0 | 0 |
| ENSMUSG00000108158 | 0 | 0 | 0 | 0 | 0 | 0 | 0 |
| ENSMUSG00000108160 | 0 | 0 | 0 | 0 | 0 | 0 | 0 |
| ENSMUSG00000108161 | 0 | 0 | 0 | 0 | 0 | 0 | 0 |
| ENSMUSG00000108162 | 0.45815691 | 0.580482024 | 0.25 | 0.396240625 | 0.568060967 | 0.301029996 | 0.224243824 |
| ENSMUSG00000108163 | 0 | 0 | 0 | 0 | 0 | 0 | 0 |

| | | | | | | | |
|---|---|---|---|---|---|---|---|
| ENSMUSG00000108164 | 0 | 0 | 0 | 0 | 0 | 0 | 0 |
| ENSMUSG00000108166 | 0.578129653 | 0.646240625 | 0.79248125 | 0.396240625 | 0.387761935 | 0.698970004 | 0.672731473 |
| ENSMUSG00000108167 | 0 | 0 | 0 | 0 | 0 | 0 | 0 |
| ENSMUSG00000108168 | 0 | 0 | 0.25 | 0 | 0 | 0 | 0 |
| ENSMUSG00000108169 | 0 | 0 | 0 | 0 | 0 | 0 | 0 |
| ENSMUSG00000108170 | 0 | 0 | 0 | 0 | 0.244650542 | 0 | 0 |
| ENSMUSG00000108171 | 0 | 0.25 | 0 | 0 | 0.244650542 | 0 | 0 |
| ENSMUSG00000108172 | 0 | 0 | 0 | 0 | 0 | 0 | 0 |
| ENSMUSG00000108173 | 1.862428343 | 1.720660762 | 1.660964047 | 1.664552871 | 1.576269619 | 1.77815125 | 1.517723536 |
| ENSMUSG00000108174 | 0 | 0 | 0 | 0 | 0 | 0 | 0 |
| ENSMUSG00000108175 | 0 | 0 | 0 | 0 | 0 | 0 | 0 |
| ENSMUSG00000108176 | 1.069666964 | 1.061981878 | 1.02186571 | 1.02186571 | 1.091002363 | 0.903089987 | 0.984950056 |
| ENSMUSG00000108177 | 0 | 0 | 0 | 0 | 0.244650542 | 0 | 0 |
| ENSMUSG00000108178 | 0 | 0 | 0 | 0 | 0 | 0 | 0 |
| ENSMUSG00000108179 | 0.289064826 | 0.701838731 | 0.25 | 0 | 0.244650542 | 0.301029996 | 0.520678036 |
| ENSMUSG00000108180 | 0 | 0 | 0.25 | 0.25 | 0 | 0.301029996 | 0 |
| ENSMUSG00000108181 | 1.205378646 | 1.214495249 | 1.302363341 | 1.26109853 | 1.212046813 | 1.342422681 | 1.054044578 |
| ENSMUSG00000108183 | 0.671187741 | 0.5 | 0.396240625 | 0 | 0.244650542 | 0 | 0.355418052 |
| ENSMUSG00000108184 | 0 | 0 | 0 | 0 | 0 | 0 | 0 |
| ENSMUSG00000108185 | 0 | 0 | 0 | 0 | 0 | 0 | 0 |
| ENSMUSG00000108186 | 0 | 0 | 0 | 0 | 0 | 0.301029996 | 0 |
| ENSMUSG00000108187 | 0 | 0 | 0 | 0 | 0 | 0 | 0 |
| ENSMUSG00000108188 | 0 | 0 | 0 | 0 | 0 | 0 | 0.224243824 |
| ENSMUSG00000108189 | 0.867194479 | 0.79248125 | 0.5 | 0.580482024 | 0.686820904 | 0.602059991 | 0.710836105 |
| ENSMUSG00000108190 | 0.45815691 | 0.5 | 0 | 0.25 | 0.244650542 | 0 | 0.355418052 |
| ENSMUSG00000108191 | 0 | 0 | 0 | 0 | 0 | 0 | 0 |
| ENSMUSG00000108192 | 0 | 0 | 0 | 0 | 0 | 0 | 0 |
| ENSMUSG00000108193 | 0 | 0 | 0.25 | 0 | 0 | 0 | 0 |
| ENSMUSG00000108194 | 0 | 0 | 0 | 0 | 0 | 0 | 0 |
| ENSMUSG00000108195 | 0.671187741 | 0.396240625 | 0 | 0 | 0.244650542 | 0.301029996 | 0.224243824 |
| ENSMUSG00000108196 | 0 | 0 | 0 | 0.25 | 0 | 0 | 0 |
| ENSMUSG00000108197 | 0 | 0 | 0.396240625 | 0 | 0 | 0.301029996 | 0.224243824 |
| ENSMUSG00000108198 | 0 | 0 | 0 | 0 | 0 | 0 | 0 |
| ENSMUSG00000108199 | 1.227926429 | 1.226722649 | 1.17510993 | 1.29248125 | 1.136121934 | 1.301029996 | 1.110947927 |

| | | | | | | | |
|---|---|---|---|---|---|---|---|
| ENSMUSG00000108200 | 0.289064826 | 0.580482024 | 0.396240625 | 0 | 0 | 0.602059991 | 0 |
| ENSMUSG00000108201 | 0 | 0 | 0 | 0 | 0 | 0 | 0 |
| ENSMUSG00000108202 | 0 | 0 | 0 | 0 | 0 | 0 | 0 |
| ENSMUSG00000108203 | 0 | 0 | 0 | 0 | 0 | 0 | 0 |
| ENSMUSG00000108204 | 0 | 0.396240625 | 0 | 0.25 | 0.387761935 | 0.477121255 | 0.355418052 |
| ENSMUSG00000108205 | 0 | 0 | 0 | 0 | 0 | 0 | 0 |
| ENSMUSG00000108206 | 0 | 0.25 | 0 | 0 | 0 | 0 | 0 |
| ENSMUSG00000108207 | 1.578129653 | 1.557204673 | 1.580482024 | 1.04248125 | 1.091002363 | 0.84509804 | 0.952571511 |
| ENSMUSG00000108208 | 0 | 0 | 0 | 0 | 0 | 0 | 0 |
| ENSMUSG00000108209 | 0 | 0 | 0 | 0 | 0 | 0 | 0 |
| ENSMUSG00000108210 | 0 | 0 | 0 | 0 | 0.244650542 | 0 | 0.224243824 |
| ENSMUSG00000108211 | 0.289064826 | 0 | 0.25 | 0.25 | 0 | 0 | 0 |
| ENSMUSG00000108212 | 0 | 0 | 0 | 0 | 0 | 0 | 0 |
| ENSMUSG00000108213 | 0 | 0 | 0 | 0 | 0 | 0 | 0 |
| ENSMUSG00000108214 | 0.289064826 | 0 | 0 | 0.25 | 0.244650542 | 0 | 0.224243824 |
| ENSMUSG00000108215 | 0 | 0 | 0 | 0 | 0 | 0 | 0.355418052 |
| ENSMUSG00000108216 | 0 | 0 | 0.25 | 0 | 0.489301084 | 0 | 0.355418052 |
| ENSMUSG00000108217 | 0 | 0.25 | 0.25 | 0.25 | 0.244650542 | 0 | 0 |
| ENSMUSG00000108218 | 1.655740331 | 1.606566189 | 1.610735874 | 1.610735874 | 1.576269619 | 1.568201724 | 1.459332757 |
| ENSMUSG00000108219 | 0 | 0 | 0 | 0 | 0 | 0 | 0 |
| ENSMUSG00000108220 | 0 | 0 | 0 | 0 | 0 | 0 | 0 |
| ENSMUSG00000108221 | 0 | 0 | 0 | 0 | 0 | 0 | 0 |
| ENSMUSG00000108222 | 0 | 0 | 0 | 0 | 0 | 0 | 0 |
| ENSMUSG00000108223 | 0 | 0 | 0 | 0 | 0 | 0 | 0 |
| ENSMUSG00000108224 | 0 | 0 | 0 | 0 | 0 | 0.301029996 | 0 |
| ENSMUSG00000108225 | 0 | 0 | 0 | 0 | 0 | 0 | 0 |
| ENSMUSG00000108226 | 0 | 0 | 0 | 0 | 0 | 0 | 0 |
| ENSMUSG00000108227 | 0.45815691 | 0.396240625 | 0 | 0 | 0 | 0 | 0.224243824 |
| ENSMUSG00000108228 | 0 | 0 | 0.396240625 | 0 | 0 | 0 | 0 |
| ENSMUSG00000108229 | 0 | 0 | 0 | 0 | 0 | 0 | 0 |
| ENSMUSG00000108230 | 0 | 0 | 0 | 0 | 0 | 0 | 0 |
| ENSMUSG00000108231 | 0 | 0 | 0 | 0 | 0.244650542 | 0 | 0 |
| ENSMUSG00000108232 | 0.45815691 | 0.580482024 | 0 | 0.25 | 0 | 0 | 0.224243824 |
| ENSMUSG00000108233 | 0 | 0 | 0 | 0 | 0 | 0 | 0.224243824 |

| | | | | | | | |
|---|---|---|---|---|---|---|---|
| ENSMUSG00000108234 | 0.289064826 | 0.646240625 | 0.5 | 0.580482024 | 0.489301084 | 0.698970004 | 0.448487648 |
| ENSMUSG00000108235 | 0 | 0 | 0 | 0 | 0 | 0 | 0 |
| ENSMUSG00000108236 | 0 | 0 | 0 | 0 | 0 | 0 | 0 |
| ENSMUSG00000108237 | 0.289064826 | 0.396240625 | 0.25 | 0.396240625 | 0.387761935 | 0 | 0.520678036 |
| ENSMUSG00000108238 | 0.45815691 | 0.396240625 | 0.25 | 0 | 0.387761935 | 0 | 0.355418052 |
| ENSMUSG00000108239 | 0.45815691 | 0.5 | 0.25 | 0.396240625 | 0.489301084 | 0 | 0.224243824 |
| ENSMUSG00000108240 | 0 | 0 | 0 | 0 | 0 | 0 | 0 |
| ENSMUSG00000108241 | 0 | 0 | 0 | 0 | 0 | 0 | 0 |
| ENSMUSG00000108242 | 0 | 0 | 0 | 0 | 0 | 0 | 0 |
| ENSMUSG00000108244 | 0 | 0 | 0 | 0 | 0 | 0 | 0 |
| ENSMUSG00000108245 | 0 | 0 | 0 | 0 | 0 | 0 | 0 |
| ENSMUSG00000108246 | 0.289064826 | 0 | 0 | 0 | 0 | 0 | 0 |
| ENSMUSG00000108247 | 0 | 0 | 0 | 0 | 0 | 0 | 0 |
| ENSMUSG00000108248 | 0 | 0 | 0 | 0 | 0 | 0 | 0 |
| ENSMUSG00000108249 | 1.269664473 | 1.146240625 | 1.146240625 | 1.214495249 | 1.176121988 | 1 | 0.853775828 |
| ENSMUSG00000108250 | 0 | 0 | 0 | 0 | 0 | 0 | 0 |
| ENSMUSG00000108251 | 0 | 0 | 0 | 0 | 0 | 0 | 0 |
| ENSMUSG00000108252 | 0 | 0 | 0 | 0 | 0 | 0 | 0 |
| ENSMUSG00000108253 | 0 | 0 | 0 | 0 | 0 | 0 | 0 |
| ENSMUSG00000108254 | 0.289064826 | 0.25 | 0.25 | 0 | 0 | 0 | 0 |
| ENSMUSG00000108255 | 0 | 0 | 0 | 0 | 0 | 0.301029996 | 0 |
| ENSMUSG00000108256 | 0.671187741 | 0.896240625 | 0.864857905 | 1.061981878 | 0.905314583 | 0.954242509 | 1.028149525 |
| ENSMUSG00000108257 | 0.578129653 | 0.25 | 0.396240625 | 0.396240625 | 0.244650542 | 0 | 0.224243824 |
| ENSMUSG00000108258 | 0 | 0.5 | 0.25 | 0.25 | 0 | 0 | 0 |
| ENSMUSG00000108259 | 0 | 0 | 0 | 0 | 0 | 0 | 0 |
| ENSMUSG00000108260 | 0.289064826 | 0 | 0 | 0 | 0 | 0 | 0 |
| ENSMUSG00000108261 | 0 | 0 | 0 | 0 | 0 | 0 | 0 |
| ENSMUSG00000108262 | 0 | 0 | 0 | 0 | 0 | 0 | 0 |
| ENSMUSG00000108263 | 0 | 0 | 0.25 | 0.25 | 0 | 0 | 0 |
| ENSMUSG00000108264 | 0 | 0 | 0 | 0 | 0 | 0 | 0 |
| ENSMUSG00000108265 | 0 | 0 | 0 | 0 | 0 | 0 | 0 |
| ENSMUSG00000108266 | 0.289064826 | 0.5 | 0.580482024 | 0.25 | 0.77552387 | 0.602059991 | 0.520678036 |
| ENSMUSG00000108267 | 0 | 0.25 | 0 | 0.25 | 0 | 0 | 0.579661877 |
| ENSMUSG00000108268 | 0 | 0.5 | 0.5 | 0 | 0.387761935 | 0 | 0.448487648 |

| | | | | | | | |
|---|---|---|---|---|---|---|---|
| ENSMUSG00000108269 | 0 | 0 | 0 | 0 | 0 | 0 | 0 |
| ENSMUSG00000108270 | 0 | 0 | 0.25 | 0.646240625 | 0 | 0.301029996 | 0.355418052 |
| ENSMUSG00000108271 | 0 | 0 | 0 | 0 | 0 | 0 | 0 |
| ENSMUSG00000108272 | 0 | 0 | 0 | 0 | 0 | 0 | 0 |
| ENSMUSG00000108273 | 0 | 0 | 0 | 0 | 0.387761935 | 0 | 0 |
| ENSMUSG00000108274 | 0 | 0 | 0 | 0 | 0 | 0 | 0 |
| ENSMUSG00000108275 | 0 | 0 | 0 | 0 | 0 | 0 | 0 |
| ENSMUSG00000108276 | 0 | 0 | 0.25 | 0 | 0 | 0 | 0.355418052 |
| ENSMUSG00000108277 | 0 | 0 | 0 | 0 | 0 | 0 | 0 |
| ENSMUSG00000108278 | 0 | 0 | 0 | 0.25 | 0.387761935 | 0 | 0 |
| ENSMUSG00000108279 | 0 | 0.25 | 0 | 0 | 0 | 0 | 0 |
| ENSMUSG00000108280 | 0 | 0 | 0 | 0 | 0.387761935 | 0 | 0.224243824 |
| ENSMUSG00000108282 | 0 | 0.25 | 0 | 0 | 0 | 0 | 0 |
| ENSMUSG00000108283 | 0.289064826 | 0 | 0.25 | 0 | 0 | 0 | 0 |
| ENSMUSG00000108284 | 0 | 0 | 0 | 0 | 0 | 0 | 0 |
| ENSMUSG00000108285 | 0 | 0 | 0 | 0 | 0 | 0 | 0 |
| ENSMUSG00000108286 | 0 | 0 | 0 | 0 | 0 | 0 | 0 |
| ENSMUSG00000108287 | 0 | 0 | 0 | 0 | 0 | 0 | 0 |
| ENSMUSG00000108288 | 0 | 0.396240625 | 0.25 | 0 | 0 | 0.301029996 | 0.355418052 |
| ENSMUSG00000108289 | 0 | 0 | 0 | 0 | 0 | 0 | 0 |
| ENSMUSG00000108290 | 0.578129653 | 0.5 | 0.25 | 0.580482024 | 0.489301084 | 0.602059991 | 0 |
| ENSMUSG00000108291 | 0.289064826 | 0.396240625 | 0.396240625 | 0 | 0.244650542 | 0.301029996 | 0.224243824 |
| ENSMUSG00000108292 | 0 | 0.25 | 0 | 0 | 0 | 0 | 0.224243824 |
| ENSMUSG00000108293 | 0 | 0.396240625 | 0.396240625 | 0 | 0 | 0 | 0 |
| ENSMUSG00000108294 | 0 | 0 | 0 | 0 | 0 | 0 | 0 |
| ENSMUSG00000108295 | 0 | 0 | 0 | 0 | 0 | 0 | 0 |
| ENSMUSG00000108296 | 0 | 0 | 0 | 0 | 0 | 0 | 0 |
| ENSMUSG00000108297 | 0 | 0 | 0 | 0 | 0 | 0 | 0.224243824 |
| ENSMUSG00000108298 | 0 | 0 | 0.25 | 0 | 0 | 0 | 0.448487648 |
| ENSMUSG00000108299 | 0 | 0 | 0 | 0 | 0 | 0 | 0 |

**Results of differential gene expression analysis**

| Gene | Fold Change | FDR-Adj P value |
|---|---|---|
| ENSMUSG00000000001 | -1.035775333 | 0.953731979 |
| ENSMUSG00000000003 | 1 | |
| ENSMUSG00000000028 | -1.052354387 | 0.830258855 |
| ENSMUSG00000000031 | -1.234652963 | 0.362063635 |
| ENSMUSG00000000037 | 1.021593671 | 0.907737577 |
| ENSMUSG00000000049 | 1.128345745 | 0.710698346 |
| ENSMUSG00000000056 | 1.004062115 | 1 |
| ENSMUSG00000000058 | 1.179392694 | 0.44650239 |
| ENSMUSG00000000078 | 1.014025705 | 1 |
| ENSMUSG00000000085 | 1.080220401 | 0.608497229 |
| ENSMUSG00000000088 | -1.138850381 | 0.167900538 |
| ENSMUSG00000000093 | -1.08282318 | 0.785584885 |
| ENSMUSG00000000094 | 1.132639128 | 0.217803313 |
| ENSMUSG00000000103 | 1 | |
| ENSMUSG00000000120 | -1.57658596 | 0.005249107 |
| ENSMUSG00000000125 | 1.023743961 | 0.938939969 |
| ENSMUSG00000000126 | -1.144168305 | 0.096614616 |
| ENSMUSG00000000127 | 1.162072284 | 0.454721467 |
| ENSMUSG00000000131 | 1.027651706 | 0.934375176 |
| ENSMUSG00000000134 | 1.037331727 | 0.884220301 |
| ENSMUSG00000000142 | -1.627367852 | 0.000830863 |
| ENSMUSG00000000148 | 1.033586555 | 0.715589161 |
| ENSMUSG00000000149 | 1.169963484 | 0.198491993 |
| ENSMUSG00000000154 | -1.120476206 | 0.583147495 |
| ENSMUSG00000000157 | 1 | |
| ENSMUSG00000000159 | 1.069068979 | 0.608497229 |
| ENSMUSG00000000167 | -1.143895024 | 0.378089842 |
| ENSMUSG00000000168 | -1.000223231 | 1 |
| ENSMUSG00000000171 | -1.067784074 | 0.79501506 |
| ENSMUSG00000000182 | 1.271344436 | 0.013758746 |
| ENSMUSG00000000183 | 1.059463094 | 0.698112512 |
| ENSMUSG00000000184 | 1.08431348 | 0.739137 |
| ENSMUSG00000000194 | 1.033548586 | 0.870020588 |
| ENSMUSG00000000197 | -1.083289272 | 0.869309206 |
| ENSMUSG00000000202 | 1 | |
| ENSMUSG00000000204 | 1.059463094 | 0.698112512 |
| ENSMUSG00000000214 | 1 | |
| ENSMUSG00000000215 | 1 | |
| ENSMUSG00000000216 | 1 | |
| ENSMUSG00000000223 | -1.243274778 | 0.012277316 |
| ENSMUSG00000000244 | 1.137168648 | 0.653415027 |
| ENSMUSG00000000247 | -1.413876141 | 0.025944658 |
| ENSMUSG00000000248 | -1.150851386 | 0.568864017 |
| ENSMUSG00000000253 | 1.039376203 | 0.698112512 |
| ENSMUSG00000000263 | 1.230075506 | 0.224243601 |
| ENSMUSG00000000266 | -1.050734348 | 0.810550732 |
| ENSMUSG00000000275 | -1.115601843 | 0.575909751 |
| ENSMUSG00000000276 | -1.098570106 | 0.604452802 |
| ENSMUSG00000000278 | 1.050169823 | 0.879199704 |
| ENSMUSG00000000282 | 1.12536979 | 0.048433844 |
| ENSMUSG00000000290 | -1.009673835 | 1 |
| ENSMUSG00000000295 | -1.042753574 | 0.814608539 |
| ENSMUSG00000000296 | 1.393877061 | 0.004205737 |
| ENSMUSG00000000301 | -1.155672657 | 0.169545598 |
| ENSMUSG00000000303 | 1.161873744 | 0.284005985 |
| ENSMUSG00000000305 | -1.03962336 | 0.698112512 |
| ENSMUSG00000000308 | -1.035049744 | 0.952057016 |
| ENSMUSG00000000317 | 1.058968218 | 0.60397857 |

| | | |
|---|---|---|
| ENSMUSG00000000318 | -1 | 1 |
| ENSMUSG00000000320 | -1.41682901 | 0.081043569 |
| ENSMUSG00000000325 | 1.123366672 | 0.216046507 |
| ENSMUSG00000000326 | 1.065701636 | 0.686477484 |
| ENSMUSG00000000339 | -1.010539648 | 0.979978576 |
| ENSMUSG00000000340 | -1.183892295 | 0.281564375 |
| ENSMUSG00000000346 | 1.002242232 | 1 |
| ENSMUSG00000000355 | 1.245795601 | 0.091449991 |
| ENSMUSG00000000359 | -1 | 1 |
| ENSMUSG00000000365 | -1.095294215 | 0.541735653 |
| ENSMUSG00000000374 | 1.015457221 | 0.998195115 |
| ENSMUSG00000000378 | 1.154719096 | 0.315845413 |
| ENSMUSG00000000381 | 1 | |
| ENSMUSG00000000384 | 1.03567216 | 0.835701362 |
| ENSMUSG00000000385 | -1.120476206 | 0.534652019 |
| ENSMUSG00000000386 | 1.024656056 | 0.972994469 |
| ENSMUSG00000000392 | -1.085651418 | 0.743622442 |
| ENSMUSG00000000394 | 1 | |
| ENSMUSG00000000399 | -1.009611918 | 1 |
| ENSMUSG00000000402 | -1.127622795 | 0.608497229 |
| ENSMUSG00000000409 | -1.012989812 | 0.977489164 |
| ENSMUSG00000000411 | 1 | |
| ENSMUSG00000000416 | 1.424293786 | 0.106322814 |
| ENSMUSG00000000420 | 1.03797761 | 0.935335058 |
| ENSMUSG00000000435 | 1 | |
| ENSMUSG00000000439 | 1.029003995 | 0.884451146 |
| ENSMUSG00000000440 | 1.286224871 | 0.004766653 |
| ENSMUSG00000000441 | 1.012713568 | 1 |
| ENSMUSG00000000486 | 1.370961114 | 0.029924974 |
| ENSMUSG00000000489 | 1.261059597 | 0.001601152 |
| ENSMUSG00000000530 | -1.495755914 | 0.008586051 |
| ENSMUSG00000000531 | -1.203804269 | 0.055425893 |
| ENSMUSG00000000532 | -1.08891182 | 0.303493034 |
| ENSMUSG00000000538 | -1.046445313 | 0.76235019 |
| ENSMUSG00000000544 | 1.158861639 | 0.507659484 |
| ENSMUSG00000000552 | -1.093696695 | 0.529596655 |
| ENSMUSG00000000555 | 1.403780737 | 0.098417569 |
| ENSMUSG00000000560 | -1.138766255 | 0.456588659 |
| ENSMUSG00000000561 | -1.064522318 | 0.819614655 |
| ENSMUSG00000000562 | -1.085651418 | 0.608497229 |
| ENSMUSG00000000563 | -1.100755642 | 0.706906489 |
| ENSMUSG00000000567 | 1.058751392 | 0.296212606 |
| ENSMUSG00000000568 | 1.110472881 | 0.700389541 |
| ENSMUSG00000000579 | -1.028752805 | 0.88972256 |
| ENSMUSG00000000581 | -1.07185699 | 0.771903504 |
| ENSMUSG00000000594 | -1.458022605 | 0.023467014 |
| ENSMUSG00000000600 | 1.064703462 | 0.767574573 |
| ENSMUSG00000000605 | 1.062604905 | 0.698112512 |
| ENSMUSG00000000606 | 1 | |
| ENSMUSG00000000617 | 1 | |
| ENSMUSG00000000627 | -1.775771491 | 2.25E-06 |
| ENSMUSG00000000628 | 1.180486104 | 0.065363053 |
| ENSMUSG00000000631 | -1.016037062 | 0.976977294 |
| ENSMUSG00000000632 | 1.05038453 | 0.858000979 |
| ENSMUSG00000000673 | -1.053549062 | 0.698112512 |
| ENSMUSG00000000682 | 1 | |
| ENSMUSG00000000686 | 1.016297569 | 1 |
| ENSMUSG00000000690 | 1.14262534 | 0.013875511 |
| ENSMUSG00000000693 | 1.382727779 | 0.002497395 |
| ENSMUSG00000000701 | 1 | |
| ENSMUSG00000000706 | -1.638428113 | 0.032896733 |
| ENSMUSG00000000708 | 1.052021594 | 0.814089245 |
| ENSMUSG00000000711 | -1.048534737 | 0.879267911 |

| | | |
|---|---|---|
| ENSMUSG00000000724 | -1.080029458 | 0.72956252 |
| ENSMUSG00000000730 | 1 | |
| ENSMUSG00000000731 | -1 | 1 |
| ENSMUSG00000000732 | -1.713840187 | 0.003226816 |
| ENSMUSG00000000738 | 1.052787884 | 0.716583464 |
| ENSMUSG00000000739 | -1.0965339 | 0.694271823 |
| ENSMUSG00000000740 | 1.035062757 | 0.887323956 |
| ENSMUSG00000000743 | 1.024530604 | 0.937921669 |
| ENSMUSG00000000751 | 1.034384897 | 0.95838544 |
| ENSMUSG00000000753 | 1.553618938 | 0.003303995 |
| ENSMUSG00000000759 | 1.158623053 | 0.438534282 |
| ENSMUSG00000000766 | 1.059463094 | 0.698112512 |
| ENSMUSG00000000776 | -1.104135453 | 0.394324991 |
| ENSMUSG00000000782 | -1.061386112 | 0.62545849 |
| ENSMUSG00000000787 | 1.015089232 | 1 |
| ENSMUSG00000000791 | -1.445935088 | 0.022042647 |
| ENSMUSG00000000792 | 1.185080234 | 0.167406338 |
| ENSMUSG00000000794 | -1.023144359 | 0.857167632 |
| ENSMUSG00000000804 | 1.074302985 | 0.779105249 |
| ENSMUSG00000000805 | 1 | |
| ENSMUSG00000000811 | 1.116790928 | 0.268067198 |
| ENSMUSG00000000817 | 1.0145205 | 1 |
| ENSMUSG00000000823 | -1.144259039 | 0.319789343 |
| ENSMUSG00000000826 | -1.120431275 | 0.65988728 |
| ENSMUSG00000000827 | -1.120949726 | 0.608497229 |
| ENSMUSG00000000838 | 1.075264509 | 0.807247723 |
| ENSMUSG00000000861 | -1.044273782 | 0.698112512 |
| ENSMUSG00000000869 | -1.180673275 | 0.017475966 |
| ENSMUSG00000000876 | -1.160106189 | 0.169970982 |
| ENSMUSG00000000881 | 1.147661782 | 0.439929645 |
| ENSMUSG00000000884 | 1.094584469 | 0.098098772 |
| ENSMUSG00000000889 | 1.059463094 | 0.698112512 |
| ENSMUSG00000000901 | 1.129905325 | 0.349076535 |
| ENSMUSG00000000902 | 1.030434291 | 0.933142264 |
| ENSMUSG00000000903 | -1 | 1 |
| ENSMUSG00000000915 | 1.066946092 | 0.608497229 |
| ENSMUSG00000000916 | 1.018826871 | 0.811372967 |
| ENSMUSG00000000934 | -1.043090831 | 0.762928508 |
| ENSMUSG00000000938 | 1.256675465 | 0.024123774 |
| ENSMUSG00000000942 | 1.173857954 | 0.05580329 |
| ENSMUSG00000000948 | -1.038926033 | 0.608497229 |
| ENSMUSG00000000957 | 1.19543658 | 0.388130187 |
| ENSMUSG00000000958 | 1.001856035 | 1 |
| ENSMUSG00000000959 | -1.122894959 | 0.537566955 |
| ENSMUSG00000000976 | 1.072199809 | 0.614616655 |
| ENSMUSG00000000982 | 1.059463094 | 0.608497229 |
| ENSMUSG00000000983 | 1.059463094 | 0.608497229 |
| ENSMUSG00000001014 | 1.102285378 | 0.693578301 |
| ENSMUSG00000001016 | -1.012458235 | 1 |
| ENSMUSG00000001017 | -1.029978476 | 0.950602516 |
| ENSMUSG00000001018 | 1.050975893 | 0.731295483 |
| ENSMUSG00000001020 | 1.374271269 | 0.058373191 |
| ENSMUSG00000001021 | -1.0529957 | 0.647290337 |
| ENSMUSG00000001023 | -1.037482356 | 0.880074775 |
| ENSMUSG00000001025 | 1.12868568 | 0.720282888 |
| ENSMUSG00000001027 | 1.069068979 | 0.608497229 |
| ENSMUSG00000001029 | -1.03962336 | 0.698112512 |
| ENSMUSG00000001034 | 1.046918547 | 0.716971703 |
| ENSMUSG00000001036 | 1.001040762 | 1 |
| ENSMUSG00000001039 | -1.012234705 | 0.94101868 |
| ENSMUSG00000001052 | -1.046410722 | 0.870238163 |
| ENSMUSG00000001053 | -1.04035458 | 0.879157979 |
| ENSMUSG00000001054 | -1.059933853 | 0.560827127 |

| | | |
|---|---|---|
| ENSMUSG00000001056 | -1.014518304 | 0.988263302 |
| ENSMUSG00000001062 | -1.040476406 | 0.472615914 |
| ENSMUSG00000001065 | -1.016670468 | 0.879253209 |
| ENSMUSG00000001076 | -1.22286951 | 0.544226414 |
| ENSMUSG00000001082 | 1.22121731 | 0.129476598 |
| ENSMUSG00000001089 | 1.043937932 | 0.914904517 |
| ENSMUSG00000001095 | 1 | |
| ENSMUSG00000001098 | 1.217880782 | 0.160089696 |
| ENSMUSG00000001100 | 1.05069086 | 0.852544847 |
| ENSMUSG00000001103 | -1 | 1 |
| ENSMUSG00000001105 | -1.025218142 | 0.908543402 |
| ENSMUSG00000001119 | 1.14204841 | 0.598655292 |
| ENSMUSG00000001120 | -1.030898687 | 0.790885744 |
| ENSMUSG00000001123 | 1.008443944 | 1 |
| ENSMUSG00000001127 | 1.082458109 | 0.652052918 |
| ENSMUSG00000001128 | 1.227223234 | 0.074452106 |
| ENSMUSG00000001131 | 1.202777104 | 0.336331834 |
| ENSMUSG00000001134 | -1.02230009 | 0.940298812 |
| ENSMUSG00000001138 | -1.040247548 | 0.698112512 |
| ENSMUSG00000001143 | -1.111308579 | 0.623278753 |
| ENSMUSG00000001150 | -1.02631152 | 0.943529281 |
| ENSMUSG00000001151 | -1.04590503 | 0.764394136 |
| ENSMUSG00000001155 | -1.044273782 | 0.878511951 |
| ENSMUSG00000001156 | 1.000372802 | 1 |
| ENSMUSG00000001157 | 1.003372545 | 1 |
| ENSMUSG00000001158 | 1.003593104 | 1 |
| ENSMUSG00000001166 | -1.636145271 | 0.001987328 |
| ENSMUSG00000001168 | 1 | |
| ENSMUSG00000001173 | 1.018206947 | 0.982838256 |
| ENSMUSG00000001175 | 1.039192051 | 0.945624631 |
| ENSMUSG00000001211 | 1.000835375 | 1 |
| ENSMUSG00000001225 | -1.023829345 | 0.96035552 |
| ENSMUSG00000001227 | 1.045296686 | 0.845683036 |
| ENSMUSG00000001228 | 1.064300212 | 0.760172055 |
| ENSMUSG00000001229 | 1.083565965 | 0.608497229 |
| ENSMUSG00000001240 | -1.729184706 | 0.013138433 |
| ENSMUSG00000001247 | -1.248463474 | 0.224238622 |
| ENSMUSG00000001248 | 1.163591036 | 0.114374577 |
| ENSMUSG00000001249 | -1.043306199 | 0.803260194 |
| ENSMUSG00000001260 | 1 | |
| ENSMUSG00000001270 | -1.168336957 | 0.453417704 |
| ENSMUSG00000001280 | -1.00007714 | 1 |
| ENSMUSG00000001281 | -1.407032877 | 0.001678863 |
| ENSMUSG00000001285 | -1.06470793 | 0.698112512 |
| ENSMUSG00000001288 | 1.090171596 | 0.596510629 |
| ENSMUSG00000001289 | -1.020455972 | 0.975938761 |
| ENSMUSG00000001300 | 1.162499106 | 0.402963237 |
| ENSMUSG00000001305 | -1.068322433 | 0.721007497 |
| ENSMUSG00000001313 | -1.110139001 | 0.378751351 |
| ENSMUSG00000001323 | -1.108634581 | 0.589610727 |
| ENSMUSG00000001333 | 1.549797127 | 0.015989143 |
| ENSMUSG00000001334 | -1.38857273 | 0.073053995 |
| ENSMUSG00000001348 | -1.265727016 | 0.361613976 |
| ENSMUSG00000001349 | 3.842476829 | 2.58639E-05 |
| ENSMUSG00000001366 | -1.07448296 | 0.608497229 |
| ENSMUSG00000001376 | 1.139197872 | 0.587631343 |
| ENSMUSG00000001380 | -1.035529452 | 0.925173733 |
| ENSMUSG00000001383 | -1.019127558 | 0.988519386 |
| ENSMUSG00000001403 | -1.126528034 | 0.608497229 |
| ENSMUSG00000001415 | 1.019570358 | 0.978557295 |
| ENSMUSG00000001416 | -1.058150351 | 0.896246183 |
| ENSMUSG00000001418 | 1.128917657 | 0.608497229 |
| ENSMUSG00000001419 | 1.080240558 | 0.553862807 |

| | | |
|---|---|---|
| ENSMUSG00000001420 | -1.011719055 | 1 |
| ENSMUSG00000001424 | 1.080029875 | 0.729059471 |
| ENSMUSG00000001435 | 1.635927703 | 0.002712715 |
| ENSMUSG00000001436 | 1.02356384 | 0.899402062 |
| ENSMUSG00000001440 | 1.035984181 | 0.972686282 |
| ENSMUSG00000001441 | 1.036222206 | 0.946935473 |
| ENSMUSG00000001444 | -1.044273782 | 0.698112512 |
| ENSMUSG00000001445 | 1.026556523 | 0.923395918 |
| ENSMUSG00000001467 | 1.034049731 | 0.94374396 |
| ENSMUSG00000001472 | -1.022228936 | 0.984432303 |
| ENSMUSG00000001473 | 1.149893061 | 0.314948458 |
| ENSMUSG00000001482 | -1.12056565 | 0.555413989 |
| ENSMUSG00000001493 | 1.11166259 | 0.686411811 |
| ENSMUSG00000001494 | -1.03962336 | 0.698112512 |
| ENSMUSG00000001496 | 1 | |
| ENSMUSG00000001497 | -1.044796633 | 0.749577885 |
| ENSMUSG00000001504 | -1.234352308 | 0.031257401 |
| ENSMUSG00000001506 | 1.494683559 | 0.051664013 |
| ENSMUSG00000001507 | 1.216102726 | 0.215991492 |
| ENSMUSG00000001508 | 1.104090066 | 0.608497229 |
| ENSMUSG00000001510 | -1.655516819 | 0.013806256 |
| ENSMUSG00000001517 | 1.04627953 | 0.852557372 |
| ENSMUSG00000001518 | -1.034232353 | 0.813485995 |
| ENSMUSG00000001520 | -1.114958973 | 0.534887579 |
| ENSMUSG00000001521 | 1.042944299 | 0.875313849 |
| ENSMUSG00000001524 | 1.014544061 | 0.961127302 |
| ENSMUSG00000001525 | 1.100532996 | 0.770958539 |
| ENSMUSG00000001542 | 1.066516515 | 0.774554935 |
| ENSMUSG00000001552 | 1.126634566 | 0.344261 |
| ENSMUSG00000001555 | 1.09048885 | 0.646760484 |
| ENSMUSG00000001558 | 1.069185486 | 0.65350336 |
| ENSMUSG00000001566 | -1.084645496 | 0.567496595 |
| ENSMUSG00000001569 | 1.010115575 | 1 |
| ENSMUSG00000001576 | 1.093594124 | 0.712544875 |
| ENSMUSG00000001583 | 1.154253937 | 0.238219732 |
| ENSMUSG00000001588 | -1.227568746 | 0.511583996 |
| ENSMUSG00000001604 | 1.06027406 | 0.608497229 |
| ENSMUSG00000001622 | -1.03962336 | 0.698112512 |
| ENSMUSG00000001627 | 1.085852688 | 0.736225935 |
| ENSMUSG00000001630 | -1.009415398 | 1 |
| ENSMUSG00000001632 | 1.108436931 | 0.287801879 |
| ENSMUSG00000001642 | 1.040716528 | 0.886380256 |
| ENSMUSG00000001655 | -1.023687458 | 0.976752232 |
| ENSMUSG00000001656 | -1.227084586 | 0.18200153 |
| ENSMUSG00000001657 | -1.509570609 | 0.001867243 |
| ENSMUSG00000001661 | -1.582705972 | 0.010341689 |
| ENSMUSG00000001663 | 1.098100892 | 0.232738517 |
| ENSMUSG00000001665 | -1.319564588 | 0.02555769 |
| ENSMUSG00000001666 | -1.099371426 | 0.30664718 |
| ENSMUSG00000001670 | 1 | |
| ENSMUSG00000001672 | -1.189068624 | 0.375655143 |
| ENSMUSG00000001674 | -1.008952061 | 1 |
| ENSMUSG00000001687 | 1.025965993 | 0.911347495 |
| ENSMUSG00000001700 | 1.017998591 | 0.895376993 |
| ENSMUSG00000001707 | -1.055524683 | 0.804810549 |
| ENSMUSG00000001729 | 1.188185592 | 0.313649348 |
| ENSMUSG00000001739 | 1.417502188 | 0.026024582 |
| ENSMUSG00000001741 | -1.025579669 | 0.975677002 |
| ENSMUSG00000001750 | -1.006051791 | 1 |
| ENSMUSG00000001751 | 1.157486703 | 0.069445601 |
| ENSMUSG00000001755 | 1.127475646 | 0.279918853 |
| ENSMUSG00000001761 | 1.062598821 | 0.783468419 |
| ENSMUSG00000001763 | -1.248721756 | 0.01536947 |

| | | |
|---|---|---|
| ENSMUSG00000001767 | -1.092129778 | 0.638420696 |
| ENSMUSG00000001768 | 1.077199673 | 0.356969956 |
| ENSMUSG00000001773 | -1.044273782 | 0.698112512 |
| ENSMUSG00000001774 | -1.087327724 | 0.717035356 |
| ENSMUSG00000001783 | -1.026926919 | 0.97573893 |
| ENSMUSG00000001785 | -1.055509147 | 0.841885126 |
| ENSMUSG00000001786 | -1.061287621 | 0.698112512 |
| ENSMUSG00000001794 | 1.019328657 | 0.9858962 |
| ENSMUSG00000001802 | -1.36994952 | 0.090064492 |
| ENSMUSG00000001804 | 1 | |
| ENSMUSG00000001815 | -1.043306199 | 0.79291651 |
| ENSMUSG00000001819 | 1 | |
| ENSMUSG00000001823 | 1 | |
| ENSMUSG00000001827 | -1.015511103 | 0.991425134 |
| ENSMUSG00000001829 | -1.029252659 | 0.87354217 |
| ENSMUSG00000001833 | 1.00243344 | 1 |
| ENSMUSG00000001844 | 1.040175815 | 0.764370878 |
| ENSMUSG00000001847 | -1.008514201 | 1 |
| ENSMUSG00000001855 | 1.040368426 | 0.853642123 |
| ENSMUSG00000001864 | 1.121425612 | 0.589051474 |
| ENSMUSG00000001865 | 1 | |
| ENSMUSG00000001870 | 1.240494595 | 0.365927614 |
| ENSMUSG00000001891 | 1.050979902 | 0.867939428 |
| ENSMUSG00000001901 | 1.161036672 | 0.251049598 |
| ENSMUSG00000001909 | 1.120365689 | 0.323395097 |
| ENSMUSG00000001910 | 1.025463364 | 0.96088958 |
| ENSMUSG00000001911 | 1.157928688 | 0.275279115 |
| ENSMUSG00000001918 | -1.072961895 | 0.577714817 |
| ENSMUSG00000001924 | 1.067460347 | 0.855779369 |
| ENSMUSG00000001930 | 1.441262801 | 0.050976163 |
| ENSMUSG00000001942 | -1.031149817 | 0.908366811 |
| ENSMUSG00000001943 | 1 | |
| ENSMUSG00000001946 | -1.003610865 | 1 |
| ENSMUSG00000001948 | -1.164289674 | 0.608497229 |
| ENSMUSG00000001962 | 1.107773743 | 0.608497229 |
| ENSMUSG00000001964 | 1.140519227 | 0.451231014 |
| ENSMUSG00000001983 | -1.055661593 | 0.378445257 |
| ENSMUSG00000001985 | 1.000185611 | 1 |
| ENSMUSG00000001986 | 1.528790946 | 0.005344566 |
| ENSMUSG00000001988 | -1.053670208 | 0.879375104 |
| ENSMUSG00000001995 | 1.287072122 | 0.000877343 |
| ENSMUSG00000001998 | -1.121684513 | 0.53003575 |
| ENSMUSG00000001999 | -1.085808333 | 0.682621073 |
| ENSMUSG00000002006 | -1.026100481 | 0.903464611 |
| ENSMUSG00000002007 | -1.160533857 | 0.608497229 |
| ENSMUSG00000002010 | 1.041695869 | 0.859839608 |
| ENSMUSG00000002012 | 1.028980088 | 0.948628145 |
| ENSMUSG00000002014 | 1.106349816 | 0.59984182 |
| ENSMUSG00000002015 | -1.057094556 | 0.866880844 |
| ENSMUSG00000002017 | 1.027671504 | 0.961096559 |
| ENSMUSG00000002020 | 1.915550368 | 0.000397241 |
| ENSMUSG00000002028 | 1.007183491 | 1 |
| ENSMUSG00000002031 | -1.035899242 | 0.911516344 |
| ENSMUSG00000002032 | -1.184085193 | 0.148284344 |
| ENSMUSG00000002033 | -1 | 1 |
| ENSMUSG00000002043 | 1.060732352 | 0.803072249 |
| ENSMUSG00000002052 | 1.005083425 | 1 |
| ENSMUSG00000002055 | -1.064395674 | 0.816422917 |
| ENSMUSG00000002057 | 1.27516987 | 0.203085735 |
| ENSMUSG00000002058 | 1.019038757 | 0.934760809 |
| ENSMUSG00000002059 | 1.099554524 | 0.650512852 |
| ENSMUSG00000002064 | 1.007375726 | 1 |
| ENSMUSG00000002068 | 1.291321546 | 0.031701113 |

| | | |
|---|---|---|
| ENSMUSG00000002076 | -1.122983117 | 0.793735215 |
| ENSMUSG00000002083 | 1.124876559 | 0.698112512 |
| ENSMUSG00000002100 | 1 | |
| ENSMUSG00000002102 | -1.098204851 | 0.69439677 |
| ENSMUSG00000002103 | -1.124062536 | 0.576689161 |
| ENSMUSG00000002104 | 1.051736333 | 0.850820147 |
| ENSMUSG00000002105 | -1.021175839 | 0.908878383 |
| ENSMUSG00000002107 | 1.237996516 | 0.185777014 |
| ENSMUSG00000002108 | -1.026566201 | 0.905432869 |
| ENSMUSG00000002109 | -1.010023173 | 0.978211686 |
| ENSMUSG00000002111 | 1.069068979 | 0.767473873 |
| ENSMUSG00000002129 | -1.041579696 | 0.888895298 |
| ENSMUSG00000002147 | 1.063213085 | 0.814079613 |
| ENSMUSG00000002190 | -1.239209048 | 0.126063768 |
| ENSMUSG00000002204 | 1.059463094 | 0.698112512 |
| ENSMUSG00000002205 | -1.055676367 | 0.586009346 |
| ENSMUSG00000002210 | 1.132296917 | 0.18200153 |
| ENSMUSG00000002221 | 1.064677817 | 0.710318196 |
| ENSMUSG00000002222 | 1.015749147 | 1 |
| ENSMUSG00000002227 | -1.008922494 | 1 |
| ENSMUSG00000002228 | 1.232312768 | 0.608497229 |
| ENSMUSG00000002233 | 1.056468053 | 0.80536789 |
| ENSMUSG00000002240 | 1 | |
| ENSMUSG00000002249 | 1.111683455 | 0.1351252 |
| ENSMUSG00000002250 | -1.008015801 | 0.99692619 |
| ENSMUSG00000002257 | 1.020469079 | 0.971410312 |
| ENSMUSG00000002265 | -2.075411374 | 0.000612262 |
| ENSMUSG00000002266 | 1.019083579 | 0.965621885 |
| ENSMUSG00000002274 | -1.31864698 | 0.088438086 |
| ENSMUSG00000002279 | 1.074391013 | 0.519847934 |
| ENSMUSG00000002280 | 1.049445564 | 0.669081593 |
| ENSMUSG00000002289 | 1.103716131 | 0.115613806 |
| ENSMUSG00000002297 | 1.091434574 | 0.710698346 |
| ENSMUSG00000002307 | 1.097946912 | 0.297198661 |
| ENSMUSG00000002308 | -1.007446581 | 1 |
| ENSMUSG00000002319 | -1.098851874 | 0.612496955 |
| ENSMUSG00000002320 | -1.113408139 | 0.572700408 |
| ENSMUSG00000002324 | 1.028323353 | 0.903464611 |
| ENSMUSG00000002325 | -1.293387598 | 0.016532413 |
| ENSMUSG00000002326 | -1.200098764 | 0.101338159 |
| ENSMUSG00000002329 | -1.153994977 | 0.181635915 |
| ENSMUSG00000002332 | -1.087944214 | 0.601666207 |
| ENSMUSG00000002341 | 1.039597911 | 0.884018396 |
| ENSMUSG00000002342 | -1.031879043 | 0.782962111 |
| ENSMUSG00000002343 | 1.102973305 | 0.511583996 |
| ENSMUSG00000002345 | 1.020563118 | 0.879026266 |
| ENSMUSG00000002346 | -1.019146785 | 0.926908514 |
| ENSMUSG00000002365 | 1.053890476 | 0.814608539 |
| ENSMUSG00000002372 | 1.031430327 | 0.896700076 |
| ENSMUSG00000002379 | 1.028449152 | 0.879253209 |
| ENSMUSG00000002384 | -1.148007082 | 0.445897268 |
| ENSMUSG00000002393 | 1.035736918 | 0.954424095 |
| ENSMUSG00000002395 | 1.088724227 | 0.420161906 |
| ENSMUSG00000002396 | 1.128652522 | 0.224336342 |
| ENSMUSG00000002409 | 1.021551943 | 0.877390802 |
| ENSMUSG00000002413 | -1.01501845 | 0.995771559 |
| ENSMUSG00000002416 | -1.094015373 | 0.552302688 |
| ENSMUSG00000002428 | -1.01629653 | 0.983659314 |
| ENSMUSG00000002455 | -1.107233325 | 0.608497229 |
| ENSMUSG00000002458 | -1.052673806 | 0.732625293 |
| ENSMUSG00000002459 | -1.175118629 | 0.078328848 |
| ENSMUSG00000002475 | -1.77087723 | 0.001540286 |
| ENSMUSG00000002477 | -1.028586843 | 0.956972448 |

| | | |
|---|---|---|
| ENSMUSG00000002486 | 1.066874216 | 0.533026432 |
| ENSMUSG00000002489 | 1.015534078 | 0.946925656 |
| ENSMUSG00000002496 | 1.00321777 | 1 |
| ENSMUSG00000002500 | 1.24122829 | 0.042914396 |
| ENSMUSG00000002504 | -1.086270667 | 0.439929645 |
| ENSMUSG00000002524 | 1.033723505 | 0.928184245 |
| ENSMUSG00000002546 | 1.086766136 | 0.685141313 |
| ENSMUSG00000002550 | 1.001581315 | 1 |
| ENSMUSG00000002565 | -1.204565254 | 0.286114407 |
| ENSMUSG00000002578 | -1.259518113 | 0.020911567 |
| ENSMUSG00000002580 | -1.036172741 | 0.876275381 |
| ENSMUSG00000002588 | 1.059463094 | 0.608497229 |
| ENSMUSG00000002602 | 1.138639097 | 0.608497229 |
| ENSMUSG00000002603 | 1.104013399 | 0.315908474 |
| ENSMUSG00000002608 | 1.081092885 | 0.329606768 |
| ENSMUSG00000002617 | 1.037275966 | 0.830872429 |
| ENSMUSG00000002625 | -1.047184693 | 0.76326282 |
| ENSMUSG00000002633 | 1 | |
| ENSMUSG00000002635 | 1.048250389 | 0.698112512 |
| ENSMUSG00000002658 | 1.021178796 | 0.956972448 |
| ENSMUSG00000002660 | 1.070499475 | 0.658430243 |
| ENSMUSG00000002661 | -1.031060065 | 0.692460578 |
| ENSMUSG00000002664 | -1.079417599 | 0.727782327 |
| ENSMUSG00000002668 | -1 | 1 |
| ENSMUSG00000002679 | 1.04461522 | 0.841409496 |
| ENSMUSG00000002688 | 1.094613735 | 0.608497229 |
| ENSMUSG00000002699 | 1.132639128 | 0.546484592 |
| ENSMUSG00000002718 | -1.091416858 | 0.770958539 |
| ENSMUSG00000002728 | -1.044440406 | 0.829826148 |
| ENSMUSG00000002731 | -1.126824503 | 0.334265395 |
| ENSMUSG00000002732 | -1.204648303 | 0.083152509 |
| ENSMUSG00000002733 | -1.026570959 | 0.917717739 |
| ENSMUSG00000002741 | 1.012876927 | 1 |
| ENSMUSG00000002748 | 1.059384577 | 0.878506664 |
| ENSMUSG00000002763 | 1.022457469 | 0.915469433 |
| ENSMUSG00000002767 | 1.042841924 | 0.634115399 |
| ENSMUSG00000002768 | -1.000034543 | 1 |
| ENSMUSG00000002769 | 1.005268644 | 1 |
| ENSMUSG00000002771 | -1.054100133 | 0.698112512 |
| ENSMUSG00000002778 | -1.013627297 | 0.995747207 |
| ENSMUSG00000002781 | -1.014269621 | 0.989729435 |
| ENSMUSG00000002791 | -1.169177327 | 0.59095666 |
| ENSMUSG00000002797 | -1.01520279 | 0.851023075 |
| ENSMUSG00000002799 | -1.182858893 | 0.486930276 |
| ENSMUSG00000002803 | 1.179969191 | 0.129583481 |
| ENSMUSG00000002804 | 1.007759927 | 1 |
| ENSMUSG00000002808 | 1.046094457 | 0.858409678 |
| ENSMUSG00000002812 | -1.023440663 | 0.959117643 |
| ENSMUSG00000002814 | -1.02175619 | 0.937535342 |
| ENSMUSG00000002820 | -1.104688754 | 0.327963287 |
| ENSMUSG00000002825 | -1.007176383 | 1 |
| ENSMUSG00000002831 | 3.58302111 | 8.37393E-05 |
| ENSMUSG00000002833 | 1.084660912 | 0.591684936 |
| ENSMUSG00000002835 | 1.038565042 | 0.893822825 |
| ENSMUSG00000002844 | 1.197870197 | 0.025381344 |
| ENSMUSG00000002845 | 1.059998216 | 0.746451077 |
| ENSMUSG00000002846 | 1.046391911 | 0.699340382 |
| ENSMUSG00000002847 | 1.059388861 | 0.532369657 |
| ENSMUSG00000002870 | 1.102846301 | 0.615405708 |
| ENSMUSG00000002871 | 1.17231366 | 0.025781546 |
| ENSMUSG00000002881 | 1.048822408 | 0.848546119 |
| ENSMUSG00000002885 | 1.182543362 | 0.150574977 |
| ENSMUSG00000002897 | -1.06185119 | 0.717035356 |

| | | |
|---|---|---|
| ENSMUSG00000002900 | 1.073703466 | 0.778208086 |
| ENSMUSG00000002908 | -1.415974268 | 0.004990486 |
| ENSMUSG00000002910 | -1.01330947 | 0.959892369 |
| ENSMUSG00000002930 | 1 | |
| ENSMUSG00000002944 | -1 | 1 |
| ENSMUSG00000002948 | 1.09694765 | 0.608863038 |
| ENSMUSG00000002949 | 1.114030008 | 0.583049259 |
| ENSMUSG00000002957 | -1.000045387 | 1 |
| ENSMUSG00000002963 | 1.083226777 | 0.234435619 |
| ENSMUSG00000002968 | 1.04763706 | 0.716896135 |
| ENSMUSG00000002980 | -1.469286253 | 0.010366876 |
| ENSMUSG00000002981 | -1.032176649 | 0.914922209 |
| ENSMUSG00000002983 | -1.049702476 | 0.608497229 |
| ENSMUSG00000002984 | 1.108667295 | 0.376977845 |
| ENSMUSG00000002985 | 1.14613064 | 0.608497229 |
| ENSMUSG00000002992 | 1 | |
| ENSMUSG00000002996 | 1.189622327 | 0.319987608 |
| ENSMUSG00000002997 | 1.184316312 | 0.440395661 |
| ENSMUSG00000003031 | 1.120771558 | 0.543095973 |
| ENSMUSG00000003032 | 1.403180058 | 0.013676301 |
| ENSMUSG00000003033 | -1.013893389 | 0.990807767 |
| ENSMUSG00000003037 | 1.016501538 | 0.972980766 |
| ENSMUSG00000003038 | -1.094851144 | 0.51470764 |
| ENSMUSG00000003039 | 1.006576493 | 1 |
| ENSMUSG00000003051 | 1.229528955 | 0.532369657 |
| ENSMUSG00000003053 | 1.019083579 | 0.968937442 |
| ENSMUSG00000003062 | 1.034972559 | 0.870778586 |
| ENSMUSG00000003068 | 1.042150344 | 0.744351452 |
| ENSMUSG00000003070 | -1.251219546 | 0.021719773 |
| ENSMUSG00000003072 | 1.068810557 | 0.698112512 |
| ENSMUSG00000003099 | 1.014504241 | 0.968055611 |
| ENSMUSG00000003119 | -1.051124945 | 0.848456999 |
| ENSMUSG00000003123 | -1.113513003 | 0.306976682 |
| ENSMUSG00000003131 | -1.03380281 | 0.959892369 |
| ENSMUSG00000003134 | -1.170473566 | 0.013060973 |
| ENSMUSG00000003135 | -1.027402234 | 0.944833562 |
| ENSMUSG00000003153 | 1.02179513 | 0.972980766 |
| ENSMUSG00000003154 | 1.057131158 | 0.698112512 |
| ENSMUSG00000003161 | 1.001539484 | 1 |
| ENSMUSG00000003166 | -1.007622179 | 1 |
| ENSMUSG00000003184 | 1.055747551 | 0.608497229 |
| ENSMUSG00000003190 | 1.130576566 | 0.449109628 |
| ENSMUSG00000003198 | 1.017903565 | 0.949812939 |
| ENSMUSG00000003199 | 1.177190646 | 0.208516141 |
| ENSMUSG00000003200 | 1.109026789 | 0.56794807 |
| ENSMUSG00000003206 | 1 | |
| ENSMUSG00000003208 | -1.052712823 | 0.65063484 |
| ENSMUSG00000003226 | 1.013215641 | 1 |
| ENSMUSG00000003227 | -1.187941777 | 0.511163311 |
| ENSMUSG00000003228 | 1.015382893 | 0.972980766 |
| ENSMUSG00000003233 | 1.140137666 | 0.110354258 |
| ENSMUSG00000003234 | -1.023010875 | 0.944649069 |
| ENSMUSG00000003235 | 1.038614888 | 0.890958598 |
| ENSMUSG00000003269 | -1.026640491 | 0.926294398 |
| ENSMUSG00000003271 | -1.736429833 | 0.009543463 |
| ENSMUSG00000003273 | 1.109578381 | 0.752398528 |
| ENSMUSG00000003279 | 1.472211935 | 0.019901782 |
| ENSMUSG00000003282 | 1.2564162 | 0.009011209 |
| ENSMUSG00000003283 | -1.062336276 | 0.698112512 |
| ENSMUSG00000003299 | -1.15858283 | 0.232855843 |
| ENSMUSG00000003308 | -1.021848217 | 0.828564184 |
| ENSMUSG00000003309 | 1 | |
| ENSMUSG00000003316 | -1.016035812 | 1 |

| | | |
|---|---|---|
| ENSMUSG00000003341 | 1.84962203 | 0.003320656 |
| ENSMUSG00000003344 | -1.05342329 | 0.698112512 |
| ENSMUSG00000003345 | 1.072653957 | 0.608497229 |
| ENSMUSG00000003346 | -1.000710724 | 1 |
| ENSMUSG00000003348 | 1.053714291 | 0.736523033 |
| ENSMUSG00000003352 | 1.074327013 | 0.608497229 |
| ENSMUSG00000003354 | 1.061522547 | 0.832587504 |
| ENSMUSG00000003355 | 1.041803022 | 0.697557233 |
| ENSMUSG00000003360 | 1.065813071 | 0.757482614 |
| ENSMUSG00000003363 | 1.08967127 | 0.50546101 |
| ENSMUSG00000003378 | 1.050924724 | 0.620398673 |
| ENSMUSG00000003379 | 1.012830131 | 1 |
| ENSMUSG00000003380 | 1.092480021 | 0.606157471 |
| ENSMUSG00000003382 | -1.014482541 | 0.970025232 |
| ENSMUSG00000003402 | 1.006367917 | 1 |
| ENSMUSG00000003410 | 1.061340802 | 0.856456807 |
| ENSMUSG00000003411 | -1.127006078 | 0.26504751 |
| ENSMUSG00000003418 | -1.381882995 | 0.007325542 |
| ENSMUSG00000003420 | -1.102297663 | 0.440837715 |
| ENSMUSG00000003421 | -1.006652003 | 1 |
| ENSMUSG00000003423 | 1.066856265 | 0.608497229 |
| ENSMUSG00000003429 | 1.041509421 | 0.931480186 |
| ENSMUSG00000003435 | 1.002635995 | 1 |
| ENSMUSG00000003436 | -1.015511103 | 0.991425134 |
| ENSMUSG00000003437 | -1.034634432 | 0.86146627 |
| ENSMUSG00000003438 | 1.005092736 | 1 |
| ENSMUSG00000003444 | -1.045814345 | 0.586769798 |
| ENSMUSG00000003452 | -1.084422318 | 0.608497229 |
| ENSMUSG00000003458 | -1.044213538 | 0.89654333 |
| ENSMUSG00000003464 | -1.072814837 | 0.750635131 |
| ENSMUSG00000003469 | -1.20776913 | 0.272787379 |
| ENSMUSG00000003476 | 1 | |
| ENSMUSG00000003477 | 1 | |
| ENSMUSG00000003484 | 1 | |
| ENSMUSG00000003500 | 1.017213638 | 0.929295642 |
| ENSMUSG00000003505 | 1 | |
| ENSMUSG00000003518 | -1.118219431 | 0.462017587 |
| ENSMUSG00000003526 | -1.242461535 | 0.050370861 |
| ENSMUSG00000003527 | 1.002009765 | 1 |
| ENSMUSG00000003528 | -1.077689625 | 0.608497229 |
| ENSMUSG00000003531 | -1.012977094 | 0.912140254 |
| ENSMUSG00000003534 | 1.317620926 | 0.035709006 |
| ENSMUSG00000003541 | 1.151394482 | 0.127885078 |
| ENSMUSG00000003545 | -1.038642806 | 0.930202553 |
| ENSMUSG00000003546 | 1.043147989 | 0.757125368 |
| ENSMUSG00000003549 | 1.126445892 | 0.024207926 |
| ENSMUSG00000003555 | -1 | 1 |
| ENSMUSG00000003559 | 1.064895386 | 0.787803345 |
| ENSMUSG00000003573 | 1.109455091 | 0.320408152 |
| ENSMUSG00000003575 | 1.007319774 | 0.982961337 |
| ENSMUSG00000003581 | -1.02123211 | 0.84360497 |
| ENSMUSG00000003585 | -1.421334627 | 0.013244138 |
| ENSMUSG00000003604 | -1.185238048 | 0.268545131 |
| ENSMUSG00000003617 | -1.305155365 | 0.082408466 |
| ENSMUSG00000003623 | 1.001783323 | 1 |
| ENSMUSG00000003644 | -1.192219513 | 0.025339655 |
| ENSMUSG00000003657 | 1 | |
| ENSMUSG00000003660 | -1.037023421 | 0.947248675 |
| ENSMUSG00000003662 | -1.144336813 | 0.347405668 |
| ENSMUSG00000003665 | 1.060925547 | 0.783694537 |
| ENSMUSG00000003680 | 1.107694988 | 0.441354969 |
| ENSMUSG00000003721 | 1.022027888 | 0.961353875 |
| ENSMUSG00000003731 | -1.066916287 | 0.812652035 |

| | | |
|---|---|---|
| ENSMUSG00000003746 | -1.086087094 | 0.608497229 |
| ENSMUSG00000003752 | 1.048970015 | 0.69605651 |
| ENSMUSG00000003762 | 1.094270071 | 0.557169918 |
| ENSMUSG00000003778 | 1.023910646 | 0.973205114 |
| ENSMUSG00000003779 | 1.028532643 | 0.969506516 |
| ENSMUSG00000003808 | -1.09688674 | 0.608497229 |
| ENSMUSG00000003809 | -1.054841971 | 0.756949808 |
| ENSMUSG00000003810 | 1.08943023 | 0.698112512 |
| ENSMUSG00000003812 | 1.197495966 | 0.060326805 |
| ENSMUSG00000003813 | 1.042193615 | 0.797105186 |
| ENSMUSG00000003814 | -1.017803295 | 1 |
| ENSMUSG00000003824 | -1.039006258 | 0.698112512 |
| ENSMUSG00000003847 | -1.022895374 | 0.993169601 |
| ENSMUSG00000003848 | 1.02534725 | 0.90514546 |
| ENSMUSG00000003849 | 1.381054459 | 0.015688343 |
| ENSMUSG00000003863 | -1.005335632 | 1 |
| ENSMUSG00000003865 | 1.009121129 | 1 |
| ENSMUSG00000003868 | 1.037801608 | 0.786691536 |
| ENSMUSG00000003872 | -1.073696587 | 0.725290225 |
| ENSMUSG00000003873 | 1.21182598 | 0.046774529 |
| ENSMUSG00000003882 | 1 | |
| ENSMUSG00000003923 | -1.001389582 | 1 |
| ENSMUSG00000003929 | 1.054251786 | 0.781492355 |
| ENSMUSG00000003934 | -1.10249302 | 0.388491505 |
| ENSMUSG00000003948 | 1.209509772 | 0.255077148 |
| ENSMUSG00000003949 | -1.303938278 | 0.068396557 |
| ENSMUSG00000003955 | 1.062832322 | 0.698112512 |
| ENSMUSG00000003970 | 1.063070808 | 0.834464052 |
| ENSMUSG00000003974 | -1.018998401 | 0.986389736 |
| ENSMUSG00000003992 | -1.001380469 | 1 |
| ENSMUSG00000004018 | 1.038762283 | 0.803641195 |
| ENSMUSG00000004031 | 1.019083579 | 0.949812939 |
| ENSMUSG00000004032 | -1.275035678 | 0.171401407 |
| ENSMUSG00000004035 | -1.183195919 | 0.44691802 |
| ENSMUSG00000004038 | 1 | |
| ENSMUSG00000004040 | 1.038949241 | 0.883615907 |
| ENSMUSG00000004043 | 1.090212664 | 0.300948082 |
| ENSMUSG00000004044 | 1.069068625 | 0.810530624 |
| ENSMUSG00000004054 | 1.074059325 | 0.69183734 |
| ENSMUSG00000004056 | 1.063406993 | 0.698112512 |
| ENSMUSG00000004069 | 1.042977816 | 0.821145247 |
| ENSMUSG00000004070 | -1.098125894 | 0.608497229 |
| ENSMUSG00000004071 | 1.000855581 | 1 |
| ENSMUSG00000004085 | -1.064894951 | 0.772179799 |
| ENSMUSG00000004096 | -1.090700427 | 0.728022344 |
| ENSMUSG00000004098 | -1.460971345 | 0.004054559 |
| ENSMUSG00000004099 | -1.040758753 | 0.924061737 |
| ENSMUSG00000004100 | 1.075895059 | 0.565559141 |
| ENSMUSG00000004105 | 1.431010261 | 0.015462827 |
| ENSMUSG00000004110 | -1.191243581 | 0.456459883 |
| ENSMUSG00000004113 | -1.063792678 | 0.71950142 |
| ENSMUSG00000004127 | -1.142983494 | 0.431683218 |
| ENSMUSG00000004151 | 1.066876115 | 0.716562452 |
| ENSMUSG00000004187 | -1.0266575 | 0.968623752 |
| ENSMUSG00000004207 | 1.156264564 | 0.611443851 |
| ENSMUSG00000004221 | 1.184483257 | 0.382981515 |
| ENSMUSG00000004231 | -1.312461824 | 0.363271112 |
| ENSMUSG00000004233 | -1.035009718 | 0.828133867 |
| ENSMUSG00000004263 | 1.076795819 | 0.608497229 |
| ENSMUSG00000004264 | 1.053100909 | 0.819222912 |
| ENSMUSG00000004266 | 1.178159053 | 0.052692618 |
| ENSMUSG00000004267 | 1.185507928 | 0.071499573 |
| ENSMUSG00000004268 | -1.004972656 | 1 |

| | | |
|---|---|---|
| ENSMUSG00000004270 | -1.017892718 | 0.978259071 |
| ENSMUSG00000004285 | 1.102757238 | 0.415659839 |
| ENSMUSG00000004296 | -1.044273782 | 0.698112512 |
| ENSMUSG00000004317 | 1.021282476 | 0.982541766 |
| ENSMUSG00000004319 | -1.098004898 | 0.698112512 |
| ENSMUSG00000004328 | -1.043306199 | 0.698112512 |
| ENSMUSG00000004341 | 1 | |
| ENSMUSG00000004344 | 1 | |
| ENSMUSG00000004347 | -1.049876361 | 0.683065028 |
| ENSMUSG00000004356 | 1.038260903 | 0.941387596 |
| ENSMUSG00000004359 | -1.074132086 | 0.698112512 |
| ENSMUSG00000004360 | 1.681502415 | 0.003129069 |
| ENSMUSG00000004364 | 1.015036432 | 1 |
| ENSMUSG00000004366 | 1 | |
| ENSMUSG00000004371 | 1.211752532 | 0.394258871 |
| ENSMUSG00000004383 | 1.125060224 | 0.575027867 |
| ENSMUSG00000004393 | -1.024686417 | 0.939477533 |
| ENSMUSG00000004394 | -1.005782267 | 1 |
| ENSMUSG00000004415 | 1.03159717 | 0.973126641 |
| ENSMUSG00000004446 | 1.115041484 | 0.608497229 |
| ENSMUSG00000004451 | 1.094171517 | 0.632015291 |
| ENSMUSG00000004455 | 1.036483797 | 0.950120678 |
| ENSMUSG00000004460 | 1.011609127 | 1 |
| ENSMUSG00000004462 | 1.096597813 | 0.403010702 |
| ENSMUSG00000004473 | -1.223915092 | 0.240114463 |
| ENSMUSG00000004500 | 1.005755373 | 0.999180249 |
| ENSMUSG00000004508 | 1.037903429 | 0.804390383 |
| ENSMUSG00000004530 | 1.234659982 | 0.250477583 |
| ENSMUSG00000004535 | 1.039206123 | 0.924324458 |
| ENSMUSG00000004540 | -1.002329314 | 1 |
| ENSMUSG00000004542 | -1.059043112 | 0.842500885 |
| ENSMUSG00000004552 | 1 | |
| ENSMUSG00000004558 | -1.224425797 | 0.080928024 |
| ENSMUSG00000004561 | -1.080194044 | 0.48990233 |
| ENSMUSG00000004562 | -1.032519472 | 0.846346254 |
| ENSMUSG00000004565 | 1.183748292 | 0.123116889 |
| ENSMUSG00000004567 | 1.326149416 | 0.055300829 |
| ENSMUSG00000004568 | 1.210307451 | 0.2399089 |
| ENSMUSG00000004591 | 1.021045459 | 0.990051834 |
| ENSMUSG00000004609 | -1.280138841 | 0.284069684 |
| ENSMUSG00000004610 | 1.115319982 | 0.36644281 |
| ENSMUSG00000004612 | 1 | |
| ENSMUSG00000004613 | 1 | |
| ENSMUSG00000004626 | 1.030026509 | 0.832128735 |
| ENSMUSG00000004630 | 1.059463094 | 0.752398528 |
| ENSMUSG00000004631 | 1.110474288 | 0.608497229 |
| ENSMUSG00000004633 | -1.578044517 | 0.007137365 |
| ENSMUSG00000004637 | -1.112828613 | 0.088040386 |
| ENSMUSG00000004642 | -1.030711705 | 0.951551605 |
| ENSMUSG00000004651 | 1.00391473 | 1 |
| ENSMUSG00000004654 | 1.132639128 | 0.217803313 |
| ENSMUSG00000004655 | 1.675641634 | 0.000182192 |
| ENSMUSG00000004661 | -1.007131586 | 0.959583351 |
| ENSMUSG00000004665 | 1.265503835 | 0.17934439 |
| ENSMUSG00000004667 | 1.069492917 | 0.667576553 |
| ENSMUSG00000004668 | -1.043306199 | 0.798487595 |
| ENSMUSG00000004677 | 1.101064501 | 0.614071182 |
| ENSMUSG00000004698 | 1.667057796 | 0.014320264 |
| ENSMUSG00000004707 | -1.276951634 | 0.013985847 |
| ENSMUSG00000004709 | 1 | |
| ENSMUSG00000004730 | 1 | |
| ENSMUSG00000004748 | 1.002918503 | 1 |
| ENSMUSG00000004768 | 1.041730615 | 0.858049519 |

| | | |
|---|---|---|
| ENSMUSG00000004771 | -1.060136054 | 0.805033642 |
| ENSMUSG00000004788 | 1.142507795 | 0.208647437 |
| ENSMUSG00000004789 | 1.104242704 | 0.698112512 |
| ENSMUSG00000004791 | 1.583016372 | 0.001400491 |
| ENSMUSG00000004798 | -1.076812864 | 0.723432738 |
| ENSMUSG00000004814 | 1 | |
| ENSMUSG00000004815 | 1.09214219 | 0.314088903 |
| ENSMUSG00000004821 | 1 | |
| ENSMUSG00000004837 | -1.178184847 | 0.608497229 |
| ENSMUSG00000004842 | -1.190772645 | 0.367237552 |
| ENSMUSG00000004843 | 1.029638576 | 0.932792747 |
| ENSMUSG00000004846 | 1.014699025 | 0.988424085 |
| ENSMUSG00000004849 | -1.021144867 | 0.971782973 |
| ENSMUSG00000004864 | -1.06216425 | 0.698112512 |
| ENSMUSG00000004865 | 1.064472488 | 0.81152628 |
| ENSMUSG00000004872 | -1.056599098 | 0.698112512 |
| ENSMUSG00000004880 | -1.034406391 | 0.910155457 |
| ENSMUSG00000004885 | 1.305940885 | 0.003094517 |
| ENSMUSG00000004891 | -1.013192705 | 1 |
| ENSMUSG00000004892 | -1.183317168 | 0.519338787 |
| ENSMUSG00000004894 | 1.019083579 | 0.949812939 |
| ENSMUSG00000004895 | -1.033680777 | 0.837784532 |
| ENSMUSG00000004896 | -1.011750723 | 0.961127302 |
| ENSMUSG00000004897 | 1.05900028 | 0.872412742 |
| ENSMUSG00000004902 | -1.038463447 | 0.962290304 |
| ENSMUSG00000004929 | -1.080161769 | 0.569629019 |
| ENSMUSG00000004931 | 1.036560823 | 0.899940304 |
| ENSMUSG00000004933 | -1.078531255 | 0.617786513 |
| ENSMUSG00000004934 | 1.094393389 | 0.325381273 |
| ENSMUSG00000004936 | -1.027868783 | 0.94695034 |
| ENSMUSG00000004937 | 1.026740838 | 0.853246308 |
| ENSMUSG00000004939 | -1.002064566 | 1 |
| ENSMUSG00000004945 | -1.010720485 | 0.998981421 |
| ENSMUSG00000004947 | 1.119977956 | 0.158121745 |
| ENSMUSG00000004948 | -1.054169715 | 0.813485995 |
| ENSMUSG00000004951 | -1.03465509 | 0.7938895 |
| ENSMUSG00000004952 | 1.126008167 | 0.103917091 |
| ENSMUSG00000004961 | -1.086092137 | 0.699340382 |
| ENSMUSG00000004980 | 1.042893028 | 0.956977788 |
| ENSMUSG00000004988 | 1.045283284 | 0.935109793 |
| ENSMUSG00000004994 | 1.043232048 | 0.608497229 |
| ENSMUSG00000004996 | 1.074443242 | 0.217803313 |
| ENSMUSG00000005034 | 1.016820614 | 0.99660387 |
| ENSMUSG00000005043 | 1.020009336 | 0.860341518 |
| ENSMUSG00000005045 | 1.214928407 | 0.049968879 |
| ENSMUSG00000005054 | 1.050542892 | 0.818144938 |
| ENSMUSG00000005057 | 1.061818929 | 0.698112512 |
| ENSMUSG00000005069 | 1.082372243 | 0.608497229 |
| ENSMUSG00000005078 | 1.11438504 | 0.415214532 |
| ENSMUSG00000005087 | -1.044746405 | 0.918280975 |
| ENSMUSG00000005089 | -1.122480399 | 0.357169145 |
| ENSMUSG00000005102 | -1.097766389 | 0.574350373 |
| ENSMUSG00000005103 | 1.26910077 | 0.369930124 |
| ENSMUSG00000005107 | 1.098658766 | 0.422263747 |
| ENSMUSG00000005124 | 1.141763383 | 0.652382402 |
| ENSMUSG00000005125 | 1.21491495 | 0.25953157 |
| ENSMUSG00000005131 | -1.065852599 | 0.803343809 |
| ENSMUSG00000005142 | -1.065704687 | 0.698112512 |
| ENSMUSG00000005148 | -1.135884004 | 0.037404406 |
| ENSMUSG00000005150 | -1.101178599 | 0.290098581 |
| ENSMUSG00000005161 | -1.012974956 | 1 |
| ENSMUSG00000005198 | -1.089995128 | 0.790851299 |
| ENSMUSG00000005202 | -1.053549062 | 0.778372272 |

| | | |
|---|---|---|
| ENSMUSG00000005204 | -1.054129756 | 0.790753571 |
| ENSMUSG00000005220 | -1.176660386 | 0.317733967 |
| ENSMUSG00000005225 | -1.00219595 | 1 |
| ENSMUSG00000005232 | -1 | 1 |
| ENSMUSG00000005233 | -1.11500082 | 0.60490228 |
| ENSMUSG00000005237 | 1.141849885 | 0.698112512 |
| ENSMUSG00000005251 | -2.23526722 | 0.000812907 |
| ENSMUSG00000005262 | -1.053879145 | 0.817152639 |
| ENSMUSG00000005267 | -1.104218192 | 0.457528841 |
| ENSMUSG00000005268 | 1.178345123 | 0.518775336 |
| ENSMUSG00000005299 | -1.001080218 | 1 |
| ENSMUSG00000005312 | -1.049078539 | 0.880698868 |
| ENSMUSG00000005320 | -1.305601963 | 0.18211237 |
| ENSMUSG00000005338 | -1.633236109 | 0.000101355 |
| ENSMUSG00000005339 | 1 | |
| ENSMUSG00000005354 | 1.064651548 | 0.748682651 |
| ENSMUSG00000005355 | -1.076770764 | 0.867068766 |
| ENSMUSG00000005357 | -1.385171563 | 0.080861974 |
| ENSMUSG00000005360 | 1.495577996 | 0.000484809 |
| ENSMUSG00000005362 | 1.05482897 | 0.773487331 |
| ENSMUSG00000005364 | 1.003940447 | 1 |
| ENSMUSG00000005370 | 1.050787096 | 0.87354792 |
| ENSMUSG00000005371 | 1.077383767 | 0.764664548 |
| ENSMUSG00000005373 | -1.484896381 | 0.080496557 |
| ENSMUSG00000005374 | 1.020282342 | 0.957288273 |
| ENSMUSG00000005378 | -1.00829403 | 0.998378474 |
| ENSMUSG00000005397 | 1.104273303 | 0.72956252 |
| ENSMUSG00000005410 | -1.009206897 | 1 |
| ENSMUSG00000005413 | 1.257515953 | 0.064245831 |
| ENSMUSG00000005417 | -1.036825905 | 0.947618587 |
| ENSMUSG00000005442 | 1.221683434 | 0.067294168 |
| ENSMUSG00000005447 | 1.108622386 | 0.423856307 |
| ENSMUSG00000005465 | -1.198452191 | 0.245568722 |
| ENSMUSG00000005469 | -1.051895244 | 0.871647276 |
| ENSMUSG00000005470 | -1.164225566 | 0.477203336 |
| ENSMUSG00000005474 | 1.212410478 | 0.121862618 |
| ENSMUSG00000005481 | 1.047778031 | 0.877593058 |
| ENSMUSG00000005483 | -1.048015816 | 0.836520228 |
| ENSMUSG00000005493 | 1.213163396 | 0.534944201 |
| ENSMUSG00000005501 | -1.029407297 | 0.934760809 |
| ENSMUSG00000005503 | 1 | |
| ENSMUSG00000005505 | -1.183298953 | 0.217849559 |
| ENSMUSG00000005506 | -1.102410897 | 0.719181945 |
| ENSMUSG00000005510 | -1.248700082 | 0.149426766 |
| ENSMUSG00000005514 | 1.201081023 | 0.139677126 |
| ENSMUSG00000005533 | 1.177160669 | 0.075650942 |
| ENSMUSG00000005534 | 1.237875133 | 0.202428693 |
| ENSMUSG00000005540 | -1.05143605 | 0.803072249 |
| ENSMUSG00000005547 | 1 | |
| ENSMUSG00000005553 | 1.026321584 | 0.962058001 |
| ENSMUSG00000005566 | 1.084067079 | 0.750067671 |
| ENSMUSG00000005575 | 1.048143776 | 0.841614159 |
| ENSMUSG00000005580 | -1.005835707 | 1 |
| ENSMUSG00000005583 | 1.098770239 | 0.369232813 |
| ENSMUSG00000005609 | -1.006686562 | 1 |
| ENSMUSG00000005610 | 1.049220719 | 0.946099281 |
| ENSMUSG00000005611 | 1.297259178 | 0.269530391 |
| ENSMUSG00000005615 | -1.017980109 | 0.982838256 |
| ENSMUSG00000005621 | 1.063406361 | 0.698112512 |
| ENSMUSG00000005625 | -1.007037304 | 1 |
| ENSMUSG00000005628 | 1.091523016 | 0.736695141 |
| ENSMUSG00000005640 | -1.069502966 | 0.744868995 |
| ENSMUSG00000005649 | -1.053549062 | 0.698112512 |

| | | |
|---|---|---|
| ENSMUSG00000005656 | 1.015678996 | 1 |
| ENSMUSG00000005667 | 1.048016995 | 0.875804019 |
| ENSMUSG00000005672 | 2.516680781 | 0.000381466 |
| ENSMUSG00000005674 | 1.041612903 | 0.617510841 |
| ENSMUSG00000005677 | -1.019108526 | 0.979258908 |
| ENSMUSG00000005681 | -1.251408229 | 0.031516897 |
| ENSMUSG00000005682 | -1.043832955 | 0.842184576 |
| ENSMUSG00000005683 | 1.010466144 | 1 |
| ENSMUSG00000005686 | 1.160977087 | 0.074225911 |
| ENSMUSG00000005687 | 1.007863838 | 1 |
| ENSMUSG00000005696 | -1.063525418 | 0.751979636 |
| ENSMUSG00000005698 | -1.075287525 | 0.776021306 |
| ENSMUSG00000005699 | 1.066004108 | 0.698112512 |
| ENSMUSG00000005705 | 1.099895318 | 0.397756797 |
| ENSMUSG00000005716 | 1 | |
| ENSMUSG00000005718 | 1.140633989 | 0.125254383 |
| ENSMUSG00000005732 | 1.024981546 | 0.961487493 |
| ENSMUSG00000005763 | 1.24122829 | 0.181635915 |
| ENSMUSG00000005774 | -1.119574413 | 0.262806495 |
| ENSMUSG00000005779 | -1.078592551 | 0.772656514 |
| ENSMUSG00000005800 | 1 | |
| ENSMUSG00000005802 | -1.012608093 | 1 |
| ENSMUSG00000005803 | -1.11363513 | 0.431748912 |
| ENSMUSG00000005804 | 1.057201505 | 0.728027496 |
| ENSMUSG00000005813 | -1.069216967 | 0.784381159 |
| ENSMUSG00000005823 | 1.019846485 | 0.949218699 |
| ENSMUSG00000005824 | 1 | |
| ENSMUSG00000005836 | -1.409737295 | 0.004558819 |
| ENSMUSG00000005846 | 1.011910328 | 1 |
| ENSMUSG00000005864 | -1.043306199 | 0.698112512 |
| ENSMUSG00000005871 | 1.007038098 | 1 |
| ENSMUSG00000005873 | -1.037064903 | 0.937719236 |
| ENSMUSG00000005881 | 1.00172148 | 1 |
| ENSMUSG00000005882 | -1.152276757 | 0.416883739 |
| ENSMUSG00000005883 | -1.219624708 | 0.224566943 |
| ENSMUSG00000005886 | 1.049581929 | 0.815954183 |
| ENSMUSG00000005891 | 1 | |
| ENSMUSG00000005892 | 1 | |
| ENSMUSG00000005893 | 1.076161968 | 0.716161214 |
| ENSMUSG00000005897 | 1.063079278 | 0.665697605 |
| ENSMUSG00000005899 | 1.008425303 | 1 |
| ENSMUSG00000005907 | 1.038389931 | 0.790885744 |
| ENSMUSG00000005917 | -1.053686075 | 0.642639207 |
| ENSMUSG00000005936 | -1.013184119 | 1 |
| ENSMUSG00000005947 | -1.074881228 | 0.701687166 |
| ENSMUSG00000005949 | 1.224027391 | 0.066445442 |
| ENSMUSG00000005950 | -1.218468733 | 0.00466733 |
| ENSMUSG00000005951 | -1.114005386 | 0.536673416 |
| ENSMUSG00000005952 | -1 | 1 |
| ENSMUSG00000005958 | 1.317787375 | 0.020246535 |
| ENSMUSG00000005968 | 1.091230894 | 0.608497229 |
| ENSMUSG00000005973 | -1.041677864 | 0.905701209 |
| ENSMUSG00000005980 | -1.055160832 | 0.717987279 |
| ENSMUSG00000005981 | 1.035856672 | 0.914862692 |
| ENSMUSG00000005982 | 1.110880749 | 0.235064476 |
| ENSMUSG00000005983 | 1.061917942 | 0.741893019 |
| ENSMUSG00000005986 | -1.149103513 | 0.385302577 |
| ENSMUSG00000005994 | 1 | |
| ENSMUSG00000006005 | 1.002046855 | 1 |
| ENSMUSG00000006007 | 1 | |
| ENSMUSG00000006010 | -1.017021009 | 0.985623907 |
| ENSMUSG00000006014 | -1.218341794 | 0.185602567 |
| ENSMUSG00000006019 | 1.145505917 | 0.289929426 |

| | | |
|---|---|---|
| ENSMUSG00000006021 | -1.042320107 | 0.759784795 |
| ENSMUSG00000006024 | 1.081972281 | 0.688874325 |
| ENSMUSG00000006050 | 1.042335557 | 0.832767487 |
| ENSMUSG00000006056 | -1.22915166 | 0.343739781 |
| ENSMUSG00000006057 | 1.042438806 | 0.832803619 |
| ENSMUSG00000006058 | -1.026292008 | 0.956374899 |
| ENSMUSG00000006095 | 1.0134651 | 0.969592305 |
| ENSMUSG00000006127 | 1.032770729 | 0.779445807 |
| ENSMUSG00000006134 | 1.009555806 | 1 |
| ENSMUSG00000006143 | 1.141411496 | 0.608497229 |
| ENSMUSG00000006154 | -1 | 1 |
| ENSMUSG00000006169 | 1.005451928 | 1 |
| ENSMUSG00000006179 | -2.135403701 | 0.014320264 |
| ENSMUSG00000006191 | -1.023758389 | 0.911521144 |
| ENSMUSG00000006200 | 1 | |
| ENSMUSG00000006204 | -1.142727245 | 0.202428693 |
| ENSMUSG00000006205 | -1.058520671 | 0.812228807 |
| ENSMUSG00000006215 | 1.059227756 | 0.686538971 |
| ENSMUSG00000006216 | 1.229451956 | 0.396246591 |
| ENSMUSG00000006218 | -1.110395831 | 0.70299173 |
| ENSMUSG00000006219 | 1.357723523 | 0.061839295 |
| ENSMUSG00000006221 | 1.66355728 | 0.025137317 |
| ENSMUSG00000006235 | 1.001663721 | 1 |
| ENSMUSG00000006241 | -1.043266368 | 0.698112512 |
| ENSMUSG00000006262 | 1.103700319 | 0.698112512 |
| ENSMUSG00000006269 | -1.13161737 | 0.685708044 |
| ENSMUSG00000006270 | 1 | |
| ENSMUSG00000006273 | 1.125434887 | 0.608497229 |
| ENSMUSG00000006276 | -1.008895238 | 1 |
| ENSMUSG00000006281 | 1.002311757 | 1 |
| ENSMUSG00000006288 | -1.150684558 | 0.149977043 |
| ENSMUSG00000006289 | -1.045562841 | 0.803852736 |
| ENSMUSG00000006299 | 1.005617131 | 1 |
| ENSMUSG00000006301 | 1.026830256 | 0.950684488 |
| ENSMUSG00000006304 | 1.040271336 | 0.93957055 |
| ENSMUSG00000006307 | 1.113607924 | 0.348477203 |
| ENSMUSG00000006310 | -1.072460681 | 0.857167632 |
| ENSMUSG00000006311 | 1.04424822 | 0.814523317 |
| ENSMUSG00000006313 | 1.359142746 | 0.200917987 |
| ENSMUSG00000006315 | -1.040931472 | 0.816416314 |
| ENSMUSG00000006333 | 1.096477429 | 0.693579485 |
| ENSMUSG00000006335 | -1.056055834 | 0.610486174 |
| ENSMUSG00000006342 | 1.079756444 | 0.698112512 |
| ENSMUSG00000006344 | 1.059463094 | 0.750182067 |
| ENSMUSG00000006345 | 1.150421103 | 0.608497229 |
| ENSMUSG00000006356 | 1.259404211 | 0.081382929 |
| ENSMUSG00000006360 | 1.180816183 | 0.048809518 |
| ENSMUSG00000006362 | -1.24359105 | 0.108172814 |
| ENSMUSG00000006369 | 1.045335916 | 0.698112512 |
| ENSMUSG00000006373 | 1.068332311 | 0.790723371 |
| ENSMUSG00000006378 | 1.017002935 | 0.951744672 |
| ENSMUSG00000006386 | 1.331891231 | 0.16125008 |
| ENSMUSG00000006389 | 1 | |
| ENSMUSG00000006390 | 1.053487383 | 0.764429998 |
| ENSMUSG00000006392 | 1.017308288 | 0.972980766 |
| ENSMUSG00000006395 | 1.046574362 | 0.608497229 |
| ENSMUSG00000006398 | 1.032738153 | 0.913105899 |
| ENSMUSG00000006403 | -1.237261395 | 0.0045402 |
| ENSMUSG00000006411 | -1.054495217 | 0.915158215 |
| ENSMUSG00000006412 | -1.088672079 | 0.624448448 |
| ENSMUSG00000006418 | -1.117044903 | 0.56488333 |
| ENSMUSG00000006423 | 1.022796029 | 0.978109404 |
| ENSMUSG00000006435 | 1.162632799 | 0.026024582 |

| | | |
|---|---|---|
| ENSMUSG00000006442 | 1.019004452 | 0.971162438 |
| ENSMUSG00000006445 | -1.039267301 | 0.845175363 |
| ENSMUSG00000006456 | 1.044904373 | 0.736737342 |
| ENSMUSG00000006457 | 1.78678193 | 0.099116846 |
| ENSMUSG00000006462 | 1.104390967 | 0.714200725 |
| ENSMUSG00000006463 | -1.01099163 | 0.968937442 |
| ENSMUSG00000006464 | -1.04831269 | 0.629824986 |
| ENSMUSG00000006469 | 1 | |
| ENSMUSG00000006471 | 1.087733646 | 0.562856014 |
| ENSMUSG00000006476 | -1.033521174 | 0.828564184 |
| ENSMUSG00000006488 | 1.051463648 | 0.940197654 |
| ENSMUSG00000006490 | 1 | |
| ENSMUSG00000006494 | -1.20759129 | 0.138124791 |
| ENSMUSG00000006498 | 1.111110616 | 0.608497229 |
| ENSMUSG00000006517 | -1.010178377 | 0.970025232 |
| ENSMUSG00000006519 | -2.128226373 | 0.000610536 |
| ENSMUSG00000006522 | 1.267400994 | 0.159066803 |
| ENSMUSG00000006526 | -1.110410119 | 0.317011706 |
| ENSMUSG00000006527 | -1.120819874 | 0.532369657 |
| ENSMUSG00000006529 | 1 | |
| ENSMUSG00000006538 | -1.069502966 | 0.699340382 |
| ENSMUSG00000006542 | -1.169530994 | 0.245670602 |
| ENSMUSG00000006546 | 1 | |
| ENSMUSG00000006567 | -1.199442126 | 0.608497229 |
| ENSMUSG00000006570 | 1 | |
| ENSMUSG00000006574 | 1 | |
| ENSMUSG00000006575 | 1.000863844 | 1 |
| ENSMUSG00000006576 | 1.067755309 | 0.793146717 |
| ENSMUSG00000006585 | 1.050225497 | 0.698112512 |
| ENSMUSG00000006586 | 1.038592733 | 0.798186535 |
| ENSMUSG00000006587 | -1.49076128 | 0.027656649 |
| ENSMUSG00000006589 | 1.031056384 | 0.90453963 |
| ENSMUSG00000006599 | 1.154042721 | 0.374710101 |
| ENSMUSG00000006611 | -1.288854693 | 0.049065949 |
| ENSMUSG00000006638 | -1.042917739 | 0.836532337 |
| ENSMUSG00000006641 | 1.08371113 | 0.413623796 |
| ENSMUSG00000006642 | 1.069068979 | 0.767473873 |
| ENSMUSG00000006649 | -1.004895262 | 1 |
| ENSMUSG00000006651 | -1.109853765 | 0.367040972 |
| ENSMUSG00000006673 | -1.001098702 | 1 |
| ENSMUSG00000006675 | 1.00653617 | 1 |
| ENSMUSG00000006676 | 1.037236666 | 0.856915299 |
| ENSMUSG00000006678 | -1.002488069 | 1 |
| ENSMUSG00000006699 | -1.001482688 | 1 |
| ENSMUSG00000006705 | -1.01254837 | 0.990316106 |
| ENSMUSG00000006711 | -1.083478679 | 0.608497229 |
| ENSMUSG00000006715 | -1.130616747 | 0.592265042 |
| ENSMUSG00000006717 | 1.090600699 | 0.560827127 |
| ENSMUSG00000006720 | -1.079282873 | 0.315920721 |
| ENSMUSG00000006724 | -1.096299593 | 0.768407406 |
| ENSMUSG00000006728 | 1.1233601 | 0.698112512 |
| ENSMUSG00000006731 | -1.178944003 | 0.124353172 |
| ENSMUSG00000006732 | -1.040797295 | 0.741283521 |
| ENSMUSG00000006736 | -1.078163248 | 0.698112512 |
| ENSMUSG00000006740 | 1.060236781 | 0.874775588 |
| ENSMUSG00000006763 | 1.034822513 | 0.832617993 |
| ENSMUSG00000006764 | 1.055416288 | 0.848848499 |
| ENSMUSG00000006777 | 1 | |
| ENSMUSG00000006782 | -1.328016435 | 0.016272817 |
| ENSMUSG00000006784 | 1.094162905 | 0.687710355 |
| ENSMUSG00000006800 | 1.212784725 | 0.125155544 |
| ENSMUSG00000006818 | -1.063952039 | 0.754389693 |
| ENSMUSG00000006850 | 1.134408909 | 0.117245227 |

| | | |
|---|---|---|
| ENSMUSG00000006906 | 1.001650434 | 1 |
| ENSMUSG00000006920 | -1.003062478 | 1 |
| ENSMUSG00000006930 | -1.569637869 | 0.001867243 |
| ENSMUSG00000006931 | 1.161205817 | 0.293309442 |
| ENSMUSG00000006932 | 1.086509113 | 0.802320762 |
| ENSMUSG00000006941 | -1.037739083 | 0.839420872 |
| ENSMUSG00000006948 | 1 | |
| ENSMUSG00000006958 | -1.183744126 | 0.241584492 |
| ENSMUSG00000006998 | -1.005647454 | 1 |
| ENSMUSG00000007021 | -1.551459334 | 0.00218008 |
| ENSMUSG00000007029 | 1.126613533 | 0.534944201 |
| ENSMUSG00000007030 | 1.011969043 | 1 |
| ENSMUSG00000007033 | 1.050568897 | 0.719653179 |
| ENSMUSG00000007034 | -1.113515092 | 0.608497229 |
| ENSMUSG00000007035 | 1.069280848 | 0.638390533 |
| ENSMUSG00000007036 | -1.006726521 | 1 |
| ENSMUSG00000007038 | 1.165454482 | 0.147255843 |
| ENSMUSG00000007039 | 1.212345204 | 0.106715962 |
| ENSMUSG00000007041 | 1.145980079 | 0.608497229 |
| ENSMUSG00000007050 | 1.043164559 | 0.837704961 |
| ENSMUSG00000007080 | 1.028308957 | 0.968158081 |
| ENSMUSG00000007097 | -1.038858794 | 0.934024342 |
| ENSMUSG00000007107 | 1 | |
| ENSMUSG00000007122 | -1.231157139 | 0.562711584 |
| ENSMUSG00000007207 | -1.042554426 | 0.724904908 |
| ENSMUSG00000007209 | 1 | |
| ENSMUSG00000007216 | 1.076206287 | 0.253355411 |
| ENSMUSG00000007279 | -1.369857598 | 0.034053148 |
| ENSMUSG00000007338 | -1.041828019 | 0.893388118 |
| ENSMUSG00000007379 | -1.026240883 | 0.899894987 |
| ENSMUSG00000007411 | 1.041014945 | 0.878551492 |
| ENSMUSG00000007415 | 1.040923069 | 0.902754356 |
| ENSMUSG00000007440 | -1 | 1 |
| ENSMUSG00000007457 | 1 | |
| ENSMUSG00000007458 | 1.02184695 | 0.979814083 |
| ENSMUSG00000007476 | 1.127735804 | 0.188909523 |
| ENSMUSG00000007480 | 1.11166259 | 0.608497229 |
| ENSMUSG00000007564 | 1.062639603 | 0.849630524 |
| ENSMUSG00000007570 | 1.13254951 | 0.262993046 |
| ENSMUSG00000007589 | -1.16558068 | 0.120016484 |
| ENSMUSG00000007591 | 1.02238451 | 0.885184589 |
| ENSMUSG00000007594 | 1.115885423 | 0.283607343 |
| ENSMUSG00000007603 | 1.0051971 | 1 |
| ENSMUSG00000007610 | 1.013277621 | 0.950378392 |
| ENSMUSG00000007613 | 1.199944683 | 0.369721892 |
| ENSMUSG00000007617 | -1.112222137 | 0.604452802 |
| ENSMUSG00000007646 | -1.073479641 | 0.70299173 |
| ENSMUSG00000007653 | 1.069068979 | 0.608497229 |
| ENSMUSG00000007655 | 1.07886389 | 0.809950986 |
| ENSMUSG00000007656 | -1.046993082 | 0.908543402 |
| ENSMUSG00000007659 | 1.04726606 | 0.695921515 |
| ENSMUSG00000007670 | 1.102611968 | 0.608497229 |
| ENSMUSG00000007682 | -1.053549062 | 0.807892349 |
| ENSMUSG00000007721 | 1.068027117 | 0.73343521 |
| ENSMUSG00000007739 | -1.079787938 | 0.817330349 |
| ENSMUSG00000007777 | -1.006289115 | 1 |
| ENSMUSG00000007783 | 1.080879236 | 0.492813798 |
| ENSMUSG00000007805 | 1.006629624 | 0.933031586 |
| ENSMUSG00000007812 | 1.051913116 | 0.834794437 |
| ENSMUSG00000007815 | -1.00091728 | 1 |
| ENSMUSG00000007817 | 1.078705161 | 0.64269949 |
| ENSMUSG00000007827 | 1.038361574 | 0.879253209 |
| ENSMUSG00000007833 | 1.124516595 | 0.513603252 |

| | | |
|---|---|---|
| ENSMUSG00000007836 | -1.041548794 | 0.926178824 |
| ENSMUSG00000007837 | 1.152597189 | 0.248095211 |
| ENSMUSG00000007850 | 1.002353116 | 1 |
| ENSMUSG00000007867 | 1.13676345 | 0.205858736 |
| ENSMUSG00000007872 | 1.445932853 | 0.012944412 |
| ENSMUSG00000007877 | 1.183690384 | 0.417931834 |
| ENSMUSG00000007880 | 1.143556957 | 0.485348239 |
| ENSMUSG00000007888 | 1.214779252 | 0.074402564 |
| ENSMUSG00000007891 | 1.201038801 | 0.43994505 |
| ENSMUSG00000007892 | 1.164415576 | 0.44650239 |
| ENSMUSG00000007907 | 1 | |
| ENSMUSG00000007908 | 1.754202794 | 0.002119905 |
| ENSMUSG00000007944 | 1 | |
| ENSMUSG00000007946 | 1.024693402 | 0.933031586 |
| ENSMUSG00000007950 | 1.145433283 | 0.433678407 |
| ENSMUSG00000007987 | -1.025777519 | 0.835367905 |
| ENSMUSG00000007989 | -1.137256632 | 0.292973656 |
| ENSMUSG00000008028 | -1.085651418 | 0.532369657 |
| ENSMUSG00000008035 | 1.407056638 | 0.00038977 |
| ENSMUSG00000008036 | -1.00394748 | 1 |
| ENSMUSG00000008090 | 1.156760223 | 0.188651469 |
| ENSMUSG00000008129 | -1.165615518 | 0.305230222 |
| ENSMUSG00000008136 | 1.36529092 | 0.033934076 |
| ENSMUSG00000008140 | 1.150402625 | 0.183504366 |
| ENSMUSG00000008153 | -1.331926223 | 0.33717311 |
| ENSMUSG00000008167 | 1.029168933 | 0.834580969 |
| ENSMUSG00000008193 | -1.180656251 | 0.433615344 |
| ENSMUSG00000008200 | -1.111188606 | 0.608497229 |
| ENSMUSG00000008206 | 1.275925073 | 0.025691028 |
| ENSMUSG00000008226 | -1.100690685 | 0.608497229 |
| ENSMUSG00000008301 | 1.021823208 | 0.943705943 |
| ENSMUSG00000008305 | 1.138205395 | 0.087589235 |
| ENSMUSG00000008307 | -1.016269825 | 0.918009323 |
| ENSMUSG00000008318 | 1.137379028 | 0.034835992 |
| ENSMUSG00000008333 | -1.090562576 | 0.698112512 |
| ENSMUSG00000008348 | 1.076954681 | 0.734674233 |
| ENSMUSG00000008373 | 1.004315427 | 1 |
| ENSMUSG00000008384 | 1.085855623 | 0.300750932 |
| ENSMUSG00000008393 | -1.011775781 | 1 |
| ENSMUSG00000008398 | 1.120365399 | 0.579219991 |
| ENSMUSG00000008429 | 1.004581272 | 1 |
| ENSMUSG00000008435 | 1.106517474 | 0.373661522 |
| ENSMUSG00000008438 | 1.154397872 | 0.698112512 |
| ENSMUSG00000008450 | -1.031316199 | 0.918280975 |
| ENSMUSG00000008461 | 1.161036672 | 0.251049598 |
| ENSMUSG00000008475 | 1.20025218 | 0.478706958 |
| ENSMUSG00000008482 | -1.005854108 | 1 |
| ENSMUSG00000008489 | 1.112282092 | 0.624676024 |
| ENSMUSG00000008496 | -1.13409489 | 0.613963375 |
| ENSMUSG00000008540 | 1.726303584 | 0.000658499 |
| ENSMUSG00000008575 | 1.061281411 | 0.823681172 |
| ENSMUSG00000008590 | 1.069068979 | 0.608497229 |
| ENSMUSG00000008601 | -1.001430011 | 1 |
| ENSMUSG00000008604 | 1.001903278 | 1 |
| ENSMUSG00000008658 | 1.844425718 | 0.000658313 |
| ENSMUSG00000008668 | 1.08130461 | 0.740381823 |
| ENSMUSG00000008682 | 1.090052659 | 0.698112512 |
| ENSMUSG00000008683 | -1.003521765 | 1 |
| ENSMUSG00000008690 | 1.048979359 | 0.8730672 |
| ENSMUSG00000008730 | 1.05903733 | 0.807247723 |
| ENSMUSG00000008734 | 1.017332338 | 0.870683145 |
| ENSMUSG00000008763 | -1.001181035 | 1 |
| ENSMUSG00000008789 | 1 | |

| | | |
|---|---|---|
| ENSMUSG00000008813 | 1 | |
| ENSMUSG00000008822 | 1.051548787 | 0.658044065 |
| ENSMUSG00000008843 | 1 | |
| ENSMUSG00000008845 | 1.178139999 | 0.33317946 |
| ENSMUSG00000008855 | 1.198863146 | 0.134244013 |
| ENSMUSG00000008859 | -1.022129548 | 0.977730887 |
| ENSMUSG00000008874 | 1 | |
| ENSMUSG00000008892 | -1.019591614 | 0.992917761 |
| ENSMUSG00000008932 | -1.023154242 | 0.990891819 |
| ENSMUSG00000008958 | 1.031183732 | 0.780124168 |
| ENSMUSG00000008976 | 1.033586684 | 0.933895276 |
| ENSMUSG00000008999 | -1.037934057 | 0.934760809 |
| ENSMUSG00000009013 | -1.00786486 | 1 |
| ENSMUSG00000009030 | 1.049975777 | 0.822116066 |
| ENSMUSG00000009035 | 1.09530902 | 0.608497229 |
| ENSMUSG00000009047 | 1 | |
| ENSMUSG00000009070 | -1.063525418 | 0.698112512 |
| ENSMUSG00000009073 | -1.026041075 | 0.956714001 |
| ENSMUSG00000009075 | 1.112843644 | 0.609147349 |
| ENSMUSG00000009076 | -1.068434669 | 0.141932891 |
| ENSMUSG00000009079 | 1.085788464 | 0.710097421 |
| ENSMUSG00000009090 | -1.034060681 | 0.902767556 |
| ENSMUSG00000009092 | 1.03460784 | 0.969044948 |
| ENSMUSG00000009093 | -1.131875116 | 0.640646424 |
| ENSMUSG00000009097 | -1.084645496 | 0.608497229 |
| ENSMUSG00000009108 | 1.181259493 | 0.045301689 |
| ENSMUSG00000009112 | -1.08748533 | 0.698112512 |
| ENSMUSG00000009114 | 1.05038453 | 0.828564184 |
| ENSMUSG00000009115 | 1 | |
| ENSMUSG00000009145 | 1.012194823 | 0.984982552 |
| ENSMUSG00000009185 | -1.003834322 | 1 |
| ENSMUSG00000009207 | -1.055580112 | 0.778372272 |
| ENSMUSG00000009210 | 1 | |
| ENSMUSG00000009214 | 1 | |
| ENSMUSG00000009216 | 1 | |
| ENSMUSG00000009246 | 1.183193407 | 0.509571857 |
| ENSMUSG00000009248 | 1 | |
| ENSMUSG00000009281 | 1.095872691 | 0.608497229 |
| ENSMUSG00000009291 | -1.111719572 | 0.698112512 |
| ENSMUSG00000009292 | 1.112539128 | 0.193727098 |
| ENSMUSG00000009293 | -1.008496016 | 1 |
| ENSMUSG00000009350 | -1.085651418 | 0.615267463 |
| ENSMUSG00000009356 | -1.238472058 | 0.139593229 |
| ENSMUSG00000009376 | 1.110038742 | 0.698112512 |
| ENSMUSG00000009378 | 1.069068979 | 0.608497229 |
| ENSMUSG00000009394 | -1.324001414 | 0.151654465 |
| ENSMUSG00000009406 | 1.173366232 | 0.215991492 |
| ENSMUSG00000009418 | 1.038498991 | 0.89107594 |
| ENSMUSG00000009470 | -1.018525764 | 1 |
| ENSMUSG00000009471 | 1 | |
| ENSMUSG00000009487 | 1.059463094 | 0.608497229 |
| ENSMUSG00000009535 | 1.023560971 | 0.968593113 |
| ENSMUSG00000009545 | -1.085651418 | 0.689423135 |
| ENSMUSG00000009549 | -1.095907998 | 0.624040338 |
| ENSMUSG00000009551 | -1.038858794 | 0.912425817 |
| ENSMUSG00000009555 | 1.1184644 | 0.308708274 |
| ENSMUSG00000009563 | -1.016791712 | 0.934938369 |
| ENSMUSG00000009566 | -1.025341179 | 0.823038846 |
| ENSMUSG00000009569 | 1.126798322 | 0.56853419 |
| ENSMUSG00000009575 | 1.045108857 | 0.949024451 |
| ENSMUSG00000009580 | 1 | |
| ENSMUSG00000009585 | -1.041590234 | 0.844771174 |
| ENSMUSG00000009588 | 1 | |

| | | |
|---|---|---|
| ENSMUSG00000009596 | -1.161107412 | 0.492689376 |
| ENSMUSG00000009614 | 1.412992057 | 0.000556642 |
| ENSMUSG00000009621 | 1.06627489 | 0.608497229 |
| ENSMUSG00000009628 | 1.065818135 | 0.608497229 |
| ENSMUSG00000009630 | 1.007957134 | 1 |
| ENSMUSG00000009633 | 1.045261892 | 0.933031586 |
| ENSMUSG00000009640 | -1.043234854 | 0.764317155 |
| ENSMUSG00000009646 | 1 | |
| ENSMUSG00000009647 | 1.049270772 | 0.787206313 |
| ENSMUSG00000009654 | 1.151140418 | 0.398767938 |
| ENSMUSG00000009670 | -1.03962336 | 0.698112512 |
| ENSMUSG00000009681 | -1.002122302 | 1 |
| ENSMUSG00000009687 | -1.015089955 | 0.968423939 |
| ENSMUSG00000009731 | 1.174353622 | 0.012831121 |
| ENSMUSG00000009733 | 1.024325798 | 0.912464846 |
| ENSMUSG00000009734 | 1 | |
| ENSMUSG00000009739 | -1.059331303 | 0.693468357 |
| ENSMUSG00000009741 | 1.016638547 | 0.987419045 |
| ENSMUSG00000009772 | 1.064222753 | 0.608497229 |
| ENSMUSG00000009828 | 1.004486734 | 1 |
| ENSMUSG00000009863 | 1.022728355 | 0.949679283 |
| ENSMUSG00000009876 | 1.059463094 | 0.822158059 |
| ENSMUSG00000009894 | -1.081269626 | 0.698112512 |
| ENSMUSG00000009900 | 1.05038453 | 0.828564184 |
| ENSMUSG00000009905 | 1.196647922 | 0.091438359 |
| ENSMUSG00000009907 | 1.009441164 | 1 |
| ENSMUSG00000009927 | -1.036047935 | 0.953011516 |
| ENSMUSG00000009941 | -1.28593513 | 0.330381384 |
| ENSMUSG00000009995 | 1.029440478 | 0.904050368 |
| ENSMUSG00000010021 | -1.53309594 | 0.058947589 |
| ENSMUSG00000010025 | -1.102229578 | 0.612478337 |
| ENSMUSG00000010044 | -1.030062723 | 0.951554277 |
| ENSMUSG00000010045 | -1.006224374 | 1 |
| ENSMUSG00000010047 | 1.100389964 | 0.352652488 |
| ENSMUSG00000010048 | -1.02315398 | 0.847902259 |
| ENSMUSG00000010051 | 1.021589722 | 0.904364508 |
| ENSMUSG00000010054 | -1.015865273 | 0.950565815 |
| ENSMUSG00000010057 | -1.071628514 | 0.499503564 |
| ENSMUSG00000010064 | -1.790662441 | 0.001393029 |
| ENSMUSG00000010066 | -1.274926712 | 0.250477583 |
| ENSMUSG00000010067 | 1.01624695 | 0.918370464 |
| ENSMUSG00000010080 | -1.026302926 | 0.79126147 |
| ENSMUSG00000010086 | -1.186706136 | 0.608497229 |
| ENSMUSG00000010095 | 1.08726959 | 0.698112512 |
| ENSMUSG00000010097 | 1.000520032 | 1 |
| ENSMUSG00000010110 | 1.031381208 | 0.907350511 |
| ENSMUSG00000010122 | -1.187111528 | 0.558213369 |
| ENSMUSG00000010136 | 1 | |
| ENSMUSG00000010142 | 1 | |
| ENSMUSG00000010154 | -1.377646069 | 0.005216743 |
| ENSMUSG00000010175 | -1.517031203 | 0.037470738 |
| ENSMUSG00000010205 | -1.056927281 | 0.744237314 |
| ENSMUSG00000010277 | -1.079430267 | 0.814524472 |
| ENSMUSG00000010290 | 1.003470663 | 1 |
| ENSMUSG00000010307 | 1.045239838 | 0.608497229 |
| ENSMUSG00000010311 | 2.028144359 | 0.002932067 |
| ENSMUSG00000010342 | -1.241880425 | 0.400452977 |
| ENSMUSG00000010358 | -1.15693551 | 0.365279648 |
| ENSMUSG00000010362 | -1.091193386 | 0.251396564 |
| ENSMUSG00000010376 | -1.142875238 | 0.608497229 |
| ENSMUSG00000010392 | -1.041061627 | 0.899219824 |
| ENSMUSG00000010406 | -1.11262169 | 0.563698099 |
| ENSMUSG00000010435 | 1 | |

| | | |
|---|---|---|
| ENSMUSG00000010453 | 1.001833584 | 1 |
| ENSMUSG00000010461 | -1.001067386 | 1 |
| ENSMUSG00000010476 | -1.109923838 | 0.569885354 |
| ENSMUSG00000010492 | -1.208421059 | 0.388781089 |
| ENSMUSG00000010505 | -1.043306199 | 0.79291651 |
| ENSMUSG00000010517 | 1.065062871 | 0.799327301 |
| ENSMUSG00000010529 | -1 | 1 |
| ENSMUSG00000010538 | 1.056291762 | 0.760867819 |
| ENSMUSG00000010554 | 1.00693132 | 1 |
| ENSMUSG00000010592 | -1.044273782 | 0.698112512 |
| ENSMUSG00000010601 | 1.142553173 | 0.121862618 |
| ENSMUSG00000010607 | 1.107696111 | 0.608497229 |
| ENSMUSG00000010608 | 1.095750195 | 0.730246345 |
| ENSMUSG00000010609 | -1.107918807 | 0.357395634 |
| ENSMUSG00000010651 | 1.05922735 | 0.848820161 |
| ENSMUSG00000010660 | -1.019517226 | 0.829918229 |
| ENSMUSG00000010663 | -1.012620156 | 1 |
| ENSMUSG00000010721 | -1.118015633 | 0.397645313 |
| ENSMUSG00000010751 | -1.678409063 | 0.004481625 |
| ENSMUSG00000010755 | 1.088125882 | 0.696497392 |
| ENSMUSG00000010760 | 1.019083579 | 0.949812939 |
| ENSMUSG00000010796 | 1 | |
| ENSMUSG00000010797 | -1.210934121 | 0.535052277 |
| ENSMUSG00000010803 | -1.023769022 | 0.934760809 |
| ENSMUSG00000010825 | -1.362965792 | 0.032833606 |
| ENSMUSG00000010830 | 1.089541901 | 0.698112512 |
| ENSMUSG00000010841 | -1 | 1 |
| ENSMUSG00000010911 | -1.227206462 | 0.097580606 |
| ENSMUSG00000010914 | -1.147028664 | 0.345777646 |
| ENSMUSG00000010936 | 1.113129413 | 0.49062471 |
| ENSMUSG00000011008 | 1.692005538 | 0.006411 |
| ENSMUSG00000011034 | -1.044273782 | 0.834556513 |
| ENSMUSG00000011052 | 1.059463094 | 0.608497229 |
| ENSMUSG00000011096 | 1.089203712 | 0.381214697 |
| ENSMUSG00000011114 | -1.062141672 | 0.778150345 |
| ENSMUSG00000011118 | 1.11166259 | 0.649807331 |
| ENSMUSG00000011148 | 1.829786913 | 0.005662914 |
| ENSMUSG00000011154 | 1 | |
| ENSMUSG00000011158 | 1.079547763 | 0.441129618 |
| ENSMUSG00000011171 | -1.223947136 | 0.016189057 |
| ENSMUSG00000011179 | 1.010883372 | 1 |
| ENSMUSG00000011254 | -1.121205327 | 0.434482395 |
| ENSMUSG00000011256 | 1.175569326 | 0.192663913 |
| ENSMUSG00000011257 | 1.120840567 | 0.535052277 |
| ENSMUSG00000011263 | -1.08081673 | 0.698112512 |
| ENSMUSG00000011267 | 1.102978678 | 0.698112512 |
| ENSMUSG00000011305 | -1.021715738 | 1 |
| ENSMUSG00000011306 | -1.029411108 | 0.857327881 |
| ENSMUSG00000011349 | 1 | |
| ENSMUSG00000011350 | 1 | |
| ENSMUSG00000011382 | -1.205944819 | 0.122036074 |
| ENSMUSG00000011427 | -1.056310415 | 0.608497229 |
| ENSMUSG00000011463 | 1 | |
| ENSMUSG00000011486 | -1.168999771 | 0.208940434 |
| ENSMUSG00000011589 | 1.099507187 | 0.169545598 |
| ENSMUSG00000011632 | -1.010960635 | 1 |
| ENSMUSG00000011658 | 1.062197776 | 0.309889276 |
| ENSMUSG00000011751 | 1.124753308 | 0.746451077 |
| ENSMUSG00000011752 | 1.024181541 | 0.986785919 |
| ENSMUSG00000011831 | 1.103623385 | 0.68735438 |
| ENSMUSG00000011832 | 1.06112927 | 0.642725233 |
| ENSMUSG00000011837 | 1.232318359 | 0.022567224 |
| ENSMUSG00000011877 | 1.267739644 | 0.048221774 |

| | | |
|---|---|---|
| ENSMUSG00000011884 | -1.018592044 | 0.97004457 |
| ENSMUSG00000011958 | 1.007752083 | 1 |
| ENSMUSG00000011960 | 1.023812122 | 0.974064994 |
| ENSMUSG00000012017 | 1.285106452 | 0.043387702 |
| ENSMUSG00000012042 | 1 | |
| ENSMUSG00000012076 | 1.04613287 | 0.849464079 |
| ENSMUSG00000012114 | 1.080870632 | 0.593009252 |
| ENSMUSG00000012117 | -1.062209551 | 0.746451077 |
| ENSMUSG00000012123 | -1.122495963 | 0.641039836 |
| ENSMUSG00000012126 | 1.046533325 | 0.892754266 |
| ENSMUSG00000012187 | -1.03962336 | 0.698112512 |
| ENSMUSG00000012211 | 1.059463094 | 0.608497229 |
| ENSMUSG00000012282 | 1 | |
| ENSMUSG00000012296 | 1.114734972 | 0.292023541 |
| ENSMUSG00000012350 | 1 | |
| ENSMUSG00000012396 | 1.123690815 | 0.246522583 |
| ENSMUSG00000012405 | -1.107295714 | 0.684413983 |
| ENSMUSG00000012422 | 1.008635637 | 1 |
| ENSMUSG00000012428 | -1.270052294 | 0.166044784 |
| ENSMUSG00000012429 | -1.052138655 | 0.722165151 |
| ENSMUSG00000012443 | -1.034867963 | 0.956752624 |
| ENSMUSG00000012483 | 1.090390877 | 0.54453778 |
| ENSMUSG00000012519 | 1.084217415 | 0.641039836 |
| ENSMUSG00000012520 | 1 | |
| ENSMUSG00000012535 | 1.021414437 | 0.982800873 |
| ENSMUSG00000012609 | 1.050328964 | 0.766730405 |
| ENSMUSG00000012640 | 1.058962281 | 0.669894357 |
| ENSMUSG00000012705 | -1.089497311 | 0.722951066 |
| ENSMUSG00000012777 | 1 | |
| ENSMUSG00000012819 | 1.291789487 | 0.075826548 |
| ENSMUSG00000012848 | 1.111371845 | 0.634336751 |
| ENSMUSG00000012889 | 1.109265699 | 0.1540272 |
| ENSMUSG00000013033 | 1.073043977 | 0.532369657 |
| ENSMUSG00000013076 | 1.043881607 | 0.879253209 |
| ENSMUSG00000013083 | 1 | |
| ENSMUSG00000013089 | -1.337651097 | 0.037054918 |
| ENSMUSG00000013091 | -1.002437625 | 1 |
| ENSMUSG00000013150 | 1.022020407 | 0.828564184 |
| ENSMUSG00000013155 | -1.031740793 | 0.862843183 |
| ENSMUSG00000013158 | -1.131439726 | 0.671149171 |
| ENSMUSG00000013160 | 1.065986509 | 0.775199822 |
| ENSMUSG00000013236 | 1.074923596 | 0.608497229 |
| ENSMUSG00000013275 | -1.010046084 | 0.994371982 |
| ENSMUSG00000013338 | -1.095294215 | 0.608497229 |
| ENSMUSG00000013353 | 1 | |
| ENSMUSG00000013367 | 1.168488695 | 0.074136375 |
| ENSMUSG00000013415 | 1.067782181 | 0.723401948 |
| ENSMUSG00000013418 | -1.004597703 | 1 |
| ENSMUSG00000013419 | 1.340975935 | 0.002716748 |
| ENSMUSG00000013465 | 1.035617739 | 0.893604387 |
| ENSMUSG00000013483 | -1.024718486 | 0.951399668 |
| ENSMUSG00000013495 | 1.083770809 | 0.466407623 |
| ENSMUSG00000013523 | -1.043306199 | 0.79291651 |
| ENSMUSG00000013539 | 1.016455641 | 0.898307357 |
| ENSMUSG00000013562 | 1.059463094 | 0.608497229 |
| ENSMUSG00000013584 | -1.136679298 | 0.608497229 |
| ENSMUSG00000013593 | -1.073727817 | 0.790851299 |
| ENSMUSG00000013611 | -1.012480761 | 1 |
| ENSMUSG00000013622 | 1.030279511 | 0.9154415 |
| ENSMUSG00000013629 | 1.111042652 | 0.597079148 |
| ENSMUSG00000013643 | 1 | |
| ENSMUSG00000013646 | -1.072934526 | 0.608497229 |
| ENSMUSG00000013653 | -1.193320102 | 0.000306852 |

| | | |
|---|---|---|
| ENSMUSG00000013662 | 1.070560655 | 0.804276221 |
| ENSMUSG00000013663 | 1.124216773 | 0.698112512 |
| ENSMUSG00000013668 | 1 | |
| ENSMUSG00000013698 | -1.055894301 | 0.87071239 |
| ENSMUSG00000013701 | -1.15360627 | 0.353990524 |
| ENSMUSG00000013707 | -1.143452648 | 0.698112512 |
| ENSMUSG00000013736 | 1.052397915 | 0.814089245 |
| ENSMUSG00000013766 | 1 | |
| ENSMUSG00000013787 | 1.063748011 | 0.706029849 |
| ENSMUSG00000013822 | -1.043700557 | 0.810407267 |
| ENSMUSG00000013833 | 1.111082017 | 0.433678407 |
| ENSMUSG00000013846 | 1.543317969 | 0.002629721 |
| ENSMUSG00000013858 | 1.072270194 | 0.608497229 |
| ENSMUSG00000013878 | 1.00048322 | 1 |
| ENSMUSG00000013921 | 1.036783811 | 0.791103495 |
| ENSMUSG00000013928 | -1.04278615 | 0.924766459 |
| ENSMUSG00000013936 | -1.126773846 | 0.608497229 |
| ENSMUSG00000013973 | 1.008558033 | 0.97479937 |
| ENSMUSG00000013974 | 1.059463094 | 0.608497229 |
| ENSMUSG00000013997 | -1.077900609 | 0.649821813 |
| ENSMUSG00000014030 | 1 | |
| ENSMUSG00000014039 | 1.063827375 | 0.558034973 |
| ENSMUSG00000014074 | 1.033644609 | 0.915070045 |
| ENSMUSG00000014075 | 1.020114434 | 0.968102702 |
| ENSMUSG00000014077 | -1.17994906 | 0.556125043 |
| ENSMUSG00000014104 | 1 | |
| ENSMUSG00000014158 | -1.024874196 | 0.87310927 |
| ENSMUSG00000014164 | -1.41143169 | 0.090390754 |
| ENSMUSG00000014177 | -1.036621026 | 0.886850795 |
| ENSMUSG00000014195 | -1.011781173 | 1 |
| ENSMUSG00000014198 | -1.205281846 | 0.200917987 |
| ENSMUSG00000014226 | -1.004226732 | 1 |
| ENSMUSG00000014232 | -1.090888369 | 0.501821225 |
| ENSMUSG00000014243 | -1.041559487 | 0.766916917 |
| ENSMUSG00000014245 | -1.028096626 | 0.835111579 |
| ENSMUSG00000014294 | 1.073883212 | 0.608497229 |
| ENSMUSG00000014301 | 1.045719478 | 0.698112512 |
| ENSMUSG00000014303 | 1.01997845 | 0.939446078 |
| ENSMUSG00000014313 | 1.073856742 | 0.746921078 |
| ENSMUSG00000014329 | 1.04062438 | 0.81152628 |
| ENSMUSG00000014349 | -1.008995114 | 1 |
| ENSMUSG00000014351 | 1.095872691 | 0.731173125 |
| ENSMUSG00000014353 | -1.179612404 | 0.314012257 |
| ENSMUSG00000014355 | -1.070999926 | 0.842981083 |
| ENSMUSG00000014361 | 1.027993807 | 0.762425598 |
| ENSMUSG00000014402 | 1.026494001 | 0.949812939 |
| ENSMUSG00000014418 | -1.023979162 | 0.930877134 |
| ENSMUSG00000014426 | 1.081677997 | 0.608497229 |
| ENSMUSG00000014444 | 1.120036611 | 0.52997988 |
| ENSMUSG00000014453 | 1.069068979 | 0.608497229 |
| ENSMUSG00000014470 | 1.076254959 | 0.608497229 |
| ENSMUSG00000014496 | -1.123511286 | 0.614071182 |
| ENSMUSG00000014498 | -1.025202538 | 0.959747602 |
| ENSMUSG00000014503 | 1.120177528 | 0.217803313 |
| ENSMUSG00000014504 | 1.00325595 | 1 |
| ENSMUSG00000014529 | 1 | |
| ENSMUSG00000014542 | 1 | |
| ENSMUSG00000014543 | 1 | |
| ENSMUSG00000014547 | -1.063173119 | 0.546997193 |
| ENSMUSG00000014550 | 1.020011765 | 0.970734719 |
| ENSMUSG00000014551 | -1.005891639 | 1 |
| ENSMUSG00000014554 | -1.033429353 | 0.782883548 |
| ENSMUSG00000014592 | -1.020234461 | 0.80586932 |

| | | |
|---|---|---|
| ENSMUSG00000014599 | 1.1088942 | 0.713688844 |
| ENSMUSG00000014601 | -1.024349569 | 0.944186417 |
| ENSMUSG00000014602 | 2.036322766 | 7.77839E-05 |
| ENSMUSG00000014603 | -1.507928386 | 0.002964254 |
| ENSMUSG00000014606 | 1.012301858 | 1 |
| ENSMUSG00000014609 | -1.03962336 | 0.698112512 |
| ENSMUSG00000014633 | -1.046676382 | 0.698112512 |
| ENSMUSG00000014668 | 1.045141429 | 0.819817237 |
| ENSMUSG00000014686 | 1 | |
| ENSMUSG00000014704 | 1.085933706 | 0.509007402 |
| ENSMUSG00000014725 | -1.024551908 | 0.949679283 |
| ENSMUSG00000014747 | -1.124996039 | 0.197131782 |
| ENSMUSG00000014748 | -1.026824087 | 0.955101438 |
| ENSMUSG00000014763 | 1.016740305 | 0.977857637 |
| ENSMUSG00000014767 | 1.025517376 | 0.954497929 |
| ENSMUSG00000014769 | -1.002165352 | 1 |
| ENSMUSG00000014771 | 1 | |
| ENSMUSG00000014773 | 1.01889686 | 0.982838256 |
| ENSMUSG00000014776 | -1.102394498 | 0.364739674 |
| ENSMUSG00000014778 | 1.052477785 | 0.634661859 |
| ENSMUSG00000014782 | -1.009340299 | 0.993993248 |
| ENSMUSG00000014786 | 1.038043425 | 0.724838192 |
| ENSMUSG00000014791 | 1.074393997 | 0.608497229 |
| ENSMUSG00000014813 | 1.116567227 | 0.662840906 |
| ENSMUSG00000014837 | -1.049654514 | 0.608497229 |
| ENSMUSG00000014846 | -1.169294698 | 0.286002013 |
| ENSMUSG00000014850 | -1.15739353 | 0.476737035 |
| ENSMUSG00000014852 | -1.043306199 | 0.79291651 |
| ENSMUSG00000014856 | 1.075864707 | 0.604782134 |
| ENSMUSG00000014859 | -1.013708754 | 0.997279592 |
| ENSMUSG00000014867 | 1.069997049 | 0.813989817 |
| ENSMUSG00000014873 | 1.066350749 | 0.698112512 |
| ENSMUSG00000014905 | 1.083775545 | 0.678185531 |
| ENSMUSG00000014907 | 1.019830505 | 0.943391303 |
| ENSMUSG00000014932 | -1.007086893 | 1 |
| ENSMUSG00000014956 | 1.074239355 | 0.828564184 |
| ENSMUSG00000014959 | -1.04019819 | 0.887323956 |
| ENSMUSG00000014980 | -1.030114177 | 0.898763027 |
| ENSMUSG00000015001 | -1.053549062 | 0.698112512 |
| ENSMUSG00000015002 | -1.006520162 | 1 |
| ENSMUSG00000015013 | -1.063640936 | 0.700990092 |
| ENSMUSG00000015016 | -1.011837336 | 0.971527799 |
| ENSMUSG00000015023 | -1.027339567 | 0.934007289 |
| ENSMUSG00000015027 | -1.071515739 | 0.515309246 |
| ENSMUSG00000015053 | 1.111389603 | 0.26714897 |
| ENSMUSG00000015083 | 1.060996588 | 0.644980656 |
| ENSMUSG00000015085 | -1.149170004 | 0.608497229 |
| ENSMUSG00000015087 | -1.035467348 | 0.889126023 |
| ENSMUSG00000015090 | -1.024885547 | 0.975677002 |
| ENSMUSG00000015092 | 1.108764316 | 0.335753246 |
| ENSMUSG00000015093 | -1.063582803 | 0.6996346 |
| ENSMUSG00000015094 | -1.074975692 | 0.608497229 |
| ENSMUSG00000015095 | 1.022272925 | 0.913438732 |
| ENSMUSG00000015112 | 1.185316215 | 0.374471212 |
| ENSMUSG00000015120 | 1.078163222 | 0.783310558 |
| ENSMUSG00000015126 | 1.012536948 | 0.949693072 |
| ENSMUSG00000015127 | 1.015880199 | 0.813485995 |
| ENSMUSG00000015133 | 1.158465948 | 0.413752343 |
| ENSMUSG00000015134 | 1.279876688 | 0.001315883 |
| ENSMUSG00000015143 | 1.466543927 | 0.103173815 |
| ENSMUSG00000015149 | 1.058150889 | 0.654371356 |
| ENSMUSG00000015165 | 1.035094207 | 0.952207945 |
| ENSMUSG00000015176 | 1.040386693 | 0.917928314 |

| | | |
|---|---|---|
| ENSMUSG00000015189 | 1.181779086 | 0.36135224 |
| ENSMUSG00000015202 | 1.083689114 | 0.389746121 |
| ENSMUSG00000015214 | -1.122262042 | 0.579642083 |
| ENSMUSG00000015217 | 1.118722823 | 0.608497229 |
| ENSMUSG00000015222 | 1.427744632 | 0.020353969 |
| ENSMUSG00000015224 | 1.204639063 | 0.544838788 |
| ENSMUSG00000015242 | 1 | |
| ENSMUSG00000015243 | -1.046725989 | 0.698112512 |
| ENSMUSG00000015247 | 1.040294945 | 0.860027857 |
| ENSMUSG00000015289 | -1.051421132 | 0.797105186 |
| ENSMUSG00000015290 | -1.099376519 | 0.624750046 |
| ENSMUSG00000015291 | -1.056774984 | 0.861137762 |
| ENSMUSG00000015305 | -1.192259093 | 0.148622601 |
| ENSMUSG00000015312 | -1.118344763 | 0.380217203 |
| ENSMUSG00000015314 | -1.069502966 | 0.744868995 |
| ENSMUSG00000015316 | 1.117670847 | 0.631765716 |
| ENSMUSG00000015335 | 1.114019665 | 0.508515749 |
| ENSMUSG00000015337 | 1.124305062 | 0.745842121 |
| ENSMUSG00000015340 | -1 | 1 |
| ENSMUSG00000015341 | 1.023437773 | 0.964995983 |
| ENSMUSG00000015342 | 1.062918645 | 0.533633568 |
| ENSMUSG00000015354 | 1.04087931 | 0.839420872 |
| ENSMUSG00000015355 | 1 | |
| ENSMUSG00000015357 | -1.113562193 | 0.608497229 |
| ENSMUSG00000015363 | 1.062558966 | 0.68491446 |
| ENSMUSG00000015365 | -1.044273782 | 0.801443616 |
| ENSMUSG00000015377 | 1.034935198 | 0.731295483 |
| ENSMUSG00000015396 | -1.420155979 | 0.031194528 |
| ENSMUSG00000015401 | 1 | |
| ENSMUSG00000015405 | -1 | 1 |
| ENSMUSG00000015437 | -1.03962336 | 0.698112512 |
| ENSMUSG00000015441 | 1 | |
| ENSMUSG00000015443 | 1 | |
| ENSMUSG00000015451 | -1.17600201 | 0.224521642 |
| ENSMUSG00000015452 | -1.204372898 | 0.33285921 |
| ENSMUSG00000015461 | -1.025356272 | 0.895588956 |
| ENSMUSG00000015467 | 1.094846634 | 0.845618448 |
| ENSMUSG00000015468 | -1.145684921 | 0.642210022 |
| ENSMUSG00000015474 | -1.135603813 | 0.297399155 |
| ENSMUSG00000015476 | 1.039816326 | 0.88325758 |
| ENSMUSG00000015478 | -1.013138555 | 0.936995747 |
| ENSMUSG00000015484 | -1.044273782 | 0.698112512 |
| ENSMUSG00000015488 | -1.00665901 | 1 |
| ENSMUSG00000015501 | -1.079089583 | 0.608497229 |
| ENSMUSG00000015519 | 1 | |
| ENSMUSG00000015522 | -1.051707674 | 0.827713826 |
| ENSMUSG00000015533 | 1.187528713 | 0.216833706 |
| ENSMUSG00000015536 | -1.152820844 | 0.534652019 |
| ENSMUSG00000015542 | 1.046775048 | 0.499572952 |
| ENSMUSG00000015568 | 1.856994954 | 0.000248898 |
| ENSMUSG00000015575 | -1.043040468 | 0.896902223 |
| ENSMUSG00000015579 | 1.013940071 | 1 |
| ENSMUSG00000015597 | -1.020210406 | 0.97609342 |
| ENSMUSG00000015599 | -1.336402403 | 0.098590033 |
| ENSMUSG00000015605 | 1.246858802 | 0.044140978 |
| ENSMUSG00000015619 | 1.090987776 | 0.327126149 |
| ENSMUSG00000015627 | -1.089497311 | 0.532369657 |
| ENSMUSG00000015647 | 1.374979628 | 0.03743915 |
| ENSMUSG00000015652 | -1.000356739 | 1 |
| ENSMUSG00000015653 | -1.003515747 | 1 |
| ENSMUSG00000015656 | -1.029735959 | 0.981456085 |
| ENSMUSG00000015659 | 1.198345563 | 0.045045157 |
| ENSMUSG00000015665 | 1 | |

| | | |
|---|---|---|
| ENSMUSG00000015668 | 1.05242126 | 0.830868034 |
| ENSMUSG00000015671 | -1.076388713 | 0.819222912 |
| ENSMUSG00000015672 | -1.09241951 | 0.609589476 |
| ENSMUSG00000015697 | -1.008155121 | 1 |
| ENSMUSG00000015702 | 1.110669913 | 0.308488402 |
| ENSMUSG00000015709 | 1.167078938 | 0.34067437 |
| ENSMUSG00000015711 | -1.046998761 | 0.819994892 |
| ENSMUSG00000015714 | -1.060838081 | 0.829460784 |
| ENSMUSG00000015721 | -1.227002046 | 0.295479694 |
| ENSMUSG00000015733 | 1.123171454 | 0.698112512 |
| ENSMUSG00000015745 | 1.34341041 | 0.004029711 |
| ENSMUSG00000015747 | -1.024408434 | 0.946232149 |
| ENSMUSG00000015748 | 1.013827093 | 0.994172456 |
| ENSMUSG00000015749 | -1.069261579 | 0.835531958 |
| ENSMUSG00000015750 | -1.010254771 | 0.988519386 |
| ENSMUSG00000015755 | 1.104593547 | 0.698112512 |
| ENSMUSG00000015757 | -1.047389935 | 0.84360497 |
| ENSMUSG00000015759 | -1.124769368 | 0.608497229 |
| ENSMUSG00000015766 | 1.002068072 | 1 |
| ENSMUSG00000015776 | 1.036389905 | 0.867969972 |
| ENSMUSG00000015787 | 1 | |
| ENSMUSG00000015790 | 1.05374338 | 0.694886902 |
| ENSMUSG00000015804 | 1.06052281 | 0.737214232 |
| ENSMUSG00000015806 | -1.053671059 | 0.698112512 |
| ENSMUSG00000015812 | -1.148257651 | 0.385303452 |
| ENSMUSG00000015829 | -1.005317348 | 1 |
| ENSMUSG00000015837 | 1.195297934 | 0.43994505 |
| ENSMUSG00000015839 | -1.095879198 | 0.633631542 |
| ENSMUSG00000015843 | -1.085651418 | 0.532369657 |
| ENSMUSG00000015846 | 1.187658894 | 0.074010245 |
| ENSMUSG00000015850 | 1.007733515 | 0.995215546 |
| ENSMUSG00000015852 | 1 | |
| ENSMUSG00000015854 | 1 | |
| ENSMUSG00000015869 | -1.025500835 | 0.934760809 |
| ENSMUSG00000015879 | 1.048104613 | 0.76331024 |
| ENSMUSG00000015880 | 1.006289885 | 1 |
| ENSMUSG00000015882 | 1.103707686 | 0.608497229 |
| ENSMUSG00000015889 | -1.040853923 | 0.914913657 |
| ENSMUSG00000015890 | 1.029490685 | 0.938939969 |
| ENSMUSG00000015932 | 1.301023998 | 0.316089517 |
| ENSMUSG00000015937 | -1.161493982 | 0.565559141 |
| ENSMUSG00000015942 | 1.094660237 | 0.145247026 |
| ENSMUSG00000015943 | 1.054484596 | 0.890342946 |
| ENSMUSG00000015944 | -1.007359275 | 1 |
| ENSMUSG00000015947 | 1.059463094 | 0.608497229 |
| ENSMUSG00000015950 | 1.135228563 | 0.608497229 |
| ENSMUSG00000015957 | -1.282409918 | 0.276523052 |
| ENSMUSG00000015961 | -1.022539982 | 0.980731794 |
| ENSMUSG00000015962 | -1.04278615 | 0.952086196 |
| ENSMUSG00000015966 | 1.184376802 | 0.337583151 |
| ENSMUSG00000015968 | -1.084645496 | 0.532369657 |
| ENSMUSG00000015970 | 1 | |
| ENSMUSG00000015971 | -1.132745372 | 0.443913715 |
| ENSMUSG00000015980 | -1.058250928 | 0.608497229 |
| ENSMUSG00000015981 | -1.591619809 | 0.001867243 |
| ENSMUSG00000015994 | -1.050491523 | 0.870427333 |
| ENSMUSG00000016018 | -1.106638481 | 0.697178241 |
| ENSMUSG00000016024 | -1.051143849 | 0.698112512 |
| ENSMUSG00000016028 | -1.188419743 | 0.083371725 |
| ENSMUSG00000016087 | -2.066148757 | 0.001559423 |
| ENSMUSG00000016128 | 1.0615094 | 0.717035356 |
| ENSMUSG00000016150 | 1.132639128 | 0.568304986 |
| ENSMUSG00000016179 | 1.059463094 | 0.699340382 |

| | | |
|---|---|---|
| ENSMUSG00000016181 | 1.006469298 | 1 |
| ENSMUSG00000016194 | 1 | |
| ENSMUSG00000016200 | -3.056085732 | 0.000207005 |
| ENSMUSG00000016206 | 1.256111609 | 0.009708135 |
| ENSMUSG00000016239 | 1.31316866 | 0.001540286 |
| ENSMUSG00000016252 | 1.010215005 | 0.983170063 |
| ENSMUSG00000016253 | -1.075569055 | 0.608497229 |
| ENSMUSG00000016255 | 1 | |
| ENSMUSG00000016256 | -1.132926337 | 0.37859055 |
| ENSMUSG00000016257 | -1.050963032 | 0.821016426 |
| ENSMUSG00000016262 | 1.193518348 | 0.413631177 |
| ENSMUSG00000016283 | -1.075542744 | 0.793033871 |
| ENSMUSG00000016308 | -1.036573151 | 0.903448666 |
| ENSMUSG00000016319 | 1.032346177 | 0.949679283 |
| ENSMUSG00000016327 | 1.059463094 | 0.608497229 |
| ENSMUSG00000016344 | 1.006528649 | 0.981214754 |
| ENSMUSG00000016346 | 1.139151239 | 0.532369657 |
| ENSMUSG00000016349 | -1.188446785 | 0.219185116 |
| ENSMUSG00000016356 | 1.025408388 | 0.908007394 |
| ENSMUSG00000016382 | 1.353921938 | 0.210876111 |
| ENSMUSG00000016386 | 1.011010231 | 0.963565014 |
| ENSMUSG00000016409 | -1.051966208 | 0.804170272 |
| ENSMUSG00000016427 | 1.076630171 | 0.633309501 |
| ENSMUSG00000016458 | -1.185786166 | 0.481515508 |
| ENSMUSG00000016477 | 1.00700611 | 1 |
| ENSMUSG00000016481 | -1.008598829 | 1 |
| ENSMUSG00000016487 | 1.118104776 | 0.698112512 |
| ENSMUSG00000016493 | -1.035645199 | 0.932570845 |
| ENSMUSG00000016494 | -2.123922128 | 0.001127478 |
| ENSMUSG00000016495 | -1.000478969 | 1 |
| ENSMUSG00000016496 | -1.092365645 | 0.375668528 |
| ENSMUSG00000016498 | 1.059463094 | 0.608497229 |
| ENSMUSG00000016503 | 1.05016837 | 0.456459883 |
| ENSMUSG00000016510 | -1.028885504 | 0.859429731 |
| ENSMUSG00000016520 | 1.191983847 | 0.02024466 |
| ENSMUSG00000016524 | 1.059463094 | 0.699340382 |
| ENSMUSG00000016526 | -1.121934941 | 0.28456385 |
| ENSMUSG00000016528 | 1.103141734 | 0.668750825 |
| ENSMUSG00000016529 | 1 | |
| ENSMUSG00000016534 | 1.206873619 | 0.566497027 |
| ENSMUSG00000016541 | 1.01045864 | 1 |
| ENSMUSG00000016552 | -1.307826063 | 0.004528705 |
| ENSMUSG00000016554 | 1.036096313 | 0.930202553 |
| ENSMUSG00000016559 | -1.00338746 | 1 |
| ENSMUSG00000016619 | 1.031532797 | 0.951098417 |
| ENSMUSG00000016624 | 1.06873985 | 0.698112512 |
| ENSMUSG00000016626 | 1 | |
| ENSMUSG00000016637 | 1.019576298 | 0.923181046 |
| ENSMUSG00000016664 | 1.047940742 | 0.857820356 |
| ENSMUSG00000016756 | 1.140809559 | 0.6100587 |
| ENSMUSG00000016757 | 1.14516123 | 0.475169715 |
| ENSMUSG00000016758 | 1.765998247 | 0.027606265 |
| ENSMUSG00000016763 | -1.529701384 | 0.044184368 |
| ENSMUSG00000016831 | -1.21420964 | 0.291044503 |
| ENSMUSG00000016833 | -1.032295534 | 0.911642049 |
| ENSMUSG00000016918 | 1.953735399 | 0.001032697 |
| ENSMUSG00000016921 | -1.099227915 | 0.738123855 |
| ENSMUSG00000016933 | 1.065511295 | 0.755320109 |
| ENSMUSG00000016940 | 1.09305002 | 0.173614736 |
| ENSMUSG00000016942 | -1.290882438 | 0.102003417 |
| ENSMUSG00000016946 | 1.16903829 | 0.436400681 |
| ENSMUSG00000016982 | 1.05530209 | 0.812255606 |
| ENSMUSG00000016984 | -1.058888846 | 0.698112512 |

| | | |
|---|---|---|
| ENSMUSG00000016995 | -1.053549062 | 0.745625104 |
| ENSMUSG00000016998 | 1 | |
| ENSMUSG00000017000 | 1 | |
| ENSMUSG00000017002 | -1.064334191 | 0.860853691 |
| ENSMUSG00000017003 | 1 | |
| ENSMUSG00000017004 | 1 | |
| ENSMUSG00000017007 | -1.084645496 | 0.532369657 |
| ENSMUSG00000017009 | -1.040164902 | 0.881084157 |
| ENSMUSG00000017049 | 1 | |
| ENSMUSG00000017057 | 1.006505128 | 1 |
| ENSMUSG00000017064 | 1 | |
| ENSMUSG00000017119 | -1.029242275 | 0.958303165 |
| ENSMUSG00000017132 | -1.084601888 | 0.369728829 |
| ENSMUSG00000017144 | 1.004157397 | 1 |
| ENSMUSG00000017146 | -1.054798215 | 0.860992632 |
| ENSMUSG00000017165 | 1.095872691 | 0.608497229 |
| ENSMUSG00000017167 | 1.091825893 | 0.263945222 |
| ENSMUSG00000017176 | -1.028930362 | 0.921490876 |
| ENSMUSG00000017188 | 1.077404995 | 0.628392912 |
| ENSMUSG00000017195 | -1.203254079 | 0.236724318 |
| ENSMUSG00000017204 | -1.071716244 | 0.807743274 |
| ENSMUSG00000017210 | 1.057325133 | 0.747974227 |
| ENSMUSG00000017211 | -1.044273782 | 0.698112512 |
| ENSMUSG00000017221 | 1.028988203 | 0.933031586 |
| ENSMUSG00000017264 | -1.006797932 | 1 |
| ENSMUSG00000017286 | -1.061115896 | 0.762976518 |
| ENSMUSG00000017288 | -1.026957478 | 0.94200166 |
| ENSMUSG00000017291 | 1.033909173 | 0.957300905 |
| ENSMUSG00000017299 | -1.10035398 | 0.371852779 |
| ENSMUSG00000017300 | -1.053549062 | 0.769745556 |
| ENSMUSG00000017307 | -1.14360514 | 0.115310535 |
| ENSMUSG00000017309 | -1.03962336 | 0.88732209 |
| ENSMUSG00000017310 | 1 | |
| ENSMUSG00000017311 | 1 | |
| ENSMUSG00000017314 | -1.073233655 | 0.472093503 |
| ENSMUSG00000017316 | 1 | |
| ENSMUSG00000017344 | -1.043306199 | 0.79291651 |
| ENSMUSG00000017376 | 1.028203143 | 0.940277706 |
| ENSMUSG00000017386 | 1.183752834 | 0.201035112 |
| ENSMUSG00000017390 | -1.099705413 | 0.608497229 |
| ENSMUSG00000017400 | 1.093686086 | 0.258650556 |
| ENSMUSG00000017404 | 1.050989463 | 0.872815894 |
| ENSMUSG00000017405 | 1.042706635 | 0.698112512 |
| ENSMUSG00000017412 | 1.15238629 | 0.240189729 |
| ENSMUSG00000017417 | -1.162311484 | 0.576169279 |
| ENSMUSG00000017418 | -1.000292591 | 1 |
| ENSMUSG00000017421 | 1.024268244 | 0.984782014 |
| ENSMUSG00000017428 | -1.004664311 | 1 |
| ENSMUSG00000017446 | 1.144893838 | 0.278288653 |
| ENSMUSG00000017453 | 1.178139999 | 0.486216685 |
| ENSMUSG00000017466 | 1.10269304 | 0.786423254 |
| ENSMUSG00000017478 | 1.019080339 | 0.962045523 |
| ENSMUSG00000017485 | -1.083406296 | 0.777175622 |
| ENSMUSG00000017491 | -1.062788411 | 0.480458517 |
| ENSMUSG00000017493 | 1.169961341 | 0.010046599 |
| ENSMUSG00000017499 | -1.054260884 | 0.82784586 |
| ENSMUSG00000017548 | -1.027053348 | 0.972980766 |
| ENSMUSG00000017550 | -1.059752867 | 0.834621543 |
| ENSMUSG00000017561 | -1.006214105 | 1 |
| ENSMUSG00000017588 | 1 | |
| ENSMUSG00000017607 | -1.143815032 | 0.56288192 |
| ENSMUSG00000017615 | 1.01480395 | 1 |
| ENSMUSG00000017631 | 1.078420859 | 0.745713519 |

| | | |
|---|---|---|
| ENSMUSG00000017639 | -1.370446168 | 0.014736529 |
| ENSMUSG00000017652 | -1.331362794 | 0.078014703 |
| ENSMUSG00000017664 | 1.051818031 | 0.508515749 |
| ENSMUSG00000017667 | -1.013591096 | 0.972143455 |
| ENSMUSG00000017670 | -1.024510462 | 0.931480186 |
| ENSMUSG00000017677 | 1.013917744 | 1 |
| ENSMUSG00000017679 | -1.11151983 | 0.608497229 |
| ENSMUSG00000017686 | -1.035019706 | 0.93818309 |
| ENSMUSG00000017688 | 1 | |
| ENSMUSG00000017692 | -1.124605566 | 0.291122014 |
| ENSMUSG00000017697 | -1.306112189 | 0.026297047 |
| ENSMUSG00000017707 | -1.096243397 | 0.698112512 |
| ENSMUSG00000017713 | -1.004744859 | 1 |
| ENSMUSG00000017715 | 1.086269719 | 0.608497229 |
| ENSMUSG00000017716 | -1.053276301 | 0.856857802 |
| ENSMUSG00000017718 | 1.172935598 | 0.12958272 |
| ENSMUSG00000017720 | -1.120056821 | 0.678985493 |
| ENSMUSG00000017721 | -1.032589493 | 0.872815393 |
| ENSMUSG00000017723 | 1.033391092 | 0.971314913 |
| ENSMUSG00000017724 | -1.175768761 | 0.215991492 |
| ENSMUSG00000017733 | 1 | |
| ENSMUSG00000017734 | 1.013223956 | 1 |
| ENSMUSG00000017737 | -1.279012826 | 0.068433456 |
| ENSMUSG00000017740 | -1.346937406 | 0.005884452 |
| ENSMUSG00000017747 | -1.016305986 | 0.94592202 |
| ENSMUSG00000017754 | 1.103684221 | 0.080458117 |
| ENSMUSG00000017756 | 1.014023578 | 0.984094159 |
| ENSMUSG00000017760 | 1.148263479 | 0.482403115 |
| ENSMUSG00000017764 | -1.120461276 | 0.082138282 |
| ENSMUSG00000017765 | 1.043690584 | 0.707311809 |
| ENSMUSG00000017767 | 1.015486245 | 0.984826086 |
| ENSMUSG00000017774 | 1.028962008 | 0.961861013 |
| ENSMUSG00000017776 | 1.049199575 | 0.907920686 |
| ENSMUSG00000017778 | 1.022412003 | 0.904050368 |
| ENSMUSG00000017781 | 1.02771158 | 0.971869901 |
| ENSMUSG00000017801 | 1.01275773 | 0.993466371 |
| ENSMUSG00000017802 | 1.033507219 | 0.905574572 |
| ENSMUSG00000017817 | 3.86597043 | 0.000071721 |
| ENSMUSG00000017830 | -1.167999866 | 0.092667548 |
| ENSMUSG00000017831 | 1.073968286 | 0.759038956 |
| ENSMUSG00000017832 | -1.164283619 | 0.458934581 |
| ENSMUSG00000017837 | 1.066422046 | 0.472312635 |
| ENSMUSG00000017843 | 1.07986294 | 0.718265321 |
| ENSMUSG00000017858 | -1.098628449 | 0.608497229 |
| ENSMUSG00000017861 | -1.066859337 | 0.661640678 |
| ENSMUSG00000017868 | -1.048277865 | 0.87354792 |
| ENSMUSG00000017897 | -1.205220788 | 0.250477583 |
| ENSMUSG00000017922 | 1 | |
| ENSMUSG00000017929 | 1.035982452 | 0.736446356 |
| ENSMUSG00000017943 | -1.411407033 | 0.029512753 |
| ENSMUSG00000017950 | 1 | |
| ENSMUSG00000017969 | 1.062574622 | 0.698112512 |
| ENSMUSG00000017978 | -1.063525418 | 0.698112512 |
| ENSMUSG00000017999 | -1.035601298 | 0.864480156 |
| ENSMUSG00000018001 | 1.025693281 | 0.941803539 |
| ENSMUSG00000018008 | -1.216643792 | 0.56288192 |
| ENSMUSG00000018012 | 1.714736576 | 0.000759427 |
| ENSMUSG00000018040 | -1.028683461 | 0.91188534 |
| ENSMUSG00000018042 | 1.102001867 | 0.698112512 |
| ENSMUSG00000018068 | -1.008782222 | 1 |
| ENSMUSG00000018076 | 1.186384395 | 0.27459474 |
| ENSMUSG00000018102 | -1.438821233 | 0.020817525 |
| ENSMUSG00000018126 | -1.28670943 | 0.304090886 |

| | | |
|---|---|---|
| ENSMUSG00000018143 | -1.001456853 | 1 |
| ENSMUSG00000018160 | 1.026125036 | 0.956026696 |
| ENSMUSG00000018166 | -1.391497401 | 0.000658313 |
| ENSMUSG00000018167 | 1.02852759 | 0.88325758 |
| ENSMUSG00000018168 | -1.03962336 | 0.698112512 |
| ENSMUSG00000018169 | 1.030415139 | 0.972980766 |
| ENSMUSG00000018171 | 1.268149914 | 0.294286866 |
| ENSMUSG00000018189 | 1.005561985 | 1 |
| ENSMUSG00000018196 | 1.030112004 | 0.88732209 |
| ENSMUSG00000018199 | -1.087968981 | 0.726686255 |
| ENSMUSG00000018209 | -1.086609785 | 0.698112512 |
| ENSMUSG00000018211 | 1 | |
| ENSMUSG00000018217 | 1.011968661 | 1 |
| ENSMUSG00000018238 | -1.17334783 | 0.353239233 |
| ENSMUSG00000018239 | -1.149947003 | 0.239349513 |
| ENSMUSG00000018259 | 1 | |
| ENSMUSG00000018263 | 1.05038453 | 0.877529894 |
| ENSMUSG00000018286 | -1.045967623 | 0.876279035 |
| ENSMUSG00000018287 | 1.028864559 | 0.905695722 |
| ENSMUSG00000018293 | 1.139268893 | 0.632746776 |
| ENSMUSG00000018322 | -1.083267792 | 0.608497229 |
| ENSMUSG00000018326 | -1.061468151 | 0.872692845 |
| ENSMUSG00000018334 | 1.124849467 | 0.378522977 |
| ENSMUSG00000018339 | -1.159504733 | 0.638899001 |
| ENSMUSG00000018340 | 1.164668253 | 0.608497229 |
| ENSMUSG00000018341 | -1.517781328 | 0.01921251 |
| ENSMUSG00000018347 | -1.129436765 | 0.442420587 |
| ENSMUSG00000018362 | -1.015966501 | 1 |
| ENSMUSG00000018363 | 1.008165649 | 1 |
| ENSMUSG00000018372 | 1.013279523 | 0.969704392 |
| ENSMUSG00000018377 | 1.028430016 | 0.951748628 |
| ENSMUSG00000018378 | 1.106403435 | 0.35640372 |
| ENSMUSG00000018379 | -1.000166533 | 1 |
| ENSMUSG00000018381 | 1.132639128 | 0.217803313 |
| ENSMUSG00000018387 | -1.245040514 | 0.004795301 |
| ENSMUSG00000018395 | -1.102311886 | 0.644513855 |
| ENSMUSG00000018398 | -1.095768992 | 0.698112512 |
| ENSMUSG00000018401 | 1.019937514 | 0.984094159 |
| ENSMUSG00000018405 | 1.024996399 | 0.818468929 |
| ENSMUSG00000018411 | -1.455358217 | 0.113934609 |
| ENSMUSG00000018412 | 1.021822721 | 0.967438206 |
| ENSMUSG00000018415 | -1.109746809 | 0.608497229 |
| ENSMUSG00000018417 | 1.170346566 | 0.43122333 |
| ENSMUSG00000018425 | 1.075397227 | 0.745625104 |
| ENSMUSG00000018427 | 1.034275183 | 0.390586581 |
| ENSMUSG00000018428 | 1.089680786 | 0.66521153 |
| ENSMUSG00000018433 | -1.050378776 | 0.871242925 |
| ENSMUSG00000018442 | 1.015040746 | 0.987819411 |
| ENSMUSG00000018446 | 1.006758712 | 1 |
| ENSMUSG00000018449 | 1.014923594 | 0.981741913 |
| ENSMUSG00000018451 | 1.067475584 | 0.698112512 |
| ENSMUSG00000018459 | 1.023743961 | 0.938939969 |
| ENSMUSG00000018470 | -1.210553891 | 0.026141714 |
| ENSMUSG00000018474 | -1.008257265 | 1 |
| ENSMUSG00000018476 | 1.132625405 | 0.145051274 |
| ENSMUSG00000018479 | 1.034245398 | 0.981307697 |
| ENSMUSG00000018481 | -1.002869379 | 1 |
| ENSMUSG00000018486 | 1 | |
| ENSMUSG00000018500 | 1.294330849 | 0.048954387 |
| ENSMUSG00000018501 | -1.084785888 | 0.773508225 |
| ENSMUSG00000018507 | 1.314809267 | 0.001011241 |
| ENSMUSG00000018509 | -1.339253662 | 0.002797687 |
| ENSMUSG00000018537 | 1.050550414 | 0.612786983 |

| | | |
|---|---|---|
| ENSMUSG00000018541 | -1.03398529 | 0.834621543 |
| ENSMUSG00000018543 | 1.058806668 | 0.917953029 |
| ENSMUSG00000018547 | -1.041112403 | 0.894558701 |
| ENSMUSG00000018548 | 1.057487707 | 0.850478054 |
| ENSMUSG00000018554 | -1.255625001 | 0.43627839 |
| ENSMUSG00000018559 | 1.068894241 | 0.751979636 |
| ENSMUSG00000018565 | 1.001309418 | 1 |
| ENSMUSG00000018566 | -1.068411596 | 0.65908139 |
| ENSMUSG00000018567 | 1.16803921 | 0.608497229 |
| ENSMUSG00000018569 | -1.084645496 | 0.70299173 |
| ENSMUSG00000018570 | 1.054908119 | 0.790294437 |
| ENSMUSG00000018572 | 1.186020552 | 0.249180157 |
| ENSMUSG00000018574 | -1.060762505 | 0.770017269 |
| ENSMUSG00000018581 | 1.116201826 | 0.662474108 |
| ENSMUSG00000018583 | -1.031662699 | 0.959492944 |
| ENSMUSG00000018585 | -1.092377496 | 0.608497229 |
| ENSMUSG00000018589 | -1 | 1 |
| ENSMUSG00000018593 | 1.012154128 | 1 |
| ENSMUSG00000018595 | -1.170395304 | 0.515417737 |
| ENSMUSG00000018599 | 1.013731483 | 0.925075932 |
| ENSMUSG00000018604 | -1.312430239 | 0.199329505 |
| ENSMUSG00000018620 | 1 | |
| ENSMUSG00000018623 | 1 | |
| ENSMUSG00000018634 | -1.03962336 | 0.698112512 |
| ENSMUSG00000018648 | -1.165757423 | 0.148771925 |
| ENSMUSG00000018651 | 1.006837167 | 1 |
| ENSMUSG00000018654 | -1.085651418 | 0.625496737 |
| ENSMUSG00000018656 | -1.053549062 | 0.840310372 |
| ENSMUSG00000018659 | -1.139386443 | 0.337410766 |
| ENSMUSG00000018661 | 1.016065411 | 0.956148535 |
| ENSMUSG00000018666 | 1.008071768 | 1 |
| ENSMUSG00000018669 | 1.043288256 | 0.775673129 |
| ENSMUSG00000018672 | 1.07874005 | 0.608497229 |
| ENSMUSG00000018677 | 1.107154298 | 0.541735653 |
| ENSMUSG00000018678 | 1.029925538 | 0.701811773 |
| ENSMUSG00000018697 | 1.059463328 | 0.769272045 |
| ENSMUSG00000018698 | -1.643583227 | 0.006041499 |
| ENSMUSG00000018707 | 1.137230831 | 0.63907663 |
| ENSMUSG00000018727 | 1.07550552 | 0.698112512 |
| ENSMUSG00000018733 | 1.038813144 | 0.700181211 |
| ENSMUSG00000018736 | 1.022547121 | 0.958303165 |
| ENSMUSG00000018740 | -1.101701862 | 0.608497229 |
| ENSMUSG00000018750 | -1.169370524 | 0.115685577 |
| ENSMUSG00000018752 | -1.140075771 | 0.527384609 |
| ENSMUSG00000018761 | 1.009234626 | 1 |
| ENSMUSG00000018765 | -1.004160409 | 1 |
| ENSMUSG00000018770 | -1.075096258 | 0.770958539 |
| ENSMUSG00000018774 | 1.08542374 | 0.675418318 |
| ENSMUSG00000018776 | -1.029065166 | 0.982838256 |
| ENSMUSG00000018796 | 1.078310219 | 0.698112512 |
| ENSMUSG00000018800 | -1.068352082 | 0.74677467 |
| ENSMUSG00000018809 | 1.096311288 | 0.583431551 |
| ENSMUSG00000018819 | 1.61113614 | 0.001601152 |
| ENSMUSG00000018820 | -1.044096196 | 0.770122654 |
| ENSMUSG00000018821 | 1.032665182 | 0.868154326 |
| ENSMUSG00000018822 | -1.170083925 | 0.206620396 |
| ENSMUSG00000018830 | 1.15867274 | 0.531052031 |
| ENSMUSG00000018841 | 1.102593621 | 0.392118296 |
| ENSMUSG00000018844 | -1.008983169 | 1 |
| ENSMUSG00000018845 | -1.03962336 | 0.698112512 |
| ENSMUSG00000018846 | -1.059023263 | 0.860645152 |
| ENSMUSG00000018848 | -1.058174237 | 0.870427333 |
| ENSMUSG00000018849 | -1.14464001 | 0.375655143 |

| | | |
|---|---|---|
| ENSMUSG00000018858 | -1.086411779 | 0.618238982 |
| ENSMUSG00000018861 | 1.066764922 | 0.66817972 |
| ENSMUSG00000018862 | 1 | |
| ENSMUSG00000018865 | -2.458711299 | 0.001032697 |
| ENSMUSG00000018868 | 1 | |
| ENSMUSG00000018882 | -1.008538341 | 1 |
| ENSMUSG00000018893 | 1.059463094 | 0.608497229 |
| ENSMUSG00000018899 | -1.28304634 | 0.205169472 |
| ENSMUSG00000018900 | -1.207153099 | 0.079015056 |
| ENSMUSG00000018906 | 1.021801441 | 0.95819506 |
| ENSMUSG00000018907 | -1.109582661 | 0.549347972 |
| ENSMUSG00000018909 | -1.049239005 | 0.769165946 |
| ENSMUSG00000018914 | 1 | |
| ENSMUSG00000018916 | -1.237299691 | 0.201977401 |
| ENSMUSG00000018919 | 1.132639128 | 0.217803313 |
| ENSMUSG00000018920 | -1.488176459 | 0.00077513 |
| ENSMUSG00000018921 | 1.079891489 | 0.597779413 |
| ENSMUSG00000018923 | -1.032718392 | 0.793735215 |
| ENSMUSG00000018924 | -1.134462398 | 0.643072778 |
| ENSMUSG00000018925 | -1.111456291 | 0.717035356 |
| ENSMUSG00000018927 | 1.211797127 | 0.250726313 |
| ENSMUSG00000018930 | 1.059463094 | 0.608497229 |
| ENSMUSG00000018931 | -1.087211296 | 0.614307028 |
| ENSMUSG00000018932 | -1.075493882 | 0.734053159 |
| ENSMUSG00000018965 | 1.069142768 | 0.812183377 |
| ENSMUSG00000018973 | 1.059463094 | 0.608497229 |
| ENSMUSG00000018974 | 1.115788311 | 0.501910794 |
| ENSMUSG00000018983 | -1.100879638 | 0.32848288 |
| ENSMUSG00000018986 | 1 | |
| ENSMUSG00000018995 | 1.011333898 | 0.972684567 |
| ENSMUSG00000018999 | 1.101529758 | 0.608497229 |
| ENSMUSG00000019027 | -1.155321732 | 0.381530795 |
| ENSMUSG00000019039 | -1.018674623 | 0.951399668 |
| ENSMUSG00000019054 | 1.078536469 | 0.628147978 |
| ENSMUSG00000019055 | -1.03228262 | 0.900651611 |
| ENSMUSG00000019066 | -1.222597679 | 0.018796561 |
| ENSMUSG00000019080 | 1.002692185 | 1 |
| ENSMUSG00000019082 | 1.095994829 | 0.393187622 |
| ENSMUSG00000019087 | 1.055397843 | 0.870427333 |
| ENSMUSG00000019088 | 1.137416678 | 0.286393177 |
| ENSMUSG00000019102 | 2.013742868 | 0.000290807 |
| ENSMUSG00000019122 | 1.960010728 | 0.000417999 |
| ENSMUSG00000019124 | -1.020256844 | 0.970208206 |
| ENSMUSG00000019132 | -1.044888796 | 0.910882003 |
| ENSMUSG00000019139 | -1.008282878 | 1 |
| ENSMUSG00000019143 | 1.003742646 | 1 |
| ENSMUSG00000019146 | 1.059463094 | 0.608497229 |
| ENSMUSG00000019158 | -1.012775076 | 1 |
| ENSMUSG00000019173 | 1.019926535 | 0.963110173 |
| ENSMUSG00000019178 | -1.15904038 | 0.633643463 |
| ENSMUSG00000019179 | 1.053719023 | 0.87457737 |
| ENSMUSG00000019188 | -1.005129528 | 1 |
| ENSMUSG00000019189 | -1.031436053 | 0.938708567 |
| ENSMUSG00000019194 | 1.203887863 | 0.206114416 |
| ENSMUSG00000019210 | 1.080135879 | 0.710698346 |
| ENSMUSG00000019214 | -1.002677215 | 1 |
| ENSMUSG00000019230 | 1.573164252 | 0.000573666 |
| ENSMUSG00000019232 | 1.05038453 | 0.828564184 |
| ENSMUSG00000019235 | -1.029404462 | 0.963965793 |
| ENSMUSG00000019254 | 1.043996776 | 0.645756877 |
| ENSMUSG00000019256 | 1.036723885 | 0.855488553 |
| ENSMUSG00000019261 | 1.146507318 | 0.670045943 |
| ENSMUSG00000019278 | 1 | |

| | | |
|---|---|---|
| ENSMUSG00000019295 | 1.025919261 | 0.906750223 |
| ENSMUSG00000019297 | -1.160735988 | 0.328718061 |
| ENSMUSG00000019301 | 1.22226954 | 0.273601178 |
| ENSMUSG00000019302 | 1.089402931 | 0.634968525 |
| ENSMUSG00000019303 | 1.051742203 | 0.57595853 |
| ENSMUSG00000019312 | -1.15462599 | 0.330039993 |
| ENSMUSG00000019320 | -1.0181342 | 1 |
| ENSMUSG00000019326 | 2.145161037 | 0.0001547 |
| ENSMUSG00000019338 | 1.040655981 | 0.807176388 |
| ENSMUSG00000019359 | 1.155013505 | 0.648983806 |
| ENSMUSG00000019362 | 1.089216277 | 0.452689651 |
| ENSMUSG00000019368 | 1.047976605 | 0.849421383 |
| ENSMUSG00000019370 | 1.008042183 | 1 |
| ENSMUSG00000019373 | -1.074652574 | 0.763242941 |
| ENSMUSG00000019428 | 1.058238952 | 0.743396505 |
| ENSMUSG00000019429 | 1 | |
| ENSMUSG00000019432 | 1.02085812 | 0.986389736 |
| ENSMUSG00000019433 | -1.000069585 | 1 |
| ENSMUSG00000019437 | 1.078854089 | 0.608497229 |
| ENSMUSG00000019461 | 1.029105385 | 0.885561194 |
| ENSMUSG00000019464 | 1.114654071 | 0.211170208 |
| ENSMUSG00000019467 | 1.132000436 | 0.282101888 |
| ENSMUSG00000019470 | 1.189245792 | 0.056310079 |
| ENSMUSG00000019471 | -1.05096184 | 0.793891026 |
| ENSMUSG00000019478 | -1.161478895 | 0.071318353 |
| ENSMUSG00000019487 | 1.065808917 | 0.608497229 |
| ENSMUSG00000019489 | -1.095294215 | 0.541735653 |
| ENSMUSG00000019494 | -1.035613599 | 0.935937536 |
| ENSMUSG00000019505 | -1.170457382 | 0.462692733 |
| ENSMUSG00000019518 | 1.026063068 | 0.869548547 |
| ENSMUSG00000019528 | -1.048397937 | 0.858124279 |
| ENSMUSG00000019539 | 1.217967263 | 0.05764497 |
| ENSMUSG00000019558 | 1.284806804 | 0.099764565 |
| ENSMUSG00000019564 | 1.05857729 | 0.426006517 |
| ENSMUSG00000019577 | 1.523742912 | 0.017650728 |
| ENSMUSG00000019578 | 1.074573894 | 0.608497229 |
| ENSMUSG00000019579 | 1.054431263 | 0.704913902 |
| ENSMUSG00000019590 | -1.24422 | 0.009056704 |
| ENSMUSG00000019647 | -1.386444923 | 0.005344566 |
| ENSMUSG00000019659 | -1.016920509 | 0.954497929 |
| ENSMUSG00000019689 | 1.076521425 | 0.453483686 |
| ENSMUSG00000019699 | 1.25284748 | 0.154089282 |
| ENSMUSG00000019710 | 1.018839144 | 0.959264294 |
| ENSMUSG00000019715 | 1.00041389 | 1 |
| ENSMUSG00000019718 | 1.511506396 | 0.002245208 |
| ENSMUSG00000019726 | 1.135252893 | 0.531723579 |
| ENSMUSG00000019731 | -1.025343208 | 0.956972448 |
| ENSMUSG00000019732 | -1.075169862 | 0.210605021 |
| ENSMUSG00000019734 | 1.217479547 | 0.019808022 |
| ENSMUSG00000019737 | 1.048062564 | 0.943663271 |
| ENSMUSG00000019738 | 1.048995711 | 0.760563995 |
| ENSMUSG00000019756 | -1.021242786 | 0.989603176 |
| ENSMUSG00000019761 | 1.109542761 | 0.205926548 |
| ENSMUSG00000019762 | -1 | 1 |
| ENSMUSG00000019763 | 1.056027842 | 0.713643114 |
| ENSMUSG00000019767 | 1.009005271 | 1 |
| ENSMUSG00000019768 | -1.045888114 | 0.779616759 |
| ENSMUSG00000019772 | 1 | |
| ENSMUSG00000019773 | 1.029484994 | 0.923833811 |
| ENSMUSG00000019774 | -1.113314819 | 0.412837113 |
| ENSMUSG00000019775 | 1.327117887 | 0.005110127 |
| ENSMUSG00000019777 | 1.049307963 | 0.88972256 |
| ENSMUSG00000019779 | 2.159038398 | 0.00039634 |

| | | |
|---|---|---|
| ENSMUSG00000019782 | -1.054240513 | 0.8349433 |
| ENSMUSG00000019785 | -1.043306199 | 0.79291651 |
| ENSMUSG00000019787 | -1 | 1 |
| ENSMUSG00000019789 | 1.731784909 | 0.017786486 |
| ENSMUSG00000019790 | -1.02422188 | 0.953731979 |
| ENSMUSG00000019791 | 1.026754683 | 0.77920725 |
| ENSMUSG00000019792 | -1.018792235 | 0.960158467 |
| ENSMUSG00000019794 | -1.018880837 | 0.971782973 |
| ENSMUSG00000019795 | -1.011178899 | 1 |
| ENSMUSG00000019796 | -1.265740376 | 0.046977175 |
| ENSMUSG00000019797 | 1.004959098 | 1 |
| ENSMUSG00000019802 | -1.006609347 | 1 |
| ENSMUSG00000019803 | -1.053549062 | 0.769745556 |
| ENSMUSG00000019804 | -1.003695049 | 1 |
| ENSMUSG00000019806 | -1.050285484 | 0.734803948 |
| ENSMUSG00000019808 | -1.122628762 | 0.272029197 |
| ENSMUSG00000019809 | -1.022063116 | 0.96035552 |
| ENSMUSG00000019810 | 1.098547304 | 0.625876002 |
| ENSMUSG00000019813 | 1.01685749 | 0.954497929 |
| ENSMUSG00000019814 | -1.054034946 | 0.82683189 |
| ENSMUSG00000019815 | -1.043306199 | 0.698112512 |
| ENSMUSG00000019817 | 1.097808382 | 0.45721653 |
| ENSMUSG00000019818 | 1.018865446 | 1 |
| ENSMUSG00000019820 | 1.11289764 | 0.692325103 |
| ENSMUSG00000019822 | -1.043324239 | 0.763700143 |
| ENSMUSG00000019823 | 1.246173788 | 0.014825903 |
| ENSMUSG00000019826 | -1.016824185 | 0.899648217 |
| ENSMUSG00000019828 | 1.210869356 | 0.327655754 |
| ENSMUSG00000019831 | -1.07358905 | 0.624676024 |
| ENSMUSG00000019832 | -1.003282735 | 1 |
| ENSMUSG00000019834 | 1 | |
| ENSMUSG00000019836 | -1.024530911 | 0.949679283 |
| ENSMUSG00000019837 | -1.108288838 | 0.608497229 |
| ENSMUSG00000019838 | 1.014517577 | 0.979002724 |
| ENSMUSG00000019841 | -1.016719952 | 0.997582482 |
| ENSMUSG00000019842 | 1.006604729 | 0.991890136 |
| ENSMUSG00000019843 | 1.269667837 | 0.099764565 |
| ENSMUSG00000019845 | -1.045998436 | 0.737954782 |
| ENSMUSG00000019846 | 1.751324703 | 0.002684214 |
| ENSMUSG00000019848 | -1.023769022 | 0.9543611 |
| ENSMUSG00000019849 | 1.041714645 | 0.894828773 |
| ENSMUSG00000019850 | 1.17160841 | 0.37007882 |
| ENSMUSG00000019851 | -1.081207712 | 0.698112512 |
| ENSMUSG00000019852 | -1.179203865 | 0.481210734 |
| ENSMUSG00000019853 | -1.060180246 | 0.734340101 |
| ENSMUSG00000019854 | 1.038080211 | 0.88051937 |
| ENSMUSG00000019856 | -1.244347583 | 0.050235993 |
| ENSMUSG00000019857 | -1.032318753 | 0.911321264 |
| ENSMUSG00000019861 | 1.013598251 | 0.998632737 |
| ENSMUSG00000019863 | -1.010329399 | 0.992217862 |
| ENSMUSG00000019864 | -1.009521116 | 1 |
| ENSMUSG00000019865 | 1.022951384 | 0.998118754 |
| ENSMUSG00000019866 | -1.057637235 | 0.811775815 |
| ENSMUSG00000019867 | 1.06585783 | 0.826758538 |
| ENSMUSG00000019868 | 1.022946959 | 0.958181659 |
| ENSMUSG00000019872 | 1.412663661 | 0.00777869 |
| ENSMUSG00000019873 | 1.019701774 | 0.991944074 |
| ENSMUSG00000019874 | 1.475033422 | 0.0407346 |
| ENSMUSG00000019876 | 1.13507402 | 0.553525456 |
| ENSMUSG00000019877 | 1.005010917 | 1 |
| ENSMUSG00000019878 | 1.015949472 | 0.988424085 |
| ENSMUSG00000019880 | 1.099601477 | 0.59120845 |
| ENSMUSG00000019883 | 1.063154744 | 0.608497229 |

| | | |
|---|---|---|
| ENSMUSG00000019888 | 1.132639128 | 0.428732467 |
| ENSMUSG00000019889 | 1.059605962 | 0.84019351 |
| ENSMUSG00000019890 | -1.043306199 | 0.768421979 |
| ENSMUSG00000019891 | -1.055437867 | 0.698112512 |
| ENSMUSG00000019892 | -1.059527646 | 0.812570482 |
| ENSMUSG00000019893 | -1.053549062 | 0.698112512 |
| ENSMUSG00000019894 | 1.744123372 | 0.001039002 |
| ENSMUSG00000019897 | -1.012744396 | 0.996876674 |
| ENSMUSG00000019899 | -1.48324693 | 0.056847573 |
| ENSMUSG00000019900 | -1.099174267 | 0.535052277 |
| ENSMUSG00000019905 | 1 | |
| ENSMUSG00000019906 | -1.502206401 | 0.004795301 |
| ENSMUSG00000019907 | 1.086180936 | 0.721403818 |
| ENSMUSG00000019909 | 1 | |
| ENSMUSG00000019913 | 1.107565762 | 0.698112512 |
| ENSMUSG00000019916 | 1.065659332 | 0.823104511 |
| ENSMUSG00000019917 | 1.07346491 | 0.70299173 |
| ENSMUSG00000019920 | 1.088977334 | 0.768101821 |
| ENSMUSG00000019923 | 1.01192835 | 1 |
| ENSMUSG00000019927 | 1.005513341 | 1 |
| ENSMUSG00000019929 | 1.088584791 | 0.509386492 |
| ENSMUSG00000019932 | 1 | |
| ENSMUSG00000019933 | -1.044273782 | 0.698112512 |
| ENSMUSG00000019935 | 1.267707981 | 0.06782433 |
| ENSMUSG00000019936 | 1 | |
| ENSMUSG00000019942 | -1.035142069 | 0.957940455 |
| ENSMUSG00000019943 | 1.057925446 | 0.860554063 |
| ENSMUSG00000019944 | 1.158686465 | 0.146736063 |
| ENSMUSG00000019945 | -1 | 1 |
| ENSMUSG00000019947 | 1.187993526 | 0.122533263 |
| ENSMUSG00000019948 | -1.084305963 | 0.657060635 |
| ENSMUSG00000019951 | 1.010488224 | 1 |
| ENSMUSG00000019952 | -1.083905264 | 0.67230273 |
| ENSMUSG00000019960 | -1.099077138 | 0.33505564 |
| ENSMUSG00000019961 | 1.010716697 | 1 |
| ENSMUSG00000019966 | 1.11860722 | 0.378017455 |
| ENSMUSG00000019969 | 1.060737003 | 0.792690927 |
| ENSMUSG00000019970 | 1.023558753 | 0.914954453 |
| ENSMUSG00000019971 | -1.030792706 | 0.893431198 |
| ENSMUSG00000019975 | 1.058692145 | 0.797105186 |
| ENSMUSG00000019977 | -1.027366418 | 0.95554732 |
| ENSMUSG00000019978 | 1.027088709 | 0.978182137 |
| ENSMUSG00000019979 | 1.03142585 | 0.93692717 |
| ENSMUSG00000019982 | -1.032813602 | 0.74782372 |
| ENSMUSG00000019984 | 1.024268876 | 0.956756676 |
| ENSMUSG00000019986 | -1.092062511 | 0.638991399 |
| ENSMUSG00000019987 | 1.11759186 | 0.636090674 |
| ENSMUSG00000019988 | -1.021637425 | 0.98012623 |
| ENSMUSG00000019989 | 1.095872691 | 0.753905984 |
| ENSMUSG00000019990 | 1.089470641 | 0.740841499 |
| ENSMUSG00000019992 | -1.04085568 | 0.827942378 |
| ENSMUSG00000019996 | -1.531287511 | 0.153716858 |
| ENSMUSG00000019997 | 1.241160336 | 0.355608366 |
| ENSMUSG00000019998 | -1.009332627 | 1 |
| ENSMUSG00000020000 | -1.389988124 | 0.021070191 |
| ENSMUSG00000020003 | -1.147958189 | 0.189746534 |
| ENSMUSG00000020007 | -1.043306199 | 0.698112512 |
| ENSMUSG00000020009 | -1.022312405 | 0.958181659 |
| ENSMUSG00000020010 | 1 | |
| ENSMUSG00000020014 | 1.242499785 | 0.501577656 |
| ENSMUSG00000020015 | 1.049638284 | 0.823018955 |
| ENSMUSG00000020017 | 1.163870837 | 0.003083671 |
| ENSMUSG00000020018 | -1.070338766 | 0.698112512 |

| | | |
|---|---|---|
| ENSMUSG00000020019 | -1.117244544 | 0.103846634 |
| ENSMUSG00000020020 | -1.188148209 | 0.02536973 |
| ENSMUSG00000020021 | -1.098710535 | 0.608497229 |
| ENSMUSG00000020022 | -1.150568113 | 0.350426777 |
| ENSMUSG00000020023 | -1.059046596 | 0.751653251 |
| ENSMUSG00000020024 | 1.02444191 | 0.936794036 |
| ENSMUSG00000020027 | 1.189021517 | 0.112340347 |
| ENSMUSG00000020029 | 1.197983802 | 0.481065348 |
| ENSMUSG00000020032 | 1.027563099 | 0.897751034 |
| ENSMUSG00000020033 | -1.089497311 | 0.532369657 |
| ENSMUSG00000020034 | -1.044509461 | 0.583147495 |
| ENSMUSG00000020037 | 1 | |
| ENSMUSG00000020038 | 1.003912131 | 1 |
| ENSMUSG00000020042 | 1.032528575 | 0.765233309 |
| ENSMUSG00000020044 | 1.358586797 | 0.1631694 |
| ENSMUSG00000020048 | 1.020932 | 1 |
| ENSMUSG00000020051 | 1 | |
| ENSMUSG00000020052 | 1 | |
| ENSMUSG00000020053 | -1.297650954 | 0.113640702 |
| ENSMUSG00000020056 | -1.014350412 | 0.98226424 |
| ENSMUSG00000020057 | 1.005411517 | 1 |
| ENSMUSG00000020059 | -1.133488037 | 0.679913326 |
| ENSMUSG00000020061 | -1.009476377 | 1 |
| ENSMUSG00000020062 | -1.012829633 | 1 |
| ENSMUSG00000020063 | -1.115961902 | 0.6097568 |
| ENSMUSG00000020064 | 1.024423088 | 0.968410203 |
| ENSMUSG00000020067 | 1.290681034 | 0.028809274 |
| ENSMUSG00000020069 | -1.049125667 | 0.819614655 |
| ENSMUSG00000020070 | -1.072030325 | 0.698112512 |
| ENSMUSG00000020072 | 1.25124303 | 0.143272335 |
| ENSMUSG00000020074 | 1.012804154 | 1 |
| ENSMUSG00000020075 | -1.017418928 | 1 |
| ENSMUSG00000020076 | 1.025092616 | 0.968937442 |
| ENSMUSG00000020077 | -1.114877987 | 0.608497229 |
| ENSMUSG00000020078 | 1.06294343 | 0.819614655 |
| ENSMUSG00000020079 | -1.062446115 | 0.749958629 |
| ENSMUSG00000020080 | -1.001348012 | 1 |
| ENSMUSG00000020081 | -1.003834322 | 1 |
| ENSMUSG00000020083 | -1.325290096 | 0.031082053 |
| ENSMUSG00000020085 | -1.14625461 | 0.303261282 |
| ENSMUSG00000020086 | -1.063862762 | 0.698112512 |
| ENSMUSG00000020087 | -1.092014782 | 0.195814049 |
| ENSMUSG00000020088 | 1.028177958 | 0.974807466 |
| ENSMUSG00000020089 | -1.097892935 | 0.710224291 |
| ENSMUSG00000020090 | -1.139982257 | 0.624040338 |
| ENSMUSG00000020091 | 1.016384347 | 0.979814083 |
| ENSMUSG00000020092 | -1.051401601 | 0.707507155 |
| ENSMUSG00000020096 | -1.007690427 | 1 |
| ENSMUSG00000020097 | 1.053366283 | 0.884942525 |
| ENSMUSG00000020098 | -1.42346169 | 0.004739776 |
| ENSMUSG00000020099 | 1.204324809 | 0.216046507 |
| ENSMUSG00000020100 | -1.016833269 | 0.96797322 |
| ENSMUSG00000020101 | 1.31196953 | 0.004906305 |
| ENSMUSG00000020102 | 1.481815013 | 0.091827709 |
| ENSMUSG00000020105 | 1.084309179 | 0.601666207 |
| ENSMUSG00000020107 | 1.003831661 | 1 |
| ENSMUSG00000020108 | 1.301679665 | 0.011391085 |
| ENSMUSG00000020109 | 1.02022375 | 0.924587329 |
| ENSMUSG00000020111 | 1.055973839 | 0.82134515 |
| ENSMUSG00000020114 | 1.028844791 | 0.975275648 |
| ENSMUSG00000020115 | 1.045482732 | 0.851171651 |
| ENSMUSG00000020116 | -1.056458393 | 0.807247723 |
| ENSMUSG00000020120 | -1.175244992 | 0.559196817 |

| | | |
|---|---|---|
| ENSMUSG00000020121 | 1.862809608 | 2.50441E-05 |
| ENSMUSG00000020122 | 1.246466788 | 0.06980995 |
| ENSMUSG00000020123 | -1.112319663 | 0.777583275 |
| ENSMUSG00000020124 | -1.035309908 | 0.945279098 |
| ENSMUSG00000020125 | 1 | |
| ENSMUSG00000020128 | -1.060766456 | 0.828564184 |
| ENSMUSG00000020130 | -1.003699694 | 1 |
| ENSMUSG00000020131 | 1.196838278 | 0.608497229 |
| ENSMUSG00000020132 | 1.024703474 | 0.950034218 |
| ENSMUSG00000020133 | 1.028405865 | 0.825645788 |
| ENSMUSG00000020134 | -1.066429581 | 0.744412315 |
| ENSMUSG00000020135 | 1.151577582 | 0.357169145 |
| ENSMUSG00000020137 | 1.087082263 | 0.608497229 |
| ENSMUSG00000020140 | -1.701172101 | 0.00466733 |
| ENSMUSG00000020142 | 1.054359322 | 0.743300655 |
| ENSMUSG00000020143 | 1 | |
| ENSMUSG00000020149 | -1.033711325 | 0.957616514 |
| ENSMUSG00000020150 | 1.122150871 | 0.220181381 |
| ENSMUSG00000020151 | 1.10776854 | 0.625496737 |
| ENSMUSG00000020152 | 1.014358025 | 1 |
| ENSMUSG00000020153 | 1.026394257 | 0.945299421 |
| ENSMUSG00000020154 | 1.323994223 | 0.049798568 |
| ENSMUSG00000020155 | 1.276746305 | 0.250477583 |
| ENSMUSG00000020156 | 1.03312776 | 0.828425932 |
| ENSMUSG00000020159 | 1 | |
| ENSMUSG00000020160 | 1.286829245 | 0.070968316 |
| ENSMUSG00000020163 | 1.059350075 | 0.731295483 |
| ENSMUSG00000020164 | 1 | |
| ENSMUSG00000020166 | -1.050795976 | 0.847428179 |
| ENSMUSG00000020167 | 1.139324683 | 0.283605457 |
| ENSMUSG00000020168 | 1 | |
| ENSMUSG00000020169 | -1.088313104 | 0.783112516 |
| ENSMUSG00000020170 | 1.059925972 | 0.820789565 |
| ENSMUSG00000020171 | -1.044527213 | 0.795609888 |
| ENSMUSG00000020173 | 1.365634518 | 0.001160262 |
| ENSMUSG00000020175 | -1.007349771 | 0.991432966 |
| ENSMUSG00000020176 | 1.358849315 | 0.198900518 |
| ENSMUSG00000020177 | 1.059463094 | 0.608497229 |
| ENSMUSG00000020178 | 1.323631064 | 0.140595679 |
| ENSMUSG00000020180 | -1.021879679 | 0.984602435 |
| ENSMUSG00000020181 | 1.300867859 | 0.008595113 |
| ENSMUSG00000020182 | -1.110426459 | 0.70299173 |
| ENSMUSG00000020183 | -1.703522166 | 0.00314868 |
| ENSMUSG00000020184 | 1.125314804 | 0.61231779 |
| ENSMUSG00000020185 | 1.017403641 | 0.987241494 |
| ENSMUSG00000020186 | 1.000365779 | 1 |
| ENSMUSG00000020189 | -1.036267192 | 0.922391825 |
| ENSMUSG00000020190 | 1.038665669 | 0.846399261 |
| ENSMUSG00000020191 | 1.019083579 | 0.968937442 |
| ENSMUSG00000020193 | 1.069293585 | 0.815807184 |
| ENSMUSG00000020196 | 1.045482936 | 0.835382855 |
| ENSMUSG00000020198 | 1.067578789 | 0.749908606 |
| ENSMUSG00000020205 | 1.218641409 | 0.035744898 |
| ENSMUSG00000020211 | 1.062271898 | 0.6926778 |
| ENSMUSG00000020212 | -1.010590016 | 0.985522133 |
| ENSMUSG00000020213 | -1.043306199 | 0.698112512 |
| ENSMUSG00000020214 | 1 | |
| ENSMUSG00000020216 | 1.079681442 | 0.705088559 |
| ENSMUSG00000020218 | -1.834767191 | 0.019657416 |
| ENSMUSG00000020219 | 1.072221144 | 0.698112512 |
| ENSMUSG00000020220 | 1.030602482 | 0.925982957 |
| ENSMUSG00000020224 | -1.064763806 | 0.708068647 |
| ENSMUSG00000020225 | -1.135260754 | 0.500630907 |

| | | |
|---|---|---|
| ENSMUSG00000020226 | 1.099850986 | 0.803764946 |
| ENSMUSG00000020227 | -1.028709819 | 0.859429731 |
| ENSMUSG00000020228 | -1.02806875 | 0.86902396 |
| ENSMUSG00000020229 | -1.053549062 | 0.769745556 |
| ENSMUSG00000020230 | 1.01278939 | 0.982838256 |
| ENSMUSG00000020231 | -1.03177771 | 0.910895334 |
| ENSMUSG00000020232 | 1.076664978 | 0.342989506 |
| ENSMUSG00000020234 | 1.035015266 | 0.941994554 |
| ENSMUSG00000020235 | 1.066725119 | 0.644606499 |
| ENSMUSG00000020238 | 1.072736162 | 0.670176133 |
| ENSMUSG00000020241 | 1.343344641 | 0.023252559 |
| ENSMUSG00000020246 | 1.016442156 | 0.988424085 |
| ENSMUSG00000020248 | 1.019008674 | 0.972980766 |
| ENSMUSG00000020250 | 1.072440439 | 0.815601444 |
| ENSMUSG00000020251 | 1.13309924 | 0.133674794 |
| ENSMUSG00000020253 | -1.024325149 | 0.913163672 |
| ENSMUSG00000020255 | -1.032142888 | 0.935353013 |
| ENSMUSG00000020256 | 1.29169847 | 0.077817735 |
| ENSMUSG00000020257 | -1.055738059 | 0.86119789 |
| ENSMUSG00000020258 | 1.054200691 | 0.627656175 |
| ENSMUSG00000020260 | 1.15300458 | 0.313158244 |
| ENSMUSG00000020261 | 1.099106111 | 0.530378324 |
| ENSMUSG00000020262 | -1.06412119 | 0.457213896 |
| ENSMUSG00000020263 | -1.051245185 | 0.698112512 |
| ENSMUSG00000020264 | 1 | |
| ENSMUSG00000020265 | -1.054376118 | 0.877257217 |
| ENSMUSG00000020267 | -1.023048387 | 0.977563419 |
| ENSMUSG00000020268 | -1.01589915 | 0.942757921 |
| ENSMUSG00000020270 | 1 | |
| ENSMUSG00000020271 | -1.00501288 | 1 |
| ENSMUSG00000020272 | 1.082058497 | 0.572727047 |
| ENSMUSG00000020273 | -1.044560149 | 0.835839899 |
| ENSMUSG00000020275 | -1.126165138 | 0.572781054 |
| ENSMUSG00000020277 | -1.00965589 | 1 |
| ENSMUSG00000020279 | 1.024097153 | 0.982534887 |
| ENSMUSG00000020280 | -1.162601109 | 0.532369657 |
| ENSMUSG00000020282 | 1.046817937 | 0.746258638 |
| ENSMUSG00000020283 | -1.098611489 | 0.612610841 |
| ENSMUSG00000020284 | -1.033670224 | 0.699340382 |
| ENSMUSG00000020286 | -1.043306199 | 0.803260194 |
| ENSMUSG00000020287 | -1.09735033 | 0.178255456 |
| ENSMUSG00000020288 | -1.11706902 | 0.486060044 |
| ENSMUSG00000020289 | 1.040072018 | 0.781240509 |
| ENSMUSG00000020290 | -1.023522731 | 0.98923405 |
| ENSMUSG00000020295 | 1 | |
| ENSMUSG00000020297 | -1.044273782 | 0.698112512 |
| ENSMUSG00000020299 | 1 | |
| ENSMUSG00000020300 | -1.079617964 | 0.698112512 |
| ENSMUSG00000020303 | 1.320466353 | 0.288134604 |
| ENSMUSG00000020305 | -1.012630427 | 1 |
| ENSMUSG00000020307 | 1.075226229 | 0.537225049 |
| ENSMUSG00000020308 | -1.049817585 | 0.946772604 |
| ENSMUSG00000020309 | -1.390372759 | 0.041401294 |
| ENSMUSG00000020310 | 1.000305362 | 1 |
| ENSMUSG00000020311 | -1.044153828 | 0.855528455 |
| ENSMUSG00000020312 | -1.158409335 | 0.240858255 |
| ENSMUSG00000020315 | 1.043055237 | 0.937575412 |
| ENSMUSG00000020317 | 1 | |
| ENSMUSG00000020319 | -1.046375771 | 0.808606444 |
| ENSMUSG00000020321 | -1.059035904 | 0.865845831 |
| ENSMUSG00000020323 | 1.132639128 | 0.217803313 |
| ENSMUSG00000020325 | 1.180805986 | 0.67232595 |
| ENSMUSG00000020326 | 1.163842257 | 0.608497229 |

| | | |
|---|---|---|
| ENSMUSG00000020327 | 1.064985481 | 0.826197304 |
| ENSMUSG00000020328 | -1.020580739 | 0.971848081 |
| ENSMUSG00000020329 | 1.077835627 | 0.698112512 |
| ENSMUSG00000020330 | -1.140507067 | 0.615267463 |
| ENSMUSG00000020331 | -1.195112548 | 0.038104479 |
| ENSMUSG00000020332 | 1 | |
| ENSMUSG00000020333 | -1.635773224 | 0.003298632 |
| ENSMUSG00000020334 | -1.230209667 | 0.039155055 |
| ENSMUSG00000020335 | -1.084760444 | 0.79291651 |
| ENSMUSG00000020340 | -1.877157413 | 0.000534462 |
| ENSMUSG00000020346 | 1.066839996 | 0.612292929 |
| ENSMUSG00000020349 | -1.074677684 | 0.840253751 |
| ENSMUSG00000020354 | -1.188446785 | 0.39980163 |
| ENSMUSG00000020357 | -1.458750548 | 0.001238515 |
| ENSMUSG00000020358 | -1.084412443 | 0.784232093 |
| ENSMUSG00000020359 | -1.160214222 | 0.217803313 |
| ENSMUSG00000020361 | -1.062463781 | 0.886841005 |
| ENSMUSG00000020362 | -1.115934458 | 0.710698346 |
| ENSMUSG00000020363 | -1.846946033 | 0.001610049 |
| ENSMUSG00000020364 | -1.09436251 | 0.36904104 |
| ENSMUSG00000020366 | -1.144408395 | 0.588635667 |
| ENSMUSG00000020368 | -1.006756883 | 1 |
| ENSMUSG00000020372 | 1.022814853 | 0.988424085 |
| ENSMUSG00000020374 | -1.043306199 | 0.79291651 |
| ENSMUSG00000020375 | -1.075230107 | 0.674022895 |
| ENSMUSG00000020376 | -1.571314213 | 0.003935818 |
| ENSMUSG00000020377 | -1.052304694 | 0.807247723 |
| ENSMUSG00000020380 | -1.059568659 | 0.79137734 |
| ENSMUSG00000020381 | -1.130950124 | 0.017287546 |
| ENSMUSG00000020383 | 1.024693402 | 0.933031586 |
| ENSMUSG00000020385 | -1.008095692 | 1 |
| ENSMUSG00000020386 | -1.054931632 | 0.787977591 |
| ENSMUSG00000020387 | 1.035734795 | 0.764129089 |
| ENSMUSG00000020388 | -1.398275297 | 0.01222575 |
| ENSMUSG00000020389 | -1.075853643 | 0.572892557 |
| ENSMUSG00000020390 | -1.026714221 | 0.961940234 |
| ENSMUSG00000020392 | -1.086385306 | 0.650560874 |
| ENSMUSG00000020393 | -1.000148361 | 1 |
| ENSMUSG00000020395 | -1.044273782 | 0.698112512 |
| ENSMUSG00000020396 | -1.077167372 | 0.869434134 |
| ENSMUSG00000020397 | -1.05721799 | 0.608497229 |
| ENSMUSG00000020399 | -1.089821813 | 0.511163311 |
| ENSMUSG00000020400 | -1.070487277 | 0.671073601 |
| ENSMUSG00000020401 | 1.095872691 | 0.608497229 |
| ENSMUSG00000020402 | -1.122430546 | 0.698112512 |
| ENSMUSG00000020405 | 1 | |
| ENSMUSG00000020407 | 1.023002031 | 0.810722095 |
| ENSMUSG00000020409 | -1.090530191 | 0.698112512 |
| ENSMUSG00000020411 | 1.294965863 | 0.014289981 |
| ENSMUSG00000020412 | -1.025789239 | 0.950565815 |
| ENSMUSG00000020413 | 1.007429809 | 1 |
| ENSMUSG00000020415 | -1.087239266 | 0.723412916 |
| ENSMUSG00000020419 | 1.138000172 | 0.376011909 |
| ENSMUSG00000020420 | -1.029405093 | 0.83141203 |
| ENSMUSG00000020422 | 1.109916663 | 0.653331397 |
| ENSMUSG00000020423 | 1.070893216 | 0.62717574 |
| ENSMUSG00000020424 | 1.049761731 | 0.59120845 |
| ENSMUSG00000020427 | 2.450701421 | 0.000822888 |
| ENSMUSG00000020428 | 1 | |
| ENSMUSG00000020429 | 1 | |
| ENSMUSG00000020430 | 1.024930112 | 0.965872389 |
| ENSMUSG00000020431 | 1.442937499 | 0.001490182 |
| ENSMUSG00000020432 | 1.258282068 | 0.01965858 |

| | | |
|---|---|---|
| ENSMUSG00000020434 | 1.044859725 | 0.850820147 |
| ENSMUSG00000020435 | -1.567359276 | 0.039761862 |
| ENSMUSG00000020436 | 1 | |
| ENSMUSG00000020437 | 1.059592506 | 0.778372272 |
| ENSMUSG00000020439 | 1.168432794 | 0.159480741 |
| ENSMUSG00000020440 | 1.056637506 | 0.675967223 |
| ENSMUSG00000020441 | -1.142427841 | 0.134585334 |
| ENSMUSG00000020444 | -1.075620156 | 0.608497229 |
| ENSMUSG00000020447 | 1.015486245 | 0.986542466 |
| ENSMUSG00000020448 | 1.061980313 | 0.701154921 |
| ENSMUSG00000020451 | 1.078275109 | 0.662474108 |
| ENSMUSG00000020453 | 1.182057613 | 0.057291025 |
| ENSMUSG00000020454 | 1.0469442 | 0.87177448 |
| ENSMUSG00000020455 | 1.002958341 | 1 |
| ENSMUSG00000020456 | -1.047237977 | 0.884451146 |
| ENSMUSG00000020457 | -1.025916224 | 0.974508422 |
| ENSMUSG00000020458 | 1.042093362 | 0.940197654 |
| ENSMUSG00000020459 | -1.038655398 | 0.845511995 |
| ENSMUSG00000020460 | -1.057972756 | 0.876422702 |
| ENSMUSG00000020461 | -1.033762817 | 0.982175268 |
| ENSMUSG00000020462 | -1.151469999 | 0.328811959 |
| ENSMUSG00000020463 | -1.108004932 | 0.698112512 |
| ENSMUSG00000020464 | -1.10687805 | 0.608497229 |
| ENSMUSG00000020467 | -2.678876775 | 0.000135858 |
| ENSMUSG00000020469 | 1.049411284 | 0.91218384 |
| ENSMUSG00000020471 | 1.058500403 | 0.694492485 |
| ENSMUSG00000020472 | -1.030079956 | 0.865583959 |
| ENSMUSG00000020473 | 1.440260276 | 0.013838802 |
| ENSMUSG00000020474 | 1.015876371 | 0.916565695 |
| ENSMUSG00000020475 | -1.048406857 | 0.819474306 |
| ENSMUSG00000020476 | 1.024767942 | 0.919215198 |
| ENSMUSG00000020477 | -1.006869822 | 0.960836234 |
| ENSMUSG00000020481 | 1 | |
| ENSMUSG00000020482 | -1.046632147 | 0.827202174 |
| ENSMUSG00000020483 | 1.088720557 | 0.698112512 |
| ENSMUSG00000020484 | 1.072156727 | 0.70443552 |
| ENSMUSG00000020485 | 1.106898761 | 0.608497229 |
| ENSMUSG00000020486 | -1.364883319 | 0.08119166 |
| ENSMUSG00000020490 | -1.044273782 | 0.698112512 |
| ENSMUSG00000020491 | -1.202405083 | 0.144820051 |
| ENSMUSG00000020492 | 1.05671324 | 0.824532064 |
| ENSMUSG00000020493 | -1.087000923 | 0.70299173 |
| ENSMUSG00000020495 | -1.008389203 | 1 |
| ENSMUSG00000020496 | -1.073532349 | 0.698112512 |
| ENSMUSG00000020513 | -1.02494561 | 0.892084693 |
| ENSMUSG00000020514 | -1.084599122 | 0.608497229 |
| ENSMUSG00000020515 | -1.021667052 | 0.974593713 |
| ENSMUSG00000020516 | -1.004343706 | 1 |
| ENSMUSG00000020519 | -1.063348955 | 0.608497229 |
| ENSMUSG00000020520 | -1.046334365 | 0.811272147 |
| ENSMUSG00000020521 | 1.062753565 | 0.790885744 |
| ENSMUSG00000020522 | -1.074718334 | 0.725290225 |
| ENSMUSG00000020523 | -1.050400755 | 0.842923096 |
| ENSMUSG00000020524 | 1.069068979 | 0.608497229 |
| ENSMUSG00000020525 | 1.008195675 | 1 |
| ENSMUSG00000020526 | 1.007215461 | 1 |
| ENSMUSG00000020527 | 1.022561158 | 0.944099922 |
| ENSMUSG00000020528 | -1.30489377 | 0.053002168 |
| ENSMUSG00000020530 | 1.038749412 | 0.920955554 |
| ENSMUSG00000020532 | -1.037578019 | 0.935237716 |
| ENSMUSG00000020534 | -1.027018502 | 0.914288146 |
| ENSMUSG00000020536 | 1.037385768 | 0.817630189 |
| ENSMUSG00000020537 | -1.135416401 | 0.602100972 |

| | | |
|---|---|---|
| ENSMUSG00000020538 | -1.026359596 | 0.895588956 |
| ENSMUSG00000020541 | 1.06395194 | 0.722577651 |
| ENSMUSG00000020542 | -1.03420529 | 0.908354926 |
| ENSMUSG00000020544 | 1.059879739 | 0.752411447 |
| ENSMUSG00000020545 | 1 | |
| ENSMUSG00000020546 | -1.015848849 | 0.986359381 |
| ENSMUSG00000020547 | 1.152858502 | 0.476544766 |
| ENSMUSG00000020548 | -1 | 1 |
| ENSMUSG00000020549 | -1.055182882 | 0.763831324 |
| ENSMUSG00000020553 | -1.282346786 | 0.094918259 |
| ENSMUSG00000020561 | 1.041722285 | 0.84122575 |
| ENSMUSG00000020562 | -1.053549062 | 0.769745556 |
| ENSMUSG00000020564 | 1.07271744 | 0.608497229 |
| ENSMUSG00000020566 | -1.920152459 | 0.000957737 |
| ENSMUSG00000020570 | 1.128666007 | 0.612674163 |
| ENSMUSG00000020571 | 1.016610163 | 1 |
| ENSMUSG00000020572 | 1.129685863 | 0.608497229 |
| ENSMUSG00000020573 | 1 | |
| ENSMUSG00000020576 | -1.00689904 | 1 |
| ENSMUSG00000020577 | -1.133087379 | 0.43764772 |
| ENSMUSG00000020580 | 1.09638934 | 0.730851014 |
| ENSMUSG00000020581 | 1 | |
| ENSMUSG00000020583 | 1.010515549 | 0.984130305 |
| ENSMUSG00000020585 | 1.091579776 | 0.729018288 |
| ENSMUSG00000020589 | -1.963673527 | 0.00117079 |
| ENSMUSG00000020590 | 1.110614595 | 0.616230752 |
| ENSMUSG00000020591 | 1 | |
| ENSMUSG00000020592 | 1.273878732 | 0.024123774 |
| ENSMUSG00000020593 | 1.000503256 | 1 |
| ENSMUSG00000020594 | 1.0739246 | 0.811563366 |
| ENSMUSG00000020598 | 1.156777198 | 0.698112512 |
| ENSMUSG00000020599 | -1.169608628 | 0.086033075 |
| ENSMUSG00000020600 | 1.059463094 | 0.608497229 |
| ENSMUSG00000020601 | 1.816520004 | 8.03973E-05 |
| ENSMUSG00000020604 | 1.162496706 | 0.17294368 |
| ENSMUSG00000020605 | 1.213655206 | 0.093115569 |
| ENSMUSG00000020607 | 1.215983299 | 0.541926749 |
| ENSMUSG00000020608 | 1.048550496 | 0.886195057 |
| ENSMUSG00000020609 | 1.547858696 | 0.01205988 |
| ENSMUSG00000020610 | 1.013839883 | 0.991658036 |
| ENSMUSG00000020611 | 1.039338258 | 0.924402067 |
| ENSMUSG00000020612 | 1.022154333 | 0.990051834 |
| ENSMUSG00000020614 | 1.778631925 | 0.000711519 |
| ENSMUSG00000020617 | 1.059463094 | 0.608497229 |
| ENSMUSG00000020620 | 1.293500849 | 0.008515961 |
| ENSMUSG00000020621 | 1.005862218 | 1 |
| ENSMUSG00000020622 | -1.024718486 | 0.931480186 |
| ENSMUSG00000020623 | 1.057148618 | 0.70299173 |
| ENSMUSG00000020624 | 1 | |
| ENSMUSG00000020627 | -1.247789791 | 0.154205805 |
| ENSMUSG00000020628 | 1.041557226 | 0.725117221 |
| ENSMUSG00000020629 | -1.085478577 | 0.679217411 |
| ENSMUSG00000020630 | 1.158159653 | 0.010275103 |
| ENSMUSG00000020633 | 1.12896782 | 0.244537877 |
| ENSMUSG00000020634 | -1.14963601 | 0.376308441 |
| ENSMUSG00000020635 | -1.172388578 | 0.062348119 |
| ENSMUSG00000020636 | 1 | |
| ENSMUSG00000020638 | -1.211989554 | 0.174472625 |
| ENSMUSG00000020639 | -1.153706825 | 0.611728136 |
| ENSMUSG00000020640 | -1.020767764 | 0.977563419 |
| ENSMUSG00000020641 | -1.159108002 | 0.608497229 |
| ENSMUSG00000020642 | -1.002759711 | 1 |
| ENSMUSG00000020644 | 1.265302865 | 0.091760234 |

| | | |
|---|---|---|
| ENSMUSG00000020646 | -1.120467906 | 0.572947856 |
| ENSMUSG00000020647 | -1.00202493 | 1 |
| ENSMUSG00000020648 | 1.106400534 | 0.184953023 |
| ENSMUSG00000020649 | -1.009989353 | 1 |
| ENSMUSG00000020650 | 1.144278929 | 0.391351306 |
| ENSMUSG00000020651 | -1 | 1 |
| ENSMUSG00000020652 | -1.080659786 | 0.631570736 |
| ENSMUSG00000020653 | 1.154803589 | 0.026038425 |
| ENSMUSG00000020654 | -1.068195669 | 0.518298719 |
| ENSMUSG00000020656 | -1.04090986 | 0.801199277 |
| ENSMUSG00000020657 | 1.040264859 | 0.80659229 |
| ENSMUSG00000020658 | -1.214532628 | 0.014748278 |
| ENSMUSG00000020659 | 1.061221887 | 0.713964595 |
| ENSMUSG00000020660 | -1.084645496 | 0.532369657 |
| ENSMUSG00000020661 | 1.045962282 | 0.747082435 |
| ENSMUSG00000020664 | 1.067826439 | 0.814089245 |
| ENSMUSG00000020668 | 1.17352108 | 0.131758166 |
| ENSMUSG00000020669 | 1.10698386 | 0.08867248 |
| ENSMUSG00000020671 | 1.105670789 | 0.698112512 |
| ENSMUSG00000020672 | 1 | |
| ENSMUSG00000020673 | 1 | |
| ENSMUSG00000020674 | 1.073405318 | 0.795079697 |
| ENSMUSG00000020676 | 1.069068979 | 0.721840456 |
| ENSMUSG00000020677 | -1.015129804 | 1 |
| ENSMUSG00000020679 | -1.118291056 | 0.655005967 |
| ENSMUSG00000020680 | 1.168380302 | 0.53463993 |
| ENSMUSG00000020681 | -1.152349418 | 0.608497229 |
| ENSMUSG00000020682 | 1.310608616 | 0.007194115 |
| ENSMUSG00000020684 | -1.162576303 | 0.249042326 |
| ENSMUSG00000020686 | 1 | |
| ENSMUSG00000020687 | -1.006032749 | 1 |
| ENSMUSG00000020689 | 1.865342615 | 5.68991E-05 |
| ENSMUSG00000020690 | 1 | |
| ENSMUSG00000020691 | -1.020746958 | 0.956374899 |
| ENSMUSG00000020692 | 1.18924891 | 0.20691966 |
| ENSMUSG00000020694 | 1.035903475 | 0.920928417 |
| ENSMUSG00000020695 | 1.153357294 | 0.287437444 |
| ENSMUSG00000020696 | 1.035312467 | 0.711140135 |
| ENSMUSG00000020697 | 1.021250079 | 0.96035552 |
| ENSMUSG00000020698 | -1.012668111 | 1 |
| ENSMUSG00000020700 | 1.070125756 | 0.778372272 |
| ENSMUSG00000020701 | 1 | |
| ENSMUSG00000020702 | 1 | |
| ENSMUSG00000020703 | -1.095294215 | 0.608497229 |
| ENSMUSG00000020704 | 1 | |
| ENSMUSG00000020705 | 1.002674643 | 1 |
| ENSMUSG00000020706 | -1.010416589 | 1 |
| ENSMUSG00000020707 | 1.049276648 | 0.457213896 |
| ENSMUSG00000020708 | 1.006796414 | 1 |
| ENSMUSG00000020709 | -1.306791625 | 0.086747947 |
| ENSMUSG00000020712 | -1.002163121 | 1 |
| ENSMUSG00000020713 | 1 | |
| ENSMUSG00000020715 | -1.140317088 | 0.534944201 |
| ENSMUSG00000020716 | 1.062261819 | 0.812228807 |
| ENSMUSG00000020717 | 1.064528607 | 0.84360497 |
| ENSMUSG00000020718 | 1.02969426 | 0.717686641 |
| ENSMUSG00000020719 | 1.003986299 | 1 |
| ENSMUSG00000020720 | -1.030057009 | 0.961255441 |
| ENSMUSG00000020721 | -1.04529032 | 0.872980877 |
| ENSMUSG00000020722 | 1 | |
| ENSMUSG00000020723 | 1.111925027 | 0.698112512 |
| ENSMUSG00000020728 | -1.001533762 | 1 |
| ENSMUSG00000020732 | 1.138070563 | 0.635422995 |

| | | |
|---|---|---|
| ENSMUSG00000020733 | 1.084029032 | 0.553779197 |
| ENSMUSG00000020734 | 1 | |
| ENSMUSG00000020736 | -1.047846174 | 0.681952832 |
| ENSMUSG00000020737 | 1.021888447 | 0.961487493 |
| ENSMUSG00000020738 | 1.004166963 | 1 |
| ENSMUSG00000020739 | 1.034521379 | 0.941510974 |
| ENSMUSG00000020740 | 1.013340594 | 0.986542466 |
| ENSMUSG00000020741 | 1.096831012 | 0.608497229 |
| ENSMUSG00000020743 | -1.034625042 | 0.648538976 |
| ENSMUSG00000020744 | -1.011658456 | 0.93376866 |
| ENSMUSG00000020745 | 1.017286614 | 1 |
| ENSMUSG00000020747 | 1.074520595 | 0.626099661 |
| ENSMUSG00000020752 | 1.059506827 | 0.532369657 |
| ENSMUSG00000020755 | 1.028387485 | 0.927169398 |
| ENSMUSG00000020758 | 1.074046278 | 0.410139679 |
| ENSMUSG00000020766 | 1.094232059 | 0.608497229 |
| ENSMUSG00000020770 | 1.081639628 | 0.563092624 |
| ENSMUSG00000020773 | 1.192222373 | 0.131010602 |
| ENSMUSG00000020774 | -1.013223975 | 1 |
| ENSMUSG00000020775 | -1.050800061 | 0.718280665 |
| ENSMUSG00000020776 | 1.098795481 | 0.378924067 |
| ENSMUSG00000020777 | 1.050572546 | 0.861137762 |
| ENSMUSG00000020778 | 1.059518474 | 0.686605292 |
| ENSMUSG00000020780 | -1.001983922 | 1 |
| ENSMUSG00000020781 | -1.007569047 | 1 |
| ENSMUSG00000020782 | -1.218964938 | 0.232747283 |
| ENSMUSG00000020783 | -1.007421201 | 1 |
| ENSMUSG00000020785 | 1.054526888 | 0.608497229 |
| ENSMUSG00000020787 | 1.015486245 | 0.972980766 |
| ENSMUSG00000020788 | -1.308383954 | 0.148537072 |
| ENSMUSG00000020790 | 1.033192741 | 0.954310937 |
| ENSMUSG00000020792 | 1.020699536 | 0.92584202 |
| ENSMUSG00000020793 | -1.10093155 | 0.587902215 |
| ENSMUSG00000020794 | 1.112992384 | 0.645756877 |
| ENSMUSG00000020798 | -1.274249452 | 0.143586578 |
| ENSMUSG00000020799 | 1 | |
| ENSMUSG00000020801 | 1.042362529 | 0.843451697 |
| ENSMUSG00000020802 | -1.009944119 | 1 |
| ENSMUSG00000020803 | -1.001753138 | 1 |
| ENSMUSG00000020804 | 1.189706618 | 0.216046507 |
| ENSMUSG00000020805 | -1.561285275 | 0.003047978 |
| ENSMUSG00000020806 | 1.005538704 | 1 |
| ENSMUSG00000020807 | 1.052663543 | 0.72715392 |
| ENSMUSG00000020808 | -1.000667914 | 1 |
| ENSMUSG00000020810 | -1.962218159 | 0.001851304 |
| ENSMUSG00000020811 | -1 | 1 |
| ENSMUSG00000020812 | -1.008098962 | 0.995747207 |
| ENSMUSG00000020814 | -1.033465681 | 0.902080215 |
| ENSMUSG00000020817 | 1.048465623 | 0.890792876 |
| ENSMUSG00000020818 | 1.141094148 | 0.489706317 |
| ENSMUSG00000020821 | 1.141619581 | 0.52710437 |
| ENSMUSG00000020823 | 1.025141701 | 0.954310937 |
| ENSMUSG00000020826 | 1.302151187 | 0.008802103 |
| ENSMUSG00000020827 | 1.053223217 | 0.745840049 |
| ENSMUSG00000020828 | 1.079372156 | 0.608497229 |
| ENSMUSG00000020829 | -1.345778489 | 0.013027074 |
| ENSMUSG00000020830 | 1.059463094 | 0.608497229 |
| ENSMUSG00000020831 | 1.017046454 | 0.989614069 |
| ENSMUSG00000020832 | -1.029613739 | 0.842184576 |
| ENSMUSG00000020834 | -1.057080793 | 0.650252997 |
| ENSMUSG00000020836 | 1.659969647 | 0.000894866 |
| ENSMUSG00000020838 | -1.252812038 | 0.300750932 |
| ENSMUSG00000020839 | 1 | |

| | | |
|---|---|---|
| ENSMUSG00000020840 | -1.019858471 | 0.984239005 |
| ENSMUSG00000020841 | -1.078283813 | 0.816842594 |
| ENSMUSG00000020843 | -1.015989766 | 0.978259071 |
| ENSMUSG00000020844 | 1.083930422 | 0.609323856 |
| ENSMUSG00000020846 | 1.067870364 | 0.698112512 |
| ENSMUSG00000020847 | -1.108189796 | 0.587480427 |
| ENSMUSG00000020848 | -1.106138521 | 0.202613226 |
| ENSMUSG00000020849 | -1.016340191 | 1 |
| ENSMUSG00000020850 | 1.045398695 | 0.940197654 |
| ENSMUSG00000020857 | -1.071115453 | 0.833851312 |
| ENSMUSG00000020859 | 1.066233582 | 0.820430396 |
| ENSMUSG00000020863 | 1.004445151 | 1 |
| ENSMUSG00000020864 | 1.034589169 | 0.925214847 |
| ENSMUSG00000020865 | -1.01869336 | 0.853859685 |
| ENSMUSG00000020866 | 1.369299353 | 0.013831763 |
| ENSMUSG00000020867 | 1.34694251 | 0.063399406 |
| ENSMUSG00000020868 | -1.048463846 | 0.698112512 |
| ENSMUSG00000020869 | 1.061039971 | 0.848601737 |
| ENSMUSG00000020870 | 1.019245206 | 1 |
| ENSMUSG00000020871 | -1.572511037 | 0.012403517 |
| ENSMUSG00000020872 | -1.146775327 | 0.564325372 |
| ENSMUSG00000020873 | -1.01277912 | 1 |
| ENSMUSG00000020875 | 1.045517023 | 0.891597525 |
| ENSMUSG00000020876 | 1.06489959 | 0.608497229 |
| ENSMUSG00000020877 | -1.015757879 | 0.951360958 |
| ENSMUSG00000020878 | -1.077445042 | 0.680599207 |
| ENSMUSG00000020882 | 1.006425043 | 1 |
| ENSMUSG00000020883 | 1.082219307 | 0.683415331 |
| ENSMUSG00000020884 | 1.060273769 | 0.814384306 |
| ENSMUSG00000020886 | 1.063429062 | 0.572947856 |
| ENSMUSG00000020887 | -1.090950423 | 0.770429489 |
| ENSMUSG00000020888 | 1.223899899 | 0.027986873 |
| ENSMUSG00000020889 | 1.330923161 | 0.010415162 |
| ENSMUSG00000020890 | -1.327548748 | 0.002998607 |
| ENSMUSG00000020891 | -1.160697821 | 0.554866155 |
| ENSMUSG00000020892 | -1.17645022 | 0.18851436 |
| ENSMUSG00000020893 | -1.079190124 | 0.52201333 |
| ENSMUSG00000020894 | -1.118914678 | 0.608497229 |
| ENSMUSG00000020895 | -1.029071247 | 0.698112512 |
| ENSMUSG00000020897 | -1.110617813 | 0.608497229 |
| ENSMUSG00000020898 | 1.017669331 | 0.977592554 |
| ENSMUSG00000020899 | -1.017189892 | 0.958403618 |
| ENSMUSG00000020900 | 1.142524721 | 0.620311605 |
| ENSMUSG00000020901 | 1.027300522 | 0.790851299 |
| ENSMUSG00000020902 | 1.088482436 | 0.608497229 |
| ENSMUSG00000020903 | -1.074484588 | 0.694267185 |
| ENSMUSG00000020904 | -1.595072085 | 0.003006398 |
| ENSMUSG00000020905 | 1.033981137 | 0.681952832 |
| ENSMUSG00000020907 | 1 | |
| ENSMUSG00000020908 | -1.044273782 | 0.698112512 |
| ENSMUSG00000020910 | -1.105831469 | 0.469713876 |
| ENSMUSG00000020911 | 1.00314539 | 1 |
| ENSMUSG00000020912 | 1 | |
| ENSMUSG00000020913 | 1 | |
| ENSMUSG00000020914 | -1.040058635 | 0.964542328 |
| ENSMUSG00000020916 | 1 | |
| ENSMUSG00000020917 | -1.031572583 | 0.961394002 |
| ENSMUSG00000020918 | -1.007281845 | 1 |
| ENSMUSG00000020919 | 1.107670033 | 0.467119761 |
| ENSMUSG00000020921 | -1.104595729 | 0.369149486 |
| ENSMUSG00000020922 | 1.004585711 | 1 |
| ENSMUSG00000020923 | 1.07793132 | 0.698909395 |
| ENSMUSG00000020925 | -1.081883713 | 0.710698346 |

| | | |
|---|---|---|
| ENSMUSG00000020926 | 1.114658125 | 0.72073896 |
| ENSMUSG00000020928 | 1.018670051 | 0.996913409 |
| ENSMUSG00000020929 | 1.003157915 | 1 |
| ENSMUSG00000020930 | 1.139251684 | 0.385303452 |
| ENSMUSG00000020932 | -1.049404962 | 0.949812939 |
| ENSMUSG00000020935 | 1.022119118 | 0.916178207 |
| ENSMUSG00000020936 | -1.01121792 | 1 |
| ENSMUSG00000020937 | 1.098290076 | 0.498086706 |
| ENSMUSG00000020940 | 1.09600655 | 0.628392912 |
| ENSMUSG00000020941 | 1.038189474 | 0.783619477 |
| ENSMUSG00000020945 | 1 | |
| ENSMUSG00000020946 | -1.019117316 | 0.991021717 |
| ENSMUSG00000020948 | 1.036319849 | 0.734674233 |
| ENSMUSG00000020949 | 1.08139513 | 0.624722543 |
| ENSMUSG00000020950 | 1.246703517 | 0.147531835 |
| ENSMUSG00000020952 | 1.03733166 | 0.915376822 |
| ENSMUSG00000020953 | -1.371964802 | 0.034323846 |
| ENSMUSG00000020954 | 1.088219939 | 0.717035356 |
| ENSMUSG00000020955 | 1.178044568 | 0.078328848 |
| ENSMUSG00000020956 | 1.045013254 | 0.736292585 |
| ENSMUSG00000020961 | -1.335474711 | 0.010415162 |
| ENSMUSG00000020962 | 1.195409836 | 0.146899474 |
| ENSMUSG00000020963 | 1.234951516 | 0.177605433 |
| ENSMUSG00000020964 | 1.058398389 | 0.834386993 |
| ENSMUSG00000020973 | 1.037407901 | 0.797105186 |
| ENSMUSG00000020974 | 1.026197141 | 0.950378392 |
| ENSMUSG00000020978 | 1.098285247 | 0.686477484 |
| ENSMUSG00000020982 | 1.066764693 | 0.765233309 |
| ENSMUSG00000020986 | 1.115864368 | 0.660622868 |
| ENSMUSG00000020988 | -1.01414324 | 0.985869746 |
| ENSMUSG00000020990 | 1.446100479 | 0.024867053 |
| ENSMUSG00000020992 | -1.009091449 | 1 |
| ENSMUSG00000020993 | 1.090275006 | 0.608497229 |
| ENSMUSG00000020994 | 1.080064024 | 0.741546126 |
| ENSMUSG00000021000 | 1.107516728 | 0.619226796 |
| ENSMUSG00000021003 | 1.037749767 | 0.87643689 |
| ENSMUSG00000021007 | -1.008576908 | 1 |
| ENSMUSG00000021009 | 1.076887251 | 0.698112512 |
| ENSMUSG00000021010 | 1.469969796 | 0.089742081 |
| ENSMUSG00000021012 | 1.043416133 | 0.89915331 |
| ENSMUSG00000021013 | 1.096850243 | 0.594181956 |
| ENSMUSG00000021018 | 1.046221929 | 0.782883548 |
| ENSMUSG00000021022 | 1.038102077 | 0.835701362 |
| ENSMUSG00000021023 | 1.06489221 | 0.687444744 |
| ENSMUSG00000021024 | 1.055732881 | 0.874206421 |
| ENSMUSG00000021025 | 1.083802679 | 0.608497229 |
| ENSMUSG00000021027 | 1.044816559 | 0.874662201 |
| ENSMUSG00000021028 | 1.064302085 | 0.698112512 |
| ENSMUSG00000021032 | -1.06136117 | 0.608497229 |
| ENSMUSG00000021033 | 1.050116127 | 0.804767157 |
| ENSMUSG00000021036 | 1.075803102 | 0.750584613 |
| ENSMUSG00000021037 | 1.060389664 | 0.802790558 |
| ENSMUSG00000021038 | 1.055876734 | 0.768665349 |
| ENSMUSG00000021039 | 1.022217486 | 0.960666316 |
| ENSMUSG00000021040 | 1.056284453 | 0.700124141 |
| ENSMUSG00000021044 | 1.265234676 | 0.014774446 |
| ENSMUSG00000021047 | 1.353111107 | 0.021374944 |
| ENSMUSG00000021048 | 1.078599047 | 0.698112512 |
| ENSMUSG00000021051 | 1.103204708 | 0.614265194 |
| ENSMUSG00000021054 | -1.007663365 | 1 |
| ENSMUSG00000021055 | 1.117955055 | 0.616779951 |
| ENSMUSG00000021056 | 1 | |
| ENSMUSG00000021057 | -1.698019761 | 0.000986973 |

| | | |
|---|---|---|
| ENSMUSG00000021061 | -1.085226118 | 0.476895694 |
| ENSMUSG00000021062 | -2.271855661 | 0.000387682 |
| ENSMUSG00000021065 | 1.275105973 | 0.054295624 |
| ENSMUSG00000021066 | 1.19472689 | 0.064831556 |
| ENSMUSG00000021067 | 1.103784971 | 0.629904463 |
| ENSMUSG00000021068 | 1.09332364 | 0.669808588 |
| ENSMUSG00000021069 | -1.801368051 | 0.019253091 |
| ENSMUSG00000021070 | -1.634716211 | 0.044140978 |
| ENSMUSG00000021071 | 1 | |
| ENSMUSG00000021072 | 1.081132392 | 0.754841426 |
| ENSMUSG00000021076 | 1.023799784 | 0.976562349 |
| ENSMUSG00000021078 | -1.03962336 | 0.803072249 |
| ENSMUSG00000021079 | 1.083227181 | 0.545124422 |
| ENSMUSG00000021081 | 1 | |
| ENSMUSG00000021086 | 1 | |
| ENSMUSG00000021087 | -1.230405735 | 0.240114463 |
| ENSMUSG00000021090 | -1.063525418 | 0.698961838 |
| ENSMUSG00000021091 | 1.29454102 | 0.208689436 |
| ENSMUSG00000021094 | 1.026488215 | 0.835701362 |
| ENSMUSG00000021095 | -1.063525418 | 0.698112512 |
| ENSMUSG00000021096 | 1.172698224 | 0.348316817 |
| ENSMUSG00000021097 | -1.467178194 | 0.003253473 |
| ENSMUSG00000021098 | -1.103445398 | 0.724003054 |
| ENSMUSG00000021099 | 1 | |
| ENSMUSG00000021101 | 1 | |
| ENSMUSG00000021102 | 1.042775116 | 0.61897445 |
| ENSMUSG00000021103 | 1.063768536 | 0.70299173 |
| ENSMUSG00000021108 | -1.113728968 | 0.608497229 |
| ENSMUSG00000021109 | 1.210443186 | 0.475208338 |
| ENSMUSG00000021111 | 1.059740678 | 0.862524901 |
| ENSMUSG00000021112 | 1.150805468 | 0.524392493 |
| ENSMUSG00000021113 | 1.129471197 | 0.428190934 |
| ENSMUSG00000021114 | 1.161261732 | 0.497425906 |
| ENSMUSG00000021115 | 1.027093464 | 0.936800157 |
| ENSMUSG00000021116 | 1.090088991 | 0.73130425 |
| ENSMUSG00000021118 | -1.258370585 | 0.01967661 |
| ENSMUSG00000021120 | -1.092022721 | 0.341402872 |
| ENSMUSG00000021123 | -1.201718201 | 0.477029641 |
| ENSMUSG00000021124 | 1.232455705 | 0.256911606 |
| ENSMUSG00000021125 | 1.073525476 | 0.608497229 |
| ENSMUSG00000021127 | 1.232847215 | 0.04416297 |
| ENSMUSG00000021130 | -1.212247213 | 0.155985055 |
| ENSMUSG00000021131 | 1.088354241 | 0.718265321 |
| ENSMUSG00000021133 | 1.19058554 | 0.264627385 |
| ENSMUSG00000021134 | 1.019343071 | 0.993466371 |
| ENSMUSG00000021135 | 1.009100337 | 1 |
| ENSMUSG00000021136 | 3.170098302 | 0.000416272 |
| ENSMUSG00000021139 | 1.144511219 | 0.597925656 |
| ENSMUSG00000021140 | 1.055498967 | 0.751346992 |
| ENSMUSG00000021143 | 1.163079504 | 0.216046507 |
| ENSMUSG00000021144 | 1.133499909 | 0.332304017 |
| ENSMUSG00000021147 | -1.026784323 | 0.942848464 |
| ENSMUSG00000021148 | 1 | |
| ENSMUSG00000021149 | -1.021882119 | 0.982039556 |
| ENSMUSG00000021156 | -1.024255775 | 0.970020309 |
| ENSMUSG00000021171 | 1.207548391 | 0.418367117 |
| ENSMUSG00000021175 | 1.126495366 | 0.260812761 |
| ENSMUSG00000021176 | 1.105269777 | 0.339943743 |
| ENSMUSG00000021177 | 1.071176414 | 0.608497229 |
| ENSMUSG00000021178 | 1.03471733 | 0.94497866 |
| ENSMUSG00000021179 | 1.057655857 | 0.608497229 |
| ENSMUSG00000021180 | 1.046432142 | 0.705231684 |
| ENSMUSG00000021182 | 1.312754793 | 0.028031065 |

| | | |
|---|---|---|
| ENSMUSG00000021185 | 1.991104207 | 0.00684332 |
| ENSMUSG00000021186 | 1.373509505 | 0.075890251 |
| ENSMUSG00000021187 | -1.044534224 | 0.929923764 |
| ENSMUSG00000021188 | 1.038434309 | 0.899484505 |
| ENSMUSG00000021189 | 1.136755085 | 0.413517796 |
| ENSMUSG00000021190 | 1.166351371 | 0.404380039 |
| ENSMUSG00000021192 | 1.087923993 | 0.608497229 |
| ENSMUSG00000021193 | 1.070701904 | 0.817623473 |
| ENSMUSG00000021194 | -1.13161737 | 0.630379139 |
| ENSMUSG00000021196 | -1.098189582 | 0.608497229 |
| ENSMUSG00000021198 | -1.02564575 | 0.979358161 |
| ENSMUSG00000021200 | -1.023769022 | 0.934760809 |
| ENSMUSG00000021203 | 1.030073267 | 0.866289429 |
| ENSMUSG00000021207 | -1 | 1 |
| ENSMUSG00000021208 | -1.03962336 | 0.698112512 |
| ENSMUSG00000021209 | -1.753986387 | 0.035878863 |
| ENSMUSG00000021210 | 1 | |
| ENSMUSG00000021211 | -1 | 1 |
| ENSMUSG00000021213 | 1.14011644 | 0.608497229 |
| ENSMUSG00000021214 | 1 | |
| ENSMUSG00000021215 | 1.038944813 | 0.895227243 |
| ENSMUSG00000021216 | 1.122462048 | 0.601164141 |
| ENSMUSG00000021217 | 1.149950446 | 0.081652295 |
| ENSMUSG00000021218 | -1.03063323 | 0.979002724 |
| ENSMUSG00000021219 | 1.161036672 | 0.251049598 |
| ENSMUSG00000021221 | -1.132666855 | 0.18851436 |
| ENSMUSG00000021222 | -1.002366863 | 1 |
| ENSMUSG00000021223 | 1.015486245 | 0.985522133 |
| ENSMUSG00000021224 | 1.071258635 | 0.641404094 |
| ENSMUSG00000021226 | 1.17813667 | 0.024722503 |
| ENSMUSG00000021228 | 1 | |
| ENSMUSG00000021234 | 1.010667723 | 0.953922475 |
| ENSMUSG00000021235 | 1.092475116 | 0.589600948 |
| ENSMUSG00000021236 | -1.087819393 | 0.623140373 |
| ENSMUSG00000021238 | 1.018770477 | 0.949812939 |
| ENSMUSG00000021239 | -1.044273782 | 0.790851299 |
| ENSMUSG00000021240 | 1.201487118 | 0.06812145 |
| ENSMUSG00000021241 | 1.005916004 | 1 |
| ENSMUSG00000021242 | 1.220586757 | 0.371928606 |
| ENSMUSG00000021243 | 1.005578124 | 1 |
| ENSMUSG00000021244 | 1.059765712 | 0.814089245 |
| ENSMUSG00000021245 | -1.029229694 | 0.887044308 |
| ENSMUSG00000021248 | 1.064033123 | 0.825410965 |
| ENSMUSG00000021250 | 1.050876024 | 0.730297627 |
| ENSMUSG00000021252 | 1.060167166 | 0.717035356 |
| ENSMUSG00000021253 | 1.288003631 | 0.076392989 |
| ENSMUSG00000021254 | 1.069649353 | 0.566175333 |
| ENSMUSG00000021255 | -1.690403397 | 0.056497717 |
| ENSMUSG00000021256 | 1.038222522 | 0.901216307 |
| ENSMUSG00000021257 | -1.217315647 | 0.058032144 |
| ENSMUSG00000021258 | 1.05991303 | 0.70299173 |
| ENSMUSG00000021259 | 1.014192793 | 0.964236174 |
| ENSMUSG00000021260 | 1.117138436 | 0.275279115 |
| ENSMUSG00000021262 | 1.084361582 | 0.6035234 |
| ENSMUSG00000021263 | 1.206036109 | 0.093784839 |
| ENSMUSG00000021264 | 1.06725603 | 0.794948027 |
| ENSMUSG00000021265 | -1.067892025 | 0.608497229 |
| ENSMUSG00000021266 | 1.145591013 | 0.395872943 |
| ENSMUSG00000021268 | 3.090604934 | 0.000231334 |
| ENSMUSG00000021270 | 1.04360213 | 0.949007391 |
| ENSMUSG00000021271 | 1.081858288 | 0.375972387 |
| ENSMUSG00000021273 | -1.09022058 | 0.698112512 |
| ENSMUSG00000021275 | 1.061086328 | 0.700389541 |

| | | |
|---|---|---|
| ENSMUSG00000021276 | 1.04764467 | 0.825086771 |
| ENSMUSG00000021277 | 1.095164353 | 0.608497229 |
| ENSMUSG00000021278 | -1.089497311 | 0.614307028 |
| ENSMUSG00000021279 | 1.137865335 | 0.378445257 |
| ENSMUSG00000021280 | 1.380313412 | 0.122036074 |
| ENSMUSG00000021281 | 1.163787728 | 0.301441914 |
| ENSMUSG00000021282 | 1.070814794 | 0.847803402 |
| ENSMUSG00000021285 | 1.089986805 | 0.602166181 |
| ENSMUSG00000021286 | 1.169982639 | 0.05296589 |
| ENSMUSG00000021287 | 1.157460826 | 0.044245893 |
| ENSMUSG00000021288 | 1.053765484 | 0.779557148 |
| ENSMUSG00000021290 | 1.058438037 | 0.805025542 |
| ENSMUSG00000021294 | -1.457438995 | 0.124648722 |
| ENSMUSG00000021298 | 1.059463094 | 0.608497229 |
| ENSMUSG00000021301 | 1.874620787 | 0.000429559 |
| ENSMUSG00000021302 | -1.023945442 | 0.943121172 |
| ENSMUSG00000021303 | -1.323821193 | 0.27203762 |
| ENSMUSG00000021306 | 1.17031114 | 0.30668853 |
| ENSMUSG00000021311 | -1.071316796 | 0.672261937 |
| ENSMUSG00000021313 | 1.085624843 | 0.759442175 |
| ENSMUSG00000021314 | -1.665744257 | 0.010548076 |
| ENSMUSG00000021318 | -1.037329674 | 0.895711764 |
| ENSMUSG00000021319 | 1.460027715 | 0.023911565 |
| ENSMUSG00000021322 | 1.059463094 | 0.699340382 |
| ENSMUSG00000021326 | -1.018514068 | 0.982541766 |
| ENSMUSG00000021327 | 1.153036568 | 0.481568621 |
| ENSMUSG00000021335 | 1 | |
| ENSMUSG00000021336 | 1 | |
| ENSMUSG00000021337 | 1 | |
| ENSMUSG00000021338 | -1.073883117 | 0.632326652 |
| ENSMUSG00000021339 | -1.03235617 | 0.886618881 |
| ENSMUSG00000021340 | -1.140896477 | 0.18177052 |
| ENSMUSG00000021342 | 1.059463094 | 0.608497229 |
| ENSMUSG00000021345 | 1 | |
| ENSMUSG00000021346 | 1 | |
| ENSMUSG00000021347 | 1 | |
| ENSMUSG00000021348 | 1 | |
| ENSMUSG00000021356 | -1.05143605 | 0.803072249 |
| ENSMUSG00000021357 | -1.023538847 | 0.958299667 |
| ENSMUSG00000021359 | 1.004013864 | 1 |
| ENSMUSG00000021360 | 1.382375645 | 0.004756452 |
| ENSMUSG00000021361 | -1.05061596 | 0.802991244 |
| ENSMUSG00000021362 | 1 | |
| ENSMUSG00000021363 | -1.330010175 | 0.010165101 |
| ENSMUSG00000021364 | -1.42835792 | 0.063557551 |
| ENSMUSG00000021365 | 1.021740087 | 0.962883651 |
| ENSMUSG00000021366 | 1.152799705 | 0.145583863 |
| ENSMUSG00000021367 | 1.416655221 | 0.009685294 |
| ENSMUSG00000021368 | -1.085632312 | 0.613693314 |
| ENSMUSG00000021371 | -1.119576619 | 0.580694717 |
| ENSMUSG00000021373 | 1.266423439 | 0.006411 |
| ENSMUSG00000021374 | -1.014481811 | 1 |
| ENSMUSG00000021375 | 1.010336449 | 1 |
| ENSMUSG00000021376 | -1.015529456 | 0.97004457 |
| ENSMUSG00000021377 | -1.076611807 | 0.818891358 |
| ENSMUSG00000021379 | 1.075540045 | 0.710698346 |
| ENSMUSG00000021381 | -1.345791402 | 0.005279442 |
| ENSMUSG00000021384 | -1.228574869 | 0.573272886 |
| ENSMUSG00000021385 | -1.035285881 | 0.861653133 |
| ENSMUSG00000021388 | 1.272358639 | 0.159466751 |
| ENSMUSG00000021390 | 1.517705945 | 0.012153229 |
| ENSMUSG00000021391 | -1.054765606 | 0.779972988 |
| ENSMUSG00000021392 | -1.019815466 | 0.975957032 |

| | | |
|---|---|---|
| ENSMUSG00000021395 | 1.033866407 | 0.936103707 |
| ENSMUSG00000021396 | 1 | |
| ENSMUSG00000021400 | -1.016415697 | 0.982838256 |
| ENSMUSG00000021403 | 1.262336976 | 0.041452314 |
| ENSMUSG00000021404 | 1.529752851 | 0.022525722 |
| ENSMUSG00000021408 | -1.061288441 | 0.784686694 |
| ENSMUSG00000021411 | -1.007961725 | 0.986126951 |
| ENSMUSG00000021413 | -1.000717307 | 1 |
| ENSMUSG00000021414 | -1.085651418 | 0.692881216 |
| ENSMUSG00000021415 | 1.059463094 | 0.608497229 |
| ENSMUSG00000021416 | -1.097906049 | 0.686194514 |
| ENSMUSG00000021417 | -1.065584002 | 0.747479869 |
| ENSMUSG00000021418 | 1.024878652 | 0.846518693 |
| ENSMUSG00000021420 | -1.046050828 | 0.755673717 |
| ENSMUSG00000021423 | -1.044273782 | 0.698112512 |
| ENSMUSG00000021427 | -1.045254021 | 0.933031586 |
| ENSMUSG00000021428 | -1.066045943 | 0.76052861 |
| ENSMUSG00000021431 | -1.064299064 | 0.608497229 |
| ENSMUSG00000021432 | -1.084513689 | 0.608497229 |
| ENSMUSG00000021439 | 1.132639128 | 0.608497229 |
| ENSMUSG00000021440 | 1 | |
| ENSMUSG00000021441 | 1 | |
| ENSMUSG00000021448 | -1.213118765 | 0.243993149 |
| ENSMUSG00000021451 | -1.260403857 | 0.017849136 |
| ENSMUSG00000021453 | 1.145994843 | 0.518394456 |
| ENSMUSG00000021456 | 1 | |
| ENSMUSG00000021457 | 1.160206296 | 0.334859726 |
| ENSMUSG00000021458 | -1.078431952 | 0.686194514 |
| ENSMUSG00000021460 | -1.033219573 | 0.835701362 |
| ENSMUSG00000021461 | -1.069538207 | 0.698112512 |
| ENSMUSG00000021464 | -1.316053574 | 0.014865686 |
| ENSMUSG00000021466 | 1.061007378 | 0.698112512 |
| ENSMUSG00000021468 | -1.06015611 | 0.792578392 |
| ENSMUSG00000021469 | 1.886830575 | 0.000367459 |
| ENSMUSG00000021470 | -1.117611542 | 0.608497229 |
| ENSMUSG00000021474 | -1.144949166 | 0.626820546 |
| ENSMUSG00000021476 | -1.191478929 | 0.233412931 |
| ENSMUSG00000021477 | 1.066807177 | 0.839924047 |
| ENSMUSG00000021478 | -1.181718351 | 0.305952299 |
| ENSMUSG00000021481 | -1.063592644 | 0.715156234 |
| ENSMUSG00000021482 | -1.021618966 | 0.850403297 |
| ENSMUSG00000021483 | 1.075317244 | 0.378431826 |
| ENSMUSG00000021484 | -1.122404906 | 0.608497229 |
| ENSMUSG00000021485 | 1.073570331 | 0.608497229 |
| ENSMUSG00000021486 | -1.054267818 | 0.818987862 |
| ENSMUSG00000021488 | -1.066490492 | 0.833884989 |
| ENSMUSG00000021490 | 1 | |
| ENSMUSG00000021492 | 1 | |
| ENSMUSG00000021493 | 1.284720106 | 0.03278599 |
| ENSMUSG00000021494 | -1.05903813 | 0.722170219 |
| ENSMUSG00000021495 | -1.014953972 | 0.955452914 |
| ENSMUSG00000021496 | -1.041814734 | 0.730618472 |
| ENSMUSG00000021497 | -1.147837781 | 0.464009618 |
| ENSMUSG00000021499 | -1.117536118 | 0.422735733 |
| ENSMUSG00000021500 | -1.063924379 | 0.825517832 |
| ENSMUSG00000021501 | -1.148568955 | 0.13028877 |
| ENSMUSG00000021504 | -1.047138321 | 0.651518688 |
| ENSMUSG00000021506 | -1 | 1 |
| ENSMUSG00000021508 | -1.222972786 | 0.259344963 |
| ENSMUSG00000021509 | -1.859654284 | 0.002994134 |
| ENSMUSG00000021510 | -1.063119323 | 0.66920741 |
| ENSMUSG00000021514 | -1.039614323 | 0.792761139 |
| ENSMUSG00000021518 | -1.107805768 | 0.698112512 |

| | | |
|---|---|---|
| ENSMUSG00000021519 | -1.041260921 | 0.847985364 |
| ENSMUSG00000021520 | -1.161491801 | 0.457225492 |
| ENSMUSG00000021532 | -1.00318469 | 1 |
| ENSMUSG00000021534 | -1.258580471 | 0.261338893 |
| ENSMUSG00000021536 | 1 | |
| ENSMUSG00000021537 | -1.060229142 | 0.819052063 |
| ENSMUSG00000021538 | 1 | |
| ENSMUSG00000021539 | 1 | 1 |
| ENSMUSG00000021540 | -1.027655391 | 0.969560962 |
| ENSMUSG00000021541 | 1 | |
| ENSMUSG00000021545 | 1 | |
| ENSMUSG00000021546 | 1.015180233 | 1 |
| ENSMUSG00000021548 | 1.001487451 | 1 |
| ENSMUSG00000021549 | 1.122183886 | 0.638545116 |
| ENSMUSG00000021550 | -2.516332698 | 0.000296212 |
| ENSMUSG00000021552 | -1.190471317 | 0.326250872 |
| ENSMUSG00000021553 | 1.050330666 | 0.814132742 |
| ENSMUSG00000021555 | -1.058554279 | 0.834839709 |
| ENSMUSG00000021556 | -1.227811749 | 0.375579698 |
| ENSMUSG00000021557 | -1.079541995 | 0.712130748 |
| ENSMUSG00000021559 | -1.591953545 | 0.001583273 |
| ENSMUSG00000021565 | -1.179226843 | 0.297578032 |
| ENSMUSG00000021566 | 1 | |
| ENSMUSG00000021567 | -1.460432292 | 0.003226816 |
| ENSMUSG00000021569 | -1.088350363 | 0.705231684 |
| ENSMUSG00000021572 | -1.15894568 | 0.140192832 |
| ENSMUSG00000021573 | -1.074998225 | 0.698112512 |
| ENSMUSG00000021575 | -1.868881769 | 0.001384356 |
| ENSMUSG00000021576 | -1.11027883 | 0.623156021 |
| ENSMUSG00000021577 | -1.086918659 | 0.785077984 |
| ENSMUSG00000021578 | -1.052465011 | 0.841492919 |
| ENSMUSG00000021579 | 1 | |
| ENSMUSG00000021583 | -1.288872888 | 0.212776118 |
| ENSMUSG00000021585 | -1.048024606 | 0.888895298 |
| ENSMUSG00000021587 | -1.127622795 | 0.581397416 |
| ENSMUSG00000021589 | 1.007461423 | 1 |
| ENSMUSG00000021590 | -1.160646446 | 0.608497229 |
| ENSMUSG00000021591 | 1.103605088 | 0.608497229 |
| ENSMUSG00000021592 | -1.056508536 | 0.751713062 |
| ENSMUSG00000021594 | -1.166791496 | 0.0681917 |
| ENSMUSG00000021595 | -1.023406021 | 0.98122995 |
| ENSMUSG00000021596 | 1.227369395 | 0.322554204 |
| ENSMUSG00000021597 | -1.10666188 | 0.608497229 |
| ENSMUSG00000021598 | 1 | |
| ENSMUSG00000021600 | 1 | |
| ENSMUSG00000021604 | 1 | |
| ENSMUSG00000021606 | -1.055186017 | 0.698112512 |
| ENSMUSG00000021607 | -1.098990931 | 0.434676351 |
| ENSMUSG00000021608 | -1.133335524 | 0.561097031 |
| ENSMUSG00000021609 | -1.339855477 | 0.052553784 |
| ENSMUSG00000021610 | -1.066503923 | 0.818306581 |
| ENSMUSG00000021611 | -1.751606782 | 0.008750044 |
| ENSMUSG00000021612 | 1.026062929 | 0.871020532 |
| ENSMUSG00000021613 | 1 | |
| ENSMUSG00000021614 | 1.270112535 | 0.195607801 |
| ENSMUSG00000021615 | -1.072147106 | 0.690952475 |
| ENSMUSG00000021619 | 1.037118462 | 0.765218754 |
| ENSMUSG00000021620 | -1.043306199 | 0.698112512 |
| ENSMUSG00000021621 | -1.043379918 | 0.793885968 |
| ENSMUSG00000021622 | 1 | |
| ENSMUSG00000021624 | -1.010960635 | 1 |
| ENSMUSG00000021629 | -1.063809431 | 0.79291651 |
| ENSMUSG00000021631 | 1 | |

| | | |
|---|---|---|
| ENSMUSG00000021635 | -1.005945227 | 1 |
| ENSMUSG00000021636 | 1.095872691 | 0.608497229 |
| ENSMUSG00000021638 | -1.628830302 | 0.004430069 |
| ENSMUSG00000021639 | -1.154854986 | 0.449109628 |
| ENSMUSG00000021640 | 1 | |
| ENSMUSG00000021643 | 1.022441056 | 0.889575988 |
| ENSMUSG00000021645 | -1.110177399 | 0.534861373 |
| ENSMUSG00000021646 | -1.123870125 | 0.313099907 |
| ENSMUSG00000021647 | 1 | |
| ENSMUSG00000021650 | -1.078487006 | 0.671987971 |
| ENSMUSG00000021660 | -1.101207879 | 0.76326282 |
| ENSMUSG00000021661 | -1.023528508 | 0.884182493 |
| ENSMUSG00000021662 | 1.039801782 | 0.839150023 |
| ENSMUSG00000021665 | 1.005451538 | 1 |
| ENSMUSG00000021666 | -1.0726816 | 0.698112512 |
| ENSMUSG00000021668 | 1.010941104 | 0.999068789 |
| ENSMUSG00000021669 | -1.084446529 | 0.754044495 |
| ENSMUSG00000021670 | -1.084714818 | 0.726628453 |
| ENSMUSG00000021671 | -1.10831656 | 0.534700732 |
| ENSMUSG00000021675 | -1.191452974 | 0.15169538 |
| ENSMUSG00000021676 | -1.774138748 | 0.000701466 |
| ENSMUSG00000021678 | -2.419324288 | 0.000371886 |
| ENSMUSG00000021679 | 1 | |
| ENSMUSG00000021680 | -1.517423219 | 0.037370544 |
| ENSMUSG00000021681 | -1.082168161 | 0.698112512 |
| ENSMUSG00000021684 | 1.17340989 | 0.115565352 |
| ENSMUSG00000021685 | 1.015486245 | 0.972980766 |
| ENSMUSG00000021686 | -1.016760589 | 0.992213082 |
| ENSMUSG00000021687 | -1.10400767 | 0.653471423 |
| ENSMUSG00000021690 | -1.070382028 | 0.717035356 |
| ENSMUSG00000021692 | -1.090913876 | 0.608497229 |
| ENSMUSG00000021693 | -1.047128767 | 0.886665438 |
| ENSMUSG00000021694 | -1.086926747 | 0.56051496 |
| ENSMUSG00000021696 | -1.133406866 | 0.308958388 |
| ENSMUSG00000021697 | 1.019899821 | 0.923395918 |
| ENSMUSG00000021699 | 1.492986051 | 0.000522056 |
| ENSMUSG00000021700 | 1.195276439 | 0.455529882 |
| ENSMUSG00000021701 | -1.132172296 | 0.673736989 |
| ENSMUSG00000021702 | -1.245586948 | 0.534652019 |
| ENSMUSG00000021703 | -1.216830729 | 0.268545131 |
| ENSMUSG00000021704 | -1.122988126 | 0.497309743 |
| ENSMUSG00000021706 | -1.047580318 | 0.857281639 |
| ENSMUSG00000021707 | -1.151243139 | 0.608497229 |
| ENSMUSG00000021708 | 1.109371301 | 0.744141915 |
| ENSMUSG00000021709 | 1.053666184 | 0.894591069 |
| ENSMUSG00000021710 | -1.013095418 | 1 |
| ENSMUSG00000021711 | -1.078573802 | 0.74773997 |
| ENSMUSG00000021712 | -1.073950879 | 0.713705885 |
| ENSMUSG00000021713 | -1.1000226 | 0.647538556 |
| ENSMUSG00000021714 | -1.173094723 | 0.286114407 |
| ENSMUSG00000021715 | -1.00913923 | 1 |
| ENSMUSG00000021716 | -1.052261456 | 0.754795782 |
| ENSMUSG00000021718 | 1 | |
| ENSMUSG00000021719 | -1.053549062 | 0.769745556 |
| ENSMUSG00000021720 | -1.067063048 | 0.828564184 |
| ENSMUSG00000021721 | 1 | |
| ENSMUSG00000021725 | 1.021790363 | 0.956972448 |
| ENSMUSG00000021728 | 1.081855706 | 0.698112512 |
| ENSMUSG00000021730 | 1.045395528 | 0.930788965 |
| ENSMUSG00000021731 | -1.073914563 | 0.70299173 |
| ENSMUSG00000021732 | 1.145695167 | 0.032054822 |
| ENSMUSG00000021733 | 1.014002334 | 1 |
| ENSMUSG00000021737 | -1.189738349 | 0.437701473 |

| | | |
|---|---|---|
| ENSMUSG00000021738 | -1.094666253 | 0.608497229 |
| ENSMUSG00000021741 | 1.019083579 | 0.949812939 |
| ENSMUSG00000021743 | 1 | |
| ENSMUSG00000021745 | -1.030640959 | 0.85864022 |
| ENSMUSG00000021747 | 1.013972085 | 0.956148535 |
| ENSMUSG00000021748 | -1.185848237 | 0.396083088 |
| ENSMUSG00000021749 | -1 | 1 |
| ENSMUSG00000021750 | -1.286250659 | 0.153133942 |
| ENSMUSG00000021751 | -1.053173184 | 0.87989052 |
| ENSMUSG00000021752 | -1.193893129 | 0.147046443 |
| ENSMUSG00000021754 | -1.177171617 | 0.085389756 |
| ENSMUSG00000021756 | -1.200672344 | 0.56853419 |
| ENSMUSG00000021758 | -1.049272231 | 0.854777354 |
| ENSMUSG00000021759 | -1.030415877 | 0.899132938 |
| ENSMUSG00000021760 | -1.098036982 | 0.684251204 |
| ENSMUSG00000021763 | -2.168731375 | 0.00254139 |
| ENSMUSG00000021764 | -1.09531905 | 0.698112512 |
| ENSMUSG00000021765 | 1.30443671 | 0.005064467 |
| ENSMUSG00000021767 | -1.000150072 | 1 |
| ENSMUSG00000021768 | 1.093819733 | 0.729157734 |
| ENSMUSG00000021770 | -1.122750465 | 0.608497229 |
| ENSMUSG00000021771 | -1.049157662 | 0.90550818 |
| ENSMUSG00000021772 | -1.177170674 | 0.171144596 |
| ENSMUSG00000021773 | 1.148103262 | 0.698112512 |
| ENSMUSG00000021774 | -1.190104666 | 0.065183251 |
| ENSMUSG00000021775 | -1.025290837 | 0.958181659 |
| ENSMUSG00000021779 | 1.078198701 | 0.675657574 |
| ENSMUSG00000021782 | -1.121323967 | 0.589600824 |
| ENSMUSG00000021785 | -1.169691315 | 0.40358341 |
| ENSMUSG00000021786 | -1.186250849 | 0.080928024 |
| ENSMUSG00000021789 | 1 | |
| ENSMUSG00000021790 | 1 | |
| ENSMUSG00000021791 | 1 | |
| ENSMUSG00000021792 | -1.028075365 | 0.84360497 |
| ENSMUSG00000021794 | -1.148189226 | 0.608497229 |
| ENSMUSG00000021795 | -1.043306199 | 0.698112512 |
| ENSMUSG00000021796 | -1.099542684 | 0.717035356 |
| ENSMUSG00000021797 | 1 | |
| ENSMUSG00000021798 | 1.557595953 | 0.004838378 |
| ENSMUSG00000021799 | 1 | |
| ENSMUSG00000021803 | 1.149620357 | 0.494986942 |
| ENSMUSG00000021804 | 1 | |
| ENSMUSG00000021806 | 1.041175366 | 0.814089245 |
| ENSMUSG00000021807 | -1.11648355 | 0.608497229 |
| ENSMUSG00000021809 | -1.215273344 | 0.010744336 |
| ENSMUSG00000021810 | -1.190797216 | 0.327640301 |
| ENSMUSG00000021811 | -1.21042637 | 0.364739674 |
| ENSMUSG00000021814 | -1.138796815 | 0.608497229 |
| ENSMUSG00000021815 | -1.253156043 | 0.113372157 |
| ENSMUSG00000021816 | -1.163774022 | 0.592544392 |
| ENSMUSG00000021819 | 1.009342808 | 1 |
| ENSMUSG00000021820 | 1.026230137 | 0.949812939 |
| ENSMUSG00000021822 | 1.276537948 | 0.121902592 |
| ENSMUSG00000021823 | 1.068290959 | 0.812796034 |
| ENSMUSG00000021824 | -1.135261488 | 0.596750715 |
| ENSMUSG00000021830 | -1.019913264 | 0.948149998 |
| ENSMUSG00000021831 | -1.1375462 | 0.612263934 |
| ENSMUSG00000021832 | -1.185907523 | 0.576129149 |
| ENSMUSG00000021835 | 1.065727839 | 0.473906907 |
| ENSMUSG00000021838 | -1.049418609 | 0.773362331 |
| ENSMUSG00000021840 | -1.083459212 | 0.772846594 |
| ENSMUSG00000021843 | -1.119266579 | 0.698112512 |
| ENSMUSG00000021846 | 1.176120837 | 0.013748391 |

| | | |
|---|---|---|
| ENSMUSG00000021848 | -1.044273782 | 0.698112512 |
| ENSMUSG00000021850 | 1 | |
| ENSMUSG00000021852 | 1 | |
| ENSMUSG00000021866 | -1.217230916 | 0.036394414 |
| ENSMUSG00000021867 | 1 | |
| ENSMUSG00000021868 | -1.146113475 | 0.209767231 |
| ENSMUSG00000021870 | -1.104108222 | 0.691285587 |
| ENSMUSG00000021871 | -1.069572565 | 0.700963448 |
| ENSMUSG00000021872 | 1 | |
| ENSMUSG00000021874 | -1.14619023 | 0.362989791 |
| ENSMUSG00000021876 | -1.12178515 | 0.43627839 |
| ENSMUSG00000021877 | -1.061131168 | 0.863200874 |
| ENSMUSG00000021879 | 1.121243997 | 0.656277822 |
| ENSMUSG00000021880 | 1 | |
| ENSMUSG00000021884 | -1.127911613 | 0.212650564 |
| ENSMUSG00000021886 | 1.030415139 | 0.949812939 |
| ENSMUSG00000021890 | -1.157467621 | 0.535214235 |
| ENSMUSG00000021891 | -1.120348999 | 0.532369657 |
| ENSMUSG00000021892 | -1.370124714 | 0.025443439 |
| ENSMUSG00000021893 | -1.241101876 | 0.271104421 |
| ENSMUSG00000021895 | -1.257351566 | 0.087823048 |
| ENSMUSG00000021898 | -1.248514591 | 0.580520513 |
| ENSMUSG00000021900 | -1.130625303 | 0.195148393 |
| ENSMUSG00000021901 | -1.126162334 | 0.570717076 |
| ENSMUSG00000021902 | -1.146730254 | 0.245273718 |
| ENSMUSG00000021903 | 1.023702974 | 0.871797708 |
| ENSMUSG00000021904 | -1.253648128 | 0.18810426 |
| ENSMUSG00000021905 | -1.129728654 | 0.428732467 |
| ENSMUSG00000021906 | -1.180284771 | 0.240669724 |
| ENSMUSG00000021907 | 1 | |
| ENSMUSG00000021908 | -1.089497311 | 0.608497229 |
| ENSMUSG00000021910 | -1.095241777 | 0.698112512 |
| ENSMUSG00000021911 | -1.13427059 | 0.581669442 |
| ENSMUSG00000021913 | 1.530074342 | 3.77621E-05 |
| ENSMUSG00000021916 | -1.067355225 | 0.70141105 |
| ENSMUSG00000021917 | -1.181870974 | 0.488763836 |
| ENSMUSG00000021918 | -1.150387769 | 0.317323566 |
| ENSMUSG00000021919 | -1.053549062 | 0.769745556 |
| ENSMUSG00000021922 | 1.024074931 | 0.998224129 |
| ENSMUSG00000021927 | -1.030976176 | 0.963734554 |
| ENSMUSG00000021928 | -1.218763838 | 0.104432727 |
| ENSMUSG00000021929 | -1.05852163 | 0.852845061 |
| ENSMUSG00000021930 | -1.148290133 | 0.405927158 |
| ENSMUSG00000021932 | -1.187380391 | 0.323036219 |
| ENSMUSG00000021933 | 1.059463094 | 0.608497229 |
| ENSMUSG00000021936 | -1.1405651 | 0.547016119 |
| ENSMUSG00000021938 | -1.205998831 | 0.36554144 |
| ENSMUSG00000021939 | 1.046846034 | 0.912425817 |
| ENSMUSG00000021940 | 1 | |
| ENSMUSG00000021943 | 1 | |
| ENSMUSG00000021944 | 1.005300174 | 0.999675223 |
| ENSMUSG00000021945 | -1.082014843 | 0.713864779 |
| ENSMUSG00000021947 | -1.195063523 | 0.034879046 |
| ENSMUSG00000021948 | -1.018357545 | 0.974807466 |
| ENSMUSG00000021950 | 1.310810065 | 0.040580159 |
| ENSMUSG00000021951 | -1.289804461 | 0.080934479 |
| ENSMUSG00000021952 | -1.169980976 | 0.517192194 |
| ENSMUSG00000021953 | 1 | |
| ENSMUSG00000021957 | -1.092815372 | 0.608497229 |
| ENSMUSG00000021958 | -1.15414089 | 0.244035142 |
| ENSMUSG00000021959 | -1.157762931 | 0.418326994 |
| ENSMUSG00000021961 | 1 | |
| ENSMUSG00000021962 | -1.13287689 | 0.477644658 |

| | | |
|---|---|---|
| ENSMUSG00000021963 | -1.183936457 | 0.387880421 |
| ENSMUSG00000021965 | -1.139425559 | 0.515271234 |
| ENSMUSG00000021966 | -1 | 1 |
| ENSMUSG00000021967 | -1.113319054 | 0.469213979 |
| ENSMUSG00000021969 | -1.187309268 | 0.48994642 |
| ENSMUSG00000021972 | -1.133907931 | 0.527262102 |
| ENSMUSG00000021973 | -1.153090167 | 0.603061081 |
| ENSMUSG00000021974 | 1.019083579 | 0.968937442 |
| ENSMUSG00000021975 | -1.161938919 | 0.397645313 |
| ENSMUSG00000021977 | 1 | |
| ENSMUSG00000021978 | -1.068105942 | 0.698112512 |
| ENSMUSG00000021981 | -1.163864689 | 0.503351299 |
| ENSMUSG00000021982 | -1.202906992 | 0.103413017 |
| ENSMUSG00000021983 | -1.375331672 | 0.098085953 |
| ENSMUSG00000021986 | -1.049075043 | 0.829371346 |
| ENSMUSG00000021987 | -1.13356994 | 0.556118084 |
| ENSMUSG00000021990 | 1.17807771 | 0.026120635 |
| ENSMUSG00000021991 | -1.041574599 | 0.886379277 |
| ENSMUSG00000021993 | -1.202160122 | 0.172195613 |
| ENSMUSG00000021994 | 1.388531289 | 0.003378766 |
| ENSMUSG00000021996 | -1.086648714 | 0.764491299 |
| ENSMUSG00000021997 | 1 | |
| ENSMUSG00000021998 | -1.124174706 | 0.410389737 |
| ENSMUSG00000021999 | 1.069068979 | 0.608497229 |
| ENSMUSG00000022000 | -1.113912732 | 0.608497229 |
| ENSMUSG00000022002 | 1 | |
| ENSMUSG00000022003 | -1.034813657 | 0.854777354 |
| ENSMUSG00000022008 | -1.185223096 | 0.358940839 |
| ENSMUSG00000022009 | -1.15469596 | 0.415175995 |
| ENSMUSG00000022010 | 1.104504754 | 0.685593205 |
| ENSMUSG00000022012 | -1.357118498 | 0.067346977 |
| ENSMUSG00000022013 | -1.036597996 | 0.807725855 |
| ENSMUSG00000022014 | -1.206338534 | 0.246763095 |
| ENSMUSG00000022015 | -1.30460298 | 0.15540061 |
| ENSMUSG00000022016 | -1.130267068 | 0.63154731 |
| ENSMUSG00000022018 | -1.658354899 | 0.006815155 |
| ENSMUSG00000022019 | -1.05219756 | 0.728908992 |
| ENSMUSG00000022020 | -1.172556655 | 0.234218025 |
| ENSMUSG00000022021 | -1.153655739 | 0.608497229 |
| ENSMUSG00000022022 | -1.483929074 | 0.025806845 |
| ENSMUSG00000022023 | -1.070351599 | 0.698112512 |
| ENSMUSG00000022024 | -1.145473937 | 0.608497229 |
| ENSMUSG00000022025 | 1 | |
| ENSMUSG00000022026 | 1 | |
| ENSMUSG00000022031 | -1.090776867 | 0.698112512 |
| ENSMUSG00000022032 | -1.397660083 | 0.032980279 |
| ENSMUSG00000022033 | -1.127242739 | 0.608497229 |
| ENSMUSG00000022034 | -1.253095643 | 0.24276078 |
| ENSMUSG00000022035 | -1.174415787 | 0.302032096 |
| ENSMUSG00000022037 | -1.072952317 | 0.608497229 |
| ENSMUSG00000022039 | -1.071075483 | 0.698112512 |
| ENSMUSG00000022040 | -2.683433922 | 0.000411964 |
| ENSMUSG00000022041 | -1.089497311 | 0.532369657 |
| ENSMUSG00000022043 | -1.053849777 | 0.853296211 |
| ENSMUSG00000022044 | -1.29784857 | 0.225946641 |
| ENSMUSG00000022048 | 1.070313777 | 0.698112512 |
| ENSMUSG00000022051 | -1.168307793 | 0.503351299 |
| ENSMUSG00000022052 | -1.123552221 | 0.636056232 |
| ENSMUSG00000022053 | -1.893100627 | 0.000519315 |
| ENSMUSG00000022054 | -1.176686249 | 0.362226215 |
| ENSMUSG00000022055 | -1.089497311 | 0.608497229 |
| ENSMUSG00000022056 | 1 | |
| ENSMUSG00000022057 | 1 | |

| | | |
|---|---|---|
| ENSMUSG00000022061 | -1.133717313 | 0.608497229 |
| ENSMUSG00000022064 | -1.108930648 | 0.451541924 |
| ENSMUSG00000022066 | 1 | |
| ENSMUSG00000022070 | -1.172509142 | 0.251049598 |
| ENSMUSG00000022074 | -1.11455147 | 0.60397857 |
| ENSMUSG00000022075 | -1.085073262 | 0.608497229 |
| ENSMUSG00000022076 | 1.059463094 | 0.608497229 |
| ENSMUSG00000022085 | 1.005613439 | 1 |
| ENSMUSG00000022089 | -1.171591481 | 0.226178334 |
| ENSMUSG00000022090 | -1.035639849 | 0.78287223 |
| ENSMUSG00000022091 | -1.124941733 | 0.472052821 |
| ENSMUSG00000022092 | -1.152201602 | 0.449109628 |
| ENSMUSG00000022094 | -1.092714253 | 0.641264745 |
| ENSMUSG00000022095 | -1.079293201 | 0.441887419 |
| ENSMUSG00000022096 | -1.695599257 | 0.001460821 |
| ENSMUSG00000022097 | -1 | 1 |
| ENSMUSG00000022098 | -1.025231513 | 0.947248675 |
| ENSMUSG00000022099 | -1.117545322 | 0.480690111 |
| ENSMUSG00000022100 | -1.130613522 | 0.647290337 |
| ENSMUSG00000022101 | -1.043306199 | 0.698112512 |
| ENSMUSG00000022102 | -1.005511336 | 1 |
| ENSMUSG00000022103 | -1.151396308 | 0.172075112 |
| ENSMUSG00000022105 | -1.039945412 | 0.853941024 |
| ENSMUSG00000022106 | -1.013039118 | 0.988043098 |
| ENSMUSG00000022108 | -1.047074188 | 0.899219824 |
| ENSMUSG00000022109 | -1.137174313 | 0.524534614 |
| ENSMUSG00000022110 | -1.112744042 | 0.652467077 |
| ENSMUSG00000022111 | -1.164512363 | 0.377229 |
| ENSMUSG00000022112 | 1.019083579 | 0.949812939 |
| ENSMUSG00000022113 | -1.123630124 | 0.698112512 |
| ENSMUSG00000022114 | -1.130680869 | 0.391570427 |
| ENSMUSG00000022116 | 1 | |
| ENSMUSG00000022119 | -1.099132528 | 0.698112512 |
| ENSMUSG00000022120 | -1.076411871 | 0.679448217 |
| ENSMUSG00000022122 | -1.043306199 | 0.803260194 |
| ENSMUSG00000022123 | -1.292076169 | 0.210508637 |
| ENSMUSG00000022124 | -1.144660954 | 0.592572342 |
| ENSMUSG00000022125 | -1.039062334 | 0.864480156 |
| ENSMUSG00000022126 | -1 | 1 |
| ENSMUSG00000022129 | 1 | |
| ENSMUSG00000022130 | -1.068513811 | 0.67362935 |
| ENSMUSG00000022131 | -1.054314605 | 0.698112512 |
| ENSMUSG00000022132 | -1.159782692 | 0.631106957 |
| ENSMUSG00000022136 | -1.210234693 | 0.428314043 |
| ENSMUSG00000022139 | -1.032539298 | 0.947600344 |
| ENSMUSG00000022141 | 1.03643316 | 0.945694084 |
| ENSMUSG00000022142 | 1.026378697 | 0.979978576 |
| ENSMUSG00000022144 | -1.082773488 | 0.698112512 |
| ENSMUSG00000022146 | 1.065262211 | 0.729591121 |
| ENSMUSG00000022148 | -1.143828558 | 0.698112512 |
| ENSMUSG00000022149 | -1 | 1 |
| ENSMUSG00000022150 | -1.224744138 | 0.417843828 |
| ENSMUSG00000022151 | 1.019073172 | 0.965872389 |
| ENSMUSG00000022155 | 1 | |
| ENSMUSG00000022156 | -1.03962336 | 0.803072249 |
| ENSMUSG00000022157 | 1.142033317 | 0.564682496 |
| ENSMUSG00000022159 | -1.18672389 | 0.160676302 |
| ENSMUSG00000022160 | -1.105267276 | 0.543453165 |
| ENSMUSG00000022174 | -1.132640033 | 0.532586 |
| ENSMUSG00000022175 | -1.063174917 | 0.702897175 |
| ENSMUSG00000022176 | 1.469927553 | 0.010969662 |
| ENSMUSG00000022177 | -1.04854681 | 0.702897175 |
| ENSMUSG00000022178 | -1.190764414 | 0.374471212 |

| | | |
|---|---|---|
| ENSMUSG00000022179 | -1.017683206 | 0.869548547 |
| ENSMUSG00000022180 | -2.167468177 | 0.000315676 |
| ENSMUSG00000022181 | -1.071075483 | 0.743300655 |
| ENSMUSG00000022184 | 1.042159469 | 0.814608539 |
| ENSMUSG00000022185 | -1.021496909 | 0.973756648 |
| ENSMUSG00000022186 | 1.12710213 | 0.698112512 |
| ENSMUSG00000022187 | -1.137305458 | 0.702425678 |
| ENSMUSG00000022191 | 1.060460376 | 0.830120378 |
| ENSMUSG00000022193 | -1.150745846 | 0.307003105 |
| ENSMUSG00000022194 | -1.06128186 | 0.608497229 |
| ENSMUSG00000022195 | 1.057917514 | 0.768188119 |
| ENSMUSG00000022197 | -1.354653885 | 0.04182893 |
| ENSMUSG00000022199 | -1.56675666 | 0.003666852 |
| ENSMUSG00000022200 | 1.000612733 | 1 |
| ENSMUSG00000022201 | 1.051987097 | 0.879917523 |
| ENSMUSG00000022203 | 1.380887155 | 0.122922169 |
| ENSMUSG00000022204 | -1.184531489 | 0.331568086 |
| ENSMUSG00000022205 | -1.083640248 | 0.745135758 |
| ENSMUSG00000022206 | 1.322540593 | 0.110552384 |
| ENSMUSG00000022208 | -1.630866819 | 0.024388617 |
| ENSMUSG00000022209 | 1 | |
| ENSMUSG00000022210 | -1.21447191 | 0.009277859 |
| ENSMUSG00000022211 | -1.288425638 | 0.549713136 |
| ENSMUSG00000022212 | 1 | |
| ENSMUSG00000022214 | -1.225684453 | 0.115727957 |
| ENSMUSG00000022215 | 1 | |
| ENSMUSG00000022216 | -1.346360729 | 0.02013427 |
| ENSMUSG00000022217 | -1.076642378 | 0.43649425 |
| ENSMUSG00000022218 | -1.190389413 | 0.435634716 |
| ENSMUSG00000022219 | -1.194507178 | 0.386955663 |
| ENSMUSG00000022220 | -1.275940027 | 0.210047602 |
| ENSMUSG00000022221 | -1.394605458 | 0.043367921 |
| ENSMUSG00000022223 | -1.098219594 | 0.173059112 |
| ENSMUSG00000022225 | 1 | |
| ENSMUSG00000022226 | 1 | |
| ENSMUSG00000022227 | 1 | |
| ENSMUSG00000022228 | -1.019251765 | 0.965532475 |
| ENSMUSG00000022229 | 1 | |
| ENSMUSG00000022231 | -1.251723638 | 0.111366442 |
| ENSMUSG00000022234 | 1.034668737 | 0.970833538 |
| ENSMUSG00000022235 | -1.136192609 | 0.568864017 |
| ENSMUSG00000022236 | 1.207714447 | 0.129473223 |
| ENSMUSG00000022237 | -1.460317537 | 0.025332826 |
| ENSMUSG00000022240 | 1.639778369 | 0.014851689 |
| ENSMUSG00000022241 | 1.084839071 | 0.734674233 |
| ENSMUSG00000022243 | 1 | |
| ENSMUSG00000022244 | 1.131938661 | 0.091124716 |
| ENSMUSG00000022245 | -1.085651418 | 0.532369657 |
| ENSMUSG00000022246 | 1.211908285 | 0.343789847 |
| ENSMUSG00000022247 | 1.033739213 | 0.941225684 |
| ENSMUSG00000022248 | 1.103178498 | 0.503351299 |
| ENSMUSG00000022249 | 1 | |
| ENSMUSG00000022253 | 1.017251017 | 0.981964814 |
| ENSMUSG00000022255 | 1.02690604 | 0.976928427 |
| ENSMUSG00000022257 | 1.090847175 | 0.646341738 |
| ENSMUSG00000022261 | 1.120017677 | 0.680567652 |
| ENSMUSG00000022262 | -1.032403253 | 0.955775173 |
| ENSMUSG00000022263 | 1.184188448 | 0.389456347 |
| ENSMUSG00000022265 | 1.484158618 | 0.016879104 |
| ENSMUSG00000022269 | 1 | |
| ENSMUSG00000022270 | -1.224770523 | 0.317618146 |
| ENSMUSG00000022272 | 1.099972588 | 0.656547319 |
| ENSMUSG00000022280 | 1.041627889 | 0.881850101 |

| | | |
|---|---|---|
| ENSMUSG00000022283 | 1.051488368 | 0.934760809 |
| ENSMUSG00000022285 | 1.023478736 | 0.996913409 |
| ENSMUSG00000022286 | 1.139434012 | 0.600772569 |
| ENSMUSG00000022288 | 1 | |
| ENSMUSG00000022292 | -1.037975615 | 0.897684198 |
| ENSMUSG00000022295 | 1.125389569 | 0.630015435 |
| ENSMUSG00000022296 | 1.020842118 | 0.819222912 |
| ENSMUSG00000022297 | -1.014851771 | 1 |
| ENSMUSG00000022299 | 1.075506201 | 0.698112512 |
| ENSMUSG00000022300 | 1.034748832 | 0.934760809 |
| ENSMUSG00000022303 | 1 | |
| ENSMUSG00000022304 | -1.019108526 | 0.979258908 |
| ENSMUSG00000022305 | 1.075326135 | 0.710698346 |
| ENSMUSG00000022306 | 1.004117171 | 1 |
| ENSMUSG00000022307 | 1.045561492 | 0.898763027 |
| ENSMUSG00000022309 | 1.338730339 | 0.024295713 |
| ENSMUSG00000022311 | -1.109582661 | 0.605986581 |
| ENSMUSG00000022312 | 1.025553722 | 0.982968316 |
| ENSMUSG00000022313 | 1.082784732 | 0.608497229 |
| ENSMUSG00000022314 | 1.03494458 | 0.969895686 |
| ENSMUSG00000022315 | 1.015486245 | 0.972980766 |
| ENSMUSG00000022321 | 1.691838596 | 0.002730903 |
| ENSMUSG00000022322 | 1.030005756 | 0.953731979 |
| ENSMUSG00000022323 | 1.053731605 | 0.725839797 |
| ENSMUSG00000022324 | 1.631935708 | 0.004326525 |
| ENSMUSG00000022325 | 1.010349088 | 1 |
| ENSMUSG00000022329 | -1.029555198 | 0.947189871 |
| ENSMUSG00000022330 | -1.374059416 | 0.113867265 |
| ENSMUSG00000022332 | 1.10679017 | 0.56072388 |
| ENSMUSG00000022335 | -1.039846639 | 0.686432851 |
| ENSMUSG00000022336 | 1.04266783 | 0.940222871 |
| ENSMUSG00000022337 | 1.026439212 | 0.95314795 |
| ENSMUSG00000022338 | 1.046440816 | 0.863449094 |
| ENSMUSG00000022339 | 1.00945314 | 1 |
| ENSMUSG00000022340 | -1.055043849 | 0.777977356 |
| ENSMUSG00000022342 | 1 | |
| ENSMUSG00000022346 | 1.032093582 | 0.809842777 |
| ENSMUSG00000022347 | 1 | |
| ENSMUSG00000022350 | 1.208968236 | 0.486216685 |
| ENSMUSG00000022351 | -1.021899156 | 0.981137599 |
| ENSMUSG00000022353 | -1.208786795 | 0.104898224 |
| ENSMUSG00000022354 | 1.003760753 | 1 |
| ENSMUSG00000022357 | -1 | 1 |
| ENSMUSG00000022358 | 1.136284232 | 0.274420574 |
| ENSMUSG00000022359 | 1.176517877 | 0.141293109 |
| ENSMUSG00000022360 | 1.061454878 | 0.877639675 |
| ENSMUSG00000022361 | -1.414485454 | 0.069329311 |
| ENSMUSG00000022362 | 1.132639128 | 0.505928387 |
| ENSMUSG00000022364 | 1.007765945 | 1 |
| ENSMUSG00000022365 | 1.089484009 | 0.698112512 |
| ENSMUSG00000022366 | 1 | |
| ENSMUSG00000022367 | 1.285256995 | 0.087671702 |
| ENSMUSG00000022369 | -1.035761596 | 0.924629556 |
| ENSMUSG00000022370 | -1.008297367 | 1 |
| ENSMUSG00000022371 | 1.293051269 | 0.075032805 |
| ENSMUSG00000022372 | 1.281317005 | 0.004868138 |
| ENSMUSG00000022375 | 1 | |
| ENSMUSG00000022376 | 1.200241468 | 0.350223743 |
| ENSMUSG00000022377 | 1.201599049 | 0.482252735 |
| ENSMUSG00000022378 | 1.06406811 | 0.809081946 |
| ENSMUSG00000022382 | -1.859935665 | 0.007857726 |
| ENSMUSG00000022383 | -1.266522641 | 0.151833616 |
| ENSMUSG00000022385 | 1.105718826 | 0.461619314 |

| | | |
|---|---|---|
| ENSMUSG00000022386 | -1.0563689 | 0.608497229 |
| ENSMUSG00000022387 | 1.125498325 | 0.474488116 |
| ENSMUSG00000022388 | 1 | |
| ENSMUSG00000022389 | 1.164994379 | 0.482433619 |
| ENSMUSG00000022390 | -1.040769735 | 0.883615907 |
| ENSMUSG00000022391 | 1.004123057 | 1 |
| ENSMUSG00000022394 | -1.012163497 | 0.997869436 |
| ENSMUSG00000022400 | 1.008248766 | 1 |
| ENSMUSG00000022401 | 1.014721884 | 0.988665808 |
| ENSMUSG00000022403 | 1.097564751 | 0.764370878 |
| ENSMUSG00000022404 | 1.007103357 | 1 |
| ENSMUSG00000022407 | 1.09181939 | 0.608497229 |
| ENSMUSG00000022408 | 1 | |
| ENSMUSG00000022412 | 1.05681891 | 0.770128273 |
| ENSMUSG00000022414 | 1.052561233 | 0.698112512 |
| ENSMUSG00000022415 | -1.233771924 | 0.067552132 |
| ENSMUSG00000022416 | -1.11080631 | 0.796433064 |
| ENSMUSG00000022419 | 1.235299793 | 0.065240532 |
| ENSMUSG00000022420 | 1.101764427 | 0.316200725 |
| ENSMUSG00000022421 | -1.169810833 | 0.4498175 |
| ENSMUSG00000022422 | -1.050814739 | 0.723401948 |
| ENSMUSG00000022425 | -1.37158507 | 0.014245646 |
| ENSMUSG00000022426 | -1.008073017 | 1 |
| ENSMUSG00000022427 | -1.01032852 | 1 |
| ENSMUSG00000022428 | -1.00322409 | 1 |
| ENSMUSG00000022429 | -1 | 1 |
| ENSMUSG00000022431 | -1.206338534 | 0.283365781 |
| ENSMUSG00000022432 | 1.113441446 | 0.763339717 |
| ENSMUSG00000022433 | 1.126064909 | 0.541094967 |
| ENSMUSG00000022434 | 1.12871929 | 0.608497229 |
| ENSMUSG00000022435 | 1 | |
| ENSMUSG00000022436 | -1.038726621 | 0.784772061 |
| ENSMUSG00000022437 | 1.051257722 | 0.801042518 |
| ENSMUSG00000022438 | 1.137437301 | 0.530301279 |
| ENSMUSG00000022439 | 1.20243393 | 0.289843376 |
| ENSMUSG00000022440 | -1.053781901 | 0.71045278 |
| ENSMUSG00000022441 | 1 | |
| ENSMUSG00000022442 | 1.030826681 | 0.871840478 |
| ENSMUSG00000022443 | 1.296487134 | 0.25531461 |
| ENSMUSG00000022445 | 1 | |
| ENSMUSG00000022449 | -1.575508232 | 0.000573666 |
| ENSMUSG00000022450 | 1.028077184 | 0.793836508 |
| ENSMUSG00000022451 | -1.011757962 | 1 |
| ENSMUSG00000022452 | 1.038419153 | 0.817101562 |
| ENSMUSG00000022453 | 1.081815333 | 0.476806427 |
| ENSMUSG00000022454 | -1.132996488 | 0.608497229 |
| ENSMUSG00000022455 | 1 | |
| ENSMUSG00000022456 | -1.156835561 | 0.119464073 |
| ENSMUSG00000022462 | 1.071342955 | 0.841812929 |
| ENSMUSG00000022463 | 1.009191848 | 1 |
| ENSMUSG00000022464 | -1.950443512 | 0.005370097 |
| ENSMUSG00000022466 | -1.042056362 | 0.828289041 |
| ENSMUSG00000022468 | 1.059463094 | 0.776953697 |
| ENSMUSG00000022469 | 1.074196149 | 0.681952832 |
| ENSMUSG00000022471 | -1.036496793 | 0.933031586 |
| ENSMUSG00000022472 | -1.077007838 | 0.766407898 |
| ENSMUSG00000022474 | 1.013108558 | 0.979212363 |
| ENSMUSG00000022475 | 1.115759462 | 0.5192873 |
| ENSMUSG00000022476 | -1.017807784 | 0.952049609 |
| ENSMUSG00000022477 | 1.033673626 | 0.945425517 |
| ENSMUSG00000022479 | -1.127193002 | 0.518302063 |
| ENSMUSG00000022483 | 1.753340362 | 0.002092992 |
| ENSMUSG00000022484 | -1.454556284 | 0.05698665 |

| | | |
|---|---|---|
| ENSMUSG00000022485 | -1.597326255 | 0.013148037 |
| ENSMUSG00000022487 | 1.415195754 | 0.098384366 |
| ENSMUSG00000022488 | 1.395943963 | 0.215991492 |
| ENSMUSG00000022489 | 1.324287474 | 0.02024466 |
| ENSMUSG00000022490 | 1.128312813 | 0.698112512 |
| ENSMUSG00000022491 | 1 | |
| ENSMUSG00000022494 | 1 | |
| ENSMUSG00000022496 | 1 | |
| ENSMUSG00000022498 | -1.023752851 | 0.920477428 |
| ENSMUSG00000022500 | 1.122420732 | 0.608497229 |
| ENSMUSG00000022501 | 1.161036672 | 0.251049598 |
| ENSMUSG00000022503 | 1.01808268 | 0.940975268 |
| ENSMUSG00000022504 | 1.091194986 | 0.710698346 |
| ENSMUSG00000022505 | 1.181228258 | 0.406675037 |
| ENSMUSG00000022507 | 1.107137006 | 0.608497229 |
| ENSMUSG00000022508 | 1.048928829 | 0.608497229 |
| ENSMUSG00000022510 | 1.393094558 | 0.001408702 |
| ENSMUSG00000022512 | -1.274634948 | 0.00367642 |
| ENSMUSG00000022514 | -1.047855229 | 0.713861573 |
| ENSMUSG00000022515 | 1.009562958 | 0.989648415 |
| ENSMUSG00000022516 | -1.025571678 | 0.864123949 |
| ENSMUSG00000022517 | 1.114201024 | 0.526180599 |
| ENSMUSG00000022518 | 1.239354098 | 0.430635163 |
| ENSMUSG00000022519 | 1.1022699 | 0.608497229 |
| ENSMUSG00000022521 | 1.038684657 | 0.809624788 |
| ENSMUSG00000022523 | -1.043306199 | 0.698112512 |
| ENSMUSG00000022525 | 1.069068979 | 0.608497229 |
| ENSMUSG00000022526 | 1.020919048 | 0.924804062 |
| ENSMUSG00000022528 | 1.333169497 | 0.002917302 |
| ENSMUSG00000022529 | -1.020924435 | 0.94814388 |
| ENSMUSG00000022533 | 1.141681018 | 0.644981762 |
| ENSMUSG00000022534 | -1.043306199 | 0.698112512 |
| ENSMUSG00000022536 | 1.009906137 | 1 |
| ENSMUSG00000022537 | 1.386244589 | 0.001198045 |
| ENSMUSG00000022538 | 1.009198822 | 1 |
| ENSMUSG00000022540 | -1.09696185 | 0.454968457 |
| ENSMUSG00000022542 | -1.044273782 | 0.698112512 |
| ENSMUSG00000022543 | -1.076999483 | 0.59469635 |
| ENSMUSG00000022544 | 1.009067857 | 0.986542466 |
| ENSMUSG00000022545 | -1.028349716 | 0.769745556 |
| ENSMUSG00000022546 | -1.011195466 | 0.977489164 |
| ENSMUSG00000022548 | -1.129231493 | 0.708329262 |
| ENSMUSG00000022550 | 1.022152283 | 0.933592873 |
| ENSMUSG00000022551 | -1.019977986 | 0.986542466 |
| ENSMUSG00000022552 | -1.023418748 | 0.910497937 |
| ENSMUSG00000022553 | 1.047567376 | 0.794786733 |
| ENSMUSG00000022554 | -1.001582011 | 1 |
| ENSMUSG00000022555 | -1.056088373 | 0.670305632 |
| ENSMUSG00000022556 | 1.029785627 | 0.884400885 |
| ENSMUSG00000022557 | 1.003483321 | 1 |
| ENSMUSG00000022558 | 1.263381585 | 0.059597632 |
| ENSMUSG00000022559 | 1.080849121 | 0.618607897 |
| ENSMUSG00000022560 | 1.100460011 | 0.505560817 |
| ENSMUSG00000022561 | 1.081981304 | 0.629688349 |
| ENSMUSG00000022562 | 1.174323051 | 0.065015108 |
| ENSMUSG00000022564 | 1.06371772 | 0.761233604 |
| ENSMUSG00000022565 | 1.165315052 | 0.382029378 |
| ENSMUSG00000022568 | 1.127068757 | 0.50840255 |
| ENSMUSG00000022570 | 1.017633801 | 0.953168721 |
| ENSMUSG00000022571 | -1.005588266 | 1 |
| ENSMUSG00000022574 | 1.119712002 | 0.308712585 |
| ENSMUSG00000022575 | -1.321074183 | 0.038158125 |
| ENSMUSG00000022577 | -1.170083925 | 0.206620396 |

| | | |
|---|---|---|
| ENSMUSG00000022579 | 1 | |
| ENSMUSG00000022580 | -1.280550493 | 0.131640936 |
| ENSMUSG00000022582 | -1 | 1 |
| ENSMUSG00000022583 | -1.041110033 | 0.934760809 |
| ENSMUSG00000022584 | 1.213944366 | 0.452349814 |
| ENSMUSG00000022586 | 1.24188384 | 0.248461741 |
| ENSMUSG00000022587 | 1.208531671 | 0.151784997 |
| ENSMUSG00000022589 | 1 | |
| ENSMUSG00000022591 | 1 | |
| ENSMUSG00000022594 | -1.141760603 | 0.251315265 |
| ENSMUSG00000022595 | 1.069437807 | 0.729206866 |
| ENSMUSG00000022596 | 1.059463094 | 0.608497229 |
| ENSMUSG00000022598 | -1.065852599 | 0.860712821 |
| ENSMUSG00000022601 | -1.061364514 | 0.743170465 |
| ENSMUSG00000022602 | 1.602673618 | 0.00088997 |
| ENSMUSG00000022603 | 1.059463094 | 0.608497229 |
| ENSMUSG00000022604 | 1.122940368 | 0.425025602 |
| ENSMUSG00000022607 | 1.071332122 | 0.801659949 |
| ENSMUSG00000022610 | 1.133838405 | 0.373031949 |
| ENSMUSG00000022613 | 1 | |
| ENSMUSG00000022614 | 1.14824745 | 0.246522583 |
| ENSMUSG00000022615 | 1.130670957 | 0.698112512 |
| ENSMUSG00000022617 | 1.034734916 | 0.834376699 |
| ENSMUSG00000022619 | -1.023769022 | 0.934760809 |
| ENSMUSG00000022620 | 1.145106109 | 0.095339876 |
| ENSMUSG00000022621 | -1.081206501 | 0.608497229 |
| ENSMUSG00000022622 | 1 | |
| ENSMUSG00000022623 | -1.65544107 | 0.019490879 |
| ENSMUSG00000022629 | 1.165636519 | 0.335377108 |
| ENSMUSG00000022634 | 1.069887041 | 0.698112512 |
| ENSMUSG00000022635 | 1.023062596 | 0.961127302 |
| ENSMUSG00000022636 | -1.070640143 | 0.698112512 |
| ENSMUSG00000022637 | 1.255477428 | 0.115685577 |
| ENSMUSG00000022639 | 1.099948845 | 0.331568086 |
| ENSMUSG00000022641 | -1.001268691 | 1 |
| ENSMUSG00000022650 | -1.043306199 | 0.698112512 |
| ENSMUSG00000022651 | 1 | |
| ENSMUSG00000022652 | 1 | |
| ENSMUSG00000022656 | 1.19374753 | 0.315720798 |
| ENSMUSG00000022657 | 1 | |
| ENSMUSG00000022658 | 1.290838689 | 0.144429723 |
| ENSMUSG00000022659 | 1 | |
| ENSMUSG00000022661 | -1.001348012 | 1 |
| ENSMUSG00000022663 | -1.013994949 | 0.996314864 |
| ENSMUSG00000022664 | -1.029503933 | 0.912782976 |
| ENSMUSG00000022665 | 1.002773123 | 1 |
| ENSMUSG00000022667 | 1.019083579 | 0.971519533 |
| ENSMUSG00000022668 | 1.02384394 | 0.941782235 |
| ENSMUSG00000022671 | 1.073215485 | 0.459632837 |
| ENSMUSG00000022672 | 1.037819499 | 0.841710814 |
| ENSMUSG00000022673 | 1.048207708 | 0.908149431 |
| ENSMUSG00000022674 | -1.011415828 | 1 |
| ENSMUSG00000022676 | 1.098732472 | 0.521644523 |
| ENSMUSG00000022677 | -1.011262139 | 1 |
| ENSMUSG00000022678 | 1.050293106 | 0.81831173 |
| ENSMUSG00000022679 | 1.060287104 | 0.608497229 |
| ENSMUSG00000022680 | 1.075748692 | 0.748682651 |
| ENSMUSG00000022681 | -1.069689138 | 0.698112512 |
| ENSMUSG00000022682 | 1.037865652 | 0.9209431 |
| ENSMUSG00000022683 | -1.086681236 | 0.693199918 |
| ENSMUSG00000022684 | -1.01669635 | 0.975727707 |
| ENSMUSG00000022685 | 1.018437726 | 0.972895772 |
| ENSMUSG00000022686 | -1.780599996 | 0.026780517 |

| | | |
|---|---|---|
| ENSMUSG00000022687 | -1.100220879 | 0.049280906 |
| ENSMUSG00000022696 | -1.043306199 | 0.698112512 |
| ENSMUSG00000022698 | 1.063850193 | 0.84515778 |
| ENSMUSG00000022701 | -1.027550805 | 0.874206421 |
| ENSMUSG00000022702 | 1.070396026 | 0.462961827 |
| ENSMUSG00000022704 | 1.022297008 | 0.961127302 |
| ENSMUSG00000022705 | 1 | |
| ENSMUSG00000022706 | -1.031229509 | 0.894873464 |
| ENSMUSG00000022707 | -1.106326564 | 0.531058358 |
| ENSMUSG00000022708 | 1.212708359 | 0.267657341 |
| ENSMUSG00000022710 | 1.011751344 | 1 |
| ENSMUSG00000022711 | 1.009839541 | 1 |
| ENSMUSG00000022715 | 1 | |
| ENSMUSG00000022718 | 1.009653716 | 1 |
| ENSMUSG00000022721 | 1.062892805 | 0.685325398 |
| ENSMUSG00000022722 | 1.025160899 | 0.907626515 |
| ENSMUSG00000022723 | -1.021595148 | 0.95838544 |
| ENSMUSG00000022724 | 1.0046128 | 1 |
| ENSMUSG00000022738 | 1 | |
| ENSMUSG00000022742 | -1.006018075 | 1 |
| ENSMUSG00000022744 | -1.03991362 | 0.89855495 |
| ENSMUSG00000022747 | -1.34007759 | 0.005222649 |
| ENSMUSG00000022748 | -1.156425396 | 0.071318353 |
| ENSMUSG00000022749 | 1.013944673 | 1 |
| ENSMUSG00000022750 | 1.142844968 | 0.430728007 |
| ENSMUSG00000022751 | -1.009992355 | 1 |
| ENSMUSG00000022752 | 1.018827079 | 0.998847216 |
| ENSMUSG00000022753 | 1.161952327 | 0.414312229 |
| ENSMUSG00000022754 | 1.097674266 | 0.535052277 |
| ENSMUSG00000022755 | 1 | |
| ENSMUSG00000022756 | 1.145055457 | 0.643264293 |
| ENSMUSG00000022757 | 1.081766985 | 0.703849495 |
| ENSMUSG00000022758 | 1.397108018 | 0.003114197 |
| ENSMUSG00000022759 | 1.023884886 | 0.998452655 |
| ENSMUSG00000022760 | 1.193367796 | 0.138602115 |
| ENSMUSG00000022761 | 1.088306505 | 0.629902583 |
| ENSMUSG00000022762 | -1.043306199 | 0.698112512 |
| ENSMUSG00000022763 | -1.041999587 | 0.875804019 |
| ENSMUSG00000022765 | -1.053338137 | 0.766837546 |
| ENSMUSG00000022766 | -1.117459691 | 0.608497229 |
| ENSMUSG00000022768 | -1.086032573 | 0.66920741 |
| ENSMUSG00000022769 | -1.06031316 | 0.698310385 |
| ENSMUSG00000022770 | 1.032631014 | 0.949526329 |
| ENSMUSG00000022771 | -1.051131317 | 0.814089245 |
| ENSMUSG00000022772 | 1.103612749 | 0.678985493 |
| ENSMUSG00000022773 | -1.004532801 | 1 |
| ENSMUSG00000022774 | 1.002883361 | 1 |
| ENSMUSG00000022779 | 1.041256177 | 0.859989151 |
| ENSMUSG00000022780 | -1.03962336 | 0.698112512 |
| ENSMUSG00000022781 | 1.032491463 | 0.954497929 |
| ENSMUSG00000022783 | -1.043306199 | 0.698112512 |
| ENSMUSG00000022787 | -1.001561592 | 1 |
| ENSMUSG00000022788 | -1.062990925 | 0.690651532 |
| ENSMUSG00000022789 | 1.002441742 | 1 |
| ENSMUSG00000022790 | 1.800923063 | 0.065153614 |
| ENSMUSG00000022791 | 1.133361884 | 0.257408696 |
| ENSMUSG00000022792 | 1.001925135 | 1 |
| ENSMUSG00000022793 | -1.17322043 | 0.189614251 |
| ENSMUSG00000022797 | -1.061919599 | 0.879253209 |
| ENSMUSG00000022798 | 1 | |
| ENSMUSG00000022799 | 1.084365248 | 0.698112512 |
| ENSMUSG00000022800 | 1.004559626 | 1 |
| ENSMUSG00000022801 | 1.01002778 | 1 |

| | | |
|---|---|---|
| ENSMUSG00000022802 | 1.065171212 | 0.585570411 |
| ENSMUSG00000022803 | 1.267042412 | 0.435797088 |
| ENSMUSG00000022805 | 1.20838637 | 0.493659452 |
| ENSMUSG00000022807 | 1.025808559 | 0.925214847 |
| ENSMUSG00000022808 | -1.029376152 | 0.952057016 |
| ENSMUSG00000022809 | -1 | 1 |
| ENSMUSG00000022811 | 1.000025728 | 1 |
| ENSMUSG00000022812 | 1.141331084 | 0.608497229 |
| ENSMUSG00000022814 | 1.015063942 | 0.996048848 |
| ENSMUSG00000022816 | 1.124825389 | 0.72715392 |
| ENSMUSG00000022817 | 1.245172958 | 0.123569983 |
| ENSMUSG00000022818 | 1 | |
| ENSMUSG00000022820 | 1.108229259 | 0.030673254 |
| ENSMUSG00000022821 | 1.080907075 | 0.797220014 |
| ENSMUSG00000022822 | 1.053667435 | 0.814557105 |
| ENSMUSG00000022824 | 1.122462048 | 0.541735653 |
| ENSMUSG00000022827 | -1.01053501 | 0.993803785 |
| ENSMUSG00000022828 | 1.072206718 | 0.698112512 |
| ENSMUSG00000022829 | -1.063525418 | 0.698112512 |
| ENSMUSG00000022831 | 1 | |
| ENSMUSG00000022832 | 1 | |
| ENSMUSG00000022833 | 1.063010488 | 0.698112512 |
| ENSMUSG00000022836 | 1.221853339 | 0.002527181 |
| ENSMUSG00000022837 | 1.000759473 | 1 |
| ENSMUSG00000022838 | -1.173690514 | 0.066778109 |
| ENSMUSG00000022840 | -1.034989968 | 0.949812939 |
| ENSMUSG00000022841 | -1.04765811 | 0.913105899 |
| ENSMUSG00000022842 | 1.07751676 | 0.696137525 |
| ENSMUSG00000022843 | -1.121611316 | 0.048686166 |
| ENSMUSG00000022844 | 1.253088684 | 0.063399406 |
| ENSMUSG00000022847 | -1.049364285 | 0.874672445 |
| ENSMUSG00000022848 | 1.019193664 | 0.929591952 |
| ENSMUSG00000022849 | -1.030042702 | 0.816836779 |
| ENSMUSG00000022853 | 1.009855508 | 1 |
| ENSMUSG00000022855 | -1.008207425 | 1 |
| ENSMUSG00000022856 | 1.088617593 | 0.556783372 |
| ENSMUSG00000022857 | 1 | |
| ENSMUSG00000022858 | -1.009937122 | 1 |
| ENSMUSG00000022860 | 1.042210396 | 0.738263421 |
| ENSMUSG00000022861 | -1.22825078 | 0.13439304 |
| ENSMUSG00000022863 | 1.010888325 | 0.979002724 |
| ENSMUSG00000022864 | -1.018070239 | 0.924766459 |
| ENSMUSG00000022865 | -1.317985913 | 0.117564123 |
| ENSMUSG00000022867 | -1.073098193 | 0.749854154 |
| ENSMUSG00000022868 | 1 | |
| ENSMUSG00000022871 | 1.133029889 | 0.608497229 |
| ENSMUSG00000022875 | 1 | |
| ENSMUSG00000022876 | -1.053549062 | 0.698112512 |
| ENSMUSG00000022877 | 1 | |
| ENSMUSG00000022878 | 1.059463094 | 0.608497229 |
| ENSMUSG00000022881 | -1.021344686 | 0.969506516 |
| ENSMUSG00000022883 | -1.174465101 | 0.16872433 |
| ENSMUSG00000022884 | 1.028671599 | 0.978211686 |
| ENSMUSG00000022885 | 1.452732035 | 0.003301399 |
| ENSMUSG00000022886 | 1 | |
| ENSMUSG00000022887 | -1.066901323 | 0.714200725 |
| ENSMUSG00000022889 | -1.055719123 | 0.717035356 |
| ENSMUSG00000022890 | 1.022311717 | 0.969506516 |
| ENSMUSG00000022892 | 1.130327424 | 0.628315739 |
| ENSMUSG00000022893 | 1.227543572 | 0.225402295 |
| ENSMUSG00000022894 | 1.664079776 | 0.004772284 |
| ENSMUSG00000022895 | 1.334380333 | 0.058015231 |
| ENSMUSG00000022897 | 1.103440935 | 0.623341196 |

| | | |
|---|---|---|
| ENSMUSG00000022898 | 1.114866029 | 0.608497229 |
| ENSMUSG00000022899 | 1.11166259 | 0.731295483 |
| ENSMUSG00000022900 | 1.059463094 | 0.698112512 |
| ENSMUSG00000022901 | 1 | |
| ENSMUSG00000022902 | 1 | |
| ENSMUSG00000022905 | 1.006424338 | 1 |
| ENSMUSG00000022906 | -1.391852988 | 0.063704667 |
| ENSMUSG00000022911 | 1.006796851 | 1 |
| ENSMUSG00000022912 | -1.069477857 | 0.717891577 |
| ENSMUSG00000022913 | 1.026864482 | 0.841678369 |
| ENSMUSG00000022914 | 1.01524375 | 1 |
| ENSMUSG00000022915 | 1 | |
| ENSMUSG00000022931 | 1 | |
| ENSMUSG00000022935 | -1.069502966 | 0.79291651 |
| ENSMUSG00000022938 | 1 | |
| ENSMUSG00000022940 | 1.096126534 | 0.145630941 |
| ENSMUSG00000022941 | -1.029054171 | 0.938939969 |
| ENSMUSG00000022945 | 1.117067396 | 0.512505963 |
| ENSMUSG00000022946 | -1.206149962 | 0.06505161 |
| ENSMUSG00000022947 | -1.583479899 | 0.001540286 |
| ENSMUSG00000022948 | 1.023161215 | 0.824542865 |
| ENSMUSG00000022949 | -1.193320102 | 0.265112993 |
| ENSMUSG00000022951 | -1.013808138 | 0.964539279 |
| ENSMUSG00000022952 | -1.14718427 | 0.280829377 |
| ENSMUSG00000022956 | -1.015633737 | 0.995747207 |
| ENSMUSG00000022957 | -1.028424143 | 0.93938607 |
| ENSMUSG00000022960 | -1.071808064 | 0.624225454 |
| ENSMUSG00000022961 | 1.025336694 | 0.983339718 |
| ENSMUSG00000022962 | 1.052383973 | 0.818908221 |
| ENSMUSG00000022964 | -1.104613192 | 0.485491618 |
| ENSMUSG00000022965 | -1.050970277 | 0.78965915 |
| ENSMUSG00000022967 | 1.042442269 | 0.865272535 |
| ENSMUSG00000022969 | 1.051767432 | 0.757988445 |
| ENSMUSG00000022971 | -1.036753149 | 0.803072249 |
| ENSMUSG00000022972 | 1.004266873 | 1 |
| ENSMUSG00000022973 | -1.011925577 | 1 |
| ENSMUSG00000022974 | 1.059478568 | 0.756295996 |
| ENSMUSG00000022978 | -1.01919518 | 0.943661908 |
| ENSMUSG00000022982 | -1.022190385 | 0.969544524 |
| ENSMUSG00000022983 | 1.062362575 | 0.689164262 |
| ENSMUSG00000022986 | 1.019083579 | 0.949812939 |
| ENSMUSG00000022987 | -1.173333974 | 0.048734589 |
| ENSMUSG00000022991 | 1 | |
| ENSMUSG00000022992 | 1.042991231 | 0.890958598 |
| ENSMUSG00000022993 | 1.13078166 | 0.698112512 |
| ENSMUSG00000022994 | 1.107397695 | 0.548958635 |
| ENSMUSG00000022995 | -1.056779719 | 0.844612733 |
| ENSMUSG00000022996 | -1.012955974 | 0.970822886 |
| ENSMUSG00000022997 | 1.116785864 | 0.70299173 |
| ENSMUSG00000022999 | -1.008288858 | 0.982838256 |
| ENSMUSG00000023000 | -1.428207648 | 0.199581726 |
| ENSMUSG00000023004 | 1.059450353 | 0.83997236 |
| ENSMUSG00000023007 | 1.087559467 | 0.239747126 |
| ENSMUSG00000023008 | 1.139156863 | 0.543724675 |
| ENSMUSG00000023009 | 1.117209637 | 0.608497229 |
| ENSMUSG00000023010 | 1.033967474 | 0.921490876 |
| ENSMUSG00000023011 | 1.699036688 | 0.001966665 |
| ENSMUSG00000023013 | 1.024693402 | 0.933031586 |
| ENSMUSG00000023015 | 1.010249726 | 1 |
| ENSMUSG00000023017 | 1.128722721 | 0.189170051 |
| ENSMUSG00000023018 | 1.007304279 | 1 |
| ENSMUSG00000023019 | 1.217178939 | 0.386153341 |
| ENSMUSG00000023020 | 1.162690873 | 0.106611663 |

| | | |
|---|---|---|
| ENSMUSG00000023021 | -1.065357103 | 0.841806844 |
| ENSMUSG00000023022 | 1.287263502 | 0.219185116 |
| ENSMUSG00000023025 | 1.009611863 | 1 |
| ENSMUSG00000023026 | 1.109468821 | 0.608497229 |
| ENSMUSG00000023027 | 1.020841317 | 0.97393994 |
| ENSMUSG00000023030 | 1.032170494 | 0.925881237 |
| ENSMUSG00000023031 | -1.064295887 | 0.608497229 |
| ENSMUSG00000023032 | 1.020943217 | 0.924330849 |
| ENSMUSG00000023033 | -1.426212131 | 0.118917727 |
| ENSMUSG00000023034 | 1.08310889 | 0.608497229 |
| ENSMUSG00000023036 | 1.069210072 | 0.128303257 |
| ENSMUSG00000023039 | 2.189725411 | 0.000522905 |
| ENSMUSG00000023041 | 1 | |
| ENSMUSG00000023043 | -1.596046585 | 0.001393029 |
| ENSMUSG00000023044 | -1.004650579 | 1 |
| ENSMUSG00000023045 | 1.120063048 | 0.698112512 |
| ENSMUSG00000023046 | -1.151758426 | 0.675127885 |
| ENSMUSG00000023047 | 1.102358556 | 0.698112512 |
| ENSMUSG00000023048 | -1.141948721 | 0.456679024 |
| ENSMUSG00000023050 | 1.083810391 | 0.456160368 |
| ENSMUSG00000023051 | 1.055202393 | 0.698112512 |
| ENSMUSG00000023052 | -1.164168724 | 0.179450399 |
| ENSMUSG00000023055 | 1.030339285 | 0.831816848 |
| ENSMUSG00000023057 | 1 | |
| ENSMUSG00000023064 | 1.24122829 | 0.03679259 |
| ENSMUSG00000023066 | 1.034271105 | 0.892023203 |
| ENSMUSG00000023067 | 1.734574222 | 0.000433572 |
| ENSMUSG00000023068 | -1.004638922 | 1 |
| ENSMUSG00000023070 | -1 | 1 |
| ENSMUSG00000023072 | -1.016038913 | 0.937182478 |
| ENSMUSG00000023073 | 1.014545335 | 0.979002724 |
| ENSMUSG00000023074 | 1.091536009 | 0.608497229 |
| ENSMUSG00000023075 | -1.034902804 | 0.907767688 |
| ENSMUSG00000023078 | 1.059463094 | 0.608497229 |
| ENSMUSG00000023079 | 1.104679428 | 0.60397857 |
| ENSMUSG00000023083 | -1.198963835 | 0.460496269 |
| ENSMUSG00000023084 | -1.058065387 | 0.824967755 |
| ENSMUSG00000023087 | 1.015295587 | 0.998812725 |
| ENSMUSG00000023088 | 1.118207564 | 0.618829386 |
| ENSMUSG00000023089 | -1.003754673 | 1 |
| ENSMUSG00000023092 | 1.836105304 | 0.006094015 |
| ENSMUSG00000023093 | 1 | |
| ENSMUSG00000023094 | 1.036973027 | 0.794274354 |
| ENSMUSG00000023104 | -1.00240285 | 1 |
| ENSMUSG00000023106 | 1.066475002 | 0.809746302 |
| ENSMUSG00000023110 | -1.181707011 | 0.483795883 |
| ENSMUSG00000023118 | 1.074576342 | 0.64374886 |
| ENSMUSG00000023120 | 1 | |
| ENSMUSG00000023122 | -1 | 1 |
| ENSMUSG00000023132 | 1 | |
| ENSMUSG00000023140 | -1.137443589 | 0.532369657 |
| ENSMUSG00000023143 | 1.160826649 | 0.10969528 |
| ENSMUSG00000023147 | 1.301509592 | 0.064293809 |
| ENSMUSG00000023150 | 1.034560213 | 0.94894091 |
| ENSMUSG00000023151 | 1.059463094 | 0.608497229 |
| ENSMUSG00000023153 | -1.084645496 | 0.532369657 |
| ENSMUSG00000023156 | -1.101739752 | 0.629902583 |
| ENSMUSG00000023159 | 1 | |
| ENSMUSG00000023165 | 1 | |
| ENSMUSG00000023169 | 1.14393835 | 0.608497229 |
| ENSMUSG00000023170 | 1.030279598 | 0.790851299 |
| ENSMUSG00000023175 | 1.032130481 | 0.949812939 |
| ENSMUSG00000023176 | -1 | 1 |

| | | |
|---|---|---|
| ENSMUSG00000023185 | 1 | |
| ENSMUSG00000023186 | -1.150660092 | 0.472138496 |
| ENSMUSG00000023191 | 1.099447697 | 0.283173741 |
| ENSMUSG00000023192 | -1.089497311 | 0.608497229 |
| ENSMUSG00000023206 | -1.433657467 | 0.001752613 |
| ENSMUSG00000023210 | 1 | |
| ENSMUSG00000023216 | 1.196463402 | 0.608497229 |
| ENSMUSG00000023224 | 1.267059297 | 0.415929354 |
| ENSMUSG00000023232 | 1.118013968 | 0.114261689 |
| ENSMUSG00000023235 | 1.220508543 | 0.019122348 |
| ENSMUSG00000023236 | -1.051577863 | 0.888246981 |
| ENSMUSG00000023243 | -2.832919572 | 0.000398998 |
| ENSMUSG00000023247 | 1 | |
| ENSMUSG00000023249 | 1.059959235 | 0.694333885 |
| ENSMUSG00000023257 | 1 | |
| ENSMUSG00000023259 | -1.037453252 | 0.637096543 |
| ENSMUSG00000023262 | 1.013272621 | 0.954533008 |
| ENSMUSG00000023263 | 1 | |
| ENSMUSG00000023266 | 1.223953372 | 0.545359962 |
| ENSMUSG00000023267 | -1.099174267 | 0.535052277 |
| ENSMUSG00000023272 | -1.005236221 | 1 |
| ENSMUSG00000023274 | 1 | |
| ENSMUSG00000023277 | -1.071474043 | 0.567125573 |
| ENSMUSG00000023279 | -1 | 1 |
| ENSMUSG00000023284 | 1.107127697 | 0.224521642 |
| ENSMUSG00000023286 | -1.028082814 | 0.86336752 |
| ENSMUSG00000023289 | 1 | |
| ENSMUSG00000023307 | 1.005918086 | 1 |
| ENSMUSG00000023328 | 1.033906491 | 0.872376523 |
| ENSMUSG00000023330 | 1.275725502 | 0.072336855 |
| ENSMUSG00000023333 | 1 | |
| ENSMUSG00000023336 | 1 | |
| ENSMUSG00000023341 | -1.297904569 | 0.221797188 |
| ENSMUSG00000023345 | -1.091095944 | 0.400463057 |
| ENSMUSG00000023348 | 1.104920682 | 0.475771979 |
| ENSMUSG00000023349 | 1 | |
| ENSMUSG00000023350 | -1.03962336 | 0.698112512 |
| ENSMUSG00000023353 | 1.174212092 | 0.138802306 |
| ENSMUSG00000023367 | -1.038135377 | 0.868791781 |
| ENSMUSG00000023387 | 1 | |
| ENSMUSG00000023391 | 1.114108048 | 0.608497229 |
| ENSMUSG00000023393 | 1.162552257 | 0.428966059 |
| ENSMUSG00000023403 | 1.059463094 | 0.608497229 |
| ENSMUSG00000023411 | -1.064222489 | 0.608497229 |
| ENSMUSG00000023433 | 1 | |
| ENSMUSG00000023439 | -1.019515214 | 1 |
| ENSMUSG00000023443 | -1.54234929 | 0.006427103 |
| ENSMUSG00000023452 | 1.119349492 | 0.385303452 |
| ENSMUSG00000023456 | 1.017696141 | 1 |
| ENSMUSG00000023460 | 1.004231895 | 1 |
| ENSMUSG00000023467 | 1.085585276 | 0.73275545 |
| ENSMUSG00000023473 | 1.035275751 | 0.944738166 |
| ENSMUSG00000023484 | -1.516778727 | 0.085919896 |
| ENSMUSG00000023495 | 1.104562136 | 0.495999195 |
| ENSMUSG00000023505 | -1.005857239 | 1 |
| ENSMUSG00000023571 | 1.095210875 | 0.74505857 |
| ENSMUSG00000023572 | -1.03992818 | 0.806949992 |
| ENSMUSG00000023577 | 1 | |
| ENSMUSG00000023707 | 1.121655408 | 0.299492291 |
| ENSMUSG00000023723 | 1.036305317 | 0.857163394 |
| ENSMUSG00000023737 | 1.010501973 | 1 |
| ENSMUSG00000023755 | -1.078321811 | 0.532369657 |
| ENSMUSG00000023764 | -1.007948904 | 0.997060234 |

| | | |
|---|---|---|
| ENSMUSG00000023781 | 1.189706618 | 0.216046507 |
| ENSMUSG00000023791 | 1.00988254 | 1 |
| ENSMUSG00000023795 | 1.022959243 | 0.900817819 |
| ENSMUSG00000023800 | 1.100817009 | 0.608497229 |
| ENSMUSG00000023802 | 1 | |
| ENSMUSG00000023805 | 1.418175285 | 0.004151516 |
| ENSMUSG00000023806 | -1.027151996 | 0.996069743 |
| ENSMUSG00000023809 | 1.103270219 | 0.609615508 |
| ENSMUSG00000023826 | -1.110695811 | 0.649951761 |
| ENSMUSG00000023827 | 1.118708168 | 0.412276983 |
| ENSMUSG00000023828 | 1 | |
| ENSMUSG00000023829 | -1 | 1 |
| ENSMUSG00000023830 | 1.017012316 | 0.989603176 |
| ENSMUSG00000023832 | -1.089413172 | 0.698112512 |
| ENSMUSG00000023845 | 1.104071008 | 0.697813098 |
| ENSMUSG00000023852 | 1.029965757 | 0.956804939 |
| ENSMUSG00000023861 | -1.076861591 | 0.611938648 |
| ENSMUSG00000023868 | 1.081082677 | 0.763030715 |
| ENSMUSG00000023873 | 1.099610164 | 0.732612801 |
| ENSMUSG00000023882 | 1.109536703 | 0.20280389 |
| ENSMUSG00000023883 | -1.018169604 | 0.986389736 |
| ENSMUSG00000023885 | 1.008560533 | 0.982838256 |
| ENSMUSG00000023886 | 2.008791456 | 0.000101355 |
| ENSMUSG00000023892 | 1.022165572 | 0.938939969 |
| ENSMUSG00000023902 | -1 | 1 |
| ENSMUSG00000023903 | -1.669299874 | 0.026578278 |
| ENSMUSG00000023904 | 1.285408462 | 0.017542255 |
| ENSMUSG00000023905 | 1.023060591 | 0.956659802 |
| ENSMUSG00000023906 | -1 | 1 |
| ENSMUSG00000023908 | 1.08092555 | 0.63076311 |
| ENSMUSG00000023909 | -1.05532428 | 0.608497229 |
| ENSMUSG00000023911 | 1.043861669 | 0.778372272 |
| ENSMUSG00000023912 | 1.129796946 | 0.034342216 |
| ENSMUSG00000023913 | 1.143301547 | 0.195607801 |
| ENSMUSG00000023914 | 1.303219601 | 0.325380902 |
| ENSMUSG00000023915 | -1.073920266 | 0.287097207 |
| ENSMUSG00000023918 | 1.005212439 | 1 |
| ENSMUSG00000023919 | -1.039645751 | 0.87310927 |
| ENSMUSG00000023921 | 1.04947894 | 0.833800367 |
| ENSMUSG00000023923 | 1.040601337 | 0.869548547 |
| ENSMUSG00000023926 | -1.009091449 | 1 |
| ENSMUSG00000023927 | 1.213249713 | 0.02683365 |
| ENSMUSG00000023930 | -1 | 1 |
| ENSMUSG00000023931 | -1.129667185 | 0.590149159 |
| ENSMUSG00000023932 | -1.031572463 | 0.951492347 |
| ENSMUSG00000023935 | 1.160224851 | 0.608497229 |
| ENSMUSG00000023938 | -1.05648007 | 0.698112512 |
| ENSMUSG00000023939 | 1.067696499 | 0.619527805 |
| ENSMUSG00000023940 | 1.017097687 | 0.991944074 |
| ENSMUSG00000023942 | 1.41341953 | 0.034065365 |
| ENSMUSG00000023943 | 1 | |
| ENSMUSG00000023944 | 1.030337037 | 0.993582397 |
| ENSMUSG00000023945 | 1.346553027 | 0.21967784 |
| ENSMUSG00000023947 | -1.402678738 | 0.004775376 |
| ENSMUSG00000023949 | 1 | |
| ENSMUSG00000023951 | 1.485916485 | 0.004445046 |
| ENSMUSG00000023952 | 1.030129735 | 0.870043872 |
| ENSMUSG00000023953 | 1.014121653 | 0.98820811 |
| ENSMUSG00000023959 | -1.130004659 | 0.407630951 |
| ENSMUSG00000023960 | 1.183840395 | 0.360936635 |
| ENSMUSG00000023961 | 1.028656724 | 0.935530845 |
| ENSMUSG00000023963 | 1.459962064 | 0.018349356 |
| ENSMUSG00000023964 | 1.015486245 | 0.985522133 |

| | | |
|---|---|---|
| ENSMUSG00000023965 | 1.044732659 | 0.828564184 |
| ENSMUSG00000023966 | -1.162049362 | 0.079964742 |
| ENSMUSG00000023967 | 1.015097322 | 0.973248005 |
| ENSMUSG00000023968 | -1.18980823 | 0.552336922 |
| ENSMUSG00000023971 | 1.084934884 | 0.320982015 |
| ENSMUSG00000023972 | 1.050041149 | 0.698112512 |
| ENSMUSG00000023973 | 1.041440789 | 0.749224689 |
| ENSMUSG00000023977 | 1.049622884 | 0.860992632 |
| ENSMUSG00000023978 | -1 | 1 |
| ENSMUSG00000023979 | -1.15462599 | 0.532369657 |
| ENSMUSG00000023980 | -1.023491483 | 0.919000509 |
| ENSMUSG00000023982 | -1.226276019 | 0.256544909 |
| ENSMUSG00000023984 | -1.273745414 | 0.355014525 |
| ENSMUSG00000023987 | -1 | 1 |
| ENSMUSG00000023988 | 1.027973743 | 0.897719303 |
| ENSMUSG00000023990 | 1.128652642 | 0.608497229 |
| ENSMUSG00000023991 | 1.129793506 | 0.23017929 |
| ENSMUSG00000023992 | 1.059463094 | 0.808713047 |
| ENSMUSG00000023993 | 1 | |
| ENSMUSG00000023994 | 1.027604812 | 0.944649069 |
| ENSMUSG00000023995 | -1 | 1 |
| ENSMUSG00000023999 | 1.509783998 | 0.014804417 |
| ENSMUSG00000024002 | 1.049724753 | 0.828564184 |
| ENSMUSG00000024006 | 1.042333349 | 0.901216307 |
| ENSMUSG00000024007 | 1.035078469 | 0.896090268 |
| ENSMUSG00000024008 | -1.048034918 | 0.873810332 |
| ENSMUSG00000024011 | -1.175273974 | 0.568529369 |
| ENSMUSG00000024012 | 1.08256302 | 0.734674233 |
| ENSMUSG00000024013 | -1.141394474 | 0.698112512 |
| ENSMUSG00000024014 | 1.073225983 | 0.493659452 |
| ENSMUSG00000024018 | -1.257362787 | 0.048406505 |
| ENSMUSG00000024019 | -1.055911243 | 0.593054478 |
| ENSMUSG00000024026 | -1.194214089 | 0.532369657 |
| ENSMUSG00000024027 | 1.059463094 | 0.608497229 |
| ENSMUSG00000024028 | 1 | |
| ENSMUSG00000024029 | 1 | |
| ENSMUSG00000024030 | -1.730995584 | 0.027236743 |
| ENSMUSG00000024032 | 1 | |
| ENSMUSG00000024033 | -2.036451782 | 0.003180003 |
| ENSMUSG00000024034 | 1.059463094 | 0.608497229 |
| ENSMUSG00000024036 | -2.024814626 | 0.009956803 |
| ENSMUSG00000024037 | 1.081333558 | 0.482403115 |
| ENSMUSG00000024038 | 1.03643344 | 0.771926153 |
| ENSMUSG00000024039 | -1.389892877 | 0.178527696 |
| ENSMUSG00000024041 | 1 | |
| ENSMUSG00000024042 | 1.01790025 | 0.902345046 |
| ENSMUSG00000024043 | -1.110665593 | 0.608497229 |
| ENSMUSG00000024044 | 1.114716169 | 0.698112512 |
| ENSMUSG00000024045 | 1.016646982 | 0.989801616 |
| ENSMUSG00000024048 | 1.193971984 | 0.566175333 |
| ENSMUSG00000024049 | 1.754368529 | 0.006292594 |
| ENSMUSG00000024050 | 1.070102584 | 0.608497229 |
| ENSMUSG00000024052 | 1.183617866 | 0.330594361 |
| ENSMUSG00000024053 | 1.664792197 | 0.000573666 |
| ENSMUSG00000024054 | 1.009122952 | 1 |
| ENSMUSG00000024055 | 1.045769886 | 0.754820307 |
| ENSMUSG00000024056 | 1.043780037 | 0.880365937 |
| ENSMUSG00000024059 | -1.074956284 | 0.449851954 |
| ENSMUSG00000024063 | 1.152141977 | 0.357647366 |
| ENSMUSG00000024064 | -1.571026796 | 0.002849214 |
| ENSMUSG00000024065 | 1.040253343 | 0.848803478 |
| ENSMUSG00000024066 | 2.340697706 | 0.001316084 |
| ENSMUSG00000024067 | -1.017935428 | 0.984628579 |

| | | |
|---|---|---|
| ENSMUSG00000024068 | -1.039153532 | 0.898763027 |
| ENSMUSG00000024069 | 1.00742381 | 1 |
| ENSMUSG00000024070 | 1.080906199 | 0.763439211 |
| ENSMUSG00000024072 | -1.008193383 | 1 |
| ENSMUSG00000024073 | 1.052091736 | 0.900730943 |
| ENSMUSG00000024074 | 1.175540297 | 0.559786559 |
| ENSMUSG00000024076 | -2.043700364 | 0.000960133 |
| ENSMUSG00000024077 | -1.026789737 | 0.9607713 |
| ENSMUSG00000024078 | -1.021668969 | 0.958299667 |
| ENSMUSG00000024079 | -1.108282753 | 0.698112512 |
| ENSMUSG00000024081 | -1.004290488 | 1 |
| ENSMUSG00000024082 | 1.026971553 | 0.933537875 |
| ENSMUSG00000024083 | 1.053935688 | 0.883021131 |
| ENSMUSG00000024084 | 1.587913963 | 0.002932067 |
| ENSMUSG00000024085 | 1.123162188 | 0.672115339 |
| ENSMUSG00000024087 | 1.280435186 | 0.363578301 |
| ENSMUSG00000024088 | 1 | |
| ENSMUSG00000024091 | 1.084875233 | 0.755613721 |
| ENSMUSG00000024095 | 1.020038689 | 0.988424085 |
| ENSMUSG00000024096 | 1.046781414 | 0.815347141 |
| ENSMUSG00000024097 | 1.008414119 | 1 |
| ENSMUSG00000024098 | -1.013975088 | 1 |
| ENSMUSG00000024099 | -1.041984616 | 0.902565126 |
| ENSMUSG00000024101 | 1.088755231 | 0.541442591 |
| ENSMUSG00000024104 | 1.156814158 | 0.536664112 |
| ENSMUSG00000024105 | 1 | |
| ENSMUSG00000024107 | 1 | |
| ENSMUSG00000024109 | -1 | 1 |
| ENSMUSG00000024112 | 1.562977145 | 0.019077018 |
| ENSMUSG00000024114 | 2.202467447 | 0.004728769 |
| ENSMUSG00000024116 | 1 | |
| ENSMUSG00000024118 | 1.011635745 | 0.984902895 |
| ENSMUSG00000024120 | 1.110412656 | 0.698112512 |
| ENSMUSG00000024121 | 1.103986588 | 0.050824368 |
| ENSMUSG00000024122 | 1.023979086 | 0.974055135 |
| ENSMUSG00000024124 | 1.122462048 | 0.628993873 |
| ENSMUSG00000024125 | 1 | |
| ENSMUSG00000024127 | 1.166065152 | 0.505928387 |
| ENSMUSG00000024128 | 1 | |
| ENSMUSG00000024130 | -1.04549563 | 0.83672351 |
| ENSMUSG00000024131 | -1.218081704 | 0.608497229 |
| ENSMUSG00000024132 | -1.044074865 | 0.728386343 |
| ENSMUSG00000024134 | 1.003509506 | 1 |
| ENSMUSG00000024135 | -1.028597274 | 0.943705943 |
| ENSMUSG00000024136 | 1.054414368 | 0.725290225 |
| ENSMUSG00000024137 | 1.08329921 | 0.473417727 |
| ENSMUSG00000024140 | -1.817566023 | 0.001642767 |
| ENSMUSG00000024142 | -1.104817126 | 0.608497229 |
| ENSMUSG00000024143 | 1.322280872 | 0.101447883 |
| ENSMUSG00000024145 | -1.039337438 | 0.779227741 |
| ENSMUSG00000024146 | -1.044129598 | 0.889802291 |
| ENSMUSG00000024150 | 1.04126568 | 0.870778586 |
| ENSMUSG00000024151 | 1.097186732 | 0.690669961 |
| ENSMUSG00000024154 | -1.069502966 | 0.698112512 |
| ENSMUSG00000024155 | -1.299880915 | 0.128539749 |
| ENSMUSG00000024158 | -1.141233718 | 0.373988904 |
| ENSMUSG00000024160 | 1.10615673 | 0.29527688 |
| ENSMUSG00000024163 | 1.098382757 | 0.608497229 |
| ENSMUSG00000024164 | 1.263296155 | 0.005557298 |
| ENSMUSG00000024165 | -1.027621522 | 0.978354388 |
| ENSMUSG00000024168 | -1.000509578 | 1 |
| ENSMUSG00000024169 | 1.088651271 | 0.615404603 |
| ENSMUSG00000024170 | 1.018160106 | 0.939580717 |

| | | |
|---|---|---|
| ENSMUSG00000024171 | 1 | |
| ENSMUSG00000024172 | 1 | |
| ENSMUSG00000024173 | 1.5720848 | 0.12725367 |
| ENSMUSG00000024174 | 1.045454611 | 0.746274305 |
| ENSMUSG00000024175 | -1 | 1 |
| ENSMUSG00000024176 | -1.454598606 | 0.070855636 |
| ENSMUSG00000024177 | -1.039682957 | 0.640707404 |
| ENSMUSG00000024180 | -1.213519481 | 0.011978159 |
| ENSMUSG00000024181 | 1.004085202 | 1 |
| ENSMUSG00000024182 | 1.04535093 | 0.851971494 |
| ENSMUSG00000024184 | 1 | |
| ENSMUSG00000024186 | -1.087673015 | 0.854074987 |
| ENSMUSG00000024187 | -1.052066886 | 0.790706187 |
| ENSMUSG00000024188 | 1.017786715 | 0.988424085 |
| ENSMUSG00000024190 | -1.086559598 | 0.608497229 |
| ENSMUSG00000024191 | 1.035353079 | 0.736165429 |
| ENSMUSG00000024193 | 1.041495603 | 0.608497229 |
| ENSMUSG00000024194 | 1.045188899 | 0.717035356 |
| ENSMUSG00000024197 | 1.107665134 | 0.291044503 |
| ENSMUSG00000024201 | 1.074528222 | 0.552584793 |
| ENSMUSG00000024205 | -1 | 1 |
| ENSMUSG00000024206 | -1.042130607 | 0.613129956 |
| ENSMUSG00000024207 | -1 | 1 |
| ENSMUSG00000024208 | 1.016299821 | 0.958403618 |
| ENSMUSG00000024209 | -1.456965084 | 0.009956796 |
| ENSMUSG00000024210 | 1.16164987 | 0.608497229 |
| ENSMUSG00000024211 | 1 | |
| ENSMUSG00000024212 | 1.074949083 | 0.608497229 |
| ENSMUSG00000024213 | 1.073087725 | 0.698112512 |
| ENSMUSG00000024215 | -1.094422282 | 0.698112512 |
| ENSMUSG00000024217 | -1.007545262 | 1 |
| ENSMUSG00000024218 | -1.045487799 | 0.823969666 |
| ENSMUSG00000024219 | 1.044202916 | 0.698112512 |
| ENSMUSG00000024220 | 1.054964225 | 0.731932479 |
| ENSMUSG00000024222 | -1.09442809 | 0.676441661 |
| ENSMUSG00000024223 | -1.03962336 | 0.698112512 |
| ENSMUSG00000024224 | 1 | |
| ENSMUSG00000024225 | 1 | |
| ENSMUSG00000024227 | 1 | |
| ENSMUSG00000024228 | 1.01542826 | 0.97954245 |
| ENSMUSG00000024231 | 1.014165723 | 1 |
| ENSMUSG00000024232 | 1.380519648 | 0.032195513 |
| ENSMUSG00000024233 | 1 | |
| ENSMUSG00000024234 | 1.024291833 | 0.951494707 |
| ENSMUSG00000024235 | 1.18934487 | 0.18445321 |
| ENSMUSG00000024236 | 1.332826646 | 0.071961713 |
| ENSMUSG00000024238 | 1.117896306 | 0.608497229 |
| ENSMUSG00000024240 | 1.015937613 | 0.975499304 |
| ENSMUSG00000024241 | 1.056444551 | 0.836148658 |
| ENSMUSG00000024242 | 1.034855689 | 0.893388118 |
| ENSMUSG00000024245 | -1 | 1 |
| ENSMUSG00000024246 | 1.276219553 | 0.00590229 |
| ENSMUSG00000024247 | 1.232869424 | 0.061097647 |
| ENSMUSG00000024248 | 1.131334899 | 0.607649304 |
| ENSMUSG00000024251 | -1.007280989 | 1 |
| ENSMUSG00000024253 | 1.014509049 | 0.972671752 |
| ENSMUSG00000024254 | 1.014545335 | 0.989648415 |
| ENSMUSG00000024256 | 1 | |
| ENSMUSG00000024258 | -1.013795661 | 1 |
| ENSMUSG00000024259 | 1.012660461 | 1 |
| ENSMUSG00000024260 | 1.093925238 | 0.608497229 |
| ENSMUSG00000024261 | 1 | |
| ENSMUSG00000024266 | 1 | |

| | | |
|---|---|---|
| ENSMUSG00000024268 | -1.03195846 | 0.725290225 |
| ENSMUSG00000024269 | 1.015815186 | 0.986394584 |
| ENSMUSG00000024270 | 1.157946849 | 0.487513292 |
| ENSMUSG00000024271 | 1.029105447 | 0.958653795 |
| ENSMUSG00000024273 | -1.021123357 | 0.883566187 |
| ENSMUSG00000024274 | 1.152861661 | 0.200376666 |
| ENSMUSG00000024276 | 1.084715382 | 0.698112512 |
| ENSMUSG00000024277 | -1.021590031 | 0.965252226 |
| ENSMUSG00000024283 | 1.056287232 | 0.84766519 |
| ENSMUSG00000024286 | 1.047784845 | 0.845866242 |
| ENSMUSG00000024287 | 1.002984509 | 1 |
| ENSMUSG00000024290 | 1.0341094 | 0.925151724 |
| ENSMUSG00000024292 | 1.059463094 | 0.608497229 |
| ENSMUSG00000024293 | 1.016888608 | 0.964579585 |
| ENSMUSG00000024294 | 1.037282741 | 0.940569501 |
| ENSMUSG00000024295 | -1.02398313 | 0.994915989 |
| ENSMUSG00000024298 | 1.000255311 | 1 |
| ENSMUSG00000024299 | 1.253816182 | 0.05687956 |
| ENSMUSG00000024300 | -1.043306199 | 0.803260194 |
| ENSMUSG00000024301 | 1.106850919 | 0.473417727 |
| ENSMUSG00000024302 | 1.717314233 | 0.00050352 |
| ENSMUSG00000024304 | 1.065580117 | 0.790851299 |
| ENSMUSG00000024306 | -1 | 1 |
| ENSMUSG00000024308 | -1.218816907 | 0.230325652 |
| ENSMUSG00000024309 | 1.058286764 | 0.684693086 |
| ENSMUSG00000024312 | -1.003296042 | 1 |
| ENSMUSG00000024313 | 1 | |
| ENSMUSG00000024317 | 1.013175243 | 0.986362318 |
| ENSMUSG00000024319 | -1.01655828 | 0.964379862 |
| ENSMUSG00000024325 | 1.016330534 | 0.969949129 |
| ENSMUSG00000024327 | -1.023136014 | 0.888246981 |
| ENSMUSG00000024330 | 1.243176355 | 0.251544797 |
| ENSMUSG00000024331 | 1.038117106 | 0.638758774 |
| ENSMUSG00000024334 | 1 | |
| ENSMUSG00000024335 | 1.000119023 | 1 |
| ENSMUSG00000024338 | -2.465881734 | 4.68417E-05 |
| ENSMUSG00000024339 | -1.736529118 | 0.000974735 |
| ENSMUSG00000024340 | 1 | |
| ENSMUSG00000024346 | -1.064852158 | 0.749681041 |
| ENSMUSG00000024347 | -1.092663169 | 0.717035356 |
| ENSMUSG00000024349 | -1.462609708 | 0.008577483 |
| ENSMUSG00000024350 | -1.073740974 | 0.69852739 |
| ENSMUSG00000024352 | 1.132334296 | 0.007234395 |
| ENSMUSG00000024353 | -1 | 1 |
| ENSMUSG00000024354 | 1.03971413 | 0.908007394 |
| ENSMUSG00000024357 | 1.045128295 | 0.698112512 |
| ENSMUSG00000024359 | 1.111838572 | 0.751455346 |
| ENSMUSG00000024360 | 1.058781587 | 0.875315552 |
| ENSMUSG00000024365 | -1 | 1 |
| ENSMUSG00000024366 | -1.03962336 | 0.698112512 |
| ENSMUSG00000024369 | -1.018701703 | 0.905298097 |
| ENSMUSG00000024370 | -1.039086785 | 0.914857505 |
| ENSMUSG00000024371 | -1.467038957 | 0.106792554 |
| ENSMUSG00000024376 | -1.473112434 | 0.003453741 |
| ENSMUSG00000024378 | -1.009927008 | 1 |
| ENSMUSG00000024379 | 1.221121054 | 0.080928024 |
| ENSMUSG00000024381 | 1.103091572 | 0.579232604 |
| ENSMUSG00000024382 | 1.011387755 | 1 |
| ENSMUSG00000024383 | 1.096257456 | 0.698112512 |
| ENSMUSG00000024384 | -1.002045122 | 1 |
| ENSMUSG00000024386 | 1.059463094 | 0.608497229 |
| ENSMUSG00000024387 | 1.000038213 | 1 |
| ENSMUSG00000024388 | 1 | |

| | | |
|---|---|---|
| ENSMUSG00000024391 | 1.18176657 | 0.376374334 |
| ENSMUSG00000024392 | 1.107144047 | 0.608497229 |
| ENSMUSG00000024393 | 1.132282767 | 0.577978082 |
| ENSMUSG00000024395 | 2.255961253 | 0.001888726 |
| ENSMUSG00000024397 | -1.174113566 | 0.416622982 |
| ENSMUSG00000024399 | -1.039768583 | 0.950513949 |
| ENSMUSG00000024400 | 1.0244069 | 0.921959803 |
| ENSMUSG00000024401 | -1.053549062 | 0.698112512 |
| ENSMUSG00000024402 | -1.015511103 | 0.982777255 |
| ENSMUSG00000024403 | 1.06969483 | 0.397706863 |
| ENSMUSG00000024404 | 1.020198171 | 0.976752232 |
| ENSMUSG00000024406 | 1.023743961 | 0.938939969 |
| ENSMUSG00000024409 | -1.028364146 | 0.970020309 |
| ENSMUSG00000024410 | 1.008864168 | 1 |
| ENSMUSG00000024411 | 1 | |
| ENSMUSG00000024413 | 1.098269702 | 0.608497229 |
| ENSMUSG00000024414 | 1.007268719 | 1 |
| ENSMUSG00000024420 | 2.641538187 | 4.55E-06 |
| ENSMUSG00000024421 | 1.159789297 | 0.183617727 |
| ENSMUSG00000024422 | 1.0307038 | 0.865800394 |
| ENSMUSG00000024423 | 1.137377924 | 0.567758634 |
| ENSMUSG00000024424 | 1.029627486 | 0.893252049 |
| ENSMUSG00000024425 | 1.005317175 | 1 |
| ENSMUSG00000024426 | 1.039719104 | 0.73068702 |
| ENSMUSG00000024427 | 1.034790339 | 0.824258933 |
| ENSMUSG00000024429 | 1.033253476 | 0.867939428 |
| ENSMUSG00000024430 | -1.064273502 | 0.723641964 |
| ENSMUSG00000024431 | 1.124375747 | 0.608497229 |
| ENSMUSG00000024436 | 1.051656881 | 0.699272448 |
| ENSMUSG00000024437 | 1.177765487 | 0.277823925 |
| ENSMUSG00000024440 | 1.093819733 | 0.699168433 |
| ENSMUSG00000024442 | -1.004798816 | 1 |
| ENSMUSG00000024446 | 1.13850732 | 0.286450465 |
| ENSMUSG00000024448 | 1 | |
| ENSMUSG00000024451 | 1.70624866 | 0.004730529 |
| ENSMUSG00000024454 | -1.01395221 | 1 |
| ENSMUSG00000024456 | 1.023392254 | 0.958084572 |
| ENSMUSG00000024457 | -1.022808767 | 0.937505238 |
| ENSMUSG00000024459 | 1.497870307 | 0.138222143 |
| ENSMUSG00000024462 | 1.16702293 | 0.155088532 |
| ENSMUSG00000024471 | 1 | |
| ENSMUSG00000024472 | -1.026677714 | 0.96035552 |
| ENSMUSG00000024474 | -1.045821982 | 0.90832083 |
| ENSMUSG00000024477 | -1.002983974 | 1 |
| ENSMUSG00000024479 | -1.213305182 | 0.536942078 |
| ENSMUSG00000024480 | 1.002784385 | 1 |
| ENSMUSG00000024481 | 1.282062471 | 0.087721407 |
| ENSMUSG00000024483 | 1.073371859 | 0.778239858 |
| ENSMUSG00000024485 | -1.044273782 | 0.698112512 |
| ENSMUSG00000024486 | -1.029548085 | 0.9073708 |
| ENSMUSG00000024487 | 1.057531302 | 0.787379965 |
| ENSMUSG00000024491 | 1.024332108 | 0.95838544 |
| ENSMUSG00000024493 | 1.112891822 | 0.698112512 |
| ENSMUSG00000024497 | -1.044273782 | 0.698112512 |
| ENSMUSG00000024498 | 1.022272052 | 0.976640829 |
| ENSMUSG00000024500 | 1.616897062 | 0.030651229 |
| ENSMUSG00000024501 | 1.263169783 | 0.297024201 |
| ENSMUSG00000024502 | 1.32304421 | 0.007935162 |
| ENSMUSG00000024503 | 1 | |
| ENSMUSG00000024505 | -1.104852311 | 0.180644327 |
| ENSMUSG00000024507 | 1.037163597 | 0.908007394 |
| ENSMUSG00000024510 | 1 | |
| ENSMUSG00000024511 | 1.112801622 | 0.516869627 |

| | | |
|---|---|---|
| ENSMUSG00000024512 | -1.225243136 | 0.462829509 |
| ENSMUSG00000024513 | 1.048358841 | 0.829324104 |
| ENSMUSG00000024515 | 1.098984553 | 0.683457122 |
| ENSMUSG00000024516 | -1.058099509 | 0.698112512 |
| ENSMUSG00000024517 | 1 | |
| ENSMUSG00000024518 | -1.063525418 | 0.751979636 |
| ENSMUSG00000024519 | 1 | |
| ENSMUSG00000024521 | 1.333132247 | 0.035417051 |
| ENSMUSG00000024524 | 1.000361072 | 1 |
| ENSMUSG00000024525 | 1.048437773 | 0.608497229 |
| ENSMUSG00000024526 | -1.109582661 | 0.549347972 |
| ENSMUSG00000024527 | 1.052074353 | 0.770958539 |
| ENSMUSG00000024528 | 1.100840357 | 0.440910868 |
| ENSMUSG00000024529 | 1.933042465 | 0.005738392 |
| ENSMUSG00000024530 | 1.150038027 | 0.022005475 |
| ENSMUSG00000024532 | 1.059463094 | 0.608497229 |
| ENSMUSG00000024533 | 1.086777765 | 0.617823788 |
| ENSMUSG00000024534 | -1.143787033 | 0.349938995 |
| ENSMUSG00000024535 | 1.156069503 | 0.287588797 |
| ENSMUSG00000024537 | 1.031457795 | 0.844158013 |
| ENSMUSG00000024538 | 1.015657079 | 1 |
| ENSMUSG00000024539 | -1.026806693 | 0.94733141 |
| ENSMUSG00000024542 | 1.050533953 | 0.864123949 |
| ENSMUSG00000024544 | 1.404944615 | 0.004795301 |
| ENSMUSG00000024546 | 1 | |
| ENSMUSG00000024548 | 1.453345401 | 0.003129069 |
| ENSMUSG00000024552 | 1.05410549 | 0.805157604 |
| ENSMUSG00000024553 | 1 | |
| ENSMUSG00000024556 | -1.02664725 | 0.965125182 |
| ENSMUSG00000024558 | 1.265598895 | 0.298705314 |
| ENSMUSG00000024560 | 1.112470226 | 0.232072861 |
| ENSMUSG00000024561 | 1.112097812 | 0.277459955 |
| ENSMUSG00000024563 | 1.078366212 | 0.711225772 |
| ENSMUSG00000024565 | -1.043306199 | 0.698112512 |
| ENSMUSG00000024566 | 1.048550362 | 0.774758227 |
| ENSMUSG00000024570 | 1.083407792 | 0.062507642 |
| ENSMUSG00000024571 | 1.15028436 | 0.36733642 |
| ENSMUSG00000024575 | 1.178666684 | 0.124934541 |
| ENSMUSG00000024576 | 1.081131351 | 0.796531827 |
| ENSMUSG00000024578 | 1 | |
| ENSMUSG00000024579 | 1.164362673 | 0.249020999 |
| ENSMUSG00000024580 | 1.009792828 | 1 |
| ENSMUSG00000024581 | 1.072298554 | 0.698112512 |
| ENSMUSG00000024583 | 1.021034914 | 0.9872207 |
| ENSMUSG00000024587 | 1.067486676 | 0.841437469 |
| ENSMUSG00000024588 | -1.03259622 | 0.9209431 |
| ENSMUSG00000024589 | 1.118885234 | 0.431488771 |
| ENSMUSG00000024590 | 1.041983457 | 0.908065256 |
| ENSMUSG00000024592 | 1.116904789 | 0.414627266 |
| ENSMUSG00000024593 | 3.614706188 | 7.72145E-05 |
| ENSMUSG00000024594 | 1.11152982 | 0.439772071 |
| ENSMUSG00000024597 | 1.277291087 | 0.223170554 |
| ENSMUSG00000024598 | -1.038296262 | 0.905346712 |
| ENSMUSG00000024600 | 1.502879847 | 0.051171361 |
| ENSMUSG00000024601 | -1.016292487 | 0.965978247 |
| ENSMUSG00000024603 | 1.061846297 | 0.806152724 |
| ENSMUSG00000024604 | 1.011388689 | 1 |
| ENSMUSG00000024608 | 1.088775258 | 0.72715392 |
| ENSMUSG00000024610 | -1.16855285 | 0.280944764 |
| ENSMUSG00000024613 | 1.124818295 | 0.390934291 |
| ENSMUSG00000024614 | 1.02075519 | 0.989500016 |
| ENSMUSG00000024617 | 1.121697966 | 0.381312112 |
| ENSMUSG00000024619 | 1.122462048 | 0.215991492 |

| | | |
|---|---|---|
| ENSMUSG00000024620 | 1.443888041 | 0.031701113 |
| ENSMUSG00000024621 | -1.148993699 | 0.571472291 |
| ENSMUSG00000024622 | 1.065931346 | 0.70299173 |
| ENSMUSG00000024639 | 1.112002261 | 0.64090177 |
| ENSMUSG00000024640 | 1.012133842 | 1 |
| ENSMUSG00000024642 | 1.22992506 | 0.071001672 |
| ENSMUSG00000024644 | -1.000967297 | 1 |
| ENSMUSG00000024645 | 1.024946434 | 0.896152472 |
| ENSMUSG00000024646 | 1.099631587 | 0.623851799 |
| ENSMUSG00000024647 | 1 | |
| ENSMUSG00000024650 | 1 | |
| ENSMUSG00000024653 | 1.059463094 | 0.608497229 |
| ENSMUSG00000024654 | -1.076855132 | 0.557811245 |
| ENSMUSG00000024658 | 1.196358003 | 0.55387742 |
| ENSMUSG00000024659 | 1.323448424 | 0.388977095 |
| ENSMUSG00000024660 | -1.009186098 | 1 |
| ENSMUSG00000024661 | 1.134057646 | 0.698112512 |
| ENSMUSG00000024663 | 1.26669603 | 0.013945157 |
| ENSMUSG00000024664 | 1.047258841 | 0.854127758 |
| ENSMUSG00000024665 | -1.006797234 | 1 |
| ENSMUSG00000024666 | 1.024586691 | 0.929405589 |
| ENSMUSG00000024667 | -1.023535159 | 0.934760809 |
| ENSMUSG00000024668 | -1.027170874 | 0.956784261 |
| ENSMUSG00000024669 | 1.095872691 | 0.608497229 |
| ENSMUSG00000024670 | 1.171563499 | 0.233412931 |
| ENSMUSG00000024672 | -1.052791364 | 0.890267277 |
| ENSMUSG00000024673 | -1.083520654 | 0.776953697 |
| ENSMUSG00000024675 | -1.03962336 | 0.698112512 |
| ENSMUSG00000024677 | 1 | |
| ENSMUSG00000024678 | 1.271141891 | 0.144329565 |
| ENSMUSG00000024679 | -1.084645496 | 0.532369657 |
| ENSMUSG00000024680 | -1.084645496 | 0.698112512 |
| ENSMUSG00000024681 | 1 | |
| ENSMUSG00000024682 | -1.095939461 | 0.682524927 |
| ENSMUSG00000024683 | 1.001720454 | 1 |
| ENSMUSG00000024687 | 1.03758305 | 0.927807415 |
| ENSMUSG00000024691 | 2.876064246 | 0.000102593 |
| ENSMUSG00000024694 | 1 | |
| ENSMUSG00000024695 | 1.030586759 | 0.969873524 |
| ENSMUSG00000024696 | -1.228648518 | 0.104270187 |
| ENSMUSG00000024697 | -1.288673043 | 0.115445674 |
| ENSMUSG00000024712 | -1.009561995 | 1 |
| ENSMUSG00000024713 | -2.206394638 | 1.70232E-05 |
| ENSMUSG00000024725 | 1.030252073 | 0.914862692 |
| ENSMUSG00000024726 | -1.003499565 | 1 |
| ENSMUSG00000024727 | -1.0789184 | 0.814384306 |
| ENSMUSG00000024728 | 1 | |
| ENSMUSG00000024729 | -1.044273782 | 0.698112512 |
| ENSMUSG00000024730 | 1 | |
| ENSMUSG00000024731 | -1 | 1 |
| ENSMUSG00000024732 | -1.014270638 | 0.941957289 |
| ENSMUSG00000024734 | -1.03962336 | 0.698112512 |
| ENSMUSG00000024735 | 1.038389142 | 0.943705943 |
| ENSMUSG00000024736 | 1.202400876 | 0.291424711 |
| ENSMUSG00000024737 | 1.304828368 | 0.311403906 |
| ENSMUSG00000024738 | 1.059463094 | 0.608497229 |
| ENSMUSG00000024740 | 1.085463725 | 0.849421383 |
| ENSMUSG00000024742 | 1.027122778 | 0.93979453 |
| ENSMUSG00000024743 | -1.320624849 | 0.032459896 |
| ENSMUSG00000024747 | 2.191638855 | 0.000346236 |
| ENSMUSG00000024749 | -1.044273782 | 0.698112512 |
| ENSMUSG00000024750 | 1.115203933 | 0.654902646 |
| ENSMUSG00000024754 | 1.097696186 | 0.608497229 |

| | | |
|---|---|---|
| ENSMUSG00000024757 | 1.084618945 | 0.698112512 |
| ENSMUSG00000024758 | 1.000250949 | 1 |
| ENSMUSG00000024759 | -1.003208267 | 1 |
| ENSMUSG00000024761 | 1.059463094 | 0.704170349 |
| ENSMUSG00000024764 | 1.050901336 | 0.826787852 |
| ENSMUSG00000024766 | -2.543971389 | 0.0001059 |
| ENSMUSG00000024767 | 1.026242487 | 0.95838544 |
| ENSMUSG00000024768 | 1 | |
| ENSMUSG00000024769 | 1.12232587 | 0.251544797 |
| ENSMUSG00000024770 | -1 | 1 |
| ENSMUSG00000024771 | -1.099174267 | 0.535052277 |
| ENSMUSG00000024772 | -1.024062579 | 0.953416829 |
| ENSMUSG00000024773 | 1.012195584 | 0.972980766 |
| ENSMUSG00000024774 | 1 | |
| ENSMUSG00000024776 | 1.039338894 | 0.800128289 |
| ENSMUSG00000024777 | -1.086453969 | 0.554252607 |
| ENSMUSG00000024778 | 1.408782656 | 0.017903363 |
| ENSMUSG00000024780 | 1.059311872 | 0.771903504 |
| ENSMUSG00000024781 | 1.065986379 | 0.80586932 |
| ENSMUSG00000024782 | 1.054637711 | 0.859839608 |
| ENSMUSG00000024784 | -1.090040692 | 0.837812805 |
| ENSMUSG00000024785 | 1.016358246 | 0.982175268 |
| ENSMUSG00000024786 | 1 | |
| ENSMUSG00000024787 | 1.116377746 | 0.162910566 |
| ENSMUSG00000024789 | -1.007451116 | 1 |
| ENSMUSG00000024790 | 1.067407153 | 0.59011989 |
| ENSMUSG00000024791 | -1.044875183 | 0.856445097 |
| ENSMUSG00000024792 | 1.090614416 | 0.496282519 |
| ENSMUSG00000024793 | 1.021884712 | 0.9780134 |
| ENSMUSG00000024795 | 1.042196596 | 0.925005828 |
| ENSMUSG00000024797 | 1.043702739 | 0.698112512 |
| ENSMUSG00000024798 | 1.028260703 | 0.934024342 |
| ENSMUSG00000024799 | -1.07737073 | 0.584587773 |
| ENSMUSG00000024800 | -1.062796371 | 0.803641195 |
| ENSMUSG00000024803 | 1.342222213 | 0.184749146 |
| ENSMUSG00000024805 | 1.098634458 | 0.608497229 |
| ENSMUSG00000024806 | 1.059463094 | 0.776953697 |
| ENSMUSG00000024807 | 1.073603604 | 0.546710863 |
| ENSMUSG00000024810 | 1.063819287 | 0.650046387 |
| ENSMUSG00000024811 | 1.050139172 | 0.917717739 |
| ENSMUSG00000024812 | -1.011044899 | 0.999883317 |
| ENSMUSG00000024815 | 1 | |
| ENSMUSG00000024816 | 1.080735782 | 0.511583996 |
| ENSMUSG00000024817 | 1.150185386 | 0.608497229 |
| ENSMUSG00000024818 | 1.085352749 | 0.76247781 |
| ENSMUSG00000024824 | 1.189674164 | 0.1003224 |
| ENSMUSG00000024826 | -1.053264279 | 0.816130675 |
| ENSMUSG00000024827 | -1.217872116 | 0.19964444 |
| ENSMUSG00000024829 | 1.022738217 | 0.929952088 |
| ENSMUSG00000024830 | -1.044062901 | 0.750067671 |
| ENSMUSG00000024831 | 1.069192862 | 0.527800513 |
| ENSMUSG00000024833 | -1.001946341 | 1 |
| ENSMUSG00000024835 | 1.122147752 | 0.588253297 |
| ENSMUSG00000024837 | -1.053549062 | 0.698112512 |
| ENSMUSG00000024841 | 1.086059292 | 0.698112512 |
| ENSMUSG00000024842 | 1.012540082 | 0.987291582 |
| ENSMUSG00000024843 | -1.066670038 | 0.717157933 |
| ENSMUSG00000024844 | 1.043276922 | 0.899860639 |
| ENSMUSG00000024845 | 1.060606963 | 0.706932054 |
| ENSMUSG00000024846 | 1.326491939 | 0.03679259 |
| ENSMUSG00000024847 | 1.03344258 | 0.870199809 |
| ENSMUSG00000024851 | 1.093164224 | 0.473553354 |
| ENSMUSG00000024853 | 1.035392639 | 0.941803539 |

| | | |
|---|---|---|
| ENSMUSG00000024854 | 1.104441438 | 0.110964535 |
| ENSMUSG00000024855 | -1.033390774 | 0.897684198 |
| ENSMUSG00000024856 | 1.076971942 | 0.608497229 |
| ENSMUSG00000024857 | 1 | |
| ENSMUSG00000024858 | 1.012891709 | 0.998794845 |
| ENSMUSG00000024862 | 1.028923095 | 0.859120996 |
| ENSMUSG00000024863 | 1 | |
| ENSMUSG00000024866 | -1.147035917 | 0.410661139 |
| ENSMUSG00000024867 | 1.122257272 | 0.421969918 |
| ENSMUSG00000024868 | 1.167088505 | 0.316625679 |
| ENSMUSG00000024869 | -1.064631696 | 0.686194514 |
| ENSMUSG00000024870 | -1.039461383 | 0.886090539 |
| ENSMUSG00000024871 | -1.06045819 | 0.812147631 |
| ENSMUSG00000024873 | -1.10171831 | 0.352642286 |
| ENSMUSG00000024875 | 1.101317288 | 0.549176447 |
| ENSMUSG00000024878 | -1.008303253 | 1 |
| ENSMUSG00000024883 | 1.06199244 | 0.667576553 |
| ENSMUSG00000024885 | -1.389700051 | 0.002539983 |
| ENSMUSG00000024887 | -1.018291252 | 0.971396058 |
| ENSMUSG00000024889 | -1.028222713 | 0.795446726 |
| ENSMUSG00000024891 | 1.093172558 | 0.156443748 |
| ENSMUSG00000024892 | -1.072908151 | 0.608497229 |
| ENSMUSG00000024896 | 1.05533553 | 0.793735215 |
| ENSMUSG00000024897 | 1.157855751 | 0.036626025 |
| ENSMUSG00000024899 | 1.031475528 | 0.886850795 |
| ENSMUSG00000024900 | 1.23250673 | 0.257433776 |
| ENSMUSG00000024901 | -1.014515724 | 0.978182137 |
| ENSMUSG00000024902 | -1.013211796 | 0.98408779 |
| ENSMUSG00000024903 | 1 | |
| ENSMUSG00000024905 | -1.038081156 | 0.853888273 |
| ENSMUSG00000024906 | 1.187358486 | 0.082663287 |
| ENSMUSG00000024907 | -1.217053278 | 0.608497229 |
| ENSMUSG00000024908 | 1.019865386 | 0.995955521 |
| ENSMUSG00000024909 | 1.125629824 | 0.563989684 |
| ENSMUSG00000024910 | 1.307291436 | 0.066036399 |
| ENSMUSG00000024911 | -1.009893118 | 0.99895137 |
| ENSMUSG00000024912 | -1.020905019 | 0.914446795 |
| ENSMUSG00000024913 | -1.146414248 | 0.280470274 |
| ENSMUSG00000024914 | 1.063985118 | 0.698112512 |
| ENSMUSG00000024921 | -1.124571405 | 0.584324822 |
| ENSMUSG00000024922 | 1.076553254 | 0.885192826 |
| ENSMUSG00000024924 | 2.352788354 | 0.000241554 |
| ENSMUSG00000024925 | 1.047270154 | 0.608497229 |
| ENSMUSG00000024926 | 1.039575452 | 0.794508734 |
| ENSMUSG00000024927 | 1.02408787 | 0.946232149 |
| ENSMUSG00000024935 | -1.089544887 | 0.738953156 |
| ENSMUSG00000024936 | -1.054169715 | 0.842733205 |
| ENSMUSG00000024937 | 1.276404696 | 0.056310079 |
| ENSMUSG00000024939 | 1.059463094 | 0.608497229 |
| ENSMUSG00000024940 | 1.20407859 | 0.092834572 |
| ENSMUSG00000024941 | 1.046060548 | 0.822190115 |
| ENSMUSG00000024942 | -1.032280716 | 0.883593753 |
| ENSMUSG00000024943 | 1.069676429 | 0.808053609 |
| ENSMUSG00000024944 | 1.101542221 | 0.477222649 |
| ENSMUSG00000024947 | 1.014432987 | 0.96035552 |
| ENSMUSG00000024948 | -1.099345308 | 0.552287505 |
| ENSMUSG00000024949 | 1.014751192 | 1 |
| ENSMUSG00000024952 | 1.092976283 | 0.359379869 |
| ENSMUSG00000024953 | 1.10180035 | 0.567860422 |
| ENSMUSG00000024955 | 1.152934177 | 0.405237049 |
| ENSMUSG00000024957 | 1.014545335 | 0.991944074 |
| ENSMUSG00000024958 | 1.004144822 | 1 |
| ENSMUSG00000024959 | 1.033157829 | 0.782658704 |

| | | |
|---|---|---|
| ENSMUSG00000024960 | 1.075631704 | 0.698112512 |
| ENSMUSG00000024962 | 1.236657067 | 0.012970042 |
| ENSMUSG00000024963 | 1.096539295 | 0.520525659 |
| ENSMUSG00000024965 | 1.063881788 | 0.770958539 |
| ENSMUSG00000024966 | -1.06679124 | 0.873511844 |
| ENSMUSG00000024968 | 1.128831669 | 0.224341885 |
| ENSMUSG00000024969 | 1.05418109 | 0.704892696 |
| ENSMUSG00000024970 | 1.081547163 | 0.608497229 |
| ENSMUSG00000024972 | 1 | |
| ENSMUSG00000024973 | 1 | |
| ENSMUSG00000024974 | 1.017686533 | 1 |
| ENSMUSG00000024975 | 1.016325937 | 0.978259071 |
| ENSMUSG00000024976 | 1.058966923 | 0.851465363 |
| ENSMUSG00000024978 | 1.021285329 | 0.934704406 |
| ENSMUSG00000024979 | 1 | |
| ENSMUSG00000024981 | -1.048645719 | 0.89915331 |
| ENSMUSG00000024982 | 1.029398267 | 0.947248675 |
| ENSMUSG00000024983 | 1.003785472 | 1 |
| ENSMUSG00000024985 | 1.052603903 | 0.71354136 |
| ENSMUSG00000024986 | -1.01060834 | 0.974891994 |
| ENSMUSG00000024987 | 1.12691148 | 0.608497229 |
| ENSMUSG00000024989 | -1.011757568 | 1 |
| ENSMUSG00000024990 | 1.017835844 | 0.996880111 |
| ENSMUSG00000024991 | 1.07951415 | 0.847073978 |
| ENSMUSG00000024992 | 1 | |
| ENSMUSG00000024993 | -1.006182915 | 1 |
| ENSMUSG00000024997 | -1.059817167 | 0.845289985 |
| ENSMUSG00000024998 | 1.438193114 | 0.004511839 |
| ENSMUSG00000024999 | 1.040016521 | 0.880481993 |
| ENSMUSG00000025001 | -1.003487235 | 1 |
| ENSMUSG00000025002 | -1.052119387 | 0.737739172 |
| ENSMUSG00000025003 | 1 | |
| ENSMUSG00000025004 | 1 | |
| ENSMUSG00000025006 | 1.486284218 | 0.002447579 |
| ENSMUSG00000025007 | 1.152268081 | 0.532586 |
| ENSMUSG00000025008 | 1.078336212 | 0.608497229 |
| ENSMUSG00000025010 | 1.02677457 | 0.879611226 |
| ENSMUSG00000025013 | -1.048277865 | 0.829945662 |
| ENSMUSG00000025014 | 1 | |
| ENSMUSG00000025016 | 1.010463597 | 1 |
| ENSMUSG00000025017 | -1.323363745 | 0.116410779 |
| ENSMUSG00000025019 | -1.047217528 | 0.777580731 |
| ENSMUSG00000025020 | -1.396807192 | 0.184582094 |
| ENSMUSG00000025024 | 1.017101919 | 0.995386171 |
| ENSMUSG00000025025 | -1.106999094 | 0.429887106 |
| ENSMUSG00000025026 | -1.157799355 | 0.324252919 |
| ENSMUSG00000025027 | 1.002422312 | 1 |
| ENSMUSG00000025034 | 1.10185323 | 0.379622425 |
| ENSMUSG00000025035 | -1.034943559 | 0.829303578 |
| ENSMUSG00000025036 | -1.077877868 | 0.448217312 |
| ENSMUSG00000025037 | 1.136289387 | 0.614648167 |
| ENSMUSG00000025038 | 1.089470641 | 0.657060635 |
| ENSMUSG00000025040 | 1.061199658 | 0.759891908 |
| ENSMUSG00000025041 | 1.015236037 | 0.996913409 |
| ENSMUSG00000025043 | 1 | |
| ENSMUSG00000025044 | 1.059463094 | 0.608497229 |
| ENSMUSG00000025047 | 1.032493143 | 0.926977462 |
| ENSMUSG00000025049 | -1.105656318 | 0.313693502 |
| ENSMUSG00000025050 | -1.105021889 | 0.608497229 |
| ENSMUSG00000025051 | 1 | |
| ENSMUSG00000025056 | 1 | |
| ENSMUSG00000025058 | -1.116428035 | 0.655462417 |
| ENSMUSG00000025059 | 1.055470562 | 0.785561203 |

| | | |
|---|---|---|
| ENSMUSG00000025060 | -1.017866675 | 1 |
| ENSMUSG00000025064 | 1.20697895 | 0.304375679 |
| ENSMUSG00000025066 | 1.045207057 | 0.905877432 |
| ENSMUSG00000025068 | 1.108663556 | 0.608497229 |
| ENSMUSG00000025069 | 1.329029583 | 0.072705092 |
| ENSMUSG00000025075 | 1.059463094 | 0.608497229 |
| ENSMUSG00000025076 | -1.17849427 | 0.351569054 |
| ENSMUSG00000025077 | 1.092219728 | 0.596276142 |
| ENSMUSG00000025078 | 1.05896089 | 0.738026665 |
| ENSMUSG00000025081 | -1 | 1 |
| ENSMUSG00000025082 | -1.623223524 | 0.003599328 |
| ENSMUSG00000025083 | -1.356696767 | 0.017759734 |
| ENSMUSG00000025085 | 1.063764332 | 0.770498527 |
| ENSMUSG00000025086 | 1.029856316 | 0.837038217 |
| ENSMUSG00000025089 | 1.63178712 | 0.010879816 |
| ENSMUSG00000025090 | 1 | |
| ENSMUSG00000025091 | -1 | 1 |
| ENSMUSG00000025092 | 1.225909281 | 0.012012138 |
| ENSMUSG00000025094 | 1.025358462 | 0.924804062 |
| ENSMUSG00000025102 | 1.095231521 | 0.059043319 |
| ENSMUSG00000025103 | -1.024290049 | 0.949812939 |
| ENSMUSG00000025104 | -1.014877498 | 0.993224268 |
| ENSMUSG00000025105 | -1.21672567 | 0.159807598 |
| ENSMUSG00000025127 | 1.033929369 | 0.966209176 |
| ENSMUSG00000025128 | 1 | |
| ENSMUSG00000025129 | 1.030415139 | 0.968937442 |
| ENSMUSG00000025130 | 1.07049236 | 0.870764638 |
| ENSMUSG00000025132 | 1.102984943 | 0.698361557 |
| ENSMUSG00000025133 | 1.015511502 | 0.993851713 |
| ENSMUSG00000025134 | -1.017542739 | 0.988424085 |
| ENSMUSG00000025135 | 1.121769869 | 0.294533546 |
| ENSMUSG00000025137 | -1.087634174 | 0.593695779 |
| ENSMUSG00000025138 | 1.010125233 | 0.969695497 |
| ENSMUSG00000025139 | 1.046949406 | 0.844158013 |
| ENSMUSG00000025140 | 1.245012628 | 0.029793193 |
| ENSMUSG00000025141 | -1 | 1 |
| ENSMUSG00000025142 | 1.045333638 | 0.712991717 |
| ENSMUSG00000025144 | -1.061745094 | 0.532369657 |
| ENSMUSG00000025145 | 1.06626562 | 0.642639207 |
| ENSMUSG00000025147 | 1.067952159 | 0.503647839 |
| ENSMUSG00000025150 | 2.400819385 | 0.000122514 |
| ENSMUSG00000025151 | 1.155474182 | 0.608497229 |
| ENSMUSG00000025153 | 1.110318533 | 0.608497229 |
| ENSMUSG00000025154 | -1.168559734 | 0.544317329 |
| ENSMUSG00000025155 | -1.041029837 | 0.80586932 |
| ENSMUSG00000025156 | 1.046362796 | 0.854494616 |
| ENSMUSG00000025157 | -1.0077613 | 1 |
| ENSMUSG00000025158 | -1.050851257 | 0.640707404 |
| ENSMUSG00000025159 | 1.05411525 | 0.804157993 |
| ENSMUSG00000025161 | -2.187138223 | 0.000228972 |
| ENSMUSG00000025162 | 1.068627941 | 0.774498429 |
| ENSMUSG00000025163 | 1 | |
| ENSMUSG00000025165 | -1.084645496 | 0.532369657 |
| ENSMUSG00000025169 | -1.114788422 | 0.318043984 |
| ENSMUSG00000025170 | -1.322016653 | 0.07645053 |
| ENSMUSG00000025171 | 1.230008129 | 0.116831796 |
| ENSMUSG00000025172 | 1.199989356 | 0.000021314 |
| ENSMUSG00000025173 | 1.010947565 | 1 |
| ENSMUSG00000025175 | -1.303715841 | 0.142601984 |
| ENSMUSG00000025176 | -1.083117507 | 0.368865011 |
| ENSMUSG00000025178 | -1.009759007 | 1 |
| ENSMUSG00000025184 | 1.021000059 | 0.966820366 |
| ENSMUSG00000025185 | -1.046048399 | 0.849862818 |

| | | |
|---|---|---|
| ENSMUSG00000025188 | -1.010407629 | 0.973521304 |
| ENSMUSG00000025189 | -1.200311323 | 0.191845319 |
| ENSMUSG00000025190 | 1.011193365 | 1 |
| ENSMUSG00000025192 | 1.036568241 | 0.805720656 |
| ENSMUSG00000025193 | -1.014321285 | 0.854777354 |
| ENSMUSG00000025194 | -1.003834322 | 1 |
| ENSMUSG00000025195 | 1.071622808 | 0.608497229 |
| ENSMUSG00000025196 | -1.044273782 | 0.698112512 |
| ENSMUSG00000025197 | 1.015486245 | 0.972980766 |
| ENSMUSG00000025198 | -1.080866613 | 0.769745556 |
| ENSMUSG00000025199 | 1.075652764 | 0.775269614 |
| ENSMUSG00000025200 | -1.011302963 | 0.998386582 |
| ENSMUSG00000025201 | 1 | |
| ENSMUSG00000025202 | 1.007298922 | 1 |
| ENSMUSG00000025203 | 1.02735905 | 0.988519386 |
| ENSMUSG00000025204 | -1.032087174 | 0.892812751 |
| ENSMUSG00000025207 | -1.166654542 | 0.06897581 |
| ENSMUSG00000025208 | 1.001009622 | 1 |
| ENSMUSG00000025209 | 1.122519932 | 0.370890497 |
| ENSMUSG00000025212 | 1.077317283 | 0.698112512 |
| ENSMUSG00000025213 | -1.120982322 | 0.055233089 |
| ENSMUSG00000025215 | -1.465437593 | 6.67169E-05 |
| ENSMUSG00000025216 | -1.109582661 | 0.549347972 |
| ENSMUSG00000025217 | -1.037463606 | 0.841209017 |
| ENSMUSG00000025218 | -1.029125166 | 0.785886031 |
| ENSMUSG00000025219 | -1.03962336 | 0.803072249 |
| ENSMUSG00000025220 | 1.074277578 | 0.790753571 |
| ENSMUSG00000025221 | 1.077658816 | 0.709356753 |
| ENSMUSG00000025223 | 1.188452167 | 0.183617727 |
| ENSMUSG00000025224 | 1.00477252 | 1 |
| ENSMUSG00000025225 | -1.081138678 | 0.698112512 |
| ENSMUSG00000025226 | 1.006997351 | 1 |
| ENSMUSG00000025227 | -1.069373894 | 0.70299173 |
| ENSMUSG00000025228 | 1.029213591 | 0.962886319 |
| ENSMUSG00000025229 | -1.084645496 | 0.628300894 |
| ENSMUSG00000025231 | 1.039457585 | 0.837321292 |
| ENSMUSG00000025232 | 1.355963337 | 0.059227853 |
| ENSMUSG00000025234 | 1.020623824 | 0.986542466 |
| ENSMUSG00000025235 | -1.07470012 | 0.608497229 |
| ENSMUSG00000025236 | -1.010711929 | 1 |
| ENSMUSG00000025237 | 1.02650171 | 0.890533327 |
| ENSMUSG00000025239 | 1.00962891 | 1 |
| ENSMUSG00000025240 | -1.064887431 | 0.811222342 |
| ENSMUSG00000025241 | 1.024634232 | 0.915070045 |
| ENSMUSG00000025243 | -1.111221915 | 0.608497229 |
| ENSMUSG00000025245 | -1.031309835 | 0.870231634 |
| ENSMUSG00000025246 | 1.114604242 | 0.697167887 |
| ENSMUSG00000025255 | -1.081438888 | 0.467178083 |
| ENSMUSG00000025257 | -1.126235585 | 0.659841524 |
| ENSMUSG00000025260 | -1.093462818 | 0.642256709 |
| ENSMUSG00000025261 | -1.067949461 | 0.866675229 |
| ENSMUSG00000025262 | -1.234437185 | 0.113598345 |
| ENSMUSG00000025264 | -1.168103163 | 0.375734352 |
| ENSMUSG00000025265 | -1.001978181 | 1 |
| ENSMUSG00000025266 | -1.138542077 | 0.608497229 |
| ENSMUSG00000025268 | 1.242452975 | 0.159653354 |
| ENSMUSG00000025269 | -1.075283001 | 0.686622012 |
| ENSMUSG00000025270 | 1.040557558 | 0.895127848 |
| ENSMUSG00000025271 | -1.087631644 | 0.38417754 |
| ENSMUSG00000025272 | -1.401488066 | 0.010006001 |
| ENSMUSG00000025277 | -1.178954869 | 0.251049598 |
| ENSMUSG00000025278 | 1.069971341 | 0.837704961 |
| ENSMUSG00000025279 | 1.154253937 | 0.277459955 |

| | | |
|---|---|---|
| ENSMUSG00000025280 | -1.097518285 | 0.608497229 |
| ENSMUSG00000025283 | 1.069757493 | 0.734819177 |
| ENSMUSG00000025287 | -1.142101072 | 0.608497229 |
| ENSMUSG00000025288 | 1 | |
| ENSMUSG00000025289 | -1.048252743 | 0.865607861 |
| ENSMUSG00000025290 | -1.127745 | 0.698112512 |
| ENSMUSG00000025314 | -1.088258773 | 0.608497229 |
| ENSMUSG00000025316 | 1.085566866 | 0.375254184 |
| ENSMUSG00000025317 | -1.030440399 | 0.981620431 |
| ENSMUSG00000025318 | -1.111705722 | 0.686194514 |
| ENSMUSG00000025321 | 1.516314002 | 0.002390679 |
| ENSMUSG00000025323 | 1.080715703 | 0.554762219 |
| ENSMUSG00000025324 | 1.111219213 | 0.124275535 |
| ENSMUSG00000025326 | 1.016414824 | 1 |
| ENSMUSG00000025328 | -1.018292245 | 0.979002724 |
| ENSMUSG00000025329 | -1 | 1 |
| ENSMUSG00000025330 | 1 | |
| ENSMUSG00000025332 | -1.068154693 | 0.750464541 |
| ENSMUSG00000025333 | -1.043306199 | 0.698112512 |
| ENSMUSG00000025337 | 1.094612668 | 0.608497229 |
| ENSMUSG00000025340 | 1.145671725 | 0.388393823 |
| ENSMUSG00000025347 | -1.043306199 | 0.79291651 |
| ENSMUSG00000025348 | -1.054598905 | 0.6973511 |
| ENSMUSG00000025350 | 1.250379363 | 0.008951509 |
| ENSMUSG00000025351 | 1.155866556 | 0.608497229 |
| ENSMUSG00000025352 | -1.093402178 | 0.61620788 |
| ENSMUSG00000025353 | -1.062657752 | 0.681190723 |
| ENSMUSG00000025354 | 1.000344819 | 1 |
| ENSMUSG00000025355 | 1.484099214 | 0.003735566 |
| ENSMUSG00000025357 | -1.13630173 | 0.223995462 |
| ENSMUSG00000025358 | -1.075795156 | 0.772179799 |
| ENSMUSG00000025359 | -1.147748038 | 0.385934169 |
| ENSMUSG00000025362 | 1.094032347 | 0.608556438 |
| ENSMUSG00000025364 | -1.055177895 | 0.894591069 |
| ENSMUSG00000025366 | 1.024087198 | 0.960812739 |
| ENSMUSG00000025369 | -1.036059177 | 0.924061737 |
| ENSMUSG00000025370 | 1 | |
| ENSMUSG00000025371 | 1.077439753 | 0.553156258 |
| ENSMUSG00000025372 | 1.139566729 | 0.253189308 |
| ENSMUSG00000025373 | 1.088708994 | 0.35863794 |
| ENSMUSG00000025374 | -1.106186971 | 0.573420551 |
| ENSMUSG00000025375 | 1.007462988 | 1 |
| ENSMUSG00000025377 | 1.053136768 | 0.647969257 |
| ENSMUSG00000025380 | 1.178861698 | 0.647921523 |
| ENSMUSG00000025381 | 1.036147295 | 0.889728438 |
| ENSMUSG00000025383 | -1.155222017 | 0.608497229 |
| ENSMUSG00000025384 | 1.223341368 | 0.115583265 |
| ENSMUSG00000025386 | 1.092855464 | 0.325380902 |
| ENSMUSG00000025389 | 1.019083579 | 0.968937442 |
| ENSMUSG00000025393 | -1.016363211 | 1 |
| ENSMUSG00000025395 | -1.078077615 | 0.773508225 |
| ENSMUSG00000025396 | 1 | |
| ENSMUSG00000025400 | 1 | |
| ENSMUSG00000025401 | 1.129836239 | 0.496377492 |
| ENSMUSG00000025402 | 1.278640849 | 0.008641609 |
| ENSMUSG00000025403 | 1.102572124 | 0.595042751 |
| ENSMUSG00000025404 | -1.012979718 | 0.997108523 |
| ENSMUSG00000025405 | 1 | |
| ENSMUSG00000025407 | -1.450557783 | 0.043159855 |
| ENSMUSG00000025408 | 1.325486383 | 0.007291063 |
| ENSMUSG00000025409 | 1.113629923 | 0.266019054 |
| ENSMUSG00000025410 | 1.010780113 | 1 |
| ENSMUSG00000025413 | -1.073321879 | 0.729018288 |

| | | |
|---|---|---|
| ENSMUSG00000025417 | 1.058121804 | 0.782393128 |
| ENSMUSG00000025418 | 1 | |
| ENSMUSG00000025420 | 1.003529192 | 1 |
| ENSMUSG00000025421 | 1.067110598 | 0.717035356 |
| ENSMUSG00000025422 | -1.520274025 | 0.082216669 |
| ENSMUSG00000025423 | 1.091892631 | 0.698112512 |
| ENSMUSG00000025425 | 1 | |
| ENSMUSG00000025427 | -1.530759152 | 0.024820889 |
| ENSMUSG00000025428 | 1.019084083 | 1 |
| ENSMUSG00000025429 | 1.018026194 | 0.889813301 |
| ENSMUSG00000025431 | 1 | |
| ENSMUSG00000025432 | 1.06174949 | 0.713806698 |
| ENSMUSG00000025433 | 1 | |
| ENSMUSG00000025436 | -1.069334369 | 0.35312092 |
| ENSMUSG00000025437 | -1.060596223 | 0.762748967 |
| ENSMUSG00000025439 | 1.008166644 | 1 |
| ENSMUSG00000025450 | -1.037198067 | 0.888757001 |
| ENSMUSG00000025451 | -1.093895495 | 0.717035356 |
| ENSMUSG00000025453 | 1.014428379 | 1 |
| ENSMUSG00000025461 | 1 | |
| ENSMUSG00000025464 | 1.121823735 | 0.229745553 |
| ENSMUSG00000025465 | 1.119715194 | 0.393150867 |
| ENSMUSG00000025466 | 1.495469574 | 0.003431136 |
| ENSMUSG00000025467 | 1 | |
| ENSMUSG00000025468 | 1 | |
| ENSMUSG00000025469 | -1.129231493 | 0.605619163 |
| ENSMUSG00000025470 | -1.060639728 | 0.576603197 |
| ENSMUSG00000025473 | -1.119307034 | 0.361485097 |
| ENSMUSG00000025474 | 1.094784867 | 0.608497229 |
| ENSMUSG00000025475 | 1.411556877 | 0.133076586 |
| ENSMUSG00000025477 | 1.112557155 | 0.295346574 |
| ENSMUSG00000025478 | -1.899729883 | 0.003996405 |
| ENSMUSG00000025479 | 1.007477354 | 1 |
| ENSMUSG00000025480 | 1 | |
| ENSMUSG00000025481 | -1.044273782 | 0.801443616 |
| ENSMUSG00000025482 | -1 | 1 |
| ENSMUSG00000025484 | 1.060830029 | 0.608497229 |
| ENSMUSG00000025485 | 1.042982395 | 0.705199845 |
| ENSMUSG00000025486 | 1.008436664 | 0.977192639 |
| ENSMUSG00000025487 | 1.002397608 | 1 |
| ENSMUSG00000025488 | 1 | |
| ENSMUSG00000025489 | 1.035156016 | 0.973126641 |
| ENSMUSG00000025491 | -1.484180972 | 0.002092992 |
| ENSMUSG00000025492 | 1.005593466 | 1 |
| ENSMUSG00000025494 | -1.067791758 | 0.821106 |
| ENSMUSG00000025495 | -1.075452411 | 0.608497229 |
| ENSMUSG00000025496 | -1.084645496 | 0.532369657 |
| ENSMUSG00000025497 | -1.043306199 | 0.698112512 |
| ENSMUSG00000025498 | -1.222838501 | 0.378431826 |
| ENSMUSG00000025499 | -1.032584083 | 0.856445097 |
| ENSMUSG00000025500 | -1.147502257 | 0.641410989 |
| ENSMUSG00000025503 | 1.024315214 | 0.885529952 |
| ENSMUSG00000025504 | -1.120476206 | 0.568311292 |
| ENSMUSG00000025505 | -1.046727278 | 0.451754539 |
| ENSMUSG00000025507 | 1.142822597 | 0.28024635 |
| ENSMUSG00000025508 | 1.123842568 | 0.405411757 |
| ENSMUSG00000025509 | -1.062817818 | 0.636222788 |
| ENSMUSG00000025510 | 1.133716214 | 0.471811361 |
| ENSMUSG00000025511 | 1.100747315 | 0.480007964 |
| ENSMUSG00000025512 | 1.002596581 | 1 |
| ENSMUSG00000025515 | -1.263042605 | 0.481246252 |
| ENSMUSG00000025519 | 1 | |
| ENSMUSG00000025521 | 1.017424721 | 0.964995983 |

| | | |
|---|---|---|
| ENSMUSG00000025525 | -1.05857909 | 0.705940113 |
| ENSMUSG00000025527 | -1.176660386 | 0.300576589 |
| ENSMUSG00000025528 | 1 | |
| ENSMUSG00000025529 | -1 | 1 |
| ENSMUSG00000025531 | -1.090670788 | 0.698112512 |
| ENSMUSG00000025532 | 1.019866125 | 0.954466467 |
| ENSMUSG00000025533 | -1.045194798 | 0.785059168 |
| ENSMUSG00000025534 | 1.142772049 | 0.604163646 |
| ENSMUSG00000025537 | -1.043984903 | 0.780219261 |
| ENSMUSG00000025538 | 1.175614009 | 0.026983666 |
| ENSMUSG00000025544 | -1.147469551 | 0.636035234 |
| ENSMUSG00000025545 | -1.17483195 | 0.134790095 |
| ENSMUSG00000025551 | 1 | |
| ENSMUSG00000025555 | -1.051932743 | 0.842404015 |
| ENSMUSG00000025557 | -1.224792133 | 0.374379691 |
| ENSMUSG00000025558 | -1.161081514 | 0.29974472 |
| ENSMUSG00000025571 | 1.03392644 | 0.871696426 |
| ENSMUSG00000025572 | -1.114494592 | 0.425606078 |
| ENSMUSG00000025573 | 1.059463094 | 0.698112512 |
| ENSMUSG00000025574 | 1.070053677 | 0.742727573 |
| ENSMUSG00000025575 | 1.059755047 | 0.69104961 |
| ENSMUSG00000025576 | 1 | |
| ENSMUSG00000025577 | 1.073926087 | 0.586769798 |
| ENSMUSG00000025578 | 1.112163584 | 0.180644327 |
| ENSMUSG00000025579 | 1.236027746 | 0.111576748 |
| ENSMUSG00000025580 | 1.005510497 | 1 |
| ENSMUSG00000025582 | -1.445695956 | 0.066745907 |
| ENSMUSG00000025583 | 1.071187132 | 0.698112512 |
| ENSMUSG00000025584 | -1.022088058 | 0.88617786 |
| ENSMUSG00000025586 | 1.413948396 | 0.003453741 |
| ENSMUSG00000025588 | -1.044273782 | 0.698112512 |
| ENSMUSG00000025591 | 1.025439042 | 0.885895888 |
| ENSMUSG00000025592 | 1.132639128 | 0.217803313 |
| ENSMUSG00000025597 | 1 | |
| ENSMUSG00000025602 | 1.006556238 | 0.970124054 |
| ENSMUSG00000025607 | 1.024963238 | 0.95702926 |
| ENSMUSG00000025608 | 1.092032608 | 0.453446164 |
| ENSMUSG00000025609 | -1.019475369 | 0.988924922 |
| ENSMUSG00000025610 | -1.165852623 | 0.541289429 |
| ENSMUSG00000025612 | 1.121015811 | 0.544344891 |
| ENSMUSG00000025613 | 1.012181273 | 1 |
| ENSMUSG00000025616 | 1.011931601 | 1 |
| ENSMUSG00000025626 | 1.011935284 | 1 |
| ENSMUSG00000025630 | -1.008484548 | 1 |
| ENSMUSG00000025644 | -1.066303125 | 0.827942378 |
| ENSMUSG00000025645 | 1.028577319 | 0.608497229 |
| ENSMUSG00000025646 | -1.033057782 | 0.855186894 |
| ENSMUSG00000025647 | -1.022001613 | 0.954958649 |
| ENSMUSG00000025648 | 1.016623737 | 0.954579929 |
| ENSMUSG00000025650 | 1.052466835 | 0.689572034 |
| ENSMUSG00000025651 | -1.005842285 | 1 |
| ENSMUSG00000025652 | 1 | |
| ENSMUSG00000025656 | 1.045480589 | 0.761985238 |
| ENSMUSG00000025658 | 1.416838559 | 0.033686074 |
| ENSMUSG00000025665 | -1.118761334 | 0.595536024 |
| ENSMUSG00000025666 | 1.799349593 | 0.002175591 |
| ENSMUSG00000025701 | -1.171710767 | 0.652300235 |
| ENSMUSG00000025702 | 1.178872968 | 0.372103656 |
| ENSMUSG00000025716 | 1 | |
| ENSMUSG00000025722 | 1.00265928 | 1 |
| ENSMUSG00000025723 | -1.217300946 | 0.159696685 |
| ENSMUSG00000025724 | 1.020257501 | 0.967310189 |
| ENSMUSG00000025726 | 1.059463094 | 0.608497229 |

| | | |
|---|---|---|
| ENSMUSG00000025727 | -1.149484141 | 0.61897445 |
| ENSMUSG00000025728 | -1.017896839 | 0.976815801 |
| ENSMUSG00000025730 | 1.123478515 | 0.212569102 |
| ENSMUSG00000025731 | -1.03789521 | 0.698112512 |
| ENSMUSG00000025732 | 1.046234347 | 0.734645261 |
| ENSMUSG00000025733 | -1.050744059 | 0.608497229 |
| ENSMUSG00000025735 | 1.216211692 | 0.376759062 |
| ENSMUSG00000025736 | -1.071495195 | 0.543489251 |
| ENSMUSG00000025737 | 1.023598361 | 0.827836364 |
| ENSMUSG00000025738 | 1.045774342 | 0.708359577 |
| ENSMUSG00000025739 | 1.003517909 | 1 |
| ENSMUSG00000025742 | -1.108154836 | 0.648436514 |
| ENSMUSG00000025743 | 1.453009435 | 0.008618666 |
| ENSMUSG00000025745 | 1.042518984 | 0.900965317 |
| ENSMUSG00000025746 | 1.537790943 | 0.071677212 |
| ENSMUSG00000025747 | 1.031540512 | 0.943858405 |
| ENSMUSG00000025754 | 1 | |
| ENSMUSG00000025757 | -1.030493779 | 0.959463747 |
| ENSMUSG00000025758 | -1.016662483 | 0.997227391 |
| ENSMUSG00000025759 | -1.028169411 | 0.9079117 |
| ENSMUSG00000025762 | 1.034770036 | 0.697178241 |
| ENSMUSG00000025764 | 1.391993023 | 0.044718571 |
| ENSMUSG00000025766 | -1.06319694 | 0.580870462 |
| ENSMUSG00000025774 | -1 | 1 |
| ENSMUSG00000025776 | -1.03962336 | 0.803072249 |
| ENSMUSG00000025777 | -1.043415084 | 0.69491764 |
| ENSMUSG00000025779 | 1.126036892 | 0.496647257 |
| ENSMUSG00000025780 | -1.175328236 | 0.170767215 |
| ENSMUSG00000025781 | 1.012238425 | 1 |
| ENSMUSG00000025782 | -1.020455971 | 0.948533308 |
| ENSMUSG00000025783 | 1.12189305 | 0.608497229 |
| ENSMUSG00000025784 | -1.010960635 | 1 |
| ENSMUSG00000025785 | -1.027417388 | 0.928180271 |
| ENSMUSG00000025786 | -1.030858025 | 0.954424095 |
| ENSMUSG00000025787 | 1.019826425 | 0.989636119 |
| ENSMUSG00000025789 | -1.323427299 | 0.041814519 |
| ENSMUSG00000025790 | 1.224925994 | 0.019043897 |
| ENSMUSG00000025791 | 1.032481177 | 0.902811791 |
| ENSMUSG00000025792 | 1.030071799 | 0.858911269 |
| ENSMUSG00000025793 | 1.033330053 | 0.842184576 |
| ENSMUSG00000025794 | 1.000581345 | 1 |
| ENSMUSG00000025795 | -1.074183272 | 0.66440866 |
| ENSMUSG00000025804 | 1 | |
| ENSMUSG00000025808 | 1 | |
| ENSMUSG00000025809 | 1.110178355 | 0.783637632 |
| ENSMUSG00000025810 | 1.342954081 | 0.087200971 |
| ENSMUSG00000025812 | -1.051742416 | 0.813545446 |
| ENSMUSG00000025813 | -1.673818862 | 0.004890823 |
| ENSMUSG00000025815 | 1.131687392 | 0.32059496 |
| ENSMUSG00000025816 | -1.01976622 | 0.978763926 |
| ENSMUSG00000025817 | 1.01297102 | 0.998601098 |
| ENSMUSG00000025821 | 1.010858798 | 0.970855463 |
| ENSMUSG00000025823 | 1.006660765 | 1 |
| ENSMUSG00000025825 | 1.086598264 | 0.519083776 |
| ENSMUSG00000025838 | 1.069068979 | 0.74773997 |
| ENSMUSG00000025839 | 1.059463094 | 0.698112512 |
| ENSMUSG00000025854 | 1.22272896 | 0.013667037 |
| ENSMUSG00000025855 | -1.103940444 | 0.441459973 |
| ENSMUSG00000025856 | 1.301178043 | 0.002362668 |
| ENSMUSG00000025857 | 1.06800542 | 0.439929645 |
| ENSMUSG00000025858 | 1.070222993 | 0.661883023 |
| ENSMUSG00000025860 | 1.02470713 | 0.979063432 |
| ENSMUSG00000025862 | 1.042868512 | 0.931368384 |

| | | |
|---|---|---|
| ENSMUSG00000025867 | -1.045809882 | 0.655571244 |
| ENSMUSG00000025868 | -1.093036938 | 0.432344545 |
| ENSMUSG00000025869 | -1.08353442 | 0.615683483 |
| ENSMUSG00000025870 | 1.040628003 | 0.720991126 |
| ENSMUSG00000025871 | -1.095303575 | 0.698112512 |
| ENSMUSG00000025872 | -1.132019219 | 0.59127359 |
| ENSMUSG00000025873 | -1.091637439 | 0.71324469 |
| ENSMUSG00000025875 | 1.117931033 | 0.149572521 |
| ENSMUSG00000025876 | -1.08081673 | 0.70510908 |
| ENSMUSG00000025877 | 1.059463094 | 0.608497229 |
| ENSMUSG00000025878 | -1.079014383 | 0.698112512 |
| ENSMUSG00000025880 | 1.252284396 | 0.032803185 |
| ENSMUSG00000025885 | -1.008057816 | 0.997108523 |
| ENSMUSG00000025887 | -1.108323971 | 0.651358924 |
| ENSMUSG00000025888 | -1.053549062 | 0.774498429 |
| ENSMUSG00000025889 | 1.926855615 | 0.000641345 |
| ENSMUSG00000025892 | 1.028165331 | 0.965762892 |
| ENSMUSG00000025893 | -1.007655186 | 1 |
| ENSMUSG00000025894 | -1.011768313 | 0.998352825 |
| ENSMUSG00000025898 | -1.081087333 | 0.681952832 |
| ENSMUSG00000025899 | -1.036630528 | 0.889575988 |
| ENSMUSG00000025900 | 1 | |
| ENSMUSG00000025902 | 1 | |
| ENSMUSG00000025903 | -1.101577486 | 0.698112512 |
| ENSMUSG00000025905 | 1 | |
| ENSMUSG00000025907 | 1.165852202 | 0.54014214 |
| ENSMUSG00000025909 | 1 | |
| ENSMUSG00000025911 | -1.034461609 | 0.910166397 |
| ENSMUSG00000025912 | -1.033702285 | 0.845679504 |
| ENSMUSG00000025915 | 1.001509729 | 1 |
| ENSMUSG00000025916 | -1.153946192 | 0.648096477 |
| ENSMUSG00000025917 | -1.059825914 | 0.84360497 |
| ENSMUSG00000025920 | -1.037385645 | 0.865865121 |
| ENSMUSG00000025921 | -1.069182227 | 0.655832783 |
| ENSMUSG00000025922 | 1.036587445 | 0.958979658 |
| ENSMUSG00000025925 | -1.020588011 | 0.944721414 |
| ENSMUSG00000025927 | 1.132639128 | 0.568304986 |
| ENSMUSG00000025929 | 1 | |
| ENSMUSG00000025930 | -1.065321276 | 0.814089245 |
| ENSMUSG00000025931 | 1.133576275 | 0.231320305 |
| ENSMUSG00000025932 | 1.004721789 | 1 |
| ENSMUSG00000025933 | -1.204626214 | 0.124474599 |
| ENSMUSG00000025934 | 1.188604282 | 0.608497229 |
| ENSMUSG00000025935 | -1.022880012 | 0.979212363 |
| ENSMUSG00000025936 | 1 | |
| ENSMUSG00000025937 | 1.031753635 | 0.868791781 |
| ENSMUSG00000025938 | -1.344539942 | 0.097043927 |
| ENSMUSG00000025939 | 1.02131933 | 0.969506516 |
| ENSMUSG00000025940 | 1.020565391 | 0.979002724 |
| ENSMUSG00000025945 | 1 | |
| ENSMUSG00000025946 | 1.050110043 | 0.767772211 |
| ENSMUSG00000025949 | -1.004689762 | 1 |
| ENSMUSG00000025950 | 1.026767792 | 0.950331262 |
| ENSMUSG00000025952 | 1 | |
| ENSMUSG00000025955 | -1.133717313 | 0.608497229 |
| ENSMUSG00000025956 | 1.105103217 | 0.556811875 |
| ENSMUSG00000025958 | -1.003694819 | 1 |
| ENSMUSG00000025959 | 1.335878997 | 0.009008372 |
| ENSMUSG00000025961 | -1.038741809 | 0.917717739 |
| ENSMUSG00000025962 | 1.014118861 | 0.992580079 |
| ENSMUSG00000025963 | 1.132639128 | 0.701586391 |
| ENSMUSG00000025964 | -1.005080108 | 1 |
| ENSMUSG00000025967 | 1.034101954 | 0.959757321 |

| | | |
|---|---|---|
| ENSMUSG00000025968 | -1.038523442 | 0.93400515 |
| ENSMUSG00000025969 | 1.21211675 | 0.424899423 |
| ENSMUSG00000025971 | -1.101451229 | 0.499935975 |
| ENSMUSG00000025977 | -1.013449573 | 1 |
| ENSMUSG00000025978 | 1.076064787 | 0.279927732 |
| ENSMUSG00000025979 | -1.02576358 | 0.9607713 |
| ENSMUSG00000025980 | -1.002573887 | 1 |
| ENSMUSG00000025981 | -1.057306553 | 0.776953697 |
| ENSMUSG00000025982 | 1.027588552 | 0.983829376 |
| ENSMUSG00000025983 | 1.051728433 | 0.762289856 |
| ENSMUSG00000025986 | 1.019781285 | 0.980617339 |
| ENSMUSG00000025991 | -1.085651418 | 0.625496737 |
| ENSMUSG00000025993 | 1.687367919 | 0.006294552 |
| ENSMUSG00000025995 | 1.059332213 | 0.828391261 |
| ENSMUSG00000025997 | 1.056733148 | 0.720501596 |
| ENSMUSG00000026000 | 1.076807119 | 0.684879215 |
| ENSMUSG00000026003 | 1.070757584 | 0.767574573 |
| ENSMUSG00000026004 | 1.235521075 | 0.05916082 |
| ENSMUSG00000026005 | -1.01779951 | 0.981432083 |
| ENSMUSG00000026009 | 1.058036848 | 0.86264687 |
| ENSMUSG00000026011 | -1.053549062 | 0.698112512 |
| ENSMUSG00000026012 | 1.034368904 | 0.93376866 |
| ENSMUSG00000026014 | -1.021961898 | 0.972473428 |
| ENSMUSG00000026017 | -1.083810067 | 0.608497229 |
| ENSMUSG00000026018 | -1.324162085 | 0.001127478 |
| ENSMUSG00000026019 | 1.005123845 | 1 |
| ENSMUSG00000026020 | -1.015179732 | 1 |
| ENSMUSG00000026021 | -1.045599803 | 0.898948667 |
| ENSMUSG00000026023 | 1.314368947 | 0.197152281 |
| ENSMUSG00000026024 | 1.085575653 | 0.698112512 |
| ENSMUSG00000026027 | -1.035837079 | 0.748295996 |
| ENSMUSG00000026028 | -1.050891687 | 0.854494616 |
| ENSMUSG00000026029 | -1.035807435 | 0.889802291 |
| ENSMUSG00000026031 | 1.023978314 | 0.966570161 |
| ENSMUSG00000026032 | -1.06368318 | 0.708068647 |
| ENSMUSG00000026034 | -1.034694148 | 0.91711909 |
| ENSMUSG00000026035 | -1.012752895 | 0.992095533 |
| ENSMUSG00000026036 | -1.090699233 | 0.655851508 |
| ENSMUSG00000026037 | -1.033932045 | 0.933031586 |
| ENSMUSG00000026039 | -1.073302543 | 0.691462757 |
| ENSMUSG00000026042 | -1.08226287 | 0.781435694 |
| ENSMUSG00000026043 | 2.351644489 | 0.000480694 |
| ENSMUSG00000026047 | 1.001241167 | 1 |
| ENSMUSG00000026048 | -1.002311004 | 1 |
| ENSMUSG00000026049 | -1.018036225 | 0.977413596 |
| ENSMUSG00000026051 | -1.100841795 | 0.760912081 |
| ENSMUSG00000026058 | 1.060715927 | 0.86902396 |
| ENSMUSG00000026062 | -1.485170677 | 0.020664923 |
| ENSMUSG00000026063 | 1 | |
| ENSMUSG00000026064 | 1.123596247 | 0.661501071 |
| ENSMUSG00000026065 | 1 | |
| ENSMUSG00000026068 | 1.221826405 | 0.010607881 |
| ENSMUSG00000026069 | 1.732924489 | 0.022495042 |
| ENSMUSG00000026070 | -1.145866846 | 0.690648064 |
| ENSMUSG00000026072 | 1.0398669 | 0.840860016 |
| ENSMUSG00000026073 | -1.030858705 | 0.9607713 |
| ENSMUSG00000026074 | 1.012499006 | 1 |
| ENSMUSG00000026077 | -1.233948724 | 0.397706863 |
| ENSMUSG00000026078 | 1.032897378 | 0.938931916 |
| ENSMUSG00000026080 | -1.077482697 | 0.608497229 |
| ENSMUSG00000026082 | 1.063027933 | 0.765464893 |
| ENSMUSG00000026083 | -1.02267684 | 0.983837837 |
| ENSMUSG00000026085 | 1 | |

| | | |
|---|---|---|
| ENSMUSG00000026087 | -1.018761424 | 0.981626226 |
| ENSMUSG00000026088 | -1.152422376 | 0.266163985 |
| ENSMUSG00000026090 | -1.289385313 | 0.061225644 |
| ENSMUSG00000026094 | 1.135730127 | 0.608497229 |
| ENSMUSG00000026095 | 1.016481595 | 0.97479937 |
| ENSMUSG00000026096 | -1.035222383 | 0.826758538 |
| ENSMUSG00000026097 | -1.033467982 | 0.893765795 |
| ENSMUSG00000026098 | -1.017277726 | 0.968937442 |
| ENSMUSG00000026100 | -1 | 1 |
| ENSMUSG00000026102 | -1.106597573 | 0.58386781 |
| ENSMUSG00000026103 | 1.034675778 | 0.954310937 |
| ENSMUSG00000026104 | -1.061749909 | 0.718565659 |
| ENSMUSG00000026107 | 1.143076157 | 0.381678195 |
| ENSMUSG00000026109 | 1.27365104 | 0.068119685 |
| ENSMUSG00000026110 | -1.511871474 | 0.001850311 |
| ENSMUSG00000026111 | -1.023965354 | 0.928822702 |
| ENSMUSG00000026112 | -1.005775574 | 1 |
| ENSMUSG00000026113 | -1.039565734 | 0.759784795 |
| ENSMUSG00000026114 | 1.093819733 | 0.706051768 |
| ENSMUSG00000026116 | -1.044888056 | 0.886986912 |
| ENSMUSG00000026117 | -1.11061171 | 0.547016491 |
| ENSMUSG00000026121 | 1.084029714 | 0.527231267 |
| ENSMUSG00000026123 | 1.129227964 | 0.60397857 |
| ENSMUSG00000026124 | 1 | |
| ENSMUSG00000026125 | 1 | |
| ENSMUSG00000026126 | -1.2573953 | 0.205071867 |
| ENSMUSG00000026127 | 1.005085987 | 1 |
| ENSMUSG00000026131 | -1.025928208 | 0.858124279 |
| ENSMUSG00000026134 | -1.086078018 | 0.698112512 |
| ENSMUSG00000026135 | 1.044108362 | 0.780431248 |
| ENSMUSG00000026141 | -1.083415624 | 0.706213769 |
| ENSMUSG00000026142 | -1.063122751 | 0.742268585 |
| ENSMUSG00000026147 | 1 | |
| ENSMUSG00000026149 | -1.01566718 | 1 |
| ENSMUSG00000026150 | 1.003297741 | 1 |
| ENSMUSG00000026153 | -1.121417969 | 0.608497229 |
| ENSMUSG00000026154 | 1.043789566 | 0.778376464 |
| ENSMUSG00000026155 | -1.015904975 | 0.96035552 |
| ENSMUSG00000026156 | -1.030606775 | 0.922343833 |
| ENSMUSG00000026158 | -1.226986808 | 0.132707257 |
| ENSMUSG00000026159 | 1.005345643 | 1 |
| ENSMUSG00000026162 | -1.122507534 | 0.342459489 |
| ENSMUSG00000026163 | 1.059463094 | 0.608497229 |
| ENSMUSG00000026166 | -1.352821423 | 0.146687451 |
| ENSMUSG00000026167 | 1.124952944 | 0.251445232 |
| ENSMUSG00000026170 | -1.375260084 | 0.010841524 |
| ENSMUSG00000026171 | -1.065367112 | 0.662710277 |
| ENSMUSG00000026172 | 1.034087772 | 0.816814078 |
| ENSMUSG00000026173 | 1.120639616 | 0.60397857 |
| ENSMUSG00000026174 | -1.009206152 | 1 |
| ENSMUSG00000026175 | -1.011526474 | 1 |
| ENSMUSG00000026176 | 1.096169729 | 0.595296772 |
| ENSMUSG00000026177 | 1.441876717 | 0.001922135 |
| ENSMUSG00000026179 | -1.068748441 | 0.608497229 |
| ENSMUSG00000026180 | 1.086522514 | 0.641039836 |
| ENSMUSG00000026181 | 1.025182809 | 0.96035552 |
| ENSMUSG00000026182 | 1 | |
| ENSMUSG00000026185 | -1.04526435 | 0.911630239 |
| ENSMUSG00000026187 | 1.098815226 | 0.638991399 |
| ENSMUSG00000026188 | -1.405451546 | 0.136807154 |
| ENSMUSG00000026189 | -1.111617214 | 0.15169538 |
| ENSMUSG00000026192 | -1.016545829 | 1 |
| ENSMUSG00000026193 | 1.466553426 | 0.221797188 |

| | | |
|---|---|---|
| ENSMUSG00000026196 | -1.014438521 | 0.983338279 |
| ENSMUSG00000026197 | -1.055429895 | 0.616733662 |
| ENSMUSG00000026198 | -1.04176736 | 0.783487011 |
| ENSMUSG00000026199 | 1.079034634 | 0.608497229 |
| ENSMUSG00000026200 | -1.177004976 | 0.054191461 |
| ENSMUSG00000026201 | -1.08060475 | 0.697178241 |
| ENSMUSG00000026202 | 1.056172803 | 0.736680637 |
| ENSMUSG00000026203 | 1.067644385 | 0.196672866 |
| ENSMUSG00000026204 | 1.461059537 | 0.012423383 |
| ENSMUSG00000026205 | -1.01210918 | 1 |
| ENSMUSG00000026207 | 1.292684362 | 0.004081733 |
| ENSMUSG00000026208 | 1.27425916 | 0.011713655 |
| ENSMUSG00000026209 | 1.086366507 | 0.605159739 |
| ENSMUSG00000026211 | -1.081323468 | 0.567948807 |
| ENSMUSG00000026213 | 1.027532136 | 0.837704961 |
| ENSMUSG00000026219 | -1.0129888 | 1 |
| ENSMUSG00000026220 | -1.03962336 | 0.808074244 |
| ENSMUSG00000026222 | -1.296634803 | 0.400200807 |
| ENSMUSG00000026223 | -1.052808208 | 0.807247723 |
| ENSMUSG00000026224 | -1.069502966 | 0.698112512 |
| ENSMUSG00000026226 | 1 | |
| ENSMUSG00000026227 | -1.043306199 | 0.803260194 |
| ENSMUSG00000026228 | -1.187745816 | 0.532586 |
| ENSMUSG00000026229 | -1.025620133 | 0.985522133 |
| ENSMUSG00000026234 | -1.025209326 | 0.998002628 |
| ENSMUSG00000026235 | 1.365281197 | 0.018089249 |
| ENSMUSG00000026237 | -1.043306199 | 0.798487595 |
| ENSMUSG00000026238 | 1.069810953 | 0.836150137 |
| ENSMUSG00000026239 | -1.027137739 | 0.933683189 |
| ENSMUSG00000026240 | -1.041044932 | 0.858073326 |
| ENSMUSG00000026241 | 1 | |
| ENSMUSG00000026245 | -1.079660279 | 0.731295483 |
| ENSMUSG00000026246 | 1.283884444 | 0.111305241 |
| ENSMUSG00000026247 | -1.309975156 | 0.086164458 |
| ENSMUSG00000026248 | -1.028711188 | 0.882411555 |
| ENSMUSG00000026249 | 1.858646351 | 0.000242314 |
| ENSMUSG00000026251 | 1 | |
| ENSMUSG00000026253 | -1 | 1 |
| ENSMUSG00000026254 | -1.057290789 | 0.847073978 |
| ENSMUSG00000026255 | -1.223075747 | 0.535052277 |
| ENSMUSG00000026258 | 1.059463094 | 0.608497229 |
| ENSMUSG00000026259 | -1.092297713 | 0.379433976 |
| ENSMUSG00000026260 | -1.007133397 | 1 |
| ENSMUSG00000026269 | 1.030113072 | 0.798487595 |
| ENSMUSG00000026270 | 1.019709869 | 0.903464611 |
| ENSMUSG00000026271 | 1.129920903 | 0.563360084 |
| ENSMUSG00000026272 | 1 | |
| ENSMUSG00000026273 | -1.038798777 | 0.698112512 |
| ENSMUSG00000026274 | 1.028339864 | 0.920920204 |
| ENSMUSG00000026275 | -1.103046032 | 0.634567735 |
| ENSMUSG00000026276 | 1.074321136 | 0.834839709 |
| ENSMUSG00000026277 | -1.008311907 | 1 |
| ENSMUSG00000026278 | 1.538404218 | 0.000685987 |
| ENSMUSG00000026279 | -1.009718002 | 0.978259071 |
| ENSMUSG00000026280 | -1.011321799 | 1 |
| ENSMUSG00000026281 | 1.070621204 | 0.725189744 |
| ENSMUSG00000026283 | 1.002289702 | 1 |
| ENSMUSG00000026285 | -1.03962336 | 0.698112512 |
| ENSMUSG00000026288 | -1.310622568 | 0.051335794 |
| ENSMUSG00000026289 | 1.009350622 | 1 |
| ENSMUSG00000026295 | 1 | |
| ENSMUSG00000026301 | 1.013583879 | 1 |
| ENSMUSG00000026303 | -1.035361515 | 0.961868983 |

| | | |
|---|---|---|
| ENSMUSG00000026304 | 1.019083579 | 0.979345947 |
| ENSMUSG00000026305 | 1.09353593 | 0.70299173 |
| ENSMUSG00000026307 | 1.029455573 | 0.837016778 |
| ENSMUSG00000026308 | 1.036554303 | 0.866870512 |
| ENSMUSG00000026309 | 1.060421614 | 0.762279343 |
| ENSMUSG00000026311 | -1.070578178 | 0.659252773 |
| ENSMUSG00000026312 | -1.044273782 | 0.790851299 |
| ENSMUSG00000026313 | 1.14340496 | 0.234875213 |
| ENSMUSG00000026315 | 2.417516904 | 3.77621E-05 |
| ENSMUSG00000026317 | 1.094078877 | 0.437917288 |
| ENSMUSG00000026319 | -1.054314306 | 0.814085406 |
| ENSMUSG00000026321 | 1.09115302 | 0.261204136 |
| ENSMUSG00000026322 | 1.015486245 | 0.983339718 |
| ENSMUSG00000026327 | 1.059463094 | 0.608497229 |
| ENSMUSG00000026331 | 1.019631466 | 0.987669167 |
| ENSMUSG00000026333 | -1.034201404 | 0.849793055 |
| ENSMUSG00000026335 | 1.013819544 | 1 |
| ENSMUSG00000026336 | 1 | |
| ENSMUSG00000026339 | 1.054483916 | 0.823367179 |
| ENSMUSG00000026341 | 1.125941081 | 0.698112512 |
| ENSMUSG00000026342 | 1.044707806 | 0.822562428 |
| ENSMUSG00000026343 | -1.135512723 | 0.549209743 |
| ENSMUSG00000026344 | 1.101981549 | 0.382326401 |
| ENSMUSG00000026347 | -1 | 1 |
| ENSMUSG00000026348 | -1.219327449 | 0.121862618 |
| ENSMUSG00000026349 | 1.014580568 | 1 |
| ENSMUSG00000026353 | -1.018435311 | 0.99092221 |
| ENSMUSG00000026354 | -1.151670879 | 0.559945355 |
| ENSMUSG00000026355 | 1.052028689 | 0.899219824 |
| ENSMUSG00000026356 | -1.043165627 | 0.911965446 |
| ENSMUSG00000026357 | 1 | |
| ENSMUSG00000026358 | -1 | 1 |
| ENSMUSG00000026360 | 1.281954477 | 0.18445321 |
| ENSMUSG00000026361 | -1.036633477 | 0.935661297 |
| ENSMUSG00000026365 | 1.199158641 | 0.117177979 |
| ENSMUSG00000026368 | 1.122462048 | 0.601164141 |
| ENSMUSG00000026374 | -1.006188697 | 1 |
| ENSMUSG00000026377 | -1.060485585 | 0.809553648 |
| ENSMUSG00000026380 | -1.493694181 | 0.007060227 |
| ENSMUSG00000026383 | -1.152912045 | 0.373282501 |
| ENSMUSG00000026384 | -1.129889841 | 0.608497229 |
| ENSMUSG00000026385 | 1.005766009 | 1 |
| ENSMUSG00000026387 | 1 | |
| ENSMUSG00000026388 | -1.105501728 | 0.385592008 |
| ENSMUSG00000026389 | 1.189599209 | 0.004323453 |
| ENSMUSG00000026390 | 1 | |
| ENSMUSG00000026393 | 1.118745527 | 0.666120375 |
| ENSMUSG00000026394 | 1 | |
| ENSMUSG00000026395 | -1.063525418 | 0.698961838 |
| ENSMUSG00000026398 | -1.229784166 | 0.467052335 |
| ENSMUSG00000026399 | -1.441928536 | 0.026445754 |
| ENSMUSG00000026401 | 1.059463094 | 0.608497229 |
| ENSMUSG00000026404 | -1.060971274 | 0.70072217 |
| ENSMUSG00000026405 | -1.093136706 | 0.790851299 |
| ENSMUSG00000026407 | -1 | 1 |
| ENSMUSG00000026409 | 1.069919002 | 0.561789645 |
| ENSMUSG00000026411 | 1.031598363 | 0.833851312 |
| ENSMUSG00000026413 | 1.034888984 | 0.847344209 |
| ENSMUSG00000026414 | 2.081652709 | 0.000328611 |
| ENSMUSG00000026415 | 1 | |
| ENSMUSG00000026416 | 1 | |
| ENSMUSG00000026417 | 1 | |
| ENSMUSG00000026418 | 1.460050024 | 0.013017191 |

| | | |
|---|---|---|
| ENSMUSG00000026420 | -1.400149822 | 0.000622261 |
| ENSMUSG00000026421 | 1.31627224 | 0.195814049 |
| ENSMUSG00000026424 | 1 | |
| ENSMUSG00000026425 | 1.069072484 | 0.756389254 |
| ENSMUSG00000026426 | 1.112812312 | 0.608497229 |
| ENSMUSG00000026427 | 1.017517075 | 0.96024157 |
| ENSMUSG00000026429 | 1.044750972 | 0.698112512 |
| ENSMUSG00000026430 | -1.233195481 | 0.050325876 |
| ENSMUSG00000026432 | -1.043306199 | 0.79291651 |
| ENSMUSG00000026433 | -1.132594428 | 0.269617125 |
| ENSMUSG00000026434 | -1.027879085 | 0.982838256 |
| ENSMUSG00000026435 | 1.009241052 | 0.972785312 |
| ENSMUSG00000026436 | 1.020418294 | 0.957402357 |
| ENSMUSG00000026437 | 1.03510172 | 0.846170406 |
| ENSMUSG00000026439 | -1.001530376 | 1 |
| ENSMUSG00000026442 | -1.227004466 | 0.343739781 |
| ENSMUSG00000026443 | -1.109582661 | 0.66912118 |
| ENSMUSG00000026447 | 1.116364704 | 0.229884308 |
| ENSMUSG00000026450 | -1.03962336 | 0.698112512 |
| ENSMUSG00000026452 | 1.200798598 | 0.087809131 |
| ENSMUSG00000026455 | -1.068814085 | 0.742268585 |
| ENSMUSG00000026456 | 1.079625631 | 0.686129647 |
| ENSMUSG00000026457 | -1.009216797 | 1 |
| ENSMUSG00000026458 | 1.037356286 | 0.910895334 |
| ENSMUSG00000026459 | 1 | |
| ENSMUSG00000026463 | 1.376366577 | 0.061476532 |
| ENSMUSG00000026466 | -1.027937568 | 0.958872312 |
| ENSMUSG00000026468 | -1.600447445 | 0.005521566 |
| ENSMUSG00000026469 | -1.109201344 | 0.657796687 |
| ENSMUSG00000026470 | 1.117873782 | 0.336300705 |
| ENSMUSG00000026471 | -1.482434851 | 0.004667441 |
| ENSMUSG00000026473 | -1.137915949 | 0.476305515 |
| ENSMUSG00000026475 | -1.105480765 | 0.111305241 |
| ENSMUSG00000026478 | 1.052262207 | 0.870130672 |
| ENSMUSG00000026479 | -1.0860956 | 0.43994505 |
| ENSMUSG00000026480 | -1.221070503 | 0.149977043 |
| ENSMUSG00000026482 | 1.132284696 | 0.361740791 |
| ENSMUSG00000026483 | 1.118734497 | 0.621197991 |
| ENSMUSG00000026484 | -1.039281875 | 0.910334649 |
| ENSMUSG00000026489 | -1.330871298 | 0.001448487 |
| ENSMUSG00000026490 | 1.074043953 | 0.775469341 |
| ENSMUSG00000026491 | 1.017692768 | 1 |
| ENSMUSG00000026492 | -1.111331787 | 0.339751436 |
| ENSMUSG00000026494 | 1.047294118 | 0.712130748 |
| ENSMUSG00000026495 | -1.021024951 | 0.933031586 |
| ENSMUSG00000026496 | -1.031312976 | 0.932242329 |
| ENSMUSG00000026497 | 1.12189305 | 0.585646836 |
| ENSMUSG00000026499 | 1.025316146 | 0.967123546 |
| ENSMUSG00000026500 | -1.063525418 | 0.698112512 |
| ENSMUSG00000026502 | 1.11228279 | 0.654902646 |
| ENSMUSG00000026504 | -1.000515704 | 1 |
| ENSMUSG00000026509 | 1.068108539 | 0.84234437 |
| ENSMUSG00000026510 | -1.006964692 | 1 |
| ENSMUSG00000026511 | -1.018426334 | 0.980901245 |
| ENSMUSG00000026514 | 1.084618945 | 0.73684216 |
| ENSMUSG00000026516 | -1.024406941 | 0.973152299 |
| ENSMUSG00000026519 | 1.097219791 | 0.559354144 |
| ENSMUSG00000026520 | 1.033190637 | 0.770255677 |
| ENSMUSG00000026523 | 1 | |
| ENSMUSG00000026525 | 1.031656037 | 0.749782622 |
| ENSMUSG00000026526 | -1.008012481 | 1 |
| ENSMUSG00000026527 | -1 | 1 |
| ENSMUSG00000026531 | 1 | |

| | | |
|---|---|---|
| ENSMUSG00000026532 | -3.275277624 | 0.000101355 |
| ENSMUSG00000026535 | 1.431859928 | 0.005771807 |
| ENSMUSG00000026536 | -1.1433188 | 0.378431826 |
| ENSMUSG00000026542 | 1 | |
| ENSMUSG00000026544 | -1.169451534 | 0.234005431 |
| ENSMUSG00000026546 | -1.07557985 | 0.760912081 |
| ENSMUSG00000026547 | 1.194917653 | 0.473421853 |
| ENSMUSG00000026548 | -1 | 1 |
| ENSMUSG00000026553 | -1.015969746 | 1 |
| ENSMUSG00000026554 | -1.0156039 | 0.988424085 |
| ENSMUSG00000026556 | 1.124353599 | 0.288413286 |
| ENSMUSG00000026558 | -1.008207917 | 1 |
| ENSMUSG00000026560 | 1 | |
| ENSMUSG00000026563 | -1.140755604 | 0.122969725 |
| ENSMUSG00000026564 | 1.637644294 | 0.005175804 |
| ENSMUSG00000026565 | -1.043341395 | 0.851424217 |
| ENSMUSG00000026566 | 1.041691381 | 0.837118968 |
| ENSMUSG00000026567 | 1.095467929 | 0.785561203 |
| ENSMUSG00000026568 | -1.099734991 | 0.585282823 |
| ENSMUSG00000026571 | 1.048889526 | 0.794274354 |
| ENSMUSG00000026572 | -1.044273782 | 0.790851299 |
| ENSMUSG00000026573 | 1 | |
| ENSMUSG00000026574 | 1.306851767 | 0.266336621 |
| ENSMUSG00000026575 | -1.075159357 | 0.69307409 |
| ENSMUSG00000026576 | 1.353394687 | 0.066281833 |
| ENSMUSG00000026577 | -1.042170321 | 0.816599259 |
| ENSMUSG00000026578 | -1.002766404 | 1 |
| ENSMUSG00000026579 | -1.350112323 | 0.279328609 |
| ENSMUSG00000026580 | -1.013902033 | 1 |
| ENSMUSG00000026581 | 1 | |
| ENSMUSG00000026582 | 1.059463094 | 0.698112512 |
| ENSMUSG00000026584 | -1.035614146 | 0.826009263 |
| ENSMUSG00000026585 | 1.037999053 | 0.897053041 |
| ENSMUSG00000026586 | 1.31096881 | 0.128223726 |
| ENSMUSG00000026587 | -1.069628355 | 0.786025504 |
| ENSMUSG00000026589 | 1.093527855 | 0.608497229 |
| ENSMUSG00000026592 | 1.030445456 | 0.937517551 |
| ENSMUSG00000026594 | 1.041212429 | 0.783662919 |
| ENSMUSG00000026596 | 1.015156844 | 0.999180249 |
| ENSMUSG00000026600 | 1.122858628 | 0.623872809 |
| ENSMUSG00000026601 | -1.03370111 | 0.951063333 |
| ENSMUSG00000026602 | -1.773558491 | 0.001410834 |
| ENSMUSG00000026603 | -1.070458749 | 0.691945548 |
| ENSMUSG00000026604 | 1.00802549 | 1 |
| ENSMUSG00000026605 | -1.02383166 | 0.978763926 |
| ENSMUSG00000026608 | 1.012835888 | 0.980773838 |
| ENSMUSG00000026609 | -1 | 1 |
| ENSMUSG00000026610 | -1.234043078 | 0.501554528 |
| ENSMUSG00000026611 | 1.076118789 | 0.608497229 |
| ENSMUSG00000026614 | -1.142727245 | 0.570749474 |
| ENSMUSG00000026615 | -1.014469095 | 1 |
| ENSMUSG00000026616 | 1.15842836 | 0.210955689 |
| ENSMUSG00000026617 | -1.092598363 | 0.685142053 |
| ENSMUSG00000026618 | -1.033733373 | 0.927544616 |
| ENSMUSG00000026620 | -1.071398125 | 0.608497229 |
| ENSMUSG00000026621 | 1 | |
| ENSMUSG00000026622 | 1.041923194 | 0.883138619 |
| ENSMUSG00000026623 | -1.037067196 | 0.939359524 |
| ENSMUSG00000026626 | -1.056245734 | 0.801443616 |
| ENSMUSG00000026627 | -1.029472728 | 0.869461232 |
| ENSMUSG00000026628 | -1.227253579 | 0.071386941 |
| ENSMUSG00000026630 | -1.0993412 | 0.608497229 |
| ENSMUSG00000026632 | -1.128473212 | 0.157561294 |

| | | |
|---|---|---|
| ENSMUSG00000026634 | 1.021516126 | 0.967899076 |
| ENSMUSG00000026637 | 1.306657136 | 0.001559906 |
| ENSMUSG00000026638 | -1.404015046 | 0.028001026 |
| ENSMUSG00000026639 | -1.324630979 | 0.016884263 |
| ENSMUSG00000026640 | 1.100610923 | 0.57044161 |
| ENSMUSG00000026641 | 1.073454752 | 0.606801085 |
| ENSMUSG00000026643 | -1.035549153 | 0.941524846 |
| ENSMUSG00000026644 | -1.082677361 | 0.764370878 |
| ENSMUSG00000026645 | 1 | |
| ENSMUSG00000026646 | -1.001940181 | 1 |
| ENSMUSG00000026648 | -1.009747016 | 1 |
| ENSMUSG00000026649 | 1.027267789 | 0.802773642 |
| ENSMUSG00000026650 | -1.120476206 | 0.677143478 |
| ENSMUSG00000026655 | -1.076434132 | 0.77681405 |
| ENSMUSG00000026656 | 1.11166259 | 0.608497229 |
| ENSMUSG00000026657 | 1.204709971 | 0.28224749 |
| ENSMUSG00000026659 | 1.029950139 | 0.865598094 |
| ENSMUSG00000026662 | 1.049183524 | 0.879036808 |
| ENSMUSG00000026663 | -1.020472186 | 0.98145432 |
| ENSMUSG00000026664 | 1.177574223 | 0.192770351 |
| ENSMUSG00000026667 | -1.000370498 | 1 |
| ENSMUSG00000026668 | 1 | |
| ENSMUSG00000026669 | -1.010564376 | 1 |
| ENSMUSG00000026670 | 1.060620019 | 0.796403921 |
| ENSMUSG00000026672 | 1.065209375 | 0.693692316 |
| ENSMUSG00000026674 | -1.785266541 | 0.026808199 |
| ENSMUSG00000026675 | 1.076595348 | 0.698112512 |
| ENSMUSG00000026676 | 1.142908482 | 0.698112512 |
| ENSMUSG00000026678 | 1.466227879 | 0.039256585 |
| ENSMUSG00000026679 | -1.124594559 | 0.651651813 |
| ENSMUSG00000026683 | -1.051246539 | 0.878551492 |
| ENSMUSG00000026686 | 1 | |
| ENSMUSG00000026687 | -1.028346261 | 0.931675054 |
| ENSMUSG00000026688 | 1.313021805 | 0.010491035 |
| ENSMUSG00000026691 | 1.177765487 | 0.276639872 |
| ENSMUSG00000026692 | 1 | |
| ENSMUSG00000026694 | 1.092886397 | 0.573420551 |
| ENSMUSG00000026696 | 1.004016744 | 1 |
| ENSMUSG00000026697 | 1 | |
| ENSMUSG00000026698 | 1.093491004 | 0.465687935 |
| ENSMUSG00000026700 | -1.03962336 | 0.698112512 |
| ENSMUSG00000026701 | 1.134071796 | 0.608497229 |
| ENSMUSG00000026705 | -1.041142193 | 0.851662555 |
| ENSMUSG00000026707 | 1.00974973 | 0.986542466 |
| ENSMUSG00000026708 | -1.013049251 | 0.986799592 |
| ENSMUSG00000026709 | -1.043496652 | 0.843449443 |
| ENSMUSG00000026712 | -1.192370134 | 0.342054851 |
| ENSMUSG00000026715 | -1.085254096 | 0.604750847 |
| ENSMUSG00000026718 | 1.045441838 | 0.865583959 |
| ENSMUSG00000026721 | -1.031090136 | 0.914187043 |
| ENSMUSG00000026723 | -1.206255417 | 0.144904333 |
| ENSMUSG00000026725 | 1.230850716 | 0.151271623 |
| ENSMUSG00000026726 | 1.117135044 | 0.341894005 |
| ENSMUSG00000026727 | 1.20496404 | 0.369149486 |
| ENSMUSG00000026728 | 1.20266077 | 0.605619163 |
| ENSMUSG00000026729 | 1 | |
| ENSMUSG00000026730 | 1.162829976 | 0.183748402 |
| ENSMUSG00000026734 | 1 | |
| ENSMUSG00000026735 | 1 | |
| ENSMUSG00000026736 | 1.107489364 | 0.744648261 |
| ENSMUSG00000026737 | 1.061350938 | 0.74010437 |
| ENSMUSG00000026739 | 1.113251975 | 0.619047344 |
| ENSMUSG00000026740 | 1.087926645 | 0.608497229 |

| | | |
|---|---|---|
| ENSMUSG00000026743 | 1.165452751 | 0.220604008 |
| ENSMUSG00000026748 | 1.145895989 | 0.467119761 |
| ENSMUSG00000026749 | 1.124390809 | 0.608497229 |
| ENSMUSG00000026750 | -1.071176963 | 0.794274354 |
| ENSMUSG00000026751 | 1.059463094 | 0.608497229 |
| ENSMUSG00000026753 | -1.04638076 | 0.878204365 |
| ENSMUSG00000026754 | 1.024905701 | 0.943496372 |
| ENSMUSG00000026755 | 1.028926573 | 0.944468485 |
| ENSMUSG00000026761 | 1.066613854 | 0.757419067 |
| ENSMUSG00000026764 | 1.41327616 | 0.003793294 |
| ENSMUSG00000026765 | -1.184259924 | 0.263492792 |
| ENSMUSG00000026766 | 1.006650445 | 1 |
| ENSMUSG00000026767 | 1.019973891 | 0.968055611 |
| ENSMUSG00000026768 | 1.540857758 | 0.01315387 |
| ENSMUSG00000026770 | 1.030256832 | 0.928585511 |
| ENSMUSG00000026771 | 1.050295593 | 0.773508225 |
| ENSMUSG00000026773 | 1.190153056 | 0.099525225 |
| ENSMUSG00000026774 | -1 | 1 |
| ENSMUSG00000026775 | 1.051492931 | 0.89103058 |
| ENSMUSG00000026778 | -1.004296146 | 1 |
| ENSMUSG00000026779 | 1.024186742 | 0.954761885 |
| ENSMUSG00000026781 | 1.014120563 | 0.996913409 |
| ENSMUSG00000026782 | -1.015663044 | 0.988424085 |
| ENSMUSG00000026784 | -1.03162158 | 0.85318934 |
| ENSMUSG00000026785 | 1.089099908 | 0.326561157 |
| ENSMUSG00000026786 | 1.187743643 | 0.188870702 |
| ENSMUSG00000026787 | -1.060503829 | 0.698112512 |
| ENSMUSG00000026788 | -1.00168824 | 1 |
| ENSMUSG00000026790 | 1.02866361 | 0.929344107 |
| ENSMUSG00000026791 | 1.023888092 | 0.953521521 |
| ENSMUSG00000026792 | 1.116158235 | 0.110097673 |
| ENSMUSG00000026796 | 1.256951627 | 0.136338589 |
| ENSMUSG00000026797 | -1.072773505 | 0.730317075 |
| ENSMUSG00000026798 | 1.015686564 | 0.797105186 |
| ENSMUSG00000026799 | 1.013833245 | 0.945548979 |
| ENSMUSG00000026803 | -1.005658939 | 1 |
| ENSMUSG00000026805 | 1.059463094 | 0.698112512 |
| ENSMUSG00000026806 | 1.02931693 | 0.805157604 |
| ENSMUSG00000026807 | 1.084618945 | 0.698112512 |
| ENSMUSG00000026809 | -1.169347953 | 0.698112512 |
| ENSMUSG00000026810 | 1.041444654 | 0.752411447 |
| ENSMUSG00000026811 | 1.007199546 | 1 |
| ENSMUSG00000026812 | 1.066531105 | 0.763700143 |
| ENSMUSG00000026814 | -2.72885642 | 0.000556642 |
| ENSMUSG00000026815 | -1.044273782 | 0.698112512 |
| ENSMUSG00000026816 | -1.029527269 | 0.922590208 |
| ENSMUSG00000026817 | 1.272228091 | 0.050815501 |
| ENSMUSG00000026818 | -1.068899451 | 0.698112512 |
| ENSMUSG00000026819 | -1.037386685 | 0.782165604 |
| ENSMUSG00000026820 | 1.066341591 | 0.608497229 |
| ENSMUSG00000026821 | 1.108301508 | 0.183274333 |
| ENSMUSG00000026822 | 1.309352122 | 0.195494168 |
| ENSMUSG00000026824 | 1.014545335 | 0.979002724 |
| ENSMUSG00000026825 | -1.059178247 | 0.683457122 |
| ENSMUSG00000026826 | 1.088555428 | 0.608497229 |
| ENSMUSG00000026827 | 1.075869537 | 0.73688717 |
| ENSMUSG00000026828 | 1 | |
| ENSMUSG00000026829 | 1.034095782 | 0.893303059 |
| ENSMUSG00000026830 | -1 | 1 |
| ENSMUSG00000026831 | -1.189107718 | 0.362935692 |
| ENSMUSG00000026832 | 1.607984878 | 0.006484725 |
| ENSMUSG00000026833 | 1.05535946 | 0.583037087 |
| ENSMUSG00000026834 | 1.010262388 | 1 |

| | | |
|---|---|---|
| ENSMUSG00000026835 | 1 | |
| ENSMUSG00000026836 | 1.263903946 | 0.108097704 |
| ENSMUSG00000026837 | 1.153560676 | 0.347722428 |
| ENSMUSG00000026839 | 1.047443505 | 0.863472381 |
| ENSMUSG00000026840 | -1.200301184 | 0.425108981 |
| ENSMUSG00000026841 | -1.140476302 | 0.532369657 |
| ENSMUSG00000026842 | 1.161471282 | 0.283605457 |
| ENSMUSG00000026843 | 1.056141226 | 0.806551098 |
| ENSMUSG00000026848 | 1.014674322 | 0.995703142 |
| ENSMUSG00000026849 | 1.057642681 | 0.699036146 |
| ENSMUSG00000026851 | -1.055874675 | 0.730297627 |
| ENSMUSG00000026853 | 1.000970016 | 1 |
| ENSMUSG00000026854 | 1.079529954 | 0.60397857 |
| ENSMUSG00000026856 | -1.004117585 | 1 |
| ENSMUSG00000026857 | 1.119015555 | 0.290003326 |
| ENSMUSG00000026858 | 1.021209499 | 0.91832956 |
| ENSMUSG00000026860 | 1.043992657 | 0.698112512 |
| ENSMUSG00000026864 | -1.011337866 | 1 |
| ENSMUSG00000026866 | 1 | |
| ENSMUSG00000026867 | 1.019527188 | 0.99092221 |
| ENSMUSG00000026869 | -1.006738343 | 1 |
| ENSMUSG00000026870 | 1.044013406 | 0.839354385 |
| ENSMUSG00000026872 | 1.085864665 | 0.605159739 |
| ENSMUSG00000026873 | -1.007924994 | 1 |
| ENSMUSG00000026874 | 1.030415139 | 0.960321094 |
| ENSMUSG00000026875 | -1.377594366 | 0.003785514 |
| ENSMUSG00000026878 | 1.032730374 | 0.9602503 |
| ENSMUSG00000026879 | 1.276258932 | 0.185996876 |
| ENSMUSG00000026880 | 1.023400394 | 0.956374899 |
| ENSMUSG00000026882 | 1 | |
| ENSMUSG00000026883 | 1.172650442 | 0.125949509 |
| ENSMUSG00000026885 | 1.385222241 | 0.000620329 |
| ENSMUSG00000026887 | 1.022278048 | 0.87470173 |
| ENSMUSG00000026888 | 1.849485821 | 0.0001059 |
| ENSMUSG00000026889 | 1.028298935 | 0.946048651 |
| ENSMUSG00000026890 | 1.014879514 | 0.960471166 |
| ENSMUSG00000026893 | 1.011875528 | 0.987097991 |
| ENSMUSG00000026894 | -1.085524987 | 0.495717496 |
| ENSMUSG00000026895 | 1.020220636 | 0.949812939 |
| ENSMUSG00000026896 | -1.263748281 | 0.059468055 |
| ENSMUSG00000026904 | 1 | |
| ENSMUSG00000026914 | 1.061925975 | 0.836051413 |
| ENSMUSG00000026915 | 1.129198538 | 0.462046668 |
| ENSMUSG00000026917 | 1.019778186 | 0.982838256 |
| ENSMUSG00000026918 | 1.120325658 | 0.434077492 |
| ENSMUSG00000026919 | 1 | |
| ENSMUSG00000026921 | -1.005189258 | 1 |
| ENSMUSG00000026922 | 1.022837943 | 0.830704102 |
| ENSMUSG00000026923 | 1.218433455 | 0.091167679 |
| ENSMUSG00000026924 | 1.037968816 | 0.888631178 |
| ENSMUSG00000026925 | 1.085958799 | 0.565049171 |
| ENSMUSG00000026926 | 1.01324213 | 1 |
| ENSMUSG00000026927 | 1.04444906 | 0.807247723 |
| ENSMUSG00000026928 | 1.024284648 | 0.968055611 |
| ENSMUSG00000026930 | 1.104338523 | 0.257120957 |
| ENSMUSG00000026931 | 1.069068979 | 0.608497229 |
| ENSMUSG00000026932 | 1.457220629 | 0.009067615 |
| ENSMUSG00000026933 | 1.114668416 | 0.607014477 |
| ENSMUSG00000026934 | 1.059463094 | 0.608497229 |
| ENSMUSG00000026936 | 1 | |
| ENSMUSG00000026937 | 1 | |
| ENSMUSG00000026938 | 1.170681764 | 0.541735653 |
| ENSMUSG00000026939 | -2.175180391 | 0.000619063 |

| | | |
|---|---|---|
| ENSMUSG00000026940 | 1.09394959 | 0.681259514 |
| ENSMUSG00000026941 | 1.104265736 | 0.327642404 |
| ENSMUSG00000026942 | 1.193726295 | 0.251544797 |
| ENSMUSG00000026943 | 1 | |
| ENSMUSG00000026944 | -1.10985426 | 0.470928144 |
| ENSMUSG00000026946 | -1.247766193 | 0.127490223 |
| ENSMUSG00000026950 | 1.0078955 | 1 |
| ENSMUSG00000026955 | 1.039405112 | 0.657385147 |
| ENSMUSG00000026956 | 1.368778442 | 0.018576481 |
| ENSMUSG00000026958 | 1.348208605 | 0.03311356 |
| ENSMUSG00000026959 | -1.028348525 | 0.936995747 |
| ENSMUSG00000026960 | 1.077479206 | 0.698112512 |
| ENSMUSG00000026961 | 1 | |
| ENSMUSG00000026963 | 1 | |
| ENSMUSG00000026965 | 1.051667768 | 0.846865416 |
| ENSMUSG00000026966 | 1.088595823 | 0.408387318 |
| ENSMUSG00000026969 | 1.015486245 | 0.987775262 |
| ENSMUSG00000026970 | 1.156457892 | 0.60052883 |
| ENSMUSG00000026971 | -1.14774293 | 0.608497229 |
| ENSMUSG00000026972 | -1.071867619 | 0.546994714 |
| ENSMUSG00000026974 | 1.044832496 | 0.82756824 |
| ENSMUSG00000026975 | 1.010572499 | 0.951098417 |
| ENSMUSG00000026976 | -1.020154831 | 1 |
| ENSMUSG00000026977 | 1.070132625 | 0.790558275 |
| ENSMUSG00000026979 | -1.129652002 | 0.70299173 |
| ENSMUSG00000026980 | 1.140425069 | 0.612496955 |
| ENSMUSG00000026981 | -1.191178226 | 0.608497229 |
| ENSMUSG00000026983 | 1 | |
| ENSMUSG00000026984 | 1 | |
| ENSMUSG00000026985 | 1 | |
| ENSMUSG00000026986 | 1 | |
| ENSMUSG00000026987 | 1.075657748 | 0.698112512 |
| ENSMUSG00000026988 | 1.258324761 | 0.015033335 |
| ENSMUSG00000026989 | -1.053549062 | 0.698112512 |
| ENSMUSG00000026991 | 1.045396259 | 0.839694706 |
| ENSMUSG00000026994 | 1.355404702 | 0.064233998 |
| ENSMUSG00000026999 | -1.135007696 | 0.350471673 |
| ENSMUSG00000027001 | -1.183133636 | 0.172335229 |
| ENSMUSG00000027002 | -1.035594563 | 0.951706877 |
| ENSMUSG00000027004 | -1.583683168 | 0.006560104 |
| ENSMUSG00000027006 | -1.006473706 | 1 |
| ENSMUSG00000027007 | 1.040352641 | 0.815671448 |
| ENSMUSG00000027009 | -1.02064379 | 0.964699214 |
| ENSMUSG00000027010 | -1.091345472 | 0.608497229 |
| ENSMUSG00000027011 | -1.074514625 | 0.753766685 |
| ENSMUSG00000027012 | -1.059464686 | 0.854896758 |
| ENSMUSG00000027014 | -1.116908082 | 0.608497229 |
| ENSMUSG00000027015 | -1.346167774 | 0.102900586 |
| ENSMUSG00000027016 | -1.069502966 | 0.744868995 |
| ENSMUSG00000027018 | -1.090150036 | 0.753731647 |
| ENSMUSG00000027022 | 1 | |
| ENSMUSG00000027030 | -1.199820671 | 0.179873578 |
| ENSMUSG00000027035 | 1.13493088 | 0.584587773 |
| ENSMUSG00000027048 | 1.122462048 | 0.608497229 |
| ENSMUSG00000027067 | -1.08087856 | 0.820232117 |
| ENSMUSG00000027068 | 1.15472312 | 0.289579094 |
| ENSMUSG00000027070 | -1.850579129 | 0.008272169 |
| ENSMUSG00000027071 | -1.006487559 | 1 |
| ENSMUSG00000027072 | 1 | |
| ENSMUSG00000027073 | -1 | 1 |
| ENSMUSG00000027074 | -1.464373116 | 0.008040974 |
| ENSMUSG00000027075 | 1.746093132 | 0.001987328 |
| ENSMUSG00000027076 | -1.055292932 | 0.717035356 |

| | | |
|---|---|---|
| ENSMUSG00000027077 | 1.079681442 | 0.608497229 |
| ENSMUSG00000027078 | -1.189032329 | 0.177322007 |
| ENSMUSG00000027079 | -1.076109112 | 0.671280603 |
| ENSMUSG00000027080 | -1.153481044 | 0.433413213 |
| ENSMUSG00000027082 | -2.10423962 | 0.001751748 |
| ENSMUSG00000027086 | -1.032690422 | 0.878910251 |
| ENSMUSG00000027087 | 1.022466698 | 0.989698707 |
| ENSMUSG00000027088 | -1.08188484 | 0.608497229 |
| ENSMUSG00000027091 | -1.061443577 | 0.825401367 |
| ENSMUSG00000027099 | 1.002652104 | 1 |
| ENSMUSG00000027102 | 3.198145659 | 9.23305E-05 |
| ENSMUSG00000027104 | 1.001841305 | 1 |
| ENSMUSG00000027107 | 1.005021373 | 1 |
| ENSMUSG00000027108 | -1.135218 | 0.608497229 |
| ENSMUSG00000027109 | -1.06072961 | 0.832767487 |
| ENSMUSG00000027111 | -1.09547544 | 0.658590104 |
| ENSMUSG00000027115 | -1.075124913 | 0.698112512 |
| ENSMUSG00000027120 | 1 | |
| ENSMUSG00000027122 | 1.082980559 | 0.698112512 |
| ENSMUSG00000027130 | -1.040138672 | 0.838164927 |
| ENSMUSG00000027131 | -1.158427063 | 0.325020498 |
| ENSMUSG00000027132 | -1.083001387 | 0.684694376 |
| ENSMUSG00000027133 | -1.141152676 | 0.532369657 |
| ENSMUSG00000027134 | -1.233845439 | 0.04530822 |
| ENSMUSG00000027157 | 1 | |
| ENSMUSG00000027160 | 1.039566367 | 0.828568813 |
| ENSMUSG00000027162 | -1.131161665 | 0.614071182 |
| ENSMUSG00000027163 | -1.027107385 | 0.770958539 |
| ENSMUSG00000027164 | -1.110983488 | 0.608497229 |
| ENSMUSG00000027165 | -1.164446952 | 0.271595502 |
| ENSMUSG00000027166 | -1.105765299 | 0.214418537 |
| ENSMUSG00000027167 | -1.063568157 | 0.637081332 |
| ENSMUSG00000027168 | -1.68899313 | 9.32404E-05 |
| ENSMUSG00000027170 | -1.09835544 | 0.698112512 |
| ENSMUSG00000027171 | -1.371902288 | 0.023182546 |
| ENSMUSG00000027173 | -1.162955506 | 0.193346479 |
| ENSMUSG00000027175 | -1.139151039 | 0.344960749 |
| ENSMUSG00000027176 | 1.037172456 | 0.895127848 |
| ENSMUSG00000027177 | -1.053223875 | 0.839713755 |
| ENSMUSG00000027180 | -1.15345062 | 0.541735653 |
| ENSMUSG00000027184 | -1.039329865 | 0.955137373 |
| ENSMUSG00000027185 | -1.045856441 | 0.804844905 |
| ENSMUSG00000027186 | -1.065521517 | 0.803072249 |
| ENSMUSG00000027187 | -1.105927402 | 0.581630482 |
| ENSMUSG00000027188 | -1.513909423 | 0.026425705 |
| ENSMUSG00000027189 | -1.073503776 | 0.793839591 |
| ENSMUSG00000027193 | -1.067731003 | 0.818459784 |
| ENSMUSG00000027194 | -1.040532087 | 0.824800385 |
| ENSMUSG00000027195 | -1.071852625 | 0.768649774 |
| ENSMUSG00000027196 | -1.140485656 | 0.30870479 |
| ENSMUSG00000027198 | -1.138324312 | 0.510359867 |
| ENSMUSG00000027199 | -1.894515369 | 0.001504475 |
| ENSMUSG00000027200 | 1.6592102 | 0.002796445 |
| ENSMUSG00000027201 | -1.056915116 | 0.835436989 |
| ENSMUSG00000027202 | 1.149172711 | 0.44650239 |
| ENSMUSG00000027203 | -1.123816165 | 0.608497229 |
| ENSMUSG00000027204 | -1.095644458 | 0.719087885 |
| ENSMUSG00000027206 | 1.000542643 | 1 |
| ENSMUSG00000027207 | 1.131854105 | 0.376889921 |
| ENSMUSG00000027208 | -1.432793437 | 0.019218017 |
| ENSMUSG00000027209 | 1.188604282 | 0.370262985 |
| ENSMUSG00000027210 | -1.047028013 | 0.698112512 |
| ENSMUSG00000027215 | -1.133687675 | 0.335436382 |

| | | |
|---|---|---|
| ENSMUSG00000027217 | 1.29563191 | 0.016758819 |
| ENSMUSG00000027219 | 1.059463094 | 0.608497229 |
| ENSMUSG00000027220 | -1.508698657 | 0.002572504 |
| ENSMUSG00000027221 | 1.506269636 | 0.00095885 |
| ENSMUSG00000027222 | -1.114355777 | 0.276229467 |
| ENSMUSG00000027223 | -1.341689106 | 0.003259982 |
| ENSMUSG00000027224 | 1.493179273 | 0.155919956 |
| ENSMUSG00000027225 | 1.059463094 | 0.608497229 |
| ENSMUSG00000027227 | 1.355202068 | 0.036059831 |
| ENSMUSG00000027229 | 1 | |
| ENSMUSG00000027230 | 1.194940177 | 0.085378767 |
| ENSMUSG00000027233 | 1.019083579 | 0.97045957 |
| ENSMUSG00000027236 | -1.053166065 | 0.770923502 |
| ENSMUSG00000027238 | -1.404453346 | 0.102871968 |
| ENSMUSG00000027239 | -1.802676081 | 0.001353988 |
| ENSMUSG00000027242 | -1.017907841 | 0.976254251 |
| ENSMUSG00000027243 | 1.004629615 | 1 |
| ENSMUSG00000027244 | -1.065992181 | 0.623464143 |
| ENSMUSG00000027245 | 1.016353389 | 0.93979453 |
| ENSMUSG00000027246 | 1.057478344 | 0.856857802 |
| ENSMUSG00000027247 | 1.042174584 | 0.8920795 |
| ENSMUSG00000027248 | 1.034438748 | 0.972389453 |
| ENSMUSG00000027249 | -1.085651418 | 0.692881216 |
| ENSMUSG00000027253 | -1.164689371 | 0.260949892 |
| ENSMUSG00000027254 | 1.102889536 | 0.608497229 |
| ENSMUSG00000027255 | -1.138385988 | 0.368825616 |
| ENSMUSG00000027257 | 1.002760293 | 1 |
| ENSMUSG00000027259 | 1.045289946 | 0.698112512 |
| ENSMUSG00000027261 | 1.199989356 | 0.000021314 |
| ENSMUSG00000027263 | -1.013267339 | 0.998152646 |
| ENSMUSG00000027270 | 1 | |
| ENSMUSG00000027272 | -1.035302212 | 0.924587329 |
| ENSMUSG00000027273 | -1.044273782 | 0.698112512 |
| ENSMUSG00000027274 | 1.056847553 | 0.752411447 |
| ENSMUSG00000027276 | 1.460256267 | 0.00139517 |
| ENSMUSG00000027281 | -1.073767786 | 0.68491446 |
| ENSMUSG00000027282 | -1.148252423 | 0.608497229 |
| ENSMUSG00000027284 | 1.039363471 | 0.774235108 |
| ENSMUSG00000027285 | -1.049188955 | 0.777977356 |
| ENSMUSG00000027286 | -1.013423981 | 0.949812939 |
| ENSMUSG00000027287 | 1.006407753 | 1 |
| ENSMUSG00000027288 | 1.031098215 | 0.965637841 |
| ENSMUSG00000027291 | -1.022117394 | 0.957340912 |
| ENSMUSG00000027293 | -1.306803883 | 0.102925573 |
| ENSMUSG00000027296 | -1.165681946 | 0.608497229 |
| ENSMUSG00000027297 | -1.827871811 | 0.002435143 |
| ENSMUSG00000027298 | -1.141896762 | 0.304506715 |
| ENSMUSG00000027300 | -1.026239289 | 0.698112512 |
| ENSMUSG00000027301 | -1.053549062 | 0.698112512 |
| ENSMUSG00000027303 | -1.081070263 | 0.724253808 |
| ENSMUSG00000027304 | -1.129355842 | 0.608497229 |
| ENSMUSG00000027305 | -1.157702865 | 0.238770358 |
| ENSMUSG00000027306 | -1.183367515 | 0.501683218 |
| ENSMUSG00000027309 | -1.086568017 | 0.678985493 |
| ENSMUSG00000027312 | -1.071305498 | 0.724472305 |
| ENSMUSG00000027313 | 1.408502243 | 0.00787737 |
| ENSMUSG00000027314 | -1.392935763 | 0.03505174 |
| ENSMUSG00000027315 | -1.984179661 | 0.000191318 |
| ENSMUSG00000027316 | -1.140161434 | 0.640091848 |
| ENSMUSG00000027317 | -1.043306199 | 0.698112512 |
| ENSMUSG00000027318 | 1.011607416 | 1 |
| ENSMUSG00000027322 | -1.013665834 | 1 |
| ENSMUSG00000027323 | -1.139190565 | 0.541838059 |

| | | |
|---|---|---|
| ENSMUSG00000027324 | -1.06332681 | 0.153819953 |
| ENSMUSG00000027326 | -1.142330751 | 0.57564331 |
| ENSMUSG00000027327 | -1.007584785 | 0.994172456 |
| ENSMUSG00000027329 | -1.043320275 | 0.748046931 |
| ENSMUSG00000027330 | -1.079479032 | 0.662725458 |
| ENSMUSG00000027331 | -1.134078207 | 0.608497229 |
| ENSMUSG00000027332 | -1.134609196 | 0.52201333 |
| ENSMUSG00000027333 | -1.075445457 | 0.608497229 |
| ENSMUSG00000027335 | 1.599692558 | 0.035027529 |
| ENSMUSG00000027338 | 1 | |
| ENSMUSG00000027339 | 1.245262785 | 0.028002557 |
| ENSMUSG00000027340 | -1.113650442 | 0.60048332 |
| ENSMUSG00000027341 | -1.057835023 | 0.681952832 |
| ENSMUSG00000027342 | -1.013030716 | 1 |
| ENSMUSG00000027344 | -1.132183958 | 0.563957069 |
| ENSMUSG00000027345 | 1 | |
| ENSMUSG00000027346 | -1.134243317 | 0.424948752 |
| ENSMUSG00000027347 | 1.053554232 | 0.803072249 |
| ENSMUSG00000027349 | -1.130712673 | 0.594567601 |
| ENSMUSG00000027350 | 1.006784519 | 1 |
| ENSMUSG00000027351 | 1.02880023 | 0.94612635 |
| ENSMUSG00000027353 | -1.142027549 | 0.39537357 |
| ENSMUSG00000027355 | 1 | |
| ENSMUSG00000027356 | 1.210869356 | 0.398767938 |
| ENSMUSG00000027357 | -1.104690999 | 0.608497229 |
| ENSMUSG00000027358 | -1.824712651 | 0.00115117 |
| ENSMUSG00000027359 | 1.069293585 | 0.797105186 |
| ENSMUSG00000027360 | -1.142727245 | 0.608497229 |
| ENSMUSG00000027361 | -1.006398044 | 1 |
| ENSMUSG00000027363 | -1.088871484 | 0.698112512 |
| ENSMUSG00000027364 | -1.200376151 | 0.3305975 |
| ENSMUSG00000027365 | -1.033432678 | 0.948750604 |
| ENSMUSG00000027366 | -1.098644635 | 0.698112512 |
| ENSMUSG00000027367 | -1.136783958 | 0.608497229 |
| ENSMUSG00000027368 | -1.031879585 | 0.886195057 |
| ENSMUSG00000027371 | -2.198785559 | 1.25105E-05 |
| ENSMUSG00000027374 | -1.0820317 | 0.608497229 |
| ENSMUSG00000027375 | -1.869405317 | 0.007654092 |
| ENSMUSG00000027376 | 1 | |
| ENSMUSG00000027377 | -1.277443017 | 0.062439995 |
| ENSMUSG00000027378 | -1.123996791 | 0.433678407 |
| ENSMUSG00000027379 | -1.144039271 | 0.608497229 |
| ENSMUSG00000027380 | 1.519751934 | 0.011705868 |
| ENSMUSG00000027381 | -1.021900781 | 0.913882562 |
| ENSMUSG00000027384 | -1.108757343 | 0.509048875 |
| ENSMUSG00000027386 | -1.112184417 | 0.703468074 |
| ENSMUSG00000027387 | -1.092987387 | 0.34135669 |
| ENSMUSG00000027394 | -1.006951473 | 1 |
| ENSMUSG00000027395 | -1.10042204 | 0.555413989 |
| ENSMUSG00000027397 | 1.020093696 | 0.981360815 |
| ENSMUSG00000027398 | -1 | 1 |
| ENSMUSG00000027399 | 1.011497958 | 1 |
| ENSMUSG00000027400 | 1 | |
| ENSMUSG00000027401 | 1.085624843 | 0.69305964 |
| ENSMUSG00000027403 | 1.059463094 | 0.608497229 |
| ENSMUSG00000027404 | -1.009967289 | 1 |
| ENSMUSG00000027405 | -1.034607549 | 0.938161311 |
| ENSMUSG00000027406 | -1.088806082 | 0.684413983 |
| ENSMUSG00000027408 | -1.385852134 | 0.113956233 |
| ENSMUSG00000027409 | 1.030445456 | 0.938939969 |
| ENSMUSG00000027411 | -1.034629463 | 0.894828773 |
| ENSMUSG00000027412 | 1.365206609 | 0.041151366 |
| ENSMUSG00000027416 | 1 | |

| | | |
|---|---|---|
| ENSMUSG00000027419 | 1.177752631 | 0.460987702 |
| ENSMUSG00000027420 | 1.128312813 | 0.624722543 |
| ENSMUSG00000027422 | 1.054242254 | 0.811884447 |
| ENSMUSG00000027423 | -1.020207486 | 0.988665808 |
| ENSMUSG00000027424 | 1.002523843 | 1 |
| ENSMUSG00000027425 | 1.036818829 | 0.8349433 |
| ENSMUSG00000027427 | -1.131398633 | 0.606134298 |
| ENSMUSG00000027428 | -1.109457598 | 0.601666207 |
| ENSMUSG00000027429 | -1.140619424 | 0.608497229 |
| ENSMUSG00000027430 | -1.27850007 | 0.01442917 |
| ENSMUSG00000027431 | 1 | |
| ENSMUSG00000027433 | -1.015735471 | 1 |
| ENSMUSG00000027434 | -1.053549062 | 0.698112512 |
| ENSMUSG00000027435 | 1.283536374 | 0.143908326 |
| ENSMUSG00000027438 | -1.063964357 | 0.431188813 |
| ENSMUSG00000027439 | -1.066221398 | 0.713861573 |
| ENSMUSG00000027442 | 1 | |
| ENSMUSG00000027443 | 1 | |
| ENSMUSG00000027444 | 1 | |
| ENSMUSG00000027445 | 1 | |
| ENSMUSG00000027446 | 1.059463094 | 0.608497229 |
| ENSMUSG00000027447 | 1.048712664 | 0.770958539 |
| ENSMUSG00000027452 | -1.143787033 | 0.201977401 |
| ENSMUSG00000027454 | -1.115415473 | 0.287585529 |
| ENSMUSG00000027455 | -1.094300976 | 0.689888255 |
| ENSMUSG00000027456 | 1.20068119 | 0.437095676 |
| ENSMUSG00000027457 | -1.428315613 | 0.002527181 |
| ENSMUSG00000027459 | -1.039829309 | 0.731295483 |
| ENSMUSG00000027460 | -1.044846375 | 0.855652226 |
| ENSMUSG00000027463 | -1.243341108 | 0.547111524 |
| ENSMUSG00000027465 | -1.025234678 | 0.956714001 |
| ENSMUSG00000027466 | -1.004692218 | 1 |
| ENSMUSG00000027468 | 1 | |
| ENSMUSG00000027469 | -1.046307844 | 0.911872344 |
| ENSMUSG00000027470 | 1.815753003 | 0.001601152 |
| ENSMUSG00000027472 | -1.036132588 | 0.894390869 |
| ENSMUSG00000027474 | 1 | |
| ENSMUSG00000027475 | -1.100106228 | 0.676050382 |
| ENSMUSG00000027478 | -1.23508408 | 0.004433343 |
| ENSMUSG00000027479 | -1.109151406 | 0.734200167 |
| ENSMUSG00000027480 | 1 | |
| ENSMUSG00000027481 | 1 | |
| ENSMUSG00000027482 | 1 | |
| ENSMUSG00000027483 | 1 | |
| ENSMUSG00000027484 | 1 | |
| ENSMUSG00000027485 | 1 | |
| ENSMUSG00000027487 | -1.148321108 | 0.323887839 |
| ENSMUSG00000027488 | 1.064227476 | 0.608497229 |
| ENSMUSG00000027489 | 1.056164081 | 0.775412181 |
| ENSMUSG00000027490 | -1.017327173 | 0.952867397 |
| ENSMUSG00000027495 | -1.230206186 | 0.122925715 |
| ENSMUSG00000027496 | -1.07319328 | 0.728184651 |
| ENSMUSG00000027498 | -1.002556145 | 1 |
| ENSMUSG00000027499 | 1.027493321 | 0.947862126 |
| ENSMUSG00000027500 | -1.421508151 | 0.002843163 |
| ENSMUSG00000027502 | -1.051968111 | 0.801443616 |
| ENSMUSG00000027505 | 1.060802788 | 0.75366313 |
| ENSMUSG00000027506 | -1.062958022 | 0.704170349 |
| ENSMUSG00000027508 | -1.300238789 | 0.045175971 |
| ENSMUSG00000027509 | -1.045147707 | 0.819614655 |
| ENSMUSG00000027510 | -1.036168485 | 0.834794437 |
| ENSMUSG00000027513 | 1.059463094 | 0.608497229 |
| ENSMUSG00000027514 | -1.189107718 | 0.414873034 |

| | | |
|---|---|---|
| ENSMUSG00000027517 | 1 | |
| ENSMUSG00000027518 | 1 | |
| ENSMUSG00000027519 | -1.005413873 | 1 |
| ENSMUSG00000027520 | -2.185765176 | 0.000424592 |
| ENSMUSG00000027522 | -1.031702345 | 0.845766674 |
| ENSMUSG00000027523 | 1.003679788 | 1 |
| ENSMUSG00000027524 | 1 | |
| ENSMUSG00000027525 | -1.069502966 | 0.744868995 |
| ENSMUSG00000027528 | 1 | |
| ENSMUSG00000027530 | 1 | |
| ENSMUSG00000027531 | -1.059233459 | 0.794675895 |
| ENSMUSG00000027533 | 1.128581227 | 0.608497229 |
| ENSMUSG00000027534 | 1.094649855 | 0.551892689 |
| ENSMUSG00000027536 | 1.087411066 | 0.608497229 |
| ENSMUSG00000027540 | -1.110210061 | 0.447986619 |
| ENSMUSG00000027544 | -1.201160384 | 0.271993081 |
| ENSMUSG00000027546 | -2.789450096 | 2.89E-06 |
| ENSMUSG00000027547 | 1 | |
| ENSMUSG00000027550 | 1.075969183 | 0.698112512 |
| ENSMUSG00000027551 | -1.004402746 | 1 |
| ENSMUSG00000027552 | -1.003669259 | 1 |
| ENSMUSG00000027555 | 1.165580719 | 0.369255557 |
| ENSMUSG00000027556 | 1 | |
| ENSMUSG00000027559 | -1.398609258 | 0.103613445 |
| ENSMUSG00000027560 | 1.595252984 | 0.005924207 |
| ENSMUSG00000027562 | -1.329331355 | 0.38968625 |
| ENSMUSG00000027564 | 1 | |
| ENSMUSG00000027566 | -1.058253133 | 0.838634229 |
| ENSMUSG00000027568 | -1.133231767 | 0.591151279 |
| ENSMUSG00000027569 | -1.022752347 | 0.801054439 |
| ENSMUSG00000027570 | 1.021696736 | 0.989742633 |
| ENSMUSG00000027573 | -1.054278414 | 0.74527374 |
| ENSMUSG00000027574 | -1.302591532 | 0.251049598 |
| ENSMUSG00000027575 | 1.056061654 | 0.737047064 |
| ENSMUSG00000027577 | -1.359227594 | 0.017758048 |
| ENSMUSG00000027579 | -1.095294215 | 0.541735653 |
| ENSMUSG00000027580 | -1.242991666 | 0.034639136 |
| ENSMUSG00000027581 | 1 | |
| ENSMUSG00000027582 | -1.049314478 | 0.673162184 |
| ENSMUSG00000027583 | -1.254619515 | 0.031764753 |
| ENSMUSG00000027584 | 1.037893788 | 0.944468485 |
| ENSMUSG00000027589 | -1.072621501 | 0.658879457 |
| ENSMUSG00000027593 | 1.003274027 | 1 |
| ENSMUSG00000027596 | 1 | |
| ENSMUSG00000027597 | -1.043584915 | 0.794274354 |
| ENSMUSG00000027598 | -1.075330177 | 0.756255421 |
| ENSMUSG00000027599 | -1.019824505 | 0.980378626 |
| ENSMUSG00000027601 | 1.023560463 | 0.967420999 |
| ENSMUSG00000027602 | 1.00827889 | 0.988622398 |
| ENSMUSG00000027603 | -1.102936657 | 0.782383746 |
| ENSMUSG00000027605 | -1.446066485 | 0.008751654 |
| ENSMUSG00000027606 | 1 | |
| ENSMUSG00000027610 | -1.099865345 | 0.432387542 |
| ENSMUSG00000027611 | -1.275910355 | 0.025430361 |
| ENSMUSG00000027612 | -1.423142641 | 0.015994956 |
| ENSMUSG00000027613 | -1.067038608 | 0.698112512 |
| ENSMUSG00000027615 | -1.025323111 | 0.874837619 |
| ENSMUSG00000027618 | -1.030657392 | 0.866135971 |
| ENSMUSG00000027620 | -1.04141301 | 0.933623827 |
| ENSMUSG00000027624 | -1.030428225 | 0.903464611 |
| ENSMUSG00000027628 | -1.074830218 | 0.622070988 |
| ENSMUSG00000027630 | 1.063413775 | 0.780320794 |
| ENSMUSG00000027634 | -1.155006112 | 0.588331755 |

| | | |
|---|---|---|
| ENSMUSG00000027635 | -1.038111018 | 0.883306173 |
| ENSMUSG00000027636 | -1.443212187 | 0.030910704 |
| ENSMUSG00000027637 | -1.064240237 | 0.608497229 |
| ENSMUSG00000027639 | -1.198263322 | 0.522155042 |
| ENSMUSG00000027641 | -1.028332963 | 0.948609119 |
| ENSMUSG00000027642 | -1.043069262 | 0.921950992 |
| ENSMUSG00000027643 | -1.053549062 | 0.698112512 |
| ENSMUSG00000027646 | 1.019088646 | 0.928340769 |
| ENSMUSG00000027649 | -1.105255719 | 0.461544768 |
| ENSMUSG00000027650 | -1.10836282 | 0.585657353 |
| ENSMUSG00000027651 | -1.075799301 | 0.698112512 |
| ENSMUSG00000027652 | -1.051792428 | 0.863410724 |
| ENSMUSG00000027654 | -1.084453116 | 0.596276142 |
| ENSMUSG00000027655 | -1.019040407 | 0.929847429 |
| ENSMUSG00000027656 | 1.246543209 | 0.012910322 |
| ENSMUSG00000027660 | 1.076252991 | 0.705199845 |
| ENSMUSG00000027661 | -1.009838638 | 0.984094159 |
| ENSMUSG00000027663 | 1.198627833 | 0.169990135 |
| ENSMUSG00000027665 | 1.048055585 | 0.86264687 |
| ENSMUSG00000027667 | 1.031066482 | 0.881483647 |
| ENSMUSG00000027668 | -1.089245285 | 0.698112512 |
| ENSMUSG00000027669 | -1.027577754 | 0.9389052 |
| ENSMUSG00000027670 | 1.03288904 | 0.965252226 |
| ENSMUSG00000027671 | -1.038088205 | 0.895127848 |
| ENSMUSG00000027673 | 1.015379479 | 0.991944074 |
| ENSMUSG00000027674 | 1 | |
| ENSMUSG00000027676 | -1.127182532 | 0.304885651 |
| ENSMUSG00000027677 | -1.014656272 | 0.991673305 |
| ENSMUSG00000027678 | -1.065892848 | 0.713861573 |
| ENSMUSG00000027679 | -1.009280936 | 0.981456085 |
| ENSMUSG00000027680 | -1.016969088 | 1 |
| ENSMUSG00000027684 | 1.219492296 | 0.110887279 |
| ENSMUSG00000027690 | -1 | 1 |
| ENSMUSG00000027692 | 1.438566518 | 0.050826125 |
| ENSMUSG00000027694 | -1 | 1 |
| ENSMUSG00000027695 | -1.014139043 | 0.918726272 |
| ENSMUSG00000027698 | -1.00524008 | 1 |
| ENSMUSG00000027699 | -1.08927966 | 0.770017269 |
| ENSMUSG00000027702 | 1.034095782 | 0.914904517 |
| ENSMUSG00000027703 | 1 | |
| ENSMUSG00000027706 | -1.007625514 | 1 |
| ENSMUSG00000027708 | 1.002311483 | 1 |
| ENSMUSG00000027709 | 1.032083158 | 0.863472381 |
| ENSMUSG00000027710 | -1.019413253 | 0.968623752 |
| ENSMUSG00000027712 | 1.131146632 | 0.694271823 |
| ENSMUSG00000027713 | 1.141991974 | 0.397906691 |
| ENSMUSG00000027714 | -1.03238883 | 0.914862692 |
| ENSMUSG00000027715 | -1.04403136 | 0.926703252 |
| ENSMUSG00000027716 | -1.4471682 | 0.011183142 |
| ENSMUSG00000027718 | 1 | |
| ENSMUSG00000027719 | 1 | |
| ENSMUSG00000027720 | 1.015486245 | 0.998386582 |
| ENSMUSG00000027722 | 1.036805848 | 0.879178663 |
| ENSMUSG00000027737 | 1.131936342 | 0.290003326 |
| ENSMUSG00000027739 | -1.00565751 | 1 |
| ENSMUSG00000027742 | -1.035091029 | 0.905976807 |
| ENSMUSG00000027744 | 1.171563499 | 0.233412931 |
| ENSMUSG00000027746 | 1.084582626 | 0.690985372 |
| ENSMUSG00000027748 | -1.053549062 | 0.698112512 |
| ENSMUSG00000027750 | 3.097896502 | 0.00051542 |
| ENSMUSG00000027751 | 1.062471221 | 0.754804776 |
| ENSMUSG00000027752 | -1.047809686 | 0.852081869 |
| ENSMUSG00000027761 | -1.044273782 | 0.698112512 |

| | | |
|---|---|---|
| ENSMUSG00000027762 | 1 | |
| ENSMUSG00000027763 | 1.156537764 | 0.608497229 |
| ENSMUSG00000027765 | -1.041608621 | 0.838178918 |
| ENSMUSG00000027770 | -1.027711022 | 0.971164403 |
| ENSMUSG00000027774 | 1.003951584 | 1 |
| ENSMUSG00000027775 | 1.154591292 | 0.608497229 |
| ENSMUSG00000027776 | 1 | |
| ENSMUSG00000027777 | 1.257741093 | 0.404197146 |
| ENSMUSG00000027778 | -1.038115807 | 0.90205528 |
| ENSMUSG00000027782 | 1.015066618 | 1 |
| ENSMUSG00000027784 | -1.03676924 | 0.88492938 |
| ENSMUSG00000027787 | -1.028474889 | 0.949812939 |
| ENSMUSG00000027788 | 1.132639128 | 0.217803313 |
| ENSMUSG00000027790 | -1.053549062 | 0.698112512 |
| ENSMUSG00000027792 | 1.026314363 | 0.951063333 |
| ENSMUSG00000027793 | 1.224234039 | 0.331384766 |
| ENSMUSG00000027794 | 1.091519092 | 0.726433403 |
| ENSMUSG00000027796 | -1.004815481 | 1 |
| ENSMUSG00000027797 | 1.103898331 | 0.698112512 |
| ENSMUSG00000027799 | 1.045450472 | 0.856445097 |
| ENSMUSG00000027800 | -1.0836569 | 0.698112512 |
| ENSMUSG00000027801 | 1.429699737 | 0.034373414 |
| ENSMUSG00000027803 | 1.076889094 | 0.751036024 |
| ENSMUSG00000027804 | -1.053882484 | 0.866289429 |
| ENSMUSG00000027805 | -1.126593491 | 0.608497229 |
| ENSMUSG00000027806 | 1.068879773 | 0.760615796 |
| ENSMUSG00000027808 | 1.028422234 | 0.971627344 |
| ENSMUSG00000027809 | -1.02268629 | 0.968895456 |
| ENSMUSG00000027810 | -1.03469564 | 0.933031586 |
| ENSMUSG00000027811 | -1.02227075 | 0.927513137 |
| ENSMUSG00000027820 | 1.21832028 | 0.011795574 |
| ENSMUSG00000027822 | 1.01316474 | 0.988424085 |
| ENSMUSG00000027823 | 1.009644739 | 1 |
| ENSMUSG00000027824 | -1.053549062 | 0.778372272 |
| ENSMUSG00000027827 | 1.689207518 | 0.000468075 |
| ENSMUSG00000027828 | 1.036558292 | 0.956714001 |
| ENSMUSG00000027829 | -1.017021744 | 0.987056723 |
| ENSMUSG00000027831 | 1.255041023 | 0.002673185 |
| ENSMUSG00000027832 | 1.08165912 | 0.760792946 |
| ENSMUSG00000027833 | 1.244979124 | 0.012374571 |
| ENSMUSG00000027834 | -1.245749251 | 0.087890629 |
| ENSMUSG00000027835 | 1.011508076 | 1 |
| ENSMUSG00000027840 | -1.264617664 | 0.678783707 |
| ENSMUSG00000027843 | -1.061372833 | 0.807743274 |
| ENSMUSG00000027845 | -1.07339686 | 0.608497229 |
| ENSMUSG00000027848 | 1.915875408 | 0.000144589 |
| ENSMUSG00000027849 | 1.48578256 | 0.034909854 |
| ENSMUSG00000027852 | 1.026307942 | 0.970020309 |
| ENSMUSG00000027854 | 1.024585127 | 0.934760809 |
| ENSMUSG00000027855 | -1.051236092 | 0.734408673 |
| ENSMUSG00000027857 | 1.261042287 | 0.388375021 |
| ENSMUSG00000027858 | 1.383364823 | 0.070982993 |
| ENSMUSG00000027859 | 1.063214872 | 0.571060395 |
| ENSMUSG00000027860 | 1.023571715 | 0.936995747 |
| ENSMUSG00000027861 | -1.054169715 | 0.845282019 |
| ENSMUSG00000027863 | 1 | |
| ENSMUSG00000027864 | 1.095411798 | 0.661969313 |
| ENSMUSG00000027865 | -1.038901175 | 0.867195992 |
| ENSMUSG00000027867 | 1 | |
| ENSMUSG00000027868 | -1.099935596 | 0.608497229 |
| ENSMUSG00000027869 | 1 | |
| ENSMUSG00000027870 | 1 | |
| ENSMUSG00000027871 | 1 | |

| | | |
|---|---|---|
| ENSMUSG00000027875 | -1 | 1 |
| ENSMUSG00000027876 | 1.059463094 | 0.608497229 |
| ENSMUSG00000027878 | 1.091147918 | 0.621723903 |
| ENSMUSG00000027879 | 1.020190301 | 0.977489164 |
| ENSMUSG00000027880 | 1.24122829 | 0.042914396 |
| ENSMUSG00000027881 | -1.002060119 | 1 |
| ENSMUSG00000027882 | 1.011520471 | 1 |
| ENSMUSG00000027883 | 1.025392732 | 0.963621048 |
| ENSMUSG00000027884 | -1.097770724 | 0.612309538 |
| ENSMUSG00000027886 | -1.424178325 | 0.047955893 |
| ENSMUSG00000027887 | -1.245835383 | 0.319960972 |
| ENSMUSG00000027889 | 1.015019371 | 0.979002724 |
| ENSMUSG00000027890 | -1.33431282 | 0.009277859 |
| ENSMUSG00000027893 | 1.049263066 | 0.923351678 |
| ENSMUSG00000027894 | 1.189868469 | 0.273959198 |
| ENSMUSG00000027895 | 1 | |
| ENSMUSG00000027896 | 1.000932014 | 1 |
| ENSMUSG00000027900 | -1.07879629 | 0.648490166 |
| ENSMUSG00000027901 | -1.056814847 | 0.930711391 |
| ENSMUSG00000027902 | 1 | |
| ENSMUSG00000027905 | -1.08968449 | 0.686035733 |
| ENSMUSG00000027907 | -1.05698019 | 0.894591069 |
| ENSMUSG00000027908 | 1 | |
| ENSMUSG00000027912 | -1.084645496 | 0.608497229 |
| ENSMUSG00000027913 | -1.024718486 | 0.931480186 |
| ENSMUSG00000027919 | 1 | |
| ENSMUSG00000027923 | 1 | |
| ENSMUSG00000027925 | 1.161036672 | 0.399744772 |
| ENSMUSG00000027927 | 1.199989356 | 0.000021314 |
| ENSMUSG00000027931 | 1.264110753 | 0.146932868 |
| ENSMUSG00000027932 | -1.331796581 | 0.01222575 |
| ENSMUSG00000027933 | -1.048288994 | 0.861137762 |
| ENSMUSG00000027935 | -1.028688006 | 0.864155141 |
| ENSMUSG00000027936 | -1.022753407 | 0.910897509 |
| ENSMUSG00000027937 | -1.000060015 | 1 |
| ENSMUSG00000027938 | -1.10299923 | 0.556783372 |
| ENSMUSG00000027939 | -1.189128081 | 0.001907923 |
| ENSMUSG00000027940 | -1.15125685 | 0.585646836 |
| ENSMUSG00000027942 | 1.062277167 | 0.768480212 |
| ENSMUSG00000027944 | -1.034633497 | 0.922446617 |
| ENSMUSG00000027947 | 1.065678136 | 0.608497229 |
| ENSMUSG00000027950 | 1.055095081 | 0.86498088 |
| ENSMUSG00000027951 | -1.048328071 | 0.775885777 |
| ENSMUSG00000027952 | -1.211063873 | 0.072407958 |
| ENSMUSG00000027953 | -1.002367828 | 1 |
| ENSMUSG00000027954 | -1.09330366 | 0.519338787 |
| ENSMUSG00000027955 | 1.284829609 | 0.274269417 |
| ENSMUSG00000027956 | 1.302918202 | 0.017980376 |
| ENSMUSG00000027957 | -1.128748119 | 0.608497229 |
| ENSMUSG00000027959 | -1.156579651 | 0.375822989 |
| ENSMUSG00000027961 | -1.218423298 | 0.427054117 |
| ENSMUSG00000027962 | -1.239049464 | 0.26210641 |
| ENSMUSG00000027963 | -1.038172339 | 0.874341108 |
| ENSMUSG00000027965 | 1 | |
| ENSMUSG00000027966 | -2.687062148 | 0.000129787 |
| ENSMUSG00000027967 | 1 | |
| ENSMUSG00000027968 | -1.146534203 | 0.585282823 |
| ENSMUSG00000027971 | -1.294395019 | 0.022079225 |
| ENSMUSG00000027973 | 1 | |
| ENSMUSG00000027977 | -1.059437071 | 0.677009735 |
| ENSMUSG00000027978 | 1.074476297 | 0.778372272 |
| ENSMUSG00000027981 | -1.069685023 | 0.698112512 |
| ENSMUSG00000027983 | -1.056880554 | 0.273221776 |

| | | |
|---|---|---|
| ENSMUSG00000027984 | -1.081503471 | 0.656277822 |
| ENSMUSG00000027985 | 1.037191383 | 0.698112512 |
| ENSMUSG00000027993 | -1.019755216 | 0.977625236 |
| ENSMUSG00000027994 | 1.056038687 | 0.662474108 |
| ENSMUSG00000027995 | 1.13527215 | 0.368089426 |
| ENSMUSG00000027996 | 1.189760276 | 0.014331568 |
| ENSMUSG00000027997 | -1.057725346 | 0.719170252 |
| ENSMUSG00000027998 | -1.025450902 | 0.968575228 |
| ENSMUSG00000027999 | -1.002641223 | 1 |
| ENSMUSG00000028001 | 1 | |
| ENSMUSG00000028003 | -1.085651418 | 0.608497229 |
| ENSMUSG00000028004 | -1 | 1 |
| ENSMUSG00000028005 | 1.026674953 | 0.82835102 |
| ENSMUSG00000028007 | -1.011791886 | 0.991890136 |
| ENSMUSG00000028008 | 1 | |
| ENSMUSG00000028009 | 1.24122829 | 0.042914396 |
| ENSMUSG00000028010 | -1.018066959 | 0.958600282 |
| ENSMUSG00000028011 | -1.028456669 | 0.990475194 |
| ENSMUSG00000028012 | -1.034369725 | 0.947794359 |
| ENSMUSG00000028013 | -1.007106822 | 1 |
| ENSMUSG00000028015 | -1.192362552 | 0.044301458 |
| ENSMUSG00000028016 | -1.041795341 | 0.860327141 |
| ENSMUSG00000028017 | -1.179766957 | 0.020070692 |
| ENSMUSG00000028018 | -1.044424653 | 0.856246989 |
| ENSMUSG00000028019 | 1.18890123 | 0.238348873 |
| ENSMUSG00000028020 | -1.174943658 | 0.486060044 |
| ENSMUSG00000028023 | -1.221963874 | 0.051327493 |
| ENSMUSG00000028024 | 1.189207115 | 0.313896003 |
| ENSMUSG00000028028 | -1.05220364 | 0.674977785 |
| ENSMUSG00000028029 | -1.054840945 | 0.831206498 |
| ENSMUSG00000028030 | -1.105287096 | 0.541735653 |
| ENSMUSG00000028031 | 1.428207243 | 0.080284922 |
| ENSMUSG00000028032 | -1.029645411 | 0.923395918 |
| ENSMUSG00000028033 | 1.011724659 | 0.981456085 |
| ENSMUSG00000028034 | 1.048660119 | 0.882514615 |
| ENSMUSG00000028035 | 1.108476867 | 0.608497229 |
| ENSMUSG00000028036 | 1.471869879 | 0.009992125 |
| ENSMUSG00000028037 | -1.083415624 | 0.857820356 |
| ENSMUSG00000028039 | -1.02002554 | 0.887896098 |
| ENSMUSG00000028040 | 1.10335239 | 0.236224875 |
| ENSMUSG00000028041 | -1.032537349 | 0.82683189 |
| ENSMUSG00000028042 | 1.227967056 | 0.029399381 |
| ENSMUSG00000028044 | 1.035308529 | 0.872807809 |
| ENSMUSG00000028047 | -1.026162871 | 0.884125698 |
| ENSMUSG00000028048 | 1.139734913 | 0.46330082 |
| ENSMUSG00000028049 | 1.042313865 | 0.778849035 |
| ENSMUSG00000028051 | -1.119132387 | 0.608497229 |
| ENSMUSG00000028053 | -1.01544967 | 1 |
| ENSMUSG00000028057 | -1.011304268 | 0.997667419 |
| ENSMUSG00000028059 | 1.08634612 | 0.698112512 |
| ENSMUSG00000028060 | 1.009145058 | 1 |
| ENSMUSG00000028062 | -1.000715932 | 1 |
| ENSMUSG00000028063 | 1.189870447 | 0.231448981 |
| ENSMUSG00000028064 | 1.105698266 | 0.538976184 |
| ENSMUSG00000028066 | -1.005504333 | 1 |
| ENSMUSG00000028068 | -1.042373906 | 0.828701947 |
| ENSMUSG00000028069 | -1.046657863 | 0.793821774 |
| ENSMUSG00000028070 | -1.065109712 | 0.693859529 |
| ENSMUSG00000028071 | -1 | 1 |
| ENSMUSG00000028072 | -1.247667166 | 0.311204699 |
| ENSMUSG00000028073 | -1.406487223 | 0.005765635 |
| ENSMUSG00000028076 | -1.378935039 | 0.004416753 |
| ENSMUSG00000028078 | 1.093422521 | 0.277459955 |

| | | |
|---|---|---|
| ENSMUSG00000028080 | -1.107754832 | 0.608497229 |
| ENSMUSG00000028081 | -1.024551086 | 0.988263302 |
| ENSMUSG00000028082 | -1.168171682 | 0.394093242 |
| ENSMUSG00000028085 | 1.020654018 | 0.865583959 |
| ENSMUSG00000028086 | 1.068942886 | 0.608497229 |
| ENSMUSG00000028088 | -1.126392128 | 0.232747283 |
| ENSMUSG00000028089 | -1.047113952 | 0.803684084 |
| ENSMUSG00000028093 | -1.184753793 | 0.07148774 |
| ENSMUSG00000028096 | -1.115198641 | 0.519338787 |
| ENSMUSG00000028098 | -1.043273881 | 0.88904823 |
| ENSMUSG00000028099 | -1.049934963 | 0.814608539 |
| ENSMUSG00000028100 | -1.132651343 | 0.694414714 |
| ENSMUSG00000028101 | -1.002120947 | 1 |
| ENSMUSG00000028102 | -1.053326608 | 0.740841499 |
| ENSMUSG00000028104 | -1.190647388 | 0.059761804 |
| ENSMUSG00000028106 | -1.01117196 | 1 |
| ENSMUSG00000028107 | -1.010563306 | 0.985059412 |
| ENSMUSG00000028108 | 1.176890179 | 0.216046507 |
| ENSMUSG00000028109 | 1 | |
| ENSMUSG00000028111 | -1.226278262 | 0.211778008 |
| ENSMUSG00000028114 | -1.059087186 | 0.704170349 |
| ENSMUSG00000028115 | -1.119741203 | 0.728544956 |
| ENSMUSG00000028116 | 1.038967555 | 0.952049609 |
| ENSMUSG00000028121 | 1.115559966 | 0.098925408 |
| ENSMUSG00000028124 | -1.068737568 | 0.778239858 |
| ENSMUSG00000028125 | -1.007653289 | 1 |
| ENSMUSG00000028126 | 1.019692265 | 0.970734719 |
| ENSMUSG00000028127 | -1.043010261 | 0.891020425 |
| ENSMUSG00000028128 | 1.885445097 | 0.000620276 |
| ENSMUSG00000028132 | -1.050021727 | 0.739850278 |
| ENSMUSG00000028133 | 1.007921461 | 0.988424085 |
| ENSMUSG00000028134 | 1.022609334 | 0.973528795 |
| ENSMUSG00000028136 | -1.093133111 | 0.719652885 |
| ENSMUSG00000028137 | 1.057851258 | 0.698112512 |
| ENSMUSG00000028138 | -1.070928389 | 0.781240509 |
| ENSMUSG00000028139 | -1.03752261 | 0.877273255 |
| ENSMUSG00000028140 | -1.020523658 | 0.969282824 |
| ENSMUSG00000028141 | 1.312396535 | 0.440395661 |
| ENSMUSG00000028145 | -1.029254865 | 0.875837626 |
| ENSMUSG00000028148 | 1 | |
| ENSMUSG00000028149 | -1.038163797 | 0.927576194 |
| ENSMUSG00000028150 | 1.679873981 | 0.002923873 |
| ENSMUSG00000028152 | -1.010567399 | 1 |
| ENSMUSG00000028156 | -1.116341095 | 0.698112512 |
| ENSMUSG00000028158 | -1.035478748 | 0.730297627 |
| ENSMUSG00000028159 | -1.09321836 | 0.394788021 |
| ENSMUSG00000028161 | 1.08204444 | 0.698112512 |
| ENSMUSG00000028163 | 1.016693517 | 0.988424085 |
| ENSMUSG00000028164 | 1.146217947 | 0.414805624 |
| ENSMUSG00000028165 | -1.042473862 | 0.87726305 |
| ENSMUSG00000028167 | 1.364899427 | 0.028511322 |
| ENSMUSG00000028172 | -1.03962336 | 0.698112512 |
| ENSMUSG00000028173 | -1.117662577 | 0.737649562 |
| ENSMUSG00000028174 | 1 | |
| ENSMUSG00000028175 | -1.097188338 | 0.682782733 |
| ENSMUSG00000028176 | 2.124436446 | 0.00182595 |
| ENSMUSG00000028177 | -1.115675116 | 0.70141105 |
| ENSMUSG00000028179 | 1.035836163 | 0.887835871 |
| ENSMUSG00000028180 | -1.0078879 | 1 |
| ENSMUSG00000028182 | 1.022589841 | 0.969400467 |
| ENSMUSG00000028184 | 1.232250781 | 0.163324662 |
| ENSMUSG00000028185 | 1 | |
| ENSMUSG00000028186 | -1.043306199 | 0.79291651 |

| | | |
|---|---|---|
| ENSMUSG00000028187 | -1.077728949 | 0.66875675 |
| ENSMUSG00000028188 | 1.098904155 | 0.563957069 |
| ENSMUSG00000028189 | -1.006502635 | 1 |
| ENSMUSG00000028191 | -1.069747762 | 0.710698346 |
| ENSMUSG00000028194 | -1.022983585 | 0.976289775 |
| ENSMUSG00000028195 | 1.0516483 | 0.880723094 |
| ENSMUSG00000028197 | 1.085932308 | 0.64374886 |
| ENSMUSG00000028199 | 1.054687597 | 0.721785396 |
| ENSMUSG00000028207 | -1.026003404 | 0.965125182 |
| ENSMUSG00000028211 | 1.428419169 | 0.027266023 |
| ENSMUSG00000028212 | 1.016390663 | 0.998129222 |
| ENSMUSG00000028214 | 1.360945841 | 0.007767672 |
| ENSMUSG00000028217 | -1.055048236 | 0.915646233 |
| ENSMUSG00000028218 | -1.008985569 | 1 |
| ENSMUSG00000028221 | 1.078850048 | 0.70299173 |
| ENSMUSG00000028222 | -1.073680831 | 0.698112512 |
| ENSMUSG00000028223 | 1.089010028 | 0.615267463 |
| ENSMUSG00000028224 | -1.012410444 | 1 |
| ENSMUSG00000028226 | 1.319463586 | 0.010805293 |
| ENSMUSG00000028228 | -1.009844969 | 1 |
| ENSMUSG00000028229 | -1.075724514 | 0.43857647 |
| ENSMUSG00000028232 | 1.017284855 | 0.96812207 |
| ENSMUSG00000028233 | -1.015299504 | 0.985845812 |
| ENSMUSG00000028234 | 1.107613338 | 0.66877097 |
| ENSMUSG00000028236 | 1 | |
| ENSMUSG00000028238 | 1.059463094 | 0.608497229 |
| ENSMUSG00000028240 | -1 | 1 |
| ENSMUSG00000028243 | -1.071339813 | 0.698112512 |
| ENSMUSG00000028244 | 1 | |
| ENSMUSG00000028245 | 1.01168621 | 1 |
| ENSMUSG00000028246 | 1.137385715 | 0.458420615 |
| ENSMUSG00000028247 | -1.017249344 | 0.964995983 |
| ENSMUSG00000028248 | 1.055473054 | 0.828564184 |
| ENSMUSG00000028249 | 1.024435557 | 0.982838256 |
| ENSMUSG00000028251 | 1.010157093 | 0.985522133 |
| ENSMUSG00000028252 | -1.050375148 | 0.819421998 |
| ENSMUSG00000028255 | -1.116853908 | 0.679917411 |
| ENSMUSG00000028256 | -1.047017155 | 0.829954854 |
| ENSMUSG00000028259 | 1 | |
| ENSMUSG00000028261 | 1.007717444 | 1 |
| ENSMUSG00000028262 | 1.288798261 | 0.312205997 |
| ENSMUSG00000028264 | 1 | |
| ENSMUSG00000028266 | 1.153256753 | 0.353003387 |
| ENSMUSG00000028268 | -1.475412283 | 0.011904275 |
| ENSMUSG00000028270 | -1.515762332 | 0.005329648 |
| ENSMUSG00000028271 | -1.062036512 | 0.752411447 |
| ENSMUSG00000028273 | 1.409292456 | 0.079359143 |
| ENSMUSG00000028274 | -1.019618534 | 0.964236174 |
| ENSMUSG00000028277 | -1.024612262 | 0.966465087 |
| ENSMUSG00000028278 | 1.2368373 | 0.065674478 |
| ENSMUSG00000028280 | -1.03962336 | 0.698112512 |
| ENSMUSG00000028282 | 1.028356207 | 0.950828026 |
| ENSMUSG00000028284 | -1.027368923 | 0.964707763 |
| ENSMUSG00000028287 | 1.222252407 | 0.492712594 |
| ENSMUSG00000028289 | 1.118098718 | 0.608497229 |
| ENSMUSG00000028291 | 1.027976936 | 0.938593635 |
| ENSMUSG00000028292 | -1.001955081 | 1 |
| ENSMUSG00000028293 | 1.000304256 | 1 |
| ENSMUSG00000028294 | -1.263518511 | 0.02397285 |
| ENSMUSG00000028295 | -1.01266685 | 0.94533054 |
| ENSMUSG00000028298 | 1 | |
| ENSMUSG00000028300 | -1.035239376 | 0.886993753 |
| ENSMUSG00000028307 | -1.03962336 | 0.814089245 |

| | | |
|---|---|---|
| ENSMUSG00000028309 | -1.024984223 | 0.964236174 |
| ENSMUSG00000028310 | 1 | |
| ENSMUSG00000028312 | -1.000215504 | 1 |
| ENSMUSG00000028314 | 1 | |
| ENSMUSG00000028318 | -1.109504017 | 0.486457995 |
| ENSMUSG00000028322 | 1.004278912 | 1 |
| ENSMUSG00000028327 | -1.120476206 | 0.534652019 |
| ENSMUSG00000028328 | 1.216869895 | 0.40700929 |
| ENSMUSG00000028329 | -1.139551285 | 0.126666392 |
| ENSMUSG00000028330 | -1.013553184 | 1 |
| ENSMUSG00000028331 | 1.025098295 | 0.751729496 |
| ENSMUSG00000028332 | 1 | |
| ENSMUSG00000028333 | 1.051350064 | 0.906826915 |
| ENSMUSG00000028334 | 1.027496087 | 0.929196072 |
| ENSMUSG00000028337 | -2.007498485 | 0.000242314 |
| ENSMUSG00000028339 | 2.272269907 | 0.000609185 |
| ENSMUSG00000028341 | -1.317614681 | 0.157466578 |
| ENSMUSG00000028343 | 1.010600304 | 1 |
| ENSMUSG00000028344 | 1.0224187 | 0.938939969 |
| ENSMUSG00000028345 | 1.043667019 | 0.880048186 |
| ENSMUSG00000028347 | -1.11463257 | 0.521195844 |
| ENSMUSG00000028348 | -1.261520168 | 0.113805611 |
| ENSMUSG00000028351 | 1.014545335 | 0.989648415 |
| ENSMUSG00000028354 | -2.094746034 | 0.000188484 |
| ENSMUSG00000028356 | 1 | |
| ENSMUSG00000028357 | 1 | |
| ENSMUSG00000028358 | 1.338049518 | 0.004223952 |
| ENSMUSG00000028359 | 1.171563499 | 0.233412931 |
| ENSMUSG00000028360 | -1.044273782 | 0.790851299 |
| ENSMUSG00000028362 | -1.044273782 | 0.698112512 |
| ENSMUSG00000028364 | 1.11706505 | 0.622419777 |
| ENSMUSG00000028367 | 1.023695517 | 0.981758965 |
| ENSMUSG00000028369 | 1.085586822 | 0.61572102 |
| ENSMUSG00000028370 | 2.169205365 | 0.000406474 |
| ENSMUSG00000028373 | -1.053549062 | 0.698112512 |
| ENSMUSG00000028378 | -1.088005642 | 0.608502051 |
| ENSMUSG00000028381 | -1.05473065 | 0.84515778 |
| ENSMUSG00000028382 | -1.007606912 | 1 |
| ENSMUSG00000028383 | -1.083817536 | 0.64008092 |
| ENSMUSG00000028385 | 1.086357081 | 0.608497229 |
| ENSMUSG00000028386 | -1 | 1 |
| ENSMUSG00000028389 | -1.179273376 | 0.286384775 |
| ENSMUSG00000028391 | -1.097003854 | 0.03650352 |
| ENSMUSG00000028392 | -1.080291586 | 0.698112512 |
| ENSMUSG00000028393 | 1.111315922 | 0.523553912 |
| ENSMUSG00000028394 | -1.007504141 | 1 |
| ENSMUSG00000028396 | 1 | |
| ENSMUSG00000028397 | -1.008711804 | 1 |
| ENSMUSG00000028398 | 1.041040802 | 0.608497229 |
| ENSMUSG00000028399 | -1.016858686 | 0.983829376 |
| ENSMUSG00000028402 | 1.098346871 | 0.698112512 |
| ENSMUSG00000028403 | 1.017278205 | 0.986542466 |
| ENSMUSG00000028405 | 1.007978564 | 1 |
| ENSMUSG00000028407 | 1.04360581 | 0.937505238 |
| ENSMUSG00000028409 | -1.032451018 | 0.959892369 |
| ENSMUSG00000028410 | -1.033207959 | 0.956807067 |
| ENSMUSG00000028411 | -1.003035837 | 1 |
| ENSMUSG00000028412 | -1.016918543 | 0.993643933 |
| ENSMUSG00000028413 | 1.13663646 | 0.608497229 |
| ENSMUSG00000028414 | 1.028149184 | 0.945625364 |
| ENSMUSG00000028415 | 1.199989356 | 0.000021314 |
| ENSMUSG00000028416 | -1.018760624 | 0.995295685 |
| ENSMUSG00000028417 | -1.089497311 | 0.614307028 |

| | | |
|---|---|---|
| ENSMUSG00000028419 | -1.061002482 | 0.806553232 |
| ENSMUSG00000028420 | 1.099541433 | 0.387707014 |
| ENSMUSG00000028423 | 1.031769082 | 0.938939969 |
| ENSMUSG00000028426 | 1.052958915 | 0.8920795 |
| ENSMUSG00000028427 | 1.077175441 | 0.838710806 |
| ENSMUSG00000028430 | 1.016257681 | 0.955042196 |
| ENSMUSG00000028431 | 1.057520498 | 0.813372967 |
| ENSMUSG00000028433 | 1.065174166 | 0.731295483 |
| ENSMUSG00000028434 | -1.28375406 | 0.238348873 |
| ENSMUSG00000028435 | -1.106875245 | 0.698112512 |
| ENSMUSG00000028436 | -1.05030652 | 0.857214752 |
| ENSMUSG00000028437 | 1.00396383 | 1 |
| ENSMUSG00000028438 | -1.06886504 | 0.609541771 |
| ENSMUSG00000028439 | -1.133438371 | 0.174584313 |
| ENSMUSG00000028441 | -1.227574303 | 0.335805609 |
| ENSMUSG00000028443 | 1.033852559 | 0.806526711 |
| ENSMUSG00000028444 | -1.594032973 | 0.026386967 |
| ENSMUSG00000028445 | -1.245557959 | 0.442177828 |
| ENSMUSG00000028447 | 1.008856818 | 0.979002724 |
| ENSMUSG00000028451 | 1.055095489 | 0.724087707 |
| ENSMUSG00000028452 | -1.005425863 | 1 |
| ENSMUSG00000028453 | -1.036137664 | 0.786110591 |
| ENSMUSG00000028454 | -1.001863984 | 1 |
| ENSMUSG00000028455 | -1.061866255 | 0.756778138 |
| ENSMUSG00000028456 | -1.295471196 | 0.03424578 |
| ENSMUSG00000028457 | 1.088512609 | 0.715092973 |
| ENSMUSG00000028458 | 1.051881335 | 0.657292031 |
| ENSMUSG00000028459 | -1.108850765 | 0.666909797 |
| ENSMUSG00000028460 | 1 | |
| ENSMUSG00000028461 | -1.030849616 | 0.861522196 |
| ENSMUSG00000028463 | 1.36953923 | 0.009960788 |
| ENSMUSG00000028464 | 1.555684947 | 0.022971084 |
| ENSMUSG00000028465 | 1.150122932 | 0.568304986 |
| ENSMUSG00000028466 | -1.058482659 | 0.722552422 |
| ENSMUSG00000028467 | -1.117825003 | 0.257777505 |
| ENSMUSG00000028468 | 1.023245823 | 0.940277706 |
| ENSMUSG00000028469 | -1.04338837 | 0.743842221 |
| ENSMUSG00000028470 | -1.14597657 | 0.129954829 |
| ENSMUSG00000028475 | 1.09981644 | 0.731295483 |
| ENSMUSG00000028476 | -1.108678492 | 0.430536837 |
| ENSMUSG00000028478 | 1.031082478 | 0.936995747 |
| ENSMUSG00000028479 | 1.05779397 | 0.534944201 |
| ENSMUSG00000028480 | 1.118866261 | 0.656734444 |
| ENSMUSG00000028483 | -1.001304481 | 1 |
| ENSMUSG00000028484 | 1.002957413 | 1 |
| ENSMUSG00000028487 | -1.166749852 | 0.073443065 |
| ENSMUSG00000028488 | -1.496579001 | 0.041628071 |
| ENSMUSG00000028492 | -1.03962336 | 0.698112512 |
| ENSMUSG00000028494 | 1.133904062 | 0.539959585 |
| ENSMUSG00000028495 | -1.006945192 | 1 |
| ENSMUSG00000028496 | 1.187432854 | 0.27163566 |
| ENSMUSG00000028497 | 1.084123096 | 0.698112512 |
| ENSMUSG00000028514 | -1.001789186 | 1 |
| ENSMUSG00000028517 | 1.635348151 | 0.000811218 |
| ENSMUSG00000028518 | 1.023006014 | 0.899151213 |
| ENSMUSG00000028519 | 1 | |
| ENSMUSG00000028520 | 1 | |
| ENSMUSG00000028521 | -1.046500077 | 0.784698888 |
| ENSMUSG00000028522 | 1.013979738 | 1 |
| ENSMUSG00000028523 | 1 | |
| ENSMUSG00000028524 | 1.059414397 | 0.709821953 |
| ENSMUSG00000028525 | 1.189162177 | 0.047464904 |
| ENSMUSG00000028527 | 1.301724416 | 0.000960133 |

| | | |
|---|---|---|
| ENSMUSG00000028528 | -1.161107412 | 0.37067934 |
| ENSMUSG00000028530 | 1.027508231 | 0.976562967 |
| ENSMUSG00000028532 | 1.03589275 | 0.839688843 |
| ENSMUSG00000028533 | -1.03962336 | 0.698112512 |
| ENSMUSG00000028536 | 1 | |
| ENSMUSG00000028538 | 1.021266781 | 0.843734664 |
| ENSMUSG00000028539 | -1.053342831 | 0.608497229 |
| ENSMUSG00000028540 | 1.034629952 | 0.757318144 |
| ENSMUSG00000028541 | 1.144654214 | 0.040506622 |
| ENSMUSG00000028542 | 1.439132548 | 0.003401228 |
| ENSMUSG00000028544 | 1 | |
| ENSMUSG00000028545 | -1.006005872 | 0.94651494 |
| ENSMUSG00000028546 | 1.46839725 | 0.158414349 |
| ENSMUSG00000028549 | -1.017019324 | 0.968158081 |
| ENSMUSG00000028550 | 1.036399093 | 0.709821953 |
| ENSMUSG00000028551 | 1.088817409 | 0.572892557 |
| ENSMUSG00000028552 | -1.03479618 | 0.941297739 |
| ENSMUSG00000028553 | -1.116853908 | 0.608497229 |
| ENSMUSG00000028555 | 1.168672293 | 0.018885322 |
| ENSMUSG00000028556 | 1.039264123 | 0.932242329 |
| ENSMUSG00000028557 | 1.09040582 | 0.72320398 |
| ENSMUSG00000028558 | 1.19479981 | 0.342440157 |
| ENSMUSG00000028559 | 1.013993245 | 1 |
| ENSMUSG00000028560 | 1.048779123 | 0.884638056 |
| ENSMUSG00000028563 | -1.003960031 | 1 |
| ENSMUSG00000028565 | 1.131730254 | 0.182140735 |
| ENSMUSG00000028567 | 1.02372626 | 0.963621048 |
| ENSMUSG00000028568 | 1.011116434 | 1 |
| ENSMUSG00000028571 | -1.02819776 | 0.963838183 |
| ENSMUSG00000028572 | -2.574935206 | 0.000263567 |
| ENSMUSG00000028573 | 1 | 1 |
| ENSMUSG00000028575 | -1.042426269 | 0.951254241 |
| ENSMUSG00000028576 | -1.011833616 | 1 |
| ENSMUSG00000028577 | 1.033823554 | 0.948083048 |
| ENSMUSG00000028578 | -1.011601955 | 0.993169601 |
| ENSMUSG00000028580 | 1.065178485 | 0.76318686 |
| ENSMUSG00000028581 | 1.132639128 | 0.217803313 |
| ENSMUSG00000028582 | 1.02722272 | 0.876260678 |
| ENSMUSG00000028583 | -4.214825772 | 0.000101355 |
| ENSMUSG00000028584 | 1 | |
| ENSMUSG00000028587 | 1.119513176 | 0.393883005 |
| ENSMUSG00000028589 | 1 | |
| ENSMUSG00000028590 | 1 | |
| ENSMUSG00000028591 | 1 | |
| ENSMUSG00000028593 | 1 | |
| ENSMUSG00000028597 | -2.58312232 | 2.06857E-05 |
| ENSMUSG00000028599 | 1.169756851 | 0.408022197 |
| ENSMUSG00000028600 | 1.405549744 | 0.099129119 |
| ENSMUSG00000028601 | -1.083840184 | 0.116240509 |
| ENSMUSG00000028602 | -1.047306572 | 0.894839966 |
| ENSMUSG00000028603 | -1.006105096 | 1 |
| ENSMUSG00000028607 | 1.007798436 | 1 |
| ENSMUSG00000028608 | 1.014816282 | 0.976788866 |
| ENSMUSG00000028609 | -1.014818452 | 0.998756776 |
| ENSMUSG00000028610 | 1 | |
| ENSMUSG00000028612 | 1 | |
| ENSMUSG00000028613 | -1.109577326 | 0.501683218 |
| ENSMUSG00000028614 | -1.070389568 | 0.806002111 |
| ENSMUSG00000028617 | 1.066813349 | 0.719191842 |
| ENSMUSG00000028618 | -1.011497615 | 1 |
| ENSMUSG00000028619 | 1.024234945 | 0.856047166 |
| ENSMUSG00000028621 | -1.042270266 | 0.688449757 |
| ENSMUSG00000028622 | -1.025130779 | 0.939658598 |

| | | |
|---|---|---|
| ENSMUSG00000028626 | 1.028323353 | 0.936340683 |
| ENSMUSG00000028629 | 1.10793014 | 0.3305975 |
| ENSMUSG00000028630 | 1.134355326 | 0.247298265 |
| ENSMUSG00000028631 | -1.03962336 | 0.803072249 |
| ENSMUSG00000028633 | -1.024618909 | 0.982838256 |
| ENSMUSG00000028634 | 1.048823004 | 0.608497229 |
| ENSMUSG00000028635 | 1.005212439 | 1 |
| ENSMUSG00000028636 | -1.004961522 | 1 |
| ENSMUSG00000028637 | 1.001055056 | 1 |
| ENSMUSG00000028639 | 1.082203286 | 0.787803345 |
| ENSMUSG00000028640 | 1.059463094 | 0.608497229 |
| ENSMUSG00000028641 | -1.014758101 | 0.980235982 |
| ENSMUSG00000028642 | -1.071075483 | 0.743300655 |
| ENSMUSG00000028643 | -1.035993288 | 0.642880527 |
| ENSMUSG00000028644 | 1.100395091 | 0.698112512 |
| ENSMUSG00000028645 | 1.250274549 | 0.05762786 |
| ENSMUSG00000028646 | 1.20322884 | 0.364493369 |
| ENSMUSG00000028647 | -1.134842089 | 0.29674248 |
| ENSMUSG00000028648 | 1.01206442 | 0.96035552 |
| ENSMUSG00000028649 | 1.018493728 | 1 |
| ENSMUSG00000028651 | -1.02088404 | 0.949295635 |
| ENSMUSG00000028653 | -1.002435188 | 1 |
| ENSMUSG00000028654 | 1.238280088 | 0.035388598 |
| ENSMUSG00000028655 | -1.341704078 | 0.028539747 |
| ENSMUSG00000028656 | 1.187343765 | 0.539222653 |
| ENSMUSG00000028657 | 1.098879754 | 0.629872539 |
| ENSMUSG00000028661 | -1.219624708 | 0.299600207 |
| ENSMUSG00000028664 | -1.048413511 | 0.624047623 |
| ENSMUSG00000028668 | -1.034682073 | 0.924141494 |
| ENSMUSG00000028669 | -1.061471779 | 0.70299173 |
| ENSMUSG00000028670 | 1.028262771 | 0.867016657 |
| ENSMUSG00000028671 | -1.151875929 | 0.132868001 |
| ENSMUSG00000028672 | -1.04531751 | 0.783946584 |
| ENSMUSG00000028673 | -1.093344303 | 0.630205913 |
| ENSMUSG00000028675 | -1.036552055 | 0.921185021 |
| ENSMUSG00000028676 | 1.026850524 | 0.972980766 |
| ENSMUSG00000028677 | 1.075750633 | 0.608497229 |
| ENSMUSG00000028678 | -1.023743664 | 0.970918992 |
| ENSMUSG00000028680 | -1.07203097 | 0.57224125 |
| ENSMUSG00000028681 | -1.134630705 | 0.514653872 |
| ENSMUSG00000028683 | -1.042473254 | 0.825401367 |
| ENSMUSG00000028684 | 1.011597951 | 0.979002724 |
| ENSMUSG00000028687 | -1.01625785 | 0.864550826 |
| ENSMUSG00000028688 | 1.019378476 | 0.94160226 |
| ENSMUSG00000028689 | -1.06810586 | 0.532451763 |
| ENSMUSG00000028690 | 1.058593593 | 0.685669046 |
| ENSMUSG00000028691 | 1.037380536 | 0.961940234 |
| ENSMUSG00000028692 | 1.034778235 | 0.950378392 |
| ENSMUSG00000028693 | -1.010284174 | 1 |
| ENSMUSG00000028696 | -1.037191916 | 0.879800829 |
| ENSMUSG00000028698 | -1.022603293 | 0.971347296 |
| ENSMUSG00000028699 | 1.085852928 | 0.758575113 |
| ENSMUSG00000028700 | -1.001599673 | 1 |
| ENSMUSG00000028701 | -1.101268461 | 0.252861426 |
| ENSMUSG00000028702 | -1.088223877 | 0.698112512 |
| ENSMUSG00000028703 | 1.016843531 | 0.951492347 |
| ENSMUSG00000028706 | -1.074497242 | 0.608497229 |
| ENSMUSG00000028707 | -1.34025079 | 0.26048326 |
| ENSMUSG00000028708 | -1.054401351 | 0.698112512 |
| ENSMUSG00000028709 | -1.129016048 | 0.273277814 |
| ENSMUSG00000028710 | -1.07065637 | 0.677534336 |
| ENSMUSG00000028712 | 1.05410549 | 0.805157604 |
| ENSMUSG00000028713 | 1 | |

| | | |
|---|---|---|
| ENSMUSG00000028715 | 1 | |
| ENSMUSG00000028716 | -1.043306199 | 0.698112512 |
| ENSMUSG00000028717 | -1 | 1 |
| ENSMUSG00000028718 | -1.00949191 | 1 |
| ENSMUSG00000028719 | -1.03679413 | 0.928742903 |
| ENSMUSG00000028729 | 1.047077298 | 0.867939428 |
| ENSMUSG00000028730 | -1.029113823 | 0.929923764 |
| ENSMUSG00000028736 | 1 | |
| ENSMUSG00000028737 | -1.076833236 | 0.355528294 |
| ENSMUSG00000028738 | -1.195474018 | 0.469856779 |
| ENSMUSG00000028741 | -1.029649064 | 0.939380482 |
| ENSMUSG00000028743 | -1.012170755 | 0.992464064 |
| ENSMUSG00000028744 | 1.094219556 | 0.048258714 |
| ENSMUSG00000028745 | 1.133717361 | 0.608497229 |
| ENSMUSG00000028747 | -1.043306199 | 0.698112512 |
| ENSMUSG00000028749 | 1 | |
| ENSMUSG00000028750 | 1 | |
| ENSMUSG00000028751 | 1.095872691 | 0.698112512 |
| ENSMUSG00000028753 | 1 | |
| ENSMUSG00000028755 | 1.064587874 | 0.608497229 |
| ENSMUSG00000028756 | 1.104609654 | 0.278692299 |
| ENSMUSG00000028757 | 1.03164309 | 0.952057016 |
| ENSMUSG00000028758 | -1.142095915 | 0.608497229 |
| ENSMUSG00000028759 | 1.047752547 | 0.866317664 |
| ENSMUSG00000028760 | 1.105609962 | 0.60397857 |
| ENSMUSG00000028763 | 1.277305801 | 0.165208661 |
| ENSMUSG00000028766 | 1.245425369 | 0.073253366 |
| ENSMUSG00000028771 | 1.150942139 | 0.532470425 |
| ENSMUSG00000028772 | -1.065690861 | 0.757276824 |
| ENSMUSG00000028773 | 1.406705945 | 0.00431659 |
| ENSMUSG00000028776 | 1.07921618 | 0.608497229 |
| ENSMUSG00000028777 | 1 | |
| ENSMUSG00000028778 | 1.012792449 | 1 |
| ENSMUSG00000028779 | -1.010441588 | 1 |
| ENSMUSG00000028780 | 1.175744428 | 0.395481973 |
| ENSMUSG00000028782 | -1.364262235 | 0.079090855 |
| ENSMUSG00000028784 | 1.034095782 | 0.921763911 |
| ENSMUSG00000028785 | 1.267400994 | 0.235993175 |
| ENSMUSG00000028786 | 1.005212439 | 1 |
| ENSMUSG00000028788 | 1.043866298 | 0.931317924 |
| ENSMUSG00000028789 | -1.006062683 | 1 |
| ENSMUSG00000028790 | 1.037955209 | 0.949679283 |
| ENSMUSG00000028792 | -1.02850701 | 0.934760809 |
| ENSMUSG00000028793 | -1.048978646 | 0.799720228 |
| ENSMUSG00000028794 | -1.100276215 | 0.792416038 |
| ENSMUSG00000028795 | -1.16960279 | 0.056148024 |
| ENSMUSG00000028796 | 1.12340618 | 0.372103656 |
| ENSMUSG00000028797 | 1.00648356 | 1 |
| ENSMUSG00000028798 | 1.003174522 | 1 |
| ENSMUSG00000028799 | 1.104626393 | 0.327326823 |
| ENSMUSG00000028800 | -1.067798123 | 0.794274354 |
| ENSMUSG00000028801 | -1.023769022 | 0.954761885 |
| ENSMUSG00000028803 | -1.084559373 | 0.386955663 |
| ENSMUSG00000028804 | -1.056339972 | 0.748723464 |
| ENSMUSG00000028807 | -1.004054552 | 1 |
| ENSMUSG00000028809 | 1.043276815 | 0.908543402 |
| ENSMUSG00000028811 | 1.056743841 | 0.840991792 |
| ENSMUSG00000028813 | 1.059463094 | 0.608497229 |
| ENSMUSG00000028820 | 1.042330151 | 0.936995747 |
| ENSMUSG00000028821 | -1.011751202 | 0.991156701 |
| ENSMUSG00000028822 | 1.030525284 | 0.911212167 |
| ENSMUSG00000028825 | -1.111368617 | 0.690869613 |
| ENSMUSG00000028826 | 1.020785211 | 0.970543056 |

| | | |
|---|---|---|
| ENSMUSG00000028830 | 1.014692704 | 0.979152632 |
| ENSMUSG00000028832 | 1.063492965 | 0.810597167 |
| ENSMUSG00000028833 | 1.078004374 | 0.608497229 |
| ENSMUSG00000028834 | 1.059463094 | 0.743396505 |
| ENSMUSG00000028836 | -1.043077979 | 0.816693551 |
| ENSMUSG00000028837 | -1.004537522 | 1 |
| ENSMUSG00000028838 | -1.390435093 | 0.003733909 |
| ENSMUSG00000028840 | -1.10201663 | 0.785543892 |
| ENSMUSG00000028841 | -1.157184854 | 0.70299173 |
| ENSMUSG00000028842 | -1.008304655 | 1 |
| ENSMUSG00000028843 | 1.034583379 | 0.798123081 |
| ENSMUSG00000028845 | 1.040982152 | 0.789737136 |
| ENSMUSG00000028847 | 1.011775206 | 1 |
| ENSMUSG00000028848 | 1.018697597 | 0.807247723 |
| ENSMUSG00000028849 | 1.187416802 | 0.147541255 |
| ENSMUSG00000028850 | 1.079945827 | 0.425531155 |
| ENSMUSG00000028851 | -1.008320889 | 1 |
| ENSMUSG00000028854 | 1.015665628 | 0.949519706 |
| ENSMUSG00000028857 | 1.009382823 | 1 |
| ENSMUSG00000028859 | 1.159692085 | 0.5819787 |
| ENSMUSG00000028860 | -1.123466014 | 0.608497229 |
| ENSMUSG00000028861 | -1.023367325 | 0.950565815 |
| ENSMUSG00000028862 | -1.005241824 | 1 |
| ENSMUSG00000028863 | -1.000803905 | 1 |
| ENSMUSG00000028864 | 1.992280432 | 0.000761085 |
| ENSMUSG00000028865 | 1.059463094 | 0.698112512 |
| ENSMUSG00000028868 | 1.108499901 | 0.608497229 |
| ENSMUSG00000028869 | -1.023599435 | 0.96035552 |
| ENSMUSG00000028871 | -1.322145407 | 0.105884496 |
| ENSMUSG00000028873 | -1.019821302 | 0.974047716 |
| ENSMUSG00000028874 | 1 | |
| ENSMUSG00000028876 | 1.069068979 | 0.698112512 |
| ENSMUSG00000028878 | 1.053779276 | 0.781240509 |
| ENSMUSG00000028879 | -1.000051174 | 1 |
| ENSMUSG00000028882 | 1.014356871 | 1 |
| ENSMUSG00000028883 | 1.598063964 | 0.009701675 |
| ENSMUSG00000028884 | -1.023016884 | 0.964239113 |
| ENSMUSG00000028885 | -1.003845013 | 1 |
| ENSMUSG00000028886 | -1.004448876 | 1 |
| ENSMUSG00000028889 | 1.014415059 | 0.986247701 |
| ENSMUSG00000028890 | 1.044550072 | 0.737121422 |
| ENSMUSG00000028893 | 1.233976275 | 0.007628795 |
| ENSMUSG00000028894 | -1.108497441 | 0.608497229 |
| ENSMUSG00000028896 | 1.068734385 | 0.6973511 |
| ENSMUSG00000028898 | 1.008647721 | 1 |
| ENSMUSG00000028899 | -1.04189139 | 0.866289429 |
| ENSMUSG00000028901 | 1.050875283 | 0.773508225 |
| ENSMUSG00000028902 | -1.01137506 | 1 |
| ENSMUSG00000028906 | -1.005198013 | 1 |
| ENSMUSG00000028907 | 1.016195676 | 0.982037739 |
| ENSMUSG00000028909 | -1.006771768 | 1 |
| ENSMUSG00000028910 | -1.000427237 | 1 |
| ENSMUSG00000028911 | 1.062687701 | 0.477335321 |
| ENSMUSG00000028914 | 1.048077412 | 0.632838524 |
| ENSMUSG00000028917 | 1.075929379 | 0.532369657 |
| ENSMUSG00000028919 | 1.198585471 | 0.265145519 |
| ENSMUSG00000028920 | 1.00705355 | 1 |
| ENSMUSG00000028923 | 1.012794375 | 0.960579094 |
| ENSMUSG00000028926 | 1.16084375 | 0.446657568 |
| ENSMUSG00000028927 | 1.031988779 | 0.949218699 |
| ENSMUSG00000028931 | -1.095146676 | 0.245715752 |
| ENSMUSG00000028932 | 1.001915563 | 1 |
| ENSMUSG00000028933 | 1.039184777 | 0.822013059 |

| | | |
|---|---|---|
| ENSMUSG00000028936 | 1.018381013 | 0.998924031 |
| ENSMUSG00000028937 | -1.03071323 | 0.880338841 |
| ENSMUSG00000028938 | 1 | |
| ENSMUSG00000028940 | -1 | 1 |
| ENSMUSG00000028943 | -1.378996426 | 0.068433456 |
| ENSMUSG00000028944 | 1.245898672 | 0.122875649 |
| ENSMUSG00000028945 | 1.189472035 | 0.389331183 |
| ENSMUSG00000028946 | 1.059463094 | 0.608497229 |
| ENSMUSG00000028948 | 1.006286961 | 1 |
| ENSMUSG00000028949 | 1.074171421 | 0.458934581 |
| ENSMUSG00000028950 | -1.116408704 | 0.641354847 |
| ENSMUSG00000028952 | 1.031580857 | 0.874468781 |
| ENSMUSG00000028953 | 1.019343017 | 0.987029357 |
| ENSMUSG00000028954 | -1.015514074 | 0.990316418 |
| ENSMUSG00000028955 | -1.017333341 | 0.996913409 |
| ENSMUSG00000028957 | 1.247743364 | 0.039462477 |
| ENSMUSG00000028958 | 1.148324756 | 0.38030038 |
| ENSMUSG00000028959 | 1.110957526 | 0.378751351 |
| ENSMUSG00000028960 | 1.11619743 | 0.629323727 |
| ENSMUSG00000028961 | -1.013381181 | 0.998864352 |
| ENSMUSG00000028962 | 1.147006562 | 0.315874173 |
| ENSMUSG00000028963 | -1 | 1 |
| ENSMUSG00000028964 | 1.066811129 | 0.754823263 |
| ENSMUSG00000028965 | 1.052510562 | 0.608497229 |
| ENSMUSG00000028967 | 1.143896788 | 0.481851765 |
| ENSMUSG00000028969 | 1.109121188 | 0.484584889 |
| ENSMUSG00000028970 | 1.04677989 | 0.814608539 |
| ENSMUSG00000028971 | -1.050155915 | 0.883306173 |
| ENSMUSG00000028972 | 1.015486245 | 0.972980766 |
| ENSMUSG00000028973 | 1.027623746 | 0.79291651 |
| ENSMUSG00000028974 | -1.138541128 | 0.273601178 |
| ENSMUSG00000028975 | 1.084361375 | 0.613449588 |
| ENSMUSG00000028976 | -1.018998401 | 0.968937442 |
| ENSMUSG00000028977 | -1.271281137 | 0.055789053 |
| ENSMUSG00000028978 | -1.07452894 | 0.811581858 |
| ENSMUSG00000028979 | 1.143259513 | 0.64712035 |
| ENSMUSG00000028980 | 1.018542706 | 0.955101438 |
| ENSMUSG00000028982 | -1.05780701 | 0.660336669 |
| ENSMUSG00000028986 | -1.030647675 | 0.924587329 |
| ENSMUSG00000028988 | -1.134725081 | 0.212065431 |
| ENSMUSG00000028989 | 1.019208058 | 1 |
| ENSMUSG00000028990 | 1.08792071 | 0.608497229 |
| ENSMUSG00000028991 | 1.046321491 | 0.781475119 |
| ENSMUSG00000028992 | -1.045629819 | 0.779557148 |
| ENSMUSG00000028995 | 1.003119107 | 1 |
| ENSMUSG00000028996 | -1.044273782 | 0.698112512 |
| ENSMUSG00000028998 | 1.003847177 | 1 |
| ENSMUSG00000028999 | 1.056391185 | 0.757039001 |
| ENSMUSG00000029001 | -1.067264582 | 0.468615237 |
| ENSMUSG00000029003 | -1.014732564 | 0.952789999 |
| ENSMUSG00000029004 | 1.067009606 | 0.773487331 |
| ENSMUSG00000029005 | -1.237332719 | 0.080580067 |
| ENSMUSG00000029007 | -1.126206369 | 0.335498586 |
| ENSMUSG00000029009 | 1.138241641 | 0.193512301 |
| ENSMUSG00000029012 | 1.041889058 | 0.805587306 |
| ENSMUSG00000029014 | 1.046897572 | 0.87726305 |
| ENSMUSG00000029015 | 1 | |
| ENSMUSG00000029016 | 1.036557293 | 0.789142226 |
| ENSMUSG00000029017 | -1.016228983 | 0.999215557 |
| ENSMUSG00000029019 | 1.125102401 | 0.75692508 |
| ENSMUSG00000029020 | 1.039047399 | 0.905645252 |
| ENSMUSG00000029022 | 1.121692441 | 0.174108732 |
| ENSMUSG00000029026 | 1 | |

| | | |
|---|---|---|
| ENSMUSG00000029027 | 1.020126817 | 0.900934705 |
| ENSMUSG00000029028 | 1.001952937 | 1 |
| ENSMUSG00000029029 | 1.116434449 | 0.039522277 |
| ENSMUSG00000029030 | 1.003132859 | 1 |
| ENSMUSG00000029032 | -1.375176619 | 0.367797304 |
| ENSMUSG00000029033 | 1.037014124 | 0.698112512 |
| ENSMUSG00000029034 | -1.057710773 | 0.80937145 |
| ENSMUSG00000029036 | 1.026706185 | 0.929746187 |
| ENSMUSG00000029038 | 1.004512402 | 1 |
| ENSMUSG00000029044 | 1 | |
| ENSMUSG00000029047 | -1.112465836 | 0.330381384 |
| ENSMUSG00000029048 | 1.002340843 | 1 |
| ENSMUSG00000029049 | 1.106647135 | 0.698112512 |
| ENSMUSG00000029050 | 1.083506091 | 0.613963375 |
| ENSMUSG00000029053 | -1.6074904 | 0.001365136 |
| ENSMUSG00000029054 | 1 | |
| ENSMUSG00000029055 | -1.137443589 | 0.608497229 |
| ENSMUSG00000029056 | -1.024747915 | 0.869434134 |
| ENSMUSG00000029059 | -2.523024174 | 9.50669E-05 |
| ENSMUSG00000029060 | 1.112775805 | 0.535052277 |
| ENSMUSG00000029061 | 1.907011571 | 0.00095885 |
| ENSMUSG00000029062 | 1.028897522 | 0.945377365 |
| ENSMUSG00000029063 | 1.021228932 | 0.972980766 |
| ENSMUSG00000029064 | 1.053427257 | 0.914055277 |
| ENSMUSG00000029066 | -1.030601903 | 0.888008875 |
| ENSMUSG00000029068 | -1.042794945 | 0.863085098 |
| ENSMUSG00000029070 | 1.152067442 | 0.342054851 |
| ENSMUSG00000029071 | 1.024722792 | 0.8918465 |
| ENSMUSG00000029072 | -1.349208766 | 0.176776279 |
| ENSMUSG00000029073 | -1.015764707 | 0.959892369 |
| ENSMUSG00000029074 | 1 | |
| ENSMUSG00000029075 | 1.035750049 | 0.983469255 |
| ENSMUSG00000029076 | -1.024305596 | 0.976129946 |
| ENSMUSG00000029082 | -1.181172082 | 0.381622253 |
| ENSMUSG00000029084 | -1.29380053 | 0.141875736 |
| ENSMUSG00000029086 | -1.006359949 | 1 |
| ENSMUSG00000029088 | -1.276367163 | 0.025983253 |
| ENSMUSG00000029089 | 1.008783942 | 1 |
| ENSMUSG00000029090 | 1.089393171 | 0.698112512 |
| ENSMUSG00000029092 | 1.081642974 | 0.718265321 |
| ENSMUSG00000029093 | 1.532070924 | 0.000459033 |
| ENSMUSG00000029094 | 1.124571493 | 0.46330082 |
| ENSMUSG00000029095 | -1.584868055 | 0.055180382 |
| ENSMUSG00000029096 | 1.307812889 | 0.098346026 |
| ENSMUSG00000029097 | 1.087345916 | 0.138888882 |
| ENSMUSG00000029098 | -1.005292839 | 1 |
| ENSMUSG00000029101 | 1.128911394 | 0.413557762 |
| ENSMUSG00000029102 | -1.12843057 | 0.540891969 |
| ENSMUSG00000029103 | -1.172051052 | 0.419455697 |
| ENSMUSG00000029104 | 1.030671391 | 0.945888211 |
| ENSMUSG00000029106 | 1.049658039 | 0.886777003 |
| ENSMUSG00000029108 | 1.094708793 | 0.644596644 |
| ENSMUSG00000029110 | 1.067960988 | 0.81686894 |
| ENSMUSG00000029111 | 1.04832379 | 0.814089245 |
| ENSMUSG00000029112 | 1 | |
| ENSMUSG00000029119 | 1.061963403 | 0.621010054 |
| ENSMUSG00000029120 | -1.179614078 | 0.534887579 |
| ENSMUSG00000029121 | -1.06076275 | 0.70299173 |
| ENSMUSG00000029122 | 1.108407418 | 0.350223743 |
| ENSMUSG00000029123 | 1.39448093 | 0.071386941 |
| ENSMUSG00000029125 | -1.048959004 | 0.717035356 |
| ENSMUSG00000029126 | -1.070561089 | 0.698112512 |
| ENSMUSG00000029127 | 1.129097033 | 0.294194175 |

| | | |
|---|---|---|
| ENSMUSG00000029128 | -1.003131332 | 1 |
| ENSMUSG00000029130 | -1.248475757 | 0.03287112 |
| ENSMUSG00000029131 | 1.065332906 | 0.75280299 |
| ENSMUSG00000029134 | 1.406166871 | 0.053311176 |
| ENSMUSG00000029135 | 1.16077726 | 0.394324991 |
| ENSMUSG00000029136 | -1.145659129 | 0.499865648 |
| ENSMUSG00000029138 | 1.059463094 | 0.752398528 |
| ENSMUSG00000029141 | -1.016681727 | 0.981964814 |
| ENSMUSG00000029145 | -1.037597195 | 0.820334863 |
| ENSMUSG00000029146 | 1.009077717 | 1 |
| ENSMUSG00000029147 | 1.037615988 | 0.943663271 |
| ENSMUSG00000029148 | 1.020689213 | 0.979002724 |
| ENSMUSG00000029149 | 1.115889017 | 0.153364178 |
| ENSMUSG00000029151 | -1.26217421 | 0.255363785 |
| ENSMUSG00000029152 | 1.061702224 | 0.82448954 |
| ENSMUSG00000029153 | 1.165327751 | 0.243993149 |
| ENSMUSG00000029154 | -1 | 1 |
| ENSMUSG00000029155 | 1 | |
| ENSMUSG00000029156 | -1.178438699 | 0.251544797 |
| ENSMUSG00000029158 | 1 | |
| ENSMUSG00000029161 | -1.005217126 | 1 |
| ENSMUSG00000029162 | -1.000562164 | 1 |
| ENSMUSG00000029163 | 1.437774166 | 0.012481057 |
| ENSMUSG00000029165 | 1.087542743 | 0.250726313 |
| ENSMUSG00000029166 | -1.039056945 | 0.569787565 |
| ENSMUSG00000029167 | 1.786074512 | 0.002956463 |
| ENSMUSG00000029168 | -1.661562541 | 0.001448487 |
| ENSMUSG00000029169 | 1.020907281 | 0.998847216 |
| ENSMUSG00000029171 | -1.074261903 | 0.729860853 |
| ENSMUSG00000029173 | 1.013902505 | 0.97004457 |
| ENSMUSG00000029174 | -1.017174406 | 0.981214754 |
| ENSMUSG00000029175 | 1.148319522 | 0.191104658 |
| ENSMUSG00000029176 | 1.067525409 | 0.808818132 |
| ENSMUSG00000029177 | -1.012033318 | 1 |
| ENSMUSG00000029178 | 1.161674348 | 0.499658973 |
| ENSMUSG00000029179 | 1.030459463 | 0.873622104 |
| ENSMUSG00000029182 | 1 | |
| ENSMUSG00000029184 | 1 | |
| ENSMUSG00000029185 | 1.245030988 | 0.206620396 |
| ENSMUSG00000029186 | 1.026175845 | 0.936807101 |
| ENSMUSG00000029188 | -1.03962336 | 0.698112512 |
| ENSMUSG00000029189 | 1.014346713 | 1 |
| ENSMUSG00000029190 | 1.008124415 | 1 |
| ENSMUSG00000029191 | 1.00071498 | 1 |
| ENSMUSG00000029192 | 1.003315268 | 1 |
| ENSMUSG00000029193 | -1 | 1 |
| ENSMUSG00000029195 | 1.065713358 | 0.692423811 |
| ENSMUSG00000029196 | 1.024504145 | 0.94339214 |
| ENSMUSG00000029198 | 1.039058865 | 0.882079974 |
| ENSMUSG00000029199 | 1.023340305 | 0.955137373 |
| ENSMUSG00000029201 | 1.052155433 | 0.877639675 |
| ENSMUSG00000029202 | 1.052534456 | 0.89498287 |
| ENSMUSG00000029203 | 1.013806801 | 1 |
| ENSMUSG00000029204 | -1.029113823 | 0.963209064 |
| ENSMUSG00000029205 | 1 | |
| ENSMUSG00000029206 | 1.251996008 | 0.036681922 |
| ENSMUSG00000029207 | 1.262995501 | 0.181128445 |
| ENSMUSG00000029208 | -1.017443126 | 0.97567552 |
| ENSMUSG00000029209 | 1.180515647 | 0.114183481 |
| ENSMUSG00000029211 | 1.024693402 | 0.933031586 |
| ENSMUSG00000029212 | 1.146425306 | 0.697178241 |
| ENSMUSG00000029213 | 1.124866533 | 0.568304986 |
| ENSMUSG00000029217 | -1.020974396 | 0.924587329 |

| | | |
|---|---|---|
| ENSMUSG00000029219 | -1.113515092 | 0.576169279 |
| ENSMUSG00000029221 | 1.023423906 | 0.970771742 |
| ENSMUSG00000029223 | 1.02435866 | 0.858652152 |
| ENSMUSG00000029227 | 1.015227116 | 1 |
| ENSMUSG00000029228 | 1.12861328 | 0.687333535 |
| ENSMUSG00000029229 | 1.003193262 | 1 |
| ENSMUSG00000029231 | 1.429244782 | 0.000372179 |
| ENSMUSG00000029233 | 1.139359809 | 0.319685456 |
| ENSMUSG00000029234 | 1.107814411 | 0.632945896 |
| ENSMUSG00000029235 | -1.03962336 | 0.698112512 |
| ENSMUSG00000029236 | 1.019083579 | 0.968937442 |
| ENSMUSG00000029238 | 1.053629343 | 0.851662555 |
| ENSMUSG00000029245 | -1.0795912 | 0.739981019 |
| ENSMUSG00000029246 | 1.056585918 | 0.832839286 |
| ENSMUSG00000029247 | 1.049338684 | 0.920766577 |
| ENSMUSG00000029248 | -1.03962336 | 0.698112512 |
| ENSMUSG00000029249 | 1.094152507 | 0.687636369 |
| ENSMUSG00000029250 | 1.004380375 | 1 |
| ENSMUSG00000029253 | 1.099986112 | 0.648441922 |
| ENSMUSG00000029254 | -1.080687155 | 0.869361059 |
| ENSMUSG00000029255 | 1 | |
| ENSMUSG00000029260 | 1 | |
| ENSMUSG00000029263 | 1.080410391 | 0.608497229 |
| ENSMUSG00000029265 | 1.07527416 | 0.778195329 |
| ENSMUSG00000029267 | 1.10338804 | 0.671280603 |
| ENSMUSG00000029268 | 1 | |
| ENSMUSG00000029269 | 1 | |
| ENSMUSG00000029270 | 1.165360494 | 0.250477583 |
| ENSMUSG00000029272 | 1 | |
| ENSMUSG00000029273 | -1 | 1 |
| ENSMUSG00000029275 | -1.0795912 | 0.698112512 |
| ENSMUSG00000029276 | 1.046459724 | 0.788231748 |
| ENSMUSG00000029279 | -1.091604276 | 0.297090969 |
| ENSMUSG00000029280 | 1 | |
| ENSMUSG00000029281 | 1 | |
| ENSMUSG00000029282 | 1 | |
| ENSMUSG00000029283 | 1.080075702 | 0.698112512 |
| ENSMUSG00000029286 | 1 | |
| ENSMUSG00000029287 | 1.455258789 | 0.002261078 |
| ENSMUSG00000029288 | 1 | |
| ENSMUSG00000029290 | 1.023451722 | 0.959659993 |
| ENSMUSG00000029291 | 1.035153131 | 0.861528342 |
| ENSMUSG00000029298 | -1.32672912 | 0.021977352 |
| ENSMUSG00000029299 | -1 | 1 |
| ENSMUSG00000029304 | 1.820788287 | 0.00095885 |
| ENSMUSG00000029306 | 1 | |
| ENSMUSG00000029307 | 1 | |
| ENSMUSG00000029309 | 1.069437807 | 0.735725261 |
| ENSMUSG00000029310 | 1.131603201 | 0.274420574 |
| ENSMUSG00000029311 | 1.28002966 | 0.05762786 |
| ENSMUSG00000029312 | 1.028699238 | 0.779062229 |
| ENSMUSG00000029313 | 1.229011119 | 0.316332674 |
| ENSMUSG00000029314 | -1.283433137 | 0.088704932 |
| ENSMUSG00000029319 | 1.068638002 | 0.677534336 |
| ENSMUSG00000029320 | -1 | 1 |
| ENSMUSG00000029321 | 1 | |
| ENSMUSG00000029322 | 1.322818589 | 0.129167216 |
| ENSMUSG00000029326 | 1.125312467 | 0.295058308 |
| ENSMUSG00000029328 | 1.087147583 | 0.76060068 |
| ENSMUSG00000029330 | -1.468472718 | 0.025781546 |
| ENSMUSG00000029331 | 1 | |
| ENSMUSG00000029333 | -1.280186558 | 0.006306676 |
| ENSMUSG00000029334 | 1.664355175 | 0.005339511 |

| | | |
|---|---|---|
| ENSMUSG00000029335 | -1.05536902 | 0.872430778 |
| ENSMUSG00000029337 | -2.12284257 | 0.001006347 |
| ENSMUSG00000029338 | 1.127218262 | 0.534944201 |
| ENSMUSG00000029343 | 1.132639128 | 0.398767938 |
| ENSMUSG00000029344 | 1.195651976 | 0.035594665 |
| ENSMUSG00000029345 | 1.119143745 | 0.276523052 |
| ENSMUSG00000029346 | 1.053483479 | 0.473579199 |
| ENSMUSG00000029348 | -1.549418562 | 0.009914504 |
| ENSMUSG00000029352 | -1.222692478 | 0.472623153 |
| ENSMUSG00000029359 | 1.198839533 | 0.576169279 |
| ENSMUSG00000029360 | 1.229645575 | 0.608497229 |
| ENSMUSG00000029361 | -1.126344191 | 0.656389155 |
| ENSMUSG00000029363 | 1.005828827 | 1 |
| ENSMUSG00000029364 | 1.151943911 | 0.453518554 |
| ENSMUSG00000029366 | 1.112432855 | 0.698112512 |
| ENSMUSG00000029368 | 1.059463094 | 0.608497229 |
| ENSMUSG00000029369 | 1.069068979 | 0.698112512 |
| ENSMUSG00000029370 | 1.791379108 | 0.020077189 |
| ENSMUSG00000029371 | 1.339839231 | 0.005833055 |
| ENSMUSG00000029372 | -1.080615268 | 0.663593888 |
| ENSMUSG00000029373 | -1.044273782 | 0.698112512 |
| ENSMUSG00000029375 | 1.56056632 | 0.065945481 |
| ENSMUSG00000029376 | 1.094825643 | 0.62941321 |
| ENSMUSG00000029377 | 1.281943021 | 0.009616706 |
| ENSMUSG00000029378 | 1.342640805 | 0.018218585 |
| ENSMUSG00000029379 | -1.314962749 | 0.388393823 |
| ENSMUSG00000029380 | 1.012504908 | 0.997451029 |
| ENSMUSG00000029381 | -1.136577481 | 0.234965016 |
| ENSMUSG00000029384 | -1.047306572 | 0.844827111 |
| ENSMUSG00000029385 | 1.285427406 | 0.07931917 |
| ENSMUSG00000029386 | -1.032530939 | 0.807247723 |
| ENSMUSG00000029387 | 1.02118262 | 0.954523477 |
| ENSMUSG00000029388 | -1.011497015 | 1 |
| ENSMUSG00000029389 | 1.081283246 | 0.357543507 |
| ENSMUSG00000029390 | 1.088159822 | 0.797491747 |
| ENSMUSG00000029392 | 1.095943927 | 0.193393634 |
| ENSMUSG00000029394 | 1.00518675 | 1 |
| ENSMUSG00000029397 | 1.079949211 | 0.699841912 |
| ENSMUSG00000029401 | -1.008904138 | 1 |
| ENSMUSG00000029402 | 1.018981738 | 0.920955554 |
| ENSMUSG00000029403 | -1.04426813 | 0.816599259 |
| ENSMUSG00000029404 | 1.074504067 | 0.600541744 |
| ENSMUSG00000029405 | 1.029603611 | 0.959999413 |
| ENSMUSG00000029406 | 1.218104962 | 0.054812809 |
| ENSMUSG00000029407 | 1.026914806 | 0.971248775 |
| ENSMUSG00000029408 | -1.262765668 | 0.150281045 |
| ENSMUSG00000029409 | -1.551765175 | 0.006382739 |
| ENSMUSG00000029410 | 1.145935543 | 0.608497229 |
| ENSMUSG00000029413 | -1.06015591 | 0.772347686 |
| ENSMUSG00000029414 | 1.052014337 | 0.861402041 |
| ENSMUSG00000029415 | 1.083205599 | 0.70141105 |
| ENSMUSG00000029416 | 1.186389215 | 0.162855518 |
| ENSMUSG00000029417 | -1.124067181 | 0.677534336 |
| ENSMUSG00000029419 | -1 | 1 |
| ENSMUSG00000029420 | 1.028323353 | 0.903464611 |
| ENSMUSG00000029422 | 1.041567451 | 0.910832671 |
| ENSMUSG00000029423 | 1 | |
| ENSMUSG00000029426 | 1.051767203 | 0.864936713 |
| ENSMUSG00000029427 | 1.025709777 | 0.917717739 |
| ENSMUSG00000029428 | 1.136845539 | 0.608497229 |
| ENSMUSG00000029430 | 1.108195353 | 0.739099222 |
| ENSMUSG00000029431 | 1.069068979 | 0.608497229 |
| ENSMUSG00000029432 | 1.006025704 | 1 |

| | | |
|---|---|---|
| ENSMUSG00000029433 | -1.005141861 | 1 |
| ENSMUSG00000029434 | 1.064399176 | 0.783545142 |
| ENSMUSG00000029436 | -1.01968506 | 0.941763301 |
| ENSMUSG00000029437 | 1 | |
| ENSMUSG00000029438 | 1.174277197 | 0.153295884 |
| ENSMUSG00000029439 | 1.131647616 | 0.482433619 |
| ENSMUSG00000029440 | 1.005291564 | 1 |
| ENSMUSG00000029442 | 1.054072311 | 0.860645152 |
| ENSMUSG00000029445 | 1 | |
| ENSMUSG00000029446 | 1.068711145 | 0.710224291 |
| ENSMUSG00000029447 | 1.032264817 | 0.975187808 |
| ENSMUSG00000029449 | 1.055039554 | 0.608497229 |
| ENSMUSG00000029451 | 1 | |
| ENSMUSG00000029452 | 1.092192797 | 0.698112512 |
| ENSMUSG00000029454 | 1.193923261 | 0.224935244 |
| ENSMUSG00000029455 | 1.174745028 | 0.318032671 |
| ENSMUSG00000029456 | 1.185857137 | 0.136863623 |
| ENSMUSG00000029458 | 1.06241948 | 0.763700143 |
| ENSMUSG00000029461 | 1.104030614 | 0.674686528 |
| ENSMUSG00000029462 | 1.113038086 | 0.692325103 |
| ENSMUSG00000029463 | -1.009797805 | 1 |
| ENSMUSG00000029464 | 1.046413333 | 0.754389693 |
| ENSMUSG00000029465 | 1.04379826 | 0.897684198 |
| ENSMUSG00000029466 | 1.086620377 | 0.697178241 |
| ENSMUSG00000029467 | 1.119778908 | 0.71045278 |
| ENSMUSG00000029468 | 1.191952832 | 0.216046507 |
| ENSMUSG00000029469 | 1.057831913 | 0.763234037 |
| ENSMUSG00000029470 | 1.067386637 | 0.627212815 |
| ENSMUSG00000029471 | 1.058370365 | 0.72360967 |
| ENSMUSG00000029472 | 1.051045146 | 0.904568793 |
| ENSMUSG00000029474 | 1.04769858 | 0.83500866 |
| ENSMUSG00000029475 | -1.020779896 | 0.927315966 |
| ENSMUSG00000029477 | -1.083415624 | 0.62221251 |
| ENSMUSG00000029478 | 1.288720018 | 0.058381103 |
| ENSMUSG00000029480 | 1.141969833 | 0.218897371 |
| ENSMUSG00000029482 | 1.100960225 | 0.534861373 |
| ENSMUSG00000029484 | 1.131425647 | 0.698112512 |
| ENSMUSG00000029486 | 1.020476514 | 0.95702926 |
| ENSMUSG00000029490 | 1.562588509 | 0.004923779 |
| ENSMUSG00000029491 | 1.331309708 | 0.411295631 |
| ENSMUSG00000029499 | -1.159409938 | 0.243283933 |
| ENSMUSG00000029500 | 1.05103366 | 0.734025178 |
| ENSMUSG00000029501 | 1.026865095 | 0.964821074 |
| ENSMUSG00000029502 | 1.113804658 | 0.681952832 |
| ENSMUSG00000029503 | -1.005309846 | 1 |
| ENSMUSG00000029504 | 1.054308151 | 0.742635518 |
| ENSMUSG00000029505 | 1.097800258 | 0.698112512 |
| ENSMUSG00000029507 | 1.101042648 | 0.503617425 |
| ENSMUSG00000029510 | -1.14832604 | 0.243959074 |
| ENSMUSG00000029512 | 1.167271206 | 0.080092733 |
| ENSMUSG00000029513 | 1.091035077 | 0.57702934 |
| ENSMUSG00000029516 | 1.140445991 | 0.347571293 |
| ENSMUSG00000029517 | 1.09538577 | 0.698112512 |
| ENSMUSG00000029518 | 1.089762981 | 0.676812025 |
| ENSMUSG00000029521 | 1.153603221 | 0.228055228 |
| ENSMUSG00000029522 | -1.043306199 | 0.79291651 |
| ENSMUSG00000029524 | 1.062057039 | 0.364897726 |
| ENSMUSG00000029526 | -1.08081673 | 0.698112512 |
| ENSMUSG00000029528 | 1.032810344 | 0.893388118 |
| ENSMUSG00000029530 | -1.003834322 | 1 |
| ENSMUSG00000029534 | 1.124919096 | 0.454976536 |
| ENSMUSG00000029535 | 1.024573396 | 0.874187032 |
| ENSMUSG00000029536 | 1.003092053 | 1 |

| | | |
|---|---|---|
| ENSMUSG00000029538 | 1.07923738 | 0.627085115 |
| ENSMUSG00000029541 | 1.059463094 | 0.608497229 |
| ENSMUSG00000029544 | 1.236487199 | 0.344160208 |
| ENSMUSG00000029545 | 1.202346581 | 0.064932631 |
| ENSMUSG00000029546 | 1.059463094 | 0.698112512 |
| ENSMUSG00000029547 | 1.063821927 | 0.698112512 |
| ENSMUSG00000029550 | 1.220063138 | 0.119190358 |
| ENSMUSG00000029551 | 1.001235575 | 1 |
| ENSMUSG00000029552 | 1.066957712 | 0.766902409 |
| ENSMUSG00000029553 | 1 | |
| ENSMUSG00000029554 | 1.023906578 | 0.92904411 |
| ENSMUSG00000029556 | 1.081478107 | 0.698112512 |
| ENSMUSG00000029557 | 1.05081331 | 0.608497229 |
| ENSMUSG00000029559 | 1.089746303 | 0.282221758 |
| ENSMUSG00000029560 | 1.105456674 | 0.388781089 |
| ENSMUSG00000029561 | -1.221953459 | 0.2274442 |
| ENSMUSG00000029563 | 1.69835709 | 0.000417999 |
| ENSMUSG00000029564 | -1.105673636 | 0.608497229 |
| ENSMUSG00000029569 | 1.118777067 | 0.608497229 |
| ENSMUSG00000029570 | 1.15634854 | 0.160526425 |
| ENSMUSG00000029571 | 1.174691862 | 0.552584793 |
| ENSMUSG00000029575 | 1.030019524 | 0.847428179 |
| ENSMUSG00000029576 | -1.375627947 | 0.040767591 |
| ENSMUSG00000029577 | 1.111106283 | 0.608497229 |
| ENSMUSG00000029578 | 1.01256733 | 1 |
| ENSMUSG00000029580 | -2.457700437 | 0.005939428 |
| ENSMUSG00000029581 | 1.120842848 | 0.608497229 |
| ENSMUSG00000029586 | 1.040172434 | 0.896152472 |
| ENSMUSG00000029587 | 1.042929487 | 0.773508225 |
| ENSMUSG00000029591 | 1.06811478 | 0.608497229 |
| ENSMUSG00000029592 | 1.050572648 | 0.736688878 |
| ENSMUSG00000029594 | 1.112931864 | 0.404271378 |
| ENSMUSG00000029595 | 1 | |
| ENSMUSG00000029596 | -1.068087458 | 0.893213207 |
| ENSMUSG00000029597 | 1.211752532 | 0.358873127 |
| ENSMUSG00000029598 | 1.223188323 | 0.125368251 |
| ENSMUSG00000029599 | 1.166318477 | 0.205169472 |
| ENSMUSG00000029600 | 1.158065835 | 0.338563396 |
| ENSMUSG00000029601 | 1.280321327 | 0.361740791 |
| ENSMUSG00000029602 | 1.095586318 | 0.698112512 |
| ENSMUSG00000029603 | 1.026509487 | 0.949679283 |
| ENSMUSG00000029605 | -1.478879237 | 0.067602202 |
| ENSMUSG00000029607 | 1.005112968 | 1 |
| ENSMUSG00000029608 | -1.143787033 | 0.201977401 |
| ENSMUSG00000029610 | 1.022484956 | 0.870388412 |
| ENSMUSG00000029613 | -1.057566801 | 0.802061488 |
| ENSMUSG00000029614 | 1.097286629 | 0.749276359 |
| ENSMUSG00000029616 | 1.075553432 | 0.676185514 |
| ENSMUSG00000029617 | -1.021383227 | 0.978053547 |
| ENSMUSG00000029618 | 1 | |
| ENSMUSG00000029620 | 1 | |
| ENSMUSG00000029621 | -1.002201426 | 1 |
| ENSMUSG00000029622 | 1.127400275 | 0.408685749 |
| ENSMUSG00000029623 | 1.03339732 | 0.91979387 |
| ENSMUSG00000029624 | 1.025338521 | 0.823367179 |
| ENSMUSG00000029625 | 1.066536308 | 0.667189083 |
| ENSMUSG00000029627 | 1.043239412 | 0.76659604 |
| ENSMUSG00000029629 | 1.151005335 | 0.405978588 |
| ENSMUSG00000029630 | 1 | |
| ENSMUSG00000029632 | 1.158270396 | 0.581997959 |
| ENSMUSG00000029633 | -1.071075483 | 0.750635131 |
| ENSMUSG00000029634 | 1.048021405 | 0.851353899 |
| ENSMUSG00000029635 | 1.128041344 | 0.578757926 |

| | | |
|---|---|---|
| ENSMUSG00000029636 | -1.711588673 | 0.001986739 |
| ENSMUSG00000029638 | -1.069485343 | 0.769745556 |
| ENSMUSG00000029640 | 1.068241659 | 0.729018288 |
| ENSMUSG00000029641 | 1.298146044 | 0.250477583 |
| ENSMUSG00000029642 | -1.012945155 | 0.998981421 |
| ENSMUSG00000029644 | -1 | 1 |
| ENSMUSG00000029646 | -1.043306199 | 0.698112512 |
| ENSMUSG00000029647 | 1.095282953 | 0.649847774 |
| ENSMUSG00000029648 | -1.284810584 | 0.042349611 |
| ENSMUSG00000029649 | -1.057381256 | 0.849942153 |
| ENSMUSG00000029650 | -1.118710599 | 0.126063768 |
| ENSMUSG00000029651 | 1.251301607 | 0.468615237 |
| ENSMUSG00000029655 | -1.030464282 | 0.934760809 |
| ENSMUSG00000029656 | 1 | |
| ENSMUSG00000029657 | -1.093492001 | 0.765513459 |
| ENSMUSG00000029658 | 1.059463094 | 0.608497229 |
| ENSMUSG00000029659 | 1.738043286 | 0.000522056 |
| ENSMUSG00000029660 | 1.024073662 | 0.89811529 |
| ENSMUSG00000029661 | 1.177594713 | 0.110955169 |
| ENSMUSG00000029663 | 1.059463094 | 0.608497229 |
| ENSMUSG00000029664 | -1.158708082 | 0.350957446 |
| ENSMUSG00000029669 | 1.394765269 | 0.009594118 |
| ENSMUSG00000029670 | -1.007015018 | 1 |
| ENSMUSG00000029671 | 1.616962397 | 0.005295819 |
| ENSMUSG00000029672 | 1.140631495 | 0.431683218 |
| ENSMUSG00000029673 | -1.712859296 | 2.66714E-05 |
| ENSMUSG00000029674 | -1.097547768 | 0.461030406 |
| ENSMUSG00000029675 | -1.079755021 | 0.747479869 |
| ENSMUSG00000029676 | 1.049218834 | 0.818942935 |
| ENSMUSG00000029678 | 1 | |
| ENSMUSG00000029679 | 1 | |
| ENSMUSG00000029680 | 1 | |
| ENSMUSG00000029681 | 1.103282333 | 0.423331221 |
| ENSMUSG00000029682 | 1 | |
| ENSMUSG00000029683 | 1 | |
| ENSMUSG00000029684 | 1.060856982 | 0.79291651 |
| ENSMUSG00000029685 | -1.0678334 | 0.861137762 |
| ENSMUSG00000029686 | 1.080156431 | 0.766364416 |
| ENSMUSG00000029687 | 1.090743489 | 0.698112512 |
| ENSMUSG00000029695 | -1.202288982 | 0.257684055 |
| ENSMUSG00000029697 | 1 | |
| ENSMUSG00000029699 | -1.054892908 | 0.698112512 |
| ENSMUSG00000029700 | 1.139467335 | 0.608497229 |
| ENSMUSG00000029701 | 1.004362936 | 1 |
| ENSMUSG00000029703 | -1.015395618 | 0.95702926 |
| ENSMUSG00000029705 | 1.155741601 | 0.496494186 |
| ENSMUSG00000029706 | 1 | |
| ENSMUSG00000029707 | -1.013419756 | 0.988424085 |
| ENSMUSG00000029708 | 1.065268374 | 0.608497229 |
| ENSMUSG00000029710 | 1.013224109 | 0.979002724 |
| ENSMUSG00000029711 | 1 | |
| ENSMUSG00000029712 | -1.043306199 | 0.803260194 |
| ENSMUSG00000029713 | 1.171473283 | 0.375579698 |
| ENSMUSG00000029714 | 1.232029034 | 0.037841921 |
| ENSMUSG00000029715 | 1.128811994 | 0.410470609 |
| ENSMUSG00000029716 | -1.011554623 | 1 |
| ENSMUSG00000029718 | 1.188314763 | 0.415460837 |
| ENSMUSG00000029720 | 1.039961952 | 0.842184576 |
| ENSMUSG00000029722 | 1.075503235 | 0.608497229 |
| ENSMUSG00000029723 | 1.18456734 | 0.079202887 |
| ENSMUSG00000029725 | 1.051293681 | 0.890958598 |
| ENSMUSG00000029726 | 1.052325755 | 0.698112512 |
| ENSMUSG00000029727 | -1.234867245 | 0.302625055 |

| | | |
|---|---|---|
| ENSMUSG00000029729 | -1.003909984 | 1 |
| ENSMUSG00000029730 | 1.031003734 | 0.963664162 |
| ENSMUSG00000029735 | -1.121201819 | 0.385487949 |
| ENSMUSG00000029736 | -1 | 1 |
| ENSMUSG00000029752 | 1.208256427 | 0.38325813 |
| ENSMUSG00000029754 | 1.129471935 | 0.458934581 |
| ENSMUSG00000029755 | 1.29881532 | 0.456745331 |
| ENSMUSG00000029757 | -1.004098868 | 1 |
| ENSMUSG00000029759 | 1.911699622 | 0.000923517 |
| ENSMUSG00000029761 | 1.329735802 | 0.335419479 |
| ENSMUSG00000029762 | 1.181244862 | 0.337694222 |
| ENSMUSG00000029763 | 1.061126247 | 0.801057489 |
| ENSMUSG00000029765 | 1.987743946 | 0.005689524 |
| ENSMUSG00000029766 | 1 | |
| ENSMUSG00000029767 | 1.053638514 | 0.92498148 |
| ENSMUSG00000029769 | 1.085688217 | 0.536673416 |
| ENSMUSG00000029771 | -1.076811245 | 0.282043208 |
| ENSMUSG00000029772 | -1.050607323 | 0.813331954 |
| ENSMUSG00000029775 | 1.060283588 | 0.816599259 |
| ENSMUSG00000029776 | -1.018557918 | 0.974355392 |
| ENSMUSG00000029777 | 1.088655053 | 0.776021306 |
| ENSMUSG00000029778 | 3.493203447 | 0.000381466 |
| ENSMUSG00000029780 | -1.021536852 | 0.953084201 |
| ENSMUSG00000029781 | 1.130707743 | 0.667110773 |
| ENSMUSG00000029782 | 1.001906585 | 1 |
| ENSMUSG00000029784 | -1.06568528 | 0.867355119 |
| ENSMUSG00000029787 | 1.057953914 | 0.780302499 |
| ENSMUSG00000029788 | -1 | 1 |
| ENSMUSG00000029790 | 1.029208157 | 0.775301936 |
| ENSMUSG00000029797 | 1.028323353 | 0.961565399 |
| ENSMUSG00000029798 | -1.113711354 | 0.470841398 |
| ENSMUSG00000029802 | -1.108072846 | 0.608497229 |
| ENSMUSG00000029804 | -1.128708123 | 0.473648456 |
| ENSMUSG00000029810 | -1.046455989 | 0.874491458 |
| ENSMUSG00000029811 | -1 | 1 |
| ENSMUSG00000029813 | 1 | |
| ENSMUSG00000029814 | 1.215519556 | 0.195148393 |
| ENSMUSG00000029815 | -1.030843321 | 0.839653389 |
| ENSMUSG00000029816 | 1.056601175 | 0.861653133 |
| ENSMUSG00000029817 | 1.008353807 | 1 |
| ENSMUSG00000029819 | -1.099174267 | 0.608497229 |
| ENSMUSG00000029821 | 1.12844969 | 0.242876366 |
| ENSMUSG00000029822 | 1.008002604 | 1 |
| ENSMUSG00000029823 | 1.041680735 | 0.919112514 |
| ENSMUSG00000029826 | -1.105980732 | 0.694955689 |
| ENSMUSG00000029828 | -1.173428378 | 0.414627266 |
| ENSMUSG00000029829 | 1 | |
| ENSMUSG00000029830 | -1.024718486 | 0.954310937 |
| ENSMUSG00000029831 | 1 | |
| ENSMUSG00000029832 | -1.347876783 | 0.025289387 |
| ENSMUSG00000029833 | 1.067526217 | 0.70299173 |
| ENSMUSG00000029836 | 1.094487733 | 0.746563203 |
| ENSMUSG00000029837 | 1 | |
| ENSMUSG00000029838 | 2.079887845 | 0.001039425 |
| ENSMUSG00000029840 | 1.064715604 | 0.858911269 |
| ENSMUSG00000029843 | 1.111812527 | 0.659884278 |
| ENSMUSG00000029844 | 1.070966208 | 0.491794669 |
| ENSMUSG00000029847 | -1.109582661 | 0.682183489 |
| ENSMUSG00000029848 | 1.024693402 | 0.933031586 |
| ENSMUSG00000029851 | -1.157423481 | 0.250203718 |
| ENSMUSG00000029859 | -1.28115524 | 0.005497832 |
| ENSMUSG00000029860 | 1.173967271 | 0.366138691 |
| ENSMUSG00000029861 | 1.049839036 | 0.689713779 |

| | | |
|---|---|---|
| ENSMUSG00000029862 | 1.392075188 | 0.219229721 |
| ENSMUSG00000029863 | 1.062230553 | 0.779813128 |
| ENSMUSG00000029864 | -1.160031575 | 0.608497229 |
| ENSMUSG00000029865 | 1 | |
| ENSMUSG00000029866 | 1 | |
| ENSMUSG00000029867 | 1 | |
| ENSMUSG00000029868 | 1.072550195 | 0.835701362 |
| ENSMUSG00000029869 | 1.394360413 | 0.012970042 |
| ENSMUSG00000029875 | 1.443859646 | 0.084732247 |
| ENSMUSG00000029878 | -1.066285281 | 0.793885968 |
| ENSMUSG00000029882 | -1 | 1 |
| ENSMUSG00000029883 | 1 | |
| ENSMUSG00000029885 | 1 | |
| ENSMUSG00000029909 | 1 | |
| ENSMUSG00000029910 | 1.004421623 | 1 |
| ENSMUSG00000029911 | 1.07319705 | 0.696249781 |
| ENSMUSG00000029913 | 1.092016402 | 0.472623484 |
| ENSMUSG00000029915 | 1 | |
| ENSMUSG00000029916 | 1.001342553 | 1 |
| ENSMUSG00000029917 | 1 | |
| ENSMUSG00000029918 | 1.019245029 | 0.972031343 |
| ENSMUSG00000029919 | -1.060575156 | 0.835189692 |
| ENSMUSG00000029920 | 1.103383063 | 0.654374885 |
| ENSMUSG00000029922 | -1.080365998 | 0.643361947 |
| ENSMUSG00000029923 | -1.063525418 | 0.716583464 |
| ENSMUSG00000029924 | 1.008539444 | 1 |
| ENSMUSG00000029925 | 1.023735632 | 0.986389736 |
| ENSMUSG00000029992 | 1.013539427 | 1 |
| ENSMUSG00000029993 | 1.037672738 | 0.74256555 |
| ENSMUSG00000029994 | 1.059996958 | 0.831092008 |
| ENSMUSG00000029998 | -1.044009118 | 0.895375561 |
| ENSMUSG00000029999 | -1.125882057 | 0.17554978 |
| ENSMUSG00000030000 | -1.575163813 | 0.000371886 |
| ENSMUSG00000030001 | 1 | |
| ENSMUSG00000030002 | -1.001094655 | 1 |
| ENSMUSG00000030004 | 1 | |
| ENSMUSG00000030007 | -1.0024717 | 1 |
| ENSMUSG00000030008 | 1.000218121 | 1 |
| ENSMUSG00000030016 | 1.037538011 | 0.93647054 |
| ENSMUSG00000030017 | -1 | 1 |
| ENSMUSG00000030019 | 1.030197561 | 0.835498864 |
| ENSMUSG00000030020 | 1.267839143 | 0.005658129 |
| ENSMUSG00000030022 | 1.10700805 | 0.403738027 |
| ENSMUSG00000030029 | 1.219339483 | 0.250203718 |
| ENSMUSG00000030030 | -1.118237073 | 0.528005802 |
| ENSMUSG00000030031 | -1.024599103 | 0.883448618 |
| ENSMUSG00000030032 | 1.106303795 | 0.151792895 |
| ENSMUSG00000030034 | 1.00334825 | 1 |
| ENSMUSG00000030035 | 1.124187236 | 0.313613158 |
| ENSMUSG00000030036 | 1.066694093 | 0.617912429 |
| ENSMUSG00000030037 | -1.063525418 | 0.698112512 |
| ENSMUSG00000030041 | 1.38892309 | 0.206114416 |
| ENSMUSG00000030042 | -1.020423561 | 0.972980766 |
| ENSMUSG00000030043 | -1.055749065 | 0.861034664 |
| ENSMUSG00000030045 | -1.033400298 | 0.870877911 |
| ENSMUSG00000030046 | 1 | |
| ENSMUSG00000030047 | -1.085651418 | 0.532369657 |
| ENSMUSG00000030048 | 1 | |
| ENSMUSG00000030049 | 1 | |
| ENSMUSG00000030050 | -1.03962336 | 0.698112512 |
| ENSMUSG00000030051 | -1.043689788 | 0.770461131 |
| ENSMUSG00000030054 | 1 | |
| ENSMUSG00000030055 | 1.046309239 | 0.925075932 |

| | | |
|---|---|---|
| ENSMUSG00000030056 | -1.082546604 | 0.608497229 |
| ENSMUSG00000030057 | 1.03352056 | 0.963664162 |
| ENSMUSG00000030058 | 1.079794537 | 0.773499772 |
| ENSMUSG00000030059 | 1.061489658 | 0.781832523 |
| ENSMUSG00000030060 | -1.072359719 | 0.698112512 |
| ENSMUSG00000030061 | -1.003260233 | 1 |
| ENSMUSG00000030062 | -1.011391271 | 1 |
| ENSMUSG00000030064 | -1.661393692 | 0.001540286 |
| ENSMUSG00000030067 | 1.115206476 | 0.608497229 |
| ENSMUSG00000030068 | 1.161036672 | 0.251049598 |
| ENSMUSG00000030069 | 1 | |
| ENSMUSG00000030074 | -1.267229828 | 0.258738987 |
| ENSMUSG00000030075 | -1.017813114 | 0.998601098 |
| ENSMUSG00000030077 | 1 | |
| ENSMUSG00000030079 | 1.005589914 | 1 |
| ENSMUSG00000030082 | 1.064831363 | 0.775765171 |
| ENSMUSG00000030083 | 1.049908942 | 0.608497229 |
| ENSMUSG00000030084 | 1.238486929 | 0.127006648 |
| ENSMUSG00000030086 | 1.066872804 | 0.373031949 |
| ENSMUSG00000030087 | -1.084072947 | 0.760374524 |
| ENSMUSG00000030088 | 1.385483523 | 0.060091435 |
| ENSMUSG00000030089 | 1.100613337 | 0.246549364 |
| ENSMUSG00000030091 | 1.095466151 | 0.344313417 |
| ENSMUSG00000030092 | 1 | |
| ENSMUSG00000030093 | -1.130037519 | 0.628392912 |
| ENSMUSG00000030094 | 1.052356468 | 0.667576553 |
| ENSMUSG00000030095 | 1.060390935 | 0.79291651 |
| ENSMUSG00000030096 | 1.204500287 | 0.49801899 |
| ENSMUSG00000030098 | -1.190907694 | 0.23684718 |
| ENSMUSG00000030101 | 1.044749642 | 0.829488293 |
| ENSMUSG00000030102 | 1.104470684 | 0.608497229 |
| ENSMUSG00000030103 | 1.230671383 | 0.066778109 |
| ENSMUSG00000030104 | -1.020199542 | 0.978053547 |
| ENSMUSG00000030105 | 1.116433769 | 0.667778769 |
| ENSMUSG00000030107 | -1.369678089 | 0.006886236 |
| ENSMUSG00000030108 | -1.069502966 | 0.698112512 |
| ENSMUSG00000030109 | -1.05948903 | 0.725549871 |
| ENSMUSG00000030110 | 1.332875555 | 0.07248718 |
| ENSMUSG00000030111 | 1 | |
| ENSMUSG00000030114 | 1 | |
| ENSMUSG00000030116 | -1 | 1 |
| ENSMUSG00000030117 | 1.132639128 | 0.217803313 |
| ENSMUSG00000030120 | 1.09630176 | 0.608497229 |
| ENSMUSG00000030122 | 1.272507457 | 0.067151105 |
| ENSMUSG00000030123 | 1.414482885 | 0.00496943 |
| ENSMUSG00000030124 | 1.263599722 | 0.17455173 |
| ENSMUSG00000030125 | 1.026072142 | 0.954310937 |
| ENSMUSG00000030126 | 1.007595735 | 1 |
| ENSMUSG00000030127 | 1.077373071 | 0.671280603 |
| ENSMUSG00000030131 | 1 | |
| ENSMUSG00000030134 | 1.363865407 | 0.001748121 |
| ENSMUSG00000030137 | -1.055662102 | 0.875804019 |
| ENSMUSG00000030138 | 1.030488107 | 0.939883155 |
| ENSMUSG00000030142 | 1 | |
| ENSMUSG00000030143 | 1 | |
| ENSMUSG00000030144 | 1 | |
| ENSMUSG00000030145 | 1.054207399 | 0.655408594 |
| ENSMUSG00000030147 | 1 | |
| ENSMUSG00000030148 | -1.158274388 | 0.608497229 |
| ENSMUSG00000030149 | -1 | 1 |
| ENSMUSG00000030154 | -1.142727245 | 0.36554144 |
| ENSMUSG00000030155 | -1.094422282 | 0.698112512 |
| ENSMUSG00000030156 | 1.032867246 | 0.951746255 |

| | | |
|---|---|---|
| ENSMUSG00000030157 | 1.745479617 | 0.004156852 |
| ENSMUSG00000030158 | 1 | |
| ENSMUSG00000030159 | 1 | |
| ENSMUSG00000030160 | 1 | |
| ENSMUSG00000030161 | 1.184566162 | 0.19154846 |
| ENSMUSG00000030162 | -1.043306199 | 0.880677217 |
| ENSMUSG00000030165 | -1 | 1 |
| ENSMUSG00000030166 | 1.13423068 | 0.098922723 |
| ENSMUSG00000030167 | 1 | |
| ENSMUSG00000030168 | -1.088678623 | 0.698112512 |
| ENSMUSG00000030170 | -2.158341326 | 0.004416753 |
| ENSMUSG00000030172 | -1.022167394 | 0.969531691 |
| ENSMUSG00000030173 | 1 | |
| ENSMUSG00000030177 | -1.012143616 | 0.996913409 |
| ENSMUSG00000030178 | 1 | |
| ENSMUSG00000030180 | 1.013240518 | 1 |
| ENSMUSG00000030187 | -1.025365038 | 0.9780134 |
| ENSMUSG00000030188 | 1.054046331 | 0.783468419 |
| ENSMUSG00000030189 | 1.144602308 | 0.431084768 |
| ENSMUSG00000030194 | 1 | |
| ENSMUSG00000030196 | 1 | |
| ENSMUSG00000030199 | 1.082393539 | 0.635981177 |
| ENSMUSG00000030200 | 1 | |
| ENSMUSG00000030201 | 1.075446928 | 0.771728893 |
| ENSMUSG00000030203 | -1.06723865 | 0.733645428 |
| ENSMUSG00000030204 | -1.008388874 | 1 |
| ENSMUSG00000030205 | -1.044273782 | 0.698112512 |
| ENSMUSG00000030206 | 1.107416398 | 0.687984391 |
| ENSMUSG00000030207 | -1.050986522 | 0.709654578 |
| ENSMUSG00000030208 | 1.027944255 | 0.984782014 |
| ENSMUSG00000030209 | -1.089497311 | 0.608497229 |
| ENSMUSG00000030213 | 1.090201234 | 0.619725454 |
| ENSMUSG00000030214 | -1.230294675 | 0.256283628 |
| ENSMUSG00000030216 | 1.030881327 | 0.911346192 |
| ENSMUSG00000030217 | 1 | |
| ENSMUSG00000030218 | 2.633319165 | 0.001717312 |
| ENSMUSG00000030219 | 1.059463094 | 0.608497229 |
| ENSMUSG00000030220 | -1.359445159 | 0.005175804 |
| ENSMUSG00000030222 | 1.314072384 | 0.004054559 |
| ENSMUSG00000030223 | 1 | |
| ENSMUSG00000030224 | 1.030016793 | 0.968937442 |
| ENSMUSG00000030225 | -1.060128824 | 0.662289659 |
| ENSMUSG00000030226 | -1.053549062 | 0.698112512 |
| ENSMUSG00000030228 | -1 | 1 |
| ENSMUSG00000030230 | 1 | |
| ENSMUSG00000030231 | 1.117662297 | 0.397805018 |
| ENSMUSG00000030232 | 1.086112025 | 0.734674233 |
| ENSMUSG00000030235 | 1 | |
| ENSMUSG00000030236 | 1 | |
| ENSMUSG00000030237 | -1.03962336 | 0.698112512 |
| ENSMUSG00000030243 | 1.011105626 | 1 |
| ENSMUSG00000030244 | 1 | |
| ENSMUSG00000030245 | 1.068392959 | 0.760216213 |
| ENSMUSG00000030246 | -1.191024512 | 0.256116326 |
| ENSMUSG00000030247 | -1.206338534 | 0.336943469 |
| ENSMUSG00000030249 | -1.077265267 | 0.683683209 |
| ENSMUSG00000030254 | -1.065545139 | 0.783616541 |
| ENSMUSG00000030255 | 1.337497261 | 0.048095607 |
| ENSMUSG00000030256 | 1.817181046 | 0.002167797 |
| ENSMUSG00000030257 | 1.344420653 | 0.005995788 |
| ENSMUSG00000030259 | 1.10355265 | 0.466284441 |
| ENSMUSG00000030263 | 1.24122829 | 0.142571618 |
| ENSMUSG00000030264 | 1.036424425 | 0.885463996 |

| | | |
|---|---|---|
| ENSMUSG00000030265 | 1.039741001 | 0.917086603 |
| ENSMUSG00000030268 | 1.087208372 | 0.711347805 |
| ENSMUSG00000030269 | 1.061971384 | 0.682836879 |
| ENSMUSG00000030270 | 1.015406766 | 1 |
| ENSMUSG00000030271 | 1.080201104 | 0.472188908 |
| ENSMUSG00000030272 | 1.005247 | 1 |
| ENSMUSG00000030275 | 1.032347609 | 0.953613038 |
| ENSMUSG00000030276 | 1.326994306 | 0.007835576 |
| ENSMUSG00000030278 | 1 | |
| ENSMUSG00000030279 | -1.088685503 | 0.616230752 |
| ENSMUSG00000030281 | 1.017307345 | 0.934760809 |
| ENSMUSG00000030282 | -1.041027207 | 0.812570482 |
| ENSMUSG00000030283 | -1.042179374 | 0.923090637 |
| ENSMUSG00000030284 | -1.001351468 | 1 |
| ENSMUSG00000030286 | -1.056870621 | 0.821797531 |
| ENSMUSG00000030287 | 1.052982586 | 0.608497229 |
| ENSMUSG00000030291 | 1.022535577 | 0.942691576 |
| ENSMUSG00000030292 | 1.881896354 | 0.005267835 |
| ENSMUSG00000030298 | 1.042363671 | 0.891104352 |
| ENSMUSG00000030301 | 1.047841992 | 0.700389541 |
| ENSMUSG00000030302 | -1.078590893 | 0.647384212 |
| ENSMUSG00000030303 | -1.032905808 | 0.819222912 |
| ENSMUSG00000030304 | 1.006015171 | 1 |
| ENSMUSG00000030306 | -1.653461038 | 0.031646778 |
| ENSMUSG00000030307 | -1.044273782 | 0.698112512 |
| ENSMUSG00000030309 | 1.003749532 | 1 |
| ENSMUSG00000030310 | -1.005345324 | 1 |
| ENSMUSG00000030313 | -1.076548519 | 0.683168971 |
| ENSMUSG00000030314 | 1.045535716 | 0.730049589 |
| ENSMUSG00000030315 | 1.130626707 | 0.192531722 |
| ENSMUSG00000030316 | 1.024704989 | 0.925509543 |
| ENSMUSG00000030317 | -1.043306199 | 0.858911269 |
| ENSMUSG00000030319 | 1.046802346 | 0.656201315 |
| ENSMUSG00000030321 | -1.009347007 | 1 |
| ENSMUSG00000030322 | -1.001198663 | 1 |
| ENSMUSG00000030323 | 1.05583025 | 0.66278123 |
| ENSMUSG00000030324 | 1.458137563 | 0.1245451 |
| ENSMUSG00000030325 | 1 | |
| ENSMUSG00000030327 | -1.004438293 | 1 |
| ENSMUSG00000030329 | 2.054832369 | 0.000739381 |
| ENSMUSG00000030330 | -1.147662821 | 0.327857845 |
| ENSMUSG00000030335 | -1.000588852 | 1 |
| ENSMUSG00000030336 | -1.26590368 | 0.143187387 |
| ENSMUSG00000030337 | -1.125759032 | 0.333663986 |
| ENSMUSG00000030339 | 1.124638586 | 0.408951276 |
| ENSMUSG00000030340 | 1.081557512 | 0.386294287 |
| ENSMUSG00000030341 | 1.07876789 | 0.6659463 |
| ENSMUSG00000030342 | 1.067450311 | 0.803072249 |
| ENSMUSG00000030344 | 1.069068979 | 0.74773997 |
| ENSMUSG00000030345 | -1 | 1 |
| ENSMUSG00000030346 | 1.021158705 | 0.945697001 |
| ENSMUSG00000030347 | 1.078945578 | 0.64570247 |
| ENSMUSG00000030350 | -1.043306199 | 0.698112512 |
| ENSMUSG00000030351 | 1.171563499 | 0.23546922 |
| ENSMUSG00000030352 | -1.031822483 | 0.915070045 |
| ENSMUSG00000030353 | -1.235561754 | 0.038007669 |
| ENSMUSG00000030357 | 1.021099987 | 0.97946024 |
| ENSMUSG00000030359 | -1.063525418 | 0.821376503 |
| ENSMUSG00000030361 | -1.044273782 | 0.698112512 |
| ENSMUSG00000030363 | 1 | |
| ENSMUSG00000030364 | -1.091287282 | 0.755282061 |
| ENSMUSG00000030365 | 1.015486245 | 0.972980766 |
| ENSMUSG00000030366 | 1 | |

| | | |
|---|---|---|
| ENSMUSG00000030368 | 1 | |
| ENSMUSG00000030373 | 1 | |
| ENSMUSG00000030374 | 1.028651744 | 0.872807809 |
| ENSMUSG00000030376 | -1.347442089 | 0.280829377 |
| ENSMUSG00000030378 | 1 | |
| ENSMUSG00000030380 | 1.130831453 | 0.41274756 |
| ENSMUSG00000030382 | 1.065519011 | 0.814089245 |
| ENSMUSG00000030385 | 1 | |
| ENSMUSG00000030386 | 1.073175753 | 0.478878776 |
| ENSMUSG00000030393 | 1.088290289 | 0.608497229 |
| ENSMUSG00000030397 | 1.001911217 | 1 |
| ENSMUSG00000030399 | 1 | |
| ENSMUSG00000030400 | 1.038428948 | 0.698112512 |
| ENSMUSG00000030401 | -1.000589742 | 1 |
| ENSMUSG00000030402 | 1 | |
| ENSMUSG00000030403 | 1.336818422 | 0.030124489 |
| ENSMUSG00000030406 | 1.059463094 | 0.698112512 |
| ENSMUSG00000030407 | -1.092637166 | 0.358940839 |
| ENSMUSG00000030409 | 1.30279513 | 0.014959231 |
| ENSMUSG00000030410 | 1.046964513 | 0.698112512 |
| ENSMUSG00000030411 | -1.019706254 | 1 |
| ENSMUSG00000030413 | -1.019108526 | 0.968937442 |
| ENSMUSG00000030417 | -1.047755215 | 0.788660205 |
| ENSMUSG00000030421 | 1.077570297 | 0.63568866 |
| ENSMUSG00000030423 | 1.053301249 | 0.648701353 |
| ENSMUSG00000030424 | -1.048211233 | 0.698112512 |
| ENSMUSG00000030427 | 1 | |
| ENSMUSG00000030428 | 1.041830896 | 0.870777622 |
| ENSMUSG00000030431 | 1.076926636 | 0.61015029 |
| ENSMUSG00000030432 | 1.056294469 | 0.728400252 |
| ENSMUSG00000030433 | -1.053549062 | 0.769745556 |
| ENSMUSG00000030435 | 1.048644339 | 0.835701362 |
| ENSMUSG00000030443 | -1.077371676 | 0.608497229 |
| ENSMUSG00000030446 | -1.09829499 | 0.527447007 |
| ENSMUSG00000030447 | -1.00025707 | 1 |
| ENSMUSG00000030450 | 1 | |
| ENSMUSG00000030451 | 1.053827655 | 0.833460635 |
| ENSMUSG00000030452 | 1.02976703 | 0.934760809 |
| ENSMUSG00000030463 | -1.03962336 | 0.698112512 |
| ENSMUSG00000030465 | 1.146106061 | 0.282982046 |
| ENSMUSG00000030468 | 1.014836126 | 0.939903859 |
| ENSMUSG00000030469 | -1.021017808 | 0.930780134 |
| ENSMUSG00000030470 | 1 | |
| ENSMUSG00000030471 | -1.012288999 | 1 |
| ENSMUSG00000030472 | 1 | |
| ENSMUSG00000030474 | 1.059463094 | 0.776953697 |
| ENSMUSG00000030483 | 1 | |
| ENSMUSG00000030484 | 1 | |
| ENSMUSG00000030486 | 1.086280696 | 0.698112512 |
| ENSMUSG00000030491 | -1 | 1 |
| ENSMUSG00000030492 | 1 | |
| ENSMUSG00000030493 | -1.063120574 | 0.608497229 |
| ENSMUSG00000030494 | -1.238781912 | 0.019538052 |
| ENSMUSG00000030495 | 1 | |
| ENSMUSG00000030498 | 1.360088317 | 0.001717312 |
| ENSMUSG00000030499 | 1.014545335 | 1 |
| ENSMUSG00000030500 | 1 | |
| ENSMUSG00000030505 | 1.010215681 | 1 |
| ENSMUSG00000030507 | 1 | |
| ENSMUSG00000030509 | 1.044831008 | 0.716583464 |
| ENSMUSG00000030510 | 1.097962432 | 0.742692237 |
| ENSMUSG00000030512 | 1.013446512 | 0.994207501 |
| ENSMUSG00000030513 | 1.605517122 | 0.002242306 |

| | | |
|---|---|---|
| ENSMUSG00000030515 | -1.001967749 | 1 |
| ENSMUSG00000030516 | -1.07151673 | 0.800729124 |
| ENSMUSG00000030518 | -1.156717385 | 0.584727336 |
| ENSMUSG00000030519 | -1.03962336 | 0.698112512 |
| ENSMUSG00000030521 | 1.01079301 | 1 |
| ENSMUSG00000030522 | -1.031807977 | 0.854739033 |
| ENSMUSG00000030523 | -1.043306199 | 0.698112512 |
| ENSMUSG00000030525 | -1.071075483 | 0.698112512 |
| ENSMUSG00000030527 | 1.085466656 | 0.440447402 |
| ENSMUSG00000030528 | -1.02291216 | 0.959892369 |
| ENSMUSG00000030530 | 1.19620148 | 0.062335064 |
| ENSMUSG00000030532 | 1.090655489 | 0.470974528 |
| ENSMUSG00000030533 | 1.055672225 | 0.630994898 |
| ENSMUSG00000030534 | 1.027762141 | 0.788750828 |
| ENSMUSG00000030536 | 1.148772412 | 0.608497229 |
| ENSMUSG00000030538 | -1.035440408 | 0.747846 |
| ENSMUSG00000030539 | 1.070226554 | 0.608497229 |
| ENSMUSG00000030541 | 1.197036793 | 0.324252919 |
| ENSMUSG00000030543 | -1.275508087 | 0.100749055 |
| ENSMUSG00000030544 | 1 | |
| ENSMUSG00000030545 | -1.068916965 | 0.251544797 |
| ENSMUSG00000030546 | 1.028323353 | 0.903464611 |
| ENSMUSG00000030549 | -1.103388856 | 0.608497229 |
| ENSMUSG00000030551 | 1.106351157 | 0.361182798 |
| ENSMUSG00000030553 | -1 | 1 |
| ENSMUSG00000030554 | 1.478217395 | 0.005393352 |
| ENSMUSG00000030555 | 1.081101308 | 0.481529404 |
| ENSMUSG00000030556 | 1.046152673 | 0.613255791 |
| ENSMUSG00000030557 | 1.17724925 | 0.39928706 |
| ENSMUSG00000030559 | -1.736763967 | 0.010744223 |
| ENSMUSG00000030560 | -1.14694268 | 0.479262818 |
| ENSMUSG00000030562 | 1.254962343 | 0.043041541 |
| ENSMUSG00000030577 | -1.028348525 | 0.960665692 |
| ENSMUSG00000030579 | 1 | |
| ENSMUSG00000030583 | 1.110104913 | 0.321024879 |
| ENSMUSG00000030584 | -1.024230945 | 0.872140518 |
| ENSMUSG00000030587 | 1.788915709 | 0.029735092 |
| ENSMUSG00000030588 | 1.13447699 | 0.206114416 |
| ENSMUSG00000030589 | 1.123054873 | 0.718265321 |
| ENSMUSG00000030590 | -1.128056357 | 0.24033917 |
| ENSMUSG00000030591 | -1.016532973 | 0.994003797 |
| ENSMUSG00000030592 | -1.034147473 | 0.860554063 |
| ENSMUSG00000030595 | -1.051548173 | 0.730297627 |
| ENSMUSG00000030598 | 1.011286245 | 0.942325266 |
| ENSMUSG00000030600 | -1.714126152 | 0.005521566 |
| ENSMUSG00000030602 | 1.098743807 | 0.5159274 |
| ENSMUSG00000030603 | -1.037786851 | 0.88284878 |
| ENSMUSG00000030604 | 1.038526171 | 0.815347141 |
| ENSMUSG00000030605 | 1.062054653 | 0.729973031 |
| ENSMUSG00000030606 | -1.109582661 | 0.698112512 |
| ENSMUSG00000030607 | 1.075043815 | 0.731328848 |
| ENSMUSG00000030609 | 1.142094077 | 0.22483686 |
| ENSMUSG00000030610 | 1.097337893 | 0.259152109 |
| ENSMUSG00000030611 | 1.023283947 | 0.79291651 |
| ENSMUSG00000030612 | -1.065297224 | 0.698112512 |
| ENSMUSG00000030613 | 1.089622868 | 0.608497229 |
| ENSMUSG00000030614 | 1.067496408 | 0.705199845 |
| ENSMUSG00000030615 | 1.077002521 | 0.608497229 |
| ENSMUSG00000030616 | 1.571876614 | 0.005796379 |
| ENSMUSG00000030617 | 1 | |
| ENSMUSG00000030619 | 1.035702531 | 0.894183102 |
| ENSMUSG00000030621 | 1.225262478 | 0.029569754 |
| ENSMUSG00000030623 | 1.069071129 | 0.698112512 |

| | | |
|---|---|---|
| ENSMUSG00000030629 | -1.025100181 | 0.95050544 |
| ENSMUSG00000030630 | 1.227323266 | 0.038839753 |
| ENSMUSG00000030636 | -1 | 1 |
| ENSMUSG00000030638 | -1.043757224 | 0.806865153 |
| ENSMUSG00000030641 | 1.088658055 | 0.608497229 |
| ENSMUSG00000030643 | -1.09005615 | 0.540731673 |
| ENSMUSG00000030647 | -1.019000616 | 0.917717739 |
| ENSMUSG00000030649 | -1.01894415 | 0.953416513 |
| ENSMUSG00000030650 | -1.037551163 | 0.83500866 |
| ENSMUSG00000030651 | 1 | |
| ENSMUSG00000030652 | -1.024354156 | 0.883395055 |
| ENSMUSG00000030653 | 1.315278441 | 0.062449758 |
| ENSMUSG00000030654 | 1.079957049 | 0.76409535 |
| ENSMUSG00000030655 | 1.069986032 | 0.819614655 |
| ENSMUSG00000030657 | -1.3447775 | 0.003813643 |
| ENSMUSG00000030659 | 1.037289956 | 0.734254834 |
| ENSMUSG00000030660 | 1.067541845 | 0.799888905 |
| ENSMUSG00000030662 | -1.029037505 | 0.972980766 |
| ENSMUSG00000030663 | -1.027354305 | 0.950560821 |
| ENSMUSG00000030664 | 1 | |
| ENSMUSG00000030666 | 1 | |
| ENSMUSG00000030669 | 1.020240423 | 0.964236174 |
| ENSMUSG00000030670 | 1 | |
| ENSMUSG00000030671 | 1.10187768 | 0.608497229 |
| ENSMUSG00000030672 | -1.153888367 | 0.698112512 |
| ENSMUSG00000030674 | -1.043306199 | 0.698112512 |
| ENSMUSG00000030677 | -1.020430232 | 0.965872389 |
| ENSMUSG00000030678 | 1.112043246 | 0.417142231 |
| ENSMUSG00000030680 | 1 | |
| ENSMUSG00000030681 | 1.039338164 | 0.762603775 |
| ENSMUSG00000030682 | 1.020571975 | 0.91353552 |
| ENSMUSG00000030683 | 1.22957636 | 0.406890538 |
| ENSMUSG00000030685 | 1.09686643 | 0.232204165 |
| ENSMUSG00000030688 | -1.368325168 | 0.002579592 |
| ENSMUSG00000030689 | 1.003086019 | 1 |
| ENSMUSG00000030691 | 1.077884141 | 0.340921812 |
| ENSMUSG00000030693 | 1.106485641 | 0.608497229 |
| ENSMUSG00000030695 | 1.170220785 | 0.49801899 |
| ENSMUSG00000030697 | 1.06289819 | 0.698112512 |
| ENSMUSG00000030699 | 1.138788635 | 0.608497229 |
| ENSMUSG00000030701 | -1.2364519 | 0.355811155 |
| ENSMUSG00000030703 | 1.455920536 | 0.016541591 |
| ENSMUSG00000030704 | 1.018092224 | 0.995338384 |
| ENSMUSG00000030706 | -1.105258137 | 0.608497229 |
| ENSMUSG00000030707 | -1.415376319 | 0.075028755 |
| ENSMUSG00000030708 | -1.131830234 | 0.715053961 |
| ENSMUSG00000030711 | -1.05290791 | 0.752398528 |
| ENSMUSG00000030713 | -1.072882392 | 0.778376464 |
| ENSMUSG00000030714 | 1.0269904 | 0.784279473 |
| ENSMUSG00000030717 | 1.308650597 | 0.02015623 |
| ENSMUSG00000030718 | 1.000095853 | 1 |
| ENSMUSG00000030720 | 1.376277891 | 0.003114197 |
| ENSMUSG00000030722 | -1.13085487 | 0.270289587 |
| ENSMUSG00000030724 | 1 | |
| ENSMUSG00000030725 | 1.109754328 | 0.698112512 |
| ENSMUSG00000030726 | -1.035616784 | 0.858932381 |
| ENSMUSG00000030727 | 1.114034448 | 0.420245309 |
| ENSMUSG00000030729 | -1.081439311 | 0.642132679 |
| ENSMUSG00000030730 | 1.047883597 | 0.906415966 |
| ENSMUSG00000030731 | -1.311838065 | 0.062913564 |
| ENSMUSG00000030732 | -1.053549062 | 0.698112512 |
| ENSMUSG00000030733 | 1.034327734 | 0.769745556 |
| ENSMUSG00000030735 | 1.055416288 | 0.798926662 |

| | | |
|---|---|---|
| ENSMUSG00000030737 | 1 | |
| ENSMUSG00000030738 | 1.024334684 | 0.982838256 |
| ENSMUSG00000030739 | -1.185992246 | 0.459883463 |
| ENSMUSG00000030741 | 1.158351637 | 0.251049598 |
| ENSMUSG00000030742 | 1.113974798 | 0.698360568 |
| ENSMUSG00000030744 | 1.035372267 | 0.963147621 |
| ENSMUSG00000030745 | 1.049411284 | 0.863033776 |
| ENSMUSG00000030747 | -1.003221054 | 1 |
| ENSMUSG00000030748 | 1.188088907 | 0.111579015 |
| ENSMUSG00000030750 | 1.031442564 | 0.870522488 |
| ENSMUSG00000030751 | -1.064797207 | 0.823969666 |
| ENSMUSG00000030752 | 1.053281493 | 0.608497229 |
| ENSMUSG00000030753 | 1.061221053 | 0.806949992 |
| ENSMUSG00000030754 | -1.013848271 | 1 |
| ENSMUSG00000030757 | -1.364145414 | 0.089664798 |
| ENSMUSG00000030759 | 1.096930792 | 0.717035356 |
| ENSMUSG00000030760 | -1.077811744 | 0.684150912 |
| ENSMUSG00000030761 | 1.832492357 | 0.00051542 |
| ENSMUSG00000030762 | 1 | |
| ENSMUSG00000030763 | -1.049653772 | 0.698112512 |
| ENSMUSG00000030766 | 1.053580686 | 0.650518769 |
| ENSMUSG00000030768 | 1.026783491 | 0.879624748 |
| ENSMUSG00000030769 | -1 | 1 |
| ENSMUSG00000030770 | 1.055571768 | 0.878741277 |
| ENSMUSG00000030771 | 1.176915854 | 0.281868105 |
| ENSMUSG00000030772 | -1.079117505 | 0.527649743 |
| ENSMUSG00000030774 | 1.041201016 | 0.889219349 |
| ENSMUSG00000030775 | 1 | |
| ENSMUSG00000030779 | -1.020421282 | 0.979002724 |
| ENSMUSG00000030780 | 1.141316462 | 0.07220259 |
| ENSMUSG00000030781 | 1.179016771 | 0.531424605 |
| ENSMUSG00000030782 | 1.357236841 | 0.012124546 |
| ENSMUSG00000030785 | 1.015486245 | 0.985522133 |
| ENSMUSG00000030786 | -1.000195812 | 1 |
| ENSMUSG00000030787 | 1 | |
| ENSMUSG00000030788 | -1.087751038 | 0.608497229 |
| ENSMUSG00000030789 | 1 | |
| ENSMUSG00000030790 | 1.888840714 | 0.022999073 |
| ENSMUSG00000030792 | -1.484306114 | 0.043433777 |
| ENSMUSG00000030793 | 1.025046921 | 0.97088369 |
| ENSMUSG00000030795 | 1.042995498 | 0.917495228 |
| ENSMUSG00000030796 | 1.059990988 | 0.750742698 |
| ENSMUSG00000030798 | -1.10336434 | 0.777418199 |
| ENSMUSG00000030800 | -1 | 1 |
| ENSMUSG00000030801 | 1.091299128 | 0.499865648 |
| ENSMUSG00000030802 | -1.008282621 | 1 |
| ENSMUSG00000030804 | 1 | |
| ENSMUSG00000030805 | 1.059711147 | 0.652946678 |
| ENSMUSG00000030806 | -1.732702673 | 0.022971084 |
| ENSMUSG00000030811 | 1.063991956 | 0.253249273 |
| ENSMUSG00000030814 | 1.012014522 | 0.968423939 |
| ENSMUSG00000030815 | -1.058915891 | 0.403030305 |
| ENSMUSG00000030816 | 1.019011946 | 0.937535342 |
| ENSMUSG00000030822 | 1.086697466 | 0.354640815 |
| ENSMUSG00000030823 | -1.04568923 | 0.949679283 |
| ENSMUSG00000030824 | 1.0188717 | 0.968937442 |
| ENSMUSG00000030825 | -1.096486055 | 0.764317155 |
| ENSMUSG00000030826 | 1.028489465 | 0.79291651 |
| ENSMUSG00000030827 | 1.530375083 | 0.021271808 |
| ENSMUSG00000030830 | -1.084645496 | 0.617024566 |
| ENSMUSG00000030834 | 1.040562055 | 0.947868053 |
| ENSMUSG00000030835 | -1.009765005 | 1 |
| ENSMUSG00000030838 | -1.044273782 | 0.699340382 |

| | | |
|---|---|---|
| ENSMUSG00000030839 | -1.099655028 | 0.252861426 |
| ENSMUSG00000030842 | 1.167321363 | 0.199750453 |
| ENSMUSG00000030844 | -1.231193867 | 0.070336944 |
| ENSMUSG00000030846 | -1.023641279 | 0.970576293 |
| ENSMUSG00000030847 | 1.090996373 | 0.512782961 |
| ENSMUSG00000030849 | 1.47837277 | 0.0020022 |
| ENSMUSG00000030850 | 1.154845474 | 0.428531045 |
| ENSMUSG00000030851 | 1 | |
| ENSMUSG00000030852 | 1.102097114 | 0.453905691 |
| ENSMUSG00000030854 | -1.083087117 | 0.765798257 |
| ENSMUSG00000030858 | 1.015486245 | 1 |
| ENSMUSG00000030859 | -1.069502966 | 0.698112512 |
| ENSMUSG00000030861 | 1.027739858 | 0.924934548 |
| ENSMUSG00000030862 | 1 | |
| ENSMUSG00000030865 | 1 | |
| ENSMUSG00000030866 | 1 | |
| ENSMUSG00000030867 | 1.029955894 | 0.816748888 |
| ENSMUSG00000030868 | -1.023603264 | 0.959892369 |
| ENSMUSG00000030869 | 1.017333049 | 0.972980766 |
| ENSMUSG00000030870 | 1.011061358 | 1 |
| ENSMUSG00000030871 | -1.014939844 | 0.97658187 |
| ENSMUSG00000030872 | 1.093052118 | 0.66102156 |
| ENSMUSG00000030873 | -1.03962336 | 0.698112512 |
| ENSMUSG00000030876 | 1.017511794 | 0.99362872 |
| ENSMUSG00000030877 | 1.005613439 | 1 |
| ENSMUSG00000030878 | 1.095642847 | 0.633269628 |
| ENSMUSG00000030879 | 1.055539606 | 0.698112512 |
| ENSMUSG00000030880 | 1.078269408 | 0.451546148 |
| ENSMUSG00000030881 | -1.006005978 | 1 |
| ENSMUSG00000030882 | 1.095359383 | 0.300091823 |
| ENSMUSG00000030884 | -1.017393448 | 0.996705669 |
| ENSMUSG00000030887 | 1.29454102 | 0.261984714 |
| ENSMUSG00000030888 | 1.091916076 | 0.608497229 |
| ENSMUSG00000030889 | -1.011863891 | 1 |
| ENSMUSG00000030890 | 1.514382375 | 0.066036399 |
| ENSMUSG00000030894 | 1.253481621 | 0.215991492 |
| ENSMUSG00000030895 | 1.102342851 | 0.608497229 |
| ENSMUSG00000030897 | 1.059463094 | 0.698112512 |
| ENSMUSG00000030898 | 1 | |
| ENSMUSG00000030905 | -1.069502966 | 0.744868995 |
| ENSMUSG00000030909 | 1 | |
| ENSMUSG00000030911 | 1 | |
| ENSMUSG00000030917 | -1.010470986 | 0.931153764 |
| ENSMUSG00000030921 | -1.212047621 | 0.266286552 |
| ENSMUSG00000030922 | 1.025002403 | 0.904797193 |
| ENSMUSG00000030924 | -1.005295872 | 0.998386582 |
| ENSMUSG00000030929 | -1.021763437 | 0.932501815 |
| ENSMUSG00000030930 | 1.130445178 | 0.112920219 |
| ENSMUSG00000030934 | 1.055498007 | 0.844780803 |
| ENSMUSG00000030935 | 1.075727585 | 0.91754678 |
| ENSMUSG00000030942 | 1.018144056 | 0.972684567 |
| ENSMUSG00000030945 | -1.073877404 | 0.83094827 |
| ENSMUSG00000030946 | -1.145055601 | 0.250477583 |
| ENSMUSG00000030954 | 1 | |
| ENSMUSG00000030956 | 1.073532556 | 0.608497229 |
| ENSMUSG00000030960 | -1.015684305 | 0.914954453 |
| ENSMUSG00000030963 | 1 | |
| ENSMUSG00000030965 | 1.022792872 | 0.945625364 |
| ENSMUSG00000030966 | -1.217952448 | 0.103537668 |
| ENSMUSG00000030967 | 1.031945739 | 0.865865121 |
| ENSMUSG00000030968 | 1 | |
| ENSMUSG00000030970 | 1.098733569 | 0.304680629 |
| ENSMUSG00000030972 | 1 | |

| | | |
|---|---|---|
| ENSMUSG00000030976 | 1 | |
| ENSMUSG00000030978 | 1.002548842 | 1 |
| ENSMUSG00000030979 | -1.008431928 | 1 |
| ENSMUSG00000030980 | 1.012989362 | 1 |
| ENSMUSG00000030981 | 1.005613439 | 1 |
| ENSMUSG00000030982 | 1.120686217 | 0.578706451 |
| ENSMUSG00000030983 | 1.011778467 | 0.997108523 |
| ENSMUSG00000030986 | -1.047711745 | 0.793726432 |
| ENSMUSG00000030987 | -1.09041145 | 0.524356958 |
| ENSMUSG00000030990 | 1.043408157 | 0.698112512 |
| ENSMUSG00000030994 | -1 | 1 |
| ENSMUSG00000030996 | 1 | |
| ENSMUSG00000031004 | -1.028346032 | 0.978016801 |
| ENSMUSG00000031007 | 1.167318034 | 0.576959009 |
| ENSMUSG00000031010 | 1.054364183 | 0.901920241 |
| ENSMUSG00000031012 | 1.022049644 | 0.956374899 |
| ENSMUSG00000031015 | 1.140056982 | 0.39537357 |
| ENSMUSG00000031016 | 1.13535855 | 0.250726313 |
| ENSMUSG00000031021 | -1.084739867 | 0.661640678 |
| ENSMUSG00000031022 | 1.02829818 | 0.975677002 |
| ENSMUSG00000031023 | 1.081630987 | 0.464062002 |
| ENSMUSG00000031024 | 1.190902937 | 0.123237825 |
| ENSMUSG00000031026 | 1.68089219 | 0.011824721 |
| ENSMUSG00000031027 | 1.024656056 | 0.972980766 |
| ENSMUSG00000031028 | -1.534861317 | 0.042441598 |
| ENSMUSG00000031029 | 1.029230326 | 0.951063333 |
| ENSMUSG00000031059 | 1.036307005 | 0.825297403 |
| ENSMUSG00000031060 | 1.059092163 | 0.698112512 |
| ENSMUSG00000031065 | 1.061403905 | 0.815715406 |
| ENSMUSG00000031066 | -1.015005806 | 0.989217536 |
| ENSMUSG00000031068 | 1.018436281 | 0.979002724 |
| ENSMUSG00000031070 | 1.448289245 | 0.005624846 |
| ENSMUSG00000031072 | -1.046923332 | 0.750748143 |
| ENSMUSG00000031073 | 1 | |
| ENSMUSG00000031074 | 1 | |
| ENSMUSG00000031075 | 1.313452983 | 0.033216362 |
| ENSMUSG00000031077 | -1.093403506 | 0.349205235 |
| ENSMUSG00000031078 | 1.120681456 | 0.617510841 |
| ENSMUSG00000031079 | -1.181967487 | 0.098972885 |
| ENSMUSG00000031085 | 1 | |
| ENSMUSG00000031089 | 1 | |
| ENSMUSG00000031090 | 1.034003584 | 0.927544616 |
| ENSMUSG00000031093 | 1.100668201 | 0.698112512 |
| ENSMUSG00000031095 | 1.029566511 | 0.958181659 |
| ENSMUSG00000031097 | 1.059463094 | 0.698112512 |
| ENSMUSG00000031098 | -1.084645496 | 0.532369657 |
| ENSMUSG00000031099 | -1.333764643 | 0.041709686 |
| ENSMUSG00000031101 | -1 | 1 |
| ENSMUSG00000031103 | -1.007750156 | 1 |
| ENSMUSG00000031104 | -1.160891938 | 0.670176133 |
| ENSMUSG00000031105 | 1.020883371 | 0.904124333 |
| ENSMUSG00000031107 | 1.015319522 | 0.970208206 |
| ENSMUSG00000031109 | 1.044211743 | 0.808074244 |
| ENSMUSG00000031111 | 1.041610047 | 0.746274305 |
| ENSMUSG00000031112 | 1.156528229 | 0.368144988 |
| ENSMUSG00000031118 | 1 | |
| ENSMUSG00000031119 | -1.009896233 | 0.998367147 |
| ENSMUSG00000031125 | -1 | 1 |
| ENSMUSG00000031129 | -1.057040589 | 0.698112512 |
| ENSMUSG00000031130 | 1 | |
| ENSMUSG00000031131 | 1 | |
| ENSMUSG00000031132 | 1 | |
| ENSMUSG00000031133 | -1.069974478 | 0.698112512 |

| | | |
|---|---|---|
| ENSMUSG00000031134 | -1.025744969 | 0.945251611 |
| ENSMUSG00000031137 | -1.089497311 | 0.532369657 |
| ENSMUSG00000031138 | 1 | |
| ENSMUSG00000031139 | -1.053549062 | 0.698112512 |
| ENSMUSG00000031142 | 1 | |
| ENSMUSG00000031143 | 1 | |
| ENSMUSG00000031144 | -1.199801466 | 0.378017455 |
| ENSMUSG00000031145 | -1.385115567 | 0.21880353 |
| ENSMUSG00000031146 | 1.051415904 | 0.73388071 |
| ENSMUSG00000031147 | 1.132193967 | 0.567496595 |
| ENSMUSG00000031148 | -1.010659671 | 1 |
| ENSMUSG00000031149 | 1.003697379 | 1 |
| ENSMUSG00000031150 | -1.035396833 | 0.742810723 |
| ENSMUSG00000031153 | -1.021589755 | 0.945481169 |
| ENSMUSG00000031154 | 1.022756212 | 0.944099922 |
| ENSMUSG00000031155 | -1.074231281 | 0.608497229 |
| ENSMUSG00000031156 | 1.119246621 | 0.288217114 |
| ENSMUSG00000031157 | -1.033317113 | 0.874317758 |
| ENSMUSG00000031158 | 1.074253503 | 0.492698532 |
| ENSMUSG00000031160 | 1 | |
| ENSMUSG00000031161 | -1.020191882 | 0.94733141 |
| ENSMUSG00000031162 | -1.071075483 | 0.750635131 |
| ENSMUSG00000031163 | -1 | 1 |
| ENSMUSG00000031165 | -1.130883057 | 0.629550689 |
| ENSMUSG00000031166 | 1.087569936 | 0.608497229 |
| ENSMUSG00000031167 | 1.036693655 | 0.968623752 |
| ENSMUSG00000031168 | -1.030111409 | 0.911250323 |
| ENSMUSG00000031169 | 1.037375016 | 0.721386781 |
| ENSMUSG00000031170 | -1.095294215 | 0.541735653 |
| ENSMUSG00000031171 | -1.004446312 | 1 |
| ENSMUSG00000031173 | 1 | |
| ENSMUSG00000031174 | -1.068187422 | 0.651538962 |
| ENSMUSG00000031176 | 1.101770258 | 0.674987219 |
| ENSMUSG00000031179 | 1 | |
| ENSMUSG00000031181 | 1 | |
| ENSMUSG00000031182 | 1 | |
| ENSMUSG00000031189 | 1.350161748 | 0.006020085 |
| ENSMUSG00000031194 | 1 | |
| ENSMUSG00000031196 | 1.141789161 | 0.314360782 |
| ENSMUSG00000031197 | -1.067593752 | 0.794274354 |
| ENSMUSG00000031198 | 1.00763624 | 1 |
| ENSMUSG00000031200 | -1.01833455 | 0.949812939 |
| ENSMUSG00000031201 | -1.022333065 | 0.963664162 |
| ENSMUSG00000031202 | 1.063295037 | 0.608497229 |
| ENSMUSG00000031204 | 1 | |
| ENSMUSG00000031207 | 1.294903366 | 0.404493254 |
| ENSMUSG00000031209 | 1.11166259 | 0.614424414 |
| ENSMUSG00000031210 | -1.031691022 | 0.979002724 |
| ENSMUSG00000031212 | 1.366399777 | 0.00968643 |
| ENSMUSG00000031214 | 1.038235989 | 0.775673129 |
| ENSMUSG00000031216 | -1.230762677 | 0.051762831 |
| ENSMUSG00000031217 | 1.021054349 | 0.895711764 |
| ENSMUSG00000031220 | 1.062527465 | 0.718265321 |
| ENSMUSG00000031221 | 1.020152141 | 0.978763926 |
| ENSMUSG00000031224 | -1.184871248 | 0.366653332 |
| ENSMUSG00000031226 | 1.032971829 | 0.932855261 |
| ENSMUSG00000031227 | 1.0053247 | 1 |
| ENSMUSG00000031229 | 1.020027112 | 0.999347497 |
| ENSMUSG00000031230 | -1 | 1 |
| ENSMUSG00000031231 | 1.023707826 | 0.96513 |
| ENSMUSG00000031232 | 1.070594768 | 0.807247723 |
| ENSMUSG00000031233 | 1 | |
| ENSMUSG00000031239 | -1.021515198 | 0.93979453 |

| | | |
|---|---|---|
| ENSMUSG00000031241 | -1 | 1 |
| ENSMUSG00000031242 | 1.013571797 | 0.996913409 |
| ENSMUSG00000031245 | 1.005800761 | 1 |
| ENSMUSG00000031246 | 1.377481875 | 0.17386461 |
| ENSMUSG00000031250 | 1 | |
| ENSMUSG00000031253 | -1.112010624 | 0.712485242 |
| ENSMUSG00000031255 | -1.293663126 | 0.059242486 |
| ENSMUSG00000031256 | -1.134466255 | 0.608497229 |
| ENSMUSG00000031257 | -1.086720843 | 0.805373432 |
| ENSMUSG00000031258 | 1.002505822 | 1 |
| ENSMUSG00000031262 | -1.142842442 | 0.608497229 |
| ENSMUSG00000031264 | 1.055416288 | 0.825058682 |
| ENSMUSG00000031266 | -1.008411461 | 1 |
| ENSMUSG00000031270 | 1 | |
| ENSMUSG00000031271 | 1 | |
| ENSMUSG00000031273 | 1.184821448 | 0.036775998 |
| ENSMUSG00000031274 | 1.032721054 | 0.949812939 |
| ENSMUSG00000031278 | -1.219357429 | 0.329995202 |
| ENSMUSG00000031283 | 1.083768932 | 0.454424089 |
| ENSMUSG00000031284 | 1.04839438 | 0.852557372 |
| ENSMUSG00000031285 | -1.100193663 | 0.608497229 |
| ENSMUSG00000031286 | -1.046821685 | 0.713861573 |
| ENSMUSG00000031289 | 1.161036672 | 0.608092964 |
| ENSMUSG00000031290 | 1.042961586 | 0.888871716 |
| ENSMUSG00000031292 | -1.284478347 | 0.102925573 |
| ENSMUSG00000031293 | -1.116853908 | 0.62484781 |
| ENSMUSG00000031294 | -1.175121877 | 0.490063384 |
| ENSMUSG00000031295 | -1.080638721 | 0.677534336 |
| ENSMUSG00000031297 | 1.211837414 | 0.095919352 |
| ENSMUSG00000031298 | -1.109431965 | 0.36085659 |
| ENSMUSG00000031299 | -1.045219841 | 0.890792876 |
| ENSMUSG00000031302 | -1.009116151 | 1 |
| ENSMUSG00000031303 | -1.540578392 | 0.002242306 |
| ENSMUSG00000031304 | -1.525511356 | 0.005165495 |
| ENSMUSG00000031309 | -1.028775994 | 0.958187754 |
| ENSMUSG00000031310 | 1.048737277 | 0.807743274 |
| ENSMUSG00000031311 | 1.034346677 | 0.965659325 |
| ENSMUSG00000031312 | 1.148669342 | 0.672115339 |
| ENSMUSG00000031314 | -1.01703196 | 1 |
| ENSMUSG00000031320 | 1.037262017 | 0.969891536 |
| ENSMUSG00000031323 | -1.095294215 | 0.541735653 |
| ENSMUSG00000031325 | 1.241656513 | 0.525842026 |
| ENSMUSG00000031326 | 1 | |
| ENSMUSG00000031327 | 1.083868507 | 0.608497229 |
| ENSMUSG00000031328 | 1.399362533 | 0.295833164 |
| ENSMUSG00000031329 | 1.177693733 | 0.500630907 |
| ENSMUSG00000031330 | -1 | 1 |
| ENSMUSG00000031333 | -1.072601901 | 0.737180328 |
| ENSMUSG00000031337 | -1.041832404 | 0.732127066 |
| ENSMUSG00000031340 | 1.118469536 | 0.38142723 |
| ENSMUSG00000031342 | -1.451038583 | 0.028615506 |
| ENSMUSG00000031343 | 1.767437752 | 0.013686068 |
| ENSMUSG00000031344 | -1.099174267 | 0.608497229 |
| ENSMUSG00000031347 | -1.097724094 | 0.608497229 |
| ENSMUSG00000031349 | -1.087315379 | 0.697515847 |
| ENSMUSG00000031351 | -1.217692076 | 0.112629065 |
| ENSMUSG00000031352 | -1.0435471 | 0.829826148 |
| ENSMUSG00000031353 | -1.010542744 | 1 |
| ENSMUSG00000031354 | 1 | |
| ENSMUSG00000031355 | -1.359619154 | 0.006127679 |
| ENSMUSG00000031357 | -1.153410538 | 0.525557653 |
| ENSMUSG00000031358 | -1.065681575 | 0.738145566 |
| ENSMUSG00000031360 | 1.012165918 | 1 |

| | | |
|---|---|---|
| ENSMUSG00000031362 | 1 | |
| ENSMUSG00000031364 | 1.005613439 | 1 |
| ENSMUSG00000031365 | 1.092005969 | 0.698112512 |
| ENSMUSG00000031367 | -1.01206459 | 1 |
| ENSMUSG00000031370 | -1.075104644 | 0.701318973 |
| ENSMUSG00000031371 | 1.206617709 | 0.312960909 |
| ENSMUSG00000031372 | 1 | |
| ENSMUSG00000031373 | -1.028308132 | 0.911521144 |
| ENSMUSG00000031374 | -1.043306199 | 0.698112512 |
| ENSMUSG00000031375 | -1.983244245 | 0.011103532 |
| ENSMUSG00000031376 | 1.015263614 | 1 |
| ENSMUSG00000031377 | 1 | |
| ENSMUSG00000031378 | -1.01184608 | 0.974925935 |
| ENSMUSG00000031379 | 1.009101938 | 0.988057225 |
| ENSMUSG00000031380 | -1.260564886 | 0.039249559 |
| ENSMUSG00000031381 | -1.060363274 | 0.755673717 |
| ENSMUSG00000031382 | 1 | |
| ENSMUSG00000031383 | 1.026113384 | 0.778239858 |
| ENSMUSG00000031384 | 1 | |
| ENSMUSG00000031385 | 1.154412827 | 0.657110556 |
| ENSMUSG00000031386 | 1.045806002 | 0.884088177 |
| ENSMUSG00000031387 | 1.941997433 | 0.000144027 |
| ENSMUSG00000031388 | 1.06366221 | 0.662840906 |
| ENSMUSG00000031389 | -1.082556607 | 0.698112512 |
| ENSMUSG00000031390 | -1.147838869 | 0.195607801 |
| ENSMUSG00000031391 | -1.4037272 | 0.001515144 |
| ENSMUSG00000031392 | 1.031298281 | 0.918344559 |
| ENSMUSG00000031393 | 1.045919964 | 0.883188695 |
| ENSMUSG00000031394 | -1.181718351 | 0.305952299 |
| ENSMUSG00000031397 | 1.029096068 | 0.973126641 |
| ENSMUSG00000031398 | 1.132625892 | 0.375579698 |
| ENSMUSG00000031399 | -1.028219935 | 0.794274354 |
| ENSMUSG00000031400 | -1.074173218 | 0.784039458 |
| ENSMUSG00000031402 | -1.044348547 | 0.88284878 |
| ENSMUSG00000031403 | -1.01966583 | 0.989387709 |
| ENSMUSG00000031409 | 1 | |
| ENSMUSG00000031410 | 1 | |
| ENSMUSG00000031411 | 1 | |
| ENSMUSG00000031422 | -1.014193013 | 1 |
| ENSMUSG00000031424 | 1 | |
| ENSMUSG00000031425 | -1.164940203 | 0.065517197 |
| ENSMUSG00000031428 | -1.284561257 | 0.007149705 |
| ENSMUSG00000031429 | -1.207310442 | 0.28720897 |
| ENSMUSG00000031430 | 1 | |
| ENSMUSG00000031431 | 1.071028981 | 0.608497229 |
| ENSMUSG00000031432 | -1.13958041 | 0.59223593 |
| ENSMUSG00000031433 | -1.191277884 | 0.255874194 |
| ENSMUSG00000031434 | -1.053372737 | 0.74724097 |
| ENSMUSG00000031438 | 1.013143172 | 0.972684567 |
| ENSMUSG00000031441 | -1.012871055 | 1 |
| ENSMUSG00000031442 | 1.236311556 | 0.038158125 |
| ENSMUSG00000031443 | 1 | |
| ENSMUSG00000031444 | -1.070510097 | 0.837350977 |
| ENSMUSG00000031445 | -1.157556633 | 0.487033708 |
| ENSMUSG00000031446 | 1.114109083 | 0.696025422 |
| ENSMUSG00000031447 | 1.347829321 | 0.299531359 |
| ENSMUSG00000031448 | 1 | |
| ENSMUSG00000031449 | 1.028323353 | 0.934760809 |
| ENSMUSG00000031450 | -1.085102125 | 0.628300894 |
| ENSMUSG00000031451 | -1.061181375 | 0.479111229 |
| ENSMUSG00000031452 | -1.088487825 | 0.729018288 |
| ENSMUSG00000031453 | 1.220817002 | 0.255959421 |
| ENSMUSG00000031458 | 1.198547014 | 0.166729085 |

| | | |
|---|---|---|
| ENSMUSG00000031461 | -1.068166409 | 0.608497229 |
| ENSMUSG00000031465 | 1.258516715 | 0.038638661 |
| ENSMUSG00000031467 | -1.024860887 | 0.928573362 |
| ENSMUSG00000031471 | 1 | |
| ENSMUSG00000031478 | -1.012706346 | 0.943298961 |
| ENSMUSG00000031479 | -1.045411166 | 0.858134675 |
| ENSMUSG00000031480 | 1.073605014 | 0.46324962 |
| ENSMUSG00000031481 | 1.112016081 | 0.654443366 |
| ENSMUSG00000031482 | 1.003845529 | 1 |
| ENSMUSG00000031483 | 1.080882742 | 0.735417419 |
| ENSMUSG00000031485 | 1.116819548 | 0.608497229 |
| ENSMUSG00000031486 | 1.138639994 | 0.274076373 |
| ENSMUSG00000031487 | 1.035045635 | 0.716562452 |
| ENSMUSG00000031488 | -1.134951713 | 0.327722667 |
| ENSMUSG00000031489 | -1.245749446 | 0.519289106 |
| ENSMUSG00000031490 | 1.207147539 | 0.085876598 |
| ENSMUSG00000031491 | 1 | |
| ENSMUSG00000031492 | 1 | |
| ENSMUSG00000031493 | 1.167101109 | 0.459223522 |
| ENSMUSG00000031494 | 1.069068979 | 0.608497229 |
| ENSMUSG00000031495 | 1 | |
| ENSMUSG00000031497 | 1.04898545 | 0.902822111 |
| ENSMUSG00000031502 | 1.332288944 | 0.183203649 |
| ENSMUSG00000031503 | 1.288910959 | 0.132502833 |
| ENSMUSG00000031504 | 1.220103922 | 0.009888628 |
| ENSMUSG00000031505 | 1.178011175 | 0.044230261 |
| ENSMUSG00000031506 | -1.127018601 | 0.698112512 |
| ENSMUSG00000031508 | 1.167268944 | 0.312000097 |
| ENSMUSG00000031509 | 1.014545335 | 0.979002724 |
| ENSMUSG00000031510 | 1 | |
| ENSMUSG00000031511 | 1.159531399 | 0.312970963 |
| ENSMUSG00000031512 | 1.059463094 | 0.608497229 |
| ENSMUSG00000031513 | 1.016761152 | 0.981992363 |
| ENSMUSG00000031516 | -1.042544553 | 0.773487331 |
| ENSMUSG00000031517 | -1.206338534 | 0.26979254 |
| ENSMUSG00000031518 | 1.069068979 | 0.608497229 |
| ENSMUSG00000031519 | 1.031293674 | 0.895048791 |
| ENSMUSG00000031520 | 1.052146299 | 0.486930276 |
| ENSMUSG00000031521 | 1.261564139 | 0.076584005 |
| ENSMUSG00000031523 | 1.006862001 | 1 |
| ENSMUSG00000031527 | -1.021769552 | 0.982178232 |
| ENSMUSG00000031529 | 1.056770515 | 0.839099152 |
| ENSMUSG00000031530 | -1.083052008 | 0.313474869 |
| ENSMUSG00000031532 | -1.004174719 | 1 |
| ENSMUSG00000031533 | 1.010900494 | 1 |
| ENSMUSG00000031534 | 1.098887261 | 0.453850153 |
| ENSMUSG00000031535 | 1 | |
| ENSMUSG00000031536 | -1.061144172 | 0.698112512 |
| ENSMUSG00000031537 | -1.002975598 | 1 |
| ENSMUSG00000031538 | 1.019655336 | 0.972136247 |
| ENSMUSG00000031539 | -1.013557508 | 0.989387709 |
| ENSMUSG00000031540 | 1.003054079 | 1 |
| ENSMUSG00000031543 | 1.2160479 | 0.014987351 |
| ENSMUSG00000031545 | 1.07373281 | 0.755282061 |
| ENSMUSG00000031546 | 1.010714976 | 1 |
| ENSMUSG00000031548 | -1.578994462 | 0.009764274 |
| ENSMUSG00000031549 | 1.100596556 | 0.70819406 |
| ENSMUSG00000031551 | 1.05410549 | 0.835701362 |
| ENSMUSG00000031552 | 1.069068979 | 0.608497229 |
| ENSMUSG00000031553 | 1.023959045 | 0.979002724 |
| ENSMUSG00000031554 | 1.275704208 | 0.018759913 |
| ENSMUSG00000031555 | 1.087057205 | 0.765746941 |
| ENSMUSG00000031556 | 1.002556528 | 1 |

| | | |
|---|---|---|
| ENSMUSG00000031557 | 1.064972659 | 0.789690099 |
| ENSMUSG00000031558 | 1.210193505 | 0.216046507 |
| ENSMUSG00000031559 | 1.064243295 | 0.836133893 |
| ENSMUSG00000031561 | -4.2402306 | 0.00037608 |
| ENSMUSG00000031562 | 1.133443682 | 0.202428693 |
| ENSMUSG00000031563 | 1.066929425 | 0.777145624 |
| ENSMUSG00000031565 | 1.220291077 | 0.171092221 |
| ENSMUSG00000031568 | 1.028417459 | 0.92971816 |
| ENSMUSG00000031570 | 1.024698274 | 0.928424364 |
| ENSMUSG00000031574 | -1.03846593 | 0.788152462 |
| ENSMUSG00000031575 | -1.007380651 | 1 |
| ENSMUSG00000031576 | 1.059463094 | 0.608497229 |
| ENSMUSG00000031577 | 1.024160543 | 0.902709435 |
| ENSMUSG00000031578 | 1.007525436 | 1 |
| ENSMUSG00000031583 | -1.012784909 | 1 |
| ENSMUSG00000031584 | -1.005831495 | 1 |
| ENSMUSG00000031585 | -1.025749723 | 0.959453108 |
| ENSMUSG00000031586 | -1.181790409 | 0.005581993 |
| ENSMUSG00000031590 | -1.004974508 | 1 |
| ENSMUSG00000031591 | 1.09972632 | 0.699309119 |
| ENSMUSG00000031592 | -1.008337448 | 1 |
| ENSMUSG00000031594 | 1.055439738 | 0.917030977 |
| ENSMUSG00000031595 | 1.338948271 | 0.03042531 |
| ENSMUSG00000031596 | 1.273361296 | 0.068335953 |
| ENSMUSG00000031600 | -1.025915556 | 0.953962555 |
| ENSMUSG00000031601 | -1.003077989 | 1 |
| ENSMUSG00000031603 | -1.044273782 | 0.698112512 |
| ENSMUSG00000031604 | 1.015600949 | 1 |
| ENSMUSG00000031605 | 1.089552224 | 0.667576553 |
| ENSMUSG00000031608 | -1.040818495 | 0.843683484 |
| ENSMUSG00000031609 | 1.057787697 | 0.698112512 |
| ENSMUSG00000031610 | 1 | |
| ENSMUSG00000031613 | 1.059463094 | 0.692423811 |
| ENSMUSG00000031616 | 1.089470641 | 0.733794666 |
| ENSMUSG00000031617 | -1.047726923 | 0.848096322 |
| ENSMUSG00000031618 | -1.147493521 | 0.658590644 |
| ENSMUSG00000031620 | 1 | |
| ENSMUSG00000031621 | 1 | |
| ENSMUSG00000031622 | -1.041290701 | 0.87505994 |
| ENSMUSG00000031626 | 1.57924853 | 0.003589328 |
| ENSMUSG00000031627 | -1.030504912 | 0.881823779 |
| ENSMUSG00000031628 | -1.000084562 | 1 |
| ENSMUSG00000031629 | -1.003161557 | 1 |
| ENSMUSG00000031631 | 1.000012416 | 1 |
| ENSMUSG00000031633 | 1.066439054 | 0.819614655 |
| ENSMUSG00000031634 | -1.029124493 | 0.93853737 |
| ENSMUSG00000031635 | 1.019083579 | 0.949812939 |
| ENSMUSG00000031636 | 1.294949469 | 0.1351252 |
| ENSMUSG00000031637 | 1.041274494 | 0.873626752 |
| ENSMUSG00000031639 | -1.130137881 | 0.481361055 |
| ENSMUSG00000031640 | 1 | |
| ENSMUSG00000031641 | -1.003752785 | 1 |
| ENSMUSG00000031642 | -1.003272005 | 1 |
| ENSMUSG00000031644 | 1.028475643 | 0.914868665 |
| ENSMUSG00000031645 | 1 | |
| ENSMUSG00000031647 | 1.126943177 | 0.43150353 |
| ENSMUSG00000031651 | 1 | |
| ENSMUSG00000031652 | 1.038947211 | 0.889846433 |
| ENSMUSG00000031654 | -1.347263424 | 0.120194303 |
| ENSMUSG00000031657 | -1.103009866 | 0.701672557 |
| ENSMUSG00000031659 | -1.597828918 | 0.000439578 |
| ENSMUSG00000031660 | -1.055955403 | 0.825809529 |
| ENSMUSG00000031661 | -2.100204875 | 0.001174334 |

| | | |
|---|---|---|
| ENSMUSG00000031662 | -1 | 1 |
| ENSMUSG00000031665 | 2.625746864 | 4.27E-07 |
| ENSMUSG00000031666 | -1.038884327 | 0.844897753 |
| ENSMUSG00000031667 | -1.075157833 | 0.640834198 |
| ENSMUSG00000031668 | 1.005026278 | 1 |
| ENSMUSG00000031669 | -1.054453831 | 0.758244074 |
| ENSMUSG00000031671 | -1.081531112 | 0.608497229 |
| ENSMUSG00000031672 | 1.022810097 | 0.979002724 |
| ENSMUSG00000031673 | 1.007566158 | 1 |
| ENSMUSG00000031681 | 1.102224974 | 0.477335321 |
| ENSMUSG00000031682 | -1 | 1 |
| ENSMUSG00000031683 | -1.055500482 | 0.832767487 |
| ENSMUSG00000031684 | 1.07205142 | 0.673318777 |
| ENSMUSG00000031688 | 1 | |
| ENSMUSG00000031691 | 1.083591306 | 0.72984031 |
| ENSMUSG00000031696 | 1.092082283 | 0.789694282 |
| ENSMUSG00000031697 | 1.00335148 | 1 |
| ENSMUSG00000031698 | -1.041666343 | 0.945710765 |
| ENSMUSG00000031700 | 1.171840675 | 0.296219427 |
| ENSMUSG00000031701 | 1.00441011 | 1 |
| ENSMUSG00000031703 | -1.002547065 | 1 |
| ENSMUSG00000031706 | 1.105767667 | 0.608497229 |
| ENSMUSG00000031708 | 1.013433197 | 1 |
| ENSMUSG00000031709 | -1.146263426 | 0.278630662 |
| ENSMUSG00000031710 | -1 | 1 |
| ENSMUSG00000031711 | -1.025538553 | 0.934760809 |
| ENSMUSG00000031712 | -1.096130134 | 0.401921535 |
| ENSMUSG00000031714 | 1.100316144 | 0.605619163 |
| ENSMUSG00000031715 | -1.009489931 | 1 |
| ENSMUSG00000031722 | 1.290681034 | 0.028809274 |
| ENSMUSG00000031723 | 1.00131905 | 1 |
| ENSMUSG00000031725 | -1 | 1 |
| ENSMUSG00000031727 | 1 | |
| ENSMUSG00000031728 | 1.059564547 | 0.424846573 |
| ENSMUSG00000031729 | -1.02419472 | 0.970800919 |
| ENSMUSG00000031730 | 1.019356774 | 0.897283466 |
| ENSMUSG00000031731 | -1.046633375 | 0.912454307 |
| ENSMUSG00000031732 | -1.038442965 | 0.903139568 |
| ENSMUSG00000031734 | -1.030635337 | 0.958979658 |
| ENSMUSG00000031736 | 1.0796899 | 0.567151281 |
| ENSMUSG00000031737 | -1.091752579 | 0.487401224 |
| ENSMUSG00000031738 | -1 | 1 |
| ENSMUSG00000031740 | 1.077339162 | 0.76916778 |
| ENSMUSG00000031748 | 1.572954936 | 0.000410078 |
| ENSMUSG00000031749 | 1.06804995 | 0.696603049 |
| ENSMUSG00000031750 | -1.137864077 | 0.130670603 |
| ENSMUSG00000031751 | -1.018865521 | 0.996913409 |
| ENSMUSG00000031753 | -1.001450184 | 1 |
| ENSMUSG00000031754 | -1.032766229 | 0.950444171 |
| ENSMUSG00000031755 | -1.100827848 | 0.547016491 |
| ENSMUSG00000031756 | -1.064024022 | 0.744030455 |
| ENSMUSG00000031757 | 1 | |
| ENSMUSG00000031758 | -1.00455597 | 1 |
| ENSMUSG00000031760 | 1.011607416 | 1 |
| ENSMUSG00000031762 | 1.707226441 | 0.016132785 |
| ENSMUSG00000031765 | 1.698244477 | 0.006778907 |
| ENSMUSG00000031766 | -1.043306199 | 0.698112512 |
| ENSMUSG00000031767 | 1.197173297 | 0.014804417 |
| ENSMUSG00000031770 | 1.141736388 | 0.268563805 |
| ENSMUSG00000031772 | 1.11166259 | 0.608497229 |
| ENSMUSG00000031774 | 1.042584074 | 0.84372915 |
| ENSMUSG00000031775 | -1.017708674 | 1 |
| ENSMUSG00000031776 | -1.115835644 | 0.636089827 |

| | | |
|---|---|---|
| ENSMUSG00000031778 | 1.077580786 | 0.562685409 |
| ENSMUSG00000031779 | 1 | |
| ENSMUSG00000031780 | -1.281728708 | 0.22408326 |
| ENSMUSG00000031781 | -1.114613273 | 0.608497229 |
| ENSMUSG00000031782 | -1.173782058 | 0.262929409 |
| ENSMUSG00000031783 | -1.036107871 | 0.887026138 |
| ENSMUSG00000031785 | 1.06140802 | 0.608497229 |
| ENSMUSG00000031786 | -1.053549062 | 0.778372272 |
| ENSMUSG00000031787 | 1.012742804 | 0.969695497 |
| ENSMUSG00000031788 | 1.014473722 | 0.982838256 |
| ENSMUSG00000031789 | -1.330936474 | 0.202942571 |
| ENSMUSG00000031790 | -1.144443611 | 0.472690788 |
| ENSMUSG00000031791 | 1.005386372 | 1 |
| ENSMUSG00000031792 | 1.02488707 | 0.891337812 |
| ENSMUSG00000031796 | 1.072404642 | 0.6769285 |
| ENSMUSG00000031799 | 1.160347139 | 0.68293521 |
| ENSMUSG00000031802 | -1 | 1 |
| ENSMUSG00000031803 | -1.07008665 | 0.79291651 |
| ENSMUSG00000031805 | 1.119400843 | 0.494597256 |
| ENSMUSG00000031807 | 1.092036415 | 0.495999195 |
| ENSMUSG00000031808 | 1.533203982 | 0.003737264 |
| ENSMUSG00000031809 | -1.063525418 | 0.698112512 |
| ENSMUSG00000031812 | 1.079993762 | 0.723432738 |
| ENSMUSG00000031813 | 1.00537758 | 1 |
| ENSMUSG00000031816 | -1.038603293 | 0.795285331 |
| ENSMUSG00000031818 | 1.0477774 | 0.866033961 |
| ENSMUSG00000031819 | -1.078017653 | 0.698112512 |
| ENSMUSG00000031820 | 1.043141803 | 0.698112512 |
| ENSMUSG00000031821 | 1.028053714 | 0.869468685 |
| ENSMUSG00000031822 | 1.103725955 | 0.608497229 |
| ENSMUSG00000031823 | -1.005639531 | 1 |
| ENSMUSG00000031824 | -1.195894112 | 0.210349702 |
| ENSMUSG00000031825 | 1.594417443 | 0.00435101 |
| ENSMUSG00000031826 | -1.010922745 | 0.99744561 |
| ENSMUSG00000031827 | 1.026289784 | 0.850076143 |
| ENSMUSG00000031828 | -1.506389736 | 0.098085953 |
| ENSMUSG00000031831 | -1.138693452 | 0.608497229 |
| ENSMUSG00000031832 | 1.069822811 | 0.698112512 |
| ENSMUSG00000031833 | -1.029709551 | 0.775027206 |
| ENSMUSG00000031834 | 1.009408056 | 0.996913409 |
| ENSMUSG00000031835 | -1.003398127 | 1 |
| ENSMUSG00000031837 | 1.009398673 | 1 |
| ENSMUSG00000031838 | -1.024835928 | 0.900308456 |
| ENSMUSG00000031839 | -1.065734422 | 0.776953697 |
| ENSMUSG00000031840 | -1.076956972 | 0.563742992 |
| ENSMUSG00000031841 | 1.012841092 | 0.979002724 |
| ENSMUSG00000031842 | -1.353814003 | 0.0220709 |
| ENSMUSG00000031843 | -1.092192208 | 0.608497229 |
| ENSMUSG00000031844 | 1 | |
| ENSMUSG00000031845 | 1 | |
| ENSMUSG00000031847 | -1.122151198 | 0.311631163 |
| ENSMUSG00000031848 | 1.022504517 | 0.924587329 |
| ENSMUSG00000031849 | -1.221162906 | 0.437095676 |
| ENSMUSG00000031851 | -1.228942881 | 0.002225593 |
| ENSMUSG00000031853 | -1.088487825 | 0.698112512 |
| ENSMUSG00000031858 | -1.023155984 | 0.965037112 |
| ENSMUSG00000031860 | -1.050282393 | 0.768480212 |
| ENSMUSG00000031861 | 1.034687834 | 0.811497438 |
| ENSMUSG00000031862 | 1.036688515 | 0.850244888 |
| ENSMUSG00000031864 | 1.028784268 | 0.92989327 |
| ENSMUSG00000031865 | 1.123068326 | 0.381036055 |
| ENSMUSG00000031870 | 1 | |
| ENSMUSG00000031871 | -1.177298609 | 0.221427514 |

| | | |
|---|---|---|
| ENSMUSG00000031872 | -1.485436624 | 0.033510579 |
| ENSMUSG00000031875 | 1.209511187 | 0.163186799 |
| ENSMUSG00000031876 | 1.132570123 | 0.48164954 |
| ENSMUSG00000031877 | -1.282519459 | 0.234728821 |
| ENSMUSG00000031878 | -1.014970492 | 1 |
| ENSMUSG00000031879 | 1.010027228 | 0.978259071 |
| ENSMUSG00000031880 | 1.059057565 | 0.621868642 |
| ENSMUSG00000031881 | -1.043306199 | 0.873296215 |
| ENSMUSG00000031883 | -1.03182308 | 0.909335407 |
| ENSMUSG00000031884 | -1.03962336 | 0.698112512 |
| ENSMUSG00000031885 | -1.059542342 | 0.864768883 |
| ENSMUSG00000031886 | -1.231184281 | 0.056310079 |
| ENSMUSG00000031887 | -1.044431308 | 0.698112512 |
| ENSMUSG00000031889 | -1.01949844 | 0.968152125 |
| ENSMUSG00000031891 | -1.405425847 | 0.129463386 |
| ENSMUSG00000031893 | -1.062262719 | 0.911620796 |
| ENSMUSG00000031896 | 1.012893425 | 1 |
| ENSMUSG00000031897 | -1.315631564 | 0.026808199 |
| ENSMUSG00000031898 | 1 | |
| ENSMUSG00000031901 | 1.038575362 | 0.706694652 |
| ENSMUSG00000031902 | -1.001481151 | 1 |
| ENSMUSG00000031903 | -1.015517078 | 0.958653795 |
| ENSMUSG00000031904 | -1.048780268 | 0.880677217 |
| ENSMUSG00000031906 | 1.42006464 | 0.002110532 |
| ENSMUSG00000031907 | 1.171334349 | 0.6659463 |
| ENSMUSG00000031910 | 1.036963274 | 0.78352577 |
| ENSMUSG00000031913 | -1.048321222 | 0.854163865 |
| ENSMUSG00000031916 | -1.002137491 | 1 |
| ENSMUSG00000031917 | -1.023998413 | 0.969089779 |
| ENSMUSG00000031918 | -1.066217162 | 0.822781131 |
| ENSMUSG00000031919 | 1.059463094 | 0.608497229 |
| ENSMUSG00000031921 | -1.026459116 | 0.934760809 |
| ENSMUSG00000031922 | -1.012943779 | 1 |
| ENSMUSG00000031924 | -1.016349547 | 1 |
| ENSMUSG00000031925 | 1.116686277 | 0.412997769 |
| ENSMUSG00000031927 | 1.218240674 | 0.217803313 |
| ENSMUSG00000031928 | -1.042791574 | 0.889581381 |
| ENSMUSG00000031930 | 1.051964171 | 0.793291876 |
| ENSMUSG00000031931 | -1.051558137 | 0.772196124 |
| ENSMUSG00000031932 | -1.020155149 | 0.996811331 |
| ENSMUSG00000031933 | 1 | |
| ENSMUSG00000031934 | -1.392183177 | 0.010780205 |
| ENSMUSG00000031935 | -1.029490426 | 0.922165724 |
| ENSMUSG00000031936 | -1 | 1 |
| ENSMUSG00000031937 | -1.799668476 | 0.000464325 |
| ENSMUSG00000031938 | 1.034913677 | 0.864972543 |
| ENSMUSG00000031939 | -1.059203241 | 0.778132999 |
| ENSMUSG00000031948 | -1.066826819 | 0.839420872 |
| ENSMUSG00000031949 | -1.050926748 | 0.668830453 |
| ENSMUSG00000031950 | -1.0912722 | 0.684693086 |
| ENSMUSG00000031951 | -1.224185111 | 0.059357562 |
| ENSMUSG00000031952 | -1.139982257 | 0.624040338 |
| ENSMUSG00000031953 | -1.120532643 | 0.15169538 |
| ENSMUSG00000031954 | -1.017368411 | 0.998847216 |
| ENSMUSG00000031955 | 1.167110057 | 0.229884308 |
| ENSMUSG00000031957 | 1.11166259 | 0.608497229 |
| ENSMUSG00000031958 | -1.078590893 | 0.648436514 |
| ENSMUSG00000031959 | 1.018011229 | 0.953276227 |
| ENSMUSG00000031960 | 1.099704246 | 0.717035356 |
| ENSMUSG00000031962 | -1 | 1 |
| ENSMUSG00000031963 | 1.044786753 | 0.733377791 |
| ENSMUSG00000031965 | -1.439035766 | 0.027606265 |
| ENSMUSG00000031966 | 1 | |

| | | |
|---|---|---|
| ENSMUSG00000031967 | -1.012173699 | 1 |
| ENSMUSG00000031969 | -1.017596623 | 0.949566828 |
| ENSMUSG00000031970 | -1.168357085 | 0.07721161 |
| ENSMUSG00000031971 | -1.156641824 | 0.276229467 |
| ENSMUSG00000031972 | 2.138013951 | 0.002723641 |
| ENSMUSG00000031974 | -1.058523341 | 0.731323613 |
| ENSMUSG00000031976 | -1.047171106 | 0.873409014 |
| ENSMUSG00000031979 | 1.033170512 | 0.839582974 |
| ENSMUSG00000031980 | -1.053549062 | 0.807892349 |
| ENSMUSG00000031981 | 1.132639128 | 0.398767938 |
| ENSMUSG00000031982 | -1.054105309 | 0.608497229 |
| ENSMUSG00000031983 | 1.084637936 | 0.685141313 |
| ENSMUSG00000031984 | 1.013898639 | 0.972389453 |
| ENSMUSG00000031985 | -1.00917079 | 1 |
| ENSMUSG00000031986 | -1.048521174 | 0.734674233 |
| ENSMUSG00000031987 | 1.031455906 | 0.897684198 |
| ENSMUSG00000031988 | -1.044257937 | 0.877881982 |
| ENSMUSG00000031990 | -1.089820255 | 0.608497229 |
| ENSMUSG00000031991 | 1 | |
| ENSMUSG00000031993 | -1.046497978 | 0.835701362 |
| ENSMUSG00000031994 | -1.089497311 | 0.698112512 |
| ENSMUSG00000031995 | -1.048277865 | 0.880781039 |
| ENSMUSG00000031996 | 1.040262185 | 0.948385094 |
| ENSMUSG00000031997 | 1.271344436 | 0.012381271 |
| ENSMUSG00000031998 | -1.025690504 | 0.952057016 |
| ENSMUSG00000032000 | -1.274165101 | 0.067998626 |
| ENSMUSG00000032002 | -1.049472711 | 0.854890151 |
| ENSMUSG00000032006 | -1.4372987 | 0.013783049 |
| ENSMUSG00000032009 | -1.134467807 | 0.399444438 |
| ENSMUSG00000032010 | -1.12459845 | 0.349914416 |
| ENSMUSG00000032011 | 2.280218506 | 0.000658053 |
| ENSMUSG00000032012 | 1.313578063 | 0.07425059 |
| ENSMUSG00000032013 | 1 | |
| ENSMUSG00000032014 | 1.268983061 | 0.021171782 |
| ENSMUSG00000032015 | 1 | |
| ENSMUSG00000032017 | 1.169163022 | 0.608497229 |
| ENSMUSG00000032018 | 1.02369373 | 0.963609512 |
| ENSMUSG00000032020 | -1.189149041 | 0.083676757 |
| ENSMUSG00000032021 | -1.03962336 | 0.698112512 |
| ENSMUSG00000032023 | 1 | |
| ENSMUSG00000032024 | 1.308299714 | 0.005249107 |
| ENSMUSG00000032026 | 1.102765762 | 0.609481654 |
| ENSMUSG00000032028 | -1.047731725 | 0.946232149 |
| ENSMUSG00000032030 | -1.034089664 | 0.935780763 |
| ENSMUSG00000032033 | 1 | |
| ENSMUSG00000032034 | 1 | |
| ENSMUSG00000032035 | -1.057211748 | 0.848720308 |
| ENSMUSG00000032036 | 1.114281776 | 0.037835897 |
| ENSMUSG00000032038 | 1.242219116 | 0.196259378 |
| ENSMUSG00000032040 | -1.022390399 | 0.944383136 |
| ENSMUSG00000032041 | -1.05533028 | 0.778372272 |
| ENSMUSG00000032042 | -1.076581319 | 0.775673129 |
| ENSMUSG00000032044 | -1.049854783 | 0.698112512 |
| ENSMUSG00000032046 | -1.037897254 | 0.854890151 |
| ENSMUSG00000032047 | -1.0825809 | 0.731295483 |
| ENSMUSG00000032048 | 1 | |
| ENSMUSG00000032050 | -1.032346099 | 0.962368837 |
| ENSMUSG00000032051 | -1.121037588 | 0.373282501 |
| ENSMUSG00000032053 | 1 | |
| ENSMUSG00000032056 | 1.132639128 | 0.217803313 |
| ENSMUSG00000032057 | 1.509883636 | 0.012083743 |
| ENSMUSG00000032058 | -1.000352691 | 1 |
| ENSMUSG00000032059 | -1.013823739 | 0.979629796 |

| | | |
|---|---|---|
| ENSMUSG00000032060 | 1.325511785 | 0.051171361 |
| ENSMUSG00000032062 | -1.005921439 | 1 |
| ENSMUSG00000032064 | 1.308727633 | 0.07338776 |
| ENSMUSG00000032065 | 1 | |
| ENSMUSG00000032066 | -1.207255912 | 0.159807598 |
| ENSMUSG00000032067 | -1.267213356 | 0.021751537 |
| ENSMUSG00000032068 | -1.014570171 | 0.996048848 |
| ENSMUSG00000032076 | -1.158908337 | 0.486959986 |
| ENSMUSG00000032077 | -1.126031212 | 0.388567904 |
| ENSMUSG00000032078 | -1.059829777 | 0.789448756 |
| ENSMUSG00000032079 | -1.03962336 | 0.698112512 |
| ENSMUSG00000032080 | 1 | |
| ENSMUSG00000032081 | 1 | |
| ENSMUSG00000032083 | 1.005212439 | 1 |
| ENSMUSG00000032085 | 2.208613498 | 0.00209931 |
| ENSMUSG00000032086 | -1.044872336 | 0.878551492 |
| ENSMUSG00000032087 | 1 | |
| ENSMUSG00000032089 | 1 | |
| ENSMUSG00000032091 | 1 | |
| ENSMUSG00000032092 | -1.743673556 | 0.000682614 |
| ENSMUSG00000032093 | 1 | |
| ENSMUSG00000032094 | -1.069502966 | 0.698112512 |
| ENSMUSG00000032096 | -1.04716755 | 0.931480186 |
| ENSMUSG00000032097 | -1.025723619 | 0.982838256 |
| ENSMUSG00000032098 | 1 | |
| ENSMUSG00000032099 | 1 | |
| ENSMUSG00000032101 | -2.267399544 | 0.000145902 |
| ENSMUSG00000032103 | 1 | |
| ENSMUSG00000032105 | 1 | |
| ENSMUSG00000032108 | 1 | |
| ENSMUSG00000032109 | -1.707244249 | 0.010755875 |
| ENSMUSG00000032110 | 1.096218584 | 0.515817146 |
| ENSMUSG00000032112 | -1.005304631 | 1 |
| ENSMUSG00000032113 | -1.059095595 | 0.804348174 |
| ENSMUSG00000032114 | -1.076097686 | 0.327318612 |
| ENSMUSG00000032115 | -1.114763635 | 0.69530468 |
| ENSMUSG00000032116 | -1.054086952 | 0.90849465 |
| ENSMUSG00000032118 | -1.552115035 | 0.105136419 |
| ENSMUSG00000032119 | -1.080954257 | 0.532369657 |
| ENSMUSG00000032120 | 1.006234276 | 1 |
| ENSMUSG00000032121 | -1.175568763 | 0.015464544 |
| ENSMUSG00000032122 | 1.077910959 | 0.559011327 |
| ENSMUSG00000032123 | -1.076981537 | 0.698112512 |
| ENSMUSG00000032125 | 1 | |
| ENSMUSG00000032126 | -1.022302917 | 0.943224821 |
| ENSMUSG00000032127 | 1.032873023 | 0.902013045 |
| ENSMUSG00000032128 | 1.372638437 | 0.166729085 |
| ENSMUSG00000032131 | -1.186580855 | 0.185330171 |
| ENSMUSG00000032134 | -1 | 1 |
| ENSMUSG00000032135 | 1.307393737 | 0.108119116 |
| ENSMUSG00000032171 | 1.007176406 | 1 |
| ENSMUSG00000032172 | -1.320237577 | 0.251049598 |
| ENSMUSG00000032174 | -1.257776892 | 0.607716251 |
| ENSMUSG00000032175 | 1.059199918 | 0.385772778 |
| ENSMUSG00000032177 | 1.070627947 | 0.608497229 |
| ENSMUSG00000032178 | 1.02881725 | 0.949177474 |
| ENSMUSG00000032179 | -1 | 1 |
| ENSMUSG00000032180 | 1.001924298 | 1 |
| ENSMUSG00000032181 | 1.033391092 | 0.942924901 |
| ENSMUSG00000032182 | 1.033210994 | 0.648566407 |
| ENSMUSG00000032184 | -1.112406452 | 0.097579999 |
| ENSMUSG00000032185 | -1.005218588 | 1 |
| ENSMUSG00000032186 | -1.239331559 | 0.12255265 |

| | | |
|---|---|---|
| ENSMUSG00000032187 | 1.027280316 | 0.9548419 |
| ENSMUSG00000032191 | 1.059463094 | 0.608497229 |
| ENSMUSG00000032192 | -1.000742546 | 1 |
| ENSMUSG00000032193 | -1.114381742 | 0.608497229 |
| ENSMUSG00000032194 | 1.038792567 | 0.825256294 |
| ENSMUSG00000032198 | -1.009959352 | 0.9858962 |
| ENSMUSG00000032199 | -1.049858576 | 0.905298189 |
| ENSMUSG00000032202 | -1.118536927 | 0.387036025 |
| ENSMUSG00000032204 | 1.065519011 | 0.818267146 |
| ENSMUSG00000032207 | -1 | 1 |
| ENSMUSG00000032212 | -1.026340395 | 0.953731979 |
| ENSMUSG00000032215 | -1.053097674 | 0.778057094 |
| ENSMUSG00000032216 | 1.080647485 | 0.865583959 |
| ENSMUSG00000032217 | 1.024481245 | 0.959892369 |
| ENSMUSG00000032218 | -1.10620928 | 0.661155252 |
| ENSMUSG00000032220 | 1.177731164 | 0.285204165 |
| ENSMUSG00000032221 | -1.16344559 | 0.312657959 |
| ENSMUSG00000032224 | -1.192109593 | 0.145828618 |
| ENSMUSG00000032226 | 1.007837529 | 1 |
| ENSMUSG00000032228 | 1.0426291 | 0.919744809 |
| ENSMUSG00000032231 | -1.071666274 | 0.880723094 |
| ENSMUSG00000032232 | -1.207429479 | 0.25982125 |
| ENSMUSG00000032235 | -1.048715326 | 0.803938171 |
| ENSMUSG00000032238 | -1.436159892 | 0.050815501 |
| ENSMUSG00000032239 | -1.091716634 | 0.582976361 |
| ENSMUSG00000032243 | 2.674391834 | 0.000474677 |
| ENSMUSG00000032244 | -1.015018056 | 1 |
| ENSMUSG00000032245 | -1.081084945 | 0.603483156 |
| ENSMUSG00000032246 | 1.415732338 | 0.071577777 |
| ENSMUSG00000032249 | 1.039456195 | 0.929923764 |
| ENSMUSG00000032251 | 1.065786008 | 0.658430243 |
| ENSMUSG00000032252 | 1.00112987 | 1 |
| ENSMUSG00000032253 | 1.055760889 | 0.862562879 |
| ENSMUSG00000032254 | -1.124495867 | 0.685141313 |
| ENSMUSG00000032257 | -1.037482356 | 0.89979145 |
| ENSMUSG00000032258 | -1.060696102 | 0.692782277 |
| ENSMUSG00000032259 | 1 | |
| ENSMUSG00000032261 | 1.05849035 | 0.608497229 |
| ENSMUSG00000032262 | -1.036008148 | 0.860327141 |
| ENSMUSG00000032263 | -1.070725598 | 0.698112512 |
| ENSMUSG00000032264 | -1.026042518 | 0.958690483 |
| ENSMUSG00000032265 | 1.67901117 | 0.000556642 |
| ENSMUSG00000032267 | -1.02476873 | 0.963386993 |
| ENSMUSG00000032268 | -1 | 1 |
| ENSMUSG00000032269 | 1 | |
| ENSMUSG00000032271 | 1 | |
| ENSMUSG00000032274 | -1.043306199 | 0.698112512 |
| ENSMUSG00000032278 | -1.185575082 | 0.18636028 |
| ENSMUSG00000032279 | -1.071515639 | 0.796457226 |
| ENSMUSG00000032280 | 1.060827088 | 0.637832253 |
| ENSMUSG00000032281 | 1.712064899 | 0.000143293 |
| ENSMUSG00000032285 | -1.137455181 | 0.110492512 |
| ENSMUSG00000032288 | -1.039519461 | 0.801443616 |
| ENSMUSG00000032289 | 1.4914567 | 0.190645128 |
| ENSMUSG00000032290 | -1.07857238 | 0.734408673 |
| ENSMUSG00000032291 | -1.23712605 | 0.01140441 |
| ENSMUSG00000032292 | 1 | |
| ENSMUSG00000032293 | -1.042354998 | 0.93481577 |
| ENSMUSG00000032294 | 1.064679453 | 0.87088305 |
| ENSMUSG00000032295 | 1.039526534 | 0.758608542 |
| ENSMUSG00000032297 | 1.137399458 | 0.657985991 |
| ENSMUSG00000032298 | 1.04119486 | 0.695859665 |
| ENSMUSG00000032299 | 1.101041449 | 0.454669379 |

| | | |
|---|---|---|
| ENSMUSG00000032300 | 1.173393358 | 0.0700306 |
| ENSMUSG00000032301 | -1.09042518 | 0.764370878 |
| ENSMUSG00000032303 | -1.043306199 | 0.79291651 |
| ENSMUSG00000032305 | -1.044738512 | 0.783974347 |
| ENSMUSG00000032306 | -1.109251543 | 0.608497229 |
| ENSMUSG00000032307 | -1.005566067 | 1 |
| ENSMUSG00000032308 | -1.01954596 | 0.699280765 |
| ENSMUSG00000032309 | -1.024606497 | 0.934364841 |
| ENSMUSG00000032310 | 1 | |
| ENSMUSG00000032311 | -1.03872304 | 0.92650471 |
| ENSMUSG00000032312 | 1.045721249 | 0.736686851 |
| ENSMUSG00000032313 | 1.212908519 | 0.453446164 |
| ENSMUSG00000032314 | -1.031474986 | 0.955457497 |
| ENSMUSG00000032315 | -1.224792133 | 0.38417754 |
| ENSMUSG00000032316 | -1.028887565 | 0.855328455 |
| ENSMUSG00000032318 | -1.317977811 | 0.173371035 |
| ENSMUSG00000032320 | -1.011699313 | 1 |
| ENSMUSG00000032322 | 1.03063878 | 0.972650737 |
| ENSMUSG00000032323 | 1.049411284 | 0.863033776 |
| ENSMUSG00000032324 | -1.033600802 | 0.946232149 |
| ENSMUSG00000032327 | -1.978626247 | 0.002655169 |
| ENSMUSG00000032328 | -1.019961324 | 0.994915989 |
| ENSMUSG00000032329 | -1.011614054 | 0.99339328 |
| ENSMUSG00000032330 | -1.010163896 | 1 |
| ENSMUSG00000032332 | 1.13645616 | 0.629405465 |
| ENSMUSG00000032333 | -1.004594855 | 1 |
| ENSMUSG00000032334 | 1.184992133 | 0.130204206 |
| ENSMUSG00000032336 | 1.007175641 | 1 |
| ENSMUSG00000032338 | -1.199669418 | 0.59127359 |
| ENSMUSG00000032340 | 1.228558585 | 0.398797577 |
| ENSMUSG00000032342 | -1.063890476 | 0.698112512 |
| ENSMUSG00000032343 | 1 | |
| ENSMUSG00000032344 | -1.16822064 | 0.394398328 |
| ENSMUSG00000032346 | -1.025643787 | 0.974533047 |
| ENSMUSG00000032348 | 1.032158794 | 0.863853621 |
| ENSMUSG00000032349 | -1.03001987 | 0.949679283 |
| ENSMUSG00000032350 | -1.031128201 | 0.915158215 |
| ENSMUSG00000032352 | 1.08200567 | 0.43764772 |
| ENSMUSG00000032353 | 1.103444897 | 0.572947856 |
| ENSMUSG00000032355 | -1.054169715 | 0.813485995 |
| ENSMUSG00000032356 | 1.693643831 | 0.008040974 |
| ENSMUSG00000032357 | -1 | 1 |
| ENSMUSG00000032358 | -1.29355546 | 0.038351204 |
| ENSMUSG00000032359 | -1.182325533 | 0.286691901 |
| ENSMUSG00000032360 | 1 | |
| ENSMUSG00000032363 | 1.245334178 | 0.004990486 |
| ENSMUSG00000032366 | 1.156055855 | 0.69305964 |
| ENSMUSG00000032368 | -3.653911307 | 0.000244121 |
| ENSMUSG00000032369 | -1.022684712 | 0.966355452 |
| ENSMUSG00000032370 | -1.177799561 | 0.251544797 |
| ENSMUSG00000032372 | 1.146061711 | 0.437909374 |
| ENSMUSG00000032373 | -1.404922069 | 0.071001672 |
| ENSMUSG00000032374 | -1.060780088 | 0.83233787 |
| ENSMUSG00000032375 | -1.039533355 | 0.854279112 |
| ENSMUSG00000032376 | -1.053113354 | 0.820337436 |
| ENSMUSG00000032377 | 1.161351664 | 0.211158311 |
| ENSMUSG00000032380 | -1.101207018 | 0.698112512 |
| ENSMUSG00000032381 | 1.029215535 | 0.917209568 |
| ENSMUSG00000032382 | -1.094631747 | 0.698112512 |
| ENSMUSG00000032383 | -1.015000768 | 0.968584032 |
| ENSMUSG00000032384 | -1.015608349 | 0.997227391 |
| ENSMUSG00000032386 | -1.141176647 | 0.608497229 |
| ENSMUSG00000032387 | 1.050935882 | 0.608497229 |

| | | |
|---|---|---|
| ENSMUSG00000032388 | 1.046005243 | 0.865124175 |
| ENSMUSG00000032392 | -1.00127709 | 1 |
| ENSMUSG00000032393 | -1.054693867 | 0.854968144 |
| ENSMUSG00000032394 | -1.388681643 | 0.012944412 |
| ENSMUSG00000032396 | -1.077267572 | 0.649280935 |
| ENSMUSG00000032397 | -1.081587384 | 0.769895472 |
| ENSMUSG00000032398 | -1.109940536 | 0.542536371 |
| ENSMUSG00000032399 | 1.005778609 | 1 |
| ENSMUSG00000032400 | -1.060837222 | 0.803072249 |
| ENSMUSG00000032401 | 1.44535423 | 0.013319415 |
| ENSMUSG00000032402 | 1.005340422 | 1 |
| ENSMUSG00000032403 | -1.043392332 | 0.755972525 |
| ENSMUSG00000032405 | -1.057077179 | 0.791482968 |
| ENSMUSG00000032407 | -1.023586591 | 0.979002724 |
| ENSMUSG00000032409 | -1.003966943 | 1 |
| ENSMUSG00000032410 | -1.080873604 | 0.748686313 |
| ENSMUSG00000032411 | -1.087976504 | 0.698112512 |
| ENSMUSG00000032412 | -1.001795205 | 1 |
| ENSMUSG00000032413 | 1.013832753 | 0.979928923 |
| ENSMUSG00000032415 | -1.023639002 | 0.907536891 |
| ENSMUSG00000032417 | -1.253003862 | 0.130858186 |
| ENSMUSG00000032418 | -1.025072963 | 0.977413596 |
| ENSMUSG00000032419 | 1.065208404 | 0.608497229 |
| ENSMUSG00000032420 | 1.021700516 | 0.938939969 |
| ENSMUSG00000032422 | 1.00786657 | 1 |
| ENSMUSG00000032423 | 1.032898895 | 0.963664162 |
| ENSMUSG00000032425 | -1.029119048 | 0.922064713 |
| ENSMUSG00000032431 | 1.035697309 | 0.882794734 |
| ENSMUSG00000032434 | -1.091588046 | 0.608497229 |
| ENSMUSG00000032435 | -1.088016828 | 0.704728898 |
| ENSMUSG00000032436 | 1.063855129 | 0.698112512 |
| ENSMUSG00000032437 | -1.035950647 | 0.938479652 |
| ENSMUSG00000032440 | -1.24363674 | 0.291233266 |
| ENSMUSG00000032443 | -1.093674795 | 0.608497229 |
| ENSMUSG00000032446 | -1.418307801 | 0.062398364 |
| ENSMUSG00000032449 | -1.026252484 | 0.958646994 |
| ENSMUSG00000032451 | 1 | |
| ENSMUSG00000032452 | -1.847066505 | 0.003083671 |
| ENSMUSG00000032454 | -1.063525418 | 0.698112512 |
| ENSMUSG00000032456 | 1.091955798 | 0.24033917 |
| ENSMUSG00000032458 | -1.012161959 | 1 |
| ENSMUSG00000032459 | -1.081038688 | 0.608497229 |
| ENSMUSG00000032462 | -1.142998287 | 0.608497229 |
| ENSMUSG00000032463 | -1.051590941 | 0.770255677 |
| ENSMUSG00000032468 | -1.038152951 | 0.893672909 |
| ENSMUSG00000032469 | -1.050337751 | 0.778372272 |
| ENSMUSG00000032470 | -1.060585808 | 0.775235085 |
| ENSMUSG00000032473 | -1.03962336 | 0.803072249 |
| ENSMUSG00000032475 | 1.017398559 | 0.981964814 |
| ENSMUSG00000032477 | -1.037956102 | 0.889846433 |
| ENSMUSG00000032478 | 1.031825874 | 0.88697865 |
| ENSMUSG00000032479 | 1.02160919 | 0.989895137 |
| ENSMUSG00000032480 | 1.014871447 | 0.984277851 |
| ENSMUSG00000032481 | 1.001702792 | 1 |
| ENSMUSG00000032482 | -1.060328162 | 0.744351452 |
| ENSMUSG00000032484 | -1 | 1 |
| ENSMUSG00000032485 | 1.018460075 | 0.930108107 |
| ENSMUSG00000032487 | 1.883808034 | 0.001866476 |
| ENSMUSG00000032489 | -1.14371553 | 0.473906907 |
| ENSMUSG00000032491 | 1.163612293 | 0.148031611 |
| ENSMUSG00000032492 | -1.308356867 | 0.223646364 |
| ENSMUSG00000032493 | 1 | |
| ENSMUSG00000032494 | -1 | 1 |

| | | |
|---|---|---|
| ENSMUSG00000032495 | -1.111192823 | 0.769745556 |
| ENSMUSG00000032496 | 1 | |
| ENSMUSG00000032497 | -1.046702554 | 0.837704961 |
| ENSMUSG00000032498 | -1.002935349 | 1 |
| ENSMUSG00000032500 | -1.112010624 | 0.670176133 |
| ENSMUSG00000032501 | 1.324223608 | 0.017028941 |
| ENSMUSG00000032502 | -1.043306199 | 0.717251435 |
| ENSMUSG00000032503 | -1.03962336 | 0.698112512 |
| ENSMUSG00000032504 | -1.024079847 | 0.980617339 |
| ENSMUSG00000032507 | -1.124035041 | 0.257889684 |
| ENSMUSG00000032508 | -1.165043183 | 0.220011422 |
| ENSMUSG00000032511 | -1.487965869 | 0.014346397 |
| ENSMUSG00000032512 | -1.017954537 | 0.985522133 |
| ENSMUSG00000032513 | 1.085008072 | 0.408528362 |
| ENSMUSG00000032514 | -1.086667286 | 0.833505594 |
| ENSMUSG00000032515 | -1.018239067 | 0.908543402 |
| ENSMUSG00000032517 | 1 | |
| ENSMUSG00000032518 | 1.066883188 | 0.842680844 |
| ENSMUSG00000032519 | -1.007004216 | 1 |
| ENSMUSG00000032523 | 1.393231649 | 0.010702717 |
| ENSMUSG00000032525 | 1.014246152 | 1 |
| ENSMUSG00000032526 | -1.099226898 | 0.464075945 |
| ENSMUSG00000032527 | -1.058849715 | 0.607004292 |
| ENSMUSG00000032528 | -1.030198322 | 0.968977347 |
| ENSMUSG00000032530 | 1 | |
| ENSMUSG00000032531 | 1.206459843 | 0.293999722 |
| ENSMUSG00000032532 | 1 | |
| ENSMUSG00000032534 | -1.007653741 | 1 |
| ENSMUSG00000032536 | -1.011077156 | 0.95904617 |
| ENSMUSG00000032537 | -1 | 1 |
| ENSMUSG00000032540 | -1.010953555 | 0.997620696 |
| ENSMUSG00000032547 | -1.021661188 | 0.982961337 |
| ENSMUSG00000032548 | 1.134479128 | 0.350223743 |
| ENSMUSG00000032549 | 1.206603917 | 0.02542895 |
| ENSMUSG00000032551 | -1.068839227 | 0.698112512 |
| ENSMUSG00000032553 | -1.0183625 | 0.96569197 |
| ENSMUSG00000032554 | 1.037456959 | 0.949679283 |
| ENSMUSG00000032555 | -1.026054072 | 0.979002724 |
| ENSMUSG00000032556 | 1.069068979 | 0.608497229 |
| ENSMUSG00000032557 | 1.018623755 | 0.980942914 |
| ENSMUSG00000032558 | -1.063674659 | 0.608497229 |
| ENSMUSG00000032560 | 1.056799178 | 0.881117973 |
| ENSMUSG00000032561 | -1.335693613 | 0.013224637 |
| ENSMUSG00000032562 | -1.00536531 | 1 |
| ENSMUSG00000032563 | -1.042437669 | 0.884447694 |
| ENSMUSG00000032564 | 1 | |
| ENSMUSG00000032565 | -1.064158572 | 0.608497229 |
| ENSMUSG00000032566 | 1 | |
| ENSMUSG00000032567 | 1.060383671 | 0.608497229 |
| ENSMUSG00000032570 | -1.017021381 | 0.991944074 |
| ENSMUSG00000032571 | -1.045711553 | 0.766191522 |
| ENSMUSG00000032572 | -1.536156151 | 0.033882553 |
| ENSMUSG00000032575 | -1.057577054 | 0.809396188 |
| ENSMUSG00000032577 | 1.090912504 | 0.487490682 |
| ENSMUSG00000032578 | 1.26347583 | 0.008479602 |
| ENSMUSG00000032579 | -1.051057358 | 0.437045401 |
| ENSMUSG00000032580 | -1.032027704 | 0.950042971 |
| ENSMUSG00000032582 | -1.006723534 | 1 |
| ENSMUSG00000032583 | -1.031012517 | 0.838710806 |
| ENSMUSG00000032584 | -1.246328198 | 0.570749474 |
| ENSMUSG00000032586 | -1.098629281 | 0.608497229 |
| ENSMUSG00000032589 | 1.003176717 | 1 |
| ENSMUSG00000032590 | -1.079434294 | 0.608497229 |

| | | |
|---|---|---|
| ENSMUSG00000032591 | -1.162714164 | 0.608497229 |
| ENSMUSG00000032593 | -1.118419513 | 0.311753963 |
| ENSMUSG00000032594 | 1.034767623 | 0.880101094 |
| ENSMUSG00000032595 | -1.043306199 | 0.698112512 |
| ENSMUSG00000032596 | -1.386364132 | 0.003077096 |
| ENSMUSG00000032598 | 1.05416044 | 0.698112512 |
| ENSMUSG00000032599 | 1.016532009 | 0.943121172 |
| ENSMUSG00000032601 | 1.016059419 | 1 |
| ENSMUSG00000032602 | -1.086295662 | 0.633312178 |
| ENSMUSG00000032604 | -1.041604996 | 0.911122256 |
| ENSMUSG00000032606 | -1.04448561 | 0.796102353 |
| ENSMUSG00000032607 | -1.005614775 | 0.973099033 |
| ENSMUSG00000032609 | -1.099847202 | 0.408046973 |
| ENSMUSG00000032611 | -1.032725316 | 0.982838256 |
| ENSMUSG00000032612 | -1.048274199 | 0.892026583 |
| ENSMUSG00000032615 | -1.020938751 | 0.698961838 |
| ENSMUSG00000032621 | -1.00858444 | 1 |
| ENSMUSG00000032623 | 1.095872691 | 0.608497229 |
| ENSMUSG00000032624 | 1.058862698 | 0.804680856 |
| ENSMUSG00000032625 | 1.028559049 | 0.872692845 |
| ENSMUSG00000032633 | 1.127063665 | 0.344542597 |
| ENSMUSG00000032637 | 1.094570586 | 0.608497229 |
| ENSMUSG00000032640 | 1.139173716 | 0.308657915 |
| ENSMUSG00000032641 | 1.239097495 | 0.025023351 |
| ENSMUSG00000032643 | 1.2878638 | 0.009348865 |
| ENSMUSG00000032648 | 1.140104518 | 0.451532357 |
| ENSMUSG00000032649 | -1.272444033 | 0.048441465 |
| ENSMUSG00000032652 | 1.033284186 | 0.803260194 |
| ENSMUSG00000032656 | 1.006221374 | 1 |
| ENSMUSG00000032657 | -1.714025928 | 0.026278438 |
| ENSMUSG00000032661 | -1.217243297 | 0.524476431 |
| ENSMUSG00000032666 | 1.003836325 | 1 |
| ENSMUSG00000032667 | 1.061859506 | 0.740742803 |
| ENSMUSG00000032671 | 1 | |
| ENSMUSG00000032673 | -1.288263816 | 0.043055794 |
| ENSMUSG00000032679 | -2.197130413 | 0.00070961 |
| ENSMUSG00000032680 | -1.030405406 | 0.982534887 |
| ENSMUSG00000032688 | -1.150578315 | 0.37308318 |
| ENSMUSG00000032690 | -1.177298609 | 0.565138487 |
| ENSMUSG00000032691 | -1.043306199 | 0.79291651 |
| ENSMUSG00000032698 | -1.129912829 | 0.684325094 |
| ENSMUSG00000032702 | 1.136500974 | 0.214783921 |
| ENSMUSG00000032705 | 1.143799341 | 0.202428693 |
| ENSMUSG00000032712 | 1.142222852 | 0.608497229 |
| ENSMUSG00000032714 | -1.000868694 | 1 |
| ENSMUSG00000032715 | 1.380673599 | 0.01222575 |
| ENSMUSG00000032717 | -1.588887843 | 0.042790808 |
| ENSMUSG00000032718 | -1.455384357 | 0.028714164 |
| ENSMUSG00000032719 | -1 | 1 |
| ENSMUSG00000032724 | -1.063984524 | 0.388093649 |
| ENSMUSG00000032725 | 1.02261662 | 0.966355452 |
| ENSMUSG00000032726 | -1.257211759 | 0.251049598 |
| ENSMUSG00000032727 | -1.095974144 | 0.674124002 |
| ENSMUSG00000032733 | 1.087706151 | 0.562156304 |
| ENSMUSG00000032735 | 1.204071528 | 0.032804406 |
| ENSMUSG00000032737 | 1.181427946 | 0.161261603 |
| ENSMUSG00000032739 | 1.10634245 | 0.690985372 |
| ENSMUSG00000032740 | -1.034264661 | 0.93878065 |
| ENSMUSG00000032741 | 1.025569264 | 0.852588732 |
| ENSMUSG00000032743 | 1.064973605 | 0.616740374 |
| ENSMUSG00000032744 | -1.084645496 | 0.532369657 |
| ENSMUSG00000032745 | -1.108971772 | 0.698112512 |
| ENSMUSG00000032750 | -1.157838297 | 0.615267463 |

| | | |
|---|---|---|
| ENSMUSG00000032754 | 1.132569601 | 0.216046507 |
| ENSMUSG00000032757 | 1.131718936 | 0.482433619 |
| ENSMUSG00000032758 | 1 | |
| ENSMUSG00000032763 | 1.143799435 | 0.069479432 |
| ENSMUSG00000032766 | 1.363388458 | 0.007124727 |
| ENSMUSG00000032769 | 1.024804142 | 0.956661971 |
| ENSMUSG00000032773 | -1.095849567 | 0.820469444 |
| ENSMUSG00000032776 | -1.089711575 | 0.536942078 |
| ENSMUSG00000032777 | 1.033303589 | 0.895931483 |
| ENSMUSG00000032782 | -1.068863823 | 0.608497229 |
| ENSMUSG00000032783 | 1.010437153 | 0.992355016 |
| ENSMUSG00000032786 | 1.032485398 | 0.851353899 |
| ENSMUSG00000032788 | -1.063338455 | 0.698112512 |
| ENSMUSG00000032796 | -1.834404149 | 0.003636347 |
| ENSMUSG00000032802 | -1.06609432 | 0.739850278 |
| ENSMUSG00000032803 | 1.044256804 | 0.933031586 |
| ENSMUSG00000032806 | 1.020366182 | 0.951744672 |
| ENSMUSG00000032807 | -1.317969732 | 0.071698973 |
| ENSMUSG00000032808 | 1 | |
| ENSMUSG00000032812 | 1.103962166 | 0.182129884 |
| ENSMUSG00000032815 | -1.086106155 | 0.613187574 |
| ENSMUSG00000032816 | -1.268340298 | 0.006338472 |
| ENSMUSG00000032818 | 1 | |
| ENSMUSG00000032826 | -1.013830499 | 0.933537875 |
| ENSMUSG00000032827 | 1.29193733 | 0.011767594 |
| ENSMUSG00000032834 | -1.014570636 | 0.995214337 |
| ENSMUSG00000032839 | -1.060200525 | 0.752153213 |
| ENSMUSG00000032840 | 1.088404361 | 0.608497229 |
| ENSMUSG00000032841 | 1.11960374 | 0.081282078 |
| ENSMUSG00000032842 | -1.017211117 | 0.93510318 |
| ENSMUSG00000032845 | 1.085535503 | 0.608497229 |
| ENSMUSG00000032846 | -1.271336876 | 0.068133347 |
| ENSMUSG00000032849 | -1.036821903 | 0.735098711 |
| ENSMUSG00000032850 | -1.038858696 | 0.698112512 |
| ENSMUSG00000032852 | 1.005613439 | 1 |
| ENSMUSG00000032854 | -1.665753121 | 0.006306899 |
| ENSMUSG00000032855 | 1.158264468 | 0.46774472 |
| ENSMUSG00000032860 | -1.372888358 | 0.021891251 |
| ENSMUSG00000032864 | -1.053549062 | 0.769745556 |
| ENSMUSG00000032867 | 1.054303954 | 0.837531941 |
| ENSMUSG00000032869 | -1.121282678 | 0.445632848 |
| ENSMUSG00000032870 | 1.041447996 | 0.82784586 |
| ENSMUSG00000032872 | -1.104184764 | 0.608497229 |
| ENSMUSG00000032875 | 1.217725908 | 0.096128731 |
| ENSMUSG00000032878 | 1.161036672 | 0.474472862 |
| ENSMUSG00000032883 | 1.000707275 | 1 |
| ENSMUSG00000032889 | -1.192692435 | 0.312283428 |
| ENSMUSG00000032890 | -1.704945481 | 0.000785016 |
| ENSMUSG00000032892 | -1.043306199 | 0.698112512 |
| ENSMUSG00000032894 | 1 | |
| ENSMUSG00000032897 | -1.024902679 | 0.925679199 |
| ENSMUSG00000032898 | 1.054378446 | 0.698112512 |
| ENSMUSG00000032899 | 1.34358437 | 0.03854438 |
| ENSMUSG00000032900 | 1 | |
| ENSMUSG00000032902 | -1.007699453 | 1 |
| ENSMUSG00000032905 | -1.003418999 | 1 |
| ENSMUSG00000032908 | -1.300383978 | 0.093577622 |
| ENSMUSG00000032911 | -3.683193333 | 0.000560801 |
| ENSMUSG00000032913 | -1.051846363 | 0.774235108 |
| ENSMUSG00000032915 | -1.044273782 | 0.790851299 |
| ENSMUSG00000032921 | 1.079681442 | 0.608497229 |
| ENSMUSG00000032925 | 1.014594439 | 0.992493283 |
| ENSMUSG00000032932 | 1.006233979 | 1 |

| | | |
|---|---|---|
| ENSMUSG00000032936 | -1.736150141 | 0.007814932 |
| ENSMUSG00000032937 | 1 | |
| ENSMUSG00000032939 | -1.032702729 | 0.951780193 |
| ENSMUSG00000032940 | -1.26234396 | 0.440395661 |
| ENSMUSG00000032942 | 1 | |
| ENSMUSG00000032946 | 1.042913754 | 0.915785585 |
| ENSMUSG00000032948 | 1.095872691 | 0.608497229 |
| ENSMUSG00000032952 | 1.013730291 | 0.993182545 |
| ENSMUSG00000032959 | 1.0717013 | 0.761101793 |
| ENSMUSG00000032965 | -1.014604582 | 0.978602033 |
| ENSMUSG00000032966 | -1.00888711 | 1 |
| ENSMUSG00000032968 | 1.03248156 | 0.996719799 |
| ENSMUSG00000032977 | 1.035484687 | 0.739206552 |
| ENSMUSG00000032978 | -1.03962336 | 0.698112512 |
| ENSMUSG00000032985 | 1 | |
| ENSMUSG00000032987 | 1 | |
| ENSMUSG00000032988 | 1.12264798 | 0.66674596 |
| ENSMUSG00000032997 | 1.135042897 | 0.608497229 |
| ENSMUSG00000032998 | 1.064666442 | 0.782447561 |
| ENSMUSG00000032999 | -1.084645496 | 0.608497229 |
| ENSMUSG00000033004 | -1.03738959 | 0.911972816 |
| ENSMUSG00000033006 | 1 | |
| ENSMUSG00000033007 | -1.037482356 | 0.908706036 |
| ENSMUSG00000033009 | -1.082429204 | 0.713688844 |
| ENSMUSG00000033014 | 1.001874672 | 1 |
| ENSMUSG00000033015 | 1 | |
| ENSMUSG00000033016 | 1.152136371 | 0.095890856 |
| ENSMUSG00000033020 | 1.059872599 | 0.642645794 |
| ENSMUSG00000033021 | 1.013643138 | 0.975362601 |
| ENSMUSG00000033022 | -1.368304728 | 0.014825903 |
| ENSMUSG00000033024 | 1 | |
| ENSMUSG00000033027 | 1 | |
| ENSMUSG00000033029 | 1.074623555 | 0.698112512 |
| ENSMUSG00000033031 | -1.065129758 | 0.839099152 |
| ENSMUSG00000033032 | -1.113461542 | 0.292787347 |
| ENSMUSG00000033033 | 1.073079559 | 0.608497229 |
| ENSMUSG00000033036 | -1.053331522 | 0.776640705 |
| ENSMUSG00000033039 | 1.011624904 | 0.975872952 |
| ENSMUSG00000033044 | 1 | |
| ENSMUSG00000033047 | 1.000331199 | 1 |
| ENSMUSG00000033053 | 1 | |
| ENSMUSG00000033054 | -1.022353294 | 0.982838256 |
| ENSMUSG00000033055 | -1.02914523 | 0.841369183 |
| ENSMUSG00000033059 | -1.072184016 | 0.716954584 |
| ENSMUSG00000033060 | -1.091671167 | 0.608497229 |
| ENSMUSG00000033061 | 1 | |
| ENSMUSG00000033063 | -1 | 1 |
| ENSMUSG00000033065 | -1.06070926 | 0.751613087 |
| ENSMUSG00000033066 | -1.720759344 | 0.001604705 |
| ENSMUSG00000033068 | -1.097841395 | 0.237969364 |
| ENSMUSG00000033075 | 1.024624425 | 0.973422289 |
| ENSMUSG00000033080 | -1.053549062 | 0.698112512 |
| ENSMUSG00000033082 | -1.037896228 | 0.916141522 |
| ENSMUSG00000033083 | -1.176144864 | 0.170993912 |
| ENSMUSG00000033088 | 1.034236619 | 0.814089245 |
| ENSMUSG00000033096 | -1.197795136 | 0.321451717 |
| ENSMUSG00000033099 | 1.007020238 | 1 |
| ENSMUSG00000033102 | -1.035229554 | 0.908114884 |
| ENSMUSG00000033105 | -1.085349594 | 0.684982671 |
| ENSMUSG00000033106 | -1.019284623 | 0.959175283 |
| ENSMUSG00000033107 | -1.228645998 | 0.164430272 |
| ENSMUSG00000033111 | -1.085124088 | 0.512137514 |
| ENSMUSG00000033114 | -1.004521653 | 1 |

| | | |
|---|---|---|
| ENSMUSG00000033122 | 1 | |
| ENSMUSG00000033124 | -1.080000709 | 0.608497229 |
| ENSMUSG00000033126 | -1.018911711 | 0.912464846 |
| ENSMUSG00000033128 | 1.102204977 | 0.49411169 |
| ENSMUSG00000033147 | -1.123862782 | 0.205228244 |
| ENSMUSG00000033149 | -1.0244968 | 0.976498231 |
| ENSMUSG00000033152 | -1.363146169 | 0.041796348 |
| ENSMUSG00000033156 | 1 | |
| ENSMUSG00000033157 | 1.048104262 | 0.756938573 |
| ENSMUSG00000033159 | 1.09563689 | 0.462634602 |
| ENSMUSG00000033161 | 1.079675821 | 0.792578392 |
| ENSMUSG00000033166 | -1.149039432 | 0.58386781 |
| ENSMUSG00000033170 | -1.096042154 | 0.518298719 |
| ENSMUSG00000033174 | 1.349646406 | 0.012970042 |
| ENSMUSG00000033177 | 1.015486245 | 0.984826086 |
| ENSMUSG00000033182 | -1 | 1 |
| ENSMUSG00000033184 | 1.026493118 | 0.972980766 |
| ENSMUSG00000033186 | -1.068548613 | 0.752077931 |
| ENSMUSG00000033187 | 1 | |
| ENSMUSG00000033191 | -1.115393597 | 0.662840906 |
| ENSMUSG00000033192 | 1.223310412 | 0.04351828 |
| ENSMUSG00000033196 | 1 | |
| ENSMUSG00000033200 | 1 | |
| ENSMUSG00000033207 | 1.255432534 | 0.294533094 |
| ENSMUSG00000033208 | -1.174541399 | 0.561688776 |
| ENSMUSG00000033209 | 1.281688658 | 0.044971634 |
| ENSMUSG00000033210 | 1 | |
| ENSMUSG00000033213 | 1.019083579 | 0.992310203 |
| ENSMUSG00000033214 | 1.084557277 | 0.823091034 |
| ENSMUSG00000033216 | -1.034323525 | 0.609615508 |
| ENSMUSG00000033219 | 1 | |
| ENSMUSG00000033220 | -1.066776513 | 0.698112512 |
| ENSMUSG00000033222 | -1.046188077 | 0.827942378 |
| ENSMUSG00000033227 | -1.463395269 | 0.103412426 |
| ENSMUSG00000033228 | 1.037415198 | 0.951767617 |
| ENSMUSG00000033233 | 1.106195041 | 0.185642391 |
| ENSMUSG00000033237 | 1.053256604 | 0.835590468 |
| ENSMUSG00000033249 | 1.04532682 | 0.698112512 |
| ENSMUSG00000033253 | 1.096393532 | 0.480690111 |
| ENSMUSG00000033255 | 1 | |
| ENSMUSG00000033256 | 1.194400734 | 0.069477846 |
| ENSMUSG00000033257 | -1.058322928 | 0.835645651 |
| ENSMUSG00000033268 | 1.580071334 | 0.013792637 |
| ENSMUSG00000033272 | -1.005758763 | 1 |
| ENSMUSG00000033276 | -1.140354317 | 0.080507499 |
| ENSMUSG00000033278 | 1.041007142 | 0.643085685 |
| ENSMUSG00000033282 | -1.062968538 | 0.715124238 |
| ENSMUSG00000033285 | 1.008262243 | 1 |
| ENSMUSG00000033287 | -1.02626918 | 0.87654539 |
| ENSMUSG00000033294 | 1.05925047 | 0.633129068 |
| ENSMUSG00000033295 | 1.202092623 | 0.131176568 |
| ENSMUSG00000033306 | 1.375896237 | 0.194068755 |
| ENSMUSG00000033307 | 1.131172096 | 0.436466116 |
| ENSMUSG00000033308 | -1.226546812 | 0.198211427 |
| ENSMUSG00000033313 | 1.050790343 | 0.737377137 |
| ENSMUSG00000033316 | -1 | 1 |
| ENSMUSG00000033318 | -1.014430581 | 0.984902895 |
| ENSMUSG00000033319 | 1.039872125 | 0.901216307 |
| ENSMUSG00000033323 | 1.23953299 | 0.025153728 |
| ENSMUSG00000033326 | 1.063339766 | 0.761802747 |
| ENSMUSG00000033327 | 1.018272578 | 0.903070447 |
| ENSMUSG00000033335 | -1.058763488 | 0.777123525 |
| ENSMUSG00000033342 | -1.063525418 | 0.698112512 |

| | | |
|---|---|---|
| ENSMUSG00000033343 | 1 | |
| ENSMUSG00000033350 | 1.164257997 | 0.029939521 |
| ENSMUSG00000033352 | -1.032532947 | 0.949812939 |
| ENSMUSG00000033355 | -1.410043648 | 0.085979471 |
| ENSMUSG00000033356 | 1.018890346 | 0.936702665 |
| ENSMUSG00000033361 | -1.819162795 | 0.001601152 |
| ENSMUSG00000033364 | -1.034226708 | 0.925537834 |
| ENSMUSG00000033365 | 1.087674093 | 0.608497229 |
| ENSMUSG00000033368 | 1.072886805 | 0.818593278 |
| ENSMUSG00000033373 | 1.133317609 | 0.14872159 |
| ENSMUSG00000033377 | -1.147838869 | 0.195607801 |
| ENSMUSG00000033379 | 1.184244795 | 0.206620396 |
| ENSMUSG00000033382 | 1.075360835 | 0.747886461 |
| ENSMUSG00000033383 | 1 | |
| ENSMUSG00000033386 | 1.019980393 | 0.942160514 |
| ENSMUSG00000033389 | -1.053009738 | 0.290098581 |
| ENSMUSG00000033392 | -1.002666663 | 1 |
| ENSMUSG00000033396 | 1.053587805 | 0.805587306 |
| ENSMUSG00000033400 | 1.02869148 | 0.956148535 |
| ENSMUSG00000033405 | -1.269005977 | 0.085349601 |
| ENSMUSG00000033409 | -1 | 1 |
| ENSMUSG00000033411 | 1.037980721 | 0.890893742 |
| ENSMUSG00000033416 | 1.035932919 | 0.863033776 |
| ENSMUSG00000033417 | 1.126895583 | 0.653153855 |
| ENSMUSG00000033419 | 1.115093963 | 0.704767557 |
| ENSMUSG00000033420 | 1.242255716 | 0.188113029 |
| ENSMUSG00000033423 | 1.101099665 | 0.600067315 |
| ENSMUSG00000033427 | -1.063103825 | 0.75692508 |
| ENSMUSG00000033429 | 1.052604784 | 0.70299173 |
| ENSMUSG00000033430 | -1.119515519 | 0.530797672 |
| ENSMUSG00000033434 | 1.095039003 | 0.608497229 |
| ENSMUSG00000033436 | -1.070107079 | 0.806949992 |
| ENSMUSG00000033439 | -1.045417729 | 0.746451077 |
| ENSMUSG00000033444 | 1.000292335 | 1 |
| ENSMUSG00000033446 | -1.195901324 | 0.150025983 |
| ENSMUSG00000033450 | 1.098878255 | 0.66521153 |
| ENSMUSG00000033453 | -1.078765755 | 0.851577835 |
| ENSMUSG00000033454 | 1.116396402 | 0.585646836 |
| ENSMUSG00000033458 | 1.023684143 | 0.76973744 |
| ENSMUSG00000033460 | 1.119118156 | 0.402514484 |
| ENSMUSG00000033467 | 1.271109066 | 0.480690111 |
| ENSMUSG00000033470 | 1 | |
| ENSMUSG00000033475 | 1.029796997 | 0.985988771 |
| ENSMUSG00000033478 | 1.087981411 | 0.608497229 |
| ENSMUSG00000033486 | 1.050256505 | 0.608497229 |
| ENSMUSG00000033487 | 1.014499305 | 1 |
| ENSMUSG00000033488 | 1.030318189 | 0.88492938 |
| ENSMUSG00000033491 | -1.08855149 | 0.698112512 |
| ENSMUSG00000033498 | 1.015486245 | 0.985522133 |
| ENSMUSG00000033499 | 1.025396968 | 0.972980766 |
| ENSMUSG00000033501 | 1.028323353 | 0.903464611 |
| ENSMUSG00000033502 | -1.026695582 | 0.837704961 |
| ENSMUSG00000033508 | 1.122462048 | 0.420161906 |
| ENSMUSG00000033510 | -1.347442089 | 0.239107704 |
| ENSMUSG00000033520 | -1.084645496 | 0.532369657 |
| ENSMUSG00000033526 | -1.02132091 | 0.924861084 |
| ENSMUSG00000033530 | -1.13439312 | 0.015070481 |
| ENSMUSG00000033533 | 1 | |
| ENSMUSG00000033538 | -1.291952818 | 0.068733821 |
| ENSMUSG00000033540 | 1.147287695 | 0.06812145 |
| ENSMUSG00000033542 | 1.068499943 | 0.762835214 |
| ENSMUSG00000033543 | -1.02875523 | 0.919932362 |
| ENSMUSG00000033544 | -1.132666855 | 0.375668528 |

| | | |
|---|---|---|
| ENSMUSG00000033545 | 1.105192515 | 0.49735617 |
| ENSMUSG00000033554 | -1.020756342 | 0.908044228 |
| ENSMUSG00000033557 | -1.028841649 | 0.951554277 |
| ENSMUSG00000033565 | 1.097392132 | 0.717035356 |
| ENSMUSG00000033569 | 1.009769373 | 1 |
| ENSMUSG00000033576 | 1.432973599 | 0.001572194 |
| ENSMUSG00000033577 | -1.217993291 | 0.257120957 |
| ENSMUSG00000033578 | -1.03962336 | 0.845175363 |
| ENSMUSG00000033579 | -1.151172433 | 0.539222653 |
| ENSMUSG00000033581 | 1.303474787 | 0.054812809 |
| ENSMUSG00000033585 | 1.221562964 | 0.006288332 |
| ENSMUSG00000033589 | -1.094898358 | 0.608497229 |
| ENSMUSG00000033590 | 1.772908896 | 0.000743415 |
| ENSMUSG00000033594 | -1.417480677 | 0.002261078 |
| ENSMUSG00000033595 | -1.219624708 | 0.567431692 |
| ENSMUSG00000033596 | -1.070680863 | 0.807247723 |
| ENSMUSG00000033597 | -1.133661878 | 0.506762906 |
| ENSMUSG00000033610 | 1.163084823 | 0.170218352 |
| ENSMUSG00000033615 | 1.015486245 | 0.985522133 |
| ENSMUSG00000033618 | -1.004738627 | 1 |
| ENSMUSG00000033623 | 1.123707861 | 0.635686545 |
| ENSMUSG00000033624 | -1.157305263 | 0.469213979 |
| ENSMUSG00000033628 | 1.014588311 | 0.982175268 |
| ENSMUSG00000033629 | -1.14898273 | 0.608497229 |
| ENSMUSG00000033632 | -1.084120476 | 0.608497229 |
| ENSMUSG00000033633 | -1.015511103 | 0.982777255 |
| ENSMUSG00000033634 | 1 | |
| ENSMUSG00000033644 | 1.026752671 | 0.948385094 |
| ENSMUSG00000033653 | 1.077781654 | 0.69305964 |
| ENSMUSG00000033658 | -1.072710285 | 0.653375474 |
| ENSMUSG00000033669 | -1.055732811 | 0.741713695 |
| ENSMUSG00000033671 | -1.02922027 | 0.933848047 |
| ENSMUSG00000033676 | -1.113515092 | 0.608497229 |
| ENSMUSG00000033684 | 1.068803875 | 0.746362525 |
| ENSMUSG00000033685 | 1.204578287 | 0.131428776 |
| ENSMUSG00000033688 | 1.065410325 | 0.764982341 |
| ENSMUSG00000033697 | -1.022915401 | 0.888246981 |
| ENSMUSG00000033701 | -1.040767671 | 0.7859015 |
| ENSMUSG00000033703 | -1.01132653 | 0.988424085 |
| ENSMUSG00000033705 | 1.134166212 | 0.46900134 |
| ENSMUSG00000033706 | 1.020750925 | 0.958403618 |
| ENSMUSG00000033707 | 1.049857924 | 0.66278123 |
| ENSMUSG00000033712 | -1.116132867 | 0.48348831 |
| ENSMUSG00000033713 | 1.185036411 | 0.086604839 |
| ENSMUSG00000033715 | 1.600545214 | 9.27689E-05 |
| ENSMUSG00000033717 | 1.506918678 | 0.075819483 |
| ENSMUSG00000033720 | 1.099389326 | 0.06870141 |
| ENSMUSG00000033721 | 1.05564078 | 0.608497229 |
| ENSMUSG00000033722 | 1.04703562 | 0.698112512 |
| ENSMUSG00000033726 | 1 | |
| ENSMUSG00000033728 | 1.092064503 | 0.120256583 |
| ENSMUSG00000033730 | 1.071111136 | 0.829324104 |
| ENSMUSG00000033731 | 1 | |
| ENSMUSG00000033732 | -1.001051298 | 1 |
| ENSMUSG00000033735 | 1.051253969 | 0.527384609 |
| ENSMUSG00000033737 | -1.133717313 | 0.608497229 |
| ENSMUSG00000033739 | -1.251015007 | 0.00218152 |
| ENSMUSG00000033740 | -1.071075483 | 0.775016379 |
| ENSMUSG00000033751 | -1.057033805 | 0.432383198 |
| ENSMUSG00000033752 | -1.124952311 | 0.142654234 |
| ENSMUSG00000033760 | 1.007131538 | 1 |
| ENSMUSG00000033762 | -1.073510806 | 0.520525659 |
| ENSMUSG00000033763 | 1.544114949 | 0.000743415 |

| | | |
|---|---|---|
| ENSMUSG00000033765 | 1 | |
| ENSMUSG00000033767 | 1.065196839 | 0.656900799 |
| ENSMUSG00000033768 | -1.409748423 | 0.121942907 |
| ENSMUSG00000033769 | 1.011826137 | 1 |
| ENSMUSG00000033770 | 1.059463094 | 0.608497229 |
| ENSMUSG00000033773 | 1.034115558 | 0.87756119 |
| ENSMUSG00000033774 | -1.058065387 | 0.80586932 |
| ENSMUSG00000033777 | -1.023769022 | 0.934760809 |
| ENSMUSG00000033781 | -1.036762639 | 0.818205891 |
| ENSMUSG00000033788 | 1.628808649 | 0.00105645 |
| ENSMUSG00000033790 | 1.023882536 | 0.935057992 |
| ENSMUSG00000033792 | -1.008870946 | 1 |
| ENSMUSG00000033793 | 1.080998378 | 0.732612801 |
| ENSMUSG00000033794 | 1 | |
| ENSMUSG00000033799 | -1.06877411 | 0.789636114 |
| ENSMUSG00000033805 | -1.078947844 | 0.861582022 |
| ENSMUSG00000033808 | 1.010454507 | 1 |
| ENSMUSG00000033809 | -1.016410352 | 0.959635626 |
| ENSMUSG00000033813 | -1.040509956 | 0.93818309 |
| ENSMUSG00000033819 | 1.066130219 | 0.698112512 |
| ENSMUSG00000033825 | 1 | |
| ENSMUSG00000033826 | -1.774355061 | 0.003402016 |
| ENSMUSG00000033831 | 1 | |
| ENSMUSG00000033834 | 1 | |
| ENSMUSG00000033837 | -1.081255487 | 0.772656514 |
| ENSMUSG00000033845 | -1.064149524 | 0.771440831 |
| ENSMUSG00000033847 | -1.03961979 | 0.773393951 |
| ENSMUSG00000033849 | -1.085864002 | 0.785584885 |
| ENSMUSG00000033850 | 1 | |
| ENSMUSG00000033852 | 1 | |
| ENSMUSG00000033854 | 1.024549129 | 0.897149458 |
| ENSMUSG00000033855 | 1.00540125 | 1 |
| ENSMUSG00000033857 | -1.127748371 | 0.349560553 |
| ENSMUSG00000033860 | -1 | 1 |
| ENSMUSG00000033862 | -1.065955728 | 0.681762447 |
| ENSMUSG00000033863 | 1.255006298 | 0.01536947 |
| ENSMUSG00000033871 | 1.331937626 | 0.261139089 |
| ENSMUSG00000033872 | 1.122462048 | 0.215991492 |
| ENSMUSG00000033880 | -1.912921969 | 0.000263878 |
| ENSMUSG00000033882 | -1.043306199 | 0.807247723 |
| ENSMUSG00000033883 | -1.048228297 | 0.790851299 |
| ENSMUSG00000033885 | -1.156189237 | 0.462046668 |
| ENSMUSG00000033898 | 1 | |
| ENSMUSG00000033900 | 1.055024966 | 0.698112512 |
| ENSMUSG00000033902 | -1.105971909 | 0.451532357 |
| ENSMUSG00000033904 | -1.042078595 | 0.866582371 |
| ENSMUSG00000033906 | 1.076161612 | 0.698112512 |
| ENSMUSG00000033909 | 1.054573334 | 0.783268112 |
| ENSMUSG00000033910 | 1.014545335 | 0.988902119 |
| ENSMUSG00000033916 | 1.034859771 | 0.893540818 |
| ENSMUSG00000033917 | -1.087339609 | 0.608497229 |
| ENSMUSG00000033918 | -1.04320995 | 0.842184576 |
| ENSMUSG00000033931 | -1.066594241 | 0.744030455 |
| ENSMUSG00000033933 | -1.088016601 | 0.660336669 |
| ENSMUSG00000033938 | 1.031734315 | 0.889846433 |
| ENSMUSG00000033940 | 1.044706747 | 0.894133318 |
| ENSMUSG00000033943 | -1.000057509 | 1 |
| ENSMUSG00000033948 | -1.505629032 | 0.003303995 |
| ENSMUSG00000033949 | -1.327128291 | 0.0161393 |
| ENSMUSG00000033952 | -1.018294232 | 0.992110111 |
| ENSMUSG00000033953 | -1.010743737 | 1 |
| ENSMUSG00000033955 | 1.03911842 | 0.840478821 |
| ENSMUSG00000033960 | 1.131537678 | 0.608497229 |

| | | |
|---|---|---|
| ENSMUSG00000033961 | 1.033772162 | 0.730008273 |
| ENSMUSG00000033963 | 1.04328066 | 0.906415966 |
| ENSMUSG00000033964 | -1.002055414 | 1 |
| ENSMUSG00000033965 | -1.122225267 | 0.152717869 |
| ENSMUSG00000033966 | 1.290681034 | 0.070657138 |
| ENSMUSG00000033967 | 1.084454733 | 0.623464439 |
| ENSMUSG00000033970 | 1.028100434 | 0.952049609 |
| ENSMUSG00000033972 | 1.061744425 | 0.651052209 |
| ENSMUSG00000033981 | -1.11061171 | 0.547016491 |
| ENSMUSG00000033983 | -1.096323847 | 0.516977167 |
| ENSMUSG00000033985 | -1.066251447 | 0.698112512 |
| ENSMUSG00000033987 | 1.096771181 | 0.608497229 |
| ENSMUSG00000033991 | -1.079444991 | 0.731295483 |
| ENSMUSG00000033998 | -1.198658038 | 0.375794285 |
| ENSMUSG00000034000 | 1.059463094 | 0.608497229 |
| ENSMUSG00000034006 | 1.162519372 | 0.159066803 |
| ENSMUSG00000034007 | 1.004638453 | 1 |
| ENSMUSG00000034009 | 1.089470641 | 0.657060635 |
| ENSMUSG00000034021 | 1.065890948 | 0.810597167 |
| ENSMUSG00000034022 | 1.034653567 | 0.896201698 |
| ENSMUSG00000034023 | -1.096683244 | 0.633673099 |
| ENSMUSG00000034024 | -1.038299857 | 0.961565399 |
| ENSMUSG00000034028 | 1 | |
| ENSMUSG00000034031 | 1 | |
| ENSMUSG00000034032 | -1.078046693 | 0.541962118 |
| ENSMUSG00000034035 | -1.13923135 | 0.732773395 |
| ENSMUSG00000034037 | -1.083415624 | 0.698112512 |
| ENSMUSG00000034039 | 1 | |
| ENSMUSG00000034040 | 1.290040212 | 0.019122348 |
| ENSMUSG00000034041 | -1.15462599 | 0.188125313 |
| ENSMUSG00000034042 | 1.021225347 | 0.955522519 |
| ENSMUSG00000034055 | 1.013290812 | 0.99457094 |
| ENSMUSG00000034057 | -1.1199802 | 0.320703325 |
| ENSMUSG00000034059 | 1.167933692 | 0.095579196 |
| ENSMUSG00000034063 | 1 | |
| ENSMUSG00000034064 | 1.079438198 | 0.698112512 |
| ENSMUSG00000034066 | -1.014155424 | 0.971405508 |
| ENSMUSG00000034071 | 1.09599856 | 0.238702368 |
| ENSMUSG00000034075 | -1.12892956 | 0.608497229 |
| ENSMUSG00000034083 | -1.061751917 | 0.70299173 |
| ENSMUSG00000034087 | 1 | |
| ENSMUSG00000034088 | 1.001733926 | 1 |
| ENSMUSG00000034098 | 1.059463094 | 0.608497229 |
| ENSMUSG00000034101 | 1.032369244 | 0.922261327 |
| ENSMUSG00000034105 | -1.096092896 | 0.17124452 |
| ENSMUSG00000034107 | 1.024693402 | 0.933031586 |
| ENSMUSG00000034108 | -1.026075582 | 0.883306173 |
| ENSMUSG00000034109 | 1.050931312 | 0.841678369 |
| ENSMUSG00000034110 | 1.003769316 | 1 |
| ENSMUSG00000034111 | 1.01384885 | 0.994172456 |
| ENSMUSG00000034112 | -1.043306199 | 0.803260194 |
| ENSMUSG00000034115 | -1.053549062 | 0.698112512 |
| ENSMUSG00000034116 | 1 | |
| ENSMUSG00000034117 | 1.086056362 | 0.749531064 |
| ENSMUSG00000034118 | 1.086419709 | 0.645756877 |
| ENSMUSG00000034120 | 1.022788714 | 0.979002724 |
| ENSMUSG00000034121 | -1.13088158 | 0.08283904 |
| ENSMUSG00000034122 | 1 | |
| ENSMUSG00000034126 | 1.092815131 | 0.60753527 |
| ENSMUSG00000034127 | 1.028323353 | 0.90607899 |
| ENSMUSG00000034135 | -1.051358403 | 0.811257973 |
| ENSMUSG00000034139 | 1 | |
| ENSMUSG00000034145 | -1.043331739 | 0.899903545 |

| | | |
|---|---|---|
| ENSMUSG00000034151 | 1 | |
| ENSMUSG00000034152 | -1.12965179 | 0.608497229 |
| ENSMUSG00000034154 | -1.093315103 | 0.653960635 |
| ENSMUSG00000034156 | 2.262140774 | 0.000500173 |
| ENSMUSG00000034157 | 1.103686684 | 0.608497229 |
| ENSMUSG00000034158 | 1.202739114 | 0.413417973 |
| ENSMUSG00000034159 | -1.111880267 | 0.608497229 |
| ENSMUSG00000034160 | 1.025270347 | 0.975677002 |
| ENSMUSG00000034161 | 1.125739245 | 0.181012503 |
| ENSMUSG00000034163 | 1.031215134 | 0.926666063 |
| ENSMUSG00000034164 | -1.112281452 | 0.671280603 |
| ENSMUSG00000034165 | -1.037467843 | 0.803072249 |
| ENSMUSG00000034168 | 1.18119006 | 0.076089694 |
| ENSMUSG00000034171 | -1.365315458 | 0.004426679 |
| ENSMUSG00000034173 | 1.078236493 | 0.479515459 |
| ENSMUSG00000034175 | 1.17724044 | 0.14921102 |
| ENSMUSG00000034177 | 1.969460219 | 0.00034694 |
| ENSMUSG00000034185 | 1 | |
| ENSMUSG00000034187 | -1.119428364 | 0.63149515 |
| ENSMUSG00000034189 | 1.038982167 | 0.765639272 |
| ENSMUSG00000034190 | -1.126843911 | 0.493961265 |
| ENSMUSG00000034192 | -1.072667582 | 0.715712804 |
| ENSMUSG00000034194 | -1.046814604 | 0.669959104 |
| ENSMUSG00000034201 | 1.071128331 | 0.608497229 |
| ENSMUSG00000034203 | -1.036914406 | 0.807933619 |
| ENSMUSG00000034205 | -1.137939245 | 0.696603049 |
| ENSMUSG00000034206 | -1.095489186 | 0.638218577 |
| ENSMUSG00000034209 | -1.037439001 | 0.912782976 |
| ENSMUSG00000034210 | -1.025825711 | 0.959974958 |
| ENSMUSG00000034211 | 1.064092422 | 0.709124035 |
| ENSMUSG00000034212 | -1.027799907 | 0.947248675 |
| ENSMUSG00000034216 | 1.018207318 | 0.945694084 |
| ENSMUSG00000034218 | -1.003553507 | 1 |
| ENSMUSG00000034219 | 1.177765487 | 0.577958514 |
| ENSMUSG00000034220 | 1.064748324 | 0.608497229 |
| ENSMUSG00000034224 | 1.069068979 | 0.698112512 |
| ENSMUSG00000034226 | -1.233820451 | 0.067146994 |
| ENSMUSG00000034227 | 1.201362935 | 0.608497229 |
| ENSMUSG00000034235 | -1.102172148 | 0.608497229 |
| ENSMUSG00000034239 | -1 | 1 |
| ENSMUSG00000034243 | 1.087777104 | 0.715092973 |
| ENSMUSG00000034245 | 1.190068139 | 0.066068145 |
| ENSMUSG00000034247 | 1.017463967 | 0.920717469 |
| ENSMUSG00000034248 | -1.046213983 | 0.793396162 |
| ENSMUSG00000034252 | 1.028759592 | 0.944833562 |
| ENSMUSG00000034254 | 1.014166565 | 0.974451204 |
| ENSMUSG00000034255 | 1.355944021 | 0.07510254 |
| ENSMUSG00000034258 | 1.128473984 | 0.108139585 |
| ENSMUSG00000034259 | 1.016809075 | 0.928552664 |
| ENSMUSG00000034263 | -1.098202485 | 0.608497229 |
| ENSMUSG00000034265 | -1.411183286 | 0.007628795 |
| ENSMUSG00000034266 | 1.005613439 | 1 |
| ENSMUSG00000034269 | 1.102849413 | 0.707580011 |
| ENSMUSG00000034271 | 1.216640367 | 0.022073797 |
| ENSMUSG00000034274 | 1.006980474 | 1 |
| ENSMUSG00000034275 | 1.045303362 | 0.844040246 |
| ENSMUSG00000034278 | -1.249339685 | 0.01222575 |
| ENSMUSG00000034282 | 1.012174333 | 1 |
| ENSMUSG00000034285 | -1.069028128 | 0.608497229 |
| ENSMUSG00000034290 | 1.057496472 | 0.840162455 |
| ENSMUSG00000034292 | 1.002817956 | 1 |
| ENSMUSG00000034295 | -1.07560991 | 0.66102156 |
| ENSMUSG00000034297 | 1.03504925 | 0.946925656 |

| | | |
|---|---|---|
| ENSMUSG00000034300 | -1.012346605 | 1 |
| ENSMUSG00000034303 | -1.081149209 | 0.327642404 |
| ENSMUSG00000034308 | 1.095872691 | 0.614079363 |
| ENSMUSG00000034310 | -1 | 1 |
| ENSMUSG00000034311 | -1.056997798 | 0.844771174 |
| ENSMUSG00000034312 | 1.092661406 | 0.608497229 |
| ENSMUSG00000034317 | 1.009508411 | 1 |
| ENSMUSG00000034320 | -1.085436748 | 0.608497229 |
| ENSMUSG00000034321 | 1.058491051 | 0.722165151 |
| ENSMUSG00000034324 | 1 | |
| ENSMUSG00000034327 | -1.271076246 | 0.175332035 |
| ENSMUSG00000034329 | -1.011951262 | 1 |
| ENSMUSG00000034330 | 1.107302284 | 0.260153539 |
| ENSMUSG00000034333 | 1.02492925 | 0.948628145 |
| ENSMUSG00000034334 | -1.099994168 | 0.608497229 |
| ENSMUSG00000034336 | -1.861105688 | 0.001032697 |
| ENSMUSG00000034339 | 1 | |
| ENSMUSG00000034341 | 1.035461143 | 0.906470281 |
| ENSMUSG00000034342 | -1.027189487 | 0.96035552 |
| ENSMUSG00000034343 | 1.007215753 | 1 |
| ENSMUSG00000034345 | -1.001929781 | 1 |
| ENSMUSG00000034349 | -1.045848194 | 0.933511041 |
| ENSMUSG00000034353 | -1.056662203 | 0.953521521 |
| ENSMUSG00000034354 | 1.067338735 | 0.748571751 |
| ENSMUSG00000034359 | 1 | |
| ENSMUSG00000034361 | 1.220908985 | 0.15502163 |
| ENSMUSG00000034362 | 1.122462048 | 0.420161906 |
| ENSMUSG00000034371 | 1.14210685 | 0.074387647 |
| ENSMUSG00000034377 | 1.08867492 | 0.768470445 |
| ENSMUSG00000034379 | 1.048163139 | 0.670305632 |
| ENSMUSG00000034382 | -1.241578614 | 0.036510341 |
| ENSMUSG00000034384 | -1.069502966 | 0.744868995 |
| ENSMUSG00000034387 | -1 | 1 |
| ENSMUSG00000034390 | 1.044748996 | 0.797767171 |
| ENSMUSG00000034391 | -1.009594728 | 1 |
| ENSMUSG00000034394 | 1.180741075 | 0.010217196 |
| ENSMUSG00000034401 | 1.038866488 | 0.874375366 |
| ENSMUSG00000034402 | 1.346809975 | 0.036626025 |
| ENSMUSG00000034403 | 1.036339634 | 0.825162695 |
| ENSMUSG00000034412 | 1.042552389 | 0.756303914 |
| ENSMUSG00000034413 | 1.121498546 | 0.306730283 |
| ENSMUSG00000034416 | 1 | |
| ENSMUSG00000034422 | -2.119324148 | 0.000666524 |
| ENSMUSG00000034424 | -1.058503931 | 0.806949992 |
| ENSMUSG00000034427 | -1.136728935 | 0.634019547 |
| ENSMUSG00000034429 | 1.042123298 | 0.698112512 |
| ENSMUSG00000034430 | 1.112603024 | 0.477370624 |
| ENSMUSG00000034432 | 1.036985346 | 0.925051265 |
| ENSMUSG00000034435 | -1.539532448 | 0.029735092 |
| ENSMUSG00000034437 | 1.076322097 | 0.814608539 |
| ENSMUSG00000034438 | 1 | |
| ENSMUSG00000034442 | 1.046074323 | 0.720798524 |
| ENSMUSG00000034445 | 1.150069399 | 0.124255505 |
| ENSMUSG00000034449 | 1.033989733 | 0.803260194 |
| ENSMUSG00000034450 | -1.03962336 | 0.698112512 |
| ENSMUSG00000034452 | 1 | |
| ENSMUSG00000034453 | -1.020829652 | 0.975148944 |
| ENSMUSG00000034456 | 1.059463094 | 0.608497229 |
| ENSMUSG00000034457 | 1.893564765 | 0.002497395 |
| ENSMUSG00000034459 | -1.390401997 | 0.146384185 |
| ENSMUSG00000034460 | 1.125534984 | 0.329218885 |
| ENSMUSG00000034462 | 1.149558913 | 0.562995613 |
| ENSMUSG00000034463 | 1.026873056 | 0.917717739 |

| | | |
|---|---|---|
| ENSMUSG00000034467 | -1.03962336 | 0.814089245 |
| ENSMUSG00000034471 | 1.085359306 | 0.605682526 |
| ENSMUSG00000034472 | 1.069068979 | 0.767473873 |
| ENSMUSG00000034473 | 1.016693432 | 0.968937442 |
| ENSMUSG00000034480 | -1.044682651 | 0.809010178 |
| ENSMUSG00000034482 | 1 | |
| ENSMUSG00000034484 | 1.11226283 | 0.608497229 |
| ENSMUSG00000034485 | -1.116133081 | 0.486930276 |
| ENSMUSG00000034486 | -1.331986746 | 0.039886178 |
| ENSMUSG00000034487 | 1.065047345 | 0.717686641 |
| ENSMUSG00000034488 | 1.515398696 | 0.000215953 |
| ENSMUSG00000034493 | -1.089497311 | 0.585199025 |
| ENSMUSG00000034501 | 1.105058468 | 0.565305007 |
| ENSMUSG00000034509 | 1.003206976 | 1 |
| ENSMUSG00000034518 | 1.005479507 | 1 |
| ENSMUSG00000034520 | 1.012086982 | 1 |
| ENSMUSG00000034522 | 1.081000555 | 0.312960909 |
| ENSMUSG00000034525 | 1.053403543 | 0.821942765 |
| ENSMUSG00000034528 | 1.112249711 | 0.661085946 |
| ENSMUSG00000034532 | -1.323634713 | 0.135556873 |
| ENSMUSG00000034533 | -1.053549062 | 0.698112512 |
| ENSMUSG00000034538 | 1.025392988 | 0.890258111 |
| ENSMUSG00000034543 | 1.023757886 | 0.961487493 |
| ENSMUSG00000034544 | -1.029735419 | 0.927576194 |
| ENSMUSG00000034551 | -1.320231717 | 0.022941523 |
| ENSMUSG00000034552 | 1 | |
| ENSMUSG00000034555 | 1 | |
| ENSMUSG00000034557 | -1.08285708 | 0.698112512 |
| ENSMUSG00000034560 | 1.061153962 | 0.804920765 |
| ENSMUSG00000034563 | 1.059463094 | 0.820462971 |
| ENSMUSG00000034566 | -1.018778198 | 0.99595788 |
| ENSMUSG00000034570 | -1.01579034 | 0.964333491 |
| ENSMUSG00000034573 | 1.243691704 | 0.160779489 |
| ENSMUSG00000034574 | 1.059302406 | 0.768634972 |
| ENSMUSG00000034575 | 1.028494404 | 0.893303059 |
| ENSMUSG00000034579 | 1.343219093 | 0.314058579 |
| ENSMUSG00000034583 | 1 | |
| ENSMUSG00000034584 | 1.130558969 | 0.572988678 |
| ENSMUSG00000034586 | 1.017554503 | 0.924629556 |
| ENSMUSG00000034587 | 1.02197417 | 0.852371141 |
| ENSMUSG00000034591 | -1.059400472 | 0.755105069 |
| ENSMUSG00000034593 | 1.072656917 | 0.817252008 |
| ENSMUSG00000034595 | 1.151020737 | 0.317557615 |
| ENSMUSG00000034601 | 1.108039718 | 0.48973629 |
| ENSMUSG00000034602 | 1.032079714 | 0.933999769 |
| ENSMUSG00000034607 | 1 | |
| ENSMUSG00000034610 | 1.082246822 | 0.698112512 |
| ENSMUSG00000034612 | 1.050701193 | 0.811496247 |
| ENSMUSG00000034613 | 1.206056821 | 0.03311356 |
| ENSMUSG00000034614 | 1.297395067 | 0.009957878 |
| ENSMUSG00000034616 | 1.206439489 | 0.03675998 |
| ENSMUSG00000034617 | 1.016991293 | 0.970020309 |
| ENSMUSG00000034620 | 1.038260484 | 0.887572464 |
| ENSMUSG00000034621 | -1.002000359 | 1 |
| ENSMUSG00000034623 | 1 | |
| ENSMUSG00000034634 | 1 | |
| ENSMUSG00000034636 | -1.039647174 | 0.922280769 |
| ENSMUSG00000034639 | -1.005769952 | 1 |
| ENSMUSG00000034640 | -1.077268224 | 0.704767557 |
| ENSMUSG00000034641 | 1.177765487 | 0.458872271 |
| ENSMUSG00000034645 | -1.051732208 | 0.854378996 |
| ENSMUSG00000034647 | 1.167538938 | 0.372430251 |
| ENSMUSG00000034648 | 1 | |

| | | |
|---|---|---|
| ENSMUSG00000034652 | 1.313616685 | 0.349340409 |
| ENSMUSG00000034653 | -1.015846317 | 0.979002724 |
| ENSMUSG00000034656 | 1.364686585 | 0.029355734 |
| ENSMUSG00000034659 | 1.073911398 | 0.698112512 |
| ENSMUSG00000034660 | 1.142908482 | 0.608497229 |
| ENSMUSG00000034663 | 1.176109759 | 0.369234602 |
| ENSMUSG00000034664 | 1.267292301 | 0.003171734 |
| ENSMUSG00000034667 | 1.078553264 | 0.755105483 |
| ENSMUSG00000034673 | -1.031264775 | 0.793885968 |
| ENSMUSG00000034674 | -1.141964705 | 0.505050548 |
| ENSMUSG00000034675 | -1.024316053 | 0.930216294 |
| ENSMUSG00000034677 | 1 | |
| ENSMUSG00000034681 | 1.03931682 | 0.863410724 |
| ENSMUSG00000034683 | -1.057338222 | 0.87756119 |
| ENSMUSG00000034684 | 1.270690934 | 0.017843579 |
| ENSMUSG00000034685 | 1.166048654 | 0.447353132 |
| ENSMUSG00000034686 | -1.025436797 | 1 |
| ENSMUSG00000034687 | -1.111345709 | 0.554334144 |
| ENSMUSG00000034689 | 1 | |
| ENSMUSG00000034690 | 1.014545335 | 0.979002724 |
| ENSMUSG00000034701 | -1 | 1 |
| ENSMUSG00000034706 | 1.081671374 | 0.637400576 |
| ENSMUSG00000034707 | 1.130586182 | 0.643361947 |
| ENSMUSG00000034708 | 1.363285967 | 0.087671702 |
| ENSMUSG00000034709 | 1.033356019 | 0.877083324 |
| ENSMUSG00000034714 | -1.011755345 | 0.978664718 |
| ENSMUSG00000034723 | 1.00028382 | 1 |
| ENSMUSG00000034724 | 1.085795345 | 0.724021681 |
| ENSMUSG00000034729 | -1.014417754 | 0.978211686 |
| ENSMUSG00000034730 | 1 | |
| ENSMUSG00000034731 | 1.171853892 | 0.094546471 |
| ENSMUSG00000034732 | -1 | 1 |
| ENSMUSG00000034738 | -1.095213954 | 0.698112512 |
| ENSMUSG00000034739 | 1.104427563 | 0.608497229 |
| ENSMUSG00000034744 | 1.252153405 | 0.036681922 |
| ENSMUSG00000034748 | 1.092290114 | 0.412661563 |
| ENSMUSG00000034751 | 1.191888502 | 0.216822321 |
| ENSMUSG00000034755 | 1.122933516 | 0.608497229 |
| ENSMUSG00000034757 | 1.031835084 | 0.857392806 |
| ENSMUSG00000034758 | 1.434021416 | 0.002031442 |
| ENSMUSG00000034761 | 1.144395762 | 0.59127359 |
| ENSMUSG00000034762 | 1.124001143 | 0.698112512 |
| ENSMUSG00000034764 | 1.433584912 | 0.149442298 |
| ENSMUSG00000034765 | 1.220635418 | 0.029382872 |
| ENSMUSG00000034768 | 1.199989356 | 0.200733632 |
| ENSMUSG00000034771 | 1.12986732 | 0.338256051 |
| ENSMUSG00000034773 | 1.044497211 | 0.698112512 |
| ENSMUSG00000034774 | 1 | |
| ENSMUSG00000034777 | -1.48509547 | 0.06189968 |
| ENSMUSG00000034780 | 1.059191146 | 0.744393068 |
| ENSMUSG00000034781 | -1.04481995 | 0.807247723 |
| ENSMUSG00000034783 | 1 | |
| ENSMUSG00000034785 | 1 | |
| ENSMUSG00000034786 | -1.168354076 | 0.481742107 |
| ENSMUSG00000034789 | 1.021254665 | 0.937719236 |
| ENSMUSG00000034792 | -1.044273782 | 0.699340382 |
| ENSMUSG00000034793 | 1.289248844 | 0.010383572 |
| ENSMUSG00000034795 | -1.004428454 | 1 |
| ENSMUSG00000034796 | -1.085651418 | 0.580293942 |
| ENSMUSG00000034799 | -1.115150214 | 0.217297788 |
| ENSMUSG00000034800 | -1.102234289 | 0.35680391 |
| ENSMUSG00000034801 | 1.103121983 | 0.608497229 |
| ENSMUSG00000034807 | 1.006957434 | 1 |

| | | |
|---|---|---|
| ENSMUSG00000034810 | 1.321997061 | 0.007615822 |
| ENSMUSG00000034813 | -1.247828232 | 0.151117627 |
| ENSMUSG00000034818 | -1.222264834 | 0.006382739 |
| ENSMUSG00000034820 | -1.038207503 | 0.924629556 |
| ENSMUSG00000034825 | -1.36323066 | 0.010391539 |
| ENSMUSG00000034826 | 1.087923756 | 0.698112512 |
| ENSMUSG00000034829 | -1.387862312 | 0.05649953 |
| ENSMUSG00000034832 | 1.029486743 | 0.891297795 |
| ENSMUSG00000034833 | 1 | |
| ENSMUSG00000034837 | 1.19790055 | 0.532369657 |
| ENSMUSG00000034839 | 1.167004257 | 0.015217793 |
| ENSMUSG00000034842 | -1.057338222 | 0.874017398 |
| ENSMUSG00000034845 | 1.027081781 | 0.958181659 |
| ENSMUSG00000034848 | 1.115410062 | 0.608497229 |
| ENSMUSG00000034850 | -1.067400248 | 0.75013293 |
| ENSMUSG00000034853 | 1.388846938 | 0.035175931 |
| ENSMUSG00000034854 | -1.147263646 | 0.205926548 |
| ENSMUSG00000034855 | -1.941668271 | 0.000367459 |
| ENSMUSG00000034858 | -1.072713454 | 0.601792863 |
| ENSMUSG00000034863 | 1.005829244 | 1 |
| ENSMUSG00000034867 | 1.102734818 | 0.581446386 |
| ENSMUSG00000034868 | -1.148793277 | 0.51470764 |
| ENSMUSG00000034871 | -1 | 1 |
| ENSMUSG00000034872 | -1.044273782 | 0.819994892 |
| ENSMUSG00000034875 | -1.03617669 | 0.744868995 |
| ENSMUSG00000034880 | -1.009127262 | 1 |
| ENSMUSG00000034881 | 1.054079687 | 0.925679199 |
| ENSMUSG00000034883 | 1.033124846 | 0.699340382 |
| ENSMUSG00000034889 | 1.146908546 | 0.250501816 |
| ENSMUSG00000034891 | -1.08081673 | 0.698112512 |
| ENSMUSG00000034892 | 1.192064811 | 0.169323119 |
| ENSMUSG00000034893 | -1.075236135 | 0.702070725 |
| ENSMUSG00000034898 | 1.059463094 | 0.698112512 |
| ENSMUSG00000034902 | -1.016326316 | 0.97045957 |
| ENSMUSG00000034903 | 1.147857228 | 0.476305515 |
| ENSMUSG00000034906 | -1.093893927 | 0.698112512 |
| ENSMUSG00000034908 | -1.006957959 | 1 |
| ENSMUSG00000034910 | -1.426547224 | 0.007522209 |
| ENSMUSG00000034911 | 1.014133649 | 1 |
| ENSMUSG00000034912 | 1.054898462 | 0.877680691 |
| ENSMUSG00000034913 | 1 | |
| ENSMUSG00000034917 | -1.541385044 | 0.131758166 |
| ENSMUSG00000034918 | -1.044273782 | 0.790851299 |
| ENSMUSG00000034919 | 1 | |
| ENSMUSG00000034923 | 1 | |
| ENSMUSG00000034926 | -1.066041913 | 0.698112512 |
| ENSMUSG00000034928 | 1.054899666 | 0.68459952 |
| ENSMUSG00000034930 | 1.053631095 | 0.690669961 |
| ENSMUSG00000034931 | 1.030986156 | 0.938982669 |
| ENSMUSG00000034932 | -1.054058378 | 0.698112512 |
| ENSMUSG00000034936 | 1.244633146 | 0.019038104 |
| ENSMUSG00000034940 | -1.005526326 | 1 |
| ENSMUSG00000034947 | -1.222821271 | 0.168431691 |
| ENSMUSG00000034949 | -1.042485693 | 0.760912081 |
| ENSMUSG00000034951 | -1.002349283 | 1 |
| ENSMUSG00000034957 | -1.154670789 | 0.608497229 |
| ENSMUSG00000034958 | -1 | 1 |
| ENSMUSG00000034959 | -1.033622861 | 0.748251831 |
| ENSMUSG00000034968 | 1.015486245 | 0.985522133 |
| ENSMUSG00000034973 | -1.059596512 | 0.773508225 |
| ENSMUSG00000034974 | 1.185906235 | 0.131758166 |
| ENSMUSG00000034981 | 1.468684016 | 0.000822888 |
| ENSMUSG00000034987 | 1 | |

| | | |
|---|---|---|
| ENSMUSG00000034990 | -1.03962336 | 0.803072249 |
| ENSMUSG00000034993 | 1.192405474 | 0.356170373 |
| ENSMUSG00000034994 | 1.109835851 | 0.698112512 |
| ENSMUSG00000034997 | 2.02328898 | 9.43242E-05 |
| ENSMUSG00000034998 | 1.048895821 | 0.807857063 |
| ENSMUSG00000035000 | -1.03962336 | 0.803072249 |
| ENSMUSG00000035004 | -1.010745438 | 1 |
| ENSMUSG00000035007 | -1.0458224 | 0.698112512 |
| ENSMUSG00000035011 | 1.03298193 | 0.781563957 |
| ENSMUSG00000035020 | -1.006606532 | 1 |
| ENSMUSG00000035021 | 1.059824591 | 0.827883256 |
| ENSMUSG00000035024 | -1.051210771 | 0.84200912 |
| ENSMUSG00000035027 | 1.116927279 | 0.506663701 |
| ENSMUSG00000035031 | 1 | |
| ENSMUSG00000035032 | 1.102024305 | 0.70299173 |
| ENSMUSG00000035033 | 1.217443603 | 0.3305975 |
| ENSMUSG00000035041 | -1.130226573 | 0.698112512 |
| ENSMUSG00000035042 | -1.360875531 | 0.006199711 |
| ENSMUSG00000035045 | 1.045461734 | 0.725858943 |
| ENSMUSG00000035047 | -1.081572005 | 0.351532819 |
| ENSMUSG00000035048 | -1.07500956 | 0.608497229 |
| ENSMUSG00000035049 | 1.058498585 | 0.698112512 |
| ENSMUSG00000035051 | 1.030306397 | 0.92584165 |
| ENSMUSG00000035057 | 1.019083579 | 0.949812939 |
| ENSMUSG00000035062 | 1.074202139 | 0.608497229 |
| ENSMUSG00000035064 | 1.24788395 | 0.035620703 |
| ENSMUSG00000035067 | -1.024993443 | 0.844675441 |
| ENSMUSG00000035069 | -1.060756594 | 0.608497229 |
| ENSMUSG00000035078 | -1.138418211 | 0.507659484 |
| ENSMUSG00000035085 | 1.11511412 | 0.777123525 |
| ENSMUSG00000035086 | -1.037258155 | 0.908543402 |
| ENSMUSG00000035093 | 1.051451964 | 0.80046457 |
| ENSMUSG00000035095 | -1.024303005 | 0.90529721 |
| ENSMUSG00000035104 | 1.163446329 | 0.060326805 |
| ENSMUSG00000035105 | 1.133590098 | 0.31625175 |
| ENSMUSG00000035107 | -1.044324061 | 0.909073664 |
| ENSMUSG00000035109 | 1.322192871 | 0.004223952 |
| ENSMUSG00000035112 | 1.05810344 | 0.595296772 |
| ENSMUSG00000035121 | -1.098247218 | 0.727718584 |
| ENSMUSG00000035125 | 1.02880617 | 0.877881982 |
| ENSMUSG00000035126 | 1.087736722 | 0.560722159 |
| ENSMUSG00000035129 | -1.133231767 | 0.608497229 |
| ENSMUSG00000035131 | -1.03962336 | 0.8950853 |
| ENSMUSG00000035133 | 1.10942904 | 0.698112512 |
| ENSMUSG00000035139 | -1.055069946 | 0.698112512 |
| ENSMUSG00000035142 | 1.076854329 | 0.532369657 |
| ENSMUSG00000035148 | 1 | |
| ENSMUSG00000035150 | 1.013700721 | 1 |
| ENSMUSG00000035151 | 1.004978781 | 1 |
| ENSMUSG00000035152 | -1.037669478 | 0.946232149 |
| ENSMUSG00000035158 | 1.052556487 | 0.64090177 |
| ENSMUSG00000035161 | -1.021324429 | 0.949558509 |
| ENSMUSG00000035164 | -1.01839757 | 0.945947634 |
| ENSMUSG00000035165 | -1.209492167 | 0.340254744 |
| ENSMUSG00000035168 | 1.207834236 | 0.230667008 |
| ENSMUSG00000035171 | -1.013861605 | 1 |
| ENSMUSG00000035172 | 1.274194172 | 0.067428441 |
| ENSMUSG00000035173 | -1.023301914 | 0.971164403 |
| ENSMUSG00000035177 | -1.085078338 | 0.749613696 |
| ENSMUSG00000035179 | -1.027702938 | 0.990700261 |
| ENSMUSG00000035181 | 1.173879212 | 0.452871976 |
| ENSMUSG00000035183 | -1.28247251 | 0.032953839 |
| ENSMUSG00000035184 | 1.509794206 | 0.013748391 |

| | | |
|---|---|---|
| ENSMUSG00000035186 | -1.216163975 | 0.383314592 |
| ENSMUSG00000035187 | -1.95831199 | 9.66874E-05 |
| ENSMUSG00000035189 | 1.089287663 | 0.70615525 |
| ENSMUSG00000035191 | 1.059463094 | 0.608497229 |
| ENSMUSG00000035198 | 1.028979185 | 0.913700746 |
| ENSMUSG00000035199 | -1.048801907 | 0.884088177 |
| ENSMUSG00000035200 | -1.060102335 | 0.860712821 |
| ENSMUSG00000035201 | 1.095872691 | 0.657060635 |
| ENSMUSG00000035202 | 1.072737263 | 0.79291651 |
| ENSMUSG00000035203 | 1.004444872 | 1 |
| ENSMUSG00000035206 | -1.053784162 | 0.698112512 |
| ENSMUSG00000035208 | -1.17077323 | 0.548371979 |
| ENSMUSG00000035211 | -1.077839627 | 0.846939416 |
| ENSMUSG00000035212 | 1.104014676 | 0.608497229 |
| ENSMUSG00000035215 | -1.114749405 | 0.414441821 |
| ENSMUSG00000035226 | -1.063525418 | 0.751979636 |
| ENSMUSG00000035227 | -1.067805358 | 0.769745556 |
| ENSMUSG00000035228 | 1.115127629 | 0.441129618 |
| ENSMUSG00000035232 | 1.03099657 | 0.939641924 |
| ENSMUSG00000035234 | 1.064421526 | 0.670176133 |
| ENSMUSG00000035235 | -1.113700048 | 0.364237673 |
| ENSMUSG00000035236 | -1.005635963 | 1 |
| ENSMUSG00000035237 | 1.008906265 | 0.964239113 |
| ENSMUSG00000035238 | 1 | |
| ENSMUSG00000035239 | 1.006087887 | 1 |
| ENSMUSG00000035242 | -1.005819666 | 1 |
| ENSMUSG00000035245 | 1.005171643 | 1 |
| ENSMUSG00000035246 | 1.049620936 | 0.773508225 |
| ENSMUSG00000035247 | 1.077590206 | 0.803641195 |
| ENSMUSG00000035248 | -1.041742823 | 0.879837259 |
| ENSMUSG00000035258 | 1.227288254 | 0.106805757 |
| ENSMUSG00000035262 | 1.171288978 | 0.59120845 |
| ENSMUSG00000035266 | 1.081670309 | 0.661414046 |
| ENSMUSG00000035268 | -1.028105074 | 0.897070422 |
| ENSMUSG00000035270 | -1.030130857 | 0.949218699 |
| ENSMUSG00000035273 | -2.532678379 | 0.000397241 |
| ENSMUSG00000035274 | 1.064671692 | 0.725189744 |
| ENSMUSG00000035275 | -1.501455677 | 0.040849575 |
| ENSMUSG00000035277 | -1 | 1 |
| ENSMUSG00000035278 | 1.016597165 | 0.915612762 |
| ENSMUSG00000035279 | 1.621413583 | 0.001814637 |
| ENSMUSG00000035283 | -1.153548054 | 0.608497229 |
| ENSMUSG00000035284 | 1.027835511 | 0.951698249 |
| ENSMUSG00000035285 | 1.384470083 | 0.102003417 |
| ENSMUSG00000035293 | 1.078902727 | 0.761305587 |
| ENSMUSG00000035295 | 1.057165869 | 0.881421285 |
| ENSMUSG00000035296 | 1 | |
| ENSMUSG00000035297 | 1.000947944 | 1 |
| ENSMUSG00000035298 | 1.05943716 | 0.829632807 |
| ENSMUSG00000035299 | 1.113727863 | 0.10939414 |
| ENSMUSG00000035305 | -1.241345018 | 0.023920434 |
| ENSMUSG00000035310 | 1.058784884 | 0.792578392 |
| ENSMUSG00000035311 | 1.0405514 | 0.900667245 |
| ENSMUSG00000035314 | -1.005126227 | 1 |
| ENSMUSG00000035325 | 1.224905652 | 0.404271378 |
| ENSMUSG00000035329 | -1.001510301 | 1 |
| ENSMUSG00000035337 | -1.028348525 | 0.936995747 |
| ENSMUSG00000035342 | 1.251293286 | 0.215801263 |
| ENSMUSG00000035349 | 1.272127852 | 0.364216566 |
| ENSMUSG00000035351 | 1.008943805 | 1 |
| ENSMUSG00000035352 | 1 | |
| ENSMUSG00000035354 | 1.062047523 | 0.698112512 |
| ENSMUSG00000035355 | -1.047332209 | 0.915274791 |

| | | |
|---|---|---|
| ENSMUSG00000035356 | 1.363443546 | 0.014748278 |
| ENSMUSG00000035357 | 1.380345489 | 0.000238003 |
| ENSMUSG00000035364 | 1.150017513 | 0.51470764 |
| ENSMUSG00000035365 | -1.024632733 | 0.944192075 |
| ENSMUSG00000035367 | 1.010620853 | 0.968937442 |
| ENSMUSG00000035370 | 1.099334516 | 0.608497229 |
| ENSMUSG00000035371 | 1 | |
| ENSMUSG00000035372 | 1.013902078 | 0.980912194 |
| ENSMUSG00000035373 | 1.92866953 | 0.001601152 |
| ENSMUSG00000035376 | -1.022210738 | 0.949340341 |
| ENSMUSG00000035378 | 1.089184876 | 0.388786254 |
| ENSMUSG00000035382 | -1.032869282 | 0.901903519 |
| ENSMUSG00000035383 | -1.111930034 | 0.49839667 |
| ENSMUSG00000035385 | 1.294582613 | 0.18705267 |
| ENSMUSG00000035387 | 1 | |
| ENSMUSG00000035390 | 1.084310869 | 0.228410454 |
| ENSMUSG00000035392 | 1.024957725 | 0.912742948 |
| ENSMUSG00000035394 | -1.012827107 | 1 |
| ENSMUSG00000035395 | -1.043306199 | 0.698112512 |
| ENSMUSG00000035397 | -1.045370138 | 0.773362331 |
| ENSMUSG00000035399 | -1.015915135 | 0.972980766 |
| ENSMUSG00000035401 | 1.046101006 | 0.80258014 |
| ENSMUSG00000035403 | 1.237106647 | 0.085543976 |
| ENSMUSG00000035407 | 1.189220528 | 0.608497229 |
| ENSMUSG00000035413 | 1.140248688 | 0.43764772 |
| ENSMUSG00000035420 | 1 | |
| ENSMUSG00000035427 | 1 | |
| ENSMUSG00000035429 | 1.122462048 | 0.215991492 |
| ENSMUSG00000035431 | 1.069068979 | 0.608497229 |
| ENSMUSG00000035435 | -1.014034612 | 1 |
| ENSMUSG00000035437 | 1.088503833 | 0.719652885 |
| ENSMUSG00000035439 | 1.024150253 | 0.926076285 |
| ENSMUSG00000035441 | 1.064081075 | 0.61763676 |
| ENSMUSG00000035443 | -1.074768641 | 0.676086178 |
| ENSMUSG00000035448 | 1 | |
| ENSMUSG00000035451 | 2.358192332 | 0.000370384 |
| ENSMUSG00000035454 | 1 | |
| ENSMUSG00000035455 | 1.02832752 | 0.950560821 |
| ENSMUSG00000035456 | -1.110513728 | 0.698112512 |
| ENSMUSG00000035458 | -1.136436699 | 0.693220376 |
| ENSMUSG00000035459 | 1.114060652 | 0.718265321 |
| ENSMUSG00000035469 | -1.010968217 | 1 |
| ENSMUSG00000035472 | -1.041574599 | 0.906689125 |
| ENSMUSG00000035473 | -1.035699748 | 0.884451146 |
| ENSMUSG00000035476 | 1.058626154 | 0.752745311 |
| ENSMUSG00000035478 | 1.094220332 | 0.608497229 |
| ENSMUSG00000035486 | -1.128668575 | 0.578221424 |
| ENSMUSG00000035491 | 1 | |
| ENSMUSG00000035493 | -1.059205096 | 0.751413483 |
| ENSMUSG00000035495 | 1.053898043 | 0.70357926 |
| ENSMUSG00000035498 | -1.095294215 | 0.608497229 |
| ENSMUSG00000035504 | -1.330294649 | 0.0038831 |
| ENSMUSG00000035505 | 1.145252931 | 0.14448103 |
| ENSMUSG00000035506 | -1.018624283 | 1 |
| ENSMUSG00000035509 | -1.450798191 | 0.004327743 |
| ENSMUSG00000035513 | -1.277563227 | 0.608497229 |
| ENSMUSG00000035517 | -1.043894615 | 0.835212486 |
| ENSMUSG00000035521 | 1.085492376 | 0.346160682 |
| ENSMUSG00000035522 | 1 | |
| ENSMUSG00000035523 | 1 | |
| ENSMUSG00000035528 | -1 | 1 |
| ENSMUSG00000035529 | 1.101976801 | 0.656511899 |
| ENSMUSG00000035530 | 1.091586474 | 0.771391875 |

| | | |
|---|---|---|
| ENSMUSG00000035539 | 1.005484436 | 1 |
| ENSMUSG00000035540 | 1 | |
| ENSMUSG00000035545 | 1.082014192 | 0.608497229 |
| ENSMUSG00000035547 | 1.007023705 | 0.937480258 |
| ENSMUSG00000035551 | 1.149355167 | 0.442138282 |
| ENSMUSG00000035557 | -1.03962336 | 0.698112512 |
| ENSMUSG00000035559 | 1.010105351 | 0.978002867 |
| ENSMUSG00000035560 | 1.019083579 | 0.949812939 |
| ENSMUSG00000035561 | -1.498066528 | 0.011133276 |
| ENSMUSG00000035566 | 1.270452795 | 0.25067776 |
| ENSMUSG00000035569 | 1.103246711 | 0.634617162 |
| ENSMUSG00000035572 | -1.022580389 | 0.96513 |
| ENSMUSG00000035575 | -1.066827344 | 0.819817237 |
| ENSMUSG00000035576 | 1.039597911 | 0.887094209 |
| ENSMUSG00000035578 | -1.151179296 | 0.608497229 |
| ENSMUSG00000035580 | 1 | |
| ENSMUSG00000035582 | 1 | |
| ENSMUSG00000035585 | 1.098140676 | 0.502133637 |
| ENSMUSG00000035592 | 1 | |
| ENSMUSG00000035594 | -1.452062024 | 0.024753959 |
| ENSMUSG00000035595 | 1.012919965 | 0.924804062 |
| ENSMUSG00000035596 | 1.015344938 | 0.959892369 |
| ENSMUSG00000035597 | 1.092009104 | 0.698112512 |
| ENSMUSG00000035601 | -1.03586626 | 0.817479993 |
| ENSMUSG00000035606 | 1 | |
| ENSMUSG00000035614 | 1.164145623 | 0.327642404 |
| ENSMUSG00000035615 | -1.862664742 | 0.000522056 |
| ENSMUSG00000035620 | -1.027926422 | 0.912594541 |
| ENSMUSG00000035621 | 1.144727235 | 0.175686119 |
| ENSMUSG00000035623 | -1.009143469 | 1 |
| ENSMUSG00000035626 | 1 | |
| ENSMUSG00000035629 | 1.048652268 | 0.77584181 |
| ENSMUSG00000035632 | 1.016936065 | 0.925741825 |
| ENSMUSG00000035637 | 1.012682311 | 0.945276261 |
| ENSMUSG00000035638 | -1.053549062 | 0.698112512 |
| ENSMUSG00000035640 | 1.319779579 | 0.270002824 |
| ENSMUSG00000035642 | -1.185843649 | 0.076873028 |
| ENSMUSG00000035649 | 1.016566923 | 0.984902895 |
| ENSMUSG00000035651 | 1 | |
| ENSMUSG00000035653 | 1 | |
| ENSMUSG00000035666 | 1.043740564 | 0.899760042 |
| ENSMUSG00000035671 | -1.087119 | 0.469213979 |
| ENSMUSG00000035673 | 1.068182667 | 0.629044949 |
| ENSMUSG00000035674 | -1.046240996 | 0.790872757 |
| ENSMUSG00000035678 | 1.027987163 | 0.674096026 |
| ENSMUSG00000035681 | -1.135975857 | 0.437095676 |
| ENSMUSG00000035683 | -1.091602357 | 0.70362646 |
| ENSMUSG00000035686 | -1.113515092 | 0.642593813 |
| ENSMUSG00000035692 | -1.929559971 | 0.003283306 |
| ENSMUSG00000035694 | 1.009597664 | 1 |
| ENSMUSG00000035696 | 1.097960146 | 0.654473929 |
| ENSMUSG00000035697 | -1.015523377 | 1 |
| ENSMUSG00000035699 | 1.024693402 | 0.933031586 |
| ENSMUSG00000035704 | -1.011329992 | 0.978109404 |
| ENSMUSG00000035711 | 1.068026664 | 0.894274714 |
| ENSMUSG00000035713 | 1.042052533 | 0.715058791 |
| ENSMUSG00000035722 | 1.155986185 | 0.165445236 |
| ENSMUSG00000035725 | -1.108691872 | 0.608497229 |
| ENSMUSG00000035726 | -1.115008434 | 0.698112512 |
| ENSMUSG00000035735 | 1.114252237 | 0.355608366 |
| ENSMUSG00000035745 | 1.227307986 | 0.576169279 |
| ENSMUSG00000035754 | 1.052510023 | 0.665598609 |
| ENSMUSG00000035757 | -1.048806654 | 0.698112512 |

| | | |
|---|---|---|
| ENSMUSG00000035759 | -1.095690783 | 0.55750619 |
| ENSMUSG00000035762 | -1.072262166 | 0.698112512 |
| ENSMUSG00000035764 | 1.031610191 | 0.915158215 |
| ENSMUSG00000035765 | -1.007469025 | 1 |
| ENSMUSG00000035769 | -1.115886458 | 0.047157545 |
| ENSMUSG00000035770 | 1.035105607 | 0.93853582 |
| ENSMUSG00000035772 | -1.020317914 | 0.947276435 |
| ENSMUSG00000035773 | -1.038058186 | 0.917717739 |
| ENSMUSG00000035775 | -1.019108526 | 0.968937442 |
| ENSMUSG00000035776 | -1.011925693 | 1 |
| ENSMUSG00000035778 | 1.210710485 | 0.251544797 |
| ENSMUSG00000035780 | 1 | |
| ENSMUSG00000035781 | -1.00772901 | 1 |
| ENSMUSG00000035783 | 3.539826377 | 0.000960895 |
| ENSMUSG00000035785 | 1 | |
| ENSMUSG00000035790 | -1.026650516 | 0.884018396 |
| ENSMUSG00000035798 | -1.083761965 | 0.698248693 |
| ENSMUSG00000035799 | 1.020483072 | 0.908878383 |
| ENSMUSG00000035804 | 1 | |
| ENSMUSG00000035805 | 1.059463094 | 0.608497229 |
| ENSMUSG00000035811 | 1 | |
| ENSMUSG00000035818 | -1.282934204 | 0.18851436 |
| ENSMUSG00000035824 | -1.000756474 | 1 |
| ENSMUSG00000035828 | 1.071521195 | 0.571944655 |
| ENSMUSG00000035829 | -1.132440898 | 0.216046507 |
| ENSMUSG00000035831 | 1 | |
| ENSMUSG00000035834 | -1.070012141 | 0.698112512 |
| ENSMUSG00000035835 | 1.342693114 | 0.104947513 |
| ENSMUSG00000035836 | 1 | |
| ENSMUSG00000035840 | -1.000027853 | 1 |
| ENSMUSG00000035842 | -1.020064806 | 0.963734554 |
| ENSMUSG00000035845 | 1.010005204 | 0.944985904 |
| ENSMUSG00000035847 | -1.02092709 | 0.984902895 |
| ENSMUSG00000035849 | -1.139484711 | 0.691595649 |
| ENSMUSG00000035851 | 1.072461328 | 0.770016961 |
| ENSMUSG00000035852 | 1.022736512 | 0.986542466 |
| ENSMUSG00000035860 | 1 | |
| ENSMUSG00000035861 | -1.140609708 | 0.633679938 |
| ENSMUSG00000035863 | 1.102611192 | 0.453624705 |
| ENSMUSG00000035864 | -1.033399257 | 0.762710889 |
| ENSMUSG00000035868 | -1.008162302 | 1 |
| ENSMUSG00000035873 | 1.076478816 | 0.657060635 |
| ENSMUSG00000035875 | 1.162065005 | 0.698112512 |
| ENSMUSG00000035877 | -1.026253162 | 0.890896234 |
| ENSMUSG00000035878 | -1.129071558 | 0.293999722 |
| ENSMUSG00000035885 | 1.046971828 | 0.748191299 |
| ENSMUSG00000035890 | 1.054864879 | 0.751234476 |
| ENSMUSG00000035891 | 1.063117495 | 0.608497229 |
| ENSMUSG00000035896 | 1 | |
| ENSMUSG00000035898 | 1.115809582 | 0.683942136 |
| ENSMUSG00000035900 | -1.012757021 | 0.995214337 |
| ENSMUSG00000035901 | 1.006747548 | 1 |
| ENSMUSG00000035910 | -1 | 1 |
| ENSMUSG00000035914 | -1.056433292 | 0.747904191 |
| ENSMUSG00000035916 | -1.226095182 | 0.505070892 |
| ENSMUSG00000035919 | -1.146985599 | 0.3305975 |
| ENSMUSG00000035923 | 1 | |
| ENSMUSG00000035929 | -1.763986967 | 0.00116918 |
| ENSMUSG00000035930 | 1.024656056 | 0.972980766 |
| ENSMUSG00000035932 | 1 | |
| ENSMUSG00000035933 | 1.027512724 | 0.891400877 |
| ENSMUSG00000035934 | -1.308505698 | 0.031696139 |
| ENSMUSG00000035936 | -1.229328645 | 0.128065713 |

| | | |
|---|---|---|
| ENSMUSG00000035941 | 1.035888338 | 0.924137791 |
| ENSMUSG00000035944 | -1.217524254 | 0.013470885 |
| ENSMUSG00000035946 | 1 | |
| ENSMUSG00000035948 | -1.125515849 | 0.463609146 |
| ENSMUSG00000035949 | 1.01536728 | 0.988424085 |
| ENSMUSG00000035951 | 1.031212313 | 0.969949129 |
| ENSMUSG00000035953 | 1.051504321 | 0.746674883 |
| ENSMUSG00000035954 | 1.056042732 | 0.702070725 |
| ENSMUSG00000035958 | -1.00639964 | 1 |
| ENSMUSG00000035960 | -1.043181527 | 0.698112512 |
| ENSMUSG00000035963 | 1 | |
| ENSMUSG00000035964 | -1.188753693 | 0.577204849 |
| ENSMUSG00000035967 | -1.040377318 | 0.751912162 |
| ENSMUSG00000035969 | 1.442169302 | 0.003599328 |
| ENSMUSG00000035983 | 1.144956958 | 0.101456114 |
| ENSMUSG00000035984 | -1.114804816 | 0.685379059 |
| ENSMUSG00000035992 | 1.100850875 | 0.697557233 |
| ENSMUSG00000036002 | 1.04749464 | 0.580660018 |
| ENSMUSG00000036006 | -1.141194367 | 0.399832022 |
| ENSMUSG00000036009 | -1.021333626 | 0.894828773 |
| ENSMUSG00000036013 | 1 | |
| ENSMUSG00000036019 | -1.108774434 | 0.73168857 |
| ENSMUSG00000036022 | -1.113991313 | 0.507659484 |
| ENSMUSG00000036023 | -1.237244268 | 0.199954817 |
| ENSMUSG00000036026 | 1.117592505 | 0.511036937 |
| ENSMUSG00000036027 | 1 | |
| ENSMUSG00000036030 | 1.00822916 | 1 |
| ENSMUSG00000036036 | 1.098917298 | 0.508515749 |
| ENSMUSG00000036040 | 1.238012572 | 0.510864398 |
| ENSMUSG00000036045 | 1.055416288 | 0.794274354 |
| ENSMUSG00000036046 | -1.04724822 | 0.848803478 |
| ENSMUSG00000036052 | 1.2088558 | 0.009184967 |
| ENSMUSG00000036053 | 1.013901723 | 1 |
| ENSMUSG00000036054 | 1.003269428 | 1 |
| ENSMUSG00000036057 | 1.109392324 | 0.608497229 |
| ENSMUSG00000036061 | -1.026960903 | 0.828564184 |
| ENSMUSG00000036062 | -1.524564737 | 0.007421655 |
| ENSMUSG00000036067 | -1.22710036 | 0.067808895 |
| ENSMUSG00000036073 | -1.091081008 | 0.038939734 |
| ENSMUSG00000036078 | -1.022063215 | 0.934760809 |
| ENSMUSG00000036083 | 1 | |
| ENSMUSG00000036086 | -1.027310122 | 0.912782976 |
| ENSMUSG00000036087 | 1.03411065 | 0.92584202 |
| ENSMUSG00000036091 | -1.108117771 | 0.731295483 |
| ENSMUSG00000036093 | 1.154669853 | 0.608497229 |
| ENSMUSG00000036095 | -1.043306199 | 0.698112512 |
| ENSMUSG00000036097 | 1.032488915 | 0.949812939 |
| ENSMUSG00000036098 | -1.618372745 | 0.008364329 |
| ENSMUSG00000036099 | 1.002328385 | 1 |
| ENSMUSG00000036103 | 1.241272078 | 0.224740129 |
| ENSMUSG00000036104 | -1.047047252 | 0.863882521 |
| ENSMUSG00000036106 | -1.020971458 | 0.891373394 |
| ENSMUSG00000036109 | 1.051252594 | 0.750243435 |
| ENSMUSG00000036110 | 1 | |
| ENSMUSG00000036111 | -1.443212187 | 0.010116338 |
| ENSMUSG00000036112 | -1.041743525 | 0.936761877 |
| ENSMUSG00000036114 | 1.054604101 | 0.698112512 |
| ENSMUSG00000036117 | -1.08081673 | 0.73636577 |
| ENSMUSG00000036120 | -1.032609107 | 0.837740447 |
| ENSMUSG00000036123 | -1.03962336 | 0.698112512 |
| ENSMUSG00000036131 | 1 | |
| ENSMUSG00000036136 | -1.036526021 | 0.896733108 |
| ENSMUSG00000036138 | -1.038481643 | 0.7938895 |

| | | |
|---|---|---|
| ENSMUSG00000036139 | -2.053297376 | 0.004413461 |
| ENSMUSG00000036144 | 1 | |
| ENSMUSG00000036151 | 1.074676495 | 0.717035356 |
| ENSMUSG00000036155 | -1.003256426 | 1 |
| ENSMUSG00000036158 | -1.384134742 | 0.004953893 |
| ENSMUSG00000036160 | 1.033150772 | 0.848385245 |
| ENSMUSG00000036162 | -1.043306199 | 0.698112512 |
| ENSMUSG00000036167 | -1.017760594 | 0.993549524 |
| ENSMUSG00000036168 | -1.058927046 | 0.709356753 |
| ENSMUSG00000036169 | 1 | |
| ENSMUSG00000036172 | -1.085651418 | 0.608497229 |
| ENSMUSG00000036180 | 1.068965588 | 0.690669961 |
| ENSMUSG00000036181 | -1.46060059 | 0.001167589 |
| ENSMUSG00000036185 | 1.155013505 | 0.546960642 |
| ENSMUSG00000036186 | 1.170337974 | 0.251544797 |
| ENSMUSG00000036188 | 1.055842362 | 0.76055463 |
| ENSMUSG00000036192 | -1.014110535 | 0.943705943 |
| ENSMUSG00000036196 | 1.066200696 | 0.884295887 |
| ENSMUSG00000036197 | -1.127505834 | 0.612433013 |
| ENSMUSG00000036198 | -1.4086967 | 0.003038921 |
| ENSMUSG00000036199 | 1.042492993 | 0.746451077 |
| ENSMUSG00000036202 | 1.022559269 | 0.979152632 |
| ENSMUSG00000036206 | 1.156667446 | 0.299006106 |
| ENSMUSG00000036208 | 1.05568746 | 0.747163034 |
| ENSMUSG00000036211 | -1.053549062 | 0.698112512 |
| ENSMUSG00000036214 | 1.05527757 | 0.608497229 |
| ENSMUSG00000036216 | -1.179226843 | 0.584107608 |
| ENSMUSG00000036218 | 1 | |
| ENSMUSG00000036223 | -1.007953438 | 1 |
| ENSMUSG00000036225 | 1.127301025 | 0.112575012 |
| ENSMUSG00000036231 | 1 | |
| ENSMUSG00000036241 | 1.071369381 | 0.770958539 |
| ENSMUSG00000036242 | 1 | |
| ENSMUSG00000036244 | 1.059463094 | 0.704170349 |
| ENSMUSG00000036246 | 1.002980289 | 1 |
| ENSMUSG00000036249 | -1.062190155 | 0.698112512 |
| ENSMUSG00000036251 | -1.29236123 | 0.120083872 |
| ENSMUSG00000036256 | 1.598051981 | 0.011463001 |
| ENSMUSG00000036257 | 1.104699506 | 0.561881571 |
| ENSMUSG00000036264 | -1 | 1 |
| ENSMUSG00000036270 | 1.014073483 | 0.985925873 |
| ENSMUSG00000036273 | 1.056861301 | 0.819614655 |
| ENSMUSG00000036275 | -1.070554197 | 0.813372967 |
| ENSMUSG00000036278 | 1.021131147 | 0.903356824 |
| ENSMUSG00000036281 | 1.049414436 | 0.693696067 |
| ENSMUSG00000036282 | -1.168050695 | 0.416439753 |
| ENSMUSG00000036285 | -1.009353012 | 1 |
| ENSMUSG00000036291 | -1.164329886 | 0.279947521 |
| ENSMUSG00000036292 | -1.339181344 | 0.250726313 |
| ENSMUSG00000036295 | 1.177765487 | 0.588718419 |
| ENSMUSG00000036298 | -1.001749642 | 1 |
| ENSMUSG00000036299 | 1.0321802 | 0.903753349 |
| ENSMUSG00000036304 | -1.218735929 | 0.454976536 |
| ENSMUSG00000036305 | -1.093542818 | 0.435929624 |
| ENSMUSG00000036306 | -1.306320118 | 0.072705092 |
| ENSMUSG00000036309 | -1.089195342 | 0.770923502 |
| ENSMUSG00000036315 | -1.00803255 | 1 |
| ENSMUSG00000036322 | 1 | |
| ENSMUSG00000036323 | -1.014373974 | 1 |
| ENSMUSG00000036327 | -1.025724064 | 0.895970183 |
| ENSMUSG00000036330 | -1.241950994 | 0.211115461 |
| ENSMUSG00000036333 | 1.030767768 | 0.949812939 |
| ENSMUSG00000036334 | 1.23817551 | 0.035333808 |

| | | |
|---|---|---|
| ENSMUSG00000036339 | -1.102682201 | 0.434612975 |
| ENSMUSG00000036352 | -1.027764255 | 0.942585548 |
| ENSMUSG00000036353 | 1.019083579 | 0.949812939 |
| ENSMUSG00000036356 | 1.682952687 | 0.000433572 |
| ENSMUSG00000036357 | 1 | |
| ENSMUSG00000036362 | 1 | |
| ENSMUSG00000036368 | 1.133965599 | 0.378445257 |
| ENSMUSG00000036371 | 1.045809673 | 0.908065256 |
| ENSMUSG00000036372 | 1.016448112 | 0.993795117 |
| ENSMUSG00000036376 | -1.075967541 | 0.287416182 |
| ENSMUSG00000036377 | 1.604879753 | 0.000110329 |
| ENSMUSG00000036381 | 1.01980401 | 0.979814083 |
| ENSMUSG00000036390 | 1.039290996 | 0.757039001 |
| ENSMUSG00000036391 | -1.012503669 | 1 |
| ENSMUSG00000036395 | -1.679901002 | 0.009529771 |
| ENSMUSG00000036398 | -1.037035824 | 0.903188698 |
| ENSMUSG00000036401 | 1 | |
| ENSMUSG00000036402 | 1.052370111 | 0.864629923 |
| ENSMUSG00000036403 | -1.037541674 | 0.854640873 |
| ENSMUSG00000036411 | -1.046295781 | 0.495914183 |
| ENSMUSG00000036412 | 1.090312574 | 0.608497229 |
| ENSMUSG00000036422 | -1 | 1 |
| ENSMUSG00000036427 | 1.04630954 | 0.876422702 |
| ENSMUSG00000036430 | 1.006868407 | 1 |
| ENSMUSG00000036432 | 1.026681621 | 0.923464844 |
| ENSMUSG00000036435 | 1.083374269 | 0.713861573 |
| ENSMUSG00000036437 | 1.56763841 | 0.007129528 |
| ENSMUSG00000036438 | 1.067622325 | 0.874672445 |
| ENSMUSG00000036442 | -1.020553247 | 0.927315966 |
| ENSMUSG00000036446 | -1 | 1 |
| ENSMUSG00000036449 | -1.044273782 | 0.698112512 |
| ENSMUSG00000036450 | -1.049741477 | 0.848960406 |
| ENSMUSG00000036452 | -1.121107665 | 0.174235548 |
| ENSMUSG00000036459 | 1.15541672 | 0.123449197 |
| ENSMUSG00000036461 | -1.060575854 | 0.731295483 |
| ENSMUSG00000036463 | 1 | |
| ENSMUSG00000036466 | -1.098255242 | 0.371499647 |
| ENSMUSG00000036469 | -1.116296275 | 0.554354454 |
| ENSMUSG00000036473 | 1.027886577 | 0.913228374 |
| ENSMUSG00000036478 | 1.187994019 | 0.378010412 |
| ENSMUSG00000036480 | 1.059463094 | 0.698112512 |
| ENSMUSG00000036492 | 1.104049742 | 0.770958539 |
| ENSMUSG00000036499 | 1.010913232 | 1 |
| ENSMUSG00000036500 | 1.639826543 | 0.026801377 |
| ENSMUSG00000036501 | 1.076306043 | 0.725290225 |
| ENSMUSG00000036502 | -1.083462935 | 0.744030455 |
| ENSMUSG00000036503 | 1.031336574 | 0.945469287 |
| ENSMUSG00000036504 | 1.0933259 | 0.469310516 |
| ENSMUSG00000036510 | 1 | |
| ENSMUSG00000036513 | 1.009327766 | 1 |
| ENSMUSG00000036521 | 1 | |
| ENSMUSG00000036523 | 1.101623201 | 0.550576507 |
| ENSMUSG00000036526 | 1 | |
| ENSMUSG00000036528 | 1.572266304 | 0.002126951 |
| ENSMUSG00000036529 | 1.072700454 | 0.698112512 |
| ENSMUSG00000036533 | 1.314477507 | 0.029295209 |
| ENSMUSG00000036534 | 1.298953948 | 0.022910867 |
| ENSMUSG00000036537 | 1.018914938 | 0.943705943 |
| ENSMUSG00000036545 | 2.218727786 | 2.98633E-05 |
| ENSMUSG00000036550 | -1.009107312 | 1 |
| ENSMUSG00000036551 | -1.023769022 | 0.956401771 |
| ENSMUSG00000036552 | 1.067304 | 0.678985493 |
| ENSMUSG00000036553 | 1.008850422 | 1 |

| | | |
|---|---|---|
| ENSMUSG00000036555 | -1.007208574 | 1 |
| ENSMUSG00000036557 | -1.044273782 | 0.698112512 |
| ENSMUSG00000036560 | 1.095872691 | 0.608497229 |
| ENSMUSG00000036561 | -1.087151913 | 0.541735653 |
| ENSMUSG00000036564 | 1.069558318 | 0.615267463 |
| ENSMUSG00000036565 | 1.173409527 | 0.23320787 |
| ENSMUSG00000036568 | 1.080213864 | 0.50927285 |
| ENSMUSG00000036570 | 1 | |
| ENSMUSG00000036572 | -1.016797926 | 0.975499304 |
| ENSMUSG00000036574 | 1 | |
| ENSMUSG00000036578 | 1.059463094 | 0.704170349 |
| ENSMUSG00000036580 | 1.030686286 | 0.916607312 |
| ENSMUSG00000036585 | 1.217954966 | 0.559827647 |
| ENSMUSG00000036586 | 1 | |
| ENSMUSG00000036587 | 1.00177232 | 1 |
| ENSMUSG00000036591 | 1.095936501 | 0.650566948 |
| ENSMUSG00000036594 | -1.354429562 | 0.065461774 |
| ENSMUSG00000036596 | -1.11061171 | 0.608497229 |
| ENSMUSG00000036598 | -1.168999771 | 0.208940434 |
| ENSMUSG00000036599 | 1.165981699 | 0.181009235 |
| ENSMUSG00000036602 | 3.751165344 | 0.000129787 |
| ENSMUSG00000036606 | 1.041113973 | 0.906826915 |
| ENSMUSG00000036611 | 1.093822085 | 0.237600636 |
| ENSMUSG00000036613 | 1.110150263 | 0.418131538 |
| ENSMUSG00000036615 | -1.048004191 | 0.791358811 |
| ENSMUSG00000036617 | -1.132741198 | 0.234728821 |
| ENSMUSG00000036620 | 1.15176175 | 0.420355052 |
| ENSMUSG00000036622 | -1.092261901 | 0.693682258 |
| ENSMUSG00000036632 | -1.010044125 | 1 |
| ENSMUSG00000036634 | 1.035626983 | 0.959892369 |
| ENSMUSG00000036636 | 1.132253199 | 0.534887579 |
| ENSMUSG00000036639 | 1.009877608 | 0.997250566 |
| ENSMUSG00000036641 | -1.006720492 | 1 |
| ENSMUSG00000036644 | -1.076155629 | 0.698112512 |
| ENSMUSG00000036646 | -1.004653515 | 1 |
| ENSMUSG00000036647 | 1 | |
| ENSMUSG00000036655 | 1.150659893 | 0.617912429 |
| ENSMUSG00000036658 | 1 | |
| ENSMUSG00000036661 | 1.017694963 | 0.923478466 |
| ENSMUSG00000036667 | 1.027319873 | 0.95829726 |
| ENSMUSG00000036672 | 1.007478714 | 1 |
| ENSMUSG00000036676 | -1.065427869 | 0.832252345 |
| ENSMUSG00000036678 | 1.020085681 | 0.975677002 |
| ENSMUSG00000036686 | 1.002042261 | 1 |
| ENSMUSG00000036687 | -1.029822524 | 0.958977086 |
| ENSMUSG00000036693 | 1.042579815 | 0.872903084 |
| ENSMUSG00000036698 | 1.145025506 | 0.608497229 |
| ENSMUSG00000036699 | 1.117640663 | 0.608497229 |
| ENSMUSG00000036707 | 1.001673733 | 1 |
| ENSMUSG00000036712 | -1.169704408 | 0.585646836 |
| ENSMUSG00000036718 | 1.191375678 | 0.210180454 |
| ENSMUSG00000036721 | -1.047601534 | 0.710222167 |
| ENSMUSG00000036731 | 1 | |
| ENSMUSG00000036733 | 1.005613439 | 1 |
| ENSMUSG00000036737 | -1.040693024 | 0.898042571 |
| ENSMUSG00000036743 | 1 | |
| ENSMUSG00000036744 | 1 | |
| ENSMUSG00000036745 | -1.059603523 | 0.814089245 |
| ENSMUSG00000036748 | 1.12096466 | 0.429352448 |
| ENSMUSG00000036749 | -1.003841375 | 1 |
| ENSMUSG00000036751 | -1.036110873 | 0.909211737 |
| ENSMUSG00000036752 | -1.097530301 | 0.670787984 |
| ENSMUSG00000036760 | 1 | |

| | | |
|---|---|---|
| ENSMUSG00000036764 | -1.137115887 | 0.407219461 |
| ENSMUSG00000036766 | -1.027230783 | 0.982838256 |
| ENSMUSG00000036768 | -1.090186785 | 0.698112512 |
| ENSMUSG00000036769 | 1.037305558 | 0.815540288 |
| ENSMUSG00000036770 | 1.010351027 | 1 |
| ENSMUSG00000036775 | 1.00844118 | 1 |
| ENSMUSG00000036777 | -1.041271121 | 0.940213048 |
| ENSMUSG00000036779 | -1.076279798 | 0.741009626 |
| ENSMUSG00000036781 | -1.009059038 | 1 |
| ENSMUSG00000036782 | 1.220292492 | 0.180628841 |
| ENSMUSG00000036790 | 1.126722214 | 0.623872809 |
| ENSMUSG00000036792 | 1.044520655 | 0.770958539 |
| ENSMUSG00000036800 | 1.069068979 | 0.74773997 |
| ENSMUSG00000036805 | 1 | |
| ENSMUSG00000036810 | 1.061356649 | 0.811257973 |
| ENSMUSG00000036813 | 1.059463094 | 0.704170349 |
| ENSMUSG00000036814 | -1.044273782 | 0.698112512 |
| ENSMUSG00000036815 | -1.019476438 | 0.968375666 |
| ENSMUSG00000036816 | 1 | |
| ENSMUSG00000036817 | 1.039989529 | 0.906480064 |
| ENSMUSG00000036819 | -1.091241834 | 0.590629201 |
| ENSMUSG00000036820 | 1.573515422 | 0.014562998 |
| ENSMUSG00000036822 | 1.050850641 | 0.857392806 |
| ENSMUSG00000036825 | -1.101656752 | 0.63198593 |
| ENSMUSG00000036826 | 1 | |
| ENSMUSG00000036832 | -1.015810725 | 0.946112838 |
| ENSMUSG00000036833 | 1.097669019 | 0.346184754 |
| ENSMUSG00000036834 | 1.194991315 | 0.10982693 |
| ENSMUSG00000036835 | 1.081227006 | 0.513200095 |
| ENSMUSG00000036840 | 1.06057263 | 0.698112512 |
| ENSMUSG00000036845 | 1.046204119 | 0.698112512 |
| ENSMUSG00000036850 | 1.024744029 | 0.858511334 |
| ENSMUSG00000036853 | 1.572006566 | 0.006294552 |
| ENSMUSG00000036854 | 1.465700306 | 0.007325542 |
| ENSMUSG00000036855 | 1 | |
| ENSMUSG00000036856 | -1.041293239 | 0.698112512 |
| ENSMUSG00000036858 | 1.050647003 | 0.920928249 |
| ENSMUSG00000036860 | -1.036820511 | 0.711602885 |
| ENSMUSG00000036862 | -1.076597425 | 0.541783276 |
| ENSMUSG00000036863 | -1.041362049 | 0.775575739 |
| ENSMUSG00000036864 | 1.128281756 | 0.232747283 |
| ENSMUSG00000036867 | 1.096176492 | 0.323633476 |
| ENSMUSG00000036872 | -1.143787033 | 0.377110629 |
| ENSMUSG00000036873 | -1.155984638 | 0.367780446 |
| ENSMUSG00000036875 | -1.000127937 | 1 |
| ENSMUSG00000036879 | 1.008544709 | 1 |
| ENSMUSG00000036880 | 1.097148075 | 0.608497229 |
| ENSMUSG00000036882 | 1.000408072 | 1 |
| ENSMUSG00000036885 | 1.236252133 | 0.2794251 |
| ENSMUSG00000036887 | -1 | 1 |
| ENSMUSG00000036890 | 1.065992513 | 0.537660024 |
| ENSMUSG00000036892 | 1.075295999 | 0.819474306 |
| ENSMUSG00000036893 | 1.078540797 | 0.650713852 |
| ENSMUSG00000036894 | 1.050335097 | 0.609147349 |
| ENSMUSG00000036896 | 1 | |
| ENSMUSG00000036898 | 1.052228188 | 0.762584863 |
| ENSMUSG00000036899 | -1 | 1 |
| ENSMUSG00000036902 | -2.279913205 | 0.000822888 |
| ENSMUSG00000036904 | 1.470038181 | 0.035334161 |
| ENSMUSG00000036905 | 1 | |
| ENSMUSG00000036907 | 1 | |
| ENSMUSG00000036908 | 1.175562859 | 0.263602582 |
| ENSMUSG00000036912 | 1.065410325 | 0.764982341 |

| | | |
|---|---|---|
| ENSMUSG00000036913 | -1.024718486 | 0.931480186 |
| ENSMUSG00000036915 | 1.005613439 | 1 |
| ENSMUSG00000036916 | 1.05188925 | 0.813450442 |
| ENSMUSG00000036918 | 1.080237924 | 0.155885816 |
| ENSMUSG00000036921 | 1 | |
| ENSMUSG00000036923 | -1.168316139 | 0.694083715 |
| ENSMUSG00000036924 | 1 | |
| ENSMUSG00000036925 | 1 | |
| ENSMUSG00000036928 | -1.269236409 | 0.275589689 |
| ENSMUSG00000036931 | -1.514905051 | 0.026780517 |
| ENSMUSG00000036932 | 1.06576043 | 0.807247723 |
| ENSMUSG00000036934 | -1.017744919 | 0.971396058 |
| ENSMUSG00000036938 | -1.03962336 | 0.698112512 |
| ENSMUSG00000036940 | 1.071444562 | 0.803347469 |
| ENSMUSG00000036941 | 1.019980101 | 0.927544616 |
| ENSMUSG00000036943 | -1.161245297 | 0.570229056 |
| ENSMUSG00000036944 | -1.097349941 | 0.698112512 |
| ENSMUSG00000036948 | 1.063001558 | 0.608497229 |
| ENSMUSG00000036949 | -1 | 1 |
| ENSMUSG00000036951 | 1 | |
| ENSMUSG00000036955 | -1.098713264 | 0.66875675 |
| ENSMUSG00000036957 | 1.193324062 | 0.294194175 |
| ENSMUSG00000036958 | 1 | |
| ENSMUSG00000036959 | 1.052810048 | 0.698112512 |
| ENSMUSG00000036960 | 1.649279497 | 0.000965153 |
| ENSMUSG00000036961 | 1 | |
| ENSMUSG00000036962 | -1 | 1 |
| ENSMUSG00000036964 | -1.026039891 | 0.972634204 |
| ENSMUSG00000036966 | 1.142361895 | 0.369135776 |
| ENSMUSG00000036968 | -1.059227448 | 0.79291651 |
| ENSMUSG00000036972 | -2.11010186 | 0.000309783 |
| ENSMUSG00000036975 | 1.015941424 | 0.957523075 |
| ENSMUSG00000036977 | -1.016819545 | 0.982854947 |
| ENSMUSG00000036980 | 1.023820378 | 0.942279278 |
| ENSMUSG00000036983 | 1.062028451 | 0.698112512 |
| ENSMUSG00000036985 | 1.008953471 | 1 |
| ENSMUSG00000036986 | 1.019514419 | 0.934760809 |
| ENSMUSG00000036989 | 1.187108141 | 0.061225644 |
| ENSMUSG00000036990 | 1.028789085 | 0.965353324 |
| ENSMUSG00000036992 | -1.004079766 | 1 |
| ENSMUSG00000036995 | -1.106191488 | 0.157466578 |
| ENSMUSG00000037001 | -1.266099903 | 0.0586397 |
| ENSMUSG00000037003 | 1.125953009 | 0.444934552 |
| ENSMUSG00000037005 | 1.225670344 | 0.036626025 |
| ENSMUSG00000037007 | -1.058426227 | 0.731295483 |
| ENSMUSG00000037010 | 1.054927842 | 0.698112512 |
| ENSMUSG00000037012 | 1.057211509 | 0.786025504 |
| ENSMUSG00000037013 | 1.075876866 | 0.703046429 |
| ENSMUSG00000037014 | 1 | |
| ENSMUSG00000037016 | -1.551240691 | 0.009999823 |
| ENSMUSG00000037017 | 1.022924176 | 0.944721414 |
| ENSMUSG00000037020 | 1.011772266 | 0.972794583 |
| ENSMUSG00000037022 | 1.033314435 | 0.786485853 |
| ENSMUSG00000037025 | -1.372886284 | 0.050085259 |
| ENSMUSG00000037028 | -1 | 1 |
| ENSMUSG00000037029 | 1.053938989 | 0.761235181 |
| ENSMUSG00000037031 | -1.05348294 | 0.716622726 |
| ENSMUSG00000037032 | 1.08450898 | 0.363615297 |
| ENSMUSG00000037033 | -1 | 1 |
| ENSMUSG00000037034 | 1 | |
| ENSMUSG00000037035 | 1.252257934 | 0.576246406 |
| ENSMUSG00000037049 | -1.002197036 | 1 |
| ENSMUSG00000037053 | 1 | |

| | | |
|---|---|---|
| ENSMUSG00000037058 | 1.045200878 | 0.885136117 |
| ENSMUSG00000037060 | 1.403544228 | 0.002088219 |
| ENSMUSG00000037062 | -1.005976459 | 1 |
| ENSMUSG00000037064 | 1 | |
| ENSMUSG00000037070 | 1.059399766 | 0.757848756 |
| ENSMUSG00000037071 | -1.03771168 | 0.934760809 |
| ENSMUSG00000037072 | -1.012063148 | 1 |
| ENSMUSG00000037075 | 1.023117636 | 0.964579585 |
| ENSMUSG00000037085 | -1.019092021 | 0.881483647 |
| ENSMUSG00000037086 | 1 | |
| ENSMUSG00000037089 | 1.079557363 | 0.698112512 |
| ENSMUSG00000037095 | -1.044273782 | 0.698112512 |
| ENSMUSG00000037096 | 1 | |
| ENSMUSG00000037098 | 1.034697028 | 0.849464079 |
| ENSMUSG00000037101 | -1 | 1 |
| ENSMUSG00000037103 | 1.004038264 | 1 |
| ENSMUSG00000037104 | 1.152838535 | 0.477044609 |
| ENSMUSG00000037106 | 1.393244618 | 0.001661811 |
| ENSMUSG00000037108 | -1.002797539 | 1 |
| ENSMUSG00000037110 | -1.112393597 | 0.534887579 |
| ENSMUSG00000037111 | -1.035885159 | 0.944389804 |
| ENSMUSG00000037112 | 1.047140345 | 0.778372272 |
| ENSMUSG00000037119 | 1.022057157 | 0.992424863 |
| ENSMUSG00000037124 | 1 | |
| ENSMUSG00000037126 | -1.10732223 | 0.125324229 |
| ENSMUSG00000037129 | 1.089470641 | 0.698112512 |
| ENSMUSG00000037130 | 1 | |
| ENSMUSG00000037134 | -1.000013011 | 1 |
| ENSMUSG00000037138 | -1.176919401 | 0.036046595 |
| ENSMUSG00000037139 | -1.069095149 | 0.686194514 |
| ENSMUSG00000037140 | -1.043306199 | 0.698112512 |
| ENSMUSG00000037143 | -1.302967834 | 0.145761595 |
| ENSMUSG00000037145 | 1 | |
| ENSMUSG00000037148 | -1.049873093 | 0.795446726 |
| ENSMUSG00000037149 | -1.033996268 | 0.953416513 |
| ENSMUSG00000037151 | -1.013069725 | 0.966954883 |
| ENSMUSG00000037152 | -1.044241273 | 0.698112512 |
| ENSMUSG00000037157 | -1 | 1 |
| ENSMUSG00000037159 | -1.053549062 | 0.698112512 |
| ENSMUSG00000037161 | -1.613071343 | 0.008586051 |
| ENSMUSG00000037166 | 1.668251399 | 0.004687172 |
| ENSMUSG00000037167 | 1.019083579 | 0.968937442 |
| ENSMUSG00000037169 | -1.270818951 | 0.153011055 |
| ENSMUSG00000037171 | -1.100193663 | 0.534887579 |
| ENSMUSG00000037172 | -1.176067315 | 0.193437983 |
| ENSMUSG00000037174 | -1.048579478 | 0.854777354 |
| ENSMUSG00000037185 | -1.215728231 | 0.055871312 |
| ENSMUSG00000037188 | -1.07452894 | 0.83500866 |
| ENSMUSG00000037190 | -1.013753819 | 0.879458055 |
| ENSMUSG00000037196 | 1.007241021 | 1 |
| ENSMUSG00000037197 | 1.017448497 | 0.988424085 |
| ENSMUSG00000037202 | -1 | 1 |
| ENSMUSG00000037204 | 1.039612338 | 0.71045278 |
| ENSMUSG00000037206 | -1.001171345 | 1 |
| ENSMUSG00000037210 | 1.07840711 | 0.676032103 |
| ENSMUSG00000037211 | 1.348470116 | 0.035832281 |
| ENSMUSG00000037214 | -1.013285653 | 0.937151862 |
| ENSMUSG00000037216 | -1.093646431 | 0.38414595 |
| ENSMUSG00000037217 | 1.156729919 | 0.087675226 |
| ENSMUSG00000037221 | 1.069310959 | 0.608497229 |
| ENSMUSG00000037224 | -1.396223581 | 0.036382222 |
| ENSMUSG00000037225 | 1.222814997 | 0.009436296 |
| ENSMUSG00000037234 | 1.040216149 | 0.92472818 |

| | | |
|---|---|---|
| ENSMUSG00000037235 | 1.060630077 | 0.743396505 |
| ENSMUSG00000037236 | 1.068563424 | 0.843685576 |
| ENSMUSG00000037239 | -1.04195505 | 0.713429278 |
| ENSMUSG00000037242 | 1.096775685 | 0.756679803 |
| ENSMUSG00000037243 | -1.047659459 | 0.600484108 |
| ENSMUSG00000037246 | 1 | |
| ENSMUSG00000037247 | 1.138788635 | 0.595042751 |
| ENSMUSG00000037251 | 1.046508008 | 0.72827364 |
| ENSMUSG00000037253 | 1.139274077 | 0.608497229 |
| ENSMUSG00000037254 | 1.986050886 | 0.011041188 |
| ENSMUSG00000037257 | -1.064949962 | 0.743920175 |
| ENSMUSG00000037259 | -1.541901577 | 0.002397042 |
| ENSMUSG00000037260 | -1.025194106 | 0.951697621 |
| ENSMUSG00000037262 | -1.000608054 | 1 |
| ENSMUSG00000037263 | -1.043306199 | 0.698112512 |
| ENSMUSG00000037266 | 1.054615491 | 0.820871905 |
| ENSMUSG00000037270 | -1.079916224 | 0.760042353 |
| ENSMUSG00000037275 | -1.09749326 | 0.68910337 |
| ENSMUSG00000037278 | -1.030117986 | 0.929405589 |
| ENSMUSG00000037279 | 1.059463094 | 0.608497229 |
| ENSMUSG00000037280 | -1.428504266 | 0.002964254 |
| ENSMUSG00000037286 | -1.033858287 | 0.948722639 |
| ENSMUSG00000037287 | -1.011441467 | 0.996913409 |
| ENSMUSG00000037295 | -1.143529862 | 0.234971141 |
| ENSMUSG00000037296 | -1.050170589 | 0.794597624 |
| ENSMUSG00000037300 | -1.046498934 | 0.836447677 |
| ENSMUSG00000037306 | 1.138507571 | 0.07590923 |
| ENSMUSG00000037307 | 1 | |
| ENSMUSG00000037313 | -1.01347072 | 1 |
| ENSMUSG00000037315 | 1.106457053 | 0.593770025 |
| ENSMUSG00000037316 | -1.058595379 | 0.778683365 |
| ENSMUSG00000037318 | -1.102107151 | 0.756258014 |
| ENSMUSG00000037321 | -1.561097619 | 0.000573666 |
| ENSMUSG00000037325 | -1.01185138 | 0.998924031 |
| ENSMUSG00000037326 | 1.159536742 | 0.517011747 |
| ENSMUSG00000037331 | 1.010635035 | 1 |
| ENSMUSG00000037334 | 1 | |
| ENSMUSG00000037335 | 1 | |
| ENSMUSG00000037336 | -1.178011445 | 0.543612473 |
| ENSMUSG00000037337 | 1.112757641 | 0.262993046 |
| ENSMUSG00000037339 | 1.130919143 | 0.050544467 |
| ENSMUSG00000037341 | 1.032515304 | 0.699340382 |
| ENSMUSG00000037343 | 1.02292394 | 0.975677002 |
| ENSMUSG00000037344 | 1.090214802 | 0.458862417 |
| ENSMUSG00000037346 | 1 | |
| ENSMUSG00000037347 | 1.101569786 | 0.608497229 |
| ENSMUSG00000037348 | 1.034029635 | 0.615569604 |
| ENSMUSG00000037349 | -1.054430158 | 0.678783707 |
| ENSMUSG00000037351 | -1.049052201 | 0.845084852 |
| ENSMUSG00000037353 | 1.089576448 | 0.602343211 |
| ENSMUSG00000037355 | 1.085298257 | 0.539222653 |
| ENSMUSG00000037358 | 1 | |
| ENSMUSG00000037361 | 1.018725258 | 0.978253323 |
| ENSMUSG00000037362 | 1.401934663 | 0.001082335 |
| ENSMUSG00000037363 | 1.058997255 | 0.693148633 |
| ENSMUSG00000037364 | 1.015763445 | 0.982838256 |
| ENSMUSG00000037366 | 1.104105892 | 0.424948752 |
| ENSMUSG00000037369 | 1.076507327 | 0.698112512 |
| ENSMUSG00000037370 | 1.084564184 | 0.565305007 |
| ENSMUSG00000037373 | 1.090385013 | 0.608497229 |
| ENSMUSG00000037375 | -1.006120903 | 0.989648415 |
| ENSMUSG00000037376 | -1.082396274 | 0.698112512 |
| ENSMUSG00000037379 | 1.326898861 | 0.064787973 |

| | | |
|---|---|---|
| ENSMUSG00000037386 | -1.190419644 | 0.217993289 |
| ENSMUSG00000037390 | 1.014545335 | 0.979002724 |
| ENSMUSG00000037393 | 1.059463094 | 0.608497229 |
| ENSMUSG00000037395 | 1.013945154 | 0.968575228 |
| ENSMUSG00000037400 | -1.084154555 | 0.738404029 |
| ENSMUSG00000037405 | -1.652065466 | 0.005650136 |
| ENSMUSG00000037406 | 1.116980306 | 0.677496354 |
| ENSMUSG00000037408 | -1.031642867 | 0.833490776 |
| ENSMUSG00000037410 | 1.023516375 | 0.933031586 |
| ENSMUSG00000037411 | 1.360515814 | 0.103979661 |
| ENSMUSG00000037415 | -1.017518342 | 0.977433779 |
| ENSMUSG00000037416 | 1.044724823 | 0.871609782 |
| ENSMUSG00000037418 | 1.154253937 | 0.452069807 |
| ENSMUSG00000037419 | -1.261096255 | 0.048683099 |
| ENSMUSG00000037424 | 1 | |
| ENSMUSG00000037426 | 1.059865121 | 0.747340207 |
| ENSMUSG00000037428 | 1.028323353 | 0.933031586 |
| ENSMUSG00000037432 | -1.130084716 | 0.250726313 |
| ENSMUSG00000037434 | 1.045471226 | 0.748642958 |
| ENSMUSG00000037437 | 1.052148224 | 0.939105243 |
| ENSMUSG00000037438 | 1 | |
| ENSMUSG00000037440 | -1.318302442 | 0.149078169 |
| ENSMUSG00000037443 | -1.000257934 | 1 |
| ENSMUSG00000037446 | 1.069597823 | 0.874375366 |
| ENSMUSG00000037447 | -1.290351871 | 0.009164773 |
| ENSMUSG00000037451 | 1 | |
| ENSMUSG00000037455 | -1.094906463 | 0.608497229 |
| ENSMUSG00000037458 | 1.018506855 | 1 |
| ENSMUSG00000037461 | -1.011040877 | 1 |
| ENSMUSG00000037463 | 1.019083579 | 0.979002724 |
| ENSMUSG00000037465 | 1.167404306 | 0.323355045 |
| ENSMUSG00000037466 | 1.10292408 | 0.418158341 |
| ENSMUSG00000037469 | 1 | |
| ENSMUSG00000037470 | -1.071715499 | 0.823038846 |
| ENSMUSG00000037474 | -1.05986652 | 0.828564184 |
| ENSMUSG00000037475 | 1.031155716 | 0.956374899 |
| ENSMUSG00000037477 | 1 | |
| ENSMUSG00000037482 | 1 | |
| ENSMUSG00000037486 | 1.06297352 | 0.774081197 |
| ENSMUSG00000037487 | 1.083132012 | 0.807847618 |
| ENSMUSG00000037490 | -1.34724846 | 0.107942936 |
| ENSMUSG00000037492 | -1.093674795 | 0.698112512 |
| ENSMUSG00000037493 | -1.22682635 | 0.127132146 |
| ENSMUSG00000037499 | 1.048861919 | 0.608497229 |
| ENSMUSG00000037503 | 1.081143364 | 0.789364755 |
| ENSMUSG00000037509 | -1.08840545 | 0.278380177 |
| ENSMUSG00000037514 | -1.066029135 | 0.743041027 |
| ENSMUSG00000037519 | 1.052144845 | 0.79291651 |
| ENSMUSG00000037523 | -1.078788805 | 0.681491935 |
| ENSMUSG00000037525 | 1.130579496 | 0.035301792 |
| ENSMUSG00000037526 | -1.129027603 | 0.477644658 |
| ENSMUSG00000037529 | 1 | |
| ENSMUSG00000037531 | -1.00493464 | 1 |
| ENSMUSG00000037533 | -1.074369185 | 0.755673717 |
| ENSMUSG00000037535 | -1.043306199 | 0.79291651 |
| ENSMUSG00000037536 | -1.131973566 | 0.340098181 |
| ENSMUSG00000037537 | 1 | |
| ENSMUSG00000037541 | -1.034742306 | 0.749776086 |
| ENSMUSG00000037542 | 1.059463094 | 0.776953697 |
| ENSMUSG00000037544 | -1.167971914 | 0.52201333 |
| ENSMUSG00000037548 | -1.195232488 | 0.192975124 |
| ENSMUSG00000037552 | 1.213329128 | 0.089234284 |
| ENSMUSG00000037553 | 1.011909572 | 0.96504537 |

| | | |
|---|---|---|
| ENSMUSG00000037563 | 1.062346632 | 0.770958539 |
| ENSMUSG00000037568 | 1.077497675 | 0.698112512 |
| ENSMUSG00000037570 | 1.044101629 | 0.807247723 |
| ENSMUSG00000037572 | -1.111782922 | 0.650904225 |
| ENSMUSG00000037573 | 1.157146862 | 0.310630871 |
| ENSMUSG00000037577 | -1.014872356 | 0.98680668 |
| ENSMUSG00000037578 | 1.023743961 | 0.938939969 |
| ENSMUSG00000037579 | -1.528285537 | 0.028386648 |
| ENSMUSG00000037580 | -1.170608972 | 0.002297492 |
| ENSMUSG00000037583 | 1.059463094 | 0.608497229 |
| ENSMUSG00000037593 | -1.186284108 | 0.547016491 |
| ENSMUSG00000037594 | 1.073969763 | 0.555973705 |
| ENSMUSG00000037600 | -1.412281493 | 0.080934479 |
| ENSMUSG00000037601 | 1.00417042 | 1 |
| ENSMUSG00000037603 | 1 | |
| ENSMUSG00000037605 | 1.392842753 | 0.008476176 |
| ENSMUSG00000037606 | -1.005448017 | 1 |
| ENSMUSG00000037608 | -1.051030039 | 0.901784675 |
| ENSMUSG00000037610 | 1 | |
| ENSMUSG00000037613 | 1.134100411 | 0.476917435 |
| ENSMUSG00000037617 | -1.040832455 | 0.751020285 |
| ENSMUSG00000037621 | 1.407192907 | 0.000620329 |
| ENSMUSG00000037622 | 1.006328471 | 1 |
| ENSMUSG00000037624 | 1.102397205 | 0.081108012 |
| ENSMUSG00000037625 | 1 | |
| ENSMUSG00000037627 | -1.132666855 | 0.18851436 |
| ENSMUSG00000037628 | -1.165761459 | 0.091827709 |
| ENSMUSG00000037636 | 1.419259991 | 0.094105753 |
| ENSMUSG00000037638 | 1.136522187 | 0.215991492 |
| ENSMUSG00000037640 | -1.011087684 | 0.992856012 |
| ENSMUSG00000037643 | -1.094275683 | 0.732769483 |
| ENSMUSG00000037646 | -1.020116093 | 0.979002724 |
| ENSMUSG00000037649 | -1.035309784 | 0.734408673 |
| ENSMUSG00000037652 | -1.002728226 | 1 |
| ENSMUSG00000037653 | 1 | |
| ENSMUSG00000037656 | 1.156043688 | 0.157532985 |
| ENSMUSG00000037660 | 1 | |
| ENSMUSG00000037661 | 1.039818332 | 0.924766459 |
| ENSMUSG00000037664 | -1.951925285 | 0.000416272 |
| ENSMUSG00000037669 | 1.078053537 | 0.698112512 |
| ENSMUSG00000037674 | -1.104983186 | 0.662744367 |
| ENSMUSG00000037679 | 1.288967986 | 0.064932631 |
| ENSMUSG00000037681 | -1.384053158 | 0.035827026 |
| ENSMUSG00000037683 | 1 | |
| ENSMUSG00000037685 | -1.463985136 | 0.021901731 |
| ENSMUSG00000037686 | -1.612451966 | 0.00435101 |
| ENSMUSG00000037689 | 1 | |
| ENSMUSG00000037692 | 1.33347378 | 0.086855814 |
| ENSMUSG00000037697 | -1.202879344 | 0.121201239 |
| ENSMUSG00000037703 | -1.133790107 | 0.158361632 |
| ENSMUSG00000037705 | 1.182649099 | 0.608497229 |
| ENSMUSG00000037706 | 1.246262147 | 0.191586302 |
| ENSMUSG00000037708 | 1.136719427 | 0.608497229 |
| ENSMUSG00000037709 | 1.012559717 | 0.965978247 |
| ENSMUSG00000037710 | -1.048560635 | 0.784381159 |
| ENSMUSG00000037712 | 1.033782288 | 0.946641274 |
| ENSMUSG00000037716 | -1.369857598 | 0.010960146 |
| ENSMUSG00000037720 | 1.00146463 | 1 |
| ENSMUSG00000037722 | -1.1310224 | 0.570229056 |
| ENSMUSG00000037725 | 1.095547598 | 0.698112512 |
| ENSMUSG00000037727 | 1 | |
| ENSMUSG00000037730 | -1.073332206 | 0.698112512 |
| ENSMUSG00000037731 | -1.113837159 | 0.755340787 |

| | | |
|---|---|---|
| ENSMUSG00000037735 | -1.044273782 | 0.698112512 |
| ENSMUSG00000037736 | 1.660368536 | 0.000519315 |
| ENSMUSG00000037737 | -1.311159057 | 0.026040928 |
| ENSMUSG00000037738 | 1.040172434 | 0.916960199 |
| ENSMUSG00000037740 | -1.084673665 | 0.552737712 |
| ENSMUSG00000037742 | 1.041237615 | 0.979186916 |
| ENSMUSG00000037747 | -1.149959939 | 0.375822989 |
| ENSMUSG00000037750 | 1.028311631 | 0.806547092 |
| ENSMUSG00000037752 | -1.005917089 | 1 |
| ENSMUSG00000037754 | -1.043870538 | 0.825478278 |
| ENSMUSG00000037759 | -1.044273782 | 0.698112512 |
| ENSMUSG00000037761 | -1.144216644 | 0.453698036 |
| ENSMUSG00000037762 | -1.059853245 | 0.605159739 |
| ENSMUSG00000037771 | 1 | |
| ENSMUSG00000037772 | 1.07693474 | 0.578712437 |
| ENSMUSG00000037773 | -1.003138066 | 1 |
| ENSMUSG00000037780 | 1 | |
| ENSMUSG00000037784 | 1.074481931 | 0.613963375 |
| ENSMUSG00000037787 | 1.147189374 | 0.056047616 |
| ENSMUSG00000037788 | 1.110940074 | 0.608497229 |
| ENSMUSG00000037790 | 1 | |
| ENSMUSG00000037791 | 1.092612112 | 0.567496595 |
| ENSMUSG00000037795 | 1.04426917 | 0.861493556 |
| ENSMUSG00000037797 | 1 | |
| ENSMUSG00000037798 | 1 | |
| ENSMUSG00000037801 | 1.054108365 | 0.887303181 |
| ENSMUSG00000037805 | 1.051962577 | 0.818891358 |
| ENSMUSG00000037808 | -1.025853715 | 0.960579094 |
| ENSMUSG00000037813 | -1.075830317 | 0.782801761 |
| ENSMUSG00000037815 | 1.093863473 | 0.743794828 |
| ENSMUSG00000037816 | 1.005062836 | 1 |
| ENSMUSG00000037818 | -1.128219563 | 0.02533587 |
| ENSMUSG00000037820 | 1.593812581 | 0.0009722 |
| ENSMUSG00000037822 | 1.055539512 | 0.773508225 |
| ENSMUSG00000037824 | -1.088676729 | 0.608497229 |
| ENSMUSG00000037826 | -1.164005217 | 0.123633764 |
| ENSMUSG00000037827 | 1 | |
| ENSMUSG00000037833 | -1.083415624 | 0.62221251 |
| ENSMUSG00000037843 | 1 | |
| ENSMUSG00000037845 | -1.071629663 | 0.608497229 |
| ENSMUSG00000037846 | -1.07101301 | 0.540020537 |
| ENSMUSG00000037847 | 1.158042687 | 0.162436861 |
| ENSMUSG00000037849 | 1 | |
| ENSMUSG00000037851 | -1.024927224 | 0.984858767 |
| ENSMUSG00000037852 | 1.209125137 | 0.516815689 |
| ENSMUSG00000037855 | 1.170765346 | 0.063896413 |
| ENSMUSG00000037857 | 1.039816373 | 0.906416365 |
| ENSMUSG00000037860 | -1.029302237 | 0.962931288 |
| ENSMUSG00000037868 | 1.306660064 | 0.239824023 |
| ENSMUSG00000037872 | -1.071075483 | 0.698112512 |
| ENSMUSG00000037876 | 1.001774783 | 1 |
| ENSMUSG00000037884 | 1 | |
| ENSMUSG00000037885 | -1.052066514 | 0.749353361 |
| ENSMUSG00000037887 | -1.169532323 | 0.076089694 |
| ENSMUSG00000037890 | 1.037446408 | 0.828564184 |
| ENSMUSG00000037892 | -1.003601942 | 1 |
| ENSMUSG00000037894 | -1.042170633 | 0.903868918 |
| ENSMUSG00000037896 | 1.070437164 | 0.787313076 |
| ENSMUSG00000037902 | -1.572987765 | 0.007792328 |
| ENSMUSG00000037904 | 1.098448425 | 0.559095313 |
| ENSMUSG00000037905 | 1.087602867 | 0.612919974 |
| ENSMUSG00000037907 | 1.311605935 | 0.009480098 |
| ENSMUSG00000037910 | 1 | |

| | | |
|---|---|---|
| ENSMUSG00000037913 | 1 | |
| ENSMUSG00000037916 | 1.047207009 | 0.825048065 |
| ENSMUSG00000037921 | -1.512156141 | 0.007960323 |
| ENSMUSG00000037922 | 1.024693402 | 0.952057016 |
| ENSMUSG00000037924 | 1 | |
| ENSMUSG00000037926 | -1.015630252 | 0.986542466 |
| ENSMUSG00000037933 | -1.01070008 | 1 |
| ENSMUSG00000037935 | 1.003721482 | 1 |
| ENSMUSG00000037936 | -1.002979832 | 1 |
| ENSMUSG00000037938 | -1.016462084 | 0.945892181 |
| ENSMUSG00000037940 | 1.007954995 | 1 |
| ENSMUSG00000037942 | -1.043306199 | 0.79291651 |
| ENSMUSG00000037944 | -1.044273782 | 0.801443616 |
| ENSMUSG00000037946 | 1.137454068 | 0.575782278 |
| ENSMUSG00000037949 | -1.032397821 | 0.928180271 |
| ENSMUSG00000037953 | 1.024804142 | 0.940018654 |
| ENSMUSG00000037957 | 1.058101972 | 0.628392912 |
| ENSMUSG00000037958 | -1.043407995 | 0.860286104 |
| ENSMUSG00000037960 | 1.003133955 | 1 |
| ENSMUSG00000037962 | 1.033864273 | 0.775966113 |
| ENSMUSG00000037965 | -1.02058052 | 0.966570161 |
| ENSMUSG00000037966 | -1.110789362 | 0.374710101 |
| ENSMUSG00000037971 | -1.069474143 | 0.355765483 |
| ENSMUSG00000037972 | 1.14486355 | 0.420403679 |
| ENSMUSG00000037973 | 1 | |
| ENSMUSG00000037974 | -1.03962336 | 0.698112512 |
| ENSMUSG00000037977 | 1.128706988 | 0.668428121 |
| ENSMUSG00000037979 | 1.044609522 | 0.711581415 |
| ENSMUSG00000037982 | 1 | |
| ENSMUSG00000037984 | 1 | |
| ENSMUSG00000037989 | -1.93564545 | 0.007835576 |
| ENSMUSG00000037990 | 1.328868093 | 0.000994822 |
| ENSMUSG00000037991 | -1.008249438 | 0.966843099 |
| ENSMUSG00000037992 | 1.361560356 | 0.009448316 |
| ENSMUSG00000037993 | 1.044463936 | 0.81396159 |
| ENSMUSG00000037994 | 1.122071757 | 0.39980163 |
| ENSMUSG00000037995 | 1.650489921 | 0.001790637 |
| ENSMUSG00000037996 | -1 | 1 |
| ENSMUSG00000037997 | -1.016591146 | 0.973676317 |
| ENSMUSG00000037999 | -1.185163225 | 0.073228372 |
| ENSMUSG00000038000 | 1.017362573 | 0.954197515 |
| ENSMUSG00000038002 | 1.043199333 | 0.815293592 |
| ENSMUSG00000038005 | 1.016561096 | 0.98569451 |
| ENSMUSG00000038007 | -1.238640319 | 0.01024994 |
| ENSMUSG00000038009 | 1.044339361 | 0.698249621 |
| ENSMUSG00000038010 | 1.09245362 | 0.497089027 |
| ENSMUSG00000038011 | 1.011437135 | 1 |
| ENSMUSG00000038013 | -1.004709704 | 1 |
| ENSMUSG00000038014 | 1.036088337 | 0.936805205 |
| ENSMUSG00000038015 | -1.03962336 | 0.698112512 |
| ENSMUSG00000038020 | -1.041275151 | 0.628154042 |
| ENSMUSG00000038022 | 1.055410444 | 0.608497229 |
| ENSMUSG00000038023 | 1.079676733 | 0.70219856 |
| ENSMUSG00000038024 | 1.015932449 | 0.990807767 |
| ENSMUSG00000038025 | 1.018688816 | 0.958403618 |
| ENSMUSG00000038026 | -1 | 1 |
| ENSMUSG00000038028 | 1.032530854 | 0.818024118 |
| ENSMUSG00000038034 | 1.19825978 | 0.171521817 |
| ENSMUSG00000038037 | -1.110279054 | 0.659878362 |
| ENSMUSG00000038039 | -1.061446242 | 0.812255606 |
| ENSMUSG00000038042 | 1.094257956 | 0.425405236 |
| ENSMUSG00000038044 | 1 | |
| ENSMUSG00000038045 | 1.153807559 | 0.425575324 |

| | | |
|---|---|---|
| ENSMUSG00000038046 | -1.047201831 | 0.608497229 |
| ENSMUSG00000038047 | -1.001920334 | 1 |
| ENSMUSG00000038048 | 1 | |
| ENSMUSG00000038055 | 1.016828901 | 0.933683189 |
| ENSMUSG00000038056 | 1.0525948 | 0.793885968 |
| ENSMUSG00000038057 | 1.030541003 | 0.974984587 |
| ENSMUSG00000038058 | 1.268365163 | 0.002705352 |
| ENSMUSG00000038059 | -1.077577651 | 0.663887079 |
| ENSMUSG00000038060 | -1.279081331 | 0.18851436 |
| ENSMUSG00000038064 | -1.089105072 | 0.817310293 |
| ENSMUSG00000038065 | 1.164222794 | 0.044619484 |
| ENSMUSG00000038067 | 1.074221595 | 0.790772938 |
| ENSMUSG00000038068 | -1.279690019 | 0.018220428 |
| ENSMUSG00000038069 | 1.062012306 | 0.726684101 |
| ENSMUSG00000038070 | 1.014485685 | 0.984826086 |
| ENSMUSG00000038071 | 1 | |
| ENSMUSG00000038072 | -1.07739809 | 0.675035111 |
| ENSMUSG00000038074 | 1.039747297 | 0.877826334 |
| ENSMUSG00000038077 | -1.376791404 | 0.019868827 |
| ENSMUSG00000038079 | -1.06947936 | 0.702897175 |
| ENSMUSG00000038080 | -1.186338784 | 0.223461269 |
| ENSMUSG00000038084 | 1.024151389 | 0.970771742 |
| ENSMUSG00000038085 | 1.027500113 | 0.894828773 |
| ENSMUSG00000038086 | 2.150587097 | 0.008688055 |
| ENSMUSG00000038092 | 1 | |
| ENSMUSG00000038094 | 1 | |
| ENSMUSG00000038095 | 1.093709961 | 0.750182067 |
| ENSMUSG00000038102 | -1.00882234 | 1 |
| ENSMUSG00000038112 | -1.043306199 | 0.701745693 |
| ENSMUSG00000038115 | -1.142727245 | 0.202428693 |
| ENSMUSG00000038116 | -1.024317621 | 0.95829726 |
| ENSMUSG00000038119 | -1.061993367 | 0.838318712 |
| ENSMUSG00000038121 | 1.038352995 | 0.90047549 |
| ENSMUSG00000038122 | -1.025736354 | 0.920128007 |
| ENSMUSG00000038126 | 1.042052653 | 0.881421285 |
| ENSMUSG00000038127 | 1.003481026 | 1 |
| ENSMUSG00000038128 | -1.149520448 | 0.125975833 |
| ENSMUSG00000038132 | 1.257068502 | 0.477417651 |
| ENSMUSG00000038135 | 1.015486245 | 0.972980766 |
| ENSMUSG00000038141 | -1.03338546 | 0.895127848 |
| ENSMUSG00000038143 | -2.200713622 | 0.001384356 |
| ENSMUSG00000038145 | 1.020091352 | 0.834072451 |
| ENSMUSG00000038146 | 2.100033929 | 0.000603481 |
| ENSMUSG00000038147 | 1.063532453 | 0.725189744 |
| ENSMUSG00000038148 | 1 | |
| ENSMUSG00000038150 | 1.051123029 | 0.751828376 |
| ENSMUSG00000038151 | -1.044273782 | 0.790851299 |
| ENSMUSG00000038152 | 1.257932827 | 0.228706738 |
| ENSMUSG00000038155 | -1 | 1 |
| ENSMUSG00000038156 | -1.323702596 | 0.087449305 |
| ENSMUSG00000038160 | 1.034190251 | 0.895623306 |
| ENSMUSG00000038165 | 1 | |
| ENSMUSG00000038167 | 1.332190621 | 0.214148278 |
| ENSMUSG00000038168 | -1.012227232 | 0.938939969 |
| ENSMUSG00000038170 | 1.190151465 | 0.34500751 |
| ENSMUSG00000038172 | 1.093090231 | 0.621604894 |
| ENSMUSG00000038173 | 1 | |
| ENSMUSG00000038174 | -1.051188666 | 0.801620398 |
| ENSMUSG00000038175 | -1.427965442 | 0.002225593 |
| ENSMUSG00000038178 | 1.138307609 | 0.194996487 |
| ENSMUSG00000038179 | 1 | |
| ENSMUSG00000038180 | -1.085688085 | 0.608497229 |
| ENSMUSG00000038181 | 1.067468618 | 0.506802808 |

| | | |
|---|---|---|
| ENSMUSG00000038187 | 1.128342185 | 0.493659452 |
| ENSMUSG00000038188 | 1.145838633 | 0.408631273 |
| ENSMUSG00000038192 | -1.03962336 | 0.698112512 |
| ENSMUSG00000038193 | -1.044273782 | 0.88284878 |
| ENSMUSG00000038194 | 1.028323353 | 0.936340683 |
| ENSMUSG00000038195 | -1.079083512 | 0.819541918 |
| ENSMUSG00000038199 | 1 | |
| ENSMUSG00000038201 | -1.113515092 | 0.608497229 |
| ENSMUSG00000038203 | 1.030415139 | 0.952462131 |
| ENSMUSG00000038204 | 1 | |
| ENSMUSG00000038205 | 1.181907807 | 0.242806824 |
| ENSMUSG00000038206 | -1.007064398 | 1 |
| ENSMUSG00000038208 | -1.01927584 | 0.874869207 |
| ENSMUSG00000038209 | -1 | 1 |
| ENSMUSG00000038210 | -1.08749201 | 0.525132023 |
| ENSMUSG00000038212 | -1.095660467 | 0.698112512 |
| ENSMUSG00000038213 | -1.156911809 | 0.075612848 |
| ENSMUSG00000038214 | 1.085327359 | 0.444718012 |
| ENSMUSG00000038215 | 1.025765599 | 0.920803649 |
| ENSMUSG00000038216 | 1 | |
| ENSMUSG00000038217 | -1.044433043 | 0.77122853 |
| ENSMUSG00000038218 | 1 | |
| ENSMUSG00000038224 | 1.25954694 | 0.532369657 |
| ENSMUSG00000038225 | 1.085775185 | 0.331843752 |
| ENSMUSG00000038227 | 1.509306228 | 0.004636749 |
| ENSMUSG00000038233 | -1.126595046 | 0.608497229 |
| ENSMUSG00000038235 | -1.26551636 | 0.215991492 |
| ENSMUSG00000038236 | 1.22048147 | 0.012849032 |
| ENSMUSG00000038239 | 1.046403186 | 0.924324458 |
| ENSMUSG00000038240 | -1.05510904 | 0.667576553 |
| ENSMUSG00000038241 | -1.005018446 | 1 |
| ENSMUSG00000038242 | -1.023769022 | 0.975580864 |
| ENSMUSG00000038244 | 1.530945502 | 0.015278471 |
| ENSMUSG00000038246 | 1 | |
| ENSMUSG00000038248 | -1.906247696 | 0.032862743 |
| ENSMUSG00000038250 | 1.006296769 | 1 |
| ENSMUSG00000038252 | 1.123903259 | 0.629924473 |
| ENSMUSG00000038253 | 1.253294197 | 0.044259536 |
| ENSMUSG00000038255 | 1 | |
| ENSMUSG00000038256 | 1.072288052 | 0.608497229 |
| ENSMUSG00000038257 | -1.044273782 | 0.790851299 |
| ENSMUSG00000038259 | -1.072259793 | 0.714596508 |
| ENSMUSG00000038260 | 1.145974077 | 0.381225514 |
| ENSMUSG00000038264 | -1.214029022 | 0.057574416 |
| ENSMUSG00000038267 | -1.086225005 | 0.576169279 |
| ENSMUSG00000038268 | -1.18739113 | 0.106894562 |
| ENSMUSG00000038271 | -1.048150656 | 0.630771413 |
| ENSMUSG00000038274 | 1.1714608 | 0.313140716 |
| ENSMUSG00000038276 | 1.160401087 | 0.234269835 |
| ENSMUSG00000038279 | 1.061290316 | 0.752472633 |
| ENSMUSG00000038280 | -1.009240222 | 1 |
| ENSMUSG00000038286 | -1.001821344 | 1 |
| ENSMUSG00000038290 | -1.006379664 | 1 |
| ENSMUSG00000038291 | -1.06399426 | 0.803391392 |
| ENSMUSG00000038292 | -1.130229437 | 0.708080874 |
| ENSMUSG00000038295 | 1.076415212 | 0.647299437 |
| ENSMUSG00000038296 | 1.836158156 | 0.003301399 |
| ENSMUSG00000038298 | -1.053549062 | 0.769745556 |
| ENSMUSG00000038299 | 1.042872849 | 0.89632135 |
| ENSMUSG00000038300 | 1 | |
| ENSMUSG00000038301 | 1.015287989 | 0.986431796 |
| ENSMUSG00000038302 | -1.032819695 | 0.847756749 |
| ENSMUSG00000038304 | -1.331252382 | 0.043138582 |

| | | |
|---|---|---|
| ENSMUSG00000038305 | -1.914163367 | 0.007382681 |
| ENSMUSG00000038311 | 1.132639128 | 0.608497229 |
| ENSMUSG00000038312 | -1.110561219 | 0.455417569 |
| ENSMUSG00000038319 | 2.200500073 | 0.006094015 |
| ENSMUSG00000038323 | -1.055102095 | 0.750343426 |
| ENSMUSG00000038324 | -1.098938757 | 0.698112512 |
| ENSMUSG00000038327 | 1.177765487 | 0.276639872 |
| ENSMUSG00000038330 | 1 | |
| ENSMUSG00000038331 | -1.081920898 | 0.556411689 |
| ENSMUSG00000038332 | -1.07109279 | 0.737954782 |
| ENSMUSG00000038335 | -1.025129496 | 0.962382822 |
| ENSMUSG00000038342 | 1.037996005 | 0.886747391 |
| ENSMUSG00000038344 | -1.178027902 | 0.327642404 |
| ENSMUSG00000038346 | 1.034336689 | 0.911752755 |
| ENSMUSG00000038347 | 1.03359863 | 0.724290685 |
| ENSMUSG00000038349 | 1.413369846 | 0.003466971 |
| ENSMUSG00000038351 | 1.150341531 | 0.043183444 |
| ENSMUSG00000038352 | 1.030475717 | 0.903464611 |
| ENSMUSG00000038354 | -1.739517289 | 0.019224376 |
| ENSMUSG00000038357 | 1.040172434 | 0.928585511 |
| ENSMUSG00000038365 | 1.119203485 | 0.440702663 |
| ENSMUSG00000038366 | 1.130858 | 0.645934663 |
| ENSMUSG00000038368 | 1.062947114 | 0.647277306 |
| ENSMUSG00000038369 | -1.037684811 | 0.879253209 |
| ENSMUSG00000038370 | -1.711069441 | 0.004504222 |
| ENSMUSG00000038371 | 1.144003064 | 0.455298078 |
| ENSMUSG00000038372 | -1.063122275 | 0.706014595 |
| ENSMUSG00000038374 | -1.124510919 | 0.595433188 |
| ENSMUSG00000038375 | -1.049626222 | 0.797643564 |
| ENSMUSG00000038379 | -1.090646451 | 0.698112512 |
| ENSMUSG00000038383 | -1.06923455 | 0.677534336 |
| ENSMUSG00000038384 | 1.1259088 | 0.08148224 |
| ENSMUSG00000038387 | 1.007337272 | 1 |
| ENSMUSG00000038388 | 1.09283415 | 0.658943194 |
| ENSMUSG00000038390 | -1.047162921 | 0.698112512 |
| ENSMUSG00000038393 | 1.104409583 | 0.578654776 |
| ENSMUSG00000038398 | 1.0506605 | 0.76868134 |
| ENSMUSG00000038400 | 1.261122415 | 0.085068744 |
| ENSMUSG00000038402 | -1.103981064 | 0.532451763 |
| ENSMUSG00000038403 | -1.161686509 | 0.607661442 |
| ENSMUSG00000038406 | 1.115991831 | 0.387036025 |
| ENSMUSG00000038408 | -1.038469959 | 0.838359329 |
| ENSMUSG00000038412 | -1.089336304 | 0.611938648 |
| ENSMUSG00000038415 | 1.136227452 | 0.698112512 |
| ENSMUSG00000038416 | 1.148438868 | 0.608497229 |
| ENSMUSG00000038417 | -1.054410283 | 0.828013032 |
| ENSMUSG00000038418 | 1.107256927 | 0.264062736 |
| ENSMUSG00000038421 | 1 | |
| ENSMUSG00000038422 | 1.062428844 | 0.507278558 |
| ENSMUSG00000038425 | 1.079791832 | 0.163630982 |
| ENSMUSG00000038429 | 1.042811519 | 0.87505994 |
| ENSMUSG00000038437 | 1.11157324 | 0.202881099 |
| ENSMUSG00000038446 | -1.023333161 | 0.97045957 |
| ENSMUSG00000038451 | 1.178384429 | 0.040397212 |
| ENSMUSG00000038453 | -1.334383734 | 0.029842389 |
| ENSMUSG00000038456 | 1.194147827 | 0.219598783 |
| ENSMUSG00000038457 | 1 | |
| ENSMUSG00000038459 | 1.026898647 | 0.93855667 |
| ENSMUSG00000038462 | -1.062121003 | 0.814089245 |
| ENSMUSG00000038463 | 1.731489434 | 0.001267143 |
| ENSMUSG00000038467 | -1.09431016 | 0.638758774 |
| ENSMUSG00000038473 | -1.130933812 | 0.74527374 |
| ENSMUSG00000038481 | 1.005714454 | 1 |

| | | |
|---|---|---|
| ENSMUSG00000038482 | 1.108517403 | 0.698112512 |
| ENSMUSG00000038485 | -1.046347711 | 0.830576988 |
| ENSMUSG00000038486 | -1.028716136 | 0.751234476 |
| ENSMUSG00000038489 | -1.050202042 | 0.629803844 |
| ENSMUSG00000038491 | -1 | 1 |
| ENSMUSG00000038495 | -1.040848351 | 0.901216307 |
| ENSMUSG00000038496 | -1 | 1 |
| ENSMUSG00000038497 | 1.07322443 | 0.670176133 |
| ENSMUSG00000038498 | 1 | |
| ENSMUSG00000038500 | -1.043980612 | 0.698112512 |
| ENSMUSG00000038502 | 1.171870576 | 0.162732522 |
| ENSMUSG00000038503 | 1.054793816 | 0.795837467 |
| ENSMUSG00000038506 | 1.075753796 | 0.625879938 |
| ENSMUSG00000038507 | -1.160231404 | 0.378017455 |
| ENSMUSG00000038508 | 1.398148693 | 0.038433268 |
| ENSMUSG00000038510 | -1.049773634 | 0.851353899 |
| ENSMUSG00000038515 | -1.199991616 | 0.015462827 |
| ENSMUSG00000038517 | 1.102055316 | 0.698112512 |
| ENSMUSG00000038518 | 1.096268488 | 0.608497229 |
| ENSMUSG00000038520 | 1.184230594 | 0.070586955 |
| ENSMUSG00000038521 | 1.583844087 | 0.001717312 |
| ENSMUSG00000038522 | -1 | 1 |
| ENSMUSG00000038523 | 1.347277487 | 0.077304067 |
| ENSMUSG00000038524 | 1.111976001 | 0.559354144 |
| ENSMUSG00000038525 | 1.040988281 | 0.849863703 |
| ENSMUSG00000038526 | 1.160019552 | 0.464062002 |
| ENSMUSG00000038527 | 1.299401565 | 0.001548396 |
| ENSMUSG00000038528 | 1.070429626 | 0.803072249 |
| ENSMUSG00000038530 | -1.12600168 | 0.25067776 |
| ENSMUSG00000038533 | -1.047732265 | 0.83493212 |
| ENSMUSG00000038534 | -1.086457459 | 0.556582772 |
| ENSMUSG00000038535 | -1.060238088 | 0.742249057 |
| ENSMUSG00000038537 | 1 | |
| ENSMUSG00000038538 | 1.021233392 | 0.979212363 |
| ENSMUSG00000038539 | 1.35914323 | 0.008428749 |
| ENSMUSG00000038540 | -1.000736654 | 1 |
| ENSMUSG00000038541 | -1.043306199 | 0.798487595 |
| ENSMUSG00000038542 | 1.06952902 | 0.698112512 |
| ENSMUSG00000038543 | 1.085684128 | 0.512218565 |
| ENSMUSG00000038544 | 1.038986723 | 0.837947689 |
| ENSMUSG00000038545 | 1.075858074 | 0.608497229 |
| ENSMUSG00000038546 | -1.043352463 | 0.876686173 |
| ENSMUSG00000038550 | 1.07107705 | 0.698112512 |
| ENSMUSG00000038552 | -1.021566577 | 0.902345046 |
| ENSMUSG00000038555 | 1.757067154 | 0.007695667 |
| ENSMUSG00000038560 | 1 | |
| ENSMUSG00000038563 | 1.061429316 | 0.650816989 |
| ENSMUSG00000038564 | 1.049773569 | 0.753629702 |
| ENSMUSG00000038567 | 1.095872691 | 0.657060635 |
| ENSMUSG00000038569 | -1.056928104 | 0.837516572 |
| ENSMUSG00000038570 | -1.162466371 | 0.464320068 |
| ENSMUSG00000038572 | 1 | |
| ENSMUSG00000038576 | -1.29398402 | 0.013472992 |
| ENSMUSG00000038578 | -1.16798883 | 0.215991492 |
| ENSMUSG00000038580 | -1.043306199 | 0.698112512 |
| ENSMUSG00000038582 | 1.102360962 | 0.561534005 |
| ENSMUSG00000038583 | -1.071075483 | 0.698112512 |
| ENSMUSG00000038587 | 1.113894179 | 0.634149843 |
| ENSMUSG00000038591 | 1.230234402 | 0.543409801 |
| ENSMUSG00000038593 | 1.092625694 | 0.476577776 |
| ENSMUSG00000038594 | -1.113933695 | 0.326678352 |
| ENSMUSG00000038598 | -1.028459293 | 0.978053547 |
| ENSMUSG00000038599 | 1 | |

| | | |
|---|---|---|
| ENSMUSG00000038600 | -1.242607144 | 0.233269975 |
| ENSMUSG00000038602 | -1.298318725 | 0.250477583 |
| ENSMUSG00000038604 | 1.024023544 | 0.942691576 |
| ENSMUSG00000038605 | 1.017323179 | 0.938670327 |
| ENSMUSG00000038607 | -1.101557534 | 0.721403818 |
| ENSMUSG00000038608 | -1.096137681 | 0.564249854 |
| ENSMUSG00000038611 | 1.063524366 | 0.698112512 |
| ENSMUSG00000038612 | -1.085336782 | 0.778372272 |
| ENSMUSG00000038615 | 1.143829605 | 0.608497229 |
| ENSMUSG00000038618 | 1.05762005 | 0.622624759 |
| ENSMUSG00000038619 | -1.027251898 | 0.952875614 |
| ENSMUSG00000038622 | -1.028785354 | 0.931480186 |
| ENSMUSG00000038623 | -1.208567643 | 0.109112928 |
| ENSMUSG00000038624 | 1.059463094 | 0.608497229 |
| ENSMUSG00000038628 | -1.059588719 | 0.730893269 |
| ENSMUSG00000038630 | -1.031780031 | 0.830576988 |
| ENSMUSG00000038633 | -1.02206907 | 0.983851582 |
| ENSMUSG00000038637 | -1.029274341 | 0.962368837 |
| ENSMUSG00000038641 | 1.132639128 | 0.398767938 |
| ENSMUSG00000038642 | -1.044273782 | 0.698112512 |
| ENSMUSG00000038644 | 1.152593978 | 0.36525262 |
| ENSMUSG00000038646 | 1.081888386 | 0.668054685 |
| ENSMUSG00000038648 | 1.125252345 | 0.583420921 |
| ENSMUSG00000038650 | 1.102820169 | 0.637788928 |
| ENSMUSG00000038651 | 1 | |
| ENSMUSG00000038656 | 1 | |
| ENSMUSG00000038658 | -1.002774792 | 1 |
| ENSMUSG00000038663 | 1.059463094 | 0.608497229 |
| ENSMUSG00000038664 | -1.03263676 | 0.953543356 |
| ENSMUSG00000038665 | 1.346809975 | 0.095721539 |
| ENSMUSG00000038668 | 1.077708153 | 0.608497229 |
| ENSMUSG00000038670 | -1.132337147 | 0.720282888 |
| ENSMUSG00000038671 | -1.056422189 | 0.740545161 |
| ENSMUSG00000038676 | 1 | |
| ENSMUSG00000038677 | -1.105516487 | 0.503745541 |
| ENSMUSG00000038679 | 1.10974272 | 0.608497229 |
| ENSMUSG00000038683 | -1.020878473 | 0.961788219 |
| ENSMUSG00000038685 | -1.090495829 | 0.676185514 |
| ENSMUSG00000038690 | -1.027417446 | 0.937535342 |
| ENSMUSG00000038691 | 1 | |
| ENSMUSG00000038692 | 1.227950183 | 0.005452662 |
| ENSMUSG00000038695 | 1.089354878 | 0.70299173 |
| ENSMUSG00000038696 | 1.077451861 | 0.643426194 |
| ENSMUSG00000038697 | 1.020166026 | 0.958403618 |
| ENSMUSG00000038700 | 1.05330935 | 0.608497229 |
| ENSMUSG00000038702 | 1.051000158 | 0.734674233 |
| ENSMUSG00000038704 | -1 | 1 |
| ENSMUSG00000038705 | -1.054914057 | 0.710350017 |
| ENSMUSG00000038708 | 1.030374489 | 0.933031586 |
| ENSMUSG00000038709 | 1 | |
| ENSMUSG00000038712 | -1.131435362 | 0.38227559 |
| ENSMUSG00000038717 | 1.033109741 | 0.895158182 |
| ENSMUSG00000038718 | -1.032587478 | 0.873012595 |
| ENSMUSG00000038721 | 1.158862884 | 0.122237883 |
| ENSMUSG00000038722 | -1.066477688 | 0.795285331 |
| ENSMUSG00000038725 | 1.2713256 | 0.169690282 |
| ENSMUSG00000038729 | 1.015074177 | 1 |
| ENSMUSG00000038732 | 1.043906763 | 0.866784751 |
| ENSMUSG00000038733 | -1.059645908 | 0.87131415 |
| ENSMUSG00000038736 | 1.045896231 | 0.878773892 |
| ENSMUSG00000038738 | -1.020155149 | 0.988424085 |
| ENSMUSG00000038740 | 1.092424764 | 0.534279144 |
| ENSMUSG00000038742 | 1.172324764 | 0.741226067 |

| | | |
|---|---|---|
| ENSMUSG00000038745 | 1 | |
| ENSMUSG00000038750 | 1 | |
| ENSMUSG00000038751 | 1.019083579 | 0.950378392 |
| ENSMUSG00000038754 | -1.110196018 | 0.529674452 |
| ENSMUSG00000038756 | 1.153124877 | 0.077747916 |
| ENSMUSG00000038759 | 1.050735783 | 0.902174523 |
| ENSMUSG00000038760 | -1 | 1 |
| ENSMUSG00000038762 | 1.002212466 | 1 |
| ENSMUSG00000038763 | 1.039597911 | 0.906480064 |
| ENSMUSG00000038764 | -1.008840651 | 0.994656991 |
| ENSMUSG00000038765 | -1.915682503 | 0.0119546 |
| ENSMUSG00000038766 | -1.068413728 | 0.810829358 |
| ENSMUSG00000038768 | -1.023769022 | 0.954761885 |
| ENSMUSG00000038770 | 1 | |
| ENSMUSG00000038773 | 1.03378286 | 0.918610481 |
| ENSMUSG00000038774 | 1.061585361 | 0.821942765 |
| ENSMUSG00000038775 | 1.329750096 | 0.009738651 |
| ENSMUSG00000038776 | 1.303196832 | 0.090224848 |
| ENSMUSG00000038777 | -1.276162492 | 0.014331568 |
| ENSMUSG00000038780 | 1.089748805 | 0.684519242 |
| ENSMUSG00000038781 | -1.1539626 | 0.088689988 |
| ENSMUSG00000038782 | -1.063350673 | 0.752472633 |
| ENSMUSG00000038784 | -1.026876291 | 0.939017294 |
| ENSMUSG00000038791 | 1 | |
| ENSMUSG00000038792 | 1 | |
| ENSMUSG00000038793 | -1.024700073 | 0.923467112 |
| ENSMUSG00000038797 | 1.197119339 | 0.520518871 |
| ENSMUSG00000038801 | 1 | |
| ENSMUSG00000038803 | 1.195075423 | 0.058312562 |
| ENSMUSG00000038805 | -2.372128341 | 0.000208442 |
| ENSMUSG00000038806 | 1.000113284 | 1 |
| ENSMUSG00000038807 | 1.187255544 | 0.052350161 |
| ENSMUSG00000038811 | 1.388605261 | 0.011125946 |
| ENSMUSG00000038812 | 1.022622187 | 0.801439557 |
| ENSMUSG00000038816 | -1.180533247 | 0.324261026 |
| ENSMUSG00000038822 | 1.006086319 | 1 |
| ENSMUSG00000038827 | -1.033602001 | 0.896299552 |
| ENSMUSG00000038828 | 1.071253356 | 0.666923506 |
| ENSMUSG00000038831 | -1.156734019 | 0.285915732 |
| ENSMUSG00000038836 | 1.108172945 | 0.034131735 |
| ENSMUSG00000038838 | 1.140205331 | 0.355031323 |
| ENSMUSG00000038840 | 1 | |
| ENSMUSG00000038843 | 1.483441265 | 0.001687496 |
| ENSMUSG00000038844 | 1.040113378 | 0.854356653 |
| ENSMUSG00000038845 | -1.040172891 | 0.87128347 |
| ENSMUSG00000038848 | -1.043899383 | 0.879458055 |
| ENSMUSG00000038855 | 1.005333402 | 1 |
| ENSMUSG00000038859 | -1.023849601 | 0.857392806 |
| ENSMUSG00000038860 | -1.131434675 | 0.189722845 |
| ENSMUSG00000038861 | -1.025892768 | 0.949218699 |
| ENSMUSG00000038866 | 1.042152857 | 0.805243413 |
| ENSMUSG00000038871 | 1.005153691 | 1 |
| ENSMUSG00000038872 | 1.174146118 | 0.070586955 |
| ENSMUSG00000038876 | 1.052594399 | 0.784729033 |
| ENSMUSG00000038879 | -1.065566524 | 0.665609688 |
| ENSMUSG00000038880 | -1.052552909 | 0.952119819 |
| ENSMUSG00000038883 | 1 | |
| ENSMUSG00000038884 | -1.473282812 | 0.035171269 |
| ENSMUSG00000038886 | 1.160273923 | 0.268545131 |
| ENSMUSG00000038888 | 1.064770224 | 0.117564123 |
| ENSMUSG00000038891 | 1 | |
| ENSMUSG00000038893 | -1.145975771 | 0.149583343 |
| ENSMUSG00000038894 | 1.334364536 | 0.028140705 |

| | | |
|---|---|---|
| ENSMUSG00000038895 | 1.090110638 | 0.499572952 |
| ENSMUSG00000038900 | 1.080851797 | 0.734950747 |
| ENSMUSG00000038902 | 1.021780902 | 0.965659325 |
| ENSMUSG00000038903 | 1.266455025 | 0.386428235 |
| ENSMUSG00000038909 | -1.011438481 | 1 |
| ENSMUSG00000038910 | -1.021790387 | 0.870427333 |
| ENSMUSG00000038914 | -1.012439743 | 1 |
| ENSMUSG00000038916 | 1.70045117 | 0.00574762 |
| ENSMUSG00000038917 | 1.245333044 | 0.332304017 |
| ENSMUSG00000038925 | -1.019412063 | 0.988424085 |
| ENSMUSG00000038930 | 1.148524563 | 0.103252121 |
| ENSMUSG00000038932 | -1.464241444 | 0.032461938 |
| ENSMUSG00000038936 | -1.100071286 | 0.643967866 |
| ENSMUSG00000038943 | -1.003290062 | 1 |
| ENSMUSG00000038949 | 1.022968538 | 0.91674081 |
| ENSMUSG00000038954 | 1.081205164 | 0.492010384 |
| ENSMUSG00000038957 | -1.039323952 | 0.867734795 |
| ENSMUSG00000038963 | -1.424786314 | 0.051327493 |
| ENSMUSG00000038965 | -1.056873284 | 0.832839286 |
| ENSMUSG00000038967 | -1.066937663 | 0.544351697 |
| ENSMUSG00000038968 | 1 | |
| ENSMUSG00000038970 | 1.007384471 | 1 |
| ENSMUSG00000038973 | -1 | 1 |
| ENSMUSG00000038975 | -1.00037997 | 1 |
| ENSMUSG00000038976 | 1.136434287 | 0.348326169 |
| ENSMUSG00000038980 | 1 | |
| ENSMUSG00000038982 | -1.069840836 | 0.473406495 |
| ENSMUSG00000038984 | 1.249659785 | 0.024850105 |
| ENSMUSG00000038987 | 1.104410004 | 0.133227444 |
| ENSMUSG00000038990 | -1.099106762 | 0.578221424 |
| ENSMUSG00000038991 | 1.039212418 | 0.934760809 |
| ENSMUSG00000038994 | -1.024530911 | 0.975108927 |
| ENSMUSG00000038997 | -1.014405749 | 1 |
| ENSMUSG00000039000 | 1.044072972 | 0.915514595 |
| ENSMUSG00000039001 | 1.058606895 | 0.72727536 |
| ENSMUSG00000039004 | -1.601829193 | 0.009566147 |
| ENSMUSG00000039005 | 1.084011251 | 0.608497229 |
| ENSMUSG00000039007 | 1.305640704 | 0.051956493 |
| ENSMUSG00000039013 | 1.200936955 | 0.247040254 |
| ENSMUSG00000039016 | -1.036956829 | 0.8171763 |
| ENSMUSG00000039018 | 1.072569924 | 0.608497229 |
| ENSMUSG00000039021 | -1.09378253 | 0.698112512 |
| ENSMUSG00000039031 | -1.040763231 | 0.867068766 |
| ENSMUSG00000039032 | 1.04424822 | 0.85864022 |
| ENSMUSG00000039033 | -1.017921033 | 0.949812939 |
| ENSMUSG00000039037 | -1.133231767 | 0.608497229 |
| ENSMUSG00000039041 | -1.009603259 | 0.97573893 |
| ENSMUSG00000039043 | -1.029521488 | 0.927355611 |
| ENSMUSG00000039046 | 1.01715402 | 0.985522133 |
| ENSMUSG00000039047 | -1.023712242 | 0.965978247 |
| ENSMUSG00000039048 | -1.071353956 | 0.677779462 |
| ENSMUSG00000039050 | -1.042277652 | 0.808074244 |
| ENSMUSG00000039055 | -1.018191507 | 0.956972448 |
| ENSMUSG00000039057 | -1.127175794 | 0.602343211 |
| ENSMUSG00000039058 | 1.10618753 | 0.335753246 |
| ENSMUSG00000039059 | 1 | |
| ENSMUSG00000039062 | 1.016320596 | 0.965058746 |
| ENSMUSG00000039063 | 1.09252871 | 0.042078126 |
| ENSMUSG00000039065 | 1.02590859 | 0.8461706 |
| ENSMUSG00000039067 | -1.037060679 | 0.936995747 |
| ENSMUSG00000039068 | -1.020256318 | 0.985869746 |
| ENSMUSG00000039069 | 1.036846035 | 0.424640946 |
| ENSMUSG00000039070 | -1 | 1 |

| | | |
|---|---|---|
| ENSMUSG00000039079 | 1.030415139 | 0.949812939 |
| ENSMUSG00000039081 | 1.143186414 | 0.091808658 |
| ENSMUSG00000039084 | 1.122598145 | 0.608497229 |
| ENSMUSG00000039086 | 1.005474134 | 1 |
| ENSMUSG00000039087 | 1.088686974 | 0.414925337 |
| ENSMUSG00000039089 | 1.088246001 | 0.608497229 |
| ENSMUSG00000039092 | 1 | |
| ENSMUSG00000039095 | -1.524048448 | 0.004569693 |
| ENSMUSG00000039096 | 1.061406274 | 0.467052335 |
| ENSMUSG00000039097 | -1.051577863 | 0.885184589 |
| ENSMUSG00000039098 | 1 | |
| ENSMUSG00000039099 | -1.270829139 | 0.275263998 |
| ENSMUSG00000039100 | 1.071432228 | 0.803641195 |
| ENSMUSG00000039103 | 1.184381292 | 0.308722709 |
| ENSMUSG00000039105 | 1.159126285 | 0.424109601 |
| ENSMUSG00000039106 | 1 | |
| ENSMUSG00000039108 | 1.046164382 | 0.698112512 |
| ENSMUSG00000039109 | -1 | 1 |
| ENSMUSG00000039110 | -1.495560523 | 0.024311662 |
| ENSMUSG00000039114 | 1.615714468 | 0.009361701 |
| ENSMUSG00000039115 | 1.019455993 | 0.903464611 |
| ENSMUSG00000039116 | 1.312662549 | 0.0864806 |
| ENSMUSG00000039117 | -1.035964511 | 0.822153525 |
| ENSMUSG00000039126 | 1.725158628 | 0.006778907 |
| ENSMUSG00000039128 | -1.016154805 | 0.998981421 |
| ENSMUSG00000039130 | -1.028080291 | 0.925214847 |
| ENSMUSG00000039131 | 1.382921322 | 0.012377342 |
| ENSMUSG00000039133 | -1.15957854 | 0.608497229 |
| ENSMUSG00000039137 | 1.359350702 | 0.022882015 |
| ENSMUSG00000039138 | 1 | |
| ENSMUSG00000039145 | 1.159714135 | 0.36554144 |
| ENSMUSG00000039146 | -1.044273782 | 0.790851299 |
| ENSMUSG00000039148 | 1.029282035 | 0.895965732 |
| ENSMUSG00000039153 | 1.128597663 | 0.5212581 |
| ENSMUSG00000039154 | -1.29355546 | 0.079761782 |
| ENSMUSG00000039155 | -1.437925156 | 0.001127478 |
| ENSMUSG00000039156 | 1.052380748 | 0.728667795 |
| ENSMUSG00000039157 | -1.000542087 | 1 |
| ENSMUSG00000039158 | 1.396974025 | 0.020186653 |
| ENSMUSG00000039159 | 1.106356198 | 0.608497229 |
| ENSMUSG00000039163 | -1.012031606 | 0.987056723 |
| ENSMUSG00000039164 | 1.032248053 | 0.698112512 |
| ENSMUSG00000039166 | 1.062260435 | 0.678897747 |
| ENSMUSG00000039167 | -1.186404996 | 0.486930276 |
| ENSMUSG00000039168 | 1.287101499 | 0.107860597 |
| ENSMUSG00000039174 | 1 | |
| ENSMUSG00000039176 | -1.026569755 | 0.856915299 |
| ENSMUSG00000039178 | 1.01005149 | 1 |
| ENSMUSG00000039179 | 1 | |
| ENSMUSG00000039182 | 1.106634126 | 0.37050419 |
| ENSMUSG00000039183 | 1.045894687 | 0.705141457 |
| ENSMUSG00000039187 | -1.08546473 | 0.69429846 |
| ENSMUSG00000039191 | 1.021065249 | 0.983149736 |
| ENSMUSG00000039193 | 1.116785864 | 0.578221424 |
| ENSMUSG00000039194 | -1.177546921 | 0.381329895 |
| ENSMUSG00000039195 | -1.146232647 | 0.534291186 |
| ENSMUSG00000039196 | 1 | |
| ENSMUSG00000039197 | -1.190539337 | 0.46637329 |
| ENSMUSG00000039198 | 1 | |
| ENSMUSG00000039199 | 1.002187898 | 1 |
| ENSMUSG00000039200 | -1.329230923 | 0.061185687 |
| ENSMUSG00000039201 | 1.016535073 | 0.924587329 |
| ENSMUSG00000039202 | 1.055105927 | 0.790851299 |

| | | |
|---|---|---|
| ENSMUSG00000039205 | 1.036455319 | 0.801199277 |
| ENSMUSG00000039206 | -1.03472495 | 0.756258014 |
| ENSMUSG00000039208 | 1.127063743 | 0.251544797 |
| ENSMUSG00000039209 | 1 | |
| ENSMUSG00000039210 | 1.051070027 | 0.724515237 |
| ENSMUSG00000039215 | 1 | |
| ENSMUSG00000039217 | -1.705193718 | 0.000957737 |
| ENSMUSG00000039218 | 1.057372041 | 0.895588956 |
| ENSMUSG00000039219 | -1.047658513 | 0.877634538 |
| ENSMUSG00000039220 | 1.062172629 | 0.764291191 |
| ENSMUSG00000039221 | -1.082159789 | 0.608497229 |
| ENSMUSG00000039224 | -1.044273782 | 0.790851299 |
| ENSMUSG00000039230 | 1.050162677 | 0.81862959 |
| ENSMUSG00000039231 | -1.012867872 | 1 |
| ENSMUSG00000039232 | -1.031273388 | 0.805330446 |
| ENSMUSG00000039233 | -1.062347015 | 0.691595649 |
| ENSMUSG00000039234 | 1.085374064 | 0.70299173 |
| ENSMUSG00000039236 | -1.153159499 | 0.048656322 |
| ENSMUSG00000039238 | 1.237106647 | 0.096219972 |
| ENSMUSG00000039239 | 1.071078946 | 0.712544875 |
| ENSMUSG00000039242 | -1.004127003 | 1 |
| ENSMUSG00000039244 | 1.038583266 | 0.784039458 |
| ENSMUSG00000039246 | -1.125963681 | 0.189699867 |
| ENSMUSG00000039252 | 1.00546335 | 1 |
| ENSMUSG00000039253 | -1.055630895 | 0.731173125 |
| ENSMUSG00000039254 | 1.023940283 | 0.918615861 |
| ENSMUSG00000039257 | 1 | |
| ENSMUSG00000039262 | 1.097412584 | 0.66036903 |
| ENSMUSG00000039263 | 1.165750476 | 0.116198478 |
| ENSMUSG00000039264 | 1.059463094 | 0.608497229 |
| ENSMUSG00000039269 | 1 | |
| ENSMUSG00000039270 | 1.097341352 | 0.608497229 |
| ENSMUSG00000039275 | 1.099859131 | 0.693836777 |
| ENSMUSG00000039278 | 1 | |
| ENSMUSG00000039285 | -1.02427921 | 0.952057016 |
| ENSMUSG00000039286 | 1.044219627 | 0.896753433 |
| ENSMUSG00000039294 | -1.041659282 | 0.868338014 |
| ENSMUSG00000039296 | 1.019083579 | 0.949812939 |
| ENSMUSG00000039298 | 1.024077796 | 0.970316487 |
| ENSMUSG00000039304 | 1.078839172 | 0.852088342 |
| ENSMUSG00000039307 | 1.024266565 | 0.826119354 |
| ENSMUSG00000039308 | -1.141174722 | 0.332282211 |
| ENSMUSG00000039313 | -1.147167612 | 0.608497229 |
| ENSMUSG00000039315 | 1 | |
| ENSMUSG00000039316 | -1.176532892 | 0.038839753 |
| ENSMUSG00000039318 | 1.066526519 | 0.74674257 |
| ENSMUSG00000039321 | 1 | |
| ENSMUSG00000039323 | -1.251142641 | 0.021291484 |
| ENSMUSG00000039328 | 1.262590111 | 0.010335029 |
| ENSMUSG00000039329 | -1.084645496 | 0.532369657 |
| ENSMUSG00000039330 | -1.220953771 | 0.210526653 |
| ENSMUSG00000039335 | 1 | |
| ENSMUSG00000039337 | -1.085651418 | 0.608497229 |
| ENSMUSG00000039339 | 1 | |
| ENSMUSG00000039342 | 1 | |
| ENSMUSG00000039345 | 1.030810821 | 0.797998079 |
| ENSMUSG00000039347 | -1.091064317 | 0.608497229 |
| ENSMUSG00000039349 | -1.504589823 | 0.062384746 |
| ENSMUSG00000039354 | 1.06394731 | 0.608497229 |
| ENSMUSG00000039356 | -1.068711589 | 0.734674233 |
| ENSMUSG00000039357 | -1.311628165 | 0.04182893 |
| ENSMUSG00000039358 | 1 | |
| ENSMUSG00000039361 | 1.155334127 | 0.608497229 |

| | | |
|---|---|---|
| ENSMUSG00000039364 | 1 | |
| ENSMUSG00000039367 | -1.12085211 | 0.614284192 |
| ENSMUSG00000039372 | -1.014410901 | 1 |
| ENSMUSG00000039375 | 1.004847951 | 1 |
| ENSMUSG00000039376 | -1.065261562 | 0.891020425 |
| ENSMUSG00000039377 | -1.016312467 | 0.951399668 |
| ENSMUSG00000039382 | 1.096577459 | 0.183611926 |
| ENSMUSG00000039384 | -1.057463504 | 0.642592496 |
| ENSMUSG00000039385 | -1.138693452 | 0.569479524 |
| ENSMUSG00000039391 | 1.161008251 | 0.35934689 |
| ENSMUSG00000039395 | -1.238559158 | 0.095175477 |
| ENSMUSG00000039396 | -1.048307967 | 0.72770134 |
| ENSMUSG00000039405 | 1.183595863 | 0.493659452 |
| ENSMUSG00000039410 | -1.03962336 | 0.803072249 |
| ENSMUSG00000039414 | -1.01271112 | 1 |
| ENSMUSG00000039419 | -1.03962336 | 0.88732209 |
| ENSMUSG00000039427 | 1.160273322 | 0.080376709 |
| ENSMUSG00000039428 | 1.014125498 | 0.964202376 |
| ENSMUSG00000039431 | -1.126454257 | 0.350790589 |
| ENSMUSG00000039438 | -1.044273782 | 0.801443616 |
| ENSMUSG00000039449 | 1.031381366 | 0.925679199 |
| ENSMUSG00000039450 | 1.299201813 | 0.004533546 |
| ENSMUSG00000039452 | -1.139258958 | 0.506801216 |
| ENSMUSG00000039456 | -1.001869194 | 1 |
| ENSMUSG00000039457 | -1.093695745 | 0.46097689 |
| ENSMUSG00000039458 | -1.035822216 | 0.89658255 |
| ENSMUSG00000039461 | -1.049388779 | 0.670305632 |
| ENSMUSG00000039462 | 1.433327957 | 0.115548937 |
| ENSMUSG00000039463 | -1.048726121 | 0.622774674 |
| ENSMUSG00000039470 | -1.151971935 | 0.532369657 |
| ENSMUSG00000039473 | 1.083116055 | 0.717244617 |
| ENSMUSG00000039474 | 1.02804145 | 0.870921434 |
| ENSMUSG00000039476 | 1.005526638 | 1 |
| ENSMUSG00000039477 | 1.340920643 | 0.024878456 |
| ENSMUSG00000039478 | -1.030059363 | 0.877375054 |
| ENSMUSG00000039480 | 1.000549977 | 1 |
| ENSMUSG00000039481 | -1.084645496 | 0.532369657 |
| ENSMUSG00000039483 | 1.16273035 | 0.248317717 |
| ENSMUSG00000039485 | -1.1692253 | 0.188562861 |
| ENSMUSG00000039488 | 1 | |
| ENSMUSG00000039492 | -1.034239753 | 0.927693427 |
| ENSMUSG00000039496 | 1.179779244 | 0.309681415 |
| ENSMUSG00000039497 | 1.057727244 | 0.813313058 |
| ENSMUSG00000039501 | -1.165255321 | 0.263445848 |
| ENSMUSG00000039508 | 1.191786762 | 0.646086873 |
| ENSMUSG00000039509 | -1.021857324 | 0.979002724 |
| ENSMUSG00000039512 | 1.009677387 | 1 |
| ENSMUSG00000039515 | 1.068964985 | 0.732211848 |
| ENSMUSG00000039518 | 2.139520812 | 0.000238309 |
| ENSMUSG00000039519 | -1.290182641 | 0.040284547 |
| ENSMUSG00000039521 | 1.236709358 | 0.531138876 |
| ENSMUSG00000039523 | -1.01829105 | 0.970006845 |
| ENSMUSG00000039529 | -1.347457385 | 0.135458517 |
| ENSMUSG00000039530 | 1.023947764 | 0.973445324 |
| ENSMUSG00000039531 | -1.042387491 | 0.836326939 |
| ENSMUSG00000039533 | -1.143787033 | 0.363110963 |
| ENSMUSG00000039536 | -1.084512274 | 0.71881308 |
| ENSMUSG00000039539 | 1 | |
| ENSMUSG00000039540 | 1 | |
| ENSMUSG00000039542 | 1.359226162 | 0.021819041 |
| ENSMUSG00000039543 | -1.089180169 | 0.769705171 |
| ENSMUSG00000039545 | 1.064160904 | 0.901824606 |
| ENSMUSG00000039546 | 1 | |

| | | |
|---|---|---|
| ENSMUSG00000039552 | 1.020020306 | 1 |
| ENSMUSG00000039555 | 1 | |
| ENSMUSG00000039556 | -1.199279465 | 0.570749474 |
| ENSMUSG00000039563 | 1.201176149 | 0.1546483 |
| ENSMUSG00000039568 | 1.392562619 | 0.001013549 |
| ENSMUSG00000039577 | 1.017852781 | 0.975399288 |
| ENSMUSG00000039578 | 1.195549523 | 0.075976049 |
| ENSMUSG00000039579 | -1.109233207 | 0.40026724 |
| ENSMUSG00000039585 | 1.031110403 | 0.950141087 |
| ENSMUSG00000039599 | -1.113067057 | 0.410982873 |
| ENSMUSG00000039601 | 2.044706377 | 0.000460073 |
| ENSMUSG00000039607 | -1.011297798 | 1 |
| ENSMUSG00000039608 | 1 | |
| ENSMUSG00000039611 | -1.22804468 | 0.05714568 |
| ENSMUSG00000039615 | 1.119796583 | 0.404768597 |
| ENSMUSG00000039616 | -1.169412247 | 0.12225429 |
| ENSMUSG00000039617 | 1 | |
| ENSMUSG00000039620 | -1.221251377 | 0.323795799 |
| ENSMUSG00000039621 | -1.848590775 | 0.00046824 |
| ENSMUSG00000039623 | 1.192773973 | 0.044084408 |
| ENSMUSG00000039628 | -1.000405949 | 1 |
| ENSMUSG00000039629 | -1.439996973 | 0.007060227 |
| ENSMUSG00000039630 | 1.021615933 | 0.98569451 |
| ENSMUSG00000039632 | -1.083232329 | 0.496494186 |
| ENSMUSG00000039633 | 1.280602711 | 0.017197931 |
| ENSMUSG00000039634 | -1.028542448 | 0.918762638 |
| ENSMUSG00000039637 | -1.01909096 | 0.918762638 |
| ENSMUSG00000039639 | -1.116853908 | 0.558122732 |
| ENSMUSG00000039640 | 1.036313678 | 0.823140628 |
| ENSMUSG00000039646 | 1.2931137 | 0.017722194 |
| ENSMUSG00000039648 | -1.001514034 | 1 |
| ENSMUSG00000039652 | -1.100573976 | 0.350417454 |
| ENSMUSG00000039653 | 1.06043669 | 0.879014609 |
| ENSMUSG00000039656 | 1.124939901 | 0.192084015 |
| ENSMUSG00000039660 | -1.004438131 | 1 |
| ENSMUSG00000039661 | -1 | 1 |
| ENSMUSG00000039662 | -1.03739185 | 0.872257321 |
| ENSMUSG00000039670 | 1.000774093 | 1 |
| ENSMUSG00000039671 | -1.107708978 | 0.599181336 |
| ENSMUSG00000039672 | 1.019083579 | 0.949812939 |
| ENSMUSG00000039676 | -1.03962336 | 0.698112512 |
| ENSMUSG00000039678 | 1.001780372 | 1 |
| ENSMUSG00000039680 | 1.088550352 | 0.609700862 |
| ENSMUSG00000039682 | 1.09392219 | 0.70299173 |
| ENSMUSG00000039683 | -1.03962336 | 0.698112512 |
| ENSMUSG00000039684 | 1.128427755 | 0.577332073 |
| ENSMUSG00000039686 | 1.065317192 | 0.608497229 |
| ENSMUSG00000039691 | 1 | |
| ENSMUSG00000039693 | -1.022321872 | 0.848848499 |
| ENSMUSG00000039697 | -1.022878645 | 0.936995747 |
| ENSMUSG00000039699 | -1.095294215 | 0.541735653 |
| ENSMUSG00000039701 | 1.01312685 | 0.991169083 |
| ENSMUSG00000039703 | -1.033929308 | 0.949812939 |
| ENSMUSG00000039704 | -1.065263731 | 0.773508225 |
| ENSMUSG00000039706 | -1 | 1 |
| ENSMUSG00000039710 | 1.019083579 | 0.949812939 |
| ENSMUSG00000039713 | 1.028977615 | 0.8410703 |
| ENSMUSG00000039714 | 1.132639128 | 0.21967784 |
| ENSMUSG00000039715 | 1.161447028 | 0.07721161 |
| ENSMUSG00000039716 | -1.424724305 | 0.076136188 |
| ENSMUSG00000039717 | -1 | 1 |
| ENSMUSG00000039720 | -1 | 1 |
| ENSMUSG00000039725 | -1.06087013 | 0.543724675 |

| | | |
|---|---|---|
| ENSMUSG00000039728 | 1.149113744 | 0.347926366 |
| ENSMUSG00000039735 | 1.010202102 | 1 |
| ENSMUSG00000039737 | -1.018764646 | 0.940197654 |
| ENSMUSG00000039738 | 1.023811803 | 0.942036436 |
| ENSMUSG00000039740 | 1.012867299 | 0.996880111 |
| ENSMUSG00000039741 | 1.08768692 | 0.608497229 |
| ENSMUSG00000039742 | -1.153756725 | 0.310755019 |
| ENSMUSG00000039745 | -1.09521656 | 0.502179335 |
| ENSMUSG00000039747 | 1.059403707 | 0.608497229 |
| ENSMUSG00000039748 | -1.063199186 | 0.723904259 |
| ENSMUSG00000039753 | 1.095929602 | 0.698112512 |
| ENSMUSG00000039754 | -1.046303372 | 0.603144355 |
| ENSMUSG00000039756 | -1.078315098 | 0.756258014 |
| ENSMUSG00000039759 | 1.055537069 | 0.704433991 |
| ENSMUSG00000039760 | 1 | |
| ENSMUSG00000039763 | 1.091757995 | 0.50530478 |
| ENSMUSG00000039765 | 1.358446281 | 0.00815974 |
| ENSMUSG00000039768 | -1.001996161 | 1 |
| ENSMUSG00000039770 | 1.054481421 | 0.78099076 |
| ENSMUSG00000039771 | -1.03359621 | 0.883111131 |
| ENSMUSG00000039774 | -1.135532239 | 0.268434099 |
| ENSMUSG00000039775 | 1 | |
| ENSMUSG00000039781 | 1.001982606 | 1 |
| ENSMUSG00000039782 | 1.471330173 | 0.006425644 |
| ENSMUSG00000039783 | 1.070073032 | 0.88284878 |
| ENSMUSG00000039785 | 1 | |
| ENSMUSG00000039787 | -1.052953783 | 0.400396831 |
| ENSMUSG00000039789 | -1.009748048 | 1 |
| ENSMUSG00000039795 | -1.069472953 | 0.608497229 |
| ENSMUSG00000039798 | -1.033197264 | 0.947154477 |
| ENSMUSG00000039801 | 1.020856463 | 0.984716624 |
| ENSMUSG00000039804 | -1.065140831 | 0.698112512 |
| ENSMUSG00000039809 | -1.168999771 | 0.328133476 |
| ENSMUSG00000039810 | 1.093933415 | 0.176121876 |
| ENSMUSG00000039813 | -1.29184958 | 0.132291893 |
| ENSMUSG00000039814 | 1.246631319 | 0.049128286 |
| ENSMUSG00000039824 | 1.171940457 | 0.110125768 |
| ENSMUSG00000039826 | 1.050782316 | 0.773508225 |
| ENSMUSG00000039828 | 1.042101691 | 0.827718961 |
| ENSMUSG00000039830 | 1 | |
| ENSMUSG00000039831 | -1.193343423 | 0.473417727 |
| ENSMUSG00000039834 | 1.038860676 | 0.711918425 |
| ENSMUSG00000039835 | -1.074028863 | 0.658590644 |
| ENSMUSG00000039838 | -1.125574008 | 0.623805174 |
| ENSMUSG00000039840 | 1.074692528 | 0.632387721 |
| ENSMUSG00000039841 | -1.022652245 | 0.946232149 |
| ENSMUSG00000039842 | -1.016937526 | 0.978120952 |
| ENSMUSG00000039844 | 1.094201852 | 0.608497229 |
| ENSMUSG00000039849 | 1.086443827 | 0.457410629 |
| ENSMUSG00000039850 | 1.018682926 | 0.904414482 |
| ENSMUSG00000039851 | -1.03962336 | 0.698112512 |
| ENSMUSG00000039852 | 1.117581845 | 0.586324142 |
| ENSMUSG00000039853 | -1.800246916 | 0.010456301 |
| ENSMUSG00000039860 | -1.300045102 | 0.062183567 |
| ENSMUSG00000039865 | 1.06585783 | 0.770891731 |
| ENSMUSG00000039873 | 1.374035764 | 0.061822335 |
| ENSMUSG00000039878 | -1.041315342 | 0.70299173 |
| ENSMUSG00000039879 | 1.112754719 | 0.317228168 |
| ENSMUSG00000039883 | -1.154410228 | 0.366665274 |
| ENSMUSG00000039886 | 1.108985203 | 0.526048279 |
| ENSMUSG00000039887 | -1.053331763 | 0.74978479 |
| ENSMUSG00000039891 | 1.319932437 | 0.345896056 |
| ENSMUSG00000039899 | -1.038127145 | 0.858168264 |

| | | |
|---|---|---|
| ENSMUSG00000039901 | -1.091901594 | 0.608497229 |
| ENSMUSG00000039903 | 1.255729019 | 0.041468164 |
| ENSMUSG00000039904 | 1.075070132 | 0.775366634 |
| ENSMUSG00000039908 | 1.083484256 | 0.350132957 |
| ENSMUSG00000039910 | 1.188333494 | 0.287426065 |
| ENSMUSG00000039911 | -1.054689405 | 0.286287911 |
| ENSMUSG00000039913 | -1.043306199 | 0.698112512 |
| ENSMUSG00000039914 | -1.038354344 | 0.810692433 |
| ENSMUSG00000039917 | -1.008072275 | 0.982838256 |
| ENSMUSG00000039929 | 1.027800728 | 0.890965503 |
| ENSMUSG00000039934 | 1.105139489 | 0.518298719 |
| ENSMUSG00000039936 | -1.129064181 | 0.275828085 |
| ENSMUSG00000039942 | 1.49328945 | 0.013034138 |
| ENSMUSG00000039943 | 1.5007303 | 0.001596503 |
| ENSMUSG00000039952 | -1.039596948 | 0.92200539 |
| ENSMUSG00000039953 | 1.001884186 | 1 |
| ENSMUSG00000039954 | -1.012660513 | 1 |
| ENSMUSG00000039956 | 1 | |
| ENSMUSG00000039958 | -1.145835315 | 0.454814065 |
| ENSMUSG00000039959 | 1.106394632 | 0.630541247 |
| ENSMUSG00000039960 | -1.008987041 | 1 |
| ENSMUSG00000039962 | 1 | |
| ENSMUSG00000039963 | -1.193957541 | 0.608497229 |
| ENSMUSG00000039967 | 1.03779613 | 0.895127848 |
| ENSMUSG00000039968 | 1.015315308 | 0.980773838 |
| ENSMUSG00000039976 | -1.09463563 | 0.480654938 |
| ENSMUSG00000039977 | -1 | 1 |
| ENSMUSG00000039981 | 1.009517015 | 1 |
| ENSMUSG00000039982 | 1.646578218 | 0.001667387 |
| ENSMUSG00000039983 | -1.092080378 | 0.526180599 |
| ENSMUSG00000039985 | 1.108387946 | 0.608497229 |
| ENSMUSG00000039987 | -1.060424238 | 0.698112512 |
| ENSMUSG00000039988 | -1.056367095 | 0.833435109 |
| ENSMUSG00000039989 | 1.030114664 | 0.684693086 |
| ENSMUSG00000039990 | -1.047072555 | 0.786178485 |
| ENSMUSG00000039994 | -1.025079822 | 0.946496258 |
| ENSMUSG00000039997 | 1.102873774 | 0.048683099 |
| ENSMUSG00000040003 | 1.197970091 | 0.223646364 |
| ENSMUSG00000040006 | -1.151909952 | 0.361955196 |
| ENSMUSG00000040007 | -1.070490466 | 0.558122732 |
| ENSMUSG00000040009 | -1.034844255 | 0.871609782 |
| ENSMUSG00000040010 | 1.036103686 | 0.926765911 |
| ENSMUSG00000040013 | -1.043306199 | 0.698112512 |
| ENSMUSG00000040016 | -1.120476206 | 0.608497229 |
| ENSMUSG00000040017 | 1.069293585 | 0.797105186 |
| ENSMUSG00000040018 | 1.003160667 | 1 |
| ENSMUSG00000040021 | 1.015547524 | 1 |
| ENSMUSG00000040022 | 1.088350915 | 0.608497229 |
| ENSMUSG00000040025 | 1.018035811 | 0.983393382 |
| ENSMUSG00000040026 | 1.023959045 | 0.974736184 |
| ENSMUSG00000040028 | 1.118839681 | 0.655462417 |
| ENSMUSG00000040029 | -1.001562447 | 1 |
| ENSMUSG00000040033 | -1.018286137 | 0.953416513 |
| ENSMUSG00000040034 | -1.075425832 | 0.711533702 |
| ENSMUSG00000040035 | -1.185713036 | 0.672090419 |
| ENSMUSG00000040037 | 1.143603129 | 0.323898801 |
| ENSMUSG00000040040 | -1.170958765 | 0.228467222 |
| ENSMUSG00000040043 | -1.047539123 | 0.89660383 |
| ENSMUSG00000040044 | -1.049630198 | 0.836447677 |
| ENSMUSG00000040046 | -1.039460158 | 0.938484564 |
| ENSMUSG00000040048 | -1.022204879 | 0.93079313 |
| ENSMUSG00000040054 | -1.030391079 | 0.949812939 |
| ENSMUSG00000040055 | -1.043306199 | 0.698112512 |

| | | |
|---|---|---|
| ENSMUSG00000040061 | -1.014570171 | 1 |
| ENSMUSG00000040065 | 1 | |
| ENSMUSG00000040078 | 1 | |
| ENSMUSG00000040084 | -1.115318961 | 0.6659463 |
| ENSMUSG00000040086 | 1.237186144 | 0.363888752 |
| ENSMUSG00000040093 | 1.20553924 | 0.02304608 |
| ENSMUSG00000040097 | 1.113712508 | 0.23546922 |
| ENSMUSG00000040102 | 1.031242406 | 0.86579205 |
| ENSMUSG00000040105 | -1.032331503 | 0.861857538 |
| ENSMUSG00000040111 | 1.157644666 | 0.100651404 |
| ENSMUSG00000040112 | 1.011349918 | 1 |
| ENSMUSG00000040113 | 1 | |
| ENSMUSG00000040118 | 1.158161721 | 0.466284441 |
| ENSMUSG00000040121 | 1.025576259 | 0.951744672 |
| ENSMUSG00000040123 | -1.115145912 | 0.608497229 |
| ENSMUSG00000040124 | 1.135623123 | 0.368089426 |
| ENSMUSG00000040125 | 1.045900989 | 0.924587329 |
| ENSMUSG00000040127 | 1 | |
| ENSMUSG00000040128 | 1.06404008 | 0.652464432 |
| ENSMUSG00000040132 | 1 | |
| ENSMUSG00000040133 | 1.224815271 | 0.081316192 |
| ENSMUSG00000040134 | 1 | |
| ENSMUSG00000040136 | -1.45117801 | 0.083152509 |
| ENSMUSG00000040138 | -1 | 1 |
| ENSMUSG00000040139 | -1.150807614 | 0.040767591 |
| ENSMUSG00000040140 | 1.206725422 | 0.369259262 |
| ENSMUSG00000040146 | -1.116569054 | 0.080861974 |
| ENSMUSG00000040147 | -1.171420412 | 0.247015918 |
| ENSMUSG00000040148 | 1 | |
| ENSMUSG00000040151 | -1.001573735 | 1 |
| ENSMUSG00000040152 | 1.284330099 | 0.391140841 |
| ENSMUSG00000040154 | 1 | |
| ENSMUSG00000040158 | 1.060552078 | 0.79291651 |
| ENSMUSG00000040163 | 1.049023683 | 0.907024843 |
| ENSMUSG00000040164 | 1 | |
| ENSMUSG00000040165 | 1.012783207 | 0.969026637 |
| ENSMUSG00000040167 | -1.002983647 | 1 |
| ENSMUSG00000040170 | -1 | 1 |
| ENSMUSG00000040174 | 1.021861205 | 0.932616842 |
| ENSMUSG00000040177 | -1.024188524 | 0.865462843 |
| ENSMUSG00000040181 | -1.011214121 | 0.977648042 |
| ENSMUSG00000040183 | -1.014196595 | 0.945749371 |
| ENSMUSG00000040187 | -1.068010471 | 0.47614255 |
| ENSMUSG00000040188 | -1.089373151 | 0.633631542 |
| ENSMUSG00000040189 | -1.286754139 | 0.357810995 |
| ENSMUSG00000040195 | -1.07780807 | 0.698112512 |
| ENSMUSG00000040197 | 1 | |
| ENSMUSG00000040204 | -1.109149122 | 0.706878376 |
| ENSMUSG00000040205 | 1.069068979 | 0.698112512 |
| ENSMUSG00000040209 | 1.010071167 | 1 |
| ENSMUSG00000040212 | -1.069117321 | 0.698112512 |
| ENSMUSG00000040213 | -1.212859354 | 0.037355797 |
| ENSMUSG00000040217 | 1 | |
| ENSMUSG00000040219 | 1.048597575 | 0.687333535 |
| ENSMUSG00000040220 | 1.019385256 | 0.916783974 |
| ENSMUSG00000040225 | 1.011268021 | 1 |
| ENSMUSG00000040229 | 1.122462048 | 0.654902646 |
| ENSMUSG00000040231 | -1.191395474 | 0.608497229 |
| ENSMUSG00000040234 | 1.093269951 | 0.608497229 |
| ENSMUSG00000040236 | 1.129434134 | 0.270080405 |
| ENSMUSG00000040242 | -1.003850379 | 1 |
| ENSMUSG00000040247 | -1.05081723 | 0.914954453 |
| ENSMUSG00000040249 | 1.369228937 | 0.087380382 |

| | | |
|---|---|---|
| ENSMUSG00000040250 | -1.049069156 | 0.846547294 |
| ENSMUSG00000040253 | -1.926930404 | 0.000957737 |
| ENSMUSG00000040254 | 1.289615964 | 0.042311392 |
| ENSMUSG00000040258 | -1.319992059 | 0.141447411 |
| ENSMUSG00000040260 | 1.512111909 | 0.000603481 |
| ENSMUSG00000040263 | 1.003505931 | 1 |
| ENSMUSG00000040264 | -1.294423543 | 0.066758526 |
| ENSMUSG00000040265 | -1.072502055 | 0.653048463 |
| ENSMUSG00000040268 | -1.053440193 | 0.772656514 |
| ENSMUSG00000040269 | -1.036943594 | 0.755673717 |
| ENSMUSG00000040270 | 1.040397547 | 0.797084953 |
| ENSMUSG00000040272 | -1.07736354 | 0.534652019 |
| ENSMUSG00000040274 | -1.110576153 | 0.647538847 |
| ENSMUSG00000040276 | -1.596023913 | 0.034812864 |
| ENSMUSG00000040280 | -1.053976749 | 0.834680332 |
| ENSMUSG00000040282 | 1.011984101 | 0.985522133 |
| ENSMUSG00000040283 | -1.029709545 | 0.975148944 |
| ENSMUSG00000040284 | 1 | |
| ENSMUSG00000040287 | 1.042090403 | 0.839688843 |
| ENSMUSG00000040289 | 1.370515437 | 0.066036399 |
| ENSMUSG00000040296 | -1.314016334 | 0.138402331 |
| ENSMUSG00000040297 | -1.002020772 | 1 |
| ENSMUSG00000040298 | -1.297044976 | 0.213163934 |
| ENSMUSG00000040302 | 1.032440095 | 0.847184973 |
| ENSMUSG00000040310 | -1.601225521 | 0.001090615 |
| ENSMUSG00000040312 | 1.144096199 | 0.102653366 |
| ENSMUSG00000040314 | 1 | |
| ENSMUSG00000040321 | -1.094926896 | 0.608497229 |
| ENSMUSG00000040322 | -1.029623928 | 0.959892369 |
| ENSMUSG00000040323 | -1.043306199 | 0.698112512 |
| ENSMUSG00000040325 | -1.008863986 | 1 |
| ENSMUSG00000040327 | 1.080514762 | 0.734674233 |
| ENSMUSG00000040328 | -1.261186606 | 0.020875656 |
| ENSMUSG00000040329 | -1.011207688 | 0.990831634 |
| ENSMUSG00000040331 | -1.032812737 | 0.912742948 |
| ENSMUSG00000040339 | -1.015089214 | 1 |
| ENSMUSG00000040340 | 1.059463094 | 0.608497229 |
| ENSMUSG00000040345 | -1.254062209 | 0.412770954 |
| ENSMUSG00000040350 | 1.11885342 | 0.100218491 |
| ENSMUSG00000040351 | 1.091517486 | 0.708068647 |
| ENSMUSG00000040354 | -1.026793064 | 0.96005778 |
| ENSMUSG00000040356 | -1.065681096 | 0.608497229 |
| ENSMUSG00000040359 | -1.01033156 | 1 |
| ENSMUSG00000040363 | 1.134114451 | 0.449661746 |
| ENSMUSG00000040364 | 1.118644273 | 0.608497229 |
| ENSMUSG00000040365 | -1.021740568 | 0.880365937 |
| ENSMUSG00000040367 | 1.482091975 | 0.001058101 |
| ENSMUSG00000040370 | 1.117418756 | 0.354932739 |
| ENSMUSG00000040372 | 1.06531439 | 0.685142053 |
| ENSMUSG00000040373 | 1 | |
| ENSMUSG00000040374 | 1.019232723 | 0.968423939 |
| ENSMUSG00000040380 | 1.142331634 | 0.416318876 |
| ENSMUSG00000040383 | -1.059307754 | 0.819817237 |
| ENSMUSG00000040385 | 1.009739653 | 1 |
| ENSMUSG00000040387 | -1.085780268 | 0.532586 |
| ENSMUSG00000040389 | -1.138069243 | 0.427987269 |
| ENSMUSG00000040390 | 1.03176313 | 0.777302582 |
| ENSMUSG00000040396 | 1.145238172 | 0.514838447 |
| ENSMUSG00000040405 | 1 | |
| ENSMUSG00000040407 | 1.025929536 | 0.962587881 |
| ENSMUSG00000040410 | 1.037872467 | 0.844771174 |
| ENSMUSG00000040412 | -1.351634127 | 0.00221616 |
| ENSMUSG00000040413 | 1 | |

| | | |
|---|---|---|
| ENSMUSG00000040414 | 1.053859162 | 0.619996008 |
| ENSMUSG00000040415 | 1.112420992 | 0.305174917 |
| ENSMUSG00000040420 | 1.255447745 | 0.011490798 |
| ENSMUSG00000040423 | 1.048126561 | 0.862443252 |
| ENSMUSG00000040424 | 1.134738056 | 0.608497229 |
| ENSMUSG00000040428 | -1.003101006 | 1 |
| ENSMUSG00000040429 | 1.155362954 | 0.337208144 |
| ENSMUSG00000040430 | -1.085896666 | 0.608497229 |
| ENSMUSG00000040432 | -1 | 1 |
| ENSMUSG00000040433 | 1.172989212 | 0.3862041 |
| ENSMUSG00000040434 | 1.023822295 | 0.996913409 |
| ENSMUSG00000040435 | -1.028631985 | 0.864561135 |
| ENSMUSG00000040441 | -1.04075631 | 0.855488553 |
| ENSMUSG00000040446 | 1.085955853 | 0.698112512 |
| ENSMUSG00000040447 | -1.238259602 | 0.014492139 |
| ENSMUSG00000040451 | 1.139074598 | 0.528868515 |
| ENSMUSG00000040452 | 1 | |
| ENSMUSG00000040455 | -1.086715169 | 0.71314753 |
| ENSMUSG00000040456 | 1.095872691 | 0.608497229 |
| ENSMUSG00000040459 | 1.135853446 | 0.605907311 |
| ENSMUSG00000040462 | -1.038656649 | 0.822158059 |
| ENSMUSG00000040463 | 1.056505345 | 0.877294154 |
| ENSMUSG00000040464 | 1.053205399 | 0.794156491 |
| ENSMUSG00000040466 | 1.042691705 | 0.904013471 |
| ENSMUSG00000040467 | 1.01602257 | 0.993999301 |
| ENSMUSG00000040471 | 1.075870199 | 0.823266891 |
| ENSMUSG00000040472 | -1.12274732 | 0.04288737 |
| ENSMUSG00000040473 | -1.009142876 | 1 |
| ENSMUSG00000040478 | 1 | |
| ENSMUSG00000040479 | -1.035215081 | 0.788000546 |
| ENSMUSG00000040481 | -1.00062175 | 1 |
| ENSMUSG00000040482 | 1.026077452 | 0.837365423 |
| ENSMUSG00000040483 | -1.58200709 | 0.002137749 |
| ENSMUSG00000040485 | 1 | |
| ENSMUSG00000040488 | 1.128204166 | 0.66759917 |
| ENSMUSG00000040489 | 1 | |
| ENSMUSG00000040490 | 1 | |
| ENSMUSG00000040495 | -1.343928243 | 0.064092836 |
| ENSMUSG00000040498 | 1.059463094 | 0.752398528 |
| ENSMUSG00000040502 | -1.115836957 | 0.251049598 |
| ENSMUSG00000040505 | -1.03962336 | 0.803072249 |
| ENSMUSG00000040506 | -1.064808074 | 0.705491405 |
| ENSMUSG00000040511 | 1.125839053 | 0.431683218 |
| ENSMUSG00000040514 | -1.03962336 | 0.698112512 |
| ENSMUSG00000040520 | 1.037639996 | 0.889445701 |
| ENSMUSG00000040521 | 1.014153502 | 0.909498685 |
| ENSMUSG00000040522 | 1 | |
| ENSMUSG00000040524 | -1.001521098 | 1 |
| ENSMUSG00000040525 | -1.084645496 | 0.608497229 |
| ENSMUSG00000040528 | 1.054307452 | 0.808155813 |
| ENSMUSG00000040532 | 1.015077772 | 0.93965105 |
| ENSMUSG00000040533 | 1 | |
| ENSMUSG00000040536 | 1.675008713 | 0.021089062 |
| ENSMUSG00000040537 | -1.360462153 | 0.019667107 |
| ENSMUSG00000040540 | -1.051018293 | 0.86119789 |
| ENSMUSG00000040541 | 1 | |
| ENSMUSG00000040543 | -1.131277635 | 0.608497229 |
| ENSMUSG00000040548 | -1.08951576 | 0.64090177 |
| ENSMUSG00000040549 | -1.127512479 | 0.698112512 |
| ENSMUSG00000040550 | -1.012698854 | 1 |
| ENSMUSG00000040552 | 1.078614944 | 0.572892557 |
| ENSMUSG00000040554 | 1.249158547 | 0.345889524 |
| ENSMUSG00000040557 | 1.03031684 | 0.887340389 |

| | | |
|---|---|---|
| ENSMUSG00000040560 | 1.06795957 | 0.698112512 |
| ENSMUSG00000040562 | -1.002808418 | 1 |
| ENSMUSG00000040563 | -1.091614212 | 0.381530795 |
| ENSMUSG00000040564 | 1 | |
| ENSMUSG00000040565 | 1.03997892 | 0.93376866 |
| ENSMUSG00000040569 | 1.059463094 | 0.608497229 |
| ENSMUSG00000040570 | -1.022890659 | 0.69533886 |
| ENSMUSG00000040576 | -1.044273782 | 0.698112512 |
| ENSMUSG00000040583 | 1.122462048 | 0.215991492 |
| ENSMUSG00000040584 | 1.262227314 | 0.247040254 |
| ENSMUSG00000040586 | -1.113654648 | 0.600132627 |
| ENSMUSG00000040591 | -1.155997716 | 0.26504751 |
| ENSMUSG00000040592 | 1 | |
| ENSMUSG00000040594 | 1.055076239 | 0.730297627 |
| ENSMUSG00000040596 | 1.091046817 | 0.659212991 |
| ENSMUSG00000040599 | 1.02917664 | 0.951480616 |
| ENSMUSG00000040600 | -1.044273782 | 0.790851299 |
| ENSMUSG00000040601 | 1 | |
| ENSMUSG00000040605 | -2.340077975 | 2.59386E-05 |
| ENSMUSG00000040606 | -1.775702005 | 0.033882553 |
| ENSMUSG00000040610 | -1.03962336 | 0.698112512 |
| ENSMUSG00000040612 | -1.174875402 | 0.462046668 |
| ENSMUSG00000040613 | -1.18749127 | 0.029143657 |
| ENSMUSG00000040614 | -1 | 1 |
| ENSMUSG00000040616 | -1.942235054 | 0.000684406 |
| ENSMUSG00000040618 | 1.10126364 | 0.579104867 |
| ENSMUSG00000040620 | 1.011697901 | 1 |
| ENSMUSG00000040621 | -1.109535235 | 0.250726313 |
| ENSMUSG00000040624 | -1.309760358 | 0.396191099 |
| ENSMUSG00000040627 | 1.010855137 | 1 |
| ENSMUSG00000040629 | 1 | |
| ENSMUSG00000040631 | -1.144160197 | 0.179720614 |
| ENSMUSG00000040632 | 1 | |
| ENSMUSG00000040640 | -1.121491923 | 0.325747783 |
| ENSMUSG00000040648 | 1.014879842 | 1 |
| ENSMUSG00000040649 | -1.044273782 | 0.698112512 |
| ENSMUSG00000040650 | 1 | |
| ENSMUSG00000040651 | -1.075557329 | 0.743396505 |
| ENSMUSG00000040652 | -1.123733081 | 0.605159739 |
| ENSMUSG00000040653 | -1.402077318 | 0.056310079 |
| ENSMUSG00000040657 | 1 | |
| ENSMUSG00000040658 | -1.072734565 | 0.61356039 |
| ENSMUSG00000040659 | 1.061834583 | 0.586366024 |
| ENSMUSG00000040660 | 1 | |
| ENSMUSG00000040661 | -1.005235905 | 1 |
| ENSMUSG00000040663 | 1.115354823 | 0.329069305 |
| ENSMUSG00000040666 | 2.238322377 | 0.001049921 |
| ENSMUSG00000040667 | -1.034976371 | 0.933031586 |
| ENSMUSG00000040669 | 1.083119113 | 0.582158676 |
| ENSMUSG00000040675 | 1.110501582 | 0.640410817 |
| ENSMUSG00000040680 | 1.049411284 | 0.836135545 |
| ENSMUSG00000040681 | 1.00469281 | 1 |
| ENSMUSG00000040687 | -1.153549783 | 0.144242528 |
| ENSMUSG00000040688 | -1.001144031 | 1 |
| ENSMUSG00000040690 | 1.300086142 | 0.029748741 |
| ENSMUSG00000040693 | 1 | |
| ENSMUSG00000040694 | 1.009866335 | 1 |
| ENSMUSG00000040697 | -1.027811487 | 0.875751698 |
| ENSMUSG00000040699 | 1.125513498 | 0.428988711 |
| ENSMUSG00000040701 | -1.135989891 | 0.084784558 |
| ENSMUSG00000040703 | -1.048433374 | 0.608497229 |
| ENSMUSG00000040705 | -1.129231493 | 0.608497229 |
| ENSMUSG00000040706 | 1.068904696 | 0.858577335 |

| | | |
|---|---|---|
| ENSMUSG00000040710 | -1.17459654 | 0.596419896 |
| ENSMUSG00000040711 | 1.161664755 | 0.534151267 |
| ENSMUSG00000040712 | 1.115438357 | 0.247420705 |
| ENSMUSG00000040713 | -1.082187306 | 0.608497229 |
| ENSMUSG00000040714 | -1.049612139 | 0.915424577 |
| ENSMUSG00000040715 | 1 | |
| ENSMUSG00000040717 | -1.673660606 | 0.001986739 |
| ENSMUSG00000040720 | 1.010541829 | 1 |
| ENSMUSG00000040721 | -1.244130396 | 0.03355902 |
| ENSMUSG00000040722 | -1.134848915 | 0.380866075 |
| ENSMUSG00000040723 | -1.127336939 | 0.769745556 |
| ENSMUSG00000040724 | 1.085624843 | 0.731830521 |
| ENSMUSG00000040725 | 1.045613425 | 0.856260317 |
| ENSMUSG00000040726 | 1 | |
| ENSMUSG00000040728 | -1.089497311 | 0.532369657 |
| ENSMUSG00000040729 | -1.097906708 | 0.404010222 |
| ENSMUSG00000040731 | 1.060845215 | 0.819614655 |
| ENSMUSG00000040732 | -1 | 1 |
| ENSMUSG00000040734 | 1.233400817 | 0.077622382 |
| ENSMUSG00000040738 | -1.043493064 | 0.896765664 |
| ENSMUSG00000040740 | 1.039560021 | 0.925209973 |
| ENSMUSG00000040746 | -1.023815259 | 0.838524395 |
| ENSMUSG00000040747 | 1.019083579 | 0.971519533 |
| ENSMUSG00000040749 | -1.138777087 | 0.365927614 |
| ENSMUSG00000040751 | 1.127760407 | 0.605619163 |
| ENSMUSG00000040752 | 1.059463094 | 0.608497229 |
| ENSMUSG00000040759 | 1 | |
| ENSMUSG00000040760 | -1.151223972 | 0.578885377 |
| ENSMUSG00000040761 | 1.108690183 | 0.345889524 |
| ENSMUSG00000040767 | -1.124944032 | 0.323719996 |
| ENSMUSG00000040770 | 1 | |
| ENSMUSG00000040771 | 1.043662182 | 0.803938171 |
| ENSMUSG00000040774 | -1.0048995 | 1 |
| ENSMUSG00000040775 | 1.013583879 | 1 |
| ENSMUSG00000040782 | 1.014027408 | 1 |
| ENSMUSG00000040785 | 1.098467177 | 0.743131616 |
| ENSMUSG00000040794 | -1.085651418 | 0.532369657 |
| ENSMUSG00000040795 | 1.023893899 | 0.879253209 |
| ENSMUSG00000040797 | -1 | 1 |
| ENSMUSG00000040808 | -1 | 1 |
| ENSMUSG00000040809 | 1 | |
| ENSMUSG00000040811 | 1.413297213 | 0.02620375 |
| ENSMUSG00000040812 | -1.188964619 | 0.145386674 |
| ENSMUSG00000040813 | 1.079871262 | 0.534609229 |
| ENSMUSG00000040818 | -1.183631161 | 0.373988904 |
| ENSMUSG00000040820 | 1.028220078 | 0.809525929 |
| ENSMUSG00000040822 | -1.089652227 | 0.608497229 |
| ENSMUSG00000040824 | -1.027972868 | 0.924807766 |
| ENSMUSG00000040828 | 1.055719505 | 0.823727281 |
| ENSMUSG00000040829 | -1.277381533 | 0.266335645 |
| ENSMUSG00000040836 | 1.089973721 | 0.608497229 |
| ENSMUSG00000040838 | 1 | |
| ENSMUSG00000040840 | -1.186953318 | 0.202727447 |
| ENSMUSG00000040841 | 1.14436891 | 0.109134668 |
| ENSMUSG00000040842 | 1.021292048 | 0.954883228 |
| ENSMUSG00000040843 | 1.007994268 | 1 |
| ENSMUSG00000040848 | -1.104438905 | 0.652734988 |
| ENSMUSG00000040850 | -1.060040627 | 0.870119185 |
| ENSMUSG00000040852 | -1.088212739 | 0.698112512 |
| ENSMUSG00000040855 | -1.333932918 | 0.143576805 |
| ENSMUSG00000040856 | 2.259450031 | 2.50188E-05 |
| ENSMUSG00000040857 | 1.142618593 | 0.130623274 |
| ENSMUSG00000040859 | 1.004060731 | 1 |

| | | |
|---|---|---|
| ENSMUSG00000040860 | 1.007845277 | 1 |
| ENSMUSG00000040865 | -1.051158348 | 0.835382855 |
| ENSMUSG00000040866 | 1 | |
| ENSMUSG00000040867 | 1.175415944 | 0.609615508 |
| ENSMUSG00000040875 | 1.218226496 | 0.135690167 |
| ENSMUSG00000040877 | 1.168229151 | 0.192763842 |
| ENSMUSG00000040883 | -1.017478697 | 0.93692717 |
| ENSMUSG00000040888 | -1.040713034 | 0.826863971 |
| ENSMUSG00000040891 | 1.125042872 | 0.726447749 |
| ENSMUSG00000040896 | 1 | |
| ENSMUSG00000040899 | 1 | |
| ENSMUSG00000040901 | 1 | |
| ENSMUSG00000040904 | 1.054079687 | 0.878872929 |
| ENSMUSG00000040907 | -1.176925332 | 0.594510871 |
| ENSMUSG00000040913 | 1.207122809 | 0.024707694 |
| ENSMUSG00000040918 | 1.099580507 | 0.576246406 |
| ENSMUSG00000040919 | -1 | 1 |
| ENSMUSG00000040928 | 1.049141011 | 0.807247723 |
| ENSMUSG00000040929 | 1.06797665 | 0.680567617 |
| ENSMUSG00000040935 | 1 | |
| ENSMUSG00000040936 | 1.196005014 | 0.029238051 |
| ENSMUSG00000040938 | 1.084993138 | 0.825776729 |
| ENSMUSG00000040940 | 1.081838807 | 0.601670517 |
| ENSMUSG00000040943 | -1.034865131 | 0.831000123 |
| ENSMUSG00000040945 | 1.172121744 | 0.369902276 |
| ENSMUSG00000040950 | 1 | |
| ENSMUSG00000040952 | 1.1123466 | 0.608497229 |
| ENSMUSG00000040957 | -1.177928124 | 0.109887659 |
| ENSMUSG00000040963 | 1.069068979 | 0.698112512 |
| ENSMUSG00000040964 | 1.233625896 | 0.060537421 |
| ENSMUSG00000040966 | 1 | |
| ENSMUSG00000040969 | -1.139116 | 0.608497229 |
| ENSMUSG00000040972 | 1 | |
| ENSMUSG00000040978 | -1.16749727 | 0.618966174 |
| ENSMUSG00000040985 | 1.059463094 | 0.704170349 |
| ENSMUSG00000040987 | 1.05410549 | 0.858169153 |
| ENSMUSG00000040990 | 1.074475238 | 0.780320794 |
| ENSMUSG00000040997 | -1.038398414 | 0.877639675 |
| ENSMUSG00000040998 | -1.039361153 | 0.902345046 |
| ENSMUSG00000041000 | -1.294115735 | 0.034158269 |
| ENSMUSG00000041009 | 1 | |
| ENSMUSG00000041012 | -1.536004304 | 0.014073837 |
| ENSMUSG00000041014 | 1 | |
| ENSMUSG00000041020 | -1.166190533 | 0.202649662 |
| ENSMUSG00000041025 | 1.056128695 | 0.764790611 |
| ENSMUSG00000041028 | -1.056880937 | 0.874129418 |
| ENSMUSG00000041035 | -1.077187412 | 0.608497229 |
| ENSMUSG00000041037 | 1.061753302 | 0.609147349 |
| ENSMUSG00000041040 | 1.076391997 | 0.608497229 |
| ENSMUSG00000041044 | 1 | |
| ENSMUSG00000041046 | -1.46232496 | 0.097035679 |
| ENSMUSG00000041052 | 1.059463094 | 0.608497229 |
| ENSMUSG00000041057 | 1.00146263 | 1 |
| ENSMUSG00000041058 | -1.068307068 | 0.775673129 |
| ENSMUSG00000041062 | 1.023770587 | 0.984094159 |
| ENSMUSG00000041064 | 1.009030407 | 0.978211686 |
| ENSMUSG00000041068 | 1 | |
| ENSMUSG00000041073 | 1.073253538 | 0.561786114 |
| ENSMUSG00000041075 | 1.091966039 | 0.528617036 |
| ENSMUSG00000041078 | -1.08081673 | 0.709164097 |
| ENSMUSG00000041079 | -1.076177558 | 0.608497229 |
| ENSMUSG00000041084 | -1.089003562 | 0.723528308 |
| ENSMUSG00000041096 | -1.08607207 | 0.613963375 |

| | | |
|---|---|---|
| ENSMUSG00000041112 | 1.044229618 | 0.784232424 |
| ENSMUSG00000041115 | 1.008459185 | 0.996402978 |
| ENSMUSG00000041117 | 1.06134215 | 0.807247723 |
| ENSMUSG00000041119 | -1.370156412 | 0.015208772 |
| ENSMUSG00000041120 | 1.334406912 | 0.00639805 |
| ENSMUSG00000041124 | 1.020317535 | 0.978259071 |
| ENSMUSG00000041126 | 1.132349793 | 0.358497627 |
| ENSMUSG00000041130 | 1.065586039 | 0.608497229 |
| ENSMUSG00000041132 | 1.007812346 | 1 |
| ENSMUSG00000041133 | -1.102630352 | 0.745686215 |
| ENSMUSG00000041134 | 1.270699966 | 0.159861411 |
| ENSMUSG00000041135 | 1.188671359 | 0.215991492 |
| ENSMUSG00000041138 | 1 | |
| ENSMUSG00000041141 | 1 | |
| ENSMUSG00000041143 | 2.677312901 | 0.001308875 |
| ENSMUSG00000041144 | 1.151502406 | 0.558569239 |
| ENSMUSG00000041147 | 1.005682275 | 1 |
| ENSMUSG00000041153 | 1.054168566 | 0.698112512 |
| ENSMUSG00000041161 | 1.076028474 | 0.152779044 |
| ENSMUSG00000041164 | 1.073037356 | 0.608497229 |
| ENSMUSG00000041165 | 1.112843644 | 0.690350304 |
| ENSMUSG00000041168 | 1.084978547 | 0.698112512 |
| ENSMUSG00000041180 | -1.039890457 | 0.843142861 |
| ENSMUSG00000041187 | -1.152727319 | 0.099764565 |
| ENSMUSG00000041189 | -1.523851764 | 0.000837706 |
| ENSMUSG00000041193 | 1 | |
| ENSMUSG00000041199 | -1.001941223 | 1 |
| ENSMUSG00000041202 | -1.022608177 | 0.975677002 |
| ENSMUSG00000041203 | -1.025354183 | 0.934760809 |
| ENSMUSG00000041205 | -1.076234129 | 0.449586213 |
| ENSMUSG00000041215 | 1.060679727 | 0.773508225 |
| ENSMUSG00000041216 | 1.335036686 | 0.26849418 |
| ENSMUSG00000041219 | -1.098958049 | 0.698112512 |
| ENSMUSG00000041220 | -1.201288297 | 0.399513586 |
| ENSMUSG00000041225 | 1.153916237 | 0.404886539 |
| ENSMUSG00000041229 | -1.117437871 | 0.520518871 |
| ENSMUSG00000041231 | -1.072293408 | 0.729513149 |
| ENSMUSG00000041235 | 1.094934712 | 0.18200153 |
| ENSMUSG00000041236 | 1.025135476 | 0.965252226 |
| ENSMUSG00000041237 | -1 | 1 |
| ENSMUSG00000041238 | -1.001582886 | 1 |
| ENSMUSG00000041241 | 1.069877553 | 0.467940264 |
| ENSMUSG00000041245 | -1.167376471 | 0.229884308 |
| ENSMUSG00000041247 | -1.100193663 | 0.608497229 |
| ENSMUSG00000041248 | -1 | 1 |
| ENSMUSG00000041255 | 1 | |
| ENSMUSG00000041258 | 1.091966129 | 0.608497229 |
| ENSMUSG00000041261 | 1.095428003 | 0.777123525 |
| ENSMUSG00000041263 | -1.282120294 | 0.001087491 |
| ENSMUSG00000041264 | 1.064891697 | 0.765313234 |
| ENSMUSG00000041268 | -1.101835886 | 0.608497229 |
| ENSMUSG00000041272 | 1.156292476 | 0.537510415 |
| ENSMUSG00000041278 | -1.072567107 | 0.716190882 |
| ENSMUSG00000041287 | -1.014570171 | 0.993999301 |
| ENSMUSG00000041293 | -1.053549062 | 0.698112512 |
| ENSMUSG00000041297 | -1.032084859 | 0.930652317 |
| ENSMUSG00000041298 | 1.175345234 | 0.295352792 |
| ENSMUSG00000041301 | -1.164496684 | 0.524401846 |
| ENSMUSG00000041303 | -1.011519206 | 1 |
| ENSMUSG00000041308 | 1.006685657 | 1 |
| ENSMUSG00000041309 | 1 | |
| ENSMUSG00000041313 | 1.029012828 | 0.945650161 |
| ENSMUSG00000041319 | 1.171099229 | 0.234435619 |

| | | |
|---|---|---|
| ENSMUSG00000041323 | 1.196004652 | 0.328793916 |
| ENSMUSG00000041324 | 1.210026896 | 0.0586397 |
| ENSMUSG00000041328 | 1.012478271 | 1 |
| ENSMUSG00000041329 | -1.4140759 | 0.000669214 |
| ENSMUSG00000041333 | 1 | |
| ENSMUSG00000041341 | 1.099895419 | 0.69533886 |
| ENSMUSG00000041343 | 1.111465923 | 0.164157817 |
| ENSMUSG00000041346 | -1.138135045 | 0.208940434 |
| ENSMUSG00000041347 | -1.126344191 | 0.413646914 |
| ENSMUSG00000041351 | 1.066339971 | 0.442177828 |
| ENSMUSG00000041353 | -1.246306562 | 0.053903116 |
| ENSMUSG00000041354 | 1.046074237 | 0.667778769 |
| ENSMUSG00000041355 | -1.075947737 | 0.789448756 |
| ENSMUSG00000041358 | -1.61163885 | 0.001127478 |
| ENSMUSG00000041359 | 1 | |
| ENSMUSG00000041360 | 1.081707585 | 0.717035356 |
| ENSMUSG00000041361 | -1.088341414 | 0.319853951 |
| ENSMUSG00000041362 | -2.226056382 | 0.000459033 |
| ENSMUSG00000041372 | -1.657951579 | 0.002657251 |
| ENSMUSG00000041375 | 1.027301342 | 0.808606444 |
| ENSMUSG00000041377 | -1.037896376 | 0.964617203 |
| ENSMUSG00000041378 | 1 | |
| ENSMUSG00000041380 | 1.059463094 | 0.608497229 |
| ENSMUSG00000041390 | 1.299788521 | 0.107605709 |
| ENSMUSG00000041396 | -1.025109054 | 0.87756119 |
| ENSMUSG00000041399 | 1 | |
| ENSMUSG00000041406 | -1.00678575 | 1 |
| ENSMUSG00000041408 | -1.114805724 | 0.698112512 |
| ENSMUSG00000041415 | 1.118510032 | 0.633718027 |
| ENSMUSG00000041417 | 1.023060039 | 0.9602503 |
| ENSMUSG00000041420 | -1.052181569 | 0.608497229 |
| ENSMUSG00000041423 | -1.110033095 | 0.749809031 |
| ENSMUSG00000041426 | -1.105311347 | 0.608497229 |
| ENSMUSG00000041429 | 1.073093711 | 0.541735653 |
| ENSMUSG00000041431 | -1.113461918 | 0.698112512 |
| ENSMUSG00000041438 | 1.051987627 | 0.854549452 |
| ENSMUSG00000041439 | -1.119868008 | 0.397756797 |
| ENSMUSG00000041440 | -1.159638019 | 0.241835098 |
| ENSMUSG00000041444 | -1.04630149 | 0.807247723 |
| ENSMUSG00000041445 | -1.186404996 | 0.534887579 |
| ENSMUSG00000041449 | 1.72421027 | 0.002932067 |
| ENSMUSG00000041453 | -1.007832861 | 1 |
| ENSMUSG00000041459 | -1.000151826 | 1 |
| ENSMUSG00000041460 | -1.00348316 | 1 |
| ENSMUSG00000041468 | -1 | 1 |
| ENSMUSG00000041471 | -1.092962188 | 0.608497229 |
| ENSMUSG00000041476 | 1 | |
| ENSMUSG00000041477 | -1.086900649 | 0.245089058 |
| ENSMUSG00000041479 | 1.12442171 | 0.698112512 |
| ENSMUSG00000041481 | 1.238923633 | 0.123449197 |
| ENSMUSG00000041482 | -1.014541826 | 0.964323862 |
| ENSMUSG00000041483 | 1.03462244 | 0.908763622 |
| ENSMUSG00000041488 | 1.154883083 | 0.338127691 |
| ENSMUSG00000041491 | 1.07207959 | 0.585537169 |
| ENSMUSG00000041498 | -1.083680162 | 0.698112512 |
| ENSMUSG00000041505 | 1 | |
| ENSMUSG00000041506 | 1.011117472 | 0.988424085 |
| ENSMUSG00000041512 | 1 | |
| ENSMUSG00000041515 | -1.808373465 | 0.001760797 |
| ENSMUSG00000041523 | 1 | |
| ENSMUSG00000041528 | -1.007303234 | 1 |
| ENSMUSG00000041530 | 1.004916866 | 1 |
| ENSMUSG00000041534 | 1 | |

| | | |
|---|---|---|
| ENSMUSG00000041536 | 1 | |
| ENSMUSG00000041538 | 1 | |
| ENSMUSG00000041540 | 1.753272915 | 0.003813643 |
| ENSMUSG00000041544 | -1.113515092 | 0.576169279 |
| ENSMUSG00000041548 | 1.143341128 | 0.459638654 |
| ENSMUSG00000041550 | 1.069068979 | 0.608497229 |
| ENSMUSG00000041552 | 1.413306049 | 0.217803313 |
| ENSMUSG00000041556 | -1.408440406 | 0.028717879 |
| ENSMUSG00000041559 | 2.645767541 | 0.000149712 |
| ENSMUSG00000041560 | 1.010788798 | 0.975842677 |
| ENSMUSG00000041565 | 1 | |
| ENSMUSG00000041566 | -1.072695076 | 0.815807184 |
| ENSMUSG00000041567 | 1.05410549 | 0.805157604 |
| ENSMUSG00000041570 | 1.018426554 | 0.989895137 |
| ENSMUSG00000041571 | 1.185095227 | 0.210268462 |
| ENSMUSG00000041577 | 1.54719379 | 0.004649604 |
| ENSMUSG00000041578 | 1.097320342 | 0.496313168 |
| ENSMUSG00000041583 | 1 | |
| ENSMUSG00000041592 | 1.030055135 | 0.973126641 |
| ENSMUSG00000041594 | 1.100229651 | 0.564317454 |
| ENSMUSG00000041596 | -1.753552745 | 0.004580147 |
| ENSMUSG00000041598 | 1.084838703 | 0.565460954 |
| ENSMUSG00000041605 | 1.028930704 | 0.803072249 |
| ENSMUSG00000041607 | 1.063364628 | 0.346734756 |
| ENSMUSG00000041608 | -1.93685268 | 0.001393029 |
| ENSMUSG00000041609 | 1.183962186 | 0.518775336 |
| ENSMUSG00000041616 | 1.059463094 | 0.638307772 |
| ENSMUSG00000041617 | 1.081055692 | 0.608497229 |
| ENSMUSG00000041620 | 1 | |
| ENSMUSG00000041623 | 1.067481372 | 0.608497229 |
| ENSMUSG00000041624 | 1.209217364 | 0.124731965 |
| ENSMUSG00000041625 | -1.106845227 | 0.427481696 |
| ENSMUSG00000041629 | 1.074251543 | 0.397673648 |
| ENSMUSG00000041632 | -1.079169317 | 0.585646836 |
| ENSMUSG00000041633 | -1.255937045 | 0.044893595 |
| ENSMUSG00000041638 | 1.193667268 | 0.246342767 |
| ENSMUSG00000041642 | 1.19057913 | 0.142769553 |
| ENSMUSG00000041644 | 1 | |
| ENSMUSG00000041645 | 1.062126567 | 0.797035369 |
| ENSMUSG00000041649 | -1.347794777 | 0.045377104 |
| ENSMUSG00000041650 | -1.198829851 | 0.169844415 |
| ENSMUSG00000041653 | -1.204227554 | 0.608497229 |
| ENSMUSG00000041654 | -1.038935633 | 0.799873922 |
| ENSMUSG00000041658 | 1.119099089 | 0.329995202 |
| ENSMUSG00000041660 | 1 | |
| ENSMUSG00000041669 | -1.043306199 | 0.698112512 |
| ENSMUSG00000041670 | -1.044273782 | 0.698112512 |
| ENSMUSG00000041671 | 1.055207592 | 0.50530478 |
| ENSMUSG00000041673 | 1 | |
| ENSMUSG00000041674 | -1.00571839 | 1 |
| ENSMUSG00000041679 | -1.072633126 | 0.747599307 |
| ENSMUSG00000041681 | -1.053549062 | 0.698112512 |
| ENSMUSG00000041684 | -1.000211789 | 1 |
| ENSMUSG00000041685 | -1.035444558 | 0.921768868 |
| ENSMUSG00000041688 | -1.296057825 | 0.20251405 |
| ENSMUSG00000041695 | -1.176832895 | 0.375579698 |
| ENSMUSG00000041696 | 1.748297264 | 0.000620329 |
| ENSMUSG00000041697 | 1.070765654 | 0.608497229 |
| ENSMUSG00000041698 | 1.059463094 | 0.608497229 |
| ENSMUSG00000041700 | 1 | |
| ENSMUSG00000041702 | 1.136445284 | 0.548047779 |
| ENSMUSG00000041703 | -1.548741559 | 0.013886264 |
| ENSMUSG00000041707 | 1 | |

| | | |
|---|---|---|
| ENSMUSG00000041708 | 1 | |
| ENSMUSG00000041710 | -1.089497311 | 0.532369657 |
| ENSMUSG00000041712 | 1.116651216 | 0.608497229 |
| ENSMUSG00000041716 | 1.03937513 | 0.608497229 |
| ENSMUSG00000041718 | -1.206222846 | 0.220914883 |
| ENSMUSG00000041720 | -1.013220618 | 1 |
| ENSMUSG00000041722 | 1 | |
| ENSMUSG00000041729 | 1.135796808 | 0.120639867 |
| ENSMUSG00000041730 | 1 | |
| ENSMUSG00000041731 | 1.871159282 | 0.001631078 |
| ENSMUSG00000041733 | 1.062567213 | 0.698112512 |
| ENSMUSG00000041734 | 1.154307986 | 0.608497229 |
| ENSMUSG00000041735 | 1 | |
| ENSMUSG00000041736 | 1.399395453 | 0.024753959 |
| ENSMUSG00000041737 | -1.152299099 | 0.608497229 |
| ENSMUSG00000041740 | 1.050309877 | 0.885184589 |
| ENSMUSG00000041741 | 1.161036672 | 0.378751351 |
| ENSMUSG00000041747 | -1.041143078 | 0.891597525 |
| ENSMUSG00000041750 | 1 | |
| ENSMUSG00000041754 | 1 | |
| ENSMUSG00000041757 | -1.352048121 | 0.011667475 |
| ENSMUSG00000041762 | -1.270817671 | 0.044140978 |
| ENSMUSG00000041763 | -1.001205334 | 1 |
| ENSMUSG00000041765 | -1.124045429 | 0.458168354 |
| ENSMUSG00000041769 | -1.036803242 | 0.8950853 |
| ENSMUSG00000041771 | -1.044273782 | 0.698112512 |
| ENSMUSG00000041773 | 1.198514233 | 0.298655863 |
| ENSMUSG00000041774 | -1.267593752 | 0.088689988 |
| ENSMUSG00000041775 | 1.034729352 | 0.731822847 |
| ENSMUSG00000041777 | -1.097376287 | 0.361870084 |
| ENSMUSG00000041779 | 1.011825407 | 1 |
| ENSMUSG00000041781 | 1.06944605 | 0.812659885 |
| ENSMUSG00000041782 | -1.31818988 | 0.030325379 |
| ENSMUSG00000041789 | -1.043306199 | 0.698112512 |
| ENSMUSG00000041791 | 1 | |
| ENSMUSG00000041794 | 1.034095782 | 0.936888542 |
| ENSMUSG00000041797 | -2.579804069 | 0.001055626 |
| ENSMUSG00000041798 | -1.028972438 | 0.971859054 |
| ENSMUSG00000041801 | 1.271032011 | 0.011933914 |
| ENSMUSG00000041805 | 1 | |
| ENSMUSG00000041809 | -1.068917436 | 0.704170349 |
| ENSMUSG00000041815 | 1.033497164 | 0.945276261 |
| ENSMUSG00000041817 | -1.204115404 | 0.079314923 |
| ENSMUSG00000041827 | -1.044273782 | 0.746899902 |
| ENSMUSG00000041828 | 1 | |
| ENSMUSG00000041831 | 1.049411284 | 0.863033776 |
| ENSMUSG00000041836 | -1.018257994 | 0.877390802 |
| ENSMUSG00000041837 | -1.013255514 | 0.9858962 |
| ENSMUSG00000041840 | -1.04764404 | 0.823140628 |
| ENSMUSG00000041841 | 1.101834411 | 0.608497229 |
| ENSMUSG00000041842 | 1.021508685 | 0.928987465 |
| ENSMUSG00000041845 | -1.040729034 | 0.828564184 |
| ENSMUSG00000041846 | 1.165240047 | 0.509571857 |
| ENSMUSG00000041849 | 1.046027437 | 0.663452129 |
| ENSMUSG00000041852 | 1.050244357 | 0.868451015 |
| ENSMUSG00000041857 | 1.059463094 | 0.608497229 |
| ENSMUSG00000041859 | -1.000885205 | 1 |
| ENSMUSG00000041870 | -1.047221567 | 0.871696426 |
| ENSMUSG00000041872 | 1.031783213 | 0.95247261 |
| ENSMUSG00000041878 | -1.170313042 | 0.571251845 |
| ENSMUSG00000041879 | -1.005667286 | 1 |
| ENSMUSG00000041881 | 1.103305754 | 0.38389163 |
| ENSMUSG00000041885 | 1 | |

| | | |
|---|---|---|
| ENSMUSG00000041886 | 2.127059413 | 0.001219416 |
| ENSMUSG00000041889 | 1.019321986 | 0.904050368 |
| ENSMUSG00000041890 | 1.079558114 | 0.698112512 |
| ENSMUSG00000041891 | 1.036186164 | 0.938462359 |
| ENSMUSG00000041895 | 1.056233765 | 0.698112512 |
| ENSMUSG00000041907 | -1.20574565 | 0.245089058 |
| ENSMUSG00000041911 | 1.069019894 | 0.244162929 |
| ENSMUSG00000041912 | 1.198030594 | 0.271595502 |
| ENSMUSG00000041915 | 1.046163139 | 0.847756749 |
| ENSMUSG00000041920 | 1.089466538 | 0.608497229 |
| ENSMUSG00000041921 | -1.098298543 | 0.579051778 |
| ENSMUSG00000041923 | 1.320903788 | 0.006243022 |
| ENSMUSG00000041926 | -1.006220367 | 1 |
| ENSMUSG00000041930 | -1.193337073 | 0.591151279 |
| ENSMUSG00000041935 | 1.081880381 | 0.747285017 |
| ENSMUSG00000041936 | -1.069462145 | 0.62641634 |
| ENSMUSG00000041939 | 1.094546883 | 0.449476937 |
| ENSMUSG00000041945 | -1.027154608 | 0.854767483 |
| ENSMUSG00000041949 | -1.041349494 | 0.698112512 |
| ENSMUSG00000041954 | 1.647928517 | 0.019853058 |
| ENSMUSG00000041957 | 1.06603359 | 0.750742698 |
| ENSMUSG00000041958 | -1.010715942 | 0.991425134 |
| ENSMUSG00000041959 | -1.105713658 | 0.735085405 |
| ENSMUSG00000041961 | 1.085497636 | 0.543962054 |
| ENSMUSG00000041966 | -1.148720856 | 0.251049598 |
| ENSMUSG00000041974 | 1.097813182 | 0.6659463 |
| ENSMUSG00000041975 | -1.091331653 | 0.608497229 |
| ENSMUSG00000041977 | -1.018301459 | 0.979002724 |
| ENSMUSG00000041984 | 1 | |
| ENSMUSG00000041986 | -1.713166 | 0.006752766 |
| ENSMUSG00000041991 | 1 | |
| ENSMUSG00000041992 | -1.426181377 | 0.141627783 |
| ENSMUSG00000041995 | -1.045205697 | 0.731295483 |
| ENSMUSG00000041997 | -1.039519329 | 0.905298097 |
| ENSMUSG00000042002 | 1 | |
| ENSMUSG00000042010 | 1.24484267 | 0.160676302 |
| ENSMUSG00000042015 | -1.148615587 | 0.387425609 |
| ENSMUSG00000042029 | 1.074922279 | 0.79291651 |
| ENSMUSG00000042031 | 1 | |
| ENSMUSG00000042032 | 1.029117528 | 0.942171713 |
| ENSMUSG00000042035 | 1.126333278 | 0.169600498 |
| ENSMUSG00000042041 | 1 | |
| ENSMUSG00000042042 | 1.013574198 | 1 |
| ENSMUSG00000042043 | -1.10286724 | 0.698112512 |
| ENSMUSG00000042045 | 1 | |
| ENSMUSG00000042046 | -1.015311858 | 0.98837815 |
| ENSMUSG00000042050 | 1.071959284 | 0.638354353 |
| ENSMUSG00000042055 | 1.009213848 | 1 |
| ENSMUSG00000042063 | 1.065868819 | 0.724515237 |
| ENSMUSG00000042064 | 1 | |
| ENSMUSG00000042066 | 1.162305554 | 0.066143749 |
| ENSMUSG00000042073 | 1.099019844 | 0.250477583 |
| ENSMUSG00000042078 | 1.169729363 | 0.56051496 |
| ENSMUSG00000042079 | 1.138706457 | 0.608497229 |
| ENSMUSG00000042082 | -1.033972002 | 0.850761932 |
| ENSMUSG00000042087 | 1.14385464 | 0.713964595 |
| ENSMUSG00000042092 | 1 | |
| ENSMUSG00000042096 | -1.137443589 | 0.608497229 |
| ENSMUSG00000042097 | -1.136979035 | 0.176183437 |
| ENSMUSG00000042099 | 1.099282726 | 0.416787698 |
| ENSMUSG00000042102 | -1 | 1 |
| ENSMUSG00000042104 | -1.074772281 | 0.743920175 |
| ENSMUSG00000042105 | -1.020980096 | 0.968180243 |

| | | |
|---|---|---|
| ENSMUSG00000042106 | -1.263468853 | 0.2729627 |
| ENSMUSG00000042109 | -1.140154348 | 0.025983253 |
| ENSMUSG00000042111 | -1.00263694 | 1 |
| ENSMUSG00000042115 | 1.012899766 | 0.942160514 |
| ENSMUSG00000042116 | -1.290164642 | 0.105136419 |
| ENSMUSG00000042118 | 1.093846635 | 0.608863038 |
| ENSMUSG00000042121 | 1.198117283 | 0.152214706 |
| ENSMUSG00000042124 | 1 | |
| ENSMUSG00000042129 | 1.49031071 | 0.000428111 |
| ENSMUSG00000042133 | -1.101887365 | 0.692415612 |
| ENSMUSG00000042138 | 1.011554051 | 0.984762771 |
| ENSMUSG00000042148 | -1.053892862 | 0.624676024 |
| ENSMUSG00000042155 | -1.112696145 | 0.608497229 |
| ENSMUSG00000042156 | -1.11303753 | 0.532369657 |
| ENSMUSG00000042157 | 1 | |
| ENSMUSG00000042165 | 1.029490685 | 0.943298961 |
| ENSMUSG00000042167 | 1.021271937 | 0.971248775 |
| ENSMUSG00000042178 | 1.063245999 | 0.730102564 |
| ENSMUSG00000042179 | 1.059463094 | 0.698112512 |
| ENSMUSG00000042182 | -1.010356169 | 0.978782337 |
| ENSMUSG00000042184 | 1 | |
| ENSMUSG00000042185 | -1.01698316 | 0.964665151 |
| ENSMUSG00000042189 | 1 | |
| ENSMUSG00000042190 | 1.26205757 | 0.246522583 |
| ENSMUSG00000042195 | -2.181774728 | 0.000905461 |
| ENSMUSG00000042197 | 1.013058398 | 1 |
| ENSMUSG00000042198 | -1.004339387 | 1 |
| ENSMUSG00000042200 | -1.053549062 | 0.769745556 |
| ENSMUSG00000042202 | -1.02589127 | 0.94651494 |
| ENSMUSG00000042203 | -1.010463046 | 0.972980766 |
| ENSMUSG00000042207 | 1.085468659 | 0.698112512 |
| ENSMUSG00000042208 | -1.139104905 | 0.561552957 |
| ENSMUSG00000042210 | -1.084642079 | 0.585282823 |
| ENSMUSG00000042211 | 1.019645826 | 0.981964814 |
| ENSMUSG00000042212 | 1 | |
| ENSMUSG00000042213 | 1.129973336 | 0.107737671 |
| ENSMUSG00000042215 | 1.091469632 | 0.698112512 |
| ENSMUSG00000042216 | -1.720250367 | 0.000456184 |
| ENSMUSG00000042219 | 1 | |
| ENSMUSG00000042225 | -1.154588756 | 0.484584722 |
| ENSMUSG00000042228 | -1.164821648 | 0.05952468 |
| ENSMUSG00000042229 | 1.013000664 | 0.994699331 |
| ENSMUSG00000042240 | 1 | |
| ENSMUSG00000042243 | 1 | |
| ENSMUSG00000042244 | 1 | |
| ENSMUSG00000042246 | -1.383320291 | 0.096520613 |
| ENSMUSG00000042248 | 1 | |
| ENSMUSG00000042249 | -1.116264357 | 0.587557629 |
| ENSMUSG00000042250 | 1 | |
| ENSMUSG00000042251 | -1.212040609 | 0.492382162 |
| ENSMUSG00000042254 | 1.050110043 | 0.767772211 |
| ENSMUSG00000042256 | 1.055849488 | 0.737673689 |
| ENSMUSG00000042258 | 1 | |
| ENSMUSG00000042262 | 1 | |
| ENSMUSG00000042265 | 1 | |
| ENSMUSG00000042268 | 1.015486245 | 0.985522133 |
| ENSMUSG00000042269 | -1.084645496 | 0.608497229 |
| ENSMUSG00000042271 | -1.120221943 | 0.532369657 |
| ENSMUSG00000042272 | -1.036108395 | 0.880723094 |
| ENSMUSG00000042275 | 1.008579473 | 0.978962085 |
| ENSMUSG00000042279 | 1 | |
| ENSMUSG00000042282 | 1 | |
| ENSMUSG00000042284 | 1.19724941 | 0.160676302 |

| | | |
|---|---|---|
| ENSMUSG00000042286 | -1.265450823 | 0.216046507 |
| ENSMUSG00000042289 | 1.06252042 | 0.544868163 |
| ENSMUSG00000042292 | 1.169400516 | 0.22008413 |
| ENSMUSG00000042293 | 1.083316837 | 0.781240509 |
| ENSMUSG00000042298 | -1.085565984 | 0.633481288 |
| ENSMUSG00000042302 | -1.03608023 | 0.887325696 |
| ENSMUSG00000042303 | 1.022250658 | 0.923758858 |
| ENSMUSG00000042305 | -1.034543744 | 0.938555263 |
| ENSMUSG00000042306 | -1.23994515 | 0.476917435 |
| ENSMUSG00000042308 | 1.0425259 | 0.698112512 |
| ENSMUSG00000042312 | -1.109491503 | 0.58026581 |
| ENSMUSG00000042320 | 1.043624155 | 0.629100036 |
| ENSMUSG00000042323 | -1.119950553 | 0.698112512 |
| ENSMUSG00000042328 | 1.101805279 | 0.105522106 |
| ENSMUSG00000042331 | 1.030314238 | 0.956784261 |
| ENSMUSG00000042333 | -1.292792545 | 0.215991492 |
| ENSMUSG00000042340 | 1.007403607 | 1 |
| ENSMUSG00000042345 | 1 | |
| ENSMUSG00000042348 | -1.110448884 | 0.489918982 |
| ENSMUSG00000042349 | -1.062629131 | 0.608497229 |
| ENSMUSG00000042350 | 1.155880181 | 0.415539236 |
| ENSMUSG00000042351 | -1.03962336 | 0.698112512 |
| ENSMUSG00000042353 | 1 | |
| ENSMUSG00000042354 | -1.08168428 | 0.705745105 |
| ENSMUSG00000042357 | 1.409993124 | 0.007912905 |
| ENSMUSG00000042359 | -1.387365318 | 0.01153622 |
| ENSMUSG00000042360 | -1.044273782 | 0.790851299 |
| ENSMUSG00000042363 | 1.082392442 | 0.698112512 |
| ENSMUSG00000042364 | 1.045321918 | 0.851742081 |
| ENSMUSG00000042367 | 1.242141635 | 0.016423797 |
| ENSMUSG00000042369 | -1.058079636 | 0.751234476 |
| ENSMUSG00000042371 | 1 | |
| ENSMUSG00000042372 | 1 | |
| ENSMUSG00000042377 | -1.270413791 | 0.587631343 |
| ENSMUSG00000042379 | 1.015486245 | 0.972980766 |
| ENSMUSG00000042380 | 1.085259761 | 0.369930124 |
| ENSMUSG00000042385 | 1 | |
| ENSMUSG00000042386 | 1.073854713 | 0.729018288 |
| ENSMUSG00000042388 | -1.280398796 | 0.486384474 |
| ENSMUSG00000042389 | -1.007345503 | 0.984902895 |
| ENSMUSG00000042390 | -1.002900349 | 1 |
| ENSMUSG00000042396 | -1.030836075 | 0.936653013 |
| ENSMUSG00000042401 | -1.085651418 | 0.532369657 |
| ENSMUSG00000042402 | 1 | |
| ENSMUSG00000042404 | 1.098576183 | 0.219469516 |
| ENSMUSG00000042406 | 1.120711407 | 0.661155252 |
| ENSMUSG00000042408 | -1.031290871 | 0.8909227 |
| ENSMUSG00000042410 | -1.087152191 | 0.756216134 |
| ENSMUSG00000042414 | 1 | |
| ENSMUSG00000042417 | -1.31637473 | 0.060794345 |
| ENSMUSG00000042419 | 1.055420242 | 0.807247723 |
| ENSMUSG00000042423 | 1.103842058 | 0.187611488 |
| ENSMUSG00000042425 | -1.132666855 | 0.534459797 |
| ENSMUSG00000042426 | -1.110321793 | 0.657973953 |
| ENSMUSG00000042428 | 1.032499917 | 0.880782387 |
| ENSMUSG00000042429 | 1.252671701 | 0.197925365 |
| ENSMUSG00000042433 | 1 | |
| ENSMUSG00000042436 | 1.193921123 | 0.078461318 |
| ENSMUSG00000042439 | 1.347588071 | 0.002044228 |
| ENSMUSG00000042444 | -1.018843483 | 0.951399668 |
| ENSMUSG00000042446 | 1.043811319 | 0.903464611 |
| ENSMUSG00000042447 | 1.068507025 | 0.698112512 |
| ENSMUSG00000042448 | 1.059463094 | 0.608497229 |

| | | |
|---|---|---|
| ENSMUSG00000042451 | -1 | 1 |
| ENSMUSG00000042453 | -1.028348525 | 0.912425817 |
| ENSMUSG00000042459 | 1 | |
| ENSMUSG00000042460 | 1.087898529 | 0.702117806 |
| ENSMUSG00000042462 | 1.030743852 | 0.814089245 |
| ENSMUSG00000042472 | 1.131164898 | 0.550973458 |
| ENSMUSG00000042473 | -1.015316685 | 0.997179782 |
| ENSMUSG00000042474 | 1 | |
| ENSMUSG00000042476 | -1.107899615 | 0.670045943 |
| ENSMUSG00000042477 | -1.965160491 | 0.000622261 |
| ENSMUSG00000042485 | 1.332096072 | 0.000550468 |
| ENSMUSG00000042487 | -1.06861536 | 0.698112512 |
| ENSMUSG00000042489 | -1.069644704 | 0.789142226 |
| ENSMUSG00000042492 | 1.005860799 | 1 |
| ENSMUSG00000042496 | 1.004043186 | 1 |
| ENSMUSG00000042498 | 1.08389928 | 0.481992415 |
| ENSMUSG00000042499 | -1.043306199 | 0.803260194 |
| ENSMUSG00000042500 | -1.157577433 | 0.197145184 |
| ENSMUSG00000042501 | -1.063811128 | 0.7616481 |
| ENSMUSG00000042502 | 1.024910231 | 0.917717739 |
| ENSMUSG00000042505 | 1.158741627 | 0.31620604 |
| ENSMUSG00000042506 | -1.073461509 | 0.770958539 |
| ENSMUSG00000042507 | 1.084598829 | 0.548901322 |
| ENSMUSG00000042508 | 1.046354235 | 0.867519242 |
| ENSMUSG00000042510 | 1.26319478 | 0.021481917 |
| ENSMUSG00000042514 | -1.053549062 | 0.698112512 |
| ENSMUSG00000042515 | -1.17699251 | 0.493975913 |
| ENSMUSG00000042520 | 1.031722592 | 0.956972448 |
| ENSMUSG00000042523 | 1.037114106 | 0.879615717 |
| ENSMUSG00000042524 | 1.054785727 | 0.737457739 |
| ENSMUSG00000042525 | 1 | |
| ENSMUSG00000042529 | 1.009414875 | 1 |
| ENSMUSG00000042532 | 1.059421989 | 0.792603413 |
| ENSMUSG00000042535 | -1.049526362 | 0.81743209 |
| ENSMUSG00000042540 | -1.044273782 | 0.870103307 |
| ENSMUSG00000042541 | 1.130162022 | 0.524801011 |
| ENSMUSG00000042548 | -1.051132913 | 0.848953901 |
| ENSMUSG00000042549 | -1.140855488 | 0.022264274 |
| ENSMUSG00000042554 | 1 | |
| ENSMUSG00000042557 | -1.010744174 | 1 |
| ENSMUSG00000042558 | 1.164989779 | 0.155073998 |
| ENSMUSG00000042564 | -1.013846536 | 0.93004429 |
| ENSMUSG00000042567 | -1.015511103 | 0.982777255 |
| ENSMUSG00000042569 | -1.141007299 | 0.44811946 |
| ENSMUSG00000042570 | 1.05760855 | 0.608497229 |
| ENSMUSG00000042572 | 1.017812562 | 0.987419045 |
| ENSMUSG00000042579 | 1.087655159 | 0.698112512 |
| ENSMUSG00000042581 | 1.144603821 | 0.615267463 |
| ENSMUSG00000042589 | -1.21462871 | 0.161228902 |
| ENSMUSG00000042590 | -1.100422166 | 0.698112512 |
| ENSMUSG00000042594 | 1.044894415 | 0.731295483 |
| ENSMUSG00000042595 | -1.076878514 | 0.637081332 |
| ENSMUSG00000042596 | 1 | |
| ENSMUSG00000042599 | 1.1163259 | 0.608497229 |
| ENSMUSG00000042604 | 1.048493559 | 0.698112512 |
| ENSMUSG00000042605 | 1.159375128 | 0.455393748 |
| ENSMUSG00000042606 | -1.015131597 | 0.977628292 |
| ENSMUSG00000042607 | -1.033791285 | 0.802047771 |
| ENSMUSG00000042608 | 1.090775527 | 0.293839654 |
| ENSMUSG00000042613 | 1.222434815 | 0.104947513 |
| ENSMUSG00000042616 | -1.108921206 | 0.357091279 |
| ENSMUSG00000042622 | -1.107385478 | 0.382027404 |
| ENSMUSG00000042625 | 1.061698007 | 0.698112512 |

| | | |
|---|---|---|
| ENSMUSG00000042626 | 1.026178045 | 0.949812939 |
| ENSMUSG00000042628 | 1.191952124 | 0.188464329 |
| ENSMUSG00000042631 | 1 | |
| ENSMUSG00000042632 | 1.056037657 | 0.698112512 |
| ENSMUSG00000042638 | -1.043306199 | 0.698112512 |
| ENSMUSG00000042641 | -1.12843057 | 0.540891969 |
| ENSMUSG00000042642 | 1.021041624 | 0.877008742 |
| ENSMUSG00000042644 | 1.155611455 | 0.310464099 |
| ENSMUSG00000042647 | 1.038699682 | 0.744050104 |
| ENSMUSG00000042650 | -1.004389793 | 1 |
| ENSMUSG00000042655 | -1.089497311 | 0.532369657 |
| ENSMUSG00000042659 | 1.01560033 | 0.971034177 |
| ENSMUSG00000042660 | -1.000029857 | 1 |
| ENSMUSG00000042662 | 1.237108713 | 0.225539252 |
| ENSMUSG00000042670 | -1.07322002 | 0.698112512 |
| ENSMUSG00000042671 | -1.084645496 | 0.608497229 |
| ENSMUSG00000042672 | -1.257639256 | 0.087746554 |
| ENSMUSG00000042675 | 1.103003167 | 0.311020871 |
| ENSMUSG00000042677 | 1.338031438 | 0.019383936 |
| ENSMUSG00000042678 | -1.229681378 | 0.322644814 |
| ENSMUSG00000042680 | 1.077330749 | 0.375579698 |
| ENSMUSG00000042682 | -1.051186894 | 0.822748231 |
| ENSMUSG00000042684 | -1.557931818 | 0.003640465 |
| ENSMUSG00000042686 | 1.150208536 | 0.289843159 |
| ENSMUSG00000042688 | 1.008164864 | 1 |
| ENSMUSG00000042694 | -1.067834364 | 0.695387231 |
| ENSMUSG00000042699 | -1.00121753 | 1 |
| ENSMUSG00000042700 | 1.244012509 | 0.160438321 |
| ENSMUSG00000042705 | -1.011406658 | 0.997546475 |
| ENSMUSG00000042707 | -1.115335079 | 0.698249621 |
| ENSMUSG00000042708 | -1.16450103 | 0.49839667 |
| ENSMUSG00000042709 | -1.06767446 | 0.62087778 |
| ENSMUSG00000042712 | 1.081905625 | 0.775469341 |
| ENSMUSG00000042717 | 1 | |
| ENSMUSG00000042719 | 1.048043927 | 0.867195992 |
| ENSMUSG00000042724 | 1.170031398 | 0.608497229 |
| ENSMUSG00000042726 | 1.047999139 | 0.73636577 |
| ENSMUSG00000042729 | 1.016340608 | 0.980617339 |
| ENSMUSG00000042734 | -1.005219744 | 1 |
| ENSMUSG00000042737 | 1.230649993 | 0.521578812 |
| ENSMUSG00000042742 | 1.116780595 | 0.579859214 |
| ENSMUSG00000042743 | -1.244590359 | 0.085009113 |
| ENSMUSG00000042744 | 1.092791195 | 0.607661815 |
| ENSMUSG00000042745 | 1.07075988 | 0.46360902 |
| ENSMUSG00000042747 | 1.115140109 | 0.208940434 |
| ENSMUSG00000042750 | -1.510663634 | 0.05942834 |
| ENSMUSG00000042751 | -1.310355818 | 0.004854846 |
| ENSMUSG00000042757 | -1.189107718 | 0.415214532 |
| ENSMUSG00000042759 | 1.029158142 | 0.797105186 |
| ENSMUSG00000042761 | -1.028348525 | 0.938427602 |
| ENSMUSG00000042763 | -1.729574879 | 0.003401228 |
| ENSMUSG00000042766 | 1.238588392 | 0.031435814 |
| ENSMUSG00000042770 | 1.049176972 | 0.698112512 |
| ENSMUSG00000042772 | 1.040665257 | 0.898763027 |
| ENSMUSG00000042774 | 1 | |
| ENSMUSG00000042784 | -1.208403516 | 0.182932251 |
| ENSMUSG00000042787 | -1.002631846 | 1 |
| ENSMUSG00000042788 | 1.015486245 | 0.985522133 |
| ENSMUSG00000042790 | 1.033253509 | 0.86902396 |
| ENSMUSG00000042793 | 1.118364196 | 0.347267031 |
| ENSMUSG00000042796 | 1 | |
| ENSMUSG00000042797 | -1.191179927 | 0.568273418 |
| ENSMUSG00000042800 | -1.095294215 | 0.666055997 |

| | | |
|---|---|---|
| ENSMUSG00000042801 | 1 | |
| ENSMUSG00000042804 | 1.002790278 | 1 |
| ENSMUSG00000042807 | -1.107646822 | 0.594452979 |
| ENSMUSG00000042808 | 1 | |
| ENSMUSG00000042810 | 1.046226735 | 0.70299173 |
| ENSMUSG00000042812 | -1.971429438 | 0.00186802 |
| ENSMUSG00000042814 | -1.076196025 | 0.608497229 |
| ENSMUSG00000042816 | 1.04424822 | 0.887979704 |
| ENSMUSG00000042817 | -1.343631505 | 0.032224317 |
| ENSMUSG00000042821 | 1.383530563 | 0.12438714 |
| ENSMUSG00000042826 | -1.428580801 | 0.007949434 |
| ENSMUSG00000042828 | -1.194100432 | 0.3282068 |
| ENSMUSG00000042831 | -1.080508959 | 0.608497229 |
| ENSMUSG00000042834 | 1.316635372 | 0.124724137 |
| ENSMUSG00000042842 | 1.988013827 | 0.000215953 |
| ENSMUSG00000042845 | 1 | |
| ENSMUSG00000042846 | -1.024718486 | 0.938427134 |
| ENSMUSG00000042848 | 1 | |
| ENSMUSG00000042849 | 1 | |
| ENSMUSG00000042851 | -1.066004703 | 0.608497229 |
| ENSMUSG00000042854 | -1.165456281 | 0.343988983 |
| ENSMUSG00000042857 | -1.022180074 | 0.815807523 |
| ENSMUSG00000042861 | 1 | |
| ENSMUSG00000042863 | 1 | |
| ENSMUSG00000042869 | 1 | |
| ENSMUSG00000042870 | 1.174927299 | 0.078512016 |
| ENSMUSG00000042873 | -1.705171187 | 0.000259425 |
| ENSMUSG00000042874 | 1 | |
| ENSMUSG00000042888 | 1 | |
| ENSMUSG00000042894 | 1 | |
| ENSMUSG00000042895 | -1.044273782 | 0.855488553 |
| ENSMUSG00000042901 | 1.042894033 | 0.892511436 |
| ENSMUSG00000042903 | -1.038881768 | 0.720669657 |
| ENSMUSG00000042909 | 1 | |
| ENSMUSG00000042918 | -1.022105482 | 1 |
| ENSMUSG00000042938 | 1 | |
| ENSMUSG00000042940 | 1 | |
| ENSMUSG00000042942 | -1.236056212 | 0.043625418 |
| ENSMUSG00000042961 | -1.26240934 | 0.208516141 |
| ENSMUSG00000042962 | -1.162056369 | 0.608497229 |
| ENSMUSG00000042976 | 1 | |
| ENSMUSG00000042978 | 1.047709447 | 0.251544797 |
| ENSMUSG00000042985 | -1.093381656 | 0.541482582 |
| ENSMUSG00000042988 | 1.038004503 | 0.824174175 |
| ENSMUSG00000042992 | 1.06088809 | 0.708149303 |
| ENSMUSG00000042993 | 1 | |
| ENSMUSG00000042997 | 1.157730349 | 0.200917987 |
| ENSMUSG00000043003 | -1 | 1 |
| ENSMUSG00000043004 | 1.063198274 | 0.698112512 |
| ENSMUSG00000043008 | 1 | |
| ENSMUSG00000043013 | 1 | |
| ENSMUSG00000043015 | -1.004975205 | 1 |
| ENSMUSG00000043017 | 1.043683676 | 0.801443616 |
| ENSMUSG00000043019 | -1.025535154 | 0.962906408 |
| ENSMUSG00000043020 | -1.044273782 | 0.698112512 |
| ENSMUSG00000043024 | -1.261870676 | 0.071698973 |
| ENSMUSG00000043029 | -1.084645496 | 0.69530468 |
| ENSMUSG00000043036 | -1.087394692 | 0.780939634 |
| ENSMUSG00000043050 | 1.059463094 | 0.608497229 |
| ENSMUSG00000043051 | -1.302750936 | 0.057039027 |
| ENSMUSG00000043059 | 1.132757309 | 0.178370009 |
| ENSMUSG00000043060 | 1 | |
| ENSMUSG00000043061 | 1.04083203 | 0.846939416 |

| | | |
|---|---|---|
| ENSMUSG00000043065 | 1.038568595 | 0.840138604 |
| ENSMUSG00000043066 | 1 | |
| ENSMUSG00000043067 | 1.131330826 | 0.608497229 |
| ENSMUSG00000043068 | -1.198605858 | 0.171318163 |
| ENSMUSG00000043073 | -1 | 1 |
| ENSMUSG00000043079 | 1.02115575 | 0.928524132 |
| ENSMUSG00000043085 | -1.034239753 | 0.949812939 |
| ENSMUSG00000043087 | 1 | |
| ENSMUSG00000043088 | 1.009517015 | 1 |
| ENSMUSG00000043089 | 1 | |
| ENSMUSG00000043090 | -1.037965961 | 0.862557896 |
| ENSMUSG00000043091 | -1.035372435 | 0.892173042 |
| ENSMUSG00000043099 | 1.079322017 | 0.888246981 |
| ENSMUSG00000043102 | -1.228542003 | 0.324252919 |
| ENSMUSG00000043110 | -1 | 1 |
| ENSMUSG00000043119 | 1 | |
| ENSMUSG00000043122 | -1.044273782 | 0.698112512 |
| ENSMUSG00000043126 | -1.093674795 | 0.623872809 |
| ENSMUSG00000043131 | 1.060689163 | 0.862652423 |
| ENSMUSG00000043140 | 1.037829762 | 0.751234476 |
| ENSMUSG00000043144 | 1.059463094 | 0.704170349 |
| ENSMUSG00000043145 | 1 | |
| ENSMUSG00000043153 | 1.284245588 | 0.006373649 |
| ENSMUSG00000043154 | 1.092975094 | 0.66861278 |
| ENSMUSG00000043155 | -1.031100252 | 0.946575044 |
| ENSMUSG00000043157 | 1.189207115 | 0.294533546 |
| ENSMUSG00000043162 | 1.116602361 | 0.231697989 |
| ENSMUSG00000043164 | 1 | |
| ENSMUSG00000043165 | -1.157636302 | 0.595148653 |
| ENSMUSG00000043168 | 1 | |
| ENSMUSG00000043183 | -1.072269871 | 0.726686255 |
| ENSMUSG00000043190 | -1.091544724 | 0.317332491 |
| ENSMUSG00000043192 | 1.079206093 | 0.509007402 |
| ENSMUSG00000043207 | -1.035936237 | 0.938816727 |
| ENSMUSG00000043219 | 1.261328415 | 0.10939414 |
| ENSMUSG00000043223 | -1.266892792 | 0.425178941 |
| ENSMUSG00000043226 | 1 | |
| ENSMUSG00000043230 | -1 | 1 |
| ENSMUSG00000043241 | -1.050367128 | 0.845772805 |
| ENSMUSG00000043243 | 1.333486041 | 0.066088064 |
| ENSMUSG00000043251 | -1.064186128 | 0.763242034 |
| ENSMUSG00000043252 | 1.241895498 | 0.292352581 |
| ENSMUSG00000043257 | 1.015689307 | 0.784729033 |
| ENSMUSG00000043259 | 1.576615241 | 0.002092992 |
| ENSMUSG00000043262 | -1.002872896 | 1 |
| ENSMUSG00000043263 | 1 | |
| ENSMUSG00000043267 | 1 | |
| ENSMUSG00000043274 | 1 | |
| ENSMUSG00000043279 | 1.000197762 | 1 |
| ENSMUSG00000043282 | -1.040555503 | 0.866311904 |
| ENSMUSG00000043284 | -1.020256719 | 0.933085377 |
| ENSMUSG00000043286 | -1.015248956 | 0.994207501 |
| ENSMUSG00000043289 | 1.187745429 | 0.572947856 |
| ENSMUSG00000043290 | -1.027509043 | 0.941601923 |
| ENSMUSG00000043295 | -1.043306199 | 0.698112512 |
| ENSMUSG00000043298 | 1 | |
| ENSMUSG00000043300 | -1.011826982 | 1 |
| ENSMUSG00000043301 | 1 | |
| ENSMUSG00000043308 | 1 | |
| ENSMUSG00000043310 | 1 | |
| ENSMUSG00000043311 | -1.048030652 | 0.698112512 |
| ENSMUSG00000043312 | 1 | |
| ENSMUSG00000043313 | 1.325468702 | 0.073927492 |

| | | |
|---|---|---|
| ENSMUSG00000043314 | 1 | |
| ENSMUSG00000043319 | 1 | |
| ENSMUSG00000043323 | 1.108748565 | 0.223739612 |
| ENSMUSG00000043329 | 1 | |
| ENSMUSG00000043331 | 1 | |
| ENSMUSG00000043333 | 1.024174073 | 0.952119819 |
| ENSMUSG00000043336 | 1.137013818 | 0.608497229 |
| ENSMUSG00000043340 | 1.017339835 | 0.979002724 |
| ENSMUSG00000043342 | 2.284208651 | 0.001039425 |
| ENSMUSG00000043346 | -1.044273782 | 0.698112512 |
| ENSMUSG00000043354 | 1 | |
| ENSMUSG00000043357 | 1 | |
| ENSMUSG00000043366 | 1 | |
| ENSMUSG00000043372 | 1.053979021 | 0.690669961 |
| ENSMUSG00000043383 | 1 | |
| ENSMUSG00000043384 | -1.182094341 | 0.317332491 |
| ENSMUSG00000043385 | -1.085651418 | 0.532369657 |
| ENSMUSG00000043388 | -1.556575222 | 0.004890823 |
| ENSMUSG00000043391 | -1.423306483 | 0.007015963 |
| ENSMUSG00000043398 | -1.03962336 | 0.922391825 |
| ENSMUSG00000043410 | 1 | |
| ENSMUSG00000043411 | 1.01869143 | 0.98837815 |
| ENSMUSG00000043415 | -1.195111895 | 0.030169802 |
| ENSMUSG00000043418 | 1.069068979 | 0.608497229 |
| ENSMUSG00000043419 | -1.779063511 | 0.000946345 |
| ENSMUSG00000043421 | 1.130270889 | 0.211534133 |
| ENSMUSG00000043424 | -1.091460114 | 0.608497229 |
| ENSMUSG00000043429 | 1 | |
| ENSMUSG00000043430 | 1.122462048 | 0.452329901 |
| ENSMUSG00000043432 | -1.013539839 | 0.968937442 |
| ENSMUSG00000043439 | -1.176385954 | 0.076794255 |
| ENSMUSG00000043441 | 1.386963324 | 0.036681922 |
| ENSMUSG00000043445 | -1.001400117 | 1 |
| ENSMUSG00000043448 | -1 | 1 |
| ENSMUSG00000043453 | 1 | |
| ENSMUSG00000043456 | -1.470490294 | 0.006484725 |
| ENSMUSG00000043458 | 1.059463094 | 0.698112512 |
| ENSMUSG00000043460 | 1 | |
| ENSMUSG00000043461 | -1 | 1 |
| ENSMUSG00000043463 | -1.123034904 | 0.695038206 |
| ENSMUSG00000043467 | -1.003840492 | 1 |
| ENSMUSG00000043468 | -1 | 1 |
| ENSMUSG00000043472 | 1 | |
| ENSMUSG00000043483 | 1.01504674 | 1 |
| ENSMUSG00000043484 | -1 | 1 |
| ENSMUSG00000043485 | 1 | |
| ENSMUSG00000043487 | 1.19720865 | 0.034430051 |
| ENSMUSG00000043488 | 1 | |
| ENSMUSG00000043496 | 1.217842673 | 0.323793283 |
| ENSMUSG00000043498 | 1.241732237 | 0.533368474 |
| ENSMUSG00000043501 | 1.143420006 | 0.541735653 |
| ENSMUSG00000043505 | 1 | |
| ENSMUSG00000043510 | 1.075802383 | 0.504256967 |
| ENSMUSG00000043518 | 1.101647022 | 0.509717267 |
| ENSMUSG00000043522 | -1.043306199 | 0.698112512 |
| ENSMUSG00000043529 | 1 | |
| ENSMUSG00000043531 | 1.533030882 | 0.002923873 |
| ENSMUSG00000043535 | 1.010285909 | 1 |
| ENSMUSG00000043537 | 1 | |
| ENSMUSG00000043541 | -1.038000494 | 0.956756676 |
| ENSMUSG00000043542 | 1.026032704 | 0.937505238 |
| ENSMUSG00000043549 | -1.27480329 | 0.011705868 |
| ENSMUSG00000043556 | 1.389957622 | 0.043367921 |

| | | |
|---|---|---|
| ENSMUSG00000043557 | -1.264422408 | 0.106143572 |
| ENSMUSG00000043569 | 1 | |
| ENSMUSG00000043572 | 1.076915905 | 0.349469864 |
| ENSMUSG00000043583 | 1 | |
| ENSMUSG00000043587 | 1.272903763 | 0.076658288 |
| ENSMUSG00000043592 | 1 | |
| ENSMUSG00000043602 | -1.110391601 | 0.051043821 |
| ENSMUSG00000043605 | 1 | |
| ENSMUSG00000043613 | 1.553893694 | 0.003822302 |
| ENSMUSG00000043614 | -1.067968853 | 0.726677903 |
| ENSMUSG00000043618 | 1.020130262 | 1 |
| ENSMUSG00000043621 | 1 | |
| ENSMUSG00000043623 | -1 | 1 |
| ENSMUSG00000043629 | -1.244363333 | 0.301753247 |
| ENSMUSG00000043631 | -1.075675297 | 0.613169248 |
| ENSMUSG00000043633 | 1.144045886 | 0.407736299 |
| ENSMUSG00000043635 | 1.285529602 | 0.045349245 |
| ENSMUSG00000043639 | 1.347200757 | 0.003477763 |
| ENSMUSG00000043644 | 1.124748687 | 0.502465177 |
| ENSMUSG00000043648 | -1.015974852 | 0.973257087 |
| ENSMUSG00000043659 | 1 | |
| ENSMUSG00000043661 | 1.04898545 | 0.906601967 |
| ENSMUSG00000043664 | -1.174739284 | 0.547111524 |
| ENSMUSG00000043668 | 1 | |
| ENSMUSG00000043670 | -1.239222927 | 0.387277186 |
| ENSMUSG00000043671 | 1.010471828 | 1 |
| ENSMUSG00000043673 | -1.961708702 | 0.004069495 |
| ENSMUSG00000043681 | 1 | |
| ENSMUSG00000043683 | 1.042625589 | 0.779557148 |
| ENSMUSG00000043687 | 1.233346357 | 0.48076246 |
| ENSMUSG00000043692 | 1 | |
| ENSMUSG00000043698 | 1 | |
| ENSMUSG00000043702 | -1.131628151 | 0.364943811 |
| ENSMUSG00000043705 | 1 | |
| ENSMUSG00000043715 | 1 | |
| ENSMUSG00000043716 | -1.023255487 | 0.993795117 |
| ENSMUSG00000043719 | -1.085651418 | 0.608497229 |
| ENSMUSG00000043727 | 1 | |
| ENSMUSG00000043733 | 1.057765792 | 0.872024936 |
| ENSMUSG00000043740 | 1.027599531 | 0.990051834 |
| ENSMUSG00000043747 | 1.150179455 | 0.565559141 |
| ENSMUSG00000043753 | -1.077818579 | 0.681952832 |
| ENSMUSG00000043760 | 1.059463094 | 0.804224641 |
| ENSMUSG00000043770 | 1 | |
| ENSMUSG00000043773 | -1.222435934 | 0.029633641 |
| ENSMUSG00000043782 | -1.130883057 | 0.608497229 |
| ENSMUSG00000043787 | 1 | |
| ENSMUSG00000043789 | 1.221837453 | 0.205489632 |
| ENSMUSG00000043794 | 1.059463094 | 0.608497229 |
| ENSMUSG00000043795 | 1.430962284 | 0.134683572 |
| ENSMUSG00000043800 | 1.006183815 | 1 |
| ENSMUSG00000043801 | -1.03962336 | 0.698112512 |
| ENSMUSG00000043807 | -1.07955242 | 0.797830164 |
| ENSMUSG00000043811 | 1.51591202 | 0.048809518 |
| ENSMUSG00000043822 | 1.139003906 | 0.303895263 |
| ENSMUSG00000043827 | 1 | |
| ENSMUSG00000043831 | 1.060791472 | 0.404695089 |
| ENSMUSG00000043832 | 1 | |
| ENSMUSG00000043833 | 1.144226778 | 0.75035743 |
| ENSMUSG00000043843 | -1 | 1 |
| ENSMUSG00000043850 | 1.046854624 | 0.923381751 |
| ENSMUSG00000043855 | 1 | |
| ENSMUSG00000043857 | -1.240791708 | 0.491794669 |

| | | |
|---|---|---|
| ENSMUSG00000043858 | 1.147845062 | 0.28024069 |
| ENSMUSG00000043859 | 1 | |
| ENSMUSG00000043865 | 1 | |
| ENSMUSG00000043866 | 1.039941429 | 0.739099222 |
| ENSMUSG00000043870 | 1 | |
| ENSMUSG00000043872 | 1.001990102 | 1 |
| ENSMUSG00000043873 | 1 | |
| ENSMUSG00000043880 | 1 | |
| ENSMUSG00000043881 | -1.162511791 | 0.167753086 |
| ENSMUSG00000043885 | -1.106897054 | 0.619107467 |
| ENSMUSG00000043889 | 1.420211674 | 0.035331602 |
| ENSMUSG00000043892 | 1 | |
| ENSMUSG00000043895 | -1.031657007 | 0.877881982 |
| ENSMUSG00000043897 | 1 | |
| ENSMUSG00000043903 | 1.314459943 | 0.007129528 |
| ENSMUSG00000043909 | -1.018797343 | 0.982489318 |
| ENSMUSG00000043911 | 1 | |
| ENSMUSG00000043913 | -1.138631955 | 0.626206128 |
| ENSMUSG00000043923 | -1.155468024 | 0.117669723 |
| ENSMUSG00000043924 | -1.316723162 | 0.288085994 |
| ENSMUSG00000043925 | 1.059463094 | 0.699340382 |
| ENSMUSG00000043929 | -1.04507036 | 0.814384306 |
| ENSMUSG00000043931 | 1 | |
| ENSMUSG00000043932 | 1 | |
| ENSMUSG00000043939 | 1.0141192 | 0.984902895 |
| ENSMUSG00000043940 | 1.11385418 | 0.608497229 |
| ENSMUSG00000043943 | -1.305762969 | 0.240278784 |
| ENSMUSG00000043945 | 1 | |
| ENSMUSG00000043948 | 1 | |
| ENSMUSG00000043953 | -1.02417757 | 0.968937442 |
| ENSMUSG00000043962 | 1.036714535 | 0.929536187 |
| ENSMUSG00000043964 | 1.061426556 | 0.666633965 |
| ENSMUSG00000043969 | 1.148635064 | 0.240825255 |
| ENSMUSG00000043972 | -1.147838869 | 0.195607801 |
| ENSMUSG00000043982 | 1 | |
| ENSMUSG00000043986 | -1.085651418 | 0.532369657 |
| ENSMUSG00000043987 | -1.000426099 | 1 |
| ENSMUSG00000043991 | -1.002569062 | 1 |
| ENSMUSG00000043993 | -1.139160638 | 0.208579897 |
| ENSMUSG00000043998 | 1.100102776 | 0.631959005 |
| ENSMUSG00000043999 | -1.208362341 | 0.449102754 |
| ENSMUSG00000044005 | -1.232629055 | 0.00846921 |
| ENSMUSG00000044006 | -1.037072151 | 0.96972873 |
| ENSMUSG00000044014 | 1.014545335 | 0.979002724 |
| ENSMUSG00000044017 | -1 | 1 |
| ENSMUSG00000044018 | 1.029082996 | 0.942062052 |
| ENSMUSG00000044021 | 1.132639128 | 0.398767938 |
| ENSMUSG00000044022 | 1.267778078 | 0.155181033 |
| ENSMUSG00000044024 | 1.011592316 | 1 |
| ENSMUSG00000044025 | 1 | |
| ENSMUSG00000044026 | -1.021632453 | 0.954497929 |
| ENSMUSG00000044029 | 1 | |
| ENSMUSG00000044030 | 1.034919467 | 0.81398533 |
| ENSMUSG00000044033 | -1.325547106 | 0.080254337 |
| ENSMUSG00000044034 | 1.010351027 | 1 |
| ENSMUSG00000044037 | -1.451427625 | 0.017903363 |
| ENSMUSG00000044039 | 1 | |
| ENSMUSG00000044040 | 1 | |
| ENSMUSG00000044041 | 1.72338764 | 0.002269744 |
| ENSMUSG00000044042 | 1.075249135 | 0.670176133 |
| ENSMUSG00000044043 | 1.38372869 | 0.217803313 |
| ENSMUSG00000044052 | 1.154183614 | 0.608497229 |
| ENSMUSG00000044055 | -1.003144677 | 1 |

| | | |
|---|---|---|
| ENSMUSG00000044056 | -1.114360938 | 0.597011942 |
| ENSMUSG00000044058 | 1 | |
| ENSMUSG00000044060 | 1.109898041 | 0.145512736 |
| ENSMUSG00000044061 | 1 | |
| ENSMUSG00000044062 | 1 | |
| ENSMUSG00000044065 | 1 | |
| ENSMUSG00000044066 | -1.016238623 | 0.979002724 |
| ENSMUSG00000044067 | 1.084618945 | 0.614071182 |
| ENSMUSG00000044068 | -1.193204468 | 0.103709162 |
| ENSMUSG00000044071 | 1.12691148 | 0.536504702 |
| ENSMUSG00000044072 | -1.086473025 | 0.197804466 |
| ENSMUSG00000044080 | 1.07230027 | 0.382393655 |
| ENSMUSG00000044081 | -1.346801586 | 0.11469433 |
| ENSMUSG00000044083 | 1.130984238 | 0.724413892 |
| ENSMUSG00000044084 | 1 | |
| ENSMUSG00000044086 | 1 | |
| ENSMUSG00000044092 | -1.032079652 | 0.856915299 |
| ENSMUSG00000044098 | 1.053299648 | 0.769745556 |
| ENSMUSG00000044103 | 1.043139565 | 0.912425817 |
| ENSMUSG00000044106 | 1 | |
| ENSMUSG00000044107 | 1 | |
| ENSMUSG00000044117 | 1.113727475 | 0.698112512 |
| ENSMUSG00000044120 | 1 | |
| ENSMUSG00000044121 | 1 | |
| ENSMUSG00000044122 | 1.066693896 | 0.674781559 |
| ENSMUSG00000044125 | 1 | |
| ENSMUSG00000044134 | 1.159346967 | 0.216046507 |
| ENSMUSG00000044139 | 1.208205767 | 0.608497229 |
| ENSMUSG00000044141 | 1.05038453 | 0.828564184 |
| ENSMUSG00000044145 | 1.055978921 | 0.730297627 |
| ENSMUSG00000044147 | 1.191359645 | 0.350957446 |
| ENSMUSG00000044148 | 1.038460228 | 0.863066555 |
| ENSMUSG00000044149 | -1.00208338 | 1 |
| ENSMUSG00000044150 | -1.109500359 | 0.596552735 |
| ENSMUSG00000044155 | 1.124486281 | 0.570914055 |
| ENSMUSG00000044156 | -1.144460846 | 0.608497229 |
| ENSMUSG00000044162 | 1.034887047 | 0.814608539 |
| ENSMUSG00000044164 | -1.153077078 | 0.460496269 |
| ENSMUSG00000044165 | 1.102024305 | 0.698112512 |
| ENSMUSG00000044167 | 1.165722502 | 0.302869899 |
| ENSMUSG00000044170 | -1.053549062 | 0.698112512 |
| ENSMUSG00000044172 | -1 | 1 |
| ENSMUSG00000044176 | -1.285944016 | 0.075140239 |
| ENSMUSG00000044177 | 1.061786598 | 0.862652423 |
| ENSMUSG00000044186 | 1 | |
| ENSMUSG00000044197 | 1.094156974 | 0.240943902 |
| ENSMUSG00000044199 | -1.155252261 | 0.596589772 |
| ENSMUSG00000044201 | -1.109926104 | 0.608497229 |
| ENSMUSG00000044205 | 1 | |
| ENSMUSG00000044206 | 1 | |
| ENSMUSG00000044211 | -1.071075483 | 0.698112512 |
| ENSMUSG00000044213 | 1 | |
| ENSMUSG00000044216 | 1.227557117 | 0.351201286 |
| ENSMUSG00000044217 | 1.836556961 | 0.002447094 |
| ENSMUSG00000044220 | -1.391891038 | 0.192289864 |
| ENSMUSG00000044221 | 1.048394804 | 0.886840208 |
| ENSMUSG00000044222 | 1 | |
| ENSMUSG00000044224 | -1.026469794 | 0.944721414 |
| ENSMUSG00000044227 | 1 | |
| ENSMUSG00000044229 | -1.153203351 | 0.048958103 |
| ENSMUSG00000044231 | 1.251593336 | 0.068335953 |
| ENSMUSG00000044243 | 1 | |
| ENSMUSG00000044244 | -1.031920916 | 0.92200539 |

| | | |
|---|---|---|
| ENSMUSG00000044248 | 1 | |
| ENSMUSG00000044249 | 1 | |
| ENSMUSG00000044250 | -1.676461124 | 0.001835833 |
| ENSMUSG00000044252 | 1.084545004 | 0.645934663 |
| ENSMUSG00000044254 | 1.480630969 | 0.035594665 |
| ENSMUSG00000044258 | 1.127218873 | 0.608497229 |
| ENSMUSG00000044265 | 1 | |
| ENSMUSG00000044268 | 1 | |
| ENSMUSG00000044276 | -1.198060551 | 0.255219415 |
| ENSMUSG00000044279 | -1.03962336 | 0.803072249 |
| ENSMUSG00000044285 | -1.050474242 | 0.859353499 |
| ENSMUSG00000044286 | -1.044273782 | 0.698112512 |
| ENSMUSG00000044287 | 1 | |
| ENSMUSG00000044288 | 1.165003909 | 0.511163311 |
| ENSMUSG00000044292 | 1 | |
| ENSMUSG00000044293 | 1 | |
| ENSMUSG00000044294 | 1 | |
| ENSMUSG00000044296 | -1.390477696 | 0.010784484 |
| ENSMUSG00000044303 | 1.168108918 | 0.194441974 |
| ENSMUSG00000044306 | -1 | 1 |
| ENSMUSG00000044308 | -1.146954092 | 0.608497229 |
| ENSMUSG00000044309 | -1 | 1 |
| ENSMUSG00000044312 | 1 | |
| ENSMUSG00000044313 | 1.560111316 | 0.026089338 |
| ENSMUSG00000044317 | -1.136856222 | 0.608497229 |
| ENSMUSG00000044320 | 1.073344276 | 0.811497438 |
| ENSMUSG00000044322 | 1.069068979 | 0.608497229 |
| ENSMUSG00000044328 | 1.083617806 | 0.650904225 |
| ENSMUSG00000044330 | -1 | 1 |
| ENSMUSG00000044337 | -1.102008066 | 0.608497229 |
| ENSMUSG00000044338 | 1.005613439 | 1 |
| ENSMUSG00000044339 | 1.230806994 | 0.101457656 |
| ENSMUSG00000044340 | 1.216765059 | 0.132458218 |
| ENSMUSG00000044345 | 1.077747529 | 0.69305964 |
| ENSMUSG00000044348 | -1.228848194 | 0.045280365 |
| ENSMUSG00000044349 | -1.254141656 | 0.187066125 |
| ENSMUSG00000044350 | -1.099281064 | 0.608497229 |
| ENSMUSG00000044352 | -1.073877404 | 0.801443616 |
| ENSMUSG00000044359 | 1.00177232 | 1 |
| ENSMUSG00000044361 | 1.182113745 | 0.435149618 |
| ENSMUSG00000044362 | 1.067949606 | 0.835189692 |
| ENSMUSG00000044364 | -1.157144386 | 0.608497229 |
| ENSMUSG00000044365 | 1.502207334 | 0.007129528 |
| ENSMUSG00000044367 | 1.113504685 | 0.014562998 |
| ENSMUSG00000044375 | 1 | |
| ENSMUSG00000044377 | 1.059463094 | 0.698112512 |
| ENSMUSG00000044378 | -1.044273782 | 0.698112512 |
| ENSMUSG00000044387 | 1 | |
| ENSMUSG00000044390 | -1.040507329 | 0.608497229 |
| ENSMUSG00000044393 | -1.653393105 | 0.022837869 |
| ENSMUSG00000044400 | -1.111880267 | 0.608497229 |
| ENSMUSG00000044405 | 1.015486245 | 0.985522133 |
| ENSMUSG00000044408 | 1.048484697 | 0.842619477 |
| ENSMUSG00000044424 | 1.147395286 | 0.698112512 |
| ENSMUSG00000044429 | 1 | |
| ENSMUSG00000044430 | -1.043306199 | 0.803260194 |
| ENSMUSG00000044433 | -1.218084579 | 0.214467205 |
| ENSMUSG00000044434 | -1 | 1 |
| ENSMUSG00000044441 | 1 | |
| ENSMUSG00000044442 | 1.009541516 | 1 |
| ENSMUSG00000044444 | 1 | |
| ENSMUSG00000044447 | -1.149267826 | 0.496494186 |
| ENSMUSG00000044452 | 1.020327412 | 0.930815125 |

| | | |
|---|---|---|
| ENSMUSG00000044453 | -1.03962336 | 0.814089245 |
| ENSMUSG00000044454 | 1 | |
| ENSMUSG00000044456 | -1.0669589 | 0.608497229 |
| ENSMUSG00000044457 | -1 | 1 |
| ENSMUSG00000044461 | -1.494998196 | 0.029941261 |
| ENSMUSG00000044465 | 1.089594283 | 0.47120728 |
| ENSMUSG00000044468 | -1.361081462 | 0.027828404 |
| ENSMUSG00000044469 | 1.095194131 | 0.116493951 |
| ENSMUSG00000044471 | -1.044273782 | 0.795446726 |
| ENSMUSG00000044475 | 1.048130036 | 0.698112512 |
| ENSMUSG00000044477 | 1.117654683 | 0.411678378 |
| ENSMUSG00000044485 | 1 | |
| ENSMUSG00000044487 | 1 | |
| ENSMUSG00000044496 | 1.050913498 | 0.70299173 |
| ENSMUSG00000044499 | -1.03962336 | 0.698112512 |
| ENSMUSG00000044501 | -1.058556396 | 0.698112512 |
| ENSMUSG00000044502 | -1.036799653 | 0.875037572 |
| ENSMUSG00000044505 | 1.30735004 | 0.00308688 |
| ENSMUSG00000044515 | 1 | |
| ENSMUSG00000044518 | 1 | |
| ENSMUSG00000044519 | 1.056037819 | 0.671958398 |
| ENSMUSG00000044522 | 1.406558196 | 0.056266471 |
| ENSMUSG00000044526 | 1 | |
| ENSMUSG00000044528 | 1.127555485 | 0.456072787 |
| ENSMUSG00000044533 | 1.106973515 | 0.697178241 |
| ENSMUSG00000044534 | 1.149113744 | 0.317618146 |
| ENSMUSG00000044537 | 1 | |
| ENSMUSG00000044538 | 1 | |
| ENSMUSG00000044542 | 1.069068979 | 0.608497229 |
| ENSMUSG00000044544 | -1.045151399 | 0.927760289 |
| ENSMUSG00000044548 | -1.356122117 | 0.111366442 |
| ENSMUSG00000044550 | 1.007241021 | 1 |
| ENSMUSG00000044551 | -1.07070496 | 0.576689161 |
| ENSMUSG00000044556 | -1.003834322 | 1 |
| ENSMUSG00000044560 | 1 | |
| ENSMUSG00000044562 | -1.092090125 | 0.731324614 |
| ENSMUSG00000044566 | 1.209354526 | 0.148284344 |
| ENSMUSG00000044573 | 1.063763815 | 0.801443616 |
| ENSMUSG00000044574 | 1.015486245 | 0.985522133 |
| ENSMUSG00000044576 | 1.750234697 | 0.00043397 |
| ENSMUSG00000044581 | 1.04991782 | 0.887835871 |
| ENSMUSG00000044583 | 1.008235195 | 1 |
| ENSMUSG00000044594 | 1 | |
| ENSMUSG00000044595 | 1.153579881 | 0.60831194 |
| ENSMUSG00000044597 | 1 | |
| ENSMUSG00000044600 | -1.047114426 | 0.87726305 |
| ENSMUSG00000044609 | -1.037931575 | 0.948385094 |
| ENSMUSG00000044617 | 1.031128077 | 0.846848031 |
| ENSMUSG00000044624 | 1 | |
| ENSMUSG00000044626 | 1.119464341 | 0.698112512 |
| ENSMUSG00000044627 | 1.04099903 | 0.917717739 |
| ENSMUSG00000044628 | -1.276845648 | 0.136048562 |
| ENSMUSG00000044629 | 1.080182965 | 0.321248448 |
| ENSMUSG00000044633 | 1 | |
| ENSMUSG00000044636 | 1.031623673 | 0.87385143 |
| ENSMUSG00000044639 | 1 | |
| ENSMUSG00000044641 | 1.010094994 | 0.917779954 |
| ENSMUSG00000044645 | -1 | 1 |
| ENSMUSG00000044646 | -1.087413865 | 0.158407923 |
| ENSMUSG00000044647 | -1.221618175 | 0.162732522 |
| ENSMUSG00000044649 | 1.059463094 | 0.608497229 |
| ENSMUSG00000044664 | -1.099174267 | 0.535052277 |
| ENSMUSG00000044667 | 1.248753852 | 0.333749058 |

| | | |
|---|---|---|
| ENSMUSG00000044674 | 1.14140073 | 0.272370683 |
| ENSMUSG00000044676 | -1.514001106 | 0.009335364 |
| ENSMUSG00000044678 | 1 | |
| ENSMUSG00000044681 | -1.159782692 | 0.565402456 |
| ENSMUSG00000044689 | -1 | 1 |
| ENSMUSG00000044694 | -1.080937125 | 0.736555608 |
| ENSMUSG00000044697 | 1 | |
| ENSMUSG00000044700 | 1.02434107 | 0.881850933 |
| ENSMUSG00000044701 | 1.059463094 | 0.608497229 |
| ENSMUSG00000044702 | 1.039006969 | 0.7290404 |
| ENSMUSG00000044703 | -1.038444538 | 0.876283388 |
| ENSMUSG00000044705 | 1 | |
| ENSMUSG00000044707 | -1.355648364 | 0.011278358 |
| ENSMUSG00000044708 | -1.29685533 | 0.453905691 |
| ENSMUSG00000044709 | 1.022120973 | 0.892237454 |
| ENSMUSG00000044712 | 1.083726394 | 0.608497229 |
| ENSMUSG00000044715 | 1.079986006 | 0.650620547 |
| ENSMUSG00000044716 | -2.14902283 | 0.001188467 |
| ENSMUSG00000044719 | 1.367667004 | 0.079509785 |
| ENSMUSG00000044724 | 1.095872691 | 0.671280603 |
| ENSMUSG00000044726 | 1.241489066 | 0.124350077 |
| ENSMUSG00000044730 | 1.051916803 | 0.698112512 |
| ENSMUSG00000044734 | 1.553623943 | 0.009478551 |
| ENSMUSG00000044737 | -1.043306199 | 0.698112512 |
| ENSMUSG00000044743 | 1 | |
| ENSMUSG00000044748 | 1 | |
| ENSMUSG00000044749 | -1.190907694 | 0.23684718 |
| ENSMUSG00000044751 | -1.152240054 | 0.327222203 |
| ENSMUSG00000044757 | 1.013796828 | 1 |
| ENSMUSG00000044763 | 1.034573837 | 0.857167632 |
| ENSMUSG00000044768 | -1.018103813 | 0.969282824 |
| ENSMUSG00000044770 | -1.158892898 | 0.608497229 |
| ENSMUSG00000044772 | 1 | |
| ENSMUSG00000044783 | 1.012269478 | 1 |
| ENSMUSG00000044786 | 1.112572929 | 0.217803313 |
| ENSMUSG00000044787 | -1.053549062 | 0.698112512 |
| ENSMUSG00000044788 | -1.238501037 | 0.241144788 |
| ENSMUSG00000044791 | 1.015817266 | 1 |
| ENSMUSG00000044792 | -1.103556171 | 0.532369657 |
| ENSMUSG00000044795 | -1.130401512 | 0.325837078 |
| ENSMUSG00000044798 | 1 | |
| ENSMUSG00000044801 | 1 | |
| ENSMUSG00000044807 | -1.071437823 | 0.698112512 |
| ENSMUSG00000044811 | 1 | |
| ENSMUSG00000044813 | -1.028109051 | 0.852371141 |
| ENSMUSG00000044814 | -1.07527053 | 0.803072249 |
| ENSMUSG00000044816 | -1.054169715 | 0.842733205 |
| ENSMUSG00000044819 | 1 | |
| ENSMUSG00000044820 | -1.128888655 | 0.640252619 |
| ENSMUSG00000044824 | -1 | 1 |
| ENSMUSG00000044827 | 1.588141368 | 0.001717312 |
| ENSMUSG00000044835 | -1.048597769 | 0.608497229 |
| ENSMUSG00000044847 | 1.002951025 | 1 |
| ENSMUSG00000044854 | 1.20415024 | 0.55675409 |
| ENSMUSG00000044857 | 1.161169786 | 0.080515718 |
| ENSMUSG00000044860 | -1.189907685 | 0.608497229 |
| ENSMUSG00000044863 | 1 | |
| ENSMUSG00000044864 | 1.178309443 | 0.404271334 |
| ENSMUSG00000044867 | 1 | |
| ENSMUSG00000044876 | 1.036202245 | 0.698112512 |
| ENSMUSG00000044881 | -1.034050214 | 0.692782277 |
| ENSMUSG00000044894 | 1.028849967 | 0.940197654 |
| ENSMUSG00000044897 | 1 | |

| | | |
|---|---|---|
| ENSMUSG00000044899 | 1 | |
| ENSMUSG00000044903 | 1 | |
| ENSMUSG00000044906 | -1.044219813 | 0.765610554 |
| ENSMUSG00000044912 | 1 | |
| ENSMUSG00000044916 | -1.329649732 | 0.126063768 |
| ENSMUSG00000044921 | 1.471270565 | 0.026527667 |
| ENSMUSG00000044923 | 1 | |
| ENSMUSG00000044927 | -1.038182731 | 0.956972448 |
| ENSMUSG00000044933 | 1 | |
| ENSMUSG00000044934 | -1.096422827 | 0.660858593 |
| ENSMUSG00000044937 | 1.049719516 | 0.73616715 |
| ENSMUSG00000044938 | -1 | 1 |
| ENSMUSG00000044948 | 1.353774764 | 0.02620375 |
| ENSMUSG00000044949 | -1.00540949 | 1 |
| ENSMUSG00000044950 | -1.06380919 | 0.698112512 |
| ENSMUSG00000044951 | -1.051018293 | 0.859335935 |
| ENSMUSG00000044952 | -1.010359695 | 0.979002724 |
| ENSMUSG00000044957 | -1.05948903 | 0.814608539 |
| ENSMUSG00000044966 | -1.035387404 | 0.780689972 |
| ENSMUSG00000044968 | -1.013621073 | 0.947484807 |
| ENSMUSG00000044976 | 1 | |
| ENSMUSG00000044982 | -1.117770678 | 0.334859726 |
| ENSMUSG00000044985 | 1 | |
| ENSMUSG00000044986 | 1.063328055 | 0.608497229 |
| ENSMUSG00000044988 | 1 | |
| ENSMUSG00000044991 | 1.029521675 | 0.717035356 |
| ENSMUSG00000044994 | 1 | |
| ENSMUSG00000044997 | 1 | |
| ENSMUSG00000045004 | -1.132666855 | 0.18851436 |
| ENSMUSG00000045005 | 1.273636169 | 0.003205101 |
| ENSMUSG00000045007 | 1.443307587 | 0.000411927 |
| ENSMUSG00000045008 | -1.264655168 | 0.224865446 |
| ENSMUSG00000045009 | -1.100069442 | 0.814384306 |
| ENSMUSG00000045010 | -1.111880267 | 0.672998864 |
| ENSMUSG00000045013 | 1 | |
| ENSMUSG00000045019 | -1.089497311 | 0.608497229 |
| ENSMUSG00000045022 | 1 | |
| ENSMUSG00000045027 | 1.564380863 | 0.001316576 |
| ENSMUSG00000045030 | 1 | |
| ENSMUSG00000045031 | -1.092470765 | 0.532369657 |
| ENSMUSG00000045034 | -1.223947136 | 0.005624846 |
| ENSMUSG00000045036 | 1.075324819 | 0.750742698 |
| ENSMUSG00000045038 | 1.02606001 | 0.843685576 |
| ENSMUSG00000045039 | 1.178199796 | 0.108172814 |
| ENSMUSG00000045045 | 1.399310094 | 0.090734064 |
| ENSMUSG00000045048 | 1 | |
| ENSMUSG00000045052 | 1 | |
| ENSMUSG00000045053 | 1 | |
| ENSMUSG00000045055 | 1.115114778 | 0.427797606 |
| ENSMUSG00000045062 | 1.011607416 | 1 |
| ENSMUSG00000045064 | 1.092894285 | 0.211880141 |
| ENSMUSG00000045065 | -1.053549062 | 0.698112512 |
| ENSMUSG00000045071 | 1.006728498 | 0.992580079 |
| ENSMUSG00000045075 | 1.299147809 | 0.214101814 |
| ENSMUSG00000045078 | 1.027289344 | 0.945910992 |
| ENSMUSG00000045083 | 1 | |
| ENSMUSG00000045087 | -1.097349941 | 0.677534336 |
| ENSMUSG00000045091 | 1 | |
| ENSMUSG00000045092 | 1.048320786 | 0.71714467 |
| ENSMUSG00000045094 | -1.294704847 | 0.119804732 |
| ENSMUSG00000045095 | 1.444314148 | 0.00046824 |
| ENSMUSG00000045098 | -1.01989276 | 0.982838256 |
| ENSMUSG00000045100 | 1.075537825 | 0.695062951 |

| | | |
|---|---|---|
| ENSMUSG00000045102 | 1.053619707 | 0.696560822 |
| ENSMUSG00000045103 | 1.327388464 | 0.003538091 |
| ENSMUSG00000045104 | 1 | |
| ENSMUSG00000045106 | -1.183366874 | 0.216046507 |
| ENSMUSG00000045107 | 1.082242661 | 0.17334066 |
| ENSMUSG00000045109 | 1 | |
| ENSMUSG00000045111 | 1 | |
| ENSMUSG00000045114 | 1.019083579 | 0.949812939 |
| ENSMUSG00000045126 | 1 | |
| ENSMUSG00000045128 | 1.077344692 | 0.724205129 |
| ENSMUSG00000045132 | 1 | |
| ENSMUSG00000045136 | 1.020577941 | 0.956374899 |
| ENSMUSG00000045140 | 1.028536479 | 0.898763027 |
| ENSMUSG00000045148 | 1 | |
| ENSMUSG00000045150 | 1 | |
| ENSMUSG00000045160 | -1.051380822 | 0.64883887 |
| ENSMUSG00000045165 | -1.308695861 | 0.089533569 |
| ENSMUSG00000045174 | 1 | |
| ENSMUSG00000045176 | -1.100712329 | 0.203727874 |
| ENSMUSG00000045178 | 1 | |
| ENSMUSG00000045179 | 1 | |
| ENSMUSG00000045180 | -1.714224201 | 0.005249107 |
| ENSMUSG00000045193 | 1.074845866 | 0.311631163 |
| ENSMUSG00000045201 | 1 | |
| ENSMUSG00000045202 | 1 | |
| ENSMUSG00000045204 | 1 | |
| ENSMUSG00000045205 | -1.015958596 | 1 |
| ENSMUSG00000045210 | -1.031590021 | 0.934760809 |
| ENSMUSG00000045211 | -1.185604241 | 0.054197279 |
| ENSMUSG00000045215 | -1.268755138 | 0.007788856 |
| ENSMUSG00000045216 | 1.194880331 | 0.103854589 |
| ENSMUSG00000045217 | 1 | |
| ENSMUSG00000045225 | 1 | |
| ENSMUSG00000045231 | 1.242822657 | 0.212677241 |
| ENSMUSG00000045232 | -1.043306199 | 0.79291651 |
| ENSMUSG00000045236 | 1 | |
| ENSMUSG00000045237 | 1.09010503 | 0.608497229 |
| ENSMUSG00000045238 | -1.082845284 | 0.788000546 |
| ENSMUSG00000045246 | 1 | |
| ENSMUSG00000045248 | 1.072578766 | 0.221797188 |
| ENSMUSG00000045251 | 1.13619392 | 0.067743282 |
| ENSMUSG00000045252 | 1.084251624 | 0.608497229 |
| ENSMUSG00000045257 | -1.037187459 | 0.698112512 |
| ENSMUSG00000045259 | 1 | |
| ENSMUSG00000045267 | 1 | |
| ENSMUSG00000045268 | 1.046289672 | 0.698112512 |
| ENSMUSG00000045273 | -1.075782164 | 0.615267463 |
| ENSMUSG00000045275 | 1.006381539 | 1 |
| ENSMUSG00000045281 | 1 | |
| ENSMUSG00000045282 | 1.036876862 | 0.883111131 |
| ENSMUSG00000045284 | -1 | 1 |
| ENSMUSG00000045287 | -1.209182227 | 0.31494917 |
| ENSMUSG00000045288 | -1.033820075 | 0.888246981 |
| ENSMUSG00000045291 | 1.020883393 | 0.977705508 |
| ENSMUSG00000045294 | -1.083634847 | 0.698112512 |
| ENSMUSG00000045302 | 1.032711412 | 0.904013471 |
| ENSMUSG00000045306 | 1 | |
| ENSMUSG00000045312 | 1.221327601 | 0.13274925 |
| ENSMUSG00000045314 | -1.021151408 | 1 |
| ENSMUSG00000045316 | 1.03536163 | 0.698112512 |
| ENSMUSG00000045318 | 1 | |
| ENSMUSG00000045319 | 1.007188381 | 1 |
| ENSMUSG00000045322 | 1 | |

| | | |
|---|---|---|
| ENSMUSG00000045326 | 1.021383522 | 0.966907235 |
| ENSMUSG00000045327 | -1.385319417 | 0.047843331 |
| ENSMUSG00000045328 | -1.065656277 | 0.82835102 |
| ENSMUSG00000045330 | 1.059463094 | 0.608497229 |
| ENSMUSG00000045331 | 1 | |
| ENSMUSG00000045333 | -1.302422805 | 0.414873034 |
| ENSMUSG00000045336 | 1 | |
| ENSMUSG00000045337 | 1 | |
| ENSMUSG00000045339 | -1 | 1 |
| ENSMUSG00000045340 | 1 | |
| ENSMUSG00000045341 | 1 | |
| ENSMUSG00000045345 | -1.111807928 | 0.703568899 |
| ENSMUSG00000045348 | 1.154004244 | 0.269500305 |
| ENSMUSG00000045349 | 1.169719873 | 0.414911123 |
| ENSMUSG00000045350 | -1.042235453 | 0.962587881 |
| ENSMUSG00000045362 | 1.256141864 | 0.434699874 |
| ENSMUSG00000045364 | 1.132639128 | 0.398767938 |
| ENSMUSG00000045374 | 1.283461167 | 0.015526202 |
| ENSMUSG00000045377 | -1.336883 | 0.033479622 |
| ENSMUSG00000045378 | -1.044273782 | 0.698112512 |
| ENSMUSG00000045381 | 1.11166259 | 0.608497229 |
| ENSMUSG00000045382 | 1.537503551 | 0.01536947 |
| ENSMUSG00000045391 | -1.044273782 | 0.698112512 |
| ENSMUSG00000045392 | 1 | |
| ENSMUSG00000045394 | -1.658610071 | 0.005249107 |
| ENSMUSG00000045395 | 1 | |
| ENSMUSG00000045404 | 1 | |
| ENSMUSG00000045409 | 1.009844181 | 1 |
| ENSMUSG00000045410 | -1.046727307 | 0.840860016 |
| ENSMUSG00000045411 | 1.067322377 | 0.608497229 |
| ENSMUSG00000045414 | -1.082436662 | 0.70141105 |
| ENSMUSG00000045417 | 1 | |
| ENSMUSG00000045421 | 1 | |
| ENSMUSG00000045427 | -1.097549095 | 0.699591599 |
| ENSMUSG00000045435 | 1.086288054 | 0.608497229 |
| ENSMUSG00000045438 | -1.052182502 | 0.608497229 |
| ENSMUSG00000045440 | 1 | |
| ENSMUSG00000045441 | 1.059596658 | 0.608497229 |
| ENSMUSG00000045455 | -1.034239753 | 0.900996516 |
| ENSMUSG00000045463 | 1 | |
| ENSMUSG00000045464 | 1.022856732 | 0.839099152 |
| ENSMUSG00000045466 | -1.010110367 | 0.937505238 |
| ENSMUSG00000045467 | -1.124780375 | 0.689164262 |
| ENSMUSG00000045471 | 1 | |
| ENSMUSG00000045474 | 1 | |
| ENSMUSG00000045475 | 1 | |
| ENSMUSG00000045479 | 1 | |
| ENSMUSG00000045482 | 1.001858915 | 1 |
| ENSMUSG00000045493 | 1 | |
| ENSMUSG00000045498 | 1.005613439 | 1 |
| ENSMUSG00000045502 | 1 | |
| ENSMUSG00000045503 | -1.06495146 | 0.499359518 |
| ENSMUSG00000045506 | 1.269617311 | 0.291044503 |
| ENSMUSG00000045508 | 1 | |
| ENSMUSG00000045509 | -1 | 1 |
| ENSMUSG00000045514 | 1 | |
| ENSMUSG00000045515 | 1.069068979 | 0.608497229 |
| ENSMUSG00000045518 | 1.04328066 | 0.867378212 |
| ENSMUSG00000045519 | 1.005327259 | 1 |
| ENSMUSG00000045521 | -1.139982257 | 0.606198803 |
| ENSMUSG00000045528 | 1 | |
| ENSMUSG00000045532 | -1.879844937 | 0.001661811 |
| ENSMUSG00000045534 | 1 | |

| | | |
|---|---|---|
| ENSMUSG00000045538 | -1.011150986 | 0.98455106 |
| ENSMUSG00000045539 | -1 | 1 |
| ENSMUSG00000045540 | 1 | |
| ENSMUSG00000045545 | -2.108628038 | 0.003599328 |
| ENSMUSG00000045551 | 1.18757839 | 0.621683209 |
| ENSMUSG00000045555 | 1.154696672 | 0.466340867 |
| ENSMUSG00000045559 | 1 | |
| ENSMUSG00000045566 | 1 | |
| ENSMUSG00000045569 | 1 | |
| ENSMUSG00000045573 | 1.626327254 | 0.001978744 |
| ENSMUSG00000045575 | 1 | |
| ENSMUSG00000045576 | -1.163756272 | 0.150051987 |
| ENSMUSG00000045581 | 1 | |
| ENSMUSG00000045584 | 1 | |
| ENSMUSG00000045587 | 1 | |
| ENSMUSG00000045589 | -1.416971076 | 0.022500491 |
| ENSMUSG00000045591 | 1.095872691 | 0.608497229 |
| ENSMUSG00000045594 | -1.015393821 | 0.988723196 |
| ENSMUSG00000045598 | 1.106205433 | 0.41649483 |
| ENSMUSG00000045608 | 1 | |
| ENSMUSG00000045609 | 1 | |
| ENSMUSG00000045613 | 1.04424822 | 0.835701362 |
| ENSMUSG00000045620 | 1 | |
| ENSMUSG00000045624 | 1.043830028 | 0.87629256 |
| ENSMUSG00000045625 | 1.069752935 | 0.758834014 |
| ENSMUSG00000045629 | 1.85477886 | 0.005913329 |
| ENSMUSG00000045631 | 1.059463094 | 0.608497229 |
| ENSMUSG00000045636 | 1.326479025 | 0.047622927 |
| ENSMUSG00000045639 | 1.028820866 | 0.858354949 |
| ENSMUSG00000045648 | 1 | |
| ENSMUSG00000045655 | 1 | |
| ENSMUSG00000045657 | 1.05530209 | 0.812255606 |
| ENSMUSG00000045658 | -1.533301179 | 0.005521566 |
| ENSMUSG00000045659 | -1.011327043 | 1 |
| ENSMUSG00000045662 | 1.234951516 | 0.251544797 |
| ENSMUSG00000045664 | 1.106341139 | 0.378140099 |
| ENSMUSG00000045665 | 1.135758924 | 0.256770279 |
| ENSMUSG00000045667 | -1.36575583 | 0.009616641 |
| ENSMUSG00000045671 | -1.051231001 | 0.773756337 |
| ENSMUSG00000045672 | 1.717531517 | 9.78671E-05 |
| ENSMUSG00000045678 | 1 | |
| ENSMUSG00000045679 | 1.009827248 | 1 |
| ENSMUSG00000045680 | -1.03962336 | 0.803072249 |
| ENSMUSG00000045684 | 1 | |
| ENSMUSG00000045689 | 1 | |
| ENSMUSG00000045690 | 1.111534869 | 0.380970157 |
| ENSMUSG00000045691 | -1.271508953 | 0.042007281 |
| ENSMUSG00000045693 | 1 | |
| ENSMUSG00000045694 | 1.075870199 | 0.642819974 |
| ENSMUSG00000045699 | -1.023016116 | 0.967414484 |
| ENSMUSG00000045708 | 1 | |
| ENSMUSG00000045709 | -1.093674795 | 0.608497229 |
| ENSMUSG00000045713 | 1 | |
| ENSMUSG00000045725 | -1.044273782 | 0.729018288 |
| ENSMUSG00000045730 | 1.413147139 | 0.00435101 |
| ENSMUSG00000045731 | -1.03962336 | 0.803072249 |
| ENSMUSG00000045733 | 1 | |
| ENSMUSG00000045744 | 1 | |
| ENSMUSG00000045746 | -1.22361269 | 0.207046069 |
| ENSMUSG00000045751 | 1.015432279 | 0.987897106 |
| ENSMUSG00000045752 | 1.134171691 | 0.251544797 |
| ENSMUSG00000045757 | -1.338762986 | 0.086651676 |
| ENSMUSG00000045761 | -1.372250124 | 0.184903824 |

| | | |
|---|---|---|
| ENSMUSG00000045763 | 1.057703042 | 0.827718961 |
| ENSMUSG00000045767 | -1.010645498 | 1 |
| ENSMUSG00000045775 | 1.144581873 | 0.608497229 |
| ENSMUSG00000045776 | 1.07913413 | 0.781240509 |
| ENSMUSG00000045777 | 1.124313245 | 0.532369657 |
| ENSMUSG00000045780 | 1 | |
| ENSMUSG00000045790 | -1.066985603 | 0.552584793 |
| ENSMUSG00000045792 | 1 | |
| ENSMUSG00000045794 | 1 | |
| ENSMUSG00000045795 | -1.044035049 | 0.698112512 |
| ENSMUSG00000045797 | 1 | |
| ENSMUSG00000045799 | -1.03962336 | 0.698112512 |
| ENSMUSG00000045802 | 1 | |
| ENSMUSG00000045812 | 1 | |
| ENSMUSG00000045813 | 1 | |
| ENSMUSG00000045815 | 1 | |
| ENSMUSG00000045817 | 1.218286088 | 0.385934169 |
| ENSMUSG00000045822 | -1.057246612 | 0.608497229 |
| ENSMUSG00000045824 | 1 | |
| ENSMUSG00000045826 | -1.189947723 | 0.302623721 |
| ENSMUSG00000045827 | -1.017112401 | 0.988655147 |
| ENSMUSG00000045835 | -1 | 1 |
| ENSMUSG00000045838 | -1.060117388 | 0.693220376 |
| ENSMUSG00000045842 | 1 | |
| ENSMUSG00000045854 | 1.000150228 | 1 |
| ENSMUSG00000045867 | -1.009792787 | 0.988424085 |
| ENSMUSG00000045868 | 1 | |
| ENSMUSG00000045871 | 1 | |
| ENSMUSG00000045875 | 1.059463094 | 0.704170349 |
| ENSMUSG00000045876 | -1.00571839 | 1 |
| ENSMUSG00000045877 | -1.088313104 | 0.741377649 |
| ENSMUSG00000045883 | 1 | |
| ENSMUSG00000045886 | -1 | 1 |
| ENSMUSG00000045896 | -1.033286767 | 0.933031586 |
| ENSMUSG00000045903 | 2.656891292 | 0.000829917 |
| ENSMUSG00000045912 | -1.487015997 | 0.043852518 |
| ENSMUSG00000045915 | 1 | |
| ENSMUSG00000045917 | 1.312749566 | 0.00639805 |
| ENSMUSG00000045928 | 1 | |
| ENSMUSG00000045929 | -1.043306199 | 0.698112512 |
| ENSMUSG00000045930 | 1.628431991 | 0.000669906 |
| ENSMUSG00000045932 | 1.189774833 | 0.380476832 |
| ENSMUSG00000045934 | 1.005975171 | 1 |
| ENSMUSG00000045942 | -1.07008665 | 0.812147631 |
| ENSMUSG00000045946 | -1.03962336 | 0.698112512 |
| ENSMUSG00000045948 | 1.099557803 | 0.698112512 |
| ENSMUSG00000045952 | -1.063525418 | 0.698112512 |
| ENSMUSG00000045954 | 1.457935853 | 0.03042531 |
| ENSMUSG00000045962 | 1.054776039 | 0.90583677 |
| ENSMUSG00000045967 | 1 | |
| ENSMUSG00000045968 | -1.077863338 | 0.698112512 |
| ENSMUSG00000045969 | 1.160936721 | 0.3305975 |
| ENSMUSG00000045973 | 1.040105295 | 0.821336725 |
| ENSMUSG00000045975 | 1.067139064 | 0.678549317 |
| ENSMUSG00000045980 | 1.123844789 | 0.265690989 |
| ENSMUSG00000045983 | 1.060471693 | 0.881925163 |
| ENSMUSG00000045989 | 1 | |
| ENSMUSG00000045991 | 1.25580102 | 0.077143981 |
| ENSMUSG00000045994 | 1 | |
| ENSMUSG00000045996 | -1.031007188 | 0.88325758 |
| ENSMUSG00000046000 | 1 | |
| ENSMUSG00000046005 | -1.239913898 | 0.266903707 |
| ENSMUSG00000046006 | 1 | |

| | | |
|---|---|---|
| ENSMUSG00000046008 | 1.059463094 | 0.608497229 |
| ENSMUSG00000046010 | -1.022105895 | 0.883448618 |
| ENSMUSG00000046016 | -1.043306199 | 0.698112512 |
| ENSMUSG00000046020 | 1.028121365 | 0.907007763 |
| ENSMUSG00000046027 | 1.070182428 | 0.276229467 |
| ENSMUSG00000046031 | -1 | 1 |
| ENSMUSG00000046032 | 1.043490376 | 0.879178663 |
| ENSMUSG00000046034 | 1.000527022 | 1 |
| ENSMUSG00000046041 | 1 | |
| ENSMUSG00000046049 | -1.03962336 | 0.698112512 |
| ENSMUSG00000046056 | -1.291397813 | 0.007699543 |
| ENSMUSG00000046057 | 1 | |
| ENSMUSG00000046058 | -1.015912982 | 0.80273087 |
| ENSMUSG00000046062 | 1.01816424 | 0.996705669 |
| ENSMUSG00000046070 | -1 | 1 |
| ENSMUSG00000046079 | 1.051723134 | 0.679374651 |
| ENSMUSG00000046080 | -1.03962336 | 0.814089245 |
| ENSMUSG00000046082 | 1 | |
| ENSMUSG00000046085 | -1.03962336 | 0.698112512 |
| ENSMUSG00000046088 | 1.059463094 | 0.608497229 |
| ENSMUSG00000046093 | -1.516778419 | 0.008233465 |
| ENSMUSG00000046095 | 1 | |
| ENSMUSG00000046096 | 1.008609239 | 1 |
| ENSMUSG00000046101 | 1.013706217 | 1 |
| ENSMUSG00000046108 | -1.168999771 | 0.210178841 |
| ENSMUSG00000046109 | 1 | |
| ENSMUSG00000046110 | 1 | |
| ENSMUSG00000046111 | -1.045585377 | 0.844612733 |
| ENSMUSG00000046119 | 1 | |
| ENSMUSG00000046130 | 1 | |
| ENSMUSG00000046133 | -1.147838869 | 0.518775336 |
| ENSMUSG00000046138 | 1.090609042 | 0.698112512 |
| ENSMUSG00000046139 | 1.04476382 | 0.887896098 |
| ENSMUSG00000046145 | 1 | |
| ENSMUSG00000046150 | 1 | |
| ENSMUSG00000046152 | 1.041442014 | 0.792783746 |
| ENSMUSG00000046157 | -1.478520572 | 0.008364861 |
| ENSMUSG00000046159 | -1 | 1 |
| ENSMUSG00000046160 | 1 | |
| ENSMUSG00000046167 | 1.122462048 | 0.215991492 |
| ENSMUSG00000046168 | -1.242545864 | 0.1991922 |
| ENSMUSG00000046169 | 1.096859354 | 0.592265042 |
| ENSMUSG00000046173 | 1 | |
| ENSMUSG00000046178 | -1.053549062 | 0.698112512 |
| ENSMUSG00000046179 | -1.004336676 | 1 |
| ENSMUSG00000046180 | 1 | |
| ENSMUSG00000046182 | 1.014545335 | 1 |
| ENSMUSG00000046185 | -1.039804192 | 0.698112512 |
| ENSMUSG00000046186 | 1.084075698 | 0.736982121 |
| ENSMUSG00000046191 | 1.740954796 | 0.011189217 |
| ENSMUSG00000046192 | -1.095738849 | 0.698112512 |
| ENSMUSG00000046196 | 1.004650133 | 1 |
| ENSMUSG00000046201 | 1.05522945 | 0.859420446 |
| ENSMUSG00000046203 | 1 | |
| ENSMUSG00000046204 | -1.519242254 | 0.00997435 |
| ENSMUSG00000046207 | -1.081225398 | 0.736695141 |
| ENSMUSG00000046210 | 1 | |
| ENSMUSG00000046213 | 1 | |
| ENSMUSG00000046215 | 1.059463094 | 0.608497229 |
| ENSMUSG00000046219 | 1.015498416 | 1 |
| ENSMUSG00000046223 | 1.399468589 | 0.000842901 |
| ENSMUSG00000046229 | 1.249909437 | 0.55103479 |
| ENSMUSG00000046230 | -1.004270385 | 1 |

| | | |
|---|---|---|
| ENSMUSG00000046240 | 1.069068979 | 0.698112512 |
| ENSMUSG00000046242 | 1 | |
| ENSMUSG00000046245 | 1.34165244 | 0.187448947 |
| ENSMUSG00000046248 | 1 | |
| ENSMUSG00000046258 | 1 | |
| ENSMUSG00000046259 | 1 | |
| ENSMUSG00000046262 | 1 | |
| ENSMUSG00000046269 | 1.083010578 | 0.547120169 |
| ENSMUSG00000046272 | -1 | 1 |
| ENSMUSG00000046275 | 1.014545335 | 0.998924031 |
| ENSMUSG00000046280 | -2.2814726 | 0.001674444 |
| ENSMUSG00000046282 | 1 | |
| ENSMUSG00000046287 | 1 | |
| ENSMUSG00000046295 | 1.018922024 | 0.950006093 |
| ENSMUSG00000046300 | 1 | |
| ENSMUSG00000046311 | -1.027178946 | 0.939477533 |
| ENSMUSG00000046312 | -1.137789049 | 0.187611488 |
| ENSMUSG00000046314 | 1.242955316 | 0.089193093 |
| ENSMUSG00000046317 | -1.129231493 | 0.608497229 |
| ENSMUSG00000046318 | 2.003481651 | 0.001905152 |
| ENSMUSG00000046321 | 1 | |
| ENSMUSG00000046323 | 1 | |
| ENSMUSG00000046324 | -1.066823066 | 0.807394771 |
| ENSMUSG00000046329 | -1.054498861 | 0.70299173 |
| ENSMUSG00000046330 | 1.032778755 | 0.934760809 |
| ENSMUSG00000046334 | -1.03962336 | 0.808074244 |
| ENSMUSG00000046337 | -1.147838869 | 0.576423047 |
| ENSMUSG00000046338 | -1.109582661 | 0.608497229 |
| ENSMUSG00000046341 | 1.019083579 | 0.968937442 |
| ENSMUSG00000046345 | 1.050356762 | 0.903736347 |
| ENSMUSG00000046351 | -1.029401755 | 0.906422894 |
| ENSMUSG00000046352 | 1.318280431 | 0.06812145 |
| ENSMUSG00000046354 | 1 | |
| ENSMUSG00000046364 | 1.031555363 | 0.936653013 |
| ENSMUSG00000046367 | 1 | |
| ENSMUSG00000046372 | 1.105566531 | 0.676404329 |
| ENSMUSG00000046378 | 1.028390773 | 0.982838256 |
| ENSMUSG00000046380 | 1.048737897 | 0.687984391 |
| ENSMUSG00000046387 | 1.881663892 | 0.001687496 |
| ENSMUSG00000046388 | 1 | |
| ENSMUSG00000046396 | 1 | |
| ENSMUSG00000046402 | -1.146909569 | 0.096445543 |
| ENSMUSG00000046404 | -1.039583801 | 0.883111131 |
| ENSMUSG00000046408 | -1.077425958 | 0.790424211 |
| ENSMUSG00000046410 | -1.633698061 | 0.003535405 |
| ENSMUSG00000046411 | 1 | |
| ENSMUSG00000046413 | 1.080095574 | 0.779432763 |
| ENSMUSG00000046415 | -1.007148435 | 1 |
| ENSMUSG00000046417 | 1.048854308 | 0.698112512 |
| ENSMUSG00000046431 | 1 | |
| ENSMUSG00000046432 | -1.047352727 | 0.847073978 |
| ENSMUSG00000046434 | 1.040133602 | 0.96504537 |
| ENSMUSG00000046435 | 1.019083579 | 0.968937442 |
| ENSMUSG00000046438 | 1 | |
| ENSMUSG00000046440 | 1.042472958 | 0.947149464 |
| ENSMUSG00000046441 | 1.066652655 | 0.698112512 |
| ENSMUSG00000046442 | -1.090373706 | 0.608497229 |
| ENSMUSG00000046447 | 1.054407888 | 0.6589481 |
| ENSMUSG00000046449 | -1.111295673 | 0.569198267 |
| ENSMUSG00000046450 | 1 | |
| ENSMUSG00000046456 | -1 | 1 |
| ENSMUSG00000046460 | 1 | |
| ENSMUSG00000046463 | 1.345754216 | 0.02298791 |

| | | |
|---|---|---|
| ENSMUSG00000046470 | -1.116984848 | 0.647487569 |
| ENSMUSG00000046474 | 1 | |
| ENSMUSG00000046480 | -1.010960635 | 1 |
| ENSMUSG00000046486 | 1 | |
| ENSMUSG00000046487 | -1.028348525 | 0.968937442 |
| ENSMUSG00000046490 | 1 | |
| ENSMUSG00000046491 | -1.137443589 | 0.666120375 |
| ENSMUSG00000046493 | 1 | |
| ENSMUSG00000046500 | 1 | |
| ENSMUSG00000046516 | 1.111838287 | 0.120269673 |
| ENSMUSG00000046518 | 1 | |
| ENSMUSG00000046519 | -1.09438007 | 0.626480879 |
| ENSMUSG00000046523 | -1.178737524 | 0.608497229 |
| ENSMUSG00000046532 | 1.510330797 | 0.009180765 |
| ENSMUSG00000046541 | 1.040605789 | 0.698112512 |
| ENSMUSG00000046546 | -1.431205504 | 0.006851645 |
| ENSMUSG00000046550 | -1.307702044 | 0.005249107 |
| ENSMUSG00000046556 | -1.012398291 | 0.858911269 |
| ENSMUSG00000046561 | 1.139037884 | 0.210047602 |
| ENSMUSG00000046562 | -1.027226723 | 0.931368384 |
| ENSMUSG00000046567 | -1.022177151 | 0.916878712 |
| ENSMUSG00000046572 | 1.101066493 | 0.608497229 |
| ENSMUSG00000046573 | -1.076769645 | 0.608497229 |
| ENSMUSG00000046574 | 1.217233311 | 0.13623371 |
| ENSMUSG00000046580 | 1.04424822 | 0.861137762 |
| ENSMUSG00000046585 | 1 | |
| ENSMUSG00000046589 | 1.013364677 | 0.904239821 |
| ENSMUSG00000046590 | 1 | |
| ENSMUSG00000046591 | -1.071375083 | 0.698112512 |
| ENSMUSG00000046593 | 1 | |
| ENSMUSG00000046598 | -1.900520644 | 0.000242314 |
| ENSMUSG00000046603 | 1.004323441 | 1 |
| ENSMUSG00000046605 | -1.054410253 | 0.608497229 |
| ENSMUSG00000046607 | 1.05410549 | 0.814089245 |
| ENSMUSG00000046610 | 1.168686685 | 0.303261282 |
| ENSMUSG00000046613 | -1.165258592 | 0.698112512 |
| ENSMUSG00000046615 | 1 | |
| ENSMUSG00000046618 | -1.044594346 | 0.805808218 |
| ENSMUSG00000046620 | 1 | |
| ENSMUSG00000046623 | -1.049903232 | 0.660684061 |
| ENSMUSG00000046634 | -1.060017444 | 0.837696424 |
| ENSMUSG00000046636 | 1.036882759 | 0.854777354 |
| ENSMUSG00000046637 | -1.063525418 | 0.698112512 |
| ENSMUSG00000046643 | 1 | |
| ENSMUSG00000046650 | 1 | |
| ENSMUSG00000046652 | 1 | |
| ENSMUSG00000046658 | 1.091634842 | 0.046492778 |
| ENSMUSG00000046667 | -1.375399369 | 0.041861099 |
| ENSMUSG00000046668 | 1.043120104 | 0.66549672 |
| ENSMUSG00000046671 | 1.03477838 | 0.866447245 |
| ENSMUSG00000046675 | 1.137561262 | 0.160676302 |
| ENSMUSG00000046676 | 1 | |
| ENSMUSG00000046678 | 1 | |
| ENSMUSG00000046679 | 1.101465383 | 0.216046507 |
| ENSMUSG00000046683 | 1 | |
| ENSMUSG00000046687 | -1.043306199 | 0.698112512 |
| ENSMUSG00000046688 | 1.151974132 | 0.186473233 |
| ENSMUSG00000046691 | 1.105406834 | 0.622074619 |
| ENSMUSG00000046694 | 1.110513453 | 0.181564875 |
| ENSMUSG00000046697 | 1 | |
| ENSMUSG00000046699 | -1.069502966 | 0.698112512 |
| ENSMUSG00000046707 | 1.00344224 | 1 |
| ENSMUSG00000046709 | 1.114427828 | 0.698112512 |

| | | |
|---|---|---|
| ENSMUSG00000046711 | 1.154159471 | 0.209767231 |
| ENSMUSG00000046714 | 1.245185459 | 0.015287164 |
| ENSMUSG00000046716 | 1 | |
| ENSMUSG00000046717 | 1 | |
| ENSMUSG00000046718 | -1.126368193 | 0.126931489 |
| ENSMUSG00000046719 | 1.199838533 | 0.44485243 |
| ENSMUSG00000046721 | -1.123547972 | 0.698112512 |
| ENSMUSG00000046722 | 1.029618641 | 0.93400515 |
| ENSMUSG00000046723 | 1 | |
| ENSMUSG00000046727 | -1.011587014 | 0.965659325 |
| ENSMUSG00000046731 | 1.032312329 | 0.860712821 |
| ENSMUSG00000046733 | -1.080944226 | 0.597079148 |
| ENSMUSG00000046743 | 1.625901853 | 0.00440537 |
| ENSMUSG00000046747 | 1.030445456 | 0.937517551 |
| ENSMUSG00000046748 | -1.168999771 | 0.351808646 |
| ENSMUSG00000046750 | 1 | |
| ENSMUSG00000046753 | -1.133970019 | 0.417037704 |
| ENSMUSG00000046755 | 1 | |
| ENSMUSG00000046756 | -1.048908311 | 0.79291651 |
| ENSMUSG00000046761 | -1.048822887 | 0.734674233 |
| ENSMUSG00000046764 | -1.03962336 | 0.698112512 |
| ENSMUSG00000046768 | 1.214381202 | 0.019236545 |
| ENSMUSG00000046774 | -1 | 1 |
| ENSMUSG00000046782 | 2.047858327 | 0.001052532 |
| ENSMUSG00000046785 | 1.003407476 | 1 |
| ENSMUSG00000046790 | 1 | |
| ENSMUSG00000046791 | 1.082960735 | 0.576959009 |
| ENSMUSG00000046792 | 1.231377963 | 0.22064254 |
| ENSMUSG00000046793 | -1.063525418 | 0.756517167 |
| ENSMUSG00000046794 | -1.48405991 | 0.09770324 |
| ENSMUSG00000046798 | 1.008297418 | 1 |
| ENSMUSG00000046804 | 1 | |
| ENSMUSG00000046805 | 1.380555806 | 0.153451765 |
| ENSMUSG00000046806 | 1.084289095 | 0.532369657 |
| ENSMUSG00000046807 | 1.344750628 | 0.019425435 |
| ENSMUSG00000046808 | 1.063258864 | 0.698112512 |
| ENSMUSG00000046811 | 1.059463094 | 0.608497229 |
| ENSMUSG00000046814 | -1.108106156 | 0.294533546 |
| ENSMUSG00000046818 | 1.889397786 | 0.000582624 |
| ENSMUSG00000046822 | 1.068527363 | 0.598916845 |
| ENSMUSG00000046826 | -1.177546921 | 0.373031949 |
| ENSMUSG00000046828 | 1.008034876 | 1 |
| ENSMUSG00000046834 | 1 | |
| ENSMUSG00000046836 | 1.037350231 | 0.903139568 |
| ENSMUSG00000046840 | 1 | |
| ENSMUSG00000046841 | 1.121837935 | 0.586328611 |
| ENSMUSG00000046844 | -1.158708082 | 0.250726313 |
| ENSMUSG00000046845 | 1 | |
| ENSMUSG00000046846 | 1 | |
| ENSMUSG00000046854 | -1.514789841 | 0.05441916 |
| ENSMUSG00000046856 | 1.019083579 | 0.968937442 |
| ENSMUSG00000046861 | 1.01635041 | 0.862843183 |
| ENSMUSG00000046862 | 1.065921126 | 0.683128036 |
| ENSMUSG00000046865 | -1.007969778 | 1 |
| ENSMUSG00000046873 | -1.137578205 | 0.60397857 |
| ENSMUSG00000046876 | 1.092338819 | 0.54024202 |
| ENSMUSG00000046879 | 1.042932825 | 0.841479645 |
| ENSMUSG00000046881 | 1 | |
| ENSMUSG00000046886 | -1.044273782 | 0.790851299 |
| ENSMUSG00000046897 | 1.037203129 | 0.929847429 |
| ENSMUSG00000046899 | -1.03962336 | 0.872692845 |
| ENSMUSG00000046908 | -1.03962336 | 0.76902772 |
| ENSMUSG00000046909 | 1.048798917 | 0.775199822 |

| | | |
|---|---|---|
| ENSMUSG00000046913 | 1 | |
| ENSMUSG00000046916 | 1 | |
| ENSMUSG00000046922 | 1 | |
| ENSMUSG00000046924 | 1 | |
| ENSMUSG00000046932 | 1 | |
| ENSMUSG00000046934 | 1.034520851 | 0.963965793 |
| ENSMUSG00000046942 | -1 | 1 |
| ENSMUSG00000046947 | 1.02010382 | 0.800329493 |
| ENSMUSG00000046949 | -1.069288275 | 0.714289536 |
| ENSMUSG00000046952 | 1 | |
| ENSMUSG00000046957 | 1 | |
| ENSMUSG00000046958 | -1.350826053 | 0.065933684 |
| ENSMUSG00000046959 | -1.128280603 | 0.69305964 |
| ENSMUSG00000046961 | -1.028878757 | 0.951298634 |
| ENSMUSG00000046962 | 1.066033124 | 0.464009618 |
| ENSMUSG00000046971 | 1.029632141 | 0.976752232 |
| ENSMUSG00000046974 | -1 | 1 |
| ENSMUSG00000046975 | 1 | |
| ENSMUSG00000046982 | 1.042558594 | 0.530278005 |
| ENSMUSG00000046985 | 1.002155376 | 1 |
| ENSMUSG00000046991 | 1.05410549 | 0.805157604 |
| ENSMUSG00000046993 | 1.0463724 | 0.890110403 |
| ENSMUSG00000046994 | -1.049089091 | 0.783112516 |
| ENSMUSG00000046995 | 1 | |
| ENSMUSG00000046997 | -1.093239514 | 0.392592923 |
| ENSMUSG00000046999 | -1.054169715 | 0.813485995 |
| ENSMUSG00000047002 | 1 | |
| ENSMUSG00000047003 | 1.117596427 | 0.363615297 |
| ENSMUSG00000047013 | -1.129770807 | 0.698112512 |
| ENSMUSG00000047014 | 1 | |
| ENSMUSG00000047021 | -1 | 1 |
| ENSMUSG00000047022 | 1.118972442 | 0.289327063 |
| ENSMUSG00000047025 | 1 | |
| ENSMUSG00000047026 | 1 | |
| ENSMUSG00000047030 | -1.03064052 | 0.814600022 |
| ENSMUSG00000047031 | 1 | |
| ENSMUSG00000047033 | 1.401336534 | 0.061185687 |
| ENSMUSG00000047034 | -1.044273782 | 0.698112512 |
| ENSMUSG00000047036 | -1.037096441 | 0.895376993 |
| ENSMUSG00000047037 | 1.012475685 | 0.937805719 |
| ENSMUSG00000047039 | 1 | |
| ENSMUSG00000047040 | 1.057700516 | 0.856925007 |
| ENSMUSG00000047044 | 1.027842539 | 0.942667157 |
| ENSMUSG00000047045 | -1.091091236 | 0.698112512 |
| ENSMUSG00000047046 | 1 | |
| ENSMUSG00000047048 | -1 | 1 |
| ENSMUSG00000047050 | 1 | |
| ENSMUSG00000047053 | 1 | |
| ENSMUSG00000047061 | -1.090507733 | 0.698112512 |
| ENSMUSG00000047067 | 1.082512475 | 0.469202786 |
| ENSMUSG00000047079 | 1 | |
| ENSMUSG00000047084 | -1.017704479 | 0.914474058 |
| ENSMUSG00000047085 | 1.019083579 | 0.968937442 |
| ENSMUSG00000047090 | 1.152810025 | 0.14679718 |
| ENSMUSG00000047094 | 1 | |
| ENSMUSG00000047098 | -1.043941114 | 0.686666203 |
| ENSMUSG00000047102 | 1 | |
| ENSMUSG00000047104 | 1.067913355 | 0.797105186 |
| ENSMUSG00000047108 | -1.07008665 | 0.809950986 |
| ENSMUSG00000047109 | 1 | |
| ENSMUSG00000047115 | -1.168434773 | 0.339500222 |
| ENSMUSG00000047117 | -1.287708362 | 0.013163378 |
| ENSMUSG00000047123 | 1.065120849 | 0.601307651 |

| | | |
|---|---|---|
| ENSMUSG00000047126 | 1.133731688 | 0.709330685 |
| ENSMUSG00000047129 | 1 | |
| ENSMUSG00000047139 | 1.131107519 | 0.698112512 |
| ENSMUSG00000047141 | -1.003725655 | 1 |
| ENSMUSG00000047143 | -1.18051812 | 0.310368041 |
| ENSMUSG00000047146 | -1.122670729 | 0.532586 |
| ENSMUSG00000047149 | 1 | |
| ENSMUSG00000047150 | -1.1395797 | 0.698112512 |
| ENSMUSG00000047153 | -1.095836398 | 0.36623954 |
| ENSMUSG00000047155 | 1.040912485 | 0.828026521 |
| ENSMUSG00000047161 | 1 | |
| ENSMUSG00000047168 | 1 | |
| ENSMUSG00000047171 | -1.053549062 | 0.698112512 |
| ENSMUSG00000047180 | -1.52916954 | 0.001448802 |
| ENSMUSG00000047181 | 1.116745172 | 0.356538835 |
| ENSMUSG00000047182 | 1.059463094 | 0.608497229 |
| ENSMUSG00000047187 | 1.07247212 | 0.796531827 |
| ENSMUSG00000047189 | 1 | |
| ENSMUSG00000047193 | -1.023629315 | 0.949812939 |
| ENSMUSG00000047197 | 1 | |
| ENSMUSG00000047203 | 1 | |
| ENSMUSG00000047205 | 1.201849101 | 0.124608459 |
| ENSMUSG00000047207 | 1 | |
| ENSMUSG00000047213 | -1.016227625 | 1 |
| ENSMUSG00000047215 | 1.020328939 | 1 |
| ENSMUSG00000047216 | 1 | |
| ENSMUSG00000047220 | -1.005725456 | 1 |
| ENSMUSG00000047221 | 1.016492427 | 0.949812939 |
| ENSMUSG00000047222 | 1 | |
| ENSMUSG00000047225 | 1 | |
| ENSMUSG00000047227 | 1.04002589 | 0.608497229 |
| ENSMUSG00000047228 | -1.044273782 | 0.801443616 |
| ENSMUSG00000047230 | -1.089497311 | 0.532369657 |
| ENSMUSG00000047237 | 1 | |
| ENSMUSG00000047238 | 1.187621184 | 0.005249107 |
| ENSMUSG00000047242 | 1.150158107 | 0.18695932 |
| ENSMUSG00000047246 | -1.059623787 | 0.835531958 |
| ENSMUSG00000047248 | -1.043109444 | 0.84784274 |
| ENSMUSG00000047250 | 1.780625982 | 0.001127478 |
| ENSMUSG00000047253 | 1.043646933 | 0.82127647 |
| ENSMUSG00000047257 | 1 | |
| ENSMUSG00000047259 | 1.283884444 | 0.083239876 |
| ENSMUSG00000047260 | -1.039948148 | 0.818272915 |
| ENSMUSG00000047261 | -1.21938293 | 0.12843928 |
| ENSMUSG00000047264 | 1.303775896 | 0.004416753 |
| ENSMUSG00000047281 | 1.138470316 | 0.704433991 |
| ENSMUSG00000047284 | 1.046833327 | 0.703046429 |
| ENSMUSG00000047286 | 1 | |
| ENSMUSG00000047293 | -1.095939461 | 0.688241341 |
| ENSMUSG00000047295 | 1 | |
| ENSMUSG00000047298 | 1.059463094 | 0.608497229 |
| ENSMUSG00000047307 | -1 | 1 |
| ENSMUSG00000047324 | 1 | |
| ENSMUSG00000047330 | -1 | 1 |
| ENSMUSG00000047342 | -1.487561428 | 0.009935399 |
| ENSMUSG00000047343 | -1.03649557 | 0.964202376 |
| ENSMUSG00000047344 | -1.068241341 | 0.698112512 |
| ENSMUSG00000047347 | 1.087450017 | 0.05848516 |
| ENSMUSG00000047352 | 1 | |
| ENSMUSG00000047356 | 1 | |
| ENSMUSG00000047361 | -1 | 1 |
| ENSMUSG00000047363 | 1.290681034 | 0.007752116 |
| ENSMUSG00000047368 | 1.025953282 | 0.956374899 |

| | | |
|---|---|---|
| ENSMUSG00000047369 | 1 | |
| ENSMUSG00000047370 | -1.03962336 | 0.698112512 |
| ENSMUSG00000047371 | 1.115543023 | 0.061839295 |
| ENSMUSG00000047379 | -1.133318693 | 0.214241028 |
| ENSMUSG00000047383 | 1 | |
| ENSMUSG00000047384 | -1.03962336 | 0.803072249 |
| ENSMUSG00000047388 | -1.022744334 | 0.911212167 |
| ENSMUSG00000047390 | 1 | |
| ENSMUSG00000047394 | -1.08081673 | 0.733498571 |
| ENSMUSG00000047407 | 1.290572075 | 0.106808755 |
| ENSMUSG00000047409 | 1.082394738 | 0.608497229 |
| ENSMUSG00000047412 | 1.070945749 | 0.660210733 |
| ENSMUSG00000047414 | 1.17847863 | 0.450928478 |
| ENSMUSG00000047415 | -1.086681236 | 0.783113058 |
| ENSMUSG00000047417 | 1.03148856 | 0.850278675 |
| ENSMUSG00000047419 | -1.089497311 | 0.698112512 |
| ENSMUSG00000047420 | 1.20958465 | 0.212056622 |
| ENSMUSG00000047423 | 1.13834586 | 0.239852828 |
| ENSMUSG00000047428 | 1.124270862 | 0.708747804 |
| ENSMUSG00000047434 | -1.075420112 | 0.681952856 |
| ENSMUSG00000047441 | -1 | 1 |
| ENSMUSG00000047443 | -1.007342085 | 0.996661676 |
| ENSMUSG00000047444 | 1 | |
| ENSMUSG00000047446 | 1.090446663 | 0.435925302 |
| ENSMUSG00000047454 | -1.006568331 | 1 |
| ENSMUSG00000047459 | -1.10478155 | 0.608497229 |
| ENSMUSG00000047462 | -1 | 1 |
| ENSMUSG00000047466 | 1.08739179 | 0.549176447 |
| ENSMUSG00000047473 | 1.023144139 | 0.807933619 |
| ENSMUSG00000047485 | 1.095872691 | 0.608497229 |
| ENSMUSG00000047490 | 1 | |
| ENSMUSG00000047492 | 1 | |
| ENSMUSG00000047495 | 1.346809975 | 0.012476765 |
| ENSMUSG00000047496 | -1.235270921 | 0.205105863 |
| ENSMUSG00000047497 | 1.416180103 | 0.005239078 |
| ENSMUSG00000047501 | -1.014570171 | 0.985623907 |
| ENSMUSG00000047502 | 1 | |
| ENSMUSG00000047507 | 1.195405785 | 0.480356555 |
| ENSMUSG00000047508 | 1 | |
| ENSMUSG00000047509 | 1 | |
| ENSMUSG00000047511 | 1 | |
| ENSMUSG00000047514 | -1.041915437 | 0.863882521 |
| ENSMUSG00000047515 | -1.099879851 | 0.698112512 |
| ENSMUSG00000047517 | 1 | |
| ENSMUSG00000047518 | 1 | |
| ENSMUSG00000047528 | 1.165323951 | 0.572708072 |
| ENSMUSG00000047531 | 1 | |
| ENSMUSG00000047534 | 1.100465311 | 0.608497229 |
| ENSMUSG00000047535 | 1 | |
| ENSMUSG00000047539 | -1.08234897 | 0.698112512 |
| ENSMUSG00000047544 | 1 | |
| ENSMUSG00000047545 | 1 | |
| ENSMUSG00000047547 | -1.21161446 | 0.10982693 |
| ENSMUSG00000047554 | -1.019243899 | 0.975843246 |
| ENSMUSG00000047557 | 1.170316077 | 0.283365781 |
| ENSMUSG00000047562 | -1.085651418 | 0.731932479 |
| ENSMUSG00000047564 | 1 | |
| ENSMUSG00000047565 | 1.052787084 | 0.931347167 |
| ENSMUSG00000047583 | -1.039911408 | 0.823872552 |
| ENSMUSG00000047586 | 1 | |
| ENSMUSG00000047591 | 1.019083579 | 0.949812939 |
| ENSMUSG00000047592 | 1.015486245 | 0.99457094 |
| ENSMUSG00000047594 | 1 | |

| | | |
|---|---|---|
| ENSMUSG00000047603 | 1.024409896 | 0.790112395 |
| ENSMUSG00000047604 | -1.081290579 | 0.543320162 |
| ENSMUSG00000047606 | 1.132639128 | 0.398767938 |
| ENSMUSG00000047613 | 1.012840072 | 0.938708567 |
| ENSMUSG00000047617 | -1.178715592 | 0.374710101 |
| ENSMUSG00000047619 | 1 | |
| ENSMUSG00000047626 | 1 | |
| ENSMUSG00000047631 | -1.314724024 | 0.063980123 |
| ENSMUSG00000047632 | -1.186060634 | 0.018531376 |
| ENSMUSG00000047635 | 1.090907055 | 0.45758853 |
| ENSMUSG00000047636 | 1 | |
| ENSMUSG00000047638 | -1.064113315 | 0.855682211 |
| ENSMUSG00000047641 | 1.380713072 | 0.218021596 |
| ENSMUSG00000047642 | -1.120476206 | 0.608497229 |
| ENSMUSG00000047643 | 1 | |
| ENSMUSG00000047648 | 1.046191629 | 0.877008742 |
| ENSMUSG00000047649 | -1.039511018 | 0.717035356 |
| ENSMUSG00000047652 | 1 | |
| ENSMUSG00000047654 | -1.095294215 | 0.541735653 |
| ENSMUSG00000047655 | 1 | |
| ENSMUSG00000047656 | 1.117083704 | 0.608497229 |
| ENSMUSG00000047657 | 1 | |
| ENSMUSG00000047658 | -1.193384339 | 0.608497229 |
| ENSMUSG00000047661 | 1 | |
| ENSMUSG00000047667 | 1 | |
| ENSMUSG00000047669 | 1.158409256 | 0.050235993 |
| ENSMUSG00000047671 | -1.033820075 | 0.918344559 |
| ENSMUSG00000047674 | 1 | |
| ENSMUSG00000047675 | 1.041487533 | 0.911346192 |
| ENSMUSG00000047676 | 1.206397991 | 0.60397857 |
| ENSMUSG00000047678 | -1.044273782 | 0.698112512 |
| ENSMUSG00000047686 | 1 | |
| ENSMUSG00000047692 | -1 | 1 |
| ENSMUSG00000047694 | -1.145164509 | 0.539222653 |
| ENSMUSG00000047696 | -1.411511715 | 0.063484316 |
| ENSMUSG00000047702 | 1 | |
| ENSMUSG00000047710 | 1.144950549 | 0.433678407 |
| ENSMUSG00000047712 | 1.033949541 | 0.865583959 |
| ENSMUSG00000047714 | -1.04269226 | 0.917316404 |
| ENSMUSG00000047716 | 1 | |
| ENSMUSG00000047719 | -1.058733938 | 0.698112512 |
| ENSMUSG00000047720 | 1 | |
| ENSMUSG00000047721 | 1.132823566 | 0.47916192 |
| ENSMUSG00000047728 | 1.230075506 | 0.292973656 |
| ENSMUSG00000047730 | -1.023769022 | 0.954761885 |
| ENSMUSG00000047731 | 1.073586711 | 0.698112512 |
| ENSMUSG00000047733 | 1.24122829 | 0.180765391 |
| ENSMUSG00000047735 | -1.080286536 | 0.631165014 |
| ENSMUSG00000047746 | -1 | 1 |
| ENSMUSG00000047747 | 1.677001866 | 0.000764285 |
| ENSMUSG00000047749 | 1.17835967 | 0.17702044 |
| ENSMUSG00000047751 | 1 | |
| ENSMUSG00000047757 | -1.111180685 | 0.541735653 |
| ENSMUSG00000047759 | -1.144495522 | 0.641259948 |
| ENSMUSG00000047766 | -1.101535046 | 0.628599199 |
| ENSMUSG00000047767 | -1.047414369 | 0.642138838 |
| ENSMUSG00000047773 | 1 | |
| ENSMUSG00000047777 | 1.022448519 | 0.914954453 |
| ENSMUSG00000047786 | -1.067718846 | 0.851367041 |
| ENSMUSG00000047787 | -1.232117496 | 0.016957915 |
| ENSMUSG00000047789 | -1.077025122 | 0.69533886 |
| ENSMUSG00000047793 | 1.172201819 | 0.09986558 |
| ENSMUSG00000047794 | 1 | |

| | | |
|---|---|---|
| ENSMUSG00000047797 | 1 | |
| ENSMUSG00000047798 | 1.059463094 | 0.704170349 |
| ENSMUSG00000047799 | 1 | |
| ENSMUSG00000047804 | -1.033159048 | 0.895320791 |
| ENSMUSG00000047807 | -1.208988299 | 0.59127359 |
| ENSMUSG00000047810 | 1.019579327 | 1 |
| ENSMUSG00000047819 | -1.177310696 | 0.440481392 |
| ENSMUSG00000047821 | -1.089783181 | 0.624088293 |
| ENSMUSG00000047822 | 1.019083579 | 0.968937442 |
| ENSMUSG00000047824 | 1.030233739 | 0.866289429 |
| ENSMUSG00000047828 | 1 | |
| ENSMUSG00000047832 | 1.150479303 | 0.388781089 |
| ENSMUSG00000047841 | 1 | |
| ENSMUSG00000047842 | 1.595637897 | 0.000500173 |
| ENSMUSG00000047843 | 1.196162087 | 0.132886711 |
| ENSMUSG00000047844 | -1.88416875 | 0.014214586 |
| ENSMUSG00000047854 | 1.096051716 | 0.681187387 |
| ENSMUSG00000047861 | 1 | |
| ENSMUSG00000047866 | -1.000595804 | 1 |
| ENSMUSG00000047867 | 1.102285378 | 0.698112512 |
| ENSMUSG00000047868 | 1 | |
| ENSMUSG00000047875 | -1.203567206 | 0.024753959 |
| ENSMUSG00000047878 | -1.603734682 | 0.011369917 |
| ENSMUSG00000047879 | 1.030289035 | 0.962368837 |
| ENSMUSG00000047880 | 1.289908992 | 0.103537668 |
| ENSMUSG00000047881 | 1.082423824 | 0.675439448 |
| ENSMUSG00000047884 | 1 | |
| ENSMUSG00000047888 | -1.001294495 | 1 |
| ENSMUSG00000047889 | 1 | |
| ENSMUSG00000047894 | -1.094836541 | 0.541227048 |
| ENSMUSG00000047897 | 1 | |
| ENSMUSG00000047898 | -1.010960635 | 1 |
| ENSMUSG00000047904 | -1.244128 | 0.467634171 |
| ENSMUSG00000047905 | -1.19653739 | 0.273660026 |
| ENSMUSG00000047907 | -1.043306199 | 0.798487595 |
| ENSMUSG00000047909 | 1.050997922 | 0.709997537 |
| ENSMUSG00000047910 | 2.096463638 | 0.000763004 |
| ENSMUSG00000047911 | 1.04213016 | 0.932855261 |
| ENSMUSG00000047921 | 1.06361625 | 0.608497229 |
| ENSMUSG00000047935 | 1.048462658 | 0.941060294 |
| ENSMUSG00000047938 | 1.017174474 | 0.99448341 |
| ENSMUSG00000047940 | -1.053549062 | 0.698112512 |
| ENSMUSG00000047945 | 1.103052523 | 0.643397479 |
| ENSMUSG00000047953 | 1.015486245 | 0.986542466 |
| ENSMUSG00000047959 | 1 | |
| ENSMUSG00000047960 | 1 | |
| ENSMUSG00000047963 | 1.224245079 | 0.17272194 |
| ENSMUSG00000047965 | 1.014133649 | 1 |
| ENSMUSG00000047969 | 1 | |
| ENSMUSG00000047976 | -1.089497311 | 0.532369657 |
| ENSMUSG00000047977 | 1.092382958 | 0.638239556 |
| ENSMUSG00000047980 | 1 | |
| ENSMUSG00000047986 | -1.323642555 | 0.489824652 |
| ENSMUSG00000047988 | 1.063301216 | 0.698112512 |
| ENSMUSG00000047989 | -1.101925608 | 0.594567601 |
| ENSMUSG00000047990 | 1 | |
| ENSMUSG00000047992 | -1.295129617 | 0.153295884 |
| ENSMUSG00000047995 | 1 | |
| ENSMUSG00000047996 | 1.145972546 | 0.373671631 |
| ENSMUSG00000048000 | 1.004397049 | 1 |
| ENSMUSG00000048001 | 1 | |
| ENSMUSG00000048003 | -1 | 1 |
| ENSMUSG00000048004 | -1 | 1 |

| | | |
|---|---|---|
| ENSMUSG00000048007 | -1.037091675 | 0.698112512 |
| ENSMUSG00000048012 | 1.051113457 | 0.802048461 |
| ENSMUSG00000048013 | 1.059463094 | 0.608497229 |
| ENSMUSG00000048015 | -1 | 1 |
| ENSMUSG00000048022 | 1 | |
| ENSMUSG00000048027 | 1.079319049 | 0.541726526 |
| ENSMUSG00000048029 | -1.003283529 | 1 |
| ENSMUSG00000048031 | 1 | |
| ENSMUSG00000048038 | 1 | |
| ENSMUSG00000048039 | -1.029802204 | 0.927258464 |
| ENSMUSG00000048040 | -1.174431866 | 0.161510586 |
| ENSMUSG00000048047 | 1.043198745 | 0.870427333 |
| ENSMUSG00000048058 | 1.060070715 | 0.698112512 |
| ENSMUSG00000048062 | 1 | |
| ENSMUSG00000048065 | -1.043306199 | 0.803260194 |
| ENSMUSG00000048067 | 1 | |
| ENSMUSG00000048070 | -1 | 1 |
| ENSMUSG00000048076 | -1.041146952 | 0.936995747 |
| ENSMUSG00000048077 | 1 | |
| ENSMUSG00000048078 | 1.314298984 | 0.012417057 |
| ENSMUSG00000048080 | 1 | |
| ENSMUSG00000048087 | -1.068741359 | 0.449386295 |
| ENSMUSG00000048096 | 3.714915277 | 0.000207005 |
| ENSMUSG00000048100 | -1.083511745 | 0.650252997 |
| ENSMUSG00000048101 | 1 | |
| ENSMUSG00000048106 | -1.029150153 | 0.908007394 |
| ENSMUSG00000048108 | 1.063432582 | 0.800287394 |
| ENSMUSG00000048109 | -1.013699291 | 0.996913409 |
| ENSMUSG00000048118 | 1.102036359 | 0.555973705 |
| ENSMUSG00000048120 | 1.170653107 | 0.608497229 |
| ENSMUSG00000048126 | 4.309577489 | 0.000263878 |
| ENSMUSG00000048138 | -1.453891896 | 0.016112968 |
| ENSMUSG00000048142 | -1.073812814 | 0.78287223 |
| ENSMUSG00000048148 | 1.029957303 | 0.981132618 |
| ENSMUSG00000048153 | -1 | 1 |
| ENSMUSG00000048154 | 1.129712599 | 0.374017624 |
| ENSMUSG00000048155 | 1 | |
| ENSMUSG00000048163 | 1.084751429 | 0.698112512 |
| ENSMUSG00000048164 | 1.116423141 | 0.687018628 |
| ENSMUSG00000048170 | 1.009553788 | 1 |
| ENSMUSG00000048173 | 1 | |
| ENSMUSG00000048174 | -1.065200986 | 0.478279422 |
| ENSMUSG00000048175 | 1.054335525 | 0.712130748 |
| ENSMUSG00000048185 | 1.059463094 | 0.608497229 |
| ENSMUSG00000048186 | -1.242269814 | 0.192078276 |
| ENSMUSG00000048188 | -1 | 1 |
| ENSMUSG00000048191 | -1.036610587 | 0.801154107 |
| ENSMUSG00000048197 | 1 | |
| ENSMUSG00000048200 | 1.068207746 | 0.250726313 |
| ENSMUSG00000048206 | 1 | |
| ENSMUSG00000048215 | 1 | |
| ENSMUSG00000048216 | 1.391305642 | 0.001532355 |
| ENSMUSG00000048217 | -1.044273782 | 0.855488553 |
| ENSMUSG00000048218 | -2.000857937 | 0.000968318 |
| ENSMUSG00000048222 | 1.005670682 | 1 |
| ENSMUSG00000048226 | 1 | |
| ENSMUSG00000048230 | -1.17077323 | 0.608497229 |
| ENSMUSG00000048231 | 1 | |
| ENSMUSG00000048232 | 1.030905039 | 0.875751698 |
| ENSMUSG00000048234 | -1.073289148 | 0.698112512 |
| ENSMUSG00000048236 | 1 | |
| ENSMUSG00000048240 | -1.51966666 | 0.009448316 |
| ENSMUSG00000048249 | 1.101259562 | 0.541227048 |

| | | |
|---|---|---|
| ENSMUSG00000048251 | -1.174186396 | 0.534861373 |
| ENSMUSG00000048261 | -1.03962336 | 0.865094641 |
| ENSMUSG00000048264 | 1.01772357 | 0.976574098 |
| ENSMUSG00000048271 | 1.072652804 | 0.740810916 |
| ENSMUSG00000048277 | -1.090908563 | 0.608497229 |
| ENSMUSG00000048279 | 1.004799506 | 1 |
| ENSMUSG00000048280 | -1.102384734 | 0.534887579 |
| ENSMUSG00000048281 | 1 | |
| ENSMUSG00000048284 | 1 | |
| ENSMUSG00000048285 | 1.019462741 | 0.978962085 |
| ENSMUSG00000048292 | 1.24122829 | 0.03679259 |
| ENSMUSG00000048294 | 1 | |
| ENSMUSG00000048299 | 1 | |
| ENSMUSG00000048304 | -1.03962336 | 0.803072249 |
| ENSMUSG00000048307 | 1.002354851 | 1 |
| ENSMUSG00000048310 | -1.058520209 | 0.717035356 |
| ENSMUSG00000048312 | 1 | |
| ENSMUSG00000048316 | 1 | |
| ENSMUSG00000048327 | -1.127817941 | 0.608497229 |
| ENSMUSG00000048329 | 1 | |
| ENSMUSG00000048330 | -1.0965339 | 0.609589476 |
| ENSMUSG00000048332 | 1.171578259 | 0.309330961 |
| ENSMUSG00000048334 | 1.069068979 | 0.608497229 |
| ENSMUSG00000048337 | 1.123041516 | 0.514301195 |
| ENSMUSG00000048347 | 1.647476317 | 0.009118829 |
| ENSMUSG00000048349 | -1.231184281 | 0.048002191 |
| ENSMUSG00000048351 | -1.030087064 | 0.848700716 |
| ENSMUSG00000048355 | -1.260350288 | 0.303893112 |
| ENSMUSG00000048356 | -1 | 1 |
| ENSMUSG00000048368 | 1.209240109 | 0.047527942 |
| ENSMUSG00000048371 | -1.064576608 | 0.608497229 |
| ENSMUSG00000048373 | -1.069654539 | 0.825838389 |
| ENSMUSG00000048376 | 1.134519555 | 0.608497229 |
| ENSMUSG00000048377 | 1 | |
| ENSMUSG00000048378 | 1 | |
| ENSMUSG00000048379 | -1.120758153 | 0.608497229 |
| ENSMUSG00000048385 | -1.03962336 | 0.698112512 |
| ENSMUSG00000048387 | 1.422510589 | 0.012065947 |
| ENSMUSG00000048388 | 1.056014251 | 0.753936646 |
| ENSMUSG00000048391 | 1 | |
| ENSMUSG00000048398 | 1 | |
| ENSMUSG00000048399 | 1.059463094 | 0.768693672 |
| ENSMUSG00000048400 | 1 | |
| ENSMUSG00000048402 | 1.19467157 | 0.058274403 |
| ENSMUSG00000048406 | -1.009897963 | 1 |
| ENSMUSG00000048410 | 1.014821567 | 0.98079728 |
| ENSMUSG00000048411 | 1 | |
| ENSMUSG00000048416 | 1.057185466 | 0.749776086 |
| ENSMUSG00000048424 | 1.102024305 | 0.654947926 |
| ENSMUSG00000048425 | 1 | |
| ENSMUSG00000048429 | -1.0602363 | 0.790210734 |
| ENSMUSG00000048439 | -1.00478002 | 1 |
| ENSMUSG00000048440 | 1.12540651 | 0.155387198 |
| ENSMUSG00000048442 | 1.089520423 | 0.825178237 |
| ENSMUSG00000048445 | 1.042992458 | 0.544543745 |
| ENSMUSG00000048450 | -2.634911564 | 2.25884E-05 |
| ENSMUSG00000048455 | 1 | |
| ENSMUSG00000048456 | 1 | |
| ENSMUSG00000048458 | 1.309247486 | 0.007291063 |
| ENSMUSG00000048469 | 1 | |
| ENSMUSG00000048473 | 1 | |
| ENSMUSG00000048478 | -1.195915902 | 0.449092158 |
| ENSMUSG00000048480 | 1 | |

| | | |
|---|---|---|
| ENSMUSG00000048481 | -1.025510922 | 0.836456594 |
| ENSMUSG00000048482 | 1.282496803 | 0.049065949 |
| ENSMUSG00000048483 | 1 | |
| ENSMUSG00000048484 | 1.020668955 | 0.979002724 |
| ENSMUSG00000048485 | -1.094712223 | 0.760685489 |
| ENSMUSG00000048486 | -1.132983115 | 0.293364956 |
| ENSMUSG00000048489 | 1.122462048 | 0.608497229 |
| ENSMUSG00000048490 | 1.059768334 | 0.698112512 |
| ENSMUSG00000048495 | -1.058441685 | 0.69104961 |
| ENSMUSG00000048497 | -1.040184006 | 0.776021306 |
| ENSMUSG00000048498 | 1 | |
| ENSMUSG00000048500 | 1 | |
| ENSMUSG00000048501 | 1 | |
| ENSMUSG00000048502 | 1 | |
| ENSMUSG00000048503 | -1.241327999 | 0.019418363 |
| ENSMUSG00000048516 | 1 | |
| ENSMUSG00000048520 | 1.144733348 | 0.608497229 |
| ENSMUSG00000048521 | -1.1927419 | 0.186603757 |
| ENSMUSG00000048528 | -1.132666855 | 0.18851436 |
| ENSMUSG00000048534 | -1.063525418 | 0.803072249 |
| ENSMUSG00000048537 | -1.005778678 | 1 |
| ENSMUSG00000048538 | -1 | 1 |
| ENSMUSG00000048540 | 1 | |
| ENSMUSG00000048544 | 1.059463094 | 0.698112512 |
| ENSMUSG00000048546 | 1.067509511 | 0.681952832 |
| ENSMUSG00000048550 | 1.09594632 | 0.608497229 |
| ENSMUSG00000048559 | 1 | |
| ENSMUSG00000048562 | 1.889490836 | 0.000388292 |
| ENSMUSG00000048572 | 1.462183684 | 0.102533756 |
| ENSMUSG00000048573 | 1 | |
| ENSMUSG00000048574 | 1 | |
| ENSMUSG00000048578 | 1.038483009 | 0.95526044 |
| ENSMUSG00000048581 | 1.061677032 | 0.608497229 |
| ENSMUSG00000048582 | -1.177388846 | 0.608497229 |
| ENSMUSG00000048583 | -1.719901355 | 0.021784237 |
| ENSMUSG00000048592 | -1.095294215 | 0.541735653 |
| ENSMUSG00000048600 | 1 | |
| ENSMUSG00000048602 | 1.05038453 | 0.877529894 |
| ENSMUSG00000048603 | 1.109396363 | 0.535052277 |
| ENSMUSG00000048612 | 1.138613541 | 0.698112512 |
| ENSMUSG00000048616 | -1.571681274 | 0.013001366 |
| ENSMUSG00000048617 | 1.264744915 | 0.156181644 |
| ENSMUSG00000048620 | 1 | |
| ENSMUSG00000048621 | 1.328600725 | 0.26415877 |
| ENSMUSG00000048626 | 1 | |
| ENSMUSG00000048628 | 1 | |
| ENSMUSG00000048636 | 1.122462048 | 0.420161906 |
| ENSMUSG00000048643 | 1 | |
| ENSMUSG00000048644 | 1.429164401 | 0.009478551 |
| ENSMUSG00000048647 | -1.113446377 | 0.379320311 |
| ENSMUSG00000048652 | 1 | |
| ENSMUSG00000048655 | -1.020873558 | 0.996385115 |
| ENSMUSG00000048661 | -1.074784074 | 0.687832464 |
| ENSMUSG00000048668 | 1.072520492 | 0.698112512 |
| ENSMUSG00000048677 | 1.024733634 | 0.878551492 |
| ENSMUSG00000048686 | 1 | |
| ENSMUSG00000048693 | 1 | |
| ENSMUSG00000048696 | 1.220957055 | 0.025176031 |
| ENSMUSG00000048697 | 1 | |
| ENSMUSG00000048699 | 1 | |
| ENSMUSG00000048701 | 1.128979356 | 0.601666207 |
| ENSMUSG00000048703 | 1 | |
| ENSMUSG00000048706 | -1.085966135 | 0.664129605 |

| | | |
|---|---|---|
| ENSMUSG00000048707 | -1.009992427 | 0.955107108 |
| ENSMUSG00000048709 | 1.05038453 | 0.828564184 |
| ENSMUSG00000048720 | 1.096885776 | 0.541534666 |
| ENSMUSG00000048721 | -1.458849207 | 0.196910724 |
| ENSMUSG00000048728 | -1.214556728 | 0.114056728 |
| ENSMUSG00000048731 | 1.283481648 | 0.254806036 |
| ENSMUSG00000048732 | -1.009192838 | 1 |
| ENSMUSG00000048745 | 1 | |
| ENSMUSG00000048747 | 1 | |
| ENSMUSG00000048752 | -1 | 1 |
| ENSMUSG00000048755 | 1.109219806 | 0.324227192 |
| ENSMUSG00000048756 | 1.17586405 | 0.279927732 |
| ENSMUSG00000048758 | -1.118277365 | 0.232747283 |
| ENSMUSG00000048763 | 1.108262617 | 0.062259568 |
| ENSMUSG00000048764 | -1.360799916 | 0.13435151 |
| ENSMUSG00000048766 | 1 | |
| ENSMUSG00000048772 | 1.079486647 | 0.672322256 |
| ENSMUSG00000048775 | -1.043306199 | 0.698112512 |
| ENSMUSG00000048776 | 1.009675516 | 1 |
| ENSMUSG00000048779 | 1.040172434 | 0.934760809 |
| ENSMUSG00000048782 | 1.138294359 | 0.642560959 |
| ENSMUSG00000048787 | 1.124954424 | 0.26133383 |
| ENSMUSG00000048794 | -1.049398742 | 0.96035552 |
| ENSMUSG00000048796 | -1.133156287 | 0.199114949 |
| ENSMUSG00000048799 | 1.070318398 | 0.740330308 |
| ENSMUSG00000048806 | 1 | |
| ENSMUSG00000048807 | 1.214376033 | 0.056767117 |
| ENSMUSG00000048810 | 1 | |
| ENSMUSG00000048814 | -1.679823848 | 0.003278432 |
| ENSMUSG00000048824 | 1 | |
| ENSMUSG00000048826 | 1.582411431 | 0.004416753 |
| ENSMUSG00000048827 | -1.170809949 | 0.240599737 |
| ENSMUSG00000048830 | 1 | |
| ENSMUSG00000048832 | 1.075457017 | 0.478777134 |
| ENSMUSG00000048833 | 1.019390826 | 0.979002724 |
| ENSMUSG00000048834 | -1 | 1 |
| ENSMUSG00000048852 | 1.11038323 | 0.643970135 |
| ENSMUSG00000048856 | 1.203490127 | 0.125154408 |
| ENSMUSG00000048865 | -1.104118099 | 0.71627176 |
| ENSMUSG00000048874 | 1.055754466 | 0.839713755 |
| ENSMUSG00000048875 | 1 | |
| ENSMUSG00000048878 | -1.003373179 | 1 |
| ENSMUSG00000048895 | 1.301834422 | 0.003084596 |
| ENSMUSG00000048897 | 1.068713391 | 0.282603542 |
| ENSMUSG00000048899 | -1.398004607 | 0.246493876 |
| ENSMUSG00000048904 | 1 | |
| ENSMUSG00000048905 | 1.067996206 | 0.825410965 |
| ENSMUSG00000048911 | 1.070255598 | 0.710698346 |
| ENSMUSG00000048915 | -1.01552818 | 0.925075932 |
| ENSMUSG00000048916 | 1.059463094 | 0.698112512 |
| ENSMUSG00000048919 | 1 | |
| ENSMUSG00000048920 | -1.015330782 | 0.826629613 |
| ENSMUSG00000048921 | 1.127811967 | 0.264869231 |
| ENSMUSG00000048922 | -1.152882167 | 0.456934057 |
| ENSMUSG00000048924 | -1.129677549 | 0.330594361 |
| ENSMUSG00000048930 | 1.12795316 | 0.110155681 |
| ENSMUSG00000048933 | 1 | |
| ENSMUSG00000048938 | 1.111812527 | 0.672818484 |
| ENSMUSG00000048939 | 1 | |
| ENSMUSG00000048949 | -1.004921656 | 1 |
| ENSMUSG00000048960 | 1.433364281 | 0.048604265 |
| ENSMUSG00000048965 | -1.119200995 | 0.155674728 |
| ENSMUSG00000048967 | 1.135487404 | 0.780320794 |

| | | |
|---|---|---|
| ENSMUSG00000048970 | 1.035362196 | 0.882707794 |
| ENSMUSG00000048978 | -1.510723113 | 0.003880882 |
| ENSMUSG00000048981 | 1 | |
| ENSMUSG00000048982 | 1.059463094 | 0.698112512 |
| ENSMUSG00000048988 | 1.646789169 | 0.065682824 |
| ENSMUSG00000048992 | 1.176313967 | 0.608497229 |
| ENSMUSG00000048994 | 1 | |
| ENSMUSG00000048996 | -1.043306199 | 0.698112512 |
| ENSMUSG00000048997 | 1.050489286 | 0.698112512 |
| ENSMUSG00000049001 | 1.062187475 | 0.756671188 |
| ENSMUSG00000049008 | 1 | |
| ENSMUSG00000049010 | 1 | |
| ENSMUSG00000049011 | 1 | |
| ENSMUSG00000049013 | -1.03962336 | 0.698112512 |
| ENSMUSG00000049015 | 1 | |
| ENSMUSG00000049018 | 1 | |
| ENSMUSG00000049028 | 1 | |
| ENSMUSG00000049036 | 1.214667946 | 0.608497229 |
| ENSMUSG00000049037 | -1 | 1 |
| ENSMUSG00000049038 | 1.016024146 | 0.961394002 |
| ENSMUSG00000049041 | 1 | |
| ENSMUSG00000049044 | -1.023769022 | 0.969544524 |
| ENSMUSG00000049047 | 1.021311156 | 0.978259071 |
| ENSMUSG00000049052 | 1 | |
| ENSMUSG00000049057 | 1 | |
| ENSMUSG00000049062 | 1 | |
| ENSMUSG00000049073 | 1 | |
| ENSMUSG00000049076 | 1.070900211 | 0.763030715 |
| ENSMUSG00000049086 | -1.293192562 | 0.326011938 |
| ENSMUSG00000049090 | 1.089925469 | 0.572721726 |
| ENSMUSG00000049091 | 1.006059249 | 0.996997119 |
| ENSMUSG00000049092 | -1.15769714 | 0.302127239 |
| ENSMUSG00000049093 | -1.310421488 | 0.04562023 |
| ENSMUSG00000049094 | 1 | |
| ENSMUSG00000049097 | -1.10504258 | 0.698112512 |
| ENSMUSG00000049098 | 1 | |
| ENSMUSG00000049100 | 1.081074865 | 0.822353727 |
| ENSMUSG00000049103 | 1.132639128 | 0.398767938 |
| ENSMUSG00000049106 | 1.105056191 | 0.50744656 |
| ENSMUSG00000049107 | -1.374991218 | 0.02776706 |
| ENSMUSG00000049109 | -1.142727245 | 0.202428693 |
| ENSMUSG00000049112 | -1.058507348 | 0.858701311 |
| ENSMUSG00000049115 | -1.24993587 | 0.151710067 |
| ENSMUSG00000049119 | 1.19879658 | 0.070911845 |
| ENSMUSG00000049122 | -1.084645496 | 0.532369657 |
| ENSMUSG00000049123 | 1.023645551 | 0.981626226 |
| ENSMUSG00000049124 | 1 | |
| ENSMUSG00000049128 | 1 | |
| ENSMUSG00000049130 | 1 | |
| ENSMUSG00000049133 | 1 | |
| ENSMUSG00000049134 | -1.045180484 | 0.939641924 |
| ENSMUSG00000049148 | 1 | |
| ENSMUSG00000049149 | 1 | |
| ENSMUSG00000049152 | -1.03962336 | 0.698112512 |
| ENSMUSG00000049154 | 1 | |
| ENSMUSG00000049160 | -1.071075483 | 0.775016379 |
| ENSMUSG00000049164 | -1.076584344 | 0.698249621 |
| ENSMUSG00000049168 | 1 | |
| ENSMUSG00000049173 | 1 | |
| ENSMUSG00000049176 | -1.044273782 | 0.892237454 |
| ENSMUSG00000049184 | 1.022748119 | 0.743576258 |
| ENSMUSG00000049191 | 1.334495043 | 0.004058621 |
| ENSMUSG00000049202 | -1.03962336 | 0.803072249 |

| | | |
|---|---|---|
| ENSMUSG00000049214 | -1.109582661 | 0.549347972 |
| ENSMUSG00000049217 | 1 | |
| ENSMUSG00000049225 | 1.205736487 | 0.14434203 |
| ENSMUSG00000049229 | 1 | |
| ENSMUSG00000049230 | 1.065410325 | 0.711602885 |
| ENSMUSG00000049231 | 1 | |
| ENSMUSG00000049232 | 1.111173064 | 0.460068234 |
| ENSMUSG00000049233 | -1.150809442 | 0.585163706 |
| ENSMUSG00000049235 | -1 | 1 |
| ENSMUSG00000049241 | 1 | |
| ENSMUSG00000049246 | -1.183404224 | 0.330417199 |
| ENSMUSG00000049249 | 1.035067229 | 0.952385403 |
| ENSMUSG00000049252 | -1.15462599 | 0.361740791 |
| ENSMUSG00000049265 | 1.580427059 | 0.005249107 |
| ENSMUSG00000049280 | 1 | |
| ENSMUSG00000049281 | 1.065818983 | 0.838148374 |
| ENSMUSG00000049285 | -1.00914849 | 1 |
| ENSMUSG00000049287 | -1.052376297 | 0.608497229 |
| ENSMUSG00000049288 | 1.031271205 | 0.932622418 |
| ENSMUSG00000049291 | 1.069068979 | 0.74773997 |
| ENSMUSG00000049295 | 1.022623132 | 0.953962555 |
| ENSMUSG00000049299 | -1.085395016 | 0.698112512 |
| ENSMUSG00000049300 | -1.063825578 | 0.67362935 |
| ENSMUSG00000049303 | -1.075041137 | 0.404886539 |
| ENSMUSG00000049305 | -1.025416005 | 0.84843268 |
| ENSMUSG00000049307 | 1.121815945 | 0.26075878 |
| ENSMUSG00000049313 | 1.182071427 | 0.048604265 |
| ENSMUSG00000049314 | 1 | |
| ENSMUSG00000049315 | 1 | |
| ENSMUSG00000049321 | -1.024586327 | 0.77589891 |
| ENSMUSG00000049323 | 1.046546431 | 0.874071622 |
| ENSMUSG00000049327 | 1.065850974 | 0.770498527 |
| ENSMUSG00000049334 | 1 | |
| ENSMUSG00000049336 | -1.085651418 | 0.689423135 |
| ENSMUSG00000049339 | 1.060090316 | 0.698112512 |
| ENSMUSG00000049349 | -1 | 1 |
| ENSMUSG00000049350 | -1 | 1 |
| ENSMUSG00000049353 | 1 | |
| ENSMUSG00000049354 | -1.006312581 | 1 |
| ENSMUSG00000049357 | -1.131994203 | 0.36733642 |
| ENSMUSG00000049362 | 1 | |
| ENSMUSG00000049372 | 1 | |
| ENSMUSG00000049382 | 1.490100753 | 0.044230959 |
| ENSMUSG00000049387 | -1.044273782 | 0.698112512 |
| ENSMUSG00000049396 | 1.162115381 | 0.608497229 |
| ENSMUSG00000049401 | -1.167615947 | 0.181047007 |
| ENSMUSG00000049404 | 1.480256749 | 0.03033847 |
| ENSMUSG00000049409 | -1.061476185 | 0.608497229 |
| ENSMUSG00000049410 | 1 | |
| ENSMUSG00000049411 | -1.102987079 | 0.549841702 |
| ENSMUSG00000049414 | 1 | |
| ENSMUSG00000049420 | 1.111329726 | 0.698112512 |
| ENSMUSG00000049421 | 1.066909347 | 0.758113599 |
| ENSMUSG00000049422 | 1.11535019 | 0.641910604 |
| ENSMUSG00000049427 | -1.063525418 | 0.788000546 |
| ENSMUSG00000049436 | -1.044273782 | 0.730192592 |
| ENSMUSG00000049439 | 1.028573902 | 0.942593054 |
| ENSMUSG00000049456 | 1 | |
| ENSMUSG00000049460 | 1.014545335 | 0.979002724 |
| ENSMUSG00000049470 | -1.047604458 | 0.922165724 |
| ENSMUSG00000049476 | 1 | |
| ENSMUSG00000049477 | 1 | |
| ENSMUSG00000049482 | 1.023809438 | 0.895591989 |

| | | |
|---|---|---|
| ENSMUSG00000049488 | 1.050072601 | 0.734674233 |
| ENSMUSG00000049489 | 1.046822538 | 0.664013268 |
| ENSMUSG00000049491 | 1 | |
| ENSMUSG00000049493 | -1.290392084 | 0.01745071 |
| ENSMUSG00000049498 | 1 | |
| ENSMUSG00000049502 | -1.490523016 | 0.00863072 |
| ENSMUSG00000049504 | 1.046658457 | 0.766589725 |
| ENSMUSG00000049506 | 1 | |
| ENSMUSG00000049511 | 1.248791877 | 0.016289153 |
| ENSMUSG00000049515 | 1.05038453 | 0.887340389 |
| ENSMUSG00000049516 | 1.006772027 | 1 |
| ENSMUSG00000049517 | -1.012992552 | 1 |
| ENSMUSG00000049521 | 1.07706035 | 0.654170062 |
| ENSMUSG00000049526 | 1.06980766 | 0.841285672 |
| ENSMUSG00000049528 | 1 | |
| ENSMUSG00000049530 | 1 | |
| ENSMUSG00000049532 | -1.040257266 | 0.805033642 |
| ENSMUSG00000049536 | 1.237517853 | 0.077622382 |
| ENSMUSG00000049537 | 1 | |
| ENSMUSG00000049538 | 1.269939456 | 0.220825929 |
| ENSMUSG00000049539 | -1.084814283 | 0.772349906 |
| ENSMUSG00000049548 | 1 | |
| ENSMUSG00000049550 | 1.012455176 | 1 |
| ENSMUSG00000049551 | -1.084699559 | 0.717035356 |
| ENSMUSG00000049553 | 1.055128521 | 0.82013376 |
| ENSMUSG00000049555 | -1.279797871 | 0.353239233 |
| ENSMUSG00000049556 | -1.147981165 | 0.698112512 |
| ENSMUSG00000049560 | 1 | |
| ENSMUSG00000049561 | 1 | |
| ENSMUSG00000049562 | 1.101349146 | 0.674749429 |
| ENSMUSG00000049565 | 1.327599164 | 0.054812809 |
| ENSMUSG00000049571 | -1.001666425 | 1 |
| ENSMUSG00000049573 | 1 | |
| ENSMUSG00000049576 | -1.024250957 | 0.989080729 |
| ENSMUSG00000049577 | 1.066284685 | 0.791778775 |
| ENSMUSG00000049580 | 1.118570495 | 0.135282107 |
| ENSMUSG00000049583 | 1.132639128 | 0.217803313 |
| ENSMUSG00000049588 | -1.311563065 | 0.050934077 |
| ENSMUSG00000049593 | 1 | |
| ENSMUSG00000049598 | -1 | 1 |
| ENSMUSG00000049600 | 1.008448536 | 1 |
| ENSMUSG00000049604 | -1.328774061 | 0.028064867 |
| ENSMUSG00000049605 | 1 | |
| ENSMUSG00000049606 | 1.153218739 | 0.551883896 |
| ENSMUSG00000049608 | 1.122462048 | 0.420161906 |
| ENSMUSG00000049612 | -1.089497311 | 0.532369657 |
| ENSMUSG00000049618 | 1 | |
| ENSMUSG00000049620 | 1.171563499 | 0.233412931 |
| ENSMUSG00000049624 | 1.00756134 | 1 |
| ENSMUSG00000049625 | 1.059463094 | 0.608497229 |
| ENSMUSG00000049630 | 1.613150015 | 0.006298149 |
| ENSMUSG00000049641 | -1.136679298 | 0.614265194 |
| ENSMUSG00000049643 | 1.118264122 | 0.251396564 |
| ENSMUSG00000049648 | 1 | |
| ENSMUSG00000049649 | -1.024718486 | 0.951399668 |
| ENSMUSG00000049653 | -1.037896228 | 0.895018604 |
| ENSMUSG00000049657 | 1.012053975 | 0.965637841 |
| ENSMUSG00000049658 | -1.081534115 | 0.741188109 |
| ENSMUSG00000049659 | -1.066186728 | 0.763429741 |
| ENSMUSG00000049670 | -1.048699997 | 0.766916917 |
| ENSMUSG00000049672 | 1.077667996 | 0.660013979 |
| ENSMUSG00000049674 | 1 | |
| ENSMUSG00000049676 | 1.141380229 | 0.347650061 |

| | | |
|---|---|---|
| ENSMUSG00000049680 | 1.063281309 | 0.50114325 |
| ENSMUSG00000049685 | 1 | |
| ENSMUSG00000049686 | 1.069593936 | 0.608497229 |
| ENSMUSG00000049687 | -1.119766761 | 0.698112512 |
| ENSMUSG00000049690 | 1.223679249 | 0.005805035 |
| ENSMUSG00000049691 | -1.03962336 | 0.838359329 |
| ENSMUSG00000049692 | 1 | |
| ENSMUSG00000049694 | 1 | |
| ENSMUSG00000049699 | -1.068537426 | 0.761801824 |
| ENSMUSG00000049708 | 1 | |
| ENSMUSG00000049709 | -1.000603363 | 1 |
| ENSMUSG00000049717 | 1.190756374 | 0.243378179 |
| ENSMUSG00000049719 | 1.182620148 | 0.320976708 |
| ENSMUSG00000049721 | -1 | 1 |
| ENSMUSG00000049723 | 1.079304712 | 0.549287883 |
| ENSMUSG00000049728 | 1.065659417 | 0.528693754 |
| ENSMUSG00000049734 | 1 | |
| ENSMUSG00000049737 | 1 | |
| ENSMUSG00000049739 | 1.05278043 | 0.698112512 |
| ENSMUSG00000049740 | 1 | |
| ENSMUSG00000049741 | -1.024718486 | 0.931480186 |
| ENSMUSG00000049744 | 1.059463094 | 0.608497229 |
| ENSMUSG00000049751 | 1.06655049 | 0.677534336 |
| ENSMUSG00000049755 | -1.149993507 | 0.251049598 |
| ENSMUSG00000049758 | 1 | |
| ENSMUSG00000049760 | 1.061621829 | 0.698112512 |
| ENSMUSG00000049761 | 1 | |
| ENSMUSG00000049764 | 1.051134274 | 0.775794967 |
| ENSMUSG00000049775 | 1.546118456 | 0.054140927 |
| ENSMUSG00000049791 | 1.181395488 | 0.08378739 |
| ENSMUSG00000049792 | 1.091033294 | 0.5103795 |
| ENSMUSG00000049796 | 1 | |
| ENSMUSG00000049797 | 1 | |
| ENSMUSG00000049799 | 1.109824797 | 0.748682651 |
| ENSMUSG00000049800 | 1.020914567 | 0.968937442 |
| ENSMUSG00000049804 | 1.020682779 | 0.975499304 |
| ENSMUSG00000049806 | 1 | |
| ENSMUSG00000049807 | 1.075626613 | 0.530270487 |
| ENSMUSG00000049809 | 1 | |
| ENSMUSG00000049811 | -1.030849836 | 0.769870659 |
| ENSMUSG00000049815 | 1 | |
| ENSMUSG00000049823 | 1.145499439 | 0.125324229 |
| ENSMUSG00000049832 | -1.113379095 | 0.532369657 |
| ENSMUSG00000049843 | 1 | |
| ENSMUSG00000049848 | 1 | |
| ENSMUSG00000049858 | -1.141927549 | 0.437902146 |
| ENSMUSG00000049864 | 1 | |
| ENSMUSG00000049866 | 1.058154392 | 0.821506378 |
| ENSMUSG00000049871 | 1.256047945 | 0.012417057 |
| ENSMUSG00000049872 | 1.11975298 | 0.154254071 |
| ENSMUSG00000049878 | 1.086296554 | 0.702193433 |
| ENSMUSG00000049881 | -1.039771477 | 0.87131415 |
| ENSMUSG00000049882 | 1.157447056 | 0.201542243 |
| ENSMUSG00000049891 | 1.133363755 | 0.608497229 |
| ENSMUSG00000049892 | -1.168120539 | 0.701146944 |
| ENSMUSG00000049894 | 1 | |
| ENSMUSG00000049897 | 1.031104702 | 0.807743274 |
| ENSMUSG00000049902 | 1.095872691 | 0.671280603 |
| ENSMUSG00000049904 | -1.0170893 | 0.94533054 |
| ENSMUSG00000049907 | 1.090850923 | 0.306528292 |
| ENSMUSG00000049908 | -1.084645496 | 0.628300894 |
| ENSMUSG00000049916 | -2.427695426 | 6.41603E-05 |
| ENSMUSG00000049922 | -1.096227773 | 0.518298719 |

| | | |
|---|---|---|
| ENSMUSG00000049926 | -1 | 1 |
| ENSMUSG00000049928 | 1 | |
| ENSMUSG00000049929 | 1.118467336 | 0.510936302 |
| ENSMUSG00000049932 | -1.197977758 | 0.083036058 |
| ENSMUSG00000049939 | -1.109582661 | 0.586366024 |
| ENSMUSG00000049940 | 1.064573362 | 0.814089245 |
| ENSMUSG00000049946 | 1 | |
| ENSMUSG00000049950 | 1.040265223 | 0.747599307 |
| ENSMUSG00000049957 | -1.037258522 | 0.834464052 |
| ENSMUSG00000049960 | -1.141320448 | 0.292376367 |
| ENSMUSG00000049969 | -1.008640872 | 1 |
| ENSMUSG00000049971 | 1.225157278 | 0.081543102 |
| ENSMUSG00000049972 | 1 | |
| ENSMUSG00000049985 | -1.158450149 | 0.698112512 |
| ENSMUSG00000049988 | 1 | |
| ENSMUSG00000049999 | 1.13641723 | 0.587742693 |
| ENSMUSG00000050002 | -1.250497922 | 0.021984475 |
| ENSMUSG00000050010 | -1.028348525 | 0.933934712 |
| ENSMUSG00000050014 | 1.770167662 | 0.027081381 |
| ENSMUSG00000050015 | 1 | |
| ENSMUSG00000050017 | 1.063585145 | 0.812147631 |
| ENSMUSG00000050022 | 1.34533367 | 0.279716146 |
| ENSMUSG00000050023 | 1 | |
| ENSMUSG00000050028 | 1 | |
| ENSMUSG00000050029 | 1.044197074 | 0.901823684 |
| ENSMUSG00000050030 | 1 | |
| ENSMUSG00000050035 | -1.045553977 | 0.615157698 |
| ENSMUSG00000050043 | -1.165289096 | 0.513703955 |
| ENSMUSG00000050050 | -1.38777778 | 0.149101712 |
| ENSMUSG00000050052 | -1.127958865 | 0.113590601 |
| ENSMUSG00000050058 | 1.069068979 | 0.608497229 |
| ENSMUSG00000050063 | -1.084645496 | 0.532369657 |
| ENSMUSG00000050064 | -1.019585404 | 0.823091034 |
| ENSMUSG00000050066 | 1.059463094 | 0.698112512 |
| ENSMUSG00000050069 | -1.216057254 | 0.608497229 |
| ENSMUSG00000050071 | -1.502231958 | 0.012190451 |
| ENSMUSG00000050074 | -1 | 1 |
| ENSMUSG00000050075 | 1 | |
| ENSMUSG00000050079 | -1.001258202 | 1 |
| ENSMUSG00000050085 | 1 | |
| ENSMUSG00000050087 | 1 | |
| ENSMUSG00000050088 | 1.081098491 | 0.629824986 |
| ENSMUSG00000050089 | 1 | |
| ENSMUSG00000050092 | 1.059463094 | 0.776953697 |
| ENSMUSG00000050097 | -1.018861695 | 0.970734719 |
| ENSMUSG00000050100 | -1.044273782 | 0.698112512 |
| ENSMUSG00000050102 | 1 | |
| ENSMUSG00000050103 | -1.085651418 | 0.689423135 |
| ENSMUSG00000050105 | 1 | |
| ENSMUSG00000050106 | 1.245842184 | 0.215845639 |
| ENSMUSG00000050107 | -1.073699871 | 0.698112512 |
| ENSMUSG00000050108 | -1.048277865 | 0.848292858 |
| ENSMUSG00000050114 | 1 | |
| ENSMUSG00000050121 | 1 | |
| ENSMUSG00000050122 | -1 | 1 |
| ENSMUSG00000050128 | 1 | |
| ENSMUSG00000050132 | -1.572283528 | 0.000916193 |
| ENSMUSG00000050134 | 1 | |
| ENSMUSG00000050138 | 1.122462048 | 0.215991492 |
| ENSMUSG00000050141 | 1 | |
| ENSMUSG00000050144 | 1.029825804 | 0.887981329 |
| ENSMUSG00000050147 | 1.030256832 | 0.90028574 |
| ENSMUSG00000050148 | -1.084549533 | 0.615243243 |

| | | |
|---|---|---|
| ENSMUSG00000050150 | -1.256024404 | 0.564226949 |
| ENSMUSG00000050157 | 1.014731082 | 0.988424085 |
| ENSMUSG00000050158 | 1 | |
| ENSMUSG00000050164 | -1.110196018 | 0.547016491 |
| ENSMUSG00000050174 | 1.086997618 | 0.626413412 |
| ENSMUSG00000050179 | 1 | |
| ENSMUSG00000050188 | 1.098148935 | 0.024467649 |
| ENSMUSG00000050189 | 1 | |
| ENSMUSG00000050190 | 1 | |
| ENSMUSG00000050192 | -1.018710673 | 0.979002724 |
| ENSMUSG00000050195 | 1.009658482 | 1 |
| ENSMUSG00000050197 | 1 | |
| ENSMUSG00000050198 | 1 | |
| ENSMUSG00000050199 | 1.341856988 | 0.041176293 |
| ENSMUSG00000050201 | -1.053549062 | 0.698112512 |
| ENSMUSG00000050211 | -1.189590435 | 0.293957289 |
| ENSMUSG00000050212 | -1.048408227 | 0.93868145 |
| ENSMUSG00000050213 | 1.030569708 | 0.893540818 |
| ENSMUSG00000050215 | 1 | |
| ENSMUSG00000050217 | 1 | |
| ENSMUSG00000050222 | -1.082477152 | 0.891020425 |
| ENSMUSG00000050224 | 1 | |
| ENSMUSG00000050229 | -1.030193747 | 0.921369049 |
| ENSMUSG00000050232 | 1 | |
| ENSMUSG00000050234 | -1.151199178 | 0.569188898 |
| ENSMUSG00000050239 | 1 | |
| ENSMUSG00000050240 | 1.036430161 | 0.774846076 |
| ENSMUSG00000050241 | 1.039560021 | 0.925209973 |
| ENSMUSG00000050243 | 1 | |
| ENSMUSG00000050244 | -1.016559372 | 0.991683288 |
| ENSMUSG00000050248 | 1.111962174 | 0.43769614 |
| ENSMUSG00000050251 | 1 | |
| ENSMUSG00000050266 | 1 | |
| ENSMUSG00000050270 | -1.057284864 | 0.834977335 |
| ENSMUSG00000050271 | 1.132322538 | 0.05764497 |
| ENSMUSG00000050272 | 1.373170703 | 0.003413712 |
| ENSMUSG00000050276 | 1 | |
| ENSMUSG00000050281 | 1 | |
| ENSMUSG00000050288 | 1.140236271 | 0.246824992 |
| ENSMUSG00000050295 | -1.027136772 | 0.908878383 |
| ENSMUSG00000050296 | -1.041132001 | 0.938484564 |
| ENSMUSG00000050299 | 1.036022986 | 0.80273087 |
| ENSMUSG00000050304 | -1.145536749 | 0.504310323 |
| ENSMUSG00000050310 | 1.03876836 | 0.911516344 |
| ENSMUSG00000050312 | -1.003390881 | 1 |
| ENSMUSG00000050315 | 1.978382028 | 0.000573666 |
| ENSMUSG00000050321 | -1 | 1 |
| ENSMUSG00000050323 | -1.093047414 | 0.532369657 |
| ENSMUSG00000050328 | -1.037678217 | 0.968575228 |
| ENSMUSG00000050332 | 1.065200001 | 0.68580047 |
| ENSMUSG00000050334 | -1.001781671 | 1 |
| ENSMUSG00000050335 | -1.237140035 | 0.177603723 |
| ENSMUSG00000050343 | 1 | |
| ENSMUSG00000050345 | 1.379979379 | 0.071944404 |
| ENSMUSG00000050347 | 1 | |
| ENSMUSG00000050350 | 1.009414875 | 1 |
| ENSMUSG00000050357 | -1.03663692 | 0.937314884 |
| ENSMUSG00000050359 | -1.230568952 | 0.024046175 |
| ENSMUSG00000050366 | 1 | |
| ENSMUSG00000050368 | 1 | |
| ENSMUSG00000050370 | 1.394618346 | 0.035571553 |
| ENSMUSG00000050373 | 1.004644064 | 1 |
| ENSMUSG00000050377 | 1.122462048 | 0.24351089 |

| | | |
|---|---|---|
| ENSMUSG00000050379 | -1.032815803 | 0.907626515 |
| ENSMUSG00000050382 | 1.020375393 | 0.929536187 |
| ENSMUSG00000050383 | 1 | |
| ENSMUSG00000050390 | 1.081862333 | 0.486774162 |
| ENSMUSG00000050394 | 1.202999411 | 0.014420891 |
| ENSMUSG00000050395 | -1.144392947 | 0.202301764 |
| ENSMUSG00000050397 | -1.003834322 | 1 |
| ENSMUSG00000050410 | 1.038999894 | 0.835624642 |
| ENSMUSG00000050423 | -1 | 1 |
| ENSMUSG00000050424 | -1.063127349 | 0.698112512 |
| ENSMUSG00000050425 | 1 | |
| ENSMUSG00000050428 | 1.096746286 | 0.532369657 |
| ENSMUSG00000050431 | 1 | |
| ENSMUSG00000050435 | 1 | |
| ENSMUSG00000050439 | -1.253827851 | 0.216046507 |
| ENSMUSG00000050440 | 1 | |
| ENSMUSG00000050445 | -1.044273782 | 0.698112512 |
| ENSMUSG00000050447 | 1.417406869 | 0.007109502 |
| ENSMUSG00000050463 | -1.12447299 | 0.608497229 |
| ENSMUSG00000050468 | 1 | |
| ENSMUSG00000050471 | -1.021707046 | 0.962886319 |
| ENSMUSG00000050473 | -1.109582661 | 0.549347972 |
| ENSMUSG00000050478 | 1 | |
| ENSMUSG00000050490 | -1.318098695 | 0.22986779 |
| ENSMUSG00000050493 | 1 | |
| ENSMUSG00000050503 | -1.069255364 | 0.608497229 |
| ENSMUSG00000050504 | 1 | |
| ENSMUSG00000050505 | 1.063384842 | 0.868430458 |
| ENSMUSG00000050511 | -1.200114593 | 0.1991922 |
| ENSMUSG00000050520 | 1 | |
| ENSMUSG00000050526 | 1.024693402 | 0.933031586 |
| ENSMUSG00000050530 | 1.053455663 | 0.70299173 |
| ENSMUSG00000050533 | 1 | |
| ENSMUSG00000050534 | -1.094422282 | 0.760180115 |
| ENSMUSG00000050538 | 1.155238177 | 0.054439097 |
| ENSMUSG00000050541 | 1.48801467 | 0.001516441 |
| ENSMUSG00000050545 | -1.073719964 | 0.777145624 |
| ENSMUSG00000050549 | -1.257415564 | 0.03586978 |
| ENSMUSG00000050550 | -1.095124786 | 0.425531155 |
| ENSMUSG00000050552 | 1.102323638 | 0.276229467 |
| ENSMUSG00000050553 | 1 | |
| ENSMUSG00000050555 | 1.002683655 | 1 |
| ENSMUSG00000050556 | -1.370660194 | 0.026016175 |
| ENSMUSG00000050558 | -1.062505631 | 0.881302915 |
| ENSMUSG00000050565 | -1.002692301 | 1 |
| ENSMUSG00000050567 | 1.038209651 | 0.838445044 |
| ENSMUSG00000050577 | 1 | |
| ENSMUSG00000050578 | 1.278980421 | 0.210271831 |
| ENSMUSG00000050587 | 1.605992882 | 0.021113563 |
| ENSMUSG00000050592 | -1.179174716 | 0.462086028 |
| ENSMUSG00000050600 | 1 | |
| ENSMUSG00000050603 | 1 | |
| ENSMUSG00000050605 | -1.090363055 | 0.091234806 |
| ENSMUSG00000050608 | 1.011220763 | 1 |
| ENSMUSG00000050612 | 1.016770329 | 0.970861412 |
| ENSMUSG00000050613 | 1 | |
| ENSMUSG00000050619 | -1.044393981 | 0.797105186 |
| ENSMUSG00000050621 | -1.090915939 | 0.648098775 |
| ENSMUSG00000050623 | 1.033922105 | 0.945694084 |
| ENSMUSG00000050625 | 1 | |
| ENSMUSG00000050627 | -1.082642354 | 0.648410415 |
| ENSMUSG00000050628 | 1.058194608 | 0.673154048 |
| ENSMUSG00000050635 | 1 | |

| | | |
|---|---|---|
| ENSMUSG00000050640 | -1.135720617 | 0.678783707 |
| ENSMUSG00000050641 | 1 | |
| ENSMUSG00000050645 | 1 | |
| ENSMUSG00000050650 | 1.059463094 | 0.608497229 |
| ENSMUSG00000050654 | 1 | |
| ENSMUSG00000050663 | -1 | 1 |
| ENSMUSG00000050666 | -1.053549062 | 0.698112512 |
| ENSMUSG00000050668 | -1.103120486 | 0.535052277 |
| ENSMUSG00000050671 | -1.323885833 | 0.35680391 |
| ENSMUSG00000050675 | 1.052300903 | 0.805009415 |
| ENSMUSG00000050677 | -1.039650847 | 0.906405443 |
| ENSMUSG00000050685 | -1 | 1 |
| ENSMUSG00000050697 | -1.026141536 | 0.973061065 |
| ENSMUSG00000050700 | 1 | |
| ENSMUSG00000050702 | 1.024097153 | 0.986001377 |
| ENSMUSG00000050704 | 1 | |
| ENSMUSG00000050705 | 1.055932261 | 0.646932348 |
| ENSMUSG00000050708 | 1.320772488 | 0.101059444 |
| ENSMUSG00000050711 | -1 | 1 |
| ENSMUSG00000050714 | 1.075365631 | 0.661371259 |
| ENSMUSG00000050721 | 1.257001354 | 0.046773596 |
| ENSMUSG00000050730 | 1.060447519 | 0.794274354 |
| ENSMUSG00000050732 | -1.060578546 | 0.754841426 |
| ENSMUSG00000050737 | -1.814279277 | 0.008875255 |
| ENSMUSG00000050742 | 1 | |
| ENSMUSG00000050747 | -1.012751181 | 0.967031732 |
| ENSMUSG00000050751 | -1.207114389 | 0.410391773 |
| ENSMUSG00000050756 | 1 | |
| ENSMUSG00000050761 | 1.055938913 | 0.911347495 |
| ENSMUSG00000050762 | 1.500496292 | 0.205169472 |
| ENSMUSG00000050763 | 1 | |
| ENSMUSG00000050765 | 1.059463094 | 0.608497229 |
| ENSMUSG00000050766 | 1 | |
| ENSMUSG00000050772 | 1 | |
| ENSMUSG00000050776 | 1 | |
| ENSMUSG00000050777 | 1.131172138 | 0.179930073 |
| ENSMUSG00000050781 | 1 | |
| ENSMUSG00000050783 | 1 | |
| ENSMUSG00000050786 | 1.006698954 | 1 |
| ENSMUSG00000050788 | 1 | |
| ENSMUSG00000050796 | 1.018317601 | 0.883646854 |
| ENSMUSG00000050799 | -1.084645496 | 0.608497229 |
| ENSMUSG00000050803 | 1 | |
| ENSMUSG00000050808 | 1 | |
| ENSMUSG00000050810 | 1 | |
| ENSMUSG00000050812 | 1.032717825 | 0.952957109 |
| ENSMUSG00000050813 | 1 | |
| ENSMUSG00000050815 | 1 | |
| ENSMUSG00000050818 | 1 | |
| ENSMUSG00000050821 | -1.037531542 | 0.772180723 |
| ENSMUSG00000050822 | -1.412092195 | 0.007023948 |
| ENSMUSG00000050824 | -1.084645496 | 0.532369657 |
| ENSMUSG00000050830 | 1 | |
| ENSMUSG00000050840 | 1 | |
| ENSMUSG00000050844 | 1 | |
| ENSMUSG00000050846 | 1.016998614 | 0.970206992 |
| ENSMUSG00000050853 | 1 | |
| ENSMUSG00000050854 | 1.059463094 | 0.608497229 |
| ENSMUSG00000050855 | -1.06616022 | 0.542903423 |
| ENSMUSG00000050856 | 1.015776082 | 0.970833538 |
| ENSMUSG00000050860 | -1.089497311 | 0.532369657 |
| ENSMUSG00000050865 | 1 | |
| ENSMUSG00000050866 | 1 | |

| | | |
|---|---|---|
| ENSMUSG00000050870 | 1 | |
| ENSMUSG00000050875 | 1 | |
| ENSMUSG00000050876 | 1 | |
| ENSMUSG00000050890 | 1.005090958 | 1 |
| ENSMUSG00000050891 | -1.027999238 | 0.922165724 |
| ENSMUSG00000050896 | -1 | 1 |
| ENSMUSG00000050900 | -1.043306199 | 0.698112512 |
| ENSMUSG00000050901 | 1 | |
| ENSMUSG00000050908 | 1.374134639 | 0.001438351 |
| ENSMUSG00000050910 | 1.008333846 | 1 |
| ENSMUSG00000050912 | -1.012322554 | 1 |
| ENSMUSG00000050914 | 1.110912942 | 0.418130164 |
| ENSMUSG00000050917 | 1 | |
| ENSMUSG00000050919 | 1.059463094 | 0.698112512 |
| ENSMUSG00000050921 | -1 | 1 |
| ENSMUSG00000050926 | 1 | |
| ENSMUSG00000050930 | -1.011371034 | 1 |
| ENSMUSG00000050931 | -1.156812025 | 0.394938139 |
| ENSMUSG00000050933 | 1 | |
| ENSMUSG00000050936 | 1.055026487 | 0.887044308 |
| ENSMUSG00000050944 | 1.166387872 | 0.051202187 |
| ENSMUSG00000050945 | 1.016668078 | 0.939076675 |
| ENSMUSG00000050947 | -1.328395178 | 0.02253597 |
| ENSMUSG00000050953 | 1.097894817 | 0.698112512 |
| ENSMUSG00000050954 | -1.003770769 | 1 |
| ENSMUSG00000050957 | 1.038334199 | 0.754181286 |
| ENSMUSG00000050961 | 1 | |
| ENSMUSG00000050963 | 1 | |
| ENSMUSG00000050965 | 1.152048322 | 0.555593235 |
| ENSMUSG00000050966 | 1.019083579 | 0.949812939 |
| ENSMUSG00000050967 | -1.116853908 | 0.608497229 |
| ENSMUSG00000050973 | 1.009508211 | 0.968421065 |
| ENSMUSG00000050974 | 1 | |
| ENSMUSG00000050978 | 1 | |
| ENSMUSG00000050982 | 1 | |
| ENSMUSG00000050989 | 1.068489444 | 0.709654578 |
| ENSMUSG00000050994 | -1.160925139 | 0.698112512 |
| ENSMUSG00000050996 | 1 | |
| ENSMUSG00000051000 | -1.074160095 | 0.584738699 |
| ENSMUSG00000051002 | 1 | |
| ENSMUSG00000051003 | 1 | |
| ENSMUSG00000051007 | 1.013651793 | 0.737022187 |
| ENSMUSG00000051008 | -1.094118927 | 0.724981165 |
| ENSMUSG00000051022 | -1.277443017 | 0.017886833 |
| ENSMUSG00000051029 | 1.21333449 | 0.026678184 |
| ENSMUSG00000051034 | 1.018220817 | 0.946099281 |
| ENSMUSG00000051036 | -1.03962336 | 0.698112512 |
| ENSMUSG00000051037 | 1.011333191 | 0.997440823 |
| ENSMUSG00000051038 | 1 | |
| ENSMUSG00000051041 | 1 | |
| ENSMUSG00000051043 | 1.623526926 | 0.001702184 |
| ENSMUSG00000051046 | 1 | |
| ENSMUSG00000051048 | 1.520746619 | 0.000878494 |
| ENSMUSG00000051050 | 1 | |
| ENSMUSG00000051051 | 1 | |
| ENSMUSG00000051052 | 1 | |
| ENSMUSG00000051054 | -1.190747999 | 0.608497229 |
| ENSMUSG00000051056 | -1.023769022 | 0.934760809 |
| ENSMUSG00000051062 | 1 | |
| ENSMUSG00000051065 | 1.146793057 | 0.12481557 |
| ENSMUSG00000051067 | -1.405704373 | 0.126536012 |
| ENSMUSG00000051074 | 1.064215019 | 0.847902259 |
| ENSMUSG00000051076 | -1 | 1 |

| | | |
|---|---|---|
| ENSMUSG00000051079 | 1 | |
| ENSMUSG00000051081 | 1 | |
| ENSMUSG00000051095 | 1 | |
| ENSMUSG00000051098 | -1.311248026 | 0.070855636 |
| ENSMUSG00000051107 | 1.160520798 | 0.591385536 |
| ENSMUSG00000051111 | 1.123040325 | 0.698112512 |
| ENSMUSG00000051113 | 1.020779405 | 0.980511343 |
| ENSMUSG00000051116 | 1.069068979 | 0.698112512 |
| ENSMUSG00000051118 | 1 | |
| ENSMUSG00000051124 | -1.21092864 | 0.545690002 |
| ENSMUSG00000051133 | 1 | |
| ENSMUSG00000051136 | -1 | 1 |
| ENSMUSG00000051146 | 1.146695737 | 0.330039993 |
| ENSMUSG00000051147 | 1.305566614 | 0.019990955 |
| ENSMUSG00000051149 | -1.126826176 | 0.717897145 |
| ENSMUSG00000051153 | 1 | |
| ENSMUSG00000051154 | 1.129598375 | 0.299792412 |
| ENSMUSG00000051156 | 1 | |
| ENSMUSG00000051159 | -1.120992057 | 0.632313789 |
| ENSMUSG00000051160 | 1 | |
| ENSMUSG00000051166 | -1.138708886 | 0.518733045 |
| ENSMUSG00000051169 | -1.012610788 | 0.997738265 |
| ENSMUSG00000051172 | 1 | |
| ENSMUSG00000051176 | 1 | |
| ENSMUSG00000051177 | 1.616203087 | 0.001365428 |
| ENSMUSG00000051180 | 1 | |
| ENSMUSG00000051182 | 1 | |
| ENSMUSG00000051184 | 1.181260633 | 0.437045401 |
| ENSMUSG00000051185 | 1.014899425 | 0.925214847 |
| ENSMUSG00000051190 | 1 | |
| ENSMUSG00000051198 | 1 | |
| ENSMUSG00000051200 | 1 | |
| ENSMUSG00000051207 | -1 | 1 |
| ENSMUSG00000051209 | 1 | |
| ENSMUSG00000051212 | -1.034990181 | 0.914913657 |
| ENSMUSG00000051219 | -1.053549062 | 0.698112512 |
| ENSMUSG00000051220 | 1.017937705 | 0.994917144 |
| ENSMUSG00000051223 | 1.014873765 | 1 |
| ENSMUSG00000051224 | -1.137011804 | 0.150238185 |
| ENSMUSG00000051225 | -1.083415624 | 0.706213769 |
| ENSMUSG00000051228 | 1.016770329 | 0.943025934 |
| ENSMUSG00000051232 | 1.037840819 | 0.869434134 |
| ENSMUSG00000051234 | 1.014101859 | 1 |
| ENSMUSG00000051235 | -1.023599508 | 0.931429981 |
| ENSMUSG00000051236 | 1.241990205 | 0.334449309 |
| ENSMUSG00000051237 | 1 | |
| ENSMUSG00000051238 | -1.019632821 | 0.909780721 |
| ENSMUSG00000051242 | 1.116971117 | 0.700190225 |
| ENSMUSG00000051243 | -1.353479973 | 0.26210641 |
| ENSMUSG00000051246 | -1.033638191 | 0.979160994 |
| ENSMUSG00000051251 | -1.03962336 | 0.851451345 |
| ENSMUSG00000051255 | 1.037487447 | 0.952049609 |
| ENSMUSG00000051256 | -1.003713439 | 1 |
| ENSMUSG00000051257 | 1 | |
| ENSMUSG00000051258 | 1 | |
| ENSMUSG00000051262 | -1.02819776 | 0.967637938 |
| ENSMUSG00000051276 | 1 | |
| ENSMUSG00000051278 | -1.043893208 | 0.847902259 |
| ENSMUSG00000051279 | -1.193320102 | 0.034332081 |
| ENSMUSG00000051285 | 1.046976663 | 0.809672812 |
| ENSMUSG00000051295 | -1.107299336 | 0.698112512 |
| ENSMUSG00000051297 | -1.053549062 | 0.769745556 |
| ENSMUSG00000051306 | 1.099984948 | 0.531123085 |

| | | |
|---|---|---|
| ENSMUSG00000051313 | 1 | |
| ENSMUSG00000051314 | 1 | |
| ENSMUSG00000051316 | -1.087032194 | 0.255315328 |
| ENSMUSG00000051319 | 1.057327754 | 0.644606499 |
| ENSMUSG00000051323 | -1.125622484 | 0.608497229 |
| ENSMUSG00000051329 | -1.093182647 | 0.698112512 |
| ENSMUSG00000051331 | 1.205828062 | 0.020353969 |
| ENSMUSG00000051335 | 1.063955741 | 0.608497229 |
| ENSMUSG00000051339 | 1.134151011 | 0.484245055 |
| ENSMUSG00000051340 | 1 | |
| ENSMUSG00000051341 | 1.040550832 | 0.794274354 |
| ENSMUSG00000051343 | 1.059105913 | 0.698112512 |
| ENSMUSG00000051344 | -1.002803447 | 1 |
| ENSMUSG00000051346 | -1.076856371 | 0.589702874 |
| ENSMUSG00000051351 | 1.067210334 | 0.480100596 |
| ENSMUSG00000051354 | 1 | |
| ENSMUSG00000051355 | -1.065367831 | 0.702662103 |
| ENSMUSG00000051359 | 1.177824539 | 0.38039555 |
| ENSMUSG00000051361 | 1 | |
| ENSMUSG00000051362 | 1 | |
| ENSMUSG00000051367 | 1.25061954 | 0.02674383 |
| ENSMUSG00000051373 | -1.053549062 | 0.778372272 |
| ENSMUSG00000051375 | 1.551446651 | 4.53364E-05 |
| ENSMUSG00000051378 | -1.004688674 | 1 |
| ENSMUSG00000051379 | 1.045406688 | 0.639346469 |
| ENSMUSG00000051390 | 1.050797472 | 0.698112512 |
| ENSMUSG00000051391 | 1.10467084 | 0.651257338 |
| ENSMUSG00000051392 | 1 | |
| ENSMUSG00000051396 | 1.024085206 | 0.966455074 |
| ENSMUSG00000051397 | -1.063127349 | 0.650186154 |
| ENSMUSG00000051401 | 1 | |
| ENSMUSG00000051403 | 1.043511683 | 0.769745556 |
| ENSMUSG00000051412 | 1.013170232 | 1 |
| ENSMUSG00000051413 | -1.069964177 | 0.698112512 |
| ENSMUSG00000051414 | 1 | |
| ENSMUSG00000051424 | 1 | |
| ENSMUSG00000051427 | 1.005303656 | 1 |
| ENSMUSG00000051431 | 1.059463094 | 0.608497229 |
| ENSMUSG00000051435 | -1.099174267 | 0.535052277 |
| ENSMUSG00000051437 | 1 | |
| ENSMUSG00000051439 | -1.01098892 | 0.950875062 |
| ENSMUSG00000051444 | 1.077382102 | 0.608497229 |
| ENSMUSG00000051451 | 1.075680571 | 0.698112512 |
| ENSMUSG00000051452 | -1.043306199 | 0.858911269 |
| ENSMUSG00000051455 | -1.033138005 | 0.976129946 |
| ENSMUSG00000051456 | -1.03962336 | 0.698112512 |
| ENSMUSG00000051457 | 1.010824455 | 1 |
| ENSMUSG00000051469 | -1.030447272 | 0.932605421 |
| ENSMUSG00000051481 | 1 | |
| ENSMUSG00000051483 | -1.005948154 | 1 |
| ENSMUSG00000051486 | -1 | 1 |
| ENSMUSG00000051490 | -1.03962336 | 0.698112512 |
| ENSMUSG00000051493 | 1 | |
| ENSMUSG00000051495 | 1.279933464 | 0.013307237 |
| ENSMUSG00000051497 | -1 | 1 |
| ENSMUSG00000051498 | 1.505967199 | 0.0045402 |
| ENSMUSG00000051499 | -1.015568505 | 0.921168619 |
| ENSMUSG00000051502 | -1.013891506 | 0.863453969 |
| ENSMUSG00000051503 | 1.196004652 | 0.143050247 |
| ENSMUSG00000051504 | 1.005613439 | 1 |
| ENSMUSG00000051506 | -1.085651418 | 0.532369657 |
| ENSMUSG00000051509 | 1 | |
| ENSMUSG00000051510 | 1.159826045 | 0.295833164 |

| | | |
|---|---|---|
| ENSMUSG00000051515 | 1 | |
| ENSMUSG00000051517 | 1.093028017 | 0.31494917 |
| ENSMUSG00000051518 | 1.016788216 | 0.94084956 |
| ENSMUSG00000051527 | 1.093604696 | 0.678330704 |
| ENSMUSG00000051528 | 1 | |
| ENSMUSG00000051537 | -1.215041912 | 0.473106921 |
| ENSMUSG00000051548 | 1 | |
| ENSMUSG00000051550 | -1.009847386 | 0.931480186 |
| ENSMUSG00000051557 | 1.109711588 | 0.408046973 |
| ENSMUSG00000051579 | 1.068709917 | 0.791219515 |
| ENSMUSG00000051582 | 1 | |
| ENSMUSG00000051586 | 1.101539215 | 0.410391773 |
| ENSMUSG00000051590 | 1.011607416 | 1 |
| ENSMUSG00000051591 | 1 | |
| ENSMUSG00000051592 | 1.059463094 | 0.608497229 |
| ENSMUSG00000051593 | 1 | |
| ENSMUSG00000051596 | -1.11061171 | 0.608497229 |
| ENSMUSG00000051599 | -1 | 1 |
| ENSMUSG00000051606 | 1 | |
| ENSMUSG00000051611 | 1 | |
| ENSMUSG00000051615 | 1.005227899 | 1 |
| ENSMUSG00000051616 | 1 | |
| ENSMUSG00000051617 | 1 | |
| ENSMUSG00000051618 | 1 | |
| ENSMUSG00000051627 | -1.142273209 | 0.698112512 |
| ENSMUSG00000051638 | 1 | |
| ENSMUSG00000051639 | 1 | |
| ENSMUSG00000051648 | -1.047885504 | 0.866699472 |
| ENSMUSG00000051650 | 1.117640663 | 0.732612801 |
| ENSMUSG00000051652 | -1.66110733 | 0.000129787 |
| ENSMUSG00000051663 | 1 | |
| ENSMUSG00000051669 | 1.665845404 | 0.000143267 |
| ENSMUSG00000051671 | -1.027529533 | 0.773508225 |
| ENSMUSG00000051674 | 1.074665683 | 0.681952832 |
| ENSMUSG00000051675 | 1.115763523 | 0.391804854 |
| ENSMUSG00000051678 | 1.024693402 | 0.955101438 |
| ENSMUSG00000051680 | 1 | |
| ENSMUSG00000051682 | 1 | |
| ENSMUSG00000051687 | 1 | |
| ENSMUSG00000051695 | 1.069589347 | 0.739206552 |
| ENSMUSG00000051703 | 1.077650749 | 0.85189266 |
| ENSMUSG00000051705 | 1.016022618 | 0.96797322 |
| ENSMUSG00000051706 | 1 | |
| ENSMUSG00000051716 | -1.115057621 | 0.717035356 |
| ENSMUSG00000051721 | 1.065244996 | 0.362935692 |
| ENSMUSG00000051723 | 1.010032815 | 1 |
| ENSMUSG00000051726 | 1.876601071 | 4.68417E-05 |
| ENSMUSG00000051727 | -1.060434227 | 0.844357937 |
| ENSMUSG00000051728 | 1.069437807 | 0.729206866 |
| ENSMUSG00000051729 | -1 | 1 |
| ENSMUSG00000051730 | -1.058436248 | 0.698112512 |
| ENSMUSG00000051732 | 1 | |
| ENSMUSG00000051735 | 1.227620047 | 0.25662825 |
| ENSMUSG00000051736 | -1.072788892 | 0.689423135 |
| ENSMUSG00000051747 | -1.02456515 | 0.914600981 |
| ENSMUSG00000051748 | 1 | |
| ENSMUSG00000051758 | -1.053549062 | 0.698112512 |
| ENSMUSG00000051768 | 1.028046068 | 0.803072249 |
| ENSMUSG00000051769 | 1 | |
| ENSMUSG00000051777 | 1 | |
| ENSMUSG00000051786 | 1.057322405 | 0.777702703 |
| ENSMUSG00000051788 | 1.069068979 | 0.608497229 |
| ENSMUSG00000051790 | -1.029438566 | 0.883646854 |

| | | |
|---|---|---|
| ENSMUSG00000051793 | 1 | |
| ENSMUSG00000051802 | 1 | |
| ENSMUSG00000051804 | 1 | |
| ENSMUSG00000051805 | 1.059463094 | 0.608497229 |
| ENSMUSG00000051811 | -1.237299691 | 0.295098609 |
| ENSMUSG00000051817 | 1.064056855 | 0.336618721 |
| ENSMUSG00000051827 | 1 | |
| ENSMUSG00000051832 | 1.386317608 | 0.017980376 |
| ENSMUSG00000051839 | 1 | |
| ENSMUSG00000051844 | -1.028074752 | 0.971248775 |
| ENSMUSG00000051851 | 1.070905837 | 0.775673129 |
| ENSMUSG00000051853 | 1.026905029 | 0.978147981 |
| ENSMUSG00000051855 | -1.279157405 | 0.0202616 |
| ENSMUSG00000051860 | 1 | |
| ENSMUSG00000051864 | 1.000771865 | 1 |
| ENSMUSG00000051877 | 1 | |
| ENSMUSG00000051879 | -1 | 1 |
| ENSMUSG00000051885 | 1 | |
| ENSMUSG00000051890 | 1.008470533 | 0.932157069 |
| ENSMUSG00000051896 | 1 | |
| ENSMUSG00000051900 | -1.043306199 | 0.698112512 |
| ENSMUSG00000051906 | 1 | |
| ENSMUSG00000051910 | 1.150179455 | 0.145828618 |
| ENSMUSG00000051917 | 1 | |
| ENSMUSG00000051920 | 2.269787781 | 0.000785016 |
| ENSMUSG00000051934 | -1.071694115 | 0.73584221 |
| ENSMUSG00000051936 | 1 | |
| ENSMUSG00000051940 | 1 | |
| ENSMUSG00000051950 | -1.048585894 | 0.813736612 |
| ENSMUSG00000051951 | -1.043306199 | 0.698112512 |
| ENSMUSG00000051952 | 1 | |
| ENSMUSG00000051956 | 1 | |
| ENSMUSG00000051965 | 1 | |
| ENSMUSG00000051969 | 1 | |
| ENSMUSG00000051977 | -1.023488394 | 0.818483291 |
| ENSMUSG00000051978 | 1.063482407 | 0.698112512 |
| ENSMUSG00000051980 | 1 | |
| ENSMUSG00000051984 | 1.32469638 | 0.009348865 |
| ENSMUSG00000051985 | -1 | 1 |
| ENSMUSG00000051986 | 1 | |
| ENSMUSG00000051989 | -1.049115611 | 0.732612801 |
| ENSMUSG00000051998 | 1.200036387 | 0.486616725 |
| ENSMUSG00000052005 | 1.15013545 | 0.35389306 |
| ENSMUSG00000052012 | 1 | |
| ENSMUSG00000052013 | 1 | |
| ENSMUSG00000052014 | 1 | |
| ENSMUSG00000052025 | 1.069068979 | 0.608497229 |
| ENSMUSG00000052026 | 1.68294231 | 0.017600706 |
| ENSMUSG00000052031 | -1.054206707 | 0.619581646 |
| ENSMUSG00000052033 | -1.107822813 | 0.608497229 |
| ENSMUSG00000052040 | 1.276950663 | 0.004416753 |
| ENSMUSG00000052056 | -1.00645504 | 1 |
| ENSMUSG00000052058 | 1 | |
| ENSMUSG00000052062 | -1.036445739 | 0.849539515 |
| ENSMUSG00000052075 | 1 | |
| ENSMUSG00000052085 | 1.113800713 | 0.096239765 |
| ENSMUSG00000052087 | -1.015511103 | 0.982777255 |
| ENSMUSG00000052099 | 1 | |
| ENSMUSG00000052102 | 1.227084402 | 0.079951524 |
| ENSMUSG00000052105 | 1.066458209 | 0.608497229 |
| ENSMUSG00000052117 | 1.05943716 | 0.848456999 |
| ENSMUSG00000052125 | -1.099153057 | 0.479889456 |
| ENSMUSG00000052131 | 1.132639128 | 0.428732467 |

| | | |
|---|---|---|
| ENSMUSG00000052133 | -1.024718486 | 0.952874798 |
| ENSMUSG00000052135 | 1.335328107 | 0.169545598 |
| ENSMUSG00000052137 | 1.050765295 | 0.746921078 |
| ENSMUSG00000052139 | 1.061191929 | 0.632732136 |
| ENSMUSG00000052142 | -1.053549062 | 0.698112512 |
| ENSMUSG00000052143 | -1.287727923 | 0.210651026 |
| ENSMUSG00000052144 | 1.081121492 | 0.770958539 |
| ENSMUSG00000052146 | 1.078080069 | 0.661155252 |
| ENSMUSG00000052151 | -1.010649223 | 1 |
| ENSMUSG00000052154 | 1.117607781 | 0.718295671 |
| ENSMUSG00000052155 | 1.039035877 | 0.844158013 |
| ENSMUSG00000052157 | -1.034239753 | 0.924324458 |
| ENSMUSG00000052160 | 1.065662724 | 0.788000546 |
| ENSMUSG00000052180 | 1.132639128 | 0.217803313 |
| ENSMUSG00000052182 | 1 | |
| ENSMUSG00000052187 | 1 | |
| ENSMUSG00000052188 | -1.03962336 | 0.698112512 |
| ENSMUSG00000052192 | 1 | |
| ENSMUSG00000052212 | 1 | |
| ENSMUSG00000052214 | -1.041132984 | 0.846072213 |
| ENSMUSG00000052217 | 1 | |
| ENSMUSG00000052221 | -1.190907694 | 0.350710755 |
| ENSMUSG00000052229 | 1.230075506 | 0.292973656 |
| ENSMUSG00000052234 | 1 | |
| ENSMUSG00000052236 | 1 | |
| ENSMUSG00000052241 | 1.079681442 | 0.608497229 |
| ENSMUSG00000052248 | -1.058335806 | 0.869434134 |
| ENSMUSG00000052253 | -1.018095903 | 0.964346207 |
| ENSMUSG00000052270 | -1.053549062 | 0.778372272 |
| ENSMUSG00000052271 | 1.061449073 | 0.893608405 |
| ENSMUSG00000052273 | -1 | 1 |
| ENSMUSG00000052276 | 1 | |
| ENSMUSG00000052281 | 1.059463094 | 0.698112512 |
| ENSMUSG00000052291 | -1.084884315 | 0.588281525 |
| ENSMUSG00000052293 | -1.132896509 | 0.608497229 |
| ENSMUSG00000052295 | 1.110292537 | 0.698112512 |
| ENSMUSG00000052296 | 1.041470083 | 0.790216277 |
| ENSMUSG00000052298 | -1.06091544 | 0.79291651 |
| ENSMUSG00000052299 | 1.001907589 | 1 |
| ENSMUSG00000052301 | 1 | |
| ENSMUSG00000052302 | -1.9981153 | 0.002796445 |
| ENSMUSG00000052303 | 1 | |
| ENSMUSG00000052305 | 1 | |
| ENSMUSG00000052310 | -1.032264637 | 0.916783974 |
| ENSMUSG00000052316 | 1.879355015 | 0.000266305 |
| ENSMUSG00000052331 | 1.074085385 | 0.698112512 |
| ENSMUSG00000052334 | -1.043306199 | 0.803260194 |
| ENSMUSG00000052336 | -1 | 1 |
| ENSMUSG00000052337 | 1.043244234 | 0.925136366 |
| ENSMUSG00000052353 | 1.104792003 | 0.291044503 |
| ENSMUSG00000052363 | 1 | |
| ENSMUSG00000052364 | -1.061026699 | 0.477644658 |
| ENSMUSG00000052368 | 1 | |
| ENSMUSG00000052369 | 1.132384914 | 0.419887592 |
| ENSMUSG00000052371 | 1.81595453 | 0.002109953 |
| ENSMUSG00000052372 | -1.044273782 | 0.795446726 |
| ENSMUSG00000052373 | -1.052098694 | 0.698112512 |
| ENSMUSG00000052374 | -1.06005009 | 0.889846433 |
| ENSMUSG00000052376 | 1 | |
| ENSMUSG00000052382 | 1 | |
| ENSMUSG00000052384 | -1.338827689 | 0.019017074 |
| ENSMUSG00000052387 | 1.020535114 | 0.969007252 |
| ENSMUSG00000052392 | -1.043306199 | 0.79291651 |

| | | |
|---|---|---|
| ENSMUSG00000052395 | -1.209296427 | 0.159880253 |
| ENSMUSG00000052396 | 1.638203262 | 5.29269E-05 |
| ENSMUSG00000052397 | 1.18211352 | 0.250477583 |
| ENSMUSG00000052403 | -1.044273782 | 0.790851299 |
| ENSMUSG00000052406 | 1.014561308 | 0.982838256 |
| ENSMUSG00000052407 | 1.048796868 | 0.818394445 |
| ENSMUSG00000052414 | 1 | |
| ENSMUSG00000052415 | -2.21729749 | 0.004430069 |
| ENSMUSG00000052417 | 1 | |
| ENSMUSG00000052419 | 1.198214711 | 0.339164066 |
| ENSMUSG00000052423 | 1.016489237 | 0.883605713 |
| ENSMUSG00000052426 | 1.024693402 | 0.952057016 |
| ENSMUSG00000052428 | 1.046930494 | 0.822562428 |
| ENSMUSG00000052429 | 1.022415035 | 0.979212363 |
| ENSMUSG00000052430 | -1.02093734 | 0.968937442 |
| ENSMUSG00000052435 | 1 | |
| ENSMUSG00000052446 | -1.017779007 | 0.967899076 |
| ENSMUSG00000052456 | 1.001665641 | 1 |
| ENSMUSG00000052459 | 1.073148598 | 0.805587306 |
| ENSMUSG00000052468 | 1 | |
| ENSMUSG00000052469 | 1 | |
| ENSMUSG00000052471 | -1 | 1 |
| ENSMUSG00000052477 | 1.023743961 | 0.938939969 |
| ENSMUSG00000052479 | -1.024718486 | 0.931480186 |
| ENSMUSG00000052482 | -1.003841375 | 1 |
| ENSMUSG00000052485 | 1.249832536 | 0.00435101 |
| ENSMUSG00000052488 | 1.056161223 | 0.698112512 |
| ENSMUSG00000052496 | -1.547010771 | 0.002191087 |
| ENSMUSG00000052504 | 1.209144687 | 0.547016491 |
| ENSMUSG00000052508 | -1.084645496 | 0.532369657 |
| ENSMUSG00000052512 | 1.283323502 | 0.00777869 |
| ENSMUSG00000052516 | -1.104862985 | 0.450311873 |
| ENSMUSG00000052520 | 1 | |
| ENSMUSG00000052525 | -1.106248237 | 0.11838992 |
| ENSMUSG00000052533 | 1.096033534 | 0.663952712 |
| ENSMUSG00000052534 | 1.277831692 | 0.12640454 |
| ENSMUSG00000052537 | 1 | |
| ENSMUSG00000052539 | -1.001887707 | 1 |
| ENSMUSG00000052544 | -1.389492869 | 0.007996319 |
| ENSMUSG00000052549 | -1.239699792 | 0.262929409 |
| ENSMUSG00000052551 | 1.080134941 | 0.726866981 |
| ENSMUSG00000052554 | 1 | |
| ENSMUSG00000052557 | 1.043181208 | 0.776021306 |
| ENSMUSG00000052560 | 1.202642837 | 0.434482395 |
| ENSMUSG00000052562 | 1 | |
| ENSMUSG00000052563 | -1.210588793 | 0.013431057 |
| ENSMUSG00000052565 | -1.068899451 | 0.823058019 |
| ENSMUSG00000052566 | 1.084378189 | 0.196704465 |
| ENSMUSG00000052572 | -1.244253939 | 0.350417454 |
| ENSMUSG00000052581 | 1.122462048 | 0.215991492 |
| ENSMUSG00000052584 | -1.080998457 | 0.528189006 |
| ENSMUSG00000052593 | -1.006770692 | 1 |
| ENSMUSG00000052595 | 1.059463094 | 0.608497229 |
| ENSMUSG00000052605 | -1.108816572 | 0.63175637 |
| ENSMUSG00000052609 | 1.182157372 | 0.116329897 |
| ENSMUSG00000052613 | 1.023743961 | 0.95702926 |
| ENSMUSG00000052616 | 1.055919176 | 0.869727357 |
| ENSMUSG00000052625 | 1 | |
| ENSMUSG00000052629 | 1.24122829 | 0.048604265 |
| ENSMUSG00000052631 | 1.527018206 | 0.000468075 |
| ENSMUSG00000052632 | 1.140014472 | 0.051664013 |
| ENSMUSG00000052642 | 1 | |
| ENSMUSG00000052656 | 1.020639614 | 0.961068159 |

| | | |
|---|---|---|
| ENSMUSG00000052658 | -1.032882776 | 0.897684198 |
| ENSMUSG00000052673 | 1.059463094 | 0.608497229 |
| ENSMUSG00000052675 | 1.075610413 | 0.229735675 |
| ENSMUSG00000052676 | 1.006656065 | 1 |
| ENSMUSG00000052681 | 1.017939571 | 1 |
| ENSMUSG00000052684 | 1.176218491 | 0.040242459 |
| ENSMUSG00000052688 | 1.112579614 | 0.210056079 |
| ENSMUSG00000052698 | 1.169127138 | 0.182650295 |
| ENSMUSG00000052707 | 1.065438832 | 0.793735215 |
| ENSMUSG00000052712 | 1.022185677 | 0.953807127 |
| ENSMUSG00000052713 | 1.002647739 | 1 |
| ENSMUSG00000052724 | 1.01393017 | 1 |
| ENSMUSG00000052726 | 1.241379171 | 0.119947827 |
| ENSMUSG00000052727 | 1.029061183 | 0.970833538 |
| ENSMUSG00000052730 | -1 | 1 |
| ENSMUSG00000052736 | 1 | |
| ENSMUSG00000052738 | -1.009881353 | 1 |
| ENSMUSG00000052748 | -1.008356658 | 1 |
| ENSMUSG00000052749 | 1 | |
| ENSMUSG00000052751 | 1.117602362 | 0.128881295 |
| ENSMUSG00000052752 | 1.014753122 | 0.995703142 |
| ENSMUSG00000052759 | 1 | |
| ENSMUSG00000052760 | -1.010531047 | 1 |
| ENSMUSG00000052763 | 1.038939282 | 0.698112512 |
| ENSMUSG00000052767 | 1 | |
| ENSMUSG00000052769 | 1.113371637 | 0.581851313 |
| ENSMUSG00000052776 | -1.450747519 | 0.033093318 |
| ENSMUSG00000052779 | 1 | |
| ENSMUSG00000052783 | -1.042971403 | 0.698112512 |
| ENSMUSG00000052785 | 1 | |
| ENSMUSG00000052790 | 1 | |
| ENSMUSG00000052794 | 1.035162701 | 0.698112512 |
| ENSMUSG00000052798 | -1.009986332 | 1 |
| ENSMUSG00000052812 | -1.044440015 | 0.770888795 |
| ENSMUSG00000052818 | 1 | |
| ENSMUSG00000052819 | 1.047947442 | 0.908354926 |
| ENSMUSG00000052821 | 1.028323353 | 0.903868918 |
| ENSMUSG00000052825 | -1.142095915 | 0.619107467 |
| ENSMUSG00000052833 | 1.004311807 | 1 |
| ENSMUSG00000052837 | 1.144330669 | 0.698112512 |
| ENSMUSG00000052848 | 1 | |
| ENSMUSG00000052850 | -1.044273782 | 0.790851299 |
| ENSMUSG00000052852 | -1.343552309 | 0.05702344 |
| ENSMUSG00000052854 | -1.209710897 | 0.03311356 |
| ENSMUSG00000052861 | 1.701244166 | 0.014289981 |
| ENSMUSG00000052865 | 1 | |
| ENSMUSG00000052889 | 1 | |
| ENSMUSG00000052906 | -1.128155193 | 0.344280463 |
| ENSMUSG00000052909 | 1 | |
| ENSMUSG00000052911 | 1.097650272 | 0.604452802 |
| ENSMUSG00000052912 | -1.044273782 | 0.83619593 |
| ENSMUSG00000052914 | 1.340904606 | 0.063236534 |
| ENSMUSG00000052915 | 1.040076696 | 0.897469779 |
| ENSMUSG00000052917 | 1.060304774 | 0.745979608 |
| ENSMUSG00000052920 | 1.13872573 | 0.378751351 |
| ENSMUSG00000052921 | 1.147985629 | 0.005401675 |
| ENSMUSG00000052922 | 1 | |
| ENSMUSG00000052926 | -1.095994816 | 0.608497229 |
| ENSMUSG00000052928 | 1.080188913 | 0.649922347 |
| ENSMUSG00000052934 | -1.091682695 | 0.281868105 |
| ENSMUSG00000052942 | -1.159951013 | 0.382663264 |
| ENSMUSG00000052949 | 1.240600519 | 0.025703509 |
| ENSMUSG00000052951 | -1.679751179 | 0.006770973 |

| | | |
|---|---|---|
| ENSMUSG00000052955 | 1 | |
| ENSMUSG00000052957 | 1.235267166 | 0.069369763 |
| ENSMUSG00000052962 | -1.079841752 | 0.698112512 |
| ENSMUSG00000052974 | 1 | |
| ENSMUSG00000052981 | -1.03962336 | 0.698112512 |
| ENSMUSG00000052997 | 1.022195174 | 0.982735249 |
| ENSMUSG00000053004 | -1.127980004 | 0.524392493 |
| ENSMUSG00000053007 | -1.085973934 | 0.486165967 |
| ENSMUSG00000053012 | 1.07510502 | 0.698112512 |
| ENSMUSG00000053024 | -1.065321276 | 0.792444161 |
| ENSMUSG00000053025 | -1.03962336 | 0.88732209 |
| ENSMUSG00000053030 | 1.018670051 | 0.982838256 |
| ENSMUSG00000053038 | 1.075334548 | 0.253249273 |
| ENSMUSG00000053040 | -1.272344521 | 0.041151366 |
| ENSMUSG00000053044 | 1.059463094 | 0.608497229 |
| ENSMUSG00000053046 | 1.024693402 | 0.972980766 |
| ENSMUSG00000053049 | 1.059463094 | 0.776953697 |
| ENSMUSG00000053054 | 1 | |
| ENSMUSG00000053062 | 1.191025605 | 0.195607801 |
| ENSMUSG00000053063 | 1.116785864 | 0.608497229 |
| ENSMUSG00000053070 | 1.05948324 | 0.608497229 |
| ENSMUSG00000053080 | 1.12170177 | 0.300032884 |
| ENSMUSG00000053081 | 1 | |
| ENSMUSG00000053091 | -1.013978683 | 0.952064824 |
| ENSMUSG00000053093 | -1.023769022 | 0.934760809 |
| ENSMUSG00000053094 | 1.098141225 | 0.638218577 |
| ENSMUSG00000053101 | -1.069095149 | 0.686194514 |
| ENSMUSG00000053110 | 1.028557933 | 0.961255441 |
| ENSMUSG00000053111 | 1.024779056 | 0.970006845 |
| ENSMUSG00000053113 | 1.102630845 | 0.209433875 |
| ENSMUSG00000053117 | 1.068648802 | 0.844158013 |
| ENSMUSG00000053119 | 1.050461571 | 0.862443252 |
| ENSMUSG00000053121 | 1 | |
| ENSMUSG00000053128 | -1.015092208 | 0.952049609 |
| ENSMUSG00000053129 | 1 | |
| ENSMUSG00000053134 | 1.013719713 | 0.969337248 |
| ENSMUSG00000053137 | 1.048995912 | 0.716562452 |
| ENSMUSG00000053141 | 1.09538577 | 0.766696422 |
| ENSMUSG00000053146 | 1 | |
| ENSMUSG00000053153 | 1.069068979 | 0.608497229 |
| ENSMUSG00000053158 | 1.256930222 | 0.024097603 |
| ENSMUSG00000053161 | 1.059463094 | 0.750182067 |
| ENSMUSG00000053164 | 1.045763856 | 0.967031732 |
| ENSMUSG00000053166 | -1.020155149 | 0.982838256 |
| ENSMUSG00000053168 | -1 | 1 |
| ENSMUSG00000053173 | -1.053549062 | 0.769745556 |
| ENSMUSG00000053175 | 1.192807183 | 0.180261993 |
| ENSMUSG00000053178 | 1.051792785 | 0.758560659 |
| ENSMUSG00000053181 | -1.029163572 | 0.9780134 |
| ENSMUSG00000053182 | 1.054940803 | 0.883188695 |
| ENSMUSG00000053184 | -1 | 1 |
| ENSMUSG00000053185 | 1 | |
| ENSMUSG00000053192 | 1.01911581 | 0.955712331 |
| ENSMUSG00000053194 | 1 | |
| ENSMUSG00000053198 | 1.087262434 | 0.205994883 |
| ENSMUSG00000053199 | 1.117694388 | 0.470928144 |
| ENSMUSG00000053205 | -1.226943413 | 0.136710153 |
| ENSMUSG00000053208 | -1 | 1 |
| ENSMUSG00000053211 | 1 | |
| ENSMUSG00000053214 | 1.271344436 | 0.053354336 |
| ENSMUSG00000053216 | -1.325651792 | 0.065119334 |
| ENSMUSG00000053217 | 1 | |
| ENSMUSG00000053219 | 1.038963086 | 0.672090419 |

| | | |
|---|---|---|
| ENSMUSG00000053226 | 1.003845124 | 1 |
| ENSMUSG00000053228 | -1 | 1 |
| ENSMUSG00000053251 | -1.01346401 | 1 |
| ENSMUSG00000053253 | -1.142149762 | 0.561261728 |
| ENSMUSG00000053263 | 1.120444696 | 0.499935975 |
| ENSMUSG00000053268 | 1 | |
| ENSMUSG00000053279 | 1.171403402 | 0.221007934 |
| ENSMUSG00000053286 | -1.04551022 | 0.816447433 |
| ENSMUSG00000053287 | 1 | |
| ENSMUSG00000053289 | -1.049513551 | 0.841213045 |
| ENSMUSG00000053291 | 1.014545335 | 0.998924031 |
| ENSMUSG00000053293 | 1.121207551 | 0.452069807 |
| ENSMUSG00000053297 | 1.030256832 | 0.910612556 |
| ENSMUSG00000053303 | 1 | |
| ENSMUSG00000053310 | 1.01393017 | 1 |
| ENSMUSG00000053317 | 1.025104884 | 0.793387188 |
| ENSMUSG00000053318 | -1.331703243 | 0.197904484 |
| ENSMUSG00000053329 | -1.06310058 | 0.608497229 |
| ENSMUSG00000053332 | 1.037095856 | 0.916323297 |
| ENSMUSG00000053333 | -1.107454656 | 0.331426543 |
| ENSMUSG00000053334 | 1.191819478 | 0.017036668 |
| ENSMUSG00000053338 | 1.12316944 | 0.694083715 |
| ENSMUSG00000053347 | -1.004969208 | 1 |
| ENSMUSG00000053353 | -1 | 1 |
| ENSMUSG00000053358 | 1.008889569 | 1 |
| ENSMUSG00000053360 | 1 | |
| ENSMUSG00000053367 | 1 | |
| ENSMUSG00000053368 | -1.03962336 | 0.698112512 |
| ENSMUSG00000053375 | -1 | 1 |
| ENSMUSG00000053388 | -1.117459691 | 0.563957069 |
| ENSMUSG00000053389 | 1 | |
| ENSMUSG00000053390 | -1.073645992 | 0.670176133 |
| ENSMUSG00000053391 | 1 | |
| ENSMUSG00000053395 | 1.297116521 | 0.151608108 |
| ENSMUSG00000053398 | 1.060614272 | 0.712790263 |
| ENSMUSG00000053399 | 1.999181489 | 0.000147458 |
| ENSMUSG00000053411 | -1.037951213 | 0.716153547 |
| ENSMUSG00000053414 | -2.127054292 | 0.001626258 |
| ENSMUSG00000053420 | 1 | |
| ENSMUSG00000053436 | 1.099629359 | 0.698043683 |
| ENSMUSG00000053441 | 1 | |
| ENSMUSG00000053442 | -1.033820075 | 0.905104202 |
| ENSMUSG00000053453 | -1.145584241 | 0.398710843 |
| ENSMUSG00000053460 | 1.009546131 | 1 |
| ENSMUSG00000053461 | 1.207441909 | 0.325037163 |
| ENSMUSG00000053465 | 1 | |
| ENSMUSG00000053469 | 1.180989026 | 0.242974035 |
| ENSMUSG00000053470 | 1.210281909 | 0.221965155 |
| ENSMUSG00000053472 | 1 | |
| ENSMUSG00000053475 | 1.237532788 | 0.192004524 |
| ENSMUSG00000053477 | 1.075761811 | 0.698112512 |
| ENSMUSG00000053483 | -1.00747311 | 1 |
| ENSMUSG00000053490 | 1 | |
| ENSMUSG00000053499 | 1 | |
| ENSMUSG00000053508 | 1 | |
| ENSMUSG00000053510 | 1.048436185 | 0.883306173 |
| ENSMUSG00000053519 | 1.059463094 | 0.698112512 |
| ENSMUSG00000053522 | -1 | 1 |
| ENSMUSG00000053528 | 1 | |
| ENSMUSG00000053536 | 1.046868245 | 0.881180973 |
| ENSMUSG00000053541 | -1.043306199 | 0.79291651 |
| ENSMUSG00000053545 | 1 | |
| ENSMUSG00000053550 | -1.084645496 | 0.69237817 |

| | | |
|---|---|---|
| ENSMUSG00000053552 | -1.247292666 | 0.085333778 |
| ENSMUSG00000053553 | 1.071656348 | 0.384106225 |
| ENSMUSG00000053559 | -1.901292364 | 0.002171462 |
| ENSMUSG00000053560 | 1.091798935 | 0.686129647 |
| ENSMUSG00000053565 | 1.039999817 | 0.84576478 |
| ENSMUSG00000053574 | -1.015538498 | 0.969007252 |
| ENSMUSG00000053580 | 1.283544177 | 0.045842071 |
| ENSMUSG00000053581 | -1.009208427 | 1 |
| ENSMUSG00000053588 | 1 | |
| ENSMUSG00000053593 | 1 | |
| ENSMUSG00000053600 | 1.026791946 | 0.850478054 |
| ENSMUSG00000053603 | 1.084618945 | 0.698112512 |
| ENSMUSG00000053604 | -1.04713905 | 0.698112512 |
| ENSMUSG00000053613 | 1.00760457 | 1 |
| ENSMUSG00000053615 | -1.308980822 | 0.135397497 |
| ENSMUSG00000053617 | 1.193206975 | 0.481246252 |
| ENSMUSG00000053624 | 1 | |
| ENSMUSG00000053626 | 2.121254692 | 2.79798E-05 |
| ENSMUSG00000053640 | 1 | |
| ENSMUSG00000053641 | -1.012071039 | 1 |
| ENSMUSG00000053644 | -1.140706562 | 0.291044503 |
| ENSMUSG00000053646 | 1.049709337 | 0.698112512 |
| ENSMUSG00000053647 | -1.075857308 | 0.608497229 |
| ENSMUSG00000053654 | 1 | |
| ENSMUSG00000053656 | 1 | |
| ENSMUSG00000053675 | 1.232987931 | 0.486930276 |
| ENSMUSG00000053678 | 1 | |
| ENSMUSG00000053684 | 1.057504525 | 0.389195911 |
| ENSMUSG00000053687 | 1.09439743 | 0.608497229 |
| ENSMUSG00000053693 | -1.217290427 | 0.508515749 |
| ENSMUSG00000053695 | 1 | |
| ENSMUSG00000053702 | 1.020594125 | 0.872376523 |
| ENSMUSG00000053706 | 1 | |
| ENSMUSG00000053714 | 1.030445456 | 0.938939969 |
| ENSMUSG00000053716 | 1.055297711 | 0.694941877 |
| ENSMUSG00000053719 | 1 | |
| ENSMUSG00000053720 | 1 | |
| ENSMUSG00000053729 | 1 | |
| ENSMUSG00000053730 | 1.184192582 | 0.008300939 |
| ENSMUSG00000053740 | -1.028348525 | 0.912425817 |
| ENSMUSG00000053742 | 1 | |
| ENSMUSG00000053746 | 1.295837076 | 0.004019068 |
| ENSMUSG00000053747 | 1 | |
| ENSMUSG00000053749 | 1.081017342 | 0.783694537 |
| ENSMUSG00000053754 | -1.148087077 | 0.604373376 |
| ENSMUSG00000053765 | 1 | |
| ENSMUSG00000053768 | 1.029571927 | 0.934704406 |
| ENSMUSG00000053769 | -1.042018399 | 0.730415115 |
| ENSMUSG00000053773 | -1.126138014 | 0.658044065 |
| ENSMUSG00000053774 | 1.021940434 | 0.977563419 |
| ENSMUSG00000053783 | -1.071075483 | 0.698112512 |
| ENSMUSG00000053790 | 1 | |
| ENSMUSG00000053797 | -1.198352625 | 0.287588797 |
| ENSMUSG00000053799 | -1.120798459 | 0.56794807 |
| ENSMUSG00000053801 | -1.004194668 | 1 |
| ENSMUSG00000053815 | 1 | |
| ENSMUSG00000053819 | 1.161592446 | 0.502459304 |
| ENSMUSG00000053820 | 1 | |
| ENSMUSG00000053821 | -1.003834322 | 1 |
| ENSMUSG00000053825 | -1.35729935 | 0.147196278 |
| ENSMUSG00000053830 | -1.008906735 | 1 |
| ENSMUSG00000053835 | -1.02179941 | 0.807933619 |
| ENSMUSG00000053838 | -1.002026664 | 1 |

| | | |
|---|---|---|
| ENSMUSG00000053841 | 1.030041003 | 0.91220985 |
| ENSMUSG00000053846 | -1.006801208 | 0.993517802 |
| ENSMUSG00000053852 | 1 | |
| ENSMUSG00000053856 | 1 | |
| ENSMUSG00000053861 | 1 | |
| ENSMUSG00000053862 | -1.053549062 | 0.769745556 |
| ENSMUSG00000053863 | 1 | |
| ENSMUSG00000053868 | 1 | |
| ENSMUSG00000053870 | -1.008585765 | 1 |
| ENSMUSG00000053873 | 1 | |
| ENSMUSG00000053877 | 1.023908172 | 0.954497929 |
| ENSMUSG00000053886 | 1.448636164 | 0.031427766 |
| ENSMUSG00000053889 | 1.062573223 | 0.915071338 |
| ENSMUSG00000053896 | 1 | |
| ENSMUSG00000053897 | 1.167506261 | 0.195607801 |
| ENSMUSG00000053898 | 1.097626799 | 0.375653391 |
| ENSMUSG00000053907 | 1.063264539 | 0.849234171 |
| ENSMUSG00000053909 | 1 | |
| ENSMUSG00000053914 | 1.039058384 | 0.934760809 |
| ENSMUSG00000053916 | 1.075070132 | 0.725426917 |
| ENSMUSG00000053925 | 1.076685932 | 0.608497229 |
| ENSMUSG00000053929 | -1.003173378 | 1 |
| ENSMUSG00000053930 | 1.127832095 | 0.606280786 |
| ENSMUSG00000053931 | 1.139740697 | 0.696249781 |
| ENSMUSG00000053950 | 1.130619054 | 0.470928144 |
| ENSMUSG00000053957 | -1.053549062 | 0.769745556 |
| ENSMUSG00000053961 | 1 | |
| ENSMUSG00000053963 | -1.239913898 | 0.266903707 |
| ENSMUSG00000053964 | 1.029114867 | 0.980511343 |
| ENSMUSG00000053965 | 1.270156632 | 0.01970464 |
| ENSMUSG00000053977 | -1.180378409 | 0.511978707 |
| ENSMUSG00000053979 | 1 | |
| ENSMUSG00000053980 | 1 | |
| ENSMUSG00000053985 | 1.045516644 | 0.78196579 |
| ENSMUSG00000054000 | 1.159030628 | 0.231320305 |
| ENSMUSG00000054003 | -1 | 1 |
| ENSMUSG00000054005 | -1.03962336 | 0.698112512 |
| ENSMUSG00000054006 | 1 | |
| ENSMUSG00000054008 | 1.053550009 | 0.797808656 |
| ENSMUSG00000054013 | 1.230075506 | 0.224243601 |
| ENSMUSG00000054021 | 1.013636269 | 0.932597278 |
| ENSMUSG00000054027 | 1.063539682 | 0.741911172 |
| ENSMUSG00000054031 | 1 | |
| ENSMUSG00000054033 | 1 | |
| ENSMUSG00000054034 | -1 | 1 |
| ENSMUSG00000054036 | 1 | |
| ENSMUSG00000054046 | 1 | |
| ENSMUSG00000054051 | -1.044034537 | 0.783658058 |
| ENSMUSG00000054052 | 1.122462048 | 0.215991492 |
| ENSMUSG00000054054 | 1 | |
| ENSMUSG00000054057 | 1.379207664 | 0.001993499 |
| ENSMUSG00000054061 | 1 | |
| ENSMUSG00000054065 | -1.010148234 | 1 |
| ENSMUSG00000054069 | 1 | |
| ENSMUSG00000054072 | -1.661292342 | 0.002466243 |
| ENSMUSG00000054074 | 1.297261926 | 0.048932904 |
| ENSMUSG00000054079 | -1.012636446 | 1 |
| ENSMUSG00000054083 | 1.059463094 | 0.698112512 |
| ENSMUSG00000054087 | 1 | |
| ENSMUSG00000054091 | -1.006244198 | 1 |
| ENSMUSG00000054099 | 1.073951802 | 0.698112512 |
| ENSMUSG00000054102 | 1 | |
| ENSMUSG00000054106 | 1 | |

| | | |
|---|---|---|
| ENSMUSG00000054115 | 1.013133392 | 1 |
| ENSMUSG00000054116 | 1 | |
| ENSMUSG00000054117 | 1 | |
| ENSMUSG00000054128 | -1.116853908 | 0.590417714 |
| ENSMUSG00000054134 | 1 | |
| ENSMUSG00000054135 | 1.168676713 | 0.634672972 |
| ENSMUSG00000054136 | 1.39494079 | 0.260949892 |
| ENSMUSG00000054141 | 1 | |
| ENSMUSG00000054142 | 1 | |
| ENSMUSG00000054146 | 1 | |
| ENSMUSG00000054150 | 1.072193062 | 0.470841398 |
| ENSMUSG00000054156 | 1 | |
| ENSMUSG00000054160 | 1 | |
| ENSMUSG00000054161 | 1.060131051 | 0.835701362 |
| ENSMUSG00000054162 | -1 | 1 |
| ENSMUSG00000054169 | 1 | |
| ENSMUSG00000054178 | 1.075909545 | 0.492773389 |
| ENSMUSG00000054181 | -1.133142804 | 0.695519259 |
| ENSMUSG00000054191 | 1.030415139 | 0.971519533 |
| ENSMUSG00000054196 | -4.487425546 | 0.000122514 |
| ENSMUSG00000054199 | 1.002947058 | 1 |
| ENSMUSG00000054200 | -1 | 1 |
| ENSMUSG00000054203 | 1.051093536 | 0.905627938 |
| ENSMUSG00000054204 | 1.059463094 | 0.698112512 |
| ENSMUSG00000054206 | 1.137503063 | 0.338075093 |
| ENSMUSG00000054215 | 1.536705276 | 0.028511322 |
| ENSMUSG00000054226 | -1.016922299 | 0.987029357 |
| ENSMUSG00000054236 | 1 | |
| ENSMUSG00000054237 | 1.023335777 | 0.922165724 |
| ENSMUSG00000054247 | -1.083415624 | 0.698112512 |
| ENSMUSG00000054252 | -1.372196403 | 0.005213906 |
| ENSMUSG00000054256 | -1.025056587 | 0.772971553 |
| ENSMUSG00000054258 | 1.271344436 | 0.053354336 |
| ENSMUSG00000054263 | -1.05928467 | 0.748295996 |
| ENSMUSG00000054266 | 1.111812527 | 0.672818484 |
| ENSMUSG00000054272 | 1 | |
| ENSMUSG00000054277 | 1.16346807 | 0.36566114 |
| ENSMUSG00000054280 | 1.020209325 | 0.97192334 |
| ENSMUSG00000054293 | -1.310550864 | 0.070657138 |
| ENSMUSG00000054302 | 1.033864402 | 0.850705298 |
| ENSMUSG00000054304 | 1.320383566 | 0.037163069 |
| ENSMUSG00000054309 | -1.000729582 | 1 |
| ENSMUSG00000054310 | 1 | |
| ENSMUSG00000054312 | -1.019999571 | 0.9605777 |
| ENSMUSG00000054320 | 1.059463094 | 0.698112512 |
| ENSMUSG00000054321 | -1.027968856 | 0.849863703 |
| ENSMUSG00000054325 | 1 | |
| ENSMUSG00000054342 | 1.280238812 | 0.22142317 |
| ENSMUSG00000054351 | 1 | |
| ENSMUSG00000054360 | 1 | |
| ENSMUSG00000054362 | 1 | |
| ENSMUSG00000054364 | 1.037808211 | 0.845096896 |
| ENSMUSG00000054379 | -1.03962336 | 0.698112512 |
| ENSMUSG00000054381 | 1.103098817 | 0.358637336 |
| ENSMUSG00000054383 | 1.047001457 | 0.861137762 |
| ENSMUSG00000054385 | 1.214792929 | 0.417353956 |
| ENSMUSG00000054387 | -1.045045984 | 0.892023203 |
| ENSMUSG00000054391 | 1 | |
| ENSMUSG00000054400 | -1.019361906 | 0.946232149 |
| ENSMUSG00000054404 | -1.314861329 | 0.071577777 |
| ENSMUSG00000054405 | 1.033772505 | 0.942279278 |
| ENSMUSG00000054406 | 1 | |
| ENSMUSG00000054408 | 1.025761513 | 0.97946024 |

| | | |
|---|---|---|
| ENSMUSG00000054409 | 1.35308594 | 0.190563636 |
| ENSMUSG00000054412 | 1.127832095 | 0.698112512 |
| ENSMUSG00000054414 | -1.070070662 | 0.734674233 |
| ENSMUSG00000054417 | 1 | |
| ENSMUSG00000054418 | -1.579235849 | 0.011862009 |
| ENSMUSG00000054422 | 1 | |
| ENSMUSG00000054423 | 1 | |
| ENSMUSG00000054426 | 1.045153844 | 0.833767346 |
| ENSMUSG00000054428 | 1.089937812 | 0.595296772 |
| ENSMUSG00000054431 | 1 | |
| ENSMUSG00000054434 | 1.030528345 | 0.74773997 |
| ENSMUSG00000054435 | -1.086192882 | 0.698112512 |
| ENSMUSG00000054446 | 1 | |
| ENSMUSG00000054450 | -1 | 1 |
| ENSMUSG00000054452 | 1.016549253 | 0.983393382 |
| ENSMUSG00000054453 | 2.153556247 | 0.000620329 |
| ENSMUSG00000054455 | -1.040857515 | 0.85351833 |
| ENSMUSG00000054457 | 1.059463094 | 0.698112512 |
| ENSMUSG00000054459 | 1.189809985 | 0.245135442 |
| ENSMUSG00000054469 | 1.001709221 | 1 |
| ENSMUSG00000054474 | -1.13935184 | 0.139593229 |
| ENSMUSG00000054477 | -1.04526435 | 0.877529894 |
| ENSMUSG00000054484 | -1.129138573 | 0.608497229 |
| ENSMUSG00000054488 | -1 | 1 |
| ENSMUSG00000054493 | -1 | 1 |
| ENSMUSG00000054497 | 1 | |
| ENSMUSG00000054498 | 1 | |
| ENSMUSG00000054499 | -1.046650694 | 0.771728893 |
| ENSMUSG00000054509 | -1.227925443 | 0.155992318 |
| ENSMUSG00000054510 | 1.014731082 | 0.998498701 |
| ENSMUSG00000054514 | -1.118046445 | 0.690141149 |
| ENSMUSG00000054517 | 1.186566674 | 0.027427122 |
| ENSMUSG00000054519 | -1.126338788 | 0.474507738 |
| ENSMUSG00000054520 | 1.115291844 | 0.119966047 |
| ENSMUSG00000054523 | -1 | 1 |
| ENSMUSG00000054526 | 1 | |
| ENSMUSG00000054537 | 1 | |
| ENSMUSG00000054545 | 1.019083579 | 0.968937442 |
| ENSMUSG00000054555 | 1.550218802 | 0.002157726 |
| ENSMUSG00000054556 | 1.186241827 | 0.59127359 |
| ENSMUSG00000054568 | 1.096535473 | 0.681022198 |
| ENSMUSG00000054580 | 2.154188746 | 0.000199503 |
| ENSMUSG00000054582 | -1.035015517 | 0.960670289 |
| ENSMUSG00000054589 | 1.182092011 | 0.606278677 |
| ENSMUSG00000054594 | 1.025046921 | 0.956524746 |
| ENSMUSG00000054598 | 1.085999383 | 0.726987474 |
| ENSMUSG00000054604 | 1.037553287 | 0.933031586 |
| ENSMUSG00000054611 | -1.004862126 | 1 |
| ENSMUSG00000054612 | 1.253707454 | 0.099597078 |
| ENSMUSG00000054618 | 1.019083579 | 0.949812939 |
| ENSMUSG00000054619 | 1.086824439 | 0.499289204 |
| ENSMUSG00000054622 | 1.085624843 | 0.698112512 |
| ENSMUSG00000054626 | 1 | |
| ENSMUSG00000054630 | 1 | |
| ENSMUSG00000054640 | 2.550281272 | 0.000143267 |
| ENSMUSG00000054641 | -1 | 1 |
| ENSMUSG00000054648 | -1.032291641 | 0.849628468 |
| ENSMUSG00000054659 | -1.089497311 | 0.608497229 |
| ENSMUSG00000054662 | 1.059463094 | 0.699340382 |
| ENSMUSG00000054666 | 1 | |
| ENSMUSG00000054667 | -1.109751192 | 0.698112512 |
| ENSMUSG00000054672 | 1 | |
| ENSMUSG00000054675 | 1.860715856 | 0.000383911 |

| | | |
|---|---|---|
| ENSMUSG00000054676 | -1.017179715 | 0.890936865 |
| ENSMUSG00000054679 | -1.791931526 | 0.005261797 |
| ENSMUSG00000054690 | 1 | |
| ENSMUSG00000054693 | 1.014490669 | 1 |
| ENSMUSG00000054702 | -1.153031888 | 0.259044719 |
| ENSMUSG00000054708 | 1.005839642 | 1 |
| ENSMUSG00000054715 | -1.70011822 | 0.000409422 |
| ENSMUSG00000054716 | -1.054143988 | 0.704170349 |
| ENSMUSG00000054717 | -1.015106506 | 1 |
| ENSMUSG00000054720 | 1.18183941 | 0.406781564 |
| ENSMUSG00000054723 | -1.047738009 | 0.608497229 |
| ENSMUSG00000054727 | 1 | |
| ENSMUSG00000054728 | -1.066167697 | 0.828564184 |
| ENSMUSG00000054733 | -1.044869221 | 0.649280935 |
| ENSMUSG00000054737 | -3.288353134 | 0.000136743 |
| ENSMUSG00000054745 | 1 | |
| ENSMUSG00000054746 | 1 | |
| ENSMUSG00000054752 | -1.046153764 | 0.833413645 |
| ENSMUSG00000054753 | -1.265341119 | 0.179450399 |
| ENSMUSG00000054757 | -1.083415624 | 0.62221251 |
| ENSMUSG00000054758 | 1 | |
| ENSMUSG00000054759 | 1 | |
| ENSMUSG00000054763 | -1.042811414 | 0.90576296 |
| ENSMUSG00000054764 | 1.069068979 | 0.608497229 |
| ENSMUSG00000054766 | 1.076328907 | 0.798487595 |
| ENSMUSG00000054770 | -1.01289336 | 0.978109404 |
| ENSMUSG00000054779 | 1.122933516 | 0.608497229 |
| ENSMUSG00000054792 | -1.059720319 | 0.608497229 |
| ENSMUSG00000054793 | 1.069061992 | 0.537316682 |
| ENSMUSG00000054808 | 1.227015894 | 0.351849863 |
| ENSMUSG00000054814 | 1.06057329 | 0.788000546 |
| ENSMUSG00000054822 | 1.116785864 | 0.608497229 |
| ENSMUSG00000054823 | 1.029111609 | 0.954497929 |
| ENSMUSG00000054827 | 1 | |
| ENSMUSG00000054836 | -1.085395172 | 0.448706039 |
| ENSMUSG00000054843 | 1.002159424 | 1 |
| ENSMUSG00000054850 | 1.21823446 | 0.022993558 |
| ENSMUSG00000054855 | -1.181616536 | 0.342054851 |
| ENSMUSG00000054863 | 1.284066971 | 0.011900948 |
| ENSMUSG00000054871 | 1.198565113 | 0.490063384 |
| ENSMUSG00000054874 | 1.155785316 | 0.234728821 |
| ENSMUSG00000054885 | 1 | |
| ENSMUSG00000054889 | -1.366336587 | 0.033703167 |
| ENSMUSG00000054890 | 1 | |
| ENSMUSG00000054892 | -1.060043393 | 0.873810332 |
| ENSMUSG00000054893 | -1.151173889 | 0.188041916 |
| ENSMUSG00000054894 | 1.091643608 | 0.608497229 |
| ENSMUSG00000054901 | 1.017286012 | 0.996192043 |
| ENSMUSG00000054905 | 1 | |
| ENSMUSG00000054909 | 1 | |
| ENSMUSG00000054910 | -1.053199206 | 0.816416314 |
| ENSMUSG00000054920 | 1.06452943 | 0.698112512 |
| ENSMUSG00000054931 | -1.053549062 | 0.764982341 |
| ENSMUSG00000054932 | -1.039754043 | 0.952556689 |
| ENSMUSG00000054934 | -1.284366496 | 0.01850635 |
| ENSMUSG00000054938 | 1.059463094 | 0.608497229 |
| ENSMUSG00000054939 | 1.056632167 | 0.608497229 |
| ENSMUSG00000054940 | 1 | |
| ENSMUSG00000054942 | 1.013627135 | 0.985522133 |
| ENSMUSG00000054944 | 1.059463094 | 0.608497229 |
| ENSMUSG00000054945 | 1.075583614 | 0.639646932 |
| ENSMUSG00000054951 | 1.015564799 | 0.93376866 |
| ENSMUSG00000054957 | 1 | |

| | | |
|---|---|---|
| ENSMUSG00000054958 | 1 | |
| ENSMUSG00000054966 | 1.059463094 | 0.608497229 |
| ENSMUSG00000054967 | 1.061413255 | 0.650221528 |
| ENSMUSG00000054976 | 1 | |
| ENSMUSG00000054978 | -1.023769022 | 0.934760809 |
| ENSMUSG00000054986 | 1 | |
| ENSMUSG00000054988 | 1 | |
| ENSMUSG00000054994 | 1 | |
| ENSMUSG00000054999 | 1.029374385 | 0.865583959 |
| ENSMUSG00000055003 | -1.092827399 | 0.758575113 |
| ENSMUSG00000055010 | 1 | |
| ENSMUSG00000055013 | 1.040429171 | 0.756494969 |
| ENSMUSG00000055015 | 1 | |
| ENSMUSG00000055022 | 1.048895035 | 0.742615858 |
| ENSMUSG00000055024 | 1.022734781 | 0.975318388 |
| ENSMUSG00000055026 | 1.059463094 | 0.608497229 |
| ENSMUSG00000055027 | -1.167457557 | 0.472312635 |
| ENSMUSG00000055030 | 1.11166259 | 0.608497229 |
| ENSMUSG00000055033 | 1 | |
| ENSMUSG00000055041 | 1.080559608 | 0.275053802 |
| ENSMUSG00000055044 | 1.062157368 | 0.777454752 |
| ENSMUSG00000055045 | 1 | |
| ENSMUSG00000055048 | 1.059592506 | 0.829954854 |
| ENSMUSG00000055053 | 1.100263478 | 0.585199025 |
| ENSMUSG00000055061 | -1.044273782 | 0.790851299 |
| ENSMUSG00000055065 | -1.012982756 | 1 |
| ENSMUSG00000055067 | -1.050041958 | 0.617886141 |
| ENSMUSG00000055069 | -1.288987208 | 0.235160214 |
| ENSMUSG00000055072 | 1.005552172 | 1 |
| ENSMUSG00000055078 | 1 | |
| ENSMUSG00000055088 | 1 | |
| ENSMUSG00000055093 | -1.145216302 | 0.68559795 |
| ENSMUSG00000055095 | 1 | |
| ENSMUSG00000055099 | 1 | |
| ENSMUSG00000055102 | -1.675370645 | 0.003402016 |
| ENSMUSG00000055108 | -1.03962336 | 0.698112512 |
| ENSMUSG00000055109 | 1 | |
| ENSMUSG00000055110 | 1 | |
| ENSMUSG00000055114 | -1.069502966 | 0.79291651 |
| ENSMUSG00000055116 | 1.038096855 | 0.752398528 |
| ENSMUSG00000055125 | 1 | |
| ENSMUSG00000055128 | -1.112002024 | 0.451548004 |
| ENSMUSG00000055134 | 1.010014464 | 0.989629241 |
| ENSMUSG00000055137 | -1.242209633 | 0.304680629 |
| ENSMUSG00000055138 | 1.069068979 | 0.608497229 |
| ENSMUSG00000055148 | -1.218461244 | 0.132365025 |
| ENSMUSG00000055150 | -1.204373222 | 0.36085659 |
| ENSMUSG00000055159 | -1.044273782 | 0.698112512 |
| ENSMUSG00000055170 | 1 | |
| ENSMUSG00000055172 | 1.171973719 | 0.037405901 |
| ENSMUSG00000055177 | 1.024693402 | 0.952057016 |
| ENSMUSG00000055184 | 1.010646566 | 0.947720549 |
| ENSMUSG00000055188 | -1.041813168 | 0.959383659 |
| ENSMUSG00000055193 | 1 | |
| ENSMUSG00000055194 | 1.005613439 | 1 |
| ENSMUSG00000055197 | 1 | |
| ENSMUSG00000055198 | 1 | |
| ENSMUSG00000055200 | -1.004697619 | 1 |
| ENSMUSG00000055202 | 1.139858854 | 0.160342423 |
| ENSMUSG00000055204 | 1.132778658 | 0.686477484 |
| ENSMUSG00000055210 | -1.120874705 | 0.770958539 |
| ENSMUSG00000055214 | -1.370750503 | 0.169990135 |
| ENSMUSG00000055216 | -1.039261712 | 0.908842909 |

| | | |
|---|---|---|
| ENSMUSG00000055228 | -1.0799558 | 0.696563202 |
| ENSMUSG00000055235 | -1.043306199 | 0.698112512 |
| ENSMUSG00000055239 | 1.08293692 | 0.654183172 |
| ENSMUSG00000055240 | -1.025882959 | 0.930238397 |
| ENSMUSG00000055254 | -1 | 1 |
| ENSMUSG00000055271 | -1.063103825 | 0.756949808 |
| ENSMUSG00000055296 | 1.0023215 | 1 |
| ENSMUSG00000055298 | 1 | |
| ENSMUSG00000055301 | 1.020817741 | 0.856732743 |
| ENSMUSG00000055302 | -1.02456544 | 0.987313439 |
| ENSMUSG00000055305 | 1.009398876 | 0.960337041 |
| ENSMUSG00000055312 | -1 | 1 |
| ENSMUSG00000055313 | -1.184951155 | 0.206620396 |
| ENSMUSG00000055319 | -1.010264049 | 1 |
| ENSMUSG00000055320 | 1.087881091 | 0.734415698 |
| ENSMUSG00000055322 | 1.234366435 | 0.242442975 |
| ENSMUSG00000055323 | 1.021582126 | 0.989648415 |
| ENSMUSG00000055333 | -1.095338671 | 0.605620689 |
| ENSMUSG00000055334 | -1.144525241 | 0.414805624 |
| ENSMUSG00000055341 | -1.039465692 | 0.740105843 |
| ENSMUSG00000055357 | 1.059463094 | 0.608497229 |
| ENSMUSG00000055360 | -1.018998401 | 1 |
| ENSMUSG00000055368 | -1.139124004 | 0.563092624 |
| ENSMUSG00000055370 | -1.104637702 | 0.731760295 |
| ENSMUSG00000055371 | 1.062294881 | 0.756618803 |
| ENSMUSG00000055373 | 1.554234178 | 0.002512807 |
| ENSMUSG00000055385 | 1 | |
| ENSMUSG00000055401 | 1.080048622 | 0.466407623 |
| ENSMUSG00000055403 | 1.179016771 | 0.531424605 |
| ENSMUSG00000055407 | 1.001356291 | 1 |
| ENSMUSG00000055408 | 1 | |
| ENSMUSG00000055409 | 1 | |
| ENSMUSG00000055413 | -1.784205029 | 0.003665403 |
| ENSMUSG00000055415 | 1 | |
| ENSMUSG00000055416 | 1.019083579 | 0.971519533 |
| ENSMUSG00000055421 | -1.075808387 | 0.608497229 |
| ENSMUSG00000055424 | -1.08384782 | 0.702897175 |
| ENSMUSG00000055430 | -1.322184358 | 0.128745364 |
| ENSMUSG00000055435 | 1.012878275 | 0.953597385 |
| ENSMUSG00000055436 | 1.010763777 | 1 |
| ENSMUSG00000055446 | 1 | |
| ENSMUSG00000055447 | 1.085434146 | 0.711533702 |
| ENSMUSG00000055452 | 1 | |
| ENSMUSG00000055456 | 1 | |
| ENSMUSG00000055471 | 1 | |
| ENSMUSG00000055480 | -1.128363601 | 0.340742086 |
| ENSMUSG00000055485 | -1.027578817 | 0.841285672 |
| ENSMUSG00000055489 | 1.059463094 | 0.608497229 |
| ENSMUSG00000055491 | 1.103112167 | 0.632670826 |
| ENSMUSG00000055493 | -1.010932686 | 0.982854947 |
| ENSMUSG00000055494 | -1.077191405 | 0.742583941 |
| ENSMUSG00000055497 | 1.036040116 | 0.934760809 |
| ENSMUSG00000055515 | 1 | |
| ENSMUSG00000055523 | 1 | |
| ENSMUSG00000055531 | 1.00748393 | 1 |
| ENSMUSG00000055538 | -1.076451555 | 0.763030715 |
| ENSMUSG00000055540 | 1 | |
| ENSMUSG00000055541 | -1.044273782 | 0.698112512 |
| ENSMUSG00000055546 | 1 | |
| ENSMUSG00000055547 | 1.059463094 | 0.608497229 |
| ENSMUSG00000055553 | 1.002424924 | 1 |
| ENSMUSG00000055555 | 1 | |
| ENSMUSG00000055560 | -1.010659465 | 0.996913409 |

| | | |
|---|---|---|
| ENSMUSG00000055561 | 1 | |
| ENSMUSG00000055567 | 1.09538577 | 0.773736245 |
| ENSMUSG00000055571 | 1 | |
| ENSMUSG00000055594 | 1 | |
| ENSMUSG00000055602 | 1.069068979 | 0.767473873 |
| ENSMUSG00000055609 | 1 | |
| ENSMUSG00000055610 | 1 | |
| ENSMUSG00000055612 | 1.016148014 | 0.968561948 |
| ENSMUSG00000055621 | 1 | |
| ENSMUSG00000055629 | 1.119123949 | 0.470354087 |
| ENSMUSG00000055632 | 1.637626305 | 0.090904488 |
| ENSMUSG00000055633 | 1.083408284 | 0.681952832 |
| ENSMUSG00000055639 | -1.042505207 | 0.930108107 |
| ENSMUSG00000055643 | 1 | |
| ENSMUSG00000055652 | 1.084952759 | 0.474788671 |
| ENSMUSG00000055653 | -1.166301255 | 0.55782474 |
| ENSMUSG00000055660 | -1.011878175 | 0.992580079 |
| ENSMUSG00000055670 | 1.098142018 | 0.678806909 |
| ENSMUSG00000055675 | -1.020646102 | 0.979002724 |
| ENSMUSG00000055676 | -1.044583175 | 0.846956027 |
| ENSMUSG00000055679 | 1 | |
| ENSMUSG00000055681 | 1.063936053 | 0.608497229 |
| ENSMUSG00000055691 | 1.032926687 | 0.949295635 |
| ENSMUSG00000055692 | 1.014845666 | 1 |
| ENSMUSG00000055694 | 1 | |
| ENSMUSG00000055700 | 1 | |
| ENSMUSG00000055704 | 1.124340693 | 0.594181956 |
| ENSMUSG00000055707 | 1.127364637 | 0.166621275 |
| ENSMUSG00000055717 | -2.019542953 | 0.000331899 |
| ENSMUSG00000055720 | -1.016644026 | 0.949539131 |
| ENSMUSG00000055723 | -1.063875575 | 0.778372272 |
| ENSMUSG00000055725 | -1.002388001 | 1 |
| ENSMUSG00000055730 | 1 | |
| ENSMUSG00000055732 | 1 | |
| ENSMUSG00000055733 | -1.278939889 | 0.046492778 |
| ENSMUSG00000055737 | 1.283410516 | 0.09778616 |
| ENSMUSG00000055745 | -2.17566603 | 0.000212323 |
| ENSMUSG00000055746 | 1 | |
| ENSMUSG00000055748 | 1.076975501 | 0.850250061 |
| ENSMUSG00000055760 | -1.004487412 | 1 |
| ENSMUSG00000055761 | 1.07913413 | 0.715341911 |
| ENSMUSG00000055762 | 1.004428339 | 1 |
| ENSMUSG00000055763 | 1.079681442 | 0.608497229 |
| ENSMUSG00000055771 | 1.171563499 | 0.233412931 |
| ENSMUSG00000055775 | -1.165123214 | 0.545246516 |
| ENSMUSG00000055780 | -1.109582661 | 0.608497229 |
| ENSMUSG00000055782 | 1.133427263 | 0.270556972 |
| ENSMUSG00000055789 | 1 | |
| ENSMUSG00000055795 | 1.177055296 | 0.51546804 |
| ENSMUSG00000055799 | 1.103430024 | 0.294914291 |
| ENSMUSG00000055805 | -1.613559405 | 0.006016739 |
| ENSMUSG00000055809 | -1.670616575 | 0.015929037 |
| ENSMUSG00000055811 | 1 | |
| ENSMUSG00000055817 | 1.080086297 | 0.395740825 |
| ENSMUSG00000055818 | 1.144492418 | 0.736861486 |
| ENSMUSG00000055820 | 1 | |
| ENSMUSG00000055826 | 1 | |
| ENSMUSG00000055827 | 1 | |
| ENSMUSG00000055833 | -1.023439908 | 0.984476939 |
| ENSMUSG00000055835 | -1.009367112 | 0.981032001 |
| ENSMUSG00000055838 | 1 | |
| ENSMUSG00000055839 | 1.175318882 | 0.223995462 |
| ENSMUSG00000055850 | -1.027110014 | 0.934497597 |

| | | |
|---|---|---|
| ENSMUSG00000055862 | 1.032734221 | 0.764129089 |
| ENSMUSG00000055865 | 1 | |
| ENSMUSG00000055866 | 1.093183291 | 0.32216057 |
| ENSMUSG00000055872 | -1.016017585 | 1 |
| ENSMUSG00000055874 | 1 | |
| ENSMUSG00000055882 | -1.24516087 | 0.53313872 |
| ENSMUSG00000055884 | 1.016086012 | 0.983830102 |
| ENSMUSG00000055891 | -1.023769022 | 0.934760809 |
| ENSMUSG00000055895 | 1 | |
| ENSMUSG00000055897 | 1.054596372 | 0.698112512 |
| ENSMUSG00000055900 | -1.590836272 | 0.004429915 |
| ENSMUSG00000055912 | 1.244719337 | 0.005995788 |
| ENSMUSG00000055917 | 1.057169779 | 0.734674233 |
| ENSMUSG00000055923 | 1.046541422 | 0.614071182 |
| ENSMUSG00000055926 | -1.381937027 | 0.005450731 |
| ENSMUSG00000055932 | 1.007082099 | 1 |
| ENSMUSG00000055933 | 1 | |
| ENSMUSG00000055937 | -1.044273782 | 0.790851299 |
| ENSMUSG00000055942 | 1 | |
| ENSMUSG00000055943 | -1.126617023 | 0.56288192 |
| ENSMUSG00000055944 | 1.017174474 | 1 |
| ENSMUSG00000055945 | 1.157058285 | 0.608001601 |
| ENSMUSG00000055958 | 1.107677833 | 0.789030564 |
| ENSMUSG00000055960 | -1.086092137 | 0.704170349 |
| ENSMUSG00000055961 | 1 | |
| ENSMUSG00000055963 | 1.061111884 | 0.590417714 |
| ENSMUSG00000055968 | 1 | |
| ENSMUSG00000055971 | 1 | |
| ENSMUSG00000055972 | -1.044273782 | 0.698112512 |
| ENSMUSG00000055976 | 1 | |
| ENSMUSG00000055978 | 1.132639128 | 0.398767938 |
| ENSMUSG00000055980 | -1.117227686 | 0.310231463 |
| ENSMUSG00000055991 | 1.004190101 | 1 |
| ENSMUSG00000055994 | -1.706757386 | 0.000772753 |
| ENSMUSG00000056004 | 1.421826279 | 0.004868138 |
| ENSMUSG00000056008 | 1 | |
| ENSMUSG00000056014 | -1.07704822 | 0.408689193 |
| ENSMUSG00000056018 | 1.0900642 | 0.603706485 |
| ENSMUSG00000056019 | -1.307196415 | 0.006889494 |
| ENSMUSG00000056023 | -1 | 1 |
| ENSMUSG00000056025 | 1.382319564 | 0.072705092 |
| ENSMUSG00000056031 | 1.204653091 | 0.194022145 |
| ENSMUSG00000056032 | 1.081017342 | 0.712533348 |
| ENSMUSG00000056035 | 1 | |
| ENSMUSG00000056043 | -1.042635269 | 0.799256428 |
| ENSMUSG00000056050 | -1.062751925 | 0.766831979 |
| ENSMUSG00000056054 | 1.005613439 | 1 |
| ENSMUSG00000056055 | -1.037395066 | 0.803072249 |
| ENSMUSG00000056061 | 1 | |
| ENSMUSG00000056069 | -1.361535863 | 0.010805005 |
| ENSMUSG00000056071 | 1 | |
| ENSMUSG00000056073 | 1.274424648 | 0.498478643 |
| ENSMUSG00000056076 | 1.041046513 | 0.9339505 |
| ENSMUSG00000056078 | 1.015486245 | 0.972980766 |
| ENSMUSG00000056089 | -1 | 1 |
| ENSMUSG00000056091 | 1.22020912 | 0.151696128 |
| ENSMUSG00000056106 | -1.019698456 | 1 |
| ENSMUSG00000056115 | 1 | |
| ENSMUSG00000056116 | -1.196455042 | 0.184445251 |
| ENSMUSG00000056121 | 1.25939482 | 0.287566569 |
| ENSMUSG00000056124 | 1.002082075 | 1 |
| ENSMUSG00000056128 | 1.132639128 | 0.217803313 |
| ENSMUSG00000056130 | -1.013431268 | 0.972684567 |

| | | |
|---|---|---|
| ENSMUSG00000056131 | -1.013585264 | 0.978384359 |
| ENSMUSG00000056133 | 1 | |
| ENSMUSG00000056144 | 1.12444444 | 0.48165686 |
| ENSMUSG00000056145 | -1.085651418 | 0.532369657 |
| ENSMUSG00000056148 | 1.028323353 | 0.903464611 |
| ENSMUSG00000056153 | 1.077076833 | 0.698112512 |
| ENSMUSG00000056155 | -1.078590893 | 0.698112512 |
| ENSMUSG00000056158 | 1.019083579 | 0.949812939 |
| ENSMUSG00000056162 | -1.093674795 | 0.608497229 |
| ENSMUSG00000056167 | 1.005822013 | 1 |
| ENSMUSG00000056174 | 1.047300491 | 0.629100036 |
| ENSMUSG00000056184 | 1 | |
| ENSMUSG00000056185 | 1.256100347 | 0.007814932 |
| ENSMUSG00000056197 | 1 | |
| ENSMUSG00000056201 | 1.157952421 | 0.676948744 |
| ENSMUSG00000056203 | 1 | |
| ENSMUSG00000056204 | -1.046295747 | 0.752411447 |
| ENSMUSG00000056209 | -1.049286175 | 0.806054257 |
| ENSMUSG00000056211 | 1.074962031 | 0.745625104 |
| ENSMUSG00000056214 | 1.109044169 | 0.069329311 |
| ENSMUSG00000056215 | -1.009858925 | 0.998253065 |
| ENSMUSG00000056216 | 1.096185681 | 0.615267463 |
| ENSMUSG00000056219 | 1 | |
| ENSMUSG00000056220 | 1.171131795 | 0.346633686 |
| ENSMUSG00000056222 | -1.03962336 | 0.698112512 |
| ENSMUSG00000056223 | 1 | |
| ENSMUSG00000056228 | 1.072749926 | 0.562116455 |
| ENSMUSG00000056234 | -1.151366792 | 0.320426081 |
| ENSMUSG00000056257 | 1.014545335 | 1 |
| ENSMUSG00000056258 | 1.143029712 | 0.608497229 |
| ENSMUSG00000056260 | -1.032183138 | 0.892732342 |
| ENSMUSG00000056267 | -1.034286635 | 0.754121635 |
| ENSMUSG00000056268 | -1.00051869 | 1 |
| ENSMUSG00000056270 | 1.059463094 | 0.608497229 |
| ENSMUSG00000056271 | -1.091058957 | 0.777145624 |
| ENSMUSG00000056281 | 1 | |
| ENSMUSG00000056288 | 1 | |
| ENSMUSG00000056290 | 1 | |
| ENSMUSG00000056293 | 1 | |
| ENSMUSG00000056296 | 1.004448592 | 1 |
| ENSMUSG00000056300 | -1.068484894 | 0.867452588 |
| ENSMUSG00000056305 | 1.029089271 | 0.949812939 |
| ENSMUSG00000056306 | -1.044273782 | 0.698112512 |
| ENSMUSG00000056310 | 1.068251371 | 0.608497229 |
| ENSMUSG00000056313 | 1.430897592 | 0.004341117 |
| ENSMUSG00000056316 | 1.122462048 | 0.608497229 |
| ENSMUSG00000056328 | -1.053549062 | 0.698112512 |
| ENSMUSG00000056329 | 1 | |
| ENSMUSG00000056332 | 1.069068979 | 0.608497229 |
| ENSMUSG00000056342 | -1.04234543 | 0.938610991 |
| ENSMUSG00000056350 | 1 | |
| ENSMUSG00000056359 | 1 | |
| ENSMUSG00000056366 | 1.059463094 | 0.750182067 |
| ENSMUSG00000056367 | 1.228421797 | 0.025087522 |
| ENSMUSG00000056370 | 1.132639128 | 0.398767938 |
| ENSMUSG00000056379 | 1 | |
| ENSMUSG00000056380 | -1.0889222 | 0.698112512 |
| ENSMUSG00000056383 | 1.064749431 | 0.57082804 |
| ENSMUSG00000056394 | 1.030716019 | 0.930108107 |
| ENSMUSG00000056399 | 1 | |
| ENSMUSG00000056411 | -1.067938213 | 0.629405465 |
| ENSMUSG00000056412 | 1 | |
| ENSMUSG00000056413 | 1.167176492 | 0.698112512 |

| | | |
|---|---|---|
| ENSMUSG00000056418 | 1.007689251 | 1 |
| ENSMUSG00000056423 | 1 | |
| ENSMUSG00000056426 | 1 | |
| ENSMUSG00000056427 | -1.804756471 | 0.000923517 |
| ENSMUSG00000056429 | -1.057473068 | 0.85864022 |
| ENSMUSG00000056436 | 1.059463094 | 0.608497229 |
| ENSMUSG00000056445 | 1.230648265 | 0.064293809 |
| ENSMUSG00000056457 | 1.07913413 | 0.807247723 |
| ENSMUSG00000056458 | -1.126597297 | 0.26570937 |
| ENSMUSG00000056459 | 1.134761617 | 0.216046507 |
| ENSMUSG00000056468 | 1.532087156 | 0.016501255 |
| ENSMUSG00000056476 | -1.635378735 | 0.009407544 |
| ENSMUSG00000056481 | 1.432920831 | 0.002635385 |
| ENSMUSG00000056486 | -1.243959989 | 0.123449197 |
| ENSMUSG00000056487 | 1 | |
| ENSMUSG00000056492 | -1.328334999 | 0.055949745 |
| ENSMUSG00000056493 | 1.138229542 | 0.209526842 |
| ENSMUSG00000056494 | 1 | |
| ENSMUSG00000056498 | -1.322000607 | 0.110887279 |
| ENSMUSG00000056501 | 1.080604066 | 0.21140672 |
| ENSMUSG00000056508 | -1.098933705 | 0.62358526 |
| ENSMUSG00000056509 | 1 | |
| ENSMUSG00000056515 | 1.197674861 | 0.462035952 |
| ENSMUSG00000056529 | -1.143474492 | 0.66920741 |
| ENSMUSG00000056531 | 1.072242366 | 0.559108563 |
| ENSMUSG00000056536 | -1.012664765 | 1 |
| ENSMUSG00000056537 | 1.090544112 | 0.736688878 |
| ENSMUSG00000056544 | 1 | |
| ENSMUSG00000056553 | 1 | |
| ENSMUSG00000056564 | 1.059463094 | 0.608497229 |
| ENSMUSG00000056569 | -1.054169715 | 0.911116262 |
| ENSMUSG00000056579 | 1.052397826 | 0.858134675 |
| ENSMUSG00000056586 | 1 | |
| ENSMUSG00000056592 | -1.048435653 | 0.608497229 |
| ENSMUSG00000056596 | -1.019736358 | 0.942468602 |
| ENSMUSG00000056598 | -1.180070117 | 0.107689552 |
| ENSMUSG00000056600 | 1.132639128 | 0.217803313 |
| ENSMUSG00000056602 | -1.7052284 | 0.000411964 |
| ENSMUSG00000056605 | 1 | |
| ENSMUSG00000056606 | 1.028323353 | 0.951772039 |
| ENSMUSG00000056608 | -1.038364073 | 0.908323578 |
| ENSMUSG00000056612 | 1.108096836 | 0.485491618 |
| ENSMUSG00000056617 | -1.103388856 | 0.694333885 |
| ENSMUSG00000056629 | 1.020368728 | 0.949812939 |
| ENSMUSG00000056632 | 1 | |
| ENSMUSG00000056643 | -1.161737199 | 0.466963788 |
| ENSMUSG00000056648 | 1.126641668 | 0.173371035 |
| ENSMUSG00000056656 | 1.067302329 | 0.837704961 |
| ENSMUSG00000056665 | 1.043001824 | 0.793885968 |
| ENSMUSG00000056666 | -1.049108286 | 0.782202661 |
| ENSMUSG00000056671 | -1.071648796 | 0.495452692 |
| ENSMUSG00000056673 | 1.098934312 | 0.608497229 |
| ENSMUSG00000056679 | -1.116349841 | 0.233160773 |
| ENSMUSG00000056682 | -1.003834322 | 1 |
| ENSMUSG00000056687 | -1.004099163 | 1 |
| ENSMUSG00000056692 | 1.029311583 | 0.933770056 |
| ENSMUSG00000056696 | 1 | |
| ENSMUSG00000056698 | -1.038640913 | 0.747365146 |
| ENSMUSG00000056699 | 1 | |
| ENSMUSG00000056706 | 1 | |
| ENSMUSG00000056708 | 1.14008718 | 0.236438455 |
| ENSMUSG00000056716 | 1 | |
| ENSMUSG00000056718 | 1.059784936 | 0.755283958 |

| | | |
|---|---|---|
| ENSMUSG00000056724 | -1.034040694 | 0.815807184 |
| ENSMUSG00000056728 | 1 | |
| ENSMUSG00000056735 | -1.113134339 | 0.030913243 |
| ENSMUSG00000056737 | 1.253801585 | 0.044245893 |
| ENSMUSG00000056738 | 1 | |
| ENSMUSG00000056749 | 1.000431694 | 1 |
| ENSMUSG00000056752 | 1.059463094 | 0.698112512 |
| ENSMUSG00000056753 | -1.081698408 | 0.787256751 |
| ENSMUSG00000056755 | -1.063525418 | 0.743394675 |
| ENSMUSG00000056758 | 1.522127759 | 0.03311356 |
| ENSMUSG00000056763 | -1.013610157 | 0.982838256 |
| ENSMUSG00000056770 | 1.086549178 | 0.690669961 |
| ENSMUSG00000056771 | -1.29677824 | 0.247650986 |
| ENSMUSG00000056772 | 1.069068979 | 0.698112512 |
| ENSMUSG00000056777 | -1 | 1 |
| ENSMUSG00000056782 | 1 | |
| ENSMUSG00000056797 | 1.059463094 | 0.704170349 |
| ENSMUSG00000056812 | -1.043306199 | 0.698112512 |
| ENSMUSG00000056815 | 1 | |
| ENSMUSG00000056820 | -1.175205119 | 0.386289966 |
| ENSMUSG00000056821 | 1.034865357 | 0.899648217 |
| ENSMUSG00000056822 | 1 | |
| ENSMUSG00000056824 | -1.03962336 | 0.698112512 |
| ENSMUSG00000056829 | 1 | |
| ENSMUSG00000056832 | 1.03921902 | 0.826909313 |
| ENSMUSG00000056836 | 1 | |
| ENSMUSG00000056849 | 1.065662724 | 0.808989997 |
| ENSMUSG00000056851 | 1.026601024 | 0.979002724 |
| ENSMUSG00000056853 | 1 | |
| ENSMUSG00000056854 | -1 | 1 |
| ENSMUSG00000056856 | 1.113683675 | 0.632857175 |
| ENSMUSG00000056858 | 1 | |
| ENSMUSG00000056863 | 1 | |
| ENSMUSG00000056870 | -1.071441666 | 0.75528471 |
| ENSMUSG00000056877 | 1.005205377 | 1 |
| ENSMUSG00000056880 | 1.07487334 | 0.656277822 |
| ENSMUSG00000056883 | 1 | |
| ENSMUSG00000056885 | 1 | |
| ENSMUSG00000056888 | 1.440817166 | 0.020282596 |
| ENSMUSG00000056895 | -1.151172433 | 0.606134298 |
| ENSMUSG00000056899 | 1.169058957 | 0.095111048 |
| ENSMUSG00000056900 | 1.018358366 | 0.94695034 |
| ENSMUSG00000056901 | 1 | |
| ENSMUSG00000056904 | 1.026020426 | 0.860197798 |
| ENSMUSG00000056910 | 1 | |
| ENSMUSG00000056912 | 1 | |
| ENSMUSG00000056917 | -1.050383148 | 0.70254369 |
| ENSMUSG00000056919 | -1.015022575 | 0.988047843 |
| ENSMUSG00000056921 | 1 | |
| ENSMUSG00000056926 | 1 | |
| ENSMUSG00000056938 | -1.057692235 | 0.608497229 |
| ENSMUSG00000056941 | -1.108286738 | 0.433615344 |
| ENSMUSG00000056943 | 1 | |
| ENSMUSG00000056946 | 1 | |
| ENSMUSG00000056947 | 1.484111699 | 0.126666392 |
| ENSMUSG00000056952 | 1.027136971 | 0.943072649 |
| ENSMUSG00000056959 | 1 | |
| ENSMUSG00000056961 | 1 | |
| ENSMUSG00000056962 | -1.007173943 | 1 |
| ENSMUSG00000056966 | -1.093247254 | 0.698112512 |
| ENSMUSG00000056972 | -1 | 1 |
| ENSMUSG00000056973 | 1 | |
| ENSMUSG00000056978 | 1 | |

| | | |
|---|---|---|
| ENSMUSG00000056987 | -1.044273782 | 0.698112512 |
| ENSMUSG00000056995 | 1 | |
| ENSMUSG00000056999 | 1.027031893 | 0.976498231 |
| ENSMUSG00000057000 | 1.015486245 | 0.972980766 |
| ENSMUSG00000057003 | 1 | |
| ENSMUSG00000057021 | -1 | 1 |
| ENSMUSG00000057036 | 1 | |
| ENSMUSG00000057037 | 1.059463094 | 0.608497229 |
| ENSMUSG00000057047 | -1.043306199 | 0.698112512 |
| ENSMUSG00000057050 | 1 | |
| ENSMUSG00000057054 | 1.036153619 | 0.812796034 |
| ENSMUSG00000057058 | -1.077378607 | 0.698112512 |
| ENSMUSG00000057060 | -1.051018293 | 0.893608405 |
| ENSMUSG00000057067 | 1 | |
| ENSMUSG00000057068 | 1 | |
| ENSMUSG00000057069 | -1.346344932 | 0.049437432 |
| ENSMUSG00000057072 | -1.153027569 | 0.608497229 |
| ENSMUSG00000057074 | -1.176621705 | 0.404695089 |
| ENSMUSG00000057092 | 1 | |
| ENSMUSG00000057093 | 1.003404092 | 1 |
| ENSMUSG00000057098 | 1.497328229 | 0.006288332 |
| ENSMUSG00000057101 | 1.080980764 | 0.608497229 |
| ENSMUSG00000057103 | 1.091480198 | 0.783644184 |
| ENSMUSG00000057110 | 1.007935378 | 1 |
| ENSMUSG00000057113 | -1.032779733 | 0.979629796 |
| ENSMUSG00000057116 | 1.116785864 | 0.608497229 |
| ENSMUSG00000057123 | 1.019083579 | 0.949812939 |
| ENSMUSG00000057130 | 1.079683596 | 0.612124048 |
| ENSMUSG00000057132 | -1.143649388 | 0.316966507 |
| ENSMUSG00000057133 | 1.017853984 | 0.982838256 |
| ENSMUSG00000057134 | 1.077926512 | 0.665539571 |
| ENSMUSG00000057135 | 1 | |
| ENSMUSG00000057137 | 1.153661004 | 0.124156493 |
| ENSMUSG00000057143 | -1.161662337 | 0.098922723 |
| ENSMUSG00000057147 | -1.040319194 | 0.831404582 |
| ENSMUSG00000057149 | 1 | |
| ENSMUSG00000057156 | -1.094390409 | 0.307805994 |
| ENSMUSG00000057157 | -1 | 1 |
| ENSMUSG00000057160 | 1 | |
| ENSMUSG00000057161 | 1 | |
| ENSMUSG00000057163 | 1 | |
| ENSMUSG00000057170 | 1 | |
| ENSMUSG00000057173 | -1.053549062 | 0.698112512 |
| ENSMUSG00000057174 | 1 | |
| ENSMUSG00000057176 | 1.052243889 | 0.822045875 |
| ENSMUSG00000057177 | 1.0130632 | 0.983973603 |
| ENSMUSG00000057179 | 1 | |
| ENSMUSG00000057181 | -1.097856107 | 0.649027513 |
| ENSMUSG00000057182 | -1.277349003 | 0.276229467 |
| ENSMUSG00000057191 | -1.194100432 | 0.608497229 |
| ENSMUSG00000057193 | 1.017105392 | 0.994869695 |
| ENSMUSG00000057195 | 1 | |
| ENSMUSG00000057203 | 1 | |
| ENSMUSG00000057207 | 1 | |
| ENSMUSG00000057215 | 1.060443017 | 0.846630937 |
| ENSMUSG00000057216 | 1 | |
| ENSMUSG00000057219 | -1.019366242 | 0.914904517 |
| ENSMUSG00000057223 | 1.015486245 | 0.985522133 |
| ENSMUSG00000057228 | -1.014844115 | 1 |
| ENSMUSG00000057229 | -1.034963139 | 0.726562954 |
| ENSMUSG00000057230 | 1.007512577 | 1 |
| ENSMUSG00000057234 | -1.080388256 | 0.323536946 |
| ENSMUSG00000057236 | 1.048736996 | 0.917688104 |

| | | |
|---|---|---|
| ENSMUSG00000057240 | 1 | |
| ENSMUSG00000057244 | -1.053549062 | 0.69923471 |
| ENSMUSG00000057246 | -1.03962336 | 0.698112512 |
| ENSMUSG00000057262 | -1.053549062 | 0.698112512 |
| ENSMUSG00000057265 | 1.084105502 | 0.59127359 |
| ENSMUSG00000057270 | 1 | |
| ENSMUSG00000057278 | -1.003522691 | 1 |
| ENSMUSG00000057280 | 1 | |
| ENSMUSG00000057286 | -1.413142383 | 0.012103059 |
| ENSMUSG00000057290 | -1.079873815 | 0.789529279 |
| ENSMUSG00000057315 | 1.599871036 | 0.000583614 |
| ENSMUSG00000057321 | -1.043306199 | 0.79291651 |
| ENSMUSG00000057322 | 1.030413461 | 0.946741635 |
| ENSMUSG00000057329 | 1.131659958 | 0.252937706 |
| ENSMUSG00000057335 | 1.083173271 | 0.774758227 |
| ENSMUSG00000057337 | 1.492268012 | 0.00039051 |
| ENSMUSG00000057342 | 1.070078401 | 0.698112512 |
| ENSMUSG00000057346 | -1.663791326 | 0.001911276 |
| ENSMUSG00000057349 | 1 | |
| ENSMUSG00000057351 | 1 | |
| ENSMUSG00000057359 | -1.008865818 | 1 |
| ENSMUSG00000057363 | -1.155683227 | 0.518298719 |
| ENSMUSG00000057367 | -1.076586199 | 0.698112512 |
| ENSMUSG00000057370 | 1.055042496 | 0.807892349 |
| ENSMUSG00000057375 | -1.073872758 | 0.711943039 |
| ENSMUSG00000057378 | 2.224571978 | 0.000658499 |
| ENSMUSG00000057381 | 1 | |
| ENSMUSG00000057388 | -1.007026562 | 1 |
| ENSMUSG00000057396 | -1.123723232 | 0.201035112 |
| ENSMUSG00000057400 | 1.059463094 | 0.608497229 |
| ENSMUSG00000057402 | -1.043306199 | 0.698112512 |
| ENSMUSG00000057406 | 1.065968033 | 0.851662555 |
| ENSMUSG00000057409 | 1.073313247 | 0.698112512 |
| ENSMUSG00000057411 | 1.042128679 | 0.75343312 |
| ENSMUSG00000057417 | 1 | |
| ENSMUSG00000057421 | 1.041489324 | 0.879602687 |
| ENSMUSG00000057424 | 1 | |
| ENSMUSG00000057425 | 1 | |
| ENSMUSG00000057439 | 1 | |
| ENSMUSG00000057440 | -1.101073917 | 0.608497229 |
| ENSMUSG00000057443 | 1 | |
| ENSMUSG00000057444 | 1 | |
| ENSMUSG00000057446 | 1.024693402 | 0.933031586 |
| ENSMUSG00000057447 | 1 | |
| ENSMUSG00000057454 | 1 | |
| ENSMUSG00000057455 | 1 | |
| ENSMUSG00000057457 | 1.082274033 | 0.312959918 |
| ENSMUSG00000057461 | 1 | |
| ENSMUSG00000057464 | 1 | |
| ENSMUSG00000057465 | 1.11166259 | 0.608497229 |
| ENSMUSG00000057469 | 1.160418109 | 0.334218464 |
| ENSMUSG00000057497 | -1.064844326 | 0.698112512 |
| ENSMUSG00000057499 | -1.11537573 | 0.698112512 |
| ENSMUSG00000057503 | 1 | |
| ENSMUSG00000057506 | -1.015515613 | 0.973126641 |
| ENSMUSG00000057513 | 1 | |
| ENSMUSG00000057522 | 1.118199802 | 0.645934895 |
| ENSMUSG00000057530 | 1.050113667 | 0.803999353 |
| ENSMUSG00000057531 | 1.0159206 | 0.977192639 |
| ENSMUSG00000057534 | 1 | |
| ENSMUSG00000057540 | 1 | |
| ENSMUSG00000057541 | 1.02084446 | 0.976788866 |
| ENSMUSG00000057551 | 1.027371884 | 0.93818309 |

| | | |
|---|---|---|
| ENSMUSG00000057554 | -1.092128933 | 0.698112512 |
| ENSMUSG00000057561 | 1.098994785 | 0.698112512 |
| ENSMUSG00000057564 | 1 | |
| ENSMUSG00000057572 | 1.088386827 | 0.608497229 |
| ENSMUSG00000057577 | 1.010016066 | 1 |
| ENSMUSG00000057580 | -1.282011559 | 0.212065431 |
| ENSMUSG00000057592 | 1 | |
| ENSMUSG00000057594 | -1.089701282 | 0.498779515 |
| ENSMUSG00000057596 | -1 | 1 |
| ENSMUSG00000057604 | 1.747860337 | 0.005465172 |
| ENSMUSG00000057605 | -1.049341547 | 0.942585548 |
| ENSMUSG00000057606 | -1.080029458 | 0.729059471 |
| ENSMUSG00000057609 | 1 | |
| ENSMUSG00000057612 | 1 | |
| ENSMUSG00000057614 | 1.524598221 | 0.019037078 |
| ENSMUSG00000057615 | 1 | |
| ENSMUSG00000057626 | 1 | |
| ENSMUSG00000057637 | -1.00648069 | 1 |
| ENSMUSG00000057649 | -1.148563087 | 0.541735653 |
| ENSMUSG00000057650 | 1 | |
| ENSMUSG00000057654 | 1 | |
| ENSMUSG00000057657 | 1.01393017 | 1 |
| ENSMUSG00000057666 | 1.190942954 | 0.479515459 |
| ENSMUSG00000057667 | -1.012693753 | 0.976080157 |
| ENSMUSG00000057672 | 1.108556448 | 0.182298956 |
| ENSMUSG00000057674 | 1 | |
| ENSMUSG00000057685 | 1.049411284 | 0.835701362 |
| ENSMUSG00000057691 | 1.014609712 | 0.941060294 |
| ENSMUSG00000057696 | -1 | 1 |
| ENSMUSG00000057697 | 1 | |
| ENSMUSG00000057699 | 1 | |
| ENSMUSG00000057706 | 1.37494634 | 0.000398998 |
| ENSMUSG00000057710 | 1 | |
| ENSMUSG00000057715 | 1.160809372 | 0.608497229 |
| ENSMUSG00000057716 | 1.038824817 | 0.948378604 |
| ENSMUSG00000057719 | -1.913185724 | 0.001174334 |
| ENSMUSG00000057722 | 1.05679065 | 0.887026138 |
| ENSMUSG00000057723 | 1 | |
| ENSMUSG00000057726 | 1.167094994 | 0.637975839 |
| ENSMUSG00000057729 | 1.222889447 | 0.186622057 |
| ENSMUSG00000057735 | 1 | |
| ENSMUSG00000057738 | -1.014749694 | 1 |
| ENSMUSG00000057751 | -3.416633818 | 0.000261679 |
| ENSMUSG00000057754 | 1 | |
| ENSMUSG00000057761 | 1 | |
| ENSMUSG00000057762 | -1 | 1 |
| ENSMUSG00000057766 | -1.243324574 | 0.02080008 |
| ENSMUSG00000057770 | 1 | |
| ENSMUSG00000057777 | 1 | |
| ENSMUSG00000057778 | 1.045166756 | 0.637081332 |
| ENSMUSG00000057788 | 1.054551631 | 0.698428398 |
| ENSMUSG00000057789 | -1.024989424 | 0.924108307 |
| ENSMUSG00000057799 | 1 | |
| ENSMUSG00000057801 | 1 | |
| ENSMUSG00000057802 | -1.181467149 | 0.567496595 |
| ENSMUSG00000057805 | 1 | |
| ENSMUSG00000057816 | 1.024693402 | 0.933031586 |
| ENSMUSG00000057817 | 1 | |
| ENSMUSG00000057818 | 1.075070132 | 0.725426917 |
| ENSMUSG00000057829 | 1 | |
| ENSMUSG00000057832 | 1 | |
| ENSMUSG00000057835 | -1.032346805 | 0.861027326 |
| ENSMUSG00000057836 | -1.03962336 | 0.698112512 |

| | | |
|---|---|---|
| ENSMUSG00000057841 | 1.015234234 | 1 |
| ENSMUSG00000057842 | -1.154941359 | 0.253189308 |
| ENSMUSG00000057855 | 1 | |
| ENSMUSG00000057858 | 1.050919253 | 0.794473471 |
| ENSMUSG00000057863 | 1.182871267 | 0.458642815 |
| ENSMUSG00000057880 | -2.032181761 | 0.002693688 |
| ENSMUSG00000057886 | -1.044273782 | 0.698112512 |
| ENSMUSG00000057890 | 1 | |
| ENSMUSG00000057894 | 1.091338796 | 0.544317329 |
| ENSMUSG00000057895 | -2.194715784 | 0.000247822 |
| ENSMUSG00000057897 | -1.140879418 | 0.310630871 |
| ENSMUSG00000057899 | 1.00177232 | 1 |
| ENSMUSG00000057903 | 1 | |
| ENSMUSG00000057913 | 1.059463094 | 0.794274354 |
| ENSMUSG00000057914 | 1.061225474 | 0.608497229 |
| ENSMUSG00000057924 | -1.063858042 | 0.613594304 |
| ENSMUSG00000057933 | -1 | 1 |
| ENSMUSG00000057945 | 1 | |
| ENSMUSG00000057948 | 1.141561261 | 0.244548392 |
| ENSMUSG00000057963 | 1.04532059 | 0.608497229 |
| ENSMUSG00000057967 | -1.251296474 | 0.143760215 |
| ENSMUSG00000057969 | 1.014057038 | 0.945195203 |
| ENSMUSG00000057977 | 1 | |
| ENSMUSG00000057981 | 1 | |
| ENSMUSG00000057982 | -1.07641238 | 0.535052277 |
| ENSMUSG00000057990 | 1.008034876 | 1 |
| ENSMUSG00000057997 | 1 | |
| ENSMUSG00000058005 | 1 | |
| ENSMUSG00000058006 | 1.061190992 | 0.824049093 |
| ENSMUSG00000058013 | 1.065606704 | 0.890994888 |
| ENSMUSG00000058014 | 1 | |
| ENSMUSG00000058017 | 1 | |
| ENSMUSG00000058019 | -1.023769022 | 0.971519533 |
| ENSMUSG00000058022 | -1.137017853 | 0.661501071 |
| ENSMUSG00000058028 | -1.044273782 | 0.698112512 |
| ENSMUSG00000058030 | 1 | |
| ENSMUSG00000058046 | -1.035196362 | 0.915070045 |
| ENSMUSG00000058050 | -1.168329485 | 0.148811477 |
| ENSMUSG00000058052 | 1 | |
| ENSMUSG00000058056 | 1.257289929 | 0.137973606 |
| ENSMUSG00000058057 | -1.016109348 | 1 |
| ENSMUSG00000058063 | 1 | |
| ENSMUSG00000058064 | 1.048105131 | 0.698112512 |
| ENSMUSG00000058070 | 1.382973473 | 0.025973012 |
| ENSMUSG00000058071 | 1 | |
| ENSMUSG00000058073 | 1 | |
| ENSMUSG00000058076 | -1.049604054 | 0.842619477 |
| ENSMUSG00000058078 | 1 | |
| ENSMUSG00000058084 | 1 | |
| ENSMUSG00000058093 | -1.078198383 | 0.608497229 |
| ENSMUSG00000058099 | -1.044273782 | 0.834556513 |
| ENSMUSG00000058101 | 1.019083579 | 0.982838256 |
| ENSMUSG00000058114 | 1 | |
| ENSMUSG00000058119 | 1 | |
| ENSMUSG00000058124 | 1 | |
| ENSMUSG00000058126 | 1.098399568 | 0.64090177 |
| ENSMUSG00000058132 | 1 | |
| ENSMUSG00000058135 | 1.215938257 | 0.096689828 |
| ENSMUSG00000058145 | -1.300993227 | 0.094855226 |
| ENSMUSG00000058147 | 1 | |
| ENSMUSG00000058152 | -1.347303906 | 0.010984356 |
| ENSMUSG00000058153 | 1 | |
| ENSMUSG00000058159 | 1.281182558 | 0.450580852 |

| | | |
|---|---|---|
| ENSMUSG00000058163 | -1.015793955 | 1 |
| ENSMUSG00000058172 | 1 | |
| ENSMUSG00000058173 | -1.056528821 | 0.727598186 |
| ENSMUSG00000058174 | 1.028323353 | 0.933031586 |
| ENSMUSG00000058176 | 1 | |
| ENSMUSG00000058183 | -1.056415273 | 0.780366278 |
| ENSMUSG00000058186 | 1.023743961 | 0.968735041 |
| ENSMUSG00000058188 | 1 | |
| ENSMUSG00000058192 | 1.017952234 | 0.952049609 |
| ENSMUSG00000058194 | 1 | |
| ENSMUSG00000058200 | 1 | |
| ENSMUSG00000058207 | 1.116785864 | 0.665646314 |
| ENSMUSG00000058216 | 1.069068979 | 0.698112512 |
| ENSMUSG00000058230 | 1.119873202 | 0.632326652 |
| ENSMUSG00000058239 | 1.127180666 | 0.208301876 |
| ENSMUSG00000058240 | -1.015009799 | 0.989996457 |
| ENSMUSG00000058244 | 1 | |
| ENSMUSG00000058246 | 1 | |
| ENSMUSG00000058248 | 1.082332121 | 0.816186113 |
| ENSMUSG00000058250 | 1 | |
| ENSMUSG00000058251 | 1 | |
| ENSMUSG00000058252 | -1.03962336 | 0.803072249 |
| ENSMUSG00000058254 | 1.21006292 | 0.022232253 |
| ENSMUSG00000058258 | -1.150587035 | 0.608497229 |
| ENSMUSG00000058260 | 1.414909481 | 0.053419075 |
| ENSMUSG00000058267 | -1.035348736 | 0.900667245 |
| ENSMUSG00000058270 | 1 | |
| ENSMUSG00000058275 | 1 | |
| ENSMUSG00000058281 | -1.000405949 | 1 |
| ENSMUSG00000058287 | -1.047668041 | 0.951480616 |
| ENSMUSG00000058290 | 1.011909631 | 1 |
| ENSMUSG00000058291 | 1.030810881 | 0.943006884 |
| ENSMUSG00000058295 | 1 | |
| ENSMUSG00000058297 | -1.310394753 | 0.063097135 |
| ENSMUSG00000058298 | -1.017231941 | 0.970096914 |
| ENSMUSG00000058301 | 1.046887567 | 0.755713814 |
| ENSMUSG00000058317 | 1.025587679 | 0.842500885 |
| ENSMUSG00000058318 | -1.005592947 | 1 |
| ENSMUSG00000058325 | 1.098026441 | 0.623655111 |
| ENSMUSG00000058328 | 1 | |
| ENSMUSG00000058331 | -1.11887805 | 0.124387865 |
| ENSMUSG00000058349 | 1 | |
| ENSMUSG00000058351 | -1.064098093 | 0.698112512 |
| ENSMUSG00000058354 | 1 | |
| ENSMUSG00000058355 | -1.036739371 | 0.959463747 |
| ENSMUSG00000058357 | -1.111880267 | 0.608497229 |
| ENSMUSG00000058360 | 1.072339777 | 0.50530478 |
| ENSMUSG00000058368 | 1 | |
| ENSMUSG00000058385 | -1 | 1 |
| ENSMUSG00000058388 | 1.056179587 | 0.785045707 |
| ENSMUSG00000058392 | 1.0198147 | 0.958245567 |
| ENSMUSG00000058396 | -1.196966665 | 0.60245982 |
| ENSMUSG00000058398 | -1.044273782 | 0.698112512 |
| ENSMUSG00000058399 | 1 | |
| ENSMUSG00000058400 | 1.116785864 | 0.578221424 |
| ENSMUSG00000058402 | 1.022016994 | 0.867408669 |
| ENSMUSG00000058407 | -1.051966914 | 0.865167999 |
| ENSMUSG00000058420 | 1.119720175 | 0.484584722 |
| ENSMUSG00000058427 | -1.335190589 | 0.18851436 |
| ENSMUSG00000058430 | -1 | 1 |
| ENSMUSG00000058435 | 1 | |
| ENSMUSG00000058440 | 1.006098863 | 1 |
| ENSMUSG00000058441 | -1.099493926 | 0.730102564 |

| | | |
|---|---|---|
| ENSMUSG00000058443 | -1.175523846 | 0.030989446 |
| ENSMUSG00000058444 | 1.010235439 | 1 |
| ENSMUSG00000058446 | 1.047545912 | 0.8349433 |
| ENSMUSG00000058447 | 1 | |
| ENSMUSG00000058454 | -1.182070343 | 0.16430711 |
| ENSMUSG00000058463 | 1 | |
| ENSMUSG00000058470 | 1 | |
| ENSMUSG00000058486 | 1.040206527 | 0.626061722 |
| ENSMUSG00000058488 | -1.121136286 | 0.66664952 |
| ENSMUSG00000058491 | 1 | |
| ENSMUSG00000058492 | 1.069068979 | 0.608497229 |
| ENSMUSG00000058498 | 1.084930669 | 0.866587403 |
| ENSMUSG00000058499 | 1 | |
| ENSMUSG00000058503 | -1.037120469 | 0.909141512 |
| ENSMUSG00000058508 | 1 | |
| ENSMUSG00000058513 | 1 | |
| ENSMUSG00000058515 | 1 | |
| ENSMUSG00000058521 | 1 | |
| ENSMUSG00000058523 | 1 | |
| ENSMUSG00000058537 | 1 | |
| ENSMUSG00000058542 | 1.122462048 | 0.601164141 |
| ENSMUSG00000058546 | -1.073430638 | 0.725610177 |
| ENSMUSG00000058550 | 1 | |
| ENSMUSG00000058558 | 1.128214922 | 0.695859665 |
| ENSMUSG00000058567 | 1.005851671 | 1 |
| ENSMUSG00000058568 | 1 | |
| ENSMUSG00000058569 | -1.0871657 | 0.70299173 |
| ENSMUSG00000058571 | 1.142876472 | 0.311020871 |
| ENSMUSG00000058579 | 1 | |
| ENSMUSG00000058581 | 1.059463094 | 0.698112512 |
| ENSMUSG00000058586 | -1.067526413 | 0.698112512 |
| ENSMUSG00000058587 | 1.033021064 | 0.955452914 |
| ENSMUSG00000058588 | 1 | |
| ENSMUSG00000058589 | 1.047842551 | 0.872692845 |
| ENSMUSG00000058590 | 1 | |
| ENSMUSG00000058594 | 1.0382042 | 0.87726305 |
| ENSMUSG00000058600 | 1.058286719 | 0.869055419 |
| ENSMUSG00000058603 | -1.073809062 | 0.709997537 |
| ENSMUSG00000058607 | -1.0965339 | 0.631890745 |
| ENSMUSG00000058613 | -1 | 1 |
| ENSMUSG00000058618 | 1 | |
| ENSMUSG00000058620 | -1.093674795 | 0.685046738 |
| ENSMUSG00000058622 | -1.320814303 | 0.021277269 |
| ENSMUSG00000058624 | -1.02419913 | 0.981214754 |
| ENSMUSG00000058625 | 1.103440022 | 0.795062098 |
| ENSMUSG00000058626 | -1.084645496 | 0.70299173 |
| ENSMUSG00000058628 | 1 | |
| ENSMUSG00000058631 | 1 | |
| ENSMUSG00000058638 | -1.004099528 | 1 |
| ENSMUSG00000058643 | 1.059463094 | 0.608497229 |
| ENSMUSG00000058655 | 1.083047058 | 0.814089245 |
| ENSMUSG00000058656 | 1.206951332 | 0.454244685 |
| ENSMUSG00000058659 | 1 | |
| ENSMUSG00000058662 | 1 | |
| ENSMUSG00000058665 | 1.170103677 | 0.116240509 |
| ENSMUSG00000058669 | -1.03962336 | 0.803072249 |
| ENSMUSG00000058670 | 1.005613439 | 1 |
| ENSMUSG00000058672 | -1.031412738 | 0.886454607 |
| ENSMUSG00000058685 | 1 | |
| ENSMUSG00000058690 | -1.116776812 | 0.615267463 |
| ENSMUSG00000058692 | 1 | |
| ENSMUSG00000058700 | 1.021812604 | 1 |
| ENSMUSG00000058704 | 1.103621893 | 0.615267463 |

| | | |
|---|---|---|
| ENSMUSG00000058706 | -1.00767177 | 1 |
| ENSMUSG00000058709 | 1.11727852 | 0.402963237 |
| ENSMUSG00000058715 | -1.063278095 | 0.730297627 |
| ENSMUSG00000058717 | 1 | |
| ENSMUSG00000058725 | 1 | |
| ENSMUSG00000058728 | 1 | |
| ENSMUSG00000058729 | -1.04994819 | 0.844780803 |
| ENSMUSG00000058731 | 1 | |
| ENSMUSG00000058740 | -1.282043321 | 0.109759063 |
| ENSMUSG00000058741 | -1.053549062 | 0.698112512 |
| ENSMUSG00000058743 | 1.054893018 | 0.938040364 |
| ENSMUSG00000058745 | 1 | |
| ENSMUSG00000058748 | 1.133596615 | 0.29498035 |
| ENSMUSG00000058755 | 1 | |
| ENSMUSG00000058756 | 1.116900231 | 0.339943743 |
| ENSMUSG00000058761 | 1.068488162 | 0.712653426 |
| ENSMUSG00000058773 | -1.098961918 | 0.72715392 |
| ENSMUSG00000058793 | -1.023986666 | 0.955887242 |
| ENSMUSG00000058794 | -1.009924012 | 1 |
| ENSMUSG00000058799 | 1.030101642 | 0.979002724 |
| ENSMUSG00000058802 | 1 | |
| ENSMUSG00000058806 | -1.043306199 | 0.698112512 |
| ENSMUSG00000058807 | 1 | |
| ENSMUSG00000058809 | -1.191126263 | 0.608497229 |
| ENSMUSG00000058812 | -1.03962336 | 0.698112512 |
| ENSMUSG00000058816 | 1 | |
| ENSMUSG00000058818 | 1.043130067 | 0.910832671 |
| ENSMUSG00000058820 | 1 | |
| ENSMUSG00000058831 | -1.065205545 | 0.867519242 |
| ENSMUSG00000058833 | 1.156141771 | 0.54086872 |
| ENSMUSG00000058835 | 1.07419572 | 0.746472926 |
| ENSMUSG00000058838 | 1 | |
| ENSMUSG00000058856 | 1 | |
| ENSMUSG00000058869 | 1.019083579 | 0.949812939 |
| ENSMUSG00000058873 | 1.069068979 | 0.608497229 |
| ENSMUSG00000058881 | 1.151444042 | 0.174701408 |
| ENSMUSG00000058883 | 1 | |
| ENSMUSG00000058884 | 1 | |
| ENSMUSG00000058886 | 1.047920169 | 0.469685706 |
| ENSMUSG00000058897 | -1.108293762 | 0.704170349 |
| ENSMUSG00000058900 | -1.326341096 | 0.111050955 |
| ENSMUSG00000058904 | 1 | |
| ENSMUSG00000058905 | 1.059463094 | 0.776953697 |
| ENSMUSG00000058908 | 1 | |
| ENSMUSG00000058914 | 1.000122148 | 1 |
| ENSMUSG00000058921 | -1.071075483 | 0.790851299 |
| ENSMUSG00000058922 | 1 | |
| ENSMUSG00000058923 | 1 | |
| ENSMUSG00000058925 | 1 | |
| ENSMUSG00000058927 | -1 | 1 |
| ENSMUSG00000058932 | 1 | |
| ENSMUSG00000058933 | 1.059463094 | 0.608497229 |
| ENSMUSG00000058934 | -1.043306199 | 0.79291651 |
| ENSMUSG00000058935 | 1 | |
| ENSMUSG00000058952 | 1.171563499 | 0.233412931 |
| ENSMUSG00000058953 | 1 | |
| ENSMUSG00000058966 | -1.297590421 | 0.434676351 |
| ENSMUSG00000058975 | -1.038444538 | 0.924629556 |
| ENSMUSG00000058976 | -1.003012737 | 1 |
| ENSMUSG00000058979 | -1.02699459 | 0.850218968 |
| ENSMUSG00000058981 | 1 | |
| ENSMUSG00000058997 | -1.113958516 | 0.577761644 |
| ENSMUSG00000059003 | -1.003567041 | 1 |

| | | |
|---|---|---|
| ENSMUSG00000059005 | -1.064433953 | 0.883306173 |
| ENSMUSG00000059013 | 1.049807005 | 0.862433631 |
| ENSMUSG00000059014 | 1 | |
| ENSMUSG00000059022 | -1.170128676 | 0.467093453 |
| ENSMUSG00000059023 | 1 | |
| ENSMUSG00000059027 | 1.026712557 | 0.966893814 |
| ENSMUSG00000059030 | 1 | |
| ENSMUSG00000059031 | 1 | |
| ENSMUSG00000059033 | 1.089470641 | 0.698112512 |
| ENSMUSG00000059040 | 1.180726182 | 0.464727974 |
| ENSMUSG00000059041 | 1 | |
| ENSMUSG00000059042 | 1 | |
| ENSMUSG00000059043 | 1 | |
| ENSMUSG00000059049 | 1.031646634 | 0.949812939 |
| ENSMUSG00000059058 | -1.252196876 | 0.13304549 |
| ENSMUSG00000059060 | 1.030113731 | 0.769724984 |
| ENSMUSG00000059064 | 1.05410549 | 0.805157604 |
| ENSMUSG00000059069 | 1 | |
| ENSMUSG00000059070 | 1.038135699 | 0.877334604 |
| ENSMUSG00000059077 | 1 | |
| ENSMUSG00000059087 | 1 | |
| ENSMUSG00000059089 | -1.043306199 | 0.698112512 |
| ENSMUSG00000059101 | 1 | |
| ENSMUSG00000059105 | 1 | |
| ENSMUSG00000059106 | 1 | |
| ENSMUSG00000059108 | -1 | 1 |
| ENSMUSG00000059112 | 1 | |
| ENSMUSG00000059113 | 1 | |
| ENSMUSG00000059114 | 1 | |
| ENSMUSG00000059119 | 1.037925563 | 0.899132938 |
| ENSMUSG00000059128 | 1 | |
| ENSMUSG00000059134 | 1 | |
| ENSMUSG00000059136 | 1 | |
| ENSMUSG00000059142 | 1.047130712 | 0.782203434 |
| ENSMUSG00000059146 | 2.0440807 | 0.000559529 |
| ENSMUSG00000059149 | 1.005251094 | 1 |
| ENSMUSG00000059150 | -1.044697704 | 0.89469925 |
| ENSMUSG00000059159 | 1 | |
| ENSMUSG00000059169 | 1 | |
| ENSMUSG00000059173 | -1.084480372 | 0.748484295 |
| ENSMUSG00000059179 | -1.044273782 | 0.790851299 |
| ENSMUSG00000059182 | 1.114877847 | 0.608497229 |
| ENSMUSG00000059183 | -1.042174092 | 0.769272045 |
| ENSMUSG00000059187 | -1.127622795 | 0.605400293 |
| ENSMUSG00000059189 | 1 | |
| ENSMUSG00000059195 | 1.302859051 | 0.255588649 |
| ENSMUSG00000059201 | 1 | |
| ENSMUSG00000059203 | 1 | |
| ENSMUSG00000059205 | 1 | |
| ENSMUSG00000059206 | 1 | |
| ENSMUSG00000059208 | 1.051040093 | 0.878223966 |
| ENSMUSG00000059213 | 1.14957175 | 0.105664704 |
| ENSMUSG00000059218 | 1 | |
| ENSMUSG00000059227 | 1 | |
| ENSMUSG00000059229 | 1.009112331 | 1 |
| ENSMUSG00000059230 | 1 | |
| ENSMUSG00000059237 | 1 | |
| ENSMUSG00000059244 | 1 | |
| ENSMUSG00000059246 | 1 | |
| ENSMUSG00000059248 | 1.151030867 | 0.469434161 |
| ENSMUSG00000059251 | 1 | |
| ENSMUSG00000059252 | 1.064176213 | 0.844780803 |
| ENSMUSG00000059256 | 1 | |

| | | |
|---|---|---|
| ENSMUSG00000059263 | 1.036407024 | 0.939634305 |
| ENSMUSG00000059273 | 1.100889745 | 0.537728798 |
| ENSMUSG00000059277 | 1.033816828 | 0.826909313 |
| ENSMUSG00000059278 | -1.041565239 | 0.738263421 |
| ENSMUSG00000059279 | 1 | |
| ENSMUSG00000059280 | 1 | |
| ENSMUSG00000059288 | 1.069873742 | 0.608497229 |
| ENSMUSG00000059291 | 1.00059727 | 1 |
| ENSMUSG00000059301 | 1.132639128 | 0.398767938 |
| ENSMUSG00000059303 | 1 | |
| ENSMUSG00000059305 | 1.034236077 | 0.941225684 |
| ENSMUSG00000059316 | -1.064607239 | 0.698112512 |
| ENSMUSG00000059319 | 1 | |
| ENSMUSG00000059323 | -1.029788792 | 0.925923334 |
| ENSMUSG00000059325 | 1.176530655 | 0.551136369 |
| ENSMUSG00000059326 | -1.580305212 | 0.011869806 |
| ENSMUSG00000059327 | 1.003116268 | 1 |
| ENSMUSG00000059334 | 1 | |
| ENSMUSG00000059336 | -1.089497311 | 0.608497229 |
| ENSMUSG00000059343 | -1.014836984 | 1 |
| ENSMUSG00000059352 | 1.059463094 | 0.608497229 |
| ENSMUSG00000059355 | 1.068940529 | 0.608497229 |
| ENSMUSG00000059361 | -1 | 1 |
| ENSMUSG00000059363 | -1.067084246 | 0.698112512 |
| ENSMUSG00000059366 | 1 | |
| ENSMUSG00000059371 | 1 | |
| ENSMUSG00000059375 | 1 | |
| ENSMUSG00000059379 | 1 | |
| ENSMUSG00000059382 | 1 | |
| ENSMUSG00000059383 | 1 | |
| ENSMUSG00000059395 | 1.010383044 | 1 |
| ENSMUSG00000059397 | 1 | |
| ENSMUSG00000059401 | 1.159797204 | 0.250477583 |
| ENSMUSG00000059406 | -1.009106367 | 1 |
| ENSMUSG00000059408 | -1.044273782 | 0.698112512 |
| ENSMUSG00000059409 | -1.03137338 | 0.95001542 |
| ENSMUSG00000059410 | 1 | |
| ENSMUSG00000059411 | 1 | |
| ENSMUSG00000059412 | 1 | |
| ENSMUSG00000059422 | -1.237212091 | 0.10378762 |
| ENSMUSG00000059423 | -1.075026069 | 0.691690748 |
| ENSMUSG00000059429 | -1 | 1 |
| ENSMUSG00000059430 | 5.695614414 | 0.000354604 |
| ENSMUSG00000059434 | -1.101277413 | 0.706116633 |
| ENSMUSG00000059436 | 1.2243804 | 0.137569458 |
| ENSMUSG00000059439 | 1.109332773 | 0.365835158 |
| ENSMUSG00000059447 | -1.009155316 | 1 |
| ENSMUSG00000059448 | 1 | |
| ENSMUSG00000059455 | -1.063525418 | 0.698112512 |
| ENSMUSG00000059456 | -1.487094296 | 0.008137765 |
| ENSMUSG00000059459 | 1 | |
| ENSMUSG00000059460 | 1 | |
| ENSMUSG00000059461 | -1.142372865 | 0.681952832 |
| ENSMUSG00000059463 | 1 | |
| ENSMUSG00000059473 | 1 | |
| ENSMUSG00000059474 | 1.03903248 | 0.872452681 |
| ENSMUSG00000059475 | -1.051847022 | 0.77638569 |
| ENSMUSG00000059479 | -1.245123057 | 0.605361141 |
| ENSMUSG00000059481 | 1 | |
| ENSMUSG00000059482 | 1.020253155 | 0.931930735 |
| ENSMUSG00000059483 | -1.024530911 | 0.961341674 |
| ENSMUSG00000059486 | 1.110831134 | 0.608497229 |
| ENSMUSG00000059488 | 1 | |

| | | |
|---|---|---|
| ENSMUSG00000059493 | 1.202302498 | 0.005803413 |
| ENSMUSG00000059495 | -1.02285562 | 0.979002724 |
| ENSMUSG00000059498 | -1.0795912 | 0.642639207 |
| ENSMUSG00000059503 | 1 | |
| ENSMUSG00000059504 | 1 | |
| ENSMUSG00000059511 | -1.043306199 | 0.698112512 |
| ENSMUSG00000059518 | -1.013989132 | 0.821450808 |
| ENSMUSG00000059534 | 1.051152032 | 0.698112512 |
| ENSMUSG00000059540 | -1.045817244 | 0.698112512 |
| ENSMUSG00000059547 | 1 | |
| ENSMUSG00000059552 | -1.086883631 | 0.698674506 |
| ENSMUSG00000059554 | -1.017333238 | 0.886637348 |
| ENSMUSG00000059555 | 1.021284254 | 0.641108184 |
| ENSMUSG00000059562 | -1.01865398 | 1 |
| ENSMUSG00000059565 | -1.03962336 | 0.808074244 |
| ENSMUSG00000059571 | 1.059463094 | 0.608497229 |
| ENSMUSG00000059585 | -1.132666855 | 0.375668528 |
| ENSMUSG00000059586 | 1.033868992 | 0.911832988 |
| ENSMUSG00000059588 | 1.378753879 | 0.003514048 |
| ENSMUSG00000059595 | 1 | |
| ENSMUSG00000059602 | 1.02252306 | 0.989742633 |
| ENSMUSG00000059606 | 1 | |
| ENSMUSG00000059610 | 1 | |
| ENSMUSG00000059623 | 1 | |
| ENSMUSG00000059625 | -1.03962336 | 0.803072249 |
| ENSMUSG00000059626 | 1 | |
| ENSMUSG00000059631 | -1.043306199 | 0.698112512 |
| ENSMUSG00000059632 | 1 | |
| ENSMUSG00000059639 | 1 | |
| ENSMUSG00000059645 | 1.059463094 | 0.608497229 |
| ENSMUSG00000059647 | 1 | |
| ENSMUSG00000059648 | 1 | |
| ENSMUSG00000059654 | 1 | |
| ENSMUSG00000059657 | 1 | |
| ENSMUSG00000059658 | -1.033820075 | 0.920928417 |
| ENSMUSG00000059659 | 1.060930255 | 0.616230752 |
| ENSMUSG00000059663 | 1 | |
| ENSMUSG00000059668 | 1 | |
| ENSMUSG00000059669 | 1.008473066 | 1 |
| ENSMUSG00000059674 | -1.180926054 | 0.106020265 |
| ENSMUSG00000059687 | 1 | |
| ENSMUSG00000059689 | 1.095301551 | 0.608497229 |
| ENSMUSG00000059690 | 1 | |
| ENSMUSG00000059695 | 1 | |
| ENSMUSG00000059706 | 1 | |
| ENSMUSG00000059708 | 1.105807689 | 0.541735653 |
| ENSMUSG00000059713 | -1.002305708 | 1 |
| ENSMUSG00000059714 | 1.118311771 | 0.10493663 |
| ENSMUSG00000059729 | 1 | |
| ENSMUSG00000059731 | 1 | |
| ENSMUSG00000059734 | 1.060232487 | 0.705910227 |
| ENSMUSG00000059741 | 1.095872691 | 0.657060635 |
| ENSMUSG00000059742 | 1.01798932 | 0.959892369 |
| ENSMUSG00000059743 | -1.198044872 | 0.003047978 |
| ENSMUSG00000059751 | 1.038307746 | 0.944721414 |
| ENSMUSG00000059762 | 1 | |
| ENSMUSG00000059763 | 1 | |
| ENSMUSG00000059768 | 1 | |
| ENSMUSG00000059772 | -1.034313629 | 0.67180178 |
| ENSMUSG00000059775 | 1.010351027 | 1 |
| ENSMUSG00000059776 | -1.226449446 | 0.273674424 |
| ENSMUSG00000059791 | -1.007437649 | 0.981516657 |
| ENSMUSG00000059796 | -1.042199993 | 0.951399668 |

| | | |
|---|---|---|
| ENSMUSG00000059810 | 1.142055919 | 0.014217336 |
| ENSMUSG00000059811 | 1.014758893 | 0.997582482 |
| ENSMUSG00000059816 | 1.019083579 | 0.949812939 |
| ENSMUSG00000059820 | -1.04450793 | 0.812379067 |
| ENSMUSG00000059821 | 1 | |
| ENSMUSG00000059824 | 1.444982657 | 0.003937082 |
| ENSMUSG00000059827 | 1 | |
| ENSMUSG00000059830 | 1.059463094 | 0.608497229 |
| ENSMUSG00000059832 | 1 | |
| ENSMUSG00000059834 | 1.027113836 | 0.911521144 |
| ENSMUSG00000059835 | -1.03962336 | 0.698112512 |
| ENSMUSG00000059839 | 1.016708206 | 0.946951599 |
| ENSMUSG00000059842 | -1.051967881 | 0.73684216 |
| ENSMUSG00000059845 | 1 | |
| ENSMUSG00000059851 | 1.031706768 | 0.807905456 |
| ENSMUSG00000059852 | 1 | |
| ENSMUSG00000059854 | -1.100193663 | 0.534887579 |
| ENSMUSG00000059857 | 1.102024305 | 0.635004673 |
| ENSMUSG00000059862 | 1 | |
| ENSMUSG00000059864 | 1.095872691 | 0.671280603 |
| ENSMUSG00000059866 | -1.010798937 | 0.97192334 |
| ENSMUSG00000059867 | 1 | |
| ENSMUSG00000059873 | 1 | |
| ENSMUSG00000059874 | 1 | |
| ENSMUSG00000059877 | 1 | |
| ENSMUSG00000059878 | 1.038648335 | 0.879253209 |
| ENSMUSG00000059883 | -1.01985979 | 0.894665389 |
| ENSMUSG00000059887 | 1 | |
| ENSMUSG00000059890 | -1.062022409 | 0.823872552 |
| ENSMUSG00000059891 | -1.044273782 | 0.698112512 |
| ENSMUSG00000059895 | 1.471561268 | 0.003770811 |
| ENSMUSG00000059897 | -1.03646951 | 0.794781383 |
| ENSMUSG00000059898 | 1.059463094 | 0.698112512 |
| ENSMUSG00000059900 | -1.471641239 | 0.040453793 |
| ENSMUSG00000059901 | 1.153465395 | 0.18635723 |
| ENSMUSG00000059908 | 1 | |
| ENSMUSG00000059910 | 1 | |
| ENSMUSG00000059912 | -1.075618801 | 0.797035369 |
| ENSMUSG00000059920 | 1.006354256 | 1 |
| ENSMUSG00000059921 | 1.236741243 | 0.107359862 |
| ENSMUSG00000059923 | 1.015323073 | 0.993518361 |
| ENSMUSG00000059934 | 1 | |
| ENSMUSG00000059939 | 1.002078949 | 1 |
| ENSMUSG00000059956 | 1 | |
| ENSMUSG00000059964 | 1 | |
| ENSMUSG00000059970 | 1.170967076 | 0.060970782 |
| ENSMUSG00000059972 | 1 | |
| ENSMUSG00000059974 | -1.044273782 | 0.870103307 |
| ENSMUSG00000059975 | -1.019487807 | 0.923414943 |
| ENSMUSG00000059981 | 1.0233978 | 0.88876099 |
| ENSMUSG00000059991 | 1.099348596 | 0.721785396 |
| ENSMUSG00000059994 | 1 | |
| ENSMUSG00000059995 | 1.152258813 | 0.346633686 |
| ENSMUSG00000059996 | 1 | |
| ENSMUSG00000060002 | -1.083796294 | 0.699591599 |
| ENSMUSG00000060012 | -1.181016414 | 0.181128445 |
| ENSMUSG00000060017 | 1 | |
| ENSMUSG00000060019 | 1.095929703 | 0.608497229 |
| ENSMUSG00000060024 | 1 | |
| ENSMUSG00000060029 | -1.047536087 | 0.894839966 |
| ENSMUSG00000060030 | 1 | |
| ENSMUSG00000060032 | 1.118896787 | 0.312623778 |
| ENSMUSG00000060034 | 1.194659242 | 0.534652019 |

| | | |
|---|---|---|
| ENSMUSG00000060036 | 1.121658269 | 0.699340382 |
| ENSMUSG00000060038 | 1.026778456 | 0.911907399 |
| ENSMUSG00000060044 | 1.445651738 | 0.007854617 |
| ENSMUSG00000060049 | 1 | |
| ENSMUSG00000060057 | 1 | |
| ENSMUSG00000060063 | -1.017305534 | 0.982854947 |
| ENSMUSG00000060068 | 1.009574969 | 1 |
| ENSMUSG00000060070 | 1 | |
| ENSMUSG00000060073 | 1.05137066 | 0.887325696 |
| ENSMUSG00000060081 | 1 | |
| ENSMUSG00000060084 | 1 | |
| ENSMUSG00000060090 | 1.055370406 | 0.812228807 |
| ENSMUSG00000060093 | 1 | |
| ENSMUSG00000060096 | -1.027395698 | 0.972188762 |
| ENSMUSG00000060098 | 1.020999059 | 0.956036866 |
| ENSMUSG00000060105 | 1 | |
| ENSMUSG00000060112 | 1 | |
| ENSMUSG00000060114 | 1 | |
| ENSMUSG00000060121 | 1.090469217 | 0.608497229 |
| ENSMUSG00000060126 | -1.086450787 | 0.818905441 |
| ENSMUSG00000060128 | 1.069293585 | 0.797105186 |
| ENSMUSG00000060131 | 1.616646177 | 0.020238393 |
| ENSMUSG00000060143 | -1.083542232 | 0.704170349 |
| ENSMUSG00000060147 | 1.000619402 | 1 |
| ENSMUSG00000060149 | 1.022222797 | 0.879253209 |
| ENSMUSG00000060152 | -1.122560952 | 0.50858486 |
| ENSMUSG00000060161 | 1 | |
| ENSMUSG00000060166 | 1.12965169 | 0.374499722 |
| ENSMUSG00000060170 | 1 | |
| ENSMUSG00000060176 | -1.258997045 | 0.056693403 |
| ENSMUSG00000060177 | 1 | |
| ENSMUSG00000060180 | 1 | |
| ENSMUSG00000060181 | 1.017890616 | 0.975875477 |
| ENSMUSG00000060183 | -2.111648411 | 0.00179575 |
| ENSMUSG00000060187 | -1 | 1 |
| ENSMUSG00000060188 | -1.058507348 | 0.860260228 |
| ENSMUSG00000060198 | 1.112843644 | 0.609147349 |
| ENSMUSG00000060201 | 1 | |
| ENSMUSG00000060204 | -1.08081673 | 0.698112512 |
| ENSMUSG00000060205 | -1.03962336 | 0.803072249 |
| ENSMUSG00000060206 | 1.137119499 | 0.439841781 |
| ENSMUSG00000060208 | 1 | |
| ENSMUSG00000060212 | 1.059463094 | 0.698112512 |
| ENSMUSG00000060216 | -1.051282929 | 0.705199845 |
| ENSMUSG00000060224 | 1.49918495 | 0.026646701 |
| ENSMUSG00000060227 | -1.105286526 | 0.685440298 |
| ENSMUSG00000060240 | -1 | 1 |
| ENSMUSG00000060244 | 1.064578333 | 0.549011445 |
| ENSMUSG00000060245 | 1 | |
| ENSMUSG00000060254 | 1 | |
| ENSMUSG00000060256 | 1 | |
| ENSMUSG00000060257 | 1 | |
| ENSMUSG00000060260 | 1.329379889 | 0.004854638 |
| ENSMUSG00000060261 | 1.059936641 | 0.829303578 |
| ENSMUSG00000060268 | 1.010855137 | 1 |
| ENSMUSG00000060275 | -1.485556844 | 0.064230353 |
| ENSMUSG00000060279 | 1.104383872 | 0.385188707 |
| ENSMUSG00000060284 | 1.006217587 | 1 |
| ENSMUSG00000060288 | -1.070580371 | 0.745293114 |
| ENSMUSG00000060301 | -1.070755153 | 0.769137942 |
| ENSMUSG00000060303 | 1 | |
| ENSMUSG00000060311 | 1 | |
| ENSMUSG00000060314 | -1.626584141 | 0.008761615 |

| | | |
|---|---|---|
| ENSMUSG00000060317 | 1 | |
| ENSMUSG00000060332 | 1.075324819 | 0.792444161 |
| ENSMUSG00000060335 | 1 | |
| ENSMUSG00000060336 | -1.051835231 | 0.698112512 |
| ENSMUSG00000060371 | 1.059463094 | 0.704170349 |
| ENSMUSG00000060373 | -1.09642672 | 0.757318144 |
| ENSMUSG00000060375 | 1 | |
| ENSMUSG00000060376 | 1.134208516 | 0.310267575 |
| ENSMUSG00000060377 | 1.059463094 | 0.608497229 |
| ENSMUSG00000060380 | -1.017156986 | 0.933583308 |
| ENSMUSG00000060397 | 1.049004255 | 0.608497229 |
| ENSMUSG00000060402 | -1.063525418 | 0.698112512 |
| ENSMUSG00000060404 | 1 | |
| ENSMUSG00000060407 | 1 | |
| ENSMUSG00000060411 | -1 | 1 |
| ENSMUSG00000060412 | 1 | |
| ENSMUSG00000060416 | 1 | |
| ENSMUSG00000060417 | 1 | |
| ENSMUSG00000060419 | 1 | |
| ENSMUSG00000060424 | 1 | |
| ENSMUSG00000060427 | -1.009690625 | 1 |
| ENSMUSG00000060429 | -1.065037659 | 0.84275122 |
| ENSMUSG00000060438 | 1.077582057 | 0.83493212 |
| ENSMUSG00000060441 | -1.172208875 | 0.597779413 |
| ENSMUSG00000060445 | -1.152299099 | 0.608497229 |
| ENSMUSG00000060450 | 1.03608231 | 0.911116262 |
| ENSMUSG00000060459 | 1 | |
| ENSMUSG00000060461 | 1.019083579 | 0.949812939 |
| ENSMUSG00000060467 | 1.222889447 | 0.195365519 |
| ENSMUSG00000060469 | 1 | |
| ENSMUSG00000060470 | 1.230592175 | 0.047894258 |
| ENSMUSG00000060475 | 1.036648091 | 0.931480186 |
| ENSMUSG00000060477 | 1.382794886 | 0.006100279 |
| ENSMUSG00000060480 | 1 | |
| ENSMUSG00000060487 | -1.099174267 | 0.608497229 |
| ENSMUSG00000060490 | 1 | |
| ENSMUSG00000060491 | 1 | |
| ENSMUSG00000060499 | -1 | 1 |
| ENSMUSG00000060503 | 1 | |
| ENSMUSG00000060508 | 1 | |
| ENSMUSG00000060509 | 1.024804955 | 1 |
| ENSMUSG00000060510 | 1.015637834 | 0.998367147 |
| ENSMUSG00000060512 | 1.009151095 | 1 |
| ENSMUSG00000060519 | -1.121022975 | 0.338831189 |
| ENSMUSG00000060523 | 1 | |
| ENSMUSG00000060530 | -1.043306199 | 0.698112512 |
| ENSMUSG00000060534 | -1.069502966 | 0.698112512 |
| ENSMUSG00000060538 | 1.061946889 | 0.698112512 |
| ENSMUSG00000060548 | -1.364976155 | 0.045654641 |
| ENSMUSG00000060550 | -1.802530356 | 0.000383911 |
| ENSMUSG00000060556 | 1 | |
| ENSMUSG00000060559 | 1.040172434 | 0.954748866 |
| ENSMUSG00000060560 | 1 | |
| ENSMUSG00000060565 | 1 | |
| ENSMUSG00000060566 | -1.089282607 | 0.531127798 |
| ENSMUSG00000060568 | 1.028323353 | 0.903464611 |
| ENSMUSG00000060572 | 1.015473959 | 0.979002724 |
| ENSMUSG00000060579 | -1.044273782 | 0.790851299 |
| ENSMUSG00000060583 | 1 | |
| ENSMUSG00000060586 | -1.089497311 | 0.681308543 |
| ENSMUSG00000060591 | 1.060175791 | 0.769215751 |
| ENSMUSG00000060594 | 1.074880396 | 0.515682352 |
| ENSMUSG00000060600 | 1.579817584 | 0.00747749 |

| | | |
|---|---|---|
| ENSMUSG00000060601 | 1.088606867 | 0.652535724 |
| ENSMUSG00000060613 | 1 | |
| ENSMUSG00000060615 | 1.059463094 | 0.698112512 |
| ENSMUSG00000060621 | 1.020631755 | 0.986431796 |
| ENSMUSG00000060630 | 1 | |
| ENSMUSG00000060636 | -1.154157092 | 0.537316682 |
| ENSMUSG00000060639 | -1.233191864 | 0.106919651 |
| ENSMUSG00000060647 | 1 | |
| ENSMUSG00000060657 | 1.108029573 | 0.614476325 |
| ENSMUSG00000060663 | 1 | |
| ENSMUSG00000060671 | 1.033880178 | 0.903169559 |
| ENSMUSG00000060673 | 1 | |
| ENSMUSG00000060675 | -1.193550451 | 0.148210003 |
| ENSMUSG00000060678 | 1 | |
| ENSMUSG00000060679 | -1.019311792 | 0.949812939 |
| ENSMUSG00000060680 | -1.089497311 | 0.532369657 |
| ENSMUSG00000060681 | -1.022109917 | 0.9607713 |
| ENSMUSG00000060685 | 1 | |
| ENSMUSG00000060688 | 1 | |
| ENSMUSG00000060691 | 1 | |
| ENSMUSG00000060699 | 1 | |
| ENSMUSG00000060701 | 1 | |
| ENSMUSG00000060703 | 1.162967675 | 0.125236397 |
| ENSMUSG00000060708 | -1.070651036 | 0.564553258 |
| ENSMUSG00000060715 | -1.099174267 | 0.535052277 |
| ENSMUSG00000060716 | 1.032607248 | 0.702074372 |
| ENSMUSG00000060724 | 1 | |
| ENSMUSG00000060726 | 1 | |
| ENSMUSG00000060727 | 1.032926687 | 0.879014609 |
| ENSMUSG00000060730 | -1.142260238 | 0.378191682 |
| ENSMUSG00000060733 | 1.051149493 | 0.847756749 |
| ENSMUSG00000060735 | 1.024693402 | 0.933031586 |
| ENSMUSG00000060738 | -1 | 1 |
| ENSMUSG00000060739 | -1.088455373 | 0.749192357 |
| ENSMUSG00000060742 | 1 | |
| ENSMUSG00000060743 | -1.014530325 | 0.898175526 |
| ENSMUSG00000060747 | 1 | |
| ENSMUSG00000060756 | -1.069502966 | 0.698112512 |
| ENSMUSG00000060759 | 1 | |
| ENSMUSG00000060765 | 1 | |
| ENSMUSG00000060771 | -1.058855041 | 0.698112512 |
| ENSMUSG00000060777 | -1.259193429 | 0.30034405 |
| ENSMUSG00000060780 | 1 | |
| ENSMUSG00000060787 | 1 | |
| ENSMUSG00000060791 | -1.148903398 | 0.698112512 |
| ENSMUSG00000060794 | 1 | |
| ENSMUSG00000060795 | 1 | |
| ENSMUSG00000060798 | 1.027390357 | 0.86093201 |
| ENSMUSG00000060802 | 1.042366301 | 0.926869014 |
| ENSMUSG00000060803 | 1.065228771 | 0.615544239 |
| ENSMUSG00000060807 | 1 | |
| ENSMUSG00000060808 | 1.065410325 | 0.792094459 |
| ENSMUSG00000060811 | 1 | |
| ENSMUSG00000060816 | 1 | |
| ENSMUSG00000060827 | 1 | |
| ENSMUSG00000060843 | -1.246152897 | 0.100818594 |
| ENSMUSG00000060860 | -1.01155061 | 1 |
| ENSMUSG00000060862 | 1.052281775 | 0.69923471 |
| ENSMUSG00000060863 | -1 | 1 |
| ENSMUSG00000060878 | 1.057478344 | 0.867016657 |
| ENSMUSG00000060879 | 1.024693402 | 0.933031586 |
| ENSMUSG00000060882 | -1.028348525 | 0.912425817 |
| ENSMUSG00000060888 | 1 | |

| | | |
|---|---|---|
| ENSMUSG00000060890 | 1.024693402 | 0.969087762 |
| ENSMUSG00000060904 | 1.021328584 | 0.992109211 |
| ENSMUSG00000060913 | 1 | |
| ENSMUSG00000060918 | 1 | |
| ENSMUSG00000060923 | 1.081369797 | 0.124648722 |
| ENSMUSG00000060924 | -1.085651418 | 0.615267463 |
| ENSMUSG00000060935 | 1.104075651 | 0.698112512 |
| ENSMUSG00000060938 | -1.065685771 | 0.822190115 |
| ENSMUSG00000060950 | 1.139983061 | 0.077274207 |
| ENSMUSG00000060956 | 1 | |
| ENSMUSG00000060961 | 1.101414845 | 0.615243243 |
| ENSMUSG00000060962 | -1 | 1 |
| ENSMUSG00000060967 | 1 | |
| ENSMUSG00000060969 | -1.804520844 | 0.00445975 |
| ENSMUSG00000060974 | 1 | |
| ENSMUSG00000060981 | -1.001876871 | 1 |
| ENSMUSG00000060985 | -1.429794106 | 0.024658538 |
| ENSMUSG00000060988 | 1.176455698 | 0.608497229 |
| ENSMUSG00000060989 | -1.096934747 | 0.698112512 |
| ENSMUSG00000060992 | -1.021606713 | 0.988424085 |
| ENSMUSG00000061000 | 1 | |
| ENSMUSG00000061013 | -1.032243607 | 0.755972525 |
| ENSMUSG00000061022 | 1 | |
| ENSMUSG00000061024 | 1.026499798 | 0.867408669 |
| ENSMUSG00000061028 | 1.027348225 | 0.768379691 |
| ENSMUSG00000061032 | -1.042126809 | 0.881302915 |
| ENSMUSG00000061039 | -1.099201174 | 0.698112512 |
| ENSMUSG00000061046 | 1.028681219 | 0.790836432 |
| ENSMUSG00000061048 | -1.299047926 | 0.013017191 |
| ENSMUSG00000061062 | -1.085380447 | 0.568304986 |
| ENSMUSG00000061065 | 1 | |
| ENSMUSG00000061068 | -1 | 1 |
| ENSMUSG00000061075 | 1 | |
| ENSMUSG00000061079 | -1.026161178 | 0.899906547 |
| ENSMUSG00000061080 | -1.071075483 | 0.698112512 |
| ENSMUSG00000061082 | -1.291477937 | 0.227172576 |
| ENSMUSG00000061086 | 1.108897852 | 0.43373562 |
| ENSMUSG00000061099 | 1.01968155 | 1 |
| ENSMUSG00000061100 | 1 | |
| ENSMUSG00000061104 | -1.100161397 | 0.807247723 |
| ENSMUSG00000061111 | 1.091306727 | 0.608497229 |
| ENSMUSG00000061115 | 1 | |
| ENSMUSG00000061118 | -1.07433426 | 0.532369657 |
| ENSMUSG00000061119 | -1.023308284 | 0.925151724 |
| ENSMUSG00000061126 | -1.063525418 | 0.751979636 |
| ENSMUSG00000061130 | 1.013374248 | 1 |
| ENSMUSG00000061132 | -1.231549888 | 0.45120857 |
| ENSMUSG00000061136 | 1.027533613 | 0.971164403 |
| ENSMUSG00000061143 | 1.215923748 | 0.019871675 |
| ENSMUSG00000061144 | 1 | |
| ENSMUSG00000061150 | 1 | |
| ENSMUSG00000061165 | 1 | |
| ENSMUSG00000061167 | 1 | |
| ENSMUSG00000061171 | 1.069068979 | 0.698112512 |
| ENSMUSG00000061175 | 1.183900224 | 0.358873127 |
| ENSMUSG00000061184 | 1.069068979 | 0.608497229 |
| ENSMUSG00000061186 | 1.167939418 | 0.003806532 |
| ENSMUSG00000061195 | 1 | |
| ENSMUSG00000061207 | -1.030952671 | 0.819614655 |
| ENSMUSG00000061208 | 1 | |
| ENSMUSG00000061210 | 1 | |
| ENSMUSG00000061232 | -1.43668562 | 0.054880317 |
| ENSMUSG00000061242 | -1.112010624 | 0.698112512 |

| | | |
|---|---|---|
| ENSMUSG00000061244 | -1.058012609 | 0.830906535 |
| ENSMUSG00000061246 | 1 | |
| ENSMUSG00000061259 | 1.009414875 | 1 |
| ENSMUSG00000061272 | 1 | |
| ENSMUSG00000061273 | -1.000767614 | 1 |
| ENSMUSG00000061286 | 1.106962224 | 0.554376355 |
| ENSMUSG00000061288 | 1.06222847 | 0.698112512 |
| ENSMUSG00000061292 | 1 | |
| ENSMUSG00000061295 | 1 | |
| ENSMUSG00000061296 | 1 | |
| ENSMUSG00000061298 | 1.022102852 | 0.955101438 |
| ENSMUSG00000061305 | 1 | |
| ENSMUSG00000061306 | 1.01253607 | 0.996972442 |
| ENSMUSG00000061311 | -1.043306199 | 0.698112512 |
| ENSMUSG00000061313 | -1.064446602 | 0.788549005 |
| ENSMUSG00000061315 | -1.004826936 | 1 |
| ENSMUSG00000061322 | -1.111880267 | 0.608497229 |
| ENSMUSG00000061330 | 1.068334851 | 0.79291651 |
| ENSMUSG00000061331 | 1.149495107 | 0.093799106 |
| ENSMUSG00000061336 | 1 | |
| ENSMUSG00000061342 | 1 | |
| ENSMUSG00000061353 | 1.53449947 | 0.085503298 |
| ENSMUSG00000061356 | -1.116853908 | 0.698112512 |
| ENSMUSG00000061360 | 1.013330462 | 1 |
| ENSMUSG00000061361 | 1 | |
| ENSMUSG00000061367 | 1 | |
| ENSMUSG00000061371 | 1.01260287 | 0.979486089 |
| ENSMUSG00000061374 | 1.043375696 | 0.664013268 |
| ENSMUSG00000061376 | 1 | |
| ENSMUSG00000061387 | 1 | |
| ENSMUSG00000061388 | 1 | |
| ENSMUSG00000061390 | 1.04424822 | 0.861137762 |
| ENSMUSG00000061392 | 1 | |
| ENSMUSG00000061393 | 1.125214997 | 0.650431683 |
| ENSMUSG00000061397 | -1.170083925 | 0.206620396 |
| ENSMUSG00000061410 | 1.081879831 | 0.630527041 |
| ENSMUSG00000061411 | -1.096729078 | 0.102900586 |
| ENSMUSG00000061414 | -1.043616943 | 0.929344107 |
| ENSMUSG00000061436 | 1.22038429 | 0.108466822 |
| ENSMUSG00000061451 | 1.122567075 | 0.427218243 |
| ENSMUSG00000061455 | -1.013455624 | 0.995292182 |
| ENSMUSG00000061457 | 1 | |
| ENSMUSG00000061458 | 1.035980182 | 0.887323956 |
| ENSMUSG00000061461 | 1.009324966 | 1 |
| ENSMUSG00000061462 | -1.400224148 | 0.298965563 |
| ENSMUSG00000061469 | 1 | |
| ENSMUSG00000061474 | -1.138531184 | 0.288139213 |
| ENSMUSG00000061477 | 1.073466331 | 0.78833944 |
| ENSMUSG00000061479 | 1.074883686 | 0.5742902 |
| ENSMUSG00000061482 | 1.089470641 | 0.608497229 |
| ENSMUSG00000061486 | 1.059463094 | 0.608497229 |
| ENSMUSG00000061488 | 1.171563499 | 0.233412931 |
| ENSMUSG00000061489 | 1 | |
| ENSMUSG00000061490 | 1 | |
| ENSMUSG00000061501 | 1 | |
| ENSMUSG00000061510 | -1.244151113 | 0.241127957 |
| ENSMUSG00000061517 | -1.440893229 | 0.076552197 |
| ENSMUSG00000061518 | -1.087223347 | 0.314360782 |
| ENSMUSG00000061520 | 1 | |
| ENSMUSG00000061524 | -2.05311233 | 7.61924E-05 |
| ENSMUSG00000061525 | 1.059463094 | 0.608497229 |
| ENSMUSG00000061527 | 1.028323353 | 0.903464611 |
| ENSMUSG00000061531 | 1.12691148 | 0.683683209 |

| | | |
|---|---|---|
| ENSMUSG00000061533 | 1.001036861 | 1 |
| ENSMUSG00000061535 | 1.059463094 | 0.608497229 |
| ENSMUSG00000061536 | 1.031406807 | 0.873118449 |
| ENSMUSG00000061540 | -1.085651418 | 0.532369657 |
| ENSMUSG00000061544 | -1.078485432 | 0.608497229 |
| ENSMUSG00000061549 | 1 | |
| ENSMUSG00000061559 | -1.081676901 | 0.719579506 |
| ENSMUSG00000061561 | 1 | |
| ENSMUSG00000061576 | 1.034095782 | 0.919000509 |
| ENSMUSG00000061577 | -1.000301086 | 1 |
| ENSMUSG00000061578 | -1.126344191 | 0.21967784 |
| ENSMUSG00000061584 | 1 | |
| ENSMUSG00000061589 | 1.050982838 | 0.71627176 |
| ENSMUSG00000061601 | -1.029113823 | 0.937805719 |
| ENSMUSG00000061602 | 1 | |
| ENSMUSG00000061603 | 1.251911808 | 0.002895506 |
| ENSMUSG00000061607 | 1.045391792 | 0.894246015 |
| ENSMUSG00000061613 | -1.021383848 | 0.977293219 |
| ENSMUSG00000061614 | 1 | |
| ENSMUSG00000061615 | 1 | |
| ENSMUSG00000061616 | 1 | |
| ENSMUSG00000061619 | 1 | |
| ENSMUSG00000061626 | 1 | |
| ENSMUSG00000061633 | -1.009476377 | 1 |
| ENSMUSG00000061637 | 1 | |
| ENSMUSG00000061650 | -1.127827587 | 0.203069723 |
| ENSMUSG00000061653 | 1 | |
| ENSMUSG00000061654 | 1.098695525 | 0.532369657 |
| ENSMUSG00000061665 | -1.020934513 | 0.988519386 |
| ENSMUSG00000061666 | -1.087790942 | 0.698112512 |
| ENSMUSG00000061669 | -1.089303093 | 0.608497229 |
| ENSMUSG00000061684 | -1.029876336 | 0.964202376 |
| ENSMUSG00000061689 | 1.054125681 | 0.641883501 |
| ENSMUSG00000061701 | 1 | |
| ENSMUSG00000061702 | 1.034384765 | 0.764618602 |
| ENSMUSG00000061707 | -1 | 1 |
| ENSMUSG00000061718 | -1.620859023 | 0.009478551 |
| ENSMUSG00000061723 | 1.271844968 | 0.202727447 |
| ENSMUSG00000061724 | 1 | |
| ENSMUSG00000061728 | -1 | 1 |
| ENSMUSG00000061731 | 1.272320582 | 0.246192072 |
| ENSMUSG00000061740 | -1.467137027 | 0.028539747 |
| ENSMUSG00000061742 | 1 | |
| ENSMUSG00000061751 | 1.432295272 | 0.001516441 |
| ENSMUSG00000061755 | 1.07609465 | 0.770671692 |
| ENSMUSG00000061758 | 1.105281551 | 0.484878393 |
| ENSMUSG00000061759 | 1.028547467 | 0.923602787 |
| ENSMUSG00000061762 | -1 | 1 |
| ENSMUSG00000061769 | 1.211527659 | 0.328602737 |
| ENSMUSG00000061777 | 1 | |
| ENSMUSG00000061778 | -1.128506834 | 0.608497229 |
| ENSMUSG00000061780 | -1 | 1 |
| ENSMUSG00000061787 | 1.02528487 | 0.97088369 |
| ENSMUSG00000061798 | 1 | |
| ENSMUSG00000061802 | 1 | |
| ENSMUSG00000061808 | 1 | |
| ENSMUSG00000061809 | 1 | |
| ENSMUSG00000061815 | 1.266936816 | 0.025022186 |
| ENSMUSG00000061816 | 1 | |
| ENSMUSG00000061825 | -1.084645496 | 0.532369657 |
| ENSMUSG00000061829 | 1 | |
| ENSMUSG00000061833 | 1.059463094 | 0.698112512 |
| ENSMUSG00000061838 | 1.013796668 | 1 |

| | | |
|---|---|---|
| ENSMUSG00000061843 | 1 | |
| ENSMUSG00000061845 | 1 | |
| ENSMUSG00000061847 | 1 | |
| ENSMUSG00000061848 | -1.052250106 | 0.884893906 |
| ENSMUSG00000061852 | 1 | |
| ENSMUSG00000061859 | 1.094816282 | 0.608497229 |
| ENSMUSG00000061863 | 1.069068979 | 0.698112512 |
| ENSMUSG00000061875 | 1 | |
| ENSMUSG00000061877 | 1 | |
| ENSMUSG00000061878 | -1.33869477 | 0.035634712 |
| ENSMUSG00000061882 | 1.084241799 | 0.734803948 |
| ENSMUSG00000061887 | 1.134563791 | 0.461880682 |
| ENSMUSG00000061894 | -1.005200477 | 1 |
| ENSMUSG00000061897 | 1.118530053 | 0.454669379 |
| ENSMUSG00000061898 | 1.018912243 | 0.932236811 |
| ENSMUSG00000061904 | -1.021093794 | 1 |
| ENSMUSG00000061906 | 1 | |
| ENSMUSG00000061911 | -1.044273782 | 0.698112512 |
| ENSMUSG00000061923 | 1.005613439 | 1 |
| ENSMUSG00000061928 | 1.028323353 | 0.936340683 |
| ENSMUSG00000061937 | -1.03962336 | 0.698112512 |
| ENSMUSG00000061947 | -1.044273782 | 0.698112512 |
| ENSMUSG00000061950 | 1.065848643 | 0.778150345 |
| ENSMUSG00000061952 | 1 | |
| ENSMUSG00000061958 | 1 | |
| ENSMUSG00000061959 | 1 | |
| ENSMUSG00000061961 | 1 | |
| ENSMUSG00000061969 | 1 | |
| ENSMUSG00000061972 | 1.165970392 | 0.479655048 |
| ENSMUSG00000061974 | -1 | 1 |
| ENSMUSG00000061975 | 1 | |
| ENSMUSG00000061977 | 1 | |
| ENSMUSG00000061979 | 1.095643276 | 0.332391635 |
| ENSMUSG00000061981 | 1.068631568 | 0.699340382 |
| ENSMUSG00000061983 | 1.082547846 | 0.710698346 |
| ENSMUSG00000061984 | 1 | |
| ENSMUSG00000061988 | -1.131382531 | 0.608497229 |
| ENSMUSG00000061991 | 1 | |
| ENSMUSG00000061992 | 1.071277874 | 0.860422396 |
| ENSMUSG00000062006 | -1.010492152 | 1 |
| ENSMUSG00000062007 | -1 | 1 |
| ENSMUSG00000062012 | -1.849999025 | 0.002321854 |
| ENSMUSG00000062014 | -1.184824753 | 0.562144433 |
| ENSMUSG00000062017 | 1 | |
| ENSMUSG00000062028 | 1 | |
| ENSMUSG00000062031 | 1.056080291 | 0.69237817 |
| ENSMUSG00000062036 | 1 | |
| ENSMUSG00000062037 | 1 | |
| ENSMUSG00000062038 | 1 | |
| ENSMUSG00000062040 | 1.007481265 | 1 |
| ENSMUSG00000062042 | 1 | |
| ENSMUSG00000062044 | 1.041689574 | 0.937505238 |
| ENSMUSG00000062046 | 1 | |
| ENSMUSG00000062054 | -1.019577576 | 0.911816741 |
| ENSMUSG00000062061 | 1 | |
| ENSMUSG00000062064 | -1.03962336 | 0.698112512 |
| ENSMUSG00000062070 | 1.064000287 | 0.850975923 |
| ENSMUSG00000062073 | 1.161036672 | 0.378751351 |
| ENSMUSG00000062074 | -1.087241753 | 0.619226796 |
| ENSMUSG00000062075 | -1.055741533 | 0.627944199 |
| ENSMUSG00000062077 | 1.738514572 | 0.014933732 |
| ENSMUSG00000062078 | 1.110716761 | 0.70299173 |
| ENSMUSG00000062081 | 1 | |

| | | |
|---|---|---|
| ENSMUSG00000062082 | 1.132639128 | 0.398767938 |
| ENSMUSG00000062083 | 1.006047292 | 1 |
| ENSMUSG00000062093 | -1.068039863 | 0.790851299 |
| ENSMUSG00000062098 | 1.019894964 | 0.922734941 |
| ENSMUSG00000062101 | 1.016399842 | 0.867181347 |
| ENSMUSG00000062103 | 1 | |
| ENSMUSG00000062105 | 1 | |
| ENSMUSG00000062110 | -1.067604505 | 0.608497229 |
| ENSMUSG00000062115 | 1.166472137 | 0.19899616 |
| ENSMUSG00000062116 | -1.044479718 | 0.698112512 |
| ENSMUSG00000062121 | 1 | |
| ENSMUSG00000062124 | -1 | 1 |
| ENSMUSG00000062127 | 1.029700978 | 0.87569322 |
| ENSMUSG00000062128 | 1 | |
| ENSMUSG00000062132 | -1 | 1 |
| ENSMUSG00000062142 | 1 | |
| ENSMUSG00000062148 | 1 | |
| ENSMUSG00000062151 | 1.293914 | 0.303135433 |
| ENSMUSG00000062152 | -1.124857661 | 0.223140601 |
| ENSMUSG00000062154 | 1 | |
| ENSMUSG00000062157 | -1.244714265 | 0.032575311 |
| ENSMUSG00000062162 | 1 | |
| ENSMUSG00000062165 | 1 | |
| ENSMUSG00000062168 | 1.026509487 | 0.938484564 |
| ENSMUSG00000062169 | -1.033436747 | 0.92650471 |
| ENSMUSG00000062170 | 1 | |
| ENSMUSG00000062175 | -1.155642677 | 0.518394456 |
| ENSMUSG00000062181 | 1 | |
| ENSMUSG00000062184 | -1.020369848 | 0.965120133 |
| ENSMUSG00000062186 | 1 | |
| ENSMUSG00000062190 | -1.048308163 | 0.808170084 |
| ENSMUSG00000062198 | 1.011170319 | 1 |
| ENSMUSG00000062199 | 1 | |
| ENSMUSG00000062200 | 1 | |
| ENSMUSG00000062201 | 1 | |
| ENSMUSG00000062202 | -1.031225656 | 0.896704424 |
| ENSMUSG00000062203 | 1.011509803 | 1 |
| ENSMUSG00000062204 | 1 | |
| ENSMUSG00000062209 | 1.059033181 | 0.801168497 |
| ENSMUSG00000062210 | -1.025985359 | 0.923192677 |
| ENSMUSG00000062224 | 1 | |
| ENSMUSG00000062226 | 1 | |
| ENSMUSG00000062232 | -1.059992256 | 0.755984843 |
| ENSMUSG00000062234 | 1.107386357 | 0.67362935 |
| ENSMUSG00000062235 | 1 | |
| ENSMUSG00000062245 | 1 | |
| ENSMUSG00000062248 | -1.055930736 | 0.787696557 |
| ENSMUSG00000062252 | 1 | |
| ENSMUSG00000062257 | -1.075293251 | 0.769346033 |
| ENSMUSG00000062270 | -1.022476629 | 0.993182545 |
| ENSMUSG00000062272 | 1 | |
| ENSMUSG00000062275 | 1 | |
| ENSMUSG00000062278 | 1 | |
| ENSMUSG00000062279 | 1.069068979 | 0.698112512 |
| ENSMUSG00000062280 | 1 | |
| ENSMUSG00000062282 | 1 | |
| ENSMUSG00000062284 | -1.210034041 | 0.597293005 |
| ENSMUSG00000062296 | -1.215486743 | 0.132291893 |
| ENSMUSG00000062300 | -1.110715048 | 0.147891125 |
| ENSMUSG00000062309 | 1.248168988 | 0.009616641 |
| ENSMUSG00000062310 | 1.06585783 | 0.856777856 |
| ENSMUSG00000062312 | 1.004141984 | 1 |
| ENSMUSG00000062314 | 1 | |

| | | |
|---|---|---|
| ENSMUSG00000062319 | 1 | |
| ENSMUSG00000062327 | 1.059463094 | 0.608497229 |
| ENSMUSG00000062328 | 1.010114546 | 1 |
| ENSMUSG00000062329 | 1 | |
| ENSMUSG00000062342 | 1.432250757 | 0.000851226 |
| ENSMUSG00000062345 | 1.887822283 | 0.001537274 |
| ENSMUSG00000062352 | 1.094698621 | 0.486179375 |
| ENSMUSG00000062353 | -1.093302381 | 0.698112512 |
| ENSMUSG00000062365 | 1.067861489 | 0.744030455 |
| ENSMUSG00000062369 | -1 | 1 |
| ENSMUSG00000062372 | 1 | |
| ENSMUSG00000062373 | 1.027094619 | 0.964442489 |
| ENSMUSG00000062376 | -1.09022619 | 0.301335308 |
| ENSMUSG00000062380 | 1.085196638 | 0.424254596 |
| ENSMUSG00000062382 | 1.020779405 | 0.970822886 |
| ENSMUSG00000062391 | 1 | |
| ENSMUSG00000062393 | -1.413471054 | 0.00859485 |
| ENSMUSG00000062397 | 1.051550059 | 0.905346712 |
| ENSMUSG00000062400 | 1 | |
| ENSMUSG00000062410 | 1.069293585 | 0.770958539 |
| ENSMUSG00000062421 | -1.167512382 | 0.443913715 |
| ENSMUSG00000062426 | 1 | |
| ENSMUSG00000062432 | 1.144363107 | 0.407080154 |
| ENSMUSG00000062433 | 1 | |
| ENSMUSG00000062434 | 1 | |
| ENSMUSG00000062438 | -1.132666855 | 0.391570427 |
| ENSMUSG00000062444 | -1.181812752 | 0.232325114 |
| ENSMUSG00000062456 | -1.02979833 | 0.985117569 |
| ENSMUSG00000062458 | 1 | |
| ENSMUSG00000062461 | 1 | |
| ENSMUSG00000062464 | -1.053549062 | 0.698112512 |
| ENSMUSG00000062470 | -1.021625676 | 0.88972256 |
| ENSMUSG00000062472 | -1.033820075 | 0.888246981 |
| ENSMUSG00000062477 | 1.014545335 | 0.989648415 |
| ENSMUSG00000062478 | 1 | |
| ENSMUSG00000062480 | -1.11442986 | 0.440702663 |
| ENSMUSG00000062483 | 1 | |
| ENSMUSG00000062488 | 1.142822568 | 0.247320454 |
| ENSMUSG00000062496 | 1.069068979 | 0.74773997 |
| ENSMUSG00000062497 | 1 | |
| ENSMUSG00000062510 | -1.09535258 | 0.608497229 |
| ENSMUSG00000062511 | 1.107416398 | 0.729018288 |
| ENSMUSG00000062515 | 1.42537756 | 0.113135627 |
| ENSMUSG00000062518 | 1.085624843 | 0.69305964 |
| ENSMUSG00000062519 | -1.084123241 | 0.594579756 |
| ENSMUSG00000062524 | 1 | |
| ENSMUSG00000062526 | 1.120583576 | 0.304966676 |
| ENSMUSG00000062527 | 1 | |
| ENSMUSG00000062528 | 1 | |
| ENSMUSG00000062542 | -1.380602596 | 0.020664923 |
| ENSMUSG00000062543 | -1 | 1 |
| ENSMUSG00000062545 | 1 | |
| ENSMUSG00000062546 | 1 | |
| ENSMUSG00000062551 | 1 | |
| ENSMUSG00000062553 | 1 | |
| ENSMUSG00000062554 | 1 | |
| ENSMUSG00000062556 | 1 | |
| ENSMUSG00000062561 | 1 | |
| ENSMUSG00000062563 | -1.088313104 | 0.754237006 |
| ENSMUSG00000062564 | 1 | |
| ENSMUSG00000062580 | -1.030916626 | 0.941510974 |
| ENSMUSG00000062582 | 1 | |
| ENSMUSG00000062585 | -1.1548396 | 0.490063384 |

| | | |
|---|---|---|
| ENSMUSG00000062588 | -1.193320102 | 0.056310079 |
| ENSMUSG00000062590 | 1.010778038 | 0.986394584 |
| ENSMUSG00000062591 | -1.066316765 | 0.478279422 |
| ENSMUSG00000062593 | 1 | |
| ENSMUSG00000062598 | 1 | |
| ENSMUSG00000062604 | 1.105683507 | 0.634663013 |
| ENSMUSG00000062608 | 1 | |
| ENSMUSG00000062609 | -1.049475223 | 0.890646431 |
| ENSMUSG00000062611 | -1.043306199 | 0.798487595 |
| ENSMUSG00000062619 | 1.108341926 | 0.489824652 |
| ENSMUSG00000062621 | 1 | |
| ENSMUSG00000062624 | 1 | |
| ENSMUSG00000062627 | -1.006133257 | 1 |
| ENSMUSG00000062629 | 1 | |
| ENSMUSG00000062638 | 1 | |
| ENSMUSG00000062646 | -1.041805346 | 0.608497229 |
| ENSMUSG00000062647 | 1.044112477 | 0.922790923 |
| ENSMUSG00000062649 | 1 | |
| ENSMUSG00000062651 | 1 | |
| ENSMUSG00000062661 | 1.116385659 | 0.608497229 |
| ENSMUSG00000062683 | 1.020806292 | 0.899827346 |
| ENSMUSG00000062687 | -1.073734115 | 0.812597809 |
| ENSMUSG00000062691 | -1.02382734 | 0.929284873 |
| ENSMUSG00000062694 | 1 | |
| ENSMUSG00000062704 | -1.227541538 | 0.245270161 |
| ENSMUSG00000062705 | 1 | |
| ENSMUSG00000062712 | 1 | |
| ENSMUSG00000062713 | -1.31589967 | 0.281564375 |
| ENSMUSG00000062727 | -1.043306199 | 0.698112512 |
| ENSMUSG00000062729 | -1.030407429 | 0.821783047 |
| ENSMUSG00000062732 | 1 | |
| ENSMUSG00000062737 | 1 | |
| ENSMUSG00000062742 | 1.154877441 | 0.608497229 |
| ENSMUSG00000062743 | 1.168445843 | 0.033141247 |
| ENSMUSG00000062751 | 1 | |
| ENSMUSG00000062753 | 1.320348809 | 0.107233877 |
| ENSMUSG00000062757 | 1 | |
| ENSMUSG00000062758 | -1.053549062 | 0.698112512 |
| ENSMUSG00000062760 | -1.369047528 | 0.016215951 |
| ENSMUSG00000062761 | 1.043297551 | 0.843609124 |
| ENSMUSG00000062762 | -1.036296007 | 0.92696103 |
| ENSMUSG00000062773 | 1 | |
| ENSMUSG00000062778 | 1 | |
| ENSMUSG00000062782 | 1 | |
| ENSMUSG00000062783 | 1.095872691 | 0.608497229 |
| ENSMUSG00000062785 | -1.612814461 | 0.041163273 |
| ENSMUSG00000062791 | 1 | |
| ENSMUSG00000062793 | 1 | |
| ENSMUSG00000062794 | -1.045032399 | 0.783204051 |
| ENSMUSG00000062797 | 1.009843844 | 1 |
| ENSMUSG00000062814 | 1 | |
| ENSMUSG00000062818 | 1 | |
| ENSMUSG00000062822 | 1.054150202 | 0.737905517 |
| ENSMUSG00000062825 | 1.330530131 | 0.456830077 |
| ENSMUSG00000062826 | 1.023743961 | 0.95029384 |
| ENSMUSG00000062833 | 1 | |
| ENSMUSG00000062844 | 1 | |
| ENSMUSG00000062846 | -1.053549062 | 0.698112512 |
| ENSMUSG00000062859 | -1.20296081 | 0.367760887 |
| ENSMUSG00000062861 | 1.014428934 | 0.872615712 |
| ENSMUSG00000062866 | -1.436977022 | 0.012307355 |
| ENSMUSG00000062867 | -1.029150632 | 0.955106668 |
| ENSMUSG00000062868 | 1 | |

| | | |
|---|---|---|
| ENSMUSG00000062873 | 1 | |
| ENSMUSG00000062874 | 1.17957616 | 0.006269391 |
| ENSMUSG00000062878 | 1 | |
| ENSMUSG00000062892 | 1 | |
| ENSMUSG00000062896 | 1.145662165 | 0.332484933 |
| ENSMUSG00000062901 | 1.322430655 | 0.054789251 |
| ENSMUSG00000062905 | -1.333774405 | 0.072111919 |
| ENSMUSG00000062906 | 1.025214324 | 0.879253209 |
| ENSMUSG00000062908 | 1.01213178 | 1 |
| ENSMUSG00000062914 | 1 | |
| ENSMUSG00000062929 | 1.087537743 | 0.756318264 |
| ENSMUSG00000062931 | -1.053717373 | 0.653078657 |
| ENSMUSG00000062933 | 1 | |
| ENSMUSG00000062937 | 1.023562246 | 0.987029357 |
| ENSMUSG00000062939 | 1.015486245 | 0.972980766 |
| ENSMUSG00000062944 | -1.049694502 | 0.403954009 |
| ENSMUSG00000062949 | -1.023000917 | 0.963386993 |
| ENSMUSG00000062952 | 1 | |
| ENSMUSG00000062960 | 1.320568445 | 0.143747578 |
| ENSMUSG00000062961 | -1.085651418 | 0.698112512 |
| ENSMUSG00000062962 | -1 | 1 |
| ENSMUSG00000062963 | -1.099020165 | 0.650713852 |
| ENSMUSG00000062980 | 1.579563187 | 0.000792101 |
| ENSMUSG00000062981 | -1.064551325 | 0.761101793 |
| ENSMUSG00000062987 | 1 | |
| ENSMUSG00000062991 | 1.247510269 | 0.028564893 |
| ENSMUSG00000062995 | 1.074848639 | 0.639884889 |
| ENSMUSG00000062997 | 1.027776111 | 0.929923764 |
| ENSMUSG00000063001 | 1.019083579 | 0.97045957 |
| ENSMUSG00000063011 | -1.173206643 | 0.318939382 |
| ENSMUSG00000063015 | 1.118905292 | 0.698112512 |
| ENSMUSG00000063018 | 1.224358958 | 0.027236743 |
| ENSMUSG00000063019 | -1.043972179 | 0.608497229 |
| ENSMUSG00000063020 | 1 | |
| ENSMUSG00000063021 | 1 | |
| ENSMUSG00000063047 | -1.048350319 | 0.698112512 |
| ENSMUSG00000063049 | 1.011213493 | 0.991874673 |
| ENSMUSG00000063052 | -1.042468494 | 0.879602687 |
| ENSMUSG00000063058 | 1 | |
| ENSMUSG00000063060 | 1.008707537 | 1 |
| ENSMUSG00000063063 | 1.129237035 | 0.50114325 |
| ENSMUSG00000063065 | 1.071745137 | 0.698112512 |
| ENSMUSG00000063077 | -1.003790439 | 1 |
| ENSMUSG00000063087 | 1.019965676 | 1 |
| ENSMUSG00000063089 | 1 | |
| ENSMUSG00000063106 | 1 | |
| ENSMUSG00000063108 | -1.109875444 | 0.608497229 |
| ENSMUSG00000063109 | -1.071075483 | 0.747163034 |
| ENSMUSG00000063116 | 1 | |
| ENSMUSG00000063120 | 1 | |
| ENSMUSG00000063129 | 1.019083579 | 0.968937442 |
| ENSMUSG00000063130 | 1 | |
| ENSMUSG00000063133 | 1 | |
| ENSMUSG00000063137 | 1 | |
| ENSMUSG00000063142 | -1.173142781 | 0.480458517 |
| ENSMUSG00000063145 | -1.079068845 | 0.608497229 |
| ENSMUSG00000063146 | 1.196354842 | 0.06980995 |
| ENSMUSG00000063157 | 1.014545335 | 0.979002724 |
| ENSMUSG00000063160 | 1.202698329 | 0.051664013 |
| ENSMUSG00000063163 | 1 | |
| ENSMUSG00000063166 | -1.132666855 | 0.18851436 |
| ENSMUSG00000063171 | -2.539445702 | 0.004310937 |
| ENSMUSG00000063172 | -1.005164007 | 1 |

| | | |
|---|---|---|
| ENSMUSG00000063173 | 1 | |
| ENSMUSG00000063176 | 1 | |
| ENSMUSG00000063177 | 1 | |
| ENSMUSG00000063179 | -1.004178105 | 1 |
| ENSMUSG00000063180 | 1.065410325 | 0.711602885 |
| ENSMUSG00000063188 | 1 | |
| ENSMUSG00000063193 | 1.516100548 | 0.01381213 |
| ENSMUSG00000063200 | -1.117356951 | 0.608497229 |
| ENSMUSG00000063206 | 1 | |
| ENSMUSG00000063216 | 1 | |
| ENSMUSG00000063221 | 1 | |
| ENSMUSG00000063225 | 1 | |
| ENSMUSG00000063229 | 1.113625784 | 0.70299173 |
| ENSMUSG00000063230 | 1 | |
| ENSMUSG00000063232 | 1.132639128 | 0.217803313 |
| ENSMUSG00000063234 | -1.089497311 | 0.532369657 |
| ENSMUSG00000063235 | -1.246257261 | 0.153133942 |
| ENSMUSG00000063236 | 1.041255564 | 0.698112512 |
| ENSMUSG00000063239 | -1.127622795 | 0.608497229 |
| ENSMUSG00000063240 | 1 | |
| ENSMUSG00000063245 | -1.363383671 | 0.042852019 |
| ENSMUSG00000063251 | 1 | |
| ENSMUSG00000063252 | 1 | |
| ENSMUSG00000063253 | -1.013481268 | 1 |
| ENSMUSG00000063254 | -1.004328165 | 1 |
| ENSMUSG00000063260 | 1 | |
| ENSMUSG00000063268 | -1.097260212 | 0.466407623 |
| ENSMUSG00000063273 | -1.028633386 | 0.972980766 |
| ENSMUSG00000063275 | 1.22318232 | 0.102254496 |
| ENSMUSG00000063277 | 1 | |
| ENSMUSG00000063281 | 1.004472176 | 1 |
| ENSMUSG00000063286 | 1.161036672 | 0.36974128 |
| ENSMUSG00000063296 | -1.878121203 | 0.005030788 |
| ENSMUSG00000063297 | -1.069502966 | 0.698112512 |
| ENSMUSG00000063305 | 1 | |
| ENSMUSG00000063314 | 1.036585803 | 0.917953029 |
| ENSMUSG00000063316 | 1.030584925 | 0.925403678 |
| ENSMUSG00000063317 | 1.106768731 | 0.608497229 |
| ENSMUSG00000063320 | -1.084062331 | 0.656734444 |
| ENSMUSG00000063328 | 1 | |
| ENSMUSG00000063334 | -1.014323774 | 0.999234847 |
| ENSMUSG00000063350 | 1 | |
| ENSMUSG00000063354 | -1.144713175 | 0.686194514 |
| ENSMUSG00000063358 | -1.003194485 | 1 |
| ENSMUSG00000063362 | -1.010738399 | 1 |
| ENSMUSG00000063364 | -1.079141397 | 0.759837182 |
| ENSMUSG00000063374 | 1 | |
| ENSMUSG00000063376 | 1 | |
| ENSMUSG00000063380 | 1 | |
| ENSMUSG00000063382 | 1.305632254 | 0.040133004 |
| ENSMUSG00000063383 | -1.084823522 | 0.239852828 |
| ENSMUSG00000063386 | 1 | |
| ENSMUSG00000063388 | -1.100193663 | 0.608497229 |
| ENSMUSG00000063394 | 1 | |
| ENSMUSG00000063406 | 1.000492509 | 1 |
| ENSMUSG00000063409 | 1.059463094 | 0.608497229 |
| ENSMUSG00000063410 | -1.102310763 | 0.611025891 |
| ENSMUSG00000063412 | -1 | 1 |
| ENSMUSG00000063415 | 1.335591956 | 0.000584777 |
| ENSMUSG00000063428 | 1 | |
| ENSMUSG00000063430 | -1.693782666 | 0.023767717 |
| ENSMUSG00000063434 | -1.003834322 | 1 |
| ENSMUSG00000063439 | -1.009748538 | 1 |

| | | |
|---|---|---|
| ENSMUSG00000063442 | 1.085431899 | 0.753332052 |
| ENSMUSG00000063445 | 1.027165244 | 0.883220031 |
| ENSMUSG00000063446 | -1 | 1 |
| ENSMUSG00000063447 | -1.03962336 | 0.698112512 |
| ENSMUSG00000063450 | 1.256655317 | 0.216046507 |
| ENSMUSG00000063455 | 1.060629689 | 0.698112512 |
| ENSMUSG00000063457 | 1.050220935 | 0.845772805 |
| ENSMUSG00000063458 | 1.075070132 | 0.78781961 |
| ENSMUSG00000063478 | 1 | |
| ENSMUSG00000063480 | -1.038415671 | 0.941524846 |
| ENSMUSG00000063488 | 1.050700135 | 0.730160452 |
| ENSMUSG00000063506 | -1.092822951 | 0.528416075 |
| ENSMUSG00000063507 | 1 | |
| ENSMUSG00000063511 | 1.01866177 | 0.96035552 |
| ENSMUSG00000063514 | 1.137739456 | 0.020664923 |
| ENSMUSG00000063522 | -1.036104018 | 0.778372272 |
| ENSMUSG00000063524 | 1.08972683 | 0.72715392 |
| ENSMUSG00000063529 | 1 | |
| ENSMUSG00000063531 | 1.060092027 | 0.726328988 |
| ENSMUSG00000063535 | -1.232409798 | 0.21055834 |
| ENSMUSG00000063543 | 1 | |
| ENSMUSG00000063548 | -1.044273782 | 0.801443616 |
| ENSMUSG00000063549 | 1 | |
| ENSMUSG00000063550 | 1.037476599 | 0.894679805 |
| ENSMUSG00000063556 | 1.059463094 | 0.608497229 |
| ENSMUSG00000063558 | 1.318654067 | 0.005312398 |
| ENSMUSG00000063564 | -2.261794838 | 0.001010847 |
| ENSMUSG00000063568 | -1.133671184 | 0.060093672 |
| ENSMUSG00000063576 | 1.044392744 | 0.782883548 |
| ENSMUSG00000063582 | 1 | |
| ENSMUSG00000063583 | 1 | |
| ENSMUSG00000063586 | -1.114167868 | 0.7616481 |
| ENSMUSG00000063590 | 1 | |
| ENSMUSG00000063594 | 1.21538509 | 0.587756011 |
| ENSMUSG00000063600 | 1 | |
| ENSMUSG00000063605 | 1.125537882 | 0.153011055 |
| ENSMUSG00000063611 | 1.034999297 | 0.910578003 |
| ENSMUSG00000063615 | 1 | |
| ENSMUSG00000063623 | -1.019290106 | 0.988424085 |
| ENSMUSG00000063626 | 1 | |
| ENSMUSG00000063628 | -1.053549062 | 0.900651611 |
| ENSMUSG00000063632 | 1.270921403 | 0.171373389 |
| ENSMUSG00000063646 | -1.116751362 | 0.769724984 |
| ENSMUSG00000063651 | -1.099174267 | 0.535052277 |
| ENSMUSG00000063652 | -1.124043488 | 0.433678407 |
| ENSMUSG00000063656 | 1 | |
| ENSMUSG00000063659 | 1.204208529 | 0.050060731 |
| ENSMUSG00000063660 | -1.043306199 | 0.698112512 |
| ENSMUSG00000063661 | 1 | |
| ENSMUSG00000063663 | -1.11566726 | 0.608497229 |
| ENSMUSG00000063672 | 1 | |
| ENSMUSG00000063681 | -1.220152793 | 0.25494774 |
| ENSMUSG00000063683 | -1.008943507 | 1 |
| ENSMUSG00000063684 | 1.05410549 | 0.835701362 |
| ENSMUSG00000063687 | 1.068424186 | 0.741127148 |
| ENSMUSG00000063689 | -1.043306199 | 0.698112512 |
| ENSMUSG00000063694 | -1.01724858 | 0.977563419 |
| ENSMUSG00000063696 | 1.240273757 | 0.085275739 |
| ENSMUSG00000063698 | -1.053771361 | 0.717035356 |
| ENSMUSG00000063704 | -1.3213142 | 0.297527773 |
| ENSMUSG00000063713 | 1 | |
| ENSMUSG00000063714 | -1.100038584 | 0.540020537 |
| ENSMUSG00000063715 | 1 | |

| | | |
|---|---|---|
| ENSMUSG00000063727 | 2.644890819 | 0.001885969 |
| ENSMUSG00000063728 | 1 | |
| ENSMUSG00000063730 | -1.027295343 | 0.968937442 |
| ENSMUSG00000063732 | 1 | |
| ENSMUSG00000063739 | -1.05875693 | 0.930108107 |
| ENSMUSG00000063751 | -1.182680997 | 0.601527473 |
| ENSMUSG00000063754 | -1.257663304 | 0.257120957 |
| ENSMUSG00000063757 | -1.007285535 | 1 |
| ENSMUSG00000063760 | 1.102857227 | 0.608497229 |
| ENSMUSG00000063762 | 1 | |
| ENSMUSG00000063764 | 1 | |
| ENSMUSG00000063765 | -1.078516354 | 0.766240401 |
| ENSMUSG00000063767 | -1.058750716 | 0.725290225 |
| ENSMUSG00000063777 | 1 | |
| ENSMUSG00000063779 | 1 | |
| ENSMUSG00000063780 | 1 | |
| ENSMUSG00000063785 | 1.086621519 | 0.681952832 |
| ENSMUSG00000063787 | -1.141459399 | 0.586009346 |
| ENSMUSG00000063796 | -1.03962336 | 0.698112512 |
| ENSMUSG00000063800 | -1.053738338 | 0.838070504 |
| ENSMUSG00000063801 | -1.066704888 | 0.725046589 |
| ENSMUSG00000063802 | 1.114233793 | 0.608497229 |
| ENSMUSG00000063804 | 1.110124334 | 0.804807576 |
| ENSMUSG00000063808 | -1.042708494 | 0.698112512 |
| ENSMUSG00000063810 | 1.07736569 | 0.619006139 |
| ENSMUSG00000063816 | -1.065647373 | 0.70299173 |
| ENSMUSG00000063820 | 1.036552144 | 0.96797322 |
| ENSMUSG00000063821 | 1 | |
| ENSMUSG00000063823 | 1 | |
| ENSMUSG00000063827 | 1 | |
| ENSMUSG00000063830 | -1.112424304 | 0.70299173 |
| ENSMUSG00000063838 | 1.052419246 | 0.698112512 |
| ENSMUSG00000063842 | 1 | |
| ENSMUSG00000063844 | 1 | |
| ENSMUSG00000063846 | 1 | |
| ENSMUSG00000063849 | 1.064954285 | 0.42877933 |
| ENSMUSG00000063851 | -1.205849876 | 0.532369657 |
| ENSMUSG00000063856 | 1.137733517 | 0.466407623 |
| ENSMUSG00000063867 | 1 | |
| ENSMUSG00000063870 | 1.046884846 | 0.914913657 |
| ENSMUSG00000063873 | 1.419460144 | 0.007471858 |
| ENSMUSG00000063875 | -1.147006962 | 0.608497229 |
| ENSMUSG00000063881 | 1 | |
| ENSMUSG00000063882 | -1.047868938 | 0.890065822 |
| ENSMUSG00000063884 | 1.010908067 | 1 |
| ENSMUSG00000063885 | 1.321214429 | 0.036626025 |
| ENSMUSG00000063887 | 1.143302785 | 0.560547927 |
| ENSMUSG00000063888 | -1.045827967 | 0.900730943 |
| ENSMUSG00000063889 | 1.055784736 | 0.624722543 |
| ENSMUSG00000063894 | -1.051469955 | 0.823428178 |
| ENSMUSG00000063895 | -1.114522426 | 0.619307317 |
| ENSMUSG00000063897 | 1.058831318 | 0.698112512 |
| ENSMUSG00000063900 | 1 | |
| ENSMUSG00000063902 | -1.080498498 | 0.837704961 |
| ENSMUSG00000063903 | 1.019083579 | 0.949812939 |
| ENSMUSG00000063904 | 1.021186325 | 0.967179343 |
| ENSMUSG00000063905 | 1 | |
| ENSMUSG00000063911 | -1.03962336 | 0.872692845 |
| ENSMUSG00000063919 | 1.934553027 | 0.002466243 |
| ENSMUSG00000063929 | 1 | |
| ENSMUSG00000063931 | 1.207970148 | 0.026727121 |
| ENSMUSG00000063932 | 1 | |
| ENSMUSG00000063935 | 1.103067059 | 0.750584613 |

| | | |
|---|---|---|
| ENSMUSG00000063952 | 1.046692717 | 0.71966832 |
| ENSMUSG00000063953 | -1.015417576 | 1 |
| ENSMUSG00000063954 | 1 | |
| ENSMUSG00000063971 | 1.448400711 | 0.059176789 |
| ENSMUSG00000063972 | 1.135658228 | 0.233888943 |
| ENSMUSG00000063975 | -1.229368627 | 0.045364838 |
| ENSMUSG00000063994 | 1 | |
| ENSMUSG00000063998 | 1 | |
| ENSMUSG00000064006 | 1 | |
| ENSMUSG00000064010 | 1 | |
| ENSMUSG00000064016 | 1.059014547 | 0.89411318 |
| ENSMUSG00000064023 | -1.251622541 | 0.439369489 |
| ENSMUSG00000064030 | -1.01019844 | 0.985522133 |
| ENSMUSG00000064032 | 1.055865073 | 0.709012023 |
| ENSMUSG00000064036 | -1.044273782 | 0.698112512 |
| ENSMUSG00000064037 | -1.017441435 | 0.982854947 |
| ENSMUSG00000064039 | 1 | |
| ENSMUSG00000064043 | 1.13318457 | 0.135282107 |
| ENSMUSG00000064044 | 1 | |
| ENSMUSG00000064052 | 1 | |
| ENSMUSG00000064057 | -1.044273782 | 0.698112512 |
| ENSMUSG00000064061 | -1.012971857 | 1 |
| ENSMUSG00000064063 | 1 | |
| ENSMUSG00000064065 | -1 | 1 |
| ENSMUSG00000064068 | 1.035404446 | 0.698112512 |
| ENSMUSG00000064080 | 1.218503686 | 0.251049598 |
| ENSMUSG00000064081 | 1.069068979 | 0.608497229 |
| ENSMUSG00000064084 | 1 | |
| ENSMUSG00000064090 | -1.110347957 | 0.595042751 |
| ENSMUSG00000064105 | 1.030997233 | 0.797105186 |
| ENSMUSG00000064109 | 1 | |
| ENSMUSG00000064110 | 1 | |
| ENSMUSG00000064115 | -1.410092105 | 0.05848516 |
| ENSMUSG00000064120 | 1.152361727 | 0.026120635 |
| ENSMUSG00000064121 | 1.122462048 | 0.215991492 |
| ENSMUSG00000064125 | -1.083352158 | 0.730297627 |
| ENSMUSG00000064127 | 1.005303893 | 1 |
| ENSMUSG00000064128 | -1.273229944 | 0.143050247 |
| ENSMUSG00000064129 | 1 | |
| ENSMUSG00000064137 | -1.013605297 | 1 |
| ENSMUSG00000064138 | -1.033027322 | 0.88284878 |
| ENSMUSG00000064140 | -1.03962336 | 0.803072249 |
| ENSMUSG00000064141 | 1.078187235 | 0.412406584 |
| ENSMUSG00000064145 | -1.021774495 | 0.956342203 |
| ENSMUSG00000064147 | 1 | |
| ENSMUSG00000064156 | 1 | |
| ENSMUSG00000064158 | -1.086192882 | 0.698112512 |
| ENSMUSG00000064165 | 1 | |
| ENSMUSG00000064168 | 1 | |
| ENSMUSG00000064177 | 1.062377276 | 0.778372272 |
| ENSMUSG00000064179 | -1.243844693 | 0.486930276 |
| ENSMUSG00000064181 | -1.049842332 | 0.801443616 |
| ENSMUSG00000064193 | 1.059463094 | 0.698112512 |
| ENSMUSG00000064194 | 1.003017659 | 1 |
| ENSMUSG00000064201 | 1 | |
| ENSMUSG00000064202 | 1.167586756 | 0.600288135 |
| ENSMUSG00000064208 | 1.059463094 | 0.608497229 |
| ENSMUSG00000064210 | 1.14245079 | 0.608497229 |
| ENSMUSG00000064213 | 1 | |
| ENSMUSG00000064215 | -1.679105979 | 0.001882191 |
| ENSMUSG00000064220 | 1 | |
| ENSMUSG00000064223 | 1 | |
| ENSMUSG00000064224 | 1 | |

| | | |
|---|---|---|
| ENSMUSG00000064225 | -1.557599024 | 0.017903363 |
| ENSMUSG00000064231 | 1.123955089 | 0.657709854 |
| ENSMUSG00000064232 | 1 | |
| ENSMUSG00000064246 | -1.03962336 | 0.731860565 |
| ENSMUSG00000064247 | 1.158687837 | 0.572275719 |
| ENSMUSG00000064252 | 1 | |
| ENSMUSG00000064254 | -1.046381344 | 0.698112512 |
| ENSMUSG00000064259 | 1 | |
| ENSMUSG00000064262 | -1.051018293 | 0.845175363 |
| ENSMUSG00000064263 | 1 | |
| ENSMUSG00000064264 | 1.056088622 | 0.707311809 |
| ENSMUSG00000064267 | 1.759544109 | 0.002321932 |
| ENSMUSG00000064272 | 1 | |
| ENSMUSG00000064280 | -1.066473521 | 0.819222912 |
| ENSMUSG00000064281 | 1.087063592 | 0.748191299 |
| ENSMUSG00000064284 | 1.048738907 | 0.644074123 |
| ENSMUSG00000064288 | -1.053549062 | 0.817479993 |
| ENSMUSG00000064289 | 1.022933024 | 0.968130843 |
| ENSMUSG00000064293 | 1 | |
| ENSMUSG00000064294 | 1 | |
| ENSMUSG00000064299 | 1 | |
| ENSMUSG00000064302 | 1.061070287 | 0.794763431 |
| ENSMUSG00000064307 | 1.112204142 | 0.493659452 |
| ENSMUSG00000064308 | -1.044273782 | 0.834556513 |
| ENSMUSG00000064310 | 1 | |
| ENSMUSG00000064317 | -1.03962336 | 0.872692845 |
| ENSMUSG00000064325 | -1.811001296 | 0.000107281 |
| ENSMUSG00000064326 | 1.232548541 | 0.158811406 |
| ENSMUSG00000064329 | 1.074990861 | 0.726328988 |
| ENSMUSG00000064330 | -1.042780998 | 0.8510899 |
| ENSMUSG00000064333 | 1 | |
| ENSMUSG00000064336 | 1.069068979 | 0.608497229 |
| ENSMUSG00000064337 | 1.2821573 | 0.242479982 |
| ENSMUSG00000064338 | 1.029490685 | 0.942171713 |
| ENSMUSG00000064339 | -1.012531697 | 1 |
| ENSMUSG00000064340 | -1.235966303 | 0.040939645 |
| ENSMUSG00000064341 | 1.07319423 | 0.908156606 |
| ENSMUSG00000064342 | -1.075899242 | 0.733570468 |
| ENSMUSG00000064343 | 1.115704272 | 0.548958635 |
| ENSMUSG00000064344 | -1.005970275 | 1 |
| ENSMUSG00000064345 | 1.089560068 | 0.848477314 |
| ENSMUSG00000064346 | -1.132401264 | 0.707458542 |
| ENSMUSG00000064347 | -1.090312011 | 0.501269661 |
| ENSMUSG00000064348 | -1.021934191 | 0.704457967 |
| ENSMUSG00000064349 | -1.020804638 | 0.951974561 |
| ENSMUSG00000064350 | 1.080135703 | 0.608497229 |
| ENSMUSG00000064351 | 1.023724865 | 1 |
| ENSMUSG00000064352 | 1.044950227 | 0.879253209 |
| ENSMUSG00000064353 | 1 | |
| ENSMUSG00000064354 | 1.162316572 | 0.710698346 |
| ENSMUSG00000064355 | -1.098765006 | 0.697665231 |
| ENSMUSG00000064356 | -1.036046696 | 0.934375176 |
| ENSMUSG00000064357 | 1.186481253 | 0.33242163 |
| ENSMUSG00000064358 | 1.063927547 | 0.798164432 |
| ENSMUSG00000064359 | -1.033209501 | 0.911346192 |
| ENSMUSG00000064360 | 1.158704861 | 0.370890497 |
| ENSMUSG00000064361 | -1.158096438 | 0.608497229 |
| ENSMUSG00000064363 | 1.118119939 | 0.762289856 |
| ENSMUSG00000064364 | -1.046151568 | 0.828564184 |
| ENSMUSG00000064365 | -1.27424025 | 0.561786114 |
| ENSMUSG00000064366 | -1.111764767 | 0.786711755 |
| ENSMUSG00000064367 | 1.117469308 | 0.777145624 |
| ENSMUSG00000064368 | 1.107993653 | 0.736194058 |

| | | |
|---|---|---|
| ENSMUSG00000064369 | 1.070956753 | 0.608497229 |
| ENSMUSG00000064370 | 1.083422699 | 0.880954906 |
| ENSMUSG00000064371 | -1.099949226 | 0.739716486 |
| ENSMUSG00000064372 | 1.017925207 | 0.933157611 |
| ENSMUSG00000064373 | 1.149921413 | 0.224411741 |
| ENSMUSG00000064377 | 1 | |
| ENSMUSG00000064380 | -1.061000526 | 0.875999519 |
| ENSMUSG00000064382 | 1 | |
| ENSMUSG00000064386 | 1 | |
| ENSMUSG00000064387 | 1 | |
| ENSMUSG00000064389 | -1.071075483 | 0.698112512 |
| ENSMUSG00000064390 | -1.163840271 | 0.608497229 |
| ENSMUSG00000064391 | 1 | |
| ENSMUSG00000064392 | 1 | |
| ENSMUSG00000064393 | 1 | |
| ENSMUSG00000064394 | 1 | |
| ENSMUSG00000064395 | 1 | |
| ENSMUSG00000064397 | 1 | |
| ENSMUSG00000064398 | 1.065410325 | 0.711602885 |
| ENSMUSG00000064399 | 1 | |
| ENSMUSG00000064400 | 1 | |
| ENSMUSG00000064403 | 1 | |
| ENSMUSG00000064405 | 1 | |
| ENSMUSG00000064407 | 1 | |
| ENSMUSG00000064408 | 1 | |
| ENSMUSG00000064409 | -1.085651418 | 0.532369657 |
| ENSMUSG00000064410 | 1 | |
| ENSMUSG00000064412 | 1 | |
| ENSMUSG00000064413 | 1 | |
| ENSMUSG00000064414 | 1 | |
| ENSMUSG00000064416 | 1 | |
| ENSMUSG00000064417 | 1.059463094 | 0.608497229 |
| ENSMUSG00000064419 | 1 | |
| ENSMUSG00000064422 | -1.044273782 | 0.698112512 |
| ENSMUSG00000064424 | 1 | |
| ENSMUSG00000064427 | 1.129237035 | 0.541735653 |
| ENSMUSG00000064428 | 1 | |
| ENSMUSG00000064429 | 1 | |
| ENSMUSG00000064432 | 1 | |
| ENSMUSG00000064433 | 1 | |
| ENSMUSG00000064435 | -1.037896228 | 0.916141522 |
| ENSMUSG00000064436 | 1 | |
| ENSMUSG00000064437 | -1.224815853 | 0.327408153 |
| ENSMUSG00000064440 | 1 | |
| ENSMUSG00000064441 | 1.0463724 | 0.869361059 |
| ENSMUSG00000064442 | 1 | |
| ENSMUSG00000064443 | 1 | |
| ENSMUSG00000064444 | 1 | |
| ENSMUSG00000064446 | 1 | |
| ENSMUSG00000064450 | 1 | |
| ENSMUSG00000064451 | -1.010960635 | 1 |
| ENSMUSG00000064452 | 1 | |
| ENSMUSG00000064453 | 1 | |
| ENSMUSG00000064455 | 1 | |
| ENSMUSG00000064460 | 1 | |
| ENSMUSG00000064463 | 1 | |
| ENSMUSG00000064464 | 1 | |
| ENSMUSG00000064467 | 1 | |
| ENSMUSG00000064468 | 1 | |
| ENSMUSG00000064471 | 1 | |
| ENSMUSG00000064472 | 1 | |
| ENSMUSG00000064476 | 1.059463094 | 0.608497229 |
| ENSMUSG00000064477 | 1 | |

| | | |
|---|---|---|
| ENSMUSG00000064481 | 1 | |
| ENSMUSG00000064485 | 1 | |
| ENSMUSG00000064486 | 1 | |
| ENSMUSG00000064487 | 1 | |
| ENSMUSG00000064489 | 1 | |
| ENSMUSG00000064490 | 1 | |
| ENSMUSG00000064491 | 1 | |
| ENSMUSG00000064492 | 1 | |
| ENSMUSG00000064493 | 1.053694875 | 0.924297242 |
| ENSMUSG00000064494 | 1.132639128 | 0.217803313 |
| ENSMUSG00000064495 | -1.095294215 | 0.698112512 |
| ENSMUSG00000064496 | 1 | |
| ENSMUSG00000064497 | 1 | |
| ENSMUSG00000064500 | 1 | |
| ENSMUSG00000064507 | 1 | |
| ENSMUSG00000064508 | 1 | |
| ENSMUSG00000064510 | 1 | |
| ENSMUSG00000064511 | 1 | |
| ENSMUSG00000064512 | 1 | |
| ENSMUSG00000064513 | 1 | |
| ENSMUSG00000064514 | -1.089693596 | 0.84019351 |
| ENSMUSG00000064515 | 1 | |
| ENSMUSG00000064517 | 1 | |
| ENSMUSG00000064519 | 1 | |
| ENSMUSG00000064522 | 1 | |
| ENSMUSG00000064524 | 1 | |
| ENSMUSG00000064525 | -1.109755328 | 0.749275715 |
| ENSMUSG00000064526 | 1 | |
| ENSMUSG00000064527 | 1 | |
| ENSMUSG00000064529 | 1 | |
| ENSMUSG00000064530 | 1.059463094 | 0.608497229 |
| ENSMUSG00000064532 | 1 | |
| ENSMUSG00000064536 | 1.059463094 | 0.608497229 |
| ENSMUSG00000064540 | 1 | |
| ENSMUSG00000064541 | 1 | |
| ENSMUSG00000064542 | -1.069502966 | 0.698112512 |
| ENSMUSG00000064543 | 1 | |
| ENSMUSG00000064545 | 1 | |
| ENSMUSG00000064547 | 1 | |
| ENSMUSG00000064548 | 1 | |
| ENSMUSG00000064549 | 1 | |
| ENSMUSG00000064550 | 1 | |
| ENSMUSG00000064551 | 1 | |
| ENSMUSG00000064553 | 1 | |
| ENSMUSG00000064554 | 1 | |
| ENSMUSG00000064555 | 1 | |
| ENSMUSG00000064556 | 1 | |
| ENSMUSG00000064558 | -1.03962336 | 0.803072249 |
| ENSMUSG00000064563 | 1 | |
| ENSMUSG00000064565 | 1 | |
| ENSMUSG00000064566 | -1.014920232 | 1 |
| ENSMUSG00000064567 | 1 | |
| ENSMUSG00000064569 | 1 | |
| ENSMUSG00000064570 | 1 | |
| ENSMUSG00000064572 | 1 | |
| ENSMUSG00000064573 | 1 | |
| ENSMUSG00000064574 | 1 | |
| ENSMUSG00000064575 | 1 | |
| ENSMUSG00000064577 | 1 | |
| ENSMUSG00000064579 | 1 | |
| ENSMUSG00000064580 | 1.024693402 | 0.933031586 |
| ENSMUSG00000064581 | 1 | |
| ENSMUSG00000064582 | 1 | |

| | | |
|---|---|---|
| ENSMUSG00000064584 | 1 | |
| ENSMUSG00000064585 | 1 | |
| ENSMUSG00000064586 | 1 | |
| ENSMUSG00000064587 | 1.059463094 | 0.608497229 |
| ENSMUSG00000064588 | 1 | |
| ENSMUSG00000064590 | 1.014545335 | 0.979002724 |
| ENSMUSG00000064591 | 1 | |
| ENSMUSG00000064595 | 1.128878069 | 0.608497229 |
| ENSMUSG00000064596 | 1 | |
| ENSMUSG00000064597 | -1 | 1 |
| ENSMUSG00000064600 | 1 | |
| ENSMUSG00000064601 | 1 | |
| ENSMUSG00000064602 | -1.107684718 | 0.752153213 |
| ENSMUSG00000064604 | 1 | |
| ENSMUSG00000064605 | 1 | |
| ENSMUSG00000064606 | 1 | |
| ENSMUSG00000064609 | -1 | 1 |
| ENSMUSG00000064612 | 1.051129405 | 0.878506664 |
| ENSMUSG00000064613 | 1 | |
| ENSMUSG00000064616 | 1 | |
| ENSMUSG00000064617 | 1 | |
| ENSMUSG00000064618 | 1 | |
| ENSMUSG00000064619 | 1 | |
| ENSMUSG00000064620 | -1 | 1 |
| ENSMUSG00000064621 | 1 | |
| ENSMUSG00000064622 | -1.03962336 | 0.838359329 |
| ENSMUSG00000064624 | 1 | |
| ENSMUSG00000064626 | 1 | |
| ENSMUSG00000064630 | 1 | |
| ENSMUSG00000064631 | 1 | |
| ENSMUSG00000064632 | 1 | |
| ENSMUSG00000064634 | -1.167557486 | 0.095607194 |
| ENSMUSG00000064637 | 1.089470641 | 0.713627281 |
| ENSMUSG00000064642 | 1 | |
| ENSMUSG00000064644 | 1 | |
| ENSMUSG00000064646 | -1 | 1 |
| ENSMUSG00000064647 | 1.143677979 | 0.565166554 |
| ENSMUSG00000064648 | 1.033906491 | 0.90616403 |
| ENSMUSG00000064652 | 1.014545335 | 0.979002724 |
| ENSMUSG00000064653 | 1 | |
| ENSMUSG00000064655 | 1 | |
| ENSMUSG00000064658 | -1.038656608 | 0.961183673 |
| ENSMUSG00000064659 | 1 | |
| ENSMUSG00000064663 | 1 | |
| ENSMUSG00000064664 | 1 | |
| ENSMUSG00000064665 | 1 | |
| ENSMUSG00000064666 | 1 | |
| ENSMUSG00000064669 | -1 | 1 |
| ENSMUSG00000064670 | 1 | |
| ENSMUSG00000064671 | 1 | |
| ENSMUSG00000064672 | -1.118547401 | 0.476577776 |
| ENSMUSG00000064673 | 1 | |
| ENSMUSG00000064675 | 1 | |
| ENSMUSG00000064677 | 1 | |
| ENSMUSG00000064679 | 1.069068979 | 0.608497229 |
| ENSMUSG00000064682 | 1 | |
| ENSMUSG00000064685 | 1 | |
| ENSMUSG00000064686 | 1 | |
| ENSMUSG00000064687 | 1 | |
| ENSMUSG00000064688 | 1 | |
| ENSMUSG00000064689 | 1 | |
| ENSMUSG00000064691 | 1 | |
| ENSMUSG00000064692 | -1.03962336 | 0.698112512 |

| | | |
|---|---|---|
| ENSMUSG00000064693 | 1 | |
| ENSMUSG00000064694 | 1 | |
| ENSMUSG00000064696 | 1 | |
| ENSMUSG00000064697 | 1 | |
| ENSMUSG00000064698 | 1 | |
| ENSMUSG00000064702 | 1 | |
| ENSMUSG00000064704 | 1 | |
| ENSMUSG00000064706 | 1 | |
| ENSMUSG00000064707 | 1 | |
| ENSMUSG00000064709 | 1 | |
| ENSMUSG00000064714 | 1 | |
| ENSMUSG00000064715 | 1 | |
| ENSMUSG00000064717 | 1 | |
| ENSMUSG00000064718 | 1 | |
| ENSMUSG00000064719 | 1 | |
| ENSMUSG00000064720 | 1 | |
| ENSMUSG00000064721 | 1.039960292 | 0.8618239 |
| ENSMUSG00000064722 | 1 | |
| ENSMUSG00000064724 | 1 | |
| ENSMUSG00000064725 | 1 | |
| ENSMUSG00000064726 | 1 | |
| ENSMUSG00000064727 | 1 | |
| ENSMUSG00000064728 | 1 | |
| ENSMUSG00000064731 | 1 | |
| ENSMUSG00000064732 | 1 | |
| ENSMUSG00000064734 | 1 | |
| ENSMUSG00000064735 | 1 | |
| ENSMUSG00000064737 | 1 | |
| ENSMUSG00000064738 | 1 | |
| ENSMUSG00000064739 | 1 | |
| ENSMUSG00000064740 | 1 | |
| ENSMUSG00000064741 | 1.00177232 | 1 |
| ENSMUSG00000064742 | 1 | |
| ENSMUSG00000064743 | 1 | |
| ENSMUSG00000064745 | 1 | |
| ENSMUSG00000064746 | 1 | |
| ENSMUSG00000064748 | 1 | |
| ENSMUSG00000064751 | 1 | |
| ENSMUSG00000064755 | 1 | |
| ENSMUSG00000064758 | 1 | |
| ENSMUSG00000064759 | 1 | |
| ENSMUSG00000064761 | 1 | |
| ENSMUSG00000064762 | 1 | |
| ENSMUSG00000064763 | 1 | |
| ENSMUSG00000064764 | 1 | |
| ENSMUSG00000064765 | -1.044273782 | 0.698112512 |
| ENSMUSG00000064766 | 1 | |
| ENSMUSG00000064767 | -1.07052105 | 0.842680402 |
| ENSMUSG00000064768 | 1 | |
| ENSMUSG00000064769 | 1 | |
| ENSMUSG00000064770 | 1.059463094 | 0.752398528 |
| ENSMUSG00000064772 | 1 | |
| ENSMUSG00000064775 | 1 | |
| ENSMUSG00000064776 | 1 | |
| ENSMUSG00000064777 | 1 | |
| ENSMUSG00000064778 | -1.013449573 | 1 |
| ENSMUSG00000064779 | 1 | |
| ENSMUSG00000064780 | 1 | |
| ENSMUSG00000064781 | 1 | |
| ENSMUSG00000064782 | 1 | |
| ENSMUSG00000064783 | 1 | |
| ENSMUSG00000064788 | 1 | |
| ENSMUSG00000064791 | 1 | |

| | | |
|---|---|---|
| ENSMUSG00000064792 | 1 | |
| ENSMUSG00000064793 | 1 | |
| ENSMUSG00000064795 | 1 | |
| ENSMUSG00000064796 | -1.044273782 | 0.698112512 |
| ENSMUSG00000064797 | -1.029779341 | 0.867433382 |
| ENSMUSG00000064798 | 1 | |
| ENSMUSG00000064800 | 1 | |
| ENSMUSG00000064801 | 1 | |
| ENSMUSG00000064802 | 1.033906491 | 0.911346192 |
| ENSMUSG00000064805 | 1 | |
| ENSMUSG00000064806 | 1 | |
| ENSMUSG00000064811 | 1.024693402 | 0.933031586 |
| ENSMUSG00000064814 | 1 | |
| ENSMUSG00000064816 | 1 | |
| ENSMUSG00000064818 | 1 | |
| ENSMUSG00000064822 | 1 | |
| ENSMUSG00000064823 | 1 | |
| ENSMUSG00000064824 | 1 | |
| ENSMUSG00000064829 | 1 | |
| ENSMUSG00000064830 | 1 | |
| ENSMUSG00000064831 | 1 | |
| ENSMUSG00000064833 | -1.043306199 | 0.79291651 |
| ENSMUSG00000064834 | 1 | |
| ENSMUSG00000064836 | 1 | |
| ENSMUSG00000064837 | 1 | |
| ENSMUSG00000064840 | 1 | |
| ENSMUSG00000064841 | 1 | |
| ENSMUSG00000064842 | 1 | |
| ENSMUSG00000064843 | 1 | |
| ENSMUSG00000064844 | 1.039408483 | 0.949812939 |
| ENSMUSG00000064845 | -1.058748442 | 0.84360497 |
| ENSMUSG00000064850 | 1 | |
| ENSMUSG00000064851 | 1 | |
| ENSMUSG00000064853 | -1.070757993 | 0.854127758 |
| ENSMUSG00000064854 | 1 | |
| ENSMUSG00000064855 | 1 | |
| ENSMUSG00000064856 | 1 | |
| ENSMUSG00000064857 | 1 | |
| ENSMUSG00000064858 | 1 | |
| ENSMUSG00000064859 | 1 | |
| ENSMUSG00000064860 | 1 | |
| ENSMUSG00000064862 | 1 | |
| ENSMUSG00000064867 | 1 | |
| ENSMUSG00000064871 | -1 | 1 |
| ENSMUSG00000064872 | 1 | |
| ENSMUSG00000064876 | 1 | |
| ENSMUSG00000064877 | 1 | |
| ENSMUSG00000064880 | 1 | |
| ENSMUSG00000064882 | 1 | |
| ENSMUSG00000064883 | 1 | |
| ENSMUSG00000064884 | 1 | |
| ENSMUSG00000064887 | 1 | |
| ENSMUSG00000064889 | 1 | |
| ENSMUSG00000064890 | 1 | |
| ENSMUSG00000064897 | 1 | |
| ENSMUSG00000064899 | 1 | |
| ENSMUSG00000064900 | 1 | |
| ENSMUSG00000064901 | -1.023016116 | 0.973242378 |
| ENSMUSG00000064904 | 1 | |
| ENSMUSG00000064905 | 1 | |
| ENSMUSG00000064906 | 1.069068979 | 0.698112512 |
| ENSMUSG00000064907 | 1 | |
| ENSMUSG00000064909 | 1 | |

| | | |
|---|---|---|
| ENSMUSG00000064915 | 1 | |
| ENSMUSG00000064916 | 1.019083579 | 0.968937442 |
| ENSMUSG00000064917 | -1.142727245 | 0.206719158 |
| ENSMUSG00000064918 | -1.011925048 | 1 |
| ENSMUSG00000064919 | -1.010830062 | 1 |
| ENSMUSG00000064921 | 1 | |
| ENSMUSG00000064923 | 1 | |
| ENSMUSG00000064925 | -1.043437345 | 0.908878383 |
| ENSMUSG00000064926 | 1 | |
| ENSMUSG00000064927 | -1.053549062 | 0.778372272 |
| ENSMUSG00000064928 | 1 | |
| ENSMUSG00000064930 | -1.049475223 | 0.833212373 |
| ENSMUSG00000064932 | 1 | |
| ENSMUSG00000064936 | -1.086192882 | 0.698112512 |
| ENSMUSG00000064938 | 1 | |
| ENSMUSG00000064939 | 1 | |
| ENSMUSG00000064940 | 1 | |
| ENSMUSG00000064941 | 1 | |
| ENSMUSG00000064943 | 1 | |
| ENSMUSG00000064945 | 1 | |
| ENSMUSG00000064948 | 1 | |
| ENSMUSG00000064949 | 1 | |
| ENSMUSG00000064952 | -1 | 1 |
| ENSMUSG00000064954 | 1 | |
| ENSMUSG00000064956 | 1 | |
| ENSMUSG00000064958 | 1 | |
| ENSMUSG00000064959 | 1 | |
| ENSMUSG00000064960 | -1.044273782 | 0.698112512 |
| ENSMUSG00000064961 | 1 | |
| ENSMUSG00000064964 | 1 | |
| ENSMUSG00000064966 | 1 | |
| ENSMUSG00000064967 | 1 | |
| ENSMUSG00000064968 | 1 | |
| ENSMUSG00000064970 | 1 | |
| ENSMUSG00000064971 | 1 | |
| ENSMUSG00000064972 | 1 | |
| ENSMUSG00000064977 | 1 | |
| ENSMUSG00000064981 | 1 | |
| ENSMUSG00000064982 | 1 | |
| ENSMUSG00000064983 | 1 | |
| ENSMUSG00000064984 | 1.083991456 | 0.760867819 |
| ENSMUSG00000064987 | 1 | |
| ENSMUSG00000064992 | 1 | |
| ENSMUSG00000064994 | -1.083415624 | 0.72770134 |
| ENSMUSG00000064995 | 1 | |
| ENSMUSG00000064999 | 1 | |
| ENSMUSG00000065004 | 1 | |
| ENSMUSG00000065005 | 1 | |
| ENSMUSG00000065006 | 1 | |
| ENSMUSG00000065007 | 1 | |
| ENSMUSG00000065011 | 1 | |
| ENSMUSG00000065012 | 1 | |
| ENSMUSG00000065013 | 1 | |
| ENSMUSG00000065014 | 1 | |
| ENSMUSG00000065016 | 1 | |
| ENSMUSG00000065017 | 1 | |
| ENSMUSG00000065018 | 1 | |
| ENSMUSG00000065022 | 1 | |
| ENSMUSG00000065027 | 1 | |
| ENSMUSG00000065028 | 1 | |
| ENSMUSG00000065029 | 1 | |
| ENSMUSG00000065030 | 1 | |
| ENSMUSG00000065031 | 1 | |

| | | |
|---|---|---|
| ENSMUSG00000065034 | 1 | |
| ENSMUSG00000065036 | 1.04424822 | 0.861137762 |
| ENSMUSG00000065037 | -1.232673695 | 0.608497229 |
| ENSMUSG00000065040 | 1 | |
| ENSMUSG00000065041 | -1.1106581 | 0.411217722 |
| ENSMUSG00000065042 | 1 | |
| ENSMUSG00000065044 | -1.038444538 | 0.895498153 |
| ENSMUSG00000065045 | -1 | 1 |
| ENSMUSG00000065047 | 1.059463094 | 0.704170349 |
| ENSMUSG00000065048 | 1 | |
| ENSMUSG00000065049 | 1.069068979 | 0.608497229 |
| ENSMUSG00000065050 | 1 | |
| ENSMUSG00000065053 | 1 | |
| ENSMUSG00000065055 | 1 | |
| ENSMUSG00000065057 | 1 | |
| ENSMUSG00000065061 | 1 | |
| ENSMUSG00000065068 | 1 | |
| ENSMUSG00000065070 | 1 | |
| ENSMUSG00000065072 | 1 | |
| ENSMUSG00000065074 | 1 | |
| ENSMUSG00000065078 | 1 | |
| ENSMUSG00000065079 | 1 | |
| ENSMUSG00000065080 | 1 | |
| ENSMUSG00000065083 | 1 | |
| ENSMUSG00000065084 | 1 | |
| ENSMUSG00000065086 | 1 | |
| ENSMUSG00000065087 | 1.022101861 | 0.867753156 |
| ENSMUSG00000065088 | 1 | |
| ENSMUSG00000065089 | 1.065410325 | 0.711602885 |
| ENSMUSG00000065090 | 1 | |
| ENSMUSG00000065091 | 1 | |
| ENSMUSG00000065094 | 1 | |
| ENSMUSG00000065095 | 1 | |
| ENSMUSG00000065096 | 1 | |
| ENSMUSG00000065097 | 1 | |
| ENSMUSG00000065100 | -1.044273782 | 0.698112512 |
| ENSMUSG00000065102 | 1 | |
| ENSMUSG00000065104 | 1 | |
| ENSMUSG00000065105 | 1 | |
| ENSMUSG00000065107 | 1 | |
| ENSMUSG00000065110 | 1 | |
| ENSMUSG00000065111 | 1 | |
| ENSMUSG00000065115 | 1 | |
| ENSMUSG00000065118 | -1.044273782 | 0.790851299 |
| ENSMUSG00000065122 | 1 | |
| ENSMUSG00000065124 | 1 | |
| ENSMUSG00000065126 | 1.147440272 | 0.698112512 |
| ENSMUSG00000065127 | -1 | 1 |
| ENSMUSG00000065130 | 1 | |
| ENSMUSG00000065132 | 1 | |
| ENSMUSG00000065138 | 1 | |
| ENSMUSG00000065141 | 1 | |
| ENSMUSG00000065143 | 1 | |
| ENSMUSG00000065145 | 1 | |
| ENSMUSG00000065147 | -1.131539196 | 0.200647863 |
| ENSMUSG00000065151 | 1 | |
| ENSMUSG00000065155 | 1 | |
| ENSMUSG00000065158 | 1 | |
| ENSMUSG00000065159 | 1 | |
| ENSMUSG00000065160 | 1 | |
| ENSMUSG00000065162 | -1.10188606 | 0.698112512 |
| ENSMUSG00000065164 | 1 | |
| ENSMUSG00000065165 | 1 | |

| | | |
|---|---|---|
| ENSMUSG00000065173 | 1 | |
| ENSMUSG00000065176 | 1 | |
| ENSMUSG00000065178 | 1 | |
| ENSMUSG00000065181 | 1 | |
| ENSMUSG00000065182 | -1.044273782 | 0.698112512 |
| ENSMUSG00000065185 | 1 | |
| ENSMUSG00000065187 | 1 | |
| ENSMUSG00000065188 | 1 | |
| ENSMUSG00000065191 | 1.212550913 | 0.411330377 |
| ENSMUSG00000065192 | 1 | |
| ENSMUSG00000065193 | -1.044273782 | 0.698112512 |
| ENSMUSG00000065195 | 1 | |
| ENSMUSG00000065196 | 1 | |
| ENSMUSG00000065198 | 1 | |
| ENSMUSG00000065199 | 1 | |
| ENSMUSG00000065200 | 1 | |
| ENSMUSG00000065201 | 1 | |
| ENSMUSG00000065202 | 1 | |
| ENSMUSG00000065203 | 1 | |
| ENSMUSG00000065204 | -1.053549062 | 0.698112512 |
| ENSMUSG00000065206 | 1 | |
| ENSMUSG00000065208 | -1.036055899 | 0.96035552 |
| ENSMUSG00000065211 | 1 | |
| ENSMUSG00000065212 | 1 | |
| ENSMUSG00000065215 | 1 | |
| ENSMUSG00000065219 | 1 | |
| ENSMUSG00000065220 | 1 | |
| ENSMUSG00000065221 | 1 | |
| ENSMUSG00000065223 | 1 | |
| ENSMUSG00000065226 | -1.120487809 | 0.608497229 |
| ENSMUSG00000065227 | 1 | |
| ENSMUSG00000065228 | 1 | |
| ENSMUSG00000065231 | 1 | |
| ENSMUSG00000065232 | -1 | 1 |
| ENSMUSG00000065233 | 1 | |
| ENSMUSG00000065235 | 1 | |
| ENSMUSG00000065236 | 1.048416787 | 0.891297795 |
| ENSMUSG00000065237 | 1 | |
| ENSMUSG00000065238 | 1 | |
| ENSMUSG00000065239 | 1 | |
| ENSMUSG00000065242 | 1 | |
| ENSMUSG00000065244 | 1 | |
| ENSMUSG00000065245 | 1 | |
| ENSMUSG00000065246 | 1 | |
| ENSMUSG00000065248 | 1 | |
| ENSMUSG00000065251 | 1 | |
| ENSMUSG00000065253 | 1 | |
| ENSMUSG00000065254 | -1.03962336 | 0.698112512 |
| ENSMUSG00000065255 | 1 | |
| ENSMUSG00000065256 | 1 | |
| ENSMUSG00000065258 | 1.139236427 | 0.599002265 |
| ENSMUSG00000065259 | -1.048277865 | 0.865583959 |
| ENSMUSG00000065260 | 1 | |
| ENSMUSG00000065261 | 1 | |
| ENSMUSG00000065262 | 1 | |
| ENSMUSG00000065265 | -1.0234022 | 0.927724559 |
| ENSMUSG00000065266 | 1 | |
| ENSMUSG00000065269 | 1 | |
| ENSMUSG00000065270 | 1 | |
| ENSMUSG00000065272 | 1 | |
| ENSMUSG00000065273 | 1.055416288 | 0.794274354 |
| ENSMUSG00000065274 | -1.03962336 | 0.698112512 |
| ENSMUSG00000065275 | 1 | |

| | | |
|---|---|---|
| ENSMUSG00000065277 | 1 | |
| ENSMUSG00000065279 | 1 | |
| ENSMUSG00000065280 | 1.058681077 | 0.734674233 |
| ENSMUSG00000065281 | -1.010245336 | 1 |
| ENSMUSG00000065282 | 1.021764721 | 0.924324458 |
| ENSMUSG00000065283 | 1 | |
| ENSMUSG00000065287 | 1 | |
| ENSMUSG00000065289 | 1 | |
| ENSMUSG00000065291 | 1 | |
| ENSMUSG00000065293 | 1 | |
| ENSMUSG00000065295 | 1 | |
| ENSMUSG00000065296 | -1 | 1 |
| ENSMUSG00000065297 | 1 | |
| ENSMUSG00000065298 | 1 | |
| ENSMUSG00000065299 | 1 | |
| ENSMUSG00000065301 | -1.053549062 | 0.733770982 |
| ENSMUSG00000065303 | 1 | |
| ENSMUSG00000065304 | 1.072851766 | 0.766115192 |
| ENSMUSG00000065305 | 1.015524901 | 0.982838256 |
| ENSMUSG00000065307 | 1 | |
| ENSMUSG00000065309 | 1 | |
| ENSMUSG00000065311 | 1 | |
| ENSMUSG00000065312 | 1 | |
| ENSMUSG00000065313 | 1 | |
| ENSMUSG00000065314 | 1 | |
| ENSMUSG00000065315 | -1.158708082 | 0.350957446 |
| ENSMUSG00000065319 | 1 | |
| ENSMUSG00000065320 | 1 | |
| ENSMUSG00000065321 | 1 | |
| ENSMUSG00000065322 | 1 | |
| ENSMUSG00000065327 | 1 | |
| ENSMUSG00000065328 | 1 | |
| ENSMUSG00000065329 | 1 | |
| ENSMUSG00000065330 | 1 | |
| ENSMUSG00000065331 | 1 | |
| ENSMUSG00000065333 | 1 | |
| ENSMUSG00000065336 | -1.01065523 | 1 |
| ENSMUSG00000065337 | 1 | |
| ENSMUSG00000065340 | 1 | |
| ENSMUSG00000065341 | -1.001738975 | 1 |
| ENSMUSG00000065343 | 1.100395091 | 0.698112512 |
| ENSMUSG00000065344 | 1 | |
| ENSMUSG00000065346 | 1 | |
| ENSMUSG00000065347 | 1 | |
| ENSMUSG00000065348 | 1 | |
| ENSMUSG00000065353 | 1 | |
| ENSMUSG00000065360 | 1.005613439 | 1 |
| ENSMUSG00000065361 | 1 | |
| ENSMUSG00000065362 | 1 | |
| ENSMUSG00000065367 | 1 | |
| ENSMUSG00000065368 | 1 | |
| ENSMUSG00000065371 | 1 | |
| ENSMUSG00000065372 | 1 | |
| ENSMUSG00000065373 | 1 | |
| ENSMUSG00000065374 | -1.044273782 | 0.698112512 |
| ENSMUSG00000065378 | 1.079382891 | 0.615267463 |
| ENSMUSG00000065379 | 1 | |
| ENSMUSG00000065380 | 1 | |
| ENSMUSG00000065381 | 1 | |
| ENSMUSG00000065382 | 1 | |
| ENSMUSG00000065384 | 1 | |
| ENSMUSG00000065385 | 1 | |
| ENSMUSG00000065387 | 1.069068979 | 0.608497229 |

| | | |
|---|---|---|
| ENSMUSG00000065388 | 1 | |
| ENSMUSG00000065389 | 1 | |
| ENSMUSG00000065392 | -1.051074015 | 0.680599207 |
| ENSMUSG00000065394 | 1 | |
| ENSMUSG00000065395 | 1 | |
| ENSMUSG00000065396 | 1 | |
| ENSMUSG00000065397 | 1 | |
| ENSMUSG00000065398 | 1 | |
| ENSMUSG00000065399 | -1.044273782 | 0.83619593 |
| ENSMUSG00000065400 | 1 | |
| ENSMUSG00000065401 | 1 | |
| ENSMUSG00000065402 | 1 | |
| ENSMUSG00000065403 | 1 | |
| ENSMUSG00000065404 | 1 | |
| ENSMUSG00000065405 | 1 | |
| ENSMUSG00000065406 | -1 | 1 |
| ENSMUSG00000065407 | 1 | |
| ENSMUSG00000065408 | 1 | |
| ENSMUSG00000065409 | 1 | |
| ENSMUSG00000065410 | 1 | |
| ENSMUSG00000065411 | 1 | |
| ENSMUSG00000065412 | 1 | |
| ENSMUSG00000065413 | 1 | |
| ENSMUSG00000065414 | 1 | |
| ENSMUSG00000065415 | 1 | |
| ENSMUSG00000065416 | 1 | |
| ENSMUSG00000065417 | 1 | |
| ENSMUSG00000065418 | -1 | 1 |
| ENSMUSG00000065420 | 1 | |
| ENSMUSG00000065421 | 1 | |
| ENSMUSG00000065422 | -1.053549062 | 0.769745556 |
| ENSMUSG00000065423 | 1 | |
| ENSMUSG00000065424 | 1 | |
| ENSMUSG00000065425 | 1 | |
| ENSMUSG00000065426 | 1 | |
| ENSMUSG00000065427 | 1 | |
| ENSMUSG00000065428 | 1 | |
| ENSMUSG00000065429 | 1 | |
| ENSMUSG00000065430 | -1 | 1 |
| ENSMUSG00000065431 | -1.10582475 | 0.608497229 |
| ENSMUSG00000065432 | 1 | |
| ENSMUSG00000065433 | 1 | |
| ENSMUSG00000065434 | -1.088434917 | 0.872304847 |
| ENSMUSG00000065435 | 1 | |
| ENSMUSG00000065436 | 1 | |
| ENSMUSG00000065437 | 1 | |
| ENSMUSG00000065438 | 1 | |
| ENSMUSG00000065439 | 1 | |
| ENSMUSG00000065440 | 1 | |
| ENSMUSG00000065441 | 1 | |
| ENSMUSG00000065442 | 1 | |
| ENSMUSG00000065443 | 1 | |
| ENSMUSG00000065444 | 1 | |
| ENSMUSG00000065445 | 1.059463094 | 0.698112512 |
| ENSMUSG00000065446 | 1 | |
| ENSMUSG00000065447 | 1 | |
| ENSMUSG00000065448 | 1 | |
| ENSMUSG00000065449 | 1 | |
| ENSMUSG00000065450 | 1 | |
| ENSMUSG00000065451 | 1 | |
| ENSMUSG00000065452 | 1 | |
| ENSMUSG00000065453 | 1 | |
| ENSMUSG00000065454 | 1 | |

| | | |
|---|---|---|
| ENSMUSG00000065455 | 1 | |
| ENSMUSG00000065456 | 1 | |
| ENSMUSG00000065457 | 1 | |
| ENSMUSG00000065458 | 1 | |
| ENSMUSG00000065460 | -1.043306199 | 0.698112512 |
| ENSMUSG00000065462 | 1 | |
| ENSMUSG00000065463 | 1 | |
| ENSMUSG00000065464 | 1 | |
| ENSMUSG00000065465 | 1 | |
| ENSMUSG00000065466 | 1 | |
| ENSMUSG00000065468 | 1.014545335 | 0.979002724 |
| ENSMUSG00000065469 | 1 | |
| ENSMUSG00000065470 | 1 | |
| ENSMUSG00000065471 | 1 | |
| ENSMUSG00000065472 | 1 | |
| ENSMUSG00000065473 | 1 | |
| ENSMUSG00000065474 | 1 | |
| ENSMUSG00000065475 | -1.100640286 | 0.697975969 |
| ENSMUSG00000065476 | 1 | |
| ENSMUSG00000065477 | 1 | |
| ENSMUSG00000065479 | 1 | |
| ENSMUSG00000065480 | 1 | |
| ENSMUSG00000065481 | 1 | |
| ENSMUSG00000065482 | 1 | |
| ENSMUSG00000065483 | -1.053549062 | 0.698112512 |
| ENSMUSG00000065484 | -1.043306199 | 0.698112512 |
| ENSMUSG00000065485 | 1 | |
| ENSMUSG00000065486 | 1 | |
| ENSMUSG00000065488 | -1 | 1 |
| ENSMUSG00000065489 | 1.015486245 | 0.972980766 |
| ENSMUSG00000065490 | 1 | |
| ENSMUSG00000065491 | 1 | |
| ENSMUSG00000065492 | 1 | |
| ENSMUSG00000065493 | 1 | |
| ENSMUSG00000065494 | 1 | |
| ENSMUSG00000065495 | 1 | |
| ENSMUSG00000065497 | 1 | |
| ENSMUSG00000065498 | 1 | |
| ENSMUSG00000065499 | 1 | |
| ENSMUSG00000065500 | 1 | |
| ENSMUSG00000065501 | 1 | |
| ENSMUSG00000065502 | 1 | |
| ENSMUSG00000065503 | 1.112349182 | 0.716190882 |
| ENSMUSG00000065505 | 1 | |
| ENSMUSG00000065507 | 1 | |
| ENSMUSG00000065508 | 1 | |
| ENSMUSG00000065509 | 1.122462048 | 0.608497229 |
| ENSMUSG00000065510 | 1 | |
| ENSMUSG00000065511 | 1 | |
| ENSMUSG00000065512 | 1 | |
| ENSMUSG00000065513 | 1 | |
| ENSMUSG00000065514 | -1.084645496 | 0.532369657 |
| ENSMUSG00000065515 | 1 | |
| ENSMUSG00000065516 | 1 | |
| ENSMUSG00000065518 | -1.043306199 | 0.698112512 |
| ENSMUSG00000065519 | -1.044273782 | 0.698112512 |
| ENSMUSG00000065520 | 1 | |
| ENSMUSG00000065521 | 1 | |
| ENSMUSG00000065523 | 1 | |
| ENSMUSG00000065524 | 1 | |
| ENSMUSG00000065526 | 1 | |
| ENSMUSG00000065527 | 1 | |
| ENSMUSG00000065528 | 1 | |

| | | |
|---|---|---|
| ENSMUSG00000065529 | 1 | |
| ENSMUSG00000065530 | 1 | |
| ENSMUSG00000065532 | 1 | |
| ENSMUSG00000065533 | 1 | |
| ENSMUSG00000065534 | 1 | |
| ENSMUSG00000065536 | 1 | |
| ENSMUSG00000065537 | 1 | |
| ENSMUSG00000065538 | 1 | |
| ENSMUSG00000065539 | 1 | |
| ENSMUSG00000065540 | 1 | |
| ENSMUSG00000065541 | 1 | |
| ENSMUSG00000065542 | -1.043306199 | 0.698112512 |
| ENSMUSG00000065543 | 1 | |
| ENSMUSG00000065544 | -1.126773846 | 0.608497229 |
| ENSMUSG00000065546 | 1.168628665 | 0.608497229 |
| ENSMUSG00000065547 | -1.043306199 | 0.79291651 |
| ENSMUSG00000065548 | 1 | |
| ENSMUSG00000065549 | 1 | |
| ENSMUSG00000065551 | 1 | |
| ENSMUSG00000065552 | 1 | |
| ENSMUSG00000065553 | 1 | |
| ENSMUSG00000065555 | 1 | |
| ENSMUSG00000065556 | 1 | |
| ENSMUSG00000065557 | 1 | |
| ENSMUSG00000065559 | 1 | |
| ENSMUSG00000065560 | 1 | |
| ENSMUSG00000065561 | 1 | |
| ENSMUSG00000065562 | 1 | |
| ENSMUSG00000065563 | 1 | |
| ENSMUSG00000065564 | 1 | |
| ENSMUSG00000065565 | 1 | |
| ENSMUSG00000065567 | 1 | |
| ENSMUSG00000065568 | 1 | |
| ENSMUSG00000065569 | 1 | |
| ENSMUSG00000065570 | 1 | |
| ENSMUSG00000065571 | 1 | |
| ENSMUSG00000065572 | 1 | |
| ENSMUSG00000065573 | 1 | |
| ENSMUSG00000065574 | 1 | |
| ENSMUSG00000065575 | 1.059463094 | 0.608497229 |
| ENSMUSG00000065577 | 1 | |
| ENSMUSG00000065578 | 1 | |
| ENSMUSG00000065579 | 1 | |
| ENSMUSG00000065580 | 1 | |
| ENSMUSG00000065581 | -1.009091449 | 1 |
| ENSMUSG00000065582 | 1 | |
| ENSMUSG00000065583 | 1 | |
| ENSMUSG00000065585 | 1 | |
| ENSMUSG00000065586 | 1 | |
| ENSMUSG00000065587 | 1 | |
| ENSMUSG00000065589 | -1.043306199 | 0.79291651 |
| ENSMUSG00000065590 | 1 | |
| ENSMUSG00000065591 | -1.096244537 | 0.82714964 |
| ENSMUSG00000065592 | 1.095872691 | 0.608497229 |
| ENSMUSG00000065593 | -1 | 1 |
| ENSMUSG00000065594 | 1 | |
| ENSMUSG00000065596 | 1 | |
| ENSMUSG00000065597 | 1 | |
| ENSMUSG00000065599 | -1.03962336 | 0.698112512 |
| ENSMUSG00000065600 | 1 | |
| ENSMUSG00000065601 | 1 | |
| ENSMUSG00000065602 | 1 | |
| ENSMUSG00000065603 | 1 | |

| | | |
|---|---|---|
| ENSMUSG00000065604 | 1 | |
| ENSMUSG00000065606 | 1.059463094 | 0.608497229 |
| ENSMUSG00000065607 | 1 | |
| ENSMUSG00000065608 | -1.053549062 | 0.698112512 |
| ENSMUSG00000065609 | 1 | |
| ENSMUSG00000065610 | 1 | |
| ENSMUSG00000065611 | -1.089497311 | 0.532369657 |
| ENSMUSG00000065612 | -1.043306199 | 0.698112512 |
| ENSMUSG00000065613 | 1 | |
| ENSMUSG00000065616 | 1 | |
| ENSMUSG00000065617 | 1 | |
| ENSMUSG00000065619 | 1 | |
| ENSMUSG00000065620 | 1 | |
| ENSMUSG00000065622 | 1 | |
| ENSMUSG00000065623 | -1.03962336 | 0.814089245 |
| ENSMUSG00000065625 | 1 | |
| ENSMUSG00000065627 | 1 | |
| ENSMUSG00000065628 | 1 | |
| ENSMUSG00000065629 | 1 | |
| ENSMUSG00000065630 | 1 | |
| ENSMUSG00000065634 | 1 | |
| ENSMUSG00000065635 | 1 | |
| ENSMUSG00000065636 | 1 | |
| ENSMUSG00000065637 | -1.002962915 | 1 |
| ENSMUSG00000065640 | 1 | |
| ENSMUSG00000065642 | 1.034340763 | 0.952057016 |
| ENSMUSG00000065643 | -1.024718486 | 0.972684567 |
| ENSMUSG00000065644 | 1 | |
| ENSMUSG00000065645 | 1 | |
| ENSMUSG00000065647 | 1 | |
| ENSMUSG00000065648 | 1 | |
| ENSMUSG00000065649 | 1 | |
| ENSMUSG00000065651 | -1.03962336 | 0.698112512 |
| ENSMUSG00000065653 | 1 | |
| ENSMUSG00000065656 | 1 | |
| ENSMUSG00000065657 | -1 | 1 |
| ENSMUSG00000065658 | 1 | |
| ENSMUSG00000065659 | 1 | |
| ENSMUSG00000065660 | 1 | |
| ENSMUSG00000065661 | 1 | |
| ENSMUSG00000065662 | 1.059463094 | 0.698112512 |
| ENSMUSG00000065663 | -1.034274428 | 0.857270703 |
| ENSMUSG00000065664 | 1 | |
| ENSMUSG00000065665 | 1 | |
| ENSMUSG00000065667 | -1.044273782 | 0.698112512 |
| ENSMUSG00000065669 | -1.000603363 | 1 |
| ENSMUSG00000065673 | 1.059463094 | 0.608497229 |
| ENSMUSG00000065674 | 1 | |
| ENSMUSG00000065675 | 1 | |
| ENSMUSG00000065676 | -1.010988769 | 1 |
| ENSMUSG00000065678 | 1 | |
| ENSMUSG00000065680 | 1 | |
| ENSMUSG00000065683 | 1 | |
| ENSMUSG00000065684 | 1 | |
| ENSMUSG00000065685 | 1 | |
| ENSMUSG00000065686 | 1.059463094 | 0.608497229 |
| ENSMUSG00000065687 | 1 | |
| ENSMUSG00000065689 | 1 | |
| ENSMUSG00000065691 | 1 | |
| ENSMUSG00000065692 | 1 | |
| ENSMUSG00000065694 | 1.095872691 | 0.608497229 |
| ENSMUSG00000065698 | 1 | |
| ENSMUSG00000065700 | 1 | |

| | | |
|---|---|---|
| ENSMUSG00000065701 | -1.028160286 | 0.964236174 |
| ENSMUSG00000065702 | 1 | |
| ENSMUSG00000065705 | 1 | |
| ENSMUSG00000065706 | 1 | |
| ENSMUSG00000065708 | 1 | |
| ENSMUSG00000065709 | 1 | |
| ENSMUSG00000065714 | 1 | |
| ENSMUSG00000065715 | 1.030842106 | 0.981782861 |
| ENSMUSG00000065716 | 1 | |
| ENSMUSG00000065718 | 1 | |
| ENSMUSG00000065719 | 1 | |
| ENSMUSG00000065721 | 1 | |
| ENSMUSG00000065722 | 1 | |
| ENSMUSG00000065725 | -1.044273782 | 0.790851299 |
| ENSMUSG00000065726 | 1 | |
| ENSMUSG00000065727 | 1 | |
| ENSMUSG00000065728 | -1.009476377 | 1 |
| ENSMUSG00000065729 | 1 | |
| ENSMUSG00000065732 | 1 | |
| ENSMUSG00000065733 | 1 | |
| ENSMUSG00000065734 | 1.06223195 | 0.785584885 |
| ENSMUSG00000065735 | 1 | |
| ENSMUSG00000065737 | 1 | |
| ENSMUSG00000065738 | -1.044631956 | 0.818635785 |
| ENSMUSG00000065739 | 1 | |
| ENSMUSG00000065742 | 1 | |
| ENSMUSG00000065743 | 1 | |
| ENSMUSG00000065745 | 1 | |
| ENSMUSG00000065746 | -1 | 1 |
| ENSMUSG00000065747 | 1 | |
| ENSMUSG00000065749 | 1 | |
| ENSMUSG00000065750 | 1.059463094 | 0.608497229 |
| ENSMUSG00000065751 | 1 | |
| ENSMUSG00000065752 | 1.266363693 | 0.34135669 |
| ENSMUSG00000065757 | 1 | |
| ENSMUSG00000065758 | 1 | |
| ENSMUSG00000065760 | 1 | |
| ENSMUSG00000065763 | 1 | |
| ENSMUSG00000065764 | 1 | |
| ENSMUSG00000065765 | 1.010259986 | 1 |
| ENSMUSG00000065767 | -1.01566718 | 1 |
| ENSMUSG00000065768 | 1 | |
| ENSMUSG00000065769 | 1 | |
| ENSMUSG00000065770 | 1 | |
| ENSMUSG00000065771 | 1.069068979 | 0.608497229 |
| ENSMUSG00000065773 | 1 | |
| ENSMUSG00000065778 | 1.059463094 | 0.608497229 |
| ENSMUSG00000065782 | 1.068550968 | 0.608497229 |
| ENSMUSG00000065787 | 1 | |
| ENSMUSG00000065788 | 1 | |
| ENSMUSG00000065789 | 1 | |
| ENSMUSG00000065791 | 1 | |
| ENSMUSG00000065792 | -1.03962336 | 0.698112512 |
| ENSMUSG00000065794 | 1 | |
| ENSMUSG00000065795 | 1 | |
| ENSMUSG00000065797 | 1 | |
| ENSMUSG00000065798 | 1 | |
| ENSMUSG00000065799 | 1 | |
| ENSMUSG00000065800 | 1 | |
| ENSMUSG00000065802 | 1 | |
| ENSMUSG00000065803 | 1 | |
| ENSMUSG00000065804 | 1 | |
| ENSMUSG00000065806 | 1.069068979 | 0.608497229 |

| | | |
|---|---|---|
| ENSMUSG00000065807 | 1 | |
| ENSMUSG00000065808 | -1.028160286 | 0.949812939 |
| ENSMUSG00000065811 | 1 | |
| ENSMUSG00000065812 | 1 | |
| ENSMUSG00000065813 | 1 | |
| ENSMUSG00000065815 | 1 | |
| ENSMUSG00000065817 | 1.059463094 | 0.608497229 |
| ENSMUSG00000065818 | 1.088377396 | 0.695198051 |
| ENSMUSG00000065819 | 1 | |
| ENSMUSG00000065820 | -1.095294215 | 0.541735653 |
| ENSMUSG00000065822 | -1 | 1 |
| ENSMUSG00000065823 | 1 | |
| ENSMUSG00000065824 | 1 | |
| ENSMUSG00000065825 | 1 | |
| ENSMUSG00000065828 | 1 | |
| ENSMUSG00000065836 | 1 | |
| ENSMUSG00000065837 | 1 | |
| ENSMUSG00000065840 | 1 | |
| ENSMUSG00000065843 | 1 | |
| ENSMUSG00000065845 | 1 | |
| ENSMUSG00000065847 | 1 | |
| ENSMUSG00000065851 | -1 | 1 |
| ENSMUSG00000065852 | 1 | |
| ENSMUSG00000065853 | 1 | |
| ENSMUSG00000065858 | 1 | |
| ENSMUSG00000065859 | 1 | |
| ENSMUSG00000065862 | 1 | |
| ENSMUSG00000065864 | 1 | |
| ENSMUSG00000065865 | 1 | |
| ENSMUSG00000065866 | 1 | |
| ENSMUSG00000065869 | 1 | |
| ENSMUSG00000065870 | 1 | |
| ENSMUSG00000065872 | 1 | |
| ENSMUSG00000065873 | -1.013347035 | 1 |
| ENSMUSG00000065874 | 1 | |
| ENSMUSG00000065875 | 1 | |
| ENSMUSG00000065876 | 1 | |
| ENSMUSG00000065878 | 1 | |
| ENSMUSG00000065881 | 1 | |
| ENSMUSG00000065882 | 1 | |
| ENSMUSG00000065883 | -1.063525418 | 0.716583464 |
| ENSMUSG00000065884 | 1 | |
| ENSMUSG00000065887 | 1 | |
| ENSMUSG00000065889 | 1 | |
| ENSMUSG00000065892 | 1 | |
| ENSMUSG00000065893 | 1 | |
| ENSMUSG00000065894 | 1 | |
| ENSMUSG00000065898 | 1 | |
| ENSMUSG00000065899 | 1 | |
| ENSMUSG00000065903 | 1 | |
| ENSMUSG00000065904 | 1.177765487 | 0.276639872 |
| ENSMUSG00000065905 | 1 | |
| ENSMUSG00000065906 | 1 | |
| ENSMUSG00000065907 | 1 | |
| ENSMUSG00000065911 | -1 | 1 |
| ENSMUSG00000065913 | 1 | |
| ENSMUSG00000065915 | 1 | |
| ENSMUSG00000065916 | 1 | |
| ENSMUSG00000065919 | 1 | |
| ENSMUSG00000065922 | 1 | |
| ENSMUSG00000065924 | 1 | |
| ENSMUSG00000065926 | 1 | |
| ENSMUSG00000065934 | 1.059463094 | 0.608497229 |

| | | |
|---|---|---|
| ENSMUSG00000065935 | 1 | |
| ENSMUSG00000065936 | 1 | |
| ENSMUSG00000065937 | 1 | |
| ENSMUSG00000065939 | 1.060678675 | 0.698112512 |
| ENSMUSG00000065940 | 1 | |
| ENSMUSG00000065943 | 1 | |
| ENSMUSG00000065944 | 1 | |
| ENSMUSG00000065945 | 1 | |
| ENSMUSG00000065947 | 1 | |
| ENSMUSG00000065950 | 1.019083579 | 0.971519533 |
| ENSMUSG00000065952 | 1.079681442 | 0.698112512 |
| ENSMUSG00000065954 | 1.121912503 | 0.608497229 |
| ENSMUSG00000065956 | 1 | |
| ENSMUSG00000065968 | 1.005021373 | 1 |
| ENSMUSG00000065979 | 1.016260905 | 0.951360958 |
| ENSMUSG00000065987 | 1 | |
| ENSMUSG00000065990 | -1.029227142 | 0.89924854 |
| ENSMUSG00000065999 | -1.360262186 | 0.074973338 |
| ENSMUSG00000066000 | 1.082364515 | 0.483527581 |
| ENSMUSG00000066007 | -1.03962336 | 0.698112512 |
| ENSMUSG00000066009 | 1 | |
| ENSMUSG00000066026 | 1.211000867 | 0.044893595 |
| ENSMUSG00000066027 | 1.059463094 | 0.608497229 |
| ENSMUSG00000066030 | 1 | |
| ENSMUSG00000066031 | 1 | |
| ENSMUSG00000066036 | 1.09342376 | 0.710673541 |
| ENSMUSG00000066037 | 1.021875222 | 0.983940831 |
| ENSMUSG00000066042 | 1.030671757 | 0.731109423 |
| ENSMUSG00000066043 | -1.032996819 | 0.927544616 |
| ENSMUSG00000066057 | 1.069058066 | 0.3862041 |
| ENSMUSG00000066058 | 1.064985481 | 0.794274354 |
| ENSMUSG00000066060 | 1 | |
| ENSMUSG00000066061 | 1 | |
| ENSMUSG00000066068 | -1 | 1 |
| ENSMUSG00000066071 | 1 | |
| ENSMUSG00000066072 | 1 | |
| ENSMUSG00000066090 | 1 | |
| ENSMUSG00000066097 | 1 | |
| ENSMUSG00000066100 | 1 | |
| ENSMUSG00000066101 | 1 | |
| ENSMUSG00000066107 | -1.036688446 | 0.759038956 |
| ENSMUSG00000066108 | -1.044273782 | 0.698112512 |
| ENSMUSG00000066113 | 1.199213707 | 0.275345684 |
| ENSMUSG00000066116 | 1.069068979 | 0.608497229 |
| ENSMUSG00000066122 | 1 | |
| ENSMUSG00000066129 | 1.024656056 | 0.972980766 |
| ENSMUSG00000066137 | -1.03962336 | 0.698112512 |
| ENSMUSG00000066141 | 1 | |
| ENSMUSG00000066148 | -1.0403707 | 0.893540818 |
| ENSMUSG00000066149 | 1.007592119 | 1 |
| ENSMUSG00000066150 | -1.05668879 | 0.84842598 |
| ENSMUSG00000066151 | -1.053395235 | 0.763339717 |
| ENSMUSG00000066152 | -1.05282845 | 0.698112512 |
| ENSMUSG00000066153 | 1 | |
| ENSMUSG00000066154 | 1 | |
| ENSMUSG00000066158 | -1.077304531 | 0.826652381 |
| ENSMUSG00000066170 | -1.096248562 | 0.698112512 |
| ENSMUSG00000066175 | -1.216015441 | 0.044535902 |
| ENSMUSG00000066176 | -1 | 1 |
| ENSMUSG00000066178 | 1.011290542 | 1 |
| ENSMUSG00000066180 | 1.095872691 | 0.608497229 |
| ENSMUSG00000066189 | 1 | |
| ENSMUSG00000066191 | 1.094236835 | 0.262306276 |

| | | |
|---|---|---|
| ENSMUSG00000066196 | -1.057391823 | 0.88054794 |
| ENSMUSG00000066197 | 1 | |
| ENSMUSG00000066224 | -1.492593037 | 0.004404638 |
| ENSMUSG00000066232 | 1.030708659 | 0.958403618 |
| ENSMUSG00000066233 | 1.033191963 | 0.785404965 |
| ENSMUSG00000066235 | 1.058521002 | 0.751653251 |
| ENSMUSG00000066239 | 1 | |
| ENSMUSG00000066240 | 1 | |
| ENSMUSG00000066241 | 1 | |
| ENSMUSG00000066242 | 1 | |
| ENSMUSG00000066245 | 1.21732381 | 0.481246252 |
| ENSMUSG00000066257 | 1 | |
| ENSMUSG00000066258 | -1.104793441 | 0.721269924 |
| ENSMUSG00000066262 | 1 | |
| ENSMUSG00000066263 | 1 | |
| ENSMUSG00000066268 | 1 | |
| ENSMUSG00000066269 | 1 | |
| ENSMUSG00000066270 | 1 | |
| ENSMUSG00000066272 | 1 | |
| ENSMUSG00000066273 | 1 | |
| ENSMUSG00000066278 | 1.03460885 | 0.728667795 |
| ENSMUSG00000066279 | -1.043306199 | 0.79291651 |
| ENSMUSG00000066306 | 1.139934577 | 0.507246493 |
| ENSMUSG00000066315 | 1 | |
| ENSMUSG00000066319 | -1.013996555 | 1 |
| ENSMUSG00000066324 | 1.039225958 | 0.944885275 |
| ENSMUSG00000066357 | 1.056965844 | 0.781649771 |
| ENSMUSG00000066359 | 1 | |
| ENSMUSG00000066361 | -1 | 1 |
| ENSMUSG00000066362 | -1.181480272 | 0.371954413 |
| ENSMUSG00000066363 | 1.373246581 | 0.015544186 |
| ENSMUSG00000066364 | 1 | |
| ENSMUSG00000066366 | 1 | |
| ENSMUSG00000066368 | 1 | |
| ENSMUSG00000066372 | 1 | |
| ENSMUSG00000066378 | 1 | |
| ENSMUSG00000066382 | 1.079681442 | 0.608497229 |
| ENSMUSG00000066383 | 1.122462048 | 0.608497229 |
| ENSMUSG00000066389 | -1.039686702 | 0.938986977 |
| ENSMUSG00000066392 | 1.009414875 | 1 |
| ENSMUSG00000066406 | 1.031905727 | 0.861034664 |
| ENSMUSG00000066407 | 1.055973703 | 0.926325112 |
| ENSMUSG00000066415 | 1.014904521 | 0.992856012 |
| ENSMUSG00000066438 | -1.03962336 | 0.698112512 |
| ENSMUSG00000066440 | 1.204705605 | 0.146384185 |
| ENSMUSG00000066441 | 1.208231392 | 0.283596572 |
| ENSMUSG00000066442 | -1.202692053 | 0.216046507 |
| ENSMUSG00000066443 | 1 | |
| ENSMUSG00000066456 | 1.047575411 | 0.815848303 |
| ENSMUSG00000066457 | 1.069068979 | 0.698112512 |
| ENSMUSG00000066463 | 1 | |
| ENSMUSG00000066475 | 1.032926687 | 0.87205152 |
| ENSMUSG00000066477 | 1.064870248 | 0.803260194 |
| ENSMUSG00000066478 | 1.059463094 | 0.608497229 |
| ENSMUSG00000066487 | 1.131135707 | 0.608497229 |
| ENSMUSG00000066489 | 1.016447115 | 1 |
| ENSMUSG00000066491 | -1.044273782 | 0.790851299 |
| ENSMUSG00000066500 | 1 | |
| ENSMUSG00000066510 | -1 | 1 |
| ENSMUSG00000066512 | 1 | |
| ENSMUSG00000066513 | 1 | |
| ENSMUSG00000066515 | 1 | |
| ENSMUSG00000066516 | 1 | |

| | | |
|---|---|---|
| ENSMUSG00000066537 | 1 | |
| ENSMUSG00000066538 | 1.05038453 | 0.860327141 |
| ENSMUSG00000066543 | -1.084645496 | 0.608497229 |
| ENSMUSG00000066551 | 1.023357204 | 0.982838256 |
| ENSMUSG00000066553 | 1 | |
| ENSMUSG00000066554 | 1.046551262 | 0.771926153 |
| ENSMUSG00000066568 | 1.065170742 | 0.750635131 |
| ENSMUSG00000066571 | 1.095430256 | 0.360176033 |
| ENSMUSG00000066583 | 1 | |
| ENSMUSG00000066584 | 1 | |
| ENSMUSG00000066586 | 1 | |
| ENSMUSG00000066592 | 1.067213122 | 0.900294918 |
| ENSMUSG00000066595 | -1.085940539 | 0.608497229 |
| ENSMUSG00000066607 | -1.005732744 | 1 |
| ENSMUSG00000066613 | -1.096675364 | 0.286287911 |
| ENSMUSG00000066620 | -1.03962336 | 0.803072249 |
| ENSMUSG00000066621 | 1.091196007 | 0.355837991 |
| ENSMUSG00000066626 | 1.000858205 | 1 |
| ENSMUSG00000066629 | 1.079681442 | 0.78131463 |
| ENSMUSG00000066632 | 1.083496801 | 0.776248867 |
| ENSMUSG00000066637 | -1.071603677 | 0.656375539 |
| ENSMUSG00000066640 | 1.188370613 | 0.044619484 |
| ENSMUSG00000066643 | 1.024222863 | 0.955841616 |
| ENSMUSG00000066647 | 1.097714564 | 0.04562023 |
| ENSMUSG00000066652 | -1 | 1 |
| ENSMUSG00000066671 | 1 | |
| ENSMUSG00000066672 | 1 | |
| ENSMUSG00000066677 | 1.09713609 | 0.758575113 |
| ENSMUSG00000066680 | 1 | |
| ENSMUSG00000066682 | -1.099174267 | 0.608497229 |
| ENSMUSG00000066684 | -1 | 1 |
| ENSMUSG00000066687 | -1.155193738 | 0.608497229 |
| ENSMUSG00000066688 | -1.033395294 | 0.969005636 |
| ENSMUSG00000066693 | 1 | |
| ENSMUSG00000066697 | 1.132639128 | 0.518443002 |
| ENSMUSG00000066704 | -1 | 1 |
| ENSMUSG00000066705 | -1.223283624 | 0.00440537 |
| ENSMUSG00000066714 | -1.044273782 | 0.698112512 |
| ENSMUSG00000066720 | 1.292359137 | 0.360552874 |
| ENSMUSG00000066721 | 1.068438367 | 0.757289696 |
| ENSMUSG00000066723 | 1 | |
| ENSMUSG00000066724 | -1.123073451 | 0.114871591 |
| ENSMUSG00000066735 | 1.038073296 | 0.924061737 |
| ENSMUSG00000066747 | 1 | |
| ENSMUSG00000066748 | 1 | |
| ENSMUSG00000066749 | 1.023743961 | 0.938939969 |
| ENSMUSG00000066750 | 1 | |
| ENSMUSG00000066752 | 1 | |
| ENSMUSG00000066755 | -1.119950137 | 0.502023826 |
| ENSMUSG00000066756 | 1 | |
| ENSMUSG00000066760 | -1.024557751 | 0.891017513 |
| ENSMUSG00000066767 | 1.075070132 | 0.725426917 |
| ENSMUSG00000066771 | -1.03962336 | 0.699591599 |
| ENSMUSG00000066772 | -1 | 1 |
| ENSMUSG00000066797 | 1 | |
| ENSMUSG00000066798 | 1.016918296 | 0.957397467 |
| ENSMUSG00000066800 | -1.261060087 | 0.022515869 |
| ENSMUSG00000066803 | 1 | |
| ENSMUSG00000066804 | 1 | |
| ENSMUSG00000066807 | -1 | 1 |
| ENSMUSG00000066809 | 1 | |
| ENSMUSG00000066810 | 1 | |
| ENSMUSG00000066820 | 1 | |

| | | |
|---|---|---|
| ENSMUSG00000066828 | 1 | |
| ENSMUSG00000066829 | -1.082990125 | 0.547580228 |
| ENSMUSG00000066838 | 1.064722895 | 0.478603495 |
| ENSMUSG00000066839 | 1.010110868 | 0.953731979 |
| ENSMUSG00000066842 | -1.098294818 | 0.402514484 |
| ENSMUSG00000066850 | -1.001578922 | 1 |
| ENSMUSG00000066861 | -1.03962336 | 0.698112512 |
| ENSMUSG00000066867 | 1 | |
| ENSMUSG00000066877 | 1.201961132 | 0.00873485 |
| ENSMUSG00000066878 | 1 | |
| ENSMUSG00000066880 | 1.028823 | 0.895965732 |
| ENSMUSG00000066892 | 1.007556972 | 0.976647797 |
| ENSMUSG00000066894 | 1.005927237 | 1 |
| ENSMUSG00000066896 | 1 | |
| ENSMUSG00000066897 | 1 | |
| ENSMUSG00000066899 | 1 | |
| ENSMUSG00000066900 | 1.118953374 | 0.530123536 |
| ENSMUSG00000066902 | 1 | |
| ENSMUSG00000066905 | 1 | |
| ENSMUSG00000066912 | 1 | |
| ENSMUSG00000066914 | 1 | |
| ENSMUSG00000066936 | 1 | |
| ENSMUSG00000066938 | 1 | |
| ENSMUSG00000066944 | 1.002448132 | 1 |
| ENSMUSG00000066952 | 1.067012261 | 0.519338787 |
| ENSMUSG00000066958 | 1 | |
| ENSMUSG00000066964 | 1.132639128 | 0.217803313 |
| ENSMUSG00000066975 | 1.402776436 | 0.076687772 |
| ENSMUSG00000066979 | 1.044675192 | 0.868146335 |
| ENSMUSG00000066983 | -1.03962336 | 0.88732209 |
| ENSMUSG00000066990 | 1 | |
| ENSMUSG00000067001 | 1.059463094 | 0.608497229 |
| ENSMUSG00000067006 | 1 | |
| ENSMUSG00000067010 | -1.044273782 | 0.698112512 |
| ENSMUSG00000067017 | 1.035846822 | 0.956288648 |
| ENSMUSG00000067028 | 1 | |
| ENSMUSG00000067038 | 1.06507254 | 0.680806726 |
| ENSMUSG00000067047 | 1 | |
| ENSMUSG00000067049 | -1.051018293 | 0.828564184 |
| ENSMUSG00000067058 | -1.040045394 | 0.896299552 |
| ENSMUSG00000067063 | -1 | 1 |
| ENSMUSG00000067064 | 1 | |
| ENSMUSG00000067071 | -1.015594454 | 0.865583959 |
| ENSMUSG00000067081 | -1.265886446 | 0.484584722 |
| ENSMUSG00000067085 | 1 | |
| ENSMUSG00000067101 | 1.059463094 | 0.608497229 |
| ENSMUSG00000067103 | 1 | |
| ENSMUSG00000067106 | -1.003567041 | 1 |
| ENSMUSG00000067121 | 1.172128996 | 0.455298078 |
| ENSMUSG00000067122 | 1 | |
| ENSMUSG00000067144 | -1.044273782 | 0.698112512 |
| ENSMUSG00000067147 | 1 | |
| ENSMUSG00000067148 | 1.007657836 | 1 |
| ENSMUSG00000067149 | -1.028348525 | 0.951360958 |
| ENSMUSG00000067150 | 1.0436654 | 0.908875934 |
| ENSMUSG00000067156 | -1.057588708 | 0.86372549 |
| ENSMUSG00000067158 | -1.117617762 | 0.552992863 |
| ENSMUSG00000067161 | -1.106396616 | 0.690723136 |
| ENSMUSG00000067173 | 1 | |
| ENSMUSG00000067186 | 1 | |
| ENSMUSG00000067189 | 1 | |
| ENSMUSG00000067194 | -1.149643607 | 0.608497229 |
| ENSMUSG00000067199 | -1.044273782 | 0.698112512 |

| | | |
|---|---|---|
| ENSMUSG00000067201 | 1 | |
| ENSMUSG00000067203 | -1.424250562 | 0.030155055 |
| ENSMUSG00000067206 | -1.03962336 | 0.814089245 |
| ENSMUSG00000067212 | -1.09712488 | 0.258650416 |
| ENSMUSG00000067215 | 1.267881226 | 0.425597339 |
| ENSMUSG00000067219 | 1.326878249 | 0.015575517 |
| ENSMUSG00000067220 | 1 | |
| ENSMUSG00000067224 | 1.033486948 | 0.943933955 |
| ENSMUSG00000067225 | 1 | |
| ENSMUSG00000067229 | -1 | 1 |
| ENSMUSG00000067231 | -1.126344191 | 0.217803313 |
| ENSMUSG00000067235 | -1.610161043 | 0.027242558 |
| ENSMUSG00000067242 | -1.149419985 | 0.262665776 |
| ENSMUSG00000067261 | 1 | |
| ENSMUSG00000067262 | 1 | |
| ENSMUSG00000067274 | 1.165235943 | 0.59223593 |
| ENSMUSG00000067276 | 1.263349309 | 0.288139213 |
| ENSMUSG00000067279 | 1.242515378 | 0.095604911 |
| ENSMUSG00000067285 | -1.132630578 | 0.617932717 |
| ENSMUSG00000067288 | 1.116420398 | 0.5713094 |
| ENSMUSG00000067292 | 1.005021373 | 1 |
| ENSMUSG00000067297 | 1.009859359 | 1 |
| ENSMUSG00000067299 | 1 | |
| ENSMUSG00000067321 | 1 | |
| ENSMUSG00000067331 | 1 | |
| ENSMUSG00000067336 | -1.04174844 | 0.924324458 |
| ENSMUSG00000067338 | -1 | 1 |
| ENSMUSG00000067341 | -1 | 1 |
| ENSMUSG00000067344 | -1.264892877 | 0.207803252 |
| ENSMUSG00000067351 | 1 | |
| ENSMUSG00000067352 | -1.047374079 | 0.93853737 |
| ENSMUSG00000067356 | 1.212858337 | 0.248294667 |
| ENSMUSG00000067360 | 1 | |
| ENSMUSG00000067365 | -1.053834461 | 0.803684084 |
| ENSMUSG00000067367 | 1.021118254 | 0.953731979 |
| ENSMUSG00000067369 | -1.149431221 | 0.289039526 |
| ENSMUSG00000067370 | 1.188577067 | 0.151380828 |
| ENSMUSG00000067377 | -1.198926186 | 0.17272194 |
| ENSMUSG00000067389 | 1.241211223 | 0.320426081 |
| ENSMUSG00000067399 | -1.017064415 | 0.963864477 |
| ENSMUSG00000067424 | 1.013610343 | 0.979811078 |
| ENSMUSG00000067430 | 1.012585282 | 0.984289155 |
| ENSMUSG00000067438 | 1 | |
| ENSMUSG00000067441 | 1 | |
| ENSMUSG00000067455 | -1.053092797 | 0.863033776 |
| ENSMUSG00000067482 | 1 | |
| ENSMUSG00000067513 | 1 | |
| ENSMUSG00000067519 | 1 | |
| ENSMUSG00000067520 | 1 | |
| ENSMUSG00000067522 | 1 | |
| ENSMUSG00000067524 | 1 | |
| ENSMUSG00000067525 | 1 | |
| ENSMUSG00000067526 | 1 | |
| ENSMUSG00000067528 | 1 | |
| ENSMUSG00000067529 | 1 | |
| ENSMUSG00000067541 | 1.006885039 | 1 |
| ENSMUSG00000067543 | 1 | |
| ENSMUSG00000067545 | 1 | |
| ENSMUSG00000067547 | 1 | |
| ENSMUSG00000067555 | 1 | |
| ENSMUSG00000067561 | 1 | |
| ENSMUSG00000067562 | 1 | |
| ENSMUSG00000067567 | -1.031218824 | 0.880191676 |

| | | |
|---|---|---|
| ENSMUSG00000067571 | 1.11166259 | 0.608497229 |
| ENSMUSG00000067575 | 1 | |
| ENSMUSG00000067577 | 1.010351027 | 1 |
| ENSMUSG00000067578 | 1 | |
| ENSMUSG00000067586 | 1.904059339 | 0.000641345 |
| ENSMUSG00000067591 | 1 | |
| ENSMUSG00000067594 | 1 | |
| ENSMUSG00000067596 | 1 | |
| ENSMUSG00000067597 | 1.079681442 | 0.697178241 |
| ENSMUSG00000067599 | 1 | |
| ENSMUSG00000067604 | 1 | |
| ENSMUSG00000067608 | -1.061342521 | 0.825058682 |
| ENSMUSG00000067610 | 1.059463094 | 0.608497229 |
| ENSMUSG00000067613 | 1.089470641 | 0.608497229 |
| ENSMUSG00000067614 | 1 | |
| ENSMUSG00000067615 | 1.059463094 | 0.608497229 |
| ENSMUSG00000067616 | 1 | |
| ENSMUSG00000067627 | 1.069068979 | 0.698112512 |
| ENSMUSG00000067629 | -1.047182918 | 0.698112512 |
| ENSMUSG00000067642 | -1.043306199 | 0.698112512 |
| ENSMUSG00000067649 | 1 | |
| ENSMUSG00000067653 | -1.080509632 | 0.698112512 |
| ENSMUSG00000067656 | 1 | |
| ENSMUSG00000067679 | 1 | |
| ENSMUSG00000067684 | 1 | |
| ENSMUSG00000067698 | 1 | |
| ENSMUSG00000067700 | 1 | |
| ENSMUSG00000067702 | -1 | 1 |
| ENSMUSG00000067704 | 1 | |
| ENSMUSG00000067713 | -1.037343332 | 0.805720656 |
| ENSMUSG00000067714 | 1 | |
| ENSMUSG00000067716 | 1 | |
| ENSMUSG00000067719 | 1 | |
| ENSMUSG00000067722 | 1.019724656 | 0.893093277 |
| ENSMUSG00000067724 | -1.096055317 | 0.806722321 |
| ENSMUSG00000067736 | 1.059463094 | 0.608497229 |
| ENSMUSG00000067745 | 1.049411284 | 0.835701362 |
| ENSMUSG00000067750 | 1 | |
| ENSMUSG00000067764 | 1 | |
| ENSMUSG00000067767 | 1 | |
| ENSMUSG00000067768 | 1 | |
| ENSMUSG00000067771 | 1 | |
| ENSMUSG00000067773 | 1 | |
| ENSMUSG00000067780 | 1.144216384 | 0.608497229 |
| ENSMUSG00000067786 | 1.016540971 | 0.968180243 |
| ENSMUSG00000067787 | 1.046815922 | 0.773499772 |
| ENSMUSG00000067795 | 1.043550045 | 0.913233275 |
| ENSMUSG00000067813 | 1.24122829 | 0.001720654 |
| ENSMUSG00000067818 | 2.947378383 | 5.75538E-05 |
| ENSMUSG00000067825 | 1.047386369 | 0.608497229 |
| ENSMUSG00000067835 | 1 | |
| ENSMUSG00000067847 | 1.006040857 | 1 |
| ENSMUSG00000067848 | 1 | |
| ENSMUSG00000067851 | 1.007651042 | 1 |
| ENSMUSG00000067855 | 1.079243324 | 0.752246983 |
| ENSMUSG00000067860 | -1.044273782 | 0.698112512 |
| ENSMUSG00000067869 | 1 | |
| ENSMUSG00000067870 | -1.123856526 | 0.040802057 |
| ENSMUSG00000067872 | 1.084993138 | 0.823318462 |
| ENSMUSG00000067873 | 1.034719966 | 0.917391267 |
| ENSMUSG00000067878 | -1.148217595 | 0.629012152 |
| ENSMUSG00000067879 | 1 | |
| ENSMUSG00000067882 | -1.071075483 | 0.750635131 |

| | | |
|---|---|---|
| ENSMUSG00000067889 | -1.078928203 | 0.561056194 |
| ENSMUSG00000067909 | 1 | |
| ENSMUSG00000067916 | -1.304170972 | 0.020353969 |
| ENSMUSG00000067919 | -1.095294215 | 0.698961838 |
| ENSMUSG00000067924 | -1.003538197 | 1 |
| ENSMUSG00000067925 | 1.005094615 | 1 |
| ENSMUSG00000067928 | 1.062961393 | 0.698112512 |
| ENSMUSG00000067929 | -1.015410163 | 0.997108523 |
| ENSMUSG00000067931 | -1.007236292 | 1 |
| ENSMUSG00000067941 | -1.182814301 | 0.418130164 |
| ENSMUSG00000067942 | 1.039245799 | 0.845175363 |
| ENSMUSG00000067949 | 1 | |
| ENSMUSG00000067951 | 1 | |
| ENSMUSG00000067971 | 1 | |
| ENSMUSG00000067973 | 1 | |
| ENSMUSG00000067995 | -1.16257991 | 0.251544797 |
| ENSMUSG00000067996 | 1 | |
| ENSMUSG00000067998 | 1.069068979 | 0.608497229 |
| ENSMUSG00000068008 | 1 | |
| ENSMUSG00000068009 | -1.043306199 | 0.698112512 |
| ENSMUSG00000068011 | 1.016827705 | 1 |
| ENSMUSG00000068014 | 1 | |
| ENSMUSG00000068015 | -1.006953586 | 1 |
| ENSMUSG00000068036 | 1.092356881 | 0.734000495 |
| ENSMUSG00000068037 | -1.047795021 | 0.923313048 |
| ENSMUSG00000068039 | 1.028359368 | 0.985522133 |
| ENSMUSG00000068040 | -1.152549528 | 0.496704482 |
| ENSMUSG00000068048 | 1 | |
| ENSMUSG00000068062 | 1 | |
| ENSMUSG00000068067 | 1 | |
| ENSMUSG00000068068 | 1 | |
| ENSMUSG00000068071 | 1 | |
| ENSMUSG00000068073 | 1 | |
| ENSMUSG00000068074 | 1 | |
| ENSMUSG00000068075 | 1 | |
| ENSMUSG00000068078 | 1 | |
| ENSMUSG00000068079 | -1.053549062 | 0.698112512 |
| ENSMUSG00000068082 | -1.03962336 | 0.698112512 |
| ENSMUSG00000068083 | 1 | |
| ENSMUSG00000068085 | 1 | |
| ENSMUSG00000068086 | 1 | |
| ENSMUSG00000068099 | 1 | |
| ENSMUSG00000068101 | 1.034310134 | 0.825032978 |
| ENSMUSG00000068105 | -1.251498208 | 0.455992192 |
| ENSMUSG00000068113 | 1 | |
| ENSMUSG00000068114 | -1.071681022 | 0.608497229 |
| ENSMUSG00000068115 | -1.0477729 | 0.608497229 |
| ENSMUSG00000068117 | 1.023743961 | 0.938939969 |
| ENSMUSG00000068120 | 1 | |
| ENSMUSG00000068122 | 1 | |
| ENSMUSG00000068129 | 1 | |
| ENSMUSG00000068130 | -1.030317806 | 0.924297242 |
| ENSMUSG00000068134 | -1.017729372 | 0.949812939 |
| ENSMUSG00000068141 | 1 | |
| ENSMUSG00000068149 | -1 | 1 |
| ENSMUSG00000068151 | -1.020155149 | 0.988424085 |
| ENSMUSG00000068154 | 1.059463094 | 0.698112512 |
| ENSMUSG00000068165 | 1 | |
| ENSMUSG00000068167 | 1 | |
| ENSMUSG00000068173 | 1 | |
| ENSMUSG00000068180 | 1 | |
| ENSMUSG00000068181 | 1 | |
| ENSMUSG00000068182 | 1 | |

| | | |
|---|---|---|
| ENSMUSG00000068184 | -1.092893114 | 0.470928144 |
| ENSMUSG00000068196 | 1.082721558 | 0.765968247 |
| ENSMUSG00000068200 | 1 | |
| ENSMUSG00000068205 | -1.016564563 | 0.996862752 |
| ENSMUSG00000068206 | 1.024121522 | 0.800651129 |
| ENSMUSG00000068218 | 1 | |
| ENSMUSG00000068219 | 1 | |
| ENSMUSG00000068220 | 1.115164193 | 0.775731377 |
| ENSMUSG00000068226 | 1.132639128 | 0.217803313 |
| ENSMUSG00000068227 | 1.025046921 | 0.96035552 |
| ENSMUSG00000068231 | -1 | 1 |
| ENSMUSG00000068232 | -1.053549062 | 0.698112512 |
| ENSMUSG00000068234 | -1.043306199 | 0.698112512 |
| ENSMUSG00000068240 | -1.019209636 | 0.996264991 |
| ENSMUSG00000068243 | -1.044273782 | 0.790851299 |
| ENSMUSG00000068245 | 1.074407693 | 0.752686953 |
| ENSMUSG00000068246 | -1.214413073 | 0.324252919 |
| ENSMUSG00000068250 | 1.106276776 | 0.502923851 |
| ENSMUSG00000068252 | 1.05038453 | 0.828564184 |
| ENSMUSG00000068259 | -1.033820075 | 0.943663271 |
| ENSMUSG00000068262 | -1.03962336 | 0.851451345 |
| ENSMUSG00000068263 | -1.17130192 | 0.608497229 |
| ENSMUSG00000068264 | 1.050906013 | 0.750430773 |
| ENSMUSG00000068267 | 1.00842248 | 1 |
| ENSMUSG00000068270 | 1.172797595 | 0.315720798 |
| ENSMUSG00000068284 | 1.051660788 | 0.831149671 |
| ENSMUSG00000068289 | 1 | |
| ENSMUSG00000068290 | -1.023928855 | 0.843685576 |
| ENSMUSG00000068299 | -1.027253291 | 0.898373612 |
| ENSMUSG00000068302 | 1 | |
| ENSMUSG00000068303 | 1.036532797 | 0.879300439 |
| ENSMUSG00000068314 | 1 | |
| ENSMUSG00000068323 | -1.069628355 | 0.807247723 |
| ENSMUSG00000068324 | 1.169915212 | 0.608863038 |
| ENSMUSG00000068327 | 1 | |
| ENSMUSG00000068328 | 1.086651956 | 0.608497229 |
| ENSMUSG00000068329 | 1.113101894 | 0.220238017 |
| ENSMUSG00000068335 | 1.273498627 | 0.037295921 |
| ENSMUSG00000068341 | 1 | |
| ENSMUSG00000068348 | 1.122462048 | 0.420161906 |
| ENSMUSG00000068349 | 1 | |
| ENSMUSG00000068373 | 1.196004652 | 0.562290479 |
| ENSMUSG00000068391 | 1.057683212 | 0.724290685 |
| ENSMUSG00000068392 | 1 | |
| ENSMUSG00000068394 | -1.069377556 | 0.668646266 |
| ENSMUSG00000068396 | 1 | |
| ENSMUSG00000068397 | -1.113515092 | 0.608497229 |
| ENSMUSG00000068399 | 1.015486245 | 0.972980766 |
| ENSMUSG00000068407 | 1 | |
| ENSMUSG00000068417 | -1.130787664 | 0.430031585 |
| ENSMUSG00000068426 | 1 | |
| ENSMUSG00000068428 | 1 | |
| ENSMUSG00000068431 | 1 | |
| ENSMUSG00000068437 | 1 | |
| ENSMUSG00000068452 | 1.222512274 | 0.439534962 |
| ENSMUSG00000068457 | -1.115712953 | 0.61763676 |
| ENSMUSG00000068463 | 1.132639128 | 0.217803313 |
| ENSMUSG00000068466 | -1.035469922 | 0.941510974 |
| ENSMUSG00000068479 | -1.029367163 | 0.861919723 |
| ENSMUSG00000068480 | -1.03962336 | 0.739206552 |
| ENSMUSG00000068487 | -1.043306199 | 0.698112512 |
| ENSMUSG00000068506 | 1 | |
| ENSMUSG00000068522 | -1.127097758 | 0.608497229 |

| | | |
|---|---|---|
| ENSMUSG00000068523 | 1.028952733 | 0.893496796 |
| ENSMUSG00000068535 | 1 | |
| ENSMUSG00000068536 | 1 | |
| ENSMUSG00000068537 | 1 | |
| ENSMUSG00000068547 | -1.044181653 | 0.944721414 |
| ENSMUSG00000068551 | 1.654304075 | 0.026869506 |
| ENSMUSG00000068566 | 1.041103768 | 0.926171634 |
| ENSMUSG00000068574 | 1 | |
| ENSMUSG00000068579 | -1.048103011 | 0.934760809 |
| ENSMUSG00000068580 | -1.140621726 | 0.113203853 |
| ENSMUSG00000068587 | 1.161036672 | 0.474472862 |
| ENSMUSG00000068600 | 1 | |
| ENSMUSG00000068601 | 1.003017659 | 1 |
| ENSMUSG00000068604 | -1.202139191 | 0.301441914 |
| ENSMUSG00000068606 | -1.073603785 | 0.754820307 |
| ENSMUSG00000068614 | 1.189207115 | 0.294533546 |
| ENSMUSG00000068615 | 1 | |
| ENSMUSG00000068617 | 1 | |
| ENSMUSG00000068629 | 1.07487334 | 0.698112512 |
| ENSMUSG00000068631 | 1.059463094 | 0.608497229 |
| ENSMUSG00000068638 | 1.014545335 | 0.990831634 |
| ENSMUSG00000068646 | 1 | |
| ENSMUSG00000068647 | 1 | |
| ENSMUSG00000068663 | -1.052889795 | 0.720958482 |
| ENSMUSG00000068674 | 1.3058238 | 0.290631638 |
| ENSMUSG00000068680 | 1 | |
| ENSMUSG00000068686 | 1.070688982 | 0.474488116 |
| ENSMUSG00000068689 | 1.119090523 | 0.751853033 |
| ENSMUSG00000068696 | 1.015486245 | 0.979002724 |
| ENSMUSG00000068697 | -1.044273782 | 0.855488553 |
| ENSMUSG00000068699 | 1.09466405 | 0.623872809 |
| ENSMUSG00000068706 | 1.089102065 | 0.785014311 |
| ENSMUSG00000068732 | -1.149378886 | 0.538858846 |
| ENSMUSG00000068735 | -1.16251747 | 0.119808842 |
| ENSMUSG00000068739 | 1.022146229 | 0.982838256 |
| ENSMUSG00000068740 | -2.226010022 | 6.60509E-05 |
| ENSMUSG00000068742 | -1.277286928 | 0.005538562 |
| ENSMUSG00000068744 | 1.170774757 | 0.008577483 |
| ENSMUSG00000068745 | 1 | |
| ENSMUSG00000068747 | -1.431263213 | 0.040378727 |
| ENSMUSG00000068748 | 3.025713375 | 0.000254504 |
| ENSMUSG00000068749 | -1.094420576 | 0.720432582 |
| ENSMUSG00000068758 | 1.518521521 | 0.305174917 |
| ENSMUSG00000068762 | -1.176247247 | 0.678897747 |
| ENSMUSG00000068776 | 1 | |
| ENSMUSG00000068794 | 1.295206637 | 0.062606813 |
| ENSMUSG00000068798 | -1.059895141 | 0.794274354 |
| ENSMUSG00000068806 | 1 | |
| ENSMUSG00000068808 | 1 | |
| ENSMUSG00000068809 | 1 | |
| ENSMUSG00000068810 | 1 | |
| ENSMUSG00000068814 | 1 | |
| ENSMUSG00000068815 | 1 | |
| ENSMUSG00000068816 | 1 | |
| ENSMUSG00000068817 | 1 | |
| ENSMUSG00000068818 | 1 | |
| ENSMUSG00000068819 | 1 | |
| ENSMUSG00000068823 | 1.011417497 | 1 |
| ENSMUSG00000068824 | 1 | |
| ENSMUSG00000068854 | -1.091159629 | 0.423856307 |
| ENSMUSG00000068855 | -1.044273782 | 0.698112512 |
| ENSMUSG00000068856 | 1.060194726 | 0.608497229 |
| ENSMUSG00000068859 | -1.120476206 | 0.698112512 |

| | | |
|---|---|---|
| ENSMUSG00000068860 | 1.161008251 | 0.43994505 |
| ENSMUSG00000068874 | -1.090507733 | 0.698112512 |
| ENSMUSG00000068876 | -1.314008603 | 0.044593269 |
| ENSMUSG00000068877 | -1.186901203 | 0.608497229 |
| ENSMUSG00000068879 | 1 | |
| ENSMUSG00000068882 | -1.094877158 | 0.737457739 |
| ENSMUSG00000068885 | 1 | |
| ENSMUSG00000068887 | 1 | |
| ENSMUSG00000068888 | 1 | |
| ENSMUSG00000068889 | 1 | |
| ENSMUSG00000068890 | 1 | |
| ENSMUSG00000068893 | 1 | |
| ENSMUSG00000068917 | 1.053229965 | 0.608497229 |
| ENSMUSG00000068921 | 1.001209782 | 1 |
| ENSMUSG00000068922 | 1.030742048 | 0.812255606 |
| ENSMUSG00000068923 | 1.018180155 | 0.954675041 |
| ENSMUSG00000068940 | 1 | |
| ENSMUSG00000068947 | -1 | 1 |
| ENSMUSG00000068948 | 1 | |
| ENSMUSG00000068950 | 1 | |
| ENSMUSG00000068957 | -1.053549062 | 0.698112512 |
| ENSMUSG00000068959 | 1.082660343 | 0.343739781 |
| ENSMUSG00000068962 | -1.043306199 | 0.79291651 |
| ENSMUSG00000068964 | 1 | |
| ENSMUSG00000068966 | -1.007250292 | 1 |
| ENSMUSG00000068969 | 1.086455536 | 0.723904259 |
| ENSMUSG00000068999 | 1 | |
| ENSMUSG00000069008 | 1 | |
| ENSMUSG00000069011 | -1.03962336 | 0.698112512 |
| ENSMUSG00000069014 | 1 | |
| ENSMUSG00000069020 | -1.008430149 | 0.995522634 |
| ENSMUSG00000069036 | 1 | |
| ENSMUSG00000069038 | 1 | |
| ENSMUSG00000069041 | -1.11061171 | 0.66585278 |
| ENSMUSG00000069044 | 1.059463094 | 0.608497229 |
| ENSMUSG00000069045 | -1.089497311 | 0.532369657 |
| ENSMUSG00000069049 | -1.089497311 | 0.614307028 |
| ENSMUSG00000069053 | -1.029624486 | 0.979700958 |
| ENSMUSG00000069072 | 1.015486245 | 0.973126641 |
| ENSMUSG00000069074 | 1 | |
| ENSMUSG00000069080 | 1 | |
| ENSMUSG00000069083 | -1.063525418 | 0.751979636 |
| ENSMUSG00000069085 | 1 | |
| ENSMUSG00000069089 | -1.070687153 | 0.791906178 |
| ENSMUSG00000069094 | -1.009725311 | 1 |
| ENSMUSG00000069114 | 1.027712538 | 0.892868903 |
| ENSMUSG00000069117 | 1 | |
| ENSMUSG00000069118 | 1 | |
| ENSMUSG00000069125 | -1.00002448 | 1 |
| ENSMUSG00000069132 | 1 | |
| ENSMUSG00000069135 | 1.119632525 | 0.608497229 |
| ENSMUSG00000069136 | 1.12189305 | 0.585646836 |
| ENSMUSG00000069170 | -2.033571956 | 0.000372179 |
| ENSMUSG00000069171 | 1.190367484 | 0.012556563 |
| ENSMUSG00000069184 | -1.048307986 | 0.772152553 |
| ENSMUSG00000069188 | 1 | |
| ENSMUSG00000069189 | -1.063525418 | 0.698112512 |
| ENSMUSG00000069196 | -1 | 1 |
| ENSMUSG00000069206 | -1.069209395 | 0.698112512 |
| ENSMUSG00000069208 | -1.065837683 | 0.608497229 |
| ENSMUSG00000069223 | -1.098893077 | 0.718265321 |
| ENSMUSG00000069227 | -3.002890438 | 3.10E-06 |
| ENSMUSG00000069236 | 1.228318929 | 0.543173564 |

| | | |
|---|---|---|
| ENSMUSG00000069237 | 1.003468051 | 1 |
| ENSMUSG00000069248 | 1 | |
| ENSMUSG00000069255 | 1.077450847 | 0.375185374 |
| ENSMUSG00000069258 | 1 | |
| ENSMUSG00000069259 | 1 | |
| ENSMUSG00000069265 | -1.089497311 | 0.608497229 |
| ENSMUSG00000069266 | 1.059463094 | 0.698112512 |
| ENSMUSG00000069267 | -1 | 1 |
| ENSMUSG00000069268 | 1 | |
| ENSMUSG00000069270 | 1 | |
| ENSMUSG00000069272 | 1.019083579 | 0.949812939 |
| ENSMUSG00000069273 | -1.03962336 | 0.698112512 |
| ENSMUSG00000069274 | 1.059463094 | 0.608497229 |
| ENSMUSG00000069276 | 1 | |
| ENSMUSG00000069280 | 1 | |
| ENSMUSG00000069289 | 1 | |
| ENSMUSG00000069292 | 1 | |
| ENSMUSG00000069294 | 1 | |
| ENSMUSG00000069295 | 1 | |
| ENSMUSG00000069296 | 1 | |
| ENSMUSG00000069297 | 1 | |
| ENSMUSG00000069299 | 1 | |
| ENSMUSG00000069300 | 1.05410549 | 0.841285672 |
| ENSMUSG00000069301 | 1 | |
| ENSMUSG00000069302 | 1 | |
| ENSMUSG00000069303 | 1 | |
| ENSMUSG00000069305 | 1 | |
| ENSMUSG00000069306 | 1 | |
| ENSMUSG00000069307 | 1 | |
| ENSMUSG00000069308 | -1.044273782 | 0.698112512 |
| ENSMUSG00000069309 | 1 | |
| ENSMUSG00000069310 | 1.04424822 | 0.861137762 |
| ENSMUSG00000069315 | -1 | 1 |
| ENSMUSG00000069324 | 1 | |
| ENSMUSG00000069355 | 1 | |
| ENSMUSG00000069372 | -1 | 1 |
| ENSMUSG00000069376 | 1.123364993 | 0.755354256 |
| ENSMUSG00000069378 | -1.431671266 | 0.175505164 |
| ENSMUSG00000069379 | -1.084645496 | 0.532369657 |
| ENSMUSG00000069385 | 1 | |
| ENSMUSG00000069390 | 1 | |
| ENSMUSG00000069421 | 1 | |
| ENSMUSG00000069430 | 1 | |
| ENSMUSG00000069439 | -1.05477085 | 0.855724967 |
| ENSMUSG00000069441 | 1 | |
| ENSMUSG00000069456 | -1.260219714 | 0.349914416 |
| ENSMUSG00000069475 | -1 | 1 |
| ENSMUSG00000069476 | -1 | 1 |
| ENSMUSG00000069495 | 1.059542446 | 0.793726432 |
| ENSMUSG00000069515 | -1.044273782 | 0.801443616 |
| ENSMUSG00000069516 | -1.218430774 | 0.203422347 |
| ENSMUSG00000069518 | 1.161265558 | 0.613468231 |
| ENSMUSG00000069520 | 1.048673908 | 0.805419186 |
| ENSMUSG00000069539 | -1.071438997 | 0.773487331 |
| ENSMUSG00000069540 | -1.02924683 | 0.983659314 |
| ENSMUSG00000069554 | -1.147838869 | 0.195607801 |
| ENSMUSG00000069565 | 1.032860545 | 0.83665746 |
| ENSMUSG00000069581 | -1.505963503 | 0.003083671 |
| ENSMUSG00000069582 | 1 | |
| ENSMUSG00000069583 | 1 | |
| ENSMUSG00000069584 | 1 | |
| ENSMUSG00000069588 | -1.187802519 | 0.314438145 |
| ENSMUSG00000069601 | 1.229086247 | 0.047784577 |

| | | |
|---|---|---|
| ENSMUSG00000069607 | 1 | |
| ENSMUSG00000069609 | 1 | |
| ENSMUSG00000069620 | 1 | |
| ENSMUSG00000069622 | -1.198352625 | 0.372103656 |
| ENSMUSG00000069631 | 1.026248832 | 0.890449109 |
| ENSMUSG00000069633 | 1.33571467 | 0.001174012 |
| ENSMUSG00000069662 | 1.275344388 | 0.265236008 |
| ENSMUSG00000069668 | 1 | |
| ENSMUSG00000069670 | 1.207623468 | 0.561619027 |
| ENSMUSG00000069678 | 1.014990061 | 0.884942525 |
| ENSMUSG00000069682 | 1.036484062 | 0.891297795 |
| ENSMUSG00000069706 | 1 | |
| ENSMUSG00000069707 | 1 | |
| ENSMUSG00000069708 | 1 | |
| ENSMUSG00000069712 | -1.078590893 | 0.698112512 |
| ENSMUSG00000069713 | -1.045424422 | 0.944099922 |
| ENSMUSG00000069717 | 1 | |
| ENSMUSG00000069718 | 1 | |
| ENSMUSG00000069720 | 1 | |
| ENSMUSG00000069721 | 1.059463094 | 0.608497229 |
| ENSMUSG00000069722 | 1 | |
| ENSMUSG00000069727 | -1.006923324 | 0.981214754 |
| ENSMUSG00000069729 | 1.150515001 | 0.223652394 |
| ENSMUSG00000069733 | 1 | |
| ENSMUSG00000069743 | -1.009145267 | 0.986641608 |
| ENSMUSG00000069744 | 1.028725081 | 0.924324458 |
| ENSMUSG00000069755 | 1.035301164 | 0.739980398 |
| ENSMUSG00000069763 | -1.043306199 | 0.839402676 |
| ENSMUSG00000069769 | 1.091319628 | 0.683209506 |
| ENSMUSG00000069785 | -1.020053668 | 0.990051834 |
| ENSMUSG00000069792 | 1 | |
| ENSMUSG00000069793 | 1.073557241 | 0.72387323 |
| ENSMUSG00000069796 | -1 | 1 |
| ENSMUSG00000069804 | -1.069062224 | 0.921950992 |
| ENSMUSG00000069805 | 1 | |
| ENSMUSG00000069806 | 1.121275662 | 0.29684302 |
| ENSMUSG00000069808 | -1.01071358 | 1 |
| ENSMUSG00000069814 | -1.210008472 | 0.338563396 |
| ENSMUSG00000069816 | 1 | |
| ENSMUSG00000069818 | 1 | |
| ENSMUSG00000069823 | 1 | |
| ENSMUSG00000069824 | 1 | |
| ENSMUSG00000069825 | 1.106574888 | 0.734624722 |
| ENSMUSG00000069830 | 1 | |
| ENSMUSG00000069833 | 1.021825263 | 1 |
| ENSMUSG00000069835 | 1.035027376 | 0.734674233 |
| ENSMUSG00000069844 | -1.066011617 | 0.66817972 |
| ENSMUSG00000069855 | 1 | |
| ENSMUSG00000069862 | 1.05410549 | 0.835701362 |
| ENSMUSG00000069867 | -1 | 1 |
| ENSMUSG00000069873 | 1 | |
| ENSMUSG00000069874 | -1.466655703 | 0.002778249 |
| ENSMUSG00000069890 | 1 | |
| ENSMUSG00000069892 | 1.169140771 | 0.344724767 |
| ENSMUSG00000069893 | 1.405015195 | 0.16449365 |
| ENSMUSG00000069895 | 1.005469242 | 1 |
| ENSMUSG00000069899 | -1.191763269 | 0.689299394 |
| ENSMUSG00000069910 | -1.12727234 | 0.518948132 |
| ENSMUSG00000069911 | 1 | |
| ENSMUSG00000069913 | -1.044273782 | 0.698112512 |
| ENSMUSG00000069917 | 1 | |
| ENSMUSG00000069919 | 1 | |
| ENSMUSG00000069920 | -1.104311598 | 0.181128445 |

| | | |
|---|---|---|
| ENSMUSG00000069922 | 1 | |
| ENSMUSG00000069925 | 1.132639128 | 0.217803313 |
| ENSMUSG00000069939 | 1.468160168 | 0.041033044 |
| ENSMUSG00000069962 | 1.07487334 | 0.84576478 |
| ENSMUSG00000069971 | 1 | |
| ENSMUSG00000069972 | 1 | |
| ENSMUSG00000069986 | -1.003834322 | 1 |
| ENSMUSG00000069996 | -1.03962336 | 0.803072249 |
| ENSMUSG00000069998 | 1 | |
| ENSMUSG00000070000 | -1.867052171 | 0.021950807 |
| ENSMUSG00000070002 | -1.041280523 | 0.729206866 |
| ENSMUSG00000070003 | -1.20285403 | 0.475208338 |
| ENSMUSG00000070023 | 1 | |
| ENSMUSG00000070025 | -1.03962336 | 0.808074244 |
| ENSMUSG00000070031 | -1.059245915 | 0.698112512 |
| ENSMUSG00000070034 | -1.043306199 | 0.698112512 |
| ENSMUSG00000070044 | -1.060783169 | 0.751234476 |
| ENSMUSG00000070047 | 1.210666415 | 0.467218078 |
| ENSMUSG00000070056 | 1.143260064 | 0.256011478 |
| ENSMUSG00000070063 | -1.01331511 | 0.966570161 |
| ENSMUSG00000070065 | 1 | |
| ENSMUSG00000070071 | 1 | |
| ENSMUSG00000070072 | 1 | |
| ENSMUSG00000070073 | 1 | |
| ENSMUSG00000070074 | -1.044273782 | 0.698112512 |
| ENSMUSG00000070075 | 1 | |
| ENSMUSG00000070076 | 1 | |
| ENSMUSG00000070077 | 1.059463094 | 0.608497229 |
| ENSMUSG00000070078 | 1 | |
| ENSMUSG00000070080 | 1 | |
| ENSMUSG00000070081 | 1 | |
| ENSMUSG00000070084 | 1 | |
| ENSMUSG00000070099 | 1 | |
| ENSMUSG00000070100 | 1 | |
| ENSMUSG00000070101 | 1 | |
| ENSMUSG00000070102 | 1 | |
| ENSMUSG00000070103 | 1 | |
| ENSMUSG00000070104 | 1 | |
| ENSMUSG00000070105 | 1 | |
| ENSMUSG00000070106 | 1 | |
| ENSMUSG00000070107 | 1 | |
| ENSMUSG00000070108 | 1 | |
| ENSMUSG00000070109 | 1 | |
| ENSMUSG00000070110 | 1 | |
| ENSMUSG00000070111 | -1.033575217 | 0.934760809 |
| ENSMUSG00000070126 | 1 | |
| ENSMUSG00000070127 | 1 | |
| ENSMUSG00000070128 | 1 | |
| ENSMUSG00000070129 | 1 | |
| ENSMUSG00000070130 | 1 | |
| ENSMUSG00000070133 | 1 | |
| ENSMUSG00000070136 | 1 | |
| ENSMUSG00000070138 | 1.059463094 | 0.608497229 |
| ENSMUSG00000070139 | 1 | |
| ENSMUSG00000070140 | 1 | |
| ENSMUSG00000070141 | 1 | |
| ENSMUSG00000070144 | 1 | |
| ENSMUSG00000070150 | 1 | |
| ENSMUSG00000070166 | -1 | 1 |
| ENSMUSG00000070167 | 1.114727698 | 0.516767179 |
| ENSMUSG00000070170 | 1.024693402 | 0.933031586 |
| ENSMUSG00000070173 | 1 | |
| ENSMUSG00000070175 | 1 | |

| | | |
|---|---|---|
| ENSMUSG00000070178 | 1 | |
| ENSMUSG00000070182 | 1 | |
| ENSMUSG00000070192 | 1 | |
| ENSMUSG00000070203 | 1 | |
| ENSMUSG00000070209 | 1 | |
| ENSMUSG00000070214 | 1 | |
| ENSMUSG00000070219 | 1 | |
| ENSMUSG00000070225 | -1.044273782 | 0.698112512 |
| ENSMUSG00000070227 | 1 | |
| ENSMUSG00000070235 | 1 | |
| ENSMUSG00000070236 | 1 | |
| ENSMUSG00000070240 | 1 | |
| ENSMUSG00000070249 | 1 | |
| ENSMUSG00000070258 | 1 | |
| ENSMUSG00000070263 | 1 | |
| ENSMUSG00000070271 | -1.014570171 | 0.997392688 |
| ENSMUSG00000070280 | -1.482906151 | 0.030092325 |
| ENSMUSG00000070282 | 1.000224443 | 1 |
| ENSMUSG00000070283 | -1.056520611 | 0.608497229 |
| ENSMUSG00000070284 | 1.030262694 | 0.750742698 |
| ENSMUSG00000070287 | -1.061566557 | 0.806851039 |
| ENSMUSG00000070291 | -1.364569376 | 0.041635084 |
| ENSMUSG00000070298 | 1.019083579 | 0.949812939 |
| ENSMUSG00000070304 | 1.033391092 | 0.959268555 |
| ENSMUSG00000070305 | -1.219243657 | 0.063595222 |
| ENSMUSG00000070306 | 1.085624843 | 0.699340382 |
| ENSMUSG00000070311 | 1 | |
| ENSMUSG00000070313 | -1.053549062 | 0.698112512 |
| ENSMUSG00000070315 | 1.212523994 | 0.10378762 |
| ENSMUSG00000070319 | -1.055292568 | 0.772292271 |
| ENSMUSG00000070323 | 1.084618945 | 0.614071182 |
| ENSMUSG00000070324 | 1 | |
| ENSMUSG00000070327 | -1.168614342 | 0.538858846 |
| ENSMUSG00000070330 | -1.254062209 | 0.262096583 |
| ENSMUSG00000070331 | 1.324551276 | 0.218315935 |
| ENSMUSG00000070332 | 1 | |
| ENSMUSG00000070334 | 1 | |
| ENSMUSG00000070335 | 1 | |
| ENSMUSG00000070336 | -1.133859594 | 0.399134697 |
| ENSMUSG00000070337 | -1.116449886 | 0.1631694 |
| ENSMUSG00000070342 | 1.17650053 | 0.567948807 |
| ENSMUSG00000070343 | -1.067671397 | 0.765303684 |
| ENSMUSG00000070345 | 1.22777087 | 0.576959009 |
| ENSMUSG00000070348 | -1.172111975 | 0.29645003 |
| ENSMUSG00000070354 | 1.028323353 | 0.972229988 |
| ENSMUSG00000070360 | 1 | |
| ENSMUSG00000070366 | 1 | |
| ENSMUSG00000070368 | -1 | 1 |
| ENSMUSG00000070369 | -1.044273782 | 0.790851299 |
| ENSMUSG00000070371 | 1.01863984 | 0.961565399 |
| ENSMUSG00000070372 | -1.117142115 | 0.642540037 |
| ENSMUSG00000070374 | 1 | |
| ENSMUSG00000070375 | 1 | |
| ENSMUSG00000070377 | 1 | |
| ENSMUSG00000070378 | 1 | |
| ENSMUSG00000070379 | 1 | |
| ENSMUSG00000070380 | 1 | |
| ENSMUSG00000070382 | 1 | |
| ENSMUSG00000070383 | 1 | |
| ENSMUSG00000070385 | 1.11790284 | 0.676809645 |
| ENSMUSG00000070388 | -1.053549062 | 0.807892349 |
| ENSMUSG00000070390 | 1 | |
| ENSMUSG00000070392 | -1 | 1 |

| | | |
|---|---|---|
| ENSMUSG00000070394 | 1.15902625 | 0.145828618 |
| ENSMUSG00000070402 | 1 | |
| ENSMUSG00000070407 | -1.34592457 | 0.182867484 |
| ENSMUSG00000070411 | 1 | |
| ENSMUSG00000070417 | 1 | |
| ENSMUSG00000070419 | 1.019083579 | 0.949812939 |
| ENSMUSG00000070420 | 1.099317784 | 0.396441539 |
| ENSMUSG00000070421 | 1 | |
| ENSMUSG00000070423 | 1 | |
| ENSMUSG00000070424 | 1.271344436 | 0.053354336 |
| ENSMUSG00000070425 | 1.046403186 | 0.909555516 |
| ENSMUSG00000070426 | -1.031996845 | 0.915071338 |
| ENSMUSG00000070427 | -1.039496045 | 0.698112512 |
| ENSMUSG00000070436 | 1.211013431 | 0.333490682 |
| ENSMUSG00000070438 | 1 | |
| ENSMUSG00000070443 | 1.342469832 | 0.062162093 |
| ENSMUSG00000070448 | 1 | |
| ENSMUSG00000070458 | 1 | |
| ENSMUSG00000070459 | 1 | |
| ENSMUSG00000070460 | 1 | |
| ENSMUSG00000070461 | 1.012809265 | 0.773325648 |
| ENSMUSG00000070462 | 1.052617655 | 0.637081332 |
| ENSMUSG00000070464 | 1 | |
| ENSMUSG00000070465 | 1 | |
| ENSMUSG00000070469 | -1.554519948 | 0.004385552 |
| ENSMUSG00000070471 | -1.193320102 | 0.056310079 |
| ENSMUSG00000070473 | -1.053549062 | 0.698112512 |
| ENSMUSG00000070476 | -1.042175913 | 0.753905984 |
| ENSMUSG00000070489 | 1 | |
| ENSMUSG00000070490 | 1.009353025 | 1 |
| ENSMUSG00000070493 | -1.027107859 | 0.9582181 |
| ENSMUSG00000070495 | 1.028022046 | 0.965978247 |
| ENSMUSG00000070498 | -1.03962336 | 0.803072249 |
| ENSMUSG00000070501 | -1.043306199 | 0.803260194 |
| ENSMUSG00000070504 | 1 | |
| ENSMUSG00000070509 | 1.537620203 | 0.000467239 |
| ENSMUSG00000070511 | 1 | |
| ENSMUSG00000070520 | 1.036353149 | 0.68754429 |
| ENSMUSG00000070522 | 1 | |
| ENSMUSG00000070524 | 1 | |
| ENSMUSG00000070526 | 1.123322288 | 0.303261282 |
| ENSMUSG00000070527 | 1.080871602 | 0.608497229 |
| ENSMUSG00000070529 | 1 | |
| ENSMUSG00000070530 | 1 | |
| ENSMUSG00000070531 | 1 | |
| ENSMUSG00000070532 | 1 | |
| ENSMUSG00000070533 | 1 | |
| ENSMUSG00000070544 | -1.074633675 | 0.782883548 |
| ENSMUSG00000070546 | 1 | |
| ENSMUSG00000070547 | 1.050891535 | 0.901216307 |
| ENSMUSG00000070550 | 1 | |
| ENSMUSG00000070551 | 1 | |
| ENSMUSG00000070552 | 1 | |
| ENSMUSG00000070563 | -1.009476377 | 1 |
| ENSMUSG00000070564 | -1.043306199 | 0.698112512 |
| ENSMUSG00000070565 | -1.046877172 | 0.865006766 |
| ENSMUSG00000070568 | 1 | |
| ENSMUSG00000070570 | -1.734422568 | 0.006348658 |
| ENSMUSG00000070572 | 1 | |
| ENSMUSG00000070574 | 1 | |
| ENSMUSG00000070576 | 1.437172628 | 0.014023003 |
| ENSMUSG00000070577 | 1 | |
| ENSMUSG00000070583 | 1.015642254 | 0.943341249 |

| | | |
|---|---|---|
| ENSMUSG00000070594 | -1 | 1 |
| ENSMUSG00000070601 | 1 | |
| ENSMUSG00000070604 | 1.030953648 | 0.96035552 |
| ENSMUSG00000070605 | -1.109540191 | 0.48909206 |
| ENSMUSG00000070606 | 1 | |
| ENSMUSG00000070609 | 1 | |
| ENSMUSG00000070610 | 1 | |
| ENSMUSG00000070616 | 1 | |
| ENSMUSG00000070617 | 1 | |
| ENSMUSG00000070618 | 1.019083579 | 0.982838256 |
| ENSMUSG00000070619 | -1.053549062 | 0.698112512 |
| ENSMUSG00000070621 | 1 | |
| ENSMUSG00000070632 | -1.007285479 | 1 |
| ENSMUSG00000070637 | 1 | |
| ENSMUSG00000070639 | 1.245741667 | 0.13435151 |
| ENSMUSG00000070641 | -1.044273782 | 0.790851299 |
| ENSMUSG00000070643 | -1.331911862 | 0.017095 |
| ENSMUSG00000070644 | -1.414445222 | 0.087823048 |
| ENSMUSG00000070645 | 1 | |
| ENSMUSG00000070661 | 1.059463094 | 0.608497229 |
| ENSMUSG00000070667 | -1.00571839 | 1 |
| ENSMUSG00000070677 | 1 | |
| ENSMUSG00000070683 | 1 | |
| ENSMUSG00000070686 | -1 | 1 |
| ENSMUSG00000070687 | 1.148945037 | 0.548740986 |
| ENSMUSG00000070690 | 1.04543363 | 0.856732743 |
| ENSMUSG00000070691 | 1.044871511 | 0.756970408 |
| ENSMUSG00000070692 | 1.058356913 | 0.790851299 |
| ENSMUSG00000070695 | -1.079873815 | 0.784381159 |
| ENSMUSG00000070697 | 1.067590444 | 0.7185059 |
| ENSMUSG00000070699 | 1.018042038 | 0.929280018 |
| ENSMUSG00000070702 | 1 | |
| ENSMUSG00000070704 | 1 | |
| ENSMUSG00000070705 | 1.086457488 | 0.23546922 |
| ENSMUSG00000070708 | 1 | |
| ENSMUSG00000070709 | 1.079701066 | 0.369031011 |
| ENSMUSG00000070713 | 1.032619126 | 0.830258476 |
| ENSMUSG00000070717 | 1.049411284 | 0.835701362 |
| ENSMUSG00000070719 | -1.069502966 | 0.698112512 |
| ENSMUSG00000070720 | -1.097023232 | 0.78833944 |
| ENSMUSG00000070729 | 1.019083579 | 0.982838256 |
| ENSMUSG00000070730 | -1.245726029 | 0.064130312 |
| ENSMUSG00000070732 | 1.069068979 | 0.608497229 |
| ENSMUSG00000070733 | 1.117633653 | 0.639770778 |
| ENSMUSG00000070737 | -1.016709407 | 0.910467721 |
| ENSMUSG00000070738 | 1.022153404 | 0.965872389 |
| ENSMUSG00000070777 | 1.028323353 | 0.903464611 |
| ENSMUSG00000070778 | -1.051347341 | 0.826172406 |
| ENSMUSG00000070780 | -1.017087767 | 0.896657787 |
| ENSMUSG00000070794 | 1.049137052 | 0.858134675 |
| ENSMUSG00000070796 | 1 | |
| ENSMUSG00000070797 | 1 | |
| ENSMUSG00000070798 | 1 | |
| ENSMUSG00000070799 | 1 | |
| ENSMUSG00000070802 | -2.160692856 | 2.98633E-05 |
| ENSMUSG00000070803 | -1.142727245 | 0.36554144 |
| ENSMUSG00000070806 | -1.043306199 | 0.79291651 |
| ENSMUSG00000070808 | 1.039160684 | 0.881931226 |
| ENSMUSG00000070810 | 1 | |
| ENSMUSG00000070811 | 1 | |
| ENSMUSG00000070814 | 1.069957917 | 0.160869438 |
| ENSMUSG00000070815 | 1 | |
| ENSMUSG00000070816 | 1 | |

| | | |
|---|---|---|
| ENSMUSG00000070817 | 1 | |
| ENSMUSG00000070820 | 1 | |
| ENSMUSG00000070821 | 1 | |
| ENSMUSG00000070822 | -1.038043907 | 0.905563166 |
| ENSMUSG00000070823 | 1.069068979 | 0.608497229 |
| ENSMUSG00000070828 | 1 | |
| ENSMUSG00000070834 | 1.059463094 | 0.608497229 |
| ENSMUSG00000070837 | 1 | |
| ENSMUSG00000070841 | 1 | |
| ENSMUSG00000070844 | -1 | 1 |
| ENSMUSG00000070847 | -1 | 1 |
| ENSMUSG00000070852 | 1 | |
| ENSMUSG00000070853 | 1 | |
| ENSMUSG00000070855 | 1 | |
| ENSMUSG00000070856 | 1 | |
| ENSMUSG00000070857 | 1 | |
| ENSMUSG00000070858 | 1.132080517 | 0.717035356 |
| ENSMUSG00000070866 | -1.024718486 | 0.968937442 |
| ENSMUSG00000070867 | 2.279217298 | 7.74828E-05 |
| ENSMUSG00000070868 | 1 | |
| ENSMUSG00000070870 | 1 | |
| ENSMUSG00000070871 | -1.027975527 | 0.926107971 |
| ENSMUSG00000070873 | 1 | |
| ENSMUSG00000070875 | 1 | |
| ENSMUSG00000070877 | -1.003834322 | 1 |
| ENSMUSG00000070880 | 1.015486245 | 0.972980766 |
| ENSMUSG00000070883 | -1.096112746 | 0.604059146 |
| ENSMUSG00000070890 | 1.019083579 | 0.949812939 |
| ENSMUSG00000070891 | -1.023769022 | 0.956401771 |
| ENSMUSG00000070896 | -1.108085261 | 0.752472633 |
| ENSMUSG00000070900 | 1 | |
| ENSMUSG00000070902 | 1 | |
| ENSMUSG00000070904 | 1 | |
| ENSMUSG00000070908 | 1 | |
| ENSMUSG00000070923 | 1.053084085 | 0.808074244 |
| ENSMUSG00000070933 | 1 | |
| ENSMUSG00000070934 | 1.027131762 | 0.945092216 |
| ENSMUSG00000070939 | -1.010218653 | 1 |
| ENSMUSG00000070942 | 1.030415139 | 0.958655379 |
| ENSMUSG00000070943 | 1 | |
| ENSMUSG00000070953 | 1.003501787 | 1 |
| ENSMUSG00000070960 | 1 | |
| ENSMUSG00000070972 | -1.052759904 | 0.785096927 |
| ENSMUSG00000070977 | 1 | |
| ENSMUSG00000070979 | 1.15516542 | 0.466704492 |
| ENSMUSG00000070980 | 1 | |
| ENSMUSG00000070983 | 1 | |
| ENSMUSG00000070984 | 1 | |
| ENSMUSG00000070985 | 1.015081164 | 1 |
| ENSMUSG00000070990 | -1.03962336 | 0.698112512 |
| ENSMUSG00000070995 | 1.062381344 | 0.870371236 |
| ENSMUSG00000070997 | -1.108998689 | 0.742692237 |
| ENSMUSG00000070999 | 1.088861202 | 0.698112512 |
| ENSMUSG00000071000 | 1 | |
| ENSMUSG00000071001 | 1.278125977 | 0.090633416 |
| ENSMUSG00000071005 | 1 | |
| ENSMUSG00000071014 | 1.07195509 | 0.698112512 |
| ENSMUSG00000071015 | 1 | |
| ENSMUSG00000071019 | 1 | |
| ENSMUSG00000071033 | 1 | |
| ENSMUSG00000071035 | -1.053549062 | 0.698112512 |
| ENSMUSG00000071036 | -1.095294215 | 0.608497229 |
| ENSMUSG00000071037 | -1.028531961 | 0.728434794 |

| | | |
|---|---|---|
| ENSMUSG00000071040 | 1 | |
| ENSMUSG00000071041 | -1.054741498 | 0.813450442 |
| ENSMUSG00000071042 | 2.097883617 | 0.000519289 |
| ENSMUSG00000071047 | 1.237106647 | 0.215763723 |
| ENSMUSG00000071052 | -1.03962336 | 0.698112512 |
| ENSMUSG00000071054 | 1.038190848 | 0.842184576 |
| ENSMUSG00000071064 | 1.286151019 | 0.032233622 |
| ENSMUSG00000071065 | 1 | |
| ENSMUSG00000071068 | 1 | |
| ENSMUSG00000071072 | 1.040536056 | 0.945122424 |
| ENSMUSG00000071073 | 1.019634634 | 0.952049609 |
| ENSMUSG00000071074 | 1.072338482 | 0.603144355 |
| ENSMUSG00000071076 | 1.006402152 | 1 |
| ENSMUSG00000071078 | 1.006807958 | 1 |
| ENSMUSG00000071083 | -1.01875214 | 0.968158081 |
| ENSMUSG00000071084 | 1 | |
| ENSMUSG00000071089 | 1.05038453 | 0.828564184 |
| ENSMUSG00000071102 | -1.066470541 | 0.808615646 |
| ENSMUSG00000071103 | 1.093470708 | 0.608497229 |
| ENSMUSG00000071104 | 1.049411284 | 0.863033776 |
| ENSMUSG00000071112 | 1.031950087 | 0.847418374 |
| ENSMUSG00000071113 | -1.051472827 | 0.881577029 |
| ENSMUSG00000071138 | 1 | |
| ENSMUSG00000071141 | 1.069068979 | 0.608497229 |
| ENSMUSG00000071147 | 1 | |
| ENSMUSG00000071149 | 1 | |
| ENSMUSG00000071150 | 1 | |
| ENSMUSG00000071151 | 1 | |
| ENSMUSG00000071158 | 1 | |
| ENSMUSG00000071165 | 1 | |
| ENSMUSG00000071169 | 1 | |
| ENSMUSG00000071172 | 1.007154765 | 1 |
| ENSMUSG00000071176 | 1.157975383 | 0.147235541 |
| ENSMUSG00000071177 | 1 | |
| ENSMUSG00000071178 | 1 | |
| ENSMUSG00000071179 | 1 | |
| ENSMUSG00000071180 | 1.062242161 | 0.775637715 |
| ENSMUSG00000071181 | 1.105845502 | 0.574596218 |
| ENSMUSG00000071185 | 1 | |
| ENSMUSG00000071192 | 1.099462 | 0.746451077 |
| ENSMUSG00000071193 | 1 | |
| ENSMUSG00000071195 | 1 | |
| ENSMUSG00000071202 | -1.006965774 | 1 |
| ENSMUSG00000071203 | 1.113765112 | 0.69923471 |
| ENSMUSG00000071204 | 1 | |
| ENSMUSG00000071207 | 1 | |
| ENSMUSG00000071217 | 1 | |
| ENSMUSG00000071226 | -1.352301045 | 0.037340456 |
| ENSMUSG00000071229 | -1 | 1 |
| ENSMUSG00000071230 | -1.108085261 | 0.698112512 |
| ENSMUSG00000071234 | 1 | |
| ENSMUSG00000071235 | 1 | |
| ENSMUSG00000071237 | 1 | |
| ENSMUSG00000071252 | -1.187625504 | 0.15502163 |
| ENSMUSG00000071253 | -1.049825795 | 0.800362789 |
| ENSMUSG00000071256 | 1.048510316 | 0.877244012 |
| ENSMUSG00000071262 | -1.089497311 | 0.532369657 |
| ENSMUSG00000071265 | -1 | 1 |
| ENSMUSG00000071266 | -1.028927197 | 0.900154956 |
| ENSMUSG00000071267 | 1.079790102 | 0.643397479 |
| ENSMUSG00000071273 | 1 | |
| ENSMUSG00000071275 | 1 | |
| ENSMUSG00000071276 | 1 | |

| | | |
|---|---|---|
| ENSMUSG00000071280 | 1 | |
| ENSMUSG00000071281 | -1.147042386 | 0.310191866 |
| ENSMUSG00000071291 | -1.071378718 | 0.203629745 |
| ENSMUSG00000071294 | 1 | |
| ENSMUSG00000071295 | 1.095872691 | 0.66521153 |
| ENSMUSG00000071302 | 1.022930307 | 0.939446078 |
| ENSMUSG00000071303 | 1.111528518 | 0.608497229 |
| ENSMUSG00000071311 | 1 | |
| ENSMUSG00000071317 | 1.104268979 | 0.318032671 |
| ENSMUSG00000071322 | 1 | |
| ENSMUSG00000071324 | -1.201307642 | 0.247755833 |
| ENSMUSG00000071332 | 1.150021168 | 0.608497229 |
| ENSMUSG00000071335 | 1.263963375 | 0.212065431 |
| ENSMUSG00000071337 | 1.041124408 | 0.896574035 |
| ENSMUSG00000071340 | 1.059463094 | 0.608497229 |
| ENSMUSG00000071341 | 1 | |
| ENSMUSG00000071342 | -1.044273782 | 0.801443616 |
| ENSMUSG00000071343 | 1.165716174 | 0.570450748 |
| ENSMUSG00000071347 | 1.059463094 | 0.608497229 |
| ENSMUSG00000071350 | -1.108055729 | 0.46945726 |
| ENSMUSG00000071356 | 1 | |
| ENSMUSG00000071359 | 1.009704952 | 1 |
| ENSMUSG00000071361 | -1 | 1 |
| ENSMUSG00000071362 | -1.081417919 | 0.626498859 |
| ENSMUSG00000071369 | -1.399463429 | 0.026267993 |
| ENSMUSG00000071379 | 1.126306006 | 0.329236078 |
| ENSMUSG00000071392 | -1.085651418 | 0.608497229 |
| ENSMUSG00000071398 | -1.128720443 | 0.698112512 |
| ENSMUSG00000071414 | -1.129231493 | 0.630993985 |
| ENSMUSG00000071415 | 1.031055255 | 0.968862261 |
| ENSMUSG00000071419 | 1.047614365 | 0.913882562 |
| ENSMUSG00000071424 | 1.015486245 | 0.973126641 |
| ENSMUSG00000071428 | 1 | |
| ENSMUSG00000071434 | -1 | 1 |
| ENSMUSG00000071451 | -1.019894862 | 0.913838795 |
| ENSMUSG00000071452 | 1 | |
| ENSMUSG00000071454 | -1.070437436 | 0.608497229 |
| ENSMUSG00000071456 | 1.012916683 | 0.98048847 |
| ENSMUSG00000071466 | 1 | |
| ENSMUSG00000071470 | -1.041426714 | 0.941128589 |
| ENSMUSG00000071471 | 1 | |
| ENSMUSG00000071475 | -1.069637405 | 0.608497229 |
| ENSMUSG00000071477 | 1.045044127 | 0.65415856 |
| ENSMUSG00000071478 | 1 | |
| ENSMUSG00000071481 | 1 | |
| ENSMUSG00000071489 | 1.069068979 | 0.608497229 |
| ENSMUSG00000071490 | 1 | |
| ENSMUSG00000071491 | 1 | |
| ENSMUSG00000071493 | 1 | |
| ENSMUSG00000071494 | 1.069068979 | 0.608497229 |
| ENSMUSG00000071497 | 1 | |
| ENSMUSG00000071506 | -1.037482356 | 0.918762638 |
| ENSMUSG00000071510 | 1 | |
| ENSMUSG00000071516 | 1 | |
| ENSMUSG00000071517 | 1 | |
| ENSMUSG00000071519 | 1 | |
| ENSMUSG00000071521 | 1 | |
| ENSMUSG00000071522 | 1 | |
| ENSMUSG00000071528 | -1.030105105 | 0.954192766 |
| ENSMUSG00000071531 | -1.090357695 | 0.755320109 |
| ENSMUSG00000071532 | 1 | |
| ENSMUSG00000071533 | -1.008646467 | 1 |
| ENSMUSG00000071537 | -1.2786754 | 0.025430361 |

| | | |
|---|---|---|
| ENSMUSG00000071540 | 1 | |
| ENSMUSG00000071547 | -1.019682561 | 0.949218699 |
| ENSMUSG00000071550 | -1.106271925 | 0.729018288 |
| ENSMUSG00000071551 | 1 | |
| ENSMUSG00000071552 | -1.326498669 | 0.242643886 |
| ENSMUSG00000071553 | -1.090721014 | 0.801443616 |
| ENSMUSG00000071561 | 1 | |
| ENSMUSG00000071562 | 1 | |
| ENSMUSG00000071567 | 1 | |
| ENSMUSG00000071568 | 1 | |
| ENSMUSG00000071573 | 1.132367418 | 0.075674707 |
| ENSMUSG00000071586 | -1.053549062 | 0.769745556 |
| ENSMUSG00000071591 | 1 | |
| ENSMUSG00000071604 | 1.218924411 | 0.026024582 |
| ENSMUSG00000071613 | 1 | |
| ENSMUSG00000071629 | 1 | |
| ENSMUSG00000071632 | -1.029957424 | 0.722892342 |
| ENSMUSG00000071633 | 1.059463094 | 0.769745556 |
| ENSMUSG00000071636 | -1.23938702 | 0.16655585 |
| ENSMUSG00000071637 | 1.319699206 | 0.172606008 |
| ENSMUSG00000071640 | 1.069068979 | 0.698112512 |
| ENSMUSG00000071644 | 1.112335919 | 0.714596508 |
| ENSMUSG00000071645 | 1.066994592 | 0.594957001 |
| ENSMUSG00000071646 | 1.055336377 | 0.85585109 |
| ENSMUSG00000071647 | -1.002198842 | 1 |
| ENSMUSG00000071648 | -1.119318033 | 0.15169538 |
| ENSMUSG00000071649 | 1.149398303 | 0.200777048 |
| ENSMUSG00000071650 | 1.006436236 | 1 |
| ENSMUSG00000071652 | 1.122717045 | 0.534861373 |
| ENSMUSG00000071653 | 1.050127056 | 0.950378392 |
| ENSMUSG00000071654 | 1.046145988 | 0.618592404 |
| ENSMUSG00000071655 | 1.07640071 | 0.664129605 |
| ENSMUSG00000071656 | 1.116252069 | 0.131060466 |
| ENSMUSG00000071657 | 1.044734295 | 0.608497229 |
| ENSMUSG00000071658 | 1.060884334 | 0.829324104 |
| ENSMUSG00000071659 | 1.013572097 | 1 |
| ENSMUSG00000071660 | 1.076740052 | 0.765218754 |
| ENSMUSG00000071661 | 1.105261759 | 0.214428334 |
| ENSMUSG00000071662 | 1.008561445 | 1 |
| ENSMUSG00000071665 | 1.059463094 | 0.608497229 |
| ENSMUSG00000071669 | -1.015184949 | 0.96035552 |
| ENSMUSG00000071671 | -1.128615549 | 0.608497229 |
| ENSMUSG00000071679 | 1 | |
| ENSMUSG00000071680 | 1.059463094 | 0.698112512 |
| ENSMUSG00000071686 | 1 | |
| ENSMUSG00000071691 | -1.150158336 | 0.698112512 |
| ENSMUSG00000071708 | -1.088454365 | 0.698112512 |
| ENSMUSG00000071711 | -1.028984556 | 0.88495285 |
| ENSMUSG00000071713 | 1.079681442 | 0.608497229 |
| ENSMUSG00000071714 | -1.02513447 | 0.945625364 |
| ENSMUSG00000071715 | -1.109582661 | 0.608497229 |
| ENSMUSG00000071716 | 1 | |
| ENSMUSG00000071719 | -1.044273782 | 0.698112512 |
| ENSMUSG00000071721 | 1 | |
| ENSMUSG00000071722 | -1.036853779 | 0.828564184 |
| ENSMUSG00000071723 | -1.067436589 | 0.686194514 |
| ENSMUSG00000071724 | -1.063258352 | 0.698112512 |
| ENSMUSG00000071726 | 1 | |
| ENSMUSG00000071738 | 1 | |
| ENSMUSG00000071745 | 1 | |
| ENSMUSG00000071748 | -1.125114201 | 0.58889649 |
| ENSMUSG00000071749 | 1 | |
| ENSMUSG00000071750 | 1 | |

| | | |
|---|---|---|
| ENSMUSG00000071753 | 1 | |
| ENSMUSG00000071757 | 2.142196055 | 0.002000539 |
| ENSMUSG00000071763 | 1 | |
| ENSMUSG00000071764 | 1 | |
| ENSMUSG00000071766 | 1 | |
| ENSMUSG00000071767 | 1 | |
| ENSMUSG00000071769 | 1 | |
| ENSMUSG00000071770 | 1 | |
| ENSMUSG00000071771 | 1 | |
| ENSMUSG00000071772 | 1 | |
| ENSMUSG00000071773 | 1 | |
| ENSMUSG00000071788 | 1 | |
| ENSMUSG00000071793 | 1.022025337 | 0.698690431 |
| ENSMUSG00000071796 | -1.082251934 | 0.630771413 |
| ENSMUSG00000071815 | 1 | |
| ENSMUSG00000071816 | 1 | |
| ENSMUSG00000071818 | 1 | |
| ENSMUSG00000071847 | 1.338924469 | 0.007788856 |
| ENSMUSG00000071855 | -1.095538981 | 0.554655964 |
| ENSMUSG00000071856 | 1.074312939 | 0.635590565 |
| ENSMUSG00000071858 | 1 | |
| ENSMUSG00000071860 | 1.059463094 | 0.608497229 |
| ENSMUSG00000071862 | 1.00580775 | 1 |
| ENSMUSG00000071866 | 1.028810976 | 0.980378626 |
| ENSMUSG00000071867 | -1.053549062 | 0.769745556 |
| ENSMUSG00000071890 | -1 | 1 |
| ENSMUSG00000071893 | -1.021632077 | 0.988424085 |
| ENSMUSG00000071909 | 1 | |
| ENSMUSG00000071937 | 1 | |
| ENSMUSG00000071952 | 1 | |
| ENSMUSG00000071984 | -1.088487825 | 0.698112512 |
| ENSMUSG00000072049 | 1 | |
| ENSMUSG00000072066 | 1.029222697 | 0.877593058 |
| ENSMUSG00000072068 | 1 | |
| ENSMUSG00000072082 | 1.047694858 | 0.813450442 |
| ENSMUSG00000072100 | 1 | |
| ENSMUSG00000072109 | 1.035844564 | 0.964202376 |
| ENSMUSG00000072115 | -1.471641239 | 0.028026721 |
| ENSMUSG00000072145 | 1 | |
| ENSMUSG00000072188 | 1 | |
| ENSMUSG00000072214 | 1.095250746 | 0.532369657 |
| ENSMUSG00000072235 | 1.017116975 | 1 |
| ENSMUSG00000072244 | 1.06899154 | 0.57779726 |
| ENSMUSG00000072249 | 1 | |
| ENSMUSG00000072255 | -1.075784115 | 0.801168497 |
| ENSMUSG00000072258 | -1.033406809 | 0.890893742 |
| ENSMUSG00000072259 | 1 | |
| ENSMUSG00000072294 | 1.714179989 | 0.018647491 |
| ENSMUSG00000072295 | 1 | |
| ENSMUSG00000072324 | -1.090800325 | 0.725290225 |
| ENSMUSG00000072381 | -1.043306199 | 0.698112512 |
| ENSMUSG00000072387 | 1 | |
| ENSMUSG00000072407 | 1 | |
| ENSMUSG00000072419 | -1.100193663 | 0.608497229 |
| ENSMUSG00000072421 | -1.057810276 | 0.901910426 |
| ENSMUSG00000072423 | -1.139361965 | 0.446939721 |
| ENSMUSG00000072432 | 1 | |
| ENSMUSG00000072437 | 1.001003444 | 1 |
| ENSMUSG00000072460 | -1.044273782 | 0.698112512 |
| ENSMUSG00000072473 | 1 | |
| ENSMUSG00000072476 | 1.090707387 | 0.717671093 |
| ENSMUSG00000072477 | 1 | |
| ENSMUSG00000072479 | 1 | |

| | | |
|---|---|---|
| ENSMUSG00000072487 | 1.059463094 | 0.699340382 |
| ENSMUSG00000072492 | -1.153519687 | 0.638239556 |
| ENSMUSG00000072494 | -1.020904738 | 0.968593113 |
| ENSMUSG00000072501 | 1.027894408 | 0.965872389 |
| ENSMUSG00000072511 | 1 | |
| ENSMUSG00000072541 | 1 | |
| ENSMUSG00000072553 | -1.043306199 | 0.698112512 |
| ENSMUSG00000072568 | 1.114425214 | 0.550831655 |
| ENSMUSG00000072571 | -1.027618047 | 0.935090686 |
| ENSMUSG00000072572 | -1.025565233 | 0.949812939 |
| ENSMUSG00000072573 | -1.174300461 | 0.404493254 |
| ENSMUSG00000072574 | 1 | |
| ENSMUSG00000072575 | 1 | |
| ENSMUSG00000072576 | 1.059463094 | 0.608497229 |
| ENSMUSG00000072582 | 1.014523036 | 0.991978016 |
| ENSMUSG00000072584 | 1 | |
| ENSMUSG00000072589 | 1.203080583 | 0.395692464 |
| ENSMUSG00000072591 | -1.084645496 | 0.608497229 |
| ENSMUSG00000072592 | 1.059463094 | 0.608497229 |
| ENSMUSG00000072594 | 1 | |
| ENSMUSG00000072595 | 1 | |
| ENSMUSG00000072596 | 1 | |
| ENSMUSG00000072598 | 1 | |
| ENSMUSG00000072599 | 1 | |
| ENSMUSG00000072600 | 1 | |
| ENSMUSG00000072601 | 1 | |
| ENSMUSG00000072605 | 1 | |
| ENSMUSG00000072612 | -1.03962336 | 0.698112512 |
| ENSMUSG00000072613 | 1 | |
| ENSMUSG00000072616 | 1 | |
| ENSMUSG00000072618 | -1.134282749 | 0.587756011 |
| ENSMUSG00000072620 | 1.108592941 | 0.698112512 |
| ENSMUSG00000072621 | 1.005404749 | 1 |
| ENSMUSG00000072623 | 1.082415881 | 0.698112512 |
| ENSMUSG00000072624 | -1.044273782 | 0.698112512 |
| ENSMUSG00000072625 | 1 | |
| ENSMUSG00000072640 | -1.025554962 | 0.925205197 |
| ENSMUSG00000072647 | -1.028354721 | 0.790851299 |
| ENSMUSG00000072653 | -1.13965082 | 0.521633709 |
| ENSMUSG00000072660 | -1.03347677 | 0.964236174 |
| ENSMUSG00000072662 | 1.085999383 | 0.739206552 |
| ENSMUSG00000072663 | 1.132639128 | 0.217803313 |
| ENSMUSG00000072664 | 1.049759971 | 0.806949992 |
| ENSMUSG00000072672 | -1.03962336 | 0.808074244 |
| ENSMUSG00000072673 | -1.022608177 | 0.993773455 |
| ENSMUSG00000072674 | 1 | |
| ENSMUSG00000072675 | 1 | |
| ENSMUSG00000072676 | 1 | |
| ENSMUSG00000072677 | 1 | |
| ENSMUSG00000072679 | -1.100193663 | 0.534887579 |
| ENSMUSG00000072680 | 1 | |
| ENSMUSG00000072683 | -1.044273782 | 0.698112512 |
| ENSMUSG00000072686 | 1 | |
| ENSMUSG00000072688 | -1.009303422 | 1 |
| ENSMUSG00000072692 | 1.031013335 | 0.827718961 |
| ENSMUSG00000072693 | 1 | |
| ENSMUSG00000072694 | 1.056037274 | 0.615241198 |
| ENSMUSG00000072704 | 1.057959798 | 0.787988343 |
| ENSMUSG00000072707 | 1 | |
| ENSMUSG00000072708 | 1 | |
| ENSMUSG00000072709 | 1 | |
| ENSMUSG00000072714 | 1 | |
| ENSMUSG00000072717 | 1 | |

| | | |
|---|---|---|
| ENSMUSG00000072718 | 1 | |
| ENSMUSG00000072720 | 1 | |
| ENSMUSG00000072721 | 1 | |
| ENSMUSG00000072722 | 1 | |
| ENSMUSG00000072723 | 1 | |
| ENSMUSG00000072726 | 1 | |
| ENSMUSG00000072731 | 1 | |
| ENSMUSG00000072738 | 1 | |
| ENSMUSG00000072739 | 1 | |
| ENSMUSG00000072753 | 1 | |
| ENSMUSG00000072754 | 1.059463094 | 0.698112512 |
| ENSMUSG00000072761 | 1.014651191 | 0.957935298 |
| ENSMUSG00000072762 | -1.001673583 | 1 |
| ENSMUSG00000072763 | 1.099309423 | 0.231578096 |
| ENSMUSG00000072768 | 1 | |
| ENSMUSG00000072769 | -1.34314964 | 0.223170554 |
| ENSMUSG00000072770 | 1.223823007 | 0.314592027 |
| ENSMUSG00000072772 | 1.075051549 | 0.401638687 |
| ENSMUSG00000072774 | 1.155131813 | 0.070844324 |
| ENSMUSG00000072778 | -1 | 1 |
| ENSMUSG00000072780 | 1 | |
| ENSMUSG00000072789 | 1.068081645 | 0.734674233 |
| ENSMUSG00000072791 | 1 | |
| ENSMUSG00000072809 | -1.041527706 | 0.698112512 |
| ENSMUSG00000072812 | 1.275600884 | 0.067890588 |
| ENSMUSG00000072813 | 1 | |
| ENSMUSG00000072814 | 1 | |
| ENSMUSG00000072821 | 1 | |
| ENSMUSG00000072822 | 1 | |
| ENSMUSG00000072825 | 1.145608027 | 0.177086637 |
| ENSMUSG00000072834 | 1 | |
| ENSMUSG00000072837 | 1 | |
| ENSMUSG00000072844 | -1.028575304 | 0.949218699 |
| ENSMUSG00000072845 | -1.147838869 | 0.243761388 |
| ENSMUSG00000072847 | -1.086286141 | 0.136686545 |
| ENSMUSG00000072849 | 1 | |
| ENSMUSG00000072852 | -1.132666855 | 0.375668528 |
| ENSMUSG00000072872 | 1.134881042 | 0.604149264 |
| ENSMUSG00000072874 | -1 | 1 |
| ENSMUSG00000072875 | -1.376586387 | 0.071698973 |
| ENSMUSG00000072878 | 1 | |
| ENSMUSG00000072884 | 1.207011299 | 0.250477583 |
| ENSMUSG00000072888 | 1 | |
| ENSMUSG00000072889 | 1.009481534 | 1 |
| ENSMUSG00000072893 | -1.06465492 | 0.351421966 |
| ENSMUSG00000072898 | -1 | 1 |
| ENSMUSG00000072900 | 1 | |
| ENSMUSG00000072902 | 1 | |
| ENSMUSG00000072905 | -1.03962336 | 0.698112512 |
| ENSMUSG00000072915 | -1.098659059 | 0.560230975 |
| ENSMUSG00000072919 | -1.081251904 | 0.828564184 |
| ENSMUSG00000072923 | 1 | |
| ENSMUSG00000072929 | 1 | |
| ENSMUSG00000072930 | 1 | |
| ENSMUSG00000072931 | 1 | |
| ENSMUSG00000072934 | 1.044412174 | 0.878957383 |
| ENSMUSG00000072940 | 1 | |
| ENSMUSG00000072941 | 1.38165141 | 0.006484725 |
| ENSMUSG00000072944 | -1.201379297 | 0.209798452 |
| ENSMUSG00000072945 | 1 | |
| ENSMUSG00000072946 | 1.025483889 | 0.908828093 |
| ENSMUSG00000072949 | -1.256823958 | 0.310630871 |
| ENSMUSG00000072950 | 1 | |

| | | |
|---|---|---|
| ENSMUSG00000072952 | 1 | |
| ENSMUSG00000072955 | -1.100193663 | 0.608497229 |
| ENSMUSG00000072962 | 1 | |
| ENSMUSG00000072963 | 1.000858205 | 1 |
| ENSMUSG00000072964 | -1.172287343 | 0.480993336 |
| ENSMUSG00000072966 | -1.567863862 | 0.002050034 |
| ENSMUSG00000072968 | 1 | |
| ENSMUSG00000072969 | -1.19414814 | 0.122196394 |
| ENSMUSG00000072972 | -1.063039293 | 0.900651611 |
| ENSMUSG00000072974 | 1.090410766 | 0.674977785 |
| ENSMUSG00000072978 | 1 | |
| ENSMUSG00000072980 | -1.161822899 | 0.321591536 |
| ENSMUSG00000072982 | 1 | |
| ENSMUSG00000072983 | 1 | |
| ENSMUSG00000072995 | 1 | |
| ENSMUSG00000072999 | -1.104860496 | 0.534631641 |
| ENSMUSG00000073000 | -1.094052268 | 0.752472633 |
| ENSMUSG00000073001 | 1 | |
| ENSMUSG00000073002 | 1.033177681 | 0.660807243 |
| ENSMUSG00000073006 | 1 | |
| ENSMUSG00000073007 | 1.020240423 | 0.977433779 |
| ENSMUSG00000073008 | 1.059463094 | 0.776953697 |
| ENSMUSG00000073010 | 1 | |
| ENSMUSG00000073012 | -1 | 1 |
| ENSMUSG00000073016 | 1.037420715 | 0.807933619 |
| ENSMUSG00000073018 | 1.252571842 | 0.484235702 |
| ENSMUSG00000073027 | 1 | |
| ENSMUSG00000073028 | 1 | |
| ENSMUSG00000073043 | 1 | |
| ENSMUSG00000073045 | -1 | 1 |
| ENSMUSG00000073051 | 1 | |
| ENSMUSG00000073052 | 1 | |
| ENSMUSG00000073060 | 1 | |
| ENSMUSG00000073062 | -1.004379082 | 1 |
| ENSMUSG00000073063 | 1 | |
| ENSMUSG00000073067 | 1.049411284 | 0.892085484 |
| ENSMUSG00000073069 | 1 | |
| ENSMUSG00000073072 | 1 | |
| ENSMUSG00000073073 | -1.018975854 | 0.989214798 |
| ENSMUSG00000073077 | 1.72768782 | 0.002386867 |
| ENSMUSG00000073079 | 1 | |
| ENSMUSG00000073085 | 1 | |
| ENSMUSG00000073094 | -1.043306199 | 0.858911269 |
| ENSMUSG00000073096 | 1.090727705 | 0.431532597 |
| ENSMUSG00000073102 | -1.090357695 | 0.755320109 |
| ENSMUSG00000073103 | 1 | |
| ENSMUSG00000073110 | 1 | |
| ENSMUSG00000073111 | 1 | |
| ENSMUSG00000073116 | 1 | |
| ENSMUSG00000073117 | 1 | |
| ENSMUSG00000073119 | 1 | |
| ENSMUSG00000073125 | -1.222972786 | 0.333071266 |
| ENSMUSG00000073130 | 1.157827942 | 0.526771915 |
| ENSMUSG00000073131 | -1.062029464 | 0.825478278 |
| ENSMUSG00000073139 | -1.0316285 | 0.925965582 |
| ENSMUSG00000073141 | 1 | |
| ENSMUSG00000073144 | -1.071075483 | 0.747163034 |
| ENSMUSG00000073145 | 1 | |
| ENSMUSG00000073147 | 1.008557912 | 0.984826086 |
| ENSMUSG00000073152 | 1 | |
| ENSMUSG00000073154 | -1.075618801 | 0.727204862 |
| ENSMUSG00000073155 | 1.234809346 | 0.038132245 |
| ENSMUSG00000073158 | -1.28303669 | 0.219185116 |

| | | |
|---|---|---|
| ENSMUSG00000073164 | -1.282064927 | 0.014492139 |
| ENSMUSG00000073173 | 1 | |
| ENSMUSG00000073174 | -1.161737199 | 0.397645313 |
| ENSMUSG00000073176 | 1.126394003 | 0.39954746 |
| ENSMUSG00000073177 | 1 | |
| ENSMUSG00000073179 | -1.00322009 | 1 |
| ENSMUSG00000073184 | -1.043306199 | 0.817489077 |
| ENSMUSG00000073197 | 1.041259584 | 0.69236387 |
| ENSMUSG00000073198 | 1 | |
| ENSMUSG00000073207 | -1.237138058 | 0.269379314 |
| ENSMUSG00000073208 | 1 | |
| ENSMUSG00000073209 | -1.03962336 | 0.814089245 |
| ENSMUSG00000073226 | 1.011497958 | 1 |
| ENSMUSG00000073233 | 1 | |
| ENSMUSG00000073234 | -1.122347111 | 0.339957009 |
| ENSMUSG00000073236 | -1.133522913 | 0.306648721 |
| ENSMUSG00000073242 | -1.043306199 | 0.798487595 |
| ENSMUSG00000073243 | 1 | |
| ENSMUSG00000073245 | 1 | |
| ENSMUSG00000073247 | 1 | |
| ENSMUSG00000073255 | 1 | |
| ENSMUSG00000073257 | 1 | |
| ENSMUSG00000073267 | 1 | |
| ENSMUSG00000073274 | -1.073310539 | 0.725290225 |
| ENSMUSG00000073290 | -1.111880267 | 0.572947856 |
| ENSMUSG00000073291 | 1 | |
| ENSMUSG00000073293 | -1.272995484 | 0.258738987 |
| ENSMUSG00000073294 | 1 | |
| ENSMUSG00000073295 | -1.214003634 | 0.38692062 |
| ENSMUSG00000073367 | 1 | |
| ENSMUSG00000073371 | 1.077658816 | 0.713861573 |
| ENSMUSG00000073374 | 1.058588408 | 0.698112512 |
| ENSMUSG00000073375 | 1 | |
| ENSMUSG00000073377 | 1 | |
| ENSMUSG00000073380 | -1.043306199 | 0.698112512 |
| ENSMUSG00000073386 | 1 | |
| ENSMUSG00000073388 | 1.019083579 | 0.956784261 |
| ENSMUSG00000073394 | 1.290567546 | 0.327217124 |
| ENSMUSG00000073396 | 1 | |
| ENSMUSG00000073399 | -1.03962336 | 0.865094641 |
| ENSMUSG00000073400 | -1 | 1 |
| ENSMUSG00000073402 | -1.204522173 | 0.526991558 |
| ENSMUSG00000073403 | -1.355461853 | 0.067517981 |
| ENSMUSG00000073405 | -1.183144353 | 0.364691511 |
| ENSMUSG00000073406 | -1.29086582 | 0.250726313 |
| ENSMUSG00000073407 | -1.170083925 | 0.506638168 |
| ENSMUSG00000073408 | 1.059463094 | 0.752398528 |
| ENSMUSG00000073409 | -1.694821552 | 0.001469981 |
| ENSMUSG00000073411 | -2.3022047 | 0.001410834 |
| ENSMUSG00000073412 | -1.139094357 | 0.304952879 |
| ENSMUSG00000073413 | -1 | 1 |
| ENSMUSG00000073414 | 1.122599155 | 0.608497229 |
| ENSMUSG00000073415 | 1.049197525 | 0.884451146 |
| ENSMUSG00000073418 | -1.230864329 | 0.50530478 |
| ENSMUSG00000073420 | 1 | |
| ENSMUSG00000073421 | 1.116897366 | 0.608497229 |
| ENSMUSG00000073422 | -1.132868669 | 0.561619027 |
| ENSMUSG00000073423 | 1.083863882 | 0.444208259 |
| ENSMUSG00000073424 | -1.03962336 | 0.698112512 |
| ENSMUSG00000073427 | -1.106329189 | 0.284965266 |
| ENSMUSG00000073430 | 1 | |
| ENSMUSG00000073433 | -1.06318136 | 0.647288746 |
| ENSMUSG00000073434 | -1.000974727 | 1 |

| | | |
|---|---|---|
| ENSMUSG00000073435 | 1 | |
| ENSMUSG00000073436 | -1.046877599 | 0.698112512 |
| ENSMUSG00000073437 | 1.124775701 | 0.082281453 |
| ENSMUSG00000073448 | -1.022567344 | 0.96035552 |
| ENSMUSG00000073457 | -1 | 1 |
| ENSMUSG00000073458 | 1 | |
| ENSMUSG00000073460 | -1.089553913 | 0.615243243 |
| ENSMUSG00000073468 | 1.033479017 | 0.814089245 |
| ENSMUSG00000073471 | 1.007239526 | 0.995473049 |
| ENSMUSG00000073478 | -1.041434409 | 0.698112512 |
| ENSMUSG00000073481 | -1.113862693 | 0.578238275 |
| ENSMUSG00000073482 | 1.132639128 | 0.217803313 |
| ENSMUSG00000073485 | 1.082193654 | 0.793735215 |
| ENSMUSG00000073486 | 1 | |
| ENSMUSG00000073489 | 1.130655539 | 0.336683729 |
| ENSMUSG00000073490 | 1.38229377 | 0.014950379 |
| ENSMUSG00000073491 | 1.069068979 | 0.766837546 |
| ENSMUSG00000073492 | 1.069068979 | 0.608497229 |
| ENSMUSG00000073494 | 1 | |
| ENSMUSG00000073497 | 1 | |
| ENSMUSG00000073510 | 1 | |
| ENSMUSG00000073514 | 1.2886569 | 0.185529637 |
| ENSMUSG00000073528 | 1 | |
| ENSMUSG00000073529 | 1.177765487 | 0.276639872 |
| ENSMUSG00000073530 | -1.009916536 | 1 |
| ENSMUSG00000073532 | 1 | |
| ENSMUSG00000073535 | -1.047419904 | 0.852081869 |
| ENSMUSG00000073538 | 1.115992654 | 0.767048884 |
| ENSMUSG00000073542 | 1.023036199 | 0.955262241 |
| ENSMUSG00000073551 | 1 | |
| ENSMUSG00000073554 | 1 | |
| ENSMUSG00000073555 | -1.089932089 | 0.758810982 |
| ENSMUSG00000073557 | -1.02397086 | 0.914913657 |
| ENSMUSG00000073559 | 1.10679692 | 0.745625104 |
| ENSMUSG00000073563 | 1.094717224 | 0.640707404 |
| ENSMUSG00000073565 | -1.134282749 | 0.65063484 |
| ENSMUSG00000073568 | -1.094488708 | 0.714698678 |
| ENSMUSG00000073569 | 1.059463094 | 0.698112512 |
| ENSMUSG00000073572 | 1 | |
| ENSMUSG00000073573 | 1 | |
| ENSMUSG00000073574 | 1 | |
| ENSMUSG00000073590 | 2.088998578 | 0.000547549 |
| ENSMUSG00000073591 | 1.058812025 | 0.608497229 |
| ENSMUSG00000073594 | -1 | 1 |
| ENSMUSG00000073598 | -1.113515092 | 0.642593813 |
| ENSMUSG00000073599 | -1.501775672 | 0.011041188 |
| ENSMUSG00000073600 | -1.020544934 | 0.828564184 |
| ENSMUSG00000073601 | 1 | |
| ENSMUSG00000073602 | 1 | |
| ENSMUSG00000073607 | -1.513624987 | 0.089033461 |
| ENSMUSG00000073608 | 1.167088505 | 0.319470416 |
| ENSMUSG00000073609 | 1.07361824 | 0.608497229 |
| ENSMUSG00000073610 | 1 | |
| ENSMUSG00000073616 | 1.01652602 | 0.959659993 |
| ENSMUSG00000073627 | 1.017333592 | 0.959463747 |
| ENSMUSG00000073628 | 1 | |
| ENSMUSG00000073631 | 1 | |
| ENSMUSG00000073633 | 1.02009878 | 0.93969881 |
| ENSMUSG00000073639 | 1.052627216 | 0.87452476 |
| ENSMUSG00000073640 | -1.089497311 | 0.532369657 |
| ENSMUSG00000073643 | 1.323372303 | 0.075927903 |
| ENSMUSG00000073647 | 1.01230007 | 0.950717906 |
| ENSMUSG00000073650 | 1.406049696 | 0.124608459 |

| | | |
|---|---|---|
| ENSMUSG00000073652 | -1.150867783 | 0.608497229 |
| ENSMUSG00000073656 | -1.024718486 | 0.931480186 |
| ENSMUSG00000073658 | 1 | |
| ENSMUSG00000073664 | 1.0521423 | 0.837740447 |
| ENSMUSG00000073676 | 1.016358007 | 0.997292167 |
| ENSMUSG00000073678 | -1.034612902 | 0.874019009 |
| ENSMUSG00000073679 | -1.033575217 | 0.95029384 |
| ENSMUSG00000073680 | 1.121477351 | 0.722165151 |
| ENSMUSG00000073682 | -1.07599997 | 0.840313932 |
| ENSMUSG00000073684 | 1.144099413 | 0.022503601 |
| ENSMUSG00000073686 | 1 | |
| ENSMUSG00000073699 | 1 | |
| ENSMUSG00000073700 | 1.071493007 | 0.589702874 |
| ENSMUSG00000073702 | -1.02397536 | 0.98408779 |
| ENSMUSG00000073705 | -1.000722886 | 1 |
| ENSMUSG00000073716 | 1.080726187 | 0.807892349 |
| ENSMUSG00000073721 | 1 | |
| ENSMUSG00000073722 | 1 | |
| ENSMUSG00000073723 | 1 | |
| ENSMUSG00000073724 | 1 | |
| ENSMUSG00000073725 | -1.095988423 | 0.66521153 |
| ENSMUSG00000073728 | 1.070149901 | 0.806002111 |
| ENSMUSG00000073730 | 1 | |
| ENSMUSG00000073731 | 1 | |
| ENSMUSG00000073733 | -1.0504352 | 0.833800367 |
| ENSMUSG00000073735 | 1 | |
| ENSMUSG00000073737 | 1 | |
| ENSMUSG00000073739 | -1.043616943 | 0.93689525 |
| ENSMUSG00000073747 | 1 | |
| ENSMUSG00000073752 | 1.059463094 | 0.608497229 |
| ENSMUSG00000073753 | 1.124923714 | 0.698112512 |
| ENSMUSG00000073755 | 1.267184099 | 0.000149712 |
| ENSMUSG00000073758 | -1.108719399 | 0.771960624 |
| ENSMUSG00000073759 | -1.235483129 | 0.267119406 |
| ENSMUSG00000073761 | 1.069158342 | 0.808074244 |
| ENSMUSG00000073764 | 1 | |
| ENSMUSG00000073765 | 1 | |
| ENSMUSG00000073768 | 1 | |
| ENSMUSG00000073769 | 1 | |
| ENSMUSG00000073770 | 1 | |
| ENSMUSG00000073771 | 1.135027755 | 0.24872542 |
| ENSMUSG00000073774 | 1 | |
| ENSMUSG00000073775 | 1.007743236 | 1 |
| ENSMUSG00000073778 | -1.154895927 | 0.608497229 |
| ENSMUSG00000073779 | 1.069068979 | 0.698112512 |
| ENSMUSG00000073781 | 1 | |
| ENSMUSG00000073785 | 1 | |
| ENSMUSG00000073786 | 1 | |
| ENSMUSG00000073787 | 1.172097463 | 0.698112512 |
| ENSMUSG00000073789 | 1 | |
| ENSMUSG00000073791 | 1.036301396 | 0.8920795 |
| ENSMUSG00000073792 | -1.067590045 | 0.744149026 |
| ENSMUSG00000073794 | -1.051018293 | 0.86119789 |
| ENSMUSG00000073795 | 1.014545335 | 0.979002724 |
| ENSMUSG00000073797 | 1 | |
| ENSMUSG00000073802 | 1.08205231 | 0.608497229 |
| ENSMUSG00000073803 | 1 | |
| ENSMUSG00000073804 | -1.053549062 | 0.769745556 |
| ENSMUSG00000073805 | -1.066331808 | 0.80666716 |
| ENSMUSG00000073807 | 1 | |
| ENSMUSG00000073809 | -1.043306199 | 0.698112512 |
| ENSMUSG00000073811 | 1 | |
| ENSMUSG00000073821 | -1.001517279 | 1 |

| | | |
|---|---|---|
| ENSMUSG00000073830 | 1 | |
| ENSMUSG00000073834 | 1 | |
| ENSMUSG00000073835 | 1 | |
| ENSMUSG00000073838 | -1.00643184 | 1 |
| ENSMUSG00000073840 | 1 | |
| ENSMUSG00000073842 | 1 | |
| ENSMUSG00000073843 | 1.079681442 | 0.812379067 |
| ENSMUSG00000073844 | 1 | |
| ENSMUSG00000073846 | -1.044273782 | 0.698112512 |
| ENSMUSG00000073856 | -1.09196997 | 0.608497229 |
| ENSMUSG00000073860 | 1 | |
| ENSMUSG00000073862 | 1 | |
| ENSMUSG00000073867 | -1.093117024 | 0.698112512 |
| ENSMUSG00000073876 | 1 | |
| ENSMUSG00000073877 | 1 | |
| ENSMUSG00000073878 | 1 | |
| ENSMUSG00000073879 | 1 | |
| ENSMUSG00000073888 | -1.212488444 | 0.532586 |
| ENSMUSG00000073889 | -1.074958839 | 0.608497229 |
| ENSMUSG00000073893 | 1 | |
| ENSMUSG00000073894 | 1 | |
| ENSMUSG00000073896 | 1 | |
| ENSMUSG00000073897 | 1 | |
| ENSMUSG00000073898 | 1 | |
| ENSMUSG00000073899 | 1 | |
| ENSMUSG00000073900 | 1 | |
| ENSMUSG00000073901 | 1 | |
| ENSMUSG00000073902 | 1 | |
| ENSMUSG00000073906 | -1.028348525 | 0.912425817 |
| ENSMUSG00000073907 | 1 | |
| ENSMUSG00000073908 | 1 | |
| ENSMUSG00000073909 | 1 | |
| ENSMUSG00000073910 | -1.039008338 | 0.749367459 |
| ENSMUSG00000073913 | 1 | |
| ENSMUSG00000073914 | 1 | |
| ENSMUSG00000073915 | 1 | |
| ENSMUSG00000073916 | 1 | |
| ENSMUSG00000073917 | 1 | |
| ENSMUSG00000073919 | 1 | |
| ENSMUSG00000073920 | 1 | |
| ENSMUSG00000073922 | 1 | |
| ENSMUSG00000073923 | 1 | |
| ENSMUSG00000073924 | 1 | |
| ENSMUSG00000073925 | 1 | |
| ENSMUSG00000073926 | 1 | |
| ENSMUSG00000073927 | 1 | |
| ENSMUSG00000073928 | 1 | |
| ENSMUSG00000073929 | 1 | |
| ENSMUSG00000073931 | 1 | |
| ENSMUSG00000073932 | 1 | |
| ENSMUSG00000073937 | 1 | |
| ENSMUSG00000073938 | 1 | |
| ENSMUSG00000073940 | 1 | |
| ENSMUSG00000073943 | 1 | |
| ENSMUSG00000073944 | 1 | |
| ENSMUSG00000073945 | 1 | |
| ENSMUSG00000073946 | 1 | |
| ENSMUSG00000073947 | 1 | |
| ENSMUSG00000073948 | 1 | |
| ENSMUSG00000073949 | 1 | |
| ENSMUSG00000073950 | 1 | |
| ENSMUSG00000073951 | 1 | |
| ENSMUSG00000073952 | 1 | |

| | | |
|---|---|---|
| ENSMUSG00000073953 | 1 | |
| ENSMUSG00000073954 | 1 | |
| ENSMUSG00000073955 | 1 | |
| ENSMUSG00000073956 | 1 | |
| ENSMUSG00000073959 | 1 | |
| ENSMUSG00000073960 | 1 | |
| ENSMUSG00000073961 | 1 | |
| ENSMUSG00000073962 | 1 | |
| ENSMUSG00000073963 | 1 | |
| ENSMUSG00000073964 | 1 | |
| ENSMUSG00000073965 | 1 | |
| ENSMUSG00000073966 | 1 | |
| ENSMUSG00000073967 | 1 | |
| ENSMUSG00000073968 | 1.120138103 | 0.253323351 |
| ENSMUSG00000073969 | 1 | |
| ENSMUSG00000073970 | 1 | |
| ENSMUSG00000073971 | 1 | |
| ENSMUSG00000073972 | 1 | |
| ENSMUSG00000073973 | 1 | |
| ENSMUSG00000073974 | 1 | |
| ENSMUSG00000073975 | 1 | |
| ENSMUSG00000073976 | 1 | |
| ENSMUSG00000073977 | -1.044273782 | 0.698112512 |
| ENSMUSG00000073978 | 1 | |
| ENSMUSG00000073979 | 1 | |
| ENSMUSG00000073980 | -1.03962336 | 0.851451345 |
| ENSMUSG00000073982 | 1.06154146 | 0.463933796 |
| ENSMUSG00000073985 | 1.314804881 | 0.17124452 |
| ENSMUSG00000073987 | -1.014323404 | 0.979486089 |
| ENSMUSG00000073988 | -1.178904939 | 0.237389238 |
| ENSMUSG00000073991 | 1 | |
| ENSMUSG00000073995 | -1.043722544 | 0.908842909 |
| ENSMUSG00000073997 | 1 | |
| ENSMUSG00000073998 | 1 | |
| ENSMUSG00000074001 | -1.03962336 | 0.698112512 |
| ENSMUSG00000074003 | 1 | |
| ENSMUSG00000074004 | 1.122462048 | 0.215991492 |
| ENSMUSG00000074006 | 1.086306196 | 0.709184271 |
| ENSMUSG00000074011 | 1.059463094 | 0.608497229 |
| ENSMUSG00000074024 | 1.037732448 | 0.807247723 |
| ENSMUSG00000074026 | 1 | |
| ENSMUSG00000074028 | -1.048703412 | 0.830576988 |
| ENSMUSG00000074029 | 1.059463094 | 0.608497229 |
| ENSMUSG00000074030 | 1.044669887 | 0.731295483 |
| ENSMUSG00000074034 | 1.163849123 | 0.608497229 |
| ENSMUSG00000074037 | 1.015486245 | 1 |
| ENSMUSG00000074039 | 1.174979913 | 0.256453848 |
| ENSMUSG00000074052 | -1.127486238 | 0.756938573 |
| ENSMUSG00000074059 | 1 | |
| ENSMUSG00000074060 | 1 | |
| ENSMUSG00000074061 | -1.084119349 | 0.790851299 |
| ENSMUSG00000074062 | 1 | |
| ENSMUSG00000074063 | 1.150495263 | 0.582072289 |
| ENSMUSG00000074064 | -1.030716068 | 0.79291651 |
| ENSMUSG00000074065 | 1.059463094 | 0.776953697 |
| ENSMUSG00000074067 | 1 | |
| ENSMUSG00000074071 | -1.054551959 | 0.839354385 |
| ENSMUSG00000074072 | 1 | |
| ENSMUSG00000074088 | -1.000445808 | 1 |
| ENSMUSG00000074092 | -1.044273782 | 0.698112512 |
| ENSMUSG00000074093 | 1.002240164 | 1 |
| ENSMUSG00000074102 | 1.036134279 | 0.865594818 |
| ENSMUSG00000074108 | 1.015486245 | 0.985522133 |

| | | |
|---|---|---|
| ENSMUSG00000074109 | 1 | |
| ENSMUSG00000074111 | 1 | |
| ENSMUSG00000074113 | 1 | |
| ENSMUSG00000074115 | 1 | |
| ENSMUSG00000074121 | -1.1859437 | 0.416813767 |
| ENSMUSG00000074123 | 1 | |
| ENSMUSG00000074127 | -1.029606215 | 0.983066615 |
| ENSMUSG00000074128 | -1.044273782 | 0.795446726 |
| ENSMUSG00000074129 | 1.093301693 | 0.696137525 |
| ENSMUSG00000074136 | -1.030779792 | 0.842184576 |
| ENSMUSG00000074139 | -1.03962336 | 0.731860565 |
| ENSMUSG00000074141 | 1.030223487 | 0.881456795 |
| ENSMUSG00000074146 | -1.027395698 | 0.981759143 |
| ENSMUSG00000074149 | -1 | 1 |
| ENSMUSG00000074151 | -1.274299738 | 0.002043913 |
| ENSMUSG00000074155 | 1 | |
| ENSMUSG00000074156 | 1 | |
| ENSMUSG00000074158 | 1.030205935 | 0.786640356 |
| ENSMUSG00000074164 | 1.085624843 | 0.69305964 |
| ENSMUSG00000074165 | 1.076324022 | 0.572892557 |
| ENSMUSG00000074166 | -1.056529082 | 0.736861486 |
| ENSMUSG00000074170 | 1.113260657 | 0.171177439 |
| ENSMUSG00000074171 | 1 | |
| ENSMUSG00000074178 | 1.019083579 | 0.97045957 |
| ENSMUSG00000074179 | 1.059463094 | 0.608497229 |
| ENSMUSG00000074182 | -1.130459631 | 0.420161906 |
| ENSMUSG00000074183 | 1.142870303 | 0.696594777 |
| ENSMUSG00000074184 | 1 | |
| ENSMUSG00000074194 | -1.248000717 | 0.012403517 |
| ENSMUSG00000074195 | 1 | |
| ENSMUSG00000074196 | 1 | |
| ENSMUSG00000074199 | -1.044273782 | 0.790851299 |
| ENSMUSG00000074203 | 1.119689435 | 0.698112512 |
| ENSMUSG00000074206 | 1.059463094 | 0.698112512 |
| ENSMUSG00000074207 | 1 | |
| ENSMUSG00000074210 | 1 | |
| ENSMUSG00000074211 | -1.005238492 | 1 |
| ENSMUSG00000074212 | -1.06116593 | 0.777702703 |
| ENSMUSG00000074213 | -1.063525418 | 0.698112512 |
| ENSMUSG00000074215 | -1.15908103 | 0.529171798 |
| ENSMUSG00000074217 | -1.142727245 | 0.379260898 |
| ENSMUSG00000074218 | 1.159813123 | 0.680806726 |
| ENSMUSG00000074219 | -1.073291408 | 0.698112512 |
| ENSMUSG00000074220 | 1.058901194 | 0.608497229 |
| ENSMUSG00000074221 | 1.055018086 | 0.778150345 |
| ENSMUSG00000074224 | 1 | |
| ENSMUSG00000074227 | -1.497858678 | 0.0038831 |
| ENSMUSG00000074228 | 1.059463094 | 0.776953697 |
| ENSMUSG00000074232 | -1.048116588 | 0.889802291 |
| ENSMUSG00000074235 | 1 | |
| ENSMUSG00000074236 | -1.03962336 | 0.803072249 |
| ENSMUSG00000074237 | 1 | |
| ENSMUSG00000074238 | 1.0025957 | 1 |
| ENSMUSG00000074240 | 1 | |
| ENSMUSG00000074243 | 1.041156662 | 0.922821172 |
| ENSMUSG00000074247 | 1.000930997 | 1 |
| ENSMUSG00000074248 | 1 | |
| ENSMUSG00000074250 | 1.141501657 | 0.251445232 |
| ENSMUSG00000074252 | 1 | 1 |
| ENSMUSG00000074254 | 1 | |
| ENSMUSG00000074256 | 1.161036672 | 0.378751351 |
| ENSMUSG00000074259 | -1.012248199 | 0.913942535 |
| ENSMUSG00000074261 | -1.063525418 | 0.698112512 |

| | | |
|---|---|---|
| ENSMUSG00000074264 | -1.104892015 | 0.747659585 |
| ENSMUSG00000074268 | 1.059463094 | 0.608497229 |
| ENSMUSG00000074269 | 1.051221758 | 0.814608539 |
| ENSMUSG00000074272 | 1.019676874 | 1 |
| ENSMUSG00000074274 | -1.03962336 | 0.698112512 |
| ENSMUSG00000074277 | -1.135597115 | 0.670176133 |
| ENSMUSG00000074280 | 1.199182804 | 0.127720171 |
| ENSMUSG00000074281 | -1.000342988 | 1 |
| ENSMUSG00000074282 | -1.01728477 | 0.888895298 |
| ENSMUSG00000074283 | 1.045410338 | 0.803235814 |
| ENSMUSG00000074284 | -1.012062641 | 1 |
| ENSMUSG00000074289 | 1 | |
| ENSMUSG00000074291 | 1 | |
| ENSMUSG00000074292 | 1.014930716 | 0.979002724 |
| ENSMUSG00000074300 | 1.095872691 | 0.608497229 |
| ENSMUSG00000074302 | 1 | |
| ENSMUSG00000074305 | -1.011987423 | 1 |
| ENSMUSG00000074311 | 1 | |
| ENSMUSG00000074322 | 1 | |
| ENSMUSG00000074336 | 1 | |
| ENSMUSG00000074340 | 1.121457453 | 0.437864659 |
| ENSMUSG00000074342 | -1.111252839 | 0.733770982 |
| ENSMUSG00000074343 | 1 | |
| ENSMUSG00000074344 | 1 | |
| ENSMUSG00000074345 | -1.053549062 | 0.825517832 |
| ENSMUSG00000074346 | 1 | |
| ENSMUSG00000074354 | -1.126344191 | 0.413646914 |
| ENSMUSG00000074355 | 1.194885329 | 0.122496827 |
| ENSMUSG00000074357 | -1.025996002 | 0.944460898 |
| ENSMUSG00000074358 | 1.228598638 | 0.503716121 |
| ENSMUSG00000074359 | 1.080993649 | 0.83270954 |
| ENSMUSG00000074361 | 1.064985481 | 0.757904398 |
| ENSMUSG00000074364 | 1.127142653 | 0.452380966 |
| ENSMUSG00000074365 | 1 | |
| ENSMUSG00000074366 | 1.813897108 | 0.001488665 |
| ENSMUSG00000074369 | 1 | |
| ENSMUSG00000074373 | 1 | |
| ENSMUSG00000074375 | 1 | |
| ENSMUSG00000074377 | 1 | |
| ENSMUSG00000074378 | 1 | |
| ENSMUSG00000074384 | 1.047699489 | 0.698798799 |
| ENSMUSG00000074385 | 1 | |
| ENSMUSG00000074388 | 1.061449073 | 0.840253751 |
| ENSMUSG00000074397 | -1.155252261 | 0.596589772 |
| ENSMUSG00000074398 | 1.129830974 | 0.639882806 |
| ENSMUSG00000074401 | 1 | |
| ENSMUSG00000074402 | 1 | |
| ENSMUSG00000074403 | 1 | |
| ENSMUSG00000074404 | 1 | |
| ENSMUSG00000074405 | 1.194972133 | 0.440837715 |
| ENSMUSG00000074406 | 1.184637282 | 0.42110385 |
| ENSMUSG00000074412 | -1.021592502 | 0.831364805 |
| ENSMUSG00000074415 | 1.175394274 | 0.43090216 |
| ENSMUSG00000074417 | 1 | |
| ENSMUSG00000074418 | 1 | |
| ENSMUSG00000074419 | 1 | |
| ENSMUSG00000074420 | 1 | |
| ENSMUSG00000074424 | -1 | 1 |
| ENSMUSG00000074433 | 1 | |
| ENSMUSG00000074434 | 1 | |
| ENSMUSG00000074435 | 1 | |
| ENSMUSG00000074436 | 1 | |
| ENSMUSG00000074437 | 1 | |

| | | |
|---|---|---|
| ENSMUSG00000074439 | 1 | |
| ENSMUSG00000074440 | 1 | |
| ENSMUSG00000074441 | 1 | |
| ENSMUSG00000074442 | 1 | |
| ENSMUSG00000074443 | 1 | |
| ENSMUSG00000074444 | 1 | |
| ENSMUSG00000074445 | -1 | 1 |
| ENSMUSG00000074446 | 1 | |
| ENSMUSG00000074447 | 1 | |
| ENSMUSG00000074448 | -1.038444538 | 0.874303881 |
| ENSMUSG00000074449 | -1 | 1 |
| ENSMUSG00000074452 | 1.265223817 | 0.447986619 |
| ENSMUSG00000074454 | 1 | |
| ENSMUSG00000074457 | 1.576158122 | 0.001744595 |
| ENSMUSG00000074461 | -1.090384387 | 0.648197854 |
| ENSMUSG00000074465 | 1.023586644 | 0.979814083 |
| ENSMUSG00000074466 | 1.111851062 | 0.532369657 |
| ENSMUSG00000074467 | 1 | |
| ENSMUSG00000074469 | 1 | |
| ENSMUSG00000074472 | -1.033575217 | 0.915070045 |
| ENSMUSG00000074473 | 1 | |
| ENSMUSG00000074476 | -1.127692588 | 0.313953572 |
| ENSMUSG00000074479 | -1.053549062 | 0.778372272 |
| ENSMUSG00000074480 | 1.076189945 | 0.704170349 |
| ENSMUSG00000074482 | 1.069068979 | 0.698112512 |
| ENSMUSG00000074483 | 1.034865357 | 0.931480186 |
| ENSMUSG00000074485 | 1.059463094 | 0.608497229 |
| ENSMUSG00000074486 | 1 | |
| ENSMUSG00000074489 | 1.022640321 | 0.944611559 |
| ENSMUSG00000074491 | 1 | |
| ENSMUSG00000074497 | -1.104032256 | 0.784381159 |
| ENSMUSG00000074500 | 1.130483453 | 0.540254765 |
| ENSMUSG00000074501 | 1 | |
| ENSMUSG00000074502 | 1.069293585 | 0.797105186 |
| ENSMUSG00000074505 | -1 | 1 |
| ENSMUSG00000074506 | 1 | |
| ENSMUSG00000074507 | -1.004497769 | 1 |
| ENSMUSG00000074508 | 1 | |
| ENSMUSG00000074513 | -1.012587679 | 0.935778761 |
| ENSMUSG00000074516 | -1.109582661 | 0.608497229 |
| ENSMUSG00000074517 | -1.03962336 | 0.698112512 |
| ENSMUSG00000074519 | -1.092308573 | 0.532369657 |
| ENSMUSG00000074521 | -1.109582661 | 0.608497229 |
| ENSMUSG00000074527 | -1.207417176 | 0.203228595 |
| ENSMUSG00000074529 | -1.189107718 | 0.442420587 |
| ENSMUSG00000074547 | 1.149490416 | 0.635590565 |
| ENSMUSG00000074553 | 1 | |
| ENSMUSG00000074555 | -1.044273782 | 0.790851299 |
| ENSMUSG00000074564 | 1 | |
| ENSMUSG00000074569 | -1.103069804 | 0.765233309 |
| ENSMUSG00000074570 | 1.138788635 | 0.595042751 |
| ENSMUSG00000074575 | -1.703808945 | 0.005721794 |
| ENSMUSG00000074576 | -1.119055465 | 0.03876409 |
| ENSMUSG00000074577 | -1.390471449 | 0.143499353 |
| ENSMUSG00000074578 | 1.026577157 | 0.942186096 |
| ENSMUSG00000074579 | 1.050245789 | 0.608497229 |
| ENSMUSG00000074580 | -1.219750074 | 0.251544797 |
| ENSMUSG00000074582 | -1.005665576 | 1 |
| ENSMUSG00000074589 | 1 | |
| ENSMUSG00000074591 | -1.085651418 | 0.692881216 |
| ENSMUSG00000074593 | 1 | |
| ENSMUSG00000074594 | 1 | |
| ENSMUSG00000074595 | 1 | |

| | | |
|---|---|---|
| ENSMUSG00000074596 | 1 | |
| ENSMUSG00000074603 | 1.024656056 | 0.973061065 |
| ENSMUSG00000074604 | 1.118758231 | 0.29684302 |
| ENSMUSG00000074607 | -1 | 1 |
| ENSMUSG00000074619 | -1.220755813 | 0.051171361 |
| ENSMUSG00000074620 | 1 | |
| ENSMUSG00000074622 | -1.008156464 | 1 |
| ENSMUSG00000074623 | 1 | |
| ENSMUSG00000074625 | 1.059463094 | 0.608497229 |
| ENSMUSG00000074627 | 1.019083579 | 0.968937442 |
| ENSMUSG00000074628 | -1.023769022 | 0.954761885 |
| ENSMUSG00000074629 | -1.160287354 | 0.414803203 |
| ENSMUSG00000074634 | -1.548550858 | 0.001032697 |
| ENSMUSG00000074635 | -1.116048458 | 0.456745331 |
| ENSMUSG00000074637 | -1 | 1 |
| ENSMUSG00000074639 | 1 | |
| ENSMUSG00000074643 | -1.144871996 | 0.564571928 |
| ENSMUSG00000074646 | -1.13301991 | 0.404493254 |
| ENSMUSG00000074647 | -1.15285904 | 0.697673897 |
| ENSMUSG00000074649 | -1.278617935 | 0.033504471 |
| ENSMUSG00000074650 | -1.287846914 | 0.517177088 |
| ENSMUSG00000074651 | -1.120476206 | 0.534652019 |
| ENSMUSG00000074652 | -1.363111584 | 0.063595222 |
| ENSMUSG00000074653 | -1.019108526 | 0.968937442 |
| ENSMUSG00000074655 | 1 | |
| ENSMUSG00000074656 | -1.068508555 | 0.831149671 |
| ENSMUSG00000074657 | -1.285400635 | 0.034835992 |
| ENSMUSG00000074664 | 1 | |
| ENSMUSG00000074665 | -1.03962336 | 0.698112512 |
| ENSMUSG00000074671 | -2.078020037 | 0.003934585 |
| ENSMUSG00000074673 | -1.340972605 | 0.044843448 |
| ENSMUSG00000074676 | 2.236060778 | 0.000972108 |
| ENSMUSG00000074677 | 1 | |
| ENSMUSG00000074678 | 1 | |
| ENSMUSG00000074679 | 1 | |
| ENSMUSG00000074680 | 1 | |
| ENSMUSG00000074681 | 1 | |
| ENSMUSG00000074682 | -1.075074555 | 0.347480564 |
| ENSMUSG00000074695 | 1 | |
| ENSMUSG00000074698 | -1.077622943 | 0.784381159 |
| ENSMUSG00000074704 | 1 | |
| ENSMUSG00000074715 | -1.961432702 | 0.009277859 |
| ENSMUSG00000074720 | 1 | |
| ENSMUSG00000074731 | 1.27919515 | 0.009361701 |
| ENSMUSG00000074733 | -1.018899546 | 0.925653135 |
| ENSMUSG00000074734 | 1 | |
| ENSMUSG00000074735 | -1.300967066 | 0.047958006 |
| ENSMUSG00000074736 | 1.321997061 | 0.006353019 |
| ENSMUSG00000074737 | 1 | |
| ENSMUSG00000074738 | 1 | |
| ENSMUSG00000074739 | 1.059463094 | 0.698112512 |
| ENSMUSG00000074743 | 1.752397232 | 0.000666524 |
| ENSMUSG00000074746 | 1.101444066 | 0.698112512 |
| ENSMUSG00000074748 | 1.112959895 | 0.689833241 |
| ENSMUSG00000074749 | -1.120270411 | 0.452437882 |
| ENSMUSG00000074753 | 1 | |
| ENSMUSG00000074754 | -1.020248548 | 0.778372272 |
| ENSMUSG00000074758 | 1 | |
| ENSMUSG00000074763 | 1 | |
| ENSMUSG00000074764 | 1 | |
| ENSMUSG00000074766 | -1.051018293 | 0.828564184 |
| ENSMUSG00000074768 | -1.044273782 | 0.698112512 |
| ENSMUSG00000074771 | 1.053991584 | 0.916014403 |

| | | |
|---|---|---|
| ENSMUSG00000074776 | 1 | |
| ENSMUSG00000074780 | 1.014545335 | 0.979002724 |
| ENSMUSG00000074781 | -1.046493792 | 0.916836847 |
| ENSMUSG00000074782 | -1.009083027 | 1 |
| ENSMUSG00000074783 | -1.02513447 | 0.968937442 |
| ENSMUSG00000074785 | -1.411949198 | 0.016003376 |
| ENSMUSG00000074793 | -1.191700959 | 0.094165307 |
| ENSMUSG00000074794 | -1.025627857 | 0.934760809 |
| ENSMUSG00000074796 | -1.010032031 | 0.980617339 |
| ENSMUSG00000074797 | -1.060849864 | 0.750742698 |
| ENSMUSG00000074800 | -1.007008185 | 1 |
| ENSMUSG00000074802 | 1.021746307 | 0.970020309 |
| ENSMUSG00000074804 | 1 | |
| ENSMUSG00000074805 | 1.059463094 | 0.820462971 |
| ENSMUSG00000074807 | -1.024780156 | 0.955452914 |
| ENSMUSG00000074811 | 1.065544951 | 0.602100972 |
| ENSMUSG00000074812 | 1 | |
| ENSMUSG00000074813 | -1.116535213 | 0.250137903 |
| ENSMUSG00000074817 | -1.381198941 | 0.01260323 |
| ENSMUSG00000074818 | 1.119766607 | 0.608497229 |
| ENSMUSG00000074824 | -1.084469591 | 0.670176133 |
| ENSMUSG00000074825 | -1.036378345 | 0.764343109 |
| ENSMUSG00000074826 | -1.013988437 | 0.9784582 |
| ENSMUSG00000074829 | 1.084278774 | 0.525842026 |
| ENSMUSG00000074830 | -1 | 1 |
| ENSMUSG00000074832 | -1.032914392 | 0.982838256 |
| ENSMUSG00000074837 | 1 | |
| ENSMUSG00000074841 | 1 | |
| ENSMUSG00000074846 | 1 | |
| ENSMUSG00000074847 | 1.039694307 | 0.955452914 |
| ENSMUSG00000074849 | 1 | |
| ENSMUSG00000074852 | 1 | |
| ENSMUSG00000074862 | 1.032420173 | 0.848848499 |
| ENSMUSG00000074863 | -1.058271395 | 0.656691142 |
| ENSMUSG00000074865 | -1.048017738 | 0.729018288 |
| ENSMUSG00000074867 | -1.025711459 | 0.886981862 |
| ENSMUSG00000074868 | 1.160607842 | 0.608497229 |
| ENSMUSG00000074870 | 1 | |
| ENSMUSG00000074871 | 1 | |
| ENSMUSG00000074872 | 1 | |
| ENSMUSG00000074873 | 1.042664899 | 0.608497229 |
| ENSMUSG00000074874 | 1.266416129 | 0.102101889 |
| ENSMUSG00000074876 | -1.030748997 | 0.942171713 |
| ENSMUSG00000074877 | 1 | |
| ENSMUSG00000074881 | 1.122242299 | 0.636101368 |
| ENSMUSG00000074882 | 1 | |
| ENSMUSG00000074884 | 1.105665061 | 0.420138683 |
| ENSMUSG00000074885 | 1 | |
| ENSMUSG00000074886 | -1.006454984 | 1 |
| ENSMUSG00000074887 | -1.043306199 | 0.698112512 |
| ENSMUSG00000074890 | 1.031680567 | 0.702862297 |
| ENSMUSG00000074892 | -1.117792522 | 0.532369657 |
| ENSMUSG00000074895 | -1.044273782 | 0.698112512 |
| ENSMUSG00000074896 | 1.040420874 | 0.767512509 |
| ENSMUSG00000074899 | -1.049475223 | 0.833212373 |
| ENSMUSG00000074903 | 1 | |
| ENSMUSG00000074909 | -1.019341317 | 0.976760734 |
| ENSMUSG00000074912 | -1.120476206 | 0.534652019 |
| ENSMUSG00000074913 | 1 | |
| ENSMUSG00000074916 | -1.038364089 | 0.706051768 |
| ENSMUSG00000074918 | -1.139494188 | 0.147316517 |
| ENSMUSG00000074919 | 1 | |
| ENSMUSG00000074922 | 1.061204149 | 0.657796687 |

| | | |
|---|---|---|
| ENSMUSG00000074923 | -1.114185589 | 0.615569604 |
| ENSMUSG00000074925 | 1.055377245 | 0.760180115 |
| ENSMUSG00000074928 | 1 | |
| ENSMUSG00000074930 | 1.122462048 | 0.420161906 |
| ENSMUSG00000074934 | 1.59436594 | 0.008670441 |
| ENSMUSG00000074939 | 1 | |
| ENSMUSG00000074943 | 1 | |
| ENSMUSG00000074945 | 1 | |
| ENSMUSG00000074946 | 1 | |
| ENSMUSG00000074947 | 1 | |
| ENSMUSG00000074951 | 1 | |
| ENSMUSG00000074952 | 1 | |
| ENSMUSG00000074955 | 1 | |
| ENSMUSG00000074957 | 1 | |
| ENSMUSG00000074965 | 1 | |
| ENSMUSG00000074966 | 1 | |
| ENSMUSG00000074968 | 1.562938159 | 0.004492904 |
| ENSMUSG00000074970 | 1 | |
| ENSMUSG00000074971 | 1.413306049 | 0.132613684 |
| ENSMUSG00000074973 | 1.150021168 | 0.509795474 |
| ENSMUSG00000074978 | 1.284154183 | 0.05050906 |
| ENSMUSG00000074981 | 1.019083579 | 0.97045957 |
| ENSMUSG00000074985 | 1 | |
| ENSMUSG00000074987 | -1 | 1 |
| ENSMUSG00000074991 | 1 | |
| ENSMUSG00000074994 | -1.08413501 | 0.734674233 |
| ENSMUSG00000074995 | 1 | |
| ENSMUSG00000074996 | 1 | |
| ENSMUSG00000074997 | 1 | |
| ENSMUSG00000074999 | -1.143787033 | 0.242169098 |
| ENSMUSG00000075000 | 1.007274005 | 1 |
| ENSMUSG00000075002 | 1 | |
| ENSMUSG00000075006 | 1 | |
| ENSMUSG00000075010 | -1 | 1 |
| ENSMUSG00000075012 | 1.295353108 | 0.101456114 |
| ENSMUSG00000075014 | -1.069502966 | 0.814385946 |
| ENSMUSG00000075015 | 1.028323353 | 0.933031586 |
| ENSMUSG00000075020 | 1 | |
| ENSMUSG00000075023 | 1.059463094 | 0.608497229 |
| ENSMUSG00000075024 | 1 | |
| ENSMUSG00000075025 | 1.05410549 | 0.838710806 |
| ENSMUSG00000075027 | -1.107299336 | 0.639279667 |
| ENSMUSG00000075028 | -1.091420314 | 0.169545598 |
| ENSMUSG00000075029 | 1 | |
| ENSMUSG00000075031 | -1.209492167 | 0.348596402 |
| ENSMUSG00000075033 | 1.033398041 | 0.814132742 |
| ENSMUSG00000075036 | 1 | |
| ENSMUSG00000075040 | -1.03241306 | 0.698112512 |
| ENSMUSG00000075042 | 1 | |
| ENSMUSG00000075044 | 1 | |
| ENSMUSG00000075045 | 1 | |
| ENSMUSG00000075046 | -1.043306199 | 0.698112512 |
| ENSMUSG00000075053 | 1.099513476 | 0.712991717 |
| ENSMUSG00000075054 | -1.011033779 | 0.989648415 |
| ENSMUSG00000075061 | 1 | |
| ENSMUSG00000075062 | 1 | |
| ENSMUSG00000075063 | 1 | |
| ENSMUSG00000075064 | 1 | |
| ENSMUSG00000075065 | 1 | |
| ENSMUSG00000075066 | 1 | |
| ENSMUSG00000075067 | 1 | |
| ENSMUSG00000075068 | 1 | |
| ENSMUSG00000075069 | 1 | |

| | | |
|---|---|---|
| ENSMUSG00000075070 | -1 | 1 |
| ENSMUSG00000075072 | 1 | |
| ENSMUSG00000075073 | 1 | |
| ENSMUSG00000075074 | 1 | |
| ENSMUSG00000075075 | 1 | |
| ENSMUSG00000075078 | 1 | |
| ENSMUSG00000075079 | 1 | |
| ENSMUSG00000075081 | 1 | |
| ENSMUSG00000075084 | 1 | |
| ENSMUSG00000075085 | 1 | |
| ENSMUSG00000075086 | 1 | |
| ENSMUSG00000075088 | 1 | |
| ENSMUSG00000075089 | 1 | |
| ENSMUSG00000075090 | 1 | |
| ENSMUSG00000075091 | 1 | |
| ENSMUSG00000075092 | 1 | |
| ENSMUSG00000075093 | 1 | |
| ENSMUSG00000075094 | 1 | |
| ENSMUSG00000075095 | 1 | |
| ENSMUSG00000075097 | 1 | |
| ENSMUSG00000075099 | 1 | |
| ENSMUSG00000075100 | 1 | |
| ENSMUSG00000075101 | 1 | |
| ENSMUSG00000075102 | 1 | |
| ENSMUSG00000075104 | 1 | |
| ENSMUSG00000075105 | 1 | |
| ENSMUSG00000075107 | 1 | |
| ENSMUSG00000075110 | 1 | |
| ENSMUSG00000075111 | 1 | |
| ENSMUSG00000075112 | 1 | |
| ENSMUSG00000075113 | 1 | |
| ENSMUSG00000075114 | 1 | |
| ENSMUSG00000075115 | 1 | |
| ENSMUSG00000075116 | 1 | |
| ENSMUSG00000075117 | 1 | |
| ENSMUSG00000075119 | 1 | |
| ENSMUSG00000075120 | 1 | |
| ENSMUSG00000075121 | 1 | |
| ENSMUSG00000075122 | 1.668643517 | 0.032224317 |
| ENSMUSG00000075124 | 1 | |
| ENSMUSG00000075125 | 1 | |
| ENSMUSG00000075126 | 1 | |
| ENSMUSG00000075127 | 1 | |
| ENSMUSG00000075128 | 1 | |
| ENSMUSG00000075132 | 1 | |
| ENSMUSG00000075133 | 1 | |
| ENSMUSG00000075136 | 1 | |
| ENSMUSG00000075137 | 1 | |
| ENSMUSG00000075139 | 1 | |
| ENSMUSG00000075140 | 1 | |
| ENSMUSG00000075141 | 1 | |
| ENSMUSG00000075142 | 1 | |
| ENSMUSG00000075143 | 1 | |
| ENSMUSG00000075144 | 1 | |
| ENSMUSG00000075145 | 1 | |
| ENSMUSG00000075146 | 1 | |
| ENSMUSG00000075148 | 1 | |
| ENSMUSG00000075149 | 1 | |
| ENSMUSG00000075150 | 1 | |
| ENSMUSG00000075151 | 1 | |
| ENSMUSG00000075153 | 1 | |
| ENSMUSG00000075154 | 1 | |
| ENSMUSG00000075155 | 1 | |

| | | |
|---|---|---|
| ENSMUSG00000075156 | 1 | |
| ENSMUSG00000075158 | 1 | |
| ENSMUSG00000075159 | 1 | |
| ENSMUSG00000075160 | 1 | |
| ENSMUSG00000075161 | 1 | |
| ENSMUSG00000075162 | 1 | |
| ENSMUSG00000075163 | 1 | |
| ENSMUSG00000075164 | 1 | |
| ENSMUSG00000075165 | 1 | |
| ENSMUSG00000075166 | 1 | |
| ENSMUSG00000075167 | 1 | |
| ENSMUSG00000075168 | 1 | |
| ENSMUSG00000075169 | 1 | |
| ENSMUSG00000075170 | 1 | |
| ENSMUSG00000075171 | 1 | |
| ENSMUSG00000075172 | 1 | |
| ENSMUSG00000075173 | 1 | |
| ENSMUSG00000075174 | 1 | |
| ENSMUSG00000075175 | 1 | |
| ENSMUSG00000075176 | 1 | |
| ENSMUSG00000075177 | 1 | |
| ENSMUSG00000075178 | 1 | |
| ENSMUSG00000075179 | 1 | |
| ENSMUSG00000075180 | 1 | |
| ENSMUSG00000075181 | 1 | |
| ENSMUSG00000075182 | 1 | |
| ENSMUSG00000075184 | -1 | 1 |
| ENSMUSG00000075185 | 1 | |
| ENSMUSG00000075186 | 1 | |
| ENSMUSG00000075187 | 1 | |
| ENSMUSG00000075188 | 1 | |
| ENSMUSG00000075189 | 1.079681442 | 0.697178241 |
| ENSMUSG00000075190 | 1 | |
| ENSMUSG00000075192 | 1 | |
| ENSMUSG00000075193 | 1 | |
| ENSMUSG00000075194 | 1 | |
| ENSMUSG00000075195 | 1 | |
| ENSMUSG00000075196 | 1 | |
| ENSMUSG00000075197 | 1 | |
| ENSMUSG00000075198 | 1 | |
| ENSMUSG00000075199 | 1 | |
| ENSMUSG00000075200 | 1 | |
| ENSMUSG00000075201 | 1 | |
| ENSMUSG00000075202 | 1 | |
| ENSMUSG00000075203 | 1 | |
| ENSMUSG00000075204 | 1 | |
| ENSMUSG00000075205 | 1 | |
| ENSMUSG00000075206 | 1 | |
| ENSMUSG00000075208 | 1 | |
| ENSMUSG00000075209 | 1 | |
| ENSMUSG00000075210 | 1 | |
| ENSMUSG00000075211 | 1 | |
| ENSMUSG00000075212 | 1 | |
| ENSMUSG00000075214 | 1 | |
| ENSMUSG00000075215 | 1 | |
| ENSMUSG00000075217 | 1 | |
| ENSMUSG00000075218 | 1 | |
| ENSMUSG00000075219 | 1 | |
| ENSMUSG00000075220 | 1 | |
| ENSMUSG00000075221 | 1 | |
| ENSMUSG00000075222 | 1 | |
| ENSMUSG00000075223 | 1 | |
| ENSMUSG00000075224 | -1.141886854 | 0.512702106 |

| | | |
|---|---|---|
| ENSMUSG00000075225 | -2.55538557 | 0.001427794 |
| ENSMUSG00000075227 | 1.059248427 | 0.803852736 |
| ENSMUSG00000075228 | 1 | |
| ENSMUSG00000075229 | -1.000959306 | 1 |
| ENSMUSG00000075232 | -1.048903022 | 0.845274366 |
| ENSMUSG00000075245 | 1.00822312 | 1 |
| ENSMUSG00000075249 | 1 | |
| ENSMUSG00000075254 | -1.096385917 | 0.644653062 |
| ENSMUSG00000075256 | -1.155936699 | 0.533863819 |
| ENSMUSG00000075265 | -1.000131914 | 1 |
| ENSMUSG00000075266 | -1.066444612 | 0.617912429 |
| ENSMUSG00000075267 | 1.03863466 | 0.950560821 |
| ENSMUSG00000075268 | -1 | 1 |
| ENSMUSG00000075269 | 1 | |
| ENSMUSG00000075270 | 1.045402873 | 0.923549054 |
| ENSMUSG00000075271 | -1.086205653 | 0.557851851 |
| ENSMUSG00000075272 | -1.106430219 | 0.666258087 |
| ENSMUSG00000075273 | -1.111345792 | 0.370614531 |
| ENSMUSG00000075276 | 1.059592506 | 0.776927068 |
| ENSMUSG00000075277 | 1.598430623 | 0.001723822 |
| ENSMUSG00000075279 | 1.079681442 | 0.726987474 |
| ENSMUSG00000075284 | -1.100250736 | 0.608497229 |
| ENSMUSG00000075286 | -1.044273782 | 0.698112512 |
| ENSMUSG00000075288 | 1.069068979 | 0.608497229 |
| ENSMUSG00000075289 | -1.02693459 | 0.773487331 |
| ENSMUSG00000075296 | 1 | |
| ENSMUSG00000075297 | 1.223775858 | 0.08283904 |
| ENSMUSG00000075302 | -1.382127068 | 0.224353537 |
| ENSMUSG00000075304 | -1.128668575 | 0.578712437 |
| ENSMUSG00000075307 | -1.165237207 | 0.51962022 |
| ENSMUSG00000075312 | -1 | 1 |
| ENSMUSG00000075314 | 1 | |
| ENSMUSG00000075316 | 2.049664973 | 4.11504E-05 |
| ENSMUSG00000075318 | 1.421483754 | 0.000696694 |
| ENSMUSG00000075324 | 1.128208983 | 0.143312239 |
| ENSMUSG00000075325 | -1 | 1 |
| ENSMUSG00000075327 | 1.046865665 | 0.764370878 |
| ENSMUSG00000075330 | 1 | |
| ENSMUSG00000075334 | 1.322417436 | 0.250944709 |
| ENSMUSG00000075335 | 1 | |
| ENSMUSG00000075359 | 1 | |
| ENSMUSG00000075370 | -1.043306199 | 0.79291651 |
| ENSMUSG00000075376 | 1.048777991 | 0.867016657 |
| ENSMUSG00000075377 | 1 | |
| ENSMUSG00000075378 | -1 | 1 |
| ENSMUSG00000075379 | 1 | |
| ENSMUSG00000075380 | 1 | |
| ENSMUSG00000075382 | 1 | |
| ENSMUSG00000075383 | 1 | |
| ENSMUSG00000075384 | 1 | |
| ENSMUSG00000075387 | 1 | |
| ENSMUSG00000075389 | 1.050070838 | 0.628315739 |
| ENSMUSG00000075390 | 1 | |
| ENSMUSG00000075391 | -1.457809263 | 0.005825322 |
| ENSMUSG00000075394 | -1.994383323 | 0.000050878 |
| ENSMUSG00000075395 | -1.068530808 | 0.751234476 |
| ENSMUSG00000075402 | 1 | |
| ENSMUSG00000075405 | -1.043306199 | 0.79291651 |
| ENSMUSG00000075408 | 1.012792449 | 1 |
| ENSMUSG00000075410 | -1.110907262 | 0.75177322 |
| ENSMUSG00000075415 | 1.043187187 | 0.854640873 |
| ENSMUSG00000075416 | 1.056280798 | 0.839185413 |
| ENSMUSG00000075419 | -1.01449481 | 0.976160676 |

| | | |
|---|---|---|
| ENSMUSG00000075420 | 1.03194599 | 0.97192334 |
| ENSMUSG00000075425 | -1.043306199 | 0.858911269 |
| ENSMUSG00000075427 | 1 | |
| ENSMUSG00000075431 | 1 | |
| ENSMUSG00000075437 | 1 | |
| ENSMUSG00000075463 | -1.003730133 | 1 |
| ENSMUSG00000075465 | 1 | |
| ENSMUSG00000075467 | 1.030331204 | 0.806982207 |
| ENSMUSG00000075469 | 1.049411284 | 0.892085484 |
| ENSMUSG00000075470 | 1.046038974 | 0.90404205 |
| ENSMUSG00000075478 | 1 | |
| ENSMUSG00000075480 | 1 | |
| ENSMUSG00000075485 | 1 | |
| ENSMUSG00000075486 | -1.144540181 | 0.289033137 |
| ENSMUSG00000075502 | -1.233290256 | 0.216046507 |
| ENSMUSG00000075510 | 1 | |
| ENSMUSG00000075511 | -1.048533766 | 0.806851039 |
| ENSMUSG00000075512 | 1 | |
| ENSMUSG00000075514 | 1.206042956 | 0.135595698 |
| ENSMUSG00000075517 | 1 | |
| ENSMUSG00000075520 | 1 | |
| ENSMUSG00000075524 | 1.069068979 | 0.74773997 |
| ENSMUSG00000075525 | 1 | |
| ENSMUSG00000075528 | -1.035233971 | 0.903464611 |
| ENSMUSG00000075534 | -1.053549062 | 0.698112512 |
| ENSMUSG00000075538 | 1 | |
| ENSMUSG00000075543 | 1 | |
| ENSMUSG00000075549 | 1 | |
| ENSMUSG00000075551 | 1 | |
| ENSMUSG00000075552 | 1 | |
| ENSMUSG00000075553 | -1.113515092 | 0.608497229 |
| ENSMUSG00000075555 | -1.044273782 | 0.698112512 |
| ENSMUSG00000075558 | -1.176211709 | 0.509717267 |
| ENSMUSG00000075566 | 1 | |
| ENSMUSG00000075567 | 1 | |
| ENSMUSG00000075569 | 1.110137235 | 0.710700793 |
| ENSMUSG00000075570 | -1.044273782 | 0.698112512 |
| ENSMUSG00000075571 | 1 | |
| ENSMUSG00000075572 | 1 | |
| ENSMUSG00000075573 | 1 | |
| ENSMUSG00000075574 | 1 | |
| ENSMUSG00000075576 | -1.016648059 | 0.992213082 |
| ENSMUSG00000075581 | 1.304691554 | 0.180865409 |
| ENSMUSG00000075585 | 1.341748473 | 0.008510863 |
| ENSMUSG00000075586 | 1 | |
| ENSMUSG00000075587 | -1.049961316 | 0.913105899 |
| ENSMUSG00000075588 | -1.052975067 | 0.729171813 |
| ENSMUSG00000075589 | -1.03845103 | 0.870427333 |
| ENSMUSG00000075590 | 1.049306942 | 0.608497229 |
| ENSMUSG00000075591 | -1.02522956 | 0.970707004 |
| ENSMUSG00000075592 | -1.694324953 | 0.004327743 |
| ENSMUSG00000075593 | 1.069068979 | 0.608497229 |
| ENSMUSG00000075595 | 1.008932956 | 1 |
| ENSMUSG00000075596 | -1.032140202 | 0.934760809 |
| ENSMUSG00000075598 | 1 | |
| ENSMUSG00000075599 | -1 | 1 |
| ENSMUSG00000075600 | 1.018406694 | 0.953172518 |
| ENSMUSG00000075602 | 1.642366069 | 0.004366561 |
| ENSMUSG00000075604 | 1.014545335 | 0.991944074 |
| ENSMUSG00000075605 | 1 | |
| ENSMUSG00000075609 | 1.171563499 | 0.233412931 |
| ENSMUSG00000075610 | -1.04526435 | 0.908065256 |
| ENSMUSG00000075611 | 1 | |

| | | |
|---|---|---|
| ENSMUSG00000075700 | -1.040346361 | 0.941510974 |
| ENSMUSG00000075701 | 1.00761504 | 1 |
| ENSMUSG00000075702 | 1.087968499 | 0.653471423 |
| ENSMUSG00000075703 | 1.050748551 | 0.867420842 |
| ENSMUSG00000075704 | 1.000927658 | 1 |
| ENSMUSG00000075705 | -1.051171775 | 0.836597096 |
| ENSMUSG00000075706 | 1.099807464 | 0.644157716 |
| ENSMUSG00000075707 | 1 | |
| ENSMUSG00000075719 | 1 | |
| ENSMUSG00000075726 | 1 | |
| ENSMUSG00000075760 | 1 | |
| ENSMUSG00000075770 | 1 | |
| ENSMUSG00000075826 | 1 | |
| ENSMUSG00000075829 | -1.053549062 | 0.698112512 |
| ENSMUSG00000075832 | 1 | |
| ENSMUSG00000075833 | 1 | |
| ENSMUSG00000075837 | 1.006784519 | 1 |
| ENSMUSG00000075864 | 1 | |
| ENSMUSG00000075874 | 1 | |
| ENSMUSG00000075893 | 1 | |
| ENSMUSG00000075903 | 1 | |
| ENSMUSG00000075906 | 1 | |
| ENSMUSG00000075908 | 1.015486245 | 0.972980766 |
| ENSMUSG00000075918 | 1 | |
| ENSMUSG00000075934 | 1 | |
| ENSMUSG00000075952 | 1 | |
| ENSMUSG00000075958 | 1 | |
| ENSMUSG00000075968 | 1 | |
| ENSMUSG00000075973 | 1 | |
| ENSMUSG00000075983 | 1 | |
| ENSMUSG00000076005 | 1 | |
| ENSMUSG00000076006 | 1 | |
| ENSMUSG00000076009 | 1 | |
| ENSMUSG00000076010 | 1 | |
| ENSMUSG00000076011 | 1 | |
| ENSMUSG00000076012 | 1 | |
| ENSMUSG00000076014 | 1 | |
| ENSMUSG00000076028 | 1 | |
| ENSMUSG00000076030 | 1 | |
| ENSMUSG00000076031 | 1 | |
| ENSMUSG00000076033 | 1 | |
| ENSMUSG00000076036 | 1 | |
| ENSMUSG00000076037 | 1 | |
| ENSMUSG00000076038 | 1 | |
| ENSMUSG00000076039 | -1.043306199 | 0.698112512 |
| ENSMUSG00000076041 | 1 | |
| ENSMUSG00000076042 | 1 | |
| ENSMUSG00000076049 | 1 | |
| ENSMUSG00000076050 | 1 | |
| ENSMUSG00000076051 | 1 | |
| ENSMUSG00000076052 | 1 | |
| ENSMUSG00000076060 | -1.044273782 | 0.698112512 |
| ENSMUSG00000076062 | 1 | |
| ENSMUSG00000076063 | 1 | |
| ENSMUSG00000076064 | 1 | |
| ENSMUSG00000076065 | -1 | 1 |
| ENSMUSG00000076066 | 1 | |
| ENSMUSG00000076067 | 1 | |
| ENSMUSG00000076068 | 1 | |
| ENSMUSG00000076117 | 1 | |
| ENSMUSG00000076118 | 1 | |
| ENSMUSG00000076119 | 1 | |
| ENSMUSG00000076120 | 1 | |

| | | |
|---|---|---|
| ENSMUSG00000076121 | -1.03962336 | 0.698112512 |
| ENSMUSG00000076122 | 1.095872691 | 0.608497229 |
| ENSMUSG00000076123 | 1 | |
| ENSMUSG00000076126 | 1 | |
| ENSMUSG00000076127 | 1 | |
| ENSMUSG00000076128 | 1 | |
| ENSMUSG00000076129 | 1 | |
| ENSMUSG00000076135 | -1.03962336 | 0.698112512 |
| ENSMUSG00000076136 | -1.139334642 | 0.608497229 |
| ENSMUSG00000076137 | 1.200936955 | 0.382009062 |
| ENSMUSG00000076138 | 1 | |
| ENSMUSG00000076140 | 1 | |
| ENSMUSG00000076141 | 1 | |
| ENSMUSG00000076142 | -1.044273782 | 0.698112512 |
| ENSMUSG00000076143 | 1 | |
| ENSMUSG00000076144 | 1 | |
| ENSMUSG00000076145 | 1 | |
| ENSMUSG00000076146 | 1 | |
| ENSMUSG00000076147 | 1 | |
| ENSMUSG00000076163 | 1 | |
| ENSMUSG00000076180 | 1 | |
| ENSMUSG00000076190 | 1 | |
| ENSMUSG00000076199 | 1 | |
| ENSMUSG00000076212 | 1 | |
| ENSMUSG00000076214 | 1 | |
| ENSMUSG00000076216 | 1 | |
| ENSMUSG00000076217 | 1 | |
| ENSMUSG00000076218 | 1 | |
| ENSMUSG00000076219 | 1 | |
| ENSMUSG00000076222 | 1 | |
| ENSMUSG00000076227 | 1 | |
| ENSMUSG00000076228 | 1 | |
| ENSMUSG00000076229 | -1 | 1 |
| ENSMUSG00000076230 | 1 | |
| ENSMUSG00000076231 | 1 | |
| ENSMUSG00000076232 | 1 | |
| ENSMUSG00000076233 | 1 | |
| ENSMUSG00000076234 | 1 | |
| ENSMUSG00000076236 | 1 | |
| ENSMUSG00000076237 | 1 | |
| ENSMUSG00000076238 | 1 | |
| ENSMUSG00000076240 | 1 | |
| ENSMUSG00000076241 | 1 | |
| ENSMUSG00000076246 | 1 | |
| ENSMUSG00000076251 | 1 | |
| ENSMUSG00000076252 | 1 | |
| ENSMUSG00000076253 | 1 | |
| ENSMUSG00000076255 | 1 | |
| ENSMUSG00000076256 | 1 | |
| ENSMUSG00000076258 | -1.043609611 | 0.929923764 |
| ENSMUSG00000076269 | 1 | |
| ENSMUSG00000076270 | 1 | |
| ENSMUSG00000076271 | -1.197957061 | 0.507246493 |
| ENSMUSG00000076272 | 1 | |
| ENSMUSG00000076274 | 1 | |
| ENSMUSG00000076275 | 1 | |
| ENSMUSG00000076281 | -1.03962336 | 0.698112512 |
| ENSMUSG00000076282 | 1 | |
| ENSMUSG00000076284 | 1 | |
| ENSMUSG00000076286 | 1 | |
| ENSMUSG00000076287 | 1 | |
| ENSMUSG00000076288 | 1 | |
| ENSMUSG00000076294 | 1 | |

| | | |
|---|---|---|
| ENSMUSG00000076300 | 1 | |
| ENSMUSG00000076303 | 1 | |
| ENSMUSG00000076304 | 1 | |
| ENSMUSG00000076305 | 1 | |
| ENSMUSG00000076309 | 1 | |
| ENSMUSG00000076312 | 1 | |
| ENSMUSG00000076313 | 1 | |
| ENSMUSG00000076315 | -1.099174267 | 0.608497229 |
| ENSMUSG00000076316 | 1 | |
| ENSMUSG00000076318 | 1 | |
| ENSMUSG00000076320 | 1 | |
| ENSMUSG00000076323 | 1 | |
| ENSMUSG00000076327 | 1 | |
| ENSMUSG00000076332 | 1.059463094 | 0.608497229 |
| ENSMUSG00000076333 | -1.085651418 | 0.608497229 |
| ENSMUSG00000076335 | 1 | |
| ENSMUSG00000076337 | 1 | |
| ENSMUSG00000076338 | 1 | |
| ENSMUSG00000076343 | 1 | |
| ENSMUSG00000076355 | 1 | |
| ENSMUSG00000076357 | 1 | |
| ENSMUSG00000076360 | 1 | |
| ENSMUSG00000076361 | 1 | |
| ENSMUSG00000076364 | 1 | |
| ENSMUSG00000076366 | 1 | |
| ENSMUSG00000076370 | 1 | |
| ENSMUSG00000076371 | 1 | |
| ENSMUSG00000076372 | 1 | |
| ENSMUSG00000076376 | -1.043306199 | 0.698112512 |
| ENSMUSG00000076377 | 1.059463094 | 0.608497229 |
| ENSMUSG00000076379 | 1 | |
| ENSMUSG00000076382 | 1 | |
| ENSMUSG00000076385 | 1 | |
| ENSMUSG00000076387 | 1 | |
| ENSMUSG00000076389 | 1 | |
| ENSMUSG00000076392 | 1 | |
| ENSMUSG00000076394 | -1.069502966 | 0.698112512 |
| ENSMUSG00000076395 | 1 | |
| ENSMUSG00000076396 | 1 | |
| ENSMUSG00000076398 | 1 | |
| ENSMUSG00000076399 | 1 | |
| ENSMUSG00000076407 | 1 | |
| ENSMUSG00000076424 | 1 | |
| ENSMUSG00000076430 | -1.03962336 | 0.698112512 |
| ENSMUSG00000076431 | 1.214998175 | 0.02605836 |
| ENSMUSG00000076432 | 1.022562522 | 0.988424085 |
| ENSMUSG00000076433 | 1.055220942 | 0.404493254 |
| ENSMUSG00000076434 | -1.106996765 | 0.739099222 |
| ENSMUSG00000076435 | 1.093478505 | 0.608497229 |
| ENSMUSG00000076436 | 1.001874515 | 1 |
| ENSMUSG00000076437 | 1.000872563 | 1 |
| ENSMUSG00000076438 | -1.053549062 | 0.698112512 |
| ENSMUSG00000076439 | 1.243032879 | 0.334859726 |
| ENSMUSG00000076441 | 1.340231548 | 0.010275103 |
| ENSMUSG00000076444 | 1.059463094 | 0.608497229 |
| ENSMUSG00000076445 | 1 | |
| ENSMUSG00000076448 | 1.015486245 | 0.972980766 |
| ENSMUSG00000076450 | 1 | |
| ENSMUSG00000076451 | 1 | |
| ENSMUSG00000076453 | 1 | |
| ENSMUSG00000076454 | 1 | |
| ENSMUSG00000076455 | 1 | |
| ENSMUSG00000076456 | 1 | |

| | |
|---|---|
| ENSMUSG00000076457 | 1 |
| ENSMUSG00000076458 | 1 |
| ENSMUSG00000076459 | 1 |
| ENSMUSG00000076460 | 1 |
| ENSMUSG00000076461 | 1 |
| ENSMUSG00000076462 | 1 |
| ENSMUSG00000076463 | 1 |
| ENSMUSG00000076465 | 1 |
| ENSMUSG00000076467 | 1 |
| ENSMUSG00000076469 | 1 |
| ENSMUSG00000076470 | 1 |
| ENSMUSG00000076471 | 1 |
| ENSMUSG00000076472 | 1 |
| ENSMUSG00000076473 | 1 |
| ENSMUSG00000076474 | 1 |
| ENSMUSG00000076475 | 1 |
| ENSMUSG00000076476 | 1 |
| ENSMUSG00000076477 | 1 |
| ENSMUSG00000076478 | 1 |
| ENSMUSG00000076479 | 1 |
| ENSMUSG00000076480 | 1 |
| ENSMUSG00000076481 | 1 |
| ENSMUSG00000076483 | 1 |
| ENSMUSG00000076484 | 1 |
| ENSMUSG00000076485 | 1 |
| ENSMUSG00000076486 | 1 |
| ENSMUSG00000076487 | 1 |
| ENSMUSG00000076488 | 1 |
| ENSMUSG00000076489 | 1 |
| ENSMUSG00000076490 | 1 |
| ENSMUSG00000076492 | 1 |
| ENSMUSG00000076493 | 1 |
| ENSMUSG00000076494 | 1 |
| ENSMUSG00000076495 | 1 |
| ENSMUSG00000076496 | 1 |
| ENSMUSG00000076497 | 1 |
| ENSMUSG00000076498 | 1 |
| ENSMUSG00000076499 | 1 |
| ENSMUSG00000076500 | 1 |
| ENSMUSG00000076501 | 1 |
| ENSMUSG00000076505 | 1 |
| ENSMUSG00000076508 | 1 |
| ENSMUSG00000076512 | 1 |
| ENSMUSG00000076514 | 1 |
| ENSMUSG00000076518 | 1 |
| ENSMUSG00000076522 | 1 |
| ENSMUSG00000076523 | 1 |
| ENSMUSG00000076525 | 1 |
| ENSMUSG00000076526 | 1 |
| ENSMUSG00000076530 | 1 |
| ENSMUSG00000076531 | 1 |
| ENSMUSG00000076532 | 1 |
| ENSMUSG00000076533 | 1 |
| ENSMUSG00000076534 | 1 |
| ENSMUSG00000076535 | 1 |
| ENSMUSG00000076536 | 1 |
| ENSMUSG00000076538 | 1 |
| ENSMUSG00000076539 | 1 |
| ENSMUSG00000076540 | 1 |
| ENSMUSG00000076541 | 1 |
| ENSMUSG00000076543 | 1 |
| ENSMUSG00000076545 | 1 |
| ENSMUSG00000076547 | 1 |

| | |
|---|---|
| ENSMUSG00000076548 | 1 |
| ENSMUSG00000076549 | 1 |
| ENSMUSG00000076550 | 1 |
| ENSMUSG00000076552 | 1 |
| ENSMUSG00000076555 | 1 |
| ENSMUSG00000076556 | 1 |
| ENSMUSG00000076557 | 1 |
| ENSMUSG00000076562 | 1 |
| ENSMUSG00000076563 | 1 |
| ENSMUSG00000076564 | 1 |
| ENSMUSG00000076569 | 1 |
| ENSMUSG00000076570 | 1 |
| ENSMUSG00000076571 | 1 |
| ENSMUSG00000076572 | 1 |
| ENSMUSG00000076573 | 1 |
| ENSMUSG00000076575 | 1 |
| ENSMUSG00000076576 | 1 |
| ENSMUSG00000076577 | 1 |
| ENSMUSG00000076578 | 1 |
| ENSMUSG00000076580 | 1 |
| ENSMUSG00000076581 | 1 |
| ENSMUSG00000076583 | 1 |
| ENSMUSG00000076586 | 1 |
| ENSMUSG00000076587 | 1 |
| ENSMUSG00000076589 | 1 |
| ENSMUSG00000076591 | 1 |
| ENSMUSG00000076594 | 1 |
| ENSMUSG00000076596 | 1 |
| ENSMUSG00000076598 | 1 |
| ENSMUSG00000076604 | 1 |
| ENSMUSG00000076605 | 1 |
| ENSMUSG00000076606 | 1 |
| ENSMUSG00000076607 | 1 |
| ENSMUSG00000076608 | 1 |
| ENSMUSG00000076609 | 1 |
| ENSMUSG00000076612 | 1 |
| ENSMUSG00000076613 | 1 |
| ENSMUSG00000076614 | 1 |
| ENSMUSG00000076615 | 1 |
| ENSMUSG00000076617 | 1 |
| ENSMUSG00000076618 | 1 |
| ENSMUSG00000076619 | 1 |
| ENSMUSG00000076620 | 1 |
| ENSMUSG00000076621 | 1 |
| ENSMUSG00000076630 | 1 |
| ENSMUSG00000076632 | 1 |
| ENSMUSG00000076633 | 1 |
| ENSMUSG00000076646 | 1 |
| ENSMUSG00000076652 | 1 |
| ENSMUSG00000076653 | 1 |
| ENSMUSG00000076655 | 1 |
| ENSMUSG00000076661 | 1 |
| ENSMUSG00000076665 | 1 |
| ENSMUSG00000076666 | 1 |
| ENSMUSG00000076668 | 1 |
| ENSMUSG00000076670 | 1 |
| ENSMUSG00000076671 | 1 |
| ENSMUSG00000076672 | 1 |
| ENSMUSG00000076674 | 1 |
| ENSMUSG00000076676 | 1 |
| ENSMUSG00000076677 | 1 |
| ENSMUSG00000076680 | 1 |
| ENSMUSG00000076688 | 1 |

| | | |
|---|---|---|
| ENSMUSG00000076695 | 1 | |
| ENSMUSG00000076709 | 1 | |
| ENSMUSG00000076710 | 1 | |
| ENSMUSG00000076731 | 1 | |
| ENSMUSG00000076733 | 1 | |
| ENSMUSG00000076744 | 1 | |
| ENSMUSG00000076745 | 1 | |
| ENSMUSG00000076746 | 1 | |
| ENSMUSG00000076747 | 1 | |
| ENSMUSG00000076749 | 1 | |
| ENSMUSG00000076750 | 1 | |
| ENSMUSG00000076752 | 1 | |
| ENSMUSG00000076754 | 1 | |
| ENSMUSG00000076755 | 1 | |
| ENSMUSG00000076757 | 1 | |
| ENSMUSG00000076758 | 1 | |
| ENSMUSG00000076759 | 1 | |
| ENSMUSG00000076760 | 1 | |
| ENSMUSG00000076761 | 1 | |
| ENSMUSG00000076762 | 1 | |
| ENSMUSG00000076764 | 1 | |
| ENSMUSG00000076770 | 1 | |
| ENSMUSG00000076778 | 1 | |
| ENSMUSG00000076795 | 1 | |
| ENSMUSG00000076796 | 1 | |
| ENSMUSG00000076800 | 1 | |
| ENSMUSG00000076802 | 1 | |
| ENSMUSG00000076821 | 1 | |
| ENSMUSG00000076822 | 1 | |
| ENSMUSG00000076823 | 1 | |
| ENSMUSG00000076824 | 1 | |
| ENSMUSG00000076826 | 1 | |
| ENSMUSG00000076831 | 1 | |
| ENSMUSG00000076839 | 1 | |
| ENSMUSG00000076840 | 1 | |
| ENSMUSG00000076846 | -1.085651418 | 0.532369657 |
| ENSMUSG00000076848 | 1 | |
| ENSMUSG00000076855 | 1 | |
| ENSMUSG00000076858 | 1 | |
| ENSMUSG00000076861 | 1 | |
| ENSMUSG00000076862 | 1 | |
| ENSMUSG00000076863 | 1 | |
| ENSMUSG00000076864 | 1 | |
| ENSMUSG00000076865 | 1 | |
| ENSMUSG00000076866 | 1 | |
| ENSMUSG00000076867 | 1 | |
| ENSMUSG00000076870 | 1 | |
| ENSMUSG00000076871 | 1 | |
| ENSMUSG00000076873 | 1 | |
| ENSMUSG00000076874 | 1 | |
| ENSMUSG00000076875 | 1 | |
| ENSMUSG00000076876 | 1 | |
| ENSMUSG00000076877 | 1 | |
| ENSMUSG00000076878 | 1 | |
| ENSMUSG00000076879 | 1 | |
| ENSMUSG00000076880 | 1 | |
| ENSMUSG00000076881 | 1 | |
| ENSMUSG00000076882 | 1 | |
| ENSMUSG00000076883 | 1 | |
| ENSMUSG00000076884 | 1 | |
| ENSMUSG00000076885 | 1 | |
| ENSMUSG00000076886 | 1 | |
| ENSMUSG00000076887 | 1 | |

| | | |
|---|---|---|
| ENSMUSG00000076888 | 1 | |
| ENSMUSG00000076889 | 1 | |
| ENSMUSG00000076890 | 1 | |
| ENSMUSG00000076891 | 1 | |
| ENSMUSG00000076892 | 1 | |
| ENSMUSG00000076893 | 1 | |
| ENSMUSG00000076894 | 1 | |
| ENSMUSG00000076895 | 1 | |
| ENSMUSG00000076896 | 1 | |
| ENSMUSG00000076897 | 1.210869356 | 0.327655754 |
| ENSMUSG00000076898 | 1 | |
| ENSMUSG00000076899 | 1 | |
| ENSMUSG00000076900 | 1 | |
| ENSMUSG00000076901 | 1 | |
| ENSMUSG00000076902 | 1 | |
| ENSMUSG00000076903 | 1.059463094 | 0.608497229 |
| ENSMUSG00000076904 | 1 | |
| ENSMUSG00000076905 | 1 | |
| ENSMUSG00000076906 | 1 | |
| ENSMUSG00000076907 | 1 | |
| ENSMUSG00000076908 | 1.059463094 | 0.608497229 |
| ENSMUSG00000076909 | 1.059463094 | 0.608497229 |
| ENSMUSG00000076910 | 1 | |
| ENSMUSG00000076911 | 1 | |
| ENSMUSG00000076912 | 1 | |
| ENSMUSG00000076913 | 1 | |
| ENSMUSG00000076914 | -1.043306199 | 0.698112512 |
| ENSMUSG00000076915 | 1 | |
| ENSMUSG00000076916 | 1 | |
| ENSMUSG00000076917 | 1 | |
| ENSMUSG00000076918 | 1 | |
| ENSMUSG00000076919 | 1 | |
| ENSMUSG00000076920 | 1.059463094 | 0.608497229 |
| ENSMUSG00000076921 | 1 | |
| ENSMUSG00000076922 | 1.069068979 | 0.608497229 |
| ENSMUSG00000076923 | 1 | |
| ENSMUSG00000076924 | -1 | 1 |
| ENSMUSG00000076925 | 1 | |
| ENSMUSG00000076926 | 1 | |
| ENSMUSG00000076927 | 1 | |
| ENSMUSG00000076928 | 1.435676746 | 0.007358942 |
| ENSMUSG00000076934 | 1 | |
| ENSMUSG00000076937 | 1 | |
| ENSMUSG00000076939 | 1 | |
| ENSMUSG00000076940 | 1 | |
| ENSMUSG00000076941 | 1 | |
| ENSMUSG00000076944 | 1 | |
| ENSMUSG00000076946 | 1 | |
| ENSMUSG00000076948 | 1 | |
| ENSMUSG00000076949 | 1 | |
| ENSMUSG00000076950 | 1 | |
| ENSMUSG00000076951 | 1 | |
| ENSMUSG00000076952 | 1 | |
| ENSMUSG00000076956 | 1 | |
| ENSMUSG00000076958 | 1 | |
| ENSMUSG00000076959 | 1 | |
| ENSMUSG00000076961 | -1 | 1 |
| ENSMUSG00000076964 | 1 | |
| ENSMUSG00000076966 | 1 | |
| ENSMUSG00000076972 | 1 | |
| ENSMUSG00000076976 | 1 | |
| ENSMUSG00000076977 | 1 | |
| ENSMUSG00000076983 | 1 | |

| | | |
|---|---|---|
| ENSMUSG00000076987 | 1 | |
| ENSMUSG00000076994 | 1 | |
| ENSMUSG00000076999 | 1 | |
| ENSMUSG00000077001 | 1 | |
| ENSMUSG00000077004 | 1 | |
| ENSMUSG00000077010 | 1 | |
| ENSMUSG00000077015 | 1 | |
| ENSMUSG00000077019 | 1 | |
| ENSMUSG00000077021 | 1 | |
| ENSMUSG00000077022 | 1 | |
| ENSMUSG00000077025 | 1 | |
| ENSMUSG00000077038 | 1 | |
| ENSMUSG00000077039 | 1 | |
| ENSMUSG00000077040 | 1 | |
| ENSMUSG00000077042 | 1 | |
| ENSMUSG00000077049 | 1 | |
| ENSMUSG00000077050 | 1 | |
| ENSMUSG00000077055 | 1 | |
| ENSMUSG00000077058 | 1 | |
| ENSMUSG00000077059 | 1 | |
| ENSMUSG00000077065 | 1 | |
| ENSMUSG00000077081 | 1 | |
| ENSMUSG00000077082 | 1 | |
| ENSMUSG00000077085 | 1 | |
| ENSMUSG00000077086 | 1 | |
| ENSMUSG00000077090 | 1 | |
| ENSMUSG00000077092 | 1 | |
| ENSMUSG00000077095 | 1 | |
| ENSMUSG00000077096 | 1 | |
| ENSMUSG00000077104 | 1 | |
| ENSMUSG00000077107 | 1 | |
| ENSMUSG00000077110 | 1 | |
| ENSMUSG00000077111 | 1 | |
| ENSMUSG00000077113 | 1 | |
| ENSMUSG00000077119 | 1 | |
| ENSMUSG00000077125 | 1 | |
| ENSMUSG00000077127 | 1 | |
| ENSMUSG00000077131 | 1 | |
| ENSMUSG00000077138 | 1 | |
| ENSMUSG00000077142 | 1 | |
| ENSMUSG00000077143 | 1 | |
| ENSMUSG00000077144 | 1 | |
| ENSMUSG00000077146 | 1 | |
| ENSMUSG00000077148 | -1.068104567 | 0.814608539 |
| ENSMUSG00000077149 | 1 | |
| ENSMUSG00000077150 | 1 | |
| ENSMUSG00000077152 | 1 | |
| ENSMUSG00000077153 | 1 | |
| ENSMUSG00000077155 | 1 | |
| ENSMUSG00000077156 | 1 | |
| ENSMUSG00000077158 | 1 | |
| ENSMUSG00000077160 | 1 | |
| ENSMUSG00000077161 | 1 | |
| ENSMUSG00000077163 | 1 | |
| ENSMUSG00000077166 | 1 | |
| ENSMUSG00000077167 | 1.065746409 | 0.802320762 |
| ENSMUSG00000077169 | 1 | |
| ENSMUSG00000077171 | 1 | |
| ENSMUSG00000077172 | 1 | |
| ENSMUSG00000077174 | 1 | |
| ENSMUSG00000077175 | 1 | |
| ENSMUSG00000077176 | 1 | |
| ENSMUSG00000077177 | 1 | |

| | | |
|---|---|---|
| ENSMUSG00000077178 | 1 | |
| ENSMUSG00000077180 | 1 | |
| ENSMUSG00000077181 | 1 | |
| ENSMUSG00000077184 | 1 | |
| ENSMUSG00000077185 | 1 | |
| ENSMUSG00000077188 | 1 | |
| ENSMUSG00000077190 | 1 | |
| ENSMUSG00000077191 | 1 | |
| ENSMUSG00000077192 | -1.047137823 | 0.949812939 |
| ENSMUSG00000077194 | 1 | |
| ENSMUSG00000077196 | 1 | |
| ENSMUSG00000077197 | 1 | |
| ENSMUSG00000077199 | 1 | |
| ENSMUSG00000077200 | 1 | |
| ENSMUSG00000077201 | 1 | |
| ENSMUSG00000077202 | 1.043471668 | 0.938479652 |
| ENSMUSG00000077204 | 1 | |
| ENSMUSG00000077205 | 1 | |
| ENSMUSG00000077207 | 1 | |
| ENSMUSG00000077208 | 1 | |
| ENSMUSG00000077209 | 1 | |
| ENSMUSG00000077212 | -1.090133001 | 0.627656175 |
| ENSMUSG00000077214 | 1 | |
| ENSMUSG00000077215 | 1 | |
| ENSMUSG00000077217 | 1 | |
| ENSMUSG00000077218 | 1 | |
| ENSMUSG00000077220 | 1 | |
| ENSMUSG00000077221 | 1 | |
| ENSMUSG00000077222 | 1.162081855 | 0.608497229 |
| ENSMUSG00000077223 | 1 | |
| ENSMUSG00000077224 | 1 | |
| ENSMUSG00000077227 | -1.053549062 | 0.698112512 |
| ENSMUSG00000077228 | 1 | |
| ENSMUSG00000077230 | 1 | |
| ENSMUSG00000077231 | 1 | |
| ENSMUSG00000077235 | 1 | |
| ENSMUSG00000077236 | 1 | |
| ENSMUSG00000077237 | 1 | |
| ENSMUSG00000077239 | -1.043306199 | 0.698112512 |
| ENSMUSG00000077240 | 1 | |
| ENSMUSG00000077241 | 1 | |
| ENSMUSG00000077243 | 1 | |
| ENSMUSG00000077244 | 1 | |
| ENSMUSG00000077246 | 1 | |
| ENSMUSG00000077248 | 1 | |
| ENSMUSG00000077250 | 1 | |
| ENSMUSG00000077251 | 1 | |
| ENSMUSG00000077252 | 1 | |
| ENSMUSG00000077253 | 1 | |
| ENSMUSG00000077254 | 1 | |
| ENSMUSG00000077255 | 1 | |
| ENSMUSG00000077256 | 1 | |
| ENSMUSG00000077257 | 1 | |
| ENSMUSG00000077258 | 1 | |
| ENSMUSG00000077259 | 1 | |
| ENSMUSG00000077260 | 1 | |
| ENSMUSG00000077261 | 1 | |
| ENSMUSG00000077262 | 1 | |
| ENSMUSG00000077263 | 1 | |
| ENSMUSG00000077264 | 1 | |
| ENSMUSG00000077265 | 1.095047923 | 0.765908827 |
| ENSMUSG00000077268 | 1 | |
| ENSMUSG00000077269 | 1 | |

| | | |
|---|---|---|
| ENSMUSG00000077270 | 1 | |
| ENSMUSG00000077271 | 1 | |
| ENSMUSG00000077272 | 1 | |
| ENSMUSG00000077273 | 1 | |
| ENSMUSG00000077274 | -1 | 1 |
| ENSMUSG00000077275 | 1 | |
| ENSMUSG00000077276 | 1 | |
| ENSMUSG00000077277 | 1 | |
| ENSMUSG00000077278 | 1 | |
| ENSMUSG00000077279 | 1 | |
| ENSMUSG00000077280 | 1 | |
| ENSMUSG00000077281 | 1 | |
| ENSMUSG00000077282 | 1 | |
| ENSMUSG00000077283 | 1 | |
| ENSMUSG00000077285 | 1.058951264 | 0.875315552 |
| ENSMUSG00000077287 | 1 | |
| ENSMUSG00000077288 | 1 | |
| ENSMUSG00000077289 | 1 | |
| ENSMUSG00000077290 | 1 | |
| ENSMUSG00000077291 | 1 | |
| ENSMUSG00000077292 | 1 | |
| ENSMUSG00000077293 | 1 | |
| ENSMUSG00000077294 | 1 | |
| ENSMUSG00000077295 | 1 | |
| ENSMUSG00000077298 | 1 | |
| ENSMUSG00000077299 | 1 | |
| ENSMUSG00000077302 | 1 | |
| ENSMUSG00000077303 | 1 | |
| ENSMUSG00000077304 | 1 | |
| ENSMUSG00000077305 | 1 | |
| ENSMUSG00000077306 | 1 | |
| ENSMUSG00000077309 | 1.172128996 | 0.251544797 |
| ENSMUSG00000077312 | 1 | |
| ENSMUSG00000077314 | 1 | |
| ENSMUSG00000077315 | 1 | |
| ENSMUSG00000077316 | 1 | |
| ENSMUSG00000077317 | 1 | |
| ENSMUSG00000077318 | 1.059463094 | 0.608497229 |
| ENSMUSG00000077320 | 1 | |
| ENSMUSG00000077323 | 1 | |
| ENSMUSG00000077325 | 1 | |
| ENSMUSG00000077327 | 1 | |
| ENSMUSG00000077328 | 1 | |
| ENSMUSG00000077332 | 1 | |
| ENSMUSG00000077333 | 1 | |
| ENSMUSG00000077334 | 1 | |
| ENSMUSG00000077336 | 1 | |
| ENSMUSG00000077337 | 1 | |
| ENSMUSG00000077338 | -1 | 1 |
| ENSMUSG00000077339 | 1 | |
| ENSMUSG00000077342 | 1 | |
| ENSMUSG00000077344 | 1 | |
| ENSMUSG00000077345 | -1.079755021 | 0.830576988 |
| ENSMUSG00000077347 | 1 | |
| ENSMUSG00000077348 | 1 | |
| ENSMUSG00000077349 | -1 | 1 |
| ENSMUSG00000077350 | 1.059463094 | 0.608497229 |
| ENSMUSG00000077351 | 1 | |
| ENSMUSG00000077354 | -1.03962336 | 0.698112512 |
| ENSMUSG00000077356 | 1 | |
| ENSMUSG00000077357 | 1 | |
| ENSMUSG00000077358 | 1 | |
| ENSMUSG00000077359 | 1 | |

| | | |
|---|---|---|
| ENSMUSG00000077360 | 1.014545335 | 0.979002724 |
| ENSMUSG00000077361 | 1 | |
| ENSMUSG00000077366 | 1 | |
| ENSMUSG00000077367 | -1.044273782 | 0.698112512 |
| ENSMUSG00000077368 | 1 | |
| ENSMUSG00000077369 | 1 | |
| ENSMUSG00000077376 | 1 | |
| ENSMUSG00000077377 | 1 | |
| ENSMUSG00000077378 | 1 | |
| ENSMUSG00000077379 | 1 | |
| ENSMUSG00000077380 | -1.245360772 | 0.543409801 |
| ENSMUSG00000077381 | 1 | |
| ENSMUSG00000077382 | 1.005613439 | 1 |
| ENSMUSG00000077383 | 1 | |
| ENSMUSG00000077384 | 1 | |
| ENSMUSG00000077385 | 1 | |
| ENSMUSG00000077387 | 1 | |
| ENSMUSG00000077391 | -1.037482356 | 0.902641804 |
| ENSMUSG00000077392 | 1 | |
| ENSMUSG00000077393 | 1 | |
| ENSMUSG00000077394 | -1.03962336 | 0.803072249 |
| ENSMUSG00000077396 | 1 | |
| ENSMUSG00000077397 | 1 | |
| ENSMUSG00000077398 | 1 | |
| ENSMUSG00000077399 | 1 | |
| ENSMUSG00000077400 | 1 | |
| ENSMUSG00000077401 | 1 | |
| ENSMUSG00000077402 | 1 | |
| ENSMUSG00000077403 | 1 | |
| ENSMUSG00000077404 | 1 | |
| ENSMUSG00000077406 | 1 | |
| ENSMUSG00000077407 | 1 | |
| ENSMUSG00000077408 | 1 | |
| ENSMUSG00000077409 | 1 | |
| ENSMUSG00000077410 | 1 | |
| ENSMUSG00000077411 | 1 | |
| ENSMUSG00000077413 | 1 | |
| ENSMUSG00000077415 | 1 | |
| ENSMUSG00000077416 | -1.024718486 | 0.954310937 |
| ENSMUSG00000077417 | 1 | |
| ENSMUSG00000077418 | 1 | |
| ENSMUSG00000077419 | 1 | |
| ENSMUSG00000077420 | 1 | |
| ENSMUSG00000077421 | 1 | |
| ENSMUSG00000077422 | 1 | |
| ENSMUSG00000077424 | 1 | |
| ENSMUSG00000077425 | 1 | |
| ENSMUSG00000077426 | 1.077055984 | 0.698112512 |
| ENSMUSG00000077427 | 1 | |
| ENSMUSG00000077428 | 1 | |
| ENSMUSG00000077430 | 1 | |
| ENSMUSG00000077431 | 1 | |
| ENSMUSG00000077433 | 1 | |
| ENSMUSG00000077435 | 1 | |
| ENSMUSG00000077437 | 1 | |
| ENSMUSG00000077438 | 1 | |
| ENSMUSG00000077439 | 1 | |
| ENSMUSG00000077440 | -1.183089797 | 0.250768007 |
| ENSMUSG00000077441 | 1 | |
| ENSMUSG00000077442 | 1 | |
| ENSMUSG00000077443 | 1 | |
| ENSMUSG00000077445 | 1 | |
| ENSMUSG00000077447 | 1 | |

| | | |
|---|---|---|
| ENSMUSG00000077448 | -1 | 1 |
| ENSMUSG00000077449 | -1.033820075 | 0.888246981 |
| ENSMUSG00000077450 | 1.100957794 | 0.608497229 |
| ENSMUSG00000077451 | 1 | |
| ENSMUSG00000077453 | 1 | |
| ENSMUSG00000077454 | 1.064531743 | 0.796396036 |
| ENSMUSG00000077455 | 1 | |
| ENSMUSG00000077456 | 1 | |
| ENSMUSG00000077457 | 1.088819267 | 0.747530418 |
| ENSMUSG00000077459 | 1 | |
| ENSMUSG00000077460 | 1 | |
| ENSMUSG00000077461 | 1 | |
| ENSMUSG00000077462 | 1 | |
| ENSMUSG00000077463 | 1 | |
| ENSMUSG00000077464 | 1 | |
| ENSMUSG00000077465 | 1 | |
| ENSMUSG00000077467 | 1 | |
| ENSMUSG00000077468 | 1 | |
| ENSMUSG00000077469 | 1 | |
| ENSMUSG00000077470 | -1.037482356 | 0.870427333 |
| ENSMUSG00000077471 | 1 | |
| ENSMUSG00000077472 | 1 | |
| ENSMUSG00000077473 | 1 | |
| ENSMUSG00000077474 | 1 | |
| ENSMUSG00000077476 | 1 | |
| ENSMUSG00000077477 | 1 | |
| ENSMUSG00000077480 | 1 | |
| ENSMUSG00000077481 | 1 | |
| ENSMUSG00000077483 | 1 | |
| ENSMUSG00000077484 | 1 | |
| ENSMUSG00000077485 | 1 | |
| ENSMUSG00000077487 | 1 | |
| ENSMUSG00000077488 | 1 | |
| ENSMUSG00000077492 | 1 | |
| ENSMUSG00000077493 | -1.050928782 | 0.90874738 |
| ENSMUSG00000077494 | 1 | |
| ENSMUSG00000077495 | 1 | |
| ENSMUSG00000077497 | 1 | |
| ENSMUSG00000077499 | 1 | |
| ENSMUSG00000077501 | 1 | |
| ENSMUSG00000077502 | 1 | |
| ENSMUSG00000077503 | 1 | |
| ENSMUSG00000077504 | 1 | |
| ENSMUSG00000077505 | 1 | |
| ENSMUSG00000077506 | 1.028323353 | 0.954497929 |
| ENSMUSG00000077507 | 1 | |
| ENSMUSG00000077511 | 1 | |
| ENSMUSG00000077514 | 1 | |
| ENSMUSG00000077515 | 1 | |
| ENSMUSG00000077516 | -1 | 1 |
| ENSMUSG00000077517 | 1 | |
| ENSMUSG00000077518 | 1 | |
| ENSMUSG00000077519 | 1 | |
| ENSMUSG00000077521 | 1 | |
| ENSMUSG00000077522 | 1 | |
| ENSMUSG00000077524 | 1.059463094 | 0.768693672 |
| ENSMUSG00000077526 | -1.043306199 | 0.698112512 |
| ENSMUSG00000077527 | 1.059463094 | 0.608497229 |
| ENSMUSG00000077528 | 1 | |
| ENSMUSG00000077535 | 1 | |
| ENSMUSG00000077537 | 1 | |
| ENSMUSG00000077538 | 1 | |
| ENSMUSG00000077539 | 1 | |

| | | |
|---|---|---|
| ENSMUSG00000077541 | 1 | |
| ENSMUSG00000077542 | 1 | |
| ENSMUSG00000077543 | 1 | |
| ENSMUSG00000077545 | 1 | |
| ENSMUSG00000077546 | 1 | |
| ENSMUSG00000077547 | 1 | |
| ENSMUSG00000077549 | 1.077364093 | 0.855682211 |
| ENSMUSG00000077552 | 1 | |
| ENSMUSG00000077553 | 1 | |
| ENSMUSG00000077554 | 1 | |
| ENSMUSG00000077555 | 1 | |
| ENSMUSG00000077557 | 1 | |
| ENSMUSG00000077558 | 1 | |
| ENSMUSG00000077559 | 1 | |
| ENSMUSG00000077561 | 1 | |
| ENSMUSG00000077562 | 1 | |
| ENSMUSG00000077563 | 1.123516412 | 0.044893595 |
| ENSMUSG00000077564 | -1.078335432 | 0.770958539 |
| ENSMUSG00000077565 | 1 | |
| ENSMUSG00000077567 | 1 | |
| ENSMUSG00000077569 | 1.059463094 | 0.750182067 |
| ENSMUSG00000077571 | 1 | |
| ENSMUSG00000077572 | 1 | |
| ENSMUSG00000077574 | 1 | |
| ENSMUSG00000077575 | -1.038444538 | 0.866289429 |
| ENSMUSG00000077576 | 1 | |
| ENSMUSG00000077577 | -1.043306199 | 0.698112512 |
| ENSMUSG00000077578 | 1 | |
| ENSMUSG00000077579 | 1 | |
| ENSMUSG00000077581 | 1 | |
| ENSMUSG00000077582 | 1 | |
| ENSMUSG00000077585 | 1 | |
| ENSMUSG00000077586 | 1 | |
| ENSMUSG00000077587 | 1 | |
| ENSMUSG00000077588 | 1 | |
| ENSMUSG00000077590 | 1 | |
| ENSMUSG00000077591 | 1 | |
| ENSMUSG00000077592 | 1 | |
| ENSMUSG00000077593 | 1 | |
| ENSMUSG00000077594 | 1 | |
| ENSMUSG00000077595 | 1 | |
| ENSMUSG00000077597 | 1 | |
| ENSMUSG00000077598 | 1 | |
| ENSMUSG00000077599 | 1 | |
| ENSMUSG00000077601 | 1 | |
| ENSMUSG00000077602 | 1 | |
| ENSMUSG00000077603 | 1 | |
| ENSMUSG00000077604 | 1 | |
| ENSMUSG00000077607 | -1.099174267 | 0.535052277 |
| ENSMUSG00000077608 | 1 | |
| ENSMUSG00000077609 | 1 | |
| ENSMUSG00000077611 | -1.043306199 | 0.803260194 |
| ENSMUSG00000077612 | 1 | |
| ENSMUSG00000077613 | 1 | |
| ENSMUSG00000077614 | 1.086779794 | 0.745135758 |
| ENSMUSG00000077615 | 1 | |
| ENSMUSG00000077616 | 1 | |
| ENSMUSG00000077617 | 1 | |
| ENSMUSG00000077618 | 1 | |
| ENSMUSG00000077619 | 1 | |
| ENSMUSG00000077620 | 1 | |
| ENSMUSG00000077621 | 1 | |
| ENSMUSG00000077622 | 1 | |

| | | |
|---|---|---|
| ENSMUSG00000077624 | -1.043306199 | 0.698112512 |
| ENSMUSG00000077625 | 1 | |
| ENSMUSG00000077627 | 1.059463094 | 0.608497229 |
| ENSMUSG00000077628 | 1 | |
| ENSMUSG00000077629 | 1 | |
| ENSMUSG00000077630 | 1 | |
| ENSMUSG00000077633 | 1 | |
| ENSMUSG00000077635 | 1 | |
| ENSMUSG00000077637 | -1.05091667 | 0.89811529 |
| ENSMUSG00000077639 | -1.065529003 | 0.798354947 |
| ENSMUSG00000077640 | 1 | |
| ENSMUSG00000077641 | 1 | |
| ENSMUSG00000077642 | 1 | |
| ENSMUSG00000077643 | 1 | |
| ENSMUSG00000077644 | 1 | |
| ENSMUSG00000077645 | 1 | |
| ENSMUSG00000077647 | 1 | |
| ENSMUSG00000077648 | 1 | |
| ENSMUSG00000077649 | 1 | |
| ENSMUSG00000077650 | 1 | |
| ENSMUSG00000077652 | -1.03962336 | 0.698112512 |
| ENSMUSG00000077653 | 1 | |
| ENSMUSG00000077655 | -1 | 1 |
| ENSMUSG00000077659 | 1 | |
| ENSMUSG00000077660 | 1 | |
| ENSMUSG00000077661 | 1 | |
| ENSMUSG00000077663 | 1 | |
| ENSMUSG00000077664 | 1 | |
| ENSMUSG00000077666 | 1 | |
| ENSMUSG00000077667 | 1 | |
| ENSMUSG00000077668 | 1 | |
| ENSMUSG00000077669 | 1 | |
| ENSMUSG00000077671 | 1 | |
| ENSMUSG00000077672 | 1 | |
| ENSMUSG00000077673 | 1 | |
| ENSMUSG00000077674 | 1 | |
| ENSMUSG00000077675 | 1 | |
| ENSMUSG00000077677 | 1 | |
| ENSMUSG00000077678 | 1 | |
| ENSMUSG00000077680 | 1 | |
| ENSMUSG00000077681 | 1 | |
| ENSMUSG00000077682 | 1 | |
| ENSMUSG00000077683 | 1 | |
| ENSMUSG00000077685 | 1 | |
| ENSMUSG00000077687 | 1 | |
| ENSMUSG00000077689 | 1 | |
| ENSMUSG00000077691 | 1 | |
| ENSMUSG00000077693 | 1 | |
| ENSMUSG00000077696 | 1.059463094 | 0.608497229 |
| ENSMUSG00000077697 | 1 | |
| ENSMUSG00000077698 | 1 | |
| ENSMUSG00000077700 | 1 | |
| ENSMUSG00000077701 | 1 | |
| ENSMUSG00000077702 | 1 | |
| ENSMUSG00000077703 | 1 | |
| ENSMUSG00000077704 | 1.083591082 | 0.785584885 |
| ENSMUSG00000077705 | 1 | |
| ENSMUSG00000077706 | 1 | |
| ENSMUSG00000077707 | 1 | |
| ENSMUSG00000077708 | 1 | |
| ENSMUSG00000077709 | 1 | |
| ENSMUSG00000077710 | 1 | |
| ENSMUSG00000077711 | 1.075295999 | 0.804350979 |

| | | |
|---|---|---|
| ENSMUSG00000077713 | 1 | |
| ENSMUSG00000077714 | 1.07555074 | 0.734482722 |
| ENSMUSG00000077716 | 1 | |
| ENSMUSG00000077717 | 1 | |
| ENSMUSG00000077718 | 1 | |
| ENSMUSG00000077719 | 1 | |
| ENSMUSG00000077726 | 1 | |
| ENSMUSG00000077728 | 1 | |
| ENSMUSG00000077730 | 1 | |
| ENSMUSG00000077733 | 1.059463094 | 0.608497229 |
| ENSMUSG00000077734 | 1.040646287 | 0.790885744 |
| ENSMUSG00000077735 | 1 | |
| ENSMUSG00000077736 | 1 | |
| ENSMUSG00000077737 | -1.094575682 | 0.608497229 |
| ENSMUSG00000077738 | 1 | |
| ENSMUSG00000077741 | 1 | |
| ENSMUSG00000077743 | 1 | |
| ENSMUSG00000077744 | 1 | |
| ENSMUSG00000077745 | 1 | |
| ENSMUSG00000077749 | 1 | |
| ENSMUSG00000077750 | 1 | |
| ENSMUSG00000077752 | 1 | |
| ENSMUSG00000077755 | 1 | |
| ENSMUSG00000077756 | 1 | |
| ENSMUSG00000077757 | 1 | |
| ENSMUSG00000077758 | 1 | |
| ENSMUSG00000077759 | 1 | |
| ENSMUSG00000077761 | 1 | |
| ENSMUSG00000077767 | 1 | |
| ENSMUSG00000077768 | 1 | |
| ENSMUSG00000077769 | 1 | |
| ENSMUSG00000077771 | 1 | |
| ENSMUSG00000077772 | 1 | |
| ENSMUSG00000077776 | 1 | |
| ENSMUSG00000077777 | 1 | |
| ENSMUSG00000077780 | 1 | |
| ENSMUSG00000077782 | 1 | |
| ENSMUSG00000077783 | 1 | |
| ENSMUSG00000077784 | 1 | |
| ENSMUSG00000077785 | -1.03962336 | 0.698112512 |
| ENSMUSG00000077786 | 1 | |
| ENSMUSG00000077787 | 1.05189803 | 0.856925873 |
| ENSMUSG00000077789 | 1 | |
| ENSMUSG00000077791 | 1 | |
| ENSMUSG00000077792 | 1 | |
| ENSMUSG00000077793 | 1 | |
| ENSMUSG00000077794 | 1 | |
| ENSMUSG00000077795 | -1.03962336 | 0.698112512 |
| ENSMUSG00000077796 | 1 | |
| ENSMUSG00000077797 | -1.205414839 | 0.193906963 |
| ENSMUSG00000077799 | 1 | |
| ENSMUSG00000077802 | 1 | |
| ENSMUSG00000077804 | 1 | |
| ENSMUSG00000077815 | 1 | |
| ENSMUSG00000077821 | 1 | |
| ENSMUSG00000077829 | 1 | |
| ENSMUSG00000077830 | 1 | |
| ENSMUSG00000077832 | 1 | |
| ENSMUSG00000077834 | 1 | |
| ENSMUSG00000077836 | 1 | |
| ENSMUSG00000077844 | 1 | |
| ENSMUSG00000077846 | 1 | |
| ENSMUSG00000077847 | 1 | |

| | | |
|---|---|---|
| ENSMUSG00000077851 | 1 | |
| ENSMUSG00000077857 | 1 | |
| ENSMUSG00000077858 | 1.095872691 | 0.608497229 |
| ENSMUSG00000077869 | 1 | |
| ENSMUSG00000077872 | 1 | |
| ENSMUSG00000077879 | 1 | |
| ENSMUSG00000077881 | 1 | |
| ENSMUSG00000077884 | 1 | |
| ENSMUSG00000077886 | 1 | |
| ENSMUSG00000077890 | -1.044273782 | 0.790851299 |
| ENSMUSG00000077892 | 1 | |
| ENSMUSG00000077896 | 1 | |
| ENSMUSG00000077897 | 1 | |
| ENSMUSG00000077902 | 1 | |
| ENSMUSG00000077903 | 1 | |
| ENSMUSG00000077921 | 1 | |
| ENSMUSG00000077925 | 1 | |
| ENSMUSG00000077928 | 1 | |
| ENSMUSG00000077931 | 1 | |
| ENSMUSG00000077938 | 1 | |
| ENSMUSG00000077939 | 1 | |
| ENSMUSG00000077941 | 1 | |
| ENSMUSG00000077948 | 1 | |
| ENSMUSG00000077954 | 1 | |
| ENSMUSG00000077962 | 1 | |
| ENSMUSG00000077965 | 1 | |
| ENSMUSG00000077971 | 1 | |
| ENSMUSG00000077972 | 1 | |
| ENSMUSG00000077980 | 1 | |
| ENSMUSG00000077981 | 1 | |
| ENSMUSG00000077990 | 1 | |
| ENSMUSG00000077991 | 1 | |
| ENSMUSG00000077996 | 1 | |
| ENSMUSG00000077998 | 1 | |
| ENSMUSG00000078002 | 1 | |
| ENSMUSG00000078006 | 1 | |
| ENSMUSG00000078008 | 1 | |
| ENSMUSG00000078017 | 1 | |
| ENSMUSG00000078024 | 1 | |
| ENSMUSG00000078025 | 1 | |
| ENSMUSG00000078026 | 1 | |
| ENSMUSG00000078027 | 1 | |
| ENSMUSG00000078031 | 1 | |
| ENSMUSG00000078032 | 1 | |
| ENSMUSG00000078034 | 1 | |
| ENSMUSG00000078035 | 1 | |
| ENSMUSG00000078038 | 1 | |
| ENSMUSG00000078041 | 1 | |
| ENSMUSG00000078042 | 1 | |
| ENSMUSG00000078057 | 1 | |
| ENSMUSG00000078060 | 1 | |
| ENSMUSG00000078061 | -1.053549062 | 0.698112512 |
| ENSMUSG00000078080 | 1 | |
| ENSMUSG00000078087 | 1 | |
| ENSMUSG00000078091 | 1.258918661 | 0.146932868 |
| ENSMUSG00000078097 | -1.053549062 | 0.698112512 |
| ENSMUSG00000078116 | 1 | |
| ENSMUSG00000078117 | -1 | 1 |
| ENSMUSG00000078118 | 1 | |
| ENSMUSG00000078122 | 1 | |
| ENSMUSG00000078123 | 1 | |
| ENSMUSG00000078125 | 1.081175853 | 0.793885968 |
| ENSMUSG00000078126 | 1.069068979 | 0.608497229 |

| | | |
|---|---|---|
| ENSMUSG00000078127 | 1 | |
| ENSMUSG00000078128 | 1 | |
| ENSMUSG00000078129 | 1.145465914 | 0.609541771 |
| ENSMUSG00000078130 | 1 | |
| ENSMUSG00000078131 | 1.059463094 | 0.608497229 |
| ENSMUSG00000078132 | 1 | |
| ENSMUSG00000078134 | 1.044637175 | 0.748191299 |
| ENSMUSG00000078137 | -1.024718486 | 0.951399668 |
| ENSMUSG00000078139 | -1.136140785 | 0.417843828 |
| ENSMUSG00000078141 | 1.069068979 | 0.608497229 |
| ENSMUSG00000078143 | 1 | |
| ENSMUSG00000078144 | 1.110258504 | 0.364739674 |
| ENSMUSG00000078151 | -1.043306199 | 0.698112512 |
| ENSMUSG00000078153 | 1.092517236 | 0.779502498 |
| ENSMUSG00000078154 | -1.131313978 | 0.250749195 |
| ENSMUSG00000078157 | 1.008034876 | 1 |
| ENSMUSG00000078160 | -1.102997856 | 0.608497229 |
| ENSMUSG00000078161 | -1.132666855 | 0.501269661 |
| ENSMUSG00000078162 | 1 | |
| ENSMUSG00000078173 | 1 | |
| ENSMUSG00000078177 | -1.132666855 | 0.18851436 |
| ENSMUSG00000078179 | 1 | |
| ENSMUSG00000078180 | -1.092195161 | 0.698112512 |
| ENSMUSG00000078182 | 1.023743961 | 0.938939969 |
| ENSMUSG00000078183 | -1.056640735 | 0.698112512 |
| ENSMUSG00000078184 | -1.145536749 | 0.608497229 |
| ENSMUSG00000078185 | 1.047180826 | 0.797105186 |
| ENSMUSG00000078190 | 1.077487612 | 0.46097689 |
| ENSMUSG00000078192 | -1.094134359 | 0.780113178 |
| ENSMUSG00000078193 | -1.051473659 | 0.819175987 |
| ENSMUSG00000078197 | 1 | |
| ENSMUSG00000078198 | -1 | 1 |
| ENSMUSG00000078201 | 1.187759136 | 0.310767187 |
| ENSMUSG00000078202 | 1.034245398 | 0.981307697 |
| ENSMUSG00000078206 | 1 | |
| ENSMUSG00000078208 | 1 | |
| ENSMUSG00000078210 | 1 | |
| ENSMUSG00000078212 | 1 | |
| ENSMUSG00000078213 | 1 | |
| ENSMUSG00000078214 | 1 | |
| ENSMUSG00000078218 | 1 | |
| ENSMUSG00000078234 | 1.078290557 | 0.698112512 |
| ENSMUSG00000078235 | -1.321866603 | 0.062951436 |
| ENSMUSG00000078238 | 1.019083579 | 0.949812939 |
| ENSMUSG00000078240 | 1.067830936 | 0.70299173 |
| ENSMUSG00000078247 | 1.179193068 | 0.113640702 |
| ENSMUSG00000078249 | 1.279419459 | 0.548145424 |
| ENSMUSG00000078251 | 1 | |
| ENSMUSG00000078252 | 1 | |
| ENSMUSG00000078253 | 1 | |
| ENSMUSG00000078254 | 1 | |
| ENSMUSG00000078255 | 1 | |
| ENSMUSG00000078256 | 1 | |
| ENSMUSG00000078257 | 1 | |
| ENSMUSG00000078258 | 1 | |
| ENSMUSG00000078259 | 1 | |
| ENSMUSG00000078260 | 1 | |
| ENSMUSG00000078261 | 1 | |
| ENSMUSG00000078262 | 1 | |
| ENSMUSG00000078263 | 1 | |
| ENSMUSG00000078269 | 1 | |
| ENSMUSG00000078270 | 1 | |
| ENSMUSG00000078276 | 1 | |

| | | |
|---|---|---|
| ENSMUSG00000078280 | 1 | |
| ENSMUSG00000078289 | -1.06700837 | 0.879253209 |
| ENSMUSG00000078291 | -1.009842415 | 1 |
| ENSMUSG00000078300 | -1.03962336 | 0.698112512 |
| ENSMUSG00000078302 | 1.022433551 | 0.893500268 |
| ENSMUSG00000078307 | 1.015486245 | 0.985522133 |
| ENSMUSG00000078308 | 1.122681009 | 0.27163566 |
| ENSMUSG00000078314 | 1.059463094 | 0.608497229 |
| ENSMUSG00000078315 | 1 | |
| ENSMUSG00000078317 | 1.088428615 | 0.608497229 |
| ENSMUSG00000078320 | 1 | |
| ENSMUSG00000078324 | 1 | |
| ENSMUSG00000078346 | 1 | |
| ENSMUSG00000078348 | 1.144000773 | 0.554164954 |
| ENSMUSG00000078349 | -1.329002041 | 0.331273341 |
| ENSMUSG00000078350 | 1.304662912 | 0.069231655 |
| ENSMUSG00000078354 | 1 | |
| ENSMUSG00000078355 | 1 | |
| ENSMUSG00000078365 | -1.043306199 | 0.895360578 |
| ENSMUSG00000078370 | 1.084728061 | 0.709078049 |
| ENSMUSG00000078377 | -1.023016116 | 0.973792594 |
| ENSMUSG00000078380 | 1.069624006 | 0.815671448 |
| ENSMUSG00000078384 | 1 | |
| ENSMUSG00000078420 | 1 | |
| ENSMUSG00000078427 | -1.066638959 | 0.78481421 |
| ENSMUSG00000078429 | 1.130363529 | 0.608497229 |
| ENSMUSG00000078430 | 1 | |
| ENSMUSG00000078435 | -1.02769217 | 0.802248757 |
| ENSMUSG00000078439 | 1.010326295 | 1 |
| ENSMUSG00000078440 | 1.06753953 | 0.698112512 |
| ENSMUSG00000078441 | 1.019083579 | 0.949812939 |
| ENSMUSG00000078442 | -1 | 1 |
| ENSMUSG00000078444 | 1.163441272 | 0.734674233 |
| ENSMUSG00000078451 | -1.090374545 | 0.70299173 |
| ENSMUSG00000078452 | -1.030018143 | 0.896152472 |
| ENSMUSG00000078453 | 1.14046482 | 0.570717076 |
| ENSMUSG00000078480 | -1.201027508 | 0.037440682 |
| ENSMUSG00000078481 | 1 | |
| ENSMUSG00000078484 | 1.063273744 | 0.619580945 |
| ENSMUSG00000078485 | -1.009535441 | 1 |
| ENSMUSG00000078486 | 1.01197517 | 0.980378626 |
| ENSMUSG00000078487 | -1 | 1 |
| ENSMUSG00000078488 | 1 | |
| ENSMUSG00000078489 | 1.129222406 | 0.70299173 |
| ENSMUSG00000078490 | 1.10556157 | 0.468621875 |
| ENSMUSG00000078491 | 1 | |
| ENSMUSG00000078492 | 1.069068979 | 0.608497229 |
| ENSMUSG00000078494 | -1.018054237 | 0.978664718 |
| ENSMUSG00000078495 | 1.088028043 | 0.289502183 |
| ENSMUSG00000078496 | -1.272166063 | 0.030292632 |
| ENSMUSG00000078497 | 1 | |
| ENSMUSG00000078498 | -1 | 1 |
| ENSMUSG00000078500 | 1 | |
| ENSMUSG00000078502 | -1.158169686 | 0.453843926 |
| ENSMUSG00000078503 | 1.091718931 | 0.732769483 |
| ENSMUSG00000078504 | 1 | |
| ENSMUSG00000078505 | 1 | |
| ENSMUSG00000078506 | 1 | |
| ENSMUSG00000078507 | 1 | |
| ENSMUSG00000078508 | 1 | |
| ENSMUSG00000078509 | 1 | |
| ENSMUSG00000078510 | 1 | |
| ENSMUSG00000078511 | -1.069502966 | 0.698112512 |

| | | |
|---|---|---|
| ENSMUSG00000078512 | 1 | |
| ENSMUSG00000078513 | 1 | |
| ENSMUSG00000078515 | 1.004011146 | 1 |
| ENSMUSG00000078517 | 1.027502787 | 0.881780339 |
| ENSMUSG00000078518 | 1 | |
| ENSMUSG00000078519 | 1.080158688 | 0.762874575 |
| ENSMUSG00000078520 | 1 | |
| ENSMUSG00000078521 | -1.024110384 | 0.936027951 |
| ENSMUSG00000078528 | 1.160565542 | 0.608497229 |
| ENSMUSG00000078531 | 1.028323353 | 0.903464611 |
| ENSMUSG00000078532 | 1.10784482 | 0.361740791 |
| ENSMUSG00000078537 | 1 | |
| ENSMUSG00000078546 | -1.008415329 | 0.999347497 |
| ENSMUSG00000078552 | -1.105655388 | 0.580660018 |
| ENSMUSG00000078554 | 1 | |
| ENSMUSG00000078566 | 1.0503424 | 0.75528471 |
| ENSMUSG00000078570 | 1.205165031 | 0.56288192 |
| ENSMUSG00000078572 | 1.030586645 | 0.942544372 |
| ENSMUSG00000078575 | 1 | |
| ENSMUSG00000078576 | 1 | |
| ENSMUSG00000078577 | 1 | |
| ENSMUSG00000078578 | -1.052560584 | 0.904410301 |
| ENSMUSG00000078580 | 1.127238877 | 0.552842869 |
| ENSMUSG00000078584 | -1.033490513 | 0.834556513 |
| ENSMUSG00000078586 | 1 | |
| ENSMUSG00000078588 | 1.244301514 | 0.282066625 |
| ENSMUSG00000078590 | 1 | |
| ENSMUSG00000078591 | -1 | 1 |
| ENSMUSG00000078592 | 1.122242299 | 0.609147349 |
| ENSMUSG00000078593 | 1 | |
| ENSMUSG00000078597 | 1 | |
| ENSMUSG00000078598 | -1.003834322 | 1 |
| ENSMUSG00000078599 | 1.017745225 | 0.896246617 |
| ENSMUSG00000078600 | 1 | |
| ENSMUSG00000078601 | -1.043306199 | 0.698112512 |
| ENSMUSG00000078604 | 1.014545335 | 0.979002724 |
| ENSMUSG00000078605 | 1 | |
| ENSMUSG00000078606 | 1.132639128 | 0.217803313 |
| ENSMUSG00000078607 | 1.050776495 | 0.847073978 |
| ENSMUSG00000078611 | 1 | |
| ENSMUSG00000078612 | 1.191685811 | 0.321726111 |
| ENSMUSG00000078616 | -1 | 1 |
| ENSMUSG00000078618 | 1 | |
| ENSMUSG00000078619 | -1.056145572 | 0.79421917 |
| ENSMUSG00000078620 | -1 | 1 |
| ENSMUSG00000078621 | 1 | |
| ENSMUSG00000078622 | -1.018517541 | 0.992726262 |
| ENSMUSG00000078624 | 1.013812468 | 0.954883228 |
| ENSMUSG00000078625 | 1.035449581 | 0.952049609 |
| ENSMUSG00000078626 | 1 | |
| ENSMUSG00000078627 | 1.059463094 | 0.811221598 |
| ENSMUSG00000078630 | 1.004394707 | 1 |
| ENSMUSG00000078632 | 1 | |
| ENSMUSG00000078635 | 1.122462048 | 0.420161906 |
| ENSMUSG00000078636 | 1 | |
| ENSMUSG00000078639 | 1 | |
| ENSMUSG00000078640 | 1.077658816 | 0.775199822 |
| ENSMUSG00000078648 | -1.126773846 | 0.539222653 |
| ENSMUSG00000078650 | 1 | |
| ENSMUSG00000078651 | 1.090259625 | 0.39790876 |
| ENSMUSG00000078652 | 1.01213518 | 1 |
| ENSMUSG00000078653 | -1.002571313 | 1 |
| ENSMUSG00000078655 | 1 | |

| | | |
|---|---|---|
| ENSMUSG00000078656 | -1.095864544 | 0.5819787 |
| ENSMUSG00000078657 | 1 | |
| ENSMUSG00000078664 | 1 | |
| ENSMUSG00000078668 | 1 | |
| ENSMUSG00000078670 | -1.356078793 | 0.050388327 |
| ENSMUSG00000078671 | 1.071346085 | 0.698112512 |
| ENSMUSG00000078672 | 1 | |
| ENSMUSG00000078673 | 1 | |
| ENSMUSG00000078674 | 1 | |
| ENSMUSG00000078675 | 1 | |
| ENSMUSG00000078676 | 1.005193513 | 1 |
| ENSMUSG00000078677 | 1.014545335 | 0.979002724 |
| ENSMUSG00000078680 | 1 | |
| ENSMUSG00000078681 | 1.067398563 | 0.641827556 |
| ENSMUSG00000078683 | 1 | |
| ENSMUSG00000078684 | -2.29102659 | 0.000545359 |
| ENSMUSG00000078685 | 1 | |
| ENSMUSG00000078686 | 1 | |
| ENSMUSG00000078687 | 1 | |
| ENSMUSG00000078688 | 1 | |
| ENSMUSG00000078689 | 1 | |
| ENSMUSG00000078691 | 1 | |
| ENSMUSG00000078695 | -1.093323637 | 0.315720798 |
| ENSMUSG00000078698 | 1 | |
| ENSMUSG00000078700 | -1.033514349 | 0.698112512 |
| ENSMUSG00000078706 | 1.217637756 | 0.00182836 |
| ENSMUSG00000078713 | 1.04426836 | 0.801443616 |
| ENSMUSG00000078716 | -1.08196299 | 0.679217411 |
| ENSMUSG00000078719 | 1 | |
| ENSMUSG00000078721 | 1 | |
| ENSMUSG00000078722 | 1 | |
| ENSMUSG00000078735 | 1 | |
| ENSMUSG00000078737 | 1 | |
| ENSMUSG00000078742 | -1.03962336 | 0.698112512 |
| ENSMUSG00000078746 | 1 | |
| ENSMUSG00000078747 | -1.157634469 | 0.608497229 |
| ENSMUSG00000078752 | -1 | 1 |
| ENSMUSG00000078753 | 1 | |
| ENSMUSG00000078754 | 1 | |
| ENSMUSG00000078757 | 1 | |
| ENSMUSG00000078759 | 1 | |
| ENSMUSG00000078762 | -1.029559884 | 0.903122231 |
| ENSMUSG00000078763 | 1 | |
| ENSMUSG00000078765 | -1.096766449 | 0.046329797 |
| ENSMUSG00000078768 | -1.045111244 | 0.698112512 |
| ENSMUSG00000078771 | -1.124963965 | 0.537387105 |
| ENSMUSG00000078772 | 1.160809372 | 0.594957001 |
| ENSMUSG00000078773 | -1.040763597 | 0.794117604 |
| ENSMUSG00000078776 | -1.050660127 | 0.614424414 |
| ENSMUSG00000078779 | 1.006359354 | 1 |
| ENSMUSG00000078780 | 1 | |
| ENSMUSG00000078783 | 1 | |
| ENSMUSG00000078784 | -1.01314764 | 0.982838256 |
| ENSMUSG00000078786 | 1.041141619 | 0.778372272 |
| ENSMUSG00000078787 | 1.0154594 | 1 |
| ENSMUSG00000078789 | 1.08269965 | 0.539550866 |
| ENSMUSG00000078793 | 1 | |
| ENSMUSG00000078794 | 1.236473398 | 0.117212031 |
| ENSMUSG00000078795 | 1 | |
| ENSMUSG00000078796 | -1.043306199 | 0.698112512 |
| ENSMUSG00000078798 | 1 | |
| ENSMUSG00000078799 | 1 | |
| ENSMUSG00000078800 | 1 | |

| | | |
|---|---|---|
| ENSMUSG00000078808 | 1.020110535 | 0.972980766 |
| ENSMUSG00000078810 | 1 | |
| ENSMUSG00000078812 | 1.043947871 | 0.933031586 |
| ENSMUSG00000078813 | 1.013038669 | 0.872692845 |
| ENSMUSG00000078815 | -1.063525418 | 0.698112512 |
| ENSMUSG00000078816 | 1.285212088 | 0.098922723 |
| ENSMUSG00000078817 | 1.037518982 | 0.949693072 |
| ENSMUSG00000078838 | -1.130749879 | 0.608497229 |
| ENSMUSG00000078840 | -1.044273782 | 0.790851299 |
| ENSMUSG00000078847 | 1.019083579 | 0.949812939 |
| ENSMUSG00000078850 | -1 | 1 |
| ENSMUSG00000078851 | -1.302342578 | 0.014928553 |
| ENSMUSG00000078853 | -1.344711476 | 0.00560438 |
| ENSMUSG00000078861 | -1.070044986 | 0.657060635 |
| ENSMUSG00000078862 | -1.009399328 | 1 |
| ENSMUSG00000078864 | -1.503228605 | 0.017882526 |
| ENSMUSG00000078865 | -1.03962336 | 0.803072249 |
| ENSMUSG00000078866 | -1.336341449 | 0.05762786 |
| ENSMUSG00000078867 | -1.561376656 | 0.018759913 |
| ENSMUSG00000078868 | -1.370207752 | 0.004592615 |
| ENSMUSG00000078869 | -1.147838869 | 0.265654156 |
| ENSMUSG00000078870 | -1.100349573 | 0.566490809 |
| ENSMUSG00000078872 | 1.091384972 | 0.770429489 |
| ENSMUSG00000078875 | -1.016502829 | 0.986542466 |
| ENSMUSG00000078876 | 1 | |
| ENSMUSG00000078877 | 1.308339931 | 0.438534282 |
| ENSMUSG00000078878 | -1.098972316 | 0.698112512 |
| ENSMUSG00000078879 | -1.060822484 | 0.647586132 |
| ENSMUSG00000078880 | -1.132107349 | 0.638133285 |
| ENSMUSG00000078881 | 1 | |
| ENSMUSG00000078886 | 1 | |
| ENSMUSG00000078887 | -1.088086309 | 0.495999195 |
| ENSMUSG00000078889 | 1 | |
| ENSMUSG00000078891 | 1 | |
| ENSMUSG00000078894 | 1 | |
| ENSMUSG00000078895 | 1 | |
| ENSMUSG00000078896 | 1 | |
| ENSMUSG00000078897 | 1 | |
| ENSMUSG00000078898 | -1.265372831 | 0.236925578 |
| ENSMUSG00000078899 | -1.053411784 | 0.769745556 |
| ENSMUSG00000078901 | -1.147838869 | 0.195607801 |
| ENSMUSG00000078902 | -1.002223526 | 1 |
| ENSMUSG00000078903 | -1.052575578 | 0.781488832 |
| ENSMUSG00000078905 | 1.049965527 | 0.854616424 |
| ENSMUSG00000078906 | -1.113967123 | 0.615267463 |
| ENSMUSG00000078907 | 1.05038453 | 0.878506664 |
| ENSMUSG00000078908 | 1.074611244 | 0.523675341 |
| ENSMUSG00000078912 | 1 | |
| ENSMUSG00000078915 | 1.069068979 | 0.608497229 |
| ENSMUSG00000078917 | -1.026276402 | 0.933807487 |
| ENSMUSG00000078919 | -1.124769311 | 0.27327174 |
| ENSMUSG00000078920 | -1.20094591 | 0.146197466 |
| ENSMUSG00000078921 | -1.014570171 | 0.988047236 |
| ENSMUSG00000078922 | 1.318365674 | 0.437045401 |
| ENSMUSG00000078923 | 1.132870311 | 0.635598239 |
| ENSMUSG00000078924 | 1 | |
| ENSMUSG00000078925 | 1.059463094 | 0.698112512 |
| ENSMUSG00000078926 | -1.03962336 | 0.872692845 |
| ENSMUSG00000078931 | -1.040325092 | 0.868236615 |
| ENSMUSG00000078932 | 1 | |
| ENSMUSG00000078934 | 1 | |
| ENSMUSG00000078935 | 1 | |
| ENSMUSG00000078937 | -1.025928871 | 0.997003355 |

| | | |
|---|---|---|
| ENSMUSG00000078938 | 1 | |
| ENSMUSG00000078940 | 1 | |
| ENSMUSG00000078941 | -1.051507423 | 0.814364805 |
| ENSMUSG00000078942 | -1.43530671 | 0.043944063 |
| ENSMUSG00000078945 | -1.362803181 | 0.144950944 |
| ENSMUSG00000078949 | 1 | |
| ENSMUSG00000078952 | 1.271163238 | 0.018726089 |
| ENSMUSG00000078953 | 1 | |
| ENSMUSG00000078954 | 1.203298979 | 0.012230446 |
| ENSMUSG00000078955 | 1 | |
| ENSMUSG00000078956 | 1 | |
| ENSMUSG00000078957 | 1 | |
| ENSMUSG00000078958 | 1 | |
| ENSMUSG00000078962 | 1 | |
| ENSMUSG00000078963 | 1.154060078 | 0.563756918 |
| ENSMUSG00000078964 | 1 | |
| ENSMUSG00000078965 | 1 | |
| ENSMUSG00000078967 | 1.177106795 | 0.608497229 |
| ENSMUSG00000078970 | -1.054996269 | 0.74527374 |
| ENSMUSG00000078972 | 1 | |
| ENSMUSG00000078974 | -1.117246289 | 0.572599793 |
| ENSMUSG00000078982 | 1 | |
| ENSMUSG00000078984 | -1.002064566 | 1 |
| ENSMUSG00000078994 | -1.277178656 | 0.063097135 |
| ENSMUSG00000078995 | 1.063931194 | 0.716583464 |
| ENSMUSG00000078998 | 1 | |
| ENSMUSG00000079000 | -1.03962336 | 0.698112512 |
| ENSMUSG00000079001 | 1 | |
| ENSMUSG00000079003 | 1.099325187 | 0.37982425 |
| ENSMUSG00000079005 | 1 | |
| ENSMUSG00000079006 | 1 | |
| ENSMUSG00000079007 | 1 | |
| ENSMUSG00000079008 | 1.02695495 | 0.986542466 |
| ENSMUSG00000079009 | 1.015486245 | 1 |
| ENSMUSG00000079010 | -1.025223089 | 0.997451029 |
| ENSMUSG00000079011 | -1.14222699 | 0.567985932 |
| ENSMUSG00000079012 | -1 | 1 |
| ENSMUSG00000079013 | 1 | |
| ENSMUSG00000079014 | 1.811794059 | 0.003914588 |
| ENSMUSG00000079015 | 1 | |
| ENSMUSG00000079016 | 1.015486245 | 0.972980766 |
| ENSMUSG00000079017 | 1 | |
| ENSMUSG00000079018 | 1.452312557 | 0.000684801 |
| ENSMUSG00000079019 | 1 | |
| ENSMUSG00000079020 | 1.237512743 | 0.088353712 |
| ENSMUSG00000079022 | -1.121461888 | 0.682659034 |
| ENSMUSG00000079024 | -1.048277865 | 0.860645152 |
| ENSMUSG00000079025 | 1 | |
| ENSMUSG00000079029 | 1.171724772 | 0.608497229 |
| ENSMUSG00000079030 | 1 | |
| ENSMUSG00000079031 | 1 | |
| ENSMUSG00000079033 | 1.059463094 | 0.608497229 |
| ENSMUSG00000079034 | 1.498528013 | 0.004814019 |
| ENSMUSG00000079036 | 1.119772782 | 0.424254596 |
| ENSMUSG00000079037 | 1.003361774 | 1 |
| ENSMUSG00000079038 | -1.401342546 | 0.034895153 |
| ENSMUSG00000079039 | 1 | |
| ENSMUSG00000079042 | -1 | 1 |
| ENSMUSG00000079043 | -1.061190348 | 0.753536551 |
| ENSMUSG00000079045 | -1.043306199 | 0.698112512 |
| ENSMUSG00000079048 | 1 | |
| ENSMUSG00000079049 | 1.195549877 | 0.349870173 |
| ENSMUSG00000079051 | -1.003834322 | 1 |

| | | |
|---|---|---|
| ENSMUSG00000079053 | -1.054169715 | 0.813485995 |
| ENSMUSG00000079055 | -1.436497502 | 0.078461318 |
| ENSMUSG00000079056 | -1.22711189 | 0.125368251 |
| ENSMUSG00000079057 | -1.119675565 | 0.210489399 |
| ENSMUSG00000079058 | 1 | |
| ENSMUSG00000079061 | 1 | |
| ENSMUSG00000079064 | 1 | |
| ENSMUSG00000079065 | 1.077292872 | 0.608497229 |
| ENSMUSG00000079067 | 1 | |
| ENSMUSG00000079069 | 1.111111184 | 0.730297627 |
| ENSMUSG00000079070 | 1.059463094 | 0.608497229 |
| ENSMUSG00000079071 | 1 | |
| ENSMUSG00000079076 | -1.016118664 | 1 |
| ENSMUSG00000079083 | 1.055222723 | 0.698112512 |
| ENSMUSG00000079084 | -1.044182988 | 0.750742698 |
| ENSMUSG00000079091 | 1 | |
| ENSMUSG00000079092 | 1.229687879 | 0.015031146 |
| ENSMUSG00000079093 | 1 | |
| ENSMUSG00000079101 | 1 | |
| ENSMUSG00000079103 | -1 | 1 |
| ENSMUSG00000079104 | 1.079466678 | 0.69305964 |
| ENSMUSG00000079105 | 1 | |
| ENSMUSG00000079108 | 1.015798745 | 0.954466467 |
| ENSMUSG00000079109 | -1.018044844 | 0.977412147 |
| ENSMUSG00000079110 | -1.109582661 | 0.608497229 |
| ENSMUSG00000079111 | 1.073797381 | 0.738123855 |
| ENSMUSG00000079112 | 1 | |
| ENSMUSG00000079113 | 1 | |
| ENSMUSG00000079114 | 1 | |
| ENSMUSG00000079116 | 1 | |
| ENSMUSG00000079120 | 1 | |
| ENSMUSG00000079137 | -1.109582661 | 0.698112512 |
| ENSMUSG00000079138 | -1.111880267 | 0.608497229 |
| ENSMUSG00000079139 | -1.042157572 | 0.857888386 |
| ENSMUSG00000079140 | 1 | |
| ENSMUSG00000079143 | -1.095294215 | 0.541735653 |
| ENSMUSG00000079144 | 1.060042346 | 0.328793916 |
| ENSMUSG00000079146 | -1 | 1 |
| ENSMUSG00000079156 | 1 | |
| ENSMUSG00000079157 | -1.044273782 | 0.698112512 |
| ENSMUSG00000079162 | 1.012708788 | 1 |
| ENSMUSG00000079163 | 1 | |
| ENSMUSG00000079164 | -1.141777045 | 0.681377339 |
| ENSMUSG00000079165 | -1.084645496 | 0.532369657 |
| ENSMUSG00000079168 | 1 | |
| ENSMUSG00000079169 | 1 | |
| ENSMUSG00000079170 | 1 | |
| ENSMUSG00000079171 | 1 | |
| ENSMUSG00000079173 | 1.335793524 | 0.012869031 |
| ENSMUSG00000079174 | 1 | |
| ENSMUSG00000079175 | 1 | |
| ENSMUSG00000079177 | -1.029709545 | 0.977592603 |
| ENSMUSG00000079179 | 1.078966448 | 0.605619163 |
| ENSMUSG00000079180 | 1 | |
| ENSMUSG00000079183 | 1 | |
| ENSMUSG00000079184 | -1.144781606 | 0.499289204 |
| ENSMUSG00000079186 | 1 | |
| ENSMUSG00000079190 | -1 | 1 |
| ENSMUSG00000079192 | 1 | |
| ENSMUSG00000079197 | -1.386707137 | 0.094241568 |
| ENSMUSG00000079215 | 1.129000775 | 0.619047344 |
| ENSMUSG00000079222 | 1.019083579 | 0.949812939 |
| ENSMUSG00000079223 | 1.116020789 | 0.629184378 |

| | | |
|---|---|---|
| ENSMUSG00000079224 | 1 | |
| ENSMUSG00000079225 | 1.054079687 | 0.854219068 |
| ENSMUSG00000079227 | 1.055624351 | 0.915070045 |
| ENSMUSG00000079230 | 1 | |
| ENSMUSG00000079235 | 1.419892002 | 0.030368594 |
| ENSMUSG00000079239 | 1 | |
| ENSMUSG00000079242 | -1.059036458 | 0.693468357 |
| ENSMUSG00000079243 | 1.069068979 | 0.608497229 |
| ENSMUSG00000079244 | 1 | |
| ENSMUSG00000079247 | 1 | |
| ENSMUSG00000079258 | -1.043306199 | 0.733276285 |
| ENSMUSG00000079259 | 1.229130254 | 0.030368594 |
| ENSMUSG00000079260 | -1.070950656 | 0.634912948 |
| ENSMUSG00000079261 | 1.037893788 | 0.949812939 |
| ENSMUSG00000079262 | -1.038068813 | 0.756915351 |
| ENSMUSG00000079263 | 1.059463094 | 0.608497229 |
| ENSMUSG00000079264 | 1 | |
| ENSMUSG00000079265 | -1 | 1 |
| ENSMUSG00000079266 | 1 | |
| ENSMUSG00000079267 | 1 | |
| ENSMUSG00000079269 | 1 | |
| ENSMUSG00000079271 | 1 | |
| ENSMUSG00000079272 | 1 | |
| ENSMUSG00000079273 | 1 | |
| ENSMUSG00000079276 | 1 | |
| ENSMUSG00000079277 | 1.385092438 | 0.022145461 |
| ENSMUSG00000079278 | 1 | |
| ENSMUSG00000079283 | 1.033503617 | 0.887094209 |
| ENSMUSG00000079286 | -1.044273782 | 0.795446726 |
| ENSMUSG00000079293 | 1 | |
| ENSMUSG00000079295 | 1 | |
| ENSMUSG00000079297 | 1.014731082 | 0.988424085 |
| ENSMUSG00000079298 | 1 | |
| ENSMUSG00000079299 | 1 | |
| ENSMUSG00000079304 | 1.046384699 | 0.833289367 |
| ENSMUSG00000079311 | 1.035329424 | 0.913742415 |
| ENSMUSG00000079316 | -1.050718259 | 0.796457226 |
| ENSMUSG00000079317 | -1.142076317 | 0.249734947 |
| ENSMUSG00000079323 | 1.059463094 | 0.698112512 |
| ENSMUSG00000079324 | -1.033820075 | 0.888246981 |
| ENSMUSG00000079326 | 1 | |
| ENSMUSG00000079330 | -1.059888401 | 0.732726475 |
| ENSMUSG00000079333 | 1.059463094 | 0.608497229 |
| ENSMUSG00000079334 | 1.007593483 | 1 |
| ENSMUSG00000079339 | 1.148994275 | 0.695921515 |
| ENSMUSG00000079342 | 1 | |
| ENSMUSG00000079343 | 1.18230292 | 0.56156678 |
| ENSMUSG00000079344 | -1.03962336 | 0.698112512 |
| ENSMUSG00000079346 | 1 | |
| ENSMUSG00000079349 | 1 | |
| ENSMUSG00000079350 | 1 | |
| ENSMUSG00000079353 | 1 | |
| ENSMUSG00000079355 | -1.158274388 | 0.608497229 |
| ENSMUSG00000079356 | -1.03962336 | 0.698112512 |
| ENSMUSG00000079357 | -1.218287941 | 0.452573014 |
| ENSMUSG00000079359 | 1.059463094 | 0.608497229 |
| ENSMUSG00000079362 | 1 | |
| ENSMUSG00000079363 | -1.402672013 | 0.009008372 |
| ENSMUSG00000079364 | 1 | |
| ENSMUSG00000079371 | 1 | |
| ENSMUSG00000079374 | 1 | |
| ENSMUSG00000079378 | 1 | |
| ENSMUSG00000079379 | 1 | |

| | | |
|---|---|---|
| ENSMUSG00000079380 | 1 | |
| ENSMUSG00000079382 | -1.043306199 | 0.698112512 |
| ENSMUSG00000079383 | 1 | |
| ENSMUSG00000079386 | 1 | |
| ENSMUSG00000079387 | 1 | |
| ENSMUSG00000079388 | 1 | |
| ENSMUSG00000079389 | 1 | |
| ENSMUSG00000079391 | -1.03962336 | 0.698112512 |
| ENSMUSG00000079392 | 1 | |
| ENSMUSG00000079395 | 1 | |
| ENSMUSG00000079396 | 1.099959823 | 0.698112512 |
| ENSMUSG00000079402 | 1 | |
| ENSMUSG00000079407 | -1.210265142 | 0.271454743 |
| ENSMUSG00000079409 | 1 | |
| ENSMUSG00000079410 | 1.028323353 | 0.903464611 |
| ENSMUSG00000079414 | 1.126153172 | 0.294641379 |
| ENSMUSG00000079415 | -1.052895978 | 0.608497229 |
| ENSMUSG00000079416 | 1 | |
| ENSMUSG00000079418 | -1.193800467 | 0.174363255 |
| ENSMUSG00000079419 | 1 | |
| ENSMUSG00000079421 | 1.059463094 | 0.608497229 |
| ENSMUSG00000079423 | 1 | |
| ENSMUSG00000079424 | 1 | |
| ENSMUSG00000079426 | 1.11051509 | 0.608497229 |
| ENSMUSG00000079427 | -1.057957995 | 0.730535264 |
| ENSMUSG00000079428 | -1.03962336 | 0.698112512 |
| ENSMUSG00000079429 | -1.031689327 | 0.608497229 |
| ENSMUSG00000079432 | 1 | |
| ENSMUSG00000079433 | 1 | |
| ENSMUSG00000079434 | -1.043306199 | 0.79291651 |
| ENSMUSG00000079435 | -1.095034295 | 0.717035356 |
| ENSMUSG00000079436 | -1.03962336 | 0.774810939 |
| ENSMUSG00000079438 | 1 | |
| ENSMUSG00000079439 | 1 | |
| ENSMUSG00000079440 | 1.92915943 | 0.002186224 |
| ENSMUSG00000079442 | -1.065960055 | 0.570717076 |
| ENSMUSG00000079444 | -1 | 1 |
| ENSMUSG00000079445 | -1.343531482 | 0.082540525 |
| ENSMUSG00000079450 | -1.196362677 | 0.596491107 |
| ENSMUSG00000079451 | -1.026853369 | 0.970833538 |
| ENSMUSG00000079455 | 1 | |
| ENSMUSG00000079457 | -1.022058569 | 0.979002724 |
| ENSMUSG00000079460 | 1 | |
| ENSMUSG00000079462 | 1 | |
| ENSMUSG00000079465 | 1.151225815 | 0.36275755 |
| ENSMUSG00000079466 | 1 | |
| ENSMUSG00000079467 | 1.062378903 | 0.840946951 |
| ENSMUSG00000079469 | -1.08064218 | 0.477644658 |
| ENSMUSG00000079470 | 1.008619321 | 1 |
| ENSMUSG00000079471 | 1.132639128 | 0.398767938 |
| ENSMUSG00000079476 | 1 | |
| ENSMUSG00000079477 | 1.078917743 | 0.770958539 |
| ENSMUSG00000079478 | 1.032717855 | 0.827158853 |
| ENSMUSG00000079479 | 1 | |
| ENSMUSG00000079480 | -1.034454092 | 0.831244429 |
| ENSMUSG00000079481 | 1.205109241 | 0.039226692 |
| ENSMUSG00000079484 | 1.186564563 | 0.203085735 |
| ENSMUSG00000079487 | 1.031553849 | 0.925653135 |
| ENSMUSG00000079489 | -1.208176077 | 0.549044354 |
| ENSMUSG00000079491 | -1.241549772 | 0.042078126 |
| ENSMUSG00000079492 | -1.253597615 | 0.198681831 |
| ENSMUSG00000079494 | 1 | |
| ENSMUSG00000079495 | 1.171563499 | 0.34958424 |

| | | |
|---|---|---|
| ENSMUSG00000079497 | -1 | 1 |
| ENSMUSG00000079499 | -1.388344751 | 0.028026721 |
| ENSMUSG00000079501 | 1 | |
| ENSMUSG00000079502 | -1.12906868 | 0.698112512 |
| ENSMUSG00000079505 | -1.413464219 | 0.061568005 |
| ENSMUSG00000079507 | -1.335739862 | 0.170587319 |
| ENSMUSG00000079508 | -1.049280576 | 0.69533886 |
| ENSMUSG00000079509 | -1.018308518 | 0.99092221 |
| ENSMUSG00000079511 | -1.038198631 | 0.830576988 |
| ENSMUSG00000079513 | 1 | |
| ENSMUSG00000079516 | 1 | |
| ENSMUSG00000079519 | 1 | |
| ENSMUSG00000079521 | 1 | |
| ENSMUSG00000079522 | 1 | |
| ENSMUSG00000079523 | -1.003212081 | 1 |
| ENSMUSG00000079524 | -1 | 1 |
| ENSMUSG00000079525 | 1 | |
| ENSMUSG00000079528 | 1 | |
| ENSMUSG00000079531 | 1 | |
| ENSMUSG00000079532 | 1 | |
| ENSMUSG00000079534 | 1 | |
| ENSMUSG00000079536 | 1 | |
| ENSMUSG00000079537 | 1 | |
| ENSMUSG00000079539 | 1 | |
| ENSMUSG00000079542 | 1 | |
| ENSMUSG00000079543 | 1 | |
| ENSMUSG00000079544 | 1 | |
| ENSMUSG00000079546 | 1 | |
| ENSMUSG00000079547 | -1.04091331 | 0.959583351 |
| ENSMUSG00000079550 | -1.029302237 | 0.96035552 |
| ENSMUSG00000079553 | 1.09339395 | 0.464062002 |
| ENSMUSG00000079554 | -1.02522956 | 0.970707004 |
| ENSMUSG00000079555 | 1.013351725 | 1 |
| ENSMUSG00000079557 | 1.015789169 | 0.95702926 |
| ENSMUSG00000079559 | 1.115832455 | 0.779557148 |
| ENSMUSG00000079560 | 1.253532786 | 0.024046175 |
| ENSMUSG00000079562 | 1.116117748 | 0.470841398 |
| ENSMUSG00000079563 | -1.20776913 | 0.237992347 |
| ENSMUSG00000079564 | -1.014570171 | 1 |
| ENSMUSG00000079566 | 1 | |
| ENSMUSG00000079575 | 1.077488122 | 0.476737035 |
| ENSMUSG00000079577 | 1 | |
| ENSMUSG00000079578 | 1 | |
| ENSMUSG00000079579 | 1 | |
| ENSMUSG00000079580 | -1.278261408 | 0.26073339 |
| ENSMUSG00000079583 | 1 | |
| ENSMUSG00000079584 | -1 | 1 |
| ENSMUSG00000079588 | 1 | |
| ENSMUSG00000079592 | 1 | |
| ENSMUSG00000079593 | -1.063525418 | 0.756517167 |
| ENSMUSG00000079594 | 1 | |
| ENSMUSG00000079595 | 1 | |
| ENSMUSG00000079597 | 1 | |
| ENSMUSG00000079598 | -1.341581038 | 0.147309246 |
| ENSMUSG00000079600 | 1.0948249 | 0.729018288 |
| ENSMUSG00000079602 | -1.044273782 | 0.748351609 |
| ENSMUSG00000079603 | 1 | |
| ENSMUSG00000079604 | 1 | |
| ENSMUSG00000079605 | 1 | |
| ENSMUSG00000079606 | 1 | |
| ENSMUSG00000079608 | 1.099765725 | 0.262806495 |
| ENSMUSG00000079610 | -1.019940017 | 0.92972545 |
| ENSMUSG00000079614 | -1.044292527 | 0.911882434 |

| | | |
|---|---|---|
| ENSMUSG00000079619 | 1 | |
| ENSMUSG00000079620 | -1.043306199 | 0.79291651 |
| ENSMUSG00000079625 | 1.079681442 | 0.78131463 |
| ENSMUSG00000079626 | 1 | |
| ENSMUSG00000079627 | 1 | |
| ENSMUSG00000079628 | 1.069068979 | 0.608497229 |
| ENSMUSG00000079629 | 1 | |
| ENSMUSG00000079630 | 1 | |
| ENSMUSG00000079633 | 1 | |
| ENSMUSG00000079635 | 1 | |
| ENSMUSG00000079636 | 1 | |
| ENSMUSG00000079637 | 1 | |
| ENSMUSG00000079638 | 1 | |
| ENSMUSG00000079639 | 1.014731082 | 0.988424085 |
| ENSMUSG00000079641 | 1.040617895 | 0.924629556 |
| ENSMUSG00000079642 | 1 | |
| ENSMUSG00000079644 | -1.133231767 | 0.591151279 |
| ENSMUSG00000079645 | 1.276717071 | 0.564833703 |
| ENSMUSG00000079649 | 1 | |
| ENSMUSG00000079652 | 1.009151095 | 1 |
| ENSMUSG00000079654 | -1.203561049 | 0.636056232 |
| ENSMUSG00000079655 | 1 | |
| ENSMUSG00000079657 | -1.187802519 | 0.324900293 |
| ENSMUSG00000079658 | 1.009053977 | 1 |
| ENSMUSG00000079659 | 1.015921441 | 0.838766926 |
| ENSMUSG00000079666 | -1.054169715 | 0.813485995 |
| ENSMUSG00000079669 | -1.027724852 | 0.922261327 |
| ENSMUSG00000079671 | -1.043306199 | 0.798487595 |
| ENSMUSG00000079677 | -1.054637773 | 0.698112512 |
| ENSMUSG00000079679 | 1.024693402 | 0.933031586 |
| ENSMUSG00000079681 | -1.00215463 | 1 |
| ENSMUSG00000079685 | -2.989333632 | 0.000364772 |
| ENSMUSG00000079693 | 1 | |
| ENSMUSG00000079694 | 1.054873865 | 0.793748802 |
| ENSMUSG00000079697 | -1.063525418 | 0.698961838 |
| ENSMUSG00000079698 | 1 | |
| ENSMUSG00000079699 | 1 | |
| ENSMUSG00000079700 | 1 | |
| ENSMUSG00000079701 | 1 | |
| ENSMUSG00000079702 | 1 | |
| ENSMUSG00000079703 | 1 | |
| ENSMUSG00000079704 | 1 | |
| ENSMUSG00000079705 | 1.059463094 | 0.608497229 |
| ENSMUSG00000079707 | -1.038444538 | 0.865865121 |
| ENSMUSG00000079710 | 1 | |
| ENSMUSG00000079711 | -1.03962336 | 0.698112512 |
| ENSMUSG00000079722 | 1.042979668 | 0.933031586 |
| ENSMUSG00000079737 | 1.040152536 | 0.692803919 |
| ENSMUSG00000079740 | 1 | |
| ENSMUSG00000079741 | 1 | |
| ENSMUSG00000079764 | 1 | |
| ENSMUSG00000079773 | 1 | |
| ENSMUSG00000079774 | 1 | |
| ENSMUSG00000079794 | 1 | |
| ENSMUSG00000079800 | 1 | |
| ENSMUSG00000079806 | 1 | |
| ENSMUSG00000079808 | 1.014545335 | 0.979002724 |
| ENSMUSG00000079834 | 1.152735541 | 0.346615757 |
| ENSMUSG00000079842 | 1 | |
| ENSMUSG00000079845 | 1 | |
| ENSMUSG00000079852 | 2.88049953 | 2.69E-07 |
| ENSMUSG00000079853 | 1 | |
| ENSMUSG00000079871 | -1.085651418 | 0.532369657 |

| | | |
|---|---|---|
| ENSMUSG00000079875 | 1 | |
| ENSMUSG00000079884 | 1 | |
| ENSMUSG00000079889 | 1.052130777 | 0.854777354 |
| ENSMUSG00000079890 | 1.014545335 | 0.979002724 |
| ENSMUSG00000079892 | 1 | |
| ENSMUSG00000079893 | 1 | |
| ENSMUSG00000079906 | -1.111233324 | 0.608497229 |
| ENSMUSG00000079915 | -1.013472123 | 1 |
| ENSMUSG00000079941 | 1 | |
| ENSMUSG00000079942 | -1.085651418 | 0.608497229 |
| ENSMUSG00000079962 | -1.054169715 | 0.873632939 |
| ENSMUSG00000079963 | 1.047232809 | 0.916976316 |
| ENSMUSG00000079965 | 1.010351027 | 1 |
| ENSMUSG00000079966 | 1 | |
| ENSMUSG00000079988 | -1.189473777 | 0.277170403 |
| ENSMUSG00000079993 | 1 | |
| ENSMUSG00000079998 | -1.043306199 | 0.79291651 |
| ENSMUSG00000080002 | 1.071809423 | 0.8349433 |
| ENSMUSG00000080006 | -1 | 1 |
| ENSMUSG00000080021 | -1.053613252 | 0.857392806 |
| ENSMUSG00000080046 | 1 | |
| ENSMUSG00000080050 | 1 | |
| ENSMUSG00000080051 | 1 | |
| ENSMUSG00000080054 | 1.059463094 | 0.608497229 |
| ENSMUSG00000080057 | -1.044273782 | 0.698112512 |
| ENSMUSG00000080058 | 1 | |
| ENSMUSG00000080059 | -1.181805148 | 0.173112919 |
| ENSMUSG00000080065 | 1 | |
| ENSMUSG00000080069 | 1 | |
| ENSMUSG00000080071 | 1 | |
| ENSMUSG00000080072 | 1 | |
| ENSMUSG00000080073 | 1 | |
| ENSMUSG00000080076 | 1 | |
| ENSMUSG00000080078 | -1.089497311 | 0.532369657 |
| ENSMUSG00000080115 | 1.053578591 | 0.883083574 |
| ENSMUSG00000080142 | 1.019833591 | 0.983446483 |
| ENSMUSG00000080152 | 1.069068979 | 0.698112512 |
| ENSMUSG00000080155 | 1 | |
| ENSMUSG00000080158 | 1.059463094 | 0.608497229 |
| ENSMUSG00000080175 | 1 | |
| ENSMUSG00000080180 | -1.20144032 | 0.475771979 |
| ENSMUSG00000080186 | 1.084618945 | 0.628369449 |
| ENSMUSG00000080193 | 1 | |
| ENSMUSG00000080198 | 1.310562799 | 0.15208156 |
| ENSMUSG00000080205 | -1.084645496 | 0.532369657 |
| ENSMUSG00000080222 | 1 | |
| ENSMUSG00000080223 | 1 | |
| ENSMUSG00000080227 | 1 | |
| ENSMUSG00000080237 | 1.059463094 | 0.608497229 |
| ENSMUSG00000080241 | 1 | |
| ENSMUSG00000080242 | 1.136862817 | 0.608497229 |
| ENSMUSG00000080250 | 1 | |
| ENSMUSG00000080254 | -1.03962336 | 0.698112512 |
| ENSMUSG00000080266 | 1 | |
| ENSMUSG00000080268 | -1.055613866 | 0.701679148 |
| ENSMUSG00000080283 | 1 | |
| ENSMUSG00000080298 | -1 | 1 |
| ENSMUSG00000080299 | 1 | |
| ENSMUSG00000080300 | 1 | |
| ENSMUSG00000080303 | 1.011539697 | 1 |
| ENSMUSG00000080316 | 1.116261113 | 0.608497229 |
| ENSMUSG00000080317 | 1 | |
| ENSMUSG00000080321 | 1 | |

| | | |
|---|---|---|
| ENSMUSG00000080330 | 1 | |
| ENSMUSG00000080331 | 1 | |
| ENSMUSG00000080334 | 1 | |
| ENSMUSG00000080336 | -1 | 1 |
| ENSMUSG00000080342 | 1 | |
| ENSMUSG00000080345 | 1 | |
| ENSMUSG00000080346 | -1.053549062 | 0.698112512 |
| ENSMUSG00000080347 | 1 | |
| ENSMUSG00000080348 | 1 | |
| ENSMUSG00000080349 | 1 | |
| ENSMUSG00000080350 | 1 | |
| ENSMUSG00000080352 | 1 | |
| ENSMUSG00000080356 | 1 | |
| ENSMUSG00000080358 | 1 | |
| ENSMUSG00000080359 | 1 | |
| ENSMUSG00000080360 | 1 | |
| ENSMUSG00000080363 | 1 | |
| ENSMUSG00000080364 | 1.059463094 | 0.698112512 |
| ENSMUSG00000080365 | -1.161737199 | 0.470974528 |
| ENSMUSG00000080366 | 1 | |
| ENSMUSG00000080367 | 1 | |
| ENSMUSG00000080368 | 1 | |
| ENSMUSG00000080370 | 1.081478107 | 0.698112512 |
| ENSMUSG00000080374 | 1 | |
| ENSMUSG00000080378 | 1 | |
| ENSMUSG00000080380 | 1 | |
| ENSMUSG00000080383 | 1 | |
| ENSMUSG00000080384 | 1 | |
| ENSMUSG00000080391 | 1 | |
| ENSMUSG00000080394 | 1 | |
| ENSMUSG00000080396 | 1.028323353 | 0.903464611 |
| ENSMUSG00000080401 | -1.043306199 | 0.698112512 |
| ENSMUSG00000080402 | 1 | |
| ENSMUSG00000080405 | 1 | |
| ENSMUSG00000080406 | 1 | |
| ENSMUSG00000080408 | 1 | |
| ENSMUSG00000080409 | 1 | |
| ENSMUSG00000080410 | 1 | |
| ENSMUSG00000080411 | 1 | |
| ENSMUSG00000080412 | 1 | |
| ENSMUSG00000080413 | 1 | |
| ENSMUSG00000080415 | 1 | |
| ENSMUSG00000080419 | 1 | |
| ENSMUSG00000080424 | 1 | |
| ENSMUSG00000080425 | 1 | |
| ENSMUSG00000080427 | 1 | |
| ENSMUSG00000080428 | 1 | |
| ENSMUSG00000080430 | 1 | |
| ENSMUSG00000080432 | 1 | |
| ENSMUSG00000080434 | 1 | |
| ENSMUSG00000080437 | 1 | |
| ENSMUSG00000080440 | -1 | 1 |
| ENSMUSG00000080441 | 1 | |
| ENSMUSG00000080443 | 1 | |
| ENSMUSG00000080451 | 1 | |
| ENSMUSG00000080452 | 1 | |
| ENSMUSG00000080454 | -1.107299336 | 0.639279667 |
| ENSMUSG00000080455 | 1 | |
| ENSMUSG00000080459 | 1 | |
| ENSMUSG00000080461 | 1.028323353 | 0.933031586 |
| ENSMUSG00000080463 | 1 | |
| ENSMUSG00000080465 | 1.059996698 | 0.760423805 |
| ENSMUSG00000080468 | 1 | |

| | | |
|---|---|---|
| ENSMUSG00000080469 | 1.117607781 | 0.642725233 |
| ENSMUSG00000080473 | 1 | |
| ENSMUSG00000080474 | 1 | |
| ENSMUSG00000080476 | 1 | |
| ENSMUSG00000080478 | 1 | |
| ENSMUSG00000080480 | 1 | |
| ENSMUSG00000080485 | -1.099174267 | 0.652602405 |
| ENSMUSG00000080486 | 1.061224565 | 0.775601843 |
| ENSMUSG00000080487 | 1 | |
| ENSMUSG00000080495 | 1 | |
| ENSMUSG00000080498 | 1 | |
| ENSMUSG00000080499 | 1 | |
| ENSMUSG00000080500 | 1 | |
| ENSMUSG00000080504 | 1 | |
| ENSMUSG00000080506 | 1 | |
| ENSMUSG00000080507 | 1.069068979 | 0.608497229 |
| ENSMUSG00000080508 | 1 | |
| ENSMUSG00000080511 | 1 | |
| ENSMUSG00000080515 | 1 | |
| ENSMUSG00000080516 | 1 | |
| ENSMUSG00000080517 | 1 | |
| ENSMUSG00000080518 | 1 | |
| ENSMUSG00000080519 | 1 | |
| ENSMUSG00000080522 | 1 | |
| ENSMUSG00000080524 | 1 | |
| ENSMUSG00000080526 | 1 | |
| ENSMUSG00000080527 | 1 | |
| ENSMUSG00000080529 | 1 | |
| ENSMUSG00000080530 | 1 | |
| ENSMUSG00000080533 | 1.042253985 | 0.873622104 |
| ENSMUSG00000080536 | 1 | |
| ENSMUSG00000080538 | 1 | |
| ENSMUSG00000080539 | 1 | |
| ENSMUSG00000080540 | 1 | |
| ENSMUSG00000080542 | 1 | |
| ENSMUSG00000080543 | 1 | |
| ENSMUSG00000080544 | 1 | |
| ENSMUSG00000080545 | 1 | |
| ENSMUSG00000080549 | 1 | |
| ENSMUSG00000080555 | 1 | |
| ENSMUSG00000080556 | 1 | |
| ENSMUSG00000080558 | 1 | |
| ENSMUSG00000080561 | 1 | |
| ENSMUSG00000080562 | 1 | |
| ENSMUSG00000080566 | 1 | |
| ENSMUSG00000080573 | 1.049517471 | 0.869287401 |
| ENSMUSG00000080574 | -1.063127349 | 0.737515436 |
| ENSMUSG00000080575 | 1 | |
| ENSMUSG00000080577 | 1 | |
| ENSMUSG00000080580 | 1 | |
| ENSMUSG00000080582 | 1 | |
| ENSMUSG00000080586 | 1 | |
| ENSMUSG00000080590 | 1 | |
| ENSMUSG00000080592 | 1 | |
| ENSMUSG00000080594 | 1.059463094 | 0.608497229 |
| ENSMUSG00000080595 | 1 | |
| ENSMUSG00000080596 | 1 | |
| ENSMUSG00000080597 | -1 | 1 |
| ENSMUSG00000080600 | 1 | |
| ENSMUSG00000080601 | 1 | |
| ENSMUSG00000080607 | 1 | |
| ENSMUSG00000080609 | 1 | |
| ENSMUSG00000080610 | -1.033820075 | 0.940197654 |

| | | |
|---|---|---|
| ENSMUSG00000080613 | -1.044273782 | 0.698112512 |
| ENSMUSG00000080615 | 1 | |
| ENSMUSG00000080616 | 1 | |
| ENSMUSG00000080622 | 1 | |
| ENSMUSG00000080624 | 1 | |
| ENSMUSG00000080625 | 1 | |
| ENSMUSG00000080626 | 1 | |
| ENSMUSG00000080627 | 1 | |
| ENSMUSG00000080631 | 1 | |
| ENSMUSG00000080635 | 1 | |
| ENSMUSG00000080636 | 1 | |
| ENSMUSG00000080645 | -1.084645496 | 0.532369657 |
| ENSMUSG00000080649 | 1 | |
| ENSMUSG00000080653 | 1 | |
| ENSMUSG00000080655 | 1 | |
| ENSMUSG00000080657 | 1 | |
| ENSMUSG00000080659 | 1 | |
| ENSMUSG00000080662 | 1 | |
| ENSMUSG00000080663 | 1 | |
| ENSMUSG00000080665 | 1 | |
| ENSMUSG00000080666 | -1 | 1 |
| ENSMUSG00000080668 | 1 | |
| ENSMUSG00000080669 | 1 | |
| ENSMUSG00000080671 | 1 | |
| ENSMUSG00000080673 | 1 | |
| ENSMUSG00000080676 | 1 | |
| ENSMUSG00000080678 | 1 | |
| ENSMUSG00000080680 | 1 | |
| ENSMUSG00000080681 | 1 | |
| ENSMUSG00000080683 | 1 | |
| ENSMUSG00000080684 | 1 | |
| ENSMUSG00000080686 | 1 | |
| ENSMUSG00000080687 | 1 | |
| ENSMUSG00000080689 | 1 | |
| ENSMUSG00000080694 | 1 | |
| ENSMUSG00000080695 | 1 | |
| ENSMUSG00000080696 | 1 | |
| ENSMUSG00000080697 | 1 | |
| ENSMUSG00000080698 | 1 | |
| ENSMUSG00000080700 | -1.095294215 | 0.608497229 |
| ENSMUSG00000080701 | 1.019083579 | 0.97045957 |
| ENSMUSG00000080703 | -1.023769022 | 0.934760809 |
| ENSMUSG00000080704 | 1 | |
| ENSMUSG00000080705 | 1 | |
| ENSMUSG00000080706 | 1 | |
| ENSMUSG00000080707 | 1 | |
| ENSMUSG00000080708 | 1 | |
| ENSMUSG00000080709 | -1.109826685 | 0.693148633 |
| ENSMUSG00000080710 | 1 | |
| ENSMUSG00000080711 | 1 | |
| ENSMUSG00000080712 | 1 | |
| ENSMUSG00000080713 | 1 | |
| ENSMUSG00000080714 | 1 | |
| ENSMUSG00000080715 | 1.069068979 | 0.608497229 |
| ENSMUSG00000080716 | 1 | |
| ENSMUSG00000080717 | -1.030401546 | 0.695729298 |
| ENSMUSG00000080718 | -1.044697704 | 0.870906262 |
| ENSMUSG00000080719 | 1.167673187 | 0.475124176 |
| ENSMUSG00000080720 | 1 | |
| ENSMUSG00000080722 | 1 | |
| ENSMUSG00000080723 | 1 | |
| ENSMUSG00000080725 | 1 | |
| ENSMUSG00000080726 | 1 | |

| | | |
|---|---|---|
| ENSMUSG00000080727 | -1.046215139 | 0.890449109 |
| ENSMUSG00000080728 | 1 | |
| ENSMUSG00000080729 | 1 | |
| ENSMUSG00000080730 | 1.28970965 | 0.182544212 |
| ENSMUSG00000080731 | 1.063665443 | 0.770958539 |
| ENSMUSG00000080732 | 1 | |
| ENSMUSG00000080733 | 1 | |
| ENSMUSG00000080734 | 1 | |
| ENSMUSG00000080735 | 1 | |
| ENSMUSG00000080736 | 1 | |
| ENSMUSG00000080737 | -1.068600963 | 0.698112512 |
| ENSMUSG00000080738 | 1 | |
| ENSMUSG00000080741 | 1 | |
| ENSMUSG00000080742 | 1 | |
| ENSMUSG00000080743 | 1.342495559 | 0.340244011 |
| ENSMUSG00000080744 | 1 | |
| ENSMUSG00000080746 | -1.073796839 | 0.894963534 |
| ENSMUSG00000080747 | -1.041954902 | 0.910467721 |
| ENSMUSG00000080748 | 1.014731082 | 0.997667419 |
| ENSMUSG00000080749 | 1 | |
| ENSMUSG00000080750 | 1 | |
| ENSMUSG00000080751 | 1 | |
| ENSMUSG00000080752 | 1 | |
| ENSMUSG00000080753 | 1 | |
| ENSMUSG00000080754 | 1 | |
| ENSMUSG00000080755 | 1 | |
| ENSMUSG00000080756 | 1 | |
| ENSMUSG00000080757 | 1 | |
| ENSMUSG00000080758 | 1 | |
| ENSMUSG00000080759 | -1.03962336 | 0.698112512 |
| ENSMUSG00000080760 | 1.059463094 | 0.698112512 |
| ENSMUSG00000080762 | -1.089497311 | 0.532369657 |
| ENSMUSG00000080763 | 1.059463094 | 0.608497229 |
| ENSMUSG00000080764 | 1.049411284 | 0.835701362 |
| ENSMUSG00000080765 | 1 | |
| ENSMUSG00000080766 | 1 | |
| ENSMUSG00000080767 | 1 | |
| ENSMUSG00000080768 | -1.075310537 | 0.637342941 |
| ENSMUSG00000080770 | 1 | |
| ENSMUSG00000080771 | 1 | |
| ENSMUSG00000080772 | -1.095294215 | 0.608497229 |
| ENSMUSG00000080773 | 1 | |
| ENSMUSG00000080774 | 1.07988273 | 0.698112512 |
| ENSMUSG00000080775 | -1.024069464 | 0.969005636 |
| ENSMUSG00000080776 | -1.085651418 | 0.615267463 |
| ENSMUSG00000080777 | 1.059463094 | 0.608497229 |
| ENSMUSG00000080778 | -1 | 1 |
| ENSMUSG00000080779 | 1.089470641 | 0.608497229 |
| ENSMUSG00000080780 | 1 | |
| ENSMUSG00000080781 | 1 | |
| ENSMUSG00000080782 | 1 | |
| ENSMUSG00000080783 | -1.048483131 | 0.91203619 |
| ENSMUSG00000080785 | 1 | |
| ENSMUSG00000080788 | 1 | |
| ENSMUSG00000080789 | -1 | 1 |
| ENSMUSG00000080790 | 1.204535345 | 0.3305975 |
| ENSMUSG00000080791 | -1.03962336 | 0.698112512 |
| ENSMUSG00000080792 | 1 | |
| ENSMUSG00000080793 | -1.053549062 | 0.698112512 |
| ENSMUSG00000080794 | 1 | |
| ENSMUSG00000080795 | -1.03962336 | 0.698112512 |
| ENSMUSG00000080796 | -1.053549062 | 0.698112512 |
| ENSMUSG00000080797 | 1.02439549 | 0.988270581 |

| | | |
|---|---|---|
| ENSMUSG00000080798 | 1.059463094 | 0.776953697 |
| ENSMUSG00000080799 | 1 | |
| ENSMUSG00000080800 | 1 | |
| ENSMUSG00000080801 | -1.14674712 | 0.608497229 |
| ENSMUSG00000080802 | 1 | |
| ENSMUSG00000080803 | 1 | |
| ENSMUSG00000080805 | 1 | |
| ENSMUSG00000080806 | 1.059463094 | 0.704170349 |
| ENSMUSG00000080807 | 1 | |
| ENSMUSG00000080808 | 1.010498549 | 1 |
| ENSMUSG00000080809 | 1 | |
| ENSMUSG00000080810 | 1 | |
| ENSMUSG00000080811 | -1.044273782 | 0.790851299 |
| ENSMUSG00000080812 | 1 | |
| ENSMUSG00000080813 | 1 | |
| ENSMUSG00000080814 | 1.171563499 | 0.233412931 |
| ENSMUSG00000080815 | 1.095872691 | 0.657060635 |
| ENSMUSG00000080816 | 1 | |
| ENSMUSG00000080817 | 1 | |
| ENSMUSG00000080818 | 1.015486245 | 0.985522133 |
| ENSMUSG00000080819 | 1 | |
| ENSMUSG00000080820 | 1 | |
| ENSMUSG00000080822 | 1 | |
| ENSMUSG00000080823 | -1.03962336 | 0.698112512 |
| ENSMUSG00000080824 | 1 | |
| ENSMUSG00000080825 | -1.024718486 | 0.979002724 |
| ENSMUSG00000080826 | 1 | |
| ENSMUSG00000080827 | 1 | |
| ENSMUSG00000080828 | 1 | |
| ENSMUSG00000080829 | 1.014545335 | 0.989648415 |
| ENSMUSG00000080831 | 1 | |
| ENSMUSG00000080832 | 1.070429626 | 0.767473873 |
| ENSMUSG00000080833 | 1 | |
| ENSMUSG00000080834 | 1 | |
| ENSMUSG00000080835 | 1 | |
| ENSMUSG00000080836 | 1.122462048 | 0.215991492 |
| ENSMUSG00000080839 | 1.015486245 | 0.987775262 |
| ENSMUSG00000080840 | -1.139364264 | 0.634636679 |
| ENSMUSG00000080841 | 1 | |
| ENSMUSG00000080842 | -1 | 1 |
| ENSMUSG00000080843 | 1 | |
| ENSMUSG00000080844 | 1.059463094 | 0.608497229 |
| ENSMUSG00000080845 | 1.095845865 | 0.608497229 |
| ENSMUSG00000080846 | 1.069068979 | 0.608497229 |
| ENSMUSG00000080847 | 1 | |
| ENSMUSG00000080848 | 1.030445456 | 0.938939969 |
| ENSMUSG00000080849 | -1.044273782 | 0.790851299 |
| ENSMUSG00000080850 | -1.186502317 | 0.473106921 |
| ENSMUSG00000080851 | 1.059463094 | 0.608497229 |
| ENSMUSG00000080852 | 1 | |
| ENSMUSG00000080855 | 1 | |
| ENSMUSG00000080857 | 1 | |
| ENSMUSG00000080859 | 1 | |
| ENSMUSG00000080860 | -1.143516169 | 0.546660718 |
| ENSMUSG00000080861 | 1 | |
| ENSMUSG00000080862 | 1 | |
| ENSMUSG00000080863 | 1 | |
| ENSMUSG00000080865 | 1 | |
| ENSMUSG00000080866 | 1.01716273 | 0.925214847 |
| ENSMUSG00000080868 | 1 | |
| ENSMUSG00000080869 | 1 | |
| ENSMUSG00000080870 | 1.161008251 | 0.433678407 |
| ENSMUSG00000080871 | 1 | |

| | | |
|---|---|---|
| ENSMUSG00000080872 | 1 | |
| ENSMUSG00000080873 | 1.014545335 | 0.989648415 |
| ENSMUSG00000080874 | 1 | |
| ENSMUSG00000080875 | 1.069068979 | 0.721840456 |
| ENSMUSG00000080876 | 1 | |
| ENSMUSG00000080877 | -1.18490436 | 0.165502439 |
| ENSMUSG00000080878 | 1 | |
| ENSMUSG00000080880 | 1 | |
| ENSMUSG00000080884 | 1 | |
| ENSMUSG00000080885 | 1.187085156 | 0.608497229 |
| ENSMUSG00000080886 | 1 | |
| ENSMUSG00000080887 | 1 | |
| ENSMUSG00000080888 | -1.275322407 | 0.09399025 |
| ENSMUSG00000080889 | 1 | |
| ENSMUSG00000080890 | 1 | |
| ENSMUSG00000080891 | 1 | |
| ENSMUSG00000080892 | 1 | |
| ENSMUSG00000080893 | 1.103985575 | 0.357647366 |
| ENSMUSG00000080894 | 1 | |
| ENSMUSG00000080895 | 1.918139935 | 0.005466258 |
| ENSMUSG00000080896 | 1 | |
| ENSMUSG00000080897 | 1 | |
| ENSMUSG00000080898 | 1 | |
| ENSMUSG00000080899 | 1 | |
| ENSMUSG00000080902 | 1.059352535 | 0.843480803 |
| ENSMUSG00000080903 | 1 | |
| ENSMUSG00000080904 | -1 | 1 |
| ENSMUSG00000080906 | 1 | |
| ENSMUSG00000080907 | 1 | |
| ENSMUSG00000080910 | 1 | |
| ENSMUSG00000080911 | 1 | |
| ENSMUSG00000080912 | 1.059463094 | 0.704170349 |
| ENSMUSG00000080913 | 1 | |
| ENSMUSG00000080914 | 1.199989356 | 0.354316858 |
| ENSMUSG00000080915 | 1 | |
| ENSMUSG00000080916 | 1 | |
| ENSMUSG00000080917 | 1 | |
| ENSMUSG00000080918 | 1 | |
| ENSMUSG00000080919 | 1 | |
| ENSMUSG00000080921 | -1.047306572 | 0.890958598 |
| ENSMUSG00000080922 | 1 | |
| ENSMUSG00000080923 | 1 | |
| ENSMUSG00000080924 | 1 | |
| ENSMUSG00000080925 | 1 | |
| ENSMUSG00000080927 | 1 | |
| ENSMUSG00000080928 | 1.042760588 | 0.949812939 |
| ENSMUSG00000080929 | -1.044273782 | 0.698112512 |
| ENSMUSG00000080930 | 1.059463094 | 0.608497229 |
| ENSMUSG00000080932 | -1.075051016 | 0.639439274 |
| ENSMUSG00000080933 | 1 | |
| ENSMUSG00000080934 | 1 | |
| ENSMUSG00000080935 | 1 | |
| ENSMUSG00000080936 | 1 | |
| ENSMUSG00000080937 | 1 | |
| ENSMUSG00000080938 | 1.029490685 | 0.942171713 |
| ENSMUSG00000080939 | 1 | |
| ENSMUSG00000080941 | 1 | |
| ENSMUSG00000080942 | 1 | |
| ENSMUSG00000080943 | 1 | |
| ENSMUSG00000080944 | 1.040221616 | 0.96035552 |
| ENSMUSG00000080946 | 1 | |
| ENSMUSG00000080947 | 1 | |
| ENSMUSG00000080948 | 1 | |

| | | |
|---|---|---|
| ENSMUSG00000080949 | 1 | |
| ENSMUSG00000080950 | 1.305338549 | 0.373106943 |
| ENSMUSG00000080951 | -1.018844336 | 1 |
| ENSMUSG00000080952 | 1 | |
| ENSMUSG00000080953 | 1 | |
| ENSMUSG00000080954 | 1 | |
| ENSMUSG00000080955 | 1.01393017 | 1 |
| ENSMUSG00000080956 | -1.044273782 | 0.790851299 |
| ENSMUSG00000080957 | -1.029113823 | 0.929923764 |
| ENSMUSG00000080958 | 1 | |
| ENSMUSG00000080959 | 1 | |
| ENSMUSG00000080960 | 1 | |
| ENSMUSG00000080961 | 1 | |
| ENSMUSG00000080962 | 1.059463094 | 0.698112512 |
| ENSMUSG00000080964 | 1 | |
| ENSMUSG00000080965 | 1.059463094 | 0.698112512 |
| ENSMUSG00000080966 | -1.043306199 | 0.698112512 |
| ENSMUSG00000080967 | 1 | |
| ENSMUSG00000080968 | 1.095872691 | 0.704170349 |
| ENSMUSG00000080969 | 1.065410325 | 0.805198503 |
| ENSMUSG00000080970 | 1 | |
| ENSMUSG00000080971 | 1.014545335 | 0.998616029 |
| ENSMUSG00000080972 | 1.091291226 | 0.608497229 |
| ENSMUSG00000080974 | -1.01251055 | 1 |
| ENSMUSG00000080976 | -1.044273782 | 0.698112512 |
| ENSMUSG00000080977 | 1 | |
| ENSMUSG00000080978 | 1 | |
| ENSMUSG00000080979 | 1 | |
| ENSMUSG00000080980 | 1 | |
| ENSMUSG00000080981 | 1 | |
| ENSMUSG00000080982 | 1 | |
| ENSMUSG00000080983 | 1 | |
| ENSMUSG00000080984 | 1 | |
| ENSMUSG00000080985 | -1.047306572 | 0.870103307 |
| ENSMUSG00000080986 | 1 | |
| ENSMUSG00000080987 | 1 | |
| ENSMUSG00000080989 | 1 | |
| ENSMUSG00000080990 | 1 | |
| ENSMUSG00000080992 | 1.089470641 | 0.608497229 |
| ENSMUSG00000080993 | 1 | |
| ENSMUSG00000080994 | 1.059463094 | 0.698112512 |
| ENSMUSG00000080995 | 1.059463094 | 0.608497229 |
| ENSMUSG00000080996 | -1 | 1 |
| ENSMUSG00000080997 | 1.194896481 | 0.385188707 |
| ENSMUSG00000080998 | -1.084645496 | 0.608497229 |
| ENSMUSG00000080999 | 1 | |
| ENSMUSG00000081000 | 1 | |
| ENSMUSG00000081001 | 1 | |
| ENSMUSG00000081002 | -1 | 1 |
| ENSMUSG00000081003 | 1 | |
| ENSMUSG00000081004 | 1.12189305 | 0.608497229 |
| ENSMUSG00000081005 | 1 | |
| ENSMUSG00000081006 | 1 | |
| ENSMUSG00000081007 | 1.069068979 | 0.608497229 |
| ENSMUSG00000081008 | 1 | |
| ENSMUSG00000081009 | 1 | |
| ENSMUSG00000081010 | 1.075870199 | 0.818635785 |
| ENSMUSG00000081011 | 1.016755844 | 0.920928417 |
| ENSMUSG00000081012 | 1 | |
| ENSMUSG00000081013 | 1.069068979 | 0.698112512 |
| ENSMUSG00000081014 | 1 | |
| ENSMUSG00000081016 | 1.059463094 | 0.698112512 |
| ENSMUSG00000081017 | 1 | |

| | | |
|---|---|---|
| ENSMUSG00000081018 | 1 | |
| ENSMUSG00000081020 | 1 | |
| ENSMUSG00000081021 | -1 | 1 |
| ENSMUSG00000081022 | 1 | |
| ENSMUSG00000081023 | 1.018670051 | 0.98871538 |
| ENSMUSG00000081024 | -1.03962336 | 0.698112512 |
| ENSMUSG00000081025 | 1 | |
| ENSMUSG00000081026 | 1.111812527 | 0.629725675 |
| ENSMUSG00000081027 | 1 | |
| ENSMUSG00000081029 | -1.03962336 | 0.803072249 |
| ENSMUSG00000081030 | -1.124966537 | 0.698112512 |
| ENSMUSG00000081033 | 1 | |
| ENSMUSG00000081034 | -1.044273782 | 0.698112512 |
| ENSMUSG00000081035 | 1 | |
| ENSMUSG00000081036 | 1 | |
| ENSMUSG00000081037 | 1 | |
| ENSMUSG00000081038 | 1 | |
| ENSMUSG00000081041 | -1.03962336 | 0.698112512 |
| ENSMUSG00000081042 | 1.020742201 | 0.995111383 |
| ENSMUSG00000081043 | -1.170352376 | 0.608497229 |
| ENSMUSG00000081044 | 1.059463094 | 0.608497229 |
| ENSMUSG00000081045 | 1 | |
| ENSMUSG00000081046 | -1.086192882 | 0.734674233 |
| ENSMUSG00000081047 | 1 | |
| ENSMUSG00000081048 | 1.079039843 | 0.808166155 |
| ENSMUSG00000081049 | 1.005851671 | 1 |
| ENSMUSG00000081050 | 1 | |
| ENSMUSG00000081051 | 1.081351378 | 0.79860688 |
| ENSMUSG00000081052 | -1 | 1 |
| ENSMUSG00000081053 | 1 | |
| ENSMUSG00000081054 | 1 | |
| ENSMUSG00000081055 | -1.03962336 | 0.698112512 |
| ENSMUSG00000081056 | 1 | |
| ENSMUSG00000081057 | 1 | |
| ENSMUSG00000081058 | -1.128668575 | 0.643672693 |
| ENSMUSG00000081059 | -1.042426269 | 0.873239663 |
| ENSMUSG00000081060 | 1 | |
| ENSMUSG00000081061 | 1 | |
| ENSMUSG00000081062 | 1 | |
| ENSMUSG00000081063 | 1 | |
| ENSMUSG00000081064 | 1.028323353 | 0.934760809 |
| ENSMUSG00000081065 | 1 | |
| ENSMUSG00000081066 | 1 | |
| ENSMUSG00000081067 | 1.008665561 | 1 |
| ENSMUSG00000081068 | 1.015486245 | 0.987775262 |
| ENSMUSG00000081069 | 1 | |
| ENSMUSG00000081070 | 1 | |
| ENSMUSG00000081071 | 1 | |
| ENSMUSG00000081072 | 1 | |
| ENSMUSG00000081073 | -1 | 1 |
| ENSMUSG00000081076 | 1.15984069 | 0.354925301 |
| ENSMUSG00000081077 | -1.044273782 | 0.698112512 |
| ENSMUSG00000081078 | -1.03962336 | 0.698112512 |
| ENSMUSG00000081079 | 1 | |
| ENSMUSG00000081080 | 1 | |
| ENSMUSG00000081081 | 1 | |
| ENSMUSG00000081082 | 1 | |
| ENSMUSG00000081083 | 1 | |
| ENSMUSG00000081084 | 1.055859595 | 0.856456807 |
| ENSMUSG00000081085 | 1 | |
| ENSMUSG00000081086 | -1.03962336 | 0.698112512 |
| ENSMUSG00000081087 | -1.083415624 | 0.698112512 |
| ENSMUSG00000081088 | 1 | |

| | | |
|---|---|---|
| ENSMUSG00000081089 | 1 | |
| ENSMUSG00000081092 | 1 | |
| ENSMUSG00000081093 | -1.018570571 | 0.957412647 |
| ENSMUSG00000081094 | 1.056642783 | 0.764049978 |
| ENSMUSG00000081095 | 1 | |
| ENSMUSG00000081096 | -1.044273782 | 0.698112512 |
| ENSMUSG00000081097 | 1 | |
| ENSMUSG00000081098 | -1 | 1 |
| ENSMUSG00000081099 | -1.03962336 | 0.803072249 |
| ENSMUSG00000081100 | -1.03962336 | 0.698112512 |
| ENSMUSG00000081101 | 1.059463094 | 0.608497229 |
| ENSMUSG00000081103 | 1.189936102 | 0.541735653 |
| ENSMUSG00000081104 | -1.084475226 | 0.756770525 |
| ENSMUSG00000081106 | 1 | |
| ENSMUSG00000081107 | 1 | |
| ENSMUSG00000081108 | 1 | |
| ENSMUSG00000081110 | 1 | |
| ENSMUSG00000081111 | -1.091015577 | 0.704722765 |
| ENSMUSG00000081112 | -1.084645496 | 0.532369657 |
| ENSMUSG00000081113 | 1.017434421 | 1 |
| ENSMUSG00000081114 | 1 | |
| ENSMUSG00000081115 | 1 | |
| ENSMUSG00000081116 | 1 | |
| ENSMUSG00000081117 | 1 | |
| ENSMUSG00000081118 | -1.23299257 | 0.391804854 |
| ENSMUSG00000081119 | 1 | |
| ENSMUSG00000081120 | 1.069068979 | 0.608497229 |
| ENSMUSG00000081121 | 1.014545335 | 0.991944074 |
| ENSMUSG00000081122 | 1 | |
| ENSMUSG00000081123 | 1.059463094 | 0.794274354 |
| ENSMUSG00000081124 | 1 | |
| ENSMUSG00000081125 | 1 | |
| ENSMUSG00000081126 | -1.098547616 | 0.79291651 |
| ENSMUSG00000081127 | -1.053549062 | 0.698112512 |
| ENSMUSG00000081128 | 1 | |
| ENSMUSG00000081129 | 1 | |
| ENSMUSG00000081130 | -1 | 1 |
| ENSMUSG00000081131 | 1 | |
| ENSMUSG00000081132 | 1 | |
| ENSMUSG00000081133 | 1 | |
| ENSMUSG00000081134 | 1 | |
| ENSMUSG00000081135 | 1.084618945 | 0.698112512 |
| ENSMUSG00000081136 | 1.059463094 | 0.698112512 |
| ENSMUSG00000081137 | 1.037629245 | 0.698112512 |
| ENSMUSG00000081138 | -1.028348525 | 0.956972448 |
| ENSMUSG00000081139 | -1.113967123 | 0.661044879 |
| ENSMUSG00000081141 | 1 | |
| ENSMUSG00000081142 | 1.101111052 | 0.701627111 |
| ENSMUSG00000081143 | 1.084618945 | 0.722577651 |
| ENSMUSG00000081144 | 1 | |
| ENSMUSG00000081145 | -1 | 1 |
| ENSMUSG00000081146 | 1 | |
| ENSMUSG00000081147 | 1.065410325 | 0.711602885 |
| ENSMUSG00000081148 | 1 | |
| ENSMUSG00000081150 | 1 | |
| ENSMUSG00000081151 | 1.014545335 | 0.979002724 |
| ENSMUSG00000081152 | -1.037482356 | 0.89979145 |
| ENSMUSG00000081153 | -1.095294215 | 0.608497229 |
| ENSMUSG00000081154 | 1 | |
| ENSMUSG00000081155 | 1 | |
| ENSMUSG00000081156 | 1.132639128 | 0.217803313 |
| ENSMUSG00000081157 | 1.059463094 | 0.698112512 |
| ENSMUSG00000081158 | 1 | |

| | | |
|---|---|---|
| ENSMUSG00000081159 | -1 | 1 |
| ENSMUSG00000081160 | 1 | |
| ENSMUSG00000081161 | -1.170083925 | 0.206620396 |
| ENSMUSG00000081162 | 1 | |
| ENSMUSG00000081163 | 1 | |
| ENSMUSG00000081164 | 1.059463094 | 0.698112512 |
| ENSMUSG00000081165 | 1 | |
| ENSMUSG00000081166 | 1 | |
| ENSMUSG00000081167 | 1.015486245 | 0.985522133 |
| ENSMUSG00000081168 | -1 | 1 |
| ENSMUSG00000081169 | -1.071075483 | 0.698112512 |
| ENSMUSG00000081170 | 1.064029762 | 0.865583959 |
| ENSMUSG00000081171 | 1 | |
| ENSMUSG00000081172 | 1.015486245 | 0.972980766 |
| ENSMUSG00000081173 | 1 | |
| ENSMUSG00000081174 | 1 | |
| ENSMUSG00000081175 | 1 | |
| ENSMUSG00000081176 | 1.059463094 | 0.698112512 |
| ENSMUSG00000081177 | 1 | |
| ENSMUSG00000081179 | 1.014545335 | 0.989648415 |
| ENSMUSG00000081180 | 1 | |
| ENSMUSG00000081181 | 1 | |
| ENSMUSG00000081182 | 1.024693402 | 0.933031586 |
| ENSMUSG00000081183 | 1 | |
| ENSMUSG00000081184 | 1 | |
| ENSMUSG00000081185 | 1 | |
| ENSMUSG00000081186 | 1 | |
| ENSMUSG00000081187 | -1.096120849 | 0.698112512 |
| ENSMUSG00000081188 | 1 | |
| ENSMUSG00000081189 | -1.005494935 | 1 |
| ENSMUSG00000081191 | 1.122462048 | 0.215991492 |
| ENSMUSG00000081194 | -1.099174267 | 0.535052277 |
| ENSMUSG00000081195 | 1 | |
| ENSMUSG00000081196 | 1.059463094 | 0.698112512 |
| ENSMUSG00000081197 | 1 | |
| ENSMUSG00000081199 | 1 | |
| ENSMUSG00000081200 | 1 | |
| ENSMUSG00000081201 | 1 | |
| ENSMUSG00000081202 | 1 | |
| ENSMUSG00000081205 | -1 | 1 |
| ENSMUSG00000081206 | 1.095872691 | 0.671280603 |
| ENSMUSG00000081207 | 1 | |
| ENSMUSG00000081208 | -1.029757833 | 0.886841005 |
| ENSMUSG00000081209 | -1 | 1 |
| ENSMUSG00000081210 | 1.058443389 | 0.730264189 |
| ENSMUSG00000081211 | 1 | |
| ENSMUSG00000081212 | 1 | |
| ENSMUSG00000081214 | 1 | |
| ENSMUSG00000081215 | 1 | |
| ENSMUSG00000081216 | 1 | |
| ENSMUSG00000081217 | 1.090965779 | 0.698112512 |
| ENSMUSG00000081218 | 1 | |
| ENSMUSG00000081219 | -1.124836466 | 0.627688973 |
| ENSMUSG00000081220 | 1 | |
| ENSMUSG00000081221 | 1.049411284 | 0.835701362 |
| ENSMUSG00000081222 | -1.090800325 | 0.722001393 |
| ENSMUSG00000081223 | 1.139506315 | 0.402088679 |
| ENSMUSG00000081224 | 1 | |
| ENSMUSG00000081225 | 1 | |
| ENSMUSG00000081226 | 1.059463094 | 0.608497229 |
| ENSMUSG00000081227 | -1.132666855 | 0.375668528 |
| ENSMUSG00000081228 | -1.009476377 | 1 |
| ENSMUSG00000081229 | -1 | 1 |

| | | |
|---|---|---|
| ENSMUSG00000081230 | 1 | |
| ENSMUSG00000081231 | 1 | |
| ENSMUSG00000081232 | 1.150021168 | 0.57950849 |
| ENSMUSG00000081233 | -1 | 1 |
| ENSMUSG00000081234 | 1 | |
| ENSMUSG00000081235 | -1.03962336 | 0.698112512 |
| ENSMUSG00000081236 | 1 | |
| ENSMUSG00000081237 | 1 | |
| ENSMUSG00000081238 | -1.049475223 | 0.861137762 |
| ENSMUSG00000081239 | 1.060378262 | 0.881483647 |
| ENSMUSG00000081240 | 1 | |
| ENSMUSG00000081242 | -1.043306199 | 0.698112512 |
| ENSMUSG00000081243 | 1.014545335 | 0.979002724 |
| ENSMUSG00000081244 | 1.122002463 | 0.534887579 |
| ENSMUSG00000081245 | 1 | |
| ENSMUSG00000081248 | 1 | |
| ENSMUSG00000081249 | 1.043098341 | 0.877747123 |
| ENSMUSG00000081250 | 1.014731082 | 0.988424085 |
| ENSMUSG00000081251 | 1.561869699 | 0.103081231 |
| ENSMUSG00000081252 | 1 | |
| ENSMUSG00000081253 | 1 | |
| ENSMUSG00000081254 | -1 | 1 |
| ENSMUSG00000081255 | 1.068878531 | 0.679917411 |
| ENSMUSG00000081256 | -1.089497311 | 0.614307028 |
| ENSMUSG00000081257 | 1 | |
| ENSMUSG00000081258 | 1 | |
| ENSMUSG00000081259 | 1 | |
| ENSMUSG00000081260 | 1 | |
| ENSMUSG00000081261 | 1 | |
| ENSMUSG00000081262 | 1.014545335 | 0.990831634 |
| ENSMUSG00000081263 | 1 | |
| ENSMUSG00000081264 | 1 | |
| ENSMUSG00000081265 | 1.115945356 | 0.713921439 |
| ENSMUSG00000081268 | 1 | |
| ENSMUSG00000081269 | -1.095294215 | 0.541735653 |
| ENSMUSG00000081270 | 1 | |
| ENSMUSG00000081271 | 1 | |
| ENSMUSG00000081272 | 1.07487334 | 0.770016888 |
| ENSMUSG00000081273 | 1 | |
| ENSMUSG00000081274 | -1.024445679 | 0.968575228 |
| ENSMUSG00000081275 | 1 | |
| ENSMUSG00000081276 | -1.122102401 | 0.698112512 |
| ENSMUSG00000081277 | -1.15595722 | 0.608497229 |
| ENSMUSG00000081278 | 1 | |
| ENSMUSG00000081279 | 1 | |
| ENSMUSG00000081280 | 1.059463094 | 0.608497229 |
| ENSMUSG00000081281 | -1.20574565 | 0.22672364 |
| ENSMUSG00000081282 | 1 | |
| ENSMUSG00000081283 | 1 | |
| ENSMUSG00000081284 | 1 | |
| ENSMUSG00000081285 | -1.073719964 | 0.761710922 |
| ENSMUSG00000081286 | 1 | |
| ENSMUSG00000081287 | 1 | |
| ENSMUSG00000081288 | 1 | |
| ENSMUSG00000081289 | -1.019698456 | 1 |
| ENSMUSG00000081290 | 1 | |
| ENSMUSG00000081292 | 1 | |
| ENSMUSG00000081293 | 1 | |
| ENSMUSG00000081295 | 1.059463094 | 0.698112512 |
| ENSMUSG00000081296 | 1 | |
| ENSMUSG00000081297 | 1 | |
| ENSMUSG00000081298 | 1 | |
| ENSMUSG00000081299 | 1 | |

| | | |
|---|---|---|
| ENSMUSG00000081300 | 1 | |
| ENSMUSG00000081301 | -1.182814301 | 0.316670006 |
| ENSMUSG00000081302 | -1.007466796 | 1 |
| ENSMUSG00000081303 | -1 | 1 |
| ENSMUSG00000081304 | 1 | |
| ENSMUSG00000081305 | 1.037084263 | 0.891092563 |
| ENSMUSG00000081306 | 1 | |
| ENSMUSG00000081307 | 1 | |
| ENSMUSG00000081308 | -1.104412161 | 0.608497229 |
| ENSMUSG00000081309 | 1 | |
| ENSMUSG00000081310 | 1 | |
| ENSMUSG00000081312 | 1 | |
| ENSMUSG00000081313 | 1 | |
| ENSMUSG00000081314 | 1.150021168 | 0.586857505 |
| ENSMUSG00000081315 | 1 | |
| ENSMUSG00000081316 | 1.059463094 | 0.608497229 |
| ENSMUSG00000081317 | 1.171563499 | 0.233412931 |
| ENSMUSG00000081318 | 1 | |
| ENSMUSG00000081319 | 1.055416288 | 0.82835102 |
| ENSMUSG00000081320 | -1 | 1 |
| ENSMUSG00000081321 | -1.181134615 | 0.603144355 |
| ENSMUSG00000081323 | 1 | |
| ENSMUSG00000081324 | 1 | |
| ENSMUSG00000081325 | 1 | |
| ENSMUSG00000081326 | 1 | |
| ENSMUSG00000081327 | -1.043306199 | 0.698112512 |
| ENSMUSG00000081328 | 1 | |
| ENSMUSG00000081329 | -1.163028607 | 0.624064666 |
| ENSMUSG00000081330 | 1 | |
| ENSMUSG00000081331 | -1 | 1 |
| ENSMUSG00000081332 | -1.053549062 | 0.698112512 |
| ENSMUSG00000081333 | 1 | |
| ENSMUSG00000081334 | 1 | |
| ENSMUSG00000081335 | 1 | |
| ENSMUSG00000081336 | 1 | |
| ENSMUSG00000081339 | -1.029113823 | 0.971248775 |
| ENSMUSG00000081340 | -1.128668575 | 0.608497229 |
| ENSMUSG00000081341 | -1.03962336 | 0.803072249 |
| ENSMUSG00000081342 | 1 | |
| ENSMUSG00000081343 | -1.043306199 | 0.803260194 |
| ENSMUSG00000081344 | 1 | |
| ENSMUSG00000081345 | 1 | |
| ENSMUSG00000081346 | 1 | |
| ENSMUSG00000081348 | 1 | |
| ENSMUSG00000081349 | 1 | |
| ENSMUSG00000081350 | 1.177223292 | 0.60397857 |
| ENSMUSG00000081352 | 1 | |
| ENSMUSG00000081353 | -1 | 1 |
| ENSMUSG00000081354 | 1 | |
| ENSMUSG00000081355 | 1.12535639 | 0.532369657 |
| ENSMUSG00000081357 | 1 | |
| ENSMUSG00000081358 | 1.095872691 | 0.608497229 |
| ENSMUSG00000081359 | 1.019083579 | 0.971519533 |
| ENSMUSG00000081360 | -1.145518406 | 0.532369657 |
| ENSMUSG00000081361 | 1 | |
| ENSMUSG00000081362 | 1 | |
| ENSMUSG00000081363 | 1 | |
| ENSMUSG00000081364 | 1 | |
| ENSMUSG00000081365 | 1 | |
| ENSMUSG00000081367 | 1 | |
| ENSMUSG00000081369 | 1 | |
| ENSMUSG00000081370 | 1 | |
| ENSMUSG00000081372 | -1 | 1 |

| | | |
|---|---|---|
| ENSMUSG00000081373 | -1.094213503 | 0.768777807 |
| ENSMUSG00000081374 | 1 | |
| ENSMUSG00000081375 | 1 | |
| ENSMUSG00000081376 | -1.177298609 | 0.343891923 |
| ENSMUSG00000081377 | 1 | |
| ENSMUSG00000081378 | 1.028323353 | 0.903464611 |
| ENSMUSG00000081379 | 1 | |
| ENSMUSG00000081381 | -1.043306199 | 0.858911269 |
| ENSMUSG00000081382 | -1.040045394 | 0.891655647 |
| ENSMUSG00000081383 | -1.043306199 | 0.698112512 |
| ENSMUSG00000081384 | 1 | |
| ENSMUSG00000081386 | 1 | |
| ENSMUSG00000081388 | 1 | |
| ENSMUSG00000081389 | 1 | |
| ENSMUSG00000081390 | -1 | 1 |
| ENSMUSG00000081391 | 1.069068979 | 0.698112512 |
| ENSMUSG00000081392 | 1.059463094 | 0.608497229 |
| ENSMUSG00000081394 | -1.106011338 | 0.710527393 |
| ENSMUSG00000081395 | 1 | |
| ENSMUSG00000081396 | 1 | |
| ENSMUSG00000081397 | 1 | |
| ENSMUSG00000081398 | 1 | |
| ENSMUSG00000081399 | 1 | |
| ENSMUSG00000081400 | -1.059479831 | 0.756216134 |
| ENSMUSG00000081402 | -1.06353289 | 0.842619477 |
| ENSMUSG00000081404 | -1.018784059 | 0.984958258 |
| ENSMUSG00000081405 | 1 | |
| ENSMUSG00000081406 | 1 | |
| ENSMUSG00000081407 | 1 | |
| ENSMUSG00000081408 | 1 | |
| ENSMUSG00000081409 | 1 | |
| ENSMUSG00000081410 | 1 | |
| ENSMUSG00000081412 | -1.043306199 | 0.698112512 |
| ENSMUSG00000081413 | 1 | |
| ENSMUSG00000081414 | 1 | |
| ENSMUSG00000081415 | 1 | |
| ENSMUSG00000081416 | -1.089497311 | 0.698112512 |
| ENSMUSG00000081417 | 1.189207115 | 0.395055569 |
| ENSMUSG00000081418 | -1.028972438 | 0.97128325 |
| ENSMUSG00000081420 | 1 | |
| ENSMUSG00000081421 | 1.095872691 | 0.66521153 |
| ENSMUSG00000081422 | 1 | |
| ENSMUSG00000081423 | 1.005613439 | 1 |
| ENSMUSG00000081424 | 1 | |
| ENSMUSG00000081426 | 1 | |
| ENSMUSG00000081427 | 1 | |
| ENSMUSG00000081428 | -1.023769022 | 0.934760809 |
| ENSMUSG00000081429 | -1.099174267 | 0.608497229 |
| ENSMUSG00000081431 | 1.069068979 | 0.698112512 |
| ENSMUSG00000081432 | 1.132639128 | 0.217803313 |
| ENSMUSG00000081433 | 1 | |
| ENSMUSG00000081434 | 1.14657069 | 0.079761782 |
| ENSMUSG00000081435 | 1 | |
| ENSMUSG00000081436 | 1.069068979 | 0.698112512 |
| ENSMUSG00000081437 | 1 | |
| ENSMUSG00000081438 | 1 | |
| ENSMUSG00000081439 | -1 | 1 |
| ENSMUSG00000081440 | 1 | |
| ENSMUSG00000081441 | -1.044273782 | 0.698112512 |
| ENSMUSG00000081442 | -1.053549062 | 0.698112512 |
| ENSMUSG00000081443 | -1.043306199 | 0.698112512 |
| ENSMUSG00000081444 | -1.03962336 | 0.698112512 |
| ENSMUSG00000081445 | -1.03962336 | 0.698112512 |

| | | |
|---|---|---|
| ENSMUSG00000081447 | 1.010351027 | 1 |
| ENSMUSG00000081449 | 1 | |
| ENSMUSG00000081450 | 1 | |
| ENSMUSG00000081451 | -1.03962336 | 0.698112512 |
| ENSMUSG00000081452 | 1 | |
| ENSMUSG00000081453 | 1.049269646 | 0.801439557 |
| ENSMUSG00000081455 | 1.035778838 | 0.901375104 |
| ENSMUSG00000081456 | -1.013860981 | 1 |
| ENSMUSG00000081457 | 1 | |
| ENSMUSG00000081458 | 1 | |
| ENSMUSG00000081459 | 1.024804142 | 0.985869746 |
| ENSMUSG00000081460 | 1 | |
| ENSMUSG00000081461 | 1.132179521 | 0.698112512 |
| ENSMUSG00000081463 | 1 | |
| ENSMUSG00000081464 | 1 | |
| ENSMUSG00000081465 | 1 | |
| ENSMUSG00000081466 | 1.059463094 | 0.698112512 |
| ENSMUSG00000081467 | 1.203696959 | 0.091305407 |
| ENSMUSG00000081469 | 1 | |
| ENSMUSG00000081470 | 1 | |
| ENSMUSG00000081471 | -1.014065352 | 0.950799842 |
| ENSMUSG00000081473 | -1.147838869 | 0.467497251 |
| ENSMUSG00000081474 | 1 | |
| ENSMUSG00000081475 | 1 | |
| ENSMUSG00000081476 | -1.118081306 | 0.216046507 |
| ENSMUSG00000081477 | 1 | |
| ENSMUSG00000081478 | -1.044273782 | 0.790851299 |
| ENSMUSG00000081480 | 1.069068979 | 0.78287223 |
| ENSMUSG00000081481 | 1 | |
| ENSMUSG00000081483 | -1.258326994 | 0.211190634 |
| ENSMUSG00000081484 | 1 | |
| ENSMUSG00000081485 | 1 | |
| ENSMUSG00000081486 | 1 | |
| ENSMUSG00000081487 | -1.05948903 | 0.834376699 |
| ENSMUSG00000081488 | 1 | |
| ENSMUSG00000081490 | 1.014545335 | 0.991944074 |
| ENSMUSG00000081491 | 1.05410549 | 0.805157604 |
| ENSMUSG00000081492 | 1.057478344 | 0.809553648 |
| ENSMUSG00000081494 | 1.04898545 | 0.874603826 |
| ENSMUSG00000081495 | 1 | |
| ENSMUSG00000081496 | 1 | |
| ENSMUSG00000081497 | 1.059463094 | 0.750182067 |
| ENSMUSG00000081498 | 1 | |
| ENSMUSG00000081499 | 1.023743961 | 0.959839883 |
| ENSMUSG00000081500 | 1 | |
| ENSMUSG00000081501 | 1 | |
| ENSMUSG00000081502 | 1 | |
| ENSMUSG00000081504 | 1 | |
| ENSMUSG00000081505 | 1 | |
| ENSMUSG00000081506 | 1 | |
| ENSMUSG00000081507 | 1 | |
| ENSMUSG00000081508 | 1 | |
| ENSMUSG00000081510 | 1 | |
| ENSMUSG00000081511 | -1.053549062 | 0.862599478 |
| ENSMUSG00000081512 | 1.06585783 | 0.799439325 |
| ENSMUSG00000081513 | 1 | |
| ENSMUSG00000081514 | 1 | |
| ENSMUSG00000081515 | 1 | |
| ENSMUSG00000081516 | 1.084618945 | 0.614071182 |
| ENSMUSG00000081517 | 1.095872691 | 0.657060635 |
| ENSMUSG00000081519 | 1 | |
| ENSMUSG00000081520 | 1.171563499 | 0.233412931 |
| ENSMUSG00000081522 | 1 | |

| | | |
|---|---|---|
| ENSMUSG00000081523 | 1 | |
| ENSMUSG00000081524 | 1.059463094 | 0.698112512 |
| ENSMUSG00000081525 | 1 | |
| ENSMUSG00000081526 | -1 | 1 |
| ENSMUSG00000081528 | 1 | |
| ENSMUSG00000081529 | 1 | |
| ENSMUSG00000081530 | 1 | |
| ENSMUSG00000081531 | 1 | |
| ENSMUSG00000081532 | 1 | |
| ENSMUSG00000081533 | 1 | |
| ENSMUSG00000081534 | 1.035114692 | 0.928666807 |
| ENSMUSG00000081536 | 1 | |
| ENSMUSG00000081537 | -1.028160286 | 0.949812939 |
| ENSMUSG00000081538 | 1 | |
| ENSMUSG00000081539 | 1.059463094 | 0.608497229 |
| ENSMUSG00000081540 | 1 | |
| ENSMUSG00000081541 | 1 | |
| ENSMUSG00000081542 | 1 | |
| ENSMUSG00000081543 | -1.019913324 | 0.995827459 |
| ENSMUSG00000081544 | 1 | |
| ENSMUSG00000081546 | -1 | 1 |
| ENSMUSG00000081547 | 1 | |
| ENSMUSG00000081548 | 1 | |
| ENSMUSG00000081550 | 1.165611676 | 0.329425066 |
| ENSMUSG00000081551 | -1.027105068 | 0.925644855 |
| ENSMUSG00000081552 | 1.117285345 | 0.690669422 |
| ENSMUSG00000081553 | 1.055159774 | 0.86862272 |
| ENSMUSG00000081555 | 1 | |
| ENSMUSG00000081556 | -1.03962336 | 0.814089245 |
| ENSMUSG00000081557 | 1.11961414 | 0.734674233 |
| ENSMUSG00000081559 | 1 | |
| ENSMUSG00000081560 | 1 | |
| ENSMUSG00000081561 | 1 | |
| ENSMUSG00000081562 | -1.044273782 | 0.698112512 |
| ENSMUSG00000081563 | 1 | |
| ENSMUSG00000081564 | 1.192118583 | 0.355014525 |
| ENSMUSG00000081565 | -1.037236631 | 0.951236455 |
| ENSMUSG00000081566 | 1 | |
| ENSMUSG00000081567 | 1 | |
| ENSMUSG00000081568 | 1.004411983 | 1 |
| ENSMUSG00000081569 | 1 | |
| ENSMUSG00000081570 | -1.044273782 | 0.698112512 |
| ENSMUSG00000081571 | -1 | 1 |
| ENSMUSG00000081572 | -1.133231767 | 0.643224515 |
| ENSMUSG00000081573 | 1 | |
| ENSMUSG00000081574 | -1.07939358 | 0.837531941 |
| ENSMUSG00000081575 | -1.038866093 | 0.93376866 |
| ENSMUSG00000081576 | 1 | |
| ENSMUSG00000081577 | 1 | |
| ENSMUSG00000081578 | -1.044273782 | 0.795446726 |
| ENSMUSG00000081579 | 1 | |
| ENSMUSG00000081580 | 1 | |
| ENSMUSG00000081583 | 1 | |
| ENSMUSG00000081584 | 1 | |
| ENSMUSG00000081585 | 1 | |
| ENSMUSG00000081586 | 1 | |
| ENSMUSG00000081587 | -1.12544754 | 0.608497229 |
| ENSMUSG00000081588 | 1 | |
| ENSMUSG00000081589 | 1 | |
| ENSMUSG00000081590 | 1 | |
| ENSMUSG00000081591 | 1 | |
| ENSMUSG00000081592 | 1 | |
| ENSMUSG00000081593 | 1.091519092 | 0.723501636 |

| | | |
|---|---|---|
| ENSMUSG00000081594 | 1.149113744 | 0.317618146 |
| ENSMUSG00000081595 | 1 | |
| ENSMUSG00000081596 | 1.059463094 | 0.608497229 |
| ENSMUSG00000081597 | 1.132639128 | 0.217803313 |
| ENSMUSG00000081598 | 1 | |
| ENSMUSG00000081599 | -1.043306199 | 0.698112512 |
| ENSMUSG00000081600 | 1.208772706 | 0.088486379 |
| ENSMUSG00000081601 | 1 | |
| ENSMUSG00000081602 | 1 | |
| ENSMUSG00000081603 | 1.643760407 | 0.011866184 |
| ENSMUSG00000081604 | 1 | |
| ENSMUSG00000081605 | -1.015058163 | 1 |
| ENSMUSG00000081606 | 1 | |
| ENSMUSG00000081607 | 1.085742167 | 0.750742698 |
| ENSMUSG00000081608 | -1.061098164 | 0.685046738 |
| ENSMUSG00000081609 | 1 | |
| ENSMUSG00000081610 | 1 | |
| ENSMUSG00000081611 | -1.099174267 | 0.608497229 |
| ENSMUSG00000081612 | 1 | |
| ENSMUSG00000081613 | 1.023861851 | 0.991944074 |
| ENSMUSG00000081614 | -1.043306199 | 0.803260194 |
| ENSMUSG00000081615 | -1 | 1 |
| ENSMUSG00000081616 | 1 | |
| ENSMUSG00000081617 | 1.069068979 | 0.608497229 |
| ENSMUSG00000081618 | 1 | |
| ENSMUSG00000081619 | 1.052246147 | 0.894591069 |
| ENSMUSG00000081620 | -1.03962336 | 0.698112512 |
| ENSMUSG00000081621 | 1 | |
| ENSMUSG00000081622 | 1.015486245 | 0.972980766 |
| ENSMUSG00000081623 | -1.043306199 | 0.698112512 |
| ENSMUSG00000081624 | 1 | |
| ENSMUSG00000081625 | -1.03962336 | 0.698112512 |
| ENSMUSG00000081626 | 1 | |
| ENSMUSG00000081627 | 1 | |
| ENSMUSG00000081628 | 1 | |
| ENSMUSG00000081629 | 1.059463094 | 0.608497229 |
| ENSMUSG00000081630 | 1 | |
| ENSMUSG00000081631 | -1 | 1 |
| ENSMUSG00000081632 | 1 | |
| ENSMUSG00000081633 | -1.033820075 | 0.916014403 |
| ENSMUSG00000081635 | -1 | 1 |
| ENSMUSG00000081636 | 1 | |
| ENSMUSG00000081638 | 1 | |
| ENSMUSG00000081639 | 1 | |
| ENSMUSG00000081640 | 1.146260679 | 0.086164458 |
| ENSMUSG00000081641 | -1.03962336 | 0.698112512 |
| ENSMUSG00000081642 | -1.06725841 | 0.820659432 |
| ENSMUSG00000081643 | -1.03962336 | 0.698112512 |
| ENSMUSG00000081646 | 1 | |
| ENSMUSG00000081647 | 1 | |
| ENSMUSG00000081648 | 1.014882437 | 0.983446483 |
| ENSMUSG00000081649 | 1 | |
| ENSMUSG00000081650 | -1.112287407 | 0.781447518 |
| ENSMUSG00000081651 | -1.03962336 | 0.803072249 |
| ENSMUSG00000081652 | 1 | |
| ENSMUSG00000081653 | 1.014545335 | 1 |
| ENSMUSG00000081654 | 1 | |
| ENSMUSG00000081656 | -1 | 1 |
| ENSMUSG00000081657 | -1.100925449 | 0.758764661 |
| ENSMUSG00000081658 | 1 | |
| ENSMUSG00000081659 | -1.015511103 | 0.982777255 |
| ENSMUSG00000081660 | 1 | |
| ENSMUSG00000081661 | -1.044273782 | 0.698112512 |

| | | |
|---|---|---|
| ENSMUSG00000081662 | 1.122128755 | 0.656978335 |
| ENSMUSG00000081663 | -1.043306199 | 0.698112512 |
| ENSMUSG00000081664 | -1.03962336 | 0.698112512 |
| ENSMUSG00000081665 | 1 | |
| ENSMUSG00000081666 | 1 | |
| ENSMUSG00000081667 | 1 | |
| ENSMUSG00000081668 | 1 | |
| ENSMUSG00000081669 | 1.051378421 | 0.891297795 |
| ENSMUSG00000081670 | -1.084889449 | 0.781240509 |
| ENSMUSG00000081671 | 1 | |
| ENSMUSG00000081672 | 1 | |
| ENSMUSG00000081673 | 1 | |
| ENSMUSG00000081674 | 1.015486245 | 0.972980766 |
| ENSMUSG00000081675 | 1.035626983 | 0.914705152 |
| ENSMUSG00000081676 | 1.095872691 | 0.698112512 |
| ENSMUSG00000081677 | 1 | |
| ENSMUSG00000081678 | 1 | |
| ENSMUSG00000081679 | -1.114894961 | 0.698112512 |
| ENSMUSG00000081680 | 1.084618945 | 0.698112512 |
| ENSMUSG00000081681 | 1 | |
| ENSMUSG00000081683 | -1.271754232 | 0.004890823 |
| ENSMUSG00000081684 | 1.059463094 | 0.704170349 |
| ENSMUSG00000081685 | 1 | |
| ENSMUSG00000081686 | 1 | |
| ENSMUSG00000081687 | 1 | |
| ENSMUSG00000081688 | 1 | |
| ENSMUSG00000081689 | -1 | 1 |
| ENSMUSG00000081690 | -1.053549062 | 0.698112512 |
| ENSMUSG00000081691 | -1.044273782 | 0.698112512 |
| ENSMUSG00000081692 | -1.053549062 | 0.698112512 |
| ENSMUSG00000081693 | 1 | |
| ENSMUSG00000081694 | 1 | |
| ENSMUSG00000081695 | 1.013608807 | 0.99362872 |
| ENSMUSG00000081696 | 1.122462048 | 0.215991492 |
| ENSMUSG00000081697 | 1 | |
| ENSMUSG00000081698 | 1 | |
| ENSMUSG00000081699 | 1 | |
| ENSMUSG00000081700 | -1 | 1 |
| ENSMUSG00000081701 | 1 | |
| ENSMUSG00000081703 | 1.028323353 | 0.903464611 |
| ENSMUSG00000081704 | -1 | 1 |
| ENSMUSG00000081705 | -1.053549062 | 0.698112512 |
| ENSMUSG00000081706 | 1.057478344 | 0.837014159 |
| ENSMUSG00000081707 | -1.07008665 | 0.79291651 |
| ENSMUSG00000081709 | 1 | |
| ENSMUSG00000081711 | 1 | |
| ENSMUSG00000081712 | -1.001348012 | 1 |
| ENSMUSG00000081713 | 1 | |
| ENSMUSG00000081714 | 1 | |
| ENSMUSG00000081715 | 1.059463094 | 0.608497229 |
| ENSMUSG00000081716 | 1 | |
| ENSMUSG00000081717 | 1 | |
| ENSMUSG00000081718 | 1.07487334 | 0.814138447 |
| ENSMUSG00000081719 | 1.069068979 | 0.608497229 |
| ENSMUSG00000081720 | -1.084645496 | 0.532369657 |
| ENSMUSG00000081721 | 1.101095402 | 0.730297627 |
| ENSMUSG00000081722 | -1 | 1 |
| ENSMUSG00000081723 | 1 | |
| ENSMUSG00000081724 | 1 | |
| ENSMUSG00000081728 | 1 | |
| ENSMUSG00000081729 | 1.078697592 | 0.807743274 |
| ENSMUSG00000081730 | 1.081017342 | 0.751653251 |
| ENSMUSG00000081731 | -1.084645496 | 0.698112512 |

| | | |
|---|---|---|
| ENSMUSG00000081732 | -1.08384782 | 0.686411811 |
| ENSMUSG00000081733 | -1.010426982 | 1 |
| ENSMUSG00000081734 | 1 | |
| ENSMUSG00000081735 | -1.044273782 | 0.801443616 |
| ENSMUSG00000081736 | 1 | |
| ENSMUSG00000081737 | 1 | |
| ENSMUSG00000081738 | 1.055859595 | 0.882514615 |
| ENSMUSG00000081739 | -1.011847768 | 0.999944982 |
| ENSMUSG00000081740 | 1.059463094 | 0.698112512 |
| ENSMUSG00000081741 | 1 | |
| ENSMUSG00000081742 | 1 | |
| ENSMUSG00000081743 | 1 | |
| ENSMUSG00000081744 | 1 | |
| ENSMUSG00000081745 | 1.013796828 | 1 |
| ENSMUSG00000081746 | 1 | |
| ENSMUSG00000081747 | 1.059463094 | 0.698112512 |
| ENSMUSG00000081748 | -1.085651418 | 0.532369657 |
| ENSMUSG00000081749 | 1 | |
| ENSMUSG00000081750 | 1 | |
| ENSMUSG00000081751 | -1 | 1 |
| ENSMUSG00000081752 | 1 | |
| ENSMUSG00000081755 | 1 | |
| ENSMUSG00000081756 | 1 | |
| ENSMUSG00000081757 | 1 | |
| ENSMUSG00000081758 | 1 | |
| ENSMUSG00000081761 | 1 | |
| ENSMUSG00000081762 | 1 | |
| ENSMUSG00000081763 | -1.024085706 | 0.986359381 |
| ENSMUSG00000081765 | 1 | |
| ENSMUSG00000081766 | 1 | |
| ENSMUSG00000081767 | 1 | |
| ENSMUSG00000081768 | -1 | 1 |
| ENSMUSG00000081769 | -1.25536244 | 0.12831008 |
| ENSMUSG00000081770 | 1 | |
| ENSMUSG00000081771 | 1.019083579 | 0.949812939 |
| ENSMUSG00000081772 | 1 | |
| ENSMUSG00000081773 | 1.06311014 | 0.815347141 |
| ENSMUSG00000081774 | 1 | |
| ENSMUSG00000081775 | -1.044273782 | 0.698112512 |
| ENSMUSG00000081777 | 1 | |
| ENSMUSG00000081778 | 1.059463094 | 0.608497229 |
| ENSMUSG00000081779 | 1 | |
| ENSMUSG00000081781 | -1.03962336 | 0.698112512 |
| ENSMUSG00000081782 | -1.053549062 | 0.769745556 |
| ENSMUSG00000081783 | 1.100990117 | 0.758995782 |
| ENSMUSG00000081784 | 1 | |
| ENSMUSG00000081786 | 1 | |
| ENSMUSG00000081787 | -1.069502966 | 0.698112512 |
| ENSMUSG00000081788 | -1.004508963 | 1 |
| ENSMUSG00000081789 | 1 | |
| ENSMUSG00000081791 | -1 | 1 |
| ENSMUSG00000081792 | 1.096901918 | 0.730237133 |
| ENSMUSG00000081793 | 1 | |
| ENSMUSG00000081796 | 1.059463094 | 0.608497229 |
| ENSMUSG00000081797 | 1.005212439 | 1 |
| ENSMUSG00000081798 | -1.013449573 | 1 |
| ENSMUSG00000081799 | 1 | |
| ENSMUSG00000081800 | -1 | 1 |
| ENSMUSG00000081801 | -1.053549062 | 0.774498429 |
| ENSMUSG00000081803 | 1.005613439 | 1 |
| ENSMUSG00000081804 | 1 | |
| ENSMUSG00000081805 | -1.161107412 | 0.37067934 |
| ENSMUSG00000081806 | 1 | |

| | | |
|---|---|---|
| ENSMUSG00000081807 | -1.017626803 | 1 |
| ENSMUSG00000081809 | -1.078296166 | 0.775966113 |
| ENSMUSG00000081810 | -1.11061171 | 0.608497229 |
| ENSMUSG00000081811 | 1.069068979 | 0.608497229 |
| ENSMUSG00000081812 | 1 | |
| ENSMUSG00000081813 | 1 | |
| ENSMUSG00000081814 | 1 | |
| ENSMUSG00000081816 | -1 | 1 |
| ENSMUSG00000081817 | 1.184657384 | 0.200640546 |
| ENSMUSG00000081819 | -1.089497311 | 0.532369657 |
| ENSMUSG00000081820 | -1.03962336 | 0.698112512 |
| ENSMUSG00000081821 | 1 | |
| ENSMUSG00000081822 | 1 | |
| ENSMUSG00000081823 | 1 | |
| ENSMUSG00000081824 | 1 | |
| ENSMUSG00000081827 | 1 | |
| ENSMUSG00000081828 | 1 | |
| ENSMUSG00000081829 | -1 | 1 |
| ENSMUSG00000081830 | 1 | |
| ENSMUSG00000081831 | -1.044273782 | 0.698112512 |
| ENSMUSG00000081832 | 1 | |
| ENSMUSG00000081833 | 1 | |
| ENSMUSG00000081834 | 1.069068979 | 0.608497229 |
| ENSMUSG00000081835 | 1 | |
| ENSMUSG00000081836 | 1 | |
| ENSMUSG00000081837 | 1 | |
| ENSMUSG00000081838 | 1.059463094 | 0.608497229 |
| ENSMUSG00000081839 | 1 | |
| ENSMUSG00000081840 | 1.069068979 | 0.608497229 |
| ENSMUSG00000081841 | 1 | |
| ENSMUSG00000081842 | 1.059463094 | 0.704170349 |
| ENSMUSG00000081843 | 1 | |
| ENSMUSG00000081844 | 1 | |
| ENSMUSG00000081845 | -1.03962336 | 0.698112512 |
| ENSMUSG00000081847 | 1 | |
| ENSMUSG00000081848 | 1 | |
| ENSMUSG00000081849 | 1 | |
| ENSMUSG00000081850 | 1.019083579 | 0.949812939 |
| ENSMUSG00000081851 | -1.066370674 | 0.842404015 |
| ENSMUSG00000081852 | 1 | |
| ENSMUSG00000081853 | 1 | |
| ENSMUSG00000081854 | 1 | |
| ENSMUSG00000081855 | -1.244253939 | 0.266022289 |
| ENSMUSG00000081856 | 1 | |
| ENSMUSG00000081857 | 1 | |
| ENSMUSG00000081858 | 1 | |
| ENSMUSG00000081859 | 1 | |
| ENSMUSG00000081860 | 1.059463094 | 0.608497229 |
| ENSMUSG00000081861 | 1.021173386 | 0.997389629 |
| ENSMUSG00000081862 | 1 | |
| ENSMUSG00000081863 | 1 | |
| ENSMUSG00000081864 | 1 | |
| ENSMUSG00000081865 | -1.016724842 | 0.973938676 |
| ENSMUSG00000081866 | 1 | |
| ENSMUSG00000081868 | -1 | 1 |
| ENSMUSG00000081869 | 1 | |
| ENSMUSG00000081870 | 1 | |
| ENSMUSG00000081871 | 1.07487334 | 0.69923471 |
| ENSMUSG00000081872 | 1 | |
| ENSMUSG00000081873 | 1.015486245 | 0.972980766 |
| ENSMUSG00000081874 | 1 | |
| ENSMUSG00000081875 | 1.069068979 | 0.608497229 |
| ENSMUSG00000081876 | 1 | |

| | | |
|---|---|---|
| ENSMUSG00000081877 | -1.03962336 | 0.698112512 |
| ENSMUSG00000081878 | 1 | |
| ENSMUSG00000081879 | 1 | |
| ENSMUSG00000081880 | 1 | |
| ENSMUSG00000081881 | 1 | |
| ENSMUSG00000081882 | 1.059463094 | 0.608497229 |
| ENSMUSG00000081883 | -1.138693452 | 0.608497229 |
| ENSMUSG00000081884 | 1 | |
| ENSMUSG00000081885 | -1.207791469 | 0.59984182 |
| ENSMUSG00000081886 | 1 | |
| ENSMUSG00000081887 | 1.070880472 | 0.762628275 |
| ENSMUSG00000081888 | 1.114201787 | 0.630379139 |
| ENSMUSG00000081889 | 1 | |
| ENSMUSG00000081890 | 1 | |
| ENSMUSG00000081891 | 1 | |
| ENSMUSG00000081892 | 1.002009643 | 1 |
| ENSMUSG00000081893 | 1 | |
| ENSMUSG00000081894 | 1 | |
| ENSMUSG00000081895 | -1.025148865 | 0.756618803 |
| ENSMUSG00000081896 | -1.014726102 | 1 |
| ENSMUSG00000081897 | 1 | |
| ENSMUSG00000081898 | -1.048085977 | 0.853042921 |
| ENSMUSG00000081900 | 1 | |
| ENSMUSG00000081901 | 1 | |
| ENSMUSG00000081903 | 1 | |
| ENSMUSG00000081904 | -1.044273782 | 0.698112512 |
| ENSMUSG00000081905 | 1 | |
| ENSMUSG00000081906 | -1 | 1 |
| ENSMUSG00000081907 | 1 | |
| ENSMUSG00000081908 | 1 | |
| ENSMUSG00000081909 | 1 | |
| ENSMUSG00000081910 | 1 | |
| ENSMUSG00000081911 | -1.043306199 | 0.698112512 |
| ENSMUSG00000081912 | 1 | |
| ENSMUSG00000081913 | 1 | |
| ENSMUSG00000081914 | 1 | |
| ENSMUSG00000081916 | 1 | |
| ENSMUSG00000081917 | 1 | |
| ENSMUSG00000081918 | 1 | |
| ENSMUSG00000081919 | 1 | |
| ENSMUSG00000081920 | 1 | |
| ENSMUSG00000081921 | 1 | |
| ENSMUSG00000081923 | 1 | |
| ENSMUSG00000081924 | -1.085651418 | 0.608497229 |
| ENSMUSG00000081925 | 1 | |
| ENSMUSG00000081926 | 1.05410549 | 0.835701362 |
| ENSMUSG00000081928 | 1 | |
| ENSMUSG00000081929 | 1.02288432 | 0.982838256 |
| ENSMUSG00000081931 | 1 | |
| ENSMUSG00000081932 | 1.011607416 | 1 |
| ENSMUSG00000081933 | 1.059463094 | 0.608497229 |
| ENSMUSG00000081934 | 1 | |
| ENSMUSG00000081935 | 1 | |
| ENSMUSG00000081936 | 1 | |
| ENSMUSG00000081937 | 1 | |
| ENSMUSG00000081938 | 1 | |
| ENSMUSG00000081939 | 1 | |
| ENSMUSG00000081940 | 1 | |
| ENSMUSG00000081941 | -1.102059063 | 0.722165151 |
| ENSMUSG00000081942 | 1 | |
| ENSMUSG00000081943 | 1.122462048 | 0.215991492 |
| ENSMUSG00000081944 | -1.086954847 | 0.683168971 |
| ENSMUSG00000081945 | 1 | |

| | | |
|---|---|---|
| ENSMUSG00000081946 | -1.092586173 | 0.608497229 |
| ENSMUSG00000081947 | 1 | |
| ENSMUSG00000081948 | 1 | |
| ENSMUSG00000081949 | 1 | |
| ENSMUSG00000081950 | 1.069068979 | 0.608497229 |
| ENSMUSG00000081951 | 1 | |
| ENSMUSG00000081952 | 1 | |
| ENSMUSG00000081953 | 1.069437807 | 0.771926153 |
| ENSMUSG00000081954 | -1.044273782 | 0.698112512 |
| ENSMUSG00000081955 | 1 | |
| ENSMUSG00000081957 | 1 | |
| ENSMUSG00000081958 | 1 | |
| ENSMUSG00000081959 | 1 | |
| ENSMUSG00000081960 | 1 | |
| ENSMUSG00000081961 | 1 | |
| ENSMUSG00000081962 | 1 | |
| ENSMUSG00000081964 | 1 | |
| ENSMUSG00000081965 | 1.024200778 | 0.979002724 |
| ENSMUSG00000081966 | 1 | |
| ENSMUSG00000081967 | 1.059463094 | 0.608497229 |
| ENSMUSG00000081968 | -1.029113823 | 0.929923764 |
| ENSMUSG00000081969 | 1 | |
| ENSMUSG00000081970 | 1 | |
| ENSMUSG00000081971 | 1.038393925 | 0.851446198 |
| ENSMUSG00000081972 | 1 | |
| ENSMUSG00000081973 | 1.15516542 | 0.466704492 |
| ENSMUSG00000081974 | 1.042633551 | 0.698112512 |
| ENSMUSG00000081975 | -1.014920232 | 1 |
| ENSMUSG00000081976 | 1 | |
| ENSMUSG00000081977 | 1 | |
| ENSMUSG00000081978 | 1 | |
| ENSMUSG00000081979 | -1.033820075 | 0.918344559 |
| ENSMUSG00000081980 | 1 | |
| ENSMUSG00000081982 | 1 | |
| ENSMUSG00000081984 | -1.279043084 | 0.019311157 |
| ENSMUSG00000081985 | -1 | 1 |
| ENSMUSG00000081986 | 1 | |
| ENSMUSG00000081987 | 1 | |
| ENSMUSG00000081988 | 1 | |
| ENSMUSG00000081991 | 1.095872691 | 0.608497229 |
| ENSMUSG00000081992 | -1.00093149 | 1 |
| ENSMUSG00000081993 | 1 | |
| ENSMUSG00000081994 | -1.044273782 | 0.698112512 |
| ENSMUSG00000081995 | 1 | |
| ENSMUSG00000081996 | 1.085237715 | 0.71893768 |
| ENSMUSG00000081998 | 1.118127424 | 0.449334247 |
| ENSMUSG00000081999 | -1.134282749 | 0.608497229 |
| ENSMUSG00000082000 | 1 | |
| ENSMUSG00000082001 | 1 | |
| ENSMUSG00000082002 | -1.043306199 | 0.79291651 |
| ENSMUSG00000082005 | 1 | |
| ENSMUSG00000082006 | 1.069068979 | 0.608497229 |
| ENSMUSG00000082007 | 1 | |
| ENSMUSG00000082008 | -1.100193663 | 0.649330329 |
| ENSMUSG00000082009 | 1 | |
| ENSMUSG00000082010 | 1 | |
| ENSMUSG00000082011 | -1 | 1 |
| ENSMUSG00000082012 | 1.049137052 | 0.698112512 |
| ENSMUSG00000082013 | 1.099112694 | 0.698112512 |
| ENSMUSG00000082014 | 1 | |
| ENSMUSG00000082015 | 1 | |
| ENSMUSG00000082016 | 1 | |
| ENSMUSG00000082017 | 1 | |

| | | |
|---|---|---|
| ENSMUSG00000082018 | 1.089470641 | 0.657060635 |
| ENSMUSG00000082019 | 1 | |
| ENSMUSG00000082020 | 1 | |
| ENSMUSG00000082021 | 1 | |
| ENSMUSG00000082022 | 1 | |
| ENSMUSG00000082023 | 1 | |
| ENSMUSG00000082024 | 1 | |
| ENSMUSG00000082025 | 1 | |
| ENSMUSG00000082026 | -1.151172433 | 0.608497229 |
| ENSMUSG00000082027 | 1 | |
| ENSMUSG00000082028 | -1 | 1 |
| ENSMUSG00000082029 | -1 | 1 |
| ENSMUSG00000082030 | 1 | |
| ENSMUSG00000082031 | -1.033820075 | 0.916014403 |
| ENSMUSG00000082032 | -1.003834322 | 1 |
| ENSMUSG00000082033 | 1 | |
| ENSMUSG00000082035 | 1 | |
| ENSMUSG00000082036 | 1 | |
| ENSMUSG00000082037 | 1 | |
| ENSMUSG00000082038 | 1 | |
| ENSMUSG00000082039 | 1.069068979 | 0.608497229 |
| ENSMUSG00000082040 | -1 | 1 |
| ENSMUSG00000082042 | 1 | |
| ENSMUSG00000082043 | -1.085651418 | 0.532369657 |
| ENSMUSG00000082044 | -1.028348525 | 0.954497929 |
| ENSMUSG00000082045 | 1 | |
| ENSMUSG00000082046 | -1 | 1 |
| ENSMUSG00000082047 | 1 | |
| ENSMUSG00000082048 | 1 | |
| ENSMUSG00000082049 | -1.052377551 | 0.89052268 |
| ENSMUSG00000082051 | -1.095294215 | 0.608497229 |
| ENSMUSG00000082052 | -1.028728523 | 0.981169503 |
| ENSMUSG00000082053 | 1 | |
| ENSMUSG00000082054 | 1 | |
| ENSMUSG00000082056 | 1 | |
| ENSMUSG00000082057 | -1.014371255 | 1 |
| ENSMUSG00000082058 | 1 | |
| ENSMUSG00000082059 | 1 | |
| ENSMUSG00000082060 | 1 | |
| ENSMUSG00000082061 | -1.251408229 | 0.001392472 |
| ENSMUSG00000082062 | 1.069068979 | 0.608497229 |
| ENSMUSG00000082063 | -1.111880267 | 0.608497229 |
| ENSMUSG00000082064 | 1.32607571 | 0.205913656 |
| ENSMUSG00000082065 | 1 | |
| ENSMUSG00000082066 | -1.055526437 | 0.854356653 |
| ENSMUSG00000082068 | 1 | |
| ENSMUSG00000082069 | 1 | |
| ENSMUSG00000082070 | 1.093819733 | 0.699168433 |
| ENSMUSG00000082071 | 1 | |
| ENSMUSG00000082072 | -1.005107211 | 1 |
| ENSMUSG00000082073 | 1 | |
| ENSMUSG00000082074 | 1 | |
| ENSMUSG00000082076 | -1.054741498 | 0.813450442 |
| ENSMUSG00000082077 | 1 | |
| ENSMUSG00000082078 | -1.029113823 | 0.929923764 |
| ENSMUSG00000082079 | 1 | |
| ENSMUSG00000082080 | 1 | |
| ENSMUSG00000082081 | 1 | |
| ENSMUSG00000082082 | -1.010303575 | 1 |
| ENSMUSG00000082083 | 1 | |
| ENSMUSG00000082084 | 1 | |
| ENSMUSG00000082085 | 1.069068979 | 0.608497229 |
| ENSMUSG00000082087 | -1 | 1 |

| | | |
|---|---|---|
| ENSMUSG00000082088 | -1.027395698 | 0.965343374 |
| ENSMUSG00000082089 | 1 | |
| ENSMUSG00000082090 | 1.149113744 | 0.158414349 |
| ENSMUSG00000082091 | -1.03962336 | 0.698112512 |
| ENSMUSG00000082092 | 1 | |
| ENSMUSG00000082093 | 1 | |
| ENSMUSG00000082095 | -1.132757337 | 0.608497229 |
| ENSMUSG00000082096 | 1 | |
| ENSMUSG00000082097 | 1 | |
| ENSMUSG00000082098 | 1 | |
| ENSMUSG00000082099 | 1 | |
| ENSMUSG00000082100 | 1.024693402 | 0.952057016 |
| ENSMUSG00000082101 | -1.044273782 | 0.83619593 |
| ENSMUSG00000082103 | 1 | |
| ENSMUSG00000082104 | 1 | |
| ENSMUSG00000082106 | 1 | |
| ENSMUSG00000082107 | 1.050889673 | 0.902933477 |
| ENSMUSG00000082108 | 1.122462048 | 0.420161906 |
| ENSMUSG00000082109 | 1.105162268 | 0.770888795 |
| ENSMUSG00000082110 | 1.059463094 | 0.608497229 |
| ENSMUSG00000082111 | -1 | 1 |
| ENSMUSG00000082112 | 1 | |
| ENSMUSG00000082113 | 1 | |
| ENSMUSG00000082114 | 1 | |
| ENSMUSG00000082115 | 1 | |
| ENSMUSG00000082116 | 1 | |
| ENSMUSG00000082117 | -1.053296659 | 0.751729496 |
| ENSMUSG00000082118 | -1.130720898 | 0.642593813 |
| ENSMUSG00000082119 | 1 | |
| ENSMUSG00000082120 | 1.136540403 | 0.424948752 |
| ENSMUSG00000082121 | 1.289282518 | 0.025757952 |
| ENSMUSG00000082122 | 1 | |
| ENSMUSG00000082123 | 1 | |
| ENSMUSG00000082125 | 1 | |
| ENSMUSG00000082126 | 1 | |
| ENSMUSG00000082127 | 1 | |
| ENSMUSG00000082128 | -1.109142811 | 0.624722543 |
| ENSMUSG00000082129 | 1 | |
| ENSMUSG00000082130 | 1.069068979 | 0.74773997 |
| ENSMUSG00000082132 | 1 | |
| ENSMUSG00000082134 | 1 | |
| ENSMUSG00000082135 | 1 | |
| ENSMUSG00000082136 | 1.199989356 | 0.354316858 |
| ENSMUSG00000082138 | 1 | |
| ENSMUSG00000082139 | 1 | |
| ENSMUSG00000082140 | -1.034443843 | 0.924766459 |
| ENSMUSG00000082141 | 1 | |
| ENSMUSG00000082142 | 1 | |
| ENSMUSG00000082143 | 1 | |
| ENSMUSG00000082144 | -1.028278093 | 0.975677002 |
| ENSMUSG00000082145 | -1.010966529 | 1 |
| ENSMUSG00000082146 | 1 | |
| ENSMUSG00000082147 | 1 | |
| ENSMUSG00000082148 | -1.03962336 | 0.808074244 |
| ENSMUSG00000082149 | 1.053966865 | 0.898893325 |
| ENSMUSG00000082151 | -1.095294215 | 0.608497229 |
| ENSMUSG00000082152 | 1 | |
| ENSMUSG00000082154 | 1 | |
| ENSMUSG00000082155 | 1 | |
| ENSMUSG00000082156 | 1 | |
| ENSMUSG00000082157 | 1 | |
| ENSMUSG00000082158 | 1 | |
| ENSMUSG00000082159 | 1 | |

| | | |
|---|---|---|
| ENSMUSG00000082160 | 1 | |
| ENSMUSG00000082161 | -1.03962336 | 0.698112512 |
| ENSMUSG00000082162 | 1 | |
| ENSMUSG00000082163 | -1.047456175 | 0.756618803 |
| ENSMUSG00000082164 | 1 | |
| ENSMUSG00000082165 | -1.188446785 | 0.219185116 |
| ENSMUSG00000082167 | 1 | |
| ENSMUSG00000082168 | 1 | |
| ENSMUSG00000082169 | 1 | |
| ENSMUSG00000082170 | 1 | |
| ENSMUSG00000082171 | 1.014545335 | 1 |
| ENSMUSG00000082172 | 1 | |
| ENSMUSG00000082173 | 1 | |
| ENSMUSG00000082174 | 1 | |
| ENSMUSG00000082175 | 1 | |
| ENSMUSG00000082176 | -1 | 1 |
| ENSMUSG00000082177 | 1.059463094 | 0.608497229 |
| ENSMUSG00000082178 | 1.128312813 | 0.668750825 |
| ENSMUSG00000082179 | 1.045906122 | 0.938555263 |
| ENSMUSG00000082180 | -1.02794432 | 0.98775399 |
| ENSMUSG00000082181 | 1 | |
| ENSMUSG00000082183 | -1.037482356 | 0.880262717 |
| ENSMUSG00000082185 | 1.069068979 | 0.608497229 |
| ENSMUSG00000082186 | -1.052304694 | 0.845175363 |
| ENSMUSG00000082187 | -1.03962336 | 0.698112512 |
| ENSMUSG00000082188 | 1 | |
| ENSMUSG00000082189 | 1.229666014 | 0.425561574 |
| ENSMUSG00000082190 | -1.262390404 | 0.404695089 |
| ENSMUSG00000082191 | 1 | |
| ENSMUSG00000082192 | 1 | |
| ENSMUSG00000082193 | -1.21289656 | 0.453662814 |
| ENSMUSG00000082194 | 1.065936141 | 0.653960635 |
| ENSMUSG00000082195 | 1 | |
| ENSMUSG00000082196 | -1.325165916 | 0.009663139 |
| ENSMUSG00000082197 | -1 | 1 |
| ENSMUSG00000082198 | -1.13161737 | 0.608497229 |
| ENSMUSG00000082199 | 1 | |
| ENSMUSG00000082200 | 1 | |
| ENSMUSG00000082202 | -1.03962336 | 0.698112512 |
| ENSMUSG00000082203 | 1 | |
| ENSMUSG00000082204 | -1.0965339 | 0.698112512 |
| ENSMUSG00000082205 | 1 | |
| ENSMUSG00000082207 | 1 | |
| ENSMUSG00000082208 | 1.012715511 | 1 |
| ENSMUSG00000082210 | 1 | |
| ENSMUSG00000082211 | 1 | |
| ENSMUSG00000082212 | 1 | |
| ENSMUSG00000082213 | -1.03962336 | 0.698112512 |
| ENSMUSG00000082214 | 1 | |
| ENSMUSG00000082215 | 1 | |
| ENSMUSG00000082216 | 1 | |
| ENSMUSG00000082217 | -1 | 1 |
| ENSMUSG00000082218 | 1.069437807 | 0.7938895 |
| ENSMUSG00000082220 | 1.065132083 | 0.870598295 |
| ENSMUSG00000082222 | 1 | |
| ENSMUSG00000082223 | 1 | |
| ENSMUSG00000082225 | -1 | 1 |
| ENSMUSG00000082226 | -1.03962336 | 0.698112512 |
| ENSMUSG00000082227 | 1 | |
| ENSMUSG00000082228 | 1 | |
| ENSMUSG00000082229 | -1.325454583 | 0.050544467 |
| ENSMUSG00000082233 | 1.079588507 | 0.826437915 |
| ENSMUSG00000082234 | -1 | 1 |

| | | |
|---|---|---|
| ENSMUSG00000082235 | 1.069068979 | 0.608497229 |
| ENSMUSG00000082236 | 1 | |
| ENSMUSG00000082237 | 1 | |
| ENSMUSG00000082238 | 1 | |
| ENSMUSG00000082239 | 1 | |
| ENSMUSG00000082240 | -1 | 1 |
| ENSMUSG00000082241 | 1 | |
| ENSMUSG00000082242 | 1 | |
| ENSMUSG00000082243 | -1.044273782 | 0.698112512 |
| ENSMUSG00000082244 | 1.067213122 | 0.897053041 |
| ENSMUSG00000082245 | -1 | 1 |
| ENSMUSG00000082246 | -1.094783341 | 0.750182067 |
| ENSMUSG00000082247 | -1.03962336 | 0.803072249 |
| ENSMUSG00000082248 | 1 | |
| ENSMUSG00000082249 | 1 | |
| ENSMUSG00000082250 | 1.000528147 | 1 |
| ENSMUSG00000082251 | 1 | |
| ENSMUSG00000082252 | 1 | |
| ENSMUSG00000082253 | 1.085624843 | 0.698112512 |
| ENSMUSG00000082254 | -1.03962336 | 0.872692845 |
| ENSMUSG00000082255 | 1 | |
| ENSMUSG00000082256 | 1 | |
| ENSMUSG00000082258 | -1.044273782 | 0.698112512 |
| ENSMUSG00000082259 | 1 | |
| ENSMUSG00000082260 | -1.084645496 | 0.532369657 |
| ENSMUSG00000082261 | 1 | |
| ENSMUSG00000082262 | -1.207526842 | 0.352985637 |
| ENSMUSG00000082263 | 1 | |
| ENSMUSG00000082264 | 1.005613439 | 1 |
| ENSMUSG00000082265 | -1.163987816 | 0.399317736 |
| ENSMUSG00000082266 | 1 | |
| ENSMUSG00000082267 | 1 | |
| ENSMUSG00000082268 | 1 | |
| ENSMUSG00000082269 | 1 | |
| ENSMUSG00000082270 | 1 | |
| ENSMUSG00000082271 | -1 | 1 |
| ENSMUSG00000082272 | -1.041132001 | 0.910895334 |
| ENSMUSG00000082274 | 1.059463094 | 0.699340382 |
| ENSMUSG00000082275 | 1 | |
| ENSMUSG00000082276 | -1.043306199 | 0.698112512 |
| ENSMUSG00000082277 | 1.123881404 | 0.630993985 |
| ENSMUSG00000082278 | 1 | |
| ENSMUSG00000082279 | 1.002009643 | 1 |
| ENSMUSG00000082280 | 1.059463094 | 0.608497229 |
| ENSMUSG00000082281 | 1 | |
| ENSMUSG00000082283 | -1.03962336 | 0.698112512 |
| ENSMUSG00000082284 | 1.059463094 | 0.608497229 |
| ENSMUSG00000082285 | -1.03962336 | 0.698112512 |
| ENSMUSG00000082286 | 1.07384304 | 0.608497229 |
| ENSMUSG00000082287 | 1 | |
| ENSMUSG00000082288 | 1 | |
| ENSMUSG00000082289 | -1.023769022 | 0.969544524 |
| ENSMUSG00000082291 | 1 | |
| ENSMUSG00000082292 | -1.187617818 | 0.405064266 |
| ENSMUSG00000082294 | 1 | |
| ENSMUSG00000082295 | 1 | |
| ENSMUSG00000082296 | 1.113135847 | 0.735035933 |
| ENSMUSG00000082297 | 1 | |
| ENSMUSG00000082301 | 1 | |
| ENSMUSG00000082302 | 1 | |
| ENSMUSG00000082303 | 1 | |
| ENSMUSG00000082304 | -1.000405949 | 1 |
| ENSMUSG00000082305 | 1.058326037 | 0.718616429 |

| | | |
|---|---|---|
| ENSMUSG00000082306 | 1 | |
| ENSMUSG00000082307 | -1.142876496 | 0.608497229 |
| ENSMUSG00000082308 | -1 | 1 |
| ENSMUSG00000082309 | -1.100193663 | 0.534887579 |
| ENSMUSG00000082310 | 1 | |
| ENSMUSG00000082311 | 1 | |
| ENSMUSG00000082312 | 1.059463094 | 0.608497229 |
| ENSMUSG00000082313 | -1.093474597 | 0.698112512 |
| ENSMUSG00000082314 | -1 | 1 |
| ENSMUSG00000082315 | -1.01916019 | 0.988424085 |
| ENSMUSG00000082316 | 1 | |
| ENSMUSG00000082317 | 1.07913413 | 0.73740059 |
| ENSMUSG00000082318 | -1.03962336 | 0.698112512 |
| ENSMUSG00000082319 | -1.101046843 | 0.806733779 |
| ENSMUSG00000082320 | -1 | 1 |
| ENSMUSG00000082321 | -1 | 1 |
| ENSMUSG00000082322 | 1.214559044 | 0.578221424 |
| ENSMUSG00000082323 | 1 | |
| ENSMUSG00000082324 | -1.17077323 | 0.554735246 |
| ENSMUSG00000082325 | 1.037404367 | 0.970006845 |
| ENSMUSG00000082326 | 1 | |
| ENSMUSG00000082327 | 1 | |
| ENSMUSG00000082328 | -1.03962336 | 0.803072249 |
| ENSMUSG00000082329 | 1 | |
| ENSMUSG00000082330 | 1.059463094 | 0.699340382 |
| ENSMUSG00000082332 | 1 | |
| ENSMUSG00000082333 | 1 | |
| ENSMUSG00000082334 | 1 | |
| ENSMUSG00000082336 | 1 | |
| ENSMUSG00000082337 | 1 | |
| ENSMUSG00000082338 | 1 | |
| ENSMUSG00000082339 | 1 | |
| ENSMUSG00000082340 | 1 | |
| ENSMUSG00000082341 | -1 | 1 |
| ENSMUSG00000082342 | 1.015486245 | 0.972980766 |
| ENSMUSG00000082343 | 1 | |
| ENSMUSG00000082344 | 1 | |
| ENSMUSG00000082345 | 1.117063563 | 0.738145566 |
| ENSMUSG00000082346 | 1 | |
| ENSMUSG00000082347 | 1 | |
| ENSMUSG00000082348 | 1.015486245 | 0.987775262 |
| ENSMUSG00000082350 | -1 | 1 |
| ENSMUSG00000082351 | 1 | |
| ENSMUSG00000082352 | 1 | |
| ENSMUSG00000082353 | 1 | |
| ENSMUSG00000082354 | 1 | |
| ENSMUSG00000082356 | 1 | |
| ENSMUSG00000082357 | -1.073603785 | 0.783862764 |
| ENSMUSG00000082358 | 1 | |
| ENSMUSG00000082359 | 1 | |
| ENSMUSG00000082360 | 1.019083579 | 0.968937442 |
| ENSMUSG00000082361 | 1.210926493 | 0.092041108 |
| ENSMUSG00000082362 | 1.255512191 | 0.217803313 |
| ENSMUSG00000082363 | 1 | |
| ENSMUSG00000082364 | 1 | |
| ENSMUSG00000082365 | 1 | |
| ENSMUSG00000082367 | 1 | |
| ENSMUSG00000082368 | 1 | |
| ENSMUSG00000082369 | 1 | |
| ENSMUSG00000082370 | 1 | |
| ENSMUSG00000082371 | 1 | |
| ENSMUSG00000082372 | 1 | |
| ENSMUSG00000082373 | 1 | |

| | | |
|---|---|---|
| ENSMUSG00000082374 | -1.044273782 | 0.698112512 |
| ENSMUSG00000082375 | 1.084618945 | 0.698112512 |
| ENSMUSG00000082376 | 1.034865357 | 0.910832671 |
| ENSMUSG00000082377 | -1.100193663 | 0.534887579 |
| ENSMUSG00000082378 | 1 | |
| ENSMUSG00000082379 | 1 | |
| ENSMUSG00000082380 | 1 | |
| ENSMUSG00000082381 | 1 | |
| ENSMUSG00000082382 | 1.059463094 | 0.698112512 |
| ENSMUSG00000082383 | -1.013310247 | 1 |
| ENSMUSG00000082388 | 1 | |
| ENSMUSG00000082389 | 1 | |
| ENSMUSG00000082390 | 1.008034876 | 1 |
| ENSMUSG00000082392 | 1 | |
| ENSMUSG00000082393 | 1 | |
| ENSMUSG00000082394 | -1.043306199 | 0.798487595 |
| ENSMUSG00000082395 | 1.160565542 | 0.608497229 |
| ENSMUSG00000082396 | -1.056346118 | 0.740841499 |
| ENSMUSG00000082397 | 1 | |
| ENSMUSG00000082398 | -1 | 1 |
| ENSMUSG00000082399 | 1 | |
| ENSMUSG00000082400 | -1.043306199 | 0.79291651 |
| ENSMUSG00000082402 | -1.132757337 | 0.613963375 |
| ENSMUSG00000082403 | 1 | |
| ENSMUSG00000082404 | 1.064531743 | 0.879766196 |
| ENSMUSG00000082405 | 1 | |
| ENSMUSG00000082406 | -1.028348525 | 0.912425817 |
| ENSMUSG00000082407 | 1 | |
| ENSMUSG00000082408 | 1 | |
| ENSMUSG00000082410 | 1 | |
| ENSMUSG00000082411 | 1 | |
| ENSMUSG00000082412 | 1 | |
| ENSMUSG00000082413 | 1 | |
| ENSMUSG00000082414 | 1 | |
| ENSMUSG00000082415 | 1 | |
| ENSMUSG00000082416 | 1 | |
| ENSMUSG00000082417 | 1 | |
| ENSMUSG00000082418 | -1.03962336 | 0.814089245 |
| ENSMUSG00000082419 | 1 | |
| ENSMUSG00000082420 | 1.139991353 | 0.627865861 |
| ENSMUSG00000082421 | 1 | |
| ENSMUSG00000082422 | 1 | |
| ENSMUSG00000082423 | 1.014731082 | 0.99911685 |
| ENSMUSG00000082424 | 1 | |
| ENSMUSG00000082426 | 1 | |
| ENSMUSG00000082427 | 1 | |
| ENSMUSG00000082428 | 1 | |
| ENSMUSG00000082429 | -1 | 1 |
| ENSMUSG00000082430 | 1 | |
| ENSMUSG00000082431 | 1.011497958 | 1 |
| ENSMUSG00000082432 | -1.043306199 | 0.698112512 |
| ENSMUSG00000082433 | 1 | |
| ENSMUSG00000082435 | 1 | |
| ENSMUSG00000082436 | 1.173847063 | 0.608497229 |
| ENSMUSG00000082437 | 1 | |
| ENSMUSG00000082440 | 1 | |
| ENSMUSG00000082442 | 1 | |
| ENSMUSG00000082443 | 1 | |
| ENSMUSG00000082444 | 1 | |
| ENSMUSG00000082446 | 1 | |
| ENSMUSG00000082447 | 1 | |
| ENSMUSG00000082448 | -1.03962336 | 0.698112512 |
| ENSMUSG00000082449 | 1 | |

| | | |
|---|---|---|
| ENSMUSG00000082450 | 1 | |
| ENSMUSG00000082451 | 1 | |
| ENSMUSG00000082453 | 1 | |
| ENSMUSG00000082454 | -1.039790071 | 0.890258111 |
| ENSMUSG00000082455 | -1 | 1 |
| ENSMUSG00000082456 | 1 | |
| ENSMUSG00000082457 | 1.059463094 | 0.698112512 |
| ENSMUSG00000082458 | 1.084618945 | 0.614071182 |
| ENSMUSG00000082459 | -1.044273782 | 0.698112512 |
| ENSMUSG00000082460 | 1 | |
| ENSMUSG00000082461 | 1.025046921 | 0.944839359 |
| ENSMUSG00000082462 | 1 | |
| ENSMUSG00000082463 | 1 | |
| ENSMUSG00000082464 | 1 | |
| ENSMUSG00000082465 | 1.150179455 | 0.145828618 |
| ENSMUSG00000082467 | -1.044273782 | 0.801443616 |
| ENSMUSG00000082468 | 1 | |
| ENSMUSG00000082472 | -1.053549062 | 0.698112512 |
| ENSMUSG00000082473 | 1 | |
| ENSMUSG00000082474 | -1.211513178 | 0.318915017 |
| ENSMUSG00000082475 | -1.168999771 | 0.261204136 |
| ENSMUSG00000082476 | 1.15516542 | 0.608497229 |
| ENSMUSG00000082478 | 1 | |
| ENSMUSG00000082479 | 1 | |
| ENSMUSG00000082480 | 1.049941052 | 0.84515778 |
| ENSMUSG00000082481 | 1.167088505 | 0.433678407 |
| ENSMUSG00000082482 | 1 | |
| ENSMUSG00000082483 | -1.037681082 | 0.971966027 |
| ENSMUSG00000082484 | 1.034865357 | 0.899648217 |
| ENSMUSG00000082485 | 1 | |
| ENSMUSG00000082486 | 1 | |
| ENSMUSG00000082487 | -1 | 1 |
| ENSMUSG00000082488 | 1 | |
| ENSMUSG00000082489 | 1 | |
| ENSMUSG00000082490 | 1 | |
| ENSMUSG00000082491 | -1.047699696 | 0.955522519 |
| ENSMUSG00000082492 | 1 | |
| ENSMUSG00000082494 | -1.109582661 | 0.608497229 |
| ENSMUSG00000082495 | 1 | |
| ENSMUSG00000082496 | 1 | |
| ENSMUSG00000082497 | 1 | |
| ENSMUSG00000082498 | -1.003834322 | 1 |
| ENSMUSG00000082499 | 1.028484806 | 0.974953883 |
| ENSMUSG00000082500 | 1 | |
| ENSMUSG00000082501 | 1 | |
| ENSMUSG00000082502 | 1 | |
| ENSMUSG00000082503 | 1 | |
| ENSMUSG00000082504 | 1 | |
| ENSMUSG00000082505 | 1 | |
| ENSMUSG00000082507 | -1.119233103 | 0.687624338 |
| ENSMUSG00000082508 | 1 | |
| ENSMUSG00000082509 | -1 | 1 |
| ENSMUSG00000082510 | 1.192305171 | 0.546598109 |
| ENSMUSG00000082511 | 1.059463094 | 0.608497229 |
| ENSMUSG00000082512 | -1.024718486 | 0.931480186 |
| ENSMUSG00000082513 | 1 | |
| ENSMUSG00000082514 | 1 | |
| ENSMUSG00000082515 | 1.189677495 | 0.533812934 |
| ENSMUSG00000082516 | 1 | |
| ENSMUSG00000082517 | 1 | |
| ENSMUSG00000082519 | 1 | |
| ENSMUSG00000082520 | -1 | 1 |
| ENSMUSG00000082523 | 1 | |

| | | |
|---|---|---|
| ENSMUSG00000082524 | -1.044273782 | 0.698112512 |
| ENSMUSG00000082526 | 1 | |
| ENSMUSG00000082527 | 1 | |
| ENSMUSG00000082528 | 1 | |
| ENSMUSG00000082529 | 1 | |
| ENSMUSG00000082530 | -1.03962336 | 0.698112512 |
| ENSMUSG00000082531 | 1 | |
| ENSMUSG00000082532 | -1.144460846 | 0.608497229 |
| ENSMUSG00000082533 | 1 | |
| ENSMUSG00000082534 | 1 | |
| ENSMUSG00000082535 | 1.132544736 | 0.386494097 |
| ENSMUSG00000082536 | -1.00853953 | 0.98887765 |
| ENSMUSG00000082537 | 1 | |
| ENSMUSG00000082538 | 1 | |
| ENSMUSG00000082539 | 1 | |
| ENSMUSG00000082540 | 1 | |
| ENSMUSG00000082541 | 1 | |
| ENSMUSG00000082542 | 1.019083579 | 0.949812939 |
| ENSMUSG00000082543 | 1.092206982 | 0.698112512 |
| ENSMUSG00000082544 | 1.024535939 | 0.982489318 |
| ENSMUSG00000082545 | -1.03962336 | 0.698112512 |
| ENSMUSG00000082546 | 1 | |
| ENSMUSG00000082547 | 1 | |
| ENSMUSG00000082548 | 1 | |
| ENSMUSG00000082549 | 1.133227159 | 0.698112512 |
| ENSMUSG00000082550 | -1.03962336 | 0.808074244 |
| ENSMUSG00000082551 | 1 | |
| ENSMUSG00000082552 | 1 | |
| ENSMUSG00000082553 | 1 | |
| ENSMUSG00000082554 | 1 | |
| ENSMUSG00000082555 | 1 | |
| ENSMUSG00000082556 | 1 | |
| ENSMUSG00000082558 | 1 | |
| ENSMUSG00000082560 | 1 | |
| ENSMUSG00000082561 | 1.028323353 | 0.903464611 |
| ENSMUSG00000082562 | 1 | |
| ENSMUSG00000082563 | 1 | |
| ENSMUSG00000082564 | 1 | |
| ENSMUSG00000082565 | 1 | |
| ENSMUSG00000082566 | 1 | |
| ENSMUSG00000082567 | 1 | |
| ENSMUSG00000082569 | -1 | 1 |
| ENSMUSG00000082570 | 1 | |
| ENSMUSG00000082571 | 1 | |
| ENSMUSG00000082573 | 1 | |
| ENSMUSG00000082574 | 1 | |
| ENSMUSG00000082575 | 1.068866838 | 0.680031094 |
| ENSMUSG00000082576 | 1.019083579 | 0.968937442 |
| ENSMUSG00000082577 | 1 | |
| ENSMUSG00000082578 | 1 | |
| ENSMUSG00000082579 | 1.074421866 | 0.608497229 |
| ENSMUSG00000082580 | 1.089470641 | 0.608497229 |
| ENSMUSG00000082581 | 1 | |
| ENSMUSG00000082583 | 1 | |
| ENSMUSG00000082584 | 1 | |
| ENSMUSG00000082585 | 1 | |
| ENSMUSG00000082586 | 1 | |
| ENSMUSG00000082587 | 1.006389981 | 1 |
| ENSMUSG00000082588 | 1.038283405 | 0.698112512 |
| ENSMUSG00000082589 | 1 | |
| ENSMUSG00000082590 | 1 | |
| ENSMUSG00000082591 | 1 | |
| ENSMUSG00000082592 | -1 | 1 |

| | | |
|---|---|---|
| ENSMUSG00000082593 | -1.280052201 | 0.256544909 |
| ENSMUSG00000082594 | 1 | |
| ENSMUSG00000082595 | 1 | |
| ENSMUSG00000082596 | 1.044210161 | 0.917717739 |
| ENSMUSG00000082597 | 1 | |
| ENSMUSG00000082598 | 1 | |
| ENSMUSG00000082599 | -1.157634469 | 0.544226414 |
| ENSMUSG00000082600 | 1.024693402 | 0.952057016 |
| ENSMUSG00000082601 | 1 | |
| ENSMUSG00000082602 | 1 | |
| ENSMUSG00000082604 | 1.059463094 | 0.608497229 |
| ENSMUSG00000082605 | -1.137576943 | 0.608497229 |
| ENSMUSG00000082606 | 1 | |
| ENSMUSG00000082607 | 1 | |
| ENSMUSG00000082608 | 1.075070132 | 0.698112512 |
| ENSMUSG00000082609 | 1.005613439 | 1 |
| ENSMUSG00000082610 | 1.059463094 | 0.698112512 |
| ENSMUSG00000082611 | 1 | |
| ENSMUSG00000082612 | 1 | |
| ENSMUSG00000082613 | 1 | |
| ENSMUSG00000082614 | 1 | |
| ENSMUSG00000082615 | 1 | |
| ENSMUSG00000082616 | 1 | |
| ENSMUSG00000082617 | 1 | |
| ENSMUSG00000082618 | 1.059463094 | 0.608497229 |
| ENSMUSG00000082619 | 1 | |
| ENSMUSG00000082620 | 1 | |
| ENSMUSG00000082621 | -1.003834322 | 1 |
| ENSMUSG00000082622 | 1.018424868 | 0.946951599 |
| ENSMUSG00000082623 | 1 | |
| ENSMUSG00000082624 | 1 | |
| ENSMUSG00000082625 | 1.102024305 | 0.694494111 |
| ENSMUSG00000082626 | -1.043306199 | 0.798487595 |
| ENSMUSG00000082628 | -1 | 1 |
| ENSMUSG00000082629 | 1 | |
| ENSMUSG00000082630 | 1.005059784 | 1 |
| ENSMUSG00000082632 | -1 | 1 |
| ENSMUSG00000082633 | -1.069095149 | 0.714024161 |
| ENSMUSG00000082634 | -1.044273782 | 0.698112512 |
| ENSMUSG00000082635 | 1 | |
| ENSMUSG00000082636 | 1.069068979 | 0.608497229 |
| ENSMUSG00000082637 | 1 | |
| ENSMUSG00000082638 | 1 | |
| ENSMUSG00000082639 | 1 | |
| ENSMUSG00000082641 | 1.01796141 | 0.982838256 |
| ENSMUSG00000082644 | 1 | |
| ENSMUSG00000082645 | 1 | |
| ENSMUSG00000082646 | 1.01112436 | 1 |
| ENSMUSG00000082647 | -1 | 1 |
| ENSMUSG00000082648 | 1 | |
| ENSMUSG00000082649 | 1 | |
| ENSMUSG00000082650 | 1 | |
| ENSMUSG00000082651 | -1.047306572 | 0.864480156 |
| ENSMUSG00000082652 | 1 | |
| ENSMUSG00000082653 | 1 | |
| ENSMUSG00000082655 | 1 | |
| ENSMUSG00000082656 | 1 | |
| ENSMUSG00000082657 | 1 | |
| ENSMUSG00000082658 | 1 | |
| ENSMUSG00000082660 | 1 | |
| ENSMUSG00000082661 | 1 | |
| ENSMUSG00000082662 | 1 | |
| ENSMUSG00000082663 | 1 | |

| | | |
|---|---|---|
| ENSMUSG00000082664 | 1 | |
| ENSMUSG00000082665 | -1.113109741 | 0.608497229 |
| ENSMUSG00000082666 | 1 | |
| ENSMUSG00000082667 | 1 | |
| ENSMUSG00000082669 | 1 | |
| ENSMUSG00000082670 | 1 | |
| ENSMUSG00000082671 | 1.069068979 | 0.608497229 |
| ENSMUSG00000082672 | -1 | 1 |
| ENSMUSG00000082673 | -1.044273782 | 0.795446726 |
| ENSMUSG00000082674 | 1 | |
| ENSMUSG00000082675 | 1.171563499 | 0.233412931 |
| ENSMUSG00000082676 | 1.059463094 | 0.698112512 |
| ENSMUSG00000082677 | 1 | |
| ENSMUSG00000082678 | 1.059463094 | 0.704170349 |
| ENSMUSG00000082679 | 1 | |
| ENSMUSG00000082680 | 1 | |
| ENSMUSG00000082681 | 1 | |
| ENSMUSG00000082682 | 1 | |
| ENSMUSG00000082683 | -1.03962336 | 0.803072249 |
| ENSMUSG00000082684 | -1.043306199 | 0.798487595 |
| ENSMUSG00000082686 | -1.043306199 | 0.79291651 |
| ENSMUSG00000082687 | 1 | |
| ENSMUSG00000082688 | -1.03962336 | 0.88732209 |
| ENSMUSG00000082689 | 1 | |
| ENSMUSG00000082690 | 1 | |
| ENSMUSG00000082691 | 1.028323353 | 0.903464611 |
| ENSMUSG00000082692 | -1 | 1 |
| ENSMUSG00000082693 | -1.053767744 | 0.881281964 |
| ENSMUSG00000082696 | 1 | |
| ENSMUSG00000082698 | 1 | |
| ENSMUSG00000082699 | 1.015486245 | 0.985522133 |
| ENSMUSG00000082700 | 1 | |
| ENSMUSG00000082701 | 1 | |
| ENSMUSG00000082703 | -1.03962336 | 0.808074244 |
| ENSMUSG00000082704 | 1 | |
| ENSMUSG00000082705 | -1.043306199 | 0.798487595 |
| ENSMUSG00000082706 | 1 | |
| ENSMUSG00000082707 | -1 | 1 |
| ENSMUSG00000082708 | -1 | 1 |
| ENSMUSG00000082709 | 1 | |
| ENSMUSG00000082710 | 1 | |
| ENSMUSG00000082711 | 1.059463094 | 0.698112512 |
| ENSMUSG00000082712 | 1 | |
| ENSMUSG00000082713 | 1 | |
| ENSMUSG00000082714 | 1 | |
| ENSMUSG00000082715 | 1 | |
| ENSMUSG00000082716 | 1 | |
| ENSMUSG00000082717 | 1 | |
| ENSMUSG00000082718 | 1 | |
| ENSMUSG00000082720 | 1 | |
| ENSMUSG00000082721 | 1 | |
| ENSMUSG00000082722 | 1 | |
| ENSMUSG00000082723 | 1 | |
| ENSMUSG00000082724 | 1 | |
| ENSMUSG00000082725 | 1 | |
| ENSMUSG00000082726 | 1 | |
| ENSMUSG00000082727 | -1.003834322 | 1 |
| ENSMUSG00000082728 | 1 | |
| ENSMUSG00000082729 | 1 | |
| ENSMUSG00000082730 | 1.069068979 | 0.608497229 |
| ENSMUSG00000082732 | 1 | |
| ENSMUSG00000082733 | 1 | |
| ENSMUSG00000082734 | 1 | |

| | | |
|---|---|---|
| ENSMUSG00000082735 | 1 | |
| ENSMUSG00000082736 | 1 | |
| ENSMUSG00000082737 | 1.059463094 | 0.608497229 |
| ENSMUSG00000082738 | -1.044273782 | 0.698112512 |
| ENSMUSG00000082739 | 1 | |
| ENSMUSG00000082740 | -1 | 1 |
| ENSMUSG00000082741 | -1.145518406 | 0.608497229 |
| ENSMUSG00000082743 | 1.0106701 | 1 |
| ENSMUSG00000082744 | 1 | |
| ENSMUSG00000082745 | 1 | |
| ENSMUSG00000082746 | -1.03962336 | 0.803072249 |
| ENSMUSG00000082749 | 1.069068979 | 0.608497229 |
| ENSMUSG00000082750 | 1 | |
| ENSMUSG00000082751 | 1 | |
| ENSMUSG00000082752 | 1 | |
| ENSMUSG00000082754 | 1 | |
| ENSMUSG00000082755 | 1.019083579 | 0.968937442 |
| ENSMUSG00000082757 | 1 | |
| ENSMUSG00000082758 | -1.03962336 | 0.698112512 |
| ENSMUSG00000082759 | 1 | |
| ENSMUSG00000082760 | 1 | |
| ENSMUSG00000082762 | 1.075870199 | 0.809396188 |
| ENSMUSG00000082763 | 1 | |
| ENSMUSG00000082765 | -1.119565536 | 0.753100881 |
| ENSMUSG00000082766 | 1 | |
| ENSMUSG00000082767 | 1 | |
| ENSMUSG00000082768 | -1.053549062 | 0.825517832 |
| ENSMUSG00000082769 | 1.003729568 | 1 |
| ENSMUSG00000082770 | 1 | |
| ENSMUSG00000082771 | 1 | |
| ENSMUSG00000082772 | 1 | |
| ENSMUSG00000082773 | 1 | |
| ENSMUSG00000082774 | -1 | 1 |
| ENSMUSG00000082775 | 1.023645551 | 0.97427316 |
| ENSMUSG00000082776 | -1.111880267 | 0.608497229 |
| ENSMUSG00000082777 | 1 | |
| ENSMUSG00000082778 | -1.019637694 | 0.991978016 |
| ENSMUSG00000082779 | 1 | |
| ENSMUSG00000082780 | -1 | 1 |
| ENSMUSG00000082781 | 1 | |
| ENSMUSG00000082782 | 1 | |
| ENSMUSG00000082783 | 1 | |
| ENSMUSG00000082784 | 1 | |
| ENSMUSG00000082785 | 1 | |
| ENSMUSG00000082786 | -1 | 1 |
| ENSMUSG00000082788 | 1 | |
| ENSMUSG00000082789 | -1 | 1 |
| ENSMUSG00000082790 | -1.084645496 | 0.628300894 |
| ENSMUSG00000082791 | 1.009459007 | 0.984782014 |
| ENSMUSG00000082792 | 1.053588583 | 0.648983806 |
| ENSMUSG00000082793 | 1 | |
| ENSMUSG00000082794 | 1.218572685 | 0.477644658 |
| ENSMUSG00000082795 | 1 | |
| ENSMUSG00000082796 | 1 | |
| ENSMUSG00000082797 | 1 | |
| ENSMUSG00000082798 | 1 | |
| ENSMUSG00000082799 | -1 | 1 |
| ENSMUSG00000082800 | 1 | |
| ENSMUSG00000082802 | 1 | |
| ENSMUSG00000082804 | 1.069068979 | 0.73684216 |
| ENSMUSG00000082806 | -1.03962336 | 0.808074244 |
| ENSMUSG00000082807 | 1 | |
| ENSMUSG00000082809 | 1.04806794 | 0.696316692 |

| | | |
|---|---|---|
| ENSMUSG00000082810 | 1.059463094 | 0.698112512 |
| ENSMUSG00000082812 | 1.024693402 | 0.952057016 |
| ENSMUSG00000082814 | 1 | |
| ENSMUSG00000082815 | 1.14205712 | 0.713864779 |
| ENSMUSG00000082816 | 1.169519855 | 0.145847782 |
| ENSMUSG00000082817 | 1 | |
| ENSMUSG00000082818 | 1 | |
| ENSMUSG00000082819 | 1 | |
| ENSMUSG00000082820 | -1 | 1 |
| ENSMUSG00000082821 | 1 | |
| ENSMUSG00000082822 | 1 | |
| ENSMUSG00000082823 | 1 | |
| ENSMUSG00000082824 | 1 | |
| ENSMUSG00000082825 | 1 | |
| ENSMUSG00000082826 | -1.043306199 | 0.698112512 |
| ENSMUSG00000082827 | 1 | |
| ENSMUSG00000082828 | 1 | |
| ENSMUSG00000082829 | -1.071075483 | 0.698112512 |
| ENSMUSG00000082831 | -1.105243038 | 0.776268516 |
| ENSMUSG00000082832 | 1 | |
| ENSMUSG00000082833 | 1 | |
| ENSMUSG00000082835 | -1 | 1 |
| ENSMUSG00000082836 | -1.015511103 | 1 |
| ENSMUSG00000082837 | 1 | |
| ENSMUSG00000082838 | 1 | |
| ENSMUSG00000082839 | 1 | |
| ENSMUSG00000082840 | -1 | 1 |
| ENSMUSG00000082841 | 1 | |
| ENSMUSG00000082842 | 1 | |
| ENSMUSG00000082843 | 1 | |
| ENSMUSG00000082844 | 1 | |
| ENSMUSG00000082845 | -1.044273782 | 0.698112512 |
| ENSMUSG00000082846 | 1 | |
| ENSMUSG00000082847 | 1.069068979 | 0.608497229 |
| ENSMUSG00000082848 | -1.085651418 | 0.698112512 |
| ENSMUSG00000082849 | 1.059463094 | 0.698112512 |
| ENSMUSG00000082850 | 1 | |
| ENSMUSG00000082851 | 1.005613439 | 1 |
| ENSMUSG00000082852 | 1.177765487 | 0.380970157 |
| ENSMUSG00000082853 | 1 | |
| ENSMUSG00000082855 | 1.122933516 | 0.63204289 |
| ENSMUSG00000082856 | 1 | |
| ENSMUSG00000082857 | 1 | |
| ENSMUSG00000082858 | 1 | |
| ENSMUSG00000082859 | 1 | |
| ENSMUSG00000082860 | 1.105251715 | 0.494379423 |
| ENSMUSG00000082861 | 1.015486245 | 0.985522133 |
| ENSMUSG00000082862 | 1 | |
| ENSMUSG00000082863 | 1 | |
| ENSMUSG00000082864 | 1 | |
| ENSMUSG00000082865 | -1.078590893 | 0.724197242 |
| ENSMUSG00000082867 | -1.005345324 | 1 |
| ENSMUSG00000082868 | 1 | |
| ENSMUSG00000082869 | 1 | |
| ENSMUSG00000082870 | -1.020155149 | 0.989387709 |
| ENSMUSG00000082871 | 1 | |
| ENSMUSG00000082872 | 1.16755408 | 0.608497229 |
| ENSMUSG00000082873 | 1.059463094 | 0.698112512 |
| ENSMUSG00000082874 | -1.099781752 | 0.724929955 |
| ENSMUSG00000082875 | 1 | |
| ENSMUSG00000082876 | 1.065093972 | 0.811529746 |
| ENSMUSG00000082877 | -1 | 1 |
| ENSMUSG00000082878 | -1.043533577 | 0.909060016 |

| | | |
|---|---|---|
| ENSMUSG00000082879 | -1.016511436 | 1 |
| ENSMUSG00000082882 | 1 | |
| ENSMUSG00000082883 | 1 | |
| ENSMUSG00000082884 | 1.047786406 | 0.937517551 |
| ENSMUSG00000082885 | 1 | |
| ENSMUSG00000082886 | 1 | |
| ENSMUSG00000082887 | 1 | |
| ENSMUSG00000082888 | -1 | 1 |
| ENSMUSG00000082889 | 1 | |
| ENSMUSG00000082890 | 1 | |
| ENSMUSG00000082891 | 1 | |
| ENSMUSG00000082892 | 1 | |
| ENSMUSG00000082893 | 1.005613439 | 1 |
| ENSMUSG00000082894 | 1.057757739 | 0.737159156 |
| ENSMUSG00000082895 | -1.056432493 | 0.794274354 |
| ENSMUSG00000082896 | 1.169224094 | 0.3244149 |
| ENSMUSG00000082897 | 1 | |
| ENSMUSG00000082898 | 1 | |
| ENSMUSG00000082899 | 1 | |
| ENSMUSG00000082900 | 1.133393325 | 0.720235853 |
| ENSMUSG00000082901 | -1.085651418 | 0.532369657 |
| ENSMUSG00000082902 | 1 | |
| ENSMUSG00000082903 | 1 | |
| ENSMUSG00000082905 | 1 | |
| ENSMUSG00000082906 | -1 | 1 |
| ENSMUSG00000082907 | 1 | |
| ENSMUSG00000082908 | 1 | |
| ENSMUSG00000082910 | 1 | |
| ENSMUSG00000082911 | -1 | 1 |
| ENSMUSG00000082912 | 1.069068979 | 0.608497229 |
| ENSMUSG00000082913 | 1 | |
| ENSMUSG00000082915 | 1 | |
| ENSMUSG00000082917 | -1 | 1 |
| ENSMUSG00000082918 | 1 | |
| ENSMUSG00000082919 | -1 | 1 |
| ENSMUSG00000082920 | 1.037089191 | 0.947248675 |
| ENSMUSG00000082922 | 1.03306284 | 0.93979453 |
| ENSMUSG00000082924 | -1.03962336 | 0.803072249 |
| ENSMUSG00000082925 | -1.053549062 | 0.778372272 |
| ENSMUSG00000082926 | 1.069068979 | 0.698112512 |
| ENSMUSG00000082927 | -1.022050271 | 0.995522634 |
| ENSMUSG00000082928 | 1 | |
| ENSMUSG00000082929 | -1.044273782 | 0.698112512 |
| ENSMUSG00000082930 | 1 | |
| ENSMUSG00000082931 | -1.099174267 | 0.535052277 |
| ENSMUSG00000082932 | 1 | |
| ENSMUSG00000082933 | 1 | |
| ENSMUSG00000082934 | 1 | |
| ENSMUSG00000082935 | -1.207817974 | 0.157858445 |
| ENSMUSG00000082936 | 1.295007897 | 0.316089517 |
| ENSMUSG00000082937 | 1 | |
| ENSMUSG00000082938 | 1.015486245 | 0.985522133 |
| ENSMUSG00000082939 | 1 | |
| ENSMUSG00000082940 | 1 | |
| ENSMUSG00000082941 | 1.207196766 | 0.196773466 |
| ENSMUSG00000082943 | 1 | |
| ENSMUSG00000082945 | 1 | |
| ENSMUSG00000082947 | 1.059463094 | 0.698112512 |
| ENSMUSG00000082949 | 1 | |
| ENSMUSG00000082950 | 1.014545335 | 0.979002724 |
| ENSMUSG00000082951 | -1.044273782 | 0.698112512 |
| ENSMUSG00000082952 | 1 | |
| ENSMUSG00000082953 | 1 | |

| | | |
|---|---|---|
| ENSMUSG00000082954 | -1 | 1 |
| ENSMUSG00000082955 | -1 | 1 |
| ENSMUSG00000082956 | 1 | |
| ENSMUSG00000082957 | 1 | |
| ENSMUSG00000082958 | -1.03962336 | 0.698112512 |
| ENSMUSG00000082960 | -1.043306199 | 0.698112512 |
| ENSMUSG00000082961 | -1.035298449 | 0.954523477 |
| ENSMUSG00000082962 | -1.03962336 | 0.698112512 |
| ENSMUSG00000082963 | 1 | |
| ENSMUSG00000082964 | -1.043306199 | 0.79291651 |
| ENSMUSG00000082965 | -1.069502966 | 0.79291651 |
| ENSMUSG00000082966 | 1 | |
| ENSMUSG00000082967 | 1.059463094 | 0.698112512 |
| ENSMUSG00000082968 | 1 | |
| ENSMUSG00000082969 | -1.012222786 | 1 |
| ENSMUSG00000082970 | 1 | |
| ENSMUSG00000082971 | 1.059463094 | 0.608497229 |
| ENSMUSG00000082973 | 1 | |
| ENSMUSG00000082974 | 1 | |
| ENSMUSG00000082975 | 1 | |
| ENSMUSG00000082976 | 1 | |
| ENSMUSG00000082977 | 1 | |
| ENSMUSG00000082978 | 1.131343654 | 0.698112512 |
| ENSMUSG00000082980 | 1 | |
| ENSMUSG00000082981 | -1.038444538 | 0.900396939 |
| ENSMUSG00000082982 | 1 | |
| ENSMUSG00000082983 | 1 | |
| ENSMUSG00000082985 | -1.130720898 | 0.634115399 |
| ENSMUSG00000082986 | -1.010960635 | 1 |
| ENSMUSG00000082987 | -1.03962336 | 0.803072249 |
| ENSMUSG00000082988 | 1.200241468 | 0.532369657 |
| ENSMUSG00000082989 | 1 | |
| ENSMUSG00000082990 | -1.024718486 | 0.951399668 |
| ENSMUSG00000082991 | 1 | |
| ENSMUSG00000082993 | 1 | |
| ENSMUSG00000082994 | -1 | 1 |
| ENSMUSG00000082995 | 1 | |
| ENSMUSG00000082996 | 1.069068979 | 0.608497229 |
| ENSMUSG00000082997 | 1.059463094 | 0.608497229 |
| ENSMUSG00000082998 | -1.053549062 | 0.769745556 |
| ENSMUSG00000083000 | 1 | |
| ENSMUSG00000083001 | 1 | |
| ENSMUSG00000083004 | -1.158708082 | 0.350957446 |
| ENSMUSG00000083005 | 1 | |
| ENSMUSG00000083006 | 1.177765487 | 0.380970157 |
| ENSMUSG00000083007 | 1.125934636 | 0.532369657 |
| ENSMUSG00000083008 | 1 | |
| ENSMUSG00000083009 | 1 | |
| ENSMUSG00000083010 | 1 | |
| ENSMUSG00000083011 | 1.023099031 | 0.937388947 |
| ENSMUSG00000083012 | 1.076269216 | 0.608497229 |
| ENSMUSG00000083013 | 1 | |
| ENSMUSG00000083014 | 1 | |
| ENSMUSG00000083015 | -1.043306199 | 0.698112512 |
| ENSMUSG00000083016 | -1.004241828 | 1 |
| ENSMUSG00000083017 | 1 | |
| ENSMUSG00000083018 | 1 | |
| ENSMUSG00000083019 | -1.089129277 | 0.913172334 |
| ENSMUSG00000083020 | 1 | |
| ENSMUSG00000083022 | -1.015974852 | 0.903868918 |
| ENSMUSG00000083023 | 1 | |
| ENSMUSG00000083024 | 1 | |
| ENSMUSG00000083025 | 1.059463094 | 0.704170349 |

| | | |
|---|---|---|
| ENSMUSG00000083026 | 1 | |
| ENSMUSG00000083027 | 1.006871124 | 1 |
| ENSMUSG00000083029 | 1 | |
| ENSMUSG00000083030 | 1 | |
| ENSMUSG00000083031 | 1 | |
| ENSMUSG00000083032 | 1 | |
| ENSMUSG00000083033 | -1.02709481 | 0.945529232 |
| ENSMUSG00000083034 | 1 | |
| ENSMUSG00000083036 | 1 | |
| ENSMUSG00000083037 | 1 | |
| ENSMUSG00000083038 | 1 | |
| ENSMUSG00000083039 | 1 | |
| ENSMUSG00000083041 | 1 | |
| ENSMUSG00000083042 | 1 | |
| ENSMUSG00000083043 | 1.2323813 | 0.163697151 |
| ENSMUSG00000083044 | 1.472380751 | 0.088438086 |
| ENSMUSG00000083045 | 1 | |
| ENSMUSG00000083046 | 1 | |
| ENSMUSG00000083047 | 1 | |
| ENSMUSG00000083049 | 1 | |
| ENSMUSG00000083050 | 1 | |
| ENSMUSG00000083052 | -1.069095149 | 0.724187622 |
| ENSMUSG00000083053 | 1 | |
| ENSMUSG00000083054 | 1.019083579 | 0.949812939 |
| ENSMUSG00000083055 | 1 | |
| ENSMUSG00000083057 | 1.059463094 | 0.608497229 |
| ENSMUSG00000083058 | 1 | |
| ENSMUSG00000083059 | 1 | |
| ENSMUSG00000083060 | 1 | |
| ENSMUSG00000083061 | 1 | |
| ENSMUSG00000083062 | 1 | |
| ENSMUSG00000083063 | 1 | |
| ENSMUSG00000083064 | -1.084268471 | 0.609700609 |
| ENSMUSG00000083065 | 1 | |
| ENSMUSG00000083066 | 1 | |
| ENSMUSG00000083067 | 1 | |
| ENSMUSG00000083068 | -1.095294215 | 0.608497229 |
| ENSMUSG00000083069 | 1 | |
| ENSMUSG00000083070 | 1 | |
| ENSMUSG00000083072 | 1.083521437 | 0.858911269 |
| ENSMUSG00000083073 | 1 | |
| ENSMUSG00000083074 | 1 | |
| ENSMUSG00000083075 | -1 | 1 |
| ENSMUSG00000083076 | 1.084618945 | 0.698112512 |
| ENSMUSG00000083077 | 1 | |
| ENSMUSG00000083078 | 1 | |
| ENSMUSG00000083079 | 1 | |
| ENSMUSG00000083080 | 1 | |
| ENSMUSG00000083081 | 1 | |
| ENSMUSG00000083083 | 1.018196408 | 0.999201226 |
| ENSMUSG00000083084 | 1.075070132 | 0.725426917 |
| ENSMUSG00000083085 | 1 | |
| ENSMUSG00000083086 | -1.060393457 | 0.801666546 |
| ENSMUSG00000083087 | 1.20210372 | 0.505106302 |
| ENSMUSG00000083088 | 1 | |
| ENSMUSG00000083089 | -1.089497311 | 0.532369657 |
| ENSMUSG00000083090 | 1.059463094 | 0.608497229 |
| ENSMUSG00000083091 | 1 | |
| ENSMUSG00000083093 | -1 | 1 |
| ENSMUSG00000083094 | 1.116638996 | 0.717035356 |
| ENSMUSG00000083096 | 1 | |
| ENSMUSG00000083097 | -1.035153216 | 0.968937442 |
| ENSMUSG00000083098 | -1 | 1 |

| | | |
|---|---|---|
| ENSMUSG00000083099 | 1 | |
| ENSMUSG00000083100 | -1 | 1 |
| ENSMUSG00000083101 | 1.059463094 | 0.608497229 |
| ENSMUSG00000083103 | 1 | |
| ENSMUSG00000083104 | 1.0557057 | 0.844881695 |
| ENSMUSG00000083105 | 1 | |
| ENSMUSG00000083106 | 1 | |
| ENSMUSG00000083107 | -1.346172611 | 0.139719371 |
| ENSMUSG00000083109 | 1 | |
| ENSMUSG00000083110 | 1 | |
| ENSMUSG00000083111 | -1.3978822 | 0.077822785 |
| ENSMUSG00000083112 | 1 | |
| ENSMUSG00000083113 | 1.075070132 | 0.725426917 |
| ENSMUSG00000083114 | 1 | |
| ENSMUSG00000083115 | -1 | 1 |
| ENSMUSG00000083116 | -1.064616399 | 0.698112512 |
| ENSMUSG00000083117 | 1 | |
| ENSMUSG00000083118 | 1 | |
| ENSMUSG00000083119 | 1 | |
| ENSMUSG00000083120 | 1 | |
| ENSMUSG00000083121 | 1.150179455 | 0.250477583 |
| ENSMUSG00000083122 | -1 | 1 |
| ENSMUSG00000083123 | 1 | |
| ENSMUSG00000083124 | 1 | |
| ENSMUSG00000083126 | 1 | |
| ENSMUSG00000083127 | 1 | |
| ENSMUSG00000083128 | 1 | |
| ENSMUSG00000083129 | 1 | |
| ENSMUSG00000083130 | 1 | |
| ENSMUSG00000083135 | 1 | |
| ENSMUSG00000083136 | 1 | |
| ENSMUSG00000083137 | 1 | |
| ENSMUSG00000083138 | 1 | |
| ENSMUSG00000083139 | 1 | |
| ENSMUSG00000083140 | 1 | |
| ENSMUSG00000083141 | 1 | |
| ENSMUSG00000083142 | 1 | |
| ENSMUSG00000083143 | 1 | |
| ENSMUSG00000083144 | 1 | |
| ENSMUSG00000083145 | 1.122462048 | 0.420161906 |
| ENSMUSG00000083147 | 1 | |
| ENSMUSG00000083149 | -1.129231493 | 0.608497229 |
| ENSMUSG00000083150 | 1 | |
| ENSMUSG00000083151 | -1 | 1 |
| ENSMUSG00000083152 | 1.043202648 | 0.949812939 |
| ENSMUSG00000083153 | 1 | |
| ENSMUSG00000083154 | -1 | 1 |
| ENSMUSG00000083155 | 1 | |
| ENSMUSG00000083156 | -1.044273782 | 0.698112512 |
| ENSMUSG00000083157 | 1 | |
| ENSMUSG00000083158 | 1 | |
| ENSMUSG00000083159 | -1.100193663 | 0.672413803 |
| ENSMUSG00000083160 | 1 | |
| ENSMUSG00000083161 | 1 | |
| ENSMUSG00000083162 | 1 | |
| ENSMUSG00000083163 | 1 | |
| ENSMUSG00000083165 | 1 | |
| ENSMUSG00000083166 | -1.132350732 | 0.488715018 |
| ENSMUSG00000083167 | 1 | |
| ENSMUSG00000083168 | 1 | |
| ENSMUSG00000083169 | -1.053549062 | 0.698112512 |
| ENSMUSG00000083170 | 1 | |
| ENSMUSG00000083171 | -1.043306199 | 0.698112512 |

| | | |
|---|---|---|
| ENSMUSG00000083172 | 1 | |
| ENSMUSG00000083173 | 1 | |
| ENSMUSG00000083174 | 1 | |
| ENSMUSG00000083175 | 1 | |
| ENSMUSG00000083176 | -1.043306199 | 0.698112512 |
| ENSMUSG00000083177 | -1.03962336 | 0.698112512 |
| ENSMUSG00000083178 | 1 | |
| ENSMUSG00000083179 | 1.075870199 | 0.748191299 |
| ENSMUSG00000083180 | -1.044273782 | 0.698112512 |
| ENSMUSG00000083181 | 1 | |
| ENSMUSG00000083182 | 1 | |
| ENSMUSG00000083183 | -1.049475223 | 0.8870716 |
| ENSMUSG00000083185 | 1 | |
| ENSMUSG00000083186 | -1.03962336 | 0.872692845 |
| ENSMUSG00000083188 | 1 | |
| ENSMUSG00000083190 | 1 | |
| ENSMUSG00000083191 | 1 | |
| ENSMUSG00000083192 | -1.043306199 | 0.698112512 |
| ENSMUSG00000083193 | 1.069068979 | 0.608497229 |
| ENSMUSG00000083195 | 1 | |
| ENSMUSG00000083196 | 1 | |
| ENSMUSG00000083197 | 1.059463094 | 0.608497229 |
| ENSMUSG00000083199 | -1.043306199 | 0.858911269 |
| ENSMUSG00000083200 | -1.10514284 | 0.427218243 |
| ENSMUSG00000083201 | 1.095586318 | 0.723742698 |
| ENSMUSG00000083202 | -1.043306199 | 0.698112512 |
| ENSMUSG00000083203 | 1 | |
| ENSMUSG00000083204 | -1.03962336 | 0.698112512 |
| ENSMUSG00000083205 | 1 | |
| ENSMUSG00000083206 | 1 | |
| ENSMUSG00000083207 | -1.048921239 | 0.889777136 |
| ENSMUSG00000083208 | 1 | |
| ENSMUSG00000083209 | 1.122462048 | 0.215991492 |
| ENSMUSG00000083210 | -1.044273782 | 0.698112512 |
| ENSMUSG00000083211 | 1 | |
| ENSMUSG00000083212 | 1 | |
| ENSMUSG00000083213 | 1 | |
| ENSMUSG00000083214 | 1.059463094 | 0.608497229 |
| ENSMUSG00000083215 | 1 | |
| ENSMUSG00000083216 | 1 | |
| ENSMUSG00000083218 | -1.18317927 | 0.088091226 |
| ENSMUSG00000083219 | 1.146630142 | 0.717035356 |
| ENSMUSG00000083220 | 1.158391391 | 0.593237123 |
| ENSMUSG00000083221 | 1 | |
| ENSMUSG00000083222 | 1 | |
| ENSMUSG00000083223 | -1.009360844 | 1 |
| ENSMUSG00000083224 | 1 | |
| ENSMUSG00000083225 | 1 | |
| ENSMUSG00000083226 | -1.03962336 | 0.698112512 |
| ENSMUSG00000083227 | 1 | |
| ENSMUSG00000083228 | 1 | |
| ENSMUSG00000083229 | 1 | |
| ENSMUSG00000083230 | 1 | |
| ENSMUSG00000083231 | 1 | |
| ENSMUSG00000083232 | 1 | |
| ENSMUSG00000083233 | 1 | |
| ENSMUSG00000083234 | 1 | |
| ENSMUSG00000083235 | 1 | |
| ENSMUSG00000083236 | 1.229754062 | 0.543975336 |
| ENSMUSG00000083237 | 1 | |
| ENSMUSG00000083238 | 1 | |
| ENSMUSG00000083239 | 1 | |
| ENSMUSG00000083240 | 1.059463094 | 0.608497229 |

| | | |
|---|---|---|
| ENSMUSG00000083241 | 1.015486245 | 0.972980766 |
| ENSMUSG00000083242 | 1 | |
| ENSMUSG00000083243 | 1 | |
| ENSMUSG00000083244 | 1 | |
| ENSMUSG00000083245 | 1.140982192 | 0.625094965 |
| ENSMUSG00000083246 | -1.053104934 | 0.933031586 |
| ENSMUSG00000083248 | 1 | |
| ENSMUSG00000083249 | 1 | |
| ENSMUSG00000083250 | 1.053966865 | 0.865865121 |
| ENSMUSG00000083251 | 1 | |
| ENSMUSG00000083252 | 1 | |
| ENSMUSG00000083253 | -1.118496046 | 0.717035356 |
| ENSMUSG00000083254 | 1 | |
| ENSMUSG00000083255 | 1 | |
| ENSMUSG00000083256 | 1 | |
| ENSMUSG00000083257 | 1 | |
| ENSMUSG00000083258 | 1.069437807 | 0.7938895 |
| ENSMUSG00000083260 | 1 | |
| ENSMUSG00000083261 | -1.028160286 | 0.950579028 |
| ENSMUSG00000083262 | 1 | |
| ENSMUSG00000083263 | -1 | 1 |
| ENSMUSG00000083264 | 1 | |
| ENSMUSG00000083266 | 1 | |
| ENSMUSG00000083267 | 1 | |
| ENSMUSG00000083268 | -1.054422867 | 0.876686173 |
| ENSMUSG00000083269 | 1 | |
| ENSMUSG00000083270 | -1.059004268 | 0.879253209 |
| ENSMUSG00000083271 | 1 | |
| ENSMUSG00000083272 | 1 | |
| ENSMUSG00000083274 | 1 | |
| ENSMUSG00000083275 | -1.053549062 | 0.778372272 |
| ENSMUSG00000083276 | 1 | |
| ENSMUSG00000083277 | 1.069068979 | 0.698112512 |
| ENSMUSG00000083278 | 1 | |
| ENSMUSG00000083279 | 1 | |
| ENSMUSG00000083280 | -1.03962336 | 0.698112512 |
| ENSMUSG00000083281 | -1.053549062 | 0.698112512 |
| ENSMUSG00000083282 | -1.204123184 | 0.002716623 |
| ENSMUSG00000083283 | -1.03962336 | 0.698112512 |
| ENSMUSG00000083284 | 1.016770329 | 0.943025934 |
| ENSMUSG00000083285 | 1 | |
| ENSMUSG00000083287 | 1 | |
| ENSMUSG00000083288 | -1.077797669 | 0.677534336 |
| ENSMUSG00000083289 | 1 | |
| ENSMUSG00000083290 | 1 | |
| ENSMUSG00000083291 | -1 | 1 |
| ENSMUSG00000083292 | 1 | |
| ENSMUSG00000083293 | 1 | |
| ENSMUSG00000083294 | 1 | |
| ENSMUSG00000083296 | 1 | |
| ENSMUSG00000083297 | 1 | |
| ENSMUSG00000083298 | 1 | |
| ENSMUSG00000083299 | 1 | |
| ENSMUSG00000083300 | 1.095872691 | 0.608497229 |
| ENSMUSG00000083303 | 1 | |
| ENSMUSG00000083304 | 1 | |
| ENSMUSG00000083305 | 1.059463094 | 0.698112512 |
| ENSMUSG00000083306 | 1.069068979 | 0.698112512 |
| ENSMUSG00000083307 | 1.293765693 | 0.024823555 |
| ENSMUSG00000083309 | 1 | |
| ENSMUSG00000083310 | 1 | |
| ENSMUSG00000083311 | 1.025311598 | 1 |
| ENSMUSG00000083312 | 1 | |

| | | |
|---|---|---|
| ENSMUSG00000083314 | -1.126773846 | 0.608497229 |
| ENSMUSG00000083315 | 1 | |
| ENSMUSG00000083316 | -1.053549062 | 0.698112512 |
| ENSMUSG00000083317 | 1 | |
| ENSMUSG00000083318 | 1 | |
| ENSMUSG00000083319 | -1.179744019 | 0.470878221 |
| ENSMUSG00000083320 | -1.133231767 | 0.591442961 |
| ENSMUSG00000083321 | 1 | |
| ENSMUSG00000083324 | 1 | |
| ENSMUSG00000083325 | -1.069502966 | 0.698112512 |
| ENSMUSG00000083326 | -1 | 1 |
| ENSMUSG00000083327 | 1.027703275 | 0.951410231 |
| ENSMUSG00000083328 | 1.015486245 | 0.972980766 |
| ENSMUSG00000083329 | -1 | 1 |
| ENSMUSG00000083330 | 1 | |
| ENSMUSG00000083331 | 1 | |
| ENSMUSG00000083332 | -1.038444538 | 0.865865121 |
| ENSMUSG00000083333 | 1 | |
| ENSMUSG00000083334 | 1 | |
| ENSMUSG00000083335 | 1.019083579 | 0.968937442 |
| ENSMUSG00000083336 | 1 | |
| ENSMUSG00000083337 | 1.159690735 | 0.608497229 |
| ENSMUSG00000083338 | 1 | |
| ENSMUSG00000083339 | -1.01043732 | 1 |
| ENSMUSG00000083340 | 1.122462048 | 0.420161906 |
| ENSMUSG00000083341 | 1 | |
| ENSMUSG00000083342 | 1 | |
| ENSMUSG00000083343 | -1 | 1 |
| ENSMUSG00000083344 | -1.040167329 | 0.911412228 |
| ENSMUSG00000083345 | 1 | |
| ENSMUSG00000083346 | -1 | 1 |
| ENSMUSG00000083347 | 1 | |
| ENSMUSG00000083348 | 1 | |
| ENSMUSG00000083350 | 1 | |
| ENSMUSG00000083351 | 1 | |
| ENSMUSG00000083352 | 1 | |
| ENSMUSG00000083353 | -1.137443589 | 0.608497229 |
| ENSMUSG00000083355 | -1 | 1 |
| ENSMUSG00000083356 | -1 | 1 |
| ENSMUSG00000083358 | 1 | |
| ENSMUSG00000083359 | 1.05410549 | 0.841285672 |
| ENSMUSG00000083361 | 1 | |
| ENSMUSG00000083362 | 1.018884069 | 1 |
| ENSMUSG00000083363 | 1 | |
| ENSMUSG00000083364 | 1 | |
| ENSMUSG00000083365 | 1 | |
| ENSMUSG00000083366 | 1 | |
| ENSMUSG00000083367 | 1.224879776 | 0.446142726 |
| ENSMUSG00000083368 | 1 | |
| ENSMUSG00000083369 | 1 | |
| ENSMUSG00000083371 | 1 | |
| ENSMUSG00000083372 | 1 | |
| ENSMUSG00000083373 | 1 | |
| ENSMUSG00000083374 | -1.115980712 | 0.698112512 |
| ENSMUSG00000083375 | 1.023743961 | 0.95702926 |
| ENSMUSG00000083376 | 1 | |
| ENSMUSG00000083377 | -1.003834322 | 1 |
| ENSMUSG00000083378 | -1.063127349 | 0.773756337 |
| ENSMUSG00000083379 | -1.03962336 | 0.814089245 |
| ENSMUSG00000083380 | 1 | |
| ENSMUSG00000083381 | 1 | |
| ENSMUSG00000083382 | -1.095939461 | 0.693463129 |
| ENSMUSG00000083383 | 1 | |

| | | |
|---|---|---|
| ENSMUSG00000083384 | 1 | |
| ENSMUSG00000083385 | 1.019083579 | 0.968937442 |
| ENSMUSG00000083386 | 1 | |
| ENSMUSG00000083387 | -1.044273782 | 0.698112512 |
| ENSMUSG00000083388 | -1.043306199 | 0.698112512 |
| ENSMUSG00000083389 | 1 | |
| ENSMUSG00000083390 | 1.069068979 | 0.608497229 |
| ENSMUSG00000083391 | 1.158567448 | 0.00908597 |
| ENSMUSG00000083392 | -1.090507733 | 0.698112512 |
| ENSMUSG00000083393 | 1 | |
| ENSMUSG00000083394 | -1.085651418 | 0.608497229 |
| ENSMUSG00000083395 | -1.044273782 | 0.801443616 |
| ENSMUSG00000083396 | -1.003858896 | 1 |
| ENSMUSG00000083397 | 1.07985268 | 0.752077931 |
| ENSMUSG00000083398 | 1 | |
| ENSMUSG00000083399 | 1 | |
| ENSMUSG00000083401 | 1 | |
| ENSMUSG00000083402 | 1 | |
| ENSMUSG00000083403 | 1 | |
| ENSMUSG00000083404 | 1 | |
| ENSMUSG00000083405 | 1 | |
| ENSMUSG00000083406 | 1 | |
| ENSMUSG00000083407 | -1 | 1 |
| ENSMUSG00000083409 | 1 | |
| ENSMUSG00000083410 | 1.014731082 | 0.997667419 |
| ENSMUSG00000083411 | 1 | |
| ENSMUSG00000083412 | 1 | |
| ENSMUSG00000083413 | 1 | |
| ENSMUSG00000083414 | -1.052838398 | 0.684022066 |
| ENSMUSG00000083416 | 1 | |
| ENSMUSG00000083417 | 1 | |
| ENSMUSG00000083418 | 1 | |
| ENSMUSG00000083419 | 1 | |
| ENSMUSG00000083420 | 1 | |
| ENSMUSG00000083421 | 1 | |
| ENSMUSG00000083422 | -1.036180887 | 0.928071154 |
| ENSMUSG00000083423 | 1 | |
| ENSMUSG00000083424 | 1.067678328 | 0.853042921 |
| ENSMUSG00000083425 | -1.084645496 | 0.532369657 |
| ENSMUSG00000083427 | 1 | |
| ENSMUSG00000083428 | 1 | |
| ENSMUSG00000083429 | -1.063525418 | 0.803072249 |
| ENSMUSG00000083430 | 1 | |
| ENSMUSG00000083431 | -1.105673636 | 0.608497229 |
| ENSMUSG00000083432 | 1 | |
| ENSMUSG00000083434 | -1.215713608 | 0.600132627 |
| ENSMUSG00000083435 | 1.019083579 | 0.949812939 |
| ENSMUSG00000083436 | 1 | |
| ENSMUSG00000083437 | 1 | |
| ENSMUSG00000083438 | 1 | |
| ENSMUSG00000083439 | 1 | |
| ENSMUSG00000083440 | -1.010988769 | 1 |
| ENSMUSG00000083441 | 1 | |
| ENSMUSG00000083443 | 1.008056004 | 1 |
| ENSMUSG00000083444 | 1 | |
| ENSMUSG00000083445 | 1 | |
| ENSMUSG00000083446 | 1 | |
| ENSMUSG00000083447 | 1 | |
| ENSMUSG00000083448 | 1 | |
| ENSMUSG00000083449 | 1 | |
| ENSMUSG00000083450 | 1 | |
| ENSMUSG00000083451 | 1 | |
| ENSMUSG00000083452 | 1 | |

| | | |
|---|---|---|
| ENSMUSG00000083453 | 1 | |
| ENSMUSG00000083454 | 1.132639128 | 0.428732467 |
| ENSMUSG00000083455 | 1 | |
| ENSMUSG00000083456 | 1 | |
| ENSMUSG00000083457 | 1 | |
| ENSMUSG00000083458 | 1.059463094 | 0.608497229 |
| ENSMUSG00000083460 | 1.288599435 | 0.468615237 |
| ENSMUSG00000083461 | 1 | |
| ENSMUSG00000083462 | 1 | |
| ENSMUSG00000083463 | 1 | |
| ENSMUSG00000083465 | 1.309558648 | 0.232747283 |
| ENSMUSG00000083466 | 1 | |
| ENSMUSG00000083467 | 1 | |
| ENSMUSG00000083468 | -1.038352922 | 0.959892369 |
| ENSMUSG00000083469 | 1 | |
| ENSMUSG00000083470 | -1.003834322 | 1 |
| ENSMUSG00000083471 | 1.019083579 | 0.949812939 |
| ENSMUSG00000083472 | 1 | |
| ENSMUSG00000083473 | -1 | 1 |
| ENSMUSG00000083474 | 1.092862004 | 0.815423825 |
| ENSMUSG00000083475 | 1 | |
| ENSMUSG00000083477 | -1.136679298 | 0.608497229 |
| ENSMUSG00000083478 | 1 | |
| ENSMUSG00000083481 | 1 | |
| ENSMUSG00000083482 | 1 | |
| ENSMUSG00000083483 | -1 | 1 |
| ENSMUSG00000083484 | 1 | |
| ENSMUSG00000083485 | -1 | 1 |
| ENSMUSG00000083486 | 1 | |
| ENSMUSG00000083487 | 1 | |
| ENSMUSG00000083488 | 1 | |
| ENSMUSG00000083489 | 1 | |
| ENSMUSG00000083490 | 1 | |
| ENSMUSG00000083491 | 1 | |
| ENSMUSG00000083492 | 1 | |
| ENSMUSG00000083493 | 1 | |
| ENSMUSG00000083494 | 1 | |
| ENSMUSG00000083495 | 1 | |
| ENSMUSG00000083496 | -1.033395294 | 0.98145432 |
| ENSMUSG00000083498 | -1 | 1 |
| ENSMUSG00000083499 | 1 | |
| ENSMUSG00000083500 | 1.033233219 | 0.972980766 |
| ENSMUSG00000083501 | 1 | |
| ENSMUSG00000083502 | 1 | |
| ENSMUSG00000083503 | 1.028323353 | 0.903464611 |
| ENSMUSG00000083505 | 1.002938927 | 1 |
| ENSMUSG00000083507 | 1.005613439 | 1 |
| ENSMUSG00000083508 | 1.069068979 | 0.608497229 |
| ENSMUSG00000083509 | 1 | |
| ENSMUSG00000083510 | 1 | |
| ENSMUSG00000083512 | -1.005725456 | 1 |
| ENSMUSG00000083513 | 1 | |
| ENSMUSG00000083514 | 1 | |
| ENSMUSG00000083515 | 1 | |
| ENSMUSG00000083516 | 1 | |
| ENSMUSG00000083517 | 1 | |
| ENSMUSG00000083518 | -1 | 1 |
| ENSMUSG00000083519 | 1 | |
| ENSMUSG00000083520 | 1 | |
| ENSMUSG00000083522 | 1.019083579 | 0.949812939 |
| ENSMUSG00000083524 | 1 | |
| ENSMUSG00000083527 | 1 | |
| ENSMUSG00000083528 | -1.023769022 | 0.934760809 |

| | | |
|---|---|---|
| ENSMUSG00000083530 | 1 | |
| ENSMUSG00000083533 | 1 | |
| ENSMUSG00000083534 | -1.071075483 | 0.743300655 |
| ENSMUSG00000083535 | 1 | |
| ENSMUSG00000083536 | 1 | |
| ENSMUSG00000083537 | 1 | |
| ENSMUSG00000083538 | -1.023769022 | 0.934760809 |
| ENSMUSG00000083539 | -1.009703678 | 1 |
| ENSMUSG00000083541 | 1 | |
| ENSMUSG00000083542 | 1 | |
| ENSMUSG00000083543 | 1 | |
| ENSMUSG00000083544 | 1 | |
| ENSMUSG00000083545 | 1 | |
| ENSMUSG00000083546 | 1 | |
| ENSMUSG00000083547 | -1.198658038 | 0.36135224 |
| ENSMUSG00000083548 | -1.085651418 | 0.532369657 |
| ENSMUSG00000083549 | 1 | |
| ENSMUSG00000083550 | 1 | |
| ENSMUSG00000083551 | 1.069068979 | 0.698112512 |
| ENSMUSG00000083553 | 1.165816028 | 0.432387542 |
| ENSMUSG00000083554 | -1.03962336 | 0.698112512 |
| ENSMUSG00000083555 | 1 | |
| ENSMUSG00000083556 | -1.044273782 | 0.790851299 |
| ENSMUSG00000083557 | 1 | |
| ENSMUSG00000083558 | 1 | |
| ENSMUSG00000083560 | 1 | |
| ENSMUSG00000083561 | -1.03962336 | 0.698112512 |
| ENSMUSG00000083562 | 1 | |
| ENSMUSG00000083563 | 1.027729568 | 0.812228807 |
| ENSMUSG00000083564 | 1 | |
| ENSMUSG00000083565 | 1 | |
| ENSMUSG00000083566 | 1 | |
| ENSMUSG00000083567 | -1.044273782 | 0.698112512 |
| ENSMUSG00000083568 | 1 | |
| ENSMUSG00000083569 | 1.057478344 | 0.837014159 |
| ENSMUSG00000083570 | 1 | |
| ENSMUSG00000083572 | 1 | |
| ENSMUSG00000083573 | -1 | 1 |
| ENSMUSG00000083574 | 1 | |
| ENSMUSG00000083576 | 1 | |
| ENSMUSG00000083577 | 1 | |
| ENSMUSG00000083579 | 1.095872691 | 0.657060635 |
| ENSMUSG00000083580 | 1.106093673 | 0.547016491 |
| ENSMUSG00000083581 | 1 | |
| ENSMUSG00000083582 | -1 | 1 |
| ENSMUSG00000083584 | 1 | |
| ENSMUSG00000083585 | -1.043306199 | 0.698112512 |
| ENSMUSG00000083586 | 1.25992105 | 0.215991492 |
| ENSMUSG00000083587 | -1.043306199 | 0.698112512 |
| ENSMUSG00000083588 | 1.100529502 | 0.629100036 |
| ENSMUSG00000083590 | 1 | |
| ENSMUSG00000083592 | 1.059463094 | 0.608497229 |
| ENSMUSG00000083594 | -1.01936322 | 0.950602516 |
| ENSMUSG00000083595 | -1 | 1 |
| ENSMUSG00000083596 | -1.009476377 | 1 |
| ENSMUSG00000083597 | 1 | |
| ENSMUSG00000083598 | 1 | |
| ENSMUSG00000083599 | 1 | |
| ENSMUSG00000083601 | 1 | |
| ENSMUSG00000083602 | 1 | |
| ENSMUSG00000083603 | 1.059463094 | 0.704170349 |
| ENSMUSG00000083605 | -1.043306199 | 0.698112512 |
| ENSMUSG00000083606 | -1.03962336 | 0.698112512 |

| | | |
|---|---|---|
| ENSMUSG00000083607 | 1 | |
| ENSMUSG00000083608 | 1.029400735 | 0.982838256 |
| ENSMUSG00000083609 | 1 | |
| ENSMUSG00000083610 | -1.107691188 | 0.258650416 |
| ENSMUSG00000083611 | -1.03962336 | 0.803072249 |
| ENSMUSG00000083613 | 1 | |
| ENSMUSG00000083614 | 1 | |
| ENSMUSG00000083615 | 1 | |
| ENSMUSG00000083616 | 1 | |
| ENSMUSG00000083617 | 1 | |
| ENSMUSG00000083618 | 1.122462048 | 0.215991492 |
| ENSMUSG00000083619 | 1 | |
| ENSMUSG00000083620 | 1 | |
| ENSMUSG00000083621 | -1.037482356 | 0.89979145 |
| ENSMUSG00000083622 | -1 | 1 |
| ENSMUSG00000083623 | -1.048909253 | 0.854827467 |
| ENSMUSG00000083624 | 1 | |
| ENSMUSG00000083625 | 1 | |
| ENSMUSG00000083626 | 1 | |
| ENSMUSG00000083627 | 1 | |
| ENSMUSG00000083628 | 1 | |
| ENSMUSG00000083629 | 1 | |
| ENSMUSG00000083630 | -1.128668575 | 0.578221424 |
| ENSMUSG00000083631 | 1 | |
| ENSMUSG00000083632 | 1 | |
| ENSMUSG00000083633 | 1.095658148 | 0.713921439 |
| ENSMUSG00000083635 | 1 | |
| ENSMUSG00000083636 | 1 | |
| ENSMUSG00000083637 | 1 | |
| ENSMUSG00000083640 | -1.150809442 | 0.470960596 |
| ENSMUSG00000083641 | 1 | |
| ENSMUSG00000083642 | 1 | |
| ENSMUSG00000083643 | 1 | |
| ENSMUSG00000083644 | 1 | |
| ENSMUSG00000083645 | 1 | |
| ENSMUSG00000083646 | -1.085651418 | 0.608497229 |
| ENSMUSG00000083648 | 1 | |
| ENSMUSG00000083649 | 1.105638764 | 0.782144932 |
| ENSMUSG00000083650 | 1.069633604 | 0.749276359 |
| ENSMUSG00000083651 | -1.075880679 | 0.846072213 |
| ENSMUSG00000083653 | 1 | |
| ENSMUSG00000083654 | -1 | 1 |
| ENSMUSG00000083655 | 1 | |
| ENSMUSG00000083657 | 1 | |
| ENSMUSG00000083658 | 1.127832095 | 0.608497229 |
| ENSMUSG00000083659 | -1.112433816 | 0.656734444 |
| ENSMUSG00000083660 | 1 | |
| ENSMUSG00000083661 | 1.059463094 | 0.608497229 |
| ENSMUSG00000083662 | 1 | |
| ENSMUSG00000083663 | 1 | |
| ENSMUSG00000083664 | 1 | |
| ENSMUSG00000083665 | -1 | 1 |
| ENSMUSG00000083666 | 1 | |
| ENSMUSG00000083667 | 1 | |
| ENSMUSG00000083668 | 1.122462048 | 0.420161906 |
| ENSMUSG00000083669 | 1.059463094 | 0.608497229 |
| ENSMUSG00000083670 | 1 | |
| ENSMUSG00000083671 | 1 | |
| ENSMUSG00000083672 | 1 | |
| ENSMUSG00000083673 | 1 | |
| ENSMUSG00000083674 | -1.48300577 | 0.048604265 |
| ENSMUSG00000083676 | 1.015486245 | 0.985522133 |
| ENSMUSG00000083677 | 1.059463094 | 0.698112512 |

| | | |
|---|---|---|
| ENSMUSG00000083678 | -1.145518406 | 0.532369657 |
| ENSMUSG00000083679 | -1.03962336 | 0.698112512 |
| ENSMUSG00000083680 | 1 | |
| ENSMUSG00000083681 | -1.024718486 | 0.931480186 |
| ENSMUSG00000083682 | 1 | |
| ENSMUSG00000083683 | 1 | |
| ENSMUSG00000083684 | -1 | 1 |
| ENSMUSG00000083685 | 1 | |
| ENSMUSG00000083686 | 1.059463094 | 0.698112512 |
| ENSMUSG00000083687 | 1 | |
| ENSMUSG00000083688 | -1.043306199 | 0.698112512 |
| ENSMUSG00000083689 | 1.019083579 | 0.971519533 |
| ENSMUSG00000083691 | -1 | 1 |
| ENSMUSG00000083692 | 1.030661948 | 0.960598161 |
| ENSMUSG00000083693 | -1.075365073 | 0.859576852 |
| ENSMUSG00000083695 | 1 | |
| ENSMUSG00000083697 | 1 | |
| ENSMUSG00000083698 | 1 | |
| ENSMUSG00000083699 | 1 | |
| ENSMUSG00000083700 | -1.021243822 | 0.947046509 |
| ENSMUSG00000083701 | 1 | |
| ENSMUSG00000083702 | 1 | |
| ENSMUSG00000083704 | 1 | |
| ENSMUSG00000083705 | -1.064815992 | 0.575391632 |
| ENSMUSG00000083706 | 1 | |
| ENSMUSG00000083707 | -1.053549062 | 0.698112512 |
| ENSMUSG00000083708 | 1.172010364 | 0.698112512 |
| ENSMUSG00000083709 | 1 | |
| ENSMUSG00000083710 | 1.059463094 | 0.608497229 |
| ENSMUSG00000083711 | 1 | |
| ENSMUSG00000083713 | -1.093651872 | 0.793726432 |
| ENSMUSG00000083714 | 1 | |
| ENSMUSG00000083715 | 1 | |
| ENSMUSG00000083716 | 1.054707116 | 0.802748054 |
| ENSMUSG00000083717 | 1 | |
| ENSMUSG00000083718 | -1.067187828 | 0.70299173 |
| ENSMUSG00000083719 | 1 | |
| ENSMUSG00000083720 | -1 | 1 |
| ENSMUSG00000083721 | 1 | |
| ENSMUSG00000083722 | -1 | 1 |
| ENSMUSG00000083723 | 1 | |
| ENSMUSG00000083725 | -1.099174267 | 0.535052277 |
| ENSMUSG00000083728 | 1 | |
| ENSMUSG00000083729 | 1 | |
| ENSMUSG00000083731 | -1.024530911 | 0.949679283 |
| ENSMUSG00000083732 | -1.182307728 | 0.583995634 |
| ENSMUSG00000083733 | 1 | |
| ENSMUSG00000083734 | -1.261186606 | 0.075805106 |
| ENSMUSG00000083735 | 1 | |
| ENSMUSG00000083736 | -1.00571839 | 1 |
| ENSMUSG00000083737 | 1 | |
| ENSMUSG00000083738 | -1 | 1 |
| ENSMUSG00000083740 | 1 | |
| ENSMUSG00000083741 | 1 | |
| ENSMUSG00000083742 | 1 | |
| ENSMUSG00000083743 | -1.103388856 | 0.608497229 |
| ENSMUSG00000083744 | 1 | |
| ENSMUSG00000083745 | 1.069068979 | 0.608497229 |
| ENSMUSG00000083747 | 1 | |
| ENSMUSG00000083748 | 1.26712279 | 0.125227464 |
| ENSMUSG00000083749 | 1 | |
| ENSMUSG00000083750 | 1.052248089 | 0.828707904 |
| ENSMUSG00000083751 | 1 | |

| | | |
|---|---|---|
| ENSMUSG00000083752 | 1 | |
| ENSMUSG00000083753 | 1 | |
| ENSMUSG00000083754 | 1.156216856 | 0.484875549 |
| ENSMUSG00000083755 | 1 | |
| ENSMUSG00000083757 | 1.02889509 | 0.971248775 |
| ENSMUSG00000083760 | -1 | 1 |
| ENSMUSG00000083761 | 1.038531341 | 0.949812939 |
| ENSMUSG00000083763 | 1.199989356 | 0.062913564 |
| ENSMUSG00000083764 | -1.040769408 | 0.774697482 |
| ENSMUSG00000083766 | 1 | |
| ENSMUSG00000083767 | 1.075870199 | 0.642819974 |
| ENSMUSG00000083768 | 1 | |
| ENSMUSG00000083769 | 1 | |
| ENSMUSG00000083771 | 1 | |
| ENSMUSG00000083772 | -1.065321276 | 0.79291651 |
| ENSMUSG00000083773 | 1.003468652 | 1 |
| ENSMUSG00000083774 | 1 | |
| ENSMUSG00000083775 | 1 | |
| ENSMUSG00000083776 | 1 | |
| ENSMUSG00000083777 | 1.022120194 | 0.893132339 |
| ENSMUSG00000083778 | 1 | |
| ENSMUSG00000083779 | -1.038444538 | 0.865865121 |
| ENSMUSG00000083780 | 1.152944534 | 0.296081577 |
| ENSMUSG00000083781 | 1 | |
| ENSMUSG00000083782 | 1 | |
| ENSMUSG00000083783 | -1.028348525 | 0.936995747 |
| ENSMUSG00000083784 | 1 | |
| ENSMUSG00000083785 | 1.059463094 | 0.608497229 |
| ENSMUSG00000083786 | 1 | |
| ENSMUSG00000083787 | 1 | |
| ENSMUSG00000083789 | 1 | |
| ENSMUSG00000083791 | 1 | |
| ENSMUSG00000083792 | 1 | |
| ENSMUSG00000083793 | 1.138997129 | 0.288134604 |
| ENSMUSG00000083794 | -1.084645496 | 0.532369657 |
| ENSMUSG00000083795 | 1 | |
| ENSMUSG00000083796 | 1.057700516 | 0.881811332 |
| ENSMUSG00000083798 | 1.205696885 | 0.158707421 |
| ENSMUSG00000083801 | 1 | |
| ENSMUSG00000083803 | 1.059463094 | 0.608497229 |
| ENSMUSG00000083804 | -1.044273782 | 0.698112512 |
| ENSMUSG00000083805 | 1.24122829 | 0.001720654 |
| ENSMUSG00000083806 | 1 | |
| ENSMUSG00000083807 | 1 | |
| ENSMUSG00000083808 | 1 | |
| ENSMUSG00000083809 | 1 | |
| ENSMUSG00000083810 | 1 | |
| ENSMUSG00000083811 | 1.060964082 | 0.810829358 |
| ENSMUSG00000083812 | 1.084618945 | 0.698112512 |
| ENSMUSG00000083813 | -1 | 1 |
| ENSMUSG00000083814 | 1 | |
| ENSMUSG00000083815 | -1.043306199 | 0.903448666 |
| ENSMUSG00000083816 | 1.105736066 | 0.659456205 |
| ENSMUSG00000083817 | -1.145518406 | 0.603504151 |
| ENSMUSG00000083818 | 1 | |
| ENSMUSG00000083819 | -1.082743957 | 0.775673129 |
| ENSMUSG00000083820 | -1.03962336 | 0.698112512 |
| ENSMUSG00000083821 | 1 | |
| ENSMUSG00000083822 | -1.066079037 | 0.818205891 |
| ENSMUSG00000083823 | 1.085624843 | 0.731830521 |
| ENSMUSG00000083825 | 1 | |
| ENSMUSG00000083826 | -1.044932672 | 0.956784261 |
| ENSMUSG00000083827 | 1 | |

| | | |
|---|---|---|
| ENSMUSG00000083828 | 1 | |
| ENSMUSG00000083829 | 1.071500237 | 0.819799177 |
| ENSMUSG00000083830 | 1.011607416 | 1 |
| ENSMUSG00000083831 | 1 | |
| ENSMUSG00000083832 | -1.054741498 | 0.842184576 |
| ENSMUSG00000083833 | 1 | |
| ENSMUSG00000083834 | 1.089470641 | 0.608497229 |
| ENSMUSG00000083835 | 1 | |
| ENSMUSG00000083836 | 1.018670051 | 0.998126414 |
| ENSMUSG00000083837 | -1.03962336 | 0.698112512 |
| ENSMUSG00000083838 | -1.085651418 | 0.532369657 |
| ENSMUSG00000083839 | 1 | |
| ENSMUSG00000083841 | -1.240391029 | 0.283365781 |
| ENSMUSG00000083842 | 1 | |
| ENSMUSG00000083843 | 1 | |
| ENSMUSG00000083844 | 1.052220693 | 0.692190045 |
| ENSMUSG00000083845 | -1.063525418 | 0.698112512 |
| ENSMUSG00000083846 | 1 | |
| ENSMUSG00000083847 | 1 | |
| ENSMUSG00000083848 | 1 | |
| ENSMUSG00000083849 | 1.141935216 | 0.407154251 |
| ENSMUSG00000083851 | 1 | |
| ENSMUSG00000083852 | 1.065410325 | 0.805198503 |
| ENSMUSG00000083853 | 1.036775755 | 0.803173793 |
| ENSMUSG00000083854 | 1 | |
| ENSMUSG00000083855 | 1 | |
| ENSMUSG00000083856 | -1.006948462 | 1 |
| ENSMUSG00000083857 | 1 | |
| ENSMUSG00000083858 | 1 | |
| ENSMUSG00000083859 | -1.078590893 | 0.647384212 |
| ENSMUSG00000083860 | -1.043306199 | 0.803260194 |
| ENSMUSG00000083861 | 1 | |
| ENSMUSG00000083863 | -1.149817178 | 0.645849205 |
| ENSMUSG00000083864 | -1.03962336 | 0.698112512 |
| ENSMUSG00000083866 | 1 | |
| ENSMUSG00000083867 | 1 | |
| ENSMUSG00000083868 | 1 | |
| ENSMUSG00000083869 | 1 | |
| ENSMUSG00000083870 | 1 | |
| ENSMUSG00000083871 | 1.059463094 | 0.608497229 |
| ENSMUSG00000083872 | 1 | |
| ENSMUSG00000083873 | 1 | |
| ENSMUSG00000083874 | 1 | |
| ENSMUSG00000083875 | 1 | |
| ENSMUSG00000083876 | 1 | |
| ENSMUSG00000083877 | 1.059463094 | 0.698112512 |
| ENSMUSG00000083878 | 1.019083579 | 0.97045957 |
| ENSMUSG00000083879 | 1.059463094 | 0.608497229 |
| ENSMUSG00000083880 | -1.07915597 | 0.608497229 |
| ENSMUSG00000083882 | 1 | |
| ENSMUSG00000083884 | -1.044273782 | 0.790851299 |
| ENSMUSG00000083885 | 1.019083579 | 0.949812939 |
| ENSMUSG00000083886 | 1 | |
| ENSMUSG00000083887 | 1 | |
| ENSMUSG00000083888 | 1 | |
| ENSMUSG00000083889 | 1 | |
| ENSMUSG00000083890 | -1.568102269 | 0.042658974 |
| ENSMUSG00000083892 | -1.039467461 | 0.906152436 |
| ENSMUSG00000083893 | 1 | |
| ENSMUSG00000083894 | 1 | |
| ENSMUSG00000083895 | 1 | |
| ENSMUSG00000083896 | 1 | |
| ENSMUSG00000083897 | -1.03962336 | 0.698112512 |

| | | |
|---|---|---|
| ENSMUSG00000083898 | 1 | |
| ENSMUSG00000083899 | -1.02927417 | 0.854777354 |
| ENSMUSG00000083900 | 1.112843644 | 0.66278123 |
| ENSMUSG00000083901 | 1 | |
| ENSMUSG00000083902 | -1.091374921 | 0.552584793 |
| ENSMUSG00000083903 | -1.053549062 | 0.698112512 |
| ENSMUSG00000083904 | 1 | |
| ENSMUSG00000083905 | 1.059463094 | 0.608497229 |
| ENSMUSG00000083906 | -1.053296659 | 0.891297795 |
| ENSMUSG00000083907 | 1.030641397 | 0.850218968 |
| ENSMUSG00000083908 | 1 | |
| ENSMUSG00000083909 | 1.095872691 | 0.671280603 |
| ENSMUSG00000083910 | -1.089814156 | 0.777145624 |
| ENSMUSG00000083911 | 1.12691148 | 0.608497229 |
| ENSMUSG00000083912 | 1.014545335 | 0.990831634 |
| ENSMUSG00000083913 | 1.122462048 | 0.628993873 |
| ENSMUSG00000083914 | 1.049411284 | 0.861198669 |
| ENSMUSG00000083915 | 1.12691148 | 0.688885668 |
| ENSMUSG00000083916 | 1 | |
| ENSMUSG00000083917 | -1.028348525 | 0.912425817 |
| ENSMUSG00000083918 | 1 | |
| ENSMUSG00000083919 | 1 | |
| ENSMUSG00000083920 | 1 | |
| ENSMUSG00000083921 | 1.09538577 | 0.785584885 |
| ENSMUSG00000083922 | -1.051577863 | 0.911548744 |
| ENSMUSG00000083923 | 1 | |
| ENSMUSG00000083924 | -1 | 1 |
| ENSMUSG00000083925 | 1 | |
| ENSMUSG00000083926 | 1 | |
| ENSMUSG00000083927 | 1.11166259 | 0.698112512 |
| ENSMUSG00000083928 | 1.112006103 | 0.72715392 |
| ENSMUSG00000083929 | 1 | |
| ENSMUSG00000083930 | 1 | |
| ENSMUSG00000083931 | 1 | |
| ENSMUSG00000083932 | 1 | |
| ENSMUSG00000083933 | 1.157087696 | 0.599948532 |
| ENSMUSG00000083934 | 1 | |
| ENSMUSG00000083935 | -1.012517664 | 1 |
| ENSMUSG00000083936 | 1 | |
| ENSMUSG00000083937 | 1.019083579 | 0.986542466 |
| ENSMUSG00000083938 | 1 | |
| ENSMUSG00000083939 | 1 | |
| ENSMUSG00000083940 | -1.03962336 | 0.698112512 |
| ENSMUSG00000083941 | 1 | |
| ENSMUSG00000083942 | 1 | |
| ENSMUSG00000083943 | -1.182814301 | 0.316670006 |
| ENSMUSG00000083944 | 1 | |
| ENSMUSG00000083945 | 1 | |
| ENSMUSG00000083946 | 1 | |
| ENSMUSG00000083947 | 1 | |
| ENSMUSG00000083948 | 1 | |
| ENSMUSG00000083949 | 1 | |
| ENSMUSG00000083950 | 1 | |
| ENSMUSG00000083951 | 1.015486245 | 0.987775262 |
| ENSMUSG00000083953 | 1.034095782 | 0.94160226 |
| ENSMUSG00000083954 | 1 | |
| ENSMUSG00000083955 | 1 | |
| ENSMUSG00000083956 | 1 | |
| ENSMUSG00000083957 | 1 | |
| ENSMUSG00000083958 | 1 | |
| ENSMUSG00000083960 | 1 | |
| ENSMUSG00000083961 | 1 | |
| ENSMUSG00000083962 | 1.024693402 | 0.972980766 |

| | | |
|---|---|---|
| ENSMUSG00000083963 | 1 | |
| ENSMUSG00000083964 | -1.018857002 | 0.989629241 |
| ENSMUSG00000083966 | 1 | |
| ENSMUSG00000083967 | 1 | |
| ENSMUSG00000083969 | 1 | |
| ENSMUSG00000083970 | 1 | |
| ENSMUSG00000083971 | 1.023743961 | 0.938939969 |
| ENSMUSG00000083972 | 1 | |
| ENSMUSG00000083973 | 1 | |
| ENSMUSG00000083974 | 1 | |
| ENSMUSG00000083975 | 1 | |
| ENSMUSG00000083976 | 1 | |
| ENSMUSG00000083977 | 1 | |
| ENSMUSG00000083978 | 1.019083579 | 0.949812939 |
| ENSMUSG00000083979 | 1 | |
| ENSMUSG00000083981 | 1 | |
| ENSMUSG00000083982 | 1.069003846 | 0.697178241 |
| ENSMUSG00000083983 | 1 | |
| ENSMUSG00000083985 | 1.059463094 | 0.608497229 |
| ENSMUSG00000083986 | 1 | |
| ENSMUSG00000083987 | 1 | |
| ENSMUSG00000083988 | 1.095872691 | 0.608497229 |
| ENSMUSG00000083989 | -1.099174267 | 0.608497229 |
| ENSMUSG00000083990 | 1 | |
| ENSMUSG00000083991 | 1 | |
| ENSMUSG00000083992 | -1.068169133 | 0.634024538 |
| ENSMUSG00000083993 | 1.006831432 | 1 |
| ENSMUSG00000083994 | -1.134282749 | 0.608497229 |
| ENSMUSG00000083995 | 1 | |
| ENSMUSG00000083996 | 1 | |
| ENSMUSG00000083997 | -1.08081673 | 0.73636577 |
| ENSMUSG00000083998 | -1.093674795 | 0.685046738 |
| ENSMUSG00000083999 | 1 | |
| ENSMUSG00000084000 | 1.059463094 | 0.608497229 |
| ENSMUSG00000084001 | 1 | |
| ENSMUSG00000084004 | -1.071075483 | 0.698112512 |
| ENSMUSG00000084006 | 1 | |
| ENSMUSG00000084007 | -1.16470229 | 0.609775718 |
| ENSMUSG00000084009 | 1 | |
| ENSMUSG00000084010 | 1 | |
| ENSMUSG00000084011 | -1.283574061 | 0.15199646 |
| ENSMUSG00000084012 | -1 | 1 |
| ENSMUSG00000084013 | 1 | |
| ENSMUSG00000084014 | 1 | |
| ENSMUSG00000084015 | -1.053549062 | 0.698112512 |
| ENSMUSG00000084016 | 1 | |
| ENSMUSG00000084017 | -1.059302952 | 0.872692845 |
| ENSMUSG00000084018 | 1 | |
| ENSMUSG00000084019 | 1 | |
| ENSMUSG00000084020 | 1 | |
| ENSMUSG00000084021 | -1.053549062 | 0.698112512 |
| ENSMUSG00000084022 | -1.275539202 | 0.184372783 |
| ENSMUSG00000084024 | 1 | |
| ENSMUSG00000084025 | 1.039816326 | 0.883047064 |
| ENSMUSG00000084026 | 1 | |
| ENSMUSG00000084027 | 1 | |
| ENSMUSG00000084028 | -1 | 1 |
| ENSMUSG00000084029 | 1 | |
| ENSMUSG00000084030 | 1 | |
| ENSMUSG00000084031 | 1.164605381 | 0.16140551 |
| ENSMUSG00000084032 | 1 | |
| ENSMUSG00000084033 | -1.066473979 | 0.677534336 |
| ENSMUSG00000084034 | -1 | 1 |

| | | |
|---|---|---|
| ENSMUSG00000084038 | 1 | |
| ENSMUSG00000084039 | 1 | |
| ENSMUSG00000084041 | 1 | |
| ENSMUSG00000084043 | 1 | |
| ENSMUSG00000084044 | 1 | |
| ENSMUSG00000084045 | 1 | |
| ENSMUSG00000084046 | 1 | |
| ENSMUSG00000084047 | -1.084645496 | 0.608497229 |
| ENSMUSG00000084048 | 1 | |
| ENSMUSG00000084050 | -1 | 1 |
| ENSMUSG00000084052 | -1 | 1 |
| ENSMUSG00000084053 | 1 | |
| ENSMUSG00000084054 | 1 | |
| ENSMUSG00000084055 | 1 | |
| ENSMUSG00000084056 | 1 | |
| ENSMUSG00000084057 | 1 | |
| ENSMUSG00000084058 | 1 | |
| ENSMUSG00000084059 | 1 | |
| ENSMUSG00000084060 | 1 | |
| ENSMUSG00000084061 | 1 | |
| ENSMUSG00000084063 | -1.03962336 | 0.698112512 |
| ENSMUSG00000084064 | 1 | |
| ENSMUSG00000084065 | 1 | |
| ENSMUSG00000084066 | 1 | |
| ENSMUSG00000084067 | -1.013449573 | 1 |
| ENSMUSG00000084068 | 1 | |
| ENSMUSG00000084069 | -1.03962336 | 0.803072249 |
| ENSMUSG00000084070 | 1 | |
| ENSMUSG00000084071 | 1 | |
| ENSMUSG00000084072 | 1 | |
| ENSMUSG00000084073 | -1 | 1 |
| ENSMUSG00000084074 | 1 | |
| ENSMUSG00000084076 | 1 | |
| ENSMUSG00000084077 | 1 | |
| ENSMUSG00000084078 | 1 | |
| ENSMUSG00000084079 | 1.029997013 | 0.959463747 |
| ENSMUSG00000084080 | 1 | |
| ENSMUSG00000084081 | 1.034388405 | 0.848848499 |
| ENSMUSG00000084082 | 1 | |
| ENSMUSG00000084083 | 1.014545335 | 0.989648415 |
| ENSMUSG00000084085 | 1.069068979 | 0.608497229 |
| ENSMUSG00000084086 | 1.059463094 | 0.608497229 |
| ENSMUSG00000084087 | 1.064870248 | 0.803260194 |
| ENSMUSG00000084088 | 1.202214291 | 0.356153868 |
| ENSMUSG00000084089 | 1 | |
| ENSMUSG00000084090 | 1 | |
| ENSMUSG00000084092 | 1 | |
| ENSMUSG00000084093 | 1 | |
| ENSMUSG00000084094 | -1.043306199 | 0.699340382 |
| ENSMUSG00000084095 | 1 | |
| ENSMUSG00000084096 | -1.043306199 | 0.698112512 |
| ENSMUSG00000084097 | 1 | |
| ENSMUSG00000084098 | -1.099174267 | 0.535052277 |
| ENSMUSG00000084099 | 1.059463094 | 0.608497229 |
| ENSMUSG00000084101 | -1 | 1 |
| ENSMUSG00000084102 | 1 | |
| ENSMUSG00000084103 | 1 | |
| ENSMUSG00000084104 | -1.11061171 | 0.547016491 |
| ENSMUSG00000084105 | 1.008034876 | 1 |
| ENSMUSG00000084106 | 1.133498461 | 0.608497229 |
| ENSMUSG00000084107 | 1 | |
| ENSMUSG00000084108 | 1 | |
| ENSMUSG00000084110 | 1 | |

| | | |
|---|---|---|
| ENSMUSG00000084111 | -1.003834322 | 1 |
| ENSMUSG00000084112 | 1 | |
| ENSMUSG00000084113 | -1.164894189 | 0.608497229 |
| ENSMUSG00000084114 | 1 | |
| ENSMUSG00000084115 | 1.069068979 | 0.608497229 |
| ENSMUSG00000084116 | 1 | |
| ENSMUSG00000084117 | 1 | |
| ENSMUSG00000084118 | 1 | |
| ENSMUSG00000084119 | 1.059463094 | 0.608497229 |
| ENSMUSG00000084120 | -1.095294215 | 0.608497229 |
| ENSMUSG00000084123 | 1 | |
| ENSMUSG00000084124 | -1.084645496 | 0.628300894 |
| ENSMUSG00000084125 | -1.054741498 | 0.813450442 |
| ENSMUSG00000084126 | 1 | |
| ENSMUSG00000084127 | 1 | |
| ENSMUSG00000084128 | -1.193439672 | 0.13621134 |
| ENSMUSG00000084129 | 1 | |
| ENSMUSG00000084130 | 1 | |
| ENSMUSG00000084131 | 1.224258861 | 0.52619224 |
| ENSMUSG00000084132 | -1.085651418 | 0.532369657 |
| ENSMUSG00000084133 | -1.014570171 | 1 |
| ENSMUSG00000084134 | 1 | |
| ENSMUSG00000084135 | 1 | |
| ENSMUSG00000084136 | 1 | |
| ENSMUSG00000084137 | -1.06804853 | 0.837704961 |
| ENSMUSG00000084138 | 1 | |
| ENSMUSG00000084139 | -1.018054237 | 1 |
| ENSMUSG00000084140 | 1.024284648 | 0.953521521 |
| ENSMUSG00000084142 | -1.086192882 | 0.698112512 |
| ENSMUSG00000084143 | -1 | 1 |
| ENSMUSG00000084144 | -1.014570171 | 0.996048848 |
| ENSMUSG00000084145 | -1.081225398 | 0.736695141 |
| ENSMUSG00000084146 | 1.085852928 | 0.723630549 |
| ENSMUSG00000084147 | 1 | |
| ENSMUSG00000084148 | 1.014346713 | 1 |
| ENSMUSG00000084149 | 1.025197225 | 0.979002724 |
| ENSMUSG00000084150 | 1 | |
| ENSMUSG00000084151 | -1.053549062 | 0.698112512 |
| ENSMUSG00000084152 | 1 | |
| ENSMUSG00000084153 | 1.059463094 | 0.704170349 |
| ENSMUSG00000084154 | 1 | |
| ENSMUSG00000084155 | 1 | |
| ENSMUSG00000084157 | 1.059463094 | 0.608497229 |
| ENSMUSG00000084158 | 1 | |
| ENSMUSG00000084159 | -1.08768245 | 0.532586 |
| ENSMUSG00000084160 | 1 | |
| ENSMUSG00000084161 | 1 | |
| ENSMUSG00000084162 | 1 | |
| ENSMUSG00000084163 | -1.069095149 | 0.686194514 |
| ENSMUSG00000084164 | -1.096427876 | 0.735417419 |
| ENSMUSG00000084166 | 1.132877091 | 0.608497229 |
| ENSMUSG00000084167 | 1 | |
| ENSMUSG00000084168 | 1 | |
| ENSMUSG00000084169 | 1 | |
| ENSMUSG00000084170 | 1 | |
| ENSMUSG00000084171 | 1 | |
| ENSMUSG00000084172 | 1 | |
| ENSMUSG00000084173 | 1 | |
| ENSMUSG00000084174 | 1 | |
| ENSMUSG00000084175 | 1 | |
| ENSMUSG00000084177 | -1.054169715 | 0.862822188 |
| ENSMUSG00000084178 | -1.248663974 | 0.258495519 |
| ENSMUSG00000084179 | 1 | |

| | | |
|---|---|---|
| ENSMUSG00000084180 | 1 | |
| ENSMUSG00000084181 | 1 | |
| ENSMUSG00000084182 | 1 | |
| ENSMUSG00000084183 | 1.120159739 | 0.698112512 |
| ENSMUSG00000084184 | -1.03962336 | 0.698112512 |
| ENSMUSG00000084185 | 1.069068979 | 0.608497229 |
| ENSMUSG00000084186 | 1 | |
| ENSMUSG00000084188 | 1 | |
| ENSMUSG00000084189 | 1 | |
| ENSMUSG00000084190 | -1.024304696 | 0.98026203 |
| ENSMUSG00000084191 | 1 | |
| ENSMUSG00000084192 | 1 | |
| ENSMUSG00000084193 | 1 | |
| ENSMUSG00000084195 | 1 | |
| ENSMUSG00000084196 | 1 | |
| ENSMUSG00000084197 | 1 | |
| ENSMUSG00000084198 | 1 | |
| ENSMUSG00000084199 | 1.069068979 | 0.608497229 |
| ENSMUSG00000084200 | 1 | |
| ENSMUSG00000084201 | 1 | |
| ENSMUSG00000084202 | 1 | |
| ENSMUSG00000084203 | -1.027230783 | 0.979002724 |
| ENSMUSG00000084204 | 1 | |
| ENSMUSG00000084205 | 1 | |
| ENSMUSG00000084206 | 1 | |
| ENSMUSG00000084207 | 1.059463094 | 0.608497229 |
| ENSMUSG00000084208 | 1.019083579 | 0.949812939 |
| ENSMUSG00000084209 | -1 | 1 |
| ENSMUSG00000084210 | 1 | |
| ENSMUSG00000084211 | 1.095872691 | 0.657060635 |
| ENSMUSG00000084212 | 1 | |
| ENSMUSG00000084213 | -1.044273782 | 0.801443616 |
| ENSMUSG00000084214 | 1 | |
| ENSMUSG00000084215 | -1.003834322 | 1 |
| ENSMUSG00000084216 | -1.034935882 | 0.968830305 |
| ENSMUSG00000084217 | -1.127622795 | 0.608497229 |
| ENSMUSG00000084218 | 1.059463094 | 0.704170349 |
| ENSMUSG00000084219 | 1 | |
| ENSMUSG00000084220 | -1.053549062 | 0.825517832 |
| ENSMUSG00000084221 | -1.048277865 | 0.891297795 |
| ENSMUSG00000084222 | 1 | |
| ENSMUSG00000084223 | -1.031335064 | 0.982854947 |
| ENSMUSG00000084224 | 1 | |
| ENSMUSG00000084226 | -1.044273782 | 0.698112512 |
| ENSMUSG00000084228 | 1 | |
| ENSMUSG00000084230 | 1 | |
| ENSMUSG00000084231 | -1.033059777 | 0.943705943 |
| ENSMUSG00000084234 | 1 | |
| ENSMUSG00000084235 | -1.038444538 | 0.895498153 |
| ENSMUSG00000084236 | 1 | |
| ENSMUSG00000084237 | 1 | |
| ENSMUSG00000084238 | 1 | |
| ENSMUSG00000084239 | 1 | |
| ENSMUSG00000084240 | 1 | |
| ENSMUSG00000084241 | 1 | |
| ENSMUSG00000084243 | -1.044273782 | 0.698112512 |
| ENSMUSG00000084244 | 1 | |
| ENSMUSG00000084245 | 1 | |
| ENSMUSG00000084246 | 1 | |
| ENSMUSG00000084247 | 1 | |
| ENSMUSG00000084248 | 1 | |
| ENSMUSG00000084249 | -1.017626803 | 0.994116287 |
| ENSMUSG00000084250 | 1 | |

| | | |
|---|---|---|
| ENSMUSG00000084251 | 1 | |
| ENSMUSG00000084252 | -1.084645496 | 0.69530468 |
| ENSMUSG00000084253 | 1 | |
| ENSMUSG00000084254 | 1 | |
| ENSMUSG00000084256 | 1 | |
| ENSMUSG00000084257 | 1 | |
| ENSMUSG00000084259 | 1 | |
| ENSMUSG00000084260 | 1 | |
| ENSMUSG00000084261 | 1 | |
| ENSMUSG00000084262 | 1 | |
| ENSMUSG00000084264 | -1 | 1 |
| ENSMUSG00000084265 | -1 | 1 |
| ENSMUSG00000084267 | 1 | |
| ENSMUSG00000084268 | 1 | |
| ENSMUSG00000084269 | 1 | |
| ENSMUSG00000084270 | 1 | |
| ENSMUSG00000084271 | 1.132639128 | 0.457778318 |
| ENSMUSG00000084272 | 1 | |
| ENSMUSG00000084273 | -1.044273782 | 0.698112512 |
| ENSMUSG00000084274 | 1.032577156 | 0.893865901 |
| ENSMUSG00000084275 | -1.068573373 | 0.890472112 |
| ENSMUSG00000084276 | -1.02199878 | 1 |
| ENSMUSG00000084277 | 1 | |
| ENSMUSG00000084278 | 1 | |
| ENSMUSG00000084279 | 1 | |
| ENSMUSG00000084280 | 1 | |
| ENSMUSG00000084281 | 1 | |
| ENSMUSG00000084282 | 1 | |
| ENSMUSG00000084283 | 1.070429626 | 0.767473873 |
| ENSMUSG00000084284 | -1.048558164 | 0.872692845 |
| ENSMUSG00000084285 | 1 | |
| ENSMUSG00000084286 | 1 | |
| ENSMUSG00000084287 | 1 | |
| ENSMUSG00000084288 | -1.078590893 | 0.698112512 |
| ENSMUSG00000084289 | 1.06585783 | 0.826758538 |
| ENSMUSG00000084291 | 1 | |
| ENSMUSG00000084292 | 1.059463094 | 0.608497229 |
| ENSMUSG00000084293 | 1 | |
| ENSMUSG00000084294 | 1 | |
| ENSMUSG00000084295 | 1 | |
| ENSMUSG00000084296 | 1 | |
| ENSMUSG00000084297 | 1 | |
| ENSMUSG00000084298 | 1 | |
| ENSMUSG00000084299 | -1.03962336 | 0.698112512 |
| ENSMUSG00000084300 | 1 | |
| ENSMUSG00000084301 | -1 | 1 |
| ENSMUSG00000084302 | 1.135123114 | 0.608497229 |
| ENSMUSG00000084303 | -1.063127349 | 0.698112512 |
| ENSMUSG00000084304 | -1.03962336 | 0.698112512 |
| ENSMUSG00000084305 | 1 | |
| ENSMUSG00000084306 | -1.085651418 | 0.692881216 |
| ENSMUSG00000084307 | -1.044273782 | 0.698112512 |
| ENSMUSG00000084308 | 1 | |
| ENSMUSG00000084309 | 1 | |
| ENSMUSG00000084310 | 1.059463094 | 0.608497229 |
| ENSMUSG00000084311 | 1 | |
| ENSMUSG00000084312 | 1 | |
| ENSMUSG00000084313 | 1.069068979 | 0.698112512 |
| ENSMUSG00000084314 | -1.085004452 | 0.803260194 |
| ENSMUSG00000084315 | 1.021563321 | 0.833407183 |
| ENSMUSG00000084316 | 1 | |
| ENSMUSG00000084318 | 1 | |
| ENSMUSG00000084319 | 1.07487334 | 0.656277822 |

| | | |
|---|---|---|
| ENSMUSG00000084320 | 1 | |
| ENSMUSG00000084323 | 1 | |
| ENSMUSG00000084324 | 1 | |
| ENSMUSG00000084325 | 1.010351027 | 1 |
| ENSMUSG00000084326 | -1.044273782 | 0.698112512 |
| ENSMUSG00000084327 | 1 | |
| ENSMUSG00000084328 | 1.105010143 | 0.608497229 |
| ENSMUSG00000084329 | -1.104243644 | 0.244763835 |
| ENSMUSG00000084330 | 1 | |
| ENSMUSG00000084331 | -1.044273782 | 0.698112512 |
| ENSMUSG00000084332 | 1.097516076 | 0.79291651 |
| ENSMUSG00000084333 | 1.020240423 | 0.986009441 |
| ENSMUSG00000084334 | 1 | |
| ENSMUSG00000084335 | 1 | |
| ENSMUSG00000084336 | 1 | |
| ENSMUSG00000084337 | 1 | |
| ENSMUSG00000084338 | 1 | |
| ENSMUSG00000084339 | 1.059463094 | 0.608497229 |
| ENSMUSG00000084340 | -1.044273782 | 0.801443616 |
| ENSMUSG00000084341 | 1 | |
| ENSMUSG00000084342 | 1.049411284 | 0.863033776 |
| ENSMUSG00000084343 | 1.059463094 | 0.698112512 |
| ENSMUSG00000084344 | 1 | |
| ENSMUSG00000084345 | -1.12544754 | 0.692944879 |
| ENSMUSG00000084346 | 1 | |
| ENSMUSG00000084347 | -1.008614656 | 1 |
| ENSMUSG00000084348 | 1.069068979 | 0.608497229 |
| ENSMUSG00000084349 | 1.016615281 | 0.943410634 |
| ENSMUSG00000084350 | -1.10055249 | 0.547111524 |
| ENSMUSG00000084351 | 1 | |
| ENSMUSG00000084352 | 1 | |
| ENSMUSG00000084353 | 1 | |
| ENSMUSG00000084355 | 1 | |
| ENSMUSG00000084356 | 1 | |
| ENSMUSG00000084357 | 1 | |
| ENSMUSG00000084358 | 1 | |
| ENSMUSG00000084359 | 1 | |
| ENSMUSG00000084360 | 1 | |
| ENSMUSG00000084361 | 1 | |
| ENSMUSG00000084362 | 1 | |
| ENSMUSG00000084363 | 1 | |
| ENSMUSG00000084366 | 1 | |
| ENSMUSG00000084367 | 1 | |
| ENSMUSG00000084368 | 1 | |
| ENSMUSG00000084369 | 1 | |
| ENSMUSG00000084370 | 1 | |
| ENSMUSG00000084371 | 1 | |
| ENSMUSG00000084372 | -1.019680325 | 0.981456085 |
| ENSMUSG00000084373 | 1.11349396 | 0.765184175 |
| ENSMUSG00000084374 | -1.043306199 | 0.698112512 |
| ENSMUSG00000084375 | -1.044273782 | 0.855488553 |
| ENSMUSG00000084376 | 1 | |
| ENSMUSG00000084377 | -1.053549062 | 0.698112512 |
| ENSMUSG00000084378 | 1.069987806 | 0.698112512 |
| ENSMUSG00000084379 | 1 | |
| ENSMUSG00000084380 | 1 | |
| ENSMUSG00000084381 | 1.12267039 | 0.631006723 |
| ENSMUSG00000084382 | 1 | |
| ENSMUSG00000084383 | 1 | |
| ENSMUSG00000084384 | 1.019083579 | 0.949812939 |
| ENSMUSG00000084385 | 1 | |
| ENSMUSG00000084386 | 1 | |
| ENSMUSG00000084387 | 1 | |

| | | |
|---|---|---|
| ENSMUSG00000084390 | -1 | 1 |
| ENSMUSG00000084391 | 1 | |
| ENSMUSG00000084392 | -1.043306199 | 0.698112512 |
| ENSMUSG00000084393 | 1 | |
| ENSMUSG00000084394 | 1 | |
| ENSMUSG00000084395 | 1 | |
| ENSMUSG00000084396 | 1 | |
| ENSMUSG00000084397 | 1 | |
| ENSMUSG00000084399 | 1 | |
| ENSMUSG00000084400 | 1 | |
| ENSMUSG00000084401 | 1.006784519 | 1 |
| ENSMUSG00000084403 | 1.102202777 | 0.756303785 |
| ENSMUSG00000084404 | 1.005613439 | 1 |
| ENSMUSG00000084405 | 1 | |
| ENSMUSG00000084407 | -1 | 1 |
| ENSMUSG00000084408 | 1.080254947 | 0.683415331 |
| ENSMUSG00000084411 | -1 | 1 |
| ENSMUSG00000084412 | 1 | |
| ENSMUSG00000084413 | 1.109062085 | 0.698112512 |
| ENSMUSG00000084414 | -1.043306199 | 0.698112512 |
| ENSMUSG00000084415 | 1 | |
| ENSMUSG00000084416 | -1.077967184 | 0.520333229 |
| ENSMUSG00000084417 | 1 | |
| ENSMUSG00000084418 | 1 | |
| ENSMUSG00000084419 | -1.084645496 | 0.532369657 |
| ENSMUSG00000084421 | 1.014545335 | 0.979002724 |
| ENSMUSG00000084422 | 1.162084994 | 0.608497229 |
| ENSMUSG00000084423 | 1 | |
| ENSMUSG00000084425 | 1 | |
| ENSMUSG00000084426 | -1 | 1 |
| ENSMUSG00000084428 | 1 | |
| ENSMUSG00000084431 | 1 | |
| ENSMUSG00000084432 | 1 | |
| ENSMUSG00000084433 | -1.074560108 | 0.708354476 |
| ENSMUSG00000084435 | 1 | |
| ENSMUSG00000084437 | 1 | |
| ENSMUSG00000084439 | 1 | |
| ENSMUSG00000084440 | -1.043722544 | 0.880723094 |
| ENSMUSG00000084441 | 1 | |
| ENSMUSG00000084442 | 1.069068979 | 0.608497229 |
| ENSMUSG00000084443 | 1 | |
| ENSMUSG00000084444 | 1 | |
| ENSMUSG00000084445 | 1 | |
| ENSMUSG00000084446 | 1 | |
| ENSMUSG00000084447 | 1 | |
| ENSMUSG00000084448 | 1 | |
| ENSMUSG00000084449 | 1 | |
| ENSMUSG00000084450 | 1 | |
| ENSMUSG00000084451 | 1 | |
| ENSMUSG00000084452 | 1 | |
| ENSMUSG00000084453 | -1 | 1 |
| ENSMUSG00000084454 | 1 | |
| ENSMUSG00000084456 | 1 | |
| ENSMUSG00000084457 | 1.059463094 | 0.608497229 |
| ENSMUSG00000084458 | -1.044273782 | 0.698112512 |
| ENSMUSG00000084459 | 1 | |
| ENSMUSG00000084460 | 1 | |
| ENSMUSG00000084461 | 1 | |
| ENSMUSG00000084462 | 1.059463094 | 0.608497229 |
| ENSMUSG00000084463 | 1 | |
| ENSMUSG00000084465 | 1 | |
| ENSMUSG00000084466 | 1 | |
| ENSMUSG00000084467 | 1 | |

| | | |
|---|---|---|
| ENSMUSG00000084468 | 1 | |
| ENSMUSG00000084469 | 1 | |
| ENSMUSG00000084472 | 1 | |
| ENSMUSG00000084473 | 1 | |
| ENSMUSG00000084474 | 1 | |
| ENSMUSG00000084475 | 1 | |
| ENSMUSG00000084476 | 1 | |
| ENSMUSG00000084477 | -1.044273782 | 0.698112512 |
| ENSMUSG00000084478 | 1.059463094 | 0.608497229 |
| ENSMUSG00000084479 | 1 | |
| ENSMUSG00000084480 | 1 | |
| ENSMUSG00000084484 | 1 | |
| ENSMUSG00000084485 | 1 | |
| ENSMUSG00000084487 | 1 | |
| ENSMUSG00000084492 | 1 | |
| ENSMUSG00000084493 | -1 | 1 |
| ENSMUSG00000084496 | 1.171563499 | 0.370636424 |
| ENSMUSG00000084497 | 1.171563499 | 0.34958424 |
| ENSMUSG00000084498 | 1 | |
| ENSMUSG00000084500 | 1 | |
| ENSMUSG00000084501 | 1 | |
| ENSMUSG00000084502 | 1 | |
| ENSMUSG00000084503 | 1 | |
| ENSMUSG00000084504 | 1 | |
| ENSMUSG00000084505 | 1 | |
| ENSMUSG00000084506 | 1 | |
| ENSMUSG00000084507 | 1.177765487 | 0.399055433 |
| ENSMUSG00000084508 | 1 | |
| ENSMUSG00000084511 | 1 | |
| ENSMUSG00000084512 | 1 | |
| ENSMUSG00000084513 | 1 | |
| ENSMUSG00000084514 | 1 | |
| ENSMUSG00000084515 | 1 | |
| ENSMUSG00000084516 | 1 | |
| ENSMUSG00000084517 | 1 | |
| ENSMUSG00000084518 | 1 | |
| ENSMUSG00000084519 | 1.005350682 | 1 |
| ENSMUSG00000084520 | 1 | |
| ENSMUSG00000084521 | 1 | |
| ENSMUSG00000084522 | -1.044273782 | 0.790851299 |
| ENSMUSG00000084523 | 1 | |
| ENSMUSG00000084524 | 1 | |
| ENSMUSG00000084526 | 1.014545335 | 0.979002724 |
| ENSMUSG00000084527 | 1 | |
| ENSMUSG00000084530 | 1 | |
| ENSMUSG00000084531 | 1 | |
| ENSMUSG00000084532 | 1 | |
| ENSMUSG00000084535 | 1 | |
| ENSMUSG00000084536 | 1 | |
| ENSMUSG00000084537 | 1 | |
| ENSMUSG00000084539 | 1 | |
| ENSMUSG00000084540 | 1 | |
| ENSMUSG00000084541 | 1 | |
| ENSMUSG00000084543 | -1.03962336 | 0.698112512 |
| ENSMUSG00000084544 | 1.11633269 | 0.608497229 |
| ENSMUSG00000084545 | 1.059463094 | 0.608497229 |
| ENSMUSG00000084546 | 1 | |
| ENSMUSG00000084547 | 1 | |
| ENSMUSG00000084548 | 1 | |
| ENSMUSG00000084549 | 1 | |
| ENSMUSG00000084550 | 1 | |
| ENSMUSG00000084551 | 1 | |
| ENSMUSG00000084552 | 1 | |

| | | |
|---|---|---|
| ENSMUSG00000084555 | 1 | |
| ENSMUSG00000084557 | 1 | |
| ENSMUSG00000084558 | 1 | |
| ENSMUSG00000084559 | 1 | |
| ENSMUSG00000084560 | 1 | |
| ENSMUSG00000084562 | -1.014836984 | 1 |
| ENSMUSG00000084564 | 1 | |
| ENSMUSG00000084565 | 1 | |
| ENSMUSG00000084566 | 1 | |
| ENSMUSG00000084567 | 1 | |
| ENSMUSG00000084568 | 1 | |
| ENSMUSG00000084569 | 1 | |
| ENSMUSG00000084571 | 1.014545335 | 0.979002724 |
| ENSMUSG00000084572 | 1 | |
| ENSMUSG00000084573 | 1 | |
| ENSMUSG00000084574 | 1 | |
| ENSMUSG00000084575 | 1 | |
| ENSMUSG00000084576 | 1 | |
| ENSMUSG00000084579 | 1 | |
| ENSMUSG00000084581 | 1 | |
| ENSMUSG00000084582 | 1 | |
| ENSMUSG00000084583 | 1 | |
| ENSMUSG00000084584 | 1.014545335 | 0.982132299 |
| ENSMUSG00000084587 | 1 | |
| ENSMUSG00000084588 | 1 | |
| ENSMUSG00000084589 | 1 | |
| ENSMUSG00000084590 | 1 | |
| ENSMUSG00000084591 | 1 | |
| ENSMUSG00000084592 | 1.155317502 | 0.420161906 |
| ENSMUSG00000084593 | 1 | |
| ENSMUSG00000084595 | -1.069502966 | 0.698112512 |
| ENSMUSG00000084596 | 1 | |
| ENSMUSG00000084597 | 1 | |
| ENSMUSG00000084598 | 1 | |
| ENSMUSG00000084599 | 1 | |
| ENSMUSG00000084600 | 1 | |
| ENSMUSG00000084601 | -1.03962336 | 0.698112512 |
| ENSMUSG00000084602 | 1 | |
| ENSMUSG00000084603 | -1.099894511 | 0.742727573 |
| ENSMUSG00000084604 | 1 | |
| ENSMUSG00000084605 | 1 | |
| ENSMUSG00000084606 | 1 | |
| ENSMUSG00000084607 | -1.126519467 | 0.633754375 |
| ENSMUSG00000084608 | 1 | |
| ENSMUSG00000084609 | 1 | |
| ENSMUSG00000084610 | 1 | |
| ENSMUSG00000084611 | -1 | 1 |
| ENSMUSG00000084612 | 1 | |
| ENSMUSG00000084613 | 1 | |
| ENSMUSG00000084616 | 1.12235904 | 0.698112512 |
| ENSMUSG00000084617 | 1 | |
| ENSMUSG00000084619 | 1 | |
| ENSMUSG00000084620 | 1 | |
| ENSMUSG00000084621 | 1 | |
| ENSMUSG00000084622 | 1 | |
| ENSMUSG00000084624 | -1.044273782 | 0.698112512 |
| ENSMUSG00000084625 | 1 | |
| ENSMUSG00000084627 | 1 | |
| ENSMUSG00000084628 | -1.043306199 | 0.698112512 |
| ENSMUSG00000084629 | 1 | |
| ENSMUSG00000084630 | 1 | |
| ENSMUSG00000084631 | 1 | |
| ENSMUSG00000084632 | -1.043306199 | 0.79291651 |

| | | |
|---|---|---|
| ENSMUSG00000084633 | 1 | |
| ENSMUSG00000084635 | 1 | |
| ENSMUSG00000084636 | 1 | |
| ENSMUSG00000084637 | 1 | |
| ENSMUSG00000084638 | 1 | |
| ENSMUSG00000084639 | 1 | |
| ENSMUSG00000084640 | 1 | |
| ENSMUSG00000084641 | 1 | |
| ENSMUSG00000084642 | 1 | |
| ENSMUSG00000084643 | 1 | |
| ENSMUSG00000084644 | -1.008906735 | 1 |
| ENSMUSG00000084645 | 1 | |
| ENSMUSG00000084646 | 1 | |
| ENSMUSG00000084647 | 1 | |
| ENSMUSG00000084648 | -1.003834322 | 1 |
| ENSMUSG00000084649 | 1 | |
| ENSMUSG00000084650 | 1 | |
| ENSMUSG00000084651 | 1 | |
| ENSMUSG00000084652 | 1 | |
| ENSMUSG00000084653 | 1 | |
| ENSMUSG00000084654 | 1 | |
| ENSMUSG00000084658 | 1 | |
| ENSMUSG00000084659 | 1.028323353 | 0.903464611 |
| ENSMUSG00000084660 | 1 | |
| ENSMUSG00000084661 | 1 | |
| ENSMUSG00000084663 | 1 | |
| ENSMUSG00000084664 | 1 | |
| ENSMUSG00000084665 | 1 | |
| ENSMUSG00000084666 | 1 | |
| ENSMUSG00000084667 | 1 | |
| ENSMUSG00000084668 | 1 | |
| ENSMUSG00000084669 | 1 | |
| ENSMUSG00000084671 | 1 | |
| ENSMUSG00000084673 | 1.069068979 | 0.608497229 |
| ENSMUSG00000084674 | 1.059463094 | 0.608497229 |
| ENSMUSG00000084675 | 1 | |
| ENSMUSG00000084678 | 1.069068979 | 0.698112512 |
| ENSMUSG00000084682 | 1 | |
| ENSMUSG00000084683 | 1 | |
| ENSMUSG00000084684 | 1 | |
| ENSMUSG00000084686 | -1.023769022 | 0.954761885 |
| ENSMUSG00000084688 | 1 | |
| ENSMUSG00000084689 | 1 | |
| ENSMUSG00000084690 | 1 | |
| ENSMUSG00000084691 | 1 | |
| ENSMUSG00000084693 | 1.059463094 | 0.608497229 |
| ENSMUSG00000084694 | 1.069068979 | 0.698112512 |
| ENSMUSG00000084696 | 1 | |
| ENSMUSG00000084700 | 1 | |
| ENSMUSG00000084701 | -1 | 1 |
| ENSMUSG00000084702 | 1 | |
| ENSMUSG00000084703 | 1 | |
| ENSMUSG00000084704 | 1 | |
| ENSMUSG00000084705 | 1 | |
| ENSMUSG00000084706 | 1.024693402 | 0.955101438 |
| ENSMUSG00000084708 | 1 | |
| ENSMUSG00000084710 | 1 | |
| ENSMUSG00000084711 | -1.053549062 | 0.698112512 |
| ENSMUSG00000084713 | 1 | |
| ENSMUSG00000084714 | 1 | |
| ENSMUSG00000084717 | 1 | |
| ENSMUSG00000084718 | 1 | |
| ENSMUSG00000084719 | 1 | |

| | | |
|---|---|---|
| ENSMUSG00000084722 | 1 | |
| ENSMUSG00000084725 | 1 | |
| ENSMUSG00000084728 | 1 | |
| ENSMUSG00000084729 | 1.009414875 | 1 |
| ENSMUSG00000084731 | 1 | |
| ENSMUSG00000084732 | 1 | |
| ENSMUSG00000084733 | 1 | |
| ENSMUSG00000084734 | 1 | |
| ENSMUSG00000084737 | 1 | |
| ENSMUSG00000084738 | -1.069502966 | 0.698112512 |
| ENSMUSG00000084739 | -1.043306199 | 0.698112512 |
| ENSMUSG00000084740 | 1 | |
| ENSMUSG00000084741 | 1 | |
| ENSMUSG00000084742 | 1 | |
| ENSMUSG00000084743 | 1 | |
| ENSMUSG00000084744 | 1.075870199 | 0.642819974 |
| ENSMUSG00000084745 | 1 | |
| ENSMUSG00000084746 | 1 | |
| ENSMUSG00000084747 | 1 | |
| ENSMUSG00000084748 | 1 | |
| ENSMUSG00000084749 | 1 | |
| ENSMUSG00000084750 | -1.095294215 | 0.541735653 |
| ENSMUSG00000084751 | 1 | |
| ENSMUSG00000084752 | 1 | |
| ENSMUSG00000084753 | 1 | |
| ENSMUSG00000084754 | 1.059463094 | 0.608497229 |
| ENSMUSG00000084755 | 1.036587445 | 0.948423547 |
| ENSMUSG00000084756 | -1 | 1 |
| ENSMUSG00000084757 | 1 | |
| ENSMUSG00000084758 | 1.138419748 | 0.621888633 |
| ENSMUSG00000084760 | 1 | |
| ENSMUSG00000084761 | 1 | |
| ENSMUSG00000084762 | 1 | |
| ENSMUSG00000084764 | 1 | |
| ENSMUSG00000084766 | -1.03962336 | 0.698112512 |
| ENSMUSG00000084768 | 1 | |
| ENSMUSG00000084769 | 1 | |
| ENSMUSG00000084771 | -1.034022509 | 0.949218699 |
| ENSMUSG00000084772 | 1.013977808 | 1 |
| ENSMUSG00000084773 | 1 | |
| ENSMUSG00000084774 | 1.059463094 | 0.608497229 |
| ENSMUSG00000084775 | 1 | |
| ENSMUSG00000084776 | -1.023769022 | 0.954761885 |
| ENSMUSG00000084780 | -1.010960635 | 1 |
| ENSMUSG00000084781 | -1.413876141 | 0.011991473 |
| ENSMUSG00000084782 | 1 | |
| ENSMUSG00000084783 | -1.047306572 | 0.834839709 |
| ENSMUSG00000084785 | -1.044273782 | 0.790851299 |
| ENSMUSG00000084786 | -1.070268011 | 0.753100881 |
| ENSMUSG00000084787 | 1 | |
| ENSMUSG00000084788 | 1 | |
| ENSMUSG00000084789 | -1.053549062 | 0.698112512 |
| ENSMUSG00000084790 | 1.176854992 | 0.662227282 |
| ENSMUSG00000084791 | -1.133231767 | 0.591151279 |
| ENSMUSG00000084792 | -1.092510195 | 0.608497229 |
| ENSMUSG00000084793 | -1 | 1 |
| ENSMUSG00000084794 | 1 | |
| ENSMUSG00000084795 | 1 | |
| ENSMUSG00000084796 | 1 | |
| ENSMUSG00000084797 | -1.009903943 | 1 |
| ENSMUSG00000084798 | 1.153444416 | 0.466340867 |
| ENSMUSG00000084799 | -1.098216337 | 0.164820549 |
| ENSMUSG00000084800 | -1.034990181 | 0.911643738 |

| | | |
|---|---|---|
| ENSMUSG00000084802 | 1 | |
| ENSMUSG00000084803 | -1.0965339 | 0.683821802 |
| ENSMUSG00000084804 | -1.03962336 | 0.698112512 |
| ENSMUSG00000084805 | 1 | |
| ENSMUSG00000084806 | 1.456031346 | 0.006758399 |
| ENSMUSG00000084807 | -1.064135801 | 0.698112512 |
| ENSMUSG00000084808 | -1.075390699 | 0.610080378 |
| ENSMUSG00000084809 | 1.023959045 | 0.979212363 |
| ENSMUSG00000084811 | 1 | |
| ENSMUSG00000084815 | 1 | |
| ENSMUSG00000084816 | 1 | |
| ENSMUSG00000084817 | 1.0196665 | 0.949812939 |
| ENSMUSG00000084818 | 1 | |
| ENSMUSG00000084819 | 1 | |
| ENSMUSG00000084821 | -1.063525418 | 0.751979636 |
| ENSMUSG00000084822 | -1.089497311 | 0.532369657 |
| ENSMUSG00000084823 | 1 | |
| ENSMUSG00000084824 | -1.089041054 | 0.608497229 |
| ENSMUSG00000084825 | 1 | |
| ENSMUSG00000084826 | -1.03962336 | 0.90107137 |
| ENSMUSG00000084828 | -1.024530911 | 0.949679283 |
| ENSMUSG00000084829 | 1 | |
| ENSMUSG00000084830 | 1 | |
| ENSMUSG00000084831 | 1 | |
| ENSMUSG00000084833 | 1 | |
| ENSMUSG00000084834 | 1.00478092 | 1 |
| ENSMUSG00000084835 | -1.076850214 | 0.801443616 |
| ENSMUSG00000084836 | 1 | |
| ENSMUSG00000084837 | 1 | |
| ENSMUSG00000084838 | 1 | |
| ENSMUSG00000084839 | 1 | |
| ENSMUSG00000084840 | 1 | |
| ENSMUSG00000084842 | -1.238095458 | 0.255019724 |
| ENSMUSG00000084843 | 1.122344914 | 0.698112512 |
| ENSMUSG00000084844 | -1.070154669 | 0.698112512 |
| ENSMUSG00000084845 | -1.025069235 | 0.955034299 |
| ENSMUSG00000084846 | -1.015109838 | 0.914816803 |
| ENSMUSG00000084847 | 1.059463094 | 0.608497229 |
| ENSMUSG00000084848 | 1.085624843 | 0.69305964 |
| ENSMUSG00000084849 | -1 | 1 |
| ENSMUSG00000084850 | -1.043306199 | 0.803260194 |
| ENSMUSG00000084851 | 1 | |
| ENSMUSG00000084852 | 1 | |
| ENSMUSG00000084853 | 1 | |
| ENSMUSG00000084854 | 1 | |
| ENSMUSG00000084855 | 1 | |
| ENSMUSG00000084856 | 1 | |
| ENSMUSG00000084858 | -1.100193663 | 0.534887579 |
| ENSMUSG00000084859 | 1 | |
| ENSMUSG00000084860 | 1.370042947 | 0.147938946 |
| ENSMUSG00000084861 | -1.295529661 | 0.10939414 |
| ENSMUSG00000084862 | -1.142814509 | 0.698112512 |
| ENSMUSG00000084863 | 1 | |
| ENSMUSG00000084864 | 1.117244663 | 0.608497229 |
| ENSMUSG00000084865 | 1 | |
| ENSMUSG00000084866 | -1.016691388 | 0.989207972 |
| ENSMUSG00000084869 | 1 | |
| ENSMUSG00000084870 | -1.116853908 | 0.558122732 |
| ENSMUSG00000084871 | 1 | |
| ENSMUSG00000084873 | 1 | |
| ENSMUSG00000084874 | 1 | |
| ENSMUSG00000084875 | 1 | |
| ENSMUSG00000084876 | -1.053549062 | 0.698112512 |

| | | |
|---|---|---|
| ENSMUSG00000084878 | 1.085624843 | 0.698112512 |
| ENSMUSG00000084880 | 1.165265624 | 0.61804326 |
| ENSMUSG00000084881 | -1.102820698 | 0.608497229 |
| ENSMUSG00000084882 | -1 | 1 |
| ENSMUSG00000084883 | 1.11240188 | 0.704170349 |
| ENSMUSG00000084884 | 1 | |
| ENSMUSG00000084885 | -1.243356308 | 0.009566147 |
| ENSMUSG00000084887 | 1 | |
| ENSMUSG00000084889 | 1 | |
| ENSMUSG00000084890 | 1.015486245 | 0.985522133 |
| ENSMUSG00000084891 | 1 | |
| ENSMUSG00000084892 | -1.029709545 | 0.975148944 |
| ENSMUSG00000084893 | 1.0463724 | 0.869361059 |
| ENSMUSG00000084894 | 1.059463094 | 0.608497229 |
| ENSMUSG00000084895 | 1 | |
| ENSMUSG00000084896 | 1 | |
| ENSMUSG00000084897 | 1.08475742 | 0.802748054 |
| ENSMUSG00000084898 | 1 | |
| ENSMUSG00000084899 | 1.05392877 | 0.941128589 |
| ENSMUSG00000084901 | 1 | |
| ENSMUSG00000084902 | 1 | |
| ENSMUSG00000084904 | 1.049651571 | 0.822825922 |
| ENSMUSG00000084905 | 1 | |
| ENSMUSG00000084906 | 1 | |
| ENSMUSG00000084907 | 1 | |
| ENSMUSG00000084908 | -1.010960635 | 1 |
| ENSMUSG00000084909 | 1 | |
| ENSMUSG00000084910 | 1.049302431 | 0.829324104 |
| ENSMUSG00000084911 | 1.026430948 | 0.913972125 |
| ENSMUSG00000084913 | -1.03962336 | 0.803072249 |
| ENSMUSG00000084915 | -1.156772062 | 0.608497229 |
| ENSMUSG00000084916 | 1.059463094 | 0.608497229 |
| ENSMUSG00000084917 | 1.045549562 | 0.696907273 |
| ENSMUSG00000084918 | 1 | |
| ENSMUSG00000084919 | 1 | |
| ENSMUSG00000084920 | 1 | |
| ENSMUSG00000084921 | 1 | |
| ENSMUSG00000084923 | 1 | |
| ENSMUSG00000084924 | 1 | |
| ENSMUSG00000084925 | -1.191047269 | 0.449109628 |
| ENSMUSG00000084927 | 1 | |
| ENSMUSG00000084929 | 1 | |
| ENSMUSG00000084930 | -1.064005321 | 0.636777862 |
| ENSMUSG00000084932 | 1 | |
| ENSMUSG00000084933 | -1.003942951 | 1 |
| ENSMUSG00000084934 | 1 | |
| ENSMUSG00000084935 | 1 | |
| ENSMUSG00000084936 | 1 | |
| ENSMUSG00000084937 | 1 | |
| ENSMUSG00000084938 | 1.155455492 | 0.346633686 |
| ENSMUSG00000084939 | -1.089932089 | 0.807247723 |
| ENSMUSG00000084940 | -1.03962336 | 0.698112512 |
| ENSMUSG00000084941 | 1.1058995 | 0.584549027 |
| ENSMUSG00000084942 | 1.059463094 | 0.608497229 |
| ENSMUSG00000084944 | 1 | |
| ENSMUSG00000084945 | 1.059463094 | 0.608497229 |
| ENSMUSG00000084946 | -1.101103426 | 0.510936302 |
| ENSMUSG00000084947 | 1 | |
| ENSMUSG00000084948 | -1.043306199 | 0.873170162 |
| ENSMUSG00000084949 | -1.04489665 | 0.698112512 |
| ENSMUSG00000084950 | 1.014453345 | 0.972980766 |
| ENSMUSG00000084951 | -1.03962336 | 0.698112512 |
| ENSMUSG00000084954 | 1.069068979 | 0.608497229 |

| | | |
|---|---|---|
| ENSMUSG00000084955 | 1 | |
| ENSMUSG00000084956 | 1 | |
| ENSMUSG00000084957 | 1.059319884 | 0.780242686 |
| ENSMUSG00000084959 | 1 | |
| ENSMUSG00000084960 | 1.089470641 | 0.608497229 |
| ENSMUSG00000084962 | 1 | |
| ENSMUSG00000084963 | 1 | |
| ENSMUSG00000084964 | -1.099174267 | 0.535052277 |
| ENSMUSG00000084965 | 1 | |
| ENSMUSG00000084966 | -1 | 1 |
| ENSMUSG00000084967 | -1.043306199 | 0.698112512 |
| ENSMUSG00000084968 | 1.067747658 | 0.836133893 |
| ENSMUSG00000084969 | -1.043306199 | 0.698112512 |
| ENSMUSG00000084970 | -1.163790014 | 0.193677799 |
| ENSMUSG00000084971 | 1 | |
| ENSMUSG00000084973 | -1 | 1 |
| ENSMUSG00000084974 | 1.071686201 | 0.772971553 |
| ENSMUSG00000084978 | -1.099174267 | 0.688934383 |
| ENSMUSG00000084979 | -1.160909573 | 0.543831124 |
| ENSMUSG00000084980 | 1.229255047 | 0.541735653 |
| ENSMUSG00000084981 | 1.059463094 | 0.608497229 |
| ENSMUSG00000084982 | 1 | |
| ENSMUSG00000084983 | -1 | 1 |
| ENSMUSG00000084984 | 1.098454884 | 0.578221424 |
| ENSMUSG00000084985 | -1.028897656 | 0.953434806 |
| ENSMUSG00000084986 | 1 | |
| ENSMUSG00000084987 | 1 | |
| ENSMUSG00000084989 | 1.253579507 | 0.430517842 |
| ENSMUSG00000084990 | 1 | |
| ENSMUSG00000084991 | 1 | |
| ENSMUSG00000084992 | 1 | |
| ENSMUSG00000084993 | -1.089497311 | 0.681308543 |
| ENSMUSG00000084994 | 1 | |
| ENSMUSG00000084995 | 1 | |
| ENSMUSG00000084996 | 1.070504273 | 0.748295996 |
| ENSMUSG00000084997 | 1 | |
| ENSMUSG00000084998 | 1.054357105 | 0.666468067 |
| ENSMUSG00000084999 | -1.138693452 | 0.552584793 |
| ENSMUSG00000085000 | 1 | |
| ENSMUSG00000085001 | 1 | |
| ENSMUSG00000085002 | 1 | |
| ENSMUSG00000085003 | 1 | |
| ENSMUSG00000085004 | 1 | |
| ENSMUSG00000085006 | -1.241780254 | 0.100385196 |
| ENSMUSG00000085007 | -1.03962336 | 0.872692845 |
| ENSMUSG00000085008 | -1.132949863 | 0.691565547 |
| ENSMUSG00000085009 | -1.184259924 | 0.25168922 |
| ENSMUSG00000085010 | 1 | |
| ENSMUSG00000085012 | 1.07487334 | 0.75314966 |
| ENSMUSG00000085014 | 1 | |
| ENSMUSG00000085015 | 1 | |
| ENSMUSG00000085016 | 1.09981644 | 0.770261954 |
| ENSMUSG00000085017 | 1 | |
| ENSMUSG00000085019 | 1.014731082 | 0.988424085 |
| ENSMUSG00000085020 | 1 | |
| ENSMUSG00000085021 | -1.044273782 | 0.698112512 |
| ENSMUSG00000085022 | 1.045531154 | 0.958824563 |
| ENSMUSG00000085023 | 1 | |
| ENSMUSG00000085024 | -1.151530222 | 0.359773676 |
| ENSMUSG00000085025 | 1 | |
| ENSMUSG00000085026 | -1.03962336 | 0.803072249 |
| ENSMUSG00000085027 | 1 | |
| ENSMUSG00000085028 | -1.065242351 | 0.615396291 |

| | | |
|---|---|---|
| ENSMUSG00000085029 | 1.069068979 | 0.608497229 |
| ENSMUSG00000085030 | -1.048295683 | 0.552842869 |
| ENSMUSG00000085032 | -1.03962336 | 0.698112512 |
| ENSMUSG00000085033 | -1.053549062 | 0.807247723 |
| ENSMUSG00000085034 | 1 | |
| ENSMUSG00000085035 | -1.064334191 | 0.907737577 |
| ENSMUSG00000085036 | 1 | |
| ENSMUSG00000085037 | 1.000409691 | 1 |
| ENSMUSG00000085039 | 1.003142735 | 1 |
| ENSMUSG00000085041 | 1 | |
| ENSMUSG00000085042 | 1.113135847 | 0.746899902 |
| ENSMUSG00000085043 | -1.027409007 | 0.993813312 |
| ENSMUSG00000085044 | 1 | |
| ENSMUSG00000085045 | 1 | |
| ENSMUSG00000085046 | 1 | |
| ENSMUSG00000085047 | 1 | |
| ENSMUSG00000085048 | 1 | |
| ENSMUSG00000085049 | 1 | |
| ENSMUSG00000085051 | -1.086092137 | 0.760867819 |
| ENSMUSG00000085053 | 1 | |
| ENSMUSG00000085054 | -1.021814107 | 0.966209176 |
| ENSMUSG00000085055 | 1.321997061 | 0.006353019 |
| ENSMUSG00000085057 | 1.116252217 | 0.465227872 |
| ENSMUSG00000085058 | 1.11166259 | 0.644125423 |
| ENSMUSG00000085059 | 1 | |
| ENSMUSG00000085060 | -1 | 1 |
| ENSMUSG00000085063 | -1.044273782 | 0.698112512 |
| ENSMUSG00000085066 | 1.388575658 | 0.022460487 |
| ENSMUSG00000085067 | 1.059463094 | 0.608497229 |
| ENSMUSG00000085068 | 1.156015583 | 0.357543507 |
| ENSMUSG00000085069 | 1 | |
| ENSMUSG00000085070 | 1 | |
| ENSMUSG00000085071 | 1.132639128 | 0.398767938 |
| ENSMUSG00000085072 | -1.116216152 | 0.506638168 |
| ENSMUSG00000085073 | 1 | |
| ENSMUSG00000085074 | 1 | |
| ENSMUSG00000085075 | 1 | |
| ENSMUSG00000085077 | 1 | |
| ENSMUSG00000085078 | 1.059463094 | 0.768693672 |
| ENSMUSG00000085079 | 1 | |
| ENSMUSG00000085080 | 1 | |
| ENSMUSG00000085081 | -1.116853908 | 0.608497229 |
| ENSMUSG00000085082 | 1 | |
| ENSMUSG00000085083 | 1.025046921 | 0.976160676 |
| ENSMUSG00000085084 | 1.313446885 | 0.237714495 |
| ENSMUSG00000085085 | 1.529555144 | 0.05157376 |
| ENSMUSG00000085087 | 1 | |
| ENSMUSG00000085088 | 1.129766774 | 0.207664843 |
| ENSMUSG00000085089 | 1.095872691 | 0.608497229 |
| ENSMUSG00000085090 | 1 | |
| ENSMUSG00000085091 | 1.188749468 | 0.467218078 |
| ENSMUSG00000085093 | -1.038444538 | 0.924629556 |
| ENSMUSG00000085094 | -1 | 1 |
| ENSMUSG00000085095 | 1 | |
| ENSMUSG00000085097 | 1 | |
| ENSMUSG00000085099 | -1.076127456 | 0.778239858 |
| ENSMUSG00000085100 | 1 | |
| ENSMUSG00000085101 | -1 | 1 |
| ENSMUSG00000085102 | 1 | |
| ENSMUSG00000085103 | 1 | |
| ENSMUSG00000085104 | 1 | |
| ENSMUSG00000085105 | -1.185736373 | 0.483375422 |
| ENSMUSG00000085106 | 1 | |

| | | |
|---|---|---|
| ENSMUSG00000085107 | 1 | |
| ENSMUSG00000085108 | 1 | |
| ENSMUSG00000085109 | 1.059463094 | 0.608497229 |
| ENSMUSG00000085110 | 1 | |
| ENSMUSG00000085111 | -1.087569789 | 0.608497229 |
| ENSMUSG00000085112 | 1.039968795 | 0.931480186 |
| ENSMUSG00000085113 | 1.122462048 | 0.420161906 |
| ENSMUSG00000085114 | 1.059463094 | 0.704170349 |
| ENSMUSG00000085115 | -1.043306199 | 0.698112512 |
| ENSMUSG00000085116 | -1 | 1 |
| ENSMUSG00000085118 | 1.027598766 | 0.898324888 |
| ENSMUSG00000085120 | 1 | |
| ENSMUSG00000085121 | 1 | |
| ENSMUSG00000085122 | 1 | |
| ENSMUSG00000085123 | 1 | |
| ENSMUSG00000085124 | 1.138997129 | 0.288134604 |
| ENSMUSG00000085125 | 1.12189305 | 0.633481288 |
| ENSMUSG00000085126 | 1.095872691 | 0.608497229 |
| ENSMUSG00000085127 | 1.10776854 | 0.761802747 |
| ENSMUSG00000085129 | -1 | 1 |
| ENSMUSG00000085130 | 1 | |
| ENSMUSG00000085132 | -1 | 1 |
| ENSMUSG00000085133 | -1.209417279 | 0.156131779 |
| ENSMUSG00000085134 | 1.075070132 | 0.725426917 |
| ENSMUSG00000085135 | 1.080929884 | 0.782883548 |
| ENSMUSG00000085139 | 1 | |
| ENSMUSG00000085141 | 1 | |
| ENSMUSG00000085142 | 1 | |
| ENSMUSG00000085143 | -1.024031001 | 1 |
| ENSMUSG00000085145 | -1.065321276 | 0.792716139 |
| ENSMUSG00000085146 | -1 | 1 |
| ENSMUSG00000085147 | -1.01808862 | 0.934760809 |
| ENSMUSG00000085148 | -1.01886362 | 0.94733141 |
| ENSMUSG00000085150 | 1 | |
| ENSMUSG00000085151 | -1.134282749 | 0.588487684 |
| ENSMUSG00000085152 | -1.043306199 | 0.903448666 |
| ENSMUSG00000085154 | 1.092538867 | 0.446338416 |
| ENSMUSG00000085155 | 1 | |
| ENSMUSG00000085156 | 1.170342098 | 0.017980376 |
| ENSMUSG00000085157 | 1 | |
| ENSMUSG00000085158 | 1 | |
| ENSMUSG00000085160 | 1 | |
| ENSMUSG00000085161 | 1.059463094 | 0.608497229 |
| ENSMUSG00000085162 | -1.075454467 | 0.778372272 |
| ENSMUSG00000085165 | -1.092198862 | 0.79077576 |
| ENSMUSG00000085166 | 1 | |
| ENSMUSG00000085167 | 1 | |
| ENSMUSG00000085168 | 1 | |
| ENSMUSG00000085169 | -1.150649829 | 0.589051474 |
| ENSMUSG00000085170 | -1.010960635 | 1 |
| ENSMUSG00000085171 | 1 | |
| ENSMUSG00000085172 | -1.125560317 | 0.614307028 |
| ENSMUSG00000085173 | 1 | |
| ENSMUSG00000085174 | 1.103638508 | 0.63418818 |
| ENSMUSG00000085175 | 1.135435629 | 0.423856307 |
| ENSMUSG00000085176 | 1 | |
| ENSMUSG00000085177 | 1 | |
| ENSMUSG00000085178 | 1.076818796 | 0.723432738 |
| ENSMUSG00000085179 | 1 | |
| ENSMUSG00000085180 | 1.175605891 | 0.215991492 |
| ENSMUSG00000085181 | 1 | |
| ENSMUSG00000085182 | 1 | |
| ENSMUSG00000085183 | 1.315035565 | 0.010698342 |

| | | |
|---|---|---|
| ENSMUSG00000085184 | -1.111456291 | 0.698112512 |
| ENSMUSG00000085185 | -1.057708979 | 0.815686662 |
| ENSMUSG00000085186 | 1.065410325 | 0.820462971 |
| ENSMUSG00000085187 | 1 | |
| ENSMUSG00000085188 | 1 | |
| ENSMUSG00000085189 | 1 | |
| ENSMUSG00000085192 | -1.03962336 | 0.803072249 |
| ENSMUSG00000085193 | 1 | |
| ENSMUSG00000085194 | 1 | |
| ENSMUSG00000085196 | 1.035514148 | 0.865583959 |
| ENSMUSG00000085197 | 1.218240674 | 0.294039426 |
| ENSMUSG00000085198 | 1 | |
| ENSMUSG00000085199 | 1 | |
| ENSMUSG00000085201 | 1.177765487 | 0.380970157 |
| ENSMUSG00000085202 | 1 | |
| ENSMUSG00000085203 | 1 | |
| ENSMUSG00000085204 | -1 | 1 |
| ENSMUSG00000085205 | 1.059463094 | 0.608497229 |
| ENSMUSG00000085206 | -1.053549062 | 0.698112512 |
| ENSMUSG00000085207 | -1.16224517 | 0.594567601 |
| ENSMUSG00000085208 | 1.034681251 | 0.951453535 |
| ENSMUSG00000085209 | 1 | |
| ENSMUSG00000085211 | -1.048277865 | 0.860645152 |
| ENSMUSG00000085213 | 1.1120792 | 0.464244482 |
| ENSMUSG00000085214 | -1.093511837 | 0.842347961 |
| ENSMUSG00000085217 | 1 | |
| ENSMUSG00000085218 | 1.325342194 | 0.174209262 |
| ENSMUSG00000085221 | 1 | |
| ENSMUSG00000085222 | 1 | |
| ENSMUSG00000085223 | 1 | |
| ENSMUSG00000085224 | -1.120476206 | 0.661322437 |
| ENSMUSG00000085225 | 1 | |
| ENSMUSG00000085226 | 1 | |
| ENSMUSG00000085227 | -1.175348413 | 0.030406771 |
| ENSMUSG00000085228 | 1.234951516 | 0.068335953 |
| ENSMUSG00000085229 | 1.019083579 | 0.949812939 |
| ENSMUSG00000085230 | 1 | |
| ENSMUSG00000085232 | 1 | |
| ENSMUSG00000085234 | -1.045964533 | 0.808432628 |
| ENSMUSG00000085235 | -1 | 1 |
| ENSMUSG00000085236 | 1.023743961 | 0.951410231 |
| ENSMUSG00000085237 | 1.171563499 | 0.34958424 |
| ENSMUSG00000085238 | 1 | |
| ENSMUSG00000085239 | 1 | |
| ENSMUSG00000085240 | 1 | |
| ENSMUSG00000085241 | -1.044273782 | 0.855488553 |
| ENSMUSG00000085242 | -1.10441992 | 0.768175961 |
| ENSMUSG00000085244 | 1 | |
| ENSMUSG00000085245 | 1 | |
| ENSMUSG00000085246 | 1.230850716 | 0.188127889 |
| ENSMUSG00000085247 | 1 | |
| ENSMUSG00000085248 | 1 | |
| ENSMUSG00000085249 | -1 | 1 |
| ENSMUSG00000085250 | 1.080134941 | 0.698112512 |
| ENSMUSG00000085251 | 1 | |
| ENSMUSG00000085252 | 1 | |
| ENSMUSG00000085253 | 1 | |
| ENSMUSG00000085254 | 1 | |
| ENSMUSG00000085255 | -1.074610093 | 0.813853563 |
| ENSMUSG00000085256 | -1.073533324 | 0.847853473 |
| ENSMUSG00000085257 | 1 | |
| ENSMUSG00000085260 | -1.088268782 | 0.698112512 |
| ENSMUSG00000085261 | 1.116785864 | 0.698112512 |

| | | |
|---|---|---|
| ENSMUSG00000085262 | 1.059463094 | 0.608497229 |
| ENSMUSG00000085263 | 1 | |
| ENSMUSG00000085264 | -1.043306199 | 0.698112512 |
| ENSMUSG00000085265 | 1 | |
| ENSMUSG00000085267 | 1 | |
| ENSMUSG00000085269 | 1.109392255 | 0.608497229 |
| ENSMUSG00000085271 | 1 | |
| ENSMUSG00000085272 | -1.221887966 | 0.230077154 |
| ENSMUSG00000085274 | -1.171420412 | 0.481568621 |
| ENSMUSG00000085275 | 1.142908482 | 0.615245112 |
| ENSMUSG00000085276 | 1.030661948 | 0.949007391 |
| ENSMUSG00000085277 | 1 | |
| ENSMUSG00000085278 | 1 | |
| ENSMUSG00000085279 | -1.005827504 | 1 |
| ENSMUSG00000085280 | 1 | |
| ENSMUSG00000085281 | 1 | |
| ENSMUSG00000085282 | 1.00580775 | 1 |
| ENSMUSG00000085284 | 1 | |
| ENSMUSG00000085285 | 1 | |
| ENSMUSG00000085286 | 1.084558424 | 0.793139117 |
| ENSMUSG00000085287 | 1.079407717 | 0.419079937 |
| ENSMUSG00000085288 | 1 | |
| ENSMUSG00000085289 | 1.108738655 | 0.698112512 |
| ENSMUSG00000085290 | 1 | |
| ENSMUSG00000085292 | 1 | |
| ENSMUSG00000085293 | 1 | |
| ENSMUSG00000085295 | 1.162340834 | 0.608497229 |
| ENSMUSG00000085297 | 1 | |
| ENSMUSG00000085298 | 1.019083579 | 0.982175268 |
| ENSMUSG00000085300 | 1 | |
| ENSMUSG00000085301 | 1 | |
| ENSMUSG00000085302 | 1 | |
| ENSMUSG00000085303 | 1 | |
| ENSMUSG00000085304 | 1 | |
| ENSMUSG00000085305 | 1 | |
| ENSMUSG00000085307 | -1.007366764 | 1 |
| ENSMUSG00000085310 | -1.014346856 | 1 |
| ENSMUSG00000085311 | 1.005951819 | 1 |
| ENSMUSG00000085312 | 1.111812527 | 0.699591599 |
| ENSMUSG00000085314 | 1 | |
| ENSMUSG00000085315 | 1.107156681 | 0.702500944 |
| ENSMUSG00000085316 | -1.053549062 | 0.769745556 |
| ENSMUSG00000085317 | 1 | |
| ENSMUSG00000085318 | 1 | |
| ENSMUSG00000085319 | 1 | |
| ENSMUSG00000085320 | -1 | 1 |
| ENSMUSG00000085321 | 1.014545335 | 0.979002724 |
| ENSMUSG00000085322 | 1 | |
| ENSMUSG00000085323 | -1 | 1 |
| ENSMUSG00000085324 | 1 | |
| ENSMUSG00000085325 | 1 | |
| ENSMUSG00000085326 | 1 | |
| ENSMUSG00000085327 | 1.12691148 | 0.608497229 |
| ENSMUSG00000085328 | 1.128636259 | 0.263355382 |
| ENSMUSG00000085329 | 1 | |
| ENSMUSG00000085330 | 1.24122829 | 0.03679259 |
| ENSMUSG00000085331 | -1 | 1 |
| ENSMUSG00000085333 | 1 | |
| ENSMUSG00000085334 | 1.080357207 | 0.636090674 |
| ENSMUSG00000085335 | 1.043550045 | 0.895092332 |
| ENSMUSG00000085336 | 1 | |
| ENSMUSG00000085337 | 1.098760651 | 0.716155572 |
| ENSMUSG00000085338 | 1 | |

| | | |
|---|---|---|
| ENSMUSG00000085340 | -1.069529147 | 0.837531941 |
| ENSMUSG00000085342 | -1.109582661 | 0.66912118 |
| ENSMUSG00000085345 | 1 | |
| ENSMUSG00000085347 | -1.061159997 | 0.884220301 |
| ENSMUSG00000085348 | 1 | |
| ENSMUSG00000085350 | 1.028323353 | 0.919123248 |
| ENSMUSG00000085351 | -1.111880267 | 0.572947856 |
| ENSMUSG00000085352 | 1.015486245 | 0.972980766 |
| ENSMUSG00000085353 | 1 | |
| ENSMUSG00000085354 | 1.085624843 | 0.69305964 |
| ENSMUSG00000085355 | 1.214942357 | 0.327421605 |
| ENSMUSG00000085357 | 1 | |
| ENSMUSG00000085358 | 1 | |
| ENSMUSG00000085359 | 1.059463094 | 0.608497229 |
| ENSMUSG00000085360 | 1 | |
| ENSMUSG00000085361 | -1.013347035 | 1 |
| ENSMUSG00000085363 | 1.097758242 | 0.725073475 |
| ENSMUSG00000085366 | 1 | |
| ENSMUSG00000085368 | 1 | |
| ENSMUSG00000085369 | 1 | |
| ENSMUSG00000085371 | -1 | 1 |
| ENSMUSG00000085372 | 1 | |
| ENSMUSG00000085375 | 1 | |
| ENSMUSG00000085376 | 1.133432014 | 0.698112512 |
| ENSMUSG00000085377 | 1 | |
| ENSMUSG00000085378 | 1 | |
| ENSMUSG00000085379 | -1.125047041 | 0.667576553 |
| ENSMUSG00000085380 | 1 | |
| ENSMUSG00000085382 | 1 | |
| ENSMUSG00000085383 | 1 | |
| ENSMUSG00000085385 | -1.056541372 | 0.737905517 |
| ENSMUSG00000085386 | 1 | |
| ENSMUSG00000085387 | 1 | |
| ENSMUSG00000085388 | 1 | |
| ENSMUSG00000085389 | -1.218430774 | 0.339676656 |
| ENSMUSG00000085390 | 1 | |
| ENSMUSG00000085391 | 1 | |
| ENSMUSG00000085393 | 1 | |
| ENSMUSG00000085394 | 1 | |
| ENSMUSG00000085395 | 1.056098055 | 0.713964595 |
| ENSMUSG00000085396 | 1.067100704 | 0.697178241 |
| ENSMUSG00000085397 | -1.028897656 | 0.962613939 |
| ENSMUSG00000085398 | -1 | 1 |
| ENSMUSG00000085399 | -1.197233213 | 0.079029749 |
| ENSMUSG00000085400 | 1 | |
| ENSMUSG00000085401 | 1.019083579 | 0.949812939 |
| ENSMUSG00000085402 | -1.090507733 | 0.698112512 |
| ENSMUSG00000085403 | -1.002097693 | 1 |
| ENSMUSG00000085404 | -1.044273782 | 0.698112512 |
| ENSMUSG00000085405 | 1 | |
| ENSMUSG00000085407 | 1.030256832 | 0.964821074 |
| ENSMUSG00000085408 | -1.029951268 | 0.958380135 |
| ENSMUSG00000085409 | 1 | |
| ENSMUSG00000085411 | 1.11166259 | 0.649807331 |
| ENSMUSG00000085412 | 1.205610949 | 0.608497229 |
| ENSMUSG00000085413 | 1 | |
| ENSMUSG00000085414 | 1.015486245 | 0.972980766 |
| ENSMUSG00000085415 | -1.050031875 | 0.939933736 |
| ENSMUSG00000085416 | 1 | |
| ENSMUSG00000085417 | 1 | |
| ENSMUSG00000085418 | 1 | |
| ENSMUSG00000085419 | -1.099174267 | 0.608497229 |
| ENSMUSG00000085420 | 1 | |

| | | |
|---|---|---|
| ENSMUSG00000085421 | 1.210869356 | 0.327655754 |
| ENSMUSG00000085423 | -1.089497311 | 0.532369657 |
| ENSMUSG00000085424 | 1 | |
| ENSMUSG00000085425 | 1 | |
| ENSMUSG00000085426 | 1 | |
| ENSMUSG00000085427 | 1 | |
| ENSMUSG00000085428 | 1 | |
| ENSMUSG00000085429 | -1.038444538 | 0.865865121 |
| ENSMUSG00000085430 | 1 | |
| ENSMUSG00000085431 | 1 | |
| ENSMUSG00000085433 | -1.007501943 | 1 |
| ENSMUSG00000085434 | 1.089470641 | 0.608497229 |
| ENSMUSG00000085435 | -1 | 1 |
| ENSMUSG00000085436 | -1.265978343 | 0.027909004 |
| ENSMUSG00000085437 | 1.059463094 | 0.698112512 |
| ENSMUSG00000085438 | -1.174261466 | 0.555252837 |
| ENSMUSG00000085439 | 1 | |
| ENSMUSG00000085440 | 1.352281781 | 0.018089249 |
| ENSMUSG00000085441 | 1 | |
| ENSMUSG00000085442 | -1.032942875 | 0.945470588 |
| ENSMUSG00000085443 | 1.059463094 | 0.608497229 |
| ENSMUSG00000085444 | 1.170610441 | 0.624676024 |
| ENSMUSG00000085445 | -1 | 1 |
| ENSMUSG00000085447 | -1.044273782 | 0.698112512 |
| ENSMUSG00000085448 | -1.03962336 | 0.698112512 |
| ENSMUSG00000085449 | 1 | |
| ENSMUSG00000085450 | 1 | |
| ENSMUSG00000085451 | 1.015486245 | 0.972980766 |
| ENSMUSG00000085452 | -1.044273782 | 0.698112512 |
| ENSMUSG00000085453 | 1.037748783 | 0.847073978 |
| ENSMUSG00000085454 | 1.059463094 | 0.608497229 |
| ENSMUSG00000085456 | -1 | 1 |
| ENSMUSG00000085457 | 1.141381995 | 0.299044914 |
| ENSMUSG00000085459 | 1.034095782 | 0.947248675 |
| ENSMUSG00000085463 | -1 | 1 |
| ENSMUSG00000085464 | -1 | 1 |
| ENSMUSG00000085465 | 1.092044451 | 0.732012457 |
| ENSMUSG00000085466 | 1 | |
| ENSMUSG00000085467 | -1 | 1 |
| ENSMUSG00000085468 | 1 | |
| ENSMUSG00000085469 | 1 | |
| ENSMUSG00000085470 | 1 | |
| ENSMUSG00000085471 | 1.059463094 | 0.698112512 |
| ENSMUSG00000085472 | 1.059463094 | 0.608497229 |
| ENSMUSG00000085473 | 1.023743961 | 0.938939969 |
| ENSMUSG00000085476 | 1 | |
| ENSMUSG00000085477 | 1 | |
| ENSMUSG00000085478 | 1.053886948 | 0.819977901 |
| ENSMUSG00000085479 | -1.053549062 | 0.698112512 |
| ENSMUSG00000085481 | 1 | |
| ENSMUSG00000085482 | 1.114313713 | 0.698112512 |
| ENSMUSG00000085483 | 1.069068979 | 0.608497229 |
| ENSMUSG00000085484 | 1 | |
| ENSMUSG00000085485 | 1 | |
| ENSMUSG00000085486 | 1.132639128 | 0.428732467 |
| ENSMUSG00000085487 | 1.020240423 | 0.964236174 |
| ENSMUSG00000085488 | -1.03962336 | 0.698112512 |
| ENSMUSG00000085489 | 1 | |
| ENSMUSG00000085491 | 1 | |
| ENSMUSG00000085492 | 1.062626621 | 0.858016391 |
| ENSMUSG00000085495 | -1.121744392 | 0.608497229 |
| ENSMUSG00000085496 | 1 | |
| ENSMUSG00000085497 | 1 | |

| | | |
|---|---|---|
| ENSMUSG00000085498 | -1.00603089 | 1 |
| ENSMUSG00000085499 | -1 | 1 |
| ENSMUSG00000085500 | 1 | |
| ENSMUSG00000085501 | -1.04690868 | 0.88900664 |
| ENSMUSG00000085502 | 1 | |
| ENSMUSG00000085504 | 1 | |
| ENSMUSG00000085505 | 1 | |
| ENSMUSG00000085507 | 1 | |
| ENSMUSG00000085508 | 1.059463094 | 0.750182067 |
| ENSMUSG00000085509 | 1 | |
| ENSMUSG00000085510 | 1 | |
| ENSMUSG00000085511 | -1.137443589 | 0.532369657 |
| ENSMUSG00000085512 | 1 | |
| ENSMUSG00000085513 | 1 | |
| ENSMUSG00000085514 | -1 | 1 |
| ENSMUSG00000085515 | 1 | |
| ENSMUSG00000085516 | 1 | |
| ENSMUSG00000085517 | -1.185980269 | 0.482177248 |
| ENSMUSG00000085519 | 1 | |
| ENSMUSG00000085521 | -1.088954237 | 0.790851299 |
| ENSMUSG00000085522 | 1.218628056 | 0.107860597 |
| ENSMUSG00000085523 | 1.059463094 | 0.608497229 |
| ENSMUSG00000085524 | 1 | |
| ENSMUSG00000085525 | -1.049046231 | 0.817330349 |
| ENSMUSG00000085526 | 1 | |
| ENSMUSG00000085527 | -1.090871944 | 0.698112512 |
| ENSMUSG00000085528 | 1 | |
| ENSMUSG00000085530 | 1 | |
| ENSMUSG00000085531 | 1 | |
| ENSMUSG00000085532 | 1.101584861 | 0.698112512 |
| ENSMUSG00000085533 | 1 | |
| ENSMUSG00000085534 | 1 | |
| ENSMUSG00000085536 | 1 | |
| ENSMUSG00000085538 | -1.024530911 | 0.964236174 |
| ENSMUSG00000085539 | 1.199989356 | 0.000957737 |
| ENSMUSG00000085540 | 1 | |
| ENSMUSG00000085541 | 1 | |
| ENSMUSG00000085542 | 1 | |
| ENSMUSG00000085543 | 1 | |
| ENSMUSG00000085545 | 1 | |
| ENSMUSG00000085546 | 1.065590939 | 0.850280922 |
| ENSMUSG00000085547 | 1 | |
| ENSMUSG00000085548 | -1.044273782 | 0.870103307 |
| ENSMUSG00000085549 | 1 | |
| ENSMUSG00000085550 | 1 | |
| ENSMUSG00000085551 | 1 | |
| ENSMUSG00000085552 | 1 | |
| ENSMUSG00000085553 | 1 | |
| ENSMUSG00000085555 | -1.47242402 | 0.021070191 |
| ENSMUSG00000085556 | 1 | |
| ENSMUSG00000085558 | 1.048416787 | 0.891297795 |
| ENSMUSG00000085559 | 1 | |
| ENSMUSG00000085560 | 1.122462048 | 0.437095676 |
| ENSMUSG00000085561 | 1 | |
| ENSMUSG00000085562 | 1.059463094 | 0.608497229 |
| ENSMUSG00000085563 | -1.012953039 | 0.98469591 |
| ENSMUSG00000085564 | 1.332037334 | 0.117376001 |
| ENSMUSG00000085565 | 1.015486245 | 0.985522133 |
| ENSMUSG00000085566 | 1.141907285 | 0.352500944 |
| ENSMUSG00000085568 | 1.032926687 | 0.898373612 |
| ENSMUSG00000085569 | 1.846082083 | 0.003369375 |
| ENSMUSG00000085572 | 1.117607781 | 0.608497229 |
| ENSMUSG00000085573 | -1.030926232 | 0.951492347 |

| | | |
|---|---|---|
| ENSMUSG00000085574 | 1 | |
| ENSMUSG00000085575 | 1 | |
| ENSMUSG00000085576 | -1.03962336 | 0.698112512 |
| ENSMUSG00000085577 | 1 | |
| ENSMUSG00000085578 | 1.024693402 | 0.969087762 |
| ENSMUSG00000085579 | 1 | |
| ENSMUSG00000085580 | 1 | |
| ENSMUSG00000085581 | -1.043415084 | 0.729059471 |
| ENSMUSG00000085582 | -1.028348525 | 0.940333715 |
| ENSMUSG00000085583 | -1.125405469 | 0.662289659 |
| ENSMUSG00000085584 | 1.020344501 | 0.988665808 |
| ENSMUSG00000085585 | 1 | |
| ENSMUSG00000085586 | -1.091030263 | 0.543724675 |
| ENSMUSG00000085587 | 1 | |
| ENSMUSG00000085588 | 1 | |
| ENSMUSG00000085589 | 1.3634092 | 0.102742578 |
| ENSMUSG00000085591 | 1.059463094 | 0.698112512 |
| ENSMUSG00000085592 | 1 | |
| ENSMUSG00000085593 | 1 | |
| ENSMUSG00000085594 | 1 | |
| ENSMUSG00000085595 | 1 | |
| ENSMUSG00000085596 | 1.141387599 | 0.698112512 |
| ENSMUSG00000085598 | 1 | |
| ENSMUSG00000085599 | 1 | |
| ENSMUSG00000085600 | 1 | |
| ENSMUSG00000085601 | -1 | 1 |
| ENSMUSG00000085603 | -1.137745341 | 0.443913715 |
| ENSMUSG00000085604 | 1.059463094 | 0.698112512 |
| ENSMUSG00000085606 | -1.043743086 | 0.910832671 |
| ENSMUSG00000085608 | 1 | |
| ENSMUSG00000085609 | -1.03962336 | 0.698112512 |
| ENSMUSG00000085610 | 1.162135265 | 0.48302662 |
| ENSMUSG00000085611 | 1.059463094 | 0.768693672 |
| ENSMUSG00000085612 | -1.16741823 | 0.608497229 |
| ENSMUSG00000085614 | -1.003157381 | 1 |
| ENSMUSG00000085615 | -1.053477692 | 0.698112512 |
| ENSMUSG00000085616 | 1 | |
| ENSMUSG00000085618 | 1 | |
| ENSMUSG00000085619 | 1 | |
| ENSMUSG00000085620 | 1.014545335 | 0.979002724 |
| ENSMUSG00000085622 | -1.005368966 | 1 |
| ENSMUSG00000085623 | 1.253143688 | 0.138888882 |
| ENSMUSG00000085624 | 1 | |
| ENSMUSG00000085625 | -1 | 1 |
| ENSMUSG00000085626 | 1 | |
| ENSMUSG00000085627 | 1.014731082 | 0.988424085 |
| ENSMUSG00000085628 | 1.133505795 | 0.310231137 |
| ENSMUSG00000085629 | 1 | |
| ENSMUSG00000085630 | 1 | |
| ENSMUSG00000085631 | 1 | |
| ENSMUSG00000085632 | 1 | |
| ENSMUSG00000085633 | 1.075870199 | 0.748191299 |
| ENSMUSG00000085634 | 1 | |
| ENSMUSG00000085635 | 1.087900292 | 0.408663947 |
| ENSMUSG00000085636 | -1.009753323 | 1 |
| ENSMUSG00000085637 | 1.05395905 | 0.887392155 |
| ENSMUSG00000085638 | 1 | |
| ENSMUSG00000085639 | 1.278355251 | 0.185083841 |
| ENSMUSG00000085640 | 1 | |
| ENSMUSG00000085641 | 1 | |
| ENSMUSG00000085642 | 1.050351867 | 0.642819974 |
| ENSMUSG00000085643 | 1 | |
| ENSMUSG00000085645 | 1.182700909 | 0.435009832 |

| | | |
|---|---|---|
| ENSMUSG00000085646 | 1 | |
| ENSMUSG00000085648 | 1 | |
| ENSMUSG00000085649 | 1 | |
| ENSMUSG00000085651 | 1 | |
| ENSMUSG00000085652 | 1 | |
| ENSMUSG00000085653 | 1 | |
| ENSMUSG00000085654 | 1 | |
| ENSMUSG00000085655 | 1.04276791 | 0.696276767 |
| ENSMUSG00000085656 | 1 | |
| ENSMUSG00000085657 | 1 | |
| ENSMUSG00000085658 | 1.132877091 | 0.608497229 |
| ENSMUSG00000085659 | 1 | |
| ENSMUSG00000085660 | 1 | |
| ENSMUSG00000085662 | 1 | |
| ENSMUSG00000085663 | 1.122462048 | 0.541735653 |
| ENSMUSG00000085664 | 1.094347704 | 0.68585459 |
| ENSMUSG00000085665 | 1.022831284 | 0.901047139 |
| ENSMUSG00000085666 | 1.101130984 | 0.176723455 |
| ENSMUSG00000085667 | -1.257387052 | 0.234005431 |
| ENSMUSG00000085668 | 1 | |
| ENSMUSG00000085669 | 1.092890067 | 0.790851299 |
| ENSMUSG00000085672 | 1 | |
| ENSMUSG00000085673 | 1.069068979 | 0.608497229 |
| ENSMUSG00000085675 | 1.453820295 | 0.037706518 |
| ENSMUSG00000085676 | 1 | |
| ENSMUSG00000085677 | 1 | |
| ENSMUSG00000085678 | -1.064499817 | 0.865598094 |
| ENSMUSG00000085679 | 1 | |
| ENSMUSG00000085680 | 1 | |
| ENSMUSG00000085681 | -1.053549062 | 0.774498429 |
| ENSMUSG00000085682 | -1.03962336 | 0.803072249 |
| ENSMUSG00000085683 | 1 | |
| ENSMUSG00000085684 | 1 | |
| ENSMUSG00000085686 | 1 | |
| ENSMUSG00000085687 | 1.069722773 | 0.532369657 |
| ENSMUSG00000085688 | 1 | |
| ENSMUSG00000085689 | 1.064531743 | 0.796396036 |
| ENSMUSG00000085690 | 1 | |
| ENSMUSG00000085691 | 1 | |
| ENSMUSG00000085693 | -1.02513447 | 0.945625364 |
| ENSMUSG00000085694 | 1 | |
| ENSMUSG00000085695 | -1 | 1 |
| ENSMUSG00000085696 | 1.211679254 | 0.035334161 |
| ENSMUSG00000085698 | 1.039597911 | 0.91218384 |
| ENSMUSG00000085699 | 1 | |
| ENSMUSG00000085700 | 1 | |
| ENSMUSG00000085701 | 1 | |
| ENSMUSG00000085702 | 1.134431798 | 0.608497229 |
| ENSMUSG00000085704 | -1.146839817 | 0.476212392 |
| ENSMUSG00000085705 | -1.044273782 | 0.70299173 |
| ENSMUSG00000085706 | 1 | |
| ENSMUSG00000085707 | 1.015486245 | 0.972980766 |
| ENSMUSG00000085708 | 1 | |
| ENSMUSG00000085709 | 1 | |
| ENSMUSG00000085710 | -1 | 1 |
| ENSMUSG00000085711 | -1.03962336 | 0.698112512 |
| ENSMUSG00000085712 | -1.063525418 | 0.698961838 |
| ENSMUSG00000085713 | 1 | |
| ENSMUSG00000085714 | 1 | |
| ENSMUSG00000085715 | -1.028348525 | 0.912425817 |
| ENSMUSG00000085716 | 1 | |
| ENSMUSG00000085717 | 1 | |
| ENSMUSG00000085718 | 1.043618539 | 0.932324875 |

| | | |
|---|---|---|
| ENSMUSG00000085719 | 1 | |
| ENSMUSG00000085720 | 1 | |
| ENSMUSG00000085721 | -1 | 1 |
| ENSMUSG00000085722 | 1 | |
| ENSMUSG00000085723 | -1.015511103 | 0.982777255 |
| ENSMUSG00000085724 | 1 | |
| ENSMUSG00000085725 | 1 | |
| ENSMUSG00000085728 | 1 | |
| ENSMUSG00000085730 | 1.059463094 | 0.608497229 |
| ENSMUSG00000085731 | 1.059463094 | 0.608497229 |
| ENSMUSG00000085732 | -1.043306199 | 0.698112512 |
| ENSMUSG00000085733 | 1 | |
| ENSMUSG00000085734 | 1 | |
| ENSMUSG00000085735 | 1.054929781 | 0.915070045 |
| ENSMUSG00000085737 | 1.028323353 | 0.954497929 |
| ENSMUSG00000085738 | -1.113245897 | 0.547796683 |
| ENSMUSG00000085739 | 1 | |
| ENSMUSG00000085740 | 1 | |
| ENSMUSG00000085741 | -1.012493565 | 1 |
| ENSMUSG00000085742 | 1 | |
| ENSMUSG00000085743 | 1.240949182 | 0.247083615 |
| ENSMUSG00000085744 | -1.032852743 | 0.965852017 |
| ENSMUSG00000085745 | -1 | 1 |
| ENSMUSG00000085747 | 1.015486245 | 0.998386582 |
| ENSMUSG00000085748 | 1.106766808 | 0.729354471 |
| ENSMUSG00000085749 | 1.046500801 | 0.924587329 |
| ENSMUSG00000085750 | -1.071075483 | 0.747163034 |
| ENSMUSG00000085751 | 1 | |
| ENSMUSG00000085753 | 1 | |
| ENSMUSG00000085754 | 1 | |
| ENSMUSG00000085755 | 1.019083579 | 0.968937442 |
| ENSMUSG00000085756 | 1 | |
| ENSMUSG00000085757 | 1 | |
| ENSMUSG00000085759 | 1 | |
| ENSMUSG00000085760 | -1.100841795 | 0.730620997 |
| ENSMUSG00000085761 | -1 | 1 |
| ENSMUSG00000085762 | -1.071075483 | 0.698112512 |
| ENSMUSG00000085763 | 1.075070132 | 0.698112512 |
| ENSMUSG00000085765 | -1.147838869 | 0.350417454 |
| ENSMUSG00000085766 | 1.056953672 | 0.698112512 |
| ENSMUSG00000085767 | 1.022360487 | 0.948224739 |
| ENSMUSG00000085768 | 1.069068979 | 0.698112512 |
| ENSMUSG00000085770 | 1 | |
| ENSMUSG00000085771 | 1 | |
| ENSMUSG00000085772 | 1 | |
| ENSMUSG00000085773 | -1.043306199 | 0.698112512 |
| ENSMUSG00000085774 | 1.068666371 | 0.862994256 |
| ENSMUSG00000085775 | 1 | |
| ENSMUSG00000085776 | -1.048391303 | 0.909855627 |
| ENSMUSG00000085779 | 1.089470641 | 0.608497229 |
| ENSMUSG00000085781 | 1.222769118 | 0.449109628 |
| ENSMUSG00000085782 | 1.145970788 | 0.633720309 |
| ENSMUSG00000085783 | 1 | |
| ENSMUSG00000085784 | -1.047741049 | 0.608497229 |
| ENSMUSG00000085785 | 1 | |
| ENSMUSG00000085786 | 1.218240674 | 0.251691567 |
| ENSMUSG00000085787 | -1.044273782 | 0.855488553 |
| ENSMUSG00000085788 | -1.028348525 | 0.912425817 |
| ENSMUSG00000085790 | 1.059463094 | 0.608497229 |
| ENSMUSG00000085791 | -1 | 1 |
| ENSMUSG00000085792 | 1 | |
| ENSMUSG00000085793 | 1.070848141 | 0.616603327 |
| ENSMUSG00000085794 | -1.557806049 | 0.014289981 |

| | | |
|---|---|---|
| ENSMUSG00000085795 | 1.194900894 | 0.040506622 |
| ENSMUSG00000085796 | 1 | |
| ENSMUSG00000085797 | 1 | |
| ENSMUSG00000085798 | 1 | |
| ENSMUSG00000085799 | 1.059463094 | 0.608497229 |
| ENSMUSG00000085800 | 1.014545335 | 0.991944074 |
| ENSMUSG00000085801 | 1 | |
| ENSMUSG00000085802 | 1.034738199 | 0.971519533 |
| ENSMUSG00000085803 | 1.161036672 | 0.251049598 |
| ENSMUSG00000085804 | 1 | |
| ENSMUSG00000085805 | 1 | |
| ENSMUSG00000085806 | 1.122462048 | 0.608497229 |
| ENSMUSG00000085807 | 1.079667456 | 0.807456157 |
| ENSMUSG00000085808 | -1.03962336 | 0.698112512 |
| ENSMUSG00000085810 | -1.043306199 | 0.698112512 |
| ENSMUSG00000085811 | 1.084413116 | 0.535052277 |
| ENSMUSG00000085812 | 1.014545335 | 0.991944074 |
| ENSMUSG00000085813 | 1 | |
| ENSMUSG00000085815 | -1.03962336 | 0.698112512 |
| ENSMUSG00000085816 | 1 | |
| ENSMUSG00000085817 | -1.028348525 | 0.969007252 |
| ENSMUSG00000085818 | 1.015486245 | 0.985522133 |
| ENSMUSG00000085819 | 1.038431323 | 0.968937442 |
| ENSMUSG00000085820 | 1 | |
| ENSMUSG00000085822 | -1.046037523 | 0.919950762 |
| ENSMUSG00000085823 | 1 | |
| ENSMUSG00000085824 | 1.10634245 | 0.698112512 |
| ENSMUSG00000085826 | -1.066004548 | 0.70141105 |
| ENSMUSG00000085827 | 1 | |
| ENSMUSG00000085828 | -1.193098649 | 0.532369657 |
| ENSMUSG00000085829 | 1.038975273 | 0.865583959 |
| ENSMUSG00000085830 | -1.089497311 | 0.608497229 |
| ENSMUSG00000085831 | 1 | |
| ENSMUSG00000085833 | -1.043306199 | 0.698112512 |
| ENSMUSG00000085834 | 1 | |
| ENSMUSG00000085835 | 1 | |
| ENSMUSG00000085836 | 1 | |
| ENSMUSG00000085837 | -1.100640286 | 0.644981762 |
| ENSMUSG00000085838 | 1 | |
| ENSMUSG00000085839 | 1.059463094 | 0.608497229 |
| ENSMUSG00000085840 | 1.084689441 | 0.608497229 |
| ENSMUSG00000085841 | 1 | |
| ENSMUSG00000085842 | -1.120382953 | 0.699340382 |
| ENSMUSG00000085843 | 1.006564967 | 1 |
| ENSMUSG00000085844 | 1 | |
| ENSMUSG00000085845 | 1 | |
| ENSMUSG00000085846 | 1 | |
| ENSMUSG00000085847 | 1.059463094 | 0.608497229 |
| ENSMUSG00000085851 | -1.126852886 | 0.706524102 |
| ENSMUSG00000085852 | -1.241896834 | 0.34585736 |
| ENSMUSG00000085853 | 1 | |
| ENSMUSG00000085856 | 1 | |
| ENSMUSG00000085857 | 1 | |
| ENSMUSG00000085858 | 1 | |
| ENSMUSG00000085860 | 1 | |
| ENSMUSG00000085861 | 1.091498489 | 0.789448756 |
| ENSMUSG00000085862 | 1.161252845 | 0.608497229 |
| ENSMUSG00000085863 | 1 | |
| ENSMUSG00000085864 | -1.044273782 | 0.698112512 |
| ENSMUSG00000085865 | 1 | |
| ENSMUSG00000085866 | 1 | |
| ENSMUSG00000085867 | -1 | 1 |
| ENSMUSG00000085868 | -1.162474293 | 0.608497229 |

| | | |
|---|---|---|
| ENSMUSG00000085869 | 1 | |
| ENSMUSG00000085870 | 1.069068979 | 0.698112512 |
| ENSMUSG00000085871 | 1 | |
| ENSMUSG00000085872 | 1 | |
| ENSMUSG00000085873 | 1.149737166 | 0.219185116 |
| ENSMUSG00000085875 | 1.11166259 | 0.686411811 |
| ENSMUSG00000085876 | 1 | |
| ENSMUSG00000085877 | 1 | |
| ENSMUSG00000085878 | 1.059463094 | 0.608497229 |
| ENSMUSG00000085880 | 1 | |
| ENSMUSG00000085881 | 1.087690111 | 0.837242197 |
| ENSMUSG00000085882 | -1.213971144 | 0.003144607 |
| ENSMUSG00000085883 | 1 | |
| ENSMUSG00000085884 | -1.121136286 | 0.655571244 |
| ENSMUSG00000085885 | 1 | |
| ENSMUSG00000085886 | -1.043306199 | 0.79291651 |
| ENSMUSG00000085887 | 1.095872691 | 0.698112512 |
| ENSMUSG00000085888 | -1.071075483 | 0.750635131 |
| ENSMUSG00000085889 | -1.043306199 | 0.79291651 |
| ENSMUSG00000085890 | 1.007688634 | 1 |
| ENSMUSG00000085891 | -1.172366539 | 0.608497229 |
| ENSMUSG00000085892 | 1 | |
| ENSMUSG00000085893 | 1.039705955 | 0.951098417 |
| ENSMUSG00000085894 | -1.090915903 | 0.571773179 |
| ENSMUSG00000085895 | 1 | |
| ENSMUSG00000085896 | 1 | |
| ENSMUSG00000085897 | -1.027805693 | 0.953613038 |
| ENSMUSG00000085898 | 1.03265388 | 0.725290225 |
| ENSMUSG00000085899 | 1 | |
| ENSMUSG00000085900 | -1.190907694 | 0.518298194 |
| ENSMUSG00000085902 | 1.084618945 | 0.70299173 |
| ENSMUSG00000085903 | -1 | 1 |
| ENSMUSG00000085905 | 1 | |
| ENSMUSG00000085908 | 1 | |
| ENSMUSG00000085909 | 1 | |
| ENSMUSG00000085910 | 1.019083579 | 0.949812939 |
| ENSMUSG00000085911 | 1 | |
| ENSMUSG00000085912 | 1.416283537 | 0.089418777 |
| ENSMUSG00000085913 | -1.167142061 | 0.373031949 |
| ENSMUSG00000085914 | 1.095872691 | 0.698112512 |
| ENSMUSG00000085915 | 1 | |
| ENSMUSG00000085917 | 1 | |
| ENSMUSG00000085918 | 1 | |
| ENSMUSG00000085919 | 1 | |
| ENSMUSG00000085920 | 1 | |
| ENSMUSG00000085921 | 1 | |
| ENSMUSG00000085922 | 1 | |
| ENSMUSG00000085923 | 1 | |
| ENSMUSG00000085924 | -1.043306199 | 0.698112512 |
| ENSMUSG00000085925 | 1.252219971 | 0.517160377 |
| ENSMUSG00000085926 | 1 | |
| ENSMUSG00000085927 | 1 | |
| ENSMUSG00000085928 | 1 | |
| ENSMUSG00000085929 | 1.001896181 | 1 |
| ENSMUSG00000085931 | -1.100193663 | 0.702070725 |
| ENSMUSG00000085932 | 1 | |
| ENSMUSG00000085933 | 1 | |
| ENSMUSG00000085934 | 1 | |
| ENSMUSG00000085936 | -1.060116214 | 0.748571751 |
| ENSMUSG00000085937 | -1.043306199 | 0.698112512 |
| ENSMUSG00000085938 | 1.059463094 | 0.608497229 |
| ENSMUSG00000085939 | 1.051896549 | 0.903418476 |
| ENSMUSG00000085940 | 1 | |

| | | |
|---|---|---|
| ENSMUSG00000085941 | 1.279621157 | 0.030511097 |
| ENSMUSG00000085944 | -1.053549062 | 0.698112512 |
| ENSMUSG00000085945 | 1.080134941 | 0.771267735 |
| ENSMUSG00000085946 | 1 | |
| ENSMUSG00000085949 | -1.095294215 | 0.608497229 |
| ENSMUSG00000085950 | 1.044896373 | 0.867519242 |
| ENSMUSG00000085951 | 1 | |
| ENSMUSG00000085952 | -1.044273782 | 0.698112512 |
| ENSMUSG00000085953 | 1 | |
| ENSMUSG00000085955 | -1.039056293 | 0.956401771 |
| ENSMUSG00000085956 | -1.007394476 | 1 |
| ENSMUSG00000085957 | 1.145056213 | 0.732012457 |
| ENSMUSG00000085958 | 1.089470641 | 0.657060635 |
| ENSMUSG00000085961 | 1.014545335 | 0.979002724 |
| ENSMUSG00000085962 | -1.013575326 | 1 |
| ENSMUSG00000085965 | 1 | |
| ENSMUSG00000085966 | 1 | |
| ENSMUSG00000085967 | 1 | |
| ENSMUSG00000085968 | 1 | |
| ENSMUSG00000085969 | 1 | |
| ENSMUSG00000085970 | 1 | |
| ENSMUSG00000085971 | 1.189926019 | 0.159807598 |
| ENSMUSG00000085972 | 1.11166259 | 0.698112512 |
| ENSMUSG00000085973 | 1 | |
| ENSMUSG00000085975 | -1 | 1 |
| ENSMUSG00000085976 | -1.210362854 | 0.452069807 |
| ENSMUSG00000085977 | 1 | |
| ENSMUSG00000085978 | 1 | |
| ENSMUSG00000085979 | 1 | |
| ENSMUSG00000085980 | 1 | |
| ENSMUSG00000085981 | 1 | |
| ENSMUSG00000085982 | 1.162504806 | 0.628604034 |
| ENSMUSG00000085984 | -1.203902127 | 0.503617425 |
| ENSMUSG00000085985 | 1 | |
| ENSMUSG00000085987 | -1.044273782 | 0.698112512 |
| ENSMUSG00000085989 | 1 | |
| ENSMUSG00000085990 | -1 | 1 |
| ENSMUSG00000085991 | 1 | |
| ENSMUSG00000085992 | -1.003834322 | 1 |
| ENSMUSG00000085993 | 1 | |
| ENSMUSG00000085995 | -1.471006056 | 0.009584796 |
| ENSMUSG00000085996 | 1.059463094 | 0.752398528 |
| ENSMUSG00000085998 | -1.014570171 | 1 |
| ENSMUSG00000085999 | 1 | |
| ENSMUSG00000086000 | 1 | |
| ENSMUSG00000086001 | -1.099174267 | 0.535052277 |
| ENSMUSG00000086003 | 1.173639885 | 0.455298078 |
| ENSMUSG00000086004 | -1.044273782 | 0.698112512 |
| ENSMUSG00000086006 | 1.132639128 | 0.225438217 |
| ENSMUSG00000086007 | 1 | |
| ENSMUSG00000086008 | 1.122462048 | 0.215991492 |
| ENSMUSG00000086009 | 1 | |
| ENSMUSG00000086010 | 1 | |
| ENSMUSG00000086011 | 1 | |
| ENSMUSG00000086012 | 1 | |
| ENSMUSG00000086013 | 1.133854707 | 0.182040757 |
| ENSMUSG00000086014 | -1 | 1 |
| ENSMUSG00000086015 | 1.058183296 | 0.653960635 |
| ENSMUSG00000086016 | 1.12316944 | 0.617501656 |
| ENSMUSG00000086017 | 1 | |
| ENSMUSG00000086018 | 1 | |
| ENSMUSG00000086019 | 1 | |
| ENSMUSG00000086020 | 1 | |

| | | |
|---|---|---|
| ENSMUSG00000086021 | 1 | |
| ENSMUSG00000086022 | -1.012966911 | 1 |
| ENSMUSG00000086025 | 1.089470641 | 0.657060635 |
| ENSMUSG00000086026 | 1 | |
| ENSMUSG00000086027 | 1 | |
| ENSMUSG00000086028 | 1 | |
| ENSMUSG00000086029 | -1.03962336 | 0.698112512 |
| ENSMUSG00000086030 | 1 | |
| ENSMUSG00000086031 | 1 | |
| ENSMUSG00000086032 | -1 | 1 |
| ENSMUSG00000086034 | 1 | |
| ENSMUSG00000086035 | 1.010877339 | 0.982838256 |
| ENSMUSG00000086036 | 1 | |
| ENSMUSG00000086037 | 1 | |
| ENSMUSG00000086039 | -1.044273782 | 0.790851299 |
| ENSMUSG00000086040 | -1.266113256 | 0.517477442 |
| ENSMUSG00000086041 | 1.271344436 | 0.053354336 |
| ENSMUSG00000086042 | 1 | |
| ENSMUSG00000086043 | 1.124313245 | 0.549558201 |
| ENSMUSG00000086044 | 1 | |
| ENSMUSG00000086046 | -1 | 1 |
| ENSMUSG00000086047 | 1.081588278 | 0.746504552 |
| ENSMUSG00000086048 | 1 | |
| ENSMUSG00000086049 | -1.138693452 | 0.608497229 |
| ENSMUSG00000086050 | -1.15462599 | 0.317733967 |
| ENSMUSG00000086052 | 1 | |
| ENSMUSG00000086053 | 1.014545335 | 0.991944074 |
| ENSMUSG00000086054 | 1.059463094 | 0.608497229 |
| ENSMUSG00000086055 | 1 | |
| ENSMUSG00000086056 | 1 | |
| ENSMUSG00000086057 | 1 | |
| ENSMUSG00000086058 | 1 | |
| ENSMUSG00000086060 | 1 | |
| ENSMUSG00000086061 | -1.089497311 | 0.585199025 |
| ENSMUSG00000086063 | 1 | |
| ENSMUSG00000086064 | 1 | |
| ENSMUSG00000086065 | 1 | |
| ENSMUSG00000086066 | 1 | |
| ENSMUSG00000086067 | 1.372330679 | 0.003144607 |
| ENSMUSG00000086068 | 1 | |
| ENSMUSG00000086069 | 1 | |
| ENSMUSG00000086070 | 1.095872691 | 0.749353361 |
| ENSMUSG00000086071 | 1 | |
| ENSMUSG00000086072 | 1 | |
| ENSMUSG00000086074 | -1 | 1 |
| ENSMUSG00000086075 | 1.04321977 | 0.706806763 |
| ENSMUSG00000086076 | -1.13161737 | 0.630379139 |
| ENSMUSG00000086077 | 1 | |
| ENSMUSG00000086078 | 1.132639128 | 0.414898324 |
| ENSMUSG00000086080 | 1 | |
| ENSMUSG00000086081 | 1 | |
| ENSMUSG00000086083 | -1.009476377 | 1 |
| ENSMUSG00000086087 | 1 | |
| ENSMUSG00000086089 | 1 | |
| ENSMUSG00000086090 | 1 | |
| ENSMUSG00000086092 | 1 | |
| ENSMUSG00000086093 | -1.03962336 | 0.698112512 |
| ENSMUSG00000086095 | 1.10476636 | 0.784686694 |
| ENSMUSG00000086096 | -1.03962336 | 0.698112512 |
| ENSMUSG00000086097 | 1 | |
| ENSMUSG00000086098 | -1.121136286 | 0.608497229 |
| ENSMUSG00000086103 | -1.0469665 | 0.776021306 |
| ENSMUSG00000086104 | 1 | |

| | | |
|---|---|---|
| ENSMUSG00000086105 | 1.152301238 | 0.564553258 |
| ENSMUSG00000086107 | 1 | |
| ENSMUSG00000086108 | -1.122345898 | 0.715866916 |
| ENSMUSG00000086109 | 1 | |
| ENSMUSG00000086111 | 1 | |
| ENSMUSG00000086112 | 1 | |
| ENSMUSG00000086114 | 1 | |
| ENSMUSG00000086115 | 1 | |
| ENSMUSG00000086117 | 1 | |
| ENSMUSG00000086118 | 1 | |
| ENSMUSG00000086119 | -1.129489754 | 0.47476794 |
| ENSMUSG00000086121 | -1.053549062 | 0.778372272 |
| ENSMUSG00000086122 | 1 | |
| ENSMUSG00000086123 | -1.069502966 | 0.778372272 |
| ENSMUSG00000086124 | 1 | |
| ENSMUSG00000086125 | 1.059463094 | 0.608497229 |
| ENSMUSG00000086126 | 1 | |
| ENSMUSG00000086127 | 1.05914025 | 0.813331954 |
| ENSMUSG00000086128 | 1 | |
| ENSMUSG00000086129 | -1.043306199 | 0.698112512 |
| ENSMUSG00000086130 | 1.061713892 | 0.608497229 |
| ENSMUSG00000086132 | 1 | |
| ENSMUSG00000086133 | 1 | |
| ENSMUSG00000086134 | -1.044697704 | 0.889420116 |
| ENSMUSG00000086135 | 1 | |
| ENSMUSG00000086136 | 1 | |
| ENSMUSG00000086137 | 1 | |
| ENSMUSG00000086138 | 1 | |
| ENSMUSG00000086139 | 1 | |
| ENSMUSG00000086140 | 1 | |
| ENSMUSG00000086141 | 1 | |
| ENSMUSG00000086142 | 1 | |
| ENSMUSG00000086144 | 1 | |
| ENSMUSG00000086145 | 1.059463094 | 0.608497229 |
| ENSMUSG00000086146 | -1.043306199 | 0.698112512 |
| ENSMUSG00000086147 | -1.03962336 | 0.807247723 |
| ENSMUSG00000086148 | 1 | |
| ENSMUSG00000086149 | 1.084618945 | 0.698112512 |
| ENSMUSG00000086150 | 1.167431987 | 0.608497229 |
| ENSMUSG00000086151 | 1.059463094 | 0.608497229 |
| ENSMUSG00000086152 | -1.085651418 | 0.608497229 |
| ENSMUSG00000086153 | 1 | |
| ENSMUSG00000086154 | -1.043306199 | 0.698112512 |
| ENSMUSG00000086155 | -1.019108526 | 0.979258908 |
| ENSMUSG00000086156 | 1 | |
| ENSMUSG00000086157 | 1 | |
| ENSMUSG00000086158 | 1.022258728 | 0.888882286 |
| ENSMUSG00000086159 | -1.054741498 | 0.872692845 |
| ENSMUSG00000086160 | 1 | |
| ENSMUSG00000086161 | 1.015486245 | 0.987775262 |
| ENSMUSG00000086162 | 1 | |
| ENSMUSG00000086163 | 1 | |
| ENSMUSG00000086164 | -1.040045394 | 0.89103058 |
| ENSMUSG00000086165 | 1.17792434 | 0.607014477 |
| ENSMUSG00000086166 | -1.044820185 | 0.892464004 |
| ENSMUSG00000086167 | 1 | |
| ENSMUSG00000086168 | 1 | |
| ENSMUSG00000086170 | 1 | |
| ENSMUSG00000086171 | 1.059463094 | 0.698112512 |
| ENSMUSG00000086172 | -1.199425955 | 0.440395661 |
| ENSMUSG00000086173 | -1.053549062 | 0.698112512 |
| ENSMUSG00000086174 | 1 | |
| ENSMUSG00000086175 | 1 | |

| | | |
|---|---|---|
| ENSMUSG00000086176 | 1 | |
| ENSMUSG00000086177 | 1 | |
| ENSMUSG00000086179 | 1 | |
| ENSMUSG00000086181 | 1.491434678 | 0.15009325 |
| ENSMUSG00000086184 | -1.043306199 | 0.699340382 |
| ENSMUSG00000086185 | 1 | |
| ENSMUSG00000086186 | 1.177765487 | 0.458872271 |
| ENSMUSG00000086187 | -1.044273782 | 0.698112512 |
| ENSMUSG00000086188 | -1.001791545 | 1 |
| ENSMUSG00000086189 | -1.075232152 | 0.656734444 |
| ENSMUSG00000086190 | 1 | |
| ENSMUSG00000086191 | -1.049157586 | 0.921571363 |
| ENSMUSG00000086192 | 1.034865357 | 0.899648217 |
| ENSMUSG00000086193 | 1.095872691 | 0.608497229 |
| ENSMUSG00000086194 | 1 | |
| ENSMUSG00000086195 | 1 | |
| ENSMUSG00000086196 | 1.069068979 | 0.698112512 |
| ENSMUSG00000086197 | 1 | |
| ENSMUSG00000086199 | -1.111221915 | 0.614476325 |
| ENSMUSG00000086201 | 1 | |
| ENSMUSG00000086202 | 1 | |
| ENSMUSG00000086203 | 1.019083579 | 0.968937442 |
| ENSMUSG00000086204 | 1 | |
| ENSMUSG00000086205 | 1 | |
| ENSMUSG00000086206 | 1.059463094 | 0.608497229 |
| ENSMUSG00000086207 | -1 | 1 |
| ENSMUSG00000086209 | 1 | |
| ENSMUSG00000086211 | 1 | |
| ENSMUSG00000086212 | 1.069427779 | 0.801297922 |
| ENSMUSG00000086213 | 1.269929483 | 0.288227228 |
| ENSMUSG00000086214 | 1 | |
| ENSMUSG00000086216 | 1.023743961 | 0.968375666 |
| ENSMUSG00000086219 | 1.534742129 | 0.032820657 |
| ENSMUSG00000086220 | 1 | |
| ENSMUSG00000086221 | 1 | |
| ENSMUSG00000086222 | 1 | |
| ENSMUSG00000086224 | 1.011607416 | 1 |
| ENSMUSG00000086225 | 1 | |
| ENSMUSG00000086226 | 1 | |
| ENSMUSG00000086227 | 1 | |
| ENSMUSG00000086228 | -1.027744676 | 0.985636424 |
| ENSMUSG00000086229 | 1.069293585 | 0.770958539 |
| ENSMUSG00000086231 | 1 | |
| ENSMUSG00000086232 | 1 | |
| ENSMUSG00000086233 | 1.306949949 | 0.277228509 |
| ENSMUSG00000086234 | 1 | |
| ENSMUSG00000086235 | 1 | |
| ENSMUSG00000086236 | -1.044273782 | 0.893554938 |
| ENSMUSG00000086237 | -1.006281294 | 1 |
| ENSMUSG00000086238 | -1.044273782 | 0.698112512 |
| ENSMUSG00000086239 | -1 | 1 |
| ENSMUSG00000086240 | 1.059463094 | 0.608497229 |
| ENSMUSG00000086241 | -1.252837605 | 0.47877278 |
| ENSMUSG00000086243 | 1 | |
| ENSMUSG00000086244 | 1 | |
| ENSMUSG00000086245 | 1.013443899 | 0.989571299 |
| ENSMUSG00000086246 | 1 | |
| ENSMUSG00000086247 | 1.046677512 | 0.918034349 |
| ENSMUSG00000086248 | 1 | |
| ENSMUSG00000086249 | 1 | |
| ENSMUSG00000086250 | 1 | |
| ENSMUSG00000086251 | -1.03962336 | 0.698112512 |
| ENSMUSG00000086253 | 1 | |

| | | |
|---|---|---|
| ENSMUSG00000086254 | 1.059463094 | 0.608497229 |
| ENSMUSG00000086255 | 1 | |
| ENSMUSG00000086256 | 1.059463094 | 0.608497229 |
| ENSMUSG00000086257 | 1 | |
| ENSMUSG00000086258 | -1.03962336 | 0.698112512 |
| ENSMUSG00000086259 | 1 | |
| ENSMUSG00000086260 | 1 | |
| ENSMUSG00000086261 | -1.053549062 | 0.698112512 |
| ENSMUSG00000086262 | 1 | |
| ENSMUSG00000086263 | 1 | |
| ENSMUSG00000086264 | 1.100748059 | 0.685372066 |
| ENSMUSG00000086265 | 1 | |
| ENSMUSG00000086266 | -1.053549062 | 0.698112512 |
| ENSMUSG00000086267 | 1.116567227 | 0.647072889 |
| ENSMUSG00000086268 | 1 | |
| ENSMUSG00000086269 | 1.054912313 | 0.858345219 |
| ENSMUSG00000086272 | 1.284154183 | 0.055595649 |
| ENSMUSG00000086273 | 1.222889447 | 0.333749058 |
| ENSMUSG00000086275 | 1.015486245 | 0.972980766 |
| ENSMUSG00000086277 | 1 | |
| ENSMUSG00000086278 | 1 | |
| ENSMUSG00000086279 | 1 | |
| ENSMUSG00000086282 | 1 | |
| ENSMUSG00000086283 | 1.301988128 | 0.013472992 |
| ENSMUSG00000086284 | 1 | |
| ENSMUSG00000086286 | 1.069068979 | 0.698112512 |
| ENSMUSG00000086287 | 1.059463094 | 0.608497229 |
| ENSMUSG00000086288 | -1.029935726 | 0.917472502 |
| ENSMUSG00000086289 | 1 | |
| ENSMUSG00000086290 | 1.082182126 | 0.419455697 |
| ENSMUSG00000086291 | -1.237062975 | 0.125154408 |
| ENSMUSG00000086292 | -1.039429576 | 0.90035832 |
| ENSMUSG00000086293 | 1 | |
| ENSMUSG00000086296 | -1.239528453 | 0.361182798 |
| ENSMUSG00000086297 | 1 | |
| ENSMUSG00000086298 | 1 | |
| ENSMUSG00000086302 | 1 | |
| ENSMUSG00000086306 | 1 | |
| ENSMUSG00000086308 | 1.034340763 | 0.924324458 |
| ENSMUSG00000086310 | 1 | |
| ENSMUSG00000086311 | -1.085651418 | 0.532369657 |
| ENSMUSG00000086312 | 1 | |
| ENSMUSG00000086313 | 1 | |
| ENSMUSG00000086315 | 1 | |
| ENSMUSG00000086316 | -1.061260915 | 0.69091159 |
| ENSMUSG00000086319 | 1 | |
| ENSMUSG00000086320 | 1.186419366 | 0.037163069 |
| ENSMUSG00000086321 | -1 | 1 |
| ENSMUSG00000086322 | -1.03962336 | 0.698112512 |
| ENSMUSG00000086324 | 1.077005225 | 0.424852858 |
| ENSMUSG00000086326 | 1 | |
| ENSMUSG00000086327 | 1 | |
| ENSMUSG00000086328 | 1.015336613 | 1 |
| ENSMUSG00000086329 | 1 | |
| ENSMUSG00000086330 | 1.021793092 | 0.983523687 |
| ENSMUSG00000086331 | 1.121850956 | 0.337363023 |
| ENSMUSG00000086332 | 1 | |
| ENSMUSG00000086333 | 1 | |
| ENSMUSG00000086335 | -1.043306199 | 0.803260194 |
| ENSMUSG00000086336 | 1 | |
| ENSMUSG00000086337 | -1 | 1 |
| ENSMUSG00000086338 | 1 | |
| ENSMUSG00000086339 | 1 | |

| | | |
|---|---|---|
| ENSMUSG00000086340 | 1.005851671 | 1 |
| ENSMUSG00000086341 | -1.1686065 | 0.476379342 |
| ENSMUSG00000086342 | -1.015029776 | 1 |
| ENSMUSG00000086344 | -1.143787033 | 0.201977401 |
| ENSMUSG00000086346 | 1 | |
| ENSMUSG00000086347 | 1.079681442 | 0.726987474 |
| ENSMUSG00000086348 | 1 | |
| ENSMUSG00000086349 | 1.024908686 | 0.977049036 |
| ENSMUSG00000086350 | -1.0476602 | 0.866643172 |
| ENSMUSG00000086352 | 1.059463094 | 0.608497229 |
| ENSMUSG00000086353 | 1.059463094 | 0.608497229 |
| ENSMUSG00000086354 | 1 | |
| ENSMUSG00000086355 | 1 | |
| ENSMUSG00000086356 | 1 | |
| ENSMUSG00000086358 | 1.108630979 | 0.704854542 |
| ENSMUSG00000086359 | 1.069210644 | 0.75366313 |
| ENSMUSG00000086360 | -1.044273782 | 0.790851299 |
| ENSMUSG00000086361 | 1 | |
| ENSMUSG00000086363 | 1.10141562 | 0.457437655 |
| ENSMUSG00000086364 | 1 | |
| ENSMUSG00000086365 | 1 | |
| ENSMUSG00000086366 | 1 | |
| ENSMUSG00000086367 | 1 | |
| ENSMUSG00000086368 | 1.034204701 | 0.734674233 |
| ENSMUSG00000086369 | 1 | |
| ENSMUSG00000086370 | 1.075697969 | 0.608497229 |
| ENSMUSG00000086371 | 1 | |
| ENSMUSG00000086372 | -1 | 1 |
| ENSMUSG00000086373 | 1 | |
| ENSMUSG00000086375 | -1.043306199 | 0.698112512 |
| ENSMUSG00000086376 | 1 | |
| ENSMUSG00000086377 | 1.102024305 | 0.694494111 |
| ENSMUSG00000086378 | 1 | |
| ENSMUSG00000086379 | 1 | |
| ENSMUSG00000086380 | 1 | |
| ENSMUSG00000086381 | 1 | |
| ENSMUSG00000086382 | -1.247623686 | 0.406431792 |
| ENSMUSG00000086384 | 1 | |
| ENSMUSG00000086385 | 1 | |
| ENSMUSG00000086386 | 1 | |
| ENSMUSG00000086387 | -1.053549062 | 0.774498429 |
| ENSMUSG00000086389 | -1.103038152 | 0.410032437 |
| ENSMUSG00000086390 | -1.034162213 | 0.98328394 |
| ENSMUSG00000086391 | -1.053976749 | 0.870602199 |
| ENSMUSG00000086392 | -1.036139145 | 0.935253485 |
| ENSMUSG00000086394 | 1 | |
| ENSMUSG00000086395 | -1.10627706 | 0.698112512 |
| ENSMUSG00000086396 | 1 | |
| ENSMUSG00000086399 | -1.186036456 | 0.389555558 |
| ENSMUSG00000086401 | 1.072139908 | 0.715898555 |
| ENSMUSG00000086402 | 1 | |
| ENSMUSG00000086404 | -1.203970232 | 0.19173105 |
| ENSMUSG00000086405 | 1 | |
| ENSMUSG00000086407 | -1.03962336 | 0.698112512 |
| ENSMUSG00000086408 | 1 | |
| ENSMUSG00000086409 | 1 | |
| ENSMUSG00000086410 | 1 | |
| ENSMUSG00000086411 | 1 | |
| ENSMUSG00000086413 | -1.24499826 | 0.15799366 |
| ENSMUSG00000086414 | 1.327652993 | 0.020440998 |
| ENSMUSG00000086415 | 1 | |
| ENSMUSG00000086416 | 1.344649656 | 0.011490798 |
| ENSMUSG00000086417 | -1.084645496 | 0.532369657 |

| | | |
|---|---|---|
| ENSMUSG00000086418 | 1 | |
| ENSMUSG00000086420 | -1 | 1 |
| ENSMUSG00000086421 | -1.07921746 | 0.718130029 |
| ENSMUSG00000086422 | 1 | |
| ENSMUSG00000086423 | 1 | |
| ENSMUSG00000086424 | -1.129231493 | 0.605619163 |
| ENSMUSG00000086425 | -1 | 1 |
| ENSMUSG00000086426 | 1 | |
| ENSMUSG00000086427 | 1.031568464 | 0.958600282 |
| ENSMUSG00000086428 | 1 | |
| ENSMUSG00000086429 | 1.077952292 | 0.65443598 |
| ENSMUSG00000086430 | -1.033833467 | 0.957935298 |
| ENSMUSG00000086431 | 1.069068979 | 0.608497229 |
| ENSMUSG00000086432 | 1 | |
| ENSMUSG00000086433 | 1 | |
| ENSMUSG00000086434 | 1 | |
| ENSMUSG00000086436 | -1.03962336 | 0.698112512 |
| ENSMUSG00000086437 | 1 | |
| ENSMUSG00000086438 | -1.053549062 | 0.769745556 |
| ENSMUSG00000086439 | 1 | |
| ENSMUSG00000086441 | 1 | |
| ENSMUSG00000086442 | -1.176660386 | 0.216046507 |
| ENSMUSG00000086443 | 1.042677087 | 0.915082038 |
| ENSMUSG00000086444 | 1.134523757 | 0.668750825 |
| ENSMUSG00000086445 | 1 | |
| ENSMUSG00000086446 | 1.035626983 | 0.815601444 |
| ENSMUSG00000086447 | 1 | |
| ENSMUSG00000086448 | -1.433765862 | 0.059684367 |
| ENSMUSG00000086449 | 1 | |
| ENSMUSG00000086450 | 1.105684453 | 0.608497229 |
| ENSMUSG00000086451 | -1.111880267 | 0.608497229 |
| ENSMUSG00000086452 | 1 | |
| ENSMUSG00000086453 | -1.101049618 | 0.70733922 |
| ENSMUSG00000086454 | -1.11676183 | 0.698112512 |
| ENSMUSG00000086455 | 1.068666572 | 0.905359034 |
| ENSMUSG00000086458 | 1 | |
| ENSMUSG00000086459 | -1.043306199 | 0.698112512 |
| ENSMUSG00000086460 | 1 | |
| ENSMUSG00000086462 | 1 | |
| ENSMUSG00000086464 | 1.042970129 | 0.887325696 |
| ENSMUSG00000086465 | 1 | |
| ENSMUSG00000086467 | -1.044273782 | 0.795446726 |
| ENSMUSG00000086468 | 1.014241457 | 1 |
| ENSMUSG00000086470 | -1 | 1 |
| ENSMUSG00000086472 | 1.046536713 | 0.969007252 |
| ENSMUSG00000086473 | 1.075072849 | 0.803260194 |
| ENSMUSG00000086474 | -1.143794946 | 0.641039836 |
| ENSMUSG00000086475 | -1.037921784 | 0.873118449 |
| ENSMUSG00000086477 | -1.095939461 | 0.688241341 |
| ENSMUSG00000086478 | 1 | |
| ENSMUSG00000086479 | 1 | |
| ENSMUSG00000086480 | -1.0585642 | 0.764751192 |
| ENSMUSG00000086481 | 1 | |
| ENSMUSG00000086482 | 1 | |
| ENSMUSG00000086483 | 1 | |
| ENSMUSG00000086484 | -1.043306199 | 0.798487595 |
| ENSMUSG00000086485 | 1 | |
| ENSMUSG00000086486 | -1.186377809 | 0.608497229 |
| ENSMUSG00000086487 | 1 | |
| ENSMUSG00000086489 | 1 | |
| ENSMUSG00000086490 | 1 | |
| ENSMUSG00000086491 | 1 | |
| ENSMUSG00000086493 | 1.059463094 | 0.608497229 |

| | | |
|---|---|---|
| ENSMUSG00000086494 | 1 | |
| ENSMUSG00000086495 | -1.161107412 | 0.532369657 |
| ENSMUSG00000086496 | -1.161737199 | 0.386294287 |
| ENSMUSG00000086497 | 1 | |
| ENSMUSG00000086498 | 1.122462048 | 0.215991492 |
| ENSMUSG00000086499 | 1.11166259 | 0.608497229 |
| ENSMUSG00000086500 | 1 | |
| ENSMUSG00000086501 | -1 | 1 |
| ENSMUSG00000086502 | -1.075224067 | 0.565609452 |
| ENSMUSG00000086503 | -1.04348991 | 0.737673689 |
| ENSMUSG00000086504 | 1 | |
| ENSMUSG00000086505 | 1 | |
| ENSMUSG00000086506 | -1.003507216 | 1 |
| ENSMUSG00000086507 | -1.084645496 | 0.532369657 |
| ENSMUSG00000086508 | 1 | |
| ENSMUSG00000086509 | 1 | |
| ENSMUSG00000086512 | 1 | |
| ENSMUSG00000086513 | -1 | 1 |
| ENSMUSG00000086514 | -1.038858794 | 0.897684198 |
| ENSMUSG00000086515 | -1.150235768 | 0.634365479 |
| ENSMUSG00000086517 | 1 | |
| ENSMUSG00000086518 | 1 | |
| ENSMUSG00000086519 | 1 | |
| ENSMUSG00000086521 | 1 | |
| ENSMUSG00000086522 | 1.336795758 | 0.231006646 |
| ENSMUSG00000086523 | 1 | |
| ENSMUSG00000086524 | 1 | |
| ENSMUSG00000086525 | 1 | |
| ENSMUSG00000086526 | 1.015840365 | 1 |
| ENSMUSG00000086527 | -1 | 1 |
| ENSMUSG00000086528 | 1.173879945 | 0.295078252 |
| ENSMUSG00000086529 | -1.03962336 | 0.698112512 |
| ENSMUSG00000086530 | 1 | |
| ENSMUSG00000086531 | 1.069068979 | 0.658433487 |
| ENSMUSG00000086533 | 1.047437598 | 0.836811166 |
| ENSMUSG00000086534 | -1.07008665 | 0.812147631 |
| ENSMUSG00000086536 | -1.103288263 | 0.321248448 |
| ENSMUSG00000086537 | -1.295724383 | 0.221183946 |
| ENSMUSG00000086539 | 1.020195175 | 0.9607713 |
| ENSMUSG00000086540 | -1 | 1 |
| ENSMUSG00000086541 | 1.449401007 | 0.135458517 |
| ENSMUSG00000086542 | -1 | 1 |
| ENSMUSG00000086544 | -1.028348525 | 0.9607713 |
| ENSMUSG00000086545 | 1 | |
| ENSMUSG00000086546 | 1 | |
| ENSMUSG00000086547 | 1 | |
| ENSMUSG00000086548 | 1 | |
| ENSMUSG00000086549 | 1 | |
| ENSMUSG00000086550 | 1 | |
| ENSMUSG00000086552 | -1.277443017 | 0.025895432 |
| ENSMUSG00000086554 | 1.070224878 | 0.836347433 |
| ENSMUSG00000086555 | 1 | |
| ENSMUSG00000086556 | -1 | 1 |
| ENSMUSG00000086557 | -1 | 1 |
| ENSMUSG00000086558 | 1 | |
| ENSMUSG00000086559 | 1 | |
| ENSMUSG00000086560 | 1.471092841 | 0.309553624 |
| ENSMUSG00000086561 | -1.044273782 | 0.699340382 |
| ENSMUSG00000086562 | 1 | |
| ENSMUSG00000086563 | -1.234582794 | 0.328283742 |
| ENSMUSG00000086564 | 1.11166259 | 0.608497229 |
| ENSMUSG00000086565 | 1 | |
| ENSMUSG00000086567 | -1.021746659 | 0.934760809 |

| | | |
|---|---|---|
| ENSMUSG00000086569 | 1 | |
| ENSMUSG00000086573 | 1.101438704 | 0.695095583 |
| ENSMUSG00000086575 | 1.059463094 | 0.608497229 |
| ENSMUSG00000086576 | 1 | |
| ENSMUSG00000086577 | 1 | |
| ENSMUSG00000086578 | 1.084618945 | 0.698112512 |
| ENSMUSG00000086580 | 1.061971464 | 0.819222912 |
| ENSMUSG00000086582 | -1.053549062 | 0.769745556 |
| ENSMUSG00000086583 | 1.040172434 | 0.918344559 |
| ENSMUSG00000086584 | 1.129400416 | 0.696594777 |
| ENSMUSG00000086585 | 1 | |
| ENSMUSG00000086586 | -1.033820075 | 0.888246981 |
| ENSMUSG00000086587 | 1.017174474 | 0.987056723 |
| ENSMUSG00000086588 | 1 | |
| ENSMUSG00000086589 | 1.001110674 | 1 |
| ENSMUSG00000086591 | 1 | |
| ENSMUSG00000086592 | 1 | |
| ENSMUSG00000086593 | 1 | |
| ENSMUSG00000086594 | 1 | |
| ENSMUSG00000086595 | 1 | |
| ENSMUSG00000086596 | -1.458671355 | 0.009817733 |
| ENSMUSG00000086598 | 1.072919021 | 0.80814891 |
| ENSMUSG00000086600 | -1.111880267 | 0.572947856 |
| ENSMUSG00000086601 | 1.122462048 | 0.215991492 |
| ENSMUSG00000086602 | -1.053549062 | 0.698112512 |
| ENSMUSG00000086603 | 1 | |
| ENSMUSG00000086604 | 1.033906491 | 0.927406094 |
| ENSMUSG00000086605 | -1.262106537 | 0.060812353 |
| ENSMUSG00000086606 | -1.097805537 | 0.608497229 |
| ENSMUSG00000086607 | 1.11521584 | 0.320605742 |
| ENSMUSG00000086608 | 1 | |
| ENSMUSG00000086610 | 1.033382383 | 0.969537118 |
| ENSMUSG00000086612 | 1 | |
| ENSMUSG00000086613 | 1 | |
| ENSMUSG00000086614 | 1 | |
| ENSMUSG00000086617 | 1 | |
| ENSMUSG00000086618 | 1 | |
| ENSMUSG00000086620 | -1.116853908 | 0.608497229 |
| ENSMUSG00000086621 | 1 | |
| ENSMUSG00000086624 | 1 | |
| ENSMUSG00000086625 | -1.001876871 | 1 |
| ENSMUSG00000086628 | 1 | |
| ENSMUSG00000086629 | -1.024267407 | 0.873810332 |
| ENSMUSG00000086630 | 1 | |
| ENSMUSG00000086631 | 1 | |
| ENSMUSG00000086632 | 1 | |
| ENSMUSG00000086633 | 1 | |
| ENSMUSG00000086636 | 1.001077657 | 1 |
| ENSMUSG00000086637 | 1 | |
| ENSMUSG00000086638 | -1.132874196 | 0.717035356 |
| ENSMUSG00000086641 | 1 | |
| ENSMUSG00000086643 | 1 | |
| ENSMUSG00000086644 | 1.247967366 | 0.608497229 |
| ENSMUSG00000086645 | 1 | |
| ENSMUSG00000086646 | -1.084645496 | 0.532369657 |
| ENSMUSG00000086648 | 1 | |
| ENSMUSG00000086649 | 1.059463094 | 0.608497229 |
| ENSMUSG00000086650 | 1 | |
| ENSMUSG00000086652 | 1 | |
| ENSMUSG00000086653 | 1 | |
| ENSMUSG00000086654 | -1.043306199 | 0.698112512 |
| ENSMUSG00000086655 | -1 | 1 |
| ENSMUSG00000086656 | 1.111812527 | 0.608497229 |

| | | |
|---|---|---|
| ENSMUSG00000086657 | -1.040768231 | 0.870119185 |
| ENSMUSG00000086658 | 1 | |
| ENSMUSG00000086659 | 1 | |
| ENSMUSG00000086660 | 1 | |
| ENSMUSG00000086662 | 1.154253937 | 0.261805279 |
| ENSMUSG00000086665 | 1.009414875 | 1 |
| ENSMUSG00000086666 | 1 | |
| ENSMUSG00000086667 | -1.03962336 | 0.698112512 |
| ENSMUSG00000086668 | 1 | |
| ENSMUSG00000086669 | 1 | |
| ENSMUSG00000086670 | 1 | |
| ENSMUSG00000086671 | -1.124067181 | 0.608497229 |
| ENSMUSG00000086674 | -1.003834322 | 1 |
| ENSMUSG00000086675 | 1 | |
| ENSMUSG00000086676 | -1.009842415 | 1 |
| ENSMUSG00000086677 | -1.044273782 | 0.790851299 |
| ENSMUSG00000086679 | 1 | |
| ENSMUSG00000086680 | -1.014146555 | 1 |
| ENSMUSG00000086681 | 1.124866072 | 0.698112512 |
| ENSMUSG00000086682 | 1.011142758 | 1 |
| ENSMUSG00000086683 | -1.004175918 | 1 |
| ENSMUSG00000086684 | 1.123949617 | 0.757379736 |
| ENSMUSG00000086685 | 1 | |
| ENSMUSG00000086686 | 1 | |
| ENSMUSG00000086687 | -1 | 1 |
| ENSMUSG00000086688 | -1.103242222 | 0.643148158 |
| ENSMUSG00000086689 | -1.03962336 | 0.698112512 |
| ENSMUSG00000086690 | -1.053549062 | 0.698112512 |
| ENSMUSG00000086691 | 1 | |
| ENSMUSG00000086693 | -1.038168511 | 0.772834122 |
| ENSMUSG00000086694 | 1 | |
| ENSMUSG00000086695 | 1.06043669 | 0.837465855 |
| ENSMUSG00000086696 | 1 | |
| ENSMUSG00000086697 | -1.010555784 | 1 |
| ENSMUSG00000086698 | 1 | |
| ENSMUSG00000086699 | 1.040172434 | 0.896152472 |
| ENSMUSG00000086700 | -1 | 1 |
| ENSMUSG00000086701 | 1 | |
| ENSMUSG00000086702 | 1 | |
| ENSMUSG00000086703 | 1 | |
| ENSMUSG00000086704 | 1 | |
| ENSMUSG00000086706 | 1.059463094 | 0.608497229 |
| ENSMUSG00000086707 | -1.023298643 | 1 |
| ENSMUSG00000086708 | 1 | |
| ENSMUSG00000086709 | -1 | 1 |
| ENSMUSG00000086710 | 1 | |
| ENSMUSG00000086711 | -1.085651418 | 0.608497229 |
| ENSMUSG00000086712 | 1 | |
| ENSMUSG00000086713 | 1 | |
| ENSMUSG00000086714 | -1.048718354 | 0.87458808 |
| ENSMUSG00000086715 | 1 | |
| ENSMUSG00000086716 | 1.069068979 | 0.698112512 |
| ENSMUSG00000086717 | 1 | |
| ENSMUSG00000086718 | 1.014545335 | 1 |
| ENSMUSG00000086719 | -1 | 1 |
| ENSMUSG00000086720 | 1 | |
| ENSMUSG00000086721 | 1.113371637 | 0.66877097 |
| ENSMUSG00000086722 | 1 | |
| ENSMUSG00000086724 | 1 | |
| ENSMUSG00000086725 | -1.055053695 | 0.635422995 |
| ENSMUSG00000086726 | 1 | |
| ENSMUSG00000086727 | 1.059463094 | 0.608497229 |
| ENSMUSG00000086728 | -1.044552394 | 0.844035928 |

| | | |
|---|---|---|
| ENSMUSG00000086729 | 1 | |
| ENSMUSG00000086730 | 1 | |
| ENSMUSG00000086731 | 1 | |
| ENSMUSG00000086732 | 1.059463094 | 0.608497229 |
| ENSMUSG00000086733 | 1 | |
| ENSMUSG00000086735 | 1.110340317 | 0.630771413 |
| ENSMUSG00000086736 | 1 | |
| ENSMUSG00000086737 | 1 | |
| ENSMUSG00000086738 | 1 | |
| ENSMUSG00000086739 | 1 | |
| ENSMUSG00000086740 | 1.149945284 | 0.773508225 |
| ENSMUSG00000086741 | -1.11061171 | 0.547016491 |
| ENSMUSG00000086742 | -1.038217282 | 0.872615712 |
| ENSMUSG00000086743 | 1.081929848 | 0.744331484 |
| ENSMUSG00000086745 | 1.112843644 | 0.608497229 |
| ENSMUSG00000086746 | 1.119204685 | 0.698112512 |
| ENSMUSG00000086748 | 1 | |
| ENSMUSG00000086749 | 1 | |
| ENSMUSG00000086750 | 1.059463094 | 0.608497229 |
| ENSMUSG00000086752 | 1 | |
| ENSMUSG00000086753 | -1.037024036 | 0.811125825 |
| ENSMUSG00000086754 | -1.173142781 | 0.464009618 |
| ENSMUSG00000086755 | -1.120476206 | 0.534652019 |
| ENSMUSG00000086756 | 1 | |
| ENSMUSG00000086757 | 1 | |
| ENSMUSG00000086758 | 1 | |
| ENSMUSG00000086760 | 1 | |
| ENSMUSG00000086761 | 1.132639128 | 0.398767938 |
| ENSMUSG00000086762 | 1 | |
| ENSMUSG00000086763 | 1 | |
| ENSMUSG00000086764 | 1.059463094 | 0.608497229 |
| ENSMUSG00000086765 | 1.024693402 | 0.977433779 |
| ENSMUSG00000086766 | 1 | |
| ENSMUSG00000086767 | 1.084846819 | 0.698112512 |
| ENSMUSG00000086769 | 1.059463094 | 0.608497229 |
| ENSMUSG00000086770 | 1 | |
| ENSMUSG00000086771 | 1 | |
| ENSMUSG00000086772 | 1 | |
| ENSMUSG00000086773 | 1 | |
| ENSMUSG00000086774 | 1.014545335 | 0.979002724 |
| ENSMUSG00000086775 | -1.137296885 | 0.608497229 |
| ENSMUSG00000086777 | -1.069502966 | 0.698112512 |
| ENSMUSG00000086779 | -1.105773029 | 0.784279473 |
| ENSMUSG00000086781 | 1 | |
| ENSMUSG00000086782 | -1.023608309 | 0.968584032 |
| ENSMUSG00000086783 | 1 | |
| ENSMUSG00000086784 | 1.134287314 | 0.203085735 |
| ENSMUSG00000086785 | 1 | |
| ENSMUSG00000086786 | 1.079681442 | 0.750742698 |
| ENSMUSG00000086788 | 1 | |
| ENSMUSG00000086789 | 1 | |
| ENSMUSG00000086790 | 1 | |
| ENSMUSG00000086791 | 1 | |
| ENSMUSG00000086792 | -1 | 1 |
| ENSMUSG00000086793 | 1 | |
| ENSMUSG00000086794 | -1.151375465 | 0.418145793 |
| ENSMUSG00000086795 | 1 | |
| ENSMUSG00000086796 | -1.206848132 | 0.148285088 |
| ENSMUSG00000086798 | -1.003948763 | 1 |
| ENSMUSG00000086799 | -1.009842415 | 1 |
| ENSMUSG00000086800 | 1 | |
| ENSMUSG00000086801 | -1 | 1 |
| ENSMUSG00000086804 | 1 | |

| | | |
|---|---|---|
| ENSMUSG00000086805 | -1.03962336 | 0.698112512 |
| ENSMUSG00000086806 | 1.095658148 | 0.737780712 |
| ENSMUSG00000086807 | -1.044273782 | 0.698112512 |
| ENSMUSG00000086808 | 1 | |
| ENSMUSG00000086810 | 1 | |
| ENSMUSG00000086812 | 1 | |
| ENSMUSG00000086813 | -1.03962336 | 0.698112512 |
| ENSMUSG00000086814 | 1 | |
| ENSMUSG00000086815 | 1.315035565 | 0.021070191 |
| ENSMUSG00000086816 | -1.043306199 | 0.698112512 |
| ENSMUSG00000086817 | 1 | |
| ENSMUSG00000086818 | 1.014545335 | 0.979002724 |
| ENSMUSG00000086819 | -1.010511572 | 1 |
| ENSMUSG00000086820 | 1.024284648 | 0.988424085 |
| ENSMUSG00000086821 | 1 | |
| ENSMUSG00000086822 | 1.059463094 | 0.608497229 |
| ENSMUSG00000086823 | 1 | |
| ENSMUSG00000086824 | -1 | 1 |
| ENSMUSG00000086825 | -1.083711965 | 0.600288135 |
| ENSMUSG00000086826 | 1.094656191 | 0.608497229 |
| ENSMUSG00000086827 | -1.043306199 | 0.698112512 |
| ENSMUSG00000086828 | 1 | |
| ENSMUSG00000086830 | 1 | |
| ENSMUSG00000086831 | 1 | |
| ENSMUSG00000086833 | 1 | |
| ENSMUSG00000086834 | 1 | |
| ENSMUSG00000086835 | 1 | |
| ENSMUSG00000086836 | -1.03962336 | 0.698112512 |
| ENSMUSG00000086837 | -1.049968634 | 0.914187043 |
| ENSMUSG00000086838 | -1.132666855 | 0.18851436 |
| ENSMUSG00000086839 | 1 | |
| ENSMUSG00000086840 | 1 | |
| ENSMUSG00000086841 | 1.123782436 | 0.077770173 |
| ENSMUSG00000086842 | 1.132639128 | 0.217803313 |
| ENSMUSG00000086843 | 1 | |
| ENSMUSG00000086844 | 1.059463094 | 0.752398528 |
| ENSMUSG00000086845 | -1.043306199 | 0.850892389 |
| ENSMUSG00000086846 | 1.014545335 | 0.989648415 |
| ENSMUSG00000086847 | -1.044273782 | 0.855488553 |
| ENSMUSG00000086848 | 1 | |
| ENSMUSG00000086849 | 1 | |
| ENSMUSG00000086850 | 1.005325648 | 1 |
| ENSMUSG00000086851 | -1.021612531 | 0.982838256 |
| ENSMUSG00000086852 | 1.069068979 | 0.608497229 |
| ENSMUSG00000086853 | 1 | |
| ENSMUSG00000086854 | 1 | |
| ENSMUSG00000086855 | 1 | |
| ENSMUSG00000086856 | 1 | |
| ENSMUSG00000086858 | 1 | |
| ENSMUSG00000086859 | 1.053510745 | 0.608497229 |
| ENSMUSG00000086860 | 1 | |
| ENSMUSG00000086861 | 1 | |
| ENSMUSG00000086862 | 1 | |
| ENSMUSG00000086863 | 1 | |
| ENSMUSG00000086864 | 1 | |
| ENSMUSG00000086866 | -1 | 1 |
| ENSMUSG00000086867 | -1.044273782 | 0.847756749 |
| ENSMUSG00000086868 | -1.063525418 | 0.751979636 |
| ENSMUSG00000086870 | -1.044273782 | 0.698112512 |
| ENSMUSG00000086872 | -1.043306199 | 0.698112512 |
| ENSMUSG00000086873 | 1.04107561 | 0.794274354 |
| ENSMUSG00000086874 | 1 | |
| ENSMUSG00000086875 | 1 | |

| | | |
|---|---|---|
| ENSMUSG00000086876 | 1 | |
| ENSMUSG00000086877 | 1.036620922 | 0.817352236 |
| ENSMUSG00000086879 | -1 | 1 |
| ENSMUSG00000086881 | 1.059463094 | 0.608497229 |
| ENSMUSG00000086882 | 1 | |
| ENSMUSG00000086883 | 1 | |
| ENSMUSG00000086884 | 1 | |
| ENSMUSG00000086885 | -1.053549062 | 0.698112512 |
| ENSMUSG00000086886 | 1 | |
| ENSMUSG00000086887 | 1 | |
| ENSMUSG00000086889 | -1.006274318 | 1 |
| ENSMUSG00000086890 | -1.043306199 | 0.858911269 |
| ENSMUSG00000086891 | 1.085624843 | 0.717035356 |
| ENSMUSG00000086892 | 1 | |
| ENSMUSG00000086893 | -1.0795912 | 0.698112512 |
| ENSMUSG00000086894 | 1.052768837 | 0.747886461 |
| ENSMUSG00000086896 | 1.020779405 | 0.970822886 |
| ENSMUSG00000086898 | 1 | |
| ENSMUSG00000086899 | 1.059463094 | 0.704170349 |
| ENSMUSG00000086900 | -1.218658516 | 0.261204136 |
| ENSMUSG00000086902 | -1 | 1 |
| ENSMUSG00000086903 | -1.089497311 | 0.608497229 |
| ENSMUSG00000086904 | -1 | 1 |
| ENSMUSG00000086905 | -1 | 1 |
| ENSMUSG00000086906 | -1.022608177 | 0.985127706 |
| ENSMUSG00000086907 | -1.044720008 | 0.940045629 |
| ENSMUSG00000086908 | -1 | 1 |
| ENSMUSG00000086909 | 1 | |
| ENSMUSG00000086910 | 1 | |
| ENSMUSG00000086911 | -1.043306199 | 0.698112512 |
| ENSMUSG00000086912 | 1.059463094 | 0.699340382 |
| ENSMUSG00000086913 | -1.044273782 | 0.698112512 |
| ENSMUSG00000086914 | -1.158727175 | 0.67070595 |
| ENSMUSG00000086915 | 1.197738254 | 0.044140978 |
| ENSMUSG00000086916 | 1.018129788 | 1 |
| ENSMUSG00000086917 | -1.063525418 | 0.698112512 |
| ENSMUSG00000086918 | 1.059463094 | 0.698112512 |
| ENSMUSG00000086919 | 1.04067451 | 0.903122231 |
| ENSMUSG00000086920 | 1.054076979 | 0.836456594 |
| ENSMUSG00000086921 | 1 | |
| ENSMUSG00000086922 | -1.085651418 | 0.532369657 |
| ENSMUSG00000086923 | 1 | |
| ENSMUSG00000086924 | 1.010972308 | 1 |
| ENSMUSG00000086925 | -1.059469475 | 0.865453833 |
| ENSMUSG00000086926 | 1 | |
| ENSMUSG00000086927 | 1 | |
| ENSMUSG00000086928 | 1 | |
| ENSMUSG00000086929 | -1 | 1 |
| ENSMUSG00000086930 | 1.065410325 | 0.841568489 |
| ENSMUSG00000086931 | 1 | |
| ENSMUSG00000086932 | -1.053549062 | 0.698112512 |
| ENSMUSG00000086933 | -1.044273782 | 0.698112512 |
| ENSMUSG00000086935 | -1.139678764 | 0.622715625 |
| ENSMUSG00000086936 | 1.029280487 | 0.976498231 |
| ENSMUSG00000086937 | 1.093819733 | 0.729157734 |
| ENSMUSG00000086938 | -1.145518406 | 0.608497229 |
| ENSMUSG00000086939 | 1 | |
| ENSMUSG00000086940 | -1.126319268 | 0.666323413 |
| ENSMUSG00000086941 | 1 | |
| ENSMUSG00000086942 | 1.059178908 | 0.773060207 |
| ENSMUSG00000086943 | 1.030661948 | 0.949007391 |
| ENSMUSG00000086944 | -1.03962336 | 0.774810939 |
| ENSMUSG00000086945 | -1.03962336 | 0.698112512 |

| | | |
|---|---|---|
| ENSMUSG00000086946 | -1.043306199 | 0.858911269 |
| ENSMUSG00000086947 | 1 | |
| ENSMUSG00000086948 | 1 | |
| ENSMUSG00000086949 | 1 | |
| ENSMUSG00000086951 | 1.069068979 | 0.608497229 |
| ENSMUSG00000086952 | 1 | |
| ENSMUSG00000086953 | -1.044273782 | 0.698112512 |
| ENSMUSG00000086954 | 1 | |
| ENSMUSG00000086955 | 1 | |
| ENSMUSG00000086957 | 1 | |
| ENSMUSG00000086959 | 1 | |
| ENSMUSG00000086960 | 1 | |
| ENSMUSG00000086961 | 1.049411284 | 0.867939428 |
| ENSMUSG00000086962 | 1.043471668 | 0.930693163 |
| ENSMUSG00000086963 | 1 | |
| ENSMUSG00000086964 | 1 | |
| ENSMUSG00000086965 | 1.107522869 | 0.477118677 |
| ENSMUSG00000086968 | -1.014698655 | 0.979002724 |
| ENSMUSG00000086969 | -1.073617693 | 0.840150908 |
| ENSMUSG00000086970 | 1 | |
| ENSMUSG00000086971 | 1 | |
| ENSMUSG00000086974 | 1 | |
| ENSMUSG00000086975 | 1 | |
| ENSMUSG00000086976 | 1.085624843 | 0.608497229 |
| ENSMUSG00000086978 | 1 | |
| ENSMUSG00000086980 | 1 | |
| ENSMUSG00000086981 | 1 | |
| ENSMUSG00000086984 | 1 | |
| ENSMUSG00000086986 | -1 | 1 |
| ENSMUSG00000086987 | 1.092668604 | 0.84360497 |
| ENSMUSG00000086989 | 1 | |
| ENSMUSG00000086991 | 1 | |
| ENSMUSG00000086992 | 1 | |
| ENSMUSG00000086993 | 1.005851671 | 1 |
| ENSMUSG00000086994 | 1 | |
| ENSMUSG00000086995 | 1.019083579 | 0.968937442 |
| ENSMUSG00000086996 | 1 | |
| ENSMUSG00000086997 | 1.103178675 | 0.640091848 |
| ENSMUSG00000086998 | 1 | |
| ENSMUSG00000086999 | -1.053549062 | 0.698112512 |
| ENSMUSG00000087001 | 1.14009503 | 0.661969313 |
| ENSMUSG00000087002 | 1 | |
| ENSMUSG00000087003 | 1 | |
| ENSMUSG00000087004 | 1.028323353 | 0.903464611 |
| ENSMUSG00000087006 | 1.539257324 | 0.05507961 |
| ENSMUSG00000087007 | 1 | |
| ENSMUSG00000087008 | -1.069502966 | 0.698112512 |
| ENSMUSG00000087009 | 1 | |
| ENSMUSG00000087010 | 1 | |
| ENSMUSG00000087011 | 1.059463094 | 0.608497229 |
| ENSMUSG00000087012 | 1 | |
| ENSMUSG00000087013 | -1.033575217 | 0.933906187 |
| ENSMUSG00000087014 | 1.408851827 | 0.250726313 |
| ENSMUSG00000087015 | 1 | |
| ENSMUSG00000087016 | 1 | |
| ENSMUSG00000087017 | 1.24122829 | 0.181635915 |
| ENSMUSG00000087018 | -1.072686 | 0.812442422 |
| ENSMUSG00000087020 | 1.005613439 | 1 |
| ENSMUSG00000087021 | 1 | |
| ENSMUSG00000087022 | -1.024313512 | 0.95029384 |
| ENSMUSG00000087024 | -1 | 1 |
| ENSMUSG00000087026 | -1.013082712 | 0.970208206 |
| ENSMUSG00000087027 | 1 | |

| | | |
|---|---|---|
| ENSMUSG00000087028 | -1.163035322 | 0.436466116 |
| ENSMUSG00000087029 | 1 | |
| ENSMUSG00000087030 | -1.044273782 | 0.698112512 |
| ENSMUSG00000087031 | 1 | |
| ENSMUSG00000087032 | 1 | |
| ENSMUSG00000087033 | 1 | |
| ENSMUSG00000087034 | 1 | |
| ENSMUSG00000087035 | -1.132666855 | 0.18851436 |
| ENSMUSG00000087036 | 1.089470641 | 0.657060635 |
| ENSMUSG00000087037 | 1.132639128 | 0.217803313 |
| ENSMUSG00000087038 | 1 | |
| ENSMUSG00000087039 | 1 | |
| ENSMUSG00000087040 | 1 | |
| ENSMUSG00000087041 | 1 | |
| ENSMUSG00000087042 | -1.089497311 | 0.608497229 |
| ENSMUSG00000087043 | 1 | |
| ENSMUSG00000087044 | 1 | |
| ENSMUSG00000087045 | 1 | |
| ENSMUSG00000087047 | 1 | |
| ENSMUSG00000087049 | -1 | 1 |
| ENSMUSG00000087050 | -1.25842122 | 0.272914614 |
| ENSMUSG00000087051 | 1 | |
| ENSMUSG00000087052 | 1.050330666 | 0.814089245 |
| ENSMUSG00000087053 | 1 | |
| ENSMUSG00000087054 | 1 | |
| ENSMUSG00000087055 | -1.15147561 | 0.608497229 |
| ENSMUSG00000087056 | -1.053549062 | 0.698112512 |
| ENSMUSG00000087057 | 1 | |
| ENSMUSG00000087058 | 1 | |
| ENSMUSG00000087059 | -1 | 1 |
| ENSMUSG00000087060 | 1.027123703 | 0.756255421 |
| ENSMUSG00000087061 | 1 | |
| ENSMUSG00000087062 | 1 | |
| ENSMUSG00000087063 | -1.03962336 | 0.698112512 |
| ENSMUSG00000087064 | 1.186338807 | 0.608497229 |
| ENSMUSG00000087066 | -1.023769022 | 0.948378604 |
| ENSMUSG00000087067 | 1.014545335 | 0.989648415 |
| ENSMUSG00000087069 | 1 | |
| ENSMUSG00000087070 | -1.033820075 | 0.919518522 |
| ENSMUSG00000087071 | 1.059463094 | 0.608497229 |
| ENSMUSG00000087072 | 1 | |
| ENSMUSG00000087074 | 1 | |
| ENSMUSG00000087075 | -1.043306199 | 0.698112512 |
| ENSMUSG00000087076 | -1.03962336 | 0.698112512 |
| ENSMUSG00000087077 | 1.271344436 | 0.012381271 |
| ENSMUSG00000087079 | 1.059463094 | 0.608497229 |
| ENSMUSG00000087080 | 1 | |
| ENSMUSG00000087082 | 1 | |
| ENSMUSG00000087083 | 1 | |
| ENSMUSG00000087084 | 1 | |
| ENSMUSG00000087085 | 1 | |
| ENSMUSG00000087086 | 1 | |
| ENSMUSG00000087088 | -1.013268619 | 1 |
| ENSMUSG00000087089 | 1 | |
| ENSMUSG00000087090 | 1 | |
| ENSMUSG00000087093 | -1.053549062 | 0.698112512 |
| ENSMUSG00000087094 | -1.044273782 | 0.790851299 |
| ENSMUSG00000087095 | 1.093750011 | 0.60831194 |
| ENSMUSG00000087097 | 1 | |
| ENSMUSG00000087098 | 1 | |
| ENSMUSG00000087099 | 1 | |
| ENSMUSG00000087100 | 1 | |
| ENSMUSG00000087102 | 1 | |

| | | |
|---|---|---|
| ENSMUSG00000087103 | 1 | |
| ENSMUSG00000087104 | 1 | |
| ENSMUSG00000087106 | 1.07487334 | 0.656277822 |
| ENSMUSG00000087107 | 1 | |
| ENSMUSG00000087108 | -1.000685443 | 1 |
| ENSMUSG00000087109 | 1 | |
| ENSMUSG00000087110 | 1.069068979 | 0.608497229 |
| ENSMUSG00000087111 | 1.015486245 | 0.972980766 |
| ENSMUSG00000087112 | -1.03962336 | 0.803072249 |
| ENSMUSG00000087113 | 1.295478754 | 0.191749554 |
| ENSMUSG00000087114 | 1 | |
| ENSMUSG00000087115 | 1 | |
| ENSMUSG00000087116 | 1 | |
| ENSMUSG00000087117 | 1 | |
| ENSMUSG00000087119 | -1.049475223 | 0.861137762 |
| ENSMUSG00000087120 | -1.06353289 | 0.893822825 |
| ENSMUSG00000087121 | -1.09323489 | 0.578757926 |
| ENSMUSG00000087122 | 1 | |
| ENSMUSG00000087123 | 1.085624843 | 0.608497229 |
| ENSMUSG00000087124 | 1 | |
| ENSMUSG00000087125 | -1 | 1 |
| ENSMUSG00000087127 | 1 | |
| ENSMUSG00000087128 | -1.092470714 | 0.627499942 |
| ENSMUSG00000087129 | -1.03962336 | 0.698112512 |
| ENSMUSG00000087131 | 1 | |
| ENSMUSG00000087132 | 1.160823827 | 0.628300894 |
| ENSMUSG00000087133 | 1 | |
| ENSMUSG00000087135 | 1.144363107 | 0.39821235 |
| ENSMUSG00000087136 | 1 | |
| ENSMUSG00000087137 | 1 | |
| ENSMUSG00000087138 | -1.076193699 | 0.608497229 |
| ENSMUSG00000087139 | 1 | |
| ENSMUSG00000087140 | 1 | |
| ENSMUSG00000087141 | 1.053536938 | 0.698112512 |
| ENSMUSG00000087142 | 1.127619746 | 0.488601722 |
| ENSMUSG00000087143 | 1.10274851 | 0.373988904 |
| ENSMUSG00000087144 | 1 | |
| ENSMUSG00000087145 | 1 | |
| ENSMUSG00000087146 | 1 | |
| ENSMUSG00000087147 | 1 | |
| ENSMUSG00000087148 | 1.069068979 | 0.608497229 |
| ENSMUSG00000087149 | 1.059463094 | 0.608497229 |
| ENSMUSG00000087150 | -1.239246105 | 0.381036055 |
| ENSMUSG00000087151 | 1.17571767 | 0.608497229 |
| ENSMUSG00000087152 | 1 | |
| ENSMUSG00000087153 | -1.004089708 | 1 |
| ENSMUSG00000087155 | -1.053549062 | 0.778372272 |
| ENSMUSG00000087157 | 1 | |
| ENSMUSG00000087158 | -1.043306199 | 0.698112512 |
| ENSMUSG00000087159 | 1.033000091 | 0.735035933 |
| ENSMUSG00000087160 | 1 | |
| ENSMUSG00000087162 | -1 | 1 |
| ENSMUSG00000087163 | 1.04424822 | 0.855328455 |
| ENSMUSG00000087164 | 1 | |
| ENSMUSG00000087165 | -1.139498484 | 0.282023792 |
| ENSMUSG00000087166 | 1 | |
| ENSMUSG00000087167 | 1 | |
| ENSMUSG00000087168 | 1.13246302 | 0.103173815 |
| ENSMUSG00000087170 | 1 | |
| ENSMUSG00000087171 | 1 | |
| ENSMUSG00000087172 | 1.217611191 | 0.608497229 |
| ENSMUSG00000087173 | 1 | |
| ENSMUSG00000087174 | 1.024911357 | 0.921071544 |

| | | |
|---|---|---|
| ENSMUSG00000087175 | 1 | |
| ENSMUSG00000087176 | -1.094488708 | 0.714698678 |
| ENSMUSG00000087177 | -1.013887716 | 0.93855667 |
| ENSMUSG00000087178 | -1.027561518 | 0.89553586 |
| ENSMUSG00000087179 | -1.036200926 | 0.734674233 |
| ENSMUSG00000087180 | 1.023743961 | 0.95702926 |
| ENSMUSG00000087181 | 1 | |
| ENSMUSG00000087184 | 1 | |
| ENSMUSG00000087185 | 1 | |
| ENSMUSG00000087187 | 1.540647667 | 0.002723641 |
| ENSMUSG00000087190 | 1 | |
| ENSMUSG00000087192 | 1.24122829 | 0.03679259 |
| ENSMUSG00000087193 | 1.112016081 | 0.660210733 |
| ENSMUSG00000087194 | -1.024718486 | 0.931480186 |
| ENSMUSG00000087195 | 1 | |
| ENSMUSG00000087196 | 1 | |
| ENSMUSG00000087197 | 1 | |
| ENSMUSG00000087198 | -1.06005608 | 0.886345617 |
| ENSMUSG00000087199 | 1 | |
| ENSMUSG00000087200 | -1.139361965 | 0.608497229 |
| ENSMUSG00000087201 | -1.069502966 | 0.744868995 |
| ENSMUSG00000087202 | 1 | |
| ENSMUSG00000087203 | 1.351147692 | 0.058764965 |
| ENSMUSG00000087204 | -1.114128859 | 0.667110773 |
| ENSMUSG00000087205 | 1 | |
| ENSMUSG00000087206 | 1.075870199 | 0.756255421 |
| ENSMUSG00000087207 | -1.043306199 | 0.858911269 |
| ENSMUSG00000087208 | 1 | |
| ENSMUSG00000087210 | 1 | |
| ENSMUSG00000087211 | -1.288068239 | 0.551136369 |
| ENSMUSG00000087213 | -1.048741327 | 0.710698346 |
| ENSMUSG00000087214 | 1 | |
| ENSMUSG00000087215 | 1 | |
| ENSMUSG00000087216 | 1 | |
| ENSMUSG00000087217 | -1 | 1 |
| ENSMUSG00000087219 | 1 | |
| ENSMUSG00000087220 | -1.084645496 | 0.532369657 |
| ENSMUSG00000087221 | 1.035306494 | 0.901903401 |
| ENSMUSG00000087222 | 1.059463094 | 0.698112512 |
| ENSMUSG00000087223 | 1 | |
| ENSMUSG00000087224 | -1.03962336 | 0.803072249 |
| ENSMUSG00000087225 | 1 | |
| ENSMUSG00000087226 | 1.059463094 | 0.768693672 |
| ENSMUSG00000087228 | 1 | |
| ENSMUSG00000087230 | 1.048519362 | 0.865598094 |
| ENSMUSG00000087231 | -1.068287749 | 0.698112512 |
| ENSMUSG00000087232 | 1 | |
| ENSMUSG00000087233 | 1 | |
| ENSMUSG00000087234 | 1 | |
| ENSMUSG00000087235 | 1 | |
| ENSMUSG00000087236 | -1.044273782 | 0.698112512 |
| ENSMUSG00000087238 | 1 | |
| ENSMUSG00000087239 | -1.071075483 | 0.801443616 |
| ENSMUSG00000087240 | 1.005613439 | 1 |
| ENSMUSG00000087241 | 1 | |
| ENSMUSG00000087242 | 1.137399458 | 0.6510358 |
| ENSMUSG00000087244 | 1 | |
| ENSMUSG00000087245 | 1 | |
| ENSMUSG00000087246 | -1.024718486 | 0.954310937 |
| ENSMUSG00000087247 | 2.508982694 | 0.001103396 |
| ENSMUSG00000087248 | 1 | |
| ENSMUSG00000087249 | 1.017290115 | 0.944468485 |
| ENSMUSG00000087250 | -1.053549062 | 0.698112512 |

| | | |
|---|---|---|
| ENSMUSG00000087252 | 1.009005271 | 1 |
| ENSMUSG00000087253 | 1.014545335 | 0.989648415 |
| ENSMUSG00000087254 | 1 | |
| ENSMUSG00000087255 | 1 | |
| ENSMUSG00000087256 | 1.011497958 | 1 |
| ENSMUSG00000087258 | -1.03962336 | 0.698112512 |
| ENSMUSG00000087259 | 1.257787146 | 0.005260694 |
| ENSMUSG00000087260 | -1.051130423 | 0.835679601 |
| ENSMUSG00000087261 | -1 | 1 |
| ENSMUSG00000087263 | 1.023743961 | 0.959839883 |
| ENSMUSG00000087264 | -1.214864586 | 0.519682156 |
| ENSMUSG00000087265 | 1.069068979 | 0.608497229 |
| ENSMUSG00000087266 | 1 | |
| ENSMUSG00000087267 | -1.116410383 | 0.709787982 |
| ENSMUSG00000087268 | 1 | |
| ENSMUSG00000087269 | 1.168899229 | 0.234728821 |
| ENSMUSG00000087270 | 1 | |
| ENSMUSG00000087272 | 1.283568842 | 0.34958424 |
| ENSMUSG00000087273 | 1.115660465 | 0.575592633 |
| ENSMUSG00000087276 | 1 | |
| ENSMUSG00000087277 | 1 | |
| ENSMUSG00000087278 | 1 | |
| ENSMUSG00000087279 | -1.043306199 | 0.698112512 |
| ENSMUSG00000087281 | -1.044273782 | 0.698112512 |
| ENSMUSG00000087282 | 1.112843644 | 0.609147349 |
| ENSMUSG00000087283 | -1 | 1 |
| ENSMUSG00000087286 | 1.105554153 | 0.608497229 |
| ENSMUSG00000087288 | 1 | |
| ENSMUSG00000087289 | 1 | |
| ENSMUSG00000087290 | 1 | |
| ENSMUSG00000087291 | 1.096401653 | 0.70299173 |
| ENSMUSG00000087292 | -1.099174267 | 0.535052277 |
| ENSMUSG00000087293 | -1 | 1 |
| ENSMUSG00000087294 | -1.03962336 | 0.698112512 |
| ENSMUSG00000087295 | 1 | |
| ENSMUSG00000087296 | 1 | |
| ENSMUSG00000087297 | 1 | |
| ENSMUSG00000087298 | -1.119290457 | 0.708569505 |
| ENSMUSG00000087299 | 1 | |
| ENSMUSG00000087300 | 1 | |
| ENSMUSG00000087301 | 1 | |
| ENSMUSG00000087302 | 1 | |
| ENSMUSG00000087303 | 1.123011825 | 0.629363816 |
| ENSMUSG00000087304 | 1 | |
| ENSMUSG00000087305 | 1.041498381 | 0.959892369 |
| ENSMUSG00000087306 | 1.218572685 | 0.584879768 |
| ENSMUSG00000087307 | 1.160565542 | 0.39790876 |
| ENSMUSG00000087308 | -1.044273782 | 0.801443616 |
| ENSMUSG00000087309 | 1 | |
| ENSMUSG00000087312 | 1.059463094 | 0.608497229 |
| ENSMUSG00000087314 | 1.173485669 | 0.251544797 |
| ENSMUSG00000087316 | 1 | |
| ENSMUSG00000087317 | 1 | |
| ENSMUSG00000087318 | 1 | |
| ENSMUSG00000087319 | 1.069068979 | 0.608497229 |
| ENSMUSG00000087321 | -1.023769022 | 0.934760809 |
| ENSMUSG00000087322 | -1 | 1 |
| ENSMUSG00000087323 | 1 | |
| ENSMUSG00000087324 | 1 | |
| ENSMUSG00000087325 | 1 | |
| ENSMUSG00000087326 | 1.069068979 | 0.608497229 |
| ENSMUSG00000087327 | 1.010855137 | 1 |
| ENSMUSG00000087328 | 1 | |

| | | |
|---|---|---|
| ENSMUSG00000087329 | 1 | |
| ENSMUSG00000087330 | 1 | |
| ENSMUSG00000087331 | 1.080914732 | 0.755879035 |
| ENSMUSG00000087332 | 1 | |
| ENSMUSG00000087333 | 1 | |
| ENSMUSG00000087334 | 1 | |
| ENSMUSG00000087335 | 1 | |
| ENSMUSG00000087336 | -1.135720617 | 0.693220376 |
| ENSMUSG00000087337 | 1 | |
| ENSMUSG00000087338 | -1.034990181 | 0.981214754 |
| ENSMUSG00000087339 | 1 | |
| ENSMUSG00000087340 | 2.146612045 | 0.000129787 |
| ENSMUSG00000087341 | 1.122462048 | 0.215991492 |
| ENSMUSG00000087342 | 1 | |
| ENSMUSG00000087343 | 1 | |
| ENSMUSG00000087344 | -1 | 1 |
| ENSMUSG00000087345 | 1 | |
| ENSMUSG00000087346 | 1 | |
| ENSMUSG00000087347 | -1.013496532 | 1 |
| ENSMUSG00000087348 | 1.028323353 | 0.903464611 |
| ENSMUSG00000087349 | -1.053549062 | 0.698112512 |
| ENSMUSG00000087350 | 1 | |
| ENSMUSG00000087351 | 1.005613439 | 1 |
| ENSMUSG00000087352 | 1 | |
| ENSMUSG00000087353 | 1 | |
| ENSMUSG00000087354 | -1.077510925 | 0.770958539 |
| ENSMUSG00000087355 | -1.066370674 | 0.82782338 |
| ENSMUSG00000087356 | -1.049272231 | 0.88972256 |
| ENSMUSG00000087357 | -1.03962336 | 0.698112512 |
| ENSMUSG00000087358 | -1.03962336 | 0.698112512 |
| ENSMUSG00000087360 | -1.283526117 | 0.002035148 |
| ENSMUSG00000087361 | 1.059463094 | 0.608497229 |
| ENSMUSG00000087362 | 1.016786017 | 0.982290355 |
| ENSMUSG00000087363 | 1 | |
| ENSMUSG00000087364 | -1.071075483 | 0.743300655 |
| ENSMUSG00000087365 | 1.316475139 | 0.004761018 |
| ENSMUSG00000087366 | 1.197866142 | 0.028902899 |
| ENSMUSG00000087367 | -1 | 1 |
| ENSMUSG00000087368 | -1.082261595 | 0.527982264 |
| ENSMUSG00000087369 | 1.07487334 | 0.656277822 |
| ENSMUSG00000087370 | -1.148245942 | 0.253736193 |
| ENSMUSG00000087371 | 1.086678356 | 0.698112512 |
| ENSMUSG00000087373 | -1.163170867 | 0.227658014 |
| ENSMUSG00000087374 | -1.023769022 | 0.954761885 |
| ENSMUSG00000087376 | 1 | |
| ENSMUSG00000087377 | 1.010104741 | 1 |
| ENSMUSG00000087378 | 1 | |
| ENSMUSG00000087379 | 1 | |
| ENSMUSG00000087380 | 1.07726198 | 0.740331062 |
| ENSMUSG00000087381 | 1.262704156 | 0.083235554 |
| ENSMUSG00000087382 | 1 | |
| ENSMUSG00000087383 | 1 | |
| ENSMUSG00000087384 | -1.237299691 | 0.326447608 |
| ENSMUSG00000087385 | 1.067724868 | 0.676676328 |
| ENSMUSG00000087386 | 1 | |
| ENSMUSG00000087387 | -1 | 1 |
| ENSMUSG00000087388 | 1 | |
| ENSMUSG00000087389 | -1.089497311 | 0.698112512 |
| ENSMUSG00000087390 | 1.046322894 | 0.876279035 |
| ENSMUSG00000087393 | 1 | |
| ENSMUSG00000087394 | 1.040172434 | 0.896152472 |
| ENSMUSG00000087395 | 1.069216143 | 0.879766196 |
| ENSMUSG00000087396 | 1.087360818 | 0.551295069 |

| | | |
|---|---|---|
| ENSMUSG00000087397 | 1.173639885 | 0.455298078 |
| ENSMUSG00000087398 | 1 | |
| ENSMUSG00000087399 | 1 | |
| ENSMUSG00000087400 | 1.007625832 | 1 |
| ENSMUSG00000087401 | 1 | |
| ENSMUSG00000087402 | 1 | |
| ENSMUSG00000087403 | -1.073578513 | 0.698112512 |
| ENSMUSG00000087404 | 1 | |
| ENSMUSG00000087405 | 1 | |
| ENSMUSG00000087406 | 1 | |
| ENSMUSG00000087407 | 1 | |
| ENSMUSG00000087408 | 1.077605404 | 0.541735653 |
| ENSMUSG00000087409 | 1 | |
| ENSMUSG00000087410 | 1 | |
| ENSMUSG00000087411 | 1 | |
| ENSMUSG00000087412 | -1.05451026 | 0.896765664 |
| ENSMUSG00000087413 | -1.120476206 | 0.608497229 |
| ENSMUSG00000087415 | 1.059463094 | 0.608497229 |
| ENSMUSG00000087416 | 1 | |
| ENSMUSG00000087417 | 1 | |
| ENSMUSG00000087422 | 1 | |
| ENSMUSG00000087424 | -1.046643042 | 0.915070045 |
| ENSMUSG00000087425 | -1.053549062 | 0.698112512 |
| ENSMUSG00000087428 | 1 | |
| ENSMUSG00000087429 | -1.03962336 | 0.803072249 |
| ENSMUSG00000087430 | 1.059463094 | 0.776953697 |
| ENSMUSG00000087431 | 1.199989356 | 0.045045157 |
| ENSMUSG00000087432 | 1 | |
| ENSMUSG00000087433 | 1.112006103 | 0.730441273 |
| ENSMUSG00000087434 | -1 | 1 |
| ENSMUSG00000087435 | -1.043306199 | 0.698112512 |
| ENSMUSG00000087436 | 1.101776656 | 0.727718584 |
| ENSMUSG00000087437 | 1 | |
| ENSMUSG00000087439 | -1.182754998 | 0.341391246 |
| ENSMUSG00000087440 | -1.029962984 | 0.875804019 |
| ENSMUSG00000087441 | 1 | |
| ENSMUSG00000087442 | 1.216705411 | 0.439085964 |
| ENSMUSG00000087443 | -1.016098951 | 1 |
| ENSMUSG00000087444 | -1.12236823 | 0.608497229 |
| ENSMUSG00000087445 | -1.05398434 | 0.850975923 |
| ENSMUSG00000087449 | -1 | 1 |
| ENSMUSG00000087450 | 1 | |
| ENSMUSG00000087451 | -1.084645496 | 0.608497229 |
| ENSMUSG00000087452 | 1.075070132 | 0.698112512 |
| ENSMUSG00000087454 | 1 | |
| ENSMUSG00000087455 | 1 | |
| ENSMUSG00000087456 | 1 | |
| ENSMUSG00000087457 | 1.002938927 | 1 |
| ENSMUSG00000087458 | -1.015083696 | 1 |
| ENSMUSG00000087459 | 1 | |
| ENSMUSG00000087460 | 1 | |
| ENSMUSG00000087461 | 1.319254719 | 0.034474274 |
| ENSMUSG00000087462 | 1 | |
| ENSMUSG00000087464 | 1 | |
| ENSMUSG00000087465 | 1 | |
| ENSMUSG00000087466 | 1.011497958 | 1 |
| ENSMUSG00000087467 | 1 | |
| ENSMUSG00000087468 | 1 | |
| ENSMUSG00000087469 | -1.004534381 | 1 |
| ENSMUSG00000087470 | -1.139116 | 0.608497229 |
| ENSMUSG00000087473 | 1 | |
| ENSMUSG00000087474 | 1 | |
| ENSMUSG00000087475 | 1 | |

| | | |
|---|---|---|
| ENSMUSG00000087476 | 1.122462048 | 0.215991492 |
| ENSMUSG00000087477 | 1 | |
| ENSMUSG00000087478 | -1.210030262 | 0.608497229 |
| ENSMUSG00000087479 | -1.107597544 | 0.717035356 |
| ENSMUSG00000087480 | 1.269057871 | 0.04905375 |
| ENSMUSG00000087481 | -1.003155205 | 1 |
| ENSMUSG00000087482 | -1.07008665 | 0.82993954 |
| ENSMUSG00000087484 | 1.254989012 | 0.03753054 |
| ENSMUSG00000087485 | 1 | |
| ENSMUSG00000087486 | 1 | |
| ENSMUSG00000087487 | 1 | |
| ENSMUSG00000087489 | 1.069068979 | 0.608497229 |
| ENSMUSG00000087491 | 1 | |
| ENSMUSG00000087492 | 1 | |
| ENSMUSG00000087495 | 1 | |
| ENSMUSG00000087496 | 1.139042611 | 0.68735438 |
| ENSMUSG00000087497 | -1.062089421 | 0.624363396 |
| ENSMUSG00000087500 | 1 | |
| ENSMUSG00000087501 | -1.048971517 | 0.729018288 |
| ENSMUSG00000087502 | 1.210356402 | 0.097546287 |
| ENSMUSG00000087504 | 1 | |
| ENSMUSG00000087505 | 1.059463094 | 0.608497229 |
| ENSMUSG00000087507 | 1.012301836 | 1 |
| ENSMUSG00000087508 | 1 | |
| ENSMUSG00000087510 | 1 | |
| ENSMUSG00000087512 | 1 | |
| ENSMUSG00000087513 | -1.022548043 | 1 |
| ENSMUSG00000087514 | 1 | |
| ENSMUSG00000087515 | 1 | |
| ENSMUSG00000087516 | -1.063525418 | 0.761580827 |
| ENSMUSG00000087517 | 1 | |
| ENSMUSG00000087518 | 1.059463094 | 0.608497229 |
| ENSMUSG00000087519 | 1.040263416 | 0.90258325 |
| ENSMUSG00000087520 | 1 | |
| ENSMUSG00000087521 | 1 | |
| ENSMUSG00000087522 | -1.03962336 | 0.698112512 |
| ENSMUSG00000087523 | 1.152760354 | 0.467526851 |
| ENSMUSG00000087524 | 1 | |
| ENSMUSG00000087525 | 1 | |
| ENSMUSG00000087526 | 1.069068979 | 0.698112512 |
| ENSMUSG00000087527 | 1 | |
| ENSMUSG00000087528 | -1.134785444 | 0.310368041 |
| ENSMUSG00000087530 | 1.031212313 | 0.95221036 |
| ENSMUSG00000087531 | 1 | |
| ENSMUSG00000087532 | 1.09529959 | 0.702054321 |
| ENSMUSG00000087534 | 1 | |
| ENSMUSG00000087535 | 1.116289651 | 0.702455751 |
| ENSMUSG00000087536 | 1 | |
| ENSMUSG00000087537 | 1 | |
| ENSMUSG00000087538 | 1 | |
| ENSMUSG00000087539 | 1 | |
| ENSMUSG00000087540 | 1 | |
| ENSMUSG00000087541 | 1.059463094 | 0.608497229 |
| ENSMUSG00000087542 | -1.003493695 | 1 |
| ENSMUSG00000087543 | 1.103329649 | 0.02304608 |
| ENSMUSG00000087544 | 1 | |
| ENSMUSG00000087545 | 1.14032354 | 0.629028898 |
| ENSMUSG00000087547 | -1.043306199 | 0.698112512 |
| ENSMUSG00000087549 | 1.030971948 | 0.938939969 |
| ENSMUSG00000087550 | -1.068104567 | 0.811581858 |
| ENSMUSG00000087552 | 1.13926696 | 0.698112512 |
| ENSMUSG00000087553 | 1 | |
| ENSMUSG00000087554 | 1 | |

| | | |
|---|---|---|
| ENSMUSG00000087555 | 1 | |
| ENSMUSG00000087556 | 1.035432047 | 0.922613598 |
| ENSMUSG00000087557 | 1.047947442 | 0.896902223 |
| ENSMUSG00000087559 | 1 | |
| ENSMUSG00000087560 | -1.011709035 | 1 |
| ENSMUSG00000087561 | 1 | |
| ENSMUSG00000087563 | 1 | |
| ENSMUSG00000087564 | 1 | |
| ENSMUSG00000087565 | 1 | |
| ENSMUSG00000087568 | 1 | |
| ENSMUSG00000087569 | 1 | |
| ENSMUSG00000087570 | 1.069068979 | 0.608497229 |
| ENSMUSG00000087571 | 1 | |
| ENSMUSG00000087574 | 1.174458726 | 0.219160732 |
| ENSMUSG00000087575 | 1.029175231 | 0.942160514 |
| ENSMUSG00000087577 | 1 | |
| ENSMUSG00000087578 | 1.22770107 | 0.526184828 |
| ENSMUSG00000087579 | 1.028464908 | 0.989217536 |
| ENSMUSG00000087581 | 1 | |
| ENSMUSG00000087582 | 1 | |
| ENSMUSG00000087584 | 1 | |
| ENSMUSG00000087585 | 1 | |
| ENSMUSG00000087586 | 1 | |
| ENSMUSG00000087587 | -1.053549062 | 0.774498429 |
| ENSMUSG00000087588 | 1 | |
| ENSMUSG00000087589 | -1.012966945 | 0.984190423 |
| ENSMUSG00000087590 | -1.088879573 | 0.671462128 |
| ENSMUSG00000087591 | 1 | |
| ENSMUSG00000087592 | 1 | |
| ENSMUSG00000087593 | 1.004606297 | 1 |
| ENSMUSG00000087594 | -1.004403216 | 1 |
| ENSMUSG00000087595 | -1.087416885 | 0.693115493 |
| ENSMUSG00000087596 | -1.187802519 | 0.216046507 |
| ENSMUSG00000087597 | -1.084645496 | 0.608497229 |
| ENSMUSG00000087598 | -1.017743156 | 0.977563419 |
| ENSMUSG00000087600 | 1.059463094 | 0.699340382 |
| ENSMUSG00000087601 | 1 | |
| ENSMUSG00000087602 | 1 | |
| ENSMUSG00000087603 | 1 | |
| ENSMUSG00000087604 | 1.069068979 | 0.608497229 |
| ENSMUSG00000087605 | -1.003834322 | 1 |
| ENSMUSG00000087606 | -1.144251352 | 0.278020735 |
| ENSMUSG00000087607 | 1 | |
| ENSMUSG00000087608 | 1.005613439 | 1 |
| ENSMUSG00000087609 | 1 | |
| ENSMUSG00000087610 | 1.165214928 | 0.608497229 |
| ENSMUSG00000087611 | -1.088145459 | 0.726135084 |
| ENSMUSG00000087612 | 1 | |
| ENSMUSG00000087613 | -1 | 1 |
| ENSMUSG00000087614 | 1.132639128 | 0.217803313 |
| ENSMUSG00000087615 | 1.034095782 | 0.919000509 |
| ENSMUSG00000087616 | -1.228703104 | 0.358000616 |
| ENSMUSG00000087617 | 1 | |
| ENSMUSG00000087618 | 1 | |
| ENSMUSG00000087620 | -1 | 1 |
| ENSMUSG00000087621 | -1.043306199 | 0.698112512 |
| ENSMUSG00000087622 | -1.043306199 | 0.698112512 |
| ENSMUSG00000087623 | 1.000650359 | 1 |
| ENSMUSG00000087624 | -1.199747809 | 0.59011989 |
| ENSMUSG00000087625 | 1 | |
| ENSMUSG00000087626 | 1.246209297 | 0.016466394 |
| ENSMUSG00000087627 | 1.035736682 | 0.961643545 |
| ENSMUSG00000087628 | 1 | |

| | | |
|---|---|---|
| ENSMUSG00000087629 | -1.16022982 | 0.626480879 |
| ENSMUSG00000087630 | 1.059463094 | 0.608497229 |
| ENSMUSG00000087632 | -1 | 1 |
| ENSMUSG00000087633 | 1.059463094 | 0.608497229 |
| ENSMUSG00000087635 | 1 | |
| ENSMUSG00000087636 | 1 | |
| ENSMUSG00000087637 | -1 | 1 |
| ENSMUSG00000087638 | 1.009414875 | 1 |
| ENSMUSG00000087639 | -1.173012665 | 0.698112512 |
| ENSMUSG00000087640 | -1.069502966 | 0.698112512 |
| ENSMUSG00000087641 | 1 | |
| ENSMUSG00000087642 | 1 | |
| ENSMUSG00000087643 | 1 | |
| ENSMUSG00000087644 | 1.177765487 | 0.277823925 |
| ENSMUSG00000087646 | 1 | |
| ENSMUSG00000087647 | 1 | |
| ENSMUSG00000087648 | -1.095294215 | 0.608497229 |
| ENSMUSG00000087649 | 1 | |
| ENSMUSG00000087651 | -1.872708732 | 0.001127478 |
| ENSMUSG00000087652 | 1 | |
| ENSMUSG00000087653 | 1 | |
| ENSMUSG00000087654 | 1 | |
| ENSMUSG00000087657 | 1 | |
| ENSMUSG00000087658 | 1.105588176 | 0.775673129 |
| ENSMUSG00000087659 | 1.023743961 | 0.95838544 |
| ENSMUSG00000087660 | -1.095939461 | 0.693463129 |
| ENSMUSG00000087662 | 1 | |
| ENSMUSG00000087663 | 1 | |
| ENSMUSG00000087666 | 1 | |
| ENSMUSG00000087667 | -1.139364264 | 0.608497229 |
| ENSMUSG00000087668 | 1 | |
| ENSMUSG00000087669 | 1 | |
| ENSMUSG00000087670 | -1 | 1 |
| ENSMUSG00000087672 | 1.237289901 | 0.268129698 |
| ENSMUSG00000087674 | 1 | |
| ENSMUSG00000087675 | 1 | |
| ENSMUSG00000087677 | 1 | |
| ENSMUSG00000087678 | 1.059463094 | 0.608497229 |
| ENSMUSG00000087679 | 1.168481501 | 0.120256583 |
| ENSMUSG00000087681 | 1 | |
| ENSMUSG00000087682 | 1 | |
| ENSMUSG00000087683 | 1 | |
| ENSMUSG00000087684 | 1.085624843 | 0.656734444 |
| ENSMUSG00000087685 | 1.11166259 | 0.6884821 |
| ENSMUSG00000087687 | 1.121762563 | 0.058947589 |
| ENSMUSG00000087688 | 1 | |
| ENSMUSG00000087689 | 1 | |
| ENSMUSG00000087690 | 1 | |
| ENSMUSG00000087691 | -1.03962336 | 0.803072249 |
| ENSMUSG00000087692 | 1.154696672 | 0.608497229 |
| ENSMUSG00000087693 | 1 | |
| ENSMUSG00000087694 | 1 | |
| ENSMUSG00000087695 | 1 | |
| ENSMUSG00000087698 | 1 | |
| ENSMUSG00000087700 | 1.457314772 | 0.115548937 |
| ENSMUSG00000087701 | 1.059463094 | 0.608497229 |
| ENSMUSG00000087702 | 1 | |
| ENSMUSG00000087703 | 1.019083579 | 0.971519533 |
| ENSMUSG00000087704 | 1 | |
| ENSMUSG00000087707 | 1 | |
| ENSMUSG00000087708 | 1 | |
| ENSMUSG00000087710 | 1 | |
| ENSMUSG00000087712 | 1 | |

| | | |
|---|---|---|
| ENSMUSG00000087714 | -1 | 1 |
| ENSMUSG00000087715 | 1 | |
| ENSMUSG00000087716 | 1 | |
| ENSMUSG00000087717 | 1 | |
| ENSMUSG00000087718 | 1 | |
| ENSMUSG00000087719 | -1.053549062 | 0.807247723 |
| ENSMUSG00000087720 | 1 | |
| ENSMUSG00000087721 | 1.124378687 | 0.608497229 |
| ENSMUSG00000087722 | -1.095294215 | 0.608497229 |
| ENSMUSG00000087723 | 1 | |
| ENSMUSG00000087724 | 1 | |
| ENSMUSG00000087726 | 1 | |
| ENSMUSG00000087728 | 1 | |
| ENSMUSG00000087730 | 1 | |
| ENSMUSG00000087731 | 1.069068979 | 0.608497229 |
| ENSMUSG00000087732 | 1 | |
| ENSMUSG00000087734 | 1 | |
| ENSMUSG00000087736 | 1 | |
| ENSMUSG00000087737 | 1 | |
| ENSMUSG00000087739 | -1.124404745 | 0.716583464 |
| ENSMUSG00000087741 | 1 | |
| ENSMUSG00000087742 | 1 | |
| ENSMUSG00000087743 | 1 | |
| ENSMUSG00000087744 | 1 | |
| ENSMUSG00000087746 | 1 | |
| ENSMUSG00000087747 | 1 | |
| ENSMUSG00000087748 | 1 | |
| ENSMUSG00000087750 | 1.047933867 | 0.95838544 |
| ENSMUSG00000087752 | 1 | |
| ENSMUSG00000087754 | 1 | |
| ENSMUSG00000087756 | 1 | |
| ENSMUSG00000087758 | 1 | |
| ENSMUSG00000087759 | 1 | |
| ENSMUSG00000087760 | 1 | |
| ENSMUSG00000087761 | 1 | |
| ENSMUSG00000087764 | 1 | |
| ENSMUSG00000087765 | 1 | |
| ENSMUSG00000087766 | 1 | |
| ENSMUSG00000087767 | -1.044273782 | 0.698112512 |
| ENSMUSG00000087768 | 1 | |
| ENSMUSG00000087769 | 1 | |
| ENSMUSG00000087772 | 1 | |
| ENSMUSG00000087774 | 1 | |
| ENSMUSG00000087775 | 1 | |
| ENSMUSG00000087776 | 1 | |
| ENSMUSG00000087777 | 1 | |
| ENSMUSG00000087779 | 1 | |
| ENSMUSG00000087780 | 1 | |
| ENSMUSG00000087781 | 1 | |
| ENSMUSG00000087782 | 1 | |
| ENSMUSG00000087783 | 1 | |
| ENSMUSG00000087784 | 1.150179455 | 0.608497229 |
| ENSMUSG00000087785 | 1.014545335 | 0.979002724 |
| ENSMUSG00000087786 | 1 | |
| ENSMUSG00000087789 | 1 | |
| ENSMUSG00000087790 | -1.009476377 | 1 |
| ENSMUSG00000087791 | 1 | |
| ENSMUSG00000087792 | 1 | |
| ENSMUSG00000087793 | 1 | |
| ENSMUSG00000087794 | 1.023151691 | 0.979002724 |
| ENSMUSG00000087795 | 1.005613439 | 1 |
| ENSMUSG00000087796 | 1 | |
| ENSMUSG00000087798 | 1 | |

| | | |
|---|---|---|
| ENSMUSG00000087799 | 1 | |
| ENSMUSG00000087800 | 1 | |
| ENSMUSG00000087801 | 1 | |
| ENSMUSG00000087802 | -1 | 1 |
| ENSMUSG00000087804 | 1 | |
| ENSMUSG00000087805 | 1 | |
| ENSMUSG00000087806 | 1 | |
| ENSMUSG00000087807 | 1.069068979 | 0.608497229 |
| ENSMUSG00000087808 | 1.132639128 | 0.428732467 |
| ENSMUSG00000087810 | 1 | |
| ENSMUSG00000087812 | 1 | |
| ENSMUSG00000087813 | 1 | |
| ENSMUSG00000087815 | 1 | |
| ENSMUSG00000087816 | 1 | |
| ENSMUSG00000087817 | 1 | |
| ENSMUSG00000087819 | 1.051378421 | 0.865598094 |
| ENSMUSG00000087820 | 1 | |
| ENSMUSG00000087823 | 1 | |
| ENSMUSG00000087825 | 1 | |
| ENSMUSG00000087829 | 1 | |
| ENSMUSG00000087830 | 1 | |
| ENSMUSG00000087831 | 1 | |
| ENSMUSG00000087832 | 1 | |
| ENSMUSG00000087833 | 1.059463094 | 0.608497229 |
| ENSMUSG00000087834 | 1 | |
| ENSMUSG00000087835 | 1 | |
| ENSMUSG00000087836 | 1 | |
| ENSMUSG00000087838 | -1 | 1 |
| ENSMUSG00000087839 | 1 | |
| ENSMUSG00000087841 | 1 | |
| ENSMUSG00000087842 | 1 | |
| ENSMUSG00000087843 | 1 | |
| ENSMUSG00000087844 | 1 | |
| ENSMUSG00000087845 | 1 | |
| ENSMUSG00000087846 | 1 | |
| ENSMUSG00000087847 | 1 | |
| ENSMUSG00000087848 | 1 | |
| ENSMUSG00000087850 | 1 | |
| ENSMUSG00000087852 | 1 | |
| ENSMUSG00000087853 | 1 | |
| ENSMUSG00000087854 | 1 | |
| ENSMUSG00000087855 | 1 | |
| ENSMUSG00000087857 | 1 | |
| ENSMUSG00000087858 | 1 | |
| ENSMUSG00000087859 | 1 | |
| ENSMUSG00000087860 | 1 | |
| ENSMUSG00000087861 | 1 | |
| ENSMUSG00000087862 | 1 | |
| ENSMUSG00000087863 | 1 | |
| ENSMUSG00000087867 | 1 | |
| ENSMUSG00000087869 | 1 | |
| ENSMUSG00000087870 | -1 | 1 |
| ENSMUSG00000087871 | 1 | |
| ENSMUSG00000087872 | 1 | |
| ENSMUSG00000087873 | 1 | |
| ENSMUSG00000087875 | 1 | |
| ENSMUSG00000087876 | 1.212674027 | 0.380468454 |
| ENSMUSG00000087877 | 1 | |
| ENSMUSG00000087878 | 1 | |
| ENSMUSG00000087879 | 1 | |
| ENSMUSG00000087880 | 1 | |
| ENSMUSG00000087881 | -1.084645496 | 0.617024566 |
| ENSMUSG00000087883 | 1 | |

| | | |
|---|---|---|
| ENSMUSG00000087884 | 1 | |
| ENSMUSG00000087885 | 1.059463094 | 0.608497229 |
| ENSMUSG00000087886 | 1 | |
| ENSMUSG00000087887 | -1 | 1 |
| ENSMUSG00000087890 | 1 | |
| ENSMUSG00000087891 | 1 | |
| ENSMUSG00000087892 | 1 | |
| ENSMUSG00000087893 | 1 | |
| ENSMUSG00000087894 | 1 | |
| ENSMUSG00000087895 | 1.059463094 | 0.608497229 |
| ENSMUSG00000087896 | 1 | |
| ENSMUSG00000087897 | 1 | |
| ENSMUSG00000087898 | 1 | |
| ENSMUSG00000087899 | 1.069068979 | 0.608497229 |
| ENSMUSG00000087902 | 1 | |
| ENSMUSG00000087903 | 1 | |
| ENSMUSG00000087904 | 1 | |
| ENSMUSG00000087905 | 1 | |
| ENSMUSG00000087908 | 1 | |
| ENSMUSG00000087909 | 1 | |
| ENSMUSG00000087910 | 1 | |
| ENSMUSG00000087911 | 1 | |
| ENSMUSG00000087912 | -1.243513524 | 0.28393292 |
| ENSMUSG00000087913 | 1 | |
| ENSMUSG00000087914 | 1 | |
| ENSMUSG00000087915 | 1 | |
| ENSMUSG00000087916 | 1 | |
| ENSMUSG00000087917 | 1 | |
| ENSMUSG00000087919 | 1 | |
| ENSMUSG00000087921 | 1 | |
| ENSMUSG00000087922 | 1 | |
| ENSMUSG00000087923 | 1 | |
| ENSMUSG00000087924 | 1 | |
| ENSMUSG00000087925 | 1.069068979 | 0.608497229 |
| ENSMUSG00000087926 | 1 | |
| ENSMUSG00000087929 | 1 | |
| ENSMUSG00000087930 | 1 | |
| ENSMUSG00000087932 | 1 | |
| ENSMUSG00000087933 | 1 | |
| ENSMUSG00000087935 | 1 | |
| ENSMUSG00000087936 | 1 | |
| ENSMUSG00000087937 | 1 | |
| ENSMUSG00000087938 | -1.038858794 | 0.914196315 |
| ENSMUSG00000087939 | 1 | |
| ENSMUSG00000087940 | 1 | |
| ENSMUSG00000087941 | 1 | |
| ENSMUSG00000087942 | 1 | |
| ENSMUSG00000087943 | 1 | |
| ENSMUSG00000087945 | 1 | |
| ENSMUSG00000087946 | 1 | |
| ENSMUSG00000087947 | 1 | |
| ENSMUSG00000087949 | 1 | |
| ENSMUSG00000087950 | 1 | |
| ENSMUSG00000087951 | 1 | |
| ENSMUSG00000087954 | 1 | |
| ENSMUSG00000087956 | 1 | |
| ENSMUSG00000087957 | 1 | |
| ENSMUSG00000087958 | 1 | |
| ENSMUSG00000087959 | 1 | |
| ENSMUSG00000087960 | 1 | |
| ENSMUSG00000087961 | 1 | |
| ENSMUSG00000087963 | -1.044273782 | 0.801443616 |
| ENSMUSG00000087964 | 1 | |

| | | |
|---|---|---|
| ENSMUSG00000087965 | 1 | |
| ENSMUSG00000087966 | 1.059463094 | 0.608497229 |
| ENSMUSG00000087967 | 1 | |
| ENSMUSG00000087968 | -1 | 1 |
| ENSMUSG00000087973 | 1 | |
| ENSMUSG00000087975 | 1 | |
| ENSMUSG00000087979 | 1 | |
| ENSMUSG00000087980 | 1 | |
| ENSMUSG00000087981 | 1 | |
| ENSMUSG00000087982 | 1.049411284 | 0.835701362 |
| ENSMUSG00000087983 | 1 | |
| ENSMUSG00000087985 | 1 | |
| ENSMUSG00000087987 | 1 | |
| ENSMUSG00000087989 | 1 | |
| ENSMUSG00000087990 | 1 | |
| ENSMUSG00000087991 | -1.03962336 | 0.872692845 |
| ENSMUSG00000087992 | 1 | |
| ENSMUSG00000087993 | 1 | |
| ENSMUSG00000087994 | 1 | |
| ENSMUSG00000087995 | 1 | |
| ENSMUSG00000087997 | 1 | |
| ENSMUSG00000087999 | 1 | |
| ENSMUSG00000088000 | 1 | |
| ENSMUSG00000088001 | 1 | |
| ENSMUSG00000088002 | 1 | |
| ENSMUSG00000088004 | 1 | |
| ENSMUSG00000088005 | 1 | |
| ENSMUSG00000088008 | 1.139475542 | 0.70299173 |
| ENSMUSG00000088009 | 1 | |
| ENSMUSG00000088010 | 1 | |
| ENSMUSG00000088014 | 1 | |
| ENSMUSG00000088015 | -1 | 1 |
| ENSMUSG00000088016 | 1.059463094 | 0.608497229 |
| ENSMUSG00000088017 | 1 | |
| ENSMUSG00000088018 | -1 | 1 |
| ENSMUSG00000088019 | 1 | |
| ENSMUSG00000088020 | 1 | |
| ENSMUSG00000088021 | 1 | |
| ENSMUSG00000088022 | 1 | |
| ENSMUSG00000088025 | 1 | |
| ENSMUSG00000088026 | 1 | |
| ENSMUSG00000088027 | 1 | |
| ENSMUSG00000088028 | 1 | |
| ENSMUSG00000088030 | 1.059463094 | 0.608497229 |
| ENSMUSG00000088031 | 1 | |
| ENSMUSG00000088036 | 1 | |
| ENSMUSG00000088037 | 1 | |
| ENSMUSG00000088039 | 1 | |
| ENSMUSG00000088040 | 1 | |
| ENSMUSG00000088041 | 1 | |
| ENSMUSG00000088043 | 1 | |
| ENSMUSG00000088044 | 1 | |
| ENSMUSG00000088045 | 1 | |
| ENSMUSG00000088046 | 1 | |
| ENSMUSG00000088047 | 1 | |
| ENSMUSG00000088048 | 1 | |
| ENSMUSG00000088049 | 1 | |
| ENSMUSG00000088050 | 1 | |
| ENSMUSG00000088051 | 1 | |
| ENSMUSG00000088052 | 1 | |
| ENSMUSG00000088054 | 1 | |
| ENSMUSG00000088055 | 1 | |
| ENSMUSG00000088056 | 1 | |

| | | |
|---|---|---|
| ENSMUSG00000088057 | 1 | |
| ENSMUSG00000088058 | 1 | |
| ENSMUSG00000088059 | 1 | |
| ENSMUSG00000088060 | 1 | |
| ENSMUSG00000088062 | 1 | |
| ENSMUSG00000088063 | 1 | |
| ENSMUSG00000088064 | 1 | |
| ENSMUSG00000088066 | 1.069068979 | 0.608497229 |
| ENSMUSG00000088067 | 1 | |
| ENSMUSG00000088068 | 1 | |
| ENSMUSG00000088069 | -1.128888655 | 0.608497229 |
| ENSMUSG00000088070 | 1 | |
| ENSMUSG00000088071 | 1 | |
| ENSMUSG00000088073 | 1 | |
| ENSMUSG00000088075 | -1.043306199 | 0.698112512 |
| ENSMUSG00000088076 | 1 | |
| ENSMUSG00000088077 | 1 | |
| ENSMUSG00000088083 | 1 | |
| ENSMUSG00000088084 | 1 | |
| ENSMUSG00000088085 | 1 | |
| ENSMUSG00000088086 | 1 | |
| ENSMUSG00000088088 | 1.089470641 | 0.608497229 |
| ENSMUSG00000088090 | 1 | |
| ENSMUSG00000088091 | 1 | |
| ENSMUSG00000088093 | 1 | |
| ENSMUSG00000088094 | 1 | |
| ENSMUSG00000088095 | 1 | |
| ENSMUSG00000088096 | 1 | |
| ENSMUSG00000088097 | 1 | |
| ENSMUSG00000088098 | 1 | |
| ENSMUSG00000088099 | 1 | |
| ENSMUSG00000088102 | 1 | |
| ENSMUSG00000088103 | 1 | |
| ENSMUSG00000088104 | 1 | |
| ENSMUSG00000088105 | 1 | |
| ENSMUSG00000088106 | 1 | |
| ENSMUSG00000088108 | 1.00124823 | 1 |
| ENSMUSG00000088111 | 1 | |
| ENSMUSG00000088112 | 1 | |
| ENSMUSG00000088113 | 1.059463094 | 0.608497229 |
| ENSMUSG00000088114 | 1 | |
| ENSMUSG00000088115 | 1 | |
| ENSMUSG00000088116 | 1 | |
| ENSMUSG00000088118 | 1 | |
| ENSMUSG00000088119 | 1.026828809 | 0.983542059 |
| ENSMUSG00000088121 | 1 | |
| ENSMUSG00000088122 | 1 | |
| ENSMUSG00000088123 | 1 | |
| ENSMUSG00000088124 | 1 | |
| ENSMUSG00000088125 | 1 | |
| ENSMUSG00000088126 | 1 | |
| ENSMUSG00000088128 | 1 | |
| ENSMUSG00000088132 | 1 | |
| ENSMUSG00000088133 | 1 | |
| ENSMUSG00000088134 | 1.095872691 | 0.608497229 |
| ENSMUSG00000088135 | 1 | |
| ENSMUSG00000088136 | 1 | |
| ENSMUSG00000088138 | 1 | |
| ENSMUSG00000088139 | -1 | 1 |
| ENSMUSG00000088141 | 1 | |
| ENSMUSG00000088143 | 1 | |
| ENSMUSG00000088144 | 1 | |
| ENSMUSG00000088145 | 1 | |

| | | |
|---|---|---|
| ENSMUSG00000088146 | 1 | |
| ENSMUSG00000088147 | 1 | |
| ENSMUSG00000088148 | 1 | |
| ENSMUSG00000088149 | 1 | |
| ENSMUSG00000088150 | 1 | |
| ENSMUSG00000088151 | -1.054169715 | 0.862822188 |
| ENSMUSG00000088152 | 1 | |
| ENSMUSG00000088153 | 1.030445456 | 0.931897759 |
| ENSMUSG00000088154 | 1 | |
| ENSMUSG00000088155 | 1 | |
| ENSMUSG00000088156 | 1 | |
| ENSMUSG00000088157 | 1 | |
| ENSMUSG00000088158 | -1.043306199 | 0.698112512 |
| ENSMUSG00000088159 | 1 | |
| ENSMUSG00000088160 | 1 | |
| ENSMUSG00000088161 | 1 | |
| ENSMUSG00000088162 | 1 | |
| ENSMUSG00000088164 | 1 | |
| ENSMUSG00000088165 | 1 | |
| ENSMUSG00000088166 | 1 | |
| ENSMUSG00000088167 | 1 | |
| ENSMUSG00000088168 | 1 | |
| ENSMUSG00000088169 | -1.03962336 | 0.698112512 |
| ENSMUSG00000088170 | -1.181812752 | 0.248732933 |
| ENSMUSG00000088171 | -1.043306199 | 0.698112512 |
| ENSMUSG00000088172 | 1 | |
| ENSMUSG00000088175 | 1 | |
| ENSMUSG00000088176 | 1.059463094 | 0.698112512 |
| ENSMUSG00000088177 | 1 | |
| ENSMUSG00000088179 | 1 | |
| ENSMUSG00000088180 | 1 | |
| ENSMUSG00000088181 | 1 | |
| ENSMUSG00000088182 | 1 | |
| ENSMUSG00000088183 | -1.044273782 | 0.698112512 |
| ENSMUSG00000088184 | 1 | |
| ENSMUSG00000088185 | 1.059463094 | 0.608497229 |
| ENSMUSG00000088187 | 1 | |
| ENSMUSG00000088188 | -1.085651418 | 0.532369657 |
| ENSMUSG00000088189 | 1 | |
| ENSMUSG00000088191 | 1 | |
| ENSMUSG00000088192 | 1 | |
| ENSMUSG00000088193 | 1 | |
| ENSMUSG00000088194 | 1 | |
| ENSMUSG00000088195 | 1 | |
| ENSMUSG00000088196 | 1 | |
| ENSMUSG00000088197 | 1 | |
| ENSMUSG00000088198 | 1 | |
| ENSMUSG00000088200 | 1.005613439 | 1 |
| ENSMUSG00000088201 | 1 | |
| ENSMUSG00000088203 | 1 | |
| ENSMUSG00000088204 | 1 | |
| ENSMUSG00000088205 | 1 | |
| ENSMUSG00000088207 | 1 | |
| ENSMUSG00000088208 | 1.200241468 | 0.113598345 |
| ENSMUSG00000088209 | 1 | |
| ENSMUSG00000088210 | 1 | |
| ENSMUSG00000088211 | 1 | |
| ENSMUSG00000088212 | 1 | |
| ENSMUSG00000088213 | 1.019083579 | 0.979002724 |
| ENSMUSG00000088216 | 1 | |
| ENSMUSG00000088217 | 1.019083579 | 0.971519533 |
| ENSMUSG00000088218 | 1 | |
| ENSMUSG00000088219 | 1 | |

| | | |
|---|---|---|
| ENSMUSG00000088220 | 1 | |
| ENSMUSG00000088221 | 1 | |
| ENSMUSG00000088222 | 1 | |
| ENSMUSG00000088223 | 1 | |
| ENSMUSG00000088224 | -1 | 1 |
| ENSMUSG00000088225 | 1 | |
| ENSMUSG00000088226 | 1.123410078 | 0.640486591 |
| ENSMUSG00000088227 | 1 | |
| ENSMUSG00000088228 | 1 | |
| ENSMUSG00000088229 | 1 | |
| ENSMUSG00000088230 | 1 | |
| ENSMUSG00000088231 | 1 | |
| ENSMUSG00000088232 | 1 | |
| ENSMUSG00000088233 | 1.059463094 | 0.608497229 |
| ENSMUSG00000088234 | 1 | |
| ENSMUSG00000088236 | 1 | |
| ENSMUSG00000088237 | 1 | |
| ENSMUSG00000088239 | 1 | |
| ENSMUSG00000088240 | 1 | |
| ENSMUSG00000088241 | 1 | |
| ENSMUSG00000088242 | 1 | |
| ENSMUSG00000088243 | 1 | |
| ENSMUSG00000088244 | 1 | |
| ENSMUSG00000088245 | 1 | |
| ENSMUSG00000088246 | 1 | |
| ENSMUSG00000088248 | 1 | |
| ENSMUSG00000088249 | 1.089470641 | 0.608497229 |
| ENSMUSG00000088250 | -1.03962336 | 0.698112512 |
| ENSMUSG00000088252 | 1 | |
| ENSMUSG00000088253 | 1 | |
| ENSMUSG00000088254 | 1.059463094 | 0.768693672 |
| ENSMUSG00000088255 | 1 | |
| ENSMUSG00000088257 | 1 | |
| ENSMUSG00000088258 | 1 | |
| ENSMUSG00000088259 | 1 | |
| ENSMUSG00000088260 | 1 | |
| ENSMUSG00000088261 | 1 | |
| ENSMUSG00000088264 | 1 | |
| ENSMUSG00000088265 | 1 | |
| ENSMUSG00000088266 | 1 | |
| ENSMUSG00000088268 | -1.081242712 | 0.722351115 |
| ENSMUSG00000088269 | 1 | |
| ENSMUSG00000088270 | 1 | |
| ENSMUSG00000088273 | 1 | |
| ENSMUSG00000088275 | 1 | |
| ENSMUSG00000088276 | 1 | |
| ENSMUSG00000088278 | 1 | |
| ENSMUSG00000088279 | 1 | |
| ENSMUSG00000088280 | 1 | |
| ENSMUSG00000088281 | 1 | |
| ENSMUSG00000088282 | 1 | |
| ENSMUSG00000088283 | 1 | |
| ENSMUSG00000088284 | 1 | |
| ENSMUSG00000088285 | 1 | |
| ENSMUSG00000088286 | 1 | |
| ENSMUSG00000088288 | 1 | |
| ENSMUSG00000088291 | 1 | |
| ENSMUSG00000088293 | 1 | |
| ENSMUSG00000088294 | 1 | |
| ENSMUSG00000088295 | 1 | |
| ENSMUSG00000088296 | 1 | |
| ENSMUSG00000088297 | 1 | |
| ENSMUSG00000088298 | -1.03962336 | 0.803072249 |

| | | |
|---|---|---|
| ENSMUSG00000088299 | 1 | |
| ENSMUSG00000088301 | 1 | |
| ENSMUSG00000088302 | 1 | |
| ENSMUSG00000088303 | 1 | |
| ENSMUSG00000088304 | 1 | |
| ENSMUSG00000088307 | 1 | |
| ENSMUSG00000088308 | -1.127622795 | 0.608497229 |
| ENSMUSG00000088310 | 1 | |
| ENSMUSG00000088311 | 1 | |
| ENSMUSG00000088312 | 1 | |
| ENSMUSG00000088313 | 1 | |
| ENSMUSG00000088314 | 1 | |
| ENSMUSG00000088317 | 1 | |
| ENSMUSG00000088318 | -1 | 1 |
| ENSMUSG00000088320 | 1 | |
| ENSMUSG00000088322 | 1 | |
| ENSMUSG00000088323 | 1 | |
| ENSMUSG00000088324 | 1 | |
| ENSMUSG00000088325 | -1 | 1 |
| ENSMUSG00000088326 | -1.152071125 | 0.647290337 |
| ENSMUSG00000088327 | 1 | |
| ENSMUSG00000088329 | 1 | |
| ENSMUSG00000088331 | 1 | |
| ENSMUSG00000088332 | -1.053549062 | 0.698112512 |
| ENSMUSG00000088333 | 1 | |
| ENSMUSG00000088334 | 1 | |
| ENSMUSG00000088335 | 1 | |
| ENSMUSG00000088337 | 1 | |
| ENSMUSG00000088338 | 1 | |
| ENSMUSG00000088339 | 1 | |
| ENSMUSG00000088340 | 1 | |
| ENSMUSG00000088342 | 1 | |
| ENSMUSG00000088343 | 1 | |
| ENSMUSG00000088344 | 1 | |
| ENSMUSG00000088346 | 1 | |
| ENSMUSG00000088348 | 1 | |
| ENSMUSG00000088349 | 1 | |
| ENSMUSG00000088350 | 1 | |
| ENSMUSG00000088351 | 1 | |
| ENSMUSG00000088352 | 1 | |
| ENSMUSG00000088353 | 1 | |
| ENSMUSG00000088354 | 1 | |
| ENSMUSG00000088356 | 1 | |
| ENSMUSG00000088357 | -1.044273782 | 0.790851299 |
| ENSMUSG00000088358 | 1 | |
| ENSMUSG00000088359 | 1 | |
| ENSMUSG00000088360 | 1 | |
| ENSMUSG00000088361 | 1 | |
| ENSMUSG00000088362 | 1 | |
| ENSMUSG00000088363 | 1 | |
| ENSMUSG00000088364 | 1 | |
| ENSMUSG00000088365 | 1 | |
| ENSMUSG00000088366 | 1 | |
| ENSMUSG00000088367 | 1 | |
| ENSMUSG00000088368 | 1 | |
| ENSMUSG00000088369 | 1 | |
| ENSMUSG00000088370 | 1 | |
| ENSMUSG00000088371 | 1 | |
| ENSMUSG00000088372 | 1 | |
| ENSMUSG00000088373 | 1 | |
| ENSMUSG00000088374 | 1 | |
| ENSMUSG00000088377 | 1 | |
| ENSMUSG00000088378 | 1.014545335 | 0.991944074 |

| | | |
|---|---|---|
| ENSMUSG00000088379 | 1 | |
| ENSMUSG00000088380 | 1 | |
| ENSMUSG00000088383 | 1 | |
| ENSMUSG00000088384 | 1 | |
| ENSMUSG00000088385 | -1.044273782 | 0.698112512 |
| ENSMUSG00000088386 | 1 | |
| ENSMUSG00000088387 | 1 | |
| ENSMUSG00000088389 | 1 | |
| ENSMUSG00000088391 | 1 | |
| ENSMUSG00000088392 | 1 | |
| ENSMUSG00000088393 | 1 | |
| ENSMUSG00000088394 | 1 | |
| ENSMUSG00000088395 | 1 | |
| ENSMUSG00000088397 | 1 | |
| ENSMUSG00000088398 | 1 | |
| ENSMUSG00000088400 | -1 | 1 |
| ENSMUSG00000088402 | -1.044273782 | 0.698112512 |
| ENSMUSG00000088403 | 1 | |
| ENSMUSG00000088404 | 1 | |
| ENSMUSG00000088405 | 1 | |
| ENSMUSG00000088406 | 1 | |
| ENSMUSG00000088407 | 1 | |
| ENSMUSG00000088408 | 1 | |
| ENSMUSG00000088409 | 1 | |
| ENSMUSG00000088410 | 1 | |
| ENSMUSG00000088412 | -1.113515092 | 0.608497229 |
| ENSMUSG00000088414 | -1.043306199 | 0.698112512 |
| ENSMUSG00000088415 | 1 | |
| ENSMUSG00000088416 | 1 | |
| ENSMUSG00000088417 | 1 | |
| ENSMUSG00000088418 | 1 | |
| ENSMUSG00000088419 | 1 | |
| ENSMUSG00000088420 | 1 | |
| ENSMUSG00000088421 | 1 | |
| ENSMUSG00000088422 | 1 | |
| ENSMUSG00000088423 | 1 | |
| ENSMUSG00000088425 | 1 | |
| ENSMUSG00000088426 | 1 | |
| ENSMUSG00000088428 | 1 | |
| ENSMUSG00000088429 | 1 | |
| ENSMUSG00000088432 | 1 | |
| ENSMUSG00000088433 | 1 | |
| ENSMUSG00000088435 | 1 | |
| ENSMUSG00000088436 | 1 | |
| ENSMUSG00000088437 | 1.015486245 | 0.972980766 |
| ENSMUSG00000088438 | 1 | |
| ENSMUSG00000088439 | 1 | |
| ENSMUSG00000088440 | 1 | |
| ENSMUSG00000088441 | 1 | |
| ENSMUSG00000088442 | -1 | 1 |
| ENSMUSG00000088443 | 1 | |
| ENSMUSG00000088444 | 1 | |
| ENSMUSG00000088445 | 1 | |
| ENSMUSG00000088446 | 1 | |
| ENSMUSG00000088447 | 1 | |
| ENSMUSG00000088448 | 1 | |
| ENSMUSG00000088449 | 1 | |
| ENSMUSG00000088450 | 1 | |
| ENSMUSG00000088451 | 1 | |
| ENSMUSG00000088453 | -1.044273782 | 0.790851299 |
| ENSMUSG00000088454 | 1 | |
| ENSMUSG00000088456 | 1 | |
| ENSMUSG00000088458 | 1 | |

| | | |
|---|---|---|
| ENSMUSG00000088459 | -1.03962336 | 0.698112512 |
| ENSMUSG00000088460 | 1 | |
| ENSMUSG00000088461 | 1 | |
| ENSMUSG00000088463 | 1 | |
| ENSMUSG00000088464 | -1.059527646 | 0.839785964 |
| ENSMUSG00000088465 | 1 | |
| ENSMUSG00000088468 | 1 | |
| ENSMUSG00000088469 | 1 | |
| ENSMUSG00000088470 | 1 | |
| ENSMUSG00000088471 | 1 | |
| ENSMUSG00000088472 | 1 | |
| ENSMUSG00000088473 | -1.166301255 | 0.539222653 |
| ENSMUSG00000088474 | 1 | |
| ENSMUSG00000088475 | 1.079681442 | 0.776953697 |
| ENSMUSG00000088476 | 1 | |
| ENSMUSG00000088477 | 1 | |
| ENSMUSG00000088478 | 1 | |
| ENSMUSG00000088479 | -1.043306199 | 0.698112512 |
| ENSMUSG00000088480 | 1 | |
| ENSMUSG00000088481 | 1 | |
| ENSMUSG00000088482 | 1 | |
| ENSMUSG00000088483 | 1 | |
| ENSMUSG00000088484 | 1 | |
| ENSMUSG00000088485 | 1 | |
| ENSMUSG00000088487 | 1 | |
| ENSMUSG00000088490 | -1.086681236 | 0.70439137 |
| ENSMUSG00000088491 | 1 | |
| ENSMUSG00000088493 | 1.019083579 | 0.949812939 |
| ENSMUSG00000088494 | 1.015486245 | 0.985522133 |
| ENSMUSG00000088495 | 1 | |
| ENSMUSG00000088497 | 1 | |
| ENSMUSG00000088498 | 1 | |
| ENSMUSG00000088499 | 1 | |
| ENSMUSG00000088500 | 1 | |
| ENSMUSG00000088501 | 1 | |
| ENSMUSG00000088502 | 1 | |
| ENSMUSG00000088504 | -1.109582661 | 0.608497229 |
| ENSMUSG00000088505 | 1 | |
| ENSMUSG00000088506 | 1 | |
| ENSMUSG00000088509 | 1 | |
| ENSMUSG00000088510 | 1 | |
| ENSMUSG00000088511 | 1 | |
| ENSMUSG00000088512 | 1 | |
| ENSMUSG00000088514 | 1 | |
| ENSMUSG00000088515 | 1 | |
| ENSMUSG00000088516 | 1 | |
| ENSMUSG00000088517 | 1 | |
| ENSMUSG00000088518 | 1 | |
| ENSMUSG00000088519 | 1 | |
| ENSMUSG00000088521 | 1 | |
| ENSMUSG00000088522 | 1 | |
| ENSMUSG00000088523 | 1 | |
| ENSMUSG00000088524 | 1 | |
| ENSMUSG00000088525 | 1 | |
| ENSMUSG00000088526 | 1 | |
| ENSMUSG00000088527 | 1 | |
| ENSMUSG00000088528 | 1 | |
| ENSMUSG00000088529 | 1 | |
| ENSMUSG00000088530 | 1 | |
| ENSMUSG00000088532 | 1 | |
| ENSMUSG00000088533 | 1 | |
| ENSMUSG00000088535 | 1 | |
| ENSMUSG00000088536 | 1 | |

| | | |
|---|---|---|
| ENSMUSG00000088537 | 1 | |
| ENSMUSG00000088538 | 1 | |
| ENSMUSG00000088539 | 1 | |
| ENSMUSG00000088541 | 1 | |
| ENSMUSG00000088542 | 1 | |
| ENSMUSG00000088543 | 1 | |
| ENSMUSG00000088544 | 1 | |
| ENSMUSG00000088545 | -1 | 1 |
| ENSMUSG00000088546 | 1 | |
| ENSMUSG00000088547 | 1 | |
| ENSMUSG00000088548 | 1 | |
| ENSMUSG00000088549 | 1 | |
| ENSMUSG00000088550 | 1 | |
| ENSMUSG00000088551 | 1 | |
| ENSMUSG00000088552 | 1.069068979 | 0.608497229 |
| ENSMUSG00000088553 | 1 | |
| ENSMUSG00000088554 | 1 | |
| ENSMUSG00000088555 | -1.043306199 | 0.803260194 |
| ENSMUSG00000088558 | 1 | |
| ENSMUSG00000088559 | 1 | |
| ENSMUSG00000088562 | 1 | |
| ENSMUSG00000088563 | 1 | |
| ENSMUSG00000088564 | 1 | |
| ENSMUSG00000088565 | 1 | |
| ENSMUSG00000088566 | 1 | |
| ENSMUSG00000088570 | 1 | |
| ENSMUSG00000088571 | 1 | |
| ENSMUSG00000088572 | 1 | |
| ENSMUSG00000088573 | 1 | |
| ENSMUSG00000088574 | 1 | |
| ENSMUSG00000088575 | 1 | |
| ENSMUSG00000088576 | 1 | |
| ENSMUSG00000088577 | 1 | |
| ENSMUSG00000088578 | 1 | |
| ENSMUSG00000088579 | 1 | |
| ENSMUSG00000088580 | 1 | |
| ENSMUSG00000088581 | 1 | |
| ENSMUSG00000088582 | 1 | |
| ENSMUSG00000088583 | -1.053549062 | 0.698112512 |
| ENSMUSG00000088585 | 1 | |
| ENSMUSG00000088587 | 1 | |
| ENSMUSG00000088589 | 1 | |
| ENSMUSG00000088591 | -1.060239133 | 0.825478278 |
| ENSMUSG00000088594 | 1 | |
| ENSMUSG00000088595 | -1.03962336 | 0.698112512 |
| ENSMUSG00000088596 | 1 | |
| ENSMUSG00000088597 | 1 | |
| ENSMUSG00000088598 | 1 | |
| ENSMUSG00000088600 | 1 | |
| ENSMUSG00000088601 | 1 | |
| ENSMUSG00000088602 | 1.095872691 | 0.621184073 |
| ENSMUSG00000088603 | 1 | |
| ENSMUSG00000088605 | -1.173142781 | 0.36904104 |
| ENSMUSG00000088606 | 1 | |
| ENSMUSG00000088608 | 1 | |
| ENSMUSG00000088609 | 1 | |
| ENSMUSG00000088611 | 1 | |
| ENSMUSG00000088614 | 1.059463094 | 0.608497229 |
| ENSMUSG00000088615 | 1.011607416 | 1 |
| ENSMUSG00000088616 | 1 | |
| ENSMUSG00000088617 | 1 | |
| ENSMUSG00000088618 | 1 | |
| ENSMUSG00000088620 | 1 | |

| | | |
|---|---|---|
| ENSMUSG00000088621 | 1 | |
| ENSMUSG00000088622 | 1 | |
| ENSMUSG00000088623 | -1.03962336 | 0.698112512 |
| ENSMUSG00000088624 | 1 | |
| ENSMUSG00000088625 | -1.041574599 | 0.941510974 |
| ENSMUSG00000088626 | -1.03962336 | 0.698112512 |
| ENSMUSG00000088627 | 1 | |
| ENSMUSG00000088628 | 1 | |
| ENSMUSG00000088629 | 1 | |
| ENSMUSG00000088630 | 1 | |
| ENSMUSG00000088631 | -1 | 1 |
| ENSMUSG00000088632 | 1.010838709 | 1 |
| ENSMUSG00000088633 | 1 | |
| ENSMUSG00000088634 | 1 | |
| ENSMUSG00000088635 | 1 | |
| ENSMUSG00000088637 | 1 | |
| ENSMUSG00000088638 | 1 | |
| ENSMUSG00000088639 | -1.069502966 | 0.752398528 |
| ENSMUSG00000088640 | 1 | |
| ENSMUSG00000088641 | 1 | |
| ENSMUSG00000088643 | -1.085651418 | 0.608497229 |
| ENSMUSG00000088644 | 1 | |
| ENSMUSG00000088646 | 1 | |
| ENSMUSG00000088647 | 1 | |
| ENSMUSG00000088649 | 1 | |
| ENSMUSG00000088650 | -1.084645496 | 0.532369657 |
| ENSMUSG00000088651 | 1 | |
| ENSMUSG00000088653 | 1 | |
| ENSMUSG00000088654 | 1 | |
| ENSMUSG00000088656 | 1 | |
| ENSMUSG00000088657 | 1 | |
| ENSMUSG00000088658 | 1 | |
| ENSMUSG00000088659 | 1 | |
| ENSMUSG00000088660 | 1 | |
| ENSMUSG00000088661 | 1 | |
| ENSMUSG00000088663 | 1 | |
| ENSMUSG00000088666 | 1 | |
| ENSMUSG00000088668 | 1 | |
| ENSMUSG00000088669 | 1 | |
| ENSMUSG00000088670 | 1 | |
| ENSMUSG00000088671 | 1 | |
| ENSMUSG00000088672 | 1 | |
| ENSMUSG00000088674 | -1 | 1 |
| ENSMUSG00000088675 | 1.132639128 | 0.217803313 |
| ENSMUSG00000088677 | 1 | |
| ENSMUSG00000088678 | 1 | |
| ENSMUSG00000088679 | 1 | |
| ENSMUSG00000088680 | 1 | |
| ENSMUSG00000088682 | 1 | |
| ENSMUSG00000088683 | 1 | |
| ENSMUSG00000088684 | 1 | |
| ENSMUSG00000088685 | -1 | 1 |
| ENSMUSG00000088686 | 1 | |
| ENSMUSG00000088687 | 1 | |
| ENSMUSG00000088688 | 1 | |
| ENSMUSG00000088689 | 1.059463094 | 0.698112512 |
| ENSMUSG00000088691 | 1 | |
| ENSMUSG00000088692 | 1 | |
| ENSMUSG00000088693 | 1 | |
| ENSMUSG00000088695 | 1.095872691 | 0.657060635 |
| ENSMUSG00000088696 | 1 | |
| ENSMUSG00000088697 | 1 | |
| ENSMUSG00000088699 | -1 | 1 |

| | | |
|---|---|---|
| ENSMUSG00000088700 | 1 | |
| ENSMUSG00000088701 | 1 | |
| ENSMUSG00000088702 | 1 | |
| ENSMUSG00000088703 | 1 | |
| ENSMUSG00000088704 | 1 | |
| ENSMUSG00000088705 | -1.023769022 | 0.971519533 |
| ENSMUSG00000088706 | 1 | |
| ENSMUSG00000088707 | 1 | |
| ENSMUSG00000088708 | -1.043306199 | 0.698112512 |
| ENSMUSG00000088709 | 1 | |
| ENSMUSG00000088710 | 1 | |
| ENSMUSG00000088711 | 1 | |
| ENSMUSG00000088712 | 1.001874515 | 1 |
| ENSMUSG00000088713 | 1 | |
| ENSMUSG00000088714 | 1 | |
| ENSMUSG00000088715 | 1 | |
| ENSMUSG00000088716 | 1 | |
| ENSMUSG00000088717 | 1 | |
| ENSMUSG00000088718 | 1 | |
| ENSMUSG00000088719 | 1 | |
| ENSMUSG00000088721 | 1 | |
| ENSMUSG00000088722 | 1 | |
| ENSMUSG00000088726 | 1 | |
| ENSMUSG00000088730 | 1 | |
| ENSMUSG00000088731 | 1 | |
| ENSMUSG00000088732 | 1 | |
| ENSMUSG00000088733 | 1 | |
| ENSMUSG00000088734 | 1 | |
| ENSMUSG00000088735 | 1 | |
| ENSMUSG00000088736 | 1 | |
| ENSMUSG00000088737 | 1 | |
| ENSMUSG00000088738 | 1 | |
| ENSMUSG00000088739 | 1 | |
| ENSMUSG00000088740 | 1 | |
| ENSMUSG00000088742 | 1 | |
| ENSMUSG00000088743 | 1 | |
| ENSMUSG00000088744 | 1 | |
| ENSMUSG00000088746 | 1 | |
| ENSMUSG00000088747 | 1 | |
| ENSMUSG00000088748 | 1 | |
| ENSMUSG00000088749 | 1 | |
| ENSMUSG00000088750 | 1 | |
| ENSMUSG00000088751 | 1 | |
| ENSMUSG00000088752 | 1 | |
| ENSMUSG00000088753 | 1 | |
| ENSMUSG00000088754 | 1 | |
| ENSMUSG00000088756 | 1 | |
| ENSMUSG00000088758 | 1 | |
| ENSMUSG00000088760 | 1 | |
| ENSMUSG00000088761 | 1 | |
| ENSMUSG00000088763 | -1.03962336 | 0.698112512 |
| ENSMUSG00000088764 | 1 | |
| ENSMUSG00000088766 | 1 | |
| ENSMUSG00000088767 | 1 | |
| ENSMUSG00000088768 | 1 | |
| ENSMUSG00000088769 | 1.021772483 | 1 |
| ENSMUSG00000088772 | 1 | |
| ENSMUSG00000088773 | 1 | |
| ENSMUSG00000088774 | 1 | |
| ENSMUSG00000088775 | 1 | |
| ENSMUSG00000088776 | 1 | |
| ENSMUSG00000088777 | -1.064334191 | 0.821493744 |
| ENSMUSG00000088779 | 1 | |

| | | |
|---|---|---|
| ENSMUSG00000088781 | 1 | |
| ENSMUSG00000088782 | 1 | |
| ENSMUSG00000088783 | 1 | |
| ENSMUSG00000088784 | 1 | |
| ENSMUSG00000088786 | 1 | |
| ENSMUSG00000088787 | 1 | |
| ENSMUSG00000088789 | 1.090857254 | 0.77943049 |
| ENSMUSG00000088790 | 1 | |
| ENSMUSG00000088791 | 1 | |
| ENSMUSG00000088792 | 1 | |
| ENSMUSG00000088793 | 1.059463094 | 0.698112512 |
| ENSMUSG00000088794 | 1 | |
| ENSMUSG00000088796 | 1 | |
| ENSMUSG00000088797 | 1 | |
| ENSMUSG00000088798 | 1 | |
| ENSMUSG00000088799 | 1 | |
| ENSMUSG00000088800 | 1 | |
| ENSMUSG00000088802 | 1 | |
| ENSMUSG00000088803 | 1 | |
| ENSMUSG00000088804 | 1 | |
| ENSMUSG00000088808 | 1 | |
| ENSMUSG00000088809 | 1 | |
| ENSMUSG00000088810 | 1 | |
| ENSMUSG00000088811 | 1 | |
| ENSMUSG00000088812 | 1 | |
| ENSMUSG00000088813 | 1 | |
| ENSMUSG00000088814 | 1 | |
| ENSMUSG00000088816 | 1 | |
| ENSMUSG00000088817 | 1 | |
| ENSMUSG00000088818 | 1 | |
| ENSMUSG00000088819 | -1.044273782 | 0.698112512 |
| ENSMUSG00000088820 | 1 | |
| ENSMUSG00000088822 | 1 | |
| ENSMUSG00000088823 | 1 | |
| ENSMUSG00000088824 | 1 | |
| ENSMUSG00000088825 | 1 | |
| ENSMUSG00000088829 | 1 | |
| ENSMUSG00000088830 | 1 | |
| ENSMUSG00000088832 | -1.047306572 | 0.885591365 |
| ENSMUSG00000088833 | 1 | |
| ENSMUSG00000088835 | -1.150879559 | 0.539222653 |
| ENSMUSG00000088836 | 1 | |
| ENSMUSG00000088839 | 1 | |
| ENSMUSG00000088840 | 1 | |
| ENSMUSG00000088842 | 1 | |
| ENSMUSG00000088843 | 1 | |
| ENSMUSG00000088845 | 1 | |
| ENSMUSG00000088846 | 1 | |
| ENSMUSG00000088848 | 1 | |
| ENSMUSG00000088849 | 1 | |
| ENSMUSG00000088850 | 1 | |
| ENSMUSG00000088851 | 1 | |
| ENSMUSG00000088852 | 1 | |
| ENSMUSG00000088853 | 1 | |
| ENSMUSG00000088854 | 1 | |
| ENSMUSG00000088855 | -1 | 1 |
| ENSMUSG00000088856 | -1.001876871 | 1 |
| ENSMUSG00000088857 | 1 | |
| ENSMUSG00000088858 | 1 | |
| ENSMUSG00000088860 | 1 | |
| ENSMUSG00000088861 | 1 | |
| ENSMUSG00000088862 | 1 | |
| ENSMUSG00000088863 | 1 | |

| | | |
|---|---|---|
| ENSMUSG00000088864 | 1 | |
| ENSMUSG00000088865 | 1 | |
| ENSMUSG00000088867 | 1 | |
| ENSMUSG00000088868 | 1 | |
| ENSMUSG00000088870 | 1 | |
| ENSMUSG00000088872 | 1 | |
| ENSMUSG00000088873 | 1 | |
| ENSMUSG00000088874 | 1 | |
| ENSMUSG00000088875 | 1 | |
| ENSMUSG00000088876 | 1 | |
| ENSMUSG00000088877 | 1 | |
| ENSMUSG00000088878 | 1 | |
| ENSMUSG00000088879 | 1 | |
| ENSMUSG00000088880 | 1 | |
| ENSMUSG00000088883 | 1 | |
| ENSMUSG00000088884 | 1 | |
| ENSMUSG00000088885 | 1 | |
| ENSMUSG00000088888 | 1 | |
| ENSMUSG00000088889 | 1 | |
| ENSMUSG00000088890 | 1 | |
| ENSMUSG00000088891 | -1.024718486 | 0.982777255 |
| ENSMUSG00000088892 | 1 | |
| ENSMUSG00000088894 | 1 | |
| ENSMUSG00000088895 | 1 | |
| ENSMUSG00000088896 | 1 | |
| ENSMUSG00000088897 | 1 | |
| ENSMUSG00000088898 | 1 | |
| ENSMUSG00000088900 | 1 | |
| ENSMUSG00000088901 | -1.100193663 | 0.608497229 |
| ENSMUSG00000088902 | 1 | |
| ENSMUSG00000088903 | 1 | |
| ENSMUSG00000088904 | 1 | |
| ENSMUSG00000088905 | 1 | |
| ENSMUSG00000088906 | 1 | |
| ENSMUSG00000088907 | 1 | |
| ENSMUSG00000088908 | 1 | |
| ENSMUSG00000088909 | 1 | |
| ENSMUSG00000088910 | 1 | |
| ENSMUSG00000088911 | 1 | |
| ENSMUSG00000088913 | 1.059463094 | 0.608497229 |
| ENSMUSG00000088914 | 1 | |
| ENSMUSG00000088915 | 1 | |
| ENSMUSG00000088916 | 1 | |
| ENSMUSG00000088918 | 1 | |
| ENSMUSG00000088920 | 1 | |
| ENSMUSG00000088922 | 1 | |
| ENSMUSG00000088923 | 1 | |
| ENSMUSG00000088924 | 1 | |
| ENSMUSG00000088925 | 1 | |
| ENSMUSG00000088926 | 1 | |
| ENSMUSG00000088928 | 1 | |
| ENSMUSG00000088929 | -1.151373707 | 0.227002255 |
| ENSMUSG00000088930 | 1 | |
| ENSMUSG00000088932 | 1 | |
| ENSMUSG00000088933 | 1 | |
| ENSMUSG00000088935 | 1 | |
| ENSMUSG00000088937 | 1 | |
| ENSMUSG00000088938 | 1 | |
| ENSMUSG00000088939 | -1 | 1 |
| ENSMUSG00000088940 | 1 | |
| ENSMUSG00000088941 | 1 | |
| ENSMUSG00000088942 | 1 | |
| ENSMUSG00000088943 | 1 | |

| | | |
|---|---|---|
| ENSMUSG00000088944 | 1 | |
| ENSMUSG00000088947 | 1 | |
| ENSMUSG00000088948 | 1.085624843 | 0.709353016 |
| ENSMUSG00000088949 | 1 | |
| ENSMUSG00000088950 | 1 | |
| ENSMUSG00000088951 | -1.03962336 | 0.698112512 |
| ENSMUSG00000088953 | 1 | |
| ENSMUSG00000088954 | 1 | |
| ENSMUSG00000088955 | 1 | |
| ENSMUSG00000088956 | 1 | |
| ENSMUSG00000088957 | 1 | |
| ENSMUSG00000088958 | 1.005613439 | 1 |
| ENSMUSG00000088959 | 1 | |
| ENSMUSG00000088960 | 1 | |
| ENSMUSG00000088961 | 1.07487334 | 0.797760476 |
| ENSMUSG00000088964 | 1 | |
| ENSMUSG00000088966 | 1 | |
| ENSMUSG00000088969 | 1.132639128 | 0.398767938 |
| ENSMUSG00000088970 | 1 | |
| ENSMUSG00000088971 | 1 | |
| ENSMUSG00000088972 | 1 | |
| ENSMUSG00000088973 | 1 | |
| ENSMUSG00000088975 | 1 | |
| ENSMUSG00000088976 | 1 | |
| ENSMUSG00000088978 | -1 | 1 |
| ENSMUSG00000088979 | 1 | |
| ENSMUSG00000088980 | 1 | |
| ENSMUSG00000088981 | 1 | |
| ENSMUSG00000088982 | -1.111880267 | 0.694494111 |
| ENSMUSG00000088983 | 1 | |
| ENSMUSG00000088984 | -1.023016116 | 0.979186916 |
| ENSMUSG00000088985 | 1.059463094 | 0.608497229 |
| ENSMUSG00000088986 | 1 | |
| ENSMUSG00000088987 | 1 | |
| ENSMUSG00000088988 | 1 | |
| ENSMUSG00000088989 | 1 | |
| ENSMUSG00000088990 | -1.05948903 | 0.841678369 |
| ENSMUSG00000088991 | 1 | |
| ENSMUSG00000088992 | 1 | |
| ENSMUSG00000088993 | -1.199425955 | 0.440395661 |
| ENSMUSG00000088994 | 1 | |
| ENSMUSG00000088995 | 1 | |
| ENSMUSG00000088996 | 1 | |
| ENSMUSG00000088997 | 1 | |
| ENSMUSG00000088999 | 1 | |
| ENSMUSG00000089000 | 1 | |
| ENSMUSG00000089001 | 1 | |
| ENSMUSG00000089002 | 1 | |
| ENSMUSG00000089003 | 1 | |
| ENSMUSG00000089004 | 1 | |
| ENSMUSG00000089005 | 1 | |
| ENSMUSG00000089007 | 1 | |
| ENSMUSG00000089008 | 1 | |
| ENSMUSG00000089009 | 1 | |
| ENSMUSG00000089011 | 1 | |
| ENSMUSG00000089014 | -1.044273782 | 0.790851299 |
| ENSMUSG00000089015 | -1 | 1 |
| ENSMUSG00000089016 | 1 | |
| ENSMUSG00000089017 | 1 | |
| ENSMUSG00000089018 | 1 | |
| ENSMUSG00000089019 | 1 | |
| ENSMUSG00000089021 | 1 | |
| ENSMUSG00000089023 | 1 | |

| | | |
|---|---|---|
| ENSMUSG00000089024 | 1 | |
| ENSMUSG00000089025 | 1 | |
| ENSMUSG00000089026 | 1.065410325 | 0.711602885 |
| ENSMUSG00000089027 | 1 | |
| ENSMUSG00000089028 | 1 | |
| ENSMUSG00000089029 | 1 | |
| ENSMUSG00000089030 | 1 | |
| ENSMUSG00000089033 | 1 | |
| ENSMUSG00000089034 | 1 | |
| ENSMUSG00000089035 | 1 | |
| ENSMUSG00000089036 | 1 | |
| ENSMUSG00000089037 | 1 | |
| ENSMUSG00000089038 | 1 | |
| ENSMUSG00000089039 | 1 | |
| ENSMUSG00000089040 | 1 | |
| ENSMUSG00000089041 | 1 | |
| ENSMUSG00000089043 | 1 | |
| ENSMUSG00000089044 | 1 | |
| ENSMUSG00000089045 | -1.043306199 | 0.698112512 |
| ENSMUSG00000089046 | 1 | |
| ENSMUSG00000089047 | 1 | |
| ENSMUSG00000089048 | -1.053549062 | 0.769745556 |
| ENSMUSG00000089049 | 1 | |
| ENSMUSG00000089050 | 1 | |
| ENSMUSG00000089051 | 1 | |
| ENSMUSG00000089052 | 1 | |
| ENSMUSG00000089053 | 1 | |
| ENSMUSG00000089054 | -1 | 1 |
| ENSMUSG00000089055 | 1 | |
| ENSMUSG00000089057 | 1 | |
| ENSMUSG00000089060 | 1 | |
| ENSMUSG00000089061 | 1 | |
| ENSMUSG00000089062 | 1 | |
| ENSMUSG00000089065 | 1 | |
| ENSMUSG00000089066 | 1 | |
| ENSMUSG00000089068 | 1 | |
| ENSMUSG00000089069 | 1 | |
| ENSMUSG00000089070 | 1 | |
| ENSMUSG00000089071 | 1 | |
| ENSMUSG00000089072 | -1.121163731 | 0.717035356 |
| ENSMUSG00000089073 | 1 | |
| ENSMUSG00000089074 | 1 | |
| ENSMUSG00000089075 | 1 | |
| ENSMUSG00000089077 | 1 | |
| ENSMUSG00000089078 | 1 | |
| ENSMUSG00000089079 | 1 | |
| ENSMUSG00000089080 | 1 | |
| ENSMUSG00000089081 | 1 | |
| ENSMUSG00000089082 | 1 | |
| ENSMUSG00000089083 | 1 | |
| ENSMUSG00000089084 | 1 | |
| ENSMUSG00000089085 | 1 | |
| ENSMUSG00000089087 | 1 | |
| ENSMUSG00000089088 | 1 | |
| ENSMUSG00000089090 | 1 | |
| ENSMUSG00000089091 | 1 | |
| ENSMUSG00000089092 | -1 | 1 |
| ENSMUSG00000089093 | -1 | 1 |
| ENSMUSG00000089094 | 1 | |
| ENSMUSG00000089095 | 1 | |
| ENSMUSG00000089096 | 1 | |
| ENSMUSG00000089097 | 1 | |
| ENSMUSG00000089098 | 1 | |

| | | |
|---|---|---|
| ENSMUSG00000089102 | 1 | |
| ENSMUSG00000089103 | -1 | 1 |
| ENSMUSG00000089105 | 1.05410549 | 0.805157604 |
| ENSMUSG00000089106 | 1 | |
| ENSMUSG00000089107 | 1 | |
| ENSMUSG00000089109 | 1 | |
| ENSMUSG00000089111 | 1 | |
| ENSMUSG00000089113 | 1.059463094 | 0.608497229 |
| ENSMUSG00000089114 | 1 | |
| ENSMUSG00000089115 | 1 | |
| ENSMUSG00000089116 | 1 | |
| ENSMUSG00000089117 | 1 | |
| ENSMUSG00000089118 | 1 | |
| ENSMUSG00000089119 | 1 | |
| ENSMUSG00000089120 | 1 | |
| ENSMUSG00000089123 | 1 | |
| ENSMUSG00000089124 | 1 | |
| ENSMUSG00000089125 | -1.103388856 | 0.610406338 |
| ENSMUSG00000089127 | 1 | |
| ENSMUSG00000089128 | 1 | |
| ENSMUSG00000089129 | -1.047588501 | 0.803072249 |
| ENSMUSG00000089130 | 1 | |
| ENSMUSG00000089131 | 1 | |
| ENSMUSG00000089132 | 1 | |
| ENSMUSG00000089133 | 1 | |
| ENSMUSG00000089134 | 1 | |
| ENSMUSG00000089136 | 1 | |
| ENSMUSG00000089137 | 1 | |
| ENSMUSG00000089138 | 1 | |
| ENSMUSG00000089139 | 1 | |
| ENSMUSG00000089141 | 1 | |
| ENSMUSG00000089143 | -1 | 1 |
| ENSMUSG00000089144 | 1 | |
| ENSMUSG00000089145 | 1 | |
| ENSMUSG00000089146 | -1 | 1 |
| ENSMUSG00000089147 | 1 | |
| ENSMUSG00000089148 | 1 | |
| ENSMUSG00000089149 | 1 | |
| ENSMUSG00000089151 | -1.071075483 | 0.801443616 |
| ENSMUSG00000089152 | 1 | |
| ENSMUSG00000089153 | 1 | |
| ENSMUSG00000089155 | -1.03962336 | 0.698112512 |
| ENSMUSG00000089156 | 1 | |
| ENSMUSG00000089157 | 1 | |
| ENSMUSG00000089158 | 1 | |
| ENSMUSG00000089161 | 1 | |
| ENSMUSG00000089162 | 1 | |
| ENSMUSG00000089163 | 1 | |
| ENSMUSG00000089164 | 1 | |
| ENSMUSG00000089168 | 1 | |
| ENSMUSG00000089169 | 1 | |
| ENSMUSG00000089170 | -1.033820075 | 0.920928417 |
| ENSMUSG00000089171 | 1 | |
| ENSMUSG00000089174 | 1 | |
| ENSMUSG00000089175 | 1 | |
| ENSMUSG00000089177 | -1 | 1 |
| ENSMUSG00000089178 | 1 | |
| ENSMUSG00000089179 | -1.03962336 | 0.698112512 |
| ENSMUSG00000089182 | 1 | |
| ENSMUSG00000089183 | 1 | |
| ENSMUSG00000089184 | 1 | |
| ENSMUSG00000089185 | -1.043306199 | 0.698112512 |
| ENSMUSG00000089186 | 1 | |

| | | |
|---|---|---|
| ENSMUSG00000089187 | 1.059463094 | 0.608497229 |
| ENSMUSG00000089188 | 1 | |
| ENSMUSG00000089189 | 1 | |
| ENSMUSG00000089190 | 1 | |
| ENSMUSG00000089191 | 1 | |
| ENSMUSG00000089194 | 1 | |
| ENSMUSG00000089195 | 1 | |
| ENSMUSG00000089197 | 1 | |
| ENSMUSG00000089199 | -1 | 1 |
| ENSMUSG00000089203 | 1 | |
| ENSMUSG00000089204 | 1 | |
| ENSMUSG00000089205 | 1 | |
| ENSMUSG00000089206 | 1 | |
| ENSMUSG00000089207 | 1 | |
| ENSMUSG00000089208 | 1 | |
| ENSMUSG00000089209 | 1 | |
| ENSMUSG00000089210 | 1 | |
| ENSMUSG00000089211 | 1 | |
| ENSMUSG00000089213 | 1 | |
| ENSMUSG00000089214 | 1 | |
| ENSMUSG00000089217 | 1 | |
| ENSMUSG00000089218 | 1 | |
| ENSMUSG00000089219 | 1 | |
| ENSMUSG00000089220 | 1 | |
| ENSMUSG00000089222 | 1 | |
| ENSMUSG00000089223 | 1 | |
| ENSMUSG00000089224 | 1.069068979 | 0.74773997 |
| ENSMUSG00000089226 | 1 | |
| ENSMUSG00000089227 | 1 | |
| ENSMUSG00000089228 | 1 | |
| ENSMUSG00000089229 | -1.083814013 | 0.786939259 |
| ENSMUSG00000089230 | 1 | |
| ENSMUSG00000089231 | 1 | |
| ENSMUSG00000089233 | 1 | |
| ENSMUSG00000089234 | 1 | |
| ENSMUSG00000089235 | -1.084645496 | 0.608497229 |
| ENSMUSG00000089237 | 1 | |
| ENSMUSG00000089239 | 1.095872691 | 0.608497229 |
| ENSMUSG00000089241 | 1 | |
| ENSMUSG00000089242 | 1 | |
| ENSMUSG00000089244 | 1 | |
| ENSMUSG00000089245 | 1 | |
| ENSMUSG00000089247 | 1 | |
| ENSMUSG00000089248 | -1.024718486 | 0.931480186 |
| ENSMUSG00000089250 | 1.059463094 | 0.608497229 |
| ENSMUSG00000089251 | 1 | |
| ENSMUSG00000089252 | 1 | |
| ENSMUSG00000089253 | 1.05410549 | 0.835701362 |
| ENSMUSG00000089255 | 1.016832042 | 1 |
| ENSMUSG00000089256 | 1 | |
| ENSMUSG00000089257 | 1 | |
| ENSMUSG00000089258 | 1 | |
| ENSMUSG00000089259 | 1 | |
| ENSMUSG00000089260 | -1.132666855 | 0.18851436 |
| ENSMUSG00000089261 | 1.059463094 | 0.608497229 |
| ENSMUSG00000089262 | 1 | |
| ENSMUSG00000089263 | 1 | |
| ENSMUSG00000089264 | 1 | |
| ENSMUSG00000089265 | -1 | 1 |
| ENSMUSG00000089266 | 1 | |
| ENSMUSG00000089267 | 1.059463094 | 0.608497229 |
| ENSMUSG00000089268 | 1 | |
| ENSMUSG00000089269 | 1 | |

| | | |
|---|---|---|
| ENSMUSG00000089270 | 1 | |
| ENSMUSG00000089271 | 1 | |
| ENSMUSG00000089272 | 1 | |
| ENSMUSG00000089273 | -1 | 1 |
| ENSMUSG00000089274 | 1 | |
| ENSMUSG00000089275 | 1 | |
| ENSMUSG00000089276 | 1 | |
| ENSMUSG00000089277 | 1 | |
| ENSMUSG00000089278 | 1 | |
| ENSMUSG00000089279 | 1 | |
| ENSMUSG00000089280 | 1 | |
| ENSMUSG00000089281 | -1.038444538 | 0.931480186 |
| ENSMUSG00000089282 | 1 | |
| ENSMUSG00000089284 | 1 | |
| ENSMUSG00000089285 | 1 | |
| ENSMUSG00000089286 | 1 | |
| ENSMUSG00000089287 | 1 | |
| ENSMUSG00000089288 | 1 | |
| ENSMUSG00000089289 | 1 | |
| ENSMUSG00000089290 | 1 | |
| ENSMUSG00000089291 | 1 | |
| ENSMUSG00000089292 | -1.03962336 | 0.698112512 |
| ENSMUSG00000089293 | 1 | |
| ENSMUSG00000089294 | 1 | |
| ENSMUSG00000089296 | 1.005827049 | 1 |
| ENSMUSG00000089297 | 1 | |
| ENSMUSG00000089298 | -1.044273782 | 0.698112512 |
| ENSMUSG00000089299 | 1 | |
| ENSMUSG00000089301 | 1 | |
| ENSMUSG00000089302 | 1 | |
| ENSMUSG00000089304 | 1 | |
| ENSMUSG00000089305 | 1 | |
| ENSMUSG00000089306 | 1 | |
| ENSMUSG00000089307 | 1 | |
| ENSMUSG00000089308 | -1.03962336 | 0.851451345 |
| ENSMUSG00000089309 | 1 | |
| ENSMUSG00000089310 | 1 | |
| ENSMUSG00000089311 | 1 | |
| ENSMUSG00000089312 | 1 | |
| ENSMUSG00000089313 | 1.059463094 | 0.608497229 |
| ENSMUSG00000089314 | 1 | |
| ENSMUSG00000089316 | 1 | |
| ENSMUSG00000089317 | 1.218240674 | 0.217803313 |
| ENSMUSG00000089318 | 1 | |
| ENSMUSG00000089320 | 1 | |
| ENSMUSG00000089321 | -1 | 1 |
| ENSMUSG00000089322 | 1 | |
| ENSMUSG00000089323 | 1 | |
| ENSMUSG00000089324 | 1 | |
| ENSMUSG00000089326 | 1 | |
| ENSMUSG00000089327 | 1 | |
| ENSMUSG00000089331 | 1 | |
| ENSMUSG00000089333 | 1 | |
| ENSMUSG00000089334 | 1 | |
| ENSMUSG00000089335 | 1 | |
| ENSMUSG00000089336 | 1 | |
| ENSMUSG00000089337 | -1.037482356 | 0.870427333 |
| ENSMUSG00000089338 | 1 | |
| ENSMUSG00000089341 | 1 | |
| ENSMUSG00000089342 | 1 | |
| ENSMUSG00000089343 | 1.059463094 | 0.704170349 |
| ENSMUSG00000089344 | 1 | |
| ENSMUSG00000089345 | 1 | |

| | | |
|---|---|---|
| ENSMUSG00000089346 | 1 | |
| ENSMUSG00000089348 | 1 | |
| ENSMUSG00000089349 | 1 | |
| ENSMUSG00000089350 | 1 | |
| ENSMUSG00000089351 | 1 | |
| ENSMUSG00000089352 | 1 | |
| ENSMUSG00000089353 | 1 | |
| ENSMUSG00000089357 | 1 | |
| ENSMUSG00000089358 | 1 | |
| ENSMUSG00000089359 | 1 | |
| ENSMUSG00000089363 | 1 | |
| ENSMUSG00000089364 | 1 | |
| ENSMUSG00000089365 | 1.005613439 | 1 |
| ENSMUSG00000089366 | 1 | |
| ENSMUSG00000089369 | 1 | |
| ENSMUSG00000089370 | 1 | |
| ENSMUSG00000089371 | 1 | |
| ENSMUSG00000089372 | 1 | |
| ENSMUSG00000089373 | 1 | |
| ENSMUSG00000089375 | 1 | |
| ENSMUSG00000089376 | 1 | |
| ENSMUSG00000089377 | 1 | |
| ENSMUSG00000089380 | 1 | |
| ENSMUSG00000089381 | 1.059463094 | 0.608497229 |
| ENSMUSG00000089382 | 1 | |
| ENSMUSG00000089383 | 1 | |
| ENSMUSG00000089384 | 1 | |
| ENSMUSG00000089385 | 1 | |
| ENSMUSG00000089387 | 1 | |
| ENSMUSG00000089389 | 1.069068979 | 0.608497229 |
| ENSMUSG00000089390 | 1 | |
| ENSMUSG00000089391 | 1 | |
| ENSMUSG00000089392 | 1 | |
| ENSMUSG00000089393 | 1 | |
| ENSMUSG00000089394 | 1 | |
| ENSMUSG00000089395 | 1 | |
| ENSMUSG00000089396 | -1.117792522 | 0.698112512 |
| ENSMUSG00000089397 | 1 | |
| ENSMUSG00000089398 | 1 | |
| ENSMUSG00000089399 | 1 | |
| ENSMUSG00000089401 | 1 | |
| ENSMUSG00000089404 | 1.059463094 | 0.608497229 |
| ENSMUSG00000089405 | 1 | |
| ENSMUSG00000089406 | 1 | |
| ENSMUSG00000089407 | 1 | |
| ENSMUSG00000089408 | 1 | |
| ENSMUSG00000089409 | 1 | |
| ENSMUSG00000089410 | 1 | |
| ENSMUSG00000089413 | 1 | |
| ENSMUSG00000089414 | 1 | |
| ENSMUSG00000089415 | 1 | |
| ENSMUSG00000089416 | 1 | |
| ENSMUSG00000089417 | -1.112470278 | 0.70299173 |
| ENSMUSG00000089418 | 1 | |
| ENSMUSG00000089420 | 1 | |
| ENSMUSG00000089422 | 1 | |
| ENSMUSG00000089423 | 1 | |
| ENSMUSG00000089424 | 1 | |
| ENSMUSG00000089425 | 1 | |
| ENSMUSG00000089427 | 1 | |
| ENSMUSG00000089429 | 1 | |
| ENSMUSG00000089431 | 1.023743961 | 0.938939969 |
| ENSMUSG00000089432 | 1.059463094 | 0.608497229 |

| | | |
|---|---|---|
| ENSMUSG00000089434 | 1 | |
| ENSMUSG00000089436 | 1 | |
| ENSMUSG00000089437 | -1.080560526 | 0.793387188 |
| ENSMUSG00000089438 | 1 | |
| ENSMUSG00000089439 | 1 | |
| ENSMUSG00000089441 | 1 | |
| ENSMUSG00000089442 | 1 | |
| ENSMUSG00000089443 | 1 | |
| ENSMUSG00000089444 | 1 | |
| ENSMUSG00000089445 | 1 | |
| ENSMUSG00000089446 | 1 | |
| ENSMUSG00000089447 | 1 | |
| ENSMUSG00000089450 | 1 | |
| ENSMUSG00000089451 | 1 | |
| ENSMUSG00000089452 | 1 | |
| ENSMUSG00000089453 | 1.015486245 | 0.972980766 |
| ENSMUSG00000089454 | 1 | |
| ENSMUSG00000089455 | 1 | |
| ENSMUSG00000089458 | 1 | |
| ENSMUSG00000089459 | 1 | |
| ENSMUSG00000089460 | 1 | |
| ENSMUSG00000089461 | 1 | |
| ENSMUSG00000089462 | 1 | |
| ENSMUSG00000089464 | 1 | |
| ENSMUSG00000089465 | 1 | |
| ENSMUSG00000089467 | 1 | |
| ENSMUSG00000089468 | 1 | |
| ENSMUSG00000089469 | 1 | |
| ENSMUSG00000089470 | 1 | |
| ENSMUSG00000089471 | 1 | |
| ENSMUSG00000089472 | 1 | |
| ENSMUSG00000089473 | 1 | |
| ENSMUSG00000089475 | -1.03962336 | 0.814089245 |
| ENSMUSG00000089476 | 1.182713286 | 0.327642404 |
| ENSMUSG00000089477 | 1 | |
| ENSMUSG00000089479 | 1 | |
| ENSMUSG00000089480 | 1.059463094 | 0.608497229 |
| ENSMUSG00000089481 | 1 | |
| ENSMUSG00000089482 | 1 | |
| ENSMUSG00000089484 | 1 | |
| ENSMUSG00000089485 | 1 | |
| ENSMUSG00000089486 | 1 | |
| ENSMUSG00000089487 | 1 | |
| ENSMUSG00000089488 | 1 | |
| ENSMUSG00000089489 | 1 | |
| ENSMUSG00000089490 | 1 | |
| ENSMUSG00000089491 | 1.059463094 | 0.704170349 |
| ENSMUSG00000089493 | -1.03962336 | 0.698112512 |
| ENSMUSG00000089497 | 1 | |
| ENSMUSG00000089498 | 1.015486245 | 0.972980766 |
| ENSMUSG00000089499 | 1 | |
| ENSMUSG00000089501 | 1 | |
| ENSMUSG00000089504 | 1 | |
| ENSMUSG00000089506 | 1 | |
| ENSMUSG00000089507 | 1 | |
| ENSMUSG00000089508 | 1 | |
| ENSMUSG00000089509 | 1 | |
| ENSMUSG00000089510 | 1 | |
| ENSMUSG00000089512 | 1.065410325 | 0.764982341 |
| ENSMUSG00000089513 | -1 | 1 |
| ENSMUSG00000089514 | 1 | |
| ENSMUSG00000089516 | 1 | |
| ENSMUSG00000089517 | 1 | |

| | | |
|---|---|---|
| ENSMUSG00000089520 | 1 | |
| ENSMUSG00000089522 | 1 | |
| ENSMUSG00000089524 | 1 | |
| ENSMUSG00000089525 | 1.141990419 | 0.698112512 |
| ENSMUSG00000089528 | 1 | |
| ENSMUSG00000089529 | 1 | |
| ENSMUSG00000089530 | 1 | |
| ENSMUSG00000089531 | 1 | |
| ENSMUSG00000089533 | 1 | |
| ENSMUSG00000089534 | 1 | |
| ENSMUSG00000089536 | -1 | 1 |
| ENSMUSG00000089538 | 1 | |
| ENSMUSG00000089539 | 1 | |
| ENSMUSG00000089541 | 1 | |
| ENSMUSG00000089542 | -1.119958417 | 0.732012457 |
| ENSMUSG00000089543 | 1 | |
| ENSMUSG00000089544 | 1 | |
| ENSMUSG00000089545 | -1.095294215 | 0.541735653 |
| ENSMUSG00000089546 | 1 | |
| ENSMUSG00000089548 | 1 | |
| ENSMUSG00000089549 | 1 | |
| ENSMUSG00000089550 | 1 | |
| ENSMUSG00000089551 | 1 | |
| ENSMUSG00000089552 | 1 | |
| ENSMUSG00000089553 | 1 | |
| ENSMUSG00000089555 | 1 | |
| ENSMUSG00000089556 | 1 | |
| ENSMUSG00000089558 | 1 | |
| ENSMUSG00000089560 | 1 | |
| ENSMUSG00000089561 | 1 | |
| ENSMUSG00000089562 | 1 | |
| ENSMUSG00000089563 | -1.085651418 | 0.608497229 |
| ENSMUSG00000089564 | 1.069068979 | 0.608497229 |
| ENSMUSG00000089570 | 1 | |
| ENSMUSG00000089571 | 1.122128755 | 0.688919977 |
| ENSMUSG00000089575 | 1 | |
| ENSMUSG00000089576 | 1 | |
| ENSMUSG00000089578 | 1 | |
| ENSMUSG00000089580 | 1 | |
| ENSMUSG00000089581 | 1.075230487 | 0.608497229 |
| ENSMUSG00000089583 | 1 | |
| ENSMUSG00000089584 | 1 | |
| ENSMUSG00000089585 | 1 | |
| ENSMUSG00000089586 | 1 | |
| ENSMUSG00000089587 | 1 | |
| ENSMUSG00000089588 | -1.044273782 | 0.698112512 |
| ENSMUSG00000089589 | 1 | |
| ENSMUSG00000089590 | 1 | |
| ENSMUSG00000089591 | 1 | |
| ENSMUSG00000089593 | 1 | |
| ENSMUSG00000089596 | 1 | |
| ENSMUSG00000089597 | -1 | 1 |
| ENSMUSG00000089598 | -1 | 1 |
| ENSMUSG00000089599 | 1 | |
| ENSMUSG00000089601 | 1 | |
| ENSMUSG00000089603 | 1 | |
| ENSMUSG00000089605 | 1 | |
| ENSMUSG00000089606 | 1 | |
| ENSMUSG00000089607 | 1 | |
| ENSMUSG00000089608 | 1 | |
| ENSMUSG00000089609 | 1 | |
| ENSMUSG00000089610 | 1 | |
| ENSMUSG00000089612 | 1 | |

| | | |
|---|---|---|
| ENSMUSG00000089613 | -1 | 1 |
| ENSMUSG00000089614 | 1 | |
| ENSMUSG00000089615 | 1 | |
| ENSMUSG00000089616 | 1 | |
| ENSMUSG00000089617 | 1 | |
| ENSMUSG00000089618 | 1 | |
| ENSMUSG00000089619 | 1 | |
| ENSMUSG00000089621 | 1 | |
| ENSMUSG00000089622 | 1 | |
| ENSMUSG00000089624 | 1 | |
| ENSMUSG00000089625 | 1 | |
| ENSMUSG00000089627 | 1 | |
| ENSMUSG00000089628 | 1 | |
| ENSMUSG00000089630 | -1.043306199 | 0.698112512 |
| ENSMUSG00000089631 | 1.086612475 | 0.712649653 |
| ENSMUSG00000089633 | 1.059463094 | 0.608497229 |
| ENSMUSG00000089634 | -1.084645496 | 0.628300894 |
| ENSMUSG00000089635 | 1.059463094 | 0.608497229 |
| ENSMUSG00000089636 | 1.260460361 | 0.149977043 |
| ENSMUSG00000089639 | 1.024693402 | 0.971248775 |
| ENSMUSG00000089640 | -1.03962336 | 0.814089245 |
| ENSMUSG00000089642 | 1 | |
| ENSMUSG00000089643 | -1.043306199 | 0.698112512 |
| ENSMUSG00000089645 | 1.13166109 | 0.432805863 |
| ENSMUSG00000089646 | -1.061339324 | 0.869713949 |
| ENSMUSG00000089647 | 1.045949855 | 0.865718572 |
| ENSMUSG00000089648 | 1.059463094 | 0.608497229 |
| ENSMUSG00000089651 | -1.085555638 | 0.805707168 |
| ENSMUSG00000089652 | 1 | |
| ENSMUSG00000089653 | 1 | |
| ENSMUSG00000089654 | 1 | |
| ENSMUSG00000089655 | 1 | |
| ENSMUSG00000089656 | 1 | |
| ENSMUSG00000089657 | 1.074554177 | 0.86577545 |
| ENSMUSG00000089659 | 1 | |
| ENSMUSG00000089661 | 1.059463094 | 0.698112512 |
| ENSMUSG00000089662 | -1.043306199 | 0.759248678 |
| ENSMUSG00000089663 | 1 | |
| ENSMUSG00000089664 | -1.058507348 | 0.863134264 |
| ENSMUSG00000089665 | -1.095778914 | 0.698112512 |
| ENSMUSG00000089666 | 1.07913413 | 0.731295483 |
| ENSMUSG00000089667 | 1.199989356 | 0.458778123 |
| ENSMUSG00000089669 | -1.007384839 | 1 |
| ENSMUSG00000089670 | 1.21551829 | 0.505070892 |
| ENSMUSG00000089671 | 1.133505795 | 0.224782045 |
| ENSMUSG00000089672 | 1 | |
| ENSMUSG00000089673 | 1 | |
| ENSMUSG00000089675 | 1 | |
| ENSMUSG00000089678 | 1 | |
| ENSMUSG00000089679 | 1.095872691 | 0.66521153 |
| ENSMUSG00000089680 | 1.019083579 | 0.971519533 |
| ENSMUSG00000089682 | -1.063523134 | 0.698112512 |
| ENSMUSG00000089683 | 1 | |
| ENSMUSG00000089684 | 1 | |
| ENSMUSG00000089685 | 1 | |
| ENSMUSG00000089686 | 1 | |
| ENSMUSG00000089687 | 1.144363107 | 0.255144284 |
| ENSMUSG00000089689 | 1 | |
| ENSMUSG00000089690 | -1.044273782 | 0.698112512 |
| ENSMUSG00000089694 | -1 | 1 |
| ENSMUSG00000089695 | -1.158135713 | 0.658665546 |
| ENSMUSG00000089696 | -1 | 1 |
| ENSMUSG00000089697 | 1.005851671 | 1 |

| | | |
|---|---|---|
| ENSMUSG00000089698 | 1 | |
| ENSMUSG00000089699 | 1 | |
| ENSMUSG00000089700 | 1 | |
| ENSMUSG00000089701 | 1.024693402 | 0.933031586 |
| ENSMUSG00000089702 | -1 | 1 |
| ENSMUSG00000089703 | 1 | |
| ENSMUSG00000089704 | 1 | |
| ENSMUSG00000089705 | 1 | |
| ENSMUSG00000089706 | -1.113396922 | 0.613963375 |
| ENSMUSG00000089707 | -1.043306199 | 0.698112512 |
| ENSMUSG00000089708 | -1.044273782 | 0.795446726 |
| ENSMUSG00000089709 | -1.03962336 | 0.814089245 |
| ENSMUSG00000089710 | 1 | |
| ENSMUSG00000089711 | -1.028348525 | 0.912425817 |
| ENSMUSG00000089712 | -1 | 1 |
| ENSMUSG00000089713 | 1 | |
| ENSMUSG00000089714 | 1 | |
| ENSMUSG00000089715 | 1.19044015 | 0.258010062 |
| ENSMUSG00000089716 | 1.012316313 | 1 |
| ENSMUSG00000089717 | 1 | |
| ENSMUSG00000089718 | 1 | |
| ENSMUSG00000089719 | -1.145977785 | 0.712130748 |
| ENSMUSG00000089720 | 1 | |
| ENSMUSG00000089722 | 1 | |
| ENSMUSG00000089723 | 1 | |
| ENSMUSG00000089724 | 1 | |
| ENSMUSG00000089726 | -1.086404226 | 0.608497229 |
| ENSMUSG00000089727 | 1 | |
| ENSMUSG00000089728 | 1 | |
| ENSMUSG00000089729 | 1 | |
| ENSMUSG00000089730 | -1.083352158 | 0.816059796 |
| ENSMUSG00000089731 | 1 | |
| ENSMUSG00000089732 | 1 | |
| ENSMUSG00000089733 | 1 | |
| ENSMUSG00000089735 | 1.255100778 | 0.13059051 |
| ENSMUSG00000089736 | 1.02975644 | 0.911577488 |
| ENSMUSG00000089737 | 1 | |
| ENSMUSG00000089739 | 1 | |
| ENSMUSG00000089742 | 1.001182517 | 1 |
| ENSMUSG00000089743 | 1 | |
| ENSMUSG00000089744 | -1.043306199 | 0.698112512 |
| ENSMUSG00000089745 | 1.059463094 | 0.608497229 |
| ENSMUSG00000089746 | -1.044273782 | 0.855488553 |
| ENSMUSG00000089748 | 1.059463094 | 0.608497229 |
| ENSMUSG00000089749 | 1 | |
| ENSMUSG00000089751 | 1 | |
| ENSMUSG00000089753 | 1 | |
| ENSMUSG00000089754 | 1 | |
| ENSMUSG00000089755 | 1 | |
| ENSMUSG00000089756 | -1.053437562 | 0.613255791 |
| ENSMUSG00000089757 | 1 | |
| ENSMUSG00000089759 | -1.104932855 | 0.658430243 |
| ENSMUSG00000089760 | -1.044273782 | 0.790851299 |
| ENSMUSG00000089761 | 1 | |
| ENSMUSG00000089762 | 1.354910723 | 0.130113317 |
| ENSMUSG00000089763 | 1 | |
| ENSMUSG00000089764 | 1.030192399 | 0.975094751 |
| ENSMUSG00000089766 | -1.023769022 | 0.934760809 |
| ENSMUSG00000089767 | 1 | |
| ENSMUSG00000089768 | -1.165665483 | 0.59011989 |
| ENSMUSG00000089769 | 1.181241012 | 0.667576553 |
| ENSMUSG00000089771 | 1 | |
| ENSMUSG00000089773 | 1 | |

| | | |
|---|---|---|
| ENSMUSG00000089774 | 1.187203548 | 0.390586581 |
| ENSMUSG00000089775 | 1 | |
| ENSMUSG00000089776 | 1.095872691 | 0.608497229 |
| ENSMUSG00000089779 | 1 | |
| ENSMUSG00000089780 | 1.079681442 | 0.698112512 |
| ENSMUSG00000089781 | 1 | |
| ENSMUSG00000089782 | -1.041117953 | 0.914142953 |
| ENSMUSG00000089783 | 1.059463094 | 0.608497229 |
| ENSMUSG00000089785 | -1.198352625 | 0.287588797 |
| ENSMUSG00000089787 | 1 | |
| ENSMUSG00000089788 | 1.017473449 | 0.982290355 |
| ENSMUSG00000089789 | -1.012498681 | 0.950042971 |
| ENSMUSG00000089790 | 1.113653231 | 0.744134639 |
| ENSMUSG00000089791 | 1.029372892 | 0.835382855 |
| ENSMUSG00000089792 | 1 | |
| ENSMUSG00000089793 | 1 | |
| ENSMUSG00000089794 | 1.059463094 | 0.608497229 |
| ENSMUSG00000089795 | 1 | |
| ENSMUSG00000089796 | 1.058649313 | 0.862393883 |
| ENSMUSG00000089797 | 1 | |
| ENSMUSG00000089798 | -1.124067181 | 0.608497229 |
| ENSMUSG00000089800 | 1.189706618 | 0.136197019 |
| ENSMUSG00000089802 | 1 | |
| ENSMUSG00000089803 | 1 | |
| ENSMUSG00000089804 | 1 | |
| ENSMUSG00000089806 | 1 | |
| ENSMUSG00000089807 | -1.064334191 | 0.794274354 |
| ENSMUSG00000089808 | -1.081706823 | 0.698112512 |
| ENSMUSG00000089809 | -1.089497311 | 0.532369657 |
| ENSMUSG00000089810 | 1.010974475 | 0.992095533 |
| ENSMUSG00000089811 | 1.095872691 | 0.608497229 |
| ENSMUSG00000089812 | 1.221712448 | 0.359354928 |
| ENSMUSG00000089814 | 1 | |
| ENSMUSG00000089815 | 1 | |
| ENSMUSG00000089816 | 1 | |
| ENSMUSG00000089817 | 1.193641091 | 0.028580998 |
| ENSMUSG00000089818 | 1.05045833 | 0.878204365 |
| ENSMUSG00000089819 | 1 | |
| ENSMUSG00000089820 | 1.283997419 | 0.441591007 |
| ENSMUSG00000089821 | 1 | |
| ENSMUSG00000089822 | -1 | 1 |
| ENSMUSG00000089824 | -1.054973678 | 0.717035356 |
| ENSMUSG00000089826 | 1 | |
| ENSMUSG00000089827 | 1.069068979 | 0.698112512 |
| ENSMUSG00000089828 | 1 | |
| ENSMUSG00000089829 | 1.015486245 | 1 |
| ENSMUSG00000089830 | 1 | |
| ENSMUSG00000089831 | 1 | |
| ENSMUSG00000089832 | 1.186825731 | 0.250726313 |
| ENSMUSG00000089833 | -1 | 1 |
| ENSMUSG00000089834 | 1 | |
| ENSMUSG00000089835 | 1 | |
| ENSMUSG00000089836 | 1 | |
| ENSMUSG00000089837 | 1 | |
| ENSMUSG00000089838 | 1 | |
| ENSMUSG00000089839 | -1.043306199 | 0.698112512 |
| ENSMUSG00000089840 | 1 | |
| ENSMUSG00000089842 | -1.085889729 | 0.70299173 |
| ENSMUSG00000089844 | -1.018079582 | 0.825178237 |
| ENSMUSG00000089847 | 1.026506631 | 0.900028588 |
| ENSMUSG00000089849 | -1.130958943 | 0.608497229 |
| ENSMUSG00000089851 | -1.132943025 | 0.608497229 |
| ENSMUSG00000089853 | 1.165366837 | 0.484584722 |

| | | |
|---|---|---|
| ENSMUSG00000089854 | 1.211901443 | 0.452437882 |
| ENSMUSG00000089855 | 1.061397393 | 0.608497229 |
| ENSMUSG00000089857 | -1.251879741 | 0.252944999 |
| ENSMUSG00000089859 | 1 | |
| ENSMUSG00000089861 | 1 | |
| ENSMUSG00000089862 | 1.099580023 | 0.512218565 |
| ENSMUSG00000089865 | 1 | |
| ENSMUSG00000089866 | 1 | |
| ENSMUSG00000089868 | -1.03962336 | 0.698112512 |
| ENSMUSG00000089870 | 1.024693402 | 0.955101438 |
| ENSMUSG00000089871 | 1.019083579 | 0.968937442 |
| ENSMUSG00000089872 | 1.015243267 | 0.968937442 |
| ENSMUSG00000089873 | 1 | |
| ENSMUSG00000089874 | 1 | |
| ENSMUSG00000089875 | -1.018438061 | 0.949812939 |
| ENSMUSG00000089876 | -1.31870163 | 0.071577777 |
| ENSMUSG00000089877 | -1.083415624 | 0.721058851 |
| ENSMUSG00000089879 | 1 | |
| ENSMUSG00000089881 | 1 | |
| ENSMUSG00000089883 | 1.097750827 | 0.770958539 |
| ENSMUSG00000089884 | 1 | |
| ENSMUSG00000089885 | -1.044273782 | 0.698112512 |
| ENSMUSG00000089886 | 1.24126125 | 0.291494241 |
| ENSMUSG00000089887 | 1 | |
| ENSMUSG00000089888 | 1 | |
| ENSMUSG00000089889 | -1.001188786 | 1 |
| ENSMUSG00000089890 | 1 | |
| ENSMUSG00000089891 | 1 | |
| ENSMUSG00000089892 | 1 | |
| ENSMUSG00000089893 | 1 | |
| ENSMUSG00000089894 | 1 | |
| ENSMUSG00000089898 | 1 | |
| ENSMUSG00000089900 | -1.058281046 | 0.906415966 |
| ENSMUSG00000089901 | 1.269339461 | 0.044024826 |
| ENSMUSG00000089902 | 1.122006571 | 0.608497229 |
| ENSMUSG00000089904 | 1 | |
| ENSMUSG00000089905 | 1 | |
| ENSMUSG00000089906 | 1 | |
| ENSMUSG00000089907 | -1.181718351 | 0.305952299 |
| ENSMUSG00000089908 | -1.089497311 | 0.614307028 |
| ENSMUSG00000089909 | 1 | |
| ENSMUSG00000089910 | 1 | |
| ENSMUSG00000089911 | -1.078218321 | 0.698112512 |
| ENSMUSG00000089914 | 1 | |
| ENSMUSG00000089916 | 1 | |
| ENSMUSG00000089917 | 1.06089357 | 0.294003641 |
| ENSMUSG00000089922 | 1.093426056 | 0.207046069 |
| ENSMUSG00000089924 | 1.059463094 | 0.608497229 |
| ENSMUSG00000089926 | 1 | |
| ENSMUSG00000089927 | 1 | |
| ENSMUSG00000089929 | 1 | |
| ENSMUSG00000089931 | 1 | |
| ENSMUSG00000089932 | -1 | 1 |
| ENSMUSG00000089933 | 1 | |
| ENSMUSG00000089934 | 1 | |
| ENSMUSG00000089935 | -1 | 1 |
| ENSMUSG00000089936 | 1 | |
| ENSMUSG00000089937 | 1 | |
| ENSMUSG00000089938 | 1.015486245 | 0.972980766 |
| ENSMUSG00000089940 | 1.053800798 | 0.70166846 |
| ENSMUSG00000089941 | 1 | |
| ENSMUSG00000089942 | 1 | |
| ENSMUSG00000089943 | 1 | |

| | | |
|---|---|---|
| ENSMUSG00000089945 | 1 | |
| ENSMUSG00000089947 | 1 | |
| ENSMUSG00000089948 | -1.1909253 | 0.54187854 |
| ENSMUSG00000089949 | 1 | |
| ENSMUSG00000089950 | 1 | |
| ENSMUSG00000089951 | 1.002949532 | 1 |
| ENSMUSG00000089952 | -1.109634706 | 0.657709854 |
| ENSMUSG00000089953 | 1.059463094 | 0.608497229 |
| ENSMUSG00000089956 | 1 | |
| ENSMUSG00000089957 | 1 | |
| ENSMUSG00000089958 | 1.123054873 | 0.63446288 |
| ENSMUSG00000089959 | -1.191047317 | 0.377254808 |
| ENSMUSG00000089960 | 1 | |
| ENSMUSG00000089961 | 1 | |
| ENSMUSG00000089963 | 1 | |
| ENSMUSG00000089964 | 1 | |
| ENSMUSG00000089965 | 1.116785864 | 0.60397857 |
| ENSMUSG00000089966 | 1 | |
| ENSMUSG00000089967 | 1 | |
| ENSMUSG00000089968 | -1.02365112 | 0.956659802 |
| ENSMUSG00000089972 | -1 | 1 |
| ENSMUSG00000089973 | -1.044273782 | 0.698112512 |
| ENSMUSG00000089975 | 1.150179455 | 0.564735809 |
| ENSMUSG00000089976 | 1 | |
| ENSMUSG00000089978 | 1 | |
| ENSMUSG00000089979 | -1.015490374 | 1 |
| ENSMUSG00000089982 | 1 | |
| ENSMUSG00000089983 | -1 | 1 |
| ENSMUSG00000089984 | -1.095606032 | 0.727991426 |
| ENSMUSG00000089987 | -1 | 1 |
| ENSMUSG00000089988 | 1.022154149 | 0.933031586 |
| ENSMUSG00000089989 | -1.241667427 | 0.410032437 |
| ENSMUSG00000089990 | 1 | |
| ENSMUSG00000089991 | 1 | |
| ENSMUSG00000089992 | -1.084645496 | 0.533812934 |
| ENSMUSG00000089993 | -1.002865154 | 1 |
| ENSMUSG00000089994 | 1.071598395 | 0.860260228 |
| ENSMUSG00000089995 | -1.043306199 | 0.79291651 |
| ENSMUSG00000089996 | 1 | |
| ENSMUSG00000089997 | 1 | |
| ENSMUSG00000089998 | 1.045958121 | 0.802567859 |
| ENSMUSG00000089999 | -1.084645496 | 0.608497229 |
| ENSMUSG00000090000 | 1.096263879 | 0.6926778 |
| ENSMUSG00000090001 | 1.228500694 | 0.439929645 |
| ENSMUSG00000090002 | -1.051018293 | 0.828564184 |
| ENSMUSG00000090003 | 1 | |
| ENSMUSG00000090004 | 1.034095782 | 0.923549054 |
| ENSMUSG00000090005 | 1.199989356 | 0.291044503 |
| ENSMUSG00000090006 | 1 | |
| ENSMUSG00000090007 | 1.122462048 | 0.314853927 |
| ENSMUSG00000090008 | 1.475932272 | 0.025408276 |
| ENSMUSG00000090009 | 1 | |
| ENSMUSG00000090012 | 1.059463094 | 0.704170349 |
| ENSMUSG00000090013 | 1 | |
| ENSMUSG00000090015 | 1.066305108 | 0.608497229 |
| ENSMUSG00000090016 | 1 | |
| ENSMUSG00000090017 | 1 | |
| ENSMUSG00000090018 | 1 | |
| ENSMUSG00000090019 | 1.048416787 | 0.878962673 |
| ENSMUSG00000090021 | -1.049475223 | 0.861137762 |
| ENSMUSG00000090023 | 1.018550636 | 0.97045957 |
| ENSMUSG00000090024 | 1.161252174 | 0.447817625 |
| ENSMUSG00000090026 | 1.023959045 | 0.982838256 |

| | | |
|---|---|---|
| ENSMUSG00000090027 | 1 | |
| ENSMUSG00000090030 | 1.019083579 | 0.949812939 |
| ENSMUSG00000090031 | 1.088627201 | 0.443127085 |
| ENSMUSG00000090033 | 1 | |
| ENSMUSG00000090034 | -1.024718486 | 0.931480186 |
| ENSMUSG00000090035 | -1.023833691 | 0.930693163 |
| ENSMUSG00000090036 | 1 | |
| ENSMUSG00000090038 | -1.044273782 | 0.698112512 |
| ENSMUSG00000090039 | -1.201738795 | 0.608497229 |
| ENSMUSG00000090040 | 1 | |
| ENSMUSG00000090041 | -1.073603785 | 0.808185344 |
| ENSMUSG00000090042 | 1 | |
| ENSMUSG00000090043 | -1.03962336 | 0.803072249 |
| ENSMUSG00000090044 | 1 | |
| ENSMUSG00000090047 | 1 | |
| ENSMUSG00000090048 | 1.075870199 | 0.642819974 |
| ENSMUSG00000090050 | 1 | |
| ENSMUSG00000090051 | 1 | |
| ENSMUSG00000090053 | 1.252404754 | 0.019871675 |
| ENSMUSG00000090054 | 1 | |
| ENSMUSG00000090057 | 1 | |
| ENSMUSG00000090059 | 1 | |
| ENSMUSG00000090061 | -1.028348525 | 0.938939969 |
| ENSMUSG00000090062 | 1 | |
| ENSMUSG00000090063 | -1.046978474 | 0.772834122 |
| ENSMUSG00000090066 | -1.564827864 | 0.028386648 |
| ENSMUSG00000090067 | 1.019083579 | 0.949812939 |
| ENSMUSG00000090069 | 1.01053084 | 1 |
| ENSMUSG00000090070 | 1.040453468 | 0.881655151 |
| ENSMUSG00000090071 | -1.345965228 | 0.014295069 |
| ENSMUSG00000090073 | 1 | |
| ENSMUSG00000090074 | 1 | |
| ENSMUSG00000090075 | -1 | 1 |
| ENSMUSG00000090077 | -1.069963358 | 0.89259544 |
| ENSMUSG00000090078 | 1.070429626 | 0.767473873 |
| ENSMUSG00000090079 | 1.059463094 | 0.608497229 |
| ENSMUSG00000090080 | 1 | |
| ENSMUSG00000090081 | 1 | |
| ENSMUSG00000090083 | 1.039638165 | 0.711404854 |
| ENSMUSG00000090084 | -1.044273782 | 0.738194282 |
| ENSMUSG00000090086 | 1.127019663 | 0.471308522 |
| ENSMUSG00000090087 | 1.132639128 | 0.398767938 |
| ENSMUSG00000090088 | 1 | |
| ENSMUSG00000090089 | 1 | |
| ENSMUSG00000090090 | 1 | |
| ENSMUSG00000090093 | 1.040113968 | 0.608497229 |
| ENSMUSG00000090095 | 1.069068979 | 0.608497229 |
| ENSMUSG00000090097 | 1 | |
| ENSMUSG00000090098 | 1 | |
| ENSMUSG00000090099 | 1 | |
| ENSMUSG00000090100 | 1.000016239 | 1 |
| ENSMUSG00000090101 | 1.042677087 | 0.924834342 |
| ENSMUSG00000090102 | 1 | |
| ENSMUSG00000090103 | 1 | |
| ENSMUSG00000090104 | 1.106109051 | 0.313896003 |
| ENSMUSG00000090105 | -1.03962336 | 0.698112512 |
| ENSMUSG00000090106 | 1 | |
| ENSMUSG00000090107 | 1 | |
| ENSMUSG00000090108 | 1.061225963 | 0.942039503 |
| ENSMUSG00000090109 | 1 | |
| ENSMUSG00000090110 | -1.013360151 | 0.954675041 |
| ENSMUSG00000090111 | 1 | |
| ENSMUSG00000090112 | -1.021397456 | 0.980154107 |

| | | |
|---|---|---|
| ENSMUSG00000090113 | 1.079681442 | 0.743396505 |
| ENSMUSG00000090114 | 1 | |
| ENSMUSG00000090115 | -1.039614996 | 0.819610935 |
| ENSMUSG00000090116 | 1 | |
| ENSMUSG00000090117 | 1 | |
| ENSMUSG00000090118 | 1 | |
| ENSMUSG00000090119 | 1.014545335 | 0.979002724 |
| ENSMUSG00000090120 | -1.000246834 | 1 |
| ENSMUSG00000090121 | -1 | 1 |
| ENSMUSG00000090122 | 1 | |
| ENSMUSG00000090124 | 1.069068979 | 0.698112512 |
| ENSMUSG00000090125 | -2.150817429 | 0.000200489 |
| ENSMUSG00000090126 | 1.019083579 | 0.982838256 |
| ENSMUSG00000090128 | 1 | |
| ENSMUSG00000090129 | -1.071075483 | 0.747163034 |
| ENSMUSG00000090131 | 1 | |
| ENSMUSG00000090132 | 1 | |
| ENSMUSG00000090135 | 1 | |
| ENSMUSG00000090136 | 1.072395878 | 0.78301543 |
| ENSMUSG00000090137 | 1.024005105 | 0.914705152 |
| ENSMUSG00000090138 | 1 | |
| ENSMUSG00000090141 | -1 | 1 |
| ENSMUSG00000090142 | -1.063525418 | 0.814089245 |
| ENSMUSG00000090145 | 1 | |
| ENSMUSG00000090146 | 1 | |
| ENSMUSG00000090147 | 1 | |
| ENSMUSG00000090150 | -1.016638327 | 0.981169503 |
| ENSMUSG00000090151 | 1 | |
| ENSMUSG00000090152 | 1 | |
| ENSMUSG00000090157 | 1 | |
| ENSMUSG00000090159 | 1 | |
| ENSMUSG00000090160 | -1.044273782 | 0.698112512 |
| ENSMUSG00000090162 | 1.059463094 | 0.698112512 |
| ENSMUSG00000090163 | 1 | |
| ENSMUSG00000090164 | 1 | |
| ENSMUSG00000090165 | 1 | |
| ENSMUSG00000090166 | 1 | |
| ENSMUSG00000090168 | 1 | |
| ENSMUSG00000090171 | 1 | |
| ENSMUSG00000090173 | -1.0435664 | 0.828873225 |
| ENSMUSG00000090174 | 1 | |
| ENSMUSG00000090175 | 1 | |
| ENSMUSG00000090176 | -1.084645496 | 0.734674233 |
| ENSMUSG00000090179 | 1 | |
| ENSMUSG00000090180 | 1 | |
| ENSMUSG00000090185 | 1 | |
| ENSMUSG00000090186 | 1 | |
| ENSMUSG00000090188 | -1.021810391 | 0.96035552 |
| ENSMUSG00000090191 | -1.107589204 | 0.698112512 |
| ENSMUSG00000090192 | 1.059463094 | 0.608497229 |
| ENSMUSG00000090193 | 1 | |
| ENSMUSG00000090194 | 1.092086589 | 0.751113028 |
| ENSMUSG00000090197 | -1.044273782 | 0.698112512 |
| ENSMUSG00000090199 | 1 | |
| ENSMUSG00000090200 | 1.019083579 | 0.982838256 |
| ENSMUSG00000090202 | 1 | |
| ENSMUSG00000090203 | 1 | |
| ENSMUSG00000090206 | -1.167613928 | 0.257664788 |
| ENSMUSG00000090207 | 1.069068979 | 0.608497229 |
| ENSMUSG00000090208 | 1 | |
| ENSMUSG00000090210 | -1.077308522 | 0.627831591 |
| ENSMUSG00000090211 | 1.048862481 | 0.911250323 |
| ENSMUSG00000090212 | 1 | |

| | | |
|---|---|---|
| ENSMUSG00000090213 | -1.04184112 | 0.775673129 |
| ENSMUSG00000090214 | 1 | |
| ENSMUSG00000090215 | 1.091686812 | 0.739981019 |
| ENSMUSG00000090216 | 1 | |
| ENSMUSG00000090217 | 1.148782218 | 0.603144355 |
| ENSMUSG00000090219 | 1 | |
| ENSMUSG00000090220 | -1.085651418 | 0.608497229 |
| ENSMUSG00000090222 | -1.053549062 | 0.769745556 |
| ENSMUSG00000090223 | 1 | |
| ENSMUSG00000090224 | -1.085651418 | 0.608497229 |
| ENSMUSG00000090225 | 1 | |
| ENSMUSG00000090226 | 1 | |
| ENSMUSG00000090227 | 1 | |
| ENSMUSG00000090228 | 1 | |
| ENSMUSG00000090230 | 1.075870199 | 0.793885968 |
| ENSMUSG00000090231 | -1.211487332 | 0.106919651 |
| ENSMUSG00000090232 | 1 | |
| ENSMUSG00000090235 | 1.045776193 | 0.854616424 |
| ENSMUSG00000090236 | -1.107299336 | 0.639279667 |
| ENSMUSG00000090237 | 1 | |
| ENSMUSG00000090242 | 1.059463094 | 0.608497229 |
| ENSMUSG00000090243 | 1.060620016 | 0.486216685 |
| ENSMUSG00000090244 | 1.019083579 | 0.979345947 |
| ENSMUSG00000090246 | 1 | |
| ENSMUSG00000090247 | -1.05621518 | 0.67884395 |
| ENSMUSG00000090248 | 1.142847535 | 0.678185531 |
| ENSMUSG00000090249 | 1 | |
| ENSMUSG00000090252 | 1 | |
| ENSMUSG00000090254 | 1 | |
| ENSMUSG00000090255 | 1 | |
| ENSMUSG00000090256 | 1.120359279 | 0.698112512 |
| ENSMUSG00000090257 | 1.035886516 | 0.861402041 |
| ENSMUSG00000090258 | 1.018552145 | 0.941011898 |
| ENSMUSG00000090259 | 1 | |
| ENSMUSG00000090260 | 1.089470641 | 0.608497229 |
| ENSMUSG00000090263 | 1.059463094 | 0.698112512 |
| ENSMUSG00000090264 | 1.127939888 | 0.698112512 |
| ENSMUSG00000090265 | 1 | |
| ENSMUSG00000090266 | 1.047775988 | 0.814608539 |
| ENSMUSG00000090268 | 1 | |
| ENSMUSG00000090270 | -1.111880267 | 0.608497229 |
| ENSMUSG00000090272 | 1.236368254 | 0.174363255 |
| ENSMUSG00000090273 | 1.081939842 | 0.828564184 |
| ENSMUSG00000090277 | 1 | |
| ENSMUSG00000090284 | 1.019083579 | 0.949812939 |
| ENSMUSG00000090286 | 1.145490319 | 0.418130164 |
| ENSMUSG00000090290 | 1.163954289 | 0.111243864 |
| ENSMUSG00000090291 | -1.181812752 | 0.14776651 |
| ENSMUSG00000090292 | 1 | |
| ENSMUSG00000090293 | 1.095872691 | 0.608497229 |
| ENSMUSG00000090298 | 1 | |
| ENSMUSG00000090300 | 1 | |
| ENSMUSG00000090302 | 1 | |
| ENSMUSG00000090303 | 1.069068979 | 0.608497229 |
| ENSMUSG00000090304 | 1 | |
| ENSMUSG00000090305 | -1.015248956 | 1 |
| ENSMUSG00000090306 | 1 | |
| ENSMUSG00000090307 | -1.015511103 | 0.99339328 |
| ENSMUSG00000090308 | 1 | |
| ENSMUSG00000090314 | 1 | |
| ENSMUSG00000090315 | 1 | |
| ENSMUSG00000090317 | -1.044273782 | 0.698112512 |
| ENSMUSG00000090318 | 1 | |

| | | |
|---|---|---|
| ENSMUSG00000090319 | -1.011566762 | 0.972684567 |
| ENSMUSG00000090322 | 1.059463094 | 0.752398528 |
| ENSMUSG00000090323 | 1 | |
| ENSMUSG00000090326 | 1 | |
| ENSMUSG00000090327 | 1.192142379 | 0.545359962 |
| ENSMUSG00000090328 | 1 | |
| ENSMUSG00000090329 | -1.043306199 | 0.698112512 |
| ENSMUSG00000090330 | 1.030235697 | 0.915205952 |
| ENSMUSG00000090331 | -1 | 1 |
| ENSMUSG00000090332 | 1 | |
| ENSMUSG00000090333 | 1 | |
| ENSMUSG00000090334 | 1.020344501 | 0.983178084 |
| ENSMUSG00000090336 | 1.059463094 | 0.608497229 |
| ENSMUSG00000090338 | 1 | |
| ENSMUSG00000090341 | 1 | |
| ENSMUSG00000090342 | 1 | |
| ENSMUSG00000090346 | 1 | |
| ENSMUSG00000090350 | -1 | 1 |
| ENSMUSG00000090351 | 1 | |
| ENSMUSG00000090352 | 1 | |
| ENSMUSG00000090353 | -1.037787403 | 0.828564184 |
| ENSMUSG00000090355 | 1 | |
| ENSMUSG00000090356 | 1 | |
| ENSMUSG00000090357 | -1.03962336 | 0.698112512 |
| ENSMUSG00000090358 | -1.098719945 | 0.630771413 |
| ENSMUSG00000090362 | 1 | |
| ENSMUSG00000090363 | 1.079681442 | 0.743396505 |
| ENSMUSG00000090364 | 1 | |
| ENSMUSG00000090365 | 1 | |
| ENSMUSG00000090369 | -1.062630016 | 0.769705171 |
| ENSMUSG00000090370 | 1 | |
| ENSMUSG00000090374 | 1 | |
| ENSMUSG00000090379 | 1 | |
| ENSMUSG00000090381 | -1 | 1 |
| ENSMUSG00000090383 | 1 | |
| ENSMUSG00000090385 | 1 | |
| ENSMUSG00000090386 | -1.847149084 | 0.019383936 |
| ENSMUSG00000090387 | -1.234037493 | 0.346299851 |
| ENSMUSG00000090389 | -1.131871888 | 0.682524927 |
| ENSMUSG00000090394 | 1.258531394 | 0.173498129 |
| ENSMUSG00000090395 | 1 | |
| ENSMUSG00000090396 | 1 | |
| ENSMUSG00000090397 | -1.092629659 | 0.454703142 |
| ENSMUSG00000090399 | 1.026708393 | 0.860327141 |
| ENSMUSG00000090404 | 1 | |
| ENSMUSG00000090406 | 1.35394505 | 0.141547558 |
| ENSMUSG00000090408 | -1.043306199 | 0.79291651 |
| ENSMUSG00000090411 | 1 | |
| ENSMUSG00000090417 | 1 | |
| ENSMUSG00000090427 | -1 | 1 |
| ENSMUSG00000090429 | 1 | |
| ENSMUSG00000090436 | 1 | |
| ENSMUSG00000090439 | 1 | |
| ENSMUSG00000090440 | -1.048277865 | 0.860645152 |
| ENSMUSG00000090441 | 1 | |
| ENSMUSG00000090448 | -1 | 1 |
| ENSMUSG00000090449 | 1 | |
| ENSMUSG00000090451 | -1.211618431 | 0.596419896 |
| ENSMUSG00000090455 | 1 | |
| ENSMUSG00000090457 | 1 | |
| ENSMUSG00000090458 | 1 | |
| ENSMUSG00000090459 | -1.188845164 | 0.565305007 |
| ENSMUSG00000090460 | -1.03962336 | 0.698112512 |

| | | |
|---|---|---|
| ENSMUSG00000090461 | 1 | |
| ENSMUSG00000090462 | 1 | |
| ENSMUSG00000090464 | 1 | |
| ENSMUSG00000090465 | 1 | |
| ENSMUSG00000090466 | 1 | |
| ENSMUSG00000090467 | 1 | |
| ENSMUSG00000090468 | 1 | |
| ENSMUSG00000090470 | 1 | |
| ENSMUSG00000090471 | 1 | |
| ENSMUSG00000090472 | 1 | |
| ENSMUSG00000090474 | 1 | |
| ENSMUSG00000090475 | -1.043306199 | 0.698112512 |
| ENSMUSG00000090476 | 1 | |
| ENSMUSG00000090478 | 1 | |
| ENSMUSG00000090481 | 1 | |
| ENSMUSG00000090482 | 1 | |
| ENSMUSG00000090485 | 1 | |
| ENSMUSG00000090486 | 1.316640831 | 0.024718723 |
| ENSMUSG00000090487 | 1 | |
| ENSMUSG00000090489 | 1.103335749 | 0.713078249 |
| ENSMUSG00000090494 | 1 | |
| ENSMUSG00000090497 | 1 | |
| ENSMUSG00000090500 | 1.011211697 | 1 |
| ENSMUSG00000090505 | 1 | |
| ENSMUSG00000090509 | -1.230294675 | 0.119808842 |
| ENSMUSG00000090512 | 1 | |
| ENSMUSG00000090515 | 1 | |
| ENSMUSG00000090516 | -1.071075483 | 0.743300655 |
| ENSMUSG00000090519 | 1 | |
| ENSMUSG00000090523 | 1.202606939 | 0.225312378 |
| ENSMUSG00000090527 | 1 | |
| ENSMUSG00000090533 | 1 | |
| ENSMUSG00000090534 | -1.044273782 | 0.698112512 |
| ENSMUSG00000090535 | 1 | |
| ENSMUSG00000090538 | -1.03962336 | 0.698112512 |
| ENSMUSG00000090539 | 1 | |
| ENSMUSG00000090544 | 1.14537085 | 0.608497229 |
| ENSMUSG00000090546 | 1 | |
| ENSMUSG00000090547 | 1 | |
| ENSMUSG00000090549 | 1 | |
| ENSMUSG00000090550 | 1 | |
| ENSMUSG00000090551 | 1.30383792 | 0.251512864 |
| ENSMUSG00000090552 | 1 | |
| ENSMUSG00000090553 | -1.015702406 | 0.998378474 |
| ENSMUSG00000090555 | 1 | |
| ENSMUSG00000090556 | 1 | |
| ENSMUSG00000090558 | 1 | |
| ENSMUSG00000090559 | -1.121136286 | 0.608497229 |
| ENSMUSG00000090561 | -1 | 1 |
| ENSMUSG00000090564 | 1.211350831 | 0.061707775 |
| ENSMUSG00000090565 | -1.009476377 | 1 |
| ENSMUSG00000090570 | 1 | |
| ENSMUSG00000090572 | 1 | |
| ENSMUSG00000090574 | 1 | |
| ENSMUSG00000090576 | 1.080115243 | 0.685044927 |
| ENSMUSG00000090577 | 1 | |
| ENSMUSG00000090578 | 1 | |
| ENSMUSG00000090581 | 1 | |
| ENSMUSG00000090582 | 1.040291945 | 0.94904267 |
| ENSMUSG00000090584 | 1 | |
| ENSMUSG00000090585 | 1 | |
| ENSMUSG00000090588 | 1.019083579 | 0.949812939 |
| ENSMUSG00000090589 | 1 | |

| | | |
|---|---|---|
| ENSMUSG00000090592 | -1.044273782 | 0.698112512 |
| ENSMUSG00000090594 | 1 | |
| ENSMUSG00000090598 | 1 | |
| ENSMUSG00000090602 | -1.233467038 | 0.141447411 |
| ENSMUSG00000090603 | 1 | |
| ENSMUSG00000090607 | 1 | |
| ENSMUSG00000090608 | 1.034865357 | 0.949812939 |
| ENSMUSG00000090610 | 1.028344344 | 0.987698908 |
| ENSMUSG00000090618 | 1 | |
| ENSMUSG00000090619 | 1 | |
| ENSMUSG00000090622 | 1.169511821 | 0.174012011 |
| ENSMUSG00000090623 | 1 | |
| ENSMUSG00000090624 | -1.163394565 | 0.594095902 |
| ENSMUSG00000090625 | -1 | 1 |
| ENSMUSG00000090626 | -1.053240932 | 0.698112512 |
| ENSMUSG00000090627 | 1 | |
| ENSMUSG00000090628 | 1.073175172 | 0.920203742 |
| ENSMUSG00000090629 | 1 | |
| ENSMUSG00000090631 | -1.334433578 | 0.099318501 |
| ENSMUSG00000090634 | 1 | |
| ENSMUSG00000090635 | 1 | |
| ENSMUSG00000090636 | 1 | |
| ENSMUSG00000090637 | 1 | |
| ENSMUSG00000090638 | 2.281351881 | 0.000855074 |
| ENSMUSG00000090639 | 1 | |
| ENSMUSG00000090641 | -1.049420031 | 0.78252598 |
| ENSMUSG00000090642 | 1 | |
| ENSMUSG00000090643 | 1 | |
| ENSMUSG00000090655 | 1 | |
| ENSMUSG00000090657 | 1 | |
| ENSMUSG00000090658 | 1 | |
| ENSMUSG00000090659 | -1.00032582 | 1 |
| ENSMUSG00000090660 | 1 | |
| ENSMUSG00000090662 | 1 | |
| ENSMUSG00000090665 | 1 | |
| ENSMUSG00000090666 | -1.206338534 | 0.434482395 |
| ENSMUSG00000090667 | -1 | 1 |
| ENSMUSG00000090671 | 1 | |
| ENSMUSG00000090673 | -1.001651038 | 1 |
| ENSMUSG00000090674 | -1.126793513 | 0.661668632 |
| ENSMUSG00000090675 | 1 | |
| ENSMUSG00000090681 | 1 | |
| ENSMUSG00000090682 | 1 | |
| ENSMUSG00000090684 | 1 | |
| ENSMUSG00000090685 | 1 | |
| ENSMUSG00000090688 | 1 | |
| ENSMUSG00000090690 | 1 | |
| ENSMUSG00000090691 | -1.084645496 | 0.628300894 |
| ENSMUSG00000090692 | 1 | |
| ENSMUSG00000090693 | 1.11297715 | 0.608497229 |
| ENSMUSG00000090698 | 1.37422482 | 0.016532413 |
| ENSMUSG00000090699 | 1 | |
| ENSMUSG00000090700 | 1 | |
| ENSMUSG00000090701 | 1 | |
| ENSMUSG00000090704 | 1 | |
| ENSMUSG00000090706 | -1 | 1 |
| ENSMUSG00000090707 | -1.053549062 | 0.698112512 |
| ENSMUSG00000090708 | -1.084645496 | 0.532369657 |
| ENSMUSG00000090709 | 1.210869356 | 0.327655754 |
| ENSMUSG00000090710 | 1 | |
| ENSMUSG00000090713 | 1.059463094 | 0.608497229 |
| ENSMUSG00000090714 | 1.059463094 | 0.608497229 |
| ENSMUSG00000090715 | 1 | |

| | | |
|---|---|---|
| ENSMUSG00000090716 | 1 | |
| ENSMUSG00000090718 | 1 | |
| ENSMUSG00000090722 | -1 | 1 |
| ENSMUSG00000090724 | 1 | |
| ENSMUSG00000090726 | 1 | |
| ENSMUSG00000090727 | 1 | |
| ENSMUSG00000090728 | 1 | |
| ENSMUSG00000090731 | 1 | |
| ENSMUSG00000090732 | 1 | |
| ENSMUSG00000090733 | -1.035413524 | 0.940197654 |
| ENSMUSG00000090736 | 1 | |
| ENSMUSG00000090737 | -1.03962336 | 0.698112512 |
| ENSMUSG00000090738 | 1 | |
| ENSMUSG00000090740 | 1 | |
| ENSMUSG00000090742 | 1 | |
| ENSMUSG00000090743 | 1 | |
| ENSMUSG00000090744 | -1.036381869 | 0.879267911 |
| ENSMUSG00000090747 | 1 | |
| ENSMUSG00000090750 | 1 | |
| ENSMUSG00000090751 | 1 | |
| ENSMUSG00000090753 | -1.044273782 | 0.790851299 |
| ENSMUSG00000090758 | 1 | |
| ENSMUSG00000090761 | -1.063058326 | 0.533954522 |
| ENSMUSG00000090762 | 1 | |
| ENSMUSG00000090764 | 1 | |
| ENSMUSG00000090765 | 1 | |
| ENSMUSG00000090772 | 1 | |
| ENSMUSG00000090773 | 1 | |
| ENSMUSG00000090774 | 1 | |
| ENSMUSG00000090777 | 1.023959045 | 0.979002724 |
| ENSMUSG00000090778 | 1.12189305 | 0.585646836 |
| ENSMUSG00000090779 | -1.03962336 | 0.698112512 |
| ENSMUSG00000090781 | 1 | |
| ENSMUSG00000090782 | 1 | |
| ENSMUSG00000090785 | -1.043306199 | 0.79291651 |
| ENSMUSG00000090788 | 1 | |
| ENSMUSG00000090790 | 1.059463094 | 0.608497229 |
| ENSMUSG00000090793 | 1.132639128 | 0.414898324 |
| ENSMUSG00000090794 | 1 | |
| ENSMUSG00000090795 | 1 | |
| ENSMUSG00000090796 | 1 | |
| ENSMUSG00000090798 | 1 | |
| ENSMUSG00000090799 | -1.033724264 | 0.958250633 |
| ENSMUSG00000090801 | 1 | |
| ENSMUSG00000090802 | 1 | |
| ENSMUSG00000090805 | 1 | |
| ENSMUSG00000090806 | 1 | |
| ENSMUSG00000090808 | 1 | |
| ENSMUSG00000090812 | -1.018864595 | 0.964236174 |
| ENSMUSG00000090816 | 1 | |
| ENSMUSG00000090817 | 1 | |
| ENSMUSG00000090818 | 1 | |
| ENSMUSG00000090821 | 1 | |
| ENSMUSG00000090824 | 1 | |
| ENSMUSG00000090827 | 1 | |
| ENSMUSG00000090839 | 1 | |
| ENSMUSG00000090840 | 1.105835042 | 0.678151972 |
| ENSMUSG00000090841 | 1.21229825 | 0.504256967 |
| ENSMUSG00000090843 | -1.095294215 | 0.541735653 |
| ENSMUSG00000090849 | 1 | |
| ENSMUSG00000090853 | 1 | |
| ENSMUSG00000090854 | 1 | |
| ENSMUSG00000090855 | 1 | |

| | | |
|---|---|---|
| ENSMUSG00000090860 | 1 | |
| ENSMUSG00000090861 | -1.084645496 | 0.698112512 |
| ENSMUSG00000090862 | 1.020749966 | 0.96175742 |
| ENSMUSG00000090863 | 1.037443274 | 0.949679283 |
| ENSMUSG00000090864 | 1 | |
| ENSMUSG00000090865 | 1 | |
| ENSMUSG00000090868 | 1 | |
| ENSMUSG00000090872 | 1 | |
| ENSMUSG00000090873 | 1.038778564 | 0.873111932 |
| ENSMUSG00000090874 | 1 | |
| ENSMUSG00000090877 | 1.025821829 | 0.698112512 |
| ENSMUSG00000090881 | -1.033820075 | 0.888246981 |
| ENSMUSG00000090882 | 1 | |
| ENSMUSG00000090889 | 1 | |
| ENSMUSG00000090891 | 1.019158394 | 0.969026637 |
| ENSMUSG00000090892 | 1 | |
| ENSMUSG00000090894 | 1 | |
| ENSMUSG00000090897 | 1 | |
| ENSMUSG00000090900 | 1 | |
| ENSMUSG00000090907 | -1.03962336 | 0.803072249 |
| ENSMUSG00000090919 | -1.03193593 | 0.94049193 |
| ENSMUSG00000090925 | -1.083397223 | 0.608497229 |
| ENSMUSG00000090927 | 1 | |
| ENSMUSG00000090935 | 1.022665255 | 0.9602503 |
| ENSMUSG00000090936 | 1.14449055 | 0.608497229 |
| ENSMUSG00000090939 | 1.079681442 | 0.726987474 |
| ENSMUSG00000090941 | 1.059463094 | 0.698112512 |
| ENSMUSG00000090942 | 1.198137776 | 0.422229812 |
| ENSMUSG00000090946 | 1.202322532 | 0.060236933 |
| ENSMUSG00000090947 | 1 | |
| ENSMUSG00000090949 | 1 | |
| ENSMUSG00000090951 | 1 | |
| ENSMUSG00000090952 | 1.117326996 | 0.578885377 |
| ENSMUSG00000090955 | -1.043657968 | 0.901514756 |
| ENSMUSG00000090957 | -1.053549062 | 0.778372272 |
| ENSMUSG00000090958 | 1.948856494 | 0.000378008 |
| ENSMUSG00000090961 | 1 | |
| ENSMUSG00000090963 | -1.036892351 | 0.887323956 |
| ENSMUSG00000090964 | 1.132639128 | 0.428732467 |
| ENSMUSG00000090965 | -1.040506081 | 0.924324458 |
| ENSMUSG00000090966 | 1 | |
| ENSMUSG00000090967 | 1 | |
| ENSMUSG00000090968 | 1 | |
| ENSMUSG00000090975 | 1 | |
| ENSMUSG00000090981 | 1 | |
| ENSMUSG00000090987 | 1 | |
| ENSMUSG00000090989 | 1 | |
| ENSMUSG00000090990 | 1 | |
| ENSMUSG00000090991 | 1 | |
| ENSMUSG00000090994 | 1.057478344 | 0.856857802 |
| ENSMUSG00000090996 | 1 | |
| ENSMUSG00000091002 | 1 | |
| ENSMUSG00000091003 | 1 | |
| ENSMUSG00000091004 | 1 | |
| ENSMUSG00000091005 | 1 | |
| ENSMUSG00000091006 | 1 | |
| ENSMUSG00000091007 | 1.059463094 | 0.698112512 |
| ENSMUSG00000091008 | 1 | |
| ENSMUSG00000091010 | 1 | |
| ENSMUSG00000091011 | 1 | |
| ENSMUSG00000091012 | 1 | |
| ENSMUSG00000091013 | 1.059463094 | 0.608497229 |
| ENSMUSG00000091014 | -1.071075483 | 0.743300655 |

| | | |
|---|---|---|
| ENSMUSG00000091016 | 1 | |
| ENSMUSG00000091017 | 1 | |
| ENSMUSG00000091019 | 1.086455536 | 0.680547155 |
| ENSMUSG00000091020 | -1.168914602 | 0.134244013 |
| ENSMUSG00000091021 | 1.163836832 | 0.33729677 |
| ENSMUSG00000091022 | 1 | |
| ENSMUSG00000091027 | 1 | |
| ENSMUSG00000091028 | 1 | |
| ENSMUSG00000091031 | 1 | |
| ENSMUSG00000091034 | 1 | |
| ENSMUSG00000091037 | 1 | |
| ENSMUSG00000091039 | 1 | |
| ENSMUSG00000091040 | -1.03962336 | 0.698112512 |
| ENSMUSG00000091041 | 1 | |
| ENSMUSG00000091043 | -1.171420412 | 0.346721275 |
| ENSMUSG00000091045 | 1.069877155 | 0.845175363 |
| ENSMUSG00000091048 | 1 | |
| ENSMUSG00000091049 | 1 | |
| ENSMUSG00000091050 | 1.129826644 | 0.608497229 |
| ENSMUSG00000091053 | 1 | |
| ENSMUSG00000091055 | 1 | |
| ENSMUSG00000091058 | -1.044273782 | 0.698112512 |
| ENSMUSG00000091059 | 1 | |
| ENSMUSG00000091060 | -1.044273782 | 0.801443616 |
| ENSMUSG00000091061 | 1.069068979 | 0.608497229 |
| ENSMUSG00000091071 | 1 | 1 |
| ENSMUSG00000091076 | 1 | |
| ENSMUSG00000091077 | 1 | |
| ENSMUSG00000091078 | 1 | |
| ENSMUSG00000091079 | -1 | 1 |
| ENSMUSG00000091080 | 1 | |
| ENSMUSG00000091083 | -1.116853908 | 0.608497229 |
| ENSMUSG00000091086 | 1.112169157 | 0.747886461 |
| ENSMUSG00000091087 | 1 | |
| ENSMUSG00000091089 | 1.109058584 | 0.698112512 |
| ENSMUSG00000091091 | 1 | |
| ENSMUSG00000091095 | 1 | |
| ENSMUSG00000091096 | -1 | 1 |
| ENSMUSG00000091099 | 1 | |
| ENSMUSG00000091101 | 1 | |
| ENSMUSG00000091102 | -1.079467562 | 0.33491989 |
| ENSMUSG00000091105 | 1.001429165 | 1 |
| ENSMUSG00000091109 | 1.020779405 | 0.976498231 |
| ENSMUSG00000091110 | 1 | |
| ENSMUSG00000091114 | 1 | |
| ENSMUSG00000091116 | 1 | |
| ENSMUSG00000091117 | -1.03962336 | 0.698112512 |
| ENSMUSG00000091119 | -1.049075043 | 0.889802291 |
| ENSMUSG00000091122 | 1 | |
| ENSMUSG00000091123 | 1 | |
| ENSMUSG00000091129 | 1 | |
| ENSMUSG00000091131 | 1 | |
| ENSMUSG00000091132 | 1 | |
| ENSMUSG00000091133 | -1.11311759 | 0.397756797 |
| ENSMUSG00000091135 | 1 | |
| ENSMUSG00000091140 | 1 | |
| ENSMUSG00000091142 | 1 | |
| ENSMUSG00000091144 | -1.107206966 | 0.266023891 |
| ENSMUSG00000091146 | 1 | |
| ENSMUSG00000091147 | -1.044273782 | 0.698112512 |
| ENSMUSG00000091148 | 1 | |
| ENSMUSG00000091149 | 1 | |
| ENSMUSG00000091151 | 1 | |

| | | |
|---|---|---|
| ENSMUSG00000091154 | 1.057478344 | 0.837014159 |
| ENSMUSG00000091155 | -1.351433432 | 0.025806845 |
| ENSMUSG00000091157 | 1 | |
| ENSMUSG00000091159 | 1 | |
| ENSMUSG00000091160 | 1.260450255 | 0.237706049 |
| ENSMUSG00000091163 | 1 | |
| ENSMUSG00000091165 | 1 | |
| ENSMUSG00000091172 | 1 | |
| ENSMUSG00000091174 | 1 | |
| ENSMUSG00000091175 | 1 | |
| ENSMUSG00000091177 | 1 | |
| ENSMUSG00000091183 | 1.038164627 | 0.766307541 |
| ENSMUSG00000091184 | -1.121216742 | 0.654640324 |
| ENSMUSG00000091185 | 1 | |
| ENSMUSG00000091189 | 1 | |
| ENSMUSG00000091191 | -1.251080495 | 0.27366404 |
| ENSMUSG00000091192 | 1 | |
| ENSMUSG00000091193 | 1 | |
| ENSMUSG00000091194 | 1.052275609 | 0.923549054 |
| ENSMUSG00000091195 | 1 | |
| ENSMUSG00000091198 | -1.043306199 | 0.698112512 |
| ENSMUSG00000091199 | 1 | |
| ENSMUSG00000091204 | -1.043306199 | 0.698112512 |
| ENSMUSG00000091205 | 1 | |
| ENSMUSG00000091206 | 1 | |
| ENSMUSG00000091212 | 1 | |
| ENSMUSG00000091215 | -1.063525418 | 0.814089245 |
| ENSMUSG00000091216 | 1 | |
| ENSMUSG00000091221 | 1 | |
| ENSMUSG00000091223 | -1.161243541 | 0.476098886 |
| ENSMUSG00000091227 | 1.030415139 | 0.971519533 |
| ENSMUSG00000091228 | 1 | |
| ENSMUSG00000091230 | 1.1501513 | 0.608497229 |
| ENSMUSG00000091233 | 1.059463094 | 0.608497229 |
| ENSMUSG00000091237 | 1 | |
| ENSMUSG00000091238 | 1.128312813 | 0.552754447 |
| ENSMUSG00000091239 | 1 | |
| ENSMUSG00000091243 | 1.130013759 | 0.526298164 |
| ENSMUSG00000091248 | 1 | |
| ENSMUSG00000091255 | 1 | |
| ENSMUSG00000091259 | 1 | |
| ENSMUSG00000091260 | 1.059463094 | 0.608497229 |
| ENSMUSG00000091264 | -1.017940533 | 0.976562349 |
| ENSMUSG00000091268 | 1 | |
| ENSMUSG00000091269 | 1 | |
| ENSMUSG00000091271 | 1 | |
| ENSMUSG00000091272 | 1 | |
| ENSMUSG00000091273 | 1 | |
| ENSMUSG00000091275 | 1 | |
| ENSMUSG00000091277 | -1.369310127 | 0.151833616 |
| ENSMUSG00000091280 | 1 | |
| ENSMUSG00000091281 | 1 | |
| ENSMUSG00000091283 | 1 | |
| ENSMUSG00000091285 | 1.13805685 | 0.165163153 |
| ENSMUSG00000091286 | 1 | |
| ENSMUSG00000091288 | 1 | |
| ENSMUSG00000091289 | 1 | |
| ENSMUSG00000091290 | 1 | |
| ENSMUSG00000091292 | 1 | |
| ENSMUSG00000091293 | 1.122047485 | 0.698112512 |
| ENSMUSG00000091296 | 1 | |
| ENSMUSG00000091297 | 1 | |
| ENSMUSG00000091304 | 1 | |

| | | |
|---|---|---|
| ENSMUSG00000091305 | 1 | |
| ENSMUSG00000091306 | 1 | |
| ENSMUSG00000091307 | 1 | |
| ENSMUSG00000091311 | 1 | |
| ENSMUSG00000091312 | 1 | |
| ENSMUSG00000091313 | -1.141886854 | 0.29310289 |
| ENSMUSG00000091315 | -1 | 1 |
| ENSMUSG00000091318 | 1.15209161 | 0.57564331 |
| ENSMUSG00000091325 | 1 | |
| ENSMUSG00000091326 | 1 | |
| ENSMUSG00000091329 | 1.081478107 | 0.841678369 |
| ENSMUSG00000091336 | 1.059463094 | 0.608497229 |
| ENSMUSG00000091337 | 1.085229146 | 0.698112512 |
| ENSMUSG00000091345 | -2.217589622 | 0.00314868 |
| ENSMUSG00000091347 | -1 | 1 |
| ENSMUSG00000091350 | 1 | |
| ENSMUSG00000091351 | -1.053549062 | 0.778372272 |
| ENSMUSG00000091355 | 1 | |
| ENSMUSG00000091358 | 1.059463094 | 0.608497229 |
| ENSMUSG00000091361 | -1.033820075 | 0.916014403 |
| ENSMUSG00000091363 | 1.115389061 | 0.727488094 |
| ENSMUSG00000091366 | -1.069502966 | 0.79291651 |
| ENSMUSG00000091367 | 1 | |
| ENSMUSG00000091368 | 1 | |
| ENSMUSG00000091370 | -1.043306199 | 0.698112512 |
| ENSMUSG00000091373 | -1.262272781 | 0.159939487 |
| ENSMUSG00000091375 | 1 | |
| ENSMUSG00000091376 | 1 | |
| ENSMUSG00000091378 | 1.014731082 | 0.988424085 |
| ENSMUSG00000091381 | 1 | |
| ENSMUSG00000091382 | 1 | |
| ENSMUSG00000091383 | 1 | |
| ENSMUSG00000091386 | 1 | |
| ENSMUSG00000091387 | -1.120708549 | 0.532586 |
| ENSMUSG00000091388 | 1 | |
| ENSMUSG00000091390 | 1.095586318 | 0.705199845 |
| ENSMUSG00000091393 | -1.359633602 | 0.186861854 |
| ENSMUSG00000091396 | 1 | |
| ENSMUSG00000091400 | 1 | |
| ENSMUSG00000091401 | 1.173139855 | 0.385188707 |
| ENSMUSG00000091402 | 1 | |
| ENSMUSG00000091404 | 1 | |
| ENSMUSG00000091405 | 1.150179455 | 0.612496955 |
| ENSMUSG00000091407 | 1 | |
| ENSMUSG00000091408 | -1.013017098 | 1 |
| ENSMUSG00000091410 | 1.208666617 | 0.505560817 |
| ENSMUSG00000091411 | 1 | |
| ENSMUSG00000091412 | -1 | 1 |
| ENSMUSG00000091415 | -1.095261873 | 0.730744646 |
| ENSMUSG00000091416 | 1 | |
| ENSMUSG00000091418 | 1.059463094 | 0.608497229 |
| ENSMUSG00000091419 | 1.095872691 | 0.698112512 |
| ENSMUSG00000091421 | -1 | 1 |
| ENSMUSG00000091422 | 1 | |
| ENSMUSG00000091423 | 1.189648646 | 0.608497229 |
| ENSMUSG00000091425 | 1 | |
| ENSMUSG00000091426 | 1.087252606 | 0.769111661 |
| ENSMUSG00000091429 | 1 | |
| ENSMUSG00000091430 | 1 | |
| ENSMUSG00000091435 | 1 | |
| ENSMUSG00000091438 | 1.069068979 | 0.698112512 |
| ENSMUSG00000091441 | 1 | |
| ENSMUSG00000091442 | -1.044273782 | 0.698112512 |

| | | |
|---|---|---|
| ENSMUSG00000091443 | 1.257824979 | 0.307303709 |
| ENSMUSG00000091449 | 1.10141562 | 0.729293671 |
| ENSMUSG00000091450 | 1 | |
| ENSMUSG00000091453 | 1 | |
| ENSMUSG00000091455 | -1.176723811 | 0.608497229 |
| ENSMUSG00000091457 | 1 | |
| ENSMUSG00000091459 | 1 | |
| ENSMUSG00000091460 | 1.211527659 | 0.457237932 |
| ENSMUSG00000091462 | -1.268812687 | 0.16934681 |
| ENSMUSG00000091466 | 1 | |
| ENSMUSG00000091468 | 1 | |
| ENSMUSG00000091470 | 1 | |
| ENSMUSG00000091471 | 1 | |
| ENSMUSG00000091472 | -1.085651418 | 0.608497229 |
| ENSMUSG00000091474 | 1.052952444 | 0.698112512 |
| ENSMUSG00000091475 | -1.539094244 | 0.018601523 |
| ENSMUSG00000091476 | 1.014545335 | 0.979002724 |
| ENSMUSG00000091477 | 1 | |
| ENSMUSG00000091478 | 1.014724921 | 0.954497929 |
| ENSMUSG00000091479 | 1 | |
| ENSMUSG00000091481 | 1 | |
| ENSMUSG00000091483 | 1 | |
| ENSMUSG00000091486 | 1 | |
| ENSMUSG00000091487 | 1 | |
| ENSMUSG00000091490 | 1 | |
| ENSMUSG00000091491 | 1 | |
| ENSMUSG00000091492 | 1 | |
| ENSMUSG00000091494 | 1 | |
| ENSMUSG00000091496 | 1 | |
| ENSMUSG00000091497 | 1 | |
| ENSMUSG00000091498 | 1 | |
| ENSMUSG00000091504 | 1.019083579 | 0.949812939 |
| ENSMUSG00000091507 | 1.116289651 | 0.760615796 |
| ENSMUSG00000091509 | -1.194483945 | 0.086566011 |
| ENSMUSG00000091510 | 1.11026164 | 0.608497229 |
| ENSMUSG00000091511 | -1 | 1 |
| ENSMUSG00000091512 | -1.016230791 | 0.964665151 |
| ENSMUSG00000091514 | -1.000971397 | 1 |
| ENSMUSG00000091515 | 1.00613998 | 1 |
| ENSMUSG00000091519 | 1 | |
| ENSMUSG00000091523 | -1.084645496 | 0.532369657 |
| ENSMUSG00000091526 | 1 | |
| ENSMUSG00000091528 | 1 | |
| ENSMUSG00000091530 | -1.009477408 | 1 |
| ENSMUSG00000091531 | 1.059463094 | 0.698112512 |
| ENSMUSG00000091534 | 1.199757411 | 0.067669179 |
| ENSMUSG00000091537 | -1.100517117 | 0.698112512 |
| ENSMUSG00000091539 | 1 | |
| ENSMUSG00000091541 | 1 | |
| ENSMUSG00000091542 | 1.022121746 | 0.948663192 |
| ENSMUSG00000091548 | 1 | |
| ENSMUSG00000091549 | 1.090997726 | 0.566927541 |
| ENSMUSG00000091550 | 1 | |
| ENSMUSG00000091552 | 1 | |
| ENSMUSG00000091553 | 1.663658025 | 2.59386E-05 |
| ENSMUSG00000091556 | -1.044273782 | 0.790851299 |
| ENSMUSG00000091557 | 1.059463094 | 0.608497229 |
| ENSMUSG00000091558 | -1.092661439 | 0.773508225 |
| ENSMUSG00000091561 | 1.050110043 | 0.778097229 |
| ENSMUSG00000091562 | -1.089355839 | 0.582158676 |
| ENSMUSG00000091563 | 1 | |
| ENSMUSG00000091568 | 1 | |
| ENSMUSG00000091569 | 1 | |

| | | |
|---|---|---|
| ENSMUSG00000091570 | -1 | 1 |
| ENSMUSG00000091572 | 1 | |
| ENSMUSG00000091573 | -1 | 1 |
| ENSMUSG00000091575 | -1.094488708 | 0.698135423 |
| ENSMUSG00000091577 | 1.051295273 | 0.847418374 |
| ENSMUSG00000091580 | 1.084251103 | 0.67884395 |
| ENSMUSG00000091583 | -1.044273782 | 0.698112512 |
| ENSMUSG00000091584 | 1 | |
| ENSMUSG00000091585 | 1 | |
| ENSMUSG00000091586 | 1.20776921 | 0.087972991 |
| ENSMUSG00000091587 | 1.075324819 | 0.750742698 |
| ENSMUSG00000091594 | 1.059463094 | 0.698112512 |
| ENSMUSG00000091595 | 1 | |
| ENSMUSG00000091601 | 1 | |
| ENSMUSG00000091604 | 1.005613439 | 1 |
| ENSMUSG00000091605 | 1 | |
| ENSMUSG00000091606 | 1 | |
| ENSMUSG00000091613 | -1.023769022 | 0.934760809 |
| ENSMUSG00000091614 | -1.03962336 | 0.698112512 |
| ENSMUSG00000091615 | 1 | |
| ENSMUSG00000091617 | 1 | |
| ENSMUSG00000091618 | 1.513874408 | 0.001039425 |
| ENSMUSG00000091619 | 1 | |
| ENSMUSG00000091620 | 1 | |
| ENSMUSG00000091623 | 1.315914911 | 0.221427514 |
| ENSMUSG00000091624 | 1 | |
| ENSMUSG00000091625 | -1.000407596 | 1 |
| ENSMUSG00000091627 | 1 | |
| ENSMUSG00000091631 | 1 | |
| ENSMUSG00000091635 | 1.059463094 | 0.698112512 |
| ENSMUSG00000091636 | 1 | |
| ENSMUSG00000091638 | 1 | |
| ENSMUSG00000091639 | -1.13161737 | 0.608497229 |
| ENSMUSG00000091648 | 1.302383313 | 0.282504068 |
| ENSMUSG00000091649 | -1.043306199 | 0.858911269 |
| ENSMUSG00000091650 | 1 | |
| ENSMUSG00000091651 | 1 | |
| ENSMUSG00000091652 | 1 | |
| ENSMUSG00000091655 | 1 | |
| ENSMUSG00000091656 | 1 | |
| ENSMUSG00000091657 | 1 | |
| ENSMUSG00000091659 | 1 | |
| ENSMUSG00000091661 | 1 | |
| ENSMUSG00000091662 | 1 | |
| ENSMUSG00000091665 | 1 | |
| ENSMUSG00000091668 | -1 | 1 |
| ENSMUSG00000091669 | 1 | |
| ENSMUSG00000091670 | 1 | |
| ENSMUSG00000091671 | 1 | |
| ENSMUSG00000091672 | 1 | |
| ENSMUSG00000091676 | 1 | |
| ENSMUSG00000091679 | 1 | |
| ENSMUSG00000091680 | 1 | |
| ENSMUSG00000091682 | 1 | |
| ENSMUSG00000091685 | -1 | 1 |
| ENSMUSG00000091686 | -1.043306199 | 0.698112512 |
| ENSMUSG00000091692 | 1 | |
| ENSMUSG00000091694 | 1.253163119 | 0.242341101 |
| ENSMUSG00000091695 | 1 | |
| ENSMUSG00000091697 | 1.059463094 | 0.608497229 |
| ENSMUSG00000091698 | -1 | 1 |
| ENSMUSG00000091700 | 1 | |
| ENSMUSG00000091703 | 1 | |

| | | |
|---|---|---|
| ENSMUSG00000091705 | -1.668236043 | 0.001135818 |
| ENSMUSG00000091707 | 1 | |
| ENSMUSG00000091709 | 1 | |
| ENSMUSG00000091712 | 1.021266192 | 1 |
| ENSMUSG00000091713 | 1 | |
| ENSMUSG00000091718 | 1 | |
| ENSMUSG00000091721 | 1 | |
| ENSMUSG00000091722 | 1 | |
| ENSMUSG00000091723 | 1 | |
| ENSMUSG00000091725 | -1.053549062 | 0.769745556 |
| ENSMUSG00000091730 | 1.218240674 | 0.217803313 |
| ENSMUSG00000091731 | 1 | |
| ENSMUSG00000091732 | 1.072379968 | 0.608497229 |
| ENSMUSG00000091733 | 1 | |
| ENSMUSG00000091734 | 1 | |
| ENSMUSG00000091735 | -1.110001877 | 0.698112512 |
| ENSMUSG00000091736 | 1 | |
| ENSMUSG00000091740 | 1 | |
| ENSMUSG00000091741 | 1 | |
| ENSMUSG00000091742 | -1.02513447 | 0.948467896 |
| ENSMUSG00000091743 | -1.03962336 | 0.698112512 |
| ENSMUSG00000091744 | -1.456429312 | 0.100537573 |
| ENSMUSG00000091745 | 1 | |
| ENSMUSG00000091749 | 1 | |
| ENSMUSG00000091752 | 1 | |
| ENSMUSG00000091754 | 1.09526813 | 0.773508225 |
| ENSMUSG00000091756 | 1 | |
| ENSMUSG00000091757 | 1 | |
| ENSMUSG00000091762 | 1 | |
| ENSMUSG00000091764 | 1.016551229 | 0.911116262 |
| ENSMUSG00000091768 | 1 | |
| ENSMUSG00000091771 | 1 | |
| ENSMUSG00000091775 | -1 | 1 |
| ENSMUSG00000091776 | 1 | |
| ENSMUSG00000091779 | 1 | |
| ENSMUSG00000091780 | 1 | |
| ENSMUSG00000091784 | 1.038940017 | 0.842601989 |
| ENSMUSG00000091785 | 1.019083579 | 0.949812939 |
| ENSMUSG00000091786 | 1.014201295 | 0.964323862 |
| ENSMUSG00000091788 | 1 | |
| ENSMUSG00000091792 | 1 | |
| ENSMUSG00000091794 | -1.044273782 | 0.698112512 |
| ENSMUSG00000091800 | 1 | |
| ENSMUSG00000091803 | 1.001203501 | 1 |
| ENSMUSG00000091805 | 1 | |
| ENSMUSG00000091806 | 1 | |
| ENSMUSG00000091808 | 1 | |
| ENSMUSG00000091809 | 1 | |
| ENSMUSG00000091811 | 1.022454478 | 0.794274354 |
| ENSMUSG00000091812 | 1 | |
| ENSMUSG00000091813 | 1 | |
| ENSMUSG00000091814 | 1.059463094 | 0.704170349 |
| ENSMUSG00000091815 | 1 | |
| ENSMUSG00000091816 | 1 | |
| ENSMUSG00000091817 | 1 | |
| ENSMUSG00000091818 | 1.046083503 | 0.941215236 |
| ENSMUSG00000091821 | 1 | |
| ENSMUSG00000091825 | -1.002901227 | 1 |
| ENSMUSG00000091827 | 1 | |
| ENSMUSG00000091831 | -1.040391774 | 0.972980766 |
| ENSMUSG00000091832 | -1 | 1 |
| ENSMUSG00000091834 | 1 | |
| ENSMUSG00000091842 | 1 | |

| | | |
|---|---|---|
| ENSMUSG00000091844 | -1.083910675 | 0.698112512 |
| ENSMUSG00000091845 | 1.29759408 | 0.608497229 |
| ENSMUSG00000091847 | 1 | |
| ENSMUSG00000091849 | 1 | |
| ENSMUSG00000091859 | 1 | |
| ENSMUSG00000091863 | 1 | |
| ENSMUSG00000091864 | 1.008742867 | 1 |
| ENSMUSG00000091866 | -1.216168767 | 0.221797188 |
| ENSMUSG00000091867 | 1 | |
| ENSMUSG00000091873 | 1 | |
| ENSMUSG00000091874 | 1 | |
| ENSMUSG00000091876 | 1 | |
| ENSMUSG00000091879 | 1 | |
| ENSMUSG00000091881 | -1.048992887 | 0.722351115 |
| ENSMUSG00000091882 | 1 | |
| ENSMUSG00000091887 | -1.043306199 | 0.698112512 |
| ENSMUSG00000091888 | 1 | |
| ENSMUSG00000091890 | 1 | |
| ENSMUSG00000091894 | 1 | |
| ENSMUSG00000091896 | 1.050070982 | 0.896920619 |
| ENSMUSG00000091897 | 1 | |
| ENSMUSG00000091898 | -1.079112059 | 0.698112512 |
| ENSMUSG00000091900 | -1.160739507 | 0.312162776 |
| ENSMUSG00000091902 | 1 | |
| ENSMUSG00000091903 | 1 | |
| ENSMUSG00000091905 | 1.095872691 | 0.749353361 |
| ENSMUSG00000091906 | 1.059463094 | 0.608497229 |
| ENSMUSG00000091908 | 1 | |
| ENSMUSG00000091911 | 1 | |
| ENSMUSG00000091922 | 1 | |
| ENSMUSG00000091923 | 1 | |
| ENSMUSG00000091924 | 1 | |
| ENSMUSG00000091926 | 1 | |
| ENSMUSG00000091929 | 1 | |
| ENSMUSG00000091930 | 1.069068979 | 0.608497229 |
| ENSMUSG00000091931 | 1.086705696 | 0.420161906 |
| ENSMUSG00000091933 | 1 | |
| ENSMUSG00000091936 | 1 | |
| ENSMUSG00000091937 | 1 | |
| ENSMUSG00000091938 | -1.043306199 | 0.698112512 |
| ENSMUSG00000091941 | -1.064113315 | 0.863165955 |
| ENSMUSG00000091942 | -1.095294215 | 0.541735653 |
| ENSMUSG00000091945 | 1 | |
| ENSMUSG00000091947 | -1.122102401 | 0.698112512 |
| ENSMUSG00000091950 | -1.044273782 | 0.698112512 |
| ENSMUSG00000091952 | -1.105062778 | 0.734674233 |
| ENSMUSG00000091955 | -1.01025236 | 1 |
| ENSMUSG00000091956 | 1 | |
| ENSMUSG00000091957 | -1.047959538 | 0.395255112 |
| ENSMUSG00000091960 | 1 | |
| ENSMUSG00000091961 | 1 | |
| ENSMUSG00000091962 | 1 | |
| ENSMUSG00000091967 | 1 | |
| ENSMUSG00000091968 | -1.155201718 | 0.608497229 |
| ENSMUSG00000091970 | 1 | |
| ENSMUSG00000091971 | 1.151772931 | 0.100326676 |
| ENSMUSG00000091972 | 1 | |
| ENSMUSG00000091980 | 1 | |
| ENSMUSG00000091983 | 1 | |
| ENSMUSG00000091985 | -1.101135229 | 0.563100765 |
| ENSMUSG00000091987 | 1 | |
| ENSMUSG00000091989 | 1 | |
| ENSMUSG00000091990 | 1 | |

| | | |
|---|---|---|
| ENSMUSG00000091993 | 1.153177845 | 0.403838785 |
| ENSMUSG00000091994 | 1.02393121 | 0.619047344 |
| ENSMUSG00000091995 | 1 | |
| ENSMUSG00000091996 | 1 | |
| ENSMUSG00000091997 | -1.240391029 | 0.274585817 |
| ENSMUSG00000091998 | 1 | |
| ENSMUSG00000092004 | 1 | |
| ENSMUSG00000092006 | 1 | |
| ENSMUSG00000092008 | 1 | |
| ENSMUSG00000092009 | -1.0795912 | 0.642639207 |
| ENSMUSG00000092014 | 1 | |
| ENSMUSG00000092019 | 1.069068979 | 0.698112512 |
| ENSMUSG00000092021 | -1.03962336 | 0.698112512 |
| ENSMUSG00000092027 | 1 | |
| ENSMUSG00000092029 | -1.044273782 | 0.698112512 |
| ENSMUSG00000092031 | 1 | |
| ENSMUSG00000092032 | 1 | |
| ENSMUSG00000092035 | 1.672455505 | 0.053615266 |
| ENSMUSG00000092036 | -1.085651418 | 0.532369657 |
| ENSMUSG00000092039 | 1 | |
| ENSMUSG00000092040 | 1 | |
| ENSMUSG00000092042 | 1 | |
| ENSMUSG00000092043 | 1.069068979 | 0.608497229 |
| ENSMUSG00000092045 | 1 | |
| ENSMUSG00000092047 | 1 | |
| ENSMUSG00000092048 | 1 | |
| ENSMUSG00000092049 | 1 | |
| ENSMUSG00000092051 | 1 | |
| ENSMUSG00000092054 | 1 | |
| ENSMUSG00000092055 | 1 | |
| ENSMUSG00000092056 | 1.05410549 | 0.8920795 |
| ENSMUSG00000092060 | -1.110015825 | 0.608497229 |
| ENSMUSG00000092062 | -1.052875465 | 0.844158013 |
| ENSMUSG00000092066 | 1 | |
| ENSMUSG00000092071 | 1.21327335 | 0.462450889 |
| ENSMUSG00000092072 | 1.017581977 | 1 |
| ENSMUSG00000092073 | 1 | |
| ENSMUSG00000092074 | 1.019083579 | 0.956663244 |
| ENSMUSG00000092075 | 1 | |
| ENSMUSG00000092077 | 1.189207115 | 0.294533546 |
| ENSMUSG00000092080 | -1.03962336 | 0.698112512 |
| ENSMUSG00000092083 | -1.024718486 | 0.954310937 |
| ENSMUSG00000092085 | -1.053549062 | 0.778372272 |
| ENSMUSG00000092086 | 1.015729899 | 0.971519533 |
| ENSMUSG00000092090 | 1.059463094 | 0.608497229 |
| ENSMUSG00000092092 | 1 | |
| ENSMUSG00000092093 | 1 | |
| ENSMUSG00000092094 | 1 | |
| ENSMUSG00000092097 | -1.034990181 | 0.88846936 |
| ENSMUSG00000092103 | -1.044273782 | 0.847756749 |
| ENSMUSG00000092107 | 1 | |
| ENSMUSG00000092109 | 1 | |
| ENSMUSG00000092111 | 1 | |
| ENSMUSG00000092116 | 1.005613439 | 1 |
| ENSMUSG00000092118 | 1.115922249 | 0.372534438 |
| ENSMUSG00000092120 | 1 | |
| ENSMUSG00000092123 | 1 | |
| ENSMUSG00000092124 | 1.12194735 | 0.392352483 |
| ENSMUSG00000092125 | 1 | |
| ENSMUSG00000092130 | 1.517749245 | 0.030231993 |
| ENSMUSG00000092131 | -1.063127349 | 0.698112512 |
| ENSMUSG00000092134 | 1.011497958 | 1 |
| ENSMUSG00000092137 | 1 | |

| | | |
|---|---|---|
| ENSMUSG00000092142 | -1.053549062 | 0.698112512 |
| ENSMUSG00000092144 | 1 | |
| ENSMUSG00000092147 | 1 | |
| ENSMUSG00000092148 | 1 | |
| ENSMUSG00000092152 | 1 | |
| ENSMUSG00000092157 | 1 | |
| ENSMUSG00000092162 | -1 | 1 |
| ENSMUSG00000092164 | 1 | |
| ENSMUSG00000092165 | -1 | 1 |
| ENSMUSG00000092166 | 1 | |
| ENSMUSG00000092167 | 1.199989356 | 0.000021314 |
| ENSMUSG00000092168 | 1 | |
| ENSMUSG00000092171 | 1.129504554 | 0.608497229 |
| ENSMUSG00000092172 | 1 | |
| ENSMUSG00000092173 | 1 | |
| ENSMUSG00000092174 | 1 | |
| ENSMUSG00000092175 | 1 | |
| ENSMUSG00000092176 | 1 | |
| ENSMUSG00000092178 | 1.019396403 | 0.877593058 |
| ENSMUSG00000092179 | 1 | |
| ENSMUSG00000092181 | 1 | |
| ENSMUSG00000092182 | -1 | 1 |
| ENSMUSG00000092183 | -1.152062829 | 0.427797606 |
| ENSMUSG00000092184 | -1 | 1 |
| ENSMUSG00000092185 | 1 | |
| ENSMUSG00000092186 | -1.138072347 | 0.698112512 |
| ENSMUSG00000092187 | 1.032211592 | 0.91991357 |
| ENSMUSG00000092188 | -1.044273782 | 0.795446726 |
| ENSMUSG00000092189 | -1 | 1 |
| ENSMUSG00000092190 | -1.181718351 | 0.394458958 |
| ENSMUSG00000092191 | 1 | |
| ENSMUSG00000092192 | 1 | |
| ENSMUSG00000092193 | -1 | 1 |
| ENSMUSG00000092195 | 1 | |
| ENSMUSG00000092196 | 1 | |
| ENSMUSG00000092197 | 1 | |
| ENSMUSG00000092199 | 1 | |
| ENSMUSG00000092200 | 1 | |
| ENSMUSG00000092201 | 1 | |
| ENSMUSG00000092202 | -1.046307045 | 0.906405443 |
| ENSMUSG00000092203 | 1.176195874 | 0.16875903 |
| ENSMUSG00000092204 | 1 | |
| ENSMUSG00000092208 | 1 | |
| ENSMUSG00000092210 | 1 | |
| ENSMUSG00000092211 | 1 | |
| ENSMUSG00000092212 | -1.237055826 | 0.352182712 |
| ENSMUSG00000092213 | 1 | |
| ENSMUSG00000092214 | 1 | |
| ENSMUSG00000092216 | 1 | |
| ENSMUSG00000092217 | 1.069068979 | 0.608497229 |
| ENSMUSG00000092218 | 1 | |
| ENSMUSG00000092219 | -1 | 1 |
| ENSMUSG00000092220 | 1 | |
| ENSMUSG00000092221 | 1 | |
| ENSMUSG00000092222 | 1 | |
| ENSMUSG00000092223 | -1.063525418 | 0.698112512 |
| ENSMUSG00000092224 | 1 | |
| ENSMUSG00000092225 | 1.029490685 | 0.973126641 |
| ENSMUSG00000092227 | 1 | |
| ENSMUSG00000092228 | 1.171563499 | 0.370636424 |
| ENSMUSG00000092229 | 1.122462048 | 0.420161906 |
| ENSMUSG00000092230 | -1.044273782 | 0.790851299 |
| ENSMUSG00000092231 | 1 | |

| | | |
|---|---|---|
| ENSMUSG00000092232 | 1 | |
| ENSMUSG00000092233 | -1.084645496 | 0.532369657 |
| ENSMUSG00000092237 | 1 | |
| ENSMUSG00000092238 | 1 | |
| ENSMUSG00000092239 | 1.000939184 | 1 |
| ENSMUSG00000092240 | -1.03962336 | 0.814089245 |
| ENSMUSG00000092241 | 1.161658033 | 0.608497229 |
| ENSMUSG00000092242 | 1 | |
| ENSMUSG00000092243 | 1 | |
| ENSMUSG00000092244 | 1 | |
| ENSMUSG00000092245 | 1 | |
| ENSMUSG00000092246 | 1.177765487 | 0.380970157 |
| ENSMUSG00000092247 | 1.059463094 | 0.608497229 |
| ENSMUSG00000092248 | 1.015486245 | 0.985522133 |
| ENSMUSG00000092250 | -1.003834322 | 1 |
| ENSMUSG00000092251 | 1 | |
| ENSMUSG00000092252 | 1 | |
| ENSMUSG00000092253 | 1 | |
| ENSMUSG00000092255 | 1 | |
| ENSMUSG00000092256 | 1 | |
| ENSMUSG00000092257 | 1.065662724 | 0.821500382 |
| ENSMUSG00000092258 | 1 | |
| ENSMUSG00000092259 | 1 | |
| ENSMUSG00000092260 | -1.03979259 | 0.752472633 |
| ENSMUSG00000092261 | -1.034990181 | 0.914913657 |
| ENSMUSG00000092263 | 1 | |
| ENSMUSG00000092264 | 1 | |
| ENSMUSG00000092265 | -1.100193663 | 0.534887579 |
| ENSMUSG00000092266 | 1 | |
| ENSMUSG00000092267 | 1 | |
| ENSMUSG00000092268 | 1.019083579 | 0.97045957 |
| ENSMUSG00000092269 | 1 | |
| ENSMUSG00000092270 | 1 | |
| ENSMUSG00000092271 | 1 | |
| ENSMUSG00000092272 | 1 | |
| ENSMUSG00000092274 | 1.284270934 | 0.151037288 |
| ENSMUSG00000092275 | 1.059463094 | 0.608497229 |
| ENSMUSG00000092276 | 1 | |
| ENSMUSG00000092277 | 1 | |
| ENSMUSG00000092278 | -1.075416662 | 0.767574573 |
| ENSMUSG00000092281 | -1.137634574 | 0.616130965 |
| ENSMUSG00000092283 | 1 | |
| ENSMUSG00000092284 | 1.161036672 | 0.251049598 |
| ENSMUSG00000092285 | 1 | |
| ENSMUSG00000092286 | 1.070148225 | 0.911787527 |
| ENSMUSG00000092287 | 1 | |
| ENSMUSG00000092289 | -1.053549062 | 0.698112512 |
| ENSMUSG00000092290 | 1.059463094 | 0.608497229 |
| ENSMUSG00000092292 | 1 | |
| ENSMUSG00000092293 | 1 | |
| ENSMUSG00000092294 | 1 | |
| ENSMUSG00000092295 | 1 | |
| ENSMUSG00000092296 | 1 | |
| ENSMUSG00000092297 | 1 | |
| ENSMUSG00000092298 | 1 | |
| ENSMUSG00000092300 | 1.085931761 | 0.764959211 |
| ENSMUSG00000092301 | 1.089470641 | 0.608497229 |
| ENSMUSG00000092302 | 1 | |
| ENSMUSG00000092303 | 1 | |
| ENSMUSG00000092304 | 1 | |
| ENSMUSG00000092305 | 1 | |
| ENSMUSG00000092306 | 1 | |
| ENSMUSG00000092307 | 1 | |

| | | |
|---|---|---|
| ENSMUSG00000092308 | 1 | |
| ENSMUSG00000092309 | 1 | |
| ENSMUSG00000092310 | 1 | |
| ENSMUSG00000092311 | 1 | |
| ENSMUSG00000092312 | -1.043306199 | 0.698112512 |
| ENSMUSG00000092313 | 1 | |
| ENSMUSG00000092314 | 1 | |
| ENSMUSG00000092316 | 1 | |
| ENSMUSG00000092317 | -1.03962336 | 0.698112512 |
| ENSMUSG00000092318 | 1 | |
| ENSMUSG00000092322 | 1 | |
| ENSMUSG00000092323 | 1 | |
| ENSMUSG00000092325 | -1.059802455 | 0.730441273 |
| ENSMUSG00000092326 | 1 | |
| ENSMUSG00000092328 | 1 | |
| ENSMUSG00000092329 | 1 | |
| ENSMUSG00000092330 | 1 | |
| ENSMUSG00000092331 | 1 | |
| ENSMUSG00000092332 | 1 | |
| ENSMUSG00000092335 | -1.413084758 | 0.011725879 |
| ENSMUSG00000092336 | 1 | |
| ENSMUSG00000092338 | 1 | |
| ENSMUSG00000092339 | 1 | |
| ENSMUSG00000092340 | 1 | |
| ENSMUSG00000092341 | 1.131883036 | 0.697557233 |
| ENSMUSG00000092342 | 1 | |
| ENSMUSG00000092343 | 1 | |
| ENSMUSG00000092344 | 1.049585115 | 0.879253209 |
| ENSMUSG00000092345 | 1 | |
| ENSMUSG00000092346 | 1 | |
| ENSMUSG00000092349 | -1.04676943 | 0.861653133 |
| ENSMUSG00000092350 | 1 | |
| ENSMUSG00000092351 | 1 | |
| ENSMUSG00000092352 | 1 | |
| ENSMUSG00000092353 | -1.069095149 | 0.686194514 |
| ENSMUSG00000092354 | 1.061907756 | 0.780320794 |
| ENSMUSG00000092356 | 1 | |
| ENSMUSG00000092357 | 1 | |
| ENSMUSG00000092358 | -1.101469545 | 0.698112512 |
| ENSMUSG00000092359 | 1 | |
| ENSMUSG00000092360 | 1 | |
| ENSMUSG00000092362 | 1 | |
| ENSMUSG00000092363 | 1 | |
| ENSMUSG00000092364 | -1.086192882 | 0.698112512 |
| ENSMUSG00000092365 | 1 | |
| ENSMUSG00000092366 | -1.136679298 | 0.608497229 |
| ENSMUSG00000092367 | 1 | |
| ENSMUSG00000092368 | 1 | |
| ENSMUSG00000092369 | 1 | |
| ENSMUSG00000092370 | 1 | |
| ENSMUSG00000092371 | -1.043306199 | 0.698112512 |
| ENSMUSG00000092372 | 1 | |
| ENSMUSG00000092373 | -1.02794432 | 0.967179343 |
| ENSMUSG00000092374 | 1 | |
| ENSMUSG00000092375 | 1.122462048 | 0.279947521 |
| ENSMUSG00000092376 | -1.053549062 | 0.769745556 |
| ENSMUSG00000092377 | 1 | |
| ENSMUSG00000092381 | 1.028323353 | 0.903464611 |
| ENSMUSG00000092382 | 1 | |
| ENSMUSG00000092383 | 1 | |
| ENSMUSG00000092384 | 1.125672422 | 0.770958539 |
| ENSMUSG00000092385 | -1.03962336 | 0.698112512 |
| ENSMUSG00000092386 | 1 | |

| | | |
|---|---|---|
| ENSMUSG00000092387 | 1 | |
| ENSMUSG00000092388 | 1 | |
| ENSMUSG00000092389 | 1 | |
| ENSMUSG00000092390 | -1.292150232 | 0.035744898 |
| ENSMUSG00000092392 | 1.222922263 | 0.60397857 |
| ENSMUSG00000092393 | 1 | |
| ENSMUSG00000092394 | 1.014545335 | 0.989648415 |
| ENSMUSG00000092395 | 1 | |
| ENSMUSG00000092396 | 1 | |
| ENSMUSG00000092397 | 1 | |
| ENSMUSG00000092398 | 1 | |
| ENSMUSG00000092399 | 1.095872691 | 0.685180517 |
| ENSMUSG00000092401 | 1 | |
| ENSMUSG00000092402 | 1 | |
| ENSMUSG00000092403 | 1 | |
| ENSMUSG00000092404 | 1.059463094 | 0.608497229 |
| ENSMUSG00000092405 | 1 | |
| ENSMUSG00000092406 | 1 | |
| ENSMUSG00000092408 | 1 | |
| ENSMUSG00000092409 | 1 | |
| ENSMUSG00000092410 | 1 | |
| ENSMUSG00000092411 | 1 | |
| ENSMUSG00000092412 | -1.017407094 | 1 |
| ENSMUSG00000092413 | 1 | |
| ENSMUSG00000092414 | 1 | |
| ENSMUSG00000092415 | 1 | |
| ENSMUSG00000092416 | 1.058252509 | 0.358965438 |
| ENSMUSG00000092417 | 1.02948725 | 0.530449026 |
| ENSMUSG00000092418 | -1.043306199 | 0.798487595 |
| ENSMUSG00000092419 | 1 | |
| ENSMUSG00000092422 | 1 | |
| ENSMUSG00000092425 | 1 | |
| ENSMUSG00000092426 | 1 | |
| ENSMUSG00000092427 | 1 | |
| ENSMUSG00000092428 | -1.014570171 | 0.985623907 |
| ENSMUSG00000092429 | 1 | |
| ENSMUSG00000092431 | 1 | |
| ENSMUSG00000092433 | 1 | |
| ENSMUSG00000092435 | 1 | |
| ENSMUSG00000092437 | -1.03962336 | 0.698112512 |
| ENSMUSG00000092438 | 1.075870199 | 0.642819974 |
| ENSMUSG00000092439 | 1 | |
| ENSMUSG00000092440 | 1 | |
| ENSMUSG00000092441 | 1 | |
| ENSMUSG00000092443 | -1.009476377 | 1 |
| ENSMUSG00000092444 | -1.11373934 | 0.608497229 |
| ENSMUSG00000092447 | 1 | |
| ENSMUSG00000092448 | 1.122462048 | 0.215991492 |
| ENSMUSG00000092450 | 1 | |
| ENSMUSG00000092451 | 1 | |
| ENSMUSG00000092452 | 1 | |
| ENSMUSG00000092454 | 1 | |
| ENSMUSG00000092455 | 1 | |
| ENSMUSG00000092456 | 1 | |
| ENSMUSG00000092457 | -1.44285408 | 0.005520018 |
| ENSMUSG00000092458 | -1 | 1 |
| ENSMUSG00000092460 | 1 | |
| ENSMUSG00000092461 | 1 | |
| ENSMUSG00000092462 | 1 | |
| ENSMUSG00000092463 | 1 | |
| ENSMUSG00000092464 | -1.006193779 | 1 |
| ENSMUSG00000092465 | 1.435600202 | 0.18421873 |
| ENSMUSG00000092466 | 1 | |

| | | |
|---|---|---|
| ENSMUSG00000092467 | 1 | |
| ENSMUSG00000092468 | 1 | |
| ENSMUSG00000092469 | 1 | |
| ENSMUSG00000092470 | 1.05943716 | 0.698112512 |
| ENSMUSG00000092471 | 1 | |
| ENSMUSG00000092472 | 1 | |
| ENSMUSG00000092473 | 1 | |
| ENSMUSG00000092474 | -1.173975439 | 0.58386781 |
| ENSMUSG00000092476 | 1 | |
| ENSMUSG00000092477 | 1 | |
| ENSMUSG00000092480 | 1 | |
| ENSMUSG00000092482 | 1.069068979 | 0.698112512 |
| ENSMUSG00000092483 | 1 | |
| ENSMUSG00000092485 | 1 | |
| ENSMUSG00000092486 | -1.191271501 | 0.122533263 |
| ENSMUSG00000092487 | 1 | |
| ENSMUSG00000092488 | 1 | |
| ENSMUSG00000092489 | 1 | |
| ENSMUSG00000092490 | -1.087027344 | 0.721403818 |
| ENSMUSG00000092492 | 1.015486245 | 0.972980766 |
| ENSMUSG00000092493 | 1 | |
| ENSMUSG00000092494 | 1 | |
| ENSMUSG00000092495 | 1.104868836 | 0.757552239 |
| ENSMUSG00000092496 | 1 | |
| ENSMUSG00000092499 | 1 | |
| ENSMUSG00000092500 | -1.174891231 | 0.515621021 |
| ENSMUSG00000092502 | -1 | 1 |
| ENSMUSG00000092503 | 1 | |
| ENSMUSG00000092504 | 1 | |
| ENSMUSG00000092505 | -1 | 1 |
| ENSMUSG00000092506 | 1 | |
| ENSMUSG00000092507 | 1 | |
| ENSMUSG00000092508 | 1 | |
| ENSMUSG00000092509 | -1.413962067 | 0.021163083 |
| ENSMUSG00000092510 | 1 | |
| ENSMUSG00000092511 | 1 | |
| ENSMUSG00000092512 | 1 | |
| ENSMUSG00000092513 | 1 | |
| ENSMUSG00000092514 | 1 | |
| ENSMUSG00000092515 | -1.222309803 | 0.390934291 |
| ENSMUSG00000092516 | 1 | |
| ENSMUSG00000092517 | 1 | |
| ENSMUSG00000092518 | -1.043306199 | 0.79291651 |
| ENSMUSG00000092519 | 1 | |
| ENSMUSG00000092520 | 1 | |
| ENSMUSG00000092521 | -1.043306199 | 0.698112512 |
| ENSMUSG00000092522 | 1 | |
| ENSMUSG00000092523 | 1 | |
| ENSMUSG00000092524 | 1.136863383 | 0.610093216 |
| ENSMUSG00000092525 | 1 | |
| ENSMUSG00000092526 | 1.091686812 | 0.714698678 |
| ENSMUSG00000092528 | 1 | |
| ENSMUSG00000092529 | 1 | |
| ENSMUSG00000092530 | 1 | |
| ENSMUSG00000092532 | 1 | |
| ENSMUSG00000092533 | 1 | |
| ENSMUSG00000092534 | 1 | |
| ENSMUSG00000092535 | 1 | |
| ENSMUSG00000092536 | -1.03962336 | 0.698112512 |
| ENSMUSG00000092538 | 1 | |
| ENSMUSG00000092539 | 1.070429626 | 0.763407099 |
| ENSMUSG00000092540 | -1.129231493 | 0.608497229 |
| ENSMUSG00000092541 | 1 | |

| | | |
|---|---|---|
| ENSMUSG00000092542 | 1 | |
| ENSMUSG00000092543 | 1.075870199 | 0.698112512 |
| ENSMUSG00000092544 | 1 | |
| ENSMUSG00000092545 | 1.059463094 | 0.608497229 |
| ENSMUSG00000092547 | 1 | |
| ENSMUSG00000092548 | 1 | |
| ENSMUSG00000092549 | 1 | |
| ENSMUSG00000092550 | -1.213118765 | 0.243993149 |
| ENSMUSG00000092551 | 1 | |
| ENSMUSG00000092552 | 1 | |
| ENSMUSG00000092553 | -1.044273782 | 0.801443616 |
| ENSMUSG00000092554 | 1 | |
| ENSMUSG00000092555 | 1 | |
| ENSMUSG00000092556 | 1.090076375 | 0.776953697 |
| ENSMUSG00000092557 | 1.024693402 | 0.953922475 |
| ENSMUSG00000092558 | 1.00066408 | 1 |
| ENSMUSG00000092559 | -1.044273782 | 0.698112512 |
| ENSMUSG00000092560 | 1 | |
| ENSMUSG00000092561 | 1 | |
| ENSMUSG00000092562 | 1 | |
| ENSMUSG00000092563 | -1.043306199 | 0.698112512 |
| ENSMUSG00000092564 | 1.065627985 | 0.936055893 |
| ENSMUSG00000092565 | 1 | |
| ENSMUSG00000092566 | 1 | |
| ENSMUSG00000092567 | 1 | |
| ENSMUSG00000092568 | 1 | |
| ENSMUSG00000092569 | 1.464181145 | 0.040928762 |
| ENSMUSG00000092570 | 1 | |
| ENSMUSG00000092571 | 1.019083579 | 0.979002724 |
| ENSMUSG00000092572 | -1.03962336 | 0.698112512 |
| ENSMUSG00000092573 | 1 | |
| ENSMUSG00000092574 | 1 | |
| ENSMUSG00000092575 | 1 | |
| ENSMUSG00000092576 | 1 | |
| ENSMUSG00000092577 | -1.033845382 | 0.786939259 |
| ENSMUSG00000092578 | 1 | |
| ENSMUSG00000092579 | 1 | |
| ENSMUSG00000092580 | 1 | |
| ENSMUSG00000092581 | 1 | |
| ENSMUSG00000092583 | 1 | |
| ENSMUSG00000092584 | 1 | |
| ENSMUSG00000092586 | 1.085543235 | 0.698112512 |
| ENSMUSG00000092587 | 1 | |
| ENSMUSG00000092588 | 1 | |
| ENSMUSG00000092591 | 1 | |
| ENSMUSG00000092592 | 1.1576987 | 0.184424856 |
| ENSMUSG00000092593 | 1.139475542 | 0.670233909 |
| ENSMUSG00000092595 | -1.000603363 | 1 |
| ENSMUSG00000092596 | 1 | |
| ENSMUSG00000092597 | 1 | |
| ENSMUSG00000092599 | 1 | |
| ENSMUSG00000092600 | 1.05943716 | 0.698112512 |
| ENSMUSG00000092601 | 1 | |
| ENSMUSG00000092602 | 1 | 1 |
| ENSMUSG00000092603 | 1 | |
| ENSMUSG00000092604 | 1 | |
| ENSMUSG00000092606 | 1 | |
| ENSMUSG00000092607 | -1.200393067 | 0.086872908 |
| ENSMUSG00000092608 | 1 | |
| ENSMUSG00000092609 | 1 | |
| ENSMUSG00000092610 | 1 | |
| ENSMUSG00000092611 | 1 | |
| ENSMUSG00000092612 | 1 | |

| | | |
|---|---|---|
| ENSMUSG00000092614 | 1 | |
| ENSMUSG00000092615 | 1 | |
| ENSMUSG00000092616 | 1 | |
| ENSMUSG00000092617 | -1.024718486 | 0.931480186 |
| ENSMUSG00000092618 | 1 | |
| ENSMUSG00000092619 | 1.059463094 | 0.608497229 |
| ENSMUSG00000092620 | 1 | |
| ENSMUSG00000092621 | 1 | |
| ENSMUSG00000092622 | -1 | 1 |
| ENSMUSG00000092624 | -1.053549062 | 0.698112512 |
| ENSMUSG00000092625 | 1.023276667 | 0.806526905 |
| ENSMUSG00000092626 | 1.039733268 | 0.938939969 |
| ENSMUSG00000092627 | -1.561684943 | 0.055769496 |
| ENSMUSG00000092628 | 1 | |
| ENSMUSG00000092629 | 1 | |
| ENSMUSG00000092631 | 1 | |
| ENSMUSG00000092633 | 1 | |
| ENSMUSG00000092636 | 1 | |
| ENSMUSG00000092638 | 1 | |
| ENSMUSG00000092639 | 1 | |
| ENSMUSG00000092641 | -1.044319384 | 0.952898197 |
| ENSMUSG00000092643 | 1 | |
| ENSMUSG00000092644 | -1.086452977 | 0.787009986 |
| ENSMUSG00000092647 | 1 | |
| ENSMUSG00000092648 | 1.059463094 | 0.608497229 |
| ENSMUSG00000092650 | 1 | |
| ENSMUSG00000092652 | 1 | |
| ENSMUSG00000092653 | 1 | |
| ENSMUSG00000092655 | 1 | |
| ENSMUSG00000092656 | 1 | |
| ENSMUSG00000092657 | 1.059463094 | 0.698112512 |
| ENSMUSG00000092659 | 1.059463094 | 0.608497229 |
| ENSMUSG00000092660 | -1.041354166 | 0.96035552 |
| ENSMUSG00000092661 | 1 | |
| ENSMUSG00000092664 | 1 | |
| ENSMUSG00000092666 | 1 | |
| ENSMUSG00000092668 | 1 | |
| ENSMUSG00000092671 | 1 | |
| ENSMUSG00000092673 | 1.059463094 | 0.608497229 |
| ENSMUSG00000092674 | 1 | |
| ENSMUSG00000092675 | 1 | |
| ENSMUSG00000092677 | 1 | |
| ENSMUSG00000092679 | 1 | |
| ENSMUSG00000092680 | 1 | |
| ENSMUSG00000092683 | 1 | |
| ENSMUSG00000092685 | 1 | |
| ENSMUSG00000092686 | 1 | |
| ENSMUSG00000092687 | 1 | |
| ENSMUSG00000092693 | 1 | |
| ENSMUSG00000092695 | 1 | |
| ENSMUSG00000092696 | 1 | |
| ENSMUSG00000092697 | 1 | |
| ENSMUSG00000092698 | 1 | |
| ENSMUSG00000092701 | 1 | |
| ENSMUSG00000092702 | 1 | |
| ENSMUSG00000092704 | 1 | |
| ENSMUSG00000092705 | 1 | |
| ENSMUSG00000092706 | 1 | |
| ENSMUSG00000092710 | 1 | |
| ENSMUSG00000092713 | 1.132639128 | 0.217803313 |
| ENSMUSG00000092714 | 1 | |
| ENSMUSG00000092715 | 1 | |
| ENSMUSG00000092718 | -1 | 1 |

| | | |
|---|---|---|
| ENSMUSG00000092719 | -1.043306199 | 0.698112512 |
| ENSMUSG00000092720 | 1 | |
| ENSMUSG00000092723 | 1 | |
| ENSMUSG00000092724 | 1 | |
| ENSMUSG00000092725 | 1 | |
| ENSMUSG00000092727 | -1.069502966 | 0.698112512 |
| ENSMUSG00000092730 | 1 | |
| ENSMUSG00000092731 | 1 | |
| ENSMUSG00000092732 | 1 | |
| ENSMUSG00000092734 | -1.03962336 | 0.698112512 |
| ENSMUSG00000092737 | 1 | |
| ENSMUSG00000092738 | 1 | |
| ENSMUSG00000092741 | -1.043306199 | 0.698112512 |
| ENSMUSG00000092743 | 1 | |
| ENSMUSG00000092745 | 1 | |
| ENSMUSG00000092746 | 1 | |
| ENSMUSG00000092747 | 1 | |
| ENSMUSG00000092748 | 1 | |
| ENSMUSG00000092751 | 1 | |
| ENSMUSG00000092752 | 1 | |
| ENSMUSG00000092753 | 1 | |
| ENSMUSG00000092758 | 1 | |
| ENSMUSG00000092759 | 1 | |
| ENSMUSG00000092760 | -1.08081673 | 0.733498571 |
| ENSMUSG00000092761 | 1 | |
| ENSMUSG00000092762 | 1 | |
| ENSMUSG00000092763 | 1 | |
| ENSMUSG00000092764 | 1 | |
| ENSMUSG00000092765 | 1 | |
| ENSMUSG00000092767 | 1 | |
| ENSMUSG00000092770 | 1 | |
| ENSMUSG00000092774 | 1 | |
| ENSMUSG00000092776 | 1 | |
| ENSMUSG00000092777 | 1 | |
| ENSMUSG00000092781 | 1.0106701 | 1 |
| ENSMUSG00000092784 | 1 | |
| ENSMUSG00000092787 | 1 | |
| ENSMUSG00000092790 | 1 | |
| ENSMUSG00000092792 | 1 | |
| ENSMUSG00000092793 | 1 | |
| ENSMUSG00000092794 | 1 | |
| ENSMUSG00000092795 | -1.041574599 | 0.886379277 |
| ENSMUSG00000092796 | 1 | |
| ENSMUSG00000092799 | 1 | |
| ENSMUSG00000092802 | -1.029302237 | 0.968410203 |
| ENSMUSG00000092805 | 1.07487334 | 0.698112512 |
| ENSMUSG00000092806 | 1 | |
| ENSMUSG00000092807 | 1 | |
| ENSMUSG00000092811 | 1 | |
| ENSMUSG00000092812 | 1 | |
| ENSMUSG00000092813 | 1 | |
| ENSMUSG00000092814 | 1 | |
| ENSMUSG00000092815 | 1 | |
| ENSMUSG00000092816 | -1.053549062 | 0.698112512 |
| ENSMUSG00000092818 | 1 | |
| ENSMUSG00000092819 | 1.049411284 | 0.883237063 |
| ENSMUSG00000092821 | 1 | |
| ENSMUSG00000092822 | 1 | |
| ENSMUSG00000092825 | 1 | |
| ENSMUSG00000092826 | 1 | |
| ENSMUSG00000092827 | -1.03962336 | 0.698112512 |
| ENSMUSG00000092829 | 1 | |
| ENSMUSG00000092830 | 1 | |

| | | |
|---|---|---|
| ENSMUSG00000092832 | 1 | |
| ENSMUSG00000092833 | 1 | |
| ENSMUSG00000092834 | 1 | |
| ENSMUSG00000092835 | 1 | |
| ENSMUSG00000092837 | -1.043306199 | 0.698112512 |
| ENSMUSG00000092838 | 1 | |
| ENSMUSG00000092840 | 1 | |
| ENSMUSG00000092841 | -1 | 1 |
| ENSMUSG00000092842 | 1 | |
| ENSMUSG00000092843 | 1 | |
| ENSMUSG00000092845 | 1 | |
| ENSMUSG00000092847 | 1 | |
| ENSMUSG00000092848 | 1 | |
| ENSMUSG00000092849 | 1.059463094 | 0.698112512 |
| ENSMUSG00000092850 | 1 | |
| ENSMUSG00000092851 | 1 | |
| ENSMUSG00000092852 | 1 | |
| ENSMUSG00000092853 | 1 | |
| ENSMUSG00000092855 | 1 | |
| ENSMUSG00000092856 | 1 | |
| ENSMUSG00000092862 | 1 | |
| ENSMUSG00000092866 | 1 | |
| ENSMUSG00000092867 | 1.355798509 | 0.065183251 |
| ENSMUSG00000092868 | 1 | |
| ENSMUSG00000092870 | 1 | |
| ENSMUSG00000092871 | 1 | |
| ENSMUSG00000092875 | 1 | |
| ENSMUSG00000092876 | 1 | |
| ENSMUSG00000092878 | 1 | |
| ENSMUSG00000092879 | 1 | |
| ENSMUSG00000092880 | 1 | |
| ENSMUSG00000092885 | 1 | |
| ENSMUSG00000092887 | 1.034865357 | 0.971519533 |
| ENSMUSG00000092888 | 1 | |
| ENSMUSG00000092889 | 1.039816326 | 0.897684198 |
| ENSMUSG00000092891 | 1 | |
| ENSMUSG00000092894 | 1 | |
| ENSMUSG00000092896 | 1 | |
| ENSMUSG00000092897 | 1 | |
| ENSMUSG00000092898 | 1 | |
| ENSMUSG00000092901 | 1 | |
| ENSMUSG00000092903 | -1.147838869 | 0.195607801 |
| ENSMUSG00000092905 | -1.254679684 | 0.121150119 |
| ENSMUSG00000092906 | 1 | |
| ENSMUSG00000092907 | 1 | |
| ENSMUSG00000092908 | 1 | |
| ENSMUSG00000092909 | 1.154253937 | 0.452069807 |
| ENSMUSG00000092912 | 1 | |
| ENSMUSG00000092913 | 1 | |
| ENSMUSG00000092914 | 1 | |
| ENSMUSG00000092915 | 1 | |
| ENSMUSG00000092916 | 1 | |
| ENSMUSG00000092917 | 1 | |
| ENSMUSG00000092918 | -1.063525418 | 0.756517167 |
| ENSMUSG00000092919 | 1 | |
| ENSMUSG00000092920 | 1 | |
| ENSMUSG00000092921 | 1 | |
| ENSMUSG00000092922 | 1 | |
| ENSMUSG00000092923 | 1 | |
| ENSMUSG00000092928 | 1 | |
| ENSMUSG00000092931 | 1 | |
| ENSMUSG00000092932 | 1.129030327 | 0.653331421 |
| ENSMUSG00000092934 | 1 | |

| | | |
|---|---|---|
| ENSMUSG00000092935 | 1 | |
| ENSMUSG00000092939 | 1 | |
| ENSMUSG00000092948 | 1 | |
| ENSMUSG00000092949 | 1 | |
| ENSMUSG00000092952 | 1 | |
| ENSMUSG00000092953 | 1 | |
| ENSMUSG00000092956 | 1 | |
| ENSMUSG00000092957 | 1 | |
| ENSMUSG00000092958 | 1 | |
| ENSMUSG00000092961 | 1 | |
| ENSMUSG00000092963 | 1 | |
| ENSMUSG00000092965 | 1 | |
| ENSMUSG00000092966 | 1 | |
| ENSMUSG00000092972 | 1 | |
| ENSMUSG00000092973 | 1 | |
| ENSMUSG00000092974 | 1 | |
| ENSMUSG00000092978 | 1 | |
| ENSMUSG00000092979 | 1 | |
| ENSMUSG00000092981 | 1.120616814 | 0.818942935 |
| ENSMUSG00000092985 | 1 | |
| ENSMUSG00000092986 | -1.028348525 | 0.936995747 |
| ENSMUSG00000092988 | 1 | |
| ENSMUSG00000092990 | 1.059463094 | 0.608497229 |
| ENSMUSG00000092994 | 1 | |
| ENSMUSG00000092995 | 1 | |
| ENSMUSG00000092997 | 1 | |
| ENSMUSG00000092998 | 1 | |
| ENSMUSG00000092999 | 1 | |
| ENSMUSG00000093000 | 1 | |
| ENSMUSG00000093002 | 1 | |
| ENSMUSG00000093003 | 1 | |
| ENSMUSG00000093004 | 1 | |
| ENSMUSG00000093005 | 1 | |
| ENSMUSG00000093006 | 1 | |
| ENSMUSG00000093007 | -1 | 1 |
| ENSMUSG00000093009 | 1 | |
| ENSMUSG00000093010 | 1 | |
| ENSMUSG00000093011 | 1 | |
| ENSMUSG00000093013 | 1 | |
| ENSMUSG00000093014 | 1 | |
| ENSMUSG00000093015 | 1 | |
| ENSMUSG00000093017 | 1 | |
| ENSMUSG00000093019 | 1 | |
| ENSMUSG00000093021 | 1 | |
| ENSMUSG00000093023 | 1.141931237 | 0.592727722 |
| ENSMUSG00000093024 | 1 | |
| ENSMUSG00000093025 | 1 | |
| ENSMUSG00000093026 | 1 | |
| ENSMUSG00000093029 | 1 | |
| ENSMUSG00000093030 | 1.059463094 | 0.698112512 |
| ENSMUSG00000093031 | 1 | |
| ENSMUSG00000093032 | -1.044273782 | 0.698112512 |
| ENSMUSG00000093033 | 1 | |
| ENSMUSG00000093034 | 1 | |
| ENSMUSG00000093036 | 1 | |
| ENSMUSG00000093037 | 1 | |
| ENSMUSG00000093039 | 1 | |
| ENSMUSG00000093040 | -1 | 1 |
| ENSMUSG00000093041 | 1 | |
| ENSMUSG00000093042 | 1 | |
| ENSMUSG00000093044 | 1.017174474 | 0.995295685 |
| ENSMUSG00000093046 | 1 | |
| ENSMUSG00000093051 | 1 | |

| | | |
|---|---|---|
| ENSMUSG00000093052 | 1 | |
| ENSMUSG00000093054 | 1 | |
| ENSMUSG00000093056 | 1 | |
| ENSMUSG00000093058 | 1 | |
| ENSMUSG00000093059 | 1.030394552 | 0.968575228 |
| ENSMUSG00000093060 | 1 | |
| ENSMUSG00000093061 | 1 | |
| ENSMUSG00000093063 | 1 | |
| ENSMUSG00000093064 | 1 | |
| ENSMUSG00000093065 | 1 | |
| ENSMUSG00000093067 | 1 | |
| ENSMUSG00000093070 | 1 | |
| ENSMUSG00000093071 | 1 | |
| ENSMUSG00000093072 | 1 | |
| ENSMUSG00000093073 | 1 | |
| ENSMUSG00000093074 | 1 | |
| ENSMUSG00000093075 | -1.111880267 | 0.70299173 |
| ENSMUSG00000093076 | 1 | |
| ENSMUSG00000093079 | 1 | |
| ENSMUSG00000093080 | 1.122128755 | 0.61203231 |
| ENSMUSG00000093081 | 1 | |
| ENSMUSG00000093082 | 1 | |
| ENSMUSG00000093083 | 1 | |
| ENSMUSG00000093085 | -1.028348525 | 0.940333715 |
| ENSMUSG00000093087 | 1 | |
| ENSMUSG00000093089 | 1 | |
| ENSMUSG00000093090 | 1 | |
| ENSMUSG00000093091 | 1 | |
| ENSMUSG00000093092 | 1 | |
| ENSMUSG00000093093 | 1.04328066 | 0.867378212 |
| ENSMUSG00000093096 | 1 | |
| ENSMUSG00000093097 | 1 | |
| ENSMUSG00000093100 | 1 | |
| ENSMUSG00000093101 | 1 | |
| ENSMUSG00000093102 | 1 | |
| ENSMUSG00000093105 | -1.014570171 | 0.988424085 |
| ENSMUSG00000093106 | 1 | |
| ENSMUSG00000093107 | 1 | |
| ENSMUSG00000093108 | 1 | |
| ENSMUSG00000093112 | 1 | |
| ENSMUSG00000093113 | 1 | |
| ENSMUSG00000093114 | 1 | |
| ENSMUSG00000093115 | 1 | |
| ENSMUSG00000093116 | 1 | |
| ENSMUSG00000093117 | 1 | |
| ENSMUSG00000093118 | 1 | |
| ENSMUSG00000093119 | -1.018605114 | 1 |
| ENSMUSG00000093120 | -1.03962336 | 0.698112512 |
| ENSMUSG00000093122 | 1 | |
| ENSMUSG00000093124 | 1.089470641 | 0.657060635 |
| ENSMUSG00000093125 | 1 | |
| ENSMUSG00000093128 | 1.069068979 | 0.608497229 |
| ENSMUSG00000093129 | 1 | |
| ENSMUSG00000093132 | 1 | |
| ENSMUSG00000093133 | 1 | |
| ENSMUSG00000093134 | 1.014545335 | 0.989648415 |
| ENSMUSG00000093137 | 1 | |
| ENSMUSG00000093138 | 1 | |
| ENSMUSG00000093139 | 1 | |
| ENSMUSG00000093140 | 1 | |
| ENSMUSG00000093142 | 1 | |
| ENSMUSG00000093147 | -1.095294215 | 0.541735653 |
| ENSMUSG00000093148 | 1 | |

| | | |
|---|---|---|
| ENSMUSG00000093149 | 1 | |
| ENSMUSG00000093152 | 1 | |
| ENSMUSG00000093153 | 1 | |
| ENSMUSG00000093155 | -1.113515092 | 0.576169279 |
| ENSMUSG00000093156 | -1.03962336 | 0.698112512 |
| ENSMUSG00000093157 | 1 | |
| ENSMUSG00000093158 | 1 | |
| ENSMUSG00000093159 | 1 | |
| ENSMUSG00000093160 | -1 | 1 |
| ENSMUSG00000093161 | -1.03962336 | 0.698112512 |
| ENSMUSG00000093162 | 1 | |
| ENSMUSG00000093163 | -1.044273782 | 0.698112512 |
| ENSMUSG00000093164 | -1.099174267 | 0.608497229 |
| ENSMUSG00000093167 | 1 | |
| ENSMUSG00000093168 | -1.099174267 | 0.535052277 |
| ENSMUSG00000093169 | 1 | |
| ENSMUSG00000093170 | 1 | |
| ENSMUSG00000093172 | 1 | |
| ENSMUSG00000093173 | 1 | |
| ENSMUSG00000093176 | 1.014545335 | 0.989648415 |
| ENSMUSG00000093178 | 1.008856821 | 1 |
| ENSMUSG00000093180 | 1.059463094 | 0.608497229 |
| ENSMUSG00000093183 | 1 | |
| ENSMUSG00000093184 | 1 | |
| ENSMUSG00000093185 | 1 | |
| ENSMUSG00000093187 | 1 | |
| ENSMUSG00000093189 | 1 | |
| ENSMUSG00000093190 | -1.121773109 | 0.612381584 |
| ENSMUSG00000093191 | 1 | |
| ENSMUSG00000093192 | 1 | |
| ENSMUSG00000093202 | 1 | |
| ENSMUSG00000093203 | -1.009476377 | 1 |
| ENSMUSG00000093206 | 1 | |
| ENSMUSG00000093207 | 1 | |
| ENSMUSG00000093208 | 1 | |
| ENSMUSG00000093211 | 1 | |
| ENSMUSG00000093212 | 1 | |
| ENSMUSG00000093214 | 1 | |
| ENSMUSG00000093215 | 1 | |
| ENSMUSG00000093216 | 1 | |
| ENSMUSG00000093217 | 1 | |
| ENSMUSG00000093218 | 1 | |
| ENSMUSG00000093219 | 1 | |
| ENSMUSG00000093221 | 1 | |
| ENSMUSG00000093223 | 1 | |
| ENSMUSG00000093224 | 1 | |
| ENSMUSG00000093226 | 1 | |
| ENSMUSG00000093227 | 1 | |
| ENSMUSG00000093228 | 1 | |
| ENSMUSG00000093230 | 1 | |
| ENSMUSG00000093231 | -1.017305534 | 0.98847058 |
| ENSMUSG00000093232 | 1 | |
| ENSMUSG00000093234 | 1 | |
| ENSMUSG00000093236 | 1 | |
| ENSMUSG00000093238 | 1 | |
| ENSMUSG00000093239 | 1 | |
| ENSMUSG00000093240 | 1 | |
| ENSMUSG00000093241 | 1 | |
| ENSMUSG00000093242 | 1 | |
| ENSMUSG00000093243 | 1 | |
| ENSMUSG00000093245 | 1 | |
| ENSMUSG00000093246 | 1 | |
| ENSMUSG00000093249 | -1.038444538 | 0.893582779 |

| | | |
|---|---|---|
| ENSMUSG00000093251 | 1 | |
| ENSMUSG00000093254 | 1 | |
| ENSMUSG00000093256 | -1 | 1 |
| ENSMUSG00000093258 | 1 | |
| ENSMUSG00000093260 | -1.044273782 | 0.698112512 |
| ENSMUSG00000093261 | 1 | |
| ENSMUSG00000093262 | 1 | |
| ENSMUSG00000093264 | 1.128878069 | 0.608497229 |
| ENSMUSG00000093266 | 1 | |
| ENSMUSG00000093269 | 1 | |
| ENSMUSG00000093270 | 1.059463094 | 0.608497229 |
| ENSMUSG00000093271 | 1 | |
| ENSMUSG00000093273 | 1 | |
| ENSMUSG00000093274 | 1 | |
| ENSMUSG00000093276 | -1.043306199 | 0.79291651 |
| ENSMUSG00000093277 | 1 | |
| ENSMUSG00000093278 | 1 | |
| ENSMUSG00000093279 | 1 | |
| ENSMUSG00000093280 | 1 | |
| ENSMUSG00000093281 | 1 | |
| ENSMUSG00000093282 | 1.059463094 | 0.608497229 |
| ENSMUSG00000093283 | -1 | 1 |
| ENSMUSG00000093286 | -1 | 1 |
| ENSMUSG00000093289 | -1 | 1 |
| ENSMUSG00000093290 | 1 | |
| ENSMUSG00000093291 | 1 | |
| ENSMUSG00000093292 | 1 | |
| ENSMUSG00000093293 | 1 | |
| ENSMUSG00000093294 | 1 | |
| ENSMUSG00000093296 | 1 | |
| ENSMUSG00000093298 | 1 | |
| ENSMUSG00000093300 | 1 | |
| ENSMUSG00000093301 | 1 | |
| ENSMUSG00000093302 | 1 | |
| ENSMUSG00000093303 | 1 | |
| ENSMUSG00000093306 | 1 | |
| ENSMUSG00000093311 | 1 | |
| ENSMUSG00000093314 | 1.014545335 | 0.979002724 |
| ENSMUSG00000093315 | 1 | |
| ENSMUSG00000093317 | 1 | |
| ENSMUSG00000093319 | -1.075182332 | 0.826758538 |
| ENSMUSG00000093322 | 1 | |
| ENSMUSG00000093323 | 1 | |
| ENSMUSG00000093325 | 1 | |
| ENSMUSG00000093327 | -1 | 1 |
| ENSMUSG00000093329 | 1 | |
| ENSMUSG00000093331 | 1 | |
| ENSMUSG00000093334 | 1 | |
| ENSMUSG00000093335 | 1 | |
| ENSMUSG00000093336 | 1 | |
| ENSMUSG00000093338 | 1 | |
| ENSMUSG00000093340 | 1 | |
| ENSMUSG00000093341 | 1 | |
| ENSMUSG00000093342 | 1 | |
| ENSMUSG00000093344 | 1 | |
| ENSMUSG00000093345 | -1.03962336 | 0.698112512 |
| ENSMUSG00000093346 | 1 | |
| ENSMUSG00000093348 | 1 | |
| ENSMUSG00000093351 | 1.059463094 | 0.698112512 |
| ENSMUSG00000093352 | 1 | |
| ENSMUSG00000093354 | 1 | |
| ENSMUSG00000093355 | 1 | |
| ENSMUSG00000093356 | 1 | |

| | | |
|---|---|---|
| ENSMUSG00000093358 | 1 | |
| ENSMUSG00000093359 | 1 | |
| ENSMUSG00000093362 | 1.059463094 | 0.608497229 |
| ENSMUSG00000093363 | 1.040172434 | 0.918344559 |
| ENSMUSG00000093365 | 1 | |
| ENSMUSG00000093368 | 1 | |
| ENSMUSG00000093373 | 1 | |
| ENSMUSG00000093376 | 1 | |
| ENSMUSG00000093378 | 1.067157119 | 0.735204038 |
| ENSMUSG00000093379 | 1 | |
| ENSMUSG00000093380 | 1 | |
| ENSMUSG00000093381 | 1 | |
| ENSMUSG00000093382 | 1 | |
| ENSMUSG00000093383 | 1 | |
| ENSMUSG00000093384 | -1.044273782 | 0.698112512 |
| ENSMUSG00000093385 | -1.02723833 | 0.895798715 |
| ENSMUSG00000093386 | 1 | |
| ENSMUSG00000093387 | 1 | |
| ENSMUSG00000093388 | 1.059463094 | 0.699340382 |
| ENSMUSG00000093390 | -1.084645496 | 0.608497229 |
| ENSMUSG00000093392 | 1 | |
| ENSMUSG00000093394 | -1.052598651 | 0.894788694 |
| ENSMUSG00000093395 | 1 | |
| ENSMUSG00000093397 | 1 | |
| ENSMUSG00000093398 | 1 | |
| ENSMUSG00000093400 | 1 | |
| ENSMUSG00000093401 | 1 | |
| ENSMUSG00000093402 | 1 | |
| ENSMUSG00000093403 | 1 | |
| ENSMUSG00000093404 | 1.014545335 | 0.979002724 |
| ENSMUSG00000093405 | 1 | |
| ENSMUSG00000093406 | 1 | |
| ENSMUSG00000093408 | 1 | |
| ENSMUSG00000093409 | 1.005613439 | 1 |
| ENSMUSG00000093410 | -1 | 1 |
| ENSMUSG00000093411 | 1 | |
| ENSMUSG00000093413 | 1.059463094 | 0.608497229 |
| ENSMUSG00000093414 | 1 | |
| ENSMUSG00000093416 | -1.022788389 | 1 |
| ENSMUSG00000093418 | 1.069068979 | 0.608497229 |
| ENSMUSG00000093419 | 1 | |
| ENSMUSG00000093420 | 1 | |
| ENSMUSG00000093421 | 1 | |
| ENSMUSG00000093424 | 1.009571778 | 1 |
| ENSMUSG00000093426 | 1 | |
| ENSMUSG00000093427 | 1 | |
| ENSMUSG00000093428 | 1 | |
| ENSMUSG00000093429 | 1.059463094 | 0.698112512 |
| ENSMUSG00000093433 | 1 | |
| ENSMUSG00000093434 | 1 | |
| ENSMUSG00000093436 | 1 | |
| ENSMUSG00000093437 | -1.175275973 | 0.608497229 |
| ENSMUSG00000093441 | 1 | |
| ENSMUSG00000093443 | 1 | |
| ENSMUSG00000093444 | 1 | |
| ENSMUSG00000093445 | 1.017233885 | 1 |
| ENSMUSG00000093446 | 1 | |
| ENSMUSG00000093447 | -1.075161737 | 0.698112512 |
| ENSMUSG00000093449 | 1 | |
| ENSMUSG00000093451 | 1 | |
| ENSMUSG00000093452 | 1 | |
| ENSMUSG00000093453 | 1 | |
| ENSMUSG00000093454 | 1 | |

| | | |
|---|---|---|
| ENSMUSG00000093456 | 1 | |
| ENSMUSG00000093458 | -1.113515092 | 0.576169279 |
| ENSMUSG00000093460 | -1.478331735 | 0.035541651 |
| ENSMUSG00000093461 | 1 | |
| ENSMUSG00000093465 | 1.015486245 | 0.972980766 |
| ENSMUSG00000093467 | 1 | |
| ENSMUSG00000093469 | 1 | |
| ENSMUSG00000093472 | 1 | |
| ENSMUSG00000093473 | 1 | |
| ENSMUSG00000093474 | 1 | |
| ENSMUSG00000093476 | -1.142727245 | 0.20412635 |
| ENSMUSG00000093479 | 1 | |
| ENSMUSG00000093480 | 1 | |
| ENSMUSG00000093482 | 1 | |
| ENSMUSG00000093483 | 1.005957997 | 0.997392688 |
| ENSMUSG00000093484 | -1.044273782 | 0.698112512 |
| ENSMUSG00000093485 | -1.03962336 | 0.698112512 |
| ENSMUSG00000093487 | 1 | |
| ENSMUSG00000093489 | 1 | |
| ENSMUSG00000093490 | 1 | |
| ENSMUSG00000093492 | 1 | |
| ENSMUSG00000093493 | 1 | |
| ENSMUSG00000093495 | 1.06585783 | 0.802248757 |
| ENSMUSG00000093497 | 1 | |
| ENSMUSG00000093498 | 1 | |
| ENSMUSG00000093499 | 1 | |
| ENSMUSG00000093501 | 1 | |
| ENSMUSG00000093502 | 1 | |
| ENSMUSG00000093503 | 1 | |
| ENSMUSG00000093504 | 1 | |
| ENSMUSG00000093505 | 1 | |
| ENSMUSG00000093507 | 1.122462048 | 0.693968585 |
| ENSMUSG00000093508 | 1 | |
| ENSMUSG00000093509 | 1 | |
| ENSMUSG00000093510 | 1 | |
| ENSMUSG00000093511 | 1 | |
| ENSMUSG00000093513 | 1 | |
| ENSMUSG00000093514 | 1.166591266 | 0.608497229 |
| ENSMUSG00000093515 | 1.069068979 | 0.608497229 |
| ENSMUSG00000093516 | 1 | |
| ENSMUSG00000093518 | 1 | |
| ENSMUSG00000093519 | 1 | |
| ENSMUSG00000093520 | 1 | |
| ENSMUSG00000093522 | 1 | |
| ENSMUSG00000093523 | 1 | |
| ENSMUSG00000093525 | 1 | |
| ENSMUSG00000093528 | 1 | |
| ENSMUSG00000093529 | 1 | |
| ENSMUSG00000093530 | 1 | |
| ENSMUSG00000093531 | 1 | |
| ENSMUSG00000093533 | 1 | |
| ENSMUSG00000093534 | 1 | |
| ENSMUSG00000093535 | -1.161624871 | 0.553671405 |
| ENSMUSG00000093536 | -1.189350556 | 0.447072383 |
| ENSMUSG00000093537 | 1 | |
| ENSMUSG00000093538 | -1.184259924 | 0.343197313 |
| ENSMUSG00000093539 | 1 | |
| ENSMUSG00000093540 | 1 | |
| ENSMUSG00000093541 | 1.139018652 | 0.698112512 |
| ENSMUSG00000093543 | -1.03962336 | 0.803072249 |
| ENSMUSG00000093544 | 1 | |
| ENSMUSG00000093545 | 1.005613439 | 1 |
| ENSMUSG00000093546 | 1 | |

| | | |
|---|---|---|
| ENSMUSG00000093547 | 1 | |
| ENSMUSG00000093548 | 1.230075506 | 0.292973656 |
| ENSMUSG00000093549 | 1 | |
| ENSMUSG00000093550 | 1 | |
| ENSMUSG00000093551 | 1 | |
| ENSMUSG00000093552 | -1.006541125 | 1 |
| ENSMUSG00000093553 | 1.281716071 | 0.139199954 |
| ENSMUSG00000093554 | 1 | |
| ENSMUSG00000093556 | 1 | |
| ENSMUSG00000093557 | 1 | |
| ENSMUSG00000093559 | 1.171563499 | 0.233412931 |
| ENSMUSG00000093560 | 1.014545335 | 0.989648415 |
| ENSMUSG00000093561 | -1.176660386 | 0.420817778 |
| ENSMUSG00000093562 | -1 | 1 |
| ENSMUSG00000093563 | 1 | |
| ENSMUSG00000093564 | 1 | |
| ENSMUSG00000093565 | 1.013046788 | 0.979002724 |
| ENSMUSG00000093567 | 1 | |
| ENSMUSG00000093568 | 1 | |
| ENSMUSG00000093570 | 1 | |
| ENSMUSG00000093571 | 1 | |
| ENSMUSG00000093572 | -1.03962336 | 0.698112512 |
| ENSMUSG00000093573 | -1.044273782 | 0.790851299 |
| ENSMUSG00000093574 | 1 | |
| ENSMUSG00000093575 | 1.069068979 | 0.698112512 |
| ENSMUSG00000093577 | 1.271444129 | 0.414138469 |
| ENSMUSG00000093579 | 1.023125763 | 0.887851178 |
| ENSMUSG00000093580 | 1 | |
| ENSMUSG00000093581 | 1 | |
| ENSMUSG00000093582 | 1 | |
| ENSMUSG00000093584 | 1 | |
| ENSMUSG00000093587 | -1.205839554 | 0.505106302 |
| ENSMUSG00000093588 | 1 | |
| ENSMUSG00000093589 | -1.121487286 | 0.748791066 |
| ENSMUSG00000093590 | 1 | |
| ENSMUSG00000093592 | 1 | |
| ENSMUSG00000093593 | 1.059463094 | 0.608497229 |
| ENSMUSG00000093594 | 1.037967124 | 0.83493212 |
| ENSMUSG00000093595 | 1 | |
| ENSMUSG00000093598 | 1 | |
| ENSMUSG00000093599 | -1 | 1 |
| ENSMUSG00000093601 | 1 | |
| ENSMUSG00000093602 | 1 | |
| ENSMUSG00000093606 | -1.0379631 | 0.924629556 |
| ENSMUSG00000093607 | 1 | |
| ENSMUSG00000093610 | 1 | |
| ENSMUSG00000093614 | -1.03962336 | 0.698112512 |
| ENSMUSG00000093615 | 1 | |
| ENSMUSG00000093616 | 1 | |
| ENSMUSG00000093617 | 1 | |
| ENSMUSG00000093618 | 1 | |
| ENSMUSG00000093619 | 1 | |
| ENSMUSG00000093621 | 1 | |
| ENSMUSG00000093622 | 1.059463094 | 0.704170349 |
| ENSMUSG00000093623 | 1 | |
| ENSMUSG00000093624 | 1 | |
| ENSMUSG00000093625 | 1 | |
| ENSMUSG00000093626 | 1 | |
| ENSMUSG00000093627 | 1 | |
| ENSMUSG00000093628 | 1 | |
| ENSMUSG00000093629 | 1.188570462 | 0.576169279 |
| ENSMUSG00000093632 | 1 | |
| ENSMUSG00000093634 | -1.044273782 | 0.698112512 |

| | | |
|---|---|---|
| ENSMUSG00000093635 | 1 | |
| ENSMUSG00000093637 | 1.06536899 | 0.759071586 |
| ENSMUSG00000093639 | 1 | |
| ENSMUSG00000093642 | 1 | |
| ENSMUSG00000093644 | 1 | |
| ENSMUSG00000093645 | 1 | |
| ENSMUSG00000093648 | 1 | |
| ENSMUSG00000093649 | 1.059463094 | 0.608497229 |
| ENSMUSG00000093650 | -1.013742517 | 0.969337248 |
| ENSMUSG00000093651 | 1.059463094 | 0.704170349 |
| ENSMUSG00000093652 | 1 | |
| ENSMUSG00000093656 | 1.132877091 | 0.608497229 |
| ENSMUSG00000093658 | -1 | 1 |
| ENSMUSG00000093660 | 1 | |
| ENSMUSG00000093661 | -1.106817083 | 0.522542762 |
| ENSMUSG00000093662 | 1 | |
| ENSMUSG00000093663 | 1 | |
| ENSMUSG00000093664 | 1 | |
| ENSMUSG00000093665 | 1 | |
| ENSMUSG00000093666 | 1 | |
| ENSMUSG00000093668 | 1.13926696 | 0.587480427 |
| ENSMUSG00000093669 | 1.059463094 | 0.704170349 |
| ENSMUSG00000093671 | 1 | |
| ENSMUSG00000093672 | 1.068703605 | 0.918774962 |
| ENSMUSG00000093673 | -1.071543562 | 0.710698346 |
| ENSMUSG00000093674 | -1.001943913 | 1 |
| ENSMUSG00000093675 | 1 | |
| ENSMUSG00000093677 | 1.026628836 | 0.815715406 |
| ENSMUSG00000093679 | 1 | |
| ENSMUSG00000093681 | 1 | |
| ENSMUSG00000093682 | -1.163954093 | 0.583458429 |
| ENSMUSG00000093683 | -1 | 1 |
| ENSMUSG00000093684 | 1 | |
| ENSMUSG00000093685 | 1 | |
| ENSMUSG00000093686 | -1 | 1 |
| ENSMUSG00000093687 | 1.052739823 | 0.891373394 |
| ENSMUSG00000093688 | 1 | |
| ENSMUSG00000093689 | 1 | |
| ENSMUSG00000093690 | -1 | 1 |
| ENSMUSG00000093692 | 1 | |
| ENSMUSG00000093695 | 1.102285378 | 0.693578301 |
| ENSMUSG00000093696 | 1 | |
| ENSMUSG00000093697 | 1 | |
| ENSMUSG00000093699 | 1 | |
| ENSMUSG00000093701 | 1 | |
| ENSMUSG00000093702 | 1 | |
| ENSMUSG00000093703 | 1 | |
| ENSMUSG00000093705 | 1 | |
| ENSMUSG00000093707 | 1 | |
| ENSMUSG00000093708 | 1 | |
| ENSMUSG00000093710 | 1 | |
| ENSMUSG00000093711 | 1 | |
| ENSMUSG00000093714 | 1 | |
| ENSMUSG00000093716 | 1.024656056 | 0.972980766 |
| ENSMUSG00000093719 | 1 | |
| ENSMUSG00000093720 | -1.03962336 | 0.698112512 |
| ENSMUSG00000093721 | 1 | |
| ENSMUSG00000093722 | 1 | |
| ENSMUSG00000093724 | 1 | |
| ENSMUSG00000093725 | 1 | |
| ENSMUSG00000093726 | -1.187649297 | 0.132086791 |
| ENSMUSG00000093727 | 1 | |
| ENSMUSG00000093728 | 1 | |

| | | |
|---|---|---|
| ENSMUSG00000093729 | 1 | |
| ENSMUSG00000093730 | 1 | |
| ENSMUSG00000093731 | 1 | |
| ENSMUSG00000093732 | 1 | |
| ENSMUSG00000093734 | -1.029956602 | 0.882657068 |
| ENSMUSG00000093736 | 1 | |
| ENSMUSG00000093738 | 1 | |
| ENSMUSG00000093739 | -1.089987187 | 0.803343809 |
| ENSMUSG00000093740 | 1.059463094 | 0.608497229 |
| ENSMUSG00000093741 | 1 | |
| ENSMUSG00000093743 | 1 | |
| ENSMUSG00000093744 | 1 | |
| ENSMUSG00000093745 | 1 | |
| ENSMUSG00000093746 | 1 | |
| ENSMUSG00000093747 | -1.086192882 | 0.698112512 |
| ENSMUSG00000093748 | 1 | |
| ENSMUSG00000093749 | -1.028972438 | 0.96504537 |
| ENSMUSG00000093751 | -1.02664175 | 0.990831634 |
| ENSMUSG00000093752 | 1 | |
| ENSMUSG00000093753 | 1 | |
| ENSMUSG00000093755 | 1 | |
| ENSMUSG00000093756 | 1 | |
| ENSMUSG00000093757 | 1 | |
| ENSMUSG00000093758 | -1.053549062 | 0.698112512 |
| ENSMUSG00000093759 | -1 | 1 |
| ENSMUSG00000093760 | 1.135412481 | 0.226472473 |
| ENSMUSG00000093761 | 1 | |
| ENSMUSG00000093762 | 1 | |
| ENSMUSG00000093763 | 1 | |
| ENSMUSG00000093764 | 1 | |
| ENSMUSG00000093765 | 1 | |
| ENSMUSG00000093766 | 1 | |
| ENSMUSG00000093767 | 1 | |
| ENSMUSG00000093768 | 1 | |
| ENSMUSG00000093769 | 1 | |
| ENSMUSG00000093770 | 1 | |
| ENSMUSG00000093771 | 1 | |
| ENSMUSG00000093772 | -1.044273782 | 0.855488553 |
| ENSMUSG00000093773 | 1 | |
| ENSMUSG00000093774 | 1.04328066 | 0.86119789 |
| ENSMUSG00000093775 | 1 | |
| ENSMUSG00000093776 | 1 | |
| ENSMUSG00000093779 | -1 | 1 |
| ENSMUSG00000093780 | 1.218240674 | 0.294039426 |
| ENSMUSG00000093781 | 1 | |
| ENSMUSG00000093782 | 1 | |
| ENSMUSG00000093784 | 1 | |
| ENSMUSG00000093785 | 1 | |
| ENSMUSG00000093787 | 1 | |
| ENSMUSG00000093788 | 1 | |
| ENSMUSG00000093789 | 1 | |
| ENSMUSG00000093791 | 1 | |
| ENSMUSG00000093793 | 1 | |
| ENSMUSG00000093795 | 1 | |
| ENSMUSG00000093796 | 1 | |
| ENSMUSG00000093798 | -1 | 1 |
| ENSMUSG00000093800 | -1.018570571 | 0.968483554 |
| ENSMUSG00000093801 | 1 | |
| ENSMUSG00000093802 | 1 | |
| ENSMUSG00000093803 | 1.07937145 | 0.846386256 |
| ENSMUSG00000093804 | 1 | |
| ENSMUSG00000093805 | 1.150137535 | 0.624250802 |
| ENSMUSG00000093806 | 1 | |

| | | |
|---|---|---|
| ENSMUSG00000093807 | 1 | |
| ENSMUSG00000093808 | 1 | |
| ENSMUSG00000093812 | 1 | |
| ENSMUSG00000093813 | 1 | |
| ENSMUSG00000093814 | 1 | |
| ENSMUSG00000093815 | 1 | |
| ENSMUSG00000093816 | 1 | |
| ENSMUSG00000093817 | 1 | |
| ENSMUSG00000093818 | 1 | |
| ENSMUSG00000093820 | 1 | |
| ENSMUSG00000093822 | 1 | |
| ENSMUSG00000093823 | 1 | |
| ENSMUSG00000093824 | 1 | |
| ENSMUSG00000093825 | 1 | |
| ENSMUSG00000093826 | 1 | |
| ENSMUSG00000093827 | 1 | |
| ENSMUSG00000093828 | 1 | |
| ENSMUSG00000093829 | 1 | |
| ENSMUSG00000093830 | 1 | |
| ENSMUSG00000093831 | 1 | |
| ENSMUSG00000093833 | 1 | |
| ENSMUSG00000093834 | 1 | |
| ENSMUSG00000093835 | 1 | |
| ENSMUSG00000093838 | 1 | |
| ENSMUSG00000093839 | 1 | |
| ENSMUSG00000093843 | 1.059463094 | 0.608497229 |
| ENSMUSG00000093844 | 1 | |
| ENSMUSG00000093846 | 1.29655355 | 0.085709466 |
| ENSMUSG00000093847 | 1.059463094 | 0.608497229 |
| ENSMUSG00000093848 | 1 | |
| ENSMUSG00000093849 | 1.161036672 | 0.251049598 |
| ENSMUSG00000093851 | 1 | |
| ENSMUSG00000093852 | 1 | |
| ENSMUSG00000093853 | 1 | |
| ENSMUSG00000093854 | 1 | |
| ENSMUSG00000093856 | 1 | |
| ENSMUSG00000093857 | 1 | |
| ENSMUSG00000093858 | -1.052411173 | 0.899648217 |
| ENSMUSG00000093859 | -1.139116 | 0.608497229 |
| ENSMUSG00000093860 | 1 | |
| ENSMUSG00000093861 | 1 | |
| ENSMUSG00000093862 | 1.116201826 | 0.608497229 |
| ENSMUSG00000093863 | 1 | |
| ENSMUSG00000093864 | 1 | |
| ENSMUSG00000093865 | 1.028323353 | 0.903464611 |
| ENSMUSG00000093866 | 1 | |
| ENSMUSG00000093867 | 1.06712641 | 0.761839849 |
| ENSMUSG00000093868 | 1 | |
| ENSMUSG00000093869 | 1 | |
| ENSMUSG00000093870 | 1 | |
| ENSMUSG00000093871 | 1 | |
| ENSMUSG00000093873 | 1 | |
| ENSMUSG00000093874 | 1 | |
| ENSMUSG00000093875 | 1 | |
| ENSMUSG00000093876 | 1 | |
| ENSMUSG00000093877 | 1 | |
| ENSMUSG00000093878 | 1 | |
| ENSMUSG00000093879 | 1 | |
| ENSMUSG00000093880 | -1.338337601 | 0.061632252 |
| ENSMUSG00000093883 | 1 | |
| ENSMUSG00000093884 | 1 | |
| ENSMUSG00000093885 | 1.019083579 | 0.949812939 |
| ENSMUSG00000093886 | -1.03962336 | 0.803072249 |

| | | |
|---|---|---|
| ENSMUSG00000093887 | 1 | |
| ENSMUSG00000093889 | 1 | |
| ENSMUSG00000093890 | 1 | |
| ENSMUSG00000093891 | 1 | |
| ENSMUSG00000093892 | 1 | |
| ENSMUSG00000093893 | 1 | |
| ENSMUSG00000093894 | 1 | |
| ENSMUSG00000093895 | 1 | |
| ENSMUSG00000093896 | 1 | |
| ENSMUSG00000093897 | -1.044273782 | 0.847756749 |
| ENSMUSG00000093898 | 1 | |
| ENSMUSG00000093899 | -1 | 1 |
| ENSMUSG00000093900 | 1 | |
| ENSMUSG00000093901 | 1 | |
| ENSMUSG00000093902 | 1 | |
| ENSMUSG00000093903 | 1 | |
| ENSMUSG00000093904 | 1.016090633 | 1 |
| ENSMUSG00000093905 | 1 | |
| ENSMUSG00000093906 | 1 | |
| ENSMUSG00000093907 | 1 | |
| ENSMUSG00000093908 | -1.033292015 | 0.899159787 |
| ENSMUSG00000093909 | 1 | |
| ENSMUSG00000093910 | -1.003834322 | 1 |
| ENSMUSG00000093912 | -1.099174267 | 0.535052277 |
| ENSMUSG00000093913 | 1 | |
| ENSMUSG00000093914 | 1 | |
| ENSMUSG00000093915 | 1 | |
| ENSMUSG00000093916 | 1 | |
| ENSMUSG00000093917 | 1 | |
| ENSMUSG00000093918 | 1 | |
| ENSMUSG00000093919 | 1 | |
| ENSMUSG00000093920 | 1 | |
| ENSMUSG00000093921 | 1.116373378 | 0.698112512 |
| ENSMUSG00000093922 | 1 | |
| ENSMUSG00000093923 | 1 | |
| ENSMUSG00000093924 | 1 | |
| ENSMUSG00000093926 | 1 | |
| ENSMUSG00000093927 | 1 | |
| ENSMUSG00000093928 | 1 | |
| ENSMUSG00000093929 | 1 | |
| ENSMUSG00000093930 | -1.07472484 | 0.778889847 |
| ENSMUSG00000093931 | 1 | |
| ENSMUSG00000093932 | 1 | |
| ENSMUSG00000093933 | -1.159878974 | 0.608497229 |
| ENSMUSG00000093934 | 1 | |
| ENSMUSG00000093935 | 1 | |
| ENSMUSG00000093937 | 1 | |
| ENSMUSG00000093938 | 1 | |
| ENSMUSG00000093939 | 1 | |
| ENSMUSG00000093940 | 1 | |
| ENSMUSG00000093941 | 1 | |
| ENSMUSG00000093942 | 1 | |
| ENSMUSG00000093943 | 1 | |
| ENSMUSG00000093944 | 1 | |
| ENSMUSG00000093945 | 1 | |
| ENSMUSG00000093946 | 1 | |
| ENSMUSG00000093947 | 1 | |
| ENSMUSG00000093948 | 1 | |
| ENSMUSG00000093949 | 1 | |
| ENSMUSG00000093950 | 1 | |
| ENSMUSG00000093952 | 1 | |
| ENSMUSG00000093953 | 1 | |
| ENSMUSG00000093954 | 1 | |

| | | |
|---|---|---|
| ENSMUSG00000093955 | 1 | |
| ENSMUSG00000093956 | 1 | |
| ENSMUSG00000093957 | 1 | |
| ENSMUSG00000093958 | 1 | |
| ENSMUSG00000093959 | 1 | |
| ENSMUSG00000093960 | 1 | |
| ENSMUSG00000093961 | 1 | |
| ENSMUSG00000093962 | 1 | |
| ENSMUSG00000093963 | 1 | |
| ENSMUSG00000093964 | 1 | |
| ENSMUSG00000093965 | 1 | |
| ENSMUSG00000093966 | 1 | |
| ENSMUSG00000093967 | 1.023099224 | 0.986726708 |
| ENSMUSG00000093968 | 1 | |
| ENSMUSG00000093969 | 1 | |
| ENSMUSG00000093970 | 1 | |
| ENSMUSG00000093971 | 1 | |
| ENSMUSG00000093973 | 1 | |
| ENSMUSG00000093974 | 1 | |
| ENSMUSG00000093975 | 1.059463094 | 0.608497229 |
| ENSMUSG00000093976 | 1 | |
| ENSMUSG00000093977 | 1 | |
| ENSMUSG00000093978 | 1 | |
| ENSMUSG00000093979 | 1.097728744 | 0.698112512 |
| ENSMUSG00000093980 | 1 | |
| ENSMUSG00000093981 | 1 | |
| ENSMUSG00000093983 | 1 | |
| ENSMUSG00000093984 | 1 | |
| ENSMUSG00000093985 | 1 | |
| ENSMUSG00000093986 | 1 | |
| ENSMUSG00000093987 | 1 | |
| ENSMUSG00000093988 | 1 | |
| ENSMUSG00000093989 | 1.022079841 | 1 |
| ENSMUSG00000093990 | 1 | |
| ENSMUSG00000093992 | 1 | |
| ENSMUSG00000093993 | 1 | |
| ENSMUSG00000093994 | 1 | |
| ENSMUSG00000093995 | 1 | |
| ENSMUSG00000093996 | 1 | |
| ENSMUSG00000093997 | 1 | |
| ENSMUSG00000093998 | 1 | |
| ENSMUSG00000093999 | 1 | |
| ENSMUSG00000094001 | 1 | |
| ENSMUSG00000094002 | 1 | |
| ENSMUSG00000094003 | 1 | |
| ENSMUSG00000094004 | 1 | |
| ENSMUSG00000094005 | 1 | |
| ENSMUSG00000094006 | 1 | |
| ENSMUSG00000094007 | 1 | |
| ENSMUSG00000094008 | 1 | |
| ENSMUSG00000094009 | 1 | |
| ENSMUSG00000094010 | 1 | |
| ENSMUSG00000094011 | 1 | |
| ENSMUSG00000094012 | 1 | |
| ENSMUSG00000094013 | 1 | |
| ENSMUSG00000094015 | 1 | |
| ENSMUSG00000094016 | 1 | |
| ENSMUSG00000094017 | -1.003942951 | 1 |
| ENSMUSG00000094018 | -1.044273782 | 0.698112512 |
| ENSMUSG00000094020 | 1 | |
| ENSMUSG00000094021 | 1 | |
| ENSMUSG00000094023 | 1 | |
| ENSMUSG00000094024 | 1 | |

| | | |
|---|---|---|
| ENSMUSG00000094025 | 1 | |
| ENSMUSG00000094026 | 1 | |
| ENSMUSG00000094027 | 1.059463094 | 0.608497229 |
| ENSMUSG00000094028 | 1 | |
| ENSMUSG00000094029 | 1 | |
| ENSMUSG00000094030 | 1.000804399 | 1 |
| ENSMUSG00000094031 | 1 | |
| ENSMUSG00000094032 | 1 | |
| ENSMUSG00000094033 | 1.059463094 | 0.608497229 |
| ENSMUSG00000094034 | 1 | |
| ENSMUSG00000094035 | -1.03962336 | 0.698112512 |
| ENSMUSG00000094036 | 1 | |
| ENSMUSG00000094037 | 1 | |
| ENSMUSG00000094038 | 1 | |
| ENSMUSG00000094039 | 1 | |
| ENSMUSG00000094041 | 1 | |
| ENSMUSG00000094043 | 1 | |
| ENSMUSG00000094044 | 1 | |
| ENSMUSG00000094048 | 1 | |
| ENSMUSG00000094049 | 1 | |
| ENSMUSG00000094050 | 1 | |
| ENSMUSG00000094051 | 1 | |
| ENSMUSG00000094052 | 1 | |
| ENSMUSG00000094053 | 1 | |
| ENSMUSG00000094054 | 1 | |
| ENSMUSG00000094057 | 1 | |
| ENSMUSG00000094059 | 1 | |
| ENSMUSG00000094060 | 1 | |
| ENSMUSG00000094063 | 1 | |
| ENSMUSG00000094064 | 1 | |
| ENSMUSG00000094065 | 1 | |
| ENSMUSG00000094066 | 1 | |
| ENSMUSG00000094067 | 1 | |
| ENSMUSG00000094068 | 1 | |
| ENSMUSG00000094069 | 1 | |
| ENSMUSG00000094071 | 1 | |
| ENSMUSG00000094072 | 1 | |
| ENSMUSG00000094074 | 1 | |
| ENSMUSG00000094075 | 1 | |
| ENSMUSG00000094076 | 1 | |
| ENSMUSG00000094077 | 1 | |
| ENSMUSG00000094078 | 1 | |
| ENSMUSG00000094079 | 1 | |
| ENSMUSG00000094080 | 1 | |
| ENSMUSG00000094081 | 1 | |
| ENSMUSG00000094082 | 1 | |
| ENSMUSG00000094083 | 1.059463094 | 0.608497229 |
| ENSMUSG00000094084 | 1 | |
| ENSMUSG00000094085 | 1 | |
| ENSMUSG00000094086 | -1.26007302 | 0.250726313 |
| ENSMUSG00000094087 | 1 | |
| ENSMUSG00000094088 | 1 | |
| ENSMUSG00000094090 | 1.02558623 | 0.821336725 |
| ENSMUSG00000094091 | -1.084645496 | 0.532369657 |
| ENSMUSG00000094092 | 1.009042047 | 1 |
| ENSMUSG00000094093 | 1 | |
| ENSMUSG00000094094 | 1 | |
| ENSMUSG00000094095 | 1 | |
| ENSMUSG00000094096 | 1 | |
| ENSMUSG00000094097 | 1 | |
| ENSMUSG00000094098 | -1 | 1 |
| ENSMUSG00000094099 | 1 | |
| ENSMUSG00000094101 | 1 | |

| | | |
|---|---|---|
| ENSMUSG00000094102 | 1 | |
| ENSMUSG00000094103 | 1 | |
| ENSMUSG00000094104 | 1 | |
| ENSMUSG00000094105 | 1 | |
| ENSMUSG00000094107 | 1 | |
| ENSMUSG00000094108 | 1 | |
| ENSMUSG00000094109 | 1 | |
| ENSMUSG00000094111 | 1 | |
| ENSMUSG00000094112 | 1 | |
| ENSMUSG00000094113 | 1 | |
| ENSMUSG00000094114 | -1.640646931 | 0.009956796 |
| ENSMUSG00000094115 | 1 | |
| ENSMUSG00000094116 | 1 | |
| ENSMUSG00000094117 | 1 | |
| ENSMUSG00000094118 | 1 | |
| ENSMUSG00000094119 | 1 | |
| ENSMUSG00000094120 | 1 | |
| ENSMUSG00000094121 | 1 | |
| ENSMUSG00000094122 | -1.03962336 | 0.698112512 |
| ENSMUSG00000094123 | 1 | |
| ENSMUSG00000094124 | 1 | |
| ENSMUSG00000094125 | 1 | |
| ENSMUSG00000094127 | 1 | |
| ENSMUSG00000094128 | 1 | |
| ENSMUSG00000094129 | 1 | |
| ENSMUSG00000094131 | 1 | |
| ENSMUSG00000094132 | 1 | |
| ENSMUSG00000094133 | 1 | |
| ENSMUSG00000094134 | 1 | |
| ENSMUSG00000094135 | 1 | |
| ENSMUSG00000094136 | 1 | |
| ENSMUSG00000094137 | 1 | |
| ENSMUSG00000094139 | 1 | |
| ENSMUSG00000094140 | 1 | |
| ENSMUSG00000094144 | 1 | |
| ENSMUSG00000094145 | 1 | |
| ENSMUSG00000094146 | 1 | |
| ENSMUSG00000094147 | 1 | |
| ENSMUSG00000094148 | 1 | |
| ENSMUSG00000094149 | 1 | |
| ENSMUSG00000094150 | 1 | |
| ENSMUSG00000094151 | -1 | 1 |
| ENSMUSG00000094152 | 1 | |
| ENSMUSG00000094153 | 1.059463094 | 0.704170349 |
| ENSMUSG00000094154 | 1 | |
| ENSMUSG00000094155 | 1 | |
| ENSMUSG00000094156 | 1 | |
| ENSMUSG00000094157 | 1 | |
| ENSMUSG00000094158 | 1.091568689 | 0.72301587 |
| ENSMUSG00000094159 | 1 | |
| ENSMUSG00000094160 | 1 | |
| ENSMUSG00000094161 | 1 | |
| ENSMUSG00000094162 | 1 | |
| ENSMUSG00000094163 | 1 | |
| ENSMUSG00000094164 | 1 | |
| ENSMUSG00000094165 | 1 | |
| ENSMUSG00000094167 | 1 | |
| ENSMUSG00000094168 | 1 | |
| ENSMUSG00000094170 | 1 | |
| ENSMUSG00000094172 | 1 | |
| ENSMUSG00000094173 | 1 | |
| ENSMUSG00000094174 | 1 | |
| ENSMUSG00000094175 | 1 | |

| | | |
|---|---|---|
| ENSMUSG00000094176 | 1 | |
| ENSMUSG00000094177 | 1 | |
| ENSMUSG00000094178 | 1 | |
| ENSMUSG00000094179 | 1 | |
| ENSMUSG00000094180 | 1 | |
| ENSMUSG00000094181 | 1 | |
| ENSMUSG00000094182 | 1 | |
| ENSMUSG00000094183 | 1 | |
| ENSMUSG00000094184 | 1 | |
| ENSMUSG00000094185 | 1.023743961 | 0.95838544 |
| ENSMUSG00000094186 | 1 | |
| ENSMUSG00000094187 | 1 | |
| ENSMUSG00000094188 | 1 | |
| ENSMUSG00000094189 | 1 | |
| ENSMUSG00000094190 | 1 | |
| ENSMUSG00000094191 | 1.018662952 | 1 |
| ENSMUSG00000094192 | 1 | |
| ENSMUSG00000094194 | 1 | |
| ENSMUSG00000094195 | 1 | |
| ENSMUSG00000094196 | 1 | |
| ENSMUSG00000094197 | 1 | |
| ENSMUSG00000094198 | 1 | |
| ENSMUSG00000094199 | 1 | |
| ENSMUSG00000094200 | 1 | |
| ENSMUSG00000094201 | 1 | |
| ENSMUSG00000094202 | 1 | |
| ENSMUSG00000094203 | 1 | |
| ENSMUSG00000094204 | 1 | |
| ENSMUSG00000094205 | 1 | |
| ENSMUSG00000094206 | 1 | |
| ENSMUSG00000094208 | 1 | |
| ENSMUSG00000094210 | 1 | |
| ENSMUSG00000094211 | 1 | |
| ENSMUSG00000094212 | -1.044273782 | 0.698112512 |
| ENSMUSG00000094213 | 1 | |
| ENSMUSG00000094214 | 1 | |
| ENSMUSG00000094215 | -1 | 1 |
| ENSMUSG00000094217 | 1 | |
| ENSMUSG00000094218 | 1 | |
| ENSMUSG00000094219 | 1.059463094 | 0.608497229 |
| ENSMUSG00000094220 | 1 | |
| ENSMUSG00000094221 | 1 | |
| ENSMUSG00000094222 | 1 | |
| ENSMUSG00000094226 | 1 | |
| ENSMUSG00000094227 | 1 | |
| ENSMUSG00000094228 | 1 | |
| ENSMUSG00000094229 | 1 | |
| ENSMUSG00000094230 | 1 | |
| ENSMUSG00000094231 | 1 | |
| ENSMUSG00000094232 | 1 | |
| ENSMUSG00000094233 | 1 | |
| ENSMUSG00000094235 | 1 | |
| ENSMUSG00000094236 | 1 | |
| ENSMUSG00000094237 | 1 | |
| ENSMUSG00000094238 | 1 | |
| ENSMUSG00000094239 | 1 | |
| ENSMUSG00000094240 | 1 | |
| ENSMUSG00000094241 | 1 | |
| ENSMUSG00000094242 | 1 | |
| ENSMUSG00000094243 | 1 | |
| ENSMUSG00000094244 | 1 | |
| ENSMUSG00000094245 | -1.064228315 | 0.835083879 |
| ENSMUSG00000094247 | -1 | 1 |

| | | |
|---|---|---|
| ENSMUSG00000094248 | 1 | |
| ENSMUSG00000094249 | 1 | |
| ENSMUSG00000094250 | 1.059463094 | 0.608497229 |
| ENSMUSG00000094251 | 1.095872691 | 0.608497229 |
| ENSMUSG00000094252 | 1 | |
| ENSMUSG00000094253 | -1 | 1 |
| ENSMUSG00000094254 | 1 | |
| ENSMUSG00000094255 | 1 | |
| ENSMUSG00000094256 | 1.059463094 | 0.608497229 |
| ENSMUSG00000094257 | -1.044273782 | 0.790851299 |
| ENSMUSG00000094258 | 1 | |
| ENSMUSG00000094259 | 1 | |
| ENSMUSG00000094261 | 1 | |
| ENSMUSG00000094262 | 1 | |
| ENSMUSG00000094263 | 1 | |
| ENSMUSG00000094264 | 1 | |
| ENSMUSG00000094265 | 1 | |
| ENSMUSG00000094266 | 1 | |
| ENSMUSG00000094267 | 1 | |
| ENSMUSG00000094268 | 1 | |
| ENSMUSG00000094269 | 1 | |
| ENSMUSG00000094270 | 1 | |
| ENSMUSG00000094271 | 1 | |
| ENSMUSG00000094273 | 1 | |
| ENSMUSG00000094274 | 1 | |
| ENSMUSG00000094275 | 1 | |
| ENSMUSG00000094276 | 1 | |
| ENSMUSG00000094277 | 1 | |
| ENSMUSG00000094278 | 1 | |
| ENSMUSG00000094280 | 1 | |
| ENSMUSG00000094281 | -1.246797111 | 0.340244011 |
| ENSMUSG00000094282 | -1.069502966 | 0.698112512 |
| ENSMUSG00000094283 | 1 | |
| ENSMUSG00000094284 | 1 | |
| ENSMUSG00000094285 | 1 | |
| ENSMUSG00000094286 | -1.120750893 | 0.581069969 |
| ENSMUSG00000094287 | 1 | |
| ENSMUSG00000094288 | 1 | |
| ENSMUSG00000094289 | 1 | |
| ENSMUSG00000094290 | 1 | |
| ENSMUSG00000094291 | 1 | |
| ENSMUSG00000094292 | 1 | |
| ENSMUSG00000094293 | 1 | |
| ENSMUSG00000094294 | 1 | |
| ENSMUSG00000094295 | 1 | |
| ENSMUSG00000094296 | -1.174896771 | 0.241336215 |
| ENSMUSG00000094297 | 1 | |
| ENSMUSG00000094298 | 1 | |
| ENSMUSG00000094301 | 1 | |
| ENSMUSG00000094302 | 1 | |
| ENSMUSG00000094303 | 1 | |
| ENSMUSG00000094304 | 1 | |
| ENSMUSG00000094305 | 1 | |
| ENSMUSG00000094306 | 1 | |
| ENSMUSG00000094307 | 1 | |
| ENSMUSG00000094308 | 1 | |
| ENSMUSG00000094309 | 1 | |
| ENSMUSG00000094310 | 1 | |
| ENSMUSG00000094311 | 1 | |
| ENSMUSG00000094312 | 1 | |
| ENSMUSG00000094313 | 1 | |
| ENSMUSG00000094314 | 1 | |
| ENSMUSG00000094315 | 1 | |

| | | |
|---|---|---|
| ENSMUSG00000094316 | 1 | |
| ENSMUSG00000094317 | 1 | |
| ENSMUSG00000094318 | 1 | |
| ENSMUSG00000094319 | 1 | |
| ENSMUSG00000094320 | 1 | |
| ENSMUSG00000094321 | 1 | |
| ENSMUSG00000094322 | 1 | |
| ENSMUSG00000094323 | 1 | |
| ENSMUSG00000094325 | 1 | |
| ENSMUSG00000094326 | 1 | |
| ENSMUSG00000094327 | 1 | |
| ENSMUSG00000094328 | 1 | |
| ENSMUSG00000094330 | 1 | |
| ENSMUSG00000094334 | 1.089470641 | 0.608497229 |
| ENSMUSG00000094335 | 1 | |
| ENSMUSG00000094336 | 1 | |
| ENSMUSG00000094337 | 1 | |
| ENSMUSG00000094338 | 1 | |
| ENSMUSG00000094340 | 1 | |
| ENSMUSG00000094342 | 1 | |
| ENSMUSG00000094344 | -1.111880267 | 0.608497229 |
| ENSMUSG00000094345 | 1 | |
| ENSMUSG00000094346 | 1 | |
| ENSMUSG00000094347 | 1 | |
| ENSMUSG00000094349 | 1 | |
| ENSMUSG00000094350 | 1 | |
| ENSMUSG00000094351 | 1 | |
| ENSMUSG00000094352 | 1 | |
| ENSMUSG00000094353 | 1 | |
| ENSMUSG00000094354 | 1 | |
| ENSMUSG00000094356 | 1 | |
| ENSMUSG00000094357 | 1 | |
| ENSMUSG00000094358 | 1 | |
| ENSMUSG00000094359 | 1 | |
| ENSMUSG00000094360 | 1 | |
| ENSMUSG00000094361 | 1 | |
| ENSMUSG00000094362 | 1 | |
| ENSMUSG00000094363 | 1 | |
| ENSMUSG00000094364 | 1 | |
| ENSMUSG00000094365 | 1 | |
| ENSMUSG00000094366 | 1 | |
| ENSMUSG00000094367 | 1 | |
| ENSMUSG00000094369 | 1 | |
| ENSMUSG00000094370 | 1 | |
| ENSMUSG00000094371 | 1 | |
| ENSMUSG00000094372 | 1 | |
| ENSMUSG00000094374 | 1 | |
| ENSMUSG00000094377 | 1 | |
| ENSMUSG00000094378 | 1 | |
| ENSMUSG00000094379 | 1 | |
| ENSMUSG00000094380 | 1 | |
| ENSMUSG00000094381 | 1 | |
| ENSMUSG00000094382 | -1.063525418 | 0.756517167 |
| ENSMUSG00000094383 | -1.071075483 | 0.698112512 |
| ENSMUSG00000094384 | 1 | |
| ENSMUSG00000094385 | 1 | |
| ENSMUSG00000094386 | 1 | |
| ENSMUSG00000094388 | 1 | |
| ENSMUSG00000094389 | 1 | |
| ENSMUSG00000094390 | 1 | |
| ENSMUSG00000094391 | 1 | |
| ENSMUSG00000094392 | 1 | |
| ENSMUSG00000094394 | 1 | |

| | | |
|---|---|---|
| ENSMUSG00000094396 | 1 | |
| ENSMUSG00000094399 | 1 | |
| ENSMUSG00000094400 | -1 | 1 |
| ENSMUSG00000094401 | 1 | |
| ENSMUSG00000094402 | 1 | |
| ENSMUSG00000094403 | 1 | |
| ENSMUSG00000094404 | 1 | |
| ENSMUSG00000094405 | 1 | |
| ENSMUSG00000094406 | 1 | |
| ENSMUSG00000094407 | 1 | |
| ENSMUSG00000094408 | 1 | |
| ENSMUSG00000094409 | 1 | |
| ENSMUSG00000094410 | -1.008483488 | 0.988424085 |
| ENSMUSG00000094411 | 1.029199389 | 0.949812939 |
| ENSMUSG00000094412 | 1 | |
| ENSMUSG00000094413 | 1 | |
| ENSMUSG00000094415 | 1 | |
| ENSMUSG00000094416 | 1 | |
| ENSMUSG00000094417 | 1 | |
| ENSMUSG00000094419 | 1 | |
| ENSMUSG00000094420 | 1 | |
| ENSMUSG00000094421 | 1 | |
| ENSMUSG00000094422 | 1 | |
| ENSMUSG00000094423 | 1 | |
| ENSMUSG00000094424 | 1 | |
| ENSMUSG00000094426 | 1 | |
| ENSMUSG00000094427 | 1 | |
| ENSMUSG00000094428 | 1 | |
| ENSMUSG00000094429 | 1 | |
| ENSMUSG00000094430 | 1 | |
| ENSMUSG00000094431 | 1 | |
| ENSMUSG00000094432 | 1 | |
| ENSMUSG00000094433 | 1 | |
| ENSMUSG00000094435 | -1.03962336 | 0.698112512 |
| ENSMUSG00000094436 | 1 | |
| ENSMUSG00000094437 | 1 | |
| ENSMUSG00000094438 | 1 | |
| ENSMUSG00000094439 | 1 | |
| ENSMUSG00000094440 | 1 | |
| ENSMUSG00000094441 | -1.055822652 | 0.698112512 |
| ENSMUSG00000094442 | 1 | |
| ENSMUSG00000094443 | -1.084645496 | 0.628300894 |
| ENSMUSG00000094444 | 1 | |
| ENSMUSG00000094445 | 1 | |
| ENSMUSG00000094446 | 1 | |
| ENSMUSG00000094447 | 1 | |
| ENSMUSG00000094449 | 1 | |
| ENSMUSG00000094452 | 1 | |
| ENSMUSG00000094454 | 1 | |
| ENSMUSG00000094455 | 1 | |
| ENSMUSG00000094456 | 1 | |
| ENSMUSG00000094457 | 1 | |
| ENSMUSG00000094458 | 1 | |
| ENSMUSG00000094460 | 1.059463094 | 0.608497229 |
| ENSMUSG00000094461 | 1 | |
| ENSMUSG00000094462 | 1 | |
| ENSMUSG00000094463 | 1 | |
| ENSMUSG00000094464 | 1 | |
| ENSMUSG00000094465 | 1 | |
| ENSMUSG00000094466 | 1 | |
| ENSMUSG00000094467 | -1.044273782 | 0.698112512 |
| ENSMUSG00000094468 | 1 | |
| ENSMUSG00000094470 | 1 | |

| | | |
|---|---|---|
| ENSMUSG00000094471 | 1 | |
| ENSMUSG00000094472 | 1.084369352 | 0.529596655 |
| ENSMUSG00000094473 | 1 | |
| ENSMUSG00000094474 | 1 | |
| ENSMUSG00000094475 | 1 | |
| ENSMUSG00000094476 | 1 | |
| ENSMUSG00000094477 | 1 | |
| ENSMUSG00000094478 | 1 | |
| ENSMUSG00000094481 | 1 | |
| ENSMUSG00000094482 | 1 | |
| ENSMUSG00000094483 | 1.017335965 | 1 |
| ENSMUSG00000094484 | 1 | |
| ENSMUSG00000094485 | 1 | |
| ENSMUSG00000094486 | -1.003834322 | 1 |
| ENSMUSG00000094487 | 1 | |
| ENSMUSG00000094488 | 1 | |
| ENSMUSG00000094489 | 1 | |
| ENSMUSG00000094491 | 1 | |
| ENSMUSG00000094493 | 1 | |
| ENSMUSG00000094494 | 1 | |
| ENSMUSG00000094496 | 1 | |
| ENSMUSG00000094497 | 1 | |
| ENSMUSG00000094498 | 1 | |
| ENSMUSG00000094499 | 1 | |
| ENSMUSG00000094500 | 1.049517471 | 0.892893968 |
| ENSMUSG00000094501 | 1 | |
| ENSMUSG00000094502 | 1 | |
| ENSMUSG00000094504 | 1 | |
| ENSMUSG00000094505 | 1 | |
| ENSMUSG00000094506 | 1 | |
| ENSMUSG00000094507 | 1 | |
| ENSMUSG00000094508 | 1 | |
| ENSMUSG00000094509 | 1 | |
| ENSMUSG00000094510 | 1 | |
| ENSMUSG00000094511 | 1 | |
| ENSMUSG00000094513 | 1 | |
| ENSMUSG00000094514 | 1 | |
| ENSMUSG00000094515 | 1 | |
| ENSMUSG00000094516 | 1 | |
| ENSMUSG00000094517 | 1 | |
| ENSMUSG00000094518 | 1.112524439 | 0.741162074 |
| ENSMUSG00000094519 | 1 | |
| ENSMUSG00000094520 | 1 | |
| ENSMUSG00000094521 | 1 | |
| ENSMUSG00000094522 | 1 | |
| ENSMUSG00000094523 | 1 | |
| ENSMUSG00000094524 | 1 | |
| ENSMUSG00000094525 | 1 | |
| ENSMUSG00000094526 | -1 | 1 |
| ENSMUSG00000094527 | 1 | |
| ENSMUSG00000094528 | 1 | |
| ENSMUSG00000094529 | 1 | |
| ENSMUSG00000094530 | -1.017057627 | 1 |
| ENSMUSG00000094531 | 1 | |
| ENSMUSG00000094532 | 1 | |
| ENSMUSG00000094533 | 1 | |
| ENSMUSG00000094534 | -1 | 1 |
| ENSMUSG00000094535 | 1 | |
| ENSMUSG00000094537 | 1 | |
| ENSMUSG00000094538 | 1 | |
| ENSMUSG00000094539 | 1 | |
| ENSMUSG00000094540 | 1 | |
| ENSMUSG00000094541 | 1 | |

| | | |
|---|---|---|
| ENSMUSG00000094542 | 1 | |
| ENSMUSG00000094543 | 1 | |
| ENSMUSG00000094544 | 1 | |
| ENSMUSG00000094545 | 1 | |
| ENSMUSG00000094546 | 1 | |
| ENSMUSG00000094548 | 1 | |
| ENSMUSG00000094550 | 1 | |
| ENSMUSG00000094551 | 1.02286406 | 0.908149431 |
| ENSMUSG00000094552 | 1 | |
| ENSMUSG00000094553 | 1 | |
| ENSMUSG00000094554 | 1 | |
| ENSMUSG00000094555 | 1 | |
| ENSMUSG00000094556 | 1 | |
| ENSMUSG00000094557 | -1.028348525 | 0.936995747 |
| ENSMUSG00000094558 | 1 | |
| ENSMUSG00000094559 | 1 | |
| ENSMUSG00000094560 | 1 | |
| ENSMUSG00000094561 | 1 | |
| ENSMUSG00000094562 | 1 | |
| ENSMUSG00000094563 | 1 | |
| ENSMUSG00000094564 | 1 | |
| ENSMUSG00000094565 | 1 | |
| ENSMUSG00000094566 | 1 | |
| ENSMUSG00000094567 | 1 | |
| ENSMUSG00000094568 | 1.005613439 | 1 |
| ENSMUSG00000094569 | 1 | |
| ENSMUSG00000094570 | 1 | |
| ENSMUSG00000094571 | 1 | |
| ENSMUSG00000094572 | 1 | |
| ENSMUSG00000094573 | 1 | |
| ENSMUSG00000094574 | 1 | |
| ENSMUSG00000094575 | 1 | |
| ENSMUSG00000094576 | 1 | |
| ENSMUSG00000094577 | 1 | |
| ENSMUSG00000094578 | 1 | |
| ENSMUSG00000094579 | 1 | |
| ENSMUSG00000094581 | 1 | |
| ENSMUSG00000094582 | 1 | |
| ENSMUSG00000094584 | 1 | |
| ENSMUSG00000094585 | 1 | |
| ENSMUSG00000094586 | 1 | |
| ENSMUSG00000094587 | 1 | |
| ENSMUSG00000094588 | 1 | |
| ENSMUSG00000094589 | 1 | |
| ENSMUSG00000094590 | -1.044273782 | 0.698112512 |
| ENSMUSG00000094592 | 1 | |
| ENSMUSG00000094593 | 1 | |
| ENSMUSG00000094594 | -1.053549062 | 0.778372272 |
| ENSMUSG00000094595 | -1.040304217 | 0.758615668 |
| ENSMUSG00000094596 | 1 | |
| ENSMUSG00000094597 | -1.053549062 | 0.698112512 |
| ENSMUSG00000094598 | 1 | |
| ENSMUSG00000094599 | 1 | |
| ENSMUSG00000094600 | 1 | |
| ENSMUSG00000094601 | 1 | |
| ENSMUSG00000094602 | 1 | |
| ENSMUSG00000094603 | 1.024693402 | 0.933031586 |
| ENSMUSG00000094604 | 1 | |
| ENSMUSG00000094605 | 1 | |
| ENSMUSG00000094606 | 1 | |
| ENSMUSG00000094607 | 1 | |
| ENSMUSG00000094608 | 1 | |
| ENSMUSG00000094609 | 1 | |

| | | |
|---|---|---|
| ENSMUSG00000094610 | 1 | |
| ENSMUSG00000094612 | 1 | |
| ENSMUSG00000094613 | 1 | |
| ENSMUSG00000094614 | -1.091755469 | 0.612278915 |
| ENSMUSG00000094615 | 1.100747781 | 0.777153053 |
| ENSMUSG00000094616 | 1 | |
| ENSMUSG00000094618 | 1.009517015 | 1 |
| ENSMUSG00000094619 | 1 | |
| ENSMUSG00000094620 | 1 | |
| ENSMUSG00000094621 | 1 | |
| ENSMUSG00000094622 | 1 | |
| ENSMUSG00000094623 | 1 | |
| ENSMUSG00000094624 | 1 | |
| ENSMUSG00000094625 | 1 | |
| ENSMUSG00000094626 | 1.069068979 | 0.608497229 |
| ENSMUSG00000094627 | -1.115549051 | 0.608497229 |
| ENSMUSG00000094628 | 1 | |
| ENSMUSG00000094630 | 1 | |
| ENSMUSG00000094631 | 1 | |
| ENSMUSG00000094632 | 1 | |
| ENSMUSG00000094634 | -1.041207564 | 0.921768868 |
| ENSMUSG00000094635 | 1.069068979 | 0.608497229 |
| ENSMUSG00000094636 | 1 | |
| ENSMUSG00000094637 | 1 | |
| ENSMUSG00000094638 | 1 | |
| ENSMUSG00000094639 | 1 | |
| ENSMUSG00000094641 | 1 | |
| ENSMUSG00000094642 | 1 | |
| ENSMUSG00000094643 | 1 | |
| ENSMUSG00000094647 | 1 | |
| ENSMUSG00000094648 | 1 | |
| ENSMUSG00000094649 | 1.143065142 | 0.552474963 |
| ENSMUSG00000094650 | 1.142908482 | 0.618829386 |
| ENSMUSG00000094651 | 1 | |
| ENSMUSG00000094652 | 1 | |
| ENSMUSG00000094654 | 1 | |
| ENSMUSG00000094655 | -1.044273782 | 0.698112512 |
| ENSMUSG00000094656 | 1 | |
| ENSMUSG00000094657 | 1 | |
| ENSMUSG00000094658 | 1 | |
| ENSMUSG00000094659 | 1 | |
| ENSMUSG00000094660 | 1 | |
| ENSMUSG00000094661 | 1 | |
| ENSMUSG00000094662 | 1 | |
| ENSMUSG00000094663 | 1 | |
| ENSMUSG00000094664 | 1 | |
| ENSMUSG00000094665 | 1 | |
| ENSMUSG00000094667 | 1 | |
| ENSMUSG00000094668 | 1 | |
| ENSMUSG00000094669 | 1 | |
| ENSMUSG00000094670 | 1 | |
| ENSMUSG00000094672 | 1 | |
| ENSMUSG00000094673 | 1 | |
| ENSMUSG00000094675 | 1 | |
| ENSMUSG00000094676 | -1 | 1 |
| ENSMUSG00000094677 | 1.069068979 | 0.608497229 |
| ENSMUSG00000094678 | 1 | |
| ENSMUSG00000094679 | 1 | |
| ENSMUSG00000094680 | 1 | |
| ENSMUSG00000094681 | 1 | |
| ENSMUSG00000094682 | 1 | |
| ENSMUSG00000094683 | -1.099174267 | 0.608497229 |
| ENSMUSG00000094684 | 1 | |

| | | |
|---|---|---|
| ENSMUSG00000094685 | 1.124393692 | 0.11356889 |
| ENSMUSG00000094686 | 1 | |
| ENSMUSG00000094687 | 1 | |
| ENSMUSG00000094688 | 1 | |
| ENSMUSG00000094689 | 1 | |
| ENSMUSG00000094690 | 1.142908482 | 0.608497229 |
| ENSMUSG00000094691 | 1 | |
| ENSMUSG00000094692 | 1 | |
| ENSMUSG00000094693 | 1 | |
| ENSMUSG00000094694 | 1 | |
| ENSMUSG00000094695 | 1 | |
| ENSMUSG00000094696 | 1 | |
| ENSMUSG00000094697 | 1 | |
| ENSMUSG00000094698 | 1 | |
| ENSMUSG00000094699 | 1 | |
| ENSMUSG00000094700 | 1 | |
| ENSMUSG00000094701 | 1 | |
| ENSMUSG00000094702 | 1 | |
| ENSMUSG00000094703 | 1 | |
| ENSMUSG00000094704 | 1 | |
| ENSMUSG00000094706 | 1 | |
| ENSMUSG00000094707 | 1.059463094 | 0.698112512 |
| ENSMUSG00000094708 | 1 | |
| ENSMUSG00000094709 | 1 | |
| ENSMUSG00000094710 | 1 | |
| ENSMUSG00000094712 | 1 | |
| ENSMUSG00000094713 | 1 | |
| ENSMUSG00000094714 | 1 | |
| ENSMUSG00000094715 | 1 | |
| ENSMUSG00000094717 | 1 | |
| ENSMUSG00000094718 | 1 | |
| ENSMUSG00000094719 | -1.054169715 | 0.813485995 |
| ENSMUSG00000094720 | 1 | |
| ENSMUSG00000094721 | 1 | |
| ENSMUSG00000094722 | 1 | |
| ENSMUSG00000094723 | 1 | |
| ENSMUSG00000094724 | 1.09866125 | 0.643397479 |
| ENSMUSG00000094725 | 1 | |
| ENSMUSG00000094726 | 1 | |
| ENSMUSG00000094727 | 1 | |
| ENSMUSG00000094728 | 1 | |
| ENSMUSG00000094729 | 1 | |
| ENSMUSG00000094730 | 1 | |
| ENSMUSG00000094732 | 1.069068979 | 0.698112512 |
| ENSMUSG00000094733 | 1 | |
| ENSMUSG00000094734 | 1 | |
| ENSMUSG00000094735 | 1 | |
| ENSMUSG00000094736 | 1 | |
| ENSMUSG00000094737 | 1 | |
| ENSMUSG00000094738 | 1 | |
| ENSMUSG00000094739 | 1 | |
| ENSMUSG00000094740 | 1 | |
| ENSMUSG00000094741 | 1 | |
| ENSMUSG00000094743 | 1 | |
| ENSMUSG00000094745 | 1 | |
| ENSMUSG00000094746 | 1 | |
| ENSMUSG00000094747 | 1 | |
| ENSMUSG00000094748 | 1 | |
| ENSMUSG00000094750 | 1 | |
| ENSMUSG00000094751 | 1 | |
| ENSMUSG00000094752 | 1 | |
| ENSMUSG00000094753 | 1 | |
| ENSMUSG00000094754 | 1 | |

| | | |
|---|---|---|
| ENSMUSG00000094755 | 1 | |
| ENSMUSG00000094757 | 1 | |
| ENSMUSG00000094758 | 1 | |
| ENSMUSG00000094759 | 1 | |
| ENSMUSG00000094760 | 1 | |
| ENSMUSG00000094762 | 1 | |
| ENSMUSG00000094763 | 1 | |
| ENSMUSG00000094764 | 1 | |
| ENSMUSG00000094765 | 1 | |
| ENSMUSG00000094766 | 1 | |
| ENSMUSG00000094767 | 1 | |
| ENSMUSG00000094768 | 1 | |
| ENSMUSG00000094771 | 1 | |
| ENSMUSG00000094772 | 1.133237328 | 0.591151279 |
| ENSMUSG00000094773 | 1 | |
| ENSMUSG00000094775 | 1 | |
| ENSMUSG00000094777 | 1 | |
| ENSMUSG00000094778 | 1 | |
| ENSMUSG00000094779 | 1 | |
| ENSMUSG00000094780 | 1 | |
| ENSMUSG00000094782 | 1 | |
| ENSMUSG00000094784 | 1 | |
| ENSMUSG00000094785 | 1 | |
| ENSMUSG00000094786 | -1.405136276 | 0.009529771 |
| ENSMUSG00000094787 | 1 | |
| ENSMUSG00000094789 | 1 | |
| ENSMUSG00000094790 | 1.000320512 | 1 |
| ENSMUSG00000094791 | 1 | |
| ENSMUSG00000094792 | 1 | |
| ENSMUSG00000094793 | 1 | |
| ENSMUSG00000094794 | 1 | |
| ENSMUSG00000094796 | 1 | |
| ENSMUSG00000094797 | 1 | |
| ENSMUSG00000094798 | 1 | |
| ENSMUSG00000094799 | 1 | |
| ENSMUSG00000094800 | 1.132639128 | 0.428732467 |
| ENSMUSG00000094801 | 1 | |
| ENSMUSG00000094802 | 1 | |
| ENSMUSG00000094803 | 1 | |
| ENSMUSG00000094804 | 1 | |
| ENSMUSG00000094805 | 1 | |
| ENSMUSG00000094806 | 1 | |
| ENSMUSG00000094807 | 1 | |
| ENSMUSG00000094808 | 1.059463094 | 0.608497229 |
| ENSMUSG00000094810 | 1 | |
| ENSMUSG00000094811 | 1 | |
| ENSMUSG00000094812 | 1 | |
| ENSMUSG00000094813 | 1 | |
| ENSMUSG00000094814 | 1 | |
| ENSMUSG00000094815 | 1 | |
| ENSMUSG00000094817 | 1 | |
| ENSMUSG00000094818 | 1 | |
| ENSMUSG00000094819 | 1 | |
| ENSMUSG00000094820 | -1.053549062 | 0.698112512 |
| ENSMUSG00000094821 | 1 | |
| ENSMUSG00000094822 | 1 | |
| ENSMUSG00000094823 | 1 | |
| ENSMUSG00000094824 | 1 | |
| ENSMUSG00000094825 | 1 | |
| ENSMUSG00000094826 | 1 | |
| ENSMUSG00000094827 | 1 | |
| ENSMUSG00000094828 | 1 | |
| ENSMUSG00000094829 | 1 | |

| | | |
|---|---|---|
| ENSMUSG00000094830 | 1 | |
| ENSMUSG00000094831 | 1 | |
| ENSMUSG00000094832 | 1.084618945 | 0.698112512 |
| ENSMUSG00000094833 | 1 | |
| ENSMUSG00000094834 | -1.07062035 | 0.782526702 |
| ENSMUSG00000094836 | 1 | |
| ENSMUSG00000094837 | 1 | |
| ENSMUSG00000094838 | 1 | |
| ENSMUSG00000094840 | -1.03962336 | 0.698112512 |
| ENSMUSG00000094841 | 1 | |
| ENSMUSG00000094842 | -1.043306199 | 0.79291651 |
| ENSMUSG00000094843 | 1 | |
| ENSMUSG00000094844 | -1 | 1 |
| ENSMUSG00000094845 | -1.190372793 | 0.615267463 |
| ENSMUSG00000094846 | 1 | |
| ENSMUSG00000094847 | 1 | |
| ENSMUSG00000094848 | 1 | |
| ENSMUSG00000094849 | -1.044273782 | 0.790851299 |
| ENSMUSG00000094850 | 1 | |
| ENSMUSG00000094852 | 1.059463094 | 0.608497229 |
| ENSMUSG00000094854 | 1 | |
| ENSMUSG00000094855 | 1 | |
| ENSMUSG00000094856 | -1.144020616 | 0.608497229 |
| ENSMUSG00000094858 | 1 | |
| ENSMUSG00000094860 | 1 | |
| ENSMUSG00000094861 | 1 | |
| ENSMUSG00000094862 | 1 | |
| ENSMUSG00000094864 | 1 | |
| ENSMUSG00000094865 | -1.044273782 | 0.698112512 |
| ENSMUSG00000094866 | 1 | |
| ENSMUSG00000094867 | 1 | |
| ENSMUSG00000094868 | 1 | |
| ENSMUSG00000094869 | 1 | |
| ENSMUSG00000094870 | -1.022049806 | 0.975658364 |
| ENSMUSG00000094872 | 1 | |
| ENSMUSG00000094873 | 1 | |
| ENSMUSG00000094874 | 1 | |
| ENSMUSG00000094875 | 1 | |
| ENSMUSG00000094876 | 1 | |
| ENSMUSG00000094877 | 1 | |
| ENSMUSG00000094878 | 1 | |
| ENSMUSG00000094879 | 1 | |
| ENSMUSG00000094880 | 1 | |
| ENSMUSG00000094881 | 1 | |
| ENSMUSG00000094883 | 1 | |
| ENSMUSG00000094884 | 1 | |
| ENSMUSG00000094885 | 1 | |
| ENSMUSG00000094886 | 1 | |
| ENSMUSG00000094887 | 1 | |
| ENSMUSG00000094888 | 1 | |
| ENSMUSG00000094889 | 1 | |
| ENSMUSG00000094890 | 1 | |
| ENSMUSG00000094891 | 1.069068979 | 0.608497229 |
| ENSMUSG00000094892 | 1 | |
| ENSMUSG00000094893 | -1.045151399 | 0.846956027 |
| ENSMUSG00000094894 | 1 | |
| ENSMUSG00000094895 | 1 | |
| ENSMUSG00000094896 | 1 | |
| ENSMUSG00000094897 | 1 | |
| ENSMUSG00000094898 | 1 | |
| ENSMUSG00000094901 | 1 | |
| ENSMUSG00000094902 | 1 | |
| ENSMUSG00000094903 | 1 | |

| | | |
|---|---|---|
| ENSMUSG00000094904 | 1 | |
| ENSMUSG00000094905 | 1 | |
| ENSMUSG00000094906 | 1 | |
| ENSMUSG00000094910 | -1.200114593 | 0.1991922 |
| ENSMUSG00000094911 | 1 | |
| ENSMUSG00000094912 | 1 | |
| ENSMUSG00000094913 | 1 | |
| ENSMUSG00000094914 | 1 | |
| ENSMUSG00000094915 | 1 | |
| ENSMUSG00000094916 | 1 | |
| ENSMUSG00000094917 | 1 | |
| ENSMUSG00000094918 | 1 | |
| ENSMUSG00000094919 | 1 | |
| ENSMUSG00000094920 | 1 | |
| ENSMUSG00000094921 | 1 | |
| ENSMUSG00000094922 | 1 | |
| ENSMUSG00000094923 | 1 | |
| ENSMUSG00000094925 | 1 | |
| ENSMUSG00000094926 | 1 | |
| ENSMUSG00000094927 | 1 | |
| ENSMUSG00000094928 | 1 | |
| ENSMUSG00000094929 | 1 | |
| ENSMUSG00000094930 | 1 | |
| ENSMUSG00000094931 | 1 | |
| ENSMUSG00000094932 | 1 | |
| ENSMUSG00000094933 | 1 | |
| ENSMUSG00000094934 | 1 | |
| ENSMUSG00000094935 | 1.202073302 | 0.227144703 |
| ENSMUSG00000094936 | 1.013868871 | 0.949558509 |
| ENSMUSG00000094937 | 1 | |
| ENSMUSG00000094938 | 1 | |
| ENSMUSG00000094939 | 1 | |
| ENSMUSG00000094940 | 1 | |
| ENSMUSG00000094941 | 1 | |
| ENSMUSG00000094942 | -1.032433427 | 0.817330349 |
| ENSMUSG00000094943 | 1 | |
| ENSMUSG00000094945 | 1 | |
| ENSMUSG00000094946 | 1 | |
| ENSMUSG00000094947 | 1 | |
| ENSMUSG00000094948 | 1 | |
| ENSMUSG00000094949 | 1 | |
| ENSMUSG00000094950 | 1 | |
| ENSMUSG00000094951 | 1 | |
| ENSMUSG00000094952 | 1 | |
| ENSMUSG00000094954 | 1 | |
| ENSMUSG00000094955 | 1 | |
| ENSMUSG00000094956 | 1 | |
| ENSMUSG00000094957 | 1 | |
| ENSMUSG00000094958 | -1.037482356 | 0.870778586 |
| ENSMUSG00000094959 | 1 | |
| ENSMUSG00000094960 | 1 | |
| ENSMUSG00000094962 | -1.043306199 | 0.698112512 |
| ENSMUSG00000094963 | 1 | |
| ENSMUSG00000094964 | 1 | |
| ENSMUSG00000094965 | 1 | |
| ENSMUSG00000094966 | 1 | |
| ENSMUSG00000094967 | 1 | |
| ENSMUSG00000094968 | 1 | |
| ENSMUSG00000094969 | -1.015511103 | 0.99339328 |
| ENSMUSG00000094970 | 1 | |
| ENSMUSG00000094971 | 1 | |
| ENSMUSG00000094972 | 1 | |
| ENSMUSG00000094973 | -1.153027569 | 0.608497229 |

| | | |
|---|---|---|
| ENSMUSG00000094974 | 1 | |
| ENSMUSG00000094975 | 1 | |
| ENSMUSG00000094976 | 1 | |
| ENSMUSG00000094977 | 1 | |
| ENSMUSG00000094978 | 1 | |
| ENSMUSG00000094979 | 1 | |
| ENSMUSG00000094980 | 1 | |
| ENSMUSG00000094981 | 1 | |
| ENSMUSG00000094982 | 1 | |
| ENSMUSG00000094983 | 1 | |
| ENSMUSG00000094985 | 1 | |
| ENSMUSG00000094986 | 1 | |
| ENSMUSG00000094987 | 1 | |
| ENSMUSG00000094989 | 1.059463094 | 0.608497229 |
| ENSMUSG00000094991 | 1 | |
| ENSMUSG00000094992 | 1 | |
| ENSMUSG00000094993 | 1 | |
| ENSMUSG00000094994 | 1 | |
| ENSMUSG00000094995 | 1.069068979 | 0.698112512 |
| ENSMUSG00000094997 | 1 | |
| ENSMUSG00000094998 | 1 | |
| ENSMUSG00000094999 | 1 | |
| ENSMUSG00000095000 | 1 | |
| ENSMUSG00000095002 | 1 | |
| ENSMUSG00000095003 | 1 | |
| ENSMUSG00000095005 | 1 | |
| ENSMUSG00000095006 | 1 | |
| ENSMUSG00000095007 | 1 | |
| ENSMUSG00000095008 | 1 | |
| ENSMUSG00000095009 | 1 | |
| ENSMUSG00000095010 | -1.03962336 | 0.814089245 |
| ENSMUSG00000095011 | 1 | |
| ENSMUSG00000095015 | 1 | |
| ENSMUSG00000095016 | 1 | |
| ENSMUSG00000095017 | 1 | |
| ENSMUSG00000095019 | 1 | |
| ENSMUSG00000095020 | 1 | |
| ENSMUSG00000095021 | 1 | |
| ENSMUSG00000095022 | 1 | |
| ENSMUSG00000095023 | 1 | |
| ENSMUSG00000095024 | 1 | |
| ENSMUSG00000095025 | 1 | |
| ENSMUSG00000095026 | -1.044273782 | 0.698112512 |
| ENSMUSG00000095027 | 1 | |
| ENSMUSG00000095028 | -1 | 1 |
| ENSMUSG00000095029 | 1 | |
| ENSMUSG00000095030 | 1 | |
| ENSMUSG00000095031 | 1 | |
| ENSMUSG00000095032 | 1 | |
| ENSMUSG00000095033 | 1 | |
| ENSMUSG00000095034 | 1 | |
| ENSMUSG00000095035 | 1 | |
| ENSMUSG00000095037 | 1 | |
| ENSMUSG00000095040 | 1 | |
| ENSMUSG00000095041 | 1.060648762 | 0.825357421 |
| ENSMUSG00000095042 | -1 | 1 |
| ENSMUSG00000095044 | 1 | |
| ENSMUSG00000095045 | 1 | |
| ENSMUSG00000095047 | 1.005613439 | 1 |
| ENSMUSG00000095048 | 1 | |
| ENSMUSG00000095049 | 1 | |
| ENSMUSG00000095051 | 1 | |
| ENSMUSG00000095052 | 1 | |

| | | |
|---|---|---|
| ENSMUSG00000095053 | 1 | |
| ENSMUSG00000095054 | 1.059463094 | 0.698112512 |
| ENSMUSG00000095056 | 1 | |
| ENSMUSG00000095057 | 1.014545335 | 0.979002724 |
| ENSMUSG00000095059 | 1 | |
| ENSMUSG00000095060 | 1 | |
| ENSMUSG00000095061 | 1 | |
| ENSMUSG00000095063 | 1 | |
| ENSMUSG00000095064 | 1 | |
| ENSMUSG00000095065 | 1 | |
| ENSMUSG00000095066 | 1 | |
| ENSMUSG00000095068 | 1.059463094 | 0.608497229 |
| ENSMUSG00000095069 | 1 | |
| ENSMUSG00000095070 | 1 | |
| ENSMUSG00000095071 | 1 | |
| ENSMUSG00000095072 | 1 | |
| ENSMUSG00000095073 | 1 | |
| ENSMUSG00000095074 | 1 | |
| ENSMUSG00000095075 | 1 | |
| ENSMUSG00000095076 | 1 | |
| ENSMUSG00000095078 | 1 | |
| ENSMUSG00000095079 | 1 | |
| ENSMUSG00000095081 | 1 | |
| ENSMUSG00000095082 | 1 | |
| ENSMUSG00000095085 | 1 | |
| ENSMUSG00000095086 | 1 | |
| ENSMUSG00000095087 | -1.111705722 | 0.686522899 |
| ENSMUSG00000095088 | 1 | |
| ENSMUSG00000095089 | 1 | |
| ENSMUSG00000095091 | 1 | |
| ENSMUSG00000095092 | 1 | |
| ENSMUSG00000095093 | 1 | |
| ENSMUSG00000095094 | 1 | |
| ENSMUSG00000095095 | 1 | |
| ENSMUSG00000095096 | 1 | |
| ENSMUSG00000095097 | 1 | |
| ENSMUSG00000095098 | 1.080882852 | 0.903464611 |
| ENSMUSG00000095100 | 1 | |
| ENSMUSG00000095101 | 1 | |
| ENSMUSG00000095102 | 1 | |
| ENSMUSG00000095103 | 1 | |
| ENSMUSG00000095104 | 1 | |
| ENSMUSG00000095105 | -1.45713026 | 0.004854638 |
| ENSMUSG00000095107 | 1 | |
| ENSMUSG00000095108 | 1 | |
| ENSMUSG00000095110 | 1 | |
| ENSMUSG00000095111 | 1 | |
| ENSMUSG00000095112 | 1 | |
| ENSMUSG00000095113 | 1 | |
| ENSMUSG00000095115 | 1.105708974 | 0.608497229 |
| ENSMUSG00000095116 | 1 | |
| ENSMUSG00000095117 | 1 | |
| ENSMUSG00000095118 | 1 | |
| ENSMUSG00000095119 | 1 | |
| ENSMUSG00000095121 | 1 | |
| ENSMUSG00000095122 | 1 | |
| ENSMUSG00000095123 | -1.040400647 | 0.730160452 |
| ENSMUSG00000095125 | 1 | |
| ENSMUSG00000095126 | 1 | |
| ENSMUSG00000095127 | 1 | |
| ENSMUSG00000095128 | 1 | |
| ENSMUSG00000095129 | 1 | |
| ENSMUSG00000095130 | 1 | |

| | | |
|---|---|---|
| ENSMUSG00000095131 | 1 | |
| ENSMUSG00000095132 | 1 | |
| ENSMUSG00000095133 | 1 | |
| ENSMUSG00000095134 | -1 | 1 |
| ENSMUSG00000095135 | 1 | |
| ENSMUSG00000095136 | 1 | |
| ENSMUSG00000095137 | 1 | |
| ENSMUSG00000095138 | 1 | |
| ENSMUSG00000095139 | 1.015081164 | 0.991978016 |
| ENSMUSG00000095140 | 1 | |
| ENSMUSG00000095141 | 1 | |
| ENSMUSG00000095142 | 1 | |
| ENSMUSG00000095143 | 1 | |
| ENSMUSG00000095146 | 1 | |
| ENSMUSG00000095147 | 1 | |
| ENSMUSG00000095148 | 1 | |
| ENSMUSG00000095149 | 1 | |
| ENSMUSG00000095150 | -1.065753742 | 0.777302582 |
| ENSMUSG00000095151 | 1 | |
| ENSMUSG00000095152 | 1 | |
| ENSMUSG00000095153 | 1 | |
| ENSMUSG00000095154 | 1 | |
| ENSMUSG00000095155 | 1 | |
| ENSMUSG00000095156 | 1 | |
| ENSMUSG00000095159 | -1.058211741 | 0.815502708 |
| ENSMUSG00000095160 | 1 | |
| ENSMUSG00000095161 | 1 | |
| ENSMUSG00000095163 | 1 | |
| ENSMUSG00000095165 | 1 | |
| ENSMUSG00000095168 | 1.171547077 | 0.656294028 |
| ENSMUSG00000095169 | 1 | |
| ENSMUSG00000095170 | 1 | |
| ENSMUSG00000095171 | 1 | |
| ENSMUSG00000095172 | 1 | |
| ENSMUSG00000095174 | 1 | |
| ENSMUSG00000095175 | 1 | |
| ENSMUSG00000095176 | 1 | |
| ENSMUSG00000095178 | 1.122933516 | 0.652395499 |
| ENSMUSG00000095179 | 1 | |
| ENSMUSG00000095180 | -1.091186874 | 0.722001393 |
| ENSMUSG00000095182 | 1 | |
| ENSMUSG00000095183 | 1 | |
| ENSMUSG00000095184 | 1.059463094 | 0.608497229 |
| ENSMUSG00000095185 | 1 | |
| ENSMUSG00000095186 | 1 | |
| ENSMUSG00000095187 | 1 | |
| ENSMUSG00000095188 | 1 | |
| ENSMUSG00000095189 | 1 | |
| ENSMUSG00000095190 | 1 | |
| ENSMUSG00000095191 | 1 | |
| ENSMUSG00000095192 | 1 | |
| ENSMUSG00000095193 | 1.046317916 | 0.798131311 |
| ENSMUSG00000095194 | 1 | |
| ENSMUSG00000095195 | -1.043306199 | 0.79291651 |
| ENSMUSG00000095196 | 1 | |
| ENSMUSG00000095197 | 1 | |
| ENSMUSG00000095198 | 1 | |
| ENSMUSG00000095199 | 1 | |
| ENSMUSG00000095200 | 1 | |
| ENSMUSG00000095201 | 1 | |
| ENSMUSG00000095202 | 1.059463094 | 0.608497229 |
| ENSMUSG00000095203 | 1 | |
| ENSMUSG00000095204 | 1 | |

| | | |
|---|---|---|
| ENSMUSG00000095205 | 1.10527898 | 0.538858846 |
| ENSMUSG00000095206 | 1 | |
| ENSMUSG00000095207 | 1 | |
| ENSMUSG00000095209 | 1 | |
| ENSMUSG00000095210 | 1 | |
| ENSMUSG00000095212 | 1 | |
| ENSMUSG00000095213 | 1 | |
| ENSMUSG00000095214 | 1 | |
| ENSMUSG00000095217 | -1.044273782 | 0.698112512 |
| ENSMUSG00000095218 | 1 | |
| ENSMUSG00000095220 | 1 | |
| ENSMUSG00000095221 | 1 | |
| ENSMUSG00000095222 | 1 | |
| ENSMUSG00000095224 | 1 | |
| ENSMUSG00000095226 | 1 | |
| ENSMUSG00000095228 | 1 | |
| ENSMUSG00000095229 | 1 | |
| ENSMUSG00000095231 | 1 | |
| ENSMUSG00000095232 | 1 | |
| ENSMUSG00000095233 | 1 | |
| ENSMUSG00000095234 | 1 | |
| ENSMUSG00000095236 | 1 | |
| ENSMUSG00000095238 | 1 | |
| ENSMUSG00000095239 | 1 | |
| ENSMUSG00000095240 | 1 | |
| ENSMUSG00000095241 | 1 | |
| ENSMUSG00000095242 | 1 | |
| ENSMUSG00000095243 | 1 | |
| ENSMUSG00000095245 | 1 | |
| ENSMUSG00000095246 | 1 | |
| ENSMUSG00000095247 | 1 | |
| ENSMUSG00000095248 | 1 | |
| ENSMUSG00000095249 | 1 | |
| ENSMUSG00000095250 | 1 | |
| ENSMUSG00000095251 | 1 | |
| ENSMUSG00000095253 | 1.007338094 | 1 |
| ENSMUSG00000095254 | 1 | |
| ENSMUSG00000095255 | 1.11166259 | 0.608497229 |
| ENSMUSG00000095256 | 1 | |
| ENSMUSG00000095257 | 1 | |
| ENSMUSG00000095258 | 1 | |
| ENSMUSG00000095259 | 1 | |
| ENSMUSG00000095260 | 1 | |
| ENSMUSG00000095261 | 1 | |
| ENSMUSG00000095262 | -1.043306199 | 0.698112512 |
| ENSMUSG00000095263 | 1 | |
| ENSMUSG00000095264 | 1 | |
| ENSMUSG00000095265 | 1 | |
| ENSMUSG00000095266 | 1.059463094 | 0.608497229 |
| ENSMUSG00000095267 | 1 | |
| ENSMUSG00000095269 | 1 | |
| ENSMUSG00000095270 | 1 | |
| ENSMUSG00000095272 | 1 | |
| ENSMUSG00000095273 | 1 | |
| ENSMUSG00000095274 | 1 | |
| ENSMUSG00000095275 | 1 | |
| ENSMUSG00000095276 | 1 | |
| ENSMUSG00000095277 | 1 | |
| ENSMUSG00000095278 | 1.059463094 | 0.608497229 |
| ENSMUSG00000095279 | 1 | |
| ENSMUSG00000095280 | -1 | 1 |
| ENSMUSG00000095281 | 1 | |
| ENSMUSG00000095282 | 1 | |

| | | |
|---|---|---|
| ENSMUSG00000095283 | 1 | |
| ENSMUSG00000095284 | 1 | |
| ENSMUSG00000095285 | 1 | |
| ENSMUSG00000095286 | 1 | |
| ENSMUSG00000095287 | 1 | |
| ENSMUSG00000095288 | 1 | |
| ENSMUSG00000095289 | 1 | |
| ENSMUSG00000095290 | 1 | |
| ENSMUSG00000095291 | 1 | |
| ENSMUSG00000095292 | 1 | |
| ENSMUSG00000095293 | 1 | |
| ENSMUSG00000095294 | 1 | |
| ENSMUSG00000095295 | 1 | |
| ENSMUSG00000095296 | 1 | |
| ENSMUSG00000095297 | 1 | |
| ENSMUSG00000095298 | 1 | |
| ENSMUSG00000095299 | 1 | |
| ENSMUSG00000095300 | 1 | |
| ENSMUSG00000095301 | 1 | |
| ENSMUSG00000095302 | 1 | |
| ENSMUSG00000095304 | 1 | |
| ENSMUSG00000095305 | 1 | |
| ENSMUSG00000095306 | -1.043306199 | 0.803260194 |
| ENSMUSG00000095307 | 1.069068979 | 0.608497229 |
| ENSMUSG00000095309 | 1 | |
| ENSMUSG00000095310 | 1 | |
| ENSMUSG00000095311 | -1.053549062 | 0.698112512 |
| ENSMUSG00000095312 | 1 | |
| ENSMUSG00000095314 | 1 | |
| ENSMUSG00000095315 | -1.432443895 | 0.049625918 |
| ENSMUSG00000095316 | 1 | |
| ENSMUSG00000095318 | 1 | |
| ENSMUSG00000095319 | 1 | |
| ENSMUSG00000095320 | 1 | |
| ENSMUSG00000095321 | 1 | |
| ENSMUSG00000095322 | 1 | |
| ENSMUSG00000095323 | 1 | |
| ENSMUSG00000095324 | 1 | |
| ENSMUSG00000095325 | -1.162838636 | 0.214753594 |
| ENSMUSG00000095327 | 1 | |
| ENSMUSG00000095328 | 1 | |
| ENSMUSG00000095329 | -1.043306199 | 0.698112512 |
| ENSMUSG00000095330 | 1 | |
| ENSMUSG00000095331 | 1 | |
| ENSMUSG00000095332 | -1.170609014 | 0.583037087 |
| ENSMUSG00000095333 | 1 | |
| ENSMUSG00000095334 | 1 | |
| ENSMUSG00000095335 | 1 | |
| ENSMUSG00000095336 | 1 | |
| ENSMUSG00000095338 | 1 | |
| ENSMUSG00000095339 | 1 | |
| ENSMUSG00000095340 | 1 | |
| ENSMUSG00000095341 | 1 | |
| ENSMUSG00000095342 | 1 | |
| ENSMUSG00000095344 | 1.024656056 | 0.984785517 |
| ENSMUSG00000095345 | 1 | |
| ENSMUSG00000095346 | 1 | |
| ENSMUSG00000095347 | 1 | |
| ENSMUSG00000095348 | 1 | |
| ENSMUSG00000095349 | 1 | |
| ENSMUSG00000095350 | 1 | |
| ENSMUSG00000095351 | 1 | |
| ENSMUSG00000095353 | 1 | |

| | | |
|---|---|---|
| ENSMUSG00000095356 | 1 | |
| ENSMUSG00000095357 | 1 | |
| ENSMUSG00000095358 | 1 | |
| ENSMUSG00000095360 | 1 | |
| ENSMUSG00000095361 | 1 | |
| ENSMUSG00000095362 | -1.119119717 | 0.461811948 |
| ENSMUSG00000095363 | 1 | |
| ENSMUSG00000095365 | 1 | |
| ENSMUSG00000095366 | 1 | |
| ENSMUSG00000095367 | 1 | |
| ENSMUSG00000095368 | 1 | |
| ENSMUSG00000095369 | 1 | |
| ENSMUSG00000095370 | 1 | |
| ENSMUSG00000095371 | 1 | |
| ENSMUSG00000095372 | 1 | |
| ENSMUSG00000095373 | 1 | |
| ENSMUSG00000095374 | 1 | |
| ENSMUSG00000095376 | 1 | |
| ENSMUSG00000095377 | -1.043306199 | 0.79291651 |
| ENSMUSG00000095379 | 1 | |
| ENSMUSG00000095380 | 1 | |
| ENSMUSG00000095382 | 1 | |
| ENSMUSG00000095383 | 1 | |
| ENSMUSG00000095384 | 1 | |
| ENSMUSG00000095385 | 1.059463094 | 0.698112512 |
| ENSMUSG00000095386 | 1 | |
| ENSMUSG00000095387 | 1 | |
| ENSMUSG00000095388 | 1 | |
| ENSMUSG00000095389 | 1 | |
| ENSMUSG00000095390 | 1 | |
| ENSMUSG00000095392 | 1 | |
| ENSMUSG00000095393 | 1 | |
| ENSMUSG00000095394 | 1 | |
| ENSMUSG00000095395 | 1 | |
| ENSMUSG00000095396 | -1.053549062 | 0.778372272 |
| ENSMUSG00000095398 | 1 | |
| ENSMUSG00000095399 | 1 | |
| ENSMUSG00000095401 | 1 | |
| ENSMUSG00000095402 | 1 | |
| ENSMUSG00000095403 | -1.074012757 | 0.674977785 |
| ENSMUSG00000095406 | 1.059463094 | 0.698112512 |
| ENSMUSG00000095407 | 1 | |
| ENSMUSG00000095408 | 1 | |
| ENSMUSG00000095409 | 1 | |
| ENSMUSG00000095410 | 1 | |
| ENSMUSG00000095411 | 1 | |
| ENSMUSG00000095412 | 1 | |
| ENSMUSG00000095413 | 1 | |
| ENSMUSG00000095414 | 1 | |
| ENSMUSG00000095416 | 1 | |
| ENSMUSG00000095418 | 1 | |
| ENSMUSG00000095419 | -1.03962336 | 0.698112512 |
| ENSMUSG00000095420 | 1 | |
| ENSMUSG00000095422 | 1 | |
| ENSMUSG00000095423 | 1 | |
| ENSMUSG00000095424 | 1 | |
| ENSMUSG00000095425 | 1 | |
| ENSMUSG00000095426 | 1 | |
| ENSMUSG00000095427 | -1.083415624 | 0.719087885 |
| ENSMUSG00000095428 | 1 | |
| ENSMUSG00000095429 | 1 | |
| ENSMUSG00000095430 | 1 | |
| ENSMUSG00000095431 | 1 | |

| | | |
|---|---|---|
| ENSMUSG00000095432 | -1.148638755 | 0.418492024 |
| ENSMUSG00000095433 | 1 | |
| ENSMUSG00000095434 | 1 | |
| ENSMUSG00000095435 | 1 | |
| ENSMUSG00000095436 | 1 | |
| ENSMUSG00000095437 | 1 | |
| ENSMUSG00000095438 | -1.009842415 | 1 |
| ENSMUSG00000095440 | -1.310070721 | 0.063588408 |
| ENSMUSG00000095442 | 1 | |
| ENSMUSG00000095444 | 1 | |
| ENSMUSG00000095445 | 1 | |
| ENSMUSG00000095446 | 1 | |
| ENSMUSG00000095447 | 1 | |
| ENSMUSG00000095448 | 1 | |
| ENSMUSG00000095449 | 1 | |
| ENSMUSG00000095450 | 1 | |
| ENSMUSG00000095452 | 1 | |
| ENSMUSG00000095453 | 1 | |
| ENSMUSG00000095455 | 1 | |
| ENSMUSG00000095456 | 1 | |
| ENSMUSG00000095457 | 1 | |
| ENSMUSG00000095458 | 1 | |
| ENSMUSG00000095460 | 1 | |
| ENSMUSG00000095462 | 1 | |
| ENSMUSG00000095463 | 1 | |
| ENSMUSG00000095464 | 1 | |
| ENSMUSG00000095465 | 1 | |
| ENSMUSG00000095466 | 1 | |
| ENSMUSG00000095467 | 1 | |
| ENSMUSG00000095469 | 1 | |
| ENSMUSG00000095470 | 1 | |
| ENSMUSG00000095472 | 1 | |
| ENSMUSG00000095474 | 1 | |
| ENSMUSG00000095475 | 1 | |
| ENSMUSG00000095476 | 1 | |
| ENSMUSG00000095477 | 1.019083579 | 0.949812939 |
| ENSMUSG00000095478 | -1.03962336 | 0.814089245 |
| ENSMUSG00000095479 | 1 | |
| ENSMUSG00000095482 | 1 | |
| ENSMUSG00000095483 | 1 | |
| ENSMUSG00000095484 | 1 | |
| ENSMUSG00000095486 | 1 | |
| ENSMUSG00000095487 | 1 | |
| ENSMUSG00000095489 | 1 | |
| ENSMUSG00000095490 | 1 | |
| ENSMUSG00000095491 | 1 | |
| ENSMUSG00000095492 | 1 | |
| ENSMUSG00000095493 | 1.07913413 | 0.830576988 |
| ENSMUSG00000095494 | 1 | |
| ENSMUSG00000095495 | 1 | |
| ENSMUSG00000095497 | 1 | |
| ENSMUSG00000095498 | 1 | |
| ENSMUSG00000095499 | 1 | |
| ENSMUSG00000095500 | 1 | |
| ENSMUSG00000095501 | 1 | |
| ENSMUSG00000095502 | 1 | |
| ENSMUSG00000095503 | 1 | |
| ENSMUSG00000095504 | 1 | |
| ENSMUSG00000095505 | 1 | |
| ENSMUSG00000095506 | 1 | |
| ENSMUSG00000095507 | 1 | |
| ENSMUSG00000095508 | 1 | |
| ENSMUSG00000095509 | 1 | |

| | | |
|---|---|---|
| ENSMUSG00000095510 | 1 | |
| ENSMUSG00000095511 | 1 | |
| ENSMUSG00000095512 | 1 | |
| ENSMUSG00000095513 | 1 | |
| ENSMUSG00000095514 | 1.132639128 | 0.217803313 |
| ENSMUSG00000095516 | 1 | |
| ENSMUSG00000095517 | 1 | |
| ENSMUSG00000095518 | 1 | |
| ENSMUSG00000095519 | 1 | |
| ENSMUSG00000095520 | 1 | |
| ENSMUSG00000095521 | 1 | |
| ENSMUSG00000095522 | 1 | |
| ENSMUSG00000095523 | 1 | |
| ENSMUSG00000095524 | 1.097820634 | 0.70934529 |
| ENSMUSG00000095525 | 1 | |
| ENSMUSG00000095526 | 1 | |
| ENSMUSG00000095527 | 1 | |
| ENSMUSG00000095528 | 1 | |
| ENSMUSG00000095529 | 1 | |
| ENSMUSG00000095530 | 1.014545335 | 0.979002724 |
| ENSMUSG00000095531 | 1 | |
| ENSMUSG00000095532 | 1 | |
| ENSMUSG00000095533 | 1 | |
| ENSMUSG00000095534 | 1 | |
| ENSMUSG00000095536 | 1 | |
| ENSMUSG00000095537 | 1 | |
| ENSMUSG00000095538 | 1 | |
| ENSMUSG00000095539 | 1 | |
| ENSMUSG00000095540 | 1 | |
| ENSMUSG00000095541 | 1 | |
| ENSMUSG00000095542 | 1 | |
| ENSMUSG00000095543 | 1 | |
| ENSMUSG00000095544 | 1 | |
| ENSMUSG00000095545 | 1 | |
| ENSMUSG00000095546 | 1 | |
| ENSMUSG00000095547 | 1 | |
| ENSMUSG00000095549 | 1 | |
| ENSMUSG00000095550 | 1 | |
| ENSMUSG00000095551 | 1 | |
| ENSMUSG00000095552 | 1 | |
| ENSMUSG00000095553 | 1 | |
| ENSMUSG00000095554 | 1 | |
| ENSMUSG00000095555 | 1 | |
| ENSMUSG00000095557 | 1 | |
| ENSMUSG00000095558 | 1 | |
| ENSMUSG00000095559 | 1 | |
| ENSMUSG00000095560 | 1 | |
| ENSMUSG00000095561 | 1 | |
| ENSMUSG00000095562 | -1.03962336 | 0.698112512 |
| ENSMUSG00000095563 | 1 | |
| ENSMUSG00000095564 | 1 | |
| ENSMUSG00000095565 | 1 | |
| ENSMUSG00000095567 | 1.069534027 | 0.711581415 |
| ENSMUSG00000095569 | 1 | |
| ENSMUSG00000095570 | 1 | |
| ENSMUSG00000095571 | 1 | |
| ENSMUSG00000095572 | 1 | |
| ENSMUSG00000095573 | 1 | |
| ENSMUSG00000095574 | 1 | |
| ENSMUSG00000095576 | 1 | |
| ENSMUSG00000095577 | 1 | |
| ENSMUSG00000095578 | 1 | |
| ENSMUSG00000095580 | 1 | |

| | | |
|---|---|---|
| ENSMUSG00000095581 | 1 | |
| ENSMUSG00000095583 | 1 | |
| ENSMUSG00000095584 | 1 | |
| ENSMUSG00000095585 | 1 | |
| ENSMUSG00000095586 | 1 | |
| ENSMUSG00000095587 | 1 | |
| ENSMUSG00000095588 | -1.03962336 | 0.698112512 |
| ENSMUSG00000095589 | 1 | |
| ENSMUSG00000095590 | 1 | |
| ENSMUSG00000095591 | 1 | |
| ENSMUSG00000095592 | 1 | |
| ENSMUSG00000095593 | 1 | |
| ENSMUSG00000095594 | 1 | |
| ENSMUSG00000095595 | -1.222502199 | 0.320257489 |
| ENSMUSG00000095596 | 1 | |
| ENSMUSG00000095597 | 1.47331616 | 0.009579947 |
| ENSMUSG00000095599 | 1 | |
| ENSMUSG00000095600 | -1.051018293 | 0.828564184 |
| ENSMUSG00000095601 | 1 | |
| ENSMUSG00000095602 | 1 | |
| ENSMUSG00000095603 | 1 | |
| ENSMUSG00000095604 | 1 | |
| ENSMUSG00000095605 | 1 | |
| ENSMUSG00000095606 | 1 | |
| ENSMUSG00000095607 | 1 | |
| ENSMUSG00000095608 | 1 | |
| ENSMUSG00000095609 | 1 | |
| ENSMUSG00000095610 | 1 | |
| ENSMUSG00000095612 | 1 | |
| ENSMUSG00000095614 | -1.084645496 | 0.532369657 |
| ENSMUSG00000095615 | 1 | |
| ENSMUSG00000095616 | 1 | |
| ENSMUSG00000095618 | 1 | |
| ENSMUSG00000095619 | 1 | |
| ENSMUSG00000095620 | -1.044273782 | 0.698112512 |
| ENSMUSG00000095621 | 1 | |
| ENSMUSG00000095623 | 1 | |
| ENSMUSG00000095624 | 1 | |
| ENSMUSG00000095625 | 1 | |
| ENSMUSG00000095626 | 1 | |
| ENSMUSG00000095627 | 1 | |
| ENSMUSG00000095628 | 1 | |
| ENSMUSG00000095629 | 1 | |
| ENSMUSG00000095630 | 1 | |
| ENSMUSG00000095631 | 1 | |
| ENSMUSG00000095632 | 1 | |
| ENSMUSG00000095633 | 1 | |
| ENSMUSG00000095634 | 1 | |
| ENSMUSG00000095635 | 1.059463094 | 0.608497229 |
| ENSMUSG00000095636 | 1 | |
| ENSMUSG00000095637 | 1 | |
| ENSMUSG00000095638 | 1 | |
| ENSMUSG00000095639 | 1 | |
| ENSMUSG00000095640 | 1 | |
| ENSMUSG00000095641 | 1 | |
| ENSMUSG00000095642 | 1 | |
| ENSMUSG00000095643 | 1 | |
| ENSMUSG00000095645 | 1 | |
| ENSMUSG00000095646 | 1 | |
| ENSMUSG00000095647 | 1 | |
| ENSMUSG00000095648 | 1 | |
| ENSMUSG00000095649 | 1 | |
| ENSMUSG00000095650 | 1 | |

| | | |
|---|---|---|
| ENSMUSG00000095651 | -1.075173119 | 0.532369657 |
| ENSMUSG00000095652 | 1 | |
| ENSMUSG00000095653 | 1.059463094 | 0.608497229 |
| ENSMUSG00000095654 | 1 | |
| ENSMUSG00000095655 | 1 | |
| ENSMUSG00000095656 | 1 | |
| ENSMUSG00000095657 | 1 | |
| ENSMUSG00000095658 | 1 | |
| ENSMUSG00000095659 | 1 | |
| ENSMUSG00000095660 | 1 | |
| ENSMUSG00000095661 | -1.090507733 | 0.77378344 |
| ENSMUSG00000095662 | 1 | |
| ENSMUSG00000095663 | 1 | |
| ENSMUSG00000095664 | 1 | |
| ENSMUSG00000095666 | 1 | |
| ENSMUSG00000095667 | 1 | |
| ENSMUSG00000095668 | 1 | |
| ENSMUSG00000095670 | 1 | |
| ENSMUSG00000095671 | 1 | |
| ENSMUSG00000095672 | 1 | |
| ENSMUSG00000095673 | 1 | |
| ENSMUSG00000095674 | 1 | |
| ENSMUSG00000095675 | 1 | |
| ENSMUSG00000095676 | 1 | |
| ENSMUSG00000095677 | 1.035165778 | 0.751670161 |
| ENSMUSG00000095678 | 1 | |
| ENSMUSG00000095680 | 1 | |
| ENSMUSG00000095681 | 1 | |
| ENSMUSG00000095682 | 1 | |
| ENSMUSG00000095683 | 1 | |
| ENSMUSG00000095684 | 1 | |
| ENSMUSG00000095685 | 1 | |
| ENSMUSG00000095686 | 1 | |
| ENSMUSG00000095687 | 1.073609354 | 0.702922227 |
| ENSMUSG00000095689 | 1 | |
| ENSMUSG00000095690 | 1 | |
| ENSMUSG00000095691 | 1 | |
| ENSMUSG00000095692 | 1 | |
| ENSMUSG00000095693 | 1 | |
| ENSMUSG00000095696 | 1 | |
| ENSMUSG00000095697 | 1 | |
| ENSMUSG00000095698 | 1 | |
| ENSMUSG00000095699 | 1 | |
| ENSMUSG00000095700 | 1 | |
| ENSMUSG00000095701 | 1 | |
| ENSMUSG00000095702 | 1 | |
| ENSMUSG00000095703 | 1 | |
| ENSMUSG00000095704 | 1 | 1 |
| ENSMUSG00000095705 | 1 | |
| ENSMUSG00000095706 | 1 | |
| ENSMUSG00000095709 | 1 | |
| ENSMUSG00000095710 | 1 | |
| ENSMUSG00000095711 | 1 | |
| ENSMUSG00000095712 | 1 | |
| ENSMUSG00000095713 | 1 | |
| ENSMUSG00000095714 | 1 | |
| ENSMUSG00000095716 | 1 | |
| ENSMUSG00000095717 | 1 | |
| ENSMUSG00000095718 | 1 | |
| ENSMUSG00000095720 | 1 | |
| ENSMUSG00000095721 | 1 | |
| ENSMUSG00000095722 | 1 | |
| ENSMUSG00000095723 | 1 | |

| | | |
|---|---|---|
| ENSMUSG00000095724 | -1.067882909 | 0.823091034 |
| ENSMUSG00000095725 | 1 | |
| ENSMUSG00000095727 | 1 | |
| ENSMUSG00000095728 | 1 | |
| ENSMUSG00000095729 | 1 | |
| ENSMUSG00000095730 | 1.132639128 | 0.398767938 |
| ENSMUSG00000095731 | 1 | |
| ENSMUSG00000095732 | 1 | |
| ENSMUSG00000095734 | 1 | |
| ENSMUSG00000095736 | 1 | |
| ENSMUSG00000095737 | 1 | |
| ENSMUSG00000095738 | 1 | |
| ENSMUSG00000095739 | 1 | |
| ENSMUSG00000095740 | -1.043306199 | 0.698112512 |
| ENSMUSG00000095741 | 1 | |
| ENSMUSG00000095742 | 1.023970213 | 1 |
| ENSMUSG00000095743 | 1 | |
| ENSMUSG00000095744 | 1 | |
| ENSMUSG00000095745 | 1 | |
| ENSMUSG00000095747 | 1 | |
| ENSMUSG00000095749 | 1 | |
| ENSMUSG00000095752 | -1 | 1 |
| ENSMUSG00000095753 | 1 | |
| ENSMUSG00000095754 | 1 | |
| ENSMUSG00000095755 | 1 | |
| ENSMUSG00000095756 | 1 | |
| ENSMUSG00000095757 | 1 | |
| ENSMUSG00000095758 | 1 | |
| ENSMUSG00000095759 | 1 | |
| ENSMUSG00000095761 | 1 | |
| ENSMUSG00000095762 | -1.028348525 | 0.912425817 |
| ENSMUSG00000095763 | 1 | |
| ENSMUSG00000095764 | 1 | |
| ENSMUSG00000095765 | 1 | |
| ENSMUSG00000095766 | 1 | |
| ENSMUSG00000095768 | 1 | |
| ENSMUSG00000095769 | 1 | |
| ENSMUSG00000095770 | 1 | |
| ENSMUSG00000095771 | 1 | |
| ENSMUSG00000095772 | 1 | |
| ENSMUSG00000095773 | 1 | |
| ENSMUSG00000095774 | 1 | |
| ENSMUSG00000095776 | 1 | |
| ENSMUSG00000095778 | 1 | |
| ENSMUSG00000095779 | 1 | |
| ENSMUSG00000095780 | 1.059463094 | 0.608497229 |
| ENSMUSG00000095782 | 1 | |
| ENSMUSG00000095785 | 1 | |
| ENSMUSG00000095786 | 1 | |
| ENSMUSG00000095787 | 1 | |
| ENSMUSG00000095788 | -1.122034075 | 0.713731505 |
| ENSMUSG00000095789 | -1.122027951 | 0.729018288 |
| ENSMUSG00000095791 | 1 | |
| ENSMUSG00000095792 | 1 | |
| ENSMUSG00000095793 | 1 | |
| ENSMUSG00000095794 | 1 | |
| ENSMUSG00000095796 | 1 | |
| ENSMUSG00000095797 | 1 | |
| ENSMUSG00000095798 | 1 | |
| ENSMUSG00000095799 | 1 | |
| ENSMUSG00000095802 | 1 | |
| ENSMUSG00000095803 | 1 | |
| ENSMUSG00000095804 | 1 | |

| | | |
|---|---|---|
| ENSMUSG00000095805 | 1 | |
| ENSMUSG00000095806 | 1 | |
| ENSMUSG00000095807 | 1 | |
| ENSMUSG00000095809 | 1 | |
| ENSMUSG00000095812 | -1.078590893 | 0.698112512 |
| ENSMUSG00000095813 | 1 | |
| ENSMUSG00000095814 | 1 | |
| ENSMUSG00000095816 | 1 | |
| ENSMUSG00000095817 | 1 | |
| ENSMUSG00000095818 | 1 | |
| ENSMUSG00000095820 | 1 | |
| ENSMUSG00000095821 | 1 | |
| ENSMUSG00000095822 | 1 | |
| ENSMUSG00000095824 | 1 | |
| ENSMUSG00000095825 | 1 | |
| ENSMUSG00000095826 | 1.189207115 | 0.294533546 |
| ENSMUSG00000095827 | 1 | |
| ENSMUSG00000095828 | 1 | |
| ENSMUSG00000095829 | 1 | |
| ENSMUSG00000095831 | 1 | |
| ENSMUSG00000095834 | 1 | |
| ENSMUSG00000095836 | -1.021087346 | 0.979212363 |
| ENSMUSG00000095837 | 1 | |
| ENSMUSG00000095839 | 1 | |
| ENSMUSG00000095841 | 1 | |
| ENSMUSG00000095842 | 1 | |
| ENSMUSG00000095843 | 1 | |
| ENSMUSG00000095845 | -1.032291545 | 0.917472106 |
| ENSMUSG00000095846 | 1 | |
| ENSMUSG00000095847 | -1.043722544 | 0.869658231 |
| ENSMUSG00000095849 | 1 | |
| ENSMUSG00000095850 | 1 | |
| ENSMUSG00000095851 | 1 | |
| ENSMUSG00000095852 | 1 | |
| ENSMUSG00000095853 | 1 | |
| ENSMUSG00000095854 | 1 | |
| ENSMUSG00000095856 | 1 | |
| ENSMUSG00000095857 | 1 | |
| ENSMUSG00000095858 | 1 | |
| ENSMUSG00000095859 | 1 | |
| ENSMUSG00000095860 | 1 | |
| ENSMUSG00000095862 | 1 | |
| ENSMUSG00000095863 | 1 | |
| ENSMUSG00000095864 | 1 | |
| ENSMUSG00000095865 | 1 | |
| ENSMUSG00000095866 | 1 | |
| ENSMUSG00000095867 | 1 | |
| ENSMUSG00000095868 | 1.127832095 | 0.608497229 |
| ENSMUSG00000095869 | 1 | |
| ENSMUSG00000095870 | 1 | |
| ENSMUSG00000095871 | 1 | |
| ENSMUSG00000095872 | 1 | |
| ENSMUSG00000095873 | 1 | |
| ENSMUSG00000095874 | 1 | |
| ENSMUSG00000095876 | -1 | 1 |
| ENSMUSG00000095878 | 1.059463094 | 0.608497229 |
| ENSMUSG00000095879 | 1 | |
| ENSMUSG00000095880 | 1 | |
| ENSMUSG00000095883 | 1 | |
| ENSMUSG00000095884 | 1 | |
| ENSMUSG00000095885 | 1 | |
| ENSMUSG00000095886 | 1 | |
| ENSMUSG00000095887 | 1 | |

| | | |
|---|---|---|
| ENSMUSG00000095889 | 1 | |
| ENSMUSG00000095890 | 1 | |
| ENSMUSG00000095891 | 1.059463094 | 0.608497229 |
| ENSMUSG00000095892 | 1.059463094 | 0.608497229 |
| ENSMUSG00000095893 | 1 | |
| ENSMUSG00000095894 | 1 | |
| ENSMUSG00000095895 | -1.043306199 | 0.698112512 |
| ENSMUSG00000095896 | 1 | |
| ENSMUSG00000095897 | 1 | |
| ENSMUSG00000095898 | 1 | |
| ENSMUSG00000095899 | 1 | |
| ENSMUSG00000095900 | 1 | |
| ENSMUSG00000095901 | 1 | |
| ENSMUSG00000095902 | 1 | |
| ENSMUSG00000095903 | 1 | |
| ENSMUSG00000095905 | 1 | |
| ENSMUSG00000095906 | 1 | |
| ENSMUSG00000095907 | 1 | |
| ENSMUSG00000095909 | -1.254679684 | 0.019659061 |
| ENSMUSG00000095910 | 1 | |
| ENSMUSG00000095911 | 1 | |
| ENSMUSG00000095912 | 1 | |
| ENSMUSG00000095913 | -1.223451821 | 0.425805785 |
| ENSMUSG00000095914 | 1 | |
| ENSMUSG00000095915 | 1 | |
| ENSMUSG00000095916 | 1 | |
| ENSMUSG00000095917 | 1 | |
| ENSMUSG00000095918 | 1 | |
| ENSMUSG00000095920 | -1.044273782 | 0.698112512 |
| ENSMUSG00000095922 | 1 | |
| ENSMUSG00000095923 | 1 | |
| ENSMUSG00000095924 | 1 | |
| ENSMUSG00000095925 | 1 | |
| ENSMUSG00000095926 | 1 | |
| ENSMUSG00000095927 | 1 | |
| ENSMUSG00000095928 | 1 | |
| ENSMUSG00000095929 | 1 | |
| ENSMUSG00000095930 | -1.132219452 | 0.311652447 |
| ENSMUSG00000095931 | 1 | |
| ENSMUSG00000095932 | 1 | |
| ENSMUSG00000095933 | 1 | |
| ENSMUSG00000095934 | 1 | |
| ENSMUSG00000095935 | 1 | |
| ENSMUSG00000095936 | 1 | |
| ENSMUSG00000095937 | 1 | |
| ENSMUSG00000095938 | 1 | |
| ENSMUSG00000095939 | 1 | |
| ENSMUSG00000095940 | 1.019083579 | 0.968937442 |
| ENSMUSG00000095941 | 1 | |
| ENSMUSG00000095942 | 1 | |
| ENSMUSG00000095943 | 1.05038453 | 0.887340389 |
| ENSMUSG00000095944 | 1 | |
| ENSMUSG00000095945 | 1 | |
| ENSMUSG00000095946 | 1 | |
| ENSMUSG00000095947 | 1 | |
| ENSMUSG00000095948 | 1 | |
| ENSMUSG00000095950 | 1 | |
| ENSMUSG00000095951 | 1 | |
| ENSMUSG00000095952 | 1 | |
| ENSMUSG00000095953 | 1 | |
| ENSMUSG00000095954 | 1 | |
| ENSMUSG00000095955 | 1 | |
| ENSMUSG00000095956 | 1 | |

| | | |
|---|---|---|
| ENSMUSG00000095957 | 1 | |
| ENSMUSG00000095958 | 1 | |
| ENSMUSG00000095959 | 1 | |
| ENSMUSG00000095960 | 1 | |
| ENSMUSG00000095961 | 1 | |
| ENSMUSG00000095962 | 1 | |
| ENSMUSG00000095963 | 1 | |
| ENSMUSG00000095964 | 1 | |
| ENSMUSG00000095965 | 1 | |
| ENSMUSG00000095966 | 1 | |
| ENSMUSG00000095968 | 1 | |
| ENSMUSG00000095969 | 1 | |
| ENSMUSG00000095970 | 1 | |
| ENSMUSG00000095971 | 1 | |
| ENSMUSG00000095972 | 1 | |
| ENSMUSG00000095973 | 1 | |
| ENSMUSG00000095974 | 1 | |
| ENSMUSG00000095975 | 1 | |
| ENSMUSG00000095976 | 1 | |
| ENSMUSG00000095977 | 1 | |
| ENSMUSG00000095978 | 1 | |
| ENSMUSG00000095979 | 1 | |
| ENSMUSG00000095980 | 1 | |
| ENSMUSG00000095981 | 1 | |
| ENSMUSG00000095982 | 1 | |
| ENSMUSG00000095984 | 1 | |
| ENSMUSG00000095985 | 1 | |
| ENSMUSG00000095986 | 1 | |
| ENSMUSG00000095987 | 1 | |
| ENSMUSG00000095988 | 1 | |
| ENSMUSG00000095989 | 1 | |
| ENSMUSG00000095990 | -1.004473935 | 1 |
| ENSMUSG00000095991 | 1 | |
| ENSMUSG00000095992 | 1 | |
| ENSMUSG00000095993 | -1.069095149 | 0.756109967 |
| ENSMUSG00000095994 | 1 | |
| ENSMUSG00000095996 | 1 | |
| ENSMUSG00000095997 | 1 | |
| ENSMUSG00000095998 | 1 | |
| ENSMUSG00000096000 | 1 | |
| ENSMUSG00000096001 | -1.01999916 | 0.901903401 |
| ENSMUSG00000096002 | 1 | |
| ENSMUSG00000096003 | 1 | |
| ENSMUSG00000096005 | 1 | |
| ENSMUSG00000096006 | 1.065838392 | 0.836005168 |
| ENSMUSG00000096008 | 1 | |
| ENSMUSG00000096009 | 1 | |
| ENSMUSG00000096010 | -1.000322333 | 1 |
| ENSMUSG00000096011 | 1 | |
| ENSMUSG00000096014 | 1 | |
| ENSMUSG00000096015 | 1 | |
| ENSMUSG00000096016 | 1 | |
| ENSMUSG00000096017 | 1 | |
| ENSMUSG00000096018 | 1 | |
| ENSMUSG00000096019 | 1 | |
| ENSMUSG00000096020 | 1 | |
| ENSMUSG00000096021 | 1 | |
| ENSMUSG00000096022 | 1 | |
| ENSMUSG00000096023 | 1 | |
| ENSMUSG00000096024 | 1.014731082 | 0.99911685 |
| ENSMUSG00000096025 | 1.122462048 | 0.215991492 |
| ENSMUSG00000096027 | 1 | |
| ENSMUSG00000096028 | 1 | |

| | | |
|---|---|---|
| ENSMUSG00000096029 | 1 | |
| ENSMUSG00000096030 | 1 | |
| ENSMUSG00000096031 | 1 | |
| ENSMUSG00000096033 | -1.044273782 | 0.698112512 |
| ENSMUSG00000096035 | -1.03124175 | 0.925352084 |
| ENSMUSG00000096036 | 1 | |
| ENSMUSG00000096037 | 1 | |
| ENSMUSG00000096038 | 1 | |
| ENSMUSG00000096039 | -1.18980823 | 0.449109628 |
| ENSMUSG00000096041 | 1 | |
| ENSMUSG00000096042 | 1 | |
| ENSMUSG00000096043 | 1 | |
| ENSMUSG00000096044 | -1.111880267 | 0.641410989 |
| ENSMUSG00000096045 | 1 | |
| ENSMUSG00000096046 | 1 | |
| ENSMUSG00000096047 | 1 | |
| ENSMUSG00000096048 | 1 | |
| ENSMUSG00000096049 | 1 | |
| ENSMUSG00000096051 | -1 | 1 |
| ENSMUSG00000096052 | 1.008034876 | 1 |
| ENSMUSG00000096053 | 1 | |
| ENSMUSG00000096054 | 1.122531824 | 0.160807489 |
| ENSMUSG00000096055 | 1 | |
| ENSMUSG00000096056 | -1.081819798 | 0.608497229 |
| ENSMUSG00000096057 | 1 | |
| ENSMUSG00000096061 | 1 | |
| ENSMUSG00000096065 | 1 | |
| ENSMUSG00000096066 | 1 | |
| ENSMUSG00000096067 | 1 | |
| ENSMUSG00000096068 | 1 | |
| ENSMUSG00000096069 | 1 | |
| ENSMUSG00000096070 | 1 | |
| ENSMUSG00000096071 | 1 | |
| ENSMUSG00000096072 | 1 | |
| ENSMUSG00000096073 | 1 | |
| ENSMUSG00000096074 | 1 | |
| ENSMUSG00000096075 | 1 | |
| ENSMUSG00000096076 | 1 | |
| ENSMUSG00000096078 | 1 | |
| ENSMUSG00000096079 | 1 | |
| ENSMUSG00000096080 | 1 | |
| ENSMUSG00000096081 | 1 | |
| ENSMUSG00000096082 | 1 | |
| ENSMUSG00000096083 | 1 | |
| ENSMUSG00000096084 | 1 | |
| ENSMUSG00000096085 | 1 | |
| ENSMUSG00000096086 | -1.006483203 | 1 |
| ENSMUSG00000096087 | 1 | |
| ENSMUSG00000096088 | 1 | |
| ENSMUSG00000096090 | 1 | |
| ENSMUSG00000096091 | 1 | |
| ENSMUSG00000096092 | 1 | |
| ENSMUSG00000096094 | -1.024530911 | 0.949679283 |
| ENSMUSG00000096095 | 1 | |
| ENSMUSG00000096096 | -1.044273782 | 0.698112512 |
| ENSMUSG00000096097 | 1 | |
| ENSMUSG00000096098 | 1 | |
| ENSMUSG00000096099 | 1 | |
| ENSMUSG00000096100 | 1 | |
| ENSMUSG00000096102 | -1.053549062 | 0.698112512 |
| ENSMUSG00000096105 | 1 | |
| ENSMUSG00000096106 | 1 | |
| ENSMUSG00000096107 | -1 | 1 |

| | | |
|---|---|---|
| ENSMUSG00000096108 | 1 | |
| ENSMUSG00000096109 | 1 | |
| ENSMUSG00000096110 | 1 | |
| ENSMUSG00000096111 | -1.069362274 | 0.870103307 |
| ENSMUSG00000096112 | 1 | |
| ENSMUSG00000096113 | 1 | |
| ENSMUSG00000096114 | 1 | |
| ENSMUSG00000096117 | 1 | |
| ENSMUSG00000096119 | 1 | |
| ENSMUSG00000096120 | 1 | |
| ENSMUSG00000096122 | 1 | |
| ENSMUSG00000096124 | 1 | |
| ENSMUSG00000096126 | 1 | |
| ENSMUSG00000096127 | 1 | |
| ENSMUSG00000096129 | 1 | |
| ENSMUSG00000096130 | 1 | |
| ENSMUSG00000096131 | 1 | |
| ENSMUSG00000096132 | 1 | |
| ENSMUSG00000096133 | 1 | |
| ENSMUSG00000096134 | 1 | |
| ENSMUSG00000096135 | 1 | |
| ENSMUSG00000096136 | 1 | |
| ENSMUSG00000096137 | 1 | |
| ENSMUSG00000096138 | 1 | |
| ENSMUSG00000096139 | 1 | |
| ENSMUSG00000096140 | -1.069628355 | 0.808989997 |
| ENSMUSG00000096141 | 1.059463094 | 0.822158059 |
| ENSMUSG00000096142 | 1 | |
| ENSMUSG00000096144 | 1.059463094 | 0.608497229 |
| ENSMUSG00000096145 | -1.042308121 | 0.698112512 |
| ENSMUSG00000096146 | -1.03962336 | 0.814089245 |
| ENSMUSG00000096147 | 1 | |
| ENSMUSG00000096148 | 1 | |
| ENSMUSG00000096149 | 1 | |
| ENSMUSG00000096150 | 1 | |
| ENSMUSG00000096151 | 1 | |
| ENSMUSG00000096152 | 1 | |
| ENSMUSG00000096153 | 1 | |
| ENSMUSG00000096154 | 1 | |
| ENSMUSG00000096156 | 1 | |
| ENSMUSG00000096157 | 1.028323353 | 0.903464611 |
| ENSMUSG00000096158 | 1 | |
| ENSMUSG00000096159 | 1 | |
| ENSMUSG00000096160 | -1.03962336 | 0.698112512 |
| ENSMUSG00000096161 | 1 | |
| ENSMUSG00000096163 | 1 | |
| ENSMUSG00000096164 | 1 | |
| ENSMUSG00000096166 | 1 | |
| ENSMUSG00000096167 | 1 | |
| ENSMUSG00000096168 | 1 | |
| ENSMUSG00000096169 | 1.019083579 | 0.949812939 |
| ENSMUSG00000096170 | 1 | |
| ENSMUSG00000096171 | -1.11061171 | 0.547016491 |
| ENSMUSG00000096172 | 1 | |
| ENSMUSG00000096173 | 1 | |
| ENSMUSG00000096174 | 1 | |
| ENSMUSG00000096175 | -1 | 1 |
| ENSMUSG00000096176 | 1 | |
| ENSMUSG00000096177 | 1.054881277 | 0.832803619 |
| ENSMUSG00000096178 | 1 | |
| ENSMUSG00000096179 | 1 | |
| ENSMUSG00000096180 | 1 | |
| ENSMUSG00000096181 | 1 | |

| | | |
|---|---|---|
| ENSMUSG00000096182 | 1 | |
| ENSMUSG00000096183 | 1 | |
| ENSMUSG00000096184 | 1 | |
| ENSMUSG00000096185 | 1 | |
| ENSMUSG00000096186 | 1 | |
| ENSMUSG00000096188 | 1.079037069 | 0.76525317 |
| ENSMUSG00000096189 | 1 | |
| ENSMUSG00000096190 | 1 | |
| ENSMUSG00000096191 | 1 | |
| ENSMUSG00000096192 | -1.043306199 | 0.79291651 |
| ENSMUSG00000096193 | 1 | |
| ENSMUSG00000096194 | 1 | |
| ENSMUSG00000096195 | 1.161036672 | 0.251049598 |
| ENSMUSG00000096196 | 1 | |
| ENSMUSG00000096197 | 1 | |
| ENSMUSG00000096198 | 1 | |
| ENSMUSG00000096199 | -1.032378587 | 0.698112512 |
| ENSMUSG00000096200 | 1 | |
| ENSMUSG00000096201 | 1 | |
| ENSMUSG00000096202 | 1 | |
| ENSMUSG00000096205 | 1 | |
| ENSMUSG00000096206 | 1 | |
| ENSMUSG00000096208 | 1 | |
| ENSMUSG00000096209 | 1 | |
| ENSMUSG00000096210 | -1.008269716 | 1 |
| ENSMUSG00000096211 | 1 | |
| ENSMUSG00000096212 | 1 | |
| ENSMUSG00000096213 | 1 | |
| ENSMUSG00000096214 | 1 | |
| ENSMUSG00000096215 | 1 | |
| ENSMUSG00000096216 | 1 | |
| ENSMUSG00000096217 | 1 | |
| ENSMUSG00000096218 | 1 | |
| ENSMUSG00000096219 | 1 | |
| ENSMUSG00000096220 | 1 | |
| ENSMUSG00000096221 | 1.045926746 | 0.698112512 |
| ENSMUSG00000096222 | 1 | |
| ENSMUSG00000096223 | 1 | |
| ENSMUSG00000096225 | 1 | |
| ENSMUSG00000096228 | 1 | |
| ENSMUSG00000096229 | 1 | |
| ENSMUSG00000096230 | 1 | |
| ENSMUSG00000096231 | 1 | |
| ENSMUSG00000096232 | 1 | |
| ENSMUSG00000096233 | -1 | 1 |
| ENSMUSG00000096235 | 1 | |
| ENSMUSG00000096236 | 1 | |
| ENSMUSG00000096237 | 1 | |
| ENSMUSG00000096238 | 1 | |
| ENSMUSG00000096239 | 1 | |
| ENSMUSG00000096240 | 1 | |
| ENSMUSG00000096241 | 1 | |
| ENSMUSG00000096243 | -1.03962336 | 0.698112512 |
| ENSMUSG00000096244 | 1 | |
| ENSMUSG00000096245 | 1 | |
| ENSMUSG00000096247 | 1 | |
| ENSMUSG00000096248 | 1 | |
| ENSMUSG00000096249 | 1 | |
| ENSMUSG00000096250 | 1 | |
| ENSMUSG00000096252 | 1.023743961 | 0.938939969 |
| ENSMUSG00000096254 | 1 | |
| ENSMUSG00000096255 | -1.013749217 | 1 |
| ENSMUSG00000096256 | 1 | |

| | | |
|---|---|---|
| ENSMUSG00000096257 | -1.044273782 | 0.698112512 |
| ENSMUSG00000096258 | -1 | 1 |
| ENSMUSG00000096259 | 1 | |
| ENSMUSG00000096263 | 1 | |
| ENSMUSG00000096264 | 1 | |
| ENSMUSG00000096265 | 1 | |
| ENSMUSG00000096268 | 1 | |
| ENSMUSG00000096269 | -1.03962336 | 0.698112512 |
| ENSMUSG00000096270 | 1 | |
| ENSMUSG00000096271 | 1 | |
| ENSMUSG00000096272 | 1 | |
| ENSMUSG00000096273 | 1 | |
| ENSMUSG00000096274 | 1 | |
| ENSMUSG00000096275 | 1 | |
| ENSMUSG00000096276 | 1 | |
| ENSMUSG00000096277 | 1 | |
| ENSMUSG00000096278 | 1 | |
| ENSMUSG00000096279 | 1 | |
| ENSMUSG00000096280 | 1 | |
| ENSMUSG00000096281 | 1 | |
| ENSMUSG00000096282 | 1 | |
| ENSMUSG00000096283 | 1 | |
| ENSMUSG00000096284 | 1.015486245 | 0.972980766 |
| ENSMUSG00000096285 | 1 | |
| ENSMUSG00000096286 | 1 | |
| ENSMUSG00000096287 | 1 | |
| ENSMUSG00000096288 | 1 | |
| ENSMUSG00000096289 | 1 | |
| ENSMUSG00000096290 | 1 | |
| ENSMUSG00000096291 | 1 | |
| ENSMUSG00000096292 | 1 | |
| ENSMUSG00000096293 | 1 | |
| ENSMUSG00000096294 | 1 | |
| ENSMUSG00000096295 | 1 | |
| ENSMUSG00000096296 | -1.080615268 | 0.608497229 |
| ENSMUSG00000096297 | 1 | |
| ENSMUSG00000096298 | 1 | |
| ENSMUSG00000096299 | -1.157432403 | 0.368089426 |
| ENSMUSG00000096300 | 1 | |
| ENSMUSG00000096301 | 1 | |
| ENSMUSG00000096302 | 1 | |
| ENSMUSG00000096303 | 1 | |
| ENSMUSG00000096304 | 1 | |
| ENSMUSG00000096305 | 1 | |
| ENSMUSG00000096306 | 1 | |
| ENSMUSG00000096307 | 1 | |
| ENSMUSG00000096308 | 1 | |
| ENSMUSG00000096309 | 1 | |
| ENSMUSG00000096310 | 1 | |
| ENSMUSG00000096311 | 1 | |
| ENSMUSG00000096313 | 1 | |
| ENSMUSG00000096314 | 1 | |
| ENSMUSG00000096316 | 1 | |
| ENSMUSG00000096318 | 1 | |
| ENSMUSG00000096320 | 1 | |
| ENSMUSG00000096322 | 1 | |
| ENSMUSG00000096323 | 1 | |
| ENSMUSG00000096324 | 1 | |
| ENSMUSG00000096326 | 1 | |
| ENSMUSG00000096327 | 1 | |
| ENSMUSG00000096328 | 1 | |
| ENSMUSG00000096329 | 1 | |
| ENSMUSG00000096330 | -1.132666855 | 0.18851436 |

| | | |
|---|---|---|
| ENSMUSG00000096331 | 1 | |
| ENSMUSG00000096332 | 1 | |
| ENSMUSG00000096333 | 1 | |
| ENSMUSG00000096335 | 1 | |
| ENSMUSG00000096336 | 1 | |
| ENSMUSG00000096337 | 1 | |
| ENSMUSG00000096338 | 1 | |
| ENSMUSG00000096339 | 1 | |
| ENSMUSG00000096340 | 1 | |
| ENSMUSG00000096341 | 1 | |
| ENSMUSG00000096342 | 1 | |
| ENSMUSG00000096343 | 1 | |
| ENSMUSG00000096344 | 1 | |
| ENSMUSG00000096345 | 1 | |
| ENSMUSG00000096346 | 1 | |
| ENSMUSG00000096347 | 1 | |
| ENSMUSG00000096348 | 1 | |
| ENSMUSG00000096349 | 1 | |
| ENSMUSG00000096350 | 1 | |
| ENSMUSG00000096351 | 1.182756393 | 0.428820025 |
| ENSMUSG00000096352 | 1 | |
| ENSMUSG00000096353 | 1 | |
| ENSMUSG00000096354 | 1 | |
| ENSMUSG00000096355 | 1 | |
| ENSMUSG00000096356 | 1 | |
| ENSMUSG00000096357 | 1 | |
| ENSMUSG00000096358 | 1 | |
| ENSMUSG00000096359 | 1 | |
| ENSMUSG00000096360 | 1 | |
| ENSMUSG00000096361 | 1 | |
| ENSMUSG00000096362 | 1 | |
| ENSMUSG00000096363 | 1 | |
| ENSMUSG00000096365 | 1 | |
| ENSMUSG00000096366 | 1 | |
| ENSMUSG00000096367 | 1 | |
| ENSMUSG00000096368 | 1 | |
| ENSMUSG00000096370 | -1.088245534 | 0.548809407 |
| ENSMUSG00000096371 | 1 | |
| ENSMUSG00000096372 | 1 | |
| ENSMUSG00000096373 | 1 | |
| ENSMUSG00000096374 | 1 | |
| ENSMUSG00000096375 | 1 | |
| ENSMUSG00000096376 | 1 | |
| ENSMUSG00000096377 | 1 | |
| ENSMUSG00000096379 | 1 | |
| ENSMUSG00000096380 | 1 | |
| ENSMUSG00000096381 | 1 | |
| ENSMUSG00000096383 | 1 | |
| ENSMUSG00000096384 | 1 | |
| ENSMUSG00000096385 | 1 | |
| ENSMUSG00000096386 | 1 | |
| ENSMUSG00000096387 | 1 | |
| ENSMUSG00000096388 | 1 | |
| ENSMUSG00000096390 | 1 | |
| ENSMUSG00000096391 | 1 | |
| ENSMUSG00000096392 | 1 | |
| ENSMUSG00000096393 | 1 | |
| ENSMUSG00000096396 | 1 | |
| ENSMUSG00000096397 | 1 | |
| ENSMUSG00000096398 | 1 | |
| ENSMUSG00000096399 | 1 | |
| ENSMUSG00000096400 | -1.158708082 | 0.450661175 |
| ENSMUSG00000096401 | -1.120541105 | 0.041401294 |

| | | |
|---|---|---|
| ENSMUSG00000096402 | 1 | |
| ENSMUSG00000096403 | 1.028728047 | 0.820284036 |
| ENSMUSG00000096405 | 1 | |
| ENSMUSG00000096406 | 1 | |
| ENSMUSG00000096407 | 1 | |
| ENSMUSG00000096408 | 1 | |
| ENSMUSG00000096409 | 1 | |
| ENSMUSG00000096410 | 1 | |
| ENSMUSG00000096411 | 1 | |
| ENSMUSG00000096412 | 1 | |
| ENSMUSG00000096413 | 1 | |
| ENSMUSG00000096414 | 1 | |
| ENSMUSG00000096415 | 1 | |
| ENSMUSG00000096417 | 1 | |
| ENSMUSG00000096420 | 1 | |
| ENSMUSG00000096421 | 1 | |
| ENSMUSG00000096422 | 1 | |
| ENSMUSG00000096423 | 1 | |
| ENSMUSG00000096424 | 1 | |
| ENSMUSG00000096425 | 1 | |
| ENSMUSG00000096426 | 1 | |
| ENSMUSG00000096427 | 1 | |
| ENSMUSG00000096429 | 1 | |
| ENSMUSG00000096430 | 1 | |
| ENSMUSG00000096431 | 1 | |
| ENSMUSG00000096432 | 1.059463094 | 0.608497229 |
| ENSMUSG00000096433 | 1.019263128 | 0.951410231 |
| ENSMUSG00000096434 | 1 | |
| ENSMUSG00000096435 | 1 | |
| ENSMUSG00000096436 | 1 | |
| ENSMUSG00000096437 | -1.044273782 | 0.698112512 |
| ENSMUSG00000096438 | 1 | |
| ENSMUSG00000096439 | 1 | |
| ENSMUSG00000096440 | 1.057478344 | 0.809553648 |
| ENSMUSG00000096441 | 1 | |
| ENSMUSG00000096442 | 1 | |
| ENSMUSG00000096444 | 1 | |
| ENSMUSG00000096445 | 1 | |
| ENSMUSG00000096446 | -1 | 1 |
| ENSMUSG00000096447 | 1 | |
| ENSMUSG00000096448 | 1.095872691 | 0.608497229 |
| ENSMUSG00000096449 | 1.059582998 | 0.926475461 |
| ENSMUSG00000096450 | 1 | |
| ENSMUSG00000096451 | 1 | |
| ENSMUSG00000096452 | 1 | |
| ENSMUSG00000096453 | 1 | |
| ENSMUSG00000096454 | 1 | |
| ENSMUSG00000096455 | 1 | |
| ENSMUSG00000096457 | 1 | |
| ENSMUSG00000096458 | 1 | |
| ENSMUSG00000096459 | 1 | |
| ENSMUSG00000096461 | 1 | |
| ENSMUSG00000096462 | 1 | |
| ENSMUSG00000096463 | -1.094376052 | 0.861027326 |
| ENSMUSG00000096464 | 1 | |
| ENSMUSG00000096465 | 1 | |
| ENSMUSG00000096467 | 1 | |
| ENSMUSG00000096468 | 1 | |
| ENSMUSG00000096470 | 1 | |
| ENSMUSG00000096472 | -1.049984385 | 0.698112512 |
| ENSMUSG00000096473 | 1 | |
| ENSMUSG00000096474 | -1.009476377 | 1 |
| ENSMUSG00000096475 | 1 | |

| | | |
|---|---|---|
| ENSMUSG00000096476 | 1 | |
| ENSMUSG00000096477 | 1 | |
| ENSMUSG00000096478 | 1 | |
| ENSMUSG00000096479 | 1 | |
| ENSMUSG00000096480 | 1 | |
| ENSMUSG00000096481 | 1 | |
| ENSMUSG00000096482 | 1 | |
| ENSMUSG00000096483 | 1 | |
| ENSMUSG00000096484 | 1 | |
| ENSMUSG00000096485 | 1 | |
| ENSMUSG00000096486 | 1 | |
| ENSMUSG00000096487 | 1 | |
| ENSMUSG00000096488 | 1 | |
| ENSMUSG00000096490 | 1 | |
| ENSMUSG00000096492 | 1 | |
| ENSMUSG00000096496 | 1 | |
| ENSMUSG00000096497 | 1 | |
| ENSMUSG00000096498 | 1 | |
| ENSMUSG00000096499 | 1 | |
| ENSMUSG00000096500 | 1 | |
| ENSMUSG00000096502 | 1 | |
| ENSMUSG00000096504 | 1 | |
| ENSMUSG00000096505 | 1 | |
| ENSMUSG00000096506 | 1 | |
| ENSMUSG00000096507 | 1 | |
| ENSMUSG00000096508 | 1 | |
| ENSMUSG00000096510 | 1 | |
| ENSMUSG00000096512 | 1 | |
| ENSMUSG00000096513 | 1 | |
| ENSMUSG00000096514 | 1 | |
| ENSMUSG00000096515 | 1 | |
| ENSMUSG00000096516 | 1 | |
| ENSMUSG00000096517 | 1 | |
| ENSMUSG00000096518 | 1 | |
| ENSMUSG00000096519 | 1 | |
| ENSMUSG00000096520 | 1 | |
| ENSMUSG00000096521 | 1 | |
| ENSMUSG00000096522 | 1 | |
| ENSMUSG00000096523 | 1 | |
| ENSMUSG00000096524 | 1 | |
| ENSMUSG00000096525 | 1 | |
| ENSMUSG00000096527 | 1 | |
| ENSMUSG00000096528 | -1.077189453 | 0.674209009 |
| ENSMUSG00000096529 | 1 | |
| ENSMUSG00000096530 | 1 | |
| ENSMUSG00000096531 | 1 | |
| ENSMUSG00000096532 | 1 | |
| ENSMUSG00000096533 | 1 | |
| ENSMUSG00000096534 | 1 | |
| ENSMUSG00000096535 | 1 | |
| ENSMUSG00000096536 | 1 | |
| ENSMUSG00000096537 | 1 | |
| ENSMUSG00000096538 | 1 | |
| ENSMUSG00000096539 | 1 | |
| ENSMUSG00000096541 | 1 | |
| ENSMUSG00000096544 | 1 | |
| ENSMUSG00000096546 | -1 | 1 |
| ENSMUSG00000096547 | -1.043306199 | 0.698112512 |
| ENSMUSG00000096549 | -1 | 1 |
| ENSMUSG00000096550 | 1 | |
| ENSMUSG00000096551 | 1 | |
| ENSMUSG00000096553 | 1 | |
| ENSMUSG00000096554 | 1 | |

| | | |
|---|---|---|
| ENSMUSG00000096555 | 1 | |
| ENSMUSG00000096557 | 1 | |
| ENSMUSG00000096558 | 1 | |
| ENSMUSG00000096559 | 1 | |
| ENSMUSG00000096560 | 1 | |
| ENSMUSG00000096561 | 1 | |
| ENSMUSG00000096563 | 1 | |
| ENSMUSG00000096564 | 1 | |
| ENSMUSG00000096565 | 1 | |
| ENSMUSG00000096566 | 1 | |
| ENSMUSG00000096567 | 1 | |
| ENSMUSG00000096568 | 1 | |
| ENSMUSG00000096569 | 1 | |
| ENSMUSG00000096573 | 1 | |
| ENSMUSG00000096574 | -1 | 1 |
| ENSMUSG00000096576 | 1 | |
| ENSMUSG00000096577 | 1 | |
| ENSMUSG00000096578 | 1 | |
| ENSMUSG00000096579 | 1 | |
| ENSMUSG00000096580 | 1 | |
| ENSMUSG00000096582 | 1 | |
| ENSMUSG00000096583 | 1 | |
| ENSMUSG00000096584 | 1 | |
| ENSMUSG00000096585 | 1 | |
| ENSMUSG00000096586 | 1 | |
| ENSMUSG00000096587 | 1 | |
| ENSMUSG00000096589 | 1 | |
| ENSMUSG00000096591 | -1.044273782 | 0.698112512 |
| ENSMUSG00000096592 | 1 | |
| ENSMUSG00000096593 | 1 | |
| ENSMUSG00000096594 | 1 | |
| ENSMUSG00000096595 | 1 | |
| ENSMUSG00000096596 | 1 | |
| ENSMUSG00000096597 | 1 | |
| ENSMUSG00000096598 | 1 | |
| ENSMUSG00000096599 | 1 | |
| ENSMUSG00000096600 | 1 | |
| ENSMUSG00000096601 | 1 | |
| ENSMUSG00000096604 | 1 | |
| ENSMUSG00000096605 | 1 | |
| ENSMUSG00000096606 | -1.251408229 | 0.001392472 |
| ENSMUSG00000096607 | -1.085651418 | 0.608497229 |
| ENSMUSG00000096612 | 1 | |
| ENSMUSG00000096613 | 1 | |
| ENSMUSG00000096614 | 1 | |
| ENSMUSG00000096615 | 1 | |
| ENSMUSG00000096617 | 1 | |
| ENSMUSG00000096618 | 1 | |
| ENSMUSG00000096619 | 1.059463094 | 0.608497229 |
| ENSMUSG00000096620 | 1 | |
| ENSMUSG00000096621 | 1 | |
| ENSMUSG00000096623 | -1.180566046 | 0.428732467 |
| ENSMUSG00000096624 | 1 | |
| ENSMUSG00000096626 | 1 | |
| ENSMUSG00000096627 | 1 | |
| ENSMUSG00000096628 | 1 | |
| ENSMUSG00000096629 | -1.043306199 | 0.79291651 |
| ENSMUSG00000096630 | -1 | 1 |
| ENSMUSG00000096631 | 1 | |
| ENSMUSG00000096632 | 1 | |
| ENSMUSG00000096633 | 1 | |
| ENSMUSG00000096634 | 1 | |
| ENSMUSG00000096635 | 1.034865357 | 0.899648217 |

| | | |
|---|---|---|
| ENSMUSG00000096638 | 1 | |
| ENSMUSG00000096639 | 1 | |
| ENSMUSG00000096640 | 1 | |
| ENSMUSG00000096641 | 1.095872691 | 0.671280603 |
| ENSMUSG00000096642 | 1 | |
| ENSMUSG00000096644 | 1 | |
| ENSMUSG00000096645 | 1 | |
| ENSMUSG00000096646 | 1 | |
| ENSMUSG00000096647 | -1.03962336 | 0.698112512 |
| ENSMUSG00000096648 | 1 | |
| ENSMUSG00000096649 | 1 | |
| ENSMUSG00000096650 | 1 | |
| ENSMUSG00000096651 | 1 | |
| ENSMUSG00000096652 | 1 | |
| ENSMUSG00000096653 | 1 | |
| ENSMUSG00000096654 | 1 | |
| ENSMUSG00000096655 | -1.007966273 | 1 |
| ENSMUSG00000096656 | 1 | |
| ENSMUSG00000096658 | 1 | |
| ENSMUSG00000096659 | 1 | |
| ENSMUSG00000096660 | 1 | |
| ENSMUSG00000096661 | 1 | |
| ENSMUSG00000096663 | 1 | |
| ENSMUSG00000096664 | 1 | |
| ENSMUSG00000096666 | 1 | |
| ENSMUSG00000096667 | 1 | |
| ENSMUSG00000096668 | 1 | |
| ENSMUSG00000096670 | 1 | |
| ENSMUSG00000096672 | 1 | |
| ENSMUSG00000096673 | 1 | |
| ENSMUSG00000096674 | 1 | |
| ENSMUSG00000096675 | 1 | |
| ENSMUSG00000096676 | 1 | |
| ENSMUSG00000096677 | 1 | |
| ENSMUSG00000096678 | 1 | |
| ENSMUSG00000096679 | 1 | |
| ENSMUSG00000096680 | 1 | |
| ENSMUSG00000096681 | 1 | |
| ENSMUSG00000096682 | 1 | |
| ENSMUSG00000096683 | 1 | |
| ENSMUSG00000096684 | 1 | |
| ENSMUSG00000096685 | 1 | |
| ENSMUSG00000096686 | 1 | |
| ENSMUSG00000096687 | 1.072854134 | 0.553862807 |
| ENSMUSG00000096688 | 1 | |
| ENSMUSG00000096689 | 1 | |
| ENSMUSG00000096690 | 1 | |
| ENSMUSG00000096691 | 1 | |
| ENSMUSG00000096692 | 1 | |
| ENSMUSG00000096694 | 1 | |
| ENSMUSG00000096695 | 1 | |
| ENSMUSG00000096696 | -1.043797599 | 0.704170349 |
| ENSMUSG00000096697 | 1 | |
| ENSMUSG00000096698 | 1 | |
| ENSMUSG00000096699 | -1.058323531 | 0.927169398 |
| ENSMUSG00000096700 | 1 | |
| ENSMUSG00000096701 | -1.253248205 | 0.331764656 |
| ENSMUSG00000096703 | 1 | |
| ENSMUSG00000096704 | 1 | |
| ENSMUSG00000096705 | 1 | |
| ENSMUSG00000096706 | 1 | |
| ENSMUSG00000096707 | 1 | |
| ENSMUSG00000096708 | 1 | |

| | | |
|---|---|---|
| ENSMUSG00000096709 | 1 | |
| ENSMUSG00000096710 | 1 | |
| ENSMUSG00000096711 | 1 | |
| ENSMUSG00000096712 | 1.059463094 | 0.698112512 |
| ENSMUSG00000096713 | 1 | |
| ENSMUSG00000096714 | 1 | |
| ENSMUSG00000096715 | 1 | |
| ENSMUSG00000096716 | 1 | |
| ENSMUSG00000096717 | 1 | |
| ENSMUSG00000096718 | 1.042760588 | 0.908136117 |
| ENSMUSG00000096719 | 1 | |
| ENSMUSG00000096721 | 1 | |
| ENSMUSG00000096722 | -1.113957691 | 0.608497229 |
| ENSMUSG00000096724 | 1 | |
| ENSMUSG00000096726 | -1.001371787 | 1 |
| ENSMUSG00000096727 | -2.295060511 | 0.00350191 |
| ENSMUSG00000096728 | 1 | |
| ENSMUSG00000096729 | 1 | |
| ENSMUSG00000096730 | 1 | |
| ENSMUSG00000096731 | 1 | |
| ENSMUSG00000096732 | 1 | |
| ENSMUSG00000096733 | 1 | |
| ENSMUSG00000096734 | 1 | |
| ENSMUSG00000096735 | 1 | |
| ENSMUSG00000096736 | 1 | |
| ENSMUSG00000096737 | 1 | |
| ENSMUSG00000096738 | 1 | |
| ENSMUSG00000096740 | -1.024194846 | 0.920705506 |
| ENSMUSG00000096741 | 1 | |
| ENSMUSG00000096742 | 1 | |
| ENSMUSG00000096743 | 1 | |
| ENSMUSG00000096744 | 1 | |
| ENSMUSG00000096745 | -1 | 1 |
| ENSMUSG00000096746 | 1 | |
| ENSMUSG00000096747 | 1 | |
| ENSMUSG00000096748 | 1 | |
| ENSMUSG00000096749 | 1 | |
| ENSMUSG00000096750 | 1 | |
| ENSMUSG00000096751 | -1.143787033 | 0.363110963 |
| ENSMUSG00000096753 | 1 | |
| ENSMUSG00000096755 | 1.142033317 | 0.608497229 |
| ENSMUSG00000096756 | 1 | |
| ENSMUSG00000096757 | 1 | |
| ENSMUSG00000096758 | 1 | |
| ENSMUSG00000096759 | 1 | |
| ENSMUSG00000096760 | 1 | |
| ENSMUSG00000096761 | 1 | |
| ENSMUSG00000096763 | 1 | |
| ENSMUSG00000096764 | -1.03962336 | 0.698112512 |
| ENSMUSG00000096765 | 1 | |
| ENSMUSG00000096766 | 1 | |
| ENSMUSG00000096767 | 1 | |
| ENSMUSG00000096768 | -1.157176496 | 0.292504162 |
| ENSMUSG00000096769 | 1 | |
| ENSMUSG00000096770 | 1 | |
| ENSMUSG00000096771 | 1 | |
| ENSMUSG00000096772 | 1 | |
| ENSMUSG00000096773 | 1 | |
| ENSMUSG00000096774 | 1 | |
| ENSMUSG00000096775 | -1.043306199 | 0.79291651 |
| ENSMUSG00000096776 | 1 | |
| ENSMUSG00000096777 | 1 | |
| ENSMUSG00000096779 | 1 | |

| | | |
|---|---|---|
| ENSMUSG00000096780 | 1.008456167 | 0.992095533 |
| ENSMUSG00000096782 | 1 | |
| ENSMUSG00000096784 | 1 | |
| ENSMUSG00000096785 | 1 | |
| ENSMUSG00000096786 | 1 | |
| ENSMUSG00000096788 | 1 | |
| ENSMUSG00000096789 | 1 | |
| ENSMUSG00000096790 | 1 | |
| ENSMUSG00000096791 | 1 | |
| ENSMUSG00000096792 | 1 | |
| ENSMUSG00000096793 | 1 | |
| ENSMUSG00000096794 | 1 | |
| ENSMUSG00000096795 | -1.17634621 | 0.244099696 |
| ENSMUSG00000096796 | 1 | |
| ENSMUSG00000096797 | 1 | |
| ENSMUSG00000096798 | 1 | |
| ENSMUSG00000096799 | 1 | |
| ENSMUSG00000096800 | 1 | |
| ENSMUSG00000096801 | 1 | |
| ENSMUSG00000096802 | 1 | |
| ENSMUSG00000096803 | 1.069068979 | 0.698112512 |
| ENSMUSG00000096804 | 1 | |
| ENSMUSG00000096805 | 1 | |
| ENSMUSG00000096806 | 1 | |
| ENSMUSG00000096807 | 1.095872691 | 0.608497229 |
| ENSMUSG00000096808 | 1.153588415 | 0.636968599 |
| ENSMUSG00000096809 | 1 | |
| ENSMUSG00000096810 | 1.015486245 | 0.998386582 |
| ENSMUSG00000096811 | 1 | |
| ENSMUSG00000096812 | 1 | |
| ENSMUSG00000096813 | 1 | |
| ENSMUSG00000096814 | 1 | |
| ENSMUSG00000096815 | 1 | |
| ENSMUSG00000096817 | 1 | |
| ENSMUSG00000096818 | 1 | |
| ENSMUSG00000096819 | 1 | |
| ENSMUSG00000096820 | 1 | |
| ENSMUSG00000096821 | 1 | |
| ENSMUSG00000096822 | 1 | |
| ENSMUSG00000096823 | 1 | |
| ENSMUSG00000096824 | 1 | |
| ENSMUSG00000096825 | 1 | |
| ENSMUSG00000096826 | 1 | |
| ENSMUSG00000096827 | 1 | |
| ENSMUSG00000096828 | 1 | |
| ENSMUSG00000096829 | 1 | |
| ENSMUSG00000096830 | 1 | |
| ENSMUSG00000096831 | 1 | |
| ENSMUSG00000096832 | -1 | 1 |
| ENSMUSG00000096833 | 1 | |
| ENSMUSG00000096834 | 1 | |
| ENSMUSG00000096835 | 1 | |
| ENSMUSG00000096838 | 1 | |
| ENSMUSG00000096839 | 1 | |
| ENSMUSG00000096840 | 1 | |
| ENSMUSG00000096841 | 1.005811684 | 1 |
| ENSMUSG00000096842 | 1 | |
| ENSMUSG00000096844 | 1 | |
| ENSMUSG00000096845 | 1 | |
| ENSMUSG00000096846 | 1 | |
| ENSMUSG00000096847 | -1.651400928 | 0.025782394 |
| ENSMUSG00000096848 | 1 | |
| ENSMUSG00000096849 | 1 | |

| | | |
|---|---|---|
| ENSMUSG00000096850 | 1 | |
| ENSMUSG00000096851 | 1 | |
| ENSMUSG00000096852 | 1 | |
| ENSMUSG00000096853 | 1 | |
| ENSMUSG00000096854 | 1.115223861 | 0.836597096 |
| ENSMUSG00000096855 | 1 | |
| ENSMUSG00000096856 | -1.136348423 | 0.621868642 |
| ENSMUSG00000096857 | 1 | |
| ENSMUSG00000096858 | 1 | |
| ENSMUSG00000096859 | 1 | |
| ENSMUSG00000096861 | 1 | |
| ENSMUSG00000096862 | 1 | |
| ENSMUSG00000096863 | 1 | |
| ENSMUSG00000096864 | 1 | |
| ENSMUSG00000096865 | 1 | |
| ENSMUSG00000096866 | 1 | |
| ENSMUSG00000096867 | 1.059463094 | 0.608497229 |
| ENSMUSG00000096868 | 1 | |
| ENSMUSG00000096869 | 1 | |
| ENSMUSG00000096870 | 1 | |
| ENSMUSG00000096871 | 1 | |
| ENSMUSG00000096872 | 1 | |
| ENSMUSG00000096873 | 1 | |
| ENSMUSG00000096874 | -1 | 1 |
| ENSMUSG00000096875 | 1 | |
| ENSMUSG00000096876 | 1 | |
| ENSMUSG00000096878 | 1 | |
| ENSMUSG00000096879 | 1.069068979 | 0.698112512 |
| ENSMUSG00000096880 | 1 | |
| ENSMUSG00000096881 | 1 | |
| ENSMUSG00000096882 | 1 | |
| ENSMUSG00000096883 | -1.110426459 | 0.698112512 |
| ENSMUSG00000096884 | 1 | |
| ENSMUSG00000096885 | 1 | |
| ENSMUSG00000096887 | -1.044273782 | 0.698112512 |
| ENSMUSG00000096889 | 1 | |
| ENSMUSG00000096890 | -1 | 1 |
| ENSMUSG00000096891 | 1 | |
| ENSMUSG00000096893 | -1 | 1 |
| ENSMUSG00000096894 | 1 | |
| ENSMUSG00000096895 | 1 | |
| ENSMUSG00000096896 | 1 | |
| ENSMUSG00000096897 | 1 | |
| ENSMUSG00000096898 | 1 | |
| ENSMUSG00000096899 | 1 | |
| ENSMUSG00000096900 | 1 | |
| ENSMUSG00000096901 | 1 | |
| ENSMUSG00000096902 | 1 | |
| ENSMUSG00000096903 | 1 | |
| ENSMUSG00000096904 | 1 | |
| ENSMUSG00000096905 | 1 | |
| ENSMUSG00000096906 | 1 | |
| ENSMUSG00000096907 | 1 | |
| ENSMUSG00000096908 | 1 | |
| ENSMUSG00000096910 | 1.046937282 | 0.698112512 |
| ENSMUSG00000096911 | 1 | |
| ENSMUSG00000096912 | 1 | |
| ENSMUSG00000096913 | 1 | |
| ENSMUSG00000096914 | 1.091686812 | 0.783046373 |
| ENSMUSG00000096915 | 1 | |
| ENSMUSG00000096916 | -1.044961059 | 0.825410965 |
| ENSMUSG00000096917 | 1.112984152 | 0.534944201 |
| ENSMUSG00000096918 | 1 | |

| | | |
|---|---|---|
| ENSMUSG00000096919 | 1.059463094 | 0.608497229 |
| ENSMUSG00000096920 | -1.049475223 | 0.863410724 |
| ENSMUSG00000096921 | -1.043306199 | 0.79291651 |
| ENSMUSG00000096922 | 1 | |
| ENSMUSG00000096923 | 1.076888206 | 0.698112512 |
| ENSMUSG00000096924 | 1.152292804 | 0.698112512 |
| ENSMUSG00000096926 | -1.13078477 | 0.698112512 |
| ENSMUSG00000096927 | 1 | |
| ENSMUSG00000096929 | 1.045402873 | 0.887610675 |
| ENSMUSG00000096930 | 1 | |
| ENSMUSG00000096931 | 1 | |
| ENSMUSG00000096932 | 1.089294043 | 0.745135758 |
| ENSMUSG00000096934 | 1 | |
| ENSMUSG00000096935 | -1.311950161 | 0.099764565 |
| ENSMUSG00000096936 | 1 | |
| ENSMUSG00000096937 | -1.043306199 | 0.837704961 |
| ENSMUSG00000096938 | 1.196008636 | 0.547016491 |
| ENSMUSG00000096939 | 1 | |
| ENSMUSG00000096940 | 1.132877091 | 0.608497229 |
| ENSMUSG00000096941 | 1 | |
| ENSMUSG00000096942 | 1 | |
| ENSMUSG00000096943 | 1 | |
| ENSMUSG00000096944 | 1.029639635 | 0.984732631 |
| ENSMUSG00000096945 | -1.105459774 | 0.698112512 |
| ENSMUSG00000096946 | 1.194195956 | 0.453753001 |
| ENSMUSG00000096947 | 1 | |
| ENSMUSG00000096948 | 1.047437598 | 0.794815197 |
| ENSMUSG00000096949 | 1 | |
| ENSMUSG00000096950 | 1.264336659 | 0.34335694 |
| ENSMUSG00000096953 | 1 | |
| ENSMUSG00000096954 | 1.24101031 | 0.04001886 |
| ENSMUSG00000096955 | 1 | |
| ENSMUSG00000096956 | 1.610774662 | 0.002996937 |
| ENSMUSG00000096957 | 1.037884101 | 0.884451146 |
| ENSMUSG00000096958 | 1.015486245 | 0.972980766 |
| ENSMUSG00000096959 | -1.180435034 | 0.449109628 |
| ENSMUSG00000096960 | 1.212410478 | 0.288251542 |
| ENSMUSG00000096962 | 1.019083579 | 0.949812939 |
| ENSMUSG00000096963 | 1 | |
| ENSMUSG00000096964 | -1.132666855 | 0.18851436 |
| ENSMUSG00000096965 | 1.442225561 | 0.010217196 |
| ENSMUSG00000096966 | 1 | |
| ENSMUSG00000096967 | -1.000919549 | 1 |
| ENSMUSG00000096968 | 1.067303011 | 0.869603732 |
| ENSMUSG00000096969 | 1 | |
| ENSMUSG00000096971 | 1.224848118 | 0.184121889 |
| ENSMUSG00000096972 | 1 | |
| ENSMUSG00000096973 | 1.084618945 | 0.614071182 |
| ENSMUSG00000096974 | -1.02522956 | 0.972496782 |
| ENSMUSG00000096975 | -1.05575982 | 0.692782277 |
| ENSMUSG00000096976 | 1.044210161 | 0.90416245 |
| ENSMUSG00000096977 | -1.081972557 | 0.697178241 |
| ENSMUSG00000096978 | -1.065096629 | 0.779623674 |
| ENSMUSG00000096979 | 1.47497114 | 0.048604265 |
| ENSMUSG00000096980 | 1.085624843 | 0.725611516 |
| ENSMUSG00000096981 | -1.117211114 | 0.490251676 |
| ENSMUSG00000096982 | 1 | |
| ENSMUSG00000096983 | -1.042570342 | 0.801443616 |
| ENSMUSG00000096984 | -1.028972438 | 0.972684567 |
| ENSMUSG00000096986 | -1.004497769 | 1 |
| ENSMUSG00000096987 | 1.143466909 | 0.443539027 |
| ENSMUSG00000096988 | 1.049806226 | 0.908214192 |
| ENSMUSG00000096990 | 1.129237035 | 0.608497229 |

| | | |
|---|---|---|
| ENSMUSG00000096991 | 1.12807594 | 0.200623453 |
| ENSMUSG00000096992 | 1.082902945 | 0.698112512 |
| ENSMUSG00000096993 | -1.053549062 | 0.769745556 |
| ENSMUSG00000096995 | 1.044921492 | 0.702608113 |
| ENSMUSG00000096996 | 1.093857462 | 0.608497229 |
| ENSMUSG00000096997 | 1 | |
| ENSMUSG00000096999 | 1 | |
| ENSMUSG00000097000 | 1.074115557 | 0.593594356 |
| ENSMUSG00000097001 | 1 | |
| ENSMUSG00000097002 | 1.39352436 | 0.012171525 |
| ENSMUSG00000097003 | -1.028160286 | 0.938931916 |
| ENSMUSG00000097004 | -1.042041425 | 0.916688447 |
| ENSMUSG00000097005 | 1 | |
| ENSMUSG00000097006 | 1.068105432 | 0.631334172 |
| ENSMUSG00000097007 | 1.060465576 | 0.848663214 |
| ENSMUSG00000097008 | 1.033486948 | 0.96035552 |
| ENSMUSG00000097009 | 1.132639128 | 0.385302577 |
| ENSMUSG00000097010 | -1.053549062 | 0.698112512 |
| ENSMUSG00000097011 | 1.059463094 | 0.681952832 |
| ENSMUSG00000097012 | 1 | |
| ENSMUSG00000097013 | 1.142753088 | 0.684376758 |
| ENSMUSG00000097014 | 1.019243582 | 0.996278978 |
| ENSMUSG00000097015 | -1.061647984 | 0.608497229 |
| ENSMUSG00000097016 | 1.075324819 | 0.792444161 |
| ENSMUSG00000097017 | -1.156756964 | 0.399470186 |
| ENSMUSG00000097018 | 1 | |
| ENSMUSG00000097019 | 1.069068979 | 0.608497229 |
| ENSMUSG00000097020 | -1.031704886 | 0.9089388 |
| ENSMUSG00000097021 | 1.015267009 | 1 |
| ENSMUSG00000097022 | 1.195724159 | 0.608497229 |
| ENSMUSG00000097023 | 1 | |
| ENSMUSG00000097024 | 1 | |
| ENSMUSG00000097025 | -1.03962336 | 0.803072249 |
| ENSMUSG00000097026 | 1.202331687 | 0.034373414 |
| ENSMUSG00000097027 | 1.047211891 | 0.915280515 |
| ENSMUSG00000097028 | 1.560376593 | 0.000302217 |
| ENSMUSG00000097029 | 1 | |
| ENSMUSG00000097030 | 1 | |
| ENSMUSG00000097031 | 1 | |
| ENSMUSG00000097032 | -1.056302448 | 0.931576531 |
| ENSMUSG00000097033 | 1.07854092 | 0.035230443 |
| ENSMUSG00000097035 | -1.020155149 | 0.989387709 |
| ENSMUSG00000097036 | -1.053549062 | 0.698112512 |
| ENSMUSG00000097037 | 1.019083579 | 0.949812939 |
| ENSMUSG00000097038 | 1 | |
| ENSMUSG00000097039 | 1.040631679 | 0.807173091 |
| ENSMUSG00000097040 | 1.068522702 | 0.79291651 |
| ENSMUSG00000097041 | -1 | 1 |
| ENSMUSG00000097042 | 1.026372203 | 0.794274354 |
| ENSMUSG00000097043 | 1 | |
| ENSMUSG00000097044 | 1 | |
| ENSMUSG00000097045 | -1.090507733 | 0.770016888 |
| ENSMUSG00000097047 | 1.253098857 | 0.349636986 |
| ENSMUSG00000097048 | -1.10977939 | 0.088216238 |
| ENSMUSG00000097049 | 1 | |
| ENSMUSG00000097050 | -1.084645496 | 0.720992214 |
| ENSMUSG00000097051 | 1.015323785 | 0.973205114 |
| ENSMUSG00000097052 | 1.171563499 | 0.441129618 |
| ENSMUSG00000097053 | 1 | |
| ENSMUSG00000097054 | 1 | |
| ENSMUSG00000097055 | 1.072180084 | 0.828701947 |
| ENSMUSG00000097056 | 1.178549514 | 0.316625679 |
| ENSMUSG00000097057 | -1.058391718 | 0.69307409 |

| | | |
|---|---|---|
| ENSMUSG00000097058 | 1.082193611 | 0.829727984 |
| ENSMUSG00000097059 | -1.132299951 | 0.440481392 |
| ENSMUSG00000097060 | 1 | |
| ENSMUSG00000097061 | 1.12027701 | 0.337606399 |
| ENSMUSG00000097062 | 1.096786361 | 0.553671405 |
| ENSMUSG00000097063 | 1 | |
| ENSMUSG00000097065 | 1.093333724 | 0.797643564 |
| ENSMUSG00000097066 | 1 | |
| ENSMUSG00000097067 | -1.044273782 | 0.698112512 |
| ENSMUSG00000097068 | 1 | |
| ENSMUSG00000097069 | 1.015486245 | 0.998386582 |
| ENSMUSG00000097071 | -1.117459691 | 0.563957069 |
| ENSMUSG00000097072 | -1.379503492 | 0.039761862 |
| ENSMUSG00000097073 | 1.110457118 | 0.563511137 |
| ENSMUSG00000097074 | 1 | |
| ENSMUSG00000097075 | -1.133231767 | 0.662710277 |
| ENSMUSG00000097076 | -1 | 1 |
| ENSMUSG00000097077 | -1.00303729 | 1 |
| ENSMUSG00000097078 | 1 | |
| ENSMUSG00000097079 | 1.019833591 | 0.984476939 |
| ENSMUSG00000097080 | 1.245480742 | 0.057027448 |
| ENSMUSG00000097081 | 1.345856331 | 0.232799305 |
| ENSMUSG00000097082 | 1 | |
| ENSMUSG00000097083 | -1.020688931 | 0.992932129 |
| ENSMUSG00000097084 | 1.57672072 | 0.13659965 |
| ENSMUSG00000097085 | 1.01287802 | 1 |
| ENSMUSG00000097086 | 1 | |
| ENSMUSG00000097087 | 1 | |
| ENSMUSG00000097088 | 1.085624843 | 0.717035356 |
| ENSMUSG00000097089 | 1 | |
| ENSMUSG00000097090 | 1.098000576 | 0.790851299 |
| ENSMUSG00000097091 | -1.083415624 | 0.719087885 |
| ENSMUSG00000097092 | 1 | |
| ENSMUSG00000097093 | 1.112998152 | 0.70299173 |
| ENSMUSG00000097094 | 1 | |
| ENSMUSG00000097095 | 1.020742201 | 0.987775262 |
| ENSMUSG00000097097 | 1 | |
| ENSMUSG00000097098 | 1.015486245 | 0.99457094 |
| ENSMUSG00000097099 | 1.051990305 | 0.640707404 |
| ENSMUSG00000097100 | 1.161036672 | 0.389746121 |
| ENSMUSG00000097101 | 1.074892039 | 0.678897747 |
| ENSMUSG00000097102 | -1.053549062 | 0.769745556 |
| ENSMUSG00000097103 | -1.097906514 | 0.47322914 |
| ENSMUSG00000097104 | 1.177765487 | 0.276639872 |
| ENSMUSG00000097106 | 1 | |
| ENSMUSG00000097107 | -1 | 1 |
| ENSMUSG00000097108 | -1.103040932 | 0.714200725 |
| ENSMUSG00000097109 | 1 | |
| ENSMUSG00000097110 | 1 | |
| ENSMUSG00000097111 | -1.099174267 | 0.608497229 |
| ENSMUSG00000097112 | 1 | |
| ENSMUSG00000097113 | -1.049945032 | 0.84515778 |
| ENSMUSG00000097115 | -1.120476206 | 0.608497229 |
| ENSMUSG00000097116 | 1 | |
| ENSMUSG00000097117 | 1.101669358 | 0.792783746 |
| ENSMUSG00000097118 | -1 | 1 |
| ENSMUSG00000097119 | 1.022577765 | 0.953731979 |
| ENSMUSG00000097120 | 1.069068979 | 0.608497229 |
| ENSMUSG00000097121 | 1.141359621 | 0.624039874 |
| ENSMUSG00000097122 | 1.153144412 | 0.401882202 |
| ENSMUSG00000097123 | 1.005728045 | 1 |
| ENSMUSG00000097124 | -1 | 1 |
| ENSMUSG00000097125 | -1 | 1 |

| | | |
|---|---|---|
| ENSMUSG00000097127 | 1 | |
| ENSMUSG00000097128 | 1.010353783 | 1 |
| ENSMUSG00000097129 | 1.160217575 | 0.343407129 |
| ENSMUSG00000097130 | 1.001874515 | 1 |
| ENSMUSG00000097131 | -1.058983389 | 0.363615297 |
| ENSMUSG00000097133 | 1.020455649 | 0.944468485 |
| ENSMUSG00000097134 | 1.085522042 | 0.838524395 |
| ENSMUSG00000097135 | 1 | |
| ENSMUSG00000097136 | 1.035125408 | 0.903074635 |
| ENSMUSG00000097137 | 1 | |
| ENSMUSG00000097138 | 1.040919012 | 0.940292889 |
| ENSMUSG00000097139 | -1.147232962 | 0.617786513 |
| ENSMUSG00000097140 | 1.00201892 | 1 |
| ENSMUSG00000097141 | 1.043087882 | 0.600663794 |
| ENSMUSG00000097142 | 1 | |
| ENSMUSG00000097145 | 1.20735355 | 0.07529351 |
| ENSMUSG00000097146 | 1.019083579 | 0.949812939 |
| ENSMUSG00000097147 | 1.019083579 | 0.968937442 |
| ENSMUSG00000097148 | 1.063798447 | 0.631765716 |
| ENSMUSG00000097149 | -1 | 1 |
| ENSMUSG00000097150 | 1.000489329 | 1 |
| ENSMUSG00000097151 | -1.051093505 | 0.698112512 |
| ENSMUSG00000097152 | 1.054270992 | 0.88915084 |
| ENSMUSG00000097154 | 1.327237448 | 0.177256987 |
| ENSMUSG00000097155 | -1.009476377 | 1 |
| ENSMUSG00000097156 | 1.064870248 | 0.80586932 |
| ENSMUSG00000097157 | 1 | |
| ENSMUSG00000097158 | -1.03468975 | 0.855911847 |
| ENSMUSG00000097159 | -1.028343479 | 0.977293219 |
| ENSMUSG00000097160 | -1 | 1 |
| ENSMUSG00000097162 | -1.005107211 | 1 |
| ENSMUSG00000097163 | 1.079691425 | 0.608497229 |
| ENSMUSG00000097164 | -1.169221849 | 0.225814318 |
| ENSMUSG00000097165 | 1.366172615 | 0.260191528 |
| ENSMUSG00000097166 | -1.083738315 | 0.755534368 |
| ENSMUSG00000097167 | -1.140788869 | 0.400113308 |
| ENSMUSG00000097168 | -1.063670771 | 0.816059796 |
| ENSMUSG00000097169 | 1 | |
| ENSMUSG00000097170 | -1.005430159 | 1 |
| ENSMUSG00000097171 | -1.151876726 | 0.608497229 |
| ENSMUSG00000097172 | -1.069502966 | 0.752398528 |
| ENSMUSG00000097173 | -1.044273782 | 0.698112512 |
| ENSMUSG00000097174 | 1.021758236 | 0.988263302 |
| ENSMUSG00000097175 | 1 | |
| ENSMUSG00000097176 | 1.106516153 | 0.70299173 |
| ENSMUSG00000097177 | 1.043147516 | 0.698112512 |
| ENSMUSG00000097178 | -1.038444538 | 0.865865121 |
| ENSMUSG00000097180 | -1.045851604 | 0.763407099 |
| ENSMUSG00000097181 | 1 | |
| ENSMUSG00000097182 | -1.044273782 | 0.698112512 |
| ENSMUSG00000097183 | 1.017227335 | 0.94196041 |
| ENSMUSG00000097184 | -1.093195985 | 0.698112512 |
| ENSMUSG00000097185 | 1.116785864 | 0.608497229 |
| ENSMUSG00000097186 | -1.028348525 | 0.912425817 |
| ENSMUSG00000097187 | 1.182314209 | 0.291044503 |
| ENSMUSG00000097189 | 1.074788289 | 0.807247723 |
| ENSMUSG00000097190 | 1 | |
| ENSMUSG00000097191 | 1.087085567 | 0.751338468 |
| ENSMUSG00000097193 | 1.155027827 | 0.187421921 |
| ENSMUSG00000097194 | -1.075832219 | 0.768101821 |
| ENSMUSG00000097195 | 1.059402955 | 0.722238034 |
| ENSMUSG00000097196 | -1.044697704 | 0.870906262 |
| ENSMUSG00000097197 | 1.078592542 | 0.807247723 |

| | | |
|---|---|---|
| ENSMUSG00000097199 | 1.059463094 | 0.608497229 |
| ENSMUSG00000097201 | 1 | |
| ENSMUSG00000097202 | 1 | |
| ENSMUSG00000097203 | 1.075983107 | 0.784280144 |
| ENSMUSG00000097204 | 1.023700751 | 0.828564184 |
| ENSMUSG00000097205 | -1.093674795 | 0.608497229 |
| ENSMUSG00000097206 | 1 | |
| ENSMUSG00000097207 | 1.088050946 | 0.223282095 |
| ENSMUSG00000097208 | 1.247338214 | 0.328718061 |
| ENSMUSG00000097209 | -1.168999771 | 0.208940434 |
| ENSMUSG00000097211 | -1.03962336 | 0.698112512 |
| ENSMUSG00000097213 | -1.053549062 | 0.698112512 |
| ENSMUSG00000097214 | 1.059463094 | 0.698112512 |
| ENSMUSG00000097215 | 1 | |
| ENSMUSG00000097216 | 1.028523534 | 0.98122995 |
| ENSMUSG00000097217 | -1.056174695 | 0.764429998 |
| ENSMUSG00000097218 | 1 | |
| ENSMUSG00000097219 | 1.027751948 | 0.85499844 |
| ENSMUSG00000097220 | -1.067285765 | 0.745045575 |
| ENSMUSG00000097221 | 1 | |
| ENSMUSG00000097222 | -1.199783199 | 0.383793776 |
| ENSMUSG00000097223 | -1.034990181 | 0.88846936 |
| ENSMUSG00000097224 | 1 | |
| ENSMUSG00000097226 | -1.151121934 | 0.608497229 |
| ENSMUSG00000097228 | 1 | |
| ENSMUSG00000097229 | 1 | |
| ENSMUSG00000097230 | -1.205615019 | 0.205169472 |
| ENSMUSG00000097231 | -1.137235672 | 0.239239326 |
| ENSMUSG00000097232 | 1.121764653 | 0.706116633 |
| ENSMUSG00000097233 | -1.091155551 | 0.681952832 |
| ENSMUSG00000097234 | 1.062597606 | 0.72073896 |
| ENSMUSG00000097235 | 1 | |
| ENSMUSG00000097236 | 1.183193407 | 0.53003575 |
| ENSMUSG00000097237 | 1.028323353 | 0.903464611 |
| ENSMUSG00000097239 | 1.006962856 | 1 |
| ENSMUSG00000097240 | -1.044273782 | 0.801443616 |
| ENSMUSG00000097241 | 1 | |
| ENSMUSG00000097242 | -1.090410747 | 0.532369657 |
| ENSMUSG00000097243 | -1.123535114 | 0.57778758 |
| ENSMUSG00000097244 | -1.147838869 | 0.195607801 |
| ENSMUSG00000097245 | 1.016831851 | 0.960578704 |
| ENSMUSG00000097246 | -1.131184271 | 0.608497229 |
| ENSMUSG00000097247 | -1.033820075 | 0.916014403 |
| ENSMUSG00000097248 | -1.285661387 | 0.00179575 |
| ENSMUSG00000097249 | 1 | |
| ENSMUSG00000097250 | 1.28575105 | 0.238959702 |
| ENSMUSG00000097251 | -1.262165069 | 0.125949509 |
| ENSMUSG00000097252 | -1.047082376 | 0.656734444 |
| ENSMUSG00000097253 | 1.226110247 | 0.040083424 |
| ENSMUSG00000097254 | -1.093674795 | 0.698112512 |
| ENSMUSG00000097255 | 1.059463094 | 0.608497229 |
| ENSMUSG00000097256 | 1 | |
| ENSMUSG00000097257 | 1.014347152 | 1 |
| ENSMUSG00000097258 | -1.076443052 | 0.612356812 |
| ENSMUSG00000097259 | 1.070558043 | 0.815547434 |
| ENSMUSG00000097260 | 1.005613439 | 1 |
| ENSMUSG00000097261 | -1.149910693 | 0.632838524 |
| ENSMUSG00000097262 | -1.08081673 | 0.733498571 |
| ENSMUSG00000097263 | -1.18913278 | 0.025901314 |
| ENSMUSG00000097264 | -1 | 1 |
| ENSMUSG00000097265 | 1.03683369 | 0.941803539 |
| ENSMUSG00000097266 | 1.019083579 | 0.949812939 |
| ENSMUSG00000097267 | 1.150372485 | 0.215645894 |

| | | |
|---|---|---|
| ENSMUSG00000097268 | -1.180378409 | 0.548486281 |
| ENSMUSG00000097271 | 1.220289056 | 0.158373059 |
| ENSMUSG00000097272 | 1 | |
| ENSMUSG00000097274 | 1.112843644 | 0.698112512 |
| ENSMUSG00000097275 | 1.077136475 | 0.817890998 |
| ENSMUSG00000097276 | 1.010351027 | 1 |
| ENSMUSG00000097277 | -1.009840629 | 0.988665808 |
| ENSMUSG00000097278 | -1.043306199 | 0.803260194 |
| ENSMUSG00000097279 | -1.060523703 | 0.770958539 |
| ENSMUSG00000097280 | 1.014545335 | 1 |
| ENSMUSG00000097281 | 1.019083579 | 0.949812939 |
| ENSMUSG00000097282 | 1.247458887 | 0.101863729 |
| ENSMUSG00000097283 | -1.121136286 | 0.655571244 |
| ENSMUSG00000097284 | -1.058110984 | 0.778372272 |
| ENSMUSG00000097285 | -1.073353294 | 0.866289429 |
| ENSMUSG00000097286 | 1 | |
| ENSMUSG00000097287 | 1.194446146 | 0.364929458 |
| ENSMUSG00000097288 | 1.161036672 | 0.399744772 |
| ENSMUSG00000097289 | -1.033890127 | 0.765391057 |
| ENSMUSG00000097290 | -1.039027472 | 0.698112512 |
| ENSMUSG00000097291 | 1 | |
| ENSMUSG00000097292 | 1.038031784 | 0.932652618 |
| ENSMUSG00000097293 | 1 | |
| ENSMUSG00000097294 | -1.190279052 | 0.541735653 |
| ENSMUSG00000097295 | 1.069068979 | 0.608497229 |
| ENSMUSG00000097296 | 1.075020499 | 0.430242898 |
| ENSMUSG00000097298 | -1.101237275 | 0.689572034 |
| ENSMUSG00000097299 | 1 | |
| ENSMUSG00000097300 | -1.03962336 | 0.814089245 |
| ENSMUSG00000097301 | 1 | |
| ENSMUSG00000097302 | 1 | |
| ENSMUSG00000097303 | -1.127614384 | 0.698112512 |
| ENSMUSG00000097304 | -1 | 1 |
| ENSMUSG00000097305 | 1.040172434 | 0.913882562 |
| ENSMUSG00000097307 | 1.049411284 | 0.835701362 |
| ENSMUSG00000097308 | 1.033771968 | 0.981964814 |
| ENSMUSG00000097309 | -1.196251461 | 0.60397857 |
| ENSMUSG00000097310 | -1.043306199 | 0.79291651 |
| ENSMUSG00000097311 | 1 | |
| ENSMUSG00000097312 | -1 | 1 |
| ENSMUSG00000097313 | 1.038017113 | 0.948918502 |
| ENSMUSG00000097314 | 1 | |
| ENSMUSG00000097315 | 1 | |
| ENSMUSG00000097316 | -1.14100357 | 0.219009364 |
| ENSMUSG00000097317 | 1.135459005 | 0.525842026 |
| ENSMUSG00000097318 | 1.1040945 | 0.608497229 |
| ENSMUSG00000097320 | 1.006748575 | 1 |
| ENSMUSG00000097321 | -1.353481268 | 0.036059831 |
| ENSMUSG00000097322 | -1.029709545 | 0.98105314 |
| ENSMUSG00000097323 | 1.084514683 | 0.80375069 |
| ENSMUSG00000097324 | 1.442305562 | 0.032337574 |
| ENSMUSG00000097325 | -1.14503164 | 0.608497229 |
| ENSMUSG00000097326 | 1 | |
| ENSMUSG00000097327 | -1.000439235 | 1 |
| ENSMUSG00000097328 | -1.147226143 | 0.339332567 |
| ENSMUSG00000097329 | -1.087913248 | 0.737064932 |
| ENSMUSG00000097330 | 1.277466845 | 0.473190726 |
| ENSMUSG00000097331 | 1.050640266 | 0.698112512 |
| ENSMUSG00000097332 | 1 | |
| ENSMUSG00000097333 | -1.071364056 | 0.694333885 |
| ENSMUSG00000097334 | -1.08081673 | 0.803938171 |
| ENSMUSG00000097335 | 1.053565996 | 0.822533388 |
| ENSMUSG00000097336 | -1.796321579 | 0.010645876 |

| | | |
|---|---|---|
| ENSMUSG00000097337 | 1.106334373 | 0.310630871 |
| ENSMUSG00000097338 | 1 | |
| ENSMUSG00000097339 | 1.014545335 | 0.989648415 |
| ENSMUSG00000097340 | 1.019798086 | 0.819817237 |
| ENSMUSG00000097342 | 1 | |
| ENSMUSG00000097343 | 1.052424405 | 0.77235507 |
| ENSMUSG00000097344 | 1.330341956 | 0.03901261 |
| ENSMUSG00000097345 | -1.012263771 | 1 |
| ENSMUSG00000097346 | 1 | |
| ENSMUSG00000097347 | 1.054150493 | 0.698112512 |
| ENSMUSG00000097348 | 1.089470641 | 0.657060635 |
| ENSMUSG00000097349 | 1 | |
| ENSMUSG00000097350 | 1.083133043 | 0.164411003 |
| ENSMUSG00000097352 | 1.330957397 | 0.039049511 |
| ENSMUSG00000097353 | -1.016617382 | 0.969625585 |
| ENSMUSG00000097354 | 1.341011523 | 0.197112751 |
| ENSMUSG00000097355 | 1 | |
| ENSMUSG00000097356 | 1.036120851 | 0.934760809 |
| ENSMUSG00000097357 | 1.047622966 | 0.770958539 |
| ENSMUSG00000097358 | -1.308671662 | 0.270086974 |
| ENSMUSG00000097360 | -1.193320102 | 0.297891376 |
| ENSMUSG00000097361 | 1 | |
| ENSMUSG00000097362 | 1 | |
| ENSMUSG00000097363 | 1 | |
| ENSMUSG00000097364 | 1.084618945 | 0.614071182 |
| ENSMUSG00000097365 | -1.084645496 | 0.532369657 |
| ENSMUSG00000097366 | 1.036199311 | 0.93818309 |
| ENSMUSG00000097367 | 1 | |
| ENSMUSG00000097368 | 1.049965527 | 0.883615907 |
| ENSMUSG00000097369 | -1.166301255 | 0.607021905 |
| ENSMUSG00000097370 | -1.044273782 | 0.698112512 |
| ENSMUSG00000097371 | 1 | |
| ENSMUSG00000097372 | -1.133618758 | 0.576603197 |
| ENSMUSG00000097373 | -1.032102584 | 0.933142264 |
| ENSMUSG00000097374 | 1 | |
| ENSMUSG00000097375 | -1.099593844 | 0.370893803 |
| ENSMUSG00000097376 | -1.014034612 | 1 |
| ENSMUSG00000097377 | -1.034826058 | 0.871725323 |
| ENSMUSG00000097378 | -1.08770062 | 0.698112512 |
| ENSMUSG00000097379 | 1.023019566 | 0.918280975 |
| ENSMUSG00000097380 | 1.044193845 | 0.792690927 |
| ENSMUSG00000097381 | -1.045151399 | 0.867939428 |
| ENSMUSG00000097382 | 1 | |
| ENSMUSG00000097383 | 1.069217628 | 0.764071649 |
| ENSMUSG00000097384 | 1.04328066 | 0.86119789 |
| ENSMUSG00000097385 | 1.069068979 | 0.608497229 |
| ENSMUSG00000097386 | 1.010855137 | 1 |
| ENSMUSG00000097387 | 1.059463094 | 0.608497229 |
| ENSMUSG00000097388 | -1.010349482 | 1 |
| ENSMUSG00000097390 | 1.062047058 | 0.608497229 |
| ENSMUSG00000097391 | 2.302165197 | 0.001460821 |
| ENSMUSG00000097392 | 1.117593625 | 0.158823567 |
| ENSMUSG00000097393 | 1.159158834 | 0.430273916 |
| ENSMUSG00000097396 | 1 | |
| ENSMUSG00000097397 | 1.247901611 | 0.380583826 |
| ENSMUSG00000097398 | -1.166437992 | 0.608497229 |
| ENSMUSG00000097399 | -1.236282638 | 0.116143171 |
| ENSMUSG00000097401 | 1.240724619 | 0.28488401 |
| ENSMUSG00000097402 | 1.113307429 | 0.702897175 |
| ENSMUSG00000097403 | 1.083094714 | 0.834621543 |
| ENSMUSG00000097404 | 1.073675068 | 0.698112512 |
| ENSMUSG00000097405 | -1.101105591 | 0.730441273 |
| ENSMUSG00000097406 | 1 | |

| | | |
|---|---|---|
| ENSMUSG00000097407 | -1.03962336 | 0.698112512 |
| ENSMUSG00000097408 | 1.089404266 | 0.698112512 |
| ENSMUSG00000097409 | 1 | |
| ENSMUSG00000097410 | -1.333076519 | 0.114168179 |
| ENSMUSG00000097411 | 1.101226948 | 0.698112512 |
| ENSMUSG00000097412 | -1.00644264 | 0.982854947 |
| ENSMUSG00000097413 | -1.179226843 | 0.493671621 |
| ENSMUSG00000097414 | -1.101021689 | 0.564912005 |
| ENSMUSG00000097415 | -1.012104484 | 0.996264991 |
| ENSMUSG00000097416 | -1.100228207 | 0.757998917 |
| ENSMUSG00000097417 | 1 | |
| ENSMUSG00000097418 | 1.022252994 | 0.939510904 |
| ENSMUSG00000097419 | 1.132639128 | 0.217803313 |
| ENSMUSG00000097420 | 1 | |
| ENSMUSG00000097421 | 1 | |
| ENSMUSG00000097422 | -1.089932089 | 0.799112791 |
| ENSMUSG00000097423 | 1.133237328 | 0.379223114 |
| ENSMUSG00000097424 | 1.069068979 | 0.698112512 |
| ENSMUSG00000097425 | 1 | |
| ENSMUSG00000097426 | 1.122462048 | 0.452329901 |
| ENSMUSG00000097427 | 1 | |
| ENSMUSG00000097428 | -1.125300565 | 0.618620792 |
| ENSMUSG00000097429 | 1.10141562 | 0.697515847 |
| ENSMUSG00000097430 | 1.059463094 | 0.752398528 |
| ENSMUSG00000097431 | 1.034244532 | 0.72146869 |
| ENSMUSG00000097433 | -1.177310696 | 0.536694773 |
| ENSMUSG00000097434 | -1.21051692 | 0.329995202 |
| ENSMUSG00000097435 | 1 | |
| ENSMUSG00000097437 | 1 | |
| ENSMUSG00000097438 | 1 | |
| ENSMUSG00000097439 | 1.075508317 | 0.709012023 |
| ENSMUSG00000097440 | -1.05948903 | 0.769745556 |
| ENSMUSG00000097441 | 1 | |
| ENSMUSG00000097442 | -1.044273782 | 0.801443616 |
| ENSMUSG00000097443 | 1.033783726 | 0.873652259 |
| ENSMUSG00000097444 | -1.053549062 | 0.854827467 |
| ENSMUSG00000097445 | -1.000068121 | 1 |
| ENSMUSG00000097446 | -1.037462873 | 0.947189871 |
| ENSMUSG00000097447 | -1.043306199 | 0.698112512 |
| ENSMUSG00000097448 | -1.193369735 | 0.429352448 |
| ENSMUSG00000097450 | 1 | |
| ENSMUSG00000097451 | 2.946397004 | 0.000111096 |
| ENSMUSG00000097452 | 1.037356286 | 0.939140695 |
| ENSMUSG00000097453 | 1 | |
| ENSMUSG00000097454 | 1 | |
| ENSMUSG00000097455 | -1.043630463 | 0.860128739 |
| ENSMUSG00000097456 | -1.134282749 | 0.608497229 |
| ENSMUSG00000097457 | -1.133004215 | 0.469213979 |
| ENSMUSG00000097458 | 1 | |
| ENSMUSG00000097459 | 1 | |
| ENSMUSG00000097460 | 1.010657329 | 1 |
| ENSMUSG00000097461 | 1 | |
| ENSMUSG00000097462 | -1.137301949 | 0.484875549 |
| ENSMUSG00000097463 | 1 | |
| ENSMUSG00000097464 | -1.064228315 | 0.844298111 |
| ENSMUSG00000097465 | 1 | |
| ENSMUSG00000097466 | 1 | |
| ENSMUSG00000097467 | -1.259953524 | 0.005391509 |
| ENSMUSG00000097468 | 1 | |
| ENSMUSG00000097469 | -1.241364419 | 0.532369657 |
| ENSMUSG00000097470 | -1.041360475 | 0.871840478 |
| ENSMUSG00000097471 | -1.08384782 | 0.686411811 |
| ENSMUSG00000097472 | 1.211035135 | 0.308510084 |

| | | |
|---|---|---|
| ENSMUSG00000097473 | 1 | |
| ENSMUSG00000097474 | -1.029302237 | 0.968937442 |
| ENSMUSG00000097476 | -1.041574599 | 0.922165724 |
| ENSMUSG00000097479 | 1.060064166 | 0.922261327 |
| ENSMUSG00000097480 | 1 | |
| ENSMUSG00000097481 | -1.044273782 | 0.698112512 |
| ENSMUSG00000097482 | 1 | |
| ENSMUSG00000097483 | 1 | |
| ENSMUSG00000097484 | 1 | |
| ENSMUSG00000097486 | 1 | |
| ENSMUSG00000097487 | 1.12995955 | 0.430495431 |
| ENSMUSG00000097488 | -1.020215242 | 0.944858734 |
| ENSMUSG00000097489 | 1.089470641 | 0.657060635 |
| ENSMUSG00000097491 | 1.105811708 | 0.784381159 |
| ENSMUSG00000097493 | 1.386925316 | 0.01315387 |
| ENSMUSG00000097494 | -1.011158594 | 1 |
| ENSMUSG00000097495 | 1.080103091 | 0.770958539 |
| ENSMUSG00000097496 | 1 | |
| ENSMUSG00000097497 | -1.085651418 | 0.532369657 |
| ENSMUSG00000097500 | 1 | |
| ENSMUSG00000097501 | 1.059463094 | 0.608497229 |
| ENSMUSG00000097502 | 1.100151335 | 0.744505578 |
| ENSMUSG00000097503 | 1.182342574 | 0.104581081 |
| ENSMUSG00000097504 | 1.023341599 | 0.984289155 |
| ENSMUSG00000097505 | 1 | |
| ENSMUSG00000097507 | 1.055733059 | 0.698112512 |
| ENSMUSG00000097509 | 1.059463094 | 0.699340382 |
| ENSMUSG00000097511 | 1 | |
| ENSMUSG00000097513 | -1.009476377 | 1 |
| ENSMUSG00000097514 | 1 | |
| ENSMUSG00000097515 | 1.138095746 | 0.640910749 |
| ENSMUSG00000097516 | 1 | |
| ENSMUSG00000097517 | 1 | |
| ENSMUSG00000097518 | -1.067579267 | 0.585362499 |
| ENSMUSG00000097519 | -1.045074972 | 0.798487595 |
| ENSMUSG00000097520 | 1.050165139 | 0.924587329 |
| ENSMUSG00000097522 | 1 | |
| ENSMUSG00000097523 | 1.058358024 | 0.734881876 |
| ENSMUSG00000097524 | 1 | |
| ENSMUSG00000097525 | 1.015486245 | 0.972980766 |
| ENSMUSG00000097526 | -1.102487507 | 0.698112512 |
| ENSMUSG00000097527 | -1.049645365 | 0.89661194 |
| ENSMUSG00000097529 | 1.075070132 | 0.698112512 |
| ENSMUSG00000097530 | 1.014545335 | 0.979002724 |
| ENSMUSG00000097532 | -1.127180405 | 0.354776885 |
| ENSMUSG00000097533 | -1.041132001 | 0.90718945 |
| ENSMUSG00000097534 | -1.070098701 | 0.813485995 |
| ENSMUSG00000097535 | 1.019083579 | 0.949812939 |
| ENSMUSG00000097536 | 1.01068296 | 0.982838256 |
| ENSMUSG00000097537 | -1.016260404 | 0.950239521 |
| ENSMUSG00000097538 | 1 | |
| ENSMUSG00000097539 | 1 | |
| ENSMUSG00000097540 | -1.084645496 | 0.532369657 |
| ENSMUSG00000097541 | -1.085651418 | 0.532369657 |
| ENSMUSG00000097542 | 1 | |
| ENSMUSG00000097543 | -1.038444538 | 0.895498153 |
| ENSMUSG00000097544 | 1 | |
| ENSMUSG00000097545 | 1 | |
| ENSMUSG00000097546 | -1.035049744 | 0.949812939 |
| ENSMUSG00000097547 | 1 | |
| ENSMUSG00000097548 | 1.004718404 | 1 |
| ENSMUSG00000097550 | 1 | |
| ENSMUSG00000097551 | 1 | |

| | | |
|---|---|---|
| ENSMUSG00000097552 | 1.095872691 | 0.608497229 |
| ENSMUSG00000097553 | -1 | 1 |
| ENSMUSG00000097554 | -1.015182824 | 0.989648415 |
| ENSMUSG00000097557 | 1.024779056 | 0.914474058 |
| ENSMUSG00000097558 | 1.214044268 | 0.063898214 |
| ENSMUSG00000097559 | 1.213012127 | 0.096445543 |
| ENSMUSG00000097560 | 1.165856327 | 0.444934552 |
| ENSMUSG00000097561 | -1.000697071 | 1 |
| ENSMUSG00000097562 | 1 | |
| ENSMUSG00000097563 | 1.084618945 | 0.644224767 |
| ENSMUSG00000097564 | 1.195832674 | 0.099129119 |
| ENSMUSG00000097565 | 1 | |
| ENSMUSG00000097566 | 1 | |
| ENSMUSG00000097567 | 1.127191814 | 0.608497229 |
| ENSMUSG00000097569 | 1.084618945 | 0.698112512 |
| ENSMUSG00000097571 | 1.080234431 | 0.647384212 |
| ENSMUSG00000097572 | 1.080235031 | 0.608497229 |
| ENSMUSG00000097573 | 1.021667102 | 0.927832746 |
| ENSMUSG00000097574 | 1.111361371 | 0.612309538 |
| ENSMUSG00000097575 | -1.043306199 | 0.698112512 |
| ENSMUSG00000097576 | 1.121754272 | 0.530100816 |
| ENSMUSG00000097577 | -1.176390618 | 0.13534116 |
| ENSMUSG00000097578 | 1.006651517 | 1 |
| ENSMUSG00000097579 | 1.025254253 | 0.893500268 |
| ENSMUSG00000097580 | 1.059463094 | 0.608497229 |
| ENSMUSG00000097581 | 1.367428922 | 0.112138014 |
| ENSMUSG00000097582 | 1 | |
| ENSMUSG00000097583 | -1.100064626 | 0.698112512 |
| ENSMUSG00000097584 | 1.010351027 | 1 |
| ENSMUSG00000097585 | -1.103388856 | 0.632059035 |
| ENSMUSG00000097586 | -1.044273782 | 0.790851299 |
| ENSMUSG00000097587 | 1.069068979 | 0.698112512 |
| ENSMUSG00000097589 | -1.157541621 | 0.312657959 |
| ENSMUSG00000097590 | 1 | |
| ENSMUSG00000097591 | -1.063185175 | 0.819614655 |
| ENSMUSG00000097592 | -1.324087294 | 0.126679497 |
| ENSMUSG00000097593 | 1 | |
| ENSMUSG00000097594 | 1 | |
| ENSMUSG00000097595 | -1.120476206 | 0.547016491 |
| ENSMUSG00000097596 | 1 | |
| ENSMUSG00000097597 | -1.069502966 | 0.698112512 |
| ENSMUSG00000097598 | 1 | |
| ENSMUSG00000097599 | 1 | |
| ENSMUSG00000097601 | 1.150179455 | 0.608497229 |
| ENSMUSG00000097602 | -1.044273782 | 0.801443616 |
| ENSMUSG00000097603 | 1.118644273 | 0.608497229 |
| ENSMUSG00000097604 | -1.044273782 | 0.698112512 |
| ENSMUSG00000097605 | 1 | |
| ENSMUSG00000097608 | 1 | |
| ENSMUSG00000097609 | -1.137619208 | 0.506373506 |
| ENSMUSG00000097610 | 1.030371249 | 0.914474058 |
| ENSMUSG00000097612 | -1.03962336 | 0.872692845 |
| ENSMUSG00000097613 | 1.078229685 | 0.762277843 |
| ENSMUSG00000097614 | 1 | |
| ENSMUSG00000097615 | -1.043306199 | 0.803260194 |
| ENSMUSG00000097616 | 1.262388595 | 0.091808658 |
| ENSMUSG00000097617 | 1.346123955 | 0.10131895 |
| ENSMUSG00000097618 | 1.112843644 | 0.608497229 |
| ENSMUSG00000097619 | 1.061035153 | 0.684413983 |
| ENSMUSG00000097620 | -1.111384924 | 0.608497229 |
| ENSMUSG00000097621 | 1.037681441 | 0.873810332 |
| ENSMUSG00000097622 | 1 | |
| ENSMUSG00000097623 | 1 | |

| | | |
|---|---|---|
| ENSMUSG00000097624 | 1 | |
| ENSMUSG00000097625 | -1.043306199 | 0.698112512 |
| ENSMUSG00000097626 | 1.488172062 | 0.151654465 |
| ENSMUSG00000097627 | 1 | |
| ENSMUSG00000097628 | 1.117510132 | 0.717035356 |
| ENSMUSG00000097630 | 1 | |
| ENSMUSG00000097631 | 1 | |
| ENSMUSG00000097632 | -1.043306199 | 0.698112512 |
| ENSMUSG00000097633 | -1.159180354 | 0.473406495 |
| ENSMUSG00000097634 | 1 | |
| ENSMUSG00000097635 | 1.354691064 | 0.124350077 |
| ENSMUSG00000097636 | -1.619359057 | 0.001965711 |
| ENSMUSG00000097637 | -1.041561228 | 0.936962652 |
| ENSMUSG00000097638 | -1.084645496 | 0.532369657 |
| ENSMUSG00000097639 | 1.075870199 | 0.814608539 |
| ENSMUSG00000097640 | 1.233977077 | 0.202224588 |
| ENSMUSG00000097641 | 1 | |
| ENSMUSG00000097642 | 1 | |
| ENSMUSG00000097643 | 1.154073294 | 0.608497229 |
| ENSMUSG00000097644 | 1 | |
| ENSMUSG00000097645 | -1.043722544 | 0.869995712 |
| ENSMUSG00000097646 | 1 | |
| ENSMUSG00000097648 | -1 | 1 |
| ENSMUSG00000097649 | 1.045321081 | 0.660863315 |
| ENSMUSG00000097650 | -1.069095149 | 0.77499309 |
| ENSMUSG00000097651 | 1.116079869 | 0.643361947 |
| ENSMUSG00000097652 | 1 | |
| ENSMUSG00000097653 | 1 | |
| ENSMUSG00000097654 | -1 | 1 |
| ENSMUSG00000097655 | 1 | |
| ENSMUSG00000097656 | 1 | |
| ENSMUSG00000097657 | 1.009288193 | 0.99457094 |
| ENSMUSG00000097658 | 1.049756759 | 0.81188464 |
| ENSMUSG00000097659 | 1 | |
| ENSMUSG00000097660 | 1.006972216 | 1 |
| ENSMUSG00000097661 | 1.177765487 | 0.48302662 |
| ENSMUSG00000097662 | 1.144603532 | 0.608497229 |
| ENSMUSG00000097664 | -1.098171958 | 0.417353956 |
| ENSMUSG00000097665 | -1.172299073 | 0.608497229 |
| ENSMUSG00000097666 | -1.044273782 | 0.790851299 |
| ENSMUSG00000097667 | 1.159841294 | 0.518298719 |
| ENSMUSG00000097668 | 1.059463094 | 0.698112512 |
| ENSMUSG00000097669 | -1.043306199 | 0.698112512 |
| ENSMUSG00000097670 | 1 | |
| ENSMUSG00000097671 | 1 | |
| ENSMUSG00000097672 | 1.12189305 | 0.608497229 |
| ENSMUSG00000097673 | -1.156541528 | 0.669959104 |
| ENSMUSG00000097674 | -1.054046247 | 0.861342848 |
| ENSMUSG00000097675 | 1.069068979 | 0.74773997 |
| ENSMUSG00000097676 | 1.059463094 | 0.608497229 |
| ENSMUSG00000097677 | -1.03962336 | 0.698112512 |
| ENSMUSG00000097679 | -1 | 1 |
| ENSMUSG00000097680 | -1.03068233 | 0.882166828 |
| ENSMUSG00000097681 | -1.03962336 | 0.698112512 |
| ENSMUSG00000097682 | 1 | |
| ENSMUSG00000097683 | 1.059463094 | 0.608497229 |
| ENSMUSG00000097684 | -1.05029985 | 0.800933962 |
| ENSMUSG00000097685 | -1.084645496 | 0.698112512 |
| ENSMUSG00000097686 | 1.02332983 | 0.995827459 |
| ENSMUSG00000097689 | 1.069068979 | 0.608497229 |
| ENSMUSG00000097690 | -1.03962336 | 0.698112512 |
| ENSMUSG00000097691 | 1 | |
| ENSMUSG00000097692 | 1.262545733 | 0.411582807 |

| | | |
|---|---|---|
| ENSMUSG00000097693 | 1 | |
| ENSMUSG00000097694 | -1.23372742 | 0.542351223 |
| ENSMUSG00000097695 | 1.046576576 | 0.713861573 |
| ENSMUSG00000097697 | 2.036731689 | 0.00025238 |
| ENSMUSG00000097698 | 1 | |
| ENSMUSG00000097699 | -1.201027237 | 0.179873578 |
| ENSMUSG00000097700 | 1 | |
| ENSMUSG00000097701 | 1.023743961 | 0.938939969 |
| ENSMUSG00000097702 | 1.015486245 | 0.985522133 |
| ENSMUSG00000097703 | -1.075131869 | 0.572781054 |
| ENSMUSG00000097704 | 1 | |
| ENSMUSG00000097705 | 1.020668955 | 0.969214526 |
| ENSMUSG00000097706 | 1.005613439 | 1 |
| ENSMUSG00000097707 | -1.043306199 | 0.698112512 |
| ENSMUSG00000097709 | 1.055597279 | 0.782042968 |
| ENSMUSG00000097710 | 1 | |
| ENSMUSG00000097711 | 1.139613677 | 0.159151577 |
| ENSMUSG00000097712 | 1.069068979 | 0.608497229 |
| ENSMUSG00000097713 | 1.05038453 | 0.860327141 |
| ENSMUSG00000097714 | 1 | |
| ENSMUSG00000097715 | 1.193025725 | 0.098181208 |
| ENSMUSG00000097716 | -1 | 1 |
| ENSMUSG00000097717 | 1.020344501 | 0.983178084 |
| ENSMUSG00000097718 | -1.166321578 | 0.253249273 |
| ENSMUSG00000097719 | -1.084645496 | 0.532369657 |
| ENSMUSG00000097720 | -1.240536453 | 0.272370683 |
| ENSMUSG00000097722 | -1.018978751 | 0.986431796 |
| ENSMUSG00000097723 | 1.014948756 | 0.989603176 |
| ENSMUSG00000097724 | -1.129300295 | 0.602078588 |
| ENSMUSG00000097725 | 1 | |
| ENSMUSG00000097726 | 1.014731082 | 0.997667419 |
| ENSMUSG00000097727 | 1.480107351 | 0.022564197 |
| ENSMUSG00000097728 | -1.360993624 | 0.222341018 |
| ENSMUSG00000097729 | 1.286842586 | 0.00767335 |
| ENSMUSG00000097730 | 1 | |
| ENSMUSG00000097731 | -1.020726133 | 0.996913409 |
| ENSMUSG00000097732 | 1.023959045 | 0.976752232 |
| ENSMUSG00000097734 | -1.079914346 | 0.784232424 |
| ENSMUSG00000097735 | 1.05410549 | 0.858911269 |
| ENSMUSG00000097736 | -1.085651418 | 0.532369657 |
| ENSMUSG00000097737 | 1.014195281 | 0.984902895 |
| ENSMUSG00000097738 | 1.127732926 | 0.573100862 |
| ENSMUSG00000097739 | 1.017706799 | 1 |
| ENSMUSG00000097740 | 1 | |
| ENSMUSG00000097742 | 1 | |
| ENSMUSG00000097743 | -1.105312711 | 0.449677095 |
| ENSMUSG00000097744 | 1.0301485 | 0.991978016 |
| ENSMUSG00000097745 | -1.03962336 | 0.872692845 |
| ENSMUSG00000097746 | 1.121065345 | 0.725290225 |
| ENSMUSG00000097747 | 1.142946526 | 0.087421547 |
| ENSMUSG00000097748 | -1.141886854 | 0.512702106 |
| ENSMUSG00000097749 | -1.018694905 | 0.940298812 |
| ENSMUSG00000097750 | 1.041694484 | 0.814089245 |
| ENSMUSG00000097751 | -1.021136267 | 0.843690197 |
| ENSMUSG00000097752 | 1 | |
| ENSMUSG00000097753 | 1 | |
| ENSMUSG00000097754 | -1 | 1 |
| ENSMUSG00000097755 | -1.109894823 | 0.426233042 |
| ENSMUSG00000097756 | 1.132639128 | 0.608497229 |
| ENSMUSG00000097757 | 1.065250457 | 0.59127359 |
| ENSMUSG00000097759 | 1.162329143 | 0.608497229 |
| ENSMUSG00000097760 | -1.031391793 | 0.839907907 |
| ENSMUSG00000097761 | -1.237062975 | 0.125154408 |

| | | |
|---|---|---|
| ENSMUSG00000097762 | 1.098645647 | 0.715606039 |
| ENSMUSG00000097763 | 1 | |
| ENSMUSG00000097764 | 1 | |
| ENSMUSG00000097766 | 1.069068979 | 0.698112512 |
| ENSMUSG00000097767 | -1.085651418 | 0.731932479 |
| ENSMUSG00000097768 | -1.135512723 | 0.53799277 |
| ENSMUSG00000097769 | 1.006481591 | 1 |
| ENSMUSG00000097770 | 1 | |
| ENSMUSG00000097772 | 1.149467677 | 0.251285035 |
| ENSMUSG00000097773 | -1.060379434 | 0.85351833 |
| ENSMUSG00000097774 | 1 | |
| ENSMUSG00000097775 | -1.053549062 | 0.769745556 |
| ENSMUSG00000097776 | 1 | |
| ENSMUSG00000097777 | -1.136655084 | 0.608497229 |
| ENSMUSG00000097779 | 1.069068979 | 0.608497229 |
| ENSMUSG00000097780 | 1 | |
| ENSMUSG00000097781 | 1.337888666 | 0.282066625 |
| ENSMUSG00000097783 | 1 | |
| ENSMUSG00000097784 | 1.076653244 | 0.71501825 |
| ENSMUSG00000097785 | 1.104061436 | 0.608497229 |
| ENSMUSG00000097786 | -1.043306199 | 0.798487595 |
| ENSMUSG00000097787 | 1.022013433 | 0.971878966 |
| ENSMUSG00000097788 | 1.264359055 | 0.226413509 |
| ENSMUSG00000097789 | 1.556598258 | 0.001674444 |
| ENSMUSG00000097790 | 1 | |
| ENSMUSG00000097791 | 1.023044722 | 1 |
| ENSMUSG00000097792 | -1.059683416 | 0.740073725 |
| ENSMUSG00000097793 | -1.159913159 | 0.279487784 |
| ENSMUSG00000097794 | 1 | |
| ENSMUSG00000097795 | 1.005613439 | 1 |
| ENSMUSG00000097796 | 1.090344457 | 0.256122242 |
| ENSMUSG00000097797 | -1.042689383 | 0.925075932 |
| ENSMUSG00000097798 | 1 | |
| ENSMUSG00000097799 | 1.132639128 | 0.217803313 |
| ENSMUSG00000097800 | -1.044273782 | 0.790851299 |
| ENSMUSG00000097801 | 1 | |
| ENSMUSG00000097803 | 1.015486245 | 1 |
| ENSMUSG00000097804 | -1.147331624 | 0.3938984 |
| ENSMUSG00000097805 | -1.053549062 | 0.769745556 |
| ENSMUSG00000097806 | -1.05309619 | 0.684694376 |
| ENSMUSG00000097807 | 1 | |
| ENSMUSG00000097809 | -1 | 1 |
| ENSMUSG00000097810 | 1.122462048 | 0.420161906 |
| ENSMUSG00000097811 | 1.005613439 | 1 |
| ENSMUSG00000097812 | 1.024908686 | 0.975677002 |
| ENSMUSG00000097814 | 1.0373132 | 0.934760809 |
| ENSMUSG00000097815 | 1.167872049 | 0.490248108 |
| ENSMUSG00000097816 | 1.17245429 | 0.254152111 |
| ENSMUSG00000097817 | 1 | |
| ENSMUSG00000097818 | -1 | 1 |
| ENSMUSG00000097819 | -1.037056164 | 0.918289894 |
| ENSMUSG00000097820 | 1.299212853 | 0.066196433 |
| ENSMUSG00000097821 | 1 | |
| ENSMUSG00000097822 | 1 | |
| ENSMUSG00000097823 | 1 | |
| ENSMUSG00000097824 | 1 | |
| ENSMUSG00000097825 | -1.120476206 | 0.547016491 |
| ENSMUSG00000097827 | -1.010960635 | 1 |
| ENSMUSG00000097828 | -1.164052161 | 0.422094064 |
| ENSMUSG00000097831 | -1.123736245 | 0.814092395 |
| ENSMUSG00000097832 | 1.061747899 | 0.872376523 |
| ENSMUSG00000097833 | 1.077658816 | 0.762178997 |
| ENSMUSG00000097834 | 1 | |

| | | |
|---|---|---|
| ENSMUSG00000097835 | 1.133941335 | 0.608497229 |
| ENSMUSG00000097836 | 1 | |
| ENSMUSG00000097837 | 1 | |
| ENSMUSG00000097838 | -1.063985937 | 0.856445097 |
| ENSMUSG00000097839 | -1.037304692 | 0.867008219 |
| ENSMUSG00000097840 | -1.116853908 | 0.559354144 |
| ENSMUSG00000097841 | 1 | |
| ENSMUSG00000097842 | 1.161278819 | 0.697512751 |
| ENSMUSG00000097843 | 1.069068979 | 0.698112512 |
| ENSMUSG00000097845 | -1.003834322 | 1 |
| ENSMUSG00000097847 | 1.123866579 | 0.650221528 |
| ENSMUSG00000097848 | 1 | |
| ENSMUSG00000097849 | 1 | |
| ENSMUSG00000097850 | 1.059463094 | 0.608497229 |
| ENSMUSG00000097851 | 1 | |
| ENSMUSG00000097852 | -1.008485483 | 1 |
| ENSMUSG00000097853 | 1 | |
| ENSMUSG00000097854 | 1 | |
| ENSMUSG00000097855 | 1.423458142 | 0.032175391 |
| ENSMUSG00000097857 | -1.075182332 | 0.770958539 |
| ENSMUSG00000097858 | -1.099512938 | 0.698112512 |
| ENSMUSG00000097859 | -1.088497035 | 0.803314907 |
| ENSMUSG00000097860 | -1.139116 | 0.534479221 |
| ENSMUSG00000097861 | 1 | |
| ENSMUSG00000097862 | 1 | |
| ENSMUSG00000097863 | 1.020240423 | 0.964236174 |
| ENSMUSG00000097864 | -1.11061171 | 0.608497229 |
| ENSMUSG00000097867 | -1.131016783 | 0.272023587 |
| ENSMUSG00000097868 | -1.043534012 | 0.870100853 |
| ENSMUSG00000097869 | -1.045575678 | 0.745589898 |
| ENSMUSG00000097870 | 1 | |
| ENSMUSG00000097871 | 1 | |
| ENSMUSG00000097872 | 1 | |
| ENSMUSG00000097873 | 1.086869377 | 0.606122889 |
| ENSMUSG00000097874 | 1.006564967 | 1 |
| ENSMUSG00000097875 | 1.115794926 | 0.58386781 |
| ENSMUSG00000097876 | -1.166678327 | 0.215597649 |
| ENSMUSG00000097877 | 1.080164328 | 0.835255454 |
| ENSMUSG00000097878 | 1 | |
| ENSMUSG00000097879 | -1.088743422 | 0.159049699 |
| ENSMUSG00000097881 | 1.122462048 | 0.215991492 |
| ENSMUSG00000097882 | 1.018727262 | 0.973833564 |
| ENSMUSG00000097883 | -1.033820075 | 0.888246981 |
| ENSMUSG00000097884 | 1.059463094 | 0.704170349 |
| ENSMUSG00000097885 | 1.019697539 | 0.982838256 |
| ENSMUSG00000097886 | 1 | |
| ENSMUSG00000097887 | -1.008696069 | 1 |
| ENSMUSG00000097888 | 1.220695418 | 0.249012323 |
| ENSMUSG00000097889 | -1.238058362 | 0.059080872 |
| ENSMUSG00000097890 | 1.114597197 | 0.477029641 |
| ENSMUSG00000097891 | -1.011001473 | 0.975993819 |
| ENSMUSG00000097892 | 1.015486245 | 0.99457094 |
| ENSMUSG00000097893 | -1.384621678 | 0.02414153 |
| ENSMUSG00000097895 | -1 | 1 |
| ENSMUSG00000097896 | -1 | 1 |
| ENSMUSG00000097897 | 1 | |
| ENSMUSG00000097898 | 1 | |
| ENSMUSG00000097899 | 1.075870199 | 0.818635785 |
| ENSMUSG00000097900 | 1.277473343 | 0.287312665 |
| ENSMUSG00000097901 | 1.029972857 | 0.884810702 |
| ENSMUSG00000097902 | 1.026752671 | 0.965978247 |
| ENSMUSG00000097903 | 1.069068979 | 0.608497229 |
| ENSMUSG00000097904 | -1.008583216 | 1 |

| | | |
|---|---|---|
| ENSMUSG00000097906 | 1.130634521 | 0.269617125 |
| ENSMUSG00000097907 | 1 | |
| ENSMUSG00000097908 | 1.263410061 | 0.063898214 |
| ENSMUSG00000097910 | -1 | 1 |
| ENSMUSG00000097911 | -1.004140089 | 1 |
| ENSMUSG00000097912 | -1.030696879 | 0.931949042 |
| ENSMUSG00000097913 | 1 | |
| ENSMUSG00000097914 | -1.195677205 | 0.502179335 |
| ENSMUSG00000097915 | -1.155652769 | 0.608497229 |
| ENSMUSG00000097917 | 1 | |
| ENSMUSG00000097918 | 1 | |
| ENSMUSG00000097919 | 1 | |
| ENSMUSG00000097920 | 1.011497958 | 1 |
| ENSMUSG00000097921 | 1.070429626 | 0.763407099 |
| ENSMUSG00000097922 | 1 | |
| ENSMUSG00000097923 | 1 | |
| ENSMUSG00000097924 | 1.17558988 | 0.352661382 |
| ENSMUSG00000097925 | 1.290200756 | 0.364820257 |
| ENSMUSG00000097926 | 1.023151691 | 0.989387709 |
| ENSMUSG00000097927 | 1 | |
| ENSMUSG00000097928 | 1.222743451 | 0.277795964 |
| ENSMUSG00000097929 | -1.099174267 | 0.688934383 |
| ENSMUSG00000097930 | 1.166909201 | 0.482800078 |
| ENSMUSG00000097931 | 1 | |
| ENSMUSG00000097932 | 1 | |
| ENSMUSG00000097933 | 1.069068979 | 0.698112512 |
| ENSMUSG00000097934 | 1.154253937 | 0.608497229 |
| ENSMUSG00000097935 | 1.11166259 | 0.698112512 |
| ENSMUSG00000097936 | 1.155564223 | 0.39980163 |
| ENSMUSG00000097937 | -1.03962336 | 0.803072249 |
| ENSMUSG00000097938 | 1 | |
| ENSMUSG00000097939 | -1 | 1 |
| ENSMUSG00000097940 | 1 | |
| ENSMUSG00000097941 | 1.184998242 | 0.564226949 |
| ENSMUSG00000097942 | -1 | 1 |
| ENSMUSG00000097943 | 1.019083579 | 0.991073538 |
| ENSMUSG00000097944 | 1.02767497 | 0.859338216 |
| ENSMUSG00000097945 | 1.030800959 | 0.932652618 |
| ENSMUSG00000097946 | 1.030800959 | 0.93251749 |
| ENSMUSG00000097947 | 1 | |
| ENSMUSG00000097948 | 1 | |
| ENSMUSG00000097949 | 1 | |
| ENSMUSG00000097950 | 1 | |
| ENSMUSG00000097951 | -1.051018293 | 0.828564184 |
| ENSMUSG00000097952 | 1 | |
| ENSMUSG00000097953 | 1.189207115 | 0.385934169 |
| ENSMUSG00000097954 | 1.182007788 | 0.316119178 |
| ENSMUSG00000097955 | 1 | |
| ENSMUSG00000097957 | 1 | |
| ENSMUSG00000097958 | 1.216580369 | 0.534291186 |
| ENSMUSG00000097959 | 1 | |
| ENSMUSG00000097960 | 1.285322042 | 0.007795608 |
| ENSMUSG00000097961 | 1 | |
| ENSMUSG00000097962 | 1 | |
| ENSMUSG00000097963 | 1 | |
| ENSMUSG00000097964 | 1 | |
| ENSMUSG00000097965 | -1.069095149 | 0.686194514 |
| ENSMUSG00000097966 | -1.03962336 | 0.808074244 |
| ENSMUSG00000097967 | 1.002009643 | 1 |
| ENSMUSG00000097968 | 1 | |
| ENSMUSG00000097969 | 1 | |
| ENSMUSG00000097970 | 1.156174695 | 0.662840906 |
| ENSMUSG00000097971 | 1.193696317 | 0.55369055 |

| | | |
|---|---|---|
| ENSMUSG00000097972 | 1 | |
| ENSMUSG00000097973 | 1.073839184 | 0.608497229 |
| ENSMUSG00000097974 | 1.059463094 | 0.608497229 |
| ENSMUSG00000097975 | 1.161036672 | 0.251049598 |
| ENSMUSG00000097976 | 1 | |
| ENSMUSG00000097977 | 1.205244041 | 0.630771413 |
| ENSMUSG00000097979 | 1.264494278 | 0.116410779 |
| ENSMUSG00000097980 | 1 | |
| ENSMUSG00000097981 | -1.00002448 | 1 |
| ENSMUSG00000097982 | 1 | |
| ENSMUSG00000097983 | 1.095872691 | 0.657060635 |
| ENSMUSG00000097984 | 1 | |
| ENSMUSG00000097985 | -1.014975048 | 1 |
| ENSMUSG00000097986 | 1 | |
| ENSMUSG00000097987 | 1.059463094 | 0.608497229 |
| ENSMUSG00000097988 | 1.069068979 | 0.608497229 |
| ENSMUSG00000097989 | 1.18450613 | 0.383324143 |
| ENSMUSG00000097990 | -1.029426411 | 0.905393341 |
| ENSMUSG00000097991 | 1.059463094 | 0.608497229 |
| ENSMUSG00000097992 | -1.223947136 | 0.074570023 |
| ENSMUSG00000097993 | 1.197666642 | 0.642327967 |
| ENSMUSG00000097994 | -1.026130559 | 0.760180115 |
| ENSMUSG00000097995 | -1.194616283 | 0.536694773 |
| ENSMUSG00000097996 | -1.023769022 | 0.934760809 |
| ENSMUSG00000097997 | -1 | 1 |
| ENSMUSG00000097998 | 1 | |
| ENSMUSG00000097999 | 1.150179455 | 0.145828618 |
| ENSMUSG00000098000 | 1.015486245 | 0.972980766 |
| ENSMUSG00000098001 | 1.128427755 | 0.372103656 |
| ENSMUSG00000098002 | -1.100193663 | 0.534887579 |
| ENSMUSG00000098003 | -1.043306199 | 0.79291651 |
| ENSMUSG00000098004 | 1 | |
| ENSMUSG00000098005 | 1.095872691 | 0.657060635 |
| ENSMUSG00000098006 | 1.250131101 | 0.250477583 |
| ENSMUSG00000098007 | 1 | |
| ENSMUSG00000098008 | 1 | |
| ENSMUSG00000098009 | -1.03962336 | 0.814089245 |
| ENSMUSG00000098010 | 1.059463094 | 0.608497229 |
| ENSMUSG00000098011 | 1 | |
| ENSMUSG00000098012 | 1 | |
| ENSMUSG00000098013 | 1 | |
| ENSMUSG00000098014 | -1.044273782 | 0.698112512 |
| ENSMUSG00000098015 | 1 | |
| ENSMUSG00000098016 | 1.059463094 | 0.699340382 |
| ENSMUSG00000098017 | -1 | 1 |
| ENSMUSG00000098019 | 1.087548918 | 0.698112512 |
| ENSMUSG00000098021 | 1.202277353 | 0.399217442 |
| ENSMUSG00000098022 | -1.079036265 | 0.829487702 |
| ENSMUSG00000098023 | 1 | |
| ENSMUSG00000098024 | -1.13006158 | 0.724737513 |
| ENSMUSG00000098025 | 1 | |
| ENSMUSG00000098026 | 1 | |
| ENSMUSG00000098027 | 1 | |
| ENSMUSG00000098028 | 1.232460493 | 0.400452977 |
| ENSMUSG00000098029 | 1.059463094 | 0.608497229 |
| ENSMUSG00000098030 | -1 | 1 |
| ENSMUSG00000098031 | 1.229666014 | 0.335919822 |
| ENSMUSG00000098033 | 1.059463094 | 0.608497229 |
| ENSMUSG00000098035 | -1.005951913 | 1 |
| ENSMUSG00000098036 | -1.03962336 | 0.814089245 |
| ENSMUSG00000098037 | -1 | 1 |
| ENSMUSG00000098038 | 1 | |
| ENSMUSG00000098039 | 1.059463094 | 0.608497229 |

| | | |
|---|---|---|
| ENSMUSG00000098040 | 1 | |
| ENSMUSG00000098041 | -1.008328258 | 1 |
| ENSMUSG00000098042 | -1.053549062 | 0.698112512 |
| ENSMUSG00000098043 | 1.059463094 | 0.608497229 |
| ENSMUSG00000098044 | 1 | |
| ENSMUSG00000098045 | 1 | |
| ENSMUSG00000098046 | -1.028348525 | 0.936995747 |
| ENSMUSG00000098047 | 1 | |
| ENSMUSG00000098048 | 1.04328066 | 0.867378212 |
| ENSMUSG00000098050 | 1.222889447 | 0.14566047 |
| ENSMUSG00000098051 | 1 | |
| ENSMUSG00000098052 | 1.171563499 | 0.370636424 |
| ENSMUSG00000098053 | 1 | |
| ENSMUSG00000098054 | 1 | |
| ENSMUSG00000098055 | 1.534379147 | 0.020837631 |
| ENSMUSG00000098056 | 1.024693402 | 0.952057016 |
| ENSMUSG00000098057 | 1.05943716 | 0.641618005 |
| ENSMUSG00000098058 | 1 | |
| ENSMUSG00000098059 | 1 | |
| ENSMUSG00000098060 | 1 | |
| ENSMUSG00000098061 | 1.972449013 | 0.00095885 |
| ENSMUSG00000098062 | 1.060054384 | 0.731587932 |
| ENSMUSG00000098064 | 1 | |
| ENSMUSG00000098065 | -1 | 1 |
| ENSMUSG00000098066 | -1.016564192 | 0.988424085 |
| ENSMUSG00000098067 | 1.095872691 | 0.608497229 |
| ENSMUSG00000098068 | -1 | 1 |
| ENSMUSG00000098069 | 1 | |
| ENSMUSG00000098070 | 1 | |
| ENSMUSG00000098071 | 1 | |
| ENSMUSG00000098072 | 1 | |
| ENSMUSG00000098073 | 1.090778897 | 0.732012457 |
| ENSMUSG00000098074 | 1 | |
| ENSMUSG00000098075 | 1 | |
| ENSMUSG00000098076 | -1.100448966 | 0.748944522 |
| ENSMUSG00000098077 | -1.084645496 | 0.608497229 |
| ENSMUSG00000098078 | 1 | |
| ENSMUSG00000098079 | 1.059463094 | 0.698112512 |
| ENSMUSG00000098080 | 1.069068979 | 0.698112512 |
| ENSMUSG00000098081 | 1 | |
| ENSMUSG00000098082 | 1.047347415 | 0.889802291 |
| ENSMUSG00000098083 | -1.03962336 | 0.803072249 |
| ENSMUSG00000098084 | 1 | |
| ENSMUSG00000098085 | -1 | 1 |
| ENSMUSG00000098086 | -1 | 1 |
| ENSMUSG00000098087 | 1 | |
| ENSMUSG00000098088 | 1 | |
| ENSMUSG00000098089 | 1 | |
| ENSMUSG00000098090 | 1.007565835 | 1 |
| ENSMUSG00000098091 | 1.150837682 | 0.547016491 |
| ENSMUSG00000098092 | -1.073603785 | 0.783862764 |
| ENSMUSG00000098093 | -1.03962336 | 0.803072249 |
| ENSMUSG00000098094 | 1 | |
| ENSMUSG00000098095 | 1 | |
| ENSMUSG00000098096 | 1 | |
| ENSMUSG00000098097 | 1 | |
| ENSMUSG00000098098 | 1.776596657 | 0.005039944 |
| ENSMUSG00000098099 | 1.05410549 | 0.833490776 |
| ENSMUSG00000098100 | -1.044273782 | 0.698112512 |
| ENSMUSG00000098101 | -1.230101541 | 0.379210853 |
| ENSMUSG00000098102 | -1.044273782 | 0.698112512 |
| ENSMUSG00000098103 | 1 | |
| ENSMUSG00000098104 | 1 | |

| | | |
|---|---|---|
| ENSMUSG00000098105 | 1.116785864 | 0.698112512 |
| ENSMUSG00000098106 | -1.087569789 | 0.729018288 |
| ENSMUSG00000098107 | 1 | |
| ENSMUSG00000098108 | 1 | |
| ENSMUSG00000098109 | 1.059463094 | 0.699340382 |
| ENSMUSG00000098110 | -1.03962336 | 0.698112512 |
| ENSMUSG00000098111 | 1.2104378 | 0.518775375 |
| ENSMUSG00000098112 | -1.02513447 | 0.945625364 |
| ENSMUSG00000098113 | 1.149070419 | 0.136267625 |
| ENSMUSG00000098114 | 1 | |
| ENSMUSG00000098115 | -1 | 1 |
| ENSMUSG00000098116 | 1.084618945 | 0.614071182 |
| ENSMUSG00000098118 | 1 | |
| ENSMUSG00000098119 | 1 | |
| ENSMUSG00000098120 | 1.122434572 | 0.547368903 |
| ENSMUSG00000098121 | 1 | |
| ENSMUSG00000098122 | 1 | |
| ENSMUSG00000098123 | -1.026529187 | 0.953613038 |
| ENSMUSG00000098124 | 1 | |
| ENSMUSG00000098125 | 1.161036672 | 0.378751351 |
| ENSMUSG00000098126 | 1 | |
| ENSMUSG00000098127 | -1.053549062 | 0.769745556 |
| ENSMUSG00000098128 | 1 | |
| ENSMUSG00000098129 | 1 | |
| ENSMUSG00000098130 | 1 | |
| ENSMUSG00000098131 | 1 | |
| ENSMUSG00000098132 | 1.028323353 | 0.936340683 |
| ENSMUSG00000098134 | 1 | |
| ENSMUSG00000098135 | 1.059463094 | 0.698112512 |
| ENSMUSG00000098136 | 1.059463094 | 0.698112512 |
| ENSMUSG00000098137 | 1 | |
| ENSMUSG00000098138 | 1.079681442 | 0.608497229 |
| ENSMUSG00000098139 | 1 | |
| ENSMUSG00000098140 | 1 | |
| ENSMUSG00000098141 | 1.156007461 | 0.503812942 |
| ENSMUSG00000098142 | 1.171418576 | 0.36623954 |
| ENSMUSG00000098143 | 1 | |
| ENSMUSG00000098144 | 1 | |
| ENSMUSG00000098145 | 1 | |
| ENSMUSG00000098146 | 1.10613438 | 0.771741564 |
| ENSMUSG00000098147 | 1 | |
| ENSMUSG00000098149 | 1.077988547 | 0.698112512 |
| ENSMUSG00000098150 | 1.132639128 | 0.398767938 |
| ENSMUSG00000098151 | 1 | |
| ENSMUSG00000098152 | 1 | |
| ENSMUSG00000098153 | 1 | |
| ENSMUSG00000098154 | 1.182756393 | 0.539963645 |
| ENSMUSG00000098155 | 1 | |
| ENSMUSG00000098156 | 1.099610164 | 0.790851299 |
| ENSMUSG00000098157 | 1 | |
| ENSMUSG00000098158 | -1.044273782 | 0.790851299 |
| ENSMUSG00000098159 | 1 | |
| ENSMUSG00000098160 | 1 | |
| ENSMUSG00000098161 | 1 | |
| ENSMUSG00000098162 | -1.010960635 | 1 |
| ENSMUSG00000098163 | 1 | |
| ENSMUSG00000098164 | -1.241737856 | 0.385487949 |
| ENSMUSG00000098165 | -1 | 1 |
| ENSMUSG00000098166 | 1 | |
| ENSMUSG00000098167 | 1 | |
| ENSMUSG00000098169 | -1.03962336 | 0.698112512 |
| ENSMUSG00000098170 | 1 | |
| ENSMUSG00000098171 | 1.069068979 | 0.608497229 |

| | | |
|---|---|---|
| ENSMUSG00000098172 | 1 | |
| ENSMUSG00000098173 | 1 | |
| ENSMUSG00000098174 | 1 | |
| ENSMUSG00000098175 | -1.053549062 | 0.774498429 |
| ENSMUSG00000098176 | -1.134214409 | 0.652116097 |
| ENSMUSG00000098177 | -1 | 1 |
| ENSMUSG00000098178 | 1.17348724 | 0.752472633 |
| ENSMUSG00000098179 | 1 | |
| ENSMUSG00000098180 | 1.367453556 | 0.19967677 |
| ENSMUSG00000098181 | 1.005613439 | 1 |
| ENSMUSG00000098183 | -1.011415248 | 0.970256467 |
| ENSMUSG00000098184 | 1.079681442 | 0.794909137 |
| ENSMUSG00000098185 | 1 | |
| ENSMUSG00000098186 | 1 | |
| ENSMUSG00000098187 | 1.199989356 | 0.054295624 |
| ENSMUSG00000098188 | 1 | |
| ENSMUSG00000098189 | 1 | |
| ENSMUSG00000098190 | -1.084645496 | 0.608497229 |
| ENSMUSG00000098191 | 1 | |
| ENSMUSG00000098192 | 1.112016081 | 0.697178241 |
| ENSMUSG00000098193 | 1 | |
| ENSMUSG00000098194 | -1.067718846 | 0.821592265 |
| ENSMUSG00000098195 | 1.015486245 | 0.985522133 |
| ENSMUSG00000098196 | -1.089497311 | 0.608497229 |
| ENSMUSG00000098197 | 1 | |
| ENSMUSG00000098198 | -1.044273782 | 0.870059001 |
| ENSMUSG00000098199 | 1.189666502 | 0.566649749 |
| ENSMUSG00000098200 | -1.043306199 | 0.698112512 |
| ENSMUSG00000098201 | -1.100193663 | 0.534887579 |
| ENSMUSG00000098202 | 1.059463094 | 0.608497229 |
| ENSMUSG00000098203 | 1.132639128 | 0.398767938 |
| ENSMUSG00000098204 | 1.069068979 | 0.698112512 |
| ENSMUSG00000098205 | 1 | |
| ENSMUSG00000098206 | -1.164110628 | 0.440837715 |
| ENSMUSG00000098207 | 1.069068979 | 0.608497229 |
| ENSMUSG00000098208 | 1.233010731 | 0.139593229 |
| ENSMUSG00000098209 | -1 | 1 |
| ENSMUSG00000098210 | 1.092525005 | 0.765197976 |
| ENSMUSG00000098211 | 1 | |
| ENSMUSG00000098212 | 1 | |
| ENSMUSG00000098213 | 1 | |
| ENSMUSG00000098214 | 1 | |
| ENSMUSG00000098215 | 1.059463094 | 0.608497229 |
| ENSMUSG00000098216 | 1 | |
| ENSMUSG00000098217 | 1 | |
| ENSMUSG00000098218 | 1 | |
| ENSMUSG00000098219 | 1.015486245 | 0.987775262 |
| ENSMUSG00000098221 | -1.184871248 | 0.511359722 |
| ENSMUSG00000098222 | 1.144363107 | 0.692423811 |
| ENSMUSG00000098223 | 1 | |
| ENSMUSG00000098224 | 1.097750068 | 0.698112512 |
| ENSMUSG00000098225 | 1 | |
| ENSMUSG00000098226 | 1 | |
| ENSMUSG00000098227 | -1.053549062 | 0.698112512 |
| ENSMUSG00000098228 | 1 | |
| ENSMUSG00000098229 | -1 | 1 |
| ENSMUSG00000098230 | 1 | |
| ENSMUSG00000098231 | 1.059463094 | 0.776953697 |
| ENSMUSG00000098232 | 1.034865357 | 0.899648217 |
| ENSMUSG00000098233 | 1 | |
| ENSMUSG00000098234 | 1.030327008 | 0.888903436 |
| ENSMUSG00000098235 | -1.069955956 | 0.649280935 |
| ENSMUSG00000098236 | 1.059463094 | 0.608497229 |

| | | |
|---|---|---|
| ENSMUSG00000098237 | 1 | |
| ENSMUSG00000098238 | 1.156954974 | 0.053274222 |
| ENSMUSG00000098240 | 1.069068979 | 0.698112512 |
| ENSMUSG00000098241 | 1 | |
| ENSMUSG00000098242 | 1 | |
| ENSMUSG00000098243 | -1.132666855 | 0.391570427 |
| ENSMUSG00000098244 | 1 | |
| ENSMUSG00000098245 | 1 | |
| ENSMUSG00000098246 | 1.015486245 | 0.972980766 |
| ENSMUSG00000098247 | 1 | |
| ENSMUSG00000098248 | 1 | |
| ENSMUSG00000098249 | 1 | |
| ENSMUSG00000098251 | 1 | |
| ENSMUSG00000098252 | 1 | |
| ENSMUSG00000098253 | 1 | |
| ENSMUSG00000098254 | 1 | |
| ENSMUSG00000098255 | 1 | |
| ENSMUSG00000098257 | 1 | |
| ENSMUSG00000098258 | 1 | |
| ENSMUSG00000098259 | 1.003467434 | 1 |
| ENSMUSG00000098260 | 1 | |
| ENSMUSG00000098261 | 1 | |
| ENSMUSG00000098262 | 1 | |
| ENSMUSG00000098263 | 1 | |
| ENSMUSG00000098264 | 1 | |
| ENSMUSG00000098265 | 1 | |
| ENSMUSG00000098266 | 1 | |
| ENSMUSG00000098267 | -1.003834322 | 1 |
| ENSMUSG00000098268 | 1 | |
| ENSMUSG00000098269 | 1 | |
| ENSMUSG00000098270 | 1 | |
| ENSMUSG00000098271 | 1.069068979 | 0.608497229 |
| ENSMUSG00000098272 | 1 | |
| ENSMUSG00000098273 | 1 | |
| ENSMUSG00000098274 | 1.0294299 | 0.934558611 |
| ENSMUSG00000098275 | 1.015486245 | 0.986542466 |
| ENSMUSG00000098276 | 1 | |
| ENSMUSG00000098278 | 1 | |
| ENSMUSG00000098279 | 1 | |
| ENSMUSG00000098280 | 1 | |
| ENSMUSG00000098281 | 1 | |
| ENSMUSG00000098282 | 1 | |
| ENSMUSG00000098284 | 1 | |
| ENSMUSG00000098285 | 1 | |
| ENSMUSG00000098286 | -1.069112205 | 0.506670163 |
| ENSMUSG00000098287 | 1 | |
| ENSMUSG00000098288 | 1 | |
| ENSMUSG00000098289 | 1 | |
| ENSMUSG00000098290 | 1 | |
| ENSMUSG00000098291 | -1.088487825 | 0.698112512 |
| ENSMUSG00000098292 | -1.071075483 | 0.750635131 |
| ENSMUSG00000098293 | 1.116785864 | 0.578221424 |
| ENSMUSG00000098294 | 1 | |
| ENSMUSG00000098295 | 1 | |
| ENSMUSG00000098296 | 1 | |
| ENSMUSG00000098297 | 1 | |
| ENSMUSG00000098298 | 1 | |
| ENSMUSG00000098299 | 1 | |
| ENSMUSG00000098300 | 1 | |
| ENSMUSG00000098302 | 1 | |
| ENSMUSG00000098303 | 1 | |
| ENSMUSG00000098304 | 1.095872691 | 0.608497229 |
| ENSMUSG00000098305 | 1.059463094 | 0.608497229 |

| | | |
|---|---|---|
| ENSMUSG00000098306 | 1 | |
| ENSMUSG00000098307 | 1 | |
| ENSMUSG00000098308 | 1 | |
| ENSMUSG00000098309 | 1 | |
| ENSMUSG00000098310 | 1 | |
| ENSMUSG00000098311 | 1 | |
| ENSMUSG00000098312 | 1 | |
| ENSMUSG00000098313 | 1 | |
| ENSMUSG00000098314 | 1 | |
| ENSMUSG00000098315 | 1 | |
| ENSMUSG00000098316 | 1 | |
| ENSMUSG00000098317 | 1 | |
| ENSMUSG00000098318 | 1.177064537 | 0.024548214 |
| ENSMUSG00000098319 | 1 | |
| ENSMUSG00000098320 | 1 | |
| ENSMUSG00000098322 | -1 | 1 |
| ENSMUSG00000098323 | 1 | |
| ENSMUSG00000098325 | 1 | |
| ENSMUSG00000098326 | 1 | |
| ENSMUSG00000098327 | 1 | |
| ENSMUSG00000098328 | -1.03962336 | 0.853933078 |
| ENSMUSG00000098329 | 1 | |
| ENSMUSG00000098330 | 1 | |
| ENSMUSG00000098331 | -1.03962336 | 0.814089245 |
| ENSMUSG00000098332 | 1.012400953 | 1 |
| ENSMUSG00000098333 | 1 | |
| ENSMUSG00000098334 | 1 | |
| ENSMUSG00000098335 | 1 | |
| ENSMUSG00000098336 | 1 | |
| ENSMUSG00000098338 | 1 | |
| ENSMUSG00000098339 | 1 | |
| ENSMUSG00000098340 | -1.043306199 | 0.79291651 |
| ENSMUSG00000098341 | -1.03962336 | 0.698112512 |
| ENSMUSG00000098342 | 1 | |
| ENSMUSG00000098343 | 1.059982015 | 0.763234043 |
| ENSMUSG00000098344 | 1.057478344 | 0.837014159 |
| ENSMUSG00000098345 | 1 | |
| ENSMUSG00000098346 | 1.059463094 | 0.608497229 |
| ENSMUSG00000098347 | 1 | |
| ENSMUSG00000098348 | 1 | |
| ENSMUSG00000098349 | 1 | |
| ENSMUSG00000098350 | 1 | |
| ENSMUSG00000098351 | 1 | |
| ENSMUSG00000098353 | 1 | |
| ENSMUSG00000098354 | 1.059033181 | 0.801168497 |
| ENSMUSG00000098355 | 1 | |
| ENSMUSG00000098356 | 1 | |
| ENSMUSG00000098357 | 1 | |
| ENSMUSG00000098358 | 1 | |
| ENSMUSG00000098359 | 1 | |
| ENSMUSG00000098360 | 1 | |
| ENSMUSG00000098361 | 1 | |
| ENSMUSG00000098362 | 1 | |
| ENSMUSG00000098363 | 1.058763308 | 0.895137669 |
| ENSMUSG00000098364 | 1 | |
| ENSMUSG00000098365 | -1.044273782 | 0.698112512 |
| ENSMUSG00000098366 | 1 | |
| ENSMUSG00000098367 | 1 | |
| ENSMUSG00000098368 | 1 | |
| ENSMUSG00000098369 | 1 | |
| ENSMUSG00000098370 | 1 | |
| ENSMUSG00000098371 | 1 | |
| ENSMUSG00000098373 | 1 | |

| | | |
|---|---|---|
| ENSMUSG00000098374 | -1.124880617 | 0.234936663 |
| ENSMUSG00000098376 | -1.084645496 | 0.608497229 |
| ENSMUSG00000098378 | 1 | |
| ENSMUSG00000098379 | 1 | |
| ENSMUSG00000098380 | -1 | 1 |
| ENSMUSG00000098381 | 1 | |
| ENSMUSG00000098382 | 1 | |
| ENSMUSG00000098383 | 1 | |
| ENSMUSG00000098385 | -1.043306199 | 0.698112512 |
| ENSMUSG00000098386 | 1 | |
| ENSMUSG00000098387 | 1 | |
| ENSMUSG00000098388 | 1 | |
| ENSMUSG00000098389 | 1 | |
| ENSMUSG00000098390 | 1 | |
| ENSMUSG00000098391 | 1 | |
| ENSMUSG00000098392 | 1 | |
| ENSMUSG00000098393 | 1 | |
| ENSMUSG00000098394 | 1 | |
| ENSMUSG00000098395 | 1 | |
| ENSMUSG00000098396 | 1 | |
| ENSMUSG00000098397 | -1.13161737 | 0.608497229 |
| ENSMUSG00000098399 | 1 | |
| ENSMUSG00000098400 | 1 | |
| ENSMUSG00000098401 | 1 | |
| ENSMUSG00000098402 | 1 | |
| ENSMUSG00000098403 | -1.110619514 | 0.675170087 |
| ENSMUSG00000098404 | -1 | 1 |
| ENSMUSG00000098405 | 1 | |
| ENSMUSG00000098406 | 1 | |
| ENSMUSG00000098407 | 1 | |
| ENSMUSG00000098408 | 1 | |
| ENSMUSG00000098409 | 1 | |
| ENSMUSG00000098410 | 1 | |
| ENSMUSG00000098412 | -1.144460846 | 0.608497229 |
| ENSMUSG00000098413 | 1 | |
| ENSMUSG00000098414 | 1.015486245 | 0.998386582 |
| ENSMUSG00000098415 | -1.043306199 | 0.803260194 |
| ENSMUSG00000098416 | 1 | |
| ENSMUSG00000098417 | 1 | |
| ENSMUSG00000098418 | 1 | |
| ENSMUSG00000098419 | 1.13984477 | 0.608497229 |
| ENSMUSG00000098420 | 1.069068979 | 0.608497229 |
| ENSMUSG00000098421 | 1 | |
| ENSMUSG00000098422 | 1 | |
| ENSMUSG00000098423 | 1 | |
| ENSMUSG00000098424 | 1 | |
| ENSMUSG00000098425 | 1 | |
| ENSMUSG00000098426 | 1 | |
| ENSMUSG00000098428 | 1 | |
| ENSMUSG00000098429 | 1 | |
| ENSMUSG00000098430 | 1 | |
| ENSMUSG00000098431 | 1 | |
| ENSMUSG00000098432 | 1 | |
| ENSMUSG00000098433 | 1 | |
| ENSMUSG00000098434 | -1.00002448 | 1 |
| ENSMUSG00000098435 | 1 | |
| ENSMUSG00000098436 | 1 | |
| ENSMUSG00000098437 | 1 | |
| ENSMUSG00000098438 | 1 | |
| ENSMUSG00000098439 | 1 | |
| ENSMUSG00000098441 | 1 | |
| ENSMUSG00000098442 | 1 | |
| ENSMUSG00000098443 | 1 | |

| | | |
|---|---|---|
| ENSMUSG00000098444 | 1 | |
| ENSMUSG00000098445 | -1.077571277 | 0.816821022 |
| ENSMUSG00000098446 | 1 | |
| ENSMUSG00000098447 | -1.177546921 | 0.451479855 |
| ENSMUSG00000098448 | 1 | |
| ENSMUSG00000098449 | 1.019083579 | 0.996913409 |
| ENSMUSG00000098451 | 1 | |
| ENSMUSG00000098452 | 1 | |
| ENSMUSG00000098453 | 1 | |
| ENSMUSG00000098454 | 1 | |
| ENSMUSG00000098455 | 1 | |
| ENSMUSG00000098456 | 1.049517471 | 0.929746187 |
| ENSMUSG00000098457 | 1 | |
| ENSMUSG00000098458 | 1 | |
| ENSMUSG00000098459 | 1 | |
| ENSMUSG00000098460 | 1 | |
| ENSMUSG00000098462 | 1 | |
| ENSMUSG00000098463 | 1 | |
| ENSMUSG00000098464 | 1 | |
| ENSMUSG00000098465 | 1.059463094 | 0.698112512 |
| ENSMUSG00000098466 | 1 | |
| ENSMUSG00000098467 | 1 | |
| ENSMUSG00000098468 | 1 | |
| ENSMUSG00000098469 | 1 | |
| ENSMUSG00000098470 | -1.068398149 | 0.734771161 |
| ENSMUSG00000098471 | 1 | |
| ENSMUSG00000098472 | 1 | |
| ENSMUSG00000098473 | 1 | |
| ENSMUSG00000098474 | 1 | |
| ENSMUSG00000098475 | 1 | |
| ENSMUSG00000098476 | 1 | |
| ENSMUSG00000098477 | 1 | |
| ENSMUSG00000098478 | 1 | |
| ENSMUSG00000098479 | 1 | |
| ENSMUSG00000098480 | 1 | |
| ENSMUSG00000098481 | -1 | 1 |
| ENSMUSG00000098482 | 1 | |
| ENSMUSG00000098483 | 1.177765487 | 0.276639872 |
| ENSMUSG00000098484 | 1 | |
| ENSMUSG00000098485 | 1 | |
| ENSMUSG00000098486 | 1 | |
| ENSMUSG00000098487 | 1 | |
| ENSMUSG00000098488 | -1.253041886 | 0.062916574 |
| ENSMUSG00000098489 | 1 | |
| ENSMUSG00000098490 | -1.063127349 | 0.698112512 |
| ENSMUSG00000098491 | 1 | |
| ENSMUSG00000098492 | 1.059463094 | 0.608497229 |
| ENSMUSG00000098493 | 1 | |
| ENSMUSG00000098494 | -1.084645496 | 0.70299173 |
| ENSMUSG00000098496 | 1.06499219 | 0.59677702 |
| ENSMUSG00000098497 | 1 | |
| ENSMUSG00000098498 | 1 | |
| ENSMUSG00000098499 | -1 | 1 |
| ENSMUSG00000098500 | 1 | |
| ENSMUSG00000098501 | 1 | |
| ENSMUSG00000098502 | 1 | |
| ENSMUSG00000098503 | 1 | |
| ENSMUSG00000098504 | 1 | |
| ENSMUSG00000098505 | 1 | |
| ENSMUSG00000098506 | 1.099938056 | 0.810722095 |
| ENSMUSG00000098507 | -1 | 1 |
| ENSMUSG00000098508 | 1 | |
| ENSMUSG00000098509 | 1 | |

| | | |
|---|---|---|
| ENSMUSG00000098510 | -1.132666855 | 0.18851436 |
| ENSMUSG00000098511 | 1 | |
| ENSMUSG00000098513 | -1 | 1 |
| ENSMUSG00000098514 | 1 | |
| ENSMUSG00000098515 | -1.135312311 | 0.698112512 |
| ENSMUSG00000098516 | 1.059463094 | 0.608497229 |
| ENSMUSG00000098518 | 1 | |
| ENSMUSG00000098519 | 1 | |
| ENSMUSG00000098520 | -1.014570171 | 0.985623907 |
| ENSMUSG00000098521 | 1 | |
| ENSMUSG00000098522 | 1 | |
| ENSMUSG00000098524 | 1 | |
| ENSMUSG00000098525 | 1 | |
| ENSMUSG00000098526 | 1 | |
| ENSMUSG00000098527 | 1 | |
| ENSMUSG00000098528 | 1 | |
| ENSMUSG00000098529 | 1 | |
| ENSMUSG00000098530 | 1 | |
| ENSMUSG00000098531 | -1 | 1 |
| ENSMUSG00000098532 | -1.044273782 | 0.698112512 |
| ENSMUSG00000098533 | 1 | |
| ENSMUSG00000098534 | -1.161107412 | 0.381043202 |
| ENSMUSG00000098535 | 1 | |
| ENSMUSG00000098536 | 1 | |
| ENSMUSG00000098537 | 1 | |
| ENSMUSG00000098538 | 1 | |
| ENSMUSG00000098539 | 1 | |
| ENSMUSG00000098540 | 1 | |
| ENSMUSG00000098541 | -1 | 1 |
| ENSMUSG00000098542 | 1 | |
| ENSMUSG00000098543 | 1 | |
| ENSMUSG00000098544 | 1 | |
| ENSMUSG00000098547 | -1.014570171 | 0.996048848 |
| ENSMUSG00000098548 | 1 | |
| ENSMUSG00000098549 | -1.014357059 | 1 |
| ENSMUSG00000098550 | 1 | |
| ENSMUSG00000098552 | 1 | |
| ENSMUSG00000098553 | 1 | |
| ENSMUSG00000098555 | 1 | |
| ENSMUSG00000098557 | -1.644617916 | 0.013388685 |
| ENSMUSG00000098558 | -1 | 1 |
| ENSMUSG00000098559 | -1.211309146 | 0.210508637 |
| ENSMUSG00000098560 | 1 | |
| ENSMUSG00000098561 | 1 | |
| ENSMUSG00000098562 | -1 | 1 |
| ENSMUSG00000098563 | 1 | |
| ENSMUSG00000098564 | 1 | |
| ENSMUSG00000098565 | 1 | |
| ENSMUSG00000098566 | 1 | |
| ENSMUSG00000098567 | 1 | |
| ENSMUSG00000098568 | 1 | |
| ENSMUSG00000098569 | -1 | 1 |
| ENSMUSG00000098570 | 1 | |
| ENSMUSG00000098572 | 1 | |
| ENSMUSG00000098573 | 1 | |
| ENSMUSG00000098574 | 1 | |
| ENSMUSG00000098575 | 1 | |
| ENSMUSG00000098577 | 1 | |
| ENSMUSG00000098578 | 1 | |
| ENSMUSG00000098579 | 1 | |
| ENSMUSG00000098580 | 1 | |
| ENSMUSG00000098582 | 1 | |
| ENSMUSG00000098583 | 1 | |

| | | |
|---|---|---|
| ENSMUSG00000098585 | 1 | |
| ENSMUSG00000098586 | 1 | |
| ENSMUSG00000098588 | 1 | |
| ENSMUSG00000098589 | -1.142727245 | 0.202428693 |
| ENSMUSG00000098590 | 1 | |
| ENSMUSG00000098591 | 1 | |
| ENSMUSG00000098592 | 1 | |
| ENSMUSG00000098593 | 1 | |
| ENSMUSG00000098594 | 1 | |
| ENSMUSG00000098595 | 1 | |
| ENSMUSG00000098596 | 1 | |
| ENSMUSG00000098597 | 1 | |
| ENSMUSG00000098598 | 1 | |
| ENSMUSG00000098599 | 1 | |
| ENSMUSG00000098600 | -1.023769022 | 0.956401771 |
| ENSMUSG00000098601 | 1 | |
| ENSMUSG00000098602 | 1 | |
| ENSMUSG00000098603 | 1 | |
| ENSMUSG00000098604 | 1 | |
| ENSMUSG00000098606 | 1 | |
| ENSMUSG00000098607 | 1 | |
| ENSMUSG00000098608 | 1 | |
| ENSMUSG00000098609 | -1.153765044 | 0.698112512 |
| ENSMUSG00000098611 | 1 | |
| ENSMUSG00000098612 | 1 | |
| ENSMUSG00000098613 | -1 | 1 |
| ENSMUSG00000098614 | -1.044273782 | 0.698112512 |
| ENSMUSG00000098617 | 1 | |
| ENSMUSG00000098618 | 1 | |
| ENSMUSG00000098619 | 1 | |
| ENSMUSG00000098620 | 1.089910716 | 0.815784461 |
| ENSMUSG00000098621 | 1 | |
| ENSMUSG00000098622 | 1 | |
| ENSMUSG00000098623 | 1 | |
| ENSMUSG00000098624 | 1 | |
| ENSMUSG00000098625 | 1 | |
| ENSMUSG00000098626 | 1 | |
| ENSMUSG00000098627 | 1 | |
| ENSMUSG00000098628 | 1 | |
| ENSMUSG00000098629 | 1 | |
| ENSMUSG00000098630 | 1 | |
| ENSMUSG00000098631 | 1 | |
| ENSMUSG00000098632 | 1 | |
| ENSMUSG00000098634 | 1 | |
| ENSMUSG00000098635 | 1 | |
| ENSMUSG00000098636 | 1 | |
| ENSMUSG00000098638 | 1 | |
| ENSMUSG00000098640 | 1.059463094 | 0.608497229 |
| ENSMUSG00000098641 | 1 | |
| ENSMUSG00000098642 | 1 | |
| ENSMUSG00000098644 | 1 | |
| ENSMUSG00000098645 | 1 | |
| ENSMUSG00000098646 | 1 | |
| ENSMUSG00000098647 | 1 | |
| ENSMUSG00000098648 | 1 | |
| ENSMUSG00000098649 | 1.059463094 | 0.608497229 |
| ENSMUSG00000098650 | 1 | |
| ENSMUSG00000098651 | 1 | |
| ENSMUSG00000098652 | 1 | |
| ENSMUSG00000098653 | 1 | |
| ENSMUSG00000098654 | -1 | 1 |
| ENSMUSG00000098655 | 1 | |
| ENSMUSG00000098656 | 1.069068979 | 0.608497229 |

| | | |
|---|---|---|
| ENSMUSG00000098657 | 1 | |
| ENSMUSG00000098658 | -1 | 1 |
| ENSMUSG00000098659 | 1 | |
| ENSMUSG00000098660 | 1 | |
| ENSMUSG00000098661 | 1.132639128 | 0.217803313 |
| ENSMUSG00000098662 | 1 | |
| ENSMUSG00000098663 | -1.043306199 | 0.698112512 |
| ENSMUSG00000098664 | 1 | |
| ENSMUSG00000098665 | 1 | |
| ENSMUSG00000098666 | 1.079681442 | 0.697178241 |
| ENSMUSG00000098667 | 1 | |
| ENSMUSG00000098668 | 1 | |
| ENSMUSG00000098669 | -1.058018035 | 0.818229917 |
| ENSMUSG00000098670 | 1.161036672 | 0.389746121 |
| ENSMUSG00000098671 | 1 | |
| ENSMUSG00000098672 | 1 | |
| ENSMUSG00000098674 | 1 | |
| ENSMUSG00000098675 | 1.049411284 | 0.835701362 |
| ENSMUSG00000098676 | 1 | |
| ENSMUSG00000098677 | 1 | |
| ENSMUSG00000098678 | -1.043943699 | 0.965153373 |
| ENSMUSG00000098679 | -1.283904146 | 0.276412037 |
| ENSMUSG00000098680 | 1 | |
| ENSMUSG00000098682 | 1 | |
| ENSMUSG00000098683 | -1.053549062 | 0.825517832 |
| ENSMUSG00000098684 | 1 | |
| ENSMUSG00000098685 | -1.038913334 | 0.935126512 |
| ENSMUSG00000098686 | 1 | |
| ENSMUSG00000098687 | 1 | |
| ENSMUSG00000098688 | 1 | |
| ENSMUSG00000098689 | -1 | 1 |
| ENSMUSG00000098690 | 1.040950401 | 0.949933632 |
| ENSMUSG00000098691 | 1 | |
| ENSMUSG00000098692 | -1.029113823 | 0.929923764 |
| ENSMUSG00000098693 | 1 | |
| ENSMUSG00000098694 | 1 | |
| ENSMUSG00000098695 | 1 | |
| ENSMUSG00000098696 | 1 | |
| ENSMUSG00000098697 | 1 | |
| ENSMUSG00000098698 | -1.025690504 | 0.958181659 |
| ENSMUSG00000098700 | 1 | |
| ENSMUSG00000098701 | 1 | |
| ENSMUSG00000098702 | 1.162313089 | 0.025087522 |
| ENSMUSG00000098703 | 1 | |
| ENSMUSG00000098704 | 1 | |
| ENSMUSG00000098705 | 1 | |
| ENSMUSG00000098706 | 1 | |
| ENSMUSG00000098707 | 1 | |
| ENSMUSG00000098708 | -1 | 1 |
| ENSMUSG00000098709 | 1 | |
| ENSMUSG00000098710 | 1.059463094 | 0.608497229 |
| ENSMUSG00000098711 | 1 | |
| ENSMUSG00000098712 | 1 | |
| ENSMUSG00000098713 | 1.102619452 | 0.618537911 |
| ENSMUSG00000098714 | -1.158708082 | 0.216046507 |
| ENSMUSG00000098715 | 1 | |
| ENSMUSG00000098716 | 1 | |
| ENSMUSG00000098717 | 1 | |
| ENSMUSG00000098718 | 1 | |
| ENSMUSG00000098719 | 1 | |
| ENSMUSG00000098720 | 1 | |
| ENSMUSG00000098721 | 1 | |
| ENSMUSG00000098722 | 1 | |

| | | |
|---|---|---|
| ENSMUSG00000098723 | 1 | |
| ENSMUSG00000098725 | 1 | |
| ENSMUSG00000098726 | 1 | |
| ENSMUSG00000098727 | 1 | |
| ENSMUSG00000098728 | 1 | |
| ENSMUSG00000098729 | 1 | |
| ENSMUSG00000098730 | 1 | |
| ENSMUSG00000098731 | 1.226417076 | 0.608497229 |
| ENSMUSG00000098732 | -1 | 1 |
| ENSMUSG00000098733 | 1 | |
| ENSMUSG00000098734 | 1 | |
| ENSMUSG00000098735 | 1 | |
| ENSMUSG00000098736 | 1 | |
| ENSMUSG00000098737 | 1 | |
| ENSMUSG00000098739 | 1 | |
| ENSMUSG00000098740 | 1 | |
| ENSMUSG00000098741 | 1 | |
| ENSMUSG00000098742 | 1 | |
| ENSMUSG00000098743 | 1 | |
| ENSMUSG00000098744 | 1 | |
| ENSMUSG00000098745 | 1 | |
| ENSMUSG00000098747 | 1 | |
| ENSMUSG00000098748 | 1 | |
| ENSMUSG00000098749 | 1.060443131 | 0.911299384 |
| ENSMUSG00000098750 | -1.053549062 | 0.698112512 |
| ENSMUSG00000098751 | 1 | |
| ENSMUSG00000098752 | 1 | |
| ENSMUSG00000098754 | 1 | |
| ENSMUSG00000098755 | -1.130334237 | 0.608497229 |
| ENSMUSG00000098756 | 1 | |
| ENSMUSG00000098757 | 1 | |
| ENSMUSG00000098758 | 1 | |
| ENSMUSG00000098759 | 1.014545335 | 0.989648415 |
| ENSMUSG00000098760 | -1.053549062 | 0.698112512 |
| ENSMUSG00000098761 | -1.069502966 | 0.698112512 |
| ENSMUSG00000098762 | 1 | |
| ENSMUSG00000098763 | 1 | |
| ENSMUSG00000098764 | 1 | |
| ENSMUSG00000098765 | 1 | |
| ENSMUSG00000098766 | -1.104995253 | 0.698112512 |
| ENSMUSG00000098767 | 1 | |
| ENSMUSG00000098768 | 1 | |
| ENSMUSG00000098769 | -1.03962336 | 0.698112512 |
| ENSMUSG00000098770 | 1 | |
| ENSMUSG00000098771 | 1 | |
| ENSMUSG00000098772 | 1 | |
| ENSMUSG00000098773 | 1 | |
| ENSMUSG00000098774 | 1 | |
| ENSMUSG00000098775 | 1 | |
| ENSMUSG00000098776 | 1 | |
| ENSMUSG00000098777 | 1 | |
| ENSMUSG00000098778 | 1 | |
| ENSMUSG00000098779 | 1.095872691 | 0.657060635 |
| ENSMUSG00000098780 | 1 | |
| ENSMUSG00000098781 | 1 | |
| ENSMUSG00000098782 | -1 | 1 |
| ENSMUSG00000098783 | 1 | |
| ENSMUSG00000098784 | 1 | |
| ENSMUSG00000098785 | 1 | |
| ENSMUSG00000098786 | 1.023743961 | 0.968735041 |
| ENSMUSG00000098787 | 1 | |
| ENSMUSG00000098788 | 1 | |
| ENSMUSG00000098789 | -1.195192018 | 0.592238062 |

| | | |
|---|---|---|
| ENSMUSG00000098790 | 1.083551589 | 0.734674233 |
| ENSMUSG00000098791 | 1 | |
| ENSMUSG00000098792 | 1 | |
| ENSMUSG00000098793 | 1 | |
| ENSMUSG00000098794 | 1 | |
| ENSMUSG00000098795 | 1 | |
| ENSMUSG00000098796 | 1 | |
| ENSMUSG00000098797 | 1 | |
| ENSMUSG00000098798 | 1 | |
| ENSMUSG00000098799 | 1 | |
| ENSMUSG00000098800 | 1 | |
| ENSMUSG00000098801 | 1 | |
| ENSMUSG00000098802 | 1 | |
| ENSMUSG00000098803 | -1.043306199 | 0.803260194 |
| ENSMUSG00000098805 | 1 | |
| ENSMUSG00000098807 | 1 | |
| ENSMUSG00000098808 | 1 | |
| ENSMUSG00000098809 | 1 | |
| ENSMUSG00000098811 | -1.161737199 | 0.608497229 |
| ENSMUSG00000098812 | 1.059463094 | 0.698112512 |
| ENSMUSG00000098813 | 1.03092687 | 0.944885562 |
| ENSMUSG00000098814 | 1 | |
| ENSMUSG00000098815 | 1 | |
| ENSMUSG00000098816 | 1 | |
| ENSMUSG00000098817 | 1 | |
| ENSMUSG00000098818 | 1 | |
| ENSMUSG00000098819 | 1 | |
| ENSMUSG00000098820 | -1.044273782 | 0.698112512 |
| ENSMUSG00000098821 | 1 | |
| ENSMUSG00000098822 | 1 | |
| ENSMUSG00000098823 | 1 | |
| ENSMUSG00000098824 | 1 | |
| ENSMUSG00000098825 | 1 | |
| ENSMUSG00000098826 | 1 | |
| ENSMUSG00000098827 | 1 | |
| ENSMUSG00000098828 | 1 | |
| ENSMUSG00000098829 | 1 | |
| ENSMUSG00000098830 | -1.03962336 | 0.698112512 |
| ENSMUSG00000098831 | 1 | |
| ENSMUSG00000098832 | -1.128668575 | 0.578221424 |
| ENSMUSG00000098833 | 1 | |
| ENSMUSG00000098834 | 1 | |
| ENSMUSG00000098836 | 1 | |
| ENSMUSG00000098837 | 1 | |
| ENSMUSG00000098838 | 1 | |
| ENSMUSG00000098839 | 1 | |
| ENSMUSG00000098840 | 1.184376802 | 0.422407646 |
| ENSMUSG00000098842 | 1 | |
| ENSMUSG00000098843 | -1.173214258 | 0.608497229 |
| ENSMUSG00000098844 | 1 | |
| ENSMUSG00000098845 | 1 | |
| ENSMUSG00000098846 | 1 | |
| ENSMUSG00000098847 | 1 | |
| ENSMUSG00000098848 | 1 | |
| ENSMUSG00000098849 | 1 | |
| ENSMUSG00000098850 | 1 | |
| ENSMUSG00000098851 | 1 | |
| ENSMUSG00000098852 | -1.043306199 | 0.698112512 |
| ENSMUSG00000098853 | 1 | |
| ENSMUSG00000098854 | 1 | |
| ENSMUSG00000098855 | 1 | |
| ENSMUSG00000098856 | 1 | |
| ENSMUSG00000098857 | 1 | |

| | | |
|---|---|---|
| ENSMUSG00000098858 | 1 | |
| ENSMUSG00000098859 | 1 | |
| ENSMUSG00000098860 | 1 | |
| ENSMUSG00000098861 | 1 | |
| ENSMUSG00000098862 | 1 | |
| ENSMUSG00000098863 | 1 | |
| ENSMUSG00000098864 | 1 | |
| ENSMUSG00000098865 | 1 | |
| ENSMUSG00000098866 | 1 | |
| ENSMUSG00000098867 | 1 | |
| ENSMUSG00000098868 | 1 | |
| ENSMUSG00000098869 | 1 | |
| ENSMUSG00000098871 | 1 | |
| ENSMUSG00000098873 | 1 | |
| ENSMUSG00000098874 | 1.122462048 | 0.639157886 |
| ENSMUSG00000098875 | 1 | |
| ENSMUSG00000098876 | 1 | |
| ENSMUSG00000098877 | 1 | |
| ENSMUSG00000098878 | 1 | |
| ENSMUSG00000098879 | 1 | |
| ENSMUSG00000098880 | 1 | |
| ENSMUSG00000098881 | 1 | |
| ENSMUSG00000098882 | 1 | |
| ENSMUSG00000098883 | 1 | |
| ENSMUSG00000098884 | 1 | |
| ENSMUSG00000098885 | 1 | |
| ENSMUSG00000098886 | 1 | |
| ENSMUSG00000098887 | 1 | |
| ENSMUSG00000098888 | 1 | |
| ENSMUSG00000098889 | 1.091125748 | 0.608497229 |
| ENSMUSG00000098890 | 1 | |
| ENSMUSG00000098891 | 1 | |
| ENSMUSG00000098893 | 1.04424822 | 0.892617752 |
| ENSMUSG00000098894 | 1 | |
| ENSMUSG00000098895 | 1 | |
| ENSMUSG00000098896 | 1 | |
| ENSMUSG00000098897 | 1 | |
| ENSMUSG00000098898 | 1.059463094 | 0.698112512 |
| ENSMUSG00000098899 | 1 | |
| ENSMUSG00000098900 | -1 | 1 |
| ENSMUSG00000098901 | 1 | |
| ENSMUSG00000098903 | 1 | |
| ENSMUSG00000098904 | -1.13161737 | 0.655149007 |
| ENSMUSG00000098905 | -1.080737224 | 0.585362499 |
| ENSMUSG00000098906 | 1 | |
| ENSMUSG00000098907 | -1.084645496 | 0.608497229 |
| ENSMUSG00000098908 | 1 | |
| ENSMUSG00000098909 | 1 | |
| ENSMUSG00000098910 | 1 | |
| ENSMUSG00000098912 | -1.120329932 | 0.374710101 |
| ENSMUSG00000098914 | 1 | |
| ENSMUSG00000098915 | -1.115460207 | 0.698112512 |
| ENSMUSG00000098916 | 1 | |
| ENSMUSG00000098917 | 1 | |
| ENSMUSG00000098918 | 1.049411284 | 0.895965732 |
| ENSMUSG00000098919 | 1 | |
| ENSMUSG00000098920 | -1.03962336 | 0.698112512 |
| ENSMUSG00000098922 | 1 | |
| ENSMUSG00000098923 | 1.024284648 | 0.953521521 |
| ENSMUSG00000098924 | 1 | |
| ENSMUSG00000098925 | 1 | |
| ENSMUSG00000098926 | -1.03962336 | 0.698112512 |
| ENSMUSG00000098928 | -1 | 1 |

| | | |
|---|---|---|
| ENSMUSG00000098929 | -1 | 1 |
| ENSMUSG00000098931 | 1 | |
| ENSMUSG00000098932 | 1 | |
| ENSMUSG00000098934 | 1 | |
| ENSMUSG00000098935 | 1 | |
| ENSMUSG00000098937 | 1 | |
| ENSMUSG00000098938 | 1 | |
| ENSMUSG00000098939 | 1 | |
| ENSMUSG00000098940 | 1 | |
| ENSMUSG00000098941 | 1 | |
| ENSMUSG00000098942 | 1 | |
| ENSMUSG00000098943 | 1 | |
| ENSMUSG00000098944 | 1 | |
| ENSMUSG00000098945 | 1 | |
| ENSMUSG00000098946 | 1 | |
| ENSMUSG00000098947 | 1 | |
| ENSMUSG00000098948 | 1 | |
| ENSMUSG00000098949 | 1 | |
| ENSMUSG00000098950 | 1 | |
| ENSMUSG00000098951 | 1.059463094 | 0.608497229 |
| ENSMUSG00000098952 | 1 | |
| ENSMUSG00000098953 | 1 | |
| ENSMUSG00000098954 | 1 | |
| ENSMUSG00000098955 | 1 | |
| ENSMUSG00000098956 | 1 | |
| ENSMUSG00000098957 | 1 | |
| ENSMUSG00000098958 | 1 | |
| ENSMUSG00000098959 | 1 | |
| ENSMUSG00000098960 | 1 | |
| ENSMUSG00000098961 | 1 | |
| ENSMUSG00000098962 | 1 | |
| ENSMUSG00000098963 | 1 | |
| ENSMUSG00000098964 | 1 | |
| ENSMUSG00000098965 | 1 | |
| ENSMUSG00000098966 | 1 | |
| ENSMUSG00000098967 | 1 | |
| ENSMUSG00000098968 | 1 | |
| ENSMUSG00000098969 | 1 | |
| ENSMUSG00000098970 | 1 | |
| ENSMUSG00000098972 | 1 | |
| ENSMUSG00000098973 | 1.204021701 | 0.352607274 |
| ENSMUSG00000098974 | 1 | |
| ENSMUSG00000098975 | 1 | |
| ENSMUSG00000098976 | -1.089497311 | 0.532369657 |
| ENSMUSG00000098977 | 1 | |
| ENSMUSG00000098978 | 1 | |
| ENSMUSG00000098979 | -1.044273782 | 0.698112512 |
| ENSMUSG00000098980 | -1.044273782 | 0.698112512 |
| ENSMUSG00000098981 | 1 | |
| ENSMUSG00000098982 | 1 | |
| ENSMUSG00000098983 | 1 | |
| ENSMUSG00000098984 | 1 | |
| ENSMUSG00000098985 | -1.113957691 | 0.692944879 |
| ENSMUSG00000098986 | 1 | |
| ENSMUSG00000098987 | 1 | |
| ENSMUSG00000098988 | 1 | |
| ENSMUSG00000098989 | 1.059463094 | 0.608497229 |
| ENSMUSG00000098990 | 1 | |
| ENSMUSG00000098991 | 1 | |
| ENSMUSG00000098992 | 1 | |
| ENSMUSG00000098993 | 1 | |
| ENSMUSG00000098994 | 1 | |
| ENSMUSG00000098995 | 1 | |

| | | |
|---|---|---|
| ENSMUSG00000098996 | 1 | |
| ENSMUSG00000098997 | 1 | |
| ENSMUSG00000098998 | 1 | |
| ENSMUSG00000098999 | 1 | |
| ENSMUSG00000099000 | 1 | |
| ENSMUSG00000099001 | 1 | |
| ENSMUSG00000099002 | 1 | |
| ENSMUSG00000099003 | 1 | |
| ENSMUSG00000099004 | 1.095872691 | 0.608497229 |
| ENSMUSG00000099005 | 1 | |
| ENSMUSG00000099006 | 1 | |
| ENSMUSG00000099008 | 1.023743961 | 0.938939969 |
| ENSMUSG00000099009 | 1 | |
| ENSMUSG00000099010 | 1 | |
| ENSMUSG00000099011 | 1 | |
| ENSMUSG00000099012 | 1 | |
| ENSMUSG00000099013 | 1 | |
| ENSMUSG00000099014 | 1 | |
| ENSMUSG00000099015 | 1 | |
| ENSMUSG00000099016 | 1 | |
| ENSMUSG00000099017 | 1 | |
| ENSMUSG00000099018 | 1 | |
| ENSMUSG00000099019 | -1.037896228 | 0.895018604 |
| ENSMUSG00000099020 | 1 | |
| ENSMUSG00000099021 | 1 | |
| ENSMUSG00000099022 | 1 | |
| ENSMUSG00000099023 | 1 | |
| ENSMUSG00000099024 | -1 | 1 |
| ENSMUSG00000099025 | 1 | |
| ENSMUSG00000099027 | 1 | |
| ENSMUSG00000099030 | 1 | |
| ENSMUSG00000099031 | 1 | |
| ENSMUSG00000099032 | -1.624569012 | 0.004728769 |
| ENSMUSG00000099033 | -1.068308909 | 0.809553648 |
| ENSMUSG00000099034 | 1.038026752 | 0.638758774 |
| ENSMUSG00000099035 | 1 | |
| ENSMUSG00000099036 | 1 | |
| ENSMUSG00000099037 | 1 | |
| ENSMUSG00000099039 | 1 | |
| ENSMUSG00000099041 | 1.069068979 | 0.608497229 |
| ENSMUSG00000099042 | 1 | |
| ENSMUSG00000099043 | 1 | |
| ENSMUSG00000099045 | 1 | |
| ENSMUSG00000099047 | 1 | |
| ENSMUSG00000099049 | 1 | |
| ENSMUSG00000099050 | 1 | |
| ENSMUSG00000099051 | -1.0778529 | 0.820185183 |
| ENSMUSG00000099052 | 1.059463094 | 0.608497229 |
| ENSMUSG00000099053 | 1 | |
| ENSMUSG00000099054 | 1 | |
| ENSMUSG00000099055 | -1.053549062 | 0.698112512 |
| ENSMUSG00000099056 | 1 | |
| ENSMUSG00000099057 | 1 | |
| ENSMUSG00000099058 | -1.03962336 | 0.803072249 |
| ENSMUSG00000099059 | 1 | |
| ENSMUSG00000099060 | 1.069068979 | 0.608497229 |
| ENSMUSG00000099061 | 1.070202264 | 0.835307733 |
| ENSMUSG00000099062 | 1 | |
| ENSMUSG00000099063 | -1 | 1 |
| ENSMUSG00000099064 | 1 | |
| ENSMUSG00000099066 | 1 | |
| ENSMUSG00000099068 | 1.019337424 | 1 |
| ENSMUSG00000099069 | 1 | |

| | | |
|---|---|---|
| ENSMUSG00000099070 | 1 | |
| ENSMUSG00000099071 | 1 | |
| ENSMUSG00000099072 | 1 | |
| ENSMUSG00000099073 | 1 | |
| ENSMUSG00000099074 | 1 | |
| ENSMUSG00000099075 | 1 | |
| ENSMUSG00000099076 | -1.03962336 | 0.698112512 |
| ENSMUSG00000099078 | 1 | |
| ENSMUSG00000099079 | 1 | |
| ENSMUSG00000099080 | 1 | |
| ENSMUSG00000099081 | 1 | |
| ENSMUSG00000099082 | 1 | |
| ENSMUSG00000099083 | 1.065298097 | 0.801177726 |
| ENSMUSG00000099085 | 1 | |
| ENSMUSG00000099086 | 1 | |
| ENSMUSG00000099087 | 1 | |
| ENSMUSG00000099088 | 1 | |
| ENSMUSG00000099089 | 1 | |
| ENSMUSG00000099091 | 1 | |
| ENSMUSG00000099092 | 1 | |
| ENSMUSG00000099093 | 1 | |
| ENSMUSG00000099094 | 1.052432126 | 0.70299173 |
| ENSMUSG00000099095 | 1 | |
| ENSMUSG00000099096 | 1 | |
| ENSMUSG00000099097 | 1 | |
| ENSMUSG00000099098 | -1.121136286 | 0.677781728 |
| ENSMUSG00000099099 | 1 | |
| ENSMUSG00000099101 | 1 | |
| ENSMUSG00000099102 | 1.059463094 | 0.608497229 |
| ENSMUSG00000099103 | 1 | |
| ENSMUSG00000099104 | -1.109582661 | 0.549347972 |
| ENSMUSG00000099105 | 1 | |
| ENSMUSG00000099106 | 1 | |
| ENSMUSG00000099107 | -1.00002448 | 1 |
| ENSMUSG00000099108 | 1 | |
| ENSMUSG00000099109 | 1 | |
| ENSMUSG00000099110 | 1 | |
| ENSMUSG00000099111 | 1 | |
| ENSMUSG00000099112 | 1.128982743 | 0.698112512 |
| ENSMUSG00000099114 | 1 | |
| ENSMUSG00000099115 | 1.023743961 | 0.95702926 |
| ENSMUSG00000099117 | 1 | |
| ENSMUSG00000099118 | 1 | |
| ENSMUSG00000099119 | 1 | |
| ENSMUSG00000099120 | 1 | |
| ENSMUSG00000099123 | 1.095872691 | 0.608497229 |
| ENSMUSG00000099124 | 1 | |
| ENSMUSG00000099125 | 1 | |
| ENSMUSG00000099126 | 1 | |
| ENSMUSG00000099127 | -1.053549062 | 0.698112512 |
| ENSMUSG00000099128 | 1 | |
| ENSMUSG00000099129 | 1 | |
| ENSMUSG00000099130 | 1.069068979 | 0.698112512 |
| ENSMUSG00000099131 | -1.044697704 | 0.896152472 |
| ENSMUSG00000099132 | 1 | |
| ENSMUSG00000099133 | -1.170487877 | 0.563878852 |
| ENSMUSG00000099134 | 1 | |
| ENSMUSG00000099135 | 1 | |
| ENSMUSG00000099136 | 1 | |
| ENSMUSG00000099137 | -1.052034482 | 0.934375176 |
| ENSMUSG00000099138 | 1 | |
| ENSMUSG00000099139 | 1 | |
| ENSMUSG00000099140 | 1.05410549 | 0.805157604 |

| | | |
|---|---|---|
| ENSMUSG00000099141 | 1 | |
| ENSMUSG00000099142 | 1 | |
| ENSMUSG00000099143 | 1 | |
| ENSMUSG00000099144 | 1 | |
| ENSMUSG00000099145 | 1 | |
| ENSMUSG00000099146 | 1.005613439 | 1 |
| ENSMUSG00000099147 | 1 | |
| ENSMUSG00000099148 | 1.02235921 | 0.961470381 |
| ENSMUSG00000099149 | 1 | |
| ENSMUSG00000099150 | 1 | |
| ENSMUSG00000099151 | 1 | |
| ENSMUSG00000099152 | 1 | |
| ENSMUSG00000099153 | 1 | |
| ENSMUSG00000099154 | 1 | |
| ENSMUSG00000099155 | 1 | |
| ENSMUSG00000099157 | 1 | |
| ENSMUSG00000099158 | 1 | |
| ENSMUSG00000099159 | 1 | |
| ENSMUSG00000099160 | 1 | |
| ENSMUSG00000099161 | 1 | |
| ENSMUSG00000099162 | -1.147838869 | 0.462961827 |
| ENSMUSG00000099163 | -1.044273782 | 0.795446726 |
| ENSMUSG00000099165 | 1 | |
| ENSMUSG00000099166 | 1 | |
| ENSMUSG00000099168 | 1 | |
| ENSMUSG00000099169 | 1 | |
| ENSMUSG00000099170 | -1.054741498 | 0.846939416 |
| ENSMUSG00000099172 | 1 | |
| ENSMUSG00000099173 | 1 | |
| ENSMUSG00000099174 | 1.005613439 | 1 |
| ENSMUSG00000099176 | 1 | |
| ENSMUSG00000099178 | -1.043306199 | 0.895360578 |
| ENSMUSG00000099179 | 1 | |
| ENSMUSG00000099180 | 1 | |
| ENSMUSG00000099181 | 1 | |
| ENSMUSG00000099182 | 1 | |
| ENSMUSG00000099183 | 1 | |
| ENSMUSG00000099184 | 1 | |
| ENSMUSG00000099185 | 1 | |
| ENSMUSG00000099186 | 1 | |
| ENSMUSG00000099188 | -1.03962336 | 0.698112512 |
| ENSMUSG00000099189 | 1 | |
| ENSMUSG00000099190 | -1 | 1 |
| ENSMUSG00000099191 | 1 | |
| ENSMUSG00000099192 | 1 | |
| ENSMUSG00000099193 | 1.095872691 | 0.608497229 |
| ENSMUSG00000099196 | 1.059463094 | 0.608497229 |
| ENSMUSG00000099197 | 1 | |
| ENSMUSG00000099198 | 1 | |
| ENSMUSG00000099199 | 1 | |
| ENSMUSG00000099200 | 1 | |
| ENSMUSG00000099201 | 1 | |
| ENSMUSG00000099202 | 1 | |
| ENSMUSG00000099203 | 1 | |
| ENSMUSG00000099204 | 1 | |
| ENSMUSG00000099205 | 1 | |
| ENSMUSG00000099206 | -1.044273782 | 0.698112512 |
| ENSMUSG00000099207 | 1 | |
| ENSMUSG00000099208 | 1 | |
| ENSMUSG00000099209 | 1 | |
| ENSMUSG00000099210 | 1.132639128 | 0.217803313 |
| ENSMUSG00000099211 | 1 | |
| ENSMUSG00000099212 | 1 | |

| | | |
|---|---|---|
| ENSMUSG00000099213 | 1 | |
| ENSMUSG00000099214 | 1 | |
| ENSMUSG00000099215 | 1 | |
| ENSMUSG00000099216 | 1 | |
| ENSMUSG00000099218 | 1 | |
| ENSMUSG00000099219 | 1 | |
| ENSMUSG00000099220 | 1 | |
| ENSMUSG00000099221 | 1 | |
| ENSMUSG00000099222 | 1 | |
| ENSMUSG00000099223 | 1 | |
| ENSMUSG00000099224 | 1 | |
| ENSMUSG00000099225 | 1 | |
| ENSMUSG00000099226 | 1.005613439 | 1 |
| ENSMUSG00000099227 | 1 | |
| ENSMUSG00000099228 | 1 | |
| ENSMUSG00000099229 | 1 | |
| ENSMUSG00000099230 | 1.069068979 | 0.608497229 |
| ENSMUSG00000099232 | 1 | |
| ENSMUSG00000099233 | 1 | |
| ENSMUSG00000099235 | 1 | |
| ENSMUSG00000099236 | 1 | |
| ENSMUSG00000099237 | 1 | |
| ENSMUSG00000099238 | 1.080134941 | 0.726866981 |
| ENSMUSG00000099239 | 1 | |
| ENSMUSG00000099240 | 1 | |
| ENSMUSG00000099241 | 1 | |
| ENSMUSG00000099242 | -1.043306199 | 0.698112512 |
| ENSMUSG00000099243 | 1 | |
| ENSMUSG00000099244 | 1 | |
| ENSMUSG00000099245 | 1 | |
| ENSMUSG00000099246 | -1.053549062 | 0.698112512 |
| ENSMUSG00000099247 | 1 | |
| ENSMUSG00000099248 | 1 | |
| ENSMUSG00000099250 | 1 | |
| ENSMUSG00000099252 | 1 | |
| ENSMUSG00000099253 | -1.03962336 | 0.880365937 |
| ENSMUSG00000099254 | -1.058785717 | 0.778395222 |
| ENSMUSG00000099255 | 1 | |
| ENSMUSG00000099256 | 1 | |
| ENSMUSG00000099257 | 1 | |
| ENSMUSG00000099258 | 1 | |
| ENSMUSG00000099259 | 1 | |
| ENSMUSG00000099260 | 1 | |
| ENSMUSG00000099261 | -1.037482356 | 0.870427333 |
| ENSMUSG00000099263 | 1 | |
| ENSMUSG00000099264 | 1 | |
| ENSMUSG00000099266 | 1 | |
| ENSMUSG00000099267 | 1.137709437 | 0.608497229 |
| ENSMUSG00000099268 | 1.059463094 | 0.608497229 |
| ENSMUSG00000099269 | 1 | |
| ENSMUSG00000099270 | 1 | |
| ENSMUSG00000099272 | 1 | |
| ENSMUSG00000099273 | 1 | |
| ENSMUSG00000099274 | 1 | |
| ENSMUSG00000099275 | 1 | |
| ENSMUSG00000099276 | 1 | |
| ENSMUSG00000099278 | 1 | |
| ENSMUSG00000099279 | 1 | |
| ENSMUSG00000099280 | -1.03962336 | 0.814089245 |
| ENSMUSG00000099283 | -1.084645496 | 0.532369657 |
| ENSMUSG00000099284 | 1.210795585 | 0.260191528 |
| ENSMUSG00000099285 | 1 | |
| ENSMUSG00000099286 | 1 | |

| | | |
|---|---|---|
| ENSMUSG00000099287 | 1.089470641 | 0.723528308 |
| ENSMUSG00000099289 | 1 | |
| ENSMUSG00000099290 | 1 | |
| ENSMUSG00000099291 | 1 | |
| ENSMUSG00000099292 | 1.069068979 | 0.698112512 |
| ENSMUSG00000099293 | 1 | |
| ENSMUSG00000099294 | 1.028323353 | 0.903464611 |
| ENSMUSG00000099295 | 1 | |
| ENSMUSG00000099298 | 1 | |
| ENSMUSG00000099299 | 1 | |
| ENSMUSG00000099300 | 1 | |
| ENSMUSG00000099301 | -1.03962336 | 0.803072249 |
| ENSMUSG00000099302 | 1 | |
| ENSMUSG00000099303 | -1.043306199 | 0.698112512 |
| ENSMUSG00000099304 | 1 | |
| ENSMUSG00000099307 | 1 | |
| ENSMUSG00000099308 | 1 | |
| ENSMUSG00000099309 | 1 | |
| ENSMUSG00000099310 | -1.043306199 | 0.698112512 |
| ENSMUSG00000099311 | 1 | |
| ENSMUSG00000099312 | 1 | |
| ENSMUSG00000099313 | 1.008034876 | 1 |
| ENSMUSG00000099314 | -1.053549062 | 0.698112512 |
| ENSMUSG00000099315 | 1 | |
| ENSMUSG00000099316 | 1 | |
| ENSMUSG00000099317 | 1 | |
| ENSMUSG00000099318 | 1 | |
| ENSMUSG00000099319 | 1 | |
| ENSMUSG00000099320 | -1 | 1 |
| ENSMUSG00000099321 | 1 | |
| ENSMUSG00000099322 | 1 | |
| ENSMUSG00000099323 | 1 | |
| ENSMUSG00000099324 | 1 | |
| ENSMUSG00000099325 | 1 | |
| ENSMUSG00000099326 | 1 | |
| ENSMUSG00000099328 | -1 | 1 |
| ENSMUSG00000099329 | 1 | |
| ENSMUSG00000099330 | -1 | 1 |
| ENSMUSG00000099331 | 1 | |
| ENSMUSG00000099332 | 1 | |
| ENSMUSG00000099334 | 1 | |
| ENSMUSG00000099335 | 1 | |
| ENSMUSG00000099336 | 1 | |
| ENSMUSG00000099337 | -1.043306199 | 0.79291651 |
| ENSMUSG00000099338 | 1.05410549 | 0.805157604 |
| ENSMUSG00000099339 | 1 | |
| ENSMUSG00000099340 | 1 | |
| ENSMUSG00000099341 | 1 | |
| ENSMUSG00000099342 | 1.08404915 | 0.541735653 |
| ENSMUSG00000099343 | 1 | |
| ENSMUSG00000099344 | 1 | |
| ENSMUSG00000099346 | 1 | |
| ENSMUSG00000099347 | 1 | |
| ENSMUSG00000099348 | 1 | |
| ENSMUSG00000099349 | 1 | |
| ENSMUSG00000099350 | 1 | |
| ENSMUSG00000099351 | 1 | |
| ENSMUSG00000099352 | 1 | |
| ENSMUSG00000099353 | 1 | |
| ENSMUSG00000099354 | -1.078882374 | 0.815802999 |
| ENSMUSG00000099355 | 1 | |
| ENSMUSG00000099356 | 1 | |
| ENSMUSG00000099357 | 1 | |

| | | |
|---|---|---|
| ENSMUSG00000099359 | 1 | |
| ENSMUSG00000099360 | 1.059463094 | 0.608497229 |
| ENSMUSG00000099361 | 1 | |
| ENSMUSG00000099362 | 1 | |
| ENSMUSG00000099363 | 1 | |
| ENSMUSG00000099364 | 1.069068979 | 0.608497229 |
| ENSMUSG00000099366 | 1 | |
| ENSMUSG00000099367 | 1 | |
| ENSMUSG00000099368 | 1 | |
| ENSMUSG00000099369 | 1 | |
| ENSMUSG00000099370 | 1.206606684 | 0.588281525 |
| ENSMUSG00000099371 | 1 | |
| ENSMUSG00000099372 | 1 | |
| ENSMUSG00000099373 | 1.024656056 | 0.972980766 |
| ENSMUSG00000099374 | 1 | |
| ENSMUSG00000099375 | -1.004895262 | 1 |
| ENSMUSG00000099376 | 1 | |
| ENSMUSG00000099377 | -1.084645496 | 0.736377134 |
| ENSMUSG00000099378 | 1 | |
| ENSMUSG00000099379 | 1 | |
| ENSMUSG00000099380 | 1 | |
| ENSMUSG00000099381 | 1 | |
| ENSMUSG00000099382 | 1 | |
| ENSMUSG00000099383 | 1 | |
| ENSMUSG00000099384 | 1.028323353 | 0.933031586 |
| ENSMUSG00000099385 | -1 | 1 |
| ENSMUSG00000099386 | 1 | |
| ENSMUSG00000099387 | 1 | |
| ENSMUSG00000099388 | 1 | |
| ENSMUSG00000099389 | 1 | |
| ENSMUSG00000099390 | 1 | |
| ENSMUSG00000099391 | -1.043306199 | 0.698112512 |
| ENSMUSG00000099392 | 1 | |
| ENSMUSG00000099393 | 1 | |
| ENSMUSG00000099394 | 1 | |
| ENSMUSG00000099395 | 1 | |
| ENSMUSG00000099396 | 1 | |
| ENSMUSG00000099397 | 1.123032408 | 0.204264186 |
| ENSMUSG00000099398 | 1 | |
| ENSMUSG00000099399 | 1 | |
| ENSMUSG00000099400 | 1 | |
| ENSMUSG00000099401 | 1.059463094 | 0.608497229 |
| ENSMUSG00000099403 | 1 | |
| ENSMUSG00000099404 | 1 | |
| ENSMUSG00000099405 | -1.03962336 | 0.814089245 |
| ENSMUSG00000099406 | 1.156310786 | 0.709330685 |
| ENSMUSG00000099407 | 1 | |
| ENSMUSG00000099408 | 1 | |
| ENSMUSG00000099409 | 1 | |
| ENSMUSG00000099410 | 1 | |
| ENSMUSG00000099411 | 1.059463094 | 0.608497229 |
| ENSMUSG00000099412 | 1 | |
| ENSMUSG00000099413 | -1.000768698 | 1 |
| ENSMUSG00000099414 | 1 | |
| ENSMUSG00000099415 | 1 | |
| ENSMUSG00000099416 | 1 | |
| ENSMUSG00000099417 | -1.048085977 | 0.909181325 |
| ENSMUSG00000099418 | 1.059463094 | 0.608497229 |
| ENSMUSG00000099419 | 1 | |
| ENSMUSG00000099420 | 1 | |
| ENSMUSG00000099421 | 1 | |
| ENSMUSG00000099422 | 1 | |
| ENSMUSG00000099423 | -1.171420412 | 0.32560015 |

| | | |
|---|---|---|
| ENSMUSG00000099424 | 1.018670051 | 0.996913409 |
| ENSMUSG00000099425 | 1 | |
| ENSMUSG00000099426 | 1 | |
| ENSMUSG00000099427 | 1 | |
| ENSMUSG00000099428 | 1.069068979 | 0.724838192 |
| ENSMUSG00000099429 | 1.200936955 | 0.215991492 |
| ENSMUSG00000099430 | 1 | |
| ENSMUSG00000099431 | 1 | |
| ENSMUSG00000099432 | 1 | |
| ENSMUSG00000099433 | 1 | |
| ENSMUSG00000099434 | 1 | |
| ENSMUSG00000099435 | 1 | |
| ENSMUSG00000099436 | 1 | |
| ENSMUSG00000099437 | 1 | |
| ENSMUSG00000099438 | 1 | |
| ENSMUSG00000099439 | 1 | |
| ENSMUSG00000099440 | 1 | |
| ENSMUSG00000099441 | 1 | |
| ENSMUSG00000099442 | 1 | |
| ENSMUSG00000099443 | 1 | |
| ENSMUSG00000099444 | -1.041938665 | 0.770958539 |
| ENSMUSG00000099445 | 1 | |
| ENSMUSG00000099446 | 1.139351443 | 0.698112512 |
| ENSMUSG00000099447 | 1 | |
| ENSMUSG00000099448 | 1 | |
| ENSMUSG00000099449 | 1 | |
| ENSMUSG00000099450 | 1 | |
| ENSMUSG00000099451 | 1 | |
| ENSMUSG00000099452 | 1 | |
| ENSMUSG00000099453 | 1 | |
| ENSMUSG00000099454 | 1 | |
| ENSMUSG00000099455 | 1 | |
| ENSMUSG00000099456 | 1 | |
| ENSMUSG00000099457 | 1 | |
| ENSMUSG00000099458 | 1 | |
| ENSMUSG00000099459 | 1 | |
| ENSMUSG00000099460 | 1 | |
| ENSMUSG00000099461 | 1 | |
| ENSMUSG00000099462 | 1 | |
| ENSMUSG00000099463 | 1 | |
| ENSMUSG00000099464 | 1 | |
| ENSMUSG00000099465 | 1.059463094 | 0.608497229 |
| ENSMUSG00000099466 | 1 | |
| ENSMUSG00000099467 | 1 | |
| ENSMUSG00000099468 | 1 | |
| ENSMUSG00000099469 | 1 | |
| ENSMUSG00000099470 | -1.042755471 | 0.896364347 |
| ENSMUSG00000099471 | -1.015511103 | 0.991425134 |
| ENSMUSG00000099472 | -1 | 1 |
| ENSMUSG00000099473 | 1 | |
| ENSMUSG00000099474 | 1 | |
| ENSMUSG00000099476 | 1 | |
| ENSMUSG00000099477 | 1 | |
| ENSMUSG00000099478 | 1.008122688 | 0.984809847 |
| ENSMUSG00000099479 | 1 | |
| ENSMUSG00000099480 | 1 | |
| ENSMUSG00000099481 | -1.005520707 | 1 |
| ENSMUSG00000099482 | 1 | |
| ENSMUSG00000099483 | 1 | |
| ENSMUSG00000099484 | 1 | |
| ENSMUSG00000099485 | 1 | |
| ENSMUSG00000099486 | 1 | |
| ENSMUSG00000099487 | 1 | |

| | | |
|---|---|---|
| ENSMUSG00000099488 | 1 | |
| ENSMUSG00000099489 | 1 | |
| ENSMUSG00000099490 | 1 | |
| ENSMUSG00000099491 | 1.059463094 | 0.698112512 |
| ENSMUSG00000099492 | 1.144656366 | 0.617501656 |
| ENSMUSG00000099493 | 1 | |
| ENSMUSG00000099494 | 1 | |
| ENSMUSG00000099495 | 1 | |
| ENSMUSG00000099496 | 1 | |
| ENSMUSG00000099497 | 1 | |
| ENSMUSG00000099498 | 1 | |
| ENSMUSG00000099499 | 1 | |
| ENSMUSG00000099500 | 1 | |
| ENSMUSG00000099501 | 1 | |
| ENSMUSG00000099502 | 1 | |
| ENSMUSG00000099503 | 1.059463094 | 0.608497229 |
| ENSMUSG00000099504 | 1 | |
| ENSMUSG00000099505 | 1 | |
| ENSMUSG00000099506 | 1 | |
| ENSMUSG00000099507 | 1 | |
| ENSMUSG00000099508 | 1 | |
| ENSMUSG00000099509 | 1 | |
| ENSMUSG00000099510 | 1 | |
| ENSMUSG00000099511 | 1 | |
| ENSMUSG00000099512 | 1 | |
| ENSMUSG00000099513 | 1 | |
| ENSMUSG00000099514 | 1 | |
| ENSMUSG00000099516 | -1.113515092 | 0.576169279 |
| ENSMUSG00000099517 | -1.03962336 | 0.698112512 |
| ENSMUSG00000099518 | 1 | |
| ENSMUSG00000099519 | 1 | |
| ENSMUSG00000099520 | 1 | |
| ENSMUSG00000099521 | 1 | |
| ENSMUSG00000099522 | 1 | |
| ENSMUSG00000099523 | 1 | |
| ENSMUSG00000099524 | 1 | |
| ENSMUSG00000099525 | 1 | |
| ENSMUSG00000099526 | 1 | |
| ENSMUSG00000099527 | 1 | |
| ENSMUSG00000099528 | 1 | |
| ENSMUSG00000099529 | 1 | |
| ENSMUSG00000099530 | 1 | |
| ENSMUSG00000099531 | 1 | |
| ENSMUSG00000099532 | 1 | |
| ENSMUSG00000099534 | -1 | 1 |
| ENSMUSG00000099535 | 1 | |
| ENSMUSG00000099536 | 1 | |
| ENSMUSG00000099537 | 1 | |
| ENSMUSG00000099539 | 1 | |
| ENSMUSG00000099541 | 1 | |
| ENSMUSG00000099542 | 1 | |
| ENSMUSG00000099543 | -1 | 1 |
| ENSMUSG00000099544 | 1 | |
| ENSMUSG00000099545 | 1 | |
| ENSMUSG00000099546 | 1 | |
| ENSMUSG00000099547 | 1 | |
| ENSMUSG00000099548 | 1 | |
| ENSMUSG00000099549 | 1 | |
| ENSMUSG00000099550 | 1 | |
| ENSMUSG00000099551 | 1 | |
| ENSMUSG00000099552 | -1.033693811 | 0.949812939 |
| ENSMUSG00000099553 | 1 | |
| ENSMUSG00000099554 | 1 | |

| | | |
|---|---|---|
| ENSMUSG00000099555 | 1 | |
| ENSMUSG00000099556 | -1 | 1 |
| ENSMUSG00000099558 | 1 | |
| ENSMUSG00000099559 | 1 | |
| ENSMUSG00000099560 | 1 | |
| ENSMUSG00000099561 | -1.085651418 | 0.532369657 |
| ENSMUSG00000099562 | -1.069502966 | 0.748944522 |
| ENSMUSG00000099563 | 1 | |
| ENSMUSG00000099564 | 1 | |
| ENSMUSG00000099565 | 1 | |
| ENSMUSG00000099566 | 1 | |
| ENSMUSG00000099568 | 1 | |
| ENSMUSG00000099569 | 1 | |
| ENSMUSG00000099570 | 1 | |
| ENSMUSG00000099571 | 1.055737967 | 0.896765664 |
| ENSMUSG00000099572 | 1 | |
| ENSMUSG00000099573 | 1 | |
| ENSMUSG00000099574 | -1 | 1 |
| ENSMUSG00000099575 | 1 | |
| ENSMUSG00000099576 | 1.059463094 | 0.704170349 |
| ENSMUSG00000099577 | 1 | |
| ENSMUSG00000099578 | 1 | |
| ENSMUSG00000099579 | -1.036374409 | 0.826221824 |
| ENSMUSG00000099581 | 1 | |
| ENSMUSG00000099582 | 1 | |
| ENSMUSG00000099583 | -1.105673636 | 0.642256709 |
| ENSMUSG00000099584 | 1.059463094 | 0.608497229 |
| ENSMUSG00000099585 | 1 | |
| ENSMUSG00000099586 | 1 | |
| ENSMUSG00000099587 | 1 | |
| ENSMUSG00000099588 | 1 | |
| ENSMUSG00000099589 | 1 | |
| ENSMUSG00000099590 | -1 | 1 |
| ENSMUSG00000099591 | -1.043306199 | 0.79291651 |
| ENSMUSG00000099592 | 1 | |
| ENSMUSG00000099593 | 1 | |
| ENSMUSG00000099594 | 1 | |
| ENSMUSG00000099595 | 1 | |
| ENSMUSG00000099596 | -1.085651418 | 0.625496737 |
| ENSMUSG00000099597 | 1.167715905 | 0.108297221 |
| ENSMUSG00000099598 | 1 | |
| ENSMUSG00000099599 | 1 | |
| ENSMUSG00000099600 | 1 | |
| ENSMUSG00000099601 | 1.059463094 | 0.608497229 |
| ENSMUSG00000099603 | 1 | |
| ENSMUSG00000099604 | 1 | |
| ENSMUSG00000099605 | 1 | |
| ENSMUSG00000099606 | 1 | |
| ENSMUSG00000099607 | 1 | |
| ENSMUSG00000099608 | 1.020636438 | 0.887323956 |
| ENSMUSG00000099609 | 1 | |
| ENSMUSG00000099610 | 1 | |
| ENSMUSG00000099611 | 1 | |
| ENSMUSG00000099612 | 1 | |
| ENSMUSG00000099613 | 1 | |
| ENSMUSG00000099614 | 1 | |
| ENSMUSG00000099615 | 1.128313112 | 0.27459474 |
| ENSMUSG00000099616 | 1 | |
| ENSMUSG00000099617 | 1 | |
| ENSMUSG00000099618 | -1 | 1 |
| ENSMUSG00000099619 | 1 | |
| ENSMUSG00000099620 | 1 | |
| ENSMUSG00000099621 | 1 | |

| | | |
|---|---|---|
| ENSMUSG00000099622 | 1 | |
| ENSMUSG00000099623 | 1 | |
| ENSMUSG00000099624 | -1.085241428 | 0.843734664 |
| ENSMUSG00000099625 | -1.044146242 | 0.823266891 |
| ENSMUSG00000099626 | 1 | |
| ENSMUSG00000099627 | -1 | 1 |
| ENSMUSG00000099628 | 1 | |
| ENSMUSG00000099629 | 1 | |
| ENSMUSG00000099630 | 1 | |
| ENSMUSG00000099631 | 1.00108122 | 1 |
| ENSMUSG00000099632 | 1.270334747 | 0.411952847 |
| ENSMUSG00000099633 | 1 | |
| ENSMUSG00000099634 | 1 | |
| ENSMUSG00000099635 | -1.084645496 | 0.608497229 |
| ENSMUSG00000099636 | 1 | |
| ENSMUSG00000099637 | 1 | |
| ENSMUSG00000099638 | 1 | |
| ENSMUSG00000099639 | 1 | |
| ENSMUSG00000099640 | 1 | |
| ENSMUSG00000099641 | 1 | |
| ENSMUSG00000099642 | 1 | |
| ENSMUSG00000099643 | 1 | |
| ENSMUSG00000099644 | 1 | |
| ENSMUSG00000099645 | 1 | |
| ENSMUSG00000099646 | 1 | |
| ENSMUSG00000099647 | 1 | |
| ENSMUSG00000099648 | 1 | |
| ENSMUSG00000099649 | 1 | |
| ENSMUSG00000099650 | 1 | |
| ENSMUSG00000099651 | 1 | |
| ENSMUSG00000099652 | 1 | |
| ENSMUSG00000099653 | 1 | |
| ENSMUSG00000099654 | 1 | |
| ENSMUSG00000099655 | 1.103662588 | 0.608497229 |
| ENSMUSG00000099656 | 1 | |
| ENSMUSG00000099657 | 1 | |
| ENSMUSG00000099658 | 1 | |
| ENSMUSG00000099659 | 1 | |
| ENSMUSG00000099660 | 1 | |
| ENSMUSG00000099661 | 1 | |
| ENSMUSG00000099662 | 1 | |
| ENSMUSG00000099663 | 1 | |
| ENSMUSG00000099664 | 1 | |
| ENSMUSG00000099665 | -1 | 1 |
| ENSMUSG00000099666 | 1 | |
| ENSMUSG00000099667 | 1 | |
| ENSMUSG00000099668 | 1 | |
| ENSMUSG00000099669 | 1 | |
| ENSMUSG00000099670 | 1 | |
| ENSMUSG00000099671 | 1 | |
| ENSMUSG00000099672 | 1 | |
| ENSMUSG00000099673 | 1 | |
| ENSMUSG00000099674 | 1 | |
| ENSMUSG00000099675 | 1 | |
| ENSMUSG00000099676 | 1 | |
| ENSMUSG00000099677 | 1 | |
| ENSMUSG00000099678 | 1 | |
| ENSMUSG00000099679 | 1 | |
| ENSMUSG00000099681 | 1.035051221 | 0.754130549 |
| ENSMUSG00000099682 | 1 | |
| ENSMUSG00000099683 | 1 | |
| ENSMUSG00000099684 | 1 | |
| ENSMUSG00000099685 | 1 | |

| | | |
|---|---|---|
| ENSMUSG00000099686 | 1 | |
| ENSMUSG00000099687 | 1 | |
| ENSMUSG00000099688 | 1 | |
| ENSMUSG00000099689 | 1.016671569 | 0.940197654 |
| ENSMUSG00000099690 | 1 | |
| ENSMUSG00000099691 | 1 | |
| ENSMUSG00000099692 | -1.03962336 | 0.698112512 |
| ENSMUSG00000099693 | 1 | |
| ENSMUSG00000099694 | -1.047306572 | 0.867068766 |
| ENSMUSG00000099695 | 1 | |
| ENSMUSG00000099696 | -1.043306199 | 0.698112512 |
| ENSMUSG00000099697 | 1.035749015 | 0.731295483 |
| ENSMUSG00000099698 | 1 | |
| ENSMUSG00000099699 | 1 | |
| ENSMUSG00000099700 | 1 | |
| ENSMUSG00000099701 | 1.019083579 | 0.949812939 |
| ENSMUSG00000099702 | 1 | |
| ENSMUSG00000099703 | 1.077658816 | 0.717035356 |
| ENSMUSG00000099704 | 1 | |
| ENSMUSG00000099706 | 1 | |
| ENSMUSG00000099707 | 1.220724818 | 0.570749474 |
| ENSMUSG00000099708 | 1.080933986 | 0.805009415 |
| ENSMUSG00000099709 | 1 | |
| ENSMUSG00000099711 | 1 | |
| ENSMUSG00000099712 | 1 | |
| ENSMUSG00000099713 | 1 | |
| ENSMUSG00000099714 | -1 | 1 |
| ENSMUSG00000099715 | 1 | |
| ENSMUSG00000099716 | 1 | |
| ENSMUSG00000099717 | -1 | 1 |
| ENSMUSG00000099718 | 1 | |
| ENSMUSG00000099719 | -1.088396847 | 0.757619873 |
| ENSMUSG00000099721 | 1 | |
| ENSMUSG00000099722 | 1 | |
| ENSMUSG00000099723 | 1 | |
| ENSMUSG00000099724 | 1 | |
| ENSMUSG00000099725 | 1 | |
| ENSMUSG00000099726 | 1 | |
| ENSMUSG00000099727 | 1 | |
| ENSMUSG00000099728 | 1 | |
| ENSMUSG00000099729 | 1 | |
| ENSMUSG00000099730 | 1 | |
| ENSMUSG00000099731 | 1 | |
| ENSMUSG00000099732 | 1 | |
| ENSMUSG00000099733 | 1 | |
| ENSMUSG00000099734 | 1 | |
| ENSMUSG00000099735 | 1 | |
| ENSMUSG00000099736 | 1 | |
| ENSMUSG00000099737 | 1 | |
| ENSMUSG00000099738 | 1 | |
| ENSMUSG00000099739 | 1 | |
| ENSMUSG00000099740 | 1 | |
| ENSMUSG00000099741 | -1 | 1 |
| ENSMUSG00000099742 | 1 | |
| ENSMUSG00000099743 | 1 | |
| ENSMUSG00000099744 | 1 | |
| ENSMUSG00000099745 | 1 | |
| ENSMUSG00000099746 | 1.046938251 | 0.935104155 |
| ENSMUSG00000099747 | 1 | |
| ENSMUSG00000099748 | 1 | |
| ENSMUSG00000099749 | 1.059463094 | 0.608497229 |
| ENSMUSG00000099750 | 1 | |
| ENSMUSG00000099751 | -1.03962336 | 0.698112512 |

| | | |
|---|---|---|
| ENSMUSG00000099752 | 1 | |
| ENSMUSG00000099753 | 1 | |
| ENSMUSG00000099754 | 1 | |
| ENSMUSG00000099755 | 1 | |
| ENSMUSG00000099756 | 1 | |
| ENSMUSG00000099757 | 1 | |
| ENSMUSG00000099758 | 1 | |
| ENSMUSG00000099759 | 1.049582719 | 0.794034798 |
| ENSMUSG00000099760 | -1.000322333 | 1 |
| ENSMUSG00000099761 | 1 | |
| ENSMUSG00000099762 | 1 | |
| ENSMUSG00000099763 | 1 | |
| ENSMUSG00000099764 | -1.078590893 | 0.698112512 |
| ENSMUSG00000099765 | 1 | |
| ENSMUSG00000099766 | 1 | |
| ENSMUSG00000099767 | 1 | |
| ENSMUSG00000099768 | 1 | |
| ENSMUSG00000099770 | 1 | |
| ENSMUSG00000099771 | 1 | |
| ENSMUSG00000099772 | 1 | |
| ENSMUSG00000099773 | 1.194113527 | 0.584017943 |
| ENSMUSG00000099774 | 1 | |
| ENSMUSG00000099775 | 1 | |
| ENSMUSG00000099776 | 1 | |
| ENSMUSG00000099777 | 1 | |
| ENSMUSG00000099778 | 1 | |
| ENSMUSG00000099779 | -1.04154363 | 0.744351452 |
| ENSMUSG00000099780 | 1 | |
| ENSMUSG00000099781 | 1 | |
| ENSMUSG00000099782 | 1 | |
| ENSMUSG00000099783 | 1 | |
| ENSMUSG00000099784 | 1.023743961 | 0.938939969 |
| ENSMUSG00000099785 | 1 | |
| ENSMUSG00000099786 | 1 | |
| ENSMUSG00000099787 | 1 | |
| ENSMUSG00000099788 | 1 | |
| ENSMUSG00000099789 | 1 | |
| ENSMUSG00000099790 | 1 | |
| ENSMUSG00000099791 | 1 | |
| ENSMUSG00000099792 | 1 | |
| ENSMUSG00000099793 | 1 | |
| ENSMUSG00000099794 | 1 | |
| ENSMUSG00000099795 | 1 | |
| ENSMUSG00000099796 | 1 | |
| ENSMUSG00000099797 | 1.156803262 | 0.637668756 |
| ENSMUSG00000099798 | 1 | |
| ENSMUSG00000099799 | 1 | |
| ENSMUSG00000099800 | 1 | |
| ENSMUSG00000099801 | 1 | |
| ENSMUSG00000099802 | 1 | |
| ENSMUSG00000099803 | 1 | |
| ENSMUSG00000099804 | -1.024718486 | 0.931480186 |
| ENSMUSG00000099805 | 1 | |
| ENSMUSG00000099806 | 1 | |
| ENSMUSG00000099807 | 1 | |
| ENSMUSG00000099808 | 1 | |
| ENSMUSG00000099809 | 1 | |
| ENSMUSG00000099810 | 1 | |
| ENSMUSG00000099811 | 1 | |
| ENSMUSG00000099812 | 1.023743961 | 0.938939969 |
| ENSMUSG00000099813 | 1 | |
| ENSMUSG00000099814 | 1 | |
| ENSMUSG00000099815 | 1 | |

| | | |
|---|---|---|
| ENSMUSG00000099816 | 1 | |
| ENSMUSG00000099817 | 1 | |
| ENSMUSG00000099818 | 1 | |
| ENSMUSG00000099819 | 1 | |
| ENSMUSG00000099820 | 1 | |
| ENSMUSG00000099821 | 1 | |
| ENSMUSG00000099822 | 1 | |
| ENSMUSG00000099823 | 1 | |
| ENSMUSG00000099824 | 1 | |
| ENSMUSG00000099825 | -1.079873815 | 0.784381159 |
| ENSMUSG00000099826 | 1 | |
| ENSMUSG00000099827 | -1 | 1 |
| ENSMUSG00000099828 | 1 | |
| ENSMUSG00000099829 | 1 | |
| ENSMUSG00000099831 | 1 | |
| ENSMUSG00000099832 | 1 | |
| ENSMUSG00000099833 | 1 | |
| ENSMUSG00000099834 | 1 | |
| ENSMUSG00000099835 | 1.245567577 | 0.128994536 |
| ENSMUSG00000099836 | 1 | |
| ENSMUSG00000099838 | 1 | |
| ENSMUSG00000099839 | 1 | |
| ENSMUSG00000099840 | 1 | |
| ENSMUSG00000099841 | 1 | |
| ENSMUSG00000099842 | -1.205113258 | 0.418910176 |
| ENSMUSG00000099843 | 1.196527034 | 0.493111652 |
| ENSMUSG00000099845 | 1 | |
| ENSMUSG00000099846 | 1 | |
| ENSMUSG00000099847 | -1 | 1 |
| ENSMUSG00000099848 | -1.170083925 | 0.206620396 |
| ENSMUSG00000099849 | 1 | |
| ENSMUSG00000099850 | 1.009414875 | 1 |
| ENSMUSG00000099851 | 1 | |
| ENSMUSG00000099852 | 1 | |
| ENSMUSG00000099853 | 1 | |
| ENSMUSG00000099854 | 1.069068979 | 0.698112512 |
| ENSMUSG00000099855 | 1 | |
| ENSMUSG00000099856 | 1 | |
| ENSMUSG00000099858 | -1 | 1 |
| ENSMUSG00000099860 | 1 | |
| ENSMUSG00000099861 | 1 | |
| ENSMUSG00000099862 | 1 | |
| ENSMUSG00000099863 | 1 | |
| ENSMUSG00000099864 | 1 | |
| ENSMUSG00000099865 | 1 | |
| ENSMUSG00000099866 | 1 | |
| ENSMUSG00000099867 | 1 | |
| ENSMUSG00000099868 | 1 | |
| ENSMUSG00000099869 | 1 | |
| ENSMUSG00000099870 | -1 | 1 |
| ENSMUSG00000099871 | 1 | |
| ENSMUSG00000099872 | -1.063525418 | 0.698112512 |
| ENSMUSG00000099873 | 1 | |
| ENSMUSG00000099874 | -1.053549062 | 0.698112512 |
| ENSMUSG00000099875 | 1.098077881 | 0.711727974 |
| ENSMUSG00000099876 | 1 | |
| ENSMUSG00000099877 | 1.040172434 | 0.915158215 |
| ENSMUSG00000099878 | -1.086192882 | 0.707359283 |
| ENSMUSG00000099879 | -1.135553958 | 0.572947856 |
| ENSMUSG00000099880 | 1 | |
| ENSMUSG00000099881 | 1.015185246 | 0.9599243 |
| ENSMUSG00000099882 | 1 | |
| ENSMUSG00000099883 | 1 | |

| | | |
|---|---|---|
| ENSMUSG00000099884 | 1 | |
| ENSMUSG00000099886 | 1 | |
| ENSMUSG00000099887 | 1 | |
| ENSMUSG00000099888 | 1 | |
| ENSMUSG00000099889 | 1 | |
| ENSMUSG00000099890 | -1.032066481 | 0.856857802 |
| ENSMUSG00000099891 | -1.107299336 | 0.700712295 |
| ENSMUSG00000099892 | 1 | |
| ENSMUSG00000099893 | -1.044273782 | 0.698112512 |
| ENSMUSG00000099894 | 1 | |
| ENSMUSG00000099895 | 1 | |
| ENSMUSG00000099896 | 1 | |
| ENSMUSG00000099897 | 1 | |
| ENSMUSG00000099898 | 1 | |
| ENSMUSG00000099899 | -1 | 1 |
| ENSMUSG00000099900 | 1 | |
| ENSMUSG00000099901 | 1 | |
| ENSMUSG00000099902 | 1 | |
| ENSMUSG00000099903 | 1 | |
| ENSMUSG00000099904 | 1 | |
| ENSMUSG00000099905 | 1 | |
| ENSMUSG00000099906 | 1.059463094 | 0.698112512 |
| ENSMUSG00000099907 | 1 | |
| ENSMUSG00000099908 | 1 | |
| ENSMUSG00000099909 | 1 | |
| ENSMUSG00000099910 | 1 | |
| ENSMUSG00000099911 | -1 | 1 |
| ENSMUSG00000099912 | 1 | |
| ENSMUSG00000099913 | 1.093914205 | 0.511378706 |
| ENSMUSG00000099914 | 1 | |
| ENSMUSG00000099915 | 1 | |
| ENSMUSG00000099916 | 1 | |
| ENSMUSG00000099917 | 1 | |
| ENSMUSG00000099918 | 1 | |
| ENSMUSG00000099920 | 1 | |
| ENSMUSG00000099921 | 1 | |
| ENSMUSG00000099922 | 1 | |
| ENSMUSG00000099923 | -1.126344191 | 0.312657959 |
| ENSMUSG00000099924 | 1.05530209 | 0.843433337 |
| ENSMUSG00000099925 | 1 | |
| ENSMUSG00000099926 | 1 | |
| ENSMUSG00000099927 | 1 | |
| ENSMUSG00000099928 | -1.111456291 | 0.698112512 |
| ENSMUSG00000099929 | 1 | |
| ENSMUSG00000099930 | -1.063525418 | 0.698112512 |
| ENSMUSG00000099931 | 1 | |
| ENSMUSG00000099932 | 1 | |
| ENSMUSG00000099933 | 1.059463094 | 0.608497229 |
| ENSMUSG00000099934 | -1.043306199 | 0.79291651 |
| ENSMUSG00000099935 | 1 | |
| ENSMUSG00000099936 | -1.069095149 | 0.686194514 |
| ENSMUSG00000099937 | 1 | |
| ENSMUSG00000099938 | 1 | |
| ENSMUSG00000099939 | 1 | |
| ENSMUSG00000099940 | 1 | |
| ENSMUSG00000099941 | 1 | |
| ENSMUSG00000099942 | 1 | |
| ENSMUSG00000099943 | 1 | |
| ENSMUSG00000099944 | 1 | |
| ENSMUSG00000099945 | 1 | |
| ENSMUSG00000099946 | -1.03962336 | 0.698112512 |
| ENSMUSG00000099947 | 1.059463094 | 0.608497229 |
| ENSMUSG00000099948 | 1 | |

| | | |
|---|---|---|
| ENSMUSG00000099949 | -1.033820075 | 0.940197654 |
| ENSMUSG00000099950 | 1 | |
| ENSMUSG00000099951 | 1 | |
| ENSMUSG00000099952 | 1 | |
| ENSMUSG00000099953 | 1 | |
| ENSMUSG00000099954 | -1.043306199 | 0.79291651 |
| ENSMUSG00000099955 | 1 | |
| ENSMUSG00000099956 | 1 | |
| ENSMUSG00000099957 | -1 | 1 |
| ENSMUSG00000099958 | 1.171563499 | 0.48302662 |
| ENSMUSG00000099959 | 1 | |
| ENSMUSG00000099960 | 1 | |
| ENSMUSG00000099961 | 1 | |
| ENSMUSG00000099962 | 1 | |
| ENSMUSG00000099963 | 1 | |
| ENSMUSG00000099964 | 1.044978665 | 0.884233459 |
| ENSMUSG00000099965 | 1 | |
| ENSMUSG00000099966 | -1.125543235 | 0.18851436 |
| ENSMUSG00000099967 | 1 | |
| ENSMUSG00000099968 | 1 | |
| ENSMUSG00000099969 | 1.231453793 | 0.133561111 |
| ENSMUSG00000099970 | 1 | |
| ENSMUSG00000099971 | 1 | |
| ENSMUSG00000099972 | 1 | |
| ENSMUSG00000099973 | 1 | |
| ENSMUSG00000099974 | 1 | |
| ENSMUSG00000099975 | 1 | |
| ENSMUSG00000099976 | 1 | |
| ENSMUSG00000099977 | 1 | |
| ENSMUSG00000099978 | 1 | |
| ENSMUSG00000099979 | -1.116428035 | 0.695387231 |
| ENSMUSG00000099980 | -1.113153193 | 0.744141915 |
| ENSMUSG00000099981 | 1 | |
| ENSMUSG00000099982 | 1 | |
| ENSMUSG00000099983 | 1 | |
| ENSMUSG00000099985 | 1 | |
| ENSMUSG00000099986 | 1 | |
| ENSMUSG00000099988 | 1 | |
| ENSMUSG00000099989 | 1 | |
| ENSMUSG00000099990 | 1 | |
| ENSMUSG00000099991 | 1 | |
| ENSMUSG00000099992 | 1 | |
| ENSMUSG00000099993 | 1 | |
| ENSMUSG00000099994 | 1 | |
| ENSMUSG00000099995 | 1 | |
| ENSMUSG00000099996 | 1 | |
| ENSMUSG00000099997 | -1.023053403 | 0.991944074 |
| ENSMUSG00000099998 | -1.043306199 | 0.698112512 |
| ENSMUSG00000099999 | 1 | |
| ENSMUSG00000100000 | 1 | |
| ENSMUSG00000100001 | 1 | |
| ENSMUSG00000100002 | -1.089497311 | 0.532369657 |
| ENSMUSG00000100003 | 1 | |
| ENSMUSG00000100004 | 1 | |
| ENSMUSG00000100005 | 1.271065034 | 0.250749195 |
| ENSMUSG00000100006 | 1 | |
| ENSMUSG00000100007 | -1.03962336 | 0.872692845 |
| ENSMUSG00000100008 | 1 | |
| ENSMUSG00000100009 | -1.107853039 | 0.729059471 |
| ENSMUSG00000100010 | 1.015486245 | 0.986542466 |
| ENSMUSG00000100011 | 1 | |
| ENSMUSG00000100012 | 1 | |
| ENSMUSG00000100013 | 1 | |

| | | |
|---|---|---|
| ENSMUSG00000100014 | 1 | |
| ENSMUSG00000100015 | 1 | |
| ENSMUSG00000100016 | 1 | |
| ENSMUSG00000100017 | -1.149492636 | 0.179873578 |
| ENSMUSG00000100018 | 1 | |
| ENSMUSG00000100019 | 1 | |
| ENSMUSG00000100020 | 1 | |
| ENSMUSG00000100021 | 1 | |
| ENSMUSG00000100022 | 1.059463094 | 0.698112512 |
| ENSMUSG00000100023 | 1 | |
| ENSMUSG00000100024 | 1 | |
| ENSMUSG00000100025 | -1.114200147 | 0.449851954 |
| ENSMUSG00000100026 | 1 | |
| ENSMUSG00000100027 | -1.044273782 | 0.698112512 |
| ENSMUSG00000100028 | 1 | |
| ENSMUSG00000100029 | -1 | 1 |
| ENSMUSG00000100031 | 1 | |
| ENSMUSG00000100032 | 1 | |
| ENSMUSG00000100033 | -1.179735745 | 0.527875797 |
| ENSMUSG00000100034 | 1 | |
| ENSMUSG00000100035 | 1 | |
| ENSMUSG00000100036 | 1 | |
| ENSMUSG00000100037 | 1 | |
| ENSMUSG00000100038 | 1 | |
| ENSMUSG00000100039 | 1 | |
| ENSMUSG00000100040 | 1 | |
| ENSMUSG00000100041 | -1.037896228 | 0.936702575 |
| ENSMUSG00000100042 | 1 | |
| ENSMUSG00000100043 | 1 | |
| ENSMUSG00000100044 | 1 | |
| ENSMUSG00000100045 | 1 | |
| ENSMUSG00000100046 | 1 | |
| ENSMUSG00000100047 | 1 | |
| ENSMUSG00000100048 | 1 | |
| ENSMUSG00000100049 | 1 | |
| ENSMUSG00000100050 | 1 | |
| ENSMUSG00000100051 | 1 | |
| ENSMUSG00000100052 | 1 | |
| ENSMUSG00000100053 | 1 | |
| ENSMUSG00000100054 | 1 | |
| ENSMUSG00000100055 | 1 | |
| ENSMUSG00000100056 | 1 | |
| ENSMUSG00000100057 | 1 | |
| ENSMUSG00000100058 | 1 | |
| ENSMUSG00000100059 | 1 | |
| ENSMUSG00000100060 | 1 | |
| ENSMUSG00000100061 | 1 | |
| ENSMUSG00000100062 | 1 | |
| ENSMUSG00000100063 | 1 | |
| ENSMUSG00000100064 | 1 | |
| ENSMUSG00000100065 | -1.063525418 | 0.698112512 |
| ENSMUSG00000100066 | 1 | |
| ENSMUSG00000100067 | 1 | |
| ENSMUSG00000100068 | -1 | 1 |
| ENSMUSG00000100069 | -1.03962336 | 0.698112512 |
| ENSMUSG00000100070 | 1 | |
| ENSMUSG00000100071 | -1.044273782 | 0.698112512 |
| ENSMUSG00000100072 | 1 | |
| ENSMUSG00000100073 | 1 | |
| ENSMUSG00000100074 | 1 | |
| ENSMUSG00000100075 | 1.101367525 | 0.693463129 |
| ENSMUSG00000100076 | 1 | |
| ENSMUSG00000100077 | 1 | |

| | | |
|---|---|---|
| ENSMUSG00000100078 | 1.001956816 | 1 |
| ENSMUSG00000100079 | 1 | |
| ENSMUSG00000100080 | 1 | |
| ENSMUSG00000100081 | 1 | |
| ENSMUSG00000100082 | 1 | |
| ENSMUSG00000100083 | 1 | |
| ENSMUSG00000100084 | -1.056728947 | 0.795047 |
| ENSMUSG00000100085 | 1 | |
| ENSMUSG00000100086 | 1 | |
| ENSMUSG00000100087 | 1 | |
| ENSMUSG00000100088 | 1 | |
| ENSMUSG00000100089 | 1 | |
| ENSMUSG00000100090 | -1.001876871 | 1 |
| ENSMUSG00000100091 | 1 | |
| ENSMUSG00000100092 | 1 | |
| ENSMUSG00000100093 | 1 | |
| ENSMUSG00000100094 | -1.053549062 | 0.698112512 |
| ENSMUSG00000100095 | 1 | |
| ENSMUSG00000100096 | -1.03962336 | 0.698112512 |
| ENSMUSG00000100097 | 1 | |
| ENSMUSG00000100098 | 1 | |
| ENSMUSG00000100099 | 1 | |
| ENSMUSG00000100100 | 1 | |
| ENSMUSG00000100101 | 1 | |
| ENSMUSG00000100103 | 1 | |
| ENSMUSG00000100104 | 1.12347554 | 0.698112512 |
| ENSMUSG00000100105 | 1 | |
| ENSMUSG00000100106 | -1.053549062 | 0.698112512 |
| ENSMUSG00000100107 | 1 | |
| ENSMUSG00000100108 | -1.089497311 | 0.608497229 |
| ENSMUSG00000100109 | 1 | |
| ENSMUSG00000100110 | 1 | |
| ENSMUSG00000100111 | -1 | 1 |
| ENSMUSG00000100112 | 1 | |
| ENSMUSG00000100113 | 1 | |
| ENSMUSG00000100115 | 1 | |
| ENSMUSG00000100116 | 1 | |
| ENSMUSG00000100117 | 1 | |
| ENSMUSG00000100118 | 1 | |
| ENSMUSG00000100119 | 1.059463094 | 0.608497229 |
| ENSMUSG00000100120 | 1 | |
| ENSMUSG00000100121 | 1.094830497 | 0.698112512 |
| ENSMUSG00000100122 | 1 | |
| ENSMUSG00000100123 | -1.044273782 | 0.698112512 |
| ENSMUSG00000100124 | 1 | |
| ENSMUSG00000100125 | 1 | |
| ENSMUSG00000100126 | 1 | |
| ENSMUSG00000100127 | -1.069502966 | 0.744868995 |
| ENSMUSG00000100128 | 1 | |
| ENSMUSG00000100129 | 1 | |
| ENSMUSG00000100130 | 1 | |
| ENSMUSG00000100131 | -1.071075483 | 0.743300655 |
| ENSMUSG00000100132 | 1 | |
| ENSMUSG00000100133 | 1 | |
| ENSMUSG00000100134 | 1 | |
| ENSMUSG00000100135 | 1 | |
| ENSMUSG00000100136 | 1 | |
| ENSMUSG00000100137 | 1 | |
| ENSMUSG00000100138 | 1 | |
| ENSMUSG00000100140 | 1 | |
| ENSMUSG00000100141 | -1.0889222 | 0.660651921 |
| ENSMUSG00000100142 | 1 | |
| ENSMUSG00000100143 | -1 | 1 |

| | | |
|---|---|---|
| ENSMUSG00000100144 | 1 | |
| ENSMUSG00000100145 | 1 | |
| ENSMUSG00000100146 | 1 | |
| ENSMUSG00000100147 | 1 | |
| ENSMUSG00000100148 | 1 | |
| ENSMUSG00000100149 | 1 | |
| ENSMUSG00000100150 | 1 | |
| ENSMUSG00000100151 | 1 | |
| ENSMUSG00000100152 | 1 | |
| ENSMUSG00000100153 | 1 | |
| ENSMUSG00000100154 | 1 | |
| ENSMUSG00000100155 | 1 | |
| ENSMUSG00000100156 | 1 | |
| ENSMUSG00000100157 | 1.059463094 | 0.608497229 |
| ENSMUSG00000100158 | 1.040663135 | 0.970833538 |
| ENSMUSG00000100159 | 1 | |
| ENSMUSG00000100160 | 1.019083579 | 0.979345947 |
| ENSMUSG00000100161 | 1 | |
| ENSMUSG00000100162 | 1 | |
| ENSMUSG00000100164 | -1.080893645 | 0.568304986 |
| ENSMUSG00000100165 | 1 | |
| ENSMUSG00000100166 | 1 | |
| ENSMUSG00000100167 | 1 | |
| ENSMUSG00000100168 | 1 | |
| ENSMUSG00000100169 | 1 | |
| ENSMUSG00000100170 | 1 | |
| ENSMUSG00000100171 | 1 | |
| ENSMUSG00000100172 | 1 | |
| ENSMUSG00000100174 | 1 | |
| ENSMUSG00000100175 | -1 | 1 |
| ENSMUSG00000100176 | -1.063525418 | 0.814089245 |
| ENSMUSG00000100177 | 1 | |
| ENSMUSG00000100178 | 1.059463094 | 0.608497229 |
| ENSMUSG00000100179 | 1 | |
| ENSMUSG00000100180 | 1 | |
| ENSMUSG00000100181 | -1.043306199 | 0.698112512 |
| ENSMUSG00000100182 | 1 | |
| ENSMUSG00000100183 | 1.043683819 | 0.847697385 |
| ENSMUSG00000100184 | 1 | |
| ENSMUSG00000100185 | 1 | |
| ENSMUSG00000100186 | 1 | |
| ENSMUSG00000100187 | 1 | |
| ENSMUSG00000100188 | 1.059463094 | 0.608497229 |
| ENSMUSG00000100189 | 1.059463094 | 0.698112512 |
| ENSMUSG00000100190 | -1 | 1 |
| ENSMUSG00000100191 | 1 | |
| ENSMUSG00000100192 | 1 | |
| ENSMUSG00000100193 | 1 | |
| ENSMUSG00000100194 | 1 | |
| ENSMUSG00000100195 | 1 | |
| ENSMUSG00000100196 | 1 | |
| ENSMUSG00000100197 | 1 | |
| ENSMUSG00000100198 | 1 | |
| ENSMUSG00000100199 | -1.013156088 | 0.978182137 |
| ENSMUSG00000100200 | -1.043306199 | 0.698112512 |
| ENSMUSG00000100201 | -1.028463613 | 0.905432869 |
| ENSMUSG00000100202 | 1 | |
| ENSMUSG00000100203 | 1 | |
| ENSMUSG00000100204 | 1.018670051 | 0.988326998 |
| ENSMUSG00000100205 | 1 | |
| ENSMUSG00000100206 | 1 | |
| ENSMUSG00000100207 | 1 | |
| ENSMUSG00000100208 | 1 | |

| | | |
|---|---|---|
| ENSMUSG00000100209 | -1 | 1 |
| ENSMUSG00000100210 | -1 | 1 |
| ENSMUSG00000100211 | -1.044273782 | 0.698112512 |
| ENSMUSG00000100212 | 1 | |
| ENSMUSG00000100213 | 1.211058028 | 0.235412953 |
| ENSMUSG00000100214 | 1.118006222 | 0.203826519 |
| ENSMUSG00000100215 | -1.107151815 | 0.666633965 |
| ENSMUSG00000100216 | 1 | |
| ENSMUSG00000100217 | 1 | |
| ENSMUSG00000100218 | 1 | |
| ENSMUSG00000100219 | 1 | |
| ENSMUSG00000100220 | 1 | |
| ENSMUSG00000100221 | 1 | |
| ENSMUSG00000100222 | -1 | 1 |
| ENSMUSG00000100223 | 1 | |
| ENSMUSG00000100224 | 1 | |
| ENSMUSG00000100225 | 1 | |
| ENSMUSG00000100226 | 1 | |
| ENSMUSG00000100227 | 1 | |
| ENSMUSG00000100228 | -1.065680862 | 0.854827467 |
| ENSMUSG00000100229 | 1 | |
| ENSMUSG00000100230 | 1.069068979 | 0.608497229 |
| ENSMUSG00000100231 | 1 | |
| ENSMUSG00000100232 | 1 | |
| ENSMUSG00000100233 | 1 | |
| ENSMUSG00000100234 | 1 | |
| ENSMUSG00000100235 | -1.081898318 | 0.7938895 |
| ENSMUSG00000100236 | -1 | 1 |
| ENSMUSG00000100237 | 1 | |
| ENSMUSG00000100238 | 1 | |
| ENSMUSG00000100239 | 1 | |
| ENSMUSG00000100240 | 1 | |
| ENSMUSG00000100241 | -1 | 1 |
| ENSMUSG00000100242 | 1 | |
| ENSMUSG00000100243 | 1 | |
| ENSMUSG00000100244 | 1 | |
| ENSMUSG00000100245 | 1 | |
| ENSMUSG00000100246 | 1 | |
| ENSMUSG00000100247 | 1.132611403 | 0.675170087 |
| ENSMUSG00000100248 | 1 | |
| ENSMUSG00000100249 | 1 | |
| ENSMUSG00000100250 | 1 | |
| ENSMUSG00000100251 | 1 | |
| ENSMUSG00000100252 | 1.023743961 | 0.95702926 |
| ENSMUSG00000100253 | 1 | |
| ENSMUSG00000100254 | 1 | |
| ENSMUSG00000100255 | 1.250993224 | 0.561097031 |
| ENSMUSG00000100256 | 1 | |
| ENSMUSG00000100257 | 1.069068979 | 0.698112512 |
| ENSMUSG00000100258 | 1 | |
| ENSMUSG00000100259 | 1.059463094 | 0.608497229 |
| ENSMUSG00000100260 | 1 | |
| ENSMUSG00000100261 | -1.023769022 | 0.975499304 |
| ENSMUSG00000100262 | 1 | |
| ENSMUSG00000100263 | 1 | |
| ENSMUSG00000100264 | 1 | |
| ENSMUSG00000100265 | 1 | |
| ENSMUSG00000100266 | 1 | |
| ENSMUSG00000100267 | 1 | |
| ENSMUSG00000100268 | 1 | |
| ENSMUSG00000100269 | 1 | |
| ENSMUSG00000100270 | 1 | |
| ENSMUSG00000100271 | 1 | |

| | | |
|---|---|---|
| ENSMUSG00000100272 | 1 | |
| ENSMUSG00000100273 | 1 | |
| ENSMUSG00000100274 | 1 | |
| ENSMUSG00000100275 | 1 | |
| ENSMUSG00000100276 | 1 | |
| ENSMUSG00000100277 | 1 | |
| ENSMUSG00000100278 | 1 | |
| ENSMUSG00000100279 | 1 | |
| ENSMUSG00000100280 | 1.054327468 | 0.657796687 |
| ENSMUSG00000100281 | -1.085960923 | 0.723212059 |
| ENSMUSG00000100282 | 1 | |
| ENSMUSG00000100283 | 1 | |
| ENSMUSG00000100284 | 1 | |
| ENSMUSG00000100285 | 1 | |
| ENSMUSG00000100286 | 1 | |
| ENSMUSG00000100287 | -1.05948903 | 0.769837343 |
| ENSMUSG00000100288 | 1 | |
| ENSMUSG00000100289 | 1 | |
| ENSMUSG00000100290 | 1 | |
| ENSMUSG00000100291 | 1 | |
| ENSMUSG00000100292 | 1 | |
| ENSMUSG00000100294 | 1.069068979 | 0.608497229 |
| ENSMUSG00000100295 | 1 | |
| ENSMUSG00000100296 | 1 | |
| ENSMUSG00000100297 | 1 | |
| ENSMUSG00000100298 | 1 | |
| ENSMUSG00000100299 | 1 | |
| ENSMUSG00000100300 | 1 | |
| ENSMUSG00000100301 | 1 | |
| ENSMUSG00000100302 | 1.199989356 | 0.000021314 |
| ENSMUSG00000100303 | -1.089497311 | 0.681308543 |
| ENSMUSG00000100304 | 1 | |
| ENSMUSG00000100305 | 1 | |
| ENSMUSG00000100306 | 1 | |
| ENSMUSG00000100307 | 1 | |
| ENSMUSG00000100308 | 1 | |
| ENSMUSG00000100309 | 1 | |
| ENSMUSG00000100310 | 1 | |
| ENSMUSG00000100311 | 1 | |
| ENSMUSG00000100312 | 1 | |
| ENSMUSG00000100313 | 1 | |
| ENSMUSG00000100314 | 1 | |
| ENSMUSG00000100315 | -1.044273782 | 0.698112512 |
| ENSMUSG00000100316 | 1 | |
| ENSMUSG00000100317 | 1 | |
| ENSMUSG00000100318 | -1.03962336 | 0.698112512 |
| ENSMUSG00000100319 | 1.07487334 | 0.777702703 |
| ENSMUSG00000100320 | 1.057478344 | 0.867016657 |
| ENSMUSG00000100321 | 1 | |
| ENSMUSG00000100322 | 1 | |
| ENSMUSG00000100323 | 1 | |
| ENSMUSG00000100324 | 1 | |
| ENSMUSG00000100325 | 1 | |
| ENSMUSG00000100326 | 1 | |
| ENSMUSG00000100328 | 1 | |
| ENSMUSG00000100329 | 1 | |
| ENSMUSG00000100330 | 1 | |
| ENSMUSG00000100331 | 1 | |
| ENSMUSG00000100332 | 1 | |
| ENSMUSG00000100333 | 1 | |
| ENSMUSG00000100334 | -1.037482356 | 0.904446274 |
| ENSMUSG00000100335 | 1.059463094 | 0.699340382 |
| ENSMUSG00000100336 | 1 | |

| | | |
|---|---|---|
| ENSMUSG00000100337 | -1.043306199 | 0.698112512 |
| ENSMUSG00000100338 | 1 | |
| ENSMUSG00000100339 | 1 | |
| ENSMUSG00000100340 | 1 | |
| ENSMUSG00000100341 | 1 | |
| ENSMUSG00000100343 | 1 | |
| ENSMUSG00000100344 | 1 | |
| ENSMUSG00000100345 | -1.044273782 | 0.698112512 |
| ENSMUSG00000100346 | 1 | |
| ENSMUSG00000100347 | 1.019083579 | 0.949812939 |
| ENSMUSG00000100348 | 1 | |
| ENSMUSG00000100349 | 1 | |
| ENSMUSG00000100350 | 1 | |
| ENSMUSG00000100351 | 1.105693296 | 0.698112512 |
| ENSMUSG00000100352 | 1 | |
| ENSMUSG00000100353 | 1 | |
| ENSMUSG00000100354 | -1.086192882 | 0.698112512 |
| ENSMUSG00000100355 | 1 | |
| ENSMUSG00000100356 | 1 | |
| ENSMUSG00000100357 | -1 | 1 |
| ENSMUSG00000100358 | 1 | |
| ENSMUSG00000100359 | 1 | |
| ENSMUSG00000100360 | 1 | |
| ENSMUSG00000100361 | -1.030465624 | 0.835701362 |
| ENSMUSG00000100362 | 1 | |
| ENSMUSG00000100363 | 1 | |
| ENSMUSG00000100364 | 1 | |
| ENSMUSG00000100365 | 1 | |
| ENSMUSG00000100366 | 1 | |
| ENSMUSG00000100367 | -1 | 1 |
| ENSMUSG00000100368 | 1 | |
| ENSMUSG00000100369 | 1 | |
| ENSMUSG00000100370 | 1.063703399 | 0.698112512 |
| ENSMUSG00000100371 | 1.069068979 | 0.608497229 |
| ENSMUSG00000100372 | 1 | |
| ENSMUSG00000100373 | 1 | |
| ENSMUSG00000100374 | 1 | |
| ENSMUSG00000100375 | 1 | |
| ENSMUSG00000100376 | 1 | |
| ENSMUSG00000100377 | 1 | |
| ENSMUSG00000100378 | 1 | |
| ENSMUSG00000100379 | 1 | |
| ENSMUSG00000100380 | 1 | |
| ENSMUSG00000100381 | 1 | |
| ENSMUSG00000100382 | 1.059463094 | 0.608497229 |
| ENSMUSG00000100383 | 1.184376802 | 0.357543507 |
| ENSMUSG00000100384 | 1 | |
| ENSMUSG00000100385 | 1 | |
| ENSMUSG00000100386 | 1 | |
| ENSMUSG00000100387 | 1 | |
| ENSMUSG00000100388 | 1 | |
| ENSMUSG00000100389 | 1 | |
| ENSMUSG00000100390 | 1 | |
| ENSMUSG00000100391 | 1 | |
| ENSMUSG00000100392 | 1 | |
| ENSMUSG00000100393 | 1.05038453 | 0.894963534 |
| ENSMUSG00000100394 | 1.036242784 | 0.929191194 |
| ENSMUSG00000100395 | 1 | |
| ENSMUSG00000100396 | -1.06672954 | 0.810722095 |
| ENSMUSG00000100397 | 1 | |
| ENSMUSG00000100398 | 1 | |
| ENSMUSG00000100399 | 1 | |
| ENSMUSG00000100400 | 1.059463094 | 0.608497229 |

| | | |
|---|---|---|
| ENSMUSG00000100401 | 1 | |
| ENSMUSG00000100402 | 1 | |
| ENSMUSG00000100403 | 1 | |
| ENSMUSG00000100404 | 1 | |
| ENSMUSG00000100405 | 1 | |
| ENSMUSG00000100406 | 1 | |
| ENSMUSG00000100407 | 1 | |
| ENSMUSG00000100408 | 1 | |
| ENSMUSG00000100409 | 1 | |
| ENSMUSG00000100410 | 1 | |
| ENSMUSG00000100411 | 1 | |
| ENSMUSG00000100412 | 1 | |
| ENSMUSG00000100413 | 1 | |
| ENSMUSG00000100415 | 1 | |
| ENSMUSG00000100416 | 1.024693402 | 0.952057016 |
| ENSMUSG00000100417 | 1.019833591 | 0.974898785 |
| ENSMUSG00000100418 | 1 | |
| ENSMUSG00000100420 | 1 | |
| ENSMUSG00000100421 | -1 | 1 |
| ENSMUSG00000100422 | 1 | |
| ENSMUSG00000100423 | 1 | |
| ENSMUSG00000100424 | 1 | |
| ENSMUSG00000100425 | 1 | |
| ENSMUSG00000100426 | -1.043306199 | 0.803260194 |
| ENSMUSG00000100428 | 1 | |
| ENSMUSG00000100429 | 1 | |
| ENSMUSG00000100430 | 1 | |
| ENSMUSG00000100431 | 1 | |
| ENSMUSG00000100432 | -1.03962336 | 0.698112512 |
| ENSMUSG00000100433 | 1 | |
| ENSMUSG00000100434 | -1 | 1 |
| ENSMUSG00000100435 | 1 | |
| ENSMUSG00000100436 | 1 | |
| ENSMUSG00000100437 | 1 | |
| ENSMUSG00000100438 | 1 | |
| ENSMUSG00000100439 | -1 | 1 |
| ENSMUSG00000100440 | 1 | |
| ENSMUSG00000100441 | -1.044273782 | 0.801443616 |
| ENSMUSG00000100442 | 1 | |
| ENSMUSG00000100443 | 1 | |
| ENSMUSG00000100444 | 1.34694251 | 0.021271808 |
| ENSMUSG00000100445 | 1 | |
| ENSMUSG00000100446 | 1 | |
| ENSMUSG00000100447 | 1 | |
| ENSMUSG00000100448 | 1 | |
| ENSMUSG00000100449 | 1 | |
| ENSMUSG00000100450 | 1 | |
| ENSMUSG00000100451 | 1 | |
| ENSMUSG00000100452 | 1 | |
| ENSMUSG00000100453 | 1 | |
| ENSMUSG00000100454 | 1 | |
| ENSMUSG00000100455 | -1.05861608 | 0.586328611 |
| ENSMUSG00000100456 | 1 | |
| ENSMUSG00000100457 | 1 | |
| ENSMUSG00000100458 | 1 | |
| ENSMUSG00000100459 | 1 | |
| ENSMUSG00000100460 | 1 | |
| ENSMUSG00000100461 | 1 | |
| ENSMUSG00000100462 | 1 | |
| ENSMUSG00000100463 | 1 | |
| ENSMUSG00000100464 | 1 | |
| ENSMUSG00000100465 | -1.043306199 | 0.837704961 |
| ENSMUSG00000100466 | 1 | |

| | | |
|---|---|---|
| ENSMUSG00000100467 | 1 | |
| ENSMUSG00000100468 | 1 | |
| ENSMUSG00000100469 | 1 | |
| ENSMUSG00000100470 | 1 | |
| ENSMUSG00000100471 | 1 | |
| ENSMUSG00000100472 | 1 | |
| ENSMUSG00000100473 | 1 | |
| ENSMUSG00000100474 | 1 | |
| ENSMUSG00000100475 | 1 | |
| ENSMUSG00000100476 | 1 | |
| ENSMUSG00000100477 | 1 | |
| ENSMUSG00000100478 | 1 | |
| ENSMUSG00000100479 | 1 | |
| ENSMUSG00000100480 | -1.318874941 | 0.304585116 |
| ENSMUSG00000100481 | 1.059463094 | 0.608497229 |
| ENSMUSG00000100483 | -1.028348525 | 0.956972448 |
| ENSMUSG00000100484 | 1 | |
| ENSMUSG00000100485 | 1 | |
| ENSMUSG00000100486 | 1 | |
| ENSMUSG00000100487 | 1 | |
| ENSMUSG00000100488 | 1 | |
| ENSMUSG00000100489 | 1 | |
| ENSMUSG00000100490 | -1.156072222 | 0.639135481 |
| ENSMUSG00000100491 | 1 | |
| ENSMUSG00000100492 | 1 | |
| ENSMUSG00000100494 | 1 | |
| ENSMUSG00000100495 | 1 | |
| ENSMUSG00000100496 | 1 | |
| ENSMUSG00000100497 | 1 | |
| ENSMUSG00000100499 | 1 | |
| ENSMUSG00000100500 | 1 | |
| ENSMUSG00000100501 | 1 | |
| ENSMUSG00000100502 | 1 | |
| ENSMUSG00000100503 | 1 | |
| ENSMUSG00000100504 | 1 | |
| ENSMUSG00000100505 | -1 | 1 |
| ENSMUSG00000100506 | 1 | |
| ENSMUSG00000100507 | 1 | |
| ENSMUSG00000100508 | 1 | |
| ENSMUSG00000100509 | 1 | |
| ENSMUSG00000100510 | 1.145390819 | 0.698112512 |
| ENSMUSG00000100511 | 1 | |
| ENSMUSG00000100512 | -1 | 1 |
| ENSMUSG00000100513 | 1 | |
| ENSMUSG00000100514 | 1.11166259 | 0.698112512 |
| ENSMUSG00000100515 | 1 | |
| ENSMUSG00000100516 | 1 | |
| ENSMUSG00000100517 | 1 | |
| ENSMUSG00000100518 | 1 | |
| ENSMUSG00000100519 | 1 | |
| ENSMUSG00000100520 | 1 | |
| ENSMUSG00000100521 | 1 | |
| ENSMUSG00000100522 | 1 | |
| ENSMUSG00000100523 | 1 | |
| ENSMUSG00000100524 | 1 | |
| ENSMUSG00000100525 | 1 | |
| ENSMUSG00000100526 | 1 | |
| ENSMUSG00000100527 | 1 | |
| ENSMUSG00000100528 | 1 | |
| ENSMUSG00000100529 | 1 | |
| ENSMUSG00000100530 | 1 | |
| ENSMUSG00000100531 | 1 | |
| ENSMUSG00000100532 | 1 | |

| | | |
|---|---|---|
| ENSMUSG00000100533 | 1 | |
| ENSMUSG00000100534 | 1.179016771 | 0.601307651 |
| ENSMUSG00000100535 | 1 | |
| ENSMUSG00000100536 | -1.271754232 | 0.085160795 |
| ENSMUSG00000100537 | 1 | |
| ENSMUSG00000100538 | 1 | |
| ENSMUSG00000100539 | 1 | |
| ENSMUSG00000100540 | 1 | |
| ENSMUSG00000100541 | 1 | |
| ENSMUSG00000100542 | 1 | |
| ENSMUSG00000100543 | 1 | |
| ENSMUSG00000100544 | 1 | |
| ENSMUSG00000100546 | 1 | |
| ENSMUSG00000100547 | 1 | |
| ENSMUSG00000100548 | 1.020182806 | 0.968180243 |
| ENSMUSG00000100549 | 1 | |
| ENSMUSG00000100550 | 1.099959823 | 0.624848794 |
| ENSMUSG00000100551 | -1.044273782 | 0.698112512 |
| ENSMUSG00000100552 | -1.028160286 | 0.933897165 |
| ENSMUSG00000100553 | 1 | |
| ENSMUSG00000100554 | -1.03962336 | 0.698112512 |
| ENSMUSG00000100555 | -1.014570171 | 1 |
| ENSMUSG00000100556 | 1 | |
| ENSMUSG00000100557 | 1 | |
| ENSMUSG00000100558 | 1 | |
| ENSMUSG00000100559 | 1 | |
| ENSMUSG00000100560 | 1 | |
| ENSMUSG00000100561 | 1.107972302 | 0.699196281 |
| ENSMUSG00000100562 | 1 | |
| ENSMUSG00000100563 | 1 | |
| ENSMUSG00000100564 | 1 | |
| ENSMUSG00000100565 | 1 | |
| ENSMUSG00000100566 | 1 | |
| ENSMUSG00000100567 | 1 | |
| ENSMUSG00000100568 | -1.089497311 | 0.608497229 |
| ENSMUSG00000100569 | 1 | |
| ENSMUSG00000100570 | 1 | |
| ENSMUSG00000100571 | 1 | |
| ENSMUSG00000100572 | 1.050586185 | 0.708480353 |
| ENSMUSG00000100573 | 1 | |
| ENSMUSG00000100574 | 1 | |
| ENSMUSG00000100575 | 1 | |
| ENSMUSG00000100576 | 1 | |
| ENSMUSG00000100577 | 1 | |
| ENSMUSG00000100578 | 1 | |
| ENSMUSG00000100579 | 1 | |
| ENSMUSG00000100580 | 1.14672896 | 0.423106595 |
| ENSMUSG00000100581 | 1 | |
| ENSMUSG00000100582 | 1 | |
| ENSMUSG00000100583 | 1.019083579 | 0.968937442 |
| ENSMUSG00000100584 | 1 | |
| ENSMUSG00000100585 | 1 | |
| ENSMUSG00000100586 | -1 | 1 |
| ENSMUSG00000100587 | 1 | |
| ENSMUSG00000100588 | 1 | |
| ENSMUSG00000100589 | 1 | |
| ENSMUSG00000100590 | 1 | |
| ENSMUSG00000100591 | 1 | |
| ENSMUSG00000100592 | 1 | |
| ENSMUSG00000100593 | -1.044273782 | 0.790851299 |
| ENSMUSG00000100594 | 1.090334524 | 0.698112512 |
| ENSMUSG00000100595 | -1.085651418 | 0.731932479 |
| ENSMUSG00000100596 | -1.03962336 | 0.698112512 |

| | | |
|---|---|---|
| ENSMUSG00000100597 | 1 | |
| ENSMUSG00000100598 | 1 | |
| ENSMUSG00000100599 | 1.063831093 | 0.823164089 |
| ENSMUSG00000100600 | -1.012736809 | 1 |
| ENSMUSG00000100601 | 1 | |
| ENSMUSG00000100602 | -1.019108526 | 0.992110111 |
| ENSMUSG00000100603 | 1 | |
| ENSMUSG00000100604 | 1 | |
| ENSMUSG00000100605 | -1.044273782 | 0.698112512 |
| ENSMUSG00000100606 | 1 | |
| ENSMUSG00000100607 | 1 | |
| ENSMUSG00000100608 | 1 | |
| ENSMUSG00000100609 | 1 | |
| ENSMUSG00000100610 | 1 | |
| ENSMUSG00000100611 | 1 | |
| ENSMUSG00000100612 | 1 | |
| ENSMUSG00000100613 | 1 | |
| ENSMUSG00000100614 | 1 | |
| ENSMUSG00000100615 | -1.222105224 | 0.420161906 |
| ENSMUSG00000100616 | 1 | |
| ENSMUSG00000100617 | 1 | |
| ENSMUSG00000100618 | 1 | |
| ENSMUSG00000100619 | -1.00303729 | 1 |
| ENSMUSG00000100620 | 1.025370635 | 0.988324779 |
| ENSMUSG00000100621 | 1 | |
| ENSMUSG00000100622 | -1.262114908 | 0.031133889 |
| ENSMUSG00000100623 | 1 | |
| ENSMUSG00000100624 | 1 | |
| ENSMUSG00000100625 | 1 | |
| ENSMUSG00000100626 | 1 | |
| ENSMUSG00000100627 | -1.056351261 | 0.887044308 |
| ENSMUSG00000100628 | 1 | |
| ENSMUSG00000100629 | 1.034478173 | 0.784698888 |
| ENSMUSG00000100630 | 1 | |
| ENSMUSG00000100631 | 1 | |
| ENSMUSG00000100632 | -1.160431826 | 0.362989791 |
| ENSMUSG00000100634 | 1 | |
| ENSMUSG00000100635 | -1.013310247 | 1 |
| ENSMUSG00000100636 | 1.059463094 | 0.698112512 |
| ENSMUSG00000100637 | 1 | |
| ENSMUSG00000100639 | 1 | |
| ENSMUSG00000100640 | 1 | |
| ENSMUSG00000100641 | 1 | |
| ENSMUSG00000100642 | 1.489460736 | 0.003785514 |
| ENSMUSG00000100643 | 1 | |
| ENSMUSG00000100644 | 1 | |
| ENSMUSG00000100645 | 1 | |
| ENSMUSG00000100646 | 1 | |
| ENSMUSG00000100647 | 1 | |
| ENSMUSG00000100648 | 1.005984237 | 1 |
| ENSMUSG00000100649 | 1 | |
| ENSMUSG00000100650 | 1 | |
| ENSMUSG00000100651 | 1 | |
| ENSMUSG00000100652 | 1.021105581 | 0.908543402 |
| ENSMUSG00000100653 | 1 | |
| ENSMUSG00000100654 | 1 | |
| ENSMUSG00000100655 | 1 | |
| ENSMUSG00000100656 | 1 | |
| ENSMUSG00000100657 | 1 | |
| ENSMUSG00000100658 | 1.221890797 | 0.608497229 |
| ENSMUSG00000100659 | 1 | |
| ENSMUSG00000100660 | 1 | |
| ENSMUSG00000100661 | 1 | |

| | | |
|---|---|---|
| ENSMUSG00000100662 | 1.019083579 | 0.949812939 |
| ENSMUSG00000100663 | 1 | |
| ENSMUSG00000100664 | 1 | |
| ENSMUSG00000100665 | 1 | |
| ENSMUSG00000100666 | 1 | |
| ENSMUSG00000100667 | 1 | |
| ENSMUSG00000100668 | 1 | |
| ENSMUSG00000100669 | 1 | |
| ENSMUSG00000100670 | 1 | |
| ENSMUSG00000100671 | 1 | |
| ENSMUSG00000100672 | -1.107598576 | 0.527745314 |
| ENSMUSG00000100673 | 1 | |
| ENSMUSG00000100674 | 1 | |
| ENSMUSG00000100675 | 1 | |
| ENSMUSG00000100676 | 1 | |
| ENSMUSG00000100677 | 1 | |
| ENSMUSG00000100678 | 1 | |
| ENSMUSG00000100679 | 1 | |
| ENSMUSG00000100680 | 1.134282516 | 0.536694773 |
| ENSMUSG00000100681 | 1 | |
| ENSMUSG00000100682 | 1 | |
| ENSMUSG00000100683 | 1 | |
| ENSMUSG00000100684 | 1 | |
| ENSMUSG00000100685 | -1.043306199 | 0.698112512 |
| ENSMUSG00000100687 | 1 | |
| ENSMUSG00000100689 | 1 | |
| ENSMUSG00000100690 | 1 | |
| ENSMUSG00000100691 | 1.128663607 | 0.443913715 |
| ENSMUSG00000100692 | 1 | |
| ENSMUSG00000100693 | 1 | |
| ENSMUSG00000100694 | 1 | |
| ENSMUSG00000100695 | 1 | |
| ENSMUSG00000100696 | -1.147838869 | 0.195607801 |
| ENSMUSG00000100697 | 1 | |
| ENSMUSG00000100698 | 1 | |
| ENSMUSG00000100699 | 1 | |
| ENSMUSG00000100700 | 1 | |
| ENSMUSG00000100701 | -1.132635486 | 0.120932298 |
| ENSMUSG00000100702 | 1.183443915 | 0.536673416 |
| ENSMUSG00000100703 | 1 | |
| ENSMUSG00000100704 | 1 | |
| ENSMUSG00000100705 | 1 | |
| ENSMUSG00000100706 | -1.48629873 | 0.025430361 |
| ENSMUSG00000100707 | 1 | |
| ENSMUSG00000100708 | 1 | |
| ENSMUSG00000100709 | 1 | |
| ENSMUSG00000100710 | 1 | |
| ENSMUSG00000100711 | 1 | |
| ENSMUSG00000100712 | 1 | |
| ENSMUSG00000100713 | 1 | |
| ENSMUSG00000100714 | 1.200936955 | 0.215991492 |
| ENSMUSG00000100715 | 1 | |
| ENSMUSG00000100716 | 1 | |
| ENSMUSG00000100717 | -1 | 1 |
| ENSMUSG00000100718 | 1 | |
| ENSMUSG00000100719 | 1 | |
| ENSMUSG00000100720 | -1.010960635 | 1 |
| ENSMUSG00000100721 | 1 | |
| ENSMUSG00000100722 | 1 | |
| ENSMUSG00000100723 | 1 | |
| ENSMUSG00000100724 | 1 | |
| ENSMUSG00000100725 | 1 | |
| ENSMUSG00000100726 | 1 | |

| | | |
|---|---|---|
| ENSMUSG00000100727 | 1 | |
| ENSMUSG00000100728 | 1 | |
| ENSMUSG00000100729 | 1 | |
| ENSMUSG00000100730 | 1 | |
| ENSMUSG00000100731 | 1 | |
| ENSMUSG00000100732 | 1 | |
| ENSMUSG00000100733 | 1.069068979 | 0.724838192 |
| ENSMUSG00000100734 | 1 | |
| ENSMUSG00000100736 | 1.13770279 | 0.70299173 |
| ENSMUSG00000100737 | 1 | |
| ENSMUSG00000100738 | 1 | |
| ENSMUSG00000100739 | 1 | |
| ENSMUSG00000100740 | 1 | |
| ENSMUSG00000100741 | 1 | |
| ENSMUSG00000100742 | 1 | |
| ENSMUSG00000100743 | 1 | |
| ENSMUSG00000100744 | -1.023769022 | 0.942541168 |
| ENSMUSG00000100745 | 1 | |
| ENSMUSG00000100746 | 1 | |
| ENSMUSG00000100747 | 1.236487199 | 0.328585197 |
| ENSMUSG00000100748 | 1 | |
| ENSMUSG00000100749 | 1 | |
| ENSMUSG00000100750 | 1.13933316 | 0.028839696 |
| ENSMUSG00000100751 | 1 | |
| ENSMUSG00000100752 | 1 | |
| ENSMUSG00000100753 | 1 | |
| ENSMUSG00000100754 | 1 | |
| ENSMUSG00000100755 | 1 | |
| ENSMUSG00000100756 | 1 | |
| ENSMUSG00000100757 | 1 | |
| ENSMUSG00000100758 | 1 | |
| ENSMUSG00000100759 | 1 | |
| ENSMUSG00000100760 | 1 | |
| ENSMUSG00000100761 | 1 | |
| ENSMUSG00000100762 | 1 | |
| ENSMUSG00000100763 | -1.110619514 | 0.625496737 |
| ENSMUSG00000100764 | -1.393514 | 0.063603912 |
| ENSMUSG00000100765 | 1 | |
| ENSMUSG00000100767 | 1.049411284 | 0.835701362 |
| ENSMUSG00000100768 | 1.06702496 | 0.698112512 |
| ENSMUSG00000100769 | 1.044085745 | 0.953416513 |
| ENSMUSG00000100770 | 1 | |
| ENSMUSG00000100771 | 1 | |
| ENSMUSG00000100772 | 1 | |
| ENSMUSG00000100773 | 1 | |
| ENSMUSG00000100774 | 1.015486245 | 1 |
| ENSMUSG00000100775 | -1.004394359 | 1 |
| ENSMUSG00000100776 | 1 | |
| ENSMUSG00000100777 | 1 | |
| ENSMUSG00000100778 | 1 | |
| ENSMUSG00000100779 | 1.03863466 | 0.951780193 |
| ENSMUSG00000100780 | 1 | |
| ENSMUSG00000100781 | -1.053549062 | 0.698112512 |
| ENSMUSG00000100782 | 1.132639128 | 0.568304986 |
| ENSMUSG00000100783 | 1.039597911 | 0.850281678 |
| ENSMUSG00000100784 | 1 | |
| ENSMUSG00000100785 | 1 | |
| ENSMUSG00000100786 | -1 | 1 |
| ENSMUSG00000100787 | 1 | |
| ENSMUSG00000100788 | 1 | |
| ENSMUSG00000100789 | -1.089497311 | 0.614307028 |
| ENSMUSG00000100790 | 1 | |
| ENSMUSG00000100791 | 1 | |

| | | |
|---|---|---|
| ENSMUSG00000100792 | 1 | |
| ENSMUSG00000100793 | 1 | |
| ENSMUSG00000100794 | 1.061495364 | 0.712070475 |
| ENSMUSG00000100795 | 1 | |
| ENSMUSG00000100796 | 1 | |
| ENSMUSG00000100797 | 1 | |
| ENSMUSG00000100798 | -1.304146544 | 0.182340781 |
| ENSMUSG00000100799 | 1 | |
| ENSMUSG00000100800 | 1 | |
| ENSMUSG00000100801 | 1 | |
| ENSMUSG00000100802 | 1 | |
| ENSMUSG00000100803 | 1 | |
| ENSMUSG00000100804 | 1 | |
| ENSMUSG00000100805 | 1 | |
| ENSMUSG00000100806 | 1 | |
| ENSMUSG00000100807 | 1 | |
| ENSMUSG00000100808 | 1 | |
| ENSMUSG00000100809 | 1 | |
| ENSMUSG00000100810 | 1 | |
| ENSMUSG00000100811 | 1.028323353 | 0.936340683 |
| ENSMUSG00000100812 | 1 | |
| ENSMUSG00000100813 | 1.042776804 | 0.794260077 |
| ENSMUSG00000100814 | 1 | |
| ENSMUSG00000100815 | 1 | |
| ENSMUSG00000100816 | 1.208657847 | 0.370165361 |
| ENSMUSG00000100817 | 1 | |
| ENSMUSG00000100818 | 1 | |
| ENSMUSG00000100819 | 1.069068979 | 0.698112512 |
| ENSMUSG00000100820 | 1 | |
| ENSMUSG00000100821 | 1 | |
| ENSMUSG00000100822 | 1 | |
| ENSMUSG00000100823 | 1 | |
| ENSMUSG00000100824 | 1 | |
| ENSMUSG00000100825 | -1.063525418 | 0.698112512 |
| ENSMUSG00000100826 | 1.006134895 | 1 |
| ENSMUSG00000100827 | 1 | |
| ENSMUSG00000100828 | 1 | |
| ENSMUSG00000100830 | -1.131509746 | 0.661155252 |
| ENSMUSG00000100831 | 1 | |
| ENSMUSG00000100832 | 1 | |
| ENSMUSG00000100833 | -1.105687959 | 0.773508225 |
| ENSMUSG00000100834 | 1 | |
| ENSMUSG00000100835 | 1 | |
| ENSMUSG00000100836 | 1 | |
| ENSMUSG00000100837 | -1.011410153 | 0.998567129 |
| ENSMUSG00000100838 | 1 | |
| ENSMUSG00000100839 | -1.171420412 | 0.346721275 |
| ENSMUSG00000100841 | 1 | |
| ENSMUSG00000100842 | 1 | |
| ENSMUSG00000100843 | -1 | 1 |
| ENSMUSG00000100844 | -1 | 1 |
| ENSMUSG00000100845 | 1 | |
| ENSMUSG00000100846 | 1 | |
| ENSMUSG00000100847 | 1 | |
| ENSMUSG00000100848 | 1 | |
| ENSMUSG00000100849 | 1 | |
| ENSMUSG00000100850 | 1 | |
| ENSMUSG00000100851 | 1 | |
| ENSMUSG00000100852 | 1 | |
| ENSMUSG00000100853 | 1 | |
| ENSMUSG00000100854 | 1 | |
| ENSMUSG00000100855 | 1.059463094 | 0.608497229 |
| ENSMUSG00000100856 | 1 | |

| | | |
|---|---|---|
| ENSMUSG00000100857 | 1 | |
| ENSMUSG00000100858 | 1 | |
| ENSMUSG00000100859 | 1 | |
| ENSMUSG00000100860 | 1 | |
| ENSMUSG00000100861 | 1 | |
| ENSMUSG00000100862 | -1.000559704 | 1 |
| ENSMUSG00000100863 | 1.069068979 | 0.698112512 |
| ENSMUSG00000100864 | 1 | |
| ENSMUSG00000100865 | -1.054559254 | 0.888281505 |
| ENSMUSG00000100866 | 1 | |
| ENSMUSG00000100867 | -1 | 1 |
| ENSMUSG00000100868 | 1.059463094 | 0.608497229 |
| ENSMUSG00000100869 | 1 | |
| ENSMUSG00000100870 | 1 | |
| ENSMUSG00000100871 | 1 | |
| ENSMUSG00000100872 | 1.101617271 | 0.763074704 |
| ENSMUSG00000100873 | 1 | |
| ENSMUSG00000100874 | 1 | |
| ENSMUSG00000100875 | 1 | |
| ENSMUSG00000100876 | 1.094868708 | 0.608497229 |
| ENSMUSG00000100877 | 1 | |
| ENSMUSG00000100878 | 1 | |
| ENSMUSG00000100879 | 1 | |
| ENSMUSG00000100880 | -1.043306199 | 0.698112512 |
| ENSMUSG00000100881 | 1 | |
| ENSMUSG00000100882 | 1 | |
| ENSMUSG00000100883 | 1 | |
| ENSMUSG00000100884 | 1 | |
| ENSMUSG00000100885 | 1 | |
| ENSMUSG00000100886 | 1 | |
| ENSMUSG00000100887 | 1 | |
| ENSMUSG00000100888 | 1 | |
| ENSMUSG00000100889 | 1 | |
| ENSMUSG00000100890 | 1 | |
| ENSMUSG00000100891 | -1.043306199 | 0.79291651 |
| ENSMUSG00000100892 | 1 | |
| ENSMUSG00000100893 | 1 | |
| ENSMUSG00000100894 | -1.109582661 | 0.698112512 |
| ENSMUSG00000100895 | 1 | |
| ENSMUSG00000100896 | 1 | |
| ENSMUSG00000100897 | 1 | |
| ENSMUSG00000100898 | 1 | |
| ENSMUSG00000100899 | 1 | |
| ENSMUSG00000100900 | 1 | |
| ENSMUSG00000100901 | 1 | |
| ENSMUSG00000100902 | 1 | |
| ENSMUSG00000100903 | 1 | |
| ENSMUSG00000100904 | 1 | |
| ENSMUSG00000100905 | 1 | |
| ENSMUSG00000100906 | 1 | |
| ENSMUSG00000100907 | 1 | |
| ENSMUSG00000100908 | 1 | |
| ENSMUSG00000100909 | 1 | |
| ENSMUSG00000100910 | 1 | |
| ENSMUSG00000100911 | 1 | |
| ENSMUSG00000100912 | 1 | |
| ENSMUSG00000100913 | -1.043306199 | 0.698112512 |
| ENSMUSG00000100914 | 1 | |
| ENSMUSG00000100915 | 1 | |
| ENSMUSG00000100916 | 1 | |
| ENSMUSG00000100917 | 1 | |
| ENSMUSG00000100918 | 1 | |
| ENSMUSG00000100919 | 1 | |

| | | |
|---|---|---|
| ENSMUSG00000100920 | 1 | |
| ENSMUSG00000100922 | 1.17495337 | 0.478341625 |
| ENSMUSG00000100923 | 1 | |
| ENSMUSG00000100924 | 1 | |
| ENSMUSG00000100925 | 1 | |
| ENSMUSG00000100926 | 1 | |
| ENSMUSG00000100927 | 1 | |
| ENSMUSG00000100928 | 1 | |
| ENSMUSG00000100929 | -1 | 1 |
| ENSMUSG00000100930 | 1 | |
| ENSMUSG00000100931 | 1.091388659 | 0.608497229 |
| ENSMUSG00000100932 | 1 | |
| ENSMUSG00000100933 | 1 | |
| ENSMUSG00000100935 | 1 | |
| ENSMUSG00000100936 | 1 | |
| ENSMUSG00000100937 | 1.289130977 | 0.121201239 |
| ENSMUSG00000100938 | 1 | |
| ENSMUSG00000100939 | 1 | |
| ENSMUSG00000100940 | 1 | |
| ENSMUSG00000100941 | 1 | |
| ENSMUSG00000100942 | 1 | |
| ENSMUSG00000100943 | 1 | |
| ENSMUSG00000100944 | 1 | |
| ENSMUSG00000100945 | 1 | |
| ENSMUSG00000100946 | 1 | |
| ENSMUSG00000100947 | 1 | |
| ENSMUSG00000100948 | 1 | |
| ENSMUSG00000100949 | 1 | |
| ENSMUSG00000100950 | 1 | |
| ENSMUSG00000100951 | 1 | |
| ENSMUSG00000100952 | 1 | |
| ENSMUSG00000100953 | 1 | |
| ENSMUSG00000100954 | -1.032817476 | 0.935057992 |
| ENSMUSG00000100955 | 1 | |
| ENSMUSG00000100956 | 1 | |
| ENSMUSG00000100957 | 1 | |
| ENSMUSG00000100958 | 1 | |
| ENSMUSG00000100959 | 1.089470641 | 0.70443552 |
| ENSMUSG00000100960 | 1 | |
| ENSMUSG00000100961 | 1 | |
| ENSMUSG00000100962 | 1 | |
| ENSMUSG00000100963 | 1.059463094 | 0.698112512 |
| ENSMUSG00000100964 | 1 | |
| ENSMUSG00000100965 | 1 | |
| ENSMUSG00000100966 | 1 | |
| ENSMUSG00000100967 | -1.060545185 | 0.575351077 |
| ENSMUSG00000100968 | 1 | |
| ENSMUSG00000100969 | 1.015486245 | 0.985522133 |
| ENSMUSG00000100970 | 1 | |
| ENSMUSG00000100971 | 1 | |
| ENSMUSG00000100972 | 1 | |
| ENSMUSG00000100973 | 1 | |
| ENSMUSG00000100974 | 1 | |
| ENSMUSG00000100975 | 1.055357849 | 0.683942136 |
| ENSMUSG00000100976 | 1 | |
| ENSMUSG00000100977 | 1 | |
| ENSMUSG00000100978 | 1 | |
| ENSMUSG00000100979 | 1 | |
| ENSMUSG00000100980 | 1.08230051 | 0.608497229 |
| ENSMUSG00000100981 | 1 | |
| ENSMUSG00000100982 | 1 | |
| ENSMUSG00000100983 | 1 | |
| ENSMUSG00000100984 | 1 | |

| | | |
|---|---|---|
| ENSMUSG00000100985 | -1.044273782 | 0.790851299 |
| ENSMUSG00000100986 | 1 | |
| ENSMUSG00000100987 | 1.33979607 | 0.093261127 |
| ENSMUSG00000100988 | 1 | |
| ENSMUSG00000100989 | 1 | |
| ENSMUSG00000100990 | -1.03962336 | 0.698112512 |
| ENSMUSG00000100991 | 1 | |
| ENSMUSG00000100992 | 1 | |
| ENSMUSG00000100993 | 1 | |
| ENSMUSG00000100994 | 1 | |
| ENSMUSG00000100995 | 1 | |
| ENSMUSG00000100996 | 1 | |
| ENSMUSG00000100998 | 1 | |
| ENSMUSG00000100999 | 1 | |
| ENSMUSG00000101000 | 1 | |
| ENSMUSG00000101001 | 1 | |
| ENSMUSG00000101002 | 1 | |
| ENSMUSG00000101003 | 1 | |
| ENSMUSG00000101004 | 1.005613439 | 1 |
| ENSMUSG00000101005 | 1 | |
| ENSMUSG00000101006 | 1 | |
| ENSMUSG00000101007 | 1.122462048 | 0.54644471 |
| ENSMUSG00000101008 | 1 | |
| ENSMUSG00000101009 | -1.023608309 | 0.978692019 |
| ENSMUSG00000101010 | 1 | |
| ENSMUSG00000101011 | -1.042755471 | 0.921185021 |
| ENSMUSG00000101012 | 1 | |
| ENSMUSG00000101013 | -1.004597703 | 1 |
| ENSMUSG00000101014 | 1 | |
| ENSMUSG00000101015 | 1 | |
| ENSMUSG00000101016 | 1 | |
| ENSMUSG00000101017 | 1 | |
| ENSMUSG00000101018 | 1 | |
| ENSMUSG00000101019 | 1 | |
| ENSMUSG00000101020 | 1 | |
| ENSMUSG00000101021 | 1 | |
| ENSMUSG00000101022 | 1 | |
| ENSMUSG00000101023 | 1 | |
| ENSMUSG00000101024 | 1 | |
| ENSMUSG00000101025 | 1 | |
| ENSMUSG00000101026 | 1 | |
| ENSMUSG00000101027 | 1 | |
| ENSMUSG00000101028 | -1 | 1 |
| ENSMUSG00000101029 | 1 | |
| ENSMUSG00000101030 | 1 | |
| ENSMUSG00000101031 | 1 | |
| ENSMUSG00000101032 | 1 | |
| ENSMUSG00000101033 | 1 | |
| ENSMUSG00000101035 | 1 | |
| ENSMUSG00000101036 | -1.03962336 | 0.698112512 |
| ENSMUSG00000101037 | -1.095910413 | 0.799372007 |
| ENSMUSG00000101038 | 1 | |
| ENSMUSG00000101039 | 1 | |
| ENSMUSG00000101040 | 1 | |
| ENSMUSG00000101041 | 1 | |
| ENSMUSG00000101042 | 1 | |
| ENSMUSG00000101043 | 1 | |
| ENSMUSG00000101044 | 1 | |
| ENSMUSG00000101045 | 1 | |
| ENSMUSG00000101046 | 1 | |
| ENSMUSG00000101049 | -1.043306199 | 0.698112512 |
| ENSMUSG00000101050 | 1 | |
| ENSMUSG00000101051 | 1 | |

| | | |
|---|---|---|
| ENSMUSG00000101052 | 1 | |
| ENSMUSG00000101053 | 1 | |
| ENSMUSG00000101054 | 1 | |
| ENSMUSG00000101055 | 1 | |
| ENSMUSG00000101056 | 1 | |
| ENSMUSG00000101057 | 1 | |
| ENSMUSG00000101058 | 1 | |
| ENSMUSG00000101059 | -1.08081673 | 0.698112512 |
| ENSMUSG00000101060 | 1 | |
| ENSMUSG00000101061 | -1.113493764 | 0.608497229 |
| ENSMUSG00000101062 | 1 | |
| ENSMUSG00000101063 | 1 | |
| ENSMUSG00000101064 | 1 | |
| ENSMUSG00000101065 | 1 | |
| ENSMUSG00000101066 | -1 | 1 |
| ENSMUSG00000101067 | 1 | |
| ENSMUSG00000101068 | 1 | |
| ENSMUSG00000101069 | 1 | |
| ENSMUSG00000101070 | 1 | |
| ENSMUSG00000101071 | 1 | |
| ENSMUSG00000101072 | 1 | |
| ENSMUSG00000101073 | 1 | |
| ENSMUSG00000101074 | 1 | |
| ENSMUSG00000101075 | 1 | |
| ENSMUSG00000101076 | 1 | |
| ENSMUSG00000101077 | 1.059463094 | 0.608497229 |
| ENSMUSG00000101078 | 1 | |
| ENSMUSG00000101079 | 1 | |
| ENSMUSG00000101080 | 1 | |
| ENSMUSG00000101081 | 1 | |
| ENSMUSG00000101082 | 1 | |
| ENSMUSG00000101083 | 1 | |
| ENSMUSG00000101084 | 1 | |
| ENSMUSG00000101085 | 1 | |
| ENSMUSG00000101086 | 1 | |
| ENSMUSG00000101087 | 1 | |
| ENSMUSG00000101088 | 1.024693402 | 0.933031586 |
| ENSMUSG00000101089 | -1 | 1 |
| ENSMUSG00000101090 | 1 | |
| ENSMUSG00000101091 | 1 | |
| ENSMUSG00000101092 | 1 | |
| ENSMUSG00000101093 | -1.03962336 | 0.698112512 |
| ENSMUSG00000101094 | 1 | |
| ENSMUSG00000101095 | 1 | |
| ENSMUSG00000101096 | 1 | |
| ENSMUSG00000101097 | -1.084645496 | 0.532369657 |
| ENSMUSG00000101098 | 1 | |
| ENSMUSG00000101099 | 1 | |
| ENSMUSG00000101100 | 1 | |
| ENSMUSG00000101101 | 1 | |
| ENSMUSG00000101102 | 1.059463094 | 0.608497229 |
| ENSMUSG00000101103 | 1 | |
| ENSMUSG00000101104 | 1 | |
| ENSMUSG00000101105 | 1 | |
| ENSMUSG00000101106 | 1 | |
| ENSMUSG00000101107 | -1.044273782 | 0.801443616 |
| ENSMUSG00000101108 | 1 | |
| ENSMUSG00000101109 | 1 | |
| ENSMUSG00000101110 | 1 | |
| ENSMUSG00000101111 | 1.049411284 | 0.863033776 |
| ENSMUSG00000101112 | 1 | |
| ENSMUSG00000101113 | 1 | |
| ENSMUSG00000101114 | 1 | |

| | | |
|---|---|---|
| ENSMUSG00000101115 | 1 | |
| ENSMUSG00000101116 | 1 | |
| ENSMUSG00000101117 | 1 | |
| ENSMUSG00000101118 | 1 | |
| ENSMUSG00000101119 | 1 | |
| ENSMUSG00000101120 | 1 | |
| ENSMUSG00000101122 | 1.111393559 | 0.609700609 |
| ENSMUSG00000101123 | 1 | |
| ENSMUSG00000101124 | 1 | |
| ENSMUSG00000101125 | 1 | |
| ENSMUSG00000101126 | 1 | |
| ENSMUSG00000101127 | 1 | |
| ENSMUSG00000101128 | 1 | |
| ENSMUSG00000101129 | -1.043306199 | 0.698112512 |
| ENSMUSG00000101130 | 1 | |
| ENSMUSG00000101131 | 1 | |
| ENSMUSG00000101132 | 1 | |
| ENSMUSG00000101133 | 1.199856674 | 0.578284342 |
| ENSMUSG00000101134 | 1 | |
| ENSMUSG00000101135 | 1.069068979 | 0.645934663 |
| ENSMUSG00000101136 | 1.014731082 | 1 |
| ENSMUSG00000101137 | 1 | |
| ENSMUSG00000101138 | 1 | |
| ENSMUSG00000101139 | 1 | |
| ENSMUSG00000101140 | 1 | |
| ENSMUSG00000101141 | 1 | |
| ENSMUSG00000101142 | 1 | |
| ENSMUSG00000101143 | 1 | |
| ENSMUSG00000101144 | 1.059463094 | 0.608497229 |
| ENSMUSG00000101145 | 1 | |
| ENSMUSG00000101146 | 1 | |
| ENSMUSG00000101147 | 1 | |
| ENSMUSG00000101148 | 1 | |
| ENSMUSG00000101149 | 1 | |
| ENSMUSG00000101150 | 1 | |
| ENSMUSG00000101151 | 1 | |
| ENSMUSG00000101152 | 1 | |
| ENSMUSG00000101153 | 1 | |
| ENSMUSG00000101154 | 1 | |
| ENSMUSG00000101155 | 1 | |
| ENSMUSG00000101156 | 1.005851671 | 1 |
| ENSMUSG00000101157 | 1 | |
| ENSMUSG00000101158 | 1 | |
| ENSMUSG00000101159 | 1 | |
| ENSMUSG00000101160 | 1 | |
| ENSMUSG00000101161 | 1 | |
| ENSMUSG00000101162 | 1.11166259 | 0.638758774 |
| ENSMUSG00000101163 | 1 | |
| ENSMUSG00000101164 | 1 | |
| ENSMUSG00000101166 | 1.069068979 | 0.698112512 |
| ENSMUSG00000101167 | 1.086023414 | 0.73432497 |
| ENSMUSG00000101168 | 1.05410549 | 0.805157604 |
| ENSMUSG00000101170 | 1 | |
| ENSMUSG00000101171 | -1.034990181 | 0.918975822 |
| ENSMUSG00000101172 | -1.143400433 | 0.642819974 |
| ENSMUSG00000101173 | 1 | |
| ENSMUSG00000101174 | 1.5995174 | 0.019383936 |
| ENSMUSG00000101176 | 1 | |
| ENSMUSG00000101177 | 1 | |
| ENSMUSG00000101178 | 1 | |
| ENSMUSG00000101179 | -1.20776913 | 0.412457217 |
| ENSMUSG00000101180 | 1 | |
| ENSMUSG00000101181 | 1 | |

| | | |
|---|---|---|
| ENSMUSG00000101182 | 1 | |
| ENSMUSG00000101183 | 1 | |
| ENSMUSG00000101184 | 1 | |
| ENSMUSG00000101185 | 1 | |
| ENSMUSG00000101186 | 1 | |
| ENSMUSG00000101188 | -1.17128027 | 0.575788842 |
| ENSMUSG00000101189 | 1 | |
| ENSMUSG00000101190 | 1 | |
| ENSMUSG00000101191 | -1.084257835 | 0.704553551 |
| ENSMUSG00000101192 | 1 | |
| ENSMUSG00000101193 | 1 | |
| ENSMUSG00000101194 | 1 | |
| ENSMUSG00000101195 | 1 | |
| ENSMUSG00000101196 | 1 | |
| ENSMUSG00000101197 | 1 | |
| ENSMUSG00000101198 | 1 | |
| ENSMUSG00000101199 | 1.132639128 | 0.217803313 |
| ENSMUSG00000101200 | 1 | |
| ENSMUSG00000101201 | 1 | |
| ENSMUSG00000101202 | 1 | |
| ENSMUSG00000101203 | 1 | |
| ENSMUSG00000101204 | 1 | |
| ENSMUSG00000101205 | 1 | |
| ENSMUSG00000101206 | -1.158708082 | 0.350957446 |
| ENSMUSG00000101207 | 1 | |
| ENSMUSG00000101208 | 1 | |
| ENSMUSG00000101209 | 1.019083579 | 0.979002724 |
| ENSMUSG00000101210 | -1.171269326 | 0.462095672 |
| ENSMUSG00000101211 | 1.257314573 | 0.205169472 |
| ENSMUSG00000101212 | 1 | |
| ENSMUSG00000101213 | 1 | |
| ENSMUSG00000101214 | 1 | |
| ENSMUSG00000101215 | 1 | |
| ENSMUSG00000101216 | 1 | |
| ENSMUSG00000101218 | 1 | |
| ENSMUSG00000101219 | -1.024718486 | 0.96275118 |
| ENSMUSG00000101220 | 1.069293585 | 0.797105186 |
| ENSMUSG00000101221 | 1 | |
| ENSMUSG00000101222 | 1 | |
| ENSMUSG00000101223 | -1.044273782 | 0.698112512 |
| ENSMUSG00000101224 | 1.059463094 | 0.752398528 |
| ENSMUSG00000101225 | 1.078755637 | 0.801168497 |
| ENSMUSG00000101226 | 1 | |
| ENSMUSG00000101227 | 1 | |
| ENSMUSG00000101228 | 1 | |
| ENSMUSG00000101229 | 1 | |
| ENSMUSG00000101230 | 1 | |
| ENSMUSG00000101231 | 1 | |
| ENSMUSG00000101232 | 1 | |
| ENSMUSG00000101233 | 1 | |
| ENSMUSG00000101234 | 1.326238551 | 0.167694565 |
| ENSMUSG00000101235 | 1 | |
| ENSMUSG00000101236 | 1 | |
| ENSMUSG00000101237 | 1 | |
| ENSMUSG00000101238 | 1 | |
| ENSMUSG00000101239 | 1 | |
| ENSMUSG00000101240 | 1.019083579 | 0.97045957 |
| ENSMUSG00000101241 | 1 | |
| ENSMUSG00000101242 | -1.126773846 | 0.539222653 |
| ENSMUSG00000101243 | 1 | |
| ENSMUSG00000101244 | 1 | |
| ENSMUSG00000101245 | 1 | |
| ENSMUSG00000101246 | 1 | |

| | | |
|---|---|---|
| ENSMUSG00000101247 | 1 | |
| ENSMUSG00000101248 | 1 | |
| ENSMUSG00000101249 | -1.212514029 | 0.346890143 |
| ENSMUSG00000101250 | 1 | |
| ENSMUSG00000101251 | 1 | |
| ENSMUSG00000101252 | 1 | |
| ENSMUSG00000101253 | 1 | |
| ENSMUSG00000101254 | 1 | |
| ENSMUSG00000101255 | 1 | |
| ENSMUSG00000101256 | 1 | |
| ENSMUSG00000101257 | 1.039597911 | 0.884018396 |
| ENSMUSG00000101258 | -1 | 1 |
| ENSMUSG00000101259 | 1 | |
| ENSMUSG00000101260 | 1 | |
| ENSMUSG00000101261 | 1 | |
| ENSMUSG00000101262 | 1.01251125 | 0.998924353 |
| ENSMUSG00000101263 | 1 | |
| ENSMUSG00000101264 | -1 | 1 |
| ENSMUSG00000101265 | 1 | |
| ENSMUSG00000101266 | 1 | |
| ENSMUSG00000101267 | 1 | |
| ENSMUSG00000101268 | 1.177765487 | 0.380970157 |
| ENSMUSG00000101270 | 1 | |
| ENSMUSG00000101271 | 1 | |
| ENSMUSG00000101272 | 1 | |
| ENSMUSG00000101273 | 1 | |
| ENSMUSG00000101275 | 1 | |
| ENSMUSG00000101276 | 1 | |
| ENSMUSG00000101277 | 1 | |
| ENSMUSG00000101278 | 1 | |
| ENSMUSG00000101279 | 1 | |
| ENSMUSG00000101280 | 1 | |
| ENSMUSG00000101281 | -1.044273782 | 0.790851299 |
| ENSMUSG00000101282 | 1 | |
| ENSMUSG00000101283 | 1 | |
| ENSMUSG00000101284 | 1 | |
| ENSMUSG00000101285 | 1 | |
| ENSMUSG00000101286 | 1 | |
| ENSMUSG00000101287 | 1 | |
| ENSMUSG00000101288 | 1 | |
| ENSMUSG00000101289 | 1 | |
| ENSMUSG00000101290 | 1 | |
| ENSMUSG00000101291 | 1 | |
| ENSMUSG00000101292 | 1 | |
| ENSMUSG00000101294 | 1 | |
| ENSMUSG00000101295 | 1 | |
| ENSMUSG00000101296 | 1 | |
| ENSMUSG00000101297 | 1 | |
| ENSMUSG00000101298 | 1 | |
| ENSMUSG00000101299 | 1 | |
| ENSMUSG00000101300 | 1 | |
| ENSMUSG00000101301 | 1.023743961 | 0.938939969 |
| ENSMUSG00000101302 | 1 | |
| ENSMUSG00000101303 | -1.044273782 | 0.698112512 |
| ENSMUSG00000101304 | 1 | |
| ENSMUSG00000101305 | 1 | |
| ENSMUSG00000101306 | 1 | |
| ENSMUSG00000101307 | 1 | |
| ENSMUSG00000101308 | -1.079361923 | 0.828564184 |
| ENSMUSG00000101309 | 1 | |
| ENSMUSG00000101310 | 1 | |
| ENSMUSG00000101311 | 1 | |
| ENSMUSG00000101312 | 1 | |

| | | |
|---|---|---|
| ENSMUSG00000101313 | 1 | |
| ENSMUSG00000101314 | 1 | |
| ENSMUSG00000101315 | 1 | |
| ENSMUSG00000101316 | -1.059458736 | 0.560498981 |
| ENSMUSG00000101317 | 1 | |
| ENSMUSG00000101318 | 1 | |
| ENSMUSG00000101319 | 1 | |
| ENSMUSG00000101320 | -1.113515092 | 0.576169279 |
| ENSMUSG00000101321 | 1 | |
| ENSMUSG00000101322 | 1 | |
| ENSMUSG00000101323 | 1 | |
| ENSMUSG00000101324 | 1 | |
| ENSMUSG00000101325 | 1 | |
| ENSMUSG00000101326 | 1 | |
| ENSMUSG00000101327 | 1 | |
| ENSMUSG00000101328 | 1.049411284 | 0.835701362 |
| ENSMUSG00000101329 | 1 | |
| ENSMUSG00000101330 | 1.059463094 | 0.608497229 |
| ENSMUSG00000101331 | 1 | |
| ENSMUSG00000101332 | 1 | |
| ENSMUSG00000101333 | 1 | |
| ENSMUSG00000101334 | 1 | |
| ENSMUSG00000101335 | -1.16958796 | 0.110964535 |
| ENSMUSG00000101336 | 1 | |
| ENSMUSG00000101337 | -1.048853822 | 0.865865121 |
| ENSMUSG00000101338 | 1 | |
| ENSMUSG00000101339 | 1 | |
| ENSMUSG00000101340 | 1 | |
| ENSMUSG00000101341 | 1 | |
| ENSMUSG00000101342 | 1 | |
| ENSMUSG00000101343 | 1 | |
| ENSMUSG00000101344 | 1 | |
| ENSMUSG00000101345 | 1 | |
| ENSMUSG00000101346 | 1 | |
| ENSMUSG00000101347 | 1 | |
| ENSMUSG00000101348 | 1 | |
| ENSMUSG00000101349 | 1 | |
| ENSMUSG00000101350 | 1 | |
| ENSMUSG00000101351 | 1 | |
| ENSMUSG00000101352 | 1 | |
| ENSMUSG00000101353 | 1 | |
| ENSMUSG00000101354 | 1 | |
| ENSMUSG00000101355 | 1 | |
| ENSMUSG00000101356 | -1.041670734 | 0.951767617 |
| ENSMUSG00000101357 | 1 | |
| ENSMUSG00000101358 | 1 | |
| ENSMUSG00000101359 | 1 | |
| ENSMUSG00000101360 | 1 | |
| ENSMUSG00000101361 | 1 | |
| ENSMUSG00000101362 | 1 | |
| ENSMUSG00000101363 | 1 | |
| ENSMUSG00000101364 | 1 | |
| ENSMUSG00000101365 | 1.304351766 | 0.292376367 |
| ENSMUSG00000101366 | 1 | |
| ENSMUSG00000101367 | 1 | |
| ENSMUSG00000101368 | 1 | |
| ENSMUSG00000101369 | 1 | |
| ENSMUSG00000101370 | 1 | |
| ENSMUSG00000101371 | 1 | |
| ENSMUSG00000101372 | -1.03962336 | 0.803072249 |
| ENSMUSG00000101373 | 1 | |
| ENSMUSG00000101374 | 1 | |
| ENSMUSG00000101375 | 1 | |

| | | |
|---|---|---|
| ENSMUSG00000101376 | 1 | |
| ENSMUSG00000101377 | 1 | |
| ENSMUSG00000101378 | 1 | |
| ENSMUSG00000101379 | 1 | |
| ENSMUSG00000101380 | 1 | |
| ENSMUSG00000101381 | 1 | |
| ENSMUSG00000101382 | 1 | |
| ENSMUSG00000101383 | 1 | |
| ENSMUSG00000101385 | 1 | |
| ENSMUSG00000101386 | 1.065093972 | 0.842680402 |
| ENSMUSG00000101387 | 1.002937339 | 1 |
| ENSMUSG00000101388 | 1 | |
| ENSMUSG00000101389 | 1 | |
| ENSMUSG00000101390 | 1 | |
| ENSMUSG00000101391 | 1 | |
| ENSMUSG00000101392 | 1 | |
| ENSMUSG00000101393 | 1 | |
| ENSMUSG00000101394 | -1.086192882 | 0.698112512 |
| ENSMUSG00000101395 | 1 | |
| ENSMUSG00000101396 | 1 | |
| ENSMUSG00000101397 | 1 | |
| ENSMUSG00000101398 | 1 | |
| ENSMUSG00000101399 | 1 | |
| ENSMUSG00000101400 | -1.115014359 | 0.608497229 |
| ENSMUSG00000101401 | 1 | |
| ENSMUSG00000101402 | 1 | |
| ENSMUSG00000101403 | 1 | |
| ENSMUSG00000101404 | 1 | |
| ENSMUSG00000101405 | 1 | |
| ENSMUSG00000101406 | 1 | |
| ENSMUSG00000101407 | 1 | |
| ENSMUSG00000101408 | 1 | |
| ENSMUSG00000101409 | 1 | |
| ENSMUSG00000101410 | 1 | |
| ENSMUSG00000101411 | 1 | |
| ENSMUSG00000101412 | 1 | |
| ENSMUSG00000101413 | -1 | 1 |
| ENSMUSG00000101414 | -1.004544855 | 1 |
| ENSMUSG00000101415 | 1 | |
| ENSMUSG00000101416 | 1 | |
| ENSMUSG00000101417 | 1 | |
| ENSMUSG00000101418 | 1 | |
| ENSMUSG00000101419 | 1 | |
| ENSMUSG00000101420 | 1 | |
| ENSMUSG00000101421 | 1 | |
| ENSMUSG00000101422 | 1 | |
| ENSMUSG00000101423 | 1 | |
| ENSMUSG00000101424 | 1 | |
| ENSMUSG00000101425 | 1 | |
| ENSMUSG00000101426 | 1 | |
| ENSMUSG00000101427 | 1 | |
| ENSMUSG00000101428 | 1 | |
| ENSMUSG00000101429 | -1.043306199 | 0.698112512 |
| ENSMUSG00000101431 | 1.318457764 | 0.035910623 |
| ENSMUSG00000101433 | 1 | |
| ENSMUSG00000101434 | 1 | |
| ENSMUSG00000101435 | 1 | |
| ENSMUSG00000101436 | -1 | 1 |
| ENSMUSG00000101437 | -1 | 1 |
| ENSMUSG00000101438 | 1.019083579 | 0.971519533 |
| ENSMUSG00000101439 | 1 | |
| ENSMUSG00000101440 | 1 | |
| ENSMUSG00000101441 | 1 | |

| | | |
|---|---|---|
| ENSMUSG00000101442 | 1 | |
| ENSMUSG00000101443 | 1.059463094 | 0.608497229 |
| ENSMUSG00000101444 | 1 | |
| ENSMUSG00000101445 | 1 | |
| ENSMUSG00000101446 | 1 | |
| ENSMUSG00000101447 | -1.011833254 | 1 |
| ENSMUSG00000101448 | 1 | |
| ENSMUSG00000101449 | 1.014545335 | 0.979002724 |
| ENSMUSG00000101450 | 1.004641251 | 1 |
| ENSMUSG00000101451 | 1.117436079 | 0.665646314 |
| ENSMUSG00000101452 | 1.060632214 | 0.698112512 |
| ENSMUSG00000101453 | 1 | |
| ENSMUSG00000101454 | 1 | |
| ENSMUSG00000101455 | 1 | |
| ENSMUSG00000101456 | 1 | |
| ENSMUSG00000101457 | 1 | |
| ENSMUSG00000101458 | -1.123044321 | 0.368089426 |
| ENSMUSG00000101459 | 1 | |
| ENSMUSG00000101462 | 1.272775903 | 0.121175467 |
| ENSMUSG00000101463 | 1 | |
| ENSMUSG00000101464 | 1 | |
| ENSMUSG00000101465 | 1 | |
| ENSMUSG00000101466 | 1 | |
| ENSMUSG00000101467 | 1 | |
| ENSMUSG00000101468 | 1 | |
| ENSMUSG00000101469 | 1 | |
| ENSMUSG00000101470 | 1 | |
| ENSMUSG00000101471 | 1 | |
| ENSMUSG00000101472 | 1 | |
| ENSMUSG00000101473 | 1 | |
| ENSMUSG00000101474 | 1 | |
| ENSMUSG00000101475 | 1 | |
| ENSMUSG00000101476 | -1.043306199 | 0.698112512 |
| ENSMUSG00000101477 | 1 | |
| ENSMUSG00000101478 | 1 | |
| ENSMUSG00000101479 | 1 | |
| ENSMUSG00000101480 | 1 | |
| ENSMUSG00000101481 | -1 | 1 |
| ENSMUSG00000101482 | 1 | |
| ENSMUSG00000101483 | -1.043306199 | 0.79291651 |
| ENSMUSG00000101484 | 1 | |
| ENSMUSG00000101485 | 1 | |
| ENSMUSG00000101486 | 1 | |
| ENSMUSG00000101487 | 1 | |
| ENSMUSG00000101488 | 1 | |
| ENSMUSG00000101489 | 1 | |
| ENSMUSG00000101490 | 1 | |
| ENSMUSG00000101491 | 1 | |
| ENSMUSG00000101492 | 1 | |
| ENSMUSG00000101493 | -1.092280538 | 0.577714817 |
| ENSMUSG00000101494 | 1 | |
| ENSMUSG00000101495 | 1 | |
| ENSMUSG00000101496 | 1.002938927 | 1 |
| ENSMUSG00000101497 | 1 | |
| ENSMUSG00000101498 | 1 | |
| ENSMUSG00000101499 | 1 | |
| ENSMUSG00000101500 | 1 | |
| ENSMUSG00000101501 | 1.005613439 | 1 |
| ENSMUSG00000101502 | 1.005613439 | 1 |
| ENSMUSG00000101503 | 1 | |
| ENSMUSG00000101504 | 1 | |
| ENSMUSG00000101505 | 1.005613439 | 1 |
| ENSMUSG00000101506 | 1 | |

| | | |
|---|---|---|
| ENSMUSG00000101507 | 1 | |
| ENSMUSG00000101508 | 1 | |
| ENSMUSG00000101509 | 1 | |
| ENSMUSG00000101510 | 1 | |
| ENSMUSG00000101511 | 1 | |
| ENSMUSG00000101512 | 1 | |
| ENSMUSG00000101513 | 1 | |
| ENSMUSG00000101514 | 1 | |
| ENSMUSG00000101515 | 1 | |
| ENSMUSG00000101516 | 1 | |
| ENSMUSG00000101517 | -1.113515092 | 0.608497229 |
| ENSMUSG00000101518 | 1 | |
| ENSMUSG00000101519 | -1.053549062 | 0.698112512 |
| ENSMUSG00000101520 | 1 | |
| ENSMUSG00000101521 | 1 | |
| ENSMUSG00000101522 | 1 | |
| ENSMUSG00000101523 | 1 | |
| ENSMUSG00000101525 | 1 | |
| ENSMUSG00000101526 | 1 | |
| ENSMUSG00000101527 | 1 | |
| ENSMUSG00000101528 | 1 | |
| ENSMUSG00000101529 | 1 | |
| ENSMUSG00000101530 | 1 | |
| ENSMUSG00000101531 | 1 | |
| ENSMUSG00000101532 | 1.019083579 | 0.949812939 |
| ENSMUSG00000101533 | 1 | |
| ENSMUSG00000101534 | 1 | |
| ENSMUSG00000101535 | 1 | |
| ENSMUSG00000101536 | 1 | |
| ENSMUSG00000101537 | 1 | |
| ENSMUSG00000101538 | 1 | |
| ENSMUSG00000101539 | 1 | |
| ENSMUSG00000101540 | 1.000503655 | 1 |
| ENSMUSG00000101541 | 1 | |
| ENSMUSG00000101542 | 1 | |
| ENSMUSG00000101543 | 1 | |
| ENSMUSG00000101544 | 1 | |
| ENSMUSG00000101545 | 1.02889509 | 0.965120133 |
| ENSMUSG00000101546 | 1 | |
| ENSMUSG00000101547 | 1 | |
| ENSMUSG00000101548 | 1 | |
| ENSMUSG00000101549 | -1.042990024 | 0.828564184 |
| ENSMUSG00000101550 | 1 | |
| ENSMUSG00000101551 | -1.006483203 | 1 |
| ENSMUSG00000101552 | 1 | |
| ENSMUSG00000101553 | 1 | |
| ENSMUSG00000101554 | 1 | |
| ENSMUSG00000101555 | -1.11061171 | 0.608497229 |
| ENSMUSG00000101556 | -1.126344191 | 0.413646914 |
| ENSMUSG00000101557 | 1 | |
| ENSMUSG00000101558 | 1 | |
| ENSMUSG00000101559 | 1.023743961 | 0.938939969 |
| ENSMUSG00000101560 | 1 | |
| ENSMUSG00000101562 | 1 | |
| ENSMUSG00000101563 | 1 | |
| ENSMUSG00000101564 | -1 | 1 |
| ENSMUSG00000101565 | 1 | |
| ENSMUSG00000101566 | 1 | |
| ENSMUSG00000101567 | -1.028348525 | 0.912425817 |
| ENSMUSG00000101568 | -1.010960635 | 1 |
| ENSMUSG00000101569 | 1 | |
| ENSMUSG00000101570 | 1 | |
| ENSMUSG00000101571 | 1.084618945 | 0.698112512 |

| | | |
|---|---|---|
| ENSMUSG00000101573 | 1 | |
| ENSMUSG00000101574 | 1.132639128 | 0.518443002 |
| ENSMUSG00000101575 | 1 | |
| ENSMUSG00000101576 | 1 | |
| ENSMUSG00000101578 | 1 | |
| ENSMUSG00000101579 | 1 | |
| ENSMUSG00000101580 | 1 | |
| ENSMUSG00000101581 | -1.007792392 | 1 |
| ENSMUSG00000101582 | 1 | |
| ENSMUSG00000101583 | 1 | |
| ENSMUSG00000101584 | 1 | |
| ENSMUSG00000101585 | -1.233355716 | 0.484584722 |
| ENSMUSG00000101586 | 1.11147739 | 0.245293156 |
| ENSMUSG00000101587 | 1.10141562 | 0.60397857 |
| ENSMUSG00000101588 | -1.071075483 | 0.743300655 |
| ENSMUSG00000101589 | -1.019929085 | 0.987476743 |
| ENSMUSG00000101590 | 1 | |
| ENSMUSG00000101591 | 1 | |
| ENSMUSG00000101592 | 1 | |
| ENSMUSG00000101593 | 1 | |
| ENSMUSG00000101594 | 1 | |
| ENSMUSG00000101595 | -1 | 1 |
| ENSMUSG00000101596 | 1 | |
| ENSMUSG00000101597 | 1 | |
| ENSMUSG00000101598 | 1 | |
| ENSMUSG00000101599 | -1.125043226 | 0.367108843 |
| ENSMUSG00000101600 | 1 | |
| ENSMUSG00000101601 | 1 | |
| ENSMUSG00000101602 | 1 | |
| ENSMUSG00000101603 | 1.090272455 | 0.824542865 |
| ENSMUSG00000101605 | 1 | |
| ENSMUSG00000101606 | 1 | |
| ENSMUSG00000101607 | 1 | |
| ENSMUSG00000101608 | 1.128745599 | 0.608497229 |
| ENSMUSG00000101609 | 1.032857503 | 0.895965732 |
| ENSMUSG00000101610 | -1.029302237 | 0.96035552 |
| ENSMUSG00000101611 | 1 | |
| ENSMUSG00000101612 | 1 | |
| ENSMUSG00000101613 | 1 | |
| ENSMUSG00000101614 | 1 | |
| ENSMUSG00000101615 | -1 | 1 |
| ENSMUSG00000101616 | 1 | |
| ENSMUSG00000101617 | 1 | |
| ENSMUSG00000101618 | 1 | |
| ENSMUSG00000101619 | 1 | |
| ENSMUSG00000101621 | 1 | |
| ENSMUSG00000101622 | 1 | |
| ENSMUSG00000101623 | 1 | |
| ENSMUSG00000101624 | 1 | |
| ENSMUSG00000101625 | -1.087357476 | 0.813372967 |
| ENSMUSG00000101626 | 1 | |
| ENSMUSG00000101627 | 1 | |
| ENSMUSG00000101628 | -1.03962336 | 0.698112512 |
| ENSMUSG00000101629 | 1 | |
| ENSMUSG00000101630 | 1.069068979 | 0.608497229 |
| ENSMUSG00000101631 | 1 | |
| ENSMUSG00000101632 | 1 | |
| ENSMUSG00000101633 | 1 | |
| ENSMUSG00000101634 | 1 | |
| ENSMUSG00000101636 | 1 | |
| ENSMUSG00000101637 | 1 | |
| ENSMUSG00000101638 | 1 | |
| ENSMUSG00000101639 | 1.059463094 | 0.608497229 |

| | | |
|---|---|---|
| ENSMUSG00000101640 | 1 | |
| ENSMUSG00000101641 | -1.142727245 | 0.202428693 |
| ENSMUSG00000101642 | 1 | |
| ENSMUSG00000101643 | 1 | |
| ENSMUSG00000101644 | 1 | |
| ENSMUSG00000101645 | 1 | |
| ENSMUSG00000101646 | 1 | |
| ENSMUSG00000101647 | 1 | |
| ENSMUSG00000101648 | 1 | |
| ENSMUSG00000101649 | 1 | |
| ENSMUSG00000101650 | 1 | |
| ENSMUSG00000101651 | 1 | |
| ENSMUSG00000101652 | 1.069267893 | 0.835701362 |
| ENSMUSG00000101653 | 1 | |
| ENSMUSG00000101654 | 1 | |
| ENSMUSG00000101655 | 1.069068979 | 0.698112512 |
| ENSMUSG00000101656 | 1 | |
| ENSMUSG00000101657 | 1 | |
| ENSMUSG00000101658 | 1 | |
| ENSMUSG00000101659 | 1 | |
| ENSMUSG00000101660 | 1 | |
| ENSMUSG00000101661 | -1 | 1 |
| ENSMUSG00000101662 | -1.084645496 | 0.734674233 |
| ENSMUSG00000101664 | 1 | |
| ENSMUSG00000101665 | 1 | |
| ENSMUSG00000101666 | 1 | |
| ENSMUSG00000101667 | 1 | |
| ENSMUSG00000101668 | 1 | |
| ENSMUSG00000101669 | 1 | |
| ENSMUSG00000101670 | 1 | |
| ENSMUSG00000101671 | 1 | |
| ENSMUSG00000101672 | 1 | |
| ENSMUSG00000101673 | 1 | |
| ENSMUSG00000101674 | 1 | |
| ENSMUSG00000101675 | 1 | |
| ENSMUSG00000101676 | 1 | |
| ENSMUSG00000101677 | -1.053549062 | 0.698112512 |
| ENSMUSG00000101678 | -1.029141097 | 0.972895772 |
| ENSMUSG00000101679 | 1 | |
| ENSMUSG00000101680 | 1 | |
| ENSMUSG00000101681 | 1 | |
| ENSMUSG00000101682 | 1.024844571 | 0.854640873 |
| ENSMUSG00000101683 | 1 | |
| ENSMUSG00000101684 | 1 | |
| ENSMUSG00000101685 | 1.059463094 | 0.698112512 |
| ENSMUSG00000101686 | 1 | |
| ENSMUSG00000101687 | 1 | |
| ENSMUSG00000101688 | 1 | |
| ENSMUSG00000101689 | 1 | |
| ENSMUSG00000101690 | 1 | |
| ENSMUSG00000101691 | 1 | |
| ENSMUSG00000101692 | 1 | |
| ENSMUSG00000101693 | -1 | 1 |
| ENSMUSG00000101694 | 1 | |
| ENSMUSG00000101695 | 1 | |
| ENSMUSG00000101696 | 1 | |
| ENSMUSG00000101698 | -1.033272356 | 0.770196474 |
| ENSMUSG00000101699 | 1 | |
| ENSMUSG00000101700 | -1 | 1 |
| ENSMUSG00000101701 | 1.029490685 | 0.94494564 |
| ENSMUSG00000101702 | 1 | |
| ENSMUSG00000101703 | 1 | |
| ENSMUSG00000101704 | 1 | |

| | | |
|---|---|---|
| ENSMUSG00000101705 | 1 | |
| ENSMUSG00000101706 | 1 | |
| ENSMUSG00000101707 | 1 | |
| ENSMUSG00000101708 | 1 | |
| ENSMUSG00000101709 | 1 | |
| ENSMUSG00000101710 | 1 | |
| ENSMUSG00000101711 | -1.266512616 | 0.253300743 |
| ENSMUSG00000101712 | 1 | |
| ENSMUSG00000101713 | 1 | |
| ENSMUSG00000101714 | 1 | |
| ENSMUSG00000101715 | 1 | |
| ENSMUSG00000101716 | 1.132639128 | 0.217803313 |
| ENSMUSG00000101717 | 1 | |
| ENSMUSG00000101718 | 1 | |
| ENSMUSG00000101719 | 1 | |
| ENSMUSG00000101720 | 1 | |
| ENSMUSG00000101721 | 1 | |
| ENSMUSG00000101722 | -1.084645496 | 0.628300894 |
| ENSMUSG00000101723 | 1 | |
| ENSMUSG00000101724 | 1 | |
| ENSMUSG00000101725 | 1 | |
| ENSMUSG00000101726 | 1.069068979 | 0.698112512 |
| ENSMUSG00000101727 | 1 | |
| ENSMUSG00000101728 | 1 | |
| ENSMUSG00000101729 | 1.059463094 | 0.698112512 |
| ENSMUSG00000101730 | 1.122462048 | 0.452329901 |
| ENSMUSG00000101731 | -1.065126268 | 0.832767487 |
| ENSMUSG00000101732 | 1 | |
| ENSMUSG00000101733 | 1 | |
| ENSMUSG00000101734 | 1 | |
| ENSMUSG00000101735 | -1 | 1 |
| ENSMUSG00000101736 | 1.089470641 | 0.657060635 |
| ENSMUSG00000101737 | 1 | |
| ENSMUSG00000101738 | 1 | |
| ENSMUSG00000101739 | 1 | |
| ENSMUSG00000101740 | 1 | |
| ENSMUSG00000101741 | 1 | |
| ENSMUSG00000101742 | 1 | |
| ENSMUSG00000101743 | 1 | |
| ENSMUSG00000101744 | 1 | |
| ENSMUSG00000101745 | 1 | |
| ENSMUSG00000101746 | 1.005613439 | 1 |
| ENSMUSG00000101747 | 1 | |
| ENSMUSG00000101748 | 1 | |
| ENSMUSG00000101750 | 1.059463094 | 0.608497229 |
| ENSMUSG00000101751 | -1.03962336 | 0.814089245 |
| ENSMUSG00000101752 | 1 | |
| ENSMUSG00000101753 | 1 | |
| ENSMUSG00000101754 | 1 | |
| ENSMUSG00000101755 | 1 | |
| ENSMUSG00000101756 | 1 | |
| ENSMUSG00000101757 | 1 | |
| ENSMUSG00000101758 | 1 | |
| ENSMUSG00000101759 | 1 | |
| ENSMUSG00000101760 | 1 | |
| ENSMUSG00000101761 | 1 | |
| ENSMUSG00000101762 | 1 | |
| ENSMUSG00000101763 | 1 | |
| ENSMUSG00000101764 | -1 | 1 |
| ENSMUSG00000101765 | 1 | |
| ENSMUSG00000101766 | 1 | |
| ENSMUSG00000101767 | 1 | |
| ENSMUSG00000101768 | 1.059463094 | 0.776953697 |

| | | |
|---|---|---|
| ENSMUSG00000101769 | 1 | |
| ENSMUSG00000101770 | 1 | |
| ENSMUSG00000101771 | 1 | |
| ENSMUSG00000101772 | 1 | |
| ENSMUSG00000101773 | 1 | |
| ENSMUSG00000101774 | 1 | |
| ENSMUSG00000101775 | 1 | |
| ENSMUSG00000101776 | 1.069657177 | 0.420660281 |
| ENSMUSG00000101777 | 1.069068979 | 0.608497229 |
| ENSMUSG00000101778 | -1.053549062 | 0.862599478 |
| ENSMUSG00000101779 | 1 | |
| ENSMUSG00000101780 | 1 | |
| ENSMUSG00000101781 | 1 | |
| ENSMUSG00000101782 | 1 | |
| ENSMUSG00000101783 | 1 | |
| ENSMUSG00000101784 | 1 | |
| ENSMUSG00000101785 | 1 | |
| ENSMUSG00000101786 | 1 | |
| ENSMUSG00000101787 | 1 | |
| ENSMUSG00000101788 | 1 | |
| ENSMUSG00000101789 | 1 | |
| ENSMUSG00000101790 | 1 | |
| ENSMUSG00000101791 | 1.101111538 | 0.771926153 |
| ENSMUSG00000101792 | -1.043306199 | 0.698112512 |
| ENSMUSG00000101793 | 1 | |
| ENSMUSG00000101794 | 1 | |
| ENSMUSG00000101795 | 1.131324033 | 0.630148234 |
| ENSMUSG00000101796 | 1 | |
| ENSMUSG00000101797 | 1 | |
| ENSMUSG00000101798 | 1.059463094 | 0.608497229 |
| ENSMUSG00000101799 | 1.132639128 | 0.428732467 |
| ENSMUSG00000101800 | 1 | |
| ENSMUSG00000101801 | -1 | 1 |
| ENSMUSG00000101802 | 1 | |
| ENSMUSG00000101803 | 1 | |
| ENSMUSG00000101804 | 1 | |
| ENSMUSG00000101805 | 1 | |
| ENSMUSG00000101806 | 1 | |
| ENSMUSG00000101807 | 1 | |
| ENSMUSG00000101808 | 1 | |
| ENSMUSG00000101809 | 1 | |
| ENSMUSG00000101810 | -1.03962336 | 0.865094641 |
| ENSMUSG00000101811 | 1 | |
| ENSMUSG00000101812 | 1 | |
| ENSMUSG00000101813 | 1 | |
| ENSMUSG00000101814 | 1 | |
| ENSMUSG00000101815 | 1 | |
| ENSMUSG00000101816 | 1 | |
| ENSMUSG00000101817 | 1 | |
| ENSMUSG00000101818 | 1 | |
| ENSMUSG00000101819 | 1 | |
| ENSMUSG00000101820 | 1 | |
| ENSMUSG00000101821 | 1 | |
| ENSMUSG00000101822 | 1 | |
| ENSMUSG00000101823 | 1.036778919 | 0.698112512 |
| ENSMUSG00000101824 | 1 | |
| ENSMUSG00000101825 | 1 | |
| ENSMUSG00000101826 | 1.059463094 | 0.776953697 |
| ENSMUSG00000101827 | -1.043306199 | 0.698112512 |
| ENSMUSG00000101828 | -1.136604631 | 0.757318144 |
| ENSMUSG00000101829 | 1 | |
| ENSMUSG00000101830 | 1 | |
| ENSMUSG00000101831 | 1 | |

| | | |
|---|---|---|
| ENSMUSG00000101832 | -1.03962336 | 0.698112512 |
| ENSMUSG00000101833 | 1 | |
| ENSMUSG00000101834 | 1 | |
| ENSMUSG00000101835 | 1 | |
| ENSMUSG00000101836 | 1 | |
| ENSMUSG00000101837 | 1 | |
| ENSMUSG00000101838 | 1.024656056 | 0.973061065 |
| ENSMUSG00000101839 | 1 | |
| ENSMUSG00000101840 | 1 | |
| ENSMUSG00000101841 | -1.071075483 | 0.743300655 |
| ENSMUSG00000101842 | 1 | |
| ENSMUSG00000101843 | 1 | |
| ENSMUSG00000101844 | 1 | |
| ENSMUSG00000101845 | -1.022105482 | 1 |
| ENSMUSG00000101846 | 1 | |
| ENSMUSG00000101847 | 1 | |
| ENSMUSG00000101848 | 1.122462048 | 0.654902646 |
| ENSMUSG00000101849 | 1.30736793 | 0.367760887 |
| ENSMUSG00000101850 | 1 | |
| ENSMUSG00000101851 | 1 | |
| ENSMUSG00000101852 | 1 | |
| ENSMUSG00000101853 | 1 | |
| ENSMUSG00000101854 | 1 | |
| ENSMUSG00000101855 | 1 | |
| ENSMUSG00000101856 | -1.050935112 | 0.835182707 |
| ENSMUSG00000101857 | 1 | |
| ENSMUSG00000101858 | 1 | |
| ENSMUSG00000101859 | 1.059033181 | 0.848848499 |
| ENSMUSG00000101860 | 1 | |
| ENSMUSG00000101861 | 1 | |
| ENSMUSG00000101862 | -1.044273782 | 0.698112512 |
| ENSMUSG00000101863 | 1 | |
| ENSMUSG00000101864 | 1 | |
| ENSMUSG00000101865 | 1 | |
| ENSMUSG00000101866 | 1 | |
| ENSMUSG00000101867 | 1 | |
| ENSMUSG00000101868 | 1 | |
| ENSMUSG00000101869 | -1 | 1 |
| ENSMUSG00000101870 | 1 | |
| ENSMUSG00000101871 | 1 | |
| ENSMUSG00000101872 | 1.173096674 | 0.438463556 |
| ENSMUSG00000101873 | 1 | |
| ENSMUSG00000101874 | 1 | |
| ENSMUSG00000101875 | 1.035626983 | 0.961487493 |
| ENSMUSG00000101877 | 1.059463094 | 0.608497229 |
| ENSMUSG00000101878 | 1 | |
| ENSMUSG00000101879 | 1 | |
| ENSMUSG00000101880 | -1.053549062 | 0.698112512 |
| ENSMUSG00000101881 | 1 | |
| ENSMUSG00000101882 | 1 | |
| ENSMUSG00000101883 | 1 | |
| ENSMUSG00000101884 | 1 | |
| ENSMUSG00000101885 | 1 | |
| ENSMUSG00000101886 | 1 | |
| ENSMUSG00000101887 | 1 | |
| ENSMUSG00000101888 | 1 | |
| ENSMUSG00000101890 | 1 | |
| ENSMUSG00000101891 | 1 | |
| ENSMUSG00000101892 | 1.048670452 | 0.629924473 |
| ENSMUSG00000101893 | 1 | |
| ENSMUSG00000101894 | 1 | |
| ENSMUSG00000101895 | -1.053549062 | 0.698112512 |
| ENSMUSG00000101896 | 1.225079193 | 0.132613684 |

| | | |
|---|---|---|
| ENSMUSG00000101897 | 1 | |
| ENSMUSG00000101898 | 1 | |
| ENSMUSG00000101899 | 1 | |
| ENSMUSG00000101900 | 1 | |
| ENSMUSG00000101901 | -1 | 1 |
| ENSMUSG00000101902 | 1 | |
| ENSMUSG00000101903 | 1 | |
| ENSMUSG00000101904 | -1 | 1 |
| ENSMUSG00000101905 | 1 | |
| ENSMUSG00000101906 | 1.018298777 | 0.978763926 |
| ENSMUSG00000101907 | 1 | |
| ENSMUSG00000101908 | 1 | |
| ENSMUSG00000101909 | 1 | |
| ENSMUSG00000101910 | 1 | |
| ENSMUSG00000101911 | 1 | |
| ENSMUSG00000101912 | 1 | |
| ENSMUSG00000101913 | 1 | |
| ENSMUSG00000101914 | 1 | |
| ENSMUSG00000101915 | 1 | |
| ENSMUSG00000101916 | 1 | |
| ENSMUSG00000101917 | 1 | |
| ENSMUSG00000101918 | 1 | |
| ENSMUSG00000101919 | 1 | |
| ENSMUSG00000101920 | 1 | |
| ENSMUSG00000101921 | 1 | |
| ENSMUSG00000101922 | 1 | |
| ENSMUSG00000101923 | 1 | |
| ENSMUSG00000101924 | 1 | |
| ENSMUSG00000101925 | 1.122462048 | 0.628993873 |
| ENSMUSG00000101926 | 1 | |
| ENSMUSG00000101927 | 1 | |
| ENSMUSG00000101928 | 1 | |
| ENSMUSG00000101929 | 1 | |
| ENSMUSG00000101930 | 1 | |
| ENSMUSG00000101931 | 1 | |
| ENSMUSG00000101932 | 1.059463094 | 0.608497229 |
| ENSMUSG00000101933 | 1 | |
| ENSMUSG00000101934 | 1 | |
| ENSMUSG00000101935 | 1 | |
| ENSMUSG00000101936 | 1 | |
| ENSMUSG00000101937 | -1 | 1 |
| ENSMUSG00000101938 | 1 | |
| ENSMUSG00000101939 | 1.077008891 | 0.489504781 |
| ENSMUSG00000101940 | 1 | |
| ENSMUSG00000101941 | 1 | |
| ENSMUSG00000101942 | 1 | |
| ENSMUSG00000101943 | 1 | |
| ENSMUSG00000101944 | 1 | |
| ENSMUSG00000101945 | -1 | 1 |
| ENSMUSG00000101946 | 1 | |
| ENSMUSG00000101947 | 1 | |
| ENSMUSG00000101948 | 1 | |
| ENSMUSG00000101949 | 1 | |
| ENSMUSG00000101950 | 1 | |
| ENSMUSG00000101951 | 1 | |
| ENSMUSG00000101952 | -1.132666855 | 0.18851436 |
| ENSMUSG00000101953 | 1.172621325 | 0.591115055 |
| ENSMUSG00000101954 | 1 | |
| ENSMUSG00000101955 | -1 | 1 |
| ENSMUSG00000101956 | 1 | |
| ENSMUSG00000101957 | 1 | |
| ENSMUSG00000101958 | -1.133217178 | 0.577958765 |
| ENSMUSG00000101959 | 1 | |

| | | |
|---|---|---|
| ENSMUSG00000101960 | 1 | |
| ENSMUSG00000101961 | 1 | |
| ENSMUSG00000101962 | 1 | |
| ENSMUSG00000101963 | 1.045347292 | 0.802061488 |
| ENSMUSG00000101964 | -1 | 1 |
| ENSMUSG00000101965 | 1.059463094 | 0.638307772 |
| ENSMUSG00000101966 | -1.101638832 | 0.66115023 |
| ENSMUSG00000101967 | 1 | |
| ENSMUSG00000101968 | 1 | |
| ENSMUSG00000101969 | 1.059463094 | 0.626808553 |
| ENSMUSG00000101970 | 1.169798189 | 0.113467919 |
| ENSMUSG00000101971 | 1 | |
| ENSMUSG00000101972 | 1 | |
| ENSMUSG00000101973 | 1.111812527 | 0.629725675 |
| ENSMUSG00000101974 | 1 | |
| ENSMUSG00000101975 | 1 | |
| ENSMUSG00000101976 | 1 | |
| ENSMUSG00000101977 | 1 | |
| ENSMUSG00000101978 | 1 | |
| ENSMUSG00000101980 | 1 | |
| ENSMUSG00000101981 | 1 | |
| ENSMUSG00000101982 | 1 | |
| ENSMUSG00000101983 | 1 | |
| ENSMUSG00000101984 | 1 | |
| ENSMUSG00000101985 | 1 | |
| ENSMUSG00000101986 | -1 | 1 |
| ENSMUSG00000101987 | 1 | |
| ENSMUSG00000101988 | 1 | |
| ENSMUSG00000101989 | 1 | |
| ENSMUSG00000101990 | 1 | |
| ENSMUSG00000101991 | 1 | |
| ENSMUSG00000101992 | 1 | |
| ENSMUSG00000101993 | 1 | |
| ENSMUSG00000101994 | 1 | |
| ENSMUSG00000101995 | -1.0889222 | 0.745531957 |
| ENSMUSG00000101996 | 1 | |
| ENSMUSG00000101997 | 1 | |
| ENSMUSG00000101998 | 1 | |
| ENSMUSG00000102000 | 1 | |
| ENSMUSG00000102001 | 1 | |
| ENSMUSG00000102002 | 1 | |
| ENSMUSG00000102003 | 1 | |
| ENSMUSG00000102004 | 1 | |
| ENSMUSG00000102005 | 1 | |
| ENSMUSG00000102006 | 1 | |
| ENSMUSG00000102008 | 1 | |
| ENSMUSG00000102009 | 1.033906491 | 0.90876612 |
| ENSMUSG00000102010 | 1 | |
| ENSMUSG00000102011 | 1 | |
| ENSMUSG00000102012 | -1.044273782 | 0.698112512 |
| ENSMUSG00000102013 | -1.053549062 | 0.825517832 |
| ENSMUSG00000102014 | -1.09174665 | 0.771903504 |
| ENSMUSG00000102015 | 1 | |
| ENSMUSG00000102016 | 1 | |
| ENSMUSG00000102017 | 1 | |
| ENSMUSG00000102018 | -1.330775927 | 0.008618666 |
| ENSMUSG00000102019 | 1 | |
| ENSMUSG00000102020 | 1 | |
| ENSMUSG00000102022 | 1 | |
| ENSMUSG00000102023 | 1 | |
| ENSMUSG00000102025 | 1 | |
| ENSMUSG00000102026 | 1 | |
| ENSMUSG00000102027 | 1.128878069 | 0.608497229 |

| | | |
|---|---|---|
| ENSMUSG00000102028 | 1 | |
| ENSMUSG00000102029 | 1 | |
| ENSMUSG00000102030 | -1.012475031 | 1 |
| ENSMUSG00000102031 | 1 | |
| ENSMUSG00000102032 | 1 | |
| ENSMUSG00000102033 | 1 | |
| ENSMUSG00000102035 | 1 | |
| ENSMUSG00000102036 | 1 | |
| ENSMUSG00000102037 | 1 | |
| ENSMUSG00000102038 | 1.072209197 | 0.849501316 |
| ENSMUSG00000102039 | 1 | |
| ENSMUSG00000102041 | 1 | |
| ENSMUSG00000102042 | 1 | |
| ENSMUSG00000102043 | 1 | |
| ENSMUSG00000102044 | 1 | |
| ENSMUSG00000102045 | 1 | |
| ENSMUSG00000102046 | -1.044273782 | 0.698112512 |
| ENSMUSG00000102047 | 1 | |
| ENSMUSG00000102048 | -1.034379482 | 0.966523329 |
| ENSMUSG00000102049 | 1 | |
| ENSMUSG00000102050 | 1 | |
| ENSMUSG00000102051 | 1.089470641 | 0.608497229 |
| ENSMUSG00000102052 | -1 | 1 |
| ENSMUSG00000102053 | 1 | |
| ENSMUSG00000102054 | 1 | |
| ENSMUSG00000102055 | 1.114809048 | 0.698112512 |
| ENSMUSG00000102056 | 1 | |
| ENSMUSG00000102057 | 1 | |
| ENSMUSG00000102058 | 1 | |
| ENSMUSG00000102059 | -1.001609404 | 1 |
| ENSMUSG00000102060 | 1 | |
| ENSMUSG00000102061 | 1 | |
| ENSMUSG00000102062 | 1 | |
| ENSMUSG00000102063 | 1 | |
| ENSMUSG00000102064 | 1.04328066 | 0.86119789 |
| ENSMUSG00000102065 | 1 | |
| ENSMUSG00000102067 | 1 | |
| ENSMUSG00000102068 | 1 | |
| ENSMUSG00000102069 | 1 | |
| ENSMUSG00000102070 | 1.056280798 | 0.857309537 |
| ENSMUSG00000102071 | 1 | |
| ENSMUSG00000102072 | 1 | |
| ENSMUSG00000102073 | 1 | |
| ENSMUSG00000102074 | 1 | |
| ENSMUSG00000102075 | -1 | 1 |
| ENSMUSG00000102076 | 1 | |
| ENSMUSG00000102077 | 1 | |
| ENSMUSG00000102078 | 1 | |
| ENSMUSG00000102079 | 1 | |
| ENSMUSG00000102080 | 1 | |
| ENSMUSG00000102081 | 1 | |
| ENSMUSG00000102082 | 1 | |
| ENSMUSG00000102083 | -1.044273782 | 0.698112512 |
| ENSMUSG00000102084 | 1 | |
| ENSMUSG00000102085 | -1.161737199 | 0.592803883 |
| ENSMUSG00000102087 | 1 | |
| ENSMUSG00000102088 | 1 | |
| ENSMUSG00000102090 | 1.059463094 | 0.704170349 |
| ENSMUSG00000102091 | 1 | |
| ENSMUSG00000102092 | 1 | |
| ENSMUSG00000102093 | -1 | 1 |
| ENSMUSG00000102095 | -1.089497311 | 0.608497229 |
| ENSMUSG00000102096 | 1 | |

| | | |
|---|---|---|
| ENSMUSG00000102097 | 1.090507733 | 0.777977356 |
| ENSMUSG00000102098 | 1 | |
| ENSMUSG00000102099 | 1 | |
| ENSMUSG00000102100 | 1 | |
| ENSMUSG00000102101 | 1.08131387 | 0.733978911 |
| ENSMUSG00000102102 | 1 | |
| ENSMUSG00000102103 | 1 | |
| ENSMUSG00000102104 | 1 | |
| ENSMUSG00000102105 | 1 | |
| ENSMUSG00000102106 | 1 | |
| ENSMUSG00000102107 | 1 | |
| ENSMUSG00000102108 | 1 | |
| ENSMUSG00000102109 | 1 | |
| ENSMUSG00000102110 | 1 | |
| ENSMUSG00000102111 | 1 | |
| ENSMUSG00000102112 | -1.24499826 | 0.085979471 |
| ENSMUSG00000102113 | 1 | |
| ENSMUSG00000102114 | 1 | |
| ENSMUSG00000102115 | 1 | |
| ENSMUSG00000102116 | 1 | |
| ENSMUSG00000102117 | 1.003904005 | 1 |
| ENSMUSG00000102118 | 1 | |
| ENSMUSG00000102119 | 1 | |
| ENSMUSG00000102120 | 1 | |
| ENSMUSG00000102121 | 1 | |
| ENSMUSG00000102122 | 1 | |
| ENSMUSG00000102123 | 1.059463094 | 0.608497229 |
| ENSMUSG00000102124 | 1 | |
| ENSMUSG00000102125 | 1 | |
| ENSMUSG00000102126 | 1 | |
| ENSMUSG00000102127 | 1 | |
| ENSMUSG00000102128 | 1 | |
| ENSMUSG00000102129 | 1 | |
| ENSMUSG00000102130 | 1 | |
| ENSMUSG00000102131 | 1 | |
| ENSMUSG00000102132 | 1 | |
| ENSMUSG00000102133 | 1.141544718 | 0.698112512 |
| ENSMUSG00000102134 | 1.122933516 | 0.576055627 |
| ENSMUSG00000102135 | 1.062291814 | 0.867753156 |
| ENSMUSG00000102136 | -1 | 1 |
| ENSMUSG00000102137 | 1.20697895 | 0.308657915 |
| ENSMUSG00000102138 | 1 | |
| ENSMUSG00000102139 | -1.03962336 | 0.803072249 |
| ENSMUSG00000102140 | 1.132639128 | 0.428732467 |
| ENSMUSG00000102141 | -1.03962336 | 0.698112512 |
| ENSMUSG00000102142 | -1.019108526 | 0.968937442 |
| ENSMUSG00000102143 | 1.108453197 | 0.534887579 |
| ENSMUSG00000102144 | 1 | |
| ENSMUSG00000102145 | 1.054154852 | 0.823872552 |
| ENSMUSG00000102146 | 1 | |
| ENSMUSG00000102147 | 1 | |
| ENSMUSG00000102148 | 1.059463094 | 0.608497229 |
| ENSMUSG00000102149 | 1 | |
| ENSMUSG00000102150 | 1 | |
| ENSMUSG00000102151 | -1.044273782 | 0.756135604 |
| ENSMUSG00000102152 | -1.505705698 | 0.03119947 |
| ENSMUSG00000102153 | 1.00177232 | 1 |
| ENSMUSG00000102154 | 1 | |
| ENSMUSG00000102155 | 1 | |
| ENSMUSG00000102156 | 1 | |
| ENSMUSG00000102157 | -1.000405949 | 1 |
| ENSMUSG00000102158 | 1.045402873 | 0.917209568 |
| ENSMUSG00000102159 | 1 | |

| | | |
|---|---|---|
| ENSMUSG00000102160 | 1 | |
| ENSMUSG00000102162 | -1.111221915 | 0.619554344 |
| ENSMUSG00000102163 | 1 | |
| ENSMUSG00000102164 | 1 | |
| ENSMUSG00000102166 | 1 | |
| ENSMUSG00000102167 | 1 | |
| ENSMUSG00000102168 | 1 | |
| ENSMUSG00000102169 | 1 | |
| ENSMUSG00000102170 | 1 | |
| ENSMUSG00000102171 | -1.27842502 | 0.137422973 |
| ENSMUSG00000102172 | -1.043306199 | 0.698112512 |
| ENSMUSG00000102173 | 1 | |
| ENSMUSG00000102174 | 1 | |
| ENSMUSG00000102175 | -1 | 1 |
| ENSMUSG00000102176 | 1 | |
| ENSMUSG00000102177 | 1 | |
| ENSMUSG00000102178 | 1 | |
| ENSMUSG00000102179 | -1.066969463 | 0.871479109 |
| ENSMUSG00000102180 | 1 | |
| ENSMUSG00000102181 | 1 | |
| ENSMUSG00000102182 | 1 | |
| ENSMUSG00000102183 | -1.119233103 | 0.73688717 |
| ENSMUSG00000102184 | 1 | |
| ENSMUSG00000102185 | 1 | |
| ENSMUSG00000102186 | 1.015486245 | 0.985522133 |
| ENSMUSG00000102187 | 1 | |
| ENSMUSG00000102188 | 1 | |
| ENSMUSG00000102189 | -1.22331357 | 0.385772778 |
| ENSMUSG00000102190 | 1 | |
| ENSMUSG00000102191 | -1 | 1 |
| ENSMUSG00000102192 | -1 | 1 |
| ENSMUSG00000102193 | -1.038383956 | 0.93969881 |
| ENSMUSG00000102194 | 1 | |
| ENSMUSG00000102195 | 1 | |
| ENSMUSG00000102196 | -1.043306199 | 0.698112512 |
| ENSMUSG00000102197 | -1.129231493 | 0.605619163 |
| ENSMUSG00000102198 | 1 | |
| ENSMUSG00000102199 | 1 | |
| ENSMUSG00000102200 | 1.132639128 | 0.217803313 |
| ENSMUSG00000102201 | 1 | |
| ENSMUSG00000102202 | -1.239367438 | 0.047347626 |
| ENSMUSG00000102203 | 1 | |
| ENSMUSG00000102204 | -1.130826364 | 0.709538083 |
| ENSMUSG00000102205 | 1 | |
| ENSMUSG00000102206 | 1 | |
| ENSMUSG00000102207 | 1.049517471 | 0.837704961 |
| ENSMUSG00000102208 | 1 | |
| ENSMUSG00000102211 | 1.084618945 | 0.614071182 |
| ENSMUSG00000102212 | 1.059463094 | 0.698112512 |
| ENSMUSG00000102213 | 1 | |
| ENSMUSG00000102214 | 1 | |
| ENSMUSG00000102215 | 1 | |
| ENSMUSG00000102216 | 1 | |
| ENSMUSG00000102217 | 1 | |
| ENSMUSG00000102218 | 1 | |
| ENSMUSG00000102219 | -1 | 1 |
| ENSMUSG00000102220 | -1.047558106 | 0.721007497 |
| ENSMUSG00000102221 | 1 | |
| ENSMUSG00000102222 | 1 | |
| ENSMUSG00000102223 | 1 | |
| ENSMUSG00000102224 | -1.260227528 | 0.023776852 |
| ENSMUSG00000102225 | -1.045919705 | 0.890893742 |
| ENSMUSG00000102227 | 1 | |

| | | |
|---|---|---|
| ENSMUSG00000102229 | 1 | |
| ENSMUSG00000102230 | 1 | |
| ENSMUSG00000102231 | 1 | |
| ENSMUSG00000102232 | 1 | |
| ENSMUSG00000102233 | 1 | |
| ENSMUSG00000102234 | 1.132639128 | 0.217803313 |
| ENSMUSG00000102235 | -1.033591808 | 0.964694571 |
| ENSMUSG00000102236 | 1 | |
| ENSMUSG00000102237 | 1 | |
| ENSMUSG00000102238 | 1 | |
| ENSMUSG00000102239 | 1 | |
| ENSMUSG00000102240 | 1 | |
| ENSMUSG00000102241 | 1.061592237 | 0.868022945 |
| ENSMUSG00000102242 | 1 | |
| ENSMUSG00000102243 | -1.107993947 | 0.721386781 |
| ENSMUSG00000102244 | 1.069068979 | 0.608497229 |
| ENSMUSG00000102245 | 1 | |
| ENSMUSG00000102246 | -1.144336658 | 0.608497229 |
| ENSMUSG00000102247 | 1 | |
| ENSMUSG00000102248 | 1.084618945 | 0.644224767 |
| ENSMUSG00000102249 | 1 | |
| ENSMUSG00000102250 | 1 | |
| ENSMUSG00000102251 | 1 | |
| ENSMUSG00000102252 | 1.024066607 | 0.924061737 |
| ENSMUSG00000102253 | 1 | |
| ENSMUSG00000102254 | -1.053549062 | 0.807892349 |
| ENSMUSG00000102255 | 1 | |
| ENSMUSG00000102256 | 1 | |
| ENSMUSG00000102257 | 1 | |
| ENSMUSG00000102258 | -1.036601951 | 0.854116965 |
| ENSMUSG00000102259 | -1.03962336 | 0.698112512 |
| ENSMUSG00000102260 | 1 | |
| ENSMUSG00000102261 | 1 | |
| ENSMUSG00000102262 | 1 | |
| ENSMUSG00000102263 | 1 | |
| ENSMUSG00000102264 | 1 | |
| ENSMUSG00000102265 | 1 | |
| ENSMUSG00000102266 | -1.071075483 | 0.698112512 |
| ENSMUSG00000102267 | -1.278512347 | 0.17272194 |
| ENSMUSG00000102268 | 1.063936254 | 0.802143138 |
| ENSMUSG00000102269 | 1.059463094 | 0.776953697 |
| ENSMUSG00000102270 | 1 | |
| ENSMUSG00000102271 | 1 | |
| ENSMUSG00000102272 | 1 | |
| ENSMUSG00000102273 | 1 | |
| ENSMUSG00000102274 | 1 | |
| ENSMUSG00000102275 | -1.031974021 | 0.943010905 |
| ENSMUSG00000102276 | 1 | |
| ENSMUSG00000102277 | 1 | |
| ENSMUSG00000102278 | -1.023209019 | 0.94193565 |
| ENSMUSG00000102279 | 1 | |
| ENSMUSG00000102280 | -1 | 1 |
| ENSMUSG00000102281 | 1 | |
| ENSMUSG00000102282 | -1.132666855 | 0.18851436 |
| ENSMUSG00000102283 | 1 | |
| ENSMUSG00000102284 | 1 | |
| ENSMUSG00000102285 | 1 | |
| ENSMUSG00000102286 | 1 | |
| ENSMUSG00000102287 | 1 | |
| ENSMUSG00000102288 | 1 | |
| ENSMUSG00000102289 | 1.027595792 | 0.813372967 |
| ENSMUSG00000102290 | 1 | |
| ENSMUSG00000102291 | 1 | |

| | | |
|---|---|---|
| ENSMUSG00000102293 | 1.059463094 | 0.608497229 |
| ENSMUSG00000102294 | 1 | |
| ENSMUSG00000102295 | 1 | |
| ENSMUSG00000102296 | 1 | |
| ENSMUSG00000102297 | -1.11061171 | 0.608497229 |
| ENSMUSG00000102298 | -1.008594059 | 1 |
| ENSMUSG00000102299 | 1.132639128 | 0.217803313 |
| ENSMUSG00000102300 | -1.023769022 | 0.934760809 |
| ENSMUSG00000102301 | 1 | |
| ENSMUSG00000102302 | 1.086647918 | 0.608497229 |
| ENSMUSG00000102303 | 1 | |
| ENSMUSG00000102304 | 1 | |
| ENSMUSG00000102305 | -1.065301822 | 0.608497229 |
| ENSMUSG00000102306 | 1 | |
| ENSMUSG00000102307 | 1.166665754 | 0.309330961 |
| ENSMUSG00000102308 | 1 | |
| ENSMUSG00000102309 | 1 | |
| ENSMUSG00000102310 | 1 | |
| ENSMUSG00000102311 | -1 | 1 |
| ENSMUSG00000102312 | 1 | |
| ENSMUSG00000102313 | 1 | |
| ENSMUSG00000102314 | 1 | |
| ENSMUSG00000102315 | 1 | |
| ENSMUSG00000102316 | 1 | |
| ENSMUSG00000102317 | -1.004754033 | 1 |
| ENSMUSG00000102318 | 1 | |
| ENSMUSG00000102319 | -1.064499817 | 0.867115751 |
| ENSMUSG00000102320 | 1 | |
| ENSMUSG00000102321 | -1.053549062 | 0.698112512 |
| ENSMUSG00000102322 | 1 | |
| ENSMUSG00000102323 | -1.181219193 | 0.262156296 |
| ENSMUSG00000102324 | -1.009098802 | 1 |
| ENSMUSG00000102325 | -1.024347651 | 0.966523329 |
| ENSMUSG00000102326 | -1.002523738 | 1 |
| ENSMUSG00000102327 | 1 | |
| ENSMUSG00000102328 | 1 | |
| ENSMUSG00000102329 | 1.059463094 | 0.608497229 |
| ENSMUSG00000102330 | 1.000859464 | 1 |
| ENSMUSG00000102331 | -1.177267787 | 0.433615344 |
| ENSMUSG00000102332 | -1.084645496 | 0.532369657 |
| ENSMUSG00000102333 | 1 | |
| ENSMUSG00000102334 | 1 | |
| ENSMUSG00000102335 | 1 | |
| ENSMUSG00000102336 | 1.360719908 | 0.00292437 |
| ENSMUSG00000102337 | 1 | |
| ENSMUSG00000102338 | 1 | |
| ENSMUSG00000102339 | 1 | |
| ENSMUSG00000102340 | -1.11455472 | 0.598655292 |
| ENSMUSG00000102341 | 1 | |
| ENSMUSG00000102342 | 1 | |
| ENSMUSG00000102343 | -1.028712946 | 0.914456038 |
| ENSMUSG00000102344 | 1 | |
| ENSMUSG00000102345 | 1 | |
| ENSMUSG00000102346 | 1 | |
| ENSMUSG00000102347 | 1 | |
| ENSMUSG00000102348 | 1 | |
| ENSMUSG00000102349 | 1 | |
| ENSMUSG00000102350 | 1 | |
| ENSMUSG00000102352 | 1 | |
| ENSMUSG00000102353 | -1.011244717 | 1 |
| ENSMUSG00000102354 | 1 | |
| ENSMUSG00000102355 | 1 | |
| ENSMUSG00000102356 | 1 | |

| | | |
|---|---|---|
| ENSMUSG00000102357 | -1 | 1 |
| ENSMUSG00000102358 | 1 | |
| ENSMUSG00000102359 | -1.03962336 | 0.698112512 |
| ENSMUSG00000102360 | -1.03962336 | 0.803072249 |
| ENSMUSG00000102362 | -1.03962336 | 0.698112512 |
| ENSMUSG00000102363 | 1 | |
| ENSMUSG00000102364 | 1 | |
| ENSMUSG00000102365 | 1.033084247 | 0.96035552 |
| ENSMUSG00000102366 | 1 | |
| ENSMUSG00000102367 | 1 | |
| ENSMUSG00000102368 | 1.059463094 | 0.608497229 |
| ENSMUSG00000102369 | -1.071075483 | 0.698112512 |
| ENSMUSG00000102370 | 1.117589842 | 0.631445633 |
| ENSMUSG00000102371 | -1.043306199 | 0.698112512 |
| ENSMUSG00000102372 | -1.043306199 | 0.698112512 |
| ENSMUSG00000102373 | 1 | |
| ENSMUSG00000102374 | 1.151570665 | 0.620873309 |
| ENSMUSG00000102375 | 1.019083579 | 0.982175268 |
| ENSMUSG00000102376 | -1.0795912 | 0.642639207 |
| ENSMUSG00000102377 | 1 | |
| ENSMUSG00000102378 | 1 | |
| ENSMUSG00000102379 | 1.019083579 | 0.949812939 |
| ENSMUSG00000102380 | -1.085651418 | 0.608497229 |
| ENSMUSG00000102381 | 1 | |
| ENSMUSG00000102382 | 1 | |
| ENSMUSG00000102383 | -1.010800322 | 1 |
| ENSMUSG00000102384 | -1.083312429 | 0.772049403 |
| ENSMUSG00000102385 | 1 | |
| ENSMUSG00000102386 | 1.059463094 | 0.776953697 |
| ENSMUSG00000102387 | 1 | |
| ENSMUSG00000102388 | 1 | |
| ENSMUSG00000102389 | -1.03962336 | 0.698112512 |
| ENSMUSG00000102390 | 1 | |
| ENSMUSG00000102391 | 1 | |
| ENSMUSG00000102392 | 1 | |
| ENSMUSG00000102393 | 1 | |
| ENSMUSG00000102394 | -1.089497311 | 0.532369657 |
| ENSMUSG00000102395 | 1 | |
| ENSMUSG00000102396 | 1 | |
| ENSMUSG00000102397 | 1 | |
| ENSMUSG00000102398 | -1.100193663 | 0.534887579 |
| ENSMUSG00000102399 | -1 | 1 |
| ENSMUSG00000102400 | 1 | |
| ENSMUSG00000102401 | 1.019083579 | 0.982838256 |
| ENSMUSG00000102402 | 1 | |
| ENSMUSG00000102403 | -1.051444952 | 0.874341108 |
| ENSMUSG00000102404 | 1.134692666 | 0.538079305 |
| ENSMUSG00000102405 | 1.069068979 | 0.608497229 |
| ENSMUSG00000102406 | 1 | |
| ENSMUSG00000102407 | 1 | |
| ENSMUSG00000102408 | 1 | |
| ENSMUSG00000102409 | 1 | |
| ENSMUSG00000102410 | 1 | |
| ENSMUSG00000102411 | 1.007513816 | 0.992856012 |
| ENSMUSG00000102413 | 1.069068979 | 0.608497229 |
| ENSMUSG00000102414 | -1.041213731 | 0.943663271 |
| ENSMUSG00000102415 | 1.153245452 | 0.608497229 |
| ENSMUSG00000102416 | 1 | |
| ENSMUSG00000102417 | 1 | |
| ENSMUSG00000102418 | 1.03167193 | 0.988324779 |
| ENSMUSG00000102419 | 1.069068979 | 0.608497229 |
| ENSMUSG00000102420 | 1 | |
| ENSMUSG00000102421 | 1 | |

| | | |
|---|---|---|
| ENSMUSG00000102422 | 1 | |
| ENSMUSG00000102423 | -1.071075483 | 0.743300655 |
| ENSMUSG00000102424 | 1 | |
| ENSMUSG00000102425 | -1.020379047 | 0.986394584 |
| ENSMUSG00000102426 | -1.173336674 | 0.13093628 |
| ENSMUSG00000102427 | 1 | |
| ENSMUSG00000102428 | -1.125166966 | 0.470547213 |
| ENSMUSG00000102429 | 1 | |
| ENSMUSG00000102430 | -1 | 1 |
| ENSMUSG00000102431 | 1 | |
| ENSMUSG00000102432 | 1 | |
| ENSMUSG00000102433 | 1.246784702 | 0.481515508 |
| ENSMUSG00000102434 | -1.03962336 | 0.887262126 |
| ENSMUSG00000102436 | 1.02950269 | 0.924796887 |
| ENSMUSG00000102437 | 1.339717816 | 0.059543567 |
| ENSMUSG00000102438 | 1 | |
| ENSMUSG00000102439 | 1 | |
| ENSMUSG00000102440 | 1.914280496 | 0.001014855 |
| ENSMUSG00000102441 | 1 | |
| ENSMUSG00000102442 | 1 | |
| ENSMUSG00000102443 | 1.050591337 | 0.79291651 |
| ENSMUSG00000102445 | 1 | |
| ENSMUSG00000102446 | 1 | |
| ENSMUSG00000102447 | 1.089470641 | 0.698112512 |
| ENSMUSG00000102448 | 1.085624843 | 0.608497229 |
| ENSMUSG00000102449 | 1 | |
| ENSMUSG00000102450 | 1 | |
| ENSMUSG00000102451 | -1.069502966 | 0.698112512 |
| ENSMUSG00000102452 | 1.252654605 | 0.408924423 |
| ENSMUSG00000102453 | 1 | |
| ENSMUSG00000102456 | 1 | |
| ENSMUSG00000102457 | 1.132639128 | 0.398767938 |
| ENSMUSG00000102458 | 1 | |
| ENSMUSG00000102459 | 1.014545335 | 0.979002724 |
| ENSMUSG00000102460 | 1 | |
| ENSMUSG00000102461 | 1.10141562 | 0.457437655 |
| ENSMUSG00000102462 | 1 | |
| ENSMUSG00000102463 | 1.069068979 | 0.608497229 |
| ENSMUSG00000102464 | 1 | |
| ENSMUSG00000102465 | 1.010644453 | 1 |
| ENSMUSG00000102466 | 1.11297715 | 0.711411495 |
| ENSMUSG00000102467 | 1 | |
| ENSMUSG00000102469 | 1 | |
| ENSMUSG00000102470 | 1.300370355 | 0.111540087 |
| ENSMUSG00000102471 | -1.044273782 | 0.698112512 |
| ENSMUSG00000102473 | 1 | |
| ENSMUSG00000102474 | 1.084618945 | 0.698112512 |
| ENSMUSG00000102475 | 1.069068979 | 0.608497229 |
| ENSMUSG00000102476 | 1 | |
| ENSMUSG00000102478 | 1.090730247 | 0.776953697 |
| ENSMUSG00000102479 | -1.082411773 | 0.778239858 |
| ENSMUSG00000102480 | 1.021987153 | 0.949812939 |
| ENSMUSG00000102481 | -1.043306199 | 0.803260194 |
| ENSMUSG00000102482 | -1.0965339 | 0.610568556 |
| ENSMUSG00000102483 | 1 | |
| ENSMUSG00000102484 | 1 | |
| ENSMUSG00000102485 | 1 | |
| ENSMUSG00000102486 | 1 | |
| ENSMUSG00000102487 | 1 | |
| ENSMUSG00000102488 | 1 | |
| ENSMUSG00000102489 | 1 | |
| ENSMUSG00000102490 | -1.121136286 | 0.655571244 |
| ENSMUSG00000102491 | 1 | |

| | | |
|---|---|---|
| ENSMUSG00000102492 | 1 | |
| ENSMUSG00000102493 | -1.339724199 | 0.017028941 |
| ENSMUSG00000102494 | 1.030006853 | 0.94327115 |
| ENSMUSG00000102495 | 1 | |
| ENSMUSG00000102496 | 1.124482352 | 0.576169279 |
| ENSMUSG00000102497 | 1 | |
| ENSMUSG00000102498 | 1 | |
| ENSMUSG00000102499 | 1 | |
| ENSMUSG00000102500 | 1 | |
| ENSMUSG00000102501 | 1.069752935 | 0.769745556 |
| ENSMUSG00000102502 | 1 | |
| ENSMUSG00000102503 | 1.171563499 | 0.233412931 |
| ENSMUSG00000102504 | -1.075182332 | 0.751475418 |
| ENSMUSG00000102505 | 1 | |
| ENSMUSG00000102506 | 1 | |
| ENSMUSG00000102507 | 1 | |
| ENSMUSG00000102508 | 1 | |
| ENSMUSG00000102509 | -1.028972438 | 0.96504537 |
| ENSMUSG00000102510 | 1 | |
| ENSMUSG00000102511 | 1 | |
| ENSMUSG00000102512 | 1 | |
| ENSMUSG00000102513 | 1 | |
| ENSMUSG00000102514 | -1.089497311 | 0.532369657 |
| ENSMUSG00000102515 | 1 | |
| ENSMUSG00000102516 | 1.014545335 | 0.998924031 |
| ENSMUSG00000102517 | 1.010073917 | 1 |
| ENSMUSG00000102518 | -1.022911911 | 0.987645678 |
| ENSMUSG00000102520 | 1 | |
| ENSMUSG00000102521 | 1.075070132 | 0.698112512 |
| ENSMUSG00000102522 | 1 | |
| ENSMUSG00000102523 | -1.059573666 | 0.866888163 |
| ENSMUSG00000102524 | 1 | |
| ENSMUSG00000102525 | -1.044273782 | 0.698112512 |
| ENSMUSG00000102526 | 1.023718867 | 0.954374429 |
| ENSMUSG00000102527 | 1 | |
| ENSMUSG00000102528 | 1 | |
| ENSMUSG00000102529 | 1 | |
| ENSMUSG00000102530 | 1 | |
| ENSMUSG00000102531 | 1 | |
| ENSMUSG00000102532 | 1 | |
| ENSMUSG00000102533 | 1.171042947 | 0.268643495 |
| ENSMUSG00000102534 | 1 | |
| ENSMUSG00000102535 | 1 | |
| ENSMUSG00000102536 | 1 | |
| ENSMUSG00000102537 | 1 | |
| ENSMUSG00000102538 | 1 | |
| ENSMUSG00000102539 | 1.059463094 | 0.608497229 |
| ENSMUSG00000102540 | -1.008906735 | 1 |
| ENSMUSG00000102541 | 1 | |
| ENSMUSG00000102542 | 1 | |
| ENSMUSG00000102543 | 1.464704689 | 0.000842901 |
| ENSMUSG00000102544 | 1.024693402 | 0.933031586 |
| ENSMUSG00000102545 | 1.112843644 | 0.665814242 |
| ENSMUSG00000102546 | 1.084618945 | 0.780872337 |
| ENSMUSG00000102547 | 1 | |
| ENSMUSG00000102548 | 1 | |
| ENSMUSG00000102549 | 1 | |
| ENSMUSG00000102550 | 1.05351655 | 0.793821774 |
| ENSMUSG00000102551 | 1 | |
| ENSMUSG00000102552 | 1 | |
| ENSMUSG00000102553 | 1.139135785 | 0.54086872 |
| ENSMUSG00000102554 | -1.095939461 | 0.698112512 |
| ENSMUSG00000102555 | -1.107547267 | 0.607021905 |

| | | |
|---|---|---|
| ENSMUSG00000102556 | -1.044273782 | 0.790851299 |
| ENSMUSG00000102558 | 1 | |
| ENSMUSG00000102559 | 1.07047934 | 0.61890292 |
| ENSMUSG00000102561 | 1.075870199 | 0.698112512 |
| ENSMUSG00000102562 | -1.915546125 | 0.002484312 |
| ENSMUSG00000102563 | 1 | |
| ENSMUSG00000102564 | -1 | 1 |
| ENSMUSG00000102565 | -1.043306199 | 0.698112512 |
| ENSMUSG00000102566 | 1 | |
| ENSMUSG00000102567 | 1 | |
| ENSMUSG00000102568 | 1.035337544 | 0.956374899 |
| ENSMUSG00000102569 | 1 | |
| ENSMUSG00000102570 | 1.037403635 | 0.835701362 |
| ENSMUSG00000102571 | 1.059463094 | 0.608497229 |
| ENSMUSG00000102572 | 1 | |
| ENSMUSG00000102573 | 1.122599258 | 0.521942749 |
| ENSMUSG00000102574 | 1 | |
| ENSMUSG00000102575 | 1.059463094 | 0.608497229 |
| ENSMUSG00000102576 | 1 | |
| ENSMUSG00000102577 | -1.051881429 | 0.89016096 |
| ENSMUSG00000102578 | 1.095872691 | 0.608497229 |
| ENSMUSG00000102579 | 1 | |
| ENSMUSG00000102580 | 1 | |
| ENSMUSG00000102581 | 1.059463094 | 0.608497229 |
| ENSMUSG00000102582 | 1 | |
| ENSMUSG00000102583 | 1 | |
| ENSMUSG00000102584 | 1 | |
| ENSMUSG00000102585 | 1 | |
| ENSMUSG00000102586 | 1 | |
| ENSMUSG00000102587 | 1 | |
| ENSMUSG00000102588 | -1 | 1 |
| ENSMUSG00000102589 | 1 | |
| ENSMUSG00000102590 | -1.043306199 | 0.698112512 |
| ENSMUSG00000102591 | -1.033627033 | 0.840946951 |
| ENSMUSG00000102592 | 1.019083579 | 0.949812939 |
| ENSMUSG00000102593 | 1.002741024 | 1 |
| ENSMUSG00000102594 | -1.009476377 | 1 |
| ENSMUSG00000102595 | 1 | |
| ENSMUSG00000102596 | 1 | |
| ENSMUSG00000102597 | 1 | |
| ENSMUSG00000102598 | 1 | |
| ENSMUSG00000102599 | -1.000501298 | 1 |
| ENSMUSG00000102600 | -1.043722544 | 0.869995712 |
| ENSMUSG00000102601 | 1 | |
| ENSMUSG00000102602 | 1.02164745 | 0.962368837 |
| ENSMUSG00000102603 | 1 | |
| ENSMUSG00000102605 | -1 | 1 |
| ENSMUSG00000102606 | -1.044273782 | 0.790851299 |
| ENSMUSG00000102607 | 1 | |
| ENSMUSG00000102608 | 1 | |
| ENSMUSG00000102609 | 1 | |
| ENSMUSG00000102610 | -1.07194389 | 0.698112512 |
| ENSMUSG00000102611 | 1 | |
| ENSMUSG00000102612 | 1 | |
| ENSMUSG00000102613 | 1 | |
| ENSMUSG00000102614 | -1.03962336 | 0.698112512 |
| ENSMUSG00000102615 | 1.059463094 | 0.699340382 |
| ENSMUSG00000102616 | 1 | |
| ENSMUSG00000102617 | 1 | |
| ENSMUSG00000102618 | 1 | |
| ENSMUSG00000102619 | -1.148903398 | 0.608497229 |
| ENSMUSG00000102620 | 1 | |
| ENSMUSG00000102621 | -1.059374542 | 0.812570482 |

| | | |
|---|---|---|
| ENSMUSG00000102622 | 1 | |
| ENSMUSG00000102623 | 1.076438416 | 0.80429734 |
| ENSMUSG00000102624 | 1 | |
| ENSMUSG00000102625 | 1.059463094 | 0.608497229 |
| ENSMUSG00000102626 | -1 | 1 |
| ENSMUSG00000102627 | 1 | |
| ENSMUSG00000102628 | 1 | |
| ENSMUSG00000102629 | 1.059463094 | 0.704170349 |
| ENSMUSG00000102630 | 1.047976605 | 0.874341108 |
| ENSMUSG00000102632 | -1.023016116 | 0.975094751 |
| ENSMUSG00000102633 | -1.028093353 | 0.886379277 |
| ENSMUSG00000102634 | 1 | |
| ENSMUSG00000102635 | 1.069068979 | 0.721840456 |
| ENSMUSG00000102636 | 1 | |
| ENSMUSG00000102637 | 1.059463094 | 0.608497229 |
| ENSMUSG00000102639 | 1 | |
| ENSMUSG00000102641 | 1 | |
| ENSMUSG00000102642 | -1.03962336 | 0.698112512 |
| ENSMUSG00000102643 | 1 | |
| ENSMUSG00000102644 | -1.003217924 | 1 |
| ENSMUSG00000102645 | 1 | |
| ENSMUSG00000102646 | 1 | |
| ENSMUSG00000102647 | -1.046913744 | 0.817935709 |
| ENSMUSG00000102648 | 1.079681442 | 0.608497229 |
| ENSMUSG00000102649 | -1.139932517 | 0.613865524 |
| ENSMUSG00000102650 | 1 | |
| ENSMUSG00000102651 | -1.044273782 | 0.855488553 |
| ENSMUSG00000102652 | -1.010960635 | 1 |
| ENSMUSG00000102653 | 1 | |
| ENSMUSG00000102654 | 1 | |
| ENSMUSG00000102655 | 1 | |
| ENSMUSG00000102656 | 1 | |
| ENSMUSG00000102657 | 1.192018723 | 0.661322437 |
| ENSMUSG00000102658 | 1 | |
| ENSMUSG00000102659 | -1.247698242 | 0.471396077 |
| ENSMUSG00000102660 | 1 | |
| ENSMUSG00000102661 | 1 | |
| ENSMUSG00000102662 | -1.210008472 | 0.256913989 |
| ENSMUSG00000102663 | -1.03962336 | 0.698112512 |
| ENSMUSG00000102664 | 1.125190212 | 0.653695234 |
| ENSMUSG00000102665 | -1.132757337 | 0.644779634 |
| ENSMUSG00000102666 | 1.095872691 | 0.608497229 |
| ENSMUSG00000102667 | 1 | |
| ENSMUSG00000102668 | 1 | |
| ENSMUSG00000102669 | -1 | 1 |
| ENSMUSG00000102670 | 1.030256832 | 0.93004429 |
| ENSMUSG00000102672 | 1.053398234 | 0.677534336 |
| ENSMUSG00000102673 | 1.180775596 | 0.608497229 |
| ENSMUSG00000102674 | 1.069293585 | 0.802544094 |
| ENSMUSG00000102676 | 1.019083579 | 0.949812939 |
| ENSMUSG00000102677 | 1 | |
| ENSMUSG00000102678 | 1 | |
| ENSMUSG00000102679 | 1 | |
| ENSMUSG00000102680 | -1.044273782 | 0.698112512 |
| ENSMUSG00000102681 | 1 | |
| ENSMUSG00000102682 | 1 | |
| ENSMUSG00000102683 | 1 | |
| ENSMUSG00000102684 | 1.059463094 | 0.699340382 |
| ENSMUSG00000102685 | -1.054130142 | 0.868154326 |
| ENSMUSG00000102686 | 1 | |
| ENSMUSG00000102687 | -1.053549062 | 0.698112512 |
| ENSMUSG00000102688 | 1 | |
| ENSMUSG00000102689 | 1 | |

| | | |
|---|---|---|
| ENSMUSG00000102690 | 1.051378421 | 0.934760809 |
| ENSMUSG00000102691 | -1.423539774 | 0.075008317 |
| ENSMUSG00000102692 | -1.335769008 | 0.01634912 |
| ENSMUSG00000102693 | 1 | |
| ENSMUSG00000102694 | -1.007652866 | 1 |
| ENSMUSG00000102696 | 1 | |
| ENSMUSG00000102697 | 1.014545335 | 0.991944074 |
| ENSMUSG00000102698 | -1.03962336 | 0.698112512 |
| ENSMUSG00000102699 | 1 | |
| ENSMUSG00000102700 | 1.162864188 | 0.608497229 |
| ENSMUSG00000102701 | -1.084645496 | 0.532369657 |
| ENSMUSG00000102702 | -1.12574963 | 0.698112512 |
| ENSMUSG00000102703 | 1 | |
| ENSMUSG00000102704 | 1.217880763 | 0.50840255 |
| ENSMUSG00000102705 | 1.106805166 | 0.224172237 |
| ENSMUSG00000102706 | 1.600676629 | 0.00680252 |
| ENSMUSG00000102707 | 1 | |
| ENSMUSG00000102709 | 1.084618945 | 0.73684216 |
| ENSMUSG00000102710 | 1 | |
| ENSMUSG00000102712 | 1.072667987 | 0.782828979 |
| ENSMUSG00000102713 | 1.173716832 | 0.58889649 |
| ENSMUSG00000102714 | 1.075870199 | 0.698112512 |
| ENSMUSG00000102715 | -1.139364264 | 0.64332945 |
| ENSMUSG00000102716 | 1 | |
| ENSMUSG00000102717 | -1.008592986 | 1 |
| ENSMUSG00000102718 | 1.036029124 | 0.874822203 |
| ENSMUSG00000102719 | 1.013039178 | 0.974593713 |
| ENSMUSG00000102721 | 1 | |
| ENSMUSG00000102722 | 1.171563499 | 0.233412931 |
| ENSMUSG00000102723 | 1 | |
| ENSMUSG00000102724 | 1 | |
| ENSMUSG00000102725 | 1 | |
| ENSMUSG00000102726 | 1 | |
| ENSMUSG00000102727 | 1 | |
| ENSMUSG00000102728 | 1.014545335 | 0.979002724 |
| ENSMUSG00000102729 | -1.043306199 | 0.698112512 |
| ENSMUSG00000102730 | 1 | |
| ENSMUSG00000102731 | 1 | |
| ENSMUSG00000102732 | 1.015486245 | 0.973126641 |
| ENSMUSG00000102733 | -1 | 1 |
| ENSMUSG00000102734 | 1 | |
| ENSMUSG00000102735 | 1.118362357 | 0.7530201 |
| ENSMUSG00000102737 | -1 | 1 |
| ENSMUSG00000102738 | 1 | |
| ENSMUSG00000102739 | 1 | |
| ENSMUSG00000102740 | 1 | |
| ENSMUSG00000102741 | 1 | |
| ENSMUSG00000102742 | 1.0917839 | 0.756361387 |
| ENSMUSG00000102743 | 1 | |
| ENSMUSG00000102744 | -1.03962336 | 0.698112512 |
| ENSMUSG00000102745 | -1 | 1 |
| ENSMUSG00000102746 | -1.111670129 | 0.574033444 |
| ENSMUSG00000102747 | 1 | |
| ENSMUSG00000102748 | 1.11084001 | 0.166531048 |
| ENSMUSG00000102749 | 1.059463094 | 0.608497229 |
| ENSMUSG00000102750 | -1.043306199 | 0.79291651 |
| ENSMUSG00000102751 | 1 | |
| ENSMUSG00000102752 | -1.01581828 | 0.70969478 |
| ENSMUSG00000102753 | 1.059463094 | 0.608497229 |
| ENSMUSG00000102754 | 1 | |
| ENSMUSG00000102755 | 1 | |
| ENSMUSG00000102756 | 1 | |
| ENSMUSG00000102757 | 1 | |

| | | |
|---|---|---|
| ENSMUSG00000102758 | -1.213907141 | 0.251049598 |
| ENSMUSG00000102759 | 1.165716974 | 0.608497229 |
| ENSMUSG00000102760 | 1.021186614 | 0.959583351 |
| ENSMUSG00000102761 | 1 | |
| ENSMUSG00000102762 | -1.027395698 | 0.981779194 |
| ENSMUSG00000102763 | 1 | |
| ENSMUSG00000102764 | 1.014545335 | 1 |
| ENSMUSG00000102765 | 1 | |
| ENSMUSG00000102766 | 1 | |
| ENSMUSG00000102768 | 1 | |
| ENSMUSG00000102769 | 1 | |
| ENSMUSG00000102770 | -1.043306199 | 0.79291651 |
| ENSMUSG00000102771 | 1 | |
| ENSMUSG00000102772 | 1 | |
| ENSMUSG00000102773 | 1 | |
| ENSMUSG00000102774 | 1 | |
| ENSMUSG00000102775 | -1.110907262 | 0.698112512 |
| ENSMUSG00000102776 | 1.00414412 | 1 |
| ENSMUSG00000102777 | 1 | |
| ENSMUSG00000102778 | 1 | |
| ENSMUSG00000102779 | -1.044273782 | 0.698112512 |
| ENSMUSG00000102780 | -1.029200399 | 0.783644184 |
| ENSMUSG00000102782 | 1.026615208 | 0.858354949 |
| ENSMUSG00000102783 | 1 | |
| ENSMUSG00000102784 | 1.049411284 | 0.888246981 |
| ENSMUSG00000102785 | 1 | |
| ENSMUSG00000102786 | 1 | |
| ENSMUSG00000102787 | 1 | |
| ENSMUSG00000102788 | 1 | |
| ENSMUSG00000102789 | 1 | |
| ENSMUSG00000102790 | 1 | |
| ENSMUSG00000102791 | 1 | |
| ENSMUSG00000102792 | -1.053549062 | 0.698112512 |
| ENSMUSG00000102793 | 1 | |
| ENSMUSG00000102794 | 1 | |
| ENSMUSG00000102795 | 1 | |
| ENSMUSG00000102796 | -1.03962336 | 0.698112512 |
| ENSMUSG00000102797 | 1 | |
| ENSMUSG00000102798 | -1.005239444 | 1 |
| ENSMUSG00000102799 | 1 | |
| ENSMUSG00000102800 | 1.07913413 | 0.698112512 |
| ENSMUSG00000102801 | -1 | 1 |
| ENSMUSG00000102802 | 1 | |
| ENSMUSG00000102803 | -1.113515092 | 0.608497229 |
| ENSMUSG00000102804 | 1 | |
| ENSMUSG00000102805 | -1 | 1 |
| ENSMUSG00000102806 | 1 | |
| ENSMUSG00000102807 | 1 | |
| ENSMUSG00000102808 | -1.03962336 | 0.698112512 |
| ENSMUSG00000102809 | 1.131005027 | 0.641910604 |
| ENSMUSG00000102810 | 1 | |
| ENSMUSG00000102813 | 1.147336016 | 0.028539747 |
| ENSMUSG00000102814 | -1.043306199 | 0.698112512 |
| ENSMUSG00000102815 | -1.187941777 | 0.275279115 |
| ENSMUSG00000102816 | 1 | |
| ENSMUSG00000102817 | -1.060017444 | 0.855488553 |
| ENSMUSG00000102818 | 1 | |
| ENSMUSG00000102819 | 1 | |
| ENSMUSG00000102820 | 1 | |
| ENSMUSG00000102821 | 1 | |
| ENSMUSG00000102822 | 1 | |
| ENSMUSG00000102824 | 1 | |
| ENSMUSG00000102825 | -1.053549062 | 0.774498429 |

| | | |
|---|---|---|
| ENSMUSG00000102826 | 1 | |
| ENSMUSG00000102827 | -1.03962336 | 0.698112512 |
| ENSMUSG00000102828 | 1 | |
| ENSMUSG00000102829 | 1 | |
| ENSMUSG00000102830 | -1.116428035 | 0.608497229 |
| ENSMUSG00000102831 | 1 | |
| ENSMUSG00000102832 | 1 | |
| ENSMUSG00000102833 | 1.047947442 | 0.89553586 |
| ENSMUSG00000102834 | 1 | |
| ENSMUSG00000102835 | 1 | |
| ENSMUSG00000102836 | 1 | |
| ENSMUSG00000102837 | -1.03962336 | 0.803072249 |
| ENSMUSG00000102838 | 1 | |
| ENSMUSG00000102839 | 1 | |
| ENSMUSG00000102840 | 1.139962167 | 0.479655048 |
| ENSMUSG00000102841 | -1.047306572 | 0.894839966 |
| ENSMUSG00000102842 | 1 | |
| ENSMUSG00000102843 | 1 | |
| ENSMUSG00000102844 | 1.04839098 | 0.877320429 |
| ENSMUSG00000102845 | 1 | |
| ENSMUSG00000102846 | 1 | |
| ENSMUSG00000102847 | 1 | |
| ENSMUSG00000102848 | 1 | |
| ENSMUSG00000102849 | 1 | |
| ENSMUSG00000102850 | -1.097438894 | 0.809807951 |
| ENSMUSG00000102851 | 1 | |
| ENSMUSG00000102852 | -1.113515092 | 0.576169279 |
| ENSMUSG00000102853 | 1 | |
| ENSMUSG00000102854 | -1.084328386 | 0.844117076 |
| ENSMUSG00000102855 | 1 | |
| ENSMUSG00000102856 | 1.064964203 | 0.902615861 |
| ENSMUSG00000102857 | 1 | |
| ENSMUSG00000102858 | 1.064452362 | 0.6659463 |
| ENSMUSG00000102859 | -1.009476377 | 1 |
| ENSMUSG00000102860 | 1 | |
| ENSMUSG00000102861 | -1.031738932 | 0.729293671 |
| ENSMUSG00000102862 | -1.043306199 | 0.698112512 |
| ENSMUSG00000102863 | 1.193528274 | 0.307303709 |
| ENSMUSG00000102864 | 1 | |
| ENSMUSG00000102865 | 1 | |
| ENSMUSG00000102866 | 1 | |
| ENSMUSG00000102867 | 1 | |
| ENSMUSG00000102868 | 1.014545335 | 0.985522133 |
| ENSMUSG00000102869 | -1.135136019 | 0.07619545 |
| ENSMUSG00000102870 | 1 | |
| ENSMUSG00000102871 | 1 | |
| ENSMUSG00000102872 | -1.009047667 | 1 |
| ENSMUSG00000102873 | 1 | |
| ENSMUSG00000102874 | 1.028323353 | 0.951772039 |
| ENSMUSG00000102875 | 1.037116955 | 0.954244503 |
| ENSMUSG00000102876 | 1 | |
| ENSMUSG00000102877 | 1 | |
| ENSMUSG00000102878 | 1 | |
| ENSMUSG00000102879 | 1 | |
| ENSMUSG00000102880 | 1 | |
| ENSMUSG00000102881 | -1.033575217 | 0.954289064 |
| ENSMUSG00000102882 | 1.065791798 | 0.899132938 |
| ENSMUSG00000102883 | 1 | |
| ENSMUSG00000102884 | 1 | |
| ENSMUSG00000102885 | 1 | |
| ENSMUSG00000102886 | 1 | |
| ENSMUSG00000102887 | 1 | |
| ENSMUSG00000102888 | 1 | |

| | | |
|---|---|---|
| ENSMUSG00000102889 | 1 | |
| ENSMUSG00000102890 | 1.059463094 | 0.608497229 |
| ENSMUSG00000102891 | 1 | |
| ENSMUSG00000102892 | -1.102582607 | 0.790851299 |
| ENSMUSG00000102893 | 1.062871489 | 0.839099152 |
| ENSMUSG00000102894 | 1 | |
| ENSMUSG00000102895 | -1.09953309 | 0.757289696 |
| ENSMUSG00000102896 | 1 | |
| ENSMUSG00000102897 | 1 | |
| ENSMUSG00000102898 | 1 | |
| ENSMUSG00000102899 | -1.062900008 | 0.835382855 |
| ENSMUSG00000102900 | 1 | |
| ENSMUSG00000102901 | 1 | |
| ENSMUSG00000102902 | -1.003834322 | 1 |
| ENSMUSG00000102903 | 1 | |
| ENSMUSG00000102904 | -1.033820075 | 0.888246981 |
| ENSMUSG00000102905 | 1 | |
| ENSMUSG00000102906 | 1 | |
| ENSMUSG00000102907 | 1 | |
| ENSMUSG00000102908 | 1.220866531 | 0.368089426 |
| ENSMUSG00000102909 | -1.014570171 | 0.985713389 |
| ENSMUSG00000102910 | 1 | |
| ENSMUSG00000102911 | 1 | |
| ENSMUSG00000102912 | 1.041908502 | 0.984902895 |
| ENSMUSG00000102913 | 1 | |
| ENSMUSG00000102914 | 1.069068979 | 0.608497229 |
| ENSMUSG00000102915 | 1 | |
| ENSMUSG00000102916 | 1 | |
| ENSMUSG00000102917 | -1.012397719 | 0.963843528 |
| ENSMUSG00000102918 | 1.167770926 | 0.050235993 |
| ENSMUSG00000102919 | 1.050112723 | 0.931949042 |
| ENSMUSG00000102920 | 1 | |
| ENSMUSG00000102921 | 1 | |
| ENSMUSG00000102922 | 1 | |
| ENSMUSG00000102923 | -1.084645496 | 0.532369657 |
| ENSMUSG00000102924 | 1 | |
| ENSMUSG00000102925 | 1 | |
| ENSMUSG00000102926 | 1 | |
| ENSMUSG00000102927 | -1.024087213 | 0.919984257 |
| ENSMUSG00000102928 | 1 | |
| ENSMUSG00000102929 | -1.027398866 | 0.92584202 |
| ENSMUSG00000102930 | -1.109582661 | 0.638646996 |
| ENSMUSG00000102931 | 1.190047299 | 0.449109628 |
| ENSMUSG00000102932 | 1 | |
| ENSMUSG00000102933 | 1 | |
| ENSMUSG00000102934 | 1 | |
| ENSMUSG00000102935 | 1.11038179 | 0.628596302 |
| ENSMUSG00000102936 | 1 | |
| ENSMUSG00000102937 | 1 | |
| ENSMUSG00000102938 | 1 | |
| ENSMUSG00000102939 | -1.038858794 | 0.890857745 |
| ENSMUSG00000102940 | 1.059463094 | 0.608497229 |
| ENSMUSG00000102941 | 1 | |
| ENSMUSG00000102942 | 1 | |
| ENSMUSG00000102943 | 1 | |
| ENSMUSG00000102945 | 1.069068979 | 0.608497229 |
| ENSMUSG00000102946 | 1 | |
| ENSMUSG00000102948 | 1 | |
| ENSMUSG00000102949 | 1 | |
| ENSMUSG00000102950 | -1.054169715 | 0.863907734 |
| ENSMUSG00000102951 | 1.05410549 | 0.841285672 |
| ENSMUSG00000102952 | 1 | |
| ENSMUSG00000102953 | 1.179016771 | 0.608497229 |

| | | |
|---|---|---|
| ENSMUSG00000102955 | 1 | |
| ENSMUSG00000102956 | 1.005588822 | 1 |
| ENSMUSG00000102957 | 1 | |
| ENSMUSG00000102958 | 1 | |
| ENSMUSG00000102959 | 1 | |
| ENSMUSG00000102960 | 1.003729568 | 1 |
| ENSMUSG00000102961 | 1 | |
| ENSMUSG00000102962 | 1 | |
| ENSMUSG00000102963 | -1.053549062 | 0.698112512 |
| ENSMUSG00000102964 | -1.066423912 | 0.75366313 |
| ENSMUSG00000102965 | 1 | |
| ENSMUSG00000102966 | 1 | |
| ENSMUSG00000102967 | 1.059463094 | 0.608497229 |
| ENSMUSG00000102968 | 1 | |
| ENSMUSG00000102969 | -1.183404224 | 0.283596572 |
| ENSMUSG00000102970 | 1.040146971 | 0.724366149 |
| ENSMUSG00000102971 | -1 | 1 |
| ENSMUSG00000102972 | 1 | |
| ENSMUSG00000102973 | 1 | |
| ENSMUSG00000102975 | 1 | |
| ENSMUSG00000102976 | 1.022301574 | 0.958690483 |
| ENSMUSG00000102977 | 1 | |
| ENSMUSG00000102979 | 1.079681442 | 0.608497229 |
| ENSMUSG00000102980 | 1 | |
| ENSMUSG00000102981 | 1 | |
| ENSMUSG00000102982 | -1.044259287 | 0.940277706 |
| ENSMUSG00000102983 | 1.024693402 | 0.952057016 |
| ENSMUSG00000102984 | 1 | |
| ENSMUSG00000102985 | 1.005613439 | 1 |
| ENSMUSG00000102986 | 1 | |
| ENSMUSG00000102987 | 1 | |
| ENSMUSG00000102988 | 1 | |
| ENSMUSG00000102989 | -1 | 1 |
| ENSMUSG00000102990 | -1.043306199 | 0.79291651 |
| ENSMUSG00000102991 | 1 | |
| ENSMUSG00000102992 | 1 | |
| ENSMUSG00000102993 | 1 | |
| ENSMUSG00000102994 | 1.025418628 | 0.99762469 |
| ENSMUSG00000102995 | 1 | |
| ENSMUSG00000102997 | 1.138419748 | 0.621888633 |
| ENSMUSG00000102998 | 1 | |
| ENSMUSG00000102999 | 1.023743961 | 0.95702926 |
| ENSMUSG00000103000 | -1.084949475 | 0.869966868 |
| ENSMUSG00000103001 | -1 | 1 |
| ENSMUSG00000103002 | -1.099174267 | 0.608497229 |
| ENSMUSG00000103003 | 1 | |
| ENSMUSG00000103004 | 1 | |
| ENSMUSG00000103005 | 1 | |
| ENSMUSG00000103006 | -1.04365757 | 0.959839883 |
| ENSMUSG00000103007 | 1 | |
| ENSMUSG00000103008 | 1 | |
| ENSMUSG00000103009 | 1.025851687 | 0.900996516 |
| ENSMUSG00000103010 | 1.069068979 | 0.608497229 |
| ENSMUSG00000103011 | -1.139116 | 0.625496737 |
| ENSMUSG00000103012 | 1 | |
| ENSMUSG00000103013 | 1.096862175 | 0.698112512 |
| ENSMUSG00000103015 | -1.019108526 | 0.985836448 |
| ENSMUSG00000103016 | 1 | |
| ENSMUSG00000103017 | -1.043306199 | 0.698112512 |
| ENSMUSG00000103018 | 1 | |
| ENSMUSG00000103020 | 1 | |
| ENSMUSG00000103021 | -1.043306199 | 0.698112512 |
| ENSMUSG00000103023 | 1 | |

| | | |
|---|---|---|
| ENSMUSG00000103024 | -1.016098951 | 1 |
| ENSMUSG00000103025 | 1 | |
| ENSMUSG00000103026 | 1 | |
| ENSMUSG00000103027 | 1 | |
| ENSMUSG00000103028 | 1 | |
| ENSMUSG00000103029 | 1 | |
| ENSMUSG00000103030 | 1.091660089 | 0.750026575 |
| ENSMUSG00000103031 | 1 | |
| ENSMUSG00000103032 | 1.313768723 | 0.064638418 |
| ENSMUSG00000103033 | 1 | |
| ENSMUSG00000103034 | -1.201433521 | 0.215991492 |
| ENSMUSG00000103035 | 1 | |
| ENSMUSG00000103037 | 1.076152694 | 0.819465115 |
| ENSMUSG00000103038 | 1 | |
| ENSMUSG00000103039 | 1.103588203 | 0.15569007 |
| ENSMUSG00000103040 | 1 | |
| ENSMUSG00000103041 | 1.018428621 | 0.889722436 |
| ENSMUSG00000103042 | 1 | |
| ENSMUSG00000103043 | 1 | |
| ENSMUSG00000103044 | 1 | |
| ENSMUSG00000103045 | 1 | |
| ENSMUSG00000103046 | 1.062710099 | 0.608497229 |
| ENSMUSG00000103047 | 1.269112919 | 0.174109967 |
| ENSMUSG00000103049 | 1 | |
| ENSMUSG00000103050 | 1.059463094 | 0.608497229 |
| ENSMUSG00000103051 | -1 | 1 |
| ENSMUSG00000103052 | 1 | |
| ENSMUSG00000103053 | -1.000144934 | 1 |
| ENSMUSG00000103054 | 1 | |
| ENSMUSG00000103055 | 1 | |
| ENSMUSG00000103056 | -1.105673636 | 0.642256709 |
| ENSMUSG00000103058 | 1 | |
| ENSMUSG00000103059 | 1 | |
| ENSMUSG00000103060 | 1 | |
| ENSMUSG00000103061 | 1 | |
| ENSMUSG00000103062 | -1.181389669 | 0.581311911 |
| ENSMUSG00000103063 | 1 | |
| ENSMUSG00000103064 | 1 | |
| ENSMUSG00000103065 | 1 | |
| ENSMUSG00000103066 | 1 | |
| ENSMUSG00000103067 | -1.008623876 | 1 |
| ENSMUSG00000103069 | 1 | |
| ENSMUSG00000103070 | 1.034494732 | 0.770958539 |
| ENSMUSG00000103071 | -1.051055372 | 0.777702703 |
| ENSMUSG00000103072 | 1 | |
| ENSMUSG00000103073 | 1 | |
| ENSMUSG00000103074 | 1 | |
| ENSMUSG00000103075 | 1.059463094 | 0.776953697 |
| ENSMUSG00000103076 | 1.148603887 | 0.374017624 |
| ENSMUSG00000103077 | 1 | |
| ENSMUSG00000103078 | 1 | |
| ENSMUSG00000103079 | -1 | 1 |
| ENSMUSG00000103080 | 1.013877317 | 0.998988117 |
| ENSMUSG00000103081 | 1.209465296 | 0.049951226 |
| ENSMUSG00000103082 | 1.020344501 | 0.988665808 |
| ENSMUSG00000103083 | -1.00002448 | 1 |
| ENSMUSG00000103084 | 1 | |
| ENSMUSG00000103085 | 1.061728572 | 0.929923764 |
| ENSMUSG00000103086 | 1 | |
| ENSMUSG00000103087 | 1.095872691 | 0.608497229 |
| ENSMUSG00000103088 | 1.433404069 | 0.003962533 |
| ENSMUSG00000103089 | 1.019083579 | 0.949812939 |
| ENSMUSG00000103090 | 1 | |

| | | |
|---|---|---|
| ENSMUSG00000103091 | 1.059463094 | 0.608497229 |
| ENSMUSG00000103092 | 1.059463094 | 0.608497229 |
| ENSMUSG00000103094 | 1.001933759 | 1 |
| ENSMUSG00000103095 | 1 | |
| ENSMUSG00000103096 | -1.073680831 | 0.724563674 |
| ENSMUSG00000103097 | 1 | |
| ENSMUSG00000103098 | 1.036585803 | 0.969949129 |
| ENSMUSG00000103099 | -1.065852599 | 0.797808656 |
| ENSMUSG00000103100 | 1.021186614 | 0.972142128 |
| ENSMUSG00000103101 | -1.039440247 | 0.702275094 |
| ENSMUSG00000103102 | 1.147794708 | 0.608497229 |
| ENSMUSG00000103103 | 1.414422098 | 0.004223952 |
| ENSMUSG00000103105 | 1 | |
| ENSMUSG00000103106 | 1 | |
| ENSMUSG00000103107 | -1.071635909 | 0.792716139 |
| ENSMUSG00000103108 | 1.059463094 | 0.608497229 |
| ENSMUSG00000103110 | 1 | |
| ENSMUSG00000103111 | 1 | |
| ENSMUSG00000103112 | 1 | |
| ENSMUSG00000103113 | 1 | |
| ENSMUSG00000103114 | -1.010960635 | 1 |
| ENSMUSG00000103115 | -1.056881566 | 0.865128496 |
| ENSMUSG00000103116 | -1 | 1 |
| ENSMUSG00000103118 | 1 | |
| ENSMUSG00000103119 | -1.112010624 | 0.698112512 |
| ENSMUSG00000103120 | 1 | |
| ENSMUSG00000103121 | 1 | |
| ENSMUSG00000103122 | -1.043306199 | 0.698112512 |
| ENSMUSG00000103123 | 1.067542381 | 0.842601989 |
| ENSMUSG00000103124 | 1 | |
| ENSMUSG00000103125 | 1 | |
| ENSMUSG00000103126 | 1 | |
| ENSMUSG00000103127 | 1 | |
| ENSMUSG00000103129 | 1 | |
| ENSMUSG00000103130 | -1 | 1 |
| ENSMUSG00000103131 | -1 | 1 |
| ENSMUSG00000103132 | -1.132666855 | 0.18851436 |
| ENSMUSG00000103133 | -1.067640993 | 0.709356753 |
| ENSMUSG00000103135 | 1.009232369 | 1 |
| ENSMUSG00000103136 | 1 | |
| ENSMUSG00000103137 | 1 | |
| ENSMUSG00000103138 | 1.294283985 | 0.335498586 |
| ENSMUSG00000103139 | 1.024693402 | 0.933031586 |
| ENSMUSG00000103140 | 1 | |
| ENSMUSG00000103141 | 1.059463094 | 0.608497229 |
| ENSMUSG00000103142 | 1 | |
| ENSMUSG00000103143 | 1.019083579 | 0.949812939 |
| ENSMUSG00000103144 | -1.109722906 | 0.480854229 |
| ENSMUSG00000103145 | -1.024718486 | 0.968937442 |
| ENSMUSG00000103146 | 1.006679456 | 1 |
| ENSMUSG00000103147 | 1 | |
| ENSMUSG00000103148 | 1.144572621 | 0.608497229 |
| ENSMUSG00000103149 | 1.018589762 | 0.998386582 |
| ENSMUSG00000103150 | 1 | |
| ENSMUSG00000103151 | -1.03962336 | 0.698112512 |
| ENSMUSG00000103152 | 1 | |
| ENSMUSG00000103153 | 1 | |
| ENSMUSG00000103154 | 1 | |
| ENSMUSG00000103155 | -1.053549062 | 0.817479993 |
| ENSMUSG00000103156 | 1.132639128 | 0.217803313 |
| ENSMUSG00000103157 | 1 | |
| ENSMUSG00000103158 | 1 | |
| ENSMUSG00000103159 | 1.009517015 | 1 |

| | | |
|---|---|---|
| ENSMUSG00000103160 | -1.05403356 | 0.893822825 |
| ENSMUSG00000103161 | -1.029402133 | 0.698112512 |
| ENSMUSG00000103162 | -1.028348525 | 0.938939969 |
| ENSMUSG00000103163 | -1 | 1 |
| ENSMUSG00000103164 | -1.350295376 | 0.029364702 |
| ENSMUSG00000103165 | 1 | |
| ENSMUSG00000103166 | 1.059463094 | 0.608497229 |
| ENSMUSG00000103167 | 1 | |
| ENSMUSG00000103168 | 1.142209427 | 0.057683229 |
| ENSMUSG00000103169 | 1 | |
| ENSMUSG00000103170 | 1.178139999 | 0.202649662 |
| ENSMUSG00000103172 | 1 | |
| ENSMUSG00000103174 | -1.165881203 | 0.464078131 |
| ENSMUSG00000103175 | 1.107693774 | 0.72715392 |
| ENSMUSG00000103176 | 1 | |
| ENSMUSG00000103177 | 1 | |
| ENSMUSG00000103178 | 1 | |
| ENSMUSG00000103179 | 1 | |
| ENSMUSG00000103180 | 1 | |
| ENSMUSG00000103181 | 1.059463094 | 0.608497229 |
| ENSMUSG00000103182 | -1.03183032 | 0.821953663 |
| ENSMUSG00000103183 | 1.248010545 | 0.040928762 |
| ENSMUSG00000103184 | 1 | |
| ENSMUSG00000103185 | 1 | |
| ENSMUSG00000103186 | -1.112797927 | 0.70141105 |
| ENSMUSG00000103187 | 1 | |
| ENSMUSG00000103188 | 1 | |
| ENSMUSG00000103189 | 1 | |
| ENSMUSG00000103190 | 1 | |
| ENSMUSG00000103191 | 1.111812527 | 0.608497229 |
| ENSMUSG00000103192 | 1 | |
| ENSMUSG00000103193 | -1.044273782 | 0.698112512 |
| ENSMUSG00000103194 | 1 | |
| ENSMUSG00000103195 | 1.177765487 | 0.479257008 |
| ENSMUSG00000103196 | 1 | |
| ENSMUSG00000103197 | 1 | |
| ENSMUSG00000103198 | 1 | |
| ENSMUSG00000103199 | -1.065321276 | 0.844881695 |
| ENSMUSG00000103200 | 1.030415139 | 0.960321094 |
| ENSMUSG00000103201 | 1 | |
| ENSMUSG00000103202 | 1 | |
| ENSMUSG00000103203 | 1 | |
| ENSMUSG00000103204 | 1 | |
| ENSMUSG00000103206 | 1 | |
| ENSMUSG00000103207 | -1.268471902 | 0.464579163 |
| ENSMUSG00000103208 | 1 | |
| ENSMUSG00000103209 | 1 | |
| ENSMUSG00000103210 | 1 | |
| ENSMUSG00000103211 | 1 | |
| ENSMUSG00000103213 | 1 | |
| ENSMUSG00000103214 | 1 | |
| ENSMUSG00000103215 | 1 | |
| ENSMUSG00000103216 | -1.03962336 | 0.698112512 |
| ENSMUSG00000103217 | 1 | |
| ENSMUSG00000103218 | 1 | |
| ENSMUSG00000103219 | 1 | |
| ENSMUSG00000103220 | -1.071075483 | 0.698112512 |
| ENSMUSG00000103221 | 1 | |
| ENSMUSG00000103222 | -1.131857941 | 0.532369657 |
| ENSMUSG00000103223 | 1.11166259 | 0.638758774 |
| ENSMUSG00000103224 | 1 | |
| ENSMUSG00000103225 | 1 | |
| ENSMUSG00000103226 | 1 | |

| | | |
|---|---|---|
| ENSMUSG00000103227 | -1.053231832 | 0.885529952 |
| ENSMUSG00000103228 | 1 | |
| ENSMUSG00000103230 | 1.059463094 | 0.608497229 |
| ENSMUSG00000103231 | 1 | |
| ENSMUSG00000103232 | 1.084618945 | 0.614071182 |
| ENSMUSG00000103233 | 1.064870248 | 0.84515778 |
| ENSMUSG00000103234 | 1.032822525 | 0.807247723 |
| ENSMUSG00000103236 | 1 | |
| ENSMUSG00000103237 | 1 | |
| ENSMUSG00000103238 | 1.104269878 | 0.764664548 |
| ENSMUSG00000103239 | 1 | |
| ENSMUSG00000103240 | 1 | |
| ENSMUSG00000103241 | 1.138419748 | 0.660684061 |
| ENSMUSG00000103242 | 1 | |
| ENSMUSG00000103243 | 1 | |
| ENSMUSG00000103244 | 1 | |
| ENSMUSG00000103245 | 1 | |
| ENSMUSG00000103246 | 1 | |
| ENSMUSG00000103247 | -1.188005937 | 0.344478207 |
| ENSMUSG00000103248 | 1 | |
| ENSMUSG00000103249 | 1 | |
| ENSMUSG00000103250 | 1 | |
| ENSMUSG00000103251 | 1.081478107 | 0.778509736 |
| ENSMUSG00000103252 | 1.019863356 | 0.988424085 |
| ENSMUSG00000103253 | 1 | |
| ENSMUSG00000103254 | 1 | |
| ENSMUSG00000103255 | 1 | |
| ENSMUSG00000103256 | 1 | |
| ENSMUSG00000103257 | -1.03962336 | 0.698112512 |
| ENSMUSG00000103258 | -1.08384782 | 0.69829433 |
| ENSMUSG00000103259 | 1 | |
| ENSMUSG00000103260 | -1.024826341 | 0.978053547 |
| ENSMUSG00000103261 | 1.005613439 | 1 |
| ENSMUSG00000103262 | 1 | |
| ENSMUSG00000103263 | 1.015486245 | 0.972980766 |
| ENSMUSG00000103264 | 1 | |
| ENSMUSG00000103265 | 1.059463094 | 0.608497229 |
| ENSMUSG00000103266 | 1 | |
| ENSMUSG00000103267 | 1 | |
| ENSMUSG00000103269 | 1 | |
| ENSMUSG00000103270 | -1.11061171 | 0.66585278 |
| ENSMUSG00000103271 | 1 | |
| ENSMUSG00000103272 | 1.127426054 | 0.608497229 |
| ENSMUSG00000103273 | 1 | |
| ENSMUSG00000103274 | 1 | |
| ENSMUSG00000103275 | 1 | |
| ENSMUSG00000103276 | -1.066870979 | 0.221007934 |
| ENSMUSG00000103277 | 1.039597911 | 0.887094209 |
| ENSMUSG00000103279 | 1 | |
| ENSMUSG00000103280 | -1.316431486 | 0.089930408 |
| ENSMUSG00000103281 | 1 | |
| ENSMUSG00000103282 | 1 | |
| ENSMUSG00000103283 | -1.100930858 | 0.279837771 |
| ENSMUSG00000103284 | 1 | |
| ENSMUSG00000103285 | -1.042170988 | 0.788000546 |
| ENSMUSG00000103286 | 1 | |
| ENSMUSG00000103287 | -1.044273782 | 0.698112512 |
| ENSMUSG00000103288 | 1 | |
| ENSMUSG00000103289 | 1 | |
| ENSMUSG00000103290 | 1 | |
| ENSMUSG00000103291 | -1.053549062 | 0.778372272 |
| ENSMUSG00000103292 | -1.152297514 | 0.5742902 |
| ENSMUSG00000103293 | -1.173142781 | 0.36904104 |

| | | |
|---|---|---|
| ENSMUSG00000103294 | 1.059463094 | 0.608497229 |
| ENSMUSG00000103295 | 1 | |
| ENSMUSG00000103296 | 1 | |
| ENSMUSG00000103297 | 1 | |
| ENSMUSG00000103298 | 1 | |
| ENSMUSG00000103299 | 1 | |
| ENSMUSG00000103300 | 1 | |
| ENSMUSG00000103301 | 1.059463094 | 0.704170349 |
| ENSMUSG00000103302 | 1 | |
| ENSMUSG00000103303 | 1.047976605 | 0.911250323 |
| ENSMUSG00000103304 | -1.044273782 | 0.698112512 |
| ENSMUSG00000103305 | 1 | |
| ENSMUSG00000103306 | 1 | |
| ENSMUSG00000103307 | 1 | |
| ENSMUSG00000103308 | -1.196541655 | 0.182650295 |
| ENSMUSG00000103309 | 1.314985504 | 0.201977401 |
| ENSMUSG00000103310 | 1 | |
| ENSMUSG00000103311 | 1 | |
| ENSMUSG00000103312 | -1.044273782 | 0.698112512 |
| ENSMUSG00000103313 | 1.044073474 | 0.744149026 |
| ENSMUSG00000103314 | -1.028160286 | 0.933897165 |
| ENSMUSG00000103315 | 1 | |
| ENSMUSG00000103317 | 1.095872691 | 0.628154042 |
| ENSMUSG00000103318 | 1 | |
| ENSMUSG00000103320 | 1.132639128 | 0.217803313 |
| ENSMUSG00000103321 | 1 | |
| ENSMUSG00000103322 | 1.04190726 | 0.859839502 |
| ENSMUSG00000103323 | 1.107899193 | 0.608497229 |
| ENSMUSG00000103324 | -1.053549062 | 0.698112512 |
| ENSMUSG00000103325 | 1 | |
| ENSMUSG00000103327 | 1 | |
| ENSMUSG00000103328 | 1 | |
| ENSMUSG00000103329 | 1 | |
| ENSMUSG00000103330 | 1 | |
| ENSMUSG00000103331 | -1.084645496 | 0.608497229 |
| ENSMUSG00000103332 | 1.1863949 | 0.066000062 |
| ENSMUSG00000103333 | 1 | |
| ENSMUSG00000103334 | -1 | 1 |
| ENSMUSG00000103335 | 1 | |
| ENSMUSG00000103336 | 1 | |
| ENSMUSG00000103338 | 1 | |
| ENSMUSG00000103339 | 1 | |
| ENSMUSG00000103340 | 1 | |
| ENSMUSG00000103341 | 1 | |
| ENSMUSG00000103342 | 1 | |
| ENSMUSG00000103343 | 1.059463094 | 0.698112512 |
| ENSMUSG00000103345 | 1.069068979 | 0.698112512 |
| ENSMUSG00000103346 | 1 | |
| ENSMUSG00000103347 | 1 | |
| ENSMUSG00000103348 | 1.059463094 | 0.608497229 |
| ENSMUSG00000103349 | -1.069502966 | 0.752398528 |
| ENSMUSG00000103350 | 1 | |
| ENSMUSG00000103351 | -1.03962336 | 0.814089245 |
| ENSMUSG00000103352 | 1 | |
| ENSMUSG00000103353 | 1 | |
| ENSMUSG00000103354 | 1 | |
| ENSMUSG00000103355 | 1.38968879 | 0.016219295 |
| ENSMUSG00000103356 | -1 | 1 |
| ENSMUSG00000103357 | 1 | |
| ENSMUSG00000103358 | 1 | |
| ENSMUSG00000103360 | 1.084618945 | 0.698112512 |
| ENSMUSG00000103361 | 1 | |
| ENSMUSG00000103362 | 1 | |

| | | |
|---|---|---|
| ENSMUSG00000103364 | -1.095939461 | 0.730441273 |
| ENSMUSG00000103365 | 1 | |
| ENSMUSG00000103366 | -1.044273782 | 0.834556513 |
| ENSMUSG00000103367 | 1.207145346 | 0.043800056 |
| ENSMUSG00000103368 | 1 | |
| ENSMUSG00000103369 | 1 | |
| ENSMUSG00000103370 | 1 | |
| ENSMUSG00000103371 | 1 | |
| ENSMUSG00000103372 | 1 | |
| ENSMUSG00000103373 | -1.164434015 | 0.608497229 |
| ENSMUSG00000103374 | 1 | |
| ENSMUSG00000103375 | -1.043306199 | 0.698112512 |
| ENSMUSG00000103376 | 1.044210161 | 0.924766459 |
| ENSMUSG00000103377 | 1 | |
| ENSMUSG00000103378 | -1.043306199 | 0.698112512 |
| ENSMUSG00000103379 | 1 | |
| ENSMUSG00000103380 | 1 | |
| ENSMUSG00000103382 | 1.178793349 | 0.407591381 |
| ENSMUSG00000103383 | 1.019083579 | 0.949812939 |
| ENSMUSG00000103384 | 1 | |
| ENSMUSG00000103385 | 1 | |
| ENSMUSG00000103386 | -1.03962336 | 0.698112512 |
| ENSMUSG00000103387 | 1 | |
| ENSMUSG00000103388 | -1.011722189 | 1 |
| ENSMUSG00000103389 | 1 | |
| ENSMUSG00000103390 | 1 | |
| ENSMUSG00000103391 | -1.042426269 | 0.873239663 |
| ENSMUSG00000103392 | 1 | |
| ENSMUSG00000103393 | 1 | |
| ENSMUSG00000103394 | 1 | |
| ENSMUSG00000103395 | 1 | |
| ENSMUSG00000103396 | 1 | |
| ENSMUSG00000103397 | 1 | |
| ENSMUSG00000103398 | 1.083991456 | 0.729783787 |
| ENSMUSG00000103399 | -1.069502966 | 0.698112512 |
| ENSMUSG00000103400 | -1.087602994 | 0.823693432 |
| ENSMUSG00000103401 | 1 | |
| ENSMUSG00000103402 | 1 | |
| ENSMUSG00000103403 | 1.011065285 | 1 |
| ENSMUSG00000103404 | 1.086349124 | 0.698112512 |
| ENSMUSG00000103406 | 1 | |
| ENSMUSG00000103407 | 1 | |
| ENSMUSG00000103408 | -1.032214272 | 0.967899076 |
| ENSMUSG00000103409 | -1.092139265 | 0.709775561 |
| ENSMUSG00000103410 | 1 | |
| ENSMUSG00000103411 | 1 | |
| ENSMUSG00000103412 | 1 | |
| ENSMUSG00000103413 | 1 | |
| ENSMUSG00000103415 | -1.043306199 | 0.698112512 |
| ENSMUSG00000103416 | 1 | |
| ENSMUSG00000103417 | 1 | |
| ENSMUSG00000103419 | 1.059463094 | 0.608497229 |
| ENSMUSG00000103420 | 1 | |
| ENSMUSG00000103421 | 1 | |
| ENSMUSG00000103422 | -1.203254079 | 0.502038506 |
| ENSMUSG00000103423 | -1.195571312 | 0.366576514 |
| ENSMUSG00000103424 | -1.043306199 | 0.837704961 |
| ENSMUSG00000103425 | 1 | |
| ENSMUSG00000103426 | 1 | |
| ENSMUSG00000103427 | -1.134410835 | 0.545891535 |
| ENSMUSG00000103428 | 1 | |
| ENSMUSG00000103430 | 1 | |
| ENSMUSG00000103431 | 1 | |

| | | |
|---|---|---|
| ENSMUSG00000103432 | -1.038444538 | 0.924629556 |
| ENSMUSG00000103433 | 1 | |
| ENSMUSG00000103434 | -1.069502966 | 0.698112512 |
| ENSMUSG00000103435 | 1.056280798 | 0.879839072 |
| ENSMUSG00000103436 | 1.04930469 | 0.772180723 |
| ENSMUSG00000103437 | 1 | |
| ENSMUSG00000103438 | 1 | |
| ENSMUSG00000103439 | -1.03962336 | 0.698112512 |
| ENSMUSG00000103440 | 1 | |
| ENSMUSG00000103441 | 1.059463094 | 0.608497229 |
| ENSMUSG00000103442 | 1 | |
| ENSMUSG00000103443 | -1.281847878 | 0.071467772 |
| ENSMUSG00000103444 | 1 | |
| ENSMUSG00000103445 | -1.117823397 | 0.42110385 |
| ENSMUSG00000103446 | 1 | |
| ENSMUSG00000103448 | 1 | |
| ENSMUSG00000103449 | 1 | |
| ENSMUSG00000103451 | 1 | |
| ENSMUSG00000103452 | -1 | 1 |
| ENSMUSG00000103454 | 1 | |
| ENSMUSG00000103456 | 1 | |
| ENSMUSG00000103457 | 1 | |
| ENSMUSG00000103458 | 1 | |
| ENSMUSG00000103459 | 1.049411284 | 0.867939428 |
| ENSMUSG00000103460 | 1 | |
| ENSMUSG00000103461 | 1 | |
| ENSMUSG00000103462 | 1 | |
| ENSMUSG00000103463 | 1 | |
| ENSMUSG00000103464 | 1.089199536 | 0.765197976 |
| ENSMUSG00000103465 | 1 | |
| ENSMUSG00000103466 | 1 | |
| ENSMUSG00000103467 | -1.016099579 | 0.990492329 |
| ENSMUSG00000103468 | 1 | |
| ENSMUSG00000103469 | -1 | 1 |
| ENSMUSG00000103470 | 1 | |
| ENSMUSG00000103471 | 1 | |
| ENSMUSG00000103472 | 1.938479102 | 0.013645979 |
| ENSMUSG00000103473 | -1.072615536 | 0.756946413 |
| ENSMUSG00000103474 | 1 | |
| ENSMUSG00000103475 | 1.071217535 | 0.734674233 |
| ENSMUSG00000103476 | 1 | |
| ENSMUSG00000103477 | 1.005613439 | 1 |
| ENSMUSG00000103478 | 1 | |
| ENSMUSG00000103479 | 1 | |
| ENSMUSG00000103481 | 1 | |
| ENSMUSG00000103482 | 1 | |
| ENSMUSG00000103483 | 1 | |
| ENSMUSG00000103484 | 1 | |
| ENSMUSG00000103485 | -1.03962336 | 0.803072249 |
| ENSMUSG00000103486 | -1.047419904 | 0.894828773 |
| ENSMUSG00000103487 | 1 | |
| ENSMUSG00000103488 | 1 | |
| ENSMUSG00000103489 | 1 | |
| ENSMUSG00000103491 | 1 | |
| ENSMUSG00000103492 | 1.085624843 | 0.698112512 |
| ENSMUSG00000103493 | 1.156423362 | 0.133561111 |
| ENSMUSG00000103494 | -1.105673636 | 0.608497229 |
| ENSMUSG00000103495 | -1.100193663 | 0.608497229 |
| ENSMUSG00000103496 | 1 | |
| ENSMUSG00000103497 | 1.059463094 | 0.608497229 |
| ENSMUSG00000103498 | 1 | |
| ENSMUSG00000103499 | 1 | |
| ENSMUSG00000103500 | 1 | |

| | | |
|---|---|---|
| ENSMUSG00000103501 | 1 | |
| ENSMUSG00000103502 | 1 | |
| ENSMUSG00000103503 | 1 | |
| ENSMUSG00000103504 | 1 | |
| ENSMUSG00000103505 | -1.13108607 | 0.729206866 |
| ENSMUSG00000103506 | 1.015888954 | 1 |
| ENSMUSG00000103507 | 1.017632426 | 0.979002724 |
| ENSMUSG00000103509 | 1.098829342 | 0.594095902 |
| ENSMUSG00000103510 | 1 | |
| ENSMUSG00000103511 | 1 | |
| ENSMUSG00000103512 | 1 | |
| ENSMUSG00000103513 | 1 | |
| ENSMUSG00000103514 | 1 | |
| ENSMUSG00000103515 | 1.032096017 | 0.698112512 |
| ENSMUSG00000103516 | 1 | |
| ENSMUSG00000103517 | 1 | |
| ENSMUSG00000103518 | 1 | |
| ENSMUSG00000103519 | 1.059463094 | 0.608497229 |
| ENSMUSG00000103520 | 1.145293987 | 0.608497229 |
| ENSMUSG00000103521 | 1 | |
| ENSMUSG00000103522 | 1.059463094 | 0.608497229 |
| ENSMUSG00000103523 | 1 | |
| ENSMUSG00000103524 | 1 | |
| ENSMUSG00000103525 | 1.075070132 | 0.725426917 |
| ENSMUSG00000103526 | 1 | |
| ENSMUSG00000103527 | -1 | 1 |
| ENSMUSG00000103528 | 1 | |
| ENSMUSG00000103529 | 1.222889447 | 0.297855218 |
| ENSMUSG00000103530 | 1 | |
| ENSMUSG00000103531 | 1 | |
| ENSMUSG00000103532 | -1.045366693 | 0.940298812 |
| ENSMUSG00000103533 | 1 | |
| ENSMUSG00000103534 | -1.053549062 | 0.698112512 |
| ENSMUSG00000103535 | 1 | |
| ENSMUSG00000103536 | 1 | |
| ENSMUSG00000103537 | 1 | |
| ENSMUSG00000103538 | 1 | |
| ENSMUSG00000103539 | -1.065302584 | 0.698112512 |
| ENSMUSG00000103540 | 1 | |
| ENSMUSG00000103541 | 1 | |
| ENSMUSG00000103542 | 1 | |
| ENSMUSG00000103543 | -1.139579749 | 0.342377288 |
| ENSMUSG00000103544 | -1.205013732 | 0.190110443 |
| ENSMUSG00000103545 | 1 | |
| ENSMUSG00000103546 | 1.005613439 | 1 |
| ENSMUSG00000103547 | 1 | |
| ENSMUSG00000103548 | 1 | |
| ENSMUSG00000103549 | 1 | |
| ENSMUSG00000103550 | 1 | |
| ENSMUSG00000103552 | 1.059463094 | 0.608497229 |
| ENSMUSG00000103553 | 1.184532816 | 0.585646836 |
| ENSMUSG00000103554 | 1 | |
| ENSMUSG00000103555 | 1 | |
| ENSMUSG00000103556 | 1.016074389 | 1 |
| ENSMUSG00000103557 | 1 | |
| ENSMUSG00000103558 | 1.014545335 | 0.989648415 |
| ENSMUSG00000103559 | 1 | |
| ENSMUSG00000103560 | 1.032513206 | 0.800620278 |
| ENSMUSG00000103561 | -1 | 1 |
| ENSMUSG00000103562 | -1 | 1 |
| ENSMUSG00000103563 | -1.03962336 | 0.698112512 |
| ENSMUSG00000103564 | -1 | 1 |
| ENSMUSG00000103565 | 1 | |

| | | |
|---|---|---|
| ENSMUSG00000103566 | 1 | |
| ENSMUSG00000103567 | -1.252112926 | 0.001918435 |
| ENSMUSG00000103568 | 1 | |
| ENSMUSG00000103569 | 1.114043271 | 0.80707931 |
| ENSMUSG00000103570 | 1.057478344 | 0.841812929 |
| ENSMUSG00000103571 | 1 | |
| ENSMUSG00000103572 | 1.24122829 | 0.142571618 |
| ENSMUSG00000103573 | 1 | |
| ENSMUSG00000103574 | -1.027366946 | 0.983648187 |
| ENSMUSG00000103575 | 1 | |
| ENSMUSG00000103576 | 1.014163704 | 1 |
| ENSMUSG00000103577 | 1 | |
| ENSMUSG00000103579 | 1 | |
| ENSMUSG00000103580 | -1.093674795 | 0.685046738 |
| ENSMUSG00000103581 | 1.028323353 | 0.903464611 |
| ENSMUSG00000103583 | 1 | |
| ENSMUSG00000103584 | 1 | |
| ENSMUSG00000103585 | 1.06447928 | 0.698112512 |
| ENSMUSG00000103586 | 1 | |
| ENSMUSG00000103587 | 1 | |
| ENSMUSG00000103588 | 1.207239036 | 0.559354144 |
| ENSMUSG00000103589 | 1 | |
| ENSMUSG00000103590 | 1.017997209 | 0.948407797 |
| ENSMUSG00000103591 | -1.21138205 | 0.608497229 |
| ENSMUSG00000103592 | 1.055495299 | 0.928398572 |
| ENSMUSG00000103593 | 1.224182002 | 0.526949181 |
| ENSMUSG00000103594 | 1 | |
| ENSMUSG00000103595 | 1 | |
| ENSMUSG00000103596 | -1.085651418 | 0.608497229 |
| ENSMUSG00000103597 | 1 | |
| ENSMUSG00000103598 | 1 | |
| ENSMUSG00000103599 | -1.044273782 | 0.698112512 |
| ENSMUSG00000103600 | 1 | |
| ENSMUSG00000103601 | 1 | |
| ENSMUSG00000103602 | 1 | |
| ENSMUSG00000103603 | 1 | |
| ENSMUSG00000103604 | 1 | |
| ENSMUSG00000103605 | 1 | |
| ENSMUSG00000103606 | 1 | |
| ENSMUSG00000103607 | 1 | |
| ENSMUSG00000103608 | 1 | |
| ENSMUSG00000103609 | -1.126018518 | 0.744505578 |
| ENSMUSG00000103610 | 1 | |
| ENSMUSG00000103611 | -1.00002448 | 1 |
| ENSMUSG00000103612 | 1 | |
| ENSMUSG00000103613 | -1.081519275 | 0.776344308 |
| ENSMUSG00000103614 | -1.205281846 | 0.273956478 |
| ENSMUSG00000103615 | 1 | |
| ENSMUSG00000103616 | 1.019083579 | 0.949812939 |
| ENSMUSG00000103617 | 1 | |
| ENSMUSG00000103618 | 1.069068979 | 0.608497229 |
| ENSMUSG00000103620 | -1.043306199 | 0.698112512 |
| ENSMUSG00000103621 | -1.257298785 | 0.31092952 |
| ENSMUSG00000103622 | 1.060936768 | 0.698112512 |
| ENSMUSG00000103624 | -1.028348525 | 0.936995747 |
| ENSMUSG00000103625 | 1.209392349 | 0.553156258 |
| ENSMUSG00000103626 | 1 | |
| ENSMUSG00000103628 | 1.059463094 | 0.608497229 |
| ENSMUSG00000103629 | 1 | |
| ENSMUSG00000103630 | -1.061109718 | 0.608497229 |
| ENSMUSG00000103631 | 1 | |
| ENSMUSG00000103632 | 1.047622966 | 0.88732209 |
| ENSMUSG00000103633 | 1 | |

| | | |
|---|---|---|
| ENSMUSG00000103634 | 1.081229559 | 0.785584885 |
| ENSMUSG00000103635 | 1 | |
| ENSMUSG00000103636 | 1.112552835 | 0.608497229 |
| ENSMUSG00000103637 | 1 | |
| ENSMUSG00000103638 | 1 | |
| ENSMUSG00000103639 | 1 | |
| ENSMUSG00000103640 | 1 | |
| ENSMUSG00000103641 | 1.015486245 | 0.998386582 |
| ENSMUSG00000103642 | -1 | 1 |
| ENSMUSG00000103643 | 1 | |
| ENSMUSG00000103644 | 1 | |
| ENSMUSG00000103645 | 1 | |
| ENSMUSG00000103646 | 1.031212313 | 0.971164403 |
| ENSMUSG00000103647 | 1 | |
| ENSMUSG00000103648 | 1 | |
| ENSMUSG00000103649 | -1.017198576 | 0.957447586 |
| ENSMUSG00000103650 | 1 | |
| ENSMUSG00000103651 | 1 | |
| ENSMUSG00000103652 | -1.03962336 | 0.698112512 |
| ENSMUSG00000103653 | -1.063525418 | 0.698112512 |
| ENSMUSG00000103654 | 1 | |
| ENSMUSG00000103655 | 1 | |
| ENSMUSG00000103656 | 1 | |
| ENSMUSG00000103657 | 1.051729 | 0.777151142 |
| ENSMUSG00000103658 | -1.03962336 | 0.698112512 |
| ENSMUSG00000103659 | 1 | |
| ENSMUSG00000103660 | 1 | |
| ENSMUSG00000103661 | 1 | |
| ENSMUSG00000103662 | 1 | |
| ENSMUSG00000103663 | 1 | |
| ENSMUSG00000103664 | 1 | |
| ENSMUSG00000103665 | 1 | |
| ENSMUSG00000103666 | 1 | |
| ENSMUSG00000103667 | 1 | |
| ENSMUSG00000103668 | 1.019083579 | 0.968937442 |
| ENSMUSG00000103669 | 1.065662724 | 0.832767487 |
| ENSMUSG00000103670 | 1.054958861 | 0.683942136 |
| ENSMUSG00000103672 | -1.009712554 | 0.976788866 |
| ENSMUSG00000103673 | -1 | 1 |
| ENSMUSG00000103674 | 1.069068979 | 0.608497229 |
| ENSMUSG00000103675 | 1 | |
| ENSMUSG00000103676 | 1 | |
| ENSMUSG00000103677 | 2.224447803 | 0.000523799 |
| ENSMUSG00000103678 | 1 | |
| ENSMUSG00000103679 | 1 | |
| ENSMUSG00000103680 | 1.005613439 | 1 |
| ENSMUSG00000103681 | 1 | |
| ENSMUSG00000103682 | -1.228755592 | 0.293523334 |
| ENSMUSG00000103684 | 1 | |
| ENSMUSG00000103685 | -1.05548846 | 0.841171009 |
| ENSMUSG00000103686 | 1 | |
| ENSMUSG00000103688 | 1 | |
| ENSMUSG00000103689 | 1 | |
| ENSMUSG00000103690 | 1 | |
| ENSMUSG00000103691 | 1 | |
| ENSMUSG00000103692 | 1 | |
| ENSMUSG00000103693 | 1.297926193 | 0.063905003 |
| ENSMUSG00000103694 | -1.089497311 | 0.532369657 |
| ENSMUSG00000103695 | 1 | |
| ENSMUSG00000103696 | 1.028693659 | 0.895672059 |
| ENSMUSG00000103697 | 1.086501477 | 0.521105972 |
| ENSMUSG00000103698 | 1 | |
| ENSMUSG00000103699 | 1 | |

| | | |
|---|---|---|
| ENSMUSG00000103700 | 1 | |
| ENSMUSG00000103701 | 1 | |
| ENSMUSG00000103702 | -1.044273782 | 0.698112512 |
| ENSMUSG00000103703 | -1.223462828 | 0.420161906 |
| ENSMUSG00000103704 | 1 | |
| ENSMUSG00000103705 | 1 | |
| ENSMUSG00000103706 | -1.066405009 | 0.683942136 |
| ENSMUSG00000103707 | 1 | |
| ENSMUSG00000103708 | 1 | |
| ENSMUSG00000103709 | 1 | |
| ENSMUSG00000103710 | -1.057079863 | 0.873810332 |
| ENSMUSG00000103711 | -1.038343851 | 0.868547243 |
| ENSMUSG00000103712 | 1 | |
| ENSMUSG00000103713 | -1 | 1 |
| ENSMUSG00000103714 | 1 | |
| ENSMUSG00000103715 | -1.007481581 | 1 |
| ENSMUSG00000103716 | 1 | |
| ENSMUSG00000103717 | 1.028323353 | 0.934760809 |
| ENSMUSG00000103718 | 1 | |
| ENSMUSG00000103719 | 1 | |
| ENSMUSG00000103720 | 1 | |
| ENSMUSG00000103722 | 1.009822651 | 1 |
| ENSMUSG00000103723 | 1.059463094 | 0.608497229 |
| ENSMUSG00000103724 | -1 | 1 |
| ENSMUSG00000103725 | 1 | |
| ENSMUSG00000103726 | 1.105317356 | 0.608497229 |
| ENSMUSG00000103727 | -1.03962336 | 0.698112512 |
| ENSMUSG00000103728 | -1.024530911 | 0.964236174 |
| ENSMUSG00000103729 | -1 | 1 |
| ENSMUSG00000103730 | 1 | |
| ENSMUSG00000103731 | -1.055999182 | 0.608497229 |
| ENSMUSG00000103732 | 1 | |
| ENSMUSG00000103733 | -1 | 1 |
| ENSMUSG00000103734 | -1.053549062 | 0.769745556 |
| ENSMUSG00000103735 | -1.03962336 | 0.698112512 |
| ENSMUSG00000103736 | 1 | |
| ENSMUSG00000103737 | 1 | |
| ENSMUSG00000103738 | 1 | |
| ENSMUSG00000103739 | 1.005613439 | 1 |
| ENSMUSG00000103740 | 1 | |
| ENSMUSG00000103741 | -1.063525418 | 0.698112512 |
| ENSMUSG00000103742 | 1 | |
| ENSMUSG00000103744 | 1 | |
| ENSMUSG00000103745 | 1 | |
| ENSMUSG00000103746 | -1.029548358 | 0.996880111 |
| ENSMUSG00000103747 | 1.023743961 | 0.968735041 |
| ENSMUSG00000103748 | -1.034006264 | 0.968423939 |
| ENSMUSG00000103749 | 1.045973314 | 0.865583959 |
| ENSMUSG00000103750 | 1 | |
| ENSMUSG00000103751 | -1 | 1 |
| ENSMUSG00000103752 | 1 | |
| ENSMUSG00000103753 | -1.043306199 | 0.698112512 |
| ENSMUSG00000103754 | -1.142755218 | 0.60311292 |
| ENSMUSG00000103755 | -1.067540749 | 0.809511199 |
| ENSMUSG00000103756 | 1.076358224 | 0.608497229 |
| ENSMUSG00000103757 | 1 | |
| ENSMUSG00000103758 | 1 | |
| ENSMUSG00000103759 | -1.129231493 | 0.608497229 |
| ENSMUSG00000103760 | 1.005212439 | 1 |
| ENSMUSG00000103761 | 1 | |
| ENSMUSG00000103762 | 1.132639128 | 0.398767938 |
| ENSMUSG00000103763 | -1.084014399 | 0.733276285 |
| ENSMUSG00000103764 | 1 | |

| | | |
|---|---|---|
| ENSMUSG00000103765 | 1 | |
| ENSMUSG00000103766 | -1 | 1 |
| ENSMUSG00000103767 | -1 | 1 |
| ENSMUSG00000103768 | 1 | |
| ENSMUSG00000103769 | 1 | |
| ENSMUSG00000103770 | 1 | |
| ENSMUSG00000103771 | -1.075182332 | 0.780023249 |
| ENSMUSG00000103772 | -1.044273782 | 0.698112512 |
| ENSMUSG00000103773 | 1 | |
| ENSMUSG00000103774 | 1.008911686 | 1 |
| ENSMUSG00000103775 | 1 | |
| ENSMUSG00000103776 | 1 | |
| ENSMUSG00000103777 | 1 | |
| ENSMUSG00000103778 | 1.069068979 | 0.608497229 |
| ENSMUSG00000103779 | 1.059463094 | 0.608497229 |
| ENSMUSG00000103780 | 1 | |
| ENSMUSG00000103781 | 1 | |
| ENSMUSG00000103782 | 1 | |
| ENSMUSG00000103783 | -1.200114593 | 0.287659118 |
| ENSMUSG00000103784 | 1 | |
| ENSMUSG00000103785 | -1.099174267 | 0.535052277 |
| ENSMUSG00000103786 | -1.137443589 | 0.608497229 |
| ENSMUSG00000103787 | 1 | |
| ENSMUSG00000103788 | 1 | |
| ENSMUSG00000103789 | -1.071836213 | 0.786546174 |
| ENSMUSG00000103790 | 1 | |
| ENSMUSG00000103791 | 1 | |
| ENSMUSG00000103792 | 1.05410549 | 0.869434134 |
| ENSMUSG00000103793 | -1.140133758 | 0.617823788 |
| ENSMUSG00000103794 | 1 | |
| ENSMUSG00000103795 | 1 | |
| ENSMUSG00000103796 | 1 | |
| ENSMUSG00000103797 | 1 | |
| ENSMUSG00000103798 | 1 | |
| ENSMUSG00000103799 | 1 | |
| ENSMUSG00000103800 | 1 | |
| ENSMUSG00000103801 | -1.069502966 | 0.698112512 |
| ENSMUSG00000103802 | 1 | |
| ENSMUSG00000103803 | -1.043306199 | 0.858911269 |
| ENSMUSG00000103804 | -1.142999928 | 0.381329895 |
| ENSMUSG00000103805 | 1 | |
| ENSMUSG00000103807 | 1.132639128 | 0.217803313 |
| ENSMUSG00000103808 | -1.049475223 | 0.897856602 |
| ENSMUSG00000103809 | -1 | 1 |
| ENSMUSG00000103810 | -1.355415621 | 0.138852703 |
| ENSMUSG00000103811 | -1 | 1 |
| ENSMUSG00000103812 | 1 | |
| ENSMUSG00000103813 | 1 | |
| ENSMUSG00000103814 | 1 | |
| ENSMUSG00000103815 | 1 | |
| ENSMUSG00000103816 | 1 | |
| ENSMUSG00000103817 | -1 | 1 |
| ENSMUSG00000103818 | 1.069068979 | 0.698112512 |
| ENSMUSG00000103819 | 1 | |
| ENSMUSG00000103820 | 1 | |
| ENSMUSG00000103821 | -1.070817129 | 0.698112512 |
| ENSMUSG00000103822 | -1.01266613 | 1 |
| ENSMUSG00000103823 | 1.122462048 | 0.420161906 |
| ENSMUSG00000103824 | -1 | 1 |
| ENSMUSG00000103825 | 1.007818844 | 1 |
| ENSMUSG00000103826 | 1 | |
| ENSMUSG00000103827 | -1 | 1 |
| ENSMUSG00000103828 | -1.044273782 | 0.698112512 |

| | | |
|---|---|---|
| ENSMUSG00000103829 | 1 | |
| ENSMUSG00000103830 | 1 | |
| ENSMUSG00000103831 | 1 | |
| ENSMUSG00000103832 | 1 | |
| ENSMUSG00000103833 | -1.01914567 | 0.995747207 |
| ENSMUSG00000103834 | 1 | |
| ENSMUSG00000103835 | -1.077189453 | 0.609323856 |
| ENSMUSG00000103836 | 1 | |
| ENSMUSG00000103837 | 1 | |
| ENSMUSG00000103839 | 1.063372219 | 0.608497229 |
| ENSMUSG00000103840 | 1 | |
| ENSMUSG00000103841 | 1 | |
| ENSMUSG00000103842 | 1 | |
| ENSMUSG00000103843 | 1.132639128 | 0.217803313 |
| ENSMUSG00000103844 | 1 | |
| ENSMUSG00000103845 | 1.020344501 | 0.988665808 |
| ENSMUSG00000103846 | 1 | |
| ENSMUSG00000103847 | -1.043764923 | 0.930108107 |
| ENSMUSG00000103848 | 1 | |
| ENSMUSG00000103849 | -1.03962336 | 0.803072249 |
| ENSMUSG00000103850 | 1 | |
| ENSMUSG00000103851 | -1.1686065 | 0.544838788 |
| ENSMUSG00000103852 | 1.069068979 | 0.608497229 |
| ENSMUSG00000103854 | 1.038531341 | 0.929955254 |
| ENSMUSG00000103855 | 1 | |
| ENSMUSG00000103856 | 1 | |
| ENSMUSG00000103857 | 1.001718606 | 1 |
| ENSMUSG00000103858 | 1.283884444 | 0.000429559 |
| ENSMUSG00000103859 | 1 | |
| ENSMUSG00000103860 | 1 | |
| ENSMUSG00000103861 | -1.03962336 | 0.803072249 |
| ENSMUSG00000103862 | 1 | |
| ENSMUSG00000103863 | 1 | |
| ENSMUSG00000103864 | 1.055416288 | 0.888667844 |
| ENSMUSG00000103865 | -1.158951053 | 0.191019295 |
| ENSMUSG00000103867 | -1.161107412 | 0.511036937 |
| ENSMUSG00000103869 | -1.219624708 | 0.249329047 |
| ENSMUSG00000103870 | 1 | |
| ENSMUSG00000103871 | 1 | |
| ENSMUSG00000103872 | 1 | |
| ENSMUSG00000103873 | 1 | |
| ENSMUSG00000103874 | 1 | |
| ENSMUSG00000103875 | 1 | |
| ENSMUSG00000103876 | 1 | |
| ENSMUSG00000103877 | 1.095872691 | 0.608497229 |
| ENSMUSG00000103878 | -1 | 1 |
| ENSMUSG00000103880 | 1.117720869 | 0.608497229 |
| ENSMUSG00000103881 | 1.084618945 | 0.698112512 |
| ENSMUSG00000103882 | 1.095872691 | 0.671280603 |
| ENSMUSG00000103883 | 1.0988361 | 0.110665406 |
| ENSMUSG00000103884 | 1 | |
| ENSMUSG00000103885 | -1.044273782 | 0.698112512 |
| ENSMUSG00000103886 | 1 | |
| ENSMUSG00000103887 | 1 | |
| ENSMUSG00000103888 | 1 | |
| ENSMUSG00000103889 | 1.00880322 | 1 |
| ENSMUSG00000103890 | 1 | |
| ENSMUSG00000103891 | -1.113957691 | 0.578238275 |
| ENSMUSG00000103892 | 1 | |
| ENSMUSG00000103893 | 1 | |
| ENSMUSG00000103894 | 1 | |
| ENSMUSG00000103895 | 1 | |
| ENSMUSG00000103896 | 1 | |

| | | |
|---|---|---|
| ENSMUSG00000103897 | 1.415121452 | 0.0399955 |
| ENSMUSG00000103898 | 1.177189741 | 0.430495431 |
| ENSMUSG00000103899 | -1.128790552 | 0.636056232 |
| ENSMUSG00000103900 | -1.037934057 | 0.951399668 |
| ENSMUSG00000103901 | -1.021415472 | 0.922446617 |
| ENSMUSG00000103902 | 1 | |
| ENSMUSG00000103903 | -1.085651418 | 0.532369657 |
| ENSMUSG00000103904 | -1.041460398 | 0.921959803 |
| ENSMUSG00000103905 | 1 | |
| ENSMUSG00000103906 | -1.048298977 | 0.91686299 |
| ENSMUSG00000103907 | -1.043306199 | 0.698112512 |
| ENSMUSG00000103908 | 1.019083579 | 0.949812939 |
| ENSMUSG00000103909 | 1 | |
| ENSMUSG00000103911 | -1 | 1 |
| ENSMUSG00000103912 | -1.11061171 | 0.547016491 |
| ENSMUSG00000103913 | 1.059463094 | 0.608497229 |
| ENSMUSG00000103914 | 1 | |
| ENSMUSG00000103915 | 1 | |
| ENSMUSG00000103916 | 1 | |
| ENSMUSG00000103917 | 1 | |
| ENSMUSG00000103918 | 1 | |
| ENSMUSG00000103919 | 1 | |
| ENSMUSG00000103920 | 1.075295999 | 0.780320794 |
| ENSMUSG00000103921 | 1.059463094 | 0.608497229 |
| ENSMUSG00000103922 | -1 | 1 |
| ENSMUSG00000103923 | 1.014545335 | 0.979002724 |
| ENSMUSG00000103924 | 1 | |
| ENSMUSG00000103925 | -1.094488708 | 0.698112512 |
| ENSMUSG00000103926 | -1.211487332 | 0.313649348 |
| ENSMUSG00000103927 | -1.003012737 | 1 |
| ENSMUSG00000103928 | -1.007637818 | 1 |
| ENSMUSG00000103929 | 1 | |
| ENSMUSG00000103930 | 1 | |
| ENSMUSG00000103931 | 1 | |
| ENSMUSG00000103932 | -1.188005937 | 0.247755833 |
| ENSMUSG00000103933 | 1.074082397 | 0.801154107 |
| ENSMUSG00000103934 | 1 | |
| ENSMUSG00000103935 | 1 | |
| ENSMUSG00000103936 | 1 | |
| ENSMUSG00000103937 | -1.065096629 | 0.807900897 |
| ENSMUSG00000103938 | 1 | |
| ENSMUSG00000103939 | 1 | |
| ENSMUSG00000103940 | -1.142727245 | 0.291233266 |
| ENSMUSG00000103942 | 1 | |
| ENSMUSG00000103943 | 1 | |
| ENSMUSG00000103944 | 1 | |
| ENSMUSG00000103945 | 1.150845036 | 0.516012944 |
| ENSMUSG00000103947 | 1 | |
| ENSMUSG00000103948 | 1.089470641 | 0.657060635 |
| ENSMUSG00000103949 | -1.222783585 | 0.437045401 |
| ENSMUSG00000103950 | 1 | |
| ENSMUSG00000103951 | -1.043306199 | 0.698112512 |
| ENSMUSG00000103952 | 1.14283885 | 0.698112512 |
| ENSMUSG00000103953 | -1.099174267 | 0.535052277 |
| ENSMUSG00000103954 | 1 | |
| ENSMUSG00000103955 | 1 | |
| ENSMUSG00000103956 | 1 | |
| ENSMUSG00000103957 | 1.059463094 | 0.608497229 |
| ENSMUSG00000103958 | 1 | |
| ENSMUSG00000103959 | 1.014545335 | 0.979002724 |
| ENSMUSG00000103960 | 1 | |
| ENSMUSG00000103961 | -1.03962336 | 0.698112512 |
| ENSMUSG00000103962 | 1.021436306 | 0.981761701 |

| | | |
|---|---|---|
| ENSMUSG00000103963 | 1 | |
| ENSMUSG00000103964 | 1.05410549 | 0.858169153 |
| ENSMUSG00000103965 | 1.059463094 | 0.608497229 |
| ENSMUSG00000103966 | 1.061698341 | 0.731173125 |
| ENSMUSG00000103967 | -1.151876726 | 0.639682514 |
| ENSMUSG00000103968 | 1 | |
| ENSMUSG00000103969 | 1 | |
| ENSMUSG00000103970 | -1.079794376 | 0.807247723 |
| ENSMUSG00000103971 | 1 | |
| ENSMUSG00000103972 | 1 | |
| ENSMUSG00000103973 | -1.052715633 | 0.839885707 |
| ENSMUSG00000103975 | 1 | |
| ENSMUSG00000103976 | -1.126611283 | 0.540959285 |
| ENSMUSG00000103977 | 1 | |
| ENSMUSG00000103978 | 1 | |
| ENSMUSG00000103979 | 1 | |
| ENSMUSG00000103980 | 1 | |
| ENSMUSG00000103981 | 1 | |
| ENSMUSG00000103982 | 1 | |
| ENSMUSG00000103983 | -1.096010634 | 0.608497229 |
| ENSMUSG00000103984 | 1.128286421 | 0.6428944 |
| ENSMUSG00000103985 | 1.132639128 | 0.398767938 |
| ENSMUSG00000103986 | 1.074221595 | 0.8072494 |
| ENSMUSG00000103987 | 1 | |
| ENSMUSG00000103989 | 1 | |
| ENSMUSG00000103990 | 1 | |
| ENSMUSG00000103991 | 1 | |
| ENSMUSG00000103992 | 1.034095782 | 0.919000509 |
| ENSMUSG00000103993 | 1 | |
| ENSMUSG00000103994 | 1 | |
| ENSMUSG00000103995 | 1 | |
| ENSMUSG00000103996 | 1.059463094 | 0.698112512 |
| ENSMUSG00000103997 | 1 | |
| ENSMUSG00000103998 | -1.001770549 | 1 |
| ENSMUSG00000103999 | 1.053859357 | 0.875934753 |
| ENSMUSG00000104000 | 1.069068979 | 0.698112512 |
| ENSMUSG00000104001 | 1 | |
| ENSMUSG00000104002 | 1.105166679 | 0.623140373 |
| ENSMUSG00000104003 | -1.244369178 | 0.323795799 |
| ENSMUSG00000104004 | -1.03962336 | 0.698112512 |
| ENSMUSG00000104006 | 1 | |
| ENSMUSG00000104007 | 1 | |
| ENSMUSG00000104009 | 1 | |
| ENSMUSG00000104010 | -1.143256789 | 0.608497229 |
| ENSMUSG00000104011 | -1.003012737 | 1 |
| ENSMUSG00000104012 | -1 | 1 |
| ENSMUSG00000104013 | 1 | |
| ENSMUSG00000104014 | 1 | |
| ENSMUSG00000104015 | 1.014545335 | 0.979002724 |
| ENSMUSG00000104016 | -1.001451142 | 1 |
| ENSMUSG00000104017 | 1 | |
| ENSMUSG00000104018 | 1.130785691 | 0.712130748 |
| ENSMUSG00000104020 | 1.059463094 | 0.699340382 |
| ENSMUSG00000104021 | 1 | |
| ENSMUSG00000104022 | 1.126351041 | 0.698112512 |
| ENSMUSG00000104023 | -1.053549062 | 0.769745556 |
| ENSMUSG00000104024 | 1 | |
| ENSMUSG00000104025 | 1 | |
| ENSMUSG00000104026 | -1.089497311 | 0.532369657 |
| ENSMUSG00000104027 | -1.043306199 | 0.698112512 |
| ENSMUSG00000104028 | 1 | |
| ENSMUSG00000104029 | 1 | |
| ENSMUSG00000104030 | -1.03962336 | 0.698112512 |

| | | |
|---|---|---|
| ENSMUSG00000104031 | -1.213118765 | 0.387056636 |
| ENSMUSG00000104032 | 1 | |
| ENSMUSG00000104033 | 1.059463094 | 0.698112512 |
| ENSMUSG00000104034 | 1.059463094 | 0.608497229 |
| ENSMUSG00000104035 | 1 | |
| ENSMUSG00000104036 | 1.186312313 | 0.136710699 |
| ENSMUSG00000104037 | 1.05289069 | 0.840506033 |
| ENSMUSG00000104038 | 1 | |
| ENSMUSG00000104039 | 1 | |
| ENSMUSG00000104040 | -1.043306199 | 0.79291651 |
| ENSMUSG00000104041 | 1 | |
| ENSMUSG00000104042 | 1 | |
| ENSMUSG00000104043 | 1.186622869 | 0.116506115 |
| ENSMUSG00000104044 | 1.005851671 | 1 |
| ENSMUSG00000104045 | 1.577873406 | 0.000122514 |
| ENSMUSG00000104046 | -1.000190151 | 1 |
| ENSMUSG00000104048 | 1 | |
| ENSMUSG00000104050 | 1 | |
| ENSMUSG00000104051 | -1.013101236 | 0.848502156 |
| ENSMUSG00000104052 | -1.023769022 | 0.975580864 |
| ENSMUSG00000104053 | -1.03962336 | 0.803072249 |
| ENSMUSG00000104054 | 1 | |
| ENSMUSG00000104055 | 1 | |
| ENSMUSG00000104056 | -1.086192882 | 0.698112512 |
| ENSMUSG00000104057 | -1.009091449 | 1 |
| ENSMUSG00000104058 | 1 | |
| ENSMUSG00000104059 | 1 | |
| ENSMUSG00000104060 | 1.023168148 | 0.827942378 |
| ENSMUSG00000104061 | -1.153237309 | 0.511378706 |
| ENSMUSG00000104062 | 1 | |
| ENSMUSG00000104063 | 1.288664021 | 0.265061919 |
| ENSMUSG00000104064 | 1 | |
| ENSMUSG00000104065 | -1 | 1 |
| ENSMUSG00000104066 | 1.132639128 | 0.217803313 |
| ENSMUSG00000104067 | 1 | |
| ENSMUSG00000104068 | 1 | |
| ENSMUSG00000104069 | -1.091329919 | 0.724042346 |
| ENSMUSG00000104070 | 1 | |
| ENSMUSG00000104071 | 1 | |
| ENSMUSG00000104072 | 1 | |
| ENSMUSG00000104073 | -1 | 1 |
| ENSMUSG00000104074 | 1.102700218 | 0.741916095 |
| ENSMUSG00000104075 | 1 | |
| ENSMUSG00000104076 | -1.044273782 | 0.801443616 |
| ENSMUSG00000104077 | 1 | |
| ENSMUSG00000104078 | 1.059463094 | 0.608497229 |
| ENSMUSG00000104079 | 1 | |
| ENSMUSG00000104080 | 1.066531716 | 0.756671188 |
| ENSMUSG00000104081 | -1 | 1 |
| ENSMUSG00000104082 | 1.005613439 | 1 |
| ENSMUSG00000104083 | 1.178139999 | 0.311521398 |
| ENSMUSG00000104084 | 1 | |
| ENSMUSG00000104085 | 1 | |
| ENSMUSG00000104086 | 1 | |
| ENSMUSG00000104087 | -1.053549062 | 0.698112512 |
| ENSMUSG00000104088 | 1.218828701 | 0.186329296 |
| ENSMUSG00000104089 | 1 | |
| ENSMUSG00000104090 | -1.013468747 | 1 |
| ENSMUSG00000104091 | 1.007021636 | 1 |
| ENSMUSG00000104092 | 1.005613439 | 1 |
| ENSMUSG00000104093 | 1.178777735 | 0.210424884 |
| ENSMUSG00000104094 | 1.340862131 | 0.347267031 |
| ENSMUSG00000104095 | -1.016001493 | 0.939910929 |

| | | |
|---|---|---|
| ENSMUSG00000104096 | 1 | |
| ENSMUSG00000104097 | 1.079681442 | 0.776953697 |
| ENSMUSG00000104098 | 1 | |
| ENSMUSG00000104099 | 1 | |
| ENSMUSG00000104100 | 1.014545335 | 0.991944074 |
| ENSMUSG00000104101 | -1.043306199 | 0.698112512 |
| ENSMUSG00000104102 | 1.05943716 | 0.770958539 |
| ENSMUSG00000104103 | 1.059463094 | 0.608497229 |
| ENSMUSG00000104104 | 1.05410549 | 0.835701362 |
| ENSMUSG00000104105 | 1 | |
| ENSMUSG00000104106 | -1 | 1 |
| ENSMUSG00000104107 | -1.053549062 | 0.698112512 |
| ENSMUSG00000104108 | 1.079681442 | 0.807933619 |
| ENSMUSG00000104109 | 1.020344501 | 0.983178084 |
| ENSMUSG00000104110 | 1 | |
| ENSMUSG00000104111 | 1 | |
| ENSMUSG00000104112 | 1 | |
| ENSMUSG00000104113 | 1.099338164 | 0.698112512 |
| ENSMUSG00000104114 | 1.024804142 | 0.956661971 |
| ENSMUSG00000104115 | 1 | |
| ENSMUSG00000104116 | 1.199989356 | 0.064133166 |
| ENSMUSG00000104117 | -1.053549062 | 0.769745556 |
| ENSMUSG00000104118 | -1 | 1 |
| ENSMUSG00000104120 | -1.028348525 | 0.936995747 |
| ENSMUSG00000104121 | 1.075324819 | 0.750742698 |
| ENSMUSG00000104122 | -1.024718486 | 0.970771742 |
| ENSMUSG00000104123 | 1 | |
| ENSMUSG00000104125 | 1.082332121 | 0.785919418 |
| ENSMUSG00000104126 | -1.053549062 | 0.698112512 |
| ENSMUSG00000104127 | 1 | |
| ENSMUSG00000104129 | 1 | |
| ENSMUSG00000104130 | -1.03962336 | 0.698112512 |
| ENSMUSG00000104131 | 1 | |
| ENSMUSG00000104132 | 1 | |
| ENSMUSG00000104133 | 1 | |
| ENSMUSG00000104134 | 1 | |
| ENSMUSG00000104135 | 1 | |
| ENSMUSG00000104136 | 1.115316151 | 0.565271791 |
| ENSMUSG00000104137 | 1 | |
| ENSMUSG00000104138 | -1.044273782 | 0.698112512 |
| ENSMUSG00000104140 | -1.122662044 | 0.608497229 |
| ENSMUSG00000104141 | -1.038444538 | 0.897848537 |
| ENSMUSG00000104142 | 1 | |
| ENSMUSG00000104144 | 1 | |
| ENSMUSG00000104145 | -1.081225398 | 0.769745556 |
| ENSMUSG00000104146 | 1 | |
| ENSMUSG00000104147 | -1.078590893 | 0.744030455 |
| ENSMUSG00000104148 | 1 | |
| ENSMUSG00000104149 | 1.032745674 | 0.912030056 |
| ENSMUSG00000104150 | -1 | 1 |
| ENSMUSG00000104152 | 1 | |
| ENSMUSG00000104153 | 1 | |
| ENSMUSG00000104154 | -1.008380008 | 1 |
| ENSMUSG00000104155 | 1 | |
| ENSMUSG00000104156 | 1.072593414 | 0.837321292 |
| ENSMUSG00000104157 | 1 | |
| ENSMUSG00000104158 | 1 | |
| ENSMUSG00000104159 | -1.091302857 | 0.506638168 |
| ENSMUSG00000104161 | 1 | |
| ENSMUSG00000104162 | 1 | |
| ENSMUSG00000104163 | 1 | |
| ENSMUSG00000104164 | 1.064509339 | 0.698112512 |
| ENSMUSG00000104165 | 1 | |

| | | |
|---|---|---|
| ENSMUSG00000104166 | 1.065410325 | 0.764982341 |
| ENSMUSG00000104167 | 1 | |
| ENSMUSG00000104168 | -1.102631374 | 0.776001651 |
| ENSMUSG00000104169 | 1 | |
| ENSMUSG00000104170 | 1.014545335 | 0.991944074 |
| ENSMUSG00000104171 | 1 | |
| ENSMUSG00000104172 | 1 | |
| ENSMUSG00000104173 | -1.037482356 | 0.902641804 |
| ENSMUSG00000104174 | 1 | |
| ENSMUSG00000104175 | 1 | |
| ENSMUSG00000104176 | 1 | |
| ENSMUSG00000104177 | 1 | |
| ENSMUSG00000104178 | 1 | |
| ENSMUSG00000104180 | 1 | |
| ENSMUSG00000104181 | -1.074213609 | 0.698112512 |
| ENSMUSG00000104182 | 1 | |
| ENSMUSG00000104183 | 1.20839735 | 0.313664201 |
| ENSMUSG00000104184 | 1.504179965 | 0.069826703 |
| ENSMUSG00000104185 | 1 | |
| ENSMUSG00000104186 | 1.02889509 | 0.960812739 |
| ENSMUSG00000104187 | -1.053549062 | 0.769745556 |
| ENSMUSG00000104188 | -1 | 1 |
| ENSMUSG00000104189 | 1.019083579 | 0.97045957 |
| ENSMUSG00000104190 | 1.005212439 | 1 |
| ENSMUSG00000104191 | 1 | |
| ENSMUSG00000104192 | 1 | |
| ENSMUSG00000104194 | 1 | |
| ENSMUSG00000104195 | 1.016250869 | 0.988424085 |
| ENSMUSG00000104197 | -1.044273782 | 0.801443616 |
| ENSMUSG00000104198 | 1 | |
| ENSMUSG00000104199 | 1 | |
| ENSMUSG00000104200 | 1 | |
| ENSMUSG00000104201 | 1 | |
| ENSMUSG00000104202 | 1 | |
| ENSMUSG00000104203 | 1 | |
| ENSMUSG00000104204 | 1.121015602 | 0.547817379 |
| ENSMUSG00000104205 | 1 | |
| ENSMUSG00000104206 | 1 | |
| ENSMUSG00000104207 | 1 | |
| ENSMUSG00000104208 | -1 | 1 |
| ENSMUSG00000104209 | 1 | |
| ENSMUSG00000104210 | -1.044273782 | 0.790851299 |
| ENSMUSG00000104211 | 1.088030006 | 0.486930276 |
| ENSMUSG00000104212 | -1.207114389 | 0.316089517 |
| ENSMUSG00000104213 | 1 | |
| ENSMUSG00000104214 | -1.028605469 | 0.972980766 |
| ENSMUSG00000104215 | 1 | |
| ENSMUSG00000104216 | 1 | |
| ENSMUSG00000104217 | 1 | |
| ENSMUSG00000104218 | -1 | 1 |
| ENSMUSG00000104219 | 1 | |
| ENSMUSG00000104220 | 1 | |
| ENSMUSG00000104221 | -1 | 1 |
| ENSMUSG00000104222 | -1.082001498 | 0.735723981 |
| ENSMUSG00000104223 | 1 | |
| ENSMUSG00000104224 | 1 | |
| ENSMUSG00000104225 | 1 | |
| ENSMUSG00000104226 | 1 | |
| ENSMUSG00000104227 | 1 | |
| ENSMUSG00000104228 | 1 | |
| ENSMUSG00000104229 | 1 | |
| ENSMUSG00000104230 | 1 | |
| ENSMUSG00000104231 | -1.052534886 | 0.72715392 |

| | | |
|---|---|---|
| ENSMUSG00000104232 | 1 | |
| ENSMUSG00000104233 | 1 | |
| ENSMUSG00000104234 | 1 | |
| ENSMUSG00000104235 | 1.161291605 | 0.469202786 |
| ENSMUSG00000104236 | 1 | |
| ENSMUSG00000104237 | 1.064531743 | 0.845217963 |
| ENSMUSG00000104238 | 1 | |
| ENSMUSG00000104239 | -1 | 1 |
| ENSMUSG00000104240 | 1 | |
| ENSMUSG00000104241 | -1.233467038 | 0.375845184 |
| ENSMUSG00000104242 | 1 | |
| ENSMUSG00000104243 | 1.059463094 | 0.608497229 |
| ENSMUSG00000104244 | 1.501636009 | 0.030120212 |
| ENSMUSG00000104245 | 1.123739359 | 0.075178706 |
| ENSMUSG00000104247 | 1 | |
| ENSMUSG00000104248 | -1.02513447 | 0.972980766 |
| ENSMUSG00000104249 | 1 | |
| ENSMUSG00000104251 | 1 | |
| ENSMUSG00000104252 | 1 | |
| ENSMUSG00000104253 | 1.074771674 | 0.50744656 |
| ENSMUSG00000104254 | 1 | |
| ENSMUSG00000104255 | 1 | |
| ENSMUSG00000104256 | 1 | |
| ENSMUSG00000104257 | -1.003834322 | 1 |
| ENSMUSG00000104258 | 1 | |
| ENSMUSG00000104259 | 1 | |
| ENSMUSG00000104260 | 1 | |
| ENSMUSG00000104261 | 1 | |
| ENSMUSG00000104262 | 1 | |
| ENSMUSG00000104263 | -1.016716156 | 0.938066471 |
| ENSMUSG00000104264 | 1 | |
| ENSMUSG00000104265 | 1 | |
| ENSMUSG00000104266 | 1.199989356 | 4.02961E-05 |
| ENSMUSG00000104267 | 1 | |
| ENSMUSG00000104268 | 1 | |
| ENSMUSG00000104269 | 1 | |
| ENSMUSG00000104270 | 1 | |
| ENSMUSG00000104271 | -1.176079149 | 0.578987143 |
| ENSMUSG00000104272 | -1.075182332 | 0.751475418 |
| ENSMUSG00000104273 | -1.022327813 | 0.958914121 |
| ENSMUSG00000104274 | -1.085651418 | 0.698112512 |
| ENSMUSG00000104275 | -1.043492091 | 0.913882562 |
| ENSMUSG00000104276 | 1 | |
| ENSMUSG00000104277 | -1.027562261 | 0.979224582 |
| ENSMUSG00000104278 | 1 | |
| ENSMUSG00000104279 | 1.041330357 | 0.894881355 |
| ENSMUSG00000104280 | 1.210812682 | 0.417843828 |
| ENSMUSG00000104281 | 1 | |
| ENSMUSG00000104282 | 1 | |
| ENSMUSG00000104283 | 1 | |
| ENSMUSG00000104284 | 1 | |
| ENSMUSG00000104285 | 1 | |
| ENSMUSG00000104286 | -1.03962336 | 0.698112512 |
| ENSMUSG00000104287 | 1 | |
| ENSMUSG00000104288 | 1 | |
| ENSMUSG00000104289 | 1 | |
| ENSMUSG00000104290 | 1 | |
| ENSMUSG00000104291 | 1.069068979 | 0.698112512 |
| ENSMUSG00000104292 | -1.034990181 | 0.963057282 |
| ENSMUSG00000104293 | 1.085267407 | 0.717035356 |
| ENSMUSG00000104294 | 1 | |
| ENSMUSG00000104295 | 1 | |
| ENSMUSG00000104296 | 1 | |

| | | |
|---|---|---|
| ENSMUSG00000104297 | -1.043306199 | 0.698112512 |
| ENSMUSG00000104298 | 1.005613439 | 1 |
| ENSMUSG00000104299 | -1.057338222 | 0.877688995 |
| ENSMUSG00000104301 | 1.069068979 | 0.698112512 |
| ENSMUSG00000104302 | -1 | 1 |
| ENSMUSG00000104303 | 1.057314073 | 0.693220376 |
| ENSMUSG00000104304 | 1 | |
| ENSMUSG00000104306 | -1.043306199 | 0.79291651 |
| ENSMUSG00000104307 | 1 | |
| ENSMUSG00000104308 | 1 | |
| ENSMUSG00000104309 | 1.038531341 | 0.914474058 |
| ENSMUSG00000104310 | 1 | |
| ENSMUSG00000104311 | 1 | |
| ENSMUSG00000104312 | 1.023743961 | 0.95702926 |
| ENSMUSG00000104313 | 1 | |
| ENSMUSG00000104314 | 1.059463094 | 0.608497229 |
| ENSMUSG00000104315 | 1.057383712 | 0.895137669 |
| ENSMUSG00000104316 | -1.085651418 | 0.795837467 |
| ENSMUSG00000104317 | 1 | |
| ENSMUSG00000104318 | 1 | |
| ENSMUSG00000104319 | 1 | |
| ENSMUSG00000104320 | 1 | |
| ENSMUSG00000104321 | 1 | |
| ENSMUSG00000104323 | 1 | |
| ENSMUSG00000104324 | -1.026069229 | 0.850372813 |
| ENSMUSG00000104325 | 1.004398972 | 1 |
| ENSMUSG00000104326 | 1 | |
| ENSMUSG00000104327 | 1.059463094 | 0.608497229 |
| ENSMUSG00000104328 | 1 | |
| ENSMUSG00000104329 | 1.27770457 | 0.00774897 |
| ENSMUSG00000104330 | 1.069068979 | 0.608497229 |
| ENSMUSG00000104331 | -1.089773232 | 0.857547928 |
| ENSMUSG00000104332 | -1 | 1 |
| ENSMUSG00000104333 | 1 | |
| ENSMUSG00000104334 | 1 | |
| ENSMUSG00000104335 | 1 | |
| ENSMUSG00000104336 | 1 | |
| ENSMUSG00000104337 | 1 | |
| ENSMUSG00000104338 | 1 | |
| ENSMUSG00000104339 | -1.085651418 | 0.698112512 |
| ENSMUSG00000104340 | -1.044273782 | 0.698112512 |
| ENSMUSG00000104341 | 1.06076715 | 0.943819872 |
| ENSMUSG00000104342 | -1.133587263 | 0.229824504 |
| ENSMUSG00000104343 | 1 | |
| ENSMUSG00000104344 | -1.046413774 | 0.608497229 |
| ENSMUSG00000104345 | 1 | |
| ENSMUSG00000104346 | 1.131080567 | 0.625224412 |
| ENSMUSG00000104347 | 1.069068979 | 0.698112512 |
| ENSMUSG00000104348 | 1.12289259 | 0.642819974 |
| ENSMUSG00000104349 | -1.043306199 | 0.698112512 |
| ENSMUSG00000104350 | -1.217770253 | 0.091167679 |
| ENSMUSG00000104351 | -1.06989077 | 0.698112512 |
| ENSMUSG00000104352 | 1 | |
| ENSMUSG00000104354 | 1 | |
| ENSMUSG00000104355 | 1 | |
| ENSMUSG00000104356 | 1 | |
| ENSMUSG00000104358 | -1.032544731 | 0.825401367 |
| ENSMUSG00000104360 | -1.03962336 | 0.814089245 |
| ENSMUSG00000104361 | 1.009538174 | 0.980773838 |
| ENSMUSG00000104362 | 1 | |
| ENSMUSG00000104363 | -1.116853908 | 0.558122732 |
| ENSMUSG00000104364 | 1 | |
| ENSMUSG00000104365 | -1 | 1 |

| | | |
|---|---|---|
| ENSMUSG00000104366 | 1 | |
| ENSMUSG00000104367 | -1.03962336 | 0.698112512 |
| ENSMUSG00000104368 | -1.034990181 | 0.965872389 |
| ENSMUSG00000104369 | -1.021365545 | 0.936351125 |
| ENSMUSG00000104370 | 1.084618945 | 0.614071182 |
| ENSMUSG00000104371 | 1 | |
| ENSMUSG00000104373 | 1 | |
| ENSMUSG00000104374 | 1 | |
| ENSMUSG00000104375 | 1 | |
| ENSMUSG00000104376 | 1 | |
| ENSMUSG00000104377 | 1.063055321 | 0.608497229 |
| ENSMUSG00000104378 | -1.003834322 | 1 |
| ENSMUSG00000104379 | -1.043306199 | 0.699340382 |
| ENSMUSG00000104380 | 1 | |
| ENSMUSG00000104381 | 1 | |
| ENSMUSG00000104382 | 1 | |
| ENSMUSG00000104383 | 1.344299606 | 0.161510586 |
| ENSMUSG00000104384 | -1.081352162 | 0.78301543 |
| ENSMUSG00000104385 | -1.044273782 | 0.801443616 |
| ENSMUSG00000104386 | 1 | |
| ENSMUSG00000104388 | -1.145536749 | 0.608497229 |
| ENSMUSG00000104389 | -1.317024107 | 0.014891518 |
| ENSMUSG00000104390 | 1 | |
| ENSMUSG00000104391 | 1.177875245 | 0.616916789 |
| ENSMUSG00000104392 | 1.00707884 | 1 |
| ENSMUSG00000104393 | 1 | |
| ENSMUSG00000104394 | -1.134823511 | 0.012878734 |
| ENSMUSG00000104395 | 1 | |
| ENSMUSG00000104396 | -1.095294215 | 0.541735653 |
| ENSMUSG00000104397 | 1 | |
| ENSMUSG00000104398 | -1 | 1 |
| ENSMUSG00000104399 | -1.037004768 | 0.836879252 |
| ENSMUSG00000104400 | 1.059463094 | 0.698112512 |
| ENSMUSG00000104401 | -1.03962336 | 0.698112512 |
| ENSMUSG00000104402 | -1.063957155 | 0.825942088 |
| ENSMUSG00000104403 | 1 | |
| ENSMUSG00000104404 | 1 | |
| ENSMUSG00000104405 | 1 | |
| ENSMUSG00000104406 | -1.090698382 | 0.788886948 |
| ENSMUSG00000104407 | 1 | |
| ENSMUSG00000104408 | -1.028176558 | 0.954497929 |
| ENSMUSG00000104409 | 1 | |
| ENSMUSG00000104410 | 1 | |
| ENSMUSG00000104411 | 1 | |
| ENSMUSG00000104412 | 1 | |
| ENSMUSG00000104413 | 1.175127719 | 0.608497229 |
| ENSMUSG00000104414 | 1 | |
| ENSMUSG00000104415 | 1.052663434 | 0.864629923 |
| ENSMUSG00000104416 | 1.016199469 | 1 |
| ENSMUSG00000104417 | 1 | |
| ENSMUSG00000104418 | -1.060102335 | 0.81188464 |
| ENSMUSG00000104419 | 1 | |
| ENSMUSG00000104420 | -1.07008665 | 0.851424496 |
| ENSMUSG00000104422 | 1 | |
| ENSMUSG00000104423 | 1.059463094 | 0.608497229 |
| ENSMUSG00000104424 | 1 | |
| ENSMUSG00000104425 | 1 | |
| ENSMUSG00000104426 | 1 | |
| ENSMUSG00000104427 | -1 | 1 |
| ENSMUSG00000104428 | 1 | |
| ENSMUSG00000104429 | 1 | |
| ENSMUSG00000104430 | 1 | |
| ENSMUSG00000104431 | 1 | |

| | | |
|---|---|---|
| ENSMUSG00000104432 | -1.018467515 | 1 |
| ENSMUSG00000104433 | 1.134080683 | 0.672108621 |
| ENSMUSG00000104434 | -1.072715868 | 0.756159003 |
| ENSMUSG00000104435 | 1.043122087 | 0.698112512 |
| ENSMUSG00000104436 | -1.034990181 | 0.954497929 |
| ENSMUSG00000104437 | 1 | |
| ENSMUSG00000104438 | 1 | |
| ENSMUSG00000104439 | 1 | |
| ENSMUSG00000104440 | 1 | |
| ENSMUSG00000104441 | 1 | |
| ENSMUSG00000104442 | 1 | |
| ENSMUSG00000104443 | 1.086003619 | 0.608497229 |
| ENSMUSG00000104444 | 1 | |
| ENSMUSG00000104445 | -1.084645496 | 0.532369657 |
| ENSMUSG00000104446 | 1 | |
| ENSMUSG00000104448 | 1 | |
| ENSMUSG00000104449 | 1.136780113 | 0.608497229 |
| ENSMUSG00000104450 | 1 | |
| ENSMUSG00000104451 | 1 | |
| ENSMUSG00000104452 | 1 | |
| ENSMUSG00000104453 | 1.134755104 | 0.424196581 |
| ENSMUSG00000104454 | 1.069068979 | 0.698112512 |
| ENSMUSG00000104455 | 1 | |
| ENSMUSG00000104456 | 1 | |
| ENSMUSG00000104457 | 1.059463094 | 0.608497229 |
| ENSMUSG00000104458 | 1 | |
| ENSMUSG00000104459 | 1.049742512 | 0.813405047 |
| ENSMUSG00000104460 | 1 | |
| ENSMUSG00000104461 | 1 | |
| ENSMUSG00000104462 | 1 | |
| ENSMUSG00000104464 | 1 | |
| ENSMUSG00000104465 | 1 | |
| ENSMUSG00000104466 | 1 | |
| ENSMUSG00000104467 | -1.120457584 | 0.443360404 |
| ENSMUSG00000104468 | 1 | |
| ENSMUSG00000104469 | 1.251296114 | 0.180548941 |
| ENSMUSG00000104470 | 1.079681442 | 0.698112512 |
| ENSMUSG00000104471 | 1 | |
| ENSMUSG00000104472 | 1 | |
| ENSMUSG00000104473 | 1 | |
| ENSMUSG00000104474 | 1 | |
| ENSMUSG00000104475 | -1.123259653 | 0.319960972 |
| ENSMUSG00000104476 | -1.229814271 | 0.330351517 |
| ENSMUSG00000104477 | 1 | |
| ENSMUSG00000104478 | 1 | |
| ENSMUSG00000104479 | 1.111812527 | 0.608497229 |
| ENSMUSG00000104480 | 1 | |
| ENSMUSG00000104481 | 1 | |
| ENSMUSG00000104482 | 1 | |
| ENSMUSG00000104484 | 1 | |
| ENSMUSG00000104485 | 1.161220813 | 0.425531155 |
| ENSMUSG00000104486 | 1.028323353 | 0.903464611 |
| ENSMUSG00000104487 | 1 | |
| ENSMUSG00000104488 | 1 | |
| ENSMUSG00000104489 | 1 | |
| ENSMUSG00000104490 | 1 | |
| ENSMUSG00000104491 | 1 | |
| ENSMUSG00000104493 | 1.014545335 | 0.991944074 |
| ENSMUSG00000104494 | 1.024693402 | 0.953922475 |
| ENSMUSG00000104496 | -1.048277865 | 0.829945662 |
| ENSMUSG00000104497 | 1 | |
| ENSMUSG00000104498 | 1 | |
| ENSMUSG00000104499 | 1 | |

| | | |
|---|---|---|
| ENSMUSG00000104500 | 1 | |
| ENSMUSG00000104501 | -1 | 1 |
| ENSMUSG00000104502 | 1 | |
| ENSMUSG00000104503 | 1.045500122 | 0.931480186 |
| ENSMUSG00000104504 | 1 | |
| ENSMUSG00000104505 | 1 | |
| ENSMUSG00000104506 | 1 | |
| ENSMUSG00000104507 | 1.059463094 | 0.608497229 |
| ENSMUSG00000104508 | -1.03962336 | 0.698112512 |
| ENSMUSG00000104509 | 1.069068979 | 0.608497229 |
| ENSMUSG00000104510 | 1 | |
| ENSMUSG00000104511 | 1 | |
| ENSMUSG00000104512 | -1 | 1 |
| ENSMUSG00000104513 | 1 | |
| ENSMUSG00000104514 | -1.245097715 | 0.318397362 |
| ENSMUSG00000104515 | 1.107269468 | 0.755135841 |
| ENSMUSG00000104516 | 1 | |
| ENSMUSG00000104517 | -1 | 1 |
| ENSMUSG00000104518 | 1 | |
| ENSMUSG00000104519 | 1.143529836 | 0.665607244 |
| ENSMUSG00000104520 | 1 | |
| ENSMUSG00000104521 | 1 | |
| ENSMUSG00000104522 | 1 | |
| ENSMUSG00000104523 | 1 | |
| ENSMUSG00000104524 | -1.003834322 | 1 |
| ENSMUSG00000104525 | -1.084645496 | 0.532369657 |
| ENSMUSG00000104526 | 1 | |
| ENSMUSG00000104527 | 1 | |
| ENSMUSG00000104528 | 1.173258561 | 0.442587908 |
| ENSMUSG00000104529 | -1 | 1 |
| ENSMUSG00000104530 | 1 | |
| ENSMUSG00000104531 | 1 | |
| ENSMUSG00000104532 | 1 | |
| ENSMUSG00000104533 | 1 | |
| ENSMUSG00000104534 | 1 | |
| ENSMUSG00000104535 | 1.02672102 | 0.939817611 |
| ENSMUSG00000104536 | 1 | |
| ENSMUSG00000104537 | 1 | |
| ENSMUSG00000104538 | 1 | |
| ENSMUSG00000104539 | 1 | |
| ENSMUSG00000104540 | 1 | |
| ENSMUSG00000104541 | 1 | |
| ENSMUSG00000104542 | 1 | |
| ENSMUSG00000104543 | -1.020155149 | 0.988424085 |
| ENSMUSG00000104544 | 1 | |
| ENSMUSG00000104545 | -1.135358696 | 0.608497229 |
| ENSMUSG00000104546 | -1.091530623 | 0.260565052 |
| ENSMUSG00000104547 | 1 | |
| ENSMUSG00000104548 | -1.185571636 | 0.257433776 |
| ENSMUSG00000104549 | 1 | |
| ENSMUSG00000104550 | -1.211513178 | 0.351239487 |
| ENSMUSG00000104551 | 1 | |
| ENSMUSG00000104552 | 1 | |
| ENSMUSG00000104553 | 1 | |
| ENSMUSG00000104554 | -1.177298609 | 0.221427514 |
| ENSMUSG00000104555 | -1.03962336 | 0.803072249 |
| ENSMUSG00000104556 | 1.019833591 | 0.983446483 |
| ENSMUSG00000104557 | 1 | |
| ENSMUSG00000104558 | 1 | |
| ENSMUSG00000104559 | 1 | |
| ENSMUSG00000104560 | 1.059463094 | 0.704170349 |
| ENSMUSG00000104561 | 1 | |
| ENSMUSG00000104562 | 1 | |

| | | |
|---|---|---|
| ENSMUSG00000104563 | 1.191053053 | 0.154927303 |
| ENSMUSG00000104564 | -1.053549062 | 0.698112512 |
| ENSMUSG00000104566 | 1 | |
| ENSMUSG00000104567 | -1.084645496 | 0.532369657 |
| ENSMUSG00000104568 | 1.079681442 | 0.608497229 |
| ENSMUSG00000104569 | -1.102325586 | 0.608497229 |
| ENSMUSG00000104570 | 1 | |
| ENSMUSG00000104571 | -1.043306199 | 0.698112512 |
| ENSMUSG00000104572 | 1 | |
| ENSMUSG00000104573 | 1 | |
| ENSMUSG00000104574 | 1 | |
| ENSMUSG00000104575 | 1 | |
| ENSMUSG00000104576 | 1.065410325 | 0.711602885 |
| ENSMUSG00000104577 | -1.05870646 | 0.879253209 |
| ENSMUSG00000104578 | 1 | |
| ENSMUSG00000104579 | 1 | |
| ENSMUSG00000104580 | 1 | |
| ENSMUSG00000104581 | -1.023769022 | 0.954761885 |
| ENSMUSG00000104582 | 1 | |
| ENSMUSG00000104583 | -1.206859842 | 0.200927632 |
| ENSMUSG00000104584 | 1 | |
| ENSMUSG00000104585 | 1.059463094 | 0.608497229 |
| ENSMUSG00000104586 | 1 | |
| ENSMUSG00000104587 | 1.008179265 | 1 |
| ENSMUSG00000104588 | -1.043306199 | 0.698112512 |
| ENSMUSG00000104589 | 1 | |
| ENSMUSG00000104590 | 1 | |
| ENSMUSG00000104591 | 1.167748221 | 0.61149727 |
| ENSMUSG00000104592 | -1.098124897 | 0.670336873 |
| ENSMUSG00000104593 | 1 | |
| ENSMUSG00000104594 | 1 | |
| ENSMUSG00000104595 | 1 | |
| ENSMUSG00000104596 | 1 | |
| ENSMUSG00000104597 | 1.411975949 | 0.147951988 |
| ENSMUSG00000104598 | 1 | |
| ENSMUSG00000104599 | 1 | |
| ENSMUSG00000104600 | 1 | |
| ENSMUSG00000104601 | 1 | |
| ENSMUSG00000104602 | 1 | |
| ENSMUSG00000104603 | 1 | |
| ENSMUSG00000104604 | -1.043306199 | 0.698112512 |
| ENSMUSG00000104605 | 1 | |
| ENSMUSG00000104606 | -1.212656185 | 0.215991492 |
| ENSMUSG00000104607 | 1 | |
| ENSMUSG00000104608 | 1 | |
| ENSMUSG00000104609 | 1 | |
| ENSMUSG00000104610 | -1.035320271 | 0.949295635 |
| ENSMUSG00000104612 | 1 | |
| ENSMUSG00000104613 | 1 | |
| ENSMUSG00000104614 | 1 | |
| ENSMUSG00000104615 | -1.043306199 | 0.698112512 |
| ENSMUSG00000104616 | -1.029113823 | 0.946575044 |
| ENSMUSG00000104617 | 1 | |
| ENSMUSG00000104618 | 1 | |
| ENSMUSG00000104619 | 1 | |
| ENSMUSG00000104620 | 1 | |
| ENSMUSG00000104621 | 1.201379336 | 0.035832281 |
| ENSMUSG00000104622 | -1.003942951 | 1 |
| ENSMUSG00000104623 | 1.145838205 | 0.608497229 |
| ENSMUSG00000104624 | 1 | |
| ENSMUSG00000104625 | -1.037745104 | 0.926061629 |
| ENSMUSG00000104626 | 1.128878069 | 0.596589772 |
| ENSMUSG00000104627 | 1.023743961 | 0.946213735 |

| | | |
|---|---|---|
| ENSMUSG00000104628 | 1 | |
| ENSMUSG00000104629 | 1 | |
| ENSMUSG00000104630 | 1 | |
| ENSMUSG00000104631 | 1 | |
| ENSMUSG00000104632 | -1.005214803 | 1 |
| ENSMUSG00000104633 | 1 | |
| ENSMUSG00000104634 | 1.014545335 | 0.979002724 |
| ENSMUSG00000104635 | 1 | |
| ENSMUSG00000104636 | 1 | |
| ENSMUSG00000104637 | 1 | |
| ENSMUSG00000104638 | 1 | |
| ENSMUSG00000104639 | -1.03962336 | 0.803072249 |
| ENSMUSG00000104640 | 1 | |
| ENSMUSG00000104641 | -1.057490426 | 0.902032983 |
| ENSMUSG00000104642 | 1 | |
| ENSMUSG00000104643 | 1.028323353 | 0.933031586 |
| ENSMUSG00000104644 | 1 | |
| ENSMUSG00000104645 | 1 | |
| ENSMUSG00000104646 | 1 | |
| ENSMUSG00000104647 | -1.028348525 | 0.912425817 |
| ENSMUSG00000104648 | 1 | |
| ENSMUSG00000104649 | -1.121041497 | 0.608497229 |
| ENSMUSG00000104650 | 1 | |
| ENSMUSG00000104651 | 1 | |
| ENSMUSG00000104652 | 1 | |
| ENSMUSG00000104653 | -1 | 1 |
| ENSMUSG00000104654 | -1 | 1 |
| ENSMUSG00000104655 | 1 | |
| ENSMUSG00000104656 | 1 | |
| ENSMUSG00000104657 | 1 | |
| ENSMUSG00000104658 | 1 | |
| ENSMUSG00000104659 | 1 | |
| ENSMUSG00000104660 | 1 | |
| ENSMUSG00000104661 | 1.04128401 | 0.944271488 |
| ENSMUSG00000104662 | -1.020232 | 0.991944074 |
| ENSMUSG00000104663 | -1.043306199 | 0.698112512 |
| ENSMUSG00000104664 | -1 | 1 |
| ENSMUSG00000104665 | -1.077766213 | 0.798926662 |
| ENSMUSG00000104666 | 1 | |
| ENSMUSG00000104667 | -1.085844855 | 0.74174982 |
| ENSMUSG00000104668 | 1 | |
| ENSMUSG00000104669 | 1.069437807 | 0.729206866 |
| ENSMUSG00000104670 | 1.075870199 | 0.809396188 |
| ENSMUSG00000104671 | -1.19238516 | 0.153355481 |
| ENSMUSG00000104672 | 1 | |
| ENSMUSG00000104673 | 1 | |
| ENSMUSG00000104674 | 1.077504054 | 0.760311821 |
| ENSMUSG00000104675 | -1.053549062 | 0.769745556 |
| ENSMUSG00000104676 | -1.072882392 | 0.816599259 |
| ENSMUSG00000104677 | 1.024693402 | 0.933031586 |
| ENSMUSG00000104678 | 1.095872691 | 0.608497229 |
| ENSMUSG00000104679 | 1 | |
| ENSMUSG00000104680 | -1.03962336 | 0.698112512 |
| ENSMUSG00000104681 | 1 | |
| ENSMUSG00000104682 | -1.076592925 | 0.444566747 |
| ENSMUSG00000104683 | 1 | |
| ENSMUSG00000104684 | 1.069068979 | 0.698112512 |
| ENSMUSG00000104685 | -1.034857663 | 0.87080301 |
| ENSMUSG00000104686 | 1.132885541 | 0.608497229 |
| ENSMUSG00000104687 | 1.040562055 | 0.933031586 |
| ENSMUSG00000104688 | 1 | |
| ENSMUSG00000104689 | 1 | |
| ENSMUSG00000104690 | 1 | |

| | | |
|---|---|---|
| ENSMUSG00000104691 | 1 | |
| ENSMUSG00000104692 | 1 | |
| ENSMUSG00000104693 | 1.038223822 | 0.924108307 |
| ENSMUSG00000104694 | -1.111880267 | 0.608497229 |
| ENSMUSG00000104695 | -1.008201504 | 1 |
| ENSMUSG00000104696 | -1.03962336 | 0.872692845 |
| ENSMUSG00000104697 | 1 | |
| ENSMUSG00000104698 | 1 | |
| ENSMUSG00000104699 | -1.021258981 | 0.956908483 |
| ENSMUSG00000104700 | 1 | |
| ENSMUSG00000104701 | -1 | 1 |
| ENSMUSG00000104702 | 1 | |
| ENSMUSG00000104703 | 1 | |
| ENSMUSG00000104704 | 1.074847028 | 0.731587932 |
| ENSMUSG00000104705 | 1 | |
| ENSMUSG00000104706 | 1.13438124 | 0.653695234 |
| ENSMUSG00000104707 | -1.03962336 | 0.814089245 |
| ENSMUSG00000104708 | -1.073345802 | 0.628207137 |
| ENSMUSG00000104709 | 1.059463094 | 0.840915886 |
| ENSMUSG00000104710 | 1 | |
| ENSMUSG00000104711 | 1 | |
| ENSMUSG00000104712 | 1 | |
| ENSMUSG00000104713 | -1.157634469 | 0.616230752 |
| ENSMUSG00000104714 | 1 | |
| ENSMUSG00000104715 | 1 | |
| ENSMUSG00000104716 | 1 | |
| ENSMUSG00000104717 | 1 | |
| ENSMUSG00000104718 | 1 | |
| ENSMUSG00000104719 | 1.19701422 | 0.158730308 |
| ENSMUSG00000104720 | -1.081783312 | 0.722479145 |
| ENSMUSG00000104721 | 1.057700516 | 0.842811366 |
| ENSMUSG00000104722 | 1 | |
| ENSMUSG00000104723 | 1 | |
| ENSMUSG00000104724 | 1.052283458 | 0.718265321 |
| ENSMUSG00000104725 | 1.069068979 | 0.608497229 |
| ENSMUSG00000104726 | 1 | |
| ENSMUSG00000104727 | 1 | |
| ENSMUSG00000104728 | -1.106216701 | 0.634256792 |
| ENSMUSG00000104729 | 1 | |
| ENSMUSG00000104730 | 1 | |
| ENSMUSG00000104731 | 1 | |
| ENSMUSG00000104732 | 1.059463094 | 0.608497229 |
| ENSMUSG00000104733 | 1 | |
| ENSMUSG00000104734 | 1 | |
| ENSMUSG00000104735 | -1.128668575 | 0.665496439 |
| ENSMUSG00000104736 | 1 | |
| ENSMUSG00000104737 | -1.012884804 | 1 |
| ENSMUSG00000104738 | 1 | |
| ENSMUSG00000104739 | 1 | |
| ENSMUSG00000104740 | 1 | |
| ENSMUSG00000104741 | 1 | |
| ENSMUSG00000104742 | 1 | |
| ENSMUSG00000104743 | 1 | |
| ENSMUSG00000104744 | 1 | |
| ENSMUSG00000104745 | -1.03962336 | 0.698112512 |
| ENSMUSG00000104746 | 1 | |
| ENSMUSG00000104747 | 1 | |
| ENSMUSG00000104748 | 1.019083579 | 0.986542466 |
| ENSMUSG00000104749 | 1 | |
| ENSMUSG00000104750 | 1 | |
| ENSMUSG00000104751 | 1 | |
| ENSMUSG00000104752 | 1 | |
| ENSMUSG00000104753 | 1 | |

| | | |
|---|---|---|
| ENSMUSG00000104754 | 1 | |
| ENSMUSG00000104755 | 1.014545335 | 0.990831634 |
| ENSMUSG00000104756 | 1.045476229 | 0.954102616 |
| ENSMUSG00000104757 | 1 | |
| ENSMUSG00000104758 | -1.043306199 | 0.698112512 |
| ENSMUSG00000104759 | 1 | |
| ENSMUSG00000104760 | 1 | |
| ENSMUSG00000104761 | 1.064115538 | 0.608497229 |
| ENSMUSG00000104762 | -1.043306199 | 0.79291651 |
| ENSMUSG00000104763 | -1 | 1 |
| ENSMUSG00000104764 | 1 | |
| ENSMUSG00000104765 | -1.12591365 | 0.698112512 |
| ENSMUSG00000104766 | 1.059463094 | 0.698112512 |
| ENSMUSG00000104767 | 1 | |
| ENSMUSG00000104768 | -1.029302237 | 0.96035552 |
| ENSMUSG00000104769 | 1 | |
| ENSMUSG00000104770 | 1 | |
| ENSMUSG00000104771 | 1 | |
| ENSMUSG00000104772 | 1 | |
| ENSMUSG00000104773 | 1 | |
| ENSMUSG00000104774 | 1 | |
| ENSMUSG00000104775 | 1.014545335 | 0.989648415 |
| ENSMUSG00000104776 | 1 | |
| ENSMUSG00000104777 | 1 | |
| ENSMUSG00000104778 | 1 | |
| ENSMUSG00000104779 | 1 | |
| ENSMUSG00000104780 | -1.03962336 | 0.698112512 |
| ENSMUSG00000104781 | -1.0795912 | 0.748682651 |
| ENSMUSG00000104782 | 1 | |
| ENSMUSG00000104783 | 1 | |
| ENSMUSG00000104784 | -1.100193663 | 0.534887579 |
| ENSMUSG00000104785 | 1 | |
| ENSMUSG00000104786 | -1.116223673 | 0.608497229 |
| ENSMUSG00000104787 | 1 | |
| ENSMUSG00000104788 | -1.02522956 | 0.970707004 |
| ENSMUSG00000104789 | 1 | |
| ENSMUSG00000104790 | 1 | |
| ENSMUSG00000104791 | 1 | |
| ENSMUSG00000104792 | 1 | |
| ENSMUSG00000104793 | 1.011334143 | 1 |
| ENSMUSG00000104794 | 1 | |
| ENSMUSG00000104795 | -1.155156971 | 0.698112512 |
| ENSMUSG00000104796 | 1 | |
| ENSMUSG00000104797 | 1 | |
| ENSMUSG00000104798 | 1 | |
| ENSMUSG00000104799 | 1 | |
| ENSMUSG00000104800 | 1 | |
| ENSMUSG00000104801 | -1 | 1 |
| ENSMUSG00000104802 | 1.002574398 | 1 |
| ENSMUSG00000104803 | 1 | |
| ENSMUSG00000104804 | 1 | |
| ENSMUSG00000104805 | 1 | |
| ENSMUSG00000104806 | -1.030752983 | 0.910155457 |
| ENSMUSG00000104807 | 1 | |
| ENSMUSG00000104808 | -1 | 1 |
| ENSMUSG00000104809 | 1 | |
| ENSMUSG00000104810 | 1 | |
| ENSMUSG00000104811 | 1 | |
| ENSMUSG00000104812 | 1 | |
| ENSMUSG00000104813 | 1 | |
| ENSMUSG00000104814 | 1.152004458 | 0.266148043 |
| ENSMUSG00000104815 | 1 | |
| ENSMUSG00000104816 | -1.048277865 | 0.860645152 |

| | | |
|---|---|---|
| ENSMUSG00000104817 | 1 | |
| ENSMUSG00000104818 | 1.00145941 | 1 |
| ENSMUSG00000104819 | 1 | |
| ENSMUSG00000104820 | 1 | |
| ENSMUSG00000104821 | 1.06585783 | 0.867420842 |
| ENSMUSG00000104822 | -1.073959562 | 0.817330349 |
| ENSMUSG00000104823 | 1 | |
| ENSMUSG00000104824 | 1 | |
| ENSMUSG00000104825 | 1 | |
| ENSMUSG00000104826 | 1 | |
| ENSMUSG00000104827 | 1 | |
| ENSMUSG00000104828 | 1.11166259 | 0.649807331 |
| ENSMUSG00000104829 | 1 | |
| ENSMUSG00000104830 | 1.137569893 | 0.429700116 |
| ENSMUSG00000104831 | 1 | |
| ENSMUSG00000104832 | 1 | |
| ENSMUSG00000104833 | 1 | |
| ENSMUSG00000104834 | 1 | |
| ENSMUSG00000104835 | 1 | |
| ENSMUSG00000104836 | 1 | |
| ENSMUSG00000104837 | -1.126703685 | 0.349636986 |
| ENSMUSG00000104838 | 1 | |
| ENSMUSG00000104839 | 1 | |
| ENSMUSG00000104840 | 1 | |
| ENSMUSG00000104841 | 1.059463094 | 0.698112512 |
| ENSMUSG00000104842 | 1 | |
| ENSMUSG00000104843 | -1.017191164 | 0.980958779 |
| ENSMUSG00000104844 | 1 | |
| ENSMUSG00000104845 | 1 | |
| ENSMUSG00000104846 | 1 | |
| ENSMUSG00000104847 | 1 | |
| ENSMUSG00000104848 | 1 | |
| ENSMUSG00000104849 | 1 | |
| ENSMUSG00000104850 | 1.229924362 | 0.332232753 |
| ENSMUSG00000104851 | 1 | |
| ENSMUSG00000104852 | -1.031876331 | 0.968954528 |
| ENSMUSG00000104853 | 1 | |
| ENSMUSG00000104854 | -1.044273782 | 0.698112512 |
| ENSMUSG00000104855 | 1 | |
| ENSMUSG00000104856 | 1 | |
| ENSMUSG00000104857 | 1 | |
| ENSMUSG00000104858 | 1 | |
| ENSMUSG00000104859 | 1.059033181 | 0.801168497 |
| ENSMUSG00000104860 | -1.097349941 | 0.645833191 |
| ENSMUSG00000104861 | 1.077800869 | 0.48076246 |
| ENSMUSG00000104862 | 1.064985481 | 0.757318144 |
| ENSMUSG00000104863 | 1 | |
| ENSMUSG00000104864 | 1.133661809 | 0.661155252 |
| ENSMUSG00000104865 | 1 | |
| ENSMUSG00000104866 | 1 | |
| ENSMUSG00000104867 | 1.010115111 | 1 |
| ENSMUSG00000104868 | 1 | |
| ENSMUSG00000104869 | 1.213587389 | 0.303039084 |
| ENSMUSG00000104870 | 1 | |
| ENSMUSG00000104871 | 1.079681442 | 0.776029362 |
| ENSMUSG00000104872 | 1.074421866 | 0.626104035 |
| ENSMUSG00000104873 | 1.116785864 | 0.578221424 |
| ENSMUSG00000104874 | 1.014545335 | 0.989648415 |
| ENSMUSG00000104875 | -1 | 1 |
| ENSMUSG00000104876 | -1 | 1 |
| ENSMUSG00000104877 | -1.008906735 | 1 |
| ENSMUSG00000104878 | 1 | |
| ENSMUSG00000104879 | 1 | |

| | | |
|---|---|---|
| ENSMUSG00000104880 | 1 | |
| ENSMUSG00000104881 | -1.100193663 | 0.534887579 |
| ENSMUSG00000104882 | -1.044273782 | 0.801443616 |
| ENSMUSG00000104883 | 1 | |
| ENSMUSG00000104884 | 1 | |
| ENSMUSG00000104885 | -1.014607149 | 1 |
| ENSMUSG00000104886 | 1 | |
| ENSMUSG00000104887 | 1 | |
| ENSMUSG00000104888 | -1.11061171 | 0.547016491 |
| ENSMUSG00000104889 | 1 | |
| ENSMUSG00000104890 | 1 | |
| ENSMUSG00000104891 | 1.081541652 | 0.608497229 |
| ENSMUSG00000104892 | 1.059463094 | 0.608497229 |
| ENSMUSG00000104893 | 1.000702347 | 1 |
| ENSMUSG00000104894 | 1 | |
| ENSMUSG00000104895 | -1.044273782 | 0.698112512 |
| ENSMUSG00000104896 | 1 | |
| ENSMUSG00000104897 | -1.021138931 | 0.970734719 |
| ENSMUSG00000104898 | 1 | |
| ENSMUSG00000104899 | 1 | |
| ENSMUSG00000104900 | 1 | |
| ENSMUSG00000104901 | 1 | |
| ENSMUSG00000104902 | 1 | |
| ENSMUSG00000104903 | -1.111880267 | 0.572947856 |
| ENSMUSG00000104904 | 1.149482339 | 0.608497229 |
| ENSMUSG00000104905 | 1 | |
| ENSMUSG00000104906 | -1.03962336 | 0.698112512 |
| ENSMUSG00000104908 | 1 | |
| ENSMUSG00000104909 | 1 | |
| ENSMUSG00000104910 | -1.023769022 | 0.951748628 |
| ENSMUSG00000104911 | 1 | |
| ENSMUSG00000104912 | 1.12189305 | 0.585646836 |
| ENSMUSG00000104913 | 1.048825188 | 0.812228807 |
| ENSMUSG00000104914 | -1.024530911 | 0.964236174 |
| ENSMUSG00000104915 | 1 | |
| ENSMUSG00000104916 | 1.059463094 | 0.750182067 |
| ENSMUSG00000104917 | 1.074408627 | 0.825401367 |
| ENSMUSG00000104918 | 1.07487334 | 0.698112512 |
| ENSMUSG00000104919 | 1 | |
| ENSMUSG00000104920 | 1.019083579 | 0.949812939 |
| ENSMUSG00000104921 | 1 | |
| ENSMUSG00000104922 | 1 | |
| ENSMUSG00000104923 | -1.053549062 | 0.769745556 |
| ENSMUSG00000104924 | -1.094488708 | 0.714698678 |
| ENSMUSG00000104925 | 1.068522702 | 0.86287706 |
| ENSMUSG00000104926 | -1.043306199 | 0.698112512 |
| ENSMUSG00000104927 | 1 | |
| ENSMUSG00000104928 | 1 | |
| ENSMUSG00000104929 | 1 | |
| ENSMUSG00000104930 | 1 | |
| ENSMUSG00000104931 | 1 | |
| ENSMUSG00000104932 | 1.193921123 | 0.188066027 |
| ENSMUSG00000104933 | 1 | |
| ENSMUSG00000104934 | 1 | |
| ENSMUSG00000104935 | -1 | 1 |
| ENSMUSG00000104936 | 1 | |
| ENSMUSG00000104937 | 1.171563499 | 0.339783652 |
| ENSMUSG00000104938 | -1.193320102 | 0.359354928 |
| ENSMUSG00000104939 | 1 | |
| ENSMUSG00000104940 | 1.11166259 | 0.608497229 |
| ENSMUSG00000104941 | 1 | |
| ENSMUSG00000104942 | 1 | |
| ENSMUSG00000104943 | -1.113259873 | 0.713861573 |

| | | |
|---|---|---|
| ENSMUSG00000104944 | 1 | |
| ENSMUSG00000104945 | -1.044273782 | 0.698112512 |
| ENSMUSG00000104946 | 1 | |
| ENSMUSG00000104947 | -1.044273782 | 0.801443616 |
| ENSMUSG00000104948 | 1 | |
| ENSMUSG00000104949 | 1 | |
| ENSMUSG00000104950 | 1.052378925 | 0.794274354 |
| ENSMUSG00000104951 | 1.054758081 | 0.924629556 |
| ENSMUSG00000104952 | 1 | |
| ENSMUSG00000104953 | 1 | |
| ENSMUSG00000104954 | -1 | 1 |
| ENSMUSG00000104955 | 1.028323353 | 0.903464611 |
| ENSMUSG00000104956 | 1 | |
| ENSMUSG00000104957 | 1 | |
| ENSMUSG00000104958 | 1 | |
| ENSMUSG00000104959 | 1 | |
| ENSMUSG00000104960 | 1.1326384 | 0.42275393 |
| ENSMUSG00000104961 | 1 | |
| ENSMUSG00000104962 | 1 | |
| ENSMUSG00000104963 | -1.149170004 | 0.608497229 |
| ENSMUSG00000104964 | 1 | |
| ENSMUSG00000104965 | -1.282009569 | 0.20602422 |
| ENSMUSG00000104966 | 1 | |
| ENSMUSG00000104967 | -1.014570171 | 0.996048848 |
| ENSMUSG00000104968 | 1 | |
| ENSMUSG00000104969 | 1.069187287 | 0.925075932 |
| ENSMUSG00000104970 | 1 | |
| ENSMUSG00000104971 | -1.044273782 | 0.698112512 |
| ENSMUSG00000104972 | 1.059463094 | 0.704170349 |
| ENSMUSG00000104973 | -1.112283437 | 0.608497229 |
| ENSMUSG00000104974 | -1 | 1 |
| ENSMUSG00000104975 | 1 | |
| ENSMUSG00000104976 | 1 | |
| ENSMUSG00000104978 | 1 | |
| ENSMUSG00000104979 | 1 | |
| ENSMUSG00000104980 | 1 | |
| ENSMUSG00000104981 | 1 | |
| ENSMUSG00000104982 | 1 | |
| ENSMUSG00000104983 | -1.053549062 | 0.698112512 |
| ENSMUSG00000104984 | 1 | |
| ENSMUSG00000104985 | 1 | |
| ENSMUSG00000104986 | -1.122689526 | 0.268643495 |
| ENSMUSG00000104987 | 1 | |
| ENSMUSG00000104988 | 1.007629259 | 0.989387709 |
| ENSMUSG00000104989 | 1 | |
| ENSMUSG00000104990 | 1.015486245 | 0.987775262 |
| ENSMUSG00000104991 | 1 | |
| ENSMUSG00000104992 | 1 | |
| ENSMUSG00000104993 | -1 | 1 |
| ENSMUSG00000104994 | 1 | |
| ENSMUSG00000104995 | 1 | |
| ENSMUSG00000104996 | 1 | |
| ENSMUSG00000104997 | 1 | |
| ENSMUSG00000104998 | -1 | 1 |
| ENSMUSG00000104999 | 1.05038453 | 0.828564184 |
| ENSMUSG00000105000 | 1 | |
| ENSMUSG00000105001 | 1 | |
| ENSMUSG00000105002 | 1 | |
| ENSMUSG00000105003 | 1.059463094 | 0.608497229 |
| ENSMUSG00000105004 | 1.068424331 | 0.698112512 |
| ENSMUSG00000105005 | 1 | |
| ENSMUSG00000105006 | 1.174227848 | 0.269056268 |
| ENSMUSG00000105007 | 1 | |

| | | |
|---|---|---|
| ENSMUSG00000105008 | 1.149113744 | 0.158414349 |
| ENSMUSG00000105009 | 1 | |
| ENSMUSG00000105010 | 1 | |
| ENSMUSG00000105011 | -1.113041925 | 0.660688301 |
| ENSMUSG00000105012 | 1 | |
| ENSMUSG00000105013 | 1 | |
| ENSMUSG00000105014 | 1 | |
| ENSMUSG00000105015 | -1.043306199 | 0.79291651 |
| ENSMUSG00000105016 | 1 | |
| ENSMUSG00000105017 | 1 | |
| ENSMUSG00000105018 | 1 | |
| ENSMUSG00000105019 | -1.044273782 | 0.698112512 |
| ENSMUSG00000105020 | 1 | |
| ENSMUSG00000105021 | -1.518930228 | 0.013881289 |
| ENSMUSG00000105022 | 1.045399152 | 0.805243413 |
| ENSMUSG00000105023 | 1 | |
| ENSMUSG00000105024 | 1 | |
| ENSMUSG00000105025 | 1 | |
| ENSMUSG00000105026 | 1 | |
| ENSMUSG00000105027 | 1 | |
| ENSMUSG00000105028 | 1 | |
| ENSMUSG00000105029 | 1 | |
| ENSMUSG00000105030 | 1 | |
| ENSMUSG00000105031 | -1.095294215 | 0.608497229 |
| ENSMUSG00000105032 | 1 | |
| ENSMUSG00000105033 | -1.00216679 | 1 |
| ENSMUSG00000105034 | 1 | |
| ENSMUSG00000105035 | 1 | |
| ENSMUSG00000105036 | 1.059463094 | 0.608497229 |
| ENSMUSG00000105037 | 1 | |
| ENSMUSG00000105038 | 1 | |
| ENSMUSG00000105039 | -1.11061171 | 0.547016491 |
| ENSMUSG00000105040 | -1.016098951 | 1 |
| ENSMUSG00000105041 | -1.063525418 | 0.751979636 |
| ENSMUSG00000105042 | -1.06003939 | 0.734803948 |
| ENSMUSG00000105043 | 1 | |
| ENSMUSG00000105044 | -1.111703988 | 0.608497229 |
| ENSMUSG00000105045 | -1.193444643 | 0.531127798 |
| ENSMUSG00000105046 | 1 | |
| ENSMUSG00000105047 | -1 | 1 |
| ENSMUSG00000105048 | -1 | 1 |
| ENSMUSG00000105049 | -1.053549062 | 0.698112512 |
| ENSMUSG00000105050 | -1 | 1 |
| ENSMUSG00000105051 | -1.03962336 | 0.698112512 |
| ENSMUSG00000105052 | 1 | |
| ENSMUSG00000105053 | 1.059463094 | 0.776953697 |
| ENSMUSG00000105054 | -1 | 1 |
| ENSMUSG00000105055 | 1 | |
| ENSMUSG00000105056 | 1 | |
| ENSMUSG00000105057 | 1 | |
| ENSMUSG00000105058 | 1 | |
| ENSMUSG00000105059 | 1 | |
| ENSMUSG00000105060 | 1 | |
| ENSMUSG00000105061 | 1 | |
| ENSMUSG00000105062 | 1.055159774 | 0.848720308 |
| ENSMUSG00000105063 | -1.041023137 | 0.962991083 |
| ENSMUSG00000105064 | -1.043306199 | 0.698112512 |
| ENSMUSG00000105065 | 1.090212144 | 0.209767231 |
| ENSMUSG00000105066 | 1 | |
| ENSMUSG00000105067 | 1 | |
| ENSMUSG00000105068 | 1 | |
| ENSMUSG00000105070 | 1 | |
| ENSMUSG00000105071 | 1.072658765 | 0.709012023 |

| | | |
|---|---|---|
| ENSMUSG00000105072 | 1 | |
| ENSMUSG00000105073 | 1 | |
| ENSMUSG00000105074 | 1 | |
| ENSMUSG00000105075 | 1 | |
| ENSMUSG00000105076 | -1.003012737 | 1 |
| ENSMUSG00000105077 | -1.003799607 | 1 |
| ENSMUSG00000105078 | 1 | |
| ENSMUSG00000105079 | -1.155193738 | 0.541735653 |
| ENSMUSG00000105080 | 1 | |
| ENSMUSG00000105081 | 1 | |
| ENSMUSG00000105082 | -1.034239753 | 0.924324458 |
| ENSMUSG00000105083 | 1.085852891 | 0.685652394 |
| ENSMUSG00000105084 | 1.024693402 | 0.933031586 |
| ENSMUSG00000105085 | 1.069068979 | 0.608497229 |
| ENSMUSG00000105086 | 1 | |
| ENSMUSG00000105087 | -1 | 1 |
| ENSMUSG00000105088 | 1 | |
| ENSMUSG00000105089 | 1 | |
| ENSMUSG00000105090 | 1 | |
| ENSMUSG00000105091 | 1 | |
| ENSMUSG00000105092 | 1 | |
| ENSMUSG00000105093 | -1.27480329 | 0.101525914 |
| ENSMUSG00000105094 | 1 | |
| ENSMUSG00000105095 | 1.029963228 | 0.925679199 |
| ENSMUSG00000105096 | -1 | 1 |
| ENSMUSG00000105097 | 1 | |
| ENSMUSG00000105098 | 1 | |
| ENSMUSG00000105099 | 1 | |
| ENSMUSG00000105100 | -1.023769022 | 0.934760809 |
| ENSMUSG00000105101 | 1 | |
| ENSMUSG00000105102 | -1.084645496 | 0.532369657 |
| ENSMUSG00000105103 | 1 | |
| ENSMUSG00000105104 | 1.122462048 | 0.420161906 |
| ENSMUSG00000105105 | 1 | |
| ENSMUSG00000105106 | 1 | |
| ENSMUSG00000105107 | -1.218532123 | 0.486384474 |
| ENSMUSG00000105108 | 1 | |
| ENSMUSG00000105109 | 1 | |
| ENSMUSG00000105110 | 1 | |
| ENSMUSG00000105111 | -1 | 1 |
| ENSMUSG00000105112 | 1.204410379 | 0.011965994 |
| ENSMUSG00000105113 | -1 | 1 |
| ENSMUSG00000105114 | 1 | |
| ENSMUSG00000105115 | 1 | |
| ENSMUSG00000105116 | 1 | |
| ENSMUSG00000105117 | 1 | |
| ENSMUSG00000105118 | 1 | |
| ENSMUSG00000105119 | -1.099781752 | 0.724929955 |
| ENSMUSG00000105120 | 1 | |
| ENSMUSG00000105121 | 1 | |
| ENSMUSG00000105122 | 1 | |
| ENSMUSG00000105123 | 1 | |
| ENSMUSG00000105124 | 1.095872691 | 0.698112512 |
| ENSMUSG00000105125 | 1.059463094 | 0.608497229 |
| ENSMUSG00000105126 | 1.059463094 | 0.608497229 |
| ENSMUSG00000105127 | -1.085651418 | 0.591151279 |
| ENSMUSG00000105128 | -1 | 1 |
| ENSMUSG00000105129 | 1 | |
| ENSMUSG00000105130 | 1.057302306 | 0.922581095 |
| ENSMUSG00000105131 | 1 | |
| ENSMUSG00000105132 | 1 | |
| ENSMUSG00000105133 | 1 | |
| ENSMUSG00000105134 | 1.001081606 | 1 |

| | | |
|---|---|---|
| ENSMUSG00000105135 | -1.283321476 | 0.030650703 |
| ENSMUSG00000105136 | 1 | |
| ENSMUSG00000105137 | 1.06443814 | 0.889643231 |
| ENSMUSG00000105138 | 1 | |
| ENSMUSG00000105139 | 1.037372746 | 0.715760774 |
| ENSMUSG00000105140 | -1.196552689 | 0.608497229 |
| ENSMUSG00000105141 | 1 | |
| ENSMUSG00000105142 | 1 | |
| ENSMUSG00000105143 | 1 | |
| ENSMUSG00000105144 | -1 | 1 |
| ENSMUSG00000105145 | 1 | |
| ENSMUSG00000105146 | 1 | |
| ENSMUSG00000105147 | 1 | |
| ENSMUSG00000105148 | 1 | |
| ENSMUSG00000105149 | 1 | |
| ENSMUSG00000105150 | -1.043306199 | 0.803260194 |
| ENSMUSG00000105151 | 1 | |
| ENSMUSG00000105152 | -1.043306199 | 0.881483647 |
| ENSMUSG00000105153 | -1.086192882 | 0.76439484 |
| ENSMUSG00000105154 | 1.059463094 | 0.608497229 |
| ENSMUSG00000105155 | -1.23300368 | 0.30650354 |
| ENSMUSG00000105156 | 1.097540288 | 0.608497229 |
| ENSMUSG00000105157 | 1 | |
| ENSMUSG00000105158 | 1 | |
| ENSMUSG00000105159 | 1 | |
| ENSMUSG00000105160 | -1.11061171 | 0.547016491 |
| ENSMUSG00000105161 | 1.000139703 | 1 |
| ENSMUSG00000105162 | 1.030256832 | 0.95029384 |
| ENSMUSG00000105163 | -1.003834322 | 1 |
| ENSMUSG00000105164 | 1 | |
| ENSMUSG00000105165 | 1 | |
| ENSMUSG00000105166 | 1.108429674 | 0.76182764 |
| ENSMUSG00000105167 | -1.041213731 | 0.925453232 |
| ENSMUSG00000105168 | -1 | 1 |
| ENSMUSG00000105169 | 1 | |
| ENSMUSG00000105170 | 1 | |
| ENSMUSG00000105171 | -1.19761686 | 0.119808842 |
| ENSMUSG00000105172 | -1.044146242 | 0.810884991 |
| ENSMUSG00000105173 | -1 | 1 |
| ENSMUSG00000105174 | 1 | |
| ENSMUSG00000105175 | -1.086192882 | 0.698112512 |
| ENSMUSG00000105176 | -1.167651702 | 0.482523609 |
| ENSMUSG00000105177 | 1 | |
| ENSMUSG00000105178 | 1 | |
| ENSMUSG00000105179 | -1.037482356 | 0.924739422 |
| ENSMUSG00000105180 | 1 | |
| ENSMUSG00000105181 | -1.153308664 | 0.608497229 |
| ENSMUSG00000105182 | 1 | |
| ENSMUSG00000105183 | 1.199989356 | 0.362989791 |
| ENSMUSG00000105184 | 1 | |
| ENSMUSG00000105185 | 1 | |
| ENSMUSG00000105186 | 1.086541237 | 0.810156519 |
| ENSMUSG00000105187 | 1 | |
| ENSMUSG00000105188 | -1.053549062 | 0.769745556 |
| ENSMUSG00000105189 | -1 | 1 |
| ENSMUSG00000105190 | 1 | |
| ENSMUSG00000105191 | 1 | |
| ENSMUSG00000105192 | 1 | |
| ENSMUSG00000105193 | 1 | |
| ENSMUSG00000105194 | 1 | |
| ENSMUSG00000105195 | 1 | |
| ENSMUSG00000105196 | 1 | |
| ENSMUSG00000105197 | 1 | |

| | | |
|---|---|---|
| ENSMUSG00000105198 | -1.054851488 | 0.814250774 |
| ENSMUSG00000105199 | -1.192994026 | 0.498630818 |
| ENSMUSG00000105200 | 1 | |
| ENSMUSG00000105201 | -1.132183056 | 0.581669442 |
| ENSMUSG00000105202 | 1.084846819 | 0.698112512 |
| ENSMUSG00000105203 | -1.143787033 | 0.201977401 |
| ENSMUSG00000105204 | 1 | |
| ENSMUSG00000105205 | 1 | |
| ENSMUSG00000105206 | -1.051444952 | 0.854968144 |
| ENSMUSG00000105207 | 1.070790205 | 0.698112512 |
| ENSMUSG00000105208 | 1 | |
| ENSMUSG00000105209 | 1 | |
| ENSMUSG00000105210 | 1 | |
| ENSMUSG00000105211 | 1 | |
| ENSMUSG00000105212 | 1 | |
| ENSMUSG00000105213 | 1 | |
| ENSMUSG00000105214 | 1 | |
| ENSMUSG00000105215 | 1 | |
| ENSMUSG00000105216 | 1 | |
| ENSMUSG00000105217 | -1.099174267 | 0.604452802 |
| ENSMUSG00000105218 | 1 | |
| ENSMUSG00000105219 | 1 | |
| ENSMUSG00000105220 | 1 | |
| ENSMUSG00000105221 | -1.089497311 | 0.608497229 |
| ENSMUSG00000105222 | 1 | |
| ENSMUSG00000105223 | 1 | |
| ENSMUSG00000105224 | 1 | |
| ENSMUSG00000105225 | 1 | |
| ENSMUSG00000105226 | -1.043306199 | 0.698112512 |
| ENSMUSG00000105227 | 1 | |
| ENSMUSG00000105228 | 1.000975865 | 1 |
| ENSMUSG00000105229 | 1 | |
| ENSMUSG00000105230 | -1.043306199 | 0.698112512 |
| ENSMUSG00000105231 | 1 | |
| ENSMUSG00000105232 | -1.043306199 | 0.698112512 |
| ENSMUSG00000105233 | 1.075870199 | 0.642819974 |
| ENSMUSG00000105234 | 1 | |
| ENSMUSG00000105235 | 1 | |
| ENSMUSG00000105236 | 1 | |
| ENSMUSG00000105237 | 1 | |
| ENSMUSG00000105238 | 1 | |
| ENSMUSG00000105239 | 1 | |
| ENSMUSG00000105240 | 1 | |
| ENSMUSG00000105241 | 1 | |
| ENSMUSG00000105242 | 1 | |
| ENSMUSG00000105243 | -1.03962336 | 0.814089245 |
| ENSMUSG00000105244 | -1.043306199 | 0.698112512 |
| ENSMUSG00000105245 | 1.162135265 | 0.475208338 |
| ENSMUSG00000105246 | 1 | |
| ENSMUSG00000105247 | 1.001363153 | 1 |
| ENSMUSG00000105248 | 1 | |
| ENSMUSG00000105249 | 1 | |
| ENSMUSG00000105250 | 1 | |
| ENSMUSG00000105251 | 1 | |
| ENSMUSG00000105252 | 1 | |
| ENSMUSG00000105253 | -1.174257662 | 0.211170208 |
| ENSMUSG00000105254 | 1.143882641 | 0.62953729 |
| ENSMUSG00000105255 | 1 | |
| ENSMUSG00000105256 | 1 | |
| ENSMUSG00000105257 | 1 | |
| ENSMUSG00000105258 | 1.059463094 | 0.608497229 |
| ENSMUSG00000105259 | -1.080227194 | 0.761911709 |
| ENSMUSG00000105260 | 1 | |

| | | |
|---|---|---|
| ENSMUSG00000105261 | -1.03962336 | 0.698112512 |
| ENSMUSG00000105262 | 1 | |
| ENSMUSG00000105263 | -1.053211779 | 0.823140628 |
| ENSMUSG00000105264 | -1.121975939 | 0.729206866 |
| ENSMUSG00000105265 | 1.127088412 | 0.608497229 |
| ENSMUSG00000105266 | 1 | |
| ENSMUSG00000105267 | 1 | |
| ENSMUSG00000105269 | 1 | |
| ENSMUSG00000105270 | 1.089349158 | 0.399744772 |
| ENSMUSG00000105271 | 1 | |
| ENSMUSG00000105272 | -1.002971829 | 1 |
| ENSMUSG00000105273 | 1 | |
| ENSMUSG00000105274 | 1 | |
| ENSMUSG00000105275 | -1.019108526 | 0.981356107 |
| ENSMUSG00000105276 | -1.100193663 | 0.534887579 |
| ENSMUSG00000105277 | -1 | 1 |
| ENSMUSG00000105278 | 1.018670051 | 0.99368696 |
| ENSMUSG00000105279 | 1 | |
| ENSMUSG00000105280 | 1 | |
| ENSMUSG00000105281 | 1 | |
| ENSMUSG00000105282 | 1 | |
| ENSMUSG00000105283 | 1 | |
| ENSMUSG00000105284 | 1 | |
| ENSMUSG00000105285 | 1.062415025 | 0.699611704 |
| ENSMUSG00000105286 | 1.059463094 | 0.776953697 |
| ENSMUSG00000105287 | -1.05909872 | 0.608497229 |
| ENSMUSG00000105288 | 1 | |
| ENSMUSG00000105289 | 1.059463094 | 0.698112512 |
| ENSMUSG00000105290 | 1 | |
| ENSMUSG00000105291 | -1.084645496 | 0.532369657 |
| ENSMUSG00000105292 | 1 | |
| ENSMUSG00000105293 | 1.171563499 | 0.233412931 |
| ENSMUSG00000105294 | 1.069068979 | 0.698112512 |
| ENSMUSG00000105295 | 1 | |
| ENSMUSG00000105296 | 1 | |
| ENSMUSG00000105297 | 1 | |
| ENSMUSG00000105298 | 1 | |
| ENSMUSG00000105299 | 1 | |
| ENSMUSG00000105300 | 1 | |
| ENSMUSG00000105301 | 1 | |
| ENSMUSG00000105302 | 1.039597911 | 0.91218384 |
| ENSMUSG00000105303 | 1.359154898 | 0.034712517 |
| ENSMUSG00000105304 | -1.136905997 | 0.608497229 |
| ENSMUSG00000105305 | 1 | |
| ENSMUSG00000105306 | 1 | |
| ENSMUSG00000105307 | 1 | |
| ENSMUSG00000105308 | 1 | |
| ENSMUSG00000105309 | 1 | |
| ENSMUSG00000105310 | 1 | |
| ENSMUSG00000105311 | 1 | |
| ENSMUSG00000105312 | 1 | |
| ENSMUSG00000105313 | 1 | |
| ENSMUSG00000105314 | 1 | |
| ENSMUSG00000105315 | 1 | |
| ENSMUSG00000105316 | 1 | |
| ENSMUSG00000105317 | 1 | |
| ENSMUSG00000105318 | 1 | |
| ENSMUSG00000105319 | 1 | |
| ENSMUSG00000105320 | 1 | |
| ENSMUSG00000105321 | 1 | |
| ENSMUSG00000105322 | 1 | |
| ENSMUSG00000105323 | 1 | |
| ENSMUSG00000105324 | 1 | |

| | | |
|---|---|---|
| ENSMUSG00000105325 | 1 | |
| ENSMUSG00000105326 | -1.01566718 | 1 |
| ENSMUSG00000105327 | -1.043616943 | 0.935744977 |
| ENSMUSG00000105328 | 1.171536022 | 0.52584487 |
| ENSMUSG00000105329 | 1 | |
| ENSMUSG00000105330 | 1 | |
| ENSMUSG00000105331 | 1 | |
| ENSMUSG00000105332 | 1.049411284 | 0.835701362 |
| ENSMUSG00000105333 | -1.043306199 | 0.698112512 |
| ENSMUSG00000105334 | 1 | |
| ENSMUSG00000105335 | 1 | |
| ENSMUSG00000105336 | -1.037482356 | 0.870427333 |
| ENSMUSG00000105337 | 1 | |
| ENSMUSG00000105338 | 1 | |
| ENSMUSG00000105339 | 1 | |
| ENSMUSG00000105340 | -1.018483386 | 0.986542466 |
| ENSMUSG00000105341 | 1.241489066 | 0.124350077 |
| ENSMUSG00000105342 | 1 | |
| ENSMUSG00000105343 | -1.084645496 | 0.532369657 |
| ENSMUSG00000105344 | 1 | |
| ENSMUSG00000105345 | 1.217385435 | 0.391804854 |
| ENSMUSG00000105346 | 1 | |
| ENSMUSG00000105347 | -1.034661764 | 0.976788866 |
| ENSMUSG00000105348 | 1 | |
| ENSMUSG00000105349 | 1.015707025 | 0.995295685 |
| ENSMUSG00000105350 | 1.059463094 | 0.698112512 |
| ENSMUSG00000105351 | 1 | |
| ENSMUSG00000105352 | 1.059463094 | 0.608497229 |
| ENSMUSG00000105353 | -1.114551661 | 0.381312112 |
| ENSMUSG00000105354 | 1 | |
| ENSMUSG00000105355 | 1 | |
| ENSMUSG00000105356 | -1.008721168 | 0.994172456 |
| ENSMUSG00000105357 | 1 | |
| ENSMUSG00000105358 | -1.337090582 | 0.223253407 |
| ENSMUSG00000105359 | -1.090988886 | 0.706967075 |
| ENSMUSG00000105360 | -1 | 1 |
| ENSMUSG00000105361 | 1 | |
| ENSMUSG00000105362 | 1 | |
| ENSMUSG00000105363 | 1 | |
| ENSMUSG00000105364 | -1.043306199 | 0.698112512 |
| ENSMUSG00000105365 | 1 | |
| ENSMUSG00000105366 | 1.593610934 | 0.003010378 |
| ENSMUSG00000105367 | 1 | |
| ENSMUSG00000105368 | 1.022024763 | 0.976843242 |
| ENSMUSG00000105369 | 1 | |
| ENSMUSG00000105370 | 1.04519852 | 0.698112512 |
| ENSMUSG00000105371 | 1 | |
| ENSMUSG00000105372 | 1 | |
| ENSMUSG00000105373 | 1 | |
| ENSMUSG00000105374 | 1 | |
| ENSMUSG00000105375 | 1 | |
| ENSMUSG00000105376 | 1.335184066 | 0.323247578 |
| ENSMUSG00000105377 | 1 | |
| ENSMUSG00000105378 | 1.026239213 | 0.949671033 |
| ENSMUSG00000105379 | 1.059463094 | 0.608497229 |
| ENSMUSG00000105380 | 1 | |
| ENSMUSG00000105381 | 1 | |
| ENSMUSG00000105382 | 1.036754564 | 0.783658058 |
| ENSMUSG00000105383 | -1.091372079 | 0.835701362 |
| ENSMUSG00000105384 | 1 | |
| ENSMUSG00000105385 | 1 | |
| ENSMUSG00000105386 | 1 | |
| ENSMUSG00000105387 | 1.065007221 | 0.892937692 |

| | | |
|---|---|---|
| ENSMUSG00000105388 | -1.046923187 | 0.922446617 |
| ENSMUSG00000105389 | 1.132639128 | 0.217803313 |
| ENSMUSG00000105390 | -1 | 1 |
| ENSMUSG00000105391 | 1.059463094 | 0.608497229 |
| ENSMUSG00000105392 | 1 | |
| ENSMUSG00000105393 | 1 | |
| ENSMUSG00000105394 | 1 | |
| ENSMUSG00000105395 | -1.192826061 | 0.523469825 |
| ENSMUSG00000105396 | 1.069068979 | 0.698112512 |
| ENSMUSG00000105397 | 1.121243997 | 0.70299173 |
| ENSMUSG00000105398 | 1 | |
| ENSMUSG00000105399 | 1 | |
| ENSMUSG00000105400 | 1 | |
| ENSMUSG00000105401 | 1.140144063 | 0.635394047 |
| ENSMUSG00000105402 | 1.487626555 | 0.000413551 |
| ENSMUSG00000105403 | 1.069068979 | 0.698112512 |
| ENSMUSG00000105404 | 1.199989356 | 0.045045157 |
| ENSMUSG00000105405 | 1 | |
| ENSMUSG00000105406 | -1.027126002 | 0.982838256 |
| ENSMUSG00000105407 | 1 | |
| ENSMUSG00000105408 | 1.001354487 | 1 |
| ENSMUSG00000105409 | 1 | |
| ENSMUSG00000105410 | 1 | |
| ENSMUSG00000105411 | 1 | |
| ENSMUSG00000105412 | 1.014545335 | 0.979002724 |
| ENSMUSG00000105413 | 1 | |
| ENSMUSG00000105415 | 1 | |
| ENSMUSG00000105416 | 1.092111687 | 0.608497229 |
| ENSMUSG00000105418 | 1 | |
| ENSMUSG00000105419 | -1.059097125 | 0.698112512 |
| ENSMUSG00000105420 | 1 | |
| ENSMUSG00000105421 | 1 | |
| ENSMUSG00000105422 | -1.03962336 | 0.698112512 |
| ENSMUSG00000105423 | 1 | |
| ENSMUSG00000105424 | 1 | |
| ENSMUSG00000105425 | 1.250307939 | 0.311092577 |
| ENSMUSG00000105426 | 1 | |
| ENSMUSG00000105427 | 1 | |
| ENSMUSG00000105428 | 1.059463094 | 0.699340382 |
| ENSMUSG00000105429 | -1.065904764 | 0.906158117 |
| ENSMUSG00000105430 | -1.053549062 | 0.840310372 |
| ENSMUSG00000105431 | -1.024718486 | 0.951399668 |
| ENSMUSG00000105432 | 1 | |
| ENSMUSG00000105433 | 1 | |
| ENSMUSG00000105434 | 1.057653512 | 0.914323346 |
| ENSMUSG00000105435 | 1 | |
| ENSMUSG00000105436 | 1 | |
| ENSMUSG00000105437 | 1.084618945 | 0.614071182 |
| ENSMUSG00000105438 | 1 | |
| ENSMUSG00000105439 | -1 | 1 |
| ENSMUSG00000105440 | 1.009100337 | 1 |
| ENSMUSG00000105441 | 1.039597911 | 0.850281678 |
| ENSMUSG00000105442 | 1 | |
| ENSMUSG00000105443 | 1 | |
| ENSMUSG00000105444 | 1 | |
| ENSMUSG00000105445 | 1 | |
| ENSMUSG00000105446 | 1 | |
| ENSMUSG00000105447 | -1.03962336 | 0.698112512 |
| ENSMUSG00000105448 | 1 | |
| ENSMUSG00000105449 | -1.107711699 | 0.642593813 |
| ENSMUSG00000105450 | 1 | |
| ENSMUSG00000105451 | 1 | |
| ENSMUSG00000105452 | 1.026546919 | 0.968735041 |

| | | |
|---|---|---|
| ENSMUSG00000105453 | -1.010745469 | 1 |
| ENSMUSG00000105454 | -1.019731926 | 0.954947258 |
| ENSMUSG00000105455 | 1 | |
| ENSMUSG00000105456 | 1.132639128 | 0.217803313 |
| ENSMUSG00000105457 | 1.15320799 | 0.698112512 |
| ENSMUSG00000105458 | 1 | |
| ENSMUSG00000105459 | 1 | |
| ENSMUSG00000105460 | 1.059463094 | 0.608497229 |
| ENSMUSG00000105461 | 1.337294464 | 0.019065095 |
| ENSMUSG00000105462 | 1 | |
| ENSMUSG00000105463 | 1 | |
| ENSMUSG00000105464 | 1 | |
| ENSMUSG00000105465 | 1 | |
| ENSMUSG00000105466 | -1.178477407 | 0.608497229 |
| ENSMUSG00000105467 | 1 | |
| ENSMUSG00000105468 | 1 | |
| ENSMUSG00000105469 | 1.069068979 | 0.608497229 |
| ENSMUSG00000105470 | 1 | |
| ENSMUSG00000105471 | 1.01292152 | 0.980617339 |
| ENSMUSG00000105472 | -1.089497311 | 0.608497229 |
| ENSMUSG00000105473 | 1 | |
| ENSMUSG00000105474 | 1 | |
| ENSMUSG00000105475 | 1 | |
| ENSMUSG00000105476 | -1.059054441 | 0.7530201 |
| ENSMUSG00000105477 | 1 | |
| ENSMUSG00000105478 | 1.227736972 | 0.429887106 |
| ENSMUSG00000105479 | 1 | |
| ENSMUSG00000105480 | 1 | |
| ENSMUSG00000105481 | -1.053549062 | 0.698112512 |
| ENSMUSG00000105482 | 1.059463094 | 0.608497229 |
| ENSMUSG00000105483 | -1.290811762 | 0.009914504 |
| ENSMUSG00000105484 | 1 | |
| ENSMUSG00000105485 | -1 | 1 |
| ENSMUSG00000105486 | 1 | |
| ENSMUSG00000105487 | 1 | |
| ENSMUSG00000105488 | 1 | |
| ENSMUSG00000105489 | 1 | |
| ENSMUSG00000105490 | 1 | |
| ENSMUSG00000105491 | 1 | |
| ENSMUSG00000105492 | 1 | |
| ENSMUSG00000105493 | 1 | |
| ENSMUSG00000105494 | 1 | |
| ENSMUSG00000105495 | 1 | |
| ENSMUSG00000105496 | 1 | |
| ENSMUSG00000105497 | 1 | |
| ENSMUSG00000105498 | 1 | |
| ENSMUSG00000105499 | 1 | |
| ENSMUSG00000105500 | 1 | |
| ENSMUSG00000105501 | -1.107299336 | 0.71045278 |
| ENSMUSG00000105502 | -1.085651418 | 0.625496737 |
| ENSMUSG00000105503 | 1 | |
| ENSMUSG00000105504 | -1.252316018 | 0.41274756 |
| ENSMUSG00000105505 | 1 | |
| ENSMUSG00000105506 | -1.051018293 | 0.876297532 |
| ENSMUSG00000105507 | 1 | |
| ENSMUSG00000105508 | 1 | |
| ENSMUSG00000105509 | -1.063071074 | 0.576169279 |
| ENSMUSG00000105510 | -1.198351104 | 0.532586 |
| ENSMUSG00000105511 | 1 | |
| ENSMUSG00000105512 | -1.03962336 | 0.872692845 |
| ENSMUSG00000105513 | 1 | |
| ENSMUSG00000105514 | 1.047494655 | 0.807247723 |
| ENSMUSG00000105515 | 1 | |

| | | |
|---|---|---|
| ENSMUSG00000105516 | -1.120855399 | 0.608497229 |
| ENSMUSG00000105517 | -1.192435027 | 0.658539719 |
| ENSMUSG00000105518 | 1 | |
| ENSMUSG00000105519 | -1.083415624 | 0.698112512 |
| ENSMUSG00000105520 | 1 | |
| ENSMUSG00000105521 | 1 | |
| ENSMUSG00000105522 | 1 | |
| ENSMUSG00000105523 | 1 | |
| ENSMUSG00000105524 | 1 | |
| ENSMUSG00000105525 | 1 | |
| ENSMUSG00000105526 | -1.053549062 | 0.769745556 |
| ENSMUSG00000105527 | 1 | |
| ENSMUSG00000105528 | 1.059463094 | 0.608497229 |
| ENSMUSG00000105529 | -1.168898831 | 0.522018707 |
| ENSMUSG00000105530 | -1.063525418 | 0.698112512 |
| ENSMUSG00000105531 | 1 | |
| ENSMUSG00000105532 | 1 | |
| ENSMUSG00000105533 | 1 | |
| ENSMUSG00000105534 | -1.155252261 | 0.193677799 |
| ENSMUSG00000105535 | 1 | |
| ENSMUSG00000105536 | 1.096901918 | 0.698112512 |
| ENSMUSG00000105537 | 1 | |
| ENSMUSG00000105538 | 1 | |
| ENSMUSG00000105539 | 1 | |
| ENSMUSG00000105540 | 1 | |
| ENSMUSG00000105541 | -1.17055892 | 0.540744382 |
| ENSMUSG00000105542 | -1.009476377 | 1 |
| ENSMUSG00000105543 | 1 | |
| ENSMUSG00000105544 | 1 | |
| ENSMUSG00000105545 | 1 | |
| ENSMUSG00000105546 | 1 | |
| ENSMUSG00000105547 | 1 | |
| ENSMUSG00000105548 | 1 | |
| ENSMUSG00000105549 | 1.066465917 | 0.682836879 |
| ENSMUSG00000105550 | 1 | |
| ENSMUSG00000105551 | 1 | |
| ENSMUSG00000105552 | 1.175006804 | 0.608497229 |
| ENSMUSG00000105553 | 1.043550045 | 0.869676326 |
| ENSMUSG00000105554 | 1.084091858 | 0.791778775 |
| ENSMUSG00000105555 | 1 | |
| ENSMUSG00000105556 | 1.056397698 | 0.549804687 |
| ENSMUSG00000105557 | 1 | |
| ENSMUSG00000105558 | 1.131503319 | 0.473579199 |
| ENSMUSG00000105559 | 1 | |
| ENSMUSG00000105560 | -1 | 1 |
| ENSMUSG00000105561 | 1.079180038 | 0.608497229 |
| ENSMUSG00000105562 | -1 | 1 |
| ENSMUSG00000105563 | 1 | |
| ENSMUSG00000105564 | 1 | |
| ENSMUSG00000105565 | 1.157813428 | 0.698112512 |
| ENSMUSG00000105566 | 1 | |
| ENSMUSG00000105567 | 1 | |
| ENSMUSG00000105568 | 1 | |
| ENSMUSG00000105569 | 1 | |
| ENSMUSG00000105570 | -1.220755813 | 0.123596425 |
| ENSMUSG00000105571 | 1 | |
| ENSMUSG00000105572 | 1.012283577 | 0.960776894 |
| ENSMUSG00000105573 | 1 | |
| ENSMUSG00000105574 | 1 | |
| ENSMUSG00000105575 | 1 | |
| ENSMUSG00000105576 | 1 | |
| ENSMUSG00000105577 | 1 | |
| ENSMUSG00000105578 | 1.036723625 | 0.83128424 |

| | | |
|---|---|---|
| ENSMUSG00000105579 | 1.059463094 | 0.698112512 |
| ENSMUSG00000105580 | 1 | |
| ENSMUSG00000105581 | 1 | |
| ENSMUSG00000105582 | 1 | |
| ENSMUSG00000105583 | 1 | |
| ENSMUSG00000105584 | 1 | |
| ENSMUSG00000105585 | 1 | |
| ENSMUSG00000105586 | -1.180268306 | 0.496888303 |
| ENSMUSG00000105588 | 1 | |
| ENSMUSG00000105589 | 1 | |
| ENSMUSG00000105590 | 1.069068979 | 0.608497229 |
| ENSMUSG00000105591 | 1.032516284 | 0.938931916 |
| ENSMUSG00000105592 | 1 | |
| ENSMUSG00000105593 | -1 | 1 |
| ENSMUSG00000105594 | -1.271047361 | 0.304680629 |
| ENSMUSG00000105595 | 1 | |
| ENSMUSG00000105596 | 1 | |
| ENSMUSG00000105597 | 1.049622663 | 0.887841887 |
| ENSMUSG00000105598 | 1 | |
| ENSMUSG00000105599 | 1.059463094 | 0.608497229 |
| ENSMUSG00000105600 | 1.026051401 | 0.981964814 |
| ENSMUSG00000105601 | -1.00616747 | 1 |
| ENSMUSG00000105602 | -1.132666855 | 0.18851436 |
| ENSMUSG00000105603 | 1 | |
| ENSMUSG00000105604 | 1 | |
| ENSMUSG00000105605 | -1.03962336 | 0.698112512 |
| ENSMUSG00000105606 | 1 | |
| ENSMUSG00000105607 | 1.029649077 | 0.946232149 |
| ENSMUSG00000105608 | 1 | |
| ENSMUSG00000105609 | 1.075870199 | 0.642819974 |
| ENSMUSG00000105610 | 1 | |
| ENSMUSG00000105611 | -1.059358544 | 0.886137227 |
| ENSMUSG00000105612 | 1 | |
| ENSMUSG00000105613 | -1.035428135 | 0.897053041 |
| ENSMUSG00000105614 | 1 | |
| ENSMUSG00000105615 | 1 | |
| ENSMUSG00000105616 | 1 | |
| ENSMUSG00000105617 | 1 | |
| ENSMUSG00000105618 | 1 | |
| ENSMUSG00000105619 | 1 | |
| ENSMUSG00000105620 | 1 | |
| ENSMUSG00000105621 | 1 | |
| ENSMUSG00000105622 | 1.189207115 | 0.385934169 |
| ENSMUSG00000105624 | 1.133733993 | 0.473421853 |
| ENSMUSG00000105625 | 1 | |
| ENSMUSG00000105626 | 1 | |
| ENSMUSG00000105627 | 1 | |
| ENSMUSG00000105628 | 1 | |
| ENSMUSG00000105629 | 1 | |
| ENSMUSG00000105630 | 1 | |
| ENSMUSG00000105632 | 1.038996468 | 0.941321009 |
| ENSMUSG00000105633 | 1.132639128 | 0.217803313 |
| ENSMUSG00000105634 | 1 | |
| ENSMUSG00000105635 | 1 | |
| ENSMUSG00000105636 | 1.050984995 | 0.927108331 |
| ENSMUSG00000105637 | 1.095872691 | 0.608497229 |
| ENSMUSG00000105638 | 1 | |
| ENSMUSG00000105639 | 1 | |
| ENSMUSG00000105640 | 1 | |
| ENSMUSG00000105641 | -1.113515092 | 0.608497229 |
| ENSMUSG00000105642 | -1 | 1 |
| ENSMUSG00000105643 | -1.039970084 | 0.83949549 |
| ENSMUSG00000105644 | 1.00950998 | 1 |

| | | |
|---|---|---|
| ENSMUSG00000105645 | 1 | |
| ENSMUSG00000105646 | 1 | |
| ENSMUSG00000105647 | 1 | |
| ENSMUSG00000105648 | 1 | |
| ENSMUSG00000105649 | 1 | |
| ENSMUSG00000105650 | -1.053549062 | 0.698112512 |
| ENSMUSG00000105651 | 1.059463094 | 0.698112512 |
| ENSMUSG00000105652 | 1.085624843 | 0.69305964 |
| ENSMUSG00000105653 | 1.019083579 | 0.968937442 |
| ENSMUSG00000105654 | 1 | |
| ENSMUSG00000105655 | -1.032816862 | 0.848953901 |
| ENSMUSG00000105656 | -1 | 1 |
| ENSMUSG00000105657 | 1.140501929 | 0.698112512 |
| ENSMUSG00000105658 | -1.044273782 | 0.698112512 |
| ENSMUSG00000105659 | -1.111221915 | 0.689297957 |
| ENSMUSG00000105660 | 1 | |
| ENSMUSG00000105661 | 1 | |
| ENSMUSG00000105662 | 1 | |
| ENSMUSG00000105663 | 1.188607586 | 0.608497229 |
| ENSMUSG00000105664 | -1.03962336 | 0.698112512 |
| ENSMUSG00000105665 | 1 | |
| ENSMUSG00000105666 | 1 | |
| ENSMUSG00000105667 | 1 | |
| ENSMUSG00000105668 | 1 | |
| ENSMUSG00000105669 | 1 | |
| ENSMUSG00000105670 | 1.005613439 | 1 |
| ENSMUSG00000105671 | 1 | |
| ENSMUSG00000105672 | 1 | |
| ENSMUSG00000105673 | 1 | |
| ENSMUSG00000105674 | 1 | |
| ENSMUSG00000105675 | 1 | |
| ENSMUSG00000105676 | -1.0795912 | 0.698112512 |
| ENSMUSG00000105677 | 1.096128311 | 0.711581415 |
| ENSMUSG00000105678 | -1.028348525 | 0.936995747 |
| ENSMUSG00000105679 | 1.109508202 | 0.646852072 |
| ENSMUSG00000105680 | 1 | |
| ENSMUSG00000105681 | 1.054008754 | 0.933142264 |
| ENSMUSG00000105682 | 1 | |
| ENSMUSG00000105683 | 1 | |
| ENSMUSG00000105684 | 1 | |
| ENSMUSG00000105685 | 1 | |
| ENSMUSG00000105686 | 1 | |
| ENSMUSG00000105687 | -1.03962336 | 0.698112512 |
| ENSMUSG00000105688 | 1 | |
| ENSMUSG00000105689 | -1.095294215 | 0.541735653 |
| ENSMUSG00000105690 | 1 | |
| ENSMUSG00000105691 | 1.054986959 | 0.89811529 |
| ENSMUSG00000105692 | 1.09082887 | 0.527191115 |
| ENSMUSG00000105693 | 1 | |
| ENSMUSG00000105694 | 1.095872691 | 0.608497229 |
| ENSMUSG00000105695 | 1.046923501 | 0.745814988 |
| ENSMUSG00000105696 | 1 | |
| ENSMUSG00000105697 | 1 | |
| ENSMUSG00000105698 | 1.059463094 | 0.608497229 |
| ENSMUSG00000105699 | 1.150511357 | 0.698112512 |
| ENSMUSG00000105700 | -1.003834322 | 1 |
| ENSMUSG00000105701 | -1 | 1 |
| ENSMUSG00000105702 | 1 | |
| ENSMUSG00000105703 | 1.032081991 | 0.803072249 |
| ENSMUSG00000105704 | 1 | |
| ENSMUSG00000105705 | 1 | |
| ENSMUSG00000105706 | 1.059463094 | 0.608497229 |
| ENSMUSG00000105707 | 1 | |

| | | |
|---|---|---|
| ENSMUSG00000105708 | 1.506991477 | 0.060030789 |
| ENSMUSG00000105709 | 1 | |
| ENSMUSG00000105710 | -1.005000082 | 1 |
| ENSMUSG00000105711 | 1.005021373 | 1 |
| ENSMUSG00000105712 | 1 | |
| ENSMUSG00000105713 | 1 | |
| ENSMUSG00000105714 | 1 | |
| ENSMUSG00000105715 | 1 | |
| ENSMUSG00000105716 | 1 | |
| ENSMUSG00000105717 | -1.184914433 | 0.503376297 |
| ENSMUSG00000105718 | 1 | |
| ENSMUSG00000105719 | -1.367396887 | 0.058549167 |
| ENSMUSG00000105720 | -1.043306199 | 0.79291651 |
| ENSMUSG00000105721 | 1 | |
| ENSMUSG00000105722 | -1.010988769 | 1 |
| ENSMUSG00000105723 | 1 | |
| ENSMUSG00000105724 | 1 | |
| ENSMUSG00000105725 | 1 | |
| ENSMUSG00000105726 | 1 | |
| ENSMUSG00000105727 | 1 | |
| ENSMUSG00000105728 | -1.015248956 | 0.992104397 |
| ENSMUSG00000105729 | -1 | 1 |
| ENSMUSG00000105730 | -1 | 1 |
| ENSMUSG00000105731 | -1.055477714 | 0.778405037 |
| ENSMUSG00000105732 | 1 | |
| ENSMUSG00000105733 | 1 | |
| ENSMUSG00000105734 | -1.03962336 | 0.803072249 |
| ENSMUSG00000105735 | 1 | |
| ENSMUSG00000105736 | 1 | |
| ENSMUSG00000105737 | 1 | |
| ENSMUSG00000105738 | 1 | |
| ENSMUSG00000105739 | 1 | |
| ENSMUSG00000105740 | 1.039816326 | 0.9296857 |
| ENSMUSG00000105741 | -1.034085987 | 0.806069168 |
| ENSMUSG00000105742 | 1.26290645 | 0.154458644 |
| ENSMUSG00000105743 | 1 | |
| ENSMUSG00000105744 | 1 | |
| ENSMUSG00000105745 | 1.165664656 | 0.608497229 |
| ENSMUSG00000105746 | 1.069068979 | 0.608497229 |
| ENSMUSG00000105747 | 1 | |
| ENSMUSG00000105748 | -1.232259831 | 0.498779515 |
| ENSMUSG00000105749 | 1 | |
| ENSMUSG00000105750 | 1.122462048 | 0.420161906 |
| ENSMUSG00000105751 | 1 | |
| ENSMUSG00000105752 | 1 | |
| ENSMUSG00000105753 | -1.009519625 | 1 |
| ENSMUSG00000105754 | 1 | |
| ENSMUSG00000105755 | 1.014731082 | 0.988424085 |
| ENSMUSG00000105756 | 1 | |
| ENSMUSG00000105757 | 1 | |
| ENSMUSG00000105758 | 1 | |
| ENSMUSG00000105759 | 1.059463094 | 0.608497229 |
| ENSMUSG00000105760 | 1 | |
| ENSMUSG00000105761 | 1.049517471 | 0.837704961 |
| ENSMUSG00000105762 | 1.189207115 | 0.295479694 |
| ENSMUSG00000105763 | 1 | |
| ENSMUSG00000105764 | 1 | |
| ENSMUSG00000105765 | 1 | |
| ENSMUSG00000105766 | -1.053549062 | 0.698112512 |
| ENSMUSG00000105767 | 1 | |
| ENSMUSG00000105768 | 1 | |
| ENSMUSG00000105769 | 1 | |
| ENSMUSG00000105770 | 1 | |

| | | |
|---|---|---|
| ENSMUSG00000105771 | 1.211684158 | 0.56051496 |
| ENSMUSG00000105772 | -1.143787033 | 0.363110963 |
| ENSMUSG00000105773 | 1 | |
| ENSMUSG00000105774 | 1.059463094 | 0.608497229 |
| ENSMUSG00000105775 | -1.033820075 | 0.916014403 |
| ENSMUSG00000105776 | -1.027287094 | 0.938484564 |
| ENSMUSG00000105777 | 1 | |
| ENSMUSG00000105778 | 1 | |
| ENSMUSG00000105779 | -1.043306199 | 0.79291651 |
| ENSMUSG00000105780 | 1 | |
| ENSMUSG00000105781 | 1 | |
| ENSMUSG00000105782 | 1.059463094 | 0.608497229 |
| ENSMUSG00000105783 | 1 | |
| ENSMUSG00000105784 | 1 | |
| ENSMUSG00000105785 | 1 | |
| ENSMUSG00000105786 | 1 | |
| ENSMUSG00000105787 | 1.089470641 | 0.713221609 |
| ENSMUSG00000105788 | 1 | |
| ENSMUSG00000105789 | -1.160253504 | 0.548809407 |
| ENSMUSG00000105790 | 1 | |
| ENSMUSG00000105791 | 1 | |
| ENSMUSG00000105792 | 1 | |
| ENSMUSG00000105793 | -1.028348525 | 0.940333715 |
| ENSMUSG00000105794 | 1 | |
| ENSMUSG00000105795 | 1 | |
| ENSMUSG00000105796 | 1 | |
| ENSMUSG00000105797 | 1.059463094 | 0.698112512 |
| ENSMUSG00000105798 | 1 | |
| ENSMUSG00000105799 | 1 | |
| ENSMUSG00000105800 | 1 | |
| ENSMUSG00000105801 | -1.044273782 | 0.698112512 |
| ENSMUSG00000105802 | 1.282918176 | 0.120506674 |
| ENSMUSG00000105803 | 1 | |
| ENSMUSG00000105804 | -1.134921691 | 0.608497229 |
| ENSMUSG00000105805 | 1 | |
| ENSMUSG00000105806 | 1 | |
| ENSMUSG00000105807 | 1 | |
| ENSMUSG00000105808 | 1 | |
| ENSMUSG00000105809 | 1 | |
| ENSMUSG00000105810 | -1.105673636 | 0.608497229 |
| ENSMUSG00000105811 | 1 | |
| ENSMUSG00000105812 | 1 | |
| ENSMUSG00000105813 | 1.146329683 | 0.128114038 |
| ENSMUSG00000105814 | -1.034118365 | 0.833759609 |
| ENSMUSG00000105815 | 1.005613439 | 1 |
| ENSMUSG00000105816 | 1 | |
| ENSMUSG00000105817 | 1 | |
| ENSMUSG00000105818 | -1.06568528 | 0.875063517 |
| ENSMUSG00000105819 | 1 | |
| ENSMUSG00000105820 | 1 | |
| ENSMUSG00000105821 | 1 | |
| ENSMUSG00000105822 | -1 | 1 |
| ENSMUSG00000105823 | 1 | |
| ENSMUSG00000105824 | 1.068424331 | 0.731295483 |
| ENSMUSG00000105825 | 1 | |
| ENSMUSG00000105826 | -1.010531604 | 1 |
| ENSMUSG00000105827 | 1 | |
| ENSMUSG00000105828 | 1 | |
| ENSMUSG00000105829 | -1.03962336 | 0.872692845 |
| ENSMUSG00000105830 | 1 | |
| ENSMUSG00000105831 | 1 | |
| ENSMUSG00000105832 | 1.016912276 | 0.963147621 |
| ENSMUSG00000105833 | -1.049075043 | 0.90117462 |

| | | |
|---|---|---|
| ENSMUSG00000105834 | -1.063525418 | 0.698112512 |
| ENSMUSG00000105835 | 1 | |
| ENSMUSG00000105836 | -1.038444538 | 0.865865121 |
| ENSMUSG00000105837 | -1.05948903 | 0.784340478 |
| ENSMUSG00000105838 | 1.069068979 | 0.608497229 |
| ENSMUSG00000105839 | 1 | |
| ENSMUSG00000105840 | 1 | |
| ENSMUSG00000105841 | 1 | |
| ENSMUSG00000105842 | -1.141849922 | 0.577220449 |
| ENSMUSG00000105843 | 1.095872691 | 0.698112512 |
| ENSMUSG00000105844 | 1 | |
| ENSMUSG00000105845 | -1.03962336 | 0.698112512 |
| ENSMUSG00000105846 | 1.060782284 | 0.676936722 |
| ENSMUSG00000105847 | 1 | |
| ENSMUSG00000105848 | 1.149113744 | 0.158414349 |
| ENSMUSG00000105849 | 1 | |
| ENSMUSG00000105850 | 1 | |
| ENSMUSG00000105851 | 1.171248102 | 0.321024879 |
| ENSMUSG00000105852 | 1.098894583 | 0.747744437 |
| ENSMUSG00000105853 | 1 | |
| ENSMUSG00000105854 | 1.006484741 | 1 |
| ENSMUSG00000105855 | 1.08813331 | 0.659204084 |
| ENSMUSG00000105856 | 1 | |
| ENSMUSG00000105857 | 1 | |
| ENSMUSG00000105858 | 1.255635743 | 0.479262818 |
| ENSMUSG00000105859 | 1 | |
| ENSMUSG00000105860 | -1.176660386 | 0.327642404 |
| ENSMUSG00000105861 | -1 | 1 |
| ENSMUSG00000105862 | 1 | |
| ENSMUSG00000105863 | 1 | |
| ENSMUSG00000105864 | -1.048277865 | 0.860645152 |
| ENSMUSG00000105865 | 1.014346713 | 1 |
| ENSMUSG00000105866 | -1.039056293 | 0.972684567 |
| ENSMUSG00000105867 | 1.029632141 | 0.965659325 |
| ENSMUSG00000105868 | -1.007097991 | 1 |
| ENSMUSG00000105869 | 1 | |
| ENSMUSG00000105870 | 1.161036672 | 0.399744772 |
| ENSMUSG00000105871 | 1 | |
| ENSMUSG00000105872 | 1 | |
| ENSMUSG00000105873 | 1 | |
| ENSMUSG00000105874 | 1 | |
| ENSMUSG00000105875 | 1.059463094 | 0.608497229 |
| ENSMUSG00000105876 | 1 | |
| ENSMUSG00000105877 | 1.034865357 | 0.899648217 |
| ENSMUSG00000105878 | 1 | |
| ENSMUSG00000105879 | 1.122462048 | 0.215991492 |
| ENSMUSG00000105880 | 1 | |
| ENSMUSG00000105881 | 1.297620467 | 0.193677799 |
| ENSMUSG00000105882 | 1 | |
| ENSMUSG00000105883 | 1 | |
| ENSMUSG00000105884 | 1 | |
| ENSMUSG00000105885 | 1 | |
| ENSMUSG00000105886 | 1 | |
| ENSMUSG00000105887 | 1 | |
| ENSMUSG00000105888 | 1 | |
| ENSMUSG00000105889 | 1 | |
| ENSMUSG00000105890 | 1 | |
| ENSMUSG00000105891 | 1.059463094 | 0.698112512 |
| ENSMUSG00000105892 | -1.044273782 | 0.698112512 |
| ENSMUSG00000105893 | -1.028348525 | 0.940333715 |
| ENSMUSG00000105894 | 1 | |
| ENSMUSG00000105895 | -1.22879693 | 0.397756797 |
| ENSMUSG00000105896 | -1.025874325 | 0.924273043 |

| | | |
|---|---|---|
| ENSMUSG00000105897 | 1 | |
| ENSMUSG00000105898 | 1 | |
| ENSMUSG00000105899 | 1 | |
| ENSMUSG00000105900 | 1 | |
| ENSMUSG00000105901 | 1 | |
| ENSMUSG00000105902 | 1 | |
| ENSMUSG00000105903 | 1 | |
| ENSMUSG00000105904 | 1 | |
| ENSMUSG00000105905 | -1.143787033 | 0.201977401 |
| ENSMUSG00000105906 | 1 | |
| ENSMUSG00000105907 | 1 | |
| ENSMUSG00000105908 | -1 | 1 |
| ENSMUSG00000105909 | 1.059463094 | 0.698112512 |
| ENSMUSG00000105910 | 1 | |
| ENSMUSG00000105911 | 1 | |
| ENSMUSG00000105912 | 1 | |
| ENSMUSG00000105913 | 1 | |
| ENSMUSG00000105914 | 1 | |
| ENSMUSG00000105915 | 1 | |
| ENSMUSG00000105916 | 1.014731082 | 0.997667419 |
| ENSMUSG00000105917 | 1.059463094 | 0.698112512 |
| ENSMUSG00000105918 | 1 | |
| ENSMUSG00000105919 | 1 | |
| ENSMUSG00000105920 | 1 | |
| ENSMUSG00000105921 | 1 | |
| ENSMUSG00000105922 | -1.111062562 | 0.664721995 |
| ENSMUSG00000105923 | 1 | |
| ENSMUSG00000105924 | 1 | |
| ENSMUSG00000105925 | -1.043306199 | 0.803260194 |
| ENSMUSG00000105926 | 1.069068979 | 0.608497229 |
| ENSMUSG00000105927 | 1 | |
| ENSMUSG00000105928 | 1 | |
| ENSMUSG00000105929 | 1.124843036 | 0.583194801 |
| ENSMUSG00000105930 | 1 | |
| ENSMUSG00000105931 | 1.074239611 | 0.698112512 |
| ENSMUSG00000105932 | -1.044273782 | 0.698112512 |
| ENSMUSG00000105933 | 1.143882641 | 0.608497229 |
| ENSMUSG00000105934 | 1 | |
| ENSMUSG00000105935 | -1.205806604 | 0.590417714 |
| ENSMUSG00000105936 | -1.193320102 | 0.000306852 |
| ENSMUSG00000105937 | 1 | |
| ENSMUSG00000105938 | 1 | |
| ENSMUSG00000105939 | -1.181030773 | 0.522542762 |
| ENSMUSG00000105940 | 1.194414407 | 0.487265142 |
| ENSMUSG00000105941 | -1.02522956 | 0.978911254 |
| ENSMUSG00000105942 | -1.113515092 | 0.576169279 |
| ENSMUSG00000105943 | 1 | |
| ENSMUSG00000105944 | 1.132877091 | 0.629824986 |
| ENSMUSG00000105945 | -1.168109923 | 0.206114416 |
| ENSMUSG00000105946 | 1.059463094 | 0.608497229 |
| ENSMUSG00000105947 | 1.049023683 | 0.918280975 |
| ENSMUSG00000105948 | -1.044273782 | 0.698112512 |
| ENSMUSG00000105949 | 1 | |
| ENSMUSG00000105950 | 1.036038403 | 0.613187574 |
| ENSMUSG00000105951 | 1 | |
| ENSMUSG00000105952 | 1 | |
| ENSMUSG00000105953 | 1 | |
| ENSMUSG00000105954 | 1.211527659 | 0.328602737 |
| ENSMUSG00000105955 | 1 | |
| ENSMUSG00000105956 | 1 | |
| ENSMUSG00000105957 | -1.021166308 | 0.970891309 |
| ENSMUSG00000105958 | 1.069068979 | 0.608497229 |
| ENSMUSG00000105959 | 1 | |

| | | |
|---|---|---|
| ENSMUSG00000105960 | -1.053549062 | 0.698112512 |
| ENSMUSG00000105961 | -1.053549062 | 0.698112512 |
| ENSMUSG00000105962 | -1.285710621 | 0.087009937 |
| ENSMUSG00000105963 | 1 | |
| ENSMUSG00000105964 | 1 | |
| ENSMUSG00000105965 | -1.043306199 | 0.698112512 |
| ENSMUSG00000105966 | 1 | |
| ENSMUSG00000105967 | 1 | |
| ENSMUSG00000105968 | 1 | |
| ENSMUSG00000105969 | 1 | |
| ENSMUSG00000105970 | 1.039183212 | 0.906386623 |
| ENSMUSG00000105971 | 1 | |
| ENSMUSG00000105972 | 1 | |
| ENSMUSG00000105973 | -1.105371473 | 0.698112512 |
| ENSMUSG00000105974 | 1.009430119 | 1 |
| ENSMUSG00000105975 | 1.014545335 | 0.989648415 |
| ENSMUSG00000105976 | 1.018854423 | 1 |
| ENSMUSG00000105977 | 1 | |
| ENSMUSG00000105978 | 1 | |
| ENSMUSG00000105979 | 1 | |
| ENSMUSG00000105980 | -1.010960635 | 1 |
| ENSMUSG00000105981 | -1.100271916 | 0.612894107 |
| ENSMUSG00000105982 | 1 | |
| ENSMUSG00000105983 | 1 | |
| ENSMUSG00000105984 | 1 | |
| ENSMUSG00000105985 | 1 | |
| ENSMUSG00000105986 | 1.132639128 | 0.217803313 |
| ENSMUSG00000105987 | 1.025962725 | 0.958600282 |
| ENSMUSG00000105988 | 1 | |
| ENSMUSG00000105989 | 1 | |
| ENSMUSG00000105990 | -1.176530916 | 0.516349583 |
| ENSMUSG00000105991 | -1 | 1 |
| ENSMUSG00000105992 | -1.124067181 | 0.608497229 |
| ENSMUSG00000105993 | 1 | |
| ENSMUSG00000105994 | 1 | |
| ENSMUSG00000105995 | 1 | |
| ENSMUSG00000105996 | -1.03962336 | 0.803072249 |
| ENSMUSG00000105997 | 1 | |
| ENSMUSG00000105998 | 1 | |
| ENSMUSG00000105999 | -1.084645496 | 0.532369657 |
| ENSMUSG00000106000 | 1.278658097 | 0.119831785 |
| ENSMUSG00000106001 | 1.13786318 | 0.653415027 |
| ENSMUSG00000106002 | 1 | |
| ENSMUSG00000106003 | 1 | |
| ENSMUSG00000106004 | 1 | |
| ENSMUSG00000106005 | -1.044273782 | 0.790851299 |
| ENSMUSG00000106006 | 1 | |
| ENSMUSG00000106007 | 1 | |
| ENSMUSG00000106008 | 1 | |
| ENSMUSG00000106009 | 1.217842673 | 0.372187987 |
| ENSMUSG00000106010 | 1.095872691 | 0.657060635 |
| ENSMUSG00000106011 | 1.069068979 | 0.608497229 |
| ENSMUSG00000106012 | 1.122462048 | 0.420161906 |
| ENSMUSG00000106013 | 1 | |
| ENSMUSG00000106014 | 1 | |
| ENSMUSG00000106015 | 1.069068979 | 0.608497229 |
| ENSMUSG00000106016 | 1 | |
| ENSMUSG00000106017 | 1 | |
| ENSMUSG00000106018 | 1 | |
| ENSMUSG00000106019 | 1.030856315 | 0.929485372 |
| ENSMUSG00000106020 | -1 | 1 |
| ENSMUSG00000106021 | 1 | |
| ENSMUSG00000106022 | 1.062739796 | 0.69533886 |

| | | |
|---|---|---|
| ENSMUSG00000106023 | 1 | |
| ENSMUSG00000106024 | 1 | |
| ENSMUSG00000106025 | -1.037167776 | 0.971981722 |
| ENSMUSG00000106026 | 1 | |
| ENSMUSG00000106027 | 1 | |
| ENSMUSG00000106028 | 1.079681442 | 0.776953697 |
| ENSMUSG00000106029 | 1 | |
| ENSMUSG00000106030 | -1 | 1 |
| ENSMUSG00000106031 | 1 | |
| ENSMUSG00000106032 | 1.345042495 | 0.128452773 |
| ENSMUSG00000106033 | 1 | |
| ENSMUSG00000106034 | 1 | |
| ENSMUSG00000106035 | 1.143507028 | 0.732682593 |
| ENSMUSG00000106036 | 1.199989356 | 0.000021314 |
| ENSMUSG00000106037 | 1.130667558 | 0.683447331 |
| ENSMUSG00000106038 | -1.142727245 | 0.202428693 |
| ENSMUSG00000106039 | 1 | |
| ENSMUSG00000106040 | 1 | |
| ENSMUSG00000106041 | 1 | |
| ENSMUSG00000106042 | 1.019083579 | 0.949812939 |
| ENSMUSG00000106043 | -1 | 1 |
| ENSMUSG00000106044 | 1.066017174 | 0.629863169 |
| ENSMUSG00000106045 | 1 | |
| ENSMUSG00000106046 | 1 | |
| ENSMUSG00000106047 | 1 | |
| ENSMUSG00000106048 | -1.001876871 | 1 |
| ENSMUSG00000106049 | 1 | |
| ENSMUSG00000106050 | 1 | |
| ENSMUSG00000106051 | 1 | |
| ENSMUSG00000106052 | 1.059463094 | 0.608497229 |
| ENSMUSG00000106053 | -1.094712223 | 0.698112512 |
| ENSMUSG00000106054 | 1 | |
| ENSMUSG00000106056 | 1 | |
| ENSMUSG00000106057 | 1.059463094 | 0.608497229 |
| ENSMUSG00000106058 | -1.099143506 | 0.781214355 |
| ENSMUSG00000106059 | 1 | |
| ENSMUSG00000106060 | 1.268173847 | 0.217803313 |
| ENSMUSG00000106061 | 1.122462048 | 0.601164141 |
| ENSMUSG00000106062 | 1.059463094 | 0.698112512 |
| ENSMUSG00000106063 | -1.043306199 | 0.698112512 |
| ENSMUSG00000106064 | -1.044273782 | 0.698112512 |
| ENSMUSG00000106065 | 1 | |
| ENSMUSG00000106066 | 1 | |
| ENSMUSG00000106067 | 1.009528481 | 1 |
| ENSMUSG00000106068 | -1.089497311 | 0.532369657 |
| ENSMUSG00000106069 | 1 | |
| ENSMUSG00000106070 | 1.069068979 | 0.608497229 |
| ENSMUSG00000106071 | 1 | |
| ENSMUSG00000106072 | 1 | |
| ENSMUSG00000106073 | -1.053549062 | 0.769745556 |
| ENSMUSG00000106074 | 1 | |
| ENSMUSG00000106075 | 1 | |
| ENSMUSG00000106076 | 1 | |
| ENSMUSG00000106077 | -1.142727245 | 0.202428693 |
| ENSMUSG00000106078 | -1 | 1 |
| ENSMUSG00000106079 | 1 | |
| ENSMUSG00000106080 | 1 | |
| ENSMUSG00000106081 | 1 | |
| ENSMUSG00000106082 | 1 | |
| ENSMUSG00000106083 | 1 | |
| ENSMUSG00000106084 | 1 | |
| ENSMUSG00000106085 | 1 | |
| ENSMUSG00000106086 | 1 | |

| | | |
|---|---|---|
| ENSMUSG00000106087 | 1.004718762 | 1 |
| ENSMUSG00000106088 | 1 | |
| ENSMUSG00000106089 | 1.04543363 | 0.717035356 |
| ENSMUSG00000106090 | 1.11166259 | 0.608497229 |
| ENSMUSG00000106091 | 1 | |
| ENSMUSG00000106092 | -1.084645496 | 0.532369657 |
| ENSMUSG00000106093 | -1.044273782 | 0.698112512 |
| ENSMUSG00000106094 | 1.005613439 | 1 |
| ENSMUSG00000106095 | 1 | |
| ENSMUSG00000106096 | 1 | |
| ENSMUSG00000106097 | 1 | |
| ENSMUSG00000106098 | 1 | |
| ENSMUSG00000106099 | -1.020037928 | 0.899919335 |
| ENSMUSG00000106100 | -1.073603785 | 0.754820307 |
| ENSMUSG00000106101 | 1 | |
| ENSMUSG00000106102 | 1 | |
| ENSMUSG00000106103 | -1.033477846 | 0.934760809 |
| ENSMUSG00000106104 | 1 | |
| ENSMUSG00000106105 | 1 | |
| ENSMUSG00000106106 | 1.156093919 | 0.41372966 |
| ENSMUSG00000106107 | -1.030445993 | 0.731295483 |
| ENSMUSG00000106108 | 1.066467141 | 0.70219856 |
| ENSMUSG00000106109 | 1 | |
| ENSMUSG00000106110 | 1.074127192 | 0.793255016 |
| ENSMUSG00000106111 | 1 | |
| ENSMUSG00000106112 | -1.071075483 | 0.698112512 |
| ENSMUSG00000106113 | 1 | |
| ENSMUSG00000106114 | 1 | |
| ENSMUSG00000106115 | 1.094514633 | 0.75390859 |
| ENSMUSG00000106116 | -1 | 1 |
| ENSMUSG00000106117 | -1.063525418 | 0.698112512 |
| ENSMUSG00000106118 | 1 | |
| ENSMUSG00000106119 | 1 | |
| ENSMUSG00000106120 | -1.071075483 | 0.698112512 |
| ENSMUSG00000106121 | -1.045012212 | 0.833135616 |
| ENSMUSG00000106122 | 1 | |
| ENSMUSG00000106123 | 1.05410549 | 0.835701362 |
| ENSMUSG00000106124 | 1 | |
| ENSMUSG00000106125 | 1.059463094 | 0.608497229 |
| ENSMUSG00000106126 | 1 | |
| ENSMUSG00000106127 | 1.200936955 | 0.311661464 |
| ENSMUSG00000106128 | 1 | |
| ENSMUSG00000106129 | 1 | |
| ENSMUSG00000106130 | -1.100193663 | 0.608497229 |
| ENSMUSG00000106131 | 1.019083579 | 0.949812939 |
| ENSMUSG00000106132 | 1 | |
| ENSMUSG00000106133 | 1 | |
| ENSMUSG00000106134 | -1.019108526 | 0.968937442 |
| ENSMUSG00000106135 | -1.077325161 | 0.580630774 |
| ENSMUSG00000106136 | 1 | |
| ENSMUSG00000106137 | 1 | |
| ENSMUSG00000106138 | 1 | |
| ENSMUSG00000106139 | 1.095872691 | 0.671280603 |
| ENSMUSG00000106140 | 1.059463094 | 0.608497229 |
| ENSMUSG00000106141 | 1 | |
| ENSMUSG00000106142 | 1.059463094 | 0.608497229 |
| ENSMUSG00000106143 | -1.055526437 | 0.829727984 |
| ENSMUSG00000106144 | 1 | |
| ENSMUSG00000106145 | 1 | |
| ENSMUSG00000106146 | 1 | |
| ENSMUSG00000106147 | 1 | |
| ENSMUSG00000106148 | 1 | |
| ENSMUSG00000106149 | 1.055780729 | 0.911346192 |

| | | |
|---|---|---|
| ENSMUSG00000106150 | 1.014244998 | 0.980754823 |
| ENSMUSG00000106151 | -1.044273782 | 0.698112512 |
| ENSMUSG00000106152 | 1 | |
| ENSMUSG00000106153 | -1.051018293 | 0.856915299 |
| ENSMUSG00000106154 | 1 | |
| ENSMUSG00000106155 | 1 | |
| ENSMUSG00000106156 | 1 | |
| ENSMUSG00000106157 | 1 | |
| ENSMUSG00000106158 | 1.079681442 | 0.697178241 |
| ENSMUSG00000106159 | 1 | |
| ENSMUSG00000106160 | -1 | 1 |
| ENSMUSG00000106161 | 1.005613439 | 1 |
| ENSMUSG00000106162 | 1 | |
| ENSMUSG00000106163 | 1 | |
| ENSMUSG00000106164 | 1.150179455 | 0.543714852 |
| ENSMUSG00000106165 | 1 | |
| ENSMUSG00000106166 | -1.11553986 | 0.730117907 |
| ENSMUSG00000106167 | 1 | |
| ENSMUSG00000106168 | 1 | |
| ENSMUSG00000106169 | -1.028348525 | 0.950684488 |
| ENSMUSG00000106170 | 1 | |
| ENSMUSG00000106172 | 1 | |
| ENSMUSG00000106173 | 1.023243708 | 0.870602199 |
| ENSMUSG00000106174 | 1 | |
| ENSMUSG00000106175 | 1 | |
| ENSMUSG00000106176 | 1.069068979 | 0.608497229 |
| ENSMUSG00000106177 | 1 | |
| ENSMUSG00000106178 | -1.129126757 | 0.608497229 |
| ENSMUSG00000106179 | 1 | |
| ENSMUSG00000106180 | 1 | |
| ENSMUSG00000106181 | 1.070429626 | 0.723873338 |
| ENSMUSG00000106182 | 1.024693402 | 0.933031586 |
| ENSMUSG00000106183 | 1 | |
| ENSMUSG00000106184 | 1.005588822 | 1 |
| ENSMUSG00000106185 | 1 | |
| ENSMUSG00000106186 | 1.131096998 | 0.698112512 |
| ENSMUSG00000106187 | 1 | |
| ENSMUSG00000106188 | -1.087913248 | 0.797808656 |
| ENSMUSG00000106189 | -1 | 1 |
| ENSMUSG00000106190 | -1.056768771 | 0.76984393 |
| ENSMUSG00000106191 | 1 | |
| ENSMUSG00000106192 | 1 | |
| ENSMUSG00000106193 | -1 | 1 |
| ENSMUSG00000106194 | -1 | 1 |
| ENSMUSG00000106195 | 1 | |
| ENSMUSG00000106196 | 1 | |
| ENSMUSG00000106197 | -1.036701442 | 0.961353875 |
| ENSMUSG00000106198 | 1 | |
| ENSMUSG00000106199 | 1 | |
| ENSMUSG00000106200 | 1 | |
| ENSMUSG00000106201 | 1 | |
| ENSMUSG00000106202 | -1.043306199 | 0.698112512 |
| ENSMUSG00000106203 | 1 | |
| ENSMUSG00000106204 | 1 | |
| ENSMUSG00000106205 | 1.224956403 | 0.077863434 |
| ENSMUSG00000106206 | -1 | 1 |
| ENSMUSG00000106207 | 1 | |
| ENSMUSG00000106208 | -1.121136286 | 0.686194514 |
| ENSMUSG00000106209 | 1.05189803 | 0.909335407 |
| ENSMUSG00000106210 | -1.03962336 | 0.698112512 |
| ENSMUSG00000106211 | -1.099174267 | 0.675967223 |
| ENSMUSG00000106212 | -1 | 1 |
| ENSMUSG00000106213 | 1 | |

| | | |
|---|---|---|
| ENSMUSG00000106214 | -1 | 1 |
| ENSMUSG00000106215 | 1 | |
| ENSMUSG00000106216 | 1 | |
| ENSMUSG00000106217 | -1.044273782 | 0.698112512 |
| ENSMUSG00000106218 | 1 | |
| ENSMUSG00000106219 | -1.010960635 | 1 |
| ENSMUSG00000106220 | 1 | |
| ENSMUSG00000106221 | 1 | |
| ENSMUSG00000106222 | 1 | |
| ENSMUSG00000106223 | 1 | |
| ENSMUSG00000106224 | 1.281777256 | 0.104947513 |
| ENSMUSG00000106225 | 1 | |
| ENSMUSG00000106226 | 1 | |
| ENSMUSG00000106227 | 1 | |
| ENSMUSG00000106228 | 1 | |
| ENSMUSG00000106229 | -1.021413658 | 0.982838256 |
| ENSMUSG00000106230 | 1 | |
| ENSMUSG00000106231 | 1 | |
| ENSMUSG00000106232 | 1 | |
| ENSMUSG00000106233 | 1 | |
| ENSMUSG00000106234 | 1 | |
| ENSMUSG00000106235 | 1 | |
| ENSMUSG00000106236 | 1 | |
| ENSMUSG00000106237 | 1.062990386 | 0.865583959 |
| ENSMUSG00000106238 | 1 | |
| ENSMUSG00000106239 | -1.044273782 | 0.790851299 |
| ENSMUSG00000106240 | -1.069502966 | 0.698112512 |
| ENSMUSG00000106241 | -1.003212715 | 1 |
| ENSMUSG00000106242 | 1 | |
| ENSMUSG00000106243 | 1.118345316 | 0.608497229 |
| ENSMUSG00000106244 | 1 | |
| ENSMUSG00000106245 | 1 | |
| ENSMUSG00000106246 | 1 | |
| ENSMUSG00000106247 | 1 | |
| ENSMUSG00000106248 | 1.092818258 | 0.788886948 |
| ENSMUSG00000106249 | 1 | |
| ENSMUSG00000106250 | 1 | |
| ENSMUSG00000106251 | 1.012973414 | 0.98509435 |
| ENSMUSG00000106252 | -1.044273782 | 0.790851299 |
| ENSMUSG00000106253 | 1 | |
| ENSMUSG00000106254 | -1.054513355 | 0.861653133 |
| ENSMUSG00000106255 | 1 | |
| ENSMUSG00000106256 | 1 | |
| ENSMUSG00000106257 | -1.053549062 | 0.698112512 |
| ENSMUSG00000106258 | -1.053549062 | 0.778372272 |
| ENSMUSG00000106259 | 1 | |
| ENSMUSG00000106260 | 1 | |
| ENSMUSG00000106261 | 1 | |
| ENSMUSG00000106262 | -1.089868377 | 0.608497229 |
| ENSMUSG00000106263 | 1 | |
| ENSMUSG00000106264 | 1 | |
| ENSMUSG00000106265 | -1.044273782 | 0.698112512 |
| ENSMUSG00000106266 | 1 | |
| ENSMUSG00000106267 | 1 | |
| ENSMUSG00000106268 | 1.172386343 | 0.378445257 |
| ENSMUSG00000106269 | 1.05639165 | 0.855387377 |
| ENSMUSG00000106270 | 1 | |
| ENSMUSG00000106271 | 1.0304079 | 0.959583351 |
| ENSMUSG00000106272 | 1 | |
| ENSMUSG00000106273 | 1.059463094 | 0.608497229 |
| ENSMUSG00000106274 | 1 | |
| ENSMUSG00000106275 | 1 | |
| ENSMUSG00000106276 | 1 | |

| | | |
|---|---|---|
| ENSMUSG00000106278 | 1 | |
| ENSMUSG00000106279 | 1 | |
| ENSMUSG00000106280 | 1 | |
| ENSMUSG00000106281 | 1 | |
| ENSMUSG00000106282 | -1.053549062 | 0.769745556 |
| ENSMUSG00000106283 | 1.059463094 | 0.608497229 |
| ENSMUSG00000106284 | 1 | |
| ENSMUSG00000106285 | -1.028348525 | 0.956972448 |
| ENSMUSG00000106286 | 1 | |
| ENSMUSG00000106287 | -1.132666855 | 0.391570427 |
| ENSMUSG00000106288 | 1.203063358 | 0.240114463 |
| ENSMUSG00000106289 | 1 | |
| ENSMUSG00000106290 | -1.00182565 | 1 |
| ENSMUSG00000106291 | 1.040559106 | 0.807247723 |
| ENSMUSG00000106292 | 1 | |
| ENSMUSG00000106293 | 1.064531743 | 0.796396036 |
| ENSMUSG00000106294 | 1.059463094 | 0.699340382 |
| ENSMUSG00000106295 | 1 | |
| ENSMUSG00000106296 | 1.085624843 | 0.69305964 |
| ENSMUSG00000106297 | 1 | |
| ENSMUSG00000106298 | 1 | |
| ENSMUSG00000106299 | 1 | |
| ENSMUSG00000106300 | -1.015511103 | 0.99457094 |
| ENSMUSG00000106301 | 1.059463094 | 0.698112512 |
| ENSMUSG00000106302 | 1 | |
| ENSMUSG00000106303 | 1 | |
| ENSMUSG00000106304 | 1 | |
| ENSMUSG00000106305 | 1 | |
| ENSMUSG00000106306 | 1 | |
| ENSMUSG00000106307 | 1 | |
| ENSMUSG00000106308 | 1 | |
| ENSMUSG00000106309 | -1.125123913 | 0.481742107 |
| ENSMUSG00000106310 | -1.126344191 | 0.217803313 |
| ENSMUSG00000106311 | -1.043306199 | 0.698112512 |
| ENSMUSG00000106312 | 1 | |
| ENSMUSG00000106313 | 1.132639128 | 0.398767938 |
| ENSMUSG00000106314 | 1.059463094 | 0.608497229 |
| ENSMUSG00000106315 | 1 | |
| ENSMUSG00000106316 | 1 | |
| ENSMUSG00000106317 | -1.114884925 | 0.113035845 |
| ENSMUSG00000106318 | 1 | |
| ENSMUSG00000106319 | 1 | |
| ENSMUSG00000106320 | -1.051018293 | 0.828564184 |
| ENSMUSG00000106321 | 1.010351027 | 1 |
| ENSMUSG00000106322 | -1.04375794 | 0.929536187 |
| ENSMUSG00000106323 | 1 | |
| ENSMUSG00000106324 | 1 | |
| ENSMUSG00000106325 | 1 | |
| ENSMUSG00000106326 | 1 | |
| ENSMUSG00000106327 | 1 | |
| ENSMUSG00000106328 | 1 | |
| ENSMUSG00000106329 | -1.073455303 | 0.783567045 |
| ENSMUSG00000106330 | -1.044273782 | 0.698112512 |
| ENSMUSG00000106331 | -1.043306199 | 0.79291651 |
| ENSMUSG00000106332 | 1.020482121 | 0.989877474 |
| ENSMUSG00000106333 | 1 | |
| ENSMUSG00000106334 | -1.939899305 | 0.008595644 |
| ENSMUSG00000106335 | 1 | |
| ENSMUSG00000106336 | 1 | |
| ENSMUSG00000106337 | 1.014545335 | 0.989648415 |
| ENSMUSG00000106338 | 1 | |
| ENSMUSG00000106339 | 1.059463094 | 0.608497229 |
| ENSMUSG00000106340 | 1 | |

| | | |
|---|---|---|
| ENSMUSG00000106341 | 1 | |
| ENSMUSG00000106342 | 1 | |
| ENSMUSG00000106343 | 1 | |
| ENSMUSG00000106344 | 1 | |
| ENSMUSG00000106345 | 1 | |
| ENSMUSG00000106346 | 1 | |
| ENSMUSG00000106347 | 1 | |
| ENSMUSG00000106348 | 1 | |
| ENSMUSG00000106349 | 1 | |
| ENSMUSG00000106350 | -1 | 1 |
| ENSMUSG00000106351 | 1 | |
| ENSMUSG00000106352 | 1 | |
| ENSMUSG00000106353 | -1.11061171 | 0.608497229 |
| ENSMUSG00000106354 | -1.100193663 | 0.534887579 |
| ENSMUSG00000106355 | 1 | |
| ENSMUSG00000106356 | 1 | |
| ENSMUSG00000106357 | 1 | |
| ENSMUSG00000106358 | 1.014545335 | 1 |
| ENSMUSG00000106359 | 1 | |
| ENSMUSG00000106360 | 1 | |
| ENSMUSG00000106361 | 1.014133649 | 1 |
| ENSMUSG00000106362 | 1 | |
| ENSMUSG00000106363 | 1 | |
| ENSMUSG00000106364 | -1.065521517 | 0.800755473 |
| ENSMUSG00000106365 | -1.043306199 | 0.803260194 |
| ENSMUSG00000106366 | 1 | |
| ENSMUSG00000106367 | 1 | |
| ENSMUSG00000106368 | 1.059463094 | 0.699340382 |
| ENSMUSG00000106369 | 1 | |
| ENSMUSG00000106370 | 1 | |
| ENSMUSG00000106371 | -1.044273782 | 0.790851299 |
| ENSMUSG00000106372 | 1 | |
| ENSMUSG00000106373 | 1 | |
| ENSMUSG00000106374 | -1 | 1 |
| ENSMUSG00000106375 | 1.116313489 | 0.698112512 |
| ENSMUSG00000106376 | 1 | |
| ENSMUSG00000106377 | 1 | |
| ENSMUSG00000106378 | 1 | |
| ENSMUSG00000106379 | 1 | |
| ENSMUSG00000106380 | -1.059945793 | 0.912782976 |
| ENSMUSG00000106381 | 1 | |
| ENSMUSG00000106382 | 1 | |
| ENSMUSG00000106383 | -1.233349506 | 0.145913041 |
| ENSMUSG00000106384 | 1 | |
| ENSMUSG00000106385 | 1.11026164 | 0.620010094 |
| ENSMUSG00000106386 | 1.059463094 | 0.608497229 |
| ENSMUSG00000106387 | 1 | |
| ENSMUSG00000106388 | 1 | |
| ENSMUSG00000106389 | 1 | |
| ENSMUSG00000106390 | -1 | 1 |
| ENSMUSG00000106391 | 1.106544056 | 0.608497229 |
| ENSMUSG00000106392 | 1 | |
| ENSMUSG00000106393 | 1 | |
| ENSMUSG00000106394 | 1 | |
| ENSMUSG00000106395 | 1 | |
| ENSMUSG00000106396 | -1.158708082 | 0.596178824 |
| ENSMUSG00000106397 | 1 | |
| ENSMUSG00000106398 | 1 | |
| ENSMUSG00000106399 | 1 | |
| ENSMUSG00000106400 | 1 | |
| ENSMUSG00000106401 | 1 | |
| ENSMUSG00000106402 | 1 | |
| ENSMUSG00000106403 | 1 | |

| | | |
|---|---|---|
| ENSMUSG00000106404 | 1 | |
| ENSMUSG00000106405 | 1 | |
| ENSMUSG00000106406 | 1.059463094 | 0.608497229 |
| ENSMUSG00000106407 | 1 | |
| ENSMUSG00000106408 | -1.05359949 | 0.848477314 |
| ENSMUSG00000106409 | 1 | |
| ENSMUSG00000106410 | 1 | |
| ENSMUSG00000106411 | 1.007338892 | 0.988749244 |
| ENSMUSG00000106412 | 1 | |
| ENSMUSG00000106413 | 1 | |
| ENSMUSG00000106414 | 1 | |
| ENSMUSG00000106415 | 1 | |
| ENSMUSG00000106416 | -1.280564164 | 0.002796445 |
| ENSMUSG00000106417 | -1.096067949 | 0.783837368 |
| ENSMUSG00000106418 | 1 | |
| ENSMUSG00000106419 | 1.029972857 | 0.928180271 |
| ENSMUSG00000106420 | 1 | |
| ENSMUSG00000106421 | -1.077732257 | 0.797035369 |
| ENSMUSG00000106422 | 1 | |
| ENSMUSG00000106423 | 1 | |
| ENSMUSG00000106424 | 1 | |
| ENSMUSG00000106425 | 1 | |
| ENSMUSG00000106426 | 1.22560735 | 0.056310079 |
| ENSMUSG00000106427 | 1.09313441 | 0.8159501 |
| ENSMUSG00000106428 | 1 | |
| ENSMUSG00000106429 | 1 | |
| ENSMUSG00000106430 | -1.12843057 | 0.569479524 |
| ENSMUSG00000106431 | 1 | |
| ENSMUSG00000106432 | -1.110426459 | 0.731295483 |
| ENSMUSG00000106433 | 1 | |
| ENSMUSG00000106434 | 1 | |
| ENSMUSG00000106435 | 1 | |
| ENSMUSG00000106436 | 1 | |
| ENSMUSG00000106437 | 1.078596186 | 0.807247723 |
| ENSMUSG00000106438 | -1.089497311 | 0.614307028 |
| ENSMUSG00000106439 | 1 | |
| ENSMUSG00000106440 | 1 | |
| ENSMUSG00000106441 | 1 | |
| ENSMUSG00000106442 | 1 | |
| ENSMUSG00000106443 | -1.040960083 | 0.917717739 |
| ENSMUSG00000106444 | 1 | |
| ENSMUSG00000106445 | 1 | |
| ENSMUSG00000106446 | 1.244969796 | 0.507199176 |
| ENSMUSG00000106447 | -1 | 1 |
| ENSMUSG00000106448 | 1 | |
| ENSMUSG00000106450 | 1 | |
| ENSMUSG00000106451 | -1.053549062 | 0.698112512 |
| ENSMUSG00000106452 | 1 | |
| ENSMUSG00000106453 | -1 | 1 |
| ENSMUSG00000106454 | -1.084645496 | 0.608497229 |
| ENSMUSG00000106455 | 1.060273769 | 0.783637632 |
| ENSMUSG00000106456 | 1 | |
| ENSMUSG00000106457 | 1.001264502 | 1 |
| ENSMUSG00000106458 | -1.014296711 | 1 |
| ENSMUSG00000106459 | 1 | |
| ENSMUSG00000106460 | 1 | |
| ENSMUSG00000106461 | 1 | |
| ENSMUSG00000106462 | 1 | |
| ENSMUSG00000106463 | 1.068013265 | 0.713861573 |
| ENSMUSG00000106464 | 1.219171185 | 0.04984671 |
| ENSMUSG00000106465 | 1 | |
| ENSMUSG00000106466 | 1 | |
| ENSMUSG00000106467 | 1 | |

| | | |
|---|---|---|
| ENSMUSG00000106468 | 1 | |
| ENSMUSG00000106469 | 1 | |
| ENSMUSG00000106470 | 1 | |
| ENSMUSG00000106471 | 1 | |
| ENSMUSG00000106472 | 1 | |
| ENSMUSG00000106473 | 1 | |
| ENSMUSG00000106474 | 1 | |
| ENSMUSG00000106475 | -1.086720843 | 0.754175915 |
| ENSMUSG00000106476 | 1.059463094 | 0.608497229 |
| ENSMUSG00000106477 | 1 | |
| ENSMUSG00000106478 | 1.04328066 | 0.885806913 |
| ENSMUSG00000106479 | 1 | |
| ENSMUSG00000106480 | 1 | |
| ENSMUSG00000106481 | 1 | |
| ENSMUSG00000106482 | 1 | |
| ENSMUSG00000106483 | 1 | |
| ENSMUSG00000106484 | -1.046526356 | 0.913237976 |
| ENSMUSG00000106485 | 1 | |
| ENSMUSG00000106486 | 1 | |
| ENSMUSG00000106487 | 1 | |
| ENSMUSG00000106488 | 1 | |
| ENSMUSG00000106489 | 1 | |
| ENSMUSG00000106490 | 1.132639128 | 0.428732467 |
| ENSMUSG00000106491 | -1.029709545 | 0.965871218 |
| ENSMUSG00000106492 | 1.24122829 | 0.03679259 |
| ENSMUSG00000106493 | 1 | |
| ENSMUSG00000106494 | 1 | |
| ENSMUSG00000106495 | -1.049633386 | 0.915158215 |
| ENSMUSG00000106496 | 1 | |
| ENSMUSG00000106497 | 1.05038453 | 0.828564184 |
| ENSMUSG00000106498 | 1.749415693 | 0.002681197 |
| ENSMUSG00000106499 | -1.173975439 | 0.649078495 |
| ENSMUSG00000106500 | 1.079681442 | 0.608497229 |
| ENSMUSG00000106501 | 1 | |
| ENSMUSG00000106502 | 1 | |
| ENSMUSG00000106503 | 1 | |
| ENSMUSG00000106504 | 1 | |
| ENSMUSG00000106505 | 1.069068979 | 0.608497229 |
| ENSMUSG00000106506 | 1 | |
| ENSMUSG00000106507 | -1.244007152 | 0.376592592 |
| ENSMUSG00000106508 | -1.044273782 | 0.790851299 |
| ENSMUSG00000106509 | 1 | |
| ENSMUSG00000106510 | 1 | |
| ENSMUSG00000106511 | 1 | |
| ENSMUSG00000106512 | 1.080134941 | 0.698112512 |
| ENSMUSG00000106513 | 1 | |
| ENSMUSG00000106514 | 1.091686812 | 0.714698678 |
| ENSMUSG00000106515 | 1 | |
| ENSMUSG00000106516 | 1 | |
| ENSMUSG00000106517 | 1 | |
| ENSMUSG00000106518 | 1.132639128 | 0.428732467 |
| ENSMUSG00000106519 | 1.059463094 | 0.608497229 |
| ENSMUSG00000106520 | 1 | |
| ENSMUSG00000106521 | 1 | |
| ENSMUSG00000106522 | 1 | |
| ENSMUSG00000106523 | 1 | |
| ENSMUSG00000106524 | 1 | |
| ENSMUSG00000106525 | 1 | |
| ENSMUSG00000106526 | 1.059463094 | 0.608497229 |
| ENSMUSG00000106527 | 1 | |
| ENSMUSG00000106528 | 1.059463094 | 0.608497229 |
| ENSMUSG00000106529 | -1.436482803 | 0.036626025 |
| ENSMUSG00000106530 | 1 | |

| | | |
|---|---|---|
| ENSMUSG00000106531 | -1.063525418 | 0.698112512 |
| ENSMUSG00000106532 | 1 | |
| ENSMUSG00000106533 | 1 | |
| ENSMUSG00000106534 | -1.084645496 | 0.608497229 |
| ENSMUSG00000106535 | -1.11272073 | 0.092667548 |
| ENSMUSG00000106536 | 1.177207639 | 0.248113367 |
| ENSMUSG00000106537 | 1 | |
| ENSMUSG00000106538 | 1.059463094 | 0.608497229 |
| ENSMUSG00000106539 | 1 | |
| ENSMUSG00000106540 | 1 | |
| ENSMUSG00000106541 | 1.019083579 | 0.949812939 |
| ENSMUSG00000106542 | 1 | |
| ENSMUSG00000106543 | 1.147736034 | 0.608497229 |
| ENSMUSG00000106544 | 1 | |
| ENSMUSG00000106545 | 1.069068979 | 0.608497229 |
| ENSMUSG00000106546 | 1 | |
| ENSMUSG00000106547 | -1.191339545 | 0.423028112 |
| ENSMUSG00000106548 | 1 | |
| ENSMUSG00000106549 | 1.04165229 | 0.722608713 |
| ENSMUSG00000106550 | 1.059463094 | 0.608497229 |
| ENSMUSG00000106551 | 1.059463094 | 0.608497229 |
| ENSMUSG00000106552 | 1 | |
| ENSMUSG00000106553 | -1 | 1 |
| ENSMUSG00000106554 | 1 | |
| ENSMUSG00000106555 | -1.044273782 | 0.698112512 |
| ENSMUSG00000106556 | 1 | |
| ENSMUSG00000106557 | 1 | |
| ENSMUSG00000106558 | -1 | 1 |
| ENSMUSG00000106559 | 1 | |
| ENSMUSG00000106560 | -1.17878616 | 0.561104668 |
| ENSMUSG00000106561 | 1 | |
| ENSMUSG00000106562 | 1.143512105 | 0.637660529 |
| ENSMUSG00000106563 | 1 | |
| ENSMUSG00000106564 | 1 | |
| ENSMUSG00000106565 | -1.128401027 | 0.67588172 |
| ENSMUSG00000106566 | 1 | |
| ENSMUSG00000106567 | 1 | |
| ENSMUSG00000106568 | 1.049101314 | 0.711624122 |
| ENSMUSG00000106569 | 1 | |
| ENSMUSG00000106570 | 1 | |
| ENSMUSG00000106571 | 1 | |
| ENSMUSG00000106572 | 1.059463094 | 0.608497229 |
| ENSMUSG00000106573 | 1 | |
| ENSMUSG00000106574 | 1.005613439 | 1 |
| ENSMUSG00000106575 | 1.059463094 | 0.608497229 |
| ENSMUSG00000106576 | 1.123722206 | 0.608497229 |
| ENSMUSG00000106577 | 1.07487334 | 0.846673732 |
| ENSMUSG00000106578 | 1 | |
| ENSMUSG00000106579 | -1.03962336 | 0.887262126 |
| ENSMUSG00000106580 | 1.059463094 | 0.608497229 |
| ENSMUSG00000106581 | -1.03962336 | 0.698112512 |
| ENSMUSG00000106582 | -1 | 1 |
| ENSMUSG00000106583 | 1 | |
| ENSMUSG00000106584 | 1 | |
| ENSMUSG00000106585 | 1 | |
| ENSMUSG00000106586 | 1 | |
| ENSMUSG00000106587 | 1 | |
| ENSMUSG00000106588 | 1.154253937 | 0.11477493 |
| ENSMUSG00000106589 | 1 | |
| ENSMUSG00000106590 | 1.400069781 | 0.222781245 |
| ENSMUSG00000106591 | 1 | |
| ENSMUSG00000106592 | -1.043306199 | 0.698112512 |
| ENSMUSG00000106593 | 1 | |

| | | |
|---|---|---|
| ENSMUSG00000106594 | 1 | |
| ENSMUSG00000106595 | 1 | |
| ENSMUSG00000106596 | -1.03962336 | 0.698112512 |
| ENSMUSG00000106597 | 1 | |
| ENSMUSG00000106598 | -1.085651418 | 0.532369657 |
| ENSMUSG00000106599 | 1 | |
| ENSMUSG00000106600 | -1.054169715 | 0.845282019 |
| ENSMUSG00000106601 | 1 | |
| ENSMUSG00000106602 | 1 | |
| ENSMUSG00000106603 | 1.314501944 | 0.136197019 |
| ENSMUSG00000106604 | -1 | 1 |
| ENSMUSG00000106605 | 1 | |
| ENSMUSG00000106606 | 1 | |
| ENSMUSG00000106607 | -1.068104567 | 0.794274354 |
| ENSMUSG00000106608 | 1 | |
| ENSMUSG00000106609 | 1.042253985 | 0.873622104 |
| ENSMUSG00000106610 | 1.059463094 | 0.608497229 |
| ENSMUSG00000106611 | 1 | |
| ENSMUSG00000106612 | 1.122462048 | 0.216735191 |
| ENSMUSG00000106613 | 1 | |
| ENSMUSG00000106614 | 1.059463094 | 0.608497229 |
| ENSMUSG00000106615 | 1 | |
| ENSMUSG00000106616 | 1 | |
| ENSMUSG00000106617 | -1.053654132 | 0.868504188 |
| ENSMUSG00000106618 | 1 | |
| ENSMUSG00000106619 | -1.053549062 | 0.698112512 |
| ENSMUSG00000106620 | 1 | |
| ENSMUSG00000106621 | 1 | |
| ENSMUSG00000106622 | -1.043306199 | 0.698112512 |
| ENSMUSG00000106623 | 1.074662908 | 0.769764777 |
| ENSMUSG00000106624 | 1 | |
| ENSMUSG00000106625 | 1.049411284 | 0.892085484 |
| ENSMUSG00000106626 | 1 | |
| ENSMUSG00000106627 | 1 | |
| ENSMUSG00000106628 | 1.001696467 | 1 |
| ENSMUSG00000106629 | -1.043306199 | 0.698112512 |
| ENSMUSG00000106630 | 1 | |
| ENSMUSG00000106631 | 1.039965644 | 0.912030056 |
| ENSMUSG00000106632 | 1.026340503 | 0.984826086 |
| ENSMUSG00000106633 | 1 | |
| ENSMUSG00000106634 | -1.024397398 | 0.803260194 |
| ENSMUSG00000106635 | 1 | |
| ENSMUSG00000106636 | 1.13155521 | 0.532369657 |
| ENSMUSG00000106637 | 1.02111758 | 0.987775262 |
| ENSMUSG00000106639 | 1 | |
| ENSMUSG00000106640 | 1.059463094 | 0.608497229 |
| ENSMUSG00000106641 | -1.307521622 | 0.056613241 |
| ENSMUSG00000106642 | 1 | |
| ENSMUSG00000106643 | 1.079681442 | 0.78131463 |
| ENSMUSG00000106644 | 1 | |
| ENSMUSG00000106645 | 1 | |
| ENSMUSG00000106646 | 1.025100702 | 0.979811078 |
| ENSMUSG00000106647 | 1.189706618 | 0.102900586 |
| ENSMUSG00000106648 | -1.0328151 | 0.960776894 |
| ENSMUSG00000106649 | 1 | |
| ENSMUSG00000106650 | -1.038858794 | 0.915658578 |
| ENSMUSG00000106651 | 1.084703674 | 0.844612733 |
| ENSMUSG00000106652 | 1 | |
| ENSMUSG00000106653 | 1 | |
| ENSMUSG00000106654 | -1.176660386 | 0.497425906 |
| ENSMUSG00000106655 | 1 | |
| ENSMUSG00000106656 | 1 | |
| ENSMUSG00000106657 | -1.043306199 | 0.698112512 |

| | | |
|---|---|---|
| ENSMUSG00000106658 | -1.089497311 | 0.614307028 |
| ENSMUSG00000106659 | 1 | |
| ENSMUSG00000106660 | 1 | |
| ENSMUSG00000106661 | 1 | |
| ENSMUSG00000106662 | -1.005904755 | 1 |
| ENSMUSG00000106663 | 1.11166259 | 0.608497229 |
| ENSMUSG00000106664 | 1.038495176 | 0.831944442 |
| ENSMUSG00000106665 | 1.013982319 | 1 |
| ENSMUSG00000106666 | 1 | |
| ENSMUSG00000106667 | 1 | |
| ENSMUSG00000106668 | 1 | |
| ENSMUSG00000106669 | 1 | |
| ENSMUSG00000106670 | 1 | |
| ENSMUSG00000106671 | 1.026895199 | 0.923464844 |
| ENSMUSG00000106673 | 1 | |
| ENSMUSG00000106674 | -1.089497311 | 0.532369657 |
| ENSMUSG00000106675 | 1 | |
| ENSMUSG00000106676 | 1.024693402 | 0.969087762 |
| ENSMUSG00000106677 | 1 | |
| ENSMUSG00000106678 | -1.084858259 | 0.039223927 |
| ENSMUSG00000106679 | 1 | |
| ENSMUSG00000106680 | 1.071809423 | 0.855626511 |
| ENSMUSG00000106681 | 1 | |
| ENSMUSG00000106682 | 1.324165034 | 0.214091296 |
| ENSMUSG00000106683 | -1.089497311 | 0.698112512 |
| ENSMUSG00000106686 | 1 | |
| ENSMUSG00000106688 | 1.156627404 | 0.413321652 |
| ENSMUSG00000106689 | 1 | |
| ENSMUSG00000106690 | 1 | |
| ENSMUSG00000106691 | 1.175496428 | 0.340360843 |
| ENSMUSG00000106692 | -1.023769022 | 0.954761885 |
| ENSMUSG00000106693 | 1 | |
| ENSMUSG00000106694 | 1.059463094 | 0.698112512 |
| ENSMUSG00000106696 | 1.090350493 | 0.66861278 |
| ENSMUSG00000106697 | 1 | |
| ENSMUSG00000106698 | -1.037482356 | 0.870427333 |
| ENSMUSG00000106699 | 1 | |
| ENSMUSG00000106701 | 1 | |
| ENSMUSG00000106702 | 1 | |
| ENSMUSG00000106703 | -1.053549062 | 0.769745556 |
| ENSMUSG00000106704 | 1 | |
| ENSMUSG00000106705 | 1 | |
| ENSMUSG00000106706 | -1.123879417 | 0.608497229 |
| ENSMUSG00000106707 | 1.149113744 | 0.317618146 |
| ENSMUSG00000106708 | 1 | |
| ENSMUSG00000106709 | 1.095872691 | 0.608497229 |
| ENSMUSG00000106710 | 1.07487334 | 0.656277822 |
| ENSMUSG00000106712 | 1 | |
| ENSMUSG00000106713 | 1 | |
| ENSMUSG00000106714 | 1.025955135 | 0.856283628 |
| ENSMUSG00000106715 | 1 | |
| ENSMUSG00000106717 | -1.094838675 | 0.608497229 |
| ENSMUSG00000106718 | 1 | |
| ENSMUSG00000106719 | -1.084968553 | 0.726987474 |
| ENSMUSG00000106720 | 1.085624843 | 0.608497229 |
| ENSMUSG00000106721 | 1.32695866 | 0.043046077 |
| ENSMUSG00000106724 | 1 | |
| ENSMUSG00000106725 | -1.133231767 | 0.608497229 |
| ENSMUSG00000106726 | 1 | |
| ENSMUSG00000106729 | 1 | |
| ENSMUSG00000106730 | 1 | |
| ENSMUSG00000106732 | 1.161036672 | 0.323279587 |
| ENSMUSG00000106733 | -1.010960635 | 1 |

| | | |
|---|---|---|
| ENSMUSG00000106734 | 1.011257092 | 0.94668925 |
| ENSMUSG00000106735 | 1 | |
| ENSMUSG00000106736 | 1.03862275 | 0.924587329 |
| ENSMUSG00000106737 | 1 | |
| ENSMUSG00000106738 | 1 | |
| ENSMUSG00000106739 | 1.015183877 | 1 |
| ENSMUSG00000106740 | 1 | |
| ENSMUSG00000106741 | 1 | |
| ENSMUSG00000106742 | 1 | |
| ENSMUSG00000106743 | -1.103450748 | 0.445897268 |
| ENSMUSG00000106744 | -1.000338788 | 1 |
| ENSMUSG00000106745 | 1 | |
| ENSMUSG00000106746 | 1 | |
| ENSMUSG00000106747 | 1.112811008 | 0.728027496 |
| ENSMUSG00000106748 | 1 | |
| ENSMUSG00000106749 | 1 | |
| ENSMUSG00000106750 | 1.069068979 | 0.608497229 |
| ENSMUSG00000106751 | -1.156223773 | 0.16601603 |
| ENSMUSG00000106752 | 1 | |
| ENSMUSG00000106753 | 1 | |
| ENSMUSG00000106754 | 1 | |
| ENSMUSG00000106755 | -1.054551959 | 0.839354385 |
| ENSMUSG00000106756 | 1 | |
| ENSMUSG00000106757 | 1.003370641 | 1 |
| ENSMUSG00000106758 | 1 | |
| ENSMUSG00000106759 | 1.069068979 | 0.698112512 |
| ENSMUSG00000106760 | 1 | |
| ENSMUSG00000106762 | 1 | |
| ENSMUSG00000106763 | 1 | |
| ENSMUSG00000106764 | 1 | |
| ENSMUSG00000106766 | 1 | |
| ENSMUSG00000106767 | 1.201744115 | 0.430273916 |
| ENSMUSG00000106768 | 1 | |
| ENSMUSG00000106769 | 1 | |
| ENSMUSG00000106770 | 1 | |
| ENSMUSG00000106771 | 1 | |
| ENSMUSG00000106772 | 1.069068979 | 0.698112512 |
| ENSMUSG00000106773 | -1 | 1 |
| ENSMUSG00000106775 | -1.010960635 | 1 |
| ENSMUSG00000106776 | -1.043306199 | 0.698112512 |
| ENSMUSG00000106777 | 1.001434933 | 1 |
| ENSMUSG00000106778 | -1.084645496 | 0.532369657 |
| ENSMUSG00000106779 | 1 | |
| ENSMUSG00000106780 | 1 | |
| ENSMUSG00000106781 | 1 | |
| ENSMUSG00000106782 | 1 | |
| ENSMUSG00000106783 | 1 | |
| ENSMUSG00000106784 | 1 | |
| ENSMUSG00000106785 | 1 | |
| ENSMUSG00000106786 | 1 | |
| ENSMUSG00000106788 | 1 | |
| ENSMUSG00000106790 | 1 | |
| ENSMUSG00000106791 | -1.107676363 | 0.754821906 |
| ENSMUSG00000106793 | 1.014545335 | 0.979002724 |
| ENSMUSG00000106794 | 1 | |
| ENSMUSG00000106795 | 1 | |
| ENSMUSG00000106797 | 1 | |
| ENSMUSG00000106798 | 1.059463094 | 0.608497229 |
| ENSMUSG00000106799 | -1.03962336 | 0.698112512 |
| ENSMUSG00000106800 | -1 | 1 |
| ENSMUSG00000106801 | 1 | |
| ENSMUSG00000106802 | -1.094712223 | 0.735423121 |
| ENSMUSG00000106803 | 1 | |

| | | |
|---|---|---|
| ENSMUSG00000106804 | -1 | 1 |
| ENSMUSG00000106805 | 1 | |
| ENSMUSG00000106806 | 1.069068979 | 0.608497229 |
| ENSMUSG00000106807 | -1.147838869 | 0.350417454 |
| ENSMUSG00000106808 | 1 | |
| ENSMUSG00000106810 | 1 | |
| ENSMUSG00000106811 | 1 | |
| ENSMUSG00000106812 | 1 | |
| ENSMUSG00000106813 | 1 | |
| ENSMUSG00000106814 | 1 | |
| ENSMUSG00000106815 | 1.191307385 | 0.36554144 |
| ENSMUSG00000106816 | 1 | |
| ENSMUSG00000106817 | 1 | |
| ENSMUSG00000106818 | 1.076583642 | 0.720767747 |
| ENSMUSG00000106819 | -1.003063157 | 1 |
| ENSMUSG00000106820 | 1.343450547 | 0.099813032 |
| ENSMUSG00000106821 | 1.059463094 | 0.698112512 |
| ENSMUSG00000106822 | 1.059463094 | 0.608497229 |
| ENSMUSG00000106824 | 1 | |
| ENSMUSG00000106825 | 1.187317089 | 0.101050918 |
| ENSMUSG00000106826 | -1.035225448 | 0.860654711 |
| ENSMUSG00000106827 | -1.010960635 | 1 |
| ENSMUSG00000106828 | 1.24122829 | 0.03679259 |
| ENSMUSG00000106829 | 1 | |
| ENSMUSG00000106831 | 1.059463094 | 0.608497229 |
| ENSMUSG00000106832 | 1.014545335 | 0.979002724 |
| ENSMUSG00000106833 | 1 | |
| ENSMUSG00000106834 | 1.069068979 | 0.608497229 |
| ENSMUSG00000106835 | 1 | |
| ENSMUSG00000106836 | 1 | |
| ENSMUSG00000106837 | -1 | 1 |
| ENSMUSG00000106838 | 1 | |
| ENSMUSG00000106839 | 1 | |
| ENSMUSG00000106840 | 1.075781176 | 0.822679139 |
| ENSMUSG00000106841 | 1.058455527 | 0.834990067 |
| ENSMUSG00000106843 | -1.136679298 | 0.608497229 |
| ENSMUSG00000106844 | 1 | |
| ENSMUSG00000106845 | 1 | |
| ENSMUSG00000106846 | 1 | |
| ENSMUSG00000106847 | 1.109059151 | 0.373217338 |
| ENSMUSG00000106849 | 1 | |
| ENSMUSG00000106851 | 1.069068979 | 0.608497229 |
| ENSMUSG00000106852 | 1 | |
| ENSMUSG00000106853 | 1 | |
| ENSMUSG00000106854 | 1 | |
| ENSMUSG00000106855 | 1.014545335 | 0.979002724 |
| ENSMUSG00000106856 | 1.095872691 | 0.608497229 |
| ENSMUSG00000106857 | 1 | |
| ENSMUSG00000106858 | 1.127071399 | 0.533812934 |
| ENSMUSG00000106859 | 1 | |
| ENSMUSG00000106860 | -1.179614078 | 0.535338451 |
| ENSMUSG00000106861 | 1 | |
| ENSMUSG00000106862 | 1.167673187 | 0.539963645 |
| ENSMUSG00000106863 | 1.049517471 | 0.837704961 |
| ENSMUSG00000106864 | -1.027410264 | 0.939076675 |
| ENSMUSG00000106865 | 1 | |
| ENSMUSG00000106866 | 1.196004652 | 0.378431183 |
| ENSMUSG00000106867 | 1.067975749 | 0.853042921 |
| ENSMUSG00000106868 | -1.063525418 | 0.803072249 |
| ENSMUSG00000106869 | 1.014545335 | 0.979002724 |
| ENSMUSG00000106870 | 1.181142828 | 0.022132535 |
| ENSMUSG00000106871 | 1.059463094 | 0.608497229 |
| ENSMUSG00000106872 | 1 | |

| | | |
|---|---|---|
| ENSMUSG00000106873 | 1 | |
| ENSMUSG00000106874 | 1.181131578 | 0.079015056 |
| ENSMUSG00000106875 | 1.122462048 | 0.215991492 |
| ENSMUSG00000106876 | 1 | |
| ENSMUSG00000106878 | 1.155639986 | 0.456591877 |
| ENSMUSG00000106879 | 1 | |
| ENSMUSG00000106880 | 1 | |
| ENSMUSG00000106881 | 1 | |
| ENSMUSG00000106883 | -1.132666855 | 0.18851436 |
| ENSMUSG00000106884 | 1 | |
| ENSMUSG00000106885 | -1.109992797 | 0.71627176 |
| ENSMUSG00000106887 | 1.004511908 | 1 |
| ENSMUSG00000106888 | -1.037482356 | 0.870427333 |
| ENSMUSG00000106889 | 1 | |
| ENSMUSG00000106890 | 1 | |
| ENSMUSG00000106891 | -1.108782257 | 0.773857666 |
| ENSMUSG00000106892 | 1.290107556 | 0.119190358 |
| ENSMUSG00000106893 | 1 | |
| ENSMUSG00000106894 | 1.014545335 | 0.979002724 |
| ENSMUSG00000106895 | -1.016392145 | 0.979002724 |
| ENSMUSG00000106896 | -1.03962336 | 0.698112512 |
| ENSMUSG00000106897 | -1.032140202 | 0.95559778 |
| ENSMUSG00000106898 | 1 | |
| ENSMUSG00000106899 | 1 | |
| ENSMUSG00000106900 | 1 | |
| ENSMUSG00000106902 | -1 | 1 |
| ENSMUSG00000106904 | 1.053719763 | 0.880723094 |
| ENSMUSG00000106905 | 1 | |
| ENSMUSG00000106906 | 1.069068979 | 0.608497229 |
| ENSMUSG00000106909 | 1 | |
| ENSMUSG00000106910 | 1.059463094 | 0.698112512 |
| ENSMUSG00000106912 | 1 | |
| ENSMUSG00000106913 | 1 | |
| ENSMUSG00000106914 | 1 | |
| ENSMUSG00000106915 | 1 | |
| ENSMUSG00000106916 | 1 | |
| ENSMUSG00000106917 | 1.059463094 | 0.608497229 |
| ENSMUSG00000106918 | 1.042775162 | 0.883408175 |
| ENSMUSG00000106920 | 1.079681442 | 0.705088559 |
| ENSMUSG00000106924 | 1.059463094 | 0.698112512 |
| ENSMUSG00000106925 | 1.059463094 | 0.608497229 |
| ENSMUSG00000106926 | -1.035455843 | 0.834580969 |
| ENSMUSG00000106927 | 1.101234685 | 0.788886948 |
| ENSMUSG00000106928 | -1.039583006 | 0.698112512 |
| ENSMUSG00000106930 | 1.079681442 | 0.743396505 |
| ENSMUSG00000106931 | 1 | |
| ENSMUSG00000106932 | 1 | |
| ENSMUSG00000106933 | 1.340985479 | 0.158258948 |
| ENSMUSG00000106934 | 1 | |
| ENSMUSG00000106935 | 1 | |
| ENSMUSG00000106936 | 1 | |
| ENSMUSG00000106937 | 1 | |
| ENSMUSG00000106938 | 1 | |
| ENSMUSG00000106939 | 1 | |
| ENSMUSG00000106940 | -1.089864654 | 0.683236659 |
| ENSMUSG00000106941 | 1 | |
| ENSMUSG00000106943 | -1.143787033 | 0.476577776 |
| ENSMUSG00000106944 | -1.065852599 | 0.797808656 |
| ENSMUSG00000106945 | 1 | |
| ENSMUSG00000106946 | 1.159935731 | 0.511378706 |
| ENSMUSG00000106947 | 1.069068979 | 0.608497229 |
| ENSMUSG00000106948 | 1.071421129 | 0.608497229 |
| ENSMUSG00000106950 | 1 | |

| | | |
|---|---|---|
| ENSMUSG00000106951 | 1.131910394 | 0.284325772 |
| ENSMUSG00000106952 | 1 | |
| ENSMUSG00000106954 | 1 | |
| ENSMUSG00000106955 | 1 | |
| ENSMUSG00000106956 | 1 | |
| ENSMUSG00000106957 | 1.24122829 | 0.001720654 |
| ENSMUSG00000106958 | 1 | |
| ENSMUSG00000106959 | -1.144940924 | 0.458699594 |
| ENSMUSG00000106960 | 1 | |
| ENSMUSG00000106961 | 1.263042918 | 0.021070191 |
| ENSMUSG00000106962 | -1.210306185 | 0.556118084 |
| ENSMUSG00000106963 | 1.091908159 | 0.608497229 |
| ENSMUSG00000106965 | -1.106436942 | 0.724205129 |
| ENSMUSG00000106967 | 1.069068979 | 0.608497229 |
| ENSMUSG00000106968 | 1.24122829 | 0.03679259 |
| ENSMUSG00000106969 | 1 | |
| ENSMUSG00000106970 | 1 | |
| ENSMUSG00000106971 | 1 | |
| ENSMUSG00000106973 | 1 | |
| ENSMUSG00000106974 | 1 | |
| ENSMUSG00000106975 | 1 | |
| ENSMUSG00000106976 | 1 | |
| ENSMUSG00000106978 | 1.106987703 | 0.608497229 |
| ENSMUSG00000106979 | 1 | |
| ENSMUSG00000106980 | -1.028348525 | 0.956972448 |
| ENSMUSG00000106981 | 1.02165347 | 0.861653133 |
| ENSMUSG00000106982 | 1 | |
| ENSMUSG00000106985 | 1 | |
| ENSMUSG00000106986 | 1 | |
| ENSMUSG00000106987 | -1.044273782 | 0.83619593 |
| ENSMUSG00000106988 | -1.024718486 | 0.931480186 |
| ENSMUSG00000106989 | 1 | |
| ENSMUSG00000106990 | -1.030807953 | 0.79058371 |
| ENSMUSG00000106991 | 1 | |
| ENSMUSG00000106992 | -1.033820075 | 0.940197654 |
| ENSMUSG00000106993 | 1 | |
| ENSMUSG00000106994 | 1.059463094 | 0.608497229 |
| ENSMUSG00000106995 | -1.251614011 | 0.398865597 |
| ENSMUSG00000106996 | 1 | |
| ENSMUSG00000106997 | 1.09590697 | 0.147196278 |
| ENSMUSG00000106998 | 1 | |
| ENSMUSG00000106999 | 1 | |
| ENSMUSG00000107000 | 1.06690417 | 0.077081575 |
| ENSMUSG00000107002 | 1.021612736 | 0.80446981 |
| ENSMUSG00000107004 | 1 | |
| ENSMUSG00000107005 | -1.065884893 | 0.815415874 |
| ENSMUSG00000107006 | -1.03962336 | 0.872692845 |
| ENSMUSG00000107008 | -1.095294215 | 0.541735653 |
| ENSMUSG00000107009 | 1.059463094 | 0.608497229 |
| ENSMUSG00000107010 | 1 | |
| ENSMUSG00000107011 | 1 | |
| ENSMUSG00000107012 | 1.059463094 | 0.698112512 |
| ENSMUSG00000107013 | 1 | |
| ENSMUSG00000107014 | 1 | |
| ENSMUSG00000107015 | -1.043306199 | 0.79291651 |
| ENSMUSG00000107016 | 1 | |
| ENSMUSG00000107017 | -1.037236631 | 0.947264903 |
| ENSMUSG00000107018 | -1.053549062 | 0.769745556 |
| ENSMUSG00000107019 | -1.030811902 | 0.809396188 |
| ENSMUSG00000107020 | -1.043306199 | 0.698112512 |
| ENSMUSG00000107021 | -1.043306199 | 0.698112512 |
| ENSMUSG00000107022 | 1.059463094 | 0.608497229 |
| ENSMUSG00000107023 | -1.102105418 | 0.1955798 |

| | | |
|---|---|---|
| ENSMUSG00000107024 | 1 | |
| ENSMUSG00000107026 | 1.052675685 | 0.862562879 |
| ENSMUSG00000107027 | 1 | |
| ENSMUSG00000107028 | -1 | 1 |
| ENSMUSG00000107029 | 1.04328066 | 0.868156972 |
| ENSMUSG00000107030 | 1.059463094 | 0.698112512 |
| ENSMUSG00000107031 | 1 | |
| ENSMUSG00000107032 | 1 | |
| ENSMUSG00000107033 | -1.053549062 | 0.698112512 |
| ENSMUSG00000107034 | 1 | |
| ENSMUSG00000107035 | 1.079681442 | 0.608497229 |
| ENSMUSG00000107036 | 1.059463094 | 0.704170349 |
| ENSMUSG00000107037 | 1 | |
| ENSMUSG00000107040 | 1 | |
| ENSMUSG00000107041 | -1.120484078 | 0.713371693 |
| ENSMUSG00000107042 | 1 | |
| ENSMUSG00000107043 | 1.171563499 | 0.233412931 |
| ENSMUSG00000107044 | -1 | 1 |
| ENSMUSG00000107045 | -1.12843057 | 0.540891969 |
| ENSMUSG00000107046 | 1 | |
| ENSMUSG00000107047 | 1 | |
| ENSMUSG00000107048 | -1.065852599 | 0.797808656 |
| ENSMUSG00000107049 | 1.015486245 | 0.985522133 |
| ENSMUSG00000107050 | -1.144871996 | 0.304923237 |
| ENSMUSG00000107053 | 1 | |
| ENSMUSG00000107054 | -1.054169715 | 0.813485995 |
| ENSMUSG00000107056 | 1 | |
| ENSMUSG00000107058 | 1 | |
| ENSMUSG00000107059 | 1 | |
| ENSMUSG00000107060 | 1 | |
| ENSMUSG00000107061 | 1 | |
| ENSMUSG00000107062 | 1 | |
| ENSMUSG00000107063 | -1.043306199 | 0.698112512 |
| ENSMUSG00000107064 | 1 | |
| ENSMUSG00000107065 | 1.028323353 | 0.934760809 |
| ENSMUSG00000107066 | 1 | |
| ENSMUSG00000107067 | -1 | 1 |
| ENSMUSG00000107068 | 1 | |
| ENSMUSG00000107069 | 1 | |
| ENSMUSG00000107070 | -1.048277865 | 0.862879216 |
| ENSMUSG00000107071 | 1 | |
| ENSMUSG00000107072 | 1 | |
| ENSMUSG00000107073 | 1 | |
| ENSMUSG00000107074 | 1 | |
| ENSMUSG00000107075 | 1 | |
| ENSMUSG00000107076 | -1.178134217 | 0.301170672 |
| ENSMUSG00000107078 | 1 | |
| ENSMUSG00000107079 | -1.184259924 | 0.127188505 |
| ENSMUSG00000107081 | 1 | |
| ENSMUSG00000107082 | -1 | 1 |
| ENSMUSG00000107083 | 1.104229632 | 0.734624722 |
| ENSMUSG00000107084 | 1 | |
| ENSMUSG00000107085 | 1 | |
| ENSMUSG00000107086 | -1.104919702 | 0.574033444 |
| ENSMUSG00000107087 | 1 | |
| ENSMUSG00000107089 | -1 | 1 |
| ENSMUSG00000107090 | 1 | |
| ENSMUSG00000107091 | 1.049593996 | 0.926076285 |
| ENSMUSG00000107092 | 1.050882889 | 0.841285672 |
| ENSMUSG00000107093 | 1 | |
| ENSMUSG00000107094 | 1.126732949 | 0.57513806 |
| ENSMUSG00000107096 | 1.089810863 | 0.053367026 |
| ENSMUSG00000107097 | 1 | |

| | | |
|---|---|---|
| ENSMUSG00000107098 | 1 | |
| ENSMUSG00000107100 | 1 | |
| ENSMUSG00000107102 | -1.023612535 | 0.927938496 |
| ENSMUSG00000107103 | 1.122462048 | 0.215991492 |
| ENSMUSG00000107105 | -1 | 1 |
| ENSMUSG00000107106 | 1.059463094 | 0.608497229 |
| ENSMUSG00000107107 | 1.028323353 | 0.933031586 |
| ENSMUSG00000107108 | -1.054741498 | 0.872692845 |
| ENSMUSG00000107109 | 1 | |
| ENSMUSG00000107110 | 1 | |
| ENSMUSG00000107111 | 1 | |
| ENSMUSG00000107112 | 1 | |
| ENSMUSG00000107113 | 1 | |
| ENSMUSG00000107114 | 1 | |
| ENSMUSG00000107115 | 1.059463094 | 0.608497229 |
| ENSMUSG00000107116 | 1.064347708 | 0.856915299 |
| ENSMUSG00000107117 | -1.074779048 | 0.774810939 |
| ENSMUSG00000107118 | 1.19249892 | 0.317733967 |
| ENSMUSG00000107120 | 1.078030942 | 0.563009714 |
| ENSMUSG00000107121 | 1.262865859 | 0.327642404 |
| ENSMUSG00000107122 | 1 | |
| ENSMUSG00000107123 | 1 | |
| ENSMUSG00000107125 | 1 | |
| ENSMUSG00000107126 | -1.044273782 | 0.698112512 |
| ENSMUSG00000107127 | 1 | |
| ENSMUSG00000107128 | 1 | |
| ENSMUSG00000107129 | 1 | |
| ENSMUSG00000107130 | 1 | |
| ENSMUSG00000107131 | -1 | 1 |
| ENSMUSG00000107132 | 1 | |
| ENSMUSG00000107133 | 1.095872691 | 0.608497229 |
| ENSMUSG00000107134 | 1 | |
| ENSMUSG00000107135 | 1 | |
| ENSMUSG00000107136 | 1.024693402 | 0.955101438 |
| ENSMUSG00000107139 | 1 | |
| ENSMUSG00000107140 | 1 | |
| ENSMUSG00000107141 | 1 | |
| ENSMUSG00000107142 | 1 | |
| ENSMUSG00000107143 | -1.047209109 | 0.796457226 |
| ENSMUSG00000107144 | 1 | |
| ENSMUSG00000107145 | 1.059463094 | 0.608497229 |
| ENSMUSG00000107146 | 1 | |
| ENSMUSG00000107148 | -1.044273782 | 0.790851299 |
| ENSMUSG00000107149 | -1.081225398 | 0.761839849 |
| ENSMUSG00000107151 | 1.07487334 | 0.68735438 |
| ENSMUSG00000107152 | -1.117459691 | 0.608497229 |
| ENSMUSG00000107153 | 1 | |
| ENSMUSG00000107154 | 1.059463094 | 0.608497229 |
| ENSMUSG00000107155 | 1.230592175 | 0.297260943 |
| ENSMUSG00000107157 | 1 | |
| ENSMUSG00000107158 | -1 | 1 |
| ENSMUSG00000107159 | 1 | |
| ENSMUSG00000107160 | 1.130316658 | 0.698112512 |
| ENSMUSG00000107161 | -1.063014928 | 0.905344458 |
| ENSMUSG00000107162 | 1.069068979 | 0.698112512 |
| ENSMUSG00000107163 | -1.003012737 | 1 |
| ENSMUSG00000107164 | 1.240919017 | 0.256011478 |
| ENSMUSG00000107165 | 1.200146492 | 0.508515749 |
| ENSMUSG00000107167 | -1 | 1 |
| ENSMUSG00000107168 | 1.127526494 | 0.657721628 |
| ENSMUSG00000107169 | 1 | |
| ENSMUSG00000107171 | 1.103206151 | 0.424525078 |
| ENSMUSG00000107172 | 1 | |

| | | |
|---|---|---|
| ENSMUSG00000107173 | -1.165541787 | 0.247953562 |
| ENSMUSG00000107175 | 1 | |
| ENSMUSG00000107176 | 1.070963712 | 0.698112512 |
| ENSMUSG00000107177 | 1 | |
| ENSMUSG00000107178 | 1.080076996 | 0.734415698 |
| ENSMUSG00000107179 | 1 | |
| ENSMUSG00000107180 | 1 | |
| ENSMUSG00000107181 | 1 | |
| ENSMUSG00000107182 | -1.114024259 | 0.217412861 |
| ENSMUSG00000107183 | 1 | |
| ENSMUSG00000107184 | -1.000405949 | 1 |
| ENSMUSG00000107185 | 1 | |
| ENSMUSG00000107187 | 1 | |
| ENSMUSG00000107189 | -1 | 1 |
| ENSMUSG00000107190 | 1 | |
| ENSMUSG00000107191 | 1 | |
| ENSMUSG00000107192 | 1 | |
| ENSMUSG00000107193 | 1 | |
| ENSMUSG00000107194 | -1.03962336 | 0.872692845 |
| ENSMUSG00000107195 | -1.208401079 | 0.159699022 |
| ENSMUSG00000107197 | -1.043306199 | 0.698112512 |
| ENSMUSG00000107198 | 1.030256832 | 0.97132868 |
| ENSMUSG00000107199 | 1 | |
| ENSMUSG00000107200 | 1.063141874 | 0.608497229 |
| ENSMUSG00000107201 | 1.029648082 | 0.654011048 |
| ENSMUSG00000107202 | -1 | 1 |
| ENSMUSG00000107203 | -1.090507733 | 0.728067732 |
| ENSMUSG00000107204 | -1.053549062 | 0.698112512 |
| ENSMUSG00000107205 | 1.06966519 | 0.69533886 |
| ENSMUSG00000107206 | 1.075870199 | 0.727587593 |
| ENSMUSG00000107207 | -1.084645496 | 0.532369657 |
| ENSMUSG00000107208 | 1 | |
| ENSMUSG00000107209 | 1 | |
| ENSMUSG00000107211 | 1.005613439 | 1 |
| ENSMUSG00000107212 | -1.043722544 | 0.923982284 |
| ENSMUSG00000107213 | 1 | |
| ENSMUSG00000107214 | 1 | |
| ENSMUSG00000107215 | 1.189039819 | 0.209798452 |
| ENSMUSG00000107216 | 1.111766029 | 0.698112512 |
| ENSMUSG00000107217 | 1 | |
| ENSMUSG00000107218 | 1.059463094 | 0.698112512 |
| ENSMUSG00000107219 | 1 | |
| ENSMUSG00000107220 | 1 | |
| ENSMUSG00000107222 | 1 | |
| ENSMUSG00000107223 | 1 | |
| ENSMUSG00000107224 | 1 | |
| ENSMUSG00000107225 | -1.048826362 | 0.898292375 |
| ENSMUSG00000107226 | 1 | |
| ENSMUSG00000107227 | 1.066431747 | 0.62346446 |
| ENSMUSG00000107228 | 1 | |
| ENSMUSG00000107229 | 1 | |
| ENSMUSG00000107230 | 1.028603875 | 0.809466772 |
| ENSMUSG00000107231 | 1 | |
| ENSMUSG00000107232 | 1 | |
| ENSMUSG00000107233 | 1.203191122 | 0.058274403 |
| ENSMUSG00000107234 | 1.059463094 | 0.608497229 |
| ENSMUSG00000107237 | 1 | |
| ENSMUSG00000107238 | 1 | |
| ENSMUSG00000107240 | -1.085651418 | 0.532369657 |
| ENSMUSG00000107241 | 1 | |
| ENSMUSG00000107242 | 1 | |
| ENSMUSG00000107243 | 1 | |
| ENSMUSG00000107244 | 1.053067877 | 0.825798518 |

| | | |
|---|---|---|
| ENSMUSG00000107245 | 1 | |
| ENSMUSG00000107246 | -1.014034612 | 1 |
| ENSMUSG00000107247 | 1 | |
| ENSMUSG00000107248 | 1 | |
| ENSMUSG00000107251 | 1 | |
| ENSMUSG00000107252 | 1.059463094 | 0.820462971 |
| ENSMUSG00000107253 | 1.000528147 | 1 |
| ENSMUSG00000107254 | -1.146375298 | 0.608497229 |
| ENSMUSG00000107255 | 1 | |
| ENSMUSG00000107256 | -1.014093825 | 1 |
| ENSMUSG00000107257 | 1 | |
| ENSMUSG00000107258 | 1 | |
| ENSMUSG00000107259 | -1.085651418 | 0.532369657 |
| ENSMUSG00000107260 | 1 | |
| ENSMUSG00000107261 | 1 | |
| ENSMUSG00000107262 | 1 | |
| ENSMUSG00000107263 | 1.195527562 | 0.210047602 |
| ENSMUSG00000107264 | 1 | |
| ENSMUSG00000107265 | 1 | |
| ENSMUSG00000107266 | -1.085651418 | 0.532369657 |
| ENSMUSG00000107270 | 1 | |
| ENSMUSG00000107271 | -1.177158763 | 0.508942701 |
| ENSMUSG00000107272 | -1.03962336 | 0.698112512 |
| ENSMUSG00000107273 | 1 | |
| ENSMUSG00000107276 | -1.047001856 | 0.903464611 |
| ENSMUSG00000107277 | 1.0964716 | 0.652522217 |
| ENSMUSG00000107278 | 1.07045453 | 0.630205913 |
| ENSMUSG00000107279 | 1 | |
| ENSMUSG00000107280 | 1 | |
| ENSMUSG00000107281 | 1 | |
| ENSMUSG00000107282 | 1 | |
| ENSMUSG00000107283 | -1.07789069 | 0.717035356 |
| ENSMUSG00000107284 | 1 | |
| ENSMUSG00000107285 | 1 | |
| ENSMUSG00000107286 | 1 | |
| ENSMUSG00000107287 | 1 | |
| ENSMUSG00000107288 | 1.199989356 | 0.246522583 |
| ENSMUSG00000107289 | 1 | |
| ENSMUSG00000107290 | 1 | |
| ENSMUSG00000107291 | 1 | |
| ENSMUSG00000107292 | 1 | |
| ENSMUSG00000107294 | 1 | |
| ENSMUSG00000107295 | -1.044273782 | 0.698112512 |
| ENSMUSG00000107296 | 1 | |
| ENSMUSG00000107298 | 1 | |
| ENSMUSG00000107299 | 1 | |
| ENSMUSG00000107300 | -1.153571102 | 0.608497229 |
| ENSMUSG00000107301 | 1 | |
| ENSMUSG00000107303 | 1 | |
| ENSMUSG00000107304 | -1.107299336 | 0.644942465 |
| ENSMUSG00000107305 | 1 | |
| ENSMUSG00000107306 | 1.084452299 | 0.464689145 |
| ENSMUSG00000107307 | 1.059463094 | 0.698112512 |
| ENSMUSG00000107308 | 1 | |
| ENSMUSG00000107309 | 1 | |
| ENSMUSG00000107310 | 1.029394426 | 0.82448954 |
| ENSMUSG00000107312 | 1.059463094 | 0.698112512 |
| ENSMUSG00000107313 | 1.097670742 | 0.851742081 |
| ENSMUSG00000107314 | -1.043306199 | 0.698112512 |
| ENSMUSG00000107315 | 1 | |
| ENSMUSG00000107316 | 1 | |
| ENSMUSG00000107317 | -1 | 1 |
| ENSMUSG00000107318 | 1 | |

| | | |
|---|---|---|
| ENSMUSG00000107319 | -1.156562185 | 0.46010161 |
| ENSMUSG00000107320 | -1.013838841 | 0.98079728 |
| ENSMUSG00000107322 | 1 | |
| ENSMUSG00000107323 | 1 | |
| ENSMUSG00000107324 | 1 | |
| ENSMUSG00000107325 | 1 | |
| ENSMUSG00000107326 | 1 | |
| ENSMUSG00000107327 | -1.069502966 | 0.744868995 |
| ENSMUSG00000107329 | 1 | |
| ENSMUSG00000107331 | 1.132456084 | 0.072407958 |
| ENSMUSG00000107332 | 1 | |
| ENSMUSG00000107333 | 1.173139855 | 0.307311513 |
| ENSMUSG00000107334 | 1 | |
| ENSMUSG00000107335 | -1.03299708 | 0.956401771 |
| ENSMUSG00000107336 | -1.071026843 | 0.797410621 |
| ENSMUSG00000107337 | 1.021186614 | 0.973756648 |
| ENSMUSG00000107338 | -1.048826362 | 0.751229613 |
| ENSMUSG00000107340 | 1.075070132 | 0.787977591 |
| ENSMUSG00000107341 | 1 | |
| ENSMUSG00000107344 | 1 | |
| ENSMUSG00000107345 | 1 | |
| ENSMUSG00000107346 | 1.024693402 | 0.933031586 |
| ENSMUSG00000107347 | 1 | |
| ENSMUSG00000107348 | 1 | |
| ENSMUSG00000107349 | 1.059463094 | 0.608497229 |
| ENSMUSG00000107350 | 1.319294337 | 0.159480741 |
| ENSMUSG00000107351 | 1 | |
| ENSMUSG00000107352 | 1.033058884 | 0.828564184 |
| ENSMUSG00000107353 | 1 | |
| ENSMUSG00000107354 | -1.0795912 | 0.731829998 |
| ENSMUSG00000107355 | 1.29751208 | 0.066068145 |
| ENSMUSG00000107356 | 1 | |
| ENSMUSG00000107357 | -1.043306199 | 0.698112512 |
| ENSMUSG00000107358 | -1.03962336 | 0.698112512 |
| ENSMUSG00000107359 | 1.059463094 | 0.608497229 |
| ENSMUSG00000107360 | 1 | |
| ENSMUSG00000107362 | -1.03089845 | 0.986359381 |
| ENSMUSG00000107363 | 1 | |
| ENSMUSG00000107364 | 1 | |
| ENSMUSG00000107365 | -1 | 1 |
| ENSMUSG00000107368 | 1.161036672 | 0.389746121 |
| ENSMUSG00000107369 | -1 | 1 |
| ENSMUSG00000107370 | 1.066671072 | 0.815671448 |
| ENSMUSG00000107371 | 1 | |
| ENSMUSG00000107372 | 1 | |
| ENSMUSG00000107373 | 1 | |
| ENSMUSG00000107374 | 1.24138083 | 0.022993558 |
| ENSMUSG00000107375 | 1 | |
| ENSMUSG00000107376 | 1 | |
| ENSMUSG00000107377 | -1.049075043 | 0.924141494 |
| ENSMUSG00000107378 | 1.059463094 | 0.698112512 |
| ENSMUSG00000107379 | 1 | |
| ENSMUSG00000107380 | 1 | |
| ENSMUSG00000107383 | 1.021067122 | 0.912782976 |
| ENSMUSG00000107384 | 1 | |
| ENSMUSG00000107385 | 1.007451137 | 0.993548588 |
| ENSMUSG00000107386 | 1 | |
| ENSMUSG00000107387 | 1 | |
| ENSMUSG00000107388 | -1.111221915 | 0.608497229 |
| ENSMUSG00000107389 | 1.060867173 | 0.88325758 |
| ENSMUSG00000107390 | 1.103067059 | 0.59127359 |
| ENSMUSG00000107392 | 1 | |
| ENSMUSG00000107394 | 1 | |

| | | |
|---|---|---|
| ENSMUSG00000107395 | -1.03962336 | 0.698112512 |
| ENSMUSG00000107396 | 1 | |
| ENSMUSG00000107397 | 1.133237328 | 0.511010591 |
| ENSMUSG00000107398 | 1 | |
| ENSMUSG00000107399 | 1 | |
| ENSMUSG00000107400 | 1 | |
| ENSMUSG00000107401 | 1 | |
| ENSMUSG00000107402 | 1.061579705 | 0.745531957 |
| ENSMUSG00000107403 | 1 | |
| ENSMUSG00000107404 | 1 | |
| ENSMUSG00000107405 | -1.053549062 | 0.778372272 |
| ENSMUSG00000107406 | -1.010960635 | 1 |
| ENSMUSG00000107407 | 1 | |
| ENSMUSG00000107408 | 1 | |
| ENSMUSG00000107409 | 1 | |
| ENSMUSG00000107410 | 1 | |
| ENSMUSG00000107411 | -1.033256173 | 0.961127302 |
| ENSMUSG00000107412 | 1 | |
| ENSMUSG00000107413 | 1 | |
| ENSMUSG00000107414 | 1 | |
| ENSMUSG00000107415 | 1 | |
| ENSMUSG00000107416 | 1 | |
| ENSMUSG00000107417 | 1.059463094 | 0.608497229 |
| ENSMUSG00000107418 | 1 | |
| ENSMUSG00000107419 | 1 | |
| ENSMUSG00000107420 | 1.069068979 | 0.608497229 |
| ENSMUSG00000107421 | 1 | |
| ENSMUSG00000107422 | 1 | |
| ENSMUSG00000107423 | 1 | |
| ENSMUSG00000107424 | 1 | |
| ENSMUSG00000107425 | 1 | |
| ENSMUSG00000107427 | 1 | |
| ENSMUSG00000107428 | 1.059463094 | 0.608497229 |
| ENSMUSG00000107429 | -1.012925407 | 0.977035046 |
| ENSMUSG00000107430 | -1.024718486 | 0.951399668 |
| ENSMUSG00000107431 | 1 | |
| ENSMUSG00000107432 | -1.04759745 | 0.729206866 |
| ENSMUSG00000107433 | 1 | |
| ENSMUSG00000107434 | 1.376503915 | 0.117310003 |
| ENSMUSG00000107435 | 1 | |
| ENSMUSG00000107436 | 1.010855137 | 1 |
| ENSMUSG00000107438 | 1 | |
| ENSMUSG00000107439 | 1 | |
| ENSMUSG00000107440 | -1.010531047 | 1 |
| ENSMUSG00000107441 | -1 | 1 |
| ENSMUSG00000107442 | 1 | |
| ENSMUSG00000107443 | -1.018054237 | 0.994536934 |
| ENSMUSG00000107444 | 1 | |
| ENSMUSG00000107445 | 1 | |
| ENSMUSG00000107446 | 1 | |
| ENSMUSG00000107447 | -1 | 1 |
| ENSMUSG00000107448 | 1 | |
| ENSMUSG00000107449 | 1 | |
| ENSMUSG00000107450 | 1 | |
| ENSMUSG00000107451 | 1 | |
| ENSMUSG00000107452 | 1 | |
| ENSMUSG00000107453 | 1 | |
| ENSMUSG00000107454 | 1 | |
| ENSMUSG00000107455 | -1.022608177 | 0.96035552 |
| ENSMUSG00000107456 | 1 | |
| ENSMUSG00000107457 | 1 | |
| ENSMUSG00000107458 | 1.069068979 | 0.608497229 |
| ENSMUSG00000107459 | 1.150771739 | 0.383499141 |

| | | |
|---|---|---|
| ENSMUSG00000107460 | 1.090076375 | 0.775673129 |
| ENSMUSG00000107461 | -1.316517498 | 0.010908481 |
| ENSMUSG00000107462 | 1.205401078 | 0.608497229 |
| ENSMUSG00000107463 | 1 | |
| ENSMUSG00000107464 | 1 | |
| ENSMUSG00000107465 | 1 | |
| ENSMUSG00000107466 | 1 | |
| ENSMUSG00000107467 | 1 | |
| ENSMUSG00000107468 | -1.08384782 | 0.747100886 |
| ENSMUSG00000107469 | 1 | |
| ENSMUSG00000107470 | -1.148993587 | 0.228239447 |
| ENSMUSG00000107471 | 1 | |
| ENSMUSG00000107472 | -1.03962336 | 0.698112512 |
| ENSMUSG00000107473 | 1 | |
| ENSMUSG00000107474 | 1 | |
| ENSMUSG00000107475 | 1.095872691 | 0.608497229 |
| ENSMUSG00000107476 | 1.039070409 | 0.608497229 |
| ENSMUSG00000107477 | -1.031335064 | 0.961832515 |
| ENSMUSG00000107478 | 1.109121605 | 0.772834122 |
| ENSMUSG00000107479 | 1 | |
| ENSMUSG00000107480 | -1.013419756 | 0.999412999 |
| ENSMUSG00000107481 | -1 | 1 |
| ENSMUSG00000107482 | 1 | |
| ENSMUSG00000107483 | -1.044273782 | 0.698112512 |
| ENSMUSG00000107484 | 1 | |
| ENSMUSG00000107486 | 1 | |
| ENSMUSG00000107487 | -1.03962336 | 0.698112512 |
| ENSMUSG00000107488 | 1 | |
| ENSMUSG00000107489 | 1 | |
| ENSMUSG00000107490 | 1 | |
| ENSMUSG00000107491 | -1.081225398 | 0.796568072 |
| ENSMUSG00000107492 | 1 | |
| ENSMUSG00000107493 | 1 | |
| ENSMUSG00000107494 | 1 | |
| ENSMUSG00000107495 | 1.089767314 | 0.635004673 |
| ENSMUSG00000107496 | 1 | |
| ENSMUSG00000107497 | 1.020240423 | 0.968804789 |
| ENSMUSG00000107498 | 1 | |
| ENSMUSG00000107499 | -1.245835383 | 0.319960972 |
| ENSMUSG00000107500 | 1 | |
| ENSMUSG00000107501 | 1.05410549 | 0.805157604 |
| ENSMUSG00000107502 | -1.06804853 | 0.834464052 |
| ENSMUSG00000107503 | 1 | |
| ENSMUSG00000107504 | -1.063525418 | 0.698112512 |
| ENSMUSG00000107505 | 1.132851843 | 0.620221546 |
| ENSMUSG00000107506 | 1 | |
| ENSMUSG00000107507 | 1 | |
| ENSMUSG00000107508 | 1.075070132 | 0.698112512 |
| ENSMUSG00000107509 | 1.304983415 | 0.481568621 |
| ENSMUSG00000107510 | 1 | |
| ENSMUSG00000107511 | -1.006331475 | 1 |
| ENSMUSG00000107512 | 1 | |
| ENSMUSG00000107513 | 1 | |
| ENSMUSG00000107514 | 1 | |
| ENSMUSG00000107515 | 1.132639128 | 0.217803313 |
| ENSMUSG00000107516 | 1 | |
| ENSMUSG00000107517 | -1.069502966 | 0.744868995 |
| ENSMUSG00000107518 | 1 | |
| ENSMUSG00000107519 | 1 | |
| ENSMUSG00000107520 | 1 | |
| ENSMUSG00000107521 | -1.03962336 | 0.803072249 |
| ENSMUSG00000107522 | 1 | |
| ENSMUSG00000107523 | 1 | |

| | | |
|---|---|---|
| ENSMUSG00000107524 | -1.270257935 | 0.112920219 |
| ENSMUSG00000107525 | 1 | |
| ENSMUSG00000107526 | 1.196004652 | 0.183846449 |
| ENSMUSG00000107527 | 1 | |
| ENSMUSG00000107528 | 1 | |
| ENSMUSG00000107529 | -1.085651418 | 0.608497229 |
| ENSMUSG00000107530 | 1 | |
| ENSMUSG00000107531 | 1.103296893 | 0.633643463 |
| ENSMUSG00000107532 | 1 | |
| ENSMUSG00000107533 | 1 | |
| ENSMUSG00000107534 | -1.188446785 | 0.219185116 |
| ENSMUSG00000107535 | 1 | |
| ENSMUSG00000107536 | 1 | |
| ENSMUSG00000107538 | 1 | |
| ENSMUSG00000107539 | 1 | |
| ENSMUSG00000107540 | 1 | |
| ENSMUSG00000107541 | 1 | |
| ENSMUSG00000107542 | -1 | 1 |
| ENSMUSG00000107543 | 1 | |
| ENSMUSG00000107544 | 1 | |
| ENSMUSG00000107545 | -1 | 1 |
| ENSMUSG00000107546 | -1.053549062 | 0.825517832 |
| ENSMUSG00000107548 | 1 | |
| ENSMUSG00000107549 | 1.140458245 | 0.477044609 |
| ENSMUSG00000107550 | 1 | |
| ENSMUSG00000107551 | -1.053549062 | 0.698112512 |
| ENSMUSG00000107552 | -1.183677106 | 0.608497229 |
| ENSMUSG00000107553 | 1 | |
| ENSMUSG00000107554 | 1 | |
| ENSMUSG00000107555 | 1 | |
| ENSMUSG00000107556 | 1.166591266 | 0.608497229 |
| ENSMUSG00000107557 | 1 | |
| ENSMUSG00000107558 | 1 | |
| ENSMUSG00000107559 | 1.079681442 | 0.608497229 |
| ENSMUSG00000107560 | 1.037400725 | 0.779799562 |
| ENSMUSG00000107561 | 1 | |
| ENSMUSG00000107562 | 1 | |
| ENSMUSG00000107563 | 1 | |
| ENSMUSG00000107564 | 1 | |
| ENSMUSG00000107565 | 1 | |
| ENSMUSG00000107566 | 1 | |
| ENSMUSG00000107567 | 1.145373242 | 0.609541771 |
| ENSMUSG00000107568 | -1.025242841 | 0.989742633 |
| ENSMUSG00000107569 | 1 | |
| ENSMUSG00000107570 | 1.110468775 | 0.066445442 |
| ENSMUSG00000107571 | -1 | 1 |
| ENSMUSG00000107572 | 1 | |
| ENSMUSG00000107573 | 1 | |
| ENSMUSG00000107574 | 1 | |
| ENSMUSG00000107575 | 1 | |
| ENSMUSG00000107576 | -1.14503164 | 0.608497229 |
| ENSMUSG00000107577 | 1.092863046 | 0.642098458 |
| ENSMUSG00000107578 | -1 | 1 |
| ENSMUSG00000107579 | 1 | |
| ENSMUSG00000107580 | 1 | |
| ENSMUSG00000107581 | 1 | |
| ENSMUSG00000107582 | 1 | |
| ENSMUSG00000107583 | -1.099174267 | 0.535052277 |
| ENSMUSG00000107584 | 1.121188275 | 0.686089423 |
| ENSMUSG00000107585 | 1 | |
| ENSMUSG00000107586 | 1.122462048 | 0.420161906 |
| ENSMUSG00000107587 | 1 | |
| ENSMUSG00000107588 | 1 | |

| | | |
|---|---|---|
| ENSMUSG00000107590 | 1 | |
| ENSMUSG00000107591 | -1.044273782 | 0.795446726 |
| ENSMUSG00000107592 | 1 | |
| ENSMUSG00000107593 | 1.059463094 | 0.608497229 |
| ENSMUSG00000107594 | 1 | |
| ENSMUSG00000107595 | 1.059463094 | 0.608497229 |
| ENSMUSG00000107596 | 1.059463094 | 0.776953697 |
| ENSMUSG00000107597 | 1 | |
| ENSMUSG00000107598 | 1 | |
| ENSMUSG00000107599 | 1.201168605 | 0.451532357 |
| ENSMUSG00000107600 | -1 | 1 |
| ENSMUSG00000107601 | 1 | |
| ENSMUSG00000107602 | 1 | |
| ENSMUSG00000107603 | -1.199425955 | 0.433678407 |
| ENSMUSG00000107604 | 1.122462048 | 0.27820072 |
| ENSMUSG00000107605 | 1 | |
| ENSMUSG00000107606 | 1.102024305 | 0.698112512 |
| ENSMUSG00000107608 | -1.11869529 | 0.608497229 |
| ENSMUSG00000107609 | 1 | |
| ENSMUSG00000107610 | -1.043349923 | 0.790885744 |
| ENSMUSG00000107611 | 1 | |
| ENSMUSG00000107612 | 1.028323353 | 0.933031586 |
| ENSMUSG00000107613 | 1 | |
| ENSMUSG00000107614 | 1 | |
| ENSMUSG00000107615 | 1 | |
| ENSMUSG00000107616 | 1 | |
| ENSMUSG00000107618 | 1 | |
| ENSMUSG00000107619 | 1 | |
| ENSMUSG00000107620 | -1.121722906 | 0.559354144 |
| ENSMUSG00000107622 | 1 | |
| ENSMUSG00000107623 | -1.03962336 | 0.698112512 |
| ENSMUSG00000107624 | 1 | |
| ENSMUSG00000107625 | 1 | |
| ENSMUSG00000107626 | 1 | |
| ENSMUSG00000107627 | 1 | |
| ENSMUSG00000107628 | 1.079681442 | 0.750742698 |
| ENSMUSG00000107629 | 1.059463094 | 0.608497229 |
| ENSMUSG00000107630 | 1 | |
| ENSMUSG00000107631 | 1 | |
| ENSMUSG00000107632 | 1.083871761 | 0.629902617 |
| ENSMUSG00000107633 | 1.122608106 | 0.265014636 |
| ENSMUSG00000107634 | 1.189207115 | 0.401882202 |
| ENSMUSG00000107635 | -1.043306199 | 0.803260194 |
| ENSMUSG00000107637 | 1 | |
| ENSMUSG00000107638 | 1 | |
| ENSMUSG00000107639 | -1.111221915 | 0.619554344 |
| ENSMUSG00000107640 | -1 | 1 |
| ENSMUSG00000107641 | 1 | |
| ENSMUSG00000107642 | 1 | |
| ENSMUSG00000107643 | -1 | 1 |
| ENSMUSG00000107644 | -1.084645496 | 0.532369657 |
| ENSMUSG00000107645 | 1 | |
| ENSMUSG00000107646 | 1 | |
| ENSMUSG00000107647 | 1.05038453 | 0.873426185 |
| ENSMUSG00000107648 | 1 | |
| ENSMUSG00000107649 | 1 | |
| ENSMUSG00000107650 | 1 | |
| ENSMUSG00000107651 | 1 | |
| ENSMUSG00000107652 | 1 | |
| ENSMUSG00000107653 | -1.002140654 | 1 |
| ENSMUSG00000107655 | 1.122462048 | 0.608497229 |
| ENSMUSG00000107656 | 1 | |
| ENSMUSG00000107657 | 1 | |

| | | |
|---|---|---|
| ENSMUSG00000107658 | -1.091692751 | 0.698112512 |
| ENSMUSG00000107659 | 1 | |
| ENSMUSG00000107660 | 1 | |
| ENSMUSG00000107661 | 1 | |
| ENSMUSG00000107662 | 1 | |
| ENSMUSG00000107663 | 1.069068979 | 0.608497229 |
| ENSMUSG00000107664 | 1 | |
| ENSMUSG00000107666 | 1 | |
| ENSMUSG00000107667 | 1 | |
| ENSMUSG00000107668 | 1.083937586 | 0.720669657 |
| ENSMUSG00000107669 | 1 | |
| ENSMUSG00000107670 | 1 | |
| ENSMUSG00000107671 | 1 | |
| ENSMUSG00000107672 | 1 | |
| ENSMUSG00000107673 | 1 | |
| ENSMUSG00000107674 | 1 | |
| ENSMUSG00000107675 | 1.059463094 | 0.794274354 |
| ENSMUSG00000107676 | 1.079681442 | 0.697178241 |
| ENSMUSG00000107677 | 1 | |
| ENSMUSG00000107678 | 1 | |
| ENSMUSG00000107679 | 1 | |
| ENSMUSG00000107680 | -1 | 1 |
| ENSMUSG00000107681 | 1.095872691 | 0.608497229 |
| ENSMUSG00000107682 | -1.010960635 | 1 |
| ENSMUSG00000107683 | 1 | |
| ENSMUSG00000107684 | 1 | |
| ENSMUSG00000107685 | 1.005613439 | 1 |
| ENSMUSG00000107686 | 1 | |
| ENSMUSG00000107687 | 1 | |
| ENSMUSG00000107688 | 1.07487334 | 0.802330818 |
| ENSMUSG00000107689 | -1 | 1 |
| ENSMUSG00000107690 | 1.199989356 | 0.045045157 |
| ENSMUSG00000107691 | 1 | |
| ENSMUSG00000107692 | 1 | |
| ENSMUSG00000107693 | 1 | |
| ENSMUSG00000107694 | 1 | |
| ENSMUSG00000107695 | 1 | |
| ENSMUSG00000107696 | 1 | |
| ENSMUSG00000107697 | 1 | |
| ENSMUSG00000107698 | 1.067399953 | 0.908007394 |
| ENSMUSG00000107699 | -1.557930197 | 0.003599328 |
| ENSMUSG00000107700 | 1 | |
| ENSMUSG00000107701 | 1 | |
| ENSMUSG00000107702 | 1 | |
| ENSMUSG00000107703 | -1.063525418 | 0.756517167 |
| ENSMUSG00000107704 | -1.044273782 | 0.855488553 |
| ENSMUSG00000107705 | 1 | |
| ENSMUSG00000107706 | 1 | |
| ENSMUSG00000107707 | 1 | |
| ENSMUSG00000107708 | 1.095872691 | 0.657060635 |
| ENSMUSG00000107709 | 1.054569652 | 0.85284229 |
| ENSMUSG00000107710 | 1 | |
| ENSMUSG00000107711 | 1 | |
| ENSMUSG00000107712 | 1.000946217 | 1 |
| ENSMUSG00000107713 | 1 | |
| ENSMUSG00000107714 | 1 | |
| ENSMUSG00000107715 | 1 | |
| ENSMUSG00000107716 | 1 | |
| ENSMUSG00000107718 | 1 | |
| ENSMUSG00000107719 | 1 | |
| ENSMUSG00000107720 | 1.132639128 | 0.398767938 |
| ENSMUSG00000107721 | 1 | |
| ENSMUSG00000107722 | -1.03962336 | 0.698112512 |

| | | |
|---|---|---|
| ENSMUSG00000107723 | 1 | |
| ENSMUSG00000107724 | 1 | |
| ENSMUSG00000107725 | 1 | |
| ENSMUSG00000107726 | 1.334549714 | 0.129473223 |
| ENSMUSG00000107727 | 1 | |
| ENSMUSG00000107728 | -1.188005937 | 0.344478207 |
| ENSMUSG00000107729 | 1 | |
| ENSMUSG00000107730 | 1.059463094 | 0.698112512 |
| ENSMUSG00000107731 | 1 | |
| ENSMUSG00000107732 | 1 | |
| ENSMUSG00000107733 | -1.03962336 | 0.698112512 |
| ENSMUSG00000107734 | 1.065519011 | 0.834376699 |
| ENSMUSG00000107735 | 1 | |
| ENSMUSG00000107736 | 1 | |
| ENSMUSG00000107737 | 1 | |
| ENSMUSG00000107738 | 1 | |
| ENSMUSG00000107739 | 1 | |
| ENSMUSG00000107740 | 1 | |
| ENSMUSG00000107741 | -1.103790803 | 0.193677799 |
| ENSMUSG00000107742 | -1.021023058 | 0.949812939 |
| ENSMUSG00000107743 | -1.05864359 | 0.51470764 |
| ENSMUSG00000107744 | 1 | |
| ENSMUSG00000107745 | 1.019083579 | 0.950034218 |
| ENSMUSG00000107746 | 1 | |
| ENSMUSG00000107747 | 1.059463094 | 0.699340382 |
| ENSMUSG00000107748 | 1 | |
| ENSMUSG00000107749 | 1.169135088 | 0.286480782 |
| ENSMUSG00000107750 | 1.092608849 | 0.709662609 |
| ENSMUSG00000107751 | 1.178139999 | 0.33317946 |
| ENSMUSG00000107752 | 1 | |
| ENSMUSG00000107753 | 1 | |
| ENSMUSG00000107754 | 1 | |
| ENSMUSG00000107755 | 1 | |
| ENSMUSG00000107756 | 1.037802375 | 0.867407919 |
| ENSMUSG00000107757 | 1 | |
| ENSMUSG00000107758 | -1 | 1 |
| ENSMUSG00000107759 | 1 | |
| ENSMUSG00000107760 | 1.237760021 | 0.21904039 |
| ENSMUSG00000107761 | -1 | 1 |
| ENSMUSG00000107762 | 1 | |
| ENSMUSG00000107763 | 1.094452911 | 0.79007763 |
| ENSMUSG00000107764 | -1.03962336 | 0.698112512 |
| ENSMUSG00000107765 | 1 | |
| ENSMUSG00000107766 | 1 | |
| ENSMUSG00000107767 | 1 | |
| ENSMUSG00000107768 | 1.223735389 | 0.295810793 |
| ENSMUSG00000107769 | 1 | |
| ENSMUSG00000107770 | 1 | |
| ENSMUSG00000107771 | 1.002894704 | 1 |
| ENSMUSG00000107772 | 1.014133649 | 1 |
| ENSMUSG00000107773 | 1 | |
| ENSMUSG00000107774 | 1 | |
| ENSMUSG00000107775 | -1.03962336 | 0.698112512 |
| ENSMUSG00000107776 | 1 | |
| ENSMUSG00000107777 | 1 | |
| ENSMUSG00000107778 | 1 | |
| ENSMUSG00000107779 | -1.263654089 | 0.131831322 |
| ENSMUSG00000107780 | 1 | |
| ENSMUSG00000107781 | 1 | |
| ENSMUSG00000107782 | 1 | |
| ENSMUSG00000107783 | 1 | |
| ENSMUSG00000107784 | 1 | |
| ENSMUSG00000107785 | 1.059463094 | 0.752398528 |

| | | |
|---|---|---|
| ENSMUSG00000107786 | 1 | |
| ENSMUSG00000107787 | 1 | |
| ENSMUSG00000107788 | 1.2649983 | 0.433678407 |
| ENSMUSG00000107789 | 1 | |
| ENSMUSG00000107790 | 1 | |
| ENSMUSG00000107791 | 1 | |
| ENSMUSG00000107792 | 1 | |
| ENSMUSG00000107793 | 1 | |
| ENSMUSG00000107794 | 1.069068979 | 0.698112512 |
| ENSMUSG00000107795 | -1 | 1 |
| ENSMUSG00000107796 | -1.013183986 | 1 |
| ENSMUSG00000107797 | 1 | |
| ENSMUSG00000107798 | 1 | |
| ENSMUSG00000107799 | 1 | |
| ENSMUSG00000107800 | 1 | |
| ENSMUSG00000107801 | 1.069068979 | 0.608497229 |
| ENSMUSG00000107802 | 1.218963704 | 0.331656032 |
| ENSMUSG00000107803 | 1 | |
| ENSMUSG00000107804 | 1.059463094 | 0.698112512 |
| ENSMUSG00000107805 | 1 | |
| ENSMUSG00000107806 | 1 | |
| ENSMUSG00000107808 | 1 | |
| ENSMUSG00000107809 | 1.034095782 | 0.923549054 |
| ENSMUSG00000107810 | 1.00437554 | 1 |
| ENSMUSG00000107811 | 1 | |
| ENSMUSG00000107812 | 1.033906491 | 0.934760809 |
| ENSMUSG00000107813 | -1.143787033 | 0.389301785 |
| ENSMUSG00000107814 | 1.079704071 | 0.698112512 |
| ENSMUSG00000107815 | -1 | 1 |
| ENSMUSG00000107816 | 1 | |
| ENSMUSG00000107817 | 1 | |
| ENSMUSG00000107818 | 1 | |
| ENSMUSG00000107819 | 1 | |
| ENSMUSG00000107821 | 1 | |
| ENSMUSG00000107822 | 1 | |
| ENSMUSG00000107823 | 1 | |
| ENSMUSG00000107824 | 1.335983734 | 0.249885119 |
| ENSMUSG00000107825 | 1 | |
| ENSMUSG00000107826 | 1 | |
| ENSMUSG00000107828 | 1 | |
| ENSMUSG00000107829 | 1 | |
| ENSMUSG00000107830 | 1 | |
| ENSMUSG00000107831 | 1 | |
| ENSMUSG00000107832 | 1 | |
| ENSMUSG00000107833 | 1.015486245 | 0.972980766 |
| ENSMUSG00000107834 | 1 | |
| ENSMUSG00000107835 | 1 | |
| ENSMUSG00000107836 | -1.102039641 | 0.532369657 |
| ENSMUSG00000107837 | 1 | |
| ENSMUSG00000107838 | 1.086694018 | 0.723189216 |
| ENSMUSG00000107839 | 1 | |
| ENSMUSG00000107840 | -1 | 1 |
| ENSMUSG00000107841 | 1.059463094 | 0.698112512 |
| ENSMUSG00000107842 | -1.043306199 | 0.79291651 |
| ENSMUSG00000107844 | 1 | |
| ENSMUSG00000107845 | 1.014545335 | 0.989648415 |
| ENSMUSG00000107846 | 1.010855137 | 1 |
| ENSMUSG00000107847 | 1 | |
| ENSMUSG00000107848 | 1 | |
| ENSMUSG00000107849 | 1 | |
| ENSMUSG00000107850 | 1 | |
| ENSMUSG00000107851 | 1 | |
| ENSMUSG00000107852 | 1.014545335 | 0.991944074 |

| | | |
|---|---|---|
| ENSMUSG00000107853 | -1.02564575 | 0.974143926 |
| ENSMUSG00000107854 | 1 | |
| ENSMUSG00000107855 | 1 | |
| ENSMUSG00000107856 | 1.059463094 | 0.608497229 |
| ENSMUSG00000107857 | 1 | |
| ENSMUSG00000107858 | 1 | |
| ENSMUSG00000107859 | 1 | |
| ENSMUSG00000107861 | -1.084645496 | 0.532369657 |
| ENSMUSG00000107862 | 1 | |
| ENSMUSG00000107863 | -1 | 1 |
| ENSMUSG00000107864 | -1.024718486 | 0.931480186 |
| ENSMUSG00000107865 | 1.034095782 | 0.893303059 |
| ENSMUSG00000107866 | 1 | |
| ENSMUSG00000107867 | 1 | |
| ENSMUSG00000107868 | 1.153860772 | 0.327429155 |
| ENSMUSG00000107869 | 1 | |
| ENSMUSG00000107870 | -1.291104262 | 0.381036055 |
| ENSMUSG00000107872 | 1 | |
| ENSMUSG00000107873 | 1.059463094 | 0.608497229 |
| ENSMUSG00000107874 | 1 | |
| ENSMUSG00000107875 | -1.044273782 | 0.698112512 |
| ENSMUSG00000107876 | 1 | |
| ENSMUSG00000107877 | -1.052712573 | 0.5713094 |
| ENSMUSG00000107878 | -1.043306199 | 0.698112512 |
| ENSMUSG00000107879 | -1.128758979 | 0.159131403 |
| ENSMUSG00000107880 | 1.069068979 | 0.608497229 |
| ENSMUSG00000107881 | 1.777234399 | 0.000946345 |
| ENSMUSG00000107882 | 1.06585783 | 0.799439325 |
| ENSMUSG00000107883 | 1.083961291 | 0.769272045 |
| ENSMUSG00000107884 | 1.019083579 | 0.949812939 |
| ENSMUSG00000107885 | 1 | |
| ENSMUSG00000107886 | 1.07197773 | 0.698112512 |
| ENSMUSG00000107887 | 1 | |
| ENSMUSG00000107888 | 1 | |
| ENSMUSG00000107889 | 1 | |
| ENSMUSG00000107890 | 1 | |
| ENSMUSG00000107891 | 1.028323353 | 0.959463747 |
| ENSMUSG00000107892 | -1.254679684 | 0.041103668 |
| ENSMUSG00000107893 | 1 | |
| ENSMUSG00000107894 | 1.183481569 | 0.617510841 |
| ENSMUSG00000107895 | 1 | |
| ENSMUSG00000107896 | 1 | |
| ENSMUSG00000107898 | 1 | |
| ENSMUSG00000107899 | 1 | |
| ENSMUSG00000107900 | 1 | |
| ENSMUSG00000107901 | 1 | |
| ENSMUSG00000107902 | 1.076548228 | 0.807247723 |
| ENSMUSG00000107903 | 1.069068979 | 0.608497229 |
| ENSMUSG00000107904 | 1 | |
| ENSMUSG00000107905 | 1 | |
| ENSMUSG00000107906 | -1.03962336 | 0.814089245 |
| ENSMUSG00000107907 | 1 | |
| ENSMUSG00000107908 | 1 | |
| ENSMUSG00000107909 | -1.027966894 | 0.968937442 |
| ENSMUSG00000107911 | -1.024162104 | 0.964236174 |
| ENSMUSG00000107913 | 1 | |
| ENSMUSG00000107914 | 1 | |
| ENSMUSG00000107915 | 1.030689571 | 0.982838256 |
| ENSMUSG00000107916 | 1 | |
| ENSMUSG00000107917 | 1 | |
| ENSMUSG00000107918 | -1 | 1 |
| ENSMUSG00000107919 | 1 | |
| ENSMUSG00000107920 | 1 | |

| | | |
|---|---|---|
| ENSMUSG00000107921 | 1 | |
| ENSMUSG00000107922 | 1.005613439 | 1 |
| ENSMUSG00000107923 | 1 | |
| ENSMUSG00000107924 | 1 | |
| ENSMUSG00000107925 | 1 | |
| ENSMUSG00000107926 | 1 | |
| ENSMUSG00000107927 | -1.083415624 | 0.698112512 |
| ENSMUSG00000107928 | -1.065096629 | 0.847006151 |
| ENSMUSG00000107929 | 1 | |
| ENSMUSG00000107930 | 1 | |
| ENSMUSG00000107931 | 1 | |
| ENSMUSG00000107932 | 1.121613511 | 0.652685575 |
| ENSMUSG00000107933 | 1 | |
| ENSMUSG00000107934 | 1 | |
| ENSMUSG00000107935 | 1 | |
| ENSMUSG00000107936 | 1.059463094 | 0.608497229 |
| ENSMUSG00000107937 | 1.04822434 | 0.80455706 |
| ENSMUSG00000107938 | 1 | |
| ENSMUSG00000107940 | 1 | |
| ENSMUSG00000107941 | 1 | |
| ENSMUSG00000107942 | 1.059463094 | 0.608497229 |
| ENSMUSG00000107943 | 1 | |
| ENSMUSG00000107944 | -1.119888335 | 0.717035356 |
| ENSMUSG00000107945 | -1.095294215 | 0.541735653 |
| ENSMUSG00000107946 | 1.089620613 | 0.763407099 |
| ENSMUSG00000107947 | 1 | |
| ENSMUSG00000107948 | 1 | |
| ENSMUSG00000107949 | 1.015486245 | 0.972980766 |
| ENSMUSG00000107950 | 1.270716635 | 0.109982432 |
| ENSMUSG00000107951 | -1.08226438 | 0.755839148 |
| ENSMUSG00000107952 | 1.005613439 | 1 |
| ENSMUSG00000107953 | 1 | |
| ENSMUSG00000107954 | -1.033575217 | 0.949218699 |
| ENSMUSG00000107955 | 1.011497958 | 1 |
| ENSMUSG00000107956 | 1 | |
| ENSMUSG00000107957 | 1 | |
| ENSMUSG00000107959 | 1 | |
| ENSMUSG00000107960 | 1 | |
| ENSMUSG00000107961 | 1.023743961 | 0.95702926 |
| ENSMUSG00000107962 | 1.07269031 | 0.698112512 |
| ENSMUSG00000107964 | 1 | |
| ENSMUSG00000107965 | 1 | |
| ENSMUSG00000107966 | 1 | |
| ENSMUSG00000107967 | -1.07062035 | 0.804890812 |
| ENSMUSG00000107968 | 1 | |
| ENSMUSG00000107969 | 1 | |
| ENSMUSG00000107970 | -1.035541219 | 0.949990903 |
| ENSMUSG00000107971 | 1.03893347 | 0.730297627 |
| ENSMUSG00000107972 | -1.053549062 | 0.769745556 |
| ENSMUSG00000107973 | 1 | |
| ENSMUSG00000107975 | -1 | 1 |
| ENSMUSG00000107976 | -1.23938702 | 0.111942377 |
| ENSMUSG00000107977 | -1.016549018 | 1 |
| ENSMUSG00000107978 | 1 | |
| ENSMUSG00000107979 | 1 | |
| ENSMUSG00000107980 | 1 | |
| ENSMUSG00000107981 | -1.044273782 | 0.698112512 |
| ENSMUSG00000107982 | 1 | |
| ENSMUSG00000107983 | 1 | |
| ENSMUSG00000107984 | 1.014545335 | 0.979002724 |
| ENSMUSG00000107985 | -1.316625637 | 0.127490223 |
| ENSMUSG00000107986 | 1 | |
| ENSMUSG00000107987 | 1 | |

| | | |
|---|---|---|
| ENSMUSG00000107988 | 1 | |
| ENSMUSG00000107989 | 1.059208266 | 0.937535342 |
| ENSMUSG00000107990 | -1.042780998 | 0.795609888 |
| ENSMUSG00000107991 | -1.029113823 | 0.937805719 |
| ENSMUSG00000107992 | -1 | 1 |
| ENSMUSG00000107993 | 1 | |
| ENSMUSG00000107994 | 1.231239424 | 0.461535898 |
| ENSMUSG00000107995 | -1.043306199 | 0.858911269 |
| ENSMUSG00000107996 | 1.033906491 | 0.954348776 |
| ENSMUSG00000107997 | 1.131418094 | 0.608497229 |
| ENSMUSG00000107998 | 1 | |
| ENSMUSG00000107999 | 1 | |
| ENSMUSG00000108000 | 1 | |
| ENSMUSG00000108002 | 1 | |
| ENSMUSG00000108003 | 1 | |
| ENSMUSG00000108004 | 1.049459984 | 0.881180973 |
| ENSMUSG00000108005 | 1 | |
| ENSMUSG00000108006 | 1.059463094 | 0.608497229 |
| ENSMUSG00000108007 | 1 | |
| ENSMUSG00000108008 | -1.038444538 | 0.865865121 |
| ENSMUSG00000108009 | -1.03962336 | 0.908842909 |
| ENSMUSG00000108010 | 1.233901898 | 0.328793916 |
| ENSMUSG00000108011 | 1 | |
| ENSMUSG00000108012 | 1 | |
| ENSMUSG00000108013 | 1 | |
| ENSMUSG00000108014 | 1 | |
| ENSMUSG00000108015 | 1.069068979 | 0.608497229 |
| ENSMUSG00000108016 | 1 | |
| ENSMUSG00000108017 | 1 | |
| ENSMUSG00000108018 | 1 | |
| ENSMUSG00000108019 | 1 | |
| ENSMUSG00000108020 | 1.342968208 | 0.001804015 |
| ENSMUSG00000108021 | -1 | 1 |
| ENSMUSG00000108022 | 1 | |
| ENSMUSG00000108023 | 1.059463094 | 0.698112512 |
| ENSMUSG00000108024 | 1.046403186 | 0.868236615 |
| ENSMUSG00000108025 | 1 | |
| ENSMUSG00000108026 | 1.01487975 | 0.979002724 |
| ENSMUSG00000108027 | -1.15117829 | 0.553862807 |
| ENSMUSG00000108028 | 1 | |
| ENSMUSG00000108029 | 1 | |
| ENSMUSG00000108030 | -1.016963296 | 1 |
| ENSMUSG00000108032 | 1 | |
| ENSMUSG00000108033 | 1.045900989 | 0.897690067 |
| ENSMUSG00000108034 | 1.052303466 | 0.840505213 |
| ENSMUSG00000108035 | 1 | |
| ENSMUSG00000108036 | 1 | |
| ENSMUSG00000108037 | -1.026049019 | 0.922665422 |
| ENSMUSG00000108038 | 1 | |
| ENSMUSG00000108039 | 1 | |
| ENSMUSG00000108040 | 1.11166259 | 0.608497229 |
| ENSMUSG00000108041 | 1 | |
| ENSMUSG00000108042 | -1.013597819 | 0.973599977 |
| ENSMUSG00000108043 | -1.03962336 | 0.698112512 |
| ENSMUSG00000108044 | 1 | |
| ENSMUSG00000108045 | 1.059463094 | 0.699340382 |
| ENSMUSG00000108046 | 1.011529912 | 1 |
| ENSMUSG00000108047 | 1 | |
| ENSMUSG00000108048 | -1.057338222 | 0.822702238 |
| ENSMUSG00000108049 | 1.160103591 | 0.193986794 |
| ENSMUSG00000108050 | -1.044273782 | 0.698112512 |
| ENSMUSG00000108052 | 1 | |
| ENSMUSG00000108053 | 1.131247234 | 0.368508709 |

| | | |
|---|---|---|
| ENSMUSG00000108054 | 1 | |
| ENSMUSG00000108055 | 1.120729039 | 0.746828674 |
| ENSMUSG00000108056 | -1.080029458 | 0.720662147 |
| ENSMUSG00000108057 | 1.059463094 | 0.704170349 |
| ENSMUSG00000108058 | 1 | |
| ENSMUSG00000108059 | 1.060198483 | 0.913882562 |
| ENSMUSG00000108060 | -1.044273782 | 0.790851299 |
| ENSMUSG00000108061 | 1 | |
| ENSMUSG00000108062 | 1 | |
| ENSMUSG00000108063 | 1 | |
| ENSMUSG00000108064 | -1.139090732 | 0.608497229 |
| ENSMUSG00000108065 | 1 | |
| ENSMUSG00000108066 | 1 | |
| ENSMUSG00000108067 | 1 | |
| ENSMUSG00000108068 | -1.179226843 | 0.397706863 |
| ENSMUSG00000108069 | 1 | |
| ENSMUSG00000108070 | 1 | |
| ENSMUSG00000108071 | 1 | |
| ENSMUSG00000108072 | -1.093466951 | 0.698112512 |
| ENSMUSG00000108073 | 1 | |
| ENSMUSG00000108074 | 1 | |
| ENSMUSG00000108075 | 1 | |
| ENSMUSG00000108076 | 1 | |
| ENSMUSG00000108077 | -1.03962336 | 0.698112512 |
| ENSMUSG00000108078 | -1 | 1 |
| ENSMUSG00000108079 | -1.077732257 | 0.850281678 |
| ENSMUSG00000108080 | -1.190401641 | 0.398797577 |
| ENSMUSG00000108081 | 1 | |
| ENSMUSG00000108082 | 1 | |
| ENSMUSG00000108083 | 1 | |
| ENSMUSG00000108084 | 1.065093972 | 0.842680402 |
| ENSMUSG00000108085 | -1.053549062 | 0.698112512 |
| ENSMUSG00000108086 | 1.095872691 | 0.671280603 |
| ENSMUSG00000108087 | 1 | |
| ENSMUSG00000108088 | 1 | |
| ENSMUSG00000108089 | 1.160346728 | 0.650779496 |
| ENSMUSG00000108090 | 1 | |
| ENSMUSG00000108091 | 1 | |
| ENSMUSG00000108092 | 1.068154587 | 0.812927638 |
| ENSMUSG00000108093 | 1 | |
| ENSMUSG00000108094 | 1 | |
| ENSMUSG00000108095 | 1 | |
| ENSMUSG00000108096 | 1 | |
| ENSMUSG00000108097 | 1 | |
| ENSMUSG00000108098 | 1 | |
| ENSMUSG00000108099 | 1 | |
| ENSMUSG00000108100 | -1.025896084 | 0.988622398 |
| ENSMUSG00000108101 | -1.102689795 | 0.698112512 |
| ENSMUSG00000108102 | -1.037482356 | 0.870427333 |
| ENSMUSG00000108103 | 1.161036672 | 0.378751351 |
| ENSMUSG00000108104 | 1 | |
| ENSMUSG00000108105 | 1.074221595 | 0.790772938 |
| ENSMUSG00000108106 | 1 | |
| ENSMUSG00000108107 | -1.084645496 | 0.617024566 |
| ENSMUSG00000108108 | 1.12316944 | 0.641554903 |
| ENSMUSG00000108109 | -1.010960635 | 1 |
| ENSMUSG00000108110 | 1 | |
| ENSMUSG00000108111 | 1 | |
| ENSMUSG00000108112 | 1 | |
| ENSMUSG00000108113 | 1 | |
| ENSMUSG00000108114 | 1 | |
| ENSMUSG00000108115 | -1.029323987 | 0.97088369 |
| ENSMUSG00000108117 | 1 | |

| | | |
|---|---|---|
| ENSMUSG00000108118 | 1 | |
| ENSMUSG00000108119 | -1.043306199 | 0.698112512 |
| ENSMUSG00000108120 | 1.049137052 | 0.815807184 |
| ENSMUSG00000108121 | 1 | |
| ENSMUSG00000108122 | -1.134332281 | 0.713405045 |
| ENSMUSG00000108123 | 1 | |
| ENSMUSG00000108124 | -1.090507733 | 0.698112512 |
| ENSMUSG00000108125 | 1.010259986 | 1 |
| ENSMUSG00000108126 | 1.290681034 | 0.008486769 |
| ENSMUSG00000108127 | -1 | 1 |
| ENSMUSG00000108128 | 1 | |
| ENSMUSG00000108129 | 1.10330766 | 0.761235181 |
| ENSMUSG00000108130 | 1 | |
| ENSMUSG00000108131 | 1 | |
| ENSMUSG00000108132 | 1.353963366 | 0.037790746 |
| ENSMUSG00000108133 | 1 | |
| ENSMUSG00000108134 | 1 | |
| ENSMUSG00000108135 | 1 | |
| ENSMUSG00000108136 | 1 | |
| ENSMUSG00000108137 | 1.059766981 | 0.811511816 |
| ENSMUSG00000108138 | 1 | |
| ENSMUSG00000108139 | 1 | |
| ENSMUSG00000108140 | 1 | |
| ENSMUSG00000108141 | 1.059463094 | 0.608497229 |
| ENSMUSG00000108142 | -1.11686982 | 0.698112512 |
| ENSMUSG00000108143 | -1.158106243 | 0.589226004 |
| ENSMUSG00000108144 | 1 | |
| ENSMUSG00000108145 | -1 | 1 |
| ENSMUSG00000108146 | 1 | |
| ENSMUSG00000108147 | 1 | |
| ENSMUSG00000108148 | 1 | |
| ENSMUSG00000108149 | 1 | |
| ENSMUSG00000108150 | 1 | |
| ENSMUSG00000108151 | 1 | |
| ENSMUSG00000108152 | -1.03962336 | 0.698112512 |
| ENSMUSG00000108153 | 1 | |
| ENSMUSG00000108154 | 1.192886193 | 0.449109628 |
| ENSMUSG00000108155 | -1.095360949 | 0.608497229 |
| ENSMUSG00000108156 | 1 | |
| ENSMUSG00000108157 | 1 | |
| ENSMUSG00000108158 | 1 | |
| ENSMUSG00000108160 | 1 | |
| ENSMUSG00000108161 | 1 | |
| ENSMUSG00000108162 | 1.040410207 | 0.867753156 |
| ENSMUSG00000108163 | -1 | 1 |
| ENSMUSG00000108164 | 1 | |
| ENSMUSG00000108166 | 1.096843606 | 0.608497229 |
| ENSMUSG00000108167 | 1 | |
| ENSMUSG00000108168 | 1.059463094 | 0.608497229 |
| ENSMUSG00000108169 | 1 | |
| ENSMUSG00000108170 | -1.043306199 | 0.79291651 |
| ENSMUSG00000108171 | 1.015486245 | 0.985522133 |
| ENSMUSG00000108172 | 1 | |
| ENSMUSG00000108173 | 1.082107178 | 0.583420921 |
| ENSMUSG00000108174 | 1 | |
| ENSMUSG00000108175 | 1 | |
| ENSMUSG00000108176 | 1.035942902 | 0.839007844 |
| ENSMUSG00000108177 | -1.043306199 | 0.803260194 |
| ENSMUSG00000108178 | 1 | |
| ENSMUSG00000108179 | 1.107299027 | 0.738937719 |
| ENSMUSG00000108180 | -1.038444538 | 0.895498153 |
| ENSMUSG00000108181 | 1.016311457 | 0.946360929 |
| ENSMUSG00000108183 | 1.294557435 | 0.216046507 |

| | | |
|---|---|---|
| ENSMUSG00000108184 | 1 | |
| ENSMUSG00000108185 | 1 | |
| ENSMUSG00000108186 | -1.053549062 | 0.698112512 |
| ENSMUSG00000108187 | 1 | |
| ENSMUSG00000108188 | -1.03962336 | 0.698112512 |
| ENSMUSG00000108189 | 1.053245793 | 0.803072249 |
| ENSMUSG00000108190 | 1.076888206 | 0.828391261 |
| ENSMUSG00000108191 | 1 | |
| ENSMUSG00000108192 | 1 | |
| ENSMUSG00000108193 | 1.059463094 | 0.608497229 |
| ENSMUSG00000108194 | 1 | |
| ENSMUSG00000108195 | 1.119867748 | 0.708232777 |
| ENSMUSG00000108196 | -1.044273782 | 0.790851299 |
| ENSMUSG00000108197 | 1.000528147 | 1 |
| ENSMUSG00000108198 | 1 | |
| ENSMUSG00000108199 | -1.000156392 | 1 |
| ENSMUSG00000108200 | 1.206990366 | 0.589121246 |
| ENSMUSG00000108201 | 1 | |
| ENSMUSG00000108202 | -1 | 1 |
| ENSMUSG00000108203 | 1 | |
| ENSMUSG00000108204 | -1.177310696 | 0.408897318 |
| ENSMUSG00000108205 | 1 | |
| ENSMUSG00000108206 | 1.059463094 | 0.608497229 |
| ENSMUSG00000108207 | 1.504360668 | 0.00139517 |
| ENSMUSG00000108208 | 1 | |
| ENSMUSG00000108209 | 1 | |
| ENSMUSG00000108210 | -1.084645496 | 0.617024566 |
| ENSMUSG00000108211 | 1.084618945 | 0.698112512 |
| ENSMUSG00000108212 | 1 | |
| ENSMUSG00000108213 | 1 | |
| ENSMUSG00000108214 | -1.05948903 | 0.725549871 |
| ENSMUSG00000108215 | -1.063525418 | 0.751979636 |
| ENSMUSG00000108216 | -1.092661439 | 0.733855391 |
| ENSMUSG00000108217 | 1.030256832 | 0.90028574 |
| ENSMUSG00000108218 | 1.050235211 | 0.534652019 |
| ENSMUSG00000108219 | -1 | 1 |
| ENSMUSG00000108220 | 1 | |
| ENSMUSG00000108221 | 1 | |
| ENSMUSG00000108222 | 1 | |
| ENSMUSG00000108223 | 1 | |
| ENSMUSG00000108224 | -1.053549062 | 0.698112512 |
| ENSMUSG00000108225 | 1 | |
| ENSMUSG00000108226 | 1 | |
| ENSMUSG00000108227 | 1.171809639 | 0.526467236 |
| ENSMUSG00000108228 | 1.095872691 | 0.657060635 |
| ENSMUSG00000108229 | 1 | |
| ENSMUSG00000108230 | 1 | |
| ENSMUSG00000108231 | -1.043306199 | 0.873296215 |
| ENSMUSG00000108232 | 1.170927719 | 0.608497229 |
| ENSMUSG00000108233 | -1.03962336 | 0.698112512 |
| ENSMUSG00000108234 | -1.054000126 | 0.789891887 |
| ENSMUSG00000108235 | -1 | 1 |
| ENSMUSG00000108236 | 1 | |
| ENSMUSG00000108237 | -1.010032465 | 1 |
| ENSMUSG00000108238 | 1.134720919 | 0.661328906 |
| ENSMUSG00000108239 | 1.090713796 | 0.734821651 |
| ENSMUSG00000108240 | 1 | |
| ENSMUSG00000108241 | 1 | |
| ENSMUSG00000108242 | 1 | |
| ENSMUSG00000108244 | 1 | |
| ENSMUSG00000108245 | 1 | |
| ENSMUSG00000108246 | 1.069068979 | 0.608497229 |
| ENSMUSG00000108247 | 1 | |

| | | |
|---|---|---|
| ENSMUSG00000108248 | 1 | |
| ENSMUSG00000108249 | 1.091478447 | 0.608497229 |
| ENSMUSG00000108250 | 1 | |
| ENSMUSG00000108251 | 1 | |
| ENSMUSG00000108252 | 1 | |
| ENSMUSG00000108253 | 1 | |
| ENSMUSG00000108254 | 1.199989356 | 0.17554978 |
| ENSMUSG00000108255 | -1.053549062 | 0.698112512 |
| ENSMUSG00000108256 | -1.130264193 | 0.532369657 |
| ENSMUSG00000108257 | 1.142219309 | 0.608497229 |
| ENSMUSG00000108258 | 1.138788635 | 0.595042751 |
| ENSMUSG00000108259 | 1 | |
| ENSMUSG00000108260 | 1.069068979 | 0.698112512 |
| ENSMUSG00000108261 | 1 | |
| ENSMUSG00000108262 | 1 | |
| ENSMUSG00000108263 | 1.014545335 | 0.979002724 |
| ENSMUSG00000108264 | 1 | |
| ENSMUSG00000108265 | 1 | |
| ENSMUSG00000108266 | -1.057420981 | 0.828564184 |
| ENSMUSG00000108267 | -1.089814156 | 0.781214355 |
| ENSMUSG00000108268 | 1.089956786 | 0.793885968 |
| ENSMUSG00000108269 | 1 | |
| ENSMUSG00000108270 | -1.182908789 | 0.608497229 |
| ENSMUSG00000108271 | 1 | |
| ENSMUSG00000108272 | 1 | |
| ENSMUSG00000108273 | -1.069502966 | 0.698112512 |
| ENSMUSG00000108274 | 1 | |
| ENSMUSG00000108275 | 1 | |
| ENSMUSG00000108276 | -1.003834322 | 1 |
| ENSMUSG00000108277 | 1 | |
| ENSMUSG00000108278 | -1.116853908 | 0.558122732 |
| ENSMUSG00000108279 | 1.059463094 | 0.704170349 |
| ENSMUSG00000108280 | -1.111880267 | 0.572947856 |
| ENSMUSG00000108282 | 1.059463094 | 0.776953697 |
| ENSMUSG00000108283 | 1.132639128 | 0.414898324 |
| ENSMUSG00000108284 | -1 | 1 |
| ENSMUSG00000108285 | 1 | |
| ENSMUSG00000108286 | 1 | |
| ENSMUSG00000108287 | 1 | |
| ENSMUSG00000108288 | 1.036199311 | 0.931170284 |
| ENSMUSG00000108289 | 1 | |
| ENSMUSG00000108290 | 1.017302764 | 1 |
| ENSMUSG00000108291 | 1.12352659 | 0.366138691 |
| ENSMUSG00000108292 | 1.019083579 | 0.971519533 |
| ENSMUSG00000108293 | 1.200936955 | 0.215991492 |
| ENSMUSG00000108294 | 1 | |
| ENSMUSG00000108295 | 1 | |
| ENSMUSG00000108296 | 1 | |
| ENSMUSG00000108297 | -1.03962336 | 0.698112512 |
| ENSMUSG00000108298 | -1.020155149 | 0.989387709 |
| ENSMUSG00000108299 | 1 | |
