## Supplemental table 4 for "Nuclear actin-dependent Meg3 expression suppresses metabolic genes by affecting the chromatin architecture at sites of elevated H3K27 acetylation levels"

**Supplementary table 4**

**ChIRP oligos biotinilated probes**

| | |
|---|---|
| Meg3-1 | AGCCAGGTAAGAGATTCTTC |
| Meg3-2 | AGTTGCTCTGTATGGTAGTG |
| Meg3-3 | GCACTGGTTCAAGGTTTGAA |
| Meg3-4 | CTAGCAGATGAACACGAGCA |
| Meg3-5 | TGAGAGGTTCCCAAAGGGAC |
| Meg3-6 | AAGGGACATCTCCGAGAAGG |
| Meg3-7 | TTGAGACACATGAGCCATGA |
| Meg3-8 | CAATCTCATGAGTGAGTCCG |
| Meg3-9 | GGGGAAATTGGAGGTGAGGA |
| Meg3-10 | GTGGAACCTGAGCACAACAG |
| Meg3-11 | AACGTGTTGTGCGTGAAGTC |
| Meg3-12 | GCTTCCAATCGATTTACAGT |
| Meg3-13 | AAGACAGAAAACCTGCCCAT |
| Meg3-14 | CGGAACAAGAGTCCATTTGT |
| Meg3-15 | CTCTGGGCTCTAGTGAATGG |
| Meg3-16 | ATGTTCTTTTACCACCAGAT |

**ChIP-qpcr**

| | |
|---|---|
| Meg3/GTL2 -F1 | TCCTCACCTCCAATTTCCCCT |
| Meg3/GTL2-R1 | GAGCGAGAGCCGTTCGATG |
| Hs3st3b1-F | GCGGGCATTGCTGGAGTTCCTG |
| Hs3st3b1-R | GGGTTCTGGGCATCAAGTCTCGGTAC |
| PLD5-2-F | AGCGGTGGACATCATGGGA |
| PLD5-2-R | AAGAGCGGTAAGTGAAACGGG |
| Malat1-F1 | GCCTTTTGTCACCTCACT |
| Malat1-R1 | CAAACTCACTGCAAGGTCTC |

BiotinTEG
BiotinTEG
BiotinTEG
BiotinTEG
BiotinTEG
BiotinTEG
BiotinTEG
BiotinTEG
BiotinTEG
BiotinTEG
BiotinTEG
BiotinTEG
BiotinTEG
BiotinTEG
BiotinTEG
BiotinTEG
