## Supplemental table 5 for "Nuclear actin-dependent Meg3 expression suppresses metabolic genes by affecting the chromatin architecture at sites of elevated H3K27 acetylation levels"

**Supplementary figure 5**

| seqnames | start | end | width | strand | V4 | V5 | annotation | geneChr | geneStart | geneEnd | geneLength | geneStrand | geneId | transcriptId | distanceToTS | ENSEMBL | SYMBOL | GENENAME |
|---|---|---|---|---|---|---|---|---|---|---|---|---|---|---|---|---|---|---|
| chr1 | 3138982 | 3139134 | 153 | * | NA_peak_2 | 26 | Distal Interge | 1 | 3214482 | 3671498 | 457017 | 2 | 497097 | ENSMUST00 | 532364 | ENSMUSG0 | Xkr4 | X-linked Kx blood group related 4 |
| chr1 | 4020023 | 4020192 | 170 | * | NA_peak_12 | 23 | Intron (ENSM | 1 | 4344146 | 4360314 | 16169 | 2 | 19888 | ENSMUST00 | 340122 | ENSMUSG0 | Rp1 | retinitis pigmentosa 1 (human) |
| chr1 | 4493413 | 4493743 | 331 | * | NA_peak_17 | 19 | Promoter (<= | 1 | 4492465 | 4493735 | 1271 | 2 | 20671 | ENSMUST00 | 0 | ENSMUSG0 | Sox17 | SRY (sex determining region Y)-box 17 |
| chr1 | 4559853 | 4560187 | 335 | * | NA_peak_18 | 23 | Distal Interge | 1 | 4491390 | 4497354 | 5965 | 2 | 20671 | ENSMUST00 | -62499 | ENSMUSG0 | Sox17 | SRY (sex determining region Y)-box 17 |
| chr1 | 11781528 | 11781722 | 195 | * | NA_peak_10 | 21 | Intron (ENSM | 1 | 11579306 | 11596631 | 17326 | 1 | 320492 | ENSMUST00 | 202222 | ENSMUSG0 | A830018L16 | RIKEN cDNA A830018L16 gene |
| chr1 | 14088301 | 14088609 | 309 | * | NA_peak_13 | 23 | Distal Interge | 1 | 14207926 | 14231492 | 23567 | 2 | 14048 | ENSMUST00 | 142883 | ENSMUSG0 | Eya1 | EYA transcriptional coactivator and phosphatase 1 |
| chr1 | 15082335 | 15082521 | 187 | * | NA_peak_14 | 13 | Distal Interge | 1 | 14872648 | 14918862 | 46215 | 2 | 277328 | ENSMUST00 | -163473 | ENSMUSG0 | Trpa1 | transient receptor potential cation channel, subfamily A, member 1 |
| chr1 | 17384061 | 17384529 | 469 | * | NA_peak_18 | 21 | Distal Interge | 1 | 17601901 | 17630939 | 29039 | 1 | 94227 | ENSMUST00 | -217372 | ENSMUSG0 | Pi15 | peptidase inhibitor 15 |
| chr1 | 18451231 | 18451416 | 186 | * | NA_peak_19 | 13 | Distal Interge | 1 | 18250980 | 18265138 | 14159 | 2 | 77673 | ENSMUST00 | -186093 | ENSMUSG0 | Defb41 | defensin beta 41 |
| chr1 | 19129976 | 19130334 | 359 | * | NA_peak_20 | 40 | Intron (ENSM | 1 | 19107181 | 19148367 | 41187 | 1 | 226896 | ENSMUST00 | 22795 | ENSMUSG0 | Tfap2d | transcription factor AP-2, delta |
| chr1 | 21200023 | 21200680 | 658 | * | NA_peak_22 | 36 | Distal Interge | 1 | 21218575 | 21230167 | 11593 | 1 | 75712 | ENSMUST00 | -17895 | ENSMUSG0 | Tmem14a | transmembrane protein 14A |
| chr1 | 21999141 | 21999332 | 192 | * | NA_peak_23 | 21 | Distal Interge | 1 | 21398408 | 21961942 | 563535 | 2 | 226922 | ENSMUST00 | -37199 | ENSMUSG0 | Kcnq5 | potassium voltage-gated channel, subfamily Q, member 5 |
| chr1 | 22162984 | 22163021 | 38 | * | NA_peak_23 | 24 | Distal Interge | 1 | 22286915 | 22315805 | 28891 | 2 | 116837 | ENSMUST00 | 152784 | ENSMUSG0 | Rims1 | regulating synaptic membrane exocytosis 1 |
| chr1 | 24611608 | 24616109 | 4502 | * | NA_peak_26 | 1007 | Exon (ENSMU | 1 | 24263994 | 24587472 | 323479 | 2 | 12823 | ENSMUST00 | -24136 | ENSMUSG0 | Col19a1 | collagen, type XIX, alpha 1 |
| chr1 | 27755730 | 27756058 | 329 | * | NA_peak_28 | 51 | Distal Interge | 1 | 27884046 | 27902396 | 18351 | 1 | 433281 | ENSMUST00 | -127988 | ENSMUSG0 | Gm5524 | predicted gene 5524 |
| chr1 | 28188681 | 28188967 | 287 | * | NA_peak_28 | 49 | Distal Interge | 1 | 27884046 | 27902396 | 18351 | 1 | 433281 | ENSMUST00 | 304635 | ENSMUSG0 | Gm5524 | predicted gene 5524 |
| chr1 | 29266041 | 29266331 | 291 | * | NA_peak_29 | 17 | Distal Interge | 1 | 29421488 | 29421612 | 125 | 1 | 102465174 | ENSMUST00 | -155157 | ENSMUSG0 | Mir6342 | microRNA 6342 |
| chr1 | 30165242 | 30165389 | 148 | * | NA_peak_31 | 31 | Distal Interge | 1 | 30806205 | 30809105 | 2901 | 2 | 213109 | ENSMUST00 | 643716 | ENSMUSG0 | Phf3 | PHD finger protein 3 |
| chr1 | 30307072 | 30307363 | 292 | * | NA_peak_31 | 27 | Distal Interge | 1 | 30806205 | 30809105 | 2901 | 2 | 213109 | ENSMUST00 | 501742 | ENSMUSG0 | Phf3 | PHD finger protein 3 |
| chr1 | 31075433 | 31075977 | 545 | * | NA_peak_32 | 33 | Intron (ENSM | 1 | 31019188 | 31096454 | 77267 | 2 | 102634473 | ENSMUST00 | 20477 | ENSMUSG0 | Gm29669 | predicted gene 29669 |
| chr1 | 31699848 | 31700133 | 286 | * | NA_peak_32 | 18 | Distal Interge | 1 | 32172714 | 32658568 | 485855 | 1 | 170771 | ENSMUST00 | -472581 | ENSMUSG0 | Khdrbs2 | KH domain containing, RNA binding, signal transduction associated 2 |
| chr1 | 32067153 | 32067567 | 415 | * | NA_peak_33 | 21 | Distal Interge | 1 | 32172714 | 32658568 | 485855 | 1 | 170771 | ENSMUST00 | -105147 | ENSMUSG0 | Khdrbs2 | KH domain containing, RNA binding, signal transduction associated 2 |
| chr1 | 33681650 | 33681939 | 290 | * | NA_peak_35 | 49 | Distal Interge | 1 | 33669843 | 33670712 | 870 | 1 | 67503 | ENSMUST00 | 11807 | ENSMUSG0 | 1700001G17 | RIKEN cDNA 1700001G17 gene |
| chr1 | 34417766 | 34418054 | 289 | * | NA_peak_36 | 20 | Exon (ENSMU | 1 | 34433121 | 34433199 | 79 | 2 | 100628580 | ENSMUST00 | 15145 | ENSMUSG0 | Mir5103 | microRNA 5103 |
| chr1 | 34459657 | 34460010 | 354 | * | NA_peak_36 | 36 | Promoter (<= | 1 | 34459765 | 34473784 | 14020 | 1 | 19253 | ENSMUST00 | 0 | ENSMUSG0 | Ptpn18 | protein tyrosine phosphatase, non-receptor type 18 |
| chr1 | 36521605 | 36521724 | 120 | * | NA_peak_43 | 27 | Promoter (<= | 1 | 36520761 | 36525600 | 4840 | 1 | 94218 | ENSMUST00 | 844 | ENSMUSG0 | Cnnm3 | cyclin M3 |
| chr1 | 41469491 | 41469829 | 339 | * | NA_peak_51 | 27 | Distal Interge | 1 | 41167587 | 41181252 | 13666 | 2 | 74897 | ENSMUST00 | -288239 | ENSMUSG0 | 4930448I06R | RIKEN cDNA 4930448I06 gene |
| chr1 | 42335881 | 42336315 | 435 | * | NA_peak_52 | 45 | Intron (ENSM | 1 | 42648200 | 42651679 | 3480 | 2 | 66297 | ENSMUST00 | 315364 | ENSMUSG0 | Pantr1 | POU domain, class 3, transcription factor 3 adjacent noncoding transcript 1 |
| chr1 | 43851272 | 43851571 | 300 | * | NA_peak_54 | 16 | Distal Interge | 1 | 43765007 | 43827800 | 62794 | 2 | 67883 | ENSMUST00 | -23472 | ENSMUSG0 | Uxs1 | UDP-glucuronate decarboxylase 1 |
| chr1 | 44102308 | 44102634 | 327 | * | NA_peak_55 | 20 | Promoter (<= | 1 | 44086613 | 44102391 | 15779 | 2 | 75623 | ENSMUST00 | 0 | ENSMUSG0 | Tex30 | testis expressed 30 |
| chr1 | 44284243 | 44284513 | 271 | * | NA_peak_55 | 28 | Distal Interge | 1 | 44204070 | 44218961 | 14892 | 2 | 403183 | ENSMUST00 | -65282 | ENSMUSG0 | Mettl21e | methyltransferase like 21E |
| chr1 | 44478517 | 44478649 | 133 | * | NA_peak_56 | 33 | Distal Interge | 1 | 44551511 | 44796838 | 245328 | 1 | 70676 | ENSMUST00 | -72862 | ENSMUSG0 | Gulp1 | GULP, engulfment adaptor PTB domain containing 1 |
| chr1 | 44491124 | 44491552 | 429 | * | NA_peak_56 | 21 | Distal Interge | 1 | 44551511 | 44796838 | 245328 | 1 | 70676 | ENSMUST00 | -59959 | ENSMUSG0 | Gulp1 | GULP, engulfment adaptor PTB domain containing 1 |
| chr1 | 45202372 | 45202620 | 249 | * | NA_peak_57 | 28 | Distal Interge | 1 | 45311538 | 45325018 | 13481 | 1 | 12825 | ENSMUST00 | -108918 | ENSMUSG0 | Col3a1 | collagen, type III, alpha 1 |
| chr1 | 47629437 | 47629641 | 205 | * | NA_peak_61 | 30 | Distal Interge | 1 | 47467021 | 47467125 | 105 | 2 | 102466969 | ENSMUST00 | -162312 | ENSMUSG0 | Mir6350 | microRNA 6350 |
| chr1 | 49253357 | 49253597 | 241 | * | NA_peak_62 | 23 | Intron (ENSM | 1 | 49244616 | 49340443 | 95828 | 1 | 442837 | ENSMUST00 | 8741 | ENSMUSG0 | C230029F24 | RIKEN cDNA C230029F24 gene |
| chr1 | 52851940 | 52852239 | 300 | * | NA_peak_66 | 17 | Promoter (<= | 1 | 52851768 | 52860143 | 8376 | 1 | 227095 | ENSMUST00 | 172 | ENSMUSG0 | Hibch | 3-hydroxyisobutyryl-Coenzyme A hydrolase |
| chr1 | 54410529 | 54410727 | 199 | * | NA_peak_68 | 55 | Intron (ENSM | 1 | 54420149 | 54424054 | 3906 | 2 | 98488 | ENSMUST00 | 13327 | ENSMUSG0 | Gtf3c3 | general transcription factor IIIC, polypeptide 3 |
| chr1 | 55066121 | 55066204 | 84 | * | NA_peak_70 | 20 | Intron (ENSM | 1 | 55052770 | 55072702 | 19933 | 1 | 67876 | ENSMUST00 | 13351 | ENSMUSG0 | Coq10b | coenzyme Q10B |
| chr1 | 56138266 | 56138572 | 307 | * | NA_peak_72 | 36 | Distal Interge | 1 | 56019023 | 56020137 | 1115 | 1 | 69356 | ENSMUST00 | 119243 | ENSMUSG0 | 1700003I22R | RIKEN cDNA 1700003I22 gene |
| chr1 | 58646754 | 58647075 | 322 | * | NA_peak_75 | 24 | 3' UTR | 1 | 58652641 | 58655831 | 3191 | 1 | 100504501 | ENSMUST00 | -5566 | ENSMUSG0 | Gm20257 | caspase 8 pseudogene |
| chr1 | 59955520 | 59956030 | 511 | * | NA_peak_77 | 40 | Intron (ENSM | 1 | 59970680 | 59981515 | 10836 | 1 | 72750 | ENSMUST00 | -14650 | ENSMUSG0 | Fam117b | family with sequence similarity 117, member B |
| chr1 | 62295985 | 62296297 | 313 | * | NA_peak_79 | 40 | Intron (ENSM | 1 | 62131195 | 62136109 | 4915 | 2 | 105243940 | ENSMUST00 | -159876 | ENSMUSG0 | Pard3bos2 | par-3 family cell polarity regulator beta, opposite strand 2 |
| chr1 | 64472525 | 64473043 | 519 | * | NA_peak_83 | 31 | Distal Interge | 1 | 64532645 | 64597695 | 65051 | 1 | 12912 | ENSMUST00 | -59602 | ENSMUSG0 | Creb1 | cAMP responsive element binding protein 1 |
| chr1 | 64737126 | 64737454 | 329 | * | NA_peak_84 | 35 | Promoter (<= | 1 | 64734469 | 64737749 | 3281 | 2 | 14367 | ENSMUST00 | 295 | ENSMUSG0 | Fzd5 | frizzled class receptor 5 |
| chr1 | 66578659 | 66578945 | 287 | * | NA_peak_87 | 24 | Intron (ENSM | 1 | 66679699 | 66696663 | 16965 | 1 | 329178 | ENSMUST00 | -100754 | ENSMUSG0 | Unc80 | unc-80, NALCN activator |
| chr1 | 69231728 | 69232089 | 362 | * | NA_peak_90 | 17 | Distal Interge | 1 | 68032186 | 69108059 | 1075874 | 2 | 13869 | ENSMUST00 | -123669 | ENSMUSG0 | Erbb4 | erb-b2 receptor tyrosine kinase 4 |
| chr1 | 70982036 | 70982433 | 398 | * | NA_peak_92 | 54 | Distal Interge | 1 | 71027498 | 71103146 | 75649 | 2 | 12021 | ENSMUST00 | 120713 | ENSMUSG0 | Bard1 | BRCA1 associated RING domain 1 |
| chr1 | 72410588 | 72410640 | 53 | * | NA_peak_94 | 16 | Distal Interge | 1 | 72307427 | 72394952 | 87526 | 1 | 22596 | ENSMUST00 | 103161 | ENSMUSG0 | Xrcc5 | X-ray repair complementing defective repair in Chinese hamster cells 5 |
| chr1 | 74607329 | 74607931 | 603 | * | NA_peak_10 | 17 | Intron (ENSM | 1 | 74603210 | 74611256 | 8047 | 1 | 269209 | ENSMUST00 | 4119 | ENSMUSG0 | Stk36 | serine/threonine kinase 36 |
| chr1 | 78800500 | 78800892 | 393 | * | NA_peak_10 | 34 | Intron (ENSM | 1 | 78816758 | 78820028 | 3271 | 1 | 57814 | ENSMUST00 | -15866 | ENSMUSG0 | Kcne4 | potassium voltage-gated channel, Isk-related subfamily, gene 4 |
| chr1 | 82771028 | 82771242 | 215 | * | NA_peak_11 | 16 | Promoter (2-3 | 1 | 82757402 | 82768461 | 11060 | 2 | 66261 | ENSMUST00 | -2567 | ENSMUSG0 | Tm4sf20 | transmembrane 4 L six family member 20 |
| chr1 | 83937173 | 83937284 | 112 | * | NA_peak_11 | 23 | Distal Interge | 1 | 84037926 | 84053253 | 15328 | 2 | 98496 | ENSMUST00 | 115969 | ENSMUSG0 | Pid1 | phosphotyrosine interaction domain containing 1 |
| chr1 | 83984823 | 83985081 | 259 | * | NA_peak_11 | 13 | Distal Interge | 1 | 84037926 | 84053253 | 15328 | 2 | 98496 | ENSMUST00 | 68172 | ENSMUSG0 | Pid1 | phosphotyrosine interaction domain containing 1 |
| chr1 | 84265309 | 84265368 | 60 | * | NA_peak_11 | 16 | Intron (ENSM | 1 | 84080579 | 84284423 | 203845 | 2 | 98496 | ENSMUST00 | 19055 | ENSMUSG0 | Pid1 | phosphotyrosine interaction domain containing 1 |
| chr1 | 84379020 | 84379064 | 45 | * | NA_peak_11 | 27 | Intron (ENSM | 1 | 84107794 | 84364180 | 256387 | 2 | 98496 | ENSMUST00 | -14840 | ENSMUSG0 | Pid1 | phosphotyrosine interaction domain containing 1 |
| chr1 | 84454880 | 84455038 | 159 | * | NA_peak_11 | 36 | Intron (ENSM | 1 | 84107794 | 84364180 | 256387 | 2 | 98496 | ENSMUST00 | -90700 | ENSMUSG0 | Pid1 | phosphotyrosine interaction domain containing 1 |
| chr1 | 84963918 | 84964248 | 331 | * | NA_peak_11 | 27 | Exon (ENSMU | 1 | 84905898 | 84935134 | 29237 | 2 | 71781 | ENSMUST00 | -28784 | ENSMUSG0 | Slc16a14 | solute carrier family 16 (monocarboxylic acid transporters), member 14 |
| chr1 | 84965920 | 84966297 | 378 | * | NA_peak_11 | 22 | Distal Interge | 1 | 84905898 | 84935134 | 29237 | 2 | 71781 | ENSMUST00 | -30786 | ENSMUSG0 | Slc16a14 | solute carrier family 16 (monocarboxylic acid transporters), member 14 |
| chr1 | 84974896 | 84975283 | 388 | * | NA_peak_11 | 40 | Distal Interge | 1 | 84905898 | 84935134 | 29237 | 2 | 71781 | ENSMUST00 | -39762 | ENSMUSG0 | Slc16a14 | solute carrier family 16 (monocarboxylic acid transporters), member 14 |
| chr1 | 84983402 | 84983698 | 297 | * | NA_peak_11 | 29 | Distal Interge | 1 | 84905898 | 84935134 | 29237 | 2 | 71781 | ENSMUST00 | -48268 | ENSMUSG0 | Slc16a14 | solute carrier family 16 (monocarboxylic acid transporters), member 14 |
| chr1 | 85027477 | 85027779 | 303 | * | NA_peak_11 | 69 | Intron (ENSM | 1 | 85083069 | 85109449 | 26381 | 2 | 381287 | ENSMUST00 | 81670 | ENSMUSG0 | A530032D15 | RIKEN cDNA A530032D15Rik gene |
| chr1 | 85029121 | 85029492 | 372 | * | NA_peak_11 | 56 | Intron (ENSM | 1 | 85083069 | 85109449 | 26381 | 2 | 381287 | ENSMUST00 | 79957 | ENSMUSG0 | A530032D15 | RIKEN cDNA A530032D15Rik gene |
| chr1 | 85030389 | 85030720 | 332 | * | NA_peak_11 | 21 | Intron (ENSM | 1 | 85083069 | 85109449 | 26381 | 2 | 381287 | ENSMUST00 | 78729 | ENSMUSG0 | A530032D15 | RIKEN cDNA A530032D15Rik gene |
| chr1 | 85039780 | 85040194 | 415 | * | NA_peak_11 | 42 | Intron (ENSM | 1 | 85083069 | 85109449 | 26381 | 2 | 381287 | ENSMUST00 | 69255 | ENSMUSG0 | A530032D15 | RIKEN cDNA A530032D15Rik gene |
| chr1 | 85063301 | 85063618 | 318 | * | NA_peak_11 | 21 | Intron (ENSM | 1 | 85083069 | 85109449 | 26381 | 2 | 381287 | ENSMUST00 | 45831 | ENSMUSG0 | A530032D15 | RIKEN cDNA A530032D15Rik gene |
| chr1 | 85074896 | 85075278 | 383 | * | NA_peak_11 | 45 | Exon (ENSMU | 1 | 85083069 | 85109449 | 26381 | 2 | 381287 | ENSMUST00 | 34171 | ENSMUSG0 | A530032D15 | RIKEN cDNA A530032D15Rik gene |
| chr1 | 85075561 | 85076115 | 555 | * | NA_peak_11 | 34 | Intron (ENSM | 1 | 85083069 | 85109449 | 26381 | 2 | 381287 | ENSMUST00 | 33334 | ENSMUSG0 | A530032D15 | RIKEN cDNA A530032D15Rik gene |
| chr1 | 85104392 | 85104618 | 227 | * | NA_peak_11 | 44 | Intron (ENSM | 1 | 85083069 | 85109449 | 26381 | 2 | 381287 | ENSMUST00 | 4831 | ENSMUSG0 | A530032D15 | RIKEN cDNA A530032D15Rik gene |
| chr1 | 85153294 | 85154699 | 1406 | * | NA_peak_11 | 22 | Intron (ENSM | 1 | 85088407 | 85109853 | 21447 | 2 | 381287 | ENSMUST00 | -43441 | ENSMUSG0 | A530032D15 | RIKEN cDNA A530032D15Rik gene |
| chr1 | 85158663 | 85159335 | 673 | * | NA_peak_11 | 47 | Intron (ENSM | 1 | 85199613 | 85203171 | 3559 | 1 | 665378 | ENSMUST00 | -40278 | ENSMUSG0 | Gm7609 | predicted pseudogene 7609 |
| chr1 | 85161165 | 85161637 | 473 | * | NA_peak_11 | 45 | Intron (ENSM | 1 | 85199613 | 85203171 | 3559 | 1 | 665378 | ENSMUST00 | -37976 | ENSMUSG0 | Gm7609 | predicted pseudogene 7609 |
| chr1 | 85169582 | 85169920 | 339 | * | NA_peak_11 | 79 | Intron (ENSM | 1 | 85199613 | 85203171 | 3559 | 1 | 665378 | ENSMUST00 | -29693 | ENSMUSG0 | Gm7609 | predicted pseudogene 7609 |
| chr1 | 85179343 | 85179941 | 599 | * | NA_peak_11 | 16 | Intron (ENSM | 1 | 85199613 | 85203171 | 3559 | 1 | 665378 | ENSMUST00 | -19672 | ENSMUSG0 | Gm7609 | predicted pseudogene 7609 |
| chr1 | 85181576 | 85182133 | 558 | * | NA_peak_11 | 34 | Intron (ENSM | 1 | 85199613 | 85203171 | 3559 | 1 | 665378 | ENSMUST00 | -17480 | ENSMUSG0 | Gm7609 | predicted pseudogene 7609 |
| chr1 | 85186213 | 85186617 | 405 | * | NA_peak_11 | 20 | Intron (ENSM | 1 | 85199613 | 85203171 | 3559 | 1 | 665378 | ENSMUST00 | -12996 | ENSMUSG0 | Gm7609 | predicted pseudogene 7609 |
| chr1 | 85214335 | 85214650 | 316 | * | NA_peak_11 | 24 | Intron (ENSM | 1 | 85199708 | 85213786 | 14079 | 1 | 665378 | ENSMUST00 | 14627 | ENSMUSG0 | Gm7609 | predicted pseudogene 7609 |
| chr1 | 85230969 | 85231672 | 704 | * | NA_peak_11 | 32 | Intron (ENSM | 1 | 85241286 | 85254548 | 13263 | 2 | 620078 | ENSMUST00 | 22876 | ENSMUSG0 | C130026I21F | RIKEN cDNA C130026I21 gene |
| chr1 | 85251915 | 85252212 | 298 | * | NA_peak_11 | 54 | Promoter (2-3 | 1 | 85241286 | 85254548 | 13263 | 2 | 620078 | ENSMUST00 | 2336 | ENSMUSG0 | C130026I21F | RIKEN cDNA C130026I21 gene |
| chr1 | 85252867 | 85253279 | 413 | * | NA_peak_11 | 27 | Promoter (1-2 | 1 | 85241286 | 85254548 | 13263 | 2 | 620078 | ENSMUST00 | 1269 | ENSMUSG0 | C130026I21F | RIKEN cDNA C130026I21 gene |
| chr1 | 85254864 | 85255371 | 508 | * | NA_peak_11 | 15 | Promoter (<= | 1 | 85241286 | 85254548 | 13263 | 2 | 620078 | ENSMUST00 | -316 | ENSMUSG0 | C130026I21F | RIKEN cDNA C130026I21 gene |
| chr1 | 85257134 | 85257423 | 290 | * | NA_peak_11 | 41 | Promoter (2-3 | 1 | 85241286 | 85254548 | 13263 | 2 | 620078 | ENSMUST00 | -2586 | ENSMUSG0 | C130026I21F | RIKEN cDNA C130026I21 gene |
| chr1 | 85263844 | 85264112 | 269 | * | NA_peak_11 | 39 | Intron (ENSM | 1 | 85245640 | 85270143 | 24504 | 2 | 620078 | ENSMUST00 | 6031 | ENSMUSG0 | C130026I21F | RIKEN cDNA C130026I21 gene |
| chr1 | 85269711 | 85270230 | 520 | * | NA_peak_11 | 46 | Promoter (<= | 1 | 85112267 | 85270178 | 157912 | 2 | 620078 | ENSMUST00 | 0 | ENSMUSG0 | C130026I21F | RIKEN cDNA C130026I21 gene |
| chr1 | 85285394 | 85285733 | 340 | * | NA_peak_11 | 70 | Exon (ENSMU | 1 | 85246670 | 85270566 | 23897 | 2 | 620078 | ENSMUST00 | -14828 | ENSMUSG0 | C130026I21F | RIKEN cDNA C130026I21 gene |
| chr1 | 85296065 | 85296805 | 741 | * | NA_peak_12 | 22 | Intron (ENSM | 1 | 85246670 | 85270566 | 23897 | 2 | 620078 | ENSMUST00 | -25499 | ENSMUSG0 | C130026I21F | RIKEN cDNA C130026I21 gene |
| chr1 | 85298545 | 85298902 | 358 | * | NA_peak_12 | 44 | Intron (ENSM | 1 | 85246670 | 85270566 | 23897 | 2 | 620078 | ENSMUST00 | -27979 | ENSMUSG0 | C130026I21F | RIKEN cDNA C130026I21 gene |
| chr1 | 85300381 | 85301020 | 640 | * | NA_peak_12 | 37 | Intron (ENSM | 1 | 85246670 | 85270566 | 23897 | 2 | 620078 | ENSMUST00 | -29815 | ENSMUSG0 | C130026I21F | RIKEN cDNA C130026I21 gene |
| chr1 | 85301701 | 85302681 | 981 | * | NA_peak_12 | 39 | Intron (ENSM | 1 | 85246670 | 85270566 | 23897 | 2 | 620078 | ENSMUST00 | -31135 | ENSMUSG0 | C130026I21F | RIKEN cDNA C130026I21 gene |
| chr1 | 85304013 | 85304735 | 723 | * | NA_peak_12 | 41 | Intron (ENSM | 1 | 85246670 | 85270566 | 23897 | 2 | 620078 | ENSMUST00 | -33447 | ENSMUSG0 | C130026I21F | RIKEN cDNA C130026I21 gene |
| chr1 | 85308930 | 85309274 | 345 | * | NA_peak_12 | 22 | Intron (ENSM | 1 | 85246670 | 85270566 | 23897 | 2 | 620078 | ENSMUST00 | -38364 | ENSMUSG0 | C130026I21F | RIKEN cDNA C130026I21 gene |
| chr1 | 85335774 | 85335992 | 219 | * | NA_peak_12 | 27 | Intron (ENSM | 1 | 85246670 | 85270566 | 23897 | 2 | 620078 | ENSMUST00 | -65208 | ENSMUSG0 | C130026I21F | RIKEN cDNA C130026I21 gene |
| chr1 | 85338470 | 85338805 | 336 | * | NA_peak_12 | 40 | Intron (ENSM | 1 | 85246670 | 85270566 | 23897 | 2 | 620078 | ENSMUST00 | -67904 | ENSMUSG0 | C130026I21F | RIKEN cDNA C130026I21 gene |
| chr1 | 85448335 | 85448811 | 477 | * | NA_peak_12 | 53 | Distal Interge | 1 | 85579357 | 85584278 | 4922 | 2 | 109032 | ENSMUST00 | 135467 | ENSMUSG0 | Sp110 | Sp110 nuclear body protein |
| chr1 | 85451952 | 85452513 | 562 | * | NA_peak_12 | 17 | Distal Interge | 1 | 85579357 | 85584278 | 4922 | 2 | 109032 | ENSMUST00 | 131765 | ENSMUSG0 | Sp110 | Sp110 nuclear body protein |
| chr1 | 85453622 | 85454357 | 736 | * | NA_peak_12 | 31 | Distal Interge | 1 | 85579357 | 85584278 | 4922 | 2 | 109032 | ENSMUST00 | 129921 | ENSMUSG0 | Sp110 | Sp110 nuclear body protein |
| chr1 | 85455957 | 85456369 | 413 | * | NA_peak_12 | 31 | Distal Interge | 1 | 85579357 | 85584278 | 4922 | 2 | 109032 | ENSMUST00 | 127909 | ENSMUSG0 | Sp110 | Sp110 nuclear body protein |
| chr1 | 85457128 | 85457573 | 446 | * | NA_peak_12 | 52 | Distal Interge | 1 | 85579357 | 85584278 | 4922 | 2 | 109032 | ENSMUST00 | 126705 | ENSMUSG0 | Sp110 | Sp110 nuclear body protein |
| chr1 | 85464635 | 85465104 | 470 | * | NA_peak_12 | 38 | Distal Interge | 1 | 85579357 | 85584278 | 4922 | 2 | 109032 | ENSMUST00 | 119174 | ENSMUSG0 | Sp110 | Sp110 nuclear body protein |
| chr1 | 85481532 | 85482174 | 643 | * | NA_peak_12 | 40 | Distal Interge | 1 | 85579357 | 85584278 | 4922 | 2 | 109032 | ENSMUST00 | 102104 | ENSMUSG0 | Sp110 | Sp110 nuclear body protein |
| chr1 | 85502980 | 85503247 | 268 | * | NA_peak_12 | 53 | Exon (ENSMU | 1 | 85579357 | 85584278 | 4922 | 2 | 109032 | ENSMUST00 | 81031 | ENSMUSG0 | Sp110 | Sp110 nuclear body protein |
| chr1 | 85556533 | 85556915 | 383 | * | NA_peak_12 | 100 | Distal Interge | 1 | 85579357 | 85584278 | 4922 | 2 | 109032 | ENSMUST00 | 27363 | ENSMUSG0 | Sp110 | Sp110 nuclear body protein |
| chr1 | 85557900 | 85558359 | 460 | * | NA_peak_12 | 34 | Distal Interge | 1 | 85579357 | 85584278 | 4922 | 2 | 109032 | ENSMUST00 | 25919 | ENSMUSG0 | Sp110 | Sp110 nuclear body protein |
| chr1 | 85565385 | 85565677 | 293 | * | NA_peak_12 | 30 | Distal Interge | 1 | 85579357 | 85584278 | 4922 | 2 | 109032 | ENSMUST00 | 18601 | ENSMUSG0 | Sp110 | Sp110 nuclear body protein |
| chr1 | 85566175 | 85566384 | 210 | * | NA_peak_12 | 21 | Distal Interge | 1 | 85579357 | 85584278 | 4922 | 2 | 109032 | ENSMUST00 | 17894 | ENSMUSG0 | Sp110 | Sp110 nuclear body protein |

| | | | | | | | | | | | | | | | | | | |
|---|---|---|---|---|---|---|---|---|---|---|---|---|---|---|---|---|---|---|
| chr1 | 85570785 | 85571086 | 302 | * | NA_peak_12 | 47 | Distal Interge | 1 | 85579357 | 85584278 | 4922 | 2 | 109032 | ENSMUST00 | 13192 | ENSMUSG0 | Sp110 | Sp110 nuclear body protein |
| chr1 | 85578563 | 85579063 | 501 | * | NA_peak_12 | 67 | Exon (ENSM | 1 | 85579357 | 85584278 | 4922 | 2 | 109032 | ENSMUST00 | 5215 | ENSMUSG0 | Sp110 | Sp110 nuclear body protein |
| chr1 | 85584102 | 85584417 | 316 | * | NA_peak_12 | 45 | Promoter (<= | 1 | 85579357 | 85584278 | 4922 | 2 | 109032 | ENSMUST00 | 0 | ENSMUSG0 | Sp110 | Sp110 nuclear body protein |
| chr1 | 85608786 | 85609563 | 778 | * | NA_peak_12 | 50 | Promoter (1- | 1 | 85610833 | 85630539 | 19707 | 1 | 434484 | ENSMUST00 | -1270 | ENSMUSG0 | Sp140 | Sp140 nuclear body protein |
| chr1 | 85624909 | 85625242 | 334 | * | NA_peak_12 | 94 | Intron (ENSM | 1 | 85632478 | 85644942 | 12465 | 1 | 434484 | ENSMUST00 | -7236 | ENSMUSG0 | Sp140 | Sp140 nuclear body protein |
| chr1 | 85626853 | 85627711 | 859 | * | NA_peak_12 | 129 | Intron (ENSM | 1 | 85632478 | 85644942 | 12465 | 1 | 434484 | ENSMUST00 | -4767 | ENSMUSG0 | Sp140 | Sp140 nuclear body protein |
| chr1 | 85653098 | 85653430 | 333 | * | NA_peak_12 | 49 | Intron (ENSM | 1 | 85650050 | 85709998 | 59949 | 1 | 20684 | ENSMUST00 | 3048 | ENSMUSG0 | Sp100 | nuclear antigen Sp100 |
| chr1 | 86778232 | 86778753 | 522 | * | NA_peak_12 | 15 | Intron (ENSM | 1 | 86703843 | 87050092 | 346250 | 1 | 208718 | ENSMUST00 | 74389 | ENSMUSG0 | Dis3l2 | DIS3 like 3'-5' exoribonuclease 2 |
| chr1 | 87776689 | 87777017 | 329 | * | NA_peak_13 | 27 | Promoter (1- | 1 | 87775565 | 87778958 | 3394 | 1 | 77040 | ENSMUST00 | 1124 | ENSMUSG0 | Atg16l1 | autophagy related 16-like 1 (S. cerevisiae) |
| chr1 | 88301940 | 88302067 | 128 | * | NA_peak_13 | 22 | Promoter (1- | 1 | 88303283 | 88388851 | 85569 | 1 | 171382 | ENSMUST00 | -1216 | ENSMUSG0 | Trpm8 | transient receptor potential cation channel, subfamily M, member 8 |
| chr1 | 89476585 | 89476850 | 266 | * | NA_peak_13 | 16 | Intron (ENSM | 1 | 89454874 | 89890001 | 435128 | 1 | 347722 | ENSMUST00 | 21711 | ENSMUSG0 | Agap1 | ArfGAP with GTPase domain, ankyrin repeat and PH domain 1 |
| chr1 | 91304139 | 91304282 | 144 | * | NA_peak_13 | 20 | Promoter (1- | 1 | 91302797 | 91309872 | 7076 | 1 | 50880 | ENSMUST00 | 1342 | ENSMUSG0 | Scly | selenocysteine lyase |
| chr1 | 91931365 | 91931500 | 136 | * | NA_peak_13 | 16 | 3' UTR | 1 | 91933203 | 91934864 | 1662 | 2 | 208727 | ENSMUST00 | 3364 | ENSMUSG0 | Hdac4 | histone deacetylase 4 |
| chr1 | 91982078 | 91982306 | 229 | * | NA_peak_13 | 23 | Exon (ENSM | 1 | 91955374 | 91975501 | 20128 | 2 | 208727 | ENSMUST00 | -6577 | ENSMUSG0 | Hdac4 | histone deacetylase 4 |
| chr1 | 93814725 | 93815032 | 308 | * | NA_peak_13 | 27 | Intron (ENSM | 1 | 93824909 | 93826458 | 1550 | 1 | 98314 | ENSMUST00 | -9877 | ENSMUSG0 | D2hgdh | D-2-hydroxyglutarate dehydrogenase |
| chr1 | 96192496 | 96192516 | 21 | * | NA_peak_14 | 14 | Distal Interge | 1 | 95990659 | 96021758 | 31100 | 2 | 73454 | ENSMUST00 | -170738 | ENSMUSG0 | 1700063A18 | RIKEN cDNA 1700063A18 gene |
| chr1 | 99265732 | 99265930 | 199 | * | NA_peak_14 | 50 | Distal Interge | 1 | 99772765 | 100484849 | 712085 | 1 | 241175 | ENSMUST00 | -506835 | ENSMUSG0 | Cntnap5b | contactin associated protein-like 5B |
| chr1 | 101603403 | 101603758 | 356 | * | NA_peak_14 | 27 | Distal Interge | 1 | 101155809 | 101175202 | 19394 | 1 | 100504525 | ENSMUST00 | 447594 | ENSMUSG0 | Gm20268 | predicted gene, 20268 |
| chr1 | 103117523 | 103117800 | 278 | * | NA_peak_14 | 57 | Distal Interge | 1 | 104768529 | 104995481 | 226953 | 1 | 23836 | ENSMUST00 | -1650729 | ENSMUSG0 | Cdh20 | cadherin 20 |
| chr1 | 104222317 | 104222692 | 376 | * | NA_peak_14 | 76 | Distal Interge | 1 | 104768529 | 104995481 | 226953 | 1 | 23836 | ENSMUST00 | -545837 | ENSMUSG0 | Cdh20 | cadherin 20 |
| chr1 | 108353971 | 108354404 | 434 | * | NA_peak_14 | 23 | Distal Interge | 1 | 107936709 | 108028337 | 91629 | 1 | 319596 | ENSMUST00 | 417262 | ENSMUSG0 | D830032E09 | RIKEN cDNA D830032E09 gene |
| chr1 | 108414553 | 108414645 | 93 | * | NA_peak_14 | 39 | Distal Interge | 1 | 107936709 | 108028337 | 91629 | 1 | 319596 | ENSMUST00 | 477844 | ENSMUSG0 | D830032E09 | RIKEN cDNA D830032E09 gene |
| chr1 | 108414955 | 108415096 | 142 | * | NA_peak_14 | 23 | Distal Interge | 1 | 107936709 | 108028337 | 91629 | 1 | 319596 | ENSMUST00 | 478246 | ENSMUSG0 | D830032E09 | RIKEN cDNA D830032E09 gene |
| chr1 | 110177088 | 110178247 | 1160 | * | NA_peak_14 | 93 | Distal Interge | 1 | 110099295 | 110138058 | 38764 | 1 | 241201 | ENSMUST00 | 77793 | ENSMUSG0 | Cdh7 | cadherin 7, type 2 |
| chr1 | 112215205 | 112215494 | 290 | * | NA_peak_14 | 45 | Distal Interge | 1 | 111858706 | 111864918 | 6213 | 2 | 319901 | ENSMUST00 | -350287 | ENSMUSG0 | Dsel | dermatan sulfate epimerase-like |
| chr1 | 112218710 | 112219006 | 297 | * | NA_peak_14 | 23 | Distal Interge | 1 | 111858706 | 111864918 | 6213 | 2 | 319901 | ENSMUST00 | -353792 | ENSMUSG0 | Dsel | dermatan sulfate epimerase-like |
| chr1 | 112844245 | 112844529 | 285 | * | NA_peak_15 | 27 | Distal Interge | 1 | 111858706 | 111864918 | 6213 | 2 | 319901 | ENSMUST00 | -979327 | ENSMUSG0 | Dsel | dermatan sulfate epimerase-like |
| chr1 | 112930316 | 112930697 | 382 | * | NA_peak_15 | 15 | Distal Interge | 1 | 111858706 | 111864918 | 6213 | 2 | 319901 | ENSMUST00 | -1065398 | ENSMUSG0 | Dsel | dermatan sulfate epimerase-like |
| chr1 | 115799838 | 115800096 | 259 | * | NA_peak_15 | 30 | Intron (ENSM | 1 | 115684756 | 116587323 | 902568 | 1 | 636808 | ENSMUST00 | 115082 | ENSMUSG0 | Cntnap5a | contactin associated protein-like 5A |
| chr1 | 115889135 | 115889496 | 362 | * | NA_peak_15 | 15 | Intron (ENSM | 1 | 115684756 | 116587323 | 902568 | 1 | 636808 | ENSMUST00 | 204379 | ENSMUSG0 | Cntnap5a | contactin associated protein-like 5A |
| chr1 | 116858567 | 116858765 | 199 | * | NA_peak_15 | 31 | Distal Interge | 1 | 115684756 | 116587323 | 902568 | 1 | 636808 | ENSMUST00 | 1173811 | ENSMUSG0 | Cntnap5a | contactin associated protein-like 5A |
| chr1 | 117372499 | 117372750 | 252 | * | NA_peak_15 | 23 | Distal Interge | 1 | 118300889 | 118309986 | 9098 | 2 | 22099 | ENSMUST00 | 937236 | ENSMUSG0 | Tsn | translin |
| chr1 | 118588292 | 118588561 | 270 | * | NA_peak_15 | 18 | Intron (ENSM | 1 | 118627945 | 118637766 | 9822 | 1 | 81879 | ENSMUST00 | -39384 | ENSMUSG0 | Tfcp2l1 | transcription factor CP2-like 1 |
| chr1 | 121039276 | 121039626 | 351 | * | NA_peak_15 | 36 | Distal Interge | 1 | 121087405 | 121120956 | 33552 | 1 | 98452 | ENSMUST00 | -47779 | ENSMUSG0 | Celrr | cerebellum expressed regulatory RNA |
| chr1 | 125317317 | 125317337 | 21 | * | NA_peak_15 | 16 | Distal Interge | 1 | 125393484 | 125397612 | 4129 | 2 | 74117 | ENSMUST00 | 80275 | ENSMUSG0 | Actr3 | ARP3 actin-related protein 3 |
| chr1 | 126773380 | 126773677 | 298 | * | NA_peak_15 | 33 | Intron (ENSM | 1 | 126258401 | 126738202 | 479802 | 2 | 210356 | ENSMUST00 | -35178 | ENSMUSG0 | Nckap5 | NCK-associated protein 5 |
| chr1 | 132316152 | 132316375 | 224 | * | NA_peak_16 | 34 | Promoter (<= | 1 | 132316165 | 132317736 | 1572 | 1 | 74137 | ENSMUST00 | 0 | ENSMUSG0 | Nuak2 | NUAK family, SNF1-like kinase, 2 |
| chr1 | 138610615 | 138610984 | 370 | * | NA_peak_17 | 70 | Intron (ENSM | 1 | 138484112 | 138619674 | 135563 | 2 | 59125 | ENSMUST00 | 8690 | ENSMUSG0 | Nek7 | NIMA (never in mitosis gene a)-related expressed kinase 7 |
| chr1 | 140044150 | 140044539 | 390 | * | NA_peak_17 | 71 | Intron (ENSM | 1 | 140085862 | 140147812 | 61951 | 2 | 12628 | ENSMUST00 | 103273 | ENSMUSG0 | Cfh | complement component factor h |
| chr1 | 140837603 | 140837713 | 111 | * | NA_peak_17 | 27 | Distal Interge | 1 | 140570176 | 140656404 | 86229 | 2 | 75910 | ENSMUST00 | -181199 | ENSMUSG0 | 4930590L20 | RIKEN cDNA 4930590L20 gene |
| chr1 | 141035992 | 141036449 | 458 | * | NA_peak_17 | 21 | Distal Interge | 1 | 140570176 | 140656404 | 86229 | 2 | 75910 | ENSMUST00 | -379588 | ENSMUSG0 | 4930590L20 | RIKEN cDNA 4930590L20 gene |
| chr1 | 141263574 | 141263863 | 290 | * | NA_peak_17 | 31 | Distal Interge | 1 | 140570176 | 140656404 | 86229 | 2 | 75910 | ENSMUST00 | -607170 | ENSMUSG0 | 4930590L20 | RIKEN cDNA 4930590L20 gene |
| chr1 | 141264972 | 141265005 | 34 | * | NA_peak_17 | 31 | Distal Interge | 1 | 140570176 | 140656404 | 86229 | 2 | 75910 | ENSMUST00 | -608568 | ENSMUSG0 | 4930590L20 | RIKEN cDNA 4930590L20 gene |
| chr1 | 143511782 | 143512013 | 232 | * | NA_peak_17 | 18 | Distal Interge | 1 | 143606161 | 143640581 | 34421 | 2 | 214498 | ENSMUST00 | 128568 | ENSMUSG0 | Cdc73 | cell division cycle 73, Paf1/RNA polymerase II complex component |
| chr1 | 144384315 | 144384635 | 321 | * | NA_peak_17 | 27 | Distal Interge | 1 | 144242296 | 144249242 | 6947 | 2 | 50778 | ENSMUST00 | -135073 | ENSMUSG0 | Rgs1 | regulator of G-protein signaling 1 |
| chr1 | 146123453 | 146123464 | 12 | * | NA_peak_17 | 14 | Distal Interge | 1 | 146494760 | 146902471 | 407712 | 1 | 215378 | ENSMUST00 | -371296 | ENSMUSG0 | Brinp3 | bone morphogenetic protein/retinoic acid inducible neural specific 3 |
| chr1 | 146402323 | 146402620 | 298 | * | NA_peak_17 | 17 | Distal Interge | 1 | 146494760 | 146902471 | 407712 | 1 | 215378 | ENSMUST00 | -92140 | ENSMUSG0 | Brinp3 | bone morphogenetic protein/retinoic acid inducible neural specific 3 |
| chr1 | 146446903 | 146447337 | 435 | * | NA_peak_17 | 23 | Intron (ENSM | 1 | 146494760 | 146902471 | 407712 | 1 | 215378 | ENSMUST00 | -47423 | ENSMUSG0 | Brinp3 | bone morphogenetic protein/retinoic acid inducible neural specific 3 |
| chr1 | 147499639 | 147500059 | 421 | * | NA_peak_17 | 59 | Distal Interge | 1 | 146497738 | 146517554 | 19817 | 1 | 215378 | ENSMUST00 | 1001901 | ENSMUSG0 | Brinp3 | bone morphogenetic protein/retinoic acid inducible neural specific 3 |
| chr1 | 148543492 | 148543667 | 176 | * | NA_peak_17 | 13 | Distal Interge | 1 | 149886184 | 149902199 | 16016 | 2 | 18783 | ENSMUST00 | 1358532 | ENSMUSG0 | Pla2g4a | phospholipase A2, group IVA (cytosolic, calcium-dependent) |
| chr1 | 149183976 | 149184205 | 230 | * | NA_peak_17 | 50 | Distal Interge | 1 | 149886184 | 149902199 | 16016 | 2 | 18783 | ENSMUST00 | 717994 | ENSMUSG0 | Pla2g4a | phospholipase A2, group IVA (cytosolic, calcium-dependent) |
| chr1 | 149393902 | 149394336 | 435 | * | NA_peak_17 | 14 | Distal Interge | 1 | 149886184 | 149902199 | 16016 | 2 | 18783 | ENSMUST00 | 507863 | ENSMUSG0 | Pla2g4a | phospholipase A2, group IVA (cytosolic, calcium-dependent) |
| chr1 | 149519122 | 149519511 | 390 | * | NA_peak_17 | 78 | Distal Interge | 1 | 149886184 | 149902199 | 16016 | 2 | 18783 | ENSMUST00 | 382688 | ENSMUSG0 | Pla2g4a | phospholipase A2, group IVA (cytosolic, calcium-dependent) |
| chr1 | 150232725 | 150233005 | 281 | * | NA_peak_18 | 54 | Distal Interge | 1 | 150159043 | 150164948 | 5906 | 2 | 102639566 | ENSMUST00 | -67777 | ENSMUSG0 | Ptgs2os2 | prostaglandin-endoperoxide synthase 2, opposite strand 2 |
| chr1 | 158885310 | 158885555 | 246 | * | NA_peak_18 | 50 | Intron (ENSM | 1 | 158814274 | 158845032 | 30759 | 2 | 23850 | ENSMUST00 | -40278 | ENSMUSG0 | Pappa2 | pappalysin 2 |
| chr1 | 162841780 | 162841955 | 176 | * | NA_peak_19 | 17 | Intron (ENSM | 1 | 162830050 | 162834672 | 4623 | 2 | 14261 | ENSMUST00 | -7108 | ENSMUSG0 | Fmo1 | flavin containing monooxygenase 1 |
| chr1 | 162842697 | 162843050 | 354 | * | NA_peak_19 | 31 | Intron (ENSM | 1 | 162830050 | 162834672 | 4623 | 2 | 14261 | ENSMUST00 | -8025 | ENSMUSG0 | Fmo1 | flavin containing monooxygenase 1 |
| chr1 | 164742629 | 164743001 | 373 | * | NA_peak_19 | 13 | Distal Interge | 1 | 164796644 | 164824266 | 27623 | 1 | 56429 | ENSMUST00 | -53643 | ENSMUSG0 | Dpt | dermatopontin |
| chr1 | 170116617 | 170116889 | 273 | * | NA_peak_19 | 20 | Distal Interge | 1 | 170035774 | 170110762 | 74989 | 2 | 18214 | ENSMUST00 | -5855 | ENSMUSG0 | Ddr2 | discoidin domain receptor family, member 2 |
| chr1 | 171549195 | 171549464 | 270 | * | NA_peak_20 | 15 | Distal Interge | 1 | 171559193 | 171585318 | 26126 | 1 | 18106 | ENSMUST00 | -9729 | ENSMUSG0 | Cd244a | CD244 molecule A |
| chr1 | 176542628 | 176542957 | 330 | * | NA_peak_20 | 40 | Distal Interge | 1 | 176430111 | 176527273 | 97163 | 2 | 102640488 | ENSMUST00 | -15355 | ENSMUSG0 | Gm36536 | predicted gene, 36536 |
| chr1 | 177924688 | 177924996 | 309 | * | NA_peak_20 | 40 | Intron (ENSM | 1 | 177983423 | 178149962 | 166540 | 1 | 545391 | ENSMUST00 | -58427 | ENSMUSG0 | Catspere2 | cation channel sperm associated auxiliary subunit epsilon 2 |
| chr1 | 180725733 | 180726438 | 706 | * | NA_peak_20 | 36 | Promoter (<= | 1 | 180726043 | 180754204 | 28162 | 1 | 170760 | ENSMUST00 | 0 | ENSMUSG0 | Acbd3 | acyl-Coenzyme A binding domain containing 3 |
| chr1 | 182794059 | 182794276 | 218 | * | NA_peak_21 | 13 | Intron (ENSM | 1 | 182764895 | 182896591 | 131697 | 1 | 96935 | ENSMUST00 | 29164 | ENSMUSG0 | Susd4 | sushi domain containing 4 |
| chr1 | 183565481 | 183565679 | 199 | * | NA_peak_21 | 23 | Distal Interge | 1 | 183419248 | 183436808 | 17561 | 1 | 78772 | ENSMUST00 | 146233 | ENSMUSG0 | Hhipl2 | hedgehog interacting protein-like 2 |
| chr1 | 195055669 | 195055988 | 320 | * | NA_peak_22 | 17 | Intron (ENSM | 1 | 195041835 | 195069114 | 27280 | 2 | 17221 | ENSMUST00 | 13126 | ENSMUSG0 | Cd46 | CD46 antigen, complement regulatory protein |
| chr1 | 195357675 | 195358222 | 548 | * | NA_peak_22 | 50 | Distal Interge | 1 | 195136811 | 195176716 | 39906 | 2 | 12902 | ENSMUST00 | -180959 | ENSMUSG0 | Cr2 | complement receptor 2 |
| chr10 | 3134972 | 3135315 | 344 | * | NA_peak_22 | 76 | Promoter (1- | 10 | 3125091 | 3133193 | 8103 | 2 | 619314 | ENSMUST00 | -1779 | ENSMUSG0 | B020014A21 | RIKEN cDNA B020014A21 gene |
| chr10 | 3235832 | 3236120 | 289 | * | NA_peak_22 | 36 | Distal Interge | 10 | 3257930 | 3267750 | 9821 | 2 | 670558 | ENSMUST00 | 31630 | ENSMUSG0 | H60c | histocompatibility 60c |
| chr10 | 3250049 | 3250355 | 307 | * | NA_peak_22 | 37 | Distal Interge | 10 | 3257930 | 3267750 | 9821 | 2 | 670558 | ENSMUST00 | 17395 | ENSMUSG0 | H60c | histocompatibility 60c |
| chr10 | 3251169 | 3251469 | 301 | * | NA_peak_22 | 72 | Distal Interge | 10 | 3257930 | 3267750 | 9821 | 2 | 670558 | ENSMUST00 | 16281 | ENSMUSG0 | H60c | histocompatibility 60c |
| chr10 | 3252982 | 3253241 | 260 | * | NA_peak_22 | 45 | Distal Interge | 10 | 3257930 | 3267750 | 9821 | 2 | 670558 | ENSMUST00 | 14509 | ENSMUSG0 | H60c | histocompatibility 60c |
| chr10 | 3491437 | 3491708 | 272 | * | NA_peak_22 | 16 | Distal Interge | 10 | 3540240 | 3554877 | 14638 | 1 | 70337 | ENSMUST00 | -48532 | ENSMUSG0 | Iyd | iodotyrosine deiodinase |
| chr10 | 4030773 | 4030788 | 16 | * | NA_peak_22 | 29 | Promoter (1- | 10 | 4032280 | 4048951 | 16672 | 1 | 270685 | ENSMUST00 | -1492 | ENSMUSG0 | Mthfd1l | methylenetetrahydrofolate dehydrogenase (NADP+ dependent) 1-like |
| chr10 | 4298694 | 4299123 | 430 | * | NA_peak_23 | 19 | Intron (ENSM | 10 | 4266380 | 4359470 | 93091 | 1 | 83397 | ENSMUST00 | 32314 | ENSMUSG0 | Akap12 | A kinase (PRKA) anchor protein (gravin) 12 |
| chr10 | 5597237 | 5597582 | 346 | * | NA_peak_23 | 13 | Intron (ENSM | 10 | 5593775 | 5606904 | 13130 | 1 | 68632 | ENSMUST00 | 3462 | ENSMUSG0 | Myct1 | myc target 1 |
| chr10 | 5775899 | 5775909 | 11 | * | NA_peak_24 | 15 | Distal Interge | 10 | 5802200 | 5804600 | 2401 | 2 | 67141 | ENSMUST00 | 28691 | ENSMUSG0 | Fbxo5 | F-box protein 5 |
| chr10 | 6570501 | 6571290 | 790 | * | NA_peak_24 | 55 | Distal Interge | 10 | 6758506 | 6979704 | 221199 | 1 | 18390 | ENSMUST00 | -187216 | ENSMUSG0 | Oprm1 | opioid receptor, mu 1 |
| chr10 | 7403181 | 7403342 | 162 | * | NA_peak_24 | 44 | Distal Interge | 10 | 7446183 | 7473341 | 27159 | 2 | 77777 | ENSMUST00 | 69999 | ENSMUSG0 | Ulbp1 | UL16 binding protein 1 |
| chr10 | 7485726 | 7485977 | 252 | * | NA_peak_24 | 23 | Distal Interge | 10 | 7443038 | 7473638 | 30601 | 2 | 77777 | ENSMUST00 | -12088 | ENSMUSG0 | Ulbp1 | UL16 binding protein 1 |
| chr10 | 8390426 | 8390707 | 282 | * | NA_peak_25 | 19 | Intron (ENSM | 10 | 8204756 | 8302242 | 97487 | 2 | 338362 | ENSMUST00 | -88184 | ENSMUSG0 | Ust | uronyl-2-sulfotransferase |
| chr10 | 8544809 | 8545232 | 424 | * | NA_peak_25 | 58 | Distal Interge | 10 | 8204809 | 8518825 | 314017 | 2 | 338362 | ENSMUST00 | -25984 | ENSMUSG0 | Ust | uronyl-2-sulfotransferase |
| chr10 | 8881250 | 8881396 | 147 | * | NA_peak_25 | 31 | Intron (ENSM | 10 | 8813722 | 8886052 | 72331 | 2 | 70097 | ENSMUST00 | 4656 | ENSMUSG0 | Sash1 | SAM and SH3 domain containing 1 |
| chr10 | 9904630 | 9904785 | 156 | * | NA_peak_26 | 36 | Intron (ENSM | 10 | 9759958 | 9901079 | 141122 | 2 | 78808 | ENSMUST00 | -3551 | ENSMUSG0 | Stxbp5 | syntaxin binding protein 5 (tomosyn) |
| chr10 | 10143189 | 10143515 | 327 | * | NA_peak_26 | 20 | Intron (ENSM | 10 | 9759958 | 9901079 | 141122 | 2 | 78808 | ENSMUST00 | -242110 | ENSMUSG0 | Stxbp5 | syntaxin binding protein 5 (tomosyn) |
| chr10 | 10369425 | 10369526 | 102 | * | NA_peak_26 | 26 | Intron (ENSM | 10 | 10335709 | 10411744 | 76036 | 2 | 215772 | ENSMUST00 | 42218 | ENSMUSG0 | Adgb | androglobin |
| chr10 | 10572106 | 10572128 | 23 | * | NA_peak_26 | 21 | Distal Interge | 10 | 10545002 | 10558265 | 13264 | 2 | 67844 | ENSMUST00 | -13841 | ENSMUSG0 | Rab32 | RAB32, member RAS oncogene family |
| chr10 | 10584693 | 10585325 | 633 | * | NA_peak_26 | 27 | Distal Interge | 10 | 10605945 | 10616313 | 10369 | 1 | 382450 | ENSMUST00 | -20620 | ENSMUSG0 | Gm5177 | glyceraldehyde-3-phosphate dehydrogenase pseudogene |
| chr10 | 10977189 | 10977446 | 258 | * | NA_peak_26 | 62 | Intron (ENSM | 10 | 10688801 | 11080956 | 392156 | 2 | 14816 | ENSMUST00 | 103510 | ENSMUSG0 | Grm1 | glutamate receptor, metabotropic 1 |
| chr10 | 11422157 | 11422403 | 247 | * | NA_peak_26 | 31 | Intron (ENSM | 10 | 11343453 | 11448698 | 105246 | 1 | 13853 | ENSMUST00 | 78704 | ENSMUSG0 | Epm2a | epilepsy, progressive myoclonic epilepsy, type 2 gene alpha |
| chr10 | 12175469 | 12175809 | 341 | * | NA_peak_27 | 16 | Distal Interge | 10 | 12401066 | 12403075 | 2010 | 2 | 22288 | ENSMUST00 | 227266 | ENSMUSG0 | Utrn | utrophin |
| chr10 | 12387481 | 12387776 | 296 | * | NA_peak_27 | 15 | Intron (ENSM | 10 | 12401066 | 12403075 | 2010 | 2 | 22288 | ENSMUST00 | 15299 | ENSMUSG0 | Utrn | utrophin |
| chr10 | 12718157 | 12718266 | 110 | * | NA_peak_27 | 15 | Intron (ENSM | 10 | 12666423 | 12688862 | 22440 | 2 | 22288 | ENSMUST00 | -29295 | ENSMUSG0 | Utrn | utrophin |
| chr10 | 12977684 | 12978028 | 345 | * | NA_peak_27 | 40 | Distal Interge | 10 | 12941727 | 12964298 | 22572 | 2 | 74732 | ENSMUST00 | -13386 | ENSMUSG0 | Stx11 | syntaxin 11 |
| chr10 | 13619589 | 13619866 | 278 | * | NA_peak_28 | 31 | Distal Interge | 10 | 13523842 | 13553142 | 29301 | 2 | 56535 | ENSMUST00 | -66447 | ENSMUSG0 | Pex3 | peroxisomal biogenesis factor 3 |
| chr10 | 13658064 | 13658310 | 247 | * | NA_peak_28 | 23 | Intron (ENSM | 10 | 13523842 | 13553142 | 29301 | 2 | 56535 | ENSMUST00 | -104922 | ENSMUSG0 | Pex3 | peroxisomal biogenesis factor 3 |
| chr10 | 14097533 | 14097723 | 191 | * | NA_peak_28 | 19 | Intron (ENSM | 10 | 13967024 | 14079675 | 112652 | 1 | 15273 | ENSMUST00 | 130509 | ENSMUSG0 | Hivep2 | human immunodeficiency virus type I enhancer binding protein 2 |
| chr10 | 14347402 | 14347731 | 330 | * | NA_peak_28 | 36 | Distal Interge | 10 | 14530916 | 14544995 | 14080 | 2 | 215798 | ENSMUST00 | 197264 | ENSMUSG0 | Adgrg6 | adhesion G protein-coupled receptor G6 |
| chr10 | 14584737 | 14584841 | 105 | * | NA_peak_28 | 40 | Distal Interge | 10 | 14407898 | 14545659 | 137762 | 2 | 215798 | ENSMUST00 | -39078 | ENSMUSG0 | Adgrg6 | adhesion G protein-coupled receptor G6 |
| chr10 | 15236199 | 15236825 | 627 | * | NA_peak_28 | 23 | Distal Interge | 10 | 14760219 | 14770583 | 10365 | 1 | 18101 | ENSMUST00 | 475980 | ENSMUSG0 | Nmbr | neuromedin B receptor |
| chr10 | 15339608 | 15339956 | 349 | * | NA_peak_28 | 55 | Intron (ENSM | 10 | 15791777 | 15807065 | 15289 | 1 | 77538 | ENSMUST00 | -451821 | ENSMUSG0 | C330004P14 | RIKEN cDNA C330004P14 gene |
| chr10 | 15344784 | 15345064 | 281 | * | NA_peak_28 | 24 | Intron (ENSM | 10 | 15791777 | 15807065 | 15289 | 1 | 77538 | ENSMUST00 | -446713 | ENSMUSG0 | C330004P14 | RIKEN cDNA C330004P14 gene |
| chr10 | 15379402 | 15379501 | 100 | * | NA_peak_28 | 23 | Intron (ENSM | 10 | 15791777 | 15807065 | 15289 | 1 | 77538 | ENSMUST00 | -412276 | ENSMUSG0 | C330004P14 | RIKEN cDNA C330004P14 gene |
| chr10 | 15470350 | 15470632 | 283 | * | NA_peak_29 | 40 | Distal Interge | 10 | 15791777 | 15807065 | 15289 | 1 | 77538 | ENSMUST00 | -321145 | ENSMUSG0 | C330004P14 | RIKEN cDNA C330004P14 gene |
| chr10 | 15639701 | 15640017 | 317 | * | NA_peak_29 | 28 | Intron (ENSM | 10 | 15791777 | 15807065 | 15289 | 1 | 77538 | ENSMUST00 | -151760 | ENSMUSG0 | C330004P14 | RIKEN cDNA C330004P14 gene |
| chr10 | 16901757 | 16902049 | 293 | * | NA_peak_29 | 13 | Distal Interge | 10 | 16951295 | 16963180 | 11886 | 2 | 100504235 | ENSMUST00 | 61131 | ENSMUSG0 | Gm20125 | predicted gene, 20125 |

| | | | | | | | | | | | | | | | | | |
|---|---|---|---|---|---|---|---|---|---|---|---|---|---|---|---|---|---|
| chr10 | 17159315 | 17159395 | 81 | * | NA_peak_29 | 24 | Distal Interge | 10 | 16981304 | 17095807 | 114504 | 2 | 100504235 | ENSMUST00 | -63508 | ENSMUSG0 | Gm20125 | predicted gene, 20125 |
| chr10 | 18247461 | 18247528 | 68 | * | NA_peak_30 | 20 | Distal Interge | 10 | 18214078 | 18234998 | 20921 | 2 | 215814 | ENSMUST00 | -12463 | ENSMUSG0 | Ccdc28a | coiled-coil domain containing 28A |
| chr10 | 18545916 | 18546141 | 226 | * | NA_peak_30 | 23 | Promoter (<= | 10 | 18539498 | 18546076 | 6579 | 2 | 56016 | ENSMUST00 | 0 | ENSMUSG0 | Hebp2 | heme binding protein 2 |
| chr10 | 18591626 | 18591872 | 247 | * | NA_peak_30 | 21 | Exon (ENSMU | 10 | 18539498 | 18546076 | 6579 | 2 | 56016 | ENSMUST00 | -45550 | ENSMUSG0 | Hebp2 | heme binding protein 2 |
| chr10 | 18597692 | 18597758 | 67 | * | NA_peak_30 | 13 | Exon (ENSMU | 10 | 18539498 | 18546076 | 6579 | 2 | 56016 | ENSMUST00 | -51616 | ENSMUSG0 | Hebp2 | heme binding protein 2 |
| chr10 | 18940080 | 18940129 | 50 | * | NA_peak_30 | 27 | Distal Interge | 10 | 19007267 | 19008845 | 1579 | 2 | 21929 | ENSMUST00 | 68716 | ENSMUSG0 | Tnfaip3 | tumor necrosis factor, alpha-induced protein 3 |
| chr10 | 19033235 | 19033596 | 362 | * | NA_peak_30 | 23 | Distal Interge | 10 | 19008266 | 19015657 | 7392 | 2 | 21929 | ENSMUST00 | -17578 | ENSMUSG0 | Tnfaip3 | tumor necrosis factor, alpha-induced protein 3 |
| chr10 | 19784193 | 19784473 | 281 | * | NA_peak_30 | 31 | Distal Interge | 10 | 19847916 | 19851459 | 3544 | 2 | 76157 | ENSMUST00 | 66986 | ENSMUSG0 | Slc35d3 | solute carrier family 35, member D3 |
| chr10 | 19800152 | 19800542 | 391 | * | NA_peak_30 | 37 | Distal Interge | 10 | 19847916 | 19851459 | 3544 | 2 | 76157 | ENSMUST00 | 50917 | ENSMUSG0 | Slc35d3 | solute carrier family 35, member D3 |
| chr10 | 19933715 | 19934011 | 297 | * | NA_peak_31 | 13 | Promoter (<= | 10 | 19934472 | 20142753 | 208282 | 1 | 26408 | ENSMUST00 | -461 | ENSMUSG0 | Map3k5 | mitogen-activated protein kinase kinase kinase 5 |
| chr10 | 21072089 | 21072334 | 246 | * | NA_peak_31 | 16 | Promoter (2-3 | 10 | 21069845 | 21074406 | 4562 | 1 | 52906 | ENSMUST00 | 2244 | ENSMUSG0 | Ahi1 | Abelson helper integration site 1 |
| chr10 | 21416218 | 21416329 | 112 | * | NA_peak_31 | 20 | Distal Interge | 10 | 21377347 | 21388996 | 11650 | 1 | 237320 | ENSMUST00 | 38871 | ENSMUSG0 | Aldh8a1 | aldehyde dehydrogenase 8 family, member A1 |
| chr10 | 21732409 | 21732693 | 285 | * | NA_peak_31 | 41 | Distal Interge | 10 | 21690845 | 21693978 | 3134 | 1 | 432436 | ENSMUST00 | 41564 | ENSMUSG0 | Gm5420 | predicted gene 5420 |
| chr10 | 22142536 | 22142983 | 448 | * | NA_peak_32 | 242 | Intron (ENSM | 10 | 22117362 | 22148610 | 31249 | 2 | 319887 | ENSMUST00 | 5627 | ENSMUSG0 | E030030I06 | RIKEN cDNA E030030I06 gene |
| chr10 | 22182159 | 22182368 | 210 | * | NA_peak_32 | 32 | Intron (ENSM | 10 | 22173919 | 22374139 | 200221 | 1 | 19370 | ENSMUST00 | 8240 | ENSMUSG0 | Raet1c | retinoic acid early transcript gamma |
| chr10 | 22198094 | 22198470 | 377 | * | NA_peak_32 | 53 | Intron (ENSM | 10 | 22173919 | 22374139 | 200221 | 1 | 19370 | ENSMUST00 | 24175 | ENSMUSG0 | Raet1c | retinoic acid early transcript gamma |
| chr10 | 22273975 | 22274657 | 683 | * | NA_peak_32 | 49 | Promoter (<= | 10 | 22273346 | 22288849 | 15504 | 1 | 667281 | ENSMUST00 | 629 | ENSMUSG0 | H60b | histocompatibility 60b |
| chr10 | 22329617 | 22329994 | 378 | * | NA_peak_32 | 19 | Intron (ENSM | 10 | 22307208 | 22314196 | 6989 | 1 | 606736 | ENSMUST00 | 22409 | NA | C920009B18 | RIKEN cDNA C920009B18 gene |
| chr10 | 22330313 | 22330431 | 119 | * | NA_peak_32 | 53 | Intron (ENSM | 10 | 22307208 | 22314196 | 6989 | 1 | 606736 | ENSMUST00 | 23105 | NA | C920009B18 | RIKEN cDNA C920009B18 gene |
| chr10 | 22332279 | 22332436 | 158 | * | NA_peak_32 | 15 | Intron (ENSM | 10 | 22307208 | 22314196 | 6989 | 1 | 606736 | ENSMUST00 | 25071 | NA | C920009B18 | RIKEN cDNA C920009B18 gene |
| chr10 | 22560008 | 22560227 | 220 | * | NA_peak_32 | 41 | Distal Interge | 10 | 22645011 | 22704285 | 59275 | 1 | 353169 | ENSMUST00 | -84784 | ENSMUSG0 | Slc2a12 | solute carrier family 2 (facilitated glucose transporter), member 12 |
| chr10 | 22760053 | 22760357 | 305 | * | NA_peak_32 | 30 | Distal Interge | 10 | 22705381 | 22731938 | 26558 | 2 | 237336 | ENSMUST00 | -28115 | ENSMUSG0 | Tbpl1 | TATA box binding protein-like 1 |
| chr10 | 22772200 | 22772481 | 282 | * | NA_peak_32 | 15 | Distal Interge | 10 | 22705381 | 22731938 | 26558 | 2 | 237336 | ENSMUST00 | -40262 | ENSMUSG0 | Tbpl1 | TATA box binding protein-like 1 |
| chr10 | 22940691 | 22940927 | 237 | * | NA_peak_32 | 20 | Distal Interge | 10 | 22817281 | 22820174 | 2894 | 2 | 21412 | ENSMUST00 | -120517 | ENSMUSG0 | Tcf21 | transcription factor 21 |
| chr10 | 22959795 | 22959919 | 125 | * | NA_peak_32 | 24 | Distal Interge | 10 | 22817281 | 22820174 | 2894 | 2 | 21412 | ENSMUST00 | -139621 | ENSMUSG0 | Tcf21 | transcription factor 21 |
| chr10 | 23481353 | 23481700 | 348 | * | NA_peak_33 | 40 | Distal Interge | 10 | 23104763 | 23350786 | 246024 | 2 | 14051 | ENSMUST00 | -130567 | ENSMUSG0 | Eya4 | EYA transcriptional coactivator and phosphatase 4 |
| chr10 | 24321908 | 24322268 | 361 | * | NA_peak_33 | 32 | Distal Interge | 10 | 24223545 | 24231199 | 7655 | 1 | 59012 | ENSMUST00 | 98363 | ENSMUSG0 | Moxd1 | monooxygenase, DBH-like 1 |
| chr10 | 24330465 | 24330966 | 502 | * | NA_peak_33 | 64 | Distal Interge | 10 | 24223545 | 24231199 | 7655 | 1 | 59012 | ENSMUST00 | 106920 | ENSMUSG0 | Moxd1 | monooxygenase, DBH-like 1 |
| chr10 | 24526958 | 24527104 | 147 | * | NA_peak_33 | 23 | Distal Interge | 10 | 24580048 | 24593758 | 13711 | 2 | 102638890 | ENSMUST00 | 66654 | NA | Gm15270 | predicted gene 15270 |
| chr10 | 24754658 | 24754760 | 103 | * | NA_peak_33 | 27 | Distal Interge | 10 | 24773491 | 24785005 | 11515 | 2 | 209558 | ENSMUST00 | 30245 | ENSMUSG0 | Enpp3 | ectonucleotide pyrophosphatase/phosphodiesterase 3 |
| chr10 | 25420051 | 25420466 | 416 | * | NA_peak_34 | 32 | Intron (ENSM | 10 | 25408387 | 25484280 | 75894 | 1 | 13822 | ENSMUST00 | 11664 | ENSMUSG0 | Epb41l2 | erythrocyte membrane protein band 4.1 like 2 |
| chr10 | 26168760 | 26168996 | 237 | * | NA_peak_34 | 43 | Distal Interge | 10 | 26229707 | 26260804 | 31098 | 1 | 268288 | ENSMUST00 | -60711 | ENSMUSG0 | Samd3 | sterile alpha motif domain containing 3 |
| chr10 | 26731765 | 26731922 | 158 | * | NA_peak_34 | 45 | Intron (ENSM | 10 | 26753421 | 26860732 | 107312 | 1 | 73910 | ENSMUST00 | -21499 | ENSMUSG0 | Arhgap18 | Rho GTPase activating protein 18 |
| chr10 | 27794106 | 27794479 | 374 | * | NA_peak_35 | 30 | Intron (ENSM | 10 | 27422392 | 27619758 | 197367 | 2 | 16773 | ENSMUST00 | -174348 | ENSMUSG0 | Lama2 | laminin, alpha 2 |
| chr10 | 27824622 | 27824826 | 205 | * | NA_peak_35 | 14 | Distal Interge | 10 | 27422392 | 27619758 | 197367 | 2 | 16773 | ENSMUST00 | -204864 | ENSMUSG0 | Lama2 | laminin, alpha 2 |
| chr10 | 28903166 | 28903171 | 6 | * | NA_peak_35 | 16 | Intron (ENSM | 10 | 28972366 | 28974189 | 1824 | 2 | 67719 | ENSMUST00 | 71018 | ENSMUSG0 | 2310057J18 | RIKEN cDNA 2310057J18 gene |
| chr10 | 28936901 | 28937093 | 193 | * | NA_peak_35 | 36 | Distal Interge | 10 | 28972366 | 28974189 | 1824 | 2 | 67719 | ENSMUST00 | 37096 | ENSMUSG0 | 2310057J18 | RIKEN cDNA 2310057J18 gene |
| chr10 | 29761344 | 29761657 | 314 | * | NA_peak_35 | 45 | Distal Interge | 10 | 29452418 | 29535867 | 83450 | 2 | 72780 | ENSMUST00 | -225477 | ENSMUSG0 | Rspo3 | R-spondin 3 |
| chr10 | 29772164 | 29772385 | 222 | * | NA_peak_35 | 47 | Distal Interge | 10 | 29452418 | 29535867 | 83450 | 2 | 72780 | ENSMUST00 | -236297 | ENSMUSG0 | Rspo3 | R-spondin 3 |
| chr10 | 30200153 | 30200416 | 264 | * | NA_peak_36 | 28 | Promoter (<= | 10 | 30194569 | 30200492 | 5924 | 2 | 66311 | ENSMUST00 | 76 | ENSMUSG0 | Cenpw | centromere protein W |
| chr10 | 30220841 | 30221106 | 266 | * | NA_peak_36 | 30 | Distal Interge | 10 | 30196009 | 30200560 | 4552 | 2 | 66311 | ENSMUST00 | -20281 | ENSMUSG0 | Cenpw | centromere protein W |
| chr10 | 30476292 | 30476662 | 371 | * | NA_peak_36 | 31 | Distal Interge | 10 | 30559121 | 30590109 | 30989 | 2 | 73681 | ENSMUST00 | 113447 | ENSMUSG0 | Trmt11 | tRNA methyltransferase 11 |
| chr10 | 30631800 | 30632104 | 305 | * | NA_peak_36 | 18 | Promoter (2-3 | 10 | 30608231 | 30629632 | 21402 | 2 | 66847 | ENSMUST00 | -2168 | ENSMUSG0 | Hint3 | histidine triad nucleotide binding protein 3 |
| chr10 | 30816276 | 30817193 | 918 | * | NA_peak_36 | 59 | Distal Interge | 10 | 30695891 | 30803326 | 107436 | 2 | 211329 | ENSMUST00 | -12950 | ENSMUSG0 | Ncoa7 | nuclear receptor coactivator 7 |
| chr10 | 31609469 | 31609745 | 277 | * | NA_peak_36 | 18 | Promoter (<= | 10 | 31493193 | 31609184 | 115992 | 2 | 268291 | ENSMUST00 | -285 | ENSMUSG0 | Rnf217 | ring finger protein 217 |
| chr10 | 31883640 | 31883685 | 46 | * | NA_peak_36 | 45 | Intron (ENSM | 10 | 31493193 | 31609184 | 115992 | 2 | 268291 | ENSMUST00 | -274456 | ENSMUSG0 | Rnf217 | ring finger protein 217 |
| chr10 | 31997347 | 31997726 | 380 | * | NA_peak_36 | 40 | Intron (ENSM | 10 | 31493193 | 31609184 | 115992 | 2 | 268291 | ENSMUST00 | -388163 | ENSMUSG0 | Rnf217 | ring finger protein 217 |
| chr10 | 32188013 | 32188230 | 218 | * | NA_peak_37 | 23 | Intron (ENSM | 10 | 32329765 | 32410335 | 80571 | 2 | 432450 | ENSMUST00 | 222105 | ENSMUSG0 | Nkain2 | Na+/K+ transporting ATPase interacting 2 |
| chr10 | 35410228 | 35410451 | 224 | * | NA_peak_38 | 16 | Distal Interge | 10 | 34483532 | 34606280 | 122749 | 1 | 14302 | ENSMUST00 | 926696 | ENSMUSG0 | Frk | fyn-related kinase |
| chr10 | 36331421 | 36331690 | 270 | * | NA_peak_38 | 23 | Distal Interge | 10 | 36506814 | 36833773 | 326960 | 1 | 319415 | ENSMUST00 | -175124 | ENSMUSG0 | Hs3st5 | heparan sulfate (glucosamine) 3-O-sulfotransferase 5 |
| chr10 | 36642960 | 36643392 | 433 | * | NA_peak_38 | 13 | Intron (ENSM | 10 | 36507407 | 36810038 | 302632 | 1 | 319415 | ENSMUST00 | 135553 | ENSMUSG0 | Hs3st5 | heparan sulfate (glucosamine) 3-O-sulfotransferase 5 |
| chr10 | 39115907 | 39116077 | 171 | * | NA_peak_39 | 31 | Distal Interge | 10 | 39118820 | 39122314 | 3495 | 2 | 66337 | ENSMUST00 | 6237 | ENSMUSG0 | Fam229b | family with sequence similarity 229, member B |
| chr10 | 40838101 | 40838191 | 91 | * | NA_peak_40 | 20 | Intron (ENSM | 10 | 40840488 | 40845937 | 5450 | 2 | 71713 | ENSMUST00 | 7746 | ENSMUSG0 | Cdc40 | cell division cycle 40 |
| chr10 | 41777354 | 41777448 | 95 | * | NA_peak_40 | 16 | Intron (ENSM | 10 | 41746746 | 41753913 | 7168 | 1 | 100196896 | ENSMUST00 | 30608 | NA | BC048559 | cDNA sequence BC048559 |
| chr10 | 42110457 | 42110676 | 220 | * | NA_peak_40 | 22 | Distal Interge | 10 | 41916807 | 42018442 | 101636 | 2 | 213402 | ENSMUST00 | -92015 | ENSMUSG0 | Armc2 | armadillo repeat containing 2 |
| chr10 | 42591552 | 42591890 | 339 | * | NA_peak_40 | 40 | Intron (ENSM | 10 | 42583838 | 42700492 | 116655 | 1 | 14628 | ENSMUST00 | 7714 | ENSMUSG0 | Ostm1 | osteopetrosis associated transmembrane protein 1 |
| chr10 | 43511471 | 43511729 | 259 | * | NA_peak_41 | 35 | Exon (ENSMU | 10 | 43525159 | 43525843 | 685 | 2 | 67851 | ENSMUST00 | 14114 | ENSMUSG0 | Mtres1 | mitochondrial transcription rescue factor 1 |
| chr10 | 44209666 | 44209833 | 168 | * | NA_peak_41 | 22 | Intron (ENSM | 10 | 44217029 | 44219582 | 2554 | 1 | 70365 | ENSMUST00 | -7196 | ENSMUSG0 | Speer5-ps1 | spermatogenesis associated glutamate (E)-rich protein 5, pseudogene 1 |
| chr10 | 44906552 | 44906743 | 192 | * | NA_peak_42 | 19 | Intron (ENSM | 10 | 45067203 | 45167198 | 99996 | 1 | 19072 | ENSMUST00 | -160460 | ENSMUSG0 | Prep | prolyl endopeptidase |
| chr10 | 45518345 | 45518697 | 353 | * | NA_peak_42 | 30 | Distal Interge | 10 | 45418363 | 45486395 | 68033 | 2 | 380669 | ENSMUST00 | -31950 | ENSMUSG0 | Lin28b | lin-28 homolog B (C. elegans) |
| chr10 | 45574477 | 45574709 | 233 | * | NA_peak_42 | 20 | Distal Interge | 10 | 45577829 | 45712345 | 134517 | 1 | 209462 | ENSMUST00 | -3120 | ENSMUSG0 | Hace1 | HECT domain and ankyrin repeat containing, E3 ubiquitin protein ligase 1 |
| chr10 | 46380313 | 46380693 | 381 | * | NA_peak_42 | 31 | Distal Interge | 10 | 45837880 | 45852283 | 14404 | 1 | 320363 | ENSMUST00 | 542433 | ENSMUSG0 | D030045P18 | RIKEN cDNA D030045P18 gene |
| chr10 | 46914691 | 46914959 | 269 | * | NA_peak_42 | 23 | Distal Interge | 10 | 45837880 | 45852283 | 14404 | 1 | 320363 | ENSMUST00 | 1076811 | ENSMUSG0 | D030045P18 | RIKEN cDNA D030045P18 gene |
| chr10 | 46984536 | 46984971 | 436 | * | NA_peak_42 | 40 | Distal Interge | 10 | 45837880 | 45852283 | 14404 | 1 | 320363 | ENSMUST00 | 1146656 | ENSMUSG0 | D030045P18 | RIKEN cDNA D030045P18 gene |
| chr10 | 50029288 | 50029668 | 381 | * | NA_peak_43 | 50 | Distal Interge | 10 | 49094833 | 49788766 | 693934 | 2 | 14806 | ENSMUST00 | -240522 | ENSMUSG0 | Grik2 | glutamate receptor, ionotropic, kainate 2 (beta 2) |
| chr10 | 50184533 | 50184817 | 285 | * | NA_peak_43 | 43 | Distal Interge | 10 | 49094833 | 49788766 | 693934 | 2 | 14806 | ENSMUST00 | -395767 | ENSMUSG0 | Grik2 | glutamate receptor, ionotropic, kainate 2 (beta 2) |
| chr10 | 50520106 | 50520200 | 95 | * | NA_peak_43 | 16 | Distal Interge | 10 | 50592669 | 50851485 | 258817 | 1 | 77987 | ENSMUST00 | -72469 | ENSMUSG0 | Ascc3 | activating signal cointegrator 1 complex subunit 3 |
| chr10 | 51884033 | 51884272 | 240 | * | NA_peak_43 | 35 | Intron (ENSM | 10 | 51756174 | 51757194 | 1021 | 2 | 71090 | ENSMUST00 | -126839 | ENSMUSG0 | 4933411G06 | Sumo1 pseudogene |
| chr10 | 52002222 | 52002407 | 186 | * | NA_peak_43 | 23 | Distal Interge | 10 | 52022502 | 52028471 | 5970 | 1 | 215031 | ENSMUST00 | -20095 | ENSMUSG0 | Vgll2 | vestigial like family member 2 |
| chr10 | 52767147 | 52767223 | 77 | * | NA_peak_44 | 40 | Intron (ENSM | 10 | 52690533 | 53111622 | 421090 | 1 | 215085 | ENSMUST00 | 76614 | ENSMUSG0 | Slc35f1 | solute carrier family 35, member F1 |
| chr10 | 53249372 | 53249534 | 163 | * | NA_peak_44 | 31 | Intron (ENSM | 10 | 53235187 | 53260736 | 25550 | 1 | 102639992 | ENSMUST00 | 14185 | ENSMUSG0 | Gm36173 | predicted gene, 36173 |
| chr10 | 53249843 | 53250662 | 820 | * | NA_peak_44 | 31 | Intron (ENSM | 10 | 53235187 | 53260736 | 25550 | 1 | 102639992 | ENSMUST00 | 14656 | ENSMUSG0 | Gm36173 | predicted gene, 36173 |
| chr10 | 53496681 | 53496885 | 205 | * | NA_peak_44 | 15 | Intron (ENSM | 10 | 53498443 | 53518989 | 20547 | 2 | 100502868 | ENSMUST00 | 22104 | ENSMUSG0 | Gm20597 | predicted gene, 20597 |
| chr10 | 53786906 | 53787112 | 207 | * | NA_peak_44 | 20 | Distal Interge | 10 | 53633145 | 53751123 | 117979 | 2 | 75906 | ENSMUST00 | -35783 | ENSMUSG0 | Fam184a | family with sequence similarity 184, member A |
| chr10 | 54146515 | 54146790 | 276 | * | NA_peak_44 | 19 | Distal Interge | 10 | 53907444 | 54076609 | 169166 | 2 | 17155 | ENSMUST00 | -69906 | ENSMUSG0 | Man1a | mannosidase 1, alpha |
| chr10 | 54616075 | 54616366 | 292 | * | NA_peak_44 | 19 | Distal Interge | 10 | 53907444 | 54076609 | 169166 | 2 | 17155 | ENSMUST00 | -539466 | ENSMUSG0 | Man1a | mannosidase 1, alpha |
| chr10 | 55632413 | 55632772 | 360 | * | NA_peak_45 | 50 | Distal Interge | 10 | 56106917 | 56116880 | 9964 | 1 | 73390 | ENSMUST00 | -474145 | ENSMUSG0 | Msl3l2 | MSL3 like 2 |
| chr10 | 55867155 | 55867215 | 61 | * | NA_peak_45 | 23 | Distal Interge | 10 | 56106917 | 56116880 | 9964 | 1 | 73390 | ENSMUST00 | -239702 | ENSMUSG0 | Msl3l2 | MSL3 like 2 |
| chr10 | 57766743 | 57766984 | 242 | * | NA_peak_46 | 35 | Distal Interge | 10 | 57784881 | 57788450 | 3570 | 1 | 12140 | ENSMUST00 | -17897 | ENSMUSG0 | Fabp7 | fatty acid binding protein 7, brain |
| chr10 | 58763993 | 58764277 | 285 | * | NA_peak_46 | 64 | Distal Interge | 10 | 58813359 | 59050020 | 236662 | 1 | 237353 | ENSMUST00 | -49082 | ENSMUSG0 | Sh3rf3 | SH3 domain containing ring finger 3 |
| chr10 | 59842396 | 59842740 | 345 | * | NA_peak_46 | 22 | Intron (ENSM | 10 | 59827221 | 59840367 | 13147 | 1 | 216001 | ENSMUST00 | 15175 | ENSMUSG0 | Micu1 | mitochondrial calcium uptake 1 |
| chr10 | 59923144 | 59923369 | 226 | * | NA_peak_46 | 19 | Distal Interge | 10 | 59949669 | 59951834 | 2166 | 2 | 74747 | ENSMUST00 | 28465 | ENSMUSG0 | Ddit4 | DNA-damage-inducible transcript 4 |
| chr10 | 60111726 | 60111960 | 235 | * | NA_peak_47 | 17 | Intron (ENSM | 10 | 60106468 | 60133908 | 27441 | 1 | 94214 | ENSMUST00 | 5258 | ENSMUSG0 | Spock2 | sparc/osteonectin, cwcv and kazal-like domains proteoglycan 2 |
| chr10 | 60137412 | 60137509 | 98 | * | NA_peak_47 | 21 | Distal Interge | 10 | 60126776 | 60126833 | 58 | 1 | 102465546 | ENSMUST00 | 10636 | ENSMUSG0 | Mir6906 | microRNA 6906 |
| chr10 | 61410908 | 61411277 | 370 | * | NA_peak_47 | 26 | Distal Interge | 10 | 61417972 | 61425338 | 7367 | 1 | 18119 | ENSMUST00 | -6695 | ENSMUSG0 | Nodal | nodal |
| chr10 | 61513877 | 61514068 | 192 | * | NA_peak_47 | 21 | Intron (ENSM | 10 | 61475801 | 61582791 | 106991 | 1 | 216011 | ENSMUST00 | 38076 | ENSMUSG0 | Lrrc20 | leucine rich repeat containing 20 |
| chr10 | 62132766 | 62133075 | 310 | * | NA_peak_47 | 24 | Promoter (<= | 10 | 62133090 | 62134763 | 1674 | 1 | 11925 | ENSMUST00 | -15 | ENSMUSG0 | Neurog3 | neurogenin 3 |
| chr10 | 62214969 | 62215266 | 298 | * | NA_peak_47 | 18 | Intron (ENSM | 10 | 62185396 | 62231251 | 45856 | 2 | 70423 | ENSMUST00 | 15985 | ENSMUSG0 | Tspan15 | tetraspanin 15 |
| chr10 | 62749247 | 62749477 | 231 | * | NA_peak_47 | 13 | Intron (ENSM | 10 | 62753273 | 62760349 | 7077 | 2 | 67500 | ENSMUST00 | 10872 | ENSMUSG0 | Ccar1 | cell division cycle and apoptosis regulator 1 |
| chr10 | 63448395 | 63448677 | 283 | * | NA_peak_48 | 20 | Intron (ENSM | 10 | 63457510 | 65003667 | 1546158 | 1 | 216033 | ENSMUST00 | -8833 | ENSMUSG0 | Ctnna3 | catenin (cadherin associated protein), alpha 3 |
| chr10 | 65168464 | 65168823 | 360 | * | NA_peak_48 | 66 | Distal Interge | 10 | 65119526 | 65120367 | 842 | 2 | 102632700 | ENSMUST00 | -48097 | ENSMUSG0 | Gm28881 | predicted gene 28881 |
| chr10 | 65575000 | 65575299 | 300 | * | NA_peak_48 | 37 | Distal Interge | 10 | 65875788 | 65908239 | 32452 | 2 | 102632897 | ENSMUST00 | 332940 | ENSMUSG0 | Gm30853 | predicted gene, 30853 |
| chr10 | 65642655 | 65642853 | 199 | * | NA_peak_48 | 23 | Distal Interge | 10 | 65875788 | 65908239 | 32452 | 2 | 102632897 | ENSMUST00 | 265386 | ENSMUSG0 | Gm30853 | predicted gene, 30853 |
| chr10 | 66463778 | 66464046 | 269 | * | NA_peak_49 | 96 | Distal Interge | 10 | 66120605 | 66254223 | 133619 | 2 | 69433 | ENSMUST00 | -209555 | NA | 1700023F02 | RIKEN cDNA 1700023F02 gene |
| chr10 | 66464614 | 66464906 | 293 | * | NA_peak_49 | 31 | Distal Interge | 10 | 66120605 | 66254223 | 133619 | 2 | 69433 | ENSMUST00 | -210391 | NA | 1700023F02 | RIKEN cDNA 1700023F02 gene |
| chr10 | 66468521 | 66468733 | 213 | * | NA_peak_49 | 39 | Distal Interge | 10 | 66120605 | 66254223 | 133619 | 2 | 69433 | ENSMUST00 | -214298 | NA | 1700023F02 | RIKEN cDNA 1700023F02 gene |
| chr10 | 66808194 | 66808487 | 294 | * | NA_peak_49 | 33 | Distal Interge | 10 | 66915636 | 66917739 | 2104 | 2 | 68488 | ENSMUST00 | 109252 | NA | 1110002J07 | RIKEN cDNA 1110002J07 gene |
| chr10 | 67300806 | 67301177 | 372 | * | NA_peak_49 | 22 | Intron (ENSM | 10 | 67266689 | 67285305 | 18617 | 2 | 641340 | ENSMUST00 | -15501 | ENSMUSG0 | Nrbf2 | nuclear receptor binding factor 2 |
| chr10 | 68799863 | 68800011 | 149 | * | NA_peak_50 | 27 | Distal Interge | 10 | 68723824 | 68779201 | 55378 | 1 | 327766 | ENSMUST00 | 76039 | ENSMUSG0 | Tmem26 | transmembrane protein 26 |
| chr10 | 68896724 | 68897043 | 320 | * | NA_peak_50 | 20 | Distal Interge | 10 | 69020740 | 69033888 | 13149 | 2 | 102635787 | ENSMUST00 | 136845 | ENSMUSG0 | Gm33035 | predicted gene, 33035 |
| chr10 | 69053150 | 69053382 | 233 | * | NA_peak_50 | 15 | Distal Interge | 10 | 69020740 | 69033888 | 13149 | 2 | 102635787 | ENSMUST00 | -19262 | ENSMUSG0 | Gm33035 | predicted gene, 33035 |
| chr10 | 69062664 | 69062835 | 172 | * | NA_peak_50 | 16 | Distal Interge | 10 | 69020740 | 69033888 | 13149 | 2 | 102635787 | ENSMUST00 | -28776 | ENSMUSG0 | Gm33035 | predicted gene, 33035 |
| chr10 | 69699083 | 69699316 | 234 | * | NA_peak_50 | 14 | Intron (ENSM | 10 | 69706370 | 70015631 | 309262 | 1 | 11735 | ENSMUST00 | -7054 | ENSMUSG0 | Ank3 | ankyrin 3, epithelial |
| chr10 | 70511772 | 70512169 | 398 | * | NA_peak_51 | 17 | Intron (ENSM | 10 | 70440926 | 70558731 | 117806 | 1 | 71721 | ENSMUST00 | 70846 | ENSMUSG0 | Fam13c | family with sequence similarity 13, member C |
| chr10 | 70599657 | 70600062 | 406 | * | NA_peak_51 | 31 | Promoter (<= | 10 | 70557682 | 70599291 | 41610 | 2 | 70911 | ENSMUST00 | -366 | ENSMUSG0 | Phyhipl | phytanoyl-CoA hydroxylase interacting protein-like |
| chr10 | 70819216 | 70819502 | 287 | * | NA_peak_51 | 31 | Distal Interge | 10 | 70934502 | 70948489 | 13988 | 2 | 83675 | ENSMUST00 | 128987 | ENSMUSG0 | Bicc1 | BicC family RNA binding protein 1 |

| | | | | | | | | | | | | | | | | | |
|---|---|---|---|---|---|---|---|---|---|---|---|---|---|---|---|---|---|
| chr10 | 70884801 | 70885287 | 487 | * | NA_peak_513 | 52 | Intron (ENSM | 10 | 70934502 | 70948489 | 13988 | 2 | 83675 | ENSMUST00 | 63202 | ENSMUSG00 | Bicc1 | BicC family RNA binding protein 1 |
| chr10 | 70888521 | 70888900 | 380 | * | NA_peak_513 | 30 | Intron (ENSM | 10 | 70934502 | 70948489 | 13988 | 2 | 83675 | ENSMUST00 | 59589 | ENSMUSG00 | Bicc1 | BicC family RNA binding protein 1 |
| chr10 | 71098222 | 71098614 | 393 | * | NA_peak_514 | 45 | Intron (ENSM | 10 | 70925102 | 71146860 | 221759 | 2 | 83675 | ENSMUST00 | 48246 | ENSMUSG00 | Bicc1 | BicC family RNA binding protein 1 |
| chr10 | 71842637 | 71842755 | 119 | * | NA_peak_518 | 20 | Distal Interge | 10 | 71980092 | 71980634 | 543 | 2 | 108950 | ENSMUST00 | 137879 | NA | 1700049L16 | hematological and neurological expressed 1-like pseudogene |
| chr10 | 72012883 | 72013093 | 211 | * | NA_peak_519 | 20 | Distal Interge | 10 | 71980092 | 71980634 | 543 | 2 | 108950 | ENSMUST00 | -32249 | NA | 1700049L16 | hematological and neurological expressed 1-like pseudogene |
| chr10 | 73437654 | 73438182 | 529 | * | NA_peak_524 | 31 | Intron (ENSM | 10 | 73099342 | 74317264 | 1217923 | 1 | 11994 | ENSMUST00 | 338312 | ENSMUSG00 | Pcdh15 | protocadherin 15 |
| chr10 | 73582173 | 73582773 | 601 | * | NA_peak_524 | 31 | Intron (ENSM | 10 | 73821867 | 74634765 | 812899 | 1 | 11994 | ENSMUST00 | -239094 | ENSMUSG00 | Pcdh15 | protocadherin 15 |
| chr10 | 73769782 | 73769887 | 106 | * | NA_peak_525 | 23 | Intron (ENSM | 10 | 73821867 | 74634765 | 812899 | 1 | 11994 | ENSMUST00 | -51980 | ENSMUSG00 | Pcdh15 | protocadherin 15 |
| chr10 | 74155348 | 74155627 | 280 | * | NA_peak_526 | 23 | Intron (ENSM | 10 | 74303151 | 74331050 | 27900 | 1 | 11994 | ENSMUST00 | -147524 | ENSMUSG00 | Pcdh15 | protocadherin 15 |
| chr10 | 74656098 | 74656493 | 396 | * | NA_peak_528 | 27 | Distal Interge | 10 | 74631834 | 74642364 | 10531 | 1 | 11994 | ENSMUST00 | 24264 | ENSMUSG00 | Pcdh15 | protocadherin 15 |
| chr10 | 74769797 | 74770187 | 391 | * | NA_peak_529 | 38 | Distal Interge | 10 | 74631834 | 74642364 | 10531 | 1 | 11994 | ENSMUST00 | 137963 | ENSMUSG00 | Pcdh15 | protocadherin 15 |
| chr10 | 74876942 | 74877113 | 172 | * | NA_peak_529 | 15 | Distal Interge | 10 | 74967177 | 75016907 | 49731 | 1 | 14687 | ENSMUST00 | -90064 | ENSMUSG00 | Gnaz | guanine nucleotide binding protein, alpha z subunit |
| chr10 | 76093824 | 76094197 | 374 | * | NA_peak_534 | 17 | Intron (ENSM | 10 | 76078617 | 76080502 | 1886 | 1 | 102639053 | ENSMUST00 | 15207 | ENSMUSG00 | Gm16222 | predicted gene 16222 |
| chr10 | 77258835 | 77259252 | 418 | * | NA_peak_537 | 22 | Promoter (<= | 10 | 77257776 | 77259223 | 1448 | 2 | 100169878 | ENSMUST00 | 0 | ENSMUSG00 | Gm10941 | predicted gene 10941 |
| chr10 | 78021197 | 78021532 | 336 | * | NA_peak_538 | 32 | Distal Interge | 10 | 78000680 | 78010083 | 9404 | 2 | 18641 | ENSMUST00 | -11114 | ENSMUSG00 | Pfkl | phosphofructokinase, liver, B-type |
| chr10 | 78858375 | 78858675 | 301 | * | NA_peak_542 | 16 | Distal Interge | 10 | 78846951 | 78847913 | 963 | 2 | 258305 | ENSMUST00 | -10462 | ENSMUSG00 | Olfr1356 | olfactory receptor 1356 |
| chr10 | 78940473 | 78940548 | 76 | * | NA_peak_542 | 20 | Distal Interge | 10 | 78955207 | 78956139 | 933 | 1 | 18372 | ENSMUST00 | -14659 | ENSMUSG00 | Olfr8 | olfactory receptor 8 |
| chr10 | 78988323 | 78988645 | 323 | * | NA_peak_542 | 36 | Intron (ENSM | 10 | 78981115 | 78984258 | 3144 | 1 | 259074 | ENSMUST00 | 7208 | ENSMUSG00 | Olfr1352 | olfactory receptor 1352 |
| chr10 | 79300261 | 79300399 | 139 | * | NA_peak_544 | 27 | Distal Interge | 10 | 79247692 | 79294535 | 46844 | 1 | 216144 | ENSMUST00 | 52569 | ENSMUSG00 | Vmn2r81 | vomeronasal 2, receptor 81 |
| chr10 | 79927431 | 79927736 | 306 | * | NA_peak_547 | 36 | Promoter (<= | 10 | 79927341 | 79954703 | 27363 | 1 | 13496 | ENSMUST00 | 90 | ENSMUSG00 | Arid3a | AT rich interactive domain 3A (BRIGHT-like) |
| chr10 | 80093910 | 80094041 | 132 | * | NA_peak_547 | 25 | Intron (ENSM | 10 | 80057771 | 80101187 | 43417 | 2 | 216161 | ENSMUST00 | 7146 | ENSMUSG00 | Sbno2 | strawberry notch 2 |
| chr10 | 80420862 | 80421142 | 281 | * | NA_peak_548 | 26 | Promoter (<= | 10 | 80420083 | 80421860 | 1778 | 2 | 21423 | ENSMUST00 | 718 | ENSMUSG00 | Tcf3 | transcription factor 3 |
| chr10 | 80429213 | 80429293 | 81 | * | NA_peak_548 | 20 | Promoter (1-2 | 10 | 80410054 | 80430985 | 20932 | 2 | 21423 | ENSMUST00 | 1692 | ENSMUSG00 | Tcf3 | transcription factor 3 |
| chr10 | 80820866 | 80821072 | 207 | * | NA_peak_549 | 18 | Distal Interge | 10 | 80826656 | 80829290 | 2635 | 1 | 18245 | ENSMUST00 | -5584 | ENSMUSG00 | Oaz1 | ornithine decarboxylase antizyme 1 |
| chr10 | 80905941 | 80906120 | 180 | * | NA_peak_549 | 16 | Promoter (<= | 10 | 80902798 | 80906412 | 3615 | 2 | 16907 | ENSMUST00 | 292 | ENSMUSG00 | Lmnb2 | lamin B2 |
| chr10 | 82027092 | 82027099 | 8 | * | NA_peak_553 | 23 | Distal Interge | 10 | 82048123 | 82061581 | 13459 | 1 | 408062 | ENSMUST00 | -21024 | ENSMUSG00 | Zfp873 | zinc finger protein 873 |
| chr10 | 82127546 | 82127654 | 109 | * | NA_peak_554 | 32 | Promoter (<= | 10 | 82128013 | 82153065 | 25053 | 1 | 216177 | ENSMUST00 | -359 | ENSMUSG00 | AU041133 | expressed sequence AU041133 |
| chr10 | 82131479 | 82131486 | 8 | * | NA_peak_554 | 17 | Intron (ENSM | 10 | 82128013 | 82153065 | 25053 | 1 | 216177 | ENSMUST00 | 3466 | ENSMUSG00 | AU041133 | expressed sequence AU041133 |
| chr10 | 82378813 | 82378970 | 158 | * | NA_peak_557 | 16 | Exon (ENSMU | 10 | 82486537 | 82492618 | 6082 | 2 | 432480 | ENSMUST00 | 113648 | ENSMUSG00 | Gm1553 | predicted gene 1553 |
| chr10 | 84155622 | 84155839 | 218 | * | NA_peak_563 | 33 | Distal Interge | 10 | 84036560 | 84100697 | 64138 | 2 | 102637349 | ENSMUST00 | -54925 | ENSMUSG00 | Gm34184 | predicted gene, 34184 |
| chr10 | 84947715 | 84947812 | 98 | * | NA_peak_567 | 19 | Exon (ENSMU | 10 | 84937393 | 85016209 | 78817 | 1 | 237422 | ENSMUST00 | 10322 | ENSMUSG00 | Ric8b | RIC8 guanine nucleotide exchange factor B |
| chr10 | 86195148 | 86195399 | 252 | * | NA_peak_573 | 31 | Intron (ENSM | 10 | 86095859 | 86113586 | 17728 | 1 | 78170 | ENSMUST00 | 99289 | ENSMUSG00 | 4930486F22 | RIKEN cDNA 4930486F22 gene |
| chr10 | 86248551 | 86248870 | 320 | * | NA_peak_574 | 20 | Intron (ENSM | 10 | 86300372 | 86349506 | 49135 | 1 | 21859 | ENSMUST00 | -51502 | ENSMUSG00 | Timp3 | tissue inhibitor of metalloproteinase 3 |
| chr10 | 87118045 | 87118360 | 316 | * | NA_peak_577 | 16 | Intron (ENSM | 10 | 87058046 | 87230588 | 172543 | 1 | 76640 | ENSMUST00 | 59999 | ENSMUSG00 | 1700113H08 | RIKEN cDNA 1700113H08 gene |
| chr10 | 87135660 | 87136097 | 438 | * | NA_peak_578 | 23 | Intron (ENSM | 10 | 87205979 | 87230587 | 24609 | 1 | 76640 | ENSMUST00 | -69882 | ENSMUSG00 | 1700113H08 | RIKEN cDNA 1700113H08 gene |
| chr10 | 87624140 | 87624414 | 275 | * | NA_peak_580 | 25 | Distal Interge | 10 | 87546673 | 87570332 | 23660 | 1 | 18478 | ENSMUST00 | 77467 | ENSMUSG00 | Pah | phenylalanine hydroxylase |
| chr10 | 87697360 | 87697697 | 338 | * | NA_peak_581 | 20 | Distal Interge | 10 | 87546673 | 87570332 | 23660 | 1 | 18478 | ENSMUST00 | 150687 | ENSMUSG00 | Pah | phenylalanine hydroxylase |
| chr10 | 87976471 | 87976605 | 135 | * | NA_peak_582 | 20 | Distal Interge | 10 | 87966847 | 87967761 | 915 | 1 | 22172 | ENSMUST00 | 9624 | NA | Tyms-ps | thymidylate synthase, pseudogene |
| chr10 | 88073601 | 88073939 | 339 | * | NA_peak_583 | 15 | Distal Interge | 10 | 88091072 | 88092375 | 1304 | 1 | 110312 | ENSMUST00 | -17133 | ENSMUSG00 | Pmch | pro-melanin-concentrating hormone |
| chr10 | 89261410 | 89261519 | 110 | * | NA_peak_588 | 20 | Intron (ENSM | 10 | 88948994 | 89257790 | 308797 | 2 | 320091 | ENSMUST00 | -3620 | ENSMUSG00 | Ano4 | anoctamin 4 |
| chr10 | 89720528 | 89720565 | 38 | * | NA_peak_591 | 23 | Intron (ENSM | 10 | 89726704 | 89732058 | 5355 | 2 | 67019 | ENSMUST00 | 11493 | ENSMUSG00 | Actr6 | ARP6 actin-related protein 6 |
| chr10 | 90055358 | 90055422 | 65 | * | NA_peak_592 | 27 | Intron (ENSM | 10 | 90046622 | 90071078 | 24457 | 1 | 77531 | ENSMUST00 | 8736 | ENSMUSG00 | Anks1b | ankyrin repeat and sterile alpha motif domain containing 1B |
| chr10 | 91284247 | 91284285 | 39 | * | NA_peak_598 | 20 | Distal Interge | 10 | 91208258 | 91220642 | 12385 | 2 | 102633753 | ENSMUST00 | -63605 | ENSMUSG00 | 4930534H03 | RIKEN cDNA 4930534H03 gene |
| chr10 | 91332255 | 91332409 | 155 | * | NA_peak_599 | 45 | Distal Interge | 10 | 91208258 | 91220642 | 12385 | 2 | 102633753 | ENSMUST00 | -111613 | ENSMUSG00 | 4930534H03 | RIKEN cDNA 4930534H03 gene |
| chr10 | 91779890 | 91780198 | 309 | * | NA_peak_601 | 23 | Distal Interge | 10 | 91888829 | 91953580 | 64752 | 1 | 102633867 | ENSMUST00 | -108631 | ENSMUSG00 | Gm31592 | predicted gene, 31592 |
| chr10 | 92624570 | 92624694 | 125 | * | NA_peak_603 | 16 | Distal Interge | 10 | 92700894 | 92716478 | 15585 | 2 | 17997 | ENSMUST00 | 91784 | ENSMUSG00 | Nedd1 | neural precursor cell expressed, developmentally down-regulated gene 1 |
| chr10 | 93347767 | 93348041 | 275 | * | NA_peak_606 | 24 | Intron (ENSM | 10 | 93335393 | 93348877 | 13485 | 1 | 432488 | ENSMUST00 | 12374 | NA | Gm17745 | predicted gene, 17745 |
| chr10 | 93775439 | 93775472 | 34 | * | NA_peak_608 | 22 | Distal Interge | 10 | 93831555 | 93848538 | 16984 | 1 | 327799 | ENSMUST00 | -56083 | ENSMUSG00 | Usp44 | ubiquitin specific peptidase 44 |
| chr10 | 94442071 | 94442276 | 206 | * | NA_peak_611 | 37 | Intron (ENSM | 10 | 94514857 | 94590956 | 76100 | 1 | 319880 | ENSMUST00 | -72581 | ENSMUSG00 | Tmcc3 | transmembrane and coiled coil domains 3 |
| chr10 | 95251939 | 95251985 | 47 | * | NA_peak_614 | 27 | Intron (ENSM | 10 | 95174746 | 95322965 | 148220 | 2 | 12905 | ENSMUST00 | 70980 | ENSMUSG00 | Cradd | CASP2 and RIPK1 domain containing adaptor with death domain |
| chr10 | 96157559 | 96157861 | 303 | * | NA_peak_619 | 13 | Distal Interge | 10 | 96226020 | 96248866 | 22847 | 1 | 74881 | ENSMUST00 | -68159 | ENSMUSG00 | 4930459C07 | RIKEN cDNA 4930459C07 gene |
| chr10 | 96199281 | 96199556 | 276 | * | NA_peak_620 | 31 | Distal Interge | 10 | 96226020 | 96248866 | 22847 | 1 | 74881 | ENSMUST00 | -26464 | ENSMUSG00 | 4930459C07 | RIKEN cDNA 4930459C07 gene |
| chr10 | 96529476 | 96529962 | 487 | * | NA_peak_621 | 16 | Intron (ENSM | 10 | 96617006 | 96622809 | 5804 | 1 | 12226 | ENSMUST00 | -87044 | ENSMUSG00 | Btg1 | BTG anti-proliferation factor 1 |
| chr10 | 97184329 | 97184584 | 256 | * | NA_peak_623 | 61 | Distal Interge | 10 | 97035780 | 97061049 | 25270 | 1 | 75301 | ENSMUST00 | 148549 | ENSMUSG00 | 4930556N09 | RIKEN cDNA 4930556N09 gene |
| chr10 | 98546754 | 98547069 | 316 | * | NA_peak_629 | 31 | Intron (ENSM | 10 | 98299927 | 98351874 | 51948 | 2 | 667438 | ENSMUST00 | -194880 | ENSMUSG00 | Gm8633 | predicted gene 8633 |
| chr10 | 98952468 | 98952856 | 389 | * | NA_peak_630 | 35 | Intron (ENSM | 10 | 98915472 | 98996930 | 81459 | 1 | 67972 | ENSMUST00 | 36996 | ENSMUSG00 | Atp2b1 | ATPase, Ca++ transporting, plasma membrane 1 |
| chr10 | 99040018 | 99040296 | 279 | * | NA_peak_631 | 20 | Exon (ENSMU | 10 | 99019037 | 99022933 | 3897 | 1 | 67972 | ENSMUST00 | 20981 | ENSMUSG00 | Atp2b1 | ATPase, Ca++ transporting, plasma membrane 1 |
| chr10 | 99693473 | 99693836 | 364 | * | NA_peak_634 | 16 | Distal Interge | 10 | 99638731 | 99641920 | 3190 | 2 | 100417125 | ENSMUST00 | -51553 | ENSMUSG00 | Gm18409 | mediator of cell motility 1 pseudogene |
| chr10 | 100167242 | 100167396 | 155 | * | NA_peak_636 | 24 | Distal Interge | 10 | 100088056 | 100097181 | 9126 | 1 | 17311 | ENSMUST00 | 79186 | ENSMUSG00 | Kitl | kit ligand |
| chr10 | 101569085 | 101569551 | 467 | * | NA_peak_639 | 36 | Distal Interge | 10 | 101681487 | 102013222 | 331736 | 1 | 67569 | ENSMUST00 | -111936 | ENSMUSG00 | Mgat4c | MGAT4 family, member C |
| chr10 | 101570000 | 101570295 | 296 | * | NA_peak_639 | 36 | Distal Interge | 10 | 101681487 | 102013222 | 331736 | 1 | 67569 | ENSMUST00 | -111192 | ENSMUSG00 | Mgat4c | MGAT4 family, member C |
| chr10 | 103870857 | 103871145 | 289 | * | NA_peak_644 | 40 | Distal Interge | 10 | 104142987 | 104145456 | 2470 | 1 | 637578 | ENSMUST00 | -271842 | ENSMUSG00 | Gm20765 | predicted gene, 20765 |
| chr10 | 104752940 | 104753157 | 218 | * | NA_peak_645 | 45 | Distal Interge | 10 | 105189662 | 105199704 | 10043 | 2 | 278279 | ENSMUST00 | 446547 | ENSMUSG00 | Tmtc2 | transmembrane and tetratricopeptide repeat containing 2 |
| chr10 | 106391105 | 106391428 | 324 | * | NA_peak_650 | 23 | Distal Interge | 10 | 106470339 | 106932492 | 462154 | 1 | 327814 | ENSMUST00 | -78911 | ENSMUSG00 | Ppfia2 | protein tyrosine phosphatase, receptor type, f polypeptide (PTPRF), interacting protein (liprin), alpha 2 |
| chr10 | 106454642 | 106454953 | 312 | * | NA_peak_650 | 27 | Distal Interge | 10 | 106470339 | 106932492 | 462154 | 1 | 327814 | ENSMUST00 | -15386 | ENSMUSG00 | Ppfia2 | protein tyrosine phosphatase, receptor type, f polypeptide (PTPRF), interacting protein (liprin), alpha 2 |
| chr10 | 106530986 | 106531114 | 129 | * | NA_peak_650 | 36 | Intron (ENSM | 10 | 106470346 | 106935952 | 465607 | 1 | 327814 | ENSMUST00 | 60640 | ENSMUSG00 | Ppfia2 | protein tyrosine phosphatase, receptor type, f polypeptide (PTPRF), interacting protein (liprin), alpha 2 |
| chr10 | 108096720 | 108096753 | 34 | * | NA_peak_654 | 22 | Distal Interge | 10 | 108162193 | 108277351 | 115159 | 1 | 17931 | ENSMUST00 | -65440 | ENSMUSG00 | Ppp1r12a | protein phosphatase 1, regulatory subunit 12A |
| chr10 | 109217120 | 109217289 | 170 | * | NA_peak_658 | 13 | Distal Interge | 10 | 108497652 | 109010982 | 513331 | 2 | 20979 | ENSMUST00 | -206138 | ENSMUSG00 | Syt1 | synaptotagmin I |
| chr10 | 109319147 | 109319436 | 290 | * | NA_peak_658 | 28 | Distal Interge | 10 | 108497652 | 109010982 | 513331 | 2 | 20979 | ENSMUST00 | -308165 | ENSMUSG00 | Syt1 | synaptotagmin I |
| chr10 | 109408369 | 109408728 | 360 | * | NA_peak_659 | 36 | Distal Interge | 10 | 109714302 | 109723801 | 9500 | 2 | 260315 | ENSMUST00 | 315073 | ENSMUSG00 | Nav3 | neuron navigator 3 |
| chr10 | 109774677 | 109774788 | 112 | * | NA_peak_659 | 23 | Intron (ENSM | 10 | 109755132 | 109764840 | 9709 | 2 | 260315 | ENSMUST00 | -9837 | ENSMUSG00 | Nav3 | neuron navigator 3 |
| chr10 | 110932073 | 110932398 | 326 | * | NA_peak_664 | 20 | Promoter (<= | 10 | 110931628 | 110937863 | 6236 | 1 | 13008 | ENSMUST00 | 445 | ENSMUSG00 | Csrp2 | cysteine and glycine-rich protein 2 |
| chr10 | 111184298 | 111184585 | 288 | * | NA_peak_665 | 23 | Intron (ENSM | 10 | 111164802 | 111293754 | 128953 | 1 | 237542 | ENSMUST00 | 19496 | ENSMUSG00 | Osbpl8 | oxysterol binding protein-like 8 |
| chr10 | 111553736 | 111553756 | 21 | * | NA_peak_667 | 29 | Distal Interge | 10 | 111586719 | 111594273 | 7555 | 2 | 71321 | ENSMUST00 | 40517 | ENSMUSG00 | 4933440J02 | RIKEN cDNA 4933440J02 gene |
| chr10 | 112323966 | 112324052 | 87 | * | NA_peak_670 | 31 | Intron (ENSM | 10 | 112315340 | 112465874 | 150535 | 1 | 268345 | ENSMUST00 | 8626 | ENSMUSG00 | Kcnc2 | potassium voltage gated channel, Shaw-related subfamily, member 2 |
| chr10 | 112324427 | 112324653 | 227 | * | NA_peak_670 | 19 | Intron (ENSM | 10 | 112315340 | 112465874 | 150535 | 1 | 268345 | ENSMUST00 | 9087 | ENSMUSG00 | Kcnc2 | potassium voltage gated channel, Shaw-related subfamily, member 2 |
| chr10 | 112674628 | 112675025 | 398 | * | NA_peak_672 | 39 | Distal Interge | 10 | 112726888 | 112785599 | 58712 | 1 | 69355 | ENSMUST00 | -51863 | ENSMUSG00 | 1700010J16 | RIKEN cDNA 1700010J16 gene |
| chr10 | 113424931 | 113425274 | 344 | * | NA_peak_674 | 27 | Intron (ENSM | 10 | 112925430 | 112928995 | 3566 | 2 | 382423 | ENSMUST00 | -495936 | ENSMUSG00 | Atxn7l3b | ataxin 7-like 3B |
| chr10 | 114029817 | 114029909 | 93 | * | NA_peak_676 | 16 | Distal Interge | 10 | 114398823 | 114801364 | 402542 | 2 | 237553 | ENSMUST00 | 771455 | ENSMUSG00 | Trhde | TRH-degrading enzyme |
| chr10 | 114059052 | 114059578 | 527 | * | NA_peak_676 | 20 | Distal Interge | 10 | 114398823 | 114801364 | 402542 | 2 | 237553 | ENSMUST00 | 741786 | ENSMUSG00 | Trhde | TRH-degrading enzyme |
| chr10 | 114944866 | 114944944 | 79 | * | NA_peak_679 | 45 | Distal Interge | 10 | 114566167 | 114802307 | 236141 | 2 | 237553 | ENSMUST00 | -142559 | ENSMUSG00 | Trhde | TRH-degrading enzyme |
| chr10 | 115037439 | 115038122 | 684 | * | NA_peak_679 | 45 | Distal Interge | 10 | 115174080 | 115175558 | 1479 | 2 | 216343 | ENSMUST00 | 137436 | ENSMUSG00 | Tph2 | tryptophan hydroxylase 2 |
| chr10 | 115277683 | 115278390 | 708 | * | NA_peak_680 | 22 | Distal Interge | 10 | 115197872 | 115251467 | 53596 | 2 | 66687 | ENSMUST00 | -26216 | ENSMUSG00 | Tbc1d15 | TBC1 domain family, member 15 |
| chr10 | 116029171 | 116029328 | 158 | * | NA_peak_684 | 20 | Intron (ENSM | 10 | 116018287 | 116114959 | 96673 | 1 | 19279 | ENSMUST00 | 10884 | ENSMUSG00 | Ptprr | protein tyrosine phosphatase, receptor type, R |
| chr10 | 116622486 | 116622818 | 333 | * | NA_peak_686 | 31 | Distal Interge | 10 | 116509791 | 116581511 | 71721 | 2 | 72068 | ENSMUST00 | -40975 | ENSMUSG00 | Cnot2 | CCR4-NOT transcription complex, subunit 2 |
| chr10 | 117424364 | 117424568 | 205 | * | NA_peak_690 | 21 | Distal Interge | 10 | 117446015 | 117447493 | 1479 | 2 | 102634600 | ENSMUST00 | 22925 | ENSMUSG00 | Gm32141 | predicted gene, 32141 |
| chr10 | 117744990 | 117745337 | 348 | * | NA_peak_691 | 17 | Promoter (<= | 10 | 117735900 | 117744942 | 9043 | 2 | 215436 | ENSMUST00 | -48 | ENSMUSG00 | Slc35e3 | solute carrier family 35, member E3 |
| chr10 | 117765146 | 117765257 | 112 | * | NA_peak_691 | 34 | Intron (ENSM | 10 | 117757285 | 117761357 | 4073 | 2 | 103468 | ENSMUST00 | -3789 | ENSMUSG00 | Nup107 | nucleoporin 107 |
| chr10 | 118243357 | 118243450 | 94 | * | NA_peak_693 | 34 | Distal Interge | 10 | 118204942 | 118210047 | 5106 | 1 | 50929 | ENSMUST00 | 38415 | ENSMUSG00 | Il22 | interleukin 22 |
| chr10 | 118256339 | 118256679 | 341 | * | NA_peak_693 | 21 | Distal Interge | 10 | 118289633 | 118293221 | 3589 | 2 | 116849 | ENSMUST00 | 36542 | ENSMUSG00 | Il22b | interleukin 22B |
| chr10 | 118957031 | 118957313 | 283 | * | NA_peak_695 | 31 | Distal Interge | 10 | 118920381 | 118927143 | 6763 | 2 | 216377 | ENSMUST00 | -29888 | ENSMUSG00 | Gm38403 | predicted gene, 38403 |
| chr10 | 120243225 | 120243335 | 111 | * | NA_peak_701 | 26 | Distal Interge | 10 | 120227186 | 120232071 | 4886 | 1 | 66225 | ENSMUST00 | 16039 | ENSMUSG00 | Llph | LLP homolog, long-term synaptic facilitation (Aplysia) |
| chr10 | 121566140 | 121566519 | 380 | * | NA_peak_707 | 24 | Promoter (1-2 | 10 | 121563925 | 121568481 | 4557 | 2 | 56480 | ENSMUST00 | 1962 | ENSMUSG00 | Tbk1 | TANK-binding kinase 1 |
| chr10 | 122141814 | 122142404 | 591 | * | NA_peak_710 | 40 | Distal Interge | 10 | 122119700 | 122130558 | 10859 | 1 | 100043513 | ENSMUST00 | 22114 | ENSMUSG00 | Gm4489 | predicted gene 4489 |
| chr10 | 122712341 | 122712467 | 127 | * | NA_peak_713 | 30 | Intron (ENSM | 10 | 122679315 | 122679415 | 101 | 1 | 102466884 | ENSMUST00 | 33026 | ENSMUSG00 | Mir8104 | microRNA 8104 |
| chr10 | 123472676 | 123472986 | 311 | * | NA_peak_716 | 20 | Intron (ENSM | 10 | 123264985 | 123740947 | 475963 | 1 | 268354 | ENSMUST00 | 207691 | ENSMUSG00 | Tafa2 | TAFA chemokine like family member 2 |
| chr10 | 123796073 | 123796363 | 291 | * | NA_peak_718 | 24 | Distal Interge | 10 | 123264985 | 123740947 | 475963 | 1 | 268354 | ENSMUST00 | 531088 | ENSMUSG00 | Tafa2 | TAFA chemokine like family member 2 |
| chr10 | 123861084 | 123861332 | 249 | * | NA_peak_718 | 33 | Distal Interge | 10 | 123264985 | 123740947 | 475963 | 1 | 268354 | ENSMUST00 | 596099 | ENSMUSG00 | Tafa2 | TAFA chemokine like family member 2 |
| chr10 | 124466940 | 124467286 | 347 | * | NA_peak_720 | 14 | Promoter (1-2 | 10 | 124469074 | 124524488 | 55415 | 1 | 75030 | ENSMUST00 | -1788 | ENSMUSG00 | 4930503E24 | eukaryotic translation initiation factor 3, subunit I pseudogene |
| chr10 | 125617697 | 125617801 | 105 | * | NA_peak_723 | 15 | Distal Interge | 10 | 125227485 | 125389465 | 161981 | 2 | 20503 | ENSMUST00 | -228232 | ENSMUSG00 | Slc16a7 | solute carrier family 16 (monocarboxylic acid transporters), member 7 |
| chr10 | 125816696 | 125816922 | 227 | * | NA_peak_723 | 26 | Distal Interge | 10 | 125966219 | 125998013 | 31795 | 1 | 320398 | ENSMUST00 | -149297 | ENSMUSG00 | Lrig3 | leucine-rich repeats and immunoglobulin-like domains 3 |
| chr10 | 128719890 | 128720118 | 229 | * | NA_peak_737 | 24 | 3' UTR | 10 | 128706258 | 128720238 | 13981 | 1 | 20431 | ENSMUST00 | 13632 | ENSMUSG00 | Pmel | premelanosome protein |
| chr10 | 130218517 | 130218769 | 253 | * | NA_peak_743 | 36 | Distal Interge | 10 | 130210092 | 130211149 | 1058 | 2 | 258297 | ENSMUST00 | -7368 | ENSMUSG00 | Olfr827 | olfactory receptor 827 |
| chr11 | 3101947 | 3102587 | 641 | * | NA_peak_745 | 76 | Distal Interge | 11 | 3125904 | 3129396 | 3493 | 1 | 236604 | ENSMUST00 | -23317 | NA | Pisd-ps1 | phosphatidylserine decarboxylase, pseudogene 1 |
| chr11 | 3129165 | 3129594 | 430 | * | NA_peak_745 | 77 | Promoter (<= | 11 | 3128972 | 3130146 | 1175 | 1 | 236604 | ENSMUST00 | 193 | NA | Pisd-ps1 | phosphatidylserine decarboxylase, pseudogene 1 |
| chr11 | 3129877 | 3130333 | 457 | * | NA_peak_746 | 28 | Promoter (<= | 11 | 3128972 | 3130146 | 1175 | 1 | 236604 | ENSMUST00 | 905 | NA | Pisd-ps1 | phosphatidylserine decarboxylase, pseudogene 1 |

| | | | | | | | | | | | | | | | | | | |
|---|---|---|---|---|---|---|---|---|---|---|---|---|---|---|---|---|---|---|
| chr11 | 3131943 | 3132252 | 310 | * | NA_peak_746 | 53 | Promoter (2-3 | 11 | 3128972 | 3130146 | 1175 | 1 | 236604 | ENSMUST00 | 2971 | NA | Pisd-ps1 | phosphatidylserine decarboxylase, pseudogene 1 |
| chr11 | 3132777 | 3133075 | 299 | * | NA_peak_746 | 53 | Promoter (2-3 | 11 | 3131887 | 3136007 | 4121 | 2 | 78887 | ENSMUST00 | 2932 | ENSMUSG00 | Sfi1 | Sfi1 homolog, spindle assembly associated (yeast) |
| chr11 | 3133917 | 3134488 | 572 | * | NA_peak_746 | 19 | Promoter (1-2 | 11 | 3131887 | 3136007 | 4121 | 2 | 78887 | ENSMUST00 | 1519 | ENSMUSG00 | Sfi1 | Sfi1 homolog, spindle assembly associated (yeast) |
| chr11 | 3141596 | 3141990 | 395 | * | NA_peak_746 | 57 | Promoter (<= | 11 | 3131890 | 3142378 | 10489 | 2 | 78887 | ENSMUST00 | 388 | ENSMUSG00 | Sfi1 | Sfi1 homolog, spindle assembly associated (yeast) |
| chr11 | 3149714 | 3150193 | 480 | * | NA_peak_746 | 13 | Promoter (2-3 | 11 | 3144881 | 3146722 | 1842 | 2 | 78887 | ENSMUST00 | -2992 | ENSMUSG00 | Sfi1 | Sfi1 homolog, spindle assembly associated (yeast) |
| chr11 | 3155448 | 3156107 | 660 | * | NA_peak_746 | 44 | Exon (ENSMU | 11 | 3131888 | 3164597 | 32710 | 2 | 78887 | ENSMUST00 | 8490 | ENSMUSG00 | Sfi1 | Sfi1 homolog, spindle assembly associated (yeast) |
| chr11 | 3157568 | 3159072 | 1505 | * | NA_peak_747 | 50 | Exon (ENSMU | 11 | 3131888 | 3164597 | 32710 | 2 | 78887 | ENSMUST00 | 5525 | ENSMUSG00 | Sfi1 | Sfi1 homolog, spindle assembly associated (yeast) |
| chr11 | 3162910 | 3164065 | 1156 | * | NA_peak_747 | 79 | Promoter (<= | 11 | 3131888 | 3164597 | 32710 | 2 | 78887 | ENSMUST00 | 532 | ENSMUSG00 | Sfi1 | Sfi1 homolog, spindle assembly associated (yeast) |
| chr11 | 3167576 | 3168479 | 904 | * | NA_peak_747 | 13 | Promoter (2-3 | 11 | 3156812 | 3170704 | 13893 | 2 | 78887 | ENSMUST00 | 2225 | ENSMUSG00 | Sfi1 | Sfi1 homolog, spindle assembly associated (yeast) |
| chr11 | 3170746 | 3171230 | 485 | * | NA_peak_747 | 63 | Promoter (<= | 11 | 3156812 | 3170704 | 13893 | 2 | 78887 | ENSMUST00 | -42 | ENSMUSG00 | Sfi1 | Sfi1 homolog, spindle assembly associated (yeast) |
| chr11 | 3174043 | 3175822 | 1780 | * | NA_peak_747 | 53 | Promoter (<= | 11 | 3160035 | 3176466 | 16432 | 2 | 78887 | ENSMUST00 | 644 | ENSMUSG00 | Sfi1 | Sfi1 homolog, spindle assembly associated (yeast) |
| chr11 | 3178917 | 3179550 | 634 | * | NA_peak_747 | 40 | Promoter (1-2 | 11 | 3153384 | 3177493 | 24110 | 2 | 78887 | ENSMUST00 | -1424 | ENSMUSG00 | Sfi1 | Sfi1 homolog, spindle assembly associated (yeast) |
| chr11 | 3180870 | 3181371 | 502 | * | NA_peak_747 | 18 | Intron (ENSM | 11 | 3153384 | 3177493 | 24110 | 2 | 78887 | ENSMUST00 | -3377 | ENSMUSG00 | Sfi1 | Sfi1 homolog, spindle assembly associated (yeast) |
| chr11 | 3182544 | 3183277 | 734 | * | NA_peak_748 | 73 | Intron (ENSM | 11 | 3165625 | 3187355 | 21731 | 2 | 78887 | ENSMUST00 | 4078 | ENSMUSG00 | Sfi1 | Sfi1 homolog, spindle assembly associated (yeast) |
| chr11 | 3186119 | 3186947 | 829 | * | NA_peak_748 | 108 | Promoter (<= | 11 | 3165625 | 3187355 | 21731 | 2 | 78887 | ENSMUST00 | 408 | ENSMUSG00 | Sfi1 | Sfi1 homolog, spindle assembly associated (yeast) |
| chr11 | 3188845 | 3189485 | 641 | * | NA_peak_748 | 108 | Promoter (<= | 11 | 3131850 | 3189056 | 57207 | 2 | 78887 | ENSMUST00 | 0 | ENSMUSG00 | Sfi1 | Sfi1 homolog, spindle assembly associated (yeast) |
| chr11 | 3194653 | 3195595 | 943 | * | NA_peak_748 | 52 | Promoter (<= | 11 | 3193516 | 3194263 | 748 | 2 | 78887 | ENSMUST00 | -390 | ENSMUSG00 | Sfi1 | Sfi1 homolog, spindle assembly associated (yeast) |
| chr11 | 3196690 | 3197030 | 341 | * | NA_peak_748 | 18 | Promoter (2-3 | 11 | 3193516 | 3194263 | 748 | 2 | 78887 | ENSMUST00 | -2427 | ENSMUSG00 | Sfi1 | Sfi1 homolog, spindle assembly associated (yeast) |
| chr11 | 3197507 | 3198511 | 1005 | * | NA_peak_748 | 41 | Intron (ENSM | 11 | 3193516 | 3194263 | 748 | 2 | 78887 | ENSMUST00 | -3244 | ENSMUSG00 | Sfi1 | Sfi1 homolog, spindle assembly associated (yeast) |
| chr11 | 3199030 | 3199813 | 784 | * | NA_peak_748 | 80 | Promoter (2-3 | 11 | 3202392 | 3244581 | 42190 | 1 | 74203 | ENSMUST00 | -2579 | ENSMUSG00 | Eif4enif1 | eukaryotic translation initiation factor 4E nuclear import factor 1 |
| chr11 | 4050715 | 4051012 | 298 | * | NA_peak_750 | 31 | Distal Interge | 11 | 4041737 | 4046336 | 4600 | 1 | 103655 | ENSMUST00 | 8978 | ENSMUSG00 | Sec14l4 | SEC14-like lipid binding 4 |
| chr11 | 10439283 | 10439661 | 379 | * | NA_peak_756 | 57 | Distal Interge | 11 | 11114223 | 11263526 | 149304 | 1 | 319922 | ENSMUST00 | -674562 | ENSMUSG00 | Vwc2 | von Willebrand factor C domain containing 2 |
| chr11 | 10532543 | 10532884 | 342 | * | NA_peak_756 | 20 | Distal Interge | 11 | 11114223 | 11263526 | 149304 | 1 | 319922 | ENSMUST00 | -581339 | ENSMUSG00 | Vwc2 | von Willebrand factor C domain containing 2 |
| chr11 | 15168143 | 15168359 | 217 | * | NA_peak_759 | 55 | Distal Interge | 11 | 14599276 | 14600314 | 1039 | 1 | 432536 | ENSMUST00 | 568867 | ENSMUSG00 | Pom121l12 | POM121 membrane glycoprotein-like 12 |
| chr11 | 16331110 | 16331508 | 399 | * | NA_peak_761 | 31 | Intron (ENSM | 11 | 16263218 | 16284551 | 21334 | 1 | 211739 | ENSMUST00 | 67892 | ENSMUSG00 | Vstm2a | V-set and transmembrane domain containing 2A |
| chr11 | 16391693 | 16391919 | 227 | * | NA_peak_761 | 27 | Intron (ENSM | 11 | 16501643 | 16505044 | 3402 | 2 | 20335 | ENSMUST00 | 113125 | ENSMUSG00 | Sec61g | SEC61, gamma subunit |
| chr11 | 16393698 | 16394050 | 353 | * | NA_peak_761 | 20 | Intron (ENSM | 11 | 16501643 | 16505044 | 3402 | 2 | 20335 | ENSMUST00 | 110994 | ENSMUSG00 | Sec61g | SEC61, gamma subunit |
| chr11 | 16879674 | 16879995 | 322 | * | NA_peak_764 | 25 | Promoter (1-2 | 11 | 16878150 | 16884357 | 6208 | 1 | 13649 | ENSMUST00 | 1524 | ENSMUSG00 | Egfr | epidermal growth factor receptor |
| chr11 | 17404018 | 17404328 | 311 | * | NA_peak_766 | 23 | Distal Interge | 11 | 17263621 | 17266560 | 2940 | 1 | 57316 | ENSMUST00 | 140397 | ENSMUSG00 | C1d | C1D nuclear receptor co-repressor |
| chr11 | 19027572 | 19027866 | 295 | * | NA_peak_770 | 31 | Distal Interge | 11 | 18879817 | 19018985 | 139169 | 2 | 17268 | ENSMUST00 | -8587 | ENSMUSG00 | Meis1 | Meis homeobox 1 |
| chr11 | 19569107 | 19569504 | 398 | * | NA_peak_773 | 98 | Distal Interge | 11 | 19924375 | 20024026 | 99652 | 1 | 114716 | ENSMUST00 | -354871 | ENSMUSG00 | Spred2 | sprouty-related EVH1 domain containing 2 |
| chr11 | 19779072 | 19779270 | 199 | * | NA_peak_773 | 40 | Distal Interge | 11 | 19924375 | 20024026 | 99652 | 1 | 114716 | ENSMUST00 | -145105 | ENSMUSG00 | Spred2 | sprouty-related EVH1 domain containing 2 |
| chr11 | 20864873 | 20864982 | 110 | * | NA_peak_777 | 16 | Distal Interge | 11 | 20829312 | 20831056 | 1745 | 2 | 216551 | ENSMUST00 | -33817 | ENSMUSG00 | Lgalsl | lectin, galactoside binding-like |
| chr11 | 20903130 | 20903586 | 457 | * | NA_peak_777 | 17 | Distal Interge | 11 | 20829312 | 20831056 | 1745 | 2 | 216551 | ENSMUST00 | -72074 | ENSMUSG00 | Lgalsl | lectin, galactoside binding-like |
| chr11 | 22963935 | 22964077 | 143 | * | NA_peak_787 | 21 | Intron (ENSM | 11 | 22899835 | 22956947 | 57113 | 2 | 17846 | ENSMUST00 | -6988 | ENSMUSG00 | Commd1 | COMM domain containing 1 |
| chr11 | 23021032 | 23021350 | 319 | * | NA_peak_787 | 19 | Promoter (<= | 11 | 23020556 | 23023730 | 3175 | 1 | 73873 | ENSMUST00 | 476 | ENSMUSG00 | Fam161a | family with sequence similarity 161, member A |
| chr11 | 23507031 | 23507197 | 167 | * | NA_peak_789 | 20 | Distal Interge | 11 | 23488547 | 23498030 | 9484 | 2 | 268390 | ENSMUST00 | -9001 | ENSMUSG00 | Ahsa2 | AHA1, activator of heat shock protein ATPase 2 |
| chr11 | 23924580 | 23924869 | 290 | * | NA_peak_791 | 32 | Distal Interge | 11 | 23862646 | 23895253 | 32608 | 2 | 216578 | ENSMUST00 | -29327 | ENSMUSG00 | Papolg | poly(A) polymerase gamma |
| chr11 | 24596494 | 24596893 | 400 | * | NA_peak_793 | 30 | Intron (ENSM | 11 | 24481909 | 24569952 | 88044 | 1 | 71316 | ENSMUST00 | 114585 | ENSMUSG00 | 4933430M04 | RIKEN cDNA 4933430M04 gene |
| chr11 | 25197126 | 25197255 | 130 | * | NA_peak_794 | 24 | Distal Interge | 11 | 25326107 | 25367137 | 41031 | 1 | 71234 | ENSMUST00 | -128852 | ENSMUSG00 | 4933427E13 | RIKEN cDNA 4933427E13 gene |
| chr11 | 25372118 | 25372143 | 26 | * | NA_peak_795 | 30 | Distal Interge | 11 | 25329957 | 25358187 | 28231 | 1 | 71234 | ENSMUST00 | 42161 | ENSMUSG00 | 4933427E13 | RIKEN cDNA 4933427E13 gene |
| chr11 | 27019876 | 27020048 | 173 | * | NA_peak_798 | 20 | Distal Interge | 11 | 26547848 | 26593999 | 46152 | 2 | 69922 | ENSMUST00 | -425877 | ENSMUSG00 | Vrk2 | vaccinia related kinase 2 |
| chr11 | 27686096 | 27686453 | 358 | * | NA_peak_799 | 32 | Distal Interge | 11 | 28388817 | 28434110 | 45294 | 2 | 216613 | ENSMUST00 | 747657 | ENSMUSG00 | Ccdc85a | coiled-coil domain containing 85A |
| chr11 | 27823299 | 27823616 | 318 | * | NA_peak_799 | 42 | Distal Interge | 11 | 28388817 | 28434110 | 45294 | 2 | 216613 | ENSMUST00 | 610494 | ENSMUSG00 | Ccdc85a | coiled-coil domain containing 85A |
| chr11 | 27880196 | 27880500 | 305 | * | NA_peak_800 | 31 | Distal Interge | 11 | 28388817 | 28434110 | 45294 | 2 | 216613 | ENSMUST00 | 553610 | ENSMUSG00 | Ccdc85a | coiled-coil domain containing 85A |
| chr11 | 27918719 | 27919218 | 500 | * | NA_peak_801 | 53 | Distal Interge | 11 | 28388817 | 28434110 | 45294 | 2 | 216613 | ENSMUST00 | 514892 | ENSMUSG00 | Ccdc85a | coiled-coil domain containing 85A |
| chr11 | 29331980 | 29332621 | 642 | * | NA_peak_805 | 55 | Distal Interge | 11 | 29373658 | 29510808 | 137151 | 1 | 108686 | ENSMUST00 | -41037 | ENSMUSG00 | Ccdc88a | coiled coil domain containing 88A |
| chr11 | 30040026 | 30040157 | 132 | * | NA_peak_808 | 21 | Distal Interge | 11 | 29743048 | 30026033 | 282986 | 2 | 237711 | ENSMUST00 | -13993 | ENSMUSG00 | Eml6 | echinoderm microtubule associated protein like 6 |
| chr11 | 30411071 | 30411281 | 211 | * | NA_peak_809 | 34 | Distal Interge | 11 | 30426006 | 30471808 | 45803 | 2 | 75087 | ENSMUST00 | 60527 | ENSMUSG00 | 4930505A04 | RIKEN cDNA 4930505A04 gene |
| chr11 | 31344506 | 31344768 | 263 | * | NA_peak_811 | 31 | Distal Interge | 11 | 31365441 | 31368166 | 2726 | 2 | 20856 | ENSMUST00 | 23398 | ENSMUSG00 | Stc2 | stanniocalcin 2 |
| chr11 | 31702395 | 31702558 | 164 | * | NA_peak_813 | 16 | Distal Interge | 11 | 31665181 | 31671885 | 6705 | 2 | 69556 | ENSMUST00 | -30510 | ENSMUSG00 | Bod1 | biorientation of chromosomes in cell division 1 |
| chr11 | 33081396 | 33081564 | 169 | * | NA_peak_817 | 23 | Distal Interge | 11 | 33117430 | 33146795 | 29366 | 2 | 14172 | ENSMUST00 | 65231 | ENSMUSG00 | Fgf18 | fibroblast growth factor 18 |
| chr11 | 34208259 | 34208341 | 83 | * | NA_peak_822 | 15 | Promoter (<= | 11 | 34204338 | 34208089 | 3752 | 2 | 14233 | ENSMUST00 | -170 | ENSMUSG00 | Foxi1 | forkhead box I1 |
| chr11 | 35337565 | 35337723 | 159 | * | NA_peak_826 | 36 | Intron (ENSM | 11 | 35177783 | 35510128 | 332346 | 1 | 20564 | ENSMUST00 | 159782 | ENSMUSG00 | Slit3 | slit guidance ligand 3 |
| chr11 | 35572171 | 35572387 | 217 | * | NA_peak_827 | 16 | Intron (ENSM | 11 | 35544685 | 35612636 | 67952 | 1 | 20564 | ENSMUST00 | 27486 | ENSMUSG00 | Slit3 | slit guidance ligand 3 |
| chr11 | 35973193 | 35973363 | 171 | * | NA_peak_828 | 19 | Intron (ENSM | 11 | 35838400 | 35980527 | 142128 | 2 | 211652 | ENSMUST00 | 7164 | ENSMUSG00 | Wwc1 | WW, C2 and coiled-coil domain containing 1 |
| chr11 | 36066047 | 36066432 | 386 | * | NA_peak_829 | 16 | Intron (ENSM | 11 | 35838400 | 35980527 | 142128 | 2 | 211652 | ENSMUST00 | -85520 | ENSMUSG00 | Wwc1 | WW, C2 and coiled-coil domain containing 1 |
| chr11 | 40612418 | 40612672 | 255 | * | NA_peak_835 | 27 | Distal Interge | 11 | 40680710 | 40685227 | 4518 | 2 | 108645 | ENSMUST00 | 72555 | ENSMUSG00 | Mat2b | methionine adenosyltransferase II, beta |
| chr11 | 40755146 | 40755427 | 282 | * | NA_peak_836 | 27 | Promoter (<= | 11 | 40748552 | 40755311 | 6760 | 2 | 12450 | ENSMUST00 | 0 | ENSMUSG00 | Ccng1 | cyclin G1 |
| chr11 | 41108767 | 41109403 | 637 | * | NA_peak_837 | 31 | Distal Interge | 11 | 40748552 | 40755311 | 6760 | 2 | 12450 | ENSMUST00 | -353456 | ENSMUSG00 | Ccng1 | cyclin G1 |
| chr11 | 43455482 | 43455649 | 168 | * | NA_peak_842 | 24 | Distal Interge | 11 | 43441407 | 43442632 | 1226 | 1 | 193116 | ENSMUST00 | 14075 | ENSMUSG00 | Slu7 | SLU7 splicing factor homolog (S. cerevisiae) |
| chr11 | 43528187 | 43528428 | 242 | * | NA_peak_842 | 36 | Promoter (<= | 11 | 43528784 | 43586997 | 58214 | 1 | 380694 | ENSMUST00 | -356 | ENSMUSG00 | Ccnjl | cyclin J-like |
| chr11 | 45776294 | 45776651 | 358 | * | NA_peak_852 | 26 | Distal Interge | 11 | 45808087 | 45841171 | 33085 | 1 | 100009664 | ENSMUST00 | -31436 | NA | F630206G17 | RIKEN cDNA F630206G17 gene |
| chr11 | 46008308 | 46008639 | 332 | * | NA_peak_853 | 23 | Intron (ENSM | 11 | 45980310 | 46017994 | 37685 | 1 | 214105 | ENSMUST00 | 27998 | ENSMUSG00 | Sox30 | SRY (sex determining region Y)-box 30 |
| chr11 | 46312177 | 46312625 | 449 | * | NA_peak_855 | 23 | Promoter (<= | 11 | 46193850 | 46312358 | 118509 | 2 | 76884 | ENSMUST00 | 0 | ENSMUSG00 | Cyfip2 | cytoplasmic FMR1 interacting protein 2 |
| chr11 | 47855529 | 47855859 | 331 | * | NA_peak_858 | 17 | Intron (ENSM | 11 | 47351154 | 47637469 | 286316 | 2 | 24052 | ENSMUST00 | -218060 | ENSMUSG00 | Sgcd | sarcoglycan, delta (dystrophin-associated glycoprotein) |
| chr11 | 49073647 | 49073945 | 299 | * | NA_peak_862 | 23 | Promoter (2-3 | 11 | 49076587 | 49096974 | 20388 | 1 | 15953 | ENSMUST00 | -2642 | ENSMUSG00 | Ifi47 | interferon gamma inducible protein 47 |
| chr11 | 49297226 | 49297529 | 304 | * | NA_peak_863 | 40 | Distal Interge | 11 | 49293297 | 49294314 | 1018 | 1 | 258462 | ENSMUST00 | 3929 | ENSMUSG00 | Olfr1392 | olfactory receptor 1392 |
| chr11 | 49905626 | 49906356 | 731 | * | NA_peak_866 | 40 | Intron (ENSM | 11 | 49902161 | 49980994 | 78834 | 1 | 74563 | ENSMUST00 | 3465 | ENSMUSG00 | Rasgef1c | RasGEF domain family, member 1C |
| chr11 | 49910216 | 49910270 | 55 | * | NA_peak_866 | 44 | Intron (ENSM | 11 | 49902161 | 49980994 | 78834 | 1 | 74563 | ENSMUST00 | 8055 | ENSMUSG00 | Rasgef1c | RasGEF domain family, member 1C |
| chr11 | 50128646 | 50128749 | 104 | * | NA_peak_867 | 49 | Promoter (2-3 | 11 | 50131396 | 50172030 | 40635 | 1 | 76795 | ENSMUST00 | -2647 | ENSMUSG00 | Tbc1d9b | TBC1 domain family, member 9B |
| chr11 | 51338337 | 51338483 | 147 | * | NA_peak_872 | 27 | Intron (ENSM | 11 | 51289920 | 51583918 | 293999 | 1 | 237759 | ENSMUST00 | 48417 | ENSMUSG00 | Col23a1 | collagen, type XXIII, alpha 1 |
| chr11 | 51856317 | 51856669 | 353 | * | NA_peak_874 | 13 | Promoter (<= | 11 | 51842301 | 51857125 | 14825 | 2 | 76901 | ENSMUST00 | 456 | ENSMUSG00 | Jade2 | jade family PHD finger 2 |
| chr11 | 52182410 | 52182513 | 104 | * | NA_peak_875 | 27 | Distal Interge | 11 | 52154722 | 52168159 | 13438 | 2 | 257871 | ENSMUST00 | -14251 | NA | Olfr1372-ps1 | olfactory receptor 1372, pseudogene 1 |
| chr11 | 52377607 | 52378024 | 418 | * | NA_peak_876 | 23 | Intron (ENSM | 11 | 52374263 | 52388523 | 14261 | 1 | 22333 | ENSMUST00 | 3344 | ENSMUSG00 | Vdac1 | voltage-dependent anion channel 1 |
| chr11 | 53226279 | 53226303 | 25 | * | NA_peak_879 | 20 | Distal Interge | 11 | 53262880 | 53267241 | 4362 | 2 | 15525 | ENSMUST00 | 40938 | ENSMUSG00 | Hspa4 | heat shock protein 4 |
| chr11 | 54385033 | 54385249 | 217 | * | NA_peak_884 | 55 | Intron (ENSM | 11 | 54370652 | 54426790 | 56139 | 1 | 74847 | ENSMUST00 | 14381 | ENSMUSG00 | Meikin | meiotic kinetochore factor |
| chr11 | 54388097 | 54388484 | 388 | * | NA_peak_884 | 57 | Intron (ENSM | 11 | 54370652 | 54426790 | 56139 | 1 | 74847 | ENSMUST00 | 17445 | ENSMUSG00 | Meikin | meiotic kinetochore factor |
| chr11 | 55668885 | 55669069 | 185 | * | NA_peak_888 | 23 | Distal Interge | 11 | 55514238 | 55608198 | 93961 | 2 | 14654 | ENSMUST00 | -60687 | ENSMUSG00 | Glra1 | glycine receptor, alpha 1 subunit |
| chr11 | 55749905 | 55750042 | 138 | * | NA_peak_889 | 45 | Distal Interge | 11 | 55514238 | 55608198 | 93961 | 2 | 14654 | ENSMUST00 | -141707 | ENSMUSG00 | Glra1 | glycine receptor, alpha 1 subunit |
| chr11 | 56182960 | 56183388 | 429 | * | NA_peak_892 | 50 | Distal Interge | 11 | 56024987 | 56041010 | 16024 | 2 | 216749 | ENSMUST00 | -141950 | ENSMUSG00 | Nmur2 | neuromedin U receptor 2 |
| chr11 | 56777198 | 56777278 | 81 | * | NA_peak_894 | 23 | Distal Interge | 11 | 57011387 | 57015291 | 3905 | 1 | 14799 | ENSMUST00 | -234109 | ENSMUSG00 | Gria1 | glutamate receptor, ionotropic, AMPA1 (alpha 1) |
| chr11 | 56976803 | 56976832 | 30 | * | NA_peak_895 | 13 | Distal Interge | 11 | 57011387 | 57015291 | 3905 | 1 | 14799 | ENSMUST00 | -34555 | ENSMUSG00 | Gria1 | glutamate receptor, ionotropic, AMPA1 (alpha 1) |
| chr11 | 58141407 | 58141689 | 283 | * | NA_peak_900 | 13 | Promoter (1-2 | 11 | 58136923 | 58139473 | 2551 | 2 | 216766 | ENSMUST00 | -1934 | ENSMUSG00 | Gemin5 | gem nuclear organelle associated protein 5 |
| chr11 | 58687601 | 58687688 | 88 | * | NA_peak_902 | 17 | Distal Interge | 11 | 58683875 | 58684802 | 928 | 1 | 216783 | ENSMUST00 | 3726 | ENSMUSG00 | Olfr320 | olfactory receptor 320 |
| chr11 | 59004891 | 59005283 | 393 | * | NA_peak_903 | 37 | Promoter (2-3 | 11 | 59001914 | 59002361 | 448 | 2 | 380698 | ENSMUST00 | -2530 | ENSMUSG00 | Obscn | obscurin, cytoskeletal calmodulin and titin-interacting RhoGEF |
| chr11 | 60561352 | 60561391 | 40 | * | NA_peak_907 | 27 | Distal Interge | 11 | 60537978 | 60558512 | 20535 | 1 | 268420 | ENSMUST00 | 23374 | ENSMUSG00 | Alkbh5 | alkB homolog 5, RNA demethylase |
| chr11 | 60639473 | 60639735 | 263 | * | NA_peak_907 | 23 | Distal Interge | 11 | 60699723 | 60714186 | 14464 | 1 | 16897 | ENSMUST00 | -59988 | ENSMUSG00 | Llgl1 | LLGL1 scribble cell polarity complex component |
| chr11 | 61118839 | 61119007 | 169 | * | NA_peak_908 | 27 | Distal Interge | 11 | 61126755 | 61149372 | 22618 | 1 | 57916 | ENSMUST00 | -7748 | ENSMUSG00 | Tnfrsf13b | tumor necrosis factor receptor superfamily, member 13b |
| chr11 | 61421282 | 61421973 | 692 | * | NA_peak_909 | 61 | Distal Interge | 11 | 61452263 | 61453919 | 1657 | 2 | 22671 | ENSMUST00 | 31946 | ENSMUSG00 | Rnf112 | ring finger protein 112 |
| chr11 | 62130250 | 62130381 | 132 | * | NA_peak_911 | 34 | Intron (ENSM | 11 | 62077097 | 62223013 | 145917 | 1 | 432572 | ENSMUST00 | 53153 | ENSMUSG00 | Specc1 | sperm antigen with calponin homology and coiled-coil domains 1 |
| chr11 | 62202073 | 62202522 | 450 | * | NA_peak_911 | 18 | Intron (ENSM | 11 | 62248984 | 62266453 | 17470 | 1 | 11541 | ENSMUST00 | -46462 | ENSMUSG00 | Adora2b | adenosine A2b receptor |
| chr11 | 62671678 | 62672047 | 370 | * | NA_peak_913 | 34 | Distal Interge | 11 | 62649449 | 62666354 | 16906 | 1 | 216829 | ENSMUST00 | 22229 | ENSMUSG00 | Mmgt2 | membrane magnesium transporter 2 |
| chr11 | 63117722 | 63117874 | 153 | * | NA_peak_915 | 20 | Distal Interge | 11 | 63128982 | 63159546 | 30565 | 1 | 18858 | ENSMUST00 | -11108 | ENSMUSG00 | Pmp22 | peripheral myelin protein 22 |
| chr11 | 63121631 | 63122003 | 373 | * | NA_peak_915 | 22 | Distal Interge | 11 | 63128982 | 63159546 | 30565 | 1 | 18858 | ENSMUST00 | -6979 | ENSMUSG00 | Pmp22 | peripheral myelin protein 22 |
| chr11 | 64571949 | 64572324 | 376 | * | NA_peak_921 | 27 | Distal Interge | 11 | 64435332 | 64522841 | 87510 | 1 | 15478 | ENSMUST00 | 136617 | ENSMUSG00 | Hs3st3a1 | heparan sulfate (glucosamine) 3-O-sulfotransferase 3A1 |
| chr11 | 64792987 | 64793275 | 289 | * | NA_peak_921 | 36 | Distal Interge | 11 | 64979038 | 65002069 | 23032 | 1 | 68626 | ENSMUST00 | -185763 | ENSMUSG00 | Elac2 | elaC ribonuclease Z 2 |
| chr11 | 65058822 | 65058838 | 17 | * | NA_peak_922 | 22 | Promoter (<= | 11 | 65002041 | 65059834 | 57794 | 2 | 216831 | ENSMUST00 | 996 | ENSMUSG00 | Arhgap44 | Rho GTPase activating protein 44 |
| chr11 | 65723238 | 65723461 | 224 | * | NA_peak_925 | 16 | Intron (ENSM | 11 | 65690333 | 65719603 | 29271 | 2 | 26398 | ENSMUST00 | -3635 | ENSMUSG00 | Map2k4 | mitogen-activated protein kinase kinase 4 |
| chr11 | 65814485 | 65814833 | 349 | * | NA_peak_925 | 16 | Exon (ENSMU | 11 | 65807244 | 65829239 | 21996 | 1 | 52712 | ENSMUST00 | 7241 | ENSMUSG00 | Zkscan6 | zinc finger with KRAB and SCAN domains 6 |
| chr11 | 67467517 | 67467822 | 306 | * | NA_peak_931 | 23 | Intron (ENSM | 11 | 67455724 | 67665449 | 209726 | 1 | 14457 | ENSMUST00 | 11793 | ENSMUSG00 | Gas7 | growth arrest specific 7 |
| chr11 | 69089007 | 69089127 | 121 | * | NA_peak_938 | 21 | Promoter (<= | 11 | 69088543 | 69090050 | 1508 | 1 | 22318 | ENSMUST00 | 464 | ENSMUSG00 | Vamp2 | vesicle-associated membrane protein 2 |
| chr11 | 69236183 | 69236491 | 309 | * | NA_peak_938 | 18 | Promoter (<= | 11 | 69218117 | 69237022 | 18906 | 2 | 14919 | ENSMUST00 | 531 | ENSMUSG00 | Gucy2e | guanylate cyclase 2e |
| chr11 | 69633348 | 69633558 | 211 | * | NA_peak_939 | 34 | Promoter (<= | 11 | 69632790 | 69653297 | 20308 | 1 | 23879 | ENSMUST00 | 358 | ENSMUSG00 | Fxr2 | fragile X mental retardation, autosomal homolog 2 |
| chr11 | 69966086 | 69966360 | 275 | * | NA_peak_940 | 19 | Promoter (<= | 11 | 69966918 | 69967652 | 735 | 1 | 53624 | ENSMUST00 | -558 | ENSMUSG00 | Cldn7 | claudin 7 |
| chr11 | 70080062 | 70080183 | 122 | * | NA_peak_941 | 27 | Distal Interge | 11 | 70092644 | 70106187 | 13544 | 1 | 11890 | ENSMUST00 | -12461 | ENSMUSG00 | Asgr2 | asialoglycoprotein receptor 2 |

| | | | | | | | | | | | | | | | | | | |
|---|---|---|---|---|---|---|---|---|---|---|---|---|---|---|---|---|---|---|
| chr11 | 72169938 | 72170115 | 178 | * | NA_peak_94 | 23 | Intron (ENSM | 11 | 72155393 | 72175238 | 19846 | 2 | 74477 | ENSMUST00 | 5123 | ENSMUSG0 | 4933427D14 | RIKEN cDNA 4933427D14 gene |
| chr11 | 73826796 | 73827042 | 247 | * | NA_peak_95 | 31 | Distal Interge | 11 | 73811740 | 73817049 | 5310 | 2 | 259008 | ENSMUST00 | -9747 | ENSMUSG0 | Olfr392 | olfactory receptor 392 |
| chr11 | 74369677 | 74369773 | 97 | * | NA_peak_95 | 32 | Distal Interge | 11 | 74364671 | 74365609 | 939 | 1 | 258153 | ENSMUST00 | 5006 | ENSMUSG0 | Olfr412 | olfactory receptor 412 |
| chr11 | 74470336 | 74470496 | 161 | * | NA_peak_95 | 16 | Intron (ENSM | 11 | 74436924 | 74522751 | 85828 | 2 | 380711 | ENSMUST00 | 52255 | ENSMUSG0 | Rap1gap2 | RAP1 GTPase activating protein 2 |
| chr11 | 74482028 | 74482299 | 272 | * | NA_peak_95 | 26 | Intron (ENSM | 11 | 74436924 | 74522751 | 85828 | 2 | 380711 | ENSMUST00 | 40452 | ENSMUSG0 | Rap1gap2 | RAP1 GTPase activating protein 2 |
| chr11 | 74959155 | 74959412 | 258 | * | NA_peak_95 | 21 | Intron (ENSM | 11 | 74930520 | 75164448 | 233929 | 1 | 103677 | ENSMUST00 | 28635 | ENSMUSG0 | Smg6 | Smg-6 homolog, nonsense mediated mRNA decay factor (C. elegans) |
| chr11 | 75468431 | 75469047 | 617 | * | NA_peak_96 | 27 | Promoter (<= | 11 | 75468711 | 75470203 | 1493 | 1 | 380712 | ENSMUST00 | 0 | ENSMUSG0 | Tlcd2 | TLC domain containing 2 |
| chr11 | 75668594 | 75668609 | 16 | * | NA_peak_96 | 17 | Promoter (<= | 11 | 75668094 | 75670254 | 2161 | 1 | 17913 | ENSMUST00 | 500 | ENSMUSG0 | Myo1c | myosin IC |
| chr11 | 76398425 | 76398634 | 210 | * | NA_peak_96 | 23 | Promoter (<= | 11 | 76358803 | 76399068 | 40266 | 2 | 18230 | ENSMUST00 | 434 | ENSMUSG0 | Nxn | nucleoredoxin |
| chr11 | 78252455 | 78252742 | 288 | * | NA_peak_96 | 23 | Promoter (1-2 | 11 | 78250894 | 78255117 | 4224 | 1 | 20316 | ENSMUST00 | 1561 | ENSMUSG0 | Sdf2 | stromal cell derived factor 2 |
| chr11 | 79236515 | 79236596 | 82 | * | NA_peak_97 | 27 | Distal Interge | 11 | 79240307 | 79242949 | 2643 | 2 | 78889 | ENSMUST00 | 6353 | ENSMUSG0 | Wsb1 | WD repeat and SOCS box-containing 1 |
| chr11 | 81998192 | 81998610 | 419 | * | NA_peak_98 | 19 | Intron (ENSM | 11 | 80880169 | 81968457 | 1088289 | 2 | 11418 | ENSMUST00 | -29735 | ENSMUSG0 | Asic2 | acid-sensing (proton-gated) ion channel 2 |
| chr11 | 82123580 | 82123849 | 270 | * | NA_peak_98 | 40 | Distal Interge | 11 | 82115185 | 82116799 | 1615 | 1 | 20307 | ENSMUST00 | 8395 | ENSMUSG0 | Ccl8 | chemokine (C-C motif) ligand 8 |
| chr11 | 84508867 | 84509045 | 179 | * | NA_peak_99 | 16 | Intron (ENSM | 11 | 84422855 | 84513522 | 90668 | 2 | 56321 | ENSMUST00 | 4477 | ENSMUSG0 | Aatf | apoptosis antagonizing transcription factor |
| chr11 | 85291465 | 85291782 | 318 | * | NA_peak_99 | 23 | Distal Interge | 11 | 85311244 | 85347066 | 35823 | 1 | 53892 | ENSMUST00 | -19462 | ENSMUSG0 | Ppm1d | protein phosphatase 1D magnesium-dependent, delta isoform |
| chr11 | 85439839 | 85440117 | 279 | * | NA_peak_99 | 23 | Intron (ENSM | 11 | 85466159 | 85826058 | 359900 | 1 | 192197 | ENSMUST00 | -26042 | ENSMUSG0 | Bcas3 | breast carcinoma amplified sequence 3 |
| chr11 | 86065550 | 86065656 | 107 | * | NA_peak_99 | 23 | Intron (ENSM | 11 | 86195033 | 86201164 | 6132 | 2 | 237911 | ENSMUST00 | 135508 | ENSMUSG0 | Brip1 | BRCA1 interacting protein C-terminal helicase 1 |
| chr11 | 86747750 | 86747885 | 136 | * | NA_peak_10 | 25 | Intron (ENSM | 11 | 86694655 | 86757501 | 62847 | 2 | 67300 | ENSMUST00 | 9616 | ENSMUSG0 | Cltc | clathrin, heavy polypeptide (Hc) |
| chr11 | 86758197 | 86758474 | 278 | * | NA_peak_10 | 14 | Promoter (<= | 11 | 86694351 | 86757565 | 63215 | 2 | 67300 | ENSMUST00 | -632 | ENSMUSG0 | Cltc | clathrin, heavy polypeptide (Hc) |
| chr11 | 87853350 | 87853420 | 71 | * | NA_peak_10 | 24 | Promoter (<= | 11 | 87853227 | 87859313 | 6087 | 1 | 380718 | ENSMUST00 | 123 | ENSMUSG0 | Mks1 | MKS transition zone complex subunit 1 |
| chr11 | 90311187 | 90311460 | 274 | * | NA_peak_10 | 22 | Distal Interge | 11 | 90249476 | 90278589 | 29114 | 1 | 67468 | ENSMUST00 | 61711 | ENSMUSG0 | Mmd | monocyte to macrophage differentiation-associated |
| chr11 | 91024871 | 91025136 | 266 | * | NA_peak_10 | 24 | Distal Interge | 11 | 90883345 | 90894999 | 11655 | 1 | 73922 | ENSMUST00 | 141526 | ENSMUSG0 | 4930405D11 | RIKEN cDNA 4930405D11 gene |
| chr11 | 92447474 | 92447577 | 104 | * | NA_peak_10 | 27 | Distal Interge | 11 | 93098028 | 93099866 | 1839 | 1 | 72605 | ENSMUST00 | -650451 | ENSMUSG0 | Car10 | carbonic anhydrase 10 |
| chr11 | 93967980 | 93968274 | 295 | * | NA_peak_10 | 27 | Promoter (<= | 11 | 93960668 | 93968272 | 7605 | 2 | 18102 | ENSMUST00 | 0 | ENSMUSG0 | Nme1 | NME/NM23 nucleoside diphosphate kinase 1 |
| chr11 | 94182861 | 94183252 | 392 | * | NA_peak_10 | 19 | Distal Interge | 11 | 94211454 | 94215495 | 4042 | 1 | 22057 | ENSMUST00 | -28202 | ENSMUSG0 | Tob1 | transducer of ErbB-2.1 |
| chr11 | 94600021 | 94600062 | 42 | * | NA_peak_10 | 19 | Promoter (1-2 | 11 | 94570291 | 94601769 | 31479 | 2 | 264895 | ENSMUST00 | 1707 | ENSMUSG0 | Acsf2 | acyl-CoA synthetase family member 2 |
| chr11 | 94918164 | 94918513 | 350 | * | NA_peak_10 | 23 | Distal Interge | 11 | 94936224 | 94953042 | 16819 | 1 | 12842 | ENSMUST00 | -17711 | ENSMUSG0 | Col1a1 | collagen, type I, alpha 1 |
| chr11 | 95413946 | 95414140 | 195 | * | NA_peak_10 | 14 | Promoter (<= | 11 | 95414080 | 95493406 | 79327 | 1 | 20747 | ENSMUST00 | 0 | ENSMUSG0 | Spop | speckle-type BTB/POZ protein |
| chr11 | 96314875 | 96315119 | 245 | * | NA_peak_10 | 26 | Promoter (2-3 | 11 | 96317165 | 96317274 | 110 | 1 | 723893 | ENSMUST00 | -2046 | ENSMUSG0 | Mir10a | microRNA 10a |
| chr11 | 98836271 | 98836668 | 398 | * | NA_peak_10 | 24 | Promoter (<= | 11 | 98836785 | 98853697 | 16913 | 1 | 268480 | ENSMUST00 | -117 | ENSMUSG0 | Rapgefl1 | Rap guanine nucleotide exchange factor (GEF)-like 1 |
| chr11 | 100311617 | 100311784 | 168 | * | NA_peak_10 | 14 | Distal Interge | 11 | 100319885 | 100322096 | 2212 | 1 | 20918 | ENSMUST00 | -8101 | ENSMUSG0 | Eif1 | eukaryotic translation initiation factor 1 |
| chr11 | 101542319 | 101542594 | 276 | * | NA_peak_10 | 23 | Intron (ENSM | 11 | 101532083 | 101551582 | 19500 | 2 | 12189 | ENSMUST00 | 8988 | ENSMUSG0 | Brca1 | breast cancer 1, early onset |
| chr11 | 102545826 | 102546110 | 285 | * | NA_peak_10 | 36 | Exon (ENSMU | 11 | 102536887 | 102556176 | 19290 | 2 | 237943 | ENSMUST00 | 10066 | ENSMUSG0 | Gpatch8 | G patch domain containing 8 |
| chr11 | 102649399 | 102649534 | 136 | * | NA_peak_10 | 21 | Distal Interge | 11 | 102663716 | 102675308 | 11593 | 1 | 268491 | ENSMUST00 | -14182 | ENSMUSG0 | Meioc | meiosis specific with coiled-coil domain |
| chr11 | 102819338 | 102819634 | 297 | * | NA_peak_10 | 19 | Promoter (<= | 11 | 102799579 | 102819700 | 20122 | 2 | 14615 | ENSMUST00 | 66 | ENSMUSG0 | Gjc1 | gap junction protein, gamma 1 |
| chr11 | 103084519 | 103084799 | 281 | * | NA_peak_10 | 24 | Intron (ENSM | 11 | 103070304 | 103078294 | 7991 | 2 | 72469 | ENSMUST00 | -6225 | ENSMUSG0 | Plcd3 | phospholipase C, delta 3 |
| chr11 | 103218016 | 103218303 | 288 | * | NA_peak_10 | 24 | Promoter (<= | 11 | 103208127 | 103218432 | 10306 | 2 | 328019 | ENSMUST00 | 129 | ENSMUSG0 | Spata32 | spermatogenesis associated 32 |
| chr11 | 106201951 | 106202325 | 375 | * | NA_peak_10 | 50 | Promoter (<= | 11 | 106163330 | 106202168 | 38839 | 2 | 72149 | ENSMUST00 | 0 | ENSMUSG0 | Strada | STE20-related kinase adaptor alpha |
| chr11 | 106332960 | 106332983 | 24 | * | NA_peak_10 | 24 | Intron (ENSM | 11 | 106318592 | 106349389 | 30798 | 2 | 110880 | ENSMUST00 | 16406 | ENSMUSG0 | Scn4a | sodium channel, voltage-gated, type IV, alpha |
| chr11 | 108709511 | 108709858 | 348 | * | NA_peak_10 | 31 | Promoter (<= | 11 | 108710072 | 108808706 | 98635 | 1 | 76380 | ENSMUST00 | -214 | ENSMUSG0 | Cep112 | centrosomal protein 112 |
| chr11 | 109717057 | 109717249 | 193 | * | NA_peak_10 | 23 | Intron (ENSM | 11 | 109672926 | 109722256 | 49331 | 2 | 208659 | ENSMUST00 | 5007 | ENSMUSG0 | Fam20a | FAM20A, golgi associated secretory pathway pseudokinase |
| chr11 | 109721266 | 109721588 | 323 | * | NA_peak_10 | 17 | Promoter (<= | 11 | 109672926 | 109722256 | 49331 | 2 | 208659 | ENSMUST00 | 668 | ENSMUSG0 | Fam20a | FAM20A, golgi associated secretory pathway pseudokinase |
| chr11 | 110288706 | 110289013 | 308 | * | NA_peak_10 | 27 | Intron (ENSM | 11 | 110236453 | 110251776 | 15324 | 2 | 76184 | ENSMUST00 | -36930 | ENSMUSG0 | Abca6 | ATP-binding cassette, sub-family A (ABC1), member 6 |
| chr11 | 112014844 | 112015185 | 342 | * | NA_peak_10 | 22 | Distal Interge | 11 | 112664839 | 112669398 | 4560 | 2 | 217294 | ENSMUST00 | 654213 | ENSMUSG0 | BC006965 | cDNA sequence BC006965 |
| chr11 | 112683427 | 112683759 | 333 | * | NA_peak_10 | 31 | Intron (ENSM | 11 | 112669201 | 112677220 | 8020 | 2 | 217294 | ENSMUST00 | -6207 | ENSMUSG0 | BC006965 | cDNA sequence BC006965 |
| chr11 | 114170837 | 114170942 | 106 | * | NA_peak_10 | 27 | Distal Interge | 11 | 114198258 | 114199199 | 942 | 1 | 73536 | ENSMUST00 | -27316 | ENSMUSG0 | 1700092K14 | RIKEN cDNA 1700092K14 gene |
| chr11 | 114305578 | 114305753 | 176 | * | NA_peak_10 | 45 | Distal Interge | 11 | 114198258 | 114199199 | 942 | 1 | 73536 | ENSMUST00 | 107320 | ENSMUSG0 | 1700092K14 | RIKEN cDNA 1700092K14 gene |
| chr11 | 114493070 | 114493491 | 422 | * | NA_peak_10 | 20 | Distal Interge | 11 | 114413401 | 114448860 | 35460 | 2 | 319819 | ENSMUST00 | -44210 | ENSMUSG0 | 4932435O22 | RIKEN cDNA 4932435O22 gene |
| chr11 | 114660678 | 114660991 | 314 | * | NA_peak_10 | 13 | Distal Interge | 11 | 114668524 | 114672331 | 3808 | 1 | 67671 | ENSMUST00 | -7533 | ENSMUSG0 | Rpl38 | ribosomal protein L38 |
| chr11 | 115653664 | 115653985 | 322 | * | NA_peak_10 | 31 | Intron (ENSM | 11 | 115614184 | 115628295 | 14112 | 2 | 67283 | ENSMUST00 | -25369 | ENSMUSG0 | Slc25a19 | solute carrier family 25 (mitochondrial thiamine pyrophosphate carrier), member 19 |
| chr11 | 116223026 | 116223301 | 276 | * | NA_peak_10 | 24 | Exon (ENSMU | 11 | 116216540 | 116220287 | 3748 | 1 | 69681 | ENSMUST00 | 6486 | ENSMUSG0 | Cdk3 | cyclin-dependent kinase 3 |
| chr11 | 119172371 | 119172523 | 153 | * | NA_peak_10 | 27 | Intron (ENSM | 11 | 119154076 | 119154137 | 62 | 2 | 102465563 | ENSMUST00 | -18234 | ENSMUSG0 | Mir6934 | microRNA 6934 |
| chr11 | 120566633 | 120567128 | 496 | * | NA_peak_10 | 32 | Intron (ENSM | 11 | 120560945 | 120572841 | 11897 | 2 | 18453 | ENSMUST00 | 5713 | ENSMUSG0 | P4hb | prolyl 4-hydroxylase, beta polypeptide |
| chr11 | 120672921 | 120673269 | 349 | * | NA_peak_10 | 20 | Promoter (<= | 11 | 120672973 | 120674346 | 1374 | 1 | 68938 | ENSMUST00 | 0 | ENSMUSG0 | Aspscr1 | alveolar soft part sarcoma chromosome region, candidate 1 (human) |
| chr11 | 120840639 | 120840941 | 303 | * | NA_peak_10 | 26 | Intron (ENSM | 11 | 120805846 | 120824547 | 18702 | 2 | 14104 | ENSMUST00 | -16092 | ENSMUSG0 | Fasn | fatty acid synthase |
| chr11 | 120867165 | 120867355 | 191 | * | NA_peak_10 | 13 | Intron (ENSM | 11 | 120826542 | 120873671 | 47130 | 2 | 71276 | ENSMUST00 | 6316 | ENSMUSG0 | Ccdc57 | coiled-coil domain containing 57 |
| chr11 | 120873369 | 120873656 | 288 | * | NA_peak_10 | 24 | Promoter (<= | 11 | 120826542 | 120873671 | 47130 | 2 | 71276 | ENSMUST00 | 15 | ENSMUSG0 | Ccdc57 | coiled-coil domain containing 57 |
| chr11 | 120884757 | 120885157 | 401 | * | NA_peak_10 | 31 | Promoter (<= | 11 | 120873792 | 120885911 | 12120 | 2 | 71276 | ENSMUST00 | 754 | ENSMUSG0 | Ccdc57 | coiled-coil domain containing 57 |
| chr11 | 120909539 | 120909910 | 372 | * | NA_peak_10 | 29 | Intron (ENSM | 11 | 120826529 | 120932872 | 106344 | 2 | 71276 | ENSMUST00 | 22962 | ENSMUSG0 | Ccdc57 | coiled-coil domain containing 57 |
| chr11 | 121076391 | 121076707 | 317 | * | NA_peak_10 | 26 | Intron (ENSM | 11 | 121069036 | 121081125 | 12090 | 2 | 209588 | ENSMUST00 | 4418 | ENSMUSG0 | Sectm1a | secreted and transmembrane 1A |
| chr11 | 121401523 | 121401804 | 282 | * | NA_peak_10 | 43 | Distal Interge | 11 | 121356125 | 121388251 | 32127 | 2 | 217371 | ENSMUST00 | -13272 | ENSMUSG0 | Rab40b | Rab40B, member RAS oncogene family |
| chr11 | 121883821 | 121883934 | 114 | * | NA_peak_10 | 41 | Exon (ENSMU | 11 | 121830247 | 121843423 | 13177 | 1 | 74675 | ENSMUST00 | 53574 | ENSMUSG0 | Ptchd3 | patched domain containing 3 |
| chr12 | 8909400 | 8909790 | 391 | * | NA_peak_10 | 97 | Distal Interge | 12 | 8921664 | 8938742 | 17079 | 1 | 17775 | ENSMUST00 | -11874 | ENSMUSG0 | Laptm4a | lysosomal-associated protein transmembrane 4A |
| chr12 | 9680105 | 9680376 | 272 | * | NA_peak_10 | 25 | Distal Interge | 12 | 9574441 | 9581500 | 7060 | 1 | 23967 | ENSMUST00 | 105664 | ENSMUSG0 | Osr1 | odd-skipped related transcription factor 1 |
| chr12 | 15191885 | 15192220 | 336 | * | NA_peak_10 | 45 | Distal Interge | 12 | 15040194 | 15051598 | 11405 | 2 | 73972 | ENSMUST00 | -140287 | ENSMUSG0 | 4930448C13 | RIKEN cDNA 4930448C13 gene |
| chr12 | 20196994 | 20197119 | 126 | * | NA_peak_11 | 33 | Exon (ENSMU | 12 | 20804381 | 20815779 | 11399 | 2 | 69513 | ENSMUST00 | 618660 | ENSMUSG0 | 1700030C10 | RIKEN cDNA 1700030C10 gene |
| chr12 | 20221771 | 20222000 | 230 | * | NA_peak_11 | 25 | Intron (ENSM | 12 | 20804381 | 20815779 | 11399 | 2 | 69513 | ENSMUST00 | 593779 | ENSMUSG0 | 1700030C10 | RIKEN cDNA 1700030C10 gene |
| chr12 | 20227702 | 20228043 | 342 | * | NA_peak_11 | 31 | Intron (ENSM | 12 | 20804381 | 20815779 | 11399 | 2 | 69513 | ENSMUST00 | 587736 | ENSMUSG0 | 1700030C10 | RIKEN cDNA 1700030C10 gene |
| chr12 | 20232835 | 20233375 | 541 | * | NA_peak_11 | 24 | Intron (ENSM | 12 | 20804381 | 20815779 | 11399 | 2 | 69513 | ENSMUST00 | 582404 | ENSMUSG0 | 1700030C10 | RIKEN cDNA 1700030C10 gene |
| chr12 | 20312189 | 20312421 | 233 | * | NA_peak_11 | 23 | Distal Interge | 12 | 20804381 | 20815779 | 11399 | 2 | 69513 | ENSMUST00 | 503358 | ENSMUSG0 | 1700030C10 | RIKEN cDNA 1700030C10 gene |
| chr12 | 20711637 | 20711812 | 176 | * | NA_peak_11 | 28 | Distal Interge | 12 | 20804381 | 20815779 | 11399 | 2 | 69513 | ENSMUST00 | 103967 | ENSMUSG0 | 1700030C10 | RIKEN cDNA 1700030C10 gene |
| chr12 | 23058284 | 23058675 | 392 | * | NA_peak_11 | 31 | Distal Interge | 12 | 22953999 | 23010284 | 56286 | 2 | 69732 | ENSMUST00 | -48000 | ENSMUSG0 | 2410018L13 | RIKEN cDNA 2410018L13 gene |
| chr12 | 23141340 | 23141653 | 314 | * | NA_peak_11 | 16 | Distal Interge | 12 | 22953999 | 23010284 | 56286 | 2 | 69732 | ENSMUST00 | -131056 | ENSMUSG0 | 2410018L13 | RIKEN cDNA 2410018L13 gene |
| chr12 | 23802067 | 23802400 | 334 | * | NA_peak_11 | 30 | Distal Interge | 12 | 24043202 | 24097246 | 54045 | 2 | 66808 | ENSMUST00 | 294846 | ENSMUSG0 | 9030624G23 | RIKEN cDNA 9030624G23 gene |
| chr12 | 24399997 | 24400316 | 320 | * | NA_peak_11 | 45 | Distal Interge | 12 | 24498359 | 24558539 | 60181 | 1 | 21340 | ENSMUST00 | -98043 | ENSMUSG0 | Taf1b | TATA-box binding protein associated factor, RNA polymerase I, B |
| chr12 | 25861880 | 25862174 | 295 | * | NA_peak_11 | 23 | Distal Interge | 12 | 26091962 | 26210505 | 118544 | 2 | 629481 | ENSMUST00 | 348331 | ENSMUSG0 | Gm29687 | predicted gene, 29687 |
| chr12 | 30991626 | 30992069 | 444 | * | NA_peak_11 | 31 | Distal Interge | 12 | 30912187 | 30960156 | 47970 | 1 | 24057 | ENSMUST00 | 79439 | ENSMUSG0 | Sh3yl1 | Sh3 domain YSC-like 1 |
| chr12 | 35856783 | 35857201 | 419 | * | NA_peak_11 | 19 | Distal Interge | 12 | 35925620 | 35949736 | 24117 | 1 | 403205 | ENSMUST00 | -68419 | ENSMUSG0 | Agr3 | anterior gradient 3 |
| chr12 | 38078793 | 38079216 | 424 | * | NA_peak_11 | 31 | Intron (ENSM | 12 | 38019180 | 38174661 | 155482 | 1 | 217480 | ENSMUST00 | 59613 | ENSMUSG0 | Dgkb | diacylglycerol kinase, beta |
| chr12 | 41504832 | 41505197 | 366 | * | NA_peak_11 | 48 | Intron (ENSM | 12 | 41451668 | 41486431 | 34764 | 2 | 16981 | ENSMUST00 | -18401 | ENSMUSG0 | Lrrn3 | leucine rich repeat protein 3, neuronal |
| chr12 | 48849095 | 48849353 | 259 | * | NA_peak_11 | 61 | Distal Interge | 12 | 49382660 | 49384364 | 1705 | 1 | 15228 | ENSMUST00 | -533307 | ENSMUSG0 | Foxg1 | forkhead box G1 |
| chr12 | 49261505 | 49261938 | 434 | * | NA_peak_11 | 36 | Distal Interge | 12 | 49382660 | 49384364 | 1705 | 1 | 15228 | ENSMUST00 | -120722 | ENSMUSG0 | Foxg1 | forkhead box G1 |
| chr12 | 50977603 | 50977886 | 284 | * | NA_peak_11 | 23 | Distal Interge | 12 | 51005158 | 51062543 | 57386 | 1 | 75494 | ENSMUST00 | -27272 | ENSMUSG0 | 1700008C04 | RIKEN cDNA 1700008C04 gene |
| chr12 | 52114660 | 52114821 | 162 | * | NA_peak_11 | 18 | Intron (ENSM | 12 | 52097761 | 52101083 | 3323 | 1 | 76826 | ENSMUST00 | 16899 | ENSMUSG0 | Nubpl | nucleotide binding protein-like |
| chr12 | 57604489 | 57604645 | 157 | * | NA_peak_11 | 21 | Intron (ENSM | 12 | 57585975 | 57618602 | 32628 | 2 | 78758 | ENSMUST00 | 13957 | ENSMUSG0 | 4921518K17 | RIKEN cDNA 4921518K17 gene |
| chr12 | 62178210 | 62178356 | 147 | * | NA_peak_11 | 20 | Distal Interge | 12 | 62687822 | 62688245 | 424 | 1 | 73309 | ENSMUST00 | -509466 | NA | Spanxn4 | SPANX family, member N4 |
| chr12 | 63121743 | 63121932 | 190 | * | NA_peak_11 | 36 | Distal Interge | 12 | 62687822 | 62688245 | 424 | 1 | 73309 | ENSMUST00 | 433921 | NA | Spanxn4 | SPANX family, member N4 |
| chr12 | 63697233 | 63697622 | 390 | * | NA_peak_11 | 26 | Distal Interge | 12 | 64471330 | 64474690 | 3361 | 2 | 623046 | ENSMUST00 | 777068 | ENSMUSG0 | Fscb | fibrous sheath CABYR binding protein |
| chr12 | 64603106 | 64603561 | 456 | * | NA_peak_11 | 36 | Distal Interge | 12 | 64471330 | 64474690 | 3361 | 2 | 623046 | ENSMUST00 | -128416 | ENSMUSG0 | Fscb | fibrous sheath CABYR binding protein |
| chr12 | 64887528 | 64887804 | 277 | * | NA_peak_11 | 22 | Distal Interge | 12 | 64917906 | 64924591 | 6686 | 1 | 217648 | ENSMUST00 | -30102 | ENSMUSG0 | Gm527 | predicted gene 527 |
| chr12 | 67829753 | 67829924 | 172 | * | NA_peak_11 | 31 | Distal Interge | 12 | 66466060 | 67222549 | 756490 | 2 | 320772 | ENSMUST00 | -607204 | ENSMUSG0 | Mdga2 | MAM domain containing glycosylphosphatidylinositol anchor 2 |
| chr12 | 68558639 | 68558842 | 204 | * | NA_peak_11 | 24 | Distal Interge | 12 | 69158358 | 69159054 | 697 | 2 | 20090 | ENSMUST00 | 600212 | ENSMUSG0 | Rps29 | ribosomal protein S29 |
| chr12 | 69892757 | 69892956 | 200 | * | NA_peak_11 | 16 | Promoter (<= | 12 | 69803750 | 69893009 | 89260 | 2 | 399510 | ENSMUST00 | 53 | ENSMUSG0 | Map4k5 | mitogen-activated protein kinase kinase kinase kinase 5 |
| chr12 | 72208603 | 72208960 | 358 | * | NA_peak_11 | 36 | Distal Interge | 12 | 72101300 | 72185029 | 83730 | 2 | 73936 | ENSMUST00 | -23574 | ENSMUSG0 | Ccdc175 | coiled-coil domain containing 175 |
| chr12 | 74804053 | 74804486 | 434 | * | NA_peak_11 | 27 | Distal Interge | 12 | 74897220 | 75177332 | 280113 | 2 | 238271 | ENSMUST00 | 372846 | ENSMUSG0 | Kcnh5 | potassium voltage-gated channel, subfamily H (eag-related), member 5 |
| chr12 | 76222782 | 76223163 | 382 | * | NA_peak_11 | 47 | Intron (ENSM | 12 | 76214486 | 76229464 | 14979 | 2 | 80384 | ENSMUST00 | 6301 | ENSMUSG0 | Tex21 | testis expressed gene 21 |
| chr12 | 76225115 | 76225484 | 370 | * | NA_peak_11 | 31 | Intron (ENSM | 12 | 76214486 | 76229464 | 14979 | 2 | 80384 | ENSMUST00 | 3980 | ENSMUSG0 | Tex21 | testis expressed gene 21 |
| chr12 | 81553645 | 81554005 | 361 | * | NA_peak_11 | 21 | Intron (ENSM | 12 | 81558584 | 81568474 | 9891 | 2 | 56622 | ENSMUST00 | 14469 | ENSMUSG0 | Adam21 | a disintegrin and metallopeptidase domain 21 |
| chr12 | 87881204 | 87881509 | 306 | * | NA_peak_11 | 24 | Distal Interge | 12 | 87874072 | 87877859 | 3788 | 1 | 100039042 | ENSMUST00 | 7132 | ENSMUSG0 | Eif1ad3 | eukaryotic translation initiation factor 1A domain containing 3 |
| chr12 | 87937330 | 87937645 | 316 | * | NA_peak_11 | 27 | Distal Interge | 12 | 87954080 | 87959053 | 4974 | 1 | 100039087 | ENSMUST00 | -16435 | ENSMUSG0 | Gm2042 | predicted gene 2042 |
| chr12 | 91457905 | 91458279 | 375 | * | NA_peak_11 | 36 | Intron (ENSM | 12 | 91401000 | 91522541 | 121542 | 1 | 22095 | ENSMUST00 | 56905 | ENSMUSG0 | Tshr | thyroid stimulating hormone receptor |
| chr12 | 92593789 | 92594191 | 403 | * | NA_peak_11 | 40 | Distal Interge | 12 | 91806043 | 91849157 | 43115 | 2 | 20338 | ENSMUST00 | -744632 | ENSMUSG0 | Sel1l | sel-1 suppressor of lin-12-like (C. elegans) |
| chr12 | 93077137 | 93077475 | 339 | * | NA_peak_11 | 50 | Distal Interge | 12 | 93416458 | 93444823 | 28366 | 2 | 75295 | ENSMUST00 | 367348 | ENSMUSG0 | 4930559C10 | RIKEN cDNA 4930559C10 gene |
| chr12 | 95248570 | 95248895 | 326 | * | NA_peak_11 | 45 | Distal Interge | 12 | 95692226 | 95785215 | 92990 | 1 | 399558 | ENSMUST00 | -443331 | ENSMUSG0 | Flrt2 | fibronectin leucine rich transmembrane protein 2 |
| chr12 | 95341961 | 95342843 | 883 | * | NA_peak_11 | 30 | Distal Interge | 12 | 95692226 | 95785215 | 92990 | 1 | 399558 | ENSMUST00 | -349383 | ENSMUSG0 | Flrt2 | fibronectin leucine rich transmembrane protein 2 |
| chr12 | 98513235 | 98513748 | 514 | * | NA_peak_11 | 86 | Intron (ENSM | 12 | 98523844 | 98526583 | 2740 | 1 | 105245437 | ENSMUST00 | -10096 | ENSMUSG0 | Gm40893 | predicted gene, 40893 |

| | | | | | | | | | | | | | | | | | | |
|---|---|---|---|---|---|---|---|---|---|---|---|---|---|---|---|---|---|---|
| chr12 | 101702470 | 101702722 | 253 | * | NA_peak_113 | 40 | Intron (ENSM | 12 | 101649033 | 101709536 | 60504 | 2 | 74413 | ENSMUST00 | 6814 | ENSMUSG0 | Tc2n | tandem C2 domains, nuclear |
| chr12 | 112781607 | 112782277 | 671 | * | NA_peak_113 | 35 | Exon (ENSMU | 12 | 112766544 | 112767561 | 1018 | 1 | 104759 | ENSMUST00 | 15063 | ENSMUSG0 | Pld4 | phospholipase D family, member 4 |
| chr12 | 112783797 | 112784349 | 553 | * | NA_peak_113 | 60 | Exon (ENSMU | 12 | 112766544 | 112767561 | 1018 | 1 | 104759 | ENSMUST00 | 17253 | ENSMUSG0 | Pld4 | phospholipase D family, member 4 |
| chr12 | 114884837 | 114885233 | 397 | * | NA_peak_113 | 36 | Distal Interge | 12 | 114513635 | 114513694 | 60 | 2 | 102465664 | ENSMUST00 | -371143 | ENSMUSG0 | Mir7094-2 | microRNA 7094-2 |
| chr12 | 115067794 | 115068264 | 471 | * | NA_peak_113 | 45 | Exon (ENSMU | 12 | 114513635 | 114513694 | 60 | 2 | 102465664 | ENSMUST00 | -554100 | ENSMUSG0 | Mir7094-2 | microRNA 7094-2 |
| chr12 | 115081562 | 115081823 | 262 | * | NA_peak_113 | 14 | Distal Interge | 12 | 114513635 | 114513694 | 60 | 2 | 102465664 | ENSMUST00 | -567868 | ENSMUSG0 | Mir7094-2 | microRNA 7094-2 |
| chr12 | 115182273 | 115182881 | 609 | * | NA_peak_114 | 60 | Distal Interge | 12 | 115695720 | 115695794 | 75 | 2 | 102465205 | ENSMUST00 | 512913 | ENSMUSG0 | Mir6388 | microRNA 6388 |
| chr12 | 115217744 | 115217809 | 66 | * | NA_peak_114 | 18 | Distal Interge | 12 | 115695720 | 115695794 | 75 | 2 | 102465205 | ENSMUST00 | 477985 | ENSMUSG0 | Mir6388 | microRNA 6388 |
| chr12 | 115224897 | 115225198 | 302 | * | NA_peak_114 | 50 | Exon (ENSMU | 12 | 115695720 | 115695794 | 75 | 2 | 102465205 | ENSMUST00 | 470596 | ENSMUSG0 | Mir6388 | microRNA 6388 |
| chr12 | 115226106 | 115226322 | 217 | * | NA_peak_114 | 30 | Distal Interge | 12 | 115695720 | 115695794 | 75 | 2 | 102465205 | ENSMUST00 | 469472 | ENSMUSG0 | Mir6388 | microRNA 6388 |
| chr12 | 115295895 | 115296159 | 265 | * | NA_peak_114 | 40 | Distal Interge | 12 | 115695720 | 115695794 | 75 | 2 | 102465205 | ENSMUST00 | 399635 | ENSMUSG0 | Mir6388 | microRNA 6388 |
| chr12 | 115395757 | 115396851 | 1095 | * | NA_peak_114 | 40 | Exon (ENSMU | 12 | 115695720 | 115695794 | 75 | 2 | 102465205 | ENSMUST00 | 298943 | ENSMUSG0 | Mir6388 | microRNA 6388 |
| chr12 | 115449909 | 115450271 | 363 | * | NA_peak_114 | 49 | Distal Interge | 12 | 115695720 | 115695794 | 75 | 2 | 102465205 | ENSMUST00 | 245523 | ENSMUSG0 | Mir6388 | microRNA 6388 |
| chr12 | 115506107 | 115506703 | 597 | * | NA_peak_114 | 17 | Distal Interge | 12 | 115695720 | 115695794 | 75 | 2 | 102465205 | ENSMUST00 | 189091 | ENSMUSG0 | Mir6388 | microRNA 6388 |
| chr12 | 115510094 | 115510567 | 474 | * | NA_peak_114 | 25 | Distal Interge | 12 | 115695720 | 115695794 | 75 | 2 | 102465205 | ENSMUST00 | 185227 | ENSMUSG0 | Mir6388 | microRNA 6388 |
| chr12 | 115570352 | 115571317 | 966 | * | NA_peak_114 | 52 | Distal Interge | 12 | 115695720 | 115695794 | 75 | 2 | 102465205 | ENSMUST00 | 124477 | ENSMUSG0 | Mir6388 | microRNA 6388 |
| chr12 | 115587800 | 115588098 | 299 | * | NA_peak_114 | 42 | Distal Interge | 12 | 115695720 | 115695794 | 75 | 2 | 102465205 | ENSMUST00 | 107696 | ENSMUSG0 | Mir6388 | microRNA 6388 |
| chr12 | 115601960 | 115602268 | 309 | * | NA_peak_114 | 18 | Distal Interge | 12 | 115695720 | 115695794 | 75 | 2 | 102465205 | ENSMUST00 | 93526 | ENSMUSG0 | Mir6388 | microRNA 6388 |
| chr12 | 115606546 | 115607006 | 461 | * | NA_peak_114 | 19 | Distal Interge | 12 | 115695720 | 115695794 | 75 | 2 | 102465205 | ENSMUST00 | 88788 | ENSMUSG0 | Mir6388 | microRNA 6388 |
| chr12 | 115615077 | 115615358 | 282 | * | NA_peak_114 | 46 | Distal Interge | 12 | 115695720 | 115695794 | 75 | 2 | 102465205 | ENSMUST00 | 80436 | ENSMUSG0 | Mir6388 | microRNA 6388 |
| chr12 | 115618412 | 115618635 | 224 | * | NA_peak_114 | 13 | Distal Interge | 12 | 115695720 | 115695794 | 75 | 2 | 102465205 | ENSMUST00 | 77159 | ENSMUSG0 | Mir6388 | microRNA 6388 |
| chr12 | 115622127 | 115622554 | 428 | * | NA_peak_114 | 39 | Distal Interge | 12 | 115695720 | 115695794 | 75 | 2 | 102465205 | ENSMUST00 | 73240 | ENSMUSG0 | Mir6388 | microRNA 6388 |
| chr12 | 115641200 | 115641781 | 582 | * | NA_peak_114 | 72 | Distal Interge | 12 | 115695720 | 115695794 | 75 | 2 | 102465205 | ENSMUST00 | 54013 | ENSMUSG0 | Mir6388 | microRNA 6388 |
| chr12 | 115663735 | 115664400 | 666 | * | NA_peak_114 | 39 | Distal Interge | 12 | 115695720 | 115695794 | 75 | 2 | 102465205 | ENSMUST00 | 31394 | ENSMUSG0 | Mir6388 | microRNA 6388 |
| chr12 | 115679422 | 115679467 | 46 | * | NA_peak_114 | 40 | Distal Interge | 12 | 115695720 | 115695794 | 75 | 2 | 102465205 | ENSMUST00 | 16327 | ENSMUSG0 | Mir6388 | microRNA 6388 |
| chr12 | 115682595 | 115683024 | 430 | * | NA_peak_114 | 61 | Distal Interge | 12 | 115695720 | 115695794 | 75 | 2 | 102465205 | ENSMUST00 | 12770 | ENSMUSG0 | Mir6388 | microRNA 6388 |
| chr12 | 115683348 | 115683578 | 231 | * | NA_peak_114 | 61 | Distal Interge | 12 | 115695720 | 115695794 | 75 | 2 | 102465205 | ENSMUST00 | 12216 | ENSMUSG0 | Mir6388 | microRNA 6388 |
| chr12 | 115771807 | 115772169 | 363 | * | NA_peak_114 | 19 | Distal Interge | 12 | 115695720 | 115695794 | 75 | 2 | 102465205 | ENSMUST00 | -76013 | ENSMUSG0 | Mir6388 | microRNA 6388 |
| chr12 | 115820573 | 115821266 | 694 | * | NA_peak_114 | 27 | Distal Interge | 12 | 115695720 | 115695794 | 75 | 2 | 102465205 | ENSMUST00 | -124779 | ENSMUSG0 | Mir6388 | microRNA 6388 |
| chr12 | 115837858 | 115838617 | 760 | * | NA_peak_114 | 34 | Distal Interge | 12 | 115695720 | 115695794 | 75 | 2 | 102465205 | ENSMUST00 | -142064 | ENSMUSG0 | Mir6388 | microRNA 6388 |
| chr12 | 115850063 | 115850373 | 311 | * | NA_peak_114 | 40 | Distal Interge | 12 | 115695720 | 115695794 | 75 | 2 | 102465205 | ENSMUST00 | -154269 | ENSMUSG0 | Mir6388 | microRNA 6388 |
| chr12 | 119518232 | 119518603 | 372 | * | NA_peak_114 | 50 | Distal Interge | 12 | 119443410 | 119466934 | 23525 | 1 | 238455 | ENSMUST00 | 74822 | ENSMUSG0 | Macc1 | metastasis associated in colon cancer 1 |
| chr12 | 119992553 | 119992723 | 171 | * | NA_peak_114 | 23 | Intron (ENSM | 12 | 119946570 | 120021246 | 74677 | 1 | 217951 | ENSMUST00 | 45983 | ENSMUSG0 | Tmem196 | transmembrane protein 196 |
| chr13 | 3361385 | 3361661 | 277 | * | NA_peak_115 | 29 | Exon (ENSMU | 13 | 3478230 | 3486177 | 7948 | 1 | 76937 | ENSMUST00 | -116569 | NA | 2810429I04R | RIKEN cDNA 2810429I04 gene |
| chr13 | 3372784 | 3373580 | 797 | * | NA_peak_115 | 94 | Distal Interge | 13 | 3478230 | 3486177 | 7948 | 1 | 76937 | ENSMUST00 | -104650 | NA | 2810429I04R | RIKEN cDNA 2810429I04 gene |
| chr13 | 3402428 | 3402721 | 294 | * | NA_peak_115 | 27 | Distal Interge | 13 | 3478230 | 3486177 | 7948 | 1 | 76937 | ENSMUST00 | -75509 | NA | 2810429I04R | RIKEN cDNA 2810429I04 gene |
| chr13 | 3403932 | 3404222 | 291 | * | NA_peak_115 | 27 | Distal Interge | 13 | 3478230 | 3486177 | 7948 | 1 | 76937 | ENSMUST00 | -74008 | NA | 2810429I04R | RIKEN cDNA 2810429I04 gene |
| chr13 | 3408494 | 3409004 | 511 | * | NA_peak_115 | 45 | Distal Interge | 13 | 3478230 | 3486177 | 7948 | 1 | 76937 | ENSMUST00 | -69226 | NA | 2810429I04R | RIKEN cDNA 2810429I04 gene |
| chr13 | 3616384 | 3616757 | 374 | * | NA_peak_115 | 27 | Distal Interge | 13 | 3566035 | 3611108 | 45074 | 2 | 105203 | ENSMUST00 | -5276 | ENSMUSG0 | Tasor2 | transcription activation suppressor family member 2 |
| chr13 | 3964304 | 3964642 | 339 | * | NA_peak_115 | 20 | Distal Interge | 13 | 3940688 | 3945349 | 4662 | 2 | 83428 | ENSMUST00 | -18955 | ENSMUSG0 | Ucn3 | urocortin 3 |
| chr13 | 4178149 | 4178299 | 151 | * | NA_peak_115 | 16 | Distal Interge | 13 | 4191150 | 4205596 | 14447 | 1 | 27384 | ENSMUST00 | -12851 | ENSMUSG0 | Akr1c13 | aldo-keto reductase family 1, member C13 |
| chr13 | 4339266 | 4339675 | 410 | * | NA_peak_115 | 30 | Distal Interge | 13 | 4268176 | 4279433 | 11258 | 2 | 622402 | ENSMUST00 | -59833 | ENSMUSG0 | Akr1c12 | aldo-keto reductase family 1, member C12 |
| chr13 | 5201696 | 5201863 | 168 | * | NA_peak_115 | 21 | Distal Interge | 13 | 4590757 | 4609174 | 18418 | 2 | 56043 | ENSMUST00 | -592522 | ENSMUSG0 | Akr1e1 | aldo-keto reductase family 1, member E1 |
| chr13 | 5696794 | 5696948 | 155 | * | NA_peak_115 | 36 | Intron (ENSM | 13 | 5855515 | 5857561 | 2047 | 2 | 71867 | ENSMUST00 | 160613 | ENSMUSG0 | 1700016G22 | RIKEN cDNA 1700016G22 gene |
| chr13 | 5811001 | 5811288 | 288 | * | NA_peak_116 | 31 | Distal Interge | 13 | 5855515 | 5857561 | 2047 | 2 | 71867 | ENSMUST00 | 46273 | ENSMUSG0 | 1700016G22 | RIKEN cDNA 1700016G22 gene |
| chr13 | 6730355 | 6730460 | 106 | * | NA_peak_116 | 22 | Distal Interge | 13 | 6579768 | 6648777 | 69010 | 2 | 56421 | ENSMUST00 | -81578 | ENSMUSG0 | Pfkp | phosphofructokinase, platelet |
| chr13 | 7014650 | 7014881 | 232 | * | NA_peak_116 | 20 | Intron (ENSM | 13 | 6579768 | 6648777 | 69010 | 2 | 56421 | ENSMUST00 | -365873 | ENSMUSG0 | Pfkp | phosphofructokinase, platelet |
| chr13 | 7128639 | 7128833 | 195 | * | NA_peak_116 | 21 | Distal Interge | 13 | 6579768 | 6648777 | 69010 | 2 | 56421 | ENSMUST00 | -479862 | ENSMUSG0 | Pfkp | phosphofructokinase, platelet |
| chr13 | 8666931 | 8666977 | 47 | * | NA_peak_117 | 24 | Intron (ENSM | 13 | 8835389 | 8837234 | 1846 | 2 | 207615 | ENSMUST00 | 170257 | ENSMUSG0 | Wdr37 | WD repeat domain 37 |
| chr13 | 8765219 | 8765552 | 334 | * | NA_peak_117 | 40 | Intron (ENSM | 13 | 8835389 | 8837234 | 1846 | 2 | 207615 | ENSMUST00 | 71682 | ENSMUSG0 | Wdr37 | WD repeat domain 37 |
| chr13 | 8924427 | 8924756 | 330 | * | NA_peak_117 | 15 | Distal Interge | 13 | 8952863 | 8960945 | 8083 | 1 | 320581 | ENSMUST00 | -28107 | ENSMUSG0 | Idi2 | isopentenyl-diphosphate delta isomerase 2 |
| chr13 | 8950458 | 8950851 | 394 | * | NA_peak_117 | 20 | Promoter (2-3 | 13 | 8952863 | 8960945 | 8083 | 1 | 320581 | ENSMUST00 | -2012 | ENSMUSG0 | Idi2 | isopentenyl-diphosphate delta isomerase 2 |
| chr13 | 9169337 | 9169696 | 360 | * | NA_peak_117 | 17 | Promoter (<= | 13 | 9168361 | 9173121 | 4761 | 1 | 217980 | ENSMUST00 | 976 | ENSMUSG0 | Larp4b | La ribonucleoprotein domain family, member 4B |
| chr13 | 9820052 | 9820210 | 159 | * | NA_peak_117 | 21 | Distal Interge | 13 | 9691049 | 9765330 | 74282 | 2 | 66505 | ENSMUST00 | -54722 | ENSMUSG0 | Zmynd11 | zinc finger, MYND domain containing 11 |
| chr13 | 10128112 | 10128326 | 215 | * | NA_peak_117 | 50 | Intron (ENSM | 13 | 10215694 | 10360563 | 144870 | 2 | 12671 | ENSMUST00 | 232237 | ENSMUSG0 | Chrm3 | cholinergic receptor, muscarinic 3, cardiac |
| chr13 | 10193138 | 10193493 | 356 | * | NA_peak_117 | 20 | Intron (ENSM | 13 | 10215694 | 10360563 | 144870 | 2 | 12671 | ENSMUST00 | 167070 | ENSMUSG0 | Chrm3 | cholinergic receptor, muscarinic 3, cardiac |
| chr13 | 10275031 | 10275057 | 27 | * | NA_peak_117 | 20 | Intron (ENSM | 13 | 10215694 | 10360563 | 144870 | 2 | 12671 | ENSMUST00 | 85506 | ENSMUSG0 | Chrm3 | cholinergic receptor, muscarinic 3, cardiac |
| chr13 | 11188922 | 11189188 | 267 | * | NA_peak_118 | 43 | Distal Interge | 13 | 11522052 | 11526436 | 4385 | 1 | 664793 | ENSMUST00 | -332864 | ENSMUSG0 | Zp4-ps | zona pellucida glycoprotein 4, pseudogene |
| chr13 | 12134370 | 12134483 | 114 | * | NA_peak_118 | 16 | Distal Interge | 13 | 11771040 | 12106945 | 335906 | 2 | 20191 | ENSMUST00 | -27425 | ENSMUSG0 | Ryr2 | ryanodine receptor 2, cardiac |
| chr13 | 12478367 | 12478670 | 304 | * | NA_peak_118 | 28 | Exon (ENSMU | 13 | 12439415 | 12464944 | 25530 | 2 | 56048 | ENSMUST00 | -13423 | ENSMUSG0 | Lgals8 | lectin, galactose binding, soluble 8 |
| chr13 | 12764898 | 12765265 | 368 | * | NA_peak_118 | 38 | Distal Interge | 13 | 12790826 | 12792340 | 1515 | 2 | 26421 | ENSMUST00 | 27075 | ENSMUSG0 | Prl2c4 | prolactin family 2, subfamily c, member 4 |
| chr13 | 12770366 | 12770819 | 454 | * | NA_peak_118 | 44 | Distal Interge | 13 | 12790826 | 12792340 | 1515 | 2 | 26421 | ENSMUST00 | 21521 | ENSMUSG0 | Prl2c4 | prolactin family 2, subfamily c, member 4 |
| chr13 | 12806446 | 12806776 | 331 | * | NA_peak_118 | 19 | Distal Interge | 13 | 12790816 | 12800079 | 9264 | 2 | 18812 | ENSMUST00 | -6367 | ENSMUSG0 | Prl2c3 | prolactin family 2, subfamily c, member 3 |
| chr13 | 12864690 | 12865414 | 725 | * | NA_peak_118 | 41 | Exon (ENSMU | 13 | 12790816 | 12800079 | 9264 | 2 | 18812 | ENSMUST00 | -64611 | ENSMUSG0 | Prl2c3 | prolactin family 2, subfamily c, member 3 |
| chr13 | 12886192 | 12886388 | 197 | * | NA_peak_119 | 40 | Distal Interge | 13 | 12790816 | 12800079 | 9264 | 2 | 18812 | ENSMUST00 | -86113 | ENSMUSG0 | Prl2c3 | prolactin family 2, subfamily c, member 3 |
| chr13 | 12903184 | 12903506 | 323 | * | NA_peak_119 | 19 | Distal Interge | 13 | 12996126 | 12997575 | 1450 | 2 | 18811 | ENSMUST00 | 94069 | ENSMUSG0 | Prl2c2 | prolactin family 2, subfamily c, member 2 |
| chr13 | 12919440 | 12919829 | 390 | * | NA_peak_119 | 33 | Distal Interge | 13 | 12996126 | 12997575 | 1450 | 2 | 18811 | ENSMUST00 | 77746 | ENSMUSG0 | Prl2c2 | prolactin family 2, subfamily c, member 2 |
| chr13 | 12945380 | 12945740 | 361 | * | NA_peak_119 | 22 | Distal Interge | 13 | 12996126 | 12997575 | 1450 | 2 | 18811 | ENSMUST00 | 51835 | ENSMUSG0 | Prl2c2 | prolactin family 2, subfamily c, member 2 |
| chr13 | 12966395 | 12966735 | 341 | * | NA_peak_119 | 28 | Distal Interge | 13 | 12996126 | 12997575 | 1450 | 2 | 18811 | ENSMUST00 | 30840 | ENSMUSG0 | Prl2c2 | prolactin family 2, subfamily c, member 2 |
| chr13 | 12968571 | 12968910 | 340 | * | NA_peak_119 | 40 | Distal Interge | 13 | 12996126 | 12997575 | 1450 | 2 | 18811 | ENSMUST00 | 28665 | ENSMUSG0 | Prl2c2 | prolactin family 2, subfamily c, member 2 |
| chr13 | 12995400 | 12995726 | 327 | * | NA_peak_119 | 22 | Promoter (1-2 | 13 | 12996126 | 12997575 | 1450 | 2 | 18811 | ENSMUST00 | 1849 | ENSMUSG0 | Prl2c2 | prolactin family 2, subfamily c, member 2 |
| chr13 | 13019257 | 13019509 | 253 | * | NA_peak_119 | 21 | Distal Interge | 13 | 12996125 | 13005383 | 9259 | 2 | 18811 | ENSMUST00 | -13874 | ENSMUSG0 | Prl2c2 | prolactin family 2, subfamily c, member 2 |
| chr13 | 13127958 | 13128167 | 210 | * | NA_peak_119 | 31 | Distal Interge | 13 | 13182715 | 13191923 | 9209 | 1 | 107849 | ENSMUST00 | -54548 | ENSMUSG0 | Prl2c5 | prolactin family 2, subfamily c, member 5 |
| chr13 | 13279660 | 13279873 | 214 | * | NA_peak_119 | 27 | Distal Interge | 13 | 13365003 | 13367805 | 2803 | 2 | 83924 | ENSMUST00 | 87932 | ENSMUSG0 | Gpr137b | G protein-coupled receptor 137B |
| chr13 | 13294800 | 13295011 | 212 | * | NA_peak_119 | 20 | Distal Interge | 13 | 13365003 | 13367805 | 2803 | 2 | 83924 | ENSMUST00 | 72794 | ENSMUSG0 | Gpr137b | G protein-coupled receptor 137B |
| chr13 | 15432436 | 15433021 | 586 | * | NA_peak_120 | 33 | Distal Interge | 13 | 15463235 | 15715091 | 251857 | 1 | 14634 | ENSMUST00 | -30214 | ENSMUSG0 | Gli3 | GLI-Kruppel family member GLI3 |
| chr13 | 15739929 | 15739978 | 50 | * | NA_peak_120 | 27 | Distal Interge | 13 | 15781664 | 15802630 | 20967 | 2 | 71071 | ENSMUST00 | 62652 | ENSMUSG0 | 4933412O06 | RIKEN cDNA 4933412O06 gene |
| chr13 | 15964430 | 15964898 | 469 | * | NA_peak_120 | 22 | Intron (ENSM | 13 | 15915502 | 15926667 | 11166 | 2 | 100503142 | ENSMUST00 | -37763 | ENSMUSG0 | B230303A05 | U1 small nuclear ribonucleoprotein 1C pseudogene |
| chr13 | 16523672 | 16524039 | 368 | * | NA_peak_120 | 16 | Distal Interge | 13 | 15944752 | 16023423 | 78672 | 2 | 100503142 | ENSMUST00 | -500249 | ENSMUSG0 | B230303A05 | U1 small nuclear ribonucleoprotein 1C pseudogene |
| chr13 | 16611432 | 16611893 | 462 | * | NA_peak_120 | 27 | Distal Interge | 13 | 15944752 | 16023423 | 78672 | 2 | 100503142 | ENSMUST00 | -588009 | ENSMUSG0 | B230303A05 | U1 small nuclear ribonucleoprotein 1C pseudogene |
| chr13 | 17460858 | 17460890 | 33 | * | NA_peak_120 | 20 | Intron (ENSM | 13 | 17684485 | 17694736 | 10252 | 2 | 192136 | ENSMUST00 | 233846 | ENSMUSG0 | Sugct | succinyl-CoA glutarate-CoA transferase |
| chr13 | 18998062 | 18998658 | 597 | * | NA_peak_121 | 20 | Intron (ENSM | 13 | 18948398 | 19101626 | 153229 | 1 | 218038 | ENSMUST00 | 49664 | ENSMUSG0 | Amph | amphiphysin |
| chr13 | 19681492 | 19681802 | 311 | * | NA_peak_121 | 30 | Intron (ENSM | 13 | 19645078 | 19675731 | 30654 | 2 | 73412 | ENSMUST00 | -5761 | ENSMUSG0 | Nme8 | NME/NM23 family member 8 |
| chr13 | 19786815 | 19787302 | 488 | * | NA_peak_121 | 16 | Intron (ENSM | 13 | 19752264 | 19824227 | 71964 | 2 | 353346 | ENSMUST00 | 36925 | ENSMUSG0 | Gpr141 | G protein-coupled receptor 141 |
| chr13 | 20795285 | 20795464 | 180 | * | NA_peak_122 | 16 | Promoter (1-2 | 13 | 20794135 | 20851912 | 57778 | 1 | 27052 | ENSMUST00 | 1150 | ENSMUSG0 | Aoah | acyloxyacyl hydrolase |
| chr13 | 21180642 | 21180946 | 305 | * | NA_peak_122 | 46 | Promoter (<= | 13 | 21181138 | 21192851 | 11714 | 1 | 19720 | ENSMUST00 | -192 | ENSMUSG0 | Trim27 | tripartite motif-containing 27 |
| chr13 | 21435522 | 21435712 | 191 | * | NA_peak_122 | 20 | Intron (ENSM | 13 | 21440471 | 21445934 | 5464 | 1 | 105245359 | ENSMUST00 | -4759 | NA | 4930470G03 | RIKEN cDNA 4930470G03 gene |
| chr13 | 22040826 | 22041133 | 308 | * | NA_peak_122 | 16 | Promoter (<= | 13 | 22040636 | 22041362 | 727 | 2 | 319158 | ENSMUST00 | 229 | ENSMUSG0 | H4c9 | H4 clustered histone 9 |
| chr13 | 22388825 | 22389529 | 705 | * | NA_peak_122 | 20 | Distal Interge | 13 | 22394947 | 22396009 | 1063 | 1 | 171246 | ENSMUST00 | -5418 | ENSMUSG0 | Vmn1r200 | vomeronasal 1 receptor 200 |
| chr13 | 22429209 | 22429526 | 318 | * | NA_peak_122 | 23 | Distal Interge | 13 | 22441272 | 22442164 | 893 | 1 | 171245 | ENSMUST00 | -11746 | ENSMUSG0 | Vmn1r-ps103 | vomeronasal 1 receptor, pseudogene 103 |
| chr13 | 22897888 | 22898640 | 753 | * | NA_peak_123 | 45 | Exon (ENSMU | 13 | 22883042 | 22884205 | 1164 | 2 | 171275 | ENSMUST00 | -13683 | ENSMUSG0 | Vmn1r212 | vomeronasal 1 receptor 212 |
| chr13 | 22964614 | 22965019 | 406 | * | NA_peak_123 | 16 | Distal Interge | 13 | 23011249 | 23012403 | 1155 | 1 | 171249 | ENSMUST00 | -46230 | ENSMUSG0 | Vmn1r213 | vomeronasal 1 receptor 213 |
| chr13 | 23206992 | 23207394 | 403 | * | NA_peak_123 | 43 | Distal Interge | 13 | 23217373 | 23218311 | 939 | 1 | 100312485 | ENSMUST00 | -9979 | ENSMUSG0 | Vmn1r221 | vomeronasal 1 receptor 221 |
| chr13 | 23408388 | 23408870 | 483 | * | NA_peak_123 | 28 | Distal Interge | 13 | 23418361 | 23423866 | 5506 | 2 | 30946 | ENSMUST00 | 14996 | ENSMUSG0 | Abt1 | activator of basal transcription 1 |
| chr13 | 23458234 | 23458385 | 152 | * | NA_peak_123 | 18 | 3' UTR | 13 | 23464244 | 23465409 | 1166 | 2 | 12231 | ENSMUST00 | 7024 | ENSMUSG0 | Btn1a1 | butyrophilin, subfamily 1, member A1 |
| chr13 | 24499301 | 24499562 | 262 | * | NA_peak_123 | 19 | Promoter (1-2 | 13 | 24501525 | 24705198 | 203674 | 1 | 193385 | ENSMUST00 | -1963 | ENSMUSG0 | Ripor2 | RHO family interacting cell polarization regulator 2 |
| chr13 | 25044241 | 25044454 | 214 | * | NA_peak_124 | 20 | Distal Interge | 13 | 25056004 | 25121523 | 65520 | 1 | 195208 | ENSMUST00 | -11550 | ENSMUSG0 | Dcdc2a | doublecortin domain containing 2a |
| chr13 | 25321691 | 25321982 | 292 | * | NA_peak_124 | 30 | Distal Interge | 13 | 25252040 | 25270082 | 18043 | 2 | 22360 | ENSMUST00 | -51609 | ENSMUSG0 | Nrsn1 | neurensin 1 |
| chr13 | 26052783 | 26053154 | 372 | * | NA_peak_124 | 31 | Distal Interge | 13 | 25922436 | 25986293 | 63858 | 2 | 100503530 | ENSMUST00 | -66490 | ENSMUSG0 | Gm11351 | predicted gene 11351 |
| chr13 | 27789401 | 27789723 | 323 | * | NA_peak_124 | 19 | Distal Interge | 13 | 27773495 | 27780804 | 7310 | 2 | 67505 | ENSMUST00 | -8597 | ENSMUSG0 | Prl7c1 | prolactin family 7, subfamily c, member 1 |
| chr13 | 27826916 | 27826962 | 47 | * | NA_peak_124 | 27 | Distal Interge | 13 | 27849342 | 27857788 | 8447 | 1 | 666317 | ENSMUST00 | -22380 | ENSMUSG0 | Prl2c1 | Prolactin family 2, subfamily c, member 1 |
| chr13 | 27876725 | 27877087 | 363 | * | NA_peak_125 | 40 | Distal Interge | 13 | 27849402 | 27857683 | 8282 | 1 | 666317 | ENSMUST00 | 27323 | ENSMUSG0 | Prl2c1 | Prolactin family 2, subfamily c, member 1 |
| chr13 | 28038631 | 28038781 | 151 | * | NA_peak_125 | 20 | Distal Interge | 13 | 28016212 | 28023559 | 7348 | 1 | 19110 | ENSMUST00 | 22419 | ENSMUSG0 | Prl4a1 | prolactin family 4, subfamily a, member 1 |
| chr13 | 29242615 | 29242900 | 286 | * | NA_peak_125 | 22 | Intron (ENSM | 13 | 29016708 | 29040336 | 23629 | 1 | 77920 | ENSMUST00 | 225907 | ENSMUSG0 | A330102I10R | RIKEN cDNA A330102I10 gene |
| chr13 | 29283010 | 29283037 | 28 | * | NA_peak_125 | 22 | Intron (ENSM | 13 | 29402593 | 29474607 | 72015 | 2 | 68916 | ENSMUST00 | 191570 | ENSMUSG0 | Cdkal1 | CDK5 regulatory subunit associated protein 1-like 1 |
| chr13 | 29411426 | 29411437 | 12 | * | NA_peak_125 | 27 | Intron (ENSM | 13 | 29402593 | 29474607 | 72015 | 2 | 68916 | ENSMUST00 | 63170 | ENSMUSG0 | Cdkal1 | CDK5 regulatory subunit associated protein 1-like 1 |

| | | | | | | | | | | | | | | | | | | |
|---|---|---|---|---|---|---|---|---|---|---|---|---|---|---|---|---|---|---|
| chr13 | 29440531 | 29440559 | 29 | * | NA_peak_12 | 26 | Intron (ENSM | 13 | 29402593 | 29474607 | 72015 | 2 | 68916 | ENSMUST00 | 34048 | ENSMUSG0 | Cdkal1 | CDK5 regulatory subunit associated protein 1-like 1 |
| chr13 | 29611585 | 29611607 | 23 | * | NA_peak_12 | 16 | Intron (ENSM | 13 | 29443776 | 29517454 | 73679 | 2 | 68916 | ENSMUST00 | -94131 | ENSMUSG0 | Cdkal1 | CDK5 regulatory subunit associated protein 1-like 1 |
| chr13 | 30062282 | 30062400 | 119 | * | NA_peak_12 | 16 | Intron (ENSM | 13 | 30136489 | 30163625 | 27137 | 1 | 218121 | ENSMUST00 | -74089 | ENSMUSG0 | Mboat1 | membrane bound O-acyltransferase domain containing 1 |
| chr13 | 30274429 | 30274578 | 150 | * | NA_peak_12 | 23 | Distal Interge | 13 | 30231853 | 30245715 | 13863 | 1 | 218121 | ENSMUST00 | 42576 | ENSMUSG0 | Mboat1 | membrane bound O-acyltransferase domain containing 1 |
| chr13 | 30511901 | 30512105 | 205 | * | NA_peak_12 | 36 | Distal Interge | 13 | 30540308 | 30545362 | 5055 | 2 | 66694 | ENSMUST00 | 33257 | ENSMUSG0 | Uqcrfs1 | ubiquinol-cytochrome c reductase, Rieske iron-sulfur polypeptide 1 |
| chr13 | 31237959 | 31238356 | 398 | * | NA_peak_12 | 36 | Promoter (2-3 | 13 | 31213410 | 31241086 | 27677 | 2 | 73924 | ENSMUST00 | 2730 | ENSMUSG0 | 4930401O12 | RIKEN cDNA 4930401O12 gene |
| chr13 | 31738238 | 31738375 | 138 | * | NA_peak_12 | 23 | Distal Interge | 13 | 31806633 | 31812476 | 5844 | 1 | 17300 | ENSMUST00 | -68258 | ENSMUSG0 | Foxc1 | forkhead box C1 |
| chr13 | 31784682 | 31784979 | 298 | * | NA_peak_12 | 39 | Distal Interge | 13 | 31806633 | 31812476 | 5844 | 1 | 17300 | ENSMUST00 | -21654 | ENSMUSG0 | Foxc1 | forkhead box C1 |
| chr13 | 31854062 | 31854235 | 174 | * | NA_peak_12 | 20 | Intron (ENSM | 13 | 31806633 | 31812476 | 5844 | 1 | 17300 | ENSMUST00 | 47429 | ENSMUSG0 | Foxc1 | forkhead box C1 |
| chr13 | 32920262 | 32920752 | 491 | * | NA_peak_12 | 20 | Distal Interge | 13 | 32881434 | 32898211 | 16778 | 2 | 380839 | ENSMUST00 | -22051 | ENSMUSG0 | Serpinb1c | serine (or cysteine) peptidase inhibitor, clade B, member 1c |
| chr13 | 33736686 | 33736756 | 71 | * | NA_peak_12 | 23 | Distal Interge | 13 | 33661475 | 33671529 | 10055 | 1 | 238568 | ENSMUST00 | 75211 | ENSMUSG0 | Serpinb6d | serine (or cysteine) peptidase inhibitor, clade B, member 6d |
| chr13 | 33741551 | 33741559 | 9 | * | NA_peak_12 | 27 | Intron (ENSM | 13 | 33661475 | 33671529 | 10055 | 1 | 238568 | ENSMUST00 | 80076 | ENSMUSG0 | Serpinb6d | serine (or cysteine) peptidase inhibitor, clade B, member 6d |
| chr13 | 33775440 | 33775828 | 389 | * | NA_peak_12 | 26 | Intron (ENSM | 13 | 33832345 | 33843408 | 11064 | 2 | 435350 | ENSMUST00 | 67580 | ENSMUSG0 | Serpinb6e | serine (or cysteine) peptidase inhibitor, clade B, member 6e |
| chr13 | 34103093 | 34103407 | 315 | * | NA_peak_12 | 26 | Intron (ENSM | 13 | 34084155 | 34113272 | 29118 | 2 | 432745 | ENSMUST00 | 9865 | ENSMUSG0 | 4930447K03 | RIKEN cDNA 4930447K03 gene |
| chr13 | 34471676 | 34471967 | 292 | * | NA_peak_12 | 36 | Distal Interge | 13 | 34179158 | 34345182 | 166025 | 2 | 73102 | ENSMUST00 | -126494 | ENSMUSG0 | Slc22a23 | solute carrier family 22, member 23 |
| chr13 | 34515544 | 34515850 | 307 | * | NA_peak_12 | 28 | Distal Interge | 13 | 34628868 | 34648568 | 19701 | 2 | 66895 | ENSMUST00 | 132718 | ENSMUSG0 | Pxdc1 | PX domain containing 1 |
| chr13 | 34788658 | 34788918 | 261 | * | NA_peak_12 | 20 | Intron (ENSM | 13 | 34739612 | 34747622 | 8011 | 1 | 108161 | ENSMUST00 | 49046 | ENSMUSG0 | Fam50b | family with sequence similarity 50, member B |
| chr13 | 35012834 | 35013195 | 362 | * | NA_peak_12 | 16 | Intron (ENSM | 13 | 34986067 | 35027094 | 41028 | 2 | 23986 | ENSMUST00 | 13899 | ENSMUSG0 | Eci2 | enoyl-Coenzyme A delta isomerase 2 |
| chr13 | 35495685 | 35496234 | 550 | * | NA_peak_12 | 19 | Distal Interge | 13 | 35659833 | 35874063 | 214231 | 1 | 12593 | ENSMUST00 | -163599 | ENSMUSG0 | Cdyl | chromodomain protein, Y chromosome-like |
| chr13 | 35569127 | 35569860 | 734 | * | NA_peak_12 | 66 | Distal Interge | 13 | 35659833 | 35874063 | 214231 | 1 | 12593 | ENSMUST00 | -89973 | ENSMUSG0 | Cdyl | chromodomain protein, Y chromosome-like |
| chr13 | 36185521 | 36185598 | 78 | * | NA_peak_12 | 22 | Intron (ENSM | 13 | 36232122 | 36246609 | 14488 | 1 | 69955 | ENSMUST00 | -46524 | ENSMUSG0 | Fars2 | phenylalanine-tRNA synthetase 2 (mitochondrial) |
| chr13 | 37350737 | 37351056 | 320 | * | NA_peak_12 | 13 | Intron (ENSM | 13 | 37345208 | 37419036 | 73829 | 1 | 17084 | ENSMUST00 | 5529 | ENSMUSG0 | Ly86 | lymphocyte antigen 86 |
| chr13 | 37507167 | 37507402 | 236 | * | NA_peak_12 | 22 | Exon (ENSMU | 13 | 37345208 | 37419036 | 73829 | 1 | 17084 | ENSMUST00 | 161959 | ENSMUSG0 | Ly86 | lymphocyte antigen 86 |
| chr13 | 38204837 | 38205148 | 312 | * | NA_peak_12 | 22 | Promoter (<= | 13 | 38204938 | 38215430 | 10493 | 1 | 67797 | ENSMUST00 | 0 | ENSMUSG0 | Snrnp48 | small nuclear ribonucleoprotein 48 (U11/U12) |
| chr13 | 38468043 | 38468361 | 319 | * | NA_peak_12 | 16 | Intron (ENSM | 13 | 38460256 | 38461805 | 1550 | 2 | 75854 | ENSMUST00 | -6238 | ENSMUSG0 | 4930579J19 | RIKEN cDNA 4930579J19 gene |
| chr13 | 38888800 | 38889059 | 260 | * | NA_peak_12 | 20 | Intron (ENSM | 13 | 38932719 | 38943152 | 10434 | 2 | 108652 | ENSMUST00 | 54093 | ENSMUSG0 | Slc35b3 | solute carrier family 35, member B3 |
| chr13 | 39594921 | 39595229 | 309 | * | NA_peak_13 | 36 | Intron (ENSM | 13 | 38932136 | 38960875 | 28740 | 2 | 108652 | ENSMUST00 | -634046 | ENSMUSG0 | Slc35b3 | solute carrier family 35, member B3 |
| chr13 | 39595827 | 39596147 | 321 | * | NA_peak_13 | 28 | Intron (ENSM | 13 | 38932136 | 38960875 | 28740 | 2 | 108652 | ENSMUST00 | -634952 | ENSMUSG0 | Slc35b3 | solute carrier family 35, member B3 |
| chr13 | 39597992 | 39598123 | 132 | * | NA_peak_13 | 26 | Intron (ENSM | 13 | 38932136 | 38960875 | 28740 | 2 | 108652 | ENSMUST00 | -637117 | ENSMUSG0 | Slc35b3 | solute carrier family 35, member B3 |
| chr13 | 39942531 | 39942926 | 396 | * | NA_peak_13 | 87 | Distal Interge | 13 | 40001882 | 40288011 | 286130 | 2 | 218165 | ENSMUST00 | 345085 | ENSMUSG0 | Ofcc1 | orofacial cleft 1 candidate 1 |
| chr13 | 40744606 | 40744947 | 342 | * | NA_peak_13 | 27 | Distal Interge | 13 | 40716605 | 40738376 | 21772 | 2 | 21418 | ENSMUST00 | -6230 | ENSMUSG0 | Tfap2a | transcription factor AP-2, alpha |
| chr13 | 40973045 | 40973257 | 213 | * | NA_peak_13 | 31 | Intron (ENSM | 13 | 40953415 | 40958228 | 4814 | 1 | 14538 | ENSMUST00 | 19630 | ENSMUSG0 | Gcnt2 | glucosaminyl (N-acetyl) transferase 2, I-branching enzyme |
| chr13 | 41058460 | 41058505 | 46 | * | NA_peak_13 | 23 | Intron (ENSM | 13 | 41026142 | 41070459 | 44318 | 2 | 17152 | ENSMUST00 | 11954 | ENSMUSG0 | Mak | male germ cell-associated kinase |
| chr13 | 41059367 | 41059643 | 277 | * | NA_peak_13 | 36 | Intron (ENSM | 13 | 41026142 | 41070459 | 44318 | 2 | 17152 | ENSMUST00 | 10816 | ENSMUSG0 | Mak | male germ cell-associated kinase |
| chr13 | 41440811 | 41440878 | 68 | * | NA_peak_13 | 16 | Intron (ENSM | 13 | 41337816 | 41487242 | 149427 | 2 | 18003 | ENSMUST00 | 46364 | ENSMUSG0 | Nedd9 | neural precursor cell expressed, developmentally down-regulated gene 9 |
| chr13 | 44869568 | 44869939 | 372 | * | NA_peak_13 | 460 | Intron (ENSM | 13 | 44840783 | 44896481 | 55699 | 1 | 16468 | ENSMUST00 | 28785 | ENSMUSG0 | Jarid2 | jumonji, AT rich interactive domain 2 |
| chr13 | 44878964 | 44879272 | 309 | * | NA_peak_13 | 23 | Intron (ENSM | 13 | 44840783 | 44896481 | 55699 | 1 | 16468 | ENSMUST00 | 38181 | ENSMUSG0 | Jarid2 | jumonji, AT rich interactive domain 2 |
| chr13 | 44987537 | 44987951 | 415 | * | NA_peak_13 | 48 | Intron (ENSM | 13 | 44922086 | 44982997 | 60912 | 2 | 94245 | ENSMUST00 | -4540 | ENSMUSG0 | Dtnbp1 | dystrobrevin binding protein 1 |
| chr13 | 47337706 | 47337715 | 10 | * | NA_peak_13 | 14 | Intron (ENSM | 13 | 47193825 | 47246417 | 52593 | 1 | 218215 | ENSMUST00 | 143881 | ENSMUSG0 | Rnf144b | ring finger protein 144B |
| chr13 | 49125588 | 49125833 | 246 | * | NA_peak_13 | 24 | Intron (ENSM | 13 | 49036309 | 49147232 | 110924 | 2 | 75607 | ENSMUST00 | 21399 | ENSMUSG0 | Wnk2 | WNK lysine deficient protein kinase 2 |
| chr13 | 49215825 | 49215843 | 19 | * | NA_peak_13 | 47 | Promoter (<= | 13 | 49202950 | 49215860 | 12911 | 2 | 68480 | ENSMUST00 | 17 | ENSMUSG0 | Card19 | caspase recruitment domain family, member 19 |
| chr13 | 49599809 | 49599999 | 191 | * | NA_peak_13 | 17 | Intron (ENSM | 13 | 49608046 | 49624501 | 16456 | 1 | 18295 | ENSMUST00 | -8047 | ENSMUSG0 | Ogn | osteoglycin |
| chr13 | 49746655 | 49746942 | 288 | * | NA_peak_13 | 57 | Distal Interge | 13 | 49722119 | 49723096 | 978 | 1 | 105148 | ENSMUST00 | 24536 | ENSMUSG0 | Iars | isoleucine-tRNA synthetase |
| chr13 | 49748138 | 49748439 | 302 | * | NA_peak_13 | 22 | Distal Interge | 13 | 49722119 | 49723096 | 978 | 1 | 105148 | ENSMUST00 | 26019 | ENSMUSG0 | Iars | isoleucine-tRNA synthetase |
| chr13 | 50501555 | 50501938 | 384 | * | NA_peak_13 | 22 | Distal Interge | 13 | 50467307 | 50475355 | 8049 | 1 | 328250 | ENSMUST00 | 34248 | ENSMUSG0 | Nutm2 | NUT family member 2 |
| chr13 | 50689988 | 50690283 | 296 | * | NA_peak_13 | 23 | Distal Interge | 13 | 50698312 | 50703435 | 5124 | 1 | 667693 | ENSMUST00 | -8029 | ENSMUSG0 | Gm8765 | predicted gene 8765 |
| chr13 | 50692688 | 50692813 | 126 | * | NA_peak_13 | 23 | Distal Interge | 13 | 50698312 | 50703435 | 5124 | 1 | 667693 | ENSMUST00 | -5499 | ENSMUSG0 | Gm8765 | predicted gene 8765 |
| chr13 | 50718962 | 50719112 | 151 | * | NA_peak_13 | 33 | Distal Interge | 13 | 50698312 | 50703435 | 5124 | 1 | 667693 | ENSMUST00 | 20650 | ENSMUSG0 | Gm8765 | predicted gene 8765 |
| chr13 | 50897933 | 50898234 | 302 | * | NA_peak_13 | 22 | Distal Interge | 13 | 50753391 | 50765248 | 11858 | 2 | 71654 | ENSMUST00 | -132685 | ENSMUSG0 | 4930518P08 | RIKEN cDNA 4930518P08 gene |
| chr13 | 51269430 | 51270098 | 669 | * | NA_peak_13 | 28 | Distal Interge | 13 | 51202727 | 51203065 | 339 | 2 | 667728 | ENSMUST00 | -66365 | NA | H2af-ps2 | H2A histone family, pseudogene 2 |
| chr13 | 51834018 | 51834434 | 417 | * | NA_peak_13 | 27 | Distal Interge | 13 | 51846678 | 51848468 | 1791 | 1 | 23882 | ENSMUST00 | -12244 | ENSMUSG0 | Gadd45g | growth arrest and DNA-damage-inducible 45 gamma |
| chr13 | 51986806 | 51986939 | 134 | * | NA_peak_13 | 16 | Distal Interge | 13 | 52096741 | 52124158 | 27418 | 1 | 74050 | ENSMUST00 | -109802 | ENSMUSG0 | 4921525O09 | RIKEN cDNA 4921525O09 gene |
| chr13 | 53043840 | 53044241 | 402 | * | NA_peak_13 | 34 | Distal Interge | 13 | 52967209 | 52981073 | 13865 | 2 | 18030 | ENSMUST00 | -62767 | ENSMUSG0 | Nfil3 | nuclear factor, interleukin 3, regulated |
| chr13 | 53254983 | 53254993 | 11 | * | NA_peak_13 | 46 | Intron (ENSM | 13 | 53131960 | 53275658 | 143699 | 2 | 26564 | ENSMUST00 | 20665 | ENSMUSG0 | Ror2 | receptor tyrosine kinase-like orphan receptor 2 |
| chr13 | 53462702 | 53462834 | 133 | * | NA_peak_13 | 57 | Distal Interge | 13 | 53466884 | 53473074 | 6191 | 2 | 17702 | ENSMUST00 | 10240 | ENSMUSG0 | Msx2 | msh homeobox 2 |
| chr13 | 53525630 | 53526129 | 500 | * | NA_peak_13 | 25 | Exon (ENSMU | 13 | 53466884 | 53473074 | 6191 | 2 | 17702 | ENSMUST00 | -52556 | ENSMUSG0 | Msx2 | msh homeobox 2 |
| chr13 | 53644030 | 53644166 | 137 | * | NA_peak_13 | 28 | Distal Interge | 13 | 53648585 | 53650080 | 1496 | 1 | 75252 | ENSMUST00 | -4419 | ENSMUSG0 | 4930555G21 | RIKEN cDNA 4930555G21 gene |
| chr13 | 54148339 | 54148552 | 214 | * | NA_peak_13 | 31 | Distal Interge | 13 | 54192129 | 54222432 | 30304 | 1 | 15466 | ENSMUST00 | -43577 | ENSMUSG0 | Hrh2 | histamine receptor H2 |
| chr13 | 54232597 | 54232963 | 367 | * | NA_peak_13 | 23 | Intron (ENSM | 13 | 54214007 | 54236180 | 22174 | 1 | 15466 | ENSMUST00 | 18590 | ENSMUSG0 | Hrh2 | histamine receptor H2 |
| chr13 | 54667571 | 54667865 | 295 | * | NA_peak_13 | 36 | Distal Interge | 13 | 54651441 | 54655546 | 4106 | 1 | 76577 | ENSMUST00 | 16130 | ENSMUSG0 | Faf2 | Fas associated factor family member 2 |
| chr13 | 55185524 | 55185943 | 420 | * | NA_peak_13 | 104 | Distal Interge | 13 | 55209782 | 55315755 | 105974 | 1 | 18193 | ENSMUST00 | -23839 | ENSMUSG0 | Nsd1 | nuclear receptor-binding SET-domain protein 1 |
| chr13 | 55321598 | 55321980 | 383 | * | NA_peak_13 | 19 | Promoter (<= | 13 | 55319743 | 55321946 | 2204 | 2 | 19336 | ENSMUST00 | 0 | ENSMUSG0 | Rab24 | RAB24, member RAS oncogene family |
| chr13 | 55326900 | 55327165 | 266 | * | NA_peak_13 | 28 | Promoter (2-3 | 13 | 55325712 | 55329352 | 3641 | 2 | 17121 | ENSMUST00 | 2187 | ENSMUSG0 | Mxd3 | Max dimerization protein 3 |
| chr13 | 55524526 | 55525079 | 554 | * | NA_peak_13 | 43 | Exon (ENSMU | 13 | 55523235 | 55528538 | 5304 | 2 | 27261 | ENSMUST00 | 3459 | ENSMUSG0 | Dok3 | docking protein 3 |
| chr13 | 55630872 | 55631154 | 283 | * | NA_peak_13 | 27 | Promoter (2-3 | 13 | 55628281 | 55632411 | 4131 | 1 | 12328 | ENSMUST00 | 2591 | ENSMUSG0 | Caml | calcium modulating ligand |
| chr13 | 56134634 | 56134896 | 263 | * | NA_peak_13 | 30 | Promoter (<= | 13 | 56104441 | 56134825 | 30385 | 2 | 26914 | ENSMUST00 | 0 | ENSMUSG0 | Macroh2a1 | macroH2A.1 histone |
| chr13 | 56444032 | 56444169 | 138 | * | NA_peak_13 | 22 | Intron (ENSM | 13 | 56438355 | 56474239 | 35885 | 1 | 328258 | ENSMUST00 | 5677 | ENSMUSG0 | Slc25a48 | solute carrier family 25, member 48 |
| chr13 | 56716584 | 56716822 | 239 | * | NA_peak_13 | 20 | Intron (ENSM | 13 | 56703668 | 56740304 | 36637 | 1 | 17129 | ENSMUST00 | 12916 | ENSMUSG0 | Smad5 | SMAD family member 5 |
| chr13 | 56895217 | 56895434 | 218 | * | NA_peak_13 | 14 | Promoter (<= | 13 | 56773113 | 56895737 | 122625 | 2 | 26946 | ENSMUST00 | 303 | ENSMUSG0 | Trpc7 | transient receptor potential cation channel, subfamily C, member 7 |
| chr13 | 57184180 | 57184240 | 61 | * | NA_peak_13 | 27 | Distal Interge | 13 | 56773764 | 56895993 | 122230 | 2 | 26946 | ENSMUST00 | -288187 | ENSMUSG0 | Trpc7 | transient receptor potential cation channel, subfamily C, member 7 |
| chr13 | 57207734 | 57208070 | 337 | * | NA_peak_13 | 31 | Distal Interge | 13 | 56773764 | 56895993 | 122230 | 2 | 26946 | ENSMUST00 | -311741 | ENSMUSG0 | Trpc7 | transient receptor potential cation channel, subfamily C, member 7 |
| chr13 | 57356109 | 57356295 | 187 | * | NA_peak_13 | 20 | Distal Interge | 13 | 56773764 | 56895993 | 122230 | 2 | 26946 | ENSMUST00 | -460116 | ENSMUSG0 | Trpc7 | transient receptor potential cation channel, subfamily C, member 7 |
| chr13 | 57429234 | 57429400 | 167 | * | NA_peak_13 | 23 | 3' UTR | 13 | 57429295 | 57907587 | 478293 | 2 | 20745 | ENSMUST00 | 478187 | ENSMUSG0 | Spock1 | sparc/osteonectin, cwcv and kazal-like domains proteoglycan 1 |
| chr13 | 57687117 | 57687194 | 78 | * | NA_peak_13 | 24 | Intron (ENSM | 13 | 57429295 | 57907587 | 478293 | 2 | 20745 | ENSMUST00 | 220393 | ENSMUSG0 | Spock1 | sparc/osteonectin, cwcv and kazal-like domains proteoglycan 1 |
| chr13 | 58264189 | 58264320 | 132 | * | NA_peak_13 | 23 | Promoter (<= | 13 | 58250720 | 58263978 | 13259 | 2 | 56278 | ENSMUST00 | -211 | ENSMUSG0 | Gkap1 | G kinase anchoring protein 1 |
| chr13 | 58716714 | 58717059 | 346 | * | NA_peak_13 | 21 | Distal Interge | 13 | 58741074 | 58762311 | 21238 | 2 | 67585 | ENSMUST00 | 45252 | ENSMUSG0 | 4930455J16 | RIKEN cDNA 4930455J16 gene |
| chr13 | 59047307 | 59047670 | 364 | * | NA_peak_13 | 23 | Intron (ENSM | 13 | 58871735 | 58986149 | 114415 | 1 | 18212 | ENSMUST00 | 175572 | ENSMUSG0 | Ntrk2 | neurotrophic tyrosine kinase, receptor, type 2 |
| chr13 | 59227758 | 59228052 | 295 | * | NA_peak_13 | 23 | Distal Interge | 13 | 59449894 | 59465288 | 15395 | 2 | 67269 | ENSMUST00 | 237236 | ENSMUSG0 | Agtpbp1 | ATP/GTP binding protein 1 |
| chr13 | 59234177 | 59234411 | 235 | * | NA_peak_13 | 19 | Distal Interge | 13 | 59449894 | 59465288 | 15395 | 2 | 67269 | ENSMUST00 | 230877 | ENSMUSG0 | Agtpbp1 | ATP/GTP binding protein 1 |
| chr13 | 59707591 | 59707871 | 281 | * | NA_peak_13 | 31 | Promoter (1-2 | 13 | 59702683 | 59706197 | 3515 | 2 | 72219 | ENSMUST00 | -1394 | ENSMUSG0 | Spata31d1a | spermatogenesis associated 31 subfamily D, member 1A |
| chr13 | 59870805 | 59871160 | 356 | * | NA_peak_13 | 31 | Distal Interge | 13 | 59771561 | 59823147 | 51587 | 2 | 214290 | ENSMUST00 | -47658 | ENSMUSG0 | Tut7 | terminal uridylyl transferase 7 |
| chr13 | 59947420 | 59947726 | 307 | * | NA_peak_13 | 22 | Distal Interge | 13 | 60029711 | 60116669 | 86959 | 1 | 328263 | ENSMUST00 | -81985 | ENSMUSG0 | A530065N20 | RIKEN cDNA A530046M15 gene |
| chr13 | 60213045 | 60213502 | 458 | * | NA_peak_14 | 16 | Exon (ENSMU | 13 | 60205471 | 60216023 | 10553 | 1 | 100503759 | ENSMUST00 | 7574 | ENSMUSG0 | Gm5084 | predicted gene 5084 |
| chr13 | 60391081 | 60391365 | 285 | * | NA_peak_14 | 22 | Distal Interge | 13 | 60205471 | 60216023 | 10553 | 1 | 100503759 | ENSMUST00 | 185610 | ENSMUSG0 | Gm5084 | predicted gene 5084 |
| chr13 | 60402419 | 60402695 | 277 | * | NA_peak_14 | 20 | Distal Interge | 13 | 60205471 | 60216023 | 10553 | 1 | 100503759 | ENSMUST00 | 196948 | ENSMUSG0 | Gm5084 | predicted gene 5084 |
| chr13 | 61398979 | 61399648 | 670 | * | NA_peak_14 | 55 | Distal Interge | 13 | 61353677 | 61358197 | 4521 | 2 | 56092 | ENSMUST00 | -40782 | ENSMUSG0 | Cts7 | cathepsin 7 |
| chr13 | 61402124 | 61402220 | 97 | * | NA_peak_14 | 54 | Distal Interge | 13 | 61353677 | 61358197 | 4521 | 2 | 56092 | ENSMUST00 | -43927 | ENSMUSG0 | Cts7 | cathepsin 7 |
| chr13 | 61402936 | 61403260 | 325 | * | NA_peak_14 | 30 | Distal Interge | 13 | 61353677 | 61358197 | 4521 | 2 | 56092 | ENSMUST00 | -44739 | ENSMUSG0 | Cts7 | cathepsin 7 |
| chr13 | 61762627 | 61762922 | 296 | * | NA_peak_14 | 36 | Distal Interge | 13 | 61564630 | 61570127 | 5498 | 2 | 117066 | ENSMUST00 | -192500 | ENSMUSG0 | Cts3 | cathepsin 3 |
| chr13 | 61822550 | 61822838 | 289 | * | NA_peak_14 | 40 | Intron (ENSM | 13 | 61564630 | 61570127 | 5498 | 2 | 117066 | ENSMUST00 | -252423 | ENSMUSG0 | Cts3 | cathepsin 3 |
| chr13 | 62078063 | 62078475 | 413 | * | NA_peak_14 | 30 | Intron (ENSM | 13 | 62129886 | 62172258 | 42373 | 1 | 630579 | ENSMUST00 | -51411 | ENSMUSG0 | Zfp808 | zinc finger protein 808 |
| chr13 | 62093975 | 62094318 | 344 | * | NA_peak_14 | 36 | Intron (ENSM | 13 | 62129886 | 62172258 | 42373 | 1 | 630579 | ENSMUST00 | -35568 | ENSMUSG0 | Zfp808 | zinc finger protein 808 |
| chr13 | 62115207 | 62115365 | 159 | * | NA_peak_14 | 32 | Intron (ENSM | 13 | 62129886 | 62172258 | 42373 | 1 | 630579 | ENSMUST00 | -14521 | ENSMUSG0 | Zfp808 | zinc finger protein 808 |
| chr13 | 62126094 | 62126388 | 295 | * | NA_peak_14 | 61 | Exon (ENSMU | 13 | 62129886 | 62172258 | 42373 | 1 | 630579 | ENSMUST00 | -3498 | ENSMUSG0 | Zfp808 | zinc finger protein 808 |
| chr13 | 62131906 | 62132247 | 342 | * | NA_peak_14 | 24 | Promoter (1-2 | 13 | 62129933 | 62206485 | 76553 | 1 | 630579 | ENSMUST00 | 1973 | ENSMUSG0 | Zfp808 | zinc finger protein 808 |
| chr13 | 62159393 | 62159444 | 52 | * | NA_peak_14 | 17 | Intron (ENSM | 13 | 62129933 | 62206485 | 76553 | 1 | 630579 | ENSMUST00 | 29460 | ENSMUSG0 | Zfp808 | zinc finger protein 808 |
| chr13 | 62166508 | 62166648 | 141 | * | NA_peak_14 | 38 | Intron (ENSM | 13 | 62129933 | 62206485 | 76553 | 1 | 630579 | ENSMUST00 | 36575 | ENSMUSG0 | Zfp808 | zinc finger protein 808 |
| chr13 | 62179107 | 62179304 | 198 | * | NA_peak_14 | 41 | Intron (ENSM | 13 | 62129933 | 62206485 | 76553 | 1 | 630579 | ENSMUST00 | 49174 | ENSMUSG0 | Zfp808 | zinc finger protein 808 |
| chr13 | 62187828 | 62187952 | 125 | * | NA_peak_14 | 20 | Intron (ENSM | 13 | 62129933 | 62206485 | 76553 | 1 | 630579 | ENSMUST00 | 57895 | ENSMUSG0 | Zfp808 | zinc finger protein 808 |
| chr13 | 62239569 | 62239855 | 287 | * | NA_peak_14 | 31 | Distal Interge | 13 | 62129933 | 62206485 | 76553 | 1 | 630579 | ENSMUST00 | 109636 | ENSMUSG0 | Zfp808 | zinc finger protein 808 |
| chr13 | 62509637 | 62509991 | 355 | * | NA_peak_14 | 13 | Intron (ENSM | 13 | 62517500 | 62520451 | 2952 | 2 | 77117 | ENSMUST00 | 10460 | ENSMUSG0 | Zfp934 | zinc finger protein 934 |
| chr13 | 62565233 | 62565521 | 289 | * | NA_peak_14 | 22 | Intron (ENSM | 13 | 62479827 | 62558635 | 78809 | 2 | 77117 | ENSMUST00 | -6598 | ENSMUSG0 | Zfp934 | zinc finger protein 934 |
| chr13 | 62578069 | 62578364 | 296 | * | NA_peak_14 | 41 | Distal Interge | 13 | 62479827 | 62558635 | 78809 | 2 | 77117 | ENSMUST00 | -19434 | ENSMUSG0 | Zfp934 | zinc finger protein 934 |
| chr13 | 62590460 | 62590881 | 422 | * | NA_peak_14 | 16 | Distal Interge | 13 | 62605190 | 62607499 | 2310 | 2 | 211378 | ENSMUST00 | 16618 | ENSMUSG0 | 6720489N17 | RIKEN cDNA 6720489N17 gene |
| chr13 | 62716768 | 62717045 | 278 | * | NA_peak_14 | 33 | Intron (ENSM | 13 | 62760739 | 62760846 | 108 | 1 | 751548 | ENSMUST00 | -43694 | ENSMUSG0 | Mir713 | microRNA 713 |
| chr13 | 62944720 | 62945043 | 324 | * | NA_peak_14 | 20 | Exon (ENSMU | 13 | 62964893 | 63015575 | 50683 | 1 | 72061 | ENSMUST00 | -19850 | ENSMUSG0 | Aopep | aminopeptidase O |

| | | | | | | | | | | | | | | | | | | |
|---|---|---|---|---|---|---|---|---|---|---|---|---|---|---|---|---|---|---|
| chr13 | 63581481 | 63581794 | 314 | * | NA_peak_142 | 20 | Exon (ENSMU | 13 | 63543176 | 63573598 | 30423 | 2 | 19206 | ENSMUST00 | -7883 | ENSMUSG00 | Ptch1 | patched 1 |
| chr13 | 64125567 | 64125769 | 203 | * | NA_peak_142 | 25 | Intron (ENSM | 13 | 64101278 | 64129305 | 28028 | 2 | 70484 | ENSMUST00 | 3536 | ENSMUSG00 | Slc35d2 | solute carrier family 35, member D2 |
| chr13 | 64135459 | 64135520 | 62 | * | NA_peak_142 | 36 | Exon (ENSMU | 13 | 64108401 | 64129368 | 20968 | 2 | 70484 | ENSMUST00 | -6091 | ENSMUSG00 | Slc35d2 | solute carrier family 35, member D2 |
| chr13 | 64407609 | 64408219 | 611 | * | NA_peak_142 | 40 | Distal Interge | 13 | 64432314 | 64441773 | 9460 | 1 | 105278 | ENSMUST00 | -24095 | ENSMUSG00 | Cdk20 | cyclin-dependent kinase 20 |
| chr13 | 64834145 | 64834420 | 276 | * | NA_peak_142 | 24 | Intron (ENSM | 13 | 64736182 | 64903955 | 167774 | 2 | 238680 | ENSMUST00 | 69535 | ENSMUSG00 | Cntnap3 | contactin associated protein-like 3 |
| chr13 | 64975428 | 64975531 | 104 | * | NA_peak_142 | 16 | Distal Interge | 13 | 65033058 | 65038004 | 4947 | 1 | 238683 | ENSMUST00 | -57527 | ENSMUSG00 | Spata31d1c | spermatogenesis associated 31 subfamily D, member 1C |
| chr13 | 65341546 | 65342036 | 491 | * | NA_peak_142 | 82 | Intron (ENSM | 13 | 65336222 | 65348597 | 12376 | 2 | 665738 | ENSMUST00 | 6561 | ENSMUSG00 | Gm7762 | predicted gene 7762 |
| chr13 | 65351097 | 65351389 | 293 | * | NA_peak_142 | 47 | Promoter (2-3 | 13 | 65336222 | 65348597 | 12376 | 2 | 665738 | ENSMUST00 | -2500 | ENSMUSG00 | Gm7762 | predicted gene 7762 |
| chr13 | 65357281 | 65358238 | 958 | * | NA_peak_142 | 78 | Intron (ENSM | 13 | 65336222 | 65348597 | 12376 | 2 | 665738 | ENSMUST00 | -8684 | ENSMUSG00 | Gm7762 | predicted gene 7762 |
| chr13 | 65358533 | 65358876 | 344 | * | NA_peak_143 | 18 | Intron (ENSM | 13 | 65336222 | 65348597 | 12376 | 2 | 665738 | ENSMUST00 | -9936 | ENSMUSG00 | Gm7762 | predicted gene 7762 |
| chr13 | 65365188 | 65366026 | 839 | * | NA_peak_143 | 76 | Distal Interge | 13 | 65336222 | 65348597 | 12376 | 2 | 665738 | ENSMUST00 | -16591 | ENSMUSG00 | Gm7762 | predicted gene 7762 |
| chr13 | 65367775 | 65368228 | 454 | * | NA_peak_143 | 63 | Exon (ENSMU | 13 | 65336222 | 65348597 | 12376 | 2 | 665738 | ENSMUST00 | -19178 | ENSMUSG00 | Gm7762 | predicted gene 7762 |
| chr13 | 65369195 | 65369779 | 585 | * | NA_peak_143 | 82 | Distal Interge | 13 | 65336222 | 65348597 | 12376 | 2 | 665738 | ENSMUST00 | -20598 | ENSMUSG00 | Gm7762 | predicted gene 7762 |
| chr13 | 65378255 | 65378543 | 289 | * | NA_peak_143 | 86 | Distal Interge | 13 | 65336222 | 65348597 | 12376 | 2 | 665738 | ENSMUST00 | -29658 | ENSMUSG00 | Gm7762 | predicted gene 7762 |
| chr13 | 65381848 | 65382246 | 399 | * | NA_peak_143 | 35 | Exon (ENSMU | 13 | 65336222 | 65348597 | 12376 | 2 | 665738 | ENSMUST00 | -33251 | ENSMUSG00 | Gm7762 | predicted gene 7762 |
| chr13 | 65390691 | 65391131 | 441 | * | NA_peak_143 | 32 | Distal Interge | 13 | 65336222 | 65348597 | 12376 | 2 | 665738 | ENSMUST00 | -42094 | ENSMUSG00 | Gm7762 | predicted gene 7762 |
| chr13 | 65419521 | 65419819 | 299 | * | NA_peak_143 | 28 | Distal Interge | 13 | 65336222 | 65348597 | 12376 | 2 | 665738 | ENSMUST00 | -70924 | ENSMUSG00 | Gm7762 | predicted gene 7762 |
| chr13 | 65430604 | 65431275 | 672 | * | NA_peak_143 | 22 | Intron (ENSM | 13 | 65336222 | 65348597 | 12376 | 2 | 665738 | ENSMUST00 | -82007 | ENSMUSG00 | Gm7762 | predicted gene 7762 |
| chr13 | 65454162 | 65454509 | 348 | * | NA_peak_143 | 58 | Distal Interge | 13 | 65336222 | 65348597 | 12376 | 2 | 665738 | ENSMUST00 | -105565 | ENSMUSG00 | Gm7762 | predicted gene 7762 |
| chr13 | 65455296 | 65455518 | 223 | * | NA_peak_143 | 34 | Distal Interge | 13 | 65336222 | 65348597 | 12376 | 2 | 665738 | ENSMUST00 | -106699 | ENSMUSG00 | Gm7762 | predicted gene 7762 |
| chr13 | 65457446 | 65457734 | 289 | * | NA_peak_143 | 28 | Distal Interge | 13 | 65336222 | 65348597 | 12376 | 2 | 665738 | ENSMUST00 | -108849 | ENSMUSG00 | Gm7762 | predicted gene 7762 |
| chr13 | 65464230 | 65464691 | 462 | * | NA_peak_143 | 31 | Distal Interge | 13 | 65336222 | 65348597 | 12376 | 2 | 665738 | ENSMUST00 | -115633 | ENSMUSG00 | Gm7762 | predicted gene 7762 |
| chr13 | 65471994 | 65472291 | 298 | * | NA_peak_143 | 107 | Distal Interge | 13 | 65336222 | 65348597 | 12376 | 2 | 665738 | ENSMUST00 | -123397 | ENSMUSG00 | Gm7762 | predicted gene 7762 |
| chr13 | 65479517 | 65479698 | 182 | * | NA_peak_143 | 30 | Distal Interge | 13 | 65336222 | 65348597 | 12376 | 2 | 665738 | ENSMUST00 | -130920 | ENSMUSG00 | Gm7762 | predicted gene 7762 |
| chr13 | 65482001 | 65482410 | 410 | * | NA_peak_143 | 53 | Distal Interge | 13 | 65336222 | 65348597 | 12376 | 2 | 665738 | ENSMUST00 | -133404 | ENSMUSG00 | Gm7762 | predicted gene 7762 |
| chr13 | 65498335 | 65498588 | 254 | * | NA_peak_143 | 38 | Intron (ENSM | 13 | 65336222 | 65348597 | 12376 | 2 | 665738 | ENSMUST00 | -149738 | ENSMUSG00 | Gm7762 | predicted gene 7762 |
| chr13 | 65515431 | 65516138 | 708 | * | NA_peak_143 | 151 | Intron (ENSM | 13 | 65336222 | 65348597 | 12376 | 2 | 665738 | ENSMUST00 | -166834 | ENSMUSG00 | Gm7762 | predicted gene 7762 |
| chr13 | 65522882 | 65523208 | 327 | * | NA_peak_143 | 60 | Intron (ENSM | 13 | 65336222 | 65348597 | 12376 | 2 | 665738 | ENSMUST00 | -174285 | ENSMUSG00 | Gm7762 | predicted gene 7762 |
| chr13 | 65541162 | 65541831 | 670 | * | NA_peak_143 | 37 | Intron (ENSM | 13 | 65336222 | 65348597 | 12376 | 2 | 665738 | ENSMUST00 | -192565 | ENSMUSG00 | Gm7762 | predicted gene 7762 |
| chr13 | 65542272 | 65543018 | 747 | * | NA_peak_143 | 37 | Intron (ENSM | 13 | 65336222 | 65348597 | 12376 | 2 | 665738 | ENSMUST00 | -193675 | ENSMUSG00 | Gm7762 | predicted gene 7762 |
| chr13 | 65559078 | 65559753 | 676 | * | NA_peak_143 | 40 | Exon (ENSMU | 13 | 65336222 | 65348597 | 12376 | 2 | 665738 | ENSMUST00 | -210481 | ENSMUSG00 | Gm7762 | predicted gene 7762 |
| chr13 | 65560395 | 65560682 | 288 | * | NA_peak_143 | 41 | Intron (ENSM | 13 | 65336222 | 65348597 | 12376 | 2 | 665738 | ENSMUST00 | -211798 | ENSMUSG00 | Gm7762 | predicted gene 7762 |
| chr13 | 65561329 | 65561619 | 291 | * | NA_peak_143 | 46 | Intron (ENSM | 13 | 65336222 | 65348597 | 12376 | 2 | 665738 | ENSMUST00 | -212732 | ENSMUSG00 | Gm7762 | predicted gene 7762 |
| chr13 | 65594119 | 65594484 | 366 | * | NA_peak_143 | 84 | Intron (ENSM | 13 | 65336222 | 65348597 | 12376 | 2 | 665738 | ENSMUST00 | -245522 | ENSMUSG00 | Gm7762 | predicted gene 7762 |
| chr13 | 65621168 | 65621480 | 313 | * | NA_peak_143 | 65 | Intron (ENSM | 13 | 65336222 | 65348597 | 12376 | 2 | 665738 | ENSMUST00 | -272571 | ENSMUSG00 | Gm7762 | predicted gene 7762 |
| chr13 | 65626487 | 65627102 | 616 | * | NA_peak_143 | 41 | Intron (ENSM | 13 | 65336222 | 65348597 | 12376 | 2 | 665738 | ENSMUST00 | -277890 | ENSMUSG00 | Gm7762 | predicted gene 7762 |
| chr13 | 65631573 | 65632077 | 505 | * | NA_peak_143 | 29 | Distal Interge | 13 | 65336222 | 65348597 | 12376 | 2 | 665738 | ENSMUST00 | -282976 | ENSMUSG00 | Gm7762 | predicted gene 7762 |
| chr13 | 65649633 | 65649964 | 332 | * | NA_peak_143 | 79 | Exon (ENSMU | 13 | 65336222 | 65348597 | 12376 | 2 | 665738 | ENSMUST00 | -301036 | ENSMUSG00 | Gm7762 | predicted gene 7762 |
| chr13 | 65652778 | 65653136 | 359 | * | NA_peak_143 | 14 | Distal Interge | 13 | 65336222 | 65348597 | 12376 | 2 | 665738 | ENSMUST00 | -304181 | ENSMUSG00 | Gm7762 | predicted gene 7762 |
| chr13 | 65674587 | 65675088 | 502 | * | NA_peak_143 | 45 | Distal Interge | 13 | 65336222 | 65348597 | 12376 | 2 | 665738 | ENSMUST00 | -325990 | ENSMUSG00 | Gm7762 | predicted gene 7762 |
| chr13 | 65704856 | 65705151 | 296 | * | NA_peak_143 | 35 | Intron (ENSM | 13 | 65336222 | 65348597 | 12376 | 2 | 665738 | ENSMUST00 | -356259 | ENSMUSG00 | Gm7762 | predicted gene 7762 |
| chr13 | 65705694 | 65706138 | 445 | * | NA_peak_143 | 28 | Intron (ENSM | 13 | 65336222 | 65348597 | 12376 | 2 | 665738 | ENSMUST00 | -357097 | ENSMUSG00 | Gm7762 | predicted gene 7762 |
| chr13 | 65712156 | 65712442 | 287 | * | NA_peak_143 | 40 | Intron (ENSM | 13 | 65336222 | 65348597 | 12376 | 2 | 665738 | ENSMUST00 | -363559 | ENSMUSG00 | Gm7762 | predicted gene 7762 |
| chr13 | 65753199 | 65753754 | 556 | * | NA_peak_143 | 57 | Distal Interge | 13 | 66113480 | 66120204 | 6725 | 1 | 628709 | ENSMUST00 | -359726 | ENSMUSG00 | Zfp997 | zinc finger protein 997 |
| chr13 | 65766842 | 65767529 | 688 | * | NA_peak_143 | 36 | Distal Interge | 13 | 66113480 | 66120204 | 6725 | 1 | 628709 | ENSMUST00 | -345951 | ENSMUSG00 | Zfp997 | zinc finger protein 997 |
| chr13 | 65774378 | 65774756 | 379 | * | NA_peak_143 | 78 | Distal Interge | 13 | 66113480 | 66120204 | 6725 | 1 | 628709 | ENSMUST00 | -338724 | ENSMUSG00 | Zfp997 | zinc finger protein 997 |
| chr13 | 65783008 | 65783071 | 64 | * | NA_peak_143 | 32 | Distal Interge | 13 | 66113480 | 66120204 | 6725 | 1 | 628709 | ENSMUST00 | -330409 | ENSMUSG00 | Zfp997 | zinc finger protein 997 |
| chr13 | 65788527 | 65788829 | 303 | * | NA_peak_144 | 21 | Distal Interge | 13 | 66113480 | 66120204 | 6725 | 1 | 628709 | ENSMUST00 | -324651 | ENSMUSG00 | Zfp997 | zinc finger protein 997 |
| chr13 | 65793129 | 65793513 | 385 | * | NA_peak_144 | 18 | Distal Interge | 13 | 66113480 | 66120204 | 6725 | 1 | 628709 | ENSMUST00 | -319967 | ENSMUSG00 | Zfp997 | zinc finger protein 997 |
| chr13 | 65856054 | 65856314 | 261 | * | NA_peak_144 | 42 | Distal Interge | 13 | 66113480 | 66120204 | 6725 | 1 | 628709 | ENSMUST00 | -257166 | ENSMUSG00 | Zfp997 | zinc finger protein 997 |
| chr13 | 65875471 | 65875777 | 307 | * | NA_peak_144 | 16 | Distal Interge | 13 | 66113480 | 66120204 | 6725 | 1 | 628709 | ENSMUST00 | -237703 | ENSMUSG00 | Zfp997 | zinc finger protein 997 |
| chr13 | 65880258 | 65880641 | 384 | * | NA_peak_144 | 58 | Intron (ENSM | 13 | 66113480 | 66120204 | 6725 | 1 | 628709 | ENSMUST00 | -232839 | ENSMUSG00 | Zfp997 | zinc finger protein 997 |
| chr13 | 65882925 | 65883176 | 252 | * | NA_peak_144 | 42 | Intron (ENSM | 13 | 66113480 | 66120204 | 6725 | 1 | 628709 | ENSMUST00 | -230304 | ENSMUSG00 | Zfp997 | zinc finger protein 997 |
| chr13 | 65884017 | 65884759 | 743 | * | NA_peak_144 | 42 | Intron (ENSM | 13 | 66113480 | 66120204 | 6725 | 1 | 628709 | ENSMUST00 | -228721 | ENSMUSG00 | Zfp997 | zinc finger protein 997 |
| chr13 | 65887117 | 65887590 | 474 | * | NA_peak_144 | 57 | Exon (ENSMU | 13 | 66113480 | 66120204 | 6725 | 1 | 628709 | ENSMUST00 | -225890 | ENSMUSG00 | Zfp997 | zinc finger protein 997 |
| chr13 | 65908658 | 65908887 | 230 | * | NA_peak_144 | 19 | Exon (ENSMU | 13 | 66113480 | 66120204 | 6725 | 1 | 628709 | ENSMUST00 | -204593 | ENSMUSG00 | Zfp997 | zinc finger protein 997 |
| chr13 | 65911572 | 65911836 | 265 | * | NA_peak_144 | 55 | Distal Interge | 13 | 66113480 | 66120204 | 6725 | 1 | 628709 | ENSMUST00 | -201644 | ENSMUSG00 | Zfp997 | zinc finger protein 997 |
| chr13 | 65934702 | 65934793 | 92 | * | NA_peak_144 | 29 | Distal Interge | 13 | 66113480 | 66120204 | 6725 | 1 | 628709 | ENSMUST00 | -178687 | ENSMUSG00 | Zfp997 | zinc finger protein 997 |
| chr13 | 65936262 | 65936711 | 450 | * | NA_peak_144 | 89 | Distal Interge | 13 | 66113480 | 66120204 | 6725 | 1 | 628709 | ENSMUST00 | -176769 | ENSMUSG00 | Zfp997 | zinc finger protein 997 |
| chr13 | 65946499 | 65947350 | 852 | * | NA_peak_144 | 92 | Exon (ENSMU | 13 | 66113480 | 66120204 | 6725 | 1 | 628709 | ENSMUST00 | -166130 | ENSMUSG00 | Zfp997 | zinc finger protein 997 |
| chr13 | 65973003 | 65973295 | 293 | * | NA_peak_144 | 64 | Distal Interge | 13 | 66113480 | 66120204 | 6725 | 1 | 628709 | ENSMUST00 | -140185 | ENSMUSG00 | Zfp997 | zinc finger protein 997 |
| chr13 | 66026201 | 66026522 | 322 | * | NA_peak_144 | 66 | Distal Interge | 13 | 66113480 | 66120204 | 6725 | 1 | 628709 | ENSMUST00 | -86958 | ENSMUSG00 | Zfp997 | zinc finger protein 997 |
| chr13 | 66031152 | 66031753 | 602 | * | NA_peak_144 | 95 | Distal Interge | 13 | 66113480 | 66120204 | 6725 | 1 | 628709 | ENSMUST00 | -81727 | ENSMUSG00 | Zfp997 | zinc finger protein 997 |
| chr13 | 66043239 | 66043795 | 557 | * | NA_peak_144 | 31 | Distal Interge | 13 | 66113480 | 66120204 | 6725 | 1 | 628709 | ENSMUST00 | -69685 | ENSMUSG00 | Zfp997 | zinc finger protein 997 |
| chr13 | 66057514 | 66057763 | 250 | * | NA_peak_144 | 24 | Distal Interge | 13 | 66113480 | 66120204 | 6725 | 1 | 628709 | ENSMUST00 | -55717 | ENSMUSG00 | Zfp997 | zinc finger protein 997 |
| chr13 | 66062532 | 66062922 | 391 | * | NA_peak_144 | 48 | Distal Interge | 13 | 66113480 | 66120204 | 6725 | 1 | 628709 | ENSMUST00 | -50558 | ENSMUSG00 | Zfp997 | zinc finger protein 997 |
| chr13 | 66103745 | 66104094 | 350 | * | NA_peak_144 | 48 | Distal Interge | 13 | 66113480 | 66120204 | 6725 | 1 | 628709 | ENSMUST00 | -9386 | ENSMUSG00 | Zfp997 | zinc finger protein 997 |
| chr13 | 66113509 | 66114026 | 518 | * | NA_peak_144 | 31 | Promoter (<= | 13 | 66113480 | 66122468 | 8989 | 1 | 628709 | ENSMUST00 | 29 | ENSMUSG00 | Zfp997 | zinc finger protein 997 |
| chr13 | 66122050 | 66122317 | 268 | * | NA_peak_144 | 62 | Promoter (2-3 | 13 | 66119547 | 66122836 | 3290 | 1 | 628709 | ENSMUST00 | 2503 | ENSMUSG00 | Zfp997 | zinc finger protein 997 |
| chr13 | 66128125 | 66128821 | 697 | * | NA_peak_144 | 33 | Intron (ENSM | 13 | 66119547 | 66122836 | 3290 | 1 | 628709 | ENSMUST00 | 8578 | ENSMUSG00 | Zfp997 | zinc finger protein 997 |
| chr13 | 66131757 | 66132036 | 280 | * | NA_peak_144 | 23 | Distal Interge | 13 | 66119547 | 66122836 | 3290 | 1 | 628709 | ENSMUST00 | 12210 | ENSMUSG00 | Zfp997 | zinc finger protein 997 |
| chr13 | 66159566 | 66159923 | 358 | * | NA_peak_144 | 30 | Distal Interge | 13 | 66119547 | 66122836 | 3290 | 1 | 628709 | ENSMUST00 | 40019 | ENSMUSG00 | Zfp997 | zinc finger protein 997 |
| chr13 | 66204873 | 66205287 | 415 | * | NA_peak_144 | 27 | Distal Interge | 13 | 66119547 | 66122836 | 3290 | 1 | 628709 | ENSMUST00 | 85326 | ENSMUSG00 | Zfp997 | zinc finger protein 997 |
| chr13 | 66225797 | 66225856 | 60 | * | NA_peak_144 | 42 | Distal Interge | 13 | 66119547 | 66122836 | 3290 | 1 | 628709 | ENSMUST00 | 106250 | ENSMUSG00 | Zfp997 | zinc finger protein 997 |
| chr13 | 66227579 | 66227920 | 342 | * | NA_peak_144 | 33 | Distal Interge | 13 | 66119547 | 66122836 | 3290 | 1 | 628709 | ENSMUST00 | 108032 | ENSMUSG00 | Zfp997 | zinc finger protein 997 |
| chr13 | 66243818 | 66244197 | 380 | * | NA_peak_144 | 39 | Distal Interge | 13 | 66119547 | 66122836 | 3290 | 1 | 628709 | ENSMUST00 | 124271 | ENSMUSG00 | Zfp997 | zinc finger protein 997 |
| chr13 | 66258172 | 66258680 | 509 | * | NA_peak_144 | 39 | Intron (ENSM | 13 | 66119547 | 66122836 | 3290 | 1 | 628709 | ENSMUST00 | 138625 | ENSMUSG00 | Zfp997 | zinc finger protein 997 |
| chr13 | 66279451 | 66279829 | 379 | * | NA_peak_144 | 45 | Intron (ENSM | 13 | 66418322 | 66419597 | 1276 | 2 | 76803 | ENSMUST00 | 139768 | ENSMUSG00 | Zfp998 | zinc finger protein 998 |
| chr13 | 66291067 | 66291456 | 390 | * | NA_peak_144 | 36 | Intron (ENSM | 13 | 66418322 | 66419597 | 1276 | 2 | 76803 | ENSMUST00 | 128141 | ENSMUSG00 | Zfp998 | zinc finger protein 998 |
| chr13 | 66292537 | 66292909 | 373 | * | NA_peak_144 | 14 | Intron (ENSM | 13 | 66418322 | 66419597 | 1276 | 2 | 76803 | ENSMUST00 | 126688 | ENSMUSG00 | Zfp998 | zinc finger protein 998 |
| chr13 | 66295007 | 66295523 | 517 | * | NA_peak_144 | 34 | Intron (ENSM | 13 | 66418322 | 66419597 | 1276 | 2 | 76803 | ENSMUST00 | 124074 | ENSMUSG00 | Zfp998 | zinc finger protein 998 |
| chr13 | 66299976 | 66300315 | 340 | * | NA_peak_144 | 25 | Intron (ENSM | 13 | 66418322 | 66419597 | 1276 | 2 | 76803 | ENSMUST00 | 119282 | ENSMUSG00 | Zfp998 | zinc finger protein 998 |
| chr13 | 66309547 | 66309941 | 395 | * | NA_peak_144 | 61 | Intron (ENSM | 13 | 66418322 | 66419597 | 1276 | 2 | 76803 | ENSMUST00 | 109656 | ENSMUSG00 | Zfp998 | zinc finger protein 998 |
| chr13 | 66388897 | 66389231 | 335 | * | NA_peak_144 | 29 | Intron (ENSM | 13 | 66418322 | 66419597 | 1276 | 2 | 76803 | ENSMUST00 | 30366 | ENSMUSG00 | Zfp998 | zinc finger protein 998 |
| chr13 | 66392014 | 66392493 | 480 | * | NA_peak_144 | 50 | Intron (ENSM | 13 | 66418322 | 66419597 | 1276 | 2 | 76803 | ENSMUST00 | 27104 | ENSMUSG00 | Zfp998 | zinc finger protein 998 |
| chr13 | 66394737 | 66395273 | 537 | * | NA_peak_144 | 28 | Intron (ENSM | 13 | 66418322 | 66419597 | 1276 | 2 | 76803 | ENSMUST00 | 24324 | ENSMUSG00 | Zfp998 | zinc finger protein 998 |
| chr13 | 66417514 | 66417866 | 353 | * | NA_peak_145 | 32 | Promoter (1-2 | 13 | 66418322 | 66419597 | 1276 | 2 | 76803 | ENSMUST00 | 1731 | ENSMUSG00 | Zfp998 | zinc finger protein 998 |
| chr13 | 66426674 | 66427008 | 335 | * | NA_peak_145 | 38 | Intron (ENSM | 13 | 66418245 | 66433895 | 15651 | 2 | 76803 | ENSMUST00 | 6887 | ENSMUSG00 | Zfp998 | zinc finger protein 998 |
| chr13 | 66428101 | 66428404 | 304 | * | NA_peak_145 | 18 | Intron (ENSM | 13 | 66418245 | 66433895 | 15651 | 2 | 76803 | ENSMUST00 | 5491 | ENSMUSG00 | Zfp998 | zinc finger protein 998 |
| chr13 | 66433304 | 66433642 | 339 | * | NA_peak_145 | 76 | Promoter (<= | 13 | 66418245 | 66433895 | 15651 | 2 | 76803 | ENSMUST00 | 253 | ENSMUSG00 | Zfp998 | zinc finger protein 998 |
| chr13 | 66442867 | 66443194 | 328 | * | NA_peak_145 | 28 | Promoter (1-2 | 13 | 66418114 | 66441118 | 23005 | 2 | 76803 | ENSMUST00 | -1749 | ENSMUSG00 | Zfp998 | zinc finger protein 998 |
| chr13 | 66444050 | 66444610 | 561 | * | NA_peak_145 | 29 | Promoter (2-3 | 13 | 66418114 | 66441118 | 23005 | 2 | 76803 | ENSMUST00 | -2932 | ENSMUSG00 | Zfp998 | zinc finger protein 998 |
| chr13 | 66452334 | 66452624 | 291 | * | NA_peak_145 | 80 | Distal Interge | 13 | 66418114 | 66441118 | 23005 | 2 | 76803 | ENSMUST00 | -11216 | ENSMUSG00 | Zfp998 | zinc finger protein 998 |
| chr13 | 66472410 | 66473212 | 803 | * | NA_peak_145 | 79 | Distal Interge | 13 | 66418114 | 66441118 | 23005 | 2 | 76803 | ENSMUST00 | -31292 | ENSMUSG00 | Zfp998 | zinc finger protein 998 |
| chr13 | 66480061 | 66480483 | 423 | * | NA_peak_145 | 24 | Distal Interge | 13 | 66418114 | 66441118 | 23005 | 2 | 76803 | ENSMUST00 | -38943 | ENSMUSG00 | Zfp998 | zinc finger protein 998 |
| chr13 | 66493431 | 66493715 | 285 | * | NA_peak_145 | 80 | Distal Interge | 13 | 66418114 | 66441118 | 23005 | 2 | 76803 | ENSMUST00 | -52313 | ENSMUSG00 | Zfp998 | zinc finger protein 998 |
| chr13 | 66605800 | 66606102 | 303 | * | NA_peak_145 | 32 | Distal Interge | 13 | 66418114 | 66441118 | 23005 | 2 | 76803 | ENSMUST00 | -164682 | ENSMUSG00 | Zfp998 | zinc finger protein 998 |
| chr13 | 66608562 | 66609660 | 1099 | * | NA_peak_145 | 67 | Distal Interge | 13 | 66418114 | 66441118 | 23005 | 2 | 76803 | ENSMUST00 | -167444 | ENSMUSG00 | Zfp998 | zinc finger protein 998 |
| chr13 | 66637292 | 66637724 | 433 | * | NA_peak_145 | 26 | Distal Interge | 13 | 66418114 | 66441118 | 23005 | 2 | 76803 | ENSMUST00 | -196174 | ENSMUSG00 | Zfp998 | zinc finger protein 998 |
| chr13 | 66641493 | 66641800 | 308 | * | NA_peak_145 | 22 | Distal Interge | 13 | 66418114 | 66441118 | 23005 | 2 | 76803 | ENSMUST00 | -200375 | ENSMUSG00 | Zfp998 | zinc finger protein 998 |
| chr13 | 66647675 | 66648064 | 390 | * | NA_peak_145 | 44 | Distal Interge | 13 | 66418114 | 66441118 | 23005 | 2 | 76803 | ENSMUST00 | -206557 | ENSMUSG00 | Zfp998 | zinc finger protein 998 |
| chr13 | 66655011 | 66655456 | 446 | * | NA_peak_145 | 49 | Distal Interge | 13 | 66418114 | 66441118 | 23005 | 2 | 76803 | ENSMUST00 | -213893 | ENSMUSG00 | Zfp998 | zinc finger protein 998 |
| chr13 | 66683589 | 66683814 | 226 | * | NA_peak_145 | 26 | Intron (ENSM | 13 | 66901061 | 66902786 | 1726 | 2 | 67530 | ENSMUST00 | 218972 | ENSMUSG00 | Uqcrb | ubiquinol-cytochrome c reductase binding protein |
| chr13 | 66689357 | 66689608 | 252 | * | NA_peak_145 | 40 | Intron (ENSM | 13 | 66901061 | 66902786 | 1726 | 2 | 67530 | ENSMUST00 | 213178 | ENSMUSG00 | Uqcrb | ubiquinol-cytochrome c reductase binding protein |
| chr13 | 66721531 | 66722085 | 555 | * | NA_peak_145 | 43 | Distal Interge | 13 | 66901061 | 66902786 | 1726 | 2 | 67530 | ENSMUST00 | 180701 | ENSMUSG00 | Uqcrb | ubiquinol-cytochrome c reductase binding protein |
| chr13 | 66722579 | 66723061 | 483 | * | NA_peak_145 | 43 | Distal Interge | 13 | 66901061 | 66902786 | 1726 | 2 | 67530 | ENSMUST00 | 179725 | ENSMUSG00 | Uqcrb | ubiquinol-cytochrome c reductase binding protein |

| | | | | | | | | | | | | | | | | | | |
|---|---|---|---|---|---|---|---|---|---|---|---|---|---|---|---|---|---|---|
| chr13 | 66723717 | 66723978 | 262 | * | NA_peak_145 | 43 | Distal Interge | 13 | 66901061 | 66902786 | 1726 | 2 | 67530 | ENSMUST00 | 178808 | ENSMUSG00 | Uqcrb | ubiquinol-cytochrome c reductase binding protein |
| chr13 | 66725231 | 66725814 | 584 | * | NA_peak_145 | 46 | Distal Interge | 13 | 66901061 | 66902786 | 1726 | 2 | 67530 | ENSMUST00 | 176972 | ENSMUSG00 | Uqcrb | ubiquinol-cytochrome c reductase binding protein |
| chr13 | 66739308 | 66739673 | 366 | * | NA_peak_145 | 92 | Exon (ENSMU | 13 | 66901061 | 66902786 | 1726 | 2 | 67530 | ENSMUST00 | 163113 | ENSMUSG00 | Uqcrb | ubiquinol-cytochrome c reductase binding protein |
| chr13 | 66775304 | 66775813 | 510 | * | NA_peak_145 | 41 | Distal Interge | 13 | 66901061 | 66902786 | 1726 | 2 | 67530 | ENSMUST00 | 126973 | ENSMUSG00 | Uqcrb | ubiquinol-cytochrome c reductase binding protein |
| chr13 | 66781078 | 66781594 | 517 | * | NA_peak_145 | 19 | Distal Interge | 13 | 66901061 | 66902786 | 1726 | 2 | 67530 | ENSMUST00 | 121192 | ENSMUSG00 | Uqcrb | ubiquinol-cytochrome c reductase binding protein |
| chr13 | 66790438 | 66790623 | 186 | * | NA_peak_145 | 28 | Distal Interge | 13 | 66901061 | 66902786 | 1726 | 2 | 67530 | ENSMUST00 | 112163 | ENSMUSG00 | Uqcrb | ubiquinol-cytochrome c reductase binding protein |
| chr13 | 66820929 | 66821197 | 269 | * | NA_peak_145 | 62 | Distal Interge | 13 | 66901061 | 66902786 | 1726 | 2 | 67530 | ENSMUST00 | 81589 | ENSMUSG00 | Uqcrb | ubiquinol-cytochrome c reductase binding protein |
| chr13 | 66826967 | 66827686 | 720 | * | NA_peak_145 | 77 | Distal Interge | 13 | 66901061 | 66902786 | 1726 | 2 | 67530 | ENSMUST00 | 75100 | ENSMUSG00 | Uqcrb | ubiquinol-cytochrome c reductase binding protein |
| chr13 | 66856400 | 66856887 | 488 | * | NA_peak_145 | 51 | Distal Interge | 13 | 66901061 | 66902786 | 1726 | 2 | 67530 | ENSMUST00 | 45899 | ENSMUSG00 | Uqcrb | ubiquinol-cytochrome c reductase binding protein |
| chr13 | 66858879 | 66859072 | 194 | * | NA_peak_145 | 13 | Distal Interge | 13 | 66901061 | 66902786 | 1726 | 2 | 67530 | ENSMUST00 | 43714 | ENSMUSG00 | Uqcrb | ubiquinol-cytochrome c reductase binding protein |
| chr13 | 67415110 | 67415126 | 17 | * | NA_peak_145 | 18 | Intron (ENSM | 13 | 67408464 | 67421364 | 12901 | 2 | 328274 | ENSMUST00 | 6238 | ENSMUSG00 | Zfp459 | zinc finger protein 459 |
| chr13 | 67563645 | 67563943 | 299 | * | NA_peak_146 | 45 | Exon (ENSMU | 13 | 67538641 | 67553830 | 15190 | 2 | 212276 | ENSMUST00 | -9815 | ENSMUSG00 | Zfp748 | zinc finger protein 748 |
| chr13 | 67931242 | 67931658 | 417 | * | NA_peak_146 | 40 | Distal Interge | 13 | 67863326 | 67863925 | 600 | 1 | 408058 | ENSMUST00 | 67916 | ENSMUSG00 | BC048507 | cDNA sequence BC048507 |
| chr13 | 67967950 | 67968246 | 297 | * | NA_peak_146 | 61 | Exon (ENSMU | 13 | 67863326 | 67863925 | 600 | 1 | 408058 | ENSMUST00 | 104624 | ENSMUSG00 | BC048507 | cDNA sequence BC048507 |
| chr13 | 67985872 | 67986152 | 281 | * | NA_peak_146 | 23 | Exon (ENSMU | 13 | 67863326 | 67863925 | 600 | 1 | 408058 | ENSMUST00 | 122546 | ENSMUSG00 | BC048507 | cDNA sequence BC048507 |
| chr13 | 68003890 | 68004340 | 451 | * | NA_peak_146 | 17 | Distal Interge | 13 | 67863326 | 67863925 | 600 | 1 | 408058 | ENSMUST00 | 140564 | ENSMUSG00 | BC048507 | cDNA sequence BC048507 |
| chr13 | 68010281 | 68010867 | 587 | * | NA_peak_146 | 42 | Distal Interge | 13 | 67863326 | 67863925 | 600 | 1 | 408058 | ENSMUST00 | 146955 | ENSMUSG00 | BC048507 | cDNA sequence BC048507 |
| chr13 | 68019887 | 68020171 | 285 | * | NA_peak_146 | 64 | Distal Interge | 13 | 67863326 | 67863925 | 600 | 1 | 408058 | ENSMUST00 | 156561 | ENSMUSG00 | BC048507 | cDNA sequence BC048507 |
| chr13 | 68026374 | 68026706 | 333 | * | NA_peak_146 | 26 | Distal Interge | 13 | 67863326 | 67863925 | 600 | 1 | 408058 | ENSMUST00 | 163048 | ENSMUSG00 | BC048507 | cDNA sequence BC048507 |
| chr13 | 68051818 | 68052021 | 204 | * | NA_peak_146 | 15 | Distal Interge | 13 | 67863326 | 67863925 | 600 | 1 | 408058 | ENSMUST00 | 188492 | ENSMUSG00 | BC048507 | cDNA sequence BC048507 |
| chr13 | 68062763 | 68063116 | 354 | * | NA_peak_146 | 24 | Distal Interge | 13 | 67863326 | 67863925 | 600 | 1 | 408058 | ENSMUST00 | 199437 | ENSMUSG00 | BC048507 | cDNA sequence BC048507 |
| chr13 | 68087901 | 68088049 | 149 | * | NA_peak_146 | 22 | Distal Interge | 13 | 67863326 | 67863925 | 600 | 1 | 408058 | ENSMUST00 | 224575 | ENSMUSG00 | BC048507 | cDNA sequence BC048507 |
| chr13 | 68119207 | 68119833 | 627 | * | NA_peak_146 | 24 | Distal Interge | 13 | 68353364 | 68356825 | 3462 | 2 | 666272 | ENSMUST00 | 236992 | ENSMUSG00 | Gm8016 | predicted gene 8016 |
| chr13 | 68124827 | 68125130 | 304 | * | NA_peak_146 | 20 | Distal Interge | 13 | 68353364 | 68356825 | 3462 | 2 | 666272 | ENSMUST00 | 231695 | ENSMUSG00 | Gm8016 | predicted gene 8016 |
| chr13 | 68249745 | 68250232 | 488 | * | NA_peak_146 | 31 | Distal Interge | 13 | 68353364 | 68356825 | 3462 | 2 | 666272 | ENSMUST00 | 106593 | ENSMUSG00 | Gm8016 | predicted gene 8016 |
| chr13 | 68318475 | 68318971 | 497 | * | NA_peak_146 | 39 | Distal Interge | 13 | 68353364 | 68356825 | 3462 | 2 | 666272 | ENSMUST00 | 37854 | ENSMUSG00 | Gm8016 | predicted gene 8016 |
| chr13 | 68356613 | 68356910 | 298 | * | NA_peak_146 | 39 | Promoter (<= | 13 | 68353364 | 68356825 | 3462 | 2 | 666272 | ENSMUST00 | 0 | ENSMUSG00 | Gm8016 | predicted gene 8016 |
| chr13 | 68558964 | 68559201 | 238 | * | NA_peak_146 | 20 | Distal Interge | 13 | 68564149 | 68566216 | 2068 | 2 | 210009 | ENSMUST00 | 7015 | ENSMUSG00 | Mtrr | 5-methyltetrahydrofolate-homocysteine methyltransferase reductase |
| chr13 | 70458678 | 70459047 | 370 | * | NA_peak_147 | 20 | Distal Interge | 13 | 70573241 | 70576110 | 2870 | 1 | 73584 | ENSMUST00 | -114194 | ENSMUSG00 | 1700100L14 | RIKEN cDNA 1700100L14 gene |
| chr13 | 70812766 | 70813063 | 298 | * | NA_peak_147 | 26 | Intron (ENSM | 13 | 70727810 | 70841775 | 113966 | 2 | 271127 | ENSMUST00 | 28712 | ENSMUSG00 | Adamts16 | a disintegrin-like and metallopeptidase (reprolysin type) with thrombospondin type 1 motif, 16 |
| chr13 | 71430477 | 71430819 | 343 | * | NA_peak_147 | 36 | Distal Interge | 13 | 71107089 | 71107209 | 121 | 1 | 100316804 | ENSMUST00 | 323388 | ENSMUSG00 | Mir466f-4 | microRNA 466f-4 |
| chr13 | 72681033 | 72681374 | 342 | * | NA_peak_148 | 20 | Distal Interge | 13 | 72632597 | 72633770 | 1174 | 1 | 16372 | ENSMUST00 | 48436 | ENSMUSG00 | Irx2 | Iroquois homeobox 2 |
| chr13 | 73404738 | 73404888 | 151 | * | NA_peak_148 | 35 | Distal Interge | 13 | 73344665 | 73347384 | 2720 | 2 | 432777 | ENSMUST00 | -57354 | ENSMUSG00 | Eprn | ephemeron, early developmental lncRNA |
| chr13 | 73829806 | 73829829 | 24 | * | NA_peak_148 | 19 | Intron (ENSM | 13 | 73813463 | 73814988 | 1526 | 1 | 20499 | ENSMUST00 | 16343 | ENSMUSG00 | Slc12a7 | solute carrier family 12, member 7 |
| chr13 | 73987875 | 73988199 | 325 | * | NA_peak_149 | 27 | Intron (ENSM | 13 | 74009407 | 74031371 | 21965 | 1 | 72948 | ENSMUST00 | -21208 | ENSMUSG00 | Tppp | tubulin polymerization promoting protein |
| chr13 | 74713842 | 74714085 | 244 | * | NA_peak_149 | 27 | Intron (ENSM | 13 | 74699105 | 74700885 | 1781 | 2 | 12380 | ENSMUST00 | -12957 | ENSMUSG00 | Cast | calpastatin |
| chr13 | 75653393 | 75653419 | 27 | * | NA_peak_149 | 19 | Distal Interge | 13 | 75645046 | 75645141 | 96 | 1 | 751556 | ENSMUST00 | 8347 | ENSMUSG00 | Mir682 | microRNA 682 |
| chr13 | 75708538 | 75708793 | 256 | * | NA_peak_150 | 23 | Promoter (<= | 13 | 75707711 | 75762192 | 54482 | 1 | 192657 | ENSMUST00 | 827 | ENSMUSG00 | Ell2 | elongation factor for RNA polymerase II 2 |
| chr13 | 75943565 | 75943904 | 340 | * | NA_peak_150 | 28 | Promoter (<= | 13 | 75869537 | 75943867 | 74331 | 2 | 73296 | ENSMUST00 | 0 | ENSMUSG00 | Rhobtb3 | Rho-related BTB domain containing 3 |
| chr13 | 75992677 | 75992703 | 27 | * | NA_peak_150 | 20 | Intron (ENSM | 13 | 75967747 | 75978343 | 10597 | 1 | 75571 | ENSMUST00 | 24930 | ENSMUSG00 | Spata9 | spermatogenesis associated 9 |
| chr13 | 78138596 | 78138949 | 354 | * | NA_peak_150 | 45 | Intron (ENSM | 13 | 78189872 | 78190443 | 572 | 2 | 13865 | ENSMUST00 | 51494 | ENSMUSG00 | Nr2f1 | nuclear receptor subfamily 2, group F, member 1 |
| chr13 | 79189881 | 79190185 | 305 | * | NA_peak_151 | 31 | Distal Interge | 13 | 78202553 | 78231386 | 28834 | 1 | 320174 | ENSMUST00 | 987328 | ENSMUSG00 | A830082K12 | RIKEN cDNA A830082K12 gene |
| chr13 | 80067910 | 80068011 | 102 | * | NA_peak_151 | 16 | Intron (ENSM | 13 | 80883384 | 80889643 | 6260 | 1 | 105171 | ENSMUST00 | -815373 | ENSMUSG00 | Arrdc3 | arrestin domain containing 3 |
| chr13 | 80433635 | 80434056 | 422 | * | NA_peak_151 | 27 | Distal Interge | 13 | 80883384 | 80889643 | 6260 | 1 | 105171 | ENSMUST00 | -449328 | ENSMUSG00 | Arrdc3 | arrestin domain containing 3 |
| chr13 | 81818937 | 81819403 | 467 | * | NA_peak_151 | 13 | Distal Interge | 13 | 81791977 | 81793668 | 1692 | 1 | 12626 | ENSMUST00 | 26960 | ENSMUSG00 | Cetn3 | centrin 3 |
| chr13 | 85061578 | 85061618 | 41 | * | NA_peak_152 | 16 | Distal Interge | 13 | 85189408 | 85213723 | 24316 | 1 | 66671 | ENSMUST00 | -127790 | ENSMUSG00 | Ccnh | cyclin H |
| chr13 | 85871565 | 85871601 | 37 | * | NA_peak_153 | 16 | Distal Interge | 13 | 86044821 | 86046347 | 1527 | 2 | 12867 | ENSMUST00 | 174746 | ENSMUSG00 | Cox7c | cytochrome c oxidase subunit 7C |
| chr13 | 86686975 | 86687238 | 264 | * | NA_peak_153 | 40 | Distal Interge | 13 | 86044816 | 86046904 | 2089 | 2 | 12867 | ENSMUST00 | -640071 | ENSMUSG00 | Cox7c | cytochrome c oxidase subunit 7C |
| chr13 | 86755557 | 86755905 | 349 | * | NA_peak_153 | 27 | Distal Interge | 13 | 86044816 | 86046904 | 2089 | 2 | 12867 | ENSMUST00 | -708653 | ENSMUSG00 | Cox7c | cytochrome c oxidase subunit 7C |
| chr13 | 87939371 | 87939462 | 92 | * | NA_peak_153 | 30 | Distal Interge | 13 | 88821472 | 89200538 | 379067 | 1 | 13612 | ENSMUST00 | -882010 | ENSMUSG00 | Edil3 | EGF-like repeats and discoidin I-like domains 3 |
| chr13 | 89162425 | 89162740 | 316 | * | NA_peak_153 | 27 | Intron (ENSM | 13 | 88821642 | 89323223 | 501582 | 1 | 13612 | ENSMUST00 | 340783 | ENSMUSG00 | Edil3 | EGF-like repeats and discoidin I-like domains 3 |
| chr13 | 90740707 | 90740772 | 66 | * | NA_peak_154 | 15 | Distal Interge | 13 | 90883453 | 90898824 | 15372 | 2 | 435376 | ENSMUST00 | 158052 | ENSMUSG00 | Atp6ap1l | ATPase, H+ transporting, lysosomal accessory protein 1-like |
| chr13 | 91961929 | 91962394 | 466 | * | NA_peak_154 | 36 | Intron (ENSM | 13 | 91880400 | 91987989 | 107590 | 2 | 19418 | ENSMUST00 | 25595 | ENSMUSG00 | Rasgrf2 | RAS protein-specific guanine nucleotide-releasing factor 2 |
| chr13 | 91976567 | 91976573 | 7 | * | NA_peak_155 | 20 | Intron (ENSM | 13 | 91880400 | 91987989 | 107590 | 2 | 19418 | ENSMUST00 | 11416 | ENSMUSG00 | Rasgrf2 | RAS protein-specific guanine nucleotide-releasing factor 2 |
| chr13 | 92833986 | 92834292 | 307 | * | NA_peak_155 | 31 | Distal Interge | 13 | 92844760 | 92848412 | 3653 | 1 | 382793 | ENSMUST00 | -10468 | ENSMUSG00 | Mtx3 | metaxin 3 |
| chr13 | 92835497 | 92835782 | 286 | * | NA_peak_155 | 27 | Distal Interge | 13 | 92844760 | 92848412 | 3653 | 1 | 382793 | ENSMUST00 | -8978 | ENSMUSG00 | Mtx3 | metaxin 3 |
| chr13 | 95269875 | 95270102 | 228 | * | NA_peak_156 | 16 | Distal Interge | 13 | 95024454 | 95250336 | 225883 | 2 | 218461 | ENSMUST00 | -19539 | ENSMUSG00 | Pde8b | phosphodiesterase 8B |
| chr13 | 95682412 | 95682856 | 445 | * | NA_peak_156 | 23 | Promoter (<= | 13 | 95661402 | 95682937 | 21536 | 2 | 544963 | ENSMUST00 | 81 | ENSMUSG00 | Iqgap2 | IQ motif containing GTPase activating protein 2 |
| chr13 | 96347902 | 96348187 | 286 | * | NA_peak_156 | 22 | Distal Interge | 13 | 96388294 | 96417737 | 29444 | 1 | 67463 | ENSMUST00 | -40107 | ENSMUSG00 | Poc5 | POC5 centriolar protein |
| chr13 | 96904185 | 96904294 | 110 | * | NA_peak_156 | 36 | Intron (ENSM | 13 | 96924689 | 96950906 | 26218 | 1 | 218476 | ENSMUST00 | -20395 | ENSMUSG00 | Gcnt4 | glucosaminyl (N-acetyl) transferase 4, core 2 (beta-1,6-N-acetylglucosaminyltransferase) |
| chr13 | 97365709 | 97365731 | 23 | * | NA_peak_157 | 36 | Distal Interge | 13 | 97434970 | 97497007 | 62038 | 1 | 100039691 | ENSMUST00 | -69239 | ENSMUSG00 | Lncenc1 | long non-coding RNA, embryonic stem cells expressed 1 |
| chr13 | 97620381 | 97620679 | 299 | * | NA_peak_157 | 31 | Distal Interge | 13 | 97560026 | 97577418 | 17393 | 1 | 328314 | ENSMUST00 | 60355 | ENSMUSG00 | Gm5086 | predicted gene 5086 |
| chr13 | 98466843 | 98467106 | 264 | * | NA_peak_157 | 23 | Distal Interge | 13 | 98424109 | 98435065 | 10957 | 2 | 70090 | ENSMUST00 | -31778 | ENSMUSG00 | 2310005E17 | RIKEN cDNA 2310005E17 gene |
| chr13 | 99429601 | 99429806 | 206 | * | NA_peak_157 | 28 | Exon (ENSMU | 13 | 99434301 | 99443944 | 9644 | 2 | 17755 | ENSMUST00 | 14138 | ENSMUSG00 | Map1b | microtubule-associated protein 1B |
| chr13 | 100103540 | 100103822 | 283 | * | NA_peak_158 | 34 | Promoter (<= | 13 | 100023305 | 100103838 | 80534 | 2 | 544971 | ENSMUST00 | 16 | ENSMUSG00 | Bdp1 | B double prime 1, subunit of RNA polymerase III transcription initiation factor IIIB |
| chr13 | 100391088 | 100391201 | 114 | * | NA_peak_158 | 14 | Distal Interge | 13 | 100407838 | 100446461 | 38624 | 2 | 17940 | ENSMUST00 | 55260 | ENSMUSG00 | Naip1 | NLR family, apoptosis inhibitory protein 1 |
| chr13 | 100426696 | 100427176 | 481 | * | NA_peak_158 | 28 | 3' UTR | 13 | 100407838 | 100446461 | 38624 | 2 | 17940 | ENSMUST00 | 19285 | ENSMUSG00 | Naip1 | NLR family, apoptosis inhibitory protein 1 |
| chr13 | 100444159 | 100444478 | 320 | * | NA_peak_158 | 27 | Promoter (1-2 | 13 | 100407838 | 100446461 | 38624 | 2 | 17940 | ENSMUST00 | 1983 | ENSMUSG00 | Naip1 | NLR family, apoptosis inhibitory protein 1 |
| chr13 | 100927331 | 100927722 | 392 | * | NA_peak_158 | 31 | Intron (ENSM | 13 | 100802648 | 100833427 | 30780 | 2 | 69048 | ENSMUST00 | -93904 | ENSMUSG00 | Slc30a5 | solute carrier family 30 (zinc transporter), member 5 |
| chr13 | 100953483 | 100953557 | 75 | * | NA_peak_158 | 31 | Intron (ENSM | 13 | 100802648 | 100833427 | 30780 | 2 | 69048 | ENSMUST00 | -120056 | ENSMUSG00 | Slc30a5 | solute carrier family 30 (zinc transporter), member 5 |
| chr13 | 101873222 | 101873328 | 107 | * | NA_peak_159 | 31 | Distal Interge | 13 | 101680563 | 101768217 | 87655 | 2 | 18708 | ENSMUST00 | -105005 | ENSMUSG00 | Pik3r1 | phosphoinositide-3-kinase regulatory subunit 1 |
| chr13 | 103998117 | 103998292 | 176 | * | NA_peak_160 | 64 | Distal Interge | 13 | 104024585 | 104034573 | 9989 | 2 | 75805 | ENSMUST00 | 36281 | ENSMUSG00 | Nln | neurolysin (metallopeptidase M3 family) |
| chr13 | 104062054 | 104062403 | 350 | * | NA_peak_160 | 13 | Promoter (<= | 13 | 104023222 | 104061854 | 38633 | 2 | 75805 | ENSMUST00 | -200 | ENSMUSG00 | Nln | neurolysin (metallopeptidase M3 family) |
| chr13 | 104178402 | 104178911 | 510 | * | NA_peak_160 | 23 | Promoter (<= | 13 | 104173722 | 104178439 | 4718 | 2 | 113002583 | ENSMUST00 | 0 | ENSMUSG00 | Shld3 | shieldin complex subunit 3 |
| chr13 | 104995378 | 104995635 | 258 | * | NA_peak_160 | 26 | Intron (ENSM | 13 | 104945904 | 105054930 | 109027 | 2 | 52882 | ENSMUST00 | 59295 | ENSMUSG00 | Rgs7bp | regulator of G-protein signalling 7 binding protein |
| chr13 | 105075717 | 105076076 | 360 | * | NA_peak_160 | 40 | Distal Interge | 13 | 105082122 | 105121782 | 39661 | 1 | 66763 | ENSMUST00 | -6046 | ENSMUSG00 | Nt5el | 5' nucleotidase, ecto-like |
| chr13 | 105403748 | 105404020 | 273 | * | NA_peak_160 | 20 | Distal Interge | 13 | 105443639 | 105448122 | 4484 | 1 | 15550 | ENSMUST00 | -39619 | ENSMUSG00 | Htr1a | 5-hydroxytryptamine (serotonin) receptor 1A |
| chr13 | 106377305 | 106377663 | 359 | * | NA_peak_160 | 27 | Distal Interge | 13 | 106811888 | 106836321 | 24434 | 2 | 76582 | ENSMUST00 | 458658 | ENSMUSG00 | Ipo11 | importin 11 |
| chr13 | 107942949 | 107943272 | 324 | * | NA_peak_161 | 23 | Distal Interge | 13 | 107724618 | 107890064 | 165447 | 2 | 67263 | ENSMUST00 | -52885 | ENSMUSG00 | Zswim6 | zinc finger SWIM-type containing 6 |
| chr13 | 108518313 | 108518558 | 246 | * | NA_peak_161 | 23 | Intron (ENSM | 13 | 108450009 | 108860143 | 410135 | 1 | 238871 | ENSMUST00 | 68304 | ENSMUSG00 | Pde4d | phosphodiesterase 4D, cAMP specific |
| chr13 | 108795173 | 108795518 | 346 | * | NA_peak_161 | 49 | Intron (ENSM | 13 | 108860166 | 109951344 | 1091179 | 1 | 238871 | ENSMUST00 | -64648 | ENSMUSG00 | Pde4d | phosphodiesterase 4D, cAMP specific |
| chr13 | 109372114 | 109372701 | 588 | * | NA_peak_161 | 28 | Intron (ENSM | 13 | 109324744 | 109324824 | 81 | 1 | 100124489 | ENSMUST00 | 47370 | ENSMUSG00 | Mir582 | microRNA 582 |
| chr13 | 110014136 | 110014466 | 331 | * | NA_peak_162 | 16 | Distal Interge | 13 | 109935312 | 109948348 | 13037 | 1 | 238871 | ENSMUST00 | 78824 | ENSMUSG00 | Pde4d | phosphodiesterase 4D, cAMP specific |
| chr13 | 111073747 | 111074025 | 279 | * | NA_peak_162 | 13 | Distal Interge | 13 | 111211467 | 111229431 | 17965 | 2 | 73129 | ENSMUST00 | 155406 | ENSMUSG00 | 3110015C05 | RIKEN cDNA 3110015C05 gene |
| chr13 | 111489040 | 111489230 | 191 | * | NA_peak_162 | 45 | Promoter (<= | 13 | 111425932 | 111490028 | 64097 | 2 | 73274 | ENSMUST00 | 798 | ENSMUSG00 | Gpbp1 | GC-rich promoter binding protein 1 |
| chr13 | 111577630 | 111577956 | 327 | * | NA_peak_162 | 20 | Distal Interge | 13 | 111439229 | 111549947 | 110719 | 2 | 73274 | ENSMUST00 | -27683 | ENSMUSG00 | Gpbp1 | GC-rich promoter binding protein 1 |
| chr13 | 111729087 | 111729336 | 250 | * | NA_peak_162 | 22 | Intron (ENSM | 13 | 111748793 | 111751373 | 2581 | 2 | 26401 | ENSMUST00 | 22037 | ENSMUSG00 | Map3k1 | mitogen-activated protein kinase kinase kinase 1 |
| chr13 | 112297820 | 112298121 | 302 | * | NA_peak_162 | 20 | Intron (ENSM | 13 | 112288820 | 112384002 | 95183 | 1 | 77318 | ENSMUST00 | 9000 | ENSMUSG00 | Ankrd55 | ankyrin repeat domain 55 |
| chr13 | 112553896 | 112554149 | 254 | * | NA_peak_163 | 23 | Intron (ENSM | 13 | 112523557 | 112562074 | 38518 | 2 | 218624 | ENSMUST00 | 7925 | ENSMUSG00 | Il31ra | interleukin 31 receptor A |
| chr13 | 112688029 | 112688195 | 167 | * | NA_peak_163 | 31 | Intron (ENSM | 13 | 112700229 | 112704961 | 4733 | 1 | 268706 | ENSMUST00 | -12034 | ENSMUSG00 | Slc38a9 | solute carrier family 38, member 9 |
| chr13 | 113285601 | 113286051 | 451 | * | NA_peak_163 | 19 | Distal Interge | 13 | 113171608 | 113225908 | 54301 | 2 | 14945 | ENSMUST00 | -59693 | ENSMUSG00 | Gzmk | granzyme K |
| chr13 | 113302898 | 113303566 | 669 | * | NA_peak_163 | 40 | Intron (ENSM | 13 | 113171608 | 113225908 | 54301 | 2 | 14945 | ENSMUST00 | -76990 | ENSMUSG00 | Gzmk | granzyme K |
| chr13 | 113373265 | 113373527 | 263 | * | NA_peak_163 | 28 | Intron (ENSM | 13 | 113446288 | 113502844 | 56557 | 2 | 70917 | ENSMUST00 | 129317 | ENSMUSG00 | 4921509O07 | RIKEN cDNA 4921509O07 gene |
| chr13 | 113719515 | 113719771 | 257 | * | NA_peak_163 | 46 | Distal Interge | 13 | 113662896 | 113663676 | 781 | 2 | 56534 | ENSMUST00 | -55839 | ENSMUSG00 | Hspb3 | heat shock protein 3 |
| chr13 | 113721892 | 113722182 | 291 | * | NA_peak_163 | 14 | Distal Interge | 13 | 113662896 | 113663676 | 781 | 2 | 56534 | ENSMUST00 | -58216 | ENSMUSG00 | Hspb3 | heat shock protein 3 |
| chr13 | 114037558 | 114037591 | 34 | * | NA_peak_163 | 31 | Intron (ENSM | 13 | 114124348 | 114151592 | 27245 | 2 | 319803 | ENSMUST00 | 114001 | ENSMUSG00 | A430090L17 | RIKEN cDNA A430090L17 gene |
| chr13 | 115502298 | 115502795 | 498 | * | NA_peak_164 | 30 | Distal Interge | 13 | 114953096 | 115101964 | 148869 | 2 | 109700 | ENSMUST00 | -400334 | ENSMUSG00 | Itga1 | integrin alpha 1 |
| chr13 | 116066777 | 116067094 | 318 | * | NA_peak_164 | 27 | Distal Interge | 13 | 116299395 | 116303351 | 3957 | 2 | 16392 | ENSMUST00 | 236257 | ENSMUSG00 | Isl1 | ISL1 transcription factor, LIM/homeodomain |
| chr13 | 116309974 | 116310245 | 272 | * | NA_peak_164 | 20 | Promoter (<= | 13 | 116298281 | 116309689 | 11409 | 2 | 16392 | ENSMUST00 | -285 | ENSMUSG00 | Isl1 | ISL1 transcription factor, LIM/homeodomain |
| chr13 | 119491185 | 119491926 | 742 | * | NA_peak_165 | 27 | Promoter (2-3 | 13 | 119488860 | 119610458 | 121599 | 1 | 633640 | ENSMUST00 | 2325 | ENSMUSG00 | Tmem267 | transmembrane protein 267 |
| chr13 | 119671685 | 119671944 | 260 | * | NA_peak_165 | 41 | Distal Interge | 13 | 119680040 | 119681588 | 1549 | 1 | 73480 | ENSMUST00 | -8096 | ENSMUSG00 | 1700074H08 | RIKEN cDNA 1700074H08 gene |
| chr13 | 120076089 | 120076708 | 620 | * | NA_peak_165 | 40 | Distal Interge | 13 | 120043425 | 120052194 | 8770 | 2 | 100861753 | ENSMUST00 | -23895 | ENSMUSG00 | Gm21188 | predicted gene, 21188 |
| chr13 | 120314040 | 120314321 | 282 | * | NA_peak_165 | 30 | Promoter (2-3 | 13 | 120316878 | 120318178 | 1301 | 1 | 236219 | ENSMUST00 | -2557 | ENSMUSG00 | Tcstv3 | 2-cell-stage, variable group, member 3 |
| chr14 | 3368270 | 3368581 | 312 | * | NA_peak_165 | 27 | Distal Interge | 14 | 3395093 | 3403525 | 8433 | 1 | 100041774 | ENSMUST00 | -26512 | ENSMUSG00 | Gm10413 | predicted gene 10413 |
| chr14 | 3378600 | 3379131 | 532 | * | NA_peak_165 | 40 | Distal Interge | 14 | 3395093 | 3403525 | 8433 | 1 | 100041774 | ENSMUST00 | -15962 | ENSMUSG00 | Gm10413 | predicted gene 10413 |
| chr14 | 3503750 | 3504106 | 357 | * | NA_peak_166 | 23 | Intron (ENSM | 14 | 3512378 | 3543372 | 30995 | 1 | 73549 | ENSMUST00 | -8272 | ENSMUSG00 | 1700110I01 | RIKEN cDNA 1700110I01 gene |

| | | | | | | | | | | | | | | | | | | |
|---|---|---|---|---|---|---|---|---|---|---|---|---|---|---|---|---|---|---|
| chr14 | 3619764 | 3619939 | 176 | * | NA_peak_16 | 45 | Distal Interge | 14 | 3634512 | 3642943 | 8432 | 1 | 100041774 | ENSMUST00 | -14573 | ENSMUSG00 | Gm10413 | predicted gene 10413 |
| chr14 | 3775392 | 3775775 | 384 | * | NA_peak_16 | 23 | Intron (ENSM | 14 | 3751771 | 3782764 | 30994 | 1 | 73549 | ENSMUST00 | 23621 | ENSMUSG00 | 1700110I01R | RIKEN cDNA 1700110I01 gene |
| chr14 | 6283790 | 6284194 | 405 | * | NA_peak_16 | 44 | Intron (ENSM | 14 | 6259747 | 6287250 | 27504 | 2 | 100861615 | ENSMUST00 | 3056 | ENSMUSG00 | Gm3411 | predicted gene 3411 |
| chr14 | 6550662 | 6551132 | 471 | * | NA_peak_16 | 20 | Distal Interge | 14 | 6736742 | 6741430 | 4689 | 2 | 100041151 | ENSMUST00 | 190298 | ENSMUSG00 | Gm3636 | predicted gene 3636 |
| chr14 | 6554012 | 6554199 | 188 | * | NA_peak_16 | 17 | Distal Interge | 14 | 6736742 | 6741430 | 4689 | 2 | 100041151 | ENSMUST00 | 187231 | ENSMUSG00 | Gm3636 | predicted gene 3636 |
| chr14 | 9127652 | 9127793 | 142 | * | NA_peak_16 | 24 | Distal Interge | 14 | 8666446 | 8673185 | 6740 | 1 | 74859 | ENSMUST00 | 461206 | ENSMUSG00 | 4930455B14I | RIKEN cDNA 4930455B14 gene |
| chr14 | 10018555 | 10018700 | 146 | * | NA_peak_16 | 30 | Exon (ENSMU | 14 | 9886026 | 9886629 | 604 | 1 | 100042415 | ENSMUST00 | 132529 | ENSMUSG00 | Mnd1-ps | Mnd1 retrotransposed pseudogene |
| chr14 | 10496601 | 10497058 | 458 | * | NA_peak_16 | 16 | Intron (ENSM | 14 | 9550092 | 10453458 | 903367 | 2 | 14198 | ENSMUST00 | -43143 | ENSMUSG00 | Fhit | fragile histidine triad gene |
| chr14 | 19415780 | 19416360 | 581 | * | NA_peak_16 | 331 | Exon (ENSMU | 14 | 19594139 | 19602581 | 8443 | 2 | 432825 | ENSMUST00 | 186221 | ENSMUSG00 | Gm5458 | predicted gene 5458 |
| chr14 | 19416853 | 19419590 | 2738 | * | NA_peak_16 | 1001 | Exon (ENSMU | 14 | 19594139 | 19602581 | 8443 | 2 | 432825 | ENSMUST00 | 182991 | ENSMUSG00 | Gm5458 | predicted gene 5458 |
| chr14 | 20181604 | 20181808 | 205 | * | NA_peak_16 | 27 | Promoter (<= | 14 | 20140057 | 20181809 | 41753 | 2 | 16529 | ENSMUST00 | 1 | ENSMUSG00 | Kcnk5 | potassium channel, subfamily K, member 5 |
| chr14 | 20280570 | 20280881 | 312 | * | NA_peak_16 | 38 | Distal Interge | 14 | 20262756 | 20269199 | 6444 | 2 | 74571 | ENSMUST00 | -11371 | ENSMUSG00 | Kcnk16 | potassium channel, subfamily K, member 16 |
| chr14 | 28606420 | 28606688 | 269 | * | NA_peak_16 | 15 | Distal Interge | 14 | 28511451 | 28511907 | 457 | 1 | 22418 | ENSMUST00 | 94969 | ENSMUSG00 | Wnt5a | wingless-type MMTV integration site family, member 5A |
| chr14 | 29765894 | 29766241 | 348 | * | NA_peak_16 | 64 | Distal Interge | 14 | 28904943 | 29721864 | 816922 | 2 | 12294 | ENSMUST00 | -44030 | ENSMUSG00 | Cacna2d3 | calcium channel, voltage-dependent, alpha2/delta subunit 3 |
| chr14 | 34805727 | 34806127 | 401 | * | NA_peak_17 | 26 | Distal Interge | 14 | 34820108 | 35583379 | 763272 | 1 | 14803 | ENSMUST00 | -13981 | ENSMUSG00 | Grid1 | glutamate receptor, ionotropic, delta 1 |
| chr14 | 35719906 | 35720180 | 275 | * | NA_peak_17 | 22 | Distal Interge | 14 | 35809488 | 35811978 | 2491 | 2 | 239036 | ENSMUST00 | 91798 | ENSMUSG00 | 4930596D02I | RIKEN cDNA 4930596D02 gene |
| chr14 | 37596054 | 37596396 | 343 | * | NA_peak_17 | 45 | Distal Interge | 14 | 37131398 | 37135272 | 3875 | 2 | 66092 | ENSMUST00 | -460782 | ENSMUSG00 | Ghitm | growth hormone inducible transmembrane protein |
| chr14 | 38606176 | 38606487 | 312 | * | NA_peak_17 | 36 | Intron (ENSM | 14 | 38898302 | 38958794 | 60493 | 1 | 75170 | ENSMUST00 | -291815 | ENSMUSG00 | Nrg3os | neuregulin 3, opposite strand |
| chr14 | 40662267 | 40662297 | 31 | * | NA_peak_17 | 23 | Distal Interge | 14 | 40813789 | 40853305 | 39517 | 2 | 328381 | ENSMUST00 | 191008 | ENSMUSG00 | Sh2d4b | SH2 domain containing 4B |
| chr14 | 40915204 | 40915235 | 32 | * | NA_peak_17 | 23 | Intron (ENSM | 14 | 40915241 | 40934316 | 19076 | 2 | 52588 | ENSMUST00 | 19081 | ENSMUSG00 | Tspan14 | tetraspanin 14 |
| chr14 | 41297417 | 41297736 | 320 | * | NA_peak_17 | 45 | Distal Interge | 14 | 41172214 | 41185149 | 12936 | 2 | 20390 | ENSMUST00 | -112268 | ENSMUSG00 | Sftpd | surfactant associated protein D |
| chr14 | 41343070 | 41343375 | 306 | * | NA_peak_17 | 21 | Distal Interge | 14 | 41172214 | 41185149 | 12936 | 2 | 20390 | ENSMUST00 | -157921 | ENSMUSG00 | Sftpd | surfactant associated protein D |
| chr14 | 41362678 | 41362963 | 286 | * | NA_peak_17 | 27 | Distal Interge | 14 | 41172214 | 41185149 | 12936 | 2 | 20390 | ENSMUST00 | -177529 | ENSMUSG00 | Sftpd | surfactant associated protein D |
| chr14 | 41365336 | 41365592 | 257 | * | NA_peak_17 | 36 | Distal Interge | 14 | 41172214 | 41185149 | 12936 | 2 | 20390 | ENSMUST00 | -180187 | ENSMUSG00 | Sftpd | surfactant associated protein D |
| chr14 | 41455850 | 41455881 | 32 | * | NA_peak_17 | 30 | Distal Interge | 14 | 41573211 | 41581305 | 8095 | 1 | 100041722 | ENSMUST00 | -117330 | ENSMUSG00 | 1700049E17I | tandem duplication of RIKEN cDNA 1700049E17 gene, gene 2 |
| chr14 | 41467264 | 41467715 | 452 | * | NA_peak_17 | 23 | Distal Interge | 14 | 41573211 | 41581305 | 8095 | 1 | 100041722 | ENSMUST00 | -105496 | ENSMUSG00 | 1700049E17I | tandem duplication of RIKEN cDNA 1700049E17 gene, gene 2 |
| chr14 | 41468506 | 41469277 | 772 | * | NA_peak_17 | 90 | Distal Interge | 14 | 41573211 | 41581305 | 8095 | 1 | 100041722 | ENSMUST00 | -103934 | ENSMUSG00 | 1700049E17I | tandem duplication of RIKEN cDNA 1700049E17 gene, gene 2 |
| chr14 | 41576052 | 41576279 | 228 | * | NA_peak_17 | 23 | Promoter (2-3 | 14 | 41573342 | 41576186 | 2845 | 1 | 100041722 | ENSMUST00 | 2710 | ENSMUSG00 | 1700049E17I | tandem duplication of RIKEN cDNA 1700049E17 gene, gene 2 |
| chr14 | 42140160 | 42140542 | 383 | * | NA_peak_17 | 20 | Distal Interge | 14 | 42089750 | 42095277 | 5528 | 2 | 73553 | ENSMUST00 | -44883 | ENSMUSG00 | 1700091H14I | RIKEN cDNA 1700091H14 gene |
| chr14 | 42168528 | 42168796 | 269 | * | NA_peak_17 | 21 | Distal Interge | 14 | 42089750 | 42095277 | 5528 | 2 | 73553 | ENSMUST00 | -73251 | ENSMUSG00 | 1700091H14I | RIKEN cDNA 1700091H14 gene |
| chr14 | 42380143 | 42380549 | 407 | * | NA_peak_17 | 23 | Distal Interge | 14 | 42254659 | 42260048 | 5390 | 2 | 101056136 | ENSMUST00 | -120095 | ENSMUSG00 | Gm17026 | predicted gene 17026 |
| chr14 | 42406625 | 42406801 | 177 | * | NA_peak_17 | 20 | Distal Interge | 14 | 42254659 | 42260048 | 5390 | 2 | 101056136 | ENSMUST00 | -146577 | ENSMUSG00 | Gm17026 | predicted gene 17026 |
| chr14 | 43305150 | 43305498 | 349 | * | NA_peak_17 | 40 | 3' UTR | 14 | 43298872 | 43306004 | 7133 | 1 | 108168154 | ENSMUST00 | 6278 | ENSMUSG00 | Gm9732 | predicted gene 9732 |
| chr14 | 43362930 | 43363267 | 338 | * | NA_peak_17 | 25 | Distal Interge | 14 | 43342383 | 43347811 | 5429 | 2 | 71826 | ENSMUST00 | -15119 | ENSMUSG00 | 1700001F09I | RIKEN cDNA 1700001F09 gene |
| chr14 | 43367974 | 43368433 | 460 | * | NA_peak_17 | 46 | Distal Interge | 14 | 43342383 | 43347811 | 5429 | 2 | 71826 | ENSMUST00 | -20163 | ENSMUSG00 | 1700001F09I | RIKEN cDNA 1700001F09 gene |
| chr14 | 43378839 | 43379204 | 366 | * | NA_peak_17 | 40 | Distal Interge | 14 | 43342383 | 43347811 | 5429 | 2 | 71826 | ENSMUST00 | -31028 | ENSMUSG00 | 1700001F09I | RIKEN cDNA 1700001F09 gene |
| chr14 | 43523323 | 43523879 | 557 | * | NA_peak_17 | 31 | Distal Interge | 14 | 43602627 | 43608013 | 5387 | 2 | 100042342 | ENSMUST00 | 84134 | ENSMUSG00 | Gm10375 | predicted gene 10375 |
| chr14 | 43625110 | 43625482 | 373 | * | NA_peak_17 | 36 | Distal Interge | 14 | 43602630 | 43608013 | 5384 | 2 | 100042342 | ENSMUST00 | -17097 | ENSMUSG00 | Gm10375 | predicted gene 10375 |
| chr14 | 44427392 | 44427677 | 286 | * | NA_peak_17 | 23 | Distal Interge | 14 | 44492463 | 44506345 | 13883 | 1 | 385138 | ENSMUST00 | -64786 | ENSMUSG00 | BC061237 | cDNA sequence BC061237 |
| chr14 | 44762016 | 44762419 | 404 | * | NA_peak_17 | 45 | Intron (ENSM | 14 | 44717173 | 44750340 | 33168 | 2 | 666744 | ENSMUST00 | -11676 | ENSMUSG00 | Gm8267 | predicted gene 8267 |
| chr14 | 45028662 | 45028902 | 241 | * | NA_peak_17 | 22 | Distal Interge | 14 | 44958607 | 44989699 | 31093 | 2 | 73895 | ENSMUST00 | -38963 | ENSMUSG00 | 4930431P03I | RIKEN cDNA 4930431P03 gene |
| chr14 | 50683868 | 50684109 | 242 | * | NA_peak_17 | 20 | Promoter (2-3 | 14 | 50680691 | 50681645 | 955 | 2 | 258264 | ENSMUST00 | -2223 | ENSMUSG00 | Olfr747 | olfactory receptor 747 |
| chr14 | 51300836 | 51301169 | 334 | * | NA_peak_17 | 27 | Distal Interge | 14 | 51317335 | 51345489 | 28155 | 2 | 75377 | ENSMUST00 | 44320 | ENSMUSG00 | 4930597G03 | RIKEN cDNA 4930597G03 gene |
| chr14 | 51580734 | 51581160 | 427 | * | NA_peak_17 | 15 | Intron (ENSM | 14 | 51552790 | 51662953 | 110164 | 1 | 434459 | ENSMUST00 | 27944 | ENSMUSG00 | Gm5622 | predicted gene 5622 |
| chr14 | 51595342 | 51595622 | 281 | * | NA_peak_17 | 14 | Intron (ENSM | 14 | 51552790 | 51662953 | 110164 | 1 | 434459 | ENSMUST00 | 42552 | ENSMUSG00 | Gm5622 | predicted gene 5622 |
| chr14 | 51609888 | 51610311 | 424 | * | NA_peak_17 | 26 | Exon (ENSMU | 14 | 51552790 | 51662953 | 110164 | 1 | 434459 | ENSMUST00 | 57098 | ENSMUSG00 | Gm5622 | predicted gene 5622 |
| chr14 | 51611783 | 51612084 | 302 | * | NA_peak_17 | 31 | Intron (ENSM | 14 | 51552790 | 51662953 | 110164 | 1 | 434459 | ENSMUST00 | 58993 | ENSMUSG00 | Gm5622 | predicted gene 5622 |
| chr14 | 53577589 | 53579064 | 1476 | * | NA_peak_17 | 31 | Distal Interge | 14 | 53032417 | 53039466 | 7050 | 2 | 102632039 | ENSMUST00 | -538123 | ENSMUSG00 | Gm30214 | predicted gene, 30214 |
| chr14 | 54894492 | 54894752 | 261 | * | NA_peak_17 | 17 | Promoter (<= | 14 | 54894573 | 54897557 | 2985 | 1 | 54196 | ENSMUST00 | 0 | ENSMUSG00 | Pabpn1 | poly(A) binding protein, nuclear 1 |
| chr14 | 57699691 | 57699945 | 255 | * | NA_peak_17 | 36 | Promoter (<= | 14 | 57697227 | 57699890 | 2664 | 2 | 50523 | ENSMUST00 | 0 | ENSMUSG00 | Lats2 | large tumor suppressor 2 |
| chr14 | 60684011 | 60684052 | 42 | * | NA_peak_17 | 20 | Intron (ENSM | 14 | 60691975 | 60709569 | 17595 | 1 | 219140 | ENSMUST00 | -7923 | ENSMUSG00 | Spata13 | spermatogenesis associated 13 |
| chr14 | 60960339 | 60960624 | 286 | * | NA_peak_17 | 59 | Distal Interge | 14 | 60966351 | 60975885 | 9535 | 2 | 29820 | ENSMUST00 | 15261 | ENSMUSG00 | Tnfrsf19 | tumor necrosis factor receptor superfamily, member 19 |
| chr14 | 68680812 | 68681129 | 318 | * | NA_peak_17 | 13 | Distal Interge | 14 | 68607734 | 68655842 | 48109 | 2 | 13522 | ENSMUST00 | -24970 | ENSMUSG00 | Adam28 | a disintegrin and metallopeptidase domain 28 |
| chr14 | 69162023 | 69162308 | 286 | * | NA_peak_17 | 31 | Distal Interge | 14 | 69171802 | 69175518 | 3717 | 1 | 18092 | ENSMUST00 | -9494 | ENSMUSG00 | Nkx2-6 | NK2 homeobox 6 |
| chr14 | 75445226 | 75445455 | 230 | * | NA_peak_17 | 23 | Distal Interge | 14 | 75455982 | 75529708 | 73727 | 1 | 380918 | ENSMUST00 | -10527 | ENSMUSG00 | Siah3 | siah E3 ubiquitin protein ligase family member 3 |
| chr14 | 76415788 | 76416062 | 275 | * | NA_peak_17 | 13 | Promoter (<= | 14 | 76415587 | 76507762 | 92176 | 1 | 21807 | ENSMUST00 | 201 | ENSMUSG00 | Tsc22d1 | TSC22 domain family, member 1 |
| chr14 | 79835963 | 79836091 | 129 | * | NA_peak_17 | 59 | Distal Interge | 14 | 79863074 | 79868398 | 5325 | 2 | 100038734 | ENSMUST00 | 32307 | ENSMUSG00 | Gm10845 | predicted gene 10845 |
| chr14 | 80351310 | 80351662 | 353 | * | NA_peak_17 | 31 | Distal Interge | 14 | 80000302 | 80021930 | 21629 | 1 | 380924 | ENSMUST00 | 351008 | ENSMUSG00 | Olfm4 | olfactomedin 4 |
| chr14 | 81794030 | 81794308 | 279 | * | NA_peak_17 | 66 | Distal Interge | 14 | 80000302 | 80021930 | 21629 | 1 | 380924 | ENSMUST00 | 1793728 | ENSMUSG00 | Olfm4 | olfactomedin 4 |
| chr14 | 82957792 | 82958103 | 312 | * | NA_peak_17 | 25 | Distal Interge | 14 | 84046431 | 84075290 | 28860 | 2 | 102635012 | ENSMUST00 | 1117187 | ENSMUSG00 | Gm32455 | predicted gene, 32455 |
| chr14 | 82960396 | 82960679 | 284 | * | NA_peak_17 | 40 | Distal Interge | 14 | 84046431 | 84075290 | 28860 | 2 | 102635012 | ENSMUST00 | 1114611 | ENSMUSG00 | Gm32455 | predicted gene, 32455 |
| chr14 | 84856156 | 84856543 | 388 | * | NA_peak_17 | 50 | Distal Interge | 14 | 84458724 | 84476424 | 17701 | 2 | 319903 | ENSMUST00 | -379732 | ENSMUSG00 | 9630013A20I | RIKEN cDNA 9630013A20 gene |
| chr14 | 85411442 | 85412226 | 785 | * | NA_peak_17 | 31 | Distal Interge | 14 | 86245970 | 86262042 | 16073 | 1 | 75155 | ENSMUST00 | -833744 | ENSMUSG00 | 4930529K09I | RIKEN cDNA 4930529K09 gene |
| chr14 | 92036341 | 92036684 | 344 | * | NA_peak_17 | 78 | Distal Interge | 14 | 93173729 | 93885748 | 712020 | 2 | 211712 | ENSMUST00 | 1849064 | ENSMUSG00 | Pcdh9 | protocadherin 9 |
| chr14 | 97325545 | 97325825 | 281 | * | NA_peak_17 | 34 | Distal Interge | 14 | 96102736 | 96519102 | 416367 | 2 | 93688 | ENSMUST00 | -806443 | ENSMUSG00 | Klhl1 | kelch-like 1 |
| chr14 | 100249470 | 100249843 | 374 | * | NA_peak_17 | 42 | Intron (ENSM | 14 | 100022832 | 100284484 | 261653 | 2 | 16597 | ENSMUST00 | 34641 | ENSMUSG00 | Klf12 | Kruppel-like factor 12 |
| chr14 | 100800629 | 100800829 | 201 | * | NA_peak_17 | 23 | Distal Interge | 14 | 101197690 | 101200069 | 2380 | 2 | 76627 | ENSMUST00 | 399240 | ENSMUSG00 | Prr30 | proline rich 30 |
| chr14 | 101963273 | 101963497 | 225 | * | NA_peak_17 | 20 | Distal Interge | 14 | 101929261 | 101932866 | 3606 | 1 | 380928 | ENSMUST00 | 34012 | ENSMUSG00 | Lmo7 | LIM domain only 7 |
| chr14 | 103237139 | 103237462 | 324 | * | NA_peak_17 | 16 | Intron (ENSM | 14 | 103262814 | 103275795 | 12982 | 2 | 105689 | ENSMUST00 | 38333 | ENSMUSG00 | Mycbp2 | MYC binding protein 2, E3 ubiquitin protein ligase |
| chr14 | 104110483 | 104110784 | 302 | * | NA_peak_17 | 36 | Distal Interge | 14 | 103977628 | 104027142 | 49515 | 2 | 73877 | ENSMUST00 | -83341 | ENSMUSG00 | 4930432J09F | RIKEN cDNA 4930432J09 gene |
| chr14 | 109680238 | 109681087 | 850 | * | NA_peak_17 | 40 | Distal Interge | 14 | 108908800 | 108914158 | 5359 | 2 | 76965 | ENSMUST00 | -766080 | ENSMUSG00 | Slitrk1 | SLIT and NTRK-like family, member 1 |
| chr14 | 111908715 | 111909124 | 410 | * | NA_peak_17 | 51 | Distal Interge | 14 | 111675849 | 111679342 | 3494 | 1 | 75409 | ENSMUST00 | 232866 | ENSMUSG00 | Slitrk5 | SLIT and NTRK-like family, member 5 |
| chr14 | 112336519 | 112336790 | 272 | * | NA_peak_17 | 27 | Distal Interge | 14 | 111675849 | 111679342 | 3494 | 1 | 75409 | ENSMUST00 | 660670 | ENSMUSG00 | Slitrk5 | SLIT and NTRK-like family, member 5 |
| chr14 | 112369530 | 112369687 | 158 | * | NA_peak_17 | 23 | Distal Interge | 14 | 111675849 | 111679342 | 3494 | 1 | 75409 | ENSMUST00 | 693681 | ENSMUSG00 | Slitrk5 | SLIT and NTRK-like family, member 5 |
| chr14 | 112633588 | 112633772 | 185 | * | NA_peak_17 | 21 | Distal Interge | 14 | 111675849 | 111679342 | 3494 | 1 | 75409 | ENSMUST00 | 957739 | ENSMUSG00 | Slitrk5 | SLIT and NTRK-like family, member 5 |
| chr14 | 117179833 | 117180268 | 436 | * | NA_peak_17 | 15 | Intron (ENSM | 14 | 116925936 | 117975086 | 1049151 | 1 | 23888 | ENSMUST00 | 253897 | ENSMUSG00 | Gpc6 | glypican 6 |
| chr14 | 118695171 | 118695321 | 151 | * | NA_peak_17 | 27 | Intron (ENSM | 14 | 118484333 | 118699492 | 215160 | 2 | 239273 | ENSMUST00 | 4171 | ENSMUSG00 | Abcc4 | ATP-binding cassette, sub-family C (CFTR/MRP), member 4 |
| chr14 | 119829420 | 119829587 | 168 | * | NA_peak_17 | 20 | Intron (ENSM | 14 | 119749225 | 119751564 | 2340 | 1 | 75432 | ENSMUST00 | 80195 | ENSMUSG00 | 1700006F04I | RIKEN cDNA 1700006F04 gene |
| chr14 | 121185233 | 121185295 | 63 | * | NA_peak_17 | 13 | Intron (ENSM | 14 | 121254908 | 121267921 | 13014 | 1 | 223254 | ENSMUST00 | -69613 | ENSMUSG00 | Farp1 | FERM, RhoGEF (Arhgef) and pleckstrin domain protein 1 (chondrocyte-derived) |
| chr14 | 123432674 | 123432997 | 324 | * | NA_peak_17 | 27 | Intron (ENSM | 14 | 123299686 | 123626876 | 327191 | 2 | 338370 | ENSMUST00 | 193879 | ENSMUSG00 | Nalcn | sodium leak channel, non-selective |
| chr14 | 124787651 | 124787685 | 35 | * | NA_peak_17 | 27 | Distal Interge | 14 | 123977907 | 124677127 | 699221 | 2 | 14169 | ENSMUST00 | -110524 | ENSMUSG00 | Fgf14 | fibroblast growth factor 14 |
| chr14 | 124801634 | 124802034 | 401 | * | NA_peak_17 | 65 | Distal Interge | 14 | 123977907 | 124677127 | 699221 | 2 | 14169 | ENSMUST00 | -124507 | ENSMUSG00 | Fgf14 | fibroblast growth factor 14 |
| chr15 | 3053047 | 3053450 | 404 | * | NA_peak_17 | 64 | Exon (ENSMU | 15 | 3268547 | 3277273 | 8727 | 1 | 20363 | ENSMUST00 | -215097 | ENSMUSG00 | Selenop | selenoprotein P |
| chr15 | 3054319 | 3054712 | 394 | * | NA_peak_17 | 64 | Distal Interge | 15 | 3268547 | 3277273 | 8727 | 1 | 20363 | ENSMUST00 | -213835 | ENSMUSG00 | Selenop | selenoprotein P |
| chr15 | 3068407 | 3068952 | 546 | * | NA_peak_17 | 23 | Distal Interge | 15 | 3268547 | 3277273 | 8727 | 1 | 20363 | ENSMUST00 | -199595 | ENSMUSG00 | Selenop | selenoprotein P |
| chr15 | 3121400 | 3121561 | 162 | * | NA_peak_17 | 23 | Distal Interge | 15 | 3268547 | 3277273 | 8727 | 1 | 20363 | ENSMUST00 | -146986 | ENSMUSG00 | Selenop | selenoprotein P |
| chr15 | 3128562 | 3128656 | 95 | * | NA_peak_17 | 25 | Exon (ENSMU | 15 | 3268547 | 3277273 | 8727 | 1 | 20363 | ENSMUST00 | -139891 | ENSMUSG00 | Selenop | selenoprotein P |
| chr15 | 3133319 | 3133588 | 270 | * | NA_peak_17 | 16 | Intron (ENSM | 15 | 3268547 | 3277273 | 8727 | 1 | 20363 | ENSMUST00 | -134959 | ENSMUSG00 | Selenop | selenoprotein P |
| chr15 | 3143006 | 3143093 | 88 | * | NA_peak_17 | 23 | Intron (ENSM | 15 | 3268547 | 3277273 | 8727 | 1 | 20363 | ENSMUST00 | -125454 | ENSMUSG00 | Selenop | selenoprotein P |
| chr15 | 3149319 | 3149775 | 457 | * | NA_peak_17 | 27 | Intron (ENSM | 15 | 3268547 | 3277273 | 8727 | 1 | 20363 | ENSMUST00 | -118772 | ENSMUSG00 | Selenop | selenoprotein P |
| chr15 | 3152505 | 3152700 | 196 | * | NA_peak_17 | 43 | Intron (ENSM | 15 | 3268547 | 3277273 | 8727 | 1 | 20363 | ENSMUST00 | -115847 | ENSMUSG00 | Selenop | selenoprotein P |
| chr15 | 3154035 | 3154392 | 358 | * | NA_peak_17 | 22 | Intron (ENSM | 15 | 3268547 | 3277273 | 8727 | 1 | 20363 | ENSMUST00 | -114155 | ENSMUSG00 | Selenop | selenoprotein P |
| chr15 | 3155581 | 3155620 | 40 | * | NA_peak_17 | 24 | Intron (ENSM | 15 | 3268547 | 3277273 | 8727 | 1 | 20363 | ENSMUST00 | -112927 | ENSMUSG00 | Selenop | selenoprotein P |
| chr15 | 3156975 | 3157337 | 363 | * | NA_peak_17 | 16 | Intron (ENSM | 15 | 3268547 | 3277273 | 8727 | 1 | 20363 | ENSMUST00 | -111210 | ENSMUSG00 | Selenop | selenoprotein P |
| chr15 | 3162819 | 3162904 | 86 | * | NA_peak_17 | 20 | Distal Interge | 15 | 3268547 | 3277273 | 8727 | 1 | 20363 | ENSMUST00 | -105643 | ENSMUSG00 | Selenop | selenoprotein P |
| chr15 | 3165865 | 3166360 | 496 | * | NA_peak_17 | 88 | Distal Interge | 15 | 3268547 | 3277273 | 8727 | 1 | 20363 | ENSMUST00 | -102187 | ENSMUSG00 | Selenop | selenoprotein P |
| chr15 | 3178834 | 3179089 | 256 | * | NA_peak_17 | 20 | Exon (ENSMU | 15 | 3268547 | 3277273 | 8727 | 1 | 20363 | ENSMUST00 | -89458 | ENSMUSG00 | Selenop | selenoprotein P |
| chr15 | 3184994 | 3185307 | 314 | * | NA_peak_17 | 43 | Distal Interge | 15 | 3268547 | 3277273 | 8727 | 1 | 20363 | ENSMUST00 | -83240 | ENSMUSG00 | Selenop | selenoprotein P |
| chr15 | 3212440 | 3212769 | 330 | * | NA_peak_17 | 17 | Distal Interge | 15 | 3268547 | 3277273 | 8727 | 1 | 20363 | ENSMUST00 | -55778 | ENSMUSG00 | Selenop | selenoprotein P |
| chr15 | 3217659 | 3217972 | 314 | * | NA_peak_17 | 33 | Exon (ENSMU | 15 | 3268547 | 3277273 | 8727 | 1 | 20363 | ENSMUST00 | -50575 | ENSMUSG00 | Selenop | selenoprotein P |
| chr15 | 3225968 | 3226029 | 62 | * | NA_peak_17 | 27 | Distal Interge | 15 | 3268547 | 3277273 | 8727 | 1 | 20363 | ENSMUST00 | -42518 | ENSMUSG00 | Selenop | selenoprotein P |
| chr15 | 3240766 | 3241080 | 315 | * | NA_peak_17 | 36 | Distal Interge | 15 | 3268547 | 3277273 | 8727 | 1 | 20363 | ENSMUST00 | -27467 | ENSMUSG00 | Selenop | selenoprotein P |
| chr15 | 3274695 | 3274978 | 284 | * | NA_peak_17 | 20 | Promoter (2-3 | 15 | 3272085 | 3277271 | 5187 | 1 | 20363 | ENSMUST00 | 2610 | ENSMUSG00 | Selenop | selenoprotein P |
| chr15 | 3279387 | 3279747 | 361 | * | NA_peak_17 | 27 | 3' UTR | 15 | 3272085 | 3277271 | 5187 | 1 | 20363 | ENSMUST00 | 7302 | ENSMUSG00 | Selenop | selenoprotein P |
| chr15 | 3281197 | 3281591 | 395 | * | NA_peak_17 | 23 | Intron (ENSM | 15 | 3272085 | 3277271 | 5187 | 1 | 20363 | ENSMUST00 | 9112 | ENSMUSG00 | Selenop | selenoprotein P |
| chr15 | 3287019 | 3287108 | 90 | * | NA_peak_17 | 23 | Intron (ENSM | 15 | 3280070 | 3301196 | 21127 | 2 | 100039139 | ENSMUST00 | 14088 | ENSMUSG00 | Ccdc152 | coiled-coil domain containing 152 |

| | | | | | | | | | | | | | | | | | | |
|---|---|---|---|---|---|---|---|---|---|---|---|---|---|---|---|---|---|---|
| chr15 | 3294150 | 3294519 | 370 | * | NA_peak_179 | 33 | Intron (ENSM | 15 | 3280070 | 3301196 | 21127 | 2 | 100039139 | ENSMUST00 | 6677 | ENSMUSG0 | Ccdc152 | coiled-coil domain containing 152 |
| chr15 | 3309613 | 3309894 | 282 | * | NA_peak_179 | 16 | Distal Interge | 15 | 3280627 | 3303526 | 22900 | 2 | 100039139 | ENSMUST00 | -6087 | ENSMUSG0 | Ccdc152 | coiled-coil domain containing 152 |
| chr15 | 3320739 | 3321043 | 305 | * | NA_peak_179 | 23 | Exon (ENSMU | 15 | 3280627 | 3303526 | 22900 | 2 | 100039139 | ENSMUST00 | -17213 | ENSMUSG0 | Ccdc152 | coiled-coil domain containing 152 |
| chr15 | 3332221 | 3332368 | 148 | * | NA_peak_179 | 23 | Intron (ENSM | 15 | 3280627 | 3303526 | 22900 | 2 | 100039139 | ENSMUST00 | -28695 | ENSMUSG0 | Ccdc152 | coiled-coil domain containing 152 |
| chr15 | 3334946 | 3335717 | 772 | * | NA_peak_179 | 25 | Intron (ENSM | 15 | 3280627 | 3303526 | 22900 | 2 | 100039139 | ENSMUST00 | -31420 | ENSMUSG0 | Ccdc152 | coiled-coil domain containing 152 |
| chr15 | 3376072 | 3376485 | 414 | * | NA_peak_179 | 20 | Intron (ENSM | 15 | 3280627 | 3303526 | 22900 | 2 | 100039139 | ENSMUST00 | -72546 | ENSMUSG0 | Ccdc152 | coiled-coil domain containing 152 |
| chr15 | 3382868 | 3383309 | 442 | * | NA_peak_179 | 14 | Intron (ENSM | 15 | 3280627 | 3303526 | 22900 | 2 | 100039139 | ENSMUST00 | -79342 | ENSMUSG0 | Ccdc152 | coiled-coil domain containing 152 |
| chr15 | 3392363 | 3392447 | 85 | * | NA_peak_179 | 27 | Intron (ENSM | 15 | 3327539 | 3471644 | 144106 | 2 | 14600 | ENSMUST00 | 79197 | ENSMUSG0 | Ghr | growth hormone receptor |
| chr15 | 3468715 | 3468859 | 145 | * | NA_peak_179 | 30 | Promoter (2-3 | 15 | 3327539 | 3471644 | 144106 | 2 | 14600 | ENSMUST00 | 2785 | ENSMUSG0 | Ghr | growth hormone receptor |
| chr15 | 3509451 | 3509594 | 144 | * | NA_peak_179 | 22 | Intron (ENSM | 15 | 3327539 | 3471644 | 144106 | 2 | 14600 | ENSMUST00 | -37807 | ENSMUSG0 | Ghr | growth hormone receptor |
| chr15 | 3519629 | 3520019 | 391 | * | NA_peak_179 | 43 | Intron (ENSM | 15 | 3327539 | 3471644 | 144106 | 2 | 14600 | ENSMUST00 | -47985 | ENSMUSG0 | Ghr | growth hormone receptor |
| chr15 | 3535454 | 3535749 | 296 | * | NA_peak_179 | 40 | Intron (ENSM | 15 | 3319009 | 3581842 | 262834 | 2 | 14600 | ENSMUST00 | 46093 | ENSMUSG0 | Ghr | growth hormone receptor |
| chr15 | 3538040 | 3538230 | 191 | * | NA_peak_179 | 20 | Intron (ENSM | 15 | 3319009 | 3581842 | 262834 | 2 | 14600 | ENSMUST00 | 43612 | ENSMUSG0 | Ghr | growth hormone receptor |
| chr15 | 3555095 | 3555205 | 111 | * | NA_peak_179 | 31 | Intron (ENSM | 15 | 3319009 | 3581842 | 262834 | 2 | 14600 | ENSMUST00 | 26637 | ENSMUSG0 | Ghr | growth hormone receptor |
| chr15 | 3604225 | 3604485 | 261 | * | NA_peak_179 | 13 | Distal Interge | 15 | 3384309 | 3583492 | 199184 | 2 | 14600 | ENSMUST00 | -20733 | ENSMUSG0 | Ghr | growth hormone receptor |
| chr15 | 3651574 | 3651679 | 106 | * | NA_peak_180 | 27 | Distal Interge | 15 | 3384309 | 3583492 | 199184 | 2 | 14600 | ENSMUST00 | -68082 | ENSMUSG0 | Ghr | growth hormone receptor |
| chr15 | 3652143 | 3652394 | 252 | * | NA_peak_180 | 27 | Distal Interge | 15 | 3384309 | 3583492 | 199184 | 2 | 14600 | ENSMUST00 | -68651 | ENSMUSG0 | Ghr | growth hormone receptor |
| chr15 | 3653157 | 3653344 | 188 | * | NA_peak_180 | 27 | Distal Interge | 15 | 3384309 | 3583492 | 199184 | 2 | 14600 | ENSMUST00 | -69665 | ENSMUSG0 | Ghr | growth hormone receptor |
| chr15 | 3665743 | 3666166 | 424 | * | NA_peak_180 | 32 | Distal Interge | 15 | 3384309 | 3583492 | 199184 | 2 | 14600 | ENSMUST00 | -82251 | ENSMUSG0 | Ghr | growth hormone receptor |
| chr15 | 3678087 | 3678491 | 405 | * | NA_peak_180 | 45 | Distal Interge | 15 | 3384309 | 3583492 | 199184 | 2 | 14600 | ENSMUST00 | -94595 | ENSMUSG0 | Ghr | growth hormone receptor |
| chr15 | 3687259 | 3687310 | 52 | * | NA_peak_180 | 42 | Distal Interge | 15 | 3384309 | 3583492 | 199184 | 2 | 14600 | ENSMUST00 | -103767 | ENSMUSG0 | Ghr | growth hormone receptor |
| chr15 | 3712817 | 3713595 | 779 | * | NA_peak_180 | 26 | Distal Interge | 15 | 3384309 | 3583492 | 199184 | 2 | 14600 | ENSMUST00 | -129325 | ENSMUSG0 | Ghr | growth hormone receptor |
| chr15 | 3732788 | 3733283 | 496 | * | NA_peak_180 | 66 | Distal Interge | 15 | 3384309 | 3583492 | 199184 | 2 | 14600 | ENSMUST00 | -149296 | ENSMUSG0 | Ghr | growth hormone receptor |
| chr15 | 3733587 | 3733806 | 220 | * | NA_peak_180 | 66 | Distal Interge | 15 | 3384309 | 3583492 | 199184 | 2 | 14600 | ENSMUST00 | -150095 | ENSMUSG0 | Ghr | growth hormone receptor |
| chr15 | 3762320 | 3762594 | 275 | * | NA_peak_180 | 23 | Distal Interge | 15 | 3384309 | 3583492 | 199184 | 2 | 14600 | ENSMUST00 | -178828 | ENSMUSG0 | Ghr | growth hormone receptor |
| chr15 | 3807597 | 3807879 | 283 | * | NA_peak_180 | 24 | Distal Interge | 15 | 3971284 | 3971982 | 699 | 2 | 106052 | ENSMUST00 | 164103 | ENSMUSG0 | Fbxo4 | F-box protein 4 |
| chr15 | 3816870 | 3817182 | 313 | * | NA_peak_180 | 31 | Distal Interge | 15 | 3971284 | 3971982 | 699 | 2 | 106052 | ENSMUST00 | 154800 | ENSMUSG0 | Fbxo4 | F-box protein 4 |
| chr15 | 3818066 | 3818411 | 346 | * | NA_peak_180 | 13 | Distal Interge | 15 | 3971284 | 3971982 | 699 | 2 | 106052 | ENSMUST00 | 153571 | ENSMUSG0 | Fbxo4 | F-box protein 4 |
| chr15 | 3825112 | 3825211 | 100 | * | NA_peak_180 | 24 | Distal Interge | 15 | 3971284 | 3971982 | 699 | 2 | 106052 | ENSMUST00 | 146771 | ENSMUSG0 | Fbxo4 | F-box protein 4 |
| chr15 | 3852473 | 3852520 | 48 | * | NA_peak_180 | 33 | Distal Interge | 15 | 3971284 | 3971982 | 699 | 2 | 106052 | ENSMUST00 | 119462 | ENSMUSG0 | Fbxo4 | F-box protein 4 |
| chr15 | 3877560 | 3877863 | 304 | * | NA_peak_180 | 27 | Distal Interge | 15 | 3971284 | 3971982 | 699 | 2 | 106052 | ENSMUST00 | 94119 | ENSMUSG0 | Fbxo4 | F-box protein 4 |
| chr15 | 3886529 | 3886840 | 312 | * | NA_peak_180 | 54 | Distal Interge | 15 | 3971284 | 3971982 | 699 | 2 | 106052 | ENSMUST00 | 85142 | ENSMUSG0 | Fbxo4 | F-box protein 4 |
| chr15 | 3889924 | 3889983 | 60 | * | NA_peak_180 | 23 | Distal Interge | 15 | 3971284 | 3971982 | 699 | 2 | 106052 | ENSMUST00 | 81999 | ENSMUSG0 | Fbxo4 | F-box protein 4 |
| chr15 | 3893308 | 3893582 | 275 | * | NA_peak_180 | 36 | Distal Interge | 15 | 3971284 | 3971982 | 699 | 2 | 106052 | ENSMUST00 | 78400 | ENSMUSG0 | Fbxo4 | F-box protein 4 |
| chr15 | 3928094 | 3928319 | 226 | * | NA_peak_180 | 16 | Distal Interge | 15 | 3971284 | 3971982 | 699 | 2 | 106052 | ENSMUST00 | 43663 | ENSMUSG0 | Fbxo4 | F-box protein 4 |
| chr15 | 3956388 | 3956761 | 374 | * | NA_peak_180 | 30 | Distal Interge | 15 | 3971284 | 3971982 | 699 | 2 | 106052 | ENSMUST00 | 15221 | ENSMUSG0 | Fbxo4 | F-box protein 4 |
| chr15 | 3998557 | 3998849 | 293 | * | NA_peak_180 | 16 | Promoter (2-3 | 15 | 3996039 | 4015858 | 19820 | 1 | 328489 | ENSMUST00 | 2518 | ENSMUSG0 | Gm2093 | predicted gene 2093 |
| chr15 | 4115970 | 4116421 | 452 | * | NA_peak_181 | 53 | Intron (ENSM | 15 | 4142523 | 4153980 | 11458 | 1 | 67041 | ENSMUST00 | -26102 | ENSMUSG0 | Oxct1 | 3-oxoacid CoA transferase 1 |
| chr15 | 4126651 | 4126824 | 174 | * | NA_peak_181 | 40 | Intron (ENSM | 15 | 4142523 | 4153980 | 11458 | 1 | 67041 | ENSMUST00 | -15699 | ENSMUSG0 | Oxct1 | 3-oxoacid CoA transferase 1 |
| chr15 | 4190973 | 4191424 | 452 | * | NA_peak_181 | 17 | Distal Interge | 15 | 4142812 | 4153959 | 11148 | 1 | 67041 | ENSMUST00 | 48161 | ENSMUSG0 | Oxct1 | 3-oxoacid CoA transferase 1 |
| chr15 | 4236949 | 4237484 | 536 | * | NA_peak_181 | 24 | Distal Interge | 15 | 4142812 | 4153959 | 11148 | 1 | 67041 | ENSMUST00 | 94137 | ENSMUSG0 | Oxct1 | 3-oxoacid CoA transferase 1 |
| chr15 | 4298003 | 4298188 | 186 | * | NA_peak_181 | 27 | Distal Interge | 15 | 4375491 | 4520414 | 144924 | 1 | 239318 | ENSMUST00 | -77303 | ENSMUSG0 | Plcxd3 | phosphatidylinositol-specific phospholipase C, X domain containing 3 |
| chr15 | 4307573 | 4307714 | 142 | * | NA_peak_181 | 16 | Distal Interge | 15 | 4375491 | 4520414 | 144924 | 1 | 239318 | ENSMUST00 | -67777 | ENSMUSG0 | Plcxd3 | phosphatidylinositol-specific phospholipase C, X domain containing 3 |
| chr15 | 4310812 | 4311103 | 292 | * | NA_peak_181 | 32 | Distal Interge | 15 | 4375491 | 4520414 | 144924 | 1 | 239318 | ENSMUST00 | -64388 | ENSMUSG0 | Plcxd3 | phosphatidylinositol-specific phospholipase C, X domain containing 3 |
| chr15 | 4341536 | 4341823 | 288 | * | NA_peak_181 | 40 | Distal Interge | 15 | 4375491 | 4520414 | 144924 | 1 | 239318 | ENSMUST00 | -33668 | ENSMUSG0 | Plcxd3 | phosphatidylinositol-specific phospholipase C, X domain containing 3 |
| chr15 | 4345434 | 4345896 | 463 | * | NA_peak_181 | 36 | Distal Interge | 15 | 4375491 | 4520414 | 144924 | 1 | 239318 | ENSMUST00 | -29595 | ENSMUSG0 | Plcxd3 | phosphatidylinositol-specific phospholipase C, X domain containing 3 |
| chr15 | 4346392 | 4346579 | 188 | * | NA_peak_181 | 22 | Distal Interge | 15 | 4375491 | 4520414 | 144924 | 1 | 239318 | ENSMUST00 | -28912 | ENSMUSG0 | Plcxd3 | phosphatidylinositol-specific phospholipase C, X domain containing 3 |
| chr15 | 4410048 | 4410220 | 173 | * | NA_peak_181 | 27 | Intron (ENSM | 15 | 4375504 | 4575553 | 200050 | 1 | 239318 | ENSMUST00 | 34544 | ENSMUSG0 | Plcxd3 | phosphatidylinositol-specific phospholipase C, X domain containing 3 |
| chr15 | 4419652 | 4420298 | 647 | * | NA_peak_181 | 16 | Intron (ENSM | 15 | 4375504 | 4575553 | 200050 | 1 | 239318 | ENSMUST00 | 44148 | ENSMUSG0 | Plcxd3 | phosphatidylinositol-specific phospholipase C, X domain containing 3 |
| chr15 | 4496691 | 4497097 | 407 | * | NA_peak_182 | 26 | Intron (ENSM | 15 | 4375504 | 4575553 | 200050 | 1 | 239318 | ENSMUST00 | 121187 | ENSMUSG0 | Plcxd3 | phosphatidylinositol-specific phospholipase C, X domain containing 3 |
| chr15 | 4501017 | 4501304 | 288 | * | NA_peak_182 | 23 | Intron (ENSM | 15 | 4375504 | 4575553 | 200050 | 1 | 239318 | ENSMUST00 | 125513 | ENSMUSG0 | Plcxd3 | phosphatidylinositol-specific phospholipase C, X domain containing 3 |
| chr15 | 4507575 | 4507860 | 286 | * | NA_peak_182 | 19 | Intron (ENSM | 15 | 4375504 | 4575553 | 200050 | 1 | 239318 | ENSMUST00 | 132071 | ENSMUSG0 | Plcxd3 | phosphatidylinositol-specific phospholipase C, X domain containing 3 |
| chr15 | 4530429 | 4530600 | 172 | * | NA_peak_182 | 23 | Intron (ENSM | 15 | 4375504 | 4575553 | 200050 | 1 | 239318 | ENSMUST00 | 154925 | ENSMUSG0 | Plcxd3 | phosphatidylinositol-specific phospholipase C, X domain containing 3 |
| chr15 | 4641385 | 4641665 | 281 | * | NA_peak_182 | 19 | Distal Interge | 15 | 4727175 | 4804045 | 76871 | 1 | 12274 | ENSMUST00 | -85510 | ENSMUSG0 | C6 | complement component 6 |
| chr15 | 4669076 | 4669504 | 429 | * | NA_peak_182 | 65 | Distal Interge | 15 | 4727175 | 4804045 | 76871 | 1 | 12274 | ENSMUST00 | -57671 | ENSMUSG0 | C6 | complement component 6 |
| chr15 | 4677386 | 4677804 | 419 | * | NA_peak_182 | 20 | Distal Interge | 15 | 4727175 | 4804045 | 76871 | 1 | 12274 | ENSMUST00 | -49371 | ENSMUSG0 | C6 | complement component 6 |
| chr15 | 4712983 | 4713264 | 282 | * | NA_peak_182 | 55 | Distal Interge | 15 | 4727175 | 4804045 | 76871 | 1 | 12274 | ENSMUST00 | -13911 | ENSMUSG0 | C6 | complement component 6 |
| chr15 | 4723820 | 4724100 | 281 | * | NA_peak_182 | 15 | Distal Interge | 15 | 4727175 | 4804045 | 76871 | 1 | 12274 | ENSMUST00 | -3075 | ENSMUSG0 | C6 | complement component 6 |
| chr15 | 4749083 | 4749432 | 350 | * | NA_peak_182 | 36 | Intron (ENSM | 15 | 4765945 | 4791164 | 25220 | 1 | 12274 | ENSMUST00 | -16513 | ENSMUSG0 | C6 | complement component 6 |
| chr15 | 4790557 | 4790660 | 104 | * | NA_peak_182 | 31 | Intron (ENSM | 15 | 4765945 | 4791164 | 25220 | 1 | 12274 | ENSMUST00 | 24612 | ENSMUSG0 | C6 | complement component 6 |
| chr15 | 4796438 | 4796628 | 191 | * | NA_peak_182 | 34 | Intron (ENSM | 15 | 4765945 | 4791164 | 25220 | 1 | 12274 | ENSMUST00 | 30493 | ENSMUSG0 | C6 | complement component 6 |
| chr15 | 4818433 | 4818851 | 419 | * | NA_peak_182 | 40 | Distal Interge | 15 | 4765945 | 4791164 | 25220 | 1 | 12274 | ENSMUST00 | 52488 | ENSMUSG0 | C6 | complement component 6 |
| chr15 | 4822406 | 4822427 | 22 | * | NA_peak_182 | 22 | Distal Interge | 15 | 4765945 | 4791164 | 25220 | 1 | 12274 | ENSMUST00 | 56461 | ENSMUSG0 | C6 | complement component 6 |
| chr15 | 4862106 | 4862455 | 350 | * | NA_peak_182 | 30 | Distal Interge | 15 | 4898737 | 4906310 | 7574 | 1 | 223825 | ENSMUST00 | -36282 | ENSMUSG0 | Mroh2b | maestro heat-like repeat family member 2B |
| chr15 | 4878350 | 4878666 | 317 | * | NA_peak_183 | 23 | Distal Interge | 15 | 4898737 | 4906310 | 7574 | 1 | 223825 | ENSMUST00 | -20071 | ENSMUSG0 | Mroh2b | maestro heat-like repeat family member 2B |
| chr15 | 4884891 | 4885778 | 888 | * | NA_peak_183 | 40 | Exon (ENSMU | 15 | 4898737 | 4906310 | 7574 | 1 | 223825 | ENSMUST00 | -12959 | ENSMUSG0 | Mroh2b | maestro heat-like repeat family member 2B |
| chr15 | 4886279 | 4886656 | 378 | * | NA_peak_183 | 40 | Distal Interge | 15 | 4898737 | 4906310 | 7574 | 1 | 223825 | ENSMUST00 | -12081 | ENSMUSG0 | Mroh2b | maestro heat-like repeat family member 2B |
| chr15 | 4894133 | 4894827 | 695 | * | NA_peak_183 | 38 | Distal Interge | 15 | 4898737 | 4906310 | 7574 | 1 | 223825 | ENSMUST00 | -3910 | ENSMUSG0 | Mroh2b | maestro heat-like repeat family member 2B |
| chr15 | 4899546 | 4899870 | 325 | * | NA_peak_183 | 26 | Promoter (<= | 15 | 4899142 | 4962205 | 63064 | 1 | 223825 | ENSMUST00 | 404 | ENSMUSG0 | Mroh2b | maestro heat-like repeat family member 2B |
| chr15 | 4923315 | 4923551 | 237 | * | NA_peak_183 | 27 | Intron (ENSM | 15 | 4929853 | 4962205 | 32353 | 1 | 223825 | ENSMUST00 | -6302 | ENSMUSG0 | Mroh2b | maestro heat-like repeat family member 2B |
| chr15 | 4925672 | 4925686 | 15 | * | NA_peak_183 | 13 | Exon (ENSMU | 15 | 4929853 | 4962205 | 32353 | 1 | 223825 | ENSMUST00 | -4167 | ENSMUSG0 | Mroh2b | maestro heat-like repeat family member 2B |
| chr15 | 4945052 | 4945281 | 230 | * | NA_peak_183 | 43 | Exon (ENSMU | 15 | 4948482 | 4962072 | 13591 | 1 | 223825 | ENSMUST00 | -3201 | ENSMUSG0 | Mroh2b | maestro heat-like repeat family member 2B |
| chr15 | 4959816 | 4960255 | 440 | * | NA_peak_183 | 25 | Intron (ENSM | 15 | 4948482 | 4962072 | 13591 | 1 | 223825 | ENSMUST00 | 11334 | ENSMUSG0 | Mroh2b | maestro heat-like repeat family member 2B |
| chr15 | 5039114 | 5039782 | 669 | * | NA_peak_183 | 26 | Intron (ENSM | 15 | 4988762 | 5063740 | 74979 | 2 | 109828 | ENSMUST00 | 23958 | ENSMUSG0 | C7 | complement component 7 |
| chr15 | 5063512 | 5063628 | 117 | * | NA_peak_183 | 15 | Promoter (<= | 15 | 4988762 | 5063740 | 74979 | 2 | 109828 | ENSMUST00 | 112 | ENSMUSG0 | C7 | complement component 7 |
| chr15 | 5065792 | 5066127 | 336 | * | NA_peak_183 | 33 | Promoter (2-3 | 15 | 4988762 | 5063740 | 74979 | 2 | 109828 | ENSMUST00 | -2052 | ENSMUSG0 | C7 | complement component 7 |
| chr15 | 5099772 | 5100056 | 285 | * | NA_peak_183 | 20 | Exon (ENSMU | 15 | 5105137 | 5108484 | 3348 | 2 | 239319 | ENSMUST00 | 8428 | ENSMUSG0 | Card6 | caspase recruitment domain family, member 6 |
| chr15 | 5173485 | 5173887 | 403 | * | NA_peak_183 | 23 | Intron (ENSM | 15 | 5185525 | 5218336 | 32812 | 1 | 67515 | ENSMUST00 | -11638 | ENSMUSG0 | Ttc33 | tetratricopeptide repeat domain 33 |
| chr15 | 5177981 | 5178336 | 356 | * | NA_peak_183 | 30 | Intron (ENSM | 15 | 5185525 | 5218336 | 32812 | 1 | 67515 | ENSMUST00 | -7189 | ENSMUSG0 | Ttc33 | tetratricopeptide repeat domain 33 |
| chr15 | 5258560 | 5258898 | 339 | * | NA_peak_183 | 16 | Distal Interge | 15 | 5233399 | 5244187 | 10789 | 2 | 19219 | ENSMUST00 | -14373 | ENSMUSG0 | Ptger4 | prostaglandin E receptor 4 (subtype EP4) |
| chr15 | 5266925 | 5267384 | 460 | * | NA_peak_183 | 49 | Distal Interge | 15 | 5233399 | 5244187 | 10789 | 2 | 19219 | ENSMUST00 | -22738 | ENSMUSG0 | Ptger4 | prostaglandin E receptor 4 (subtype EP4) |
| chr15 | 5269638 | 5270000 | 363 | * | NA_peak_183 | 31 | Distal Interge | 15 | 5233399 | 5244187 | 10789 | 2 | 19219 | ENSMUST00 | -25451 | ENSMUSG0 | Ptger4 | prostaglandin E receptor 4 (subtype EP4) |
| chr15 | 5308473 | 5308542 | 70 | * | NA_peak_183 | 18 | Distal Interge | 15 | 5233399 | 5244187 | 10789 | 2 | 19219 | ENSMUST00 | -64286 | ENSMUSG0 | Ptger4 | prostaglandin E receptor 4 (subtype EP4) |
| chr15 | 5329326 | 5329335 | 10 | * | NA_peak_184 | 20 | Distal Interge | 15 | 5233399 | 5244187 | 10789 | 2 | 19219 | ENSMUST00 | -85139 | ENSMUSG0 | Ptger4 | prostaglandin E receptor 4 (subtype EP4) |
| chr15 | 5409153 | 5409558 | 406 | * | NA_peak_184 | 31 | Distal Interge | 15 | 5496266 | 5594984 | 98719 | 1 | 71432 | ENSMUST00 | -86708 | ENSMUSG0 | 5430437J10R | RIKEN cDNA 5430437J10 gene |
| chr15 | 5412454 | 5412931 | 478 | * | NA_peak_184 | 19 | Distal Interge | 15 | 5496266 | 5594984 | 98719 | 1 | 71432 | ENSMUST00 | -83335 | ENSMUSG0 | 5430437J10R | RIKEN cDNA 5430437J10 gene |
| chr15 | 5453983 | 5454571 | 589 | * | NA_peak_184 | 21 | Distal Interge | 15 | 5496266 | 5594984 | 98719 | 1 | 71432 | ENSMUST00 | -41695 | ENSMUSG0 | 5430437J10R | RIKEN cDNA 5430437J10 gene |
| chr15 | 5475819 | 5475995 | 177 | * | NA_peak_184 | 27 | Distal Interge | 15 | 5496266 | 5594984 | 98719 | 1 | 71432 | ENSMUST00 | -20271 | ENSMUSG0 | 5430437J10R | RIKEN cDNA 5430437J10 gene |
| chr15 | 5505130 | 5505415 | 286 | * | NA_peak_184 | 30 | Intron (ENSM | 15 | 5496318 | 5594983 | 98666 | 1 | 71432 | ENSMUST00 | 8812 | ENSMUSG0 | 5430437J10R | RIKEN cDNA 5430437J10 gene |
| chr15 | 5507094 | 5507405 | 312 | * | NA_peak_184 | 31 | Intron (ENSM | 15 | 5496318 | 5594983 | 98666 | 1 | 71432 | ENSMUST00 | 10776 | ENSMUSG0 | 5430437J10R | RIKEN cDNA 5430437J10 gene |
| chr15 | 5519467 | 5519621 | 155 | * | NA_peak_184 | 23 | Intron (ENSM | 15 | 5496318 | 5594983 | 98666 | 1 | 71432 | ENSMUST00 | 23149 | ENSMUSG0 | 5430437J10R | RIKEN cDNA 5430437J10 gene |
| chr15 | 5520779 | 5520875 | 97 | * | NA_peak_184 | 20 | Intron (ENSM | 15 | 5496318 | 5594983 | 98666 | 1 | 71432 | ENSMUST00 | 24461 | ENSMUSG0 | 5430437J10R | RIKEN cDNA 5430437J10 gene |
| chr15 | 5543856 | 5544121 | 266 | * | NA_peak_184 | 55 | Intron (ENSM | 15 | 5496318 | 5594983 | 98666 | 1 | 71432 | ENSMUST00 | 47538 | ENSMUSG0 | 5430437J10R | RIKEN cDNA 5430437J10 gene |
| chr15 | 5561947 | 5562405 | 459 | * | NA_peak_184 | 36 | Intron (ENSM | 15 | 5496318 | 5594983 | 98666 | 1 | 71432 | ENSMUST00 | 65629 | ENSMUSG0 | 5430437J10R | RIKEN cDNA 5430437J10 gene |
| chr15 | 5566939 | 5566967 | 29 | * | NA_peak_184 | 27 | Intron (ENSM | 15 | 5496318 | 5594983 | 98666 | 1 | 71432 | ENSMUST00 | 70621 | ENSMUSG0 | 5430437J10R | RIKEN cDNA 5430437J10 gene |
| chr15 | 5589193 | 5589706 | 514 | * | NA_peak_184 | 23 | Intron (ENSM | 15 | 5496318 | 5594983 | 98666 | 1 | 71432 | ENSMUST00 | 92875 | ENSMUSG0 | 5430437J10R | RIKEN cDNA 5430437J10 gene |
| chr15 | 5714735 | 5714997 | 263 | * | NA_peak_184 | 20 | Distal Interge | 15 | 5496318 | 5594983 | 98666 | 1 | 71432 | ENSMUST00 | 218417 | ENSMUSG0 | 5430437J10R | RIKEN cDNA 5430437J10 gene |
| chr15 | 5770356 | 5770724 | 369 | * | NA_peak_184 | 22 | Distal Interge | 15 | 5496318 | 5594983 | 98666 | 1 | 71432 | ENSMUST00 | 274038 | ENSMUSG0 | 5430437J10R | RIKEN cDNA 5430437J10 gene |
| chr15 | 5792802 | 5793276 | 475 | * | NA_peak_184 | 27 | Distal Interge | 15 | 5496318 | 5594983 | 98666 | 1 | 71432 | ENSMUST00 | 296484 | ENSMUSG0 | 5430437J10R | RIKEN cDNA 5430437J10 gene |
| chr15 | 5806617 | 5806914 | 298 | * | NA_peak_185 | 19 | Distal Interge | 15 | 5496318 | 5594983 | 98666 | 1 | 71432 | ENSMUST00 | 310299 | ENSMUSG0 | 5430437J10R | RIKEN cDNA 5430437J10 gene |
| chr15 | 5810977 | 5811287 | 311 | * | NA_peak_185 | 30 | Distal Interge | 15 | 5496318 | 5594983 | 98666 | 1 | 71432 | ENSMUST00 | 314659 | ENSMUSG0 | 5430437J10R | RIKEN cDNA 5430437J10 gene |
| chr15 | 5820532 | 5820798 | 267 | * | NA_peak_185 | 16 | Distal Interge | 15 | 5496318 | 5594983 | 98666 | 1 | 71432 | ENSMUST00 | 324214 | ENSMUSG0 | 5430437J10R | RIKEN cDNA 5430437J10 gene |
| chr15 | 5884110 | 5884333 | 224 | * | NA_peak_185 | 26 | Distal Interge | 15 | 5496318 | 5594983 | 98666 | 1 | 71432 | ENSMUST00 | 387792 | ENSMUSG0 | 5430437J10R | RIKEN cDNA 5430437J10 gene |
| chr15 | 5936382 | 5936628 | 247 | * | NA_peak_185 | 16 | Distal Interge | 15 | 6299788 | 6435520 | 135733 | 1 | 13132 | ENSMUST00 | -363160 | ENSMUSG0 | Dab2 | disabled 2, mitogen-responsive phosphoprotein |
| chr15 | 5982606 | 5982623 | 18 | * | NA_peak_185 | 40 | Distal Interge | 15 | 6299788 | 6435520 | 135733 | 1 | 13132 | ENSMUST00 | -317165 | ENSMUSG0 | Dab2 | disabled 2, mitogen-responsive phosphoprotein |
| chr15 | 5983817 | 5983937 | 121 | * | NA_peak_185 | 15 | Distal Interge | 15 | 6299788 | 6435520 | 135733 | 1 | 13132 | ENSMUST00 | -315851 | ENSMUSG0 | Dab2 | disabled 2, mitogen-responsive phosphoprotein |

| | | | | | | | | | | | | | | | | | | |
|---|---|---|---|---|---|---|---|---|---|---|---|---|---|---|---|---|---|---|
| chr15 | 5987019 | 5987401 | 383 | * | NA_peak_18 | 59 | Distal Interge | 15 | 6299788 | 6435520 | 135733 | 1 | 13132 | ENSMUST00 | -312387 | ENSMUSG0 | Dab2 | disabled 2, mitogen-responsive phosphoprotein |
| chr15 | 6011882 | 6011972 | 91 | * | NA_peak_18 | 36 | Distal Interge | 15 | 6299788 | 6435520 | 135733 | 1 | 13132 | ENSMUST00 | -287816 | ENSMUSG0 | Dab2 | disabled 2, mitogen-responsive phosphoprotein |
| chr15 | 6034508 | 6034790 | 283 | * | NA_peak_18 | 20 | Distal Interge | 15 | 6299788 | 6435520 | 135733 | 1 | 13132 | ENSMUST00 | -264998 | ENSMUSG0 | Dab2 | disabled 2, mitogen-responsive phosphoprotein |
| chr15 | 6126010 | 6126219 | 210 | * | NA_peak_18 | 27 | Distal Interge | 15 | 6299788 | 6435520 | 135733 | 1 | 13132 | ENSMUST00 | -173569 | ENSMUSG0 | Dab2 | disabled 2, mitogen-responsive phosphoprotein |
| chr15 | 6160963 | 6161139 | 177 | * | NA_peak_18 | 32 | Distal Interge | 15 | 6299788 | 6435520 | 135733 | 1 | 13132 | ENSMUST00 | -138649 | ENSMUSG0 | Dab2 | disabled 2, mitogen-responsive phosphoprotein |
| chr15 | 6264766 | 6264876 | 111 | * | NA_peak_18 | 18 | Distal Interge | 15 | 6299788 | 6435520 | 135733 | 1 | 13132 | ENSMUST00 | -34912 | ENSMUSG0 | Dab2 | disabled 2, mitogen-responsive phosphoprotein |
| chr15 | 6367839 | 6367989 | 151 | * | NA_peak_18 | 31 | Intron (ENSM | 15 | 6386598 | 6440705 | 54108 | 1 | 13132 | ENSMUST00 | -18609 | ENSMUSG0 | Dab2 | disabled 2, mitogen-responsive phosphoprotein |
| chr15 | 6381478 | 6382017 | 540 | * | NA_peak_18 | 19 | Intron (ENSM | 15 | 6386598 | 6440705 | 54108 | 1 | 13132 | ENSMUST00 | -4581 | ENSMUSG0 | Dab2 | disabled 2, mitogen-responsive phosphoprotein |
| chr15 | 6388620 | 6388913 | 294 | * | NA_peak_18 | 28 | Promoter (<= | 15 | 6389050 | 6418318 | 29269 | 1 | 13132 | ENSMUST00 | -137 | ENSMUSG0 | Dab2 | disabled 2, mitogen-responsive phosphoprotein |
| chr15 | 6430753 | 6430948 | 196 | * | NA_peak_18 | 42 | Intron (ENSM | 15 | 6423162 | 6425024 | 1863 | 1 | 13132 | ENSMUST00 | 7591 | ENSMUSG0 | Dab2 | disabled 2, mitogen-responsive phosphoprotein |
| chr15 | 6497049 | 6497365 | 317 | * | NA_peak_18 | 16 | 3' UTR | 15 | 6489918 | 6498744 | 8827 | 1 | 12279 | ENSMUST00 | 7131 | ENSMUSG0 | C9 | complement component 9 |
| chr15 | 6675159 | 6675450 | 292 | * | NA_peak_18 | 19 | Distal Interge | 15 | 6660405 | 6663251 | 2847 | 1 | 23880 | ENSMUST00 | 14754 | ENSMUSG0 | Fyb | FYN binding protein |
| chr15 | 6696498 | 6696740 | 243 | * | NA_peak_18 | 18 | Distal Interge | 15 | 6708379 | 6765417 | 57039 | 1 | 78757 | ENSMUST00 | -11639 | ENSMUSG0 | Rictor | RPTOR independent companion of MTOR, complex 2 |
| chr15 | 6719492 | 6719750 | 259 | * | NA_peak_18 | 34 | Intron (ENSM | 15 | 6708383 | 6800401 | 92019 | 1 | 78757 | ENSMUST00 | 11109 | ENSMUSG0 | Rictor | RPTOR independent companion of MTOR, complex 2 |
| chr15 | 6729312 | 6729624 | 313 | * | NA_peak_18 | 34 | Intron (ENSM | 15 | 6708383 | 6800401 | 92019 | 1 | 78757 | ENSMUST00 | 20929 | ENSMUSG0 | Rictor | RPTOR independent companion of MTOR, complex 2 |
| chr15 | 6801328 | 6801660 | 333 | * | NA_peak_18 | 27 | Distal Interge | 15 | 6820637 | 6824595 | 3959 | 2 | 18414 | ENSMUST00 | 22935 | ENSMUSG0 | Osmr | oncostatin M receptor |
| chr15 | 6805789 | 6806175 | 387 | * | NA_peak_18 | 24 | Distal Interge | 15 | 6820637 | 6824595 | 3959 | 2 | 18414 | ENSMUST00 | 18420 | ENSMUSG0 | Osmr | oncostatin M receptor |
| chr15 | 7101050 | 7101285 | 236 | * | NA_peak_18 | 20 | Intron (ENSM | 15 | 7090614 | 7164566 | 73953 | 1 | 16880 | ENSMUST00 | 10436 | ENSMUSG0 | Lifr | LIF receptor alpha |
| chr15 | 7111309 | 7111569 | 261 | * | NA_peak_18 | 23 | Intron (ENSM | 15 | 7129559 | 7191743 | 62185 | 1 | 16880 | ENSMUST00 | -17990 | ENSMUSG0 | Lifr | LIF receptor alpha |
| chr15 | 7122058 | 7122519 | 462 | * | NA_peak_18 | 28 | Intron (ENSM | 15 | 7129559 | 7191743 | 62185 | 1 | 16880 | ENSMUST00 | -7040 | ENSMUSG0 | Lifr | LIF receptor alpha |
| chr15 | 7131303 | 7131696 | 394 | * | NA_peak_18 | 22 | Promoter (1-2 | 15 | 7129572 | 7185343 | 55772 | 1 | 16880 | ENSMUST00 | 1731 | ENSMUSG0 | Lifr | LIF receptor alpha |
| chr15 | 7273659 | 7274004 | 346 | * | NA_peak_18 | 55 | Intron (ENSM | 15 | 7253511 | 7254768 | 1258 | 2 | 268780 | ENSMUST00 | -18891 | ENSMUSG0 | Egflam | EGF-like, fibronectin type III and laminin G domains |
| chr15 | 7276136 | 7276150 | 15 | * | NA_peak_18 | 16 | Intron (ENSM | 15 | 7253511 | 7254768 | 1258 | 2 | 268780 | ENSMUST00 | -21368 | ENSMUSG0 | Egflam | EGF-like, fibronectin type III and laminin G domains |
| chr15 | 7276462 | 7276999 | 538 | * | NA_peak_18 | 27 | Intron (ENSM | 15 | 7253511 | 7254768 | 1258 | 2 | 268780 | ENSMUST00 | -21694 | ENSMUSG0 | Egflam | EGF-like, fibronectin type III and laminin G domains |
| chr15 | 7324804 | 7324925 | 122 | * | NA_peak_18 | 31 | Intron (ENSM | 15 | 7253511 | 7254768 | 1258 | 2 | 268780 | ENSMUST00 | -70036 | ENSMUSG0 | Egflam | EGF-like, fibronectin type III and laminin G domains |
| chr15 | 7342506 | 7342757 | 252 | * | NA_peak_18 | 23 | Exon (ENSMU | 15 | 7305070 | 7398261 | 93192 | 2 | 268780 | ENSMUST00 | 55504 | ENSMUSG0 | Egflam | EGF-like, fibronectin type III and laminin G domains |
| chr15 | 7359556 | 7359702 | 147 | * | NA_peak_18 | 39 | Intron (ENSM | 15 | 7305070 | 7398261 | 93192 | 2 | 268780 | ENSMUST00 | 38559 | ENSMUSG0 | Egflam | EGF-like, fibronectin type III and laminin G domains |
| chr15 | 7418598 | 7418644 | 47 | * | NA_peak_18 | 13 | Distal Interge | 15 | 7206583 | 7398395 | 191813 | 2 | 268780 | ENSMUST00 | -20203 | ENSMUSG0 | Egflam | EGF-like, fibronectin type III and laminin G domains |
| chr15 | 7527393 | 7527931 | 539 | * | NA_peak_18 | 30 | Distal Interge | 15 | 7206583 | 7398395 | 191813 | 2 | 268780 | ENSMUST00 | -128998 | ENSMUSG0 | Egflam | EGF-like, fibronectin type III and laminin G domains |
| chr15 | 7548163 | 7548256 | 94 | * | NA_peak_18 | 27 | Distal Interge | 15 | 7206583 | 7398395 | 191813 | 2 | 268780 | ENSMUST00 | -149768 | ENSMUSG0 | Egflam | EGF-like, fibronectin type III and laminin G domains |
| chr15 | 7555304 | 7555973 | 670 | * | NA_peak_18 | 38 | Distal Interge | 15 | 7206583 | 7398395 | 191813 | 2 | 268780 | ENSMUST00 | -156909 | ENSMUSG0 | Egflam | EGF-like, fibronectin type III and laminin G domains |
| chr15 | 7618916 | 7619316 | 401 | * | NA_peak_18 | 45 | Distal Interge | 15 | 7810846 | 7837575 | 26730 | 1 | 14573 | ENSMUST00 | -191530 | ENSMUSG0 | Gdnf | glial cell line derived neurotrophic factor |
| chr15 | 7657008 | 7657402 | 395 | * | NA_peak_18 | 45 | Distal Interge | 15 | 7810846 | 7837575 | 26730 | 1 | 14573 | ENSMUST00 | -153444 | ENSMUSG0 | Gdnf | glial cell line derived neurotrophic factor |
| chr15 | 7684478 | 7684697 | 220 | * | NA_peak_18 | 22 | Distal Interge | 15 | 7810846 | 7837575 | 26730 | 1 | 14573 | ENSMUST00 | -126149 | ENSMUSG0 | Gdnf | glial cell line derived neurotrophic factor |
| chr15 | 7698706 | 7699189 | 484 | * | NA_peak_18 | 21 | Distal Interge | 15 | 7810846 | 7837575 | 26730 | 1 | 14573 | ENSMUST00 | -111657 | ENSMUSG0 | Gdnf | glial cell line derived neurotrophic factor |
| chr15 | 7850762 | 7851049 | 288 | * | NA_peak_18 | 58 | Distal Interge | 15 | 7873055 | 7886377 | 13323 | 2 | 545085 | ENSMUST00 | 35328 | ENSMUSG0 | Wdr70 | WD repeat domain 70 |
| chr15 | 7904546 | 7904899 | 354 | * | NA_peak_18 | 29 | Intron (ENSM | 15 | 7873055 | 7886377 | 13323 | 2 | 545085 | ENSMUST00 | -18169 | ENSMUSG0 | Wdr70 | WD repeat domain 70 |
| chr15 | 7994827 | 7995043 | 217 | * | NA_peak_18 | 37 | Intron (ENSM | 15 | 7873062 | 8099191 | 226130 | 2 | 545085 | ENSMUST00 | 104148 | ENSMUSG0 | Wdr70 | WD repeat domain 70 |
| chr15 | 8145633 | 8145939 | 307 | * | NA_peak_18 | 20 | Promoter (2-3 | 15 | 8143124 | 8144572 | 1449 | 1 | 170762 | ENSMUST00 | 2509 | ENSMUSG0 | Nup155 | nucleoporin 155 |
| chr15 | 8291451 | 8292023 | 573 | * | NA_peak_18 | 27 | Exon (ENSMU | 15 | 8247103 | 8252976 | 5874 | 1 | 73692 | ENSMUST00 | 44348 | ENSMUSG0 | Cplane1 | ciliogenesis and planar polarity effector 1 |
| chr15 | 8314870 | 8314978 | 109 | * | NA_peak_18 | 20 | Intron (ENSM | 15 | 8247103 | 8252976 | 5874 | 1 | 73692 | ENSMUST00 | 67767 | ENSMUSG0 | Cplane1 | ciliogenesis and planar polarity effector 1 |
| chr15 | 8342743 | 8343064 | 322 | * | NA_peak_18 | 31 | Intron (ENSM | 15 | 8247103 | 8252976 | 5874 | 1 | 73692 | ENSMUST00 | 95640 | ENSMUSG0 | Cplane1 | ciliogenesis and planar polarity effector 1 |
| chr15 | 8348150 | 8348718 | 569 | * | NA_peak_18 | 20 | Intron (ENSM | 15 | 8290617 | 8444463 | 153847 | 2 | 71175 | ENSMUST00 | 95745 | ENSMUSG0 | Nipbl | NIPBL cohesin loading factor |
| chr15 | 8400720 | 8400962 | 243 | * | NA_peak_18 | 14 | Intron (ENSM | 15 | 8290617 | 8444463 | 153847 | 2 | 71175 | ENSMUST00 | 43501 | ENSMUSG0 | Nipbl | NIPBL cohesin loading factor |
| chr15 | 8406429 | 8406592 | 164 | * | NA_peak_18 | 23 | Intron (ENSM | 15 | 8290617 | 8444463 | 153847 | 2 | 71175 | ENSMUST00 | 37871 | ENSMUSG0 | Nipbl | NIPBL cohesin loading factor |
| chr15 | 8613586 | 8613635 | 50 | * | NA_peak_18 | 27 | Distal Interge | 15 | 8642256 | 8643247 | 992 | 2 | 20512 | ENSMUST00 | 29612 | ENSMUSG0 | Slc1a3 | solute carrier family 1 (glial high affinity glutamate transporter), member 3 |
| chr15 | 8762366 | 8762421 | 56 | * | NA_peak_18 | 29 | Distal Interge | 15 | 8634124 | 8710764 | 76641 | 2 | 20512 | ENSMUST00 | -51602 | ENSMUSG0 | Slc1a3 | solute carrier family 1 (glial high affinity glutamate transporter), member 3 |
| chr15 | 8765668 | 8766107 | 440 | * | NA_peak_18 | 22 | Distal Interge | 15 | 8634124 | 8710764 | 76641 | 2 | 20512 | ENSMUST00 | -54904 | ENSMUSG0 | Slc1a3 | solute carrier family 1 (glial high affinity glutamate transporter), member 3 |
| chr15 | 8796172 | 8796576 | 405 | * | NA_peak_18 | 23 | Distal Interge | 15 | 8634124 | 8710764 | 76641 | 2 | 20512 | ENSMUST00 | -85408 | ENSMUSG0 | Slc1a3 | solute carrier family 1 (glial high affinity glutamate transporter), member 3 |
| chr15 | 8841568 | 8841591 | 24 | * | NA_peak_18 | 27 | Distal Interge | 15 | 8967949 | 8970298 | 2350 | 1 | 223332 | ENSMUST00 | -126358 | ENSMUSG0 | Ranbp3l | RAN binding protein 3-like |
| chr15 | 8890177 | 8890180 | 4 | * | NA_peak_18 | 26 | Distal Interge | 15 | 8967949 | 8970298 | 2350 | 1 | 223332 | ENSMUST00 | -77769 | ENSMUSG0 | Ranbp3l | RAN binding protein 3-like |
| chr15 | 8970408 | 8970904 | 497 | * | NA_peak_18 | 32 | Promoter (1-2 | 15 | 8968426 | 9066551 | 98126 | 1 | 223332 | ENSMUST00 | 1982 | ENSMUSG0 | Ranbp3l | RAN binding protein 3-like |
| chr15 | 9015926 | 9016275 | 350 | * | NA_peak_18 | 13 | Intron (ENSM | 15 | 9007266 | 9067335 | 60070 | 1 | 223332 | ENSMUST00 | 8660 | ENSMUSG0 | Ranbp3l | RAN binding protein 3-like |
| chr15 | 9057225 | 9057737 | 513 | * | NA_peak_18 | 28 | Exon (ENSMU | 15 | 9071260 | 9110487 | 39228 | 1 | 68646 | ENSMUST00 | -13523 | ENSMUSG0 | Nadk2 | NAD kinase 2, mitochondrial |
| chr15 | 9099817 | 9099956 | 140 | * | NA_peak_18 | 19 | Promoter (<= | 15 | 9100156 | 9107178 | 7023 | 1 | 68646 | ENSMUST00 | -200 | ENSMUSG0 | Nadk2 | NAD kinase 2, mitochondrial |
| chr15 | 9190206 | 9190406 | 201 | * | NA_peak_19 | 16 | Intron (ENSM | 15 | 9175463 | 9182586 | 7124 | 1 | 320506 | ENSMUST00 | 14743 | ENSMUSG0 | Lmbrd2 | LMBR1 domain containing 2 |
| chr15 | 9298298 | 9298648 | 351 | * | NA_peak_19 | 37 | Intron (ENSM | 15 | 9283501 | 9292027 | 8527 | 1 | 105887 | ENSMUST00 | 14797 | ENSMUSG0 | Ugt3a1 | UDP glycosyltransferases 3 family, polypeptide A1 |
| chr15 | 9303192 | 9303716 | 525 | * | NA_peak_19 | 15 | Intron (ENSM | 15 | 9283501 | 9292027 | 8527 | 1 | 105887 | ENSMUST00 | 19691 | ENSMUSG0 | Ugt3a1 | UDP glycosyltransferases 3 family, polypeptide A1 |
| chr15 | 9304831 | 9305216 | 386 | * | NA_peak_19 | 20 | Intron (ENSM | 15 | 9283501 | 9292027 | 8527 | 1 | 105887 | ENSMUST00 | 21330 | ENSMUSG0 | Ugt3a1 | UDP glycosyltransferases 3 family, polypeptide A1 |
| chr15 | 9332489 | 9332803 | 315 | * | NA_peak_19 | 30 | Promoter (2-3 | 15 | 9335550 | 9370955 | 35406 | 1 | 223337 | ENSMUST00 | -2747 | ENSMUSG0 | Ugt3a2 | UDP glycosyltransferases 3 family, polypeptide A2 |
| chr15 | 9402593 | 9402865 | 273 | * | NA_peak_19 | 20 | Distal Interge | 15 | 9436028 | 9466038 | 30011 | 1 | 75568 | ENSMUST00 | -33163 | ENSMUSG0 | Capsl | calcyphosine-like |
| chr15 | 9404912 | 9405121 | 210 | * | NA_peak_19 | 27 | Distal Interge | 15 | 9436028 | 9466038 | 30011 | 1 | 75568 | ENSMUST00 | -30907 | ENSMUSG0 | Capsl | calcyphosine-like |
| chr15 | 9414722 | 9414864 | 143 | * | NA_peak_19 | 16 | Intron (ENSM | 15 | 9436028 | 9466038 | 30011 | 1 | 75568 | ENSMUST00 | -21164 | ENSMUSG0 | Capsl | calcyphosine-like |
| chr15 | 9417199 | 9417270 | 72 | * | NA_peak_19 | 27 | Intron (ENSM | 15 | 9436028 | 9466038 | 30011 | 1 | 75568 | ENSMUST00 | -18758 | ENSMUSG0 | Capsl | calcyphosine-like |
| chr15 | 9441833 | 9442085 | 253 | * | NA_peak_19 | 30 | Intron (ENSM | 15 | 9436028 | 9466038 | 30011 | 1 | 75568 | ENSMUST00 | 5805 | ENSMUSG0 | Capsl | calcyphosine-like |
| chr15 | 9461218 | 9462088 | 871 | * | NA_peak_19 | 36 | Exon (ENSMU | 15 | 9436028 | 9466038 | 30011 | 1 | 75568 | ENSMUST00 | 25190 | ENSMUSG0 | Capsl | calcyphosine-like |
| chr15 | 9557466 | 9557749 | 284 | * | NA_peak_19 | 23 | Distal Interge | 15 | 9516409 | 9530176 | 13768 | 2 | 16197 | ENSMUST00 | -27290 | ENSMUSG0 | Il7r | interleukin 7 receptor |
| chr15 | 9560206 | 9560445 | 240 | * | NA_peak_19 | 20 | Distal Interge | 15 | 9516409 | 9530176 | 13768 | 2 | 16197 | ENSMUST00 | -30030 | ENSMUSG0 | Il7r | interleukin 7 receptor |
| chr15 | 9570254 | 9570740 | 487 | * | NA_peak_19 | 23 | Distal Interge | 15 | 9516409 | 9530176 | 13768 | 2 | 16197 | ENSMUST00 | -40078 | ENSMUSG0 | Il7r | interleukin 7 receptor |
| chr15 | 9580661 | 9580847 | 187 | * | NA_peak_19 | 36 | Intron (ENSM | 15 | 9516409 | 9530176 | 13768 | 2 | 16197 | ENSMUST00 | -50485 | ENSMUSG0 | Il7r | interleukin 7 receptor |
| chr15 | 9583791 | 9583996 | 206 | * | NA_peak_19 | 31 | Exon (ENSMU | 15 | 9516409 | 9530176 | 13768 | 2 | 16197 | ENSMUST00 | -53615 | ENSMUSG0 | Il7r | interleukin 7 receptor |
| chr15 | 9614509 | 9615030 | 522 | * | NA_peak_19 | 20 | Intron (ENSM | 15 | 9516409 | 9530176 | 13768 | 2 | 16197 | ENSMUST00 | -84333 | ENSMUSG0 | Il7r | interleukin 7 receptor |
| chr15 | 9723748 | 9723958 | 211 | * | NA_peak_19 | 26 | Intron (ENSM | 15 | 9704312 | 9748777 | 44466 | 2 | 320277 | ENSMUST00 | 24819 | ENSMUSG0 | Spef2 | sperm flagellar 2 |
| chr15 | 9755491 | 9755794 | 304 | * | NA_peak_19 | 36 | Distal Interge | 15 | 9661546 | 9748868 | 87323 | 2 | 320277 | ENSMUST00 | -6623 | ENSMUSG0 | Spef2 | sperm flagellar 2 |
| chr15 | 9776871 | 9777227 | 357 | * | NA_peak_19 | 27 | Distal Interge | 15 | 9661546 | 9748868 | 87323 | 2 | 320277 | ENSMUST00 | -28003 | ENSMUSG0 | Spef2 | sperm flagellar 2 |
| chr15 | 9870605 | 9870788 | 184 | * | NA_peak_19 | 23 | Distal Interge | 15 | 9661546 | 9748868 | 87323 | 2 | 320277 | ENSMUST00 | -121737 | ENSMUSG0 | Spef2 | sperm flagellar 2 |
| chr15 | 9905472 | 9905863 | 392 | * | NA_peak_19 | 27 | Distal Interge | 15 | 9661546 | 9748868 | 87323 | 2 | 320277 | ENSMUST00 | -156604 | ENSMUSG0 | Spef2 | sperm flagellar 2 |
| chr15 | 10106243 | 10106525 | 283 | * | NA_peak_19 | 14 | Distal Interge | 15 | 10177238 | 10329388 | 152151 | 1 | 19116 | ENSMUST00 | -70713 | ENSMUSG0 | Prlr | prolactin receptor |
| chr15 | 10274577 | 10274919 | 343 | * | NA_peak_19 | 43 | Intron (ENSM | 15 | 10249560 | 10322492 | 72933 | 1 | 19116 | ENSMUST00 | 25017 | ENSMUSG0 | Prlr | prolactin receptor |
| chr15 | 10327506 | 10327639 | 134 | * | NA_peak_19 | 23 | Promoter (2-3 | 15 | 10325380 | 10373749 | 48370 | 1 | 19116 | ENSMUST00 | 2126 | ENSMUSG0 | Prlr | prolactin receptor |
| chr15 | 10335611 | 10335853 | 243 | * | NA_peak_19 | 18 | 3' UTR | 15 | 10325380 | 10373749 | 48370 | 1 | 19116 | ENSMUST00 | 10231 | ENSMUSG0 | Prlr | prolactin receptor |
| chr15 | 10367160 | 10367487 | 328 | * | NA_peak_19 | 19 | Intron (ENSM | 15 | 10358579 | 10409738 | 51160 | 1 | 268782 | ENSMUST00 | 8581 | ENSMUSG0 | Agxt2 | alanine-glyoxylate aminotransferase 2 |
| chr15 | 10368109 | 10368763 | 655 | * | NA_peak_19 | 26 | Intron (ENSM | 15 | 10358579 | 10409738 | 51160 | 1 | 268782 | ENSMUST00 | 9530 | ENSMUSG0 | Agxt2 | alanine-glyoxylate aminotransferase 2 |
| chr15 | 10461380 | 10461809 | 430 | * | NA_peak_19 | 45 | Promoter (<= | 15 | 10446765 | 10461407 | 14643 | 2 | 78244 | ENSMUST00 | 0 | ENSMUSG0 | Dnajc21 | DnaJ heat shock protein family (Hsp40) member C21 |
| chr15 | 10513796 | 10513843 | 48 | * | NA_peak_19 | 27 | Intron (ENSM | 15 | 10486609 | 10493674 | 7066 | 1 | 19355 | ENSMUST00 | 27187 | ENSMUSG0 | Rad1 | RAD1 checkpoint DNA exonuclease |
| chr15 | 10529497 | 10529601 | 105 | * | NA_peak_19 | 18 | Intron (ENSM | 15 | 10537626 | 10558650 | 21025 | 2 | 75777 | ENSMUST00 | 29049 | ENSMUSG0 | Ttc23l | tetratricopeptide repeat domain 23-like |
| chr15 | 10604885 | 10605422 | 538 | * | NA_peak_19 | 19 | Intron (ENSM | 15 | 10600337 | 10607491 | 7155 | 1 | 100038608 | ENSMUST00 | 4548 | ENSMUSG0 | Gm10389 | predicted gene 10389 |
| chr15 | 10606886 | 10606925 | 40 | * | NA_peak_19 | 26 | Exon (ENSMU | 15 | 10600337 | 10607491 | 7155 | 1 | 100038608 | ENSMUST00 | 6549 | ENSMUSG0 | Gm10389 | predicted gene 10389 |
| chr15 | 10652912 | 10653271 | 360 | * | NA_peak_19 | 30 | Intron (ENSM | 15 | 10600337 | 10607491 | 7155 | 1 | 100038608 | ENSMUST00 | 52575 | ENSMUSG0 | Gm10389 | predicted gene 10389 |
| chr15 | 10855139 | 10855412 | 274 | * | NA_peak_19 | 26 | Distal Interge | 15 | 10814827 | 10835214 | 20388 | 1 | 100502601 | ENSMUST00 | 40312 | NA | Gm19276 | predicted gene, 19276 |
| chr15 | 10878194 | 10878570 | 377 | * | NA_peak_19 | 28 | Distal Interge | 15 | 10814827 | 10835214 | 20388 | 1 | 100502601 | ENSMUST00 | 63367 | NA | Gm19276 | predicted gene, 19276 |
| chr15 | 10968149 | 10968392 | 244 | * | NA_peak_19 | 22 | Intron (ENSM | 15 | 10981756 | 10996626 | 14871 | 1 | 17117 | ENSMUST00 | -13364 | ENSMUSG0 | Amacr | alpha-methylacyl-CoA racemase |
| chr15 | 10990643 | 10990910 | 268 | * | NA_peak_19 | 37 | Intron (ENSM | 15 | 10983406 | 10995007 | 11602 | 1 | 17117 | ENSMUST00 | 7237 | ENSMUSG0 | Amacr | alpha-methylacyl-CoA racemase |
| chr15 | 11051690 | 11051900 | 211 | * | NA_peak_19 | 16 | Distal Interge | 15 | 11064790 | 11153010 | 88221 | 1 | 239337 | ENSMUST00 | -12890 | ENSMUSG0 | Adamts12 | a disintegrin-like and metallopeptidase (reprolysin type) with thrombospondin type 1 motif, 12 |
| chr15 | 11202166 | 11202457 | 292 | * | NA_peak_19 | 22 | Intron (ENSM | 15 | 11064818 | 11217121 | 152304 | 1 | 239337 | ENSMUST00 | 137348 | ENSMUSG0 | Adamts12 | a disintegrin-like and metallopeptidase (reprolysin type) with thrombospondin type 1 motif, 12 |
| chr15 | 11212414 | 11213521 | 1108 | * | NA_peak_19 | 53 | Intron (ENSM | 15 | 11064818 | 11217121 | 152304 | 1 | 239337 | ENSMUST00 | 147596 | ENSMUSG0 | Adamts12 | a disintegrin-like and metallopeptidase (reprolysin type) with thrombospondin type 1 motif, 12 |
| chr15 | 11217872 | 11218155 | 284 | * | NA_peak_19 | 22 | Exon (ENSMU | 15 | 11064818 | 11217121 | 152304 | 1 | 239337 | ENSMUST00 | 153054 | ENSMUSG0 | Adamts12 | a disintegrin-like and metallopeptidase (reprolysin type) with thrombospondin type 1 motif, 12 |
| chr15 | 11223535 | 11223829 | 295 | * | NA_peak_19 | 20 | Intron (ENSM | 15 | 11064818 | 11217121 | 152304 | 1 | 239337 | ENSMUST00 | 158717 | ENSMUSG0 | Adamts12 | a disintegrin-like and metallopeptidase (reprolysin type) with thrombospondin type 1 motif, 12 |
| chr15 | 11246136 | 11246403 | 268 | * | NA_peak_19 | 30 | Intron (ENSM | 15 | 11384067 | 11387345 | 3279 | 2 | 110960 | ENSMUST00 | 140942 | ENSMUSG0 | Tars | threonyl-tRNA synthetase |
| chr15 | 11255508 | 11256052 | 545 | * | NA_peak_19 | 20 | Exon (ENSMU | 15 | 11384067 | 11387345 | 3279 | 2 | 110960 | ENSMUST00 | 131293 | ENSMUSG0 | Tars | threonyl-tRNA synthetase |
| chr15 | 11277978 | 11278145 | 168 | * | NA_peak_19 | 20 | Intron (ENSM | 15 | 11384067 | 11387345 | 3279 | 2 | 110960 | ENSMUST00 | 109200 | ENSMUSG0 | Tars | threonyl-tRNA synthetase |
| chr15 | 11289351 | 11289467 | 117 | * | NA_peak_19 | 23 | Intron (ENSM | 15 | 11384067 | 11387345 | 3279 | 2 | 110960 | ENSMUST00 | 97878 | ENSMUSG0 | Tars | threonyl-tRNA synthetase |
| chr15 | 11324078 | 11324389 | 312 | * | NA_peak_19 | 27 | Intron (ENSM | 15 | 11384067 | 11387345 | 3279 | 2 | 110960 | ENSMUST00 | 62956 | ENSMUSG0 | Tars | threonyl-tRNA synthetase |
| chr15 | 11488293 | 11488533 | 241 | * | NA_peak_19 | 23 | Distal Interge | 15 | 11382301 | 11399665 | 17365 | 2 | 110960 | ENSMUST00 | -88628 | ENSMUSG0 | Tars | threonyl-tRNA synthetase |
| chr15 | 11498282 | 11498487 | 206 | * | NA_peak_19 | 39 | Distal Interge | 15 | 11382301 | 11399665 | 17365 | 2 | 110960 | ENSMUST00 | -98617 | ENSMUSG0 | Tars | threonyl-tRNA synthetase |
| chr15 | 11513535 | 11513963 | 429 | * | NA_peak_19 | 14 | Distal Interge | 15 | 11382301 | 11399665 | 17365 | 2 | 110960 | ENSMUST00 | -113870 | ENSMUSG0 | Tars | threonyl-tRNA synthetase |

| | | | | | | | | | | | | | | | | | | |
|---|---|---|---|---|---|---|---|---|---|---|---|---|---|---|---|---|---|---|
| chr15 | 11521695 | 11521989 | 295 | * | NA_peak_193 | 36 | Distal Interge | 15 | 11382301 | 11399665 | 17365 | 2 | 110960 | ENSMUST00 | -122030 | ENSMUSG0( | Tars | threonyl-tRNA synthetase |
| chr15 | 11535376 | 11535777 | 402 | * | NA_peak_193 | 23 | Distal Interge | 15 | 11382301 | 11399665 | 17365 | 2 | 110960 | ENSMUST00 | -135711 | ENSMUSG0( | Tars | threonyl-tRNA synthetase |
| chr15 | 11548505 | 11549008 | 504 | * | NA_peak_193 | 31 | Distal Interge | 15 | 11382301 | 11399665 | 17365 | 2 | 110960 | ENSMUST00 | -148840 | ENSMUSG0( | Tars | threonyl-tRNA synthetase |
| chr15 | 11553828 | 11554607 | 780 | * | NA_peak_193 | 40 | Distal Interge | 15 | 11382301 | 11399665 | 17365 | 2 | 110960 | ENSMUST00 | -154163 | ENSMUSG0( | Tars | threonyl-tRNA synthetase |
| chr15 | 11588543 | 11588748 | 206 | * | NA_peak_194 | 20 | Distal Interge | 15 | 11382301 | 11399665 | 17365 | 2 | 110960 | ENSMUST00 | -188878 | ENSMUSG0( | Tars | threonyl-tRNA synthetase |
| chr15 | 11610096 | 11610682 | 587 | * | NA_peak_194 | 13 | Distal Interge | 15 | 11382301 | 11399665 | 17365 | 2 | 110960 | ENSMUST00 | -210431 | ENSMUSG0( | Tars | threonyl-tRNA synthetase |
| chr15 | 11767057 | 11767357 | 301 | * | NA_peak_194 | 28 | Distal Interge | 15 | 11842260 | 11890988 | 48729 | 2 | 18162 | ENSMUST00 | 123631 | ENSMUSG0( | Npr3 | natriuretic peptide receptor 3 |
| chr15 | 11798500 | 11798729 | 230 | * | NA_peak_194 | 21 | Distal Interge | 15 | 11842260 | 11890988 | 48729 | 2 | 18162 | ENSMUST00 | 92259 | ENSMUSG0( | Npr3 | natriuretic peptide receptor 3 |
| chr15 | 11820627 | 11820648 | 22 | * | NA_peak_194 | 40 | Distal Interge | 15 | 11842260 | 11890988 | 48729 | 2 | 18162 | ENSMUST00 | 70340 | ENSMUSG0( | Npr3 | natriuretic peptide receptor 3 |
| chr15 | 11820943 | 11820943 | 1 | * | NA_peak_194 | 16 | Distal Interge | 15 | 11842260 | 11890988 | 48729 | 2 | 18162 | ENSMUST00 | 70045 | ENSMUSG0( | Npr3 | natriuretic peptide receptor 3 |
| chr15 | 11824966 | 11825163 | 198 | * | NA_peak_194 | 36 | Distal Interge | 15 | 11842260 | 11890988 | 48729 | 2 | 18162 | ENSMUST00 | 65825 | ENSMUSG0( | Npr3 | natriuretic peptide receptor 3 |
| chr15 | 11840917 | 11841254 | 338 | * | NA_peak_194 | 23 | 3' UTR | 15 | 11842260 | 11890988 | 48729 | 2 | 18162 | ENSMUST00 | 49734 | ENSMUSG0( | Npr3 | natriuretic peptide receptor 3 |
| chr15 | 11936934 | 11937361 | 428 | * | NA_peak_194 | 28 | Distal Interge | 15 | 11964276 | 11969474 | 5199 | 1 | 73339 | ENSMUST00 | -26915 | ENSMUSG0( | 1700047G03 | RIKEN cDNA 1700047G03 gene |
| chr15 | 12061812 | 12062120 | 309 | * | NA_peak_194 | 40 | Distal Interge | 15 | 12117831 | 12185683 | 67853 | 1 | 22763 | ENSMUST00 | -55711 | ENSMUSG0( | Zfr | zinc finger RNA binding protein |
| chr15 | 12098489 | 12098558 | 70 | * | NA_peak_194 | 18 | Distal Interge | 15 | 12117831 | 12185683 | 67853 | 1 | 22763 | ENSMUST00 | -19273 | ENSMUSG0( | Zfr | zinc finger RNA binding protein |
| chr15 | 12276706 | 12276991 | 286 | * | NA_peak_194 | 19 | Distal Interge | 15 | 12321450 | 12343430 | 21981 | 1 | 66629 | ENSMUST00 | -44459 | ENSMUSG0( | Golph3 | golgi phosphoprotein 3 |
| chr15 | 12277725 | 12277807 | 83 | * | NA_peak_194 | 21 | Distal Interge | 15 | 12321450 | 12343430 | 21981 | 1 | 66629 | ENSMUST00 | -43643 | ENSMUSG0( | Golph3 | golgi phosphoprotein 3 |
| chr15 | 12325673 | 12326012 | 340 | * | NA_peak_194 | 16 | Intron (ENSM | 15 | 12321635 | 12351591 | 29957 | 1 | 66629 | ENSMUST00 | 4038 | ENSMUSG0( | Golph3 | golgi phosphoprotein 3 |
| chr15 | 12352396 | 12352680 | 285 | * | NA_peak_195 | 29 | Distal Interge | 15 | 12321635 | 12351591 | 29957 | 1 | 66629 | ENSMUST00 | 30761 | ENSMUSG0( | Golph3 | golgi phosphoprotein 3 |
| chr15 | 12360420 | 12360728 | 309 | * | NA_peak_195 | 29 | 3' UTR | 15 | 12321635 | 12351591 | 29957 | 1 | 66629 | ENSMUST00 | 38785 | ENSMUSG0( | Golph3 | golgi phosphoprotein 3 |
| chr15 | 12443601 | 12443766 | 166 | * | NA_peak_195 | 33 | Exon (ENSMU | 15 | 12404932 | 12412380 | 7449 | 2 | 68070 | ENSMUST00 | -31221 | ENSMUSG0( | Pdzd2 | PDZ domain containing 2 |
| chr15 | 12474817 | 12474922 | 106 | * | NA_peak_195 | 35 | Intron (ENSM | 15 | 12458106 | 12493559 | 35454 | 2 | 68070 | ENSMUST00 | 18637 | ENSMUSG0( | Pdzd2 | PDZ domain containing 2 |
| chr15 | 12659292 | 12659355 | 64 | * | NA_peak_195 | 30 | Intron (ENSM | 15 | 12588515 | 12599056 | 10542 | 2 | 68070 | ENSMUST00 | -60236 | ENSMUSG0( | Pdzd2 | PDZ domain containing 2 |
| chr15 | 12712120 | 12712296 | 177 | * | NA_peak_195 | 21 | Intron (ENSM | 15 | 12443331 | 12739924 | 296594 | 2 | 68070 | ENSMUST00 | 27628 | ENSMUSG0( | Pdzd2 | PDZ domain containing 2 |
| chr15 | 12715737 | 12715880 | 144 | * | NA_peak_195 | 30 | Intron (ENSM | 15 | 12443331 | 12739924 | 296594 | 2 | 68070 | ENSMUST00 | 24044 | ENSMUSG0( | Pdzd2 | PDZ domain containing 2 |
| chr15 | 12756111 | 12756493 | 383 | * | NA_peak_195 | 19 | Distal Interge | 15 | 12443331 | 12739924 | 296594 | 2 | 68070 | ENSMUST00 | -16187 | ENSMUSG0( | Pdzd2 | PDZ domain containing 2 |
| chr15 | 12805003 | 12805267 | 265 | * | NA_peak_195 | 32 | Exon (ENSMU | 15 | 12813160 | 12817900 | 4741 | 2 | 77877 | ENSMUST00 | 12633 | ENSMUSG0( | 6030458C11I | RIKEN cDNA 6030458C11 gene |
| chr15 | 12865821 | 12866153 | 333 | * | NA_peak_195 | 26 | Intron (ENSM | 15 | 12878708 | 12885230 | 6523 | 1 | 14000 | ENSMUST00 | -12555 | ENSMUSG0( | Drosha | drosha, ribonuclease type III |
| chr15 | 12969655 | 12969997 | 343 | * | NA_peak_195 | 49 | Distal Interge | 15 | 12878708 | 12885230 | 6523 | 1 | 14000 | ENSMUST00 | 90947 | ENSMUSG0( | Drosha | drosha, ribonuclease type III |
| chr15 | 12974176 | 12974469 | 294 | * | NA_peak_195 | 24 | Distal Interge | 15 | 12878708 | 12885230 | 6523 | 1 | 14000 | ENSMUST00 | 95468 | ENSMUSG0( | Drosha | drosha, ribonuclease type III |
| chr15 | 12993611 | 12994043 | 433 | * | NA_peak_195 | 27 | Distal Interge | 15 | 12878708 | 12885230 | 6523 | 1 | 14000 | ENSMUST00 | 114903 | ENSMUSG0( | Drosha | drosha, ribonuclease type III |
| chr15 | 13004785 | 13005107 | 323 | * | NA_peak_195 | 24 | Distal Interge | 15 | 12878708 | 12885230 | 6523 | 1 | 14000 | ENSMUST00 | 126077 | ENSMUSG0( | Drosha | drosha, ribonuclease type III |
| chr15 | 13041894 | 13042265 | 372 | * | NA_peak_195 | 26 | Intron (ENSM | 15 | 13028701 | 13173675 | 144975 | 2 | 12563 | ENSMUST00 | 131410 | ENSMUSG0( | Cdh6 | cadherin 6 |
| chr15 | 13111199 | 13111659 | 461 | * | NA_peak_196 | 78 | Intron (ENSM | 15 | 13028701 | 13173675 | 144975 | 2 | 12563 | ENSMUST00 | 62016 | ENSMUSG0( | Cdh6 | cadherin 6 |
| chr15 | 13186705 | 13186814 | 110 | * | NA_peak_196 | 20 | Distal Interge | 15 | 13059251 | 13173675 | 114425 | 2 | 12563 | ENSMUST00 | -13030 | ENSMUSG0( | Cdh6 | cadherin 6 |
| chr15 | 13201450 | 13201830 | 381 | * | NA_peak_196 | 50 | Distal Interge | 15 | 13059251 | 13173675 | 114425 | 2 | 12563 | ENSMUST00 | -27775 | ENSMUSG0( | Cdh6 | cadherin 6 |
| chr15 | 13339366 | 13339666 | 301 | * | NA_peak_196 | 23 | Distal Interge | 15 | 13059251 | 13173675 | 114425 | 2 | 12563 | ENSMUST00 | -165691 | ENSMUSG0( | Cdh6 | cadherin 6 |
| chr15 | 13533242 | 13533717 | 476 | * | NA_peak_196 | 20 | Distal Interge | 15 | 13059251 | 13173675 | 114425 | 2 | 12563 | ENSMUST00 | -359567 | ENSMUSG0( | Cdh6 | cadherin 6 |
| chr15 | 13755412 | 13755659 | 248 | * | NA_peak_197 | 16 | Exon (ENSMU | 15 | 13059251 | 13173675 | 114425 | 2 | 12563 | ENSMUST00 | -581737 | ENSMUSG0( | Cdh6 | cadherin 6 |
| chr15 | 13944939 | 13945413 | 475 | * | NA_peak_197 | 31 | Distal Interge | 15 | 13059251 | 13173675 | 114425 | 2 | 12563 | ENSMUST00 | -771264 | ENSMUSG0( | Cdh6 | cadherin 6 |
| chr15 | 14536468 | 14536783 | 316 | * | NA_peak_197 | 27 | Distal Interge | 15 | 13059251 | 13173675 | 114425 | 2 | 12563 | ENSMUST00 | -1362793 | ENSMUSG0( | Cdh6 | cadherin 6 |
| chr15 | 14573414 | 14573818 | 405 | * | NA_peak_197 | 20 | Distal Interge | 15 | 13059251 | 13173675 | 114425 | 2 | 12563 | ENSMUST00 | -1399739 | ENSMUSG0( | Cdh6 | cadherin 6 |
| chr15 | 14877001 | 14877781 | 781 | * | NA_peak_198 | 49 | Distal Interge | 15 | 13059251 | 13173675 | 114425 | 2 | 12563 | ENSMUST00 | -1703326 | ENSMUSG0( | Cdh6 | cadherin 6 |
| chr15 | 15059292 | 15059659 | 368 | * | NA_peak_198 | 21 | Distal Interge | 15 | 16728756 | 16857094 | 128339 | 1 | 12565 | ENSMUST00 | -1669097 | ENSMUSG0( | Cdh9 | cadherin 9 |
| chr15 | 15265981 | 15266043 | 63 | * | NA_peak_198 | 16 | Distal Interge | 15 | 16728756 | 16857094 | 128339 | 1 | 12565 | ENSMUST00 | -1462713 | ENSMUSG0( | Cdh9 | cadherin 9 |
| chr15 | 15419089 | 15419163 | 75 | * | NA_peak_198 | 16 | Distal Interge | 15 | 16728756 | 16857094 | 128339 | 1 | 12565 | ENSMUST00 | -1309593 | ENSMUSG0( | Cdh9 | cadherin 9 |
| chr15 | 15434706 | 15435217 | 512 | * | NA_peak_198 | 31 | Distal Interge | 15 | 16728756 | 16857094 | 128339 | 1 | 12565 | ENSMUST00 | -1293539 | ENSMUSG0( | Cdh9 | cadherin 9 |
| chr15 | 15550667 | 15550706 | 40 | * | NA_peak_198 | 23 | Distal Interge | 15 | 16728756 | 16857094 | 128339 | 1 | 12565 | ENSMUST00 | -1178050 | ENSMUSG0( | Cdh9 | cadherin 9 |
| chr15 | 15810661 | 15810773 | 113 | * | NA_peak_199 | 15 | Intron (ENSM | 15 | 16728756 | 16857094 | 128339 | 1 | 12565 | ENSMUST00 | -917983 | ENSMUSG0( | Cdh9 | cadherin 9 |
| chr15 | 15916684 | 15916953 | 270 | * | NA_peak_199 | 23 | Distal Interge | 15 | 16728756 | 16857094 | 128339 | 1 | 12565 | ENSMUST00 | -811803 | ENSMUSG0( | Cdh9 | cadherin 9 |
| chr15 | 16135774 | 16135851 | 78 | * | NA_peak_199 | 31 | Distal Interge | 15 | 16728756 | 16857094 | 128339 | 1 | 12565 | ENSMUST00 | -592905 | ENSMUSG0( | Cdh9 | cadherin 9 |
| chr15 | 16180821 | 16180874 | 54 | * | NA_peak_199 | 27 | Distal Interge | 15 | 16728756 | 16857094 | 128339 | 1 | 12565 | ENSMUST00 | -547882 | ENSMUSG0( | Cdh9 | cadherin 9 |
| chr15 | 16435450 | 16435800 | 351 | * | NA_peak_199 | 23 | Distal Interge | 15 | 16728756 | 16857094 | 128339 | 1 | 12565 | ENSMUST00 | -292956 | ENSMUSG0( | Cdh9 | cadherin 9 |
| chr15 | 16460318 | 16460556 | 239 | * | NA_peak_199 | 22 | Distal Interge | 15 | 16728756 | 16857094 | 128339 | 1 | 12565 | ENSMUST00 | -268200 | ENSMUSG0( | Cdh9 | cadherin 9 |
| chr15 | 16656374 | 16656445 | 72 | * | NA_peak_199 | 23 | Distal Interge | 15 | 16728756 | 16857094 | 128339 | 1 | 12565 | ENSMUST00 | -72311 | ENSMUSG0( | Cdh9 | cadherin 9 |
| chr15 | 16865263 | 16865393 | 131 | * | NA_peak_200 | 13 | Distal Interge | 15 | 16778101 | 16856767 | 78667 | 1 | 12565 | ENSMUST00 | 87162 | ENSMUSG0( | Cdh9 | cadherin 9 |
| chr15 | 16994954 | 16995111 | 158 | * | NA_peak_200 | 23 | Distal Interge | 15 | 16778101 | 16856767 | 78667 | 1 | 12565 | ENSMUST00 | 216853 | ENSMUSG0( | Cdh9 | cadherin 9 |
| chr15 | 17034777 | 17035051 | 275 | * | NA_peak_200 | 27 | Intron (ENSM | 15 | 16778101 | 16856767 | 78667 | 1 | 12565 | ENSMUST00 | 256676 | ENSMUSG0( | Cdh9 | cadherin 9 |
| chr15 | 17104442 | 17104543 | 102 | * | NA_peak_200 | 23 | Distal Interge | 15 | 16778101 | 16856767 | 78667 | 1 | 12565 | ENSMUST00 | 326341 | ENSMUSG0( | Cdh9 | cadherin 9 |
| chr15 | 17252508 | 17252936 | 429 | * | NA_peak_200 | 36 | Distal Interge | 15 | 16778101 | 16856767 | 78667 | 1 | 12565 | ENSMUST00 | 474407 | ENSMUSG0( | Cdh9 | cadherin 9 |
| chr15 | 17456279 | 17456553 | 275 | * | NA_peak_200 | 20 | Distal Interge | 15 | 16778101 | 16856767 | 78667 | 1 | 12565 | ENSMUST00 | 678178 | ENSMUSG0( | Cdh9 | cadherin 9 |
| chr15 | 17577678 | 17578177 | 500 | * | NA_peak_200 | 20 | Distal Interge | 15 | 18078118 | 18259800 | 181683 | 2 | 70886 | ENSMUST00 | 681623 | ENSMUSG0( | 4921515E04I | RIKEN cDNA 4921515E04 gene |
| chr15 | 17580522 | 17580812 | 291 | * | NA_peak_200 | 27 | Distal Interge | 15 | 18078118 | 18259800 | 181683 | 2 | 70886 | ENSMUST00 | 678988 | ENSMUSG0( | 4921515E04I | RIKEN cDNA 4921515E04 gene |
| chr15 | 17639833 | 17640228 | 396 | * | NA_peak_200 | 36 | Distal Interge | 15 | 18078118 | 18259800 | 181683 | 2 | 70886 | ENSMUST00 | 619572 | ENSMUSG0( | 4921515E04I | RIKEN cDNA 4921515E04 gene |
| chr15 | 17643421 | 17643566 | 146 | * | NA_peak_200 | 31 | Distal Interge | 15 | 18078118 | 18259800 | 181683 | 2 | 70886 | ENSMUST00 | 616234 | ENSMUSG0( | 4921515E04I | RIKEN cDNA 4921515E04 gene |
| chr15 | 17750218 | 17750509 | 292 | * | NA_peak_200 | 23 | Distal Interge | 15 | 18078118 | 18259800 | 181683 | 2 | 70886 | ENSMUST00 | 509291 | ENSMUSG0( | 4921515E04I | RIKEN cDNA 4921515E04 gene |
| chr15 | 17750799 | 17750866 | 68 | * | NA_peak_200 | 20 | Distal Interge | 15 | 18078118 | 18259800 | 181683 | 2 | 70886 | ENSMUST00 | 508934 | ENSMUSG0( | 4921515E04I | RIKEN cDNA 4921515E04 gene |
| chr15 | 17753338 | 17753349 | 12 | * | NA_peak_200 | 27 | Distal Interge | 15 | 18078118 | 18259800 | 181683 | 2 | 70886 | ENSMUST00 | 506451 | ENSMUSG0( | 4921515E04I | RIKEN cDNA 4921515E04 gene |
| chr15 | 17813269 | 17813555 | 287 | * | NA_peak_201 | 20 | Distal Interge | 15 | 18078118 | 18259800 | 181683 | 2 | 70886 | ENSMUST00 | 446245 | ENSMUSG0( | 4921515E04I | RIKEN cDNA 4921515E04 gene |
| chr15 | 17903882 | 17904297 | 416 | * | NA_peak_201 | 22 | Distal Interge | 15 | 18078118 | 18259800 | 181683 | 2 | 70886 | ENSMUST00 | 355503 | ENSMUSG0( | 4921515E04I | RIKEN cDNA 4921515E04 gene |
| chr15 | 17918945 | 17919618 | 674 | * | NA_peak_201 | 21 | Distal Interge | 15 | 18078118 | 18259800 | 181683 | 2 | 70886 | ENSMUST00 | 340182 | ENSMUSG0( | 4921515E04I | RIKEN cDNA 4921515E04 gene |
| chr15 | 17942788 | 17943115 | 328 | * | NA_peak_201 | 40 | Distal Interge | 15 | 18078118 | 18259800 | 181683 | 2 | 70886 | ENSMUST00 | 316685 | ENSMUSG0( | 4921515E04I | RIKEN cDNA 4921515E04 gene |
| chr15 | 18112323 | 18112367 | 45 | * | NA_peak_201 | 15 | Intron (ENSM | 15 | 18078118 | 18259800 | 181683 | 2 | 70886 | ENSMUST00 | 147433 | ENSMUSG0( | 4921515E04I | RIKEN cDNA 4921515E04 gene |
| chr15 | 18267409 | 18267694 | 286 | * | NA_peak_201 | 13 | Intron (ENSM | 15 | 18078118 | 18259800 | 181683 | 2 | 70886 | ENSMUST00 | -7609 | ENSMUSG0( | 4921515E04I | RIKEN cDNA 4921515E04 gene |
| chr15 | 18368310 | 18368456 | 147 | * | NA_peak_201 | 16 | Promoter (<= | 15 | 18078131 | 18368248 | 290118 | 2 | 70886 | ENSMUST00 | -62 | ENSMUSG0( | 4921515E04I | RIKEN cDNA 4921515E04 gene |
| chr15 | 18408110 | 18408477 | 368 | * | NA_peak_201 | 36 | Distal Interge | 15 | 18078137 | 18369627 | 291491 | 2 | 70886 | ENSMUST00 | -38483 | ENSMUSG0( | 4921515E04I | RIKEN cDNA 4921515E04 gene |
| chr15 | 18409584 | 18409625 | 42 | * | NA_peak_201 | 22 | Distal Interge | 15 | 18078137 | 18369627 | 291491 | 2 | 70886 | ENSMUST00 | -39957 | ENSMUSG0( | 4921515E04I | RIKEN cDNA 4921515E04 gene |
| chr15 | 18424651 | 18424927 | 277 | * | NA_peak_201 | 27 | Distal Interge | 15 | 18078137 | 18369627 | 291491 | 2 | 70886 | ENSMUST00 | -55024 | ENSMUSG0( | 4921515E04I | RIKEN cDNA 4921515E04 gene |
| chr15 | 18544734 | 18544996 | 263 | * | NA_peak_201 | 26 | Distal Interge | 15 | 18078137 | 18369627 | 291491 | 2 | 70886 | ENSMUST00 | -175107 | ENSMUSG0( | 4921515E04I | RIKEN cDNA 4921515E04 gene |
| chr15 | 18934204 | 18934469 | 266 | * | NA_peak_202 | 14 | Intron (ENSM | 15 | 18899589 | 19013682 | 114094 | 1 | 320873 | ENSMUST00 | 34615 | ENSMUSG0( | Cdh10 | cadherin 10 |
| chr15 | 19132239 | 19132637 | 399 | * | NA_peak_202 | 21 | Distal Interge | 15 | 19185373 | 19203329 | 17957 | 1 | 102639102 | ENSMUST00 | -52736 | ENSMUSG0( | Gm35496 | predicted gene, 35496 |
| chr15 | 19513178 | 19513452 | 275 | * | NA_peak_202 | 27 | Distal Interge | 15 | 19185373 | 19203329 | 17957 | 1 | 102639102 | ENSMUST00 | 327805 | ENSMUSG0( | Gm35496 | predicted gene, 35496 |
| chr15 | 19515940 | 19516332 | 393 | * | NA_peak_202 | 27 | Distal Interge | 15 | 19185373 | 19203329 | 17957 | 1 | 102639102 | ENSMUST00 | 330567 | ENSMUSG0( | Gm35496 | predicted gene, 35496 |
| chr15 | 19637868 | 19638249 | 382 | * | NA_peak_202 | 20 | Distal Interge | 15 | 19185373 | 19203329 | 17957 | 1 | 102639102 | ENSMUST00 | 452495 | ENSMUSG0( | Gm35496 | predicted gene, 35496 |
| chr15 | 19646759 | 19647084 | 326 | * | NA_peak_202 | 20 | Distal Interge | 15 | 19185373 | 19203329 | 17957 | 1 | 102639102 | ENSMUST00 | 461386 | ENSMUSG0( | Gm35496 | predicted gene, 35496 |
| chr15 | 19893447 | 19893675 | 229 | * | NA_peak_203 | 18 | Distal Interge | 15 | 19185373 | 19203329 | 17957 | 1 | 102639102 | ENSMUST00 | 708074 | ENSMUSG0( | Gm35496 | predicted gene, 35496 |
| chr15 | 20062133 | 20062326 | 194 | * | NA_peak_203 | 36 | Distal Interge | 15 | 20665214 | 20666750 | 1537 | 2 | 64833 | ENSMUST00 | 604424 | ENSMUSG0( | Acot10 | acyl-CoA thioesterase 10 |
| chr15 | 20373194 | 20373466 | 273 | * | NA_peak_203 | 23 | Distal Interge | 15 | 20665214 | 20666750 | 1537 | 2 | 64833 | ENSMUST00 | 293284 | ENSMUSG0( | Acot10 | acyl-CoA thioesterase 10 |
| chr15 | 20405562 | 20405877 | 316 | * | NA_peak_203 | 24 | Distal Interge | 15 | 20665214 | 20666750 | 1537 | 2 | 64833 | ENSMUST00 | 260873 | ENSMUSG0( | Acot10 | acyl-CoA thioesterase 10 |
| chr15 | 20514031 | 20514138 | 108 | * | NA_peak_203 | 20 | Intron (ENSM | 15 | 20665214 | 20666750 | 1537 | 2 | 64833 | ENSMUST00 | 152612 | ENSMUSG0( | Acot10 | acyl-CoA thioesterase 10 |
| chr15 | 20629713 | 20630131 | 419 | * | NA_peak_203 | 27 | Intron (ENSM | 15 | 20665214 | 20666750 | 1537 | 2 | 64833 | ENSMUST00 | 36619 | ENSMUSG0( | Acot10 | acyl-CoA thioesterase 10 |
| chr15 | 21183169 | 21183455 | 287 | * | NA_peak_204 | 13 | Intron (ENSM | 15 | 21111452 | 21589530 | 478079 | 1 | 215654 | ENSMUST00 | 71717 | ENSMUSG0( | Cdh12 | cadherin 12 |
| chr15 | 21336730 | 21337058 | 329 | * | NA_peak_204 | 45 | Intron (ENSM | 15 | 21111452 | 21589530 | 478079 | 1 | 215654 | ENSMUST00 | 225278 | ENSMUSG0( | Cdh12 | cadherin 12 |
| chr15 | 21447752 | 21447945 | 194 | * | NA_peak_204 | 30 | Intron (ENSM | 15 | 21111452 | 21589530 | 478079 | 1 | 215654 | ENSMUST00 | 336300 | ENSMUSG0( | Cdh12 | cadherin 12 |
| chr15 | 21452818 | 21452936 | 119 | * | NA_peak_204 | 22 | Intron (ENSM | 15 | 21111452 | 21589530 | 478079 | 1 | 215654 | ENSMUST00 | 341366 | ENSMUSG0( | Cdh12 | cadherin 12 |
| chr15 | 21526282 | 21526636 | 355 | * | NA_peak_204 | 31 | Intron (ENSM | 15 | 21111452 | 21589530 | 478079 | 1 | 215654 | ENSMUST00 | 414830 | ENSMUSG0( | Cdh12 | cadherin 12 |
| chr15 | 22007643 | 22008029 | 387 | * | NA_peak_204 | 20 | Distal Interge | 15 | 22549022 | 23400567 | 851546 | 1 | 320865 | ENSMUST00 | -540993 | ENSMUSG0( | Cdh18 | cadherin 18 |
| chr15 | 22288143 | 22288331 | 189 | * | NA_peak_205 | 23 | Distal Interge | 15 | 22549022 | 23400567 | 851546 | 1 | 320865 | ENSMUST00 | -260691 | ENSMUSG0( | Cdh18 | cadherin 18 |
| chr15 | 22395837 | 22395878 | 42 | * | NA_peak_205 | 20 | Distal Interge | 15 | 22549022 | 23400567 | 851546 | 1 | 320865 | ENSMUST00 | -153144 | ENSMUSG0( | Cdh18 | cadherin 18 |
| chr15 | 22594903 | 22594993 | 91 | * | NA_peak_205 | 20 | Intron (ENSM | 15 | 22549022 | 23400567 | 851546 | 1 | 320865 | ENSMUST00 | 45881 | ENSMUSG0( | Cdh18 | cadherin 18 |
| chr15 | 22699052 | 22699372 | 321 | * | NA_peak_205 | 48 | Intron (ENSM | 15 | 22713868 | 22714830 | 963 | 2 | 545091 | ENSMUST00 | 15458 | ENSMUSG0( | Hnrnpa1l2-ps | heterogeneous nuclear ribonucleoprotein A1-like 2, pseudogene 2 |
| chr15 | 22766872 | 22767153 | 282 | * | NA_peak_205 | 23 | Intron (ENSM | 15 | 22713868 | 22714830 | 963 | 2 | 545091 | ENSMUST00 | -52042 | ENSMUSG0( | Hnrnpa1l2-ps | heterogeneous nuclear ribonucleoprotein A1-like 2, pseudogene 2 |
| chr15 | 22806167 | 22806784 | 618 | * | NA_peak_205 | 36 | Intron (ENSM | 15 | 22713868 | 22714830 | 963 | 2 | 545091 | ENSMUST00 | -91337 | ENSMUSG0( | Hnrnpa1l2-ps | heterogeneous nuclear ribonucleoprotein A1-like 2, pseudogene 2 |
| chr15 | 22872257 | 22872569 | 313 | * | NA_peak_205 | 36 | Intron (ENSM | 15 | 22713868 | 22714830 | 963 | 2 | 545091 | ENSMUST00 | -157427 | ENSMUSG0( | Hnrnpa1l2-ps | heterogeneous nuclear ribonucleoprotein A1-like 2, pseudogene 2 |
| chr15 | 23222123 | 23222236 | 114 | * | NA_peak_206 | 20 | Intron (ENSM | 15 | 23173864 | 23474029 | 300166 | 1 | 320865 | ENSMUST00 | 48259 | ENSMUSG0( | Cdh18 | cadherin 18 |
| chr15 | 23445592 | 23445686 | 95 | * | NA_peak_206 | 31 | Intron (ENSM | 15 | 23173864 | 23474029 | 300166 | 1 | 320865 | ENSMUST00 | 271728 | ENSMUSG0( | Cdh18 | cadherin 18 |

| | | | | | | | | | | | | | | | | | | |
|---|---|---|---|---|---|---|---|---|---|---|---|---|---|---|---|---|---|---|
| chr15 | 23463384 | 23463581 | 198 | * | NA_peak_206 | 23 | Intron (ENSM | 15 | 23173864 | 23474029 | 300166 | 1 | 320865 | ENSMUST00 | 289520 | ENSMUSG0 | Cdh18 | cadherin 18 |
| chr15 | 23464866 | 23465186 | 321 | * | NA_peak_206 | 31 | Intron (ENSM | 15 | 23173864 | 23474029 | 300166 | 1 | 320865 | ENSMUST00 | 291002 | ENSMUSG0 | Cdh18 | cadherin 18 |
| chr15 | 23575024 | 23575376 | 353 | * | NA_peak_206 | 59 | Distal Interge | 15 | 23173864 | 23474029 | 300166 | 1 | 320865 | ENSMUST00 | 401160 | ENSMUSG0 | Cdh18 | cadherin 18 |
| chr15 | 23776009 | 23776526 | 518 | * | NA_peak_206 | 30 | Distal Interge | 15 | 23173864 | 23474029 | 300166 | 1 | 320865 | ENSMUST00 | 602145 | ENSMUSG0 | Cdh18 | cadherin 18 |
| chr15 | 23793252 | 23793295 | 44 | * | NA_peak_206 | 20 | Distal Interge | 15 | 23173864 | 23474029 | 300166 | 1 | 320865 | ENSMUST00 | 619388 | ENSMUSG0 | Cdh18 | cadherin 18 |
| chr15 | 23825736 | 23826050 | 315 | * | NA_peak_206 | 20 | Distal Interge | 15 | 23173864 | 23474029 | 300166 | 1 | 320865 | ENSMUST00 | 651872 | ENSMUSG0 | Cdh18 | cadherin 18 |
| chr15 | 23862740 | 23863042 | 303 | * | NA_peak_206 | 26 | Distal Interge | 15 | 23173864 | 23474029 | 300166 | 1 | 320865 | ENSMUST00 | 688876 | ENSMUSG0 | Cdh18 | cadherin 18 |
| chr15 | 23929438 | 23929687 | 250 | * | NA_peak_206 | 23 | Distal Interge | 15 | 23173864 | 23474029 | 300166 | 1 | 320865 | ENSMUST00 | 755574 | ENSMUSG0 | Cdh18 | cadherin 18 |
| chr15 | 23950712 | 23951116 | 405 | * | NA_peak_206 | 31 | Distal Interge | 15 | 23173864 | 23474029 | 300166 | 1 | 320865 | ENSMUST00 | 776848 | ENSMUSG0 | Cdh18 | cadherin 18 |
| chr15 | 23958229 | 23958818 | 590 | * | NA_peak_206 | 39 | Distal Interge | 15 | 23173864 | 23474029 | 300166 | 1 | 320865 | ENSMUST00 | 784365 | ENSMUSG0 | Cdh18 | cadherin 18 |
| chr15 | 23969804 | 23970470 | 667 | * | NA_peak_206 | 20 | Distal Interge | 15 | 23173864 | 23474029 | 300166 | 1 | 320865 | ENSMUST00 | 795940 | ENSMUSG0 | Cdh18 | cadherin 18 |
| chr15 | 23974024 | 23974329 | 306 | * | NA_peak_206 | 20 | Distal Interge | 15 | 23173864 | 23474029 | 300166 | 1 | 320865 | ENSMUST00 | 800160 | ENSMUSG0 | Cdh18 | cadherin 18 |
| chr15 | 24113757 | 24114029 | 273 | * | NA_peak_207 | 40 | Distal Interge | 15 | 23173864 | 23474029 | 300166 | 1 | 320865 | ENSMUST00 | 939893 | ENSMUSG0 | Cdh18 | cadherin 18 |
| chr15 | 24351088 | 24351501 | 414 | * | NA_peak_207 | 31 | Intron (ENSM | 15 | 25364857 | 25413698 | 48842 | 2 | 70350 | ENSMUST00 | 1062197 | ENSMUSG0 | Basp1 | brain abundant, membrane attached signal protein 1 |
| chr15 | 24359653 | 24359667 | 15 | * | NA_peak_207 | 22 | Intron (ENSM | 15 | 25364857 | 25413698 | 48842 | 2 | 70350 | ENSMUST00 | 1054031 | ENSMUSG0 | Basp1 | brain abundant, membrane attached signal protein 1 |
| chr15 | 24366646 | 24366885 | 240 | * | NA_peak_207 | 21 | Intron (ENSM | 15 | 25364857 | 25413698 | 48842 | 2 | 70350 | ENSMUST00 | 1046813 | ENSMUSG0 | Basp1 | brain abundant, membrane attached signal protein 1 |
| chr15 | 24529105 | 24529185 | 81 | * | NA_peak_207 | 23 | Distal Interge | 15 | 25364857 | 25413698 | 48842 | 2 | 70350 | ENSMUST00 | 884513 | ENSMUSG0 | Basp1 | brain abundant, membrane attached signal protein 1 |
| chr15 | 24637458 | 24637756 | 299 | * | NA_peak_207 | 31 | Distal Interge | 15 | 25364857 | 25413698 | 48842 | 2 | 70350 | ENSMUST00 | 775942 | ENSMUSG0 | Basp1 | brain abundant, membrane attached signal protein 1 |
| chr15 | 24647183 | 24647473 | 291 | * | NA_peak_207 | 17 | Distal Interge | 15 | 25364857 | 25413698 | 48842 | 2 | 70350 | ENSMUST00 | 766225 | ENSMUSG0 | Basp1 | brain abundant, membrane attached signal protein 1 |
| chr15 | 24665543 | 24665543 | 1 | * | NA_peak_207 | 30 | Distal Interge | 15 | 25364857 | 25413698 | 48842 | 2 | 70350 | ENSMUST00 | 748155 | ENSMUSG0 | Basp1 | brain abundant, membrane attached signal protein 1 |
| chr15 | 24668126 | 24668142 | 17 | * | NA_peak_208 | 20 | Distal Interge | 15 | 25364857 | 25413698 | 48842 | 2 | 70350 | ENSMUST00 | 745556 | ENSMUSG0 | Basp1 | brain abundant, membrane attached signal protein 1 |
| chr15 | 24713373 | 24713675 | 303 | * | NA_peak_208 | 40 | Distal Interge | 15 | 25364857 | 25413698 | 48842 | 2 | 70350 | ENSMUST00 | 700023 | ENSMUSG0 | Basp1 | brain abundant, membrane attached signal protein 1 |
| chr15 | 24734740 | 24735105 | 366 | * | NA_peak_208 | 31 | Distal Interge | 15 | 25364857 | 25413698 | 48842 | 2 | 70350 | ENSMUST00 | 678593 | ENSMUSG0 | Basp1 | brain abundant, membrane attached signal protein 1 |
| chr15 | 24823032 | 24823043 | 12 | * | NA_peak_208 | 19 | Distal Interge | 15 | 25364857 | 25413698 | 48842 | 2 | 70350 | ENSMUST00 | 590655 | ENSMUSG0 | Basp1 | brain abundant, membrane attached signal protein 1 |
| chr15 | 24896665 | 24896937 | 273 | * | NA_peak_208 | 58 | Intron (ENSM | 15 | 25364857 | 25413698 | 48842 | 2 | 70350 | ENSMUST00 | 516761 | ENSMUSG0 | Basp1 | brain abundant, membrane attached signal protein 1 |
| chr15 | 24901940 | 24902225 | 286 | * | NA_peak_208 | 23 | Intron (ENSM | 15 | 25364857 | 25413698 | 48842 | 2 | 70350 | ENSMUST00 | 511473 | ENSMUSG0 | Basp1 | brain abundant, membrane attached signal protein 1 |
| chr15 | 24906095 | 24906463 | 369 | * | NA_peak_208 | 40 | Intron (ENSM | 15 | 25364857 | 25413698 | 48842 | 2 | 70350 | ENSMUST00 | 507235 | ENSMUSG0 | Basp1 | brain abundant, membrane attached signal protein 1 |
| chr15 | 24913059 | 24913415 | 357 | * | NA_peak_208 | 40 | Distal Interge | 15 | 25364857 | 25413698 | 48842 | 2 | 70350 | ENSMUST00 | 500283 | ENSMUSG0 | Basp1 | brain abundant, membrane attached signal protein 1 |
| chr15 | 25033193 | 25033218 | 26 | * | NA_peak_208 | 31 | Distal Interge | 15 | 25364857 | 25413698 | 48842 | 2 | 70350 | ENSMUST00 | 380480 | ENSMUSG0 | Basp1 | brain abundant, membrane attached signal protein 1 |
| chr15 | 25042142 | 25042227 | 86 | * | NA_peak_208 | 26 | Distal Interge | 15 | 25364857 | 25413698 | 48842 | 2 | 70350 | ENSMUST00 | 371471 | ENSMUSG0 | Basp1 | brain abundant, membrane attached signal protein 1 |
| chr15 | 25141160 | 25141433 | 274 | * | NA_peak_208 | 36 | Intron (ENSM | 15 | 25364857 | 25413698 | 48842 | 2 | 70350 | ENSMUST00 | 272265 | ENSMUSG0 | Basp1 | brain abundant, membrane attached signal protein 1 |
| chr15 | 25429469 | 25429628 | 160 | * | NA_peak_209 | 23 | Intron (ENSM | 15 | 25415063 | 25433520 | 18458 | 1 | 432939 | ENSMUST00 | 14406 | NA | Gm48957 | predicted gene, 48957 |
| chr15 | 25753977 | 25754322 | 346 | * | NA_peak_209 | 35 | Promoter (<= | 15 | 25752979 | 25779774 | 26796 | 1 | 17909 | ENSMUST00 | 998 | ENSMUSG0 | Myo10 | myosin X |
| chr15 | 26077173 | 26077300 | 128 | * | NA_peak_209 | 39 | Distal Interge | 15 | 26054365 | 26062438 | 8074 | 2 | 105245898 | ENSMUST00 | -14735 | ENSMUSG0 | Gm41279 | predicted gene, 41279 |
| chr15 | 26101455 | 26101799 | 345 | * | NA_peak_209 | 27 | Distal Interge | 15 | 26054365 | 26062438 | 8074 | 2 | 105245898 | ENSMUST00 | -39017 | ENSMUSG0 | Gm41279 | predicted gene, 41279 |
| chr15 | 26190892 | 26191137 | 246 | * | NA_peak_209 | 16 | Distal Interge | 15 | 26309048 | 26312371 | 3324 | 1 | 211147 | ENSMUST00 | -117911 | ENSMUSG0 | Marchf11 | membrane associated ring-CH-type finger 11 |
| chr15 | 26194365 | 26194659 | 295 | * | NA_peak_209 | 31 | Distal Interge | 15 | 26309048 | 26312371 | 3324 | 1 | 211147 | ENSMUST00 | -114389 | ENSMUSG0 | Marchf11 | membrane associated ring-CH-type finger 11 |
| chr15 | 26239591 | 26240176 | 586 | * | NA_peak_209 | 22 | Distal Interge | 15 | 26309048 | 26312371 | 3324 | 1 | 211147 | ENSMUST00 | -68872 | ENSMUSG0 | Marchf11 | membrane associated ring-CH-type finger 11 |
| chr15 | 26243421 | 26243809 | 389 | * | NA_peak_209 | 23 | Distal Interge | 15 | 26309048 | 26312371 | 3324 | 1 | 211147 | ENSMUST00 | -65239 | ENSMUSG0 | Marchf11 | membrane associated ring-CH-type finger 11 |
| chr15 | 26291636 | 26291832 | 197 | * | NA_peak_209 | 27 | Distal Interge | 15 | 26309048 | 26312371 | 3324 | 1 | 211147 | ENSMUST00 | -17216 | ENSMUSG0 | Marchf11 | membrane associated ring-CH-type finger 11 |
| chr15 | 26404521 | 26404756 | 236 | * | NA_peak_210 | 50 | Intron (ENSM | 15 | 26387719 | 26409576 | 21858 | 1 | 211147 | ENSMUST00 | 16802 | ENSMUSG0 | Marchf11 | membrane associated ring-CH-type finger 11 |
| chr15 | 26476837 | 26477046 | 210 | * | NA_peak_210 | 14 | Distal Interge | 15 | 26387719 | 26409576 | 21858 | 1 | 211147 | ENSMUST00 | 89118 | ENSMUSG0 | Marchf11 | membrane associated ring-CH-type finger 11 |
| chr15 | 26509099 | 26509102 | 4 | * | NA_peak_210 | 36 | Distal Interge | 15 | 26387719 | 26409576 | 21858 | 1 | 211147 | ENSMUST00 | 121380 | ENSMUSG0 | Marchf11 | membrane associated ring-CH-type finger 11 |
| chr15 | 26608495 | 26608573 | 79 | * | NA_peak_210 | 22 | Intron (ENSM | 15 | 26387719 | 26409576 | 21858 | 1 | 211147 | ENSMUST00 | 220776 | ENSMUSG0 | Marchf11 | membrane associated ring-CH-type finger 11 |
| chr15 | 26620476 | 26620882 | 407 | * | NA_peak_210 | 27 | Exon (ENSMU | 15 | 26387719 | 26409576 | 21858 | 1 | 211147 | ENSMUST00 | 232757 | ENSMUSG0 | Marchf11 | membrane associated ring-CH-type finger 11 |
| chr15 | 26879066 | 26879609 | 544 | * | NA_peak_210 | 23 | Intron (ENSM | 15 | 26787619 | 26895518 | 107900 | 2 | 448987 | ENSMUST00 | 15909 | ENSMUSG0 | Fbxl7 | F-box and leucine-rich repeat protein 7 |
| chr15 | 26901122 | 26901381 | 260 | * | NA_peak_210 | 20 | Distal Interge | 15 | 26540459 | 26895580 | 355122 | 2 | 448987 | ENSMUST00 | -5542 | ENSMUSG0 | Fbxl7 | F-box and leucine-rich repeat protein 7 |
| chr15 | 26905447 | 26905795 | 349 | * | NA_peak_210 | 16 | Distal Interge | 15 | 26540459 | 26895580 | 355122 | 2 | 448987 | ENSMUST00 | -9867 | ENSMUSG0 | Fbxl7 | F-box and leucine-rich repeat protein 7 |
| chr15 | 26922179 | 26922258 | 80 | * | NA_peak_210 | 27 | Distal Interge | 15 | 26540459 | 26895580 | 355122 | 2 | 448987 | ENSMUST00 | -26599 | ENSMUSG0 | Fbxl7 | F-box and leucine-rich repeat protein 7 |
| chr15 | 27057980 | 27058319 | 340 | * | NA_peak_210 | 23 | Distal Interge | 15 | 26540459 | 26895580 | 355122 | 2 | 448987 | ENSMUST00 | -162400 | ENSMUSG0 | Fbxl7 | F-box and leucine-rich repeat protein 7 |
| chr15 | 27068384 | 27068699 | 316 | * | NA_peak_210 | 14 | Distal Interge | 15 | 26540459 | 26895580 | 355122 | 2 | 448987 | ENSMUST00 | -172804 | ENSMUSG0 | Fbxl7 | F-box and leucine-rich repeat protein 7 |
| chr15 | 27089585 | 27089691 | 107 | * | NA_peak_211 | 20 | Distal Interge | 15 | 26540459 | 26895580 | 355122 | 2 | 448987 | ENSMUST00 | -194005 | ENSMUSG0 | Fbxl7 | F-box and leucine-rich repeat protein 7 |
| chr15 | 27206595 | 27206761 | 167 | * | NA_peak_211 | 22 | Distal Interge | 15 | 27466677 | 27594909 | 128233 | 1 | 11732 | ENSMUST00 | -259916 | ENSMUSG0 | Ank | progressive ankylosis |
| chr15 | 27228376 | 27228638 | 263 | * | NA_peak_211 | 30 | Distal Interge | 15 | 27466677 | 27594909 | 128233 | 1 | 11732 | ENSMUST00 | -238039 | ENSMUSG0 | Ank | progressive ankylosis |
| chr15 | 27236477 | 27236489 | 13 | * | NA_peak_211 | 16 | Distal Interge | 15 | 27466677 | 27594909 | 128233 | 1 | 11732 | ENSMUST00 | -230188 | ENSMUSG0 | Ank | progressive ankylosis |
| chr15 | 27273291 | 27273557 | 267 | * | NA_peak_211 | 40 | Distal Interge | 15 | 27466677 | 27594909 | 128233 | 1 | 11732 | ENSMUST00 | -193120 | ENSMUSG0 | Ank | progressive ankylosis |
| chr15 | 27280645 | 27280728 | 84 | * | NA_peak_211 | 23 | Distal Interge | 15 | 27466677 | 27594909 | 128233 | 1 | 11732 | ENSMUST00 | -185949 | ENSMUSG0 | Ank | progressive ankylosis |
| chr15 | 27374030 | 27374054 | 25 | * | NA_peak_211 | 28 | Distal Interge | 15 | 27466677 | 27594909 | 128233 | 1 | 11732 | ENSMUST00 | -92623 | ENSMUSG0 | Ank | progressive ankylosis |
| chr15 | 27733779 | 27733815 | 37 | * | NA_peak_211 | 34 | Promoter (2-3 | 15 | 27735050 | 27736522 | 1473 | 2 | 223435 | ENSMUST00 | 2707 | ENSMUSG0 | Trio | triple functional domain (PTPRF interacting) |
| chr15 | 28242699 | 28242805 | 107 | * | NA_peak_212 | 24 | Exon (ENSMU | 15 | 28203752 | 28472052 | 268301 | 1 | 110082 | ENSMUST00 | 38947 | ENSMUSG0 | Dnah5 | dynein, axonemal, heavy chain 5 |
| chr15 | 28244967 | 28245269 | 303 | * | NA_peak_212 | 18 | Intron (ENSM | 15 | 28203752 | 28472052 | 268301 | 1 | 110082 | ENSMUST00 | 41215 | ENSMUSG0 | Dnah5 | dynein, axonemal, heavy chain 5 |
| chr15 | 28380098 | 28380203 | 106 | * | NA_peak_212 | 35 | Intron (ENSM | 15 | 28203752 | 28472052 | 268301 | 1 | 110082 | ENSMUST00 | 176346 | ENSMUSG0 | Dnah5 | dynein, axonemal, heavy chain 5 |
| chr15 | 28405513 | 28405830 | 318 | * | NA_peak_212 | 17 | Intron (ENSM | 15 | 28203752 | 28472052 | 268301 | 1 | 110082 | ENSMUST00 | 201761 | ENSMUSG0 | Dnah5 | dynein, axonemal, heavy chain 5 |
| chr15 | 28527036 | 28527138 | 103 | * | NA_peak_212 | 36 | Distal Interge | 15 | 28203752 | 28472052 | 268301 | 1 | 110082 | ENSMUST00 | 323284 | ENSMUSG0 | Dnah5 | dynein, axonemal, heavy chain 5 |
| chr15 | 28573727 | 28574049 | 323 | * | NA_peak_212 | 27 | Distal Interge | 15 | 28203752 | 28472052 | 268301 | 1 | 110082 | ENSMUST00 | 369975 | ENSMUSG0 | Dnah5 | dynein, axonemal, heavy chain 5 |
| chr15 | 28624494 | 28624713 | 220 | * | NA_peak_212 | 23 | Distal Interge | 15 | 28203752 | 28472052 | 268301 | 1 | 110082 | ENSMUST00 | 420742 | ENSMUSG0 | Dnah5 | dynein, axonemal, heavy chain 5 |
| chr15 | 28662074 | 28662496 | 423 | * | NA_peak_212 | 16 | Distal Interge | 15 | 28203752 | 28472052 | 268301 | 1 | 110082 | ENSMUST00 | 458322 | ENSMUSG0 | Dnah5 | dynein, axonemal, heavy chain 5 |
| chr15 | 28780158 | 28780476 | 319 | * | NA_peak_212 | 27 | Distal Interge | 15 | 28203752 | 28472052 | 268301 | 1 | 110082 | ENSMUST00 | 576406 | ENSMUSG0 | Dnah5 | dynein, axonemal, heavy chain 5 |
| chr15 | 28786147 | 28786563 | 417 | * | NA_peak_212 | 31 | Distal Interge | 15 | 28203752 | 28472052 | 268301 | 1 | 110082 | ENSMUST00 | 582395 | ENSMUSG0 | Dnah5 | dynein, axonemal, heavy chain 5 |
| chr15 | 28824974 | 28825338 | 365 | * | NA_peak_213 | 50 | Distal Interge | 15 | 28203752 | 28472052 | 268301 | 1 | 110082 | ENSMUST00 | 621222 | ENSMUSG0 | Dnah5 | dynein, axonemal, heavy chain 5 |
| chr15 | 28827383 | 28827987 | 605 | * | NA_peak_213 | 31 | Distal Interge | 15 | 28203752 | 28472052 | 268301 | 1 | 110082 | ENSMUST00 | 623631 | ENSMUSG0 | Dnah5 | dynein, axonemal, heavy chain 5 |
| chr15 | 28924424 | 28924601 | 178 | * | NA_peak_213 | 31 | Distal Interge | 15 | 28203752 | 28472052 | 268301 | 1 | 110082 | ENSMUST00 | 720672 | ENSMUSG0 | Dnah5 | dynein, axonemal, heavy chain 5 |
| chr15 | 29150991 | 29151305 | 315 | * | NA_peak_213 | 23 | Distal Interge | 15 | 29713331 | 29713404 | 74 | 2 | 100628607 | ENSMUST00 | 562099 | ENSMUSG0 | Mir3964 | microRNA 3964 |
| chr15 | 29172090 | 29172470 | 381 | * | NA_peak_213 | 50 | Distal Interge | 15 | 29713331 | 29713404 | 74 | 2 | 100628607 | ENSMUST00 | 540934 | ENSMUSG0 | Mir3964 | microRNA 3964 |
| chr15 | 29253132 | 29253450 | 319 | * | NA_peak_213 | 27 | Distal Interge | 15 | 29713331 | 29713404 | 74 | 2 | 100628607 | ENSMUST00 | 459954 | ENSMUSG0 | Mir3964 | microRNA 3964 |
| chr15 | 29289478 | 29289572 | 95 | * | NA_peak_213 | 40 | Distal Interge | 15 | 29713331 | 29713404 | 74 | 2 | 100628607 | ENSMUST00 | 423832 | ENSMUSG0 | Mir3964 | microRNA 3964 |
| chr15 | 29290133 | 29290583 | 451 | * | NA_peak_213 | 40 | Distal Interge | 15 | 29713331 | 29713404 | 74 | 2 | 100628607 | ENSMUST00 | 422821 | ENSMUSG0 | Mir3964 | microRNA 3964 |
| chr15 | 29369314 | 29369810 | 497 | * | NA_peak_213 | 13 | Distal Interge | 15 | 29713331 | 29713404 | 74 | 2 | 100628607 | ENSMUST00 | 343594 | ENSMUSG0 | Mir3964 | microRNA 3964 |
| chr15 | 29371949 | 29372233 | 285 | * | NA_peak_213 | 19 | Distal Interge | 15 | 29713331 | 29713404 | 74 | 2 | 100628607 | ENSMUST00 | 341171 | ENSMUSG0 | Mir3964 | microRNA 3964 |
| chr15 | 29548224 | 29548629 | 406 | * | NA_peak_214 | 26 | Distal Interge | 15 | 29713331 | 29713404 | 74 | 2 | 100628607 | ENSMUST00 | 164775 | ENSMUSG0 | Mir3964 | microRNA 3964 |
| chr15 | 29628341 | 29628745 | 405 | * | NA_peak_214 | 20 | Distal Interge | 15 | 29713331 | 29713404 | 74 | 2 | 100628607 | ENSMUST00 | 84659 | ENSMUSG0 | Mir3964 | microRNA 3964 |
| chr15 | 29649113 | 29649174 | 62 | * | NA_peak_214 | 18 | Distal Interge | 15 | 29713331 | 29713404 | 74 | 2 | 100628607 | ENSMUST00 | 64230 | ENSMUSG0 | Mir3964 | microRNA 3964 |
| chr15 | 29683822 | 29684078 | 257 | * | NA_peak_214 | 20 | Distal Interge | 15 | 29713331 | 29713404 | 74 | 2 | 100628607 | ENSMUST00 | 29326 | ENSMUSG0 | Mir3964 | microRNA 3964 |
| chr15 | 29907396 | 29907942 | 547 | * | NA_peak_214 | 30 | Distal Interge | 15 | 29713331 | 29713404 | 74 | 2 | 100628607 | ENSMUST00 | -193992 | ENSMUSG0 | Mir3964 | microRNA 3964 |
| chr15 | 29969891 | 29970204 | 314 | * | NA_peak_214 | 50 | Distal Interge | 15 | 30172593 | 31029341 | 856749 | 1 | 18163 | ENSMUST00 | -202389 | ENSMUSG0 | Ctnnd2 | catenin (cadherin associated protein), delta 2 |
| chr15 | 29992114 | 29992376 | 263 | * | NA_peak_214 | 16 | Distal Interge | 15 | 30172593 | 31029341 | 856749 | 1 | 18163 | ENSMUST00 | -180217 | ENSMUSG0 | Ctnnd2 | catenin (cadherin associated protein), delta 2 |
| chr15 | 30221896 | 30222293 | 398 | * | NA_peak_214 | 159 | Intron (ENSM | 15 | 30173138 | 30481349 | 308212 | 1 | 18163 | ENSMUST00 | 48758 | ENSMUSG0 | Ctnnd2 | catenin (cadherin associated protein), delta 2 |
| chr15 | 30255358 | 30255539 | 182 | * | NA_peak_214 | 37 | Intron (ENSM | 15 | 30173138 | 30481349 | 308212 | 1 | 18163 | ENSMUST00 | 82220 | ENSMUSG0 | Ctnnd2 | catenin (cadherin associated protein), delta 2 |
| chr15 | 30276002 | 30276138 | 137 | * | NA_peak_214 | 19 | Intron (ENSM | 15 | 30173138 | 30481349 | 308212 | 1 | 18163 | ENSMUST00 | 102864 | ENSMUSG0 | Ctnnd2 | catenin (cadherin associated protein), delta 2 |
| chr15 | 30485697 | 30485871 | 175 | * | NA_peak_215 | 20 | Intron (ENSM | 15 | 30457788 | 30504438 | 46651 | 1 | 18163 | ENSMUST00 | 27909 | ENSMUSG0 | Ctnnd2 | catenin (cadherin associated protein), delta 2 |
| chr15 | 30614215 | 30614521 | 307 | * | NA_peak_215 | 23 | Intron (ENSM | 15 | 30646532 | 30671124 | 24593 | 1 | 18163 | ENSMUST00 | -32011 | ENSMUSG0 | Ctnnd2 | catenin (cadherin associated protein), delta 2 |
| chr15 | 30945526 | 30945830 | 305 | * | NA_peak_215 | 17 | Intron (ENSM | 15 | 31004796 | 31020776 | 15981 | 1 | 18163 | ENSMUST00 | -58966 | ENSMUSG0 | Ctnnd2 | catenin (cadherin associated protein), delta 2 |
| chr15 | 30971906 | 30971982 | 77 | * | NA_peak_216 | 36 | Intron (ENSM | 15 | 31004796 | 31020776 | 15981 | 1 | 18163 | ENSMUST00 | -32814 | ENSMUSG0 | Ctnnd2 | catenin (cadherin associated protein), delta 2 |
| chr15 | 31078563 | 31078687 | 125 | * | NA_peak_216 | 28 | Distal Interge | 15 | 31039128 | 31043331 | 4204 | 1 | 73784 | ENSMUST00 | 39435 | ENSMUSG0 | 4930430F21 | RIKEN cDNA 4930430F21 gene |
| chr15 | 31086995 | 31087265 | 271 | * | NA_peak_216 | 20 | Distal Interge | 15 | 31039128 | 31043331 | 4204 | 1 | 73784 | ENSMUST00 | 47867 | ENSMUSG0 | 4930430F21 | RIKEN cDNA 4930430F21 gene |
| chr15 | 31090814 | 31091287 | 474 | * | NA_peak_216 | 27 | Distal Interge | 15 | 31039128 | 31043331 | 4204 | 1 | 73784 | ENSMUST00 | 51686 | ENSMUSG0 | 4930430F21 | RIKEN cDNA 4930430F21 gene |
| chr15 | 31298946 | 31299214 | 269 | * | NA_peak_216 | 33 | Intron (ENSM | 15 | 31297567 | 31308194 | 10628 | 2 | 67434 | ENSMUST00 | 8980 | ENSMUSG0 | Ankrd33b | ankyrin repeat domain 33B |
| chr15 | 31312575 | 31312977 | 403 | * | NA_peak_216 | 24 | Intron (ENSM | 15 | 31297567 | 31308194 | 10628 | 2 | 67434 | ENSMUST00 | -4381 | ENSMUSG0 | Ankrd33b | ankyrin repeat domain 33B |
| chr15 | 31393844 | 31393871 | 28 | * | NA_peak_216 | 27 | Distal Interge | 15 | 31294095 | 31367726 | 73632 | 2 | 67434 | ENSMUST00 | -26118 | ENSMUSG0 | Ankrd33b | ankyrin repeat domain 33B |
| chr15 | 31469189 | 31469854 | 666 | * | NA_peak_216 | 23 | Promoter (2-3 | 15 | 31465251 | 31471991 | 6741 | 2 | 223455 | ENSMUST00 | 2137 | ENSMUSG0 | Marchf6 | membrane associated ring-CH-type finger 6 |
| chr15 | 31498175 | 31498448 | 274 | * | NA_peak_216 | 20 | Intron (ENSM | 15 | 31465251 | 31471991 | 6741 | 2 | 223455 | ENSMUST00 | -26184 | ENSMUSG0 | Marchf6 | membrane associated ring-CH-type finger 6 |
| chr15 | 31502316 | 31502687 | 372 | * | NA_peak_216 | 23 | Exon (ENSMU | 15 | 31455891 | 31531053 | 75163 | 2 | 223455 | ENSMUST00 | 28366 | ENSMUSG0 | Marchf6 | membrane associated ring-CH-type finger 6 |
| chr15 | 31507410 | 31507855 | 446 | * | NA_peak_216 | 34 | Intron (ENSM | 15 | 31455891 | 31531053 | 75163 | 2 | 223455 | ENSMUST00 | 23198 | ENSMUSG0 | Marchf6 | membrane associated ring-CH-type finger 6 |
| chr15 | 31655031 | 31655284 | 254 | * | NA_peak_216 | 29 | Exon (ENSMU | 15 | 31602118 | 31618609 | 16492 | 1 | 68073 | ENSMUST00 | 52913 | ENSMUSG0 | Atpsckmt | ATP synthase C subunit lysine N-methyltransferase |
| chr15 | 31682576 | 31682928 | 353 | * | NA_peak_216 | 31 | Distal Interge | 15 | 31602118 | 31618609 | 16492 | 1 | 68073 | ENSMUST00 | 80458 | ENSMUSG0 | Atpsckmt | ATP synthase C subunit lysine N-methyltransferase |
| chr15 | 31748026 | 31748378 | 353 | * | NA_peak_216 | 33 | Distal Interge | 15 | 31602118 | 31618609 | 16492 | 1 | 68073 | ENSMUST00 | 145908 | ENSMUSG0 | Atpsckmt | ATP synthase C subunit lysine N-methyltransferase |

| | | | | | | | | | | | | | | | | |
|---|---|---|---|---|---|---|---|---|---|---|---|---|---|---|---|---|
| chr15 | 31767470 | 31767756 | 287 | * | NA_peak_216 | 31 | Distal Interge | 15 | 31602118 | 31618609 | 16492 | 1 | 68073 | ENSMUST00 | 165352 | ENSMUSG0 | Atpsckmt | ATP synthase C subunit lysine N-methyltransferase |
| chr15 | 31772234 | 31772552 | 319 | * | NA_peak_216 | 23 | Distal Interge | 15 | 31602118 | 31618609 | 16492 | 1 | 68073 | ENSMUST00 | 170116 | ENSMUSG0 | Atpsckmt | ATP synthase C subunit lysine N-methyltransferase |
| chr15 | 31783492 | 31783734 | 243 | * | NA_peak_217 | 21 | Distal Interge | 15 | 31602118 | 31618609 | 16492 | 1 | 68073 | ENSMUST00 | 181374 | ENSMUSG0 | Atpsckmt | ATP synthase C subunit lysine N-methyltransferase |
| chr15 | 31814340 | 31814662 | 323 | * | NA_peak_217 | 20 | Distal Interge | 15 | 31602118 | 31618609 | 16492 | 1 | 68073 | ENSMUST00 | 212222 | ENSMUSG0 | Atpsckmt | ATP synthase C subunit lysine N-methyltransferase |
| chr15 | 31830877 | 31831423 | 547 | * | NA_peak_217 | 21 | Distal Interge | 15 | 31602118 | 31618609 | 16492 | 1 | 68073 | ENSMUST00 | 228759 | ENSMUSG0 | Atpsckmt | ATP synthase C subunit lysine N-methyltransferase |
| chr15 | 31837211 | 31837365 | 155 | * | NA_peak_217 | 20 | Distal Interge | 15 | 31602118 | 31618609 | 16492 | 1 | 68073 | ENSMUST00 | 235093 | ENSMUSG0 | Atpsckmt | ATP synthase C subunit lysine N-methyltransferase |
| chr15 | 31838573 | 31838880 | 308 | * | NA_peak_217 | 15 | Distal Interge | 15 | 31602118 | 31618609 | 16492 | 1 | 68073 | ENSMUST00 | 236455 | ENSMUSG0 | Atpsckmt | ATP synthase C subunit lysine N-methyltransferase |
| chr15 | 31848601 | 31849020 | 420 | * | NA_peak_217 | 40 | Distal Interge | 15 | 31602118 | 31618609 | 16492 | 1 | 68073 | ENSMUST00 | 246483 | ENSMUSG0 | Atpsckmt | ATP synthase C subunit lysine N-methyltransferase |
| chr15 | 31888423 | 31888770 | 348 | * | NA_peak_217 | 55 | Distal Interge | 15 | 31602118 | 31618609 | 16492 | 1 | 68073 | ENSMUST00 | 286305 | ENSMUSG0 | Atpsckmt | ATP synthase C subunit lysine N-methyltransferase |
| chr15 | 31931822 | 31932091 | 270 | * | NA_peak_217 | 27 | Distal Interge | 15 | 32177289 | 32178294 | 1006 | 1 | 57254 | ENSMUST00 | -245198 | ENSMUSG0 | Tas2r119 | taste receptor, type 2, member 119 |
| chr15 | 31932467 | 31932536 | 70 | * | NA_peak_217 | 43 | Distal Interge | 15 | 32177289 | 32178294 | 1006 | 1 | 57254 | ENSMUST00 | -244753 | ENSMUSG0 | Tas2r119 | taste receptor, type 2, member 119 |
| chr15 | 31942292 | 31942652 | 361 | * | NA_peak_217 | 36 | Distal Interge | 15 | 32177289 | 32178294 | 1006 | 1 | 57254 | ENSMUST00 | -234637 | ENSMUSG0 | Tas2r119 | taste receptor, type 2, member 119 |
| chr15 | 31943255 | 31943259 | 5 | * | NA_peak_217 | 36 | Distal Interge | 15 | 32177289 | 32178294 | 1006 | 1 | 57254 | ENSMUST00 | -234030 | ENSMUSG0 | Tas2r119 | taste receptor, type 2, member 119 |
| chr15 | 31943608 | 31944019 | 412 | * | NA_peak_217 | 23 | Distal Interge | 15 | 32177289 | 32178294 | 1006 | 1 | 57254 | ENSMUST00 | -233270 | ENSMUSG0 | Tas2r119 | taste receptor, type 2, member 119 |
| chr15 | 31945805 | 31946195 | 391 | * | NA_peak_217 | 23 | Distal Interge | 15 | 32177289 | 32178294 | 1006 | 1 | 57254 | ENSMUST00 | -231094 | ENSMUSG0 | Tas2r119 | taste receptor, type 2, member 119 |
| chr15 | 31982690 | 31983315 | 626 | * | NA_peak_217 | 20 | Distal Interge | 15 | 32177289 | 32178294 | 1006 | 1 | 57254 | ENSMUST00 | -193974 | ENSMUSG0 | Tas2r119 | taste receptor, type 2, member 119 |
| chr15 | 32115890 | 32116141 | 252 | * | NA_peak_217 | 31 | Distal Interge | 15 | 32177289 | 32178294 | 1006 | 1 | 57254 | ENSMUST00 | -61148 | ENSMUSG0 | Tas2r119 | taste receptor, type 2, member 119 |
| chr15 | 32132252 | 32132713 | 462 | * | NA_peak_217 | 16 | Distal Interge | 15 | 32177289 | 32178294 | 1006 | 1 | 57254 | ENSMUST00 | -44576 | ENSMUSG0 | Tas2r119 | taste receptor, type 2, member 119 |
| chr15 | 32214366 | 32214739 | 374 | * | NA_peak_217 | 15 | Distal Interge | 15 | 32241853 | 32241923 | 71 | 2 | 66835 | ENSMUST00 | 27184 | ENSMUSG0 | Snord123 | small nucleolar RNA, C/D box 123 |
| chr15 | 32215728 | 32215989 | 262 | * | NA_peak_217 | 14 | Distal Interge | 15 | 32241853 | 32241923 | 71 | 2 | 66835 | ENSMUST00 | 25934 | ENSMUSG0 | Snord123 | small nucleolar RNA, C/D box 123 |
| chr15 | 32362536 | 32362742 | 207 | * | NA_peak_218 | 20 | Intron (ENSM | 15 | 32245103 | 32678594 | 433492 | 1 | 20356 | ENSMUST00 | 117433 | ENSMUSG0 | Sema5a | sema domain, seven thrombospondin repeats (type 1 and type 1-like), transmembrane domain (TM) and short cytoplasmic domain, (semaphorin) 5A |
| chr15 | 32367570 | 32368226 | 657 | * | NA_peak_218 | 23 | Intron (ENSM | 15 | 32245103 | 32678594 | 433492 | 1 | 20356 | ENSMUST00 | 122467 | ENSMUSG0 | Sema5a | sema domain, seven thrombospondin repeats (type 1 and type 1-like), transmembrane domain (TM) and short cytoplasmic domain, (semaphorin) 5A |
| chr15 | 32410349 | 32410708 | 360 | * | NA_peak_218 | 20 | Intron (ENSM | 15 | 32245103 | 32678594 | 433492 | 1 | 20356 | ENSMUST00 | 165246 | ENSMUSG0 | Sema5a | sema domain, seven thrombospondin repeats (type 1 and type 1-like), transmembrane domain (TM) and short cytoplasmic domain, (semaphorin) 5A |
| chr15 | 32413490 | 32413670 | 181 | * | NA_peak_218 | 28 | Intron (ENSM | 15 | 32245103 | 32678594 | 433492 | 1 | 20356 | ENSMUST00 | 168387 | ENSMUSG0 | Sema5a | sema domain, seven thrombospondin repeats (type 1 and type 1-like), transmembrane domain (TM) and short cytoplasmic domain, (semaphorin) 5A |
| chr15 | 32453281 | 32453303 | 23 | * | NA_peak_218 | 13 | Intron (ENSM | 15 | 32601675 | 32689636 | 87962 | 1 | 20356 | ENSMUST00 | -148372 | ENSMUSG0 | Sema5a | sema domain, seven thrombospondin repeats (type 1 and type 1-like), transmembrane domain (TM) and short cytoplasmic domain, (semaphorin) 5A |
| chr15 | 32481485 | 32481750 | 266 | * | NA_peak_218 | 40 | Intron (ENSM | 15 | 32601675 | 32689636 | 87962 | 1 | 20356 | ENSMUST00 | -119925 | ENSMUSG0 | Sema5a | sema domain, seven thrombospondin repeats (type 1 and type 1-like), transmembrane domain (TM) and short cytoplasmic domain, (semaphorin) 5A |
| chr15 | 32556646 | 32557013 | 368 | * | NA_peak_218 | 36 | Intron (ENSM | 15 | 32601675 | 32689636 | 87962 | 1 | 20356 | ENSMUST00 | -44662 | ENSMUSG0 | Sema5a | sema domain, seven thrombospondin repeats (type 1 and type 1-like), transmembrane domain (TM) and short cytoplasmic domain, (semaphorin) 5A |
| chr15 | 32595460 | 32595735 | 276 | * | NA_peak_218 | 19 | Intron (ENSM | 15 | 32601675 | 32689636 | 87962 | 1 | 20356 | ENSMUST00 | -5940 | ENSMUSG0 | Sema5a | sema domain, seven thrombospondin repeats (type 1 and type 1-like), transmembrane domain (TM) and short cytoplasmic domain, (semaphorin) 5A |
| chr15 | 32700790 | 32701030 | 241 | * | NA_peak_218 | 18 | Distal Interge | 15 | 32757416 | 32773152 | 15737 | 2 | 105245909 | ENSMUST00 | 72122 | ENSMUSG0 | Gm41289 | predicted gene, 41289 |
| chr15 | 32756610 | 32756881 | 272 | * | NA_peak_218 | 36 | Distal Interge | 15 | 32757416 | 32773152 | 15737 | 2 | 105245909 | ENSMUST00 | 16271 | ENSMUSG0 | Gm41289 | predicted gene, 41289 |
| chr15 | 32861692 | 32862148 | 457 | * | NA_peak_218 | 21 | Distal Interge | 15 | 32920723 | 33034935 | 114213 | 1 | 15529 | ENSMUST00 | -58575 | ENSMUSG0 | Sdc2 | syndecan 2 |
| chr15 | 32903306 | 32903623 | 318 | * | NA_peak_218 | 30 | Distal Interge | 15 | 32920723 | 33034935 | 114213 | 1 | 15529 | ENSMUST00 | -17100 | ENSMUSG0 | Sdc2 | syndecan 2 |
| chr15 | 32905182 | 32905784 | 603 | * | NA_peak_218 | 28 | Distal Interge | 15 | 32920723 | 33034935 | 114213 | 1 | 15529 | ENSMUST00 | -14939 | ENSMUSG0 | Sdc2 | syndecan 2 |
| chr15 | 32914894 | 32915042 | 149 | * | NA_peak_218 | 24 | Distal Interge | 15 | 32920723 | 33034935 | 114213 | 1 | 15529 | ENSMUST00 | -5681 | ENSMUSG0 | Sdc2 | syndecan 2 |
| chr15 | 32984921 | 32985292 | 372 | * | NA_peak_218 | 52 | Intron (ENSM | 15 | 32920723 | 33034935 | 114213 | 1 | 15529 | ENSMUST00 | 64198 | ENSMUSG0 | Sdc2 | syndecan 2 |
| chr15 | 32994500 | 32994791 | 292 | * | NA_peak_218 | 20 | Intron (ENSM | 15 | 32920723 | 33034935 | 114213 | 1 | 15529 | ENSMUST00 | 73777 | ENSMUSG0 | Sdc2 | syndecan 2 |
| chr15 | 33037010 | 33037496 | 487 | * | NA_peak_219 | 23 | Distal Interge | 15 | 33083129 | 33594552 | 511424 | 1 | 54381 | ENSMUST00 | -45633 | ENSMUSG0 | Cpq | carboxypeptidase Q |
| chr15 | 33058341 | 33059142 | 802 | * | NA_peak_219 | 50 | Intron (ENSM | 15 | 33083129 | 33594552 | 511424 | 1 | 54381 | ENSMUST00 | -23987 | ENSMUSG0 | Cpq | carboxypeptidase Q |
| chr15 | 33074341 | 33074748 | 408 | * | NA_peak_219 | 28 | Distal Interge | 15 | 33083129 | 33594552 | 511424 | 1 | 54381 | ENSMUST00 | -8381 | ENSMUSG0 | Cpq | carboxypeptidase Q |
| chr15 | 33082974 | 33083376 | 403 | * | NA_peak_219 | 18 | Promoter (<= | 15 | 33083129 | 33594552 | 511424 | 1 | 54381 | ENSMUST00 | 0 | ENSMUSG0 | Cpq | carboxypeptidase Q |
| chr15 | 33164439 | 33164667 | 229 | * | NA_peak_219 | 26 | Intron (ENSM | 15 | 33090308 | 33304620 | 214313 | 1 | 54381 | ENSMUST00 | 74131 | ENSMUSG0 | Cpq | carboxypeptidase Q |
| chr15 | 33215408 | 33215702 | 295 | * | NA_peak_219 | 40 | Intron (ENSM | 15 | 33090308 | 33304620 | 214313 | 1 | 54381 | ENSMUST00 | 125100 | ENSMUSG0 | Cpq | carboxypeptidase Q |
| chr15 | 33262109 | 33262286 | 178 | * | NA_peak_219 | 19 | Intron (ENSM | 15 | 33405206 | 33405939 | 734 | 2 | 73486 | ENSMUST00 | 143653 | ENSMUSG0 | 1700084J12F | ribosomal protein L7-like 1 pseudogene |
| chr15 | 33270748 | 33271852 | 1105 | * | NA_peak_219 | 50 | Intron (ENSM | 15 | 33405206 | 33405939 | 734 | 2 | 73486 | ENSMUST00 | 134087 | ENSMUSG0 | 1700084J12F | ribosomal protein L7-like 1 pseudogene |
| chr15 | 33297919 | 33298311 | 393 | * | NA_peak_219 | 36 | Intron (ENSM | 15 | 33405206 | 33405939 | 734 | 2 | 73486 | ENSMUST00 | 107628 | ENSMUSG0 | 1700084J12F | ribosomal protein L7-like 1 pseudogene |
| chr15 | 33398499 | 33398613 | 115 | * | NA_peak_219 | 20 | Intron (ENSM | 15 | 33405206 | 33405939 | 734 | 2 | 73486 | ENSMUST00 | 7326 | ENSMUSG0 | 1700084J12F | ribosomal protein L7-like 1 pseudogene |
| chr15 | 33496860 | 33497205 | 346 | * | NA_peak_219 | 13 | Exon (ENSMU | 15 | 33403985 | 33405976 | 1992 | 2 | 73486 | ENSMUST00 | -90884 | ENSMUSG0 | 1700084J12F | ribosomal protein L7-like 1 pseudogene |
| chr15 | 33509096 | 33509515 | 420 | * | NA_peak_219 | 40 | Intron (ENSM | 15 | 33403985 | 33405976 | 1992 | 2 | 73486 | ENSMUST00 | -103120 | ENSMUSG0 | 1700084J12F | ribosomal protein L7-like 1 pseudogene |
| chr15 | 33535061 | 33535355 | 295 | * | NA_peak_219 | 28 | Intron (ENSM | 15 | 33593881 | 33653436 | 59556 | 2 | 75880 | ENSMUST00 | 118081 | ENSMUSG0 | 4930592A05I | RIKEN cDNA 4930592A05 gene |
| chr15 | 33551644 | 33551824 | 181 | * | NA_peak_219 | 16 | Intron (ENSM | 15 | 33593881 | 33653436 | 59556 | 2 | 75880 | ENSMUST00 | 101612 | ENSMUSG0 | 4930592A05I | RIKEN cDNA 4930592A05 gene |
| chr15 | 33566874 | 33566925 | 52 | * | NA_peak_219 | 18 | Intron (ENSM | 15 | 33593881 | 33653436 | 59556 | 2 | 75880 | ENSMUST00 | 86511 | ENSMUSG0 | 4930592A05I | RIKEN cDNA 4930592A05 gene |
| chr15 | 33568252 | 33568781 | 530 | * | NA_peak_219 | 24 | Intron (ENSM | 15 | 33593881 | 33653436 | 59556 | 2 | 75880 | ENSMUST00 | 84655 | ENSMUSG0 | 4930592A05I | RIKEN cDNA 4930592A05 gene |
| chr15 | 33596442 | 33596685 | 244 | * | NA_peak_219 | 27 | Intron (ENSM | 15 | 33593881 | 33653436 | 59556 | 2 | 75880 | ENSMUST00 | 56751 | ENSMUSG0 | 4930592A05I | RIKEN cDNA 4930592A05 gene |
| chr15 | 33633575 | 33633772 | 198 | * | NA_peak_220 | 22 | Intron (ENSM | 15 | 33593881 | 33653436 | 59556 | 2 | 75880 | ENSMUST00 | 19664 | ENSMUSG0 | 4930592A05I | RIKEN cDNA 4930592A05 gene |
| chr15 | 33659593 | 33659876 | 284 | * | NA_peak_220 | 40 | Distal Interge | 15 | 33593881 | 33653436 | 59556 | 2 | 75880 | ENSMUST00 | -6157 | ENSMUSG0 | 4930592A05I | RIKEN cDNA 4930592A05 gene |
| chr15 | 33733292 | 33733698 | 407 | * | NA_peak_220 | 21 | Distal Interge | 15 | 33683875 | 33687884 | 4010 | 2 | 239364 | ENSMUST00 | -45408 | ENSMUSG0 | Tspyl5 | testis-specific protein, Y-encoded-like 5 |
| chr15 | 33776847 | 33777151 | 305 | * | NA_peak_220 | 22 | Distal Interge | 15 | 33683875 | 33687884 | 4010 | 2 | 239364 | ENSMUST00 | -88963 | ENSMUSG0 | Tspyl5 | testis-specific protein, Y-encoded-like 5 |
| chr15 | 33777904 | 33778139 | 236 | * | NA_peak_220 | 23 | Distal Interge | 15 | 33683875 | 33687884 | 4010 | 2 | 239364 | ENSMUST00 | -90020 | ENSMUSG0 | Tspyl5 | testis-specific protein, Y-encoded-like 5 |
| chr15 | 33824839 | 33825237 | 399 | * | NA_peak_220 | 13 | Distal Interge | 15 | 33683875 | 33687884 | 4010 | 2 | 239364 | ENSMUST00 | -136955 | ENSMUSG0 | Tspyl5 | testis-specific protein, Y-encoded-like 5 |
| chr15 | 33832757 | 33833500 | 744 | * | NA_peak_220 | 80 | Exon (ENSMU | 15 | 33683875 | 33687884 | 4010 | 2 | 239364 | ENSMUST00 | -144873 | ENSMUSG0 | Tspyl5 | testis-specific protein, Y-encoded-like 5 |
| chr15 | 33872893 | 33872987 | 95 | * | NA_peak_220 | 28 | Distal Interge | 15 | 33683875 | 33687884 | 4010 | 2 | 239364 | ENSMUST00 | -185009 | ENSMUSG0 | Tspyl5 | testis-specific protein, Y-encoded-like 5 |
| chr15 | 33911609 | 33912062 | 454 | * | NA_peak_220 | 49 | Distal Interge | 15 | 34082694 | 34103345 | 20652 | 1 | 67154 | ENSMUST00 | -170632 | ENSMUSG0 | Mtdh | metadherin |
| chr15 | 34039186 | 34039526 | 341 | * | NA_peak_220 | 33 | Distal Interge | 15 | 34082694 | 34103345 | 20652 | 1 | 67154 | ENSMUST00 | -43168 | ENSMUSG0 | Mtdh | metadherin |
| chr15 | 34066075 | 34066219 | 145 | * | NA_peak_220 | 13 | Distal Interge | 15 | 34082694 | 34103345 | 20652 | 1 | 67154 | ENSMUST00 | -16475 | ENSMUSG0 | Mtdh | metadherin |
| chr15 | 34245473 | 34245588 | 116 | * | NA_peak_221 | 31 | Intron (ENSM | 15 | 34238167 | 34276168 | 38002 | 1 | 114128 | ENSMUST00 | 7306 | ENSMUSG0 | Laptm4b | lysosomal-associated protein transmembrane 4B |
| chr15 | 34448913 | 34449162 | 250 | * | NA_peak_221 | 31 | Distal Interge | 15 | 34453305 | 34473892 | 20588 | 1 | 239368 | ENSMUST00 | -4143 | ENSMUSG0 | Erich5 | glutamate rich 5 |
| chr15 | 34490565 | 34491316 | 752 | * | NA_peak_221 | 31 | Promoter (<= | 15 | 34484024 | 34489800 | 5777 | 2 | 15473 | ENSMUST00 | -765 | ENSMUSG0 | Rida | reactive intermediate imine deaminase A homolog |
| chr15 | 34596927 | 34597241 | 315 | * | NA_peak_221 | 23 | Intron (ENSM | 15 | 34575506 | 34582771 | 7266 | 2 | 223473 | ENSMUST00 | -14156 | ENSMUSG0 | Nipal2 | NIPA-like domain containing 2 |
| chr15 | 34685950 | 34686108 | 159 | * | NA_peak_221 | 46 | Distal Interge | 15 | 34572799 | 34679212 | 106414 | 2 | 223473 | ENSMUST00 | -6738 | ENSMUSG0 | Nipal2 | NIPA-like domain containing 2 |
| chr15 | 34709195 | 34709488 | 294 | * | NA_peak_221 | 27 | Distal Interge | 15 | 34572799 | 34679212 | 106414 | 2 | 223473 | ENSMUST00 | -29983 | ENSMUSG0 | Nipal2 | NIPA-like domain containing 2 |
| chr15 | 34728938 | 34729565 | 628 | * | NA_peak_221 | 40 | Distal Interge | 15 | 34572799 | 34679212 | 106414 | 2 | 223473 | ENSMUST00 | -49726 | ENSMUSG0 | Nipal2 | NIPA-like domain containing 2 |
| chr15 | 34741509 | 34741752 | 244 | * | NA_peak_221 | 18 | Distal Interge | 15 | 34572799 | 34679212 | 106414 | 2 | 223473 | ENSMUST00 | -62297 | ENSMUSG0 | Nipal2 | NIPA-like domain containing 2 |
| chr15 | 34944264 | 34944538 | 275 | * | NA_peak_221 | 24 | Intron (ENSM | 15 | 34958564 | 34999922 | 41359 | 2 | 56274 | ENSMUST00 | 55384 | ENSMUSG0 | Stk3 | serine/threonine kinase 3 |
| chr15 | 34992037 | 34992464 | 428 | * | NA_peak_221 | 25 | Intron (ENSM | 15 | 34958564 | 34999922 | 41359 | 2 | 56274 | ENSMUST00 | 7458 | ENSMUSG0 | Stk3 | serine/threonine kinase 3 |
| chr15 | 35093223 | 35093504 | 282 | * | NA_peak_222 | 26 | Intron (ENSM | 15 | 35072479 | 35155669 | 83191 | 2 | 56274 | ENSMUST00 | 62165 | ENSMUSG0 | Stk3 | serine/threonine kinase 3 |
| chr15 | 35107602 | 35108034 | 433 | * | NA_peak_222 | 21 | Intron (ENSM | 15 | 35072479 | 35155669 | 83191 | 2 | 56274 | ENSMUST00 | 47635 | ENSMUSG0 | Stk3 | serine/threonine kinase 3 |
| chr15 | 35119833 | 35119879 | 47 | * | NA_peak_222 | 47 | Intron (ENSM | 15 | 35072479 | 35155669 | 83191 | 2 | 56274 | ENSMUST00 | 35790 | ENSMUSG0 | Stk3 | serine/threonine kinase 3 |
| chr15 | 35122353 | 35122525 | 173 | * | NA_peak_222 | 15 | Intron (ENSM | 15 | 35072479 | 35155669 | 83191 | 2 | 56274 | ENSMUST00 | 33144 | ENSMUSG0 | Stk3 | serine/threonine kinase 3 |
| chr15 | 35136531 | 35136946 | 416 | * | NA_peak_222 | 15 | Intron (ENSM | 15 | 35072479 | 35155669 | 83191 | 2 | 56274 | ENSMUST00 | 18723 | ENSMUSG0 | Stk3 | serine/threonine kinase 3 |
| chr15 | 35162090 | 35162295 | 206 | * | NA_peak_222 | 40 | Intron (ENSM | 15 | 34875496 | 35155806 | 280311 | 2 | 56274 | ENSMUST00 | -6284 | ENSMUSG0 | Stk3 | serine/threonine kinase 3 |
| chr15 | 35174705 | 35174853 | 149 | * | NA_peak_222 | 25 | Intron (ENSM | 15 | 34875496 | 35178921 | 303426 | 2 | 56274 | ENSMUST00 | 4068 | ENSMUSG0 | Stk3 | serine/threonine kinase 3 |
| chr15 | 35203721 | 35203795 | 75 | * | NA_peak_222 | 23 | Intron (ENSM | 15 | 34875496 | 35178921 | 303426 | 2 | 56274 | ENSMUST00 | -24800 | ENSMUSG0 | Stk3 | serine/threonine kinase 3 |
| chr15 | 35207034 | 35207389 | 356 | * | NA_peak_222 | 29 | Intron (ENSM | 15 | 34875496 | 35178921 | 303426 | 2 | 56274 | ENSMUST00 | -28113 | ENSMUSG0 | Stk3 | serine/threonine kinase 3 |
| chr15 | 35398679 | 35398874 | 196 | * | NA_peak_222 | 13 | Intron (ENSM | 15 | 35371566 | 35642952 | 271387 | 1 | 666173 | ENSMUST00 | 27113 | ENSMUSG0 | Vps13b | vacuolar protein sorting 13B |
| chr15 | 35490312 | 35490762 | 451 | * | NA_peak_222 | 24 | Intron (ENSM | 15 | 35371566 | 35642952 | 271387 | 1 | 666173 | ENSMUST00 | 118746 | ENSMUSG0 | Vps13b | vacuolar protein sorting 13B |
| chr15 | 35583674 | 35583757 | 84 | * | NA_peak_222 | 36 | Intron (ENSM | 15 | 35660831 | 35660918 | 88 | 2 | 100316736 | ENSMUST00 | 77161 | ENSMUSG0 | Mir599 | microRNA 599 |
| chr15 | 35623538 | 35623562 | 25 | * | NA_peak_222 | 23 | Intron (ENSM | 15 | 35660831 | 35660918 | 88 | 2 | 100316736 | ENSMUST00 | 37356 | ENSMUSG0 | Mir599 | microRNA 599 |
| chr15 | 35626538 | 35626656 | 119 | * | NA_peak_222 | 26 | Intron (ENSM | 15 | 35660831 | 35660918 | 88 | 2 | 100316736 | ENSMUST00 | 34262 | ENSMUSG0 | Mir599 | microRNA 599 |
| chr15 | 35630400 | 35630721 | 322 | * | NA_peak_222 | 31 | Intron (ENSM | 15 | 35660831 | 35660918 | 88 | 2 | 100316736 | ENSMUST00 | 30197 | ENSMUSG0 | Mir599 | microRNA 599 |
| chr15 | 35631566 | 35631961 | 396 | * | NA_peak_222 | 31 | Intron (ENSM | 15 | 35660831 | 35660918 | 88 | 2 | 100316736 | ENSMUST00 | 28957 | ENSMUSG0 | Mir599 | microRNA 599 |
| chr15 | 35827251 | 35827507 | 257 | * | NA_peak_223 | 27 | Intron (ENSM | 15 | 35834215 | 35843534 | 9320 | 1 | 666173 | ENSMUST00 | -6708 | ENSMUSG0 | Vps13b | vacuolar protein sorting 13B |
| chr15 | 35833528 | 35833732 | 205 | * | NA_peak_223 | 31 | Promoter (<= | 15 | 35834215 | 35843534 | 9320 | 1 | 666173 | ENSMUST00 | -483 | ENSMUSG0 | Vps13b | vacuolar protein sorting 13B |
| chr15 | 35866518 | 35866882 | 365 | * | NA_peak_223 | 27 | Intron (ENSM | 15 | 35834215 | 35843534 | 9320 | 1 | 666173 | ENSMUST00 | 32303 | ENSMUSG0 | Vps13b | vacuolar protein sorting 13B |
| chr15 | 36048168 | 36048689 | 522 | * | NA_peak_223 | 40 | Intron (ENSM | 15 | 36009570 | 36040679 | 31110 | 2 | 626596 | ENSMUST00 | -7489 | ENSMUSG0 | Rgs22 | regulator of G-protein signalling 22 |
| chr15 | 36076464 | 36076624 | 161 | * | NA_peak_223 | 21 | Intron (ENSM | 15 | 36048875 | 36083741 | 34867 | 2 | 626596 | ENSMUST00 | 7117 | ENSMUSG0 | Rgs22 | regulator of G-protein signalling 22 |
| chr15 | 36115980 | 36116250 | 271 | * | NA_peak_223 | 25 | Intron (ENSM | 15 | 36138221 | 36140385 | 2165 | 2 | 626596 | ENSMUST00 | 24135 | ENSMUSG0 | Rgs22 | regulator of G-protein signalling 22 |
| chr15 | 36135387 | 36135845 | 459 | * | NA_peak_223 | 34 | Intron (ENSM | 15 | 36138221 | 36140385 | 2165 | 2 | 626596 | ENSMUST00 | 4540 | ENSMUSG0 | Rgs22 | regulator of G-protein signalling 22 |
| chr15 | 36310948 | 36311179 | 232 | * | NA_peak_223 | 25 | Distal Interge | 15 | 36239933 | 36283147 | 43215 | 2 | 30945 | ENSMUST00 | -27801 | ENSMUSG0 | Rnf19a | ring finger protein 19A |
| chr15 | 36434761 | 36435067 | 307 | * | NA_peak_223 | 37 | Intron (ENSM | 15 | 36495942 | 36496715 | 774 | 2 | 68839 | ENSMUST00 | 61648 | ENSMUSG0 | Ankrd46 | ankyrin repeat domain 46 |
| chr15 | 36619529 | 36619650 | 122 | * | NA_peak_223 | 13 | Distal Interge | 15 | 36605752 | 36609668 | 3917 | 2 | 18458 | ENSMUST00 | -9861 | ENSMUSG0 | Pabpc1 | poly(A) binding protein, cytoplasmic 1 |
| chr15 | 36662322 | 36662548 | 227 | * | NA_peak_223 | 14 | Intron (ENSM | 15 | 36605752 | 36609668 | 3917 | 2 | 18458 | ENSMUST00 | -52654 | ENSMUSG0 | Pabpc1 | poly(A) binding protein, cytoplasmic 1 |
| chr15 | 36733402 | 36733787 | 386 | * | NA_peak_224 | 14 | Distal Interge | 15 | 36790712 | 36792414 | 1703 | 2 | 22631 | ENSMUST00 | 58627 | ENSMUSG0 | Ywhaz | tyrosine 3-monooxygenase/tryptophan 5-monooxygenase activation protein, zeta polypeptide |
| chr15 | 36773504 | 36773785 | 282 | * | NA_peak_224 | 19 | Intron (ENSM | 15 | 36790712 | 36792414 | 1703 | 2 | 22631 | ENSMUST00 | 18629 | ENSMUSG0 | Ywhaz | tyrosine 3-monooxygenase/tryptophan 5-monooxygenase activation protein, zeta polypeptide |
| chr15 | 36837966 | 36838099 | 134 | * | NA_peak_224 | 25 | Distal Interge | 15 | 36793539 | 36796929 | 3391 | 2 | 22631 | ENSMUST00 | -41037 | ENSMUSG0 | Ywhaz | tyrosine 3-monooxygenase/tryptophan 5-monooxygenase activation protein, zeta polypeptide |
| chr15 | 36964098 | 36964562 | 465 | * | NA_peak_224 | 16 | Distal Interge | 15 | 37000628 | 37002035 | 1408 | 2 | 68036 | ENSMUST00 | 37473 | ENSMUSG0 | Zfp706 | zinc finger protein 706 |
| chr15 | 37027424 | 37027561 | 138 | * | NA_peak_224 | 32 | Intron (ENSM | 15 | 37000814 | 37007773 | 6960 | 2 | 68036 | ENSMUST00 | -19651 | ENSMUSG0 | Zfp706 | zinc finger protein 706 |

| | | | | | | | | | | | | | | | | | | |
|---|---|---|---|---|---|---|---|---|---|---|---|---|---|---|---|---|---|---|
| chr15 | 37119903 | 37120189 | 287 | * | NA_peak_224 | 26 | Distal Interge | 15 | 37000814 | 37007773 | 6960 | 2 | 68036 | ENSMUST00 | -112130 | ENSMUSG00 | Zfp706 | zinc finger protein 706 |
| chr15 | 37332684 | 37332971 | 288 | * | NA_peak_224 | 17 | Promoter (2-3 | 15 | 37335737 | 37360940 | 25204 | 1 | 252973 | ENSMUST00 | -2766 | ENSMUSG00 | Grhl2 | grainyhead like transcription factor 2 |
| chr15 | 37473730 | 37473941 | 212 | * | NA_peak_224 | 33 | Intron (ENSM | 15 | 37366218 | 37459360 | 93143 | 2 | 52589 | ENSMUST00 | -14370 | ENSMUSG00 | Ncald | neurocalcin delta |
| chr15 | 37519237 | 37519383 | 147 | * | NA_peak_224 | 22 | Intron (ENSM | 15 | 37366218 | 37459360 | 93143 | 2 | 52589 | ENSMUST00 | -59877 | ENSMUSG00 | Ncald | neurocalcin delta |
| chr15 | 37568220 | 37568324 | 105 | * | NA_peak_224 | 13 | Intron (ENSM | 15 | 37366218 | 37459360 | 93143 | 2 | 52589 | ENSMUST00 | -108860 | ENSMUSG00 | Ncald | neurocalcin delta |
| chr15 | 37773874 | 37774177 | 304 | * | NA_peak_225 | 29 | Intron (ENSM | 15 | 37366175 | 37792000 | 425826 | 2 | 52589 | ENSMUST00 | 17823 | ENSMUSG00 | Ncald | neurocalcin delta |
| chr15 | 37821027 | 37821278 | 252 | * | NA_peak_225 | 26 | Distal Interge | 15 | 37397171 | 37792570 | 395400 | 2 | 52589 | ENSMUST00 | -28457 | ENSMUSG00 | Ncald | neurocalcin delta |
| chr15 | 37844110 | 37844184 | 75 | * | NA_peak_225 | 21 | Distal Interge | 15 | 37397171 | 37792570 | 395400 | 2 | 52589 | ENSMUST00 | -51540 | ENSMUSG00 | Ncald | neurocalcin delta |
| chr15 | 37848862 | 37849133 | 272 | * | NA_peak_225 | 29 | Distal Interge | 15 | 37397171 | 37792570 | 395400 | 2 | 52589 | ENSMUST00 | -56292 | ENSMUSG00 | Ncald | neurocalcin delta |
| chr15 | 37891295 | 37891584 | 290 | * | NA_peak_225 | 20 | Distal Interge | 15 | 37945048 | 37955138 | 10091 | 2 | 382985 | ENSMUST00 | 63554 | ENSMUSG00 | Rrm2b | ribonucleotide reductase M2 B (TP53 inducible) |
| chr15 | 37912420 | 37912829 | 410 | * | NA_peak_225 | 21 | Distal Interge | 15 | 37945048 | 37955138 | 10091 | 2 | 382985 | ENSMUST00 | 42309 | ENSMUSG00 | Rrm2b | ribonucleotide reductase M2 B (TP53 inducible) |
| chr15 | 37993265 | 37993286 | 22 | * | NA_peak_225 | 16 | Promoter (1-2 | 15 | 37988346 | 37991637 | 3292 | 2 | 70790 | ENSMUST00 | -1628 | ENSMUSG00 | Ubr5 | ubiquitin protein ligase E3 component n-recognin 5 |
| chr15 | 38095240 | 38095276 | 37 | * | NA_peak_225 | 22 | Distal Interge | 15 | 37967859 | 38078854 | 110996 | 2 | 70790 | ENSMUST00 | -16386 | ENSMUSG00 | Ubr5 | ubiquitin protein ligase E3 component n-recognin 5 |
| chr15 | 38121614 | 38121671 | 58 | * | NA_peak_225 | 33 | Distal Interge | 15 | 37967859 | 38078854 | 110996 | 2 | 70790 | ENSMUST00 | -42760 | ENSMUSG00 | Ubr5 | ubiquitin protein ligase E3 component n-recognin 5 |
| chr15 | 38259979 | 38260374 | 396 | * | NA_peak_225 | 20 | Distal Interge | 15 | 38297295 | 38298839 | 1545 | 2 | 21847 | ENSMUST00 | 38465 | ENSMUSG00 | Klf10 | Kruppel-like factor 10 |
| chr15 | 38439788 | 38440347 | 560 | * | NA_peak_225 | 47 | Intron (ENSM | 15 | 38490473 | 38490547 | 75 | 2 | 102465574 | ENSMUST00 | 50200 | ENSMUSG00 | Mir6951 | microRNA 6951 |
| chr15 | 38530527 | 38530796 | 270 | * | NA_peak_225 | 19 | Distal Interge | 15 | 38487427 | 38519266 | 31840 | 2 | 54375 | ENSMUST00 | -11261 | ENSMUSG00 | Azin1 | antizyme inhibitor 1 |
| chr15 | 38690680 | 38690998 | 319 | * | NA_peak_226 | 15 | Intron (ENSM | 15 | 38675689 | 38679368 | 3680 | 1 | 66335 | ENSMUST00 | 14991 | ENSMUSG00 | Atp6v1c1 | ATPase, H+ transporting, lysosomal V1 subunit C1 |
| chr15 | 39131139 | 39131579 | 441 | * | NA_peak_226 | 17 | Promoter (<= | 15 | 39132253 | 39146853 | 14601 | 1 | 223499 | ENSMUST00 | -674 | ENSMUSG00 | Dcaf13 | DDB1 and CUL4 associated factor 13 |
| chr15 | 39141840 | 39141934 | 95 | * | NA_peak_226 | 17 | Intron (ENSM | 15 | 39144959 | 39146856 | 1898 | 1 | 223499 | ENSMUST00 | -3025 | ENSMUSG00 | Dcaf13 | DDB1 and CUL4 associated factor 13 |
| chr15 | 39300188 | 39300415 | 228 | * | NA_peak_226 | 20 | Intron (ENSM | 15 | 39391800 | 39452232 | 60433 | 1 | 116838 | ENSMUST00 | -91385 | ENSMUSG00 | Rims2 | regulating synaptic membrane exocytosis 2 |
| chr15 | 39622111 | 39622277 | 167 | * | NA_peak_226 | 20 | Intron (ENSM | 15 | 39586707 | 39616410 | 29704 | 1 | 116838 | ENSMUST00 | 35404 | ENSMUSG00 | Rims2 | regulating synaptic membrane exocytosis 2 |
| chr15 | 39698883 | 39699310 | 428 | * | NA_peak_226 | 27 | Distal Interge | 15 | 39659566 | 39684372 | 24807 | 1 | 116838 | ENSMUST00 | 39317 | ENSMUSG00 | Rims2 | regulating synaptic membrane exocytosis 2 |
| chr15 | 39721936 | 39722136 | 201 | * | NA_peak_227 | 21 | Distal Interge | 15 | 39745932 | 39760934 | 15003 | 1 | 75766 | ENSMUST00 | -23796 | ENSMUSG00 | Dcstamp | dendrocyte expressed seven transmembrane protein |
| chr15 | 39898775 | 39899030 | 256 | * | NA_peak_227 | 33 | Intron (ENSM | 15 | 39855877 | 39857470 | 1594 | 2 | 64705 | ENSMUST00 | -41305 | ENSMUSG00 | Dpys | dihydropyrimidinase |
| chr15 | 39947048 | 39947197 | 150 | * | NA_peak_227 | 27 | Distal Interge | 15 | 39870603 | 39943994 | 73392 | 2 | 239393 | ENSMUST00 | -3054 | ENSMUSG00 | Lrp12 | low density lipoprotein-related protein 12 |
| chr15 | 40003339 | 40003470 | 132 | * | NA_peak_227 | 13 | Distal Interge | 15 | 39962649 | 39967024 | 4376 | 1 | 105976 | ENSMUST00 | 40690 | NA | AU022793 | expressed sequence AU022793 |
| chr15 | 40099902 | 40099969 | 68 | * | NA_peak_227 | 43 | Intron (ENSM | 15 | 40134365 | 40150010 | 15646 | 1 | 328531 | ENSMUST00 | -34396 | ENSMUSG00 | 9330182O14 | RIKEN cDNA 9330182O14 gene |
| chr15 | 40155458 | 40155624 | 167 | * | NA_peak_227 | 28 | Distal Interge | 15 | 40162581 | 40167719 | 5139 | 1 | 102636151 | ENSMUST00 | -6957 | ENSMUSG00 | Gm33301 | predicted gene, 33301 |
| chr15 | 40210195 | 40210527 | 333 | * | NA_peak_227 | 31 | Distal Interge | 15 | 40162607 | 40168076 | 5470 | 1 | 102636151 | ENSMUST00 | 47588 | ENSMUSG00 | Gm33301 | predicted gene, 33301 |
| chr15 | 40241858 | 40242216 | 359 | * | NA_peak_227 | 27 | Distal Interge | 15 | 40162607 | 40168076 | 5470 | 1 | 102636151 | ENSMUST00 | 79251 | ENSMUSG00 | Gm33301 | predicted gene, 33301 |
| chr15 | 40252255 | 40252737 | 483 | * | NA_peak_227 | 26 | Distal Interge | 15 | 40162607 | 40168076 | 5470 | 1 | 102636151 | ENSMUST00 | 89648 | ENSMUSG00 | Gm33301 | predicted gene, 33301 |
| chr15 | 40298116 | 40298275 | 160 | * | NA_peak_227 | 19 | Distal Interge | 15 | 40162607 | 40168076 | 5470 | 1 | 102636151 | ENSMUST00 | 135509 | ENSMUSG00 | Gm33301 | predicted gene, 33301 |
| chr15 | 40412812 | 40413119 | 308 | * | NA_peak_227 | 17 | Distal Interge | 15 | 40511322 | 40527277 | 15956 | 2 | 432945 | ENSMUST00 | 114158 | ENSMUSG00 | Gm16294 | predicted gene 16294 |
| chr15 | 40470863 | 40470887 | 25 | * | NA_peak_227 | 20 | Distal Interge | 15 | 40511322 | 40527277 | 15956 | 2 | 432945 | ENSMUST00 | 56390 | ENSMUSG00 | Gm16294 | predicted gene 16294 |
| chr15 | 40475244 | 40475547 | 304 | * | NA_peak_227 | 22 | Distal Interge | 15 | 40511322 | 40527277 | 15956 | 2 | 432945 | ENSMUST00 | 51730 | ENSMUSG00 | Gm16294 | predicted gene 16294 |
| chr15 | 40518502 | 40518798 | 297 | * | NA_peak_227 | 26 | Intron (ENSM | 15 | 40511322 | 40527277 | 15956 | 2 | 432945 | ENSMUST00 | 8479 | ENSMUSG00 | Gm16294 | predicted gene 16294 |
| chr15 | 40522668 | 40523322 | 655 | * | NA_peak_227 | 31 | Intron (ENSM | 15 | 40511322 | 40527277 | 15956 | 2 | 432945 | ENSMUST00 | 3955 | ENSMUSG00 | Gm16294 | predicted gene 16294 |
| chr15 | 40640123 | 40640245 | 123 | * | NA_peak_228 | 18 | Distal Interge | 15 | 40655035 | 40872723 | 217689 | 1 | 22762 | ENSMUST00 | -14790 | ENSMUSG00 | Zfpm2 | zinc finger protein, multitype 2 |
| chr15 | 40681340 | 40681644 | 305 | * | NA_peak_228 | 37 | Intron (ENSM | 15 | 40664990 | 41104586 | 439597 | 1 | 22762 | ENSMUST00 | 16350 | ENSMUSG00 | Zfpm2 | zinc finger protein, multitype 2 |
| chr15 | 40694962 | 40694968 | 7 | * | NA_peak_228 | 16 | Intron (ENSM | 15 | 40664990 | 41104586 | 439597 | 1 | 22762 | ENSMUST00 | 29972 | ENSMUSG00 | Zfpm2 | zinc finger protein, multitype 2 |
| chr15 | 40888575 | 40888703 | 129 | * | NA_peak_228 | 20 | Intron (ENSM | 15 | 40664990 | 41104586 | 439597 | 1 | 22762 | ENSMUST00 | 223585 | ENSMUSG00 | Zfpm2 | zinc finger protein, multitype 2 |
| chr15 | 41145769 | 41146133 | 365 | * | NA_peak_228 | 62 | Distal Interge | 15 | 41173487 | 41173868 | 382 | 1 | 100418302 | ENSMUST00 | -27354 | NA | 4930555K19I | leucine rich repeat and sterile alpha motif containing 1 pseudogene |
| chr15 | 41245398 | 41245689 | 292 | * | NA_peak_228 | 27 | Distal Interge | 15 | 41173487 | 41173868 | 382 | 1 | 100418302 | ENSMUST00 | 71911 | NA | 4930555K19I | leucine rich repeat and sterile alpha motif containing 1 pseudogene |
| chr15 | 41272674 | 41273042 | 369 | * | NA_peak_228 | 21 | Distal Interge | 15 | 41173487 | 41173868 | 382 | 1 | 100418302 | ENSMUST00 | 99187 | NA | 4930555K19I | leucine rich repeat and sterile alpha motif containing 1 pseudogene |
| chr15 | 41341503 | 41341746 | 244 | * | NA_peak_228 | 28 | Distal Interge | 15 | 41447482 | 41861045 | 413564 | 1 | 170719 | ENSMUST00 | -105736 | ENSMUSG00 | Oxr1 | oxidation resistance 1 |
| chr15 | 41343006 | 41343151 | 146 | * | NA_peak_228 | 19 | Distal Interge | 15 | 41447482 | 41861045 | 413564 | 1 | 170719 | ENSMUST00 | -104331 | ENSMUSG00 | Oxr1 | oxidation resistance 1 |
| chr15 | 41537474 | 41537909 | 436 | * | NA_peak_229 | 21 | Intron (ENSM | 15 | 41449027 | 41861047 | 412021 | 1 | 170719 | ENSMUST00 | 88447 | ENSMUSG00 | Oxr1 | oxidation resistance 1 |
| chr15 | 41676887 | 41677182 | 296 | * | NA_peak_229 | 22 | Intron (ENSM | 15 | 41710604 | 41820380 | 109777 | 1 | 170719 | ENSMUST00 | -33422 | ENSMUSG00 | Oxr1 | oxidation resistance 1 |
| chr15 | 41699997 | 41700017 | 21 | * | NA_peak_229 | 13 | Intron (ENSM | 15 | 41710604 | 41820380 | 109777 | 1 | 170719 | ENSMUST00 | -10587 | ENSMUSG00 | Oxr1 | oxidation resistance 1 |
| chr15 | 41745094 | 41745518 | 425 | * | NA_peak_229 | 23 | Intron (ENSM | 15 | 41751698 | 41820233 | 68536 | 1 | 170719 | ENSMUST00 | -6180 | ENSMUSG00 | Oxr1 | oxidation resistance 1 |
| chr15 | 41827448 | 41827868 | 421 | * | NA_peak_229 | 23 | Intron (ENSM | 15 | 41830932 | 41859308 | 28377 | 1 | 170719 | ENSMUST00 | -3064 | ENSMUSG00 | Oxr1 | oxidation resistance 1 |
| chr15 | 41884155 | 41884350 | 196 | * | NA_peak_229 | 27 | Distal Interge | 15 | 41864076 | 41869720 | 5645 | 2 | 223513 | ENSMUST00 | -14435 | ENSMUSG00 | Abra | actin-binding Rho activating protein |
| chr15 | 41960185 | 41960333 | 149 | * | NA_peak_229 | 14 | Distal Interge | 15 | 41864076 | 41869720 | 5645 | 2 | 223513 | ENSMUST00 | -90465 | ENSMUSG00 | Abra | actin-binding Rho activating protein |
| chr15 | 41978554 | 41978768 | 215 | * | NA_peak_229 | 31 | Distal Interge | 15 | 41864076 | 41869720 | 5645 | 2 | 223513 | ENSMUST00 | -108834 | ENSMUSG00 | Abra | actin-binding Rho activating protein |
| chr15 | 41995657 | 41996077 | 421 | * | NA_peak_229 | 23 | Distal Interge | 15 | 41864076 | 41869720 | 5645 | 2 | 223513 | ENSMUST00 | -125937 | ENSMUSG00 | Abra | actin-binding Rho activating protein |
| chr15 | 42136719 | 42136916 | 198 | * | NA_peak_229 | 24 | Distal Interge | 15 | 41864076 | 41869720 | 5645 | 2 | 223513 | ENSMUST00 | -266999 | ENSMUSG00 | Abra | actin-binding Rho activating protein |
| chr15 | 42400841 | 42401098 | 258 | * | NA_peak_230 | 21 | Distal Interge | 15 | 42633306 | 42676749 | 43444 | 2 | 11600 | ENSMUST00 | 275651 | ENSMUSG00 | Angpt1 | angiopoietin 1 |
| chr15 | 42411360 | 42411988 | 629 | * | NA_peak_230 | 40 | Distal Interge | 15 | 42633306 | 42676749 | 43444 | 2 | 11600 | ENSMUST00 | 264761 | ENSMUSG00 | Angpt1 | angiopoietin 1 |
| chr15 | 42579929 | 42580257 | 329 | * | NA_peak_230 | 20 | Intron (ENSM | 15 | 42633306 | 42676749 | 43444 | 2 | 11600 | ENSMUST00 | 96492 | ENSMUSG00 | Angpt1 | angiopoietin 1 |
| chr15 | 42603021 | 42603555 | 535 | * | NA_peak_230 | 16 | Intron (ENSM | 15 | 42633306 | 42676749 | 43444 | 2 | 11600 | ENSMUST00 | 73194 | ENSMUSG00 | Angpt1 | angiopoietin 1 |
| chr15 | 42723667 | 42723772 | 106 | * | NA_peak_230 | 23 | Distal Interge | 15 | 42424723 | 42676977 | 252255 | 2 | 11600 | ENSMUST00 | -46690 | ENSMUSG00 | Angpt1 | angiopoietin 1 |
| chr15 | 43042905 | 43043202 | 298 | * | NA_peak_230 | 23 | Intron (ENSM | 15 | 43020797 | 43170818 | 150022 | 2 | 239405 | ENSMUST00 | 127616 | ENSMUSG00 | Rspo2 | R-spondin 2 |
| chr15 | 43075495 | 43075817 | 323 | * | NA_peak_230 | 20 | Exon (ENSMU | 15 | 43020797 | 43170818 | 150022 | 2 | 239405 | ENSMUST00 | 95001 | ENSMUSG00 | Rspo2 | R-spondin 2 |
| chr15 | 43229395 | 43229747 | 353 | * | NA_peak_231 | 32 | Distal Interge | 15 | 43261692 | 43268487 | 6796 | 2 | 16341 | ENSMUST00 | 38740 | ENSMUSG00 | Eif3e | eukaryotic translation initiation factor 3, subunit E |
| chr15 | 43288740 | 43289088 | 349 | * | NA_peak_231 | 50 | Distal Interge | 15 | 43250058 | 43282719 | 32662 | 2 | 16341 | ENSMUST00 | -6021 | ENSMUSG00 | Eif3e | eukaryotic translation initiation factor 3, subunit E |
| chr15 | 43345555 | 43345918 | 364 | * | NA_peak_231 | 16 | Distal Interge | 15 | 43250058 | 43282719 | 32662 | 2 | 16341 | ENSMUST00 | -62836 | ENSMUSG00 | Eif3e | eukaryotic translation initiation factor 3, subunit E |
| chr15 | 43435734 | 43435982 | 249 | * | NA_peak_231 | 23 | Intron (ENSM | 15 | 43430943 | 43477036 | 46094 | 2 | 100038728 | ENSMUST00 | 41054 | ENSMUSG00 | Gm10373 | predicted gene 10373 |
| chr15 | 43471087 | 43471406 | 320 | * | NA_peak_231 | 20 | Intron (ENSM | 15 | 43430943 | 43477036 | 46094 | 2 | 100038728 | ENSMUST00 | 5630 | ENSMUSG00 | Gm10373 | predicted gene 10373 |
| chr15 | 43497911 | 43497993 | 83 | * | NA_peak_231 | 30 | Intron (ENSM | 15 | 43477241 | 43493632 | 16392 | 1 | 66736 | ENSMUST00 | 20670 | ENSMUSG00 | Emc2 | ER membrane protein complex subunit 2 |
| chr15 | 43533791 | 43534090 | 300 | * | NA_peak_231 | 17 | Distal Interge | 15 | 43477241 | 43493632 | 16392 | 1 | 66736 | ENSMUST00 | 56550 | ENSMUSG00 | Emc2 | ER membrane protein complex subunit 2 |
| chr15 | 43587194 | 43587463 | 270 | * | NA_peak_231 | 20 | Distal Interge | 15 | 43477241 | 43493632 | 16392 | 1 | 66736 | ENSMUST00 | 109953 | ENSMUSG00 | Emc2 | ER membrane protein complex subunit 2 |
| chr15 | 43594613 | 43594909 | 297 | * | NA_peak_231 | 45 | Distal Interge | 15 | 43477241 | 43493632 | 16392 | 1 | 66736 | ENSMUST00 | 117372 | ENSMUSG00 | Emc2 | ER membrane protein complex subunit 2 |
| chr15 | 43667853 | 43668075 | 223 | * | NA_peak_231 | 36 | Distal Interge | 15 | 43477241 | 43493632 | 16392 | 1 | 66736 | ENSMUST00 | 190612 | ENSMUSG00 | Emc2 | ER membrane protein complex subunit 2 |
| chr15 | 43755932 | 43756141 | 210 | * | NA_peak_231 | 20 | Distal Interge | 15 | 43864646 | 43870036 | 5391 | 2 | 239408 | ENSMUST00 | 113895 | ENSMUSG00 | Tmem74 | transmembrane protein 74 |
| chr15 | 43759562 | 43760067 | 506 | * | NA_peak_231 | 31 | Distal Interge | 15 | 43864646 | 43870036 | 5391 | 2 | 239408 | ENSMUST00 | 109969 | ENSMUSG00 | Tmem74 | transmembrane protein 74 |
| chr15 | 43842185 | 43842586 | 402 | * | NA_peak_232 | 15 | Distal Interge | 15 | 43864646 | 43870036 | 5391 | 2 | 239408 | ENSMUST00 | 27450 | ENSMUSG00 | Tmem74 | transmembrane protein 74 |
| chr15 | 44014199 | 44014592 | 394 | * | NA_peak_232 | 16 | Distal Interge | 15 | 43864646 | 43870036 | 5391 | 2 | 239408 | ENSMUST00 | -144163 | ENSMUSG00 | Tmem74 | transmembrane protein 74 |
| chr15 | 44044193 | 44044391 | 199 | * | NA_peak_232 | 16 | Distal Interge | 15 | 44196135 | 44235544 | 39410 | 1 | 22045 | ENSMUST00 | -151744 | ENSMUSG00 | Trhr | thyrotropin releasing hormone receptor |
| chr15 | 44087070 | 44087537 | 468 | * | NA_peak_232 | 30 | Distal Interge | 15 | 44196135 | 44235544 | 39410 | 1 | 22045 | ENSMUST00 | -108598 | ENSMUSG00 | Trhr | thyrotropin releasing hormone receptor |
| chr15 | 44088331 | 44088589 | 259 | * | NA_peak_232 | 35 | Distal Interge | 15 | 44196135 | 44235544 | 39410 | 1 | 22045 | ENSMUST00 | -107546 | ENSMUSG00 | Trhr | thyrotropin releasing hormone receptor |
| chr15 | 44130831 | 44130865 | 35 | * | NA_peak_232 | 23 | Distal Interge | 15 | 44196135 | 44235544 | 39410 | 1 | 22045 | ENSMUST00 | -65270 | ENSMUSG00 | Trhr | thyrotropin releasing hormone receptor |
| chr15 | 44240265 | 44240677 | 413 | * | NA_peak_232 | 27 | Distal Interge | 15 | 44196967 | 44229201 | 32235 | 1 | 22045 | ENSMUST00 | 43298 | ENSMUSG00 | Trhr | thyrotropin releasing hormone receptor |
| chr15 | 44247855 | 44248155 | 301 | * | NA_peak_232 | 43 | Distal Interge | 15 | 44196967 | 44229201 | 32235 | 1 | 22045 | ENSMUST00 | 50888 | ENSMUSG00 | Trhr | thyrotropin releasing hormone receptor |
| chr15 | 44282314 | 44282376 | 63 | * | NA_peak_232 | 23 | Intron (ENSM | 15 | 44196967 | 44229201 | 32235 | 1 | 22045 | ENSMUST00 | 85347 | ENSMUSG00 | Trhr | thyrotropin releasing hormone receptor |
| chr15 | 44319784 | 44320067 | 284 | * | NA_peak_232 | 20 | Distal Interge | 15 | 44376368 | 44427785 | 51418 | 2 | 67429 | ENSMUST00 | 107718 | ENSMUSG00 | Nudcd1 | NudC domain containing 1 |
| chr15 | 44769265 | 44769530 | 266 | * | NA_peak_233 | 27 | Intron (ENSM | 15 | 44671856 | 44752458 | 80603 | 2 | 319613 | ENSMUST00 | -16807 | ENSMUSG00 | Sybu | syntabulin (syntaxin-interacting) |
| chr15 | 44779211 | 44779234 | 24 | * | NA_peak_233 | 16 | Intron (ENSM | 15 | 44787769 | 44805827 | 18059 | 1 | 69659 | ENSMUST00 | -8535 | ENSMUSG00 | 2310069G16 | RIKEN cDNA 2310069G16 gene |
| chr15 | 44795628 | 44796033 | 406 | * | NA_peak_233 | 46 | Intron (ENSM | 15 | 44760975 | 44788063 | 27089 | 2 | 319613 | ENSMUST00 | -7565 | ENSMUSG00 | Sybu | syntabulin (syntaxin-interacting) |
| chr15 | 44909960 | 44910505 | 546 | * | NA_peak_233 | 104 | Distal Interge | 15 | 44881419 | 44884302 | 2884 | 1 | 239410 | ENSMUST00 | 28541 | ENSMUSG00 | A930017M01 | Smg-5 homolog, nonsense mediated mRNA decay factor pseudogene |
| chr15 | 44939458 | 44939598 | 141 | * | NA_peak_233 | 27 | Distal Interge | 15 | 44881419 | 44884302 | 2884 | 1 | 239410 | ENSMUST00 | 58039 | ENSMUSG00 | A930017M01 | Smg-5 homolog, nonsense mediated mRNA decay factor pseudogene |
| chr15 | 44951140 | 44951458 | 319 | * | NA_peak_233 | 36 | Distal Interge | 15 | 44881419 | 44884302 | 2884 | 1 | 239410 | ENSMUST00 | 69721 | ENSMUSG00 | A930017M01 | Smg-5 homolog, nonsense mediated mRNA decay factor pseudogene |
| chr15 | 45067606 | 45067667 | 62 | * | NA_peak_233 | 22 | Distal Interge | 15 | 45106284 | 45114827 | 8544 | 2 | 67498 | ENSMUST00 | 47160 | ENSMUSG00 | Kcnv1 | potassium channel, subfamily V, member 1 |
| chr15 | 45163666 | 45163972 | 307 | * | NA_peak_233 | 20 | Exon (ENSMU | 15 | 45112785 | 45114920 | 2136 | 2 | 67498 | ENSMUST00 | -48746 | ENSMUSG00 | Kcnv1 | potassium channel, subfamily V, member 1 |
| chr15 | 45231076 | 45231393 | 318 | * | NA_peak_233 | 23 | Distal Interge | 15 | 45112785 | 45114920 | 2136 | 2 | 67498 | ENSMUST00 | -116156 | ENSMUSG00 | Kcnv1 | potassium channel, subfamily V, member 1 |
| chr15 | 45534785 | 45535062 | 278 | * | NA_peak_234 | 22 | Distal Interge | 15 | 45112785 | 45114920 | 2136 | 2 | 67498 | ENSMUST00 | -419865 | ENSMUSG00 | Kcnv1 | potassium channel, subfamily V, member 1 |
| chr15 | 45852400 | 45852820 | 421 | * | NA_peak_234 | 23 | Distal Interge | 15 | 46324559 | 46373858 | 49300 | 1 | 71120 | ENSMUST00 | -471739 | ENSMUSG00 | 1700022A22I | RIKEN cDNA 1700022A22 gene |
| chr15 | 45856591 | 45856950 | 360 | * | NA_peak_234 | 27 | Distal Interge | 15 | 46324559 | 46373858 | 49300 | 1 | 71120 | ENSMUST00 | -467609 | ENSMUSG00 | 1700022A22I | RIKEN cDNA 1700022A22 gene |
| chr15 | 45997286 | 45997959 | 674 | * | NA_peak_234 | 22 | Distal Interge | 15 | 46324559 | 46373858 | 49300 | 1 | 71120 | ENSMUST00 | -326600 | ENSMUSG00 | 1700022A22I | RIKEN cDNA 1700022A22 gene |
| chr15 | 46027816 | 46027991 | 176 | * | NA_peak_234 | 27 | Distal Interge | 15 | 46324559 | 46373858 | 49300 | 1 | 71120 | ENSMUST00 | -296568 | ENSMUSG00 | 1700022A22I | RIKEN cDNA 1700022A22 gene |
| chr15 | 46138500 | 46138869 | 370 | * | NA_peak_234 | 31 | Distal Interge | 15 | 46324559 | 46373858 | 49300 | 1 | 71120 | ENSMUST00 | -185690 | ENSMUSG00 | 1700022A22I | RIKEN cDNA 1700022A22 gene |
| chr15 | 46205711 | 46206505 | 795 | * | NA_peak_234 | 50 | Distal Interge | 15 | 46324559 | 46373858 | 49300 | 1 | 71120 | ENSMUST00 | -118054 | ENSMUSG00 | 1700022A22I | RIKEN cDNA 1700022A22 gene |
| chr15 | 46238447 | 46238517 | 71 | * | NA_peak_234 | 20 | Distal Interge | 15 | 46324559 | 46373858 | 49300 | 1 | 71120 | ENSMUST00 | -86042 | ENSMUSG00 | 1700022A22I | RIKEN cDNA 1700022A22 gene |
| chr15 | 46257359 | 46257677 | 319 | * | NA_peak_234 | 43 | Distal Interge | 15 | 46324559 | 46373858 | 49300 | 1 | 71120 | ENSMUST00 | -66882 | ENSMUSG00 | 1700022A22I | RIKEN cDNA 1700022A22 gene |
| chr15 | 46505091 | 46505434 | 344 | * | NA_peak_235 | 31 | Distal Interge | 15 | 46435866 | 46496181 | 60316 | 2 | 74726 | ENSMUST00 | -8910 | ENSMUSG00 | 4930523O13 | RIKEN cDNA 4930523O13 gene |
| chr15 | 46707521 | 46707685 | 165 | * | NA_peak_235 | 13 | Distal Interge | 15 | 46623305 | 46639858 | 16554 | 1 | 75281 | ENSMUST00 | 84216 | ENSMUSG00 | 4930548G14 | RIKEN cDNA 4930548G14 gene |

| | | | | | | | | | | | | | | | | | | |
|---|---|---|---|---|---|---|---|---|---|---|---|---|---|---|---|---|---|---|
| chr15 | 46950476 | 46950626 | 151 | * | NA_peak_23 | 27 | Distal Interge | 15 | 46623305 | 46639858 | 16554 | 1 | 75281 | ENSMUST00 | 327171 | ENSMUSG0 | 4930548G14 | RIKEN cDNA 4930548G14 gene |
| chr15 | 47183474 | 47183908 | 435 | * | NA_peak_23 | 130 | Distal Interge | 15 | 47581726 | 47615201 | 33476 | 2 | 239420 | ENSMUST00 | 431293 | ENSMUSG0 | Csmd3 | CUB and Sushi multiple domains 3 |
| chr15 | 47379509 | 47379704 | 196 | * | NA_peak_23 | 20 | Distal Interge | 15 | 47581726 | 47615201 | 33476 | 2 | 239420 | ENSMUST00 | 235497 | ENSMUSG0 | Csmd3 | CUB and Sushi multiple domains 3 |
| chr15 | 48020241 | 48020405 | 165 | * | NA_peak_23 | 27 | Intron (ENSM | 15 | 47581723 | 48001621 | 419899 | 2 | 239420 | ENSMUST00 | -18620 | ENSMUSG0 | Csmd3 | CUB and Sushi multiple domains 3 |
| chr15 | 48183711 | 48184060 | 350 | * | NA_peak_23 | 23 | Intron (ENSM | 15 | 47581723 | 48001621 | 419899 | 2 | 239420 | ENSMUST00 | -182090 | ENSMUSG0 | Csmd3 | CUB and Sushi multiple domains 3 |
| chr15 | 48400051 | 48400301 | 251 | * | NA_peak_23 | 19 | Intron (ENSM | 15 | 47580637 | 48791933 | 1211297 | 2 | 239420 | ENSMUST00 | 391632 | ENSMUSG0 | Csmd3 | CUB and Sushi multiple domains 3 |
| chr15 | 48697319 | 48697672 | 354 | * | NA_peak_23 | 27 | Intron (ENSM | 15 | 47580637 | 48791933 | 1211297 | 2 | 239420 | ENSMUST00 | 94261 | ENSMUSG0 | Csmd3 | CUB and Sushi multiple domains 3 |
| chr15 | 49171404 | 49171622 | 219 | * | NA_peak_23 | 13 | Distal Interge | 15 | 47580722 | 48792063 | 1211342 | 2 | 239420 | ENSMUST00 | -379341 | ENSMUSG0 | Csmd3 | CUB and Sushi multiple domains 3 |
| chr15 | 49176600 | 49177048 | 449 | * | NA_peak_23 | 45 | Distal Interge | 15 | 47580722 | 48792063 | 1211342 | 2 | 239420 | ENSMUST00 | -384537 | ENSMUSG0 | Csmd3 | CUB and Sushi multiple domains 3 |
| chr15 | 49243816 | 49243917 | 102 | * | NA_peak_23 | 36 | Distal Interge | 15 | 47580722 | 48792063 | 1211342 | 2 | 239420 | ENSMUST00 | -451753 | ENSMUSG0 | Csmd3 | CUB and Sushi multiple domains 3 |
| chr15 | 49403945 | 49404353 | 409 | * | NA_peak_23 | 36 | Distal Interge | 15 | 47580722 | 48792063 | 1211342 | 2 | 239420 | ENSMUST00 | -611882 | ENSMUSG0 | Csmd3 | CUB and Sushi multiple domains 3 |
| chr15 | 49622111 | 49622113 | 3 | * | NA_peak_23 | 20 | Distal Interge | 15 | 47580722 | 48792063 | 1211342 | 2 | 239420 | ENSMUST00 | -830048 | ENSMUSG0 | Csmd3 | CUB and Sushi multiple domains 3 |
| chr15 | 49662870 | 49662972 | 103 | * | NA_peak_23 | 22 | Distal Interge | 15 | 47580722 | 48792063 | 1211342 | 2 | 239420 | ENSMUST00 | -870807 | ENSMUSG0 | Csmd3 | CUB and Sushi multiple domains 3 |
| chr15 | 50060139 | 50060475 | 337 | * | NA_peak_23 | 28 | Distal Interge | 15 | 50819499 | 50827521 | 8023 | 2 | 83925 | ENSMUST00 | 767046 | ENSMUSG0 | Trps1 | transcriptional repressor GATA binding 1 |
| chr15 | 50364559 | 50364909 | 351 | * | NA_peak_23 | 14 | Distal Interge | 15 | 50819499 | 50827521 | 8023 | 2 | 83925 | ENSMUST00 | 462612 | ENSMUSG0 | Trps1 | transcriptional repressor GATA binding 1 |
| chr15 | 50482523 | 50482957 | 435 | * | NA_peak_23 | 31 | Distal Interge | 15 | 50819499 | 50827521 | 8023 | 2 | 83925 | ENSMUST00 | 344564 | ENSMUSG0 | Trps1 | transcriptional repressor GATA binding 1 |
| chr15 | 50491927 | 50492154 | 228 | * | NA_peak_23 | 40 | Distal Interge | 15 | 50819499 | 50827521 | 8023 | 2 | 83925 | ENSMUST00 | 335367 | ENSMUSG0 | Trps1 | transcriptional repressor GATA binding 1 |
| chr15 | 50533237 | 50533574 | 338 | * | NA_peak_23 | 48 | Distal Interge | 15 | 50819499 | 50827521 | 8023 | 2 | 83925 | ENSMUST00 | 293947 | ENSMUSG0 | Trps1 | transcriptional repressor GATA binding 1 |
| chr15 | 50558340 | 50558615 | 276 | * | NA_peak_23 | 14 | Distal Interge | 15 | 50819499 | 50827521 | 8023 | 2 | 83925 | ENSMUST00 | 268906 | ENSMUSG0 | Trps1 | transcriptional repressor GATA binding 1 |
| chr15 | 50580431 | 50580720 | 290 | * | NA_peak_23 | 16 | Distal Interge | 15 | 50819499 | 50827521 | 8023 | 2 | 83925 | ENSMUST00 | 246801 | ENSMUSG0 | Trps1 | transcriptional repressor GATA binding 1 |
| chr15 | 50643363 | 50643469 | 107 | * | NA_peak_23 | 24 | Distal Interge | 15 | 50819499 | 50827521 | 8023 | 2 | 83925 | ENSMUST00 | 184052 | ENSMUSG0 | Trps1 | transcriptional repressor GATA binding 1 |
| chr15 | 50666584 | 50666740 | 157 | * | NA_peak_23 | 18 | Intron (ENSM | 15 | 50819499 | 50827521 | 8023 | 2 | 83925 | ENSMUST00 | 160781 | ENSMUSG0 | Trps1 | transcriptional repressor GATA binding 1 |
| chr15 | 51092949 | 51093235 | 287 | * | NA_peak_23 | 41 | Distal Interge | 15 | 50846176 | 50890463 | 44288 | 2 | 83925 | ENSMUST00 | -202486 | ENSMUSG0 | Trps1 | transcriptional repressor GATA binding 1 |
| chr15 | 51401454 | 51401484 | 31 | * | NA_peak_23 | 27 | Distal Interge | 15 | 51231640 | 51395610 | 163971 | 2 | 100502643 | ENSMUST00 | -5844 | ENSMUSG0 | Gm19303 | predicted gene, 19303 |
| chr15 | 51452771 | 51453336 | 566 | * | NA_peak_23 | 40 | Distal Interge | 15 | 51231640 | 51395610 | 163971 | 2 | 100502643 | ENSMUST00 | -57161 | ENSMUSG0 | Gm19303 | predicted gene, 19303 |
| chr15 | 51476146 | 51476213 | 68 | * | NA_peak_23 | 36 | Distal Interge | 15 | 51231640 | 51395610 | 163971 | 2 | 100502643 | ENSMUST00 | -80536 | ENSMUSG0 | Gm19303 | predicted gene, 19303 |
| chr15 | 51833786 | 51834117 | 332 | * | NA_peak_23 | 16 | Promoter (2-3 | 15 | 51787969 | 51836257 | 48289 | 2 | 68135 | ENSMUST00 | 2140 | ENSMUSG0 | Eif3h | eukaryotic translation initiation factor 3, subunit H |
| chr15 | 51846814 | 51847061 | 248 | * | NA_peak_23 | 26 | Intron (ENSM | 15 | 51787969 | 51836257 | 48289 | 2 | 68135 | ENSMUST00 | -10557 | ENSMUSG0 | Eif3h | eukaryotic translation initiation factor 3, subunit H |
| chr15 | 51877413 | 51877783 | 371 | * | NA_peak_23 | 45 | Promoter (<= | 15 | 51877429 | 51884614 | 7186 | 1 | 78581 | ENSMUST00 | 0 | ENSMUSG0 | Utp23 | UTP23 small subunit processome component |
| chr15 | 52334038 | 52334175 | 138 | * | NA_peak_24 | 23 | Intron (ENSM | 15 | 52295553 | 52335798 | 40246 | 1 | 239436 | ENSMUST00 | 38485 | ENSMUSG0 | Slc30a8 | solute carrier family 30 (zinc transporter), member 8 |
| chr15 | 52426355 | 52426594 | 240 | * | NA_peak_24 | 30 | Distal Interge | 15 | 52295553 | 52335798 | 40246 | 1 | 239436 | ENSMUST00 | 130802 | ENSMUSG0 | Slc30a8 | solute carrier family 30 (zinc transporter), member 8 |
| chr15 | 52650190 | 52650482 | 293 | * | NA_peak_24 | 45 | Distal Interge | 15 | 52712408 | 52715272 | 2865 | 1 | 69790 | ENSMUST00 | -61926 | ENSMUSG0 | Med30 | mediator complex subunit 30 |
| chr15 | 52662155 | 52662526 | 372 | * | NA_peak_24 | 42 | Distal Interge | 15 | 52712408 | 52715272 | 2865 | 1 | 69790 | ENSMUST00 | -49882 | ENSMUSG0 | Med30 | mediator complex subunit 30 |
| chr15 | 52729741 | 52729773 | 33 | * | NA_peak_24 | 26 | Intron (ENSM | 15 | 52712442 | 52730564 | 18123 | 1 | 69790 | ENSMUST00 | 17299 | ENSMUSG0 | Med30 | mediator complex subunit 30 |
| chr15 | 52772286 | 52772981 | 696 | * | NA_peak_24 | 39 | Intron (ENSM | 15 | 52712442 | 52730564 | 18123 | 1 | 69790 | ENSMUST00 | 59844 | ENSMUSG0 | Med30 | mediator complex subunit 30 |
| chr15 | 52782940 | 52782940 | 1 | * | NA_peak_24 | 48 | Intron (ENSM | 15 | 52712442 | 52730564 | 18123 | 1 | 69790 | ENSMUST00 | 70498 | ENSMUSG0 | Med30 | mediator complex subunit 30 |
| chr15 | 52833777 | 52833927 | 151 | * | NA_peak_24 | 14 | Distal Interge | 15 | 52712442 | 52730564 | 18123 | 1 | 69790 | ENSMUST00 | 121335 | ENSMUSG0 | Med30 | mediator complex subunit 30 |
| chr15 | 52899402 | 52899673 | 272 | * | NA_peak_24 | 23 | Distal Interge | 15 | 52712442 | 52730564 | 18123 | 1 | 69790 | ENSMUST00 | 186960 | ENSMUSG0 | Med30 | mediator complex subunit 30 |
| chr15 | 52937305 | 52937345 | 41 | * | NA_peak_24 | 23 | Distal Interge | 15 | 53089931 | 53107868 | 17938 | 2 | 14042 | ENSMUST00 | 170523 | ENSMUSG0 | Ext1 | exostosin glycosyltransferase 1 |
| chr15 | 52937621 | 52938243 | 623 | * | NA_peak_24 | 36 | Distal Interge | 15 | 53089931 | 53107868 | 17938 | 2 | 14042 | ENSMUST00 | 169625 | ENSMUSG0 | Ext1 | exostosin glycosyltransferase 1 |
| chr15 | 52970956 | 52971288 | 333 | * | NA_peak_24 | 19 | Distal Interge | 15 | 53089931 | 53107868 | 17938 | 2 | 14042 | ENSMUST00 | 136580 | ENSMUSG0 | Ext1 | exostosin glycosyltransferase 1 |
| chr15 | 53137165 | 53137622 | 458 | * | NA_peak_24 | 31 | Intron (ENSM | 15 | 53089931 | 53107868 | 17938 | 2 | 14042 | ENSMUST00 | -29297 | ENSMUSG0 | Ext1 | exostosin glycosyltransferase 1 |
| chr15 | 53200184 | 53200569 | 386 | * | NA_peak_24 | 17 | Intron (ENSM | 15 | 53083146 | 53196255 | 113110 | 2 | 14042 | ENSMUST00 | -3929 | ENSMUSG0 | Ext1 | exostosin glycosyltransferase 1 |
| chr15 | 53379166 | 53379503 | 338 | * | NA_peak_24 | 16 | Distal Interge | 15 | 53064038 | 53346159 | 282122 | 2 | 14042 | ENSMUST00 | -33007 | ENSMUSG0 | Ext1 | exostosin glycosyltransferase 1 |
| chr15 | 53470581 | 53470886 | 306 | * | NA_peak_24 | 17 | Intron (ENSM | 15 | 53064038 | 53346159 | 282122 | 2 | 14042 | ENSMUST00 | -124422 | ENSMUSG0 | Ext1 | exostosin glycosyltransferase 1 |
| chr15 | 53537449 | 53537482 | 34 | * | NA_peak_24 | 30 | Intron (ENSM | 15 | 53614205 | 53614279 | 75 | 2 | 100628567 | ENSMUST00 | 76797 | ENSMUSG0 | Mir28c | microRNA 28c |
| chr15 | 53579372 | 53579589 | 218 | * | NA_peak_24 | 57 | Intron (ENSM | 15 | 53614205 | 53614279 | 75 | 2 | 100628567 | ENSMUST00 | 34690 | ENSMUSG0 | Mir28c | microRNA 28c |
| chr15 | 53654746 | 53654839 | 94 | * | NA_peak_24 | 26 | Intron (ENSM | 15 | 53614205 | 53614279 | 75 | 2 | 100628567 | ENSMUST00 | -40467 | ENSMUSG0 | Mir28c | microRNA 28c |
| chr15 | 53745307 | 53745862 | 556 | * | NA_peak_24 | 22 | Intron (ENSM | 15 | 53614205 | 53614279 | 75 | 2 | 100628567 | ENSMUST00 | -131028 | ENSMUSG0 | Mir28c | microRNA 28c |
| chr15 | 53853615 | 53853963 | 349 | * | NA_peak_24 | 28 | Intron (ENSM | 15 | 53462194 | 53902381 | 440188 | 2 | 320679 | ENSMUST00 | 48418 | ENSMUSG0 | Samd12 | sterile alpha motif domain containing 12 |
| chr15 | 53867482 | 53867765 | 284 | * | NA_peak_24 | 22 | Intron (ENSM | 15 | 53462194 | 53902381 | 440188 | 2 | 320679 | ENSMUST00 | 34616 | ENSMUSG0 | Samd12 | sterile alpha motif domain containing 12 |
| chr15 | 53889488 | 53889701 | 214 | * | NA_peak_24 | 17 | Intron (ENSM | 15 | 53462194 | 53902381 | 440188 | 2 | 320679 | ENSMUST00 | 12680 | ENSMUSG0 | Samd12 | sterile alpha motif domain containing 12 |
| chr15 | 53954839 | 53955001 | 163 | * | NA_peak_24 | 23 | Distal Interge | 15 | 53818794 | 53902537 | 83744 | 2 | 320679 | ENSMUST00 | -52302 | ENSMUSG0 | Samd12 | sterile alpha motif domain containing 12 |
| chr15 | 54004852 | 54005123 | 272 | * | NA_peak_24 | 26 | Distal Interge | 15 | 53818794 | 53902537 | 83744 | 2 | 320679 | ENSMUST00 | -102315 | ENSMUSG0 | Samd12 | sterile alpha motif domain containing 12 |
| chr15 | 54099854 | 54100243 | 390 | * | NA_peak_24 | 36 | Distal Interge | 15 | 54250619 | 54278484 | 27866 | 2 | 18383 | ENSMUST00 | 178241 | ENSMUSG0 | Tnfrsf11b | tumor necrosis factor receptor superfamily, member 11b (osteoprotegerin) |
| chr15 | 54135655 | 54135902 | 248 | * | NA_peak_24 | 27 | Distal Interge | 15 | 54250619 | 54278484 | 27866 | 2 | 18383 | ENSMUST00 | 142582 | ENSMUSG0 | Tnfrsf11b | tumor necrosis factor receptor superfamily, member 11b (osteoprotegerin) |
| chr15 | 54136359 | 54136511 | 153 | * | NA_peak_24 | 27 | Distal Interge | 15 | 54250619 | 54278484 | 27866 | 2 | 18383 | ENSMUST00 | 141973 | ENSMUSG0 | Tnfrsf11b | tumor necrosis factor receptor superfamily, member 11b (osteoprotegerin) |
| chr15 | 54147485 | 54147914 | 430 | * | NA_peak_24 | 31 | Distal Interge | 15 | 54250619 | 54278484 | 27866 | 2 | 18383 | ENSMUST00 | 130570 | ENSMUSG0 | Tnfrsf11b | tumor necrosis factor receptor superfamily, member 11b (osteoprotegerin) |
| chr15 | 54186799 | 54187114 | 316 | * | NA_peak_24 | 27 | Distal Interge | 15 | 54250619 | 54278484 | 27866 | 2 | 18383 | ENSMUST00 | 91370 | ENSMUSG0 | Tnfrsf11b | tumor necrosis factor receptor superfamily, member 11b (osteoprotegerin) |
| chr15 | 54190883 | 54191275 | 393 | * | NA_peak_24 | 23 | Distal Interge | 15 | 54250619 | 54278484 | 27866 | 2 | 18383 | ENSMUST00 | 87209 | ENSMUSG0 | Tnfrsf11b | tumor necrosis factor receptor superfamily, member 11b (osteoprotegerin) |
| chr15 | 54233024 | 54233522 | 499 | * | NA_peak_24 | 26 | Distal Interge | 15 | 54250619 | 54278484 | 27866 | 2 | 18383 | ENSMUST00 | 44962 | ENSMUSG0 | Tnfrsf11b | tumor necrosis factor receptor superfamily, member 11b (osteoprotegerin) |
| chr15 | 54345492 | 54345604 | 113 | * | NA_peak_24 | 27 | Distal Interge | 15 | 54410774 | 54466358 | 55585 | 1 | 239447 | ENSMUST00 | -65170 | ENSMUSG0 | Colec10 | collectin sub-family member 10 |
| chr15 | 54351987 | 54352075 | 89 | * | NA_peak_24 | 27 | Distal Interge | 15 | 54410774 | 54466358 | 55585 | 1 | 239447 | ENSMUST00 | -58699 | ENSMUSG0 | Colec10 | collectin sub-family member 10 |
| chr15 | 54419674 | 54420209 | 536 | * | NA_peak_24 | 36 | Intron (ENSM | 15 | 54410774 | 54466358 | 55585 | 1 | 239447 | ENSMUST00 | 8900 | ENSMUSG0 | Colec10 | collectin sub-family member 10 |
| chr15 | 54446416 | 54446816 | 401 | * | NA_peak_24 | 27 | Intron (ENSM | 15 | 54410774 | 54466358 | 55585 | 1 | 239447 | ENSMUST00 | 35642 | ENSMUSG0 | Colec10 | collectin sub-family member 10 |
| chr15 | 54449896 | 54450274 | 379 | * | NA_peak_24 | 50 | Intron (ENSM | 15 | 54410774 | 54466358 | 55585 | 1 | 239447 | ENSMUST00 | 39122 | ENSMUSG0 | Colec10 | collectin sub-family member 10 |
| chr15 | 54558312 | 54558396 | 85 | * | NA_peak_24 | 40 | Distal Interge | 15 | 54571192 | 54602847 | 31656 | 1 | 105853 | ENSMUST00 | -12796 | ENSMUSG0 | Mal2 | mal, T cell differentiation protein 2 |
| chr15 | 54565573 | 54565909 | 337 | * | NA_peak_24 | 16 | Distal Interge | 15 | 54571192 | 54602847 | 31656 | 1 | 105853 | ENSMUST00 | -5283 | ENSMUSG0 | Mal2 | mal, T cell differentiation protein 2 |
| chr15 | 54568368 | 54568948 | 581 | * | NA_peak_24 | 31 | Promoter (2-3 | 15 | 54571192 | 54602847 | 31656 | 1 | 105853 | ENSMUST00 | -2244 | ENSMUSG0 | Mal2 | mal, T cell differentiation protein 2 |
| chr15 | 54613177 | 54613213 | 37 | * | NA_peak_24 | 16 | Distal Interge | 15 | 54571192 | 54602847 | 31656 | 1 | 105853 | ENSMUST00 | 41985 | ENSMUSG0 | Mal2 | mal, T cell differentiation protein 2 |
| chr15 | 54697400 | 54697759 | 360 | * | NA_peak_24 | 34 | Distal Interge | 15 | 54745702 | 54754039 | 8338 | 1 | 18133 | ENSMUST00 | -47943 | ENSMUSG0 | Ccn3 | cellular communication network factor 3 |
| chr15 | 54841827 | 54842095 | 269 | * | NA_peak_24 | 42 | Intron (ENSM | 15 | 54859798 | 54862686 | 2889 | 2 | 18606 | ENSMUST00 | 20591 | ENSMUSG0 | Enpp2 | ectonucleotide pyrophosphatase/phosphodiesterase 2 |
| chr15 | 54884102 | 54884197 | 96 | * | NA_peak_24 | 20 | Intron (ENSM | 15 | 54859798 | 54862686 | 2889 | 2 | 18606 | ENSMUST00 | -21416 | ENSMUSG0 | Enpp2 | ectonucleotide pyrophosphatase/phosphodiesterase 2 |
| chr15 | 54906961 | 54907118 | 158 | * | NA_peak_24 | 43 | Intron (ENSM | 15 | 54899276 | 54914718 | 15443 | 2 | 18606 | ENSMUST00 | 7600 | ENSMUSG0 | Enpp2 | ectonucleotide pyrophosphatase/phosphodiesterase 2 |
| chr15 | 54907403 | 54907615 | 213 | * | NA_peak_24 | 55 | Intron (ENSM | 15 | 54899276 | 54914718 | 15443 | 2 | 18606 | ENSMUST00 | 7103 | ENSMUSG0 | Enpp2 | ectonucleotide pyrophosphatase/phosphodiesterase 2 |
| chr15 | 55329494 | 55329689 | 196 | * | NA_peak_24 | 16 | Intron (ENSM | 15 | 55307919 | 55520203 | 212285 | 1 | 12818 | ENSMUST00 | 21575 | ENSMUSG0 | Col14a1 | collagen, type XIV, alpha 1 |
| chr15 | 55409317 | 55409708 | 392 | * | NA_peak_24 | 19 | Intron (ENSM | 15 | 55448826 | 55519689 | 70864 | 1 | 12818 | ENSMUST00 | -39118 | ENSMUSG0 | Col14a1 | collagen, type XIV, alpha 1 |
| chr15 | 55615042 | 55615233 | 192 | * | NA_peak_24 | 15 | Promoter (2-3 | 15 | 55617587 | 55619938 | 2352 | 1 | 105837 | ENSMUST00 | -2354 | ENSMUSG0 | Mtbp | Mdm2, transformed 3T3 cell double minute p53 binding protein |
| chr15 | 55641750 | 55642060 | 311 | * | NA_peak_24 | 26 | Intron (ENSM | 15 | 55621452 | 55626421 | 4970 | 1 | 105837 | ENSMUST00 | 20298 | ENSMUSG0 | Mtbp | Mdm2, transformed 3T3 cell double minute p53 binding protein |
| chr15 | 55674556 | 55674842 | 287 | * | NA_peak_24 | 34 | Intron (ENSM | 15 | 55621452 | 55626421 | 4970 | 1 | 105837 | ENSMUST00 | 53104 | ENSMUSG0 | Mtbp | Mdm2, transformed 3T3 cell double minute p53 binding protein |
| chr15 | 55781786 | 55782070 | 285 | * | NA_peak_24 | 23 | Intron (ENSM | 15 | 55791938 | 55906310 | 114373 | 2 | 20649 | ENSMUST00 | 124240 | ENSMUSG0 | Sntb1 | syntrophin, basic 1 |
| chr15 | 55794758 | 55795168 | 411 | * | NA_peak_24 | 36 | Intron (ENSM | 15 | 55791938 | 55906310 | 114373 | 2 | 20649 | ENSMUST00 | 111142 | ENSMUSG0 | Sntb1 | syntrophin, basic 1 |
| chr15 | 55802264 | 55802548 | 285 | * | NA_peak_24 | 31 | Intron (ENSM | 15 | 55791938 | 55906310 | 114373 | 2 | 20649 | ENSMUST00 | 103762 | ENSMUSG0 | Sntb1 | syntrophin, basic 1 |
| chr15 | 55809093 | 55809461 | 369 | * | NA_peak_24 | 13 | Intron (ENSM | 15 | 55791938 | 55906310 | 114373 | 2 | 20649 | ENSMUST00 | 96849 | ENSMUSG0 | Sntb1 | syntrophin, basic 1 |
| chr15 | 55858115 | 55858398 | 284 | * | NA_peak_24 | 21 | Intron (ENSM | 15 | 55791938 | 55906310 | 114373 | 2 | 20649 | ENSMUST00 | 47912 | ENSMUSG0 | Sntb1 | syntrophin, basic 1 |
| chr15 | 55873332 | 55873534 | 203 | * | NA_peak_24 | 37 | Intron (ENSM | 15 | 55791938 | 55906310 | 114373 | 2 | 20649 | ENSMUST00 | 32776 | ENSMUSG0 | Sntb1 | syntrophin, basic 1 |
| chr15 | 56088380 | 56088407 | 28 | * | NA_peak_24 | 20 | Distal Interge | 15 | 55636388 | 55906949 | 270562 | 2 | 20649 | ENSMUST00 | -181431 | ENSMUSG0 | Sntb1 | syntrophin, basic 1 |
| chr15 | 56154926 | 56155249 | 324 | * | NA_peak_24 | 45 | Distal Interge | 15 | 55636388 | 55906949 | 270562 | 2 | 20649 | ENSMUST00 | -247977 | ENSMUSG0 | Sntb1 | syntrophin, basic 1 |
| chr15 | 56228171 | 56228449 | 279 | * | NA_peak_24 | 22 | Distal Interge | 15 | 55636388 | 55906949 | 270562 | 2 | 20649 | ENSMUST00 | -321222 | ENSMUSG0 | Sntb1 | syntrophin, basic 1 |
| chr15 | 56367422 | 56367529 | 108 | * | NA_peak_24 | 21 | Distal Interge | 15 | 56689944 | 56694460 | 4517 | 1 | 594843 | ENSMUST00 | -322415 | ENSMUSG0 | Has2os | hyaluronan synthase 2, opposite strand |
| chr15 | 56394744 | 56395094 | 351 | * | NA_peak_24 | 27 | Distal Interge | 15 | 56689944 | 56694460 | 4517 | 1 | 594843 | ENSMUST00 | -294850 | ENSMUSG0 | Has2os | hyaluronan synthase 2, opposite strand |
| chr15 | 56479530 | 56479880 | 351 | * | NA_peak_24 | 29 | Distal Interge | 15 | 56689944 | 56694460 | 4517 | 1 | 594843 | ENSMUST00 | -210064 | ENSMUSG0 | Has2os | hyaluronan synthase 2, opposite strand |
| chr15 | 56748533 | 56748766 | 234 | * | NA_peak_24 | 38 | Intron (ENSM | 15 | 56665627 | 56694539 | 28913 | 2 | 15117 | ENSMUST00 | -53994 | ENSMUSG0 | Has2 | hyaluronan synthase 2 |
| chr15 | 56756136 | 56756484 | 349 | * | NA_peak_24 | 22 | Intron (ENSM | 15 | 56665627 | 56694539 | 28913 | 2 | 15117 | ENSMUST00 | -61597 | ENSMUSG0 | Has2 | hyaluronan synthase 2 |
| chr15 | 56793380 | 56793686 | 307 | * | NA_peak_24 | 22 | Distal Interge | 15 | 56665627 | 56694539 | 28913 | 2 | 15117 | ENSMUST00 | -98841 | ENSMUSG0 | Has2 | hyaluronan synthase 2 |
| chr15 | 56804341 | 56804655 | 315 | * | NA_peak_24 | 20 | Distal Interge | 15 | 56665627 | 56694539 | 28913 | 2 | 15117 | ENSMUST00 | -109802 | ENSMUSG0 | Has2 | hyaluronan synthase 2 |
| chr15 | 56844478 | 56844662 | 185 | * | NA_peak_24 | 31 | Distal Interge | 15 | 56665627 | 56694539 | 28913 | 2 | 15117 | ENSMUST00 | -149939 | ENSMUSG0 | Has2 | hyaluronan synthase 2 |
| chr15 | 56891078 | 56891431 | 354 | * | NA_peak_24 | 30 | Intron (ENSM | 15 | 56665627 | 56694539 | 28913 | 2 | 15117 | ENSMUST00 | -196539 | ENSMUSG0 | Has2 | hyaluronan synthase 2 |
| chr15 | 56902302 | 56902595 | 294 | * | NA_peak_24 | 29 | Intron (ENSM | 15 | 56665627 | 56694539 | 28913 | 2 | 15117 | ENSMUST00 | -207763 | ENSMUSG0 | Has2 | hyaluronan synthase 2 |
| chr15 | 56926389 | 56926625 | 237 | * | NA_peak_24 | 31 | Intron (ENSM | 15 | 56665627 | 56694539 | 28913 | 2 | 15117 | ENSMUST00 | -231850 | ENSMUSG0 | Has2 | hyaluronan synthase 2 |
| chr15 | 56946482 | 56946850 | 369 | * | NA_peak_24 | 32 | Intron (ENSM | 15 | 56665627 | 56694539 | 28913 | 2 | 15117 | ENSMUST00 | -251943 | ENSMUSG0 | Has2 | hyaluronan synthase 2 |
| chr15 | 56964587 | 56964828 | 242 | * | NA_peak_24 | 32 | Intron (ENSM | 15 | 56665627 | 56694539 | 28913 | 2 | 15117 | ENSMUST00 | -270048 | ENSMUSG0 | Has2 | hyaluronan synthase 2 |
| chr15 | 56987193 | 56988043 | 851 | * | NA_peak_24 | 35 | Intron (ENSM | 15 | 57256556 | 57265137 | 8582 | 2 | 210463 | ENSMUST00 | 277094 | ENSMUSG0 | Slc22a22 | solute carrier family 22 (organic cation transporter), member 22 |
| chr15 | 57045170 | 57045310 | 141 | * | NA_peak_24 | 19 | Distal Interge | 15 | 57256556 | 57265137 | 8582 | 2 | 210463 | ENSMUST00 | 219827 | ENSMUSG0 | Slc22a22 | solute carrier family 22 (organic cation transporter), member 22 |
| chr15 | 57068296 | 57068587 | 292 | * | NA_peak_24 | 25 | Distal Interge | 15 | 57256556 | 57265137 | 8582 | 2 | 210463 | ENSMUST00 | 196550 | ENSMUSG0 | Slc22a22 | solute carrier family 22 (organic cation transporter), member 22 |
| chr15 | 57206968 | 57207067 | 100 | * | NA_peak_24 | 36 | Distal Interge | 15 | 57256556 | 57265137 | 8582 | 2 | 210463 | ENSMUST00 | 58070 | ENSMUSG0 | Slc22a22 | solute carrier family 22 (organic cation transporter), member 22 |

| | | | | | | | | | | | | | | | | | | |
|---|---|---|---|---|---|---|---|---|---|---|---|---|---|---|---|---|---|---|
| chr15 | 57246969 | 57247124 | 156 | * | NA_peak_247 | 19 | Intron (ENSM | 15 | 57256556 | 57265137 | 8582 | 2 | 210463 | ENSMUST00 | 18013 | ENSMUSG0( | Slc22a22 | solute carrier family 22 (organic cation transporter), member 22 |
| chr15 | 57280681 | 57281053 | 373 | * | NA_peak_247 | 21 | Intron (ENSM | 15 | 57243767 | 57265161 | 21395 | 2 | 210463 | ENSMUST00 | -15520 | ENSMUSG0( | Slc22a22 | solute carrier family 22 (organic cation transporter), member 22 |
| chr15 | 57335311 | 57335574 | 264 | * | NA_peak_247 | 27 | Intron (ENSM | 15 | 57243767 | 57265161 | 21395 | 2 | 210463 | ENSMUST00 | -70150 | ENSMUSG0( | Slc22a22 | solute carrier family 22 (organic cation transporter), member 22 |
| chr15 | 57415176 | 57415652 | 477 | * | NA_peak_247 | 22 | Intron (ENSM | 15 | 57243771 | 57477625 | 233855 | 2 | 210463 | ENSMUST00 | 61973 | ENSMUSG0( | Slc22a22 | solute carrier family 22 (organic cation transporter), member 22 |
| chr15 | 57441669 | 57441911 | 243 | * | NA_peak_247 | 29 | Intron (ENSM | 15 | 57243771 | 57477625 | 233855 | 2 | 210463 | ENSMUST00 | 35714 | ENSMUSG0( | Slc22a22 | solute carrier family 22 (organic cation transporter), member 22 |
| chr15 | 57481997 | 57482173 | 177 | * | NA_peak_247 | 23 | Distal Interge | 15 | 57243771 | 57477625 | 233855 | 2 | 210463 | ENSMUST00 | -4372 | ENSMUSG0( | Slc22a22 | solute carrier family 22 (organic cation transporter), member 22 |
| chr15 | 57628392 | 57628781 | 390 | * | NA_peak_247 | 37 | Distal Interge | 15 | 57694665 | 57839832 | 145168 | 1 | 387609 | ENSMUST00 | -65884 | ENSMUSG0( | Zhx2 | zinc fingers and homeoboxes 2 |
| chr15 | 57804866 | 57805110 | 245 | * | NA_peak_247 | 30 | Intron (ENSM | 15 | 57871663 | 57890394 | 18732 | 2 | 67819 | ENSMUST00 | 85284 | ENSMUSG0( | Derl1 | Der1-like domain family, member 1 |
| chr15 | 58049452 | 58049911 | 460 | * | NA_peak_247 | 43 | Intron (ENSM | 15 | 58030956 | 58034325 | 3370 | 2 | 75758 | ENSMUST00 | -15127 | ENSMUSG0( | 9130401M01 | RIKEN cDNA 9130401M01 gene |
| chr15 | 58062248 | 58062263 | 16 | * | NA_peak_247 | 36 | Intron (ENSM | 15 | 58048547 | 58076183 | 27637 | 2 | 22770 | ENSMUST00 | 13920 | ENSMUSG0( | Zhx1 | zinc fingers and homeoboxes 1 |
| chr15 | 58103227 | 58103723 | 497 | * | NA_peak_248 | 45 | Exon (ENSMU | 15 | 58094044 | 58098776 | 4733 | 2 | 70472 | ENSMUST00 | -4451 | ENSMUSG0( | Atad2 | ATPase family, AAA domain containing 2 |
| chr15 | 58425269 | 58425564 | 296 | * | NA_peak_248 | 27 | Intron (ENSM | 15 | 58432260 | 58433233 | 974 | 1 | 210998 | ENSMUST00 | -6696 | ENSMUSG0( | Fam91a1 | family with sequence similarity 91, member A1 |
| chr15 | 58783491 | 58783863 | 373 | * | NA_peak_248 | 16 | 3' UTR | 15 | 58782269 | 58823530 | 41262 | 2 | 74868 | ENSMUST00 | 39667 | ENSMUSG0( | Tmem65 | transmembrane protein 65 |
| chr15 | 58788655 | 58789225 | 571 | * | NA_peak_248 | 20 | Intron (ENSM | 15 | 58782269 | 58823530 | 41262 | 2 | 74868 | ENSMUST00 | 34305 | ENSMUSG0( | Tmem65 | transmembrane protein 65 |
| chr15 | 58918281 | 58918953 | 673 | * | NA_peak_248 | 27 | Promoter (2-3 | 15 | 58901017 | 58916178 | 15162 | 2 | 69694 | ENSMUST00 | -2103 | ENSMUSG0( | Tatdn1 | TatD DNase domain containing 1 |
| chr15 | 59420531 | 59421079 | 549 | * | NA_peak_248 | 30 | Intron (ENSM | 15 | 59374287 | 59601684 | 227398 | 1 | 68501 | ENSMUST00 | 46244 | ENSMUSG0( | Nsmce2 | NSE2/MMS21 homolog, SMC5-SMC6 complex SUMO ligase |
| chr15 | 59422592 | 59422675 | 84 | * | NA_peak_248 | 31 | Intron (ENSM | 15 | 59374287 | 59601684 | 227398 | 1 | 68501 | ENSMUST00 | 48305 | ENSMUSG0( | Nsmce2 | NSE2/MMS21 homolog, SMC5-SMC6 complex SUMO ligase |
| chr15 | 59515797 | 59515873 | 77 | * | NA_peak_248 | 34 | Intron (ENSM | 15 | 59648350 | 59657099 | 8750 | 1 | 211770 | ENSMUST00 | -132477 | ENSMUSG0( | Trib1 | tribbles pseudokinase 1 |
| chr15 | 59682705 | 59683154 | 450 | * | NA_peak_249 | 26 | Distal Interge | 15 | 59648653 | 59653689 | 5037 | 1 | 211770 | ENSMUST00 | 34052 | ENSMUSG0( | Trib1 | tribbles pseudokinase 1 |
| chr15 | 59972505 | 59972609 | 105 | * | NA_peak_249 | 14 | Distal Interge | 15 | 60014866 | 60016613 | 1748 | 2 | 71088 | ENSMUST00 | 44004 | ENSMUSG0( | 4933412E24I | RIKEN cDNA 4933412E24 gene |
| chr15 | 60022974 | 60023217 | 244 | * | NA_peak_249 | 40 | Distal Interge | 15 | 60014866 | 60016613 | 1748 | 2 | 71088 | ENSMUST00 | -6361 | ENSMUSG0( | 4933412E24I | RIKEN cDNA 4933412E24 gene |
| chr15 | 60063209 | 60063494 | 286 | * | NA_peak_249 | 39 | Distal Interge | 15 | 60014866 | 60016613 | 1748 | 2 | 71088 | ENSMUST00 | -46596 | ENSMUSG0( | 4933412E24I | RIKEN cDNA 4933412E24 gene |
| chr15 | 60152662 | 60152973 | 312 | * | NA_peak_249 | 21 | Intron (ENSM | 15 | 60014866 | 60016613 | 1748 | 2 | 71088 | ENSMUST00 | -136049 | ENSMUSG0( | 4933412E24I | RIKEN cDNA 4933412E24 gene |
| chr15 | 60174136 | 60174511 | 376 | * | NA_peak_249 | 52 | Intron (ENSM | 15 | 60014866 | 60016613 | 1748 | 2 | 71088 | ENSMUST00 | -157523 | ENSMUSG0( | 4933412E24I | RIKEN cDNA 4933412E24 gene |
| chr15 | 60184507 | 60184630 | 124 | * | NA_peak_249 | 31 | Intron (ENSM | 15 | 60014866 | 60016613 | 1748 | 2 | 71088 | ENSMUST00 | -167894 | ENSMUSG0( | 4933412E24I | RIKEN cDNA 4933412E24 gene |
| chr15 | 60192663 | 60192801 | 139 | * | NA_peak_249 | 24 | Distal Interge | 15 | 60014866 | 60016613 | 1748 | 2 | 71088 | ENSMUST00 | -176050 | ENSMUSG0( | 4933412E24I | RIKEN cDNA 4933412E24 gene |
| chr15 | 60296826 | 60297281 | 456 | * | NA_peak_249 | 36 | Exon (ENSMU | 15 | 60014866 | 60016613 | 1748 | 2 | 71088 | ENSMUST00 | -280213 | ENSMUSG0( | 4933412E24I | RIKEN cDNA 4933412E24 gene |
| chr15 | 60340595 | 60340627 | 33 | * | NA_peak_249 | 27 | Distal Interge | 15 | 60014866 | 60016613 | 1748 | 2 | 71088 | ENSMUST00 | -323982 | ENSMUSG0( | 4933412E24I | RIKEN cDNA 4933412E24 gene |
| chr15 | 60341091 | 60341191 | 101 | * | NA_peak_249 | 31 | Distal Interge | 15 | 60014866 | 60016613 | 1748 | 2 | 71088 | ENSMUST00 | -324478 | ENSMUSG0( | 4933412E24I | RIKEN cDNA 4933412E24 gene |
| chr15 | 60363248 | 60363367 | 120 | * | NA_peak_249 | 20 | Distal Interge | 15 | 60014866 | 60016613 | 1748 | 2 | 71088 | ENSMUST00 | -346635 | ENSMUSG0( | 4933412E24I | RIKEN cDNA 4933412E24 gene |
| chr15 | 60470330 | 60470622 | 293 | * | NA_peak_250 | 45 | Distal Interge | 15 | 60822271 | 60828650 | 6380 | 1 | 320469 | ENSMUST00 | -351649 | NA | 9930014A18I | RIKEN cDNA 9930014A18 gene |
| chr15 | 60484143 | 60484815 | 673 | * | NA_peak_250 | 45 | Distal Interge | 15 | 60822271 | 60828650 | 6380 | 1 | 320469 | ENSMUST00 | -337456 | NA | 9930014A18I | RIKEN cDNA 9930014A18 gene |
| chr15 | 60514031 | 60514598 | 568 | * | NA_peak_250 | 32 | Distal Interge | 15 | 60822271 | 60828650 | 6380 | 1 | 320469 | ENSMUST00 | -307673 | NA | 9930014A18I | RIKEN cDNA 9930014A18 gene |
| chr15 | 60722885 | 60723162 | 278 | * | NA_peak_250 | 22 | Distal Interge | 15 | 60822271 | 60828650 | 6380 | 1 | 320469 | ENSMUST00 | -99109 | NA | 9930014A18I | RIKEN cDNA 9930014A18 gene |
| chr15 | 60727990 | 60728146 | 157 | * | NA_peak_250 | 40 | Distal Interge | 15 | 60822271 | 60828650 | 6380 | 1 | 320469 | ENSMUST00 | -94125 | NA | 9930014A18I | RIKEN cDNA 9930014A18 gene |
| chr15 | 60789994 | 60790328 | 335 | * | NA_peak_250 | 43 | Distal Interge | 15 | 60822271 | 60828650 | 6380 | 1 | 320469 | ENSMUST00 | -31943 | NA | 9930014A18I | RIKEN cDNA 9930014A18 gene |
| chr15 | 61026175 | 61026799 | 625 | * | NA_peak_250 | 21 | Distal Interge | 15 | 60919675 | 60923012 | 3338 | 2 | 117586 | ENSMUST00 | -103163 | ENSMUSG0( | A1bg | alpha-1-B glycoprotein |
| chr15 | 61027606 | 61028060 | 455 | * | NA_peak_250 | 61 | Distal Interge | 15 | 60919675 | 60923012 | 3338 | 2 | 117586 | ENSMUST00 | -104594 | ENSMUSG0( | A1bg | alpha-1-B glycoprotein |
| chr15 | 61046025 | 61046458 | 434 | * | NA_peak_250 | 27 | Distal Interge | 15 | 60919675 | 60923012 | 3338 | 2 | 117586 | ENSMUST00 | -123013 | ENSMUSG0( | A1bg | alpha-1-B glycoprotein |
| chr15 | 61065667 | 61066012 | 346 | * | NA_peak_250 | 36 | Distal Interge | 15 | 60919675 | 60923012 | 3338 | 2 | 117586 | ENSMUST00 | -142655 | ENSMUSG0( | A1bg | alpha-1-B glycoprotein |
| chr15 | 61286346 | 61286614 | 269 | * | NA_peak_251 | 36 | Distal Interge | 15 | 60919675 | 60923012 | 3338 | 2 | 117586 | ENSMUST00 | -363334 | ENSMUSG0( | A1bg | alpha-1-B glycoprotein |
| chr15 | 61288397 | 61288758 | 362 | * | NA_peak_251 | 23 | Exon (ENSMU | 15 | 60919675 | 60923012 | 3338 | 2 | 117586 | ENSMUST00 | -365385 | ENSMUSG0( | A1bg | alpha-1-B glycoprotein |
| chr15 | 61298194 | 61298555 | 362 | * | NA_peak_251 | 31 | Intron (ENSM | 15 | 60919675 | 60923012 | 3338 | 2 | 117586 | ENSMUST00 | -375182 | ENSMUSG0( | A1bg | alpha-1-B glycoprotein |
| chr15 | 61363529 | 61364049 | 521 | * | NA_peak_251 | 28 | Intron (ENSM | 15 | 61534676 | 61745184 | 210509 | 2 | 432956 | ENSMUST00 | 381135 | ENSMUSG0( | D030024E09 | RIKEN cDNA D030024E09 gene |
| chr15 | 61374584 | 61374874 | 291 | * | NA_peak_251 | 31 | Intron (ENSM | 15 | 61534676 | 61745184 | 210509 | 2 | 432956 | ENSMUST00 | 370310 | ENSMUSG0( | D030024E09 | RIKEN cDNA D030024E09 gene |
| chr15 | 61495361 | 61495800 | 440 | * | NA_peak_251 | 21 | Distal Interge | 15 | 61534676 | 61745184 | 210509 | 2 | 432956 | ENSMUST00 | 249384 | ENSMUSG0( | D030024E09 | RIKEN cDNA D030024E09 gene |
| chr15 | 61507709 | 61508033 | 325 | * | NA_peak_251 | 26 | Distal Interge | 15 | 61534676 | 61745184 | 210509 | 2 | 432956 | ENSMUST00 | 237151 | ENSMUSG0( | D030024E09 | RIKEN cDNA D030024E09 gene |
| chr15 | 61540263 | 61540725 | 463 | * | NA_peak_251 | 36 | Intron (ENSM | 15 | 61534676 | 61745184 | 210509 | 2 | 432956 | ENSMUST00 | 204459 | ENSMUSG0( | D030024E09 | RIKEN cDNA D030024E09 gene |
| chr15 | 61831879 | 61832192 | 314 | * | NA_peak_252 | 50 | Distal Interge | 15 | 61699282 | 61774451 | 75170 | 2 | 432956 | ENSMUST00 | -57428 | ENSMUSG0( | D030024E09 | RIKEN cDNA D030024E09 gene |
| chr15 | 61876751 | 61876903 | 153 | * | NA_peak_252 | 40 | Distal Interge | 15 | 61699282 | 61774451 | 75170 | 2 | 432956 | ENSMUST00 | -102300 | ENSMUSG0( | D030024E09 | RIKEN cDNA D030024E09 gene |
| chr15 | 61879020 | 61879251 | 232 | * | NA_peak_252 | 23 | Distal Interge | 15 | 61699282 | 61774451 | 75170 | 2 | 432956 | ENSMUST00 | -104569 | ENSMUSG0( | D030024E09 | RIKEN cDNA D030024E09 gene |
| chr15 | 62150632 | 62150928 | 297 | * | NA_peak_252 | 40 | Intron (ENSM | 15 | 62178175 | 62250975 | 72801 | 1 | 19296 | ENSMUST00 | -27247 | ENSMUSG0( | Pvt1 | Pvt1 oncogene |
| chr15 | 62308280 | 62308664 | 385 | * | NA_peak_252 | 20 | Intron (ENSM | 15 | 62222603 | 62242505 | 19903 | 1 | 19296 | ENSMUST00 | 85677 | ENSMUSG0( | Pvt1 | Pvt1 oncogene |
| chr15 | 62432939 | 62433283 | 345 | * | NA_peak_252 | 30 | Distal Interge | 15 | 62222603 | 62242505 | 19903 | 1 | 19296 | ENSMUST00 | 210336 | ENSMUSG0( | Pvt1 | Pvt1 oncogene |
| chr15 | 62447675 | 62448212 | 538 | * | NA_peak_252 | 25 | Distal Interge | 15 | 62222603 | 62242505 | 19903 | 1 | 19296 | ENSMUST00 | 225072 | ENSMUSG0( | Pvt1 | Pvt1 oncogene |
| chr15 | 62482728 | 62483080 | 353 | * | NA_peak_252 | 23 | Distal Interge | 15 | 62222603 | 62242505 | 19903 | 1 | 19296 | ENSMUST00 | 260125 | ENSMUSG0( | Pvt1 | Pvt1 oncogene |
| chr15 | 62553595 | 62553952 | 358 | * | NA_peak_252 | 23 | Distal Interge | 15 | 62222603 | 62242505 | 19903 | 1 | 19296 | ENSMUST00 | 330992 | ENSMUSG0( | Pvt1 | Pvt1 oncogene |
| chr15 | 62625763 | 62626035 | 273 | * | NA_peak_253 | 24 | Intron (ENSM | 15 | 62222603 | 62242505 | 19903 | 1 | 19296 | ENSMUST00 | 403160 | ENSMUSG0( | Pvt1 | Pvt1 oncogene |
| chr15 | 62640444 | 62640446 | 3 | * | NA_peak_253 | 28 | Intron (ENSM | 15 | 62222603 | 62242505 | 19903 | 1 | 19296 | ENSMUST00 | 417841 | ENSMUSG0( | Pvt1 | Pvt1 oncogene |
| chr15 | 62646061 | 62646308 | 248 | * | NA_peak_253 | 19 | Intron (ENSM | 15 | 62222603 | 62242505 | 19903 | 1 | 19296 | ENSMUST00 | 423458 | ENSMUSG0( | Pvt1 | Pvt1 oncogene |
| chr15 | 62661642 | 62661967 | 326 | * | NA_peak_253 | 36 | Intron (ENSM | 15 | 62222603 | 62242505 | 19903 | 1 | 19296 | ENSMUST00 | 439039 | ENSMUSG0( | Pvt1 | Pvt1 oncogene |
| chr15 | 62684991 | 62685198 | 208 | * | NA_peak_253 | 27 | Intron (ENSM | 15 | 62222603 | 62242505 | 19903 | 1 | 19296 | ENSMUST00 | 462388 | ENSMUSG0( | Pvt1 | Pvt1 oncogene |
| chr15 | 62995809 | 62996225 | 417 | * | NA_peak_253 | 22 | Distal Interge | 15 | 63414400 | 63479608 | 65209 | 2 | 432958 | ENSMUST00 | 483383 | ENSMUSG0( | Gm20740 | predicted gene, 20740 |
| chr15 | 63073695 | 63074111 | 417 | * | NA_peak_253 | 36 | Intron (ENSM | 15 | 63414400 | 63479608 | 65209 | 2 | 432958 | ENSMUST00 | 405497 | ENSMUSG0( | Gm20740 | predicted gene, 20740 |
| chr15 | 63118943 | 63119232 | 290 | * | NA_peak_253 | 20 | Intron (ENSM | 15 | 63414400 | 63479608 | 65209 | 2 | 432958 | ENSMUST00 | 360376 | ENSMUSG0( | Gm20740 | predicted gene, 20740 |
| chr15 | 63192536 | 63192794 | 259 | * | NA_peak_253 | 26 | Intron (ENSM | 15 | 63414400 | 63479608 | 65209 | 2 | 432958 | ENSMUST00 | 286814 | ENSMUSG0( | Gm20740 | predicted gene, 20740 |
| chr15 | 63333247 | 63333548 | 302 | * | NA_peak_254 | 20 | Intron (ENSM | 15 | 63414400 | 63479608 | 65209 | 2 | 432958 | ENSMUST00 | 146060 | ENSMUSG0( | Gm20740 | predicted gene, 20740 |
| chr15 | 63485380 | 63485897 | 518 | * | NA_peak_254 | 40 | Intron (ENSM | 15 | 63414400 | 63479608 | 65209 | 2 | 432958 | ENSMUST00 | -5772 | ENSMUSG0( | Gm20740 | predicted gene, 20740 |
| chr15 | 63509830 | 63509955 | 126 | * | NA_peak_254 | 31 | Intron (ENSM | 15 | 63414400 | 63479608 | 65209 | 2 | 432958 | ENSMUST00 | -30222 | ENSMUSG0( | Gm20740 | predicted gene, 20740 |
| chr15 | 63589671 | 63589742 | 72 | * | NA_peak_254 | 23 | Intron (ENSM | 15 | 63414400 | 63479608 | 65209 | 2 | 432958 | ENSMUST00 | -110063 | ENSMUSG0( | Gm20740 | predicted gene, 20740 |
| chr15 | 63719566 | 63720014 | 449 | * | NA_peak_254 | 20 | Promoter (1-2 | 15 | 63176838 | 63718289 | 541452 | 2 | 432958 | ENSMUST00 | -1277 | ENSMUSG0( | Gm20740 | predicted gene, 20740 |
| chr15 | 63809445 | 63809728 | 284 | * | NA_peak_254 | 23 | Promoter (<= | 15 | 63775971 | 63808759 | 32789 | 2 | 83492 | ENSMUST00 | -686 | ENSMUSG0( | Gsdmc | gasdermin C |
| chr15 | 63810997 | 63811300 | 304 | * | NA_peak_254 | 22 | Promoter (2-3 | 15 | 63775971 | 63808759 | 32789 | 2 | 83492 | ENSMUST00 | -2238 | ENSMUSG0( | Gsdmc | gasdermin C |
| chr15 | 63838810 | 63839107 | 298 | * | NA_peak_254 | 56 | Promoter (2-3 | 15 | 63824346 | 63835949 | 11604 | 2 | 331063 | ENSMUST00 | -2861 | ENSMUSG0( | Gsdmc2 | gasdermin C2 |
| chr15 | 63883237 | 63883452 | 216 | * | NA_peak_254 | 46 | Promoter (1-2 | 15 | 63882047 | 63888765 | 6719 | 1 | 665769 | ENSMUST00 | 1190 | ENSMUSG0( | Gsdmcl2 | gasdermin C-like 2 |
| chr15 | 63890209 | 63890492 | 284 | * | NA_peak_254 | 19 | Exon (ENSMU | 15 | 63882047 | 63888765 | 6719 | 1 | 665769 | ENSMUST00 | 8162 | ENSMUSG0( | Gsdmcl2 | gasdermin C-like 2 |
| chr15 | 64089559 | 64089657 | 99 | * | NA_peak_254 | 25 | Intron (ENSM | 15 | 64094234 | 64102542 | 8309 | 2 | 13196 | ENSMUST00 | 12885 | ENSMUSG0( | Asap1 | ArfGAP with SH3 domain, ankyrin repeat and PH domain1 |
| chr15 | 64425250 | 64425494 | 245 | * | NA_peak_255 | 29 | Intron (ENSM | 15 | 64088541 | 64382919 | 294379 | 2 | 13196 | ENSMUST00 | -42331 | ENSMUSG0( | Asap1 | ArfGAP with SH3 domain, ankyrin repeat and PH domain1 |
| chr15 | 64651594 | 64651885 | 292 | * | NA_peak_255 | 20 | Distal Interge | 15 | 64088541 | 64382919 | 294379 | 2 | 13196 | ENSMUST00 | -268675 | ENSMUSG0( | Asap1 | ArfGAP with SH3 domain, ankyrin repeat and PH domain1 |
| chr15 | 64808058 | 64808118 | 61 | * | NA_peak_255 | 15 | Intron (ENSM | 15 | 64782562 | 64922271 | 139710 | 2 | 11514 | ENSMUST00 | 114153 | ENSMUSG0( | Adcy8 | adenylate cyclase 8 |
| chr15 | 64888554 | 64888810 | 257 | * | NA_peak_255 | 39 | Intron (ENSM | 15 | 64782562 | 64922271 | 139710 | 2 | 11514 | ENSMUST00 | 33461 | ENSMUSG0( | Adcy8 | adenylate cyclase 8 |
| chr15 | 65143060 | 65143697 | 638 | * | NA_peak_255 | 36 | Distal Interge | 15 | 64697084 | 64922296 | 225213 | 2 | 11514 | ENSMUST00 | -220764 | ENSMUSG0( | Adcy8 | adenylate cyclase 8 |
| chr15 | 65166177 | 65166437 | 261 | * | NA_peak_255 | 16 | Distal Interge | 15 | 64697084 | 64922296 | 225213 | 2 | 11514 | ENSMUST00 | -243881 | ENSMUSG0( | Adcy8 | adenylate cyclase 8 |
| chr15 | 65382552 | 65383176 | 625 | * | NA_peak_256 | 39 | Distal Interge | 15 | 65787034 | 65791351 | 4318 | 1 | 76740 | ENSMUST00 | -403858 | ENSMUSG0( | Efr3a | EFR3 homolog A |
| chr15 | 65652671 | 65652719 | 49 | * | NA_peak_256 | 16 | Distal Interge | 15 | 65787034 | 65791351 | 4318 | 1 | 76740 | ENSMUST00 | -134315 | ENSMUSG0( | Efr3a | EFR3 homolog A |
| chr15 | 65653736 | 65654218 | 483 | * | NA_peak_256 | 29 | Distal Interge | 15 | 65787034 | 65791351 | 4318 | 1 | 76740 | ENSMUST00 | -132816 | ENSMUSG0( | Efr3a | EFR3 homolog A |
| chr15 | 65926448 | 65926460 | 13 | * | NA_peak_257 | 14 | Intron (ENSM | 15 | 65876053 | 65912397 | 36345 | 2 | 18256 | ENSMUST00 | -14051 | ENSMUSG0( | Oc90 | otoconin 90 |
| chr15 | 65963790 | 65964134 | 345 | * | NA_peak_257 | 23 | Intron (ENSM | 15 | 65916609 | 65976804 | 60196 | 2 | 654498 | ENSMUST00 | 12670 | ENSMUSG0( | Hhla1 | HERV-H LTR-associating 1 |
| chr15 | 65992271 | 65992736 | 466 | * | NA_peak_257 | 25 | 3' UTR | 15 | 65922443 | 65976804 | 54362 | 2 | 654498 | ENSMUST00 | -15467 | ENSMUSG0( | Hhla1 | HERV-H LTR-associating 1 |
| chr15 | 66000899 | 66001232 | 334 | * | NA_peak_257 | 20 | Intron (ENSM | 15 | 65922443 | 65976804 | 54362 | 2 | 654498 | ENSMUST00 | -24095 | ENSMUSG0( | Hhla1 | HERV-H LTR-associating 1 |
| chr15 | 66041467 | 66041491 | 25 | * | NA_peak_257 | 36 | Intron (ENSM | 15 | 66025116 | 66103254 | 78139 | 2 | 110862 | ENSMUST00 | 61763 | ENSMUSG0( | Kcnq3 | potassium voltage-gated channel, subfamily Q, member 3 |
| chr15 | 66064627 | 66064899 | 273 | * | NA_peak_257 | 33 | Intron (ENSM | 15 | 66025116 | 66103254 | 78139 | 2 | 110862 | ENSMUST00 | 38355 | ENSMUSG0( | Kcnq3 | potassium voltage-gated channel, subfamily Q, member 3 |
| chr15 | 66115890 | 66116315 | 426 | * | NA_peak_257 | 15 | Intron (ENSM | 15 | 66025116 | 66103254 | 78139 | 2 | 110862 | ENSMUST00 | -12636 | ENSMUSG0( | Kcnq3 | potassium voltage-gated channel, subfamily Q, member 3 |
| chr15 | 66401925 | 66402060 | 136 | * | NA_peak_257 | 18 | Intron (ENSM | 15 | 66379858 | 66500910 | 121053 | 2 | 54562 | ENSMUST00 | 98850 | ENSMUSG0( | Lrrc6 | leucine rich repeat containing 6 (testis) |
| chr15 | 66482444 | 66482649 | 206 | * | NA_peak_257 | 16 | Intron (ENSM | 15 | 66379858 | 66500910 | 121053 | 2 | 54562 | ENSMUST00 | 18261 | ENSMUSG0( | Lrrc6 | leucine rich repeat containing 6 (testis) |
| chr15 | 66624278 | 66624582 | 305 | * | NA_peak_258 | 23 | Intron (ENSM | 15 | 66629910 | 66637032 | 7123 | 1 | 239510 | ENSMUST00 | -5328 | ENSMUSG0( | Phf20l1 | PHD finger protein 20-like 1 |
| chr15 | 66656389 | 66656736 | 348 | * | NA_peak_258 | 38 | Distal Interge | 15 | 66670753 | 66850721 | 179969 | 1 | 21819 | ENSMUST00 | -14017 | ENSMUSG0( | Tg | thyroglobulin |
| chr15 | 66658350 | 66658682 | 333 | * | NA_peak_258 | 38 | Distal Interge | 15 | 66670753 | 66850721 | 179969 | 1 | 21819 | ENSMUST00 | -12071 | ENSMUSG0( | Tg | thyroglobulin |
| chr15 | 66659689 | 66660099 | 411 | * | NA_peak_258 | 45 | Distal Interge | 15 | 66670753 | 66850721 | 179969 | 1 | 21819 | ENSMUST00 | -10654 | ENSMUSG0( | Tg | thyroglobulin |
| chr15 | 66991926 | 66992010 | 85 | * | NA_peak_258 | 23 | Intron (ENSM | 15 | 66946537 | 66969640 | 23104 | 2 | 17988 | ENSMUST00 | -22286 | ENSMUSG0( | Ndrg1 | N-myc downstream regulated gene 1 |
| chr15 | 67134851 | 67135104 | 254 | * | NA_peak_258 | 30 | Intron (ENSM | 15 | 67114238 | 67129023 | 14786 | 2 | 20442 | ENSMUST00 | -5828 | ENSMUSG0( | St3gal1 | ST3 beta-galactoside alpha-2,3-sialyltransferase 1 |
| chr15 | 67367884 | 67368499 | 616 | * | NA_peak_258 | 25 | Intron (ENSM | 15 | 67226769 | 67377094 | 150326 | 1 | 69334 | ENSMUST00 | 141115 | ENSMUSG0( | 1700012I11R | RIKEN cDNA 1700012I11 gene |
| chr15 | 67378096 | 67378225 | 130 | * | NA_peak_258 | 19 | Distal Interge | 15 | 67226769 | 67377094 | 150326 | 1 | 69334 | ENSMUST00 | 151327 | ENSMUSG0( | 1700012I11R | RIKEN cDNA 1700012I11 gene |
| chr15 | 67543228 | 67543773 | 546 | * | NA_peak_258 | 40 | Distal Interge | 15 | 67226769 | 67377094 | 150326 | 1 | 69334 | ENSMUST00 | 316459 | ENSMUSG0( | 1700012I11R | RIKEN cDNA 1700012I11 gene |
| chr15 | 67556100 | 67556408 | 309 | * | NA_peak_258 | 20 | Distal Interge | 15 | 67226769 | 67377094 | 150326 | 1 | 69334 | ENSMUST00 | 329331 | ENSMUSG0( | 1700012I11R | RIKEN cDNA 1700012I11 gene |
| chr15 | 67651903 | 67652101 | 199 | * | NA_peak_258 | 20 | Distal Interge | 15 | 67226769 | 67377094 | 150326 | 1 | 69334 | ENSMUST00 | 425134 | ENSMUSG0( | 1700012I11R | RIKEN cDNA 1700012I11 gene |

| | | | | | | | | | | | | | | | | | | |
|---|---|---|---|---|---|---|---|---|---|---|---|---|---|---|---|---|---|---|
| chr15 | 67796681 | 67796832 | 152 | * | NA_peak_25 | 24 | Distal Interge | 15 | 68084321 | 68258787 | 174467 | 2 | 380993 | ENSMUST00 | 461955 | ENSMUSG0 | Zfat | zinc finger and AT hook domain containing |
| chr15 | 68136259 | 68136318 | 60 | * | NA_peak_25 | 20 | Intron (ENSM | 15 | 68084321 | 68258787 | 174467 | 2 | 380993 | ENSMUST00 | 122469 | ENSMUSG0 | Zfat | zinc finger and AT hook domain containing |
| chr15 | 68324946 | 68324957 | 12 | * | NA_peak_25 | 14 | Exon (ENSMU | 15 | 68334715 | 68337430 | 2716 | 2 | 387226 | ENSMUST00 | 12473 | ENSMUSG0 | Mir30b | microRNA 30b |
| chr15 | 68390571 | 68390892 | 322 | * | NA_peak_25 | 20 | Distal Interge | 15 | 68341208 | 68341289 | 82 | 2 | 387228 | ENSMUST00 | -49282 | ENSMUSG0 | Mir30d | microRNA 30d |
| chr15 | 68537202 | 68537662 | 461 | * | NA_peak_25 | 29 | Distal Interge | 15 | 68341208 | 68341289 | 82 | 2 | 387228 | ENSMUST00 | -195913 | ENSMUSG0 | Mir30d | microRNA 30d |
| chr15 | 68575341 | 68575698 | 358 | * | NA_peak_25 | 20 | Distal Interge | 15 | 68341208 | 68341289 | 82 | 2 | 387228 | ENSMUST00 | -234052 | ENSMUSG0 | Mir30d | microRNA 30d |
| chr15 | 68605157 | 68605521 | 365 | * | NA_peak_25 | 31 | Distal Interge | 15 | 68341208 | 68341289 | 82 | 2 | 387228 | ENSMUST00 | -263868 | ENSMUSG0 | Mir30d | microRNA 30d |
| chr15 | 68628650 | 68628969 | 320 | * | NA_peak_25 | 55 | Distal Interge | 15 | 68878315 | 68883576 | 5262 | 1 | 105245971 | ENSMUST00 | -249346 | ENSMUSG0 | Gm41341 | predicted gene, 41341 |
| chr15 | 68863863 | 68864110 | 248 | * | NA_peak_26 | 27 | Intron (ENSM | 15 | 68878315 | 68883576 | 5262 | 1 | 105245971 | ENSMUST00 | -14205 | ENSMUSG0 | Gm41341 | predicted gene, 41341 |
| chr15 | 68875033 | 68875321 | 289 | * | NA_peak_26 | 23 | Promoter (2-3 | 15 | 68878315 | 68883576 | 5262 | 1 | 105245971 | ENSMUST00 | -2994 | ENSMUSG0 | Gm41341 | predicted gene, 41341 |
| chr15 | 69078390 | 69078725 | 336 | * | NA_peak_26 | 27 | Intron (ENSM | 15 | 68930068 | 69024765 | 94698 | 1 | 13992 | ENSMUST00 | 148322 | ENSMUSG0 | Khdrbs3 | KH domain containing, RNA binding, signal transduction associated 3 |
| chr15 | 69117333 | 69117583 | 251 | * | NA_peak_26 | 33 | Distal Interge | 15 | 69270533 | 69274720 | 4188 | 1 | 75076 | ENSMUST00 | -152950 | ENSMUSG0 | 4930504C09 | RIKEN cDNA 4930504C09 gene |
| chr15 | 69221198 | 69221528 | 331 | * | NA_peak_26 | 17 | Distal Interge | 15 | 69270533 | 69274720 | 4188 | 1 | 75076 | ENSMUST00 | -49005 | ENSMUSG0 | 4930504C09 | RIKEN cDNA 4930504C09 gene |
| chr15 | 69293923 | 69293966 | 44 | * | NA_peak_26 | 31 | Distal Interge | 15 | 69270533 | 69274720 | 4188 | 1 | 75076 | ENSMUST00 | 23390 | ENSMUSG0 | 4930504C09 | RIKEN cDNA 4930504C09 gene |
| chr15 | 69646243 | 69646699 | 457 | * | NA_peak_26 | 16 | Distal Interge | 15 | 70011623 | 70077841 | 66219 | 1 | 100503580 | ENSMUST00 | -364924 | ENSMUSG0 | Gm19782 | predicted gene, 19782 |
| chr15 | 69894194 | 69894571 | 378 | * | NA_peak_26 | 20 | Distal Interge | 15 | 70011623 | 70077841 | 66219 | 1 | 100503580 | ENSMUST00 | -117052 | ENSMUSG0 | Gm19782 | predicted gene, 19782 |
| chr15 | 70087496 | 70087766 | 271 | * | NA_peak_26 | 45 | Distal Interge | 15 | 70011623 | 70077841 | 66219 | 1 | 100503580 | ENSMUST00 | 75873 | ENSMUSG0 | Gm19782 | predicted gene, 19782 |
| chr15 | 70695010 | 70695467 | 458 | * | NA_peak_26 | 16 | Distal Interge | 15 | 70011623 | 70077841 | 66219 | 1 | 100503580 | ENSMUST00 | 683387 | ENSMUSG0 | Gm19782 | predicted gene, 19782 |
| chr15 | 70697428 | 70697538 | 111 | * | NA_peak_26 | 17 | Distal Interge | 15 | 70011623 | 70077841 | 66219 | 1 | 100503580 | ENSMUST00 | 685805 | ENSMUSG0 | Gm19782 | predicted gene, 19782 |
| chr15 | 70733480 | 70733785 | 306 | * | NA_peak_26 | 31 | Distal Interge | 15 | 70011623 | 70077841 | 66219 | 1 | 100503580 | ENSMUST00 | 721857 | ENSMUSG0 | Gm19782 | predicted gene, 19782 |
| chr15 | 70856870 | 70857090 | 221 | * | NA_peak_26 | 21 | Distal Interge | 15 | 71445676 | 71462400 | 16725 | 2 | 70363 | ENSMUST00 | 605310 | ENSMUSG0 | Fam135b | family with sequence similarity 135, member B |
| chr15 | 70873693 | 70873986 | 294 | * | NA_peak_26 | 31 | Distal Interge | 15 | 71445676 | 71462400 | 16725 | 2 | 70363 | ENSMUST00 | 588414 | ENSMUSG0 | Fam135b | family with sequence similarity 135, member B |
| chr15 | 70937944 | 70938197 | 254 | * | NA_peak_26 | 27 | Distal Interge | 15 | 71445676 | 71462400 | 16725 | 2 | 70363 | ENSMUST00 | 524203 | ENSMUSG0 | Fam135b | family with sequence similarity 135, member B |
| chr15 | 71209633 | 71209887 | 255 | * | NA_peak_26 | 22 | Distal Interge | 15 | 71445676 | 71462400 | 16725 | 2 | 70363 | ENSMUST00 | 252513 | ENSMUSG0 | Fam135b | family with sequence similarity 135, member B |
| chr15 | 71345729 | 71345850 | 122 | * | NA_peak_26 | 13 | Distal Interge | 15 | 71445676 | 71462400 | 16725 | 2 | 70363 | ENSMUST00 | 116550 | ENSMUSG0 | Fam135b | family with sequence similarity 135, member B |
| chr15 | 71406543 | 71406970 | 428 | * | NA_peak_26 | 31 | Distal Interge | 15 | 71445676 | 71462400 | 16725 | 2 | 70363 | ENSMUST00 | 55430 | ENSMUSG0 | Fam135b | family with sequence similarity 135, member B |
| chr15 | 71518336 | 71518647 | 312 | * | NA_peak_26 | 31 | Intron (ENSM | 15 | 71445676 | 71462400 | 16725 | 2 | 70363 | ENSMUST00 | -55936 | ENSMUSG0 | Fam135b | family with sequence similarity 135, member B |
| chr15 | 71561148 | 71561368 | 221 | * | NA_peak_26 | 23 | Intron (ENSM | 15 | 71445676 | 71462400 | 16725 | 2 | 70363 | ENSMUST00 | -98748 | ENSMUSG0 | Fam135b | family with sequence similarity 135, member B |
| chr15 | 71582871 | 71583078 | 208 | * | NA_peak_26 | 27 | Intron (ENSM | 15 | 71445676 | 71462400 | 16725 | 2 | 70363 | ENSMUST00 | -120471 | ENSMUSG0 | Fam135b | family with sequence similarity 135, member B |
| chr15 | 71797447 | 71797456 | 10 | * | NA_peak_26 | 26 | 3' UTR | 15 | 71487411 | 71727838 | 240428 | 2 | 70363 | ENSMUST00 | -69609 | ENSMUSG0 | Fam135b | family with sequence similarity 135, member B |
| chr15 | 71821264 | 71821421 | 158 | * | NA_peak_26 | 27 | Intron (ENSM | 15 | 71868881 | 71897616 | 28736 | 2 | 69700 | ENSMUST00 | 76195 | ENSMUSG0 | Col22a1 | collagen, type XXII, alpha 1 |
| chr15 | 71841290 | 71841680 | 391 | * | NA_peak_26 | 15 | Exon (ENSMU | 15 | 71868881 | 71897616 | 28736 | 2 | 69700 | ENSMUST00 | 55936 | ENSMUSG0 | Col22a1 | collagen, type XXII, alpha 1 |
| chr15 | 71870107 | 71870574 | 468 | * | NA_peak_26 | 43 | Exon (ENSMU | 15 | 71868881 | 71897616 | 28736 | 2 | 69700 | ENSMUST00 | 27042 | ENSMUSG0 | Col22a1 | collagen, type XXII, alpha 1 |
| chr15 | 71937718 | 71938097 | 380 | * | NA_peak_26 | 22 | Exon (ENSMU | 15 | 71904728 | 71954374 | 49647 | 2 | 69700 | ENSMUST00 | 16277 | ENSMUSG0 | Col22a1 | collagen, type XXII, alpha 1 |
| chr15 | 72002892 | 72003072 | 181 | * | NA_peak_26 | 30 | Intron (ENSM | 15 | 71795795 | 72034227 | 238433 | 2 | 69700 | ENSMUST00 | 31155 | ENSMUSG0 | Col22a1 | collagen, type XXII, alpha 1 |
| chr15 | 72004683 | 72004888 | 206 | * | NA_peak_26 | 22 | Intron (ENSM | 15 | 71795795 | 72034227 | 238433 | 2 | 69700 | ENSMUST00 | 29339 | ENSMUSG0 | Col22a1 | collagen, type XXII, alpha 1 |
| chr15 | 72084964 | 72084984 | 21 | * | NA_peak_26 | 25 | Distal Interge | 15 | 71795795 | 72034227 | 238433 | 2 | 69700 | ENSMUST00 | -50737 | ENSMUSG0 | Col22a1 | collagen, type XXII, alpha 1 |
| chr15 | 72237120 | 72237492 | 373 | * | NA_peak_26 | 17 | Intron (ENSM | 15 | 71795795 | 72034227 | 238433 | 2 | 69700 | ENSMUST00 | -202893 | ENSMUSG0 | Col22a1 | collagen, type XXII, alpha 1 |
| chr15 | 72249279 | 72249513 | 235 | * | NA_peak_26 | 37 | Intron (ENSM | 15 | 71795795 | 72034227 | 238433 | 2 | 69700 | ENSMUST00 | -215052 | ENSMUSG0 | Col22a1 | collagen, type XXII, alpha 1 |
| chr15 | 72253892 | 72254225 | 334 | * | NA_peak_26 | 15 | Intron (ENSM | 15 | 71795795 | 72034227 | 238433 | 2 | 69700 | ENSMUST00 | -219665 | ENSMUSG0 | Col22a1 | collagen, type XXII, alpha 1 |
| chr15 | 72313548 | 72313570 | 23 | * | NA_peak_26 | 18 | Distal Interge | 15 | 72501089 | 72546340 | 45252 | 2 | 223604 | ENSMUST00 | 232770 | ENSMUSG0 | Kcnk9 | potassium channel, subfamily K, member 9 |
| chr15 | 72318940 | 72319200 | 261 | * | NA_peak_26 | 28 | Distal Interge | 15 | 72501089 | 72546340 | 45252 | 2 | 223604 | ENSMUST00 | 227140 | ENSMUSG0 | Kcnk9 | potassium channel, subfamily K, member 9 |
| chr15 | 72342668 | 72342673 | 6 | * | NA_peak_26 | 40 | Distal Interge | 15 | 72501089 | 72546340 | 45252 | 2 | 223604 | ENSMUST00 | 203667 | ENSMUSG0 | Kcnk9 | potassium channel, subfamily K, member 9 |
| chr15 | 72450795 | 72451139 | 345 | * | NA_peak_26 | 38 | Distal Interge | 15 | 72501089 | 72546340 | 45252 | 2 | 223604 | ENSMUST00 | 95201 | ENSMUSG0 | Kcnk9 | potassium channel, subfamily K, member 9 |
| chr15 | 72476587 | 72476946 | 360 | * | NA_peak_26 | 40 | Distal Interge | 15 | 72501089 | 72546340 | 45252 | 2 | 223604 | ENSMUST00 | 69394 | ENSMUSG0 | Kcnk9 | potassium channel, subfamily K, member 9 |
| chr15 | 72485358 | 72485384 | 27 | * | NA_peak_26 | 31 | Distal Interge | 15 | 72501089 | 72546340 | 45252 | 2 | 223604 | ENSMUST00 | 60956 | ENSMUSG0 | Kcnk9 | potassium channel, subfamily K, member 9 |
| chr15 | 72522353 | 72522729 | 377 | * | NA_peak_26 | 29 | Intron (ENSM | 15 | 72501089 | 72546340 | 45252 | 2 | 223604 | ENSMUST00 | 23611 | ENSMUSG0 | Kcnk9 | potassium channel, subfamily K, member 9 |
| chr15 | 72630551 | 72630837 | 287 | * | NA_peak_26 | 19 | Intron (ENSM | 15 | 72501089 | 72546340 | 45252 | 2 | 223604 | ENSMUST00 | -84211 | ENSMUSG0 | Kcnk9 | potassium channel, subfamily K, member 9 |
| chr15 | 72861151 | 72861554 | 404 | * | NA_peak_26 | 42 | Intron (ENSM | 15 | 72805600 | 72810324 | 4725 | 2 | 353342 | ENSMUST00 | -50827 | ENSMUSG0 | Peg13 | paternally expressed 13 |
| chr15 | 72946683 | 72947040 | 358 | * | NA_peak_26 | 26 | Intron (ENSM | 15 | 72589620 | 73055810 | 466191 | 2 | 76510 | ENSMUST00 | 108770 | ENSMUSG0 | Trappc9 | trafficking protein particle complex 9 |
| chr15 | 73244393 | 73244646 | 254 | * | NA_peak_26 | 25 | Promoter (1-2 | 15 | 73231907 | 73243247 | 11341 | 2 | 14083 | ENSMUST00 | -1146 | ENSMUSG0 | Ptk2 | PTK2 protein tyrosine kinase 2 |
| chr15 | 73362003 | 73362174 | 172 | * | NA_peak_26 | 17 | Intron (ENSM | 15 | 73215601 | 73343327 | 127727 | 2 | 14083 | ENSMUST00 | -18676 | ENSMUSG0 | Ptk2 | PTK2 protein tyrosine kinase 2 |
| chr15 | 74049738 | 74050081 | 344 | * | NA_peak_26 | 26 | Distal Interge | 15 | 73796736 | 73839699 | 42964 | 2 | 268816 | ENSMUST00 | -210039 | ENSMUSG0 | Mroh5 | maestro heat-like repeat family member 5 |
| chr15 | 74057079 | 74057095 | 17 | * | NA_peak_26 | 18 | Distal Interge | 15 | 73796736 | 73839699 | 42964 | 2 | 268816 | ENSMUST00 | -217380 | ENSMUSG0 | Mroh5 | maestro heat-like repeat family member 5 |
| chr15 | 74126112 | 74126216 | 105 | * | NA_peak_26 | 24 | Distal Interge | 15 | 73796736 | 73839699 | 42964 | 2 | 268816 | ENSMUST00 | -286413 | ENSMUSG0 | Mroh5 | maestro heat-like repeat family member 5 |
| chr15 | 74143320 | 74143632 | 313 | * | NA_peak_26 | 31 | Distal Interge | 15 | 73796736 | 73839699 | 42964 | 2 | 268816 | ENSMUST00 | -303621 | ENSMUSG0 | Mroh5 | maestro heat-like repeat family member 5 |
| chr15 | 74859078 | 74859295 | 218 | * | NA_peak_26 | 52 | Promoter (2-3 | 15 | 74851010 | 74856318 | 5309 | 2 | 13072 | ENSMUST00 | -2760 | ENSMUSG0 | Cyp11b2 | cytochrome P450, family 11, subfamily b, polypeptide 2 |
| chr15 | 74902414 | 74902707 | 294 | * | NA_peak_26 | 23 | Distal Interge | 15 | 74876987 | 74897021 | 20035 | 2 | 67038 | ENSMUST00 | -5393 | ENSMUSG0 | Ly6m | lymphocyte antigen 6 complex, locus M |
| chr15 | 75086297 | 75086933 | 637 | * | NA_peak_26 | 178 | Distal Interge | 15 | 75108331 | 75111697 | 3367 | 2 | 100041546 | ENSMUST00 | 24764 | ENSMUSG0 | Ly6c2 | lymphocyte antigen 6 complex, locus C2 |
| chr15 | 75102421 | 75102846 | 426 | * | NA_peak_26 | 35 | Distal Interge | 15 | 75108331 | 75111697 | 3367 | 2 | 100041546 | ENSMUST00 | 8851 | ENSMUSG0 | Ly6c2 | lymphocyte antigen 6 complex, locus C2 |
| chr15 | 75216412 | 75216798 | 387 | * | NA_peak_26 | 33 | Promoter (<= | 15 | 75216596 | 75222661 | 6066 | 1 | 223631 | ENSMUST00 | 0 | ENSMUSG0 | Ly6g2 | lymphocyte antigen 6 complex, locus G2 |
| chr15 | 75242789 | 75243180 | 392 | * | NA_peak_26 | 15 | Distal Interge | 15 | 75268421 | 75272234 | 3814 | 1 | 17071 | ENSMUST00 | -25241 | ENSMUSG0 | Ly6f | lymphocyte antigen 6 complex, locus F |
| chr15 | 75321221 | 75321381 | 161 | * | NA_peak_26 | 22 | Distal Interge | 15 | 75268702 | 75272235 | 3534 | 1 | 17071 | ENSMUST00 | 52519 | ENSMUSG0 | Ly6f | lymphocyte antigen 6 complex, locus F |
| chr15 | 75333895 | 75334293 | 399 | * | NA_peak_26 | 54 | Distal Interge | 15 | 75268702 | 75272235 | 3534 | 1 | 17071 | ENSMUST00 | 65193 | ENSMUSG0 | Ly6f | lymphocyte antigen 6 complex, locus F |
| chr15 | 75366028 | 75366319 | 292 | * | NA_peak_26 | 24 | Distal Interge | 15 | 75427966 | 75431436 | 3471 | 2 | 105892 | ENSMUST00 | 65117 | ENSMUSG0 | 9030619P08 | lymphocyte antigen 6 complex pseudogene |
| chr15 | 75585368 | 75585394 | 27 | * | NA_peak_26 | 29 | Distal Interge | 15 | 75596628 | 75598213 | 1586 | 1 | 68453 | ENSMUST00 | -11234 | ENSMUSG0 | Gpihbp1 | GPI-anchored HDL-binding protein 1 |
| chr15 | 75966558 | 75966834 | 277 | * | NA_peak_26 | 27 | Promoter (2-3 | 15 | 75969123 | 75975749 | 6627 | 1 | 69020 | ENSMUST00 | -2289 | ENSMUSG0 | Zfp707 | zinc finger protein 707 |
| chr15 | 76268082 | 76268325 | 244 | * | NA_peak_26 | 25 | Promoter (<= | 15 | 76267001 | 76268267 | 1267 | 2 | 100038659 | ENSMUST00 | 0 | NA | Mirt2 | myocardial infraction associated transcript 2 |
| chr15 | 76974154 | 76974548 | 395 | * | NA_peak_26 | 24 | Distal Interge | 15 | 76948749 | 76950979 | 2231 | 1 | 117171 | ENSMUST00 | 25405 | ENSMUSG0 | 1110038F14 | RIKEN cDNA 1110038F14 gene |
| chr15 | 77306368 | 77306808 | 441 | * | NA_peak_26 | 31 | Promoter (<= | 15 | 77083802 | 77306605 | 222804 | 2 | 93686 | ENSMUST00 | 0 | ENSMUSG0 | Rbfox2 | RNA binding protein, fox-1 homolog (C. elegans) 2 |
| chr15 | 77429480 | 77429742 | 263 | * | NA_peak_26 | 24 | Exon (ENSMU | 15 | 77423730 | 77446196 | 22467 | 2 | 278679 | ENSMUST00 | 16454 | ENSMUSG0 | Apol7b | apolipoprotein L 7b |
| chr15 | 77606220 | 77606385 | 166 | * | NA_peak_26 | 19 | Distal Interge | 15 | 77583819 | 77596125 | 12307 | 2 | 328561 | ENSMUST00 | -10095 | ENSMUSG0 | Apol10b | apolipoprotein L 10B |
| chr15 | 77693982 | 77694256 | 275 | * | NA_peak_26 | 30 | Distal Interge | 15 | 77698857 | 77714634 | 15778 | 1 | 666348 | ENSMUST00 | -4601 | ENSMUSG0 | Apol7e | apolipoprotein L 7e |
| chr15 | 77705759 | 77706145 | 387 | * | NA_peak_26 | 45 | Intron (ENSM | 15 | 77698905 | 77719285 | 20381 | 1 | 666348 | ENSMUST00 | 6854 | ENSMUSG0 | Apol7e | apolipoprotein L 7e |
| chr15 | 77846832 | 77846900 | 69 | * | NA_peak_26 | 13 | Distal Interge | 15 | 77760587 | 77842175 | 81589 | 2 | 17886 | ENSMUST00 | -4657 | ENSMUSG0 | Myh9 | myosin, heavy polypeptide 9, non-muscle |
| chr15 | 78820517 | 78820856 | 340 | * | NA_peak_26 | 14 | Exon (ENSMU | 15 | 78775138 | 78803042 | 27905 | 2 | 105844 | ENSMUST00 | -17475 | ENSMUSG0 | Card10 | caspase recruitment domain family, member 10 |
| chr15 | 78990986 | 78991208 | 223 | * | NA_peak_26 | 18 | Exon (ENSMU | 15 | 78983705 | 78991438 | 7734 | 1 | 110253 | ENSMUST00 | 7281 | ENSMUSG0 | Triobp | TRIO and F-actin binding protein |
| chr15 | 79576389 | 79576495 | 107 | * | NA_peak_26 | 32 | Intron (ENSM | 15 | 79562383 | 79585250 | 22868 | 2 | 13404 | ENSMUST00 | 8755 | ENSMUSG0 | Dmc1 | DNA meiotic recombinase 1 |
| chr15 | 79667129 | 79667556 | 428 | * | NA_peak_26 | 34 | 3' UTR | 15 | 79670861 | 79673400 | 2540 | 1 | 223696 | ENSMUST00 | -3305 | ENSMUSG0 | Tomm22 | translocase of outer mitochondrial membrane 22 |
| chr15 | 80233924 | 80234050 | 127 | * | NA_peak_26 | 44 | Promoter (<= | 15 | 80234018 | 80253371 | 19354 | 1 | 239555 | ENSMUST00 | 0 | ENSMUSG0 | Mief1 | mitochondrial elongation factor 1 |
| chr15 | 80572732 | 80573039 | 308 | * | NA_peak_26 | 28 | Promoter (<= | 15 | 80572594 | 80648793 | 76200 | 1 | 17444 | ENSMUST00 | 138 | ENSMUSG0 | Grap2 | GRB2-related adaptor protein 2 |
| chr15 | 81085041 | 81085114 | 74 | * | NA_peak_26 | 20 | Intron (ENSM | 15 | 81012291 | 81105083 | 92793 | 2 | 223701 | ENSMUST00 | 19969 | ENSMUSG0 | Mrtfa | myocardin related transcription factor A |
| chr15 | 81369703 | 81369761 | 59 | * | NA_peak_26 | 22 | Intron (ENSM | 15 | 81318911 | 81360812 | 41902 | 2 | 20524 | ENSMUST00 | -8891 | ENSMUSG0 | Slc25a17 | solute carrier family 25 (mitochondrial carrier, peroxisomal membrane protein), member 17 |
| chr15 | 81403931 | 81404219 | 289 | * | NA_peak_26 | 16 | Intron (ENSM | 15 | 81400197 | 81438158 | 37962 | 1 | 321003 | ENSMUST00 | 3734 | ENSMUSG0 | Xpnpep3 | X-prolyl aminopeptidase 3, mitochondrial |
| chr15 | 81629830 | 81630237 | 408 | * | NA_peak_26 | 23 | Exon (ENSMU | 15 | 81633979 | 81640529 | 6551 | 1 | 328572 | ENSMUST00 | -3742 | ENSMUSG0 | Ep300 | E1A binding protein p300 |
| chr15 | 81811396 | 81811449 | 54 | * | NA_peak_26 | 22 | Promoter (<= | 15 | 81811414 | 81826698 | 15285 | 1 | 21685 | ENSMUST00 | 0 | ENSMUSG0 | Tef | thyrotroph embryonic factor |
| chr15 | 82401400 | 82401769 | 370 | * | NA_peak_26 | 13 | Distal Interge | 15 | 82403197 | 82405967 | 2771 | 2 | 13101 | ENSMUST00 | 4198 | ENSMUSG0 | Cyp2d10 | cytochrome P450, family 2, subfamily d, polypeptide 10 |
| chr15 | 82508559 | 82508820 | 262 | * | NA_peak_26 | 32 | Intron (ENSM | 15 | 82555072 | 82557834 | 2763 | 1 | 380997 | ENSMUST00 | -46252 | ENSMUSG0 | Cyp2d12 | cytochrome P450, family 2, subfamily d, polypeptide 12 |
| chr15 | 82574890 | 82575155 | 266 | * | NA_peak_26 | 20 | Distal Interge | 15 | 82555114 | 82559380 | 4267 | 1 | 380997 | ENSMUST00 | 19776 | ENSMUSG0 | Cyp2d12 | cytochrome P450, family 2, subfamily d, polypeptide 12 |
| chr15 | 82655612 | 82655750 | 139 | * | NA_peak_26 | 20 | Intron (ENSM | 15 | 82636187 | 82642046 | 5860 | 2 | 68444 | ENSMUST00 | -13566 | ENSMUSG0 | Cyp2d13 | cytochrome P450, family 2, subfamily d, polypeptide 13 |
| chr15 | 82671321 | 82671632 | 312 | * | NA_peak_26 | 16 | Exon (ENSMU | 15 | 82688818 | 82693001 | 4184 | 2 | 627860 | ENSMUST00 | 21369 | NA | Cyp2d37-ps | cytochrome P450, family 2, subfamily d, polypeptide 37, pseudogene |
| chr15 | 82841282 | 82841571 | 290 | * | NA_peak_26 | 23 | Intron (ENSM | 15 | 82790101 | 82794294 | 4194 | 2 | 76279 | ENSMUST00 | -46988 | ENSMUSG0 | Cyp2d26 | cytochrome P450, family 2, subfamily d, polypeptide 26 |
| chr15 | 82846122 | 82846443 | 322 | * | NA_peak_26 | 36 | Intron (ENSM | 15 | 82790101 | 82794294 | 4194 | 2 | 76279 | ENSMUST00 | -51828 | ENSMUSG0 | Cyp2d26 | cytochrome P450, family 2, subfamily d, polypeptide 26 |
| chr15 | 83196019 | 83196228 | 210 | * | NA_peak_26 | 20 | Distal Interge | 15 | 83153494 | 83172592 | 19099 | 2 | 109754 | ENSMUST00 | -23427 | ENSMUSG0 | Cyb5r3 | cytochrome b5 reductase 3 |
| chr15 | 83627881 | 83628260 | 380 | * | NA_peak_26 | 20 | Exon (ENSMU | 15 | 83606631 | 83613739 | 7109 | 2 | 64706 | ENSMUST00 | -14142 | ENSMUSG0 | Scube1 | signal peptide, CUB domain, EGF-like 1 |
| chr15 | 83967299 | 83967487 | 189 | * | NA_peak_27 | 18 | Promoter (<= | 15 | 83965198 | 83967842 | 2645 | 2 | 77627 | ENSMUST00 | 355 | ENSMUSG0 | Efcab6 | EF-hand calcium binding domain 6 |
| chr15 | 84024737 | 84024889 | 153 | * | NA_peak_27 | 40 | Intron (ENSM | 15 | 83972483 | 83988982 | 16500 | 2 | 77627 | ENSMUST00 | -35755 | ENSMUSG0 | Efcab6 | EF-hand calcium binding domain 6 |
| chr15 | 84025678 | 84026129 | 452 | * | NA_peak_27 | 24 | Intron (ENSM | 15 | 83972483 | 83988982 | 16500 | 2 | 77627 | ENSMUST00 | -36696 | ENSMUSG0 | Efcab6 | EF-hand calcium binding domain 6 |
| chr15 | 84027236 | 84027293 | 58 | * | NA_peak_27 | 23 | Intron (ENSM | 15 | 83866712 | 84065349 | 198638 | 2 | 77627 | ENSMUST00 | 38056 | ENSMUSG0 | Efcab6 | EF-hand calcium binding domain 6 |
| chr15 | 84239267 | 84239666 | 400 | * | NA_peak_27 | 23 | Intron (ENSM | 15 | 84232066 | 84271318 | 39253 | 1 | 170736 | ENSMUST00 | 7201 | ENSMUSG0 | Parvb | parvin, beta |
| chr15 | 84358797 | 84358825 | 29 | * | NA_peak_27 | 33 | Distal Interge | 15 | 84326119 | 84342978 | 16860 | 1 | 64099 | ENSMUST00 | 32678 | ENSMUSG0 | Parvg | parvin, gamma |
| chr15 | 84521128 | 84521791 | 664 | * | NA_peak_27 | 25 | Distal Interge | 15 | 84553398 | 84557823 | 4426 | 2 | 223732 | ENSMUST00 | 36032 | ENSMUSG0 | Rtl6 | retrotransposon Gag like 6 |
| chr15 | 84896129 | 84896476 | 348 | * | NA_peak_27 | 28 | Distal Interge | 15 | 84923411 | 84942963 | 19553 | 1 | 18141 | ENSMUST00 | -26935 | ENSMUSG0 | Nup50 | nucleoporin 50 |
| chr15 | 85118428 | 85118536 | 109 | * | NA_peak_27 | 28 | Intron (ENSM | 15 | 85064689 | 85131964 | 67276 | 2 | 140557 | ENSMUST00 | 13428 | ENSMUSG0 | Smc1b | structural maintenance of chromosomes 1B |
| chr15 | 85661566 | 85661643 | 78 | * | NA_peak_27 | 18 | Intron (ENSM | 15 | 85653616 | 85707524 | 53909 | 1 | 102800312 | ENSMUST00 | 7950 | NA | Lncppara | long noncoding RNA near Ppara |
| chr15 | 85661959 | 85662077 | 119 | * | NA_peak_27 | 16 | Intron (ENSM | 15 | 85653616 | 85707524 | 53909 | 1 | 102800312 | ENSMUST00 | 8343 | NA | Lncppara | long noncoding RNA near Ppara |

| | | | | | | | | | | | | | | | | | | |
|---|---|---|---|---|---|---|---|---|---|---|---|---|---|---|---|---|---|---|
| chr15 | 86185088 | 86185502 | 415 | * | NA_peak_27 | 21 | Promoter (<= | 15 | 86185728 | 86187152 | 1425 | 1 | 102638208 | ENSMUST00 | -226 | ENSMUSG0 | Gm15569 | predicted gene 15569 |
| chr15 | 86194909 | 86195316 | 408 | * | NA_peak_27 | 35 | Distal Interge | 15 | 86139129 | 86186141 | 47013 | 2 | 223753 | ENSMUST00 | -8768 | ENSMUSG0 | Cerk | ceramide kinase |
| chr15 | 86497982 | 86498255 | 274 | * | NA_peak_27 | 27 | 3' UTR | 15 | 86235505 | 86292162 | 56658 | 1 | 223754 | ENSMUST00 | 262477 | ENSMUSG0 | Tbc1d22a | TBC1 domain family, member 22a |
| chr15 | 86789479 | 86789853 | 375 | * | NA_peak_27 | 43 | Distal Interge | 15 | 86235505 | 86292162 | 56658 | 1 | 223754 | ENSMUST00 | 553974 | ENSMUSG0 | Tbc1d22a | TBC1 domain family, member 22a |
| chr15 | 86848445 | 86848688 | 244 | * | NA_peak_27 | 28 | Distal Interge | 15 | 86235505 | 86292162 | 56658 | 1 | 223754 | ENSMUST00 | 612940 | ENSMUSG0 | Tbc1d22a | TBC1 domain family, member 22a |
| chr15 | 86909331 | 86909563 | 233 | * | NA_peak_27 | 18 | Distal Interge | 15 | 87544299 | 87759362 | 215064 | 1 | 106014 | ENSMUST00 | -634736 | ENSMUSG0 | Tafa5 | TAFA chemokine like family member 5 |
| chr15 | 86952194 | 86952641 | 448 | * | NA_peak_27 | 40 | Distal Interge | 15 | 87544299 | 87759362 | 215064 | 1 | 106014 | ENSMUST00 | -591658 | ENSMUSG0 | Tafa5 | TAFA chemokine like family member 5 |
| chr15 | 87018488 | 87018638 | 151 | * | NA_peak_27 | 31 | Distal Interge | 15 | 87544299 | 87759362 | 215064 | 1 | 106014 | ENSMUST00 | -525661 | ENSMUSG0 | Tafa5 | TAFA chemokine like family member 5 |
| chr15 | 87038828 | 87039225 | 398 | * | NA_peak_27 | 31 | Distal Interge | 15 | 87544299 | 87759362 | 215064 | 1 | 106014 | ENSMUST00 | -505074 | ENSMUSG0 | Tafa5 | TAFA chemokine like family member 5 |
| chr15 | 87091593 | 87091887 | 295 | * | NA_peak_27 | 45 | Distal Interge | 15 | 87544299 | 87759362 | 215064 | 1 | 106014 | ENSMUST00 | -452412 | ENSMUSG0 | Tafa5 | TAFA chemokine like family member 5 |
| chr15 | 87213280 | 87213478 | 199 | * | NA_peak_27 | 36 | Distal Interge | 15 | 87544299 | 87759362 | 215064 | 1 | 106014 | ENSMUST00 | -330821 | ENSMUSG0 | Tafa5 | TAFA chemokine like family member 5 |
| chr15 | 87219445 | 87219705 | 261 | * | NA_peak_27 | 13 | Distal Interge | 15 | 87544299 | 87759362 | 215064 | 1 | 106014 | ENSMUST00 | -324594 | ENSMUSG0 | Tafa5 | TAFA chemokine like family member 5 |
| chr15 | 87224566 | 87224855 | 290 | * | NA_peak_27 | 20 | Distal Interge | 15 | 87544299 | 87759362 | 215064 | 1 | 106014 | ENSMUST00 | -319444 | ENSMUSG0 | Tafa5 | TAFA chemokine like family member 5 |
| chr15 | 87259381 | 87259648 | 268 | * | NA_peak_27 | 31 | Distal Interge | 15 | 87544299 | 87759362 | 215064 | 1 | 106014 | ENSMUST00 | -284651 | ENSMUSG0 | Tafa5 | TAFA chemokine like family member 5 |
| chr15 | 87294900 | 87295273 | 374 | * | NA_peak_27 | 21 | Distal Interge | 15 | 87544299 | 87759362 | 215064 | 1 | 106014 | ENSMUST00 | -249026 | ENSMUSG0 | Tafa5 | TAFA chemokine like family member 5 |
| chr15 | 87321146 | 87321357 | 212 | * | NA_peak_27 | 45 | Distal Interge | 15 | 87544299 | 87759362 | 215064 | 1 | 106014 | ENSMUST00 | -222942 | ENSMUSG0 | Tafa5 | TAFA chemokine like family member 5 |
| chr15 | 87336635 | 87336983 | 349 | * | NA_peak_27 | 15 | Distal Interge | 15 | 87544299 | 87759362 | 215064 | 1 | 106014 | ENSMUST00 | -207316 | ENSMUSG0 | Tafa5 | TAFA chemokine like family member 5 |
| chr15 | 87417982 | 87418036 | 55 | * | NA_peak_27 | 25 | Distal Interge | 15 | 87544299 | 87759362 | 215064 | 1 | 106014 | ENSMUST00 | -126263 | ENSMUSG0 | Tafa5 | TAFA chemokine like family member 5 |
| chr15 | 87421971 | 87422385 | 415 | * | NA_peak_27 | 26 | Distal Interge | 15 | 87544299 | 87759362 | 215064 | 1 | 106014 | ENSMUST00 | -121914 | ENSMUSG0 | Tafa5 | TAFA chemokine like family member 5 |
| chr15 | 87470335 | 87470550 | 216 | * | NA_peak_27 | 20 | Distal Interge | 15 | 87544299 | 87759362 | 215064 | 1 | 106014 | ENSMUST00 | -73749 | ENSMUSG0 | Tafa5 | TAFA chemokine like family member 5 |
| chr15 | 87726862 | 87727224 | 363 | * | NA_peak_27 | 13 | Intron (ENSM | 15 | 87795850 | 87795943 | 94 | 2 | 102465209 | ENSMUST00 | 68719 | ENSMUSG0 | Mir6393 | microRNA 6393 |
| chr15 | 87747662 | 87747948 | 287 | * | NA_peak_27 | 29 | Intron (ENSM | 15 | 87795850 | 87795943 | 94 | 2 | 102465209 | ENSMUST00 | 47995 | ENSMUSG0 | Mir6393 | microRNA 6393 |
| chr15 | 87847563 | 87847686 | 124 | * | NA_peak_27 | 27 | Distal Interge | 15 | 87795850 | 87795943 | 94 | 2 | 102465209 | ENSMUST00 | -51620 | ENSMUSG0 | Mir6393 | microRNA 6393 |
| chr15 | 87935678 | 87935906 | 229 | * | NA_peak_27 | 24 | Distal Interge | 15 | 87795850 | 87795943 | 94 | 2 | 102465209 | ENSMUST00 | -139735 | ENSMUSG0 | Mir6393 | microRNA 6393 |
| chr15 | 87966777 | 87966778 | 2 | * | NA_peak_27 | 19 | Distal Interge | 15 | 87795850 | 87795943 | 94 | 2 | 102465209 | ENSMUST00 | -170834 | ENSMUSG0 | Mir6393 | microRNA 6393 |
| chr15 | 87987016 | 87987068 | 53 | * | NA_peak_27 | 23 | Intron (ENSM | 15 | 87795850 | 87795943 | 94 | 2 | 102465209 | ENSMUST00 | -191073 | ENSMUSG0 | Mir6393 | microRNA 6393 |
| chr15 | 88095236 | 88095440 | 205 | * | NA_peak_27 | 22 | Distal Interge | 15 | 88314874 | 88372653 | 57780 | 1 | 328579 | ENSMUST00 | -219434 | ENSMUSG0 | B230214G05 | RIKEN cDNA B230214G05 gene |
| chr15 | 88141243 | 88141346 | 104 | * | NA_peak_27 | 29 | Distal Interge | 15 | 88314874 | 88372653 | 57780 | 1 | 328579 | ENSMUST00 | -173528 | ENSMUSG0 | B230214G05 | RIKEN cDNA B230214G05 gene |
| chr15 | 88218215 | 88218430 | 216 | * | NA_peak_27 | 39 | Distal Interge | 15 | 88314874 | 88372653 | 57780 | 1 | 328579 | ENSMUST00 | -96444 | ENSMUSG0 | B230214G05 | RIKEN cDNA B230214G05 gene |
| chr15 | 88464446 | 88464824 | 379 | * | NA_peak_27 | 22 | Distal Interge | 15 | 88600302 | 88601669 | 1368 | 1 | 70073 | ENSMUST00 | -135478 | ENSMUSG0 | Zdhhc25 | zinc finger, DHHC domain containing 25 |
| chr15 | 88467915 | 88468269 | 355 | * | NA_peak_27 | 15 | Distal Interge | 15 | 88600302 | 88601669 | 1368 | 1 | 70073 | ENSMUST00 | -132033 | ENSMUSG0 | Zdhhc25 | zinc finger, DHHC domain containing 25 |
| chr15 | 88483803 | 88484049 | 247 | * | NA_peak_27 | 24 | Distal Interge | 15 | 88600302 | 88601669 | 1368 | 1 | 70073 | ENSMUST00 | -116253 | ENSMUSG0 | Zdhhc25 | zinc finger, DHHC domain containing 25 |
| chr15 | 88592772 | 88593136 | 365 | * | NA_peak_27 | 23 | Distal Interge | 15 | 88600302 | 88601669 | 1368 | 1 | 70073 | ENSMUST00 | -7166 | ENSMUSG0 | Zdhhc25 | zinc finger, DHHC domain containing 25 |
| chr15 | 88773423 | 88773714 | 292 | * | NA_peak_27 | 21 | Intron (ENSM | 15 | 88751760 | 88770876 | 19117 | 1 | 223773 | ENSMUST00 | 21663 | ENSMUSG0 | Zbed4 | zinc finger, BED type containing 4 |
| chr15 | 88805175 | 88805540 | 366 | * | NA_peak_27 | 28 | 3' UTR | 15 | 88805244 | 88816171 | 10928 | 2 | 223774 | ENSMUST00 | 10631 | ENSMUSG0 | Alg12 | asparagine-linked glycosylation 12 (alpha-1,6-mannosyltransferase) |
| chr15 | 88902587 | 88902796 | 210 | * | NA_peak_27 | 24 | Intron (ENSM | 15 | 88915669 | 88919868 | 4200 | 2 | 239591 | ENSMUST00 | 17072 | ENSMUSG0 | Ttll8 | tubulin tyrosine ligase-like family, member 8 |
| chr15 | 89398340 | 89398540 | 201 | * | NA_peak_27 | 19 | Intron (ENSM | 15 | 89386891 | 89388708 | 1818 | 1 | 546648 | ENSMUST00 | 11449 | ENSMUSG0 | Klhdc7b | kelch domain containing 7B |
| chr15 | 89620128 | 89620371 | 244 | * | NA_peak_27 | 33 | Distal Interge | 15 | 89582533 | 89591923 | 9391 | 2 | 68708 | ENSMUST00 | -28205 | ENSMUSG0 | Rabl2 | RAB, member RAS oncogene family-like 2 |
| chr15 | 89816988 | 89817159 | 172 | * | NA_peak_27 | 27 | Intron (ENSM | 15 | 89782393 | 89841860 | 59468 | 2 | 54526 | ENSMUST00 | 24701 | ENSMUSG0 | Syt10 | synaptotagmin X |
| chr15 | 89841521 | 89842138 | 618 | * | NA_peak_27 | 27 | Promoter (<= | 15 | 89782393 | 89841860 | 59468 | 2 | 54526 | ENSMUST00 | 0 | ENSMUSG0 | Syt10 | synaptotagmin X |
| chr15 | 89950869 | 89951219 | 351 | * | NA_peak_27 | 21 | Distal Interge | 15 | 89782393 | 89841860 | 59468 | 2 | 54526 | ENSMUST00 | -109009 | ENSMUSG0 | Syt10 | synaptotagmin X |
| chr15 | 90020401 | 90020555 | 155 | * | NA_peak_27 | 23 | Distal Interge | 15 | 89782393 | 89841860 | 59468 | 2 | 54526 | ENSMUST00 | -178541 | ENSMUSG0 | Syt10 | synaptotagmin X |
| chr15 | 90141973 | 90142176 | 204 | * | NA_peak_27 | 20 | Distal Interge | 15 | 90224311 | 90233471 | 9161 | 1 | 380959 | ENSMUST00 | -82135 | ENSMUSG0 | Alg10b | asparagine-linked glycosylation 10B (alpha-1,2-glucosyltransferase) |
| chr15 | 90411329 | 90411731 | 403 | * | NA_peak_27 | 29 | Distal Interge | 15 | 90353706 | 90368444 | 14739 | 1 | 102640415 | ENSMUST00 | 57623 | ENSMUSG0 | Gm36480 | predicted gene, 36480 |
| chr15 | 90641387 | 90641663 | 277 | * | NA_peak_27 | 21 | Intron (ENSM | 15 | 90643329 | 90649438 | 6110 | 2 | 66871 | ENSMUST00 | 7775 | ENSMUSG0 | Cpne8 | copine VIII |
| chr15 | 90706326 | 90706619 | 294 | * | NA_peak_27 | 36 | Distal Interge | 15 | 90487483 | 90679432 | 191950 | 2 | 66871 | ENSMUST00 | -26894 | ENSMUSG0 | Cpne8 | copine VIII |
| chr15 | 91082335 | 91082713 | 379 | * | NA_peak_27 | 42 | Distal Interge | 15 | 90933276 | 91049948 | 116673 | 2 | 16564 | ENSMUST00 | -32387 | ENSMUSG0 | Kif21a | kinesin family member 21A |
| chr15 | 91154157 | 91154367 | 211 | * | NA_peak_27 | 34 | Intron (ENSM | 15 | 91159111 | 91172185 | 13075 | 2 | 26874 | ENSMUST00 | 17818 | ENSMUSG0 | Abcd2 | ATP-binding cassette, sub-family D (ALD), member 2 |
| chr15 | 91166441 | 91166671 | 231 | * | NA_peak_27 | 26 | Intron (ENSM | 15 | 91159111 | 91172185 | 13075 | 2 | 26874 | ENSMUST00 | 5514 | ENSMUSG0 | Abcd2 | ATP-binding cassette, sub-family D (ALD), member 2 |
| chr15 | 91371001 | 91371357 | 357 | * | NA_peak_27 | 23 | Intron (ENSM | 15 | 91396832 | 91412217 | 15386 | 2 | 239606 | ENSMUST00 | 40860 | ENSMUSG0 | Slc2a13 | solute carrier family 2 (facilitated glucose transporter), member 13 |
| chr15 | 91495231 | 91495545 | 315 | * | NA_peak_27 | 31 | Intron (ENSM | 15 | 91542225 | 91573194 | 30970 | 2 | 239606 | ENSMUST00 | 77649 | ENSMUSG0 | Slc2a13 | solute carrier family 2 (facilitated glucose transporter), member 13 |
| chr15 | 91586746 | 91587035 | 290 | * | NA_peak_27 | 28 | Distal Interge | 15 | 91267696 | 91573261 | 305566 | 2 | 239606 | ENSMUST00 | -13485 | ENSMUSG0 | Slc2a13 | solute carrier family 2 (facilitated glucose transporter), member 13 |
| chr15 | 91737341 | 91737363 | 23 | * | NA_peak_27 | 23 | Promoter (1-2 | 15 | 91738948 | 91789110 | 50163 | 1 | 66725 | ENSMUST00 | -1585 | ENSMUSG0 | Lrrk2 | leucine-rich repeat kinase 2 |
| chr15 | 92018516 | 92019354 | 839 | * | NA_peak_27 | 21 | Distal Interge | 15 | 92051165 | 92249451 | 198287 | 1 | 12805 | ENSMUST00 | -31811 | ENSMUSG0 | Cntn1 | contactin 1 |
| chr15 | 92085470 | 92085926 | 457 | * | NA_peak_27 | 20 | Intron (ENSM | 15 | 92051165 | 92341967 | 290803 | 1 | 12805 | ENSMUST00 | 34305 | ENSMUSG0 | Cntn1 | contactin 1 |
| chr15 | 92359096 | 92359112 | 17 | * | NA_peak_27 | 31 | Intron (ENSM | 15 | 92344359 | 92378103 | 33745 | 1 | 105246020 | ENSMUST00 | 14737 | ENSMUSG0 | Gm26760 | predicted gene, 26760 |
| chr15 | 92402300 | 92402641 | 342 | * | NA_peak_27 | 31 | Intron (ENSM | 15 | 92396881 | 92771036 | 374156 | 1 | 239618 | ENSMUST00 | 5419 | ENSMUSG0 | Pdzrn4 | PDZ domain containing RING finger 4 |
| chr15 | 92431632 | 92431642 | 11 | * | NA_peak_27 | 22 | Intron (ENSM | 15 | 92396881 | 92771036 | 374156 | 1 | 239618 | ENSMUST00 | 34751 | ENSMUSG0 | Pdzrn4 | PDZ domain containing RING finger 4 |
| chr15 | 92543144 | 92543389 | 246 | * | NA_peak_27 | 40 | Intron (ENSM | 15 | 92597126 | 92771819 | 174694 | 1 | 239618 | ENSMUST00 | -53737 | ENSMUSG0 | Pdzrn4 | PDZ domain containing RING finger 4 |
| chr15 | 92683514 | 92683539 | 26 | * | NA_peak_27 | 23 | Intron (ENSM | 15 | 92743643 | 92746699 | 3057 | 1 | 239618 | ENSMUST00 | -60104 | ENSMUSG0 | Pdzrn4 | PDZ domain containing RING finger 4 |
| chr15 | 92685932 | 92685961 | 30 | * | NA_peak_27 | 36 | Intron (ENSM | 15 | 92743643 | 92746699 | 3057 | 1 | 239618 | ENSMUST00 | -57682 | ENSMUSG0 | Pdzrn4 | PDZ domain containing RING finger 4 |
| chr15 | 92841821 | 92842235 | 415 | * | NA_peak_27 | 20 | Distal Interge | 15 | 92743643 | 92746699 | 3057 | 1 | 239618 | ENSMUST00 | 98178 | ENSMUSG0 | Pdzrn4 | PDZ domain containing RING finger 4 |
| chr15 | 92893543 | 92893746 | 204 | * | NA_peak_27 | 22 | Distal Interge | 15 | 92743643 | 92746699 | 3057 | 1 | 239618 | ENSMUST00 | 149900 | ENSMUSG0 | Pdzrn4 | PDZ domain containing RING finger 4 |
| chr15 | 93057198 | 93057459 | 262 | * | NA_peak_27 | 16 | Distal Interge | 15 | 93244455 | 93275151 | 30697 | 2 | 223827 | ENSMUST00 | 217692 | ENSMUSG0 | Gxylt1 | glucoside xylosyltransferase 1 |
| chr15 | 93064760 | 93065080 | 321 | * | NA_peak_27 | 22 | Distal Interge | 15 | 93244455 | 93275151 | 30697 | 2 | 223827 | ENSMUST00 | 210071 | ENSMUSG0 | Gxylt1 | glucoside xylosyltransferase 1 |
| chr15 | 93136537 | 93137199 | 663 | * | NA_peak_27 | 13 | Exon (ENSMU | 15 | 93244455 | 93275151 | 30697 | 2 | 223827 | ENSMUST00 | 137952 | ENSMUSG0 | Gxylt1 | glucoside xylosyltransferase 1 |
| chr15 | 93153745 | 93154008 | 264 | * | NA_peak_27 | 30 | Distal Interge | 15 | 93244455 | 93275151 | 30697 | 2 | 223827 | ENSMUST00 | 121143 | ENSMUSG0 | Gxylt1 | glucoside xylosyltransferase 1 |
| chr15 | 93172578 | 93172840 | 263 | * | NA_peak_27 | 24 | Distal Interge | 15 | 93244455 | 93275151 | 30697 | 2 | 223827 | ENSMUST00 | 102311 | ENSMUSG0 | Gxylt1 | glucoside xylosyltransferase 1 |
| chr15 | 93178286 | 93178319 | 34 | * | NA_peak_27 | 40 | Distal Interge | 15 | 93244455 | 93275151 | 30697 | 2 | 223827 | ENSMUST00 | 96832 | ENSMUSG0 | Gxylt1 | glucoside xylosyltransferase 1 |
| chr15 | 93238990 | 93239310 | 321 | * | NA_peak_27 | 33 | Distal Interge | 15 | 93244455 | 93275151 | 30697 | 2 | 223827 | ENSMUST00 | 35841 | ENSMUSG0 | Gxylt1 | glucoside xylosyltransferase 1 |
| chr15 | 93372665 | 93373090 | 426 | * | NA_peak_27 | 13 | Distal Interge | 15 | 93387617 | 93388144 | 528 | 2 | 67197 | ENSMUST00 | 15054 | ENSMUSG0 | Zcrb1 | zinc finger CCHC-type and RNA binding motif 1 |
| chr15 | 93866923 | 93867183 | 261 | * | NA_peak_27 | 20 | Distal Interge | 15 | 93904856 | 93927111 | 22256 | 2 | 77248 | ENSMUST00 | 59928 | ENSMUSG0 | 9430014N10 | RIKEN cDNA 9430014N10 gene |
| chr15 | 94061071 | 94061421 | 351 | * | NA_peak_27 | 19 | Distal Interge | 15 | 93904856 | 93927111 | 22256 | 2 | 77248 | ENSMUST00 | -133960 | ENSMUSG0 | 9430014N10 | RIKEN cDNA 9430014N10 gene |
| chr15 | 94125328 | 94125696 | 369 | * | NA_peak_28 | 36 | Distal Interge | 15 | 93904856 | 93927111 | 22256 | 2 | 77248 | ENSMUST00 | -198217 | ENSMUSG0 | 9430014N10 | RIKEN cDNA 9430014N10 gene |
| chr15 | 94139387 | 94139606 | 220 | * | NA_peak_28 | 23 | Distal Interge | 15 | 93904856 | 93927111 | 22256 | 2 | 77248 | ENSMUST00 | -212276 | ENSMUSG0 | 9430014N10 | RIKEN cDNA 9430014N10 gene |
| chr15 | 94185692 | 94185994 | 303 | * | NA_peak_28 | 29 | Distal Interge | 15 | 94270163 | 94404258 | 134096 | 2 | 223838 | ENSMUST00 | 218264 | ENSMUSG0 | Adamts20 | a disintegrin-like and metallopeptidase (reprolysin type) with thrombospondin type 1 motif, 20 |
| chr15 | 94298949 | 94299151 | 203 | * | NA_peak_28 | 31 | Intron (ENSM | 15 | 94270163 | 94404258 | 134096 | 2 | 223838 | ENSMUST00 | 105107 | ENSMUSG0 | Adamts20 | a disintegrin-like and metallopeptidase (reprolysin type) with thrombospondin type 1 motif, 20 |
| chr15 | 94344757 | 94345038 | 282 | * | NA_peak_28 | 33 | Intron (ENSM | 15 | 94270163 | 94404258 | 134096 | 2 | 223838 | ENSMUST00 | 59220 | ENSMUSG0 | Adamts20 | a disintegrin-like and metallopeptidase (reprolysin type) with thrombospondin type 1 motif, 20 |
| chr15 | 94507156 | 94507564 | 409 | * | NA_peak_28 | 24 | Distal Interge | 15 | 94540735 | 94543486 | 2752 | 2 | 78895 | ENSMUST00 | 35922 | ENSMUSG0 | Pus7l | pseudouridylate synthase 7-like |
| chr15 | 94538407 | 94538492 | 86 | * | NA_peak_28 | 28 | Intron (ENSM | 15 | 94540735 | 94543486 | 2752 | 2 | 78895 | ENSMUST00 | 4994 | ENSMUSG0 | Pus7l | pseudouridylate synthase 7-like |
| chr15 | 94630815 | 94631102 | 288 | * | NA_peak_28 | 37 | Promoter (1-2 | 15 | 94629250 | 95096098 | 466849 | 1 | 320709 | ENSMUST00 | 1565 | ENSMUSG0 | Tmem117 | transmembrane protein 117 |
| chr15 | 95006684 | 95007015 | 332 | * | NA_peak_28 | 26 | Intron (ENSM | 15 | 95097616 | 95149318 | 51703 | 2 | 100043315 | ENSMUST00 | 142303 | ENSMUSG0 | LOC1000433 | uncharacterized LOC100043315 |
| chr15 | 95101437 | 95102421 | 985 | * | NA_peak_28 | 31 | Intron (ENSM | 15 | 95097616 | 95149318 | 51703 | 2 | 100043315 | ENSMUST00 | 46897 | ENSMUSG0 | LOC1000433 | uncharacterized LOC100043315 |
| chr15 | 95116402 | 95116669 | 268 | * | NA_peak_28 | 27 | Intron (ENSM | 15 | 95097616 | 95149318 | 51703 | 2 | 100043315 | ENSMUST00 | 32649 | ENSMUSG0 | LOC1000433 | uncharacterized LOC100043315 |
| chr15 | 95172105 | 95172336 | 232 | * | NA_peak_28 | 16 | Intron (ENSM | 15 | 95097542 | 95162394 | 64853 | 2 | 100043315 | ENSMUST00 | -9711 | ENSMUSG0 | LOC1000433 | uncharacterized LOC100043315 |
| chr15 | 95205481 | 95205771 | 291 | * | NA_peak_28 | 31 | Intron (ENSM | 15 | 95097542 | 95162394 | 64853 | 2 | 100043315 | ENSMUST00 | -43087 | ENSMUSG0 | LOC1000433 | uncharacterized LOC100043315 |
| chr15 | 95235626 | 95235871 | 246 | * | NA_peak_28 | 19 | Intron (ENSM | 15 | 95097542 | 95162394 | 64853 | 2 | 100043315 | ENSMUST00 | -73232 | ENSMUSG0 | LOC1000433 | uncharacterized LOC100043315 |
| chr15 | 95248620 | 95248703 | 84 | * | NA_peak_28 | 22 | Intron (ENSM | 15 | 95097542 | 95162394 | 64853 | 2 | 100043315 | ENSMUST00 | -86226 | ENSMUSG0 | LOC1000433 | uncharacterized LOC100043315 |
| chr15 | 95251440 | 95251678 | 239 | * | NA_peak_28 | 27 | Intron (ENSM | 15 | 95097542 | 95162394 | 64853 | 2 | 100043315 | ENSMUST00 | -89046 | ENSMUSG0 | LOC1000433 | uncharacterized LOC100043315 |
| chr15 | 95289997 | 95290184 | 188 | * | NA_peak_28 | 22 | Intron (ENSM | 15 | 95097542 | 95162394 | 64853 | 2 | 100043315 | ENSMUST00 | -127603 | ENSMUSG0 | LOC1000433 | uncharacterized LOC100043315 |
| chr15 | 95295137 | 95295452 | 316 | * | NA_peak_28 | 16 | Intron (ENSM | 15 | 95097542 | 95162394 | 64853 | 2 | 100043315 | ENSMUST00 | -132743 | ENSMUSG0 | LOC1000433 | uncharacterized LOC100043315 |
| chr15 | 95391017 | 95391350 | 334 | * | NA_peak_28 | 13 | Intron (ENSM | 15 | 95186706 | 95528178 | 341473 | 2 | 54003 | ENSMUST00 | 136828 | ENSMUSG0 | Nell2 | NEL-like 2 |
| chr15 | 95406091 | 95406192 | 102 | * | NA_peak_28 | 20 | Intron (ENSM | 15 | 95186706 | 95528178 | 341473 | 2 | 54003 | ENSMUST00 | 121986 | ENSMUSG0 | Nell2 | NEL-like 2 |
| chr15 | 95565481 | 95565709 | 229 | * | NA_peak_28 | 23 | Distal Interge | 15 | 95219450 | 95528559 | 309110 | 2 | 54003 | ENSMUST00 | -36922 | ENSMUSG0 | Nell2 | NEL-like 2 |
| chr15 | 95585652 | 95585951 | 300 | * | NA_peak_28 | 21 | Distal Interge | 15 | 95624738 | 95632828 | 8091 | 2 | 223843 | ENSMUST00 | 46877 | ENSMUSG0 | Dbx2 | developing brain homeobox 2 |
| chr15 | 96065730 | 96066206 | 477 | * | NA_peak_28 | 17 | Intron (ENSM | 15 | 96051603 | 96054825 | 3223 | 2 | 402773 | ENSMUST00 | -10905 | ENSMUSG0 | D030018L15 | nuclear receptor coactivator 2 pseudogene |
| chr15 | 96137882 | 96137918 | 37 | * | NA_peak_28 | 27 | Intron (ENSM | 15 | 96073338 | 96079553 | 6216 | 2 | 402773 | ENSMUST00 | -58329 | ENSMUSG0 | D030018L15 | nuclear receptor coactivator 2 pseudogene |
| chr15 | 96208785 | 96209003 | 219 | * | NA_peak_28 | 22 | Intron (ENSM | 15 | 96231870 | 96241789 | 9920 | 2 | 75768 | ENSMUST00 | 32786 | ENSMUSG0 | 4833422M21 | RIKEN cDNA 4833422M21 gene |
| chr15 | 96248517 | 96248625 | 109 | * | NA_peak_28 | 34 | Intron (ENSM | 15 | 96248958 | 96254616 | 5659 | 2 | 75768 | ENSMUST00 | 5991 | ENSMUSG0 | 4833422M21 | RIKEN cDNA 4833422M21 gene |
| chr15 | 96281014 | 96281113 | 100 | * | NA_peak_28 | 20 | Promoter (2-3 | 15 | 96134819 | 96283608 | 148790 | 2 | 70040 | ENSMUST00 | 2495 | NA | 2610037D02 | RIKEN cDNA 2610037D02 gene |
| chr15 | 96293458 | 96293515 | 58 | * | NA_peak_28 | 16 | Intron (ENSM | 15 | 96287573 | 96404992 | 117420 | 1 | 77044 | ENSMUST00 | 5885 | ENSMUSG0 | Arid2 | AT rich interactive domain 2 (ARID, RFX-like) |
| chr15 | 96357384 | 96357506 | 123 | * | NA_peak_28 | 36 | Promoter (<= | 15 | 96356840 | 96359783 | 2944 | 1 | 77044 | ENSMUST00 | 544 | ENSMUSG0 | Arid2 | AT rich interactive domain 2 (ARID, RFX-like) |
| chr15 | 96635916 | 96635941 | 26 | * | NA_peak_28 | 24 | Intron (ENSM | 15 | 96575987 | 96641962 | 65976 | 2 | 105727 | ENSMUST00 | 6021 | ENSMUSG0 | Slc38a1 | solute carrier family 38, member 1 |
| chr15 | 96650062 | 96650470 | 409 | * | NA_peak_28 | 28 | Distal Interge | 15 | 96571418 | 96642913 | 71496 | 2 | 105727 | ENSMUST00 | -7149 | ENSMUSG0 | Slc38a1 | solute carrier family 38, member 1 |
| chr15 | 96652593 | 96653168 | 576 | * | NA_peak_28 | 36 | Distal Interge | 15 | 96571418 | 96642913 | 71496 | 2 | 105727 | ENSMUST00 | -9680 | ENSMUSG0 | Slc38a1 | solute carrier family 38, member 1 |
| chr15 | 96703943 | 96704250 | 308 | * | NA_peak_28 | 21 | Distal Interge | 15 | 96687392 | 96699730 | 12339 | 2 | 67760 | ENSMUST00 | -4213 | ENSMUSG0 | Slc38a2 | solute carrier family 38, member 2 |

| | | | | | | | | | | | | | | | | | | |
|---|---|---|---|---|---|---|---|---|---|---|---|---|---|---|---|---|---|---|
| chr15 | 97400802 | 97400864 | 63 | * | NA_peak_283 | 13 | Distal Interge | 15 | 97361217 | 97384349 | 23133 | 1 | 239647 | ENSMUST00 | 39585 | ENSMUSG00 | Pced1b | PC-esterase domain containing 1B |
| chr15 | 97426170 | 97426492 | 323 | * | NA_peak_283 | 30 | Distal Interge | 15 | 97361217 | 97384349 | 23133 | 1 | 239647 | ENSMUST00 | 64953 | ENSMUSG00 | Pced1b | PC-esterase domain containing 1B |
| chr15 | 97485204 | 97485738 | 535 | * | NA_peak_283 | 28 | Distal Interge | 15 | 97361217 | 97384349 | 23133 | 1 | 239647 | ENSMUST00 | 123987 | ENSMUSG00 | Pced1b | PC-esterase domain containing 1B |
| chr15 | 97660481 | 97660858 | 378 | * | NA_peak_283 | 13 | Distal Interge | 15 | 97677793 | 97680353 | 2561 | 2 | 71919 | ENSMUST00 | 19495 | ENSMUSG00 | Rpap3 | RNA polymerase II associated protein 3 |
| chr15 | 97681363 | 97681711 | 349 | * | NA_peak_283 | 25 | Promoter (<= | 15 | 97680379 | 97682199 | 1821 | 2 | 71919 | ENSMUST00 | 488 | ENSMUSG00 | Rpap3 | RNA polymerase II associated protein 3 |
| chr15 | 100366209 | 100366464 | 256 | * | NA_peak_284 | 18 | Promoter (1-2 | 15 | 100364545 | 100383964 | 19420 | 1 | 380975 | ENSMUST00 | 1664 | ENSMUSG00 | Higd1c | HIG1 domain family, member 1C |
| chr15 | 100440921 | 100441244 | 324 | * | NA_peak_284 | 21 | Distal Interge | 15 | 100406306 | 100425072 | 18767 | 2 | 18174 | ENSMUST00 | -15849 | ENSMUSG00 | Slc11a2 | solute carrier family 11 (proton-coupled divalent metal ion transporters), member 2 |
| chr15 | 100636523 | 100636812 | 290 | * | NA_peak_284 | 22 | Promoter (<= | 15 | 100621362 | 100636691 | 15330 | 2 | 207818 | ENSMUST00 | 0 | ENSMUSG00 | Smagp | small cell adhesion glycoprotein |
| chr15 | 101135794 | 101136140 | 347 | * | NA_peak_284 | 18 | Promoter (<= | 15 | 101135657 | 101137032 | 1376 | 1 | 11482 | ENSMUST00 | 137 | ENSMUSG00 | Acvrl1 | activin A receptor, type II-like 1 |
| chr15 | 101646998 | 101647279 | 282 | * | NA_peak_284 | 17 | Promoter (<= | 15 | 101643020 | 101647380 | 4361 | 2 | 432987 | ENSMUST00 | 101 | ENSMUSG00 | Gm5478 | predicted pseudogene 5478 |
| chr15 | 101688122 | 101688397 | 276 | * | NA_peak_284 | 55 | Distal Interge | 15 | 101689928 | 101694279 | 4352 | 2 | 16687 | ENSMUST00 | 5882 | ENSMUSG00 | Krt6a | keratin 6A |
| chr15 | 102358121 | 102358406 | 286 | * | NA_peak_284 | 44 | 3' UTR | 15 | 102350827 | 102355139 | 4313 | 1 | 102640093 | ENSMUST00 | 7294 | ENSMUSG00 | Gm36246 | predicted gene, 36246 |
| chr15 | 102485937 | 102486223 | 287 | * | NA_peak_284 | 21 | Promoter (<= | 15 | 102485979 | 102489330 | 3352 | 1 | 18521 | ENSMUST00 | 0 | ENSMUSG00 | Pcbp2 | poly(rC) binding protein 2 |
| chr15 | 102873175 | 102873420 | 246 | * | NA_peak_285 | 17 | Distal Interge | 15 | 102921103 | 102928814 | 7712 | 1 | 15422 | ENSMUST00 | -47683 | ENSMUSG00 | Hoxc13 | homeobox C13 |
| chr15 | 103027772 | 103027928 | 157 | * | NA_peak_285 | 23 | Intron (ENSM | 15 | 103034395 | 103036852 | 2458 | 1 | 15423 | ENSMUST00 | -6467 | ENSMUSG00 | Hoxc4 | homeobox C4 |
| chr15 | 103217545 | 103217804 | 260 | * | NA_peak_285 | 28 | Promoter (2-3 | 15 | 103213082 | 103215366 | 2285 | 2 | 12419 | ENSMUST00 | -2179 | ENSMUSG00 | Cbx5 | chromobox 5 |
| chr15 | 103333979 | 103334113 | 135 | * | NA_peak_285 | 21 | Promoter (<= | 15 | 103313895 | 103334327 | 20433 | 2 | 29813 | ENSMUST00 | 214 | ENSMUSG00 | Zfp385a | zinc finger protein 385A |
| chr15 | 103631741 | 103632073 | 333 | * | NA_peak_285 | 22 | Distal Interge | 15 | 103562762 | 103565081 | 2320 | 2 | 14663 | ENSMUST00 | -66660 | ENSMUSG00 | Glycam1 | glycosylation dependent cell adhesion molecule 1 |
| chr15 | 103643434 | 103643720 | 287 | * | NA_peak_285 | 23 | Distal Interge | 15 | 103562762 | 103565081 | 2320 | 2 | 14663 | ENSMUST00 | -78353 | ENSMUSG00 | Glycam1 | glycosylation dependent cell adhesion molecule 1 |
| chr15 | 103720705 | 103721024 | 320 | * | NA_peak_285 | 19 | Distal Interge | 15 | 103752063 | 103755439 | 3377 | 2 | 20771 | ENSMUST00 | 34415 | ENSMUSG00 | Mucl1 | mucin-like 1 |
| chr15 | 103736705 | 103737009 | 305 | * | NA_peak_285 | 54 | Distal Interge | 15 | 103752063 | 103755439 | 3377 | 2 | 20771 | ENSMUST00 | 18430 | ENSMUSG00 | Mucl1 | mucin-like 1 |
| chr15 | 103757239 | 103757556 | 318 | * | NA_peak_285 | 31 | Promoter (1-2 | 15 | 103751919 | 103755469 | 3551 | 2 | 20771 | ENSMUST00 | -1770 | ENSMUSG00 | Mucl1 | mucin-like 1 |
| chr15 | 103814948 | 103815226 | 279 | * | NA_peak_285 | 39 | Distal Interge | 15 | 103751919 | 103755469 | 3551 | 2 | 20771 | ENSMUST00 | -59479 | ENSMUSG00 | Mucl1 | mucin-like 1 |
| chr15 | 103847759 | 103847937 | 179 | * | NA_peak_285 | 14 | Distal Interge | 15 | 103896916 | 103899290 | 2375 | 2 | 20770 | ENSMUST00 | 51353 | ENSMUSG00 | Mucl2 | mucin-like 2 |
| chr15 | 103890515 | 103890974 | 460 | * | NA_peak_285 | 24 | Distal Interge | 15 | 103896916 | 103899290 | 2375 | 2 | 20770 | ENSMUST00 | 8316 | ENSMUSG00 | Mucl2 | mucin-like 2 |
| chr15 | 103916787 | 103917053 | 267 | * | NA_peak_285 | 24 | Distal Interge | 15 | 103895855 | 103899312 | 3458 | 2 | 20770 | ENSMUST00 | -17475 | ENSMUSG00 | Mucl2 | mucin-like 2 |
| chr16 | 3099875 | 3099911 | 37 | * | NA_peak_285 | 16 | Distal Interge | 16 | 3592398 | 3593339 | 942 | 1 | 258859 | ENSMUST00 | -492487 | ENSMUSG00 | Olfr161 | olfactory receptor 161 |
| chr16 | 3105236 | 3105510 | 275 | * | NA_peak_286 | 31 | Distal Interge | 16 | 3592398 | 3593339 | 942 | 1 | 258859 | ENSMUST00 | -486888 | ENSMUSG00 | Olfr161 | olfactory receptor 161 |
| chr16 | 3116223 | 3116572 | 350 | * | NA_peak_286 | 331 | Distal Interge | 16 | 3592398 | 3593339 | 942 | 1 | 258859 | ENSMUST00 | -475826 | ENSMUSG00 | Olfr161 | olfactory receptor 161 |
| chr16 | 3139143 | 3140073 | 931 | * | NA_peak_286 | 85 | Distal Interge | 16 | 3592398 | 3593339 | 942 | 1 | 258859 | ENSMUST00 | -452325 | ENSMUSG00 | Olfr161 | olfactory receptor 161 |
| chr16 | 3149077 | 3149370 | 294 | * | NA_peak_286 | 30 | Distal Interge | 16 | 3592398 | 3593339 | 942 | 1 | 258859 | ENSMUST00 | -443028 | ENSMUSG00 | Olfr161 | olfactory receptor 161 |
| chr16 | 3150447 | 3150716 | 270 | * | NA_peak_286 | 61 | Distal Interge | 16 | 3592398 | 3593339 | 942 | 1 | 258859 | ENSMUST00 | -441682 | ENSMUSG00 | Olfr161 | olfactory receptor 161 |
| chr16 | 3151049 | 3151361 | 313 | * | NA_peak_286 | 61 | Distal Interge | 16 | 3592398 | 3593339 | 942 | 1 | 258859 | ENSMUST00 | -441037 | ENSMUSG00 | Olfr161 | olfactory receptor 161 |
| chr16 | 3173586 | 3173714 | 129 | * | NA_peak_286 | 37 | Distal Interge | 16 | 3592398 | 3593339 | 942 | 1 | 258859 | ENSMUST00 | -418684 | ENSMUSG00 | Olfr161 | olfactory receptor 161 |
| chr16 | 3244398 | 3244814 | 417 | * | NA_peak_286 | 15 | Distal Interge | 16 | 3592398 | 3593339 | 942 | 1 | 258859 | ENSMUST00 | -347584 | ENSMUSG00 | Olfr161 | olfactory receptor 161 |
| chr16 | 3246268 | 3247758 | 1491 | * | NA_peak_286 | 74 | Distal Interge | 16 | 3592398 | 3593339 | 942 | 1 | 258859 | ENSMUST00 | -344640 | ENSMUSG00 | Olfr161 | olfactory receptor 161 |
| chr16 | 3254968 | 3255422 | 455 | * | NA_peak_286 | 34 | Distal Interge | 16 | 3592398 | 3593339 | 942 | 1 | 258859 | ENSMUST00 | -336976 | ENSMUSG00 | Olfr161 | olfactory receptor 161 |
| chr16 | 3258316 | 3258622 | 307 | * | NA_peak_286 | 39 | Distal Interge | 16 | 3592398 | 3593339 | 942 | 1 | 258859 | ENSMUST00 | -333776 | ENSMUSG00 | Olfr161 | olfactory receptor 161 |
| chr16 | 3269298 | 3269751 | 454 | * | NA_peak_286 | 37 | Distal Interge | 16 | 3592398 | 3593339 | 942 | 1 | 258859 | ENSMUST00 | -322647 | ENSMUSG00 | Olfr161 | olfactory receptor 161 |
| chr16 | 3278623 | 3278978 | 356 | * | NA_peak_286 | 28 | Distal Interge | 16 | 3592398 | 3593339 | 942 | 1 | 258859 | ENSMUST00 | -313420 | ENSMUSG00 | Olfr161 | olfactory receptor 161 |
| chr16 | 3280842 | 3281327 | 486 | * | NA_peak_286 | 35 | Distal Interge | 16 | 3592398 | 3593339 | 942 | 1 | 258859 | ENSMUST00 | -311071 | ENSMUSG00 | Olfr161 | olfactory receptor 161 |
| chr16 | 3303081 | 3304275 | 1195 | * | NA_peak_286 | 58 | Distal Interge | 16 | 3592398 | 3593339 | 942 | 1 | 258859 | ENSMUST00 | -288123 | ENSMUSG00 | Olfr161 | olfactory receptor 161 |
| chr16 | 3325891 | 3326383 | 493 | * | NA_peak_286 | 73 | Distal Interge | 16 | 3592398 | 3593339 | 942 | 1 | 258859 | ENSMUST00 | -266015 | ENSMUSG00 | Olfr161 | olfactory receptor 161 |
| chr16 | 3345599 | 3346043 | 445 | * | NA_peak_286 | 48 | Distal Interge | 16 | 3592398 | 3593339 | 942 | 1 | 258859 | ENSMUST00 | -246355 | ENSMUSG00 | Olfr161 | olfactory receptor 161 |
| chr16 | 3349917 | 3350446 | 530 | * | NA_peak_286 | 26 | Distal Interge | 16 | 3592398 | 3593339 | 942 | 1 | 258859 | ENSMUST00 | -241952 | ENSMUSG00 | Olfr161 | olfactory receptor 161 |
| chr16 | 3351043 | 3351444 | 402 | * | NA_peak_286 | 15 | Distal Interge | 16 | 3592398 | 3593339 | 942 | 1 | 258859 | ENSMUST00 | -240954 | ENSMUSG00 | Olfr161 | olfactory receptor 161 |
| chr16 | 4886214 | 4886498 | 285 | * | NA_peak_286 | 23 | Promoter (<= | 16 | 4886249 | 4938295 | 52047 | 1 | 17237 | ENSMUST00 | 0 | ENSMUSG00 | Mgrn1 | mahogunin, ring finger 1 |
| chr16 | 8084492 | 8084626 | 135 | * | NA_peak_287 | 26 | Distal Interge | 16 | 8409275 | 8425136 | 15862 | 2 | 74720 | ENSMUST00 | 340510 | ENSMUSG00 | Tmem114 | transmembrane protein 114 |
| chr16 | 11864996 | 11865216 | 221 | * | NA_peak_287 | 16 | Intron (ENSM | 16 | 11901363 | 11930600 | 29238 | 1 | 791359 | ENSMUST00 | -36147 | ENSMUSG00 | Gm9961 | predicted gene 9961 |
| chr16 | 12626652 | 12626967 | 316 | * | NA_peak_287 | 36 | Distal Interge | 16 | 12800579 | 12818298 | 17720 | 1 | 75493 | ENSMUST00 | -173612 | ENSMUSG00 | 1700003L19 | RIKEN cDNA 1700003L19 gene |
| chr16 | 12829743 | 12829919 | 177 | * | NA_peak_288 | 13 | Intron (ENSM | 16 | 12811390 | 12848653 | 37264 | 1 | 75493 | ENSMUST00 | 18353 | ENSMUSG00 | 1700003L19 | RIKEN cDNA 1700003L19 gene |
| chr16 | 13061294 | 13061458 | 165 | * | NA_peak_288 | 13 | Distal Interge | 16 | 13109684 | 13148734 | 39051 | 1 | 50505 | ENSMUST00 | -48226 | ENSMUSG00 | Ercc4 | excision repair cross-complementing rodent repair deficiency, complementation group 4 |
| chr16 | 13750843 | 13751134 | 292 | * | NA_peak_288 | 30 | Distal Interge | 16 | 13754566 | 13758537 | 3972 | 1 | 432999 | ENSMUST00 | -3432 | ENSMUSG00 | A930007A09 | RIKEN cDNA A930007A09 gene |
| chr16 | 14572354 | 14572609 | 256 | * | NA_peak_288 | 27 | Intron (ENSM | 16 | 14580221 | 14621300 | 41080 | 1 | 545280 | ENSMUST00 | -7612 | NA | A630010A05 | RIKEN cDNA A630010A05 gene |
| chr16 | 14691284 | 14691574 | 291 | * | NA_peak_288 | 55 | Distal Interge | 16 | 14705852 | 14709385 | 3534 | 1 | 20583 | ENSMUST00 | -14278 | ENSMUSG00 | Snai2 | snail family zinc finger 2 |
| chr16 | 14813442 | 14813805 | 364 | * | NA_peak_288 | 37 | Distal Interge | 16 | 14906606 | 14938120 | 31515 | 1 | 66793 | ENSMUST00 | -92801 | ENSMUSG00 | Efcab1 | EF-hand calcium binding domain 1 |
| chr16 | 14874572 | 14874901 | 330 | * | NA_peak_288 | 23 | Distal Interge | 16 | 14906606 | 14938120 | 31515 | 1 | 66793 | ENSMUST00 | -31705 | ENSMUSG00 | Efcab1 | EF-hand calcium binding domain 1 |
| chr16 | 16507415 | 16507980 | 566 | * | NA_peak_288 | 16 | Intron (ENSM | 16 | 16435706 | 16504222 | 68517 | 2 | 224014 | ENSMUST00 | -3193 | ENSMUSG00 | Fgd4 | FYVE, RhoGEF and PH domain containing 4 |
| chr16 | 16696012 | 16696185 | 174 | * | NA_peak_288 | 28 | Distal Interge | 16 | 16707614 | 16718096 | 10483 | 2 | 74066 | ENSMUST00 | 21911 | ENSMUSG00 | 4933404G15 | RIKEN cDNA 4933404G15 gene |
| chr16 | 17127217 | 17127327 | 111 | * | NA_peak_288 | 20 | Promoter (2-3 | 16 | 17121226 | 17125167 | 3942 | 2 | 70458 | ENSMUST00 | -2050 | ENSMUSG00 | 2610318N02 | RIKEN cDNA 2610318N02 gene |
| chr16 | 19596449 | 19596850 | 402 | * | NA_peak_289 | 39 | Distal Interge | 16 | 19605696 | 19606703 | 1008 | 2 | 258959 | ENSMUST00 | 9853 | ENSMUSG00 | Olfr170 | olfactory receptor 170 |
| chr16 | 20622194 | 20622353 | 160 | * | NA_peak_289 | 16 | Promoter (<= | 16 | 20619215 | 20621287 | 2073 | 2 | 73047 | ENSMUST00 | -907 | ENSMUSG00 | Camk2n2 | calcium/calmodulin-dependent protein kinase II inhibitor 2 |
| chr16 | 21107095 | 21107526 | 432 | * | NA_peak_289 | 23 | Distal Interge | 16 | 21062213 | 21063094 | 882 | 2 | 625123 | ENSMUST00 | -44001 | NA | Gm6557 | predicted gene 6557 |
| chr16 | 21545809 | 21545907 | 99 | * | NA_peak_289 | 16 | Intron (ENSM | 16 | 21533112 | 21568434 | 35323 | 1 | 209018 | ENSMUST00 | 12697 | ENSMUSG00 | Vps8 | VPS8 CORVET complex subunit |
| chr16 | 24556570 | 24556863 | 294 | * | NA_peak_290 | 65 | Intron (ENSM | 16 | 24529394 | 24530362 | 969 | 1 | 624582 | ENSMUST00 | 27176 | NA | Morf4l1-ps1 | mortality factor 4 like 1, pseudogene 1 |
| chr16 | 25480689 | 25480783 | 95 | * | NA_peak_290 | 16 | Distal Interge | 16 | 25286821 | 25318235 | 31415 | 1 | 71338 | ENSMUST00 | 193868 | ENSMUSG00 | Tprg | transformation related protein 63 regulated |
| chr16 | 26227744 | 26227972 | 229 | * | NA_peak_290 | 31 | Distal Interge | 16 | 25959288 | 26105784 | 146497 | 2 | 210530 | ENSMUST00 | -121960 | ENSMUSG00 | P3h2 | prolyl 3-hydroxylase 2 |
| chr16 | 26260661 | 26260851 | 191 | * | NA_peak_290 | 26 | Distal Interge | 16 | 26356642 | 26371841 | 15200 | 2 | 12737 | ENSMUST00 | 110990 | ENSMUSG00 | Cldn1 | claudin 1 |
| chr16 | 27423742 | 27424075 | 334 | * | NA_peak_290 | 21 | Intron (ENSM | 16 | 27389936 | 27438349 | 48414 | 1 | 67501 | ENSMUST00 | 33806 | ENSMUSG00 | Ccdc50 | coiled-coil domain containing 50 |
| chr16 | 28294951 | 28295323 | 373 | * | NA_peak_290 | 23 | Intron (ENSM | 16 | 28189435 | 28291751 | 102317 | 2 | 14167 | ENSMUST00 | -3200 | ENSMUSG00 | Fgf12 | fibroblast growth factor 12 |
| chr16 | 30599911 | 30600055 | 145 | * | NA_peak_291 | 31 | Promoter (<= | 16 | 30599723 | 30602797 | 3075 | 1 | 224093 | ENSMUST00 | 188 | ENSMUSG00 | Fam43a | family with sequence similarity 43, member A |
| chr16 | 31663870 | 31664197 | 328 | * | NA_peak_291 | 23 | Promoter (<= | 16 | 31663909 | 31871846 | 207938 | 1 | 13383 | ENSMUST00 | 0 | ENSMUSG00 | Dlg1 | discs large MAGUK scaffold protein 1 |
| chr16 | 36298763 | 36299365 | 603 | * | NA_peak_291 | 75 | Distal Interge | 16 | 36277147 | 36285373 | 8227 | 1 | 20861 | ENSMUST00 | 21616 | ENSMUSG00 | Stfa1 | stefin A1 |
| chr16 | 37120207 | 37120557 | 351 | * | NA_peak_291 | 31 | Intron (ENSM | 16 | 37085915 | 37095417 | 9503 | 1 | 77782 | ENSMUST00 | 34292 | ENSMUSG00 | Polq | polymerase (DNA directed), theta |
| chr16 | 37738480 | 37738698 | 219 | * | NA_peak_291 | 36 | Distal Interge | 16 | 37776873 | 37836514 | 59642 | 1 | 14314 | ENSMUST00 | -38175 | ENSMUSG00 | Fstl1 | follistatin-like 1 |
| chr16 | 41057585 | 41057919 | 335 | * | NA_peak_292 | 46 | Intron (ENSM | 16 | 41532858 | 42146213 | 613356 | 1 | 268890 | ENSMUST00 | -474939 | ENSMUSG00 | Lsamp | limbic system-associated membrane protein |
| chr16 | 42843134 | 42843480 | 347 | * | NA_peak_292 | 43 | Intron (ENSM | 16 | 42725702 | 42875587 | 149886 | 2 | 75722 | ENSMUST00 | 32107 | ENSMUSG00 | 4932412D23 | RIKEN cDNA 4932412D23 gene |
| chr16 | 43768842 | 43769652 | 811 | * | NA_peak_292 | 52 | Intron (ENSM | 16 | 43762221 | 43822932 | 60712 | 1 | 13490 | ENSMUST00 | 6621 | ENSMUSG00 | Drd3 | dopamine receptor D3 |
| chr16 | 46277663 | 46277979 | 317 | * | NA_peak_292 | 36 | Distal Interge | 16 | 46152985 | 46155077 | 2093 | 2 | 11615 | ENSMUST00 | -122586 | ENSMUSG00 | Gm4737 | predicted gene 4737 |
| chr16 | 46642751 | 46643077 | 327 | * | NA_peak_292 | 23 | Distal Interge | 16 | 46458852 | 46498525 | 39674 | 2 | 58998 | ENSMUST00 | -144226 | ENSMUSG00 | Nectin3 | nectin cell adhesion molecule 3 |
| chr16 | 48857704 | 48857974 | 271 | * | NA_peak_293 | 39 | Distal Interge | 16 | 48872621 | 48874496 | 1876 | 1 | 245195 | ENSMUST00 | -14647 | ENSMUSG00 | Retnlg | resistin like gamma |
| chr16 | 48860466 | 48860718 | 253 | * | NA_peak_293 | 33 | Distal Interge | 16 | 48872621 | 48874496 | 1876 | 1 | 245195 | ENSMUST00 | -11903 | ENSMUSG00 | Retnlg | resistin like gamma |
| chr16 | 49030018 | 49030429 | 412 | * | NA_peak_293 | 27 | Distal Interge | 16 | 49015837 | 49019709 | 3873 | 1 | 224171 | ENSMUST00 | 14181 | ENSMUSG00 | Cip2a | cell proliferation regulating inhibitor of protein phosphatase 2A |
| chr16 | 50994310 | 50994548 | 239 | * | NA_peak_293 | 54 | Distal Interge | 16 | 50783478 | 50784065 | 588 | 1 | 78639 | ENSMUST00 | 210832 | ENSMUSG00 | 1700116B05 | RIKEN cDNA 1700116B05 gene |
| chr16 | 51274219 | 51274602 | 384 | * | NA_peak_293 | 31 | Distal Interge | 16 | 50783478 | 50784065 | 588 | 1 | 78639 | ENSMUST00 | 490741 | ENSMUSG00 | 1700116B05 | RIKEN cDNA 1700116B05 gene |
| chr16 | 52031456 | 52031736 | 281 | * | NA_peak_293 | 20 | Promoter (<= | 16 | 52031563 | 52205384 | 173822 | 1 | 208650 | ENSMUST00 | 0 | ENSMUSG00 | Cblb | Casitas B-lineage lymphoma b |
| chr16 | 53349330 | 53349614 | 285 | * | NA_peak_293 | 19 | Distal Interge | 16 | 52799508 | 52815744 | 16237 | 2 | 73820 | ENSMUST00 | -533586 | ENSMUSG00 | 4930404A05 | RIKEN cDNA 4930404A05 gene |
| chr16 | 54302004 | 54302164 | 161 | * | NA_peak_293 | 27 | Distal Interge | 16 | 55225175 | 55283237 | 58063 | 2 | 239852 | ENSMUST00 | 981073 | ENSMUSG00 | Zpld1 | zona pellucida like domain containing 1 |
| chr16 | 54432141 | 54432389 | 249 | * | NA_peak_293 | 22 | Distal Interge | 16 | 55225175 | 55283237 | 58063 | 2 | 239852 | ENSMUST00 | 850848 | ENSMUSG00 | Zpld1 | zona pellucida like domain containing 1 |
| chr16 | 55324396 | 55324773 | 378 | * | NA_peak_293 | 64 | Distal Interge | 16 | 55225175 | 55297986 | 72812 | 2 | 239852 | ENSMUST00 | -26410 | ENSMUSG00 | Zpld1 | zona pellucida like domain containing 1 |
| chr16 | 55429724 | 55430084 | 361 | * | NA_peak_293 | 26 | Distal Interge | 16 | 55494731 | 55494797 | 67 | 1 | 100628589 | ENSMUST00 | -64647 | ENSMUSG00 | Mir5118 | microRNA 5118 |
| chr16 | 57391224 | 57391621 | 398 | * | NA_peak_293 | 541 | Intron (ENSM | 16 | 57353277 | 57515690 | 162414 | 1 | 78749 | ENSMUST00 | 37947 | ENSMUSG00 | Filip1l | filamin A interacting protein 1-like |
| chr16 | 58374056 | 58374364 | 309 | * | NA_peak_294 | 27 | Distal Interge | 16 | 58408443 | 58469727 | 61285 | 1 | 73379 | ENSMUST00 | -34079 | ENSMUSG00 | Dcbld2 | discoidin, CUB and LCCL domain containing 2 |
| chr16 | 60773916 | 60774243 | 328 | * | NA_peak_294 | 23 | Distal Interge | 16 | 60424262 | 60605531 | 181270 | 2 | 13840 | ENSMUST00 | -168385 | ENSMUSG00 | Epha6 | Eph receptor A6 |
| chr16 | 60776707 | 60776949 | 243 | * | NA_peak_294 | 20 | Distal Interge | 16 | 60424262 | 60605531 | 181270 | 2 | 13840 | ENSMUST00 | -171176 | ENSMUSG00 | Epha6 | Eph receptor A6 |
| chr16 | 63442380 | 63442773 | 394 | * | NA_peak_294 | 55 | Distal Interge | 16 | 63611023 | 63863413 | 252391 | 2 | 13837 | ENSMUST00 | 420640 | ENSMUSG00 | Epha3 | Eph receptor A3 |
| chr16 | 63726562 | 63726872 | 311 | * | NA_peak_294 | 31 | Intron (ENSM | 16 | 63611023 | 63863413 | 252391 | 2 | 13837 | ENSMUST00 | 136541 | ENSMUSG00 | Epha3 | Eph receptor A3 |
| chr16 | 64806595 | 64806942 | 348 | * | NA_peak_294 | 31 | Intron (ENSM | 16 | 64780351 | 64786321 | 5971 | 2 | 72020 | ENSMUST00 | -20274 | ENSMUSG00 | Zfp654 | zinc finger protein 654 |
| chr16 | 64966260 | 64966555 | 296 | * | NA_peak_294 | 31 | Intron (ENSM | 16 | 64852702 | 64855986 | 3285 | 1 | 106143 | ENSMUST00 | 113558 | ENSMUSG00 | Cggbp1 | CGG triplet repeat binding protein 1 |
| chr16 | 64999102 | 64999316 | 215 | * | NA_peak_294 | 28 | Intron (ENSM | 16 | 64924729 | 65105854 | 181126 | 2 | 15557 | ENSMUST00 | 106538 | ENSMUSG00 | Htr1f | 5-hydroxytryptamine (serotonin) receptor 1F |
| chr16 | 65056289 | 65056551 | 263 | * | NA_peak_294 | 31 | Intron (ENSM | 16 | 64924729 | 65105854 | 181126 | 2 | 15557 | ENSMUST00 | 49303 | ENSMUSG00 | Htr1f | 5-hydroxytryptamine (serotonin) receptor 1F |
| chr16 | 66524183 | 66524568 | 386 | * | NA_peak_294 | 40 | Distal Interge | 16 | 66657118 | 66664626 | 7509 | 1 | 75501 | ENSMUST00 | -132550 | ENSMUSG00 | 1700010K23 | RIKEN cDNA 1700010K23 gene |
| chr16 | 69002556 | 69002919 | 364 | * | NA_peak_294 | 21 | Distal Interge | 16 | 69796135 | 69800381 | 4247 | 2 | 78110 | ENSMUST00 | 797462 | ENSMUSG00 | 4930428D20 | RIKEN cDNA 4930428D20 gene |
| chr16 | 69894885 | 69895184 | 300 | * | NA_peak_294 | 34 | Distal Interge | 16 | 69856874 | 69863744 | 6871 | 2 | 224318 | ENSMUST00 | -31141 | ENSMUSG00 | Speer2 | spermatogenesis associated glutamate (E)-rich protein 2 |
| chr16 | 73548368 | 73548656 | 289 | * | NA_peak_294 | 48 | Distal Interge | 16 | 73894664 | 73899885 | 5222 | 2 | 268902 | ENSMUST00 | 351229 | ENSMUSG00 | Robo2 | roundabout guidance receptor 2 |
| chr16 | 74973415 | 74973417 | 3 | * | NA_peak_295 | 16 | Distal Interge | 16 | 74349795 | 74411825 | 62031 | 2 | 268902 | ENSMUST00 | -561590 | ENSMUSG00 | Robo2 | roundabout guidance receptor 2 |

| | | | | | | | | | | | | | | | | | | |
|---|---|---|---|---|---|---|---|---|---|---|---|---|---|---|---|---|---|---|
| chr16 | 77156788 | 77157108 | 321 | * | NA_peak_295 | 25 | Distal Interge | 16 | 77083900 | 77108205 | 24306 | 1 | 30940 | ENSMUST00 | 72888 | ENSMUSG00 | Usp25 | ubiquitin specific peptidase 25 |
| chr16 | 78610345 | 78610515 | 171 | * | NA_peak_295 | 21 | Distal Interge | 16 | 78547706 | 78576708 | 29003 | 2 | 67102 | ENSMUST00 | -33637 | ENSMUSG00 | D16Ertd472e | DNA segment, Chr 16, ERATO Doi 472, expressed |
| chr16 | 79043857 | 79044065 | 209 | * | NA_peak_295 | 45 | Intron (ENSM | 16 | 78953008 | 79091092 | 138085 | 2 | 19146 | ENSMUST00 | 47027 | ENSMUSG00 | Tmprss15 | transmembrane protease, serine 15 |
| chr16 | 79370850 | 79371178 | 329 | * | NA_peak_295 | 30 | Distal Interge | 16 | 78953464 | 79091097 | 137634 | 2 | 19146 | ENSMUST00 | -279753 | ENSMUSG00 | Tmprss15 | transmembrane protease, serine 15 |
| chr16 | 79944101 | 79944453 | 353 | * | NA_peak_295 | 36 | Distal Interge | 16 | 79999376 | 80000546 | 1171 | 1 | 78654 | ENSMUST00 | -54923 | ENSMUSG00 | 1700066C05 | RIKEN cDNA 1700066C05 gene |
| chr16 | 81863812 | 81863909 | 98 | * | NA_peak_295 | 40 | Distal Interge | 16 | 81465623 | 81518030 | 52408 | 1 | 17968 | ENSMUST00 | 398189 | ENSMUSG00 | Ncam2 | neural cell adhesion molecule 2 |
| chr16 | 84942271 | 84942748 | 478 | * | NA_peak_295 | 21 | Distal Interge | 16 | 84954438 | 84963381 | 8944 | 2 | 11820 | ENSMUST00 | 20633 | ENSMUSG00 | App | amyloid beta (A4) precursor protein |
| chr16 | 87116599 | 87116911 | 313 | * | NA_peak_296 | 31 | Distal Interge | 16 | 87125036 | 87128439 | 3404 | 2 | 102635567 | ENSMUST00 | 11528 | ENSMUSG00 | Gm32865 | predicted gene, 32865 |
| chr16 | 87301593 | 87301911 | 319 | * | NA_peak_296 | 50 | Distal Interge | 16 | 87354185 | 87368742 | 14558 | 1 | 67768 | ENSMUST00 | -52274 | ENSMUSG00 | N6amt1 | N-6 adenine-specific DNA methyltransferase 1 (putative) |
| chr16 | 88029137 | 88029684 | 548 | * | NA_peak_296 | 19 | Intron (ENSM | 16 | 87912421 | 88056176 | 143756 | 2 | 14805 | ENSMUST00 | 26492 | ENSMUSG00 | Grik1 | glutamate receptor, ionotropic, kainate 1 |
| chr16 | 89194017 | 89194277 | 261 | * | NA_peak_296 | 13 | Distal Interge | 16 | 89205861 | 89206394 | 534 | 1 | 622935 | ENSMUST00 | -11584 | ENSMUSG00 | Krtap20-2 | keratin associated protein 20-2 |
| chr16 | 89998157 | 89998392 | 236 | * | NA_peak_296 | 23 | Intron (ENSM | 16 | 89955965 | 89980080 | 24116 | 2 | 21844 | ENSMUST00 | -18077 | ENSMUSG00 | Tiam1 | T cell lymphoma invasion and metastasis 1 |
| chr16 | 90345082 | 90345445 | 364 | * | NA_peak_296 | 87 | Intron (ENSM | 16 | 90386013 | 90499553 | 113541 | 1 | 26559 | ENSMUST00 | -40568 | ENSMUSG00 | Hunk | hormonally upregulated Neu-associated kinase |
| chr16 | 91249948 | 91250573 | 626 | * | NA_peak_296 | 33 | Distal Interge | 16 | 91269772 | 91271933 | 2162 | 1 | 50914 | ENSMUST00 | -19199 | ENSMUSG00 | Olig1 | oligodendrocyte transcription factor 1 |
| chr16 | 91255131 | 91255472 | 342 | * | NA_peak_296 | 45 | Distal Interge | 16 | 91269772 | 91271933 | 2162 | 1 | 50914 | ENSMUST00 | -14300 | ENSMUSG00 | Olig1 | oligodendrocyte transcription factor 1 |
| chr16 | 93617224 | 93617330 | 107 | * | NA_peak_297 | 23 | Distal Interge | 16 | 93607889 | 93609917 | 2029 | 1 | 12408 | ENSMUST00 | 9335 | ENSMUSG00 | Cbr1 | carbonyl reductase 1 |
| chr16 | 94481998 | 94482181 | 184 | * | NA_peak_297 | 42 | Distal Interge | 16 | 94466522 | 94468217 | 1696 | 1 | 22129 | ENSMUST00 | 15476 | ENSMUSG00 | Ttc3 | tetratricopeptide repeat domain 3 |
| chr16 | 95870511 | 95870529 | 19 | * | NA_peak_297 | 15 | Intron (ENSM | 16 | 95831117 | 95845770 | 14654 | 2 | 69776 | ENSMUST00 | -24741 | ENSMUSG00 | 1600002D24 | RIKEN cDNA 1600002D24 gene |
| chr17 | 3001215 | 3001531 | 317 | * | NA_peak_297 | 15 | Distal Interge | 17 | 3077738 | 3084040 | 6303 | 2 | 328734 | ENSMUST00 | 82509 | NA | Pisd-ps2 | phosphatidylserine decarboxylase, pseudogene 2 |
| chr17 | 3004410 | 3004717 | 308 | * | NA_peak_297 | 16 | Distal Interge | 17 | 3077738 | 3084040 | 6303 | 2 | 328734 | ENSMUST00 | 79323 | NA | Pisd-ps2 | phosphatidylserine decarboxylase, pseudogene 2 |
| chr17 | 3077579 | 3077980 | 402 | * | NA_peak_297 | 60 | Exon (ENSMU | 17 | 3077738 | 3084040 | 6303 | 2 | 328734 | ENSMUST00 | 6060 | NA | Pisd-ps2 | phosphatidylserine decarboxylase, pseudogene 2 |
| chr17 | 3080314 | 3080761 | 448 | * | NA_peak_297 | 45 | Intron (ENSM | 17 | 3077738 | 3084040 | 6303 | 2 | 328734 | ENSMUST00 | 3279 | NA | Pisd-ps2 | phosphatidylserine decarboxylase, pseudogene 2 |
| chr17 | 3114787 | 3115344 | 558 | * | NA_peak_297 | 30 | Promoter (<= | 17 | 3114972 | 3151719 | 36748 | 1 | 106583 | ENSMUST00 | 0 | ENSMUSG00 | Scaf8 | SR-related CTD-associated factor 8 |
| chr17 | 3271394 | 3271616 | 223 | * | NA_peak_297 | 27 | Intron (ENSM | 17 | 3326573 | 3518979 | 192407 | 1 | 24001 | ENSMUST00 | -54957 | ENSMUSG00 | Tiam2 | T cell lymphoma invasion and metastasis 2 |
| chr17 | 3964148 | 3964431 | 284 | * | NA_peak_298 | 20 | Distal Interge | 17 | 4044658 | 4082995 | 38338 | 2 | 75815 | ENSMUST00 | 118564 | ENSMUSG00 | 4930470H14 | RIKEN cDNA 4930470H14 gene |
| chr17 | 4205323 | 4205616 | 294 | * | NA_peak_298 | 53 | Distal Interge | 17 | 4120217 | 4122113 | 1897 | 2 | 78043 | ENSMUST00 | -83210 | ENSMUSG00 | 4930548J01 | RIKEN cDNA 4930548J01 gene |
| chr17 | 5620412 | 5620583 | 172 | * | NA_peak_299 | 27 | Intron (ENSM | 17 | 5492557 | 5753811 | 261255 | 1 | 224454 | ENSMUST00 | 127855 | ENSMUSG00 | Zdhhc14 | zinc finger, DHHC domain containing 14 |
| chr17 | 6368152 | 6368323 | 172 | * | NA_peak_299 | 31 | Intron (ENSM | 17 | 6339604 | 6361848 | 22245 | 2 | 101055731 | ENSMUST00 | -6304 | ENSMUSG00 | Gm29719 | synaptotagmin-like 3 pseudogene |
| chr17 | 6379629 | 6379807 | 179 | * | NA_peak_299 | 17 | Intron (ENSM | 17 | 6339604 | 6361848 | 22245 | 2 | 101055731 | ENSMUST00 | -17781 | ENSMUSG00 | Gm29719 | synaptotagmin-like 3 pseudogene |
| chr17 | 6397711 | 6398097 | 387 | * | NA_peak_299 | 28 | Intron (ENSM | 17 | 6369017 | 6411531 | 42515 | 2 | 101055731 | ENSMUST00 | 13434 | ENSMUSG00 | Gm29719 | synaptotagmin-like 3 pseudogene |
| chr17 | 6539687 | 6539825 | 139 | * | NA_peak_299 | 33 | Intron (ENSM | 17 | 6601671 | 6609679 | 8009 | 1 | 100040531 | ENSMUST00 | -61846 | ENSMUSG00 | Dynlt1f | dynein light chain Tctex-type 1F |
| chr17 | 6544344 | 6544677 | 334 | * | NA_peak_299 | 36 | Intron (ENSM | 17 | 6601671 | 6609679 | 8009 | 1 | 100040531 | ENSMUST00 | -56994 | ENSMUSG00 | Dynlt1f | dynein light chain Tctex-type 1F |
| chr17 | 6608546 | 6608843 | 298 | * | NA_peak_299 | 24 | 3' UTR | 17 | 6601804 | 6607549 | 5746 | 1 | 100040563 | ENSMUST00 | 6742 | ENSMUSG00 | Dynlt1c | dynein light chain Tctex-type 1C |
| chr17 | 6630501 | 6630704 | 204 | * | NA_peak_299 | 33 | Intron (ENSM | 17 | 6613753 | 6639790 | 26038 | 2 | 100040525 | ENSMUST00 | 9086 | NA | Tmem181c-p | transmembrane protein 181C, pseudogene |
| chr17 | 6681279 | 6682079 | 801 | * | NA_peak_299 | 52 | 3' UTR | 17 | 6675921 | 6687400 | 11480 | 1 | 83672 | ENSMUST00 | 5358 | ENSMUSG00 | Sytl3 | synaptotagmin-like 3 |
| chr17 | 6690051 | 6690441 | 391 | * | NA_peak_299 | 18 | Promoter (<= | 17 | 6689072 | 6738042 | 48971 | 1 | 83672 | ENSMUST00 | 979 | ENSMUSG00 | Sytl3 | synaptotagmin-like 3 |
| chr17 | 6699646 | 6699958 | 313 | * | NA_peak_299 | 16 | Promoter (2-3 | 17 | 6702952 | 6737836 | 34885 | 1 | 83672 | ENSMUST00 | -2994 | ENSMUSG00 | Sytl3 | synaptotagmin-like 3 |
| chr17 | 6841883 | 6842202 | 320 | * | NA_peak_299 | 22 | Distal Interge | 17 | 6895229 | 6895337 | 109 | 2 | 751529 | ENSMUST00 | 53135 | ENSMUSG00 | Mir692-1 | microRNA 692-1 |
| chr17 | 6924531 | 6925364 | 834 | * | NA_peak_300 | 43 | Intron (ENSM | 17 | 6920869 | 6948281 | 27413 | 2 | 100037282 | ENSMUST00 | 22917 | ENSMUSG00 | Rsph3b | radial spoke 3B homolog (Chlamydomonas) |
| chr17 | 7675381 | 7675562 | 182 | * | NA_peak_300 | 23 | Distal Interge | 17 | 7746641 | 7761015 | 14375 | 2 | 68655 | ENSMUST00 | 85453 | ENSMUSG00 | Fndc1 | fibronectin type III domain containing 1 |
| chr17 | 7969952 | 7970201 | 250 | * | NA_peak_300 | 23 | Promoter (<= | 17 | 7970222 | 7979405 | 9184 | 1 | 66832 | ENSMUST00 | -21 | ENSMUSG00 | Rsph3a | radial spoke 3A homolog (Chlamydomonas) |
| chr17 | 7992550 | 7992862 | 313 | * | NA_peak_300 | 56 | Intron (ENSM | 17 | 7975887 | 7979504 | 3618 | 1 | 66832 | ENSMUST00 | 16663 | ENSMUSG00 | Rsph3a | radial spoke 3A homolog (Chlamydomonas) |
| chr17 | 8434341 | 8434556 | 216 | * | NA_peak_300 | 25 | Promoter (<= | 17 | 8434423 | 8442496 | 8074 | 1 | 20997 | ENSMUST00 | 0 | ENSMUSG00 | T | brachyury, T-box transcription factor T |
| chr17 | 8525265 | 8525503 | 239 | * | NA_peak_300 | 50 | Promoter (<= | 17 | 8525372 | 8981671 | 456300 | 1 | 23984 | ENSMUST00 | 0 | ENSMUSG00 | Pde10a | phosphodiesterase 10A |
| chr17 | 8606485 | 8606571 | 87 | * | NA_peak_300 | 23 | Intron (ENSM | 17 | 8527026 | 8798731 | 271706 | 1 | 23984 | ENSMUST00 | 79459 | ENSMUSG00 | Pde10a | phosphodiesterase 10A |
| chr17 | 8610650 | 8610813 | 164 | * | NA_peak_300 | 23 | Intron (ENSM | 17 | 8527026 | 8798731 | 271706 | 1 | 23984 | ENSMUST00 | 83624 | ENSMUSG00 | Pde10a | phosphodiesterase 10A |
| chr17 | 8717912 | 8718247 | 336 | * | NA_peak_300 | 38 | Intron (ENSM | 17 | 8757018 | 8899193 | 142176 | 1 | 23984 | ENSMUST00 | -38771 | ENSMUSG00 | Pde10a | phosphodiesterase 10A |
| chr17 | 10020055 | 10020151 | 97 | * | NA_peak_301 | 40 | Distal Interge | 17 | 10209093 | 10213574 | 4482 | 2 | 19317 | ENSMUST00 | 193423 | ENSMUSG00 | Qk | quaking, KH domain containing RNA binding |
| chr17 | 12016119 | 12016461 | 343 | * | NA_peak_302 | 48 | Intron (ENSM | 17 | 12029916 | 12063358 | 33443 | 1 | 50873 | ENSMUST00 | -13455 | ENSMUSG00 | Prkn | parkin RBR E3 ubiquitin protein ligase |
| chr17 | 12016770 | 12016894 | 125 | * | NA_peak_302 | 48 | Intron (ENSM | 17 | 12029916 | 12063358 | 33443 | 1 | 50873 | ENSMUST00 | -13022 | ENSMUSG00 | Prkn | parkin RBR E3 ubiquitin protein ligase |
| chr17 | 13090374 | 13090691 | 318 | * | NA_peak_303 | 24 | Distal Interge | 17 | 13096679 | 13098472 | 1794 | 2 | 625850 | ENSMUST00 | 7781 | ENSMUSG00 | Gm11166 | predicted gene 11166 |
| chr17 | 13493313 | 13493350 | 38 | * | NA_peak_303 | 29 | Intron (ENSM | 17 | 13482553 | 13500336 | 17784 | 2 | 72578 | ENSMUST00 | 6986 | ENSMUSG00 | 2700054A10 | RIKEN cDNA 2700054A10 gene |
| chr17 | 13527587 | 13528062 | 476 | * | NA_peak_303 | 22 | Exon (ENSMU | 17 | 13521456 | 13528433 | 6978 | 1 | 272667 | ENSMUST00 | 6131 | ENSMUSG00 | Smok4a | sperm motility kinase 4A |
| chr17 | 14426940 | 14427405 | 466 | * | NA_peak_303 | 45 | Distal Interge | 17 | 14400786 | 14403587 | 2802 | 1 | 64074 | ENSMUST00 | 26154 | ENSMUSG00 | Smoc2 | SPARC related modular calcium binding 2 |
| chr17 | 14585821 | 14586101 | 281 | * | NA_peak_304 | 30 | Distal Interge | 17 | 14667119 | 14670866 | 3748 | 2 | 21826 | ENSMUST00 | 84765 | ENSMUSG00 | Thbs2 | thrombospondin 2 |
| chr17 | 14806413 | 14806782 | 370 | * | NA_peak_304 | 31 | Distal Interge | 17 | 14829331 | 14909519 | 80189 | 2 | 71682 | ENSMUST00 | 102737 | ENSMUSG00 | Wdr27 | WD repeat domain 27 |
| chr17 | 14865797 | 14866101 | 305 | * | NA_peak_304 | 32 | Intron (ENSM | 17 | 14829331 | 14909519 | 80189 | 2 | 71682 | ENSMUST00 | 43418 | ENSMUSG00 | Wdr27 | WD repeat domain 27 |
| chr17 | 14914567 | 14914689 | 123 | * | NA_peak_304 | 16 | Intron (ENSM | 17 | 14829331 | 14909519 | 80189 | 2 | 71682 | ENSMUST00 | -5048 | ENSMUSG00 | Wdr27 | WD repeat domain 27 |
| chr17 | 15607845 | 15608119 | 275 | * | NA_peak_304 | 20 | Distal Interge | 17 | 15631763 | 15634681 | 2919 | 1 | 71008 | ENSMUST00 | -23644 | ENSMUSG00 | 4933401D09 | RIKEN cDNA 4933401D09 gene |
| chr17 | 15608897 | 15609144 | 248 | * | NA_peak_304 | 39 | Distal Interge | 17 | 15631763 | 15634681 | 2919 | 1 | 71008 | ENSMUST00 | -22619 | ENSMUSG00 | 4933401D09 | RIKEN cDNA 4933401D09 gene |
| chr17 | 15782319 | 15782319 | 1 | * | NA_peak_304 | 16 | Distal Interge | 17 | 15756765 | 15761335 | 4571 | 1 | 12648 | ENSMUST00 | 25554 | ENSMUSG00 | Chd1 | chromodomain helicase DNA binding protein 1 |
| chr17 | 16127481 | 16128155 | 675 | * | NA_peak_304 | 72 | Distal Interge | 17 | 15821041 | 15830885 | 9845 | 2 | 68799 | ENSMUST00 | -296596 | ENSMUSG00 | Rgmb | repulsive guidance molecule family member B |
| chr17 | 16429956 | 16430273 | 318 | * | NA_peak_304 | 20 | Distal Interge | 17 | 15821041 | 15830885 | 9845 | 2 | 68799 | ENSMUST00 | -599071 | ENSMUSG00 | Rgmb | repulsive guidance molecule family member B |
| chr17 | 16625053 | 16625379 | 327 | * | NA_peak_305 | 13 | Distal Interge | 17 | 17064113 | 17089628 | 25516 | 1 | 449000 | ENSMUST00 | -438734 | ENSMUSG00 | Zfp960 | zinc finger protein 960 |
| chr17 | 16671844 | 16671900 | 57 | * | NA_peak_305 | 18 | Distal Interge | 17 | 17064113 | 17089628 | 25516 | 1 | 449000 | ENSMUST00 | -392213 | ENSMUSG00 | Zfp960 | zinc finger protein 960 |
| chr17 | 16875283 | 16875684 | 402 | * | NA_peak_305 | 22 | Distal Interge | 17 | 17064113 | 17089628 | 25516 | 1 | 449000 | ENSMUST00 | -188429 | ENSMUSG00 | Zfp960 | zinc finger protein 960 |
| chr17 | 17056285 | 17056781 | 497 | * | NA_peak_305 | 24 | Distal Interge | 17 | 17064113 | 17089628 | 25516 | 1 | 449000 | ENSMUST00 | -7332 | ENSMUSG00 | Zfp960 | zinc finger protein 960 |
| chr17 | 17057545 | 17057655 | 111 | * | NA_peak_305 | 48 | Distal Interge | 17 | 17064113 | 17089628 | 25516 | 1 | 449000 | ENSMUST00 | -6458 | ENSMUSG00 | Zfp960 | zinc finger protein 960 |
| chr17 | 17083262 | 17083599 | 338 | * | NA_peak_305 | 36 | Intron (ENSM | 17 | 17064113 | 17089628 | 25516 | 1 | 449000 | ENSMUST00 | 19149 | ENSMUSG00 | Zfp960 | zinc finger protein 960 |
| chr17 | 17106798 | 17107101 | 304 | * | NA_peak_305 | 26 | Distal Interge | 17 | 17121415 | 17146831 | 25417 | 1 | 22759 | ENSMUST00 | -14314 | ENSMUSG00 | Zfp97 | zinc finger protein 97 |
| chr17 | 17127091 | 17127426 | 336 | * | NA_peak_305 | 27 | Intron (ENSM | 17 | 17121415 | 17146831 | 25417 | 1 | 22759 | ENSMUST00 | 5676 | ENSMUSG00 | Zfp97 | zinc finger protein 97 |
| chr17 | 17139867 | 17140136 | 270 | * | NA_peak_305 | 15 | Intron (ENSM | 17 | 17121415 | 17146831 | 25417 | 1 | 22759 | ENSMUST00 | 18452 | ENSMUSG00 | Zfp97 | zinc finger protein 97 |
| chr17 | 17221816 | 17222394 | 579 | * | NA_peak_305 | 14 | Distal Interge | 17 | 17234872 | 17266063 | 31192 | 1 | 102640673 | ENSMUST00 | -12478 | ENSMUSG00 | Gm5165 | predicted gene 5165 |
| chr17 | 17223479 | 17223959 | 481 | * | NA_peak_305 | 42 | Distal Interge | 17 | 17234872 | 17266063 | 31192 | 1 | 102640673 | ENSMUST00 | -10913 | ENSMUSG00 | Gm5165 | predicted gene 5165 |
| chr17 | 17224277 | 17224546 | 270 | * | NA_peak_305 | 42 | Distal Interge | 17 | 17234872 | 17266063 | 31192 | 1 | 102640673 | ENSMUST00 | -10326 | ENSMUSG00 | Gm5165 | predicted gene 5165 |
| chr17 | 17226613 | 17226969 | 357 | * | NA_peak_305 | 15 | Distal Interge | 17 | 17234872 | 17266063 | 31192 | 1 | 102640673 | ENSMUST00 | -7903 | ENSMUSG00 | Gm5165 | predicted gene 5165 |
| chr17 | 17248012 | 17248426 | 415 | * | NA_peak_305 | 27 | Intron (ENSM | 17 | 17234872 | 17266063 | 31192 | 1 | 102640673 | ENSMUST00 | 13140 | ENSMUSG00 | Gm5165 | predicted gene 5165 |
| chr17 | 17624612 | 17624989 | 378 | * | NA_peak_305 | 27 | Promoter (<= | 17 | 17552598 | 17624641 | 72044 | 2 | 240028 | ENSMUST00 | 0 | ENSMUSG00 | Lnpep | leucyl/cystinyl aminopeptidase |
| chr17 | 17671373 | 17671377 | 5 | * | NA_peak_305 | 16 | Distal Interge | 17 | 17703747 | 17735156 | 31410 | 1 | 626942 | ENSMUST00 | -32370 | ENSMUSG00 | Vmn2r90 | vomeronasal 2, receptor 90 |
| chr17 | 17778601 | 17778920 | 320 | * | NA_peak_305 | 32 | Distal Interge | 17 | 17827158 | 17839071 | 11914 | 1 | 75202 | ENSMUST00 | -48238 | ENSMUSG00 | Spaca6 | sperm acrosome associated 6 |
| chr17 | 18792203 | 18792418 | 216 | * | NA_peak_306 | 27 | Distal Interge | 17 | 18914300 | 18933667 | 19368 | 1 | 627367 | ENSMUST00 | -121882 | ENSMUSG00 | Vmn2r97 | vomeronasal 2, receptor 97 |
| chr17 | 18994758 | 18995119 | 362 | * | NA_peak_306 | 64 | Distal Interge | 17 | 19053460 | 19082411 | 28952 | 1 | 224552 | ENSMUST00 | -58341 | ENSMUSG00 | Vmn2r98 | vomeronasal 2, receptor 98 |
| chr17 | 20166755 | 20166910 | 156 | * | NA_peak_306 | 16 | Distal Interge | 17 | 20182078 | 20183097 | 1020 | 2 | 321020 | ENSMUST00 | 16187 | ENSMUSG00 | Fpr-rs6 | formyl peptide receptor, related sequence 6 |
| chr17 | 20706639 | 20707150 | 512 | * | NA_peak_306 | 26 | Distal Interge | 17 | 20687508 | 20688404 | 897 | 1 | 171225 | ENSMUST00 | 19131 | ENSMUSG00 | Vmn1r226 | vomeronasal 1 receptor 226 |
| chr17 | 20983352 | 20983355 | 4 | * | NA_peak_306 | 42 | Distal Interge | 17 | 20991647 | 20996464 | 4818 | 2 | 171236 | ENSMUST00 | 13109 | ENSMUSG00 | Vmn1r233 | vomeronasal 1 receptor 233 |
| chr17 | 21150718 | 21150767 | 50 | * | NA_peak_307 | 18 | Distal Interge | 17 | 21100261 | 21102455 | 2195 | 1 | 328766 | ENSMUST00 | 50457 | ENSMUSG00 | Gm5092 | predicted gene 5092 |
| chr17 | 21418807 | 21419124 | 318 | * | NA_peak_307 | 23 | Distal Interge | 17 | 21423227 | 21435640 | 12414 | 1 | 22712 | ENSMUST00 | -4103 | ENSMUSG00 | Zfp54 | zinc finger protein 54 |
| chr17 | 21419865 | 21420173 | 309 | * | NA_peak_307 | 20 | Distal Interge | 17 | 21423227 | 21435640 | 12414 | 1 | 22712 | ENSMUST00 | -3054 | ENSMUSG00 | Zfp54 | zinc finger protein 54 |
| chr17 | 21489051 | 21489446 | 396 | * | NA_peak_307 | 15 | Promoter (<= | 17 | 21488998 | 21510473 | 21476 | 1 | 24132 | ENSMUST00 | 53 | ENSMUSG00 | Zfp53 | zinc finger protein 53 |
| chr17 | 21677398 | 21677680 | 283 | * | NA_peak_307 | 23 | Distal Interge | 17 | 21657582 | 21662540 | 4959 | 1 | 73229 | ENSMUST00 | 19816 | ENSMUSG00 | Zfp983 | zinc finger protein 983 |
| chr17 | 21918448 | 21918754 | 307 | * | NA_peak_307 | 28 | Distal Interge | 17 | 21879570 | 21909926 | 30357 | 2 | 70081 | ENSMUST00 | -8522 | ENSMUSG00 | Zfp995 | zinc finger protein 995 |
| chr17 | 22071199 | 22071537 | 339 | * | NA_peak_307 | 26 | Downstream | 17 | 22144359 | 22166190 | 21832 | 2 | 210853 | ENSMUST00 | 94653 | ENSMUSG00 | Zfp947 | zinc finger protein 947 |
| chr17 | 22394616 | 22394911 | 296 | * | NA_peak_307 | 50 | Distal Interge | 17 | 22424222 | 22447416 | 23195 | 1 | 74149 | ENSMUST00 | -29311 | ENSMUSG00 | Zfp946 | zinc finger protein 946 |
| chr17 | 22630049 | 22630511 | 463 | * | NA_peak_308 | 28 | Distal Interge | 17 | 22600951 | 22620842 | 19892 | 1 | 628185 | ENSMUST00 | 29098 | ENSMUSG00 | Vmn2r112 | vomeronasal 2, receptor 112 |
| chr17 | 22637660 | 22638044 | 385 | * | NA_peak_308 | 34 | Distal Interge | 17 | 22600951 | 22620842 | 19892 | 1 | 628185 | ENSMUST00 | 36709 | ENSMUSG00 | Vmn2r112 | vomeronasal 2, receptor 112 |
| chr17 | 22966246 | 22966436 | 191 | * | NA_peak_308 | 36 | Distal Interge | 17 | 22942984 | 22965658 | 22675 | 1 | 434701 | ENSMUST00 | 23262 | ENSMUSG00 | Vmn2r113 | vomeronasal 2, receptor 113 |
| chr17 | 23080384 | 23080610 | 227 | * | NA_peak_308 | 20 | 3' UTR | 17 | 23174876 | 23178539 | 3664 | 2 | 22700 | ENSMUST00 | 97929 | ENSMUSG00 | Zfp40 | zinc finger protein 40 |
| chr17 | 23291445 | 23291629 | 185 | * | NA_peak_308 | 39 | 3' UTR | 17 | 23285392 | 23312428 | 27037 | 2 | 666002 | ENSMUST00 | 20799 | ENSMUSG00 | Vmn2r114 | vomeronasal 2, receptor 114 |
| chr17 | 23293906 | 23294344 | 439 | * | NA_peak_308 | 26 | Intron (ENSM | 17 | 23285392 | 23312428 | 27037 | 2 | 666002 | ENSMUST00 | 18084 | ENSMUSG00 | Vmn2r114 | vomeronasal 2, receptor 114 |
| chr17 | 23310882 | 23311225 | 344 | * | NA_peak_308 | 14 | Promoter (1-2 | 17 | 23285392 | 23312428 | 27037 | 2 | 666002 | ENSMUST00 | 1203 | ENSMUSG00 | Vmn2r114 | vomeronasal 2, receptor 114 |
| chr17 | 23329318 | 23329801 | 484 | * | NA_peak_308 | 46 | Distal Interge | 17 | 23343741 | 23362538 | 18798 | 1 | 638102 | ENSMUST00 | -13940 | ENSMUSG00 | Vmn2r115 | vomeronasal 2, receptor 115 |
| chr17 | 23346163 | 23346397 | 235 | * | NA_peak_308 | 30 | Promoter (2-3 | 17 | 23343741 | 23362538 | 18798 | 1 | 638102 | ENSMUST00 | 2422 | ENSMUSG00 | Vmn2r115 | vomeronasal 2, receptor 115 |
| chr17 | 23347634 | 23348185 | 552 | * | NA_peak_308 | 50 | Exon (ENSMU | 17 | 23343741 | 23362538 | 18798 | 1 | 638102 | ENSMUST00 | 3893 | ENSMUSG00 | Vmn2r115 | vomeronasal 2, receptor 115 |
| chr17 | 23359170 | 23359573 | 404 | * | NA_peak_308 | 131 | 3' UTR | 17 | 23343741 | 23362538 | 18798 | 1 | 638102 | ENSMUST00 | 15429 | ENSMUSG00 | Vmn2r115 | vomeronasal 2, receptor 115 |
| chr17 | 23460688 | 23461263 | 576 | * | NA_peak_308 | 45 | Intron (ENSM | 17 | 23455173 | 23479699 | 24527 | 2 | 619788 | ENSMUST00 | 18436 | ENSMUSG00 | Vmn2r117 | vomeronasal 2, receptor 117 |
| chr17 | 23512881 | 23513208 | 328 | * | NA_peak_308 | 51 | Distal Interge | 17 | 23455173 | 23479699 | 24527 | 2 | 619788 | ENSMUST00 | -33182 | ENSMUSG00 | Vmn2r117 | vomeronasal 2, receptor 117 |

| | | | | | | | | | | | | | | | | | | |
|---|---|---|---|---|---|---|---|---|---|---|---|---|---|---|---|---|---|---|
| chr17 | 23533236 | 23533561 | 326 | * | NA_peak_308 | 53 | Distal Interge | 17 | 23550799 | 23565291 | 14493 | 1 | 320020 | ENSMUST00 | -17238 | ENSMUSG0 | 6330415G19 | RIKEN cDNA 6330415G19 gene |
| chr17 | 23535662 | 23536045 | 384 | * | NA_peak_308 | 44 | Distal Interge | 17 | 23550799 | 23565291 | 14493 | 1 | 320020 | ENSMUST00 | -14754 | ENSMUSG0 | 6330415G19 | RIKEN cDNA 6330415G19 gene |
| chr17 | 23622387 | 23622694 | 308 | * | NA_peak_308 | 18 | Distal Interge | 17 | 23607490 | 23609697 | 2208 | 1 | 332221 | ENSMUST00 | 14897 | ENSMUSG0 | Zscan10 | zinc finger and SCAN domain containing 10 |
| chr17 | 23803542 | 23803886 | 345 | * | NA_peak_308 | 31 | Promoter (<= | 17 | 23803234 | 23810028 | 6795 | 1 | 75956 | ENSMUST00 | 308 | ENSMUSG0 | Srrm2 | serine/arginine repetitive matrix 2 |
| chr17 | 24090958 | 24091064 | 107 | * | NA_peak_309 | 16 | Promoter (2-3 | 17 | 24088181 | 24093798 | 5618 | 2 | 18607 | ENSMUST00 | 2734 | ENSMUSG0 | Pdpk1 | 3-phosphoinositide dependent protein kinase 1 |
| chr17 | 26165142 | 26165414 | 273 | * | NA_peak_309 | 16 | Intron (ENSM | 17 | 26143347 | 26184314 | 40968 | 1 | 12005 | ENSMUST00 | 21795 | ENSMUSG0 | Axin1 | axin 1 |
| chr17 | 26455232 | 26455534 | 303 | * | NA_peak_309 | 22 | Intron (ENSM | 17 | 26414965 | 26446349 | 31385 | 1 | 240055 | ENSMUST00 | 40267 | ENSMUSG0 | Neurl1b | neuralized E3 ubiquitin protein ligase 1B |
| chr17 | 26564627 | 26565023 | 397 | * | NA_peak_309 | 15 | Promoter (2-3 | 17 | 26505591 | 26562128 | 56538 | 2 | 19252 | ENSMUST00 | -2499 | ENSMUSG0 | Dusp1 | dual specificity phosphatase 1 |
| chr17 | 27948056 | 27948128 | 73 | * | NA_peak_310 | 21 | Intron (ENSM | 17 | 27901126 | 27909844 | 8719 | 2 | 68776 | ENSMUST00 | -38212 | ENSMUSG0 | Taf11 | TATA-box binding protein associated factor 11 |
| chr17 | 28294129 | 28294352 | 224 | * | NA_peak_310 | 30 | Exon (ENSMU | 17 | 28313519 | 28326572 | 13054 | 1 | 72775 | ENSMUST00 | -19167 | ENSMUSG0 | Fance | Fanconi anemia, complementation group E |
| chr17 | 28298587 | 28299018 | 432 | * | NA_peak_310 | 21 | Exon (ENSMU | 17 | 28313519 | 28326572 | 13054 | 1 | 72775 | ENSMUST00 | -14501 | ENSMUSG0 | Fance | Fanconi anemia, complementation group E |
| chr17 | 29896282 | 29896556 | 275 | * | NA_peak_310 | 29 | Exon (ENSMU | 17 | 29850515 | 29889960 | 39446 | 2 | 74762 | ENSMUST00 | -6322 | ENSMUSG0 | Mdga1 | MAM domain containing glycosylphosphatidylinositol anchor 1 |
| chr17 | 30634642 | 30634863 | 222 | * | NA_peak_310 | 18 | Promoter (<= | 17 | 30635756 | 30875264 | 239509 | 1 | 13417 | ENSMUST00 | -893 | ENSMUSG0 | Dnah8 | dynein, axonemal, heavy chain 8 |
| chr17 | 31386395 | 31386654 | 260 | * | NA_peak_311 | 25 | Promoter (<= | 17 | 31386294 | 31455122 | 68829 | 1 | 18585 | ENSMUST00 | 101 | ENSMUSG0 | Pde9a | phosphodiesterase 9A |
| chr17 | 32316132 | 32316438 | 307 | * | NA_peak_311 | 26 | Promoter (<= | 17 | 32313854 | 32316390 | 2537 | 2 | 56399 | ENSMUST00 | 0 | ENSMUSG0 | Akap8 | A kinase (PRKA) anchor protein 8 |
| chr17 | 32369628 | 32369966 | 339 | * | NA_peak_311 | 23 | Promoter (<= | 17 | 32361814 | 32369452 | 7639 | 2 | 22404 | ENSMUST00 | -176 | ENSMUSG0 | Wiz | widely-interspaced zinc finger motifs |
| chr17 | 32717186 | 32717535 | 350 | * | NA_peak_311 | 36 | Distal Interge | 17 | 32685679 | 32703352 | 17674 | 1 | 106648 | ENSMUST00 | 31507 | ENSMUSG0 | Cyp4f15 | cytochrome P450, family 4, subfamily f, polypeptide 15 |
| chr17 | 32860105 | 32860438 | 334 | * | NA_peak_311 | 23 | Intron (ENSM | 17 | 32879228 | 32886083 | 6856 | 2 | 240066 | ENSMUST00 | 25645 | ENSMUSG0 | Zfp870 | zinc finger protein 870 |
| chr17 | 33050874 | 33051262 | 389 | * | NA_peak_311 | 39 | Distal Interge | 17 | 33065100 | 33067826 | 2727 | 2 | 74042 | ENSMUST00 | 16564 | ENSMUSG0 | Phf8-ps | PHD finger protein 8, pseudogene |
| chr17 | 33161287 | 33161552 | 266 | * | NA_peak_311 | 44 | Distal Interge | 17 | 33176416 | 33177363 | 948 | 1 | 100038859 | ENSMUST00 | -14864 | ENSMUSG0 | Olfr55 | olfactory receptor 55 |
| chr17 | 33170595 | 33170740 | 146 | * | NA_peak_311 | 26 | Intron (ENSM | 17 | 33176416 | 33177363 | 948 | 1 | 100038859 | ENSMUST00 | -5676 | ENSMUSG0 | Olfr55 | olfactory receptor 55 |
| chr17 | 33287638 | 33288099 | 462 | * | NA_peak_312 | 27 | Promoter (1-2 | 17 | 33289544 | 33307241 | 17698 | 1 | 100043468 | ENSMUST00 | -1445 | ENSMUSG0 | Zfp955b | zinc finger protein 955B |
| chr17 | 33929247 | 33929685 | 439 | * | NA_peak_312 | 25 | Promoter (<= | 17 | 33929543 | 33930495 | 953 | 1 | 19732 | ENSMUST00 | 0 | ENSMUSG0 | Rgl2 | ral guanine nucleotide dissociation stimulator-like 2 |
| chr17 | 34687147 | 34687431 | 285 | * | NA_peak_312 | 39 | Intron (ENSM | 17 | 34670535 | 34719815 | 49281 | 1 | 81877 | ENSMUST00 | 16612 | ENSMUSG0 | Tnxb | tenascin XB |
| chr17 | 34765237 | 34765533 | 297 | * | NA_peak_312 | 20 | Distal Interge | 17 | 34728380 | 34743882 | 15503 | 2 | 12268 | ENSMUST00 | -21355 | ENSMUSG0 | C4b | complement component 4B (Chido blood group) |
| chr17 | 34940191 | 34940534 | 344 | * | NA_peak_312 | 28 | Distal Interge | 17 | 34931863 | 34932765 | 903 | 1 | 18010 | ENSMUST00 | 8328 | ENSMUSG0 | Neu1 | neuraminidase 1 |
| chr17 | 34971395 | 34971869 | 475 | * | NA_peak_312 | 13 | Promoter (<= | 17 | 34969190 | 34972156 | 2967 | 2 | 193740 | ENSMUST00 | 287 | ENSMUSG0 | Hspa1a | heat shock protein 1A |
| chr17 | 35931856 | 35932200 | 345 | * | NA_peak_312 | 20 | 3' UTR | 17 | 35926494 | 35928329 | 1836 | 1 | 52040 | ENSMUST00 | 5362 | ENSMUSG0 | Ppp1r10 | protein phosphatase 1, regulatory subunit 10 |
| chr17 | 36138664 | 36139152 | 489 | * | NA_peak_312 | 14 | Exon (ENSMU | 17 | 36145018 | 36162374 | 17357 | 1 | 675325 | ENSMUST00 | -5866 | ENSMUSG0 | 2410017I17R | RIKEN cDNA 2410017I17 gene |
| chr17 | 36295185 | 36295410 | 226 | * | NA_peak_312 | 36 | Distal Interge | 17 | 36284227 | 36286490 | 2264 | 2 | 333715 | ENSMUST00 | -8695 | ENSMUSG0 | H2-M10.2 | histocompatibility 2, M region locus 10.2 |
| chr17 | 36380652 | 36381007 | 356 | * | NA_peak_313 | 37 | Distal Interge | 17 | 36365003 | 36368417 | 3415 | 2 | 110696 | ENSMUST00 | -12235 | ENSMUSG0 | H2-M10.3 | histocompatibility 2, M region locus 10.3 |
| chr17 | 37119617 | 37119902 | 286 | * | NA_peak_313 | 19 | 5' UTR | 17 | 37111042 | 37111980 | 939 | 2 | 258448 | ENSMUST00 | -7637 | ENSMUSG0 | Olfr92 | olfactory receptor 92 |
| chr17 | 37232098 | 37232370 | 273 | * | NA_peak_313 | 27 | Promoter (<= | 17 | 37231436 | 37232368 | 933 | 2 | 258505 | ENSMUST00 | 0 | ENSMUSG0 | Olfr97 | olfactory receptor 97 |
| chr17 | 37560926 | 37561921 | 996 | * | NA_peak_313 | 52 | 3' UTR | 17 | 37561668 | 37567094 | 5427 | 2 | 258096 | ENSMUST00 | 5173 | ENSMUSG0 | Olfr112 | olfactory receptor 112 |
| chr17 | 38562707 | 38563011 | 305 | * | NA_peak_314 | 26 | Distal Interge | 17 | 38554192 | 38560621 | 6430 | 1 | 100126773 | ENSMUST00 | 8515 | ENSMUSG0 | Esp34 | exocrine gland secreted peptide 34 |
| chr17 | 38569061 | 38569440 | 380 | * | NA_peak_314 | 26 | Distal Interge | 17 | 38554192 | 38560621 | 6430 | 1 | 100126773 | ENSMUST00 | 14869 | ENSMUSG0 | Esp34 | exocrine gland secreted peptide 34 |
| chr17 | 38617289 | 38617639 | 351 | * | NA_peak_314 | 39 | Distal Interge | 17 | 38639447 | 38645653 | 6207 | 1 | 100126768 | ENSMUST00 | -21808 | ENSMUSG0 | Esp31 | exocrine gland secreted peptide 31 |
| chr17 | 38642496 | 38642833 | 338 | * | NA_peak_314 | 30 | Intron (ENSM | 17 | 38639447 | 38645653 | 6207 | 1 | 100126768 | ENSMUST00 | 3049 | ENSMUSG0 | Esp31 | exocrine gland secreted peptide 31 |
| chr17 | 38652764 | 38653088 | 325 | * | NA_peak_314 | 50 | Distal Interge | 17 | 38639447 | 38645653 | 6207 | 1 | 100126768 | ENSMUST00 | 13317 | ENSMUSG0 | Esp31 | exocrine gland secreted peptide 31 |
| chr17 | 38653405 | 38653955 | 551 | * | NA_peak_314 | 50 | Distal Interge | 17 | 38639447 | 38645653 | 6207 | 1 | 100126768 | ENSMUST00 | 13958 | ENSMUSG0 | Esp31 | exocrine gland secreted peptide 31 |
| chr17 | 38761383 | 38761592 | 210 | * | NA_peak_314 | 36 | Distal Interge | 17 | 38639447 | 38645653 | 6207 | 1 | 100126768 | ENSMUST00 | 121936 | ENSMUSG0 | Esp31 | exocrine gland secreted peptide 31 |
| chr17 | 38782082 | 38782843 | 762 | * | NA_peak_314 | 31 | Distal Interge | 17 | 38639447 | 38645653 | 6207 | 1 | 100126768 | ENSMUST00 | 142635 | ENSMUSG0 | Esp31 | exocrine gland secreted peptide 31 |
| chr17 | 39544335 | 39544409 | 75 | * | NA_peak_314 | 14 | Distal Interge | 17 | 39536142 | 39540847 | 4706 | 1 | 100043300 | ENSMUST00 | 8193 | ENSMUSG0 | Esp16 | exocrine gland secreted peptide 16 |
| chr17 | 39848277 | 39848695 | 419 | * | NA_peak_314 | 153 | Promoter (1-2 | 17 | 39846958 | 39848788 | 1831 | 1 | 100861531 | ENSMUST00 | 1319 | NA | Rn45s | 45S pre-ribosomal RNA |
| chr17 | 39843119 | 39847974 | 4856 | * | NA_peak_314 | 153 | Promoter (<= | 17 | 39846958 | 39848788 | 1831 | 1 | 100861531 | ENSMUST00 | 0 | NA | Rn45s | 45S pre-ribosomal RNA |
| chr17 | 40123440 | 40123782 | 343 | * | NA_peak_314 | 18 | Distal Interge | 17 | 40207015 | 40208640 | 1626 | 2 | 18663 | ENSMUST00 | 84858 | ENSMUSG0 | Pgk2 | phosphoglycerate kinase 2 |
| chr17 | 40125160 | 40125410 | 251 | * | NA_peak_314 | 34 | Distal Interge | 17 | 40207015 | 40208640 | 1626 | 2 | 18663 | ENSMUST00 | 83230 | ENSMUSG0 | Pgk2 | phosphoglycerate kinase 2 |
| chr17 | 42086788 | 42086920 | 133 | * | NA_peak_315 | 36 | Distal Interge | 17 | 42315947 | 42505077 | 189131 | 1 | 627626 | ENSMUST00 | -229027 | ENSMUSG0 | Ptchd4 | patched domain containing 4 |
| chr17 | 42263893 | 42264339 | 447 | * | NA_peak_315 | 20 | Distal Interge | 17 | 42315947 | 42505077 | 189131 | 1 | 627626 | ENSMUST00 | -51608 | ENSMUSG0 | Ptchd4 | patched domain containing 4 |
| chr17 | 43854854 | 43855054 | 201 | * | NA_peak_316 | 16 | Intron (ENSM | 17 | 43804003 | 44039512 | 235510 | 1 | 53901 | ENSMUST00 | 50851 | ENSMUSG0 | Rcan2 | regulator of calcineurin 2 |
| chr17 | 44109872 | 44110149 | 278 | * | NA_peak_316 | 37 | Distal Interge | 17 | 44096308 | 44105809 | 9502 | 2 | 224794 | ENSMUST00 | -4063 | ENSMUSG0 | Enpp4 | ectonucleotide pyrophosphatase/phosphodiesterase 4 |
| chr17 | 44198837 | 44199115 | 279 | * | NA_peak_316 | 16 | Intron (ENSM | 17 | 44188584 | 44280172 | 91589 | 1 | 224796 | ENSMUST00 | 10253 | ENSMUSG0 | Clic5 | chloride intracellular channel 5 |
| chr17 | 44464303 | 44464688 | 386 | * | NA_peak_316 | 16 | Distal Interge | 17 | 44237106 | 44251719 | 14614 | 1 | 224796 | ENSMUST00 | 227197 | ENSMUSG0 | Clic5 | chloride intracellular channel 5 |
| chr17 | 44734530 | 44734753 | 224 | * | NA_peak_316 | 38 | Promoter (<= | 17 | 44639681 | 44734693 | 95013 | 2 | 12393 | ENSMUST00 | 0 | ENSMUSG0 | Runx2 | runt related transcription factor 2 |
| chr17 | 45965218 | 45965361 | 144 | * | NA_peak_316 | 19 | Distal Interge | 17 | 46018712 | 46021372 | 2661 | 2 | 22339 | ENSMUST00 | 56011 | ENSMUSG0 | Vegfa | vascular endothelial growth factor A |
| chr17 | 48446446 | 48446605 | 160 | * | NA_peak_317 | 36 | Distal Interge | 17 | 48448568 | 48450070 | 1503 | 2 | 70026 | ENSMUST00 | 3465 | ENSMUSG0 | Tspo2 | translocator protein 2 |
| chr17 | 48489639 | 48490114 | 476 | * | NA_peak_317 | 31 | Intron (ENSM | 17 | 48462404 | 48539607 | 77204 | 1 | 76589 | ENSMUST00 | 27235 | ENSMUSG0 | Unc5cl | unc-5 family C-terminal like |
| chr17 | 50315282 | 50315527 | 246 | * | NA_peak_318 | 23 | Distal Interge | 17 | 50279397 | 50293636 | 14240 | 2 | 13164 | ENSMUST00 | -21646 | ENSMUSG0 | Dazl | deleted in azoospermia-like |
| chr17 | 50625402 | 50625640 | 239 | * | NA_peak_318 | 23 | Intron (ENSM | 17 | 50606994 | 50615113 | 8120 | 1 | 224860 | ENSMUST00 | 18408 | ENSMUSG0 | Plcl2 | phospholipase C-like 2 |
| chr17 | 51037883 | 51038033 | 151 | * | NA_peak_318 | 31 | Intron (ENSM | 17 | 50953993 | 50984626 | 30634 | 2 | 72238 | ENSMUST00 | -53257 | ENSMUSG0 | Tbc1d5 | TBC1 domain family, member 5 |
| chr17 | 51866587 | 51866739 | 153 | * | NA_peak_319 | 16 | Distal Interge | 17 | 51882727 | 51965257 | 82531 | 1 | 100504169 | ENSMUST00 | -15988 | ENSMUSG0 | Gm20098 | predicted gene, 20098 |
| chr17 | 53122951 | 53123025 | 75 | * | NA_peak_319 | 17 | Distal Interge | 17 | 52950060 | 52978379 | 28320 | 1 | 211468 | ENSMUST00 | 172891 | ENSMUSG0 | Kcnh8 | potassium voltage-gated channel, subfamily H (eag-related), member 8 |
| chr17 | 53355665 | 53356019 | 355 | * | NA_peak_319 | 31 | Distal Interge | 17 | 53399148 | 53463306 | 64159 | 2 | 211482 | ENSMUST00 | 107287 | ENSMUSG0 | Efhb | EF hand domain family, member B |
| chr17 | 55904410 | 55904481 | 72 | * | NA_peak_320 | 23 | Distal Interge | 17 | 55892468 | 55897279 | 4812 | 1 | 224893 | ENSMUST00 | 11942 | ENSMUSG0 | Zfp959 | zinc finger protein 959 |
| chr17 | 56779727 | 56779991 | 265 | * | NA_peak_321 | 13 | Exon (ENSMU | 17 | 56781867 | 56783704 | 1838 | 2 | 19725 | ENSMUST00 | 3713 | ENSMUSG0 | Rfx2 | regulatory factor X, 2 (influences HLA class II expression) |
| chr17 | 57433722 | 57434144 | 423 | * | NA_peak_321 | 26 | Intron (ENSM | 17 | 57358714 | 57483523 | 124810 | 1 | 13733 | ENSMUST00 | 75008 | ENSMUSG0 | Adgre1 | adhesion G protein-coupled receptor E1 |
| chr17 | 57452836 | 57453284 | 449 | * | NA_peak_321 | 20 | Intron (ENSM | 17 | 57506453 | 57545505 | 39053 | 2 | 224916 | ENSMUST00 | 92221 | ENSMUSG0 | Vmn2r120 | vomeronasal 2, receptor 120 |
| chr17 | 57572580 | 57572880 | 301 | * | NA_peak_321 | 31 | Distal Interge | 17 | 57506453 | 57545505 | 39053 | 2 | 224916 | ENSMUST00 | -27075 | ENSMUSG0 | Vmn2r120 | vomeronasal 2, receptor 120 |
| chr17 | 58553246 | 58553533 | 288 | * | NA_peak_321 | 16 | Distal Interge | 17 | 58878808 | 58884133 | 5326 | 2 | 69239 | ENSMUST00 | 330600 | ENSMUSG0 | Pdzph1 | PDZ and pleckstrin homology domains 1 |
| chr17 | 58845504 | 58845801 | 298 | * | NA_peak_322 | 36 | Distal Interge | 17 | 58878808 | 58884133 | 5326 | 2 | 69239 | ENSMUST00 | 38332 | ENSMUSG0 | Pdzph1 | PDZ and pleckstrin homology domains 1 |
| chr17 | 59582817 | 59582882 | 66 | * | NA_peak_322 | 20 | Distal Interge | 17 | 59017314 | 59024729 | 7416 | 1 | 75039 | ENSMUST00 | 565503 | ENSMUSG0 | Nudt12os | nudix (nucleoside diphosphate linked moiety X)-type motif 12, opposite strand |
| chr17 | 59887292 | 59887561 | 270 | * | NA_peak_322 | 16 | Intron (ENSM | 17 | 59017314 | 59024729 | 7416 | 1 | 75039 | ENSMUST00 | 869978 | ENSMUSG0 | Nudt12os | nudix (nucleoside diphosphate linked moiety X)-type motif 12, opposite strand |
| chr17 | 60306383 | 60306946 | 564 | * | NA_peak_322 | 20 | Distal Interge | 17 | 59017314 | 59024729 | 7416 | 1 | 75039 | ENSMUST00 | 1289069 | ENSMUSG0 | Nudt12os | nudix (nucleoside diphosphate linked moiety X)-type motif 12, opposite strand |
| chr17 | 60442459 | 60442504 | 46 | * | NA_peak_322 | 35 | Distal Interge | 17 | 59017314 | 59024729 | 7416 | 1 | 75039 | ENSMUST00 | 1425145 | ENSMUSG0 | Nudt12os | nudix (nucleoside diphosphate linked moiety X)-type motif 12, opposite strand |
| chr17 | 62514925 | 62515091 | 167 | * | NA_peak_322 | 27 | Distal Interge | 17 | 62607319 | 62881144 | 273826 | 2 | 13640 | ENSMUST00 | 366053 | ENSMUSG0 | Efna5 | ephrin A5 |
| chr17 | 62612281 | 62612639 | 359 | * | NA_peak_322 | 20 | Intron (ENSM | 17 | 62607319 | 62881144 | 273826 | 2 | 13640 | ENSMUST00 | 268505 | ENSMUSG0 | Efna5 | ephrin A5 |
| chr17 | 63762673 | 63762847 | 175 | * | NA_peak_323 | 28 | Distal Interge | 17 | 63863062 | 63923397 | 60336 | 1 | 14158 | ENSMUST00 | -100215 | ENSMUSG0 | Fer | fer (fms/fps related) protein kinase |
| chr17 | 63777865 | 63778109 | 245 | * | NA_peak_323 | 27 | Distal Interge | 17 | 63863062 | 63923397 | 60336 | 1 | 14158 | ENSMUST00 | -84953 | ENSMUSG0 | Fer | fer (fms/fps related) protein kinase |
| chr17 | 64254335 | 64254526 | 192 | * | NA_peak_323 | 20 | Distal Interge | 17 | 64281006 | 64313031 | 32026 | 2 | 224938 | ENSMUST00 | 58505 | ENSMUSG0 | Pja2 | praja ring finger ubiquitin ligase 2 |
| chr17 | 64311913 | 64312401 | 489 | * | NA_peak_323 | 27 | Promoter (<= | 17 | 64281006 | 64313031 | 32026 | 2 | 224938 | ENSMUST00 | 630 | ENSMUSG0 | Pja2 | praja ring finger ubiquitin ligase 2 |
| chr17 | 64437039 | 64437165 | 127 | * | NA_peak_323 | 27 | Distal Interge | 17 | 64440806 | 64440907 | 102 | 1 | 102465228 | ENSMUST00 | -3641 | ENSMUSG0 | Mir6420 | microRNA 6420 |
| chr17 | 65570403 | 65570405 | 3 | * | NA_peak_324 | 28 | Distal Interge | 17 | 65580420 | 65594970 | 14551 | 2 | 30960 | ENSMUST00 | 24565 | ENSMUSG0 | Vapa | vesicle-associated membrane protein, associated protein A |
| chr17 | 66225635 | 66226009 | 375 | * | NA_peak_324 | 23 | Distal Interge | 17 | 66143709 | 66148917 | 5209 | 1 | 320209 | ENSMUST00 | 81926 | ENSMUSG0 | Ddx11 | DEAD/H box helicase 11 |
| chr17 | 66298343 | 66298791 | 449 | * | NA_peak_324 | 31 | Distal Interge | 17 | 66343946 | 66346545 | 2600 | 2 | 68617 | ENSMUST00 | 47754 | ENSMUSG0 | Mtcl1 | microtubule crosslinking factor 1 |
| chr17 | 66655858 | 66656227 | 370 | * | NA_peak_324 | 15 | Distal Interge | 17 | 66681487 | 66694033 | 12547 | 2 | 19274 | ENSMUST00 | 37806 | ENSMUSG0 | Ptprm | protein tyrosine phosphatase, receptor type, M |
| chr17 | 67655139 | 67655280 | 142 | * | NA_peak_325 | 23 | Distal Interge | 17 | 67630964 | 67632723 | 1760 | 2 | 240131 | ENSMUST00 | -22416 | ENSMUSG0 | Lrrc30 | leucine rich repeat containing 30 |
| chr17 | 68311767 | 68312116 | 350 | * | NA_peak_325 | 21 | Intron (ENSM | 17 | 68273797 | 68777961 | 504165 | 1 | 320858 | ENSMUST00 | 37970 | ENSMUSG0 | L3mbtl4 | L3MBTL4 histone methyl-lysine binding protein |
| chr17 | 68318458 | 68318521 | 64 | * | NA_peak_325 | 31 | Intron (ENSM | 17 | 68273797 | 68777961 | 504165 | 1 | 320858 | ENSMUST00 | 44661 | ENSMUSG0 | L3mbtl4 | L3MBTL4 histone methyl-lysine binding protein |
| chr17 | 69540613 | 69541055 | 443 | * | NA_peak_326 | 19 | Distal Interge | 17 | 69439834 | 69489231 | 49398 | 1 | 320722 | ENSMUST00 | 100779 | ENSMUSG0 | Akain1 | A kinase (PRKA) anchor inhibitor 1 |
| chr17 | 70501940 | 70501948 | 9 | * | NA_peak_326 | 15 | Intron (ENSM | 17 | 70516022 | 70818234 | 302213 | 1 | 224997 | ENSMUST00 | -14074 | ENSMUSG0 | Dlgap1 | DLG associated protein 1 |
| chr17 | 70790688 | 70791040 | 353 | * | NA_peak_326 | 34 | Intron (ENSM | 17 | 70809072 | 70819401 | 10330 | 1 | 224997 | ENSMUST00 | -18032 | ENSMUSG0 | Dlgap1 | DLG associated protein 1 |
| chr17 | 71373758 | 71374073 | 316 | * | NA_peak_326 | 22 | Promoter (2-3 | 17 | 71375983 | 71377030 | 1048 | 2 | 74355 | ENSMUST00 | 2957 | ENSMUSG0 | Smchd1 | SMC hinge domain containing 1 |
| chr17 | 72575760 | 72576043 | 284 | * | NA_peak_327 | 19 | Intron (ENSM | 17 | 71868972 | 72603827 | 734856 | 2 | 11682 | ENSMUST00 | 27784 | ENSMUSG0 | Alk | anaplastic lymphoma kinase |
| chr17 | 72687152 | 72687437 | 286 | * | NA_peak_327 | 23 | Distal Interge | 17 | 72204979 | 72604627 | 399649 | 2 | 11682 | ENSMUST00 | -82525 | ENSMUSG0 | Alk | anaplastic lymphoma kinase |
| chr17 | 74110614 | 74110996 | 383 | * | NA_peak_328 | 23 | Distal Interge | 17 | 74026998 | 74079046 | 52049 | 2 | 94224 | ENSMUST00 | -31568 | ENSMUSG0 | Srd5a2 | steroid 5 alpha-reductase 2 |
| chr17 | 74247338 | 74247520 | 183 | * | NA_peak_328 | 27 | Intron (ENSM | 17 | 74201116 | 74230501 | 29386 | 2 | 76890 | ENSMUST00 | -16837 | ENSMUSG0 | Memo1 | mediator of cell motility 1 |
| chr17 | 74447600 | 74447604 | 5 | * | NA_peak_328 | 16 | Promoter (<= | 17 | 74426828 | 74448206 | 21379 | 2 | 268973 | ENSMUST00 | 602 | ENSMUSG0 | Nlrc4 | NLR family, CARD domain containing 4 |
| chr17 | 75251629 | 75251724 | 96 | * | NA_peak_328 | 24 | Intron (ENSM | 17 | 75289976 | 75294369 | 4394 | 1 | 268977 | ENSMUST00 | -38252 | ENSMUSG0 | Ltbp1 | latent transforming growth factor beta binding protein 1 |
| chr17 | 75425090 | 75425648 | 559 | * | NA_peak_328 | 37 | Distal Interge | 17 | 75435896 | 75491790 | 55895 | 1 | 240168 | ENSMUST00 | -10248 | ENSMUSG0 | Rasgrp3 | RAS, guanyl releasing protein 3 |
| chr17 | 77302177 | 77302486 | 310 | * | NA_peak_329 | 38 | Distal Interge | 17 | 78200248 | 78376592 | 176345 | 1 | 50766 | ENSMUST00 | -897762 | ENSMUSG0 | Crim1 | cysteine rich transmembrane BMP regulator 1 (chordin like) |
| chr17 | 78074284 | 78074715 | 432 | * | NA_peak_329 | 31 | Distal Interge | 17 | 78200248 | 78376592 | 176345 | 1 | 50766 | ENSMUST00 | -125533 | ENSMUSG0 | Crim1 | cysteine rich transmembrane BMP regulator 1 (chordin like) |
| chr17 | 78345950 | 78346206 | 257 | * | NA_peak_330 | 24 | Exon (ENSMU | 17 | 78350889 | 78374178 | 23290 | 1 | 50766 | ENSMUST00 | -4683 | ENSMUSG0 | Crim1 | cysteine rich transmembrane BMP regulator 1 (chordin like) |
| chr17 | 78384230 | 78384342 | 113 | * | NA_peak_330 | 14 | Intron (ENSM | 17 | 78377886 | 78401430 | 23545 | 2 | 225020 | ENSMUST00 | 17088 | ENSMUSG0 | Fez2 | fasciculation and elongation protein zeta 2 (zygin II) |
| chr17 | 78491001 | 78491456 | 456 | * | NA_peak_330 | 15 | Distal Interge | 17 | 78508064 | 78627398 | 119335 | 1 | 74199 | ENSMUST00 | -16608 | ENSMUSG0 | Vit | vitrin |
| chr17 | 78736866 | 78736946 | 81 | * | NA_peak_330 | 28 | Promoter (<= | 17 | 78649913 | 78737196 | 87284 | 2 | 268980 | ENSMUST00 | 250 | ENSMUSG0 | Strn | striatin, calmodulin binding protein |
| chr17 | 78863269 | 78863435 | 167 | * | NA_peak_330 | 17 | Intron (ENSM | 17 | 78855354 | 78858980 | 3627 | 2 | 19106 | ENSMUST00 | -4289 | ENSMUSG0 | Eif2ak2 | eukaryotic translation initiation factor 2-alpha kinase 2 |

| | | | | | | | | | | | | | | | | | | |
|---|---|---|---|---|---|---|---|---|---|---|---|---|---|---|---|---|---|---|
| chr17 | 79676758 | 79677053 | 296 | * | NA_peak_33 | 27 | Intron (ENSM | 17 | 79672400 | 79693192 | 20793 | 1 | 381110 | ENSMUST00 | 4358 | ENSMUSG0 | Rmdn2 | regulator of microtubule dynamics 2 |
| chr17 | 80287434 | 80287885 | 452 | * | NA_peak_33 | 40 | Promoter (1-2 | 17 | 80252565 | 80289874 | 37310 | 2 | 106794 | ENSMUST00 | 1989 | ENSMUSG0 | Dhx57 | DEAH (Asp-Glu-Ala-Asp/His) box polypeptide 57 |
| chr17 | 80332303 | 80332497 | 195 | * | NA_peak_33 | 23 | Intron (ENSM | 17 | 80347540 | 80370404 | 22865 | 1 | 381112 | ENSMUST00 | -15043 | ENSMUSG0 | Arhgef33 | Rho guanine nucleotide exchange factor (GEF) 33 |
| chr17 | 80931251 | 80931575 | 325 | * | NA_peak_33 | 16 | Intron (ENSM | 17 | 80944632 | 81001816 | 57185 | 1 | 68027 | ENSMUST00 | -13057 | ENSMUSG0 | Tmem178 | transmembrane protein 178 |
| chr17 | 80940013 | 80940311 | 299 | * | NA_peak_33 | 16 | Distal Interge | 17 | 80944632 | 81001816 | 57185 | 1 | 68027 | ENSMUST00 | -4321 | ENSMUSG0 | Tmem178 | transmembrane protein 178 |
| chr17 | 81348477 | 81348935 | 459 | * | NA_peak_33 | 55 | Distal Interge | 17 | 81026332 | 81065082 | 38751 | 2 | 72167 | ENSMUST00 | -283395 | ENSMUSG0 | Thumpd2 | THUMP domain containing 2 |
| chr17 | 82802439 | 82802717 | 279 | * | NA_peak_33 | 40 | Distal Interge | 17 | 83033594 | 83078225 | 44632 | 2 | 100503434 | ENSMUST00 | 275508 | ENSMUSG0 | Gm19689 | predicted gene, 19689 |
| chr17 | 82803747 | 82804048 | 302 | * | NA_peak_33 | 30 | Distal Interge | 17 | 83033594 | 83078225 | 44632 | 2 | 100503434 | ENSMUST00 | 274177 | ENSMUSG0 | Gm19689 | predicted gene, 19689 |
| chr17 | 82852392 | 82852822 | 431 | * | NA_peak_33 | 37 | Distal Interge | 17 | 83033594 | 83078225 | 44632 | 2 | 100503434 | ENSMUST00 | 225403 | ENSMUSG0 | Gm19689 | predicted gene, 19689 |
| chr17 | 85540305 | 85540575 | 271 | * | NA_peak_33 | 23 | Distal Interge | 17 | 85613608 | 85625222 | 11615 | 1 | 20473 | ENSMUST00 | -73033 | ENSMUSG0 | Six3 | sine oculis-related homeobox 3 |
| chr17 | 85631009 | 85631033 | 25 | * | NA_peak_33 | 20 | Distal Interge | 17 | 85621031 | 85629302 | 8272 | 1 | 20473 | ENSMUST00 | 9978 | ENSMUSG0 | Six3 | sine oculis-related homeobox 3 |
| chr17 | 86058877 | 86059071 | 195 | * | NA_peak_33 | 19 | Promoter (1-2 | 17 | 86047975 | 86057700 | 9726 | 2 | 78586 | ENSMUST00 | -1177 | ENSMUSG0 | Srbd1 | S1 RNA binding domain 1 |
| chr17 | 86275336 | 86275482 | 147 | * | NA_peak_33 | 20 | Intron (ENSM | 17 | 86285164 | 86287178 | 2015 | 2 | 72104 | ENSMUST00 | 11696 | ENSMUSG0 | 2010106C02 | RIKEN cDNA 2010106C02 gene |
| chr17 | 88725313 | 88725963 | 651 | * | NA_peak_33 | 29 | Intron (ENSM | 17 | 88716481 | 88774056 | 57576 | 2 | 16867 | ENSMUST00 | 48093 | ENSMUSG0 | Lhcgr | luteinizing hormone/choriogonadotropin receptor |
| chr17 | 89259745 | 89260025 | 281 | * | NA_peak_33 | 35 | Distal Interge | 17 | 88984952 | 89200675 | 215724 | 2 | 14309 | ENSMUST00 | -59070 | ENSMUSG0 | Fshr | follicle stimulating hormone receptor |
| chr17 | 90694711 | 90695019 | 309 | * | NA_peak_33 | 22 | Intron (ENSM | 17 | 90590194 | 90700774 | 110581 | 2 | 18189 | ENSMUST00 | 5755 | ENSMUSG0 | Nrxn1 | neurexin I |
| chr17 | 91645702 | 91646004 | 303 | * | NA_peak_33 | 45 | Distal Interge | 17 | 90033644 | 91093071 | 1059428 | 2 | 18189 | ENSMUST00 | -552631 | ENSMUSG0 | Nrxn1 | neurexin I |
| chr17 | 91646818 | 91646894 | 77 | * | NA_peak_33 | 16 | Distal Interge | 17 | 90033644 | 91093071 | 1059428 | 2 | 18189 | ENSMUST00 | -553747 | ENSMUSG0 | Nrxn1 | neurexin I |
| chr17 | 94301025 | 94301222 | 198 | * | NA_peak_33 | 20 | Distal Interge | 17 | 94722590 | 94749892 | 27303 | 2 | 76781 | ENSMUST00 | 448670 | ENSMUSG0 | Mettl4 | methyltransferase like 4 |
| chr17 | 94669945 | 94670235 | 291 | * | NA_peak_33 | 24 | Distal Interge | 17 | 94722590 | 94749892 | 27303 | 2 | 76781 | ENSMUST00 | 79657 | ENSMUSG0 | Mettl4 | methyltransferase like 4 |
| chr18 | 3004785 | 3005163 | 379 | * | NA_peak_33 | 128 | Distal Interge | 18 | 3122492 | 3123412 | 921 | 2 | 100312476 | ENSMUST00 | 118249 | ENSMUSG0 | Vmn1r238 | vomeronasal 1 receptor, 238 |
| chr18 | 3005610 | 3005924 | 315 | * | NA_peak_33 | 488 | Distal Interge | 18 | 3122492 | 3123412 | 921 | 2 | 100312476 | ENSMUST00 | 117488 | ENSMUSG0 | Vmn1r238 | vomeronasal 1 receptor, 238 |
| chr18 | 3010320 | 3010618 | 299 | * | NA_peak_33 | 125 | Distal Interge | 18 | 3122492 | 3123412 | 921 | 2 | 100312476 | ENSMUST00 | 112794 | ENSMUSG0 | Vmn1r238 | vomeronasal 1 receptor, 238 |
| chr18 | 3020132 | 3020435 | 304 | * | NA_peak_33 | 23 | Distal Interge | 18 | 3122492 | 3123412 | 921 | 2 | 100312476 | ENSMUST00 | 102977 | ENSMUSG0 | Vmn1r238 | vomeronasal 1 receptor, 238 |
| chr18 | 3020726 | 3020778 | 53 | * | NA_peak_33 | 31 | Distal Interge | 18 | 3122492 | 3123412 | 921 | 2 | 100312476 | ENSMUST00 | 102634 | ENSMUSG0 | Vmn1r238 | vomeronasal 1 receptor, 238 |
| chr18 | 3031536 | 3031860 | 325 | * | NA_peak_33 | 28 | Distal Interge | 18 | 3122492 | 3123412 | 921 | 2 | 100312476 | ENSMUST00 | 91552 | ENSMUSG0 | Vmn1r238 | vomeronasal 1 receptor, 238 |
| chr18 | 3048369 | 3048664 | 296 | * | NA_peak_33 | 37 | Distal Interge | 18 | 3122492 | 3123412 | 921 | 2 | 100312476 | ENSMUST00 | 74748 | ENSMUSG0 | Vmn1r238 | vomeronasal 1 receptor, 238 |
| chr18 | 3064063 | 3064411 | 349 | * | NA_peak_33 | 31 | Distal Interge | 18 | 3122492 | 3123412 | 921 | 2 | 100312476 | ENSMUST00 | 59001 | ENSMUSG0 | Vmn1r238 | vomeronasal 1 receptor, 238 |
| chr18 | 3067024 | 3067430 | 407 | * | NA_peak_33 | 20 | Distal Interge | 18 | 3122492 | 3123412 | 921 | 2 | 100312476 | ENSMUST00 | 55982 | ENSMUSG0 | Vmn1r238 | vomeronasal 1 receptor, 238 |
| chr18 | 3091388 | 3091684 | 297 | * | NA_peak_33 | 22 | Distal Interge | 18 | 3122492 | 3123412 | 921 | 2 | 100312476 | ENSMUST00 | 31728 | ENSMUSG0 | Vmn1r238 | vomeronasal 1 receptor, 238 |
| chr18 | 3115632 | 3115993 | 362 | * | NA_peak_33 | 26 | Distal Interge | 18 | 3122492 | 3123412 | 921 | 2 | 100312476 | ENSMUST00 | 7419 | ENSMUSG0 | Vmn1r238 | vomeronasal 1 receptor, 238 |
| chr18 | 4023000 | 4023286 | 287 | * | NA_peak_33 | 23 | Distal Interge | 18 | 4165832 | 4182232 | 16401 | 1 | 67328 | ENSMUST00 | -142546 | ENSMUSG0 | Lyzl1 | lysozyme-like 1 |
| chr18 | 4024871 | 4025032 | 162 | * | NA_peak_33 | 49 | Distal Interge | 18 | 4165832 | 4182232 | 16401 | 1 | 67328 | ENSMUST00 | -140800 | ENSMUSG0 | Lyzl1 | lysozyme-like 1 |
| chr18 | 4196937 | 4197631 | 695 | * | NA_peak_33 | 50 | Distal Interge | 18 | 4165862 | 4182232 | 16371 | 1 | 67328 | ENSMUST00 | 31075 | ENSMUSG0 | Lyzl1 | lysozyme-like 1 |
| chr18 | 5755179 | 5755293 | 115 | * | NA_peak_33 | 21 | Intron (ENSM | 18 | 5731081 | 5772804 | 41724 | 1 | 21417 | ENSMUST00 | 24098 | ENSMUSG0 | Zeb1 | zinc finger E-box binding homeobox 1 |
| chr18 | 6372482 | 6372521 | 40 | * | NA_peak_33 | 16 | Distal Interge | 18 | 6439592 | 6441894 | 2303 | 2 | 13831 | ENSMUST00 | 69373 | ENSMUSG0 | Epc1 | enhancer of polycomb homolog 1 |
| chr18 | 12622592 | 12622823 | 232 | * | NA_peak_33 | 27 | Intron (ENSM | 18 | 12643344 | 12689445 | 46102 | 1 | 72747 | ENSMUST00 | -20521 | ENSMUSG0 | Ttc39c | tetratricopeptide repeat domain 39C |
| chr18 | 13040176 | 13040488 | 313 | * | NA_peak_33 | 55 | Distal Interge | 18 | 13007037 | 13021823 | 14787 | 1 | 225192 | ENSMUST00 | 33139 | ENSMUSG0 | Hrh4 | histamine receptor H4 |
| chr18 | 16962939 | 16963377 | 439 | * | NA_peak_33 | 137 | Distal Interge | 18 | 17019051 | 17137559 | 118509 | 1 | 75433 | ENSMUST00 | -55674 | ENSMUSG0 | 1700001G01 | RIKEN cDNA 1700001G01 gene |
| chr18 | 17350972 | 17351074 | 103 | * | NA_peak_33 | 22 | Distal Interge | 18 | 17234484 | 17235453 | 970 | 2 | 664619 | ENSMUST00 | -115519 | ENSMUSG0 | 4921533I20R | PRP18 pre-mRNA processing factor 18 homolog pseudogene |
| chr18 | 17663075 | 17663397 | 323 | * | NA_peak_33 | 36 | Distal Interge | 18 | 17568746 | 17580990 | 12245 | 2 | 75162 | ENSMUST00 | -82085 | ENSMUSG0 | 4930545E07 | RIKEN cDNA 4930545E07 gene |
| chr18 | 19284627 | 19284865 | 239 | * | NA_peak_33 | 32 | Distal Interge | 18 | 19962970 | 20001955 | 38986 | 2 | 13507 | ENSMUST00 | 717090 | ENSMUSG0 | Dsc3 | desmocollin 3 |
| chr18 | 22298428 | 22298882 | 455 | * | NA_peak_33 | 55 | Distal Interge | 18 | 22344883 | 22378950 | 34068 | 1 | 211961 | ENSMUST00 | -46001 | ENSMUSG0 | Asxl3 | ASXL transcriptional regulator 3 |
| chr18 | 24035309 | 24035627 | 319 | * | NA_peak_33 | 28 | Distal Interge | 18 | 24009702 | 24020810 | 11109 | 2 | 59057 | ENSMUST00 | -14499 | ENSMUSG0 | Zfp24 | zinc finger protein 24 |
| chr18 | 26701725 | 26702040 | 316 | * | NA_peak_33 | 25 | Distal Interge | 18 | 25479115 | 25754157 | 275043 | 2 | 108013 | ENSMUST00 | -947568 | ENSMUSG0 | Celf4 | CUGBP, Elav-like family member 4 |
| chr18 | 27234242 | 27234457 | 216 | * | NA_peak_33 | 40 | Distal Interge | 18 | 25479115 | 25754157 | 275043 | 2 | 108013 | ENSMUST00 | -1480085 | ENSMUSG0 | Celf4 | CUGBP, Elav-like family member 4 |
| chr18 | 27460322 | 27460734 | 413 | * | NA_peak_33 | 22 | Distal Interge | 18 | 28862850 | 28998176 | 135327 | 1 | 320624 | ENSMUST00 | -1402116 | ENSMUSG0 | 4930474G06 | RIKEN cDNA 4930474G06 gene |
| chr18 | 28749954 | 28750246 | 293 | * | NA_peak_33 | 32 | Distal Interge | 18 | 28862850 | 28998176 | 135327 | 1 | 320624 | ENSMUST00 | -112604 | ENSMUSG0 | 4930474G06 | RIKEN cDNA 4930474G06 gene |
| chr18 | 36479691 | 36480212 | 522 | * | NA_peak_33 | 28 | Distal Interge | 18 | 36403676 | 36454524 | 50849 | 2 | 67199 | ENSMUST00 | -25167 | ENSMUSG0 | Pfdn1 | prefoldin 1 |
| chr18 | 40606162 | 40606393 | 232 | * | NA_peak_33 | 40 | Distal Interge | 18 | 40523099 | 40531179 | 8081 | 1 | 383348 | ENSMUST00 | 83063 | ENSMUSG0 | Kctd16 | potassium channel tetramerisation domain containing 16 |
| chr18 | 42476520 | 42476817 | 298 | * | NA_peak_33 | 16 | Distal Interge | 18 | 42511487 | 42575793 | 64307 | 1 | 56070 | ENSMUST00 | -34670 | ENSMUSG0 | Tcerg1 | transcription elongation regulator 1 (CA150) |
| chr18 | 44737005 | 44737209 | 205 | * | NA_peak_33 | 28 | Intron (ENSM | 18 | 44711435 | 44724658 | 13224 | 2 | 328949 | ENSMUST00 | -12347 | ENSMUSG0 | Mcc | mutated in colorectal cancers |
| chr18 | 47740016 | 47740115 | 100 | * | NA_peak_33 | 15 | Intron (ENSM | 18 | 47761089 | 47789533 | 28445 | 2 | 100042982 | ENSMUST00 | 49418 | ENSMUSG0 | Gm4146 | predicted gene 4146 |
| chr18 | 50266383 | 50266834 | 452 | * | NA_peak_33 | 96 | Distal Interge | 18 | 50278369 | 50283014 | 4646 | 1 | 225497 | ENSMUST00 | -11535 | ENSMUSG0 | Fam170a | family with sequence similarity 170, member A |
| chr18 | 55318889 | 55318995 | 107 | * | NA_peak_33 | 15 | Distal Interge | 18 | 54888723 | 54992555 | 103833 | 2 | 269023 | ENSMUST00 | -326334 | ENSMUSG0 | Zfp608 | zinc finger protein 608 |
| chr18 | 57795734 | 57796199 | 466 | * | NA_peak_33 | 22 | Distal Interge | 18 | 57688646 | 57792360 | 103715 | 2 | 100503033 | ENSMUST00 | -3374 | ENSMUSG0 | Gm19519 | predicted gene, 19519 |
| chr18 | 59120737 | 59121023 | 287 | * | NA_peak_33 | 15 | Distal Interge | 18 | 59175401 | 59182021 | 6621 | 1 | 78923 | ENSMUST00 | -54378 | ENSMUSG0 | Chsy3 | chondroitin sulfate synthase 3 |
| chr18 | 59401726 | 59401929 | 204 | * | NA_peak_33 | 26 | Intron (ENSM | 18 | 59579630 | 59579735 | 106 | 1 | 102465182 | ENSMUST00 | -177701 | ENSMUSG0 | Mir6355 | microRNA 6355 |
| chr18 | 60446947 | 60447402 | 456 | * | NA_peak_33 | 19 | Distal Interge | 18 | 60474191 | 60501987 | 27797 | 2 | 106878 | ENSMUST00 | 54585 | ENSMUSG0 | Smim3 | small integral membrane protein 3 |
| chr18 | 63912111 | 63912449 | 339 | * | NA_peak_33 | 45 | Intron (ENSM | 18 | 63777972 | 63865290 | 87319 | 1 | 104082 | ENSMUST00 | 134139 | ENSMUSG0 | Wdr7 | WD repeat domain 7 |
| chr18 | 72512221 | 72512229 | 9 | * | NA_peak_33 | 16 | Distal Interge | 18 | 71253634 | 72351069 | 1097436 | 2 | 13176 | ENSMUST00 | -161152 | ENSMUSG0 | Dcc | deleted in colorectal carcinoma |
| chr18 | 76364262 | 76364617 | 356 | * | NA_peak_33 | 30 | Distal Interge | 18 | 76262492 | 76302642 | 40151 | 1 | 17126 | ENSMUST00 | 101770 | ENSMUSG0 | Smad2 | SMAD family member 2 |
| chr18 | 76790360 | 76790768 | 409 | * | NA_peak_33 | 55 | Distal Interge | 18 | 76856405 | 76900342 | 43938 | 1 | 664805 | ENSMUST00 | -65637 | ENSMUSG0 | Skor2 | SKI family transcriptional corepressor 2 |
| chr18 | 85079559 | 85079949 | 391 | * | NA_peak_33 | 20 | Intron (ENSM | 18 | 84965510 | 84981472 | 15963 | 1 | 50764 | ENSMUST00 | 114049 | ENSMUSG0 | Fbxo15 | F-box protein 15 |
| chr18 | 87779921 | 87780238 | 318 | * | NA_peak_33 | 17 | Distal Interge | 18 | 86713048 | 86718283 | 5236 | 1 | 12405 | ENSMUST00 | 1066873 | ENSMUSG0 | Cbln2 | cerebellin 2 precursor protein |
| chr18 | 89649752 | 89650151 | 400 | * | NA_peak_33 | 40 | Intron (ENSM | 18 | 89470148 | 89560069 | 89922 | 2 | 623279 | ENSMUST00 | -89683 | ENSMUSG0 | Dok6 | docking protein 6 |
| chr18 | 90275627 | 90275878 | 252 | * | NA_peak_33 | 40 | Distal Interge | 18 | 90510154 | 90543267 | 33114 | 1 | 67988 | ENSMUST00 | -234276 | ENSMUSG0 | Tmx3 | thioredoxin-related transmembrane protein 3 |
| chr19 | 3078885 | 3079302 | 418 | * | NA_peak_33 | 31 | Intron (ENSM | 19 | 3145946 | 3160181 | 14236 | 2 | 70011 | ENSMUST00 | 80879 | ENSMUSG0 | 1700030N03 | RIKEN cDNA 1700030N03 gene |
| chr19 | 3079838 | 3079904 | 67 | * | NA_peak_33 | 36 | Intron (ENSM | 19 | 3145946 | 3160181 | 14236 | 2 | 70011 | ENSMUST00 | 80277 | ENSMUSG0 | 1700030N03 | RIKEN cDNA 1700030N03 gene |
| chr19 | 3104211 | 3104296 | 86 | * | NA_peak_33 | 43 | Intron (ENSM | 19 | 3145946 | 3160181 | 14236 | 2 | 70011 | ENSMUST00 | 55885 | ENSMUSG0 | 1700030N03 | RIKEN cDNA 1700030N03 gene |
| chr19 | 3113815 | 3114192 | 378 | * | NA_peak_33 | 82 | Intron (ENSM | 19 | 3145946 | 3160181 | 14236 | 2 | 70011 | ENSMUST00 | 45989 | ENSMUSG0 | 1700030N03 | RIKEN cDNA 1700030N03 gene |
| chr19 | 3173264 | 3173719 | 456 | * | NA_peak_33 | 23 | Intron (ENSM | 19 | 3145946 | 3160181 | 14236 | 2 | 70011 | ENSMUST00 | -13083 | ENSMUSG0 | 1700030N03 | RIKEN cDNA 1700030N03 gene |
| chr19 | 3547980 | 3548162 | 183 | * | NA_peak_33 | 24 | Intron (ENSM | 19 | 3511495 | 3521747 | 10253 | 2 | 52036 | ENSMUST00 | -26233 | ENSMUSG0 | Ppp6r3 | protein phosphatase 6, regulatory subunit 3 |
| chr19 | 3787927 | 3788151 | 225 | * | NA_peak_33 | 13 | Intron (ENSM | 19 | 3791720 | 3799549 | 7830 | 1 | 225888 | ENSMUST00 | -3569 | ENSMUSG0 | Kmt5b | lysine methyltransferase 5B |
| chr19 | 4483967 | 4484205 | 239 | * | NA_peak_33 | 22 | Distal Interge | 19 | 4445908 | 4477447 | 31540 | 2 | 171180 | ENSMUST00 | -6520 | ENSMUSG0 | Syt12 | synaptotagmin XII |
| chr19 | 5024017 | 5024295 | 279 | * | NA_peak_33 | 15 | Promoter (<= | 19 | 5024071 | 5031969 | 7899 | 1 | 13340 | ENSMUST00 | 0 | ENSMUSG0 | Slc29a2 | solute carrier family 29 (nucleoside transporters), member 2 |
| chr19 | 5264889 | 5265154 | 266 | * | NA_peak_33 | 19 | Intron (ENSM | 19 | 5160400 | 5272707 | 112308 | 2 | 107975 | ENSMUST00 | 7553 | ENSMUSG0 | Pacs1 | phosphofurin acidic cluster sorting protein 1 |
| chr19 | 5457201 | 5457419 | 219 | * | NA_peak_33 | 35 | Promoter (<= | 19 | 5453185 | 5457504 | 4320 | 2 | 240514 | ENSMUST00 | 85 | ENSMUSG0 | Ccdc85b | coiled-coil domain containing 85B |
| chr19 | 5786624 | 5786863 | 240 | * | NA_peak_33 | 20 | Distal Interge | 19 | 5799000 | 5799085 | 86 | 2 | 72289 | ENSMUST00 | 12222 | ENSMUSG0 | Malat1 | metastasis associated lung adenocarcinoma transcript 1 (non-coding RNA) |
| chr19 | 6042101 | 6042380 | 280 | * | NA_peak_33 | 24 | Distal Interge | 19 | 6046576 | 6053712 | 7137 | 1 | 74126 | ENSMUST00 | -4196 | ENSMUSG0 | Syvn1 | synovial apoptosis inhibitor 1, synoviolin |
| chr19 | 6413225 | 6413371 | 147 | * | NA_peak_33 | 28 | 3' UTR | 19 | 6408405 | 6415211 | 6807 | 1 | 19395 | ENSMUST00 | 4820 | ENSMUSG0 | Rasgrp2 | RAS, guanyl releasing protein 2 |
| chr19 | 6888001 | 6888306 | 306 | * | NA_peak_34 | 38 | Distal Interge | 19 | 6906750 | 6908301 | 1552 | 2 | 54683 | ENSMUST00 | 19995 | ENSMUSG0 | Prdx5 | peroxiredoxin 5 |
| chr19 | 6980133 | 6980269 | 137 | * | NA_peak_34 | 15 | Promoter (<= | 19 | 6977747 | 6980250 | 2504 | 2 | 114841036 | ENSMUST00 | 0 | ENSMUSG0 | Gm51400 | predicted gene, 51400 |
| chr19 | 7464670 | 7465113 | 444 | * | NA_peak_34 | 23 | Intron (ENSM | 19 | 7425901 | 7483202 | 57302 | 2 | 20168 | ENSMUST00 | 18089 | ENSMUSG0 | Rtn3 | reticulon 3 |
| chr19 | 7490079 | 7490469 | 391 | * | NA_peak_34 | 25 | Distal Interge | 19 | 7494040 | 7539674 | 45635 | 1 | 109168 | ENSMUST00 | -3571 | ENSMUSG0 | Atl3 | atlastin GTPase 3 |
| chr19 | 7579600 | 7579874 | 275 | * | NA_peak_34 | 17 | 3' UTR | 19 | 7557485 | 7586711 | 29227 | 1 | 225845 | ENSMUST00 | 22115 | ENSMUSG0 | Plaat3 | phospholipase A and acyltransferase 3 |
| chr19 | 7631240 | 7631754 | 515 | * | NA_peak_34 | 36 | Intron (ENSM | 19 | 7612605 | 7639642 | 27038 | 1 | 66727 | ENSMUST00 | 18635 | ENSMUSG0 | Plaat5 | phospholipase A and acyltransferase 5 |
| chr19 | 7663777 | 7663851 | 75 | * | NA_peak_34 | 23 | Distal Interge | 19 | 7682277 | 7711279 | 29003 | 2 | 207151 | ENSMUST00 | 47428 | ENSMUSG0 | Slc22a19 | solute carrier family 22 (organic anion transporter), member 19 |
| chr19 | 7721334 | 7721640 | 307 | * | NA_peak_34 | 27 | Distal Interge | 19 | 7706682 | 7711310 | 4629 | 2 | 207151 | ENSMUST00 | -10024 | ENSMUSG0 | Slc22a19 | solute carrier family 22 (organic anion transporter), member 19 |
| chr19 | 7848562 | 7848768 | 207 | * | NA_peak_34 | 23 | Exon (ENSMU | 19 | 7781980 | 7802667 | 20688 | 2 | 236149 | ENSMUST00 | -45895 | ENSMUSG0 | Slc22a26 | solute carrier family 22 (organic cation transporter), member 26 |
| chr19 | 7858184 | 7858651 | 468 | * | NA_peak_34 | 23 | Distal Interge | 19 | 7781980 | 7802667 | 20688 | 2 | 236149 | ENSMUST00 | -55517 | ENSMUSG0 | Slc22a26 | solute carrier family 22 (organic cation transporter), member 26 |
| chr19 | 7915402 | 7915878 | 477 | * | NA_peak_34 | 15 | Intron (ENSM | 19 | 7864388 | 7966000 | 101613 | 2 | 171405 | ENSMUST00 | 50122 | ENSMUSG0 | Slc22a27 | solute carrier family 22, member 27 |
| chr19 | 7939562 | 7939591 | 30 | * | NA_peak_34 | 31 | Intron (ENSM | 19 | 7864388 | 7966000 | 101613 | 2 | 171405 | ENSMUST00 | 26409 | ENSMUSG0 | Slc22a27 | solute carrier family 22, member 27 |
| chr19 | 7989063 | 7989232 | 170 | * | NA_peak_34 | 40 | Distal Interge | 19 | 7864388 | 7966027 | 101640 | 2 | 171405 | ENSMUST00 | -23036 | ENSMUSG0 | Slc22a27 | solute carrier family 22, member 27 |
| chr19 | 7993134 | 7993508 | 375 | * | NA_peak_34 | 30 | Distal Interge | 19 | 7864388 | 7966027 | 101640 | 2 | 171405 | ENSMUST00 | -27107 | ENSMUSG0 | Slc22a27 | solute carrier family 22, member 27 |
| chr19 | 8027056 | 8027572 | 517 | * | NA_peak_34 | 31 | Distal Interge | 19 | 7864388 | 7966027 | 101640 | 2 | 171405 | ENSMUST00 | -61029 | ENSMUSG0 | Slc22a27 | solute carrier family 22, member 27 |
| chr19 | 8047898 | 8048970 | 1073 | * | NA_peak_34 | 28 | Distal Interge | 19 | 7864388 | 7966027 | 101640 | 2 | 171405 | ENSMUST00 | -81871 | ENSMUSG0 | Slc22a27 | solute carrier family 22, member 27 |
| chr19 | 8050048 | 8050230 | 183 | * | NA_peak_34 | 24 | Distal Interge | 19 | 8062209 | 8131982 | 69774 | 2 | 434674 | ENSMUST00 | 81752 | ENSMUSG0 | Slc22a28 | solute carrier family 22, member 28 |
| chr19 | 8058244 | 8058527 | 284 | * | NA_peak_34 | 28 | Distal Interge | 19 | 8062209 | 8131982 | 69774 | 2 | 434674 | ENSMUST00 | 73455 | ENSMUSG0 | Slc22a28 | solute carrier family 22, member 28 |
| chr19 | 8059595 | 8059893 | 299 | * | NA_peak_34 | 36 | Distal Interge | 19 | 8062209 | 8131982 | 69774 | 2 | 434674 | ENSMUST00 | 72089 | ENSMUSG0 | Slc22a28 | solute carrier family 22, member 28 |
| chr19 | 8144202 | 8144589 | 388 | * | NA_peak_34 | 27 | Distal Interge | 19 | 8062209 | 8131982 | 69774 | 2 | 434674 | ENSMUST00 | -12220 | ENSMUSG0 | Slc22a28 | solute carrier family 22, member 28 |
| chr19 | 8152836 | 8153216 | 381 | * | NA_peak_34 | 46 | Distal Interge | 19 | 8160294 | 8170045 | 9752 | 2 | 236293 | ENSMUST00 | 16829 | ENSMUSG0 | Slc22a29 | solute carrier family 22. member 29 |
| chr19 | 8163451 | 8163694 | 244 | * | NA_peak_34 | 23 | Intron (ENSM | 19 | 8160294 | 8170045 | 9752 | 2 | 236293 | ENSMUST00 | 6351 | ENSMUSG0 | Slc22a29 | solute carrier family 22. member 29 |
| chr19 | 8185759 | 8186036 | 278 | * | NA_peak_34 | 28 | Intron (ENSM | 19 | 8160294 | 8170045 | 9752 | 2 | 236293 | ENSMUST00 | -15714 | ENSMUSG0 | Slc22a29 | solute carrier family 22. member 29 |
| chr19 | 8204281 | 8204676 | 396 | * | NA_peak_34 | 34 | Intron (ENSM | 19 | 8161550 | 8218673 | 57124 | 2 | 236293 | ENSMUST00 | 13997 | ENSMUSG0 | Slc22a29 | solute carrier family 22. member 29 |
| chr19 | 8243368 | 8243722 | 355 | * | NA_peak_34 | 19 | Distal Interge | 19 | 8160304 | 8218900 | 58597 | 2 | 236293 | ENSMUST00 | -24468 | ENSMUSG0 | Slc22a29 | solute carrier family 22. member 29 |

| | | | | | | | | | | | | | | | | | | |
|---|---|---|---|---|---|---|---|---|---|---|---|---|---|---|---|---|---|---|
| chr19 | 8260531 | 8260736 | 206 | * | NA_peak_34 | 31 | Intron (ENSM | 19 | 8160304 | 8218900 | 58597 | 2 | 236293 | ENSMUST00 | -41631 | ENSMUSG0 | Slc22a29 | solute carrier family 22. member 29 |
| chr19 | 8274324 | 8274454 | 131 | * | NA_peak_34 | 16 | Exon (ENSMU | 19 | 8160304 | 8218900 | 58597 | 2 | 236293 | ENSMUST00 | -55424 | ENSMUSG0 | Slc22a29 | solute carrier family 22. member 29 |
| chr19 | 8286200 | 8286514 | 315 | * | NA_peak_34 | 28 | Intron (ENSM | 19 | 8160304 | 8218900 | 58597 | 2 | 236293 | ENSMUST00 | -67300 | ENSMUSG0 | Slc22a29 | solute carrier family 22. member 29 |
| chr19 | 8350218 | 8350408 | 191 | * | NA_peak_34 | 16 | Intron (ENSM | 19 | 8335374 | 8405060 | 69687 | 2 | 319800 | ENSMUST00 | 54652 | ENSMUSG0 | Slc22a30 | solute carrier family 22, member 30 |
| chr19 | 8355437 | 8355858 | 422 | * | NA_peak_34 | 20 | Intron (ENSM | 19 | 8335374 | 8405060 | 69687 | 2 | 319800 | ENSMUST00 | 49202 | ENSMUSG0 | Slc22a30 | solute carrier family 22, member 30 |
| chr19 | 8487898 | 8488193 | 296 | * | NA_peak_34 | 31 | Intron (ENSM | 19 | 8335371 | 8405111 | 69741 | 2 | 319800 | ENSMUST00 | -82787 | ENSMUSG0 | Slc22a30 | solute carrier family 22, member 30 |
| chr19 | 8491732 | 8492289 | 558 | * | NA_peak_34 | 20 | Exon (ENSMU | 19 | 8335371 | 8405111 | 69741 | 2 | 319800 | ENSMUST00 | -86621 | ENSMUSG0 | Slc22a30 | solute carrier family 22, member 30 |
| chr19 | 8537829 | 8537882 | 54 | * | NA_peak_34 | 17 | Intron (ENSM | 19 | 8591254 | 8593242 | 1989 | 1 | 19879 | ENSMUST00 | -53372 | ENSMUSG0 | Slc22a8 | solute carrier family 22 (organic anion transporter), member 8 |
| chr19 | 9143876 | 9144511 | 636 | * | NA_peak_34 | 31 | Intron (ENSM | 19 | 9109868 | 9135636 | 25769 | 2 | 66514 | ENSMUST00 | -8240 | ENSMUSG0 | Asrgl1 | asparaginase like 1 |
| chr19 | 9150772 | 9150910 | 139 | * | NA_peak_34 | 29 | Intron (ENSM | 19 | 9109868 | 9135636 | 25769 | 2 | 66514 | ENSMUST00 | -15136 | ENSMUSG0 | Asrgl1 | asparaginase like 1 |
| chr19 | 9447810 | 9447960 | 151 | * | NA_peak_34 | 15 | Distal Interge | 19 | 9557792 | 9559183 | 1392 | 2 | 619371 | ENSMUST00 | 111223 | NA | Stxbp3-ps | syntaxin-binding protein 3, pseudogene |
| chr19 | 9559106 | 9559363 | 258 | * | NA_peak_34 | 36 | Promoter (<= | 19 | 9557792 | 9559183 | 1392 | 2 | 619371 | ENSMUST00 | 0 | NA | Stxbp3-ps | syntaxin-binding protein 3, pseudogene |
| chr19 | 9577470 | 9577511 | 42 | * | NA_peak_34 | 27 | Distal Interge | 19 | 9557792 | 9559183 | 1392 | 2 | 619371 | ENSMUST00 | -18287 | NA | Stxbp3-ps | syntaxin-binding protein 3, pseudogene |
| chr19 | 9640158 | 9640448 | 291 | * | NA_peak_34 | 40 | Distal Interge | 19 | 9557792 | 9559183 | 1392 | 2 | 619371 | ENSMUST00 | -80975 | NA | Stxbp3-ps | syntaxin-binding protein 3, pseudogene |
| chr19 | 9671966 | 9672538 | 573 | * | NA_peak_34 | 27 | Distal Interge | 19 | 9557792 | 9559183 | 1392 | 2 | 619371 | ENSMUST00 | -112783 | NA | Stxbp3-ps | syntaxin-binding protein 3, pseudogene |
| chr19 | 9714371 | 9714689 | 319 | * | NA_peak_34 | 21 | Distal Interge | 19 | 9557792 | 9559183 | 1392 | 2 | 619371 | ENSMUST00 | -155188 | NA | Stxbp3-ps | syntaxin-binding protein 3, pseudogene |
| chr19 | 9719806 | 9719987 | 182 | * | NA_peak_34 | 36 | Distal Interge | 19 | 9872342 | 9874923 | 2582 | 2 | 16319 | ENSMUST00 | 154936 | ENSMUSG0 | Incenp | inner centromere protein |
| chr19 | 9774431 | 9774623 | 193 | * | NA_peak_34 | 54 | Distal Interge | 19 | 9872342 | 9874923 | 2582 | 2 | 16319 | ENSMUST00 | 100300 | ENSMUSG0 | Incenp | inner centromere protein |
| chr19 | 9789132 | 9789301 | 170 | * | NA_peak_34 | 28 | Distal Interge | 19 | 9872342 | 9874923 | 2582 | 2 | 16319 | ENSMUST00 | 85622 | ENSMUSG0 | Incenp | inner centromere protein |
| chr19 | 9834226 | 9834856 | 631 | * | NA_peak_34 | 72 | Intron (ENSM | 19 | 9872342 | 9874923 | 2582 | 2 | 16319 | ENSMUST00 | 40067 | ENSMUSG0 | Incenp | inner centromere protein |
| chr19 | 10533269 | 10533543 | 275 | * | NA_peak_34 | 20 | Promoter (<= | 19 | 10532389 | 10533372 | 984 | 1 | 269061 | ENSMUST00 | 880 | ENSMUSG0 | Cpsf7 | cleavage and polyadenylation specific factor 7 |
| chr19 | 10536053 | 10536386 | 334 | * | NA_peak_34 | 19 | Intron (ENSM | 19 | 10532389 | 10533372 | 984 | 1 | 269061 | ENSMUST00 | 3664 | ENSMUSG0 | Cpsf7 | cleavage and polyadenylation specific factor 7 |
| chr19 | 10618009 | 10618266 | 258 | * | NA_peak_34 | 37 | Promoter (<= | 19 | 10618994 | 10622145 | 3152 | 1 | 13194 | ENSMUST00 | -728 | ENSMUSG0 | Ddb1 | damage specific DNA binding protein 1 |
| chr19 | 11008138 | 11008413 | 276 | * | NA_peak_34 | 22 | Intron (ENSM | 19 | 10997025 | 11018031 | 21007 | 2 | 76002 | ENSMUST00 | 9618 | ENSMUSG0 | Ms4a18 | membrane-spanning 4-domains, subfamily A, member 18 |
| chr19 | 11029426 | 11029624 | 199 | * | NA_peak_34 | 42 | Distal Interge | 19 | 10997025 | 11018031 | 21007 | 2 | 76002 | ENSMUST00 | -11395 | ENSMUSG0 | Ms4a18 | membrane-spanning 4-domains, subfamily A, member 18 |
| chr19 | 11070149 | 11070361 | 213 | * | NA_peak_34 | 23 | Intron (ENSM | 19 | 11067471 | 11081102 | 13632 | 2 | 64381 | ENSMUST00 | 10741 | ENSMUSG0 | Ms4a8a | membrane-spanning 4-domains, subfamily A, member 8A |
| chr19 | 11238048 | 11238301 | 254 | * | NA_peak_34 | 19 | Intron (ENSM | 19 | 11250877 | 11254684 | 3808 | 2 | 12482 | ENSMUST00 | 16383 | ENSMUSG0 | Ms4a1 | membrane-spanning 4-domains, subfamily A, member 1 |
| chr19 | 11248590 | 11248830 | 241 | * | NA_peak_34 | 27 | Intron (ENSM | 19 | 11250877 | 11254684 | 3808 | 2 | 12482 | ENSMUST00 | 5854 | ENSMUSG0 | Ms4a1 | membrane-spanning 4-domains, subfamily A, member 1 |
| chr19 | 11334199 | 11334378 | 180 | * | NA_peak_34 | 30 | Promoter (<= | 19 | 11321039 | 11334695 | 13657 | 2 | 109225 | ENSMUST00 | 317 | ENSMUSG0 | Ms4a7 | membrane-spanning 4-domains, subfamily A, member 7 |
| chr19 | 11476626 | 11476730 | 105 | * | NA_peak_34 | 20 | Intron (ENSM | 19 | 11469425 | 11472736 | 3312 | 1 | 73656 | ENSMUST00 | 7201 | ENSMUSG0 | Ms4a6c | membrane-spanning 4-domains, subfamily A, member 6C |
| chr19 | 11490822 | 11491021 | 200 | * | NA_peak_34 | 32 | Promoter (1-2 | 19 | 11492038 | 11511708 | 19671 | 1 | 666926 | ENSMUST00 | -1017 | ENSMUSG0 | Gm8369 | predicted gene 8369 |
| chr19 | 11715741 | 11715990 | 250 | * | NA_peak_34 | 19 | Distal Interge | 19 | 11697078 | 11711874 | 14797 | 1 | 225923 | ENSMUST00 | 18663 | ENSMUSG0 | Oosp3 | oocyte secreted protein 3 |
| chr19 | 11739431 | 11739585 | 155 | * | NA_peak_34 | 41 | Distal Interge | 19 | 11747554 | 11763447 | 15894 | 1 | 14603 | ENSMUST00 | -7969 | ENSMUSG0 | Cblif | cobalamin binding intrinsic factor |
| chr19 | 11895655 | 11896141 | 487 | * | NA_peak_34 | 27 | Promoter (<= | 19 | 11895607 | 11897050 | 1444 | 1 | 258405 | ENSMUST00 | 48 | ENSMUSG0 | Olfr1420 | olfactory receptor 1420 |
| chr19 | 12058940 | 12059195 | 256 | * | NA_peak_34 | 31 | Promoter (<= | 19 | 12058809 | 12059750 | 942 | 2 | 258676 | ENSMUST00 | 555 | ENSMUSG0 | Olfr1424 | olfactory receptor 1424 |
| chr19 | 12087561 | 12087819 | 259 | * | NA_peak_34 | 15 | Promoter (2-3 | 19 | 12086746 | 12090442 | 3697 | 2 | 258805 | ENSMUST00 | 2623 | ENSMUSG0 | Olfr1426 | olfactory receptor 1426 |
| chr19 | 12137413 | 12137786 | 374 | * | NA_peak_34 | 19 | Distal Interge | 19 | 12119653 | 12120700 | 1048 | 2 | 258677 | ENSMUST00 | -16713 | ENSMUSG0 | Olfr76 | olfactory receptor 76 |
| chr19 | 12151649 | 12151893 | 245 | * | NA_peak_34 | 34 | Distal Interge | 19 | 12119653 | 12120700 | 1048 | 2 | 258677 | ENSMUST00 | -30949 | ENSMUSG0 | Olfr76 | olfactory receptor 76 |
| chr19 | 12160443 | 12160810 | 368 | * | NA_peak_34 | 43 | Distal Interge | 19 | 12119653 | 12120700 | 1048 | 2 | 258677 | ENSMUST00 | -39743 | ENSMUSG0 | Olfr76 | olfactory receptor 76 |
| chr19 | 12166987 | 12167161 | 175 | * | NA_peak_34 | 20 | 5' UTR | 19 | 12209568 | 12210506 | 939 | 1 | 258409 | ENSMUST00 | -42407 | ENSMUSG0 | Olfr1431 | olfactory receptor 1431 |
| chr19 | 12292215 | 12292424 | 210 | * | NA_peak_34 | 20 | Distal Interge | 19 | 12298183 | 12299130 | 948 | 2 | 258682 | ENSMUST00 | 6706 | ENSMUSG0 | Olfr1436 | olfactory receptor 1436 |
| chr19 | 12429359 | 12429692 | 334 | * | NA_peak_34 | 23 | Promoter (1-2 | 19 | 12427905 | 12432110 | 4206 | 1 | 56093 | ENSMUST00 | 1454 | ENSMUSG0 | Pfpl | pore forming protein-like |
| chr19 | 12598830 | 12598941 | 112 | * | NA_peak_34 | 16 | Promoter (1-2 | 19 | 12599974 | 12626873 | 26900 | 1 | 240549 | ENSMUST00 | -1033 | ENSMUSG0 | Gm4952 | predicted gene 4952 |
| chr19 | 12662558 | 12662950 | 393 | * | NA_peak_34 | 13 | Distal Interge | 19 | 12674179 | 12675243 | 1065 | 1 | 258692 | ENSMUST00 | -11229 | ENSMUSG0 | Olfr1442 | olfactory receptor 1442 |
| chr19 | 12767873 | 12768220 | 348 | * | NA_peak_34 | 22 | Promoter (2-3 | 19 | 12763660 | 12765632 | 1973 | 2 | 12803 | ENSMUST00 | -2241 | ENSMUSG0 | Cntf | ciliary neurotrophic factor |
| chr19 | 12826100 | 12826446 | 347 | * | NA_peak_34 | 36 | Intron (ENSM | 19 | 12798597 | 12833807 | 35211 | 1 | 107321 | ENSMUST00 | 27503 | ENSMUSG0 | Lpxn | leupaxin |
| chr19 | 13053173 | 13053474 | 302 | * | NA_peak_34 | 18 | Distal Interge | 19 | 13063413 | 13064336 | 924 | 1 | 258687 | ENSMUST00 | -9939 | ENSMUSG0 | Olfr1454 | olfactory receptor 1454 |
| chr19 | 13074381 | 13074662 | 282 | * | NA_peak_34 | 31 | Distal Interge | 19 | 13063413 | 13064336 | 924 | 1 | 258687 | ENSMUST00 | 10968 | ENSMUSG0 | Olfr1454 | olfactory receptor 1454 |
| chr19 | 13078034 | 13078405 | 372 | * | NA_peak_34 | 32 | Distal Interge | 19 | 13063413 | 13064336 | 924 | 1 | 258687 | ENSMUST00 | 14621 | ENSMUSG0 | Olfr1454 | olfactory receptor 1454 |
| chr19 | 13193556 | 13193589 | 34 | * | NA_peak_34 | 23 | Promoter (2-3 | 19 | 13190669 | 13191592 | 924 | 1 | 258688 | ENSMUST00 | 2887 | ENSMUSG0 | Olfr1462 | olfactory receptor 1462 |
| chr19 | 13243864 | 13244252 | 389 | * | NA_peak_34 | 53 | Distal Interge | 19 | 13234252 | 13235184 | 933 | 1 | 258120 | ENSMUST00 | 9612 | ENSMUSG0 | Olfr1463 | olfactory receptor 1463 |
| chr19 | 13323864 | 13324123 | 260 | * | NA_peak_34 | 23 | Distal Interge | 19 | 13313360 | 13314283 | 924 | 2 | 258121 | ENSMUST00 | -9581 | ENSMUSG0 | Olfr1465 | olfactory receptor 1465 |
| chr19 | 13405335 | 13405711 | 377 | * | NA_peak_34 | 26 | Distal Interge | 19 | 13410571 | 13411500 | 930 | 1 | 258690 | ENSMUST00 | -4860 | ENSMUSG0 | Olfr1469 | olfactory receptor 1469 |
| chr19 | 13757434 | 13757708 | 275 | * | NA_peak_34 | 27 | Distal Interge | 19 | 13749108 | 13750055 | 948 | 1 | 258992 | ENSMUST00 | 8326 | ENSMUSG0 | Olfr1494 | olfactory receptor 1494 |
| chr19 | 13798853 | 13798857 | 5 | * | NA_peak_34 | 20 | Distal Interge | 19 | 13794665 | 13795609 | 945 | 2 | 258736 | ENSMUST00 | -3244 | ENSMUSG0 | Olfr1497 | olfactory receptor 1497 |
| chr19 | 13801223 | 13801519 | 297 | * | NA_peak_34 | 27 | Distal Interge | 19 | 13794665 | 13795609 | 945 | 2 | 258736 | ENSMUST00 | -5614 | ENSMUSG0 | Olfr1497 | olfactory receptor 1497 |
| chr19 | 13902141 | 13902570 | 430 | * | NA_peak_34 | 27 | Distal Interge | 19 | 13887221 | 13888305 | 1085 | 2 | 258627 | ENSMUST00 | -13836 | ENSMUSG0 | Olfr1504 | olfactory receptor 1504 |
| chr19 | 13941791 | 13942237 | 447 | * | NA_peak_34 | 23 | Distal Interge | 19 | 13918942 | 13920044 | 1103 | 1 | 258151 | ENSMUST00 | 22849 | ENSMUSG0 | Olfr1505 | olfactory receptor 1505 |
| chr19 | 13994040 | 13994287 | 248 | * | NA_peak_34 | 21 | Distal Interge | 19 | 13918942 | 13920044 | 1103 | 1 | 258151 | ENSMUST00 | 75098 | ENSMUSG0 | Olfr1505 | olfactory receptor 1505 |
| chr19 | 14001040 | 14001331 | 292 | * | NA_peak_34 | 40 | Distal Interge | 19 | 13918942 | 13920044 | 1103 | 1 | 258151 | ENSMUST00 | 82098 | ENSMUSG0 | Olfr1505 | olfactory receptor 1505 |
| chr19 | 14004598 | 14004914 | 317 | * | NA_peak_34 | 40 | Distal Interge | 19 | 13918942 | 13920044 | 1103 | 1 | 258151 | ENSMUST00 | 85656 | ENSMUSG0 | Olfr1505 | olfactory receptor 1505 |
| chr19 | 14024574 | 14024580 | 7 | * | NA_peak_34 | 31 | Distal Interge | 19 | 13918942 | 13920044 | 1103 | 1 | 258151 | ENSMUST00 | 105632 | ENSMUSG0 | Olfr1505 | olfactory receptor 1505 |
| chr19 | 14085368 | 14085900 | 533 | * | NA_peak_34 | 17 | Distal Interge | 19 | 13918942 | 13920044 | 1103 | 1 | 258151 | ENSMUST00 | 166426 | ENSMUSG0 | Olfr1505 | olfactory receptor 1505 |
| chr19 | 14112157 | 14112468 | 312 | * | NA_peak_34 | 13 | Distal Interge | 19 | 13918942 | 13920044 | 1103 | 1 | 258151 | ENSMUST00 | 193215 | ENSMUSG0 | Olfr1505 | olfactory receptor 1505 |
| chr19 | 14153561 | 14153975 | 415 | * | NA_peak_34 | 20 | Distal Interge | 19 | 13918942 | 13920044 | 1103 | 1 | 258151 | ENSMUST00 | 234619 | ENSMUSG0 | Olfr1505 | olfactory receptor 1505 |
| chr19 | 14179937 | 14180397 | 461 | * | NA_peak_34 | 36 | Distal Interge | 19 | 13918942 | 13920044 | 1103 | 1 | 258151 | ENSMUST00 | 260995 | ENSMUSG0 | Olfr1505 | olfactory receptor 1505 |
| chr19 | 14345809 | 14346107 | 299 | * | NA_peak_34 | 16 | Distal Interge | 19 | 14543726 | 14595539 | 51814 | 2 | 21888 | ENSMUST00 | 249432 | ENSMUSG0 | Tle4 | transducin-like enhancer of split 4 |
| chr19 | 14491023 | 14491224 | 202 | * | NA_peak_34 | 21 | Intron (ENSM | 19 | 14543726 | 14595539 | 51814 | 2 | 21888 | ENSMUST00 | 104315 | ENSMUSG0 | Tle4 | transducin-like enhancer of split 4 |
| chr19 | 14556985 | 14557070 | 86 | * | NA_peak_34 | 23 | Intron (ENSM | 19 | 14543726 | 14595539 | 51814 | 2 | 21888 | ENSMUST00 | 38469 | ENSMUSG0 | Tle4 | transducin-like enhancer of split 4 |
| chr19 | 14557787 | 14557866 | 80 | * | NA_peak_34 | 16 | Intron (ENSM | 19 | 14543726 | 14595539 | 51814 | 2 | 21888 | ENSMUST00 | 37673 | ENSMUSG0 | Tle4 | transducin-like enhancer of split 4 |
| chr19 | 14565621 | 14565784 | 164 | * | NA_peak_34 | 20 | Intron (ENSM | 19 | 14543726 | 14595539 | 51814 | 2 | 21888 | ENSMUST00 | 29755 | ENSMUSG0 | Tle4 | transducin-like enhancer of split 4 |
| chr19 | 14649843 | 14650101 | 259 | * | NA_peak_34 | 21 | Distal Interge | 19 | 14448150 | 14598051 | 149902 | 2 | 21888 | ENSMUST00 | -51792 | ENSMUSG0 | Tle4 | transducin-like enhancer of split 4 |
| chr19 | 14672520 | 14673160 | 641 | * | NA_peak_34 | 23 | Distal Interge | 19 | 14448150 | 14598051 | 149902 | 2 | 21888 | ENSMUST00 | -74469 | ENSMUSG0 | Tle4 | transducin-like enhancer of split 4 |
| chr19 | 14697865 | 14698155 | 291 | * | NA_peak_34 | 27 | Distal Interge | 19 | 14448150 | 14598051 | 149902 | 2 | 21888 | ENSMUST00 | -99814 | ENSMUSG0 | Tle4 | transducin-like enhancer of split 4 |
| chr19 | 14845409 | 14845636 | 228 | * | NA_peak_34 | 26 | Distal Interge | 19 | 14448150 | 14598051 | 149902 | 2 | 21888 | ENSMUST00 | -247358 | ENSMUSG0 | Tle4 | transducin-like enhancer of split 4 |
| chr19 | 14855598 | 14855860 | 263 | * | NA_peak_34 | 26 | Distal Interge | 19 | 14448150 | 14598051 | 149902 | 2 | 21888 | ENSMUST00 | -257547 | ENSMUSG0 | Tle4 | transducin-like enhancer of split 4 |
| chr19 | 14924893 | 14925347 | 455 | * | NA_peak_34 | 31 | Distal Interge | 19 | 14448150 | 14598051 | 149902 | 2 | 21888 | ENSMUST00 | -326842 | ENSMUSG0 | Tle4 | transducin-like enhancer of split 4 |
| chr19 | 14998953 | 14999345 | 393 | * | NA_peak_34 | 22 | Distal Interge | 19 | 14448150 | 14598051 | 149902 | 2 | 21888 | ENSMUST00 | -400902 | ENSMUSG0 | Tle4 | transducin-like enhancer of split 4 |
| chr19 | 15258395 | 15258859 | 465 | * | NA_peak_34 | 27 | Distal Interge | 19 | 15905130 | 15909712 | 4583 | 2 | 107272 | ENSMUST00 | 650853 | ENSMUSG0 | Psat1 | phosphoserine aminotransferase 1 |
| chr19 | 15274796 | 15274810 | 15 | * | NA_peak_34 | 27 | Distal Interge | 19 | 15905130 | 15909712 | 4583 | 2 | 107272 | ENSMUST00 | 634902 | ENSMUSG0 | Psat1 | phosphoserine aminotransferase 1 |
| chr19 | 15430213 | 15431001 | 789 | * | NA_peak_34 | 31 | Distal Interge | 19 | 15905130 | 15909712 | 4583 | 2 | 107272 | ENSMUST00 | 478711 | ENSMUSG0 | Psat1 | phosphoserine aminotransferase 1 |
| chr19 | 15437065 | 15437670 | 606 | * | NA_peak_34 | 26 | Distal Interge | 19 | 15905130 | 15909712 | 4583 | 2 | 107272 | ENSMUST00 | 472042 | ENSMUSG0 | Psat1 | phosphoserine aminotransferase 1 |
| chr19 | 15549640 | 15549968 | 329 | * | NA_peak_34 | 23 | Distal Interge | 19 | 15905130 | 15909712 | 4583 | 2 | 107272 | ENSMUST00 | 359744 | ENSMUSG0 | Psat1 | phosphoserine aminotransferase 1 |
| chr19 | 15561675 | 15561935 | 261 | * | NA_peak_34 | 16 | Distal Interge | 19 | 15905130 | 15909712 | 4583 | 2 | 107272 | ENSMUST00 | 347777 | ENSMUSG0 | Psat1 | phosphoserine aminotransferase 1 |
| chr19 | 15765710 | 15765833 | 124 | * | NA_peak_34 | 20 | Distal Interge | 19 | 15905130 | 15909712 | 4583 | 2 | 107272 | ENSMUST00 | 143879 | ENSMUSG0 | Psat1 | phosphoserine aminotransferase 1 |
| chr19 | 15984748 | 15985194 | 447 | * | NA_peak_34 | 18 | Promoter (<= | 19 | 15985078 | 16010696 | 25619 | 1 | 100502653 | ENSMUST00 | 0 | ENSMUSG0 | C130060C02 | RIKEN cDNA C130060C02 gene |
| chr19 | 16031378 | 16031525 | 148 | * | NA_peak_34 | 23 | Distal Interge | 19 | 15985078 | 16010696 | 25619 | 1 | 100502653 | ENSMUST00 | 46300 | ENSMUSG0 | C130060C02 | RIKEN cDNA C130060C02 gene |
| chr19 | 16038722 | 16038958 | 237 | * | NA_peak_34 | 31 | Distal Interge | 19 | 15985078 | 16010696 | 25619 | 1 | 100502653 | ENSMUST00 | 53644 | ENSMUSG0 | C130060C02 | RIKEN cDNA C130060C02 gene |
| chr19 | 16181595 | 16181978 | 384 | * | NA_peak_34 | 40 | Intron (ENSM | 19 | 16219632 | 16317274 | 97643 | 1 | 14682 | ENSMUST00 | -37654 | ENSMUSG0 | Gnaq | guanine nucleotide binding protein, alpha q polypeptide |
| chr19 | 16333844 | 16334180 | 337 | * | NA_peak_34 | 23 | Intron (ENSM | 19 | 16314892 | 16315004 | 113 | 2 | 102465218 | ENSMUST00 | -18840 | ENSMUSG0 | Mir496b | microRNA 496b |
| chr19 | 16440446 | 16440652 | 207 | * | NA_peak_34 | 28 | Promoter (1-2 | 19 | 16439300 | 16609819 | 170520 | 1 | 14675 | ENSMUST00 | 1146 | ENSMUSG0 | Gna14 | guanine nucleotide binding protein, alpha 14 |
| chr19 | 16475457 | 16475681 | 225 | * | NA_peak_34 | 28 | Intron (ENSM | 19 | 16439300 | 16609819 | 170520 | 1 | 14675 | ENSMUST00 | 36157 | ENSMUSG0 | Gna14 | guanine nucleotide binding protein, alpha 14 |
| chr19 | 16565110 | 16565777 | 668 | * | NA_peak_35 | 34 | Intron (ENSM | 19 | 16615366 | 16631347 | 15982 | 2 | 271564 | ENSMUST00 | 65570 | ENSMUSG0 | Vps13a | vacuolar protein sorting 13A |
| chr19 | 16566188 | 16566297 | 110 | * | NA_peak_35 | 22 | Intron (ENSM | 19 | 16615366 | 16631347 | 15982 | 2 | 271564 | ENSMUST00 | 65050 | ENSMUSG0 | Vps13a | vacuolar protein sorting 13A |
| chr19 | 16567627 | 16567915 | 289 | * | NA_peak_35 | 27 | Intron (ENSM | 19 | 16615366 | 16631347 | 15982 | 2 | 271564 | ENSMUST00 | 63432 | ENSMUSG0 | Vps13a | vacuolar protein sorting 13A |
| chr19 | 16646348 | 16646749 | 402 | * | NA_peak_35 | 23 | Intron (ENSM | 19 | 16632556 | 16640618 | 8063 | 2 | 271564 | ENSMUST00 | -5730 | ENSMUSG0 | Vps13a | vacuolar protein sorting 13A |
| chr19 | 16867131 | 16867173 | 43 | * | NA_peak_35 | 31 | Distal Interge | 19 | 16872319 | 16873805 | 1487 | 2 | 14240 | ENSMUST00 | 6632 | ENSMUSG0 | Foxb2 | forkhead box B2 |
| chr19 | 16908140 | 16908698 | 559 | * | NA_peak_35 | 23 | Intron (ENSM | 19 | 16872319 | 16873805 | 1487 | 2 | 14240 | ENSMUST00 | -34335 | ENSMUSG0 | Foxb2 | forkhead box B2 |
| chr19 | 16961669 | 16961787 | 119 | * | NA_peak_35 | 31 | Intron (ENSM | 19 | 16956118 | 17223932 | 267815 | 1 | 353211 | ENSMUST00 | 5551 | ENSMUSG0 | Prune2 | prune homolog 2 |
| chr19 | 17138791 | 17138913 | 123 | * | NA_peak_35 | 25 | Promoter (<= | 19 | 17138739 | 17221194 | 82456 | 1 | 353211 | ENSMUST00 | 52 | ENSMUSG0 | Prune2 | prune homolog 2 |
| chr19 | 17275524 | 17275857 | 334 | * | NA_peak_35 | 16 | Distal Interge | 19 | 17328909 | 17335436 | 6528 | 2 | 14537 | ENSMUST00 | 59579 | ENSMUSG0 | Gcnt1 | glucosaminyl (N-acetyl) transferase 1, core 2 |
| chr19 | 17301200 | 17301607 | 408 | * | NA_peak_35 | 27 | Distal Interge | 19 | 17328909 | 17335436 | 6528 | 2 | 14537 | ENSMUST00 | 33829 | ENSMUSG0 | Gcnt1 | glucosaminyl (N-acetyl) transferase 1, core 2 |
| chr19 | 17789577 | 17789853 | 277 | * | NA_peak_35 | 20 | Intron (ENSM | 19 | 17673452 | 17837617 | 164166 | 2 | 18552 | ENSMUST00 | 47764 | ENSMUSG0 | Pcsk5 | proprotein convertase subtilisin/kexin type 5 |
| chr19 | 17889207 | 17889514 | 308 | * | NA_peak_35 | 23 | Intron (ENSM | 19 | 17670420 | 17837632 | 167213 | 2 | 18552 | ENSMUST00 | -51575 | ENSMUSG0 | Pcsk5 | proprotein convertase subtilisin/kexin type 5 |
| chr19 | 17939547 | 17939817 | 271 | * | NA_peak_35 | 33 | Distal Interge | 19 | 17670420 | 17837632 | 167213 | 2 | 18552 | ENSMUST00 | -101915 | ENSMUSG0 | Pcsk5 | proprotein convertase subtilisin/kexin type 5 |
| chr19 | 17941080 | 17941423 | 344 | * | NA_peak_35 | 24 | Distal Interge | 19 | 17670420 | 17837632 | 167213 | 2 | 18552 | ENSMUST00 | -103448 | ENSMUSG0 | Pcsk5 | proprotein convertase subtilisin/kexin type 5 |

| | | | | | | | | | | | | | | | | | | |
|---|---|---|---|---|---|---|---|---|---|---|---|---|---|---|---|---|---|---|
| chr19 | 18061811 | 18062121 | 311 | * | NA_peak_351 | 19 | Distal Interge | 19 | 17670420 | 17837632 | 167213 | 2 | 18552 | ENSMUST00 | -224179 | ENSMUSG0 | Pcsk5 | proprotein convertase subtilisin/kexin type 5 |
| chr19 | 18063224 | 18063526 | 303 | * | NA_peak_351 | 23 | Distal Interge | 19 | 17670420 | 17837632 | 167213 | 2 | 18552 | ENSMUST00 | -225592 | ENSMUSG0 | Pcsk5 | proprotein convertase subtilisin/kexin type 5 |
| chr19 | 18097772 | 18098308 | 537 | * | NA_peak_351 | 29 | Distal Interge | 19 | 17670420 | 17837632 | 167213 | 2 | 18552 | ENSMUST00 | -260140 | ENSMUSG0 | Pcsk5 | proprotein convertase subtilisin/kexin type 5 |
| chr19 | 18188218 | 18188230 | 13 | * | NA_peak_351 | 18 | Distal Interge | 19 | 17670420 | 17837632 | 167213 | 2 | 18552 | ENSMUST00 | -350586 | ENSMUSG0 | Pcsk5 | proprotein convertase subtilisin/kexin type 5 |
| chr19 | 18803968 | 18804358 | 391 | * | NA_peak_352 | 22 | Intron (ENSM | 19 | 18840637 | 18866239 | 25603 | 1 | 225997 | ENSMUST00 | -36279 | ENSMUSG0 | Trpm6 | transient receptor potential cation channel, subfamily M, member 6 |
| chr19 | 18838198 | 18838586 | 389 | * | NA_peak_352 | 22 | Promoter (2-3 | 19 | 18840637 | 18866239 | 25603 | 1 | 225997 | ENSMUST00 | -2051 | ENSMUSG0 | Trpm6 | transient receptor potential cation channel, subfamily M, member 6 |
| chr19 | 18900338 | 18900634 | 297 | * | NA_peak_352 | 22 | Distal Interge | 19 | 18840637 | 18866239 | 25603 | 1 | 225997 | ENSMUST00 | 59701 | ENSMUSG0 | Trpm6 | transient receptor potential cation channel, subfamily M, member 6 |
| chr19 | 18911968 | 18912271 | 304 | * | NA_peak_352 | 36 | Distal Interge | 19 | 18840637 | 18866239 | 25603 | 1 | 225997 | ENSMUST00 | 71331 | ENSMUSG0 | Trpm6 | transient receptor potential cation channel, subfamily M, member 6 |
| chr19 | 18943773 | 18943990 | 218 | * | NA_peak_352 | 30 | Intron (ENSM | 19 | 18975962 | 19001100 | 25139 | 2 | 225998 | ENSMUST00 | 57110 | ENSMUSG0 | Rorb | RAR-related orphan receptor beta |
| chr19 | 18960453 | 18960982 | 530 | * | NA_peak_352 | 36 | Exon (ENSMU | 19 | 18975962 | 19001100 | 25139 | 2 | 225998 | ENSMUST00 | 40118 | ENSMUSG0 | Rorb | RAR-related orphan receptor beta |
| chr19 | 18993501 | 18993801 | 301 | * | NA_peak_352 | 24 | Intron (ENSM | 19 | 18975962 | 19001100 | 25139 | 2 | 225998 | ENSMUST00 | 7299 | ENSMUSG0 | Rorb | RAR-related orphan receptor beta |
| chr19 | 19187163 | 19187446 | 284 | * | NA_peak_352 | 22 | Distal Interge | 19 | 18930605 | 19111196 | 180592 | 2 | 225998 | ENSMUST00 | -75967 | ENSMUSG0 | Rorb | RAR-related orphan receptor beta |
| chr19 | 19363699 | 19363976 | 278 | * | NA_peak_352 | 21 | Distal Interge | 19 | 18930605 | 19111196 | 180592 | 2 | 225998 | ENSMUST00 | -252503 | ENSMUSG0 | Rorb | RAR-related orphan receptor beta |
| chr19 | 19464655 | 19464736 | 82 | * | NA_peak_353 | 24 | Distal Interge | 19 | 18930605 | 19111196 | 180592 | 2 | 225998 | ENSMUST00 | -353459 | ENSMUSG0 | Rorb | RAR-related orphan receptor beta |
| chr19 | 19546654 | 19547049 | 396 | * | NA_peak_353 | 36 | Distal Interge | 19 | 18930605 | 19111196 | 180592 | 2 | 225998 | ENSMUST00 | -435458 | ENSMUSG0 | Rorb | RAR-related orphan receptor beta |
| chr19 | 19595203 | 19595424 | 222 | * | NA_peak_353 | 20 | Distal Interge | 19 | 18930605 | 19111196 | 180592 | 2 | 225998 | ENSMUST00 | -484007 | ENSMUSG0 | Rorb | RAR-related orphan receptor beta |
| chr19 | 19663300 | 19663533 | 234 | * | NA_peak_353 | 36 | Distal Interge | 19 | 18930605 | 19111196 | 180592 | 2 | 225998 | ENSMUST00 | -552104 | ENSMUSG0 | Rorb | RAR-related orphan receptor beta |
| chr19 | 19706094 | 19706751 | 658 | * | NA_peak_353 | 23 | Distal Interge | 19 | 18930605 | 19111196 | 180592 | 2 | 225998 | ENSMUST00 | -594898 | ENSMUSG0 | Rorb | RAR-related orphan receptor beta |
| chr19 | 19730910 | 19731080 | 171 | * | NA_peak_353 | 20 | Distal Interge | 19 | 20324856 | 20440923 | 116068 | 1 | 68994 | ENSMUST00 | -593776 | ENSMUSG0 | 1500015L24 | RIKEN cDNA 1500015L24 gene |
| chr19 | 19744318 | 19744364 | 47 | * | NA_peak_353 | 20 | Distal Interge | 19 | 20324856 | 20440923 | 116068 | 1 | 68994 | ENSMUST00 | -580492 | ENSMUSG0 | 1500015L24 | RIKEN cDNA 1500015L24 gene |
| chr19 | 19998526 | 19998817 | 292 | * | NA_peak_353 | 28 | Distal Interge | 19 | 20324856 | 20440923 | 116068 | 1 | 68994 | ENSMUST00 | -326039 | ENSMUSG0 | 1500015L24 | RIKEN cDNA 1500015L24 gene |
| chr19 | 20185045 | 20185371 | 327 | * | NA_peak_353 | 16 | Distal Interge | 19 | 20324856 | 20440923 | 116068 | 1 | 68994 | ENSMUST00 | -139485 | ENSMUSG0 | 1500015L24 | RIKEN cDNA 1500015L24 gene |
| chr19 | 20260117 | 20260353 | 237 | * | NA_peak_354 | 16 | Distal Interge | 19 | 20324856 | 20440923 | 116068 | 1 | 68994 | ENSMUST00 | -64503 | ENSMUSG0 | 1500015L24 | RIKEN cDNA 1500015L24 gene |
| chr19 | 20526732 | 20527084 | 353 | * | NA_peak_354 | 20 | Intron (ENSM | 19 | 20420817 | 20538950 | 118134 | 2 | 320155 | ENSMUST00 | 11866 | ENSMUSG0 | C730002L08 | RIKEN cDNA C730002L08 gene |
| chr19 | 20560964 | 20561250 | 287 | * | NA_peak_354 | 20 | Intron (ENSM | 19 | 20534527 | 20552330 | 17804 | 2 | 320155 | ENSMUST00 | -8634 | ENSMUSG0 | C730002L08 | RIKEN cDNA C730002L08 gene |
| chr19 | 20737723 | 20737983 | 261 | * | NA_peak_354 | 20 | Distal Interge | 19 | 20692953 | 20727562 | 34610 | 2 | 26358 | ENSMUST00 | -10161 | ENSMUSG0 | Aldh1a7 | aldehyde dehydrogenase family 1, subfamily A7 |
| chr19 | 20831609 | 20831649 | 41 | * | NA_peak_354 | 23 | Promoter (<= | 19 | 20824478 | 20832375 | 7898 | 2 | 13409 | ENSMUST00 | 726 | ENSMUSG0 | Tmc1 | transmembrane channel-like gene family 1 |
| chr19 | 20984548 | 20984586 | 39 | * | NA_peak_354 | 15 | Distal Interge | 19 | 20783458 | 20954202 | 170745 | 2 | 13409 | ENSMUST00 | -30346 | ENSMUSG0 | Tmc1 | transmembrane channel-like gene family 1 |
| chr19 | 21471911 | 21472476 | 566 | * | NA_peak_355 | 22 | Promoter (<= | 19 | 21418725 | 21472588 | 53864 | 2 | 14544 | ENSMUST00 | 112 | ENSMUSG0 | Gda | guanine deaminase |
| chr19 | 21507868 | 21508282 | 415 | * | NA_peak_355 | 16 | Distal Interge | 19 | 21394038 | 21473445 | 79408 | 2 | 14544 | ENSMUST00 | -34423 | ENSMUSG0 | Gda | guanine deaminase |
| chr19 | 21857462 | 21857802 | 341 | * | NA_peak_355 | 39 | 3' UTR | 19 | 21778388 | 21807902 | 29515 | 1 | 83921 | ENSMUST00 | 79074 | ENSMUSG0 | Cemip2 | cell migration inducing hyaluronidase 2 |
| chr19 | 21886307 | 21886591 | 285 | * | NA_peak_355 | 24 | Distal Interge | 19 | 21778388 | 21807902 | 29515 | 1 | 83921 | ENSMUST00 | 107919 | ENSMUSG0 | Cemip2 | cell migration inducing hyaluronidase 2 |
| chr19 | 22056817 | 22057100 | 284 | * | NA_peak_355 | 16 | Distal Interge | 19 | 22139046 | 22995410 | 856365 | 1 | 226025 | ENSMUST00 | -81946 | ENSMUSG0 | Trpm3 | transient receptor potential cation channel, subfamily M, member 3 |
| chr19 | 22098250 | 22098727 | 478 | * | NA_peak_355 | 18 | Distal Interge | 19 | 22139046 | 22995410 | 856365 | 1 | 226025 | ENSMUST00 | -40319 | ENSMUSG0 | Trpm3 | transient receptor potential cation channel, subfamily M, member 3 |
| chr19 | 22245108 | 22245477 | 370 | * | NA_peak_356 | 26 | Intron (ENSM | 19 | 22139117 | 22988332 | 849216 | 1 | 226025 | ENSMUST00 | 105991 | ENSMUSG0 | Trpm3 | transient receptor potential cation channel, subfamily M, member 3 |
| chr19 | 22301000 | 22301271 | 272 | * | NA_peak_356 | 23 | Intron (ENSM | 19 | 22448183 | 22988332 | 540150 | 1 | 226025 | ENSMUST00 | -146912 | ENSMUSG0 | Trpm3 | transient receptor potential cation channel, subfamily M, member 3 |
| chr19 | 22770759 | 22770967 | 209 | * | NA_peak_356 | 23 | Intron (ENSM | 19 | 22766634 | 22989884 | 223251 | 1 | 226025 | ENSMUST00 | 4125 | ENSMUSG0 | Trpm3 | transient receptor potential cation channel, subfamily M, member 3 |
| chr19 | 23025668 | 23025830 | 163 | * | NA_peak_356 | 28 | Intron (ENSM | 19 | 23141226 | 23168134 | 26909 | 1 | 16601 | ENSMUST00 | -115396 | ENSMUSG0 | Klf9 | Kruppel-like factor 9 |
| chr19 | 23058474 | 23058666 | 193 | * | NA_peak_356 | 34 | Intron (ENSM | 19 | 23141226 | 23168134 | 26909 | 1 | 16601 | ENSMUST00 | -82560 | ENSMUSG0 | Klf9 | Kruppel-like factor 9 |
| chr19 | 23071751 | 23072069 | 319 | * | NA_peak_356 | 27 | Intron (ENSM | 19 | 23141226 | 23168134 | 26909 | 1 | 16601 | ENSMUST00 | -69157 | ENSMUSG0 | Klf9 | Kruppel-like factor 9 |
| chr19 | 23082079 | 23082593 | 515 | * | NA_peak_356 | 30 | Distal Interge | 19 | 23141226 | 23168134 | 26909 | 1 | 16601 | ENSMUST00 | -58633 | ENSMUSG0 | Klf9 | Kruppel-like factor 9 |
| chr19 | 23361247 | 23361464 | 218 | * | NA_peak_357 | 21 | Intron (ENSM | 19 | 23302609 | 23448398 | 145790 | 2 | 71738 | ENSMUST00 | 86934 | ENSMUSG0 | Mamdc2 | MAM domain containing 2 |
| chr19 | 23457439 | 23457610 | 172 | * | NA_peak_357 | 32 | Distal Interge | 19 | 23302745 | 23448442 | 145698 | 2 | 71738 | ENSMUST00 | -8997 | ENSMUSG0 | Mamdc2 | MAM domain containing 2 |
| chr19 | 23466206 | 23466465 | 260 | * | NA_peak_357 | 16 | Distal Interge | 19 | 23302745 | 23448442 | 145698 | 2 | 71738 | ENSMUST00 | -17764 | ENSMUSG0 | Mamdc2 | MAM domain containing 2 |
| chr19 | 23562603 | 23562890 | 288 | * | NA_peak_357 | 23 | Intron (ENSM | 19 | 23558753 | 23652812 | 94060 | 2 | 67483 | ENSMUST00 | 89922 | ENSMUSG0 | 1700028P14 | RIKEN cDNA 1700028P14 gene |
| chr19 | 23570941 | 23571056 | 116 | * | NA_peak_357 | 20 | Intron (ENSM | 19 | 23558753 | 23652812 | 94060 | 2 | 67483 | ENSMUST00 | 81756 | ENSMUSG0 | 1700028P14 | RIKEN cDNA 1700028P14 gene |
| chr19 | 23686955 | 23687461 | 507 | * | NA_peak_357 | 55 | Promoter (<= | 19 | 23687429 | 23731668 | 44240 | 1 | 72351 | ENSMUST00 | 0 | ENSMUSG0 | Ptar1 | protein prenyltransferase alpha subunit repeat containing 1 |
| chr19 | 23840471 | 23840498 | 28 | * | NA_peak_357 | 27 | Intron (ENSM | 19 | 23904457 | 23947376 | 42920 | 1 | 319924 | ENSMUST00 | -63959 | ENSMUSG0 | Apba1 | amyloid beta (A4) precursor protein binding, family A, member 1 |
| chr19 | 23870464 | 23870920 | 457 | * | NA_peak_357 | 49 | Intron (ENSM | 19 | 23904457 | 23947376 | 42920 | 1 | 319924 | ENSMUST00 | -33537 | ENSMUSG0 | Apba1 | amyloid beta (A4) precursor protein binding, family A, member 1 |
| chr19 | 24060674 | 24061038 | 365 | * | NA_peak_357 | 31 | Distal Interge | 19 | 23972750 | 24031019 | 58270 | 2 | 381217 | ENSMUST00 | -29655 | ENSMUSG0 | Fam189a2 | family with sequence similarity 189, member A2 |
| chr19 | 24074846 | 24075057 | 212 | * | NA_peak_357 | 16 | Distal Interge | 19 | 23972750 | 24031019 | 58270 | 2 | 381217 | ENSMUST00 | -43827 | ENSMUSG0 | Fam189a2 | family with sequence similarity 189, member A2 |
| chr19 | 24380517 | 24380739 | 223 | * | NA_peak_357 | 23 | Intron (ENSM | 19 | 24295397 | 24317342 | 21946 | 2 | 18719 | ENSMUST00 | -63175 | ENSMUSG0 | Pip5k1b | phosphatidylinositol-4-phosphate 5-kinase, type 1 beta |
| chr19 | 24472256 | 24472591 | 336 | * | NA_peak_357 | 26 | Intron (ENSM | 19 | 24476502 | 24477356 | 855 | 2 | 68034 | ENSMUST00 | 4765 | ENSMUSG0 | Fam122a | family with sequence similarity 122, member A |
| chr19 | 24700975 | 24701277 | 303 | * | NA_peak_358 | 45 | Intron (ENSM | 19 | 24674008 | 24682233 | 8226 | 1 | 226040 | ENSMUST00 | 26967 | ENSMUSG0 | Tmem252 | transmembrane protein 252 |
| chr19 | 24755566 | 24755668 | 103 | * | NA_peak_358 | 16 | Intron (ENSM | 19 | 24684551 | 24763078 | 78528 | 2 | 226041 | ENSMUST00 | 7410 | ENSMUSG0 | Pgm5 | phosphoglucomutase 5 |
| chr19 | 24793563 | 24793637 | 75 | * | NA_peak_358 | 13 | Intron (ENSM | 19 | 24684551 | 24763078 | 78528 | 2 | 226041 | ENSMUST00 | -30485 | ENSMUSG0 | Pgm5 | phosphoglucomutase 5 |
| chr19 | 24874728 | 24874889 | 162 | * | NA_peak_358 | 22 | Distal Interge | 19 | 24683016 | 24861855 | 178840 | 2 | 226041 | ENSMUST00 | -12873 | ENSMUSG0 | Pgm5 | phosphoglucomutase 5 |
| chr19 | 25101401 | 25101561 | 161 | * | NA_peak_358 | 39 | Intron (ENSM | 19 | 25077914 | 25079902 | 1989 | 1 | 76088 | ENSMUST00 | 23487 | ENSMUSG0 | Dock8 | dedicator of cytokinesis 8 |
| chr19 | 25217269 | 25217368 | 100 | * | NA_peak_358 | 24 | Distal Interge | 19 | 25236975 | 25434496 | 197522 | 1 | 107351 | ENSMUST00 | -19607 | ENSMUSG0 | Kank1 | KN motif and ankyrin repeat domains 1 |
| chr19 | 25428657 | 25429020 | 364 | * | NA_peak_358 | 55 | Intron (ENSM | 19 | 25422239 | 25424508 | 2270 | 1 | 107351 | ENSMUST00 | 6418 | ENSMUSG0 | Kank1 | KN motif and ankyrin repeat domains 1 |
| chr19 | 25584570 | 25584761 | 192 | * | NA_peak_358 | 21 | Intron (ENSM | 19 | 25610537 | 25623921 | 13385 | 1 | 240590 | ENSMUST00 | -25776 | ENSMUSG0 | Dmrt3 | doublesex and mab-3 related transcription factor 3 |
| chr19 | 25601432 | 25601596 | 165 | * | NA_peak_358 | 13 | Intron (ENSM | 19 | 25610537 | 25623921 | 13385 | 1 | 240590 | ENSMUST00 | -8941 | ENSMUSG0 | Dmrt3 | doublesex and mab-3 related transcription factor 3 |
| chr19 | 25615600 | 25615917 | 318 | * | NA_peak_358 | 24 | Intron (ENSM | 19 | 25610537 | 25623921 | 13385 | 1 | 240590 | ENSMUST00 | 5063 | ENSMUSG0 | Dmrt3 | doublesex and mab-3 related transcription factor 3 |
| chr19 | 25742537 | 25742908 | 372 | * | NA_peak_358 | 23 | Distal Interge | 19 | 25672411 | 25678991 | 6581 | 1 | 226049 | ENSMUST00 | 70126 | ENSMUSG0 | Dmrt2 | doublesex and mab-3 related transcription factor 2 |
| chr19 | 25747203 | 25747523 | 321 | * | NA_peak_358 | 36 | Distal Interge | 19 | 25672411 | 25678991 | 6581 | 1 | 226049 | ENSMUST00 | 74792 | ENSMUSG0 | Dmrt2 | doublesex and mab-3 related transcription factor 2 |
| chr19 | 25822316 | 25822380 | 65 | * | NA_peak_359 | 23 | Distal Interge | 19 | 25672411 | 25678991 | 6581 | 1 | 226049 | ENSMUST00 | 149905 | ENSMUSG0 | Dmrt2 | doublesex and mab-3 related transcription factor 2 |
| chr19 | 25879171 | 25879510 | 340 | * | NA_peak_359 | 31 | Distal Interge | 19 | 25672411 | 25678991 | 6581 | 1 | 226049 | ENSMUST00 | 206760 | ENSMUSG0 | Dmrt2 | doublesex and mab-3 related transcription factor 2 |
| chr19 | 26064452 | 26064474 | 23 | * | NA_peak_359 | 16 | Distal Interge | 19 | 25672411 | 25678991 | 6581 | 1 | 226049 | ENSMUST00 | 392041 | ENSMUSG0 | Dmrt2 | doublesex and mab-3 related transcription factor 2 |
| chr19 | 26299933 | 26300018 | 86 | * | NA_peak_359 | 19 | Intron (ENSM | 19 | 26605050 | 26778275 | 173226 | 1 | 67155 | ENSMUST00 | -305032 | ENSMUSG0 | Smarca2 | SWI/SNF related, matrix associated, actin dependent regulator of chromatin, subfamily a, member 2 |
| chr19 | 26552025 | 26552688 | 664 | * | NA_peak_359 | 55 | Distal Interge | 19 | 26605050 | 26778275 | 173226 | 1 | 67155 | ENSMUST00 | -52362 | ENSMUSG0 | Smarca2 | SWI/SNF related, matrix associated, actin dependent regulator of chromatin, subfamily a, member 2 |
| chr19 | 26594381 | 26594536 | 156 | * | NA_peak_360 | 20 | Distal Interge | 19 | 26605050 | 26778275 | 173226 | 1 | 67155 | ENSMUST00 | -10514 | ENSMUSG0 | Smarca2 | SWI/SNF related, matrix associated, actin dependent regulator of chromatin, subfamily a, member 2 |
| chr19 | 26661852 | 26662244 | 393 | * | NA_peak_360 | 23 | Intron (ENSM | 19 | 26671098 | 26678577 | 7480 | 1 | 67155 | ENSMUST00 | -8854 | ENSMUSG0 | Smarca2 | SWI/SNF related, matrix associated, actin dependent regulator of chromatin, subfamily a, member 2 |
| chr19 | 26800725 | 26800915 | 191 | * | NA_peak_360 | 24 | Intron (ENSM | 19 | 26768801 | 26823907 | 55107 | 2 | 70942 | ENSMUST00 | 22992 | ENSMUSG0 | 4931403E22 | RIKEN cDNA 4931403E22 gene |
| chr19 | 26829247 | 26829541 | 295 | * | NA_peak_360 | 36 | Distal Interge | 19 | 26768801 | 26823907 | 55107 | 2 | 70942 | ENSMUST00 | -5340 | ENSMUSG0 | 4931403E22 | RIKEN cDNA 4931403E22 gene |
| chr19 | 27198826 | 27199435 | 610 | * | NA_peak_360 | 31 | Distal Interge | 19 | 27216484 | 27249755 | 33272 | 1 | 22359 | ENSMUST00 | -17049 | ENSMUSG0 | Vldlr | very low density lipoprotein receptor |
| chr19 | 27201868 | 27202162 | 295 | * | NA_peak_360 | 23 | Distal Interge | 19 | 27216484 | 27249755 | 33272 | 1 | 22359 | ENSMUST00 | -14322 | ENSMUSG0 | Vldlr | very low density lipoprotein receptor |
| chr19 | 27354064 | 27354416 | 353 | * | NA_peak_360 | 24 | Intron (ENSM | 19 | 27322588 | 27337179 | 14592 | 1 | 240595 | ENSMUST00 | 31476 | ENSMUSG0 | Kcnv2 | potassium channel, subfamily V, member 2 |
| chr19 | 27434544 | 27434947 | 404 | * | NA_peak_360 | 23 | Distal Interge | 19 | 27388698 | 27429825 | 41128 | 2 | 52874 | ENSMUST00 | -4719 | ENSMUSG0 | Pum3 | pumilio RNA-binding family member 3 |
| chr19 | 27524217 | 27524231 | 15 | * | NA_peak_360 | 36 | Distal Interge | 19 | 27388698 | 27429825 | 41128 | 2 | 52874 | ENSMUST00 | -94392 | ENSMUSG0 | Pum3 | pumilio RNA-binding family member 3 |
| chr19 | 27553589 | 27553894 | 306 | * | NA_peak_360 | 13 | Distal Interge | 19 | 27388698 | 27429825 | 41128 | 2 | 52874 | ENSMUST00 | -123764 | ENSMUSG0 | Pum3 | pumilio RNA-binding family member 3 |
| chr19 | 27590735 | 27591246 | 512 | * | NA_peak_360 | 16 | Distal Interge | 19 | 27388698 | 27429825 | 41128 | 2 | 52874 | ENSMUST00 | -160910 | ENSMUSG0 | Pum3 | pumilio RNA-binding family member 3 |
| chr19 | 27611110 | 27611215 | 106 | * | NA_peak_360 | 19 | Distal Interge | 19 | 27388698 | 27429825 | 41128 | 2 | 52874 | ENSMUST00 | -181285 | ENSMUSG0 | Pum3 | pumilio RNA-binding family member 3 |
| chr19 | 27664327 | 27664648 | 322 | * | NA_peak_361 | 16 | Distal Interge | 19 | 27768141 | 27836846 | 68706 | 2 | 19726 | ENSMUST00 | 172198 | ENSMUSG0 | Rfx3 | regulatory factor X, 3 (influences HLA class II expression) |
| chr19 | 27756811 | 27757537 | 727 | * | NA_peak_361 | 23 | Distal Interge | 19 | 27768141 | 27836846 | 68706 | 2 | 19726 | ENSMUST00 | 79309 | ENSMUSG0 | Rfx3 | regulatory factor X, 3 (influences HLA class II expression) |
| chr19 | 27863471 | 27863834 | 364 | * | NA_peak_361 | 36 | Intron (ENSM | 19 | 27768141 | 27836846 | 68706 | 2 | 19726 | ENSMUST00 | -26625 | ENSMUSG0 | Rfx3 | regulatory factor X, 3 (influences HLA class II expression) |
| chr19 | 27972022 | 27972177 | 156 | * | NA_peak_361 | 32 | Intron (ENSM | 19 | 27761721 | 27982948 | 221228 | 2 | 19726 | ENSMUST00 | 10771 | ENSMUSG0 | Rfx3 | regulatory factor X, 3 (influences HLA class II expression) |
| chr19 | 27973240 | 27973537 | 298 | * | NA_peak_361 | 36 | Intron (ENSM | 19 | 27761721 | 27982948 | 221228 | 2 | 19726 | ENSMUST00 | 9411 | ENSMUSG0 | Rfx3 | regulatory factor X, 3 (influences HLA class II expression) |
| chr19 | 27979120 | 27979399 | 280 | * | NA_peak_361 | 27 | Intron (ENSM | 19 | 27761721 | 27982948 | 221228 | 2 | 19726 | ENSMUST00 | 3549 | ENSMUSG0 | Rfx3 | regulatory factor X, 3 (influences HLA class II expression) |
| chr19 | 28148279 | 28148842 | 564 | * | NA_peak_361 | 17 | Distal Interge | 19 | 28263951 | 28283906 | 19956 | 2 | 226075 | ENSMUST00 | 135064 | ENSMUSG0 | Glis3 | GLIS family zinc finger 3 |
| chr19 | 28381683 | 28381978 | 296 | * | NA_peak_361 | 16 | Intron (ENSM | 19 | 28262347 | 28326006 | 63660 | 2 | 226075 | ENSMUST00 | -55677 | ENSMUSG0 | Glis3 | GLIS family zinc finger 3 |
| chr19 | 28426028 | 28426275 | 248 | * | NA_peak_361 | 31 | Intron (ENSM | 19 | 28262347 | 28326006 | 63660 | 2 | 226075 | ENSMUST00 | -100022 | ENSMUSG0 | Glis3 | GLIS family zinc finger 3 |
| chr19 | 28438316 | 28438325 | 10 | * | NA_peak_361 | 32 | Intron (ENSM | 19 | 28262973 | 28540420 | 277448 | 2 | 226075 | ENSMUST00 | 102095 | ENSMUSG0 | Glis3 | GLIS family zinc finger 3 |
| chr19 | 28504537 | 28504606 | 70 | * | NA_peak_361 | 27 | Intron (ENSM | 19 | 28262973 | 28540420 | 277448 | 2 | 226075 | ENSMUST00 | 35814 | ENSMUSG0 | Glis3 | GLIS family zinc finger 3 |
| chr19 | 28545593 | 28546342 | 750 | * | NA_peak_361 | 26 | Intron (ENSM | 19 | 28262973 | 28540420 | 277448 | 2 | 226075 | ENSMUST00 | -5173 | ENSMUSG0 | Glis3 | GLIS family zinc finger 3 |
| chr19 | 29002271 | 29002645 | 375 | * | NA_peak_362 | 16 | Promoter (2-3 | 19 | 28999317 | 29012186 | 12870 | 1 | 67072 | ENSMUST00 | 2954 | ENSMUSG0 | Cdc37l1 | cell division cycle 37-like 1 |
| chr19 | 29075395 | 29075439 | 45 | * | NA_peak_362 | 17 | Distal Interge | 19 | 29067347 | 29069503 | 2157 | 1 | 100503337 | ENSMUST00 | 8048 | ENSMUSG0 | Gm9895 | predicted gene 9895 |
| chr19 | 29077658 | 29077995 | 338 | * | NA_peak_362 | 15 | Distal Interge | 19 | 29067347 | 29069503 | 2157 | 1 | 100503337 | ENSMUST00 | 10311 | ENSMUSG0 | Gm9895 | predicted gene 9895 |
| chr19 | 29296036 | 29296416 | 381 | * | NA_peak_362 | 24 | Intron (ENSM | 19 | 29321344 | 29325356 | 4013 | 2 | 27356 | ENSMUST00 | 28940 | ENSMUSG0 | Insl6 | insulin-like 6 |
| chr19 | 29553790 | 29554047 | 258 | * | NA_peak_362 | 26 | Intron (ENSM | 19 | 29533236 | 29604200 | 70965 | 1 | 226089 | ENSMUST00 | 20554 | ENSMUSG0 | Ric1 | RAB6A GEF complex partner 1 |
| chr19 | 29705232 | 29705530 | 299 | * | NA_peak_362 | 24 | Intron (ENSM | 19 | 29697934 | 29708426 | 10493 | 1 | 77836 | ENSMUST00 | 7298 | ENSMUSG0 | Mlana | melan-A |
| chr19 | 29942531 | 29943088 | 558 | * | NA_peak_363 | 31 | Promoter (2-3 | 19 | 29945790 | 29960704 | 14915 | 1 | 77125 | ENSMUST00 | -2702 | ENSMUSG0 | Il33 | interleukin 33 |
| chr19 | 29961966 | 29962260 | 295 | * | NA_peak_363 | 25 | Distal Interge | 19 | 29951816 | 29952813 | 998 | 1 | 77125 | ENSMUST00 | 10150 | ENSMUSG0 | Il33 | interleukin 33 |
| chr19 | 30379192 | 30379253 | 62 | * | NA_peak_363 | 30 | Distal Interge | 19 | 30232942 | 30239687 | 6746 | 1 | 17195 | ENSMUST00 | 146250 | ENSMUSG0 | Mbl2 | mannose-binding lectin (protein C) 2 |
| chr19 | 30566876 | 30567241 | 366 | * | NA_peak_363 | 45 | 3' UTR | 19 | 30568961 | 30574155 | 5195 | 2 | 19091 | ENSMUST00 | 6914 | ENSMUSG0 | Prkg1 | protein kinase, cGMP-dependent, type I |
| chr19 | 30670436 | 30670958 | 523 | * | NA_peak_363 | 25 | Intron (ENSM | 19 | 30619371 | 30625491 | 6121 | 2 | 19091 | ENSMUST00 | -44945 | ENSMUSG0 | Prkg1 | protein kinase, cGMP-dependent, type I |
| chr19 | 30817382 | 30817560 | 179 | * | NA_peak_363 | 36 | Intron (ENSM | 19 | 30708885 | 30781452 | 72568 | 2 | 19091 | ENSMUST00 | -35930 | ENSMUSG0 | Prkg1 | protein kinase, cGMP-dependent, type I |

| | | | | | | | | | | | | | | | | | |
|---|---|---|---|---|---|---|---|---|---|---|---|---|---|---|---|---|---|
| chr19 | 31018571 | 31018954 | 384 | * | NA_peak_364 | 23 | Intron (ENSM | 19 | 31082837 | 31087069 | 4233 | 1 | 83410 | ENSMUST00 | -63883 | ENSMUSG0( | Cstf2t | cleavage stimulation factor, 3' pre-RNA subunit 2, tau |
| chr19 | 31089031 | 31089299 | 269 | * | NA_peak_364 | 25 | Intron (ENSM | 19 | 31082837 | 31087069 | 4233 | 1 | 83410 | ENSMUST00 | 6194 | ENSMUSG0( | Cstf2t | cleavage stimulation factor, 3' pre-RNA subunit 2, tau |
| chr19 | 31195218 | 31195386 | 169 | * | NA_peak_364 | 17 | Intron (ENSM | 19 | 31082837 | 31087069 | 4233 | 1 | 83410 | ENSMUST00 | 112381 | ENSMUSG0( | Cstf2t | cleavage stimulation factor, 3' pre-RNA subunit 2, tau |
| chr19 | 31417473 | 31417476 | 4 | * | NA_peak_364 | 30 | Intron (ENSM | 19 | 30568547 | 31664370 | 1095824 | 2 | 19091 | ENSMUST00 | 246894 | ENSMUSG0( | Prkg1 | protein kinase, cGMP-dependent, type I |
| chr19 | 31525871 | 31526077 | 207 | * | NA_peak_364 | 13 | Intron (ENSM | 19 | 30568547 | 31664370 | 1095824 | 2 | 19091 | ENSMUST00 | 138293 | ENSMUSG0( | Prkg1 | protein kinase, cGMP-dependent, type I |
| chr19 | 31652541 | 31652772 | 232 | * | NA_peak_364 | 16 | Intron (ENSM | 19 | 30568547 | 31664370 | 1095824 | 2 | 19091 | ENSMUST00 | 11598 | ENSMUSG0( | Prkg1 | protein kinase, cGMP-dependent, type I |
| chr19 | 31949272 | 31950095 | 824 | * | NA_peak_365 | 49 | Exon (ENSMU | 19 | 32012463 | 32016804 | 4342 | 2 | 54447 | ENSMUST00 | 66709 | ENSMUSG0( | Asah2 | N-acylsphingosine amidohydrolase 2 |
| chr19 | 32000944 | 32001202 | 259 | * | NA_peak_365 | 38 | Intron (ENSM | 19 | 32012463 | 32016804 | 4342 | 2 | 54447 | ENSMUST00 | 15602 | ENSMUSG0( | Asah2 | N-acylsphingosine amidohydrolase 2 |
| chr19 | 32039111 | 32039459 | 349 | * | NA_peak_365 | 40 | Intron (ENSM | 19 | 31984654 | 32061469 | 76816 | 2 | 54447 | ENSMUST00 | 22010 | ENSMUSG0( | Asah2 | N-acylsphingosine amidohydrolase 2 |
| chr19 | 32101978 | 32102281 | 304 | * | NA_peak_365 | 50 | Promoter (<= | 19 | 31986524 | 32103124 | 116601 | 2 | 54447 | ENSMUST00 | 843 | ENSMUSG0( | Asah2 | N-acylsphingosine amidohydrolase 2 |
| chr19 | 32210604 | 32210757 | 154 | * | NA_peak_365 | 41 | Promoter (<= | 19 | 32122728 | 32211013 | 88286 | 2 | 208449 | ENSMUST00 | 256 | ENSMUSG0( | Sgms1 | sphingomyelin synthase 1 |
| chr19 | 32552152 | 32552477 | 326 | * | NA_peak_365 | 43 | Intron (ENSM | 19 | 32595790 | 32665710 | 69921 | 1 | 23972 | ENSMUST00 | -43313 | ENSMUSG0( | Papss2 | 3'-phosphoadenosine 5'-phosphosulfate synthase 2 |
| chr19 | 32571754 | 32572041 | 288 | * | NA_peak_365 | 21 | Distal Interge | 19 | 32595790 | 32665710 | 69921 | 1 | 23972 | ENSMUST00 | -23749 | ENSMUSG0( | Papss2 | 3'-phosphoadenosine 5'-phosphosulfate synthase 2 |
| chr19 | 32885754 | 32886109 | 356 | * | NA_peak_365 | 17 | Distal Interge | 19 | 32811225 | 32817977 | 6753 | 1 | 19211 | ENSMUST00 | 74529 | ENSMUSG0( | Pten | phosphatase and tensin homolog |
| chr19 | 32995811 | 32996227 | 417 | * | NA_peak_365 | 31 | Distal Interge | 19 | 33157112 | 33157201 | 90 | 1 | 100628575 | ENSMUST00 | -160885 | ENSMUSG0( | Mir3970 | microRNA 3970 |
| chr19 | 33118655 | 33119014 | 360 | * | NA_peak_365 | 27 | Distal Interge | 19 | 33157112 | 33157201 | 90 | 1 | 100628575 | ENSMUST00 | -38098 | ENSMUSG0( | Mir3970 | microRNA 3970 |
| chr19 | 33197593 | 33197772 | 180 | * | NA_peak_366 | 36 | Intron (ENSM | 19 | 33157112 | 33157201 | 90 | 1 | 100628575 | ENSMUST00 | 40481 | ENSMUSG0( | Mir3970 | microRNA 3970 |
| chr19 | 33439503 | 33439695 | 193 | * | NA_peak_366 | 32 | Distal Interge | 19 | 33137747 | 33392295 | 254549 | 2 | 67795 | ENSMUST00 | -47208 | ENSMUSG0( | Rnls | renalase, FAD-dependent amine oxidase |
| chr19 | 33516050 | 33516338 | 289 | * | NA_peak_366 | 23 | Promoter (1-2 | 19 | 33498037 | 33517780 | 19744 | 2 | 628236 | ENSMUST00 | 1442 | ENSMUSG0( | Lipo4 | lipase, member O4 |
| chr19 | 33534581 | 33534732 | 152 | * | NA_peak_366 | 13 | Distal Interge | 19 | 33498037 | 33517780 | 19744 | 2 | 628236 | ENSMUST00 | -16801 | ENSMUSG0( | Lipo4 | lipase, member O4 |
| chr19 | 33603807 | 33604331 | 525 | * | NA_peak_366 | 31 | Intron (ENSM | 19 | 33555160 | 33590308 | 35149 | 2 | 381236 | ENSMUST00 | -13499 | ENSMUSG0( | Lipo3 | lipase, member O3 |
| chr19 | 33881640 | 33881790 | 151 | * | NA_peak_367 | 22 | Distal Interge | 19 | 33961187 | 33976810 | 15624 | 1 | 67717 | ENSMUST00 | -79397 | ENSMUSG0( | Lipf | lipase, gastric |
| chr19 | 33910876 | 33911025 | 150 | * | NA_peak_367 | 36 | Distal Interge | 19 | 33961187 | 33976810 | 15624 | 1 | 67717 | ENSMUST00 | -50162 | ENSMUSG0( | Lipf | lipase, gastric |
| chr19 | 33949311 | 33949553 | 243 | * | NA_peak_367 | 20 | Distal Interge | 19 | 33961187 | 33976810 | 15624 | 1 | 67717 | ENSMUST00 | -11634 | ENSMUSG0( | Lipf | lipase, gastric |
| chr19 | 34074082 | 34074259 | 178 | * | NA_peak_367 | 22 | Intron (ENSM | 19 | 34067529 | 34081158 | 13630 | 1 | 70166 | ENSMUST00 | 6553 | ENSMUSG0( | Lipn | lipase, family member N |
| chr19 | 34085768 | 34085997 | 230 | * | NA_peak_367 | 34 | Distal Interge | 19 | 34100933 | 34122124 | 21192 | 1 | 78753 | ENSMUST00 | -14936 | ENSMUSG0( | Lipm | lipase, family member M |
| chr19 | 34108265 | 34108576 | 312 | * | NA_peak_367 | 16 | Intron (ENSM | 19 | 34100940 | 34112995 | 12056 | 1 | 78753 | ENSMUST00 | 7325 | ENSMUSG0( | Lipm | lipase, family member M |
| chr19 | 34109575 | 34109933 | 359 | * | NA_peak_367 | 33 | Exon (ENSMU | 19 | 34100940 | 34112995 | 12056 | 1 | 78753 | ENSMUST00 | 8635 | ENSMUSG0( | Lipm | lipase, family member M |
| chr19 | 34133411 | 34133839 | 429 | * | NA_peak_367 | 16 | Intron (ENSM | 19 | 34122925 | 34154917 | 31993 | 2 | 52024 | ENSMUST00 | 21078 | ENSMUSG0( | Ankrd22 | ankyrin repeat domain 22 |
| chr19 | 34225641 | 34225681 | 41 | * | NA_peak_367 | 17 | Intron (ENSM | 19 | 34232706 | 34234299 | 1594 | 1 | 76630 | ENSMUST00 | -7025 | ENSMUSG0( | Stambpl1 | STAM binding protein like 1 |
| chr19 | 34368553 | 34368663 | 111 | * | NA_peak_367 | 28 | Distal Interge | 19 | 34300075 | 34316677 | 16603 | 1 | 14102 | ENSMUST00 | 68478 | ENSMUSG0( | Fas | Fas (TNF receptor superfamily member 6) |
| chr19 | 34374221 | 34374268 | 48 | * | NA_peak_367 | 26 | Distal Interge | 19 | 34300075 | 34316677 | 16603 | 1 | 14102 | ENSMUST00 | 74146 | ENSMUSG0( | Fas | Fas (TNF receptor superfamily member 6) |
| chr19 | 34569415 | 34569553 | 139 | * | NA_peak_367 | 36 | Intron (ENSM | 19 | 34583531 | 34588731 | 5201 | 1 | 15959 | ENSMUST00 | -13978 | ENSMUSG0( | Ifit3 | interferon-induced protein with tetratricopeptide repeats 3 |
| chr19 | 34629666 | 34629944 | 279 | * | NA_peak_367 | 23 | 5' UTR | 19 | 34617049 | 34640743 | 23695 | 2 | 112419 | ENSMUST00 | 10799 | ENSMUSG0( | Ifit1bl2 | interferon induced protein with tetratricopeptide repeats 1B like 2 |
| chr19 | 34637582 | 34637639 | 58 | * | NA_peak_367 | 28 | Intron (ENSM | 19 | 34617049 | 34640743 | 23695 | 2 | 112419 | ENSMUST00 | 3104 | ENSMUSG0( | Ifit1bl2 | interferon induced protein with tetratricopeptide repeats 1B like 2 |
| chr19 | 34791938 | 34792184 | 247 | * | NA_peak_367 | 16 | Distal Interge | 19 | 34820687 | 34820773 | 87 | 2 | 723826 | ENSMUST00 | 28589 | ENSMUSG0( | Mir107 | microRNA 107 |
| chr19 | 34937336 | 34937634 | 299 | * | NA_peak_368 | 23 | Promoter (1-2 | 19 | 34935686 | 34975743 | 40058 | 1 | 240641 | ENSMUST00 | 1650 | ENSMUSG0( | Kif20b | kinesin family member 20B |
| chr19 | 35426512 | 35426969 | 458 | * | NA_peak_368 | 32 | Distal Interge | 19 | 34972729 | 34975743 | 3015 | 1 | 240641 | ENSMUST00 | 453783 | ENSMUSG0( | Kif20b | kinesin family member 20B |
| chr19 | 35430995 | 35431103 | 109 | * | NA_peak_368 | 14 | Distal Interge | 19 | 34972729 | 34975743 | 3015 | 1 | 240641 | ENSMUST00 | 458266 | ENSMUSG0( | Kif20b | kinesin family member 20B |
| chr19 | 35549318 | 35549647 | 330 | * | NA_peak_368 | 36 | Distal Interge | 19 | 35836469 | 35923544 | 87076 | 2 | 329056 | ENSMUST00 | 373897 | ENSMUSG0( | A830019P07 | RIKEN cDNA A830019P07 gene |
| chr19 | 35594216 | 35594710 | 495 | * | NA_peak_368 | 66 | Distal Interge | 19 | 35836469 | 35923544 | 87076 | 2 | 329056 | ENSMUST00 | 328834 | ENSMUSG0( | A830019P07 | RIKEN cDNA A830019P07 gene |
| chr19 | 35783526 | 35783577 | 52 | * | NA_peak_368 | 27 | Distal Interge | 19 | 35836469 | 35923544 | 87076 | 2 | 329056 | ENSMUST00 | 139967 | ENSMUSG0( | A830019P07 | RIKEN cDNA A830019P07 gene |
| chr19 | 35784024 | 35784064 | 41 | * | NA_peak_368 | 27 | Distal Interge | 19 | 35836469 | 35923544 | 87076 | 2 | 329056 | ENSMUST00 | 139480 | ENSMUSG0( | A830019P07 | RIKEN cDNA A830019P07 gene |
| chr19 | 35943572 | 35943780 | 209 | * | NA_peak_369 | 27 | Distal Interge | 19 | 35835221 | 35924578 | 89358 | 2 | 329056 | ENSMUST00 | -18994 | ENSMUSG0( | A830019P07 | RIKEN cDNA A830019P07 gene |
| chr19 | 35989890 | 35990179 | 290 | * | NA_peak_369 | 25 | Intron (ENSM | 19 | 35835221 | 35924578 | 89358 | 2 | 329056 | ENSMUST00 | -65312 | ENSMUSG0( | A830019P07 | RIKEN cDNA A830019P07 gene |
| chr19 | 36249087 | 36249282 | 196 | * | NA_peak_369 | 23 | Intron (ENSM | 19 | 36348309 | 36460970 | 112662 | 1 | 76073 | ENSMUST00 | -99027 | ENSMUSG0( | Pcgf5 | polycomb group ring finger 5 |
| chr19 | 36381783 | 36382137 | 355 | * | NA_peak_369 | 24 | Promoter (2-3 | 19 | 36379106 | 36455943 | 76838 | 1 | 76073 | ENSMUST00 | 2677 | ENSMUSG0( | Pcgf5 | polycomb group ring finger 5 |
| chr19 | 36648691 | 36649087 | 397 | * | NA_peak_369 | 23 | Intron (ENSM | 19 | 36643546 | 36658584 | 15039 | 2 | 668215 | ENSMUST00 | 9497 | ENSMUSG0( | Hectd2os | Hectd2, opposite strand |
| chr19 | 36878365 | 36878757 | 393 | * | NA_peak_369 | 20 | Promoter (<= | 19 | 36878507 | 36883081 | 4575 | 1 | 74493 | ENSMUST00 | 0 | ENSMUSG0( | Tnks2 | tankyrase, TRF1-interacting ankyrin-related ADP-ribose polymerase 2 |
| chr19 | 37136700 | 37136996 | 297 | * | NA_peak_369 | 25 | Intron (ENSM | 19 | 37173843 | 37196541 | 22699 | 1 | 319931 | ENSMUST00 | -36847 | ENSMUSG0( | A330032B11 | RIKEN cDNA A330032B11 gene |
| chr19 | 37292934 | 37293015 | 82 | * | NA_peak_370 | 25 | Promoter (2-3 | 19 | 37280584 | 37290878 | 10295 | 2 | 15925 | ENSMUST00 | -2056 | ENSMUSG0( | Ide | insulin degrading enzyme |
| chr19 | 37295754 | 37296255 | 502 | * | NA_peak_370 | 26 | Exon (ENSMU | 19 | 37280584 | 37290878 | 10295 | 2 | 15925 | ENSMUST00 | -4876 | ENSMUSG0( | Ide | insulin degrading enzyme |
| chr19 | 37550579 | 37550953 | 375 | * | NA_peak_370 | 13 | Promoter (<= | 19 | 37550459 | 37683255 | 132797 | 1 | 107371 | ENSMUST00 | 120 | ENSMUSG0( | Exoc6 | exocyst complex component 6 |
| chr19 | 37573086 | 37573401 | 316 | * | NA_peak_370 | 40 | Intron (ENSM | 19 | 37550459 | 37683255 | 132797 | 1 | 107371 | ENSMUST00 | 22627 | ENSMUSG0( | Exoc6 | exocyst complex component 6 |
| chr19 | 37802099 | 37802449 | 351 | * | NA_peak_370 | 14 | Distal Interge | 19 | 37899468 | 37901503 | 2036 | 2 | 226101 | ENSMUST00 | 99054 | ENSMUSG0( | Myof | myoferlin |
| chr19 | 37826010 | 37826302 | 293 | * | NA_peak_370 | 15 | Distal Interge | 19 | 37899468 | 37901503 | 2036 | 2 | 226101 | ENSMUST00 | 75201 | ENSMUSG0( | Myof | myoferlin |
| chr19 | 37891329 | 37891335 | 7 | * | NA_peak_370 | 25 | Distal Interge | 19 | 37899468 | 37901503 | 2036 | 2 | 226101 | ENSMUST00 | 10168 | ENSMUSG0( | Myof | myoferlin |
| chr19 | 38076164 | 38076477 | 314 | * | NA_peak_370 | 14 | Distal Interge | 19 | 38097071 | 38114263 | 17193 | 1 | 107221 | ENSMUST00 | -20594 | ENSMUSG0( | Ffar4 | free fatty acid receptor 4 |
| chr19 | 38097234 | 38097684 | 451 | * | NA_peak_370 | 52 | Promoter (<= | 19 | 38097071 | 38114263 | 17193 | 1 | 107221 | ENSMUST00 | 163 | ENSMUSG0( | Ffar4 | free fatty acid receptor 4 |
| chr19 | 38192081 | 38192300 | 220 | * | NA_peak_370 | 36 | Intron (ENSM | 19 | 38188481 | 38224138 | 35658 | 2 | 70567 | ENSMUST00 | 31838 | ENSMUSG0( | Fra10ac1 | FRA10AC1 homolog (human) |
| chr19 | 38211413 | 38211666 | 254 | * | NA_peak_370 | 14 | Intron (ENSM | 19 | 38188481 | 38224138 | 35658 | 2 | 70567 | ENSMUST00 | 12472 | ENSMUSG0( | Fra10ac1 | FRA10AC1 homolog (human) |
| chr19 | 38587940 | 38588308 | 369 | * | NA_peak_371 | 23 | Intron (ENSM | 19 | 38583368 | 38652007 | 68640 | 1 | 74055 | ENSMUST00 | 4572 | ENSMUSG0( | Plce1 | phospholipase C, epsilon 1 |
| chr19 | 38703992 | 38704156 | 165 | * | NA_peak_371 | 21 | Exon (ENSMU | 19 | 38722370 | 38783588 | 61219 | 1 | 74055 | ENSMUST00 | -18214 | ENSMUSG0( | Plce1 | phospholipase C, epsilon 1 |
| chr19 | 38795252 | 38795306 | 55 | * | NA_peak_371 | 13 | Intron (ENSM | 19 | 38788128 | 38819237 | 31110 | 2 | 57753 | ENSMUST00 | 23931 | ENSMUSG0( | Noc3l | NOC3 like DNA replication regulator |
| chr19 | 38909264 | 38909554 | 291 | * | NA_peak_371 | 50 | Intron (ENSM | 19 | 38930915 | 38971051 | 40137 | 1 | 15201 | ENSMUST00 | -21361 | ENSMUSG0( | Hells | helicase, lymphoid specific |
| chr19 | 39021200 | 39021238 | 39 | * | NA_peak_371 | 19 | Intron (ENSM | 19 | 39007019 | 39042693 | 35675 | 1 | 72082 | ENSMUST00 | 14181 | ENSMUSG0( | Cyp2c55 | cytochrome P450, family 2, subfamily c, polypeptide 55 |
| chr19 | 39095692 | 39096259 | 568 | * | NA_peak_371 | 36 | Distal Interge | 19 | 39113898 | 39186756 | 72859 | 1 | 69888 | ENSMUST00 | -17639 | ENSMUSG0( | Cyp2c66 | cytochrome P450, family 2, subfamily c, polypeptide 66 |
| chr19 | 39182984 | 39183260 | 277 | * | NA_peak_371 | 23 | Intron (ENSM | 19 | 39229254 | 39274737 | 45484 | 1 | 638988 | ENSMUST00 | -45994 | ENSMUSG0( | Cyp2c53-ps | cytochrome P450, family 2, subfamily c, polypeptide 53, pseudogene |
| chr19 | 39189666 | 39189994 | 329 | * | NA_peak_371 | 20 | Distal Interge | 19 | 39229254 | 39274737 | 45484 | 1 | 638988 | ENSMUST00 | -39260 | ENSMUSG0( | Cyp2c53-ps | cytochrome P450, family 2, subfamily c, polypeptide 53, pseudogene |
| chr19 | 39235486 | 39235568 | 83 | * | NA_peak_371 | 23 | Intron (ENSM | 19 | 39229284 | 39274694 | 45411 | 1 | 638988 | ENSMUST00 | 6202 | ENSMUSG0( | Cyp2c53-ps | cytochrome P450, family 2, subfamily c, polypeptide 53, pseudogene |
| chr19 | 39266541 | 39267066 | 526 | * | NA_peak_372 | 55 | Promoter (2-3 | 19 | 39269405 | 39330648 | 61244 | 1 | 13095 | ENSMUST00 | -2339 | ENSMUSG0( | Cyp2c29 | cytochrome P450, family 2, subfamily c, polypeptide 29 |
| chr19 | 39328501 | 39328688 | 188 | * | NA_peak_372 | 31 | Intron (ENSM | 19 | 39287074 | 39330713 | 43640 | 1 | 13095 | ENSMUST00 | 41427 | ENSMUSG0( | Cyp2c29 | cytochrome P450, family 2, subfamily c, polypeptide 29 |
| chr19 | 39453640 | 39453925 | 286 | * | NA_peak_372 | 27 | Intron (ENSM | 19 | 39389556 | 39463075 | 73520 | 2 | 13097 | ENSMUST00 | 9150 | ENSMUSG0( | Cyp2c38 | cytochrome P450, family 2, subfamily c, polypeptide 38 |
| chr19 | 39470839 | 39471391 | 553 | * | NA_peak_372 | 13 | Distal Interge | 19 | 39389556 | 39463075 | 73520 | 2 | 13097 | ENSMUST00 | -7764 | ENSMUSG0( | Cyp2c38 | cytochrome P450, family 2, subfamily c, polypeptide 38 |
| chr19 | 39535158 | 39535633 | 476 | * | NA_peak_372 | 20 | Intron (ENSM | 19 | 39510862 | 39568529 | 57668 | 1 | 13098 | ENSMUST00 | 24296 | ENSMUSG0( | Cyp2c39 | cytochrome P450, family 2, subfamily c, polypeptide 39 |
| chr19 | 39566601 | 39566875 | 275 | * | NA_peak_372 | 43 | Intron (ENSM | 19 | 39510862 | 39568529 | 57668 | 1 | 13098 | ENSMUST00 | 55739 | ENSMUSG0( | Cyp2c39 | cytochrome P450, family 2, subfamily c, polypeptide 39 |
| chr19 | 39574870 | 39575010 | 141 | * | NA_peak_372 | 28 | Distal Interge | 19 | 39510862 | 39568529 | 57668 | 1 | 13098 | ENSMUST00 | 64008 | ENSMUSG0( | Cyp2c39 | cytochrome P450, family 2, subfamily c, polypeptide 39 |
| chr19 | 39644109 | 39644401 | 293 | * | NA_peak_372 | 40 | Intron (ENSM | 19 | 39608842 | 39649051 | 40210 | 2 | 545288 | ENSMUST00 | 4650 | ENSMUSG0( | Cyp2c67 | cytochrome P450, family 2, subfamily c, polypeptide 67 |
| chr19 | 39717138 | 39717425 | 288 | * | NA_peak_372 | 34 | Intron (ENSM | 19 | 39688834 | 39741054 | 52221 | 2 | 433247 | ENSMUST00 | 23629 | ENSMUSG0( | Cyp2c68 | cytochrome P450, family 2, subfamily c, polypeptide 68 |
| chr19 | 39729222 | 39729485 | 264 | * | NA_peak_372 | 23 | Intron (ENSM | 19 | 39688834 | 39741054 | 52221 | 2 | 433247 | ENSMUST00 | 11569 | ENSMUSG0( | Cyp2c68 | cytochrome P450, family 2, subfamily c, polypeptide 68 |
| chr19 | 39826503 | 39826513 | 11 | * | NA_peak_372 | 40 | Distal Interge | 19 | 39767227 | 39812814 | 45588 | 2 | 13099 | ENSMUST00 | -13689 | ENSMUSG0( | Cyp2c40 | cytochrome P450, family 2, subfamily c, polypeptide 40 |
| chr19 | 39840478 | 39840613 | 136 | * | NA_peak_372 | 18 | Distal Interge | 19 | 39767227 | 39812814 | 45588 | 2 | 13099 | ENSMUST00 | -27664 | ENSMUSG0( | Cyp2c40 | cytochrome P450, family 2, subfamily c, polypeptide 40 |
| chr19 | 40048102 | 40048451 | 350 | * | NA_peak_373 | 23 | Intron (ENSM | 19 | 40037940 | 40073827 | 35888 | 2 | 404195 | ENSMUST00 | 25376 | ENSMUSG0( | Cyp2c54 | cytochrome P450, family 2, subfamily c, polypeptide 54 |
| chr19 | 40056806 | 40056914 | 109 | * | NA_peak_373 | 20 | Intron (ENSM | 19 | 40037940 | 40073827 | 35888 | 2 | 404195 | ENSMUST00 | 16913 | ENSMUSG0( | Cyp2c54 | cytochrome P450, family 2, subfamily c, polypeptide 54 |
| chr19 | 40079175 | 40079564 | 390 | * | NA_peak_373 | 19 | Distal Interge | 19 | 40037940 | 40073827 | 35888 | 2 | 404195 | ENSMUST00 | -5348 | ENSMUSG0( | Cyp2c54 | cytochrome P450, family 2, subfamily c, polypeptide 54 |
| chr19 | 40109082 | 40109326 | 245 | * | NA_peak_373 | 14 | Intron (ENSM | 19 | 40089735 | 40092322 | 2588 | 1 | 107141 | ENSMUST00 | 19347 | ENSMUSG0( | Cyp2c50 | cytochrome P450, family 2, subfamily c, polypeptide 50 |
| chr19 | 40146907 | 40147237 | 331 | * | NA_peak_373 | 21 | Distal Interge | 19 | 40153353 | 40187286 | 33934 | 2 | 226105 | ENSMUST00 | 40049 | ENSMUSG0( | Cyp2c70 | cytochrome P450, family 2, subfamily c, polypeptide 70 |
| chr19 | 40169940 | 40170258 | 319 | * | NA_peak_373 | 13 | Intron (ENSM | 19 | 40153353 | 40187286 | 33934 | 2 | 226105 | ENSMUST00 | 17028 | ENSMUSG0( | Cyp2c70 | cytochrome P450, family 2, subfamily c, polypeptide 70 |
| chr19 | 40260793 | 40261063 | 271 | * | NA_peak_373 | 25 | Intron (ENSM | 19 | 40221173 | 40271616 | 50444 | 2 | 54132 | ENSMUST00 | 10553 | ENSMUSG0( | Pdlim1 | PDZ and LIM domain 1 (elfin) |
| chr19 | 40265613 | 40265963 | 351 | * | NA_peak_373 | 30 | Intron (ENSM | 19 | 40221173 | 40271616 | 50444 | 2 | 54132 | ENSMUST00 | 5653 | ENSMUSG0( | Pdlim1 | PDZ and LIM domain 1 (elfin) |
| chr19 | 40348789 | 40348911 | 123 | * | NA_peak_373 | 25 | Intron (ENSM | 19 | 40334032 | 40340594 | 6563 | 2 | 20411 | ENSMUST00 | -8195 | ENSMUSG0( | Sorbs1 | sorbin and SH3 domain containing 1 |
| chr19 | 40401059 | 40401373 | 315 | * | NA_peak_373 | 24 | Promoter (<= | 19 | 40295049 | 40402267 | 107219 | 2 | 20411 | ENSMUST00 | 894 | ENSMUSG0( | Sorbs1 | sorbin and SH3 domain containing 1 |
| chr19 | 40648476 | 40648496 | 21 | * | NA_peak_373 | 23 | Intron (ENSM | 19 | 40659770 | 40741602 | 81833 | 1 | 12495 | ENSMUST00 | -11274 | ENSMUSG0( | Entpd1 | ectonucleoside triphosphate diphosphohydrolase 1 |
| chr19 | 40648839 | 40648906 | 68 | * | NA_peak_373 | 22 | Intron (ENSM | 19 | 40659770 | 40741602 | 81833 | 1 | 12495 | ENSMUST00 | -10864 | ENSMUSG0( | Entpd1 | ectonucleoside triphosphate diphosphohydrolase 1 |
| chr19 | 40682273 | 40682639 | 367 | * | NA_peak_373 | 32 | Promoter (<= | 19 | 40681667 | 40723193 | 41527 | 1 | 12495 | ENSMUST00 | 606 | ENSMUSG0( | Entpd1 | ectonucleoside triphosphate diphosphohydrolase 1 |
| chr19 | 40821101 | 40821366 | 266 | * | NA_peak_373 | 22 | Intron (ENSM | 19 | 40831279 | 40848572 | 17294 | 1 | 240665 | ENSMUST00 | -9913 | ENSMUSG0( | Ccnj | cyclin J |
| chr19 | 40914917 | 40915047 | 131 | * | NA_peak_373 | 20 | Exon (ENSMU | 19 | 40908159 | 40917943 | 9785 | 1 | 72672 | ENSMUST00 | 6758 | ENSMUSG0( | Zfp518a | zinc finger protein 518A |
| chr19 | 40935834 | 40936200 | 367 | * | NA_peak_373 | 23 | Intron (ENSM | 19 | 40908159 | 40917943 | 9785 | 1 | 72672 | ENSMUST00 | 27675 | ENSMUSG0( | Zfp518a | zinc finger protein 518A |
| chr19 | 41074013 | 41074142 | 130 | * | NA_peak_373 | 27 | Promoter (2-3 | 19 | 41062473 | 41077113 | 14641 | 2 | 226115 | ENSMUST00 | 2971 | ENSMUSG0( | Opalin | oligodendrocytic myelin paranodal and inner loop protein |
| chr19 | 41566474 | 41566886 | 413 | * | NA_peak_374 | 20 | Intron (ENSM | 19 | 41575867 | 41586536 | 10670 | 1 | 212391 | ENSMUST00 | -8981 | ENSMUSG0( | Lcor | ligand dependent nuclear receptor corepressor |
| chr19 | 42141254 | 42141352 | 99 | * | NA_peak_374 | 20 | Distal Interge | 19 | 42147400 | 42151703 | 4304 | 1 | 277010 | ENSMUST00 | -6048 | ENSMUSG0( | Marveld1 | MARVEL (membrane-associating) domain containing 1 |
| chr19 | 42158434 | 42158521 | 88 | * | NA_peak_374 | 30 | Distal Interge | 19 | 42163951 | 42177726 | 13776 | 1 | 319740 | ENSMUST00 | -5430 | ENSMUSG0( | Zfyve27 | zinc finger, FYVE domain containing 27 |
| chr19 | 42389781 | 42390075 | 295 | * | NA_peak_374 | 20 | Intron (ENSM | 19 | 42283035 | 42431783 | 148749 | 2 | 72832 | ENSMUST00 | 41708 | ENSMUSG0( | Crtac1 | cartilage acidic protein 1 |
| chr19 | 43006571 | 43007095 | 525 | * | NA_peak_374 | 23 | Intron (ENSM | 19 | 42908584 | 43223689 | 315106 | 2 | 545291 | ENSMUST00 | 216594 | ENSMUSG0( | Hpse2 | heparanase 2 |
| chr19 | 43048066 | 43048409 | 344 | * | NA_peak_374 | 36 | Intron (ENSM | 19 | 42908584 | 43223689 | 315106 | 2 | 545291 | ENSMUST00 | 175280 | ENSMUSG0( | Hpse2 | heparanase 2 |
| chr19 | 43060843 | 43061166 | 324 | * | NA_peak_375 | 23 | Intron (ENSM | 19 | 42908584 | 43223689 | 315106 | 2 | 545291 | ENSMUST00 | 162523 | ENSMUSG0( | Hpse2 | heparanase 2 |

| | | | | | | | | | | | | | | | | | | |
|---|---|---|---|---|---|---|---|---|---|---|---|---|---|---|---|---|---|---|
| chr19 | 43076730 | 43077099 | 370 | * | NA_peak_375 | 31 | Intron (ENSM | 19 | 42908584 | 43223689 | 315106 | 2 | 545291 | ENSMUST00 | 146590 | ENSMUSG0( | Hpse2 | heparanase 2 |
| chr19 | 43110120 | 43110478 | 359 | * | NA_peak_375 | 23 | Intron (ENSM | 19 | 42908584 | 43223689 | 315106 | 2 | 545291 | ENSMUST00 | 113211 | ENSMUSG0( | Hpse2 | heparanase 2 |
| chr19 | 43118914 | 43119350 | 437 | * | NA_peak_375 | 61 | Intron (ENSM | 19 | 42908584 | 43223689 | 315106 | 2 | 545291 | ENSMUST00 | 104339 | ENSMUSG0( | Hpse2 | heparanase 2 |
| chr19 | 43214248 | 43214428 | 181 | * | NA_peak_375 | 13 | Intron (ENSM | 19 | 42908584 | 43223689 | 315106 | 2 | 545291 | ENSMUST00 | 9261 | ENSMUSG0( | Hpse2 | heparanase 2 |
| chr19 | 43268143 | 43268510 | 368 | * | NA_peak_375 | 30 | Intron (ENSM | 19 | 42908584 | 43223689 | 315106 | 2 | 545291 | ENSMUST00 | -44454 | ENSMUSG0( | Hpse2 | heparanase 2 |
| chr19 | 43291831 | 43292104 | 274 | * | NA_peak_375 | 31 | Intron (ENSM | 19 | 42908584 | 43223689 | 315106 | 2 | 545291 | ENSMUST00 | -68142 | ENSMUSG0( | Hpse2 | heparanase 2 |
| chr19 | 43318183 | 43318427 | 245 | * | NA_peak_375 | 23 | Intron (ENSM | 19 | 43294201 | 43388254 | 94054 | 2 | 545291 | ENSMUST00 | 69827 | ENSMUSG0( | Hpse2 | heparanase 2 |
| chr19 | 43573183 | 43573467 | 285 | * | NA_peak_375 | 20 | Exon (ENSMU | 19 | 43612325 | 43615892 | 3568 | 1 | 18089 | ENSMUST00 | -38858 | ENSMUSG0( | Nkx2-3 | NK2 homeobox 3 |
| chr19 | 43574041 | 43574167 | 127 | * | NA_peak_375 | 47 | Distal Interge | 19 | 43612325 | 43615892 | 3568 | 1 | 18089 | ENSMUST00 | -38158 | ENSMUSG0( | Nkx2-3 | NK2 homeobox 3 |
| chr19 | 43602198 | 43602455 | 258 | * | NA_peak_375 | 24 | Distal Interge | 19 | 43612325 | 43615892 | 3568 | 1 | 18089 | ENSMUST00 | -9870 | ENSMUSG0( | Nkx2-3 | NK2 homeobox 3 |
| chr19 | 44139339 | 44139463 | 125 | * | NA_peak_375 | 16 | 3' UTR | 19 | 44139606 | 44142578 | 2973 | 2 | 73689 | ENSMUST00 | 3115 | ENSMUSG0( | Bloc1s2 | biogenesis of lysosomal organelles complex-1, subunit 2 |
| chr19 | 44203007 | 44203425 | 419 | * | NA_peak_375 | 20 | Promoter (<= | 19 | 44203269 | 44244016 | 40748 | 1 | 30049 | ENSMUST00 | 0 | ENSMUSG0( | Scd3 | stearoyl-coenzyme A desaturase 3 |
| chr19 | 45194806 | 45195115 | 310 | * | NA_peak_376 | 27 | Distal Interge | 19 | 45232684 | 45235812 | 3129 | 2 | 16814 | ENSMUST00 | 40697 | ENSMUSG0( | Lbx1 | ladybird homeobox 1 |
| chr19 | 46486320 | 46486618 | 299 | * | NA_peak_376 | 17 | 3' UTR | 19 | 46483097 | 46485982 | 2886 | 1 | 24069 | ENSMUST00 | 3223 | ENSMUSG0( | Sufu | SUFU negative regulator of hedgehog signaling |
| chr19 | 46801635 | 46801891 | 257 | * | NA_peak_376 | 17 | Intron (ENSM | 19 | 46761605 | 46854052 | 92448 | 1 | 94219 | ENSMUST00 | 40030 | ENSMUSG0( | Cnnm2 | cyclin M2 |
| chr19 | 47351629 | 47351924 | 296 | * | NA_peak_377 | 22 | Intron (ENSM | 19 | 47266913 | 47314751 | 47839 | 2 | 14218 | ENSMUST00 | -36878 | ENSMUSG0( | Sh3pxd2a | SH3 and PX domains 2A |
| chr19 | 47578474 | 47578706 | 233 | * | NA_peak_377 | 16 | Promoter (<= | 19 | 47579678 | 47645246 | 65569 | 1 | 20874 | ENSMUST00 | -972 | ENSMUSG0( | Slk | STE20-like kinase |
| chr19 | 47754576 | 47754877 | 302 | * | NA_peak_377 | 24 | Promoter (<= | 19 | 47755022 | 47755157 | 136 | 2 | 102465883 | ENSMUST00 | 280 | ENSMUSG0( | Mir8090 | microRNA 8090 |
| chr19 | 47954853 | 47955296 | 444 | * | NA_peak_377 | 21 | Exon (ENSMU | 19 | 47937712 | 48035387 | 97676 | 1 | 381229 | ENSMUST00 | 17141 | ENSMUSG0( | Cfap58 | cilia and flagella associated protein 58 |
| chr19 | 47976131 | 47976165 | 35 | * | NA_peak_377 | 26 | Intron (ENSM | 19 | 47937712 | 48035387 | 97676 | 1 | 381229 | ENSMUST00 | 38419 | ENSMUSG0( | Cfap58 | cilia and flagella associated protein 58 |
| chr19 | 48014675 | 48015036 | 362 | * | NA_peak_377 | 48 | Intron (ENSM | 19 | 47937712 | 48035387 | 97676 | 1 | 381229 | ENSMUST00 | 76963 | ENSMUSG0( | Cfap58 | cilia and flagella associated protein 58 |
| chr19 | 48124322 | 48124478 | 157 | * | NA_peak_377 | 26 | Intron (ENSM | 19 | 48206025 | 48805505 | 599481 | 1 | 66673 | ENSMUST00 | -81547 | ENSMUSG0( | Sorcs3 | sortilin-related VPS10 domain containing receptor 3 |
| chr19 | 48189954 | 48190435 | 482 | * | NA_peak_377 | 42 | Distal Interge | 19 | 48206025 | 48805505 | 599481 | 1 | 66673 | ENSMUST00 | -15590 | ENSMUSG0( | Sorcs3 | sortilin-related VPS10 domain containing receptor 3 |
| chr19 | 48216833 | 48216991 | 159 | * | NA_peak_377 | 20 | Intron (ENSM | 19 | 48206025 | 48805505 | 599481 | 1 | 66673 | ENSMUST00 | 10808 | ENSMUSG0( | Sorcs3 | sortilin-related VPS10 domain containing receptor 3 |
| chr19 | 48275759 | 48275788 | 30 | * | NA_peak_377 | 16 | Intron (ENSM | 19 | 48206025 | 48805505 | 599481 | 1 | 66673 | ENSMUST00 | 69734 | ENSMUSG0( | Sorcs3 | sortilin-related VPS10 domain containing receptor 3 |
| chr19 | 48477152 | 48477379 | 228 | * | NA_peak_377 | 20 | Intron (ENSM | 19 | 48429960 | 48655310 | 225351 | 1 | 66673 | ENSMUST00 | 47192 | ENSMUSG0( | Sorcs3 | sortilin-related VPS10 domain containing receptor 3 |
| chr19 | 48613763 | 48614037 | 275 | * | NA_peak_378 | 16 | Intron (ENSM | 19 | 48704216 | 48804537 | 100322 | 1 | 66673 | ENSMUST00 | -90179 | ENSMUSG0( | Sorcs3 | sortilin-related VPS10 domain containing receptor 3 |
| chr19 | 48978758 | 48979216 | 459 | * | NA_peak_378 | 13 | Distal Interge | 19 | 48704216 | 48804537 | 100322 | 1 | 66673 | ENSMUST00 | 274542 | ENSMUSG0( | Sorcs3 | sortilin-related VPS10 domain containing receptor 3 |
| chr19 | 48979920 | 48980403 | 484 | * | NA_peak_378 | 33 | Distal Interge | 19 | 48704216 | 48804537 | 100322 | 1 | 66673 | ENSMUST00 | 275704 | ENSMUSG0( | Sorcs3 | sortilin-related VPS10 domain containing receptor 3 |
| chr19 | 49070376 | 49070424 | 49 | * | NA_peak_378 | 28 | Distal Interge | 19 | 48704216 | 48804537 | 100322 | 1 | 66673 | ENSMUST00 | 366160 | ENSMUSG0( | Sorcs3 | sortilin-related VPS10 domain containing receptor 3 |
| chr19 | 49385299 | 49385368 | 70 | * | NA_peak_378 | 27 | Distal Interge | 19 | 48704216 | 48804537 | 100322 | 1 | 66673 | ENSMUST00 | 681083 | ENSMUSG0( | Sorcs3 | sortilin-related VPS10 domain containing receptor 3 |
| chr19 | 49408404 | 49408794 | 391 | * | NA_peak_378 | 23 | Distal Interge | 19 | 48704216 | 48804537 | 100322 | 1 | 66673 | ENSMUST00 | 704188 | ENSMUSG0( | Sorcs3 | sortilin-related VPS10 domain containing receptor 3 |
| chr19 | 49665704 | 49665805 | 102 | * | NA_peak_379 | 26 | Distal Interge | 19 | 50150967 | 50252678 | 101712 | 2 | 58178 | ENSMUST00 | 586873 | ENSMUSG0( | Sorcs1 | sortilin-related VPS10 domain containing receptor 1 |
| chr19 | 49679550 | 49679835 | 286 | * | NA_peak_379 | 31 | Distal Interge | 19 | 50150967 | 50252678 | 101712 | 2 | 58178 | ENSMUST00 | 572843 | ENSMUSG0( | Sorcs1 | sortilin-related VPS10 domain containing receptor 1 |
| chr19 | 49685695 | 49686082 | 388 | * | NA_peak_379 | 31 | Distal Interge | 19 | 50150967 | 50252678 | 101712 | 2 | 58178 | ENSMUST00 | 566596 | ENSMUSG0( | Sorcs1 | sortilin-related VPS10 domain containing receptor 1 |
| chr19 | 50052992 | 50053536 | 545 | * | NA_peak_379 | 16 | Distal Interge | 19 | 50150967 | 50252678 | 101712 | 2 | 58178 | ENSMUST00 | 199142 | ENSMUSG0( | Sorcs1 | sortilin-related VPS10 domain containing receptor 1 |
| chr19 | 50410632 | 50410715 | 84 | * | NA_peak_379 | 22 | Intron (ENSM | 19 | 50150967 | 50252678 | 101712 | 2 | 58178 | ENSMUST00 | -157954 | ENSMUSG0( | Sorcs1 | sortilin-related VPS10 domain containing receptor 1 |
| chr19 | 50782054 | 50782331 | 278 | * | NA_peak_379 | 20 | Distal Interge | 19 | 50147660 | 50678646 | 530987 | 2 | 58178 | ENSMUST00 | -103408 | ENSMUSG0( | Sorcs1 | sortilin-related VPS10 domain containing receptor 1 |
| chr19 | 51341370 | 51341408 | 39 | * | NA_peak_380 | 23 | Distal Interge | 19 | 50147660 | 50678646 | 530987 | 2 | 58178 | ENSMUST00 | -662724 | ENSMUSG0( | Sorcs1 | sortilin-related VPS10 domain containing receptor 1 |
| chr19 | 51432888 | 51433262 | 375 | * | NA_peak_380 | 27 | Distal Interge | 19 | 50147660 | 50678646 | 530987 | 2 | 58178 | ENSMUST00 | -754242 | ENSMUSG0( | Sorcs1 | sortilin-related VPS10 domain containing receptor 1 |
| chr19 | 51451625 | 51451933 | 309 | * | NA_peak_380 | 23 | Distal Interge | 19 | 50147660 | 50678646 | 530987 | 2 | 58178 | ENSMUST00 | -772979 | ENSMUSG0( | Sorcs1 | sortilin-related VPS10 domain containing receptor 1 |
| chr19 | 51459930 | 51460211 | 282 | * | NA_peak_380 | 20 | Distal Interge | 19 | 50147660 | 50678646 | 530987 | 2 | 58178 | ENSMUST00 | -781284 | ENSMUSG0( | Sorcs1 | sortilin-related VPS10 domain containing receptor 1 |
| chr19 | 51532680 | 51532750 | 71 | * | NA_peak_380 | 40 | Distal Interge | 19 | 52264297 | 52265476 | 1180 | 1 | 16333 | ENSMUST00 | -731547 | ENSMUSG0( | Ins1 | insulin I |
| chr19 | 51536515 | 51536702 | 188 | * | NA_peak_380 | 31 | Distal Interge | 19 | 52264297 | 52265476 | 1180 | 1 | 16333 | ENSMUST00 | -727595 | ENSMUSG0( | Ins1 | insulin I |
| chr19 | 51549953 | 51550033 | 81 | * | NA_peak_380 | 16 | Distal Interge | 19 | 52264297 | 52265476 | 1180 | 1 | 16333 | ENSMUST00 | -714264 | ENSMUSG0( | Ins1 | insulin I |
| chr19 | 51596826 | 51596863 | 38 | * | NA_peak_380 | 27 | Distal Interge | 19 | 52264297 | 52265476 | 1180 | 1 | 16333 | ENSMUST00 | -667434 | ENSMUSG0( | Ins1 | insulin I |
| chr19 | 51651573 | 51651759 | 187 | * | NA_peak_380 | 26 | Distal Interge | 19 | 52264297 | 52265476 | 1180 | 1 | 16333 | ENSMUST00 | -612538 | ENSMUSG0( | Ins1 | insulin I |
| chr19 | 51895306 | 51895830 | 525 | * | NA_peak_380 | 60 | Intron (ENSM | 19 | 52264297 | 52265476 | 1180 | 1 | 16333 | ENSMUST00 | -368467 | ENSMUSG0( | Ins1 | insulin I |
| chr19 | 51914896 | 51914947 | 52 | * | NA_peak_380 | 45 | Intron (ENSM | 19 | 52264297 | 52265476 | 1180 | 1 | 16333 | ENSMUST00 | -349350 | ENSMUSG0( | Ins1 | insulin I |
| chr19 | 52028474 | 52028875 | 402 | * | NA_peak_380 | 17 | Distal Interge | 19 | 52264297 | 52265476 | 1180 | 1 | 16333 | ENSMUST00 | -235422 | ENSMUSG0( | Ins1 | insulin I |
| chr19 | 52225042 | 52225075 | 34 | * | NA_peak_381 | 40 | Distal Interge | 19 | 52264297 | 52265476 | 1180 | 1 | 16333 | ENSMUST00 | -39222 | ENSMUSG0( | Ins1 | insulin I |
| chr19 | 52282726 | 52283023 | 298 | * | NA_peak_381 | 31 | Distal Interge | 19 | 52264297 | 52265476 | 1180 | 1 | 16333 | ENSMUST00 | 18429 | ENSMUSG0( | Ins1 | insulin I |
| chr19 | 52361919 | 52362290 | 372 | * | NA_peak_381 | 19 | Distal Interge | 19 | 52264297 | 52265476 | 1180 | 1 | 16333 | ENSMUST00 | 97622 | ENSMUSG0( | Ins1 | insulin I |
| chr19 | 52411765 | 52412126 | 362 | * | NA_peak_381 | 23 | Distal Interge | 19 | 52264297 | 52265476 | 1180 | 1 | 16333 | ENSMUST00 | 147468 | ENSMUSG0( | Ins1 | insulin I |
| chr19 | 52670775 | 52671152 | 378 | * | NA_peak_381 | 16 | Distal Interge | 19 | 52923181 | 52926931 | 3751 | 1 | 100043450 | ENSMUST00 | -252029 | ENSMUSG0( | AA387883 | expressed sequence AA387883 |
| chr19 | 52867260 | 52867551 | 292 | * | NA_peak_381 | 19 | Distal Interge | 19 | 52923181 | 52926931 | 3751 | 1 | 100043450 | ENSMUST00 | -55630 | ENSMUSG0( | AA387883 | expressed sequence AA387883 |
| chr19 | 53140727 | 53140814 | 88 | * | NA_peak_382 | 32 | Promoter (<= | 19 | 53140517 | 53245896 | 105380 | 1 | 27360 | ENSMUST00 | 210 | ENSMUSG0( | Add3 | adducin 3 (gamma) |
| chr19 | 53156293 | 53156423 | 131 | * | NA_peak_382 | 13 | Intron (ENSM | 19 | 53143997 | 53247399 | 103403 | 1 | 27360 | ENSMUST00 | 12296 | ENSMUSG0( | Add3 | adducin 3 (gamma) |
| chr19 | 53925464 | 53925671 | 208 | * | NA_peak_382 | 19 | Intron (ENSM | 19 | 53930960 | 53944180 | 13221 | 2 | 100503572 | ENSMUST00 | 18509 | ENSMUSG0( | Bbip1 | BBSome interacting protein 1 |
| chr19 | 53997595 | 53997858 | 264 | * | NA_peak_382 | 16 | Intron (ENSM | 19 | 54003086 | 54029981 | 26896 | 1 | 56392 | ENSMUST00 | -5228 | ENSMUSG0( | Shoc2 | Shoc2, leucine rich repeat scaffold protein |
| chr19 | 54224866 | 54225051 | 186 | * | NA_peak_382 | 19 | Distal Interge | 19 | 54044151 | 54048982 | 4832 | 1 | 11551 | ENSMUST00 | 180715 | ENSMUSG0( | Adra2a | adrenergic receptor, alpha 2a |
| chr19 | 54227891 | 54228266 | 376 | * | NA_peak_382 | 33 | Distal Interge | 19 | 54044151 | 54048982 | 4832 | 1 | 11551 | ENSMUST00 | 183740 | ENSMUSG0( | Adra2a | adrenergic receptor, alpha 2a |
| chr19 | 54244939 | 54245247 | 309 | * | NA_peak_382 | 36 | Distal Interge | 19 | 54044151 | 54048982 | 4832 | 1 | 11551 | ENSMUST00 | 200788 | ENSMUSG0( | Adra2a | adrenergic receptor, alpha 2a |
| chr19 | 54275013 | 54275415 | 403 | * | NA_peak_382 | 14 | Distal Interge | 19 | 54044151 | 54048982 | 4832 | 1 | 11551 | ENSMUST00 | 230862 | ENSMUSG0( | Adra2a | adrenergic receptor, alpha 2a |
| chr19 | 54289691 | 54289894 | 204 | * | NA_peak_382 | 21 | Distal Interge | 19 | 54044151 | 54048982 | 4832 | 1 | 11551 | ENSMUST00 | 245540 | ENSMUSG0( | Adra2a | adrenergic receptor, alpha 2a |
| chr19 | 54384436 | 54384719 | 284 | * | NA_peak_383 | 31 | Distal Interge | 19 | 54044151 | 54048982 | 4832 | 1 | 11551 | ENSMUST00 | 340285 | ENSMUSG0( | Adra2a | adrenergic receptor, alpha 2a |
| chr19 | 54434411 | 54434687 | 277 | * | NA_peak_383 | 35 | Distal Interge | 19 | 54044151 | 54048982 | 4832 | 1 | 11551 | ENSMUST00 | 390260 | ENSMUSG0( | Adra2a | adrenergic receptor, alpha 2a |
| chr19 | 54554017 | 54554380 | 364 | * | NA_peak_383 | 26 | Distal Interge | 19 | 54044151 | 54048982 | 4832 | 1 | 11551 | ENSMUST00 | 509866 | ENSMUSG0( | Adra2a | adrenergic receptor, alpha 2a |
| chr19 | 54576249 | 54576871 | 623 | * | NA_peak_383 | 36 | Distal Interge | 19 | 55069734 | 55099451 | 29718 | 2 | 14732 | ENSMUST00 | 522580 | ENSMUSG0( | Gpam | glycerol-3-phosphate acyltransferase, mitochondrial |
| chr19 | 54625258 | 54625529 | 272 | * | NA_peak_383 | 27 | Distal Interge | 19 | 55069734 | 55099451 | 29718 | 2 | 14732 | ENSMUST00 | 473922 | ENSMUSG0( | Gpam | glycerol-3-phosphate acyltransferase, mitochondrial |
| chr19 | 54769869 | 54770214 | 346 | * | NA_peak_383 | 20 | Distal Interge | 19 | 55069734 | 55099451 | 29718 | 2 | 14732 | ENSMUST00 | 329237 | ENSMUSG0( | Gpam | glycerol-3-phosphate acyltransferase, mitochondrial |
| chr19 | 54804661 | 54804936 | 276 | * | NA_peak_383 | 14 | Distal Interge | 19 | 55069734 | 55099451 | 29718 | 2 | 14732 | ENSMUST00 | 294515 | ENSMUSG0( | Gpam | glycerol-3-phosphate acyltransferase, mitochondrial |
| chr19 | 54805330 | 54805396 | 67 | * | NA_peak_383 | 44 | Distal Interge | 19 | 55069734 | 55099451 | 29718 | 2 | 14732 | ENSMUST00 | 294055 | ENSMUSG0( | Gpam | glycerol-3-phosphate acyltransferase, mitochondrial |
| chr19 | 54884157 | 54884424 | 268 | * | NA_peak_383 | 48 | Distal Interge | 19 | 55069734 | 55099451 | 29718 | 2 | 14732 | ENSMUST00 | 215027 | ENSMUSG0( | Gpam | glycerol-3-phosphate acyltransferase, mitochondrial |
| chr19 | 54964355 | 54964575 | 221 | * | NA_peak_383 | 37 | Distal Interge | 19 | 55069734 | 55099451 | 29718 | 2 | 14732 | ENSMUST00 | 134876 | ENSMUSG0( | Gpam | glycerol-3-phosphate acyltransferase, mitochondrial |
| chr19 | 55041621 | 55041735 | 115 | * | NA_peak_383 | 30 | Distal Interge | 19 | 55069734 | 55099451 | 29718 | 2 | 14732 | ENSMUST00 | 57716 | ENSMUSG0( | Gpam | glycerol-3-phosphate acyltransferase, mitochondrial |
| chr19 | 55348720 | 55349017 | 298 | * | NA_peak_383 | 34 | Intron (ENSM | 19 | 55381014 | 55446979 | 65966 | 1 | 53611 | ENSMUST00 | -31997 | ENSMUSG0( | Vti1a | vesicle transport through interaction with t-SNAREs 1A |
| chr19 | 56692959 | 56693300 | 342 | * | NA_peak_384 | 40 | Distal Interge | 19 | 56722199 | 56725150 | 2952 | 1 | 11554 | ENSMUST00 | -28899 | ENSMUSG0( | Adrb1 | adrenergic receptor, beta 1 |
| chr19 | 57462656 | 57462970 | 315 | * | NA_peak_384 | 36 | Intron (ENSM | 19 | 57452957 | 57483626 | 30670 | 1 | 72133 | ENSMUST00 | 9699 | ENSMUSG0( | Trub1 | TruB pseudouridine (psi) synthase family member 1 |
| chr19 | 57880130 | 57880471 | 342 | * | NA_peak_385 | 23 | Intron (ENSM | 19 | 58051167 | 58051236 | 70 | 1 | 100885835 | ENSMUST00 | -170696 | ENSMUSG0( | Mir5623 | microRNA 5623 |
| chr19 | 57924117 | 57924325 | 209 | * | NA_peak_385 | 24 | Intron (ENSM | 19 | 58051167 | 58051236 | 70 | 1 | 100885835 | ENSMUST00 | -126842 | ENSMUSG0( | Mir5623 | microRNA 5623 |
| chr19 | 58157399 | 58157610 | 212 | * | NA_peak_385 | 18 | Distal Interge | 19 | 58051167 | 58051236 | 70 | 1 | 100885835 | ENSMUST00 | 106232 | ENSMUSG0( | Mir5623 | microRNA 5623 |
| chr19 | 58643113 | 58643411 | 299 | * | NA_peak_385 | 16 | Distal Interge | 19 | 58652498 | 58671735 | 19238 | 1 | 69060 | ENSMUST00 | -9087 | ENSMUSG0( | Pnlip | pancreatic lipase |
| chr19 | 58700618 | 58700653 | 36 | * | NA_peak_385 | 13 | Promoter (1-2 | 19 | 58699082 | 58715770 | 16689 | 1 | 69166 | ENSMUST00 | 1536 | ENSMUSG0( | 1810018F18I | RIKEN cDNA 1810018F18 gene |
| chr19 | 58739656 | 58739783 | 128 | * | NA_peak_385 | 24 | Promoter (<= | 19 | 58740436 | 58741438 | 1003 | 1 | 18946 | ENSMUST00 | -653 | ENSMUSG0( | Pnliprp1 | pancreatic lipase related protein 1 |
| chr19 | 58822268 | 58822679 | 412 | * | NA_peak_385 | 21 | Promoter (<= | 19 | 58799682 | 58822121 | 22440 | 2 | 73442 | ENSMUST00 | -147 | ENSMUSG0( | Hspa12a | heat shock protein 12A |
| chr19 | 59093336 | 59093858 | 523 | * | NA_peak_385 | 33 | Distal Interge | 19 | 58973496 | 59076100 | 102605 | 2 | 71653 | ENSMUST00 | -17236 | ENSMUSG0( | Shtn1 | shootin 1 |
| chr19 | 59151314 | 59151552 | 239 | * | NA_peak_386 | 22 | Distal Interge | 19 | 59166598 | 59166980 | 383 | 2 | 22326 | ENSMUST00 | 15428 | ENSMUSG0( | Vax1 | ventral anterior homeobox 1 |
| chr19 | 59196383 | 59196746 | 364 | * | NA_peak_386 | 24 | Distal Interge | 19 | 59219214 | 59237374 | 18161 | 1 | 332396 | ENSMUST00 | -22468 | ENSMUSG0( | Kcnk18 | potassium channel, subfamily K, member 18 |
| chr19 | 59199425 | 59199652 | 228 | * | NA_peak_386 | 31 | Distal Interge | 19 | 59219214 | 59237374 | 18161 | 1 | 332396 | ENSMUST00 | -19562 | ENSMUSG0( | Kcnk18 | potassium channel, subfamily K, member 18 |
| chr19 | 59221207 | 59221592 | 386 | * | NA_peak_386 | 16 | Promoter (1-2 | 19 | 59219214 | 59237374 | 18161 | 1 | 332396 | ENSMUST00 | 1993 | ENSMUSG0( | Kcnk18 | potassium channel, subfamily K, member 18 |
| chr19 | 59276526 | 59276650 | 125 | * | NA_peak_386 | 33 | Exon (ENSMU | 19 | 59260959 | 59295243 | 34285 | 1 | 214084 | ENSMUST00 | 15567 | ENSMUSG0( | Slc18a2 | solute carrier family 18 (vesicular monoamine), member 2 |
| chr19 | 59667012 | 59667342 | 331 | * | NA_peak_386 | 30 | Intron (ENSM | 19 | 59493136 | 59559391 | 66256 | 1 | 72647 | ENSMUST00 | 173876 | ENSMUSG0( | 2700089I24R | RIKEN cDNA 2700089I24 gene |
| chr19 | 59960810 | 59961116 | 307 | * | NA_peak_386 | 24 | Distal Interge | 19 | 59902884 | 59943654 | 40771 | 2 | 74998 | ENSMUST00 | -17156 | ENSMUSG0( | Rab11fip2 | RAB11 family interacting protein 2 (class I) |
| chr19 | 60304228 | 60304547 | 320 | * | NA_peak_387 | 30 | Distal Interge | 19 | 60199435 | 60226701 | 27267 | 2 | 76539 | ENSMUST00 | -77527 | ENSMUSG0( | Fam204a | family with sequence similarity 204, member A |
| chr19 | 60346778 | 60347092 | 315 | * | NA_peak_387 | 27 | Distal Interge | 19 | 60199435 | 60226701 | 27267 | 2 | 76539 | ENSMUST00 | -120077 | ENSMUSG0( | Fam204a | family with sequence similarity 204, member A |
| chr19 | 60401877 | 60402113 | 237 | * | NA_peak_387 | 24 | Distal Interge | 19 | 60466732 | 60468304 | 1573 | 2 | 226278 | ENSMUST00 | 66191 | ENSMUSG0( | Prlhr | prolactin releasing hormone receptor |
| chr19 | 60489417 | 60489460 | 44 | * | NA_peak_387 | 20 | Distal Interge | 19 | 60466732 | 60468304 | 1573 | 2 | 226278 | ENSMUST00 | -21113 | ENSMUSG0( | Prlhr | prolactin releasing hormone receptor |
| chr19 | 60558692 | 60558972 | 281 | * | NA_peak_387 | 23 | Intron (ENSM | 19 | 60543036 | 60546022 | 2987 | 2 | 78832 | ENSMUST00 | -12670 | ENSMUSG0( | Cacul1 | CDK2 associated, cullin domain 1 |
| chr19 | 60763665 | 60764210 | 546 | * | NA_peak_387 | 36 | Exon (ENSMU | 19 | 60755987 | 60759914 | 3928 | 1 | 332397 | ENSMUST00 | 7678 | ENSMUSG0( | Nanos1 | nanos C2HC-type zinc finger 1 |
| chr19 | 60769240 | 60769259 | 20 | * | NA_peak_387 | 24 | Promoter (2-3 | 19 | 60770786 | 60772151 | 1366 | 2 | 13669 | ENSMUST00 | 2892 | ENSMUSG0( | Eif3a | eukaryotic translation initiation factor 3, subunit A |
| chr19 | 60853708 | 60854092 | 385 | * | NA_peak_387 | 22 | 3' UTR | 19 | 60838758 | 60861424 | 22667 | 2 | 94281 | ENSMUST00 | 7332 | ENSMUSG0( | Sfxn4 | sideroflexin 4 |
| chr19 | 61264093 | 61264455 | 363 | * | NA_peak_388 | 13 | Distal Interge | 19 | 61226843 | 61228429 | 1587 | 2 | 12982 | ENSMUST00 | -35664 | ENSMUSG0( | Csf2ra | colony stimulating factor 2 receptor, alpha, low-affinity (granulocyte-macrophage) |
| chr19 | 61266560 | 61266926 | 367 | * | NA_peak_388 | 58 | Distal Interge | 19 | 61226843 | 61228429 | 1587 | 2 | 12982 | ENSMUST00 | -38131 | ENSMUSG0( | Csf2ra | colony stimulating factor 2 receptor, alpha, low-affinity (granulocyte-macrophage) |
| chr19 | 61290025 | 61290394 | 370 | * | NA_peak_388 | 27 | Distal Interge | 19 | 61226843 | 61228429 | 1587 | 2 | 12982 | ENSMUST00 | -61596 | ENSMUSG0( | Csf2ra | colony stimulating factor 2 receptor, alpha, low-affinity (granulocyte-macrophage) |
| chr19 | 61320787 | 61321127 | 341 | * | NA_peak_388 | 32 | Distal Interge | 19 | 61226843 | 61228429 | 1587 | 2 | 12982 | ENSMUST00 | -92358 | ENSMUSG0( | Csf2ra | colony stimulating factor 2 receptor, alpha, low-affinity (granulocyte-macrophage) |

| | | | | | | | | | | | | | | | | | | |
|---|---|---|---|---|---|---|---|---|---|---|---|---|---|---|---|---|---|---|
| chr2 | 6840269 | 6840877 | 609 | * | NA_peak_38 | 19 | Intron (ENSM | 2 | 6624704 | 6871972 | 247269 | 2 | 14007 | ENSMUST00 | 31095 | ENSMUSG0( | Celf2 | CUGBP, Elav-like family member 2 |
| chr2 | 8185719 | 8186447 | 729 | * | NA_peak_38 | 34 | Distal Interge | 2 | 7394076 | 7509563 | 115488 | 2 | 14007 | ENSMUST00 | -676156 | ENSMUSG0( | Celf2 | CUGBP, Elav-like family member 2 |
| chr2 | 11977751 | 11978053 | 303 | * | NA_peak_38 | 23 | Distal Interge | 2 | 12109070 | 12132552 | 23483 | 2 | 241226 | ENSMUST00 | 154499 | ENSMUSG0( | Itga8 | integrin alpha 8 |
| chr2 | 12043302 | 12043638 | 337 | * | NA_peak_38 | 30 | Distal Interge | 2 | 12109070 | 12132552 | 23483 | 2 | 241226 | ENSMUST00 | 88914 | ENSMUSG0( | Itga8 | integrin alpha 8 |
| chr2 | 12642101 | 12642306 | 206 | * | NA_peak_38 | 31 | Distal Interge | 2 | 12347263 | 12419470 | 72208 | 2 | 66960 | ENSMUST00 | -222631 | ENSMUSG0( | Mindy3 | MINDY lysine 48 deubiquitinase 3 |
| chr2 | 13184777 | 13185173 | 397 | * | NA_peak_38 | 15 | Intron (ENSM | 2 | 13218593 | 13271221 | 52629 | 2 | 20163 | ENSMUST00 | 86048 | ENSMUSG0( | Rsu1 | Ras suppressor protein 1 |
| chr2 | 13251663 | 13252001 | 339 | * | NA_peak_38 | 26 | Intron (ENSM | 2 | 13218593 | 13271221 | 52629 | 2 | 20163 | ENSMUST00 | 19220 | ENSMUSG0( | Rsu1 | Ras suppressor protein 1 |
| chr2 | 13397727 | 13398161 | 435 | * | NA_peak_38 | 23 | Intron (ENSM | 2 | 13422029 | 13455627 | 33599 | 2 | 65969 | ENSMUST00 | 57466 | ENSMUSG0( | Cubn | cubilin (intrinsic factor-cobalamin receptor) |
| chr2 | 13716517 | 13716798 | 282 | * | NA_peak_39 | 20 | Intron (ENSM | 2 | 13657154 | 13793293 | 136140 | 2 | 241230 | ENSMUST00 | 76495 | ENSMUSG0( | St8sia6 | ST8 alpha-N-acetyl-neuraminide alpha-2,8-sialyltransferase 6 |
| chr2 | 14086462 | 14086631 | 170 | * | NA_peak_39 | 19 | Intron (ENSM | 2 | 14074427 | 14129073 | 54647 | 1 | 20844 | ENSMUST00 | 12035 | ENSMUSG0( | Stam | signal transducing adaptor molecule (SH3 domain and ITAM motif) 1 |
| chr2 | 14123521 | 14123708 | 188 | * | NA_peak_39 | 23 | Intron (ENSM | 2 | 14128337 | 14129168 | 832 | 1 | 20844 | ENSMUST00 | -4629 | ENSMUSG0( | Stam | signal transducing adaptor molecule (SH3 domain and ITAM motif) 1 |
| chr2 | 15120862 | 15121133 | 272 | * | NA_peak_39 | 16 | Distal Interge | 2 | 15074999 | 15082452 | 7454 | 1 | 75869 | ENSMUST00 | 45863 | ENSMUSG0( | Arl5b | ADP-ribosylation factor-like 5B |
| chr2 | 15171135 | 15171473 | 339 | * | NA_peak_39 | 16 | Distal Interge | 2 | 15074999 | 15082452 | 7454 | 1 | 75869 | ENSMUST00 | 96136 | ENSMUSG0( | Arl5b | ADP-ribosylation factor-like 5B |
| chr2 | 17475136 | 17475286 | 151 | * | NA_peak_39 | 36 | Intron (ENSM | 2 | 17392906 | 17462789 | 69884 | 2 | 74103 | ENSMUST00 | -12347 | ENSMUSG0( | Nebl | nebulette |
| chr2 | 18556403 | 18556702 | 300 | * | NA_peak_39 | 20 | Distal Interge | 2 | 18672384 | 18676230 | 3847 | 1 | 12238 | ENSMUST00 | -115682 | ENSMUSG0( | Commd3 | COMM domain containing 3 |
| chr2 | 21301633 | 21302089 | 457 | * | NA_peak_39 | 23 | Distal Interge | 2 | 21367542 | 21830547 | 463006 | 1 | 241263 | ENSMUST00 | -65453 | ENSMUSG0( | Gpr158 | G protein-coupled receptor 158 |
| chr2 | 22077897 | 22078193 | 297 | * | NA_peak_39 | 23 | Distal Interge | 2 | 22227503 | 22618252 | 390750 | 1 | 667663 | ENSMUST00 | -149310 | ENSMUSG0( | Myo3a | myosin IIIA |
| chr2 | 22587371 | 22587905 | 535 | * | NA_peak_39 | 174 | Exon (ENSMU | 2 | 22622205 | 22693874 | 71670 | 1 | 14417 | ENSMUST00 | -34300 | ENSMUSG0( | Gad2 | glutamic acid decarboxylase 2 |
| chr2 | 22589616 | 22589919 | 304 | * | NA_peak_39 | 98 | Exon (ENSMU | 2 | 22622205 | 22693874 | 71670 | 1 | 14417 | ENSMUST00 | -32286 | ENSMUSG0( | Gad2 | glutamic acid decarboxylase 2 |
| chr2 | 23339960 | 23340364 | 405 | * | NA_peak_39 | 16 | Intron (ENSM | 2 | 23321246 | 23401973 | 80728 | 1 | 18232 | ENSMUST00 | 18714 | ENSMUSG0( | Nxph2 | neurexophilin 2 |
| chr2 | 25578394 | 25578792 | 399 | * | NA_peak_39 | 13 | Promoter (1-2 | 2 | 25576973 | 25579897 | 2925 | 2 | 381353 | ENSMUST00 | 1105 | ENSMUSG0( | Ajm1 | apical junction component 1 |
| chr2 | 26132864 | 26133175 | 312 | * | NA_peak_39 | 20 | Distal Interge | 2 | 26136814 | 26140521 | 3708 | 2 | 109299 | ENSMUST00 | 7346 | ENSMUSG0( | Tmem250-ps | transmembrane protein 250, pseudogene |
| chr2 | 26334047 | 26334112 | 66 | * | NA_peak_39 | 40 | Intron (ENSM | 2 | 26330434 | 26340531 | 10098 | 1 | 67839 | ENSMUST00 | 3613 | ENSMUSG0( | Gpsm1 | G-protein signalling modulator 1 (AGS3-like, C. elegans) |
| chr2 | 26392709 | 26392848 | 140 | * | NA_peak_39 | 27 | Promoter (1-2 | 2 | 26390927 | 26392398 | 1472 | 1 | 66865 | ENSMUST00 | 1782 | ENSMUSG0( | Pmpca | peptidase (mitochondrial processing) alpha |
| chr2 | 26714768 | 26714993 | 226 | * | NA_peak_39 | 20 | Distal Interge | 2 | 26667674 | 26671282 | 3609 | 2 | 16821 | ENSMUST00 | -43486 | ENSMUSG0( | Lcn4 | lipocalin 4 |
| chr2 | 29827373 | 29827817 | 445 | * | NA_peak_39 | 20 | Promoter (<= | 2 | 29827395 | 29833636 | 6242 | 1 | 68205 | ENSMUST00 | 0 | ENSMUSG0( | Urm1 | ubiquitin related modifier 1 |
| chr2 | 30635248 | 30635421 | 174 | * | NA_peak_39 | 29 | Intron (ENSM | 2 | 30605217 | 30608217 | 3001 | 1 | 78617 | ENSMUST00 | 30031 | ENSMUSG0( | Cstad | CSA-conditional, T cell activation-dependent protein |
| chr2 | 30947387 | 30947592 | 206 | * | NA_peak_39 | 19 | Distal Interge | 2 | 30952959 | 30959015 | 6057 | 1 | 30934 | ENSMUST00 | -5367 | ENSMUSG0( | Tor1b | torsin family 1, member B |
| chr2 | 31410800 | 31411156 | 357 | * | NA_peak_39 | 29 | Intron (ENSM | 2 | 31400277 | 31405814 | 5538 | 1 | 665700 | ENSMUST00 | 10523 | ENSMUSG0( | Hmcn2 | hemicentin 2 |
| chr2 | 32907905 | 32908373 | 469 | * | NA_peak_39 | 23 | Promoter (2-3 | 2 | 32910908 | 32912587 | 1680 | 1 | 227737 | ENSMUST00 | -2535 | ENSMUSG0( | Niban2 | niban apoptosis regulator 2 |
| chr2 | 34930587 | 34930785 | 199 | * | NA_peak_39 | 35 | Distal Interge | 2 | 34898969 | 34914026 | 15058 | 2 | 74016 | ENSMUST00 | -16561 | ENSMUSG0( | Phf19 | PHD finger protein 19 |
| chr2 | 35577856 | 35577960 | 105 | * | NA_peak_39 | 23 | Intron (ENSM | 2 | 35581060 | 35730992 | 149933 | 1 | 69601 | ENSMUST00 | -3100 | ENSMUSG0( | Dab2ip | disabled 2 interacting protein |
| chr2 | 35719971 | 35720058 | 88 | * | NA_peak_39 | 20 | 3' UTR | 2 | 35710327 | 35730994 | 20668 | 1 | 69601 | ENSMUST00 | 9644 | ENSMUSG0( | Dab2ip | disabled 2 interacting protein |
| chr2 | 36442255 | 36442697 | 443 | * | NA_peak_39 | 54 | Distal Interge | 2 | 36452587 | 36453525 | 939 | 1 | 258953 | ENSMUST00 | -9890 | ENSMUSG0( | Olfr340 | olfactory receptor 340 |
| chr2 | 36443221 | 36443316 | 96 | * | NA_peak_39 | 54 | Distal Interge | 2 | 36452587 | 36453525 | 939 | 1 | 258953 | ENSMUST00 | -9271 | ENSMUSG0( | Olfr340 | olfactory receptor 340 |
| chr2 | 38058561 | 38059399 | 839 | * | NA_peak_39 | 40 | Intron (ENSM | 2 | 38021266 | 38166081 | 144816 | 2 | 227801 | ENSMUST00 | 106682 | ENSMUSG0( | Dennd1a | DENN/MADD domain containing 1A |
| chr2 | 39362501 | 39362870 | 370 | * | NA_peak_39 | 40 | Distal Interge | 2 | 39194354 | 39226451 | 32098 | 2 | 67857 | ENSMUST00 | -136050 | ENSMUSG0( | Ppp6c | protein phosphatase 6, catalytic subunit |
| chr2 | 40236490 | 40236970 | 481 | * | NA_peak_39 | 40 | Distal Interge | 2 | 40595246 | 40615251 | 20006 | 2 | 94217 | ENSMUST00 | 378281 | ENSMUSG0( | Lrp1b | low density lipoprotein-related protein 1B |
| chr2 | 41779312 | 41779597 | 286 | * | NA_peak_39 | 27 | Intron (ENSM | 2 | 40801393 | 41789078 | 987686 | 2 | 94217 | ENSMUST00 | 9481 | ENSMUSG0( | Lrp1b | low density lipoprotein-related protein 1B |
| chr2 | 41780354 | 41781206 | 853 | * | NA_peak_39 | 48 | Intron (ENSM | 2 | 40801393 | 41789078 | 987686 | 2 | 94217 | ENSMUST00 | 7872 | ENSMUSG0( | Lrp1b | low density lipoprotein-related protein 1B |
| chr2 | 45682499 | 45682791 | 293 | * | NA_peak_39 | 40 | Distal Interge | 2 | 45696605 | 45698447 | 1843 | 1 | 69405 | ENSMUST00 | -13814 | ENSMUSG0( | 1700019E08I | RIKEN cDNA 1700019E08 gene |
| chr2 | 46578215 | 46578272 | 58 | * | NA_peak_39 | 25 | Distal Interge | 2 | 45696800 | 45698067 | 1268 | 1 | 69405 | ENSMUST00 | 881415 | ENSMUSG0( | 1700019E08I | RIKEN cDNA 1700019E08 gene |
| chr2 | 48055236 | 48055497 | 262 | * | NA_peak_39 | 16 | Distal Interge | 2 | 48814109 | 48903269 | 89161 | 1 | 11480 | ENSMUST00 | -758612 | ENSMUSG0( | Acvr2a | activin receptor IIA |
| chr2 | 49478838 | 49479123 | 286 | * | NA_peak_39 | 20 | Intron (ENSM | 2 | 49451486 | 49551948 | 100463 | 1 | 227867 | ENSMUST00 | 27352 | ENSMUSG0( | Epc2 | enhancer of polycomb homolog 2 |
| chr2 | 55198402 | 55198634 | 233 | * | NA_peak_39 | 27 | Distal Interge | 2 | 55435970 | 55598145 | 162176 | 1 | 16519 | ENSMUST00 | -237336 | ENSMUSG0( | Kcnj3 | potassium inwardly-rectifying channel, subfamily J, member 3 |
| chr2 | 55347576 | 55347752 | 177 | * | NA_peak_39 | 31 | Distal Interge | 2 | 55435970 | 55598145 | 162176 | 1 | 16519 | ENSMUST00 | -88218 | ENSMUSG0( | Kcnj3 | potassium inwardly-rectifying channel, subfamily J, member 3 |
| chr2 | 55628818 | 55628842 | 25 | * | NA_peak_39 | 16 | Distal Interge | 2 | 55437165 | 55598140 | 160976 | 1 | 16519 | ENSMUST00 | 191653 | ENSMUSG0( | Kcnj3 | potassium inwardly-rectifying channel, subfamily J, member 3 |
| chr2 | 55674290 | 55674660 | 371 | * | NA_peak_39 | 31 | Distal Interge | 2 | 55437165 | 55598140 | 160976 | 1 | 16519 | ENSMUST00 | 237125 | ENSMUSG0( | Kcnj3 | potassium inwardly-rectifying channel, subfamily J, member 3 |
| chr2 | 55871693 | 55872028 | 336 | * | NA_peak_39 | 36 | Distal Interge | 2 | 55437165 | 55598140 | 160976 | 1 | 16519 | ENSMUST00 | 434528 | ENSMUSG0( | Kcnj3 | potassium inwardly-rectifying channel, subfamily J, member 3 |
| chr2 | 56068516 | 56068832 | 317 | * | NA_peak_39 | 26 | Distal Interge | 2 | 55437165 | 55598140 | 160976 | 1 | 16519 | ENSMUST00 | 631351 | ENSMUSG0( | Kcnj3 | potassium inwardly-rectifying channel, subfamily J, member 3 |
| chr2 | 56932565 | 56932954 | 390 | * | NA_peak_39 | 23 | Distal Interge | 2 | 56785811 | 56785907 | 97 | 1 | 102465169 | ENSMUST00 | 146754 | ENSMUSG0( | Mir195b | microRNA 195b |
| chr2 | 57554542 | 57555062 | 521 | * | NA_peak_39 | 40 | Distal Interge | 2 | 57267377 | 57268108 | 732 | 1 | 14571 | ENSMUST00 | 287165 | ENSMUSG0( | Gpd2 | glycerol phosphate dehydrogenase 2, mitochondrial |
| chr2 | 58256657 | 58256992 | 336 | * | NA_peak_39 | 27 | Distal Interge | 2 | 58276779 | 58286902 | 10124 | 1 | 100502932 | ENSMUST00 | -19787 | ENSMUSG0( | Gm13544 | predicted gene 13544 |
| chr2 | 58649052 | 58649442 | 391 | * | NA_peak_39 | 39 | Intron (ENSM | 2 | 58567387 | 58792971 | 225585 | 1 | 76654 | ENSMUST00 | 81665 | ENSMUSG0( | Upp2 | uridine phosphorylase 2 |
| chr2 | 58674120 | 58674399 | 280 | * | NA_peak_39 | 31 | Intron (ENSM | 2 | 58754942 | 58791242 | 36301 | 1 | 76654 | ENSMUST00 | -80543 | ENSMUSG0( | Upp2 | uridine phosphorylase 2 |
| chr2 | 61444586 | 61444856 | 271 | * | NA_peak_39 | 19 | Distal Interge | 2 | 61578585 | 61654171 | 75587 | 1 | 21353 | ENSMUST00 | -133729 | ENSMUSG0( | Tank | TRAF family member-associated Nf-kappa B activator |
| chr2 | 63399228 | 63399426 | 199 | * | NA_peak_39 | 27 | Distal Interge | 2 | 62786281 | 63184287 | 398007 | 2 | 170738 | ENSMUST00 | -214941 | ENSMUSG0( | Kcnh7 | potassium voltage-gated channel, subfamily H (eag-related), member 7 |
| chr2 | 63838918 | 63839232 | 315 | * | NA_peak_39 | 20 | Distal Interge | 2 | 63980447 | 64097988 | 117542 | 2 | 60344 | ENSMUST00 | 258756 | ENSMUSG0( | Fign | fidgetin |
| chr2 | 64199857 | 64200242 | 386 | * | NA_peak_39 | 22 | Distal Interge | 2 | 63977337 | 64098038 | 120702 | 2 | 60344 | ENSMUST00 | -101819 | ENSMUSG0( | Fign | fidgetin |
| chr2 | 64631826 | 64632215 | 390 | * | NA_peak_39 | 35 | Distal Interge | 2 | 64912476 | 64915019 | 2544 | 2 | 50915 | ENSMUST00 | 282804 | ENSMUSG0( | Grb14 | growth factor receptor bound protein 14 |
| chr2 | 65755505 | 65756163 | 659 | * | NA_peak_39 | 33 | Intron (ENSM | 2 | 65717576 | 65728392 | 10817 | 1 | 110876 | ENSMUST00 | 37929 | ENSMUSG0( | Scn2a | sodium channel, voltage-gated, type II, alpha |
| chr2 | 66429829 | 66430032 | 204 | * | NA_peak_39 | 22 | Intron (ENSM | 2 | 66348364 | 66440814 | 92451 | 2 | 20265 | ENSMUST00 | 10782 | ENSMUSG0( | Scn1a | sodium channel, voltage-gated, type I, alpha |
| chr2 | 70076601 | 70076788 | 188 | * | NA_peak_39 | 46 | Intron (ENSM | 2 | 70083074 | 70429198 | 346125 | 1 | 329421 | ENSMUST00 | -6286 | ENSMUSG0( | Myo3b | myosin IIIB |
| chr2 | 74681892 | 74682095 | 204 | * | NA_peak_39 | 20 | Promoter (<= | 2 | 74682323 | 74684168 | 1846 | 1 | 15431 | ENSMUST00 | -228 | ENSMUSG0( | Hoxd11 | homeobox D11 |
| chr2 | 76208493 | 76208913 | 421 | * | NA_peak_39 | 42 | Intron (ENSM | 2 | 76027665 | 76215394 | 187730 | 2 | 241489 | ENSMUST00 | 6481 | ENSMUSG0( | Pde11a | phosphodiesterase 11A |
| chr2 | 78621458 | 78621761 | 304 | * | NA_peak_39 | 21 | Distal Interge | 2 | 78868124 | 78870607 | 2484 | 1 | 22193 | ENSMUST00 | -246363 | ENSMUSG0( | Ube2e3 | ubiquitin-conjugating enzyme E2E 3 |
| chr2 | 80832060 | 80832314 | 255 | * | NA_peak_39 | 13 | Distal Interge | 2 | 80644204 | 80653414 | 9211 | 1 | 69482 | ENSMUST00 | 187856 | ENSMUSG0( | Nup35 | nucleoporin 35 |
| chr2 | 84734173 | 84734187 | 15 | * | NA_peak_39 | 13 | Promoter (<= | 2 | 84734227 | 84738649 | 4423 | 1 | 241525 | ENSMUST00 | -40 | ENSMUSG0( | Ypel4 | yippee like 4 |
| chr2 | 84926935 | 84927264 | 330 | * | NA_peak_39 | 78 | Distal Interge | 2 | 84936579 | 84937834 | 1256 | 1 | 58207 | ENSMUST00 | -9315 | ENSMUSG0( | Slc43a3 | solute carrier family 43, member 3 |
| chr2 | 89899024 | 89899344 | 321 | * | NA_peak_39 | 28 | Distal Interge | 2 | 89880794 | 89881809 | 1016 | 1 | 258984 | ENSMUST00 | 18230 | ENSMUSG0( | Olfr1257 | olfactory receptor 1257 |
| chr2 | 90101094 | 90101290 | 197 | * | NA_peak_39 | 27 | Distal Interge | 2 | 90118600 | 90119617 | 1018 | 2 | 258339 | ENSMUST00 | 18327 | ENSMUSG0( | Olfr1269 | olfactory receptor 1269 |
| chr2 | 90248626 | 90249006 | 381 | * | NA_peak_39 | 19 | Distal Interge | 2 | 90252069 | 90252986 | 918 | 2 | 406186 | ENSMUST00 | 3980 | ENSMUSG0( | Olfr142 | olfactory receptor 142 |
| chr2 | 90890123 | 90890474 | 352 | * | NA_peak_39 | 13 | 3' UTR | 2 | 90885860 | 90890525 | 4666 | 1 | 67445 | ENSMUST00 | 4263 | ENSMUSG0( | C1qtnf4 | C1q and tumor necrosis factor related protein 4 |
| chr2 | 91097040 | 91097321 | 282 | * | NA_peak_39 | 19 | Promoter (<= | 2 | 91096780 | 91113315 | 16536 | 1 | 20375 | ENSMUST00 | 260 | ENSMUSG0( | Spi1 | spleen focus forming virus (SFFV) proviral integration oncogene |
| chr2 | 94802187 | 94802624 | 438 | * | NA_peak_39 | 17 | Distal Interge | 2 | 94422336 | 94438136 | 15801 | 2 | 11800 | ENSMUST00 | -364051 | ENSMUSG0( | Api5 | apoptosis inhibitor 5 |
| chr2 | 95419952 | 95420374 | 423 | * | NA_peak_39 | 24 | Distal Interge | 2 | 96318169 | 97631666 | 1313498 | 1 | 241568 | ENSMUST00 | -897795 | ENSMUSG0( | Lrrc4c | leucine rich repeat containing 4C |
| chr2 | 97838186 | 97838625 | 440 | * | NA_peak_39 | 18 | Distal Interge | 2 | 97468089 | 97629063 | 160975 | 1 | 241568 | ENSMUST00 | 370097 | ENSMUSG0( | Lrrc4c | leucine rich repeat containing 4C |
| chr2 | 97839550 | 97839696 | 147 | * | NA_peak_39 | 55 | Distal Interge | 2 | 97468089 | 97629063 | 160975 | 1 | 241568 | ENSMUST00 | 371461 | ENSMUSG0( | Lrrc4c | leucine rich repeat containing 4C |
| chr2 | 98634135 | 98634440 | 306 | * | NA_peak_39 | 48 | Distal Interge | 2 | 97468089 | 97629063 | 160975 | 1 | 241568 | ENSMUST00 | 1166046 | ENSMUSG0( | Lrrc4c | leucine rich repeat containing 4C |
| chr2 | 98662247 | 98662762 | 516 | * | NA_peak_40 | 394 | Exon (ENSMU | 2 | 97468089 | 97629063 | 160975 | 1 | 241568 | ENSMUST00 | 1194158 | ENSMUSG0( | Lrrc4c | leucine rich repeat containing 4C |
| chr2 | 98666259 | 98667315 | 1057 | * | NA_peak_40 | 1476 | Exon (ENSMU | 2 | 97468089 | 97629063 | 160975 | 1 | 241568 | ENSMUST00 | 1198170 | ENSMUSG0( | Lrrc4c | leucine rich repeat containing 4C |
| chr2 | 100547301 | 100547573 | 273 | * | NA_peak_40 | 22 | Distal Interge | 2 | 101562125 | 101576125 | 14001 | 2 | 68170 | ENSMUST00 | 1028552 | ENSMUSG0( | Iftap | intraflagellar transport associated protein |
| chr2 | 100606324 | 100606620 | 297 | * | NA_peak_40 | 45 | Distal Interge | 2 | 101562125 | 101576125 | 14001 | 2 | 68170 | ENSMUST00 | 969505 | ENSMUSG0( | Iftap | intraflagellar transport associated protein |
| chr2 | 104014233 | 104014657 | 425 | * | NA_peak_40 | 23 | Intron (ENSM | 2 | 104027721 | 104063240 | 35520 | 1 | 57443 | ENSMUST00 | -13064 | ENSMUSG0( | Fbxo3 | F-box protein 3 |
| chr2 | 108951542 | 108951632 | 91 | * | NA_peak_40 | 23 | Distal Interge | 2 | 109092300 | 109278280 | 185981 | 2 | 76894 | ENSMUST00 | 326648 | ENSMUSG0( | Mettl15 | methyltransferase like 15 |
| chr2 | 110125621 | 110125892 | 272 | * | NA_peak_40 | 31 | Intron (ENSM | 2 | 110031251 | 110050479 | 19229 | 2 | 75517 | ENSMUST00 | -75142 | ENSMUSG0( | Ccdc34os | coiled-coil domain containing 34, opposite strand |
| chr2 | 116269890 | 116270188 | 299 | * | NA_peak_40 | 16 | Distal Interge | 2 | 116075165 | 116076096 | 932 | 2 | 69974 | ENSMUST00 | -193794 | NA | 2810405F15I | RIKEN cDNA 2810405F15 gene |
| chr2 | 120638195 | 120638635 | 441 | * | NA_peak_40 | 49 | Intron (ENSM | 2 | 120629126 | 120716082 | 86957 | 1 | 668880 | ENSMUST00 | 9069 | ENSMUSG0( | Stard9 | START domain containing 9 |
| chr2 | 122681383 | 122682235 | 853 | * | NA_peak_40 | 24 | 3' UTR | 2 | 122681233 | 122702639 | 21407 | 2 | 22785 | ENSMUST00 | 20404 | ENSMUSG0( | Slc30a4 | solute carrier family 30 (zinc transporter), member 4 |
| chr2 | 122950657 | 122950917 | 261 | * | NA_peak_40 | 31 | Distal Interge | 2 | 122799762 | 122809559 | 9798 | 1 | 59010 | ENSMUST00 | 150895 | ENSMUSG0( | Sqor | sulfide quinone oxidoreductase |
| chr2 | 124255334 | 124255647 | 314 | * | NA_peak_40 | 26 | Intron (ENSM | 2 | 124217679 | 124222331 | 4653 | 2 | 75946 | ENSMUST00 | -33003 | ENSMUSG0( | 4930583P06I | RIKEN cDNA 4930583P06 gene |
| chr2 | 125411215 | 125411466 | 252 | * | NA_peak_40 | 16 | Intron (ENSM | 2 | 125389756 | 125506072 | 116317 | 2 | 14118 | ENSMUST00 | 94606 | ENSMUSG0( | Fbn1 | fibrillin 1 |
| chr2 | 129613006 | 129613327 | 322 | * | NA_peak_40 | 45 | Intron (ENSM | 2 | 129620685 | 129628044 | 7360 | 1 | 19261 | ENSMUST00 | -7358 | ENSMUSG0( | Sirpa | signal-regulatory protein alpha |
| chr2 | 131492366 | 131492452 | 87 | * | NA_peak_40 | 20 | Promoter (<= | 2 | 131492264 | 131512177 | 19914 | 1 | 228608 | ENSMUST00 | 102 | ENSMUSG0( | Smox | spermine oxidase |
| chr2 | 131562526 | 131562690 | 165 | * | NA_peak_40 | 41 | Promoter (<= | 2 | 131545850 | 131562283 | 16434 | 2 | 11550 | ENSMUST00 | -243 | ENSMUSG0( | Adra1d | adrenergic receptor, alpha 1d |
| chr2 | 133072521 | 133072907 | 387 | * | NA_peak_40 | 21 | Distal Interge | 2 | 132904389 | 132945906 | 41518 | 2 | 241639 | ENSMUST00 | -126615 | ENSMUSG0( | Fermt1 | fermitin family member 1 |
| chr2 | 133810830 | 133811061 | 232 | * | NA_peak_40 | 19 | Distal Interge | 2 | 133564708 | 133564794 | 87 | 1 | 100526500 | ENSMUST00 | 246122 | ENSMUSG0( | Mir3090 | microRNA 3090 |
| chr2 | 133913810 | 133914129 | 320 | * | NA_peak_40 | 26 | Distal Interge | 2 | 133564708 | 133564794 | 87 | 1 | 100526500 | ENSMUST00 | 349102 | ENSMUSG0( | Mir3090 | microRNA 3090 |
| chr2 | 133970198 | 133970755 | 558 | * | NA_peak_40 | 61 | Distal Interge | 2 | 133564708 | 133564794 | 87 | 1 | 100526500 | ENSMUST00 | 405490 | ENSMUSG0( | Mir3090 | microRNA 3090 |
| chr2 | 134315170 | 134315475 | 306 | * | NA_peak_40 | 36 | Distal Interge | 2 | 134530760 | 134554307 | 23548 | 2 | 15112 | ENSMUST00 | 238832 | ENSMUSG0( | Hao1 | hydroxyacid oxidase 1, liver |
| chr2 | 139067483 | 139067667 | 185 | * | NA_peak_40 | 23 | Distal Interge | 2 | 138819929 | 138820007 | 79 | 1 | 100316816 | ENSMUST00 | 247554 | ENSMUSG0( | Mir1952 | microRNA 1952 |
| chr2 | 140131464 | 140131722 | 259 | * | NA_peak_40 | 31 | Intron (ENSM | 2 | 140148810 | 140155305 | 6496 | 2 | 66580 | ENSMUST00 | 23583 | ENSMUSG0( | Esf1 | ESF1 nucleolar pre-rRNA processing protein homolog |
| chr2 | 141905327 | 141905522 | 196 | * | NA_peak_40 | 55 | Intron (ENSM | 2 | 142176607 | 142234375 | 57769 | 1 | 72899 | ENSMUST00 | -271085 | ENSMUSG0( | Macrod2 | mono-ADP ribosylhydrolase 2 |
| chr2 | 144155959 | 144156312 | 354 | * | NA_peak_40 | 32 | Intron (ENSM | 2 | 144174315 | 144179086 | 4772 | 2 | 433481 | ENSMUST00 | 22774 | ENSMUSG0( | Gm5535 | predicted gene 5535 |
| chr2 | 144344928 | 144345248 | 321 | * | NA_peak_40 | 21 | Distal Interge | 2 | 144305175 | 144332146 | 26972 | 2 | 107586 | ENSMUST00 | -12782 | ENSMUSG0( | Ovol2 | ovo like zinc finger 2 |
| chr2 | 148411656 | 148411983 | 328 | * | NA_peak_40 | 26 | Distal Interge | 2 | 148404466 | 148408188 | 3723 | 2 | 21824 | ENSMUST00 | -3468 | ENSMUSG0( | Thbd | thrombomodulin |

| | | | | | | | | | | | | | | | | | | |
|---|---|---|---|---|---|---|---|---|---|---|---|---|---|---|---|---|---|---|
| chr2 | 148619449 | 148619856 | 408 | * | NA_peak_404 | 23 | Distal Interge | 2 | 148672601 | 148676027 | 3427 | 1 | 56488 | ENSMUST00 | -52745 | ENSMUSG0( | Nxt1 | NTF2-related export protein 1 |
| chr2 | 151162008 | 151162537 | 530 | * | NA_peak_404 | 36 | Distal Interge | 2 | 151100608 | 151103041 | 2434 | 1 | 433486 | ENSMUST00 | 61400 | ENSMUSG0( | Gm14151 | predicted gene 14151 |
| chr2 | 153038025 | 153038081 | 57 | * | NA_peak_404 | 31 | Intron (ENSM | 2 | 153031852 | 153055775 | 23924 | 1 | 228787 | ENSMUST00 | 6173 | ENSMUSG0( | Xkr7 | X-linked Kx blood group related 7 |
| chr2 | 153826231 | 153826791 | 561 | * | NA_peak_404 | 15 | Exon (ENSMU | 2 | 153842981 | 153844745 | 1765 | 1 | 100504221 | ENSMUST00 | -16190 | ENSMUSG0( | Efcab8 | EF-hand calcium binding domain 8 |
| chr2 | 154143439 | 154143778 | 340 | * | NA_peak_404 | 15 | Promoter (<= | 2 | 154144303 | 154147700 | 3398 | 1 | 18843 | ENSMUST00 | -525 | ENSMUSG0( | Bpifa1 | BPI fold containing family A, member 1 |
| chr2 | 154481899 | 154482251 | 353 | * | NA_peak_405 | 21 | Intron (ENSM | 2 | 154510536 | 154539356 | 28821 | 1 | 12396 | ENSMUST00 | -28285 | ENSMUSG0( | Cbfa2t2 | CBFA2/RUNX1 translocation partner 2 |
| chr2 | 154851071 | 154851346 | 276 | * | NA_peak_405 | 13 | Intron (ENSM | 2 | 154871418 | 154892782 | 21365 | 2 | 67204 | ENSMUST00 | 41436 | ENSMUSG0( | Eif2s2 | eukaryotic translation initiation factor 2, subunit 2 (beta) |
| chr2 | 154853646 | 154853967 | 322 | * | NA_peak_405 | 20 | Intron (ENSM | 2 | 154871418 | 154892782 | 21365 | 2 | 67204 | ENSMUST00 | 38815 | ENSMUSG0( | Eif2s2 | eukaryotic translation initiation factor 2, subunit 2 (beta) |
| chr2 | 154892383 | 154892832 | 450 | * | NA_peak_405 | 55 | Promoter (<= | 2 | 154871418 | 154892782 | 21365 | 2 | 67204 | ENSMUST00 | 0 | ENSMUSG0( | Eif2s2 | eukaryotic translation initiation factor 2, subunit 2 (beta) |
| chr2 | 155178338 | 155178426 | 89 | * | NA_peak_405 | 27 | Intron (ENSM | 2 | 155133561 | 155212594 | 79034 | 1 | 16396 | ENSMUST00 | 44777 | ENSMUSG0( | Itch | itchy, E3 ubiquitin protein ligase |
| chr2 | 155944783 | 155945096 | 314 | * | NA_peak_405 | 27 | Promoter (<= | 2 | 155941023 | 155945367 | 4345 | 2 | 14563 | ENSMUST00 | 271 | ENSMUSG0( | Gdf5 | growth differentiation factor 5 |
| chr2 | 156187242 | 156187589 | 348 | * | NA_peak_405 | 31 | Distal Interge | 2 | 156174774 | 156180238 | 5465 | 2 | 170791 | ENSMUST00 | -7004 | ENSMUSG0( | Rbm39 | RNA binding motif protein 39 |
| chr2 | 156312254 | 156312398 | 145 | * | NA_peak_405 | 16 | Promoter (<= | 2 | 156312299 | 156375635 | 63337 | 1 | 70873 | ENSMUST00 | 0 | ENSMUSG0( | Cnbd2 | cyclic nucleotide binding domain containing 2 |
| chr2 | 161375214 | 161375276 | 63 | * | NA_peak_406 | 20 | Distal Interge | 2 | 161008034 | 161109075 | 101042 | 2 | 71389 | ENSMUST00 | -266139 | ENSMUSG0( | Chd6 | chromodomain helicase DNA binding protein 6 |
| chr2 | 162597710 | 162598055 | 346 | * | NA_peak_406 | 50 | Intron (ENSM | 2 | 161526357 | 162661094 | 1134738 | 2 | 19281 | ENSMUST00 | 63039 | ENSMUSG0( | Ptprt | protein tyrosine phosphatase, receptor type, T |
| chr2 | 163406640 | 163407002 | 363 | * | NA_peak_406 | 27 | Exon (ENSMU | 2 | 163406586 | 163411665 | 5080 | 2 | 66680 | ENSMUST00 | 4663 | ENSMUSG0( | Oser1 | oxidative stress responsive serine rich 1 |
| chr2 | 168692973 | 168693286 | 314 | * | NA_peak_407 | 23 | Intron (ENSM | 2 | 168705160 | 168712913 | 7754 | 2 | 11981 | ENSMUST00 | 19627 | ENSMUSG0( | Atp9a | ATPase, class II, type 9A |
| chr2 | 173462743 | 173462957 | 215 | * | NA_peak_407 | 27 | Distal Interge | 2 | 173522586 | 173528501 | 5916 | 1 | 72221 | ENSMUST00 | -59629 | ENSMUSG0( | 1700021F07I | RIKEN cDNA 1700021F07 gene |
| chr2 | 177321177 | 177321532 | 356 | * | NA_peak_408 | 45 | Intron (ENSM | 2 | 177464733 | 177478129 | 13397 | 1 | 628308 | ENSMUST00 | -143201 | ENSMUSG0( | Zfp970 | zinc finger protein 970 |
| chr2 | 181861735 | 181861838 | 104 | * | NA_peak_408 | 26 | Promoter (2-3 | 2 | 181864337 | 181870830 | 6494 | 1 | 67005 | ENSMUST00 | -2499 | ENSMUSG0( | Polr3k | polymerase (RNA) III (DNA directed) polypeptide K |
| chr2 | 181862156 | 181862302 | 147 | * | NA_peak_408 | 36 | Promoter (2-3 | 2 | 181864337 | 181870830 | 6494 | 1 | 67005 | ENSMUST00 | -2035 | ENSMUSG0( | Polr3k | polymerase (RNA) III (DNA directed) polypeptide K |
| chr2 | 181930291 | 181930941 | 651 | * | NA_peak_408 | 280 | Distal Interge | 2 | 181991129 | 182003861 | 12733 | 1 | 672125 | ENSMUST00 | -60188 | ENSMUSG0( | Gm14496 | predicted gene 14496 |
| chr3 | 3055949 | 3056227 | 279 | * | NA_peak_408 | 20 | Distal Interge | 3 | 3508030 | 3657844 | 149815 | 1 | 30942 | ENSMUST00 | -451803 | ENSMUSG0( | Hnf4g | hepatocyte nuclear factor 4, gamma |
| chr3 | 3142192 | 3142488 | 297 | * | NA_peak_408 | 40 | Distal Interge | 3 | 3508030 | 3657844 | 149815 | 1 | 30942 | ENSMUST00 | -365542 | ENSMUSG0( | Hnf4g | hepatocyte nuclear factor 4, gamma |
| chr3 | 6603111 | 6603455 | 345 | * | NA_peak_409 | 23 | Distal Interge | 3 | 6615418 | 6620443 | 5026 | 2 | 69347 | ENSMUST00 | 16988 | ENSMUSG0( | 1700008P02I | RIKEN cDNA 1700008P02 gene |
| chr3 | 8245951 | 8246365 | 415 | * | NA_peak_409 | 67 | Distal Interge | 3 | 8462228 | 8463035 | 808 | 2 | 620966 | ENSMUST00 | 216670 | NA | Gm6194 | transmembrane protein 189 pseudogene |
| chr3 | 8656168 | 8656503 | 336 | * | NA_peak_409 | 53 | Distal Interge | 3 | 8664796 | 8666625 | 1830 | 2 | 15213 | ENSMUST00 | 10122 | ENSMUSG0( | Hey1 | hairy/enhancer-of-split related with YRPW motif 1 |
| chr3 | 8923643 | 8923790 | 148 | * | NA_peak_409 | 38 | Promoter (<= | 3 | 8802146 | 8923918 | 121773 | 2 | 66230 | ENSMUST00 | 128 | ENSMUSG0( | Mrps28 | mitochondrial ribosomal protein S28 |
| chr3 | 12251820 | 12251886 | 67 | * | NA_peak_410 | 20 | Distal Interge | 3 | 11823229 | 11844359 | 21131 | 2 | 100038545 | ENSMUST00 | -407461 | ENSMUSG0( | Gm10745 | predicted gene 10745 |
| chr3 | 12837649 | 12837952 | 304 | * | NA_peak_410 | 19 | Distal Interge | 3 | 13471655 | 13777103 | 305449 | 1 | 76897 | ENSMUST00 | -633703 | ENSMUSG0( | Ralyl | RALY RNA binding protein-like |
| chr3 | 14721186 | 14721476 | 291 | * | NA_peak_410 | 26 | Distal Interge | 3 | 14766217 | 14778459 | 12243 | 2 | 12346 | ENSMUST00 | 56983 | ENSMUSG0( | Car1 | carbonic anhydrase 1 |
| chr3 | 15527408 | 15527848 | 441 | * | NA_peak_410 | 23 | Intron (ENSM | 3 | 15495751 | 15574985 | 79235 | 2 | 668101 | ENSMUST00 | 47137 | ENSMUSG0( | Sirpb1b | signal-regulatory protein beta 1B |
| chr3 | 17297549 | 17297848 | 300 | * | NA_peak_410 | 27 | Distal Interge | 3 | 17789921 | 17800073 | 10153 | 1 | 70441 | ENSMUST00 | -492073 | ENSMUSG0( | Mir124-2hg | Mir124-2 host gene (non-protein coding) |
| chr3 | 17905638 | 17905756 | 119 | * | NA_peak_410 | 27 | Distal Interge | 3 | 17948410 | 17948959 | 550 | 1 | 75439 | ENSMUST00 | -42654 | ENSMUSG0( | Cypt12 | cysteine-rich perinuclear theca 12 |
| chr3 | 19850161 | 19850579 | 419 | * | NA_peak_411 | 40 | Distal Interge | 3 | 19894886 | 19901033 | 6148 | 1 | 70808 | ENSMUST00 | -44307 | NA | 4632415L05I | RRS1 ribosome biogenesis regulator homolog pseudogene |
| chr3 | 21192235 | 21192523 | 289 | * | NA_peak_411 | 18 | Distal Interge | 3 | 20314473 | 20367177 | 52705 | 2 | 11608 | ENSMUST00 | -825058 | ENSMUSG0( | Agtr1b | angiotensin II receptor, type 1b |
| chr3 | 22673995 | 22674348 | 354 | * | NA_peak_411 | 28 | Distal Interge | 3 | 22251374 | 22251663 | 290 | 1 | 19784 | ENSMUST00 | 422621 | NA | Rprl2 | ribonuclease P RNA-like 2 |
| chr3 | 22718172 | 22718446 | 275 | * | NA_peak_411 | 16 | Distal Interge | 3 | 22251374 | 22251663 | 290 | 1 | 19784 | ENSMUST00 | 466798 | NA | Rprl2 | ribonuclease P RNA-like 2 |
| chr3 | 24090619 | 24090850 | 232 | * | NA_peak_411 | 16 | Intron (ENSM | 3 | 23804985 | 23972185 | 167201 | 2 | 635702 | ENSMUST00 | -118434 | ENSMUSG0( | Naaladl2 | N-acetylated alpha-linked acidic dipeptidase-like 2 |
| chr3 | 25036808 | 25037062 | 255 | * | NA_peak_411 | 31 | Intron (ENSM | 3 | 24551210 | 25144261 | 593052 | 2 | 635702 | ENSMUST00 | 107199 | ENSMUSG0( | Naaladl2 | N-acetylated alpha-linked acidic dipeptidase-like 2 |
| chr3 | 26090699 | 26091066 | 368 | * | NA_peak_411 | 45 | Intron (ENSM | 3 | 25431811 | 26133734 | 701924 | 2 | 192167 | ENSMUST00 | 42668 | ENSMUSG0( | Nlgn1 | neuroligin 1 |
| chr3 | 26108619 | 26108897 | 279 | * | NA_peak_411 | 24 | Intron (ENSM | 3 | 25431811 | 26133734 | 701924 | 2 | 192167 | ENSMUST00 | 24837 | ENSMUSG0( | Nlgn1 | neuroligin 1 |
| chr3 | 27153711 | 27154038 | 328 | * | NA_peak_411 | 17 | Promoter (<= | 3 | 27097222 | 27153861 | 56640 | 2 | 13605 | ENSMUST00 | 0 | ENSMUSG0( | Ect2 | ect2 oncogene |
| chr3 | 28877906 | 28878196 | 291 | * | NA_peak_412 | 21 | Distal Interge | 3 | 28892617 | 28926724 | 34108 | 1 | 385263 | ENSMUST00 | -14421 | ENSMUSG0( | Gm1527 | predicted gene 1527 |
| chr3 | 29521630 | 29522049 | 420 | * | NA_peak_412 | 36 | Intron (ENSM | 3 | 29513906 | 29670734 | 156829 | 1 | 75740 | ENSMUST00 | 7724 | ENSMUSG0( | Egfem1 | EGF-like and EMI domain containing 1 |
| chr3 | 30030185 | 30030480 | 296 | * | NA_peak_412 | 30 | Intron (ENSM | 3 | 29951306 | 30013505 | 62200 | 2 | 14013 | ENSMUST00 | -16680 | ENSMUSG0( | Mecom | MDS1 and EVI1 complex locus |
| chr3 | 30356374 | 30356537 | 164 | * | NA_peak_412 | 20 | Intron (ENSM | 3 | 29952567 | 30238162 | 285596 | 2 | 14013 | ENSMUST00 | -118212 | ENSMUSG0( | Mecom | MDS1 and EVI1 complex locus |
| chr3 | 32163050 | 32163343 | 294 | * | NA_peak_412 | 43 | Intron (ENSM | 3 | 32156114 | 32198356 | 42243 | 1 | 72413 | ENSMUST00 | 6936 | ENSMUSG0( | Kcnmb2 | potassium large conductance calcium-activated channel, subfamily M, beta member 2 |
| chr3 | 32168157 | 32168306 | 150 | * | NA_peak_412 | 45 | Intron (ENSM | 3 | 32156114 | 32198356 | 42243 | 1 | 72413 | ENSMUST00 | 12043 | ENSMUSG0( | Kcnmb2 | potassium large conductance calcium-activated channel, subfamily M, beta member 2 |
| chr3 | 32624387 | 32624522 | 136 | * | NA_peak_412 | 26 | Distal Interge | 3 | 32580332 | 32616585 | 36254 | 2 | 14696 | ENSMUST00 | -7802 | ENSMUSG0( | Gnb4 | guanine nucleotide binding protein (G protein), beta 4 |
| chr3 | 35118895 | 35119440 | 546 | * | NA_peak_413 | 27 | Distal Interge | 3 | 34922545 | 34922651 | 107 | 2 | 102466155 | ENSMUST00 | -196244 | ENSMUSG0( | Mir6378 | microRNA 6378 |
| chr3 | 35170398 | 35170994 | 597 | * | NA_peak_413 | 40 | Distal Interge | 3 | 34922545 | 34922651 | 107 | 2 | 102466155 | ENSMUST00 | -247747 | ENSMUSG0( | Mir6378 | microRNA 6378 |
| chr3 | 36521380 | 36521547 | 168 | * | NA_peak_413 | 36 | Promoter (<= | 3 | 36515057 | 36521813 | 6757 | 2 | 100503068 | ENSMUST00 | 266 | ENSMUSG0( | Smim43 | small integral membrane protein 43 |
| chr3 | 37254675 | 37255050 | 376 | * | NA_peak_413 | 20 | Distal Interge | 3 | 37222759 | 37232618 | 9860 | 2 | 60505 | ENSMUST00 | -22057 | ENSMUSG0( | Il21 | interleukin 21 |
| chr3 | 38195114 | 38195217 | 104 | * | NA_peak_413 | 26 | Distal Interge | 3 | 38203151 | 38208119 | 4969 | 2 | 71342 | ENSMUST00 | 12902 | ENSMUSG0( | 5430434I15R | RIKEN cDNA 5430434I15 gene |
| chr3 | 38431198 | 38431484 | 287 | * | NA_peak_413 | 27 | Intron (ENSM | 3 | 38452079 | 38455557 | 3479 | 2 | 99696 | ENSMUST00 | 24073 | ENSMUSG0( | Ankrd50 | ankyrin repeat domain 50 |
| chr3 | 41341109 | 41341474 | 366 | * | NA_peak_414 | 27 | Distal Interge | 3 | 41487679 | 41493190 | 5512 | 2 | 78706 | ENSMUST00 | 151716 | ENSMUSG0( | Platr4 | pluripotency associated transcript 4 |
| chr3 | 43010970 | 43011295 | 326 | * | NA_peak_414 | 50 | Distal Interge | 3 | 41751196 | 41756492 | 5297 | 1 | 73852 | ENSMUST00 | 1259774 | ENSMUSG0( | D3Ertd751e | DNA segment, Chr 3, ERATO Doi 751, expressed |
| chr3 | 43858657 | 43858698 | 42 | * | NA_peak_414 | 27 | Distal Interge | 3 | 45281350 | 45364670 | 83321 | 2 | 72511 | ENSMUST00 | 1505972 | ENSMUSG0( | 2610316D01I | RIKEN cDNA 2610316D01 gene |
| chr3 | 44007662 | 44007816 | 155 | * | NA_peak_414 | 22 | Distal Interge | 3 | 45281350 | 45364670 | 83321 | 2 | 72511 | ENSMUST00 | 1356854 | ENSMUSG0( | 2610316D01I | RIKEN cDNA 2610316D01 gene |
| chr3 | 44624250 | 44624559 | 310 | * | NA_peak_414 | 45 | Distal Interge | 3 | 45281350 | 45364670 | 83321 | 2 | 72511 | ENSMUST00 | 740111 | ENSMUSG0( | 2610316D01I | RIKEN cDNA 2610316D01 gene |
| chr3 | 46098729 | 46098771 | 43 | * | NA_peak_414 | 20 | Distal Interge | 3 | 46447056 | 46447920 | 865 | 2 | 241989 | ENSMUST00 | 349149 | ENSMUSG0( | Pabpc4l | poly(A) binding protein, cytoplasmic 4-like |
| chr3 | 46141659 | 46141962 | 304 | * | NA_peak_414 | 13 | Distal Interge | 3 | 46447056 | 46447920 | 865 | 2 | 241989 | ENSMUST00 | 305958 | ENSMUSG0( | Pabpc4l | poly(A) binding protein, cytoplasmic 4-like |
| chr3 | 46921094 | 46921381 | 288 | * | NA_peak_414 | 23 | Distal Interge | 3 | 46900668 | 46900782 | 115 | 2 | 102465199 | ENSMUST00 | -20312 | ENSMUSG0( | Mir6379 | microRNA 6379 |
| chr3 | 48959142 | 48959667 | 526 | * | NA_peak_414 | 94 | Distal Interge | 3 | 49752421 | 49755464 | 3044 | 2 | 73173 | ENSMUST00 | 795797 | ENSMUSG0( | Pcdh18 | protocadherin 18 |
| chr3 | 49478881 | 49479194 | 314 | * | NA_peak_414 | 27 | Distal Interge | 3 | 49752421 | 49755464 | 3044 | 2 | 73173 | ENSMUST00 | 276270 | ENSMUSG0( | Pcdh18 | protocadherin 18 |
| chr3 | 51484125 | 51484306 | 182 | * | NA_peak_415 | 20 | Promoter (<= | 3 | 51483966 | 51496232 | 12267 | 1 | 19338 | ENSMUST00 | 159 | ENSMUSG0( | Rab33b | RAB33B, member RAS oncogene family |
| chr3 | 53187435 | 53187714 | 280 | * | NA_peak_415 | 24 | Intron (ENSM | 3 | 53041583 | 53261679 | 220097 | 1 | 108927 | ENSMUST00 | 145852 | ENSMUSG0( | Lhfp | lipoma HMGIC fusion partner |
| chr3 | 55585594 | 55585927 | 334 | * | NA_peak_415 | 30 | Distal Interge | 3 | 55527879 | 55534312 | 6434 | 1 | 13175 | ENSMUST00 | 57715 | ENSMUSG0( | Dclk1 | doublecortin-like kinase 1 |
| chr3 | 57325093 | 57325591 | 499 | * | NA_peak_416 | 50 | Distal Interge | 3 | 57287484 | 57301988 | 14505 | 2 | 17112 | ENSMUST00 | -23105 | ENSMUSG0( | Tm4sf1 | transmembrane 4 superfamily member 1 |
| chr3 | 58766455 | 58766937 | 483 | * | NA_peak_416 | 27 | Intron (ENSM | 3 | 58732830 | 58797331 | 64502 | 1 | 71301 | ENSMUST00 | 33625 | ENSMUSG0( | 4930593A02I | RIKEN cDNA 4930593A02 gene |
| chr3 | 58909083 | 58909427 | 345 | * | NA_peak_416 | 43 | Distal Interge | 3 | 58845981 | 58885340 | 39360 | 2 | 229320 | ENSMUST00 | -23743 | ENSMUSG0( | Clrn1 | clarin 1 |
| chr3 | 60436354 | 60436626 | 273 | * | NA_peak_416 | 27 | Distal Interge | 3 | 60472830 | 60595764 | 122935 | 1 | 56758 | ENSMUST00 | -36204 | ENSMUSG0( | Mbnl1 | muscleblind like splicing factor 1 |
| chr3 | 61002383 | 61002574 | 192 | * | NA_peak_416 | 14 | Promoter (<= | 3 | 61002795 | 61008979 | 6185 | 1 | 18441 | ENSMUST00 | -221 | ENSMUSG0( | P2ry1 | purinergic receptor P2Y, G-protein coupled 1 |
| chr3 | 61720788 | 61721007 | 220 | * | NA_peak_416 | 52 | Distal Interge | 3 | 61361638 | 61368430 | 6793 | 1 | 74012 | ENSMUST00 | 359150 | ENSMUSG0( | Rap2b | RAP2B, member of RAS oncogene family |
| chr3 | 62122774 | 62122987 | 214 | * | NA_peak_416 | 39 | Distal Interge | 3 | 62338344 | 62462221 | 123878 | 1 | 622434 | ENSMUST00 | -215357 | ENSMUSG0( | Arhgef26 | Rho guanine nucleotide exchange factor (GEF) 26 |
| chr3 | 64239533 | 64239970 | 438 | * | NA_peak_416 | 16 | Distal Interge | 3 | 64258961 | 64287417 | 28457 | 2 | 637004 | ENSMUST00 | 47447 | ENSMUSG0( | Vmn2r3 | vomeronasal 2, receptor 3 |
| chr3 | 66973395 | 66973697 | 303 | * | NA_peak_417 | 31 | 3' UTR | 3 | 66972780 | 66978447 | 5668 | 2 | 20429 | ENSMUST00 | 4750 | ENSMUSG0( | Shox2 | short stature homeobox 2 |
| chr3 | 67294600 | 67294811 | 212 | * | NA_peak_417 | 43 | Intron (ENSM | 3 | 67374097 | 67399998 | 25902 | 1 | 17349 | ENSMUST00 | -79286 | ENSMUSG0( | Mlf1 | myeloid leukemia factor 1 |
| chr3 | 67410169 | 67410189 | 21 | * | NA_peak_417 | 27 | Distal Interge | 3 | 67430096 | 67476529 | 46434 | 1 | 28030 | ENSMUST00 | -19907 | ENSMUSG0( | Gfm1 | G elongation factor, mitochondrial 1 |
| chr3 | 69643950 | 69644030 | 81 | * | NA_peak_417 | 20 | Distal Interge | 3 | 69574158 | 69598960 | 24803 | 2 | 26879 | ENSMUST00 | -44990 | ENSMUSG0( | B3galnt1 | UDP-GalNAc:betaGlcNAc beta 1,3-galactosaminyltransferase, polypeptide 1 |
| chr3 | 71446437 | 71446669 | 233 | * | NA_peak_417 | 50 | Distal Interge | 3 | 70007613 | 70028708 | 21096 | 1 | 229389 | ENSMUST00 | 1438824 | ENSMUSG0( | Otol1 | otolin 1 |
| chr3 | 73737123 | 73737389 | 267 | * | NA_peak_418 | 17 | Distal Interge | 3 | 73635808 | 73708415 | 72608 | 2 | 12038 | ENSMUST00 | -28708 | ENSMUSG0( | Bche | butyrylcholinesterase |
| chr3 | 76557035 | 76557478 | 444 | * | NA_peak_418 | 26 | Intron (ENSM | 3 | 76593550 | 76707549 | 114000 | 1 | 213262 | ENSMUST00 | -36072 | ENSMUSG0( | Fstl5 | follistatin-like 5 |
| chr3 | 77602447 | 77602783 | 337 | * | NA_peak_418 | 30 | Distal Interge | 3 | 76593550 | 76707549 | 114000 | 1 | 213262 | ENSMUST00 | 1008897 | ENSMUSG0( | Fstl5 | follistatin-like 5 |
| chr3 | 78823693 | 78824028 | 336 | * | NA_peak_418 | 19 | Distal Interge | 3 | 79062516 | 79144977 | 82462 | 2 | 76089 | ENSMUST00 | 320949 | ENSMUSG0( | Rapgef2 | Rap guanine nucleotide exchange factor (GEF) 2 |
| chr3 | 80014354 | 80014676 | 323 | * | NA_peak_418 | 38 | Distal Interge | 3 | 79885930 | 79946278 | 60349 | 1 | 68659 | ENSMUST00 | 128424 | ENSMUSG0( | Gask1b | golgi associated kinase 1B |
| chr3 | 80747273 | 80747560 | 288 | * | NA_peak_418 | 27 | Intron (ENSM | 3 | 80688040 | 80802579 | 114540 | 2 | 14800 | ENSMUST00 | 55019 | ENSMUSG0( | Gria2 | glutamate receptor, ionotropic, AMPA2 (alpha 2) |
| chr3 | 82487032 | 82487257 | 226 | * | NA_peak_418 | 27 | Distal Interge | 3 | 82539949 | 82547960 | 8012 | 2 | 18167 | ENSMUST00 | 60703 | ENSMUSG0( | Npy2r | neuropeptide Y receptor Y2 |
| chr3 | 82589373 | 82589761 | 389 | * | NA_peak_418 | 40 | Distal Interge | 3 | 82538383 | 82548084 | 9702 | 2 | 18167 | ENSMUST00 | -41289 | ENSMUSG0( | Npy2r | neuropeptide Y receptor Y2 |
| chr3 | 83295008 | 83295202 | 195 | * | NA_peak_418 | 42 | Intron (ENSM | 3 | 83125276 | 83129246 | 3971 | 2 | 100038599 | ENSMUST00 | -165762 | ENSMUSG0( | Gm10710 | predicted gene 10710 |
| chr3 | 83902674 | 83902878 | 205 | * | NA_peak_418 | 23 | Intron (ENSM | 3 | 83900332 | 83907579 | 7248 | 2 | 229473 | ENSMUST00 | 4701 | ENSMUSG0( | Tmem131l | transmembrane 131 like |
| chr3 | 86335697 | 86335991 | 295 | * | NA_peak_419 | 20 | Intron (ENSM | 3 | 86225289 | 86542711 | 317423 | 1 | 80877 | ENSMUST00 | 110408 | ENSMUSG0( | Lrba | LPS-responsive beige-like anchor |
| chr3 | 92232418 | 92232796 | 379 | * | NA_peak_420 | 33 | Intron (ENSM | 3 | 92215920 | 92254564 | 38645 | 1 | 20755 | ENSMUST00 | 16498 | ENSMUSG0( | Sprr2a1 | small proline-rich protein 2A1 |
| chr3 | 92234943 | 92235331 | 389 | * | NA_peak_420 | 15 | Intron (ENSM | 3 | 92250747 | 92256724 | 5978 | 1 | 20755 | ENSMUST00 | -15416 | ENSMUSG0( | Sprr2a1 | small proline-rich protein 2A1 |
| chr3 | 92548104 | 92548478 | 375 | * | NA_peak_420 | 76 | Distal Interge | 3 | 92570902 | 92573735 | 2834 | 2 | 16447 | ENSMUST00 | 25257 | ENSMUSG0( | Ivl | involucrin |
| chr3 | 93280219 | 93280522 | 304 | * | NA_peak_420 | 24 | Exon (ENSMU | 3 | 93319749 | 93333570 | 13822 | 1 | 68723 | ENSMUST00 | -39227 | ENSMUSG0( | Hrnr | hornerin |
| chr3 | 94310066 | 94310329 | 264 | * | NA_peak_420 | 14 | Promoter (<= | 3 | 94310089 | 94332540 | 22452 | 1 | 75778 | ENSMUST00 | 0 | ENSMUSG0( | Them4 | thioesterase superfamily member 4 |
| chr3 | 95687660 | 95687953 | 294 | * | NA_peak_420 | 16 | Promoter (<= | 3 | 95676201 | 95687917 | 11717 | 2 | 229595 | ENSMUST00 | 0 | ENSMUSG0( | Adamtsl4 | ADAMTS-like 4 |
| chr3 | 96368194 | 96368563 | 370 | * | NA_peak_421 | 56 | Distal Interge | 3 | 96348627 | 96357094 | 8468 | 2 | 102632400 | ENSMUST00 | -11100 | NA | Gm26689 | predicted gene, 26689 |
| chr3 | 96596382 | 96596644 | 263 | * | NA_peak_421 | 38 | Promoter (<= | 3 | 96596636 | 96599775 | 3140 | 1 | 545554 | ENSMUST00 | 0 | ENSMUSG0( | Ankrd34a | ankyrin repeat domain 34A |
| chr3 | 104041898 | 104042218 | 321 | * | NA_peak_422 | 16 | Intron (ENSM | 3 | 103996420 | 103997565 | 1146 | 1 | 18685 | ENSMUST00 | 45478 | ENSMUSG0( | Phtf1 | putative homeodomain transcription factor 1 |
| chr3 | 104051378 | 104051706 | 329 | * | NA_peak_422 | 20 | Exon (ENSMU | 3 | 103996420 | 103997565 | 1146 | 1 | 18685 | ENSMUST00 | 54958 | ENSMUSG0( | Phtf1 | putative homeodomain transcription factor 1 |
| chr3 | 104308067 | 104308085 | 19 | * | NA_peak_422 | 21 | Distal Interge | 3 | 104013259 | 104220374 | 207116 | 2 | 99470 | ENSMUST00 | -87693 | ENSMUSG0( | Magi3 | membrane associated guanylate kinase, WW and PDZ domain containing 3 |
| chr3 | 104838375 | 104838509 | 135 | * | NA_peak_422 | 36 | Intron (ENSM | 3 | 104839888 | 104842961 | 3074 | 2 | 12340 | ENSMUST00 | 4452 | ENSMUSG0( | Capza1 | capping protein (actin filament) muscle Z-line, alpha 1 |
| chr3 | 104924547 | 104924985 | 439 | * | NA_peak_422 | 20 | Intron (ENSM | 3 | 104888958 | 104895062 | 6105 | 1 | 229681 | ENSMUST00 | 35589 | ENSMUSG0( | St7l | suppression of tumorigenicity 7-like |

| | | | | | | | | | | | | | | | | | | |
|---|---|---|---|---|---|---|---|---|---|---|---|---|---|---|---|---|---|---|
| chr3 | 104976925 | 104977220 | 296 | * | NA_peak_42 | 22 | Distal Interge | 3 | 104945272 | 104961921 | 16650 | 2 | 22414 | ENSMUST00 | -15004 | ENSMUSG0( | Wnt2b | wingless-type MMTV integration site family, member 2B |
| chr3 | 105250181 | 105250414 | 234 | * | NA_peak_42 | 27 | Distal Interge | 3 | 105066844 | 105078806 | 11963 | 1 | 75269 | ENSMUST00 | 183337 | ENSMUSG0( | 4930564D02I | RIKEN cDNA 4930564D02 gene |
| chr3 | 105766724 | 105767034 | 311 | * | NA_peak_42 | 27 | Exon (ENSMU | 3 | 105728314 | 105750328 | 22015 | 2 | 109905 | ENSMUST00 | -16396 | ENSMUSG0( | Rap1a | RAS-related protein 1a |
| chr3 | 106204728 | 106204941 | 214 | * | NA_peak_42 | 27 | Intron (ENSM | 3 | 106201490 | 106219507 | 18018 | 2 | 104183 | ENSMUST00 | 14566 | ENSMUSG0( | Chil4 | chitinase-like 4 |
| chr3 | 106362895 | 106363196 | 302 | * | NA_peak_42 | 15 | Distal Interge | 3 | 106387384 | 106406182 | 18799 | 2 | 229688 | ENSMUST00 | 42986 | ENSMUSG0( | Chil6 | chitinase-like 6 |
| chr3 | 106779047 | 106779363 | 317 | * | NA_peak_42 | 31 | Intron (ENSM | 3 | 106759921 | 106790149 | 30229 | 2 | 12508 | ENSMUST00 | 10786 | ENSMUSG0( | Cd53 | CD53 antigen |
| chr3 | 107332619 | 107332920 | 302 | * | NA_peak_42 | 33 | Promoter (<= | 3 | 107325421 | 107333673 | 8253 | 2 | 229700 | ENSMUST00 | 753 | ENSMUSG0( | Rbm15 | RNA binding motif protein 15 |
| chr3 | 107368509 | 107368683 | 175 | * | NA_peak_42 | 25 | Distal Interge | 3 | 107325421 | 107333673 | 8253 | 2 | 229700 | ENSMUST00 | -34836 | ENSMUSG0( | Rbm15 | RNA binding motif protein 15 |
| chr3 | 107643343 | 107643729 | 387 | * | NA_peak_42 | 31 | Distal Interge | 3 | 107620452 | 107631694 | 11243 | 2 | 229707 | ENSMUST00 | -11649 | ENSMUSG0( | Strip1 | striatin interacting protein 1 |
| chr3 | 108134845 | 108135099 | 255 | * | NA_peak_42 | 40 | Intron (ENSM | 3 | 108107280 | 108146146 | 38867 | 2 | 14679 | ENSMUST00 | 11047 | ENSMUSG0( | Gnai3 | guanine nucleotide binding protein (G protein), alpha inhibiting 3 |
| chr3 | 108158309 | 108158626 | 318 | * | NA_peak_42 | 21 | Distal Interge | 3 | 108148657 | 108154882 | 6226 | 2 | 229714 | ENSMUST00 | -3427 | ENSMUSG0( | Gpr61 | G protein-coupled receptor 61 |
| chr3 | 108245882 | 108246172 | 291 | * | NA_peak_42 | 22 | Distal Interge | 3 | 108256926 | 108279974 | 23049 | 1 | 26442 | ENSMUST00 | -10754 | ENSMUSG0( | Psma5 | proteasome subunit alpha 5 |
| chr3 | 108530480 | 108530766 | 287 | * | NA_peak_42 | 16 | Intron (ENSM | 3 | 108491932 | 108536459 | 44528 | 2 | 229722 | ENSMUST00 | 5693 | ENSMUSG0( | Elapor1 | endosome-lysosome associated apoptosis and autophagy regulator 1 |
| chr3 | 108589201 | 108589525 | 325 | * | NA_peak_42 | 39 | Promoter (1-2 | 3 | 108591279 | 108645719 | 54441 | 1 | 99512 | ENSMUST00 | -1754 | ENSMUSG0( | Wdr47 | WD repeat domain 47 |
| chr3 | 109223770 | 109224134 | 365 | * | NA_peak_42 | 23 | Distal Interge | 3 | 109123162 | 109158540 | 35379 | 1 | 229731 | ENSMUST00 | 100608 | ENSMUSG0( | Slc25a24 | solute carrier family 25 (mitochondrial carrier, phosphate carrier), member 24 |
| chr3 | 109398176 | 109398396 | 221 | * | NA_peak_42 | 13 | Intron (ENSM | 3 | 109341106 | 109442212 | 101107 | 1 | 57257 | ENSMUST00 | 57070 | ENSMUSG0( | Vav3 | vav 3 oncogene |
| chr3 | 110490991 | 110491348 | 358 | * | NA_peak_42 | 53 | Distal Interge | 3 | 110248549 | 110250998 | 2450 | 2 | 99890 | ENSMUST00 | -239993 | ENSMUSG0( | Prmt6 | protein arginine N-methyltransferase 6 |
| chr3 | 112285378 | 112285662 | 285 | * | NA_peak_42 | 45 | Distal Interge | 3 | 111881903 | 112082001 | 200099 | 1 | 625585 | ENSMUST00 | 403475 | ENSMUSG0( | Gm6602 | predicted gene 6602 |
| chr3 | 112400718 | 112400742 | 25 | * | NA_peak_42 | 31 | Distal Interge | 3 | 111881903 | 112082001 | 200099 | 1 | 625585 | ENSMUST00 | 518815 | ENSMUSG0( | Gm6602 | predicted gene 6602 |
| chr3 | 113313408 | 113313547 | 140 | * | NA_peak_42 | 23 | 3' UTR | 3 | 113312455 | 113324052 | 11598 | 2 | 100043686 | ENSMUST00 | 10505 | ENSMUSG0( | Amy2a3 | amylase 2a3 |
| chr3 | 113365793 | 113366198 | 406 | * | NA_peak_42 | 23 | Distal Interge | 3 | 113345055 | 113356658 | 11604 | 2 | 100043684 | ENSMUST00 | -9135 | ENSMUSG0( | Amy2a4 | amylase 2a4 |
| chr3 | 114312126 | 114312333 | 208 | * | NA_peak_42 | 13 | Distal Interge | 3 | 114165278 | 114219317 | 54040 | 1 | 12814 | ENSMUST00 | 146848 | ENSMUSG0( | Col11a1 | collagen, type XI, alpha 1 |
| chr3 | 114944299 | 114944608 | 310 | * | NA_peak_42 | 45 | Intron (ENSM | 3 | 114904635 | 115120974 | 216340 | 1 | 229759 | ENSMUST00 | 39664 | ENSMUSG0( | Olfm3 | olfactomedin 3 |
| chr3 | 115039431 | 115039455 | 25 | * | NA_peak_42 | 16 | Intron (ENSM | 3 | 115080965 | 115125256 | 44292 | 1 | 229759 | ENSMUST00 | -41510 | ENSMUSG0( | Olfm3 | olfactomedin 3 |
| chr3 | 115916082 | 115916478 | 397 | * | NA_peak_42 | 45 | Intron (ENSM | 3 | 115888195 | 115892990 | 4796 | 1 | 69740 | ENSMUST00 | 27887 | ENSMUSG0( | Dph5 | diphthamide biosynthesis 5 |
| chr3 | 116035866 | 116036100 | 235 | * | NA_peak_42 | 47 | Distal Interge | 3 | 116008220 | 116028960 | 20741 | 1 | 58193 | ENSMUST00 | 27646 | ENSMUSG0( | Extl2 | exostosin-like glycosyltransferase 2 |
| chr3 | 116347035 | 116347156 | 122 | * | NA_peak_42 | 36 | Intron (ENSM | 3 | 116322210 | 116422649 | 100440 | 2 | 229776 | ENSMUST00 | 75493 | ENSMUSG0( | Cdc14a | CDC14 cell division cycle 14A |
| chr3 | 116683137 | 116683323 | 187 | * | NA_peak_42 | 16 | Promoter (1-2 | 3 | 116641902 | 116681260 | 39359 | 2 | 15247 | ENSMUST00 | -1877 | ENSMUSG0( | Mfsd14a | major facilitator superfamily domain containing 14A |
| chr3 | 116856268 | 116856960 | 693 | * | NA_peak_42 | 23 | Promoter (2-3 | 3 | 116859464 | 116903478 | 44015 | 1 | 20321 | ENSMUST00 | -2504 | ENSMUSG0( | Frrs1 | ferric-chelate reductase 1 |
| chr3 | 116878490 | 116878808 | 319 | * | NA_peak_42 | 16 | Promoter (<= | 3 | 116878227 | 116908177 | 29951 | 1 | 20321 | ENSMUST00 | 263 | ENSMUSG0( | Frrs1 | ferric-chelate reductase 1 |
| chr3 | 116896655 | 116896941 | 287 | * | NA_peak_42 | 23 | Promoter (<= | 3 | 116896710 | 116902438 | 5729 | 1 | 20321 | ENSMUST00 | 0 | ENSMUSG0( | Frrs1 | ferric-chelate reductase 1 |
| chr3 | 116968478 | 116968672 | 195 | * | NA_peak_42 | 22 | Promoter (<= | 3 | 116921858 | 116968858 | 47001 | 2 | 114301 | ENSMUST00 | 186 | ENSMUSG0( | Palmd | palmdelphin |
| chr3 | 117920810 | 117920874 | 65 | * | NA_peak_42 | 16 | Distal Interge | 3 | 117781647 | 117868936 | 87290 | 2 | 76561 | ENSMUST00 | -51874 | ENSMUSG0( | Snx7 | sorting nexin 7 |
| chr3 | 118716544 | 118717565 | 1022 | * | NA_peak_42 | 57 | Intron (ENSM | 3 | 118811222 | 118899831 | 88610 | 1 | 99586 | ENSMUST00 | -93657 | ENSMUSG0( | Dpyd | dihydropyrimidine dehydrogenase |
| chr3 | 120827535 | 120827926 | 392 | * | NA_peak_42 | 45 | Intron (ENSM | 3 | 120772730 | 120886691 | 113962 | 2 | 320739 | ENSMUST00 | 58765 | NA | 6530403H02I | RIKEN cDNA 6530403H02 gene |
| chr3 | 122322027 | 122322379 | 353 | * | NA_peak_42 | 22 | Intron (ENSM | 3 | 122294136 | 122427148 | 133013 | 1 | 29815 | ENSMUST00 | 27891 | ENSMUSG0( | Bcar3 | breast cancer anti-estrogen resistance 3 |
| chr3 | 122643441 | 122643877 | 437 | * | NA_peak_42 | 27 | Distal Interge | 3 | 122538725 | 122619715 | 80991 | 2 | 214459 | ENSMUST00 | -23726 | ENSMUSG0( | Fnbp1l | formin binding protein 1-like |
| chr3 | 123195005 | 123195393 | 389 | * | NA_peak_42 | 27 | Intron (ENSM | 3 | 123076519 | 123236096 | 159578 | 2 | 118449 | ENSMUST00 | 40703 | ENSMUSG0( | Synpo2 | synaptopodin 2 |
| chr3 | 129214528 | 129214881 | 354 | * | NA_peak_43 | 32 | Promoter (<= | 3 | 129214275 | 129216624 | 2350 | 1 | 18741 | ENSMUST00 | 253 | ENSMUSG0( | Pitx2 | paired-like homeodomain transcription factor 2 |
| chr3 | 129218366 | 129218518 | 153 | * | NA_peak_43 | 18 | 3' UTR | 3 | 129214275 | 129216624 | 2350 | 1 | 18741 | ENSMUST00 | 4091 | ENSMUSG0( | Pitx2 | paired-like homeodomain transcription factor 2 |
| chr3 | 129477192 | 129477390 | 199 | * | NA_peak_43 | 23 | Distal Interge | 3 | 129433673 | 129434715 | 1043 | 1 | 435755 | ENSMUST00 | 43519 | ENSMUSG0( | Pou5f1-rs4 | POU domain, class 5, transcription factor 1, related sequence 4 |
| chr3 | 129893351 | 129893496 | 146 | * | NA_peak_43 | 29 | Exon (ENSMU | 3 | 129901425 | 129914103 | 12679 | 1 | 12368 | ENSMUST00 | -7929 | ENSMUSG0( | Casp6 | caspase 6 |
| chr3 | 129969628 | 129970054 | 427 | * | NA_peak_43 | 18 | Promoter (<= | 3 | 129917080 | 129969989 | 52910 | 2 | 66815 | ENSMUST00 | 0 | ENSMUSG0( | Mcub | mitochondrial calcium uniporter dominant negative beta subunit |
| chr3 | 130739141 | 130739376 | 236 | * | NA_peak_43 | 31 | Intron (ENSM | 3 | 130726839 | 130730398 | 3560 | 2 | 68436 | ENSMUST00 | -8743 | ENSMUSG0( | Rpl34 | ribosomal protein L34 |
| chr3 | 130919065 | 130919456 | 392 | * | NA_peak_43 | 40 | Intron (ENSM | 3 | 130726839 | 130730398 | 3560 | 2 | 68436 | ENSMUST00 | -188667 | ENSMUSG0( | Rpl34 | ribosomal protein L34 |
| chr3 | 131303032 | 131303332 | 301 | * | NA_peak_43 | 22 | Promoter (<= | 3 | 131288441 | 131303224 | 14784 | 2 | 71519 | ENSMUST00 | 0 | ENSMUSG0( | Cyp2u1 | cytochrome P450, family 2, subfamily u, polypeptide 1 |
| chr3 | 132625833 | 132626285 | 453 | * | NA_peak_43 | 27 | Distal Interge | 3 | 132629820 | 132645348 | 15529 | 1 | 433653 | ENSMUST00 | -3535 | ENSMUSG0( | Gimd1 | GIMAP family P-loop NTPase domain containing 1 |
| chr3 | 132827457 | 132827767 | 311 | * | NA_peak_43 | 22 | Intron (ENSM | 3 | 132786910 | 132838506 | 51597 | 1 | 271981 | ENSMUST00 | 40547 | ENSMUSG0( | Tbck | TBC1 domain containing kinase |
| chr3 | 133124402 | 133124886 | 485 | * | NA_peak_43 | 60 | Intron (ENSM | 3 | 133116432 | 133137302 | 20871 | 2 | 77669 | ENSMUST00 | 12416 | ENSMUSG0( | Arhgef38 | Rho guanine nucleotide exchange factor (GEF) 38 |
| chr3 | 133251766 | 133252251 | 486 | * | NA_peak_43 | 31 | Distal Interge | 3 | 133172880 | 133234949 | 62070 | 2 | 77669 | ENSMUST00 | -16817 | ENSMUSG0( | Arhgef38 | Rho guanine nucleotide exchange factor (GEF) 38 |
| chr3 | 133297961 | 133298053 | 93 | * | NA_peak_43 | 27 | Distal Interge | 3 | 133310110 | 133378078 | 67969 | 1 | 74776 | ENSMUST00 | -12057 | ENSMUSG0( | Ppa2 | pyrophosphatase (inorganic) 2 |
| chr3 | 133573591 | 133573699 | 109 | * | NA_peak_43 | 36 | Intron (ENSM | 3 | 133488535 | 133545139 | 56605 | 2 | 214133 | ENSMUST00 | -28452 | ENSMUSG0( | Tet2 | tet methylcytosine dioxygenase 2 |
| chr3 | 133619209 | 133619273 | 65 | * | NA_peak_43 | 16 | Distal Interge | 3 | 133488535 | 133545139 | 56605 | 2 | 214133 | ENSMUST00 | -74070 | ENSMUSG0( | Tet2 | tet methylcytosine dioxygenase 2 |
| chr3 | 134449316 | 134449628 | 313 | * | NA_peak_43 | 36 | Distal Interge | 3 | 134404143 | 134423635 | 19493 | 2 | 75208 | ENSMUST00 | -25681 | ENSMUSG0( | 4930539C22I | RIKEN cDNA 4930539C22 gene |
| chr3 | 134549378 | 134549607 | 230 | * | NA_peak_43 | 22 | Intron (ENSM | 3 | 134404143 | 134423635 | 19493 | 2 | 75208 | ENSMUST00 | -125743 | ENSMUSG0( | 4930539C22I | RIKEN cDNA 4930539C22 gene |
| chr3 | 135296828 | 135297603 | 776 | * | NA_peak_43 | 40 | Exon (ENSMU | 3 | 135288658 | 135304423 | 15766 | 1 | 69772 | ENSMUST00 | 8170 | ENSMUSG0( | Bdh2 | 3-hydroxybutyrate dehydrogenase, type 2 |
| chr3 | 135331259 | 135331437 | 179 | * | NA_peak_43 | 17 | Promoter (<= | 3 | 135330306 | 135342767 | 12462 | 1 | 97086 | ENSMUST00 | 953 | ENSMUSG0( | Slc9b2 | solute carrier family 9, subfamily B (NHA2, cation proton antiporter 2), member 2 |
| chr3 | 135392262 | 135392338 | 77 | * | NA_peak_43 | 27 | Promoter (1-2 | 3 | 135394004 | 135397824 | 3821 | 1 | 74446 | ENSMUST00 | -1666 | ENSMUSG0( | Slc9b1 | solute carrier family 9, subfamily B (NHA1, cation proton antiporter 1), member 1 |
| chr3 | 135437821 | 135438216 | 396 | * | NA_peak_43 | 30 | Promoter (<= | 3 | 135438149 | 135465237 | 27089 | 1 | 66105 | ENSMUST00 | 0 | ENSMUSG0( | Ube2d3 | ubiquitin-conjugating enzyme E2D 3 |
| chr3 | 136971937 | 136972019 | 83 | * | NA_peak_43 | 23 | Distal Interge | 3 | 136903149 | 136905659 | 2511 | 1 | 19055 | ENSMUST00 | 68788 | ENSMUSG0( | Ppp3ca | protein phosphatase 3, catalytic subunit, alpha isoform |
| chr3 | 137603867 | 137604388 | 522 | * | NA_peak_43 | 40 | Distal Interge | 3 | 137621612 | 137625399 | 3788 | 1 | 73284 | ENSMUST00 | -17224 | ENSMUSG0( | Ddit4l | DNA-damage-inducible transcript 4-like |
| chr3 | 138143583 | 138143867 | 285 | * | NA_peak_43 | 27 | Promoter (<= | 3 | 138143799 | 138148362 | 4564 | 1 | 108943 | ENSMUST00 | 0 | ENSMUSG0( | Trmt10a | tRNA methyltransferase 10A |
| chr3 | 138585134 | 138585682 | 549 | * | NA_peak_43 | 45 | Distal Interge | 3 | 138550855 | 138553721 | 2867 | 1 | 13684 | ENSMUST00 | 34279 | ENSMUSG0( | Eif4e | eukaryotic translation initiation factor 4E |
| chr3 | 138616174 | 138616216 | 43 | * | NA_peak_43 | 20 | Distal Interge | 3 | 138550855 | 138553721 | 2867 | 1 | 13684 | ENSMUST00 | 65319 | ENSMUSG0( | Eif4e | eukaryotic translation initiation factor 4E |
| chr3 | 139420017 | 139420376 | 360 | * | NA_peak_43 | 27 | Intron (ENSM | 3 | 139205893 | 139710299 | 504407 | 1 | 381476 | ENSMUST00 | 214124 | ENSMUSG0( | Stpg2 | sperm tail PG rich repeat containing 2 |
| chr3 | 141665878 | 141666270 | 393 | * | NA_peak_43 | 34 | Intron (ENSM | 3 | 141789852 | 141791986 | 2135 | 1 | 22253 | ENSMUST00 | -123582 | ENSMUSG0( | Unc5c | unc-5 netrin receptor C |
| chr3 | 142364520 | 142364981 | 462 | * | NA_peak_43 | 19 | Intron (ENSM | 3 | 142242680 | 142391808 | 149129 | 2 | 56376 | ENSMUST00 | 26827 | ENSMUSG0( | Pdlim5 | PDZ and LIM domain 5 |
| chr3 | 143868465 | 143868518 | 54 | * | NA_peak_43 | 30 | Distal Interge | 3 | 143694189 | 143712210 | 18022 | 2 | 621809 | ENSMUST00 | -156255 | ENSMUSG0( | Gm6260 | predicted gene 6260 |
| chr3 | 144030221 | 144030525 | 305 | * | NA_peak_43 | 27 | Distal Interge | 3 | 144189062 | 144194818 | 5757 | 2 | 16911 | ENSMUST00 | 164293 | ENSMUSG0( | Lmo4 | LIM domain only 4 |
| chr3 | 144125070 | 144125354 | 285 | * | NA_peak_43 | 23 | Intron (ENSM | 3 | 144189062 | 144194818 | 5757 | 2 | 16911 | ENSMUST00 | 69464 | ENSMUSG0( | Lmo4 | LIM domain only 4 |
| chr3 | 144201682 | 144201759 | 78 | * | NA_peak_43 | 19 | Promoter (<= | 3 | 144201623 | 144202300 | 678 | 2 | 16911 | ENSMUST00 | 541 | ENSMUSG0( | Lmo4 | LIM domain only 4 |
| chr3 | 144550685 | 144550955 | 271 | * | NA_peak_43 | 23 | Intron (ENSM | 3 | 144453963 | 144569881 | 115919 | 2 | 23908 | ENSMUST00 | 18926 | ENSMUSG0( | Hs2st1 | heparan sulfate 2-O-sulfotransferase 1 |
| chr3 | 144587004 | 144587132 | 129 | * | NA_peak_43 | 25 | Intron (ENSM | 3 | 144577652 | 144595432 | 17781 | 1 | 93684 | ENSMUST00 | 9352 | ENSMUSG0( | Selenof | selenoprotein F |
| chr3 | 144654513 | 144654794 | 282 | * | NA_peak_43 | 45 | Distal Interge | 3 | 144684895 | 144697467 | 12573 | 2 | 54673 | ENSMUST00 | 42673 | ENSMUSG0( | Sh3glb1 | SH3-domain GRB2-like B1 (endophilin) |
| chr3 | 144655094 | 144655384 | 291 | * | NA_peak_43 | 45 | Distal Interge | 3 | 144684895 | 144697467 | 12573 | 2 | 54673 | ENSMUST00 | 42083 | ENSMUSG0( | Sh3glb1 | SH3-domain GRB2-like B1 (endophilin) |
| chr3 | 144720169 | 144720522 | 354 | * | NA_peak_43 | 19 | Promoter (<= | 3 | 144686911 | 144720323 | 33413 | 2 | 54673 | ENSMUST00 | 0 | ENSMUSG0( | Sh3glb1 | SH3-domain GRB2-like B1 (endophilin) |
| chr3 | 144828419 | 144828476 | 58 | * | NA_peak_43 | 26 | Intron (ENSM | 3 | 144796559 | 144819494 | 22936 | 2 | 80797 | ENSMUST00 | -8925 | ENSMUSG0( | Clca3a2 | chloride channel accessory 3A2 |
| chr3 | 144856690 | 144857004 | 315 | * | NA_peak_43 | 31 | Distal Interge | 3 | 144822623 | 144849357 | 26735 | 2 | 229927 | ENSMUST00 | -7333 | ENSMUSG0( | Clca3b | chloride channel accessory 3B |
| chr3 | 144942198 | 144942481 | 284 | * | NA_peak_43 | 27 | Distal Interge | 3 | 144910921 | 144932529 | 21609 | 2 | 99709 | ENSMUST00 | -9669 | ENSMUSG0( | Clca4b | chloride channel accessory 4B |
| chr3 | 147009961 | 147010274 | 314 | * | NA_peak_43 | 23 | Distal Interge | 3 | 146943920 | 146981920 | 38001 | 1 | 70892 | ENSMUST00 | 66041 | ENSMUSG0( | Ttll7 | tubulin tyrosine ligase-like family, member 7 |
| chr3 | 147216255 | 147216448 | 194 | * | NA_peak_43 | 27 | Distal Interge | 3 | 146943920 | 146981920 | 38001 | 1 | 70892 | ENSMUST00 | 272335 | ENSMUSG0( | Ttll7 | tubulin tyrosine ligase-like family, member 7 |
| chr3 | 147222731 | 147223072 | 342 | * | NA_peak_43 | 23 | Distal Interge | 3 | 146943920 | 146981920 | 38001 | 1 | 70892 | ENSMUST00 | 278811 | ENSMUSG0( | Ttll7 | tubulin tyrosine ligase-like family, member 7 |
| chr3 | 149052532 | 149052848 | 317 | * | NA_peak_43 | 28 | Distal Interge | 3 | 148816093 | 148990555 | 174463 | 2 | 99633 | ENSMUST00 | -61977 | ENSMUSG0( | Adgrl2 | adhesion G protein-coupled receptor L2 |
| chr3 | 150162981 | 150163435 | 455 | * | NA_peak_43 | 27 | Distal Interge | 3 | 151023792 | 151023862 | 71 | 2 | 100628568 | ENSMUST00 | 860427 | ENSMUSG0( | Mir3963 | microRNA 3963 |
| chr3 | 150163826 | 150163836 | 11 | * | NA_peak_43 | 27 | Distal Interge | 3 | 151023792 | 151023862 | 71 | 2 | 100628568 | ENSMUST00 | 860026 | ENSMUSG0( | Mir3963 | microRNA 3963 |
| chr3 | 151962232 | 151962505 | 274 | * | NA_peak_43 | 61 | Distal Interge | 3 | 151796502 | 151837630 | 41129 | 2 | 19220 | ENSMUST00 | -124602 | ENSMUSG0( | Ptgfr | prostaglandin F receptor |
| chr3 | 153115486 | 153115903 | 418 | * | NA_peak_44 | 27 | Distal Interge | 3 | 153205366 | 153225783 | 20418 | 2 | 20447 | ENSMUST00 | 109880 | ENSMUSG0( | St6galnac3 | ST6 (alpha-N-acetyl-neuraminyl-2,3-beta-galactosyl-1,3)-N-acetylgalactosaminide alpha-2,6-sialyltransferase 3 |
| chr3 | 153158568 | 153158961 | 394 | * | NA_peak_44 | 16 | Distal Interge | 3 | 153205366 | 153225783 | 20418 | 2 | 20447 | ENSMUST00 | 66822 | ENSMUSG0( | St6galnac3 | ST6 (alpha-N-acetyl-neuraminyl-2,3-beta-galactosyl-1,3)-N-acetylgalactosaminide alpha-2,6-sialyltransferase 3 |
| chr3 | 153500713 | 153500940 | 228 | * | NA_peak_44 | 30 | Intron (ENSM | 3 | 153296504 | 153298649 | 2146 | 2 | 75489 | ENSMUST00 | -202064 | ENSMUSG0( | 1700012D16I | RIKEN cDNA 1700012D16 gene |
| chr3 | 154332172 | 154332479 | 308 | * | NA_peak_44 | 27 | Promoter (1-2 | 3 | 154330821 | 154334254 | 3434 | 1 | 99686 | ENSMUST00 | 1351 | NA | AI606473 | expressed sequence AI606473 |
| chr3 | 154881118 | 154881423 | 306 | * | NA_peak_44 | 20 | Intron (ENSM | 3 | 154711865 | 154714083 | 2219 | 1 | 209601 | ENSMUST00 | 169253 | ENSMUSG0( | Erich3 | glutamate rich 3 |
| chr3 | 155149299 | 155149722 | 424 | * | NA_peak_44 | 43 | Intron (ENSM | 3 | 155093588 | 155137748 | 44161 | 1 | 74435 | ENSMUST00 | 55711 | ENSMUSG0( | Lrriq3 | leucine-rich repeats and IQ motif containing 3 |
| chr3 | 155151961 | 155152282 | 322 | * | NA_peak_44 | 27 | Intron (ENSM | 3 | 155093588 | 155137748 | 44161 | 1 | 74435 | ENSMUST00 | 58373 | ENSMUSG0( | Lrriq3 | leucine-rich repeats and IQ motif containing 3 |
| chr3 | 156261386 | 156261815 | 430 | * | NA_peak_44 | 39 | Distal Interge | 3 | 155976354 | 156008592 | 32239 | 1 | 329780 | ENSMUST00 | 285032 | ENSMUSG0( | 9330178D15I | RIKEN cDNA 9330178D15 gene |
| chr3 | 157572240 | 157572521 | 282 | * | NA_peak_44 | 31 | Intron (ENSM | 3 | 157566900 | 157644756 | 77857 | 1 | 19218 | ENSMUST00 | 5340 | ENSMUSG0( | Ptger3 | prostaglandin E receptor 3 (subtype EP3) |
| chr3 | 157647048 | 157647357 | 310 | * | NA_peak_44 | 56 | Distal Interge | 3 | 157566900 | 157644756 | 77857 | 1 | 19218 | ENSMUST00 | 80148 | ENSMUSG0( | Ptger3 | prostaglandin E receptor 3 (subtype EP3) |
| chr3 | 158573949 | 158574217 | 269 | * | NA_peak_44 | 33 | Distal Interge | 3 | 158084439 | 158562221 | 477783 | 2 | 242274 | ENSMUST00 | -11728 | ENSMUSG0( | Lrrc7 | leucine rich repeat containing 7 |
| chr3 | 158628921 | 158629178 | 258 | * | NA_peak_44 | 56 | Distal Interge | 3 | 158084439 | 158562221 | 477783 | 2 | 242274 | ENSMUST00 | -66700 | ENSMUSG0( | Lrrc7 | leucine rich repeat containing 7 |
| chr3 | 159339580 | 159339871 | 292 | * | NA_peak_44 | 16 | Distal Interge | 3 | 159495433 | 159529955 | 34523 | 1 | 76131 | ENSMUST00 | -155562 | ENSMUSG0( | Depdc1a | DEP domain containing 1a |
| chr3 | 159642006 | 159642700 | 695 | * | NA_peak_44 | 36 | Distal Interge | 3 | 159599233 | 159624799 | 25567 | 1 | 19892 | ENSMUST00 | 42773 | ENSMUSG0( | Rpe65 | retinal pigment epithelium 65 |
| chr3 | 159697379 | 159697792 | 414 | * | NA_peak_44 | 34 | Distal Interge | 3 | 159599233 | 159624799 | 25567 | 1 | 19892 | ENSMUST00 | 98146 | ENSMUSG0( | Rpe65 | retinal pigment epithelium 65 |
| chr3 | 159816398 | 159816673 | 276 | * | NA_peak_44 | 22 | Distal Interge | 3 | 159839672 | 159936023 | 96352 | 1 | 68151 | ENSMUST00 | -22999 | ENSMUSG0( | Wls | wntless WNT ligand secretion mediator |
| chr4 | 3086580 | 3086896 | 317 | * | NA_peak_44 | 25 | Distal Interge | 4 | 3172083 | 3173003 | 921 | 1 | 100312470 | ENSMUST00 | -85187 | ENSMUSG0( | Vmn1r2 | vomeronasal 1 receptor 2 |
| chr4 | 3091001 | 3091492 | 492 | * | NA_peak_44 | 81 | Distal Interge | 4 | 3172083 | 3173003 | 921 | 1 | 100312470 | ENSMUST00 | -80591 | ENSMUSG0( | Vmn1r2 | vomeronasal 1 receptor 2 |
| chr4 | 3224679 | 3225200 | 522 | * | NA_peak_44 | 41 | Distal Interge | 4 | 3184348 | 3185358 | 1011 | 2 | 100312471 | ENSMUST00 | -39321 | ENSMUSG0( | Vmn1r3 | vomeronasal 1 receptor 3 |
| chr4 | 3239597 | 3239923 | 327 | * | NA_peak_44 | 17 | Distal Interge | 4 | 3184348 | 3185358 | 1011 | 2 | 100312471 | ENSMUST00 | -54239 | ENSMUSG0( | Vmn1r3 | vomeronasal 1 receptor 3 |

| | | | | | | | | | | | | | | | | | | |
|---|---|---|---|---|---|---|---|---|---|---|---|---|---|---|---|---|---|---|
| chr4 | 3249758 | 3249892 | 135 | * | NA_peak_443 | 23 | Intron (ENSM | 4 | 3184348 | 3185358 | 1011 | 2 | 100312471 | ENSMUST00 | -64400 | ENSMUSG00 | Vmn1r3 | vomeronasal 1 receptor 3 |
| chr4 | 11181175 | 11181726 | 552 | * | NA_peak_444 | 48 | Distal Interge | 4 | 11191351 | 11193961 | 2611 | 1 | 12448 | ENSMUST00 | -9625 | ENSMUSG00 | Ccne2 | cyclin E2 |
| chr4 | 12848117 | 12848411 | 295 | * | NA_peak_444 | 61 | Distal Interge | 4 | 12906838 | 12980873 | 74036 | 1 | 208820 | ENSMUST00 | -58427 | ENSMUSG00 | Triqk | triple QxxK/R motif containing |
| chr4 | 14641177 | 14641745 | 569 | * | NA_peak_444 | 52 | Intron (ENSM | 4 | 14502430 | 14621805 | 119376 | 2 | 208890 | ENSMUST00 | -19372 | ENSMUSG00 | Slc26a7 | solute carrier family 26, member 7 |
| chr4 | 14730202 | 14730203 | 2 | * | NA_peak_444 | 23 | Intron (ENSM | 4 | 14665754 | 14756845 | 91092 | 2 | 73314 | ENSMUST00 | 26642 | ENSMUSG00 | Lrrc69 | leucine rich repeat containing 69 |
| chr4 | 15048346 | 15048696 | 351 | * | NA_peak_444 | 20 | Intron (ENSM | 4 | 15110962 | 15148961 | 38000 | 2 | 69352 | ENSMUST00 | 100265 | ENSMUSG00 | Necab1 | N-terminal EF-hand calcium binding protein 1 |
| chr4 | 15055971 | 15056266 | 296 | * | NA_peak_444 | 40 | Intron (ENSM | 4 | 15110962 | 15148961 | 38000 | 2 | 69352 | ENSMUST00 | 92695 | ENSMUSG00 | Necab1 | N-terminal EF-hand calcium binding protein 1 |
| chr4 | 15241962 | 15242316 | 355 | * | NA_peak_444 | 31 | Distal Interge | 4 | 15265831 | 15286753 | 20923 | 1 | 100201 | ENSMUST00 | -23515 | ENSMUSG00 | Tmem64 | transmembrane protein 64 |
| chr4 | 17450072 | 17450369 | 298 | * | NA_peak_444 | 36 | Distal Interge | 4 | 17852893 | 18117457 | 264565 | 1 | 17389 | ENSMUST00 | -402524 | ENSMUSG00 | Mmp16 | matrix metallopeptidase 16 |
| chr4 | 20291615 | 20291879 | 265 | * | NA_peak_444 | 31 | Intron (ENSM | 4 | 20042052 | 20066750 | 24699 | 1 | 14590 | ENSMUST00 | 249563 | ENSMUSG00 | Ggh | gamma-glutamyl hydrolase |
| chr4 | 22540779 | 22541353 | 575 | * | NA_peak_445 | 45 | Distal Interge | 4 | 22490548 | 22493126 | 2579 | 1 | 105500046 | ENSMUST00 | 50231 | ENSMUSG00 | Gm30731 | predicted gene, 30731 |
| chr4 | 22853653 | 22854030 | 378 | * | NA_peak_445 | 23 | Distal Interge | 4 | 22490548 | 22493126 | 2579 | 1 | 105500046 | ENSMUST00 | 363105 | ENSMUSG00 | Gm30731 | predicted gene, 30731 |
| chr4 | 23545243 | 23545512 | 270 | * | NA_peak_445 | 23 | Distal Interge | 4 | 24496451 | 24581338 | 84888 | 1 | 212377 | ENSMUST00 | -950939 | ENSMUSG00 | Mms22l | MMS22-like, DNA repair protein |
| chr4 | 25186669 | 25187148 | 480 | * | NA_peak_445 | 41 | Intron (ENSM | 4 | 25199908 | 25242876 | 42969 | 2 | 57756 | ENSMUST00 | 55728 | ENSMUSG00 | Fhl5 | four and a half LIM domains 5 |
| chr4 | 25729059 | 25729364 | 306 | * | NA_peak_445 | 45 | Intron (ENSM | 4 | 25797703 | 25799855 | 2153 | 2 | 14348 | ENSMUST00 | 70491 | ENSMUSG00 | Fut9 | fucosyltransferase 9 |
| chr4 | 28467126 | 28467379 | 254 | * | NA_peak_445 | 20 | Distal Interge | 4 | 28813131 | 28947047 | 133917 | 1 | 13841 | ENSMUST00 | -345752 | ENSMUSG00 | Epha7 | Eph receptor A7 |
| chr4 | 29714657 | 29714943 | 287 | * | NA_peak_445 | 20 | Distal Interge | 4 | 29068079 | 29088811 | 20733 | 1 | 102634169 | ENSMUST00 | 646578 | NA | Gm11917 | predicted gene 11917 |
| chr4 | 31528003 | 31528476 | 474 | * | NA_peak_445 | 22 | Distal Interge | 4 | 31964097 | 32023467 | 59371 | 1 | 26409 | ENSMUST00 | -435621 | ENSMUSG00 | Map3k7 | mitogen-activated protein kinase kinase kinase 7 |
| chr4 | 34343418 | 34343691 | 274 | * | NA_peak_445 | 36 | Distal Interge | 4 | 34550937 | 34566908 | 15972 | 1 | 433693 | ENSMUST00 | -207246 | ENSMUSG00 | Akirin2 | akirin 2 |
| chr4 | 34504427 | 34504726 | 300 | * | NA_peak_445 | 23 | Distal Interge | 4 | 34550937 | 34566908 | 15972 | 1 | 433693 | ENSMUST00 | -46211 | ENSMUSG00 | Akirin2 | akirin 2 |
| chr4 | 34540588 | 34541122 | 535 | * | NA_peak_445 | 55 | Distal Interge | 4 | 34550937 | 34566908 | 15972 | 1 | 433693 | ENSMUST00 | -9815 | ENSMUSG00 | Akirin2 | akirin 2 |
| chr4 | 35047370 | 35047585 | 216 | * | NA_peak_446 | 36 | Intron (ENSM | 4 | 35152056 | 35154005 | 1950 | 1 | 387510 | ENSMUST00 | -104471 | ENSMUSG00 | Ifnk | interferon kappa |
| chr4 | 35071420 | 35071429 | 10 | * | NA_peak_446 | 30 | Intron (ENSM | 4 | 35152056 | 35154005 | 1950 | 1 | 387510 | ENSMUST00 | -80627 | ENSMUSG00 | Ifnk | interferon kappa |
| chr4 | 35426357 | 35426768 | 412 | * | NA_peak_446 | 36 | Distal Interge | 4 | 35212423 | 35226175 | 13753 | 2 | 73205 | ENSMUST00 | -200182 | ENSMUSG00 | C9orf72 | C9orf72, member of C9orf72-SMCR8 complex |
| chr4 | 36611526 | 36611845 | 320 | * | NA_peak_446 | 142 | Intron (ENSM | 4 | 36645373 | 36645453 | 81 | 2 | 100124458 | ENSMUST00 | 33608 | ENSMUSG00 | Mir876 | microRNA 876 |
| chr4 | 38319522 | 38319833 | 312 | * | NA_peak_446 | 23 | Distal Interge | 4 | 38571667 | 38855483 | 283817 | 2 | 102636968 | ENSMUST00 | 535650 | ENSMUSG00 | Gm12381 | predicted gene 12381 |
| chr4 | 40595579 | 40595807 | 229 | * | NA_peak_446 | 42 | Distal Interge | 4 | 40482840 | 40525548 | 42709 | 2 | 71293 | ENSMUST00 | -70031 | ENSMUSG00 | 4933428C19 | RIKEN cDNA 4933428C19 gene |
| chr4 | 44063422 | 44063780 | 359 | * | NA_peak_446 | 23 | Promoter (<= | 4 | 44055305 | 44063173 | 7869 | 2 | 50798 | ENSMUST00 | -249 | ENSMUSG00 | Gne | glucosamine (UDP-N-acetyl)-2-epimerase/N-acetylmannosamine kinase |
| chr4 | 49561772 | 49562119 | 348 | * | NA_peak_447 | 45 | Distal Interge | 4 | 49535995 | 49549546 | 13552 | 2 | 230163 | ENSMUST00 | -12226 | ENSMUSG00 | Aldob | aldolase B, fructose-bisphosphate |
| chr4 | 51650147 | 51650459 | 313 | * | NA_peak_447 | 50 | Distal Interge | 4 | 51968094 | 52051110 | 83017 | 2 | 329844 | ENSMUST00 | 400651 | ENSMUSG00 | C630028M04 | RIKEN cDNA C630028M04 gene |
| chr4 | 51654123 | 51654597 | 475 | * | NA_peak_447 | 62 | Distal Interge | 4 | 51968094 | 52051110 | 83017 | 2 | 329844 | ENSMUST00 | 396513 | ENSMUSG00 | C630028M04 | RIKEN cDNA C630028M04 gene |
| chr4 | 52190186 | 52190638 | 453 | * | NA_peak_447 | 100 | Distal Interge | 4 | 51968094 | 52051110 | 83017 | 2 | 329844 | ENSMUST00 | -139076 | ENSMUSG00 | C630028M04 | RIKEN cDNA C630028M04 gene |
| chr4 | 53946221 | 53946702 | 482 | * | NA_peak_448 | 27 | Distal Interge | 4 | 53840470 | 53854332 | 13863 | 1 | 52076 | ENSMUST00 | 105751 | ENSMUSG00 | Tmem38b | transmembrane protein 38B |
| chr4 | 56865121 | 56865473 | 353 | * | NA_peak_448 | 30 | Promoter (<= | 4 | 56810935 | 56865188 | 54254 | 2 | 54366 | ENSMUST00 | 0 | ENSMUSG00 | Ctnnal1 | catenin (cadherin associated protein), alpha-like 1 |
| chr4 | 61571761 | 61572016 | 256 | * | NA_peak_448 | 47 | Distal Interge | 4 | 61591929 | 61595871 | 3943 | 2 | 100039206 | ENSMUST00 | 23855 | ENSMUSG00 | Mup17 | major urinary protein 17 |
| chr4 | 61681563 | 61681908 | 346 | * | NA_peak_448 | 46 | Distal Interge | 4 | 61670177 | 61674136 | 3960 | 2 | 100048884 | ENSMUST00 | -7427 | ENSMUSG00 | Mup18 | major urinary protein 18 |
| chr4 | 62523193 | 62523653 | 461 | * | NA_peak_448 | 22 | Promoter (1-2 | 4 | 62524335 | 62525038 | 704 | 2 | 59001 | ENSMUST00 | 1385 | ENSMUSG00 | Pole3 | polymerase (DNA directed), epsilon 3 (p17 subunit) |
| chr4 | 64288708 | 64288908 | 201 | * | NA_peak_449 | 66 | Distal Interge | 4 | 64254053 | 64276603 | 22551 | 2 | 102633436 | ENSMUST00 | -12105 | ENSMUSG00 | Gm11217 | predicted gene 11217 |
| chr4 | 67940491 | 67940869 | 379 | * | NA_peak_449 | 107 | Distal Interge | 4 | 68312657 | 68321961 | 9305 | 2 | 102633975 | ENSMUST00 | 381092 | ENSMUSG00 | Gm12911 | predicted gene 12911 |
| chr4 | 68697794 | 68698076 | 283 | * | NA_peak_449 | 40 | Distal Interge | 4 | 68826581 | 68829358 | 2778 | 2 | 56710 | ENSMUST00 | 131282 | ENSMUSG00 | Brinp1 | bone morphogenic protein/retinoic acid inducible neural specific 1 |
| chr4 | 69711821 | 69712116 | 296 | * | NA_peak_449 | 70 | Distal Interge | 4 | 70242471 | 70243852 | 1382 | 2 | 214444 | ENSMUST00 | 531736 | ENSMUSG00 | Cdk5rap2 | CDK5 regulatory subunit associated protein 2 |
| chr4 | 70595026 | 70595157 | 132 | * | NA_peak_449 | 20 | Distal Interge | 4 | 70461826 | 70534995 | 73170 | 2 | 230316 | ENSMUST00 | -60031 | ENSMUSG00 | Megf9 | multiple EGF-like-domains 9 |
| chr4 | 71436972 | 71437254 | 283 | * | NA_peak_449 | 23 | Distal Interge | 4 | 72118220 | 72120734 | 2515 | 2 | 21885 | ENSMUST00 | 683480 | ENSMUSG00 | Tle1 | transducin-like enhancer of split 1 |
| chr4 | 71965225 | 71965324 | 100 | * | NA_peak_449 | 30 | Distal Interge | 4 | 72118220 | 72120734 | 2515 | 2 | 21885 | ENSMUST00 | 155410 | ENSMUSG00 | Tle1 | transducin-like enhancer of split 1 |
| chr4 | 72077845 | 72078099 | 255 | * | NA_peak_449 | 27 | Distal Interge | 4 | 72118220 | 72120734 | 2515 | 2 | 21885 | ENSMUST00 | 42635 | ENSMUSG00 | Tle1 | transducin-like enhancer of split 1 |
| chr4 | 72408589 | 72408756 | 168 | * | NA_peak_449 | 30 | Distal Interge | 4 | 72117142 | 72200919 | 83778 | 2 | 21885 | ENSMUST00 | -207670 | ENSMUSG00 | Tle1 | transducin-like enhancer of split 1 |
| chr4 | 74502627 | 74502998 | 372 | * | NA_peak_450 | 20 | Distal Interge | 4 | 74262810 | 74273659 | 10850 | 1 | 76804 | ENSMUST00 | 239817 | ENSMUSG00 | Kdm4c | lysine (K)-specific demethylase 4C |
| chr4 | 75072851 | 75073141 | 291 | * | NA_peak_450 | 31 | Distal Interge | 4 | 75277354 | 75278305 | 952 | 2 | 66928 | ENSMUST00 | 205164 | ENSMUSG00 | Dmac1 | distal membrane arm assembly complex 1 |
| chr4 | 75509992 | 75510271 | 280 | * | NA_peak_450 | 61 | Distal Interge | 4 | 75277354 | 75278305 | 952 | 2 | 66928 | ENSMUST00 | -231687 | ENSMUSG00 | Dmac1 | distal membrane arm assembly complex 1 |
| chr4 | 76213031 | 76213311 | 281 | * | NA_peak_450 | 26 | Intron (ENSM | 4 | 76133158 | 76134143 | 986 | 2 | 19266 | ENSMUST00 | -78888 | ENSMUSG00 | Ptprd | protein tyrosine phosphatase, receptor type, D |
| chr4 | 78478387 | 78478575 | 189 | * | NA_peak_450 | 20 | Distal Interge | 4 | 77351290 | 78211961 | 860672 | 2 | 19266 | ENSMUST00 | -266426 | ENSMUSG00 | Ptprd | protein tyrosine phosphatase, receptor type, D |
| chr4 | 78513198 | 78513623 | 426 | * | NA_peak_450 | 54 | Distal Interge | 4 | 77351290 | 78211961 | 860672 | 2 | 19266 | ENSMUST00 | -301237 | ENSMUSG00 | Ptprd | protein tyrosine phosphatase, receptor type, D |
| chr4 | 79260378 | 79260471 | 94 | * | NA_peak_450 | 13 | Distal Interge | 4 | 77351290 | 78211961 | 860672 | 2 | 19266 | ENSMUST00 | -1048417 | ENSMUSG00 | Ptprd | protein tyrosine phosphatase, receptor type, D |
| chr4 | 80190543 | 80190769 | 227 | * | NA_peak_450 | 28 | Distal Interge | 4 | 80834123 | 80837684 | 3562 | 1 | 22178 | ENSMUST00 | -643354 | ENSMUSG00 | Tyrp1 | tyrosinase-related protein 1 |
| chr4 | 80807693 | 80808182 | 490 | * | NA_peak_450 | 14 | Distal Interge | 4 | 80834123 | 80837684 | 3562 | 1 | 22178 | ENSMUST00 | -25941 | ENSMUSG00 | Tyrp1 | tyrosinase-related protein 1 |
| chr4 | 81703689 | 81704563 | 875 | * | NA_peak_450 | 45 | Distal Interge | 4 | 81416540 | 81442815 | 26276 | 2 | 17475 | ENSMUST00 | -260874 | ENSMUSG00 | Mpdz | multiple PDZ domain crumbs cell polarity complex component |
| chr4 | 82193167 | 82193428 | 262 | * | NA_peak_450 | 26 | Distal Interge | 4 | 82065957 | 82102673 | 36717 | 1 | 545641 | ENSMUST00 | 127210 | ENSMUSG00 | Gm5860 | predicted gene 5860 |
| chr4 | 83749010 | 83749442 | 433 | * | NA_peak_450 | 27 | Intron (ENSM | 4 | 83695955 | 83798676 | 102722 | 1 | 320226 | ENSMUST00 | 53055 | ENSMUSG00 | Ccdc171 | coiled-coil domain containing 171 |
| chr4 | 88056063 | 88056390 | 328 | * | NA_peak_451 | 30 | Distal Interge | 4 | 87769925 | 88033364 | 263440 | 2 | 70122 | ENSMUST00 | -22699 | ENSMUSG00 | Mllt3 | myeloid/lymphoid or mixed-lineage leukemia; translocated to, 3 |
| chr4 | 89791509 | 89791808 | 300 | * | NA_peak_451 | 20 | Distal Interge | 4 | 89688198 | 89694772 | 6575 | 1 | 242523 | ENSMUST00 | 103311 | ENSMUSG00 | Dmrta1 | doublesex and mab-3 related transcription factor like family A1 |
| chr4 | 90975322 | 90975610 | 289 | * | NA_peak_451 | 16 | Distal Interge | 4 | 91253080 | 91254582 | 1503 | 2 | 15569 | ENSMUST00 | 278972 | ENSMUSG00 | Elavl2 | ELAV like RNA binding protein 1 |
| chr4 | 93704186 | 93704543 | 358 | * | NA_peak_451 | 24 | Distal Interge | 4 | 93334138 | 93335511 | 1374 | 2 | 69136 | ENSMUST00 | -368675 | ENSMUSG00 | Tusc1 | tumor suppressor candidate 1 |
| chr4 | 99732222 | 99732504 | 283 | * | NA_peak_451 | 36 | Intron (ENSM | 4 | 99719152 | 99739473 | 20322 | 1 | 320438 | ENSMUST00 | 13070 | ENSMUSG00 | Alg6 | asparagine-linked glycosylation 6 (alpha-1,3,-glucosyltransferase) |
| chr4 | 103278525 | 103278813 | 289 | * | NA_peak_452 | 37 | Intron (ENSM | 4 | 103230445 | 103290863 | 60419 | 2 | 70941 | ENSMUST00 | 12050 | ENSMUSG00 | 4921539E11 | RIKEN cDNA 4921539E11 gene |
| chr4 | 111572567 | 111573078 | 512 | * | NA_peak_453 | 55 | Intron (ENSM | 4 | 111566725 | 111657507 | 90783 | 1 | 78933 | ENSMUST00 | 5842 | ENSMUSG00 | Agbl4 | ATP/GTP binding protein-like 4 |
| chr4 | 116276356 | 116277063 | 708 | * | NA_peak_453 | 43 | Intron (ENSM | 4 | 116308008 | 116309205 | 1198 | 2 | 17776 | ENSMUST00 | 32142 | ENSMUSG00 | Mast2 | microtubule associated serine/threonine kinase 2 |
| chr4 | 120033344 | 120033393 | 50 | * | NA_peak_453 | 20 | Promoter (<= | 4 | 120032766 | 120065063 | 32298 | 1 | 16656 | ENSMUST00 | 578 | ENSMUSG00 | Hivep3 | human immunodeficiency virus type I enhancer binding protein 3 |
| chr4 | 120267264 | 120267578 | 315 | * | NA_peak_453 | 27 | 3' UTR | 4 | 120267079 | 120287349 | 20271 | 2 | 329934 | ENSMUST00 | 19771 | ENSMUSG00 | Foxo6 | forkhead box O6 |
| chr4 | 131896175 | 131896622 | 448 | * | NA_peak_455 | 52 | Intron (ENSM | 4 | 131885208 | 131901706 | 16499 | 1 | 57317 | ENSMUST00 | 10967 | ENSMUSG00 | Srsf4 | serine and arginine-rich splicing factor 4 |
| chr4 | 132048504 | 132048852 | 349 | * | NA_peak_455 | 45 | Promoter (<= | 4 | 131923429 | 132049078 | 125650 | 2 | 269587 | ENSMUST00 | 226 | ENSMUSG00 | Epb41 | erythrocyte membrane protein band 4.1 |
| chr4 | 133063805 | 133064043 | 239 | * | NA_peak_455 | 30 | Promoter (2-3 | 4 | 133066249 | 133072063 | 5815 | 1 | 230793 | ENSMUST00 | -2206 | ENSMUSG00 | Ahdc1 | AT hook, DNA binding motif, containing 1 |
| chr4 | 135037914 | 135038227 | 314 | * | NA_peak_455 | 22 | Distal Interge | 4 | 135120652 | 135177990 | 57339 | 1 | 12399 | ENSMUST00 | -82425 | ENSMUSG00 | Runx3 | runt related transcription factor 3 |
| chr4 | 135458789 | 135458918 | 130 | * | NA_peak_455 | 36 | Intron (ENSM | 4 | 135452267 | 135455335 | 3069 | 2 | 74552 | ENSMUST00 | -3454 | ENSMUSG00 | Nipal3 | NIPA-like domain containing 3 |
| chr4 | 138331173 | 138331356 | 184 | * | NA_peak_456 | 27 | Distal Interge | 4 | 138313412 | 138326307 | 12896 | 2 | 68943 | ENSMUST00 | -4866 | ENSMUSG00 | Pink1 | PTEN induced putative kinase 1 |
| chr4 | 138634260 | 138634571 | 312 | * | NA_peak_456 | 13 | Promoter (1-2 | 4 | 138581630 | 138635884 | 54255 | 2 | 75718 | ENSMUST00 | 1313 | ENSMUSG00 | Vwa5b1 | von Willebrand factor A domain containing 5B1 |
| chr4 | 139574965 | 139575240 | 276 | * | NA_peak_456 | 23 | Promoter (<= | 4 | 139574720 | 139620382 | 45663 | 1 | 212632 | ENSMUST00 | 245 | ENSMUSG00 | Iffo2 | intermediate filament family orphan 2 |
| chr4 | 139957781 | 139958026 | 246 | * | NA_peak_456 | 36 | Distal Interge | 4 | 139967526 | 139967620 | 95 | 2 | 100316727 | ENSMUST00 | 9594 | ENSMUSG00 | Mir2139 | microRNA 2139 |
| chr4 | 141967859 | 141968119 | 261 | * | NA_peak_456 | 16 | Intron (ENSM | 4 | 141932783 | 141954194 | 21412 | 2 | 329977 | ENSMUST00 | -13665 | ENSMUSG00 | Fhad1 | forkhead-associated (FHA) phosphopeptide binding domain 1 |
| chr4 | 142160746 | 142160798 | 53 | * | NA_peak_456 | 27 | Promoter (1-2 | 4 | 142118233 | 142159054 | 40822 | 2 | 71529 | ENSMUST00 | -1692 | ENSMUSG00 | Kazn | kazrin, periplakin interacting protein |
| chr4 | 143533811 | 143534137 | 327 | * | NA_peak_456 | 23 | Promoter (1-2 | 4 | 143535954 | 143542471 | 6518 | 1 | 100040861 | ENSMUST00 | -1817 | ENSMUSG00 | Gm13057 | predicted gene 13057 |
| chr4 | 143645873 | 143646170 | 298 | * | NA_peak_456 | 22 | Distal Interge | 4 | 143653760 | 143657246 | 3487 | 2 | 277668 | ENSMUST00 | 11076 | ENSMUSG00 | Pramel22 | PRAME like 22 |
| chr4 | 145522129 | 145522262 | 134 | * | NA_peak_457 | 36 | Intron (ENSM | 4 | 145514884 | 145539188 | 24305 | 1 | 101056073 | ENSMUST00 | 7245 | ENSMUSG00 | Zfp990 | zinc finger protein 990 |
| chr4 | 145526169 | 145526601 | 433 | * | NA_peak_457 | 21 | Intron (ENSM | 4 | 145514884 | 145539188 | 24305 | 1 | 101056073 | ENSMUST00 | 11285 | ENSMUSG00 | Zfp990 | zinc finger protein 990 |
| chr4 | 145537497 | 145537859 | 363 | * | NA_peak_457 | 43 | Exon (ENSMU | 4 | 145514884 | 145539188 | 24305 | 1 | 101056073 | ENSMUST00 | 22613 | ENSMUSG00 | Zfp990 | zinc finger protein 990 |
| chr4 | 145566967 | 145567282 | 316 | * | NA_peak_457 | 15 | Distal Interge | 4 | 145585166 | 145625345 | 40180 | 1 | 433801 | ENSMUST00 | -17884 | ENSMUSG00 | Zfp268 | zinc finger protein 268 |
| chr4 | 145604765 | 145605002 | 238 | * | NA_peak_457 | 31 | Intron (ENSM | 4 | 145617146 | 145623215 | 6070 | 1 | 433801 | ENSMUST00 | -12144 | ENSMUSG00 | Zfp268 | zinc finger protein 268 |
| chr4 | 145628961 | 145629074 | 114 | * | NA_peak_457 | 30 | Distal Interge | 4 | 145617146 | 145623215 | 6070 | 1 | 433801 | ENSMUST00 | 11815 | ENSMUSG00 | Zfp268 | zinc finger protein 268 |
| chr4 | 145663996 | 145664359 | 364 | * | NA_peak_457 | 45 | Distal Interge | 4 | 145670685 | 145704441 | 33757 | 1 | 100041379 | ENSMUST00 | -6326 | ENSMUSG00 | Zfp980 | zinc finger protein 980 |
| chr4 | 145665468 | 145665964 | 497 | * | NA_peak_457 | 19 | Distal Interge | 4 | 145670685 | 145704441 | 33757 | 1 | 100041379 | ENSMUST00 | -4721 | ENSMUSG00 | Zfp980 | zinc finger protein 980 |
| chr4 | 145684715 | 145685381 | 667 | * | NA_peak_457 | 49 | Intron (ENSM | 4 | 145696161 | 145702640 | 6480 | 1 | 100041379 | ENSMUST00 | -10780 | ENSMUSG00 | Zfp980 | zinc finger protein 980 |
| chr4 | 145687113 | 145687688 | 576 | * | NA_peak_457 | 33 | Intron (ENSM | 4 | 145696161 | 145702640 | 6480 | 1 | 100041379 | ENSMUST00 | -8473 | ENSMUSG00 | Zfp980 | zinc finger protein 980 |
| chr4 | 145688225 | 145688607 | 383 | * | NA_peak_457 | 53 | Intron (ENSM | 4 | 145696161 | 145702640 | 6480 | 1 | 100041379 | ENSMUST00 | -7554 | ENSMUSG00 | Zfp980 | zinc finger protein 980 |
| chr4 | 145699583 | 145699802 | 220 | * | NA_peak_457 | 22 | Intron (ENSM | 4 | 145696161 | 145702640 | 6480 | 1 | 100041379 | ENSMUST00 | 3422 | ENSMUSG00 | Zfp980 | zinc finger protein 980 |
| chr4 | 145704670 | 145704937 | 268 | * | NA_peak_457 | 53 | Downstream | 4 | 145696161 | 145702640 | 6480 | 1 | 100041379 | ENSMUST00 | 8509 | ENSMUSG00 | Zfp980 | zinc finger protein 980 |
| chr4 | 145744966 | 145745167 | 202 | * | NA_peak_457 | 33 | Distal Interge | 4 | 145696161 | 145702640 | 6480 | 1 | 100041379 | ENSMUST00 | 48805 | ENSMUSG00 | Zfp980 | zinc finger protein 980 |
| chr4 | 145764917 | 145765196 | 280 | * | NA_peak_457 | 38 | Exon (ENSMU | 4 | 145825237 | 145829942 | 4706 | 1 | 70005 | ENSMUST00 | -60041 | NA | Znf41-ps | ZNF41, pseudogene |
| chr4 | 145767812 | 145768496 | 685 | * | NA_peak_457 | 55 | Distal Interge | 4 | 145825237 | 145829942 | 4706 | 1 | 70005 | ENSMUST00 | -56741 | NA | Znf41-ps | ZNF41, pseudogene |
| chr4 | 145815235 | 145815561 | 327 | * | NA_peak_457 | 30 | Distal Interge | 4 | 145825237 | 145829942 | 4706 | 1 | 70005 | ENSMUST00 | -9676 | NA | Znf41-ps | ZNF41, pseudogene |
| chr4 | 145821305 | 145821668 | 364 | * | NA_peak_457 | 41 | Distal Interge | 4 | 145825237 | 145829942 | 4706 | 1 | 70005 | ENSMUST00 | -3569 | NA | Znf41-ps | ZNF41, pseudogene |
| chr4 | 145875582 | 145875966 | 385 | * | NA_peak_457 | 15 | Intron (ENSM | 4 | 145837219 | 145837998 | 780 | 1 | 236069 | ENSMUST00 | 38363 | ENSMUSG00 | Gm13238 | carnitine deficiency-associated gene expressed in ventricle 3 pseudogene |
| chr4 | 146167295 | 146167584 | 290 | * | NA_peak_457 | 69 | Exon (ENSMU | 4 | 146161909 | 146195169 | 33261 | 1 | 667666 | ENSMUST00 | 5386 | ENSMUSG00 | Zfp600 | zinc finger protein 600 |
| chr4 | 146177954 | 146178529 | 576 | * | NA_peak_457 | 32 | Exon (ENSMU | 4 | 146161909 | 146195169 | 33261 | 1 | 667666 | ENSMUST00 | 16045 | ENSMUSG00 | Zfp600 | zinc finger protein 600 |
| chr4 | 146192839 | 146193248 | 410 | * | NA_peak_457 | 45 | Intron (ENSM | 4 | 146161909 | 146195169 | 33261 | 1 | 667666 | ENSMUST00 | 30930 | ENSMUSG00 | Zfp600 | zinc finger protein 600 |
| chr4 | 146202332 | 146202667 | 336 | * | NA_peak_457 | 52 | Intron (ENSM | 4 | 146161909 | 146195169 | 33261 | 1 | 667666 | ENSMUST00 | 40423 | ENSMUSG00 | Zfp600 | zinc finger protein 600 |
| chr4 | 146226291 | 146226784 | 494 | * | NA_peak_457 | 44 | Distal Interge | 4 | 146252173 | 146272975 | 20803 | 1 | 666532 | ENSMUST00 | -25389 | ENSMUSG00 | Zfp991 | zinc finger protein 991 |
| chr4 | 146259780 | 146260075 | 296 | * | NA_peak_457 | 30 | Intron (ENSM | 4 | 146252173 | 146272975 | 20803 | 1 | 666532 | ENSMUST00 | 7607 | ENSMUSG00 | Zfp991 | zinc finger protein 991 |

| | | | | | | | | | | | | | | | | | | |
|---|---|---|---|---|---|---|---|---|---|---|---|---|---|---|---|---|---|---|
| chr4 | 146264630 | 146265023 | 394 | * | NA_peak_45 | 39 | Exon (ENSMU | 4 | 146270331 | 146278378 | 8048 | 1 | 666532 | ENSMUST00 | -5308 | ENSMUSG0 | Zfp991 | zinc finger protein 991 |
| chr4 | 146274696 | 146275020 | 325 | * | NA_peak_45 | 16 | Intron (ENSM | 4 | 146270372 | 146278434 | 8063 | 1 | 666532 | ENSMUST00 | 4324 | ENSMUSG0 | Zfp991 | zinc finger protein 991 |
| chr4 | 146280685 | 146280911 | 227 | * | NA_peak_45 | 82 | Distal Interge | 4 | 146270372 | 146278434 | 8063 | 1 | 666532 | ENSMUST00 | 10313 | ENSMUSG0 | Zfp991 | zinc finger protein 991 |
| chr4 | 146425463 | 146425728 | 266 | * | NA_peak_45 | 16 | Distal Interge | 4 | 146449023 | 146470292 | 21270 | 1 | 433791 | ENSMUST00 | -23295 | ENSMUSG0 | Zfp992 | zinc finger protein 992 |
| chr4 | 146475503 | 146475811 | 309 | * | NA_peak_45 | 35 | Exon (ENSMU | 4 | 146502027 | 146539392 | 37366 | 1 | 100041433 | ENSMUST00 | -26216 | ENSMUSG0 | Zfp981 | zinc finger protein 981 |
| chr4 | 146478560 | 146478861 | 302 | * | NA_peak_45 | 35 | Distal Interge | 4 | 146502027 | 146539392 | 37366 | 1 | 100041433 | ENSMUST00 | -23166 | ENSMUSG0 | Zfp981 | zinc finger protein 981 |
| chr4 | 146483151 | 146483568 | 418 | * | NA_peak_45 | 28 | Distal Interge | 4 | 146502027 | 146539392 | 37366 | 1 | 100041433 | ENSMUST00 | -18459 | ENSMUSG0 | Zfp981 | zinc finger protein 981 |
| chr4 | 146493601 | 146494177 | 577 | * | NA_peak_45 | 25 | Distal Interge | 4 | 146502027 | 146539392 | 37366 | 1 | 100041433 | ENSMUST00 | -7850 | ENSMUSG0 | Zfp981 | zinc finger protein 981 |
| chr4 | 146501460 | 146501533 | 74 | * | NA_peak_45 | 60 | Promoter (<= | 4 | 146502027 | 146539392 | 37366 | 1 | 100041433 | ENSMUST00 | -494 | ENSMUSG0 | Zfp981 | zinc finger protein 981 |
| chr4 | 146515167 | 146515175 | 9 | * | NA_peak_45 | 21 | Promoter (<= | 4 | 146514920 | 146539395 | 24476 | 1 | 100041433 | ENSMUST00 | 247 | ENSMUSG0 | Zfp981 | zinc finger protein 981 |
| chr4 | 146537464 | 146537777 | 314 | * | NA_peak_45 | 18 | Exon (ENSMU | 4 | 146519039 | 146546873 | 27835 | 2 | 67524 | ENSMUST00 | 9096 | ENSMUSG0 | 1700095A21 | RIKEN cDNA 1700095A21 gene |
| chr4 | 146539626 | 146540084 | 459 | * | NA_peak_45 | 33 | Intron (ENSM | 4 | 146519039 | 146546873 | 27835 | 2 | 67524 | ENSMUST00 | 6789 | ENSMUSG0 | 1700095A21 | RIKEN cDNA 1700095A21 gene |
| chr4 | 146571730 | 146572042 | 313 | * | NA_peak_45 | 26 | Distal Interge | 4 | 146519039 | 146546873 | 27835 | 2 | 67524 | ENSMUST00 | -24857 | ENSMUSG0 | 1700095A21 | RIKEN cDNA 1700095A21 gene |
| chr4 | 146593280 | 146593586 | 307 | * | NA_peak_45 | 27 | Distal Interge | 4 | 146519039 | 146546873 | 27835 | 2 | 67524 | ENSMUST00 | -46407 | ENSMUSG0 | 1700095A21 | RIKEN cDNA 1700095A21 gene |
| chr4 | 146607615 | 146608081 | 467 | * | NA_peak_45 | 40 | Distal Interge | 4 | 146519039 | 146546873 | 27835 | 2 | 67524 | ENSMUST00 | -60742 | ENSMUSG0 | 1700095A21 | RIKEN cDNA 1700095A21 gene |
| chr4 | 146609220 | 146609577 | 358 | * | NA_peak_45 | 40 | Distal Interge | 4 | 146519039 | 146546873 | 27835 | 2 | 67524 | ENSMUST00 | -62347 | ENSMUSG0 | 1700095A21 | RIKEN cDNA 1700095A21 gene |
| chr4 | 146651849 | 146652139 | 291 | * | NA_peak_45 | 41 | Intron (ENSM | 4 | 146519039 | 146546873 | 27835 | 2 | 67524 | ENSMUST00 | -104976 | ENSMUSG0 | 1700095A21 | RIKEN cDNA 1700095A21 gene |
| chr4 | 146710501 | 146710914 | 414 | * | NA_peak_45 | 66 | Distal Interge | 4 | 146519039 | 146546873 | 27835 | 2 | 67524 | ENSMUST00 | -163628 | ENSMUSG0 | 1700095A21 | RIKEN cDNA 1700095A21 gene |
| chr4 | 146716740 | 146717019 | 280 | * | NA_peak_45 | 95 | Distal Interge | 4 | 146519039 | 146546873 | 27835 | 2 | 67524 | ENSMUST00 | -169867 | ENSMUSG0 | 1700095A21 | RIKEN cDNA 1700095A21 gene |
| chr4 | 146891807 | 146892116 | 310 | * | NA_peak_45 | 77 | Exon (ENSMU | 4 | 147021850 | 147060794 | 38945 | 1 | 100043034 | ENSMUST00 | -129734 | ENSMUSG0 | Rex2 | reduced expression 2 |
| chr4 | 146952996 | 146953275 | 280 | * | NA_peak_45 | 34 | Distal Interge | 4 | 147021850 | 147060794 | 38945 | 1 | 100043034 | ENSMUST00 | -68575 | ENSMUSG0 | Rex2 | reduced expression 2 |
| chr4 | 147019448 | 147019876 | 429 | * | NA_peak_45 | 21 | Promoter (1-2 | 4 | 147021850 | 147060794 | 38945 | 1 | 100043034 | ENSMUST00 | -1974 | ENSMUSG0 | Rex2 | reduced expression 2 |
| chr4 | 147022839 | 147023131 | 293 | * | NA_peak_45 | 25 | Promoter (<= | 4 | 147021878 | 147023811 | 1934 | 1 | 100043034 | ENSMUST00 | 961 | ENSMUSG0 | Rex2 | reduced expression 2 |
| chr4 | 147024922 | 147025263 | 342 | * | NA_peak_45 | 13 | Intron (ENSM | 4 | 147021878 | 147023811 | 1934 | 1 | 100043034 | ENSMUST00 | 3044 | ENSMUSG0 | Rex2 | reduced expression 2 |
| chr4 | 147033510 | 147033946 | 437 | * | NA_peak_45 | 39 | Intron (ENSM | 4 | 147021878 | 147023811 | 1934 | 1 | 100043034 | ENSMUST00 | 11632 | ENSMUSG0 | Rex2 | reduced expression 2 |
| chr4 | 147042588 | 147042909 | 322 | * | NA_peak_45 | 13 | Intron (ENSM | 4 | 147021878 | 147023811 | 1934 | 1 | 100043034 | ENSMUST00 | 20710 | ENSMUSG0 | Rex2 | reduced expression 2 |
| chr4 | 147043710 | 147043997 | 288 | * | NA_peak_45 | 20 | Intron (ENSM | 4 | 147021878 | 147023811 | 1934 | 1 | 100043034 | ENSMUST00 | 21832 | ENSMUSG0 | Rex2 | reduced expression 2 |
| chr4 | 147053120 | 147053327 | 208 | * | NA_peak_45 | 36 | Intron (ENSM | 4 | 147063366 | 147068436 | 5071 | 2 | 71270 | ENSMUST00 | 15109 | ENSMUSG0 | 4933438K21 | RIKEN cDNA 4933438K21 gene |
| chr4 | 147112037 | 147112106 | 70 | * | NA_peak_45 | 27 | Distal Interge | 4 | 147063366 | 147068436 | 5071 | 2 | 71270 | ENSMUST00 | -43601 | ENSMUSG0 | 4933438K21 | RIKEN cDNA 4933438K21 gene |
| chr4 | 147125792 | 147126048 | 257 | * | NA_peak_45 | 22 | Exon (ENSMU | 4 | 147063366 | 147068436 | 5071 | 2 | 71270 | ENSMUST00 | -57356 | ENSMUSG0 | 4933438K21 | RIKEN cDNA 4933438K21 gene |
| chr4 | 147196642 | 147196946 | 305 | * | NA_peak_45 | 50 | Distal Interge | 4 | 147063366 | 147068436 | 5071 | 2 | 71270 | ENSMUST00 | -128206 | ENSMUSG0 | 4933438K21 | RIKEN cDNA 4933438K21 gene |
| chr4 | 147217551 | 147218209 | 659 | * | NA_peak_45 | 34 | Distal Interge | 4 | 147063366 | 147068436 | 5071 | 2 | 71270 | ENSMUST00 | -149115 | ENSMUSG0 | 4933438K21 | RIKEN cDNA 4933438K21 gene |
| chr4 | 147248001 | 147248538 | 538 | * | NA_peak_45 | 40 | Intron (ENSM | 4 | 147063366 | 147068436 | 5071 | 2 | 71270 | ENSMUST00 | -179565 | ENSMUSG0 | 4933438K21 | RIKEN cDNA 4933438K21 gene |
| chr4 | 147249816 | 147249842 | 27 | * | NA_peak_45 | 18 | Intron (ENSM | 4 | 147063366 | 147068436 | 5071 | 2 | 71270 | ENSMUST00 | -181380 | ENSMUSG0 | 4933438K21 | RIKEN cDNA 4933438K21 gene |
| chr4 | 147295094 | 147295382 | 289 | * | NA_peak_45 | 22 | Distal Interge | 4 | 147492417 | 147513486 | 21070 | 1 | 195531 | ENSMUST00 | -197035 | ENSMUSG0 | Zfp982 | zinc finger protein 982 |
| chr4 | 147312695 | 147312983 | 289 | * | NA_peak_45 | 19 | Intron (ENSM | 4 | 147492417 | 147513486 | 21070 | 1 | 195531 | ENSMUST00 | -179434 | ENSMUSG0 | Zfp982 | zinc finger protein 982 |
| chr4 | 147319147 | 147319534 | 388 | * | NA_peak_45 | 24 | Intron (ENSM | 4 | 147492417 | 147513486 | 21070 | 1 | 195531 | ENSMUST00 | -172883 | ENSMUSG0 | Zfp982 | zinc finger protein 982 |
| chr4 | 147354694 | 147354900 | 207 | * | NA_peak_45 | 16 | Distal Interge | 4 | 147492417 | 147513486 | 21070 | 1 | 195531 | ENSMUST00 | -137517 | ENSMUSG0 | Zfp982 | zinc finger protein 982 |
| chr4 | 147367515 | 147367772 | 258 | * | NA_peak_45 | 37 | Intron (ENSM | 4 | 147492417 | 147513486 | 21070 | 1 | 195531 | ENSMUST00 | -124645 | ENSMUSG0 | Zfp982 | zinc finger protein 982 |
| chr4 | 147370248 | 147370484 | 237 | * | NA_peak_45 | 51 | Intron (ENSM | 4 | 147492417 | 147513486 | 21070 | 1 | 195531 | ENSMUST00 | -121933 | ENSMUSG0 | Zfp982 | zinc finger protein 982 |
| chr4 | 147401946 | 147402298 | 353 | * | NA_peak_45 | 18 | Distal Interge | 4 | 147492417 | 147513486 | 21070 | 1 | 195531 | ENSMUST00 | -90119 | ENSMUSG0 | Zfp982 | zinc finger protein 982 |
| chr4 | 147438314 | 147439164 | 851 | * | NA_peak_45 | 60 | Distal Interge | 4 | 147492417 | 147513486 | 21070 | 1 | 195531 | ENSMUST00 | -53253 | ENSMUSG0 | Zfp982 | zinc finger protein 982 |
| chr4 | 147452578 | 147452774 | 197 | * | NA_peak_45 | 25 | Distal Interge | 4 | 147492417 | 147513486 | 21070 | 1 | 195531 | ENSMUST00 | -39643 | ENSMUSG0 | Zfp982 | zinc finger protein 982 |
| chr4 | 147455452 | 147455879 | 428 | * | NA_peak_45 | 18 | Distal Interge | 4 | 147492417 | 147513486 | 21070 | 1 | 195531 | ENSMUST00 | -36538 | ENSMUSG0 | Zfp982 | zinc finger protein 982 |
| chr4 | 147497075 | 147497545 | 471 | * | NA_peak_45 | 29 | Intron (ENSM | 4 | 147492417 | 147513486 | 21070 | 1 | 195531 | ENSMUST00 | 4658 | ENSMUSG0 | Zfp982 | zinc finger protein 982 |
| chr4 | 147497834 | 147498171 | 338 | * | NA_peak_45 | 29 | Intron (ENSM | 4 | 147492417 | 147513486 | 21070 | 1 | 195531 | ENSMUST00 | 5417 | ENSMUSG0 | Zfp982 | zinc finger protein 982 |
| chr4 | 147507759 | 147508097 | 339 | * | NA_peak_45 | 45 | Promoter (<= | 4 | 147507600 | 147513420 | 5821 | 1 | 195531 | ENSMUST00 | 159 | ENSMUSG0 | Zfp982 | zinc finger protein 982 |
| chr4 | 147518355 | 147518641 | 287 | * | NA_peak_45 | 53 | Distal Interge | 4 | 147507600 | 147513420 | 5821 | 1 | 195531 | ENSMUST00 | 10755 | ENSMUSG0 | Zfp982 | zinc finger protein 982 |
| chr4 | 147551490 | 147551740 | 251 | * | NA_peak_45 | 27 | Promoter (1-2 | 4 | 147553277 | 147583126 | 29850 | 1 | 433804 | ENSMUST00 | -1537 | ENSMUSG0 | Zfp985 | zinc finger protein 985 |
| chr4 | 147688116 | 147688532 | 417 | * | NA_peak_45 | 40 | Intron (ENSM | 4 | 147673502 | 147702553 | 29052 | 2 | 100503584 | ENSMUST00 | 14021 | ENSMUSG0 | Zfp534 | zinc finger protein 534 |
| chr4 | 147707483 | 147707790 | 308 | * | NA_peak_45 | 21 | Distal Interge | 4 | 147673502 | 147702553 | 29052 | 2 | 100503584 | ENSMUST00 | -4930 | ENSMUSG0 | Zfp534 | zinc finger protein 534 |
| chr4 | 147726934 | 147727224 | 291 | * | NA_peak_45 | 19 | Distal Interge | 4 | 147673502 | 147702553 | 29052 | 2 | 100503584 | ENSMUST00 | -24381 | ENSMUSG0 | Zfp534 | zinc finger protein 534 |
| chr4 | 147817983 | 147818260 | 278 | * | NA_peak_45 | 28 | Distal Interge | 4 | 147821777 | 147822202 | 426 | 2 | 277692 | ENSMUST00 | 3942 | ENSMUSG0 | Rps19-ps3 | ribosomal protein S19, pseudogene 3 |
| chr4 | 148139799 | 148140150 | 352 | * | NA_peak_46 | 17 | Promoter (<= | 4 | 148140182 | 148145121 | 4940 | 1 | 71890 | ENSMUST00 | -32 | ENSMUSG0 | Mad2l2 | MAD2 mitotic arrest deficient-like 2 |
| chr4 | 149631998 | 149632220 | 223 | * | NA_peak_46 | 16 | Exon (ENSMU | 4 | 149627130 | 149632173 | 5044 | 1 | 65945 | ENSMUST00 | 4868 | ENSMUSG0 | Clstn1 | calsyntenin 1 |
| chr4 | 155681263 | 155681557 | 295 | * | NA_peak_46 | 50 | Distal Interge | 4 | 155654682 | 155669198 | 14517 | 2 | 76580 | ENSMUST00 | -12065 | ENSMUSG0 | Mib2 | mindbomb E3 ubiquitin protein ligase 2 |
| chr4 | 155705342 | 155705520 | 179 | * | NA_peak_46 | 27 | Promoter (<= | 4 | 155705135 | 155732189 | 27055 | 1 | 68991 | ENSMUST00 | 207 | ENSMUSG0 | Ssu72 | Ssu72 RNA polymerase II CTD phosphatase homolog (yeast) |
| chr4 | 156307341 | 156307674 | 334 | * | NA_peak_46 | 31 | Distal Interge | 4 | 156331103 | 156339496 | 8394 | 1 | 22314 | ENSMUST00 | -23429 | ENSMUSG0 | Vmn2r129 | vomeronasal 2, receptor 129 |
| chr5 | 3012534 | 3013301 | 768 | * | NA_peak_46 | 26 | Distal Interge | 5 | 3341485 | 3344453 | 2969 | 1 | 12571 | ENSMUST00 | -328184 | ENSMUSG0 | Cdk6 | cyclin-dependent kinase 6 |
| chr5 | 3092419 | 3092700 | 282 | * | NA_peak_46 | 36 | Intron (ENSM | 5 | 3341485 | 3344453 | 2969 | 1 | 12571 | ENSMUST00 | -248785 | ENSMUSG0 | Cdk6 | cyclin-dependent kinase 6 |
| chr5 | 3987860 | 3988088 | 229 | * | NA_peak_46 | 32 | Intron (ENSM | 5 | 3975616 | 3976531 | 916 | 1 | 100986 | ENSMUST00 | 12244 | ENSMUSG0 | Akap9 | A kinase (PRKA) anchor protein (yotiao) 9 |
| chr5 | 4039798 | 4040020 | 223 | * | NA_peak_46 | 27 | Intron (ENSM | 5 | 4023415 | 4032904 | 9490 | 1 | 100986 | ENSMUST00 | 16383 | ENSMUSG0 | Akap9 | A kinase (PRKA) anchor protein (yotiao) 9 |
| chr5 | 4178789 | 4178938 | 150 | * | NA_peak_46 | 16 | Distal Interge | 5 | 4192367 | 4197651 | 5285 | 1 | 208595 | ENSMUST00 | -13429 | ENSMUSG0 | Mterf1b | mitochondrial transcription termination factor 1b |
| chr5 | 4240254 | 4240569 | 316 | * | NA_peak_46 | 40 | Distal Interge | 5 | 4192367 | 4197651 | 5285 | 1 | 208595 | ENSMUST00 | 47887 | ENSMUSG0 | Mterf1b | mitochondrial transcription termination factor 1b |
| chr5 | 4314317 | 4314625 | 309 | * | NA_peak_46 | 23 | Distal Interge | 5 | 4192367 | 4197651 | 5285 | 1 | 208595 | ENSMUST00 | 121950 | ENSMUSG0 | Mterf1b | mitochondrial transcription termination factor 1b |
| chr5 | 4376869 | 4377119 | 251 | * | NA_peak_46 | 23 | Distal Interge | 5 | 4192367 | 4197651 | 5285 | 1 | 208595 | ENSMUST00 | 184502 | ENSMUSG0 | Mterf1b | mitochondrial transcription termination factor 1b |
| chr5 | 4602248 | 4602987 | 740 | * | NA_peak_46 | 36 | Distal Interge | 5 | 4753839 | 4758035 | 4197 | 2 | 14362 | ENSMUST00 | 155048 | ENSMUSG0 | Fzd1 | frizzled class receptor 1 |
| chr5 | 4938255 | 4938727 | 473 | * | NA_peak_46 | 40 | Intron (ENSM | 5 | 4931690 | 5036516 | 104827 | 2 | 18647 | ENSMUST00 | 97789 | ENSMUSG0 | Cdk14 | cyclin-dependent kinase 14 |
| chr5 | 4956320 | 4956392 | 73 | * | NA_peak_46 | 16 | Intron (ENSM | 5 | 4931690 | 5036516 | 104827 | 2 | 18647 | ENSMUST00 | 80124 | ENSMUSG0 | Cdk14 | cyclin-dependent kinase 14 |
| chr5 | 5100248 | 5100564 | 317 | * | NA_peak_46 | 20 | Intron (ENSM | 5 | 4931690 | 5036516 | 104827 | 2 | 18647 | ENSMUST00 | -63732 | ENSMUSG0 | Cdk14 | cyclin-dependent kinase 14 |
| chr5 | 5713583 | 5713750 | 168 | * | NA_peak_46 | 20 | Distal Interge | 5 | 5664829 | 5694578 | 29750 | 2 | 74051 | ENSMUST00 | -19005 | ENSMUSG0 | Steap2 | six transmembrane epithelial antigen of prostate 2 |
| chr5 | 6085405 | 6085709 | 305 | * | NA_peak_46 | 20 | Distal Interge | 5 | 5736317 | 5749326 | 13010 | 2 | 70358 | ENSMUST00 | -336079 | ENSMUSG0 | Steap1 | six transmembrane epithelial antigen of the prostate 1 |
| chr5 | 6564164 | 6564461 | 298 | * | NA_peak_46 | 27 | Distal Interge | 5 | 6769030 | 6876523 | 107494 | 2 | 207618 | ENSMUST00 | 312062 | ENSMUSG0 | Zfp804b | zinc finger protein 804B |
| chr5 | 6737805 | 6738070 | 266 | * | NA_peak_46 | 16 | Distal Interge | 5 | 6769030 | 6876523 | 107494 | 2 | 207618 | ENSMUST00 | 138453 | ENSMUSG0 | Zfp804b | zinc finger protein 804B |
| chr5 | 7541950 | 7542249 | 300 | * | NA_peak_46 | 16 | Distal Interge | 5 | 6769010 | 7344756 | 575747 | 2 | 207618 | ENSMUST00 | -197194 | ENSMUSG0 | Zfp804b | zinc finger protein 804B |
| chr5 | 8019904 | 8020190 | 287 | * | NA_peak_46 | 27 | Distal Interge | 5 | 8046078 | 8069379 | 23302 | 1 | 109552 | ENSMUST00 | -25888 | ENSMUSG0 | Sri | sorcin |
| chr5 | 8711097 | 8711362 | 266 | * | NA_peak_46 | 16 | Intron (ENSM | 5 | 8719819 | 8748570 | 28752 | 1 | 18671 | ENSMUST00 | -8457 | ENSMUSG0 | Abcb1a | ATP-binding cassette, sub-family B (MDR/TAP), member 1A |
| chr5 | 9307761 | 9308224 | 464 | * | NA_peak_46 | 22 | Intron (ENSM | 5 | 9266175 | 9461486 | 195312 | 1 | 231014 | ENSMUST00 | 41586 | ENSMUSG0 | Elapor2 | endosome-lysosome associated apoptosis and autophagy regulator family member 2 |
| chr5 | 11307416 | 11307738 | 323 | * | NA_peak_46 | 31 | Distal Interge | 5 | 11341516 | 11346286 | 4771 | 1 | 70896 | ENSMUST00 | -33778 | ENSMUSG0 | Speer1 | spermatogenesis associated glutamate (E)-rich protein 1 |
| chr5 | 12519904 | 12520190 | 287 | * | NA_peak_46 | 31 | Intron (ENSM | 5 | 12484955 | 12508522 | 23568 | 1 | 108151 | ENSMUST00 | 34949 | ENSMUSG0 | Sema3d | sema domain, immunoglobulin domain (Ig), short basic domain, secreted, (semaphorin) 3D |
| chr5 | 13664282 | 13664612 | 331 | * | NA_peak_46 | 45 | Distal Interge | 5 | 13791619 | 13796820 | 5202 | 1 | 71026 | ENSMUST00 | -127007 | ENSMUSG0 | Speer3 | spermatogenesis associated glutamate (E)-rich protein 3 |
| chr5 | 13698368 | 13698739 | 372 | * | NA_peak_46 | 14 | Distal Interge | 5 | 13791619 | 13796820 | 5202 | 1 | 71026 | ENSMUST00 | -92880 | ENSMUSG0 | Speer3 | spermatogenesis associated glutamate (E)-rich protein 3 |
| chr5 | 13885812 | 13886143 | 332 | * | NA_peak_46 | 50 | Distal Interge | 5 | 13791619 | 13796820 | 5202 | 1 | 71026 | ENSMUST00 | 94193 | ENSMUSG0 | Speer3 | spermatogenesis associated glutamate (E)-rich protein 3 |
| chr5 | 14906327 | 14906742 | 416 | * | NA_peak_46 | 60 | Distal Interge | 5 | 14913397 | 14914853 | 1457 | 2 | 381714 | ENSMUST00 | 8111 | ENSMUSG0 | Gm9758 | predicted gene 9758 |
| chr5 | 14910549 | 14911038 | 490 | * | NA_peak_46 | 22 | Exon (ENSMU | 5 | 14913397 | 14914853 | 1457 | 2 | 381714 | ENSMUST00 | 3815 | ENSMUSG0 | Gm9758 | predicted gene 9758 |
| chr5 | 14917919 | 14918209 | 291 | * | NA_peak_46 | 53 | Distal Interge | 5 | 14910122 | 14914899 | 4778 | 2 | 381714 | ENSMUST00 | -3020 | ENSMUSG0 | Gm9758 | predicted gene 9758 |
| chr5 | 14933913 | 14934002 | 90 | * | NA_peak_46 | 39 | 3' UTR | 5 | 14933221 | 14938429 | 5209 | 2 | 624245 | ENSMUST00 | 4427 | ENSMUSG0 | Speer4e | spermatogenesis associated glutamate (E)-rich protein 4e |
| chr5 | 14942085 | 14942413 | 329 | * | NA_peak_46 | 118 | Distal Interge | 5 | 14933221 | 14938429 | 5209 | 2 | 624245 | ENSMUST00 | -3656 | ENSMUSG0 | Speer4e | spermatogenesis associated glutamate (E)-rich protein 4e |
| chr5 | 14944082 | 14944434 | 353 | * | NA_peak_46 | 24 | Distal Interge | 5 | 14933221 | 14938429 | 5209 | 2 | 624245 | ENSMUST00 | -5653 | ENSMUSG0 | Speer4e | spermatogenesis associated glutamate (E)-rich protein 4e |
| chr5 | 14946878 | 14947409 | 532 | * | NA_peak_46 | 58 | Distal Interge | 5 | 14933221 | 14938429 | 5209 | 2 | 624245 | ENSMUST00 | -8449 | ENSMUSG0 | Speer4e | spermatogenesis associated glutamate (E)-rich protein 4e |
| chr5 | 14948820 | 14949335 | 516 | * | NA_peak_46 | 36 | Distal Interge | 5 | 14933221 | 14938429 | 5209 | 2 | 624245 | ENSMUST00 | -10391 | ENSMUSG0 | Speer4e | spermatogenesis associated glutamate (E)-rich protein 4e |
| chr5 | 14955189 | 14955430 | 242 | * | NA_peak_46 | 34 | Distal Interge | 5 | 14933221 | 14938429 | 5209 | 2 | 624245 | ENSMUST00 | -16760 | ENSMUSG0 | Speer4e | spermatogenesis associated glutamate (E)-rich protein 4e |
| chr5 | 14957134 | 14957436 | 303 | * | NA_peak_46 | 92 | Distal Interge | 5 | 14933221 | 14938429 | 5209 | 2 | 624245 | ENSMUST00 | -18705 | ENSMUSG0 | Speer4e | spermatogenesis associated glutamate (E)-rich protein 4e |
| chr5 | 14961559 | 14961817 | 259 | * | NA_peak_46 | 72 | Distal Interge | 5 | 14974113 | 14978935 | 4823 | 2 | 100503549 | ENSMUST00 | 17118 | ENSMUSG0 | Gm10354 | predicted gene 10354 |
| chr5 | 14962132 | 14962583 | 452 | * | NA_peak_46 | 72 | Distal Interge | 5 | 14974113 | 14978935 | 4823 | 2 | 100503549 | ENSMUST00 | 16352 | ENSMUSG0 | Gm10354 | predicted gene 10354 |
| chr5 | 14973476 | 14973858 | 383 | * | NA_peak_46 | 16 | Distal Interge | 5 | 14974113 | 14978935 | 4823 | 2 | 100503549 | ENSMUST00 | 5077 | ENSMUSG0 | Gm10354 | predicted gene 10354 |
| chr5 | 14978100 | 14978477 | 378 | * | NA_peak_46 | 18 | Promoter (<= | 5 | 14974113 | 14978935 | 4823 | 2 | 100503549 | ENSMUST00 | 458 | ENSMUSG0 | Gm10354 | predicted gene 10354 |
| chr5 | 14980473 | 14980705 | 233 | * | NA_peak_46 | 16 | Promoter (1-2 | 5 | 14974113 | 14978935 | 4823 | 2 | 100503549 | ENSMUST00 | -1538 | ENSMUSG0 | Gm10354 | predicted gene 10354 |
| chr5 | 14987891 | 14988249 | 359 | * | NA_peak_46 | 47 | Distal Interge | 5 | 14974113 | 14978935 | 4823 | 2 | 100503549 | ENSMUST00 | -8956 | ENSMUSG0 | Gm10354 | predicted gene 10354 |
| chr5 | 14994209 | 14994680 | 472 | * | NA_peak_46 | 94 | Distal Interge | 5 | 14974113 | 14978935 | 4823 | 2 | 100503549 | ENSMUST00 | -15274 | ENSMUSG0 | Gm10354 | predicted gene 10354 |
| chr5 | 15019436 | 15019831 | 396 | * | NA_peak_46 | 76 | Distal Interge | 5 | 15028950 | 15032998 | 4049 | 2 | 66773 | ENSMUST00 | 13167 | ENSMUSG0 | Gm17019 | predicted gene 17019 |
| chr5 | 15034929 | 15035339 | 411 | * | NA_peak_46 | 30 | Promoter (1-2 | 5 | 15028950 | 15032998 | 4049 | 2 | 66773 | ENSMUST00 | -1931 | ENSMUSG0 | Gm17019 | predicted gene 17019 |
| chr5 | 15037635 | 15037899 | 265 | * | NA_peak_46 | 37 | Distal Interge | 5 | 15028950 | 15032998 | 4049 | 2 | 66773 | ENSMUST00 | -4637 | ENSMUSG0 | Gm17019 | predicted gene 17019 |
| chr5 | 15038717 | 15038993 | 277 | * | NA_peak_46 | 35 | Distal Interge | 5 | 15028950 | 15032998 | 4049 | 2 | 66773 | ENSMUST00 | -5719 | ENSMUSG0 | Gm17019 | predicted gene 17019 |
| chr5 | 15042561 | 15043211 | 651 | * | NA_peak_46 | 53 | Distal Interge | 5 | 15028950 | 15032998 | 4049 | 2 | 66773 | ENSMUST00 | -9563 | ENSMUSG0 | Gm17019 | predicted gene 17019 |
| chr5 | 15060894 | 15061129 | 236 | * | NA_peak_46 | 41 | Distal Interge | 5 | 15028950 | 15032998 | 4049 | 2 | 66773 | ENSMUST00 | -27896 | ENSMUSG0 | Gm17019 | predicted gene 17019 |

| | | | | | | | | | | | | | | | | | | |
|---|---|---|---|---|---|---|---|---|---|---|---|---|---|---|---|---|---|---|
| chr5 | 15462632 | 15463162 | 531 | * | NA_peak_465 | 27 | Distal Interge | 5 | 15524456 | 15529222 | 4767 | 2 | 100861755 | ENSMUST00 | 66060 | ENSMUSG0( | Gm21190 | predicted gene, 21190 |
| chr5 | 15470110 | 15470567 | 458 | * | NA_peak_465 | 65 | Exon (ENSMU | 5 | 15524456 | 15529222 | 4767 | 2 | 100861755 | ENSMUST00 | 58655 | ENSMUSG0( | Gm21190 | predicted gene, 21190 |
| chr5 | 15473075 | 15473667 | 593 | * | NA_peak_465 | 33 | Exon (ENSMU | 5 | 15524456 | 15529222 | 4767 | 2 | 100861755 | ENSMUST00 | 55555 | ENSMUSG0( | Gm21190 | predicted gene, 21190 |
| chr5 | 15480068 | 15480597 | 530 | * | NA_peak_465 | 62 | Distal Interge | 5 | 15524456 | 15529222 | 4767 | 2 | 100861755 | ENSMUST00 | 48625 | ENSMUSG0( | Gm21190 | predicted gene, 21190 |
| chr5 | 15483492 | 15483950 | 459 | * | NA_peak_465 | 45 | Distal Interge | 5 | 15524456 | 15529222 | 4767 | 2 | 100861755 | ENSMUST00 | 45272 | ENSMUSG0( | Gm21190 | predicted gene, 21190 |
| chr5 | 15511839 | 15512314 | 476 | * | NA_peak_465 | 52 | Distal Interge | 5 | 15524456 | 15529222 | 4767 | 2 | 100861755 | ENSMUST00 | 16908 | ENSMUSG0( | Gm21190 | predicted gene, 21190 |
| chr5 | 15516737 | 15517024 | 288 | * | NA_peak_465 | 58 | Intron (ENSM | 5 | 15524456 | 15529222 | 4767 | 2 | 100861755 | ENSMUST00 | 12198 | ENSMUSG0( | Gm21190 | predicted gene, 21190 |
| chr5 | 15543830 | 15544218 | 389 | * | NA_peak_466 | 23 | Intron (ENSM | 5 | 15524456 | 15529222 | 4767 | 2 | 100861755 | ENSMUST00 | -14608 | ENSMUSG0( | Gm21190 | predicted gene, 21190 |
| chr5 | 15560902 | 15561207 | 306 | * | NA_peak_466 | 16 | Intron (ENSM | 5 | 15577333 | 15582078 | 4746 | 1 | 105242399 | ENSMUST00 | -16126 | ENSMUSG0( | Gm21083 | predicted gene, 21083 |
| chr5 | 15575345 | 15576058 | 714 | * | NA_peak_466 | 85 | Promoter (1-2 | 5 | 15577333 | 15582078 | 4746 | 1 | 105242399 | ENSMUST00 | -1275 | ENSMUSG0( | Gm21083 | predicted gene, 21083 |
| chr5 | 15596346 | 15596727 | 382 | * | NA_peak_466 | 56 | Intron (ENSM | 5 | 15577333 | 15582078 | 4746 | 1 | 105242399 | ENSMUST00 | 19013 | ENSMUSG0( | Gm21083 | predicted gene, 21083 |
| chr5 | 15597910 | 15598354 | 445 | * | NA_peak_466 | 41 | Intron (ENSM | 5 | 15577333 | 15582078 | 4746 | 1 | 105242399 | ENSMUST00 | 20577 | ENSMUSG0( | Gm21083 | predicted gene, 21083 |
| chr5 | 15598698 | 15598888 | 191 | * | NA_peak_466 | 33 | Intron (ENSM | 5 | 15619064 | 15623864 | 4801 | 1 | 360220 | ENSMUST00 | -20176 | ENSMUSG0( | Speer4d | spermatogenesis associated glutamate (E)-rich protein 4D |
| chr5 | 15604630 | 15605000 | 371 | * | NA_peak_466 | 34 | Exon (ENSMU | 5 | 15619064 | 15623864 | 4801 | 1 | 360220 | ENSMUST00 | -14064 | ENSMUSG0( | Speer4d | spermatogenesis associated glutamate (E)-rich protein 4D |
| chr5 | 15620663 | 15620897 | 235 | * | NA_peak_466 | 35 | Promoter (<= | 5 | 15620162 | 15623611 | 3450 | 1 | 360220 | ENSMUST00 | 501 | ENSMUSG0( | Speer4d | spermatogenesis associated glutamate (E)-rich protein 4D |
| chr5 | 15661349 | 15661750 | 402 | * | NA_peak_466 | 52 | Distal Interge | 5 | 15652286 | 15657059 | 4774 | 2 | 381715 | ENSMUST00 | -4290 | ENSMUSG0( | 4930572O03 | spermatogenesis associated glutamate (E)-rich protein pseudogene |
| chr5 | 15679713 | 15680341 | 629 | * | NA_peak_466 | 35 | Promoter (<= | 5 | 15680710 | 15714596 | 33887 | 1 | 75858 | ENSMUST00 | -369 | ENSMUSG0( | Speer4cos | spermatogenesis associated glutamate (E)-rich protein 4C, opposite strand transcript |
| chr5 | 15680741 | 15681160 | 420 | * | NA_peak_466 | 49 | Promoter (<= | 5 | 15680710 | 15714596 | 33887 | 1 | 75858 | ENSMUST00 | 31 | ENSMUSG0( | Speer4cos | spermatogenesis associated glutamate (E)-rich protein 4C, opposite strand transcript |
| chr5 | 15689842 | 15690814 | 973 | * | NA_peak_466 | 90 | Intron (ENSM | 5 | 15680710 | 15714596 | 33887 | 1 | 75858 | ENSMUST00 | 9132 | ENSMUSG0( | Speer4cos | spermatogenesis associated glutamate (E)-rich protein 4C, opposite strand transcript |
| chr5 | 15696190 | 15696551 | 362 | * | NA_peak_466 | 35 | Intron (ENSM | 5 | 15680710 | 15714596 | 33887 | 1 | 75858 | ENSMUST00 | 15480 | ENSMUSG0( | Speer4cos | spermatogenesis associated glutamate (E)-rich protein 4C, opposite strand transcript |
| chr5 | 15715207 | 15715517 | 311 | * | NA_peak_466 | 71 | Promoter (<= | 5 | 15711557 | 15714271 | 2715 | 2 | 100502967 | ENSMUST00 | -936 | ENSMUSG0( | Speer4c | spermatogenesis associated glutamate (E)-rich protein 4C |
| chr5 | 17048880 | 17049071 | 192 | * | NA_peak_466 | 39 | Distal Interge | 5 | 17476098 | 17480936 | 4839 | 1 | 70935 | ENSMUST00 | -427027 | ENSMUSG0( | Speer4f1 | spermatogenesis associated glutamate (E)-rich protein 4F1 |
| chr5 | 17383668 | 17383822 | 155 | * | NA_peak_466 | 27 | Distal Interge | 5 | 17476098 | 17480936 | 4839 | 1 | 70935 | ENSMUST00 | -92276 | ENSMUSG0( | Speer4f1 | spermatogenesis associated glutamate (E)-rich protein 4F1 |
| chr5 | 17920571 | 17921105 | 535 | * | NA_peak_466 | 30 | Distal Interge | 5 | 17814675 | 17888801 | 74127 | 2 | 12491 | ENSMUST00 | -31770 | ENSMUSG0( | Cd36 | CD36 molecule |
| chr5 | 18398077 | 18398343 | 267 | * | NA_peak_467 | 27 | Distal Interge | 5 | 18265135 | 18360355 | 95221 | 2 | 14677 | ENSMUST00 | -37722 | ENSMUSG0( | Gnai1 | guanine nucleotide binding protein (G protein), alpha inhibiting 1 |
| chr5 | 18734957 | 18735177 | 221 | * | NA_peak_467 | 31 | Distal Interge | 5 | 18265135 | 18360355 | 95221 | 2 | 14677 | ENSMUST00 | -374602 | ENSMUSG0( | Gnai1 | guanine nucleotide binding protein (G protein), alpha inhibiting 1 |
| chr5 | 19477137 | 19477530 | 394 | * | NA_peak_467 | 17 | Intron (ENSM | 5 | 19227055 | 20704773 | 1477719 | 1 | 50791 | ENSMUST00 | 250082 | ENSMUSG0( | Magi2 | membrane associated guanylate kinase, WW and PDZ domain containing 2 |
| chr5 | 20124104 | 20124394 | 291 | * | NA_peak_467 | 30 | Intron (ENSM | 5 | 20202569 | 20465613 | 263045 | 1 | 50791 | ENSMUST00 | -78175 | ENSMUSG0( | Magi2 | membrane associated guanylate kinase, WW and PDZ domain containing 2 |
| chr5 | 20425971 | 20426049 | 79 | * | NA_peak_467 | 28 | Intron (ENSM | 5 | 20533500 | 20600810 | 67311 | 1 | 50791 | ENSMUST00 | -107451 | ENSMUSG0( | Magi2 | membrane associated guanylate kinase, WW and PDZ domain containing 2 |
| chr5 | 21241174 | 21241405 | 232 | * | NA_peak_467 | 44 | Intron (ENSM | 5 | 21251159 | 21258283 | 7125 | 1 | 212167 | ENSMUST00 | -9754 | ENSMUSG0( | Gsap | gamma-secretase activating protein |
| chr5 | 21786393 | 21786726 | 334 | * | NA_peak_468 | 15 | Promoter (1-2 | 5 | 21785283 | 21803787 | 18505 | 1 | 19181 | ENSMUST00 | 1110 | ENSMUSG0( | Psmc2 | proteasome (prosome, macropain) 26S subunit, ATPase 2 |
| chr5 | 22215436 | 22215526 | 91 | * | NA_peak_468 | 19 | Intron (ENSM | 5 | 22060313 | 22134669 | 74357 | 2 | 19699 | ENSMUST00 | -80767 | ENSMUSG0( | Reln | reelin |
| chr5 | 22308379 | 22308430 | 52 | * | NA_peak_468 | 23 | Intron (ENSM | 5 | 21885250 | 22344690 | 459441 | 2 | 19699 | ENSMUST00 | 36260 | ENSMUSG0( | Reln | reelin |
| chr5 | 22371682 | 22372034 | 353 | * | NA_peak_468 | 27 | Distal Interge | 5 | 22342071 | 22344702 | 2632 | 2 | 19699 | ENSMUST00 | -26980 | ENSMUSG0( | Reln | reelin |
| chr5 | 22589681 | 22590082 | 402 | * | NA_peak_468 | 23 | Intron (ENSM | 5 | 22563957 | 22564252 | 296 | 1 | 105247048 | ENSMUST00 | 25724 | NA | Gm9057 | ubiquitin-conjugating enzyme E2 L3 pseudogene |
| chr5 | 23429726 | 23430013 | 288 | * | NA_peak_468 | 16 | Intron (ENSM | 5 | 23431810 | 23433919 | 2110 | 2 | 269630 | ENSMUST00 | 3906 | ENSMUSG0( | 5031425E22I | RIKEN cDNA 5031425E22 gene |
| chr5 | 23728109 | 23728453 | 345 | * | NA_peak_468 | 96 | Distal Interge | 5 | 23709996 | 23712667 | 2672 | 2 | 433855 | ENSMUST00 | -15442 | ENSMUSG0( | AI506816 | expressed sequence AI506816 |
| chr5 | 24364668 | 24364965 | 298 | * | NA_peak_468 | 21 | Promoter (<= | 5 | 24364810 | 24384327 | 19518 | 1 | 18127 | ENSMUST00 | 0 | ENSMUSG0( | Nos3 | nitric oxide synthase 3, endothelial cell |
| chr5 | 25957584 | 25957696 | 113 | * | NA_peak_469 | 16 | Exon (ENSMU | 5 | 25948253 | 25954417 | 6165 | 2 | 100862359 | ENSMUST00 | -3167 | ENSMUSG0( | Gm21671 | predicted gene, 21671 |
| chr5 | 25958423 | 25958677 | 255 | * | NA_peak_469 | 31 | Distal Interge | 5 | 25948253 | 25954417 | 6165 | 2 | 100862359 | ENSMUST00 | -4006 | ENSMUSG0( | Gm21671 | predicted gene, 21671 |
| chr5 | 25961575 | 25961667 | 93 | * | NA_peak_469 | 22 | Distal Interge | 5 | 25948253 | 25954417 | 6165 | 2 | 100862359 | ENSMUST00 | -7158 | ENSMUSG0( | Gm21671 | predicted gene, 21671 |
| chr5 | 26006892 | 26007168 | 277 | * | NA_peak_469 | 27 | Promoter (2-3 | 5 | 25999022 | 26004798 | 5777 | 2 | 100038949 | ENSMUST00 | -2094 | ENSMUSG0( | Gm1979 | predicted gene 1979 |
| chr5 | 26032615 | 26033225 | 611 | * | NA_peak_469 | 17 | 3' UTR | 5 | 26032846 | 26039558 | 6713 | 2 | 75657 | ENSMUST00 | 6333 | ENSMUSG0( | Speer4a | spermatogenesis associated glutamate (E)-rich protein 4A |
| chr5 | 26103335 | 26103745 | 411 | * | NA_peak_469 | 30 | Promoter (1-2 | 5 | 26098668 | 26105362 | 6695 | 2 | 381622 | ENSMUST00 | 1617 | ENSMUSG0( | 5031410I06R | RIKEN cDNA 5031410I06 gene |
| chr5 | 26106355 | 26106724 | 370 | * | NA_peak_469 | 19 | Promoter (<= | 5 | 26098668 | 26105362 | 6695 | 2 | 381622 | ENSMUST00 | -993 | ENSMUSG0( | 5031410I06R | RIKEN cDNA 5031410I06 gene |
| chr5 | 26119657 | 26119705 | 49 | * | NA_peak_469 | 33 | Promoter (1-2 | 5 | 26114764 | 26121421 | 6658 | 2 | 434689 | ENSMUST00 | 1716 | ENSMUSG0( | Gm10220 | predicted gene 10220 |
| chr5 | 26381565 | 26382147 | 583 | * | NA_peak_469 | 34 | Distal Interge | 5 | 26460074 | 26493278 | 33205 | 2 | 75942 | ENSMUST00 | 111131 | ENSMUSG0( | 4930584F24I | RIKEN cDNA 4930584F24 gene |
| chr5 | 26434515 | 26434730 | 216 | * | NA_peak_469 | 15 | Distal Interge | 5 | 26460074 | 26493278 | 33205 | 2 | 75942 | ENSMUST00 | 58548 | ENSMUSG0( | 4930584F24I | RIKEN cDNA 4930584F24 gene |
| chr5 | 27771932 | 27772013 | 82 | * | NA_peak_470 | 13 | Exon (ENSMU | 5 | 27759732 | 27765737 | 6006 | 2 | 55982 | ENSMUST00 | -6195 | ENSMUSG0( | Paxip1 | PAX interacting (with transcription-activation domain) protein 1 |
| chr5 | 29504934 | 29505024 | 91 | * | NA_peak_471 | 23 | Distal Interge | 5 | 29473261 | 29478470 | 5210 | 2 | 15285 | ENSMUST00 | -26464 | ENSMUSG0( | Mnx1 | motor neuron and pancreas homeobox 1 |
| chr5 | 30529560 | 30529581 | 22 | * | NA_peak_471 | 36 | Exon (ENSMU | 5 | 30485583 | 30545836 | 60254 | 2 | 73259 | ENSMUST00 | 16255 | ENSMUSG0( | Cib4 | calcium and integrin binding family member 4 |
| chr5 | 31098896 | 31099171 | 276 | * | NA_peak_472 | 23 | Intron (ENSM | 5 | 31089341 | 31094801 | 5461 | 2 | 22784 | ENSMUST00 | -4095 | ENSMUSG0( | Slc30a3 | solute carrier family 30 (zinc transporter), member 3 |
| chr5 | 31173519 | 31173890 | 372 | * | NA_peak_472 | 44 | Promoter (<= | 5 | 31173014 | 31174501 | 1488 | 2 | 71752 | ENSMUST00 | 611 | ENSMUSG0( | Gtf3c2 | general transcription factor IIIC, polypeptide 2, beta |
| chr5 | 31677688 | 31677919 | 232 | * | NA_peak_472 | 17 | Intron (ENSM | 5 | 31660045 | 31686093 | 26049 | 2 | 71336 | ENSMUST00 | 8174 | ENSMUSG0( | Rbks | ribokinase |
| chr5 | 32136730 | 32136961 | 232 | * | NA_peak_472 | 23 | Promoter (<= | 5 | 32135801 | 32157842 | 22042 | 1 | 14284 | ENSMUST00 | 929 | ENSMUSG0( | Fosl2 | fos-like antigen 2 |
| chr5 | 32630520 | 32630750 | 231 | * | NA_peak_472 | 36 | Exon (ENSMU | 5 | 32639531 | 32653183 | 13653 | 1 | 22612 | ENSMUST00 | -8781 | ENSMUSG0( | Yes1 | YES proto-oncogene 1, Src family tyrosine kinase |
| chr5 | 34043146 | 34043655 | 510 | * | NA_peak_473 | 22 | Intron (ENSM | 5 | 34007189 | 34051117 | 43929 | 2 | 272158 | ENSMUST00 | 7462 | ENSMUSG0( | Poln | DNA polymerase N |
| chr5 | 34580126 | 34580255 | 130 | * | NA_peak_473 | 13 | Intron (ENSM | 5 | 34574151 | 34574213 | 63 | 1 | 102466999 | ENSMUST00 | 5975 | ENSMUSG0( | Mir7036b | microRNA 7036b |
| chr5 | 38983356 | 38983453 | 98 | * | NA_peak_474 | 30 | Distal Interge | 5 | 38706462 | 38876812 | 170351 | 2 | 27278 | ENSMUST00 | -106544 | ENSMUSG0( | Clnk | cytokine-dependent hematopoietic cell linker |
| chr5 | 39038000 | 39038116 | 117 | * | NA_peak_474 | 13 | Distal Interge | 5 | 38706462 | 38876812 | 170351 | 2 | 27278 | ENSMUST00 | -161188 | ENSMUSG0( | Clnk | cytokine-dependent hematopoietic cell linker |
| chr5 | 39644239 | 39644578 | 340 | * | NA_peak_474 | 23 | Promoter (<= | 5 | 39615074 | 39644597 | 29524 | 2 | 15476 | ENSMUST00 | 19 | ENSMUSG0( | Hs3st1 | heparan sulfate (glucosamine) 3-O-sulfotransferase 1 |
| chr5 | 40547969 | 40548211 | 243 | * | NA_peak_474 | 36 | Distal Interge | 5 | 39613935 | 39755475 | 141541 | 2 | 15476 | ENSMUST00 | -792494 | ENSMUSG0( | Hs3st1 | heparan sulfate (glucosamine) 3-O-sulfotransferase 1 |
| chr5 | 40548530 | 40548861 | 332 | * | NA_peak_474 | 22 | Distal Interge | 5 | 39613935 | 39755475 | 141541 | 2 | 15476 | ENSMUST00 | -793055 | ENSMUSG0( | Hs3st1 | heparan sulfate (glucosamine) 3-O-sulfotransferase 1 |
| chr5 | 42377879 | 42378118 | 240 | * | NA_peak_475 | 20 | Distal Interge | 5 | 42067960 | 42216798 | 148839 | 1 | 433882 | ENSMUST00 | 309919 | ENSMUSG0( | Gm16223 | predicted gene 16223 |
| chr5 | 42378443 | 42378466 | 24 | * | NA_peak_475 | 23 | Distal Interge | 5 | 42067960 | 42216798 | 148839 | 1 | 433882 | ENSMUST00 | 310483 | ENSMUSG0( | Gm16223 | predicted gene 16223 |
| chr5 | 42611357 | 42611639 | 283 | * | NA_peak_475 | 31 | Distal Interge | 5 | 42067960 | 42216798 | 148839 | 1 | 433882 | ENSMUST00 | 543397 | ENSMUSG0( | Gm16223 | predicted gene 16223 |
| chr5 | 43407933 | 43408275 | 343 | * | NA_peak_475 | 23 | Distal Interge | 5 | 43515538 | 43618803 | 103266 | 1 | 109323 | ENSMUST00 | -107263 | ENSMUSG0( | C1qtnf7 | C1q and tumor necrosis factor related protein 7 |
| chr5 | 45427982 | 45428228 | 247 | * | NA_peak_476 | 31 | Distal Interge | 5 | 45009171 | 45424402 | 415232 | 2 | 52401 | ENSMUST00 | -3580 | ENSMUSG0( | D5Ertd615e | DNA segment, Chr 5, ERATO Doi 615, expressed |
| chr5 | 47250554 | 47250778 | 225 | * | NA_peak_476 | 16 | Distal Interge | 5 | 47983138 | 48307733 | 324596 | 1 | 20563 | ENSMUST00 | -732360 | ENSMUSG0( | Slit2 | slit guidance ligand 2 |
| chr5 | 48732809 | 48733103 | 295 | * | NA_peak_476 | 20 | Intron (ENSM | 5 | 48396034 | 48754564 | 358531 | 2 | 80334 | ENSMUST00 | 21461 | ENSMUSG0( | Kcnip4 | Kv channel interacting protein 4 |
| chr5 | 51973434 | 51974023 | 590 | * | NA_peak_477 | 16 | Distal Interge | 5 | 52135902 | 52136017 | 116 | 2 | 102466648 | ENSMUST00 | 161994 | ENSMUSG0( | Mir6417 | microRNA 6417 |
| chr5 | 52368644 | 52368740 | 97 | * | NA_peak_477 | 17 | 3' UTR | 5 | 52363791 | 52371418 | 7628 | 1 | 20657 | ENSMUST00 | 4853 | ENSMUSG0( | Sod3 | superoxide dismutase 3, extracellular |
| chr5 | 52405966 | 52406327 | 362 | * | NA_peak_477 | 23 | Promoter (<= | 5 | 52376027 | 52405122 | 29096 | 2 | 100503884 | ENSMUST00 | -844 | ENSMUSG0( | Ccdc149 | coiled-coil domain containing 149 |
| chr5 | 53088089 | 53088344 | 256 | * | NA_peak_478 | 23 | Distal Interge | 5 | 53049353 | 53066300 | 16948 | 1 | 20531 | ENSMUST00 | 38736 | ENSMUSG0( | Slc34a2 | solute carrier family 34 (sodium phosphate), member 2 |
| chr5 | 57019877 | 57020203 | 327 | * | NA_peak_479 | 50 | Distal Interge | 5 | 57570082 | 57717920 | 147839 | 2 | 319216 | ENSMUST00 | 697717 | ENSMUSG0( | 4932441J04F | RIKEN cDNA 4932441J04 gene |
| chr5 | 57656704 | 57657238 | 535 | * | NA_peak_479 | 27 | Intron (ENSM | 5 | 57570082 | 57717920 | 147839 | 2 | 319216 | ENSMUST00 | 60682 | ENSMUSG0( | 4932441J04F | RIKEN cDNA 4932441J04 gene |
| chr5 | 57997570 | 57997891 | 322 | * | NA_peak_479 | 27 | Intron (ENSM | 5 | 57722209 | 58129454 | 407246 | 1 | 54216 | ENSMUST00 | 275361 | ENSMUSG0( | Pcdh7 | protocadherin 7 |
| chr5 | 58404491 | 58404993 | 503 | * | NA_peak_479 | 28 | Distal Interge | 5 | 58443239 | 58452999 | 9761 | 2 | 78163 | ENSMUST00 | 48006 | ENSMUSG0( | 4930459L07F | RIKEN cDNA 4930459L07 gene |
| chr5 | 59311409 | 59311692 | 284 | * | NA_peak_479 | 71 | Distal Interge | 5 | 59963124 | 59985998 | 22875 | 2 | 71057 | ENSMUST00 | 674306 | ENSMUSG0( | 4933402J10F | RIKEN cDNA 4933402J10 gene |
| chr5 | 60069781 | 60070066 | 286 | * | NA_peak_479 | 66 | Distal Interge | 5 | 59963119 | 59986004 | 22886 | 2 | 71057 | ENSMUST00 | -83777 | ENSMUSG0( | 4933402J10F | RIKEN cDNA 4933402J10 gene |
| chr5 | 61919591 | 61919731 | 141 | * | NA_peak_479 | 34 | Distal Interge | 5 | 61808816 | 61811163 | 2348 | 1 | 14380 | ENSMUST00 | 110775 | ENSMUSG0( | G6pd2 | glucose-6-phosphate dehydrogenase 2 |
| chr5 | 64894921 | 64895076 | 156 | * | NA_peak_480 | 21 | Distal Interge | 5 | 64924680 | 64932761 | 8082 | 2 | 21897 | ENSMUST00 | 37685 | ENSMUSG0( | Tlr1 | toll-like receptor 1 |
| chr5 | 65296420 | 65296702 | 283 | * | NA_peak_480 | 23 | Intron (ENSM | 5 | 65264246 | 65273667 | 9422 | 2 | 19687 | ENSMUST00 | -22753 | ENSMUSG0( | Rfc1 | replication factor C (activator 1) 1 |
| chr5 | 65482543 | 65482794 | 252 | * | NA_peak_480 | 45 | Intron (ENSM | 5 | 65427427 | 65492833 | 65407 | 2 | 68552 | ENSMUST00 | 10039 | ENSMUSG0( | Smim14 | small integral membrane protein 14 |
| chr5 | 67915124 | 67915530 | 407 | * | NA_peak_481 | 105 | Distal Interge | 5 | 67941669 | 67977070 | 35402 | 1 | 330097 | ENSMUST00 | -26139 | ENSMUSG0( | Gm5108 | predicted gene 5108 |
| chr5 | 68100873 | 68101089 | 217 | * | NA_peak_481 | 36 | Intron (ENSM | 5 | 68031835 | 68166398 | 134564 | 1 | 433899 | ENSMUST00 | 69038 | ENSMUSG0( | Grxcr1 | glutaredoxin, cysteine rich 1 |
| chr5 | 68137574 | 68137901 | 328 | * | NA_peak_481 | 24 | Intron (ENSM | 5 | 68031835 | 68166398 | 134564 | 1 | 433899 | ENSMUST00 | 105739 | ENSMUSG0( | Grxcr1 | glutaredoxin, cysteine rich 1 |
| chr5 | 70425702 | 70426045 | 344 | * | NA_peak_482 | 13 | Distal Interge | 5 | 70751775 | 70842204 | 90430 | 2 | 14405 | ENSMUST00 | 416159 | ENSMUSG0( | Gabrg1 | gamma-aminobutyric acid (GABA) A receptor, subunit gamma 1 |
| chr5 | 71717685 | 71717764 | 80 | * | NA_peak_482 | 20 | Intron (ENSM | 5 | 71699932 | 72149037 | 449106 | 1 | 14400 | ENSMUST00 | 17753 | ENSMUSG0( | Gabrb1 | gamma-aminobutyric acid (GABA) A receptor, subunit beta 1 |
| chr5 | 71979677 | 71979778 | 102 | * | NA_peak_482 | 36 | Intron (ENSM | 5 | 72159011 | 72161138 | 2128 | 2 | 27784 | ENSMUST00 | 181360 | ENSMUSG0( | Commd8 | COMM domain containing 8 |
| chr5 | 72125597 | 72125963 | 367 | * | NA_peak_482 | 22 | Intron (ENSM | 5 | 72159011 | 72161138 | 2128 | 2 | 27784 | ENSMUST00 | 35175 | ENSMUSG0( | Commd8 | COMM domain containing 8 |
| chr5 | 72545178 | 72545521 | 344 | * | NA_peak_482 | 45 | Intron (ENSM | 5 | 72550222 | 72559412 | 9191 | 2 | 100978 | ENSMUST00 | 13891 | ENSMUSG0( | Nfxl1 | nuclear transcription factor, X-box binding-like 1 |
| chr5 | 72581137 | 72581147 | 11 | * | NA_peak_482 | 23 | Promoter (<= | 5 | 72577114 | 72581254 | 4141 | 2 | 317755 | ENSMUST00 | 107 | ENSMUSG0( | Zar1 | zygote arrest 1 |
| chr5 | 72737907 | 72738189 | 283 | * | NA_peak_482 | 20 | Promoter (1-2 | 5 | 72695978 | 72736602 | 40625 | 2 | 22165 | ENSMUST00 | -1305 | ENSMUSG0( | Txk | TXK tyrosine kinase |
| chr5 | 73967730 | 73968411 | 682 | * | NA_peak_482 | 31 | Distal Interge | 5 | 74026293 | 74029368 | 3076 | 2 | 69727 | ENSMUST00 | 60957 | ENSMUSG0( | Usp46 | ubiquitin specific peptidase 46 |
| chr5 | 74524721 | 74525006 | 286 | * | NA_peak_483 | 25 | Intron (ENSM | 5 | 74447780 | 74531729 | 83950 | 2 | 212986 | ENSMUST00 | 6723 | ENSMUSG0( | Scfd2 | Sec1 family domain containing 2 |
| chr5 | 75085507 | 75085535 | 29 | * | NA_peak_483 | 31 | Distal Interge | 5 | 75075602 | 75077625 | 2024 | 1 | 14843 | ENSMUST00 | 9905 | ENSMUSG0( | Gsx2 | GS homeobox 2 |
| chr5 | 75368574 | 75368847 | 274 | * | NA_peak_483 | 16 | Distal Interge | 5 | 75176831 | 75176904 | 74 | 1 | 102465620 | ENSMUST00 | 191743 | ENSMUSG0( | Mir7025 | microRNA 7025 |
| chr5 | 75561460 | 75561736 | 277 | * | NA_peak_483 | 16 | Distal Interge | 5 | 75574916 | 75656722 | 81807 | 1 | 16590 | ENSMUST00 | -13180 | ENSMUSG0( | Kit | KIT proto-oncogene receptor tyrosine kinase |
| chr5 | 76223991 | 76224303 | 313 | * | NA_peak_483 | 37 | Intron (ENSM | 5 | 76204120 | 76206033 | 1914 | 1 | 21982 | ENSMUST00 | 19871 | ENSMUSG0( | Tmem165 | transmembrane protein 165 |
| chr5 | 76452605 | 76452881 | 277 | * | NA_peak_483 | 20 | Distal Interge | 5 | 76484040 | 76516649 | 32610 | 1 | 639545 | ENSMUST00 | -31159 | ENSMUSG0( | Gm7271 | predicted gene 7271 |
| chr5 | 76505719 | 76505987 | 269 | * | NA_peak_483 | 36 | Intron (ENSM | 5 | 76484331 | 76516584 | 32254 | 1 | 639545 | ENSMUST00 | 21388 | ENSMUSG0( | Gm7271 | predicted gene 7271 |
| chr5 | 77182194 | 77182215 | 22 | * | NA_peak_484 | 16 | Distal Interge | 5 | 77205112 | 77211205 | 6094 | 2 | 69982 | ENSMUST00 | 28990 | ENSMUSG0( | Spink2 | serine peptidase inhibitor, Kazal type 2 |
| chr5 | 77406249 | 77406637 | 389 | * | NA_peak_484 | 123 | Promoter (1-2 | 5 | 77349248 | 77408040 | 58793 | 2 | 29817 | ENSMUST00 | 1403 | ENSMUSG0( | Igfbp7 | insulin-like growth factor binding protein 7 |
| chr5 | 77413213 | 77413642 | 430 | * | NA_peak_484 | 31 | Distal Interge | 5 | 77349240 | 77408041 | 58802 | 2 | 29817 | ENSMUST00 | -5172 | ENSMUSG0( | Igfbp7 | insulin-like growth factor binding protein 7 |
| chr5 | 77623496 | 77623626 | 131 | * | NA_peak_484 | 22 | Distal Interge | 5 | 77465488 | 77468851 | 3364 | 1 | 69370 | ENSMUST00 | 158008 | ENSMUSG0( | 1700017L05F | RIKEN cDNA 1700017L05 gene |

| | | | | | | | | | | | | | | | | | |
|---|---|---|---|---|---|---|---|---|---|---|---|---|---|---|---|---|---|
| chr5 | 77631951 | 77632246 | 296 | * | NA_peak_484 | 24 | Distal Interge | 5 | 77465488 | 77468851 | 3364 | 1 | 69370 | ENSMUST00 | 166463 | ENSMUSG00 | 1700017L05R | RIKEN cDNA 1700017L05 gene |
| chr5 | 77964705 | 77965107 | 403 | * | NA_peak_484 | 51 | Distal Interge | 5 | 77465488 | 77468851 | 3364 | 1 | 69370 | ENSMUST00 | 499217 | ENSMUSG00 | 1700017L05R | RIKEN cDNA 1700017L05 gene |
| chr5 | 79419246 | 79419560 | 315 | * | NA_peak_484 | 110 | Distal Interge | 5 | 81020138 | 81388784 | 368647 | 1 | 319387 | ENSMUST00 | -1600578 | ENSMUSG00 | Adgrl3 | adhesion G protein-coupled receptor L3 |
| chr5 | 79437031 | 79437400 | 370 | * | NA_peak_484 | 14 | Distal Interge | 5 | 81020138 | 81388784 | 368647 | 1 | 319387 | ENSMUST00 | -1582738 | ENSMUSG00 | Adgrl3 | adhesion G protein-coupled receptor L3 |
| chr5 | 81707886 | 81708166 | 281 | * | NA_peak_485 | 36 | Intron (ENSM | 5 | 81660303 | 81796634 | 136332 | 1 | 319387 | ENSMUST00 | 47583 | ENSMUSG00 | Adgrl3 | adhesion G protein-coupled receptor L3 |
| chr5 | 82408716 | 82408918 | 203 | * | NA_peak_485 | 22 | Distal Interge | 5 | 82122427 | 82124713 | 2287 | 2 | 67327 | ENSMUST00 | -284003 | ENSMUSG00 | 1700031L13R | RIKEN cDNA 1700031L13 gene |
| chr5 | 83141667 | 83142018 | 352 | * | NA_peak_485 | 22 | Distal Interge | 5 | 83289996 | 83354956 | 64961 | 2 | 243078 | ENSMUST00 | 212938 | ENSMUSG00 | Tecrl | trans-2,3-enoyl-CoA reductase-like |
| chr5 | 83552713 | 83552843 | 131 | * | NA_peak_485 | 31 | Distal Interge | 5 | 83278145 | 83355195 | 77051 | 2 | 243078 | ENSMUST00 | -197518 | ENSMUSG00 | Tecrl | trans-2,3-enoyl-CoA reductase-like |
| chr5 | 84375994 | 84376413 | 420 | * | NA_peak_485 | 45 | Intron (ENSM | 5 | 84057791 | 84416806 | 359016 | 2 | 13839 | ENSMUST00 | 40393 | ENSMUSG00 | Epha5 | Eph receptor A5 |
| chr5 | 84833168 | 84833383 | 216 | * | NA_peak_485 | 22 | Distal Interge | 5 | 84330775 | 84417382 | 86608 | 2 | 13839 | ENSMUST00 | -415786 | ENSMUSG00 | Epha5 | Eph receptor A5 |
| chr5 | 85169220 | 85169519 | 300 | * | NA_peak_485 | 45 | Distal Interge | 5 | 84330775 | 84417382 | 86608 | 2 | 13839 | ENSMUST00 | -751838 | ENSMUSG00 | Epha5 | Eph receptor A5 |
| chr5 | 85864659 | 85864946 | 288 | * | NA_peak_485 | 31 | Distal Interge | 5 | 86057116 | 86065522 | 8407 | 2 | 12617 | ENSMUST00 | 200576 | ENSMUSG00 | Cenpc1 | centromere protein C1 |
| chr5 | 86367516 | 86367762 | 247 | * | NA_peak_485 | 43 | Intron (ENSM | 5 | 86302217 | 86373420 | 71204 | 2 | 231382 | ENSMUST00 | 5658 | ENSMUSG00 | Tmprss11d | transmembrane protease, serine 11d |
| chr5 | 87545794 | 87546154 | 361 | * | NA_peak_486 | 19 | Distal Interge | 5 | 87513339 | 87538195 | 24857 | 2 | 56362 | ENSMUST00 | -7599 | ENSMUSG00 | Sult1b1 | sulfotransferase family 1B, member 1 |
| chr5 | 88163522 | 88163893 | 372 | * | NA_peak_486 | 26 | Distal Interge | 5 | 88086556 | 88109053 | 22498 | 1 | 20600 | ENSMUST00 | 76966 | ENSMUSG00 | Smr2 | submaxillary gland androgen regulated protein 2 |
| chr5 | 88165085 | 88165386 | 302 | * | NA_peak_486 | 88 | Distal Interge | 5 | 88086556 | 88109053 | 22498 | 1 | 20600 | ENSMUST00 | 78529 | ENSMUSG00 | Smr2 | submaxillary gland androgen regulated protein 2 |
| chr5 | 88165708 | 88167022 | 1315 | * | NA_peak_486 | 88 | Distal Interge | 5 | 88086556 | 88109053 | 22498 | 1 | 20600 | ENSMUST00 | 79152 | ENSMUSG00 | Smr2 | submaxillary gland androgen regulated protein 2 |
| chr5 | 88259359 | 88259641 | 283 | * | NA_peak_486 | 48 | Distal Interge | 5 | 88268919 | 88282835 | 13917 | 1 | 665615 | ENSMUST00 | -9278 | ENSMUSG00 | Smr2l | submaxillary gland androgen regulated protein 2 like |
| chr5 | 88547235 | 88547380 | 146 | * | NA_peak_486 | 24 | Distal Interge | 5 | 88554462 | 88556090 | 1629 | 1 | 65961 | ENSMUST00 | -7082 | ENSMUSG00 | Utp3 | UTP3 small subunit processome component |
| chr5 | 88765241 | 88765367 | 127 | * | NA_peak_486 | 39 | Promoter (<= | 5 | 88764996 | 88783281 | 18286 | 1 | 13178 | ENSMUST00 | 245 | ENSMUSG00 | Dck | deoxycytidine kinase |
| chr5 | 89112096 | 89112419 | 324 | * | NA_peak_486 | 13 | Intron (ENSM | 5 | 89028092 | 89122453 | 94362 | 1 | 54403 | ENSMUST00 | 84004 | ENSMUSG00 | Slc4a4 | solute carrier family 4 (anion exchanger), member 4 |
| chr5 | 89205069 | 89206096 | 1028 | * | NA_peak_486 | 39 | Intron (ENSM | 5 | 89028092 | 89122453 | 94362 | 1 | 54403 | ENSMUST00 | 176977 | ENSMUSG00 | Slc4a4 | solute carrier family 4 (anion exchanger), member 4 |
| chr5 | 89207976 | 89208276 | 301 | * | NA_peak_486 | 15 | Intron (ENSM | 5 | 89028092 | 89122453 | 94362 | 1 | 54403 | ENSMUST00 | 179884 | ENSMUSG00 | Slc4a4 | solute carrier family 4 (anion exchanger), member 4 |
| chr5 | 90097268 | 90097609 | 342 | * | NA_peak_486 | 43 | Distal Interge | 5 | 90214727 | 90215581 | 855 | 2 | 231430 | ENSMUST00 | 117972 | ENSMUSG00 | Cox18 | cytochrome c oxidase assembly protein 18 |
| chr5 | 91250592 | 91250889 | 298 | * | NA_peak_487 | 27 | Distal Interge | 5 | 91257571 | 91283421 | 25851 | 2 | 100503265 | ENSMUST00 | 32532 | ENSMUSG00 | Gm19619 | predicted gene, 19619 |
| chr5 | 91411907 | 91412171 | 265 | * | NA_peak_487 | 27 | Distal Interge | 5 | 91357261 | 91402994 | 45734 | 2 | 12223 | ENSMUST00 | -8913 | ENSMUSG00 | Btc | betacellulin, epidermal growth factor family member |
| chr5 | 92150712 | 92150998 | 287 | * | NA_peak_487 | 27 | Intron (ENSM | 5 | 92137966 | 92202796 | 64831 | 1 | 56041 | ENSMUST00 | 12746 | ENSMUSG00 | Uso1 | USO1 vesicle docking factor |
| chr5 | 92615725 | 92615838 | 114 | * | NA_peak_487 | 24 | Intron (ENSM | 5 | 92607954 | 92612503 | 4550 | 2 | 320696 | ENSMUST00 | -3222 | ENSMUSG00 | Ccdc158 | coiled-coil domain containing 158 |
| chr5 | 93381838 | 93382137 | 300 | * | NA_peak_488 | 28 | Distal Interge | 5 | 93268708 | 93273485 | 4778 | 1 | 12452 | ENSMUST00 | 113130 | ENSMUSG00 | Ccng2 | cyclin G2 |
| chr5 | 93479684 | 93479959 | 276 | * | NA_peak_488 | 50 | Distal Interge | 5 | 93636335 | 93638052 | 1718 | 2 | 381654 | ENSMUST00 | 158093 | ENSMUSG00 | Pramel34 | PRAME like 34 |
| chr5 | 93621681 | 93621748 | 68 | * | NA_peak_488 | 20 | Distal Interge | 5 | 93636335 | 93638052 | 1718 | 2 | 381654 | ENSMUST00 | 16304 | ENSMUSG00 | Pramel34 | PRAME like 34 |
| chr5 | 93643621 | 93643867 | 247 | * | NA_peak_488 | 27 | Intron (ENSM | 5 | 93636335 | 93638052 | 1718 | 2 | 381654 | ENSMUST00 | -5569 | ENSMUSG00 | Pramel34 | PRAME like 34 |
| chr5 | 93807724 | 93808366 | 643 | * | NA_peak_488 | 77 | Distal Interge | 5 | 93634993 | 93671551 | 36559 | 2 | 381654 | ENSMUST00 | -136173 | ENSMUSG00 | Pramel34 | PRAME like 34 |
| chr5 | 93891351 | 93891505 | 155 | * | NA_peak_488 | 23 | Distal Interge | 5 | 94066151 | 94070545 | 4395 | 2 | 666203 | ENSMUST00 | 179040 | ENSMUSG00 | Pramel50 | PRAME like 50 |
| chr5 | 93931684 | 93931908 | 225 | * | NA_peak_488 | 22 | Distal Interge | 5 | 94066151 | 94070545 | 4395 | 2 | 666203 | ENSMUST00 | 138637 | ENSMUSG00 | Pramel50 | PRAME like 50 |
| chr5 | 94143629 | 94143977 | 349 | * | NA_peak_488 | 36 | Distal Interge | 5 | 94066151 | 94070545 | 4395 | 2 | 666203 | ENSMUST00 | -73084 | ENSMUSG00 | Pramel50 | PRAME like 50 |
| chr5 | 94241669 | 94242226 | 558 | * | NA_peak_488 | 22 | Distal Interge | 5 | 94302519 | 94306796 | 4278 | 2 | 331195 | ENSMUST00 | 64570 | ENSMUSG00 | Pramel39 | PRAME like 39 |
| chr5 | 94333314 | 94333593 | 280 | * | NA_peak_488 | 18 | Distal Interge | 5 | 94339749 | 94385010 | 45262 | 1 | 100554 | ENSMUST00 | -6156 | ENSMUSG00 | AA792892 | expressed sequence AA792892 |
| chr5 | 94366628 | 94366824 | 197 | * | NA_peak_488 | 31 | Intron (ENSM | 5 | 94377317 | 94385079 | 7763 | 1 | 100554 | ENSMUST00 | -10493 | ENSMUSG00 | AA792892 | expressed sequence AA792892 |
| chr5 | 94601939 | 94602062 | 124 | * | NA_peak_489 | 44 | Distal Interge | 5 | 94534849 | 94537928 | 3080 | 1 | 100041102 | ENSMUST00 | 67090 | ENSMUSG00 | Pramel42 | PRAME like 42 |
| chr5 | 94619040 | 94619330 | 291 | * | NA_peak_489 | 45 | Distal Interge | 5 | 94534849 | 94537928 | 3080 | 1 | 100041102 | ENSMUST00 | 84191 | ENSMUSG00 | Pramel42 | PRAME like 42 |
| chr5 | 94633558 | 94633850 | 293 | * | NA_peak_489 | 18 | Distal Interge | 5 | 94534849 | 94537928 | 3080 | 1 | 100041102 | ENSMUST00 | 98709 | ENSMUSG00 | Pramel42 | PRAME like 42 |
| chr5 | 94648317 | 94648620 | 304 | * | NA_peak_489 | 33 | 5' UTR | 5 | 94534849 | 94537928 | 3080 | 1 | 100041102 | ENSMUST00 | 113468 | ENSMUSG00 | Pramel42 | PRAME like 42 |
| chr5 | 94766496 | 94767037 | 542 | * | NA_peak_489 | 16 | Distal Interge | 5 | 94865436 | 94885388 | 19953 | 1 | 100041253 | ENSMUST00 | -98399 | ENSMUSG00 | Gm16513 | predicted gene, Gm16513 |
| chr5 | 94775421 | 94775872 | 452 | * | NA_peak_489 | 18 | Distal Interge | 5 | 94865436 | 94885388 | 19953 | 1 | 100041253 | ENSMUST00 | -89564 | ENSMUSG00 | Gm16513 | predicted gene, Gm16513 |
| chr5 | 94788673 | 94789053 | 381 | * | NA_peak_489 | 25 | Distal Interge | 5 | 94865436 | 94885388 | 19953 | 1 | 100041253 | ENSMUST00 | -76383 | ENSMUSG00 | Gm16513 | predicted gene, Gm16513 |
| chr5 | 94797111 | 94797266 | 156 | * | NA_peak_489 | 45 | Distal Interge | 5 | 94865436 | 94885388 | 19953 | 1 | 100041253 | ENSMUST00 | -68170 | ENSMUSG00 | Gm16513 | predicted gene, Gm16513 |
| chr5 | 94810627 | 94810861 | 235 | * | NA_peak_489 | 29 | Distal Interge | 5 | 94865436 | 94885388 | 19953 | 1 | 100041253 | ENSMUST00 | -54575 | ENSMUSG00 | Gm16513 | predicted gene, Gm16513 |
| chr5 | 94842059 | 94842409 | 351 | * | NA_peak_489 | 23 | Distal Interge | 5 | 94865436 | 94885388 | 19953 | 1 | 100041253 | ENSMUST00 | -23027 | ENSMUSG00 | Gm16513 | predicted gene, Gm16513 |
| chr5 | 94876045 | 94876525 | 481 | * | NA_peak_489 | 35 | Promoter (2-3 | 5 | 94873797 | 94876892 | 3096 | 1 | 545762 | ENSMUST00 | 2248 | ENSMUSG00 | Gm16367 | predicted gene 16367 |
| chr5 | 94914338 | 94915106 | 769 | * | NA_peak_489 | 34 | Distal Interge | 5 | 94951537 | 94954134 | 2598 | 2 | 331195 | ENSMUST00 | 39028 | ENSMUSG00 | Pramel39 | PRAME like 39 |
| chr5 | 94996015 | 94996041 | 27 | * | NA_peak_489 | 23 | Intron (ENSM | 5 | 94988626 | 95026536 | 37911 | 1 | 622894 | ENSMUST00 | 7389 | ENSMUSG00 | Gm6367 | predicted gene 6367 |
| chr5 | 95072684 | 95072762 | 79 | * | NA_peak_490 | 37 | Distal Interge | 5 | 94988626 | 95026536 | 37911 | 1 | 622894 | ENSMUST00 | 84058 | ENSMUSG00 | Gm6367 | predicted gene 6367 |
| chr5 | 95110978 | 95111289 | 312 | * | NA_peak_490 | 30 | 3' UTR | 5 | 95182624 | 95194966 | 12343 | 1 | 100041253 | ENSMUST00 | -71335 | ENSMUSG00 | Gm16513 | predicted gene, Gm16513 |
| chr5 | 95123032 | 95123055 | 24 | * | NA_peak_490 | 20 | Distal Interge | 5 | 95182624 | 95194966 | 12343 | 1 | 100041253 | ENSMUST00 | -59569 | ENSMUSG00 | Gm16513 | predicted gene, Gm16513 |
| chr5 | 95155306 | 95155598 | 293 | * | NA_peak_490 | 18 | Distal Interge | 5 | 95182624 | 95194966 | 12343 | 1 | 100041253 | ENSMUST00 | -27026 | ENSMUSG00 | Gm16513 | predicted gene, Gm16513 |
| chr5 | 95230456 | 95230988 | 533 | * | NA_peak_490 | 35 | Distal Interge | 5 | 95190874 | 95195342 | 4469 | 1 | 100041253 | ENSMUST00 | 39582 | ENSMUSG00 | Gm16513 | predicted gene, Gm16513 |
| chr5 | 95231305 | 95231502 | 198 | * | NA_peak_490 | 20 | Distal Interge | 5 | 95190874 | 95195342 | 4469 | 1 | 100041253 | ENSMUST00 | 40431 | ENSMUSG00 | Gm16513 | predicted gene, Gm16513 |
| chr5 | 95470680 | 95471000 | 321 | * | NA_peak_490 | 23 | Promoter (2-3 | 5 | 95473553 | 95485587 | 12035 | 1 | 320549 | ENSMUST00 | -2553 | ENSMUSG00 | Pramel48 | PRAME like 48 |
| chr5 | 96273932 | 96274247 | 316 | * | NA_peak_490 | 63 | Distal Interge | 5 | 96210144 | 96239684 | 29541 | 1 | 94061 | ENSMUST00 | 63788 | ENSMUSG00 | Mrpl1 | mitochondrial ribosomal protein L1 |
| chr5 | 96293257 | 96293682 | 426 | * | NA_peak_491 | 29 | Distal Interge | 5 | 96373955 | 96784728 | 410774 | 1 | 231470 | ENSMUST00 | -80273 | ENSMUSG00 | Fras1 | Fraser extracellular matrix complex subunit 1 |
| chr5 | 98043587 | 98043690 | 104 | * | NA_peak_491 | 40 | Distal Interge | 5 | 97915373 | 98031043 | 115671 | 2 | 71914 | ENSMUST00 | -12544 | ENSMUSG00 | Antxr2 | anthrax toxin receptor 2 |
| chr5 | 100842386 | 100842413 | 28 | * | NA_peak_492 | 23 | Intron (ENSM | 5 | 100845713 | 100846810 | 1098 | 1 | 231510 | ENSMUST00 | -3300 | ENSMUSG00 | Gpat3 | glycerol-3-phosphate acyltransferase 3 |
| chr5 | 101547699 | 101547940 | 242 | * | NA_peak_492 | 27 | Distal Interge | 5 | 101658139 | 101664996 | 6858 | 2 | 18096 | ENSMUST00 | 117056 | ENSMUSG00 | Nkx6-1 | NK6 homeobox 1 |
| chr5 | 102412013 | 102412336 | 324 | * | NA_peak_492 | 25 | Distal Interge | 5 | 102481391 | 102897936 | 416546 | 1 | 231532 | ENSMUST00 | -69055 | ENSMUSG00 | Arhgap24 | Rho GTPase activating protein 24 |
| chr5 | 102651420 | 102651785 | 366 | * | NA_peak_493 | 17 | Intron (ENSM | 5 | 102724973 | 102897937 | 172965 | 1 | 231532 | ENSMUST00 | -73188 | ENSMUSG00 | Arhgap24 | Rho GTPase activating protein 24 |
| chr5 | 102700990 | 102701247 | 258 | * | NA_peak_493 | 37 | Intron (ENSM | 5 | 102724973 | 102897937 | 172965 | 1 | 231532 | ENSMUST00 | -23726 | ENSMUSG00 | Arhgap24 | Rho GTPase activating protein 24 |
| chr5 | 103413149 | 103413635 | 487 | * | NA_peak_493 | 23 | Distal Interge | 5 | 103425192 | 103541434 | 116243 | 1 | 19249 | ENSMUST00 | -11557 | ENSMUSG00 | Ptpn13 | protein tyrosine phosphatase, non-receptor type 13 |
| chr5 | 103655599 | 103656020 | 422 | * | NA_peak_493 | 31 | Promoter (<= | 5 | 103648039 | 103656010 | 7972 | 2 | 74218 | ENSMUST00 | 0 | ENSMUSG00 | 1700016H13R | RIKEN cDNA 1700016H13 gene |
| chr5 | 104362741 | 104362807 | 67 | * | NA_peak_493 | 32 | Distal Interge | 5 | 104325329 | 104338611 | 13283 | 1 | 94111 | ENSMUST00 | 37412 | ENSMUSG00 | Mepe | matrix extracellular phosphoglycoprotein with ASARM motif (bone) |
| chr5 | 104363699 | 104363980 | 282 | * | NA_peak_493 | 32 | Distal Interge | 5 | 104325329 | 104338611 | 13283 | 1 | 94111 | ENSMUST00 | 38370 | ENSMUSG00 | Mepe | matrix extracellular phosphoglycoprotein with ASARM motif (bone) |
| chr5 | 104448654 | 104448825 | 172 | * | NA_peak_493 | 20 | Distal Interge | 5 | 104459450 | 104505819 | 46370 | 1 | 18764 | ENSMUST00 | -10625 | ENSMUSG00 | Pkd2 | polycystin 2, transient receptor potential cation channel |
| chr5 | 105087605 | 105088018 | 414 | * | NA_peak_493 | 55 | Intron (ENSM | 5 | 105080136 | 105082763 | 2628 | 2 | 236573 | ENSMUST00 | -4842 | ENSMUSG00 | Gbp9 | guanylate-binding protein 9 |
| chr5 | 105360539 | 105360940 | 402 | * | NA_peak_493 | 21 | Distal Interge | 5 | 105323042 | 105346472 | 23431 | 2 | 634650 | ENSMUST00 | -14067 | ENSMUSG00 | Gbp11 | guanylate binding protein 11 |
| chr5 | 105366747 | 105367485 | 739 | * | NA_peak_493 | 125 | Distal Interge | 5 | 105323042 | 105346472 | 23431 | 2 | 634650 | ENSMUST00 | -20275 | ENSMUSG00 | Gbp11 | guanylate binding protein 11 |
| chr5 | 106425723 | 106425966 | 244 | * | NA_peak_494 | 13 | Distal Interge | 5 | 106452523 | 106458166 | 5644 | 2 | 104382 | ENSMUST00 | 32200 | ENSMUSG00 | Barhl2 | BarH like homeobox 2 |
| chr5 | 108761919 | 108762171 | 253 | * | NA_peak_494 | 27 | Promoter (1-2 | 5 | 108763842 | 108768914 | 5073 | 1 | 73295 | ENSMUST00 | -1671 | ENSMUSG00 | 1700047L14R | RIKEN cDNA 1700047L14 gene |
| chr5 | 108816965 | 108816973 | 9 | * | NA_peak_494 | 16 | Distal Interge | 5 | 108791843 | 108808850 | 17008 | 2 | 627479 | ENSMUST00 | -8115 | ENSMUSG00 | Vmn2r8 | vomeronasal 2, receptor 8 |
| chr5 | 109726375 | 109726788 | 414 | * | NA_peak_495 | 25 | Distal Interge | 5 | 109833501 | 109834145 | 645 | 2 | 667213 | ENSMUST00 | 107357 | ENSMUSG00 | Gm10416 | zinc finger protein pseudogene |
| chr5 | 109914804 | 109915118 | 315 | * | NA_peak_495 | 23 | Distal Interge | 5 | 109923400 | 109943936 | 20537 | 1 | 100642166 | ENSMUST00 | -8282 | ENSMUSG00 | Gm15446 | predicted gene 15446 |
| chr5 | 110000046 | 110000464 | 419 | * | NA_peak_495 | 23 | Intron (ENSM | 5 | 109996563 | 110003844 | 7282 | 1 | 69504 | ENSMUST00 | 3483 | ENSMUSG00 | Zfp932 | zinc finger protein 932 |
| chr5 | 111689995 | 111690068 | 74 | * | NA_peak_495 | 24 | Distal Interge | 5 | 111734410 | 111761700 | 27291 | 2 | 269683 | ENSMUST00 | 71632 | ENSMUSG00 | E130006D01 | RIKEN cDNA E130006D01 gene |
| chr5 | 112695112 | 112695380 | 269 | * | NA_peak_496 | 16 | Intron (ENSM | 5 | 112688878 | 112765931 | 77054 | 2 | 74376 | ENSMUST00 | 70551 | ENSMUSG00 | Myo18b | myosin XVIIIb |
| chr5 | 113244815 | 113245125 | 311 | * | NA_peak_496 | 30 | 3' UTR | 5 | 113226909 | 113239263 | 12355 | 1 | 333048 | ENSMUST00 | 17906 | ENSMUSG00 | Tmem211 | transmembrane protein 211 |
| chr5 | 113771517 | 113771918 | 402 | * | NA_peak_496 | 65 | Promoter (<= | 5 | 113742448 | 113771638 | 29191 | 2 | 53890 | ENSMUST00 | 0 | ENSMUSG00 | Sart3 | squamous cell carcinoma antigen recognized by T cells 3 |
| chr5 | 114679324 | 114679556 | 233 | * | NA_peak_496 | 31 | Intron (ENSM | 5 | 114670228 | 114690908 | 20681 | 2 | 56356 | ENSMUST00 | 11352 | ENSMUSG00 | Gltp | glycolipid transfer protein |
| chr5 | 115272577 | 115272637 | 61 | * | NA_peak_496 | 30 | Promoter (<= | 5 | 115241770 | 115272892 | 31123 | 2 | 50849 | ENSMUST00 | 255 | ENSMUSG00 | Rnf10 | ring finger protein 10 |
| chr5 | 115514965 | 115515076 | 112 | * | NA_peak_496 | 20 | Promoter (1-2 | 5 | 115516355 | 115544883 | 28529 | 1 | 19303 | ENSMUST00 | -1279 | ENSMUSG00 | Pxn | paxillin |
| chr5 | 116425534 | 116425872 | 339 | * | NA_peak_497 | 24 | Promoter (2-3 | 5 | 116408491 | 116422864 | 14374 | 2 | 80888 | ENSMUST00 | -2670 | ENSMUSG00 | Hspb8 | heat shock protein 8 |
| chr5 | 116600005 | 116600185 | 181 | * | NA_peak_497 | 16 | Distal Interge | 5 | 116439276 | 116591817 | 152542 | 2 | 68955 | ENSMUST00 | -8188 | ENSMUSG00 | Srrm4 | serine/arginine repetitive matrix 4 |
| chr5 | 116681632 | 116681923 | 292 | * | NA_peak_497 | 27 | Intron (ENSM | 5 | 116439276 | 116591817 | 152542 | 2 | 68955 | ENSMUST00 | -89815 | ENSMUSG00 | Srrm4 | serine/arginine repetitive matrix 4 |
| chr5 | 117223302 | 117223485 | 184 | * | NA_peak_497 | 31 | Intron (ENSM | 5 | 117240460 | 117273867 | 33408 | 1 | 330177 | ENSMUST00 | -16975 | ENSMUSG00 | Taok3 | TAO kinase 3 |
| chr5 | 117369909 | 117370319 | 411 | * | NA_peak_497 | 20 | Intron (ENSM | 5 | 117375652 | 117377480 | 1829 | 1 | 59043 | ENSMUST00 | -5333 | ENSMUSG00 | Wsb2 | WD repeat and SOCS box-containing 2 |
| chr5 | 117898818 | 117899158 | 341 | * | NA_peak_497 | 13 | Intron (ENSM | 5 | 117919175 | 117925731 | 6557 | 1 | 18125 | ENSMUST00 | -20017 | ENSMUSG00 | Nos1 | nitric oxide synthase 1, neuronal |
| chr5 | 118574928 | 118574974 | 47 | * | NA_peak_498 | 13 | Intron (ENSM | 5 | 118561575 | 118593539 | 31965 | 1 | 76199 | ENSMUST00 | 13353 | ENSMUSG00 | Med13l | mediator complex subunit 13-like |
| chr5 | 119831446 | 119831767 | 322 | * | NA_peak_498 | 20 | Promoter (<= | 5 | 119832668 | 119840886 | 8219 | 1 | 21388 | ENSMUST00 | -901 | ENSMUSG00 | Tbx5 | T-box 5 |
| chr5 | 120430780 | 120431328 | 549 | * | NA_peak_498 | 26 | Promoter (<= | 5 | 120431699 | 120441223 | 9525 | 1 | 16873 | ENSMUST00 | -371 | ENSMUSG00 | Lhx5 | LIM homeobox protein 5 |
| chr5 | 120839601 | 120839897 | 297 | * | NA_peak_498 | 26 | Distal Interge | 5 | 120847367 | 120857986 | 10620 | 1 | 243262 | ENSMUST00 | -7470 | ENSMUSG00 | Oas1f | 2'-5' oligoadenylate synthetase 1F |
| chr5 | 121077854 | 121078140 | 287 | * | NA_peak_498 | 18 | Distal Interge | 5 | 121133840 | 121143635 | 9796 | 2 | 19247 | ENSMUST00 | 65495 | ENSMUSG00 | Ptpn11 | protein tyrosine phosphatase, non-receptor type 11 |
| chr5 | 121434177 | 121434517 | 341 | * | NA_peak_498 | 20 | Intron (ENSM | 5 | 121439418 | 121444378 | 4961 | 1 | 231713 | ENSMUST00 | -4901 | ENSMUSG00 | Naa25 | N(alpha)-acetyltransferase 25, NatB auxiliary subunit |
| chr5 | 122427912 | 122428008 | 97 | * | NA_peak_499 | 13 | Intron (ENSM | 5 | 122422481 | 122443434 | 20954 | 1 | 56317 | ENSMUST00 | 5431 | ENSMUSG00 | Anapc7 | anaphase promoting complex subunit 7 |
| chr5 | 123543826 | 123544043 | 218 | * | NA_peak_499 | 16 | Intron (ENSM | 5 | 123554765 | 123559019 | 4255 | 2 | 77573 | ENSMUST00 | 14976 | ENSMUSG00 | Vps33a | VPS33A CORVET/HOPS core subunit |
| chr5 | 124425195 | 124425629 | 435 | * | NA_peak_499 | 27 | Promoter (<= | 5 | 124368708 | 124425610 | 56903 | 2 | 243272 | ENSMUST00 | 0 | ENSMUSG00 | Sbno1 | strawberry notch 1 |
| chr5 | 124880537 | 124880786 | 250 | * | NA_peak_499 | 13 | Intron (ENSM | 5 | 124884569 | 124888630 | 4062 | 1 | 269704 | ENSMUST00 | -3783 | ENSMUSG00 | Zfp664 | zinc finger protein 664 |
| chr5 | 125387658 | 125387982 | 325 | * | NA_peak_499 | 23 | Promoter (1-2 | 5 | 125387659 | 125389186 | 1528 | 2 | 22190 | ENSMUST00 | 1204 | ENSMUSG00 | Ubc | ubiquitin C |
| chr5 | 125567156 | 125567517 | 362 | * | NA_peak_499 | 23 | Intron (ENSM | 5 | 125532387 | 125792583 | 260197 | 1 | 208151 | ENSMUST00 | 34769 | ENSMUSG00 | Tmem132b | transmembrane protein 132B |

| | | | | | | | | | | | | | | | | | |
|---|---|---|---|---|---|---|---|---|---|---|---|---|---|---|---|---|---|
| chr5 | 125630242 | 125630605 | 364 | * | NA_peak_49 | 33 | Intron (ENSM | 5 | 125622488 | 125698773 | 76286 | 1 | 208151 | ENSMUST00 | 7754 | ENSMUSG0 | Tmem132b | transmembrane protein 132B |
| chr5 | 126122761 | 126122892 | 132 | * | NA_peak_50 | 23 | Distal Interge | 5 | 125622488 | 125698773 | 76286 | 1 | 208151 | ENSMUST00 | 500273 | ENSMUSG0 | Tmem132b | transmembrane protein 132B |
| chr5 | 127010071 | 127010166 | 96 | * | NA_peak_50 | 21 | Distal Interge | 5 | 127030783 | 127088529 | 57747 | 1 | 71247 | ENSMUST00 | -20617 | ENSMUSG0 | 4933438B17 | RIKEN cDNA 4933438B17 gene |
| chr5 | 127012394 | 127012759 | 366 | * | NA_peak_50 | 40 | Distal Interge | 5 | 127030783 | 127088529 | 57747 | 1 | 71247 | ENSMUST00 | -18024 | ENSMUSG0 | 4933438B17 | RIKEN cDNA 4933438B17 gene |
| chr5 | 127430152 | 127430502 | 351 | * | NA_peak_50 | 24 | Intron (ENSM | 5 | 127538260 | 127564866 | 26607 | 1 | 208213 | ENSMUST00 | -107758 | ENSMUSG0 | Tmem132c | transmembrane protein 132C |
| chr5 | 128824142 | 128824552 | 411 | * | NA_peak_50 | 23 | Intron (ENSM | 5 | 128757791 | 128820069 | 62279 | 2 | 231760 | ENSMUST00 | -4073 | ENSMUSG0 | Rimbp2 | RIMS binding protein 2 |
| chr5 | 129012494 | 129012777 | 284 | * | NA_peak_50 | 22 | Distal Interge | 5 | 128984557 | 129008574 | 24018 | 2 | 13852 | ENSMUST00 | -3920 | ENSMUSG0 | Stx2 | syntaxin 2 |
| chr5 | 129108738 | 129108922 | 185 | * | NA_peak_50 | 18 | Intron (ENSM | 5 | 129096757 | 129203680 | 106924 | 1 | 243277 | ENSMUST00 | 11981 | ENSMUSG0 | Adgrd1 | adhesion G protein-coupled receptor D1 |
| chr5 | 129467496 | 129467835 | 340 | * | NA_peak_50 | 23 | Distal Interge | 5 | 129501221 | 129571384 | 70164 | 1 | 231769 | ENSMUST00 | -33386 | ENSMUSG0 | Sfswap | splicing factor SWAP |
| chr5 | 131447375 | 131447584 | 210 | * | NA_peak_50 | 14 | Intron (ENSM | 5 | 131445424 | 131455862 | 10439 | 2 | 319974 | ENSMUST00 | 8278 | ENSMUSG0 | Auts2 | autism susceptibility candidate 2 |
| chr5 | 132811766 | 132812089 | 324 | * | NA_peak_50 | 40 | Distal Interge | 5 | 131437333 | 132543344 | 1106012 | 2 | 319974 | ENSMUST00 | -268422 | ENSMUSG0 | Auts2 | autism susceptibility candidate 2 |
| chr5 | 134368867 | 134369101 | 235 | * | NA_peak_50 | 23 | Intron (ENSM | 5 | 134357663 | 134363276 | 5614 | 2 | 57080 | ENSMUST00 | -5591 | ENSMUSG0 | Gtf2ird1 | general transcription factor II I repeat domain-containing 1 |
| chr5 | 134773203 | 134773277 | 75 | * | NA_peak_50 | 23 | Intron (ENSM | 5 | 134702593 | 134747323 | 44731 | 2 | 13717 | ENSMUST00 | -25880 | ENSMUSG0 | Eln | elastin |
| chr5 | 135561071 | 135561351 | 281 | * | NA_peak_50 | 26 | Promoter (2-3 | 5 | 135560448 | 135563569 | 3122 | 2 | 541307 | ENSMUST00 | 2218 | ENSMUSG0 | Ccl26 | chemokine (C-C motif) ligand 26 |
| chr5 | 138807124 | 138807285 | 162 | * | NA_peak_50 | 17 | Promoter (<= | 5 | 138807190 | 138809405 | 2216 | 1 | 80752 | ENSMUST00 | 0 | ENSMUSG0 | Fam20c | FAM20C, golgi associated secretory pathway kinase |
| chr5 | 138807700 | 138808004 | 305 | * | NA_peak_50 | 33 | Promoter (<= | 5 | 138807190 | 138809405 | 2216 | 1 | 80752 | ENSMUST00 | 510 | ENSMUSG0 | Fam20c | FAM20C, golgi associated secretory pathway kinase |
| chr5 | 142059085 | 142059126 | 42 | * | NA_peak_50 | 23 | Intron (ENSM | 5 | 141856787 | 142213194 | 356408 | 1 | 330222 | ENSMUST00 | 202298 | ENSMUSG0 | Sdk1 | sidekick cell adhesion molecule 1 |
| chr5 | 142816890 | 142817196 | 307 | * | NA_peak_50 | 34 | Promoter (<= | 5 | 142793005 | 142817380 | 24376 | 2 | 231861 | ENSMUST00 | 184 | ENSMUSG0 | Tnrc18 | trinucleotide repeat containing 18 |
| chr5 | 143200684 | 143200990 | 307 | * | NA_peak_50 | 21 | Intron (ENSM | 5 | 143194131 | 143204213 | 10083 | 1 | 330228 | ENSMUST00 | 6553 | ENSMUSG0 | Spdye4b | speedy/RINGO cell cycle regulator family, member E4B |
| chr5 | 144099084 | 144099338 | 255 | * | NA_peak_50 | 26 | Promoter (1-2 | 5 | 144100436 | 144188204 | 87769 | 1 | 231876 | ENSMUST00 | -1098 | ENSMUSG0 | Lmtk2 | lemur tyrosine kinase 2 |
| chr5 | 145053302 | 145053480 | 179 | * | NA_peak_50 | 20 | Intron (ENSM | 5 | 145042990 | 145045681 | 2692 | 1 | 69396 | ENSMUST00 | 10312 | ENSMUSG0 | 1700018F24 | RIKEN cDNA 1700018F24 gene |
| chr5 | 145233648 | 145233740 | 93 | * | NA_peak_50 | 22 | Promoter (1-2 | 5 | 145231757 | 145237282 | 5526 | 1 | 72611 | ENSMUST00 | 1891 | ENSMUSG0 | Zfp655 | zinc finger protein 655 |
| chr5 | 145444806 | 145445093 | 288 | * | NA_peak_50 | 23 | Intron (ENSM | 5 | 145436309 | 145469723 | 33415 | 2 | 13114 | ENSMUST00 | 24630 | ENSMUSG0 | Cyp3a16 | cytochrome P450, family 3, subfamily a, polypeptide 16 |
| chr5 | 145468778 | 145469141 | 364 | * | NA_peak_50 | 20 | Promoter (<= | 5 | 145436309 | 145469723 | 33415 | 2 | 13114 | ENSMUST00 | 582 | ENSMUSG0 | Cyp3a16 | cytochrome P450, family 3, subfamily a, polypeptide 16 |
| chr5 | 145575471 | 145575755 | 285 | * | NA_peak_50 | 37 | Intron (ENSM | 5 | 145558664 | 145584730 | 26067 | 2 | 100041375 | ENSMUST00 | 8975 | ENSMUSG0 | Cyp3a41b | cytochrome P450, family 3, subfamily a, polypeptide 41B |
| chr5 | 145599922 | 145600485 | 564 | * | NA_peak_50 | 36 | Distal Interge | 5 | 145558664 | 145584730 | 26067 | 2 | 100041375 | ENSMUST00 | -15192 | ENSMUSG0 | Cyp3a41b | cytochrome P450, family 3, subfamily a, polypeptide 41B |
| chr5 | 145671629 | 145671898 | 270 | * | NA_peak_50 | 16 | Distal Interge | 5 | 145694049 | 145720136 | 26088 | 2 | 53973 | ENSMUST00 | 48238 | ENSMUSG0 | Cyp3a41a | cytochrome P450, family 3, subfamily a, polypeptide 41A |
| chr5 | 145726205 | 145727217 | 1013 | * | NA_peak_50 | 31 | Distal Interge | 5 | 145694049 | 145720136 | 26088 | 2 | 53973 | ENSMUST00 | -6069 | ENSMUSG0 | Cyp3a41a | cytochrome P450, family 3, subfamily a, polypeptide 41A |
| chr5 | 146698236 | 146698285 | 50 | * | NA_peak_50 | 13 | Distal Interge | 5 | 146734839 | 146758448 | 23610 | 2 | 22217 | ENSMUST00 | 60163 | ENSMUSG0 | Usp12 | ubiquitin specific peptidase 12 |
| chr5 | 147586483 | 147586822 | 340 | * | NA_peak_50 | 26 | Intron (ENSM | 5 | 147614756 | 147615242 | 487 | 2 | 14254 | ENSMUST00 | 28420 | ENSMUSG0 | Flt1 | FMS-like tyrosine kinase 1 |
| chr5 | 147809702 | 147809990 | 289 | * | NA_peak_50 | 13 | Distal Interge | 5 | 147860461 | 147876456 | 15996 | 1 | 66537 | ENSMUST00 | -50471 | ENSMUSG0 | Pomp | proteasome maturation protein |
| chr5 | 148231395 | 148231470 | 76 | * | NA_peak_50 | 31 | Intron (ENSM | 5 | 148265217 | 148314071 | 48855 | 1 | 77521 | ENSMUST00 | -33747 | ENSMUSG0 | Mtus2 | microtubule associated tumor suppressor candidate 2 |
| chr5 | 148865272 | 148865566 | 295 | * | NA_peak_50 | 20 | Distal Interge | 5 | 148875683 | 148879079 | 3397 | 2 | 231912 | ENSMUST00 | 13513 | ENSMUSG0 | Katnal1 | katanin p60 subunit A-like 1 |
| chr5 | 149366263 | 149366586 | 324 | * | NA_peak_50 | 36 | Promoter (1-2 | 5 | 149368476 | 149430682 | 62207 | 1 | 70717 | ENSMUST00 | -1890 | ENSMUSG0 | Medag | mesenteric estrogen dependent adipogenesis |
| chr5 | 149489630 | 149489747 | 118 | * | NA_peak_50 | 16 | Exon (ENSMU | 5 | 149528679 | 149611894 | 83216 | 1 | 381693 | ENSMUST00 | -38932 | ENSMUSG0 | Wdr95 | WD40 repeat domain 95 |
| chr5 | 150592467 | 150592956 | 490 | * | NA_peak_50 | 23 | Promoter (1-2 | 5 | 150572665 | 150594462 | 21798 | 2 | 100637 | ENSMUST00 | 1506 | ENSMUSG0 | N4bp2l1 | NEDD4 binding protein 2-like 1 |
| chr5 | 151067720 | 151067905 | 186 | * | NA_peak_50 | 27 | Intron (ENSM | 5 | 151063543 | 151089262 | 25720 | 2 | 243362 | ENSMUST00 | 21357 | ENSMUSG0 | Stard13 | StAR-related lipid transfer (START) domain containing 13 |
| chr5 | 151357329 | 151357630 | 302 | * | NA_peak_50 | 16 | Intron (ENSM | 5 | 151368675 | 151382183 | 13509 | 1 | 625175 | ENSMUST00 | -11045 | ENSMUSG0 | 1700028E10 | RIKEN cDNA 1700028E10 gene |
| chr5 | 151456498 | 151456858 | 361 | * | NA_peak_50 | 15 | Intron (ENSM | 5 | 151063546 | 151428163 | 364618 | 2 | 243362 | ENSMUST00 | -28335 | ENSMUSG0 | Stard13 | StAR-related lipid transfer (START) domain containing 13 |
| chr5 | 151475181 | 151475460 | 280 | * | NA_peak_50 | 39 | Distal Interge | 5 | 151063546 | 151428163 | 364618 | 2 | 243362 | ENSMUST00 | -47018 | ENSMUSG0 | Stard13 | StAR-related lipid transfer (START) domain containing 13 |
| chr5 | 151528656 | 151529018 | 363 | * | NA_peak_50 | 55 | Intron (ENSM | 5 | 151561661 | 151586924 | 25264 | 2 | 632671 | ENSMUST00 | 57906 | ENSMUSG0 | Vmn2r18 | vomeronasal 2, receptor 18 |
| chr6 | 3192476 | 3193198 | 723 | * | NA_peak_50 | 39 | Distal Interge | 6 | 3288519 | 3317019 | 28501 | 1 | 667335 | ENSMUST00 | -95321 | ENSMUSG0 | Gm8579 | spermatogenesis associated glutamate (E)-rich protein 4a pseudogene |
| chr6 | 5612946 | 5613484 | 539 | * | NA_peak_50 | 36 | Distal Interge | 6 | 5725639 | 6028039 | 302401 | 1 | 13426 | ENSMUST00 | -112155 | ENSMUSG0 | Dync1i1 | dynein cytoplasmic 1 intermediate chain 1 |
| chr6 | 10147429 | 10147468 | 40 | * | NA_peak_50 | 16 | Distal Interge | 6 | 10972058 | 10974378 | 2321 | 2 | 57866 | ENSMUST00 | 826910 | NA | AA545190 | EST AA545190 |
| chr6 | 12622240 | 12622513 | 274 | * | NA_peak_50 | 47 | Intron (ENSM | 6 | 12465280 | 12555113 | 89834 | 2 | 330267 | ENSMUST00 | -67127 | ENSMUSG0 | Thsd7a | thrombospondin, type I, domain containing 7A |
| chr6 | 12870632 | 12871342 | 711 | * | NA_peak_50 | 53 | Distal Interge | 6 | 12317087 | 12749410 | 432324 | 2 | 330267 | ENSMUST00 | -121222 | ENSMUSG0 | Thsd7a | thrombospondin, type I, domain containing 7A |
| chr6 | 14797380 | 14797452 | 73 | * | NA_peak_50 | 14 | Distal Interge | 6 | 14713977 | 14755274 | 41298 | 2 | 140491 | ENSMUST00 | -42106 | ENSMUSG0 | Ppp1r3a | protein phosphatase 1, regulatory subunit 3A |
| chr6 | 17463889 | 17464188 | 300 | * | NA_peak_51 | 23 | Promoter (<= | 6 | 17464038 | 17572229 | 108192 | 1 | 17295 | ENSMUST00 | 0 | ENSMUSG0 | Met | met proto-oncogene |
| chr6 | 18064799 | 18065038 | 240 | * | NA_peak_51 | 26 | Intron (ENSM | 6 | 18073804 | 18075107 | 1304 | 2 | 74068 | ENSMUST00 | 10069 | ENSMUSG0 | Asz1 | ankyrin repeat, SAM and basic leucine zipper domain containing 1 |
| chr6 | 19071665 | 19072327 | 663 | * | NA_peak_51 | 50 | Distal Interge | 6 | 18866350 | 18879582 | 13233 | 1 | 75196 | ENSMUST00 | 205315 | ENSMUSG0 | Ankrd7 | ankyrin repeat domain 7 |
| chr6 | 21014863 | 21015000 | 138 | * | NA_peak_51 | 36 | Distal Interge | 6 | 21215503 | 21729805 | 514303 | 1 | 16508 | ENSMUST00 | -200503 | ENSMUSG0 | Kcnd2 | potassium voltage-gated channel, Shal-related family, member 2 |
| chr6 | 22126482 | 22126747 | 266 | * | NA_peak_51 | 50 | Intron (ENSM | 6 | 22135546 | 22145551 | 10006 | 1 | 214642 | ENSMUST00 | -8799 | ENSMUSG0 | Cped1 | cadherin-like and PC-esterase domain containing 1 |
| chr6 | 23066192 | 23066526 | 335 | * | NA_peak_51 | 27 | Distal Interge | 6 | 23092283 | 23127580 | 35298 | 2 | 30956 | ENSMUST00 | 61054 | ENSMUSG0 | Aass | aminoadipate-semialdehyde synthase |
| chr6 | 23480236 | 23480527 | 292 | * | NA_peak_51 | 65 | Intron (ENSM | 6 | 23263508 | 23587424 | 323917 | 2 | 320405 | ENSMUST00 | 106897 | ENSMUSG0 | Cadps2 | Ca2+-dependent activator protein for secretion 2 |
| chr6 | 23672953 | 23673272 | 320 | * | NA_peak_51 | 40 | Intron (ENSM | 6 | 23653898 | 23655136 | 1239 | 2 | 71300 | ENSMUST00 | -17817 | ENSMUSG0 | Rnf148 | ring finger protein 148 |
| chr6 | 24281246 | 24281589 | 344 | * | NA_peak_51 | 40 | Distal Interge | 6 | 24088283 | 24168092 | 79810 | 2 | 55961 | ENSMUST00 | -113154 | ENSMUSG0 | Slc13a1 | solute carrier family 13 (sodium/sulfate symporters), member 1 |
| chr6 | 24673457 | 24673777 | 321 | * | NA_peak_51 | 36 | Distal Interge | 6 | 24613805 | 24665009 | 51205 | 2 | 73178 | ENSMUST00 | -8448 | ENSMUSG0 | Wasl | WASP like actin nucleation promoting factor |
| chr6 | 25555525 | 25555544 | 20 | * | NA_peak_51 | 21 | Distal Interge | 6 | 25665878 | 25689792 | 23915 | 2 | 14763 | ENSMUST00 | 134248 | ENSMUSG0 | Gpr37 | G protein-coupled receptor 37 |
| chr6 | 26402979 | 26403265 | 287 | * | NA_peak_51 | 51 | Distal Interge | 6 | 25954485 | 25958159 | 3675 | 1 | 626904 | ENSMUST00 | 448494 | ENSMUSG0 | Gm20756 | predicted gene, 20756 |
| chr6 | 26879055 | 26879636 | 582 | * | NA_peak_51 | 53 | Distal Interge | 6 | 25954485 | 25958159 | 3675 | 1 | 626904 | ENSMUST00 | 924570 | ENSMUSG0 | Gm20756 | predicted gene, 20756 |
| chr6 | 31853188 | 31853492 | 305 | * | NA_peak_51 | 54 | Intron (ENSM | 6 | 31662220 | 31666131 | 3912 | 2 | 434003 | ENSMUST00 | -187057 | ENSMUSG0 | Gm13848 | predicted gene 13848 |
| chr6 | 32094196 | 32095006 | 811 | * | NA_peak_51 | 32 | Distal Interge | 6 | 32058141 | 32058621 | 481 | 1 | 76382 | ENSMUST00 | 36055 | ENSMUSG0 | 1700012A03 | RIKEN cDNA 1700012A03 gene |
| chr6 | 35488586 | 35488868 | 283 | * | NA_peak_51 | 31 | Distal Interge | 6 | 35521929 | 35539799 | 17871 | 2 | 14489 | ENSMUST00 | 50931 | ENSMUSG0 | Mtpn | myotrophin |
| chr6 | 36085268 | 36085725 | 458 | * | NA_peak_51 | 48 | Distal Interge | 6 | 36209620 | 36387851 | 178232 | 2 | 319472 | ENSMUST00 | 302126 | ENSMUSG0 | 9330158H04 | RIKEN cDNA 9330158H04 gene |
| chr6 | 37366120 | 37366387 | 268 | * | NA_peak_51 | 20 | Intron (ENSM | 6 | 37334292 | 37336535 | 2244 | 2 | 208647 | ENSMUST00 | -29585 | ENSMUSG0 | Creb3l2 | cAMP responsive element binding protein 3-like 2 |
| chr6 | 40541452 | 40541799 | 348 | * | NA_peak_51 | 26 | Distal Interge | 6 | 40544036 | 40544977 | 942 | 2 | 258380 | ENSMUST00 | 3178 | ENSMUSG0 | Olfr461 | olfactory receptor 461 |
| chr6 | 40991471 | 40991894 | 424 | * | NA_peak_51 | 36 | Distal Interge | 6 | 40995830 | 40999479 | 3650 | 2 | 100040208 | ENSMUST00 | 7585 | ENSMUSG0 | Gm2663 | predicted gene 2663 |
| chr6 | 41995507 | 41995597 | 91 | * | NA_peak_51 | 27 | Distal Interge | 6 | 41968140 | 41973090 | 4951 | 1 | 387564 | ENSMUST00 | 27367 | ENSMUSG0 | Sval3 | seminal vesicle antigen-like 3 |
| chr6 | 42498566 | 42498967 | 402 | * | NA_peak_51 | 21 | Distal Interge | 6 | 42486163 | 42487214 | 1052 | 2 | 259144 | ENSMUST00 | -11352 | ENSMUSG0 | Olfr456 | olfactory receptor 456 |
| chr6 | 44692129 | 44692338 | 210 | * | NA_peak_51 | 23 | Distal Interge | 6 | 45059357 | 45715945 | 656589 | 1 | 66797 | ENSMUST00 | -367019 | ENSMUSG0 | Cntnap2 | contactin associated protein-like 2 |
| chr6 | 44726177 | 44726505 | 329 | * | NA_peak_51 | 23 | Distal Interge | 6 | 45059357 | 45715945 | 656589 | 1 | 66797 | ENSMUST00 | -332852 | ENSMUSG0 | Cntnap2 | contactin associated protein-like 2 |
| chr6 | 46582149 | 46582182 | 34 | * | NA_peak_51 | 19 | Intron (ENSM | 6 | 46170215 | 46435576 | 265362 | 1 | 66797 | ENSMUST00 | 411934 | ENSMUSG0 | Cntnap2 | contactin associated protein-like 2 |
| chr6 | 47740595 | 47740853 | 259 | * | NA_peak_51 | 17 | Distal Interge | 6 | 47803576 | 47803652 | 77 | 2 | 735289 | ENSMUST00 | 62799 | ENSMUSG0 | Mir704 | microRNA 704 |
| chr6 | 47765777 | 47766257 | 481 | * | NA_peak_51 | 43 | Distal Interge | 6 | 47803576 | 47803652 | 77 | 2 | 735289 | ENSMUST00 | 37395 | ENSMUSG0 | Mir704 | microRNA 704 |
| chr6 | 47771427 | 47771727 | 301 | * | NA_peak_51 | 20 | Distal Interge | 6 | 47803576 | 47803652 | 77 | 2 | 735289 | ENSMUST00 | 31925 | ENSMUSG0 | Mir704 | microRNA 704 |
| chr6 | 48233305 | 48233624 | 320 | * | NA_peak_51 | 31 | Distal Interge | 6 | 48130906 | 48159280 | 28375 | 2 | 100503622 | ENSMUST00 | -74025 | ENSMUSG0 | Gm16630 | predicted gene, 16630 |
| chr6 | 49047884 | 49048055 | 172 | * | NA_peak_51 | 16 | Promoter (<= | 6 | 49047424 | 49048351 | 928 | 1 | 93695 | ENSMUST00 | 460 | ENSMUSG0 | Gpnmb | glycoprotein (transmembrane) nmb |
| chr6 | 49210569 | 49210575 | 7 | * | NA_peak_51 | 16 | Intron (ENSM | 6 | 49093111 | 49214161 | 121051 | 2 | 140488 | ENSMUST00 | 3586 | ENSMUSG0 | Igf2bp3 | insulin-like growth factor 2 mRNA binding protein 3 |
| chr6 | 51232848 | 51232920 | 73 | * | NA_peak_51 | 20 | Distal Interge | 6 | 51269812 | 51269910 | 99 | 2 | 387166 | ENSMUST00 | 36990 | ENSMUSG0 | Mir148a | microRNA 148a |
| chr6 | 55122074 | 55122559 | 486 | * | NA_peak_51 | 27 | Intron (ENSM | 6 | 55091186 | 55117985 | 26800 | 2 | 12922 | ENSMUST00 | -4089 | ENSMUSG0 | Crhr2 | corticotropin releasing hormone receptor 2 |
| chr6 | 57288119 | 57288282 | 164 | * | NA_peak_51 | 45 | Distal Interge | 6 | 57257952 | 57259972 | 2021 | 1 | 113863 | ENSMUST00 | 30167 | ENSMUSG0 | Vmn1r15 | vomeronasal 1 receptor 15 |
| chr6 | 58069237 | 58069600 | 364 | * | NA_peak_51 | 26 | Distal Interge | 6 | 58008183 | 58009202 | 1020 | 2 | 171190 | ENSMUST00 | -60035 | ENSMUSG0 | Vmn1r26 | vomeronasal 1 receptor 26 |
| chr6 | 59201795 | 59202083 | 289 | * | NA_peak_51 | 17 | Distal Interge | 6 | 59208870 | 59212033 | 3164 | 1 | 68140 | ENSMUST00 | -6787 | ENSMUSG0 | Tigd2 | tigger transposable element derived 2 |
| chr6 | 59288943 | 59289262 | 320 | * | NA_peak_51 | 31 | Distal Interge | 6 | 59208870 | 59212033 | 3164 | 1 | 68140 | ENSMUST00 | 80073 | ENSMUSG0 | Tigd2 | tigger transposable element derived 2 |
| chr6 | 59587566 | 59587768 | 203 | * | NA_peak_51 | 50 | Distal Interge | 6 | 59347226 | 59426294 | 79069 | 2 | 243385 | ENSMUST00 | -161272 | ENSMUSG0 | Gprin3 | GPRIN family member 3 |
| chr6 | 63175769 | 63176091 | 323 | * | NA_peak_51 | 40 | Distal Interge | 6 | 63255876 | 63314071 | 58196 | 1 | 14804 | ENSMUST00 | -79785 | ENSMUSG0 | Grid2 | glutamate receptor, ionotropic, delta 2 |
| chr6 | 63672031 | 63672270 | 240 | * | NA_peak_51 | 31 | Intron (ENSM | 6 | 64076792 | 64094748 | 17957 | 1 | 14804 | ENSMUST00 | -404522 | ENSMUSG0 | Grid2 | glutamate receptor, ionotropic, delta 2 |
| chr6 | 66769695 | 66770022 | 328 | * | NA_peak_51 | 27 | Distal Interge | 6 | 66776222 | 66777130 | 909 | 2 | 171186 | ENSMUST00 | 7108 | ENSMUSG0 | Vmn1r38 | vomeronasal 1 receptor 38 |
| chr6 | 67581857 | 67582129 | 273 | * | NA_peak_51 | 23 | Distal Interge | 6 | 67565191 | 67566446 | 1256 | 2 | 75070 | ENSMUST00 | -15411 | NA | 4930515G16 | myc induced nuclear antigen pseudogene |
| chr6 | 67690115 | 67691437 | 1323 | * | NA_peak_51 | 48 | Distal Interge | 6 | 67565191 | 67566446 | 1256 | 2 | 75070 | ENSMUST00 | -123669 | NA | 4930515G16 | myc induced nuclear antigen pseudogene |
| chr6 | 67693002 | 67693464 | 463 | * | NA_peak_51 | 38 | Distal Interge | 6 | 67565191 | 67566446 | 1256 | 2 | 75070 | ENSMUST00 | -126556 | NA | 4930515G16 | myc induced nuclear antigen pseudogene |
| chr6 | 71831414 | 71831739 | 326 | * | NA_peak_51 | 23 | Promoter (<= | 6 | 71831418 | 71875250 | 43833 | 1 | 76614 | ENSMUST00 | 0 | ENSMUSG0 | Immt | inner membrane protein, mitochondrial |
| chr6 | 78027203 | 78027489 | 287 | * | NA_peak_51 | 31 | Distal Interge | 6 | 76881637 | 77979699 | 1098063 | 2 | 12386 | ENSMUST00 | -47504 | ENSMUSG0 | Ctnna2 | catenin (cadherin associated protein), alpha 2 |
| chr6 | 78040249 | 78040554 | 306 | * | NA_peak_51 | 39 | Distal Interge | 6 | 76881637 | 77979699 | 1098063 | 2 | 12386 | ENSMUST00 | -60550 | ENSMUSG0 | Ctnna2 | catenin (cadherin associated protein), alpha 2 |
| chr6 | 78155900 | 78156224 | 325 | * | NA_peak_51 | 68 | Distal Interge | 6 | 76881637 | 77979699 | 1098063 | 2 | 12386 | ENSMUST00 | -176201 | ENSMUSG0 | Ctnna2 | catenin (cadherin associated protein), alpha 2 |
| chr6 | 79522068 | 79522394 | 327 | * | NA_peak_51 | 24 | Distal Interge | 6 | 79817916 | 79818364 | 449 | 1 | 100463512 | ENSMUST00 | -295522 | ENSMUSG0 | Gm20594 | predicted gene, 20594 |
| chr6 | 82344917 | 82345203 | 287 | * | NA_peak_51 | 31 | Distal Interge | 6 | 82402334 | 82405364 | 3031 | 1 | 21336 | ENSMUST00 | -57131 | ENSMUSG0 | Tacr1 | tachykinin receptor 1 |
| chr6 | 82626616 | 82626746 | 131 | * | NA_peak_51 | 36 | Intron (ENSM | 6 | 82647843 | 82652518 | 4676 | 2 | 66979 | ENSMUST00 | 25772 | ENSMUSG0 | Pole4 | polymerase (DNA-directed), epsilon 4 (p12 subunit) |
| chr6 | 82748852 | 82749266 | 415 | * | NA_peak_51 | 20 | Exon (ENSMU | 6 | 82760117 | 82772988 | 12872 | 2 | 15277 | ENSMUST00 | 23722 | ENSMUSG0 | Hk2 | hexokinase 2 |
| chr6 | 85875556 | 85875861 | 306 | * | NA_peak_51 | 27 | Distal Interge | 6 | 85865422 | 85869158 | 3737 | 2 | 93673 | ENSMUST00 | -6398 | ENSMUSG0 | Nat8f2 | N-acetyltransferase 8 (GCN5-related) family member 2 |
| chr6 | 86732780 | 86733023 | 244 | * | NA_peak_51 | 40 | Promoter (<= | 6 | 86720210 | 86733240 | 13031 | 2 | 23885 | ENSMUST00 | 217 | ENSMUSG0 | Gmcl1 | germ cell-less, spermatogenesis associated 1 |
| chr6 | 87402448 | 87402707 | 260 | * | NA_peak_51 | 26 | Distal Interge | 6 | 87383256 | 87388935 | 5680 | 2 | 68888 | ENSMUST00 | -13513 | ENSMUSG0 | Gkn3 | gastrokine 3 |
| chr6 | 87850450 | 87850845 | 396 | * | NA_peak_51 | 20 | Promoter (<= | 6 | 87843939 | 87851022 | 7084 | 2 | 12785 | ENSMUST00 | 177 | ENSMUSG0 | Cnbp | cellular nucleic acid binding protein |
| chr6 | 90012733 | 90013025 | 293 | * | NA_peak_51 | 23 | Distal Interge | 6 | 90021888 | 90022820 | 933 | 1 | 113846 | ENSMUST00 | -8863 | ENSMUSG0 | Vmn1r47 | vomeronasal 1 receptor 47 |
| chr6 | 90031793 | 90032086 | 294 | * | NA_peak_51 | 23 | Distal Interge | 6 | 90035933 | 90036841 | 909 | 2 | 113845 | ENSMUST00 | 4755 | ENSMUSG0 | Vmn1r48 | vomeronasal 1 receptor 48 |
| chr6 | 95214685 | 95215009 | 325 | * | NA_peak_51 | 56 | Distal Interge | 6 | 95117984 | 95121797 | 3814 | 1 | 243574 | ENSMUST00 | 96701 | ENSMUSG0 | Kbtbd8 | kelch repeat and BTB (POZ) domain containing 8 |

| | | | | | | | | | | | | | | | | | | |
|---|---|---|---|---|---|---|---|---|---|---|---|---|---|---|---|---|---|---|
| chr6 | 95659844 | 95659859 | 16 | * | NA_peak_518 | 30 | Promoter (2-3 | 6 | 95594683 | 95657468 | 62786 | 2 | 20917 | ENSMUST00 | -2376 | ENSMUSG0 | Suclg2 | succinate-Coenzyme A ligase, GDP-forming, beta subunit |
| chr6 | 95796482 | 95797031 | 550 | * | NA_peak_518 | 27 | Distal Intergei | 6 | 95654751 | 95718800 | 64050 | 2 | 20917 | ENSMUST00 | -77682 | ENSMUSG0 | Suclg2 | succinate-Coenzyme A ligase, GDP-forming, beta subunit |
| chr6 | 98823443 | 98823849 | 407 | * | NA_peak_518 | 28 | Distal Intergei | 6 | 98930171 | 98935545 | 5375 | 2 | 108655 | ENSMUST00 | 111696 | ENSMUSG0 | Foxp1 | forkhead box P1 |
| chr6 | 101783605 | 101783834 | 230 | * | NA_peak_518 | 50 | Promoter (1-2 | 6 | 101782539 | 101801815 | 19277 | 1 | 207157 | ENSMUST00 | 1066 | ENSMUSG0 | Gm9871 | predicted gene 9871 |
| chr6 | 109659814 | 109660068 | 255 | * | NA_peak_519 | 16 | Distal Intergei | 6 | 108852391 | 108855525 | 3135 | 1 | 192193 | ENSMUST00 | 807423 | ENSMUSG0 | Edem1 | ER degradation enhancer, mannosidase alpha-like 1 |
| chr6 | 110849117 | 110849412 | 296 | * | NA_peak_519 | 59 | Intron (ENSM | 6 | 110645582 | 111567230 | 921649 | 1 | 108073 | ENSMUST00 | 203535 | ENSMUSG0 | Grm7 | glutamate receptor, metabotropic 7 |
| chr6 | 117398280 | 117398574 | 295 | * | NA_peak_520 | 16 | Distal Intergei | 6 | 117495235 | 117500077 | 4843 | 1 | 100126203 | ENSMUST00 | -96661 | ENSMUSG0 | 6820426E19 | RIKEN cDNA 6820426E19 gene |
| chr6 | 117609314 | 117609587 | 274 | * | NA_peak_520 | 20 | Intron (ENSM | 6 | 117646694 | 117717263 | 70570 | 2 | 72263 | ENSMUST00 | 107676 | ENSMUSG0 | 1700030F04 | RIKEN cDNA 1700030F04 gene |
| chr6 | 118209759 | 118210084 | 326 | * | NA_peak_520 | 61 | Distal Intergei | 6 | 118154566 | 118197718 | 43153 | 2 | 19713 | ENSMUST00 | -12041 | ENSMUSG0 | Ret | ret proto-oncogene |
| chr6 | 122915340 | 122915536 | 197 | * | NA_peak_520 | 17 | Distal Intergei | 6 | 122921848 | 122934619 | 12772 | 1 | 269799 | ENSMUST00 | -6312 | ENSMUSG0 | Clec4a1 | C-type lectin domain family 4, member a1 |
| chr6 | 124914022 | 124914050 | 29 | * | NA_peak_521 | 23 | Promoter (<= | 6 | 124913688 | 124914674 | 987 | 2 | 69202 | ENSMUST00 | 624 | ENSMUSG0 | Ptms | parathymosin |
| chr6 | 131018122 | 131018406 | 285 | * | NA_peak_521 | 16 | Distal Intergei | 6 | 131219223 | 131247362 | 28140 | 2 | 16633 | ENSMUST00 | 228956 | ENSMUSG0 | Klra2 | killer cell lectin-like receptor, subfamily A, member 2 |
| chr6 | 133191811 | 133192031 | 221 | * | NA_peak_521 | 16 | Exon (ENSMU | 6 | 133105492 | 133107429 | 1938 | 1 | 381820 | ENSMUST00 | 86319 | ENSMUSG0 | Smim10l1 | small integral membrane protein 10 like 1 |
| chr6 | 133508193 | 133508513 | 321 | * | NA_peak_521 | 23 | Distal Intergei | 6 | 133529189 | 133532762 | 3574 | 1 | 545884 | ENSMUST00 | -20676 | ENSMUSG0 | Gm5885 | predicted gene 5885 |
| chr6 | 133684391 | 133684535 | 145 | * | NA_peak_521 | 16 | Distal Intergei | 6 | 133763999 | 133767403 | 3405 | 2 | 545886 | ENSMUST00 | 82868 | ENSMUSG0 | Gm5886 | predicted gene 5886 |
| chr6 | 136324421 | 136324705 | 285 | * | NA_peak_522 | 20 | Promoter (2-3 | 6 | 136327539 | 136329983 | 2445 | 1 | 668137 | ENSMUST00 | -2834 | ENSMUSG0 | Eif4a3l1 | eukaryotic translation initiation factor 4A3 like 1 |
| chr6 | 136330367 | 136330847 | 481 | * | NA_peak_522 | 36 | Promoter (2-3 | 6 | 136327539 | 136329983 | 2445 | 1 | 668137 | ENSMUST00 | 2828 | ENSMUSG0 | Eif4a3l1 | eukaryotic translation initiation factor 4A3 like 1 |
| chr6 | 138025678 | 138025920 | 243 | * | NA_peak_522 | 31 | Intron (ENSM | 6 | 138017385 | 138036903 | 19519 | 2 | 277898 | ENSMUST00 | 10983 | ENSMUSG0 | Slc15a5 | solute carrier family 15, member 5 |
| chr6 | 138098997 | 138099084 | 88 | * | NA_peak_522 | 27 | Distal Intergei | 6 | 137983590 | 138079916 | 96327 | 2 | 277898 | ENSMUST00 | -19081 | ENSMUSG0 | Slc15a5 | solute carrier family 15, member 5 |
| chr6 | 140459356 | 140459637 | 282 | * | NA_peak_522 | 49 | Intron (ENSM | 6 | 140432989 | 140553029 | 120041 | 1 | 109135 | ENSMUST00 | 26367 | ENSMUSG0 | Plekha5 | pleckstrin homology domain containing, family A member 5 |
| chr6 | 144438976 | 144439271 | 296 | * | NA_peak_522 | 55 | Intron (ENSM | 6 | 144154829 | 144209568 | 54740 | 2 | 20678 | ENSMUST00 | -229408 | ENSMUSG0 | Sox5 | SRY (sex determining region Y)-box 5 |
| chr6 | 145995216 | 145995849 | 634 | * | NA_peak_522 | 15 | Distal Intergei | 6 | 145953429 | 145965223 | 11795 | 1 | 16651 | ENSMUST00 | 41787 | ENSMUSG0 | Sspn | sarcospan |
| chr6 | 149309563 | 149309820 | 258 | * | NA_peak_523 | 31 | Promoter (<= | 6 | 149309573 | 149310455 | 883 | 1 | 67246 | ENSMUST00 | 0 | ENSMUSG0 | Resf1 | retroelement silencing factor 1 |
| chr7 | 5709726 | 5709856 | 131 | * | NA_peak_523 | 40 | Distal Intergei | 7 | 5802345 | 5805445 | 3101 | 2 | 81017 | ENSMUST00 | 95589 | ENSMUSG0 | Vmn1r63 | vomeronasal 1 receptor 63 |
| chr7 | 5863182 | 5863337 | 156 | * | NA_peak_523 | 23 | Distal Intergei | 7 | 5883580 | 5884542 | 963 | 2 | 404285 | ENSMUST00 | 21205 | ENSMUSG0 | Vmn1r64 | vomeronasal 1 receptor 64 |
| chr7 | 7377964 | 7378106 | 143 | * | NA_peak_523 | 23 | Distal Intergei | 7 | 7334149 | 7374573 | 40425 | 2 | 22306 | ENSMUST00 | -3391 | ENSMUSG0 | Vmn2r30 | vomeronasal 2, receptor 30 |
| chr7 | 7424986 | 7425305 | 320 | * | NA_peak_523 | 36 | Intron (ENSM | 7 | 7378251 | 7399703 | 21453 | 2 | 100042591 | ENSMUST00 | -25283 | ENSMUSG0 | Vmn2r31 | vomeronasal 2, receptor 31 |
| chr7 | 7447383 | 7447817 | 435 | * | NA_peak_523 | 36 | Distal Intergei | 7 | 7462096 | 7480018 | 17923 | 2 | 22311 | ENSMUST00 | 32201 | ENSMUSG0 | Vmn2r32 | vomeronasal 2, receptor 32 |
| chr7 | 7457378 | 7457752 | 375 | * | NA_peak_523 | 70 | Distal Intergei | 7 | 7462096 | 7480018 | 17923 | 2 | 22311 | ENSMUST00 | 22266 | ENSMUSG0 | Vmn2r32 | vomeronasal 2, receptor 32 |
| chr7 | 7459497 | 7459648 | 152 | * | NA_peak_523 | 24 | Distal Intergei | 7 | 7462096 | 7480018 | 17923 | 2 | 22311 | ENSMUST00 | 20370 | ENSMUSG0 | Vmn2r32 | vomeronasal 2, receptor 32 |
| chr7 | 7459999 | 7460218 | 220 | * | NA_peak_523 | 24 | Distal Intergei | 7 | 7462096 | 7480018 | 17923 | 2 | 22311 | ENSMUST00 | 19800 | ENSMUSG0 | Vmn2r32 | vomeronasal 2, receptor 32 |
| chr7 | 7467630 | 7467946 | 317 | * | NA_peak_523 | 22 | Intron (ENSM | 7 | 7462096 | 7480018 | 17923 | 2 | 22311 | ENSMUST00 | 12072 | ENSMUSG0 | Vmn2r32 | vomeronasal 2, receptor 32 |
| chr7 | 7483788 | 7484297 | 510 | * | NA_peak_523 | 50 | Distal Intergei | 7 | 7462096 | 7480018 | 17923 | 2 | 22311 | ENSMUST00 | -3770 | ENSMUSG0 | Vmn2r32 | vomeronasal 2, receptor 32 |
| chr7 | 7486980 | 7487322 | 343 | * | NA_peak_523 | 46 | Distal Intergei | 7 | 7462096 | 7480018 | 17923 | 2 | 22311 | ENSMUST00 | -6962 | ENSMUSG0 | Vmn2r32 | vomeronasal 2, receptor 32 |
| chr7 | 7495367 | 7495759 | 393 | * | NA_peak_523 | 53 | Intron (ENSM | 7 | 7462096 | 7480018 | 17923 | 2 | 22311 | ENSMUST00 | -15349 | ENSMUSG0 | Vmn2r32 | vomeronasal 2, receptor 32 |
| chr7 | 7507921 | 7508229 | 309 | * | NA_peak_523 | 39 | Distal Intergei | 7 | 7462096 | 7480018 | 17923 | 2 | 22311 | ENSMUST00 | -27903 | ENSMUSG0 | Vmn2r32 | vomeronasal 2, receptor 32 |
| chr7 | 7512541 | 7512849 | 309 | * | NA_peak_523 | 18 | Distal Intergei | 7 | 7462096 | 7480018 | 17923 | 2 | 22311 | ENSMUST00 | -32523 | ENSMUSG0 | Vmn2r32 | vomeronasal 2, receptor 32 |
| chr7 | 7521530 | 7521600 | 71 | * | NA_peak_523 | 42 | Distal Intergei | 7 | 7462096 | 7480018 | 17923 | 2 | 22311 | ENSMUST00 | -41512 | ENSMUSG0 | Vmn2r32 | vomeronasal 2, receptor 32 |
| chr7 | 7522381 | 7522781 | 401 | * | NA_peak_523 | 42 | Distal Intergei | 7 | 7462096 | 7480018 | 17923 | 2 | 22311 | ENSMUST00 | -42363 | ENSMUSG0 | Vmn2r32 | vomeronasal 2, receptor 32 |
| chr7 | 8089825 | 8090134 | 310 | * | NA_peak_523 | 50 | Promoter (2-3 | 7 | 8086387 | 8092838 | 6452 | 2 | 72716 | ENSMUST00 | 2704 | ENSMUSG0 | Gm45351 | predicted gene 45351 |
| chr7 | 8099565 | 8099940 | 376 | * | NA_peak_523 | 27 | Distal Intergei | 7 | 8086326 | 8093041 | 6716 | 2 | 72716 | ENSMUST00 | -6524 | ENSMUSG0 | Gm45351 | predicted gene 45351 |
| chr7 | 8114516 | 8114849 | 334 | * | NA_peak_523 | 50 | Intron (ENSM | 7 | 8086326 | 8093041 | 6716 | 2 | 72716 | ENSMUST00 | -21475 | ENSMUSG0 | Gm45351 | predicted gene 45351 |
| chr7 | 8230043 | 8230565 | 523 | * | NA_peak_523 | 20 | Distal Intergei | 7 | 8182372 | 8200320 | 17949 | 2 | 22310 | ENSMUST00 | -29723 | ENSMUSG0 | Vmn2r42 | vomeronasal 2, receptor 42 |
| chr7 | 8369140 | 8369446 | 307 | * | NA_peak_524 | 15 | Exon (ENSMU | 7 | 8366164 | 8383333 | 17170 | 2 | 434113 | ENSMUST00 | 13887 | ENSMUSG0 | Vmn2r44 | vomeronasal 2, receptor 44 |
| chr7 | 8375328 | 8375570 | 243 | * | NA_peak_524 | 17 | Intron (ENSM | 7 | 8366164 | 8383333 | 17170 | 2 | 434113 | ENSMUST00 | 7763 | ENSMUSG0 | Vmn2r44 | vomeronasal 2, receptor 44 |
| chr7 | 8426901 | 8426905 | 5 | * | NA_peak_524 | 49 | Distal Intergei | 7 | 8366164 | 8383333 | 17170 | 2 | 434113 | ENSMUST00 | -43568 | ENSMUSG0 | Vmn2r44 | vomeronasal 2, receptor 44 |
| chr7 | 8869381 | 8869556 | 176 | * | NA_peak_524 | 24 | Distal Intergei | 7 | 8855847 | 8862851 | 7005 | 2 | 72716 | ENSMUST00 | -6530 | ENSMUSG0 | Gm45351 | predicted gene 45351 |
| chr7 | 8882480 | 8883038 | 559 | * | NA_peak_524 | 36 | Exon (ENSMU | 7 | 8855847 | 8862851 | 7005 | 2 | 72716 | ENSMUST00 | -19629 | ENSMUSG0 | Gm45351 | predicted gene 45351 |
| chr7 | 9062173 | 9062451 | 279 | * | NA_peak_524 | 27 | Intron (ENSM | 7 | 9013806 | 9030798 | 16993 | 2 | 545909 | ENSMUST00 | -31375 | ENSMUSG0 | Vmn2r39 | vomeronasal 2, receptor 39 |
| chr7 | 9227809 | 9228140 | 332 | * | NA_peak_524 | 45 | Distal Intergei | 7 | 9205546 | 9223653 | 18108 | 2 | 22305 | ENSMUST00 | -4156 | ENSMUSG0 | Vmn2r37 | vomeronasal 2, receptor 37 |
| chr7 | 9895678 | 9895884 | 207 | * | NA_peak_524 | 23 | Intron (ENSM | 7 | 9868880 | 9876966 | 8087 | 2 | 72716 | ENSMUST00 | -18712 | ENSMUSG0 | Gm45351 | predicted gene 45351 |
| chr7 | 9962069 | 9962407 | 339 | * | NA_peak_524 | 19 | Distal Intergei | 7 | 9927624 | 9953585 | 25962 | 2 | 625580 | ENSMUST00 | -8484 | ENSMUSG0 | Vmn2r48 | vomeronasal 2, receptor 48 |
| chr7 | 9966135 | 9966652 | 518 | * | NA_peak_524 | 36 | Exon (ENSMU | 7 | 9927624 | 9953585 | 25962 | 2 | 625580 | ENSMUST00 | -12550 | ENSMUSG0 | Vmn2r48 | vomeronasal 2, receptor 48 |
| chr7 | 9970769 | 9970970 | 202 | * | NA_peak_524 | 65 | Distal Intergei | 7 | 9927624 | 9953585 | 25962 | 2 | 625580 | ENSMUST00 | -17184 | ENSMUSG0 | Vmn2r48 | vomeronasal 2, receptor 48 |
| chr7 | 9986799 | 9987618 | 820 | * | NA_peak_524 | 40 | Exon (ENSMU | 7 | 9976245 | 9992257 | 16013 | 2 | 625605 | ENSMUST00 | 4639 | ENSMUSG0 | Vmn2r49 | vomeronasal 2, receptor 49 |
| chr7 | 9998402 | 9998745 | 344 | * | NA_peak_524 | 29 | Distal Intergei | 7 | 9976245 | 9992257 | 16013 | 2 | 625605 | ENSMUST00 | -6145 | ENSMUSG0 | Vmn2r49 | vomeronasal 2, receptor 49 |
| chr7 | 10003964 | 10004277 | 314 | * | NA_peak_524 | 35 | Distal Intergei | 7 | 9976245 | 9992257 | 16013 | 2 | 625605 | ENSMUST00 | -11707 | ENSMUSG0 | Vmn2r49 | vomeronasal 2, receptor 49 |
| chr7 | 10005385 | 10005659 | 275 | * | NA_peak_524 | 24 | Distal Intergei | 7 | 9976245 | 9992257 | 16013 | 2 | 625605 | ENSMUST00 | -13128 | ENSMUSG0 | Vmn2r49 | vomeronasal 2, receptor 49 |
| chr7 | 10021029 | 10021329 | 301 | * | NA_peak_524 | 52 | Distal Intergei | 7 | 9976245 | 9992257 | 16013 | 2 | 625605 | ENSMUST00 | -28772 | ENSMUSG0 | Vmn2r49 | vomeronasal 2, receptor 49 |
| chr7 | 10023901 | 10024117 | 217 | * | NA_peak_524 | 24 | Distal Intergei | 7 | 10035836 | 10053296 | 17461 | 2 | 434117 | ENSMUST00 | 29179 | ENSMUSG0 | Vmn2r50 | vomeronasal 2, receptor 50 |
| chr7 | 10063825 | 10064126 | 302 | * | NA_peak_524 | 28 | Distal Intergei | 7 | 10035836 | 10053296 | 17461 | 2 | 434117 | ENSMUST00 | -10529 | ENSMUSG0 | Vmn2r50 | vomeronasal 2, receptor 50 |
| chr7 | 12043442 | 12043614 | 173 | * | NA_peak_524 | 23 | Promoter (2-3 | 7 | 12041299 | 12042219 | 921 | 1 | 628946 | ENSMUST00 | 2143 | ENSMUSG0 | Vmn1r77 | vomeronasal 1 receptor 77 |
| chr7 | 12178386 | 12178699 | 314 | * | NA_peak_524 | 55 | Promoter (2-3 | 7 | 12176193 | 12177113 | 921 | 1 | 100042437 | ENSMUST00 | 2193 | ENSMUSG0 | Vmn1r79 | vomeronasal 1 receptor 79 |
| chr7 | 12676297 | 12676518 | 222 | * | NA_peak_524 | 22 | Intron (ENSM | 7 | 12665463 | 12684991 | 19529 | 2 | 100042499 | ENSMUST00 | 8473 | ENSMUSG0 | Vmn2r55 | vomeronasal 2, receptor 55 |
| chr7 | 13593739 | 13594067 | 329 | * | NA_peak_524 | 26 | Distal Intergei | 7 | 13554866 | 13571067 | 16202 | 2 | 77684 | ENSMUST00 | -22672 | ENSMUSG0 | Bsph2 | binder of sperm protein homolog 2 |
| chr7 | 14822165 | 14822608 | 444 | * | NA_peak_524 | 36 | Distal Intergei | 7 | 14659745 | 14664574 | 4830 | 1 | 194588 | ENSMUST00 | 162420 | ENSMUSG0 | Obox7 | oocyte specific homeobox 7 |
| chr7 | 15182496 | 15182891 | 396 | * | NA_peak_525 | 83 | Distal Intergei | 7 | 15388721 | 15398545 | 9825 | 1 | 246792 | ENSMUST00 | -205830 | ENSMUSG0 | Obox2 | oocyte specific homeobox 2 |
| chr7 | 15252127 | 15252230 | 104 | * | NA_peak_525 | 47 | Distal Intergei | 7 | 15388721 | 15398545 | 9825 | 1 | 246792 | ENSMUST00 | -136491 | ENSMUSG0 | Obox2 | oocyte specific homeobox 2 |
| chr7 | 15431778 | 15432076 | 299 | * | NA_peak_525 | 35 | Exon (ENSMU | 7 | 15390176 | 15398129 | 7954 | 1 | 246792 | ENSMUST00 | 41602 | ENSMUSG0 | Obox2 | oocyte specific homeobox 2 |
| chr7 | 15438712 | 15439368 | 657 | * | NA_peak_525 | 53 | Intron (ENSM | 7 | 15390176 | 15398129 | 7954 | 1 | 246792 | ENSMUST00 | 48536 | ENSMUSG0 | Obox2 | oocyte specific homeobox 2 |
| chr7 | 15452137 | 15452288 | 152 | * | NA_peak_525 | 50 | Intron (ENSM | 7 | 15512340 | 15556446 | 44107 | 1 | 71468 | ENSMUST00 | -60052 | ENSMUSG0 | Obox1 | oocyte specific homeobox 1 |
| chr7 | 15463113 | 15463402 | 290 | * | NA_peak_525 | 31 | Intron (ENSM | 7 | 15512340 | 15556446 | 44107 | 1 | 71468 | ENSMUST00 | -48938 | ENSMUSG0 | Obox1 | oocyte specific homeobox 1 |
| chr7 | 21405269 | 21405610 | 342 | * | NA_peak_525 | 27 | Distal Intergei | 7 | 21360368 | 21361291 | 924 | 2 | 621510 | ENSMUST00 | -43978 | ENSMUSG0 | Vmn1r129 | vomeronasal 1 receptor 129 |
| chr7 | 24105352 | 24105794 | 443 | * | NA_peak_525 | 27 | 3' UTR | 7 | 24112314 | 24127952 | 15639 | 1 | 57745 | ENSMUST00 | -6520 | ENSMUSG0 | Zfp112 | zinc finger protein 112 |
| chr7 | 28658168 | 28658473 | 306 | * | NA_peak_525 | 45 | Intron (ENSM | 7 | 28629524 | 28631332 | 1809 | 2 | 101744 | ENSMUST00 | -26836 | ENSMUSG0 | Acp7 | acid phosphatase 7, tartrate resistant |
| chr7 | 29664498 | 29664798 | 301 | * | NA_peak_525 | 31 | Distal Intergei | 7 | 29697691 | 29725250 | 27560 | 2 | 76718 | ENSMUST00 | 60452 | ENSMUSG0 | Catsperg2 | cation channel sperm associated auxiliary subunit gamma 2 |
| chr7 | 30933449 | 30933785 | 337 | * | NA_peak_525 | 49 | Intron (ENSM | 7 | 30922395 | 30924681 | 2287 | 2 | 66438 | ENSMUST00 | -8768 | ENSMUSG0 | Hamp2 | hepcidin antimicrobial peptide 2 |
| chr7 | 31277116 | 31277283 | 168 | * | NA_peak_525 | 22 | Distal Intergei | 7 | 31290508 | 31304977 | 14470 | 1 | 381970 | ENSMUST00 | -13225 | ENSMUSG0 | Scgb2b2 | secretoglobin, family 2B, member 2 |
| chr7 | 35934138 | 35934143 | 6 | * | NA_peak_525 | 16 | Distal Intergei | 7 | 35772343 | 35802989 | 30647 | 2 | 668501 | ENSMUST00 | -131149 | ENSMUSG0 | Zfp507 | zinc finger protein 507 |
| chr7 | 35934514 | 35934575 | 62 | * | NA_peak_525 | 40 | Distal Intergei | 7 | 35772343 | 35802989 | 30647 | 2 | 668501 | ENSMUST00 | -131525 | ENSMUSG0 | Zfp507 | zinc finger protein 507 |
| chr7 | 38176930 | 38177315 | 386 | * | NA_peak_525 | 39 | Distal Intergei | 7 | 38183217 | 38197563 | 14347 | 1 | 72244 | ENSMUST00 | -5902 | ENSMUSG0 | 1600014C10 | RIKEN cDNA 1600014C10 gene |
| chr7 | 38189414 | 38190006 | 593 | * | NA_peak_525 | 19 | Intron (ENSM | 7 | 38186271 | 38195266 | 8996 | 1 | 72244 | ENSMUST00 | 3143 | ENSMUSG0 | 1600014C10 | RIKEN cDNA 1600014C10 gene |
| chr7 | 41029028 | 41029315 | 288 | * | NA_peak_526 | 28 | Intron (ENSM | 7 | 41032719 | 41045302 | 12584 | 1 | 233164 | ENSMUST00 | -3404 | ENSMUSG0 | Gm4884 | predicted gene 4884 |
| chr7 | 41032306 | 41032444 | 139 | * | NA_peak_526 | 29 | Promoter (<= | 7 | 41032719 | 41045302 | 12584 | 1 | 233164 | ENSMUST00 | -275 | ENSMUSG0 | Gm4884 | predicted gene 4884 |
| chr7 | 41430052 | 41430344 | 293 | * | NA_peak_526 | 31 | Intron (ENSM | 7 | 41395945 | 41448696 | 52752 | 2 | 269902 | ENSMUST00 | 18352 | ENSMUSG0 | Vmn2r57 | vomeronasal 2, receptor 57 |
| chr7 | 41432581 | 41432604 | 24 | * | NA_peak_526 | 15 | Intron (ENSM | 7 | 41395945 | 41448696 | 52752 | 2 | 269902 | ENSMUST00 | 16092 | ENSMUSG0 | Vmn2r57 | vomeronasal 2, receptor 57 |
| chr7 | 41847659 | 41847897 | 239 | * | NA_peak_526 | 16 | Intron (ENSM | 7 | 41835814 | 41872712 | 36899 | 2 | 628422 | ENSMUST00 | 24815 | ENSMUSG0 | Vmn2r58 | vomeronasal 2, receptor 58 |
| chr7 | 47176239 | 47176524 | 286 | * | NA_peak_526 | 39 | Distal Intergei | 7 | 47077795 | 47133684 | 55890 | 2 | 19259 | ENSMUST00 | -42555 | ENSMUSG0 | Ptpn5 | protein tyrosine phosphatase, non-receptor type 5 |
| chr7 | 50954635 | 50954907 | 273 | * | NA_peak_526 | 21 | Distal Intergei | 7 | 50599190 | 50600530 | 1341 | 1 | 243996 | ENSMUST00 | 355445 | ENSMUSG0 | 4933405O20 | RIKEN cDNA 4933405O20 gene |
| chr7 | 50955858 | 50956316 | 459 | * | NA_peak_526 | 104 | Distal Intergei | 7 | 50599190 | 50600530 | 1341 | 1 | 243996 | ENSMUST00 | 356668 | ENSMUSG0 | 4933405O20 | RIKEN cDNA 4933405O20 gene |
| chr7 | 50978379 | 50978787 | 409 | * | NA_peak_526 | 35 | Distal Intergei | 7 | 50599190 | 50600530 | 1341 | 1 | 243996 | ENSMUST00 | 379189 | ENSMUSG0 | 4933405O20 | RIKEN cDNA 4933405O20 gene |
| chr7 | 52816256 | 52816521 | 266 | * | NA_peak_526 | 45 | Distal Intergei | 7 | 52011683 | 52015728 | 4046 | 2 | 100503036 | ENSMUST00 | -800528 | ENSMUSG0 | 1700015G11 | RIKEN cDNA 1700015G11 gene |
| chr7 | 59364690 | 59365123 | 434 | * | NA_peak_526 | 29 | Distal Intergei | 7 | 59347337 | 59347424 | 88 | 2 | 102642160 | ENSMUST00 | -17266 | ENSMUSG0 | Gm22496 | predicted gene, 22496 |
| chr7 | 59521229 | 59521455 | 227 | * | NA_peak_526 | 61 | Distal Intergei | 7 | 59445703 | 59453239 | 7537 | 2 | 52480 | ENSMUST00 | -67990 | NA | Snhg14 | small nucleolar RNA host gene 14 |
| chr7 | 59529118 | 59529419 | 302 | * | NA_peak_526 | 36 | Distal Intergei | 7 | 59445703 | 59453239 | 7537 | 2 | 52480 | ENSMUST00 | -75879 | NA | Snhg14 | small nucleolar RNA host gene 14 |
| chr7 | 59600158 | 59600276 | 119 | * | NA_peak_526 | 27 | Distal Intergei | 7 | 59675988 | 59676081 | 94 | 2 | 64244 | ENSMUST00 | 75805 | ENSMUSG0 | Snord116l1 | small nucleolar RNA, C/D box 116-like 1 |
| chr7 | 59681851 | 59682406 | 556 | * | NA_peak_526 | 42 | Promoter (1-2 | 7 | 59683586 | 59683679 | 94 | 2 | 64244 | ENSMUST00 | 1273 | ENSMUSG0 | Snord116l1 | small nucleolar RNA, C/D box 116-like 1 |
| chr7 | 59760070 | 59760346 | 277 | * | NA_peak_527 | 49 | Intron (ENSM | 7 | 59751489 | 59751582 | 94 | 2 | 64243 | ENSMUST00 | -8488 | ENSMUSG0 | Gm26504 | predicted gene, 26504 |
| chr7 | 59768455 | 59768754 | 300 | * | NA_peak_527 | 48 | Intron (ENSM | 7 | 59751489 | 59751582 | 94 | 2 | 64243 | ENSMUST00 | -16873 | ENSMUSG0 | Gm26504 | predicted gene, 26504 |
| chr7 | 59891718 | 59892224 | 507 | * | NA_peak_527 | 40 | Exon (ENSMU | 7 | 59711359 | 59904686 | 193328 | 2 | 52480 | ENSMUST00 | 12462 | NA | Snhg14 | small nucleolar RNA host gene 14 |
| chr7 | 61350420 | 61350498 | 79 | * | NA_peak_527 | 20 | Distal Intergei | 7 | 61535241 | 61546155 | 10915 | 2 | 319752 | ENSMUST00 | 195657 | NA | B230209E15 | RIKEN cDNA B230209E15 gene |
| chr7 | 62535831 | 62536124 | 294 | * | NA_peak_527 | 42 | Distal Intergei | 7 | 62461871 | 62464510 | 2640 | 2 | 27412 | ENSMUST00 | -71321 | ENSMUSG0 | Peg12 | paternally expressed 12 |
| chr7 | 63374079 | 63374439 | 361 | * | NA_peak_527 | 47 | Intron (ENSM | 7 | 63344690 | 63444529 | 99840 | 2 | 77631 | ENSMUST00 | 70090 | ENSMUSG0 | 4930554H23 | RIKEN cDNA 4930554H23 gene |
| chr7 | 64474457 | 64474649 | 193 | * | NA_peak_527 | 13 | Distal Intergei | 7 | 64501706 | 64753878 | 252173 | 1 | 11784 | ENSMUST00 | -27057 | ENSMUSG0 | Apba2 | amyloid beta (A4) precursor protein-binding, family A, member 2 |
| chr7 | 71269832 | 71269873 | 42 | * | NA_peak_527 | 27 | Intron (ENSM | 7 | 71909693 | 71939191 | 29499 | 1 | 78156 | ENSMUST00 | -639820 | ENSMUSG0 | 4930441H08 | RIKEN cDNA 4930441H08 gene |
| chr7 | 72863180 | 72863464 | 285 | * | NA_peak_527 | 42 | Distal Intergei | 7 | 73097749 | 73102068 | 4320 | 2 | 625243 | ENSMUST00 | 238604 | ENSMUSG0 | Gm6567 | predicted gene 6567 |

| | | | | | | | | | | | | | | | | | | |
|---|---|---|---|---|---|---|---|---|---|---|---|---|---|---|---|---|---|---|
| chr7 | 72885397 | 72885762 | 366 | * | NA_peak_52 | 28 | Distal Interge | 7 | 73097749 | 73102068 | 4320 | 2 | 625243 | ENSMUST00 | 216306 | ENSMUSG0( | Gm6567 | predicted gene 6567 |
| chr7 | 74743040 | 74743326 | 287 | * | NA_peak_52 | 31 | Distal Interge | 7 | 74295371 | 74554780 | 259410 | 2 | 108116 | ENSMUST00 | -188260 | ENSMUSG0( | Slco3a1 | solute carrier organic anion transporter family, member 3a1 |
| chr7 | 74799969 | 74800305 | 337 | * | NA_peak_52 | 20 | Distal Interge | 7 | 75030194 | 75058162 | 27969 | 1 | 75205 | ENSMUST00 | -229889 | ENSMUSG0( | 4930533N22 | RIKEN cDNA 4930533N22 gene |
| chr7 | 75811289 | 75811638 | 350 | * | NA_peak_52 | 23 | Distal Interge | 7 | 75848310 | 75874131 | 25822 | 1 | 207952 | ENSMUST00 | -36672 | ENSMUSG0( | Klhl25 | kelch-like 25 |
| chr7 | 78418841 | 78419119 | 279 | * | NA_peak_52 | 27 | Intron (ENSM | 7 | 78356095 | 78452123 | 96029 | 2 | 18213 | ENSMUST00 | 33004 | ENSMUSG0( | Ntrk3 | neurotrophic tyrosine kinase, receptor, type 3 |
| chr7 | 82215386 | 82215575 | 190 | * | NA_peak_52 | 27 | Intron (ENSM | 7 | 82175156 | 82284158 | 109003 | 1 | 20408 | ENSMUST00 | 40230 | ENSMUSG0( | Sh3gl3 | SH3-domain GRB2-like 3 |
| chr7 | 85762975 | 85763346 | 372 | * | NA_peak_52 | 40 | Distal Interge | 7 | 85735940 | 85755097 | 19158 | 2 | 244114 | ENSMUST00 | -7878 | ENSMUSG0( | Vmn2r72 | vomeronasal 2, receptor 72 |
| chr7 | 95407163 | 95407445 | 283 | * | NA_peak_52 | 31 | Distal Interge | 7 | 96171246 | 96908569 | 737324 | 1 | 23966 | ENSMUST00 | -763801 | ENSMUSG0( | Tenm4 | teneurin transmembrane protein 4 |
| chr7 | 97274125 | 97274408 | 284 | * | NA_peak_52 | 40 | Intron (ENSM | 7 | 97312473 | 97315025 | 2553 | 2 | 244144 | ENSMUST00 | 40617 | ENSMUSG0( | Usp35 | ubiquitin specific peptidase 35 |
| chr7 | 102838895 | 102839192 | 298 | * | NA_peak_52 | 14 | Distal Interge | 7 | 102855337 | 102858755 | 3419 | 2 | 258168 | ENSMUST00 | 19563 | ENSMUSG0( | Olfr566 | olfactory receptor 566 |
| chr7 | 103112372 | 103112660 | 289 | * | NA_peak_52 | 39 | Distal Interge | 7 | 103121829 | 103122782 | 954 | 2 | 259115 | ENSMUST00 | 10122 | ENSMUSG0( | Olfr586 | olfactory receptor 586 |
| chr7 | 103288435 | 103288719 | 285 | * | NA_peak_52 | 59 | Distal Interge | 7 | 103309723 | 103310661 | 939 | 1 | 100041187 | ENSMUST00 | -21004 | ENSMUSG0( | Olfr596 | olfactory receptor 596 |
| chr7 | 107471381 | 107471683 | 303 | * | NA_peak_52 | 17 | Intron (ENSM | 7 | 107504062 | 107548653 | 44592 | 1 | 60510 | ENSMUST00 | -32379 | ENSMUSG0( | Syt9 | synaptotagmin IX |
| chr7 | 108828572 | 108828687 | 116 | * | NA_peak_52 | 13 | Promoter (2-3 | 7 | 108825065 | 108825997 | 933 | 2 | 258721 | ENSMUST00 | -2575 | ENSMUSG0( | Olfr514 | olfactory receptor 514 |
| chr7 | 117663727 | 117664359 | 633 | * | NA_peak_52 | 24 | Intron (ENSM | 7 | 117475492 | 117549359 | 73868 | 1 | 233781 | ENSMUST00 | 188235 | ENSMUSG0( | Xylt1 | xylosyltransferase 1 |
| chr7 | 122914423 | 122914879 | 457 | * | NA_peak_52 | 49 | Distal Interge | 7 | 122965686 | 122976008 | 10323 | 1 | 19647 | ENSMUST00 | -50807 | ENSMUSG0( | Rbbp6 | retinoblastoma binding protein 6, ubiquitin ligase |
| chr7 | 122915996 | 122916766 | 771 | * | NA_peak_52 | 52 | Distal Interge | 7 | 122965686 | 122976008 | 10323 | 1 | 19647 | ENSMUST00 | -48920 | ENSMUSG0( | Rbbp6 | retinoblastoma binding protein 6, ubiquitin ligase |
| chr7 | 125011078 | 125011495 | 418 | * | NA_peak_52 | 22 | Distal Interge | 7 | 124625670 | 124708935 | 83266 | 2 | 73076 | ENSMUST00 | -302143 | ENSMUSG0( | 3100003L05 | RIKEN cDNA 3100003L05 gene |
| chr7 | 138262963 | 138263305 | 343 | * | NA_peak_52 | 20 | Intron (ENSM | 7 | 138209717 | 138271329 | 61613 | 2 | 70571 | ENSMUST00 | 8024 | ENSMUSG0( | Tcerg1l | transcription elongation regulator 1-like |
| chr8 | 3132614 | 3132719 | 106 | * | NA_peak_52 | 23 | Intron (ENSM | 8 | 3122061 | 3159555 | 37495 | 2 | 16337 | ENSMUST00 | 26836 | ENSMUSG0( | Insr | insulin receptor |
| chr8 | 3306577 | 3306855 | 279 | * | NA_peak_52 | 33 | Distal Interge | 8 | 3150922 | 3279617 | 128696 | 2 | 16337 | ENSMUST00 | -26960 | ENSMUSG0( | Insr | insulin receptor |
| chr8 | 3588025 | 3588192 | 168 | * | NA_peak_52 | 30 | Promoter (<= | 8 | 3588051 | 3598927 | 10877 | 1 | 69697 | ENSMUST00 | 0 | ENSMUSG0( | Camsap3 | calmodulin regulated spectrin-associated protein family, member 3 |
| chr8 | 3634421 | 3634630 | 210 | * | NA_peak_52 | 17 | Promoter (<= | 8 | 3635416 | 3635719 | 304 | 1 | 20911 | ENSMUST00 | -786 | ENSMUSG0( | Stxbp2 | syntaxin binding protein 2 |
| chr8 | 3904584 | 3904585 | 2 | * | NA_peak_52 | 27 | Distal Interge | 8 | 3918677 | 3926813 | 8137 | 2 | 69165 | ENSMUST00 | 22228 | ENSMUSG0( | Cd209b | CD209b antigen |
| chr8 | 4135489 | 4135732 | 244 | * | NA_peak_52 | 59 | Promoter (<= | 8 | 4134652 | 4137707 | 3056 | 1 | 70192 | ENSMUST00 | 837 | ENSMUSG0( | Cd209g | CD209g antigen |
| chr8 | 4286896 | 4287212 | 317 | * | NA_peak_52 | 23 | 3' UTR | 8 | 4259731 | 4275913 | 16183 | 2 | 21856 | ENSMUST00 | -10983 | ENSMUSG0( | Timm44 | translocase of inner mitochondrial membrane 44 |
| chr8 | 4870724 | 4870808 | 85 | * | NA_peak_53 | 27 | Distal Interge | 8 | 4763468 | 4779567 | 16100 | 2 | 20419 | ENSMUST00 | -91157 | ENSMUSG0( | Shcbp1 | Shc SH2-domain binding protein 1 |
| chr8 | 7538693 | 7538899 | 207 | * | NA_peak_53 | 15 | Intron (ENSM | 8 | 8620597 | 8639363 | 18767 | 2 | 13642 | ENSMUST00 | 1100464 | ENSMUSG0( | Efnb2 | ephrin B2 |
| chr8 | 7542011 | 7542740 | 730 | * | NA_peak_53 | 27 | Intron (ENSM | 8 | 8620597 | 8639363 | 18767 | 2 | 13642 | ENSMUST00 | 1096623 | ENSMUSG0( | Efnb2 | ephrin B2 |
| chr8 | 7546202 | 7546484 | 283 | * | NA_peak_53 | 27 | Intron (ENSM | 8 | 8620597 | 8639363 | 18767 | 2 | 13642 | ENSMUST00 | 1092879 | ENSMUSG0( | Efnb2 | ephrin B2 |
| chr8 | 8393145 | 8393272 | 128 | * | NA_peak_53 | 16 | Intron (ENSM | 8 | 8620597 | 8639363 | 18767 | 2 | 13642 | ENSMUST00 | 246091 | ENSMUSG0( | Efnb2 | ephrin B2 |
| chr8 | 8532231 | 8532464 | 234 | * | NA_peak_53 | 31 | Distal Interge | 8 | 8620597 | 8639363 | 18767 | 2 | 13642 | ENSMUST00 | 106899 | ENSMUSG0( | Efnb2 | ephrin B2 |
| chr8 | 8711330 | 8711749 | 420 | * | NA_peak_53 | 24 | Exon (ENSMU | 8 | 8665075 | 8690521 | 25447 | 2 | 234023 | ENSMUST00 | -20809 | ENSMUSG0( | Arglu1 | arginine and glutamate rich 1 |
| chr8 | 9513323 | 9513540 | 218 | * | NA_peak_53 | 27 | Intron (ENSM | 8 | 9207238 | 9323711 | 116474 | 2 | 270028 | ENSMUST00 | -189612 | ENSMUSG0( | Fam155a | family with sequence similarity 155, member A |
| chr8 | 9514305 | 9514808 | 504 | * | NA_peak_53 | 36 | Intron (ENSM | 8 | 9207238 | 9323711 | 116474 | 2 | 270028 | ENSMUST00 | -190594 | ENSMUSG0( | Fam155a | family with sequence similarity 155, member A |
| chr8 | 10729278 | 10729578 | 301 | * | NA_peak_53 | 27 | Distal Interge | 8 | 10871509 | 10882435 | 10927 | 2 | 320014 | ENSMUST00 | 152857 | ENSMUSG0( | B930025P03 | RIKEN cDNA B930025P03 gene |
| chr8 | 10915234 | 10915432 | 199 | * | NA_peak_53 | 24 | Distal Interge | 8 | 10926917 | 10928444 | 1528 | 2 | 665306 | ENSMUST00 | 13012 | ENSMUSG0( | 3930402G23 | RIKEN cDNA 3930402G23 gene |
| chr8 | 11206101 | 11206324 | 224 | * | NA_peak_53 | 25 | Exon (ENSMU | 8 | 11199519 | 11210831 | 11313 | 2 | 12826 | ENSMUST00 | 4507 | ENSMUSG0( | Col4a1 | collagen, type IV, alpha 1 |
| chr8 | 12004662 | 12004848 | 187 | * | NA_peak_53 | 25 | Distal Interge | 8 | 11842061 | 11855757 | 13697 | 1 | 75528 | ENSMUST00 | 162601 | ENSMUSG0( | Tex29 | testis expressed 29 |
| chr8 | 12148928 | 12149115 | 188 | * | NA_peak_53 | 20 | Distal Interge | 8 | 12385771 | 12436729 | 50959 | 1 | 434280 | ENSMUST00 | -236656 | ENSMUSG0( | Sox1ot | Sox1 overlapping transcript |
| chr8 | 13221647 | 13221839 | 193 | * | NA_peak_53 | 13 | Exon (ENSMU | 8 | 13175162 | 13200620 | 25459 | 2 | 66790 | ENSMUST00 | -21027 | ENSMUSG0( | Grtp1 | GH regulated TBC protein 1 |
| chr8 | 14732537 | 14732865 | 329 | * | NA_peak_53 | 23 | Intron (ENSM | 8 | 14742753 | 14760902 | 18150 | 1 | 244310 | ENSMUST00 | -9888 | ENSMUSG0( | Dlgap2 | DLG associated protein 2 |
| chr8 | 15623860 | 15623893 | 34 | * | NA_peak_53 | 27 | Intron (ENSM | 8 | 15892545 | 15984812 | 92268 | 2 | 94109 | ENSMUST00 | 360919 | ENSMUSG0( | Csmd1 | CUB and Sushi multiple domains 1 |
| chr8 | 16362666 | 16363029 | 364 | * | NA_peak_53 | 27 | Intron (ENSM | 8 | 16168757 | 16168838 | 82 | 2 | 100526464 | ENSMUST00 | -193828 | ENSMUSG0( | Mir3106 | microRNA 3106 |
| chr8 | 16794045 | 16794322 | 278 | * | NA_peak_53 | 20 | Intron (ENSM | 8 | 16168757 | 16168838 | 82 | 2 | 100526464 | ENSMUST00 | -625207 | ENSMUSG0( | Mir3106 | microRNA 3106 |
| chr8 | 17458519 | 17458828 | 310 | * | NA_peak_53 | 20 | Intron (ENSM | 8 | 15892537 | 17535281 | 1642745 | 2 | 94109 | ENSMUST00 | 76453 | ENSMUSG0( | Csmd1 | CUB and Sushi multiple domains 1 |
| chr8 | 18882886 | 18883128 | 243 | * | NA_peak_53 | 23 | 3' UTR | 8 | 18846279 | 18883421 | 37143 | 1 | 52123 | ENSMUST00 | 36607 | ENSMUSG0( | Agpat5 | 1-acylglycerol-3-phosphate O-acyltransferase 5 (lysophosphatidic acid acyltransferase, epsilon) |
| chr8 | 19100461 | 19101081 | 621 | * | NA_peak_53 | 45 | Distal Interge | 8 | 19111931 | 19114839 | 2909 | 2 | 77674 | ENSMUST00 | 13758 | ENSMUSG0( | Defb12 | defensin beta 12 |
| chr8 | 19705340 | 19705545 | 206 | * | NA_peak_53 | 40 | Distal Interge | 8 | 19729576 | 19753527 | 23952 | 1 | 626415 | ENSMUST00 | -24031 | ENSMUSG0( | 4930467E23 | RIKEN cDNA 4930467E23 gene |
| chr8 | 19735249 | 19735979 | 731 | * | NA_peak_53 | 30 | Exon (ENSMU | 8 | 19729576 | 19753527 | 23952 | 1 | 626415 | ENSMUST00 | 5673 | ENSMUSG0( | 4930467E23 | RIKEN cDNA 4930467E23 gene |
| chr8 | 19737190 | 19737449 | 260 | * | NA_peak_53 | 46 | Intron (ENSM | 8 | 19729576 | 19753527 | 23952 | 1 | 626415 | ENSMUST00 | 7614 | ENSMUSG0( | 4930467E23 | RIKEN cDNA 4930467E23 gene |
| chr8 | 19755688 | 19756000 | 313 | * | NA_peak_53 | 65 | Distal Interge | 8 | 19729576 | 19753527 | 23952 | 1 | 626415 | ENSMUST00 | 26112 | ENSMUSG0( | 4930467E23 | RIKEN cDNA 4930467E23 gene |
| chr8 | 19773724 | 19774108 | 385 | * | NA_peak_53 | 60 | Distal Interge | 8 | 19729576 | 19753527 | 23952 | 1 | 626415 | ENSMUST00 | 44148 | ENSMUSG0( | 4930467E23 | RIKEN cDNA 4930467E23 gene |
| chr8 | 19911333 | 19911680 | 348 | * | NA_peak_53 | 43 | Intron (ENSM | 8 | 19729576 | 19753527 | 23952 | 1 | 626415 | ENSMUST00 | 181757 | ENSMUSG0( | 4930467E23 | RIKEN cDNA 4930467E23 gene |
| chr8 | 19919268 | 19919908 | 641 | * | NA_peak_53 | 18 | Intron (ENSM | 8 | 19729576 | 19753527 | 23952 | 1 | 626415 | ENSMUST00 | 189692 | ENSMUSG0( | 4930467E23 | RIKEN cDNA 4930467E23 gene |
| chr8 | 19933801 | 19934596 | 796 | * | NA_peak_53 | 201 | Distal Interge | 8 | 19729576 | 19753527 | 23952 | 1 | 626415 | ENSMUST00 | 204225 | ENSMUSG0( | 4930467E23 | RIKEN cDNA 4930467E23 gene |
| chr8 | 19966548 | 19966840 | 293 | * | NA_peak_53 | 42 | Distal Interge | 8 | 19729576 | 19753527 | 23952 | 1 | 626415 | ENSMUST00 | 236972 | ENSMUSG0( | 4930467E23 | RIKEN cDNA 4930467E23 gene |
| chr8 | 20006316 | 20006772 | 457 | * | NA_peak_53 | 39 | Intron (ENSM | 8 | 20262613 | 20283266 | 20654 | 2 | 547150 | ENSMUST00 | 276494 | ENSMUSG0( | 6820431F20 | cadherin 11 pseudogene |
| chr8 | 20026154 | 20026442 | 289 | * | NA_peak_53 | 63 | Distal Interge | 8 | 20262613 | 20283266 | 20654 | 2 | 547150 | ENSMUST00 | 256824 | ENSMUSG0( | 6820431F20 | cadherin 11 pseudogene |
| chr8 | 20047836 | 20048564 | 729 | * | NA_peak_53 | 49 | Distal Interge | 8 | 20262613 | 20283266 | 20654 | 2 | 547150 | ENSMUST00 | 234702 | ENSMUSG0( | 6820431F20 | cadherin 11 pseudogene |
| chr8 | 20049032 | 20049665 | 634 | * | NA_peak_53 | 81 | Distal Interge | 8 | 20262613 | 20283266 | 20654 | 2 | 547150 | ENSMUST00 | 233601 | ENSMUSG0( | 6820431F20 | cadherin 11 pseudogene |
| chr8 | 20074404 | 20074720 | 317 | * | NA_peak_53 | 45 | Distal Interge | 8 | 20262613 | 20283266 | 20654 | 2 | 547150 | ENSMUST00 | 208546 | ENSMUSG0( | 6820431F20 | cadherin 11 pseudogene |
| chr8 | 20077993 | 20078274 | 282 | * | NA_peak_53 | 16 | Distal Interge | 8 | 20262613 | 20283266 | 20654 | 2 | 547150 | ENSMUST00 | 204992 | ENSMUSG0( | 6820431F20 | cadherin 11 pseudogene |
| chr8 | 20085680 | 20085812 | 133 | * | NA_peak_53 | 41 | Distal Interge | 8 | 20262613 | 20283266 | 20654 | 2 | 547150 | ENSMUST00 | 197454 | ENSMUSG0( | 6820431F20 | cadherin 11 pseudogene |
| chr8 | 20098191 | 20098476 | 286 | * | NA_peak_53 | 73 | Distal Interge | 8 | 20262613 | 20283266 | 20654 | 2 | 547150 | ENSMUST00 | 184790 | ENSMUSG0( | 6820431F20 | cadherin 11 pseudogene |
| chr8 | 20099854 | 20100414 | 561 | * | NA_peak_53 | 27 | Distal Interge | 8 | 20262613 | 20283266 | 20654 | 2 | 547150 | ENSMUST00 | 182852 | ENSMUSG0( | 6820431F20 | cadherin 11 pseudogene |
| chr8 | 20101964 | 20102859 | 896 | * | NA_peak_53 | 110 | Distal Interge | 8 | 20262613 | 20283266 | 20654 | 2 | 547150 | ENSMUST00 | 180407 | ENSMUSG0( | 6820431F20 | cadherin 11 pseudogene |
| chr8 | 20106895 | 20107415 | 521 | * | NA_peak_53 | 37 | Distal Interge | 8 | 20262613 | 20283266 | 20654 | 2 | 547150 | ENSMUST00 | 175851 | ENSMUSG0( | 6820431F20 | cadherin 11 pseudogene |
| chr8 | 20116959 | 20117329 | 371 | * | NA_peak_53 | 70 | Distal Interge | 8 | 20262613 | 20283266 | 20654 | 2 | 547150 | ENSMUST00 | 165937 | ENSMUSG0( | 6820431F20 | cadherin 11 pseudogene |
| chr8 | 20119201 | 20119598 | 398 | * | NA_peak_53 | 55 | Distal Interge | 8 | 20262613 | 20283266 | 20654 | 2 | 547150 | ENSMUST00 | 163668 | ENSMUSG0( | 6820431F20 | cadherin 11 pseudogene |
| chr8 | 20125954 | 20126200 | 247 | * | NA_peak_53 | 36 | Distal Interge | 8 | 20262613 | 20283266 | 20654 | 2 | 547150 | ENSMUST00 | 157066 | ENSMUSG0( | 6820431F20 | cadherin 11 pseudogene |
| chr8 | 20137862 | 20139263 | 1402 | * | NA_peak_53 | 66 | Intron (ENSM | 8 | 20262613 | 20283266 | 20654 | 2 | 547150 | ENSMUST00 | 144003 | ENSMUSG0( | 6820431F20 | cadherin 11 pseudogene |
| chr8 | 20140429 | 20140781 | 353 | * | NA_peak_53 | 52 | Intron (ENSM | 8 | 20262613 | 20283266 | 20654 | 2 | 547150 | ENSMUST00 | 142485 | ENSMUSG0( | 6820431F20 | cadherin 11 pseudogene |
| chr8 | 20142616 | 20143105 | 490 | * | NA_peak_53 | 31 | Intron (ENSM | 8 | 20262613 | 20283266 | 20654 | 2 | 547150 | ENSMUST00 | 140161 | ENSMUSG0( | 6820431F20 | cadherin 11 pseudogene |
| chr8 | 20145393 | 20146335 | 943 | * | NA_peak_53 | 49 | Exon (ENSMU | 8 | 20262613 | 20283266 | 20654 | 2 | 547150 | ENSMUST00 | 136931 | ENSMUSG0( | 6820431F20 | cadherin 11 pseudogene |
| chr8 | 20147400 | 20148249 | 850 | * | NA_peak_53 | 124 | Exon (ENSMU | 8 | 20262613 | 20283266 | 20654 | 2 | 547150 | ENSMUST00 | 135017 | ENSMUSG0( | 6820431F20 | cadherin 11 pseudogene |
| chr8 | 20273731 | 20274440 | 710 | * | NA_peak_53 | 125 | Exon (ENSMU | 8 | 20262613 | 20283266 | 20654 | 2 | 547150 | ENSMUST00 | 8826 | ENSMUSG0( | 6820431F20 | cadherin 11 pseudogene |
| chr8 | 20275318 | 20275642 | 325 | * | NA_peak_53 | 97 | Intron (ENSM | 8 | 20262613 | 20283266 | 20654 | 2 | 547150 | ENSMUST00 | 7624 | ENSMUSG0( | 6820431F20 | cadherin 11 pseudogene |
| chr8 | 20279942 | 20280523 | 582 | * | NA_peak_53 | 37 | Promoter (2-3 | 8 | 20262613 | 20283266 | 20654 | 2 | 547150 | ENSMUST00 | 2743 | ENSMUSG0( | 6820431F20 | cadherin 11 pseudogene |
| chr8 | 20281964 | 20282322 | 359 | * | NA_peak_53 | 83 | Promoter (<= | 8 | 20262613 | 20283266 | 20654 | 2 | 547150 | ENSMUST00 | 944 | ENSMUSG0( | 6820431F20 | cadherin 11 pseudogene |
| chr8 | 20283718 | 20284091 | 374 | * | NA_peak_53 | 26 | Promoter (<= | 8 | 20262613 | 20283266 | 20654 | 2 | 547150 | ENSMUST00 | -452 | ENSMUSG0( | 6820431F20 | cadherin 11 pseudogene |
| chr8 | 20284925 | 20285262 | 338 | * | NA_peak_53 | 41 | Promoter (1-2 | 8 | 20262613 | 20283266 | 20654 | 2 | 547150 | ENSMUST00 | -1659 | ENSMUSG0( | 6820431F20 | cadherin 11 pseudogene |
| chr8 | 20296117 | 20296487 | 371 | * | NA_peak_53 | 44 | Promoter (<= | 8 | 20268286 | 20297432 | 29147 | 2 | 547150 | ENSMUST00 | 945 | ENSMUSG0( | 6820431F20 | cadherin 11 pseudogene |
| chr8 | 20302766 | 20303059 | 294 | * | NA_peak_53 | 38 | Distal Interge | 8 | 20268286 | 20297432 | 29147 | 2 | 547150 | ENSMUST00 | -5334 | ENSMUSG0( | 6820431F20 | cadherin 11 pseudogene |
| chr8 | 20323507 | 20323818 | 312 | * | NA_peak_53 | 47 | Distal Interge | 8 | 20268286 | 20297432 | 29147 | 2 | 547150 | ENSMUST00 | -26075 | ENSMUSG0( | 6820431F20 | cadherin 11 pseudogene |
| chr8 | 20331955 | 20332160 | 206 | * | NA_peak_53 | 44 | Distal Interge | 8 | 20337664 | 20363210 | 25547 | 2 | 100040599 | ENSMUST00 | 31050 | ENSMUSG0( | Gm15319 | predicted gene 15319 |
| chr8 | 20334116 | 20334555 | 440 | * | NA_peak_53 | 41 | Distal Interge | 8 | 20337664 | 20363210 | 25547 | 2 | 100040599 | ENSMUST00 | 28655 | ENSMUSG0( | Gm15319 | predicted gene 15319 |
| chr8 | 20346235 | 20346460 | 226 | * | NA_peak_53 | 54 | Intron (ENSM | 8 | 20337664 | 20363210 | 25547 | 2 | 100040599 | ENSMUST00 | 16750 | ENSMUSG0( | Gm15319 | predicted gene 15319 |
| chr8 | 20357635 | 20357952 | 318 | * | NA_peak_53 | 31 | Intron (ENSM | 8 | 20337664 | 20363210 | 25547 | 2 | 100040599 | ENSMUST00 | 5258 | ENSMUSG0( | Gm15319 | predicted gene 15319 |
| chr8 | 20379993 | 20380329 | 337 | * | NA_peak_53 | 13 | Distal Interge | 8 | 20337664 | 20363210 | 25547 | 2 | 100040599 | ENSMUST00 | -16783 | ENSMUSG0( | Gm15319 | predicted gene 15319 |
| chr8 | 20389350 | 20389626 | 277 | * | NA_peak_53 | 70 | Intron (ENSM | 8 | 20395392 | 20410565 | 15174 | 2 | 381598 | ENSMUST00 | 20939 | ENSMUSG0( | 2610005L07 | cadherin 11 pseudogene |
| chr8 | 20399900 | 20400265 | 366 | * | NA_peak_53 | 15 | Intron (ENSM | 8 | 20395392 | 20410565 | 15174 | 2 | 381598 | ENSMUST00 | 10300 | ENSMUSG0( | 2610005L07 | cadherin 11 pseudogene |
| chr8 | 20407499 | 20408214 | 716 | * | NA_peak_53 | 41 | Promoter (2-3 | 8 | 20395392 | 20410565 | 15174 | 2 | 381598 | ENSMUST00 | 2351 | ENSMUSG0( | 2610005L07 | cadherin 11 pseudogene |
| chr8 | 20418846 | 20419096 | 251 | * | NA_peak_53 | 49 | Intron (ENSM | 8 | 20385782 | 20424814 | 39033 | 2 | 381598 | ENSMUST00 | 5718 | ENSMUSG0( | 2610005L07 | cadherin 11 pseudogene |
| chr8 | 20419918 | 20420195 | 278 | * | NA_peak_53 | 47 | Intron (ENSM | 8 | 20385782 | 20424814 | 39033 | 2 | 381598 | ENSMUST00 | 4619 | ENSMUSG0( | 2610005L07 | cadherin 11 pseudogene |
| chr8 | 20441930 | 20442227 | 298 | * | NA_peak_53 | 24 | Distal Interge | 8 | 20385782 | 20424814 | 39033 | 2 | 381598 | ENSMUST00 | -17116 | ENSMUSG0( | 2610005L07 | cadherin 11 pseudogene |
| chr8 | 20493716 | 20494267 | 552 | * | NA_peak_53 | 55 | Distal Interge | 8 | 20385782 | 20424814 | 39033 | 2 | 381598 | ENSMUST00 | -68902 | ENSMUSG0( | 2610005L07 | cadherin 11 pseudogene |
| chr8 | 20495577 | 20495996 | 420 | * | NA_peak_53 | 65 | Distal Interge | 8 | 20385782 | 20424814 | 39033 | 2 | 381598 | ENSMUST00 | -70763 | ENSMUSG0( | 2610005L07 | cadherin 11 pseudogene |
| chr8 | 20498129 | 20498929 | 801 | * | NA_peak_53 | 62 | Distal Interge | 8 | 20385782 | 20424814 | 39033 | 2 | 381598 | ENSMUST00 | -73315 | ENSMUSG0( | 2610005L07 | cadherin 11 pseudogene |
| chr8 | 20503687 | 20504057 | 371 | * | NA_peak_53 | 39 | Distal Interge | 8 | 20385782 | 20424814 | 39033 | 2 | 381598 | ENSMUST00 | -78873 | ENSMUSG0( | 2610005L07 | cadherin 11 pseudogene |
| chr8 | 20520357 | 20520739 | 383 | * | NA_peak_53 | 166 | Distal Interge | 8 | 20614919 | 20640703 | 25785 | 1 | 100861668 | ENSMUST00 | -94180 | ENSMUSG0( | Gm21119 | predicted gene, 21119 |
| chr8 | 20526059 | 20526397 | 339 | * | NA_peak_53 | 84 | Distal Interge | 8 | 20614919 | 20640703 | 25785 | 1 | 100861668 | ENSMUST00 | -88522 | ENSMUSG0( | Gm21119 | predicted gene, 21119 |
| chr8 | 20530192 | 20530573 | 382 | * | NA_peak_53 | 88 | Distal Interge | 8 | 20614919 | 20640703 | 25785 | 1 | 100861668 | ENSMUST00 | -84346 | ENSMUSG0( | Gm21119 | predicted gene, 21119 |
| chr8 | 20532119 | 20532720 | 602 | * | NA_peak_53 | 33 | Distal Interge | 8 | 20614919 | 20640703 | 25785 | 1 | 100861668 | ENSMUST00 | -82199 | ENSMUSG0( | Gm21119 | predicted gene, 21119 |
| chr8 | 20537056 | 20537301 | 246 | * | NA_peak_53 | 39 | Distal Interge | 8 | 20614919 | 20640703 | 25785 | 1 | 100861668 | ENSMUST00 | -77618 | ENSMUSG0( | Gm21119 | predicted gene, 21119 |
| chr8 | 20539705 | 20540161 | 457 | * | NA_peak_53 | 47 | Distal Interge | 8 | 20614919 | 20640703 | 25785 | 1 | 100861668 | ENSMUST00 | -74758 | ENSMUSG0( | Gm21119 | predicted gene, 21119 |

| | | | | | | | | | | | | | | | | | | |
|---|---|---|---|---|---|---|---|---|---|---|---|---|---|---|---|---|---|---|
| chr8 | 20562134 | 20562493 | 360 | * | NA_peak_537 | 83 | Distal Interge | 8 | 20614919 | 20640703 | 25785 | 1 | 100861668 | ENSMUST00 | -52426 | ENSMUSG0( | Gm21119 | predicted gene, 21119 |
| chr8 | 20575597 | 20576356 | 760 | * | NA_peak_537 | 30 | Intron (ENSM | 8 | 20614919 | 20640703 | 25785 | 1 | 100861668 | ENSMUST00 | -38563 | ENSMUSG0( | Gm21119 | predicted gene, 21119 |
| chr8 | 20579033 | 20579403 | 371 | * | NA_peak_537 | 51 | Intron (ENSM | 8 | 20614919 | 20640703 | 25785 | 1 | 100861668 | ENSMUST00 | -35516 | ENSMUSG0( | Gm21119 | predicted gene, 21119 |
| chr8 | 20578119 | 20578490 | 372 | * | NA_peak_537 | 51 | Intron (ENSM | 8 | 20614919 | 20640703 | 25785 | 1 | 100861668 | ENSMUST00 | -36429 | ENSMUSG0( | Gm21119 | predicted gene, 21119 |
| chr8 | 20581461 | 20581843 | 383 | * | NA_peak_537 | 51 | Intron (ENSM | 8 | 20614919 | 20640703 | 25785 | 1 | 100861668 | ENSMUST00 | -33076 | ENSMUSG0( | Gm21119 | predicted gene, 21119 |
| chr8 | 20582312 | 20582694 | 383 | * | NA_peak_537 | 27 | Exon (ENSMU | 8 | 20614919 | 20640703 | 25785 | 1 | 100861668 | ENSMUST00 | -32225 | ENSMUSG0( | Gm21119 | predicted gene, 21119 |
| chr8 | 20583408 | 20583749 | 342 | * | NA_peak_537 | 19 | Distal Interge | 8 | 20614919 | 20640703 | 25785 | 1 | 100861668 | ENSMUST00 | -31170 | ENSMUSG0( | Gm21119 | predicted gene, 21119 |
| chr8 | 20601578 | 20601877 | 300 | * | NA_peak_537 | 26 | Distal Interge | 8 | 20614919 | 20640703 | 25785 | 1 | 100861668 | ENSMUST00 | -13042 | ENSMUSG0( | Gm21119 | predicted gene, 21119 |
| chr8 | 20605423 | 20605761 | 339 | * | NA_peak_537 | 60 | Distal Interge | 8 | 20614919 | 20640703 | 25785 | 1 | 100861668 | ENSMUST00 | -9158 | ENSMUSG0( | Gm21119 | predicted gene, 21119 |
| chr8 | 20608231 | 20609081 | 851 | * | NA_peak_537 | 65 | Distal Interge | 8 | 20614919 | 20640703 | 25785 | 1 | 100861668 | ENSMUST00 | -5838 | ENSMUSG0( | Gm21119 | predicted gene, 21119 |
| chr8 | 20611674 | 20612304 | 631 | * | NA_peak_537 | 34 | Promoter (2-3 | 8 | 20614919 | 20640703 | 25785 | 1 | 100861668 | ENSMUST00 | -2615 | ENSMUSG0( | Gm21119 | predicted gene, 21119 |
| chr8 | 20616017 | 20616879 | 863 | * | NA_peak_537 | 37 | Promoter (1-2 | 8 | 20614919 | 20640703 | 25785 | 1 | 100861668 | ENSMUST00 | 1098 | ENSMUSG0( | Gm21119 | predicted gene, 21119 |
| chr8 | 20624880 | 20625188 | 309 | * | NA_peak_537 | 56 | Intron (ENSM | 8 | 20614919 | 20640703 | 25785 | 1 | 100861668 | ENSMUST00 | 9961 | ENSMUSG0( | Gm21119 | predicted gene, 21119 |
| chr8 | 20626094 | 20626380 | 287 | * | NA_peak_537 | 31 | Intron (ENSM | 8 | 20614919 | 20640703 | 25785 | 1 | 100861668 | ENSMUST00 | 11175 | ENSMUSG0( | Gm21119 | predicted gene, 21119 |
| chr8 | 20631533 | 20632093 | 561 | * | NA_peak_538 | 38 | Intron (ENSM | 8 | 20614919 | 20640703 | 25785 | 1 | 100861668 | ENSMUST00 | 16614 | ENSMUSG0( | Gm21119 | predicted gene, 21119 |
| chr8 | 20641064 | 20641870 | 807 | * | NA_peak_538 | 41 | Distal Interge | 8 | 20614919 | 20640703 | 25785 | 1 | 100861668 | ENSMUST00 | 26145 | ENSMUSG0( | Gm21119 | predicted gene, 21119 |
| chr8 | 20654969 | 20655369 | 401 | * | NA_peak_538 | 45 | Distal Interge | 8 | 20614919 | 20640703 | 25785 | 1 | 100861668 | ENSMUST00 | 40050 | ENSMUSG0( | Gm21119 | predicted gene, 21119 |
| chr8 | 20674021 | 20674625 | 605 | * | NA_peak_538 | 73 | Distal Interge | 8 | 20614919 | 20640703 | 25785 | 1 | 100861668 | ENSMUST00 | 59102 | ENSMUSG0( | Gm21119 | predicted gene, 21119 |
| chr8 | 20698044 | 20698440 | 397 | * | NA_peak_538 | 20 | Distal Interge | 8 | 20614919 | 20640703 | 25785 | 1 | 100861668 | ENSMUST00 | 83125 | ENSMUSG0( | Gm21119 | predicted gene, 21119 |
| chr8 | 20813557 | 20814203 | 647 | * | NA_peak_538 | 74 | Distal Interge | 8 | 20892669 | 20897723 | 5055 | 1 | 654453 | ENSMUST00 | -78466 | ENSMUSG0( | Defb33 | defensin beta 33 |
| chr8 | 20828516 | 20828995 | 480 | * | NA_peak_538 | 31 | Intron (ENSM | 8 | 20892669 | 20897723 | 5055 | 1 | 654453 | ENSMUST00 | -63674 | ENSMUSG0( | Defb33 | defensin beta 33 |
| chr8 | 20991244 | 20991535 | 292 | * | NA_peak_538 | 51 | Distal Interge | 8 | 21025545 | 21026516 | 972 | 1 | 66298 | ENSMUST00 | -34010 | ENSMUSG0( | Defa21 | defensin, alpha, 21 |
| chr8 | 20993315 | 20993739 | 425 | * | NA_peak_538 | 33 | Distal Interge | 8 | 21025545 | 21026516 | 972 | 1 | 66298 | ENSMUST00 | -31806 | ENSMUSG0( | Defa21 | defensin, alpha, 21 |
| chr8 | 21292000 | 21292117 | 118 | * | NA_peak_538 | 21 | Distal Interge | 8 | 21287409 | 21288377 | 969 | 1 | 13237 | ENSMUST00 | 4591 | ENSMUSG0( | Defa3 | defensin, alpha, 3 |
| chr8 | 21293924 | 21294239 | 316 | * | NA_peak_538 | 16 | Distal Interge | 8 | 21297439 | 21298375 | 937 | 1 | 13239 | ENSMUST00 | -3200 | ENSMUSG0( | Defa5 | defensin, alpha, 5 |
| chr8 | 21647018 | 21647128 | 111 | * | NA_peak_538 | 20 | Distal Interge | 8 | 21655780 | 21656735 | 956 | 1 | 23855 | ENSMUST00 | -8652 | ENSMUSG0( | Defa17 | defensin, alpha, 17 |
| chr8 | 21677970 | 21678312 | 343 | * | NA_peak_538 | 20 | Distal Interge | 8 | 21665797 | 21666732 | 936 | 1 | 100041952 | ENSMUST00 | 12173 | ENSMUSG0( | Defa34 | defensin, alpha, 34 |
| chr8 | 21710855 | 21711455 | 601 | * | NA_peak_538 | 21 | Distal Interge | 8 | 21702519 | 21703645 | 1127 | 2 | 382000 | ENSMUST00 | -7210 | ENSMUSG0( | Defa39 | defensin, alpha, 39 |
| chr8 | 21987109 | 21987411 | 303 | * | NA_peak_538 | 13 | Distal Interge | 8 | 21970596 | 21974041 | 3446 | 1 | 67929 | ENSMUST00 | 16513 | ENSMUSG0( | Ccdc70 | coiled-coil domain containing 70 |
| chr8 | 22159575 | 22159925 | 351 | * | NA_peak_538 | 23 | Promoter (2-3 | 8 | 22128286 | 22162494 | 34209 | 2 | 23954 | ENSMUST00 | 2569 | ENSMUSG0( | Nek3 | NIMA (never in mitosis gene a)-related expressed kinase 3 |
| chr8 | 22295577 | 22295950 | 374 | * | NA_peak_538 | 36 | Intron (ENSM | 8 | 22283441 | 22371418 | 87978 | 1 | 234129 | ENSMUST00 | 12136 | ENSMUSG0( | Tpte | transmembrane phosphatase with tensin homology |
| chr8 | 22582106 | 22582401 | 296 | * | NA_peak_539 | 19 | Exon (ENSMU | 8 | 22581919 | 22593711 | 11793 | 2 | 22335 | ENSMUST00 | 11310 | ENSMUSG0( | Vdac3 | voltage-dependent anion channel 3 |
| chr8 | 22723601 | 22723796 | 196 | * | NA_peak_539 | 26 | Distal Interge | 8 | 22659768 | 22706589 | 46822 | 2 | 16150 | ENSMUST00 | -17012 | ENSMUSG0( | Ikbkb | inhibitor of kappaB kinase beta |
| chr8 | 24819748 | 24820122 | 375 | * | NA_peak_540 | 23 | Intron (ENSM | 8 | 24727202 | 24823985 | 96784 | 2 | 11499 | ENSMUST00 | 3863 | ENSMUSG0( | Adam5 | a disintegrin and metallopeptidase domain 5 |
| chr8 | 25454099 | 25454456 | 358 | * | NA_peak_540 | 21 | Exon (ENSMU | 8 | 25513654 | 25555568 | 41915 | 1 | 14182 | ENSMUST00 | -59198 | ENSMUSG0( | Fgfr1 | fibroblast growth factor receptor 1 |
| chr8 | 25601576 | 25602189 | 614 | * | NA_peak_540 | 39 | Promoter (<= | 8 | 25601601 | 25677214 | 75614 | 1 | 234135 | ENSMUST00 | 0 | ENSMUSG0( | Nsd3 | nuclear receptor binding SET domain protein 3 |
| chr8 | 26815971 | 26816165 | 195 | * | NA_peak_541 | 18 | Intron (ENSM | 8 | 26814593 | 26824423 | 9831 | 2 | 72316 | ENSMUST00 | 8258 | ENSMUSG0( | 2310008N11 | RIKEN cDNA 2310008N11 gene |
| chr8 | 26922267 | 26922444 | 178 | * | NA_peak_541 | 23 | Distal Interge | 8 | 26814593 | 26880582 | 65990 | 2 | 72316 | ENSMUST00 | -41685 | ENSMUSG0( | 2310008N11 | RIKEN cDNA 2310008N11 gene |
| chr8 | 26933062 | 26933333 | 272 | * | NA_peak_541 | 20 | Distal Interge | 8 | 26977325 | 26978672 | 1348 | 1 | 353310 | ENSMUST00 | -43992 | ENSMUSG0( | Zfp703 | zinc finger protein 703 |
| chr8 | 26976838 | 26977122 | 285 | * | NA_peak_541 | 19 | Promoter (<= | 8 | 26977325 | 26978672 | 1348 | 1 | 353310 | ENSMUST00 | -203 | ENSMUSG0( | Zfp703 | zinc finger protein 703 |
| chr8 | 26994394 | 26994704 | 311 | * | NA_peak_541 | 21 | Distal Interge | 8 | 26978185 | 26978692 | 508 | 1 | 353310 | ENSMUST00 | 16209 | ENSMUSG0( | Zfp703 | zinc finger protein 703 |
| chr8 | 27363867 | 27364138 | 272 | * | NA_peak_541 | 27 | Distal Interge | 8 | 27368711 | 27399730 | 31020 | 1 | 108043 | ENSMUST00 | -4573 | ENSMUSG0( | Chrnb3 | cholinergic receptor, nicotinic, beta polypeptide 3 |
| chr8 | 27686036 | 27686162 | 127 | * | NA_peak_541 | 23 | Distal Interge | 8 | 27447670 | 27495172 | 47503 | 1 | 70952 | ENSMUST00 | 238366 | ENSMUSG0( | Poteg | POTE ankyrin domain family, member G |
| chr8 | 28250392 | 28250782 | 391 | * | NA_peak_541 | 55 | Distal Interge | 8 | 28083330 | 28084920 | 1591 | 2 | 666422 | ENSMUST00 | -165472 | ENSMUSG0( | Gm8096 | 3-phosphoglycerate dehydrogenase pseudogene |
| chr8 | 28530170 | 28530404 | 235 | * | NA_peak_541 | 36 | Distal Interge | 8 | 28083330 | 28084920 | 1591 | 2 | 666422 | ENSMUST00 | -445250 | ENSMUSG0( | Gm8096 | 3-phosphoglycerate dehydrogenase pseudogene |
| chr8 | 28619255 | 28619588 | 334 | * | NA_peak_542 | 40 | Distal Interge | 8 | 28083330 | 28084920 | 1591 | 2 | 666422 | ENSMUST00 | -534335 | ENSMUSG0( | Gm8096 | 3-phosphoglycerate dehydrogenase pseudogene |
| chr8 | 28905597 | 28906238 | 642 | * | NA_peak_542 | 13 | Intron (ENSM | 8 | 28652425 | 29219427 | 567003 | 2 | 210801 | ENSMUST00 | 313189 | ENSMUSG0( | Unc5d | unc-5 netrin receptor D |
| chr8 | 29195309 | 29195548 | 240 | * | NA_peak_542 | 54 | Intron (ENSM | 8 | 28652425 | 29219427 | 567003 | 2 | 210801 | ENSMUST00 | 23879 | ENSMUSG0( | Unc5d | unc-5 netrin receptor D |
| chr8 | 29410598 | 29411103 | 506 | * | NA_peak_542 | 16 | Distal Interge | 8 | 28646717 | 29219636 | 572920 | 2 | 210801 | ENSMUST00 | -190962 | ENSMUSG0( | Unc5d | unc-5 netrin receptor D |
| chr8 | 29415736 | 29416173 | 438 | * | NA_peak_542 | 36 | Distal Interge | 8 | 28646717 | 29219636 | 572920 | 2 | 210801 | ENSMUST00 | -196100 | ENSMUSG0( | Unc5d | unc-5 netrin receptor D |
| chr8 | 31095706 | 31095878 | 173 | * | NA_peak_542 | 27 | Promoter (1-2 | 8 | 31094008 | 31096591 | 2584 | 1 | 66959 | ENSMUST00 | 1698 | ENSMUSG0( | Dusp26 | dual specificity phosphatase 26 (putative) |
| chr8 | 31357893 | 31358005 | 113 | * | NA_peak_542 | 22 | Distal Interge | 8 | 31378813 | 31382439 | 3627 | 1 | 74517 | ENSMUST00 | -20808 | ENSMUSG0( | 7420700N18 | RIKEN cDNA 7420700N18 gene |
| chr8 | 32438038 | 32438294 | 257 | * | NA_peak_543 | 23 | Intron (ENSM | 8 | 31849311 | 32292913 | 443603 | 2 | 211323 | ENSMUST00 | -145125 | ENSMUSG0( | Nrg1 | neuregulin 1 |
| chr8 | 32883988 | 32884116 | 129 | * | NA_peak_543 | 20 | Promoter (<= | 8 | 31837604 | 32883862 | 1046259 | 2 | 211323 | ENSMUST00 | -126 | ENSMUSG0( | Nrg1 | neuregulin 1 |
| chr8 | 33312372 | 33312727 | 356 | * | NA_peak_543 | 30 | Intron (ENSM | 8 | 33235684 | 33330550 | 94867 | 2 | 22427 | ENSMUST00 | 17823 | ENSMUSG0( | Wrn | Werner syndrome RecQ like helicase |
| chr8 | 33468645 | 33468814 | 170 | * | NA_peak_543 | 20 | Distal Interge | 8 | 33484196 | 33485167 | 972 | 1 | 628431 | ENSMUST00 | -15382 | NA | Hmgb1-rs17 | high mobility group box 1, related sequence 17 |
| chr8 | 33649760 | 33649792 | 33 | * | NA_peak_543 | 28 | Promoter (2-3 | 8 | 33652523 | 33674199 | 21677 | 1 | 14782 | ENSMUST00 | -2731 | ENSMUSG0( | Gsr | glutathione reductase |
| chr8 | 34122752 | 34122928 | 177 | * | NA_peak_543 | 31 | Intron (ENSM | 8 | 34115030 | 34125182 | 10153 | 1 | 234155 | ENSMUST00 | 7722 | ENSMUSG0( | Mboat4 | membrane bound O-acyltransferase domain containing 4 |
| chr8 | 34154374 | 34154641 | 268 | * | NA_peak_543 | 22 | Promoter (<= | 8 | 34154575 | 34155413 | 839 | 1 | 67887 | ENSMUST00 | 0 | ENSMUSG0( | Saraf | store-operated calcium entry-associated regulatory factor |
| chr8 | 35753063 | 35753409 | 347 | * | NA_peak_544 | 16 | Intron (ENSM | 8 | 35765790 | 35818047 | 52258 | 1 | 100502846 | ENSMUST00 | -12381 | ENSMUSG0( | Gm19410 | predicted gene, 19410 |
| chr8 | 35887983 | 35888041 | 59 | * | NA_peak_544 | 31 | Distal Interge | 8 | 35864085 | 35882957 | 18873 | 2 | 330737 | ENSMUST00 | -5026 | ENSMUSG0( | 5430403N17 | RIKEN cDNA 5430403N17 gene |
| chr8 | 36142613 | 36142699 | 87 | * | NA_peak_544 | 20 | Intron (ENSM | 8 | 36156010 | 36161034 | 5025 | 1 | 330738 | ENSMUST00 | -13311 | ENSMUSG0( | Gm38414 | predicted gene, 38414 |
| chr8 | 36952490 | 36952665 | 176 | * | NA_peak_545 | 31 | Promoter (<= | 8 | 36570249 | 36952442 | 382194 | 2 | 50768 | ENSMUST00 | -48 | ENSMUSG0( | Dlc1 | deleted in liver cancer 1 |
| chr8 | 37480417 | 37480477 | 61 | * | NA_peak_545 | 16 | Distal Interge | 8 | 36985573 | 37024238 | 38666 | 2 | 414128 | ENSMUST00 | -456179 | ENSMUSG0( | G630064G18 | RIKEN cDNA G630064G18 gene |
| chr8 | 37570118 | 37570307 | 190 | * | NA_peak_545 | 30 | Intron (ENSM | 8 | 36985573 | 37024238 | 38666 | 2 | 414128 | ENSMUST00 | -545880 | ENSMUSG0( | G630064G18 | RIKEN cDNA G630064G18 gene |
| chr8 | 37998201 | 37998503 | 303 | * | NA_peak_545 | 40 | Intron (ENSM | 8 | 38252133 | 38252202 | 70 | 2 | 723860 | ENSMUST00 | 253699 | ENSMUSG0( | Mir383 | microRNA 383 |
| chr8 | 39385094 | 39385344 | 251 | * | NA_peak_545 | 31 | Distal Interge | 8 | 39297980 | 39368268 | 70289 | 1 | 621335 | ENSMUST00 | 87114 | ENSMUSG0( | Gm6213 | predicted gene 6213 |
| chr8 | 40511859 | 40512137 | 279 | * | NA_peak_545 | 33 | Promoter (<= | 8 | 40511868 | 40544041 | 32174 | 1 | 52348 | ENSMUST00 | 0 | ENSMUSG0( | Vps37a | vacuolar protein sorting 37A |
| chr8 | 40955722 | 40956021 | 300 | * | NA_peak_545 | 31 | Intron (ENSM | 8 | 40926233 | 40990785 | 64553 | 1 | 68797 | ENSMUST00 | 29489 | ENSMUSG0( | Pdgfrl | platelet-derived growth factor receptor-like |
| chr8 | 41138668 | 41138750 | 83 | * | NA_peak_546 | 31 | Distal Interge | 8 | 41050100 | 41133726 | 83627 | 2 | 102103 | ENSMUST00 | -4942 | ENSMUSG0( | Mtus1 | mitochondrial tumor suppressor 1 |
| chr8 | 41411470 | 41411545 | 76 | * | NA_peak_546 | 31 | Promoter (1-2 | 8 | 41411237 | 41412922 | 1686 | 2 | 14300 | ENSMUST00 | 1377 | ENSMUSG0( | Frg1 | FSHD region gene 1 |
| chr8 | 41513470 | 41513904 | 435 | * | NA_peak_546 | 45 | Distal Interge | 8 | 41397459 | 41417074 | 19616 | 2 | 14300 | ENSMUST00 | -96396 | ENSMUSG0( | Frg1 | FSHD region gene 1 |
| chr8 | 41659124 | 41659425 | 302 | * | NA_peak_546 | 20 | Distal Interge | 8 | 41827267 | 42046202 | 218936 | 1 | 69966 | ENSMUST00 | -167842 | ENSMUSG0( | 2810404M03 | RIKEN cDNA 2810404M03 gene |
| chr8 | 42112139 | 42112278 | 140 | * | NA_peak_546 | 28 | Distal Interge | 8 | 41827267 | 42046202 | 218936 | 1 | 69966 | ENSMUST00 | 284872 | ENSMUSG0( | 2810404M03 | RIKEN cDNA 2810404M03 gene |
| chr8 | 44858205 | 44858705 | 501 | * | NA_peak_547 | 40 | Distal Interge | 8 | 44935447 | 45052257 | 116811 | 1 | 14107 | ENSMUST00 | -76742 | ENSMUSG0( | Fat1 | FAT atypical cadherin 1 |
| chr8 | 45961363 | 45961812 | 450 | * | NA_peak_547 | 36 | Intron (ENSM | 8 | 45953606 | 45975252 | 21647 | 2 | 69479 | ENSMUST00 | 13440 | ENSMUSG0( | 1700029J07 | RIKEN cDNA 1700029J07 gene |
| chr8 | 46693231 | 46693672 | 442 | * | NA_peak_547 | 16 | Intron (ENSM | 8 | 46668808 | 46739508 | 70701 | 2 | 100503498 | ENSMUST00 | 45836 | ENSMUSG0( | Gm16675 | predicted gene, 16675 |
| chr8 | 47477329 | 47477597 | 269 | * | NA_peak_548 | 19 | Distal Interge | 8 | 47504245 | 47511348 | 7104 | 2 | 320714 | ENSMUST00 | 33751 | ENSMUSG0( | Trappc11 | trafficking protein particle complex 11 |
| chr8 | 48072599 | 48072780 | 182 | * | NA_peak_548 | 23 | Distal Interge | 8 | 48099092 | 48140609 | 41518 | 1 | 320685 | ENSMUST00 | -26312 | ENSMUSG0( | Dctd | dCMP deaminase |
| chr8 | 48736357 | 48736639 | 283 | * | NA_peak_549 | 31 | Intron (ENSM | 8 | 48227682 | 48674690 | 447009 | 2 | 23965 | ENSMUST00 | -61667 | ENSMUSG0( | Tenm3 | teneurin transmembrane protein 3 |
| chr8 | 48789645 | 48790049 | 405 | * | NA_peak_549 | 17 | Intron (ENSM | 8 | 48646261 | 48843951 | 197691 | 2 | 23965 | ENSMUST00 | 53902 | ENSMUSG0( | Tenm3 | teneurin transmembrane protein 3 |
| chr8 | 49078998 | 49079287 | 290 | * | NA_peak_549 | 38 | Distal Interge | 8 | 48646261 | 48843951 | 197691 | 2 | 23965 | ENSMUST00 | -235047 | ENSMUSG0( | Tenm3 | teneurin transmembrane protein 3 |
| chr8 | 49835517 | 49835909 | 393 | * | NA_peak_549 | 16 | Intron (ENSM | 8 | 50368112 | 50416088 | 47977 | 1 | 100039954 | ENSMUST00 | -532203 | ENSMUSG0( | Gm2516 | predicted gene 2516 |
| chr8 | 49983118 | 49983443 | 326 | * | NA_peak_549 | 27 | Intron (ENSM | 8 | 50368112 | 50416088 | 47977 | 1 | 100039954 | ENSMUST00 | -384669 | ENSMUSG0( | Gm2516 | predicted gene 2516 |
| chr8 | 51662804 | 51663146 | 343 | * | NA_peak_549 | 16 | Distal Interge | 8 | 52356611 | 52403339 | 46729 | 1 | 330767 | ENSMUST00 | -693465 | ENSMUSG0( | 1190028D05 | RIKEN cDNA 1190028D05 gene |
| chr8 | 51734642 | 51735048 | 407 | * | NA_peak_549 | 35 | Distal Interge | 8 | 52356611 | 52403339 | 46729 | 1 | 330767 | ENSMUST00 | -621563 | ENSMUSG0( | 1190028D05 | RIKEN cDNA 1190028D05 gene |
| chr8 | 52755943 | 52756085 | 143 | * | NA_peak_550 | 24 | Distal Interge | 8 | 52356611 | 52403339 | 46729 | 1 | 330767 | ENSMUST00 | 399332 | ENSMUSG0( | 1190028D05 | RIKEN cDNA 1190028D05 gene |
| chr8 | 53265998 | 53266060 | 63 | * | NA_peak_550 | 26 | Distal Interge | 8 | 53511727 | 53523334 | 11608 | 1 | 11593 | ENSMUST00 | -245667 | ENSMUSG0( | Aga | aspartylglucosaminidase |
| chr8 | 53455243 | 53455567 | 325 | * | NA_peak_550 | 20 | Intron (ENSM | 8 | 53511727 | 53523334 | 11608 | 1 | 11593 | ENSMUST00 | -56160 | ENSMUSG0( | Aga | aspartylglucosaminidase |
| chr8 | 54729770 | 54730063 | 294 | * | NA_peak_550 | 20 | Intron (ENSM | 8 | 54629247 | 54724504 | 95258 | 2 | 244484 | ENSMUST00 | -5266 | ENSMUSG0( | Wdr17 | WD repeat domain 17 |
| chr8 | 55118916 | 55119382 | 467 | * | NA_peak_550 | 29 | Distal Interge | 8 | 54954843 | 55060871 | 106029 | 1 | 234267 | ENSMUST00 | 164073 | ENSMUSG0( | Gpm6a | glycoprotein m6a |
| chr8 | 55121990 | 55123493 | 1504 | * | NA_peak_550 | 78 | Distal Interge | 8 | 54954843 | 55060871 | 106029 | 1 | 234267 | ENSMUST00 | 167147 | ENSMUSG0( | Gpm6a | glycoprotein m6a |
| chr8 | 55354216 | 55354561 | 346 | * | NA_peak_550 | 49 | Distal Interge | 8 | 54954843 | 55060871 | 106029 | 1 | 234267 | ENSMUST00 | 399373 | ENSMUSG0( | Gpm6a | glycoprotein m6a |
| chr8 | 55374032 | 55374314 | 283 | * | NA_peak_550 | 47 | Distal Interge | 8 | 54954843 | 55060871 | 106029 | 1 | 234267 | ENSMUST00 | 419189 | ENSMUSG0( | Gpm6a | glycoprotein m6a |
| chr8 | 55381709 | 55382003 | 295 | * | NA_peak_550 | 43 | Distal Interge | 8 | 54954843 | 55060871 | 106029 | 1 | 234267 | ENSMUST00 | 426866 | ENSMUSG0( | Gpm6a | glycoprotein m6a |
| chr8 | 57103101 | 57103365 | 265 | * | NA_peak_551 | 30 | Distal Interge | 8 | 57304271 | 57305358 | 1088 | 2 | 102636514 | ENSMUST00 | 201993 | NA | Hand2os1 | Hand2, opposite strand 1 |
| chr8 | 57190512 | 57190575 | 64 | * | NA_peak_551 | 15 | Distal Interge | 8 | 57304271 | 57305358 | 1088 | 2 | 102636514 | ENSMUST00 | 114783 | NA | Hand2os1 | Hand2, opposite strand 1 |
| chr8 | 57218321 | 57218541 | 221 | * | NA_peak_551 | 16 | Distal Interge | 8 | 57304271 | 57305358 | 1088 | 2 | 102636514 | ENSMUST00 | 86817 | NA | Hand2os1 | Hand2, opposite strand 1 |
| chr8 | 57531390 | 57531600 | 211 | * | NA_peak_551 | 20 | Intron (ENSM | 8 | 57526024 | 57538453 | 12430 | 2 | 108150 | ENSMUST00 | 6853 | ENSMUSG0( | Galnt7 | polypeptide N-acetylgalactosaminyltransferase 7 |
| chr8 | 57571427 | 57571983 | 557 | * | NA_peak_551 | 43 | Intron (ENSM | 8 | 57526024 | 57538453 | 12430 | 2 | 108150 | ENSMUST00 | -32974 | ENSMUSG0( | Galnt7 | polypeptide N-acetylgalactosaminyltransferase 7 |
| chr8 | 59839142 | 59839893 | 752 | * | NA_peak_552 | 27 | Intron (ENSM | 8 | 59825649 | 59844723 | 19075 | 1 | 75017 | ENSMUST00 | 13493 | ENSMUSG0( | 4930470O06 | RIKEN cDNA 4930470O06 gene |
| chr8 | 60323378 | 60323722 | 345 | * | NA_peak_552 | 116 | Distal Interge | 8 | 60505932 | 60545677 | 39746 | 1 | 23923 | ENSMUST00 | -182210 | ENSMUSG0( | Aadat | aminoadipate aminotransferase |
| chr8 | 60767561 | 60767779 | 219 | * | NA_peak_552 | 16 | Distal Interge | 8 | 60655540 | 60672362 | 16823 | 1 | 71306 | ENSMUST00 | 112021 | ENSMUSG0( | Mfap3l | microfibrillar-associated protein 3-like |
| chr8 | 60850887 | 60851239 | 353 | * | NA_peak_552 | 31 | Distal Interge | 8 | 60890418 | 60907575 | 17158 | 1 | 72612 | ENSMUST00 | -39179 | ENSMUSG0( | Hpf1 | histone PARylation factor 1 |
| chr8 | 61952299 | 61952571 | 273 | * | NA_peak_553 | 50 | Intron (ENSM | 8 | 61928089 | 62038244 | 110156 | 1 | 234311 | ENSMUST00 | 24210 | ENSMUSG0( | Ddx60 | DExD/H box helicase 60 |
| chr8 | 62160882 | 62161324 | 443 | * | NA_peak_553 | 98 | Distal Interge | 8 | 62063744 | 62123150 | 59407 | 2 | 26359 | ENSMUST00 | -37732 | ENSMUSG0( | Anxa10 | annexin A10 |

| | | | | | | | | | | | | | | | | | | |
|---|---|---|---|---|---|---|---|---|---|---|---|---|---|---|---|---|---|---|
| chr8 | 62651913 | 62652080 | 168 | * | NA_peak_553 | 50 | Intron (ENSM | 8 | 62951009 | 63356852 | 405844 | 1 | 72902 | ENSMUST00 | -298929 | ENSMUSG0 | Spock3 | sparc/osteonectin, cwcv and kazal-like domains proteoglycan 3 |
| chr8 | 63674541 | 63674825 | 285 | * | NA_peak_553 | 23 | Distal Interge | 8 | 63924694 | 63952170 | 27477 | 2 | 244495 | ENSMUST00 | 277345 | ENSMUSG0 | Sgo2b | shugoshin 2B |
| chr8 | 64288609 | 64288894 | 286 | * | NA_peak_553 | 40 | Distal Interge | 8 | 64084326 | 64206008 | 121683 | 2 | 21892 | ENSMUST00 | -82601 | ENSMUSG0 | Tll1 | tolloid-like |
| chr8 | 64947454 | 64947664 | 211 | * | NA_peak_554 | 26 | Promoter (<= | 8 | 64947254 | 64969031 | 21778 | 1 | 73067 | ENSMUST00 | 200 | ENSMUSG0 | Tmem192 | transmembrane protein 192 |
| chr8 | 65185709 | 65186020 | 312 | * | NA_peak_554 | 48 | Distal Interge | 8 | 65085613 | 65129563 | 43951 | 2 | 407795 | ENSMUST00 | -56146 | ENSMUSG0 | Smim31 | small integral membrane protein 31 |
| chr8 | 65549779 | 65550089 | 311 | * | NA_peak_554 | 23 | Distal Interge | 8 | 65617900 | 66469960 | 852061 | 1 | 72925 | ENSMUST00 | -67811 | ENSMUSG0 | Marchf1 | membrane associated ring-CH-type finger 1 |
| chr8 | 65709375 | 65709651 | 277 | * | NA_peak_554 | 36 | Intron (ENSM | 8 | 65618525 | 66470427 | 851903 | 1 | 72925 | ENSMUST00 | 90850 | ENSMUSG0 | Marchf1 | membrane associated ring-CH-type finger 1 |
| chr8 | 66554846 | 66554889 | 44 | * | NA_peak_554 | 23 | Distal Interge | 8 | 66511759 | 66518335 | 6577 | 1 | 74419 | ENSMUST00 | 43087 | ENSMUSG0 | Tktl2 | transketolase-like 2 |
| chr8 | 66579844 | 66580091 | 248 | * | NA_peak_554 | 27 | Distal Interge | 8 | 66511759 | 66518335 | 6577 | 1 | 74419 | ENSMUST00 | 68085 | ENSMUSG0 | Tktl2 | transketolase-like 2 |
| chr8 | 66953566 | 66954087 | 522 | * | NA_peak_554 | 17 | Distal Interge | 8 | 66860957 | 66883597 | 22641 | 1 | 234344 | ENSMUST00 | 92609 | ENSMUSG0 | Naf1 | nuclear assembly factor 1 ribonucleoprotein |
| chr8 | 66956171 | 66956790 | 620 | * | NA_peak_554 | 31 | Distal Interge | 8 | 66860957 | 66883597 | 22641 | 1 | 234344 | ENSMUST00 | 95214 | ENSMUSG0 | Naf1 | nuclear assembly factor 1 ribonucleoprotein |
| chr8 | 67037814 | 67038049 | 236 | * | NA_peak_555 | 20 | Distal Interge | 8 | 66860957 | 66883597 | 22641 | 1 | 234344 | ENSMUST00 | 176857 | ENSMUSG0 | Naf1 | nuclear assembly factor 1 ribonucleoprotein |
| chr8 | 67221860 | 67222032 | 173 | * | NA_peak_555 | 16 | Distal Interge | 8 | 67490758 | 67492104 | 1347 | 1 | 17960 | ENSMUST00 | -268726 | ENSMUSG0 | Nat1 | N-acetyl transferase 1 |
| chr8 | 67412849 | 67413058 | 210 | * | NA_peak_555 | 31 | Distal Interge | 8 | 67490758 | 67492104 | 1347 | 1 | 17960 | ENSMUST00 | -77700 | ENSMUSG0 | Nat1 | N-acetyl transferase 1 |
| chr8 | 68832666 | 68832790 | 125 | * | NA_peak_555 | 20 | Distal Interge | 8 | 68820686 | 68827501 | 6816 | 1 | 70885 | ENSMUST00 | 11980 | ENSMUSG0 | Ints10 | integrator complex subunit 10 |
| chr8 | 69633581 | 69633785 | 205 | * | NA_peak_556 | 21 | Distal Interge | 8 | 69610857 | 69625548 | 14692 | 2 | 234362 | ENSMUST00 | -8033 | ENSMUSG0 | Zfp868 | zinc finger protein 868 |
| chr8 | 70863427 | 70863746 | 320 | * | NA_peak_556 | 28 | Promoter (<= | 8 | 70843906 | 70863258 | 19353 | 2 | 84036 | ENSMUST00 | -169 | ENSMUSG0 | Kcnn1 | potassium intermediate/small conductance calcium-activated channel, subfamily N, member 1 |
| chr8 | 71026521 | 71026726 | 206 | * | NA_peak_556 | 20 | Promoter (<= | 8 | 71006727 | 71026755 | 20029 | 2 | 69419 | ENSMUST00 | 29 | ENSMUSG0 | 1700026F02 | RIKEN cDNA 1700026F02 gene |
| chr8 | 71291437 | 71291831 | 395 | * | NA_peak_556 | 14 | Intron (ENSM | 8 | 71251125 | 71272934 | 21810 | 2 | 76478 | ENSMUST00 | -18503 | ENSMUSG0 | Haus8 | 4HAUS augmin-like complex, subunit 8 |
| chr8 | 71809941 | 71810258 | 318 | * | NA_peak_556 | 36 | Distal Interge | 8 | 71882019 | 71895727 | 13709 | 1 | 236193 | ENSMUST00 | -71761 | ENSMUSG0 | Zfp709 | zinc finger protein 709 |
| chr8 | 71811275 | 71811359 | 85 | * | NA_peak_556 | 37 | Distal Interge | 8 | 71882019 | 71895727 | 13709 | 1 | 236193 | ENSMUST00 | -70660 | ENSMUSG0 | Zfp709 | zinc finger protein 709 |
| chr8 | 72085329 | 72085638 | 310 | * | NA_peak_556 | 60 | Intron (ENSM | 8 | 72099762 | 72100706 | 945 | 1 | 258532 | ENSMUST00 | -14124 | ENSMUSG0 | Olfr373 | olfactory receptor 373 |
| chr8 | 73255730 | 73256186 | 457 | * | NA_peak_557 | 36 | Intron (ENSM | 8 | 72814599 | 73352556 | 537958 | 2 | 16795 | ENSMUST00 | 96370 | ENSMUSG0 | Large1 | LARGE xylosyl- and glucuronyltransferase 1 |
| chr8 | 73372675 | 73372956 | 282 | * | NA_peak_557 | 20 | Distal Interge | 8 | 72814599 | 73353540 | 538942 | 2 | 16795 | ENSMUST00 | -19135 | ENSMUSG0 | Large1 | LARGE xylosyl- and glucuronyltransferase 1 |
| chr8 | 73394827 | 73395109 | 283 | * | NA_peak_557 | 27 | Distal Interge | 8 | 72814599 | 73353540 | 538942 | 2 | 16795 | ENSMUST00 | -41287 | ENSMUSG0 | Large1 | LARGE xylosyl- and glucuronyltransferase 1 |
| chr8 | 74799858 | 74799858 | 1 | * | NA_peak_558 | 36 | Intron (ENSM | 8 | 74873173 | 74893506 | 20334 | 1 | 71597 | ENSMUST00 | -73315 | ENSMUSG0 | Isx | intestine specific homeobox |
| chr8 | 74899184 | 74899362 | 179 | * | NA_peak_558 | 20 | Distal Interge | 8 | 74873540 | 74893147 | 19608 | 1 | 71597 | ENSMUST00 | 25644 | ENSMUSG0 | Isx | intestine specific homeobox |
| chr8 | 75158635 | 75158968 | 334 | * | NA_peak_558 | 25 | Distal Interge | 8 | 75123067 | 75125228 | 2162 | 1 | 17218 | ENSMUST00 | 35568 | ENSMUSG0 | Mcm5 | minichromosome maintenance complex component 5 |
| chr8 | 75872574 | 75872901 | 328 | * | NA_peak_558 | 28 | Intron (ENSM | 8 | 75448983 | 75984503 | 535521 | 1 | 71831 | ENSMUST00 | 423591 | ENSMUSG0 | Iqcm | IQ motif containing M |
| chr8 | 76064180 | 76064217 | 38 | * | NA_peak_558 | 20 | Intron (ENSM | 8 | 75448983 | 75984503 | 535521 | 1 | 71831 | ENSMUST00 | 615197 | ENSMUSG0 | Iqcm | IQ motif containing M |
| chr8 | 77138311 | 77138791 | 481 | * | NA_peak_559 | 48 | Intron (ENSM | 8 | 77128013 | 77185872 | 57860 | 1 | 110784 | ENSMUST00 | 10298 | ENSMUSG0 | Nr3c2 | nuclear receptor subfamily 3, group C, member 2 |
| chr8 | 77356294 | 77356597 | 304 | * | NA_peak_559 | 20 | Intron (ENSM | 8 | 77517056 | 77518579 | 1524 | 1 | 70345 | ENSMUST00 | -160459 | ENSMUSG0 | 0610038B21 | RIKEN cDNA 0610038B21 gene |
| chr8 | 77582386 | 77582701 | 316 | * | NA_peak_559 | 23 | Distal Interge | 8 | 77563662 | 77565093 | 1432 | 1 | 102182 | ENSMUST00 | 18724 | ENSMUSG0 | Prmt9 | protein arginine methyltransferase 9 |
| chr8 | 77586034 | 77586268 | 235 | * | NA_peak_559 | 20 | Distal Interge | 8 | 77563662 | 77565093 | 1432 | 1 | 102182 | ENSMUST00 | 22372 | ENSMUSG0 | Prmt9 | protein arginine methyltransferase 9 |
| chr8 | 77622775 | 77623113 | 339 | * | NA_peak_559 | 16 | Promoter (1-2 | 8 | 77624172 | 77628957 | 4786 | 1 | 67357 | ENSMUST00 | -1059 | ENSMUSG0 | 1700092C02 | RIKEN cDNA 1700092C02 gene |
| chr8 | 77886564 | 77886728 | 165 | * | NA_peak_559 | 28 | Distal Interge | 8 | 77669694 | 77724464 | 54771 | 2 | 13617 | ENSMUST00 | -162100 | ENSMUSG0 | Ednra | endothelin receptor type A |
| chr8 | 79765761 | 79766134 | 374 | * | NA_peak_560 | 45 | Promoter (<= | 8 | 79765212 | 79775342 | 10131 | 1 | 68999 | ENSMUST00 | 549 | ENSMUSG0 | Anapc10 | anaphase promoting complex subunit 10 |
| chr8 | 79824381 | 79824662 | 282 | * | NA_peak_560 | 38 | Distal Interge | 8 | 79765212 | 79775342 | 10131 | 1 | 68999 | ENSMUST00 | 59169 | ENSMUSG0 | Anapc10 | anaphase promoting complex subunit 10 |
| chr8 | 79843309 | 79843570 | 262 | * | NA_peak_560 | 23 | Distal Interge | 8 | 79765212 | 79775342 | 10131 | 1 | 68999 | ENSMUST00 | 78097 | ENSMUSG0 | Anapc10 | anaphase promoting complex subunit 10 |
| chr8 | 80349499 | 80349820 | 322 | * | NA_peak_561 | 50 | Distal Interge | 8 | 80493781 | 80510542 | 16762 | 1 | 14934 | ENSMUST00 | -143961 | ENSMUSG0 | Gypa | glycophorin A |
| chr8 | 80365302 | 80365801 | 500 | * | NA_peak_561 | 20 | Distal Interge | 8 | 80493781 | 80510542 | 16762 | 1 | 14934 | ENSMUST00 | -127980 | ENSMUSG0 | Gypa | glycophorin A |
| chr8 | 80395585 | 80395939 | 355 | * | NA_peak_561 | 19 | Distal Interge | 8 | 80493781 | 80510542 | 16762 | 1 | 14934 | ENSMUST00 | -97842 | ENSMUSG0 | Gypa | glycophorin A |
| chr8 | 80643621 | 80643924 | 304 | * | NA_peak_561 | 28 | Intron (ENSM | 8 | 80611080 | 80695356 | 84277 | 1 | 333315 | ENSMUST00 | 32541 | ENSMUSG0 | Frem3 | Fras1 related extracellular matrix protein 3 |
| chr8 | 81014277 | 81014550 | 274 | * | NA_peak_561 | 52 | Promoter (<= | 8 | 80998213 | 81014913 | 16701 | 2 | 74841 | ENSMUST00 | 363 | ENSMUSG0 | Usp38 | ubiquitin specific peptidase 38 |
| chr8 | 83621049 | 83621403 | 355 | * | NA_peak_562 | 19 | Distal Interge | 8 | 83608553 | 83612653 | 4101 | 1 | 81489 | ENSMUST00 | 12496 | ENSMUSG0 | Dnajb1 | DnaJ heat shock protein family (Hsp40) member B1 |
| chr8 | 83657007 | 83657159 | 153 | * | NA_peak_562 | 42 | 5' UTR | 8 | 83652698 | 83661187 | 8490 | 1 | 67903 | ENSMUST00 | 4309 | ENSMUSG0 | Gipc1 | GIPC PDZ domain containing family, member 1 |
| chr8 | 83857729 | 83857874 | 146 | * | NA_peak_562 | 20 | Distal Interge | 8 | 83900105 | 83941954 | 41850 | 1 | 330814 | ENSMUST00 | -42231 | ENSMUSG0 | Adgrl1 | adhesion G protein-coupled receptor L1 |
| chr8 | 83875647 | 83875765 | 119 | * | NA_peak_562 | 17 | Distal Interge | 8 | 83900105 | 83941954 | 41850 | 1 | 330814 | ENSMUST00 | -24340 | ENSMUSG0 | Adgrl1 | adhesion G protein-coupled receptor L1 |
| chr8 | 84306807 | 84307066 | 260 | * | NA_peak_562 | 27 | Distal Interge | 8 | 84338639 | 84640246 | 301608 | 1 | 12286 | ENSMUST00 | -31573 | ENSMUSG0 | Cacna1a | calcium channel, voltage-dependent, P/Q type, alpha 1A subunit |
| chr8 | 84329106 | 84329251 | 146 | * | NA_peak_562 | 13 | Distal Interge | 8 | 84338639 | 84640246 | 301608 | 1 | 12286 | ENSMUST00 | -9388 | ENSMUSG0 | Cacna1a | calcium channel, voltage-dependent, P/Q type, alpha 1A subunit |
| chr8 | 84687580 | 84687975 | 396 | * | NA_peak_562 | 21 | Promoter (<= | 8 | 84670479 | 84687902 | 17424 | 2 | 66830 | ENSMUST00 | 0 | ENSMUSG0 | Nacc1 | nucleus accumbens associated 1, BEN and BTB (POZ) domain containing |
| chr8 | 84872149 | 84872392 | 244 | * | NA_peak_562 | 47 | Promoter (<= | 8 | 84872193 | 84887456 | 15264 | 1 | 71846 | ENSMUST00 | 0 | ENSMUSG0 | Syce2 | synaptonemal complex central element protein 2 |
| chr8 | 85420019 | 85420157 | 139 | * | NA_peak_562 | 22 | Intron (ENSM | 8 | 85412243 | 85432814 | 20572 | 2 | 66714 | ENSMUST00 | 12657 | ENSMUSG0 | 4921524J17 | RIKEN cDNA 4921524J17 gene |
| chr8 | 85840577 | 85840898 | 322 | * | NA_peak_563 | 13 | Promoter (<= | 8 | 85717578 | 85840921 | 123344 | 2 | 71927 | ENSMUST00 | 23 | ENSMUSG0 | Itfg1 | integrin alpha FG-GAP repeat containing 1 |
| chr8 | 86101327 | 86101669 | 343 | * | NA_peak_563 | 13 | Distal Interge | 8 | 86035646 | 86060641 | 24996 | 1 | 102093 | ENSMUST00 | 65681 | ENSMUSG0 | Phkb | phosphorylase kinase beta |
| chr8 | 86241125 | 86241481 | 357 | * | NA_peak_563 | 26 | Distal Interge | 8 | 86035646 | 86060641 | 24996 | 1 | 102093 | ENSMUST00 | 205479 | ENSMUSG0 | Phkb | phosphorylase kinase beta |
| chr8 | 86336304 | 86336497 | 194 | * | NA_peak_563 | 13 | Distal Interge | 8 | 86504583 | 86534747 | 30165 | 2 | 244562 | ENSMUST00 | 198250 | ENSMUSG0 | Abcc12 | ATP-binding cassette, sub-family C (CFTR/MRP), member 12 |
| chr8 | 86752016 | 86752453 | 438 | * | NA_peak_563 | 25 | Distal Interge | 8 | 86724007 | 86745934 | 21928 | 2 | 20437 | ENSMUST00 | -6082 | ENSMUSG0 | Siah1a | siah E3 ubiquitin protein ligase 1A |
| chr8 | 87915812 | 87916244 | 433 | * | NA_peak_564 | 36 | Intron (ENSM | 8 | 87661812 | 87895735 | 233924 | 2 | 94187 | ENSMUST00 | -20077 | ENSMUSG0 | Zfp423 | zinc finger protein 423 |
| chr8 | 88179863 | 88180180 | 318 | * | NA_peak_564 | 15 | Distal Interge | 8 | 88164482 | 88167045 | 2564 | 1 | 234549 | ENSMUST00 | 15381 | ENSMUSG0 | Heatr3 | HEAT repeat containing 3 |
| chr8 | 88398574 | 88398580 | 7 | * | NA_peak_564 | 35 | Distal Interge | 8 | 88331059 | 88362194 | 31136 | 2 | 26992 | ENSMUST00 | -36380 | ENSMUSG0 | Brd7 | bromodomain containing 7 |
| chr8 | 88789856 | 88789883 | 28 | * | NA_peak_564 | 24 | Distal Interge | 8 | 88729558 | 88737240 | 7683 | 1 | 74256 | ENSMUST00 | 60298 | ENSMUSG0 | Cyld | CYLD lysine 63 deubiquitinase |
| chr8 | 88924224 | 88924551 | 328 | * | NA_peak_564 | 16 | Distal Interge | 8 | 89024735 | 89024831 | 97 | 1 | 102465899 | ENSMUST00 | -100184 | ENSMUSG0 | Mir8110 | microRNA 8110 |
| chr8 | 88927491 | 88927656 | 166 | * | NA_peak_564 | 13 | Distal Interge | 8 | 89024735 | 89024831 | 97 | 1 | 102465899 | ENSMUST00 | -97079 | ENSMUSG0 | Mir8110 | microRNA 8110 |
| chr8 | 90314826 | 90315143 | 318 | * | NA_peak_565 | 19 | Intron (ENSM | 8 | 90257946 | 90348126 | 90181 | 2 | 244579 | ENSMUST00 | 32983 | ENSMUSG0 | Tox3 | TOX high mobility group box family member 3 |
| chr8 | 90505888 | 90506500 | 613 | * | NA_peak_565 | 20 | Distal Interge | 8 | 90247040 | 90348343 | 101304 | 2 | 244579 | ENSMUST00 | -157545 | ENSMUSG0 | Tox3 | TOX high mobility group box family member 3 |
| chr8 | 90591331 | 90591606 | 276 | * | NA_peak_565 | 36 | Distal Interge | 8 | 90816145 | 90817393 | 1249 | 2 | 100503868 | ENSMUST00 | 225787 | ENSMUSG0 | Gm19935 | predicted gene, 19935 |
| chr8 | 90857590 | 90857969 | 380 | * | NA_peak_565 | 23 | Intron (ENSM | 8 | 90849667 | 90872090 | 22424 | 1 | 109151 | ENSMUST00 | 7923 | ENSMUSG0 | Chd9 | chromodomain helicase DNA binding protein 9 |
| chr8 | 91050533 | 91050658 | 126 | * | NA_peak_565 | 16 | Intron (ENSM | 8 | 91070057 | 91123844 | 53788 | 1 | 19651 | ENSMUST00 | -19399 | ENSMUSG0 | Rbl2 | RB transcriptional corepressor like 2 |
| chr8 | 91551527 | 91551630 | 104 | * | NA_peak_565 | 24 | Intron (ENSM | 8 | 91467685 | 91666946 | 199262 | 1 | 26383 | ENSMUST00 | 83842 | ENSMUSG0 | Fto | fat mass and obesity associated |
| chr8 | 91749295 | 91749511 | 217 | * | NA_peak_565 | 31 | Distal Interge | 8 | 91798525 | 91801915 | 3391 | 2 | 16373 | ENSMUST00 | 52404 | ENSMUSG0 | Irx3 | Iroquois related homeobox 3 |
| chr8 | 91882519 | 91882609 | 91 | * | NA_peak_566 | 21 | Distal Interge | 8 | 91798525 | 91802067 | 3543 | 2 | 16373 | ENSMUST00 | -80452 | ENSMUSG0 | Irx3 | Iroquois related homeobox 3 |
| chr8 | 92064049 | 92064147 | 99 | * | NA_peak_566 | 26 | Distal Interge | 8 | 91798525 | 91802067 | 3543 | 2 | 16373 | ENSMUST00 | -261982 | ENSMUSG0 | Irx3 | Iroquois related homeobox 3 |
| chr8 | 92388079 | 92388530 | 452 | * | NA_peak_566 | 30 | Distal Interge | 8 | 92357749 | 92361456 | 3708 | 1 | 54352 | ENSMUST00 | 30330 | ENSMUSG0 | Irx5 | Iroquois homeobox 5 |
| chr8 | 92673590 | 92673631 | 42 | * | NA_peak_566 | 13 | Promoter (<= | 8 | 92674288 | 92680956 | 6669 | 1 | 64379 | ENSMUST00 | -657 | ENSMUSG0 | Irx6 | Iroquois homeobox 6 |
| chr8 | 92826332 | 92826597 | 266 | * | NA_peak_566 | 40 | Promoter (<= | 8 | 92827291 | 92853420 | 26130 | 1 | 17390 | ENSMUST00 | -694 | ENSMUSG0 | Mmp2 | matrix metallopeptidase 2 |
| chr8 | 92995811 | 92996173 | 363 | * | NA_peak_566 | 33 | Exon (ENSMU | 8 | 92960933 | 93001667 | 40735 | 1 | 20538 | ENSMUST00 | 34878 | ENSMUSG0 | Slc6a2 | solute carrier family 6 (neurotransmitter transporter, noradrenalin), member 2 |
| chr8 | 93150331 | 93150725 | 395 | * | NA_peak_566 | 61 | Intron (ENSM | 8 | 93107077 | 93131283 | 24207 | 2 | 13884 | ENSMUST00 | -19048 | ENSMUSG0 | Ces1c | carboxylesterase 1C |
| chr8 | 93876144 | 93876494 | 351 | * | NA_peak_567 | 26 | Intron (ENSM | 8 | 93896231 | 93968163 | 71933 | 1 | 14681 | ENSMUST00 | -19737 | ENSMUSG0 | Gnao1 | guanine nucleotide binding protein, alpha O |
| chr8 | 94196801 | 94197152 | 352 | * | NA_peak_567 | 30 | Distal Interge | 8 | 94214564 | 94314860 | 100297 | 1 | 71805 | ENSMUST00 | -17412 | ENSMUSG0 | Nup93 | nucleoporin 93 |
| chr8 | 94616915 | 94617341 | 427 | * | NA_peak_567 | 30 | Intron (ENSM | 8 | 94628602 | 94632180 | 3579 | 1 | 67610 | ENSMUST00 | -11261 | ENSMUSG0 | Rspry1 | ring finger and SPRY domain containing 1 |
| chr8 | 94628143 | 94628706 | 564 | * | NA_peak_567 | 36 | Promoter (<= | 8 | 94628602 | 94632180 | 3579 | 1 | 67610 | ENSMUST00 | 0 | ENSMUSG0 | Rspry1 | ring finger and SPRY domain containing 1 |
| chr8 | 94961989 | 94962071 | 83 | * | NA_peak_567 | 16 | Distal Interge | 8 | 94974751 | 95005804 | 31054 | 1 | 14766 | ENSMUST00 | -12680 | ENSMUSG0 | Adgrg1 | adhesion G protein-coupled receptor G1 |
| chr8 | 95184380 | 95184427 | 48 | * | NA_peak_567 | 23 | Promoter (1-2 | 8 | 95110249 | 95182791 | 72543 | 2 | 16582 | ENSMUST00 | -1589 | ENSMUSG0 | Kifc3 | kinesin family member C3 |
| chr8 | 95972316 | 95972618 | 303 | * | NA_peak_567 | 16 | Distal Interge | 8 | 95864134 | 95888547 | 24414 | 2 | 14719 | ENSMUST00 | -83769 | ENSMUSG0 | Got2 | glutamatic-oxaloacetic transaminase 2, mitochondrial |
| chr8 | 96418066 | 96418368 | 303 | * | NA_peak_568 | 50 | Distal Interge | 8 | 95864134 | 95888547 | 24414 | 2 | 14719 | ENSMUST00 | -529519 | ENSMUSG0 | Got2 | glutamatic-oxaloacetic transaminase 2, mitochondrial |
| chr8 | 96669336 | 96669607 | 272 | * | NA_peak_568 | 15 | Distal Interge | 8 | 95864134 | 95888547 | 24414 | 2 | 14719 | ENSMUST00 | -780789 | ENSMUSG0 | Got2 | glutamatic-oxaloacetic transaminase 2, mitochondrial |
| chr8 | 97114690 | 97115129 | 440 | * | NA_peak_568 | 78 | Distal Interge | 8 | 95864134 | 95888547 | 24414 | 2 | 14719 | ENSMUST00 | -1226143 | ENSMUSG0 | Got2 | glutamatic-oxaloacetic transaminase 2, mitochondrial |
| chr8 | 98034160 | 98034433 | 274 | * | NA_peak_568 | 31 | Distal Interge | 8 | 99011886 | 99032740 | 20855 | 1 | 100873092 | ENSMUST00 | -977453 | ENSMUSG0 | Gm15679 | predicted gene 15679 |
| chr8 | 98698524 | 98698923 | 400 | * | NA_peak_568 | 14 | Distal Interge | 8 | 99011886 | 99032740 | 20855 | 1 | 100873092 | ENSMUST00 | -312963 | ENSMUSG0 | Gm15679 | predicted gene 15679 |
| chr8 | 100279848 | 100280153 | 306 | * | NA_peak_568 | 53 | Distal Interge | 8 | 99416674 | 99733438 | 316765 | 1 | 234624 | ENSMUST00 | 863174 | ENSMUSG0 | A330008L17 | RIKEN cDNA A330008L17 gene |
| chr8 | 101066261 | 101066381 | 121 | * | NA_peak_568 | 20 | Distal Interge | 8 | 102634604 | 102637319 | 2716 | 2 | 12552 | ENSMUST00 | 1570938 | ENSMUSG0 | Cdh11 | cadherin 11 |
| chr8 | 101208805 | 101208870 | 66 | * | NA_peak_569 | 27 | Distal Interge | 8 | 102634604 | 102637319 | 2716 | 2 | 12552 | ENSMUST00 | 1428449 | ENSMUSG0 | Cdh11 | cadherin 11 |
| chr8 | 103040705 | 103040874 | 170 | * | NA_peak_569 | 27 | Distal Interge | 8 | 102661487 | 102785642 | 124156 | 2 | 12552 | ENSMUST00 | -255063 | ENSMUSG0 | Cdh11 | cadherin 11 |
| chr8 | 103688901 | 103689192 | 292 | * | NA_peak_569 | 20 | Distal Interge | 8 | 103603799 | 103680384 | 76586 | 1 | 71058 | ENSMUST00 | 85102 | ENSMUSG0 | 4933400L20 | RIKEN cDNA 4933400L20 gene |
| chr8 | 104271509 | 104271915 | 407 | * | NA_peak_569 | 38 | Distal Interge | 8 | 104262147 | 104263444 | 1298 | 1 | 75458 | ENSMUST00 | 9362 | ENSMUSG0 | Cklf | chemokine-like factor |
| chr8 | 104839421 | 104839678 | 258 | * | NA_peak_569 | 20 | 3' UTR | 8 | 104847068 | 104854483 | 7416 | 1 | 234671 | ENSMUST00 | -7390 | ENSMUSG0 | Ces2c | carboxylesterase 2C |
| chr8 | 105006774 | 105007869 | 1096 | * | NA_peak_569 | 24 | Intron (ENSM | 8 | 105000853 | 105021178 | 20326 | 1 | 436059 | ENSMUST00 | 5921 | ENSMUSG0 | Ces2h | carboxylesterase 2H |
| chr8 | 105162560 | 105162637 | 78 | * | NA_peak_570 | 20 | Distal Interge | 8 | 105170674 | 105217989 | 47316 | 1 | 12400 | ENSMUST00 | -8037 | ENSMUSG0 | Cbfb | core binding factor beta |
| chr8 | 105841772 | 105842006 | 235 | * | NA_peak_570 | 28 | Exon (ENSMU | 8 | 105845074 | 105846700 | 1627 | 2 | 320394 | ENSMUST00 | 4694 | ENSMUSG0 | Cenpt | centromere protein T |
| chr8 | 105860975 | 105861114 | 140 | * | NA_peak_570 | 15 | Promoter (<= | 8 | 105860983 | 105878979 | 17997 | 1 | 68051 | ENSMUST00 | 0 | ENSMUSG0 | Nutf2 | nuclear transport factor 2 |
| chr8 | 105861495 | 105861535 | 41 | * | NA_peak_570 | 20 | Promoter (<= | 8 | 105860983 | 105878979 | 17997 | 1 | 68051 | ENSMUST00 | 512 | ENSMUSG0 | Nutf2 | nuclear transport factor 2 |
| chr8 | 106622738 | 106622805 | 68 | * | NA_peak_570 | 20 | Intron (ENSM | 8 | 106604143 | 106653919 | 49777 | 1 | 12550 | ENSMUST00 | 18595 | ENSMUSG0 | Cdh1 | cadherin 1 |
| chr8 | 107244067 | 107244090 | 24 | * | NA_peak_570 | 16 | Distal Interge | 8 | 107293470 | 107329307 | 35838 | 1 | 54446 | ENSMUST00 | -49380 | ENSMUSG0 | Nfat5 | nuclear factor of activated T cells 5 |
| chr8 | 107390117 | 107390425 | 309 | * | NA_peak_570 | 23 | Intron (ENSM | 8 | 107388225 | 107403206 | 14982 | 2 | 18104 | ENSMUST00 | 12781 | ENSMUSG0 | Nqo1 | NAD(P)H dehydrogenase, quinone 1 |
| chr8 | 107878353 | 107878894 | 542 | * | NA_peak_571 | 20 | Intron (ENSM | 8 | 107942644 | 108961630 | 1018987 | 1 | 11906 | ENSMUST00 | -63750 | ENSMUSG0 | Zfhx3 | zinc finger homeobox 3 |

| | | | | | | | | | | | | | | | | | | |
|---|---|---|---|---|---|---|---|---|---|---|---|---|---|---|---|---|---|---|
| chr8 | 108747992 | 108748200 | 209 | * | NA_peak_57 | 20 | Intron (ENSM | 8 | 108714644 | 108961630 | 246987 | 1 | 11906 | ENSMUST00 | 33348 | ENSMUSG0( | Zfhx3 | zinc finger homeobox 3 |
| chr8 | 108852915 | 108853150 | 236 | * | NA_peak_57 | 26 | Intron (ENSM | 8 | 108936860 | 108936938 | 79 | 1 | 100526511 | ENSMUST00 | -83710 | ENSMUSG0( | Mir3108 | microRNA 3108 |
| chr8 | 108870379 | 108870700 | 322 | * | NA_peak_57 | 21 | Intron (ENSM | 8 | 108936860 | 108936938 | 79 | 1 | 100526511 | ENSMUST00 | -66160 | ENSMUSG0( | Mir3108 | microRNA 3108 |
| chr8 | 109721432 | 109721673 | 242 | * | NA_peak_57 | 19 | Intron (ENSM | 8 | 109708914 | 109724932 | 16019 | 1 | 75871 | ENSMUST00 | 12518 | ENSMUSG0( | Zfp821 | zinc finger protein 821 |
| chr8 | 110422801 | 110423266 | 466 | * | NA_peak_57 | 50 | Intron (ENSM | 8 | 110444489 | 110447518 | 3030 | 1 | 244653 | ENSMUST00 | -21223 | ENSMUSG0( | Hydin | HYDIN, axonemal central pair apparatus protein |
| chr8 | 110820434 | 110820705 | 272 | * | NA_peak_57 | 16 | Intron (ENSM | 8 | 110755525 | 110805924 | 50400 | 2 | 76527 | ENSMUST00 | -14510 | ENSMUSG0( | Il34 | interleukin 34 |
| chr8 | 111446141 | 111446226 | 86 | * | NA_peak_57 | 19 | Distal Interge | 8 | 111345141 | 111393824 | 48684 | 2 | 338521 | ENSMUST00 | -52317 | ENSMUSG0( | Fa2h | fatty acid 2-hydroxylase |
| chr8 | 112651104 | 112651351 | 248 | * | NA_peak_57 | 21 | Intron (ENSM | 8 | 112731399 | 112733519 | 2121 | 1 | 170571 | ENSMUST00 | -80048 | ENSMUSG0( | Cntnap4 | contactin associated protein-like 4 |
| chr8 | 113576215 | 113576474 | 260 | * | NA_peak_57 | 14 | Distal Interge | 8 | 113635587 | 113643806 | 8220 | 1 | 270096 | ENSMUST00 | -59113 | ENSMUSG0( | Mon1b | MON1 homolog B, secretory traffciking associated |
| chr8 | 114393356 | 114393775 | 420 | * | NA_peak_57 | 35 | Distal Interge | 8 | 114418086 | 114425846 | 7761 | 1 | 403395 | ENSMUST00 | -24311 | ENSMUSG0( | Clec3a | C-type lectin domain family 3, member a |
| chr8 | 114812329 | 114812838 | 510 | * | NA_peak_57 | 20 | Intron (ENSM | 8 | 114712167 | 115352143 | 639977 | 1 | 80707 | ENSMUST00 | 100162 | ENSMUSG0( | Wwox | WW domain-containing oxidoreductase |
| chr8 | 115257651 | 115257890 | 240 | * | NA_peak_57 | 36 | Intron (ENSM | 8 | 115701435 | 115707794 | 6360 | 2 | 17132 | ENSMUST00 | 449904 | ENSMUSG0( | Maf | avian musculoaponeurotic fibrosarcoma oncogene homolog |
| chr8 | 116792024 | 116792072 | 49 | * | NA_peak_57 | 23 | Distal Interge | 8 | 116574400 | 116732991 | 158592 | 2 | 75796 | ENSMUST00 | -59033 | ENSMUSG0( | Cdyl2 | chromodomain protein, Y chromosome-like 2 |
| chr8 | 117646479 | 117646546 | 68 | * | NA_peak_57 | 16 | Distal Interge | 8 | 117661399 | 117671547 | 10149 | 2 | 74032 | ENSMUST00 | 25001 | ENSMUSG0( | Sdr42e1 | short chain dehydrogenase/reductase family 42E, member 1 |
| chr8 | 118617150 | 118617346 | 197 | * | NA_peak_57 | 17 | Intron (ENSM | 8 | 118314413 | 118856206 | 541794 | 1 | 12554 | ENSMUST00 | 302737 | ENSMUSG0( | Cdh13 | cadherin 13 |
| chr8 | 118820803 | 118821046 | 244 | * | NA_peak_57 | 22 | Intron (ENSM | 8 | 119151465 | 119242356 | 90892 | 1 | 12554 | ENSMUST00 | -330419 | ENSMUSG0( | Cdh13 | cadherin 13 |
| chr8 | 119335451 | 119335832 | 382 | * | NA_peak_57 | 38 | Distal Interge | 8 | 119344488 | 119348922 | 4435 | 1 | 68196 | ENSMUST00 | -8656 | ENSMUSG0( | Hsbp1 | heat shock factor binding protein 1 |
| chr8 | 119403216 | 119403501 | 286 | * | NA_peak_57 | 25 | Intron (ENSM | 8 | 119394898 | 119411087 | 16190 | 1 | 56690 | ENSMUST00 | 8318 | ENSMUSG0( | Mlycd | malonyl-CoA decarboxylase |
| chr8 | 120101139 | 120101503 | 365 | * | NA_peak_57 | 18 | Promoter (<= | 8 | 120080890 | 120101482 | 20593 | 2 | 102193 | ENSMUST00 | 0 | ENSMUSG0( | Zdhhc7 | zinc finger, DHHC domain containing 7 |
| chr8 | 120594413 | 120594717 | 305 | * | NA_peak_57 | 18 | Intron (ENSM | 8 | 120578633 | 120589304 | 10672 | 2 | 272551 | ENSMUST00 | -5109 | ENSMUSG0( | Gins2 | GINS complex subunit 2 (Psf2 homolog) |
| chr8 | 120750033 | 120750261 | 229 | * | NA_peak_57 | 27 | Promoter (1-2 | 8 | 120752209 | 120753711 | 1503 | 1 | 15900 | ENSMUST00 | -1948 | ENSMUSG0( | Irf8 | interferon regulatory factor 8 |
| chr8 | 121538510 | 121538983 | 474 | * | NA_peak_57 | 21 | Promoter (2-3 | 8 | 121530828 | 121541869 | 11042 | 2 | 76405 | ENSMUST00 | 2886 | ENSMUSG0( | 1700018B08 | RIKEN cDNA 1700018B08 gene |
| chr8 | 122014333 | 122014490 | 158 | * | NA_peak_57 | 30 | Intron (ENSM | 8 | 122000847 | 122007140 | 6294 | 1 | 53325 | ENSMUST00 | 13486 | ENSMUSG0( | Banp | BTG3 associated nuclear protein |
| chr8 | 123864355 | 123864722 | 368 | * | NA_peak_57 | 20 | Intron (ENSM | 8 | 123840844 | 123860209 | 19366 | 2 | 73420 | ENSMUST00 | -4146 | ENSMUSG0( | Ccsap | centriole, cilia and spindle associated protein |
| chr8 | 124721635 | 124722348 | 714 | * | NA_peak_57 | 20 | Promoter (<= | 8 | 124671332 | 124721964 | 50633 | 2 | 234875 | ENSMUST00 | 0 | ENSMUSG0( | Ttc13 | tetratricopeptide repeat domain 13 |
| chr8 | 125684622 | 125685001 | 380 | * | NA_peak_57 | 13 | Distal Interge | 8 | 125669818 | 125673359 | 3542 | 1 | 74393 | ENSMUST00 | 14804 | ENSMUSG0( | Map10 | microtubule-associated protein 10 |
| chr8 | 126276212 | 126276432 | 221 | * | NA_peak_57 | 21 | Intron (ENSM | 8 | 126298558 | 126395482 | 96925 | 1 | 210027 | ENSMUST00 | -22126 | ENSMUSG0( | Slc35f3 | solute carrier family 35, member F3 |
| chr8 | 126895078 | 126895389 | 312 | * | NA_peak_57 | 28 | Intron (ENSM | 8 | 126930667 | 126945840 | 15174 | 2 | 67952 | ENSMUST00 | 50451 | ENSMUSG0( | Tomm20 | translocase of outer mitochondrial membrane 20 |
| chr8 | 127619392 | 127619685 | 294 | * | NA_peak_57 | 20 | Distal Interge | 8 | 127579724 | 127610848 | 31125 | 1 | 93742 | ENSMUST00 | 39668 | ENSMUSG0( | Pard3 | par-3 family cell polarity regulator |
| chr8 | 128746663 | 128747049 | 387 | * | NA_peak_57 | 22 | Intron (ENSM | 8 | 128724333 | 128732389 | 8057 | 1 | 16412 | ENSMUST00 | 22330 | ENSMUSG0( | Itgb1 | integrin beta 1 (fibronectin receptor beta) |
| chr8 | 128937198 | 128937507 | 310 | * | NA_peak_57 | 65 | Intron (ENSM | 8 | 128980101 | 129065488 | 85388 | 2 | 74703 | ENSMUST00 | 127981 | ENSMUSG0( | Ccdc7a | coiled-coil domain containing 7A |
| chr8 | 129122719 | 129122923 | 205 | * | NA_peak_57 | 23 | Intron (ENSM | 8 | 129118041 | 129183732 | 65692 | 1 | 75453 | ENSMUST00 | 4678 | ENSMUSG0( | Ccdc7b | coiled-coil domain containing 7B |
| chr8 | 129213110 | 129213612 | 503 | * | NA_peak_57 | 32 | Distal Interge | 8 | 129217724 | 129221950 | 4227 | 2 | 72139 | ENSMUST00 | 8338 | ENSMUSG0( | 2610044O15 | RIKEN cDNA 2610044O15 gene |
| chr9 | 3000067 | 3004866 | 4800 | * | NA_peak_57 | 558 | Exon (ENSMU | 9 | 3038669 | 3038743 | 75 | 2 | 100628572 | ENSMUST00 | 33877 | ENSMUSG0( | Mir101c | microRNA 101c |
| chr9 | 3006244 | 3008727 | 2484 | * | NA_peak_57 | 611 | Exon (ENSMU | 9 | 3038669 | 3038743 | 75 | 2 | 100628572 | ENSMUST00 | 30016 | ENSMUSG0( | Mir101c | microRNA 101c |
| chr9 | 3009297 | 3010373 | 1077 | * | NA_peak_57 | 283 | Distal Interge | 9 | 3038669 | 3038743 | 75 | 2 | 100628572 | ENSMUST00 | 28370 | ENSMUSG0( | Mir101c | microRNA 101c |
| chr9 | 3010795 | 3011141 | 347 | * | NA_peak_57 | 521 | Distal Interge | 9 | 3038669 | 3038743 | 75 | 2 | 100628572 | ENSMUST00 | 27602 | ENSMUSG0( | Mir101c | microRNA 101c |
| chr9 | 3012108 | 3012410 | 303 | * | NA_peak_57 | 373 | Distal Interge | 9 | 3038669 | 3038743 | 75 | 2 | 100628572 | ENSMUST00 | 26333 | ENSMUSG0( | Mir101c | microRNA 101c |
| chr9 | 3012739 | 3017251 | 4513 | * | NA_peak_57 | 494 | Exon (ENSMU | 9 | 3038669 | 3038743 | 75 | 2 | 100628572 | ENSMUST00 | 21492 | ENSMUSG0( | Mir101c | microRNA 101c |
| chr9 | 3017775 | 3019354 | 1580 | * | NA_peak_57 | 525 | Exon (ENSMU | 9 | 3038669 | 3038743 | 75 | 2 | 100628572 | ENSMUST00 | 19389 | ENSMUSG0( | Mir101c | microRNA 101c |
| chr9 | 3019829 | 3022621 | 2793 | * | NA_peak_57 | 429 | Exon (ENSMU | 9 | 3038669 | 3038743 | 75 | 2 | 100628572 | ENSMUST00 | 16122 | ENSMUSG0( | Mir101c | microRNA 101c |
| chr9 | 3023387 | 3024673 | 1287 | * | NA_peak_57 | 855 | Exon (ENSMU | 9 | 3038669 | 3038743 | 75 | 2 | 100628572 | ENSMUST00 | 14070 | ENSMUSG0( | Mir101c | microRNA 101c |
| chr9 | 3024995 | 3026024 | 1030 | * | NA_peak_57 | 361 | Exon (ENSMU | 9 | 3038669 | 3038743 | 75 | 2 | 100628572 | ENSMUST00 | 12719 | ENSMUSG0( | Mir101c | microRNA 101c |
| chr9 | 3026300 | 3026767 | 468 | * | NA_peak_57 | 369 | Exon (ENSMU | 9 | 3038669 | 3038743 | 75 | 2 | 100628572 | ENSMUST00 | 11976 | ENSMUSG0( | Mir101c | microRNA 101c |
| chr9 | 3027150 | 3028903 | 1754 | * | NA_peak_57 | 338 | Exon (ENSMU | 9 | 3038669 | 3038743 | 75 | 2 | 100628572 | ENSMUST00 | 9840 | ENSMUSG0( | Mir101c | microRNA 101c |
| chr9 | 3029339 | 3029875 | 537 | * | NA_peak_57 | 650 | Exon (ENSMU | 9 | 3038669 | 3038743 | 75 | 2 | 100628572 | ENSMUST00 | 8868 | ENSMUSG0( | Mir101c | microRNA 101c |
| chr9 | 3030501 | 3035322 | 4822 | * | NA_peak_57 | 269 | Exon (ENSMU | 9 | 3038669 | 3038743 | 75 | 2 | 100628572 | ENSMUST00 | 3421 | ENSMUSG0( | Mir101c | microRNA 101c |
| chr9 | 3035669 | 3036368 | 700 | * | NA_peak_57 | 349 | Promoter (2-3 | 9 | 3038669 | 3038743 | 75 | 2 | 100628572 | ENSMUST00 | 2375 | ENSMUSG0( | Mir101c | microRNA 101c |
| chr9 | 3036832 | 3037494 | 663 | * | NA_peak_57 | 408 | Promoter (1-2 | 9 | 3038669 | 3038743 | 75 | 2 | 100628572 | ENSMUST00 | 1249 | ENSMUSG0( | Mir101c | microRNA 101c |
| chr9 | 3037782 | 3038240 | 459 | * | NA_peak_57 | 428 | Promoter (<= | 9 | 3038669 | 3038743 | 75 | 2 | 100628572 | ENSMUST00 | 503 | ENSMUSG0( | Mir101c | microRNA 101c |
| chr9 | 3080391 | 3080857 | 467 | * | NA_peak_57 | 31 | Distal Interge | 9 | 3038669 | 3038743 | 75 | 2 | 100628572 | ENSMUST00 | -41648 | ENSMUSG0( | Mir101c | microRNA 101c |
| chr9 | 3311876 | 3312292 | 417 | * | NA_peak_57 | 74 | Distal Interge | 9 | 3335140 | 3386175 | 51036 | 1 | 67667 | ENSMUST00 | -22848 | ENSMUSG0( | Alkbh8 | alkB homolog 8, tRNA methyltransferase |
| chr9 | 3659521 | 3659878 | 358 | * | NA_peak_57 | 74 | Intron (ENSM | 9 | 3532888 | 3761177 | 228290 | 1 | 234889 | ENSMUST00 | 126633 | ENSMUSG0( | Gucy1a2 | guanylate cyclase 1, soluble, alpha 2 |
| chr9 | 3948222 | 3948520 | 299 | * | NA_peak_57 | 27 | Distal Interge | 9 | 4294793 | 4309467 | 14675 | 2 | 67618 | ENSMUST00 | 360947 | ENSMUSG0( | Aasdhppt | aminoadipate-semialdehyde dehydrogenase-phosphopantetheinyl transferase |
| chr9 | 4244456 | 4244763 | 308 | * | NA_peak_57 | 19 | Distal Interge | 9 | 4294793 | 4309467 | 14675 | 2 | 67618 | ENSMUST00 | 64704 | ENSMUSG0( | Aasdhppt | aminoadipate-semialdehyde dehydrogenase-phosphopantetheinyl transferase |
| chr9 | 4771589 | 4771595 | 7 | * | NA_peak_57 | 16 | Intron (ENSM | 9 | 4417899 | 4795519 | 377621 | 2 | 14802 | ENSMUST00 | 23924 | ENSMUSG0( | Gria4 | glutamate receptor, ionotropic, AMPA4 (alpha 4) |
| chr9 | 7373821 | 7374175 | 355 | * | NA_peak_57 | 13 | Intron (ENSM | 9 | 7368239 | 7388047 | 19809 | 2 | 83996 | ENSMUST00 | 13872 | ENSMUSG0( | Mmp1b | matrix metallopeptidase 1b (interstitial collagenase) |
| chr9 | 7799078 | 7799439 | 362 | * | NA_peak_57 | 112 | Distal Interge | 9 | 7784390 | 7784440 | 51 | 1 | 100885830 | ENSMUST00 | 14688 | ENSMUSG0( | Mir5618 | microRNA 5618 |
| chr9 | 7837030 | 7837267 | 238 | * | NA_peak_57 | 31 | Promoter (<= | 9 | 7818227 | 7837057 | 18831 | 2 | 11797 | ENSMUST00 | 0 | ENSMUSG0( | Birc2 | baculoviral IAP repeat-containing 2 |
| chr9 | 8277177 | 8277501 | 325 | * | NA_peak_57 | 20 | Distal Interge | 9 | 8246786 | 8246881 | 96 | 1 | 100316772 | ENSMUST00 | 30391 | ENSMUSG0( | Mir1899 | microRNA 1899 |
| chr9 | 8860273 | 8860564 | 292 | * | NA_peak_57 | 24 | Distal Interge | 9 | 8899833 | 8968611 | 68779 | 1 | 18667 | ENSMUST00 | -39269 | ENSMUSG0( | Pgr | progesterone receptor |
| chr9 | 9294651 | 9294949 | 299 | * | NA_peak_57 | 35 | Distal Interge | 9 | 9011238 | 9239101 | 227864 | 2 | 71544 | ENSMUST00 | -55550 | ENSMUSG0( | Arhgap42 | Rho GTPase activating protein 42 |
| chr9 | 10969591 | 10969744 | 154 | * | NA_peak_57 | 31 | Distal Interge | 9 | 9660969 | 10904775 | 1243807 | 2 | 244682 | ENSMUST00 | -64816 | ENSMUSG0( | Cntn5 | contactin 5 |
| chr9 | 11088158 | 11088589 | 432 | * | NA_peak_57 | 23 | Distal Interge | 9 | 9660969 | 10904775 | 1243807 | 2 | 244682 | ENSMUST00 | -183383 | ENSMUSG0( | Cntn5 | contactin 5 |
| chr9 | 11554247 | 11554530 | 284 | * | NA_peak_57 | 40 | Intron (ENSM | 9 | 9660969 | 10904775 | 1243807 | 2 | 244682 | ENSMUST00 | -649472 | ENSMUSG0( | Cntn5 | contactin 5 |
| chr9 | 11555027 | 11555318 | 292 | * | NA_peak_57 | 40 | Intron (ENSM | 9 | 9660969 | 10904775 | 1243807 | 2 | 244682 | ENSMUST00 | -650252 | ENSMUSG0( | Cntn5 | contactin 5 |
| chr9 | 11725452 | 11725780 | 329 | * | NA_peak_57 | 61 | Distal Interge | 9 | 9660969 | 10904775 | 1243807 | 2 | 244682 | ENSMUST00 | -820677 | ENSMUSG0( | Cntn5 | contactin 5 |
| chr9 | 11825136 | 11825466 | 331 | * | NA_peak_57 | 45 | Distal Interge | 9 | 9660969 | 10904775 | 1243807 | 2 | 244682 | ENSMUST00 | -920361 | ENSMUSG0( | Cntn5 | contactin 5 |
| chr9 | 12284865 | 12285228 | 364 | * | NA_peak_57 | 23 | Distal Interge | 9 | 13091267 | 13105855 | 14589 | 2 | 78219 | ENSMUST00 | 820627 | ENSMUSG0( | 4930568E12 | RIKEN cDNA 4930568E12 gene |
| chr9 | 17059281 | 17059543 | 263 | * | NA_peak_57 | 31 | Distal Interge | 9 | 15910189 | 16501285 | 591097 | 2 | 270120 | ENSMUST00 | -557996 | ENSMUSG0( | Fat3 | FAT atypical cadherin 3 |
| chr9 | 17061009 | 17061100 | 92 | * | NA_peak_57 | 22 | Distal Interge | 9 | 15910189 | 16501285 | 591097 | 2 | 270120 | ENSMUST00 | -559724 | ENSMUSG0( | Fat3 | FAT atypical cadherin 3 |
| chr9 | 18992815 | 18993064 | 250 | * | NA_peak_57 | 30 | Distal Interge | 9 | 18987990 | 18988928 | 939 | 1 | 258074 | ENSMUST00 | 4825 | ENSMUSG0( | Olfr834 | olfactory receptor 834 |
| chr9 | 19957696 | 19957977 | 282 | * | NA_peak_57 | 31 | Distal Interge | 9 | 19916865 | 19929124 | 12260 | 1 | 258336 | ENSMUST00 | 40831 | ENSMUSG0( | Olfr77 | olfactory receptor 77 |
| chr9 | 19958267 | 19958681 | 415 | * | NA_peak_57 | 48 | Distal Interge | 9 | 19916865 | 19929124 | 12260 | 1 | 258336 | ENSMUST00 | 41402 | ENSMUSG0( | Olfr77 | olfactory receptor 77 |
| chr9 | 20066984 | 20067020 | 37 | * | NA_peak_57 | 23 | Distal Interge | 9 | 20054466 | 20055461 | 996 | 2 | 257898 | ENSMUST00 | -11523 | ENSMUSG0( | Olfr867 | olfactory receptor 867 |
| chr9 | 20371041 | 20371404 | 364 | * | NA_peak_58 | 46 | Intron (ENSM | 9 | 20345136 | 20385177 | 40042 | 2 | 434377 | ENSMUST00 | 13773 | ENSMUSG0( | Zfp560 | zinc finger protein 560 |
| chr9 | 21880413 | 21880728 | 316 | * | NA_peak_58 | 16 | Intron (ENSM | 9 | 21800180 | 21852635 | 52456 | 2 | 319899 | ENSMUST00 | -27778 | ENSMUSG0( | Dock6 | dedicator of cytokinesis 6 |
| chr9 | 22036755 | 22036935 | 181 | * | NA_peak_58 | 36 | 3' UTR | 9 | 22016461 | 22026374 | 9914 | 2 | 15571 | ENSMUST00 | -10381 | ENSMUSG0( | Elavl3 | ELAV like RNA binding protein 3 |
| chr9 | 23184957 | 23185232 | 276 | * | NA_peak_58 | 31 | Distal Interge | 9 | 23223076 | 23485200 | 262125 | 1 | 73230 | ENSMUST00 | -37844 | ENSMUSG0( | Bmper | BMP-binding endothelial regulator |
| chr9 | 26102072 | 26102248 | 177 | * | NA_peak_58 | 26 | Distal Interge | 9 | 25482867 | 25487313 | 4447 | 1 | 67484 | ENSMUST00 | 619205 | ENSMUSG0( | Eepd1 | endonuclease/exonuclease/phosphatase family domain containing 1 |
| chr9 | 26403280 | 26403652 | 373 | * | NA_peak_58 | 40 | Distal Interge | 9 | 26733728 | 26761350 | 27623 | 1 | 76898 | ENSMUST00 | -330076 | ENSMUSG0( | B3gat1 | beta-1,3-glucuronyltransferase 1 (glucuronosyltransferase P) |
| chr9 | 27980029 | 27980097 | 69 | * | NA_peak_58 | 28 | Intron (ENSM | 9 | 27790947 | 28925410 | 1134464 | 1 | 330908 | ENSMUST00 | 189082 | ENSMUSG0( | Opcml | opioid binding protein/cell adhesion molecule-like |
| chr9 | 31186726 | 31187062 | 337 | * | NA_peak_58 | 13 | Intron (ENSM | 9 | 31168419 | 31170574 | 2156 | 2 | 11804 | ENSMUST00 | -16152 | ENSMUSG0( | Aplp2 | amyloid beta (A4) precursor-like protein 2 |
| chr9 | 31533665 | 31533882 | 218 | * | NA_peak_58 | 23 | Distal Interge | 9 | 31426196 | 31464462 | 38267 | 2 | 235135 | ENSMUST00 | -69203 | ENSMUSG0( | Tmem45b | transmembrane protein 45b |
| chr9 | 32081294 | 32081663 | 370 | * | NA_peak_58 | 28 | Distal Interge | 9 | 32116136 | 32268446 | 152311 | 1 | 330914 | ENSMUST00 | -34473 | ENSMUSG0( | Arhgap32 | Rho GTPase activating protein 32 |
| chr9 | 35305269 | 35305563 | 295 | * | NA_peak_58 | 549 | Intron (ENSM | 9 | 35281105 | 35285423 | 4319 | 1 | 71133 | ENSMUST00 | 24164 | ENSMUSG0( | 4933422A05 | RIKEN cDNA 4933422A05 gene |
| chr9 | 36595710 | 36596145 | 436 | * | NA_peak_58 | 23 | Distal Interge | 9 | 36581278 | 36582635 | 1358 | 2 | 100312948 | ENSMUST00 | -13075 | ENSMUSG0( | Pate8 | prostate and testis expressed 8 |
| chr9 | 37885582 | 37885701 | 120 | * | NA_peak_58 | 27 | Distal Interge | 9 | 37896724 | 37903519 | 6796 | 1 | 258310 | ENSMUST00 | -11023 | ENSMUSG0( | Olfr145 | olfactory receptor 145 |
| chr9 | 38242356 | 38242387 | 32 | * | NA_peak_58 | 31 | Distal Interge | 9 | 38253419 | 38254360 | 942 | 1 | 258802 | ENSMUST00 | -11032 | ENSMUSG0( | Olfr143 | olfactory receptor 143 |
| chr9 | 43210168 | 43210438 | 271 | * | NA_peak_58 | 36 | Promoter (<= | 9 | 43123939 | 43210369 | 86431 | 2 | 18988 | ENSMUST00 | 0 | ENSMUSG0( | Pou2f3 | POU domain, class 2, transcription factor 3 |
| chr9 | 46410047 | 46410339 | 293 | * | NA_peak_58 | 31 | Distal Interge | 9 | 46303361 | 46319986 | 16626 | 2 | 74361 | ENSMUST00 | -90061 | ENSMUSG0( | 4931429L15 | RIKEN cDNA 4931429L15 gene |
| chr9 | 47267059 | 47267619 | 561 | * | NA_peak_58 | 24 | Intron (ENSM | 9 | 47530173 | 47853385 | 323213 | 1 | 54725 | ENSMUST00 | -262554 | ENSMUSG0( | Cadm1 | cell adhesion molecule 1 |
| chr9 | 51425400 | 51425509 | 110 | * | NA_peak_58 | 48 | Distal Interge | 9 | 51289686 | 51328924 | 39239 | 2 | 69809 | ENSMUST00 | -96476 | ENSMUSG0( | 1810046K07 | RIKEN cDNA 1810046K07 gene |
| chr9 | 52614379 | 52614506 | 128 | * | NA_peak_58 | 20 | Distal Interge | 9 | 52673044 | 52679429 | 6386 | 2 | 330941 | ENSMUST00 | 64923 | ENSMUSG0( | AI593442 | expressed sequence AI593442 |
| chr9 | 55879980 | 55880194 | 215 | * | NA_peak_58 | 20 | Intron (ENSM | 9 | 55549883 | 55919605 | 369723 | 2 | 244891 | ENSMUST00 | 39411 | ENSMUSG0( | Scaper | S phase cyclin A-associated protein in the ER |
| chr9 | 56223573 | 56223958 | 386 | * | NA_peak_58 | 45 | Exon (ENSMU | 9 | 56201126 | 56244065 | 42940 | 2 | 244895 | ENSMUST00 | 20107 | ENSMUSG0( | Peak1 | pseudopodium-enriched atypical kinase 1 |
| chr9 | 57083363 | 57083631 | 269 | * | NA_peak_58 | 16 | Promoter (2-3 | 9 | 57081158 | 57128366 | 47209 | 1 | 20466 | ENSMUST00 | 2205 | ENSMUSG0( | Sin3a | transcriptional regulator, SIN3A (yeast) |
| chr9 | 57408528 | 57408814 | 287 | * | NA_peak_58 | 31 | 3' UTR | 9 | 57415166 | 57421318 | 6153 | 2 | 66812 | ENSMUST00 | 12504 | ENSMUSG0( | Ppcdc | phosphopantothenoylcysteine decarboxylase |
| chr9 | 57605654 | 57606128 | 475 | * | NA_peak_58 | 14 | Promoter (<= | 9 | 57602124 | 57606236 | 4113 | 2 | 235415 | ENSMUST00 | 108 | ENSMUSG0( | Cplx3 | complexin 3 |
| chr9 | 62600222 | 62600484 | 263 | * | NA_peak_58 | 31 | Distal Interge | 9 | 62419492 | 62537044 | 117553 | 2 | 235431 | ENSMUST00 | -63178 | ENSMUSG0( | Coro2b | coronin, actin binding protein, 2B |
| chr9 | 64413752 | 64413841 | 90 | * | NA_peak_58 | 31 | Intron (ENSM | 9 | 64385686 | 64709205 | 323520 | 1 | 214058 | ENSMUST00 | 28066 | ENSMUSG0( | Megf11 | multiple EGF-like-domains 11 |
| chr9 | 65146050 | 65146306 | 257 | * | NA_peak_58 | 16 | Intron (ENSM | 9 | 65141212 | 65185872 | 44661 | 1 | 19289 | ENSMUST00 | 4838 | ENSMUSG0( | Igdcc3 | immunoglobulin superfamily, DCC subclass, member 3 |
| chr9 | 71781816 | 71782204 | 389 | * | NA_peak_58 | 53 | Distal Interge | 9 | 71626509 | 71771602 | 145094 | 2 | 68178 | ENSMUST00 | -10214 | ENSMUSG0( | Cgnl1 | cingulin-like 1 |
| chr9 | 73935892 | 73936172 | 281 | * | NA_peak_58 | 40 | Promoter (2-3 | 9 | 73480870 | 73933567 | 452698 | 2 | 208898 | ENSMUST00 | -2325 | ENSMUSG0( | Unc13c | unc-13 homolog C |
| chr9 | 73938886 | 73939689 | 804 | * | NA_peak_58 | 40 | Intron (ENSM | 9 | 73480870 | 73933567 | 452698 | 2 | 208898 | ENSMUST00 | -5319 | ENSMUSG0( | Unc13c | unc-13 homolog C |
| chr9 | 74629337 | 74629691 | 355 | * | NA_peak_58 | 13 | Distal Interge | 9 | 74847983 | 74852873 | 4891 | 1 | 100503019 | ENSMUST00 | -218292 | ENSMUSG0( | Gm16551 | predicted gene 16551 |

| | | | | | | | | | | | | | | | | | | |
|---|---|---|---|---|---|---|---|---|---|---|---|---|---|---|---|---|---|---|
| chr9 | 75537924 | 75538144 | 221 | * | NA_peak_587 | 26 | Intron (ENSM | 9 | 75516558 | 75541707 | 25150 | 2 | 50875 | ENSMUST00 | 3563 | ENSMUSG00 | Tmod3 | tropomodulin 3 |
| chr9 | 75922870 | 75923024 | 155 | * | NA_peak_587 | 21 | Intron (ENSM | 9 | 76014855 | 76136350 | 121496 | 1 | 208982 | ENSMUST00 | -91831 | ENSMUSG00 | Hmgcll1 | 3-hydroxymethyl-3-methylglutaryl-Coenzyme A lyase-like 1 |
| chr9 | 76813153 | 76813445 | 293 | * | NA_peak_587 | 27 | Intron (ENSM | 9 | 76951693 | 77045749 | 94057 | 2 | 26944 | ENSMUST00 | 232304 | ENSMUSG00 | Tinag | tubulointerstitial nephritis antigen |
| chr9 | 78395757 | 78395955 | 199 | * | NA_peak_587 | 20 | Promoter (<= | 9 | 78395777 | 78423587 | 27811 | 1 | 100048658 | ENSMUST00 | 0 | ENSMUSG00 | Ddx43 | DEAD box helicase 43 |
| chr9 | 79908473 | 79908804 | 332 | * | NA_peak_587 | 13 | Intron (ENSM | 9 | 79820661 | 79962033 | 141373 | 2 | 70598 | ENSMUST00 | 53229 | ENSMUSG00 | Filip1 | filamin A interacting protein 1 |
| chr9 | 81617801 | 81618045 | 245 | * | NA_peak_588 | 50 | Distal Interge | 9 | 81631551 | 81644629 | 13079 | 1 | 319405 | ENSMUST00 | -13506 | ENSMUSG00 | D430036J16I | RIKEN cDNA D430036J16 gene |
| chr9 | 81619293 | 81619574 | 282 | * | NA_peak_588 | 61 | Distal Interge | 9 | 81631551 | 81644629 | 13079 | 1 | 319405 | ENSMUST00 | -11977 | ENSMUSG00 | D430036J16I | RIKEN cDNA D430036J16 gene |
| chr9 | 82568160 | 82568471 | 312 | * | NA_peak_588 | 40 | Distal Interge | 9 | 82829540 | 82847687 | 18148 | 1 | 65099 | ENSMUST00 | -261069 | ENSMUSG00 | Irak1bp1 | interleukin-1 receptor-associated kinase 1 binding protein 1 |
| chr9 | 84887666 | 84888094 | 429 | * | NA_peak_588 | 31 | Distal Interge | 9 | 84774512 | 84826183 | 51672 | 1 | 75253 | ENSMUST00 | 113154 | ENSMUSG00 | 4930554C24I | RIKEN cDNA 4930554C24 gene |
| chr9 | 86458929 | 86459407 | 479 | * | NA_peak_588 | 47 | Intron (ENSM | 9 | 86307306 | 86464950 | 157645 | 2 | 70348 | ENSMUST00 | 5543 | ENSMUSG00 | Ube2cbp | ubiquitin-conjugating enzyme E2C binding protein |
| chr9 | 92740529 | 92740775 | 247 | * | NA_peak_589 | 50 | Intron (ENSM | 9 | 92718716 | 92795711 | 76996 | 1 | 331003 | ENSMUST00 | 21813 | ENSMUSG00 | 4933400C23I | RIKEN cDNA 4933400C23 gene |
| chr9 | 93648576 | 93648840 | 265 | * | NA_peak_589 | 32 | Distal Interge | 9 | 92979995 | 93005739 | 25745 | 2 | 78349 | ENSMUST00 | -642837 | ENSMUSG00 | 1700034K08I | RIKEN cDNA 1700034K08 gene |
| chr9 | 93911216 | 93911574 | 359 | * | NA_peak_589 | 19 | Distal Interge | 9 | 94517864 | 94538081 | 20218 | 2 | 68861 | ENSMUST00 | 626507 | ENSMUSG00 | Dipk2a | divergent protein kinase domain 2A |
| chr9 | 94641347 | 94641752 | 406 | * | NA_peak_589 | 48 | Distal Interge | 9 | 94669909 | 95230445 | 560537 | 1 | 331004 | ENSMUST00 | -28157 | ENSMUSG00 | Slc9a9 | solute carrier family 9 (sodium/hydrogen exchanger), member 9 |
| chr9 | 96697791 | 96698117 | 327 | * | NA_peak_589 | 26 | Intron (ENSM | 9 | 96688477 | 96719565 | 31089 | 2 | 245007 | ENSMUST00 | 21448 | ENSMUSG00 | Zbtb38 | zinc finger and BTB domain containing 38 |
| chr9 | 97111032 | 97111335 | 304 | * | NA_peak_589 | 23 | Promoter (<= | 9 | 97074961 | 97111157 | 36197 | 2 | 192287 | ENSMUST00 | 0 | ENSMUSG00 | Slc25a36 | solute carrier family 25, member 36 |
| chr9 | 98432497 | 98432883 | 387 | * | NA_peak_589 | 16 | Intron (ENSM | 9 | 98422961 | 98446575 | 23615 | 1 | 19659 | ENSMUST00 | 9536 | ENSMUSG00 | Rbp1 | retinol binding protein 1, cellular |
| chr9 | 101129032 | 101129383 | 352 | * | NA_peak_590 | 36 | Intron (ENSM | 9 | 101107883 | 101125125 | 17243 | 2 | 235542 | ENSMUST00 | -3907 | ENSMUSG00 | Ppp2r3a | protein phosphatase 2, regulatory subunit B'', alpha |
| chr9 | 102612875 | 102613188 | 314 | * | NA_peak_590 | 23 | Intron (ENSM | 9 | 102601415 | 102618848 | 17434 | 2 | 28135 | ENSMUST00 | 5660 | ENSMUSG00 | Cep63 | centrosomal protein 63 |
| chr9 | 102625776 | 102626190 | 415 | * | NA_peak_590 | 21 | Promoter (<= | 9 | 102586578 | 102626124 | 39547 | 2 | 28135 | ENSMUST00 | 0 | ENSMUSG00 | Cep63 | centrosomal protein 63 |
| chr9 | 105087793 | 105088103 | 311 | * | NA_peak_590 | 17 | Exon (ENSMU | 9 | 105070859 | 105074117 | 3259 | 1 | 94062 | ENSMUST00 | 16934 | ENSMUSG00 | Mrpl3 | mitochondrial ribosomal protein L3 |
| chr9 | 107563051 | 107563277 | 227 | * | NA_peak_590 | 23 | Promoter (<= | 9 | 107563255 | 107566112 | 2858 | 1 | 80385 | ENSMUST00 | 0 | ENSMUSG00 | Tusc2 | tumor suppressor 2, mitochondrial calcium regulator |
| chr9 | 108236698 | 108237045 | 348 | * | NA_peak_590 | 23 | Intron (ENSM | 9 | 108218216 | 108263709 | 45494 | 2 | 13138 | ENSMUST00 | 26664 | ENSMUSG00 | Dag1 | dystroglycan 1 |
| chr9 | 109273061 | 109273458 | 398 | * | NA_peak_591 | 33 | Promoter (2-3 | 9 | 109270790 | 109276126 | 5337 | 2 | 50757 | ENSMUST00 | 2668 | ENSMUSG00 | Fbxw14 | F-box and WD-40 domain protein 14 |
| chr9 | 109328404 | 109328490 | 87 | * | NA_peak_591 | 45 | Intron (ENSM | 9 | 109330842 | 109339642 | 8801 | 2 | 668758 | ENSMUST00 | 11152 | ENSMUSG00 | Fbxw28 | F-box and WD-40 domain protein 28 |
| chr9 | 114146176 | 114146490 | 315 | * | NA_peak_591 | 50 | Intron (ENSM | 9 | 114057140 | 114098728 | 41589 | 1 | 382111 | ENSMUST00 | 89036 | ENSMUSG00 | Susd5 | sushi domain containing 5 |
| chr9 | 114408816 | 114409137 | 322 | * | NA_peak_591 | 15 | Intron (ENSM | 9 | 114401108 | 114474898 | 73791 | 1 | 12091 | ENSMUST00 | 7708 | ENSMUSG00 | Glb1 | galactosidase, beta 1 |
| chr9 | 114512974 | 114513284 | 311 | * | NA_peak_591 | 19 | Exon (ENSMU | 9 | 114492900 | 114504916 | 12017 | 2 | 12773 | ENSMUST00 | -8058 | ENSMUSG00 | Ccr4 | chemokine (C-C motif) receptor 4 |
| chr9 | 115081956 | 115082097 | 142 | * | NA_peak_591 | 23 | Intron (ENSM | 9 | 115088841 | 115218418 | 129578 | 1 | 74486 | ENSMUST00 | -6744 | ENSMUSG00 | Osbpl10 | oxysterol binding protein-like 10 |
| chr9 | 117292943 | 117293037 | 95 | * | NA_peak_592 | 20 | Intron (ENSM | 9 | 116578515 | 117252483 | 673969 | 2 | 207181 | ENSMUST00 | -40460 | ENSMUSG00 | Rbms3 | RNA binding motif, single stranded interacting protein |
| chr9 | 119484106 | 119484181 | 76 | * | NA_peak_592 | 16 | 3' UTR | 9 | 119444987 | 119464239 | 19253 | 1 | 208194 | ENSMUST00 | 39119 | ENSMUSG00 | Exog | endo/exonuclease (5'-3'), endonuclease G-like |
| chr9 | 119867485 | 119867723 | 239 | * | NA_peak_592 | 23 | Distal Interge | 9 | 119894878 | 119926586 | 31709 | 1 | 67561 | ENSMUST00 | -27155 | ENSMUSG00 | Wdr48 | WD repeat domain 48 |
| chr9 | 120639716 | 120640175 | 460 | * | NA_peak_592 | 13 | Distal Interge | 9 | 120577346 | 120578072 | 727 | 1 | 76100 | ENSMUST00 | 62370 | ENSMUSG00 | 5830454E08I | RIKEN cDNA 5830454E08 gene |
| chr9 | 121412166 | 121412398 | 233 | * | NA_peak_592 | 13 | Intron (ENSM | 9 | 121415975 | 121462610 | 46636 | 1 | 67095 | ENSMUST00 | -3577 | ENSMUSG00 | Trak1 | trafficking protein, kinesin binding 1 |
| chr9 | 121857577 | 121857581 | 5 | * | NA_peak_592 | 26 | Promoter (<= | 9 | 121850166 | 121857567 | 7402 | 2 | 56295 | ENSMUST00 | -10 | ENSMUSG00 | Higd1a | HIG1 domain family, member 1A |
| chr9 | 122299757 | 122300123 | 367 | * | NA_peak_592 | 50 | Intron (ENSM | 9 | 122175874 | 122294423 | 118550 | 2 | 102566 | ENSMUST00 | -5334 | ENSMUSG00 | Ano10 | anoctamin 10 |
| chrX | 3084156 | 3084521 | 366 | * | NA_peak_593 | 16 | Distal Interge | 20 | 3076875 | 3078817 | 1943 | 1 | 100861966 | ENSMUST00 | 7281 | ENSMUSG00 | Btbd35f23 | BTB domain containing 35, family member 23 |
| chrX | 3085243 | 3085621 | 379 | * | NA_peak_593 | 28 | Distal Interge | 20 | 3076875 | 3078817 | 1943 | 1 | 100861966 | ENSMUST00 | 8368 | ENSMUSG00 | Btbd35f23 | BTB domain containing 35, family member 23 |
| chrX | 3218675 | 3220290 | 1616 | * | NA_peak_593 | 48 | Distal Interge | 20 | 3076875 | 3078817 | 1943 | 1 | 100861966 | ENSMUST00 | 141800 | ENSMUSG00 | Btbd35f23 | BTB domain containing 35, family member 23 |
| chrX | 3228688 | 3228911 | 224 | * | NA_peak_593 | 43 | Intron (ENSM | 20 | 3076875 | 3078817 | 1943 | 1 | 100861966 | ENSMUST00 | 151813 | ENSMUSG00 | Btbd35f23 | BTB domain containing 35, family member 23 |
| chrX | 3229292 | 3229605 | 314 | * | NA_peak_593 | 43 | Intron (ENSM | 20 | 3076875 | 3078817 | 1943 | 1 | 100861966 | ENSMUST00 | 152417 | ENSMUSG00 | Btbd35f23 | BTB domain containing 35, family member 23 |
| chrX | 3232268 | 3232988 | 721 | * | NA_peak_593 | 61 | Distal Interge | 20 | 3076875 | 3078817 | 1943 | 1 | 100861966 | ENSMUST00 | 155393 | ENSMUSG00 | Btbd35f23 | BTB domain containing 35, family member 23 |
| chrX | 3248350 | 3248676 | 327 | * | NA_peak_593 | 39 | Distal Interge | 20 | 3076875 | 3078817 | 1943 | 1 | 100861966 | ENSMUST00 | 171475 | ENSMUSG00 | Btbd35f23 | BTB domain containing 35, family member 23 |
| chrX | 3251448 | 3251755 | 308 | * | NA_peak_593 | 32 | Distal Interge | 20 | 3076875 | 3078817 | 1943 | 1 | 100861966 | ENSMUST00 | 174573 | ENSMUSG00 | Btbd35f23 | BTB domain containing 35, family member 23 |
| chrX | 3324703 | 3325035 | 333 | * | NA_peak_593 | 31 | Distal Interge | 20 | 3441731 | 3443690 | 1960 | 2 | 630022 | ENSMUST00 | 118655 | ENSMUSG00 | Btbd35f18 | BTB domain containing 35, family member 18 |
| chrX | 3327024 | 3327353 | 330 | * | NA_peak_593 | 46 | Distal Interge | 20 | 3441731 | 3443690 | 1960 | 2 | 630022 | ENSMUST00 | 116337 | ENSMUSG00 | Btbd35f18 | BTB domain containing 35, family member 18 |
| chrX | 3341913 | 3342301 | 389 | * | NA_peak_593 | 31 | Distal Interge | 20 | 3441731 | 3443690 | 1960 | 2 | 630022 | ENSMUST00 | 101389 | ENSMUSG00 | Btbd35f18 | BTB domain containing 35, family member 18 |
| chrX | 3356777 | 3357229 | 453 | * | NA_peak_593 | 24 | Distal Interge | 20 | 3441731 | 3443690 | 1960 | 2 | 630022 | ENSMUST00 | 86461 | ENSMUSG00 | Btbd35f18 | BTB domain containing 35, family member 18 |
| chrX | 3425318 | 3425712 | 395 | * | NA_peak_593 | 15 | Distal Interge | 20 | 3441731 | 3443690 | 1960 | 2 | 630022 | ENSMUST00 | 17978 | ENSMUSG00 | Btbd35f18 | BTB domain containing 35, family member 18 |
| chrX | 3708268 | 3708594 | 327 | * | NA_peak_593 | 34 | Distal Interge | 20 | 3700233 | 3702192 | 1960 | 1 | 630022 | ENSMUST00 | 8035 | ENSMUSG00 | Btbd35f18 | BTB domain containing 35, family member 18 |
| chrX | 3775038 | 3775539 | 502 | * | NA_peak_593 | 55 | Distal Interge | 20 | 3750934 | 3752885 | 1952 | 2 | 668960 | ENSMUST00 | -22153 | ENSMUSG00 | Btbd35f10 | BTB domain containing 35, family member 10 |
| chrX | 3784403 | 3784567 | 165 | * | NA_peak_593 | 20 | Intron (ENSM | 20 | 3750934 | 3752885 | 1952 | 2 | 668960 | ENSMUST00 | -31518 | ENSMUSG00 | Btbd35f10 | BTB domain containing 35, family member 10 |
| chrX | 3789479 | 3789671 | 193 | * | NA_peak_593 | 19 | Distal Interge | 20 | 3750934 | 3752885 | 1952 | 2 | 668960 | ENSMUST00 | -36594 | ENSMUSG00 | Btbd35f10 | BTB domain containing 35, family member 10 |
| chrX | 4075806 | 4076086 | 281 | * | NA_peak_593 | 40 | Distal Interge | 20 | 4037674 | 4039616 | 1943 | 1 | 100042254 | ENSMUST00 | 38132 | ENSMUSG00 | Btbd35f4 | BTB domain containing 35, family member 4 |
| chrX | 4079448 | 4079924 | 477 | * | NA_peak_593 | 49 | Distal Interge | 20 | 4037674 | 4039616 | 1943 | 1 | 100042254 | ENSMUST00 | 41774 | ENSMUSG00 | Btbd35f4 | BTB domain containing 35, family member 4 |
| chrX | 4102553 | 4102871 | 319 | * | NA_peak_593 | 23 | Distal Interge | 20 | 4037674 | 4039616 | 1943 | 1 | 100042254 | ENSMUST00 | 64879 | ENSMUSG00 | Btbd35f4 | BTB domain containing 35, family member 4 |
| chrX | 4132070 | 4132394 | 325 | * | NA_peak_594 | 31 | Distal Interge | 20 | 4196576 | 4198535 | 1960 | 1 | 627264 | ENSMUST00 | -64182 | ENSMUSG00 | Btbd35f28 | BTB domain containing 35, family member 28 |
| chrX | 4530482 | 4530869 | 388 | * | NA_peak_594 | 18 | Distal Interge | 20 | 4515674 | 4517170 | 1497 | 1 | 546185 | ENSMUST00 | 14808 | NA | Gm5925 | germ cell-less homolog 1 family pseudogene |
| chrX | 5087981 | 5088492 | 512 | * | NA_peak_594 | 20 | Distal Interge | 20 | 4952135 | 4954077 | 1943 | 1 | 100042275 | ENSMUST00 | 135846 | ENSMUSG00 | Btbd35f27 | BTB domain containing 35, family member 27 |
| chrX | 5098815 | 5099138 | 324 | * | NA_peak_594 | 26 | Distal Interge | 20 | 4952135 | 4954077 | 1943 | 1 | 100042275 | ENSMUST00 | 146680 | ENSMUSG00 | Btbd35f27 | BTB domain containing 35, family member 27 |
| chrX | 5100926 | 5101886 | 961 | * | NA_peak_594 | 57 | Distal Interge | 20 | 4952135 | 4954077 | 1943 | 1 | 100042275 | ENSMUST00 | 148791 | ENSMUSG00 | Btbd35f27 | BTB domain containing 35, family member 27 |
| chrX | 5113554 | 5113851 | 298 | * | NA_peak_594 | 46 | Distal Interge | 20 | 4952135 | 4954077 | 1943 | 1 | 100042275 | ENSMUST00 | 161419 | ENSMUSG00 | Btbd35f27 | BTB domain containing 35, family member 27 |
| chrX | 5176928 | 5177316 | 389 | * | NA_peak_594 | 20 | Distal Interge | 20 | 4952135 | 4954077 | 1943 | 1 | 100042275 | ENSMUST00 | 224793 | ENSMUSG00 | Btbd35f27 | BTB domain containing 35, family member 27 |
| chrX | 5319209 | 5319574 | 366 | * | NA_peak_594 | 15 | Distal Interge | 20 | 5466898 | 5469265 | 2368 | 2 | 17870 | ENSMUST00 | 149691 | ENSMUSG00 | Mycs | myc-like oncogene, s-myc protein |
| chrX | 5320819 | 5321799 | 981 | * | NA_peak_594 | 62 | Distal Interge | 20 | 5466898 | 5469265 | 2368 | 2 | 17870 | ENSMUST00 | 147466 | ENSMUSG00 | Mycs | myc-like oncogene, s-myc protein |
| chrX | 5322098 | 5322383 | 286 | * | NA_peak_594 | 20 | Distal Interge | 20 | 5466898 | 5469265 | 2368 | 2 | 17870 | ENSMUST00 | 146882 | ENSMUSG00 | Mycs | myc-like oncogene, s-myc protein |
| chrX | 5369528 | 5369670 | 143 | * | NA_peak_594 | 21 | Distal Interge | 20 | 5466898 | 5469265 | 2368 | 2 | 17870 | ENSMUST00 | 99595 | ENSMUSG00 | Mycs | myc-like oncogene, s-myc protein |
| chrX | 15967339 | 15967725 | 387 | * | NA_peak_594 | 46 | Distal Interge | 20 | 16522869 | 16523689 | 821 | 1 | 66742 | ENSMUST00 | -555144 | ENSMUSG00 | Cypt1 | cysteine-rich perinuclear theca 1 |
| chrX | 17476720 | 17477454 | 735 | * | NA_peak_594 | 31 | Distal Interge | 20 | 17557898 | 17571624 | 13727 | 2 | 72018 | ENSMUST00 | 94170 | ENSMUSG00 | Fundc1 | FUN14 domain containing 1 |
| chrX | 17500347 | 17500657 | 311 | * | NA_peak_594 | 46 | Distal Interge | 20 | 17557898 | 17571624 | 13727 | 2 | 72018 | ENSMUST00 | 70967 | ENSMUSG00 | Fundc1 | FUN14 domain containing 1 |
| chrX | 18538673 | 18539514 | 842 | * | NA_peak_595 | 36 | Distal Interge | 20 | 18414881 | 18461397 | 46517 | 2 | 75905 | ENSMUST00 | -77276 | ENSMUSG00 | Dipk2b | divergent protein kinase domain 2B |
| chrX | 18722341 | 18722569 | 229 | * | NA_peak_595 | 16 | Distal Interge | 20 | 18414881 | 18461397 | 46517 | 2 | 75905 | ENSMUST00 | -260944 | ENSMUSG00 | Dipk2b | divergent protein kinase domain 2B |
| chrX | 18801655 | 18801783 | 129 | * | NA_peak_595 | 23 | Distal Interge | 20 | 18414881 | 18461397 | 46517 | 2 | 75905 | ENSMUST00 | -340258 | ENSMUSG00 | Dipk2b | divergent protein kinase domain 2B |
| chrX | 19191827 | 19192174 | 348 | * | NA_peak_595 | 16 | Distal Interge | 20 | 19146893 | 19146971 | 79 | 2 | 723828 | ENSMUST00 | -44856 | ENSMUSG00 | Mir222 | microRNA 222 |
| chrX | 19651861 | 19652242 | 382 | * | NA_peak_595 | 36 | Distal Interge | 20 | 19479420 | 19481319 | 1900 | 2 | 278255 | ENSMUST00 | -170542 | ENSMUSG00 | Tex13c3 | TEX13 family member C3 |
| chrX | 20862730 | 20862860 | 131 | * | NA_peak_595 | 16 | Promoter (<= | 20 | 20863126 | 20863182 | 57 | 2 | 100885829 | ENSMUST00 | 322 | ENSMUSG00 | Mir5617 | microRNA 5617 |
| chrX | 21760614 | 21760815 | 202 | * | NA_peak_595 | 39 | Intron (ENSM | 20 | 21743440 | 21745097 | 1658 | 2 | 100043180 | ENSMUST00 | -15517 | ENSMUSG00 | Gm28269 | predicted gene 28269 |
| chrX | 22880250 | 22880525 | 276 | * | NA_peak_595 | 29 | Distal Interge | 20 | 23219286 | 23266802 | 47517 | 2 | 67455 | ENSMUST00 | 386277 | ENSMUSG00 | Klhl13 | kelch-like 13 |
| chrX | 29735372 | 29735781 | 410 | * | NA_peak_595 | 55 | Distal Interge | 20 | 29882560 | 29915946 | 33387 | 2 | 108168459 | ENSMUST00 | 180165 | ENSMUSG00 | Gm2101 | predicted gene 2101 |
| chrX | 31158178 | 31158710 | 533 | * | NA_peak_595 | 39 | Distal Interge | 20 | 31336401 | 31338101 | 1701 | 1 | 668960 | ENSMUST00 | -177691 | ENSMUSG00 | Btbd35f10 | BTB domain containing 35, family member 10 |
| chrX | 31159729 | 31160042 | 314 | * | NA_peak_595 | 27 | Distal Interge | 20 | 31336401 | 31338101 | 1701 | 1 | 668960 | ENSMUST00 | -176359 | ENSMUSG00 | Btbd35f10 | BTB domain containing 35, family member 10 |
| chrX | 31160582 | 31160957 | 376 | * | NA_peak_595 | 46 | Distal Interge | 20 | 31336401 | 31338101 | 1701 | 1 | 668960 | ENSMUST00 | -175444 | ENSMUSG00 | Btbd35f10 | BTB domain containing 35, family member 10 |
| chrX | 31167334 | 31167916 | 583 | * | NA_peak_595 | 49 | Distal Interge | 20 | 31336401 | 31338101 | 1701 | 1 | 668960 | ENSMUST00 | -168485 | ENSMUSG00 | Btbd35f10 | BTB domain containing 35, family member 10 |
| chrX | 31169248 | 31169673 | 426 | * | NA_peak_595 | 45 | Distal Interge | 20 | 31336401 | 31338101 | 1701 | 1 | 668960 | ENSMUST00 | -166728 | ENSMUSG00 | Btbd35f10 | BTB domain containing 35, family member 10 |
| chrX | 31170365 | 31171004 | 640 | * | NA_peak_595 | 42 | Distal Interge | 20 | 31336401 | 31338101 | 1701 | 1 | 668960 | ENSMUST00 | -165397 | ENSMUSG00 | Btbd35f10 | BTB domain containing 35, family member 10 |
| chrX | 31195832 | 31196420 | 589 | * | NA_peak_595 | 42 | Distal Interge | 20 | 31336401 | 31338101 | 1701 | 1 | 668960 | ENSMUST00 | -139981 | ENSMUSG00 | Btbd35f10 | BTB domain containing 35, family member 10 |
| chrX | 31202558 | 31202878 | 321 | * | NA_peak_595 | 38 | Distal Interge | 20 | 31336401 | 31338101 | 1701 | 1 | 668960 | ENSMUST00 | -133523 | ENSMUSG00 | Btbd35f10 | BTB domain containing 35, family member 10 |
| chrX | 31203788 | 31204228 | 441 | * | NA_peak_595 | 35 | Distal Interge | 20 | 31336401 | 31338101 | 1701 | 1 | 668960 | ENSMUST00 | -132173 | ENSMUSG00 | Btbd35f10 | BTB domain containing 35, family member 10 |
| chrX | 31223687 | 31224123 | 437 | * | NA_peak_595 | 29 | Distal Interge | 20 | 31336401 | 31338101 | 1701 | 1 | 668960 | ENSMUST00 | -112278 | ENSMUSG00 | Btbd35f10 | BTB domain containing 35, family member 10 |
| chrX | 31228478 | 31229210 | 733 | * | NA_peak_595 | 69 | Distal Interge | 20 | 31336401 | 31338101 | 1701 | 1 | 668960 | ENSMUST00 | -107191 | ENSMUSG00 | Btbd35f10 | BTB domain containing 35, family member 10 |
| chrX | 31229696 | 31230171 | 476 | * | NA_peak_595 | 29 | Distal Interge | 20 | 31336401 | 31338101 | 1701 | 1 | 668960 | ENSMUST00 | -106230 | ENSMUSG00 | Btbd35f10 | BTB domain containing 35, family member 10 |
| chrX | 31231783 | 31232146 | 364 | * | NA_peak_595 | 50 | Distal Interge | 20 | 31336401 | 31338101 | 1701 | 1 | 668960 | ENSMUST00 | -104255 | ENSMUSG00 | Btbd35f10 | BTB domain containing 35, family member 10 |
| chrX | 31235299 | 31235715 | 417 | * | NA_peak_595 | 15 | Distal Interge | 20 | 31336401 | 31338101 | 1701 | 1 | 668960 | ENSMUST00 | -100686 | ENSMUSG00 | Btbd35f10 | BTB domain containing 35, family member 10 |
| chrX | 31244518 | 31244982 | 465 | * | NA_peak_595 | 45 | Distal Interge | 20 | 31336401 | 31338101 | 1701 | 1 | 668960 | ENSMUST00 | -91419 | ENSMUSG00 | Btbd35f10 | BTB domain containing 35, family member 10 |
| chrX | 31250458 | 31250658 | 201 | * | NA_peak_595 | 25 | Distal Interge | 20 | 31336401 | 31338101 | 1701 | 1 | 668960 | ENSMUST00 | -85743 | ENSMUSG00 | Btbd35f10 | BTB domain containing 35, family member 10 |
| chrX | 31253311 | 31253806 | 496 | * | NA_peak_595 | 19 | 5' UTR | 20 | 31336401 | 31338101 | 1701 | 1 | 668960 | ENSMUST00 | -82595 | ENSMUSG00 | Btbd35f10 | BTB domain containing 35, family member 10 |
| chrX | 31256869 | 31257182 | 314 | * | NA_peak_595 | 31 | Distal Interge | 20 | 31336401 | 31338101 | 1701 | 1 | 668960 | ENSMUST00 | -79219 | ENSMUSG00 | Btbd35f10 | BTB domain containing 35, family member 10 |
| chrX | 31268062 | 31268788 | 727 | * | NA_peak_595 | 55 | Distal Interge | 20 | 31336401 | 31338101 | 1701 | 1 | 668960 | ENSMUST00 | -67613 | ENSMUSG00 | Btbd35f10 | BTB domain containing 35, family member 10 |
| chrX | 31272968 | 31273261 | 294 | * | NA_peak_595 | 28 | Distal Interge | 20 | 31336401 | 31338101 | 1701 | 1 | 668960 | ENSMUST00 | -63140 | ENSMUSG00 | Btbd35f10 | BTB domain containing 35, family member 10 |
| chrX | 31673073 | 31673443 | 371 | * | NA_peak_596 | 55 | Distal Interge | 20 | 31962044 | 31962821 | 778 | 1 | 74851 | ENSMUST00 | -288601 | ENSMUSG00 | Spin2-ps1 | spindlin family, member 2, pseudogene 1 |
| chrX | 31729924 | 31730594 | 671 | * | NA_peak_596 | 27 | Distal Interge | 20 | 31962044 | 31962821 | 778 | 1 | 74851 | ENSMUST00 | -231450 | ENSMUSG00 | Spin2-ps1 | spindlin family, member 2, pseudogene 1 |
| chrX | 31737754 | 31737888 | 135 | * | NA_peak_596 | 40 | Distal Interge | 20 | 31962044 | 31962821 | 778 | 1 | 74851 | ENSMUST00 | -224156 | ENSMUSG00 | Spin2-ps1 | spindlin family, member 2, pseudogene 1 |
| chrX | 31747657 | 31748020 | 364 | * | NA_peak_596 | 30 | Distal Interge | 20 | 31962044 | 31962821 | 778 | 1 | 74851 | ENSMUST00 | -214024 | ENSMUSG00 | Spin2-ps1 | spindlin family, member 2, pseudogene 1 |
| chrX | 31750318 | 31750378 | 61 | * | NA_peak_596 | 23 | Distal Interge | 20 | 31962044 | 31962821 | 778 | 1 | 74851 | ENSMUST00 | -211666 | ENSMUSG00 | Spin2-ps1 | spindlin family, member 2, pseudogene 1 |
| chrX | 31759546 | 31759852 | 307 | * | NA_peak_596 | 22 | Distal Interge | 20 | 31962044 | 31962821 | 778 | 1 | 74851 | ENSMUST00 | -202192 | ENSMUSG00 | Spin2-ps1 | spindlin family, member 2, pseudogene 1 |
| chrX | 31761816 | 31762283 | 468 | * | NA_peak_596 | 31 | Distal Interge | 20 | 31962044 | 31962821 | 778 | 1 | 74851 | ENSMUST00 | -199761 | ENSMUSG00 | Spin2-ps1 | spindlin family, member 2, pseudogene 1 |

| | | | | | | | | | | | | | | | | | | |
|---|---|---|---|---|---|---|---|---|---|---|---|---|---|---|---|---|---|---|
| chrX | 31775125 | 31775408 | 284 | * | NA_peak_59 | 36 | Distal Interge | 20 | 31962044 | 31962821 | 778 | 1 | 74851 | ENSMUST00 | -186636 | ENSMUSG0 | Spin2-ps1 | spindlin family, member 2, pseudogene 1 |
| chrX | 31775995 | 31776281 | 287 | * | NA_peak_59 | 28 | Distal Interge | 20 | 31962044 | 31962821 | 778 | 1 | 74851 | ENSMUST00 | -185763 | ENSMUSG0 | Spin2-ps1 | spindlin family, member 2, pseudogene 1 |
| chrX | 31778592 | 31779190 | 599 | * | NA_peak_59 | 31 | Distal Interge | 20 | 31962044 | 31962821 | 778 | 1 | 74851 | ENSMUST00 | -182854 | ENSMUSG0 | Spin2-ps1 | spindlin family, member 2, pseudogene 1 |
| chrX | 31920478 | 31920997 | 520 | * | NA_peak_59 | 41 | Distal Interge | 20 | 31962044 | 31962821 | 778 | 1 | 74851 | ENSMUST00 | -41047 | ENSMUSG0 | Spin2-ps1 | spindlin family, member 2, pseudogene 1 |
| chrX | 31931846 | 31932122 | 277 | * | NA_peak_59 | 31 | Distal Interge | 20 | 31962044 | 31962821 | 778 | 1 | 74851 | ENSMUST00 | -29922 | ENSMUSG0 | Spin2-ps1 | spindlin family, member 2, pseudogene 1 |
| chrX | 31952802 | 31953112 | 311 | * | NA_peak_59 | 27 | Distal Interge | 20 | 31962044 | 31962821 | 778 | 1 | 74851 | ENSMUST00 | -8932 | ENSMUSG0 | Spin2-ps1 | spindlin family, member 2, pseudogene 1 |
| chrX | 32058501 | 32058794 | 294 | * | NA_peak_59 | 20 | Distal Interge | 20 | 31962044 | 31962821 | 778 | 1 | 74851 | ENSMUST00 | 96457 | ENSMUSG0 | Spin2-ps1 | spindlin family, member 2, pseudogene 1 |
| chrX | 32153983 | 32154308 | 326 | * | NA_peak_59 | 27 | Distal Interge | 20 | 32278435 | 32280341 | 1907 | 1 | 100040482 | ENSMUST00 | -124127 | ENSMUSG0 | Btbd35f2 | BTB domain containing 35, family member 2 |
| chrX | 32310436 | 32310684 | 249 | * | NA_peak_59 | 16 | Distal Interge | 20 | 32278435 | 32280341 | 1907 | 1 | 100040482 | ENSMUST00 | 32001 | ENSMUSG0 | Btbd35f2 | BTB domain containing 35, family member 2 |
| chrX | 32311792 | 32312278 | 487 | * | NA_peak_59 | 24 | Distal Interge | 20 | 32278435 | 32280341 | 1907 | 1 | 100040482 | ENSMUST00 | 33357 | ENSMUSG0 | Btbd35f2 | BTB domain containing 35, family member 2 |
| chrX | 32450333 | 32450683 | 351 | * | NA_peak_59 | 21 | Distal Interge | 20 | 32560096 | 32562003 | 1908 | 1 | 100040482 | ENSMUST00 | -109413 | ENSMUSG0 | Btbd35f2 | BTB domain containing 35, family member 2 |
| chrX | 32559896 | 32560329 | 434 | * | NA_peak_59 | 31 | Promoter (<= | 20 | 32560096 | 32562003 | 1908 | 1 | 100040482 | ENSMUST00 | 0 | ENSMUSG0 | Btbd35f2 | BTB domain containing 35, family member 2 |
| chrX | 32584155 | 32584465 | 311 | * | NA_peak_59 | 42 | Distal Interge | 20 | 32560096 | 32562003 | 1908 | 1 | 100040482 | ENSMUST00 | 24059 | ENSMUSG0 | Btbd35f2 | BTB domain containing 35, family member 2 |
| chrX | 32596970 | 32597316 | 347 | * | NA_peak_59 | 31 | Distal Interge | 20 | 32560096 | 32562003 | 1908 | 1 | 100040482 | ENSMUST00 | 36874 | ENSMUSG0 | Btbd35f2 | BTB domain containing 35, family member 2 |
| chrX | 32700240 | 32700538 | 299 | * | NA_peak_59 | 16 | Distal Interge | 20 | 32726109 | 32728051 | 1943 | 1 | 100862329 | ENSMUST00 | -25571 | ENSMUSG0 | Btbd35f21 | BTB domain containing 35, family member 21 |
| chrX | 32719295 | 32719638 | 344 | * | NA_peak_59 | 23 | Distal Interge | 20 | 32726109 | 32728051 | 1943 | 1 | 100862329 | ENSMUST00 | -6471 | ENSMUSG0 | Btbd35f21 | BTB domain containing 35, family member 21 |
| chrX | 32915880 | 32916163 | 284 | * | NA_peak_59 | 20 | Distal Interge | 20 | 32973938 | 32975845 | 1908 | 1 | 100040533 | ENSMUST00 | -57775 | ENSMUSG0 | Btbd35f5 | BTB domain containing 35, family member 5 |
| chrX | 32925525 | 32925883 | 359 | * | NA_peak_59 | 45 | Distal Interge | 20 | 32973938 | 32975845 | 1908 | 1 | 100040533 | ENSMUST00 | -48055 | ENSMUSG0 | Btbd35f5 | BTB domain containing 35, family member 5 |
| chrX | 32927138 | 32927482 | 345 | * | NA_peak_59 | 55 | Distal Interge | 20 | 32973938 | 32975845 | 1908 | 1 | 100040533 | ENSMUST00 | -46456 | ENSMUSG0 | Btbd35f5 | BTB domain containing 35, family member 5 |
| chrX | 32995300 | 32995571 | 272 | * | NA_peak_59 | 31 | Distal Interge | 20 | 32973938 | 32975845 | 1908 | 1 | 100040533 | ENSMUST00 | 21362 | ENSMUSG0 | Btbd35f5 | BTB domain containing 35, family member 5 |
| chrX | 33066242 | 33066575 | 334 | * | NA_peak_59 | 18 | Distal Interge | 20 | 32973938 | 32975845 | 1908 | 1 | 100040533 | ENSMUST00 | 92304 | ENSMUSG0 | Btbd35f5 | BTB domain containing 35, family member 5 |
| chrX | 33096113 | 33096458 | 346 | * | NA_peak_59 | 27 | Distal Interge | 20 | 32973938 | 32975845 | 1908 | 1 | 100040533 | ENSMUST00 | 122175 | ENSMUSG0 | Btbd35f5 | BTB domain containing 35, family member 5 |
| chrX | 33100112 | 33100256 | 145 | * | NA_peak_59 | 31 | Distal Interge | 20 | 32973938 | 32975845 | 1908 | 1 | 100040533 | ENSMUST00 | 126174 | ENSMUSG0 | Btbd35f5 | BTB domain containing 35, family member 5 |
| chrX | 33110231 | 33110428 | 198 | * | NA_peak_59 | 24 | Distal Interge | 20 | 32973938 | 32975845 | 1908 | 1 | 100040533 | ENSMUST00 | 136293 | ENSMUSG0 | Btbd35f5 | BTB domain containing 35, family member 5 |
| chrX | 33217233 | 33217341 | 109 | * | NA_peak_59 | 16 | Distal Interge | 20 | 33313379 | 33315286 | 1908 | 1 | 100040606 | ENSMUST00 | -96038 | ENSMUSG0 | Btbd35f12 | BTB domain containing 35, family member 12 |
| chrX | 33357093 | 33357379 | 287 | * | NA_peak_59 | 35 | Distal Interge | 20 | 33313379 | 33315286 | 1908 | 1 | 100040606 | ENSMUST00 | 43714 | ENSMUSG0 | Btbd35f12 | BTB domain containing 35, family member 12 |
| chrX | 33362319 | 33362578 | 260 | * | NA_peak_59 | 30 | Distal Interge | 20 | 33313379 | 33315286 | 1908 | 1 | 100040606 | ENSMUST00 | 48940 | ENSMUSG0 | Btbd35f12 | BTB domain containing 35, family member 12 |
| chrX | 33557574 | 33558045 | 472 | * | NA_peak_59 | 43 | Distal Interge | 20 | 33575398 | 33577314 | 1917 | 2 | 668960 | ENSMUST00 | 19269 | ENSMUSG0 | Btbd35f10 | BTB domain containing 35, family member 10 |
| chrX | 33920891 | 33921404 | 514 | * | NA_peak_59 | 19 | Distal Interge | 20 | 33700649 | 33702349 | 1701 | 2 | 100040732 | ENSMUST00 | -218542 | ENSMUSG0 | Btbd35f14 | BTB domain containing 35, family member 14 |
| chrX | 33921847 | 33922126 | 280 | * | NA_peak_59 | 38 | Distal Interge | 20 | 33700649 | 33702349 | 1701 | 2 | 100040732 | ENSMUST00 | -219498 | ENSMUSG0 | Btbd35f14 | BTB domain containing 35, family member 14 |
| chrX | 33922832 | 33923151 | 320 | * | NA_peak_59 | 28 | Distal Interge | 20 | 33700649 | 33702349 | 1701 | 2 | 100040732 | ENSMUST00 | -220483 | ENSMUSG0 | Btbd35f14 | BTB domain containing 35, family member 14 |
| chrX | 33926658 | 33926910 | 253 | * | NA_peak_59 | 41 | Distal Interge | 20 | 33700649 | 33702349 | 1701 | 2 | 100040732 | ENSMUST00 | -224309 | ENSMUSG0 | Btbd35f14 | BTB domain containing 35, family member 14 |
| chrX | 33936193 | 33936786 | 594 | * | NA_peak_59 | 46 | Distal Interge | 20 | 34159636 | 34161564 | 1929 | 2 | 668960 | ENSMUST00 | 224778 | ENSMUSG0 | Btbd35f10 | BTB domain containing 35, family member 10 |
| chrX | 33937821 | 33938114 | 294 | * | NA_peak_59 | 51 | Distal Interge | 20 | 34159636 | 34161564 | 1929 | 2 | 668960 | ENSMUST00 | 223450 | ENSMUSG0 | Btbd35f10 | BTB domain containing 35, family member 10 |
| chrX | 33939434 | 33939757 | 324 | * | NA_peak_59 | 33 | Distal Interge | 20 | 34159636 | 34161564 | 1929 | 2 | 668960 | ENSMUST00 | 221807 | ENSMUSG0 | Btbd35f10 | BTB domain containing 35, family member 10 |
| chrX | 33977424 | 33977840 | 417 | * | NA_peak_59 | 30 | Distal Interge | 20 | 34159636 | 34161564 | 1929 | 2 | 668960 | ENSMUST00 | 183724 | ENSMUSG0 | Btbd35f10 | BTB domain containing 35, family member 10 |
| chrX | 33988302 | 33988775 | 474 | * | NA_peak_59 | 32 | Distal Interge | 20 | 34159636 | 34161564 | 1929 | 2 | 668960 | ENSMUST00 | 172789 | ENSMUSG0 | Btbd35f10 | BTB domain containing 35, family member 10 |
| chrX | 34009330 | 34009367 | 38 | * | NA_peak_59 | 30 | Distal Interge | 20 | 34159636 | 34161564 | 1929 | 2 | 668960 | ENSMUST00 | 152197 | ENSMUSG0 | Btbd35f10 | BTB domain containing 35, family member 10 |
| chrX | 34192382 | 34192789 | 408 | * | NA_peak_59 | 32 | Distal Interge | 20 | 34159636 | 34161564 | 1929 | 2 | 668960 | ENSMUST00 | -30818 | ENSMUSG0 | Btbd35f10 | BTB domain containing 35, family member 10 |
| chrX | 34210428 | 34210756 | 329 | * | NA_peak_59 | 55 | Distal Interge | 20 | 34159636 | 34161564 | 1929 | 2 | 668960 | ENSMUST00 | -48864 | ENSMUSG0 | Btbd35f10 | BTB domain containing 35, family member 10 |
| chrX | 34228699 | 34228853 | 155 | * | NA_peak_59 | 16 | Distal Interge | 20 | 34159636 | 34161564 | 1929 | 2 | 668960 | ENSMUST00 | -67135 | ENSMUSG0 | Btbd35f10 | BTB domain containing 35, family member 10 |
| chrX | 34322342 | 34322641 | 300 | * | NA_peak_59 | 50 | Distal Interge | 20 | 34159636 | 34161564 | 1929 | 2 | 668960 | ENSMUST00 | -160778 | ENSMUSG0 | Btbd35f10 | BTB domain containing 35, family member 10 |
| chrX | 34330729 | 34331179 | 451 | * | NA_peak_59 | 29 | Distal Interge | 20 | 34159636 | 34161564 | 1929 | 2 | 668960 | ENSMUST00 | -169165 | ENSMUSG0 | Btbd35f10 | BTB domain containing 35, family member 10 |
| chrX | 34332033 | 34332386 | 354 | * | NA_peak_59 | 56 | Distal Interge | 20 | 34159636 | 34161564 | 1929 | 2 | 668960 | ENSMUST00 | -170469 | ENSMUSG0 | Btbd35f10 | BTB domain containing 35, family member 10 |
| chrX | 34468285 | 34468797 | 513 | * | NA_peak_59 | 20 | Distal Interge | 20 | 34159636 | 34161564 | 1929 | 2 | 668960 | ENSMUST00 | -306721 | ENSMUSG0 | Btbd35f10 | BTB domain containing 35, family member 10 |
| chrX | 34578374 | 34578579 | 206 | * | NA_peak_59 | 23 | Distal Interge | 20 | 34159636 | 34161564 | 1929 | 2 | 668960 | ENSMUST00 | -416810 | ENSMUSG0 | Btbd35f10 | BTB domain containing 35, family member 10 |
| chrX | 39665658 | 39666265 | 608 | * | NA_peak_59 | 50 | Distal Interge | 20 | 39862919 | 39863604 | 686 | 2 | 100504400 | ENSMUST00 | 197339 | ENSMUSG0 | Cypt14 | cysteine-rich perinuclear theca 14 |
| chrX | 41177630 | 41178014 | 385 | * | NA_peak_59 | 27 | Distal Interge | 20 | 41400854 | 41613099 | 212246 | 1 | 53623 | ENSMUST00 | -222840 | ENSMUSG0 | Gria3 | glutamate receptor, ionotropic, AMPA3 (alpha 3) |
| chrX | 41309013 | 41309269 | 257 | * | NA_peak_59 | 22 | Distal Interge | 20 | 41400854 | 41613099 | 212246 | 1 | 53623 | ENSMUST00 | -91585 | ENSMUSG0 | Gria3 | glutamate receptor, ionotropic, AMPA3 (alpha 3) |
| chrX | 41641870 | 41641928 | 59 | * | NA_peak_59 | 31 | Intron (ENSM | 20 | 41610197 | 41621262 | 11066 | 1 | 53623 | ENSMUST00 | 31673 | ENSMUSG0 | Gria3 | glutamate receptor, ionotropic, AMPA3 (alpha 3) |
| chrX | 47360753 | 47361127 | 375 | * | NA_peak_59 | 45 | Distal Interge | 20 | 47841930 | 47892396 | 50467 | 2 | 93761 | ENSMUST00 | 531269 | ENSMUSG0 | Smarca1 | SWI/SNF related, matrix associated, actin dependent regulator of chromatin, subfamily a, member 1 |
| chrX | 47738072 | 47738736 | 665 | * | NA_peak_59 | 30 | Distal Interge | 20 | 47841930 | 47892396 | 50467 | 2 | 93761 | ENSMUST00 | 153660 | ENSMUSG0 | Smarca1 | SWI/SNF related, matrix associated, actin dependent regulator of chromatin, subfamily a, member 1 |
| chrX | 47969114 | 47969508 | 395 | * | NA_peak_59 | 36 | Distal Interge | 20 | 47960289 | 47965763 | 5475 | 1 | 320634 | ENSMUST00 | 8825 | ENSMUSG0 | Ocrl | OCRL, inositol polyphosphate-5-phosphatase |
| chrX | 50000337 | 50000497 | 161 | * | NA_peak_59 | 36 | Distal Interge | 20 | 50009306 | 50010235 | 930 | 2 | 258385 | ENSMUST00 | 9738 | ENSMUSG0 | Olfr1323 | olfactory receptor 1323 |
| chrX | 54732473 | 54732781 | 309 | * | NA_peak_59 | 32 | Distal Interge | 20 | 54714790 | 54729932 | 15143 | 1 | 666122 | ENSMUST00 | 17683 | ENSMUSG0 | Gm14595 | predicted gene 14595 |
| chrX | 54744602 | 54744995 | 394 | * | NA_peak_59 | 31 | Distal Interge | 20 | 54714790 | 54729932 | 15143 | 1 | 666122 | ENSMUST00 | 29812 | ENSMUSG0 | Gm14595 | predicted gene 14595 |
| chrX | 55537735 | 55538108 | 374 | * | NA_peak_59 | 70 | Distal Interge | 20 | 55553147 | 55568358 | 15212 | 1 | 100039055 | ENSMUST00 | -15039 | ENSMUSG0 | Gm16404 | predicted gene 16404 |
| chrX | 57603311 | 57603877 | 567 | * | NA_peak_59 | 34 | Distal Interge | 20 | 57496668 | 57503757 | 7090 | 2 | 245424 | ENSMUST00 | -99554 | ENSMUSG0 | Gpr101 | G protein-coupled receptor 101 |
| chrX | 57604693 | 57605135 | 443 | * | NA_peak_59 | 42 | Distal Interge | 20 | 57496668 | 57503757 | 7090 | 2 | 245424 | ENSMUST00 | -100936 | ENSMUSG0 | Gpr101 | G protein-coupled receptor 101 |
| chrX | 64127075 | 64127561 | 487 | * | NA_peak_59 | 16 | Distal Interge | 20 | 64151405 | 64152198 | 794 | 2 | 627470 | ENSMUST00 | 24637 | ENSMUSG0 | Gm6760 | predicted gene 6760 |
| chrX | 64888458 | 64888526 | 69 | * | NA_peak_59 | 31 | Distal Interge | 20 | 65047644 | 65049014 | 1371 | 1 | 70062 | ENSMUST00 | -159118 | ENSMUSG0 | Ctag2 | cancer/testis antigen 2 |
| chrX | 65707985 | 65708279 | 295 | * | NA_peak_59 | 61 | Distal Interge | 20 | 65047644 | 65049014 | 1371 | 1 | 70062 | ENSMUST00 | 660341 | ENSMUSG0 | Ctag2 | cancer/testis antigen 2 |
| chrX | 66474767 | 66475130 | 364 | * | NA_peak_59 | 16 | Distal Interge | 20 | 66649318 | 66661393 | 12076 | 1 | 245450 | ENSMUST00 | -174188 | ENSMUSG0 | Slitrk2 | SLIT and NTRK-like family, member 2 |
| chrX | 68521388 | 68521721 | 334 | * | NA_peak_59 | 21 | Distal Interge | 20 | 68678541 | 68717963 | 39423 | 1 | 14265 | ENSMUST00 | -156820 | ENSMUSG0 | Fmr1 | FMRP translational regulator 1 |
| chrX | 69424226 | 69424410 | 185 | * | NA_peak_59 | 31 | Intron (ENSM | 20 | 69360700 | 69785176 | 424477 | 1 | 14266 | ENSMUST00 | 63526 | ENSMUSG0 | Aff2 | AF4/FMR2 family, member 2 |
| chrX | 71685177 | 71685480 | 304 | * | NA_peak_59 | 22 | Distal Interge | 20 | 71663966 | 71667392 | 3427 | 1 | 14765 | ENSMUST00 | 21211 | ENSMUSG0 | Gpr50 | G-protein-coupled receptor 50 |
| chrX | 71686127 | 71686402 | 276 | * | NA_peak_59 | 31 | Distal Interge | 20 | 71663966 | 71667392 | 3427 | 1 | 14765 | ENSMUST00 | 22161 | ENSMUSG0 | Gpr50 | G-protein-coupled receptor 50 |
| chrX | 72943123 | 72943612 | 490 | * | NA_peak_59 | 22 | Intron (ENSM | 20 | 72929752 | 72953656 | 23905 | 1 | 18194 | ENSMUST00 | 13371 | ENSMUSG0 | Nsdhl | NAD(P) dependent steroid dehydrogenase-like |
| chrX | 74620892 | 74621477 | 586 | * | NA_peak_59 | 40 | Distal Interge | 20 | 74639119 | 74645635 | 6517 | 2 | 434797 | ENSMUST00 | 24158 | ENSMUSG0 | Pwwp4c | PWWP domain containing 4C |
| chrX | 75199940 | 75199978 | 39 | * | NA_peak_59 | 27 | Intron (ENSM | 20 | 75149121 | 75163770 | 14650 | 2 | 434800 | ENSMUST00 | -36170 | ENSMUSG0 | Smim9 | small integral membrane protein 9 |
| chrX | 75712154 | 75712483 | 330 | * | NA_peak_59 | 27 | Intron (ENSM | 20 | 75731068 | 75755600 | 24533 | 2 | 74396 | ENSMUST00 | 43117 | ENSMUSG0 | 4933407K13 | RIKEN cDNA 4933407K13 gene |
| chrX | 76147945 | 76148411 | 467 | * | NA_peak_59 | 40 | Distal Interge | 20 | 76264304 | 76267738 | 3435 | 1 | 238829 | ENSMUST00 | -115893 | ENSMUSG0 | Cldn34b3 | claudin 34B3 |
| chrX | 76598685 | 76599029 | 345 | * | NA_peak_59 | 147 | Intron (ENSM | 20 | 76582610 | 76602924 | 20315 | 2 | 74987 | ENSMUST00 | 3895 | ENSMUSG0 | Cldn34d | claudin 34D |
| chrX | 78235753 | 78236070 | 318 | * | NA_peak_59 | 37 | Distal Interge | 20 | 78240462 | 78245921 | 5460 | 2 | 236874 | ENSMUST00 | 9851 | ENSMUSG0 | Gm14743 | predicted gene 14743 |
| chrX | 81866128 | 81866411 | 284 | * | NA_peak_59 | 55 | Distal Interge | 20 | 81419575 | 81458121 | 38547 | 2 | 245492 | ENSMUST00 | -408007 | ENSMUSG0 | 4930595M18 | RIKEN cDNA 4930595M18 gene |
| chrX | 82267151 | 82267388 | 238 | * | NA_peak_59 | 26 | Distal Interge | 20 | 82948870 | 83492739 | 543870 | 1 | 13405 | ENSMUST00 | -681482 | ENSMUSG0 | Dmd | dystrophin, muscular dystrophy |
| chrX | 82608340 | 82608550 | 211 | * | NA_peak_59 | 16 | Distal Interge | 20 | 82948870 | 83492739 | 543870 | 1 | 13405 | ENSMUST00 | -340320 | ENSMUSG0 | Dmd | dystrophin, muscular dystrophy |
| chrX | 84154634 | 84154924 | 291 | * | NA_peak_59 | 55 | Intron (ENSM | 20 | 84076649 | 84309894 | 233246 | 1 | 13405 | ENSMUST00 | 77985 | ENSMUSG0 | Dmd | dystrophin, muscular dystrophy |
| chrX | 84509743 | 84510101 | 359 | * | NA_peak_59 | 32 | Intron (ENSM | 20 | 84599201 | 85144413 | 545213 | 1 | 13405 | ENSMUST00 | -89100 | ENSMUSG0 | Dmd | dystrophin, muscular dystrophy |
| chrX | 85967327 | 85968010 | 684 | * | NA_peak_59 | 35 | Distal Interge | 20 | 86044199 | 86047313 | 3115 | 1 | 73495 | ENSMUST00 | -76189 | ENSMUSG0 | Samt3 | spermatogenesis associated multipass transmembrane protein 3 |
| chrX | 86221972 | 86222450 | 479 | * | NA_peak_59 | 45 | Distal Interge | 20 | 86191764 | 86195947 | 4184 | 1 | 11614 | ENSMUST00 | 30208 | ENSMUSG0 | Nr0b1 | nuclear receptor subfamily 0, group B, member 1 |
| chrX | 86897344 | 86897699 | 356 | * | NA_peak_59 | 46 | Intron (ENSM | 20 | 86250256 | 86305093 | 54838 | 2 | 434903 | ENSMUST00 | -592251 | ENSMUSG0 | Mageb4 | MAGE family member B4 |
| chrX | 92273235 | 92273471 | 237 | * | NA_peak_60 | 31 | Intron (ENSM | 20 | 92489949 | 92490681 | 733 | 1 | 546347 | ENSMUST00 | -216478 | ENSMUSG0 | Gm5941 | predicted gene 5941 |
| chrX | 92802379 | 92802775 | 397 | * | NA_peak_60 | 61 | Distal Interge | 20 | 92118879 | 92599572 | 480694 | 2 | 215641 | ENSMUST00 | -202807 | ENSMUSG0 | Mageb18 | MAGE family member B18 |
| chrX | 94878958 | 94879527 | 570 | * | NA_peak_60 | 104 | Distal Interge | 20 | 94795563 | 94798183 | 2621 | 2 | 668171 | ENSMUST00 | -80775 | NA | Zxda | zinc finger, X-linked, duplicated A |
| chrX | 94883252 | 94883786 | 535 | * | NA_peak_60 | 46 | Distal Interge | 20 | 94795563 | 94798183 | 2621 | 2 | 668171 | ENSMUST00 | -85069 | NA | Zxda | zinc finger, X-linked, duplicated A |
| chrX | 94897522 | 94897916 | 395 | * | NA_peak_60 | 45 | Distal Interge | 20 | 94795563 | 94798183 | 2621 | 2 | 668171 | ENSMUST00 | -99339 | NA | Zxda | zinc finger, X-linked, duplicated A |
| chrX | 94905034 | 94905960 | 927 | * | NA_peak_60 | 32 | Distal Interge | 20 | 94795563 | 94798183 | 2621 | 2 | 668171 | ENSMUST00 | -106851 | NA | Zxda | zinc finger, X-linked, duplicated A |
| chrX | 94907776 | 94908145 | 370 | * | NA_peak_60 | 34 | Distal Interge | 20 | 94795563 | 94798183 | 2621 | 2 | 668171 | ENSMUST00 | -109593 | NA | Zxda | zinc finger, X-linked, duplicated A |
| chrX | 94910217 | 94910578 | 362 | * | NA_peak_60 | 20 | Distal Interge | 20 | 94795563 | 94798183 | 2621 | 2 | 668171 | ENSMUST00 | -112034 | NA | Zxda | zinc finger, X-linked, duplicated A |
| chrX | 94914309 | 94915548 | 1240 | * | NA_peak_60 | 48 | Distal Interge | 20 | 95022510 | 95026682 | 4173 | 2 | 270624 | ENSMUST00 | 111134 | ENSMUSG0 | Spin4 | spindlin family, member 4 |
| chrX | 95745251 | 95745552 | 302 | * | NA_peak_60 | 59 | Intron (ENSM | 20 | 95732586 | 95733265 | 680 | 2 | 76386 | ENSMUST00 | -11986 | ENSMUSG0 | 1700010D01 | RIKEN cDNA 1700010D01 gene |
| chrX | 101713944 | 101714279 | 336 | * | NA_peak_60 | 24 | Intron (ENSM | 20 | 101731535 | 101734269 | 2735 | 2 | 12766 | ENSMUST00 | 19990 | ENSMUSG0 | Cxcr3 | chemokine (C-X-C motif) receptor 3 |
| chrX | 105615891 | 105616315 | 425 | * | NA_peak_60 | 76 | Distal Interge | 20 | 105499772 | 105500639 | 868 | 1 | 245566 | ENSMUST00 | 116119 | ENSMUSG0 | Cypt2 | cysteine-rich perinuclear theca 2 |
| chrX | 107052568 | 107052847 | 280 | * | NA_peak_60 | 13 | Distal Interge | 20 | 107088492 | 107104970 | 16479 | 1 | 78826 | ENSMUST00 | -35645 | ENSMUSG0 | P2ry10 | purinergic receptor P2Y, G-protein coupled 10 |
| chrX | 107453336 | 107453735 | 400 | * | NA_peak_60 | 46 | Distal Interge | 20 | 107397099 | 107403376 | 6278 | 2 | 16431 | ENSMUST00 | -49960 | ENSMUSG0 | Itm2a | integral membrane protein 2A |
| chrX | 107454507 | 107455298 | 792 | * | NA_peak_60 | 36 | Distal Interge | 20 | 107397099 | 107403376 | 6278 | 2 | 16431 | ENSMUST00 | -51131 | ENSMUSG0 | Itm2a | integral membrane protein 2A |
| chrX | 107457123 | 107457628 | 506 | * | NA_peak_60 | 32 | Distal Interge | 20 | 107397099 | 107403376 | 6278 | 2 | 16431 | ENSMUST00 | -53747 | ENSMUSG0 | Itm2a | integral membrane protein 2A |
| chrX | 107734666 | 107734967 | 302 | * | NA_peak_60 | 32 | Distal Interge | 20 | 107679015 | 107687622 | 8608 | 1 | 245572 | ENSMUST00 | 55651 | ENSMUSG0 | Tbx22 | T-box 22 |
| chrX | 107981864 | 107982175 | 312 | * | NA_peak_60 | 39 | Distal Interge | 20 | 107946876 | 107948465 | 1590 | 2 | 213450 | ENSMUST00 | -33399 | ENSMUSG0 | Wmp | WAVE homology in membrane protrusions |
| chrX | 108075632 | 108075821 | 190 | * | NA_peak_60 | 34 | Distal Interge | 20 | 107946876 | 107948465 | 1590 | 2 | 213450 | ENSMUST00 | -127167 | ENSMUSG0 | Wmp | WAVE homology in membrane protrusions |
| chrX | 110092083 | 110092091 | 9 | * | NA_peak_60 | 16 | Distal Interge | 20 | 110372753 | 110375158 | 2406 | 1 | 634340 | ENSMUST00 | -280662 | ENSMUSG0 | Gm7134 | kinesin family member 18B pseudogene |
| chrX | 110882992 | 110883141 | 150 | * | NA_peak_60 | 23 | Distal Interge | 20 | 110814280 | 110817207 | 2928 | 1 | 18994 | ENSMUST00 | 68712 | ENSMUSG0 | Pou3f4 | POU domain, class 3, transcription factor 4 |
| chrX | 110887312 | 110887582 | 271 | * | NA_peak_60 | 32 | Distal Interge | 20 | 110814280 | 110817207 | 2928 | 1 | 18994 | ENSMUST00 | 73032 | ENSMUSG0 | Pou3f4 | POU domain, class 3, transcription factor 4 |
| chrX | 112370854 | 112371155 | 302 | * | NA_peak_60 | 127 | Intron (ENSM | 20 | 112384303 | 112405822 | 21520 | 2 | 73809 | ENSMUST00 | 34667 | ENSMUSG0 | Satl1 | spermidine/spermine N1-acetyl transferase-like 1 |
| chrX | 114140620 | 114141204 | 585 | * | NA_peak_60 | 22 | Distal Interge | 20 | 114474333 | 114560829 | 86497 | 1 | 237010 | ENSMUST00 | -333129 | ENSMUSG0 | Klhl4 | kelch-like 4 |

| | | | | | | | | | | | | | | | | | | |
|---|---|---|---|---|---|---|---|---|---|---|---|---|---|---|---|---|---|---|
| chrX | 114267077 | 114267386 | 310 | * | NA_peak_60 | 15 | Distal Interge | 20 | 114474333 | 114560829 | 86497 | 1 | 237010 | ENSMUST00 | -206947 | ENSMUSG0( | Klhl4 | kelch-like 4 |
| chrX | 115386520 | 115386892 | 373 | * | NA_peak_60 | 49 | Distal Interge | 20 | 114980970 | 114981317 | 348 | 2 | 664969 | ENSMUST00 | -405203 | ENSMUSG0( | Gm7429 | predicted pseudogene 7429 |
| chrX | 115388287 | 115388617 | 331 | * | NA_peak_60 | 74 | Distal Interge | 20 | 114980970 | 114981317 | 348 | 2 | 664969 | ENSMUST00 | -406970 | ENSMUSG0( | Gm7429 | predicted pseudogene 7429 |
| chrX | 117088489 | 117088563 | 75 | * | NA_peak_60 | 25 | Distal Interge | 20 | 117014757 | 117015104 | 348 | 1 | 100039319 | ENSMUST00 | 73732 | ENSMUSG0( | H2ab1 | H2A.B variant histone 1 |
| chrX | 118205762 | 118206037 | 276 | * | NA_peak_60 | 40 | Distal Interge | 20 | 118427227 | 118428256 | 1030 | 1 | 100039551 | ENSMUST00 | -221190 | ENSMUSG0( | Tgif2lx2 | TGFB-induced factor homeobox 2-like, X-linked 2 |
| chrX | 119502888 | 119503133 | 246 | * | NA_peak_60 | 22 | Distal Interge | 20 | 119927196 | 119930165 | 2970 | 1 | 93728 | ENSMUST00 | -424063 | ENSMUSG0( | Pabpc5 | poly(A) binding protein, cytoplasmic 5 |
| chrX | 119505510 | 119505807 | 298 | * | NA_peak_60 | 45 | Distal Interge | 20 | 119927196 | 119930165 | 2970 | 1 | 93728 | ENSMUST00 | -421389 | ENSMUSG0( | Pabpc5 | poly(A) binding protein, cytoplasmic 5 |
| chrX | 120961964 | 120962309 | 346 | * | NA_peak_60 | 39 | Distal Interge | 20 | 120401364 | 120906344 | 504981 | 1 | 245578 | ENSMUST00 | 560600 | ENSMUSG0( | Pcdh11x | protocadherin 11 X-linked |
| chrX | 121444286 | 121444910 | 625 | * | NA_peak_60 | 45 | Distal Interge | 20 | 122394565 | 122397401 | 2837 | 2 | 54561 | ENSMUST00 | 952491 | ENSMUSG0( | Nap1l3 | nucleosome assembly protein 1-like 3 |
| chrX | 121591125 | 121591436 | 312 | * | NA_peak_60 | 24 | Distal Interge | 20 | 122394565 | 122397401 | 2837 | 2 | 54561 | ENSMUST00 | 805965 | ENSMUSG0( | Nap1l3 | nucleosome assembly protein 1-like 3 |
| chrX | 123038407 | 123038702 | 296 | * | NA_peak_60 | 123 | Distal Interge | 20 | 123103495 | 123145651 | 42157 | 1 | 73061 | ENSMUST00 | -64793 | ENSMUSG0( | Cldn34c1 | claudin 34C1 |
| chrX | 123044113 | 123044248 | 136 | * | NA_peak_60 | 27 | Distal Interge | 20 | 123103495 | 123145651 | 42157 | 1 | 73061 | ENSMUST00 | -59247 | ENSMUSG0( | Cldn34c1 | claudin 34C1 |
| chrX | 123045371 | 123045788 | 418 | * | NA_peak_60 | 75 | Distal Interge | 20 | 123103495 | 123145651 | 42157 | 1 | 73061 | ENSMUST00 | -57707 | ENSMUSG0( | Cldn34c1 | claudin 34C1 |
| chrX | 123076608 | 123076911 | 304 | * | NA_peak_60 | 64 | Distal Interge | 20 | 123103495 | 123145651 | 42157 | 1 | 73061 | ENSMUST00 | -26584 | ENSMUSG0( | Cldn34c1 | claudin 34C1 |
| chrX | 124119777 | 124120317 | 541 | * | NA_peak_60 | 14 | Distal Interge | 20 | 124124961 | 124135952 | 10992 | 2 | 100038941 | ENSMUST00 | 15635 | ENSMUSG0( | Vmn2r121 | vomeronasal 2, receptor 121 |
| chrX | 124227810 | 124228187 | 378 | * | NA_peak_60 | 36 | Distal Interge | 20 | 124124961 | 124135952 | 10992 | 2 | 100038941 | ENSMUST00 | -91858 | ENSMUSG0( | Vmn2r121 | vomeronasal 2, receptor 121 |
| chrX | 127959484 | 127960050 | 567 | * | NA_peak_60 | 40 | Distal Interge | 20 | 127721182 | 127736554 | 15373 | 2 | 73934 | ENSMUST00 | -222930 | ENSMUSG0( | Cldn34c4 | claudin 34C4 |
| chrX | 128234279 | 128234555 | 277 | * | NA_peak_60 | 31 | Distal Interge | 20 | 127721182 | 127736554 | 15373 | 2 | 73934 | ENSMUST00 | -497725 | ENSMUSG0( | Cldn34c4 | claudin 34C4 |
| chrX | 129444660 | 129445016 | 357 | * | NA_peak_60 | 28 | Distal Interge | 20 | 129749742 | 130180471 | 430730 | 1 | 54004 | ENSMUST00 | -304726 | ENSMUSG0( | Diaph2 | diaphanous related formin 2 |
| chrX | 133881156 | 133881479 | 324 | * | NA_peak_60 | 20 | Distal Interge | 20 | 133896718 | 133898113 | 1396 | 2 | 56496 | ENSMUST00 | 16634 | ENSMUSG0( | Tspan6 | tetraspanin 6 |
| chrX | 138513810 | 138513839 | 30 | * | NA_peak_60 | 31 | Intron (ENSM | 20 | 138208165 | 138209580 | 1416 | 2 | 67944 | ENSMUST00 | -304230 | ENSMUSG0( | Tex13a | testis expressed 13A |
| chrX | 140896415 | 140896673 | 259 | * | NA_peak_60 | 39 | Distal Interge | 20 | 140907608 | 140939472 | 31865 | 1 | 78789 | ENSMUST00 | -10935 | ENSMUSG0( | Vsig1 | V-set and immunoglobulin domain containing 1 |
| chrX | 141826492 | 141826838 | 347 | * | NA_peak_60 | 16 | Distal Interge | 20 | 141710998 | 141725263 | 14266 | 2 | 16370 | ENSMUST00 | -101229 | ENSMUSG0( | Irs4 | insulin receptor substrate 4 |
| chrX | 141892866 | 141893200 | 335 | * | NA_peak_60 | 17 | Distal Interge | 20 | 141710998 | 141725263 | 14266 | 2 | 16370 | ENSMUST00 | -167603 | ENSMUSG0( | Irs4 | insulin receptor substrate 4 |
| chrX | 146357630 | 146357822 | 193 | * | NA_peak_60 | 20 | Distal Interge | 20 | 146962513 | 146989328 | 26816 | 1 | 15560 | ENSMUST00 | -604691 | ENSMUSG0( | Htr2c | 5-hydroxytryptamine (serotonin) receptor 2C |
| chrX | 146358239 | 146358592 | 354 | * | NA_peak_60 | 20 | Distal Interge | 20 | 146962513 | 146989328 | 26816 | 1 | 15560 | ENSMUST00 | -603921 | ENSMUSG0( | Htr2c | 5-hydroxytryptamine (serotonin) receptor 2C |
| chrX | 147968925 | 147969440 | 516 | * | NA_peak_60 | 23 | Distal Interge | 20 | 147992993 | 148033174 | 40182 | 1 | 666184 | ENSMUST00 | -23553 | ENSMUSG0( | Gm15080 | predicted gene 15080 |
| chrX | 151743162 | 151743804 | 643 | * | NA_peak_602 | 27 | Distal Interge | 20 | 151800883 | 151804511 | 3629 | 1 | 59026 | ENSMUST00 | -57079 | ENSMUSG0( | Huwe1 | HECT, UBA and WWE domain containing 1 |
| chrX | 152527023 | 152527360 | 338 | * | NA_peak_602 | 20 | Distal Interge | 20 | 152489661 | 152491860 | 2200 | 2 | 207318 | ENSMUST00 | -35163 | ENSMUSG0( | Gm4750 | actin, beta pseudogene |
| chrX | 156069857 | 156069876 | 20 | * | NA_peak_602 | 16 | Intron (ENSM | 20 | 155586808 | 155623814 | 37007 | 2 | 211612 | ENSMUST00 | -446043 | ENSMUSG0( | Ptchd1 | patched domain containing 1 |
| chrX | 156372217 | 156372537 | 321 | * | NA_peak_602 | 39 | Distal Interge | 20 | 155586808 | 155623814 | 37007 | 2 | 211612 | ENSMUST00 | -748403 | ENSMUSG0( | Ptchd1 | patched domain containing 1 |
| chrX | 161546973 | 161547102 | 130 | * | NA_peak_602 | 19 | Distal Interge | 20 | 161717069 | 161779494 | 62426 | 1 | 24004 | ENSMUST00 | -169967 | ENSMUSG0( | Rai2 | retinoic acid induced 2 |
| chrX | 165646191 | 165646446 | 256 | * | NA_peak_602 | 76 | Distal Interge | 20 | 165129017 | 165327393 | 198377 | 2 | 237213 | ENSMUST00 | -318798 | ENSMUSG0( | Glra2 | glycine receptor, alpha 2 subunit |
| chrX | 170019614 | 170020204 | 591 | * | NA_peak_602 | 23 | Distal Interge | 20 | 169979451 | 169990798 | 11348 | 1 | 17318 | ENSMUST00 | 40163 | ENSMUSG0( | Mid1 | midline 1 |
| chrY | 90729061 | 90729279 | 219 | * | NA_peak_602 | 14 | Distal Interge | 21 | 90752427 | 90755467 | 3041 | 2 | 654820 | ENSMUST00 | 26188 | ENSMUSG0( | G530011O06 | RIKEN cDNA G530011O06 gene |
| chrY | 90783370 | 90783840 | 471 | * | NA_peak_602 | 25 | Promoter (<= | 21 | 90784738 | 90816447 | 31710 | 1 | 170942 | ENSMUST00 | -898 | ENSMUSG0( | Erdr1 | erythroid differentiation regulator 1 |
| chrY | 90793661 | 90794002 | 342 | * | NA_peak_602 | 55 | Exon (ENSMU | 21 | 90790451 | 90816465 | 26015 | 1 | 170942 | ENSMUST00 | 3210 | ENSMUSG0( | Erdr1 | erythroid differentiation regulator 1 |
| chrY | 90800450 | 90800759 | 310 | * | NA_peak_602 | 24 | Intron (ENSM | 21 | 90790451 | 90816465 | 26015 | 1 | 170942 | ENSMUST00 | 9999 | ENSMUSG0( | Erdr1 | erythroid differentiation regulator 1 |
| chrY | 90810263 | 90810855 | 593 | * | NA_peak_602 | 34 | Intron (ENSM | 21 | 90790451 | 90816465 | 26015 | 1 | 170942 | ENSMUST00 | 19812 | ENSMUSG0( | Erdr1 | erythroid differentiation regulator 1 |

| seqnames | start | end | width | strand | V4 | V5 | annotation | geneChr | geneStart | geneEnd | geneLength | geneStrand | geneId | transcriptId | distanceToTS | ENSEMBL | SYMBOL | GENENAME |
|---|---|---|---|---|---|---|---|---|---|---|---|---|---|---|---|---|---|---|
| chr1 | 4020037 | 4020389 | 353 | * | NA_peak_2 | 30 | Intron (ENSM | 1 | 4344146 | 4360314 | 16169 | 2 | 19888 | ENSMUST0 | 339925 | ENSMUSG0 | Rp1 | retinitis pigmentosa 1 (human) |
| chr1 | 9546170 | 9546172 | 3 | * | NA_peak_8 | 14 | Promoter (<= | 1 | 9545408 | 9547455 | 2048 | 1 | 59014 | ENSMUST0 | 762 | ENSMUSG0 | Rrs1 | ribosome biogenesis regulator 1 |
| chr1 | 10885111 | 10885415 | 305 | * | NA_peak_12 | 26 | Distal Interge | 1 | 10993465 | 11303681 | 310217 | 1 | 109294 | ENSMUST0 | -108050 | ENSMUSG0 | Prex2 | phosphatidylinositol-3,4,5-trisphosphate-dependent Rac exchange factor 2 |
| chr1 | 12219358 | 12219592 | 235 | * | NA_peak_19 | 19 | Distal Interge | 1 | 12425986 | 12426106 | 121 | 1 | 102465961 | ENSMUST0 | -206394 | ENSMUSG0 | Mir6341 | microRNA 6341 |
| chr1 | 19373290 | 19373857 | 568 | * | NA_peak_39 | 70 | Distal Interge | 1 | 19213879 | 19219253 | 5375 | 1 | 21419 | ENSMUST0 | 159411 | ENSMUSG0 | Tfap2b | transcription factor AP-2 beta |
| chr1 | 19435828 | 19436125 | 298 | * | NA_peak_40 | 22 | Distal Interge | 1 | 19213879 | 19219253 | 5375 | 1 | 21419 | ENSMUST0 | 221949 | ENSMUSG0 | Tfap2b | transcription factor AP-2 beta |
| chr1 | 20319977 | 20320351 | 375 | * | NA_peak_43 | 26 | Intron (ENSM | 1 | 20356452 | 20392300 | 35849 | 1 | 75028 | ENSMUST0 | -36101 | ENSMUSG0 | 4930486I03R | RIKEN cDNA 4930486I03 gene |
| chr1 | 20707917 | 20708275 | 359 | * | NA_peak_47 | 30 | Distal Interge | 1 | 20730905 | 20734496 | 3592 | 1 | 16171 | ENSMUST0 | -22630 | ENSMUSG0 | Il17a | interleukin 17A |
| chr1 | 21199957 | 21200610 | 654 | * | NA_peak_50 | 86 | Distal Interge | 1 | 21218575 | 21230167 | 11593 | 1 | 75712 | ENSMUST0 | -17965 | ENSMUSG0 | Tmem14a | transmembrane protein 14A |
| chr1 | 24611606 | 24616106 | 4501 | * | NA_peak_58 | 720 | Exon (ENSM | 1 | 24263994 | 24587472 | 323479 | 2 | 12823 | ENSMUST0 | -24134 | ENSMUSG0 | Col19a1 | collagen, type XIX, alpha 1 |
| chr1 | 27755728 | 27756115 | 388 | * | NA_peak_64 | 70 | Distal Interge | 1 | 27884046 | 27902396 | 18351 | 1 | 433281 | ENSMUST0 | -127931 | ENSMUSG0 | Gm5524 | predicted gene 5524 |
| chr1 | 28306976 | 28307388 | 413 | * | NA_peak_65 | 44 | Distal Interge | 1 | 27884046 | 27902396 | 18351 | 1 | 433281 | ENSMUST0 | 422930 | ENSMUSG0 | Gm5524 | predicted gene 5524 |
| chr1 | 29252180 | 29252479 | 300 | * | NA_peak_67 | 28 | Distal Interge | 1 | 29421488 | 29421612 | 125 | 1 | 102465174 | ENSMUST0 | -169009 | ENSMUSG0 | Mir6342 | microRNA 6342 |
| chr1 | 34080853 | 34080996 | 144 | * | NA_peak_90 | 30 | Intron (ENSM | 1 | 34077227 | 34135157 | 57931 | 1 | 13518 | ENSMUST0 | 3626 | ENSMUSG0 | Dst | dystonin |
| chr1 | 34604445 | 34604930 | 486 | * | NA_peak_93 | 30 | Distal Interge | 1 | 34579657 | 34590944 | 11288 | 1 | 211383 | ENSMUST0 | 24788 | ENSMUSG0 | Amer3 | APC membrane recruitment 3 |
| chr1 | 35413434 | 35413732 | 299 | * | NA_peak_95 | 31 | Distal Interge | 1 | 34851837 | 34863475 | 11639 | 1 | 226971 | ENSMUST0 | 561597 | ENSMUSG0 | Plekhb2 | pleckstrin homology domain containing, family B (evectins) member 2 |
| chr1 | 41472376 | 41472746 | 371 | * | NA_peak_12 | 44 | Distal Interge | 1 | 41167587 | 41181252 | 13666 | 2 | 74897 | ENSMUST0 | -291124 | ENSMUSG0 | 4930448I06R | RIKEN cDNA 4930448I06 gene |
| chr1 | 43931383 | 43931705 | 323 | * | NA_peak_13 | 26 | Promoter (1- | 1 | 43933647 | 43949405 | 15759 | 1 | 22019 | ENSMUST0 | -1942 | ENSMUSG0 | Tpp2 | tripeptidyl peptidase II |
| chr1 | 44148113 | 44148386 | 274 | * | NA_peak_13 | 23 | Promoter (<= | 1 | 44147932 | 44158255 | 10324 | 1 | 22592 | ENSMUST0 | 181 | ENSMUSG0 | Ercc5 | excision repair cross-complementing rodent repair deficiency, complementation group 5 |
| chr1 | 44284195 | 44284618 | 424 | * | NA_peak_14 | 95 | Distal Interge | 1 | 44204070 | 44218961 | 14892 | 2 | 403183 | ENSMUST0 | -65234 | ENSMUSG0 | Mettl21e | methyltransferase like 21E |
| chr1 | 44604069 | 44604443 | 375 | * | NA_peak_14 | 23 | Intron (ENSM | 1 | 44552597 | 44708632 | 156036 | 1 | 70676 | ENSMUST0 | 51472 | ENSMUSG0 | Gulp1 | GULP, engulfment adaptor PTB domain containing 1 |
| chr1 | 45204259 | 45204387 | 129 | * | NA_peak_15 | 26 | Distal Interge | 1 | 45311538 | 45325018 | 13481 | 1 | 12825 | ENSMUST0 | -107151 | ENSMUSG0 | Col3a1 | collagen, type III, alpha 1 |
| chr1 | 45558203 | 45558529 | 327 | * | NA_peak_15 | 33 | Distal Interge | 1 | 45414199 | 45503282 | 89084 | 2 | 12832 | ENSMUST0 | -54921 | ENSMUSG0 | Col5a2 | collagen, type V, alpha 2 |
| chr1 | 47629261 | 47629596 | 336 | * | NA_peak_16 | 34 | Distal Interge | 1 | 47467021 | 47467125 | 105 | 2 | 102466969 | ENSMUST0 | -162136 | ENSMUSG0 | Mir6350 | microRNA 6350 |
| chr1 | 58406051 | 58406302 | 252 | * | NA_peak_20 | 13 | 3' UTR | 1 | 58412000 | 58412861 | 862 | 2 | 12747 | ENSMUST0 | 6559 | ENSMUSG0 | Clk1 | CDC-like kinase 1 |
| chr1 | 59678675 | 59678941 | 267 | * | NA_peak_22 | 23 | Distal Interge | 1 | 59684971 | 59692666 | 7696 | 1 | 55989 | ENSMUST0 | -6030 | ENSMUSG0 | Nop58 | NOP58 ribonucleoprotein |
| chr1 | 61499340 | 61499547 | 208 | * | NA_peak_24 | 20 | Distal Interge | 1 | 61378432 | 61395904 | 17473 | 1 | 77341 | ENSMUST0 | 120908 | ENSMUSG0 | 9530026F06I | RIKEN cDNA 9530026F06 gene |
| chr1 | 64314913 | 64315132 | 220 | * | NA_peak_25 | 16 | Distal Interge | 1 | 64079103 | 64122282 | 43180 | 2 | 93691 | ENSMUST0 | -192631 | ENSMUSG0 | Klf7 | Kruppel-like factor 7 (ubiquitous) |
| chr1 | 67280503 | 67280905 | 403 | * | NA_peak_27 | 54 | Intron (ENSM | 1 | 67123026 | 67231259 | 108234 | 1 | 227231 | ENSMUST0 | 157477 | ENSMUSG0 | Cps1 | carbamoyl-phosphate synthetase 1 |
| chr1 | 72212089 | 72212329 | 241 | * | NA_peak_29 | 39 | Promoter (<= | 1 | 72159442 | 72212307 | 52866 | 2 | 381269 | ENSMUST0 | 0 | ENSMUSG0 | Mreg | melanoregulin |
| chr1 | 73322623 | 73323177 | 555 | * | NA_peak_30 | 26 | Distal Interge | 1 | 73398968 | 73430790 | 31823 | 1 | 78718 | ENSMUST0 | -75791 | ENSMUSG0 | D530049I02F | RIKEN cDNA D530049I02 gene |
| chr1 | 74263707 | 74263823 | 117 | * | NA_peak_31 | 19 | Exon (ENSM | 1 | 74278550 | 74279624 | 1075 | 1 | 227289 | ENSMUST0 | -14727 | ENSMUSG0 | Gpbar1 | G protein-coupled bile acid receptor 1 |
| chr1 | 74558094 | 74558168 | 75 | * | NA_peak_32 | 16 | Promoter (2- | 1 | 74560427 | 74565301 | 4875 | 1 | 18802 | ENSMUST0 | -2259 | ENSMUSG0 | Plcd4 | phospholipase C, delta 4 |
| chr1 | 75440789 | 75441217 | 429 | * | NA_peak_33 | 19 | Promoter (<= | 1 | 75441620 | 75442932 | 1313 | 1 | 69080 | ENSMUST0 | -403 | ENSMUSG0 | Gmppa | GDP-mannose pyrophosphorylase A |
| chr1 | 78246619 | 78246923 | 305 | * | NA_peak_34 | 23 | Distal Interge | 1 | 78101270 | 78197134 | 95865 | 2 | 18505 | ENSMUST0 | -49485 | ENSMUSG0 | Pax3 | paired box 3 |
| chr1 | 79384063 | 79384403 | 341 | * | NA_peak_34 | 34 | Distal Interge | 1 | 79434669 | 79440042 | 5374 | 2 | 20254 | ENSMUST0 | 55639 | ENSMUSG0 | Scg2 | secretogranin II |
| chr1 | 79689463 | 79689763 | 301 | * | NA_peak_34 | 44 | Distal Interge | 1 | 79606876 | 79671972 | 65097 | 2 | 252903 | ENSMUST0 | -17491 | ENSMUSG0 | Ap1s3 | adaptor-related protein complex AP-1, sigma 3 |
| chr1 | 81569962 | 81570277 | 316 | * | NA_peak_35 | 17 | Distal Interge | 1 | 81077583 | 81341764 | 264182 | 1 | 241134 | ENSMUST0 | 492379 | ENSMUSG0 | Nyap2 | neuronal tyrosine-phophorylated phosphoinositide 3-kinase adaptor 2 |
| chr1 | 82705245 | 82705604 | 360 | * | NA_peak_35 | 35 | Intron (ENSM | 1 | 82715304 | 82718884 | 3581 | 1 | 12828 | ENSMUST0 | -9700 | ENSMUSG0 | Col4a3 | collagen, type IV, alpha 3 |
| chr1 | 84966151 | 84966525 | 375 | * | NA_peak_36 | 37 | Distal Interge | 1 | 84905898 | 84935134 | 29237 | 2 | 71781 | ENSMUST0 | -31017 | ENSMUSG0 | Slc16a14 | solute carrier family 16 (monocarboxylic acid transporters), member 14 |
| chr1 | 84972106 | 84972423 | 318 | * | NA_peak_36 | 47 | Distal Interge | 1 | 84905898 | 84935134 | 29237 | 2 | 71781 | ENSMUST0 | -36972 | ENSMUSG0 | Slc16a14 | solute carrier family 16 (monocarboxylic acid transporters), member 14 |
| chr1 | 85004240 | 85005607 | 1368 | * | NA_peak_36 | 37 | Intron (ENSM | 1 | 84905898 | 84935134 | 29237 | 2 | 71781 | ENSMUST0 | -69106 | ENSMUSG0 | Slc16a14 | solute carrier family 16 (monocarboxylic acid transporters), member 14 |
| chr1 | 85011194 | 85011804 | 611 | * | NA_peak_37 | 48 | Intron (ENSM | 1 | 84905898 | 84935134 | 29237 | 2 | 71781 | ENSMUST0 | -76060 | ENSMUSG0 | Slc16a14 | solute carrier family 16 (monocarboxylic acid transporters), member 14 |
| chr1 | 85071092 | 85071435 | 344 | * | NA_peak_37 | 24 | Intron (ENSM | 1 | 85083069 | 85109449 | 26381 | 2 | 381287 | ENSMUST0 | 38014 | ENSMUSG0 | A530032D15 | RIKEN cDNA A530032D15Rik gene |
| chr1 | 85233407 | 85233852 | 446 | * | NA_peak_37 | 35 | Intron (ENSM | 1 | 85241286 | 85254548 | 13263 | 2 | 620078 | ENSMUST0 | 20696 | ENSMUSG0 | C130026I21F | RIKEN cDNA C130026I21 gene |
| chr1 | 85299091 | 85299223 | 133 | * | NA_peak_37 | 21 | Intron (ENSM | 1 | 85246670 | 85270566 | 23897 | 2 | 620078 | ENSMUST0 | -28525 | ENSMUSG0 | C130026I21F | RIKEN cDNA C130026I21 gene |
| chr1 | 85315637 | 85315921 | 285 | * | NA_peak_37 | 43 | Intron (ENSM | 1 | 85246670 | 85270566 | 23897 | 2 | 620078 | ENSMUST0 | -45071 | ENSMUSG0 | C130026I21F | RIKEN cDNA C130026I21 gene |
| chr1 | 85329948 | 85330274 | 327 | * | NA_peak_38 | 21 | Intron (ENSM | 1 | 85246670 | 85270566 | 23897 | 2 | 620078 | ENSMUST0 | -59382 | ENSMUSG0 | C130026I21F | RIKEN cDNA C130026I21 gene |
| chr1 | 85332792 | 85333024 | 233 | * | NA_peak_38 | 16 | Exon (ENSM | 1 | 85246670 | 85270566 | 23897 | 2 | 620078 | ENSMUST0 | -62226 | ENSMUSG0 | C130026I21F | RIKEN cDNA C130026I21 gene |
| chr1 | 85333333 | 85333596 | 264 | * | NA_peak_38 | 16 | Intron (ENSM | 1 | 85246670 | 85270566 | 23897 | 2 | 620078 | ENSMUST0 | -62767 | ENSMUSG0 | C130026I21F | RIKEN cDNA C130026I21 gene |
| chr1 | 85338391 | 85338721 | 331 | * | NA_peak_38 | 54 | Intron (ENSM | 1 | 85246670 | 85270566 | 23897 | 2 | 620078 | ENSMUST0 | -67825 | ENSMUSG0 | C130026I21F | RIKEN cDNA C130026I21 gene |
| chr1 | 85345015 | 85345197 | 183 | * | NA_peak_38 | 23 | Distal Interge | 1 | 85246670 | 85270566 | 23897 | 2 | 620078 | ENSMUST0 | -74449 | ENSMUSG0 | C130026I21F | RIKEN cDNA C130026I21 gene |
| chr1 | 85346364 | 85346954 | 591 | * | NA_peak_38 | 27 | Distal Interge | 1 | 85246670 | 85270566 | 23897 | 2 | 620078 | ENSMUST0 | -75798 | ENSMUSG0 | C130026I21F | RIKEN cDNA C130026I21 gene |
| chr1 | 85453787 | 85454249 | 463 | * | NA_peak_38 | 89 | Distal Interge | 1 | 85579357 | 85584278 | 4922 | 2 | 109032 | ENSMUST0 | 130029 | ENSMUSG0 | Sp110 | Sp110 nuclear body protein |
| chr1 | 85463406 | 85463649 | 244 | * | NA_peak_38 | 20 | Distal Interge | 1 | 85579357 | 85584278 | 4922 | 2 | 109032 | ENSMUST0 | 120629 | ENSMUSG0 | Sp110 | Sp110 nuclear body protein |
| chr1 | 85464644 | 85465100 | 457 | * | NA_peak_38 | 56 | Distal Interge | 1 | 85579357 | 85584278 | 4922 | 2 | 109032 | ENSMUST0 | 119178 | ENSMUSG0 | Sp110 | Sp110 nuclear body protein |
| chr1 | 85479004 | 85479710 | 707 | * | NA_peak_38 | 80 | Distal Interge | 1 | 85579357 | 85584278 | 4922 | 2 | 109032 | ENSMUST0 | 104568 | ENSMUSG0 | Sp110 | Sp110 nuclear body protein |
| chr1 | 85480709 | 85481008 | 300 | * | NA_peak_39 | 39 | Distal Interge | 1 | 85579357 | 85584278 | 4922 | 2 | 109032 | ENSMUST0 | 103270 | ENSMUSG0 | Sp110 | Sp110 nuclear body protein |
| chr1 | 85556538 | 85556951 | 414 | * | NA_peak_39 | 48 | Distal Interge | 1 | 85579357 | 85584278 | 4922 | 2 | 109032 | ENSMUST0 | 27327 | ENSMUSG0 | Sp110 | Sp110 nuclear body protein |
| chr1 | 85584173 | 85584419 | 247 | * | NA_peak_39 | 39 | Promoter (<= | 1 | 85579357 | 85584278 | 4922 | 2 | 109032 | ENSMUST0 | 0 | ENSMUSG0 | Sp110 | Sp110 nuclear body protein |
| chr1 | 85650574 | 85650860 | 287 | * | NA_peak_39 | 26 | Promoter (<= | 1 | 85650050 | 85709998 | 59949 | 1 | 20684 | ENSMUST0 | 524 | ENSMUSG0 | Sp100 | nuclear antigen Sp100 |
| chr1 | 85653124 | 85653417 | 294 | * | NA_peak_39 | 51 | Intron (ENSM | 1 | 85650050 | 85709998 | 59949 | 1 | 20684 | ENSMUST0 | 3074 | ENSMUSG0 | Sp100 | nuclear antigen Sp100 |
| chr1 | 86018366 | 86018396 | 31 | * | NA_peak_40 | 17 | Promoter (<= | 1 | 86018068 | 86029248 | 11181 | 1 | 70060 | ENSMUST0 | 298 | ENSMUSG0 | Spata3 | spermatogenesis associated 3 |
| chr1 | 86154758 | 86154943 | 186 | * | NA_peak_40 | 26 | Promoter (<= | 1 | 86154780 | 86246326 | 91547 | 1 | 78795 | ENSMUST0 | 0 | ENSMUSG0 | Armc9 | armadillo repeat containing 9 |
| chr1 | 86472669 | 86473023 | 355 | * | NA_peak_40 | 16 | Distal Interge | 1 | 86426329 | 86428052 | 1724 | 1 | 71863 | ENSMUST0 | 46340 | ENSMUSG0 | Tex44 | testis expressed 44 |
| chr1 | 86487228 | 86487813 | 586 | * | NA_peak_41 | 22 | Distal Interge | 1 | 86526726 | 86530712 | 3987 | 1 | 19231 | ENSMUST0 | -38913 | ENSMUSG0 | Ptma | prothymosin alpha |
| chr1 | 86504265 | 86504572 | 308 | * | NA_peak_41 | 26 | Distal Interge | 1 | 86526726 | 86530712 | 3987 | 1 | 19231 | ENSMUST0 | -22154 | ENSMUSG0 | Ptma | prothymosin alpha |
| chr1 | 88289955 | 88290333 | 379 | * | NA_peak_43 | 27 | Intron (ENSM | 1 | 88277661 | 88308073 | 30413 | 1 | 171382 | ENSMUST0 | 12294 | ENSMUSG0 | Trpm8 | transient receptor potential cation channel, subfamily M, member 8 |
| chr1 | 89580847 | 89581081 | 235 | * | NA_peak_45 | 41 | Promoter (<= | 1 | 89580236 | 89895277 | 315042 | 1 | 347722 | ENSMUST0 | 611 | ENSMUSG0 | Agap1 | ArfGAP with GTPase domain, ankyrin repeat and PH domain 1 |
| chr1 | 89886839 | 89887171 | 333 | * | NA_peak_45 | 17 | Intron (ENSM | 1 | 89927956 | 89931179 | 3224 | 2 | 14472 | ENSMUST0 | 44008 | ENSMUSG0 | Gbx2 | gastrulation brain homeobox 2 |
| chr1 | 89930835 | 89931316 | 482 | * | NA_peak_45 | 21 | Promoter (<= | 1 | 89927956 | 89931179 | 3224 | 2 | 14472 | ENSMUST0 | 0 | ENSMUSG0 | Gbx2 | gastrulation brain homeobox 2 |
| chr1 | 91433209 | 91433297 | 89 | * | NA_peak_48 | 30 | Intron (ENSM | 1 | 91447722 | 91449865 | 2144 | 2 | 18627 | ENSMUST0 | 16568 | ENSMUSG0 | Per2 | period circadian clock 2 |
| chr1 | 97420127 | 97420288 | 162 | * | NA_peak_52 | 27 | Intron (ENSM | 1 | 97643903 | 97661676 | 17774 | 2 | 52392 | ENSMUST0 | 241388 | ENSMUSG0 | Macir | macrophage immunometabolism regulator |
| chr1 | 98602684 | 98603023 | 340 | * | NA_peak_52 | 39 | Intron (ENSM | 1 | 98499889 | 98509383 | 9495 | 1 | 70866 | ENSMUST0 | 102795 | ENSMUSG0 | Slco6d1 | solute carrier organic anion transporter family, member 6d1 |
| chr1 | 98885960 | 98886269 | 310 | * | NA_peak_52 | 24 | Distal Interge | 1 | 98499889 | 98509383 | 9495 | 1 | 70866 | ENSMUST0 | 386071 | ENSMUSG0 | Slco6d1 | solute carrier organic anion transporter family, member 6d1 |
| chr1 | 101305178 | 101305394 | 217 | * | NA_peak_53 | 19 | Distal Interge | 1 | 101155809 | 101175202 | 19394 | 1 | 100504525 | ENSMUST0 | 149369 | ENSMUSG0 | Gm20268 | predicted gene, 20268 |
| chr1 | 103210991 | 103211267 | 277 | * | NA_peak_53 | 23 | Intron (ENSM | 1 | 104768529 | 104995481 | 226953 | 1 | 23836 | ENSMUST0 | -1557262 | ENSMUSG0 | Cdh20 | cadherin 20 |
| chr1 | 104222379 | 104222663 | 285 | * | NA_peak_54 | 36 | Distal Interge | 1 | 104768529 | 104995481 | 226953 | 1 | 23836 | ENSMUST0 | -545866 | ENSMUSG0 | Cdh20 | cadherin 20 |
| chr1 | 105837077 | 105837399 | 323 | * | NA_peak_54 | 26 | Intron (ENSM | 1 | 105819870 | 105825401 | 5532 | 1 | 21934 | ENSMUST0 | 17207 | ENSMUSG0 | Tnfrsf11a | tumor necrosis factor receptor superfamily, member 11a, NFKB activator |
| chr1 | 110330001 | 110330398 | 398 | * | NA_peak_56 | 46 | Distal Interge | 1 | 110099295 | 110138058 | 38764 | 1 | 241201 | ENSMUST0 | 230706 | ENSMUSG0 | Cdh7 | cadherin 7, type 2 |
| chr1 | 112839571 | 112840015 | 445 | * | NA_peak_57 | 23 | Distal Interge | 1 | 111858706 | 111864918 | 6213 | 2 | 319901 | ENSMUST0 | -974653 | ENSMUSG0 | Dsel | dermatan sulfate epimerase-like |
| chr1 | 113403312 | 113403640 | 329 | * | NA_peak_57 | 30 | Distal Interge | 1 | 111858706 | 111864918 | 6213 | 2 | 319901 | ENSMUST0 | -1538394 | ENSMUSG0 | Dsel | dermatan sulfate epimerase-like |
| chr1 | 114875184 | 114875494 | 311 | * | NA_peak_58 | 19 | Distal Interge | 1 | 115684756 | 116587323 | 902568 | 1 | 636808 | ENSMUST0 | -809262 | ENSMUSG0 | Cntnap5a | contactin associated protein-like 5A |
| chr1 | 116858494 | 116858908 | 415 | * | NA_peak_58 | 65 | Distal Interge | 1 | 115684756 | 116587323 | 902568 | 1 | 636808 | ENSMUST0 | 1173738 | ENSMUSG0 | Cntnap5a | contactin associated protein-like 5A |
| chr1 | 116860276 | 116860672 | 397 | * | NA_peak_58 | 24 | Distal Interge | 1 | 115684756 | 116587323 | 902568 | 1 | 636808 | ENSMUST0 | 1175520 | ENSMUSG0 | Cntnap5a | contactin associated protein-like 5A |
| chr1 | 117484796 | 117484919 | 124 | * | NA_peak_59 | 23 | Distal Interge | 1 | 118300889 | 118309986 | 9098 | 2 | 22099 | ENSMUST0 | 825067 | ENSMUSG0 | Tsn | translin |
| chr1 | 119525776 | 119526363 | 588 | * | NA_peak_60 | 40 | Promoter (<= | 1 | 119526160 | 119528983 | 2824 | 1 | 226351 | ENSMUST0 | 0 | ENSMUSG0 | Tmem185b | transmembrane protein 185B |
| chr1 | 119837097 | 119837667 | 571 | * | NA_peak_60 | 30 | Promoter (<= | 1 | 119652467 | 119837613 | 185147 | 2 | 19258 | ENSMUST0 | 0 | ENSMUSG0 | Ptpn4 | protein tyrosine phosphatase, non-receptor type 4 |
| chr1 | 124091188 | 124091510 | 323 | * | NA_peak_62 | 15 | Intron (ENSM | 1 | 123321471 | 124045559 | 724089 | 2 | 269109 | ENSMUST0 | -45629 | ENSMUSG0 | Dpp10 | dipeptidylpeptidase 10 |
| chr1 | 124967794 | 124968150 | 357 | * | NA_peak_62 | 183 | Distal Interge | 1 | 123332142 | 124846039 | 1513898 | 2 | 269109 | ENSMUST0 | -121755 | ENSMUSG0 | Dpp10 | dipeptidylpeptidase 10 |
| chr1 | 131562346 | 131562392 | 47 | * | NA_peak_64 | 14 | Distal Interge | 1 | 131527903 | 131539872 | 11970 | 1 | 108900 | ENSMUST0 | 34443 | ENSMUSG0 | Fam72a | family with sequence similarity 72, member A |
| chr1 | 131999715 | 132000397 | 683 | * | NA_peak_64 | 35 | Intron (ENSM | 1 | 132007607 | 132026448 | 18842 | 1 | 13714 | ENSMUST0 | -7210 | ENSMUSG0 | Elk4 | ELK4, member of ETS oncogene family |
| chr1 | 132098443 | 132098884 | 442 | * | NA_peak_65 | 23 | Distal Interge | 1 | 132114934 | 132116255 | 1322 | 2 | 18557 | ENSMUST0 | 17371 | ENSMUSG0 | Cdk18 | cyclin-dependent kinase 18 |
| chr1 | 133131411 | 133131940 | 530 | * | NA_peak_65 | 30 | Promoter (<= | 1 | 133131344 | 133139783 | 8440 | 1 | 108954 | ENSMUST0 | 67 | ENSMUSG0 | Ppp1r15b | protein phosphatase 1, regulatory subunit 15B |
| chr1 | 134233728 | 134234086 | 359 | * | NA_peak_67 | 23 | Promoter (<= | 1 | 134199223 | 134234578 | 35356 | 2 | 11539 | ENSMUST0 | 492 | ENSMUSG0 | Adora1 | adenosine A1 receptor |
| chr1 | 134398454 | 134398701 | 248 | * | NA_peak_67 | 16 | Distal Interge | 1 | 134405559 | 134407990 | 2432 | 1 | 72017 | ENSMUST0 | -6858 | ENSMUSG0 | Cyb5r1 | cytochrome b5 reductase 1 |
| chr1 | 141194728 | 141195044 | 317 | * | NA_peak_72 | 30 | Distal Interge | 1 | 140570176 | 140656404 | 86229 | 2 | 75910 | ENSMUST0 | -538324 | ENSMUSG0 | 4930590L20F | RIKEN cDNA 4930590L20 gene |
| chr1 | 143241434 | 143241636 | 203 | * | NA_peak_73 | 15 | Distal Interge | 1 | 143606161 | 143640581 | 34421 | 2 | 214498 | ENSMUST0 | 398945 | ENSMUSG0 | Cdc73 | cell division cycle 73, Paf1/RNA polymerase II complex component |
| chr1 | 144856743 | 144856778 | 36 | * | NA_peak_74 | 23 | Distal Interge | 1 | 144752683 | 144775435 | 22753 | 2 | 64214 | ENSMUST0 | -81308 | ENSMUSG0 | Rgs18 | regulator of G-protein signaling 18 |
| chr1 | 146192882 | 146193353 | 472 | * | NA_peak_74 | 39 | Distal Interge | 1 | 146494760 | 146902471 | 407712 | 1 | 215378 | ENSMUST0 | -301407 | ENSMUSG0 | Brinp3 | bone morphogenetic protein/retinoic acid inducible neural specific 3 |
| chr1 | 146606783 | 146607329 | 547 | * | NA_peak_75 | 39 | Intron (ENSM | 1 | 146497738 | 146517554 | 19817 | 1 | 215378 | ENSMUST0 | 109045 | ENSMUSG0 | Brinp3 | bone morphogenetic protein/retinoic acid inducible neural specific 3 |
| chr1 | 147499687 | 147500024 | 338 | * | NA_peak_75 | 24 | Distal Interge | 1 | 146497738 | 146517554 | 19817 | 1 | 215378 | ENSMUST0 | 1001949 | ENSMUSG0 | Brinp3 | bone morphogenetic protein/retinoic acid inducible neural specific 3 |
| chr1 | 149394383 | 149394579 | 197 | * | NA_peak_75 | 23 | Distal Interge | 1 | 149886184 | 149902199 | 16016 | 2 | 18783 | ENSMUST0 | 507620 | ENSMUSG0 | Pla2g4a | phospholipase A2, group IVA (cytosolic, calcium-dependent) |
| chr1 | 150342779 | 150343073 | 295 | * | NA_peak_76 | 59 | Intron (ENSM | 1 | 150320681 | 150346771 | 26091 | 2 | 75584 | ENSMUST0 | 3698 | ENSMUSG0 | 2310030A07I | RIKEN cDNA 2310030A07 gene |
| chr1 | 151201299 | 151201605 | 307 | * | NA_peak_76 | 34 | Distal Interge | 1 | 151138034 | 151142345 | 4312 | 1 | 402734 | ENSMUST0 | 63265 | NA | C730036E19 | RIKEN cDNA C730036E19 gene |
| chr1 | 152657842 | 152658233 | 392 | * | NA_peak_77 | 16 | Intron (ENSM | 1 | 152516951 | 152625217 | 108267 | 2 | 19731 | ENSMUST0 | -32625 | ENSMUSG0 | Rgl1 | ral guanine nucleotide dissociation stimulator,-like 1 |
| chr1 | 153133023 | 153133420 | 398 | * | NA_peak_78 | 21 | Intron (ENSM | 1 | 153124173 | 153127234 | 3062 | 2 | 16782 | ENSMUST0 | -5789 | ENSMUSG0 | Lamc2 | laminin, gamma 2 |
| chr1 | 153425275 | 153425667 | 393 | * | NA_peak_78 | 17 | Promoter (<= | 1 | 153425172 | 153439937 | 14766 | 1 | 71836 | ENSMUST0 | 103 | ENSMUSG0 | Shcbp1l | Shc SH2-domain binding protein 1-like |
| chr1 | 154455059 | 154455100 | 42 | * | NA_peak_79 | 44 | Intron (ENSM | 1 | 154469052 | 154476696 | 7645 | 2 | 12290 | ENSMUST0 | 21596 | ENSMUSG0 | Cacna1e | calcium channel, voltage-dependent, R type, alpha 1E subunit |
| chr1 | 155098369 | 155098572 | 204 | * | NA_peak_79 | 28 | Promoter (1- | 1 | 155096361 | 155099636 | 3276 | 2 | 15939 | ENSMUST0 | 1064 | ENSMUSG0 | Ier5 | immediate early response 5 |
| chr1 | 155099030 | 155099363 | 334 | * | NA_peak_79 | 18 | Promoter (<= | 1 | 155096361 | 155099636 | 3276 | 2 | 15939 | ENSMUST0 | 273 | ENSMUSG0 | Ier5 | immediate early response 5 |
| chr1 | 156740982 | 156741086 | 105 | * | NA_peak_81 | 30 | Distal Interge | 1 | 156678571 | 156719086 | 40516 | 2 | 215015 | ENSMUST0 | -21896 | ENSMUSG0 | Fam20b | FAM20B, glycosaminoglycan xylosylkinase |
| chr1 | 162971448 | 162972027 | 580 | * | NA_peak_84 | 20 | Intron (ENSM | 1 | 162953800 | 162984527 | 30728 | 2 | 14262 | ENSMUST0 | 12500 | ENSMUSG0 | Fmo3 | flavin containing monooxygenase 3 |
| chr1 | 163811079 | 163811432 | 354 | * | NA_peak_85 | 17 | Intron (ENSM | 1 | 163779583 | 163917109 | 137527 | 1 | 16579 | ENSMUST0 | 31496 | ENSMUSG0 | Kifap3 | kinesin-associated protein 3 |
| chr1 | 165193898 | 165194371 | 474 | * | NA_peak_87 | 29 | Promoter (<= | 1 | 165175871 | 165194321 | 18451 | 2 | 108735 | ENSMUST0 | 0 | ENSMUSG0 | Sft2d2 | SFT2 domain containing 2 |
| chr1 | 165450699 | 165450787 | 89 | * | NA_peak_88 | 19 | Intron (ENSM | 1 | 165328698 | 165460465 | 131768 | 2 | 74106 | ENSMUST0 | 9678 | ENSMUSG0 | Dcaf6 | DDB1 and CUL4 associated factor 6 |
| chr1 | 167784923 | 167785275 | 353 | * | NA_peak_89 | 30 | Intron (ENSM | 1 | 167689557 | 167848741 | 159185 | 1 | 110648 | ENSMUST0 | 95366 | ENSMUSG0 | Lmx1a | LIM homeobox transcription factor 1 alpha |
| chr1 | 170098215 | 170098368 | 154 | * | NA_peak_90 | 36 | Promoter (<= | 1 | 170083632 | 170097328 | 13697 | 2 | 18214 | ENSMUST0 | -887 | ENSMUSG0 | Ddr2 | discoidin domain receptor family, member 2 |
| chr1 | 172376773 | 172377184 | 412 | * | NA_peak_93 | 23 | Promoter (<= | 1 | 172376546 | 172384099 | 7554 | 1 | 67556 | ENSMUST0 | 227 | ENSMUSG0 | Pigm | phosphatidylinositol glycan anchor biosynthesis, class M |
| chr1 | 173847463 | 173847465 | 3 | * | NA_peak_93 | 15 | Distal Interge | 1 | 173849126 | 173862499 | 13374 | 2 | 100040462 | ENSMUST0 | 15034 | ENSMUSG0 | Mndal | myeloid nuclear differentiation antigen like |
| chr1 | 173911573 | 173911932 | 360 | * | NA_peak_94 | 21 | Promoter (1- | 1 | 173896345 | 173913046 | 16702 | 2 | 381308 | ENSMUST0 | 1114 | ENSMUSG0 | Ifi211 | interferon activated gene 211 |
| chr1 | 173916621 | 173917107 | 487 | * | NA_peak_94 | 27 | Distal Interge | 1 | 173896345 | 173913046 | 16702 | 2 | 381308 | ENSMUST0 | -3575 | ENSMUSG0 | Ifi211 | interferon activated gene 211 |
| chr1 | 173948250 | 173948709 | 460 | * | NA_peak_94 | 38 | Distal Interge | 1 | 173920407 | 173942672 | 22266 | 2 | 15950 | ENSMUST0 | -5578 | ENSMUSG0 | Ifi203 | interferon activated gene 203 |
| chr1 | 173951269 | 173951633 | 365 | * | NA_peak_94 | 17 | Distal Interge | 1 | 173920407 | 173942672 | 22266 | 2 | 15950 | ENSMUST0 | -8597 | ENSMUSG0 | Ifi203 | interferon activated gene 203 |
| chr1 | 175797870 | 175798255 | 386 | * | NA_peak_95 | 18 | Intron (ENSM | 1 | 175767038 | 175815731 | 48694 | 1 | 75820 | ENSMUST0 | 30832 | ENSMUSG0 | Wdr64 | WD repeat domain 64 |
| chr1 | 177924656 | 177924961 | 306 | * | NA_peak_96 | 22 | Intron (ENSM | 1 | 177983423 | 178149962 | 166540 | 1 | 545391 | ENSMUST0 | -58462 | ENSMUSG0 | Catspere2 | cation channel sperm associated auxiliary subunit epsilon 2 |
| chr1 | 178008984 | 178009192 | 209 | * | NA_peak_97 | 39 | Intron (ENSM | 1 | 177991436 | 178172704 | 181269 | 1 | 545391 | ENSMUST0 | 17548 | ENSMUSG0 | Catspere2 | cation channel sperm associated auxiliary subunit epsilon 2 |
| chr1 | 178527905 | 178528125 | 221 | * | NA_peak_97 | 26 | Promoter (<= | 1 | 178529125 | 178716500 | 187376 | 1 | 269152 | ENSMUST0 | -1000 | ENSMUSG0 | Kif26b | kinesin family member 26B |
| chr1 | 179791515 | 179791790 | 276 | * | NA_peak_98 | 23 | Intron (ENSM | 1 | 179798847 | 179803606 | 4760 | 2 | 226747 | ENSMUST0 | 11816 | ENSMUSG0 | Ahctf1 | AT hook containing transcription factor 1 |
| chr1 | 180850918 | 180850930 | 13 | * | NA_peak_10 | 26 | Promoter (<= | 1 | 180851127 | 180868113 | 16987 | 1 | 208768 | ENSMUST0 | -197 | ENSMUSG0 | Sde2 | SDE2 telomere maintenance homolog (S. pombe) |
| chr1 | 181493048 | 181493422 | 375 | * | NA_peak_10 | 26 | Distal Interge | 1 | 181509633 | 181511451 | 1819 | 2 | 403180 | ENSMUST0 | 18029 | ENSMUSG0 | Ccdc121 | coiled-coil domain containing 121 |
| chr1 | 181871146 | 181871474 | 329 | * | NA_peak_10 | 17 | Distal Interge | 1 | 181836058 | 181843046 | 6989 | 2 | 98386 | ENSMUST0 | -28100 | ENSMUSG0 | Lbr | lamin B receptor |
| chr1 | 183553332 | 183553768 | 437 | * | NA_peak_10 | 26 | Distal Interge | 1 | 183419248 | 183436808 | 17561 | 1 | 78772 | ENSMUST0 | 134084 | ENSMUSG0 | Hhipl2 | hedgehog interacting protein-like 2 |
| chr1 | 185626194 | 185626436 | 243 | * | NA_peak_10 | 23 | Distal Interge | 1 | 185497388 | 185498017 | 630 | 1 | 114668 | ENSMUST0 | 128806 | ENSMUSG0 | 5033404E19I | NSA2 ribosome biogenesis homolog pseudogene |
| chr1 | 186875218 | 186875540 | 323 | * | NA_peak_10 | 26 | Distal Interge | 1 | 186967416 | 186970627 | 3212 | 1 | 110957 | ENSMUST0 | -91876 | ENSMUSG0 | D1Pas1 | DNA segment, Chr 1, Pasteur Institute 1 |
| chr1 | 188342014 | 188342350 | 337 | * | NA_peak_10 | 20 | Intron (ENSM | 1 | 188263034 | 188965041 | 702008 | 1 | 22283 | ENSMUST0 | 78980 | ENSMUSG0 | Ush2a | usherin |
| chr1 | 189973389 | 189973725 | 337 | * | NA_peak_10 | 23 | Distal Interge | 1 | 189880492 | 189922363 | 41872 | 2 | 226830 | ENSMUST0 | -51026 | ENSMUSG0 | Smyd2 | SET and MYND domain containing 2 |
| chr1 | 190978915 | 190979331 | 417 | * | NA_peak_10 | 19 | Promoter (<= | 1 | 190947646 | 190979296 | 31651 | 2 | 226841 | ENSMUST0 | 0 | ENSMUSG0 | Vash2 | vasohibin 2 |
| chr1 | 191090994 | 191091265 | 272 | * | NA_peak_10 | 19 | Distal Interge | 1 | 191097847 | 191108486 | 10640 | 1 | 381319 | ENSMUST0 | -6582 | ENSMUSG0 | Batf3 | basic leucine zipper transcription factor, ATF-like 3 |
| chr1 | 192257062 | 192257350 | 289 | * | NA_peak_10 | 19 | Intron (ENSM | 1 | 192190774 | 192510159 | 319386 | 1 | 16510 | ENSMUST0 | 66288 | ENSMUSG0 | Kcnh1 | potassium voltage-gated channel, subfamily H (eag-related), member 1 |
| chr1 | 195280192 | 195280563 | 372 | * | NA_peak_10 | 39 | Distal Interge | 1 | 195136811 | 195176716 | 39906 | 2 | 12902 | ENSMUST0 | -103476 | ENSMUSG0 | Cr2 | complement receptor 2 |
| chr10 | 3221952 | 3222256 | 305 | * | NA_peak_10 | 16 | Intron (ENSM | 10 | 3257930 | 3267750 | 9821 | 2 | 670558 | ENSMUST0 | 45494 | ENSMUSG0 | H60c | histocompatibility 60c |
| chr10 | 3366537 | 3366873 | 337 | * | NA_peak_10 | 18 | Promoter (<= | 10 | 3366544 | 3464975 | 98432 | 1 | 76142 | ENSMUST0 | 0 | ENSMUSG0 | Ppp1r14c | protein phosphatase 1, regulatory inhibitor subunit 14C |
| chr10 | 4552984 | 4553146 | 163 | * | NA_peak_11 | 19 | Intron (ENSM | 10 | 4532995 | 4541701 | 8707 | 1 | 100504234 | ENSMUST0 | 19989 | ENSMUSG0 | Ccdc170 | coiled-coil domain containing 170 |
| chr10 | 5881414 | 5881561 | 148 | * | NA_peak_11 | 26 | Intron (ENSM | 10 | 5832169 | 5922366 | 90198 | 2 | 56533 | ENSMUST0 | 40805 | ENSMUSG0 | Rgs17 | regulator of G-protein signaling 17 |
| chr10 | 6399516 | 6399975 | 460 | * | NA_peak_11 | 24 | Distal Interge | 10 | 6758506 | 6979704 | 221199 | 1 | 18390 | ENSMUST0 | -358531 | ENSMUSG0 | Oprm1 | opioid receptor, mu 1 |
| chr10 | 6400356 | 6400461 | 106 | * | NA_peak_11 | 24 | Distal Interge | 10 | 6758506 | 6979704 | 221199 | 1 | 18390 | ENSMUST0 | -358045 | ENSMUSG0 | Oprm1 | opioid receptor, mu 1 |
| chr10 | 7392292 | 7392375 | 84 | * | NA_peak_11 | 19 | Distal Interge | 10 | 7446183 | 7473341 | 27159 | 2 | 77777 | ENSMUST0 | 80966 | ENSMUSG0 | Ulbp1 | UL16 binding protein 1 |
| chr10 | 7403194 | 7403427 | 234 | * | NA_peak_11 | 15 | Distal Interge | 10 | 7446183 | 7473341 | 27159 | 2 | 77777 | ENSMUST0 | 69914 | ENSMUSG0 | Ulbp1 | UL16 binding protein 1 |
| chr10 | 7479217 | 7479579 | 363 | * | NA_peak_11 | 37 | Distal Interge | 10 | 7443038 | 7473638 | 30601 | 2 | 77777 | ENSMUST0 | -5579 | ENSMUSG0 | Ulbp1 | UL16 binding protein 1 |
| chr10 | 7726088 | 7726249 | 162 | * | NA_peak_11 | 39 | Promoter (<= | 10 | 7726008 | 7752749 | 26742 | 1 | 23924 | ENSMUST0 | 80 | ENSMUSG0 | Katna1 | katanin p60 (ATPase-containing) subunit A1 |
| chr10 | 7956403 | 7956748 | 346 | * | NA_peak_11 | 44 | Promoter (<= | 10 | 7905653 | 7956230 | 50578 | 2 | 68652 | ENSMUST0 | -173 | ENSMUSG0 | Tab2 | TGF-beta activated kinase 1/MAP3K7 binding protein 2 |
| chr10 | 8454370 | 8454623 | 254 | * | NA_peak_11 | 70 | Intron (ENSM | 10 | 8204809 | 8518825 | 314017 | 2 | 338362 | ENSMUST0 | 64202 | ENSMUSG0 | Ust | uronyl-2-sulfotransferase |
| chr10 | 8549367 | 8549749 | 383 | * | NA_peak_11 | 40 | Distal Interge | 10 | 8204809 | 8518825 | 314017 | 2 | 338362 | ENSMUST0 | -30542 | ENSMUSG0 | Ust | uronyl-2-sulfotransferase |
| chr10 | 10571784 | 10572127 | 344 | * | NA_peak_11 | 20 | Distal Interge | 10 | 10545002 | 10558265 | 13264 | 2 | 67844 | ENSMUST0 | -13519 | ENSMUSG0 | Rab32 | RAB32, member RAS oncogene family |
| chr10 | 11373248 | 11373871 | 624 | * | NA_peak_11 | 22 | Intron (ENSM | 10 | 11343453 | 11448698 | 105246 | 1 | 13853 | ENSMUST0 | 29795 | ENSMUSG0 | Epm2a | epilepsy, progressive myoclonic epilepsy, type 2 gene alpha |
| chr10 | 11375287 | 11375671 | 385 | * | NA_peak_11 | 20 | Intron (ENSM | 10 | 11343453 | 11448698 | 105246 | 1 | 13853 | ENSMUST0 | 31834 | ENSMUSG0 | Epm2a | epilepsy, progressive myoclonic epilepsy, type 2 gene alpha |
| chr10 | 12387579 | 12387874 | 296 | * | NA_peak_11 | 30 | Intron (ENSM | 10 | 12401066 | 12403075 | 2010 | 2 | 22288 | ENSMUST0 | 15201 | ENSMUSG0 | Utrn | utrophin |
| chr10 | 12854944 | 12855119 | 176 | * | NA_peak_11 | 37 | Intron (ENSM | 10 | 12382188 | 12861735 | 479548 | 2 | 22288 | ENSMUST0 | 6616 | ENSMUSG0 | Utrn | utrophin |
| chr10 | 13008367 | 13008584 | 218 | * | NA_peak_11 | 23 | 5' UTR | 10 | 13005343 | 13009183 | 3841 | 1 | 66125 | ENSMUST0 | 3024 | ENSMUSG0 | Sf3b5 | splicing factor 3b, subunit 5 |
| chr10 | 14579222 | 14579423 | 202 | * | NA_peak_11 | 54 | Distal Interge | 10 | 14407898 | 14545659 | 137762 | 2 | 215798 | ENSMUST0 | -33563 | ENSMUSG0 | Adgrg6 | adhesion G protein-coupled receptor G6 |
| chr10 | 15385475 | 15385859 | 385 | * | NA_peak_12 | 23 | Intron (ENSM | 10 | 15791777 | 15807065 | 15289 | 1 | 77538 | ENSMUST0 | -405918 | ENSMUSG0 | C330004P14 | RIKEN cDNA C330004P14 gene |
| chr10 | 15469216 | 15469497 | 282 | * | NA_peak_12 | 20 | Distal Interge | 10 | 15791777 | 15807065 | 15289 | 1 | 77538 | ENSMUST0 | -322280 | ENSMUSG0 | C330004P14 | RIKEN cDNA C330004P14 gene |
| chr10 | 16821425 | 16821826 | 402 | * | NA_peak_12 | 54 | Distal Interge | 10 | 16951295 | 16963180 | 11886 | 2 | 100504235 | ENSMUST0 | 141354 | ENSMUSG0 | Gm20125 | predicted gene, 20125 |
| chr10 | 16970847 | 16971263 | 417 | * | NA_peak_12 | 20 | Intron (ENSM | 10 | 16951293 | 16977148 | 25856 | 2 | 100504235 | ENSMUST0 | 5885 | ENSMUSG0 | Gm20125 | predicted gene, 20125 |
| chr10 | 18139252 | 18139504 | 253 | * | NA_peak_12 | 20 | Promoter (<= | 10 | 18128903 | 18139114 | 10212 | 2 | 100045792 | ENSMUST0 | -138 | ENSMUSG0 | Ect2l | epithelial cell transforming sequence 2 oncogene-like |
| chr10 | 18239781 | 18240289 | 509 | * | NA_peak_12 | 24 | Intron (ENSM | 10 | 18214078 | 18234998 | 20921 | 2 | 215814 | ENSMUST0 | -4783 | ENSMUSG0 | Ccdc28a | coiled-coil domain containing 28A |
| chr10 | 20570688 | 20570787 | 100 | * | NA_peak_12 | 16 | Intron (ENSM | 10 | 20407087 | 20548470 | 141384 | 2 | 29863 | ENSMUST0 | -22218 | ENSMUSG0 | Pde7b | phosphodiesterase 7B |
| chr10 | 21926399 | 21926697 | 299 | * | NA_peak_12 | 15 | Promoter (1- | 10 | 21928470 | 21990402 | 61933 | 1 | 20393 | ENSMUST0 | -1773 | ENSMUSG0 | Sgk1 | serum/glucocorticoid regulated kinase 1 |
| chr10 | 21994209 | 21994573 | 365 | * | NA_peak_12 | 40 | Promoter (<= | 10 | 21994666 | 21999903 | 5238 | 1 | 20393 | ENSMUST0 | -93 | ENSMUSG0 | Sgk1 | serum/glucocorticoid regulated kinase 1 |
| chr10 | 22174301 | 22174707 | 407 | * | NA_peak_12 | 70 | Promoter (<= | 10 | 22173919 | 22374139 | 200221 | 1 | 19370 | ENSMUST0 | 382 | ENSMUSG0 | Raet1c | retinoic acid early transcript gamma |
| chr10 | 22176099 | 22176416 | 318 | * | NA_peak_12 | 28 | Promoter (2- | 10 | 22173919 | 22374139 | 200221 | 1 | 19370 | ENSMUST0 | 2180 | ENSMUSG0 | Raet1c | retinoic acid early transcript gamma |
| chr10 | 22425616 | 22425749 | 134 | * | NA_peak_12 | 19 | Intron (ENSM | 10 | 22402883 | 22407470 | 4588 | 1 | 100038752 | ENSMUST0 | 22733 | ENSMUSG0 | Gm10825 | predicted gene 10825 |
| chr10 | 22550395 | 22550728 | 334 | * | NA_peak_12 | 26 | Distal Interge | 10 | 22645011 | 22704285 | 59275 | 1 | 353169 | ENSMUST0 | -94283 | ENSMUSG0 | Slc2a12 | solute carrier family 2 (facilitated glucose transporter), member 12 |
| chr10 | 22557850 | 22558323 | 474 | * | NA_peak_12 | 25 | Distal Interge | 10 | 22645011 | 22704285 | 59275 | 1 | 353169 | ENSMUST0 | -86688 | ENSMUSG0 | Slc2a12 | solute carrier family 2 (facilitated glucose transporter), member 12 |
| chr10 | 22755674 | 22755850 | 177 | * | NA_peak_12 | 23 | Distal Interge | 10 | 22705381 | 22731938 | 26558 | 2 | 237336 | ENSMUST0 | -23736 | ENSMUSG0 | Tbpl1 | TATA box binding protein-like 1 |
| chr10 | 24330496 | 24330760 | 265 | * | NA_peak_13 | 27 | Distal Interge | 10 | 24223545 | 24231199 | 7655 | 1 | 59012 | ENSMUST0 | 106951 | ENSMUSG0 | Moxd1 | monooxygenase, DBH-like 1 |
| chr10 | 24566642 | 24566800 | 159 | * | NA_peak_13 | 26 | Intron (ENSM | 10 | 24580048 | 24593758 | 13711 | 2 | 102638890 | ENSMUST0 | 26958 | NA | Gm15270 | predicted gene 15270 |
| chr10 | 25310335 | 25310680 | 346 | * | NA_peak_13 | 31 | Promoter (2- | 10 | 25284024 | 25307870 | 23847 | 2 | 432442 | ENSMUST0 | -2465 | ENSMUSG0 | Akap7 | A kinase (PRKA) anchor protein 7 |
| chr10 | 25969381 | 25969657 | 277 | * | NA_peak_13 | 20 | Intron (ENSM | 10 | 25993658 | 26017543 | 23886 | 2 | 77220 | ENSMUST0 | 47886 | ENSMUSG0 | Tmem200a | transmembrane protein 200A |
| chr10 | 26593120 | 26593147 | 28 | * | NA_peak_13 | 19 | Distal Interge | 10 | 26673601 | 26674612 | 1012 | 2 | 667572 | ENSMUST0 | 81465 | ENSMUSG0 | Gm8709 | glyceraldehyde-3-phosphate dehydrogenase pseudogene |
| chr10 | 26927199 | 26927391 | 193 | * | NA_peak_13 | 42 | Distal Interge | 10 | 26980036 | 26988514 | 8479 | 2 | 16773 | ENSMUST0 | 61123 | ENSMUSG0 | Lama2 | laminin, alpha 2 |
| chr10 | 27255386 | 27255746 | 361 | * | NA_peak_13 | 49 | Intron (ENSM | 10 | 27204801 | 27212254 | 7454 | 2 | 16773 | ENSMUST0 | -43132 | ENSMUSG0 | Lama2 | laminin, alpha 2 |
| chr10 | 27556827 | 27557221 | 395 | * | NA_peak_13 | 35 | Intron (ENSM | 10 | 27187548 | 27616796 | 429249 | 2 | 16773 | ENSMUST0 | 59575 | ENSMUSG0 | Lama2 | laminin, alpha 2 |
| chr10 | 28158190 | 28158214 | 25 | * | NA_peak_13 | 23 | Intron (ENSM | 10 | 28075010 | 28264673 | 189664 | 1 | 19272 | ENSMUST0 | 83180 | ENSMUSG0 | Ptprk | protein tyrosine phosphatase, receptor type, K |
| chr10 | 30564929 | 30564964 | 36 | * | NA_peak_14 | 19 | Intron (ENSM | 10 | 30559121 | 30590109 | 30989 | 2 | 73681 | ENSMUST0 | 25145 | ENSMUSG0 | Trmt11 | tRNA methyltransferase 11 |
| chr10 | 30750915 | 30751306 | 392 | * | NA_peak_14 | 20 | Intron (ENSM | 10 | 30647388 | 30771840 | 124453 | 2 | 211329 | ENSMUST0 | 20534 | ENSMUSG0 | Ncoa7 | nuclear receptor coactivator 7 |
| chr10 | 31081011 | 31081331 | 321 | * | NA_peak_14 | 25 | Distal Interge | 10 | 31313383 | 31328204 | 14822 | 1 | 69692 | ENSMUST0 | -232052 | ENSMUSG0 | Hddc2 | HD domain containing 2 |

| | | | | | | | | | | | | | | | | | | |
|---|---|---|---|---|---|---|---|---|---|---|---|---|---|---|---|---|---|---|
| chr10 | 31406550 | 31406856 | 307 | * | NA_peak_14 | 32 | Intron (ENSM | 10 | 31413815 | 31414165 | 351 | 2 | 100041294 | ENSMUST0( | 7309 | NA | Supt4b | SPT4B, DSIF elongation factor subunit |
| chr10 | 33641064 | 33641431 | 368 | * | NA_peak_14 | 16 | Intron (ENSM | 10 | 33512286 | 33624769 | 112484 | 2 | 215890 | ENSMUST0( | -16295 | ENSMUSG0( | Clvs2 | clavesin 2 |
| chr10 | 33776350 | 33776510 | 161 | * | NA_peak_14 | 29 | Intron (ENSM | 10 | 33766424 | 33782115 | 15692 | 2 | 215895 | ENSMUST0( | 5605 | ENSMUSG0( | Sult3a2 | sulfotransferase family 3A, member 2 |
| chr10 | 34134784 | 34135143 | 360 | * | NA_peak_14 | 24 | Distal Interge | 10 | 34126067 | 34127984 | 1918 | 2 | 215900 | ENSMUST0( | -6800 | ENSMUSG0( | Calhm6 | calcium homeostasis modulator family member 6 |
| chr10 | 34348186 | 34348391 | 206 | * | NA_peak_14 | 19 | Intron (ENSM | 10 | 34288288 | 34313683 | 25396 | 2 | 319638 | ENSMUST0( | -34503 | ENSMUSG0( | Nt5dc1 | 5'-nucleotidase domain containing 1 |
| chr10 | 37238637 | 37238938 | 302 | * | NA_peak_14 | 13 | Distal Interge | 10 | 37133375 | 37138920 | 5546 | 2 | 17118 | ENSMUST0( | -99717 | ENSMUSG0( | Marcks | myristoylated alanine rich protein kinase C substrate |
| chr10 | 38845674 | 38845913 | 240 | * | NA_peak_15 | 19 | Distal Interge | 10 | 38820541 | 38821779 | 1239 | 2 | 215919 | ENSMUST0( | -23895 | ENSMUSG0( | Rfpl4b | ret finger protein-like 4B |
| chr10 | 39261988 | 39262054 | 67 | * | NA_peak_15 | 17 | Distal Interge | 10 | 39150970 | 39163794 | 12825 | 2 | 327743 | ENSMUST0( | -98194 | ENSMUSG0( | Ccn6 | cellular communication network factor 6 |
| chr10 | 40263885 | 40264253 | 369 | * | NA_peak_15 | 23 | Distal Interge | 10 | 40258467 | 40258680 | 214 | 1 | 19850 | ENSMUST0( | 5418 | ENSMUSG0( | Snord3a | small nucleolar RNA, C/D box 3A |
| chr10 | 40593889 | 40594057 | 169 | * | NA_peak_15 | 24 | Intron (ENSM | 10 | 40570613 | 40604128 | 33516 | 1 | 70840 | ENSMUST0( | 23276 | ENSMUSG0( | Slc22a16 | solute carrier family 22 (organic cation transporter), member 16 |
| chr10 | 41779719 | 41779785 | 67 | * | NA_peak_15 | 17 | Intron (ENSM | 10 | 41718846 | 41809607 | 90762 | 2 | 103268 | ENSMUST0( | 29822 | ENSMUSG0( | Cep57l1 | centrosomal protein 57-like 1 |
| chr10 | 42275239 | 42275367 | 129 | * | NA_peak_15 | 32 | Promoter (1- | 10 | 42196268 | 42276696 | 80429 | 2 | 56484 | ENSMUST0( | 1329 | ENSMUSG0( | Foxo3 | forkhead box O3 |
| chr10 | 42501544 | 42502043 | 500 | * | NA_peak_15 | 26 | Promoter (<= | 10 | 42502030 | 42534787 | 32758 | 1 | 54198 | ENSMUST0( | 0 | ENSMUSG0( | Snx3 | sorting nexin 3 |
| chr10 | 42769895 | 42770007 | 113 | * | NA_peak_15 | 26 | Intron (ENSM | 10 | 42761806 | 42812742 | 50937 | 1 | 140740 | ENSMUST0( | 8089 | ENSMUSG0( | Sec63 | SEC63-like (S. cerevisiae) |
| chr10 | 42901378 | 42901436 | 59 | * | NA_peak_15 | 23 | Promoter (1- | 10 | 42902473 | 42935139 | 32667 | 1 | 268297 | ENSMUST0( | -1037 | ENSMUSG0( | Scml4 | Scm polycomb group protein like 4 |
| chr10 | 44123968 | 44124071 | 104 | * | NA_peak_16 | 19 | Intron (ENSM | 10 | 43950636 | 44148853 | 198218 | 2 | 11630 | ENSMUST0( | 24782 | ENSMUSG0( | Crybg1 | crystallin beta-gamma domain containing 1 |
| chr10 | 45518801 | 45518932 | 132 | * | NA_peak_16 | 44 | Distal Interge | 10 | 45418363 | 45486395 | 68033 | 2 | 380669 | ENSMUST0( | -32406 | ENSMUSG0( | Lin28b | lin-28 homolog B (C. elegans) |
| chr10 | 45567921 | 45568322 | 402 | * | NA_peak_16 | 20 | Distal Interge | 10 | 45577829 | 45712345 | 134517 | 1 | 209462 | ENSMUST0( | -9507 | ENSMUSG0( | Hace1 | HECT domain and ankyrin repeat containing, E3 ubiquitin protein ligase 1 |
| chr10 | 46584743 | 46584768 | 26 | * | NA_peak_16 | 22 | Distal Interge | 10 | 45837880 | 45852283 | 14404 | 1 | 320363 | ENSMUST0( | 746863 | ENSMUSG0( | D030045P18 | RIKEN cDNA D030045P18 gene |
| chr10 | 48731522 | 48731932 | 411 | * | NA_peak_16 | 46 | Distal Interge | 10 | 49101128 | 49103931 | 2804 | 2 | 14806 | ENSMUST0( | 371999 | ENSMUSG0( | Grik2 | glutamate receptor, ionotropic, kainate 2 (beta 2) |
| chr10 | 48787168 | 48787401 | 234 | * | NA_peak_16 | 23 | Distal Interge | 10 | 49101128 | 49103931 | 2804 | 2 | 14806 | ENSMUST0( | 316530 | ENSMUSG0( | Grik2 | glutamate receptor, ionotropic, kainate 2 (beta 2) |
| chr10 | 52109055 | 52109272 | 218 | * | NA_peak_16 | 22 | Intron (ENSM | 10 | 52045721 | 52194956 | 149236 | 2 | 19886 | ENSMUST0( | 85684 | ENSMUSG0( | Ros1 | Ros1 proto-oncogene |
| chr10 | 53250309 | 53250840 | 532 | * | NA_peak_16 | 53 | Intron (ENSM | 10 | 53235187 | 53260736 | 25550 | 1 | 102639992 | ENSMUST0( | 15122 | ENSMUSG0( | Gm36173 | predicted gene, 36173 |
| chr10 | 53572849 | 53573201 | 353 | * | NA_peak_16 | 32 | Exon (ENSM | 10 | 53536329 | 53593161 | 56833 | 2 | 71567 | ENSMUST0( | 19960 | ENSMUSG0( | Mcm9 | minichromosome maintenance 9 homologous recombination repair factor |
| chr10 | 56619755 | 56619797 | 43 | * | NA_peak_16 | 23 | Distal Interge | 10 | 56431981 | 56475967 | 43987 | 1 | 102640560 | ENSMUST0( | 187774 | ENSMUSG0( | Gm36595 | predicted gene, 36595 |
| chr10 | 56704070 | 56704326 | 257 | * | NA_peak_16 | 35 | Distal Interge | 10 | 56431981 | 56475967 | 43987 | 1 | 102640560 | ENSMUST0( | 272089 | ENSMUSG0( | Gm36595 | predicted gene, 36595 |
| chr10 | 58447192 | 58447503 | 312 | * | NA_peak_17 | 28 | Promoter (<= | 10 | 58447195 | 58460973 | 13779 | 1 | 19386 | ENSMUST0( | 0 | ENSMUSG0( | Ranbp2 | RAN binding protein 2 |
| chr10 | 58720734 | 58721418 | 685 | * | NA_peak_17 | 23 | Distal Interge | 10 | 58696745 | 58704277 | 7533 | 1 | 103505 | ENSMUST0( | 23989 | ENSMUSG0( | BB019430 | expressed sequence BB019430 |
| chr10 | 58780577 | 58780617 | 41 | * | NA_peak_17 | 29 | Distal Interge | 10 | 58813359 | 59050020 | 236662 | 1 | 237353 | ENSMUST0( | -32742 | ENSMUSG0( | Sh3rf3 | SH3 domain containing ring finger 3 |
| chr10 | 59616389 | 59616717 | 329 | * | NA_peak_17 | 16 | Promoter (<= | 10 | 59446984 | 59616692 | 169709 | 2 | 215999 | ENSMUST0( | 0 | ENSMUSG0( | Mcu | mitochondrial calcium uniporter |
| chr10 | 59646317 | 59646513 | 197 | * | NA_peak_17 | 26 | Distal Interge | 10 | 59446984 | 59616692 | 169709 | 2 | 215999 | ENSMUST0( | -29625 | ENSMUSG0( | Mcu | mitochondrial calcium uniporter |
| chr10 | 61018580 | 61018884 | 305 | * | NA_peak_17 | 17 | Distal Interge | 10 | 61031303 | 61038080 | 6778 | 1 | 327760 | ENSMUST0( | -12419 | ENSMUSG0( | Gm20611 | predicted gene 20611 |
| chr10 | 61408375 | 61408881 | 507 | * | NA_peak_17 | 16 | Distal Interge | 10 | 61417972 | 61425338 | 7367 | 1 | 18119 | ENSMUST0( | -9091 | ENSMUSG0( | Nodal | nodal |
| chr10 | 61507087 | 61507447 | 361 | * | NA_peak_17 | 19 | Intron (ENSM | 10 | 61475801 | 61582791 | 106991 | 1 | 216011 | ENSMUST0( | 31286 | ENSMUSG0( | Lrrc20 | leucine rich repeat containing 20 |
| chr10 | 62369049 | 62369408 | 360 | * | NA_peak_17 | 26 | Intron (ENSM | 10 | 62315476 | 62375100 | 59625 | 2 | 15275 | ENSMUST0( | 5692 | ENSMUSG0( | Hk1 | hexokinase 1 |
| chr10 | 62870497 | 62870633 | 137 | * | NA_peak_17 | 13 | Intron (ENSM | 10 | 62839097 | 62865304 | 26208 | 2 | 52463 | ENSMUST0( | -5193 | ENSMUSG0( | Tet1 | tet methylcytosine dioxygenase 1 |
| chr10 | 62885524 | 62885781 | 258 | * | NA_peak_17 | 16 | Promoter (1- | 10 | 62879873 | 62887324 | 7452 | 2 | 52463 | ENSMUST0( | 1543 | ENSMUSG0( | Tet1 | tet methylcytosine dioxygenase 1 |
| chr10 | 63214663 | 63214677 | 15 | * | NA_peak_18 | 22 | Distal Interge | 10 | 63115795 | 63203952 | 88158 | 2 | 68802 | ENSMUST0( | -10711 | ENSMUSG0( | Mypn | myopalladin |
| chr10 | 65659652 | 65660227 | 576 | * | NA_peak_18 | 26 | Distal Interge | 10 | 65875788 | 65908239 | 32452 | 2 | 102632897 | ENSMUST0( | 248012 | ENSMUSG0( | Gm30853 | predicted gene, 30853 |
| chr10 | 66523725 | 66524048 | 324 | * | NA_peak_18 | 22 | Distal Interge | 10 | 66120605 | 66254223 | 133619 | 2 | 69433 | ENSMUST0( | -269502 | NA | 1700023F02I | RIKEN cDNA 1700023F02 gene |
| chr10 | 66645561 | 66645712 | 152 | * | NA_peak_18 | 30 | Distal Interge | 10 | 66915636 | 66917739 | 2104 | 2 | 68488 | ENSMUST0( | 272027 | NA | 1110002J07f | RIKEN cDNA 1110002J07 gene |
| chr10 | 66673255 | 66673453 | 199 | * | NA_peak_18 | 13 | Distal Interge | 10 | 66915636 | 66917739 | 2104 | 2 | 68488 | ENSMUST0( | 244286 | NA | 1110002J07f | RIKEN cDNA 1110002J07 gene |
| chr10 | 66835142 | 66835853 | 712 | * | NA_peak_18 | 43 | Distal Interge | 10 | 66915636 | 66917739 | 2104 | 2 | 68488 | ENSMUST0( | 81886 | NA | 1110002J07f | RIKEN cDNA 1110002J07 gene |
| chr10 | 67830011 | 67830022 | 12 | * | NA_peak_18 | 23 | Intron (ENSM | 10 | 67889953 | 67899007 | 9055 | 2 | 216049 | ENSMUST0( | 68985 | ENSMUSG0( | Zfp365 | zinc finger protein 365 |
| chr10 | 67979498 | 67979938 | 441 | * | NA_peak_18 | 31 | Promoter (<= | 10 | 67979570 | 68043868 | 64299 | 1 | 170799 | ENSMUST0( | 0 | ENSMUSG0( | Rtkn2 | rhotekin 2 |
| chr10 | 69357123 | 69357454 | 332 | * | NA_peak_18 | 23 | Distal Interge | 10 | 69335146 | 69352938 | 17793 | 2 | 12534 | ENSMUST0( | -4185 | ENSMUSG0( | Cdk1 | cyclin-dependent kinase 1 |
| chr10 | 69626461 | 69626899 | 439 | * | NA_peak_18 | 26 | Intron (ENSM | 10 | 69706370 | 70015631 | 309262 | 1 | 11735 | ENSMUST0( | -79471 | ENSMUSG0( | Ank3 | ankyrin 3, epithelial |
| chr10 | 70884797 | 70885080 | 284 | * | NA_peak_18 | 43 | Intron (ENSM | 10 | 70934502 | 70948489 | 13988 | 2 | 83675 | ENSMUST0( | 63409 | ENSMUSG0( | Bicc1 | BicC family RNA binding protein 1 |
| chr10 | 71344593 | 71344850 | 258 | * | NA_peak_18 | 19 | Promoter (<= | 10 | 71330480 | 71344954 | 14475 | 2 | 52637 | ENSMUST0( | 104 | ENSMUSG0( | Cisd1 | CDGSH iron sulfur domain 1 |
| chr10 | 71560407 | 71560761 | 355 | * | NA_peak_18 | 13 | Intron (ENSM | 10 | 71347843 | 71363880 | 16038 | 1 | 69718 | ENSMUST0( | 212564 | ENSMUSG0( | Ipmk | inositol polyphosphate multikinase |
| chr10 | 72239791 | 72240086 | 296 | * | NA_peak_18 | 49 | Distal Interge | 10 | 71980092 | 71980634 | 543 | 2 | 108950 | ENSMUST0( | -259157 | NA | 1700049L16f | hematological and neurological expressed 1-like pseudogene |
| chr10 | 73580677 | 73580998 | 322 | * | NA_peak_18 | 50 | Intron (ENSM | 10 | 73821867 | 74634765 | 812899 | 1 | 11994 | ENSMUST0( | -240869 | ENSMUSG0( | Pcdh15 | protocadherin 15 |
| chr10 | 74769761 | 74770322 | 562 | * | NA_peak_18 | 49 | Distal Interge | 10 | 74631834 | 74642364 | 10531 | 1 | 11994 | ENSMUST0( | 137927 | ENSMUSG0( | Pcdh15 | protocadherin 15 |
| chr10 | 74878944 | 74879173 | 230 | * | NA_peak_18 | 23 | Distal Interge | 10 | 74967177 | 75016907 | 49731 | 1 | 14687 | ENSMUST0( | -88004 | ENSMUSG0( | Gnaz | guanine nucleotide binding protein, alpha z subunit |
| chr10 | 75032712 | 75032978 | 267 | * | NA_peak_18 | 18 | Promoter (<= | 10 | 75024513 | 75032586 | 8074 | 2 | 71236 | ENSMUST0( | -126 | ENSMUSG0( | Rsph14 | radial spoke head homolog 14 (Chlamydomonas) |
| chr10 | 75107350 | 75107725 | 376 | * | NA_peak_18 | 19 | Intron (ENSM | 10 | 75060592 | 75184921 | 124330 | 1 | 110279 | ENSMUST0( | 46758 | ENSMUSG0( | Bcr | BCR activator of RhoGEF and GTPase |
| chr10 | 75767850 | 75768001 | 152 | * | NA_peak_19 | 16 | Distal Interge | 10 | 75646112 | 75764316 | 118205 | 2 | 104248 | ENSMUST0( | -3534 | ENSMUSG0( | Cabin1 | calcineurin binding protein 1 |
| chr10 | 75833106 | 75833122 | 17 | * | NA_peak_19 | 17 | Promoter (<= | 10 | 75832059 | 75833136 | 1078 | 2 | 14872 | ENSMUST0( | 14 | ENSMUSG0( | Gstt2 | glutathione S-transferase, theta 2 |
| chr10 | 75865044 | 75865665 | 622 | * | NA_peak_19 | 54 | Distal Interge | 10 | 75859353 | 75860240 | 888 | 2 | 17319 | ENSMUST0( | -4804 | ENSMUSG0( | Mif | macrophage migration inhibitory factor (glycosylation-inhibiting factor) |
| chr10 | 76237904 | 76238267 | 364 | * | NA_peak_19 | 30 | Promoter (<= | 10 | 76207222 | 76237865 | 30644 | 2 | 15468 | ENSMUST0( | -39 | ENSMUSG0( | Prmt2 | protein arginine N-methyltransferase 2 |
| chr10 | 76277786 | 76277979 | 194 | * | NA_peak_19 | 20 | Intron (ENSM | 10 | 76264672 | 76270571 | 5900 | 2 | 64451 | ENSMUST0( | -7215 | ENSMUSG0( | Dip2a | disco interacting protein 2 homolog A |
| chr10 | 76463096 | 76463410 | 315 | * | NA_peak_19 | 17 | 3' UTR | 10 | 76468927 | 76515857 | 46931 | 1 | 54387 | ENSMUST0( | -5517 | ENSMUSG0( | Mcm3ap | minichromosome maintenance complex component 3 associated protein |
| chr10 | 76523658 | 76523910 | 253 | * | NA_peak_19 | 15 | Distal Interge | 10 | 76531588 | 76535568 | 3981 | 1 | 16987 | ENSMUST0( | -7678 | ENSMUSG0( | Lss | lanosterol synthase |
| chr10 | 76524973 | 76525308 | 336 | * | NA_peak_19 | 31 | Distal Interge | 10 | 76531588 | 76535568 | 3981 | 1 | 16987 | ENSMUST0( | -6280 | ENSMUSG0( | Lss | lanosterol synthase |
| chr10 | 77126108 | 77126374 | 267 | * | NA_peak_19 | 23 | Intron (ENSM | 10 | 77053194 | 77113946 | 60753 | 2 | 12822 | ENSMUST0( | -12162 | ENSMUSG0( | Col18a1 | collagen, type XVIII, alpha 1 |
| chr10 | 77581575 | 77582094 | 520 | * | NA_peak_19 | 20 | Promoter (<= | 10 | 77581720 | 77598732 | 17013 | 1 | 108705 | ENSMUST0( | 0 | ENSMUSG0( | Pttg1ip | pituitary tumor-transforming 1 interacting protein |
| chr10 | 78818338 | 78818674 | 337 | * | NA_peak_19 | 24 | Distal Interge | 10 | 78846951 | 78847913 | 963 | 2 | 258305 | ENSMUST0( | 29239 | ENSMUSG0( | Olfr1356 | olfactory receptor 1356 |
| chr10 | 78941145 | 78941925 | 781 | * | NA_peak_19 | 142 | Distal Interge | 10 | 78955207 | 78956139 | 933 | 1 | 18372 | ENSMUST0( | -13282 | ENSMUSG0( | Olfr8 | olfactory receptor 8 |
| chr10 | 79674904 | 79675494 | 591 | * | NA_peak_20 | 32 | Exon (ENSM | 10 | 79669913 | 79675448 | 5536 | 1 | 110012 | ENSMUST0( | 4991 | ENSMUSG0( | Tpgs1 | tubulin polyglutamylase complex subunit 1 |
| chr10 | 79988413 | 79988704 | 292 | * | NA_peak_20 | 40 | Promoter (<= | 10 | 79988584 | 79996062 | 7479 | 1 | 12798 | ENSMUST0( | 0 | ENSMUSG0( | Cnn2 | calponin 2 |
| chr10 | 80036101 | 80036444 | 344 | * | NA_peak_20 | 15 | Promoter (<= | 10 | 80035949 | 80037185 | 1237 | 2 | 66420 | ENSMUST0( | 741 | ENSMUSG0( | Polr2e | polymerase (RNA) II (DNA directed) polypeptide E |
| chr10 | 80053440 | 80054294 | 855 | * | NA_peak_20 | 25 | Promoter (<= | 10 | 80053488 | 80056439 | 2952 | 1 | 625249 | ENSMUST0( | 0 | ENSMUSG0( | Gpx4 | glutathione peroxidase 4 |
| chr10 | 80096369 | 80096676 | 308 | * | NA_peak_20 | 39 | Intron (ENSM | 10 | 80057771 | 80101187 | 43417 | 2 | 216161 | ENSMUST0( | 4511 | ENSMUSG0( | Sbno2 | strawberry notch 2 |
| chr10 | 80149688 | 80149975 | 288 | * | NA_peak_20 | 51 | Promoter (<= | 10 | 80149886 | 80153687 | 3802 | 1 | 59090 | ENSMUST0( | 0 | ENSMUSG0( | Midn | midnolin |
| chr10 | 80150835 | 80150944 | 110 | * | NA_peak_20 | 51 | Promoter (<= | 10 | 80151154 | 80154472 | 3319 | 1 | 59090 | ENSMUST0( | -210 | ENSMUSG0( | Midn | midnolin |
| chr10 | 80152290 | 80152542 | 253 | * | NA_peak_20 | 24 | Promoter (1- | 10 | 80151238 | 80153696 | 2459 | 1 | 59090 | ENSMUST0( | 1052 | ENSMUSG0( | Midn | midnolin |
| chr10 | 80167548 | 80168360 | 813 | * | NA_peak_20 | 43 | Promoter (<= | 10 | 80167778 | 80171655 | 3878 | 1 | 12696 | ENSMUST0( | 0 | ENSMUSG0( | Cirbp | cold inducible RNA binding protein |
| chr10 | 80329334 | 80329643 | 310 | * | NA_peak_20 | 27 | Promoter (<= | 10 | 80321283 | 80329453 | 8171 | 2 | 18551 | ENSMUST0( | 0 | ENSMUSG0( | Pcsk4 | proprotein convertase subtilisin/kexin type 4 |
| chr10 | 80341639 | 80341651 | 13 | * | NA_peak_20 | 28 | Exon (ENSM | 10 | 80343006 | 80345238 | 2233 | 2 | 66548 | ENSMUST0( | 3587 | ENSMUSG0( | Adamtsl5 | ADAMTS-like 5 |
| chr10 | 80399036 | 80399074 | 39 | * | NA_peak_20 | 27 | Promoter (<= | 10 | 80392839 | 80399422 | 6584 | 2 | 17192 | ENSMUST0( | 348 | ENSMUSG0( | Mbd3 | methyl-CpG binding domain protein 3 |
| chr10 | 80415207 | 80415582 | 376 | * | NA_peak_20 | 25 | Promoter (<= | 10 | 80412627 | 80415487 | 2861 | 2 | 21423 | ENSMUST0( | 0 | ENSMUSG0( | Tcf3 | transcription factor 3 |
| chr10 | 80508840 | 80508865 | 26 | * | NA_peak_20 | 26 | Exon (ENSM | 10 | 80494835 | 80517276 | 22442 | 1 | 246086 | ENSMUST0( | 14005 | ENSMUSG0( | Onecut3 | one cut domain, family member 3 |
| chr10 | 80568860 | 80569255 | 396 | * | NA_peak_20 | 30 | 3' UTR | 10 | 80540925 | 80561560 | 20636 | 2 | 66932 | ENSMUST0( | -7300 | ENSMUSG0( | Rexo1 | REX1, RNA exonuclease 1 |
| chr10 | 80577030 | 80577160 | 131 | * | NA_peak_20 | 17 | Promoter (<= | 10 | 80567124 | 80577321 | 10198 | 2 | 118445 | ENSMUST0( | 161 | ENSMUSG0( | Klf16 | Kruppel-like factor 16 |
| chr10 | 80590152 | 80590434 | 283 | * | NA_peak_20 | 26 | Promoter (<= | 10 | 80583654 | 80590341 | 6688 | 2 | 216169 | ENSMUST0( | 0 | ENSMUSG0( | Abhd17a | abhydrolase domain containing 17A |
| chr10 | 80650272 | 80650591 | 320 | * | NA_peak_20 | 17 | Promoter (2- | 10 | 80644131 | 80647710 | 3580 | 2 | 208198 | ENSMUST0( | -2562 | ENSMUSG0( | Btbd2 | BTB (POZ) domain containing 2 |
| chr10 | 80732617 | 80732990 | 374 | * | NA_peak_20 | 15 | Exon (ENSM | 10 | 80722913 | 80741971 | 19059 | 2 | 11776 | ENSMUST0( | 8981 | ENSMUSG0( | Ap3d1 | adaptor-related protein complex 3, delta 1 subunit |
| chr10 | 80736711 | 80737107 | 397 | * | NA_peak_20 | 29 | Intron (ENSM | 10 | 80722913 | 80741971 | 19059 | 2 | 11776 | ENSMUST0( | 4864 | ENSMUSG0( | Ap3d1 | adaptor-related protein complex 3, delta 1 subunit |
| chr10 | 80750343 | 80750378 | 36 | * | NA_peak_20 | 30 | Distal Interge | 10 | 80755206 | 80795461 | 40256 | 1 | 208266 | ENSMUST0( | -4828 | ENSMUSG0( | Dot1l | DOT1-like, histone H3 methyltransferase (S. cerevisiae) |
| chr10 | 80754882 | 80755121 | 240 | * | NA_peak_20 | 31 | Promoter (<= | 10 | 80755206 | 80795461 | 40256 | 1 | 208266 | ENSMUST0( | -85 | ENSMUSG0( | Dot1l | DOT1-like, histone H3 methyltransferase (S. cerevisiae) |
| chr10 | 80825628 | 80826117 | 490 | * | NA_peak_20 | 26 | Promoter (<= | 10 | 80826656 | 80829290 | 2635 | 1 | 18245 | ENSMUST0( | -539 | ENSMUSG0( | Oaz1 | ornithine decarboxylase antizyme 1 |
| chr10 | 80928615 | 80928963 | 349 | * | NA_peak_20 | 39 | Promoter (1- | 10 | 80930073 | 80932204 | 2132 | 1 | 17873 | ENSMUST0( | -1110 | ENSMUSG0( | Gadd45b | growth arrest and DNA-damage-inducible 45 beta |
| chr10 | 81069791 | 81070259 | 469 | * | NA_peak_20 | 55 | Promoter (<= | 10 | 81070035 | 81082559 | 12525 | 1 | 50492 | ENSMUST0( | 0 | ENSMUSG0( | Thop1 | thimet oligopeptidase 1 |
| chr10 | 81147159 | 81147303 | 145 | * | NA_peak_20 | 25 | Intron (ENSM | 10 | 81154235 | 81156124 | 1890 | 2 | 59004 | ENSMUST0( | 8821 | ENSMUSG0( | Pias4 | protein inhibitor of activated STAT 4 |
| chr10 | 81281387 | 81281733 | 347 | * | NA_peak_21 | 26 | Promoter (1- | 10 | 81278624 | 81280117 | 1494 | 2 | 27375 | ENSMUST0( | -1270 | ENSMUSG0( | Tjp3 | tight junction protein 3 |
| chr10 | 81343023 | 81343489 | 467 | * | NA_peak_21 | 30 | Promoter (<= | 10 | 81335265 | 81343266 | 8002 | 2 | 209047 | ENSMUST0( | 0 | ENSMUSG0( | Gipc3 | GIPC PDZ domain containing family, member 3 |
| chr10 | 81350363 | 81350738 | 376 | * | NA_peak_21 | 44 | Promoter (<= | 10 | 81346048 | 81350480 | 4433 | 2 | 15353 | ENSMUST0( | 0 | ENSMUSG0( | Hmg20b | high mobility group 20B |
| chr10 | 81357283 | 81357404 | 122 | * | NA_peak_21 | 24 | Promoter (<= | 10 | 81357491 | 81366225 | 8735 | 1 | 73822 | ENSMUST0( | -87 | ENSMUSG0( | Mfsd12 | major facilitator superfamily domain containing 12 |
| chr10 | 82178302 | 82178484 | 183 | * | NA_peak_21 | 50 | Exon (ENSM | 10 | 82138253 | 82152011 | 13759 | 1 | 216177 | ENSMUST0( | 40049 | ENSMUSG0( | AU041133 | expressed sequence AU041133 |
| chr10 | 83516472 | 83517174 | 703 | * | NA_peak_21 | 23 | Promoter (2- | 10 | 83503992 | 83513720 | 9729 | 2 | 216188 | ENSMUST0( | -2752 | ENSMUSG0( | Aldh1l2 | aldehyde dehydrogenase 1 family, member L2 |
| chr10 | 85857009 | 85857172 | 164 | * | NA_peak_21 | 15 | Intron (ENSM | 10 | 85871831 | 85889096 | 17266 | 1 | 103136 | ENSMUST0( | -14659 | ENSMUSG0( | Pwp1 | PWP1 homolog, endonuclein |
| chr10 | 85957547 | 85957863 | 317 | * | NA_peak_21 | 39 | Promoter (<= | 10 | 85938637 | 85957823 | 19187 | 2 | 28088 | ENSMUST0( | 0 | ENSMUSG0( | Rtcb | RNA 2',3'-cyclic phosphate and 5'-OH ligase |
| chr10 | 86238129 | 86238576 | 448 | * | NA_peak_21 | 38 | Intron (ENSM | 10 | 86300372 | 86349506 | 49135 | 1 | 21859 | ENSMUST0( | -61796 | ENSMUSG0( | Timp3 | tissue inhibitor of metalloproteinase 3 |
| chr10 | 86676356 | 86676676 | 321 | * | NA_peak_21 | 30 | Distal Interge | 10 | 86685525 | 86689954 | 4430 | 1 | 66263 | ENSMUST0( | -8849 | ENSMUSG0( | 1810014B01I | RIKEN cDNA 1810014B01 gene |
| chr10 | 90328198 | 90328528 | 331 | * | NA_peak_21 | 44 | Intron (ENSM | 10 | 90575727 | 90973300 | 397574 | 1 | 77531 | ENSMUST0( | -247199 | ENSMUSG0( | Anks1b | ankyrin repeat and sterile alpha motif domain containing 1B |
| chr10 | 91123202 | 91123758 | 557 | * | NA_peak_21 | 39 | Promoter (<= | 10 | 91116808 | 91123955 | 7148 | 2 | 18674 | ENSMUST0( | 197 | ENSMUSG0( | Slc25a3 | solute carrier family 25 (mitochondrial carrier, phosphate carrier), member 3 |
| chr10 | 91256656 | 91257052 | 397 | * | NA_peak_22 | 24 | Intron (ENSM | 10 | 91208258 | 91220642 | 12385 | 2 | 102633753 | ENSMUST0( | -36014 | ENSMUSG0( | 4930534H03 | RIKEN cDNA 4930534H03 gene |
| chr10 | 91337913 | 91338346 | 434 | * | NA_peak_22 | 26 | Distal Interge | 10 | 91208258 | 91220642 | 12385 | 2 | 102633753 | ENSMUST0( | -117271 | ENSMUSG0( | 4930534H03 | RIKEN cDNA 4930534H03 gene |
| chr10 | 92807356 | 92807708 | 353 | * | NA_peak_22 | 23 | Intron (ENSM | 10 | 92796753 | 92836346 | 39594 | 2 | 380654 | ENSMUST0( | 28638 | ENSMUSG0( | Cfap54 | cilia and flagella associated protein 54 |
| chr10 | 94198917 | 94199262 | 346 | * | NA_peak_22 | 38 | Promoter (<= | 10 | 94198955 | 94221443 | 22489 | 1 | 66414 | ENSMUST0( | 0 | ENSMUSG0( | Ndufa12 | NADH:ubiquinone oxidoreductase subunit A12 |
| chr10 | 98547052 | 98547086 | 35 | * | NA_peak_22 | 16 | Intron (ENSM | 10 | 98299927 | 98351874 | 51948 | 2 | 667438 | ENSMUST0( | -195178 | ENSMUSG0( | Gm8633 | predicted gene 8633 |
| chr10 | 99723876 | 99724217 | 342 | * | NA_peak_22 | 26 | Exon (ENSM | 10 | 99759506 | 99830562 | 71057 | 1 | 102638712 | ENSMUST0( | -35289 | ENSMUSG0( | Gm35206 | predicted gene, 35206 |
| chr10 | 106082830 | 106083209 | 380 | * | NA_peak_22 | 19 | Distal Interge | 10 | 105841504 | 105842982 | 1479 | 1 | 52713 | ENSMUST0( | 241326 | ENSMUSG0( | Ccdc59 | coiled-coil domain containing 59 |
| chr10 | 108161930 | 108162150 | 221 | * | NA_peak_22 | 21 | Promoter (<= | 10 | 108162193 | 108277351 | 115159 | 1 | 17931 | ENSMUST0( | -43 | ENSMUSG0( | Ppp1r12a | protein phosphatase 1, regulatory subunit 12A |
| chr10 | 109412383 | 109412591 | 209 | * | NA_peak_22 | 35 | Distal Interge | 10 | 109714302 | 109723801 | 9500 | 2 | 260315 | ENSMUST0( | 311210 | ENSMUSG0( | Nav3 | neuron navigator 3 |
| chr10 | 114536996 | 114537435 | 440 | * | NA_peak_22 | 44 | Intron (ENSM | 10 | 114398823 | 114801364 | 402542 | 2 | 237553 | ENSMUST0( | 263929 | ENSMUSG0( | Trhde | TRH-degrading enzyme |
| chr10 | 117195074 | 117195433 | 360 | * | NA_peak_23 | 18 | Distal Interge | 10 | 117215428 | 117217358 | 1931 | 2 | 64050 | ENSMUST0( | 21925 | ENSMUSG0( | Yeats4 | YEATS domain containing 4 |
| chr10 | 118333977 | 118334381 | 405 | * | NA_peak_23 | 238 | Distal Interge | 10 | 118289629 | 118295038 | 5410 | 2 | 116849 | ENSMUST0( | -38939 | ENSMUSG0( | Il22b | interleukin 22B |
| chr10 | 118695764 | 118696053 | 290 | * | NA_peak_23 | 43 | Distal Interge | 10 | 118714481 | 118715726 | 1246 | 2 | 102636202 | ENSMUST0( | 19673 | ENSMUSG0( | Gm33337 | predicted gene, 33337 |
| chr10 | 119287213 | 119287460 | 248 | * | NA_peak_23 | 39 | Distal Interge | 10 | 119320234 | 119331862 | 11629 | 1 | 74905 | ENSMUST0( | -32774 | ENSMUSG0( | 4930477N07 | RIKEN cDNA 4930477N07 gene |
| chr10 | 119768475 | 119768794 | 320 | * | NA_peak_23 | 34 | Intron (ENSM | 10 | 119746063 | 119761699 | 15637 | 2 | 75321 | ENSMUST0( | -6776 | ENSMUSG0( | Grip1os2 | glutamate receptor interacting protein 1, opposite strand 2 |
| chr10 | 122391977 | 122392317 | 341 | * | NA_peak_23 | 13 | Intron (ENSM | 10 | 122448499 | 122453452 | 4954 | 1 | 54140 | ENSMUST0( | -56182 | ENSMUSG0( | Avpr1a | arginine vasopressin receptor 1A |
| chr10 | 122393292 | 122393501 | 210 | * | NA_peak_23 | 22 | Intron (ENSM | 10 | 122448499 | 122453452 | 4954 | 1 | 54140 | ENSMUST0( | -54998 | ENSMUSG0( | Avpr1a | arginine vasopressin receptor 1A |
| chr10 | 122837144 | 122837342 | 199 | * | NA_peak_23 | 13 | Intron (ENSM | 10 | 122811447 | 122886293 | 74847 | 1 | 319468 | ENSMUST0( | 25697 | ENSMUSG0( | Ppm1h | protein phosphatase 1H (PP2C domain containing) |
| chr10 | 122928279 | 122928628 | 350 | * | NA_peak_23 | 17 | Intron (ENSM | 10 | 122895368 | 122941449 | 46082 | 1 | 319468 | ENSMUST0( | 32911 | ENSMUSG0( | Ppm1h | protein phosphatase 1H (PP2C domain containing) |
| chr10 | 125822437 | 125822784 | 348 | * | NA_peak_23 | 27 | Distal Interge | 10 | 125966219 | 125998013 | 31795 | 1 | 320398 | ENSMUST0( | -143435 | ENSMUSG0( | Lrig3 | leucine-rich repeats and immunoglobulin-like domains 3 |
| chr10 | 126896782 | 126897129 | 348 | * | NA_peak_23 | 23 | Intron (ENSM | 10 | 126887249 | 126901348 | 14100 | 2 | 68876 | ENSMUST0( | 4219 | ENSMUSG0( | Atp23 | ATP23 metallopeptidase and ATP synthase assembly factor homolog |
| chr10 | 126928233 | 126928588 | 356 | * | NA_peak_23 | 26 | Distal Interge | 10 | 126868427 | 126901355 | 32929 | 2 | 68876 | ENSMUST0( | -26878 | ENSMUSG0( | Atp23 | ATP23 metallopeptidase and ATP synthase assembly factor homolog |
| chr10 | 127165225 | 127165514 | 290 | * | NA_peak_23 | 18 | Promoter (<= | 10 | 127165225 | 127172330 | 7106 | 1 | 14421 | ENSMUST0( | 0 | ENSMUSG0( | B4galnt1 | beta-1,4-N-acetyl-galactosaminyl transferase 1 |
| chr10 | 127178563 | 127178894 | 332 | * | NA_peak_23 | 22 | Promoter (1- | 10 | 127174785 | 127176989 | 2205 | 2 | 216441 | ENSMUST0( | -1574 | ENSMUSG0( | Slc26a10 | solute carrier family 26, member 10 |
| chr10 | 127459190 | 127459438 | 249 | * | NA_peak_23 | 22 | Promoter (<= | 10 | 127459465 | 127465182 | 5718 | 1 | 71750 | ENSMUST0( | -27 | ENSMUSG0( | R3hdm2 | R3H domain containing 2 |
| chr10 | 127676943 | 127677254 | 312 | * | NA_peak_23 | 13 | Promoter (<= | 10 | 127677064 | 127695168 | 18105 | 1 | 210035 | ENSMUST0( | 0 | ENSMUSG0( | Nemp1 | nuclear envelope integral membrane protein 1 |
| chr10 | 128056180 | 128056493 | 314 | * | NA_peak_23 | 26 | Promoter (2- | 10 | 128058954 | 128077272 | 18319 | 1 | 56351 | ENSMUST0( | -2461 | ENSMUSG0( | Ptges3 | prostaglandin E synthase 3 |
| chr10 | 128058739 | 128059329 | 591 | * | NA_peak_23 | 26 | Promoter (<= | 10 | 128058954 | 128077272 | 18319 | 1 | 56351 | ENSMUST0( | 0 | ENSMUSG0( | Ptges3 | prostaglandin E synthase 3 |
| chr10 | 128188209 | 128188585 | 377 | * | NA_peak_24 | 26 | Distal Interge | 10 | 128194457 | 128209939 | 15483 | 1 | 216456 | ENSMUST0( | -5872 | ENSMUSG0( | Gls2 | glutaminase 2 (liver, mitochondrial) |
| chr10 | 128232071 | 128232408 | 338 | * | NA_peak_24 | 24 | Promoter (<= | 10 | 128232080 | 128252932 | 20853 | 1 | 21853 | ENSMUST0( | 0 | ENSMUSG0( | Timeless | timeless circadian clock 1 |
| chr10 | 128242340 | 128242474 | 135 | * | NA_peak_24 | 14 | Promoter (<= | 10 | 128242408 | 128243930 | 1523 | 1 | 21853 | ENSMUST0( | 0 | ENSMUSG0( | Timeless | timeless circadian clock 1 |
| chr10 | 128337784 | 128338054 | 271 | * | NA_peak_24 | 32 | Promoter (<= | 10 | 128337734 | 128362479 | 24746 | 1 | 12974 | ENSMUST0( | 50 | ENSMUSG0( | Cs | citrate synthase |
| chr10 | 128909727 | 128910144 | 418 | * | NA_peak_24 | 18 | Promoter (<= | 10 | 128909699 | 128912815 | 3117 | 1 | 12512 | ENSMUST0( | 28 | ENSMUSG0( | Cd63 | CD63 antigen |
| chr10 | 129809017 | 129809083 | 67 | * | NA_peak_24 | 19 | Distal Interge | 10 | 129801564 | 129802523 | 960 | 2 | 258545 | ENSMUST0( | -6494 | ENSMUSG0( | Olfr811 | olfactory receptor 811 |
| chr10 | 130223345 | 130223760 | 416 | * | NA_peak_24 | 23 | Distal Interge | 10 | 130210092 | 130211149 | 1058 | 2 | 258297 | ENSMUST0( | -12196 | ENSMUSG0( | Olfr827 | olfactory receptor 827 |
| chr11 | 3100225 | 3100540 | 316 | * | NA_peak_24 | 24 | Distal Interge | 11 | 3125904 | 3129396 | 3493 | 1 | 236604 | ENSMUST0( | -25364 | NA | Pisd-ps1 | phosphatidylserine decarboxylase, pseudogene 1 |
| chr11 | 3102001 | 3102379 | 379 | * | NA_peak_24 | 13 | Distal Interge | 11 | 3125904 | 3129396 | 3493 | 1 | 236604 | ENSMUST0( | -23525 | NA | Pisd-ps1 | phosphatidylserine decarboxylase, pseudogene 1 |
| chr11 | 3112299 | 3112683 | 385 | * | NA_peak_24 | 21 | Distal Interge | 11 | 3125904 | 3129396 | 3493 | 1 | 236604 | ENSMUST0( | -13221 | NA | Pisd-ps1 | phosphatidylserine decarboxylase, pseudogene 1 |
| chr11 | 4579995 | 4580031 | 37 | * | NA_peak_24 | 23 | Intron (ENSM | 11 | 4480868 | 4594815 | 113948 | 2 | 74302 | ENSMUST0( | 14784 | ENSMUSG0( | Mtmr3 | myotubularin related protein 3 |
| chr11 | 5852288 | 5852617 | 330 | * | NA_peak_25 | 34 | Distal Interge | 11 | 5861947 | 5872088 | 10142 | 1 | 11568 | ENSMUST0( | -9330 | ENSMUSG0( | Aebp1 | AE binding protein 1 |
| chr11 | 7923633 | 7924365 | 733 | * | NA_peak_25 | 20 | Distal Interge | 11 | 8451070 | 8468675 | 17606 | 2 | 319939 | ENSMUST0( | 544310 | ENSMUSG0( | Tns3 | tensin 3 |
| chr11 | 20436059 | 20436458 | 400 | * | NA_peak_25 | 35 | Distal Interge | 11 | 20302180 | 20332713 | 30534 | 2 | 55963 | ENSMUST0( | -103346 | ENSMUSG0( | Slc1a4 | solute carrier family 1 (glutamate/neutral amino acid transporter), member 4 |
| chr11 | 20542576 | 20542849 | 274 | * | NA_peak_25 | 32 | Promoter (<= | 11 | 20543253 | 20597020 | 53768 | 1 | 58172 | ENSMUST0( | -404 | ENSMUSG0( | Sertad2 | SERTA domain containing 2 |
| chr11 | 20830646 | 20831127 | 482 | * | NA_peak_25 | 26 | Promoter (<= | 11 | 20823576 | 20831056 | 7481 | 2 | 216551 | ENSMUST0( | 0 | ENSMUSG0( | Lgalsl | lectin, galactoside binding-like |
| chr11 | 21128697 | 21128933 | 237 | * | NA_peak_25 | 13 | Intron (ENSM | 11 | 21135469 | 21150323 | 14855 | 1 | 67245 | ENSMUST0( | -6536 | ENSMUSG0( | Peli1 | pellino 1 |
| chr11 | 23223135 | 23223275 | 141 | * | NA_peak_26 | 35 | Distal Interge | 11 | 23256041 | 23297597 | 41557 | 1 | 103573 | ENSMUST0( | -32766 | ENSMUSG0( | Xpo1 | exportin 1 |
| chr11 | 23255467 | 23255876 | 410 | * | NA_peak_26 | 22 | Promoter (<= | 11 | 23256041 | 23297597 | 41557 | 1 | 103573 | ENSMUST0( | -165 | ENSMUSG0( | Xpo1 | exportin 1 |
| chr11 | 23771021 | 23771219 | 199 | * | NA_peak_26 | 23 | Promoter (<= | 11 | 23736847 | 23770970 | 34124 | 2 | 19696 | ENSMUST0( | -51 | ENSMUSG0( | Rel | reticuloendotheliosis oncogene |
| chr11 | 23839936 | 23840335 | 400 | * | NA_peak_26 | 26 | Distal Interge | 11 | 23865244 | 23868470 | 3227 | 2 | 216578 | ENSMUST0( | 28135 | ENSMUSG0( | Papolg | poly(A) polymerase gamma |
| chr11 | 25201765 | 25202172 | 408 | * | NA_peak_26 | 35 | Distal Interge | 11 | 25326107 | 25367137 | 41031 | 1 | 71234 | ENSMUST0( | -123935 | ENSMUSG0( | 4933427E13I | RIKEN cDNA 4933427E13 gene |
| chr11 | 27687179 | 27687475 | 297 | * | NA_peak_26 | 26 | Distal Interge | 11 | 28388817 | 28434110 | 45294 | 2 | 216613 | ENSMUST0( | 746635 | ENSMUSG0( | Ccdc85a | coiled-coil domain containing 85A |
| chr11 | 29693021 | 29693334 | 314 | * | NA_peak_26 | 13 | Promoter (<= | 11 | 29692948 | 29742111 | 49164 | 1 | 68585 | ENSMUST0( | 73 | ENSMUSG0( | Rtn4 | reticulon 4 |
| chr11 | 32221902 | 32222037 | 136 | * | NA_peak_26 | 19 | Promoter (<= | 11 | 32215068 | 32222244 | 7177 | 2 | 13650 | ENSMUST0( | 207 | ENSMUSG0( | Rhbdf1 | rhomboid 5 homolog 1 |
| chr11 | 32418135 | 32418443 | 309 | * | NA_peak_26 | 23 | Intron (ENSM | 11 | 32455370 | 32516687 | 61318 | 1 | 327900 | ENSMUST0( | -36927 | ENSMUSG0( | Ubtd2 | ubiquitin domain containing 2 |
| chr11 | 32454884 | 32455288 | 405 | * | NA_peak_26 | 20 | Promoter (<= | 11 | 32455370 | 32516687 | 61318 | 1 | 327900 | ENSMUST0( | -82 | ENSMUSG0( | Ubtd2 | ubiquitin domain containing 2 |
| chr11 | 32611797 | 32612108 | 312 | * | NA_peak_26 | 13 | Intron (ENSM | 11 | 32588595 | 32598528 | 9934 | 1 | 20868 | ENSMUST0( | 23202 | ENSMUSG0( | Stk10 | serine/threonine kinase 10 |
| chr11 | 33147398 | 33147553 | 156 | * | NA_peak_26 | 22 | Promoter (<= | 11 | 33117430 | 33147400 | 29971 | 2 | 14172 | ENSMUST0( | 0 | ENSMUSG0( | Fgf18 | fibroblast growth factor 18 |
| chr11 | 33200889 | 33201114 | 226 | * | NA_peak_26 | 34 | Promoter (2- | 11 | 33200752 | 33203589 | 2838 | 2 | 27140 | ENSMUST0( | 2475 | ENSMUSG0( | Tlx3 | T cell leukemia, homeobox 3 |
| chr11 | 35793988 | 35794241 | 254 | * | NA_peak_26 | 35 | Distal Interge | 11 | 35797382 | 35798884 | 1503 | 2 | 237730 | ENSMUST0( | 4643 | ENSMUSG0( | Fbll1 | fibrillarin-like 1 |
| chr11 | 40235269 | 40235564 | 296 | * | NA_peak_26 | 35 | Intron (ENSM | 11 | 40680710 | 40685227 | 4518 | 2 | 108645 | ENSMUST0( | 449663 | ENSMUSG0( | Mat2b | methionine adenosyltransferase II, beta |
| chr11 | 42685203 | 42685555 | 353 | * | NA_peak_26 | 35 | Distal Interge | 11 | 42421350 | 42626891 | 205542 | 1 | 14401 | ENSMUST0( | 263853 | ENSMUSG0( | Gabrb2 | gamma-aminobutyric acid (GABA) A receptor, subunit beta 2 |
| chr11 | 44360959 | 44361166 | 208 | * | NA_peak_26 | 18 | Distal Interge | 11 | 44400063 | 44414010 | 13948 | 1 | 16160 | ENSMUST0( | -38897 | ENSMUSG0( | Il12b | interleukin 12b |
| chr11 | 46012735 | 46012866 | 132 | * | NA_peak_26 | 39 | Intron (ENSM | 11 | 45980310 | 46017994 | 37685 | 1 | 214105 | ENSMUST0( | 32425 | ENSMUSG0( | Sox30 | SRY (sex determining region Y)-box 30 |
| chr11 | 48319270 | 48319557 | 288 | * | NA_peak_27 | 19 | Distal Interge | 11 | 48800332 | 48804974 | 4643 | 1 | 14694 | ENSMUST0( | -480775 | ENSMUSG0( | Rack1 | receptor for activated C kinase 1 |
| chr11 | 48468721 | 48469033 | 313 | * | NA_peak_27 | 35 | Distal Interge | 11 | 48800332 | 48804974 | 4643 | 1 | 14694 | ENSMUST0( | -331299 | ENSMUSG0( | Rack1 | receptor for activated C kinase 1 |
| chr11 | 48826224 | 48826591 | 368 | * | NA_peak_27 | 30 | Promoter (<= | 11 | 48826140 | 48851187 | 25048 | 1 | 94089 | ENSMUST0( | 84 | ENSMUSG0( | Trim7 | tripartite motif-containing 7 |
| chr11 | 49367636 | 49367920 | 285 | * | NA_peak_27 | 24 | Distal Interge | 11 | 49340534 | 49341469 | 936 | 1 | 259068 | ENSMUST0( | 27102 | ENSMUSG0( | Olfr1390 | olfactory receptor 1390 |
| chr11 | 50210694 | 50211099 | 406 | * | NA_peak_27 | 23 | Promoter (<= | 11 | 50210890 | 50231099 | 20210 | 1 | 103534 | ENSMUST0( | 0 | ENSMUSG0( | Mgat4b | mannoside acetylglucosaminyltransferase 4, isoenzyme B |
| chr11 | 51756476 | 51756705 | 230 | * | NA_peak_27 | 26 | Promoter (<= | 11 | 51692264 | 51756618 | 64355 | 2 | 77371 | ENSMUST0( | 0 | ENSMUSG0( | Sec24a | Sec24 related gene family, member A (S. cerevisiae) |
| chr11 | 51992032 | 51992377 | 346 | * | NA_peak_27 | 26 | Intron (ENSM | 11 | 51997570 | 52000479 | 2910 | 2 | 22210 | ENSMUST0( | 8102 | ENSMUSG0( | Ube2b | ubiquitin-conjugating enzyme E2B |
| chr11 | 53927056 | 53927430 | 375 | * | NA_peak_27 | 36 | Distal Interge | 11 | 53950563 | 53953146 | 2584 | 2 | 56517 | ENSMUST0( | 25716 | ENSMUSG0( | Slc22a21 | solute carrier family 22 (organic cation transporter), member 21 |
| chr11 | 53984857 | 53985252 | 396 | * | NA_peak_27 | 16 | Intron (ENSM | 11 | 53949965 | 53980332 | 30368 | 2 | 56517 | ENSMUST0( | -4525 | ENSMUSG0( | Slc22a21 | solute carrier family 22 (organic cation transporter), member 21 |
| chr11 | 54530545 | 54530704 | 160 | * | NA_peak_27 | 19 | Intron (ENSM | 11 | 54522946 | 54696344 | 173399 | 1 | 192786 | ENSMUST0( | 7599 | ENSMUSG0( | Rapgef6 | Rap guanine nucleotide exchange factor (GEF) 6 |
| chr11 | 55173962 | 55174166 | 205 | * | NA_peak_27 | 23 | Intron (ENSM | 11 | 55158470 | 55185077 | 26608 | 2 | 246049 | ENSMUST0( | 10911 | ENSMUSG0( | Slc36a2 | solute carrier family 36 (proton/amino acid symporter), member 2 |
| chr11 | 56182977 | 56183343 | 367 | * | NA_peak_27 | 54 | Distal Interge | 11 | 56024987 | 56041010 | 16024 | 2 | 216749 | ENSMUST0( | -141967 | ENSMUSG0( | Nmur2 | neuromedin U receptor 2 |
| chr11 | 56847626 | 56847707 | 82 | * | NA_peak_27 | 24 | Distal Interge | 11 | 57011387 | 57015291 | 3905 | 1 | 14799 | ENSMUST0( | -163680 | ENSMUSG0( | Gria1 | glutamate receptor, ionotropic, AMPA1 (alpha 1) |
| chr11 | 59006022 | 59006439 | 418 | * | NA_peak_28 | 44 | Exon (ENSM | 11 | 59001914 | 59002361 | 448 | 2 | 380698 | ENSMUST0( | -3661 | ENSMUSG0( | Obscn | obscurin, cytoskeletal calmodulin and titin-interacting RhoGEF |
| chr11 | 59454844 | 59455103 | 260 | * | NA_peak_28 | 29 | 3' UTR | 11 | 59438449 | 59451186 | 12738 | 2 | 67826 | ENSMUST0( | -3658 | ENSMUSG0( | Snap47 | synaptosomal-associated protein, 47 |
| chr11 | 59564661 | 59564843 | 183 | * | NA_peak_28 | 23 | Intron (ENSM | 11 | 59570699 | 59571813 | 1115 | 2 | 257962 | ENSMUST0( | 6970 | ENSMUSG0( | Olfr222 | olfactory receptor 222 |
| chr11 | 60186556 | 60186891 | 336 | * | NA_peak_28 | 28 | Exon (ENSM | 11 | 60175879 | 60199197 | 23319 | 1 | 19377 | ENSMUST0( | 10677 | ENSMUSG0( | Rai1 | retinoic acid induced 1 |
| chr11 | 60575451 | 60575909 | 459 | * | NA_peak_28 | 19 | Exon (ENSM | 11 | 60537978 | 60558512 | 20535 | 1 | 268420 | ENSMUST0( | 37473 | ENSMUSG0( | Alkbh5 | alkB homolog 5, RNA demethylase |
| chr11 | 61187673 | 61188035 | 363 | * | NA_peak_28 | 23 | Distal Interge | 11 | 61151785 | 61175055 | 23271 | 2 | 216825 | ENSMUST0( | -12618 | ENSMUSG0( | Usp22 | ubiquitin specific peptidase 22 |
| chr11 | 61367679 | 61367780 | 102 | * | NA_peak_28 | 35 | Promoter (2- | 11 | 61362709 | 61365652 | 2944 | 2 | 67473 | ENSMUST0( | -2027 | ENSMUSG0( | Slc47a1 | solute carrier family 47, member 1 |

| | | | | | | | | | | | | | | | | | | |
|---|---|---|---|---|---|---|---|---|---|---|---|---|---|---|---|---|---|---|
| chr11 | 62430393 | 62430722 | 330 | * | NA_peak_28 | 30 | Intron (ENSM | 11 | 62317797 | 62438481 | 120685 | 2 | 20185 | ENSMUST00 | 7759 | ENSMUSG0 | Ncor1 | nuclear receptor co-repressor 1 |
| chr11 | 62457008 | 62457339 | 332 | * | NA_peak_28 | 16 | Promoter (<= | 11 | 62316426 | 62457332 | 140907 | 2 | 20185 | ENSMUST00 | 0 | ENSMUSG0 | Ncor1 | nuclear receptor co-repressor 1 |
| chr11 | 62561173 | 62561515 | 343 | * | NA_peak_28 | 26 | Distal Interge | 11 | 62551676 | 62552853 | 1178 | 1 | 22187 | ENSMUST00 | 9497 | ENSMUSG0 | Ubb | ubiquitin B |
| chr11 | 68692486 | 68692536 | 51 | * | NA_peak_28 | 26 | Promoter (<= | 11 | 68692316 | 68816624 | 124309 | 1 | 77579 | ENSMUST00 | 170 | ENSMUSG0 | Myh10 | myosin, heavy polypeptide 10, non-muscle |
| chr11 | 69068526 | 69068949 | 424 | * | NA_peak_29 | 23 | Promoter (1- | 11 | 69070806 | 69073030 | 2225 | 1 | 66910 | ENSMUST00 | -1857 | ENSMUSG0 | Tmem107 | transmembrane protein 107 |
| chr11 | 69106910 | 69106974 | 65 | * | NA_peak_29 | 20 | Exon (ENSM | 11 | 69100522 | 69109960 | 9439 | 1 | 18626 | ENSMUST00 | 6388 | ENSMUSG0 | Per1 | period circadian clock 1 |
| chr11 | 69188452 | 69188761 | 310 | * | NA_peak_29 | 35 | Exon (ENSM | 11 | 69191537 | 69197384 | 5848 | 2 | 11688 | ENSMUST00 | 8623 | ENSMUSG0 | Alox8 | arachidonate 8-lipoxygenase |
| chr11 | 69662945 | 69663146 | 202 | * | NA_peak_29 | 15 | Promoter (<= | 11 | 69663043 | 69663125 | 83 | 1 | 100316840 | ENSMUST00 | 0 | ENSMUSG0 | Mir1934 | microRNA 1934 |
| chr11 | 69977004 | 69977026 | 23 | * | NA_peak_29 | 18 | Intron (ENSM | 11 | 69969241 | 69980061 | 10821 | 2 | 54351 | ENSMUST00 | 3035 | ENSMUSG0 | Elp5 | elongator acetyltransferase complex subunit 5 |
| chr11 | 70562559 | 70562980 | 422 | * | NA_peak_29 | 34 | Promoter (<= | 11 | 70562881 | 70614482 | 51602 | 1 | 50932 | ENSMUST00 | 0 | ENSMUSG0 | Mink1 | misshapen-like kinase 1 (zebrafish) |
| chr11 | 70669808 | 70670224 | 417 | * | NA_peak_29 | 25 | Promoter (<= | 11 | 70663771 | 70669416 | 5646 | 2 | 216873 | ENSMUST00 | -392 | ENSMUSG0 | Spag7 | sperm associated antigen 7 |
| chr11 | 72690083 | 72690514 | 432 | * | NA_peak_29 | 26 | Promoter (<= | 11 | 72690006 | 72772146 | 82141 | 1 | 11736 | ENSMUST00 | 77 | ENSMUSG0 | Ankfy1 | ankyrin repeat and FYVE domain containing 1 |
| chr11 | 72796302 | 72796624 | 323 | * | NA_peak_29 | 36 | Promoter (<= | 11 | 72796254 | 72927118 | 130865 | 1 | 195018 | ENSMUST00 | 48 | ENSMUSG0 | Zzef1 | zinc finger, ZZ-type with EF hand domain 1 |
| chr11 | 74224038 | 74224355 | 318 | * | NA_peak_29 | 18 | Distal Interge | 11 | 74206198 | 74207289 | 1092 | 2 | 258706 | ENSMUST00 | -16749 | ENSMUSG0 | Olfr43 | olfactory receptor 43 |
| chr11 | 74770845 | 74771214 | 370 | * | NA_peak_29 | 23 | Promoter (<= | 11 | 74770916 | 74773621 | 2706 | 1 | 67493 | ENSMUST00 | 0 | ENSMUSG0 | Mettl16 | methyltransferase like 16 |
| chr11 | 74832011 | 74832445 | 435 | * | NA_peak_29 | 24 | Promoter (1- | 11 | 74830920 | 74845725 | 14806 | 1 | 17428 | ENSMUST00 | 1091 | ENSMUSG0 | Mnt | max binding protein |
| chr11 | 74925490 | 74925936 | 447 | * | NA_peak_30 | 22 | Promoter (<= | 11 | 74925823 | 75164448 | 238626 | 1 | 103677 | ENSMUST00 | 0 | ENSMUSG0 | Smg6 | Smg-6 homolog, nonsense mediated mRNA decay factor (C. elegans) |
| chr11 | 75166737 | 75167044 | 308 | * | NA_peak_30 | 13 | Promoter (1- | 11 | 75167153 | 75168146 | 994 | 2 | 15248 | ENSMUST00 | 1102 | ENSMUSG0 | Hic1 | hypermethylated in cancer 1 |
| chr11 | 75194291 | 75194468 | 178 | * | NA_peak_30 | 15 | Promoter (<= | 11 | 75193783 | 75267769 | 73987 | 1 | 237847 | ENSMUST00 | 508 | ENSMUSG0 | Rtn4rl1 | reticulon 4 receptor-like 1 |
| chr11 | 75454560 | 75454620 | 61 | * | NA_peak_30 | 27 | Promoter (<= | 11 | 75440944 | 75454717 | 13774 | 2 | 192652 | ENSMUST00 | 97 | ENSMUSG0 | Wdr81 | WD repeat domain 81 |
| chr11 | 75479377 | 75479745 | 369 | * | NA_peak_30 | 21 | Distal Interge | 11 | 75486816 | 75509447 | 22632 | 1 | 192159 | ENSMUST00 | -7071 | ENSMUSG0 | Prpf8 | pre-mRNA processing factor 8 |
| chr11 | 75518642 | 75518945 | 304 | * | NA_peak_30 | 23 | Intron (ENSM | 11 | 75513575 | 75526578 | 13004 | 1 | 380713 | ENSMUST00 | 5067 | ENSMUSG0 | Scarf1 | scavenger receptor class F, member 1 |
| chr11 | 75655910 | 75656253 | 344 | * | NA_peak_30 | 49 | Promoter (<= | 11 | 75656149 | 75669587 | 13439 | 1 | 17913 | ENSMUST00 | 0 | ENSMUSG0 | Myo1c | myosin IC |
| chr11 | 76207720 | 76207878 | 159 | * | NA_peak_30 | 23 | Intron (ENSM | 11 | 76202101 | 76209274 | 7174 | 1 | 116972 | ENSMUST00 | 5619 | ENSMUSG0 | Tlcd3a | TLC domain containing 3A |
| chr11 | 76672705 | 76673487 | 783 | * | NA_peak_30 | 23 | Promoter (<= | 11 | 76672470 | 76673676 | 1207 | 1 | 320522 | ENSMUST00 | 235 | ENSMUSG0 | Bhlha9 | basic helix-loop-helix family, member a9 |
| chr11 | 77293711 | 77293713 | 3 | * | NA_peak_30 | 23 | Exon (ENSM | 11 | 77348280 | 77413635 | 65356 | 1 | 237860 | ENSMUST00 | -54567 | ENSMUSG0 | Ssh2 | slingshot protein phosphatase 2 |
| chr11 | 78188975 | 78189281 | 307 | * | NA_peak_30 | 35 | Promoter (<= | 11 | 78189240 | 78190306 | 1067 | 1 | 19376 | ENSMUST00 | 0 | ENSMUSG0 | Rab34 | RAB34, member RAS oncogene family |
| chr11 | 78261731 | 78261814 | 84 | * | NA_peak_30 | 15 | Promoter (<= | 11 | 78261752 | 78264032 | 2281 | 1 | 72503 | ENSMUST00 | 0 | ENSMUSG0 | 2610507B11 | RIKEN cDNA 2610507B11 gene |
| chr11 | 78550843 | 78551220 | 378 | * | NA_peak_30 | 26 | Promoter (<= | 11 | 78541817 | 78550777 | 8961 | 2 | 69071 | ENSMUST00 | -66 | ENSMUSG0 | Tmem97 | transmembrane protein 97 |
| chr11 | 78695929 | 78696171 | 243 | * | NA_peak_30 | 39 | Promoter (<= | 11 | 78617272 | 78696494 | 79223 | 2 | 18099 | ENSMUST00 | 323 | ENSMUSG0 | Nlk | nemo like kinase |
| chr11 | 79146135 | 79146365 | 231 | * | NA_peak_30 | 47 | Promoter (<= | 11 | 79013440 | 79146407 | 132968 | 2 | 16706 | ENSMUST00 | 42 | ENSMUSG0 | Ksr1 | kinase suppressor of ras 1 |
| chr11 | 79234258 | 79234764 | 507 | * | NA_peak_30 | 28 | Distal Interge | 11 | 79240307 | 79242949 | 2643 | 2 | 78889 | ENSMUST00 | 8185 | ENSMUSG0 | Wsb1 | WD repeat and SOCS box-containing 1 |
| chr11 | 79339960 | 79340117 | 158 | * | NA_peak_30 | 17 | Promoter (<= | 11 | 79339887 | 79430788 | 90902 | 1 | 18015 | ENSMUST00 | 73 | ENSMUSG0 | Nf1 | neurofibromin 1 |
| chr11 | 80300980 | 80301097 | 118 | * | NA_peak_31 | 44 | Promoter (<= | 11 | 80300912 | 80355955 | 55044 | 1 | 246104 | ENSMUST00 | 68 | ENSMUSG0 | Rhbdl3 | rhomboid like 3 |
| chr11 | 82123577 | 82123866 | 290 | * | NA_peak_31 | 26 | Distal Interge | 11 | 82115185 | 82116799 | 1615 | 1 | 20307 | ENSMUST00 | 8392 | ENSMUSG0 | Ccl8 | chemokine (C-C motif) ligand 8 |
| chr11 | 86134408 | 86134710 | 303 | * | NA_peak_31 | 23 | Intron (ENSM | 11 | 86195033 | 86201164 | 6132 | 2 | 237911 | ENSMUST00 | 66454 | ENSMUSG0 | Brip1 | BRCA1 interacting protein C-terminal helicase 1 |
| chr11 | 87616919 | 87617236 | 318 | * | NA_peak_31 | 16 | Promoter (<= | 11 | 87617164 | 87659542 | 42379 | 1 | 327992 | ENSMUST00 | 0 | ENSMUSG0 | Hsf5 | heat shock transcription factor family member 5 |
| chr11 | 91447137 | 91447404 | 268 | * | NA_peak_31 | 16 | Distal Interge | 11 | 91575315 | 91577558 | 2244 | 2 | 73470 | ENSMUST00 | 130154 | ENSMUSG0 | Kif2b | kinesin family member 2B |
| chr11 | 92598816 | 92599130 | 315 | * | NA_peak_31 | 44 | Distal Interge | 11 | 93098028 | 93099866 | 1839 | 1 | 72605 | ENSMUST00 | -498898 | ENSMUSG0 | Car10 | carbonic anhydrase 10 |
| chr11 | 94992234 | 94992531 | 298 | * | NA_peak_32 | 25 | Promoter (<= | 11 | 94992880 | 95001326 | 8447 | 1 | 217124 | ENSMUST00 | -349 | ENSMUSG0 | Ppp1r9b | protein phosphatase 1, regulatory subunit 9B |
| chr11 | 95076587 | 95076945 | 359 | * | NA_peak_32 | 14 | Promoter (<= | 11 | 95044474 | 95076714 | 32241 | 2 | 16400 | ENSMUST00 | 0 | ENSMUSG0 | Itga3 | integrin alpha 3 |
| chr11 | 96004862 | 96005479 | 618 | * | NA_peak_32 | 26 | Promoter (<= | 11 | 95957163 | 96005940 | 48778 | 2 | 140486 | ENSMUST00 | 461 | ENSMUSG0 | Igf2bp1 | insulin-like growth factor 2 mRNA binding protein 1 |
| chr11 | 96006019 | 96006226 | 208 | * | NA_peak_32 | 14 | Promoter (<= | 11 | 95957163 | 96005940 | 48778 | 2 | 140486 | ENSMUST00 | -79 | ENSMUSG0 | Igf2bp1 | insulin-like growth factor 2 mRNA binding protein 1 |
| chr11 | 96124377 | 96124606 | 230 | * | NA_peak_32 | 13 | Distal Interge | 11 | 96133786 | 96165451 | 31666 | 1 | 237930 | ENSMUST00 | -9180 | ENSMUSG0 | Ttll6 | tubulin tyrosine ligase-like family, member 6 |
| chr11 | 96270944 | 96271328 | 385 | * | NA_peak_32 | 30 | Promoter (<= | 11 | 96271457 | 96276595 | 5139 | 1 | 15417 | ENSMUST00 | -129 | ENSMUSG0 | Hoxb9 | homeobox B9 |
| chr11 | 96303721 | 96304252 | 532 | * | NA_peak_32 | 43 | Promoter (<= | 11 | 96303512 | 96306120 | 2609 | 1 | 15413 | ENSMUST00 | 209 | ENSMUSG0 | Hoxb5 | homeobox B5 |
| chr11 | 96320228 | 96320260 | 33 | * | NA_peak_32 | 22 | Promoter (1- | 11 | 96318267 | 96321638 | 3372 | 1 | 15412 | ENSMUST00 | 1961 | ENSMUSG0 | Hoxb4 | homeobox B4 |
| chr11 | 96764416 | 96764699 | 284 | * | NA_peak_32 | 23 | Distal Interge | 11 | 96770098 | 96771050 | 953 | 2 | 74479 | ENSMUST00 | 6351 | ENSMUSG0 | Snx11 | sorting nexin 11 |
| chr11 | 96804562 | 96804605 | 44 | * | NA_peak_32 | 16 | Intron (ENSM | 11 | 96789249 | 96808540 | 19292 | 1 | 12412 | ENSMUST00 | 15313 | ENSMUSG0 | Cbx1 | chromobox 1 |
| chr11 | 96806349 | 96806862 | 514 | * | NA_peak_32 | 23 | 3' UTR | 11 | 96821346 | 96822725 | 1380 | 2 | 18023 | ENSMUST00 | 15863 | ENSMUSG0 | Nfe2l1 | nuclear factor, erythroid derived 2,-like 1 |
| chr11 | 96836245 | 96836445 | 201 | * | NA_peak_32 | 22 | Distal Interge | 11 | 96817414 | 96829968 | 12555 | 2 | 18023 | ENSMUST00 | -6277 | ENSMUSG0 | Nfe2l1 | nuclear factor, erythroid derived 2,-like 1 |
| chr11 | 97021422 | 97021810 | 389 | * | NA_peak_32 | 30 | Promoter (2- | 11 | 97018733 | 97024731 | 5999 | 1 | 83395 | ENSMUST00 | 2689 | ENSMUSG0 | Sp6 | trans-acting transcription factor 6 |
| chr11 | 97022463 | 97022743 | 281 | * | NA_peak_32 | 29 | 3' UTR | 11 | 97020508 | 97026248 | 5741 | 2 | 100127433 | ENSMUST00 | 3505 | ENSMUSG0 | Gm11532 | predicted gene 11532 |
| chr11 | 97441714 | 97442010 | 297 | * | NA_peak_32 | 20 | Intron (ENSM | 11 | 97450160 | 97502400 | 52241 | 1 | 58996 | ENSMUST00 | -8150 | ENSMUSG0 | Arhgap23 | Rho GTPase activating protein 23 |
| chr11 | 97466501 | 97466800 | 300 | * | NA_peak_33 | 30 | Exon (ENSM | 11 | 97452130 | 97502402 | 50273 | 1 | 58996 | ENSMUST00 | 14371 | ENSMUSG0 | Arhgap23 | Rho GTPase activating protein 23 |
| chr11 | 97546433 | 97546875 | 443 | * | NA_peak_33 | 24 | Intron (ENSM | 11 | 97509340 | 97536945 | 27606 | 2 | 56013 | ENSMUST00 | -9488 | ENSMUSG0 | Srcin1 | SRC kinase signaling inhibitor 1 |
| chr11 | 97800155 | 97800348 | 194 | * | NA_peak_33 | 43 | Promoter (<= | 11 | 97800171 | 97833563 | 33393 | 1 | 16796 | ENSMUST00 | 0 | ENSMUSG0 | Lasp1 | LIM and SH3 protein 1 |
| chr11 | 98682488 | 98682886 | 399 | * | NA_peak_33 | 24 | Promoter (<= | 11 | 98682554 | 98695979 | 13426 | 1 | 22123 | ENSMUST00 | 0 | ENSMUSG0 | Psmd3 | proteasome (prosome, macropain) 26S subunit, non-ATPase, 3 |
| chr11 | 98770316 | 98770688 | 373 | * | NA_peak_33 | 34 | Promoter (<= | 11 | 98769347 | 98770426 | 1080 | 2 | 217166 | ENSMUST00 | 0 | ENSMUSG0 | Nr1d1 | nuclear receptor subfamily 1, group D, member 1 |
| chr11 | 98809978 | 98810375 | 398 | * | NA_peak_33 | 13 | Promoter (<= | 11 | 98797707 | 98810087 | 12381 | 2 | 100037262 | ENSMUST00 | 0 | ENSMUSG0 | Gm12359 | predicted gene 12359 |
| chr11 | 99264440 | 99264475 | 36 | * | NA_peak_33 | 26 | Distal Interge | 11 | 99233890 | 99244085 | 10196 | 2 | 268481 | ENSMUST00 | -20355 | ENSMUSG0 | Krt222 | keratin 222 |
| chr11 | 100663546 | 100664209 | 664 | * | NA_peak_33 | 30 | Intron (ENSM | 11 | 100630720 | 100657622 | 26903 | 2 | 278304 | ENSMUST00 | -5924 | ENSMUSG0 | Zfp385c | zinc finger protein 385C |
| chr11 | 100664855 | 100665130 | 276 | * | NA_peak_33 | 65 | Intron (ENSM | 11 | 100630720 | 100657622 | 26903 | 2 | 278304 | ENSMUST00 | -7233 | ENSMUSG0 | Zfp385c | zinc finger protein 385C |
| chr11 | 100692175 | 100692458 | 284 | * | NA_peak_33 | 15 | 5' UTR | 11 | 100695063 | 100696620 | 1558 | 2 | 80861 | ENSMUST00 | 4162 | ENSMUSG0 | Dhx58 | DEXH (Asp-Glu-X-His) box polypeptide 58 |
| chr11 | 100696338 | 100696567 | 230 | * | NA_peak_33 | 26 | Promoter (<= | 11 | 100695063 | 100696620 | 1558 | 2 | 80861 | ENSMUST00 | 53 | ENSMUSG0 | Dhx58 | DEXH (Asp-Glu-X-His) box polypeptide 58 |
| chr11 | 101144170 | 101144455 | 286 | * | NA_peak_33 | 23 | Distal Interge | 11 | 101155907 | 101161787 | 5881 | 1 | 103768 | ENSMUST00 | -11452 | ENSMUSG0 | Tubg2 | tubulin, gamma 2 |
| chr11 | 101167043 | 101167179 | 137 | * | NA_peak_33 | 23 | Promoter (1- | 11 | 101162679 | 101165620 | 2942 | 2 | 217198 | ENSMUST00 | -1423 | ENSMUSG0 | Plekhh3 | pleckstrin homology domain containing, family H (with MyTH4 domain) member 3 |
| chr11 | 101167975 | 101168374 | 400 | * | NA_peak_33 | 19 | Promoter (1- | 11 | 101167474 | 101170354 | 2881 | 2 | 217198 | ENSMUST00 | 1980 | ENSMUSG0 | Plekhh3 | pleckstrin homology domain containing, family H (with MyTH4 domain) member 3 |
| chr11 | 101260721 | 101261029 | 309 | * | NA_peak_33 | 30 | Promoter (<= | 11 | 101260655 | 101277274 | 16620 | 1 | 69847 | ENSMUST00 | 66 | ENSMUSG0 | Wnk4 | WNK lysine deficient protein kinase 4 |
| chr11 | 103088081 | 103088265 | 185 | * | NA_peak_34 | 21 | Intron (ENSM | 11 | 103070304 | 103078294 | 7991 | 2 | 72469 | ENSMUST00 | -9787 | ENSMUSG0 | Plcd3 | phospholipase C, delta 3 |
| chr11 | 103112496 | 103112543 | 48 | * | NA_peak_34 | 22 | Downstream | 11 | 103116231 | 103119725 | 3495 | 1 | 192231 | ENSMUST00 | -3688 | ENSMUSG0 | Hexim1 | hexamethylene bis-acetamide inducible 1 |
| chr11 | 105165818 | 105166345 | 528 | * | NA_peak_34 | 30 | Exon (ENSM | 11 | 105178807 | 105283959 | 105153 | 1 | 24086 | ENSMUST00 | -12462 | ENSMUSG0 | Tlk2 | tousled-like kinase 2 (Arabidopsis) |
| chr11 | 105638307 | 105638811 | 505 | * | NA_peak_34 | 26 | Intron (ENSM | 11 | 105625038 | 105715553 | 90516 | 1 | 77097 | ENSMUST00 | 13269 | ENSMUSG0 | Tanc2 | tetratricopeptide repeat, ankyrin repeat and coiled-coil containing 2 |
| chr11 | 108279792 | 108280056 | 265 | * | NA_peak_34 | 23 | Intron (ENSM | 11 | 108343354 | 108395905 | 52552 | 1 | 11818 | ENSMUST00 | -63298 | ENSMUSG0 | Apoh | apolipoprotein H |
| chr11 | 108343022 | 108343293 | 272 | * | NA_peak_34 | 16 | Promoter (<= | 11 | 108343354 | 108395905 | 52552 | 1 | 11818 | ENSMUST00 | -61 | ENSMUSG0 | Apoh | apolipoprotein H |
| chr11 | 108534736 | 108535033 | 298 | * | NA_peak_34 | 54 | Intron (ENSM | 11 | 108531406 | 108570827 | 39422 | 1 | 76380 | ENSMUST00 | 3330 | ENSMUSG0 | Cep112 | centrosomal protein 112 |
| chr11 | 112432716 | 112432748 | 33 | * | NA_peak_35 | 16 | Distal Interge | 11 | 112664839 | 112669398 | 4560 | 2 | 217294 | ENSMUST00 | 236650 | ENSMUSG0 | BC006965 | cDNA sequence BC006965 |
| chr11 | 113216843 | 113217066 | 224 | * | NA_peak_35 | 19 | Distal Interge | 11 | 113203315 | 113209669 | 6355 | 2 | 319871 | ENSMUST00 | -7174 | ENSMUSG0 | 4732490B19 | RIKEN cDNA 4732490B19 gene |
| chr11 | 113619007 | 113619449 | 443 | * | NA_peak_35 | 30 | Promoter (<= | 11 | 113619342 | 113626028 | 6687 | 1 | 20606 | ENSMUST00 | 0 | ENSMUSG0 | Sstr2 | somatostatin receptor 2 |
| chr11 | 113683576 | 113683992 | 417 | * | NA_peak_35 | 19 | Promoter (<= | 11 | 113661319 | 113684155 | 22837 | 2 | 28081 | ENSMUST00 | 163 | ENSMUSG0 | Fam104a | family with sequence similarity 104, member A |
| chr11 | 115541173 | 115541509 | 337 | * | NA_peak_35 | 24 | Distal Interge | 11 | 115523102 | 115536276 | 13175 | 2 | 170930 | ENSMUST00 | -4897 | ENSMUSG0 | Sumo2 | small ubiquitin-like modifier 2 |
| chr11 | 115859514 | 115859788 | 275 | * | NA_peak_35 | 17 | Exon (ENSM | 11 | 115855032 | 115855780 | 749 | 1 | 217325 | ENSMUST00 | 4482 | ENSMUSG0 | Llgl2 | LLGL2 scribble cell polarity complex component |
| chr11 | 116194569 | 116194893 | 325 | * | NA_peak_35 | 30 | Intron (ENSM | 11 | 116189029 | 116198729 | 9701 | 2 | 11430 | ENSMUST00 | 3836 | ENSMUSG0 | Acox1 | acyl-Coenzyme A oxidase 1, palmitoyl |
| chr11 | 116195351 | 116195513 | 163 | * | NA_peak_35 | 30 | Intron (ENSM | 11 | 116189029 | 116198729 | 9701 | 2 | 11430 | ENSMUST00 | 3216 | ENSMUSG0 | Acox1 | acyl-Coenzyme A oxidase 1, palmitoyl |
| chr11 | 116201949 | 116202366 | 418 | * | NA_peak_35 | 14 | Promoter (2- | 11 | 116171888 | 116199045 | 27158 | 2 | 11430 | ENSMUST00 | -2904 | ENSMUSG0 | Acox1 | acyl-Coenzyme A oxidase 1, palmitoyl |
| chr11 | 116623920 | 116624349 | 430 | * | NA_peak_36 | 23 | Promoter (<= | 11 | 116598165 | 116624200 | 26036 | 2 | 217344 | ENSMUST00 | 0 | ENSMUSG0 | Rhbdf2 | rhomboid 5 homolog 2 |
| chr11 | 116843262 | 116843371 | 110 | * | NA_peak_36 | 16 | Promoter (<= | 11 | 116837432 | 116843281 | 5850 | 2 | 107817 | ENSMUST00 | 0 | ENSMUSG0 | Jmjd6 | jumonji domain containing 6 |
| chr11 | 117667705 | 117668098 | 394 | * | NA_peak_36 | 26 | Intron (ENSM | 11 | 117654813 | 117763439 | 108627 | 1 | 217351 | ENSMUST00 | 12892 | ENSMUSG0 | Tnrc6c | trinucleotide repeat containing 6C |
| chr11 | 117736238 | 117736574 | 337 | * | NA_peak_36 | 15 | Intron (ENSM | 11 | 117742501 | 117749301 | 6801 | 1 | 217351 | ENSMUST00 | -5927 | ENSMUSG0 | Tnrc6c | trinucleotide repeat containing 6C |
| chr11 | 117754943 | 117755227 | 285 | * | NA_peak_36 | 18 | Intron (ENSM | 11 | 117749231 | 117751308 | 2078 | 1 | 217351 | ENSMUST00 | 5712 | ENSMUSG0 | Tnrc6c | trinucleotide repeat containing 6C |
| chr11 | 117849129 | 117849693 | 565 | * | NA_peak_36 | 34 | Promoter (<= | 11 | 117849251 | 117855732 | 6482 | 1 | 11799 | ENSMUST00 | 0 | ENSMUSG0 | Birc5 | baculoviral IAP repeat-containing 5 |
| chr11 | 118209456 | 118209745 | 290 | * | NA_peak_36 | 20 | Promoter (2- | 11 | 118164166 | 118212742 | 48577 | 2 | 19157 | ENSMUST00 | 2997 | ENSMUSG0 | Cyth1 | cytohesin 1 |
| chr11 | 118386425 | 118387416 | 992 | * | NA_peak_36 | 34 | Distal Interge | 11 | 118392751 | 118398151 | 5401 | 2 | 19039 | ENSMUST00 | 10735 | ENSMUSG0 | Lgals3bp | lectin, galactoside-binding, soluble, 3 binding protein |
| chr11 | 118417971 | 118418204 | 234 | * | NA_peak_36 | 25 | Promoter (<= | 11 | 118406290 | 118419023 | 12734 | 2 | 76025 | ENSMUST00 | 819 | ENSMUSG0 | Cant1 | calcium activated nucleotidase 1 |
| chr11 | 119020509 | 119021173 | 665 | * | NA_peak_36 | 23 | Promoter (1- | 11 | 119022962 | 119031270 | 8309 | 1 | 12416 | ENSMUST00 | -1789 | ENSMUSG0 | Cbx2 | chromobox 2 |
| chr11 | 119023515 | 119023762 | 248 | * | NA_peak_36 | 32 | Promoter (<= | 11 | 119023725 | 119028038 | 4314 | 1 | 12416 | ENSMUST00 | 0 | ENSMUSG0 | Cbx2 | chromobox 2 |
| chr11 | 119706589 | 119707065 | 477 | * | NA_peak_36 | 13 | Intron (ENSM | 11 | 119655944 | 119865810 | 209867 | 1 | 74370 | ENSMUST00 | 50645 | ENSMUSG0 | Rptor | regulatory associated protein of MTOR, complex 1 |
| chr11 | 120236908 | 120238014 | 1107 | * | NA_peak_36 | 55 | Promoter (<= | 11 | 120236699 | 120292282 | 55584 | 1 | 268515 | ENSMUST00 | 209 | ENSMUSG0 | Bahcc1 | BAH domain and coiled-coil containing 1 |
| chr11 | 120528969 | 120529463 | 495 | * | NA_peak_37 | 25 | Promoter (1- | 11 | 120530699 | 120538984 | 8286 | 1 | 14527 | ENSMUST00 | -1236 | ENSMUSG0 | Gcgr | glucagon receptor |
| chr11 | 120572992 | 120573108 | 117 | * | NA_peak_37 | 38 | Promoter (<= | 11 | 120560298 | 120573253 | 12956 | 2 | 18453 | ENSMUST00 | 145 | ENSMUSG0 | P4hb | prolyl 4-hydroxylase, beta polypeptide |
| chr11 | 120573406 | 120573623 | 218 | * | NA_peak_37 | 38 | Promoter (<= | 11 | 120560298 | 120573253 | 12956 | 2 | 18453 | ENSMUST00 | -153 | ENSMUSG0 | P4hb | prolyl 4-hydroxylase, beta polypeptide |
| chr11 | 120578569 | 120578793 | 225 | * | NA_peak_37 | 25 | Promoter (2- | 11 | 120578109 | 120581380 | 3272 | 2 | 192662 | ENSMUST00 | 2587 | ENSMUSG0 | Arhgdia | Rho GDP dissociation inhibitor (GDI) alpha |
| chr11 | 120659898 | 120660230 | 333 | * | NA_peak_37 | 33 | Promoter (<= | 11 | 120653788 | 120660606 | 6819 | 2 | 77583 | ENSMUST00 | 376 | ENSMUSG0 | Notum | notum palmitoleoyl-protein carboxylesterase |
| chr11 | 120666545 | 120667063 | 519 | * | NA_peak_37 | 23 | Distal Interge | 11 | 120657563 | 120661175 | 3613 | 2 | 77583 | ENSMUST00 | -5370 | ENSMUSG0 | Notum | notum palmitoleoyl-protein carboxylesterase |
| chr11 | 120783792 | 120784223 | 432 | * | NA_peak_37 | 14 | Promoter (<= | 11 | 120780746 | 120784185 | 3440 | 2 | 19719 | ENSMUST00 | 0 | ENSMUSG0 | Rfng | RFNG O-fucosylpeptide 3-beta-N-acetylglucosaminyltransferase |
| chr11 | 120824156 | 120824843 | 688 | * | NA_peak_37 | 14 | Promoter (<= | 11 | 120805846 | 120824547 | 18702 | 2 | 14104 | ENSMUST00 | 0 | ENSMUSG0 | Fasn | fatty acid synthase |
| chr11 | 120982169 | 120982331 | 163 | * | NA_peak_37 | 13 | Intron (ENSM | 11 | 120972472 | 120990871 | 18400 | 2 | 104318 | ENSMUST00 | 8540 | ENSMUSG0 | Csnk1d | casein kinase 1, delta |
| chr11 | 121156349 | 121156988 | 640 | * | NA_peak_37 | 46 | Distal Interge | 11 | 121160271 | 121161973 | 1703 | 1 | 217369 | ENSMUST00 | -3283 | ENSMUSG0 | Uts2r | urotensin 2 receptor |
| chr11 | 121297832 | 121298051 | 220 | * | NA_peak_37 | 21 | Promoter (<= | 11 | 121298516 | 121299850 | 1335 | 1 | 68837 | ENSMUST00 | -465 | ENSMUSG0 | Foxk2 | forkhead box K2 |
| chr11 | 121318245 | 121318595 | 351 | * | NA_peak_37 | 76 | Distal Interge | 11 | 121298516 | 121299850 | 1335 | 1 | 68837 | ENSMUST00 | 19729 | ENSMUSG0 | Foxk2 | forkhead box K2 |
| chr11 | 121549950 | 121550317 | 368 | * | NA_peak_37 | 22 | Promoter (<= | 11 | 121550111 | 121590383 | 40273 | 1 | 108903 | ENSMUST00 | 0 | ENSMUSG0 | Tbcd | tubulin-specific chaperone d |
| chr11 | 121730998 | 121731395 | 398 | * | NA_peak_37 | 36 | Distal Interge | 11 | 121702763 | 121715829 | 13067 | 1 | 210029 | ENSMUST00 | 28235 | ENSMUSG0 | Metrnl | meteorin, glial cell differentiation regulator-like |
| chr11 | 121899994 | 121900060 | 67 | * | NA_peak_37 | 27 | Distal Interge | 11 | 121830247 | 121843423 | 13177 | 1 | 74675 | ENSMUST00 | 69747 | ENSMUSG0 | Ptchd3 | patched domain containing 3 |
| chr11 | 121960740 | 121961011 | 272 | * | NA_peak_37 | 26 | Distal Interge | 11 | 121830247 | 121843423 | 13177 | 1 | 74675 | ENSMUST00 | 130493 | ENSMUSG0 | Ptchd3 | patched domain containing 3 |
| chr12 | 3002745 | 3003080 | 336 | * | NA_peak_37 | 42 | Distal Interge | 12 | 3235504 | 3241410 | 5907 | 1 | 74173 | ENSMUST00 | -232424 | ENSMUSG0 | Rab10os | RAB10, member RAS oncogene family, opposite strand |
| chr12 | 3184289 | 3184777 | 489 | * | NA_peak_37 | 23 | Intron (ENSM | 12 | 3235504 | 3241410 | 5907 | 1 | 74173 | ENSMUST00 | -50727 | ENSMUSG0 | Rab10os | RAB10, member RAS oncogene family, opposite strand |
| chr12 | 3666475 | 3666622 | 148 | * | NA_peak_37 | 21 | Intron (ENSM | 12 | 3686756 | 3718968 | 32213 | 1 | 13528 | ENSMUST00 | -20134 | ENSMUSG0 | Dtnb | dystrobrevin, beta |
| chr12 | 3741245 | 3741663 | 419 | * | NA_peak_37 | 65 | Intron (ENSM | 12 | 3749289 | 3781142 | 31854 | 1 | 13528 | ENSMUST00 | -7626 | ENSMUSG0 | Dtnb | dystrobrevin, beta |
| chr12 | 3955038 | 3955357 | 320 | * | NA_peak_37 | 14 | Promoter (<= | 12 | 3954970 | 3958260 | 3291 | 1 | 18976 | ENSMUST00 | 68 | ENSMUSG0 | Pomc | pro-opiomelanocortin-alpha |
| chr12 | 4402137 | 4402303 | 167 | * | NA_peak_38 | 17 | Intron (ENSM | 12 | 4249451 | 4416796 | 167346 | 2 | 17977 | ENSMUST00 | 14493 | ENSMUSG0 | Ncoa1 | nuclear receptor coactivator 1 |
| chr12 | 4459442 | 4460088 | 647 | * | NA_peak_38 | 23 | Intron (ENSM | 12 | 4247362 | 4477182 | 229821 | 2 | 17977 | ENSMUST00 | 17094 | ENSMUSG0 | Ncoa1 | nuclear receptor coactivator 1 |
| chr12 | 4499662 | 4500001 | 340 | * | NA_peak_38 | 18 | Distal Interge | 12 | 4247362 | 4477182 | 229821 | 2 | 17977 | ENSMUST00 | -22480 | ENSMUSG0 | Ncoa1 | nuclear receptor coactivator 1 |
| chr12 | 4655559 | 4655581 | 23 | * | NA_peak_38 | 30 | Intron (ENSM | 12 | 4664535 | 4674167 | 9633 | 1 | 20403 | ENSMUST00 | -8954 | ENSMUSG0 | Itsn2 | intersectin 2 |
| chr12 | 4911790 | 4912403 | 614 | * | NA_peak_38 | 23 | Distal Interge | 12 | 4879032 | 4907705 | 28674 | 2 | 217379 | ENSMUST00 | -4085 | ENSMUSG0 | Ubxn2a | UBX domain protein 2A |
| chr12 | 5055206 | 5055590 | 385 | * | NA_peak_38 | 28 | Distal Interge | 12 | 4982052 | 4993758 | 11707 | 1 | 320817 | ENSMUST00 | 73154 | ENSMUSG0 | Atad2b | ATPase family, AAA domain containing 2B |
| chr12 | 5831478 | 5831635 | 158 | * | NA_peak_38 | 26 | Intron (ENSM | 12 | 5411641 | 5416631 | 4991 | 1 | 72669 | ENSMUST00 | 419837 | ENSMUSG0 | 2810032G03 | RIKEN cDNA 2810032G03 gene |
| chr12 | 7802301 | 7802579 | 279 | * | NA_peak_38 | 39 | Intron (ENSM | 12 | 7977648 | 7987818 | 10171 | 1 | 238055 | ENSMUST00 | -175069 | ENSMUSG0 | Apob | apolipoprotein B |
| chr12 | 9680082 | 9680383 | 302 | * | NA_peak_38 | 23 | Distal Interge | 12 | 9574441 | 9581500 | 7060 | 1 | 23967 | ENSMUST00 | 105641 | ENSMUSG0 | Osr1 | odd-skipped related transcription factor 1 |
| chr12 | 15426003 | 15426292 | 290 | * | NA_peak_38 | 26 | Distal Interge | 12 | 15040194 | 15051598 | 11405 | 2 | 73972 | ENSMUST00 | -374405 | ENSMUSG0 | 4930448C13 | RIKEN cDNA 4930448C13 gene |
| chr12 | 17961037 | 17961352 | 316 | * | NA_peak_38 | 26 | Intron (ENSM | 12 | 17728226 | 17791126 | 62901 | 1 | 53602 | ENSMUST00 | 232811 | ENSMUSG0 | Hpcal1 | hippocalcin-like 1 |
| chr12 | 18262530 | 18263122 | 593 | * | NA_peak_38 | 23 | Distal Interge | 12 | 18514510 | 18535254 | 20745 | 1 | 236366 | ENSMUST00 | -251388 | ENSMUSG0 | 5730507C01 | RIKEN cDNA 5730507C01 gene |
| chr12 | 18940071 | 18940075 | 5 | * | NA_peak_38 | 22 | Distal Interge | 12 | 18514546 | 18535251 | 20706 | 1 | 236366 | ENSMUST00 | 425525 | ENSMUSG0 | 5730507C01 | RIKEN cDNA 5730507C01 gene |
| chr12 | 19522349 | 19522780 | 432 | * | NA_peak_38 | 30 | Distal Interge | 12 | 18514546 | 18535251 | 20706 | 1 | 236366 | ENSMUST00 | 1007803 | ENSMUSG0 | 5730507C01 | RIKEN cDNA 5730507C01 gene |
| chr12 | 19572843 | 19573012 | 170 | * | NA_peak_38 | 29 | Distal Interge | 12 | 18514546 | 18535251 | 20706 | 1 | 236366 | ENSMUST00 | 1058297 | ENSMUSG0 | 5730507C01 | RIKEN cDNA 5730507C01 gene |
| chr12 | 19575700 | 19576058 | 359 | * | NA_peak_38 | 42 | Distal Interge | 12 | 18514546 | 18535251 | 20706 | 1 | 236366 | ENSMUST00 | 1061154 | ENSMUSG0 | 5730507C01 | RIKEN cDNA 5730507C01 gene |
| chr12 | 20196909 | 20197153 | 245 | * | NA_peak_39 | 26 | Exon (ENSM | 12 | 20804381 | 20815779 | 11399 | 2 | 69513 | ENSMUST00 | 618626 | ENSMUSG0 | 1700030C10 | RIKEN cDNA 1700030C10 gene |
| chr12 | 20246787 | 20247673 | 887 | * | NA_peak_39 | 50 | Intron (ENSM | 12 | 20804381 | 20815779 | 11399 | 2 | 69513 | ENSMUST00 | 568106 | ENSMUSG0 | 1700030C10 | RIKEN cDNA 1700030C10 gene |
| chr12 | 20251811 | 20251904 | 94 | * | NA_peak_39 | 31 | Intron (ENSM | 12 | 20804381 | 20815779 | 11399 | 2 | 69513 | ENSMUST00 | 563875 | ENSMUSG0 | 1700030C10 | RIKEN cDNA 1700030C10 gene |
| chr12 | 22064836 | 22064943 | 108 | * | NA_peak_39 | 26 | Distal Interge | 12 | 22115756 | 22156646 | 40891 | 2 | 100043416 | ENSMUST00 | 91703 | ENSMUSG0 | Gm4425 | KRAB-domain protein pseudogene |
| chr12 | 23078996 | 23079281 | 286 | * | NA_peak_39 | 28 | Distal Interge | 12 | 22953999 | 23010284 | 56286 | 2 | 69732 | ENSMUST00 | -68712 | ENSMUSG0 | 2410018L13 | RIKEN cDNA 2410018L13 gene |
| chr12 | 23200014 | 23200184 | 171 | * | NA_peak_39 | 23 | Distal Interge | 12 | 22953999 | 23010284 | 56286 | 2 | 69732 | ENSMUST00 | -189730 | ENSMUSG0 | 2410018L13 | RIKEN cDNA 2410018L13 gene |
| chr12 | 23275824 | 23276172 | 349 | * | NA_peak_39 | 54 | Intron (ENSM | 12 | 22953999 | 23010284 | 56286 | 2 | 69732 | ENSMUST00 | -265540 | ENSMUSG0 | 2410018L13 | RIKEN cDNA 2410018L13 gene |
| chr12 | 23310272 | 23310542 | 271 | * | NA_peak_39 | 22 | Intron (ENSM | 12 | 22953999 | 23010284 | 56286 | 2 | 69732 | ENSMUST00 | -299988 | ENSMUSG0 | 2410018L13 | RIKEN cDNA 2410018L13 gene |
| chr12 | 32029030 | 32029275 | 246 | * | NA_peak_39 | 24 | Intron (ENSM | 12 | 31958480 | 32060589 | 102110 | 2 | 19088 | ENSMUST00 | 31314 | ENSMUSG0 | Prkar2b | protein kinase, cAMP dependent regulatory, type II beta |
| chr12 | 32904579 | 32904734 | 156 | * | NA_peak_39 | 26 | Distal Interge | 12 | 32924957 | 32953783 | 28827 | 2 | 74076 | ENSMUST00 | 49049 | ENSMUSG0 | 4933406C10 | RIKEN cDNA 4933406C10 gene |
| chr12 | 35809191 | 35809988 | 798 | * | NA_peak_39 | 43 | Distal Interge | 12 | 35672380 | 35695118 | 22739 | 2 | 71581 | ENSMUST00 | -114073 | ENSMUSG0 | 9130015A21 | RIKEN cDNA 9130015A21 gene |
| chr12 | 35811325 | 35811608 | 284 | * | NA_peak_39 | 30 | Distal Interge | 12 | 35925620 | 35949736 | 24117 | 1 | 403205 | ENSMUST00 | -114012 | ENSMUSG0 | Agr3 | anterior gradient 3 |
| chr12 | 38078648 | 38079240 | 593 | * | NA_peak_39 | 30 | Intron (ENSM | 12 | 38019180 | 38174661 | 155482 | 1 | 217480 | ENSMUST00 | 59468 | ENSMUSG0 | Dgkb | diacylglycerol kinase, beta |
| chr12 | 43753544 | 43753798 | 255 | * | NA_peak_39 | 34 | Exon (ENSM | 12 | 44208148 | 44210068 | 1921 | 2 | 27362 | ENSMUST00 | 456270 | ENSMUSG0 | Dnajb9 | DnaJ heat shock protein family (Hsp40) member B9 |
| chr12 | 44232680 | 44232984 | 305 | * | NA_peak_39 | 39 | Intron (ENSM | 12 | 44221745 | 44223373 | 1629 | 1 | 100039062 | ENSMUST00 | 10935 | NA | Gm2027 | nuclear assembly factor 1 homolog pseudogene |
| chr12 | 50980001 | 50980422 | 422 | * | NA_peak_39 | 34 | Distal Interge | 12 | 51005158 | 51062543 | 57386 | 1 | 75494 | ENSMUST00 | -24736 | ENSMUSG0 | 1700008C04 | RIKEN cDNA 1700008C04 gene |
| chr12 | 57604440 | 57604906 | 467 | * | NA_peak_39 | 58 | Intron (ENSM | 12 | 57585975 | 57618602 | 32628 | 2 | 78758 | ENSMUST00 | 13696 | ENSMUSG0 | 4921518K17 | RIKEN cDNA 4921518K17 gene |
| chr12 | 59223275 | 59223465 | 191 | * | NA_peak_39 | 16 | Distal Interge | 12 | 59200655 | 59219725 | 19071 | 2 | 70611 | ENSMUST00 | -3550 | ENSMUSG0 | Fbxo33 | F-box protein 33 |
| chr12 | 59274096 | 59274506 | 411 | * | NA_peak_39 | 49 | Distal Interge | 12 | 59200655 | 59219725 | 19071 | 2 | 70611 | ENSMUST00 | -54371 | ENSMUSG0 | Fbxo33 | F-box protein 33 |
| chr12 | 60469082 | 60469452 | 371 | * | NA_peak_39 | 30 | Distal Interge | 12 | 61499660 | 61521820 | 22161 | 2 | 100504101 | ENSMUST00 | 1052368 | ENSMUSG0 | Gm20063 | predicted gene, 20063 |
| chr12 | 61816049 | 61816354 | 306 | * | NA_peak_39 | 39 | Intron (ENSM | 12 | 61523948 | 61857928 | 333981 | 1 | 238205 | ENSMUST00 | 292101 | ENSMUSG0 | Lrfn5 | leucine rich repeat and fibronectin type III domain containing 5 |
| chr12 | 62177147 | 62177576 | 430 | * | NA_peak_40 | 54 | Distal Interge | 12 | 62687822 | 62688245 | 424 | 1 | 73309 | ENSMUST00 | -510246 | NA | Spanxn4 | SPANX family, member N4 |
| chr12 | 62986093 | 62986251 | 159 | * | NA_peak_40 | 38 | Distal Interge | 12 | 62687822 | 62688245 | 424 | 1 | 73309 | ENSMUST00 | 298271 | NA | Spanxn4 | SPANX family, member N4 |
| chr12 | 63121590 | 63121967 | 378 | * | NA_peak_40 | 39 | Distal Interge | 12 | 62687822 | 62688245 | 424 | 1 | 73309 | ENSMUST00 | 433768 | NA | Spanxn4 | SPANX family, member N4 |
| chr12 | 70480822 | 70481155 | 334 | * | NA_peak_40 | 36 | Intron (ENSM | 12 | 70459841 | 70460576 | 736 | 1 | 72736 | ENSMUST00 | 20981 | ENSMUSG0 | Tmx1 | thioredoxin-related transmembrane protein 1 |
| chr12 | 70666847 | 70667185 | 339 | * | NA_peak_40 | 31 | Intron (ENSM | 12 | 70825514 | 70902234 | 76721 | 1 | 319710 | ENSMUST00 | -158329 | ENSMUSG0 | Frmd6 | FERM domain containing 6 |
| chr12 | 76225374 | 76225659 | 286 | * | NA_peak_40 | 46 | Intron (ENSM | 12 | 76214486 | 76229464 | 14979 | 2 | 80384 | ENSMUST00 | 3805 | ENSMUSG0 | Tex21 | testis expressed gene 21 |
| chr12 | 77553168 | 77553483 | 316 | * | NA_peak_40 | 13 | Exon (ENSM | 12 | 77410141 | 77475989 | 65849 | 1 | 53618 | ENSMUST00 | 143027 | ENSMUSG0 | Fut8 | fucosyltransferase 8 |
| chr12 | 87976963 | 87977461 | 499 | * | NA_peak_40 | 26 | Distal Interge | 12 | 87954548 | 87958845 | 4298 | 1 | 100039087 | ENSMUST00 | 22415 | ENSMUSG0 | Gm2042 | predicted gene 2042 |
| chr12 | 93510554 | 93510680 | 127 | * | NA_peak_40 | 23 | Distal Interge | 12 | 93416458 | 93444823 | 28366 | 2 | 75295 | ENSMUST00 | -65731 | ENSMUSG0 | 4930559C10 | RIKEN cDNA 4930559C10 gene |
| chr12 | 93588923 | 93589231 | 309 | * | NA_peak_40 | 35 | Distal Interge | 12 | 93416458 | 93444823 | 28366 | 2 | 75295 | ENSMUST00 | -144100 | ENSMUSG0 | 4930559C10 | RIKEN cDNA 4930559C10 gene |
| chr12 | 95342084 | 95342384 | 301 | * | NA_peak_40 | 33 | Distal Interge | 12 | 95692226 | 95785215 | 92990 | 1 | 399558 | ENSMUST00 | -349842 | ENSMUSG0 | Flrt2 | fibronectin leucine rich transmembrane protein 2 |
| chr12 | 109934256 | 109934634 | 379 | * | NA_peak_40 | 26 | Distal Interge | 12 | 109747964 | 109749458 | 1495 | 1 | 100040724 | ENSMUST00 | 186292 | ENSMUSG0 | Mirg | miRNA containing gene |
| chr12 | 112776768 | 112776998 | 231 | * | NA_peak_40 | 44 | Exon (ENSM | 12 | 112766544 | 112767561 | 1018 | 1 | 104759 | ENSMUST00 | 10224 | ENSMUSG0 | Pld4 | phospholipase D family, member 4 |
| chr12 | 113718017 | 113718280 | 264 | * | NA_peak_40 | 19 | Distal Interge | 12 | 113543908 | 113546465 | 2558 | 1 | 238406 | ENSMUST00 | 174109 | ENSMUSG0 | Adam6a | a disintegrin and metallopeptidase domain 6A |
| chr12 | 113772599 | 113773151 | 553 | * | NA_peak_40 | 67 | Distal Interge | 12 | 113543908 | 113546465 | 2558 | 1 | 238406 | ENSMUST00 | 228691 | ENSMUSG0 | Adam6a | a disintegrin and metallopeptidase domain 6A |
| chr12 | 114480222 | 114480429 | 208 | * | NA_peak_40 | 23 | Distal Interge | 12 | 114469407 | 114469466 | 60 | 2 | 102465663 | ENSMUST00 | -10756 | ENSMUSG0 | Mir7094-1 | microRNA 7094-1 |
| chr12 | 114820071 | 114820521 | 451 | * | NA_peak_40 | 30 | Distal Interge | 12 | 114513635 | 114513694 | 60 | 2 | 102465664 | ENSMUST00 | -306377 | ENSMUSG0 | Mir7094-2 | microRNA 7094-2 |
| chr12 | 114832041 | 114832350 | 310 | * | NA_peak_40 | 23 | Distal Interge | 12 | 114513635 | 114513694 | 60 | 2 | 102465664 | ENSMUST00 | -318347 | ENSMUSG0 | Mir7094-2 | microRNA 7094-2 |
| chr12 | 115178268 | 115178472 | 205 | * | NA_peak_40 | 44 | Distal Interge | 12 | 115695720 | 115695794 | 75 | 2 | 102465205 | ENSMUST00 | 517322 | ENSMUSG0 | Mir6388 | microRNA 6388 |
| chr12 | 115205883 | 115206208 | 326 | * | NA_peak_40 | 25 | Distal Interge | 12 | 115695720 | 115695794 | 75 | 2 | 102465205 | ENSMUST00 | 489586 | ENSMUSG0 | Mir6388 | microRNA 6388 |
| chr12 | 115214327 | 115214814 | 488 | * | NA_peak_40 | 13 | Distal Interge | 12 | 115695720 | 115695794 | 75 | 2 | 102465205 | ENSMUST00 | 480980 | ENSMUSG0 | Mir6388 | microRNA 6388 |
| chr12 | 115215773 | 115216024 | 252 | * | NA_peak_40 | 42 | Distal Interge | 12 | 115695720 | 115695794 | 75 | 2 | 102465205 | ENSMUST00 | 479770 | ENSMUSG0 | Mir6388 | microRNA 6388 |
| chr12 | 115571253 | 115571813 | 561 | * | NA_peak_40 | 16 | Distal Interge | 12 | 115695720 | 115695794 | 75 | 2 | 102465205 | ENSMUST00 | 123981 | ENSMUSG0 | Mir6388 | microRNA 6388 |
| chr12 | 115586500 | 115586863 | 364 | * | NA_peak_40 | 59 | Distal Interge | 12 | 115695720 | 115695794 | 75 | 2 | 102465205 | ENSMUST00 | 108931 | ENSMUSG0 | Mir6388 | microRNA 6388 |
| chr12 | 115606315 | 115606839 | 525 | * | NA_peak_40 | 30 | Distal Interge | 12 | 115695720 | 115695794 | 75 | 2 | 102465205 | ENSMUST00 | 88955 | ENSMUSG0 | Mir6388 | microRNA 6388 |
| chr12 | 115684194 | 115684523 | 330 | * | NA_peak_40 | 21 | Distal Interge | 12 | 115695720 | 115695794 | 75 | 2 | 102465205 | ENSMUST00 | 11271 | ENSMUSG0 | Mir6388 | microRNA 6388 |
| chr12 | 115788660 | 115788870 | 211 | * | NA_peak_40 | 30 | Distal Interge | 12 | 115695720 | 115695794 | 75 | 2 | 102465205 | ENSMUST00 | -92866 | ENSMUSG0 | Mir6388 | microRNA 6388 |
| chr12 | 115851099 | 115851377 | 279 | * | NA_peak_40 | 15 | Distal Interge | 12 | 115695720 | 115695794 | 75 | 2 | 102465205 | ENSMUST00 | -155305 | ENSMUSG0 | Mir6388 | microRNA 6388 |
| chr12 | 119595450 | 119595596 | 147 | * | NA_peak_40 | 23 | Distal Interge | 12 | 119443410 | 119466934 | 23525 | 1 | 238455 | ENSMUST00 | 152040 | ENSMUSG0 | Macc1 | metastasis associated in colon cancer 1 |
| chr13 | 3372845 | 3373558 | 714 | * | NA_peak_40 | 29 | Distal Interge | 13 | 3478230 | 3486177 | 7948 | 1 | 76937 | ENSMUST00 | -104672 | NA | 2810429I04 | RIKEN cDNA 2810429I04 gene |
| chr13 | 6784463 | 6784848 | 386 | * | NA_peak_41 | 57 | Distal Interge | 13 | 6579768 | 6648777 | 69010 | 2 | 56421 | ENSMUST00 | -135686 | ENSMUSG0 | Pfkp | phosphofructokinase, platelet |
| chr13 | 7277602 | 7278160 | 559 | * | NA_peak_41 | 38 | Distal Interge | 13 | 6579768 | 6648777 | 69010 | 2 | 56421 | ENSMUST00 | -628825 | ENSMUSG0 | Pfkp | phosphofructokinase, platelet |
| chr13 | 9046709 | 9047072 | 364 | * | NA_peak_41 | 39 | Intron (ENSM | 13 | 9093881 | 9173988 | 80108 | 1 | 217980 | ENSMUST00 | -46809 | ENSMUSG0 | Larp4b | La ribonucleoprotein domain family, member 4B |

| | | | | | | | | | | | | | | | | | | |
|---|---|---|---|---|---|---|---|---|---|---|---|---|---|---|---|---|---|---|
| chr13 | 10305901 | 10306231 | 331 | * | NA_peak_41 | 26 | Intron (ENSM | 13 | 10215694 | 10360563 | 144870 | 2 | 12671 | ENSMUST0 | 54332 | ENSMUSG0 | Chrm3 | cholinergic receptor, muscarinic 3, cardiac |
| chr13 | 12788110 | 12788757 | 648 | * | NA_peak_41 | 25 | Distal Interge | 13 | 12790826 | 12792340 | 1515 | 2 | 26421 | ENSMUST0 | 3583 | ENSMUSG0 | Prl2c4 | prolactin family 2, subfamily c, member 4 |
| chr13 | 12962901 | 12963199 | 299 | * | NA_peak_41 | 51 | Distal Interge | 13 | 12996126 | 12997575 | 1450 | 2 | 18811 | ENSMUST0 | 34376 | ENSMUSG0 | Prl2c2 | prolactin family 2, subfamily c, member 2 |
| chr13 | 12977279 | 12977627 | 349 | * | NA_peak_41 | 20 | Distal Interge | 13 | 12996126 | 12997575 | 1450 | 2 | 18811 | ENSMUST0 | 19948 | ENSMUSG0 | Prl2c2 | prolactin family 2, subfamily c, member 2 |
| chr13 | 12978379 | 12978655 | 277 | * | NA_peak_41 | 20 | Distal Interge | 13 | 12996126 | 12997575 | 1450 | 2 | 18811 | ENSMUST0 | 18920 | ENSMUSG0 | Prl2c2 | prolactin family 2, subfamily c, member 2 |
| chr13 | 12981140 | 12981356 | 217 | * | NA_peak_41 | 35 | Distal Interge | 13 | 12996126 | 12997575 | 1450 | 2 | 18811 | ENSMUST0 | 16219 | ENSMUSG0 | Prl2c2 | prolactin family 2, subfamily c, member 2 |
| chr13 | 12996933 | 12997224 | 292 | * | NA_peak_41 | 38 | Promoter (<= | 13 | 12996126 | 12997575 | 1450 | 2 | 18811 | ENSMUST0 | 351 | ENSMUSG0 | Prl2c2 | prolactin family 2, subfamily c, member 2 |
| chr13 | 13018030 | 13018157 | 128 | * | NA_peak_41 | 24 | Distal Interge | 13 | 12996125 | 13005383 | 9259 | 2 | 18811 | ENSMUST0 | -12647 | ENSMUSG0 | Prl2c2 | prolactin family 2, subfamily c, member 2 |
| chr13 | 13309940 | 13310265 | 326 | * | NA_peak_41 | 44 | Distal Interge | 13 | 13365003 | 13367805 | 2803 | 2 | 83924 | ENSMUST0 | 57540 | ENSMUSG0 | Gpr137b | G protein-coupled receptor 137B |
| chr13 | 13599805 | 13599978 | 174 | * | NA_peak_41 | 23 | Intron (ENSM | 13 | 13590409 | 13650117 | 59709 | 1 | 17101 | ENSMUST0 | 9396 | ENSMUSG0 | Lyst | lysosomal trafficking regulator |
| chr13 | 15832124 | 15832558 | 435 | * | NA_peak_41 | 29 | Intron (ENSM | 13 | 15807243 | 15827537 | 20295 | 1 | 328190 | ENSMUST0 | 24881 | ENSMUSG0 | A530046M15 | RIKEN cDNA A530046M15 gene |
| chr13 | 16248990 | 16249256 | 267 | * | NA_peak_41 | 23 | Distal Interge | 13 | 15944752 | 16023423 | 78672 | 2 | 100503142 | ENSMUST0 | -225567 | ENSMUSG0 | B230303A05 | U1 small nuclear ribonucleoprotein 1C pseudogene |
| chr13 | 17444222 | 17444595 | 374 | * | NA_peak_41 | 39 | Intron (ENSM | 13 | 17684485 | 17694736 | 10252 | 2 | 192136 | ENSMUST0 | 250141 | ENSMUSG0 | Sugct | succinyl-CoA glutarate-CoA transferase |
| chr13 | 17465294 | 17465608 | 315 | * | NA_peak_42 | 13 | Intron (ENSM | 13 | 17684485 | 17694736 | 10252 | 2 | 192136 | ENSMUST0 | 229128 | ENSMUSG0 | Sugct | succinyl-CoA glutarate-CoA transferase |
| chr13 | 17803686 | 17804213 | 528 | * | NA_peak_42 | 39 | Promoter (<= | 13 | 17766233 | 17803468 | 37236 | 2 | 69562 | ENSMUST0 | -218 | ENSMUSG0 | Cdk13 | cyclin-dependent kinase 13 |
| chr13 | 18933536 | 18933869 | 334 | * | NA_peak_42 | 23 | Distal Interge | 13 | 18948205 | 19150921 | 202717 | 1 | 218038 | ENSMUST0 | -14336 | ENSMUSG0 | Amph | amphiphysin |
| chr13 | 18937153 | 18937484 | 332 | * | NA_peak_42 | 37 | Distal Interge | 13 | 18948205 | 19150921 | 202717 | 1 | 218038 | ENSMUST0 | -10721 | ENSMUSG0 | Amph | amphiphysin |
| chr13 | 18963413 | 18964044 | 632 | * | NA_peak_42 | 26 | Intron (ENSM | 13 | 18948398 | 19101626 | 153229 | 1 | 218038 | ENSMUST0 | 15015 | ENSMUSG0 | Amph | amphiphysin |
| chr13 | 20091008 | 20091552 | 545 | * | NA_peak_42 | 24 | Promoter (<= | 13 | 20090619 | 20608353 | 517735 | 1 | 140580 | ENSMUST0 | 389 | ENSMUSG0 | Elmo1 | engulfment and cell motility 1 |
| chr13 | 20672792 | 20673526 | 735 | * | NA_peak_42 | 29 | Distal Interge | 13 | 20794113 | 21024252 | 230140 | 1 | 27052 | ENSMUST0 | -120587 | ENSMUSG0 | Aoah | acyloxyacyl hydrolase |
| chr13 | 21717691 | 21717974 | 284 | * | NA_peak_42 | 30 | Promoter (<= | 13 | 21717659 | 21718069 | 411 | 1 | 319152 | ENSMUST0 | 32 | ENSMUSG0 | H3c10 | H3 clustered histone 10 |
| chr13 | 21764585 | 21765041 | 457 | * | NA_peak_42 | 24 | Distal Interge | 13 | 21754123 | 21754503 | 381 | 1 | 319187 | ENSMUST0 | 10462 | ENSMUSG0 | H2bc15 | H2B clustered histone 15 |
| chr13 | 22599425 | 22599577 | 153 | * | NA_peak_42 | 44 | Distal Interge | 13 | 22591900 | 22593000 | 1101 | 2 | 171251 | ENSMUST0 | -6425 | ENSMUSG0 | Vmn1r205 | vomeronasal 1 receptor 205 |
| chr13 | 22971352 | 22971646 | 295 | * | NA_peak_42 | 16 | Distal Interge | 13 | 23011249 | 23012403 | 1155 | 1 | 171249 | ENSMUST0 | -39603 | ENSMUSG0 | Vmn1r213 | vomeronasal 1 receptor 213 |
| chr13 | 24777189 | 24777247 | 59 | * | NA_peak_42 | 26 | Distal Interge | 13 | 24751845 | 24761923 | 10079 | 2 | 57441 | ENSMUST0 | -15266 | ENSMUSG0 | Gmnn | geminin |
| chr13 | 24832010 | 24832305 | 296 | * | NA_peak_42 | 17 | Promoter (<= | 13 | 24831944 | 24842152 | 10209 | 1 | 56196 | ENSMUST0 | 66 | ENSMUSG0 | Tdp2 | tyrosyl-DNA phosphodiesterase 2 |
| chr13 | 25009504 | 25009765 | 262 | * | NA_peak_42 | 44 | Intron (ENSM | 13 | 24998794 | 25001800 | 3007 | 2 | 380836 | ENSMUST0 | -7704 | ENSMUSG0 | Mrs2 | MRS2 magnesium transporter |
| chr13 | 26833868 | 26834242 | 375 | * | NA_peak_42 | 26 | Distal Interge | 13 | 26768173 | 26770165 | 1993 | 2 | 15192 | ENSMUST0 | -63703 | ENSMUSG0 | Hdgfl1 | HDGF like 1 |
| chr13 | 28961890 | 28962033 | 144 | * | NA_peak_42 | 23 | Distal Interge | 13 | 28948919 | 28953713 | 4795 | 2 | 20677 | ENSMUST0 | -8177 | ENSMUSG0 | Sox4 | SRY (sex determining region Y)-box 4 |
| chr13 | 30263555 | 30263926 | 372 | * | NA_peak_43 | 14 | Distal Interge | 13 | 30231853 | 30245715 | 13863 | 1 | 218121 | ENSMUST0 | 31702 | ENSMUSG0 | Mboat1 | membrane bound O-acyltransferase domain containing 1 |
| chr13 | 31012578 | 31012859 | 282 | * | NA_peak_43 | 43 | Distal Interge | 13 | 30923656 | 30974093 | 50438 | 2 | 66482 | ENSMUST0 | -38485 | ENSMUSG0 | Exoc2 | exocyst complex component 2 |
| chr13 | 31755011 | 31755356 | 346 | * | NA_peak_43 | 15 | Distal Interge | 13 | 31806633 | 31812476 | 5844 | 1 | 17300 | ENSMUST0 | -51277 | ENSMUSG0 | Foxc1 | forkhead box C1 |
| chr13 | 33775501 | 33775604 | 104 | * | NA_peak_43 | 30 | Intron (ENSM | 13 | 33832345 | 33843408 | 11064 | 2 | 435350 | ENSMUST0 | 67804 | ENSMUSG0 | Serpinb6e | serine (or cysteine) peptidase inhibitor, clade B, member 6e |
| chr13 | 35707270 | 35707568 | 299 | * | NA_peak_43 | 43 | Intron (ENSM | 13 | 35741313 | 35874058 | 132746 | 1 | 12593 | ENSMUST0 | -33745 | ENSMUSG0 | Cdyl | chromodomain protein, Y chromosome-like |
| chr13 | 36185338 | 36186385 | 1048 | * | NA_peak_43 | 38 | Intron (ENSM | 13 | 36232122 | 36246609 | 14488 | 1 | 69955 | ENSMUST0 | -45737 | ENSMUSG0 | Fars2 | phenylalanine-tRNA synthetase 2 (mitochondrial) |
| chr13 | 36189463 | 36189614 | 152 | * | NA_peak_43 | 29 | Exon (ENSM | 13 | 36232122 | 36246609 | 14488 | 1 | 69955 | ENSMUST0 | -42508 | ENSMUSG0 | Fars2 | phenylalanine-tRNA synthetase 2 (mitochondrial) |
| chr13 | 37037715 | 37037803 | 89 | * | NA_peak_43 | 23 | Intron (ENSM | 13 | 36938916 | 37049187 | 110272 | 2 | 74145 | ENSMUST0 | 11384 | ENSMUSG0 | F13a1 | coagulation factor XIII, A1 subunit |
| chr13 | 38658951 | 38659266 | 316 | * | NA_peak_43 | 30 | Promoter (<= | 13 | 38644207 | 38659058 | 14852 | 2 | 66143 | ENSMUST0 | 0 | ENSMUSG0 | Eef1e1 | eukaryotic translation elongation factor 1 epsilon 1 |
| chr13 | 39053498 | 39053843 | 346 | * | NA_peak_43 | 39 | Distal Interge | 13 | 38932136 | 38960875 | 28740 | 2 | 108652 | ENSMUST0 | -92623 | ENSMUSG0 | Slc35b3 | solute carrier family 35, member B3 |
| chr13 | 39942579 | 39942932 | 354 | * | NA_peak_43 | 56 | Distal Interge | 13 | 40001882 | 40288011 | 286130 | 2 | 218165 | ENSMUST0 | 345079 | ENSMUSG0 | Ofcc1 | orofacial cleft 1 candidate 1 |
| chr13 | 40588942 | 40589278 | 337 | * | NA_peak_43 | 17 | Intron (ENSM | 13 | 40727808 | 40735162 | 7355 | 1 | 102633805 | ENSMUST0 | -138530 | ENSMUSG0 | Gm26688 | predicted gene, 26688 |
| chr13 | 40592086 | 40592429 | 344 | * | NA_peak_43 | 24 | Intron (ENSM | 13 | 40727808 | 40735162 | 7355 | 1 | 102633805 | ENSMUST0 | -135379 | ENSMUSG0 | Gm26688 | predicted gene, 26688 |
| chr13 | 43615786 | 43615867 | 82 | * | NA_peak_43 | 17 | Promoter (<= | 13 | 43615710 | 43670945 | 55236 | 1 | 328234 | ENSMUST0 | 76 | ENSMUSG0 | Rnf182 | ring finger protein 182 |
| chr13 | 44869576 | 44869906 | 331 | * | NA_peak_43 | 93 | Intron (ENSM | 13 | 44840783 | 44896481 | 55699 | 1 | 16468 | ENSMUST0 | 28793 | ENSMUSG0 | Jarid2 | jumonji, AT rich interactive domain 2 |
| chr13 | 49144420 | 49144589 | 170 | * | NA_peak_44 | 19 | Promoter (2- | 13 | 49036309 | 49147232 | 110924 | 2 | 75607 | ENSMUST0 | 2643 | ENSMUSG0 | Wnk2 | WNK lysine deficient protein kinase 2 |
| chr13 | 49446496 | 49446749 | 254 | * | NA_peak_44 | 17 | 3' UTR | 13 | 49421614 | 49462901 | 41288 | 1 | 75678 | ENSMUST0 | 24882 | ENSMUSG0 | Ippk | inositol 1,3,4,5,6-pentakisphosphate 2-kinase |
| chr13 | 50807430 | 50807874 | 445 | * | NA_peak_44 | 27 | Distal Interge | 13 | 50753391 | 50765248 | 11858 | 2 | 71654 | ENSMUST0 | -42182 | ENSMUSG0 | 4930518P08 | RIKEN cDNA 4930518P08 gene |
| chr13 | 51938884 | 51939170 | 287 | * | NA_peak_44 | 20 | Distal Interge | 13 | 51846744 | 51848224 | 1481 | 1 | 23882 | ENSMUST0 | 92140 | ENSMUSG0 | Gadd45g | growth arrest and DNA-damage-inducible 45 gamma |
| chr13 | 53522754 | 53522922 | 169 | * | NA_peak_44 | 23 | Intron (ENSM | 13 | 53466884 | 53473074 | 6191 | 2 | 17702 | ENSMUST0 | -49680 | ENSMUSG0 | Msx2 | msh homeobox 2 |
| chr13 | 55472052 | 55472282 | 231 | * | NA_peak_44 | 15 | Exon (ENSM | 13 | 55477681 | 55477776 | 96 | 2 | 102465569 | ENSMUST0 | 5494 | ENSMUSG0 | Mir6944 | microRNA 6944 |
| chr13 | 55474903 | 55475410 | 508 | * | NA_peak_44 | 42 | Promoter (2- | 13 | 55477681 | 55477776 | 96 | 2 | 102465569 | ENSMUST0 | 2366 | ENSMUSG0 | Mir6944 | microRNA 6944 |
| chr13 | 56770737 | 56770976 | 240 | * | NA_peak_44 | 24 | Distal Interge | 13 | 56733017 | 56738993 | 5977 | 1 | 17129 | ENSMUST0 | 37720 | ENSMUSG0 | Smad5 | SMAD family member 5 |
| chr13 | 59769524 | 59769700 | 177 | * | NA_peak_45 | 19 | Promoter (<= | 13 | 59769497 | 59771408 | 1912 | 1 | 13996 | ENSMUST0 | 27 | NA | Etohd2 | ethanol decreased 2 |
| chr13 | 60978693 | 60979041 | 349 | * | NA_peak_45 | 19 | Distal Interge | 13 | 61000278 | 61005911 | 5634 | 2 | 26898 | ENSMUST0 | 26870 | ENSMUSG0 | Ctsj | cathepsin J |
| chr13 | 62079569 | 62079929 | 361 | * | NA_peak_45 | 27 | Intron (ENSM | 13 | 62129886 | 62172258 | 42373 | 1 | 630579 | ENSMUST0 | -49957 | ENSMUSG0 | Zfp808 | zinc finger protein 808 |
| chr13 | 62081533 | 62081842 | 310 | * | NA_peak_45 | 58 | Intron (ENSM | 13 | 62129886 | 62172258 | 42373 | 1 | 630579 | ENSMUST0 | -48044 | ENSMUSG0 | Zfp808 | zinc finger protein 808 |
| chr13 | 62093971 | 62094831 | 861 | * | NA_peak_45 | 37 | Intron (ENSM | 13 | 62129886 | 62172258 | 42373 | 1 | 630579 | ENSMUST0 | -35055 | ENSMUSG0 | Zfp808 | zinc finger protein 808 |
| chr13 | 62099600 | 62100017 | 418 | * | NA_peak_45 | 47 | Intron (ENSM | 13 | 62129886 | 62172258 | 42373 | 1 | 630579 | ENSMUST0 | -29869 | ENSMUSG0 | Zfp808 | zinc finger protein 808 |
| chr13 | 62109886 | 62109974 | 89 | * | NA_peak_45 | 28 | Intron (ENSM | 13 | 62129886 | 62172258 | 42373 | 1 | 630579 | ENSMUST0 | -19912 | ENSMUSG0 | Zfp808 | zinc finger protein 808 |
| chr13 | 62117583 | 62117840 | 258 | * | NA_peak_45 | 29 | Intron (ENSM | 13 | 62129886 | 62172258 | 42373 | 1 | 630579 | ENSMUST0 | -12046 | ENSMUSG0 | Zfp808 | zinc finger protein 808 |
| chr13 | 62419240 | 62419263 | 24 | * | NA_peak_45 | 22 | Distal Interge | 13 | 62368328 | 62383177 | 14850 | 2 | 100041979 | ENSMUST0 | -36063 | ENSMUSG0 | Gm3604 | predicted gene 3604 |
| chr13 | 62436185 | 62436657 | 473 | * | NA_peak_45 | 32 | Distal Interge | 13 | 62454913 | 62466793 | 11881 | 2 | 71508 | ENSMUST0 | 30136 | ENSMUSG0 | Zfp935 | zinc finger protein 935 |
| chr13 | 62508348 | 62508755 | 408 | * | NA_peak_45 | 72 | Intron (ENSM | 13 | 62517500 | 62520451 | 2952 | 2 | 77117 | ENSMUST0 | 11696 | ENSMUSG0 | Zfp934 | zinc finger protein 934 |
| chr13 | 62509575 | 62510015 | 441 | * | NA_peak_45 | 52 | Intron (ENSM | 13 | 62517500 | 62520451 | 2952 | 2 | 77117 | ENSMUST0 | 10436 | ENSMUSG0 | Zfp934 | zinc finger protein 934 |
| chr13 | 62544275 | 62544570 | 296 | * | NA_peak_45 | 17 | Intron (ENSM | 13 | 62453715 | 62558511 | 104797 | 2 | 71508 | ENSMUST0 | 13941 | ENSMUSG0 | Zfp935 | zinc finger protein 935 |
| chr13 | 62590674 | 62590902 | 229 | * | NA_peak_45 | 28 | Distal Interge | 13 | 62605190 | 62607499 | 2310 | 2 | 211378 | ENSMUST0 | 16597 | ENSMUSG0 | 6720489N17 | RIKEN cDNA 6720489N17 gene |
| chr13 | 63564565 | 63564735 | 171 | * | NA_peak_45 | 23 | Promoter (<= | 13 | 63508336 | 63564535 | 56200 | 2 | 19206 | ENSMUST0 | -30 | ENSMUSG0 | Ptch1 | patched 1 |
| chr13 | 65337876 | 65338012 | 137 | * | NA_peak_45 | 29 | Intron (ENSM | 13 | 65336222 | 65348597 | 12376 | 2 | 665738 | ENSMUST0 | 10585 | ENSMUSG0 | Gm7762 | predicted gene 7762 |
| chr13 | 65351183 | 65351815 | 633 | * | NA_peak_45 | 26 | Promoter (2- | 13 | 65336222 | 65348597 | 12376 | 2 | 665738 | ENSMUST0 | -2586 | ENSMUSG0 | Gm7762 | predicted gene 7762 |
| chr13 | 65358335 | 65358854 | 520 | * | NA_peak_45 | 35 | Intron (ENSM | 13 | 65336222 | 65348597 | 12376 | 2 | 665738 | ENSMUST0 | -9738 | ENSMUSG0 | Gm7762 | predicted gene 7762 |
| chr13 | 65369237 | 65370277 | 1041 | * | NA_peak_45 | 27 | Distal Interge | 13 | 65336222 | 65348597 | 12376 | 2 | 665738 | ENSMUST0 | -20640 | ENSMUSG0 | Gm7762 | predicted gene 7762 |
| chr13 | 65390651 | 65391180 | 530 | * | NA_peak_45 | 52 | Distal Interge | 13 | 65336222 | 65348597 | 12376 | 2 | 665738 | ENSMUST0 | -42054 | ENSMUSG0 | Gm7762 | predicted gene 7762 |
| chr13 | 65414826 | 65415237 | 412 | * | NA_peak_45 | 37 | Distal Interge | 13 | 65336222 | 65348597 | 12376 | 2 | 665738 | ENSMUST0 | -66229 | ENSMUSG0 | Gm7762 | predicted gene 7762 |
| chr13 | 65429274 | 65430705 | 1432 | * | NA_peak_45 | 27 | Intron (ENSM | 13 | 65336222 | 65348597 | 12376 | 2 | 665738 | ENSMUST0 | -80677 | ENSMUSG0 | Gm7762 | predicted gene 7762 |
| chr13 | 65436123 | 65436564 | 442 | * | NA_peak_45 | 21 | Exon (ENSM | 13 | 65336222 | 65348597 | 12376 | 2 | 665738 | ENSMUST0 | -87526 | ENSMUSG0 | Gm7762 | predicted gene 7762 |
| chr13 | 65468753 | 65469184 | 432 | * | NA_peak_45 | 33 | Distal Interge | 13 | 65336222 | 65348597 | 12376 | 2 | 665738 | ENSMUST0 | -120156 | ENSMUSG0 | Gm7762 | predicted gene 7762 |
| chr13 | 65484353 | 65484755 | 403 | * | NA_peak_45 | 37 | Distal Interge | 13 | 65336222 | 65348597 | 12376 | 2 | 665738 | ENSMUST0 | -135756 | ENSMUSG0 | Gm7762 | predicted gene 7762 |
| chr13 | 65514710 | 65515050 | 341 | * | NA_peak_45 | 17 | Exon (ENSM | 13 | 65336222 | 65348597 | 12376 | 2 | 665738 | ENSMUST0 | -166113 | ENSMUSG0 | Gm7762 | predicted gene 7762 |
| chr13 | 65515410 | 65515763 | 354 | * | NA_peak_45 | 49 | Intron (ENSM | 13 | 65336222 | 65348597 | 12376 | 2 | 665738 | ENSMUST0 | -166813 | ENSMUSG0 | Gm7762 | predicted gene 7762 |
| chr13 | 65522882 | 65523134 | 253 | * | NA_peak_45 | 26 | Intron (ENSM | 13 | 65336222 | 65348597 | 12376 | 2 | 665738 | ENSMUST0 | -174285 | ENSMUSG0 | Gm7762 | predicted gene 7762 |
| chr13 | 65560014 | 65560475 | 462 | * | NA_peak_45 | 40 | Intron (ENSM | 13 | 65336222 | 65348597 | 12376 | 2 | 665738 | ENSMUST0 | -211417 | ENSMUSG0 | Gm7762 | predicted gene 7762 |
| chr13 | 65569540 | 65569851 | 312 | * | NA_peak_45 | 28 | Intron (ENSM | 13 | 65336222 | 65348597 | 12376 | 2 | 665738 | ENSMUST0 | -220943 | ENSMUSG0 | Gm7762 | predicted gene 7762 |
| chr13 | 65585237 | 65585547 | 311 | * | NA_peak_45 | 18 | Intron (ENSM | 13 | 65336222 | 65348597 | 12376 | 2 | 665738 | ENSMUST0 | -236640 | ENSMUSG0 | Gm7762 | predicted gene 7762 |
| chr13 | 65607060 | 65607359 | 300 | * | NA_peak_45 | 24 | Intron (ENSM | 13 | 65336222 | 65348597 | 12376 | 2 | 665738 | ENSMUST0 | -258463 | ENSMUSG0 | Gm7762 | predicted gene 7762 |
| chr13 | 65621102 | 65621514 | 413 | * | NA_peak_45 | 85 | Intron (ENSM | 13 | 65336222 | 65348597 | 12376 | 2 | 665738 | ENSMUST0 | -272505 | ENSMUSG0 | Gm7762 | predicted gene 7762 |
| chr13 | 65624289 | 65624604 | 316 | * | NA_peak_45 | 21 | Intron (ENSM | 13 | 65336222 | 65348597 | 12376 | 2 | 665738 | ENSMUST0 | -275692 | ENSMUSG0 | Gm7762 | predicted gene 7762 |
| chr13 | 65626453 | 65626621 | 169 | * | NA_peak_45 | 17 | Intron (ENSM | 13 | 65336222 | 65348597 | 12376 | 2 | 665738 | ENSMUST0 | -277856 | ENSMUSG0 | Gm7762 | predicted gene 7762 |
| chr13 | 65630437 | 65631019 | 583 | * | NA_peak_45 | 54 | Exon (ENSM | 13 | 65336222 | 65348597 | 12376 | 2 | 665738 | ENSMUST0 | -281840 | ENSMUSG0 | Gm7762 | predicted gene 7762 |
| chr13 | 65631460 | 65631963 | 504 | * | NA_peak_45 | 41 | Distal Interge | 13 | 65336222 | 65348597 | 12376 | 2 | 665738 | ENSMUST0 | -282863 | ENSMUSG0 | Gm7762 | predicted gene 7762 |
| chr13 | 65641691 | 65642105 | 415 | * | NA_peak_45 | 16 | Distal Interge | 13 | 65336222 | 65348597 | 12376 | 2 | 665738 | ENSMUST0 | -293094 | ENSMUSG0 | Gm7762 | predicted gene 7762 |
| chr13 | 65649631 | 65649971 | 341 | * | NA_peak_45 | 57 | Exon (ENSM | 13 | 65336222 | 65348597 | 12376 | 2 | 665738 | ENSMUST0 | -301034 | ENSMUSG0 | Gm7762 | predicted gene 7762 |
| chr13 | 65650395 | 65650727 | 333 | * | NA_peak_46 | 59 | Distal Interge | 13 | 65336222 | 65348597 | 12376 | 2 | 665738 | ENSMUST0 | -301798 | ENSMUSG0 | Gm7762 | predicted gene 7762 |
| chr13 | 65657368 | 65657633 | 266 | * | NA_peak_46 | 38 | Distal Interge | 13 | 65336222 | 65348597 | 12376 | 2 | 665738 | ENSMUST0 | -308771 | ENSMUSG0 | Gm7762 | predicted gene 7762 |
| chr13 | 65658775 | 65659258 | 484 | * | NA_peak_46 | 21 | Distal Interge | 13 | 65336222 | 65348597 | 12376 | 2 | 665738 | ENSMUST0 | -310178 | ENSMUSG0 | Gm7762 | predicted gene 7762 |
| chr13 | 65682323 | 65682690 | 368 | * | NA_peak_46 | 40 | Distal Interge | 13 | 65336222 | 65348597 | 12376 | 2 | 665738 | ENSMUST0 | -333726 | ENSMUSG0 | Gm7762 | predicted gene 7762 |
| chr13 | 65711821 | 65712270 | 450 | * | NA_peak_46 | 39 | Intron (ENSM | 13 | 65336222 | 65348597 | 12376 | 2 | 665738 | ENSMUST0 | -363224 | ENSMUSG0 | Gm7762 | predicted gene 7762 |
| chr13 | 65750114 | 65750581 | 468 | * | NA_peak_46 | 28 | Distal Interge | 13 | 66113480 | 66120204 | 6725 | 1 | 628709 | ENSMUST0 | -362899 | ENSMUSG0 | Zfp997 | zinc finger protein 997 |
| chr13 | 65754744 | 65755120 | 377 | * | NA_peak_46 | 88 | Distal Interge | 13 | 66113480 | 66120204 | 6725 | 1 | 628709 | ENSMUST0 | -358360 | ENSMUSG0 | Zfp997 | zinc finger protein 997 |
| chr13 | 65793064 | 65793489 | 426 | * | NA_peak_46 | 28 | Distal Interge | 13 | 66113480 | 66120204 | 6725 | 1 | 628709 | ENSMUST0 | -319991 | ENSMUSG0 | Zfp997 | zinc finger protein 997 |
| chr13 | 65805261 | 65805842 | 582 | * | NA_peak_46 | 46 | Distal Interge | 13 | 66113480 | 66120204 | 6725 | 1 | 628709 | ENSMUST0 | -307638 | ENSMUSG0 | Zfp997 | zinc finger protein 997 |
| chr13 | 65858310 | 65858968 | 659 | * | NA_peak_46 | 37 | Distal Interge | 13 | 66113480 | 66120204 | 6725 | 1 | 628709 | ENSMUST0 | -254512 | ENSMUSG0 | Zfp997 | zinc finger protein 997 |
| chr13 | 65870434 | 65870969 | 536 | * | NA_peak_46 | 41 | Distal Interge | 13 | 66113480 | 66120204 | 6725 | 1 | 628709 | ENSMUST0 | -242511 | ENSMUSG0 | Zfp997 | zinc finger protein 997 |
| chr13 | 65874047 | 65874550 | 504 | * | NA_peak_46 | 20 | Exon (ENSM | 13 | 66113480 | 66120204 | 6725 | 1 | 628709 | ENSMUST0 | -238930 | ENSMUSG0 | Zfp997 | zinc finger protein 997 |
| chr13 | 65884390 | 65884700 | 311 | * | NA_peak_46 | 36 | Intron (ENSM | 13 | 66113480 | 66120204 | 6725 | 1 | 628709 | ENSMUST0 | -228780 | ENSMUSG0 | Zfp997 | zinc finger protein 997 |
| chr13 | 65886004 | 65886283 | 280 | * | NA_peak_46 | 53 | Exon (ENSM | 13 | 66113480 | 66120204 | 6725 | 1 | 628709 | ENSMUST0 | -227197 | ENSMUSG0 | Zfp997 | zinc finger protein 997 |
| chr13 | 65887125 | 65888526 | 1402 | * | NA_peak_46 | 35 | Exon (ENSM | 13 | 66113480 | 66120204 | 6725 | 1 | 628709 | ENSMUST0 | -224954 | ENSMUSG0 | Zfp997 | zinc finger protein 997 |
| chr13 | 65901253 | 65901760 | 508 | * | NA_peak_46 | 37 | Distal Interge | 13 | 66113480 | 66120204 | 6725 | 1 | 628709 | ENSMUST0 | -211720 | ENSMUSG0 | Zfp997 | zinc finger protein 997 |
| chr13 | 65914681 | 65915010 | 330 | * | NA_peak_46 | 17 | Distal Interge | 13 | 66113480 | 66120204 | 6725 | 1 | 628709 | ENSMUST0 | -198470 | ENSMUSG0 | Zfp997 | zinc finger protein 997 |
| chr13 | 65936264 | 65936806 | 543 | * | NA_peak_46 | 61 | Distal Interge | 13 | 66113480 | 66120204 | 6725 | 1 | 628709 | ENSMUST0 | -176674 | ENSMUSG0 | Zfp997 | zinc finger protein 997 |
| chr13 | 65941287 | 65941682 | 396 | * | NA_peak_46 | 15 | Distal Interge | 13 | 66113480 | 66120204 | 6725 | 1 | 628709 | ENSMUST0 | -171798 | ENSMUSG0 | Zfp997 | zinc finger protein 997 |
| chr13 | 65977548 | 65977921 | 374 | * | NA_peak_46 | 68 | Intron (ENSM | 13 | 66113480 | 66120204 | 6725 | 1 | 628709 | ENSMUST0 | -135559 | ENSMUSG0 | Zfp997 | zinc finger protein 997 |
| chr13 | 65982737 | 65983497 | 761 | * | NA_peak_46 | 43 | Exon (ENSM | 13 | 66113480 | 66120204 | 6725 | 1 | 628709 | ENSMUST0 | -129983 | ENSMUSG0 | Zfp997 | zinc finger protein 997 |
| chr13 | 65985349 | 65985851 | 503 | * | NA_peak_46 | 53 | Distal Interge | 13 | 66113480 | 66120204 | 6725 | 1 | 628709 | ENSMUST0 | -127629 | ENSMUSG0 | Zfp997 | zinc finger protein 997 |
| chr13 | 66002912 | 66003404 | 493 | * | NA_peak_46 | 13 | Distal Interge | 13 | 66113480 | 66120204 | 6725 | 1 | 628709 | ENSMUST0 | -110076 | ENSMUSG0 | Zfp997 | zinc finger protein 997 |
| chr13 | 66025953 | 66026337 | 385 | * | NA_peak_46 | 51 | Distal Interge | 13 | 66113480 | 66120204 | 6725 | 1 | 628709 | ENSMUST0 | -87143 | ENSMUSG0 | Zfp997 | zinc finger protein 997 |
| chr13 | 66041626 | 66041928 | 303 | * | NA_peak_46 | 50 | Distal Interge | 13 | 66113480 | 66120204 | 6725 | 1 | 628709 | ENSMUST0 | -71552 | ENSMUSG0 | Zfp997 | zinc finger protein 997 |
| chr13 | 66043015 | 66043412 | 398 | * | NA_peak_46 | 33 | Distal Interge | 13 | 66113480 | 66120204 | 6725 | 1 | 628709 | ENSMUST0 | -70068 | ENSMUSG0 | Zfp997 | zinc finger protein 997 |
| chr13 | 66062633 | 66062940 | 308 | * | NA_peak_46 | 13 | Distal Interge | 13 | 66113480 | 66120204 | 6725 | 1 | 628709 | ENSMUST0 | -50540 | ENSMUSG0 | Zfp997 | zinc finger protein 997 |
| chr13 | 66121915 | 66122228 | 314 | * | NA_peak_46 | 21 | Promoter (2- | 13 | 66119547 | 66122836 | 3290 | 1 | 628709 | ENSMUST0 | 2368 | ENSMUSG0 | Zfp997 | zinc finger protein 997 |
| chr13 | 66149708 | 66150130 | 423 | * | NA_peak_46 | 26 | Distal Interge | 13 | 66119547 | 66122836 | 3290 | 1 | 628709 | ENSMUST0 | 30161 | ENSMUSG0 | Zfp997 | zinc finger protein 997 |
| chr13 | 66157625 | 66158005 | 381 | * | NA_peak_46 | 17 | Distal Interge | 13 | 66119547 | 66122836 | 3290 | 1 | 628709 | ENSMUST0 | 38078 | ENSMUSG0 | Zfp997 | zinc finger protein 997 |
| chr13 | 66185518 | 66185833 | 316 | * | NA_peak_46 | 49 | Distal Interge | 13 | 66119547 | 66122836 | 3290 | 1 | 628709 | ENSMUST0 | 65971 | ENSMUSG0 | Zfp997 | zinc finger protein 997 |
| chr13 | 66243265 | 66244179 | 915 | * | NA_peak_46 | 63 | Distal Interge | 13 | 66119547 | 66122836 | 3290 | 1 | 628709 | ENSMUST0 | 123718 | ENSMUSG0 | Zfp997 | zinc finger protein 997 |
| chr13 | 66270981 | 66271295 | 315 | * | NA_peak_46 | 38 | Intron (ENSM | 13 | 66418322 | 66419597 | 1276 | 2 | 76803 | ENSMUST0 | 148302 | ENSMUSG0 | Zfp998 | zinc finger protein 998 |
| chr13 | 66298726 | 66299078 | 353 | * | NA_peak_46 | 39 | Intron (ENSM | 13 | 66418322 | 66419597 | 1276 | 2 | 76803 | ENSMUST0 | 120519 | ENSMUSG0 | Zfp998 | zinc finger protein 998 |
| chr13 | 66306596 | 66306999 | 404 | * | NA_peak_46 | 65 | Intron (ENSM | 13 | 66418322 | 66419597 | 1276 | 2 | 76803 | ENSMUST0 | 112598 | ENSMUSG0 | Zfp998 | zinc finger protein 998 |
| chr13 | 66364700 | 66365012 | 313 | * | NA_peak_46 | 46 | Intron (ENSM | 13 | 66418322 | 66419597 | 1276 | 2 | 76803 | ENSMUST0 | 54585 | ENSMUSG0 | Zfp998 | zinc finger protein 998 |
| chr13 | 66412383 | 66413104 | 722 | * | NA_peak_46 | 24 | Exon (ENSM | 13 | 66418322 | 66419597 | 1276 | 2 | 76803 | ENSMUST0 | 6493 | ENSMUSG0 | Zfp998 | zinc finger protein 998 |
| chr13 | 66415760 | 66416116 | 357 | * | NA_peak_46 | 20 | Intron (ENSM | 13 | 66418322 | 66419597 | 1276 | 2 | 76803 | ENSMUST0 | 3481 | ENSMUSG0 | Zfp998 | zinc finger protein 998 |
| chr13 | 66431124 | 66431451 | 328 | * | NA_peak_46 | 26 | Promoter (2- | 13 | 66418245 | 66433895 | 15651 | 2 | 76803 | ENSMUST0 | 2444 | ENSMUSG0 | Zfp998 | zinc finger protein 998 |
| chr13 | 66434457 | 66434793 | 337 | * | NA_peak_46 | 18 | Promoter (<= | 13 | 66418245 | 66433895 | 15651 | 2 | 76803 | ENSMUST0 | -562 | ENSMUSG0 | Zfp998 | zinc finger protein 998 |
| chr13 | 66469152 | 66469424 | 273 | * | NA_peak_46 | 30 | Distal Interge | 13 | 66418114 | 66441118 | 23005 | 2 | 76803 | ENSMUST0 | -28034 | ENSMUSG0 | Zfp998 | zinc finger protein 998 |
| chr13 | 66495714 | 66495924 | 211 | * | NA_peak_46 | 21 | Distal Interge | 13 | 66418114 | 66441118 | 23005 | 2 | 76803 | ENSMUST0 | -54596 | ENSMUSG0 | Zfp998 | zinc finger protein 998 |
| chr13 | 66594518 | 66594854 | 337 | * | NA_peak_46 | 33 | Distal Interge | 13 | 66418114 | 66441118 | 23005 | 2 | 76803 | ENSMUST0 | -153400 | ENSMUSG0 | Zfp998 | zinc finger protein 998 |
| chr13 | 66621265 | 66621761 | 497 | * | NA_peak_46 | 15 | Exon (ENSM | 13 | 66418114 | 66441118 | 23005 | 2 | 76803 | ENSMUST0 | -180147 | ENSMUSG0 | Zfp998 | zinc finger protein 998 |
| chr13 | 66623051 | 66623476 | 426 | * | NA_peak_46 | 18 | Distal Interge | 13 | 66418114 | 66441118 | 23005 | 2 | 76803 | ENSMUST0 | -181933 | ENSMUSG0 | Zfp998 | zinc finger protein 998 |
| chr13 | 66626175 | 66626364 | 190 | * | NA_peak_46 | 18 | Distal Interge | 13 | 66418114 | 66441118 | 23005 | 2 | 76803 | ENSMUST0 | -185057 | ENSMUSG0 | Zfp998 | zinc finger protein 998 |
| chr13 | 66630775 | 66631100 | 326 | * | NA_peak_46 | 15 | Distal Interge | 13 | 66418114 | 66441118 | 23005 | 2 | 76803 | ENSMUST0 | -189657 | ENSMUSG0 | Zfp998 | zinc finger protein 998 |
| chr13 | 66692175 | 66692477 | 303 | * | NA_peak_46 | 106 | Intron (ENSM | 13 | 66901061 | 66902786 | 1726 | 2 | 67530 | ENSMUST0 | 210309 | ENSMUSG0 | Uqcrb | ubiquinol-cytochrome c reductase binding protein |
| chr13 | 66710529 | 66710819 | 291 | * | NA_peak_46 | 67 | Distal Interge | 13 | 66901061 | 66902786 | 1726 | 2 | 67530 | ENSMUST0 | 191967 | ENSMUSG0 | Uqcrb | ubiquinol-cytochrome c reductase binding protein |
| chr13 | 66725894 | 66726415 | 522 | * | NA_peak_46 | 17 | Distal Interge | 13 | 66901061 | 66902786 | 1726 | 2 | 67530 | ENSMUST0 | 176371 | ENSMUSG0 | Uqcrb | ubiquinol-cytochrome c reductase binding protein |
| chr13 | 66773424 | 66773745 | 322 | * | NA_peak_46 | 20 | Distal Interge | 13 | 66901061 | 66902786 | 1726 | 2 | 67530 | ENSMUST0 | 129041 | ENSMUSG0 | Uqcrb | ubiquinol-cytochrome c reductase binding protein |
| chr13 | 66775393 | 66775531 | 139 | * | NA_peak_46 | 32 | Distal Interge | 13 | 66901061 | 66902786 | 1726 | 2 | 67530 | ENSMUST0 | 127255 | ENSMUSG0 | Uqcrb | ubiquinol-cytochrome c reductase binding protein |
| chr13 | 66776476 | 66776803 | 328 | * | NA_peak_46 | 14 | Distal Interge | 13 | 66901061 | 66902786 | 1726 | 2 | 67530 | ENSMUST0 | 125983 | ENSMUSG0 | Uqcrb | ubiquinol-cytochrome c reductase binding protein |
| chr13 | 66820977 | 66821833 | 857 | * | NA_peak_46 | 35 | Distal Interge | 13 | 66901061 | 66902786 | 1726 | 2 | 67530 | ENSMUST0 | 80953 | ENSMUSG0 | Uqcrb | ubiquinol-cytochrome c reductase binding protein |
| chr13 | 66858709 | 66859058 | 350 | * | NA_peak_46 | 67 | Distal Interge | 13 | 66901061 | 66902786 | 1726 | 2 | 67530 | ENSMUST0 | 43728 | ENSMUSG0 | Uqcrb | ubiquinol-cytochrome c reductase binding protein |
| chr13 | 68010268 | 68010691 | 424 | * | NA_peak_46 | 22 | Distal Interge | 13 | 67863326 | 67863925 | 600 | 1 | 408058 | ENSMUST0 | 146942 | ENSMUSG0 | BC048507 | cDNA sequence BC048507 |
| chr13 | 68079569 | 68079628 | 60 | * | NA_peak_46 | 16 | Distal Interge | 13 | 67863326 | 67863925 | 600 | 1 | 408058 | ENSMUST0 | 216243 | ENSMUSG0 | BC048507 | cDNA sequence BC048507 |
| chr13 | 68086444 | 68086877 | 434 | * | NA_peak_46 | 18 | Distal Interge | 13 | 67863326 | 67863925 | 600 | 1 | 408058 | ENSMUST0 | 223118 | ENSMUSG0 | BC048507 | cDNA sequence BC048507 |
| chr13 | 68087428 | 68087820 | 393 | * | NA_peak_46 | 40 | Distal Interge | 13 | 67863326 | 67863925 | 600 | 1 | 408058 | ENSMUST0 | 224102 | ENSMUSG0 | BC048507 | cDNA sequence BC048507 |
| chr13 | 68166122 | 68166459 | 338 | * | NA_peak_47 | 64 | Distal Interge | 13 | 68353364 | 68356825 | 3462 | 2 | 666272 | ENSMUST0 | 190366 | ENSMUSG0 | Gm8016 | predicted gene 8016 |
| chr13 | 68293468 | 68293756 | 289 | * | NA_peak_47 | 65 | Distal Interge | 13 | 68353364 | 68356825 | 3462 | 2 | 666272 | ENSMUST0 | 63069 | ENSMUSG0 | Gm8016 | predicted gene 8016 |
| chr13 | 68305089 | 68305434 | 346 | * | NA_peak_47 | 19 | Distal Interge | 13 | 68353364 | 68356825 | 3462 | 2 | 666272 | ENSMUST0 | 51391 | ENSMUSG0 | Gm8016 | predicted gene 8016 |
| chr13 | 68427687 | 68428421 | 735 | * | NA_peak_47 | 35 | Distal Interge | 13 | 68353364 | 68356825 | 3462 | 2 | 666272 | ENSMUST0 | -70862 | ENSMUSG0 | Gm8016 | predicted gene 8016 |
| chr13 | 72037511 | 72037870 | 360 | * | NA_peak_47 | 26 | Distal Interge | 13 | 71957921 | 71963723 | 5803 | 2 | 16371 | ENSMUST0 | -73788 | ENSMUSG0 | Irx1 | Iroquois homeobox 1 |
| chr13 | 72391010 | 72391233 | 224 | * | NA_peak_47 | 20 | Distal Interge | 13 | 72626213 | 72628564 | 2352 | 2 | 328287 | ENSMUST0 | 237331 | NA | Gm20554 | predicted gene, 20554 |
| chr13 | 73128320 | 73128639 | 320 | * | NA_peak_47 | 40 | Distal Interge | 13 | 73260479 | 73269608 | 9130 | 1 | 50916 | ENSMUST0 | -131840 | ENSMUSG0 | Irx4 | Iroquois homeobox 4 |
| chr13 | 73467134 | 73467364 | 231 | * | NA_peak_47 | 17 | Promoter (<= | 13 | 73467197 | 73516422 | 49226 | 1 | 210992 | ENSMUST0 | 0 | ENSMUSG0 | Lpcat1 | lysophosphatidylcholine acyltransferase 1 |
| chr13 | 74356722 | 74356953 | 232 | * | NA_peak_47 | 30 | Exon (ENSM | 13 | 74350365 | 74357048 | 6684 | 1 | 67433 | ENSMUST0 | 6357 | ENSMUSG0 | Ccdc127 | coiled-coil domain containing 127 |
| chr13 | 74544893 | 74544987 | 95 | * | NA_peak_47 | 14 | Distal Interge | 13 | 74480057 | 74493992 | 13936 | 2 | 235956 | ENSMUST0 | -50901 | ENSMUSG0 | Zfp825 | zinc finger protein 825 |
| chr13 | 78262440 | 78262958 | 519 | * | NA_peak_47 | 23 | Intron (ENSM | 13 | 78202553 | 78231386 | 28834 | 1 | 320174 | ENSMUST0 | 59887 | ENSMUSG0 | A830082K12 | RIKEN cDNA A830082K12 gene |
| chr13 | 79455422 | 79455809 | 388 | * | NA_peak_47 | 22 | Distal Interge | 13 | 78202553 | 78231386 | 28834 | 1 | 320174 | ENSMUST0 | 1252869 | ENSMUSG0 | A830082K12 | RIKEN cDNA A830082K12 gene |
| chr13 | 79565530 | 79565921 | 392 | * | NA_peak_47 | 26 | Distal Interge | 13 | 80883384 | 80889643 | 6260 | 1 | 105171 | ENSMUST0 | -1317463 | ENSMUSG0 | Arrdc3 | arrestin domain containing 3 |
| chr13 | 84441516 | 84441635 | 120 | * | NA_peak_47 | 39 | Distal Interge | 13 | 84251271 | 84286738 | 35468 | 1 | 72745 | ENSMUST0 | 190245 | ENSMUSG0 | Tmem161b | transmembrane protein 161B |
| chr13 | 92623886 | 92623919 | 34 | * | NA_peak_48 | 27 | Intron (ENSM | 13 | 92611174 | 92686193 | 75020 | 1 | 218442 | ENSMUST0 | 12712 | ENSMUSG0 | Serinc5 | serine incorporator 5 |
| chr13 | 94057753 | 94057792 | 40 | * | NA_peak_48 | 23 | Promoter (<= | 13 | 94057796 | 94195409 | 137614 | 1 | 218454 | ENSMUST0 | -4 | ENSMUSG0 | Lhfpl2 | lipoma HMGIC fusion partner-like 2 |
| chr13 | 95479515 | 95479820 | 306 | * | NA_peak_48 | 20 | Promoter (<= | 13 | 95477268 | 95478721 | 1454 | 2 | 268686 | ENSMUST0 | -794 | ENSMUSG0 | S100z | S100 calcium binding protein, zeta |
| chr13 | 96542403 | 96542795 | 393 | * | NA_peak_48 | 13 | Promoter (<= | 13 | 96480690 | 96542485 | 61796 | 2 | 27015 | ENSMUST0 | 0 | ENSMUSG0 | Polk | polymerase (DNA directed), kappa |
| chr13 | 98450599 | 98450753 | 155 | * | NA_peak_48 | 26 | Distal Interge | 13 | 98424109 | 98435065 | 10957 | 2 | 70090 | ENSMUST0 | -15534 | ENSMUSG0 | 2310005E17 | RIKEN cDNA 2310005E17 gene |
| chr13 | 98613687 | 98613767 | 81 | * | NA_peak_48 | 19 | Exon (ENSM | 13 | 98634978 | 98637410 | 2433 | 2 | 68344 | ENSMUST0 | 23643 | ENSMUSG0 | Tmem174 | transmembrane protein 174 |
| chr13 | 100364100 | 100364698 | 599 | * | NA_peak_48 | 26 | Intron (ENSM | 13 | 100281121 | 100317674 | 36554 | 2 | 17952 | ENSMUST0 | -46426 | ENSMUSG0 | Naip6 | NLR family, apoptosis inhibitory protein 6 |
| chr13 | 100652336 | 100652708 | 373 | * | NA_peak_48 | 18 | Promoter (<= | 13 | 100651626 | 100656060 | 4435 | 1 | 108143 | ENSMUST0 | 710 | ENSMUSG0 | Taf9 | TATA-box binding protein associated factor 9 |
| chr13 | 108844477 | 108844816 | 340 | * | NA_peak_49 | 57 | Intron (ENSM | 13 | 108860166 | 109951344 | 1091179 | 1 | 238871 | ENSMUST0 | -15350 | ENSMUSG0 | Pde4d | phosphodiesterase 4D, cAMP specific |
| chr13 | 109413330 | 109413586 | 257 | * | NA_peak_49 | 30 | Intron (ENSM | 13 | 109442183 | 109951364 | 509182 | 1 | 238871 | ENSMUST0 | -28597 | ENSMUSG0 | Pde4d | phosphodiesterase 4D, cAMP specific |
| chr13 | 109853821 | 109854175 | 355 | * | NA_peak_49 | 35 | Intron (ENSM | 13 | 109903102 | 109951387 | 48286 | 1 | 238871 | ENSMUST0 | -48927 | ENSMUSG0 | Pde4d | phosphodiesterase 4D, cAMP specific |
| chr13 | 109870459 | 109870492 | 34 | * | NA_peak_49 | 26 | Intron (ENSM | 13 | 109903102 | 109951387 | 48286 | 1 | 238871 | ENSMUST0 | -32610 | ENSMUSG0 | Pde4d | phosphodiesterase 4D, cAMP specific |
| chr13 | 109902568 | 109902880 | 313 | * | NA_peak_49 | 30 | Promoter (<= | 13 | 109903102 | 109951387 | 48286 | 1 | 238871 | ENSMUST0 | -222 | ENSMUSG0 | Pde4d | phosphodiesterase 4D, cAMP specific |
| chr13 | 111514945 | 111515459 | 515 | * | NA_peak_49 | 30 | Intron (ENSM | 13 | 111425680 | 111490111 | 64432 | 2 | 73274 | ENSMUST0 | -24834 | ENSMUSG0 | Gpbp1 | GC-rich promoter binding protein 1 |
| chr13 | 113285307 | 113285507 | 201 | * | NA_peak_49 | 23 | Distal Interge | 13 | 113171608 | 113225908 | 54301 | 2 | 14945 | ENSMUST0 | -59399 | ENSMUSG0 | Gzmk | granzyme K |
| chr13 | 113474438 | 113474737 | 300 | * | NA_peak_49 | 39 | Intron (ENSM | 13 | 113446288 | 113502844 | 56557 | 2 | 70917 | ENSMUST0 | 28107 | ENSMUSG0 | 4921509O07 | RIKEN cDNA 4921509O07 gene |
| chr13 | 113720997 | 113721329 | 333 | * | NA_peak_49 | 22 | Distal Interge | 13 | 113662896 | 113663676 | 781 | 2 | 56534 | ENSMUST0 | -57321 | ENSMUSG0 | Hspb3 | heat shock protein 3 |
| chr13 | 113722535 | 113722825 | 291 | * | NA_peak_49 | 31 | Distal Interge | 13 | 113662896 | 113663676 | 781 | 2 | 56534 | ENSMUST0 | -58859 | ENSMUSG0 | Hspb3 | heat shock protein 3 |
| chr13 | 114542807 | 114542898 | 92 | * | NA_peak_49 | 18 | Distal Interge | 13 | 114586513 | 114607084 | 20572 | 2 | 74909 | ENSMUST0 | 64186 | ENSMUSG0 | 4930467J12 | RIKEN cDNA 4930467J12 gene |
| chr13 | 114931709 | 114932034 | 326 | * | NA_peak_49 | 23 | Promoter (<= | 13 | 114833081 | 114932100 | 99020 | 2 | 16398 | ENSMUST0 | 66 | ENSMUSG0 | Itga2 | integrin alpha 2 |
| chr13 | 117625353 | 117625796 | 444 | * | NA_peak_49 | 26 | Intron (ENSM | 13 | 117602480 | 117987418 | 384939 | 1 | 15165 | ENSMUST0 | 22873 | ENSMUSG0 | Hcn1 | hyperpolarization activated cyclic nucleotide gated potassium channel 1 |
| chr13 | 119671740 | 119671964 | 225 | * | NA_peak_49 | 20 | Distal Interge | 13 | 119680040 | 119681588 | 1549 | 1 | 73480 | ENSMUST0 | -8076 | ENSMUSG0 | 1700074H08 | RIKEN cDNA 1700074H08 gene |
| chr14 | 4097290 | 4097542 | 253 | * | NA_peak_50 | 24 | Intron (ENSM | 14 | 4023941 | 4041370 | 17430 | 1 | 545007 | ENSMUST0 | 73349 | ENSMUSG0 | Gm5796 | predicted gene 5796 |
| chr14 | 5637685 | 5637880 | 196 | * | NA_peak_50 | 23 | Intron (ENSM | 14 | 5723156 | 5725943 | 2788 | 2 | 100041678 | ENSMUST0 | 88063 | ENSMUSG0 | Gm3500 | predicted gene 3500 |
| chr14 | 10113217 | 10113540 | 324 | * | NA_peak_50 | 26 | Intron (ENSM | 14 | 9886026 | 9886629 | 604 | 1 | 100042415 | ENSMUST0 | 227191 | ENSMUSG0 | Mnd1-ps | Mnd1 retrotransposed pseudogene |

| | | | | | | | | | | | | | | | | | | |
|---|---|---|---|---|---|---|---|---|---|---|---|---|---|---|---|---|---|---|
| chr14 | 14626050 | 14626438 | 389 | * | NA_peak_50 | 39 | Distal Interge | 14 | 14702279 | 14799940 | 97662 | 1 | 218756 | ENSMUST0( | -75841 | ENSMUSG0( | Slc4a7 | solute carrier family 4, sodium bicarbonate cotransporter, member 7 |
| chr14 | 17380611 | 17380911 | 301 | * | NA_peak_50 | 30 | Distal Interge | 14 | 17105561 | 17105876 | 316 | 1 | 100417973 | ENSMUST0( | 275050 | NA | Pou5f1-rs6 | POU domain, class 5, transcription factor 1, related sequence 6 |
| chr14 | 19415847 | 19416333 | 487 | * | NA_peak_50 | 170 | Exon (ENSM | 14 | 19594139 | 19602581 | 8443 | 2 | 432825 | ENSMUST0( | 186248 | ENSMUSG0( | Gm5458 | predicted gene 5458 |
| chr14 | 19416900 | 19418269 | 1370 | * | NA_peak_50 | 155 | Exon (ENSM | 14 | 19594139 | 19602581 | 8443 | 2 | 432825 | ENSMUST0( | 184312 | ENSMUSG0( | Gm5458 | predicted gene 5458 |
| chr14 | 19418613 | 19419500 | 888 | * | NA_peak_50 | 60 | Exon (ENSM | 14 | 19594139 | 19602581 | 8443 | 2 | 432825 | ENSMUST0( | 183081 | ENSMUSG0( | Gm5458 | predicted gene 5458 |
| chr14 | 28607404 | 28608006 | 603 | * | NA_peak_50 | 30 | Distal Interge | 14 | 28511451 | 28511907 | 457 | 1 | 22418 | ENSMUST0( | 95953 | ENSMUSG0( | Wnt5a | wingless-type MMTV integration site family, member 5A |
| chr14 | 28620701 | 28620900 | 200 | * | NA_peak_50 | 22 | Distal Interge | 14 | 28511451 | 28511907 | 457 | 1 | 22418 | ENSMUST0( | 109250 | ENSMUSG0( | Wnt5a | wingless-type MMTV integration site family, member 5A |
| chr14 | 32399608 | 32399908 | 301 | * | NA_peak_50 | 15 | Distal Interge | 14 | 32376068 | 32388391 | 12324 | 2 | 67085 | ENSMUST0( | -11217 | ENSMUSG0( | 1700024G13 | RIKEN cDNA 1700024G13 gene |
| chr14 | 33981426 | 33981538 | 113 | * | NA_peak_50 | 26 | Promoter (2- | 14 | 33967070 | 33978764 | 11695 | 2 | 382867 | ENSMUST0( | -2662 | ENSMUSG0( | Zfp488 | zinc finger protein 488 |
| chr14 | 38606215 | 38606485 | 271 | * | NA_peak_50 | 35 | Intron (ENSM | 14 | 38898302 | 38958794 | 60493 | 1 | 75170 | ENSMUST0( | -291817 | ENSMUSG0( | Nrg3os | neuregulin 3, opposite strand |
| chr14 | 40552238 | 40552830 | 593 | * | NA_peak_50 | 54 | Distal Interge | 14 | 40813789 | 40853305 | 39517 | 2 | 328381 | ENSMUST0( | 300475 | ENSMUSG0( | Sh2d4b | SH2 domain containing 4B |
| chr14 | 41297301 | 41297536 | 236 | * | NA_peak_50 | 49 | Distal Interge | 14 | 41172214 | 41185149 | 12936 | 2 | 20390 | ENSMUST0( | -112152 | ENSMUSG0( | Sftpd | surfactant associated protein D |
| chr14 | 41468761 | 41469125 | 365 | * | NA_peak_50 | 20 | Distal Interge | 14 | 41573211 | 41581305 | 8095 | 1 | 100041722 | ENSMUST0( | -104086 | ENSMUSG0( | 1700049E17 | tandem duplication of RIKEN cDNA 1700049E17 gene, gene 2 |
| chr14 | 43254558 | 43255191 | 634 | * | NA_peak_50 | 39 | Intron (ENSM | 14 | 43275036 | 43289618 | 14583 | 1 | 108168154 | ENSMUST0( | -19845 | ENSMUSG0( | Gm9732 | predicted gene 9732 |
| chr14 | 43262162 | 43262400 | 239 | * | NA_peak_50 | 33 | Intron (ENSM | 14 | 43275036 | 43289618 | 14583 | 1 | 108168154 | ENSMUST0( | -12636 | ENSMUSG0( | Gm9732 | predicted gene 9732 |
| chr14 | 43263324 | 43263725 | 402 | * | NA_peak_50 | 22 | Intron (ENSM | 14 | 43275036 | 43289618 | 14583 | 1 | 108168154 | ENSMUST0( | -11311 | ENSMUSG0( | Gm9732 | predicted gene 9732 |
| chr14 | 43274374 | 43274733 | 360 | * | NA_peak_50 | 48 | Promoter (<= | 14 | 43275036 | 43289618 | 14583 | 1 | 108168154 | ENSMUST0( | -303 | ENSMUSG0( | Gm9732 | predicted gene 9732 |
| chr14 | 43317049 | 43317217 | 169 | * | NA_peak_50 | 24 | Distal Interge | 14 | 43298872 | 43306004 | 7133 | 1 | 108168154 | ENSMUST0( | 18177 | ENSMUSG0( | Gm9732 | predicted gene 9732 |
| chr14 | 43330756 | 43330891 | 136 | * | NA_peak_50 | 16 | Distal Interge | 14 | 43342383 | 43347811 | 5429 | 2 | 71826 | ENSMUST0( | 16920 | ENSMUSG0( | 1700001F09 | RIKEN cDNA 1700001F09 gene |
| chr14 | 43331435 | 43331875 | 441 | * | NA_peak_50 | 16 | Distal Interge | 14 | 43342383 | 43347811 | 5429 | 2 | 71826 | ENSMUST0( | 15936 | ENSMUSG0( | 1700001F09 | RIKEN cDNA 1700001F09 gene |
| chr14 | 43346152 | 43346526 | 375 | * | NA_peak_50 | 21 | Promoter (1- | 14 | 43342383 | 43347811 | 5429 | 2 | 71826 | ENSMUST0( | 1285 | ENSMUSG0( | 1700001F09 | RIKEN cDNA 1700001F09 gene |
| chr14 | 43362013 | 43362883 | 871 | * | NA_peak_50 | 33 | Distal Interge | 14 | 43342383 | 43347811 | 5429 | 2 | 71826 | ENSMUST0( | -14202 | ENSMUSG0( | 1700001F09 | RIKEN cDNA 1700001F09 gene |
| chr14 | 43364567 | 43365163 | 597 | * | NA_peak_50 | 25 | Distal Interge | 14 | 43342383 | 43347811 | 5429 | 2 | 71826 | ENSMUST0( | -16756 | ENSMUSG0( | 1700001F09 | RIKEN cDNA 1700001F09 gene |
| chr14 | 43393496 | 43393814 | 319 | * | NA_peak_50 | 23 | Exon (ENSM | 14 | 43342383 | 43347811 | 5429 | 2 | 71826 | ENSMUST0( | -45685 | ENSMUSG0( | 1700001F09 | RIKEN cDNA 1700001F09 gene |
| chr14 | 43404993 | 43405485 | 493 | * | NA_peak_50 | 27 | Distal Interge | 14 | 43342383 | 43347811 | 5429 | 2 | 71826 | ENSMUST0( | -57182 | ENSMUSG0( | 1700001F09 | RIKEN cDNA 1700001F09 gene |
| chr14 | 44277676 | 44278044 | 369 | * | NA_peak_50 | 30 | Intron (ENSM | 14 | 44170306 | 44213254 | 42949 | 2 | 74328 | ENSMUST0( | -64422 | ENSMUSG0( | 1700047E10 | RIKEN cDNA 1700047E10 gene |
| chr14 | 45019565 | 45020079 | 515 | * | NA_peak_51 | 49 | Distal Interge | 14 | 44958607 | 44989699 | 31093 | 2 | 73895 | ENSMUST0( | -29866 | ENSMUSG0( | 4930431P03 | RIKEN cDNA 4930431P03 gene |
| chr14 | 51580700 | 51581006 | 307 | * | NA_peak_51 | 34 | Intron (ENSM | 14 | 51552790 | 51662953 | 110164 | 1 | 434459 | ENSMUST0( | 27910 | ENSMUSG0( | Gm5622 | predicted gene 5622 |
| chr14 | 51594536 | 51594790 | 255 | * | NA_peak_51 | 25 | Intron (ENSM | 14 | 51552790 | 51662953 | 110164 | 1 | 434459 | ENSMUST0( | 41746 | ENSMUSG0( | Gm5622 | predicted gene 5622 |
| chr14 | 51693520 | 51693912 | 393 | * | NA_peak_51 | 81 | Intron (ENSM | 14 | 51711643 | 51717183 | 5541 | 2 | 545047 | ENSMUST0( | 23271 | ENSMUSG0( | Gm5800 | predicted gene 5800 |
| chr14 | 56199507 | 56199965 | 459 | * | NA_peak_51 | 54 | Intron (ENSM | 14 | 56205263 | 56211417 | 6155 | 2 | 14943 | ENSMUST0( | 11452 | ENSMUSG0( | Gzmf | granzyme F |
| chr14 | 58326178 | 58326600 | 423 | * | NA_peak_51 | 98 | Intron (ENSM | 14 | 58076507 | 58109392 | 32886 | 1 | 14180 | ENSMUST0( | 249671 | ENSMUSG0( | Fgf9 | fibroblast growth factor 9 |
| chr14 | 60509175 | 60509563 | 389 | * | NA_peak_51 | 40 | Intron (ENSM | 14 | 60634001 | 60691367 | 57367 | 1 | 219140 | ENSMUST0( | -124438 | ENSMUSG0( | Spata13 | spermatogenesis associated 13 |
| chr14 | 69162038 | 69162434 | 397 | * | NA_peak_51 | 16 | Distal Interge | 14 | 69171802 | 69175518 | 3717 | 1 | 18092 | ENSMUST0( | -9368 | ENSMUSG0( | Nkx2-6 | NK2 homeobox 6 |
| chr14 | 69750919 | 69751169 | 251 | * | NA_peak_51 | 40 | Distal Interge | 14 | 69767472 | 69784411 | 16940 | 1 | 21933 | ENSMUST0( | -16303 | ENSMUSG0( | Tnfrsf10b | tumor necrosis factor receptor superfamily, member 10b |
| chr14 | 75439228 | 75439567 | 340 | * | NA_peak_51 | 24 | Distal Interge | 14 | 75455982 | 75529708 | 73727 | 1 | 380918 | ENSMUST0( | -16415 | ENSMUSG0( | Siah3 | siah E3 ubiquitin protein ligase family member 3 |
| chr14 | 83700151 | 83700380 | 230 | * | NA_peak_51 | 44 | Distal Interge | 14 | 84046431 | 84075290 | 28860 | 2 | 102635012 | ENSMUST0( | 374910 | ENSMUSG0( | Gm32455 | predicted gene, 32455 |
| chr14 | 84939409 | 84939608 | 200 | * | NA_peak_51 | 14 | Distal Interge | 14 | 84458724 | 84476424 | 17701 | 2 | 319903 | ENSMUST0( | -462985 | ENSMUSG0( | 9630013A20 | RIKEN cDNA 9630013A20 gene |
| chr14 | 85953402 | 85953686 | 285 | * | NA_peak_51 | 38 | Distal Interge | 14 | 86245970 | 86262042 | 16073 | 1 | 75155 | ENSMUST0( | -292284 | ENSMUSG0( | 4930529K09 | RIKEN cDNA 4930529K09 gene |
| chr14 | 100385731 | 100386099 | 369 | * | NA_peak_51 | 64 | Distal Interge | 14 | 99870632 | 100284679 | 414048 | 2 | 16597 | ENSMUST0( | -101052 | ENSMUSG0( | Klf12 | Kruppel-like factor 12 |
| chr14 | 102146533 | 102146728 | 196 | * | NA_peak_51 | 19 | Distal Interge | 14 | 101929261 | 101932866 | 3606 | 1 | 380928 | ENSMUST0( | 217272 | ENSMUSG0( | Lmo7 | LIM domain only 7 |
| chr14 | 103636225 | 103636769 | 545 | * | NA_peak_51 | 36 | Distal Interge | 14 | 103650228 | 103704907 | 54680 | 1 | 105439 | ENSMUST0( | -13459 | ENSMUSG0( | Slain1 | SLAIN motif family, member 1 |
| chr14 | 103863632 | 103864018 | 387 | * | NA_peak_51 | 23 | Distal Interge | 14 | 103814625 | 103844402 | 29778 | 2 | 13618 | ENSMUST0( | -19230 | ENSMUSG0( | Ednrb | endothelin receptor type B |
| chr14 | 106967607 | 106967883 | 277 | * | NA_peak_51 | 39 | Distal Interge | 14 | 106417187 | 106486520 | 69334 | 2 | 73631 | ENSMUST0( | -481087 | ENSMUSG0( | 1700128A07 | RIKEN cDNA 1700128A07 gene |
| chr14 | 109680777 | 109681090 | 314 | * | NA_peak_51 | 39 | Distal Interge | 14 | 108908800 | 108914158 | 5359 | 2 | 76965 | ENSMUST0( | -766619 | ENSMUSG0( | Slitrk1 | SLIT and NTRK-like family, member 1 |
| chr14 | 111157441 | 111158111 | 671 | * | NA_peak_51 | 30 | Distal Interge | 14 | 110748580 | 110755149 | 6570 | 2 | 239250 | ENSMUST0( | -402292 | ENSMUSG0( | Slitrk6 | SLIT and NTRK-like family, member 6 |
| chr14 | 112333789 | 112334115 | 327 | * | NA_peak_51 | 35 | Distal Interge | 14 | 111675849 | 111679342 | 3494 | 1 | 75409 | ENSMUST0( | 657940 | ENSMUSG0( | Slitrk5 | SLIT and NTRK-like family, member 5 |
| chr14 | 112369220 | 112369627 | 408 | * | NA_peak_51 | 32 | Distal Interge | 14 | 111675849 | 111679342 | 3494 | 1 | 75409 | ENSMUST0( | 693371 | ENSMUSG0( | Slitrk5 | SLIT and NTRK-like family, member 5 |
| chr14 | 113813115 | 113813537 | 423 | * | NA_peak_51 | 23 | Distal Interge | 14 | 114831965 | 114861327 | 29363 | 1 | 75822 | ENSMUST0( | -1018428 | ENSMUSG0( | 4930524C18 | RIKEN cDNA 4930524C18 gene |
| chr14 | 114506674 | 114507134 | 461 | * | NA_peak_51 | 23 | Distal Interge | 14 | 114831965 | 114861327 | 29363 | 1 | 75822 | ENSMUST0( | -324831 | ENSMUSG0( | 4930524C18 | RIKEN cDNA 4930524C18 gene |
| chr14 | 116233525 | 116233893 | 369 | * | NA_peak_51 | 26 | Intron (ENSM | 14 | 116925315 | 117627198 | 701884 | 1 | 23888 | ENSMUST0( | -691422 | ENSMUSG0( | Gpc6 | glypican 6 |
| chr14 | 120064161 | 120064480 | 320 | * | NA_peak_51 | 26 | Distal Interge | 14 | 120019585 | 120042435 | 22851 | 2 | 239283 | ENSMUST0( | -21726 | ENSMUSG0( | Oxgr1 | oxoglutarate (alpha-ketoglutarate) receptor 1 |
| chr14 | 123724800 | 123725077 | 278 | * | NA_peak_52 | 23 | Intron (ENSM | 14 | 123661424 | 123975618 | 314195 | 1 | 223272 | ENSMUST0( | 63376 | ENSMUSG0( | Itgbl1 | integrin, beta-like 1 |
| chr14 | 123731068 | 123731336 | 269 | * | NA_peak_52 | 30 | Intron (ENSM | 14 | 123661424 | 123975618 | 314195 | 1 | 223272 | ENSMUST0( | 69644 | ENSMUSG0( | Itgbl1 | integrin, beta-like 1 |
| chr15 | 3166978 | 3167214 | 237 | * | NA_peak_52 | 22 | Distal Interge | 15 | 3268547 | 3277273 | 8727 | 1 | 20363 | ENSMUST0( | -101333 | ENSMUSG0( | Selenop | selenoprotein P |
| chr15 | 3246103 | 3246423 | 321 | * | NA_peak_52 | 34 | Distal Interge | 15 | 3268547 | 3277273 | 8727 | 1 | 20363 | ENSMUST0( | -22124 | ENSMUSG0( | Selenop | selenoprotein P |
| chr15 | 3613181 | 3613249 | 69 | * | NA_peak_52 | 13 | Distal Interge | 15 | 3384309 | 3583492 | 199184 | 2 | 14600 | ENSMUST0( | -29689 | ENSMUSG0( | Ghr | growth hormone receptor |
| chr15 | 3676063 | 3676174 | 112 | * | NA_peak_52 | 19 | Distal Interge | 15 | 3384309 | 3583492 | 199184 | 2 | 14600 | ENSMUST0( | -92571 | ENSMUSG0( | Ghr | growth hormone receptor |
| chr15 | 3732874 | 3732957 | 84 | * | NA_peak_52 | 32 | Distal Interge | 15 | 3384309 | 3583492 | 199184 | 2 | 14600 | ENSMUST0( | -149382 | ENSMUSG0( | Ghr | growth hormone receptor |
| chr15 | 3733658 | 3733679 | 22 | * | NA_peak_52 | 49 | Distal Interge | 15 | 3384309 | 3583492 | 199184 | 2 | 14600 | ENSMUST0( | -150166 | ENSMUSG0( | Ghr | growth hormone receptor |
| chr15 | 3734204 | 3734317 | 114 | * | NA_peak_52 | 49 | Distal Interge | 15 | 3384309 | 3583492 | 199184 | 2 | 14600 | ENSMUST0( | -150712 | ENSMUSG0( | Ghr | growth hormone receptor |
| chr15 | 3783715 | 3784260 | 546 | * | NA_peak_52 | 30 | Distal Interge | 15 | 3971284 | 3971982 | 699 | 2 | 106052 | ENSMUST0( | 187722 | ENSMUSG0( | Fbxo4 | F-box protein 4 |
| chr15 | 3812578 | 3812627 | 50 | * | NA_peak_52 | 33 | Distal Interge | 15 | 3971284 | 3971982 | 699 | 2 | 106052 | ENSMUST0( | 159355 | ENSMUSG0( | Fbxo4 | F-box protein 4 |
| chr15 | 3817848 | 3817923 | 76 | * | NA_peak_52 | 20 | Distal Interge | 15 | 3971284 | 3971982 | 699 | 2 | 106052 | ENSMUST0( | 154059 | ENSMUSG0( | Fbxo4 | F-box protein 4 |
| chr15 | 3979138 | 3979525 | 388 | * | NA_peak_52 | 19 | Promoter (<= | 15 | 3965445 | 3979573 | 14129 | 2 | 106052 | ENSMUST0( | 48 | ENSMUSG0( | Fbxo4 | F-box protein 4 |
| chr15 | 4075291 | 4075577 | 287 | * | NA_peak_52 | 17 | Intron (ENSM | 15 | 4047259 | 4092493 | 45235 | 1 | 67041 | ENSMUST0( | 28032 | ENSMUSG0( | Oxct1 | 3-oxoacid CoA transferase 1 |
| chr15 | 4109067 | 4109639 | 573 | * | NA_peak_52 | 34 | Intron (ENSM | 15 | 4142523 | 4153980 | 11458 | 1 | 67041 | ENSMUST0( | -32884 | ENSMUSG0( | Oxct1 | 3-oxoacid CoA transferase 1 |
| chr15 | 4175883 | 4176290 | 408 | * | NA_peak_52 | 26 | Distal Interge | 15 | 4142812 | 4153959 | 11148 | 1 | 67041 | ENSMUST0( | 33071 | ENSMUSG0( | Oxct1 | 3-oxoacid CoA transferase 1 |
| chr15 | 4284569 | 4285066 | 498 | * | NA_peak_52 | 37 | Distal Interge | 15 | 4375491 | 4520414 | 144924 | 1 | 239318 | ENSMUST0( | -90425 | ENSMUSG0( | Plcxd3 | phosphatidylinositol-specific phospholipase C, X domain containing 3 |
| chr15 | 4669244 | 4669481 | 238 | * | NA_peak_52 | 35 | Distal Interge | 15 | 4727175 | 4804045 | 76871 | 1 | 12274 | ENSMUST0( | -57694 | ENSMUSG0( | C6 | complement component 6 |
| chr15 | 4670227 | 4670601 | 375 | * | NA_peak_52 | 19 | Distal Interge | 15 | 4727175 | 4804045 | 76871 | 1 | 12274 | ENSMUST0( | -56574 | ENSMUSG0( | C6 | complement component 6 |
| chr15 | 4862179 | 4862457 | 279 | * | NA_peak_52 | 21 | Distal Interge | 15 | 4898737 | 4906310 | 7574 | 1 | 223825 | ENSMUST0( | -36280 | ENSMUSG0( | Mroh2b | maestro heat-like repeat family member 2B |
| chr15 | 5105094 | 5105251 | 158 | * | NA_peak_52 | 26 | Exon (ENSM | 15 | 5105137 | 5108484 | 3348 | 2 | 239319 | ENSMUST0( | 3233 | ENSMUSG0( | Card6 | caspase recruitment domain family, member 6 |
| chr15 | 5183449 | 5183774 | 326 | * | NA_peak_52 | 29 | Promoter (1- | 15 | 5185525 | 5218336 | 32812 | 1 | 67515 | ENSMUST0( | -1751 | ENSMUSG0( | Ttc33 | tetratricopeptide repeat domain 33 |
| chr15 | 5268456 | 5269221 | 766 | * | NA_peak_53 | 13 | Distal Interge | 15 | 5233399 | 5244187 | 10789 | 2 | 19219 | ENSMUST0( | -24269 | ENSMUSG0( | Ptger4 | prostaglandin E receptor 4 (subtype EP4) |
| chr15 | 5379562 | 5380075 | 514 | * | NA_peak_53 | 17 | Distal Interge | 15 | 5496266 | 5594984 | 98719 | 1 | 71432 | ENSMUST0( | -116191 | ENSMUSG0( | 5430437J10 | RIKEN cDNA 5430437J10 gene |
| chr15 | 5621043 | 5621419 | 377 | * | NA_peak_53 | 26 | Distal Interge | 15 | 5496318 | 5594983 | 98666 | 1 | 71432 | ENSMUST0( | 124725 | ENSMUSG0( | 5430437J10 | RIKEN cDNA 5430437J10 gene |
| chr15 | 5830011 | 5830124 | 114 | * | NA_peak_53 | 18 | Distal Interge | 15 | 5496318 | 5594983 | 98666 | 1 | 71432 | ENSMUST0( | 333693 | ENSMUSG0( | 5430437J10 | RIKEN cDNA 5430437J10 gene |
| chr15 | 5925468 | 5925785 | 318 | * | NA_peak_53 | 19 | Distal Interge | 15 | 6299788 | 6435520 | 135733 | 1 | 13132 | ENSMUST0( | -374003 | ENSMUSG0( | Dab2 | disabled 2, mitogen-responsive phosphoprotein |
| chr15 | 6467175 | 6467490 | 316 | * | NA_peak_53 | 30 | 3' UTR | 15 | 6445372 | 6467409 | 22038 | 1 | 12279 | ENSMUST0( | 21803 | ENSMUSG0( | C9 | complement component 9 |
| chr15 | 6697648 | 6698207 | 560 | * | NA_peak_53 | 16 | Distal Interge | 15 | 6708379 | 6765417 | 57039 | 1 | 78757 | ENSMUST0( | -10172 | ENSMUSG0( | Rictor | RPTOR independent companion of MTOR, complex 2 |
| chr15 | 6746496 | 6746645 | 150 | * | NA_peak_53 | 23 | Intron (ENSM | 15 | 6708383 | 6800401 | 92019 | 1 | 78757 | ENSMUST0( | 38113 | ENSMUSG0( | Rictor | RPTOR independent companion of MTOR, complex 2 |
| chr15 | 6956480 | 6956742 | 263 | * | NA_peak_53 | 18 | Distal Interge | 15 | 6815037 | 6874969 | 59933 | 2 | 18414 | ENSMUST0( | -81511 | ENSMUSG0( | Osmr | oncostatin M receptor |
| chr15 | 7293081 | 7293470 | 390 | * | NA_peak_53 | 23 | Intron (ENSM | 15 | 7253511 | 7254768 | 1258 | 2 | 268780 | ENSMUST0( | -38313 | ENSMUSG0( | Egflam | EGF-like, fibronectin type III and laminin G domains |
| chr15 | 7364838 | 7365177 | 340 | * | NA_peak_53 | 14 | Intron (ENSM | 15 | 7305070 | 7398261 | 93192 | 2 | 268780 | ENSMUST0( | 33084 | ENSMUSG0( | Egflam | EGF-like, fibronectin type III and laminin G domains |
| chr15 | 7546422 | 7546818 | 397 | * | NA_peak_53 | 17 | Distal Interge | 15 | 7206583 | 7398395 | 191813 | 2 | 268780 | ENSMUST0( | -148027 | ENSMUSG0( | Egflam | EGF-like, fibronectin type III and laminin G domains |
| chr15 | 7847620 | 7847952 | 333 | * | NA_peak_53 | 30 | Distal Interge | 15 | 7810846 | 7837575 | 26730 | 1 | 14573 | ENSMUST0( | 36774 | ENSMUSG0( | Gdnf | glial cell line derived neurotrophic factor |
| chr15 | 7850759 | 7851119 | 361 | * | NA_peak_53 | 40 | Distal Interge | 15 | 7873055 | 7886377 | 13323 | 2 | 545085 | ENSMUST0( | 35258 | ENSMUSG0( | Wdr70 | WD repeat domain 70 |
| chr15 | 7937404 | 7937852 | 449 | * | NA_peak_53 | 23 | Intron (ENSM | 15 | 7873055 | 7886377 | 13323 | 2 | 545085 | ENSMUST0( | -51027 | ENSMUSG0( | Wdr70 | WD repeat domain 70 |
| chr15 | 7950363 | 7950578 | 216 | * | NA_peak_53 | 26 | Intron (ENSM | 15 | 7873055 | 7886377 | 13323 | 2 | 545085 | ENSMUST0( | -63986 | ENSMUSG0( | Wdr70 | WD repeat domain 70 |
| chr15 | 7978287 | 7978765 | 479 | * | NA_peak_54 | 17 | Intron (ENSM | 15 | 7873055 | 7886377 | 13323 | 2 | 545085 | ENSMUST0( | -91910 | ENSMUSG0( | Wdr70 | WD repeat domain 70 |
| chr15 | 8045418 | 8045647 | 230 | * | NA_peak_54 | 24 | Intron (ENSM | 15 | 7873062 | 8099191 | 226130 | 2 | 545085 | ENSMUST0( | 53544 | ENSMUSG0( | Wdr70 | WD repeat domain 70 |
| chr15 | 8050331 | 8050363 | 33 | * | NA_peak_54 | 18 | Intron (ENSM | 15 | 7873062 | 8099191 | 226130 | 2 | 545085 | ENSMUST0( | 48828 | ENSMUSG0( | Wdr70 | WD repeat domain 70 |
| chr15 | 8192420 | 8192482 | 63 | * | NA_peak_54 | 35 | Promoter (1- | 15 | 8194301 | 8202606 | 8306 | 1 | 73692 | ENSMUST0( | -1819 | ENSMUSG0( | Cplane1 | ciliogenesis and planar polarity effector 1 |
| chr15 | 8236308 | 8236743 | 436 | * | NA_peak_54 | 23 | Intron (ENSM | 15 | 8247103 | 8252976 | 5874 | 1 | 73692 | ENSMUST0( | -10360 | ENSMUSG0( | Cplane1 | ciliogenesis and planar polarity effector 1 |
| chr15 | 8243153 | 8243381 | 229 | * | NA_peak_54 | 26 | Intron (ENSM | 15 | 8247103 | 8252976 | 5874 | 1 | 73692 | ENSMUST0( | -3722 | ENSMUSG0( | Cplane1 | ciliogenesis and planar polarity effector 1 |
| chr15 | 8832899 | 8833188 | 290 | * | NA_peak_54 | 21 | Distal Interge | 15 | 8634124 | 8710764 | 76641 | 2 | 20512 | ENSMUST0( | -122135 | ENSMUSG0( | Slc1a3 | solute carrier family 1 (glial high affinity glutamate transporter), member 3 |
| chr15 | 8907342 | 8907870 | 529 | * | NA_peak_54 | 26 | Distal Interge | 15 | 8967949 | 8970298 | 2350 | 1 | 223332 | ENSMUST0( | -60079 | ENSMUSG0( | Ranbp3l | RAN binding protein 3-like |
| chr15 | 9127860 | 9127880 | 21 | * | NA_peak_54 | 36 | Exon (ENSM | 15 | 9127048 | 9140352 | 13305 | 2 | 27401 | ENSMUST0( | 12472 | ENSMUSG0( | Skp2 | S-phase kinase-associated protein 2 |
| chr15 | 9298077 | 9298419 | 343 | * | NA_peak_54 | 23 | Intron (ENSM | 15 | 9283501 | 9292027 | 8527 | 1 | 105887 | ENSMUST0( | 14576 | ENSMUSG0( | Ugt3a1 | UDP glycosyltransferases 3 family, polypeptide A1 |
| chr15 | 9344347 | 9344444 | 98 | * | NA_peak_54 | 19 | Intron (ENSM | 15 | 9335589 | 9362755 | 27167 | 1 | 223337 | ENSMUST0( | 8758 | ENSMUSG0( | Ugt3a2 | UDP glycosyltransferases 3 family, polypeptide A2 |
| chr15 | 9964138 | 9965106 | 969 | * | NA_peak_54 | 19 | Distal Interge | 15 | 10177238 | 10329388 | 152151 | 1 | 19116 | ENSMUST0( | -212132 | ENSMUSG0( | Prlr | prolactin receptor |
| chr15 | 10448707 | 10448774 | 68 | * | NA_peak_54 | 34 | Promoter (1- | 15 | 10446766 | 10449992 | 3227 | 2 | 78244 | ENSMUST0( | 1218 | ENSMUSG0( | Dnajc21 | DnaJ heat shock protein family (Hsp40) member C21 |
| chr15 | 10469653 | 10470104 | 452 | * | NA_peak_54 | 17 | Promoter (<= | 15 | 10446756 | 10470516 | 23761 | 2 | 78244 | ENSMUST0( | 412 | ENSMUSG0( | Dnajc21 | DnaJ heat shock protein family (Hsp40) member C21 |
| chr15 | 10519948 | 10520183 | 236 | * | NA_peak_54 | 23 | Intron (ENSM | 15 | 10486609 | 10493674 | 7066 | 1 | 19355 | ENSMUST0( | 33339 | ENSMUSG0( | Rad1 | RAD1 checkpoint DNA exonuclease |
| chr15 | 10771572 | 10771921 | 350 | * | NA_peak_54 | 26 | Intron (ENSM | 15 | 10814792 | 10835214 | 20423 | 1 | 100502601 | ENSMUST0( | -42871 | NA | Gm19276 | predicted gene, 19276 |
| chr15 | 11173122 | 11173301 | 180 | * | NA_peak_54 | 26 | Intron (ENSM | 15 | 11064818 | 11217121 | 152304 | 1 | 239337 | ENSMUST0( | 108304 | ENSMUSG0( | Adamts12 | a disintegrin-like and metallopeptidase (reprolysin type) with thrombospondin type 1 motif, 12 |
| chr15 | 11209461 | 11209876 | 416 | * | NA_peak_54 | 32 | Intron (ENSM | 15 | 11064818 | 11217121 | 152304 | 1 | 239337 | ENSMUST0( | 144643 | ENSMUSG0( | Adamts12 | a disintegrin-like and metallopeptidase (reprolysin type) with thrombospondin type 1 motif, 12 |
| chr15 | 11224376 | 11224641 | 266 | * | NA_peak_54 | 36 | Intron (ENSM | 15 | 11064818 | 11217121 | 152304 | 1 | 239337 | ENSMUST0( | 159558 | ENSMUSG0( | Adamts12 | a disintegrin-like and metallopeptidase (reprolysin type) with thrombospondin type 1 motif, 12 |
| chr15 | 11399460 | 11399668 | 209 | * | NA_peak_54 | 27 | Promoter (<= | 15 | 11382301 | 11399601 | 17301 | 2 | 110960 | ENSMUST0( | 0 | ENSMUSG0( | Tars | threonyl-tRNA synthetase |
| chr15 | 11550411 | 11550473 | 63 | * | NA_peak_55 | 18 | Distal Interge | 15 | 11382301 | 11399665 | 17365 | 2 | 110960 | ENSMUST0( | -150746 | ENSMUSG0( | Tars | threonyl-tRNA synthetase |
| chr15 | 11922384 | 11922682 | 299 | * | NA_peak_55 | 33 | Distal Interge | 15 | 11844383 | 11907287 | 62905 | 2 | 18162 | ENSMUST0( | -15097 | ENSMUSG0( | Npr3 | natriuretic peptide receptor 3 |
| chr15 | 14276632 | 14276721 | 90 | * | NA_peak_55 | 16 | Distal Interge | 15 | 13059251 | 13173675 | 114425 | 2 | 12563 | ENSMUST0( | -1102957 | ENSMUSG0( | Cdh6 | cadherin 6 |
| chr15 | 15640753 | 15640937 | 185 | * | NA_peak_55 | 26 | Distal Interge | 15 | 16728756 | 16857094 | 128339 | 1 | 12565 | ENSMUST0( | -1087819 | ENSMUSG0( | Cdh9 | cadherin 9 |
| chr15 | 15966615 | 15966889 | 275 | * | NA_peak_55 | 39 | Distal Interge | 15 | 16728756 | 16857094 | 128339 | 1 | 12565 | ENSMUST0( | -761867 | ENSMUSG0( | Cdh9 | cadherin 9 |
| chr15 | 16386617 | 16386843 | 227 | * | NA_peak_55 | 23 | Distal Interge | 15 | 16728756 | 16857094 | 128339 | 1 | 12565 | ENSMUST0( | -341913 | ENSMUSG0( | Cdh9 | cadherin 9 |
| chr15 | 17168075 | 17168357 | 283 | * | NA_peak_55 | 44 | Distal Interge | 15 | 16778101 | 16856767 | 78667 | 1 | 12565 | ENSMUST0( | 389974 | ENSMUSG0( | Cdh9 | cadherin 9 |
| chr15 | 17751593 | 17752079 | 487 | * | NA_peak_55 | 53 | Distal Interge | 15 | 18078118 | 18259800 | 181683 | 2 | 70886 | ENSMUST0( | 507721 | ENSMUSG0( | 4921515E04 | RIKEN cDNA 4921515E04 gene |
| chr15 | 18267586 | 18268002 | 417 | * | NA_peak_56 | 33 | Intron (ENSM | 15 | 18078118 | 18259800 | 181683 | 2 | 70886 | ENSMUST0( | -7786 | ENSMUSG0( | 4921515E04 | RIKEN cDNA 4921515E04 gene |
| chr15 | 18268354 | 18268402 | 49 | * | NA_peak_56 | 33 | Intron (ENSM | 15 | 18078118 | 18259800 | 181683 | 2 | 70886 | ENSMUST0( | -8554 | ENSMUSG0( | 4921515E04 | RIKEN cDNA 4921515E04 gene |
| chr15 | 18269371 | 18269597 | 227 | * | NA_peak_56 | 20 | Intron (ENSM | 15 | 18078118 | 18259800 | 181683 | 2 | 70886 | ENSMUST0( | -9571 | ENSMUSG0( | 4921515E04 | RIKEN cDNA 4921515E04 gene |
| chr15 | 18573836 | 18574144 | 309 | * | NA_peak_56 | 26 | Distal Interge | 15 | 18078137 | 18369627 | 291491 | 2 | 70886 | ENSMUST0( | -204209 | ENSMUSG0( | 4921515E04 | RIKEN cDNA 4921515E04 gene |
| chr15 | 18665421 | 18665816 | 396 | * | NA_peak_56 | 41 | Distal Interge | 15 | 18818947 | 19014236 | 195290 | 1 | 320873 | ENSMUST0( | -153131 | ENSMUSG0( | Cdh10 | cadherin 10 |
| chr15 | 20159558 | 20159667 | 110 | * | NA_peak_56 | 25 | Distal Interge | 15 | 20665214 | 20666750 | 1537 | 2 | 64833 | ENSMUST0( | 507083 | ENSMUSG0( | Acot10 | acyl-CoA thioesterase 10 |
| chr15 | 20350397 | 20351041 | 645 | * | NA_peak_56 | 15 | Distal Interge | 15 | 20665214 | 20666750 | 1537 | 2 | 64833 | ENSMUST0( | 315709 | ENSMUSG0( | Acot10 | acyl-CoA thioesterase 10 |
| chr15 | 20394459 | 20394581 | 123 | * | NA_peak_56 | 23 | Distal Interge | 15 | 20665214 | 20666750 | 1537 | 2 | 64833 | ENSMUST0( | 272169 | ENSMUSG0( | Acot10 | acyl-CoA thioesterase 10 |
| chr15 | 20536857 | 20537136 | 280 | * | NA_peak_56 | 19 | Intron (ENSM | 15 | 20665214 | 20666750 | 1537 | 2 | 64833 | ENSMUST0( | 129614 | ENSMUSG0( | Acot10 | acyl-CoA thioesterase 10 |
| chr15 | 20915101 | 20915412 | 312 | * | NA_peak_56 | 30 | Intron (ENSM | 15 | 21110125 | 21589533 | 479409 | 1 | 215654 | ENSMUST0( | -194713 | ENSMUSG0( | Cdh12 | cadherin 12 |
| chr15 | 21262079 | 21262329 | 251 | * | NA_peak_56 | 43 | Intron (ENSM | 15 | 21111452 | 21589530 | 478079 | 1 | 215654 | ENSMUST0( | 150627 | ENSMUSG0( | Cdh12 | cadherin 12 |
| chr15 | 21446923 | 21447393 | 471 | * | NA_peak_56 | 16 | Intron (ENSM | 15 | 21111452 | 21589530 | 478079 | 1 | 215654 | ENSMUST0( | 335471 | ENSMUSG0( | Cdh12 | cadherin 12 |
| chr15 | 21874550 | 21874804 | 255 | * | NA_peak_56 | 19 | Distal Interge | 15 | 22549022 | 23400567 | 851546 | 1 | 320865 | ENSMUST0( | -674218 | ENSMUSG0( | Cdh18 | cadherin 18 |
| chr15 | 22013433 | 22013552 | 120 | * | NA_peak_56 | 26 | Distal Interge | 15 | 22549022 | 23400567 | 851546 | 1 | 320865 | ENSMUST0( | -535470 | ENSMUSG0( | Cdh18 | cadherin 18 |
| chr15 | 22270832 | 22271138 | 307 | * | NA_peak_56 | 34 | Distal Interge | 15 | 22549022 | 23400567 | 851546 | 1 | 320865 | ENSMUST0( | -277884 | ENSMUSG0( | Cdh18 | cadherin 18 |
| chr15 | 22364741 | 22364988 | 248 | * | NA_peak_56 | 26 | Distal Interge | 15 | 22549022 | 23400567 | 851546 | 1 | 320865 | ENSMUST0( | -184034 | ENSMUSG0( | Cdh18 | cadherin 18 |
| chr15 | 22366921 | 22367275 | 355 | * | NA_peak_56 | 36 | Distal Interge | 15 | 22549022 | 23400567 | 851546 | 1 | 320865 | ENSMUST0( | -181747 | ENSMUSG0( | Cdh18 | cadherin 18 |
| chr15 | 22696503 | 22696526 | 24 | * | NA_peak_56 | 23 | Intron (ENSM | 15 | 22713868 | 22714830 | 963 | 2 | 545091 | ENSMUST0( | 18304 | ENSMUSG0( | Hnrnpa1l2-p | heterogeneous nuclear ribonucleoprotein A1-like 2, pseudogene 2 |
| chr15 | 23479636 | 23479969 | 334 | * | NA_peak_56 | 35 | Distal Interge | 15 | 23173864 | 23474029 | 300166 | 1 | 320865 | ENSMUST0( | 305772 | ENSMUSG0( | Cdh18 | cadherin 18 |
| chr15 | 23575085 | 23575436 | 352 | * | NA_peak_56 | 40 | Distal Interge | 15 | 23173864 | 23474029 | 300166 | 1 | 320865 | ENSMUST0( | 401221 | ENSMUSG0( | Cdh18 | cadherin 18 |
| chr15 | 23701467 | 23701815 | 349 | * | NA_peak_56 | 40 | Distal Interge | 15 | 23173864 | 23474029 | 300166 | 1 | 320865 | ENSMUST0( | 527603 | ENSMUSG0( | Cdh18 | cadherin 18 |
| chr15 | 23776476 | 23776569 | 94 | * | NA_peak_56 | 19 | Distal Interge | 15 | 23173864 | 23474029 | 300166 | 1 | 320865 | ENSMUST0( | 602612 | ENSMUSG0( | Cdh18 | cadherin 18 |
| chr15 | 24065465 | 24065987 | 523 | * | NA_peak_56 | 20 | Distal Interge | 15 | 23173864 | 23474029 | 300166 | 1 | 320865 | ENSMUST0( | 891601 | ENSMUSG0( | Cdh18 | cadherin 18 |
| chr15 | 24181689 | 24181808 | 120 | * | NA_peak_57 | 16 | Distal Interge | 15 | 23173864 | 23474029 | 300166 | 1 | 320865 | ENSMUST0( | 1007825 | ENSMUSG0( | Cdh18 | cadherin 18 |
| chr15 | 24370948 | 24371868 | 921 | * | NA_peak_57 | 44 | Intron (ENSM | 15 | 25364857 | 25413698 | 48842 | 2 | 70350 | ENSMUST0( | 1041830 | ENSMUSG0( | Basp1 | brain abundant, membrane attached signal protein 1 |
| chr15 | 24412376 | 24412492 | 117 | * | NA_peak_57 | 24 | Distal Interge | 15 | 25364857 | 25413698 | 48842 | 2 | 70350 | ENSMUST0( | 1001206 | ENSMUSG0( | Basp1 | brain abundant, membrane attached signal protein 1 |
| chr15 | 24728993 | 24729474 | 482 | * | NA_peak_57 | 35 | Distal Interge | 15 | 25364857 | 25413698 | 48842 | 2 | 70350 | ENSMUST0( | 684224 | ENSMUSG0( | Basp1 | brain abundant, membrane attached signal protein 1 |
| chr15 | 24762004 | 24762283 | 280 | * | NA_peak_57 | 30 | Distal Interge | 15 | 25364857 | 25413698 | 48842 | 2 | 70350 | ENSMUST0( | 651415 | ENSMUSG0( | Basp1 | brain abundant, membrane attached signal protein 1 |
| chr15 | 25246868 | 25247044 | 177 | * | NA_peak_57 | 54 | Distal Interge | 15 | 25364857 | 25413698 | 48842 | 2 | 70350 | ENSMUST0( | 166654 | ENSMUSG0( | Basp1 | brain abundant, membrane attached signal protein 1 |
| chr15 | 25412935 | 25413521 | 587 | * | NA_peak_57 | 28 | Promoter (<= | 15 | 25364857 | 25413698 | 48842 | 2 | 70350 | ENSMUST0( | 177 | ENSMUSG0( | Basp1 | brain abundant, membrane attached signal protein 1 |
| chr15 | 25605909 | 25606172 | 264 | * | NA_peak_57 | 17 | Distal Interge | 15 | 25622525 | 25813673 | 191149 | 1 | 17909 | ENSMUST0( | -16353 | ENSMUSG0( | Myo10 | myosin X |
| chr15 | 25669903 | 25670113 | 211 | * | NA_peak_57 | 19 | Intron (ENSM | 15 | 25674510 | 25723950 | 49441 | 1 | 17909 | ENSMUST0( | -4397 | ENSMUSG0( | Myo10 | myosin X |
| chr15 | 25842991 | 25843508 | 518 | * | NA_peak_57 | 13 | Promoter (<= | 15 | 25843180 | 25973687 | 130508 | 1 | 66270 | ENSMUST0( | 0 | ENSMUSG0( | Retreg1 | reticulophagy regulator 1 |
| chr15 | 25975654 | 25975841 | 188 | * | NA_peak_57 | 30 | Distal Interge | 15 | 25984366 | 25990464 | 6099 | 1 | 52521 | ENSMUST0( | -8525 | ENSMUSG0( | Zfp622 | zinc finger protein 622 |
| chr15 | 26482187 | 26482257 | 71 | * | NA_peak_57 | 16 | Distal Interge | 15 | 26387719 | 26409576 | 21858 | 1 | 211147 | ENSMUST0( | 94468 | ENSMUSG0( | Marchf11 | membrane associated ring-CH-type finger 11 |
| chr15 | 27920033 | 27920042 | 10 | * | NA_peak_57 | 22 | Promoter (<= | 15 | 27770536 | 27919271 | 148736 | 2 | 223435 | ENSMUST0( | -762 | ENSMUSG0( | Trio | triple functional domain (PTPRF interacting) |
| chr15 | 28983690 | 28984038 | 349 | * | NA_peak_58 | 18 | Distal Interge | 15 | 29713331 | 29713404 | 74 | 2 | 100628607 | ENSMUST0( | 729366 | ENSMUSG0( | Mir3964 | microRNA 3964 |
| chr15 | 29141127 | 29141380 | 254 | * | NA_peak_58 | 23 | Distal Interge | 15 | 29713331 | 29713404 | 74 | 2 | 100628607 | ENSMUST0( | 572024 | ENSMUSG0( | Mir3964 | microRNA 3964 |
| chr15 | 29302858 | 29302981 | 124 | * | NA_peak_58 | 23 | Distal Interge | 15 | 29713331 | 29713404 | 74 | 2 | 100628607 | ENSMUST0( | 410423 | ENSMUSG0( | Mir3964 | microRNA 3964 |
| chr15 | 29469789 | 29470109 | 321 | * | NA_peak_58 | 70 | Distal Interge | 15 | 29713331 | 29713404 | 74 | 2 | 100628607 | ENSMUST0( | 243295 | ENSMUSG0( | Mir3964 | microRNA 3964 |
| chr15 | 29548284 | 29548630 | 347 | * | NA_peak_58 | 16 | Distal Interge | 15 | 29713331 | 29713404 | 74 | 2 | 100628607 | ENSMUST0( | 164774 | ENSMUSG0( | Mir3964 | microRNA 3964 |
| chr15 | 29597558 | 29598193 | 636 | * | NA_peak_58 | 23 | Distal Interge | 15 | 29713331 | 29713404 | 74 | 2 | 100628607 | ENSMUST0( | 115211 | ENSMUSG0( | Mir3964 | microRNA 3964 |
| chr15 | 29985785 | 29986141 | 357 | * | NA_peak_58 | 23 | Distal Interge | 15 | 30172593 | 31029341 | 856749 | 1 | 18163 | ENSMUST0( | -186452 | ENSMUSG0( | Ctnnd2 | catenin (cadherin associated protein), delta 2 |
| chr15 | 30221842 | 30222249 | 408 | * | NA_peak_58 | 161 | Intron (ENSM | 15 | 30173138 | 30481349 | 308212 | 1 | 18163 | ENSMUST0( | 48704 | ENSMUSG0( | Ctnnd2 | catenin (cadherin associated protein), delta 2 |
| chr15 | 30350436 | 30350469 | 34 | * | NA_peak_58 | 16 | Intron (ENSM | 15 | 30457673 | 30480904 | 23232 | 1 | 18163 | ENSMUST0( | -107204 | ENSMUSG0( | Ctnnd2 | catenin (cadherin associated protein), delta 2 |
| chr15 | 30945516 | 30945737 | 222 | * | NA_peak_58 | 35 | Intron (ENSM | 15 | 31004796 | 31020776 | 15981 | 1 | 18163 | ENSMUST0( | -59059 | ENSMUSG0( | Ctnnd2 | catenin (cadherin associated protein), delta 2 |
| chr15 | 31069100 | 31069404 | 305 | * | NA_peak_58 | 23 | Distal Interge | 15 | 31039128 | 31043331 | 4204 | 1 | 73784 | ENSMUST0( | 29972 | ENSMUSG0( | 4930430F21 | RIKEN cDNA 4930430F21 gene |
| chr15 | 31631524 | 31631807 | 284 | * | NA_peak_58 | 14 | Distal Interge | 15 | 31602118 | 31618609 | 16492 | 1 | 68073 | ENSMUST0( | 29406 | ENSMUSG0( | Atpsckmt | ATP synthase C subunit lysine N-methyltransferase |
| chr15 | 31893122 | 31893378 | 257 | * | NA_peak_58 | 32 | Distal Interge | 15 | 32177289 | 32178294 | 1006 | 1 | 57254 | ENSMUST0( | -283911 | ENSMUSG0( | Tas2r119 | taste receptor, type 2, member 119 |
| chr15 | 32479477 | 32479778 | 302 | * | NA_peak_58 | 23 | Intron (ENSM | 15 | 32601675 | 32689636 | 87962 | 1 | 20356 | ENSMUST0( | -121897 | ENSMUSG0( | Sema5a | sema domain, seven thrombospondin repeats (type 1 and type 1-like), transmembrane domain (TM) and short cytoplasmic domain, (semaphorin) |
| chr15 | 32483402 | 32483753 | 352 | * | NA_peak_58 | 21 | Intron (ENSM | 15 | 32601675 | 32689636 | 87962 | 1 | 20356 | ENSMUST0( | -117922 | ENSMUSG0( | Sema5a | sema domain, seven thrombospondin repeats (type 1 and type 1-like), transmembrane domain (TM) and short cytoplasmic domain, (semaphorin) |
| chr15 | 32622582 | 32622644 | 63 | * | NA_peak_58 | 29 | Intron (ENSM | 15 | 32601675 | 32689636 | 87962 | 1 | 20356 | ENSMUST0( | 20907 | ENSMUSG0( | Sema5a | sema domain, seven thrombospondin repeats (type 1 and type 1-like), transmembrane domain (TM) and short cytoplasmic domain, (semaphorin) |
| chr15 | 32668549 | 32668892 | 344 | * | NA_peak_58 | 20 | Intron (ENSM | 15 | 32601675 | 32689636 | 87962 | 1 | 20356 | ENSMUST0( | 66874 | ENSMUSG0( | Sema5a | sema domain, seven thrombospondin repeats (type 1 and type 1-like), transmembrane domain (TM) and short cytoplasmic domain, (semaphorin) |
| chr15 | 32862515 | 32862562 | 48 | * | NA_peak_58 | 17 | Distal Interge | 15 | 32920723 | 33034935 | 114213 | 1 | 15529 | ENSMUST0( | -58161 | ENSMUSG0( | Sdc2 | syndecan 2 |
| chr15 | 32878926 | 32879254 | 329 | * | NA_peak_58 | 14 | Distal Interge | 15 | 32920723 | 33034935 | 114213 | 1 | 15529 | ENSMUST0( | -41469 | ENSMUSG0( | Sdc2 | syndecan 2 |
| chr15 | 32905553 | 32905780 | 228 | * | NA_peak_58 | 19 | Distal Interge | 15 | 32920723 | 33034935 | 114213 | 1 | 15529 | ENSMUST0( | -14943 | ENSMUSG0( | Sdc2 | syndecan 2 |
| chr15 | 32953711 | 32953939 | 229 | * | NA_peak_58 | 13 | Intron (ENSM | 15 | 32920723 | 33034935 | 114213 | 1 | 15529 | ENSMUST0( | 32988 | ENSMUSG0( | Sdc2 | syndecan 2 |
| chr15 | 33031024 | 33031422 | 399 | * | NA_peak_58 | 24 | Intron (ENSM | 15 | 33083129 | 33594552 | 511424 | 1 | 54381 | ENSMUST0( | -51707 | ENSMUSG0( | Cpq | carboxypeptidase Q |
| chr15 | 33203733 | 33204043 | 311 | * | NA_peak_59 | 21 | Intron (ENSM | 15 | 33090308 | 33304620 | 214313 | 1 | 54381 | ENSMUST0( | 113425 | ENSMUSG0( | Cpq | carboxypeptidase Q |
| chr15 | 33270699 | 33271168 | 470 | * | NA_peak_59 | 84 | Intron (ENSM | 15 | 33405206 | 33405939 | 734 | 2 | 73486 | ENSMUST0( | 134771 | ENSMUSG0( | 1700084J12 | ribosomal protein L7-like 1 pseudogene |
| chr15 | 33612019 | 33613020 | 1002 | * | NA_peak_59 | 32 | Intron (ENSM | 15 | 33593881 | 33653436 | 59556 | 2 | 75880 | ENSMUST0( | 40416 | ENSMUSG0( | 4930592A05 | RIKEN cDNA 4930592A05 gene |
| chr15 | 33753137 | 33753449 | 313 | * | NA_peak_59 | 13 | Distal Interge | 15 | 33683875 | 33687884 | 4010 | 2 | 239364 | ENSMUST0( | -65253 | ENSMUSG0( | Tspyl5 | testis-specific protein, Y-encoded-like 5 |
| chr15 | 33834469 | 33834751 | 283 | * | NA_peak_59 | 60 | Distal Interge | 15 | 33683875 | 33687884 | 4010 | 2 | 239364 | ENSMUST0( | -146585 | ENSMUSG0( | Tspyl5 | testis-specific protein, Y-encoded-like 5 |
| chr15 | 33835368 | 33835855 | 488 | * | NA_peak_59 | 60 | Distal Interge | 15 | 33683875 | 33687884 | 4010 | 2 | 239364 | ENSMUST0( | -147484 | ENSMUSG0( | Tspyl5 | testis-specific protein, Y-encoded-like 5 |
| chr15 | 33846884 | 33846910 | 27 | * | NA_peak_59 | 20 | Distal Interge | 15 | 33683875 | 33687884 | 4010 | 2 | 239364 | ENSMUST0( | -159000 | ENSMUSG0( | Tspyl5 | testis-specific protein, Y-encoded-like 5 |
| chr15 | 33870214 | 33870539 | 326 | * | NA_peak_59 | 25 | Distal Interge | 15 | 33683875 | 33687884 | 4010 | 2 | 239364 | ENSMUST0( | -182330 | ENSMUSG0( | Tspyl5 | testis-specific protein, Y-encoded-like 5 |
| chr15 | 33966255 | 33966624 | 370 | * | NA_peak_59 | 41 | Distal Interge | 15 | 34082694 | 34103345 | 20652 | 1 | 67154 | ENSMUST0( | -116070 | ENSMUSG0( | Mtdh | metadherin |
| chr15 | 34076697 | 34076721 | 25 | * | NA_peak_59 | 17 | Distal Interge | 15 | 34082694 | 34103345 | 20652 | 1 | 67154 | ENSMUST0( | -5973 | ENSMUSG0( | Mtdh | metadherin |
| chr15 | 34148119 | 34148473 | 355 | * | NA_peak_59 | 35 | Distal Interge | 15 | 34133490 | 34141283 | 7794 | 1 | 67154 | ENSMUST0( | 14629 | ENSMUSG0( | Mtdh | metadherin |
| chr15 | 34243185 | 34243598 | 414 | * | NA_peak_59 | 19 | Intron (ENSM | 15 | 34238167 | 34276168 | 38002 | 1 | 114128 | ENSMUST0( | 5018 | ENSMUSG0( | Laptm4b | lysosomal-associated protein transmembrane 4B |
| chr15 | 34409202 | 34409396 | 195 | * | NA_peak_59 | 18 | Intron (ENSM | 15 | 34440575 | 34441665 | 1091 | 2 | 666899 | ENSMUST0( | 32269 | NA | Gm12191 | ribosomal protein L30 pseudogene |

| | | | | | | | | | | | | | | | | | | |
|---|---|---|---|---|---|---|---|---|---|---|---|---|---|---|---|---|---|---|
| chr15 | 34488607 | 34488660 | 54 | * | NA_peak_59 | 16 | Promoter (1- | 15 | 34485604 | 34489686 | 4083 | 2 | 15473 | ENSMUST0 | 1026 | ENSMUSG0 | Rida | reactive intermediate imine deaminase A homolog |
| chr15 | 34488964 | 34489349 | 386 | * | NA_peak_59 | 16 | Promoter (<= | 15 | 34485604 | 34489686 | 4083 | 2 | 15473 | ENSMUST0 | 337 | ENSMUSG0 | Rida | reactive intermediate imine deaminase A homolog |
| chr15 | 35004610 | 35004809 | 200 | * | NA_peak_59 | 17 | Intron (ENSM | 15 | 34958564 | 34999922 | 41359 | 2 | 56274 | ENSMUST0 | -4688 | ENSMUSG0 | Stk3 | serine/threonine kinase 3 |
| chr15 | 35060753 | 35060822 | 70 | * | NA_peak_59 | 16 | Intron (ENSM | 15 | 34958564 | 34999922 | 41359 | 2 | 56274 | ENSMUST0 | -60831 | ENSMUSG0 | Stk3 | serine/threonine kinase 3 |
| chr15 | 35155421 | 35155832 | 412 | * | NA_peak_60 | 31 | Promoter (<= | 15 | 34875496 | 35155806 | 280311 | 2 | 56274 | ENSMUST0 | 0 | ENSMUSG0 | Stk3 | serine/threonine kinase 3 |
| chr15 | 35461935 | 35462289 | 355 | * | NA_peak_60 | 26 | Intron (ENSM | 15 | 35371566 | 35642952 | 271387 | 1 | 666173 | ENSMUST0 | 90369 | ENSMUSG0 | Vps13b | vacuolar protein sorting 13B |
| chr15 | 35988266 | 35988549 | 284 | * | NA_peak_60 | 28 | Distal Interge | 15 | 35931976 | 35938246 | 6271 | 2 | 12864 | ENSMUST0 | -50020 | ENSMUSG0 | Cox6c | cytochrome c oxidase subunit 6C |
| chr15 | 36135125 | 36135482 | 358 | * | NA_peak_60 | 30 | Intron (ENSM | 15 | 36138221 | 36140385 | 2165 | 2 | 626596 | ENSMUST0 | 4903 | ENSMUSG0 | Rgs22 | regulator of G-protein signalling 22 |
| chr15 | 36263284 | 36263722 | 439 | * | NA_peak_60 | 15 | Intron (ENSM | 15 | 36240351 | 36283098 | 42748 | 2 | 30945 | ENSMUST0 | 19376 | ENSMUSG0 | Rnf19a | ring finger protein 19A |
| chr15 | 36676605 | 36676755 | 151 | * | NA_peak_60 | 38 | Intron (ENSM | 15 | 36605752 | 36609668 | 3917 | 2 | 18458 | ENSMUST0 | -66937 | ENSMUSG0 | Pabpc1 | poly(A) binding protein, cytoplasmic 1 |
| chr15 | 37077934 | 37078271 | 338 | * | NA_peak_60 | 27 | Distal Interge | 15 | 37000814 | 37007773 | 6960 | 2 | 68036 | ENSMUST0 | -70161 | ENSMUSG0 | Zfp706 | zinc finger protein 706 |
| chr15 | 38253279 | 38253675 | 397 | * | NA_peak_60 | 19 | Intron (ENSM | 15 | 38219203 | 38226735 | 7533 | 1 | 18285 | ENSMUST0 | 34076 | ENSMUSG0 | Odf1 | outer dense fiber of sperm tails 1 |
| chr15 | 38530588 | 38530754 | 167 | * | NA_peak_60 | 17 | Distal Interge | 15 | 38487427 | 38519266 | 31840 | 2 | 54375 | ENSMUST0 | -11322 | ENSMUSG0 | Azin1 | antizyme inhibitor 1 |
| chr15 | 38731673 | 38731863 | 191 | * | NA_peak_61 | 29 | Distal Interge | 15 | 38713483 | 38728516 | 15034 | 2 | 105463124 | ENSMUST0 | -3157 | ENSMUSG0 | Gm45924 | predicted gene, 45924 |
| chr15 | 38884189 | 38884555 | 367 | * | NA_peak_61 | 23 | Distal Interge | 15 | 38933144 | 38951264 | 18121 | 1 | 118452 | ENSMUST0 | -48589 | ENSMUSG0 | Baalc | brain and acute leukemia, cytoplasmic |
| chr15 | 39068449 | 39068797 | 349 | * | NA_peak_61 | 26 | Distal Interge | 15 | 39076932 | 39087121 | 10190 | 1 | 68588 | ENSMUST0 | -8135 | ENSMUSG0 | Cthrc1 | collagen triple helix repeat containing 1 |
| chr15 | 39246390 | 39246652 | 263 | * | NA_peak_61 | 26 | Intron (ENSM | 15 | 39198363 | 39681764 | 483402 | 1 | 116838 | ENSMUST0 | 48027 | ENSMUSG0 | Rims2 | regulating synaptic membrane exocytosis 2 |
| chr15 | 39280671 | 39280733 | 63 | * | NA_peak_61 | 23 | Intron (ENSM | 15 | 39198363 | 39681764 | 483402 | 1 | 116838 | ENSMUST0 | 82308 | ENSMUSG0 | Rims2 | regulating synaptic membrane exocytosis 2 |
| chr15 | 39338785 | 39339004 | 220 | * | NA_peak_61 | 29 | Intron (ENSM | 15 | 39391800 | 39452232 | 60433 | 1 | 116838 | ENSMUST0 | -52796 | ENSMUSG0 | Rims2 | regulating synaptic membrane exocytosis 2 |
| chr15 | 39339425 | 39339663 | 239 | * | NA_peak_61 | 13 | Intron (ENSM | 15 | 39391800 | 39452232 | 60433 | 1 | 116838 | ENSMUST0 | -52137 | ENSMUSG0 | Rims2 | regulating synaptic membrane exocytosis 2 |
| chr15 | 40051164 | 40051516 | 353 | * | NA_peak_61 | 19 | Intron (ENSM | 15 | 40045191 | 40128982 | 83792 | 1 | 102636079 | ENSMUST0 | 5973 | ENSMUSG0 | Gm33251 | predicted gene, 33251 |
| chr15 | 40086500 | 40086887 | 388 | * | NA_peak_61 | 13 | Intron (ENSM | 15 | 40045191 | 40128982 | 83792 | 1 | 102636079 | ENSMUST0 | 41309 | ENSMUSG0 | Gm33251 | predicted gene, 33251 |
| chr15 | 40095176 | 40095517 | 342 | * | NA_peak_61 | 27 | Intron (ENSM | 15 | 40134365 | 40150010 | 15646 | 1 | 328531 | ENSMUST0 | -38848 | ENSMUSG0 | 9330182O14 | RIKEN cDNA 9330182O14 gene |
| chr15 | 40323526 | 40323587 | 62 | * | NA_peak_61 | 19 | Distal Interge | 15 | 40162607 | 40168076 | 5470 | 1 | 102636151 | ENSMUST0 | 160919 | ENSMUSG0 | Gm33301 | predicted gene, 33301 |
| chr15 | 40444678 | 40445059 | 382 | * | NA_peak_61 | 65 | Distal Interge | 15 | 40511322 | 40527277 | 15956 | 2 | 432945 | ENSMUST0 | 82218 | ENSMUSG0 | Gm16294 | predicted gene 16294 |
| chr15 | 40446024 | 40446345 | 322 | * | NA_peak_61 | 23 | Distal Interge | 15 | 40511322 | 40527277 | 15956 | 2 | 432945 | ENSMUST0 | 80932 | ENSMUSG0 | Gm16294 | predicted gene 16294 |
| chr15 | 40446970 | 40447431 | 462 | * | NA_peak_61 | 44 | Distal Interge | 15 | 40511322 | 40527277 | 15956 | 2 | 432945 | ENSMUST0 | 79846 | ENSMUSG0 | Gm16294 | predicted gene 16294 |
| chr15 | 40621693 | 40621713 | 21 | * | NA_peak_61 | 18 | Distal Interge | 15 | 40655035 | 40872723 | 217689 | 1 | 22762 | ENSMUST0 | -33322 | ENSMUSG0 | Zfpm2 | zinc finger protein, multitype 2 |
| chr15 | 40976253 | 40976302 | 50 | * | NA_peak_61 | 23 | Intron (ENSM | 15 | 41173487 | 41173868 | 382 | 1 | 100418302 | ENSMUST0 | -197185 | NA | 4930555K19 | leucine rich repeat and sterile alpha motif containing 1 pseudogene |
| chr15 | 41473595 | 41473855 | 261 | * | NA_peak_61 | 33 | Intron (ENSM | 15 | 41449027 | 41861047 | 412021 | 1 | 170719 | ENSMUST0 | 24568 | ENSMUSG0 | Oxr1 | oxidation resistance 1 |
| chr15 | 41530275 | 41530378 | 104 | * | NA_peak_61 | 19 | Intron (ENSM | 15 | 41449027 | 41861047 | 412021 | 1 | 170719 | ENSMUST0 | 81248 | ENSMUSG0 | Oxr1 | oxidation resistance 1 |
| chr15 | 41624986 | 41625290 | 305 | * | NA_peak_61 | 20 | Intron (ENSM | 15 | 41710604 | 41820380 | 109777 | 1 | 170719 | ENSMUST0 | -85314 | ENSMUSG0 | Oxr1 | oxidation resistance 1 |
| chr15 | 41788878 | 41789188 | 311 | * | NA_peak_62 | 13 | Promoter (<= | 15 | 41788941 | 41861047 | 72107 | 1 | 170719 | ENSMUST0 | 0 | ENSMUSG0 | Oxr1 | oxidation resistance 1 |
| chr15 | 42359195 | 42359807 | 613 | * | NA_peak_62 | 28 | Distal Interge | 15 | 42633306 | 42676749 | 43444 | 2 | 11600 | ENSMUST0 | 316942 | ENSMUSG0 | Angpt1 | angiopoietin 1 |
| chr15 | 42888556 | 42888562 | 7 | * | NA_peak_62 | 26 | Distal Interge | 15 | 42424723 | 42676977 | 252255 | 2 | 11600 | ENSMUST0 | -211579 | ENSMUSG0 | Angpt1 | angiopoietin 1 |
| chr15 | 42930655 | 42930941 | 287 | * | NA_peak_62 | 21 | Distal Interge | 15 | 43020797 | 43170818 | 150022 | 2 | 239405 | ENSMUST0 | 239877 | ENSMUSG0 | Rspo2 | R-spondin 2 |
| chr15 | 43182920 | 43183126 | 207 | * | NA_peak_62 | 23 | Distal Interge | 15 | 43020794 | 43170826 | 150033 | 2 | 239405 | ENSMUST0 | -12094 | ENSMUSG0 | Rspo2 | R-spondin 2 |
| chr15 | 43187441 | 43187684 | 244 | * | NA_peak_62 | 19 | Distal Interge | 15 | 43020794 | 43170826 | 150033 | 2 | 239405 | ENSMUST0 | -16615 | ENSMUSG0 | Rspo2 | R-spondin 2 |
| chr15 | 43243014 | 43243187 | 174 | * | NA_peak_62 | 13 | Distal Interge | 15 | 43261692 | 43268487 | 6796 | 2 | 16341 | ENSMUST0 | 25300 | ENSMUSG0 | Eif3e | eukaryotic translation initiation factor 3, subunit E |
| chr15 | 43287954 | 43289060 | 1107 | * | NA_peak_62 | 46 | Distal Interge | 15 | 43250058 | 43282719 | 32662 | 2 | 16341 | ENSMUST0 | -5235 | ENSMUSG0 | Eif3e | eukaryotic translation initiation factor 3, subunit E |
| chr15 | 43289747 | 43290385 | 639 | * | NA_peak_62 | 15 | Distal Interge | 15 | 43250058 | 43282719 | 32662 | 2 | 16341 | ENSMUST0 | -7028 | ENSMUSG0 | Eif3e | eukaryotic translation initiation factor 3, subunit E |
| chr15 | 43326748 | 43326891 | 144 | * | NA_peak_62 | 23 | Distal Interge | 15 | 43250058 | 43282719 | 32662 | 2 | 16341 | ENSMUST0 | -44029 | ENSMUSG0 | Eif3e | eukaryotic translation initiation factor 3, subunit E |
| chr15 | 43687757 | 43688211 | 455 | * | NA_peak_62 | 38 | Distal Interge | 15 | 43864646 | 43870036 | 5391 | 2 | 239408 | ENSMUST0 | 181825 | ENSMUSG0 | Tmem74 | transmembrane protein 74 |
| chr15 | 43742711 | 43743134 | 424 | * | NA_peak_62 | 17 | Distal Interge | 15 | 43864646 | 43870036 | 5391 | 2 | 239408 | ENSMUST0 | 126902 | ENSMUSG0 | Tmem74 | transmembrane protein 74 |
| chr15 | 43746117 | 43746492 | 376 | * | NA_peak_62 | 29 | Distal Interge | 15 | 43864646 | 43870036 | 5391 | 2 | 239408 | ENSMUST0 | 123544 | ENSMUSG0 | Tmem74 | transmembrane protein 74 |
| chr15 | 43882379 | 43882612 | 234 | * | NA_peak_62 | 19 | Distal Interge | 15 | 43864646 | 43870036 | 5391 | 2 | 239408 | ENSMUST0 | -12343 | ENSMUSG0 | Tmem74 | transmembrane protein 74 |
| chr15 | 44014271 | 44014644 | 374 | * | NA_peak_62 | 35 | Distal Interge | 15 | 43864646 | 43870036 | 5391 | 2 | 239408 | ENSMUST0 | -144235 | ENSMUSG0 | Tmem74 | transmembrane protein 74 |
| chr15 | 44183912 | 44184249 | 338 | * | NA_peak_62 | 23 | Distal Interge | 15 | 44196135 | 44235544 | 39410 | 1 | 22045 | ENSMUST0 | -11886 | ENSMUSG0 | Trhr | thyrotropin releasing hormone receptor |
| chr15 | 44298549 | 44298837 | 289 | * | NA_peak_62 | 23 | Distal Interge | 15 | 44196967 | 44229201 | 32235 | 1 | 22045 | ENSMUST0 | 101582 | ENSMUSG0 | Trhr | thyrotropin releasing hormone receptor |
| chr15 | 44427961 | 44428414 | 454 | * | NA_peak_62 | 28 | Promoter (<= | 15 | 44428042 | 44433008 | 4967 | 1 | 223527 | ENSMUST0 | 0 | ENSMUSG0 | Eny2 | ENY2 transcription and export complex 2 subunit |
| chr15 | 44791027 | 44791568 | 542 | * | NA_peak_62 | 58 | Promoter (2- | 15 | 44760975 | 44788063 | 27089 | 2 | 319613 | ENSMUST0 | -2964 | ENSMUSG0 | Sybu | syntabulin (syntaxin-interacting) |
| chr15 | 44795536 | 44796060 | 525 | * | NA_peak_62 | 13 | Intron (ENSM | 15 | 44760975 | 44788063 | 27089 | 2 | 319613 | ENSMUST0 | -7473 | ENSMUSG0 | Sybu | syntabulin (syntaxin-interacting) |
| chr15 | 44851079 | 44851187 | 109 | * | NA_peak_62 | 23 | Intron (ENSM | 15 | 44881419 | 44884302 | 2884 | 1 | 239410 | ENSMUST0 | -30232 | ENSMUSG0 | A930017M01 | Smg-5 homolog, nonsense mediated mRNA decay factor pseudogene |
| chr15 | 44952230 | 44952694 | 465 | * | NA_peak_63 | 19 | Distal Interge | 15 | 44881419 | 44884302 | 2884 | 1 | 239410 | ENSMUST0 | 70811 | ENSMUSG0 | A930017M01 | Smg-5 homolog, nonsense mediated mRNA decay factor pseudogene |
| chr15 | 44979019 | 44979374 | 356 | * | NA_peak_63 | 44 | Distal Interge | 15 | 44881419 | 44884302 | 2884 | 1 | 239410 | ENSMUST0 | 97600 | ENSMUSG0 | A930017M01 | Smg-5 homolog, nonsense mediated mRNA decay factor pseudogene |
| chr15 | 45234967 | 45235658 | 692 | * | NA_peak_63 | 30 | Distal Interge | 15 | 45112785 | 45114920 | 2136 | 2 | 67498 | ENSMUST0 | -120047 | ENSMUSG0 | Kcnv1 | potassium channel, subfamily V, member 1 |
| chr15 | 45619910 | 45620348 | 439 | * | NA_peak_63 | 26 | Distal Interge | 15 | 45112785 | 45114920 | 2136 | 2 | 67498 | ENSMUST0 | -504990 | ENSMUSG0 | Kcnv1 | potassium channel, subfamily V, member 1 |
| chr15 | 45850961 | 45851255 | 295 | * | NA_peak_63 | 19 | Distal Interge | 15 | 46324559 | 46373858 | 49300 | 1 | 71120 | ENSMUST0 | -473304 | ENSMUSG0 | 1700022A22 | RIKEN cDNA 1700022A22 gene |
| chr15 | 46211361 | 46211639 | 279 | * | NA_peak_63 | 86 | Distal Interge | 15 | 46324559 | 46373858 | 49300 | 1 | 71120 | ENSMUST0 | -112920 | ENSMUSG0 | 1700022A22 | RIKEN cDNA 1700022A22 gene |
| chr15 | 47052071 | 47052384 | 314 | * | NA_peak_63 | 19 | Distal Interge | 15 | 46623305 | 46639858 | 16554 | 1 | 75281 | ENSMUST0 | 428766 | ENSMUSG0 | 4930548G14 | RIKEN cDNA 4930548G14 gene |
| chr15 | 47220719 | 47220749 | 31 | * | NA_peak_63 | 23 | Distal Interge | 15 | 47581726 | 47615201 | 33476 | 2 | 239420 | ENSMUST0 | 394452 | ENSMUSG0 | Csmd3 | CUB and Sushi multiple domains 3 |
| chr15 | 48905399 | 48905737 | 339 | * | NA_peak_63 | 43 | Distal Interge | 15 | 47580722 | 48792063 | 1211342 | 2 | 239420 | ENSMUST0 | -113336 | ENSMUSG0 | Csmd3 | CUB and Sushi multiple domains 3 |
| chr15 | 49194625 | 49194926 | 302 | * | NA_peak_63 | 22 | Exon (ENSM | 15 | 47580722 | 48792063 | 1211342 | 2 | 239420 | ENSMUST0 | -402562 | ENSMUSG0 | Csmd3 | CUB and Sushi multiple domains 3 |
| chr15 | 49623955 | 49624277 | 323 | * | NA_peak_63 | 13 | Distal Interge | 15 | 47580722 | 48792063 | 1211342 | 2 | 239420 | ENSMUST0 | -831892 | ENSMUSG0 | Csmd3 | CUB and Sushi multiple domains 3 |
| chr15 | 49994979 | 49995442 | 464 | * | NA_peak_63 | 19 | Distal Interge | 15 | 50819499 | 50827521 | 8023 | 2 | 83925 | ENSMUST0 | 832079 | ENSMUSG0 | Trps1 | transcriptional repressor GATA binding 1 |
| chr15 | 51248210 | 51248443 | 234 | * | NA_peak_64 | 40 | Intron (ENSM | 15 | 51231640 | 51395610 | 163971 | 2 | 100502643 | ENSMUST0 | 147167 | ENSMUSG0 | Gm19303 | predicted gene, 19303 |
| chr15 | 51393058 | 51393408 | 351 | * | NA_peak_64 | 26 | Promoter (2- | 15 | 51231640 | 51395610 | 163971 | 2 | 100502643 | ENSMUST0 | 2202 | ENSMUSG0 | Gm19303 | predicted gene, 19303 |
| chr15 | 51474722 | 51475139 | 418 | * | NA_peak_64 | 39 | Distal Interge | 15 | 51231640 | 51395610 | 163971 | 2 | 100502643 | ENSMUST0 | -79112 | ENSMUSG0 | Gm19303 | predicted gene, 19303 |
| chr15 | 51608869 | 51608984 | 116 | * | NA_peak_64 | 25 | Intron (ENSM | 15 | 51786682 | 51790549 | 3868 | 2 | 68135 | ENSMUST0 | 181565 | ENSMUSG0 | Eif3h | eukaryotic translation initiation factor 3, subunit H |
| chr15 | 51752412 | 51752615 | 204 | * | NA_peak_64 | 16 | Distal Interge | 15 | 51786682 | 51790549 | 3868 | 2 | 68135 | ENSMUST0 | 37934 | ENSMUSG0 | Eif3h | eukaryotic translation initiation factor 3, subunit H |
| chr15 | 51867273 | 51867637 | 365 | * | NA_peak_64 | 24 | Promoter (1- | 15 | 51786558 | 51865523 | 78966 | 2 | 68135 | ENSMUST0 | -1750 | ENSMUSG0 | Eif3h | eukaryotic translation initiation factor 3, subunit H |
| chr15 | 52047036 | 52047085 | 50 | * | NA_peak_64 | 13 | Distal Interge | 15 | 52040107 | 52045722 | 5616 | 1 | 239435 | ENSMUST0 | 6929 | ENSMUSG0 | Aard | alanine and arginine rich domain containing protein |
| chr15 | 52206754 | 52206792 | 39 | * | NA_peak_64 | 26 | Distal Interge | 15 | 52295553 | 52335798 | 40246 | 1 | 239436 | ENSMUST0 | -88761 | ENSMUSG0 | Slc30a8 | solute carrier family 30 (zinc transporter), member 8 |
| chr15 | 52632971 | 52633126 | 156 | * | NA_peak_64 | 32 | Distal Interge | 15 | 52712408 | 52715272 | 2865 | 1 | 69790 | ENSMUST0 | -79282 | ENSMUSG0 | Med30 | mediator complex subunit 30 |
| chr15 | 52686329 | 52686819 | 491 | * | NA_peak_64 | 20 | Distal Interge | 15 | 52712408 | 52715272 | 2865 | 1 | 69790 | ENSMUST0 | -25589 | ENSMUSG0 | Med30 | mediator complex subunit 30 |
| chr15 | 52829858 | 52830121 | 264 | * | NA_peak_64 | 22 | Distal Interge | 15 | 52712442 | 52730564 | 18123 | 1 | 69790 | ENSMUST0 | 117416 | ENSMUSG0 | Med30 | mediator complex subunit 30 |
| chr15 | 52848994 | 52849393 | 400 | * | NA_peak_64 | 24 | Distal Interge | 15 | 52712442 | 52730564 | 18123 | 1 | 69790 | ENSMUST0 | 136552 | ENSMUSG0 | Med30 | mediator complex subunit 30 |
| chr15 | 52874088 | 52874287 | 200 | * | NA_peak_64 | 13 | Distal Interge | 15 | 52712442 | 52730564 | 18123 | 1 | 69790 | ENSMUST0 | 161646 | ENSMUSG0 | Med30 | mediator complex subunit 30 |
| chr15 | 53155175 | 53155497 | 323 | * | NA_peak_64 | 13 | Intron (ENSM | 15 | 53083146 | 53196255 | 113110 | 2 | 14042 | ENSMUST0 | 40758 | ENSMUSG0 | Ext1 | exostosin glycosyltransferase 1 |
| chr15 | 53422529 | 53422829 | 301 | * | NA_peak_65 | 39 | Distal Interge | 15 | 53064038 | 53346159 | 282122 | 2 | 14042 | ENSMUST0 | -76370 | ENSMUSG0 | Ext1 | exostosin glycosyltransferase 1 |
| chr15 | 53435112 | 53435463 | 352 | * | NA_peak_65 | 22 | Distal Interge | 15 | 53064038 | 53346159 | 282122 | 2 | 14042 | ENSMUST0 | -88953 | ENSMUSG0 | Ext1 | exostosin glycosyltransferase 1 |
| chr15 | 53476567 | 53477003 | 437 | * | NA_peak_65 | 17 | Intron (ENSM | 15 | 53064038 | 53346159 | 282122 | 2 | 14042 | ENSMUST0 | -130408 | ENSMUSG0 | Ext1 | exostosin glycosyltransferase 1 |
| chr15 | 53574267 | 53574584 | 318 | * | NA_peak_65 | 36 | Intron (ENSM | 15 | 53614205 | 53614279 | 75 | 2 | 100628567 | ENSMUST0 | 39695 | ENSMUSG0 | Mir28c | microRNA 28c |
| chr15 | 53713019 | 53713368 | 350 | * | NA_peak_65 | 19 | Intron (ENSM | 15 | 53614205 | 53614279 | 75 | 2 | 100628567 | ENSMUST0 | -98740 | ENSMUSG0 | Mir28c | microRNA 28c |
| chr15 | 53714610 | 53714748 | 139 | * | NA_peak_65 | 22 | Intron (ENSM | 15 | 53614205 | 53614279 | 75 | 2 | 100628567 | ENSMUST0 | -100331 | ENSMUSG0 | Mir28c | microRNA 28c |
| chr15 | 53871565 | 53871769 | 205 | * | NA_peak_65 | 44 | Intron (ENSM | 15 | 53462194 | 53902381 | 440188 | 2 | 320679 | ENSMUST0 | 30612 | ENSMUSG0 | Samd12 | sterile alpha motif domain containing 12 |
| chr15 | 53918900 | 53919218 | 319 | * | NA_peak_65 | 38 | Distal Interge | 15 | 53818794 | 53902537 | 83744 | 2 | 320679 | ENSMUST0 | -16363 | ENSMUSG0 | Samd12 | sterile alpha motif domain containing 12 |
| chr15 | 53930800 | 53931251 | 452 | * | NA_peak_65 | 29 | Distal Interge | 15 | 53818794 | 53902537 | 83744 | 2 | 320679 | ENSMUST0 | -28263 | ENSMUSG0 | Samd12 | sterile alpha motif domain containing 12 |
| chr15 | 54052069 | 54052415 | 347 | * | NA_peak_65 | 26 | Distal Interge | 15 | 53818794 | 53902537 | 83744 | 2 | 320679 | ENSMUST0 | -149532 | ENSMUSG0 | Samd12 | sterile alpha motif domain containing 12 |
| chr15 | 54137835 | 54138505 | 671 | * | NA_peak_65 | 31 | Distal Interge | 15 | 54250619 | 54278484 | 27866 | 2 | 18383 | ENSMUST0 | 139979 | ENSMUSG0 | Tnfrsf11b | tumor necrosis factor receptor superfamily, member 11b (osteoprotegerin) |
| chr15 | 54309158 | 54309352 | 195 | * | NA_peak_65 | 22 | Distal Interge | 15 | 54250619 | 54278484 | 27866 | 2 | 18383 | ENSMUST0 | -30674 | ENSMUSG0 | Tnfrsf11b | tumor necrosis factor receptor superfamily, member 11b (osteoprotegerin) |
| chr15 | 54640730 | 54641509 | 780 | * | NA_peak_65 | 21 | Distal Interge | 15 | 54571192 | 54602847 | 31656 | 1 | 105853 | ENSMUST0 | 69538 | ENSMUSG0 | Mal2 | mal, T cell differentiation protein 2 |
| chr15 | 54670923 | 54671067 | 145 | * | NA_peak_65 | 18 | Distal Interge | 15 | 54745702 | 54754039 | 8338 | 1 | 18133 | ENSMUST0 | -74635 | ENSMUSG0 | Ccn3 | cellular communication network factor 3 |
| chr15 | 54688136 | 54688375 | 240 | * | NA_peak_65 | 16 | Distal Interge | 15 | 54745702 | 54754039 | 8338 | 1 | 18133 | ENSMUST0 | -57327 | ENSMUSG0 | Ccn3 | cellular communication network factor 3 |
| chr15 | 54720624 | 54720935 | 312 | * | NA_peak_65 | 23 | Distal Interge | 15 | 54745702 | 54754039 | 8338 | 1 | 18133 | ENSMUST0 | -24767 | ENSMUSG0 | Ccn3 | cellular communication network factor 3 |
| chr15 | 55050394 | 55050511 | 118 | * | NA_peak_65 | 16 | Promoter (1- | 15 | 55049166 | 55052421 | 3256 | 2 | 319944 | ENSMUST0 | 1910 | ENSMUSG0 | Taf2 | TATA-box binding protein associated factor 2 |
| chr15 | 55068897 | 55069205 | 309 | * | NA_peak_65 | 13 | Promoter (2- | 15 | 55045120 | 55071899 | 26780 | 2 | 319944 | ENSMUST0 | 2694 | ENSMUSG0 | Taf2 | TATA-box binding protein associated factor 2 |
| chr15 | 55201340 | 55201632 | 293 | * | NA_peak_65 | 22 | Intron (ENSM | 15 | 55133453 | 55208885 | 75433 | 1 | 97998 | ENSMUST0 | 67887 | ENSMUSG0 | Deptor | DEP domain containing MTOR-interacting protein |
| chr15 | 55213094 | 55213421 | 328 | * | NA_peak_65 | 18 | Intron (ENSM | 15 | 55133453 | 55208885 | 75433 | 1 | 97998 | ENSMUST0 | 79641 | ENSMUSG0 | Deptor | DEP domain containing MTOR-interacting protein |
| chr15 | 55289286 | 55289494 | 209 | * | NA_peak_65 | 28 | Distal Interge | 15 | 55307750 | 55520803 | 213054 | 1 | 12818 | ENSMUST0 | -18256 | ENSMUSG0 | Col14a1 | collagen, type XIV, alpha 1 |
| chr15 | 55410518 | 55410756 | 239 | * | NA_peak_65 | 24 | Intron (ENSM | 15 | 55448826 | 55519689 | 70864 | 1 | 12818 | ENSMUST0 | -38070 | ENSMUSG0 | Col14a1 | collagen, type XIV, alpha 1 |
| chr15 | 55462371 | 55462954 | 584 | * | NA_peak_65 | 17 | Intron (ENSM | 15 | 55448826 | 55519689 | 70864 | 1 | 12818 | ENSMUST0 | 13545 | ENSMUSG0 | Col14a1 | collagen, type XIV, alpha 1 |
| chr15 | 55529306 | 55529512 | 207 | * | NA_peak_65 | 18 | Distal Interge | 15 | 55538636 | 55548164 | 9529 | 2 | 68537 | ENSMUST0 | 18652 | ENSMUSG0 | Mrpl13 | mitochondrial ribosomal protein L13 |
| chr15 | 55714912 | 55715349 | 438 | * | NA_peak_65 | 23 | Intron (ENSM | 15 | 55621452 | 55626421 | 4970 | 1 | 105837 | ENSMUST0 | 93460 | ENSMUSG0 | Mtbp | Mdm2, transformed 3T3 cell double minute p53 binding protein |
| chr15 | 55858439 | 55858682 | 244 | * | NA_peak_65 | 30 | Intron (ENSM | 15 | 55791938 | 55906310 | 114373 | 2 | 20649 | ENSMUST0 | 47628 | ENSMUSG0 | Sntb1 | syntrophin, basic 1 |
| chr15 | 55873626 | 55873986 | 361 | * | NA_peak_65 | 51 | Intron (ENSM | 15 | 55791938 | 55906310 | 114373 | 2 | 20649 | ENSMUST0 | 32324 | ENSMUSG0 | Sntb1 | syntrophin, basic 1 |
| chr15 | 55992246 | 55992591 | 346 | * | NA_peak_65 | 15 | Distal Interge | 15 | 55636388 | 55906949 | 270562 | 2 | 20649 | ENSMUST0 | -85297 | ENSMUSG0 | Sntb1 | syntrophin, basic 1 |
| chr15 | 55993257 | 55993438 | 182 | * | NA_peak_65 | 22 | Distal Interge | 15 | 55636388 | 55906949 | 270562 | 2 | 20649 | ENSMUST0 | -86308 | ENSMUSG0 | Sntb1 | syntrophin, basic 1 |
| chr15 | 56400825 | 56401188 | 364 | * | NA_peak_66 | 46 | Distal Interge | 15 | 56689944 | 56694460 | 4517 | 1 | 594843 | ENSMUST0 | -288756 | ENSMUSG0 | Has2os | hyaluronan synthase 2, opposite strand |
| chr15 | 56538619 | 56538981 | 363 | * | NA_peak_66 | 24 | Distal Interge | 15 | 56689944 | 56694460 | 4517 | 1 | 594843 | ENSMUST0 | -150963 | ENSMUSG0 | Has2os | hyaluronan synthase 2, opposite strand |
| chr15 | 56568978 | 56569321 | 344 | * | NA_peak_66 | 24 | Distal Interge | 15 | 56689944 | 56694460 | 4517 | 1 | 594843 | ENSMUST0 | -120623 | ENSMUSG0 | Has2os | hyaluronan synthase 2, opposite strand |
| chr15 | 56778141 | 56778232 | 92 | * | NA_peak_66 | 19 | Exon (ENSM | 15 | 56665627 | 56694539 | 28913 | 2 | 15117 | ENSMUST0 | -83602 | ENSMUSG0 | Has2 | hyaluronan synthase 2 |
| chr15 | 56827992 | 56828313 | 322 | * | NA_peak_66 | 35 | Distal Interge | 15 | 56665627 | 56694539 | 28913 | 2 | 15117 | ENSMUST0 | -133453 | ENSMUSG0 | Has2 | hyaluronan synthase 2 |
| chr15 | 56885633 | 56886252 | 620 | * | NA_peak_66 | 50 | Intron (ENSM | 15 | 56665627 | 56694539 | 28913 | 2 | 15117 | ENSMUST0 | -191094 | ENSMUSG0 | Has2 | hyaluronan synthase 2 |
| chr15 | 56955130 | 56955468 | 339 | * | NA_peak_66 | 27 | Intron (ENSM | 15 | 56665627 | 56694539 | 28913 | 2 | 15117 | ENSMUST0 | -260591 | ENSMUSG0 | Has2 | hyaluronan synthase 2 |
| chr15 | 56962176 | 56962549 | 374 | * | NA_peak_66 | 26 | Intron (ENSM | 15 | 56665627 | 56694539 | 28913 | 2 | 15117 | ENSMUST0 | -267637 | ENSMUSG0 | Has2 | hyaluronan synthase 2 |
| chr15 | 57000048 | 57000136 | 89 | * | NA_peak_66 | 23 | Intron (ENSM | 15 | 57256556 | 57265137 | 8582 | 2 | 210463 | ENSMUST0 | 265001 | ENSMUSG0 | Slc22a22 | solute carrier family 22 (organic cation transporter), member 22 |
| chr15 | 57679281 | 57679777 | 497 | * | NA_peak_66 | 26 | Distal Interge | 15 | 57694665 | 57839832 | 145168 | 1 | 387609 | ENSMUST0 | -14888 | ENSMUSG0 | Zhx2 | zinc fingers and homeoboxes 2 |
| chr15 | 57690631 | 57691037 | 407 | * | NA_peak_66 | 30 | Distal Interge | 15 | 57694665 | 57839832 | 145168 | 1 | 387609 | ENSMUST0 | -3628 | ENSMUSG0 | Zhx2 | zinc fingers and homeoboxes 2 |
| chr15 | 57695681 | 57695992 | 312 | * | NA_peak_66 | 25 | Promoter (<= | 15 | 57695729 | 57821110 | 125382 | 1 | 387609 | ENSMUST0 | 0 | ENSMUSG0 | Zhx2 | zinc fingers and homeoboxes 2 |
| chr15 | 58343354 | 58343478 | 125 | * | NA_peak_66 | 16 | Intron (ENSM | 15 | 58314573 | 58324169 | 9597 | 2 | 268807 | ENSMUST0 | -19185 | ENSMUSG0 | Klhl38 | kelch-like 38 |
| chr15 | 58363301 | 58363412 | 112 | * | NA_peak_66 | 13 | Promoter (<= | 15 | 58341874 | 58364148 | 22275 | 2 | 69787 | ENSMUST0 | 736 | ENSMUSG0 | Anxa13 | annexin A13 |
| chr15 | 58491936 | 58492232 | 297 | * | NA_peak_66 | 23 | Distal Interge | 15 | 58510048 | 58662933 | 152886 | 1 | 631797 | ENSMUST0 | -17816 | ENSMUSG0 | Fer1l6 | fer-1-like 6 (C. elegans) |
| chr15 | 58837544 | 58837992 | 449 | * | NA_peak_66 | 27 | Distal Interge | 15 | 58782269 | 58823530 | 41262 | 2 | 74868 | ENSMUST0 | -14014 | ENSMUSG0 | Tmem65 | transmembrane protein 65 |
| chr15 | 58838313 | 58838332 | 20 | * | NA_peak_66 | 27 | Distal Interge | 15 | 58782269 | 58823530 | 41262 | 2 | 74868 | ENSMUST0 | -14783 | ENSMUSG0 | Tmem65 | transmembrane protein 65 |
| chr15 | 58916954 | 58917295 | 342 | * | NA_peak_66 | 40 | Promoter (<= | 15 | 58901017 | 58916178 | 15162 | 2 | 69694 | ENSMUST0 | -776 | ENSMUSG0 | Tatdn1 | TatD DNase domain containing 1 |
| chr15 | 59300208 | 59300658 | 451 | * | NA_peak_67 | 24 | Distal Interge | 15 | 59306948 | 59312013 | 5066 | 1 | 100416830 | ENSMUST0 | -6290 | ENSMUSG0 | Zfp572 | zinc finger protein 572 |
| chr15 | 59392222 | 59392503 | 282 | * | NA_peak_67 | 24 | Intron (ENSM | 15 | 59374287 | 59601684 | 227398 | 1 | 68501 | ENSMUST0 | 17935 | ENSMUSG0 | Nsmce2 | NSE2/MMS21 homolog, SMC5-SMC6 complex SUMO ligase |
| chr15 | 59456129 | 59456361 | 233 | * | NA_peak_67 | 46 | Intron (ENSM | 15 | 59374287 | 59601684 | 227398 | 1 | 68501 | ENSMUST0 | 81842 | ENSMUSG0 | Nsmce2 | NSE2/MMS21 homolog, SMC5-SMC6 complex SUMO ligase |
| chr15 | 59474060 | 59474359 | 300 | * | NA_peak_67 | 16 | Intron (ENSM | 15 | 59374287 | 59601684 | 227398 | 1 | 68501 | ENSMUST0 | 99773 | ENSMUSG0 | Nsmce2 | NSE2/MMS21 homolog, SMC5-SMC6 complex SUMO ligase |
| chr15 | 59703376 | 59704008 | 633 | * | NA_peak_67 | 28 | Distal Interge | 15 | 59648653 | 59653689 | 5037 | 1 | 211770 | ENSMUST0 | 54723 | ENSMUSG0 | Trib1 | tribbles pseudokinase 1 |
| chr15 | 60060667 | 60060831 | 165 | * | NA_peak_67 | 23 | Distal Interge | 15 | 60014866 | 60016613 | 1748 | 2 | 71088 | ENSMUST0 | -44054 | ENSMUSG0 | 4933412E24 | RIKEN cDNA 4933412E24 gene |
| chr15 | 60444529 | 60444877 | 349 | * | NA_peak_67 | 18 | Distal Interge | 15 | 60822271 | 60828650 | 6380 | 1 | 320469 | ENSMUST0 | -377394 | NA | 9930014A18 | RIKEN cDNA 9930014A18 gene |
| chr15 | 60586675 | 60586734 | 60 | * | NA_peak_67 | 18 | Distal Interge | 15 | 60822271 | 60828650 | 6380 | 1 | 320469 | ENSMUST0 | -235537 | NA | 9930014A18 | RIKEN cDNA 9930014A18 gene |
| chr15 | 60670423 | 60670557 | 135 | * | NA_peak_67 | 16 | Intron (ENSM | 15 | 60822271 | 60828650 | 6380 | 1 | 320469 | ENSMUST0 | -151714 | NA | 9930014A18 | RIKEN cDNA 9930014A18 gene |
| chr15 | 60744677 | 60745125 | 449 | * | NA_peak_67 | 31 | Intron (ENSM | 15 | 60822271 | 60828650 | 6380 | 1 | 320469 | ENSMUST0 | -77146 | NA | 9930014A18 | RIKEN cDNA 9930014A18 gene |
| chr15 | 60802787 | 60803011 | 225 | * | NA_peak_67 | 17 | Distal Interge | 15 | 60822271 | 60828650 | 6380 | 1 | 320469 | ENSMUST0 | -19260 | NA | 9930014A18 | RIKEN cDNA 9930014A18 gene |
| chr15 | 61230222 | 61230555 | 334 | * | NA_peak_67 | 36 | Intron (ENSM | 15 | 60919675 | 60923012 | 3338 | 2 | 117586 | ENSMUST0 | -307210 | ENSMUSG0 | A1bg | alpha-1-B glycoprotein |
| chr15 | 61837552 | 61837658 | 107 | * | NA_peak_67 | 34 | Distal Interge | 15 | 61699282 | 61774451 | 75170 | 2 | 432956 | ENSMUST0 | -63101 | ENSMUSG0 | D030024E09 | RIKEN cDNA D030024E09 gene |
| chr15 | 61985008 | 61985480 | 473 | * | NA_peak_68 | 36 | Promoter (<= | 15 | 61985391 | 61990350 | 4960 | 1 | 17869 | ENSMUST0 | 0 | ENSMUSG0 | Myc | myelocytomatosis oncogene |
| chr15 | 62356539 | 62356730 | 192 | * | NA_peak_68 | 19 | Intron (ENSM | 15 | 62222603 | 62242505 | 19903 | 1 | 19296 | ENSMUST0 | 133936 | ENSMUSG0 | Pvt1 | Pvt1 oncogene |
| chr15 | 62434043 | 62434373 | 331 | * | NA_peak_68 | 23 | Distal Interge | 15 | 62222603 | 62242505 | 19903 | 1 | 19296 | ENSMUST0 | 211440 | ENSMUSG0 | Pvt1 | Pvt1 oncogene |
| chr15 | 62523339 | 62523742 | 404 | * | NA_peak_68 | 19 | Distal Interge | 15 | 62222603 | 62242505 | 19903 | 1 | 19296 | ENSMUST0 | 300736 | ENSMUSG0 | Pvt1 | Pvt1 oncogene |
| chr15 | 62552496 | 62552821 | 326 | * | NA_peak_68 | 34 | Distal Interge | 15 | 62222603 | 62242505 | 19903 | 1 | 19296 | ENSMUST0 | 329893 | ENSMUSG0 | Pvt1 | Pvt1 oncogene |
| chr15 | 62991313 | 62991483 | 171 | * | NA_peak_68 | 21 | Distal Interge | 15 | 63414400 | 63479608 | 65209 | 2 | 432958 | ENSMUST0 | 488125 | ENSMUSG0 | Gm20740 | predicted gene, 20740 |
| chr15 | 63183204 | 63183285 | 82 | * | NA_peak_68 | 35 | Intron (ENSM | 15 | 63414400 | 63479608 | 65209 | 2 | 432958 | ENSMUST0 | 296323 | ENSMUSG0 | Gm20740 | predicted gene, 20740 |
| chr15 | 63244724 | 63244863 | 140 | * | NA_peak_68 | 23 | Intron (ENSM | 15 | 63414400 | 63479608 | 65209 | 2 | 432958 | ENSMUST0 | 234745 | ENSMUSG0 | Gm20740 | predicted gene, 20740 |
| chr15 | 63563625 | 63563856 | 232 | * | NA_peak_68 | 27 | Intron (ENSM | 15 | 63414400 | 63479608 | 65209 | 2 | 432958 | ENSMUST0 | -84017 | ENSMUSG0 | Gm20740 | predicted gene, 20740 |
| chr15 | 63627876 | 63628136 | 261 | * | NA_peak_68 | 30 | Intron (ENSM | 15 | 63176838 | 63718289 | 541452 | 2 | 432958 | ENSMUST0 | 90153 | ENSMUSG0 | Gm20740 | predicted gene, 20740 |
| chr15 | 63638670 | 63638901 | 232 | * | NA_peak_68 | 23 | Intron (ENSM | 15 | 63176838 | 63718289 | 541452 | 2 | 432958 | ENSMUST0 | 79388 | ENSMUSG0 | Gm20740 | predicted gene, 20740 |
| chr15 | 63715115 | 63715319 | 205 | * | NA_peak_68 | 19 | Promoter (2- | 15 | 63176838 | 63718289 | 541452 | 2 | 432958 | ENSMUST0 | 2970 | ENSMUSG0 | Gm20740 | predicted gene, 20740 |
| chr15 | 63786902 | 63787442 | 541 | * | NA_peak_68 | 35 | Intron (ENSM | 15 | 63775968 | 63808759 | 32792 | 2 | 83492 | ENSMUST0 | 21317 | ENSMUSG0 | Gsdmc | gasdermin C |
| chr15 | 63943252 | 63943403 | 152 | * | NA_peak_68 | 39 | Exon (ENSM | 15 | 63922277 | 63923197 | 921 | 1 | 74460 | ENSMUST0 | 20975 | ENSMUSG0 | Gsdmcl-ps | gasdermin C-like, pseudogene |
| chr15 | 63989457 | 63989504 | 48 | * | NA_peak_68 | 26 | Intron (ENSM | 15 | 63930942 | 63997969 | 67028 | 2 | 223601 | ENSMUST0 | 8465 | ENSMUSG0 | Cyrib | CYFIP related Rac1 interactor B |
| chr15 | 64149133 | 64149673 | 541 | * | NA_peak_68 | 20 | Exon (ENSM | 15 | 64120091 | 64123806 | 3716 | 2 | 13196 | ENSMUST0 | -25327 | ENSMUSG0 | Asap1 | ArfGAP with SH3 domain, ankyrin repeat and PH domain1 |
| chr15 | 64230974 | 64231288 | 315 | * | NA_peak_68 | 21 | Intron (ENSM | 15 | 64158939 | 64208491 | 49553 | 2 | 13196 | ENSMUST0 | -22483 | ENSMUSG0 | Asap1 | ArfGAP with SH3 domain, ankyrin repeat and PH domain1 |
| chr15 | 64644064 | 64644196 | 133 | * | NA_peak_69 | 19 | Distal Interge | 15 | 64088541 | 64382919 | 294379 | 2 | 13196 | ENSMUST0 | -261145 | ENSMUSG0 | Asap1 | ArfGAP with SH3 domain, ankyrin repeat and PH domain1 |
| chr15 | 65155301 | 65155708 | 408 | * | NA_peak_69 | 22 | Distal Interge | 15 | 64697084 | 64922296 | 225213 | 2 | 11514 | ENSMUST0 | -233005 | ENSMUSG0 | Adcy8 | adenylate cyclase 8 |
| chr15 | 65257609 | 65257849 | 241 | * | NA_peak_69 | 20 | Distal Interge | 15 | 64697084 | 64922296 | 225213 | 2 | 11514 | ENSMUST0 | -335313 | ENSMUSG0 | Adcy8 | adenylate cyclase 8 |
| chr15 | 65714138 | 65714276 | 139 | * | NA_peak_69 | 22 | Distal Interge | 15 | 65787034 | 65791351 | 4318 | 1 | 76740 | ENSMUST0 | -72758 | ENSMUSG0 | Efr3a | EFR3 homolog A |
| chr15 | 65878346 | 65878885 | 540 | * | NA_peak_69 | 22 | Exon (ENSM | 15 | 65876520 | 65889638 | 13119 | 2 | 18256 | ENSMUST0 | 10753 | ENSMUSG0 | Oc90 | otoconin 90 |
| chr15 | 66502472 | 66502656 | 185 | * | NA_peak_69 | 16 | Promoter (1- | 15 | 66379858 | 66500910 | 121053 | 2 | 54562 | ENSMUST0 | -1562 | ENSMUSG0 | Lrrc6 | leucine rich repeat containing 6 (testis) |
| chr15 | 66540455 | 66541005 | 551 | * | NA_peak_69 | 26 | Intron (ENSM | 15 | 66526212 | 66561103 | 34892 | 2 | 213068 | ENSMUST0 | 20098 | ENSMUSG0 | Tmem71 | transmembrane protein 71 |
| chr15 | 66577308 | 66578137 | 830 | * | NA_peak_69 | 33 | Promoter (<= | 15 | 66577560 | 66611616 | 34057 | 1 | 239510 | ENSMUST0 | 0 | ENSMUSG0 | Phf20l1 | PHD finger protein 20-like 1 |
| chr15 | 67170513 | 67170847 | 335 | * | NA_peak_69 | 29 | Intron (ENSM | 15 | 67106285 | 67176712 | 70428 | 2 | 20442 | ENSMUST0 | 5865 | ENSMUSG0 | St3gal1 | ST3 beta-galactoside alpha-2,3-sialyltransferase 1 |
| chr15 | 67330665 | 67330939 | 275 | * | NA_peak_69 | 32 | Intron (ENSM | 15 | 67226769 | 67377094 | 150326 | 1 | 69334 | ENSMUST0 | 103896 | ENSMUSG0 | 1700012I11 | RIKEN cDNA 1700012I11 gene |
| chr15 | 67667904 | 67668138 | 235 | * | NA_peak_69 | 22 | Distal Interge | 15 | 67226769 | 67377094 | 150326 | 1 | 69334 | ENSMUST0 | 441135 | ENSMUSG0 | 1700012I11 | RIKEN cDNA 1700012I11 gene |
| chr15 | 68032368 | 68032730 | 363 | * | NA_peak_69 | 26 | Distal Interge | 15 | 68084321 | 68258787 | 174467 | 2 | 380993 | ENSMUST0 | 226057 | ENSMUSG0 | Zfat | zinc finger and AT hook domain containing |
| chr15 | 68340447 | 68340678 | 232 | * | NA_peak_69 | 18 | Promoter (<= | 15 | 68338295 | 68341216 | 2922 | 2 | 387228 | ENSMUST0 | 538 | ENSMUSG0 | Mir30d | microRNA 30d |
| chr15 | 68363464 | 68363765 | 302 | * | NA_peak_69 | 47 | Distal Interge | 15 | 68341208 | 68341289 | 82 | 2 | 387228 | ENSMUST0 | -22175 | ENSMUSG0 | Mir30d | microRNA 30d |
| chr15 | 69144437 | 69144480 | 44 | * | NA_peak_69 | 30 | Distal Interge | 15 | 69270533 | 69274720 | 4188 | 1 | 75076 | ENSMUST0 | -126053 | ENSMUSG0 | 4930504C09 | RIKEN cDNA 4930504C09 gene |
| chr15 | 69646344 | 69646653 | 310 | * | NA_peak_69 | 61 | Distal Interge | 15 | 70011623 | 70077841 | 66219 | 1 | 100503580 | ENSMUST0 | -364970 | ENSMUSG0 | Gm19782 | predicted gene, 19782 |
| chr15 | 71585634 | 71586168 | 535 | * | NA_peak_70 | 18 | Intron (ENSM | 15 | 71445676 | 71462400 | 16725 | 2 | 70363 | ENSMUST0 | -123234 | ENSMUSG0 | Fam135b | family with sequence similarity 135, member B |
| chr15 | 71826333 | 71826633 | 301 | * | NA_peak_70 | 26 | Intron (ENSM | 15 | 71868881 | 71897616 | 28736 | 2 | 69700 | ENSMUST0 | 70983 | ENSMUSG0 | Col22a1 | collagen, type XXII, alpha 1 |
| chr15 | 72249278 | 72249655 | 378 | * | NA_peak_70 | 30 | Intron (ENSM | 15 | 71795795 | 72034227 | 238433 | 2 | 69700 | ENSMUST0 | -215051 | ENSMUSG0 | Col22a1 | collagen, type XXII, alpha 1 |
| chr15 | 72482838 | 72483111 | 274 | * | NA_peak_70 | 19 | Distal Interge | 15 | 72501089 | 72546340 | 45252 | 2 | 223604 | ENSMUST0 | 63229 | ENSMUSG0 | Kcnk9 | potassium channel, subfamily K, member 9 |
| chr15 | 72882776 | 72882780 | 5 | * | NA_peak_70 | 20 | Intron (ENSM | 15 | 72805600 | 72810324 | 4725 | 2 | 353342 | ENSMUST0 | -72452 | ENSMUSG0 | Peg13 | paternally expressed 13 |
| chr15 | 72939896 | 72940203 | 308 | * | NA_peak_70 | 22 | Intron (ENSM | 15 | 72589620 | 73055810 | 466191 | 2 | 76510 | ENSMUST0 | 115607 | ENSMUSG0 | Trappc9 | trafficking protein particle complex 9 |
| chr15 | 73003907 | 73004061 | 155 | * | NA_peak_70 | 13 | Intron (ENSM | 15 | 72589620 | 73055810 | 466191 | 2 | 76510 | ENSMUST0 | 51749 | ENSMUSG0 | Trappc9 | trafficking protein particle complex 9 |
| chr15 | 73192703 | 73193080 | 378 | * | NA_peak_70 | 26 | Distal Interge | 15 | 73095844 | 73184935 | 89092 | 2 | 239528 | ENSMUST0 | -7768 | ENSMUSG0 | Ago2 | argonaute RISC catalytic subunit 2 |
| chr15 | 73283562 | 73283864 | 303 | * | NA_peak_70 | 34 | Intron (ENSM | 15 | 73254815 | 73254882 | 68 | 2 | 387169 | ENSMUST0 | -28680 | ENSMUSG0 | Mir151 | microRNA 151 |
| chr15 | 73341927 | 73342022 | 96 | * | NA_peak_70 | 25 | Promoter (1- | 15 | 73215557 | 73343327 | 127771 | 2 | 14083 | ENSMUST0 | 1305 | ENSMUSG0 | Ptk2 | PTK2 protein tyrosine kinase 2 |
| chr15 | 73503765 | 73504135 | 371 | * | NA_peak_70 | 41 | Distal Interge | 15 | 73512560 | 73572242 | 59683 | 1 | 105841 | ENSMUST0 | -8425 | ENSMUSG0 | Dennd3 | DENN/MADD domain containing 3 |
| chr15 | 74814704 | 74814853 | 150 | * | NA_peak_70 | 21 | Promoter (2- | 15 | 74813452 | 74817473 | 4022 | 2 | 625599 | ENSMUST0 | 2620 | ENSMUSG0 | Gml | glycosylphosphatidylinositol anchored molecule like |
| chr15 | 74815433 | 74815597 | 165 | * | NA_peak_70 | 21 | Promoter (1- | 15 | 74813452 | 74817473 | 4022 | 2 | 625599 | ENSMUST0 | 1876 | ENSMUSG0 | Gml | glycosylphosphatidylinositol anchored molecule like |
| chr15 | 74830701 | 74831052 | 352 | * | NA_peak_70 | 26 | Intron (ENSM | 15 | 74823065 | 74824436 | 1372 | 1 | 15202 | ENSMUST0 | 7636 | ENSMUSG0 | Gml2 | glycosylphosphatidylinositol anchored molecule like 2 |
| chr15 | 75333941 | 75334290 | 350 | * | NA_peak_70 | 40 | Distal Interge | 15 | 75268702 | 75272235 | 3534 | 1 | 17071 | ENSMUST0 | 65239 | ENSMUSG0 | Ly6f | lymphocyte antigen 6 complex, locus F |
| chr15 | 75494439 | 75494550 | 112 | * | NA_peak_70 | 17 | Distal Interge | 15 | 75449457 | 75451406 | 1950 | 1 | 102637705 | ENSMUST0 | 44982 | ENSMUSG0 | Ly6l | lymphocyte antigen 6 complex, locus L |
| chr15 | 75612256 | 75612483 | 228 | * | NA_peak_71 | 22 | Distal Interge | 15 | 75616679 | 75625300 | 8622 | 1 | 22701 | ENSMUST0 | -4196 | ENSMUSG0 | Zfp41 | zinc finger protein 41 |

| | | | | | | | | | | | | | | | | | | |
|---|---|---|---|---|---|---|---|---|---|---|---|---|---|---|---|---|---|---|
| chr15 | 75616149 | 75616407 | 259 | * | NA_peak_71 | 25 | Promoter (<= | 15 | 75616679 | 75625300 | 8622 | 1 | 22701 | ENSMUST00 | -272 | ENSMUSG0 | Zfp41 | zinc finger protein 41 |
| chr15 | 75969067 | 75969412 | 346 | * | NA_peak_71 | 23 | Promoter (<= | 15 | 75969123 | 75975749 | 6627 | 1 | 69020 | ENSMUST00 | 0 | ENSMUSG0 | Zfp707 | zinc finger protein 707 |
| chr15 | 76091096 | 76091505 | 410 | * | NA_peak_71 | 30 | Promoter (<= | 15 | 76085518 | 76091505 | 5988 | 2 | 223649 | ENSMUST00 | 0 | ENSMUSG0 | Nrbp2 | nuclear receptor binding protein 2 |
| chr15 | 76257936 | 76258384 | 449 | * | NA_peak_71 | 16 | Distal Interge | 15 | 76268089 | 76292572 | 24484 | 1 | 74281 | ENSMUST00 | -9705 | ENSMUSG0 | Spatc1 | spermatogenesis and centriole associated 1 |
| chr15 | 76259022 | 76259522 | 501 | * | NA_peak_71 | 35 | Distal Interge | 15 | 76268089 | 76292572 | 24484 | 1 | 74281 | ENSMUST00 | -8567 | ENSMUSG0 | Spatc1 | spermatogenesis and centriole associated 1 |
| chr15 | 76282137 | 76282287 | 151 | * | NA_peak_71 | 16 | Intron (ENSM | 15 | 76290879 | 76290968 | 90 | 2 | 100526548 | ENSMUST00 | 8681 | ENSMUSG0 | Mir3079 | microRNA 3079 |
| chr15 | 76286231 | 76286302 | 72 | * | NA_peak_71 | 19 | Intron (ENSM | 15 | 76290879 | 76290968 | 90 | 2 | 100526548 | ENSMUST00 | 4666 | ENSMUSG0 | Mir3079 | microRNA 3079 |
| chr15 | 76287882 | 76288088 | 207 | * | NA_peak_71 | 30 | Promoter (2- | 15 | 76290879 | 76290968 | 90 | 2 | 100526548 | ENSMUST00 | 2880 | ENSMUSG0 | Mir3079 | microRNA 3079 |
| chr15 | 76320148 | 76320375 | 228 | * | NA_peak_71 | 19 | Intron (ENSM | 15 | 76327397 | 76330677 | 3281 | 1 | 109075 | ENSMUST00 | -7022 | ENSMUSG0 | Exosc4 | exosome component 4 |
| chr15 | 76511667 | 76512030 | 364 | * | NA_peak_71 | 22 | Promoter (<= | 15 | 76502015 | 76511951 | 9937 | 2 | 13350 | ENSMUST00 | 0 | ENSMUSG0 | Dgat1 | diacylglycerol O-acyltransferase 1 |
| chr15 | 76530456 | 76530621 | 166 | * | NA_peak_71 | 22 | Distal Interge | 15 | 76537219 | 76537986 | 768 | 2 | 30840 | ENSMUST00 | 7365 | ENSMUSG0 | Fbxl6 | F-box and leucine-rich repeat protein 6 |
| chr15 | 76554476 | 76554816 | 341 | * | NA_peak_71 | 52 | Distal Interge | 15 | 76538964 | 76542130 | 3167 | 1 | 52710 | ENSMUST00 | 15512 | ENSMUSG0 | Slc52a2 | solute carrier protein 52, member 2 |
| chr15 | 76613453 | 76613900 | 448 | * | NA_peak_71 | 29 | Promoter (<= | 15 | 76613817 | 76614306 | 490 | 2 | 72027 | ENSMUST00 | 406 | ENSMUSG0 | Slc39a4 | solute carrier family 39 (zinc transporter), member 4 |
| chr15 | 76668822 | 76668917 | 96 | * | NA_peak_71 | 19 | Promoter (<= | 15 | 76668925 | 76669802 | 878 | 2 | 14106 | ENSMUST00 | 885 | ENSMUSG0 | Foxh1 | forkhead box H1 |
| chr15 | 76773818 | 76774068 | 251 | * | NA_peak_71 | 30 | Intron (ENSM | 15 | 76751677 | 76794825 | 43149 | 2 | 223666 | ENSMUST00 | 20757 | ENSMUSG0 | Arhgap39 | Rho GTPase activating protein 39 |
| chr15 | 76795464 | 76795728 | 265 | * | NA_peak_71 | 14 | Promoter (<= | 15 | 76751677 | 76794825 | 43149 | 2 | 223666 | ENSMUST00 | -639 | ENSMUSG0 | Arhgap39 | Rho GTPase activating protein 39 |
| chr15 | 76949183 | 76949290 | 108 | * | NA_peak_71 | 13 | Promoter (<= | 15 | 76948749 | 76950979 | 2231 | 1 | 117171 | ENSMUST00 | 434 | ENSMUSG0 | 1110038F14 | RIKEN cDNA 1110038F14 gene |
| chr15 | 77056987 | 77056991 | 5 | * | NA_peak_71 | 20 | 3' UTR | 15 | 77052751 | 77056044 | 3294 | 1 | 71939 | ENSMUST00 | 4236 | ENSMUSG0 | Apol6 | apolipoprotein L 6 |
| chr15 | 77065836 | 77065995 | 160 | * | NA_peak_71 | 27 | Distal Interge | 15 | 77052751 | 77056044 | 3294 | 1 | 71939 | ENSMUST00 | 13085 | ENSMUSG0 | Apol6 | apolipoprotein L 6 |
| chr15 | 77167603 | 77167918 | 316 | * | NA_peak_71 | 16 | Intron (ENSM | 15 | 77078991 | 77153812 | 74822 | 2 | 93686 | ENSMUST00 | -13791 | ENSMUSG0 | Rbfox2 | RNA binding protein, fox-1 homolog (C. elegans) 2 |
| chr15 | 77175326 | 77175554 | 229 | * | NA_peak_71 | 24 | Intron (ENSM | 15 | 77078991 | 77153812 | 74822 | 2 | 93686 | ENSMUST00 | -21514 | ENSMUSG0 | Rbfox2 | RNA binding protein, fox-1 homolog (C. elegans) 2 |
| chr15 | 77332105 | 77332208 | 104 | * | NA_peak_71 | 41 | Distal Interge | 15 | 77113606 | 77307004 | 193399 | 2 | 93686 | ENSMUST00 | -25101 | ENSMUSG0 | Rbfox2 | RNA binding protein, fox-1 homolog (C. elegans) 2 |
| chr15 | 77557116 | 77557456 | 341 | * | NA_peak_71 | 22 | Distal Interge | 15 | 77558239 | 77561892 | 3654 | 2 | 76409 | ENSMUST00 | 4436 | ENSMUSG0 | 1700025B11 | RIKEN cDNA 1700025B11 gene |
| chr15 | 77682523 | 77683274 | 752 | * | NA_peak_71 | 20 | Distal Interge | 15 | 77698857 | 77714634 | 15778 | 1 | 666348 | ENSMUST00 | -15583 | ENSMUSG0 | Apol7e | apolipoprotein L 7e |
| chr15 | 77698811 | 77699158 | 348 | * | NA_peak_71 | 35 | Promoter (<= | 15 | 77698857 | 77714634 | 15778 | 1 | 666348 | ENSMUST00 | 0 | ENSMUSG0 | Apol7e | apolipoprotein L 7e |
| chr15 | 77710882 | 77711273 | 392 | * | NA_peak_71 | 16 | Intron (ENSM | 15 | 77698905 | 77719285 | 20381 | 1 | 666348 | ENSMUST00 | 11977 | ENSMUSG0 | Apol7e | apolipoprotein L 7e |
| chr15 | 77841123 | 77841716 | 594 | * | NA_peak_71 | 13 | Promoter (<= | 15 | 77790741 | 77842047 | 51307 | 2 | 17886 | ENSMUST00 | 331 | ENSMUSG0 | Myh9 | myosin, heavy polypeptide 9, non-muscle |
| chr15 | 77881158 | 77881443 | 286 | * | NA_peak_71 | 25 | Distal Interge | 15 | 77760587 | 77842175 | 81589 | 2 | 17886 | ENSMUST00 | -38983 | ENSMUSG0 | Myh9 | myosin, heavy polypeptide 9, non-muscle |
| chr15 | 78318199 | 78318506 | 308 | * | NA_peak_72 | 14 | Distal Interge | 15 | 78325752 | 78351001 | 25250 | 1 | 12983 | ENSMUST00 | -7246 | ENSMUSG0 | Csf2rb | colony stimulating factor 2 receptor, beta, low-affinity (granulocyte-macrophage) |
| chr15 | 78857690 | 78857940 | 251 | * | NA_peak_72 | 38 | Promoter (2- | 15 | 78850860 | 78855529 | 4670 | 2 | 107753 | ENSMUST00 | -2161 | ENSMUSG0 | Lgals2 | lectin, galactose-binding, soluble 2 |
| chr15 | 78916512 | 78916874 | 363 | * | NA_peak_72 | 35 | Promoter (2- | 15 | 78913919 | 78919517 | 5599 | 1 | 57028 | ENSMUST00 | 2593 | ENSMUSG0 | Pdxp | pyridoxal (pyridoxine, vitamin B6) phosphatase |
| chr15 | 78939271 | 78939629 | 359 | * | NA_peak_72 | 41 | Intron (ENSM | 15 | 78935208 | 78936581 | 1374 | 1 | 97961 | ENSMUST00 | 4063 | ENSMUSG0 | Nol12 | nucleolar protein 12 |
| chr15 | 78953310 | 78953618 | 309 | * | NA_peak_72 | 31 | Intron (ENSM | 15 | 78948245 | 78953255 | 5011 | 1 | 110253 | ENSMUST00 | 5065 | ENSMUSG0 | Triobp | TRIO and F-actin binding protein |
| chr15 | 79180505 | 79180974 | 470 | * | NA_peak_72 | 18 | Intron (ENSM | 15 | 79172720 | 79216497 | 43778 | 1 | 100041655 | ENSMUST00 | 7785 | ENSMUSG0 | Gm10863 | predicted gene 10863 |
| chr15 | 79409733 | 79410059 | 327 | * | NA_peak_72 | 31 | Distal Interge | 15 | 79360684 | 79403569 | 42886 | 2 | 223693 | ENSMUST00 | -6164 | ENSMUSG0 | Tmem184b | transmembrane protein 184b |
| chr15 | 79537167 | 79537469 | 303 | * | NA_peak_72 | 26 | Exon (ENSM | 15 | 79531353 | 79546555 | 15203 | 2 | 67040 | ENSMUST00 | 9086 | ENSMUSG0 | Ddx17 | DEAD box helicase 17 |
| chr15 | 79698150 | 79698488 | 339 | * | NA_peak_72 | 31 | Intron (ENSM | 15 | 79694180 | 79712173 | 17994 | 1 | 14904 | ENSMUST00 | 3970 | ENSMUSG0 | Gtpbp1 | GTP binding protein 1 |
| chr15 | 79885770 | 79885844 | 75 | * | NA_peak_72 | 21 | Distal Interge | 15 | 79891659 | 79905547 | 13889 | 1 | 80287 | ENSMUST00 | -5815 | ENSMUSG0 | Apobec3 | apolipoprotein B mRNA editing enzyme, catalytic polypeptide 3 |
| chr15 | 80230977 | 80231229 | 253 | * | NA_peak_72 | 20 | Promoter (2- | 15 | 80234018 | 80253371 | 19354 | 1 | 239555 | ENSMUST00 | -2789 | ENSMUSG0 | Mief1 | mitochondrial elongation factor 1 |
| chr15 | 80725743 | 80725864 | 122 | * | NA_peak_72 | 16 | Intron (ENSM | 15 | 80711313 | 80941085 | 229773 | 1 | 213988 | ENSMUST00 | 14430 | ENSMUSG0 | Tnrc6b | trinucleotide repeat containing 6b |
| chr15 | 80729328 | 80729589 | 262 | * | NA_peak_72 | 19 | Intron (ENSM | 15 | 80711313 | 80941085 | 229773 | 1 | 213988 | ENSMUST00 | 18015 | ENSMUSG0 | Tnrc6b | trinucleotide repeat containing 6b |
| chr15 | 80794413 | 80794822 | 410 | * | NA_peak_72 | 44 | Intron (ENSM | 15 | 80798642 | 80881488 | 82847 | 1 | 213988 | ENSMUST00 | -3820 | ENSMUSG0 | Tnrc6b | trinucleotide repeat containing 6b |
| chr15 | 80856868 | 80856990 | 123 | * | NA_peak_72 | 22 | Intron (ENSM | 15 | 80880698 | 80930008 | 49311 | 1 | 213988 | ENSMUST00 | -23708 | ENSMUSG0 | Tnrc6b | trinucleotide repeat containing 6b |
| chr15 | 80912531 | 80912840 | 310 | * | NA_peak_72 | 27 | Intron (ENSM | 15 | 80923064 | 80930032 | 6969 | 1 | 213988 | ENSMUST00 | -10224 | ENSMUSG0 | Tnrc6b | trinucleotide repeat containing 6b |
| chr15 | 80933346 | 80933429 | 84 | * | NA_peak_72 | 15 | 3' UTR | 15 | 80940039 | 80940120 | 82 | 1 | 100628612 | ENSMUST00 | -6610 | ENSMUSG0 | Mir5113 | microRNA 5113 |
| chr15 | 80980017 | 80980409 | 393 | * | NA_peak_72 | 27 | Promoter (1- | 15 | 80978057 | 81006532 | 28476 | 1 | 105835 | ENSMUST00 | 1960 | ENSMUSG0 | Sgsm3 | small G protein signaling modulator 3 |
| chr15 | 81744998 | 81745292 | 295 | * | NA_peak_73 | 13 | Promoter (<= | 15 | 81745057 | 81784444 | 39388 | 1 | 20286 | ENSMUST00 | 0 | ENSMUSG0 | Zc3h7b | zinc finger CCCH type containing 7B |
| chr15 | 81754043 | 81754051 | 9 | * | NA_peak_73 | 30 | Intron (ENSM | 15 | 81718790 | 81745530 | 26741 | 2 | 19387 | ENSMUST00 | -8513 | ENSMUSG0 | Rangap1 | RAN GTPase activating protein 1 |
| chr15 | 81826255 | 81826491 | 237 | * | NA_peak_73 | 16 | 3' UTR | 15 | 81820943 | 81823779 | 2837 | 1 | 21685 | ENSMUST00 | 5312 | ENSMUSG0 | Tef | thyrotroph embryonic factor |
| chr15 | 81858010 | 81858275 | 266 | * | NA_peak_73 | 47 | Promoter (<= | 15 | 81848272 | 81858485 | 10214 | 2 | 57259 | ENSMUST00 | 210 | ENSMUSG0 | Tob2 | transducer of ERBB2, 2 |
| chr15 | 81858774 | 81859429 | 656 | * | NA_peak_73 | 35 | Promoter (<= | 15 | 81851633 | 81858522 | 6890 | 2 | 57259 | ENSMUST00 | -252 | ENSMUSG0 | Tob2 | transducer of ERBB2, 2 |
| chr15 | 82097535 | 82097927 | 393 | * | NA_peak_73 | 27 | Intron (ENSM | 15 | 82083163 | 82126736 | 43574 | 1 | 74369 | ENSMUST00 | 14372 | ENSMUSG0 | Mei1 | meiotic double-stranded break formation protein 1 |
| chr15 | 82263896 | 82264137 | 242 | * | NA_peak_73 | 25 | Distal Interge | 15 | 82268802 | 82294574 | 25773 | 1 | 24050 | ENSMUST00 | -4665 | ENSMUSG0 | Septin3 | septin 3 |
| chr15 | 82538132 | 82538373 | 242 | * | NA_peak_73 | 30 | Exon (ENSM | 15 | 82555072 | 82557834 | 2763 | 1 | 380997 | ENSMUST00 | -16699 | ENSMUSG0 | Cyp2d12 | cytochrome P450, family 2, subfamily d, polypeptide 12 |
| chr15 | 82550567 | 82550645 | 79 | * | NA_peak_73 | 31 | Distal Interge | 15 | 82555072 | 82557834 | 2763 | 1 | 380997 | ENSMUST00 | -4427 | ENSMUSG0 | Cyp2d12 | cytochrome P450, family 2, subfamily d, polypeptide 12 |
| chr15 | 82892887 | 82893204 | 318 | * | NA_peak_73 | 22 | Intron (ENSM | 15 | 82899080 | 82905818 | 6739 | 1 | 100504637 | ENSMUST00 | -5876 | ENSMUSG0 | Gm20324 | predicted gene, 20324 |
| chr15 | 83157425 | 83157735 | 311 | * | NA_peak_73 | 17 | 3' UTR | 15 | 83149644 | 83149794 | 151 | 1 | 104307 | ENSMUST00 | 7781 | ENSMUSG0 | Rnu12 | RNA U12, small nuclear |
| chr15 | 83180927 | 83181249 | 323 | * | NA_peak_73 | 25 | Distal Interge | 15 | 83153494 | 83172592 | 19099 | 2 | 109754 | ENSMUST00 | -8335 | ENSMUSG0 | Cyb5r3 | cytochrome b5 reductase 3 |
| chr15 | 83202134 | 83202473 | 340 | * | NA_peak_73 | 18 | Distal Interge | 15 | 83153494 | 83172592 | 19099 | 2 | 109754 | ENSMUST00 | -29542 | ENSMUSG0 | Cyb5r3 | cytochrome b5 reductase 3 |
| chr15 | 83202944 | 83203490 | 547 | * | NA_peak_73 | 18 | Distal Interge | 15 | 83153494 | 83172592 | 19099 | 2 | 109754 | ENSMUST00 | -30352 | ENSMUSG0 | Cyb5r3 | cytochrome b5 reductase 3 |
| chr15 | 83402053 | 83402390 | 338 | * | NA_peak_73 | 18 | Intron (ENSM | 15 | 83377578 | 83388602 | 11025 | 2 | 23970 | ENSMUST00 | -13451 | ENSMUSG0 | Pacsin2 | protein kinase C and casein kinase substrate in neurons 2 |
| chr15 | 83560318 | 83560354 | 37 | * | NA_peak_73 | 23 | Intron (ENSM | 15 | 83563592 | 83574203 | 10612 | 1 | 12257 | ENSMUST00 | -3238 | ENSMUSG0 | Tspo | translocator protein |
| chr15 | 84373184 | 84373552 | 369 | * | NA_peak_73 | 26 | Distal Interge | 15 | 84326119 | 84342978 | 16860 | 1 | 64099 | ENSMUST00 | 47065 | ENSMUSG0 | Parvg | parvin, gamma |
| chr15 | 85017200 | 85017543 | 344 | * | NA_peak_74 | 18 | Promoter (<= | 15 | 85017141 | 85022547 | 5407 | 1 | 22270 | ENSMUST00 | 59 | ENSMUSG0 | Upk3a | uroplakin 3A |
| chr15 | 85066110 | 85066196 | 87 | * | NA_peak_74 | 30 | Intron (ENSM | 15 | 85053697 | 85061749 | 8053 | 1 | 73225 | ENSMUST00 | 12413 | ENSMUSG0 | Fam118a | family with sequence similarity 118, member A |
| chr15 | 85408896 | 85409101 | 206 | * | NA_peak_74 | 20 | Intron (ENSM | 15 | 85336245 | 85463212 | 126968 | 1 | 54138 | ENSMUST00 | 72651 | ENSMUSG0 | Atxn10 | ataxin 10 |
| chr15 | 86162385 | 86162388 | 4 | * | NA_peak_74 | 26 | Promoter (2- | 15 | 86139128 | 86159468 | 20341 | 2 | 223753 | ENSMUST00 | -2917 | ENSMUSG0 | Cerk | ceramide kinase |
| chr15 | 86194679 | 86195317 | 639 | * | NA_peak_74 | 131 | Distal Interge | 15 | 86139129 | 86186141 | 47013 | 2 | 223753 | ENSMUST00 | -8538 | ENSMUSG0 | Cerk | ceramide kinase |
| chr15 | 86662670 | 86662912 | 243 | * | NA_peak_74 | 30 | Distal Interge | 15 | 86235505 | 86292162 | 56658 | 1 | 223754 | ENSMUST00 | 427165 | ENSMUSG0 | Tbc1d22a | TBC1 domain family, member 22a |
| chr15 | 87929778 | 87929997 | 220 | * | NA_peak_74 | 30 | Distal Interge | 15 | 87795850 | 87795943 | 94 | 2 | 102465209 | ENSMUST00 | -133835 | ENSMUSG0 | Mir6393 | microRNA 6393 |
| chr15 | 88729640 | 88729937 | 298 | * | NA_peak_74 | 27 | Promoter (<= | 15 | 88688296 | 88730771 | 42476 | 2 | 223770 | ENSMUST00 | 834 | ENSMUSG0 | Brd1 | bromodomain containing 1 |
| chr15 | 89004524 | 89004828 | 305 | * | NA_peak_74 | 22 | Intron (ENSM | 15 | 89020421 | 89023621 | 3201 | 1 | 83456 | ENSMUST00 | -15593 | ENSMUSG0 | Mov10l1 | Mov10 like RISC complex RNA helicase 1 |
| chr15 | 89204461 | 89204841 | 381 | * | NA_peak_74 | 18 | Distal Interge | 15 | 89211553 | 89253069 | 41517 | 1 | 71474 | ENSMUST00 | -6712 | ENSMUSG0 | Ppp6r2 | protein phosphatase 6, regulatory subunit 2 |
| chr15 | 89473245 | 89473563 | 319 | * | NA_peak_74 | 14 | Promoter (<= | 15 | 89473240 | 89474271 | 1032 | 2 | 11883 | ENSMUST00 | 708 | ENSMUSG0 | Arsa | arylsulfatase A |
| chr15 | 89599123 | 89599391 | 269 | * | NA_peak_74 | 21 | Distal Interge | 15 | 89582533 | 89591923 | 9391 | 2 | 68708 | ENSMUST00 | -7200 | ENSMUSG0 | Rabl2 | RAB, member RAS oncogene family-like 2 |
| chr15 | 89621708 | 89622298 | 591 | * | NA_peak_74 | 24 | Distal Interge | 15 | 89582533 | 89591923 | 9391 | 2 | 68708 | ENSMUST00 | -29785 | ENSMUSG0 | Rabl2 | RAB, member RAS oncogene family-like 2 |
| chr15 | 89826417 | 89826806 | 390 | * | NA_peak_74 | 35 | Intron (ENSM | 15 | 89782393 | 89841860 | 59468 | 2 | 54526 | ENSMUST00 | 15054 | ENSMUSG0 | Syt10 | synaptotagmin X |
| chr15 | 90411746 | 90412263 | 518 | * | NA_peak_74 | 16 | Distal Interge | 15 | 90353706 | 90368444 | 14739 | 1 | 102640415 | ENSMUST00 | 58040 | ENSMUSG0 | Gm36480 | predicted gene, 36480 |
| chr15 | 90488415 | 90488606 | 192 | * | NA_peak_74 | 20 | 3' UTR | 15 | 90353706 | 90368444 | 14739 | 1 | 102640415 | ENSMUST00 | 134709 | ENSMUSG0 | Gm36480 | predicted gene, 36480 |
| chr15 | 90525340 | 90525552 | 213 | * | NA_peak_74 | 50 | Intron (ENSM | 15 | 90643329 | 90649438 | 6110 | 2 | 66871 | ENSMUST00 | 123886 | ENSMUSG0 | Cpne8 | copine VIII |
| chr15 | 90707358 | 90707641 | 284 | * | NA_peak_75 | 31 | Distal Interge | 15 | 90487483 | 90679432 | 191950 | 2 | 66871 | ENSMUST00 | -27926 | ENSMUSG0 | Cpne8 | copine VIII |
| chr15 | 90897431 | 90897734 | 304 | * | NA_peak_75 | 23 | Promoter (<= | 15 | 90891402 | 90898428 | 7027 | 2 | 20738 | ENSMUST00 | 694 | ENSMUSG0 | Spn-ps | sialophorin, pseudogene |
| chr15 | 91118675 | 91119359 | 685 | * | NA_peak_75 | 29 | Distal Interge | 15 | 91159111 | 91172185 | 13075 | 2 | 26874 | ENSMUST00 | 52826 | ENSMUSG0 | Abcd2 | ATP-binding cassette, sub-family D (ALD), member 2 |
| chr15 | 91874990 | 91875302 | 313 | * | NA_peak_75 | 26 | Intron (ENSM | 15 | 91847915 | 91861438 | 13524 | 1 | 223809 | ENSMUST00 | 27075 | ENSMUSG0 | Smgc | submandibular gland protein C |
| chr15 | 93425078 | 93425423 | 346 | * | NA_peak_75 | 26 | Intron (ENSM | 15 | 93432945 | 93445015 | 12071 | 1 | 223828 | ENSMUST00 | -7522 | ENSMUSG0 | Pphln1 | periphilin 1 |
| chr15 | 93869455 | 93869742 | 288 | * | NA_peak_75 | 20 | Distal Interge | 15 | 93904856 | 93927111 | 22256 | 2 | 77248 | ENSMUST00 | 57369 | ENSMUSG0 | 9430014N10 | RIKEN cDNA 9430014N10 gene |
| chr15 | 94475677 | 94475737 | 61 | * | NA_peak_75 | 19 | Distal Interge | 15 | 94394734 | 94465418 | 70685 | 2 | 223838 | ENSMUST00 | -10259 | ENSMUSG0 | Adamts20 | a disintegrin-like and metallopeptidase (reprolysin type) with thrombospondin type 1 motif, 20 |
| chr15 | 95217948 | 95218381 | 434 | * | NA_peak_75 | 40 | Intron (ENSM | 15 | 95097542 | 95162394 | 64853 | 2 | 100043315 | ENSMUST00 | -55554 | ENSMUSG0 | LOC1000433 | uncharacterized LOC100043315 |
| chr15 | 95835994 | 95836264 | 271 | * | NA_peak_76 | 34 | Intron (ENSM | 15 | 95824458 | 95830072 | 5615 | 2 | 102634389 | ENSMUST00 | -5922 | ENSMUSG0 | Gm17546 | predicted gene, 17546 |
| chr15 | 96213547 | 96213653 | 107 | * | NA_peak_76 | 16 | Intron (ENSM | 15 | 96231870 | 96241789 | 9920 | 2 | 75768 | ENSMUST00 | 28136 | ENSMUSG0 | 4833422M21 | RIKEN cDNA 4833422M21 gene |
| chr15 | 96268371 | 96268372 | 2 | * | NA_peak_76 | 21 | Intron (ENSM | 15 | 96231870 | 96254684 | 22815 | 2 | 75768 | ENSMUST00 | -13687 | ENSMUSG0 | 4833422M21 | RIKEN cDNA 4833422M21 gene |
| chr15 | 96287755 | 96287991 | 237 | * | NA_peak_76 | 16 | Promoter (<= | 15 | 96287573 | 96404992 | 117420 | 1 | 77044 | ENSMUST00 | 182 | ENSMUSG0 | Arid2 | AT rich interactive domain 2 (ARID, RFX-like) |
| chr15 | 96425356 | 96425655 | 300 | * | NA_peak_76 | 19 | Intron (ENSM | 15 | 96411699 | 96418692 | 6994 | 2 | 72193 | ENSMUST00 | -6664 | ENSMUSG0 | Scaf11 | SR-related CTD-associated factor 11 |
| chr15 | 96485106 | 96485171 | 66 | * | NA_peak_76 | 43 | Distal Interge | 15 | 96413656 | 96460843 | 47188 | 2 | 72193 | ENSMUST00 | -24263 | ENSMUSG0 | Scaf11 | SR-related CTD-associated factor 11 |
| chr15 | 96488414 | 96488560 | 147 | * | NA_peak_76 | 13 | Distal Interge | 15 | 96413656 | 96460843 | 47188 | 2 | 72193 | ENSMUST00 | -27571 | ENSMUSG0 | Scaf11 | SR-related CTD-associated factor 11 |
| chr15 | 96796445 | 96796492 | 48 | * | NA_peak_76 | 17 | Distal Interge | 15 | 96687392 | 96699730 | 12339 | 2 | 67760 | ENSMUST00 | -96715 | ENSMUSG0 | Slc38a2 | solute carrier family 38, member 2 |
| chr15 | 97082656 | 97083050 | 395 | * | NA_peak_76 | 17 | Distal Interge | 15 | 96994823 | 97055956 | 61134 | 2 | 69354 | ENSMUST00 | -26700 | ENSMUSG0 | Slc38a4 | solute carrier family 38, member 4 |
| chr15 | 97801141 | 97801144 | 4 | * | NA_peak_76 | 22 | Exon (ENSM | 15 | 97799609 | 97807633 | 8025 | 2 | 56233 | ENSMUST00 | 6489 | ENSMUSG0 | Hdac7 | histone deacetylase 7 |
| chr15 | 98021932 | 98022261 | 330 | * | NA_peak_76 | 29 | Distal Interge | 15 | 97975602 | 98004695 | 29094 | 2 | 12824 | ENSMUST00 | -17237 | ENSMUSG0 | Col2a1 | collagen, type II, alpha 1 |
| chr15 | 98255723 | 98255813 | 91 | * | NA_peak_76 | 36 | Promoter (1- | 15 | 98255986 | 98257293 | 1308 | 2 | 70069 | ENSMUST00 | 1480 | ENSMUSG0 | H1f7 | H1.7 linker histone |
| chr15 | 98342118 | 98342425 | 308 | * | NA_peak_76 | 33 | Promoter (1- | 15 | 98340022 | 98340987 | 966 | 2 | 258278 | ENSMUST00 | -1131 | ENSMUSG0 | Olfr284 | olfactory receptor 284 |
| chr15 | 98532669 | 98532918 | 250 | * | NA_peak_77 | 13 | Promoter (1- | 15 | 98518808 | 98534244 | 15437 | 2 | 69612 | ENSMUST00 | 1326 | ENSMUSG0 | Kansl2 | KAT8 regulatory NSL complex subunit 2 |
| chr15 | 98726573 | 98727036 | 464 | * | NA_peak_77 | 36 | Promoter (1- | 15 | 98724366 | 98728152 | 3787 | 2 | 66120 | ENSMUST00 | 1116 | ENSMUSG0 | Fkbp11 | FK506 binding protein 11 |
| chr15 | 98881341 | 98881362 | 22 | * | NA_peak_77 | 20 | Promoter (<= | 15 | 98878154 | 98881334 | 3181 | 2 | 69159 | ENSMUST00 | -7 | ENSMUSG0 | Rhebl1 | Ras homolog enriched in brain like 1 |
| chr15 | 98917983 | 98918364 | 382 | * | NA_peak_77 | 44 | Promoter (<= | 15 | 98903917 | 98918231 | 14315 | 2 | 74775 | ENSMUST00 | 0 | ENSMUSG0 | Lmbr1l | limb region 1 like |
| chr15 | 99062599 | 99063081 | 483 | * | NA_peak_77 | 46 | Distal Interge | 15 | 99055970 | 99056874 | 905 | 1 | 19132 | ENSMUST00 | 6629 | ENSMUSG0 | Prph | peripherin |
| chr15 | 99065459 | 99065614 | 156 | * | NA_peak_77 | 21 | Distal Interge | 15 | 99074575 | 99083409 | 8835 | 1 | 78733 | ENSMUST00 | -8961 | ENSMUSG0 | Troap | trophinin associated protein |
| chr15 | 99242188 | 99242561 | 374 | * | NA_peak_77 | 27 | Promoter (1- | 15 | 99243207 | 99244248 | 1042 | 2 | 51812 | ENSMUST00 | 1687 | ENSMUSG0 | Mcrs1 | microspherule protein 1 |
| chr15 | 99426463 | 99426893 | 431 | * | NA_peak_77 | 34 | Promoter (1- | 15 | 99423615 | 99424537 | 923 | 2 | 380969 | ENSMUST00 | -1926 | ENSMUSG0 | Nckap5l | NCK-associated protein 5-like |
| chr15 | 99634038 | 99634485 | 448 | * | NA_peak_77 | 21 | Promoter (<= | 15 | 99632159 | 99634774 | 2616 | 2 | 26934 | ENSMUST00 | 289 | ENSMUSG0 | Racgap1 | Rac GTPase-activating protein 1 |
| chr15 | 99736458 | 99736512 | 55 | * | NA_peak_77 | 25 | Intron (ENSM | 15 | 99737335 | 99747070 | 9736 | 2 | 71949 | ENSMUST00 | 10558 | ENSMUSG0 | Cers5 | ceramide synthase 5 |
| chr15 | 99740462 | 99740782 | 321 | * | NA_peak_77 | 25 | Intron (ENSM | 15 | 99737335 | 99747070 | 9736 | 2 | 71949 | ENSMUST00 | 6288 | ENSMUSG0 | Cers5 | ceramide synthase 5 |
| chr15 | 99914940 | 99915238 | 299 | * | NA_peak_77 | 18 | Distal Interge | 15 | 99918348 | 99931796 | 13449 | 2 | 72277 | ENSMUST00 | 16558 | ENSMUSG0 | Fam186a | family with sequence similarity 186, member A |
| chr15 | 99973156 | 99973467 | 312 | * | NA_peak_77 | 19 | Promoter (<= | 15 | 99972158 | 99973287 | 1130 | 2 | 70281 | ENSMUST00 | 0 | ENSMUSG0 | 2310068J16 | RIKEN cDNA 2310068J16 gene |
| chr15 | 100008907 | 100009277 | 371 | * | NA_peak_77 | 34 | Intron (ENSM | 15 | 100003257 | 100011943 | 8687 | 1 | 207214 | ENSMUST00 | 5650 | ENSMUSG0 | Larp4 | La ribonucleoprotein domain family, member 4 |
| chr15 | 100228566 | 100228917 | 352 | * | NA_peak_77 | 17 | Promoter (<= | 15 | 100228263 | 100252075 | 23813 | 1 | 11908 | ENSMUST00 | 303 | ENSMUSG0 | Atf1 | activating transcription factor 1 |
| chr15 | 100422739 | 100423067 | 329 | * | NA_peak_77 | 24 | Promoter (<= | 15 | 100387898 | 100423055 | 35158 | 2 | 18174 | ENSMUST00 | 0 | ENSMUSG0 | Slc11a2 | solute carrier family 11 (proton-coupled divalent metal ion transporters), member 2 |
| chr15 | 100870066 | 100870331 | 266 | * | NA_peak_77 | 30 | Promoter (<= | 15 | 100869858 | 100936327 | 66470 | 1 | 20273 | ENSMUST00 | 208 | ENSMUSG0 | Scn8a | sodium channel, voltage-gated, type VIII, alpha |
| chr15 | 101110959 | 101110972 | 14 | * | NA_peak_77 | 54 | Distal Interge | 15 | 101115755 | 101120040 | 4286 | 1 | 208258 | ENSMUST00 | -4783 | ENSMUSG0 | Ankrd33 | ankyrin repeat domain 33 |
| chr15 | 101369813 | 101369989 | 177 | * | NA_peak_77 | 22 | Promoter (<= | 15 | 101348315 | 101370162 | 21848 | 2 | 74127 | ENSMUST00 | 173 | ENSMUSG0 | Krt80 | keratin 80 |
| chr15 | 101656690 | 101657082 | 393 | * | NA_peak_78 | 23 | Distal Interge | 15 | 101643024 | 101647380 | 4357 | 2 | 432987 | ENSMUST00 | -9310 | ENSMUSG0 | Gm5478 | predicted pseudogene 5478 |
| chr15 | 102037415 | 102037736 | 322 | * | NA_peak_78 | 44 | Distal Interge | 15 | 102028180 | 102032027 | 3848 | 1 | 16668 | ENSMUST00 | 9235 | ENSMUSG0 | Krt18 | keratin 18 |
| chr15 | 102100724 | 102101080 | 357 | * | NA_peak_78 | 56 | Promoter (<= | 15 | 102100413 | 102115996 | 15584 | 1 | 209039 | ENSMUST00 | 311 | ENSMUSG0 | Tns2 | tensin 2 |
| chr15 | 102138846 | 102139213 | 368 | * | NA_peak_78 | 32 | Promoter (2- | 15 | 102116528 | 102136234 | 19707 | 2 | 223918 | ENSMUST00 | -2612 | ENSMUSG0 | Spryd3 | SPRY domain containing 3 |
| chr15 | 102195121 | 102195299 | 179 | * | NA_peak_78 | 45 | Intron (ENSM | 15 | 102187566 | 102192059 | 4494 | 2 | 246277 | ENSMUST00 | -3062 | ENSMUSG0 | Csad | cysteine sulfinic acid decarboxylase |
| chr15 | 102203419 | 102203709 | 291 | * | NA_peak_78 | 27 | Promoter (<= | 15 | 102187660 | 102203431 | 15772 | 2 | 246277 | ENSMUST00 | 0 | ENSMUSG0 | Csad | cysteine sulfinic acid decarboxylase |
| chr15 | 102279358 | 102279611 | 254 | * | NA_peak_78 | 46 | Promoter (<= | 15 | 102279454 | 102281754 | 2301 | 1 | 106073 | ENSMUST00 | 0 | ENSMUSG0 | Mfsd5 | major facilitator superfamily domain containing 5 |
| chr15 | 102303939 | 102304383 | 445 | * | NA_peak_78 | 25 | Exon (ENSM | 15 | 102296305 | 102298399 | 2095 | 1 | 105988 | ENSMUST00 | 7634 | ENSMUSG0 | Espl1 | extra spindle pole bodies 1, separase |
| chr15 | 102306347 | 102306532 | 186 | * | NA_peak_78 | 23 | Exon (ENSM | 15 | 102316485 | 102317076 | 592 | 1 | 105988 | ENSMUST00 | -9953 | ENSMUSG0 | Espl1 | extra spindle pole bodies 1, separase |
| chr15 | 102422960 | 102423409 | 450 | * | NA_peak_78 | 29 | Intron (ENSM | 15 | 102407588 | 102408385 | 798 | 1 | 20683 | ENSMUST00 | 15372 | ENSMUSG0 | Sp1 | trans-acting transcription factor 1 |
| chr15 | 102423818 | 102423961 | 144 | * | NA_peak_78 | 29 | Intron (ENSM | 15 | 102407588 | 102408385 | 798 | 1 | 20683 | ENSMUST00 | 16230 | ENSMUSG0 | Sp1 | trans-acting transcription factor 1 |
| chr15 | 102435437 | 102435535 | 99 | * | NA_peak_78 | 13 | 3' UTR | 15 | 102445367 | 102454630 | 9264 | 1 | 110542 | ENSMUST00 | -9832 | ENSMUSG0 | Amhr2 | anti-Mullerian hormone type 2 receptor |
| chr15 | 102454054 | 102454684 | 631 | * | NA_peak_78 | 24 | Promoter (1- | 15 | 102452323 | 102452871 | 549 | 1 | 110542 | ENSMUST00 | 1731 | ENSMUSG0 | Amhr2 | anti-Mullerian hormone type 2 receptor |
| chr15 | 102505308 | 102505392 | 85 | * | NA_peak_78 | 30 | Exon (ENSM | 15 | 102498776 | 102510700 | 11925 | 2 | 26404 | ENSMUST00 | 5308 | ENSMUSG0 | Map3k12 | mitogen-activated protein kinase kinase kinase 12 |
| chr15 | 102596866 | 102597152 | 287 | * | NA_peak_78 | 59 | Intron (ENSM | 15 | 102547827 | 102624922 | 77096 | 2 | 223922 | ENSMUST00 | 27770 | ENSMUSG0 | Atf7 | activating transcription factor 7 |
| chr15 | 102676366 | 102676734 | 369 | * | NA_peak_78 | 36 | Distal Interge | 15 | 102665801 | 102672352 | 6552 | 2 | 67942 | ENSMUST00 | -4014 | ENSMUSG0 | Atp5g2 | ATP synthase, H+ transporting, mitochondrial F0 complex, subunit C2 (subunit 9) |
| chr15 | 102682683 | 102682764 | 82 | * | NA_peak_78 | 20 | Distal Interge | 15 | 102665801 | 102672352 | 6552 | 2 | 67942 | ENSMUST00 | -10331 | ENSMUSG0 | Atp5g2 | ATP synthase, H+ transporting, mitochondrial F0 complex, subunit C2 (subunit 9) |
| chr15 | 102692840 | 102693226 | 387 | * | NA_peak_78 | 65 | Distal Interge | 15 | 102706972 | 102709902 | 2931 | 2 | 67488 | ENSMUST00 | 16676 | ENSMUSG0 | Calcoco1 | calcium binding and coiled coil domain 1 |
| chr15 | 102954657 | 102955017 | 361 | * | NA_peak_78 | 40 | Promoter (<= | 15 | 102954427 | 102957708 | 3282 | 1 | 109663 | ENSMUST00 | 230 | ENSMUSG0 | Hoxc11 | homeobox C11 |
| chr15 | 102960082 | 102960292 | 211 | * | NA_peak_78 | 23 | Distal Interge | 15 | 102954427 | 102957708 | 3282 | 1 | 109663 | ENSMUST00 | 5655 | ENSMUSG0 | Hoxc11 | homeobox C11 |
| chr15 | 102963599 | 102964009 | 411 | * | NA_peak_78 | 20 | Promoter (2- | 15 | 102966796 | 102971893 | 5098 | 1 | 209448 | ENSMUST00 | -2787 | ENSMUSG0 | Hoxc10 | homeobox C10 |
| chr15 | 102975799 | 102976406 | 608 | * | NA_peak_78 | 25 | Promoter (<= | 15 | 102976439 | 102981669 | 5231 | 1 | 15427 | ENSMUST00 | -33 | ENSMUSG0 | Hoxc9 | homeobox C9 |
| chr15 | 103236519 | 103236575 | 57 | * | NA_peak_78 | 33 | Intron (ENSM | 15 | 103191544 | 103239816 | 48273 | 2 | 12419 | ENSMUST00 | 3241 | ENSMUSG0 | Cbx5 | chromobox 5 |
| chr15 | 103270067 | 103270739 | 673 | * | NA_peak_78 | 28 | Promoter (1- | 15 | 103272714 | 103296850 | 24137 | 1 | 56447 | ENSMUST00 | -1975 | ENSMUSG0 | Copz1 | coatomer protein complex, subunit zeta 1 |
| chr15 | 103407767 | 103408266 | 500 | * | NA_peak_78 | 36 | Intron (ENSM | 15 | 103402458 | 103428500 | 26043 | 2 | 74174 | ENSMUST00 | 20234 | ENSMUSG0 | Gtsf1 | gametocyte specific factor 1 |
| chr15 | 103768167 | 103768580 | 414 | * | NA_peak_78 | 17 | Distal Interge | 15 | 103751919 | 103755469 | 3551 | 2 | 20771 | ENSMUST00 | -12698 | ENSMUSG0 | Mucl1 | mucin-like 1 |
| chr15 | 103879306 | 103879744 | 439 | * | NA_peak_78 | 34 | Distal Interge | 15 | 103896916 | 103899290 | 2375 | 2 | 20770 | ENSMUST00 | 19546 | ENSMUSG0 | Mucl2 | mucin-like 2 |
| chr16 | 3048882 | 3049227 | 346 | * | NA_peak_79 | 30 | Distal Interge | 16 | 3592398 | 3593339 | 942 | 1 | 258859 | ENSMUST00 | -543171 | ENSMUSG0 | Olfr161 | olfactory receptor 161 |
| chr16 | 3116238 | 3116552 | 315 | * | NA_peak_79 | 180 | Distal Interge | 16 | 3592398 | 3593339 | 942 | 1 | 258859 | ENSMUST00 | -475846 | ENSMUSG0 | Olfr161 | olfactory receptor 161 |
| chr16 | 3139265 | 3139590 | 326 | * | NA_peak_79 | 16 | Distal Interge | 16 | 3592398 | 3593339 | 942 | 1 | 258859 | ENSMUST00 | -452808 | ENSMUSG0 | Olfr161 | olfactory receptor 161 |
| chr16 | 3141034 | 3141771 | 738 | * | NA_peak_79 | 21 | Distal Interge | 16 | 3592398 | 3593339 | 942 | 1 | 258859 | ENSMUST00 | -450627 | ENSMUSG0 | Olfr161 | olfactory receptor 161 |
| chr16 | 3242703 | 3243118 | 416 | * | NA_peak_79 | 64 | Distal Interge | 16 | 3592398 | 3593339 | 942 | 1 | 258859 | ENSMUST00 | -349280 | ENSMUSG0 | Olfr161 | olfactory receptor 161 |
| chr16 | 3268582 | 3268999 | 418 | * | NA_peak_79 | 19 | Distal Interge | 16 | 3592398 | 3593339 | 942 | 1 | 258859 | ENSMUST00 | -323399 | ENSMUSG0 | Olfr161 | olfactory receptor 161 |
| chr16 | 3269328 | 3269661 | 334 | * | NA_peak_79 | 15 | Distal Interge | 16 | 3592398 | 3593339 | 942 | 1 | 258859 | ENSMUST00 | -322737 | ENSMUSG0 | Olfr161 | olfactory receptor 161 |
| chr16 | 3271598 | 3271692 | 95 | * | NA_peak_79 | 35 | Distal Interge | 16 | 3592398 | 3593339 | 942 | 1 | 258859 | ENSMUST00 | -320706 | ENSMUSG0 | Olfr161 | olfactory receptor 161 |
| chr16 | 3338801 | 3338984 | 184 | * | NA_peak_79 | 41 | Distal Interge | 16 | 3592398 | 3593339 | 942 | 1 | 258859 | ENSMUST00 | -253414 | ENSMUSG0 | Olfr161 | olfactory receptor 161 |
| chr16 | 3341170 | 3341513 | 344 | * | NA_peak_79 | 47 | Distal Interge | 16 | 3592398 | 3593339 | 942 | 1 | 258859 | ENSMUST00 | -250885 | ENSMUSG0 | Olfr161 | olfactory receptor 161 |
| chr16 | 3349927 | 3350477 | 551 | * | NA_peak_79 | 102 | Distal Interge | 16 | 3592398 | 3593339 | 942 | 1 | 258859 | ENSMUST00 | -241921 | ENSMUSG0 | Olfr161 | olfactory receptor 161 |
| chr16 | 3351030 | 3351946 | 917 | * | NA_peak_79 | 104 | Distal Interge | 16 | 3592398 | 3593339 | 942 | 1 | 258859 | ENSMUST00 | -240452 | ENSMUSG0 | Olfr161 | olfactory receptor 161 |
| chr16 | 3447151 | 3447506 | 356 | * | NA_peak_79 | 100 | Distal Interge | 16 | 3592398 | 3593339 | 942 | 1 | 258859 | ENSMUST00 | -144892 | ENSMUSG0 | Olfr161 | olfactory receptor 161 |
| chr16 | 3453596 | 3454181 | 586 | * | NA_peak_79 | 22 | Intron (ENSM | 16 | 3592398 | 3593339 | 942 | 1 | 258859 | ENSMUST00 | -138217 | ENSMUSG0 | Olfr161 | olfactory receptor 161 |
| chr16 | 3744087 | 3744570 | 484 | * | NA_peak_79 | 22 | Promoter (<= | 16 | 3744093 | 3750781 | 6689 | 1 | 74120 | ENSMUST00 | 0 | ENSMUSG0 | Zfp263 | zinc finger protein 263 |
| chr16 | 4017826 | 4018204 | 379 | * | NA_peak_79 | 35 | Distal Interge | 16 | 3979123 | 4001680 | 22558 | 2 | 52864 | ENSMUST00 | -16146 | ENSMUSG0 | Slx4 | SLX4 structure-specific endonuclease subunit homolog (S. cerevisiae) |
| chr16 | 4161168 | 4161490 | 323 | * | NA_peak_79 | 26 | Intron (ENSM | 16 | 4091839 | 4128621 | 36783 | 2 | 12914 | ENSMUST00 | -32547 | ENSMUSG0 | Crebbp | CREB binding protein |
| chr16 | 4264297 | 4264374 | 78 | * | NA_peak_79 | 24 | Distal Interge | 16 | 4230301 | 4231099 | 799 | 1 | 436332 | ENSMUST00 | 33996 | ENSMUSG0 | Gm5766 | ribosomal protein L7a pseudogene |
| chr16 | 5049887 | 5050192 | 306 | * | NA_peak_79 | 26 | Promoter (<= | 16 | 5050057 | 5086289 | 36233 | 1 | 170644 | ENSMUST00 | 0 | ENSMUSG0 | Ubn1 | ubinuclein 1 |
| chr16 | 5050614 | 5050657 | 44 | * | NA_peak_79 | 26 | Promoter (<= | 16 | 5050734 | 5086289 | 35556 | 1 | 170644 | ENSMUST00 | -77 | ENSMUSG0 | Ubn1 | ubinuclein 1 |
| chr16 | 8084575 | 8084775 | 201 | * | NA_peak_80 | 22 | Distal Interge | 16 | 8409275 | 8425136 | 15862 | 2 | 74720 | ENSMUST00 | 340361 | ENSMUSG0 | Tmem114 | transmembrane protein 114 |
| chr16 | 8737492 | 8737946 | 455 | * | NA_peak_80 | 33 | Promoter (<= | 16 | 8690146 | 8738342 | 48197 | 2 | 252870 | ENSMUST00 | 396 | ENSMUSG0 | Usp7 | ubiquitin specific peptidase 7 |
| chr16 | 9065454 | 9065470 | 17 | * | NA_peak_80 | 32 | Distal Interge | 16 | 8831321 | 8844131 | 12811 | 1 | 69053 | ENSMUST00 | 234133 | ENSMUSG0 | 1810013L24 | RIKEN cDNA 1810013L24 gene |
| chr16 | 9068567 | 9069688 | 1122 | * | NA_peak_80 | 55 | Distal Interge | 16 | 8831321 | 8844131 | 12811 | 1 | 69053 | ENSMUST00 | 237246 | ENSMUSG0 | 1810013L24 | RIKEN cDNA 1810013L24 gene |
| chr16 | 9245564 | 9245853 | 290 | * | NA_peak_80 | 20 | Distal Interge | 16 | 8831321 | 8844131 | 12811 | 1 | 69053 | ENSMUST00 | 414243 | ENSMUSG0 | 1810013L24 | RIKEN cDNA 1810013L24 gene |
| chr16 | 10192723 | 10193163 | 441 | * | NA_peak_80 | 23 | Promoter (<= | 16 | 10192906 | 10237288 | 44383 | 1 | 75329 | ENSMUST00 | 0 | ENSMUSG0 | Atf7ip2 | activating transcription factor 7 interacting protein 2 |
| chr16 | 10542712 | 10543180 | 469 | * | NA_peak_80 | 14 | Promoter (<= | 16 | 10502531 | 10543054 | 40524 | 2 | 58239 | ENSMUST00 | 0 | ENSMUSG0 | Dexi | dexamethasone-induced transcript |
| chr16 | 10545329 | 10545732 | 404 | * | NA_peak_80 | 14 | Promoter (<= | 16 | 10545339 | 10744878 | 199540 | 1 | 74374 | ENSMUST00 | 0 | ENSMUSG0 | Clec16a | C-type lectin domain family 16, member A |
| chr16 | 11471493 | 11471883 | 391 | * | NA_peak_80 | 35 | Intron (ENSM | 16 | 11447308 | 11661496 | 214189 | 1 | 74478 | ENSMUST00 | 24185 | ENSMUSG0 | Snx29 | sorting nexin 29 |
| chr16 | 16936164 | 16936195 | 32 | * | NA_peak_80 | 19 | Distal Interge | 16 | 16896574 | 16915194 | 18621 | 1 | 68606 | ENSMUST00 | 39590 | ENSMUSG0 | Ppm1f | protein phosphatase 1F (PP2C domain containing) |
| chr16 | 20466260 | 20466565 | 306 | * | NA_peak_81 | 16 | Distal Interge | 16 | 20498817 | 20509323 | 10507 | 1 | 224045 | ENSMUST00 | -32252 | ENSMUSG0 | Eif2b5 | eukaryotic translation initiation factor 2B, subunit 5 epsilon |
| chr16 | 22009507 | 22009988 | 482 | * | NA_peak_81 | 49 | Promoter (<= | 16 | 22009512 | 22046866 | 37355 | 1 | 75826 | ENSMUST00 | 0 | ENSMUSG0 | Senp2 | SUMO/sentrin specific peptidase 2 |
| chr16 | 22161527 | 22161810 | 284 | * | NA_peak_81 | 30 | Promoter (<= | 16 | 22059460 | 22161450 | 101991 | 2 | 319765 | ENSMUST00 | -77 | ENSMUSG0 | Igf2bp2 | insulin-like growth factor 2 mRNA binding protein 2 |
| chr16 | 22278486 | 22278822 | 337 | * | NA_peak_81 | 18 | Distal Interge | 16 | 22244549 | 22266005 | 21457 | 2 | 20462 | ENSMUST00 | -12481 | ENSMUSG0 | Tra2b | transformer 2 beta |
| chr16 | 22439539 | 22439881 | 343 | * | NA_peak_81 | 31 | Promoter (<= | 16 | 22381309 | 22439719 | 58411 | 2 | 104156 | ENSMUST00 | 0 | ENSMUSG0 | Etv5 | ets variant 5 |
| chr16 | 22923745 | 22924335 | 591 | * | NA_peak_81 | 27 | Promoter (1- | 16 | 22921745 | 22939757 | 18013 | 1 | 59083 | ENSMUST00 | 2000 | ENSMUSG0 | Fetub | fetuin beta |
| chr16 | 23004770 | 23005082 | 313 | * | NA_peak_81 | 23 | Intron (ENSM | 16 | 22985874 | 22999959 | 14086 | 2 | 385643 | ENSMUST00 | -4811 | ENSMUSG0 | Kng2 | kininogen 2 |
| chr16 | 23322933 | 23323240 | 308 | * | NA_peak_81 | 13 | Intron (ENSM | 16 | 23290470 | 23358595 | 68126 | 1 | 20440 | ENSMUST00 | 32463 | ENSMUSG0 | St6gal1 | beta galactoside alpha 2,6 sialyltransferase 1 |
| chr16 | 23520512 | 23520791 | 280 | * | NA_peak_81 | 24 | Promoter (<= | 16 | 23468363 | 23520650 | 52288 | 2 | 17174 | ENSMUST00 | 0 | ENSMUSG0 | Masp1 | mannan-binding lectin serine peptidase 1 |
| chr16 | 23969978 | 23970187 | 210 | * | NA_peak_82 | 13 | Promoter (2- | 16 | 23967694 | 23972380 | 4687 | 2 | 12053 | ENSMUST00 | 2193 | ENSMUSG0 | Bcl6 | B cell leukemia/lymphoma 6 |
| chr16 | 23981128 | 23981446 | 319 | * | NA_peak_82 | 13 | Intron (ENSM | 16 | 23965052 | 23988852 | 23801 | 2 | 12053 | ENSMUST00 | 7406 | ENSMUSG0 | Bcl6 | B cell leukemia/lymphoma 6 |
| chr16 | 25518266 | 25518716 | 451 | * | NA_peak_82 | 27 | Distal Interge | 16 | 25683763 | 25821318 | 137556 | 1 | 22061 | ENSMUST00 | -165047 | ENSMUSG0 | Trp63 | transformation related protein 63 |

| | | | | | | | | | | | | | | | | | | |
|---|---|---|---|---|---|---|---|---|---|---|---|---|---|---|---|---|---|---|
| chr16 | 26658017 | 26658463 | 447 | * | NA_peak_82 | 35 | Intron (ENSM | 16 | 26624156 | 26715068 | 90913 | 1 | 16180 | ENSMUST0 | 33861 | ENSMUSG0 | Il1rap | interleukin 1 receptor accessory protein |
| chr16 | 27081931 | 27082235 | 305 | * | NA_peak_82 | 43 | Distal Interge | 16 | 26959105 | 26991652 | 32548 | 2 | 239789 | ENSMUST0 | -90279 | ENSMUSG0 | Gmnc | geminin coiled-coil domain containing |
| chr16 | 27668109 | 27668136 | 28 | * | NA_peak_82 | 16 | Distal Interge | 16 | 27849930 | 27926128 | 76199 | 1 | 100038623 | ENSMUST0 | -181794 | ENSMUSG0 | Gm10823 | predicted gene 10823 |
| chr16 | 27738912 | 27739173 | 262 | * | NA_peak_82 | 16 | Distal Interge | 16 | 27849930 | 27926128 | 76199 | 1 | 100038623 | ENSMUST0 | -110757 | ENSMUSG0 | Gm10823 | predicted gene 10823 |
| chr16 | 28193022 | 28193108 | 87 | * | NA_peak_82 | 16 | Promoter (<= | 16 | 28160111 | 28192075 | 31965 | 2 | 14167 | ENSMUST0 | -947 | ENSMUSG0 | Fgf12 | fibroblast growth factor 12 |
| chr16 | 29992380 | 29992725 | 346 | * | NA_peak_82 | 15 | Distal Interge | 16 | 29957927 | 29979184 | 21258 | 2 | 328657 | ENSMUST0 | -13196 | ENSMUSG0 | Gm1968 | predicted gene 1968 |
| chr16 | 30251413 | 30251591 | 179 | * | NA_peak_83 | 15 | Distal Interge | 16 | 30256378 | 30267499 | 11122 | 2 | 71756 | ENSMUST0 | 15908 | ENSMUSG0 | Cpn2 | carboxypeptidase N, polypeptide 2 |
| chr16 | 30331796 | 30332055 | 260 | * | NA_peak_83 | 20 | Promoter (1- | 16 | 30315560 | 30333848 | 18289 | 2 | 224088 | ENSMUST0 | 1793 | ENSMUSG0 | Atp13a3 | ATPase type 13A3 |
| chr16 | 30457362 | 30457887 | 526 | * | NA_peak_83 | 25 | Distal Interge | 16 | 30362579 | 30405975 | 43397 | 2 | 224088 | ENSMUST0 | -51387 | ENSMUSG0 | Atp13a3 | ATPase type 13A3 |
| chr16 | 30618151 | 30618222 | 72 | * | NA_peak_83 | 32 | Distal Interge | 16 | 30599723 | 30602797 | 3075 | 1 | 224093 | ENSMUST0 | 18428 | ENSMUSG0 | Fam43a | family with sequence similarity 43, member A |
| chr16 | 31274732 | 31275328 | 597 | * | NA_peak_83 | 34 | Promoter (<= | 16 | 31251537 | 31275277 | 23741 | 2 | 66849 | ENSMUST0 | 0 | ENSMUSG0 | Ppp1r2 | protein phosphatase 1, regulatory inhibitor subunit 2 |
| chr16 | 31663891 | 31664311 | 421 | * | NA_peak_83 | 56 | Promoter (<= | 16 | 31663909 | 31871846 | 207938 | 1 | 13383 | ENSMUST0 | 0 | ENSMUSG0 | Dlg1 | discs large MAGUK scaffold protein 1 |
| chr16 | 31678466 | 31679047 | 582 | * | NA_peak_83 | 22 | Intron (ENSM | 16 | 31684266 | 31773787 | 89522 | 1 | 13383 | ENSMUST0 | -5219 | ENSMUSG0 | Dlg1 | discs large MAGUK scaffold protein 1 |
| chr16 | 32143792 | 32144333 | 542 | * | NA_peak_83 | 17 | Exon (ENSM | 16 | 32142795 | 32147881 | 5087 | 2 | 224109 | ENSMUST0 | 3548 | ENSMUSG0 | Nrros | negative regulator of reactive oxygen species |
| chr16 | 32272863 | 32273188 | 326 | * | NA_peak_83 | 22 | Promoter (1- | 16 | 32271468 | 32274781 | 3314 | 1 | 69576 | ENSMUST0 | 1395 | ENSMUSG0 | Smco1 | single-pass membrane protein with coiled-coil domains 1 |
| chr16 | 32317106 | 32317155 | 50 | * | NA_peak_83 | 30 | Distal Interge | 16 | 32332257 | 32353063 | 20807 | 1 | 224111 | ENSMUST0 | -15102 | ENSMUSG0 | Ubxn7 | UBX domain protein 7 |
| chr16 | 32395392 | 32395699 | 308 | * | NA_peak_83 | 23 | Distal Interge | 16 | 32400506 | 32408227 | 7722 | 1 | 277203 | ENSMUST0 | -4807 | ENSMUSG0 | Tm4sf19 | transmembrane 4 L six family member 19 |
| chr16 | 32576046 | 32576111 | 66 | * | NA_peak_84 | 39 | Distal Interge | 16 | 32608920 | 32632794 | 23875 | 1 | 22042 | ENSMUST0 | -32809 | ENSMUSG0 | Tfrc | transferrin receptor |
| chr16 | 32877726 | 32877838 | 113 | * | NA_peak_84 | 21 | Promoter (<= | 16 | 32823859 | 32877766 | 53908 | 2 | 100502698 | ENSMUST0 | 0 | ENSMUSG0 | Rubcn | RUN domain and cysteine-rich domain containing, Beclin 1-interacting protein |
| chr16 | 32967758 | 32967975 | 218 | * | NA_peak_84 | 20 | Intron (ENSM | 16 | 32979397 | 32998578 | 19182 | 1 | 70144 | ENSMUST0 | -11422 | ENSMUSG0 | Lrch3 | leucine-rich repeats and calponin homology (CH) domain containing 3 |
| chr16 | 33041331 | 33041458 | 128 | * | NA_peak_84 | 13 | Intron (ENSM | 16 | 33016620 | 33049870 | 33251 | 2 | 69707 | ENSMUST0 | 8412 | ENSMUSG0 | Iqcg | IQ motif containing G |
| chr16 | 33861364 | 33861681 | 318 | * | NA_peak_84 | 24 | Intron (ENSM | 16 | 33885026 | 33910585 | 25560 | 1 | 16419 | ENSMUST0 | -23345 | ENSMUSG0 | Itgb5 | integrin beta 5 |
| chr16 | 38020293 | 38020604 | 312 | * | NA_peak_85 | 23 | Distal Interge | 16 | 38089001 | 38246084 | 157084 | 1 | 56637 | ENSMUST0 | -68397 | ENSMUSG0 | Gsk3b | glycogen synthase kinase 3 beta |
| chr16 | 40611694 | 40612227 | 534 | * | NA_peak_85 | 35 | Intron (ENSM | 16 | 39984595 | 40657886 | 673292 | 1 | 268890 | ENSMUST0 | 627099 | ENSMUSG0 | Lsamp | limbic system-associated membrane protein |
| chr16 | 41057545 | 41057877 | 333 | * | NA_peak_85 | 59 | Intron (ENSM | 16 | 41532858 | 42146213 | 613356 | 1 | 268890 | ENSMUST0 | -474981 | ENSMUSG0 | Lsamp | limbic system-associated membrane protein |
| chr16 | 44724581 | 44724845 | 265 | * | NA_peak_85 | 26 | Promoter (<= | 16 | 44724357 | 44736573 | 12217 | 1 | 212547 | ENSMUST0 | 224 | ENSMUSG0 | Nepro | nucleolus and neural progenitor protein |
| chr16 | 44893577 | 44894036 | 460 | * | NA_peak_85 | 14 | Intron (ENSM | 16 | 44867097 | 44915840 | 48744 | 1 | 271375 | ENSMUST0 | 26480 | ENSMUSG0 | Cd200r2 | Cd200 receptor 2 |
| chr16 | 45042810 | 45043041 | 232 | * | NA_peak_85 | 19 | Distal Interge | 16 | 45093454 | 45094904 | 1451 | 1 | 67896 | ENSMUST0 | -50413 | ENSMUSG0 | Ccdc80 | coiled-coil domain containing 80 |
| chr16 | 45057753 | 45058207 | 455 | * | NA_peak_85 | 26 | Distal Interge | 16 | 45093454 | 45094904 | 1451 | 1 | 67896 | ENSMUST0 | -35247 | ENSMUSG0 | Ccdc80 | coiled-coil domain containing 80 |
| chr16 | 45146824 | 45146988 | 165 | * | NA_peak_85 | 23 | Intron (ENSM | 16 | 45144027 | 45151921 | 7895 | 2 | 74102 | ENSMUST0 | 4933 | ENSMUSG0 | Slc35a5 | solute carrier family 35, member A5 |
| chr16 | 45807273 | 45807454 | 182 | * | NA_peak_85 | 16 | Intron (ENSM | 16 | 45747160 | 45801571 | 54412 | 2 | 208177 | ENSMUST0 | -5702 | ENSMUSG0 | Phldb2 | pleckstrin homology like domain, family B, member 2 |
| chr16 | 46080956 | 46081247 | 292 | * | NA_peak_85 | 24 | Intron (ENSM | 16 | 46035657 | 46120251 | 84595 | 2 | 84544 | ENSMUST0 | 39004 | ENSMUSG0 | Cd96 | CD96 antigen |
| chr16 | 46364774 | 46365022 | 249 | * | NA_peak_85 | 23 | Distal Interge | 16 | 46394858 | 46411169 | 16312 | 2 | 58998 | ENSMUST0 | 46147 | ENSMUSG0 | Nectin3 | nectin cell adhesion molecule 3 |
| chr16 | 48860470 | 48860767 | 298 | * | NA_peak_86 | 48 | Distal Interge | 16 | 48872621 | 48874496 | 1876 | 1 | 245195 | ENSMUST0 | -11854 | ENSMUSG0 | Retnlg | resistin like gamma |
| chr16 | 49466638 | 49466952 | 315 | * | NA_peak_86 | 17 | Distal Interge | 16 | 49699233 | 49765126 | 65894 | 1 | 73916 | ENSMUST0 | -232281 | ENSMUSG0 | Ift57 | intraflagellar transport 57 |
| chr16 | 49555184 | 49555186 | 3 | * | NA_peak_86 | 16 | Distal Interge | 16 | 49699233 | 49765126 | 65894 | 1 | 73916 | ENSMUST0 | -144047 | ENSMUSG0 | Ift57 | intraflagellar transport 57 |
| chr16 | 50910904 | 50911031 | 128 | * | NA_peak_86 | 23 | Distal Interge | 16 | 50783478 | 50784065 | 588 | 1 | 78639 | ENSMUST0 | 127426 | ENSMUSG0 | 1700116B05 | RIKEN cDNA 1700116B05 gene |
| chr16 | 51952597 | 51952663 | 67 | * | NA_peak_86 | 35 | Distal Interge | 16 | 51982484 | 51985785 | 3302 | 2 | 629091 | ENSMUST0 | 33122 | NA | Gm29686 | predicted gene, 29686 |
| chr16 | 55649588 | 55650433 | 846 | * | NA_peak_86 | 32 | Distal Interge | 16 | 55494731 | 55494797 | 67 | 1 | 100628589 | ENSMUST0 | 154857 | ENSMUSG0 | Mir5118 | microRNA 5118 |
| chr16 | 55652068 | 55652366 | 299 | * | NA_peak_86 | 35 | Distal Interge | 16 | 55494731 | 55494797 | 67 | 1 | 100628589 | ENSMUST0 | 157337 | ENSMUSG0 | Mir5118 | microRNA 5118 |
| chr16 | 57391285 | 57391596 | 312 | * | NA_peak_86 | 274 | Intron (ENSM | 16 | 57353277 | 57515690 | 162414 | 1 | 78749 | ENSMUST0 | 38008 | ENSMUSG0 | Filip1l | filamin A interacting protein 1-like |
| chr16 | 62664383 | 62664992 | 610 | * | NA_peak_86 | 15 | Distal Interge | 16 | 62735161 | 62767123 | 31963 | 2 | 106338 | ENSMUST0 | 102131 | ENSMUSG0 | Nsun3 | NOL1/NOP2/Sun domain family member 3 |
| chr16 | 62709266 | 62709590 | 325 | * | NA_peak_86 | 44 | Distal Interge | 16 | 62735161 | 62767123 | 31963 | 2 | 106338 | ENSMUST0 | 57533 | ENSMUSG0 | Nsun3 | NOL1/NOP2/Sun domain family member 3 |
| chr16 | 64399840 | 64400320 | 481 | * | NA_peak_86 | 27 | Distal Interge | 16 | 64477810 | 64602408 | 124599 | 2 | 224291 | ENSMUST0 | 202088 | ENSMUSG0 | Csnka2ip | casein kinase 2, alpha prime interacting protein |
| chr16 | 64454976 | 64455339 | 364 | * | NA_peak_86 | 27 | Distal Interge | 16 | 64477810 | 64602408 | 124599 | 2 | 224291 | ENSMUST0 | 147069 | ENSMUSG0 | Csnka2ip | casein kinase 2, alpha prime interacting protein |
| chr16 | 69894945 | 69895121 | 177 | * | NA_peak_86 | 29 | Distal Interge | 16 | 69856874 | 69863744 | 6871 | 2 | 224318 | ENSMUST0 | -31201 | ENSMUSG0 | Speer2 | spermatogenesis associated glutamate (E)-rich protein 2 |
| chr16 | 70272415 | 70272719 | 305 | * | NA_peak_86 | 23 | Distal Interge | 16 | 70313949 | 70569716 | 255768 | 1 | 74185 | ENSMUST0 | -41230 | ENSMUSG0 | Gbe1 | glucan (1,4-alpha-), branching enzyme 1 |
| chr16 | 72916812 | 72917208 | 397 | * | NA_peak_86 | 34 | Intron (ENSM | 16 | 72976349 | 72979218 | 2870 | 1 | 19876 | ENSMUST0 | -59141 | ENSMUSG0 | Robo1 | roundabout guidance receptor 1 |
| chr16 | 77158006 | 77158287 | 282 | * | NA_peak_86 | 26 | Distal Interge | 16 | 77083900 | 77108205 | 24306 | 1 | 30940 | ENSMUST0 | 74106 | ENSMUSG0 | Usp25 | ubiquitin specific peptidase 25 |
| chr16 | 81035501 | 81035787 | 287 | * | NA_peak_86 | 26 | Distal Interge | 16 | 81200697 | 81597805 | 397109 | 1 | 17968 | ENSMUST0 | -164910 | ENSMUSG0 | Ncam2 | neural cell adhesion molecule 2 |
| chr16 | 83264097 | 83264436 | 340 | * | NA_peak_86 | 44 | Distal Interge | 16 | 84371143 | 84375505 | 4363 | 1 | 75247 | ENSMUST0 | -1106707 | ENSMUSG0 | 4930553E22 | RIKEN cDNA 4930553E22 gene |
| chr16 | 83320369 | 83320669 | 301 | * | NA_peak_86 | 51 | Distal Interge | 16 | 84371143 | 84375505 | 4363 | 1 | 75247 | ENSMUST0 | -1050474 | ENSMUSG0 | 4930553E22 | RIKEN cDNA 4930553E22 gene |
| chr16 | 83485618 | 83485797 | 180 | * | NA_peak_86 | 23 | Distal Interge | 16 | 84371143 | 84375505 | 4363 | 1 | 75247 | ENSMUST0 | -885346 | ENSMUSG0 | 4930553E22 | RIKEN cDNA 4930553E22 gene |
| chr16 | 88109303 | 88109434 | 132 | * | NA_peak_87 | 24 | Intron (ENSM | 16 | 87912421 | 88056176 | 143756 | 2 | 14805 | ENSMUST0 | -53127 | ENSMUSG0 | Grik1 | glutamate receptor, ionotropic, kainate 1 |
| chr16 | 88527596 | 88527967 | 372 | * | NA_peak_87 | 39 | Distal Interge | 16 | 88505807 | 88506978 | 1172 | 2 | 239931 | ENSMUST0 | -20618 | ENSMUSG0 | Cldn17 | claudin 17 |
| chr16 | 91647129 | 91647393 | 265 | * | NA_peak_87 | 23 | Promoter (<= | 16 | 91647506 | 91679207 | 31702 | 1 | 20658 | ENSMUST0 | -113 | ENSMUSG0 | Son | Son DNA binding protein |
| chr16 | 95783193 | 95783575 | 383 | * | NA_peak_87 | 45 | Intron (ENSM | 16 | 95831117 | 95845770 | 14654 | 2 | 69776 | ENSMUST0 | 62195 | ENSMUSG0 | 1600002D24 | RIKEN cDNA 1600002D24 gene |
| chr16 | 97763163 | 97763357 | 195 | * | NA_peak_87 | 23 | Promoter (<= | 16 | 97741933 | 97763787 | 21855 | 2 | 72388 | ENSMUST0 | 430 | ENSMUSG0 | Ripk4 | receptor-interacting serine-threonine kinase 4 |
| chr16 | 98095078 | 98095468 | 391 | * | NA_peak_87 | 22 | Distal Interge | 16 | 98065444 | 98082450 | 17007 | 2 | 320586 | ENSMUST0 | -12628 | ENSMUSG0 | A630089N07 | RIKEN cDNA A630089N07 gene |
| chr17 | 3048699 | 3048724 | 26 | * | NA_peak_87 | 23 | Distal Interge | 17 | 3077738 | 3084040 | 6303 | 2 | 328734 | ENSMUST0 | 35316 | NA | Pisd-ps2 | phosphatidylserine decarboxylase, pseudogene 2 |
| chr17 | 3563830 | 3564140 | 311 | * | NA_peak_87 | 22 | Distal Interge | 17 | 3557869 | 3559862 | 1994 | 1 | 68230 | ENSMUST0 | 5961 | ENSMUSG0 | 1700102H20 | RIKEN cDNA 1700102H20 gene |
| chr17 | 3643316 | 3643594 | 279 | * | NA_peak_87 | 13 | Intron (ENSM | 17 | 3635240 | 3696261 | 61022 | 2 | 224480 | ENSMUST0 | 52667 | ENSMUSG0 | Nox3 | NADPH oxidase 3 |
| chr17 | 5159259 | 5159570 | 312 | * | NA_peak_87 | 30 | Intron (ENSM | 17 | 5040578 | 5127640 | 87063 | 1 | 239985 | ENSMUST0 | 118681 | ENSMUSG0 | Arid1b | AT rich interactive domain 1B (SWI-like) |
| chr17 | 5423457 | 5423669 | 213 | * | NA_peak_87 | 43 | Intron (ENSM | 17 | 5417323 | 5418767 | 1445 | 2 | 106557 | ENSMUST0 | -4690 | ENSMUSG0 | Ldhal6b | lactate dehydrogenase A-like 6B |
| chr17 | 6419457 | 6419720 | 264 | * | NA_peak_88 | 41 | Distal Interge | 17 | 6356813 | 6411834 | 55022 | 2 | 101055731 | ENSMUST0 | -7623 | ENSMUSG0 | Gm29719 | synaptotagmin-like 3 pseudogene |
| chr17 | 6571895 | 6572166 | 272 | * | NA_peak_88 | 49 | Intron (ENSM | 17 | 6601671 | 6609679 | 8009 | 1 | 100040531 | ENSMUST0 | -29505 | ENSMUSG0 | Dynlt1f | dynein light chain Tctex-type 1F |
| chr17 | 6672247 | 6672552 | 306 | * | NA_peak_88 | 72 | Promoter (1- | 17 | 6673583 | 6715600 | 42018 | 1 | 83672 | ENSMUST0 | -1031 | ENSMUSG0 | Sytl3 | synaptotagmin-like 3 |
| chr17 | 6698943 | 6699229 | 287 | * | NA_peak_88 | 40 | Intron (ENSM | 17 | 6702952 | 6737836 | 34885 | 1 | 83672 | ENSMUST0 | -3723 | ENSMUSG0 | Sytl3 | synaptotagmin-like 3 |
| chr17 | 6699654 | 6700305 | 652 | * | NA_peak_88 | 37 | Promoter (2- | 17 | 6702952 | 6737836 | 34885 | 1 | 83672 | ENSMUST0 | -2647 | ENSMUSG0 | Sytl3 | synaptotagmin-like 3 |
| chr17 | 6923865 | 6924448 | 584 | * | NA_peak_88 | 23 | Intron (ENSM | 17 | 6920869 | 6948281 | 27413 | 2 | 100037282 | ENSMUST0 | 23833 | ENSMUSG0 | Rsph3b | radial spoke 3B homolog (Chlamydomonas) |
| chr17 | 7158596 | 7158714 | 119 | * | NA_peak_88 | 16 | Distal Interge | 17 | 7170115 | 7303315 | 133201 | 1 | 20112 | ENSMUST0 | -11401 | ENSMUSG0 | Rps6ka2 | ribosomal protein S6 kinase, polypeptide 2 |
| chr17 | 8165848 | 8165913 | 66 | * | NA_peak_88 | 28 | Promoter (<= | 17 | 8165540 | 8196475 | 30936 | 1 | 75296 | ENSMUST0 | 308 | ENSMUSG0 | Cep43 | centrosomal protein 43 |
| chr17 | 9806688 | 9807043 | 356 | * | NA_peak_88 | 23 | Distal Interge | 17 | 9774217 | 9775867 | 1651 | 2 | 71092 | ENSMUST0 | -30821 | ENSMUSG0 | 4930452A19 | RIKEN cDNA 4930452A19 gene |
| chr17 | 10107428 | 10107920 | 493 | * | NA_peak_88 | 26 | Distal Interge | 17 | 10209093 | 10213574 | 4482 | 2 | 19317 | ENSMUST0 | 105654 | ENSMUSG0 | Qk | quaking, KH domain containing RNA binding |
| chr17 | 10952772 | 10952965 | 194 | * | NA_peak_88 | 16 | Intron (ENSM | 17 | 10840470 | 12061678 | 1221209 | 1 | 50873 | ENSMUST0 | 112302 | ENSMUSG0 | Prkn | parkin RBR E3 ubiquitin protein ligase |
| chr17 | 11164056 | 11164366 | 311 | * | NA_peak_88 | 26 | Intron (ENSM | 17 | 11067125 | 11635446 | 568322 | 1 | 50873 | ENSMUST0 | 96931 | ENSMUSG0 | Prkn | parkin RBR E3 ubiquitin protein ligase |
| chr17 | 11758834 | 11759128 | 295 | * | NA_peak_89 | 23 | Intron (ENSM | 17 | 11838602 | 11867561 | 28960 | 1 | 50873 | ENSMUST0 | -79474 | ENSMUSG0 | Prkn | parkin RBR E3 ubiquitin protein ligase |
| chr17 | 12555955 | 12556276 | 322 | * | NA_peak_89 | 23 | Distal Interge | 17 | 12584132 | 12628488 | 44357 | 1 | 20518 | ENSMUST0 | -27856 | ENSMUSG0 | Slc22a2 | solute carrier family 22 (organic cation transporter), member 2 |
| chr17 | 13037163 | 13037487 | 325 | * | NA_peak_89 | 20 | Promoter (2- | 17 | 13034976 | 13037398 | 2423 | 1 | 102639918 | ENSMUST0 | 2187 | NA | Gm36117 | predicted gene, 36117 |
| chr17 | 13761145 | 13761660 | 516 | * | NA_peak_89 | 35 | Promoter (<= | 17 | 13761605 | 13854225 | 92621 | 1 | 17356 | ENSMUST0 | 0 | ENSMUSG0 | Afdn | afadin, adherens junction formation factor |
| chr17 | 14504326 | 14504624 | 299 | * | NA_peak_89 | 17 | Distal Interge | 17 | 14400786 | 14403587 | 2802 | 1 | 64074 | ENSMUST0 | 103540 | ENSMUSG0 | Smoc2 | SPARC related modular calcium binding 2 |
| chr17 | 14962901 | 14963456 | 556 | * | NA_peak_89 | 37 | Promoter (1- | 17 | 14947738 | 14961273 | 13536 | 2 | 72057 | ENSMUST0 | -1628 | ENSMUSG0 | Phf10 | PHD finger protein 10 |
| chr17 | 14965549 | 14966085 | 537 | * | NA_peak_89 | 38 | Intron (ENSM | 17 | 14947738 | 14961273 | 13536 | 2 | 72057 | ENSMUST0 | -4276 | ENSMUSG0 | Phf10 | PHD finger protein 10 |
| chr17 | 14992222 | 14992707 | 486 | * | NA_peak_89 | 35 | Distal Interge | 17 | 14979499 | 14989374 | 9876 | 1 | 100041574 | ENSMUST0 | 12723 | ENSMUSG0 | 9030025P20 | RIKEN cDNA 9030025P20 gene |
| chr17 | 15029629 | 15030065 | 437 | * | NA_peak_89 | 45 | Intron (ENSM | 17 | 15041208 | 15063850 | 22643 | 1 | 381062 | ENSMUST0 | -11143 | ENSMUSG0 | Ermard | ER membrane associated RNA degradation |
| chr17 | 15195374 | 15195404 | 31 | * | NA_peak_89 | 24 | Distal Interge | 17 | 15231585 | 15240783 | 9199 | 2 | 328747 | ENSMUST0 | 45379 | ENSMUSG0 | Gm5091 | predicted gene 5091 |
| chr17 | 15276388 | 15276490 | 103 | * | NA_peak_89 | 17 | Intron (ENSM | 17 | 15231581 | 15247917 | 16337 | 2 | 328747 | ENSMUST0 | -28471 | ENSMUSG0 | Gm5091 | predicted gene 5091 |
| chr17 | 15635704 | 15635898 | 195 | * | NA_peak_89 | 23 | Intron (ENSM | 17 | 15631788 | 15641104 | 9317 | 1 | 71008 | ENSMUST0 | 3916 | ENSMUSG0 | 4933401D09 | RIKEN cDNA 4933401D09 gene |
| chr17 | 15705012 | 15705423 | 412 | * | NA_peak_89 | 19 | Promoter (<= | 17 | 15705018 | 15731908 | 26891 | 1 | 12648 | ENSMUST0 | 0 | ENSMUSG0 | Chd1 | chromodomain helicase DNA binding protein 1 |
| chr17 | 16122919 | 16123354 | 436 | * | NA_peak_90 | 43 | Distal Interge | 17 | 15821041 | 15830885 | 9845 | 2 | 68799 | ENSMUST0 | -292034 | ENSMUSG0 | Rgmb | repulsive guidance molecule family member B |
| chr17 | 16127784 | 16128116 | 333 | * | NA_peak_90 | 47 | Distal Interge | 17 | 15821041 | 15830885 | 9845 | 2 | 68799 | ENSMUST0 | -296899 | ENSMUSG0 | Rgmb | repulsive guidance molecule family member B |
| chr17 | 16430724 | 16431047 | 324 | * | NA_peak_90 | 23 | Distal Interge | 17 | 15821041 | 15830885 | 9845 | 2 | 68799 | ENSMUST0 | -599839 | ENSMUSG0 | Rgmb | repulsive guidance molecule family member B |
| chr17 | 17045255 | 17045876 | 622 | * | NA_peak_90 | 15 | Distal Interge | 17 | 17064113 | 17089628 | 25516 | 1 | 449000 | ENSMUST0 | -18237 | ENSMUSG0 | Zfp960 | zinc finger protein 960 |
| chr17 | 17047581 | 17047666 | 86 | * | NA_peak_90 | 27 | Distal Interge | 17 | 17064113 | 17089628 | 25516 | 1 | 449000 | ENSMUST0 | -16447 | ENSMUSG0 | Zfp960 | zinc finger protein 960 |
| chr17 | 17049909 | 17050230 | 322 | * | NA_peak_90 | 19 | Distal Interge | 17 | 17064113 | 17089628 | 25516 | 1 | 449000 | ENSMUST0 | -13883 | ENSMUSG0 | Zfp960 | zinc finger protein 960 |
| chr17 | 17157903 | 17157928 | 26 | * | NA_peak_90 | 39 | Distal Interge | 17 | 17121415 | 17146831 | 25417 | 1 | 22759 | ENSMUST0 | 36488 | ENSMUSG0 | Zfp97 | zinc finger protein 97 |
| chr17 | 17210648 | 17210867 | 220 | * | NA_peak_90 | 23 | Distal Interge | 17 | 17234872 | 17266063 | 31192 | 1 | 102640673 | ENSMUST0 | -24005 | ENSMUSG0 | Gm5165 | predicted gene 5165 |
| chr17 | 17214482 | 17215008 | 527 | * | NA_peak_90 | 18 | Distal Interge | 17 | 17234872 | 17266063 | 31192 | 1 | 102640673 | ENSMUST0 | -19864 | ENSMUSG0 | Gm5165 | predicted gene 5165 |
| chr17 | 17623841 | 17624192 | 352 | * | NA_peak_90 | 26 | Promoter (<= | 17 | 17552118 | 17624489 | 72372 | 2 | 240028 | ENSMUST0 | 297 | ENSMUSG0 | Lnpep | leucyl/cystinyl aminopeptidase |
| chr17 | 20988547 | 20988947 | 401 | * | NA_peak_90 | 18 | Distal Interge | 17 | 20991647 | 20996464 | 4818 | 2 | 171236 | ENSMUST0 | 7517 | ENSMUSG0 | Vmn1r233 | vomeronasal 1 receptor 233 |
| chr17 | 21383705 | 21384101 | 397 | * | NA_peak_90 | 16 | Promoter (<= | 17 | 21383748 | 21399262 | 15515 | 1 | 210503 | ENSMUST0 | 0 | ENSMUSG0 | Zfp677 | zinc finger protein 677 |
| chr17 | 21643252 | 21643279 | 28 | * | NA_peak_90 | 16 | Intron (ENSM | 17 | 21650624 | 21664966 | 14343 | 1 | 73229 | ENSMUST0 | -7345 | ENSMUSG0 | Zfp983 | zinc finger protein 983 |
| chr17 | 21702595 | 21703009 | 415 | * | NA_peak_90 | 22 | Distal Interge | 17 | 21707741 | 21725636 | 17896 | 1 | 240034 | ENSMUST0 | -4732 | ENSMUSG0 | Zfp760 | zinc finger protein 760 |
| chr17 | 22595976 | 22596404 | 429 | * | NA_peak_91 | 20 | Distal Interge | 17 | 22600951 | 22613183 | 12233 | 1 | 628185 | ENSMUST0 | -4547 | ENSMUSG0 | Vmn2r112 | vomeronasal 2, receptor 112 |
| chr17 | 22644029 | 22644512 | 484 | * | NA_peak_91 | 16 | Distal Interge | 17 | 22600951 | 22620842 | 19892 | 1 | 628185 | ENSMUST0 | 43078 | ENSMUSG0 | Vmn2r112 | vomeronasal 2, receptor 112 |
| chr17 | 23477784 | 23478049 | 266 | * | NA_peak_91 | 34 | Promoter (1- | 17 | 23455173 | 23479699 | 24527 | 2 | 619788 | ENSMUST0 | 1650 | ENSMUSG0 | Vmn2r117 | vomeronasal 2, receptor 117 |
| chr17 | 23533264 | 23533561 | 298 | * | NA_peak_91 | 39 | Distal Interge | 17 | 23550799 | 23565291 | 14493 | 1 | 320020 | ENSMUST0 | -17238 | ENSMUSG0 | 6330415G19 | RIKEN cDNA 6330415G19 gene |
| chr17 | 23660528 | 23660986 | 459 | * | NA_peak_91 | 24 | Promoter (<= | 17 | 23660506 | 23668621 | 8116 | 1 | 212733 | ENSMUST0 | 22 | ENSMUSG0 | Bicdl2 | BICD family like cargo adaptor 2 |
| chr17 | 23736311 | 23736715 | 405 | * | NA_peak_91 | 24 | Promoter (1- | 17 | 23734727 | 23736735 | 2009 | 1 | 268930 | ENSMUST0 | 1584 | ENSMUSG0 | Pkmyt1 | protein kinase, membrane associated tyrosine/threonine 1 |
| chr17 | 23779638 | 23779960 | 323 | * | NA_peak_91 | 17 | Promoter (<= | 17 | 23776959 | 23779791 | 2833 | 2 | 76917 | ENSMUST0 | 0 | ENSMUSG0 | Flywch2 | FLYWCH family member 2 |
| chr17 | 23803366 | 23803646 | 281 | * | NA_peak_91 | 35 | Promoter (<= | 17 | 23803234 | 23810028 | 6795 | 1 | 75956 | ENSMUST0 | 132 | ENSMUSG0 | Srrm2 | serine/arginine repetitive matrix 2 |
| chr17 | 23902012 | 23902344 | 333 | * | NA_peak_91 | 15 | Intron (ENSM | 17 | 23898734 | 23900788 | 2055 | 1 | 630537 | ENSMUST0 | 3278 | ENSMUSG0 | Dcpp2 | demilune cell and parotid protein 2 |
| chr17 | 23903146 | 23903379 | 234 | * | NA_peak_91 | 27 | Intron (ENSM | 17 | 23898734 | 23900788 | 2055 | 1 | 630537 | ENSMUST0 | 4412 | ENSMUSG0 | Dcpp2 | demilune cell and parotid protein 2 |
| chr17 | 23951475 | 23951715 | 241 | * | NA_peak_91 | 21 | Distal Interge | 17 | 23953085 | 23954919 | 1835 | 2 | 638345 | ENSMUST0 | 3204 | ENSMUSG0 | Sbpl | spermine binding protein-like |
| chr17 | 24115093 | 24115140 | 48 | * | NA_peak_91 | 17 | Intron (ENSM | 17 | 24106887 | 24120794 | 13908 | 2 | 18607 | ENSMUST0 | 5654 | ENSMUSG0 | Pdpk1 | 3-phosphoinositide dependent protein kinase 1 |
| chr17 | 24124556 | 24124742 | 187 | * | NA_peak_91 | 23 | Intron (ENSM | 17 | 24106887 | 24120794 | 13908 | 2 | 18607 | ENSMUST0 | -3762 | ENSMUSG0 | Pdpk1 | 3-phosphoinositide dependent protein kinase 1 |
| chr17 | 24317424 | 24317710 | 287 | * | NA_peak_91 | 18 | Intron (ENSM | 17 | 24326469 | 24328532 | 2064 | 2 | 381072 | ENSMUST0 | 10822 | ENSMUSG0 | Abca17 | ATP-binding cassette, sub-family A (ABC1), member 17 |
| chr17 | 24543714 | 24544016 | 303 | * | NA_peak_91 | 18 | Intron (ENSM | 17 | 24549950 | 24596508 | 46559 | 1 | 18763 | ENSMUST0 | -5934 | ENSMUSG0 | Pkd1 | polycystin 1, transient receptor potential channel interacting |
| chr17 | 24549810 | 24550063 | 254 | * | NA_peak_91 | 30 | Promoter (<= | 17 | 24549950 | 24596508 | 46559 | 1 | 18763 | ENSMUST0 | 0 | ENSMUSG0 | Pkd1 | polycystin 1, transient receptor potential channel interacting |
| chr17 | 24657824 | 24658183 | 360 | * | NA_peak_91 | 34 | Promoter (<= | 17 | 24657405 | 24658425 | 1021 | 2 | 381073 | ENSMUST0 | 242 | ENSMUSG0 | Npw | neuropeptide W |
| chr17 | 24665679 | 24665859 | 181 | * | NA_peak_91 | 28 | Distal Interge | 17 | 24669687 | 24682016 | 12330 | 1 | 213753 | ENSMUST0 | -3828 | ENSMUSG0 | Zfp598 | zinc finger protein 598 |
| chr17 | 24730145 | 24730368 | 224 | * | NA_peak_91 | 23 | Promoter (1- | 17 | 24728174 | 24733364 | 5191 | 1 | 66211 | ENSMUST0 | 1971 | ENSMUSG0 | Rpl3l | ribosomal protein L3-like |
| chr17 | 24911196 | 24911585 | 390 | * | NA_peak_91 | 17 | Intron (ENSM | 17 | 24905180 | 24906798 | 1619 | 2 | 30957 | ENSMUST0 | -4398 | ENSMUSG0 | Mapk8ip3 | mitogen-activated protein kinase 8 interacting protein 3 |
| chr17 | 25015317 | 25015683 | 367 | * | NA_peak_91 | 34 | Promoter (<= | 17 | 24961226 | 25015230 | 54005 | 2 | 57354 | ENSMUST0 | -87 | ENSMUSG0 | Cramp1 | cramped chromatin regulator 1 |
| chr17 | 25756736 | 25756744 | 9 | * | NA_peak_92 | 22 | Promoter (2- | 17 | 25748615 | 25754378 | 5764 | 2 | 56047 | ENSMUST0 | -2358 | ENSMUSG0 | Msln | mesothelin |
| chr17 | 26095439 | 26095535 | 97 | * | NA_peak_92 | 19 | Promoter (<= | 17 | 26091736 | 26095473 | 3738 | 2 | 56520 | ENSMUST0 | 0 | ENSMUSG0 | Nme4 | NME/NM23 nucleoside diphosphate kinase 4 |
| chr17 | 26793165 | 26793293 | 129 | * | NA_peak_92 | 24 | Promoter (2- | 17 | 26790775 | 26792565 | 1791 | 1 | 224630 | ENSMUST0 | 2390 | ENSMUSG0 | Bnip1 | BCL2/adenovirus E1B interacting protein 1 |
| chr17 | 27203528 | 27203851 | 324 | * | NA_peak_92 | 36 | Promoter (<= | 17 | 27189950 | 27203676 | 13727 | 2 | 224640 | ENSMUST0 | 0 | ENSMUSG0 | Lemd2 | LEM domain containing 2 |
| chr17 | 27529030 | 27529354 | 325 | * | NA_peak_92 | 20 | Distal Interge | 17 | 27425907 | 27521403 | 95497 | 2 | 268934 | ENSMUST0 | -7627 | ENSMUSG0 | Grm4 | glutamate receptor, metabotropic 4 |
| chr17 | 27820538 | 27820832 | 295 | * | NA_peak_92 | 31 | Promoter (<= | 17 | 27751235 | 27820625 | 69391 | 2 | 224647 | ENSMUST0 | 0 | ENSMUSG0 | Ilrun | inflammation and lipid regulator with UBA-like and NBR1-like domains |
| chr17 | 27942350 | 27942396 | 47 | * | NA_peak_92 | 22 | Intron (ENSM | 17 | 27901126 | 27909844 | 8719 | 2 | 68776 | ENSMUST0 | -32506 | ENSMUSG0 | Taf11 | TATA-box binding protein associated factor 11 |
| chr17 | 28142445 | 28142559 | 115 | * | NA_peak_92 | 27 | Promoter (<= | 17 | 28142526 | 28170145 | 27620 | 1 | 268935 | ENSMUST0 | 0 | ENSMUSG0 | Scube3 | signal peptide, CUB domain, EGF-like 3 |
| chr17 | 28165750 | 28165759 | 10 | * | NA_peak_92 | 25 | Promoter (<= | 17 | 28165809 | 28167651 | 1843 | 1 | 268935 | ENSMUST0 | -50 | ENSMUSG0 | Scube3 | signal peptide, CUB domain, EGF-like 3 |
| chr17 | 28333173 | 28333488 | 316 | * | NA_peak_92 | 22 | Promoter (1- | 17 | 28333506 | 28335415 | 1910 | 2 | 21678 | ENSMUST0 | 1927 | ENSMUSG0 | Tead3 | TEA domain family member 3 |
| chr17 | 28447635 | 28447939 | 305 | * | NA_peak_92 | 29 | Intron (ENSM | 17 | 28415961 | 28441722 | 25762 | 2 | 14229 | ENSMUST0 | -5913 | ENSMUSG0 | Fkbp5 | FK506 binding protein 5 |
| chr17 | 28485440 | 28485771 | 332 | * | NA_peak_92 | 49 | Promoter (<= | 17 | 28400638 | 28486093 | 85456 | 2 | 14229 | ENSMUST0 | 322 | ENSMUSG0 | Fkbp5 | FK506 binding protein 5 |
| chr17 | 28563645 | 28563799 | 155 | * | NA_peak_92 | 26 | Promoter (2- | 17 | 28558149 | 28560766 | 2618 | 2 | 109791 | ENSMUST0 | -2879 | ENSMUSG0 | Clps | colipase, pancreatic |
| chr17 | 28889077 | 28889369 | 293 | * | NA_peak_92 | 39 | 3' UTR | 17 | 28905265 | 28906771 | 1507 | 1 | 74179 | ENSMUST0 | -15896 | ENSMUSG0 | 1700030A11 | RIKEN cDNA 1700030A11 gene |
| chr17 | 29022485 | 29023134 | 650 | * | NA_peak_92 | 27 | Intron (ENSM | 17 | 29032628 | 29043373 | 10746 | 1 | 20383 | ENSMUST0 | -9494 | ENSMUSG0 | Srsf3 | serine and arginine-rich splicing factor 3 |
| chr17 | 29200155 | 29200489 | 335 | * | NA_peak_92 | 22 | Intron (ENSM | 17 | 29157916 | 29184103 | 26188 | 2 | 240058 | ENSMUST0 | -16052 | ENSMUSG0 | Cpne5 | copine V |
| chr17 | 29201535 | 29201868 | 334 | * | NA_peak_92 | 15 | Intron (ENSM | 17 | 29157916 | 29184103 | 26188 | 2 | 240058 | ENSMUST0 | -17432 | ENSMUSG0 | Cpne5 | copine V |
| chr17 | 29268772 | 29269436 | 665 | * | NA_peak_92 | 52 | Promoter (<= | 17 | 29268788 | 29302890 | 34103 | 1 | 80748 | ENSMUST0 | 0 | ENSMUSG0 | BC004004 | cDNA sequence BC004004 |
| chr17 | 29424203 | 29424544 | 342 | * | NA_peak_92 | 23 | Distal Interge | 17 | 29376501 | 29378452 | 1952 | 1 | 26382 | ENSMUST0 | 47702 | ENSMUSG0 | Fgd2 | FYVE, RhoGEF and PH domain containing 2 |
| chr17 | 29437390 | 29437517 | 128 | * | NA_peak_92 | 23 | Exon (ENSM | 17 | 29490753 | 29496112 | 5360 | 1 | 18712 | ENSMUST0 | -53236 | ENSMUSG0 | Pim1 | proviral integration site 1 |
| chr17 | 29497198 | 29497391 | 194 | * | NA_peak_92 | 20 | Distal Interge | 17 | 29493407 | 29494964 | 1558 | 1 | 18712 | ENSMUST0 | 3791 | ENSMUSG0 | Pim1 | proviral integration site 1 |
| chr17 | 29663868 | 29664113 | 246 | * | NA_peak_92 | 17 | Intron (ENSM | 17 | 29660807 | 29676146 | 15340 | 1 | 74157 | ENSMUST0 | 3061 | ENSMUSG0 | Cmtr1 | cap methyltransferase 1 |
| chr17 | 30162214 | 30162522 | 309 | * | NA_peak_93 | 35 | Intron (ENSM | 17 | 30136190 | 30210020 | 73831 | 1 | 21769 | ENSMUST0 | 26024 | ENSMUSG0 | Zfand3 | zinc finger, AN1-type domain 3 |
| chr17 | 32174703 | 32174989 | 287 | * | NA_peak_93 | 13 | Distal Interge | 17 | 32120820 | 32166852 | 46033 | 2 | 18131 | ENSMUST0 | -7851 | ENSMUSG0 | Notch3 | notch 3 |
| chr17 | 32218765 | 32219158 | 394 | * | NA_peak_93 | 28 | Intron (ENSM | 17 | 32198470 | 32201008 | 2539 | 2 | 57261 | ENSMUST0 | -17757 | ENSMUSG0 | Brd4 | bromodomain containing 4 |
| chr17 | 32287555 | 32288040 | 486 | * | NA_peak_93 | 29 | Promoter (2- | 17 | 32213610 | 32284722 | 71113 | 2 | 57261 | ENSMUST0 | -2833 | ENSMUSG0 | Brd4 | bromodomain containing 4 |
| chr17 | 32384247 | 32384344 | 98 | * | NA_peak_93 | 24 | Intron (ENSM | 17 | 32377894 | 32387688 | 9795 | 2 | 22404 | ENSMUST0 | 3344 | ENSMUSG0 | Wiz | widely-interspaced zinc finger motifs |
| chr17 | 32414187 | 32414192 | 6 | * | NA_peak_93 | 37 | Intron (ENSM | 17 | 32413361 | 32420975 | 7615 | 2 | 57757 | ENSMUST0 | 6783 | ENSMUSG0 | Pglyrp2 | peptidoglycan recognition protein 2 |
| chr17 | 33050782 | 33051311 | 530 | * | NA_peak_93 | 14 | Distal Interge | 17 | 33065100 | 33067826 | 2727 | 2 | 74042 | ENSMUST0 | 16515 | ENSMUSG0 | Phf8-ps | PHD finger protein 8, pseudogene |
| chr17 | 33313074 | 33313550 | 477 | * | NA_peak_93 | 35 | Distal Interge | 17 | 33289544 | 33307241 | 17698 | 1 | 100043468 | ENSMUST0 | 23530 | ENSMUSG0 | Zfp955b | zinc finger protein 955B |
| chr17 | 33408609 | 33408621 | 13 | * | NA_peak_93 | 44 | Distal Interge | 17 | 33380255 | 33394641 | 14387 | 2 | 22643 | ENSMUST0 | -13968 | ENSMUSG0 | Zfp101 | zinc finger protein 101 |
| chr17 | 33649409 | 33650225 | 817 | * | NA_peak_93 | 29 | Promoter (<= | 17 | 33647828 | 33649687 | 1860 | 2 | 76936 | ENSMUST0 | 0 | ENSMUSG0 | Hnrnpm | heterogeneous nuclear ribonucleoprotein M |
| chr17 | 33718477 | 33718785 | 309 | * | NA_peak_93 | 15 | Promoter (<= | 17 | 33685692 | 33718670 | 32979 | 2 | 224703 | ENSMUST0 | 0 | ENSMUSG0 | Marchf2 | membrane associated ring-CH-type finger 2 |
| chr17 | 33898851 | 33899134 | 284 | * | NA_peak_93 | 22 | Distal Interge | 17 | 33905135 | 33906675 | 1541 | 1 | 474160 | ENSMUST0 | -6001 | ENSMUSG0 | Smim40 | small integral membrane protein 40 |
| chr17 | 33919766 | 33919969 | 204 | * | NA_peak_93 | 25 | Promoter (<= | 17 | 33919828 | 33920392 | 565 | 1 | 21356 | ENSMUST0 | 0 | ENSMUSG0 | Tapbp | TAP binding protein |
| chr17 | 34118627 | 34119873 | 1247 | * | NA_peak_94 | 44 | Promoter (<= | 17 | 34119541 | 34135412 | 15872 | 1 | 14998 | ENSMUST0 | 0 | ENSMUSG0 | H2-DMa | histocompatibility 2, class II, locus DMa |
| chr17 | 34248571 | 34248868 | 298 | * | NA_peak_94 | 27 | 3' UTR | 17 | 34242081 | 34244451 | 2371 | 1 | 15002 | ENSMUST0 | 6490 | ENSMUSG0 | H2-Ob | histocompatibility 2, O region beta locus |
| chr17 | 34459404 | 34459417 | 14 | * | NA_peak_94 | 16 | Intron (ENSM | 17 | 34454394 | 34460421 | 6028 | 1 | 407788 | ENSMUST0 | 5010 | ENSMUSG0 | BC051142 | cDNA sequence BC051142 |
| chr17 | 34765176 | 34765583 | 408 | * | NA_peak_94 | 18 | Distal Interge | 17 | 34728380 | 34743882 | 15503 | 2 | 12268 | ENSMUST0 | -21294 | ENSMUSG0 | C4b | complement component 4B (Chido blood group) |
| chr17 | 34766632 | 34766807 | 176 | * | NA_peak_94 | 31 | Distal Interge | 17 | 34728380 | 34743882 | 15503 | 2 | 12268 | ENSMUST0 | -22750 | ENSMUSG0 | C4b | complement component 4B (Chido blood group) |
| chr17 | 34938790 | 34939204 | 415 | * | NA_peak_94 | 32 | Distal Interge | 17 | 34931863 | 34932765 | 903 | 1 | 18010 | ENSMUST0 | 6927 | ENSMUSG0 | Neu1 | neuraminidase 1 |
| chr17 | 34977563 | 34977763 | 201 | * | NA_peak_94 | 21 | Exon (ENSM | 17 | 34981862 | 34983981 | 2120 | 1 | 27756 | ENSMUST0 | -4099 | ENSMUSG0 | Lsm2 | LSM2 homolog, U6 small nuclear RNA and mRNA degradation associated |
| chr17 | 35135067 | 35135369 | 303 | * | NA_peak_94 | 35 | Promoter (<= | 17 | 35135178 | 35147305 | 12128 | 1 | 224727 | ENSMUST0 | 0 | ENSMUSG0 | Bag6 | BCL2-associated athanogene 6 |
| chr17 | 35205012 | 35205289 | 278 | * | NA_peak_94 | 17 | Promoter (<= | 17 | 35203165 | 35205351 | 2187 | 2 | 16992 | ENSMUST0 | 62 | ENSMUSG0 | Lta | lymphotoxin A |
| chr17 | 35265693 | 35266086 | 394 | * | NA_peak_94 | 58 | Promoter (<= | 17 | 35265477 | 35266319 | 843 | 1 | 14980 | ENSMUST0 | 216 | NA | H2-L | histocompatibility 2, D region locus L |
| chr17 | 35575677 | 35576137 | 461 | * | NA_peak_94 | 17 | Distal Interge | 17 | 35567450 | 35568952 | 1503 | 1 | 69542 | ENSMUST0 | 8227 | ENSMUSG0 | 2300002M23 | RIKEN cDNA 2300002M23 gene |
| chr17 | 35682831 | 35683082 | 252 | * | NA_peak_94 | 44 | Intron (ENSM | 17 | 35667730 | 35673739 | 6010 | 2 | 14885 | ENSMUST0 | -9092 | ENSMUSG0 | Gtf2h4 | general transcription factor II H, polypeptide 4 |
| chr17 | 35858451 | 35858592 | 142 | * | NA_peak_94 | 19 | Promoter (2- | 17 | 35861318 | 35865402 | 4085 | 1 | 106582 | ENSMUST0 | -2726 | ENSMUSG0 | Nrm | nurim (nuclear envelope membrane protein) |
| chr17 | 35896820 | 35897365 | 546 | * | NA_peak_94 | 13 | Promoter (<= | 17 | 35892677 | 35897120 | 4444 | 2 | 69662 | ENSMUST0 | 0 | ENSMUSG0 | 2310061I04 | RIKEN cDNA 2310061I04 gene |
| chr17 | 35916498 | 35917044 | 547 | * | NA_peak_94 | 21 | Promoter (<= | 17 | 35916566 | 35932283 | 15718 | 1 | 52040 | ENSMUST0 | 0 | ENSMUSG0 | Ppp1r10 | protein phosphatase 1, regulatory subunit 10 |
| chr17 | 35969487 | 35970140 | 654 | * | NA_peak_94 | 32 | Promoter (<= | 17 | 35956819 | 35969761 | 12943 | 2 | 224742 | ENSMUST0 | 0 | ENSMUSG0 | Abcf1 | ATP-binding cassette, sub-family F (GCN20), member 1 |
| chr17 | 35972405 | 35972509 | 105 | * | NA_peak_94 | 15 | Promoter (2- | 17 | 35964254 | 35969761 | 5508 | 2 | 224742 | ENSMUST0 | -2644 | ENSMUSG0 | Abcf1 | ATP-binding cassette, sub-family F (GCN20), member 1 |
| chr17 | 35988614 | 35988915 | 302 | * | NA_peak_94 | 25 | 3' UTR | 17 | 35994453 | 36038435 | 43983 | 1 | 77810 | ENSMUST0 | -5538 | NA | A930015D03 | RIKEN cDNA A930015D03 gene |
| chr17 | 36745367 | 36745416 | 50 | * | NA_peak_94 | 16 | Distal Interge | 17 | 36772910 | 36776234 | 3325 | 1 | 224761 | ENSMUST0 | -27494 | ENSMUSG0 | H2-M10.5 | histocompatibility 2, M region locus 10.5 |
| chr17 | 38513215 | 38513689 | 475 | * | NA_peak_95 | 42 | Distal Interge | 17 | 38554192 | 38560621 | 6430 | 1 | 100126773 | ENSMUST0 | -40503 | ENSMUSG0 | Esp34 | exocrine gland secreted peptide 34 |
| chr17 | 38615608 | 38615715 | 108 | * | NA_peak_95 | 25 | Distal Interge | 17 | 38639447 | 38645653 | 6207 | 1 | 100126768 | ENSMUST0 | -23732 | ENSMUSG0 | Esp31 | exocrine gland secreted peptide 31 |
| chr17 | 38644785 | 38645230 | 446 | * | NA_peak_95 | 14 | 3' UTR | 17 | 38639447 | 38645653 | 6207 | 1 | 100126768 | ENSMUST0 | 5338 | ENSMUSG0 | Esp31 | exocrine gland secreted peptide 31 |
| chr17 | 39846954 | 39847618 | 665 | * | NA_peak_95 | 32 | Promoter (<= | 17 | 39846958 | 39848788 | 1831 | 1 | 100861531 | ENSMUST0 | 0 | NA | Rn45s | 45S pre-ribosomal RNA |
| chr17 | 40125798 | 40126081 | 284 | * | NA_peak_95 | 41 | Distal Interge | 17 | 40207015 | 40208640 | 1626 | 2 | 18663 | ENSMUST0 | 82559 | ENSMUSG0 | Pgk2 | phosphoglycerate kinase 2 |
| chr17 | 40644722 | 40645000 | 279 | * | NA_peak_95 | 46 | Distal Interge | 17 | 40632075 | 40637060 | 4986 | 1 | 100126766 | ENSMUST0 | 12647 | ENSMUSG0 | Esp3 | exocrine gland secreted peptide 3 |
| chr17 | 42878776 | 42879244 | 469 | * | NA_peak_95 | 24 | Promoter (2- | 17 | 42816264 | 42876665 | 60402 | 2 | 12488 | ENSMUST0 | -2111 | ENSMUSG0 | Cd2ap | CD2-associated protein |
| chr17 | 44595919 | 44596242 | 324 | * | NA_peak_95 | 30 | Intron (ENSM | 17 | 44639681 | 44734693 | 95013 | 2 | 12393 | ENSMUST0 | 138451 | ENSMUSG0 | Runx2 | runt related transcription factor 2 |
| chr17 | 44649346 | 44649784 | 439 | * | NA_peak_95 | 22 | Intron (ENSM | 17 | 44639681 | 44734693 | 95013 | 2 | 12393 | ENSMUST0 | 84909 | ENSMUSG0 | Runx2 | runt related transcription factor 2 |
| chr17 | 45108745 | 45109302 | 558 | * | NA_peak_96 | 34 | Intron (ENSM | 17 | 44879770 | 44880295 | 526 | 2 | 75341 | ENSMUST0 | -228450 | NA | 4930564C03 | RIKEN cDNA 4930564C03 gene |
| chr17 | 45478586 | 45478809 | 224 | * | NA_peak_96 | 13 | Distal Interge | 17 | 45440578 | 45475415 | 34838 | 2 | 71020 | ENSMUST0 | -3171 | ENSMUSG0 | Spats1 | spermatogenesis associated, serine-rich 1 |

| | | | | | | | | | | | | | | | | | | |
|---|---|---|---|---|---|---|---|---|---|---|---|---|---|---|---|---|---|---|
| chr17 | 45634384 | 45634683 | 300 | * | NA_peak_96 | 13 | Intron (ENSM | 17 | 45642912 | 45659297 | 16386 | 2 | 268958 | ENSMUST0 | 24614 | ENSMUSG0 | Capn11 | calpain 11 |
| chr17 | 46100923 | 46100927 | 5 | * | NA_peak_96 | 26 | Distal Interge | 17 | 46110986 | 46128910 | 17925 | 1 | 68565 | ENSMUST0 | -10059 | ENSMUSG0 | Mrps18a | mitochondrial ribosomal protein S18A |
| chr17 | 46110937 | 46111283 | 347 | * | NA_peak_96 | 43 | Promoter (<= | 17 | 46110986 | 46128910 | 17925 | 1 | 68565 | ENSMUST0 | 0 | ENSMUSG0 | Mrps18a | mitochondrial ribosomal protein S18A |
| chr17 | 46178510 | 46178828 | 319 | * | NA_peak_96 | 49 | Intron (ENSM | 17 | 46167636 | 46168410 | 775 | 1 | 56055 | ENSMUST0 | 10874 | ENSMUSG0 | Gtpbp2 | GTP binding protein 2 |
| chr17 | 46247745 | 46247991 | 247 | * | NA_peak_96 | 48 | Promoter (<= | 17 | 46244524 | 46247932 | 3409 | 2 | 20016 | ENSMUST0 | 0 | ENSMUSG0 | Polr1c | polymerase (RNA) I polypeptide C |
| chr17 | 46346301 | 46346672 | 372 | * | NA_peak_96 | 45 | Distal Interge | 17 | 46324557 | 46328352 | 3796 | 2 | 224814 | ENSMUST0 | -17949 | ENSMUSG0 | Abcc10 | ATP-binding cassette, sub-family C (CFTR/MRP), member 10 |
| chr17 | 46354951 | 46355264 | 314 | * | NA_peak_96 | 13 | Distal Interge | 17 | 46324557 | 46328352 | 3796 | 2 | 224814 | ENSMUST0 | -26599 | ENSMUSG0 | Abcc10 | ATP-binding cassette, sub-family C (CFTR/MRP), member 10 |
| chr17 | 46895826 | 46896136 | 311 | * | NA_peak_96 | 26 | Distal Interge | 17 | 46890684 | 46891847 | 1164 | 1 | 72726 | ENSMUST0 | 5142 | ENSMUSG0 | Tbcc | tubulin-specific chaperone C |
| chr17 | 46950249 | 46950639 | 391 | * | NA_peak_96 | 16 | Intron (ENSM | 17 | 46941104 | 46945369 | 4266 | 2 | 224826 | ENSMUST0 | -4880 | ENSMUSG0 | Ubr2 | ubiquitin protein ligase E3 component n-recognin 2 |
| chr17 | 47052118 | 47052266 | 149 | * | NA_peak_96 | 23 | Distal Interge | 17 | 46956625 | 47010556 | 53932 | 2 | 224826 | ENSMUST0 | -41562 | ENSMUSG0 | Ubr2 | ubiquitin protein ligase E3 component n-recognin 2 |
| chr17 | 47083938 | 47084134 | 197 | * | NA_peak_96 | 31 | Distal Interge | 17 | 47140875 | 47329536 | 188662 | 1 | 224829 | ENSMUST0 | -56741 | ENSMUSG0 | Trerf1 | transcriptional regulating factor 1 |
| chr17 | 47481350 | 47481545 | 196 | * | NA_peak_96 | 20 | Distal Interge | 17 | 47501146 | 47502272 | 1127 | 2 | 63856 | ENSMUST0 | 20727 | ENSMUSG0 | Taf8 | TATA-box binding protein associated factor 8 |
| chr17 | 47593317 | 47593628 | 312 | * | NA_peak_96 | 29 | Promoter (<= | 17 | 47593436 | 47598174 | 4739 | 1 | 12445 | ENSMUST0 | 0 | ENSMUSG0 | Ccnd3 | cyclin D3 |
| chr17 | 47611151 | 47611279 | 129 | * | NA_peak_96 | 20 | Promoter (<= | 17 | 47599331 | 47611492 | 12162 | 2 | 53414 | ENSMUST0 | 213 | ENSMUSG0 | Bysl | bystin-like |
| chr17 | 47670142 | 47670383 | 242 | * | NA_peak_96 | 35 | Promoter (2- | 17 | 47672720 | 47681056 | 8337 | 1 | 112694759 | ENSMUST0 | -2337 | ENSMUSG0 | Gm20517 | predicted gene 20517 |
| chr17 | 47672314 | 47672658 | 345 | * | NA_peak_97 | 29 | Promoter (<= | 17 | 47672720 | 47681056 | 8337 | 1 | 112694759 | ENSMUST0 | -62 | ENSMUSG0 | Gm20517 | predicted gene 20517 |
| chr17 | 47735775 | 47735791 | 17 | * | NA_peak_97 | 27 | Promoter (1- | 17 | 47737030 | 47792419 | 55390 | 1 | 21425 | ENSMUST0 | -1239 | ENSMUSG0 | Tfeb | transcription factor EB |
| chr17 | 49451894 | 49451953 | 60 | * | NA_peak_97 | 15 | Promoter (<= | 17 | 49452284 | 49455435 | 3152 | 1 | 56738 | ENSMUST0 | -331 | ENSMUSG0 | Mocs1 | molybdenum cofactor synthesis 1 |
| chr17 | 50156974 | 50157228 | 255 | * | NA_peak_97 | 17 | Intron (ENSM | 17 | 50036623 | 50190497 | 153875 | 2 | 76438 | ENSMUST0 | 33269 | ENSMUSG0 | Rftn1 | raftlin lipid raft linker 1 |
| chr17 | 50870652 | 50870753 | 102 | * | NA_peak_97 | 23 | Intron (ENSM | 17 | 50953993 | 50984626 | 30634 | 2 | 72238 | ENSMUST0 | 113873 | ENSMUSG0 | Tbc1d5 | TBC1 domain family, member 5 |
| chr17 | 52988589 | 52988891 | 303 | * | NA_peak_97 | 26 | Distal Interge | 17 | 52950060 | 52978379 | 28320 | 1 | 211468 | ENSMUST0 | 38529 | ENSMUSG0 | Kcnh8 | potassium voltage-gated channel, subfamily H (eag-related), member 8 |
| chr17 | 53479403 | 53480042 | 640 | * | NA_peak_97 | 38 | Promoter (<= | 17 | 53479234 | 53507680 | 28447 | 1 | 271457 | ENSMUST0 | 169 | ENSMUSG0 | Rab5a | RAB5A, member RAS oncogene family |
| chr17 | 53551747 | 53552265 | 519 | * | NA_peak_97 | 23 | Distal Interge | 17 | 53507460 | 53539451 | 31992 | 2 | 110332 | ENSMUST0 | -12296 | ENSMUSG0 | Pp2d1 | protein phosphatase 2C-like domain containing 1 |
| chr17 | 55445847 | 55445912 | 66 | * | NA_peak_97 | 13 | Promoter (<= | 17 | 55445550 | 55514581 | 69032 | 1 | 240119 | ENSMUST0 | 297 | ENSMUSG0 | St6gal2 | beta galactoside alpha 2,6 sialyltransferase 2 |
| chr17 | 55840422 | 55840743 | 322 | * | NA_peak_97 | 53 | Intron (ENSM | 17 | 55864892 | 55878930 | 14039 | 2 | 104349 | ENSMUST0 | 38187 | ENSMUSG0 | Zfp119a | zinc finger protein 119a |
| chr17 | 56034774 | 56035107 | 334 | * | NA_peak_98 | 13 | Promoter (1- | 17 | 56017264 | 56036576 | 19313 | 2 | 20405 | ENSMUST0 | 1469 | ENSMUSG0 | Sh3gl1 | SH3-domain GRB2-like 1 |
| chr17 | 56036282 | 56036621 | 340 | * | NA_peak_98 | 34 | Promoter (<= | 17 | 56017264 | 56036576 | 19313 | 2 | 20405 | ENSMUST0 | 0 | ENSMUSG0 | Sh3gl1 | SH3-domain GRB2-like 1 |
| chr17 | 56476431 | 56476662 | 232 | * | NA_peak_98 | 28 | Promoter (<= | 17 | 56412426 | 56476439 | 64014 | 2 | 19280 | ENSMUST0 | 0 | ENSMUSG0 | Ptprs | protein tyrosine phosphatase, receptor type, S |
| chr17 | 56496436 | 56496906 | 471 | * | NA_peak_98 | 34 | Distal Interge | 17 | 56511248 | 56512431 | 1184 | 2 | 20834 | ENSMUST0 | 15525 | ENSMUSG0 | Znrf4 | zinc and ring finger 4 |
| chr17 | 57182405 | 57182730 | 326 | * | NA_peak_98 | 25 | Distal Interge | 17 | 57189492 | 57194177 | 4686 | 2 | 50930 | ENSMUST0 | 11447 | ENSMUSG0 | Tnfsf14 | tumor necrosis factor (ligand) superfamily, member 14 |
| chr17 | 57374363 | 57374388 | 26 | * | NA_peak_98 | 27 | Intron (ENSM | 17 | 57358714 | 57483523 | 124810 | 1 | 13733 | ENSMUST0 | 15649 | ENSMUSG0 | Adgre1 | adhesion G protein-coupled receptor E1 |
| chr17 | 57428183 | 57428518 | 336 | * | NA_peak_98 | 47 | Intron (ENSM | 17 | 57358714 | 57483523 | 124810 | 1 | 13733 | ENSMUST0 | 69469 | ENSMUSG0 | Adgre1 | adhesion G protein-coupled receptor E1 |
| chr17 | 57656635 | 57657002 | 368 | * | NA_peak_98 | 44 | Distal Interge | 17 | 57506453 | 57545505 | 39053 | 2 | 224916 | ENSMUST0 | -111130 | ENSMUSG0 | Vmn2r120 | vomeronasal 2, receptor 120 |
| chr17 | 58729446 | 58729664 | 219 | * | NA_peak_98 | 43 | Intron (ENSM | 17 | 58878808 | 58884133 | 5326 | 2 | 69239 | ENSMUST0 | 154469 | ENSMUSG0 | Pdzph1 | PDZ and pleckstrin homology domains 1 |
| chr17 | 58763574 | 58764177 | 604 | * | NA_peak_98 | 26 | Intron (ENSM | 17 | 58878808 | 58884133 | 5326 | 2 | 69239 | ENSMUST0 | 119956 | ENSMUSG0 | Pdzph1 | PDZ and pleckstrin homology domains 1 |
| chr17 | 58958580 | 58958744 | 165 | * | NA_peak_98 | 16 | Intron (ENSM | 17 | 58878808 | 58991375 | 112568 | 2 | 69239 | ENSMUST0 | 32631 | ENSMUSG0 | Pdzph1 | PDZ and pleckstrin homology domains 1 |
| chr17 | 60419861 | 60419940 | 80 | * | NA_peak_98 | 16 | Distal Interge | 17 | 59017314 | 59024729 | 7416 | 1 | 75039 | ENSMUST0 | 1402547 | ENSMUSG0 | Nudt12os | nudix (nucleoside diphosphate linked moiety X)-type motif 12, opposite strand |
| chr17 | 61760276 | 61760546 | 271 | * | NA_peak_98 | 23 | Distal Interge | 17 | 62607319 | 62881144 | 273826 | 2 | 13640 | ENSMUST0 | 1120598 | ENSMUSG0 | Efna5 | ephrin A5 |
| chr17 | 64105598 | 64105866 | 269 | * | NA_peak_99 | 30 | Intron (ENSM | 17 | 63937972 | 64139197 | 201226 | 1 | 14158 | ENSMUST0 | 167626 | ENSMUSG0 | Fer | fer (fms/fps related) protein kinase |
| chr17 | 64382930 | 64383395 | 466 | * | NA_peak_99 | 23 | Distal Interge | 17 | 64281005 | 64331916 | 50912 | 2 | 224938 | ENSMUST0 | -51014 | ENSMUSG0 | Pja2 | praja ring finger ubiquitin ligase 2 |
| chr17 | 64730209 | 64730313 | 105 | * | NA_peak_99 | 24 | Intron (ENSM | 17 | 64713619 | 64754838 | 41220 | 1 | 17158 | ENSMUST0 | 16590 | ENSMUSG0 | Man2a1 | mannosidase 2, alpha 1 |
| chr17 | 65442272 | 65442590 | 319 | * | NA_peak_99 | 26 | Intron (ENSM | 17 | 65378996 | 65436395 | 57400 | 2 | 381107 | ENSMUST0 | -5877 | ENSMUSG0 | Tmem232 | transmembrane protein 232 |
| chr17 | 65757098 | 65757298 | 201 | * | NA_peak_99 | 24 | Intron (ENSM | 17 | 65721266 | 65772640 | 51375 | 2 | 106572 | ENSMUST0 | 15342 | ENSMUSG0 | Rab31 | RAB31, member RAS oncogene family |
| chr17 | 66332379 | 66332676 | 298 | * | NA_peak_99 | 30 | Distal Interge | 17 | 66343946 | 66346545 | 2600 | 2 | 68617 | ENSMUST0 | 13869 | ENSMUSG0 | Mtcl1 | microtubule crosslinking factor 1 |
| chr17 | 67429568 | 67429906 | 339 | * | NA_peak_10 | 40 | Distal Interge | 17 | 66761075 | 67354457 | 593383 | 2 | 19274 | ENSMUST0 | -75111 | ENSMUSG0 | Ptprm | protein tyrosine phosphatase, receptor type, M |
| chr17 | 68004147 | 68004187 | 41 | * | NA_peak_10 | 28 | Promoter (<= | 17 | 67842713 | 68004120 | 161408 | 2 | 268970 | ENSMUST0 | -27 | ENSMUSG0 | Arhgap28 | Rho GTPase activating protein 28 |
| chr17 | 69393073 | 69393165 | 93 | * | NA_peak_10 | 18 | Distal Interge | 17 | 69383840 | 69391199 | 7360 | 1 | 22666 | ENSMUST0 | 9233 | ENSMUSG0 | Zbtb14 | zinc finger and BTB domain containing 14 |
| chr17 | 70843576 | 70843800 | 225 | * | NA_peak_10 | 24 | Promoter (2- | 17 | 70844869 | 70846326 | 1458 | 2 | 21815 | ENSMUST0 | 2526 | ENSMUSG0 | Tgif1 | TGFB-induced factor homeobox 1 |
| chr17 | 71772987 | 71773346 | 360 | * | NA_peak_10 | 44 | Intron (ENSM | 17 | 71769743 | 71850163 | 80421 | 1 | 78785 | ENSMUST0 | 3244 | ENSMUSG0 | Clip4 | CAP-GLY domain containing linker protein family, member 4 |
| chr17 | 71943650 | 71944040 | 391 | * | NA_peak_10 | 19 | Intron (ENSM | 17 | 71856286 | 71864273 | 7988 | 1 | 78785 | ENSMUST0 | 87364 | ENSMUSG0 | Clip4 | CAP-GLY domain containing linker protein family, member 4 |
| chr17 | 72075799 | 72075870 | 72 | * | NA_peak_10 | 23 | Intron (ENSM | 17 | 71856286 | 71864273 | 7988 | 1 | 78785 | ENSMUST0 | 219513 | ENSMUSG0 | Clip4 | CAP-GLY domain containing linker protein family, member 4 |
| chr17 | 72515790 | 72516135 | 346 | * | NA_peak_10 | 23 | Intron (ENSM | 17 | 71868972 | 72603827 | 734856 | 2 | 11682 | ENSMUST0 | 87692 | ENSMUSG0 | Alk | anaplastic lymphoma kinase |
| chr17 | 72833470 | 72833780 | 311 | * | NA_peak_10 | 20 | Promoter (2- | 17 | 72836453 | 72851195 | 14743 | 1 | 383295 | ENSMUST0 | -2673 | ENSMUSG0 | Ypel5 | yippee like 5 |
| chr17 | 73107978 | 73108211 | 234 | * | NA_peak_10 | 23 | Promoter (<= | 17 | 73107985 | 73161991 | 54007 | 1 | 225010 | ENSMUST0 | 0 | ENSMUSG0 | Lclat1 | lysocardiolipin acyltransferase 1 |
| chr17 | 73180756 | 73181061 | 306 | * | NA_peak_10 | 24 | Intron (ENSM | 17 | 73162989 | 73239908 | 76920 | 1 | 225010 | ENSMUST0 | 17767 | ENSMUSG0 | Lclat1 | lysocardiolipin acyltransferase 1 |
| chr17 | 73386287 | 73386528 | 242 | * | NA_peak_10 | 19 | Intron (ENSM | 17 | 73306464 | 73399296 | 92833 | 2 | 381122 | ENSMUST0 | 12768 | ENSMUSG0 | Capn13 | calpain 13 |
| chr17 | 73794700 | 73794817 | 118 | * | NA_peak_10 | 13 | Distal Interge | 17 | 73804841 | 73832094 | 27254 | 1 | 57440 | ENSMUST0 | -10024 | ENSMUSG0 | Ehd3 | EH-domain containing 3 |
| chr17 | 74128585 | 74128981 | 397 | * | NA_peak_10 | 26 | Distal Interge | 17 | 74026998 | 74079046 | 52049 | 2 | 94224 | ENSMUST0 | -49539 | ENSMUSG0 | Srd5a2 | steroid 5 alpha-reductase 2 |
| chr17 | 74350370 | 74350642 | 273 | * | NA_peak_10 | 21 | Intron (ENSM | 17 | 74339136 | 74375644 | 36509 | 1 | 50850 | ENSMUST0 | 11234 | ENSMUSG0 | Spast | spastin |
| chr17 | 74527570 | 74528255 | 686 | * | NA_peak_10 | 27 | Promoter (<= | 17 | 74528295 | 74703356 | 175062 | 1 | 12211 | ENSMUST0 | -40 | ENSMUSG0 | Birc6 | baculoviral IAP repeat-containing 6 |
| chr17 | 74662310 | 74662650 | 341 | * | NA_peak_10 | 39 | Exon (ENSM | 17 | 74659146 | 74668813 | 9668 | 1 | 12211 | ENSMUST0 | 3164 | ENSMUSG0 | Birc6 | baculoviral IAP repeat-containing 6 |
| chr17 | 74778061 | 74778390 | 330 | * | NA_peak_10 | 43 | Intron (ENSM | 17 | 74770066 | 74780855 | 10790 | 1 | 74196 | ENSMUST0 | 7995 | ENSMUSG0 | Ttc27 | tetratricopeptide repeat domain 27 |
| chr17 | 74896618 | 74896778 | 161 | * | NA_peak_10 | 22 | Distal Interge | 17 | 74861921 | 74863570 | 1650 | 1 | 74196 | ENSMUST0 | 34697 | ENSMUSG0 | Ttc27 | tetratricopeptide repeat domain 27 |
| chr17 | 74980849 | 74981037 | 189 | * | NA_peak_10 | 20 | Distal Interge | 17 | 75005568 | 75391328 | 385761 | 1 | 268977 | ENSMUST0 | -24531 | ENSMUSG0 | Ltbp1 | latent transforming growth factor beta binding protein 1 |
| chr17 | 75109321 | 75109637 | 317 | * | NA_peak_10 | 17 | Intron (ENSM | 17 | 75178810 | 75348933 | 170124 | 1 | 268977 | ENSMUST0 | -69173 | ENSMUSG0 | Ltbp1 | latent transforming growth factor beta binding protein 1 |
| chr17 | 75280213 | 75280604 | 392 | * | NA_peak_10 | 13 | Intron (ENSM | 17 | 75289976 | 75294369 | 4394 | 1 | 268977 | ENSMUST0 | -9372 | ENSMUSG0 | Ltbp1 | latent transforming growth factor beta binding protein 1 |
| chr17 | 75425357 | 75426228 | 872 | * | NA_peak_10 | 92 | Distal Interge | 17 | 75435896 | 75491790 | 55895 | 1 | 240168 | ENSMUST0 | -9668 | ENSMUSG0 | Rasgrp3 | RAS, guanyl releasing protein 3 |
| chr17 | 75670475 | 75670510 | 36 | * | NA_peak_10 | 22 | Distal Interge | 17 | 75537086 | 75551946 | 14861 | 2 | 72722 | ENSMUST0 | -118529 | ENSMUSG0 | Fam98a | family with sequence similarity 98, member A |
| chr17 | 75973596 | 75973647 | 52 | * | NA_peak_10 | 17 | Distal Interge | 17 | 75861023 | 75889039 | 28017 | 1 | 100043885 | ENSMUST0 | 112573 | ENSMUSG0 | Gm4710 | predicted gene 4710 |
| chr17 | 76666059 | 76666120 | 62 | * | NA_peak_10 | 26 | Distal Interge | 17 | 75861023 | 75889039 | 28017 | 1 | 100043885 | ENSMUST0 | 805036 | ENSMUSG0 | Gm4710 | predicted gene 4710 |
| chr17 | 78130075 | 78130263 | 189 | * | NA_peak_10 | 19 | Distal Interge | 17 | 78200248 | 78376592 | 176345 | 1 | 50766 | ENSMUST0 | -69985 | ENSMUSG0 | Crim1 | cysteine rich transmembrane BMP regulator 1 (chordin like) |
| chr17 | 78864974 | 78865183 | 210 | * | NA_peak_10 | 13 | Intron (ENSM | 17 | 78855354 | 78858980 | 3627 | 2 | 19106 | ENSMUST0 | -5994 | ENSMUSG0 | Eif2ak2 | eukaryotic translation initiation factor 2-alpha kinase 2 |
| chr17 | 78936225 | 78936564 | 340 | * | NA_peak_10 | 26 | Promoter (<= | 17 | 78935217 | 78937037 | 1821 | 2 | 12607 | ENSMUST0 | 473 | ENSMUSG0 | Cebpz | CCAAT/enhancer binding protein zeta |
| chr17 | 79316425 | 79316758 | 334 | * | NA_peak_10 | 20 | Distal Interge | 17 | 79335197 | 79353007 | 17811 | 2 | 260409 | ENSMUST0 | 36249 | ENSMUSG0 | Cdc42ep3 | CDC42 effector protein (Rho GTPase binding) 3 |
| chr17 | 79395239 | 79395317 | 79 | * | NA_peak_10 | 13 | Distal Interge | 17 | 79333727 | 79355091 | 21365 | 2 | 260409 | ENSMUST0 | -40148 | ENSMUSG0 | Cdc42ep3 | CDC42 effector protein (Rho GTPase binding) 3 |
| chr17 | 79615320 | 79615723 | 404 | * | NA_peak_10 | 38 | Promoter (<= | 17 | 79615062 | 79682249 | 67188 | 1 | 381110 | ENSMUST0 | 258 | ENSMUSG0 | Rmdn2 | regulator of microtubule dynamics 2 |
| chr17 | 79733156 | 79733677 | 522 | * | NA_peak_10 | 20 | Distal Interge | 17 | 79706894 | 79715061 | 8168 | 2 | 13078 | ENSMUST0 | -18095 | ENSMUSG0 | Cyp1b1 | cytochrome P450, family 1, subfamily b, polypeptide 1 |
| chr17 | 80083574 | 80083777 | 204 | * | NA_peak_10 | 27 | Distal Interge | 17 | 80029487 | 80062334 | 32848 | 2 | 72692 | ENSMUST0 | -21240 | ENSMUSG0 | Hnrnpll | heterogeneous nuclear ribonucleoprotein L-like |
| chr17 | 80275728 | 80275929 | 202 | * | NA_peak_10 | 28 | Exon (ENSM | 17 | 80265075 | 80270583 | 5509 | 2 | 106794 | ENSMUST0 | -5145 | ENSMUSG0 | Dhx57 | DEAH (Asp-Glu-Ala-Asp/His) box polypeptide 57 |
| chr17 | 80572399 | 80572644 | 246 | * | NA_peak_10 | 24 | Intron (ENSM | 17 | 80524533 | 80577484 | 52952 | 2 | 381113 | ENSMUST0 | 4840 | ENSMUSG0 | Cdkl4 | cyclin-dependent kinase-like 4 |
| chr17 | 81348483 | 81348811 | 329 | * | NA_peak_10 | 27 | Distal Interge | 17 | 81026332 | 81065082 | 38751 | 2 | 72167 | ENSMUST0 | -283401 | ENSMUSG0 | Thumpd2 | THUMP domain containing 2 |
| chr17 | 82610626 | 82610837 | 212 | * | NA_peak_10 | 17 | Distal Interge | 17 | 83033594 | 83078225 | 44632 | 2 | 100503434 | ENSMUST0 | 467388 | ENSMUSG0 | Gm19689 | predicted gene, 19689 |
| chr17 | 83256313 | 83256388 | 76 | * | NA_peak_10 | 26 | Distal Interge | 17 | 83221035 | 83224575 | 3541 | 1 | 106522 | ENSMUST0 | 35278 | ENSMUSG0 | Pkdcc | protein kinase domain containing, cytoplasmic |
| chr17 | 83473081 | 83473374 | 294 | * | NA_peak_10 | 16 | Exon (ENSM | 17 | 83456301 | 83480357 | 24057 | 1 | 78798 | ENSMUST0 | 16780 | ENSMUSG0 | Eml4 | echinoderm microtubule associated protein like 4 |
| chr17 | 83767573 | 83767889 | 317 | * | NA_peak_10 | 23 | Intron (ENSM | 17 | 83755634 | 83800240 | 44607 | 1 | 116871 | ENSMUST0 | 11939 | ENSMUSG0 | Mta3 | metastasis associated 3 |
| chr17 | 83861587 | 83861771 | 185 | * | NA_peak_10 | 30 | Distal Interge | 17 | 83851751 | 83854418 | 2668 | 2 | 320021 | ENSMUST0 | -7169 | ENSMUSG0 | C430042M11 | RIKEN cDNA C430042M11 gene |
| chr17 | 83864927 | 83865375 | 449 | * | NA_peak_10 | 27 | Distal Interge | 17 | 83851751 | 83854418 | 2668 | 2 | 320021 | ENSMUST0 | -10509 | ENSMUSG0 | C430042M11 | RIKEN cDNA C430042M11 gene |
| chr17 | 84077514 | 84077872 | 359 | * | NA_peak_10 | 34 | Promoter (<= | 17 | 84078660 | 84089875 | 11216 | 1 | 74473 | ENSMUST0 | -788 | ENSMUSG0 | 4933433H22 | RIKEN cDNA 4933433H22 gene |
| chr17 | 84323093 | 84323391 | 299 | * | NA_peak_10 | 16 | Intron (ENSM | 17 | 84309626 | 84313679 | 4054 | 2 | 240174 | ENSMUST0 | -9414 | ENSMUSG0 | Thada | thyroid adenoma associated |
| chr17 | 84514914 | 84515116 | 203 | * | NA_peak_10 | 34 | Promoter (2- | 17 | 84511923 | 84572076 | 60154 | 1 | 213556 | ENSMUST0 | 2991 | ENSMUSG0 | Plekhh2 | pleckstrin homology domain containing, family H (with MyTH4 domain) member 2 |
| chr17 | 84940069 | 84940347 | 279 | * | NA_peak_10 | 19 | Intron (ENSM | 17 | 84956741 | 85023991 | 67251 | 1 | 19043 | ENSMUST0 | -16394 | ENSMUSG0 | Ppm1b | protein phosphatase 1B, magnesium dependent, beta isoform |
| chr17 | 85051444 | 85051663 | 220 | * | NA_peak_10 | 23 | Intron (ENSM | 17 | 85028376 | 85064236 | 35861 | 1 | 20532 | ENSMUST0 | 23068 | ENSMUSG0 | Slc3a1 | solute carrier family 3, member 1 |
| chr17 | 85115576 | 85115682 | 107 | * | NA_peak_10 | 24 | Intron (ENSM | 17 | 85092083 | 85166508 | 74426 | 1 | 73582 | ENSMUST0 | 23493 | ENSMUSG0 | Camkmt | calmodulin-lysine N-methyltransferase |
| chr17 | 86766812 | 86766948 | 137 | * | NA_peak_10 | 16 | Intron (ENSM | 17 | 86753700 | 86833410 | 79711 | 1 | 13819 | ENSMUST0 | 13112 | ENSMUSG0 | Epas1 | endothelial PAS domain protein 1 |
| chr17 | 87149584 | 87149995 | 412 | * | NA_peak_10 | 24 | Distal Interge | 17 | 87107858 | 87122990 | 15133 | 1 | 56468 | ENSMUST0 | 41726 | ENSMUSG0 | Socs5 | suppressor of cytokine signaling 5 |
| chr17 | 87152357 | 87152662 | 306 | * | NA_peak_10 | 27 | Distal Interge | 17 | 87192216 | 87194942 | 2727 | 2 | 399626 | ENSMUST0 | 42280 | ENSMUSG0 | C330024C12 | RIKEN cDNA C330024C12 gene |
| chr17 | 87220207 | 87220575 | 369 | * | NA_peak_10 | 22 | Distal Interge | 17 | 87192216 | 87194942 | 2727 | 2 | 399626 | ENSMUST0 | -25265 | ENSMUSG0 | C330024C12 | RIKEN cDNA C330024C12 gene |
| chr17 | 87407767 | 87408082 | 316 | * | NA_peak_10 | 23 | Intron (ENSM | 17 | 87417490 | 87427635 | 10146 | 2 | 75467 | ENSMUST0 | 19553 | ENSMUSG0 | Stpg4 | sperm tail PG rich repeat containing 4 |
| chr17 | 87427313 | 87427348 | 36 | * | NA_peak_10 | 16 | Promoter (<= | 17 | 87417490 | 87427635 | 10146 | 2 | 75467 | ENSMUST0 | 287 | ENSMUSG0 | Stpg4 | sperm tail PG rich repeat containing 4 |
| chr17 | 87446807 | 87447201 | 395 | * | NA_peak_10 | 17 | Promoter (<= | 17 | 87433412 | 87446935 | 13524 | 2 | 12314 | ENSMUST0 | 0 | ENSMUSG0 | Calm2 | calmodulin 2 |
| chr17 | 87592187 | 87592743 | 557 | * | NA_peak_10 | 28 | Distal Interge | 17 | 87635979 | 87639964 | 3986 | 1 | 17075 | ENSMUST0 | -43236 | ENSMUSG0 | Epcam | epithelial cell adhesion molecule |
| chr17 | 87916294 | 87916478 | 185 | * | NA_peak_10 | 28 | Distal Interge | 17 | 87975062 | 87990892 | 15831 | 1 | 17688 | ENSMUST0 | -58584 | ENSMUSG0 | Msh6 | mutS homolog 6 |
| chr17 | 88064431 | 88064796 | 366 | * | NA_peak_10 | 17 | Promoter (<= | 17 | 88015595 | 88064887 | 49293 | 2 | 225055 | ENSMUST0 | 91 | ENSMUSG0 | Fbxo11 | F-box protein 11 |
| chr17 | 88530130 | 88530407 | 278 | * | NA_peak_10 | 24 | Promoter (<= | 17 | 88530216 | 88543436 | 13221 | 1 | 73825 | ENSMUST0 | 0 | ENSMUSG0 | Ppp1r21 | protein phosphatase 1, regulatory subunit 21 |
| chr17 | 89481670 | 89482294 | 625 | * | NA_peak_10 | 33 | Distal Interge | 17 | 89379241 | 89384993 | 5753 | 1 | 100043904 | ENSMUST0 | 102429 | ENSMUSG0 | Gm4719 | predicted gene 4719 |
| chr17 | 89839883 | 89840196 | 314 | * | NA_peak_10 | 39 | Distal Interge | 17 | 90036853 | 90088140 | 51288 | 2 | 18189 | ENSMUST0 | 247944 | ENSMUSG0 | Nrxn1 | neurexin I |
| chr17 | 91220076 | 91220377 | 302 | * | NA_peak_10 | 26 | Distal Interge | 17 | 90033644 | 91093071 | 1059428 | 2 | 18189 | ENSMUST0 | -127005 | ENSMUSG0 | Nrxn1 | neurexin I |
| chr17 | 91643505 | 91643591 | 87 | * | NA_peak_10 | 23 | Distal Interge | 17 | 90033644 | 91093071 | 1059428 | 2 | 18189 | ENSMUST0 | -550434 | ENSMUSG0 | Nrxn1 | neurexin I |
| chr17 | 92182585 | 92183007 | 423 | * | NA_peak_10 | 30 | Distal Interge | 17 | 93199017 | 93200096 | 1080 | 1 | 11516 | ENSMUST0 | -1016010 | ENSMUSG0 | Adcyap1 | adenylate cyclase activating polypeptide 1 |
| chr17 | 92201602 | 92201989 | 388 | * | NA_peak_10 | 13 | Distal Interge | 17 | 93199017 | 93200096 | 1080 | 1 | 11516 | ENSMUST0 | -997028 | ENSMUSG0 | Adcyap1 | adenylate cyclase activating polypeptide 1 |
| chr17 | 92988646 | 92988884 | 239 | * | NA_peak_10 | 18 | Distal Interge | 17 | 93199017 | 93200096 | 1080 | 1 | 11516 | ENSMUST0 | -210133 | ENSMUSG0 | Adcyap1 | adenylate cyclase activating polypeptide 1 |
| chr17 | 93285093 | 93286055 | 963 | * | NA_peak_10 | 41 | Distal Interge | 17 | 93202076 | 93205563 | 3488 | 1 | 11516 | ENSMUST0 | 83017 | ENSMUSG0 | Adcyap1 | adenylate cyclase activating polypeptide 1 |
| chr17 | 94786858 | 94787039 | 182 | * | NA_peak_10 | 17 | Intron (ENSM | 17 | 94761250 | 94764541 | 3292 | 2 | 100038746 | ENSMUST0 | -22317 | ENSMUSG0 | Gm1976 | predicted gene 1976 |
| chr18 | 3038744 | 3038993 | 250 | * | NA_peak_10 | 22 | Distal Interge | 18 | 3122492 | 3123412 | 921 | 2 | 100312476 | ENSMUST0 | 84419 | ENSMUSG0 | Vmn1r238 | vomeronasal 1 receptor, 238 |
| chr18 | 3040764 | 3040951 | 188 | * | NA_peak_10 | 17 | Distal Interge | 18 | 3122492 | 3123412 | 921 | 2 | 100312476 | ENSMUST0 | 82461 | ENSMUSG0 | Vmn1r238 | vomeronasal 1 receptor, 238 |
| chr18 | 3711083 | 3711390 | 308 | * | NA_peak_10 | 49 | Distal Interge | 18 | 3507923 | 3516404 | 8482 | 1 | 68010 | ENSMUST0 | 203160 | ENSMUSG0 | Bambi | BMP and activin membrane-bound inhibitor |
| chr18 | 3783968 | 3784040 | 73 | * | NA_peak_10 | 23 | Distal Interge | 18 | 3507923 | 3516404 | 8482 | 1 | 68010 | ENSMUST0 | 276045 | ENSMUSG0 | Bambi | BMP and activin membrane-bound inhibitor |
| chr18 | 5115080 | 5115174 | 95 | * | NA_peak_10 | 23 | Promoter (<= | 18 | 5114253 | 5119218 | 4966 | 1 | 225115 | ENSMUST0 | 827 | ENSMUSG0 | Svil | supervillin |
| chr18 | 9913144 | 9913672 | 529 | * | NA_peak_10 | 59 | Distal Interge | 18 | 9874679 | 9877353 | 2675 | 1 | 140792 | ENSMUST0 | 38465 | ENSMUSG0 | Colec12 | collectin sub-family member 12 |
| chr18 | 13581354 | 13581774 | 421 | * | NA_peak_10 | 44 | Intron (ENSM | 18 | 13816872 | 13843832 | 26961 | 2 | 225207 | ENSMUST0 | 262058 | ENSMUSG0 | Zfp521 | zinc finger protein 521 |
| chr18 | 14738482 | 14738663 | 182 | * | NA_peak_10 | 26 | Intron (ENSM | 18 | 14726546 | 14731471 | 4926 | 1 | 73677 | ENSMUST0 | 11936 | ENSMUSG0 | Psma8 | proteasome subunit alpha 8 |
| chr18 | 16962927 | 16963350 | 424 | * | NA_peak_10 | 95 | Distal Interge | 18 | 17019051 | 17137559 | 118509 | 1 | 75433 | ENSMUST0 | -55701 | ENSMUSG0 | 1700001G01 | RIKEN cDNA 1700001G01 gene |
| chr18 | 17438202 | 17438730 | 529 | * | NA_peak_10 | 40 | Distal Interge | 18 | 17568746 | 17580990 | 12245 | 2 | 75162 | ENSMUST0 | 142260 | ENSMUSG0 | 4930545E07 | RIKEN cDNA 4930545E07 gene |
| chr18 | 19613495 | 19613895 | 401 | * | NA_peak_10 | 53 | Distal Interge | 18 | 19962970 | 20001955 | 38986 | 2 | 13507 | ENSMUST0 | 388060 | ENSMUSG0 | Dsc3 | desmocollin 3 |
| chr18 | 19864238 | 19864611 | 374 | * | NA_peak_10 | 64 | Distal Interge | 18 | 19962970 | 20001955 | 38986 | 2 | 13507 | ENSMUST0 | 137344 | ENSMUSG0 | Dsc3 | desmocollin 3 |
| chr18 | 24035536 | 24035979 | 444 | * | NA_peak_10 | 19 | Distal Interge | 18 | 24009702 | 24020810 | 11109 | 2 | 59057 | ENSMUST0 | -14726 | ENSMUSG0 | Zfp24 | zinc finger protein 24 |
| chr18 | 24538012 | 24538187 | 176 | * | NA_peak_10 | 23 | Distal Interge | 18 | 24484960 | 24530204 | 45245 | 2 | 225283 | ENSMUST0 | -7808 | ENSMUSG0 | Rprd1a | regulation of nuclear pre-mRNA domain containing 1A |
| chr18 | 25046330 | 25046529 | 200 | * | NA_peak_10 | 19 | Intron (ENSM | 18 | 25124361 | 25130694 | 6334 | 1 | 225288 | ENSMUST0 | -77832 | ENSMUSG0 | Fhod3 | formin homology 2 domain containing 3 |
| chr18 | 33034780 | 33035070 | 291 | * | NA_peak_10 | 30 | Intron (ENSM | 18 | 32939158 | 33186270 | 247113 | 1 | 12326 | ENSMUST0 | 95622 | ENSMUSG0 | Camk4 | calcium/calmodulin-dependent protein kinase IV |
| chr18 | 37630329 | 37630638 | 310 | * | NA_peak_10 | 44 | Intron (ENSM | 18 | 37637377 | 37638745 | 1369 | 2 | 83885 | ENSMUST0 | 8107 | ENSMUSG0 | Slc25a2 | solute carrier family 25 (mitochondrial carrier, ornithine transporter) member 2 |
| chr18 | 40649274 | 40649704 | 431 | * | NA_peak_10 | 16 | Distal Interge | 18 | 40523099 | 40531179 | 8081 | 1 | 383348 | ENSMUST0 | 126175 | ENSMUSG0 | Kctd16 | potassium channel tetramerisation domain containing 16 |
| chr18 | 43476297 | 43476633 | 337 | * | NA_peak_10 | 23 | 3' UTR | 18 | 43320984 | 43438286 | 117303 | 2 | 22240 | ENSMUST0 | -38011 | ENSMUSG0 | Dpysl3 | dihydropyrimidinase-like 3 |
| chr18 | 48711774 | 48712002 | 229 | * | NA_peak_10 | 44 | Distal Interge | 18 | 48045335 | 48048412 | 3078 | 1 | 433182 | ENSMUST0 | 666439 | ENSMUSG0 | Eno1b | enolase 1B, retrotransposed |
| chr18 | 48980479 | 48980510 | 32 | * | NA_peak_10 | 33 | Distal Interge | 18 | 49523289 | 49524440 | 1152 | 1 | 67321 | ENSMUST0 | -542779 | ENSMUSG0 | 1700044K03 | RIKEN cDNA 1700044K03 gene |
| chr18 | 50256033 | 50256335 | 303 | * | NA_peak_10 | 25 | Distal Interge | 18 | 50278369 | 50283014 | 4646 | 1 | 225497 | ENSMUST0 | -22034 | ENSMUSG0 | Fam170a | family with sequence similarity 170, member A |
| chr18 | 52452045 | 52452419 | 375 | * | NA_peak_10 | 23 | Distal Interge | 18 | 52465693 | 52487603 | 21911 | 1 | 67222 | ENSMUST0 | -13274 | ENSMUSG0 | Srfbp1 | serum response factor binding protein 1 |
| chr18 | 59731360 | 59731767 | 408 | * | NA_peak_10 | 13 | Distal Interge | 18 | 59579630 | 59579735 | 106 | 1 | 102465182 | ENSMUST0 | 151730 | ENSMUSG0 | Mir6355 | microRNA 6355 |
| chr18 | 70778678 | 70779021 | 344 | * | NA_peak_10 | 35 | Distal Interge | 18 | 70617773 | 70625777 | 8005 | 1 | 17191 | ENSMUST0 | 160905 | ENSMUSG0 | Mbd2 | methyl-CpG binding domain protein 2 |
| chr18 | 72512442 | 72512494 | 53 | * | NA_peak_10 | 13 | Distal Interge | 18 | 71253634 | 72351069 | 1097436 | 2 | 13176 | ENSMUST0 | -161373 | ENSMUSG0 | Dcc | deleted in colorectal carcinoma |
| chr18 | 73262802 | 73263095 | 294 | * | NA_peak_10 | 18 | Distal Interge | 18 | 73573044 | 73592575 | 19532 | 1 | 240396 | ENSMUST0 | -309949 | ENSMUSG0 | Mex3c | mex3 RNA binding family member C |
| chr18 | 85323542 | 85323907 | 366 | * | NA_peak_10 | 14 | Distal Interge | 18 | 84965510 | 84981472 | 15963 | 1 | 50764 | ENSMUST0 | 358032 | ENSMUSG0 | Fbxo15 | F-box protein 15 |
| chr18 | 89166774 | 89166974 | 201 | * | NA_peak_10 | 23 | Distal Interge | 18 | 89176967 | 89270201 | 93235 | 1 | 225825 | ENSMUST0 | -9993 | ENSMUSG0 | Cd226 | CD226 antigen |
| chr19 | 3046277 | 3046491 | 215 | * | NA_peak_10 | 16 | Distal Interge | 19 | 3145946 | 3160181 | 14236 | 2 | 70011 | ENSMUST0 | 113690 | ENSMUSG0 | 1700030N03 | RIKEN cDNA 1700030N03 gene |
| chr19 | 3390503 | 3390643 | 141 | * | NA_peak_10 | 16 | Promoter (1- | 19 | 3389379 | 3407785 | 18407 | 1 | 17771 | ENSMUST0 | 1124 | ENSMUSG0 | Tesmin | testis expressed metallothionein like |
| chr19 | 3740838 | 3741234 | 397 | * | NA_peak_10 | 17 | Intron (ENSM | 19 | 3767421 | 3808829 | 41409 | 1 | 225888 | ENSMUST0 | -26187 | ENSMUSG0 | Kmt5b | lysine methyltransferase 5B |
| chr19 | 4073634 | 4074031 | 398 | * | NA_peak_10 | 24 | Distal Interge | 19 | 4081578 | 4085676 | 4099 | 1 | 29866 | ENSMUST0 | -7547 | ENSMUSG0 | Cabp2 | calcium binding protein 2 |
| chr19 | 4125947 | 4126215 | 269 | * | NA_peak_10 | 28 | Promoter (<= | 19 | 4125959 | 4131444 | 5486 | 1 | 66990 | ENSMUST0 | 0 | ENSMUSG0 | Tmem134 | transmembrane protein 134 |
| chr19 | 4222230 | 4222541 | 312 | * | NA_peak_10 | 27 | 3' UTR | 19 | 4214594 | 4222513 | 7920 | 1 | 56708 | ENSMUST0 | 7636 | ENSMUSG0 | Clcf1 | cardiotrophin-like cytokine factor 1 |
| chr19 | 4354070 | 4354117 | 48 | * | NA_peak_10 | 13 | Intron (ENSM | 19 | 4350186 | 4362887 | 12702 | 2 | 225876 | ENSMUST0 | 8770 | ENSMUSG0 | Kdm2a | lysine (K)-specific demethylase 2A |
| chr19 | 4375964 | 4376161 | 198 | * | NA_peak_10 | 35 | Intron (ENSM | 19 | 4350186 | 4362887 | 12702 | 2 | 225876 | ENSMUST0 | -13077 | ENSMUSG0 | Kdm2a | lysine (K)-specific demethylase 2A |
| chr19 | 4425811 | 4426132 | 322 | * | NA_peak_10 | 22 | Promoter (<= | 19 | 4426227 | 4426555 | 329 | 2 | 11854 | ENSMUST0 | 423 | ENSMUSG0 | Rhod | ras homolog family member D |
| chr19 | 4485692 | 4485866 | 175 | * | NA_peak_10 | 14 | Distal Interge | 19 | 4445908 | 4477447 | 31540 | 2 | 171180 | ENSMUST0 | -8245 | ENSMUSG0 | Syt12 | synaptotagmin XII |
| chr19 | 4765529 | 4765888 | 360 | * | NA_peak_10 | 27 | 3' UTR | 19 | 4756905 | 4762419 | 5515 | 1 | 66704 | ENSMUST0 | 8624 | ENSMUSG0 | Rbm4b | RNA binding motif protein 4B |
| chr19 | 4811251 | 4811430 | 180 | * | NA_peak_10 | 16 | Promoter (<= | 19 | 4785153 | 4811212 | 26060 | 2 | 102902673 | ENSMUST0 | -39 | ENSMUSG0 | Gm21992 | predicted gene 21992 |
| chr19 | 4971992 | 4972332 | 341 | * | NA_peak_10 | 17 | Distal Interge | 19 | 4962374 | 4963557 | 1184 | 1 | 66419 | ENSMUST0 | 9618 | ENSMUSG0 | Mrpl11 | mitochondrial ribosomal protein L11 |
| chr19 | 5414505 | 5415010 | 506 | * | NA_peak_10 | 26 | Exon (ENSM | 19 | 5406886 | 5422209 | 15324 | 1 | 74931 | ENSMUST0 | 7619 | ENSMUSG0 | 4930481A15 | RIKEN cDNA 4930481A15 gene |
| chr19 | 5450367 | 5450502 | 136 | * | NA_peak_10 | 18 | Promoter (2- | 19 | 5447921 | 5450520 | 2600 | 1 | 14283 | ENSMUST0 | 2446 | ENSMUSG0 | Fosl1 | fos-like antigen 1 |
| chr19 | 5490209 | 5491367 | 1159 | * | NA_peak_10 | 36 | Promoter (<= | 19 | 5490455 | 5495201 | 4747 | 1 | 12631 | ENSMUST0 | 0 | ENSMUSG0 | Cfl1 | cofilin 1, non-muscle |
| chr19 | 5511043 | 5511298 | 256 | * | NA_peak_10 | 15 | Promoter (<= | 19 | 5495274 | 5510538 | 15265 | 2 | 225861 | ENSMUST0 | -505 | ENSMUSG0 | Snx32 | sorting nexin 32 |
| chr19 | 5741705 | 5742043 | 339 | * | NA_peak_10 | 43 | Promoter (<= | 19 | 5742357 | 5745883 | 3527 | 1 | 16998 | ENSMUST0 | -314 | ENSMUSG0 | Ltbp3 | latent transforming growth factor beta binding protein 3 |
| chr19 | 5781535 | 5781744 | 210 | * | NA_peak_10 | 20 | Distal Interge | 19 | 5758351 | 5771419 | 13069 | 2 | 78891 | ENSMUST0 | -10116 | ENSMUSG0 | Scyl1 | SCY1-like 1 (S. cerevisiae) |
| chr19 | 5803190 | 5803360 | 171 | * | NA_peak_10 | 19 | Promoter (<= | 19 | 5795690 | 5802672 | 6983 | 2 | 72289 | ENSMUST0 | -518 | ENSMUSG0 | Malat1 | metastasis associated lung adenocarcinoma transcript 1 (non-coding RNA) |
| chr19 | 5817918 | 5818289 | 372 | * | NA_peak_10 | 20 | Distal Interge | 19 | 5795690 | 5802672 | 6983 | 2 | 72289 | ENSMUST0 | -15246 | ENSMUSG0 | Malat1 | metastasis associated lung adenocarcinoma transcript 1 (non-coding RNA) |
| chr19 | 5848228 | 5848450 | 223 | * | NA_peak_10 | 37 | Promoter (2- | 19 | 5824708 | 5845478 | 20771 | 2 | 66961 | ENSMUST0 | -2750 | ENSMUSG0 | Neat1 | nuclear paraspeckle assembly transcript 1 (non-protein coding) |
| chr19 | 6061154 | 6061467 | 314 | * | NA_peak_10 | 22 | Promoter (<= | 19 | 6061192 | 6062472 | 1281 | 1 | 29805 | ENSMUST0 | 0 | ENSMUSG0 | Znhit2 | zinc finger, HIT domain containing 2 |
| chr19 | 6227234 | 6227536 | 303 | * | NA_peak_10 | 20 | Promoter (<= | 19 | 6226388 | 6227768 | 1381 | 1 | 170458 | ENSMUST0 | 846 | ENSMUSG0 | Gpha2 | glycoprotein hormone alpha 2 |
| chr19 | 6389586 | 6390236 | 651 | * | NA_peak_10 | 21 | Promoter (2- | 19 | 6392711 | 6394184 | 1474 | 1 | 19309 | ENSMUST0 | -2475 | ENSMUSG0 | Pygm | muscle glycogen phosphorylase |
| chr19 | 6647075 | 6647164 | 90 | * | NA_peak_10 | 26 | Exon (ENSM | 19 | 6535845 | 6543032 | 7188 | 2 | 20521 | ENSMUST0 | -104043 | ENSMUSG0 | Slc22a12 | solute carrier family 22 (organic anion/cation transporter), member 12 |
| chr19 | 6651202 | 6651715 | 514 | * | NA_peak_10 | 30 | Intron (ENSM | 19 | 6535845 | 6543032 | 7188 | 2 | 20521 | ENSMUST0 | -108170 | ENSMUSG0 | Slc22a12 | solute carrier family 22 (organic anion/cation transporter), member 12 |
| chr19 | 6814129 | 6814342 | 214 | * | NA_peak_10 | 21 | Distal Interge | 19 | 6829886 | 6829944 | 59 | 2 | 100628576 | ENSMUST0 | 15602 | ENSMUSG0 | Mir5046 | microRNA 5046 |
| chr19 | 6866168 | 6866359 | 192 | * | NA_peak_10 | 15 | Distal Interge | 19 | 6844623 | 6858211 | 13589 | 2 | 78317 | ENSMUST0 | -7957 | ENSMUSG0 | Ccdc88b | coiled-coil domain containing 88B |
| chr19 | 6888855 | 6889413 | 559 | * | NA_peak_10 | 33 | Distal Interge | 19 | 6906750 | 6908301 | 1552 | 2 | 54683 | ENSMUST0 | 18888 | ENSMUSG0 | Prdx5 | peroxiredoxin 5 |
| chr19 | 6970279 | 6970830 | 552 | * | NA_peak_10 | 34 | Promoter (<= | 19 | 6953713 | 6969814 | 16102 | 2 | 18797 | ENSMUST0 | -465 | ENSMUSG0 | Plcb3 | phospholipase C, beta 3 |
| chr19 | 7028104 | 7028288 | 185 | * | NA_peak_10 | 20 | Intron (ENSM | 19 | 6998958 | 7019469 | 20512 | 2 | 108101 | ENSMUST0 | -8635 | ENSMUSG0 | Fermt3 | fermitin family member 3 |
| chr19 | 7253296 | 7253945 | 650 | * | NA_peak_10 | 31 | Distal Interge | 19 | 7225673 | 7241183 | 15511 | 2 | 70999 | ENSMUST0 | -12113 | ENSMUSG0 | Naa40 | N(alpha)-acetyltransferase 40, NatD catalytic subunit |
| chr19 | 7269531 | 7269904 | 374 | * | NA_peak_10 | 41 | Promoter (<= | 19 | 7269764 | 7275224 | 5461 | 1 | 104383 | ENSMUST0 | 0 | ENSMUSG0 | Rcor2 | REST corepressor 2 |
| chr19 | 7370842 | 7371264 | 423 | * | NA_peak_10 | 23 | Intron (ENSM | 19 | 7357686 | 7374337 | 16652 | 2 | 68229 | ENSMUST0 | 3073 | ENSMUSG0 | Spindoc | spindlin interactor and repressor of chromatin binding |
| chr19 | 7406535 | 7406830 | 296 | * | NA_peak_10 | 18 | Distal Interge | 19 | 7417625 | 7425904 | 8280 | 1 | 108899 | ENSMUST0 | -10795 | ENSMUSG0 | Zfta | zinc finger translocation associated |
| chr19 | 7423230 | 7423771 | 542 | * | NA_peak_10 | 22 | Promoter (<= | 19 | 7422573 | 7422642 | 70 | 1 | 102466781 | ENSMUST0 | 657 | ENSMUSG0 | Mir6991 | microRNA 6991 |
| chr19 | 7455002 | 7455208 | 207 | * | NA_peak_10 | 20 | Intron (ENSM | 19 | 7425901 | 7483202 | 57302 | 2 | 20168 | ENSMUST0 | 27994 | ENSMUSG0 | Rtn3 | reticulon 3 |
| chr19 | 7482805 | 7482983 | 179 | * | NA_peak_10 | 22 | Promoter (<= | 19 | 7425901 | 7483202 | 57302 | 2 | 20168 | ENSMUST0 | 219 | ENSMUSG0 | Rtn3 | reticulon 3 |
| chr19 | 7485016 | 7485176 | 161 | * | NA_peak_10 | 17 | Promoter (1- | 19 | 7425901 | 7483281 | 57381 | 2 | 20168 | ENSMUST0 | -1735 | ENSMUSG0 | Rtn3 | reticulon 3 |
| chr19 | 7492103 | 7492562 | 460 | * | NA_peak_10 | 13 | Promoter (1- | 19 | 7494040 | 7539674 | 45635 | 1 | 109168 | ENSMUST0 | -1478 | ENSMUSG0 | Atl3 | atlastin GTPase 3 |
| chr19 | 7494654 | 7495149 | 496 | * | NA_peak_10 | 21 | Promoter (<= | 19 | 7494535 | 7534200 | 39666 | 1 | 109168 | ENSMUST0 | 119 | ENSMUSG0 | Atl3 | atlastin GTPase 3 |
| chr19 | 7627217 | 7627504 | 288 | * | NA_peak_10 | 38 | Intron (ENSM | 19 | 7612605 | 7639642 | 27038 | 1 | 66727 | ENSMUST0 | 14612 | ENSMUSG0 | Plaat5 | phospholipase A and acyltransferase 5 |
| chr19 | 7641794 | 7641799 | 6 | * | NA_peak_10 | 22 | Distal Interge | 19 | 7612605 | 7639642 | 27038 | 1 | 66727 | ENSMUST0 | 29189 | ENSMUSG0 | Plaat5 | phospholipase A and acyltransferase 5 |
| chr19 | 7651610 | 7651790 | 181 | * | NA_peak_10 | 22 | Distal Interge | 19 | 7612605 | 7639642 | 27038 | 1 | 66727 | ENSMUST0 | 39005 | ENSMUSG0 | Plaat5 | phospholipase A and acyltransferase 5 |
| chr19 | 7668261 | 7668544 | 284 | * | NA_peak_10 | 23 | Exon (ENSM | 19 | 7682277 | 7711279 | 29003 | 2 | 207151 | ENSMUST0 | 42735 | ENSMUSG0 | Slc22a19 | solute carrier family 22 (organic anion transporter), member 19 |
| chr19 | 7730784 | 7731178 | 395 | * | NA_peak_10 | 24 | Distal Interge | 19 | 7706682 | 7711310 | 4629 | 2 | 207151 | ENSMUST0 | -19474 | ENSMUSG0 | Slc22a19 | solute carrier family 22 (organic anion transporter), member 19 |

| | | | | | | | | | | | | | | | | | | |
|---|---|---|---|---|---|---|---|---|---|---|---|---|---|---|---|---|---|---|
| chr19 | 7887720 | 7887955 | 236 | * | NA_peak_10 | 13 | Intron (ENSM | 19 | 7864388 | 7966000 | 101613 | 2 | 171405 | ENSMUST00 | 78045 | ENSMUSG0 | Slc22a27 | solute carrier family 22, member 27 |
| chr19 | 8243493 | 8243829 | 337 | * | NA_peak_10 | 22 | Distal Interge | 19 | 8160304 | 8218900 | 58597 | 2 | 236293 | ENSMUST00 | -24593 | ENSMUSG0 | Slc22a29 | solute carrier family 22. member 29 |
| chr19 | 8329936 | 8330712 | 777 | * | NA_peak_10 | 23 | Distal Interge | 19 | 8335374 | 8405060 | 69687 | 2 | 319800 | ENSMUST00 | 74348 | ENSMUSG0 | Slc22a30 | solute carrier family 22, member 30 |
| chr19 | 8516243 | 8516651 | 409 | * | NA_peak_10 | 49 | Intron (ENSM | 19 | 8591254 | 8593242 | 1989 | 1 | 19879 | ENSMUST00 | -74603 | ENSMUSG0 | Slc22a8 | solute carrier family 22 (organic anion transporter), member 8 |
| chr19 | 8528267 | 8528647 | 381 | * | NA_peak_10 | 19 | Intron (ENSM | 19 | 8591254 | 8593242 | 1989 | 1 | 19879 | ENSMUST00 | -62607 | ENSMUSG0 | Slc22a8 | solute carrier family 22 (organic anion transporter), member 8 |
| chr19 | 8722893 | 8723595 | 703 | * | NA_peak_11 | 34 | Promoter (<= | 19 | 8706882 | 8723369 | 16488 | 2 | 17254 | ENSMUST00 | 0 | ENSMUSG0 | Slc3a2 | solute carrier family 3 (activators of dibasic and neutral amino acid transport), member 2 |
| chr19 | 8735397 | 8736148 | 752 | * | NA_peak_11 | 16 | Promoter (<= | 19 | 8735827 | 8740618 | 4792 | 1 | 107071 | ENSMUST00 | 0 | ENSMUSG0 | Wdr74 | WD repeat domain 74 |
| chr19 | 8769435 | 8769906 | 472 | * | NA_peak_11 | 27 | Promoter (<= | 19 | 8770092 | 8770921 | 830 | 1 | 53319 | ENSMUST00 | -186 | ENSMUSG0 | Nxf1 | nuclear RNA export factor 1 |
| chr19 | 8774669 | 8775191 | 523 | * | NA_peak_11 | 42 | Promoter (<= | 19 | 8772552 | 8774467 | 1916 | 2 | 67706 | ENSMUST00 | -202 | ENSMUSG0 | Tmem179b | transmembrane protein 179B |
| chr19 | 8819870 | 8820932 | 1063 | * | NA_peak_11 | 28 | Promoter (<= | 19 | 8820010 | 8834143 | 14134 | 1 | 68693 | ENSMUST00 | 0 | ENSMUSG0 | Hnrnpul2 | heterogeneous nuclear ribonucleoprotein U-like 2 |
| chr19 | 8853555 | 8853638 | 84 | * | NA_peak_11 | 27 | Promoter (2- | 19 | 8850785 | 8853909 | 3125 | 1 | 68852 | ENSMUST00 | 2770 | ENSMUSG0 | Lrrn4cl | LRRN4 C-terminal like |
| chr19 | 8951797 | 8951826 | 30 | * | NA_peak_11 | 24 | Promoter (2- | 19 | 8953847 | 8966210 | 12364 | 1 | 70044 | ENSMUST00 | -2021 | ENSMUSG0 | Tut1 | terminal uridylyl transferase 1, U6 snRNA-specific |
| chr19 | 8989134 | 8989343 | 210 | * | NA_peak_11 | 29 | Promoter (<= | 19 | 8989284 | 9019196 | 29913 | 1 | 66395 | ENSMUST00 | 0 | ENSMUSG0 | Ahnak | AHNAK nucleoprotein (desmoyokin) |
| chr19 | 9050100 | 9050367 | 268 | * | NA_peak_11 | 25 | Intron (ENSM | 19 | 9083636 | 9087958 | 4323 | 2 | 22287 | ENSMUST00 | 37591 | ENSMUSG0 | Scgb1a1 | secretoglobin, family 1A, member 1 (uteroglobin) |
| chr19 | 9082597 | 9082911 | 315 | * | NA_peak_11 | 22 | Distal Interge | 19 | 9083636 | 9087958 | 4323 | 2 | 22287 | ENSMUST00 | 5047 | ENSMUSG0 | Scgb1a1 | secretoglobin, family 1A, member 1 (uteroglobin) |
| chr19 | 9125112 | 9125410 | 299 | * | NA_peak_11 | 39 | Intron (ENSM | 19 | 9132459 | 9135531 | 3073 | 2 | 66514 | ENSMUST00 | 10121 | ENSMUSG0 | Asrgl1 | asparaginase like 1 |
| chr19 | 9551625 | 9551902 | 278 | * | NA_peak_11 | 26 | Distal Interge | 19 | 9557792 | 9559183 | 1392 | 2 | 619371 | ENSMUST00 | 7281 | NA | Stxbp3-ps | syntaxin-binding protein 3, pseudogene |
| chr19 | 9555037 | 9555367 | 331 | * | NA_peak_11 | 30 | Distal Interge | 19 | 9557792 | 9559183 | 1392 | 2 | 619371 | ENSMUST00 | 3816 | NA | Stxbp3-ps | syntaxin-binding protein 3, pseudogene |
| chr19 | 9607117 | 9607463 | 347 | * | NA_peak_11 | 34 | Exon (ENSM | 19 | 9557792 | 9559183 | 1392 | 2 | 619371 | ENSMUST00 | -47934 | NA | Stxbp3-ps | syntaxin-binding protein 3, pseudogene |
| chr19 | 9721287 | 9721426 | 140 | * | NA_peak_11 | 16 | Distal Interge | 19 | 9872342 | 9874923 | 2582 | 2 | 16319 | ENSMUST00 | 153497 | ENSMUSG0 | Incenp | inner centromere protein |
| chr19 | 9723963 | 9724007 | 45 | * | NA_peak_11 | 13 | Distal Interge | 19 | 9872342 | 9874923 | 2582 | 2 | 16319 | ENSMUST00 | 150916 | ENSMUSG0 | Incenp | inner centromere protein |
| chr19 | 9798606 | 9798674 | 69 | * | NA_peak_11 | 45 | Distal Interge | 19 | 9872342 | 9874923 | 2582 | 2 | 16319 | ENSMUST00 | 76249 | ENSMUSG0 | Incenp | inner centromere protein |
| chr19 | 9854577 | 9854878 | 302 | * | NA_peak_11 | 52 | Distal Interge | 19 | 9872342 | 9874923 | 2582 | 2 | 16319 | ENSMUST00 | 20045 | ENSMUSG0 | Incenp | inner centromere protein |
| chr19 | 10530124 | 10530552 | 429 | * | NA_peak_11 | 17 | Promoter (1- | 19 | 10531819 | 10535247 | 3429 | 1 | 269061 | ENSMUST00 | -1267 | ENSMUSG0 | Cpsf7 | cleavage and polyadenylation specific factor 7 |
| chr19 | 10605724 | 10605865 | 142 | * | NA_peak_11 | 31 | Promoter (<= | 19 | 10605710 | 10609960 | 4251 | 1 | 13194 | ENSMUST00 | 14 | ENSMUSG0 | Ddb1 | damage specific DNA binding protein 1 |
| chr19 | 10675878 | 10676215 | 338 | * | NA_peak_11 | 20 | Promoter (<= | 19 | 10671789 | 10676817 | 5029 | 2 | 58803 | ENSMUST00 | 602 | ENSMUSG0 | Pga5 | pepsinogen 5, group I |
| chr19 | 11005493 | 11005767 | 275 | * | NA_peak_11 | 19 | Intron (ENSM | 19 | 10978307 | 10993250 | 14944 | 2 | 545279 | ENSMUST00 | -12243 | ENSMUSG0 | Ms4a15 | membrane-spanning 4-domains, subfamily A, member 15 |
| chr19 | 11202941 | 11203172 | 232 | * | NA_peak_11 | 25 | Intron (ENSM | 19 | 11196817 | 11299886 | 103070 | 1 | 75138 | ENSMUST00 | 6124 | ENSMUSG0 | 4930526L06 | RIKEN cDNA 4930526L06 gene |
| chr19 | 11342698 | 11343069 | 372 | * | NA_peak_11 | 18 | Distal Interge | 19 | 11321505 | 11336146 | 14642 | 2 | 109225 | ENSMUST00 | -6552 | ENSMUSG0 | Ms4a7 | membrane-spanning 4-domains, subfamily A, member 7 |
| chr19 | 11424058 | 11424363 | 306 | * | NA_peak_11 | 27 | Intron (ENSM | 19 | 11414841 | 11426468 | 11628 | 1 | 64380 | ENSMUST00 | 9217 | ENSMUSG0 | Ms4a4c | membrane-spanning 4-domains, subfamily A, member 4C |
| chr19 | 12036050 | 12036143 | 94 | * | NA_peak_11 | 17 | Promoter (<= | 19 | 12035808 | 12036740 | 933 | 2 | 258675 | ENSMUST00 | 597 | ENSMUSG0 | Olfr1423 | olfactory receptor 1423 |
| chr19 | 12158872 | 12159255 | 384 | * | NA_peak_11 | 17 | Distal Interge | 19 | 12119653 | 12120700 | 1048 | 2 | 258677 | ENSMUST00 | -38172 | ENSMUSG0 | Olfr76 | olfactory receptor 76 |
| chr19 | 12769333 | 12769637 | 305 | * | NA_peak_11 | 23 | 3' UTR | 19 | 12763660 | 12765632 | 1973 | 2 | 12803 | ENSMUST00 | -3701 | ENSMUSG0 | Cntf | ciliary neurotrophic factor |
| chr19 | 13902650 | 13903077 | 428 | * | NA_peak_11 | 38 | Distal Interge | 19 | 13887221 | 13888305 | 1085 | 2 | 258627 | ENSMUST00 | -14345 | ENSMUSG0 | Olfr1504 | olfactory receptor 1504 |
| chr19 | 14245314 | 14245445 | 132 | * | NA_peak_11 | 19 | Distal Interge | 19 | 13918942 | 13920044 | 1103 | 1 | 258151 | ENSMUST00 | 326372 | ENSMUSG0 | Olfr1505 | olfactory receptor 1505 |
| chr19 | 15429616 | 15429748 | 133 | * | NA_peak_11 | 17 | Distal Interge | 19 | 15905130 | 15909712 | 4583 | 2 | 107272 | ENSMUST00 | 479964 | ENSMUSG0 | Psat1 | phosphoserine aminotransferase 1 |
| chr19 | 15699926 | 15700242 | 317 | * | NA_peak_11 | 19 | Distal Interge | 19 | 15905130 | 15909712 | 4583 | 2 | 107272 | ENSMUST00 | 209470 | ENSMUSG0 | Psat1 | phosphoserine aminotransferase 1 |
| chr19 | 15716876 | 15716898 | 23 | * | NA_peak_11 | 30 | Distal Interge | 19 | 15905130 | 15909712 | 4583 | 2 | 107272 | ENSMUST00 | 192814 | ENSMUSG0 | Psat1 | phosphoserine aminotransferase 1 |
| chr19 | 15776475 | 15776872 | 398 | * | NA_peak_11 | 24 | Distal Interge | 19 | 15905130 | 15909712 | 4583 | 2 | 107272 | ENSMUST00 | 132840 | ENSMUSG0 | Psat1 | phosphoserine aminotransferase 1 |
| chr19 | 15777635 | 15777971 | 337 | * | NA_peak_11 | 19 | Distal Interge | 19 | 15905130 | 15909712 | 4583 | 2 | 107272 | ENSMUST00 | 131741 | ENSMUSG0 | Psat1 | phosphoserine aminotransferase 1 |
| chr19 | 15956849 | 15957041 | 193 | * | NA_peak_11 | 25 | Intron (ENSM | 19 | 15915009 | 15947337 | 32329 | 2 | 107272 | ENSMUST00 | -9512 | ENSMUSG0 | Psat1 | phosphoserine aminotransferase 1 |
| chr19 | 16132249 | 16132708 | 460 | * | NA_peak_11 | 48 | Promoter (<= | 19 | 16132831 | 16387463 | 254633 | 1 | 14682 | ENSMUST00 | -123 | ENSMUSG0 | Gnaq | guanine nucleotide binding protein, alpha q polypeptide |
| chr19 | 16477712 | 16477788 | 77 | * | NA_peak_11 | 16 | Intron (ENSM | 19 | 16439300 | 16609819 | 170520 | 1 | 14675 | ENSMUST00 | 38412 | ENSMUSG0 | Gna14 | guanine nucleotide binding protein, alpha 14 |
| chr19 | 16666564 | 16666872 | 309 | * | NA_peak_11 | 30 | Promoter (1- | 19 | 16635207 | 16664648 | 29442 | 2 | 271564 | ENSMUST00 | -1916 | ENSMUSG0 | Vps13a | vacuolar protein sorting 13A |
| chr19 | 16860741 | 16861150 | 410 | * | NA_peak_11 | 23 | Distal Interge | 19 | 16872319 | 16873805 | 1487 | 2 | 14240 | ENSMUST00 | 12655 | ENSMUSG0 | Foxb2 | forkhead box B2 |
| chr19 | 17892681 | 17893025 | 345 | * | NA_peak_11 | 17 | Intron (ENSM | 19 | 17670420 | 17837632 | 167213 | 2 | 18552 | ENSMUST00 | -55049 | ENSMUSG0 | Pcsk5 | proprotein convertase subtilisin/kexin type 5 |
| chr19 | 18372988 | 18373275 | 288 | * | NA_peak_11 | 30 | Distal Interge | 19 | 18593601 | 18631718 | 38118 | 2 | 20409 | ENSMUST00 | 258443 | ENSMUSG0 | Ostf1 | osteoclast stimulating factor 1 |
| chr19 | 18570851 | 18570933 | 83 | * | NA_peak_11 | 33 | Intron (ENSM | 19 | 18593601 | 18631718 | 38118 | 2 | 20409 | ENSMUST00 | 60785 | ENSMUSG0 | Ostf1 | osteoclast stimulating factor 1 |
| chr19 | 18670493 | 18670665 | 173 | * | NA_peak_11 | 22 | Promoter (<= | 19 | 18670764 | 18707200 | 36437 | 1 | 67383 | ENSMUST00 | -99 | ENSMUSG0 | Carnmt1 | carnosine N-methyltransferase 1 |
| chr19 | 18712982 | 18713476 | 495 | * | NA_peak_11 | 17 | Promoter (<= | 19 | 18713193 | 18718428 | 5236 | 1 | 225995 | ENSMUST00 | 0 | ENSMUSG0 | D030056L22 | RIKEN cDNA D030056L22 gene |
| chr19 | 18912097 | 18912301 | 205 | * | NA_peak_11 | 19 | Distal Interge | 19 | 18840637 | 18866239 | 25603 | 1 | 225997 | ENSMUST00 | 71460 | ENSMUSG0 | Trpm6 | transient receptor potential cation channel, subfamily M, member 6 |
| chr19 | 18977842 | 18977974 | 133 | * | NA_peak_11 | 19 | Exon (ENSM | 19 | 18975962 | 19001100 | 25139 | 2 | 225998 | ENSMUST00 | 23126 | ENSMUSG0 | Rorb | RAR-related orphan receptor beta |
| chr19 | 20119217 | 20119607 | 391 | * | NA_peak_11 | 23 | Intron (ENSM | 19 | 20324856 | 20440923 | 116068 | 1 | 68994 | ENSMUST00 | -205249 | ENSMUSG0 | 1500015L24 | RIKEN cDNA 1500015L24 gene |
| chr19 | 20705169 | 20705423 | 255 | * | NA_peak_11 | 23 | Intron (ENSM | 19 | 20692953 | 20727562 | 34610 | 2 | 26358 | ENSMUST00 | 22139 | ENSMUSG0 | Aldh1a7 | aldehyde dehydrogenase family 1, subfamily A7 |
| chr19 | 21230617 | 21230985 | 369 | * | NA_peak_11 | 24 | Intron (ENSM | 19 | 21272278 | 21286840 | 14563 | 1 | 22682 | ENSMUST00 | -41293 | ENSMUSG0 | Zfand5 | zinc finger, AN1-type domain 5 |
| chr19 | 21318402 | 21318493 | 92 | * | NA_peak_11 | 30 | Distal Interge | 19 | 21276040 | 21280971 | 4932 | 1 | 22682 | ENSMUST00 | 42362 | ENSMUSG0 | Zfand5 | zinc finger, AN1-type domain 5 |
| chr19 | 21761523 | 21761731 | 209 | * | NA_peak_11 | 17 | Distal Interge | 19 | 21778342 | 21858327 | 79986 | 1 | 83921 | ENSMUST00 | -16611 | ENSMUSG0 | Cemip2 | cell migration inducing hyaluronidase 2 |
| chr19 | 21799302 | 21799617 | 316 | * | NA_peak_11 | 20 | Exon (ENSM | 19 | 21778388 | 21807902 | 29515 | 1 | 83921 | ENSMUST00 | 20914 | ENSMUSG0 | Cemip2 | cell migration inducing hyaluronidase 2 |
| chr19 | 23010540 | 23010928 | 389 | * | NA_peak_11 | 17 | Intron (ENSM | 19 | 23141226 | 23168134 | 26909 | 1 | 16601 | ENSMUST00 | -130298 | ENSMUSG0 | Klf9 | Kruppel-like factor 9 |
| chr19 | 23145401 | 23145749 | 349 | * | NA_peak_11 | 31 | Promoter (1- | 19 | 23143644 | 23165471 | 21828 | 1 | 16601 | ENSMUST00 | 1757 | ENSMUSG0 | Klf9 | Kruppel-like factor 9 |
| chr19 | 23147962 | 23148266 | 305 | * | NA_peak_11 | 17 | Promoter (1- | 19 | 23149431 | 23149551 | 121 | 1 | 100316672 | ENSMUST00 | -1165 | ENSMUSG0 | Mir1192 | microRNA 1192 |
| chr19 | 23186348 | 23186598 | 251 | * | NA_peak_11 | 18 | Distal Interge | 19 | 23213755 | 23214941 | 1187 | 2 | 226026 | ENSMUST00 | 28343 | ENSMUSG0 | Smc5 | structural maintenance of chromosomes 5 |
| chr19 | 23445449 | 23445715 | 267 | * | NA_peak_11 | 26 | Promoter (2- | 19 | 23302609 | 23448398 | 145790 | 2 | 71738 | ENSMUST00 | 2683 | ENSMUSG0 | Mamdc2 | MAM domain containing 2 |
| chr19 | 23704996 | 23705296 | 301 | * | NA_peak_11 | 23 | Intron (ENSM | 19 | 23687744 | 23721012 | 33269 | 1 | 72351 | ENSMUST00 | 17252 | ENSMUSG0 | Ptar1 | protein prenyltransferase alpha subunit repeat containing 1 |
| chr19 | 24251459 | 24251827 | 369 | * | NA_peak_11 | 19 | Distal Interge | 19 | 24261862 | 24278171 | 16310 | 2 | 14297 | ENSMUST00 | 26344 | ENSMUSG0 | Fxn | frataxin |
| chr19 | 24848642 | 24848796 | 155 | * | NA_peak_11 | 14 | Intron (ENSM | 19 | 24816231 | 24861652 | 45422 | 2 | 226041 | ENSMUST00 | 12856 | ENSMUSG0 | Pgm5 | phosphoglucomutase 5 |
| chr19 | 25102887 | 25103087 | 201 | * | NA_peak_11 | 13 | Intron (ENSM | 19 | 25077914 | 25079902 | 1989 | 1 | 76088 | ENSMUST00 | 24973 | ENSMUSG0 | Dock8 | dedicator of cytokinesis 8 |
| chr19 | 25198554 | 25198969 | 416 | * | NA_peak_11 | 20 | Intron (ENSM | 19 | 25236975 | 25434496 | 197522 | 1 | 107351 | ENSMUST00 | -38006 | ENSMUSG0 | Kank1 | KN motif and ankyrin repeat domains 1 |
| chr19 | 25255475 | 25255873 | 399 | * | NA_peak_11 | 30 | Intron (ENSM | 19 | 25236975 | 25434496 | 197522 | 1 | 107351 | ENSMUST00 | 18500 | ENSMUSG0 | Kank1 | KN motif and ankyrin repeat domains 1 |
| chr19 | 25358812 | 25359149 | 338 | * | NA_peak_11 | 21 | Intron (ENSM | 19 | 25395303 | 25409772 | 14470 | 1 | 107351 | ENSMUST00 | -36154 | ENSMUSG0 | Kank1 | KN motif and ankyrin repeat domains 1 |
| chr19 | 25500454 | 25500655 | 202 | * | NA_peak_11 | 23 | Distal Interge | 19 | 25505618 | 25604329 | 98712 | 1 | 50796 | ENSMUST00 | -4963 | ENSMUSG0 | Dmrt1 | doublesex and mab-3 related transcription factor 1 |
| chr19 | 25695172 | 25695456 | 285 | * | NA_peak_11 | 19 | Distal Interge | 19 | 25672411 | 25678991 | 6581 | 1 | 226049 | ENSMUST00 | 22761 | ENSMUSG0 | Dmrt2 | doublesex and mab-3 related transcription factor 2 |
| chr19 | 25961848 | 25962312 | 465 | * | NA_peak_11 | 26 | Distal Interge | 19 | 25672411 | 25678991 | 6581 | 1 | 226049 | ENSMUST00 | 289437 | ENSMUSG0 | Dmrt2 | doublesex and mab-3 related transcription factor 2 |
| chr19 | 26232030 | 26232205 | 176 | * | NA_peak_11 | 19 | Distal Interge | 19 | 26605050 | 26778275 | 173226 | 1 | 67155 | ENSMUST00 | -372845 | ENSMUSG0 | Smarca2 | SWI/SNF related, matrix associated, actin dependent regulator of chromatin, subfamily a, member 2 |
| chr19 | 26237530 | 26237896 | 367 | * | NA_peak_11 | 35 | Distal Interge | 19 | 26605050 | 26778275 | 173226 | 1 | 67155 | ENSMUST00 | -367154 | ENSMUSG0 | Smarca2 | SWI/SNF related, matrix associated, actin dependent regulator of chromatin, subfamily a, member 2 |
| chr19 | 26845438 | 26845569 | 132 | * | NA_peak_11 | 23 | Distal Interge | 19 | 26768801 | 26823907 | 55107 | 2 | 70942 | ENSMUST00 | -21531 | ENSMUSG0 | 4931403E22 | RIKEN cDNA 4931403E22 gene |
| chr19 | 27277541 | 27277908 | 368 | * | NA_peak_11 | 23 | Distal Interge | 19 | 27238774 | 27243828 | 5055 | 1 | 22359 | ENSMUST00 | 38767 | ENSMUSG0 | Vldlr | very low density lipoprotein receptor |
| chr19 | 27279608 | 27279762 | 155 | * | NA_peak_11 | 19 | Distal Interge | 19 | 27238774 | 27243828 | 5055 | 1 | 22359 | ENSMUST00 | 40834 | ENSMUSG0 | Vldlr | very low density lipoprotein receptor |
| chr19 | 27406202 | 27406366 | 165 | * | NA_peak_11 | 26 | Intron (ENSM | 19 | 27390061 | 27412264 | 22204 | 2 | 52874 | ENSMUST00 | 5898 | ENSMUSG0 | Pum3 | pumilio RNA-binding family member 3 |
| chr19 | 27417189 | 27417477 | 289 | * | NA_peak_11 | 26 | Promoter (2- | 19 | 27418418 | 27420099 | 1682 | 2 | 52874 | ENSMUST00 | 2622 | ENSMUSG0 | Pum3 | pumilio RNA-binding family member 3 |
| chr19 | 27493709 | 27494178 | 470 | * | NA_peak_11 | 13 | Distal Interge | 19 | 27388698 | 27429825 | 41128 | 2 | 52874 | ENSMUST00 | -63884 | ENSMUSG0 | Pum3 | pumilio RNA-binding family member 3 |
| chr19 | 27732087 | 27732443 | 357 | * | NA_peak_11 | 27 | Distal Interge | 19 | 27768141 | 27836846 | 68706 | 2 | 19726 | ENSMUST00 | 104403 | ENSMUSG0 | Rfx3 | regulatory factor X, 3 (influences HLA class II expression) |
| chr19 | 28010694 | 28010978 | 285 | * | NA_peak_11 | 43 | Promoter (<= | 19 | 27867515 | 28010790 | 143276 | 2 | 19726 | ENSMUST00 | 0 | ENSMUSG0 | Rfx3 | regulatory factor X, 3 (influences HLA class II expression) |
| chr19 | 28230492 | 28231094 | 603 | * | NA_peak_11 | 23 | Distal Interge | 19 | 28263951 | 28283906 | 19956 | 2 | 226075 | ENSMUST00 | 52812 | ENSMUSG0 | Glis3 | GLIS family zinc finger 3 |
| chr19 | 28326995 | 28327022 | 28 | * | NA_peak_11 | 23 | Promoter (<= | 19 | 28262347 | 28326006 | 63660 | 2 | 226075 | ENSMUST00 | -989 | ENSMUSG0 | Glis3 | GLIS family zinc finger 3 |
| chr19 | 28683121 | 28683597 | 477 | * | NA_peak_11 | 18 | Intron (ENSM | 19 | 28349573 | 28680077 | 330505 | 2 | 226075 | ENSMUST00 | -3044 | ENSMUSG0 | Glis3 | GLIS family zinc finger 3 |
| chr19 | 28964151 | 28964621 | 471 | * | NA_peak_11 | 16 | Promoter (<= | 19 | 28923064 | 28964154 | 41091 | 2 | 381218 | ENSMUST00 | 0 | ENSMUSG0 | Spata6l | spermatogenesis associated 6 like |
| chr19 | 28978814 | 28979095 | 282 | * | NA_peak_11 | 15 | Distal Interge | 19 | 28923530 | 28967800 | 44271 | 2 | 381218 | ENSMUST00 | -11014 | ENSMUSG0 | Spata6l | spermatogenesis associated 6 like |
| chr19 | 29245136 | 29245402 | 267 | * | NA_peak_11 | 20 | Distal Interge | 19 | 29251828 | 29313080 | 61253 | 1 | 16452 | ENSMUST00 | -6426 | ENSMUSG0 | Jak2 | Janus kinase 2 |
| chr19 | 29325066 | 29325557 | 492 | * | NA_peak_11 | 44 | Promoter (<= | 19 | 29321344 | 29325356 | 4013 | 2 | 27356 | ENSMUST00 | 0 | ENSMUSG0 | Insl6 | insulin-like 6 |
| chr19 | 29393686 | 29393726 | 41 | * | NA_peak_11 | 23 | Distal Interge | 19 | 29410919 | 29471161 | 60243 | 1 | 58205 | ENSMUST00 | -17193 | ENSMUSG0 | Pdcd1lg2 | programmed cell death 1 ligand 2 |
| chr19 | 29542871 | 29543227 | 357 | * | NA_peak_11 | 26 | Intron (ENSM | 19 | 29533236 | 29604200 | 70965 | 1 | 226089 | ENSMUST00 | 9635 | ENSMUSG0 | Ric1 | RAB6A GEF complex partner 1 |
| chr19 | 29556324 | 29557054 | 731 | * | NA_peak_11 | 61 | Intron (ENSM | 19 | 29533236 | 29604200 | 70965 | 1 | 226089 | ENSMUST00 | 23088 | ENSMUSG0 | Ric1 | RAB6A GEF complex partner 1 |
| chr19 | 29936369 | 29936709 | 341 | * | NA_peak_11 | 17 | Intron (ENSM | 19 | 29945790 | 29960704 | 14915 | 1 | 77125 | ENSMUST00 | -9081 | ENSMUSG0 | Il33 | interleukin 33 |
| chr19 | 30010734 | 30011133 | 400 | * | NA_peak_11 | 19 | Distal Interge | 19 | 30003790 | 30006018 | 2229 | 1 | 66745 | ENSMUST00 | 6944 | ENSMUSG0 | Trpd52l3 | tumor protein D52-like 3 |
| chr19 | 30057322 | 30057392 | 71 | * | NA_peak_11 | 22 | Exon (ENSM | 19 | 30073783 | 30075521 | 1739 | 1 | 109113 | ENSMUST00 | -16391 | ENSMUSG0 | Uhrf2 | ubiquitin-like, containing PHD and RING finger domains 2 |
| chr19 | 30379029 | 30379313 | 285 | * | NA_peak_11 | 17 | Distal Interge | 19 | 30232942 | 30239687 | 6746 | 1 | 17195 | ENSMUST00 | 146087 | ENSMUSG0 | Mbl2 | mannose-binding lectin (protein C) 2 |
| chr19 | 30395848 | 30396179 | 332 | * | NA_peak_11 | 19 | Distal Interge | 19 | 30538928 | 30539545 | 618 | 2 | 546723 | ENSMUST00 | 143366 | NA | Ppp1r2-ps3 | protein phosphatase 1, regulatory (inhibitor) subunit 2, pseudogene 3 |
| chr19 | 30965724 | 30966000 | 277 | * | NA_peak_11 | 17 | Intron (ENSM | 19 | 31082837 | 31087069 | 4233 | 1 | 83410 | ENSMUST00 | -116837 | ENSMUSG0 | Cstf2t | cleavage stimulation factor, 3' pre-RNA subunit 2, tau |
| chr19 | 31723286 | 31723696 | 411 | * | NA_peak_11 | 26 | Intron (ENSM | 19 | 31742114 | 31764973 | 22860 | 2 | 19091 | ENSMUST00 | 41277 | ENSMUSG0 | Prkg1 | protein kinase, cGMP-dependent, type I |
| chr19 | 31724823 | 31725107 | 285 | * | NA_peak_11 | 23 | Intron (ENSM | 19 | 31742114 | 31764973 | 22860 | 2 | 19091 | ENSMUST00 | 39866 | ENSMUSG0 | Prkg1 | protein kinase, cGMP-dependent, type I |
| chr19 | 31764579 | 31764597 | 19 | * | NA_peak_11 | 17 | Promoter (<= | 19 | 31742114 | 31764973 | 22860 | 2 | 19091 | ENSMUST00 | 376 | ENSMUSG0 | Prkg1 | protein kinase, cGMP-dependent, type I |
| chr19 | 31937335 | 31937573 | 239 | * | NA_peak_11 | 27 | Intron (ENSM | 19 | 31868781 | 31911443 | 42663 | 1 | 69865 | ENSMUST00 | 68554 | ENSMUSG0 | A1cf | APOBEC1 complementation factor |
| chr19 | 32033374 | 32033696 | 323 | * | NA_peak_11 | 47 | Intron (ENSM | 19 | 32012463 | 32016804 | 4342 | 2 | 54447 | ENSMUST00 | -16570 | ENSMUSG0 | Asah2 | N-acylsphingosine amidohydrolase 2 |
| chr19 | 32094026 | 32094338 | 313 | * | NA_peak_11 | 19 | Intron (ENSM | 19 | 31986524 | 32103124 | 116601 | 2 | 54447 | ENSMUST00 | 8786 | ENSMUSG0 | Asah2 | N-acylsphingosine amidohydrolase 2 |
| chr19 | 32153797 | 32154448 | 652 | * | NA_peak_11 | 34 | Intron (ENSM | 19 | 32124147 | 32160223 | 36077 | 2 | 208449 | ENSMUST00 | 5775 | ENSMUSG0 | Sgms1 | sphingomyelin synthase 1 |
| chr19 | 32297454 | 32297801 | 348 | * | NA_peak_11 | 21 | Intron (ENSM | 19 | 32220587 | 32289579 | 68993 | 2 | 208449 | ENSMUST00 | -7875 | ENSMUSG0 | Sgms1 | sphingomyelin synthase 1 |
| chr19 | 32368779 | 32369137 | 359 | * | NA_peak_11 | 29 | Intron (ENSM | 19 | 32289279 | 32388463 | 99185 | 2 | 208449 | ENSMUST00 | 19326 | ENSMUSG0 | Sgms1 | sphingomyelin synthase 1 |
| chr19 | 32804785 | 32804902 | 118 | * | NA_peak_11 | 23 | Intron (ENSM | 19 | 32811225 | 32817977 | 6753 | 1 | 19211 | ENSMUST00 | -6323 | ENSMUSG0 | Pten | phosphatase and tensin homolog |
| chr19 | 32956956 | 32957315 | 360 | * | NA_peak_11 | 17 | Distal Interge | 19 | 32811225 | 32817977 | 6753 | 1 | 19211 | ENSMUST00 | 145731 | ENSMUSG0 | Pten | phosphatase and tensin homolog |
| chr19 | 33483795 | 33484142 | 348 | * | NA_peak_11 | 23 | Distal Interge | 19 | 33498037 | 33517780 | 19744 | 2 | 628236 | ENSMUST00 | 33638 | ENSMUSG0 | Lipo4 | lipase, member O4 |
| chr19 | 33707242 | 33707505 | 264 | * | NA_peak_11 | 20 | Intron (ENSM | 19 | 33719670 | 33730856 | 11187 | 2 | 101055671 | ENSMUST00 | 23351 | ENSMUSG0 | Lipo2 | lipase, member O2 |
| chr19 | 33749999 | 33750454 | 456 | * | NA_peak_11 | 18 | Promoter (<= | 19 | 33719799 | 33749517 | 29719 | 2 | 101055671 | ENSMUST00 | -482 | ENSMUSG0 | Lipo2 | lipase, member O2 |
| chr19 | 33955416 | 33955783 | 368 | * | NA_peak_11 | 35 | Distal Interge | 19 | 33961187 | 33976810 | 15624 | 1 | 67717 | ENSMUST00 | -5404 | ENSMUSG0 | Lipf | lipase, gastric |
| chr19 | 34192162 | 34192666 | 505 | * | NA_peak_11 | 18 | Promoter (<= | 19 | 34192229 | 34240333 | 48105 | 1 | 76630 | ENSMUST00 | 0 | ENSMUSG0 | Stambpl1 | STAM binding protein like 1 |
| chr19 | 34219288 | 34219309 | 22 | * | NA_peak_11 | 23 | Promoter (1- | 19 | 34217624 | 34240331 | 22708 | 1 | 76630 | ENSMUST00 | 1664 | ENSMUSG0 | Stambpl1 | STAM binding protein like 1 |
| chr19 | 34281049 | 34281260 | 212 | * | NA_peak_11 | 20 | Distal Interge | 19 | 34290666 | 34327772 | 37107 | 1 | 14102 | ENSMUST00 | -9406 | ENSMUSG0 | Fas | Fas (TNF receptor superfamily member 6) |
| chr19 | 34501876 | 34502172 | 297 | * | NA_peak_11 | 33 | Promoter (<= | 19 | 34494046 | 34501853 | 7808 | 2 | 16889 | ENSMUST00 | -23 | ENSMUSG0 | Lipa | lysosomal acid lipase A |
| chr19 | 34566866 | 34567260 | 395 | * | NA_peak_11 | 30 | Intron (ENSM | 19 | 34550710 | 34576419 | 25710 | 1 | 15958 | ENSMUST00 | 16156 | ENSMUSG0 | Ifit2 | interferon-induced protein with tetratricopeptide repeats 2 |
| chr19 | 34746957 | 34747231 | 275 | * | NA_peak_11 | 26 | Promoter (<= | 19 | 34668403 | 34747289 | 78887 | 2 | 240638 | ENSMUST00 | 58 | ENSMUSG0 | Slc16a12 | solute carrier family 16 (monocarboxylic acid transporters), member 12 |
| chr19 | 34849196 | 34849453 | 258 | * | NA_peak_11 | 33 | Intron (ENSM | 19 | 34840136 | 34877849 | 37714 | 2 | 75735 | ENSMUST00 | 28396 | ENSMUSG0 | Pank1 | pantothenate kinase 1 |
| chr19 | 35237258 | 35237546 | 289 | * | NA_peak_11 | 39 | Distal Interge | 19 | 34972729 | 34975743 | 3015 | 1 | 240641 | ENSMUST00 | 264529 | ENSMUSG0 | Kif20b | kinesin family member 20B |
| chr19 | 36204442 | 36204656 | 215 | * | NA_peak_11 | 19 | Intron (ENSM | 19 | 36111965 | 36120099 | 8135 | 2 | 107765 | ENSMUST00 | -84343 | ENSMUSG0 | Ankrd1 | ankyrin repeat domain 1 (cardiac muscle) |
| chr19 | 36349389 | 36349390 | 2 | * | NA_peak_11 | 26 | Promoter (<= | 19 | 36348537 | 36355613 | 7077 | 1 | 76073 | ENSMUST00 | 852 | ENSMUSG0 | Pcgf5 | polycomb group ring finger 5 |
| chr19 | 36548930 | 36549232 | 303 | * | NA_peak_11 | 13 | Distal Interge | 19 | 36554639 | 36621132 | 66494 | 1 | 226098 | ENSMUST00 | -5407 | ENSMUSG0 | Hectd2 | HECT domain E3 ubiquitin protein ligase 2 |
| chr19 | 36553758 | 36554073 | 316 | * | NA_peak_11 | 22 | Promoter (<= | 19 | 36554639 | 36621132 | 66494 | 1 | 226098 | ENSMUST00 | -566 | ENSMUSG0 | Hectd2 | HECT domain E3 ubiquitin protein ligase 2 |
| chr19 | 36897656 | 36897749 | 94 | * | NA_peak_11 | 16 | Distal Interge | 19 | 36880573 | 36887373 | 6801 | 1 | 74493 | ENSMUST00 | 17083 | ENSMUSG0 | Tnks2 | tankyrase, TRF1-interacting ankyrin-related ADP-ribose polymerase 2 |
| chr19 | 36906234 | 36906382 | 149 | * | NA_peak_11 | 14 | Distal Interge | 19 | 36917550 | 36919599 | 2050 | 2 | 72514 | ENSMUST00 | 13217 | ENSMUSG0 | Fgfbp3 | fibroblast growth factor binding protein 3 |
| chr19 | 36962458 | 36963068 | 611 | * | NA_peak_11 | 36 | Promoter (1- | 19 | 36961315 | 36965832 | 4518 | 1 | 107182 | ENSMUST00 | 1143 | ENSMUSG0 | Btaf1 | B-TFIID TATA-box binding protein associated factor 1 |
| chr19 | 36984314 | 36984457 | 144 | * | NA_peak_11 | 23 | Intron (ENSM | 19 | 36961315 | 36965832 | 4518 | 1 | 107182 | ENSMUST00 | 22999 | ENSMUSG0 | Btaf1 | B-TFIID TATA-box binding protein associated factor 1 |
| chr19 | 37018451 | 37018700 | 250 | * | NA_peak_11 | 29 | Distal Interge | 19 | 36961315 | 36965832 | 4518 | 1 | 107182 | ENSMUST00 | 57136 | ENSMUSG0 | Btaf1 | B-TFIID TATA-box binding protein associated factor 1 |
| chr19 | 37115216 | 37115513 | 298 | * | NA_peak_11 | 35 | Intron (ENSM | 19 | 37173843 | 37196541 | 22699 | 1 | 319931 | ENSMUST00 | -58330 | ENSMUSG0 | A330032B11 | RIKEN cDNA A330032B11 gene |
| chr19 | 37128653 | 37128965 | 313 | * | NA_peak_11 | 23 | Intron (ENSM | 19 | 37173843 | 37196541 | 22699 | 1 | 319931 | ENSMUST00 | -44878 | ENSMUSG0 | A330032B11 | RIKEN cDNA A330032B11 gene |
| chr19 | 37358135 | 37358471 | 337 | * | NA_peak_11 | 14 | Distal Interge | 19 | 37376403 | 37421859 | 45457 | 1 | 16551 | ENSMUST00 | -17932 | ENSMUSG0 | Kif11 | kinesin family member 11 |
| chr19 | 37451759 | 37451768 | 10 | * | NA_peak_11 | 26 | Distal Interge | 19 | 37465254 | 37471827 | 6574 | 1 | 78640 | ENSMUST00 | -13486 | ENSMUSG0 | 1700122C19 | RIKEN cDNA 1700122C19 gene |
| chr19 | 37577810 | 37578548 | 739 | * | NA_peak_11 | 26 | Exon (ENSM | 19 | 37550459 | 37683255 | 132797 | 1 | 107371 | ENSMUST00 | 27351 | ENSMUSG0 | Exoc6 | exocyst complex component 6 |
| chr19 | 37621465 | 37621615 | 151 | * | NA_peak_11 | 16 | Intron (ENSM | 19 | 37685581 | 37693398 | 7818 | 1 | 546726 | ENSMUST00 | -63966 | ENSMUSG0 | Cyp26c1 | cytochrome P450, family 26, subfamily c, polypeptide 1 |
| chr19 | 37752274 | 37752583 | 310 | * | NA_peak_11 | 19 | Distal Interge | 19 | 37698085 | 37699008 | 924 | 1 | 13082 | ENSMUST00 | 54189 | ENSMUSG0 | Cyp26a1 | cytochrome P450, family 26, subfamily a, polypeptide 1 |
| chr19 | 37790824 | 37791030 | 207 | * | NA_peak_11 | 14 | Distal Interge | 19 | 37698085 | 37699008 | 924 | 1 | 13082 | ENSMUST00 | 92739 | ENSMUSG0 | Cyp26a1 | cytochrome P450, family 26, subfamily a, polypeptide 1 |
| chr19 | 37795414 | 37795806 | 393 | * | NA_peak_11 | 20 | Distal Interge | 19 | 37698085 | 37699008 | 924 | 1 | 13082 | ENSMUST00 | 97329 | ENSMUSG0 | Cyp26a1 | cytochrome P450, family 26, subfamily a, polypeptide 1 |
| chr19 | 38033131 | 38033210 | 80 | * | NA_peak_11 | 29 | Intron (ENSM | 19 | 37899310 | 38043351 | 144042 | 2 | 226101 | ENSMUST00 | 10141 | ENSMUSG0 | Myof | myoferlin |
| chr19 | 38124531 | 38125066 | 536 | * | NA_peak_11 | 24 | Promoter (<= | 19 | 38116630 | 38124828 | 8199 | 2 | 19662 | ENSMUST00 | 0 | ENSMUSG0 | Rbp4 | retinol binding protein 4, plasma |
| chr19 | 38342204 | 38342412 | 209 | * | NA_peak_11 | 17 | Exon (ENSM | 19 | 38395980 | 38405610 | 9631 | 1 | 240660 | ENSMUST00 | -53568 | ENSMUSG0 | Slc35g1 | solute carrier family 35, member G1 |
| chr19 | 38873383 | 38873388 | 6 | * | NA_peak_11 | 13 | Intron (ENSM | 19 | 38836677 | 38919438 | 82762 | 1 | 209478 | ENSMUST00 | 36706 | ENSMUSG0 | Tbc1d12 | TBC1D12: TBC1 domain family, member 12 |
| chr19 | 38873808 | 38874072 | 265 | * | NA_peak_11 | 13 | Intron (ENSM | 19 | 38836677 | 38919438 | 82762 | 1 | 209478 | ENSMUST00 | 37131 | ENSMUSG0 | Tbc1d12 | TBC1D12: TBC1 domain family, member 12 |
| chr19 | 38983421 | 38983629 | 209 | * | NA_peak_11 | 15 | Intron (ENSM | 19 | 39007019 | 39042693 | 35675 | 1 | 72082 | ENSMUST00 | -23390 | ENSMUSG0 | Cyp2c55 | cytochrome P450, family 2, subfamily c, polypeptide 55 |
| chr19 | 39046877 | 39047362 | 486 | * | NA_peak_11 | 23 | Distal Interge | 19 | 39061015 | 39093944 | 32930 | 1 | 72303 | ENSMUST00 | -13653 | ENSMUSG0 | Cyp2c65 | cytochrome P450, family 2, subfamily c, polypeptide 65 |
| chr19 | 40148616 | 40148826 | 211 | * | NA_peak_11 | 43 | Distal Interge | 19 | 40153353 | 40187286 | 33934 | 2 | 226105 | ENSMUST00 | 38460 | ENSMUSG0 | Cyp2c70 | cytochrome P450, family 2, subfamily c, polypeptide 70 |
| chr19 | 40197788 | 40198000 | 213 | * | NA_peak_11 | 14 | Distal Interge | 19 | 40180144 | 40187333 | 7190 | 2 | 226105 | ENSMUST00 | -10455 | ENSMUSG0 | Cyp2c70 | cytochrome P450, family 2, subfamily c, polypeptide 70 |
| chr19 | 40224328 | 40224736 | 409 | * | NA_peak_11 | 22 | Promoter (<= | 19 | 40222363 | 40223356 | 994 | 2 | 54132 | ENSMUST00 | -972 | ENSMUSG0 | Pdlim1 | PDZ and LIM domain 1 (elfin) |
| chr19 | 41086038 | 41086192 | 155 | * | NA_peak_11 | 21 | Intron (ENSM | 19 | 41062473 | 41077113 | 14641 | 2 | 226115 | ENSMUST00 | -8925 | ENSMUSG0 | Opalin | oligodendrocytic myelin paranodal and inner loop protein |
| chr19 | 41250630 | 41251015 | 386 | * | NA_peak_11 | 15 | Intron (ENSM | 19 | 41213792 | 41263980 | 50189 | 2 | 107358 | ENSMUST00 | 12965 | ENSMUSG0 | Tm9sf3 | transmembrane 9 superfamily member 3 |
| chr19 | 41906766 | 41906870 | 105 | * | NA_peak_11 | 38 | Distal Interge | 19 | 41911923 | 41918660 | 6738 | 1 | 18648 | ENSMUST00 | -5053 | ENSMUSG0 | Pgam1 | phosphoglycerate mutase 1 |
| chr19 | 41907492 | 41907503 | 12 | * | NA_peak_11 | 38 | Distal Interge | 19 | 41911923 | 41918660 | 6738 | 1 | 18648 | ENSMUST00 | -4420 | ENSMUSG0 | Pgam1 | phosphoglycerate mutase 1 |
| chr19 | 41971575 | 41971804 | 230 | * | NA_peak_11 | 34 | Promoter (1- | 19 | 41962988 | 41969741 | 6754 | 2 | 72199 | ENSMUST00 | -1834 | ENSMUSG0 | Mms19 | MMS19 cytosolic iron-sulfur assembly component |
| chr19 | 42003784 | 42003946 | 163 | * | NA_peak_11 | 25 | Intron (ENSM | 19 | 42017149 | 42033695 | 16547 | 1 | 226122 | ENSMUST00 | -13203 | ENSMUSG0 | Ubtd1 | ubiquitin domain containing 1 |
| chr19 | 42021885 | 42022221 | 337 | * | NA_peak_11 | 21 | Intron (ENSM | 19 | 42017149 | 42033695 | 16547 | 1 | 226122 | ENSMUST00 | 4736 | ENSMUSG0 | Ubtd1 | ubiquitin domain containing 1 |
| chr19 | 42624846 | 42625353 | 508 | * | NA_peak_11 | 34 | Distal Interge | 19 | 42595106 | 42612813 | 17708 | 2 | 67573 | ENSMUST00 | -12033 | ENSMUSG0 | Loxl4 | lysyl oxidase-like 4 |
| chr19 | 43476930 | 43477250 | 321 | * | NA_peak_12 | 19 | Exon (ENSM | 19 | 43471674 | 43477576 | 5903 | 1 | 83674 | ENSMUST00 | 5256 | ENSMUSG0 | Cnnm1 | cyclin M1 |
| chr19 | 43689546 | 43690008 | 463 | * | NA_peak_12 | 22 | Promoter (<= | 19 | 43689672 | 43733697 | 44026 | 1 | 93685 | ENSMUST00 | 0 | ENSMUSG0 | Entpd7 | ectonucleoside triphosphate diphosphohydrolase 7 |
| chr19 | 43753668 | 43753871 | 204 | * | NA_peak_12 | 19 | Promoter (<= | 19 | 43753137 | 43768238 | 15102 | 1 | 66388 | ENSMUST00 | 531 | ENSMUSG0 | Cutc | cutC copper transporter |
| chr19 | 43990225 | 43990379 | 155 | * | NA_peak_12 | 20 | Distal Interge | 19 | 43956307 | 43986556 | 30250 | 2 | 93721 | ENSMUST00 | -3669 | ENSMUSG0 | Cpn1 | carboxypeptidase N, polypeptide 1 |
| chr19 | 44193988 | 44194275 | 288 | * | NA_peak_12 | 21 | Promoter (1- | 19 | 44147638 | 44192442 | 44805 | 2 | 329064 | ENSMUST00 | -1546 | ENSMUSG0 | Pkd2l1 | polycystic kidney disease 2-like 1 |
| chr19 | 44239378 | 44239808 | 431 | * | NA_peak_12 | 47 | Promoter (1- | 19 | 44237815 | 44242018 | 4204 | 1 | 30049 | ENSMUST00 | 1563 | ENSMUSG0 | Scd3 | stearoyl-coenzyme A desaturase 3 |
| chr19 | 44274614 | 44275046 | 433 | * | NA_peak_12 | 36 | Intron (ENSM | 19 | 44293674 | 44306864 | 13191 | 1 | 20250 | ENSMUST00 | -18628 | ENSMUSG0 | Scd2 | stearoyl-Coenzyme A desaturase 2 |
| chr19 | 44393724 | 44394083 | 360 | * | NA_peak_12 | 27 | Distal Interge | 19 | 44397993 | 44406690 | 8698 | 2 | 20249 | ENSMUST00 | 12607 | ENSMUSG0 | Scd1 | stearoyl-Coenzyme A desaturase 1 |
| chr19 | 44790717 | 44791019 | 303 | * | NA_peak_12 | 18 | Exon (ENSM | 19 | 44808406 | 44808529 | 124 | 1 | 102466645 | ENSMUST00 | -17387 | ENSMUSG0 | Mir6405 | microRNA 6405 |
| chr19 | 44823657 | 44823947 | 291 | * | NA_peak_12 | 29 | Intron (ENSM | 19 | 44828493 | 44835938 | 7446 | 2 | 73322 | ENSMUST00 | 11991 | ENSMUSG0 | 1700039E22 | RIKEN cDNA 1700039E22 gene |
| chr19 | 44825607 | 44825937 | 331 | * | NA_peak_12 | 42 | Intron (ENSM | 19 | 44828493 | 44835938 | 7446 | 2 | 73322 | ENSMUST00 | 10001 | ENSMUSG0 | 1700039E22 | RIKEN cDNA 1700039E22 gene |
| chr19 | 44845222 | 44845707 | 486 | * | NA_peak_12 | 19 | Distal Interge | 19 | 44828493 | 44835938 | 7446 | 2 | 73322 | ENSMUST00 | -9284 | ENSMUSG0 | 1700039E22 | RIKEN cDNA 1700039E22 gene |
| chr19 | 44914239 | 44914688 | 450 | * | NA_peak_12 | 15 | Distal Interge | 19 | 44931151 | 44983188 | 52038 | 1 | 226151 | ENSMUST00 | -16463 | ENSMUSG0 | Slf2 | SMC5-SMC6 complex localization factor 2 |
| chr19 | 44931442 | 44931782 | 341 | * | NA_peak_12 | 36 | Promoter (<= | 19 | 44931160 | 44982615 | 51456 | 1 | 226151 | ENSMUST00 | 282 | ENSMUSG0 | Slf2 | SMC5-SMC6 complex localization factor 2 |
| chr19 | 45200376 | 45200467 | 92 | * | NA_peak_12 | 16 | Distal Interge | 19 | 45232684 | 45235812 | 3129 | 2 | 16814 | ENSMUST00 | 35345 | ENSMUSG0 | Lbx1 | ladybird homeobox 1 |
| chr19 | 45293566 | 45293683 | 118 | * | NA_peak_12 | 14 | Distal Interge | 19 | 45232684 | 45235812 | 3129 | 2 | 16814 | ENSMUST00 | -57754 | ENSMUSG0 | Lbx1 | ladybird homeobox 1 |
| chr19 | 45462470 | 45462672 | 203 | * | NA_peak_12 | 26 | Intron (ENSM | 19 | 45445508 | 45529293 | 83786 | 1 | 12234 | ENSMUST00 | 16962 | ENSMUSG0 | Btrc | beta-transducin repeat containing protein |
| chr19 | 45737169 | 45737229 | 61 | * | NA_peak_12 | 13 | Exon (ENSM | 19 | 45736971 | 45742684 | 5714 | 2 | 14179 | ENSMUST00 | 5455 | ENSMUSG0 | Fgf8 | fibroblast growth factor 8 |
| chr19 | 45917508 | 45917955 | 448 | * | NA_peak_12 | 18 | Intron (ENSM | 19 | 45921273 | 45936429 | 15157 | 2 | 71617 | ENSMUST00 | 18474 | ENSMUSG0 | Armh3 | armadillo-like helical domain containing 3 |
| chr19 | 45983321 | 45983832 | 512 | * | NA_peak_12 | 17 | Intron (ENSM | 19 | 45998137 | 46013386 | 15250 | 1 | 100174921 | ENSMUST00 | -14305 | ENSMUSG0 | 4930505N22 | RIKEN cDNA 4930505N22 gene |
| chr19 | 45997039 | 45997400 | 362 | * | NA_peak_12 | 23 | Promoter (<= | 19 | 45998137 | 46013386 | 15250 | 1 | 100174921 | ENSMUST00 | -737 | ENSMUSG0 | 4930505N22 | RIKEN cDNA 4930505N22 gene |
| chr19 | 46025074 | 46025410 | 337 | * | NA_peak_12 | 23 | Distal Interge | 19 | 46033745 | 46034878 | 1134 | 2 | 16825 | ENSMUST00 | 9468 | ENSMUSG0 | Ldb1 | LIM domain binding 1 |
| chr19 | 46033117 | 46033407 | 291 | * | NA_peak_12 | 23 | Promoter (1- | 19 | 46033745 | 46034878 | 1134 | 2 | 16825 | ENSMUST00 | 1471 | ENSMUSG0 | Ldb1 | LIM domain binding 1 |
| chr19 | 46039030 | 46039473 | 444 | * | NA_peak_12 | 43 | Promoter (<= | 19 | 46033425 | 46039481 | 6057 | 2 | 16825 | ENSMUST00 | 8 | ENSMUSG0 | Ldb1 | LIM domain binding 1 |
| chr19 | 46044855 | 46045377 | 523 | * | NA_peak_12 | 28 | Promoter (<= | 19 | 46031570 | 46045210 | 13641 | 2 | 16825 | ENSMUST00 | 0 | ENSMUSG0 | Ldb1 | LIM domain binding 1 |
| chr19 | 46226457 | 46226744 | 288 | * | NA_peak_12 | 25 | Intron (ENSM | 19 | 46246088 | 46286509 | 40422 | 1 | 107338 | ENSMUST00 | -19344 | ENSMUSG0 | Gbf1 | golgi-specific brefeldin A-resistance factor 1 |
| chr19 | 46340847 | 46341254 | 408 | * | NA_peak_12 | 23 | Promoter (<= | 19 | 46341121 | 46375252 | 34132 | 1 | 75146 | ENSMUST00 | 0 | ENSMUSG0 | Mfsd13a | major facilitator superfamily domain containing 13a |
| chr19 | 46414613 | 46414660 | 48 | * | NA_peak_12 | 20 | Intron (ENSM | 19 | 46401124 | 46454043 | 52920 | 1 | 24069 | ENSMUST00 | 13489 | ENSMUSG0 | Sufu | SUFU negative regulator of hedgehog signaling |
| chr19 | 46502068 | 46502205 | 138 | * | NA_peak_12 | 22 | Promoter (<= | 19 | 46501702 | 46516848 | 15147 | 1 | 93679 | ENSMUST00 | 366 | ENSMUSG0 | Trim8 | tripartite motif-containing 8 |
| chr19 | 46556132 | 46556549 | 418 | * | NA_peak_12 | 18 | Promoter (1- | 19 | 46531118 | 46558392 | 27275 | 2 | 56350 | ENSMUST00 | 1843 | ENSMUSG0 | Arl3 | ADP-ribosylation factor-like 3 |
| chr19 | 47314204 | 47314564 | 361 | * | NA_peak_12 | 23 | Promoter (<= | 19 | 47266913 | 47314751 | 47839 | 2 | 14218 | ENSMUST00 | 187 | ENSMUSG0 | Sh3pxd2a | SH3 and PX domains 2A |

| | | | | | | | | | | | | | | | | | | |
|---|---|---|---|---|---|---|---|---|---|---|---|---|---|---|---|---|---|---|
| chr19 | 48177368 | 48177578 | 211 | * | NA_peak_12 | 33 | Distal Interge | 19 | 48206025 | 48805505 | 599481 | 1 | 66673 | ENSMUST0 | -28447 | ENSMUSG0 | Sorcs3 | sortilin-related VPS10 domain containing receptor 3 |
| chr19 | 48361077 | 48361401 | 325 | * | NA_peak_12 | 23 | Intron (ENSM | 19 | 48429960 | 48655310 | 225351 | 1 | 66673 | ENSMUST0 | -68559 | ENSMUSG0 | Sorcs3 | sortilin-related VPS10 domain containing receptor 3 |
| chr19 | 49749813 | 49750105 | 293 | * | NA_peak_12 | 17 | Distal Interge | 19 | 50150967 | 50252678 | 101712 | 2 | 58178 | ENSMUST0 | 502573 | ENSMUSG0 | Sorcs1 | sortilin-related VPS10 domain containing receptor 1 |
| chr19 | 51699994 | 51700336 | 343 | * | NA_peak_12 | 35 | Distal Interge | 19 | 52264297 | 52265476 | 1180 | 1 | 16333 | ENSMUST0 | -563961 | ENSMUSG0 | Ins1 | insulin I |
| chr19 | 51891916 | 51892274 | 359 | * | NA_peak_12 | 38 | Exon (ENSM | 19 | 52264297 | 52265476 | 1180 | 1 | 16333 | ENSMUST0 | -372023 | ENSMUSG0 | Ins1 | insulin I |
| chr19 | 52435084 | 52435367 | 284 | * | NA_peak_12 | 23 | Distal Interge | 19 | 52264297 | 52265476 | 1180 | 1 | 16333 | ENSMUST0 | 170787 | ENSMUSG0 | Ins1 | insulin I |
| chr19 | 52798299 | 52798606 | 308 | * | NA_peak_12 | 34 | Distal Interge | 19 | 52923181 | 52926931 | 3751 | 1 | 100043450 | ENSMUST0 | -124575 | ENSMUSG0 | AA387883 | expressed sequence AA387883 |
| chr19 | 52931206 | 52931589 | 384 | * | NA_peak_12 | 18 | Intron (ENSM | 19 | 52923242 | 52941209 | 17968 | 1 | 100043450 | ENSMUST0 | 7964 | ENSMUSG0 | AA387883 | expressed sequence AA387883 |
| chr19 | 53051510 | 53051693 | 184 | * | NA_peak_12 | 24 | Intron (ENSM | 19 | 52991183 | 53040214 | 49032 | 2 | 170750 | ENSMUST0 | -11296 | ENSMUSG0 | Xpnpep1 | X-prolyl aminopeptidase (aminopeptidase P) 1, soluble |
| chr19 | 53310656 | 53311182 | 527 | * | NA_peak_12 | 24 | Promoter (<= | 19 | 53310518 | 53373293 | 62776 | 1 | 17859 | ENSMUST0 | 138 | ENSMUSG0 | Mxi1 | MAX interactor 1, dimerization protein |
| chr19 | 53323276 | 53323731 | 456 | * | NA_peak_12 | 22 | Exon (ENSM | 19 | 53329413 | 53373154 | 43742 | 1 | 17859 | ENSMUST0 | -5682 | ENSMUSG0 | Mxi1 | MAX interactor 1, dimerization protein |
| chr19 | 53582073 | 53582210 | 138 | * | NA_peak_12 | 35 | Distal Interge | 19 | 53600398 | 53645833 | 45436 | 1 | 13006 | ENSMUST0 | -18188 | ENSMUSG0 | Smc3 | structural maintenance of chromosomes 3 |
| chr19 | 53660520 | 53660682 | 163 | * | NA_peak_12 | 33 | Distal Interge | 19 | 53677306 | 53865380 | 188075 | 1 | 73713 | ENSMUST0 | -16624 | ENSMUSG0 | Rbm20 | RNA binding motif protein 20 |
| chr19 | 53690819 | 53691213 | 395 | * | NA_peak_12 | 23 | Intron (ENSM | 19 | 53677306 | 53865380 | 188075 | 1 | 73713 | ENSMUST0 | 13513 | ENSMUSG0 | Rbm20 | RNA binding motif protein 20 |
| chr19 | 53704362 | 53704649 | 288 | * | NA_peak_12 | 28 | Intron (ENSM | 19 | 53677306 | 53865380 | 188075 | 1 | 73713 | ENSMUST0 | 27056 | ENSMUSG0 | Rbm20 | RNA binding motif protein 20 |
| chr19 | 53730977 | 53731098 | 122 | * | NA_peak_12 | 17 | Intron (ENSM | 19 | 53677306 | 53865380 | 188075 | 1 | 73713 | ENSMUST0 | 53671 | ENSMUSG0 | Rbm20 | RNA binding motif protein 20 |
| chr19 | 53774028 | 53774481 | 454 | * | NA_peak_12 | 20 | Intron (ENSM | 19 | 53793308 | 53813573 | 20266 | 1 | 73713 | ENSMUST0 | -18827 | ENSMUSG0 | Rbm20 | RNA binding motif protein 20 |
| chr19 | 54576518 | 54576794 | 277 | * | NA_peak_12 | 46 | Distal Interge | 19 | 55069734 | 55099451 | 29718 | 2 | 14732 | ENSMUST0 | 522657 | ENSMUSG0 | Gpam | glycerol-3-phosphate acyltransferase, mitochondrial |
| chr19 | 54726832 | 54727354 | 523 | * | NA_peak_12 | 22 | Distal Interge | 19 | 55069734 | 55099451 | 29718 | 2 | 14732 | ENSMUST0 | 372097 | ENSMUSG0 | Gpam | glycerol-3-phosphate acyltransferase, mitochondrial |
| chr19 | 55286565 | 55286848 | 284 | * | NA_peak_12 | 20 | Promoter (2- | 19 | 55284334 | 55288823 | 4490 | 1 | 433256 | ENSMUST0 | 2231 | ENSMUSG0 | Acsl5 | acyl-CoA synthetase long-chain family member 5 |
| chr19 | 55758939 | 55759316 | 378 | * | NA_peak_12 | 15 | Intron (ENSM | 19 | 55742845 | 55910681 | 167837 | 1 | 21416 | ENSMUST0 | 16094 | ENSMUSG0 | Tcf7l2 | transcription factor 7 like 2, T cell specific, HMG box |
| chr19 | 56410381 | 56410771 | 391 | * | NA_peak_12 | 34 | Intron (ENSM | 19 | 56397278 | 56408721 | 11444 | 1 | 12369 | ENSMUST0 | 13103 | ENSMUSG0 | Casp7 | caspase 7 |
| chr19 | 56470877 | 56471128 | 252 | * | NA_peak_12 | 25 | Exon (ENSM | 19 | 56478901 | 56479864 | 964 | 1 | 226245 | ENSMUST0 | -7773 | ENSMUSG0 | Plekhs1 | pleckstrin homology domain containing, family S member 1 |
| chr19 | 56577319 | 56577568 | 250 | * | NA_peak_12 | 39 | Intron (ENSM | 19 | 56570797 | 56591644 | 20848 | 1 | 66866 | ENSMUST0 | 6522 | ENSMUSG0 | Nhlrc2 | NHL repeat containing 2 |
| chr19 | 56803993 | 56804280 | 288 | * | NA_peak_12 | 26 | Intron (ENSM | 19 | 56797349 | 56800986 | 3638 | 2 | 213993 | ENSMUST0 | -3007 | ENSMUSG0 | Ccdc186 | coiled-coil domain containing 186 |
| chr19 | 57334417 | 57334723 | 307 | * | NA_peak_12 | 20 | Intron (ENSM | 19 | 57037251 | 57314919 | 277669 | 2 | 226251 | ENSMUST0 | -19498 | ENSMUSG0 | Ablim1 | actin-binding LIM protein 1 |
| chr19 | 58073235 | 58073605 | 371 | * | NA_peak_12 | 20 | Intron (ENSM | 19 | 58051167 | 58051236 | 70 | 1 | 100885835 | ENSMUST0 | 22068 | ENSMUSG0 | Mir5623 | microRNA 5623 |
| chr19 | 58539975 | 58540680 | 706 | * | NA_peak_12 | 33 | Intron (ENSM | 19 | 58512022 | 58553028 | 41007 | 1 | 75645 | ENSMUST0 | 27953 | ENSMUSG0 | Ccdc172 | coiled-coil domain containing 172 |
| chr19 | 58922387 | 58922456 | 70 | * | NA_peak_12 | 22 | Intron (ENSM | 19 | 58911673 | 58911759 | 87 | 2 | 100526480 | ENSMUST0 | -10628 | ENSMUSG0 | Mir3086 | microRNA 3086 |
| chr19 | 59220780 | 59220796 | 17 | * | NA_peak_12 | 15 | Promoter (1- | 19 | 59219214 | 59237374 | 18161 | 1 | 332396 | ENSMUST0 | 1566 | ENSMUSG0 | Kcnk18 | potassium channel, subfamily K, member 18 |
| chr19 | 59224255 | 59224558 | 304 | * | NA_peak_12 | 24 | Intron (ENSM | 19 | 59219214 | 59237374 | 18161 | 1 | 332396 | ENSMUST0 | 5041 | ENSMUSG0 | Kcnk18 | potassium channel, subfamily K, member 18 |
| chr19 | 59322127 | 59322233 | 107 | * | NA_peak_12 | 38 | Intron (ENSM | 19 | 59297178 | 59345780 | 48603 | 2 | 107368 | ENSMUST0 | 23547 | ENSMUSG0 | Pdzd8 | PDZ domain containing 8 |
| chr19 | 59345928 | 59346003 | 76 | * | NA_peak_12 | 43 | Promoter (<= | 19 | 59297178 | 59345780 | 48603 | 2 | 107368 | ENSMUST0 | -148 | ENSMUSG0 | Pdzd8 | PDZ domain containing 8 |
| chr19 | 59427079 | 59427241 | 163 | * | NA_peak_12 | 17 | Exon (ENSM | 19 | 59458372 | 59465357 | 6986 | 1 | 13797 | ENSMUST0 | -31131 | ENSMUSG0 | Emx2 | empty spiracles homeobox 2 |
| chr19 | 59648349 | 59648663 | 315 | * | NA_peak_12 | 13 | Distal Interge | 19 | 59493136 | 59559391 | 66256 | 1 | 72647 | ENSMUST0 | 155213 | ENSMUSG0 | 2700089I24R | RIKEN cDNA 2700089I24 gene |
| chr19 | 60268756 | 60269178 | 423 | * | NA_peak_12 | 26 | Distal Interge | 19 | 60199435 | 60226701 | 27267 | 2 | 76539 | ENSMUST0 | -42055 | ENSMUSG0 | Fam204a | family with sequence similarity 204, member A |
| chr19 | 60380177 | 60380488 | 312 | * | NA_peak_12 | 22 | Distal Interge | 19 | 60466732 | 60468304 | 1573 | 2 | 226278 | ENSMUST0 | 87816 | ENSMUSG0 | Prlhr | prolactin releasing hormone receptor |
| chr19 | 60580407 | 60580618 | 212 | * | NA_peak_12 | 19 | Promoter (<= | 19 | 60527328 | 60580955 | 53628 | 2 | 78832 | ENSMUST0 | 337 | ENSMUSG0 | Cacul1 | CDK2 associated, cullin domain 1 |
| chr19 | 60634279 | 60634659 | 381 | * | NA_peak_12 | 19 | Distal Interge | 19 | 60524696 | 60581025 | 56330 | 2 | 78832 | ENSMUST0 | -53254 | ENSMUSG0 | Cacul1 | CDK2 associated, cullin domain 1 |
| chr19 | 60644290 | 60644565 | 276 | * | NA_peak_12 | 39 | Distal Interge | 19 | 60524696 | 60581025 | 56330 | 2 | 78832 | ENSMUST0 | -63265 | ENSMUSG0 | Cacul1 | CDK2 associated, cullin domain 1 |
| chr19 | 61137304 | 61137619 | 316 | * | NA_peak_12 | 35 | Intron (ENSM | 19 | 61118131 | 61140762 | 22632 | 2 | 414758 | ENSMUST0 | 3143 | ENSMUSG0 | Zfp950 | zinc finger protein 950 |
| chr19 | 61239776 | 61240144 | 369 | * | NA_peak_12 | 21 | Distal Interge | 19 | 61226843 | 61228429 | 1587 | 2 | 12982 | ENSMUST0 | -11347 | ENSMUSG0 | Csf2ra | colony stimulating factor 2 receptor, alpha, low-affinity (granulocyte-macrophage) |
| chr19 | 61257617 | 61257881 | 265 | * | NA_peak_12 | 19 | Exon (ENSM | 19 | 61226843 | 61228429 | 1587 | 2 | 12982 | ENSMUST0 | -29188 | ENSMUSG0 | Csf2ra | colony stimulating factor 2 receptor, alpha, low-affinity (granulocyte-macrophage) |
| chr19 | 61266553 | 61267134 | 582 | * | NA_peak_12 | 59 | Distal Interge | 19 | 61226843 | 61228429 | 1587 | 2 | 12982 | ENSMUST0 | -38124 | ENSMUSG0 | Csf2ra | colony stimulating factor 2 receptor, alpha, low-affinity (granulocyte-macrophage) |
| chr2 | 3845240 | 3845504 | 265 | * | NA_peak_12 | 28 | Distal Interge | 2 | 3773946 | 3779625 | 5680 | 1 | 66540 | ENSMUST0 | 71294 | ENSMUSG0 | Fam107b | family with sequence similarity 107, member B |
| chr2 | 5951642 | 5951797 | 156 | * | NA_peak_12 | 40 | Promoter (<= | 2 | 5951469 | 6056703 | 105235 | 1 | 326622 | ENSMUST0 | 173 | ENSMUSG0 | Upf2 | UPF2 regulator of nonsense transcripts homolog (yeast) |
| chr2 | 7921775 | 7922067 | 293 | * | NA_peak_12 | 35 | Distal Interge | 2 | 7394076 | 7509563 | 115488 | 2 | 14007 | ENSMUST0 | -412212 | ENSMUSG0 | Celf2 | CUGBP, Elav-like family member 2 |
| chr2 | 8185670 | 8186576 | 907 | * | NA_peak_12 | 33 | Distal Interge | 2 | 7394076 | 7509563 | 115488 | 2 | 14007 | ENSMUST0 | -676107 | ENSMUSG0 | Celf2 | CUGBP, Elav-like family member 2 |
| chr2 | 13615999 | 13616573 | 575 | * | NA_peak_12 | 57 | Distal Interge | 2 | 13579369 | 13582478 | 3110 | 1 | 22352 | ENSMUST0 | 36630 | ENSMUSG0 | Vim | vimentin |
| chr2 | 15023920 | 15024129 | 210 | * | NA_peak_12 | 15 | Intron (ENSM | 2 | 15047221 | 15049279 | 2059 | 2 | 74455 | ENSMUST0 | 25150 | ENSMUSG0 | Nsun6 | NOL1/NOP2/Sun domain family member 6 |
| chr2 | 15377981 | 15378396 | 416 | * | NA_peak_12 | 43 | Distal Interge | 2 | 15526479 | 16255555 | 729077 | 1 | 102635496 | ENSMUST0 | -148083 | ENSMUSG0 | Malrd1 | MAM and LDL receptor class A domain containing 1 |
| chr2 | 15433895 | 15434095 | 201 | * | NA_peak_12 | 20 | Distal Interge | 2 | 15526479 | 16255555 | 729077 | 1 | 102635496 | ENSMUST0 | -92384 | ENSMUSG0 | Malrd1 | MAM and LDL receptor class A domain containing 1 |
| chr2 | 20387870 | 20388177 | 308 | * | NA_peak_12 | 40 | Intron (ENSM | 2 | 20339824 | 20661918 | 322095 | 1 | 208618 | ENSMUST0 | 48046 | ENSMUSG0 | Etl4 | enhancer trap locus 4 |
| chr2 | 22077763 | 22078179 | 417 | * | NA_peak_12 | 57 | Distal Interge | 2 | 22227503 | 22618252 | 390750 | 1 | 667663 | ENSMUST0 | -149324 | ENSMUSG0 | Myo3a | myosin IIIA |
| chr2 | 22587305 | 22588115 | 811 | * | NA_peak_12 | 157 | Exon (ENSM | 2 | 22622205 | 22693874 | 71670 | 1 | 14417 | ENSMUST0 | -34090 | ENSMUSG0 | Gad2 | glutamic acid decarboxylase 2 |
| chr2 | 23786169 | 23786280 | 112 | * | NA_peak_12 | 23 | Distal Interge | 2 | 23543243 | 23572106 | 28864 | 2 | 76857 | ENSMUST0 | -214063 | ENSMUSG0 | Spopl | speckle-type BTB/POZ protein-like |
| chr2 | 27803580 | 27803726 | 147 | * | NA_peak_12 | 33 | Distal Interge | 2 | 27740201 | 27748671 | 8471 | 1 | 20181 | ENSMUST0 | 63379 | ENSMUSG0 | Rxra | retinoid X receptor alpha |
| chr2 | 30048653 | 30049143 | 491 | * | NA_peak_12 | 23 | Promoter (<= | 2 | 30031546 | 30048881 | 17336 | 2 | 71820 | ENSMUST0 | 0 | ENSMUSG0 | Dync2i2 | dynein 2 intermediate chain 2 |
| chr2 | 30237906 | 30238116 | 211 | * | NA_peak_12 | 29 | Promoter (<= | 2 | 30236795 | 30238223 | 1429 | 2 | 67350 | ENSMUST0 | 107 | NA | 1700084E18I | RIKEN cDNA 1700084E18 gene |
| chr2 | 30600177 | 30600310 | 134 | * | NA_peak_12 | 32 | Intron (ENSM | 2 | 30605217 | 30608217 | 3001 | 1 | 78617 | ENSMUST0 | -4907 | ENSMUSG0 | Cstad | CSA-conditional, T cell activation-dependent protein |
| chr2 | 31136034 | 31136064 | 31 | * | NA_peak_12 | 26 | Intron (ENSM | 2 | 31026206 | 31141802 | 115597 | 2 | 14269 | ENSMUST0 | 5738 | ENSMUSG0 | Fnbp1 | formin binding protein 1 |
| chr2 | 32535733 | 32535994 | 262 | * | NA_peak_12 | 26 | Promoter (<= | 2 | 32535733 | 32558101 | 22369 | 1 | 98952 | ENSMUST0 | 0 | ENSMUSG0 | Fam102a | family with sequence similarity 102, member A |
| chr2 | 32981340 | 32981534 | 195 | * | NA_peak_12 | 23 | Promoter (<= | 2 | 32973585 | 32982017 | 8433 | 2 | 56017 | ENSMUST0 | 483 | ENSMUSG0 | Slc2a8 | solute carrier family 2, (facilitated glucose transporter), member 8 |
| chr2 | 33130316 | 33130436 | 121 | * | NA_peak_12 | 20 | Promoter (<= | 2 | 33054216 | 33130608 | 76393 | 2 | 99326 | ENSMUST0 | 172 | ENSMUSG0 | Garnl3 | GTPase activating RANGAP domain-like 3 |
| chr2 | 35256212 | 35256766 | 555 | * | NA_peak_12 | 49 | Promoter (<= | 2 | 35256380 | 35307890 | 51511 | 1 | 227753 | ENSMUST0 | 0 | ENSMUSG0 | Gsn | gelsolin |
| chr2 | 35544633 | 35544862 | 230 | * | NA_peak_12 | 21 | Distal Interge | 2 | 35558266 | 35730994 | 172729 | 1 | 69601 | ENSMUST0 | -13404 | ENSMUSG0 | Dab2ip | disabled 2 interacting protein |
| chr2 | 35711747 | 35712076 | 330 | * | NA_peak_12 | 17 | Promoter (1- | 2 | 35710327 | 35730994 | 20668 | 1 | 69601 | ENSMUST0 | 1420 | ENSMUSG0 | Dab2ip | disabled 2 interacting protein |
| chr2 | 35979502 | 35979666 | 165 | * | NA_peak_12 | 13 | Promoter (<= | 2 | 35751241 | 35979624 | 228384 | 2 | 74410 | ENSMUST0 | 0 | ENSMUSG0 | Ttll11 | tubulin tyrosine ligase-like family, member 11 |
| chr2 | 36442187 | 36442567 | 381 | * | NA_peak_12 | 39 | Distal Interge | 2 | 36452587 | 36453525 | 939 | 1 | 258953 | ENSMUST0 | -10020 | ENSMUSG0 | Olfr340 | olfactory receptor 340 |
| chr2 | 38768622 | 38768890 | 269 | * | NA_peak_12 | 23 | Intron (ENSM | 2 | 38742957 | 38763765 | 20809 | 2 | 14536 | ENSMUST0 | -4857 | ENSMUSG0 | Nr6a1 | nuclear receptor subfamily 6, group A, member 1 |
| chr2 | 40236300 | 40236963 | 664 | * | NA_peak_12 | 70 | Distal Interge | 2 | 40595246 | 40615251 | 20006 | 2 | 94217 | ENSMUST0 | 378288 | ENSMUSG0 | Lrp1b | low density lipoprotein-related protein 1B |
| chr2 | 40790698 | 40791000 | 303 | * | NA_peak_12 | 17 | Intron (ENSM | 2 | 40595534 | 40702800 | 107267 | 2 | 94217 | ENSMUST0 | -87898 | ENSMUSG0 | Lrp1b | low density lipoprotein-related protein 1B |
| chr2 | 42214351 | 42214855 | 505 | * | NA_peak_12 | 31 | Intron (ENSM | 2 | 42447439 | 42447568 | 130 | 2 | 102465171 | ENSMUST0 | 232713 | ENSMUSG0 | Mir6336 | microRNA 6336 |
| chr2 | 42952222 | 42952585 | 364 | * | NA_peak_12 | 40 | Distal Interge | 2 | 40595248 | 42653624 | 2058377 | 2 | 94217 | ENSMUST0 | -298598 | ENSMUSG0 | Lrp1b | low density lipoprotein-related protein 1B |
| chr2 | 43409564 | 43409791 | 228 | * | NA_peak_12 | 46 | Distal Interge | 2 | 43555329 | 43563496 | 8168 | 1 | 70789 | ENSMUST0 | -145538 | ENSMUSG0 | Kynu | kynureninase |
| chr2 | 44861786 | 44862141 | 356 | * | NA_peak_12 | 36 | Promoter (<= | 2 | 44591910 | 44862275 | 270366 | 2 | 227835 | ENSMUST0 | 134 | ENSMUSG0 | Gtdc1 | glycosyltransferase-like domain containing 1 |
| chr2 | 48814672 | 48814955 | 284 | * | NA_peak_12 | 30 | Promoter (<= | 2 | 48814109 | 48903269 | 89161 | 1 | 11480 | ENSMUST0 | 563 | ENSMUSG0 | Acvr2a | activin receptor IIA |
| chr2 | 49477952 | 49478325 | 374 | * | NA_peak_12 | 65 | Intron (ENSM | 2 | 49451486 | 49551948 | 100463 | 1 | 227867 | ENSMUST0 | 26466 | ENSMUSG0 | Epc2 | enhancer of polycomb homolog 2 |
| chr2 | 49478849 | 49479076 | 228 | * | NA_peak_12 | 65 | Intron (ENSM | 2 | 49451486 | 49551948 | 100463 | 1 | 227867 | ENSMUST0 | 27363 | ENSMUSG0 | Epc2 | enhancer of polycomb homolog 2 |
| chr2 | 50602339 | 50602523 | 185 | * | NA_peak_12 | 17 | Distal Interge | 2 | 50380400 | 50412610 | 32211 | 1 | 102634280 | ENSMUST0 | 221939 | NA | Gm48908 | predicted gene 48908 |
| chr2 | 53507930 | 53508116 | 187 | * | NA_peak_12 | 26 | Distal Interge | 2 | 53192088 | 53219220 | 27133 | 1 | 65103 | ENSMUST0 | 315842 | ENSMUSG0 | Arl6ip6 | ADP-ribosylation factor-like 6 interacting protein 6 |
| chr2 | 54887842 | 54887919 | 78 | * | NA_peak_12 | 26 | Intron (ENSM | 2 | 54516540 | 55112961 | 596422 | 1 | 271786 | ENSMUST0 | 371302 | ENSMUSG0 | Galnt13 | polypeptide N-acetylgalactosaminyltransferase 13 |
| chr2 | 59479181 | 59479210 | 30 | * | NA_peak_12 | 13 | Distal Interge | 2 | 59484653 | 59505020 | 20368 | 1 | 76747 | ENSMUST0 | -5443 | ENSMUSG0 | Dapl1 | death associated protein-like 1 |
| chr2 | 59585368 | 59585634 | 267 | * | NA_peak_12 | 13 | Distal Interge | 2 | 59612042 | 59845717 | 233676 | 1 | 66860 | ENSMUST0 | -26408 | ENSMUSG0 | Tanc1 | tetratricopeptide repeat, ankyrin repeat and coiled-coil containing 1 |
| chr2 | 61572047 | 61572180 | 134 | * | NA_peak_12 | 26 | Distal Interge | 2 | 61578585 | 61654171 | 75587 | 1 | 21353 | ENSMUST0 | -6405 | ENSMUSG0 | Tank | TRAF family member-associated Nf-kappa B activator |
| chr2 | 62195921 | 62196246 | 326 | * | NA_peak_12 | 57 | Intron (ENSM | 2 | 62147862 | 62158283 | 10422 | 1 | 94229 | ENSMUST0 | 48059 | ENSMUSG0 | Slc4a10 | solute carrier family 4, sodium bicarbonate cotransporter-like, member 10 |
| chr2 | 63645447 | 63645502 | 56 | * | NA_peak_12 | 22 | Distal Interge | 2 | 63980447 | 64097988 | 117542 | 2 | 60344 | ENSMUST0 | 452486 | ENSMUSG0 | Fign | fidgetin |
| chr2 | 64199861 | 64199957 | 97 | * | NA_peak_12 | 26 | Distal Interge | 2 | 63977337 | 64098038 | 120702 | 2 | 60344 | ENSMUST0 | -101823 | ENSMUSG0 | Fign | fidgetin |
| chr2 | 64313114 | 64313255 | 142 | * | NA_peak_12 | 20 | Distal Interge | 2 | 63977337 | 64098038 | 120702 | 2 | 60344 | ENSMUST0 | -215076 | ENSMUSG0 | Fign | fidgetin |
| chr2 | 66745513 | 66746075 | 563 | * | NA_peak_13 | 44 | Intron (ENSM | 2 | 66673425 | 66784914 | 111490 | 2 | 20272 | ENSMUST0 | 38839 | ENSMUSG0 | Scn7a | sodium channel, voltage-gated, type VII, alpha |
| chr2 | 67941350 | 67941499 | 150 | * | NA_peak_13 | 31 | Intron (ENSM | 2 | 67781043 | 67781140 | 98 | 1 | 102466627 | ENSMUST0 | 160307 | ENSMUSG0 | Mir6335 | microRNA 6335 |
| chr2 | 74370354 | 74370654 | 301 | * | NA_peak_13 | 23 | Distal Interge | 2 | 74298940 | 74307717 | 8778 | 2 | 73962 | ENSMUST0 | -62637 | ENSMUSG0 | 4930441J16R | RIKEN cDNA 4930441J16 gene |
| chr2 | 74586415 | 74586501 | 87 | * | NA_peak_13 | 16 | Distal Interge | 2 | 74548024 | 74579435 | 31412 | 2 | 69605 | ENSMUST0 | -6980 | ENSMUSG0 | Lnpk | lunapark, ER junction formation factor |
| chr2 | 80816439 | 80816794 | 356 | * | NA_peak_13 | 22 | Distal Interge | 2 | 80644204 | 80653414 | 9211 | 1 | 69482 | ENSMUST0 | 172235 | ENSMUSG0 | Nup35 | nucleoporin 35 |
| chr2 | 82489467 | 82489874 | 408 | * | NA_peak_13 | 43 | Distal Interge | 2 | 82375372 | 82392202 | 16831 | 2 | 228116 | ENSMUST0 | -97265 | ENSMUSG0 | A130030D18 | RIKEN cDNA A130030D18 gene |
| chr2 | 82491866 | 82492073 | 208 | * | NA_peak_13 | 43 | Distal Interge | 2 | 82375372 | 82392202 | 16831 | 2 | 228116 | ENSMUST0 | -99664 | ENSMUSG0 | A130030D18 | RIKEN cDNA A130030D18 gene |
| chr2 | 82919078 | 82919428 | 351 | * | NA_peak_13 | 13 | Distal Interge | 2 | 82943634 | 83008937 | 65304 | 1 | 241516 | ENSMUST0 | -24206 | ENSMUSG0 | Fsip2 | fibrous sheath-interacting protein 2 |
| chr2 | 84872182 | 84872494 | 313 | * | NA_peak_13 | 22 | Exon (ENSM | 2 | 84880523 | 84886476 | 5954 | 2 | 269295 | ENSMUST0 | 13982 | ENSMUSG0 | Rtn4rl2 | reticulon 4 receptor-like 2 |
| chr2 | 89898020 | 89898227 | 208 | * | NA_peak_13 | 43 | Distal Interge | 2 | 89880794 | 89881809 | 1016 | 1 | 258984 | ENSMUST0 | 17226 | ENSMUSG0 | Olfr1257 | olfactory receptor 1257 |
| chr2 | 92914856 | 92915116 | 261 | * | NA_peak_13 | 23 | Promoter (<= | 2 | 92915098 | 92956058 | 40961 | 1 | 80976 | ENSMUST0 | 0 | ENSMUSG0 | Syt13 | synaptotagmin XIII |
| chr2 | 94885767 | 94885906 | 140 | * | NA_peak_13 | 35 | Distal Interge | 2 | 94422336 | 94438136 | 15801 | 2 | 11800 | ENSMUST0 | -447631 | ENSMUSG0 | Api5 | apoptosis inhibitor 5 |
| chr2 | 98633935 | 98634386 | 452 | * | NA_peak_13 | 31 | Distal Interge | 2 | 97468089 | 97629063 | 160975 | 1 | 241568 | ENSMUST0 | 1165846 | ENSMUSG0 | Lrrc4c | leucine rich repeat containing 4C |
| chr2 | 98666847 | 98667170 | 324 | * | NA_peak_13 | 14 | Exon (ENSM | 2 | 97468089 | 97629063 | 160975 | 1 | 241568 | ENSMUST0 | 1198758 | ENSMUSG0 | Lrrc4c | leucine rich repeat containing 4C |
| chr2 | 105862813 | 105863146 | 334 | * | NA_peak_13 | 16 | Promoter (2- | 2 | 105731448 | 105865982 | 134535 | 2 | 77766 | ENSMUST0 | 2836 | ENSMUSG0 | Elp4 | elongator acetyltransferase complex subunit 4 |
| chr2 | 108121320 | 108121514 | 195 | * | NA_peak_13 | 29 | Distal Interge | 2 | 107291395 | 107294472 | 3078 | 1 | 16492 | ENSMUST0 | 829925 | ENSMUSG0 | Kcna4 | potassium voltage-gated channel, shaker-related subfamily, member 4 |
| chr2 | 110125521 | 110125875 | 355 | * | NA_peak_13 | 44 | Intron (ENSM | 2 | 110031251 | 110050479 | 19229 | 2 | 75517 | ENSMUST0 | -75042 | ENSMUSG0 | Ccdc34os | coiled-coil domain containing 34, opposite strand |
| chr2 | 110126360 | 110126649 | 290 | * | NA_peak_13 | 39 | Intron (ENSM | 2 | 110031251 | 110050479 | 19229 | 2 | 75517 | ENSMUST0 | -75881 | ENSMUSG0 | Ccdc34os | coiled-coil domain containing 34, opposite strand |
| chr2 | 110157382 | 110157707 | 326 | * | NA_peak_13 | 65 | Intron (ENSM | 2 | 110031251 | 110050479 | 19229 | 2 | 75517 | ENSMUST0 | -106903 | ENSMUSG0 | Ccdc34os | coiled-coil domain containing 34, opposite strand |
| chr2 | 111720373 | 111720832 | 460 | * | NA_peak_13 | 29 | Distal Interge | 2 | 111754251 | 111755189 | 939 | 1 | 258889 | ENSMUST0 | -33419 | ENSMUSG0 | Olfr1301 | olfactory receptor 1301 |
| chr2 | 116454920 | 116455262 | 343 | * | NA_peak_13 | 18 | Distal Interge | 2 | 116075165 | 116076096 | 932 | 2 | 69974 | ENSMUST0 | -378824 | NA | 2810405F15I | RIKEN cDNA 2810405F15 gene |
| chr2 | 120563651 | 120563868 | 218 | * | NA_peak_13 | 44 | Promoter (<= | 2 | 120506820 | 120563807 | 56988 | 2 | 20402 | ENSMUST0 | 0 | ENSMUSG0 | Zfp106 | zinc finger protein 106 |
| chr2 | 122028641 | 122029004 | 364 | * | NA_peak_13 | 44 | Promoter (<= | 2 | 122028652 | 122029328 | 677 | 1 | 100042807 | ENSMUST0 | 0 | ENSMUSG0 | Eif3j2 | eukaryotic translation initiation factor 3, subunit J2 |
| chr2 | 122110207 | 122110567 | 361 | * | NA_peak_13 | 18 | Intron (ENSM | 2 | 122093540 | 122118296 | 24757 | 2 | 214585 | ENSMUST0 | 7729 | ENSMUSG0 | Spg11 | SPG11, spatacsin vesicle trafficking associated |
| chr2 | 122127386 | 122127630 | 245 | * | NA_peak_13 | 21 | Exon (ENSM | 2 | 122053520 | 122118386 | 64867 | 2 | 214585 | ENSMUST0 | -9000 | ENSMUSG0 | Spg11 | SPG11, spatacsin vesicle trafficking associated |
| chr2 | 122366163 | 122366493 | 331 | * | NA_peak_13 | 18 | Promoter (<= | 2 | 122348892 | 122366828 | 17937 | 2 | 435684 | ENSMUST0 | 335 | ENSMUSG0 | Shf | Src homology 2 domain containing F |
| chr2 | 126240650 | 126240982 | 333 | * | NA_peak_13 | 29 | Distal Interge | 2 | 126154541 | 126165279 | 10739 | 1 | 69185 | ENSMUST0 | 86109 | ENSMUSG0 | Dtwd1 | DTW domain containing 1 |
| chr2 | 126244048 | 126244089 | 42 | * | NA_peak_13 | 14 | Distal Interge | 2 | 126154541 | 126165279 | 10739 | 1 | 69185 | ENSMUST0 | 89507 | ENSMUSG0 | Dtwd1 | DTW domain containing 1 |
| chr2 | 126244541 | 126244847 | 307 | * | NA_peak_13 | 14 | Distal Interge | 2 | 126154541 | 126165279 | 10739 | 1 | 69185 | ENSMUST0 | 90000 | ENSMUSG0 | Dtwd1 | DTW domain containing 1 |
| chr2 | 127674514 | 127674936 | 423 | * | NA_peak_13 | 13 | Distal Interge | 2 | 127633226 | 127656695 | 23470 | 2 | 17153 | ENSMUST0 | -17819 | ENSMUSG0 | Mal | myelin and lymphocyte protein, T cell differentiation protein |
| chr2 | 128052673 | 128053082 | 410 | * | NA_peak_13 | 38 | Intron (ENSM | 2 | 128126038 | 128130015 | 3978 | 1 | 12125 | ENSMUST0 | -72956 | ENSMUSG0 | Bcl2l11 | BCL2-like 11 (apoptosis facilitator) |
| chr2 | 128134995 | 128135283 | 289 | * | NA_peak_13 | 26 | Intron (ENSM | 2 | 128128131 | 128128737 | 607 | 1 | 12125 | ENSMUST0 | 6864 | ENSMUSG0 | Bcl2l11 | BCL2-like 11 (apoptosis facilitator) |
| chr2 | 131919158 | 131919465 | 308 | * | NA_peak_13 | 34 | Intron (ENSM | 2 | 131909957 | 131956130 | 46174 | 1 | 111368 | ENSMUST0 | 9201 | ENSMUSG0 | Prn | prion protein readthrough transcript |
| chr2 | 133969938 | 133970166 | 229 | * | NA_peak_13 | 39 | Distal Interge | 2 | 133564708 | 133564794 | 87 | 1 | 100526500 | ENSMUST0 | 405230 | ENSMUSG0 | Mir3090 | microRNA 3090 |
| chr2 | 134507951 | 134508233 | 283 | * | NA_peak_13 | 49 | Intron (ENSM | 2 | 134530760 | 134554307 | 23548 | 2 | 15112 | ENSMUST0 | 46074 | ENSMUSG0 | Hao1 | hydroxyacid oxidase 1, liver |
| chr2 | 134664108 | 134664300 | 193 | * | NA_peak_13 | 23 | Distal Interge | 2 | 134594185 | 134644145 | 49961 | 2 | 52837 | ENSMUST0 | -19963 | ENSMUSG0 | Tmx4 | thioredoxin-related transmembrane protein 4 |
| chr2 | 135516234 | 135516503 | 270 | * | NA_peak_13 | 22 | Intron (ENSM | 2 | 135580530 | 135583296 | 2767 | 2 | 320684 | ENSMUST0 | 66793 | ENSMUSG0 | 9630028H03 | RIKEN cDNA 9630028H03 gene |
| chr2 | 139403132 | 139403632 | 501 | * | NA_peak_13 | 18 | Intron (ENSM | 2 | 139493913 | 139637056 | 143144 | 1 | 228677 | ENSMUST0 | -90281 | ENSMUSG0 | Sptlc3 | serine palmitoyltransferase, long chain base subunit 3 |
| chr2 | 139472226 | 139472361 | 136 | * | NA_peak_13 | 22 | Intron (ENSM | 2 | 139493913 | 139637056 | 143144 | 1 | 228677 | ENSMUST0 | -21552 | ENSMUSG0 | Sptlc3 | serine palmitoyltransferase, long chain base subunit 3 |
| chr2 | 139473407 | 139473667 | 261 | * | NA_peak_13 | 26 | Intron (ENSM | 2 | 139493913 | 139637056 | 143144 | 1 | 228677 | ENSMUST0 | -20246 | ENSMUSG0 | Sptlc3 | serine palmitoyltransferase, long chain base subunit 3 |
| chr2 | 142901354 | 142901550 | 197 | * | NA_peak_13 | 18 | Promoter (<= | 2 | 142617474 | 142901519 | 284046 | 2 | 16558 | ENSMUST0 | 0 | ENSMUSG0 | Kif16b | kinesin family member 16B |
| chr2 | 143429126 | 143429481 | 356 | * | NA_peak_13 | 24 | Distal Interge | 2 | 143537970 | 143545977 | 8008 | 2 | 378776 | ENSMUST0 | 116496 | ENSMUSG0 | Pcsk2os1 | proprotein convertase subtilisin/kexin type 2, opposite strand 1 |
| chr2 | 144149292 | 144149490 | 199 | * | NA_peak_13 | 26 | Intron (ENSM | 2 | 144174315 | 144179086 | 4772 | 2 | 433481 | ENSMUST0 | 29596 | ENSMUSG0 | Gm5535 | predicted gene 5535 |
| chr2 | 144154835 | 144155134 | 300 | * | NA_peak_13 | 27 | Intron (ENSM | 2 | 144174315 | 144179086 | 4772 | 2 | 433481 | ENSMUST0 | 23952 | ENSMUSG0 | Gm5535 | predicted gene 5535 |
| chr2 | 145140213 | 145140543 | 331 | * | NA_peak_13 | 38 | Distal Interge | 2 | 145167754 | 145642166 | 474413 | 1 | 94249 | ENSMUST0 | -27211 | ENSMUSG0 | Slc24a3 | solute carrier family 24 (sodium/potassium/calcium exchanger), member 3 |
| chr2 | 146160827 | 146161141 | 315 | * | NA_peak_13 | 18 | Intron (ENSM | 2 | 146152344 | 146214960 | 62617 | 1 | 78774 | ENSMUST0 | 8483 | ENSMUSG0 | Cfap61 | cilia and flagella associated protein 61 |
| chr2 | 148411378 | 148411779 | 402 | * | NA_peak_13 | 13 | Distal Interge | 2 | 148404466 | 148408188 | 3723 | 2 | 21824 | ENSMUST0 | -3190 | ENSMUSG0 | Thbd | thrombomodulin |
| chr2 | 150443719 | 150444238 | 520 | * | NA_peak_13 | 13 | Intron (ENSM | 2 | 150407141 | 150451486 | 44346 | 2 | 668923 | ENSMUST0 | 7248 | ENSMUSG0 | Zfp442 | zinc finger protein 442 |
| chr2 | 152414796 | 152414970 | 175 | * | NA_peak_13 | 36 | Promoter (<= | 2 | 152411955 | 152415027 | 3073 | 2 | 67917 | ENSMUST0 | 57 | ENSMUSG0 | Zcchc3 | zinc finger, CCHC domain containing 3 |
| chr2 | 153140980 | 153141308 | 329 | * | NA_peak_13 | 13 | Intron (ENSM | 2 | 153161303 | 153204862 | 43560 | 1 | 99237 | ENSMUST0 | -19995 | ENSMUSG0 | Tm9sf4 | transmembrane 9 superfamily member 4 |
| chr2 | 153649765 | 153649935 | 171 | * | NA_peak_13 | 24 | Promoter (<= | 2 | 153649617 | 153687316 | 37700 | 1 | 13436 | ENSMUST0 | 148 | ENSMUSG0 | Dnmt3b | DNA methyltransferase 3B |
| chr2 | 155918085 | 155918210 | 126 | * | NA_peak_13 | 19 | Intron (ENSM | 2 | 155887119 | 155910604 | 23486 | 2 | 56046 | ENSMUST0 | -7481 | ENSMUSG0 | Uqcc1 | ubiquinol-cytochrome c reductase complex assembly factor 1 |
| chr2 | 155936047 | 155936317 | 271 | * | NA_peak_13 | 44 | Distal Interge | 2 | 155848259 | 155930310 | 82052 | 2 | 56046 | ENSMUST0 | -5737 | ENSMUSG0 | Uqcc1 | ubiquinol-cytochrome c reductase complex assembly factor 1 |
| chr2 | 156144115 | 156144392 | 278 | * | NA_peak_13 | 14 | Promoter (<= | 2 | 156123639 | 156144186 | 20548 | 2 | 18041 | ENSMUST0 | 0 | ENSMUSG0 | Nfs1 | nitrogen fixation gene 1 (S. cerevisiae) |
| chr2 | 156765469 | 156765847 | 379 | * | NA_peak_13 | 16 | Distal Interge | 2 | 156759945 | 156762779 | 2835 | 1 | 228836 | ENSMUST0 | 5524 | ENSMUSG0 | Dlgap4 | DLG associated protein 4 |
| chr2 | 156991869 | 156992188 | 320 | * | NA_peak_13 | 14 | Promoter (<= | 2 | 156927345 | 156992043 | 64699 | 2 | 29812 | ENSMUST0 | 0 | ENSMUSG0 | Ndrg3 | N-myc downstream regulated gene 3 |
| chr2 | 157349353 | 157349706 | 354 | * | NA_peak_13 | 30 | Promoter (1- | 2 | 157329497 | 157347506 | 18010 | 2 | 14601 | ENSMUST0 | -1847 | ENSMUSG0 | Ghrh | growth hormone releasing hormone |
| chr2 | 158115883 | 158116199 | 317 | * | NA_peak_13 | 16 | Intron (ENSM | 2 | 158134322 | 158139736 | 5415 | 2 | 21817 | ENSMUST0 | 23537 | ENSMUSG0 | Tgm2 | transglutaminase 2, C polypeptide |
| chr2 | 158611057 | 158611389 | 333 | * | NA_peak_13 | 18 | Promoter (<= | 2 | 158610767 | 158615748 | 4982 | 1 | 22348 | ENSMUST0 | 290 | ENSMUSG0 | Slc32a1 | solute carrier family 32 (GABA vesicular transporter), member 1 |
| chr2 | 160908029 | 160908396 | 368 | * | NA_peak_13 | 19 | Exon (ENSM | 2 | 160906437 | 160912317 | 5881 | 2 | 280635 | ENSMUST0 | 3921 | ENSMUSG0 | Emilin3 | elastin microfibril interfacer 3 |
| chr2 | 161649579 | 161649971 | 393 | * | NA_peak_13 | 65 | Intron (ENSM | 2 | 161698009 | 161800155 | 102147 | 2 | 19281 | ENSMUST0 | 150184 | ENSMUSG0 | Ptprt | protein tyrosine phosphatase, receptor type, T |
| chr2 | 162846094 | 162846375 | 282 | * | NA_peak_13 | 27 | Distal Interge | 2 | 162931528 | 162935208 | 3681 | 1 | 67996 | ENSMUST0 | -85153 | ENSMUSG0 | Srsf6 | serine and arginine-rich splicing factor 6 |
| chr2 | 163644874 | 163645148 | 275 | * | NA_peak_13 | 23 | Promoter (<= | 2 | 163623272 | 163645131 | 21860 | 2 | 26943 | ENSMUST0 | 0 | ENSMUSG0 | Serinc3 | serine incorporator 3 |
| chr2 | 164461020 | 164461430 | 411 | * | NA_peak_13 | 20 | Promoter (<= | 2 | 164461053 | 164479638 | 18586 | 1 | 66460 | ENSMUST0 | 0 | ENSMUSG0 | Sys1 | SYS1 Golgi-localized integral membrane protein homolog (S. cerevisiae) |
| chr2 | 164729476 | 164729691 | 216 | * | NA_peak_13 | 26 | Distal Interge | 2 | 164731180 | 164743312 | 12133 | 2 | 71856 | ENSMUST0 | 13621 | ENSMUSG0 | Wfdc3 | WAP four-disulfide core domain 3 |
| chr2 | 164745814 | 164746108 | 295 | * | NA_peak_13 | 26 | Promoter (<= | 2 | 164731180 | 164745931 | 14752 | 2 | 71856 | ENSMUST0 | 0 | ENSMUSG0 | Wfdc3 | WAP four-disulfide core domain 3 |
| chr2 | 164846923 | 164847232 | 310 | * | NA_peak_13 | 19 | Intron (ENSM | 2 | 164846813 | 164856796 | 9984 | 2 | 18830 | ENSMUST0 | 9564 | ENSMUSG0 | Pltp | phospholipid transfer protein |
| chr2 | 164997636 | 164997774 | 139 | * | NA_peak_13 | 23 | Promoter (1- | 2 | 164996591 | 164999722 | 3132 | 1 | 57138 | ENSMUST0 | 1045 | ENSMUSG0 | Slc12a5 | solute carrier family 12, member 5 |
| chr2 | 164998232 | 164998442 | 211 | * | NA_peak_13 | 20 | Promoter (1- | 2 | 164996591 | 164999722 | 3132 | 1 | 57138 | ENSMUST0 | 1641 | ENSMUSG0 | Slc12a5 | solute carrier family 12, member 5 |
| chr2 | 165361422 | 165361549 | 128 | * | NA_peak_13 | 23 | Promoter (1- | 2 | 165351298 | 165360136 | 8839 | 2 | 381405 | ENSMUST0 | -1286 | ENSMUSG0 | Zfp663 | zinc finger protein 663 |
| chr2 | 165429932 | 165430242 | 311 | * | NA_peak_13 | 26 | Exon (ENSM | 2 | 165411884 | 165420293 | 8410 | 2 | 114644 | ENSMUST0 | -9639 | ENSMUSG0 | Slc13a3 | solute carrier family 13 (sodium-dependent dicarboxylate transporter), member 3 |
| chr2 | 166679052 | 166679288 | 237 | * | NA_peak_13 | 16 | Intron (ENSM | 2 | 166633423 | 166703594 | 70172 | 2 | 277360 | ENSMUST0 | 24306 | ENSMUSG0 | Prex1 | phosphatidylinositol-3,4,5-trisphosphate-dependent Rac exchange factor 1 |
| chr2 | 167484221 | 167484253 | 33 | * | NA_peak_13 | 19 | Promoter (1- | 2 | 167484520 | 167485830 | 1311 | 2 | 263876 | ENSMUST0 | 1577 | ENSMUSG0 | Spata2 | spermatogenesis associated 2 |
| chr2 | 167895905 | 167896270 | 366 | * | NA_peak_13 | 21 | Distal Interge | 2 | 167932057 | 167974737 | 42681 | 1 | 19246 | ENSMUST0 | -35787 | ENSMUSG0 | Ptpn1 | protein tyrosine phosphatase, non-receptor type 1 |
| chr2 | 167932172 | 167932688 | 517 | * | NA_peak_13 | 26 | Promoter (<= | 2 | 167932327 | 167979385 | 47059 | 1 | 19246 | ENSMUST0 | 0 | ENSMUSG0 | Ptpn1 | protein tyrosine phosphatase, non-receptor type 1 |
| chr2 | 168007931 | 168008178 | 248 | * | NA_peak_13 | 27 | Promoter (2- | 2 | 167980164 | 168010618 | 30455 | 2 | 69553 | ENSMUST0 | 2440 | ENSMUSG0 | Ripor3 | RIPOR family member 3 |
| chr2 | 168230999 | 168231425 | 427 | * | NA_peak_13 | 13 | Promoter (<= | 2 | 168230622 | 168232594 | 1973 | 1 | 69372 | ENSMUST0 | 377 | ENSMUSG0 | Mocs3 | molybdenum cofactor synthesis 3 |
| chr2 | 170320725 | 170320878 | 154 | * | NA_peak_13 | 23 | Distal Interge | 2 | 170347931 | 170349740 | 1810 | 2 | 76960 | ENSMUST0 | 28862 | ENSMUSG0 | Bcas1 | breast carcinoma amplified sequence 1 |
| chr2 | 174299138 | 174299543 | 406 | * | NA_peak_13 | 23 | Promoter (1- | 2 | 174297859 | 174346744 | 48886 | 1 | 14683 | ENSMUST0 | 1279 | ENSMUSG0 | Gnas | GNAS (guanine nucleotide binding protein, alpha stimulating) complex locus |
| chr2 | 177252195 | 177252478 | 284 | * | NA_peak_13 | 38 | Distal Interge | 2 | 177464733 | 177478129 | 13397 | 1 | 628308 | ENSMUST0 | -212255 | ENSMUSG0 | Zfp970 | zinc finger protein 970 |
| chr2 | 177314563 | 177315081 | 519 | * | NA_peak_13 | 50 | Exon (ENSM | 2 | 177464733 | 177478129 | 13397 | 1 | 628308 | ENSMUST0 | -149652 | ENSMUSG0 | Zfp970 | zinc finger protein 970 |
| chr2 | 179310264 | 179310344 | 81 | * | NA_peak_13 | 48 | Distal Interge | 2 | 179414936 | 179416882 | 1947 | 1 | 68175 | ENSMUST0 | -104592 | ENSMUSG0 | 4930591A17I | RIKEN cDNA 4930591A17 gene |
| chr2 | 179613394 | 179613759 | 366 | * | NA_peak_13 | 35 | Exon (ENSM | 2 | 179777163 | 179797511 | 20349 | 1 | 12561 | ENSMUST0 | -163404 | ENSMUSG0 | Cdh4 | cadherin 4 |
| chr2 | 180025340 | 180025552 | 213 | * | NA_peak_13 | 23 | Promoter (<= | 2 | 180024987 | 180035461 | 10475 | 1 | 241846 | ENSMUST0 | 353 | ENSMUSG0 | Lsm14b | LSM family member 14B |
| chr2 | 180119272 | 180119530 | 259 | * | NA_peak_13 | 29 | Promoter (<= | 2 | 180119306 | 180162680 | 43375 | 1 | 228983 | ENSMUST0 | 0 | ENSMUSG0 | Osbpl2 | oxysterol binding protein-like 2 |
| chr2 | 181930292 | 181930924 | 633 | * | NA_peak_13 | 193 | Distal Interge | 2 | 181991129 | 182003861 | 12733 | 1 | 672125 | ENSMUST0 | -60205 | ENSMUSG0 | Gm14496 | predicted gene 14496 |
| chr3 | 4162265 | 4162661 | 397 | * | NA_peak_13 | 14 | Distal Interge | 3 | 3634150 | 3658052 | 23903 | 1 | 30942 | ENSMUST0 | 528115 | ENSMUSG0 | Hnf4g | hepatocyte nuclear factor 4, gamma |
| chr3 | 4310008 | 4310056 | 49 | * | NA_peak_13 | 20 | Distal Interge | 3 | 4798708 | 4814911 | 16204 | 1 | 100503859 | ENSMUST0 | -488652 | ENSMUSG0 | 1110015O18 | RIKEN cDNA 1110015O18 gene |
| chr3 | 6477581 | 6477731 | 151 | * | NA_peak_13 | 30 | Distal Interge | 3 | 6615418 | 6620443 | 5026 | 2 | 69347 | ENSMUST0 | 142712 | ENSMUSG0 | 1700008P02I | RIKEN cDNA 1700008P02 gene |
| chr3 | 7221742 | 7222040 | 299 | * | NA_peak_13 | 24 | Distal Interge | 3 | 7366669 | 7437515 | 70847 | 1 | 18767 | ENSMUST0 | -144629 | ENSMUSG0 | Pkia | protein kinase inhibitor, alpha |
| chr3 | 7790801 | 7791183 | 383 | * | NA_peak_13 | 54 | Distal Interge | 3 | 7925303 | 7954882 | 29580 | 2 | 75476 | ENSMUST0 | 163699 | ENSMUSG0 | 1700010I02R | RIKEN cDNA 1700010I02 gene |
| chr3 | 10235521 | 10235548 | 28 | * | NA_peak_13 | 22 | Distal Interge | 3 | 10245918 | 10254225 | 8308 | 2 | 75497 | ENSMUST0 | 18677 | ENSMUSG0 | Fabp12 | fatty acid binding protein 12 |
| chr3 | 10965904 | 10966318 | 415 | * | NA_peak_13 | 30 | Distal Interge | 3 | 10417817 | 10440102 | 22286 | 2 | 74718 | ENSMUST0 | -525802 | ENSMUSG0 | Snx16 | sorting nexin 16 |
| chr3 | 12469139 | 12469397 | 259 | * | NA_peak_13 | 15 | Distal Interge | 3 | 11823229 | 11844359 | 21131 | 2 | 100038545 | ENSMUST0 | -624780 | ENSMUSG0 | Gm10745 | predicted gene 10745 |
| chr3 | 14470054 | 14470350 | 297 | * | NA_peak_13 | 22 | Distal Interge | 3 | 14480697 | 14505830 | 25134 | 1 | 140918 | ENSMUST0 | -10347 | ENSMUSG0 | Slc7a12 | solute carrier family 7 (cationic amino acid transporter, y+ system), member 12 |
| chr3 | 14706566 | 14706744 | 179 | * | NA_peak_13 | 37 | Distal Interge | 3 | 14641727 | 14663002 | 21276 | 1 | 71934 | ENSMUST0 | 64839 | ENSMUSG0 | Car13 | carbonic anhydrase 13 |
| chr3 | 14707083 | 14707588 | 506 | * | NA_peak_13 | 37 | Distal Interge | 3 | 14641727 | 14663002 | 21276 | 1 | 71934 | ENSMUST0 | 65356 | ENSMUSG0 | Car13 | carbonic anhydrase 13 |
| chr3 | 15993013 | 15993047 | 35 | * | NA_peak_13 | 26 | Intron (ENSM | 3 | 15948070 | 16006332 | 58263 | 2 | 381484 | ENSMUST0 | 13285 | ENSMUSG0 | Gm5150 | predicted gene 5150 |
| chr3 | 19357005 | 19357077 | 73 | * | NA_peak_13 | 18 | Distal Interge | 3 | 19223108 | 19311322 | 88215 | 2 | 18583 | ENSMUST0 | -45683 | ENSMUSG0 | Pde7a | phosphodiesterase 7A |
| chr3 | 19357529 | 19358540 | 1012 | * | NA_peak_13 | 49 | Distal Interge | 3 | 19223108 | 19311322 | 88215 | 2 | 18583 | ENSMUST0 | -46207 | ENSMUSG0 | Pde7a | phosphodiesterase 7A |

| | | | | | | | | | | | | | | | | | | |
|---|---|---|---|---|---|---|---|---|---|---|---|---|---|---|---|---|---|---|
| chr3 | 19358847 | 19359092 | 246 | * | NA_peak_13 | 49 | Distal Interge | 3 | 19223108 | 19311322 | 88215 | 2 | 18583 | ENSMUST0( | -47525 | ENSMUSG0( | Pde7a | phosphodiesterase 7A |
| chr3 | 20050442 | 20050448 | 7 | * | NA_peak_13 | 19 | Distal Interge | 3 | 20057811 | 20117281 | 59471 | 1 | 20585 | ENSMUST0( | -7363 | ENSMUSG0( | Hltf | helicase-like transcription factor |
| chr3 | 20618140 | 20618428 | 289 | * | NA_peak_13 | 19 | Distal Interge | 3 | 20314473 | 20367177 | 52705 | 2 | 11608 | ENSMUST0( | -250963 | ENSMUSG0( | Agtr1b | angiotensin II receptor, type 1b |
| chr3 | 20742474 | 20742843 | 370 | * | NA_peak_13 | 22 | Distal Interge | 3 | 20314473 | 20367177 | 52705 | 2 | 11608 | ENSMUST0( | -375297 | ENSMUSG0( | Agtr1b | angiotensin II receptor, type 1b |
| chr3 | 21321255 | 21321571 | 317 | * | NA_peak_13 | 23 | Distal Interge | 3 | 22076652 | 22216594 | 139943 | 1 | 81004 | ENSMUST0( | -755081 | ENSMUSG0( | Tbl1xr1 | transducin (beta)-like 1X-linked receptor 1 |
| chr3 | 22365545 | 22365818 | 274 | * | NA_peak_13 | 35 | Distal Interge | 3 | 22251374 | 22251663 | 290 | 1 | 19784 | ENSMUST0( | 114171 | NA | Rprl2 | ribonuclease P RNA-like 2 |
| chr3 | 22459716 | 22459837 | 122 | * | NA_peak_13 | 43 | Distal Interge | 3 | 22251374 | 22251663 | 290 | 1 | 19784 | ENSMUST0( | 208342 | NA | Rprl2 | ribonuclease P RNA-like 2 |
| chr3 | 22497152 | 22497452 | 301 | * | NA_peak_13 | 16 | Distal Interge | 3 | 22251374 | 22251663 | 290 | 1 | 19784 | ENSMUST0( | 245778 | NA | Rprl2 | ribonuclease P RNA-like 2 |
| chr3 | 22672176 | 22672561 | 386 | * | NA_peak_13 | 13 | Distal Interge | 3 | 22251374 | 22251663 | 290 | 1 | 19784 | ENSMUST0( | 420802 | NA | Rprl2 | ribonuclease P RNA-like 2 |
| chr3 | 22734681 | 22734860 | 180 | * | NA_peak_13 | 17 | Distal Interge | 3 | 22251374 | 22251663 | 290 | 1 | 19784 | ENSMUST0( | 483307 | NA | Rprl2 | ribonuclease P RNA-like 2 |
| chr3 | 23011913 | 23011916 | 4 | * | NA_peak_13 | 30 | Distal Interge | 3 | 22251374 | 22251663 | 290 | 1 | 19784 | ENSMUST0( | 760539 | NA | Rprl2 | ribonuclease P RNA-like 2 |
| chr3 | 23312613 | 23312840 | 228 | * | NA_peak_13 | 29 | Distal Interge | 3 | 23804335 | 23939477 | 135143 | 2 | 635702 | ENSMUST0( | 626637 | ENSMUSG0( | Naaladl2 | N-acetylated alpha-linked acidic dipeptidase-like 2 |
| chr3 | 23355144 | 23355453 | 310 | * | NA_peak_13 | 24 | Distal Interge | 3 | 23804335 | 23939477 | 135143 | 2 | 635702 | ENSMUST0( | 584024 | ENSMUSG0( | Naaladl2 | N-acetylated alpha-linked acidic dipeptidase-like 2 |
| chr3 | 24191347 | 24191698 | 352 | * | NA_peak_13 | 68 | Intron (ENSM | 3 | 23804985 | 23972185 | 167201 | 2 | 635702 | ENSMUST0( | -219162 | ENSMUSG0( | Naaladl2 | N-acetylated alpha-linked acidic dipeptidase-like 2 |
| chr3 | 27659559 | 27659855 | 297 | * | NA_peak_13 | 26 | Intron (ENSM | 3 | 27416162 | 27710398 | 294237 | 2 | 72007 | ENSMUST0( | 50543 | ENSMUSG0( | Fndc3b | fibronectin type III domain containing 3B |
| chr3 | 27995437 | 27995543 | 107 | * | NA_peak_13 | 17 | Intron (ENSM | 3 | 27984173 | 28132050 | 147878 | 1 | 18805 | ENSMUST0( | 11264 | ENSMUSG0( | Pld1 | phospholipase D1 |
| chr3 | 28609301 | 28609639 | 339 | * | NA_peak_13 | 22 | Intron (ENSM | 3 | 28629872 | 28638469 | 8598 | 1 | 665113 | ENSMUST0( | -20233 | ENSMUSG0( | Tnik | TRAF2 and NCK interacting kinase |
| chr3 | 28877915 | 28878094 | 180 | * | NA_peak_13 | 15 | Distal Interge | 3 | 28892617 | 28926724 | 34108 | 1 | 385263 | ENSMUST0( | -14523 | ENSMUSG0( | Gm1527 | predicted gene 1527 |
| chr3 | 29521596 | 29522183 | 588 | * | NA_peak_13 | 44 | Intron (ENSM | 3 | 29513906 | 29670734 | 156829 | 1 | 75740 | ENSMUST0( | 7690 | ENSMUSG0( | Egfem1 | EGF-like and EMI domain containing 1 |
| chr3 | 29784039 | 29784258 | 220 | * | NA_peak_13 | 18 | Intron (ENSM | 3 | 29891014 | 29924191 | 33178 | 1 | 102636083 | ENSMUST0( | -106756 | ENSMUSG0( | Mannr | Mecom adjacent non-protein coding RNA |
| chr3 | 30084415 | 30084443 | 29 | * | NA_peak_13 | 13 | Intron (ENSM | 3 | 29951307 | 30140423 | 189117 | 2 | 14013 | ENSMUST0( | 55980 | ENSMUSG0( | Mecom | MDS1 and EVI1 complex locus |
| chr3 | 30622620 | 30622812 | 193 | * | NA_peak_13 | 19 | Distal Interge | 3 | 30613511 | 30616795 | 3285 | 1 | 80732 | ENSMUST0( | 9109 | ENSMUSG0( | Mynn | myoneurin |
| chr3 | 30793759 | 30794096 | 338 | * | NA_peak_13 | 14 | Promoter (<= | 3 | 30793336 | 30801541 | 8206 | 1 | 69276 | ENSMUST0( | 423 | ENSMUSG0( | Sec62 | SEC62 homolog (S. cerevisiae) |
| chr3 | 30851689 | 30851872 | 184 | * | NA_peak_13 | 27 | Distal Interge | 3 | 30855950 | 30879461 | 23512 | 1 | 71862 | ENSMUST0( | -4078 | ENSMUSG0( | Gpr160 | G protein-coupled receptor 160 |
| chr3 | 31164559 | 31164919 | 361 | * | NA_peak_13 | 44 | Downstream | 3 | 31149920 | 31164324 | 14405 | 1 | 18417 | ENSMUST0( | 14639 | ENSMUSG0( | Cldn11 | claudin 11 |
| chr3 | 31588777 | 31589144 | 368 | * | NA_peak_13 | 19 | Distal Interge | 3 | 31202858 | 31310378 | 107521 | 2 | 241919 | ENSMUST0( | -278399 | ENSMUSG0( | Slc7a14 | solute carrier family 7 (cationic amino acid transporter, y+ system), member 14 |
| chr3 | 32163040 | 32163364 | 325 | * | NA_peak_13 | 144 | Intron (ENSM | 3 | 32156114 | 32198356 | 42243 | 1 | 72413 | ENSMUST0( | 6926 | ENSMUSG0( | Kcnmb2 | potassium large conductance calcium-activated channel, subfamily M, beta member 2 |
| chr3 | 32240745 | 32240968 | 224 | * | NA_peak_13 | 21 | Distal Interge | 3 | 32156114 | 32198356 | 42243 | 1 | 72413 | ENSMUST0( | 84631 | ENSMUSG0( | Kcnmb2 | potassium large conductance calcium-activated channel, subfamily M, beta member 2 |
| chr3 | 32243586 | 32243699 | 114 | * | NA_peak_13 | 30 | Distal Interge | 3 | 32156114 | 32198356 | 42243 | 1 | 72413 | ENSMUST0( | 87472 | ENSMUSG0( | Kcnmb2 | potassium large conductance calcium-activated channel, subfamily M, beta member 2 |
| chr3 | 32522832 | 32522922 | 91 | * | NA_peak_13 | 16 | Distal Interge | 3 | 32518799 | 32519560 | 762 | 1 | 67778 | ENSMUST0( | 4033 | ENSMUSG0( | Zfp639 | zinc finger protein 639 |
| chr3 | 33323422 | 33323804 | 383 | * | NA_peak_14 | 19 | Distal Interge | 3 | 32954337 | 33143247 | 188911 | 2 | 58869 | ENSMUST0( | -180175 | ENSMUSG0( | Pex5l | peroxisomal biogenesis factor 5-like |
| chr3 | 33621057 | 33621628 | 572 | * | NA_peak_14 | 53 | Exon (ENSM | 3 | 33799832 | 33814860 | 15029 | 1 | 67120 | ENSMUST0( | -178204 | ENSMUSG0( | Ttc14 | tetratricopeptide repeat domain 14 |
| chr3 | 34364459 | 34364782 | 324 | * | NA_peak_14 | 26 | Intron (ENSM | 3 | 34196751 | 34638483 | 441733 | 1 | 320478 | ENSMUST0( | 167708 | ENSMUSG0( | Sox2ot | SOX2 overlapping transcript (non-protein coding) |
| chr3 | 35178915 | 35179223 | 309 | * | NA_peak_14 | 13 | Distal Interge | 3 | 34922545 | 34922651 | 107 | 2 | 102466155 | ENSMUST0( | -256264 | ENSMUSG0( | Mir6378 | microRNA 6378 |
| chr3 | 35415056 | 35415525 | 470 | * | NA_peak_14 | 30 | Distal Interge | 3 | 35618814 | 35630964 | 12151 | 2 | 625963 | ENSMUST0( | 215439 | ENSMUSG0( | Gm6639 | predicted gene 6639 |
| chr3 | 35417070 | 35417691 | 622 | * | NA_peak_14 | 27 | Distal Interge | 3 | 35618814 | 35630964 | 12151 | 2 | 625963 | ENSMUST0( | 213273 | ENSMUSG0( | Gm6639 | predicted gene 6639 |
| chr3 | 35822357 | 35822662 | 306 | * | NA_peak_14 | 26 | Intron (ENSM | 3 | 35827050 | 35829189 | 2140 | 1 | 76295 | ENSMUST0( | -4388 | ENSMUSG0( | Atp11b | ATPase, class VI, type 11B |
| chr3 | 36066078 | 36066375 | 298 | * | NA_peak_14 | 23 | Promoter (<= | 3 | 36066072 | 36091135 | 25064 | 1 | 229211 | ENSMUST0( | 6 | ENSMUSG0( | Acad9 | acyl-Coenzyme A dehydrogenase family, member 9 |
| chr3 | 36086414 | 36086784 | 371 | * | NA_peak_14 | 16 | Promoter (2-: | 3 | 36088859 | 36091446 | 2588 | 1 | 229211 | ENSMUST0( | -2075 | ENSMUSG0( | Acad9 | acyl-Coenzyme A dehydrogenase family, member 9 |
| chr3 | 36107980 | 36108045 | 66 | * | NA_peak_14 | 24 | Distal Interge | 3 | 36089837 | 36091372 | 1536 | 1 | 229211 | ENSMUST0( | 18143 | ENSMUSG0( | Acad9 | acyl-Coenzyme A dehydrogenase family, member 9 |
| chr3 | 36521376 | 36521677 | 302 | * | NA_peak_14 | 23 | Promoter (<= | 3 | 36515057 | 36521813 | 6757 | 2 | 100503068 | ENSMUST0( | 136 | ENSMUSG0( | Smim43 | small integral membrane protein 43 |
| chr3 | 36627092 | 36627219 | 128 | * | NA_peak_14 | 19 | Intron (ENSM | 3 | 36638356 | 36640620 | 2265 | 2 | 22065 | ENSMUST0( | 13401 | ENSMUSG0( | Trpc3 | transient receptor potential cation channel, subfamily C, member 3 |
| chr3 | 40351105 | 40351169 | 65 | * | NA_peak_14 | 16 | Distal Interge | 3 | 40508430 | 40509774 | 1345 | 2 | 100040882 | ENSMUST0( | 158605 | ENSMUSG0( | 1700017G19 | RIKEN cDNA 1700017G19 gene |
| chr3 | 40415550 | 40415619 | 70 | * | NA_peak_14 | 19 | Distal Interge | 3 | 40508430 | 40509774 | 1345 | 2 | 100040882 | ENSMUST0( | 94155 | ENSMUSG0( | 1700017G19 | RIKEN cDNA 1700017G19 gene |
| chr3 | 40863300 | 40863474 | 175 | * | NA_peak_14 | 39 | Promoter (1-: | 3 | 40862165 | 40869962 | 7798 | 1 | 100039027 | ENSMUST0( | 1135 | ENSMUSG0( | Gm2011 | predicted gene 2011 |
| chr3 | 41014751 | 41015097 | 347 | * | NA_peak_14 | 26 | Intron (ENSM | 3 | 40971637 | 40977376 | 5740 | 1 | 214048 | ENSMUST0( | 43114 | ENSMUSG0( | Larp1b | La ribonucleoprotein domain family, member 1B |
| chr3 | 41186671 | 41187045 | 375 | * | NA_peak_14 | 39 | Distal Interge | 3 | 41066326 | 41083046 | 16721 | 2 | 70804 | ENSMUST0( | -103625 | ENSMUSG0( | Pgrmc2 | progesterone receptor membrane component 2 |
| chr3 | 41545077 | 41545261 | 185 | * | NA_peak_14 | 30 | Distal Interge | 3 | 41555731 | 41616864 | 61134 | 1 | 269424 | ENSMUST0( | -10470 | ENSMUSG0( | Jade1 | jade family PHD finger 1 |
| chr3 | 41557340 | 41557677 | 338 | * | NA_peak_14 | 26 | Promoter (1-: | 3 | 41555760 | 41596620 | 40861 | 1 | 269424 | ENSMUST0( | 1580 | ENSMUSG0( | Jade1 | jade family PHD finger 1 |
| chr3 | 41565409 | 41565875 | 467 | * | NA_peak_14 | 21 | Promoter (<= | 3 | 41564920 | 41606695 | 41776 | 1 | 269424 | ENSMUST0( | 489 | ENSMUSG0( | Jade1 | jade family PHD finger 1 |
| chr3 | 41769463 | 41770104 | 642 | * | NA_peak_14 | 24 | Intron (ENSM | 3 | 41751196 | 41756492 | 5297 | 1 | 73852 | ENSMUST0( | 18267 | ENSMUSG0( | D3Ertd751e | DNA segment, Chr 3, ERATO Doi 751, expressed |
| chr3 | 41792125 | 41792425 | 301 | * | NA_peak_14 | 38 | Intron (ENSM | 3 | 41751196 | 41756492 | 5297 | 1 | 73852 | ENSMUST0( | 40929 | ENSMUSG0( | D3Ertd751e | DNA segment, Chr 3, ERATO Doi 751, expressed |
| chr3 | 42612216 | 42612295 | 80 | * | NA_peak_14 | 16 | Distal Interge | 3 | 41751196 | 41756492 | 5297 | 1 | 73852 | ENSMUST0( | 861020 | ENSMUSG0( | D3Ertd751e | DNA segment, Chr 3, ERATO Doi 751, expressed |
| chr3 | 42824927 | 42825357 | 431 | * | NA_peak_14 | 48 | Distal Interge | 3 | 41751196 | 41756492 | 5297 | 1 | 73852 | ENSMUST0( | 1073731 | ENSMUSG0( | D3Ertd751e | DNA segment, Chr 3, ERATO Doi 751, expressed |
| chr3 | 43487454 | 43487785 | 332 | * | NA_peak_14 | 16 | Distal Interge | 3 | 41751196 | 41756492 | 5297 | 1 | 73852 | ENSMUST0( | 1736258 | ENSMUSG0( | D3Ertd751e | DNA segment, Chr 3, ERATO Doi 751, expressed |
| chr3 | 44192426 | 44192440 | 15 | * | NA_peak_14 | 23 | Distal Interge | 3 | 45281350 | 45364670 | 83321 | 2 | 72511 | ENSMUST0( | 1172230 | ENSMUSG0( | 2610316D01 | RIKEN cDNA 2610316D01 gene |
| chr3 | 44243793 | 44244114 | 322 | * | NA_peak_14 | 16 | Distal Interge | 3 | 45281350 | 45364670 | 83321 | 2 | 72511 | ENSMUST0( | 1120556 | ENSMUSG0( | 2610316D01 | RIKEN cDNA 2610316D01 gene |
| chr3 | 45024552 | 45024867 | 316 | * | NA_peak_14 | 37 | Distal Interge | 3 | 45281350 | 45364670 | 83321 | 2 | 72511 | ENSMUST0( | 339803 | ENSMUSG0( | 2610316D01 | RIKEN cDNA 2610316D01 gene |
| chr3 | 46922682 | 46923001 | 320 | * | NA_peak_14 | 19 | Distal Interge | 3 | 46900668 | 46900782 | 115 | 2 | 102465199 | ENSMUST0( | -21900 | ENSMUSG0( | Mir6379 | microRNA 6379 |
| chr3 | 50243437 | 50243560 | 124 | * | NA_peak_14 | 16 | Intron (ENSM | 3 | 50364938 | 50443611 | 78674 | 2 | 26570 | ENSMUST0( | 200051 | ENSMUSG0( | Slc7a11 | solute carrier family 7 (cationic amino acid transporter, y+ system), member 11 |
| chr3 | 52268213 | 52268747 | 535 | * | NA_peak_14 | 30 | Promoter (<= | 3 | 52268336 | 52353221 | 84886 | 1 | 56458 | ENSMUST0( | 0 | ENSMUSG0( | Foxo1 | forkhead box O1 |
| chr3 | 53119632 | 53120292 | 661 | * | NA_peak_14 | 14 | Intron (ENSM | 3 | 53041583 | 53261679 | 220097 | 1 | 108927 | ENSMUST0( | 78049 | ENSMUSG0( | Lhfp | lipoma HMGIC fusion partner |
| chr3 | 53136136 | 53136482 | 347 | * | NA_peak_14 | 18 | Intron (ENSM | 3 | 53041583 | 53261679 | 220097 | 1 | 108927 | ENSMUST0( | 94553 | ENSMUSG0( | Lhfp | lipoma HMGIC fusion partner |
| chr3 | 55077456 | 55077835 | 380 | * | NA_peak_14 | 18 | Intron (ENSM | 3 | 55045478 | 55055501 | 10024 | 2 | 12427 | ENSMUST0( | -21955 | ENSMUSG0( | Ccna1 | cyclin A1 |
| chr3 | 55196705 | 55197023 | 319 | * | NA_peak_14 | 23 | Exon (ENSM | 3 | 55182028 | 55209957 | 27930 | 1 | 74434 | ENSMUST0( | 14677 | ENSMUSG0( | Sohlh2 | spermatogenesis and oogenesis specific basic helix-loop-helix 2 |
| chr3 | 55329243 | 55329718 | 476 | * | NA_peak_14 | 20 | Intron (ENSM | 3 | 55354134 | 55477919 | 123786 | 1 | 13175 | ENSMUST0( | -24416 | ENSMUSG0( | Dclk1 | doublecortin-like kinase 1 |
| chr3 | 55555700 | 55556031 | 332 | * | NA_peak_14 | 19 | Distal Interge | 3 | 55527879 | 55534312 | 6434 | 1 | 13175 | ENSMUST0( | 27821 | ENSMUSG0( | Dclk1 | doublecortin-like kinase 1 |
| chr3 | 55726333 | 55726615 | 283 | * | NA_peak_14 | 16 | Promoter (2-: | 3 | 55696360 | 55723947 | 27588 | 2 | 26422 | ENSMUST0( | -2386 | ENSMUSG0( | Nbea | neurobeachin |
| chr3 | 55917153 | 55917224 | 72 | * | NA_peak_14 | 23 | Intron (ENSM | 3 | 55847264 | 55848400 | 1137 | 1 | 71182 | ENSMUST0( | 69889 | ENSMUSG0( | 4933417G07 | SEC13 homolog pseudogene |
| chr3 | 56154448 | 56154610 | 163 | * | NA_peak_14 | 13 | Intron (ENSM | 3 | 55625195 | 56183701 | 558507 | 2 | 26422 | ENSMUST0( | 29091 | ENSMUSG0( | Nbea | neurobeachin |
| chr3 | 56975662 | 56975890 | 229 | * | NA_peak_14 | 14 | Distal Interge | 3 | 57292819 | 57293703 | 885 | 2 | 17112 | ENSMUST0( | 317813 | ENSMUSG0( | Tm4sf1 | transmembrane 4 superfamily member 1 |
| chr3 | 58025465 | 58025708 | 244 | * | NA_peak_14 | 22 | Distal Interge | 3 | 58141707 | 58163807 | 22101 | 2 | 67850 | ENSMUST0( | 138099 | ENSMUSG0( | 1700007F19I | RIKEN cDNA 1700007F19 gene |
| chr3 | 58228672 | 58228954 | 283 | * | NA_peak_14 | 28 | Distal Interge | 3 | 58141707 | 58163807 | 22101 | 2 | 67850 | ENSMUST0( | -64865 | ENSMUSG0( | 1700007F19I | RIKEN cDNA 1700007F19 gene |
| chr3 | 58271594 | 58271829 | 236 | * | NA_peak_14 | 23 | Distal Interge | 3 | 58141707 | 58163807 | 22101 | 2 | 67850 | ENSMUST0( | -107787 | ENSMUSG0( | 1700007F19I | RIKEN cDNA 1700007F19 gene |
| chr3 | 58371550 | 58371772 | 223 | * | NA_peak_14 | 13 | Distal Interge | 3 | 58414715 | 58466783 | 52069 | 1 | 72033 | ENSMUST0( | -42943 | ENSMUSG0( | Tsc22d2 | TSC22 domain family, member 2 |
| chr3 | 58601126 | 58601362 | 237 | * | NA_peak_14 | 30 | Distal Interge | 3 | 58576698 | 58588812 | 12115 | 1 | 69227 | ENSMUST0( | 24428 | ENSMUSG0( | Selenot | selenoprotein T |
| chr3 | 60640233 | 60640262 | 30 | * | NA_peak_14 | 17 | Distal Interge | 3 | 60623278 | 60626511 | 3234 | 1 | 56758 | ENSMUST0( | 16955 | ENSMUSG0( | Mbnl1 | muscleblind like splicing factor 1 |
| chr3 | 61446611 | 61446625 | 15 | * | NA_peak_14 | 19 | Distal Interge | 3 | 61361638 | 61368430 | 6793 | 1 | 74012 | ENSMUST0( | 84973 | ENSMUSG0( | Rap2b | RAP2B, member of RAS oncogene family |
| chr3 | 61513757 | 61513973 | 217 | * | NA_peak_14 | 35 | Distal Interge | 3 | 61361638 | 61368430 | 6793 | 1 | 74012 | ENSMUST0( | 152119 | ENSMUSG0( | Rap2b | RAP2B, member of RAS oncogene family |
| chr3 | 62122699 | 62123061 | 363 | * | NA_peak_14 | 86 | Distal Interge | 3 | 62338344 | 62462221 | 123878 | 1 | 622434 | ENSMUST0( | -215283 | ENSMUSG0( | Arhgef26 | Rho guanine nucleotide exchange factor (GEF) 26 |
| chr3 | 62209016 | 62209257 | 242 | * | NA_peak_14 | 16 | Distal Interge | 3 | 62338344 | 62462221 | 123878 | 1 | 622434 | ENSMUST0( | -129087 | ENSMUSG0( | Arhgef26 | Rho guanine nucleotide exchange factor (GEF) 26 |
| chr3 | 62877363 | 62878049 | 687 | * | NA_peak_14 | 39 | Distal Interge | 3 | 62529077 | 62605140 | 76064 | 2 | 229357 | ENSMUST0( | -272223 | ENSMUSG0( | Gpr149 | G protein-coupled receptor 149 |
| chr3 | 63964271 | 63964561 | 291 | * | NA_peak_14 | 37 | Promoter (<= | 3 | 63942414 | 63964372 | 21959 | 2 | 11416 | ENSMUST0( | 0 | ENSMUSG0( | Slc33a1 | solute carrier family 33 (acetyl-CoA transporter), member 1 |
| chr3 | 64960029 | 64960333 | 305 | * | NA_peak_14 | 19 | Intron (ENSM | 3 | 64949223 | 65223701 | 274479 | 1 | 16497 | ENSMUST0( | 10806 | ENSMUSG0( | Kcnab1 | potassium voltage-gated channel, shaker-related subfamily, beta member 1 |
| chr3 | 65339003 | 65339174 | 172 | * | NA_peak_14 | 22 | Intron (ENSM | 3 | 65324776 | 65349429 | 24654 | 2 | 319798 | ENSMUST0( | 10255 | NA | A730090N16 | RIKEN cDNA A730090N16 gene |
| chr3 | 65614018 | 65614142 | 125 | * | NA_peak_14 | 48 | Exon (ENSM | 3 | 65659288 | 65659426 | 139 | 1 | 102465905 | ENSMUST0( | -45146 | ENSMUSG0( | Mir8120 | microRNA 8120 |
| chr3 | 65735229 | 65735391 | 163 | * | NA_peak_14 | 30 | Intron (ENSM | 3 | 65666317 | 65728551 | 62235 | 1 | 624866 | ENSMUST0( | 68912 | ENSMUSG0( | Lekr1 | leucine, glutamate and lysine rich 1 |
| chr3 | 65974857 | 65974988 | 132 | * | NA_peak_14 | 30 | Distal Interge | 3 | 65946151 | 65958249 | 12099 | 2 | 56706 | ENSMUST0( | -16608 | ENSMUSG0( | Ccnl1 | cyclin L1 |
| chr3 | 66708470 | 66708925 | 456 | * | NA_peak_14 | 36 | Distal Interge | 3 | 66972780 | 66978447 | 5668 | 2 | 20429 | ENSMUST0( | 269522 | ENSMUSG0( | Shox2 | short stature homeobox 2 |
| chr3 | 67681150 | 67681482 | 333 | * | NA_peak_14 | 39 | Distal Interge | 3 | 67584746 | 67596208 | 11463 | 1 | 66868 | ENSMUST0( | 96404 | ENSMUSG0( | Mfsd1 | major facilitator superfamily domain containing 1 |
| chr3 | 68523541 | 68523656 | 116 | * | NA_peak_14 | 29 | Intron (ENSM | 3 | 68494208 | 68626463 | 132256 | 1 | 30953 | ENSMUST0( | 29333 | ENSMUSG0( | Schip1 | schwannomin interacting protein 1 |
| chr3 | 68524637 | 68524784 | 148 | * | NA_peak_14 | 15 | Intron (ENSM | 3 | 68494208 | 68626463 | 132256 | 1 | 30953 | ENSMUST0( | 30429 | ENSMUSG0( | Schip1 | schwannomin interacting protein 1 |
| chr3 | 68741913 | 68741914 | 2 | * | NA_peak_14 | 16 | Distal Interge | 3 | 68761495 | 68762667 | 1173 | 1 | 102639225 | ENSMUST0( | -19581 | ENSMUSG0( | Gm35584 | predicted gene, 35584 |
| chr3 | 69188717 | 69189010 | 294 | * | NA_peak_14 | 32 | Distal Interge | 3 | 69222419 | 69223618 | 1200 | 1 | 71619 | ENSMUST0( | -33409 | ENSMUSG0( | Arl14 | ADP-ribosylation factor-like 14 |
| chr3 | 73867850 | 73868306 | 457 | * | NA_peak_14 | 13 | Distal Interge | 3 | 74009217 | 74017277 | 8061 | 2 | 100504692 | ENSMUST0( | 148971 | ENSMUSG0( | Gm20356 | predicted gene, 20356 |
| chr3 | 75100993 | 75101331 | 339 | * | NA_peak_14 | 30 | Intron (ENSM | 3 | 75061361 | 75143703 | 82343 | 2 | 213234 | ENSMUST0( | 42372 | ENSMUSG0( | Zbbx | zinc finger, B-box domain containing |
| chr3 | 76092307 | 76092683 | 377 | * | NA_peak_14 | 44 | Intron (ENSM | 3 | 76075583 | 76708636 | 633054 | 1 | 213262 | ENSMUST0( | 16724 | ENSMUSG0( | Fstl5 | follistatin-like 5 |
| chr3 | 78497674 | 78497741 | 68 | * | NA_peak_14 | 39 | Distal Interge | 3 | 79062516 | 79144977 | 82462 | 2 | 76089 | ENSMUST0( | 647236 | ENSMUSG0( | Rapgef2 | Rap guanine nucleotide exchange factor (GEF) 2 |
| chr3 | 78814003 | 78814113 | 111 | * | NA_peak_14 | 22 | Distal Interge | 3 | 79062516 | 79144977 | 82462 | 2 | 76089 | ENSMUST0( | 330864 | ENSMUSG0( | Rapgef2 | Rap guanine nucleotide exchange factor (GEF) 2 |
| chr3 | 79066476 | 79066779 | 304 | * | NA_peak_14 | 26 | Exon (ENSM | 3 | 79062516 | 79144977 | 82462 | 2 | 76089 | ENSMUST0( | 78198 | ENSMUSG0( | Rapgef2 | Rap guanine nucleotide exchange factor (GEF) 2 |
| chr3 | 79286426 | 79286617 | 192 | * | NA_peak_14 | 33 | Promoter (<= | 3 | 79062531 | 79286517 | 223987 | 2 | 76089 | ENSMUST0( | 0 | ENSMUSG0( | Rapgef2 | Rap guanine nucleotide exchange factor (GEF) 2 |
| chr3 | 79844778 | 79844934 | 157 | * | NA_peak_14 | 35 | Promoter (1-: | 3 | 79839269 | 79842912 | 3644 | 2 | 70652 | ENSMUST0( | -1866 | ENSMUSG0( | Tmem144 | transmembrane protein 144 |
| chr3 | 79963445 | 79963733 | 289 | * | NA_peak_14 | 30 | Distal Interge | 3 | 79885930 | 79946278 | 60349 | 1 | 68659 | ENSMUST0( | 77515 | ENSMUSG0( | Gask1b | golgi associated kinase 1B |
| chr3 | 80014359 | 80014643 | 285 | * | NA_peak_14 | 26 | Distal Interge | 3 | 79885930 | 79946278 | 60349 | 1 | 68659 | ENSMUST0( | 128429 | ENSMUSG0( | Gask1b | golgi associated kinase 1B |
| chr3 | 82544721 | 82545094 | 374 | * | NA_peak_14 | 18 | Promoter (2-: | 3 | 82539949 | 82547960 | 8012 | 2 | 18167 | ENSMUST0( | 2866 | ENSMUSG0( | Npy2r | neuropeptide Y receptor Y2 |
| chr3 | 83055345 | 83055683 | 339 | * | NA_peak_14 | 34 | Promoter (<= | 3 | 83055522 | 83072355 | 16834 | 1 | 53317 | ENSMUST0( | 0 | ENSMUSG0( | Plrg1 | pleiotropic regulator 1 |
| chr3 | 83603059 | 83603165 | 107 | * | NA_peak_14 | 13 | Distal Interge | 3 | 83555363 | 83574122 | 18760 | 2 | 69460 | ENSMUST0( | -28937 | ENSMUSG0( | 1700028M03 | RIKEN cDNA 1700028M03 gene |
| chr3 | 83675059 | 83675208 | 150 | * | NA_peak_14 | 16 | Distal Interge | 3 | 83766321 | 83774316 | 7996 | 1 | 20319 | ENSMUST0( | -91113 | ENSMUSG0( | Sfrp2 | secreted frizzled-related protein 2 |
| chr3 | 83765880 | 83766307 | 428 | * | NA_peak_14 | 16 | Promoter (<= | 3 | 83766321 | 83774316 | 7996 | 1 | 20319 | ENSMUST0( | -14 | ENSMUSG0( | Sfrp2 | secreted frizzled-related protein 2 |
| chr3 | 83824552 | 83824912 | 361 | * | NA_peak_14 | 13 | Intron (ENSM | 3 | 83836272 | 83841767 | 5496 | 2 | 24088 | ENSMUST0( | 16855 | ENSMUSG0( | Tlr2 | toll-like receptor 2 |
| chr3 | 83861568 | 83861856 | 289 | * | NA_peak_14 | 20 | Distal Interge | 3 | 83836272 | 83841767 | 5496 | 2 | 24088 | ENSMUST0( | -19801 | ENSMUSG0( | Tlr2 | toll-like receptor 2 |
| chr3 | 84259200 | 84259384 | 185 | * | NA_peak_14 | 14 | Promoter (<= | 3 | 84193359 | 84259369 | 66011 | 2 | 80890 | ENSMUST0( | 0 | ENSMUSG0( | Trim2 | tripartite motif-containing 2 |
| chr3 | 84517578 | 84517768 | 191 | * | NA_peak_14 | 26 | Intron (ENSM | 3 | 84442198 | 84480429 | 38232 | 2 | 229474 | ENSMUST0( | -37149 | ENSMUSG0( | Fhdc1 | FH2 domain containing 1 |
| chr3 | 84760199 | 84760515 | 317 | * | NA_peak_14 | 30 | Intron (ENSM | 3 | 84815268 | 84979198 | 163931 | 1 | 50754 | ENSMUST0( | -54753 | ENSMUSG0( | Fbxw7 | F-box and WD-40 domain protein 7 |
| chr3 | 85826624 | 85826894 | 271 | * | NA_peak_14 | 35 | Intron (ENSM | 3 | 85660063 | 85817291 | 157229 | 2 | 229488 | ENSMUST0( | -9333 | ENSMUSG0( | Fam160a1 | family with sequence similarity 160, member A1 |
| chr3 | 87174619 | 87174932 | 314 | * | NA_peak_14 | 16 | Promoter (<= | 3 | 87078593 | 87174700 | 96108 | 2 | 170643 | ENSMUST0( | 0 | ENSMUSG0( | Kirrel | kirre like nephrin family adhesion molecule 1 |
| chr3 | 88142442 | 88142565 | 124 | * | NA_peak_14 | 32 | Promoter (<= | 3 | 88142572 | 88168655 | 26084 | 1 | 17261 | ENSMUST0( | -7 | ENSMUSG0( | Mef2d | myocyte enhancer factor 2D |
| chr3 | 89245711 | 89245756 | 46 | * | NA_peak_14 | 30 | Promoter (<= | 3 | 89234868 | 89245872 | 11005 | 2 | 360213 | ENSMUST0( | 116 | ENSMUSG0( | Trim46 | tripartite motif-containing 46 |
| chr3 | 89320824 | 89321239 | 416 | * | NA_peak_14 | 13 | Promoter (<= | 3 | 89315859 | 89321631 | 5773 | 2 | 13638 | ENSMUST0( | 392 | ENSMUSG0( | Efna3 | ephrin A3 |
| chr3 | 89367210 | 89367582 | 373 | * | NA_peak_14 | 26 | Promoter (1-: | 3 | 89350219 | 89365253 | 15035 | 2 | 77772 | ENSMUST0( | -1957 | ENSMUSG0( | Dcst1 | DC-STAMP domain containing 1 |
| chr3 | 89386853 | 89387143 | 291 | * | NA_peak_14 | 47 | Promoter (<= | 3 | 89377644 | 89387189 | 9546 | 2 | 22724 | ENSMUST0( | 46 | ENSMUSG0( | Zbtb7b | zinc finger and BTB domain containing 7B |
| chr3 | 90231634 | 90232095 | 462 | * | NA_peak_14 | 26 | Promoter (<= | 3 | 90231634 | 90235419 | 3786 | 1 | 23922 | ENSMUST0( | 0 | ENSMUSG0( | Jtb | jumping translocation breakpoint |
| chr3 | 90389042 | 90389559 | 518 | * | NA_peak_14 | 26 | Promoter (<= | 3 | 90387435 | 90388981 | 1547 | 2 | 26568 | ENSMUST0( | -61 | ENSMUSG0( | Slc27a3 | solute carrier family 27 (fatty acid transporter), member 3 |
| chr3 | 90530104 | 90530620 | 517 | * | NA_peak_14 | 35 | Distal Interge | 3 | 90526872 | 90528826 | 1955 | 1 | 66166 | ENSMUST0( | 3232 | ENSMUSG0( | S100a14 | S100 calcium binding protein A14 |
| chr3 | 91589099 | 91589375 | 277 | * | NA_peak_14 | 22 | Distal Interge | 3 | 92014583 | 92031582 | 17000 | 1 | 242100 | ENSMUST0( | -425208 | ENSMUSG0( | Pglyrp3 | peptidoglycan recognition protein 3 |
| chr3 | 92548114 | 92548689 | 576 | * | NA_peak_14 | 46 | Distal Interge | 3 | 92570902 | 92573735 | 2834 | 2 | 16447 | ENSMUST0( | 25046 | ENSMUSG0( | Ivl | involucrin |
| chr3 | 93077073 | 93077140 | 68 | * | NA_peak_14 | 19 | Distal Interge | 3 | 93073567 | 93074633 | 1067 | 1 | 229571 | ENSMUST0( | 3506 | ENSMUSG0( | Tdpoz8 | TD and POZ domain containing 8 |
| chr3 | 93809242 | 93809420 | 179 | * | NA_peak_14 | 39 | Distal Interge | 3 | 93796398 | 93797510 | 1113 | 1 | 399675 | ENSMUST0( | 12844 | ENSMUSG0( | Tdpoz4 | TD and POZ domain containing 4 |
| chr3 | 93919801 | 93919948 | 148 | * | NA_peak_14 | 20 | Distal Interge | 3 | 93960292 | 93961314 | 1023 | 2 | 399676 | ENSMUST0( | 41366 | ENSMUSG0( | Tdpoz5 | TD and POZ domain containing 5 |
| chr3 | 94582138 | 94582814 | 677 | * | NA_peak_14 | 34 | Promoter (<= | 3 | 94497544 | 94582716 | 85173 | 2 | 76742 | ENSMUST0( | 0 | ENSMUSG0( | Snx27 | sorting nexin family member 27 |
| chr3 | 94843142 | 94843417 | 276 | * | NA_peak_14 | 23 | Intron (ENSM | 3 | 94837713 | 94882324 | 44612 | 1 | 229584 | ENSMUST0( | 5429 | ENSMUSG0( | Pogz | pogo transposable element with ZNF domain |
| chr3 | 95093931 | 95094009 | 79 | * | NA_peak_14 | 23 | Intron (ENSM | 3 | 95078105 | 95083232 | 5128 | 2 | 18720 | ENSMUST0( | -10699 | ENSMUSG0( | Pip5k1a | phosphatidylinositol-4-phosphate 5-kinase, type 1 alpha |
| chr3 | 95099622 | 95099966 | 345 | * | NA_peak_14 | 35 | Intron (ENSM | 3 | 95058569 | 95106779 | 48211 | 2 | 18720 | ENSMUST0( | 6813 | ENSMUSG0( | Pip5k1a | phosphatidylinositol-4-phosphate 5-kinase, type 1 alpha |
| chr3 | 95106252 | 95107020 | 769 | * | NA_peak_14 | 23 | Promoter (<= | 3 | 95058530 | 95106875 | 48346 | 2 | 18720 | ENSMUST0( | 0 | ENSMUSG0( | Pip5k1a | phosphatidylinositol-4-phosphate 5-kinase, type 1 alpha |
| chr3 | 95160505 | 95161511 | 1007 | * | NA_peak_14 | 40 | Promoter (<= | 3 | 95160505 | 95174024 | 13520 | 1 | 20360 | ENSMUST0( | 0 | ENSMUSG0( | Sema6c | sema domain, transmembrane domain (TM), and cytoplasmic domain, (semaphorin) 6C |
| chr3 | 95203970 | 95204701 | 732 | * | NA_peak_14 | 32 | Exon (ENSM | 3 | 95181773 | 95208937 | 27165 | 2 | 213054 | ENSMUST0( | 4236 | ENSMUSG0( | Gabpb2 | GA repeat binding protein, beta 2 |
| chr3 | 95217256 | 95217639 | 384 | * | NA_peak_14 | 24 | Promoter (<= | 3 | 95217457 | 95226658 | 9202 | 1 | 100504029 | ENSMUST0( | 0 | ENSMUSG0( | Gm16740 | predicted gene, 16740 |
| chr3 | 95240252 | 95240597 | 346 | * | NA_peak_14 | 31 | Promoter (1-: | 3 | 95236920 | 95241598 | 4679 | 2 | 229588 | ENSMUST0( | 1001 | ENSMUSG0( | Gm128 | predicted gene 128 |
| chr3 | 95659321 | 95659636 | 316 | * | NA_peak_14 | 38 | Promoter (<= | 3 | 95658857 | 95660823 | 1967 | 1 | 17210 | ENSMUST0( | 464 | ENSMUSG0( | Mcl1 | myeloid cell leukemia sequence 1 |
| chr3 | 95818909 | 95819256 | 348 | * | NA_peak_14 | 21 | Promoter (<= | 3 | 95783968 | 95818863 | 34896 | 2 | 75137 | ENSMUST0( | -46 | ENSMUSG0( | Rprd2 | regulation of nuclear pre-mRNA domain containing 2 |
| chr3 | 96176639 | 96177120 | 482 | * | NA_peak_14 | 22 | Exon (ENSM | 3 | 96181151 | 96195521 | 14371 | 1 | 64051 | ENSMUST0( | -4031 | ENSMUSG0( | Sv2a | synaptic vesicle glycoprotein 2 a |
| chr3 | 96207291 | 96207663 | 373 | * | NA_peak_14 | 39 | Intron (ENSM | 3 | 96196588 | 96198489 | 1902 | 2 | 69168 | ENSMUST0( | -8802 | ENSMUSG0( | Bola1 | bolA-like 1 (E. coli) |
| chr3 | 96390513 | 96390852 | 340 | * | NA_peak_14 | 21 | Distal Interge | 3 | 96414437 | 96414833 | 397 | 2 | 21748 | ENSMUST0( | 23981 | ENSMUSG0( | Terc | telomerase RNA component |
| chr3 | 96557689 | 96557998 | 310 | * | NA_peak_14 | 47 | Promoter (<= | 3 | 96555765 | 96557815 | 2051 | 2 | 100038464 | ENSMUST0( | 0 | ENSMUSG0( | Gm15441 | predicted gene 15441 |
| chr3 | 96598261 | 96598714 | 454 | * | NA_peak_14 | 55 | Promoter (1-: | 3 | 96596636 | 96599775 | 3140 | 1 | 545554 | ENSMUST0( | 1625 | ENSMUSG0( | Ankrd34a | ankyrin repeat domain 34A |
| chr3 | 96628932 | 96629085 | 154 | * | NA_peak_14 | 18 | Promoter (<= | 3 | 96606662 | 96629819 | 23158 | 2 | 229613 | ENSMUST0( | 734 | ENSMUSG0( | 6330549D23 | RIKEN cDNA 6330549D23 gene |
| chr3 | 96696347 | 96696524 | 178 | * | NA_peak_14 | 15 | Promoter (<= | 3 | 96696384 | 96705962 | 9579 | 1 | 229615 | ENSMUST0( | 0 | ENSMUSG0( | Pias3 | protein inhibitor of activated STAT 3 |
| chr3 | 97148228 | 97149007 | 780 | * | NA_peak_14 | 34 | Distal Interge | 3 | 97158777 | 97176574 | 17798 | 1 | 66659 | ENSMUST0( | -9770 | ENSMUSG0( | Acp6 | acid phosphatase 6, lysophosphatidic |
| chr3 | 97944243 | 97944480 | 238 | * | NA_peak_15 | 23 | Intron (ENSM | 3 | 97930173 | 97967018 | 36846 | 1 | 433632 | ENSMUST0( | 14070 | ENSMUSG0( | Gm5544 | predicted gene 5544 |
| chr3 | 98013220 | 98013504 | 285 | * | NA_peak_15 | 31 | Promoter (<= | 3 | 98013527 | 98100659 | 87133 | 1 | 18129 | ENSMUST0( | -23 | ENSMUSG0( | Notch2 | notch 2 |
| chr3 | 98274373 | 98274453 | 81 | * | NA_peak_15 | 19 | Distal Interge | 3 | 98280435 | 98310732 | 30298 | 1 | 15360 | ENSMUST0( | -5982 | ENSMUSG0( | Hmgcs2 | 3-hydroxy-3-methylglutaryl-Coenzyme A synthase 2 |
| chr3 | 98642804 | 98643083 | 280 | * | NA_peak_15 | 54 | Intron (ENSM | 3 | 98618634 | 98630252 | 11619 | 2 | 15496 | ENSMUST0( | -12552 | ENSMUSG0( | Hsd3b5 | hydroxy-delta-5-steroid dehydrogenase, 3 beta- and steroid delta-isomerase 5 |
| chr3 | 99906165 | 99906610 | 446 | * | NA_peak_15 | 32 | Intron (ENSM | 3 | 99885417 | 100143322 | 257906 | 1 | 74362 | ENSMUST0( | 20748 | ENSMUSG0( | Spag17 | sperm associated antigen 17 |
| chr3 | 100630050 | 100630368 | 319 | * | NA_peak_15 | 26 | Exon (ENSM | 3 | 100587910 | 100652592 | 64683 | 2 | 17156 | ENSMUST0( | 22224 | ENSMUSG0( | Man1a2 | mannosidase, alpha, class 1A, member 2 |
| chr3 | 100744989 | 100745121 | 133 | * | NA_peak_15 | 23 | Distal Interge | 3 | 100562208 | 100685503 | 123296 | 2 | 17156 | ENSMUST0( | -59486 | ENSMUSG0( | Man1a2 | mannosidase, alpha, class 1A, member 2 |
| chr3 | 100852851 | 100852901 | 51 | * | NA_peak_15 | 23 | Intron (ENSM | 3 | 100825459 | 100896922 | 71464 | 1 | 242122 | ENSMUST0( | 27392 | ENSMUSG0( | Vtcn1 | V-set domain containing T cell activation inhibitor 1 |
| chr3 | 100880187 | 100880492 | 306 | * | NA_peak_15 | 22 | Intron (ENSM | 3 | 100922202 | 100931702 | 9501 | 1 | 229644 | ENSMUST0( | -41710 | ENSMUSG0( | Trim45 | tripartite motif-containing 45 |
| chr3 | 101110142 | 101110456 | 315 | * | NA_peak_15 | 37 | Promoter (<= | 3 | 101040232 | 101110278 | 70047 | 2 | 19221 | ENSMUST0( | 0 | ENSMUSG0( | Ptgfrn | prostaglandin F2 receptor negative regulator |
| chr3 | 101369940 | 101370069 | 130 | * | NA_peak_15 | 21 | Distal Interge | 3 | 101377083 | 101379540 | 2458 | 1 | 78908 | ENSMUST0( | -7014 | ENSMUSG0( | Igsf3 | immunoglobulin superfamily, member 3 |
| chr3 | 102421256 | 102421741 | 486 | * | NA_peak_15 | 30 | Exon (ENSM | 3 | 102469919 | 102520848 | 50930 | 1 | 18049 | ENSMUST0( | -48178 | ENSMUSG0( | Ngf | nerve growth factor |
| chr3 | 103229053 | 103229204 | 152 | * | NA_peak_15 | 23 | Distal Interge | 3 | 103279293 | 103358775 | 79483 | 1 | 94093 | ENSMUST0( | -50089 | ENSMUSG0( | Trim33 | tripartite motif-containing 33 |
| chr3 | 103272677 | 103272777 | 101 | * | NA_peak_15 | 26 | Distal Interge | 3 | 103279293 | 103358775 | 79483 | 1 | 94093 | ENSMUST0( | -6516 | ENSMUSG0( | Trim33 | tripartite motif-containing 33 |
| chr3 | 103649910 | 103650252 | 343 | * | NA_peak_15 | 17 | Distal Interge | 3 | 103576142 | 103645569 | 69428 | 1 | 54524 | ENSMUST0( | 73768 | ENSMUSG0( | Syt6 | synaptotagmin VI |
| chr3 | 103858154 | 103858279 | 126 | * | NA_peak_15 | 16 | Promoter (1-: | 3 | 103859795 | 103912244 | 52450 | 1 | 19260 | ENSMUST0( | -1516 | ENSMUSG0( | Ptpn22 | protein tyrosine phosphatase, non-receptor type 22 (lymphoid) |
| chr3 | 103860099 | 103860234 | 136 | * | NA_peak_15 | 22 | Promoter (<= | 3 | 103860277 | 103912247 | 51971 | 1 | 19260 | ENSMUST0( | -43 | ENSMUSG0( | Ptpn22 | protein tyrosine phosphatase, non-receptor type 22 (lymphoid) |
| chr3 | 103973583 | 103973797 | 215 | * | NA_peak_15 | 23 | Intron (ENSM | 3 | 103968925 | 103997521 | 28597 | 1 | 18685 | ENSMUST0( | 4658 | ENSMUSG0( | Phtf1 | putative homeodomain transcription factor 1 |
| chr3 | 104122153 | 104122342 | 190 | * | NA_peak_15 | 39 | Intron (ENSM | 3 | 104013625 | 104220103 | 206479 | 2 | 99470 | ENSMUST0( | 97761 | ENSMUSG0( | Magi3 | membrane associated guanylate kinase, WW and PDZ domain containing 3 |
| chr3 | 104173016 | 104173227 | 212 | * | NA_peak_15 | 16 | Intron (ENSM | 3 | 104013625 | 104220103 | 206479 | 2 | 99470 | ENSMUST0( | 46876 | ENSMUSG0( | Magi3 | membrane associated guanylate kinase, WW and PDZ domain containing 3 |
| chr3 | 104227589 | 104227936 | 348 | * | NA_peak_15 | 19 | Distal Interge | 3 | 104013259 | 104220374 | 207116 | 2 | 99470 | ENSMUST0( | -7215 | ENSMUSG0( | Magi3 | membrane associated guanylate kinase, WW and PDZ domain containing 3 |
| chr3 | 104237162 | 104237535 | 374 | * | NA_peak_15 | 23 | Distal Interge | 3 | 104013259 | 104220374 | 207116 | 2 | 99470 | ENSMUST0( | -16788 | ENSMUSG0( | Magi3 | membrane associated guanylate kinase, WW and PDZ domain containing 3 |
| chr3 | 104239039 | 104239343 | 305 | * | NA_peak_15 | 33 | Distal Interge | 3 | 104013259 | 104220374 | 207116 | 2 | 99470 | ENSMUST0( | -18665 | ENSMUSG0( | Magi3 | membrane associated guanylate kinase, WW and PDZ domain containing 3 |
| chr3 | 104511166 | 104511627 | 462 | * | NA_peak_15 | 26 | Promoter (<= | 3 | 104481275 | 104511661 | 30387 | 2 | 269473 | ENSMUST0( | 34 | ENSMUSG0( | Lrig2 | leucine-rich repeats and immunoglobulin-like domains 2 |
| chr3 | 104567405 | 104567662 | 258 | * | NA_peak_15 | 15 | Distal Interge | 3 | 104457346 | 104511918 | 54573 | 2 | 269473 | ENSMUST0( | -55487 | ENSMUSG0( | Lrig2 | leucine-rich repeats and immunoglobulin-like domains 2 |
| chr3 | 104741271 | 104741454 | 184 | * | NA_peak_15 | 18 | Distal Interge | 3 | 104753123 | 104773270 | 20148 | 2 | 329731 | ENSMUST0( | 31816 | ENSMUSG0( | Tafa3 | TAFA chemokine like family member 3 |
| chr3 | 104817482 | 104817952 | 471 | * | NA_peak_15 | 34 | Promoter (<= | 3 | 104804550 | 104817354 | 12805 | 2 | 17454 | ENSMUST0( | -128 | ENSMUSG0( | Mov10 | Mov10 RISC complex RNA helicase |
| chr3 | 105212718 | 105212955 | 238 | * | NA_peak_15 | 22 | Distal Interge | 3 | 105066844 | 105078806 | 11963 | 1 | 75269 | ENSMUST0( | 145874 | ENSMUSG0( | 4930564D02 | RIKEN cDNA 4930564D02 gene |
| chr3 | 105249846 | 105250242 | 397 | * | NA_peak_15 | 30 | Distal Interge | 3 | 105066844 | 105078806 | 11963 | 1 | 75269 | ENSMUST0( | 183002 | ENSMUSG0( | 4930564D02 | RIKEN cDNA 4930564D02 gene |
| chr3 | 105671047 | 105671295 | 249 | * | NA_peak_15 | 22 | 3' UTR | 3 | 105681489 | 105687444 | 5956 | 2 | 53975 | ENSMUST0( | 16149 | ENSMUSG0( | Ddx20 | DEAD box helicase 20 |
| chr3 | 105687687 | 105687889 | 203 | * | NA_peak_15 | 23 | Promoter (<= | 3 | 105678270 | 105687574 | 9305 | 2 | 53975 | ENSMUST0( | -113 | ENSMUSG0( | Ddx20 | DEAD box helicase 20 |
| chr3 | 106016525 | 106016804 | 280 | * | NA_peak_15 | 13 | Promoter (1-: | 3 | 105996957 | 106014646 | 17690 | 2 | 100503311 | ENSMUST0( | -1879 | ENSMUSG0( | Pifo | primary cilia formation |
| chr3 | 106082226 | 106082542 | 317 | * | NA_peak_15 | 18 | Distal Interge | 3 | 106113229 | 106132120 | 18892 | 1 | 81600 | ENSMUST0( | -30687 | ENSMUSG0( | Chia1 | chitinase, acidic 1 |
| chr3 | 107643244 | 107643490 | 247 | * | NA_peak_15 | 20 | Distal Interge | 3 | 107620452 | 107631694 | 11243 | 2 | 229707 | ENSMUST0( | -11550 | ENSMUSG0( | Strip1 | striatin interacting protein 1 |
| chr3 | 107644075 | 107644321 | 247 | * | NA_peak_15 | 20 | Distal Interge | 3 | 107620452 | 107631694 | 11243 | 2 | 229707 | ENSMUST0( | -12381 | ENSMUSG0( | Strip1 | striatin interacting protein 1 |
| chr3 | 107750894 | 107750974 | 81 | * | NA_peak_15 | 28 | Promoter (<= | 3 | 107743014 | 107750979 | 7966 | 2 | 12977 | ENSMUST0( | 5 | ENSMUSG0( | Csf1 | colony stimulating factor 1 (macrophage) |
| chr3 | 107782943 | 107783205 | 263 | * | NA_peak_15 | 34 | Distal Interge | 3 | 107741048 | 107760469 | 19422 | 2 | 12977 | ENSMUST0( | -22474 | ENSMUSG0( | Csf1 | colony stimulating factor 1 (macrophage) |
| chr3 | 108214203 | 108214503 | 301 | * | NA_peak_15 | 26 | 3' UTR | 3 | 108202222 | 108210934 | 8713 | 2 | 72522 | ENSMUST0( | -3269 | ENSMUSG0( | Atxn7l2 | ataxin 7-like 2 |
| chr3 | 108280703 | 108281070 | 368 | * | NA_peak_15 | 20 | Distal Interge | 3 | 108284082 | 108361443 | 77362 | 1 | 20661 | ENSMUST0( | -3012 | ENSMUSG0( | Sort1 | sortilin 1 |
| chr3 | 108434108 | 108434128 | 21 | * | NA_peak_15 | 19 | Exon (ENSM | 3 | 108428231 | 108445143 | 16913 | 2 | 20226 | ENSMUST0( | 11015 | ENSMUSG0( | Sars | seryl-aminoacyl-tRNA synthetase |
| chr3 | 108554241 | 108554759 | 519 | * | NA_peak_15 | 23 | Promoter (<= | 3 | 108554338 | 108554751 | 414 | 2 | 100217438 | ENSMUST0( | 0 | NA | Scarna2 | small Cajal body-specific RNA 2 |

| | | | | | | | | | | | | | | | | | |
|---|---|---|---|---|---|---|---|---|---|---|---|---|---|---|---|---|---|
| chr3 | 108604311 | 108604590 | 280 | * | NA_peak_15 | 44 | Intron (ENSM | 3 | 108591389 | 108614960 | 23572 | 1 | 99512 | ENSMUST0( | 12922 | ENSMUSG0( | Wdr47 | WD repeat domain 47 |
| chr3 | 108681729 | 108682016 | 288 | * | NA_peak_15 | 16 | Exon (ENSM | 3 | 108667851 | 108668833 | 983 | 1 | 229725 | ENSMUST0( | 13878 | ENSMUSG0( | Clcc1 | chloride channel CLIC-like 1 |
| chr3 | 108722040 | 108722141 | 102 | * | NA_peak_15 | 40 | Promoter (<= | 3 | 108702976 | 108722116 | 19141 | 2 | 76123 | ENSMUST0( | 0 | ENSMUSG0( | Gpsm2 | G-protein signalling modulator 2 (AGS3-like, C. elegans) |
| chr3 | 108858179 | 108858517 | 339 | * | NA_peak_15 | 24 | Promoter (<= | 3 | 108856088 | 108858609 | 2522 | 2 | 320181 | ENSMUST0( | 92 | ENSMUSG0( | Fndc7 | fibronectin type III domain containing 7 |
| chr3 | 109038308 | 109038649 | 342 | * | NA_peak_15 | 21 | Distal Interge | 3 | 108970997 | 109027607 | 56611 | 2 | 329739 | ENSMUST0( | -10701 | ENSMUSG0( | Fam102b | family with sequence similarity 102, member B |
| chr3 | 109144693 | 109144705 | 13 | * | NA_peak_15 | 23 | Intron (ENSM | 3 | 109123162 | 109158540 | 35379 | 1 | 229731 | ENSMUST0( | 21531 | ENSMUSG0( | Slc25a24 | solute carrier family 25 (mitochondrial carrier, phosphate carrier), member 24 |
| chr3 | 109145079 | 109145399 | 321 | * | NA_peak_15 | 44 | Intron (ENSM | 3 | 109123162 | 109158540 | 35379 | 1 | 229731 | ENSMUST0( | 21917 | ENSMUSG0( | Slc25a24 | solute carrier family 25 (mitochondrial carrier, phosphate carrier), member 24 |
| chr3 | 109147012 | 109147803 | 792 | * | NA_peak_15 | 18 | Intron (ENSM | 3 | 109123162 | 109158540 | 35379 | 1 | 229731 | ENSMUST0( | 23850 | ENSMUSG0( | Slc25a24 | solute carrier family 25 (mitochondrial carrier, phosphate carrier), member 24 |
| chr3 | 109659781 | 109660035 | 255 | * | NA_peak_15 | 16 | Intron (ENSM | 3 | 109647524 | 109682903 | 35380 | 1 | 57257 | ENSMUST0( | 12257 | ENSMUSG0( | Vav3 | vav 3 oncogene |
| chr3 | 112281194 | 112281313 | 120 | * | NA_peak_15 | 39 | Distal Interge | 3 | 111881903 | 112082001 | 200099 | 1 | 625585 | ENSMUST0( | 399291 | ENSMUSG0( | Gm6602 | predicted gene 6602 |
| chr3 | 113270391 | 113270772 | 382 | * | NA_peak_15 | 27 | Distal Interge | 3 | 113247178 | 113258837 | 11660 | 2 | 109959 | ENSMUST0( | -11554 | ENSMUSG0( | Amy2a5 | amylase 2a5 |
| chr3 | 113984825 | 113985182 | 358 | * | NA_peak_15 | 54 | Distal Interge | 3 | 114030540 | 114108324 | 77785 | 1 | 12814 | ENSMUST0( | -45358 | ENSMUSG0( | Col11a1 | collagen, type XI, alpha 1 |
| chr3 | 117722215 | 117722282 | 68 | * | NA_peak_15 | 13 | Distal Interge | 3 | 117782173 | 117846708 | 64536 | 2 | 76561 | ENSMUST0( | 124426 | ENSMUSG0( | Snx7 | sorting nexin 7 |
| chr3 | 119775048 | 119775313 | 266 | * | NA_peak_15 | 39 | Intron (ENSM | 3 | 119720320 | 119783234 | 62915 | 2 | 56195 | ENSMUST0( | 7921 | ENSMUSG0( | Ptbp2 | polypyrimidine tract binding protein 2 |
| chr3 | 121426653 | 121427313 | 661 | * | NA_peak_15 | 32 | Promoter (<= | 3 | 121426770 | 121457811 | 31042 | 1 | 71994 | ENSMUST0( | 0 | ENSMUSG0( | Cnn3 | calponin 3, acidic |
| chr3 | 122396341 | 122396403 | 63 | * | NA_peak_15 | 29 | Intron (ENSM | 3 | 122419757 | 122426866 | 7110 | 1 | 29815 | ENSMUST0( | -23354 | ENSMUSG0( | Bcar3 | breast cancer anti-estrogen resistance 3 |
| chr3 | 122482737 | 122483093 | 357 | * | NA_peak_15 | 36 | Promoter (<= | 3 | 122483723 | 122508448 | 24726 | 1 | 29815 | ENSMUST0( | -630 | ENSMUSG0( | Bcar3 | breast cancer anti-estrogen resistance 3 |
| chr3 | 123012714 | 123012729 | 16 | * | NA_peak_15 | 22 | Intron (ENSM | 3 | 123006517 | 123034844 | 28328 | 2 | 59006 | ENSMUST0( | 22115 | ENSMUSG0( | Myoz2 | myozenin 2 |
| chr3 | 124235565 | 124235832 | 268 | * | NA_peak_15 | 23 | Distal Interge | 3 | 124320855 | 124324743 | 3889 | 1 | 229801 | ENSMUST0( | -85023 | ENSMUSG0( | Tram1l1 | translocation associated membrane protein 1-like 1 |
| chr3 | 125353593 | 125353881 | 289 | * | NA_peak_15 | 23 | Distal Interge | 3 | 125404076 | 125728899 | 324824 | 1 | 64580 | ENSMUST0( | -50195 | ENSMUSG0( | Ndst4 | N-deacetylase/N-sulfotransferase (heparin glucosaminyl) 4 |
| chr3 | 126504423 | 126504429 | 7 | * | NA_peak_15 | 23 | Distal Interge | 3 | 126596302 | 126684688 | 88387 | 1 | 108058 | ENSMUST0( | -91873 | ENSMUSG0( | Camk2d | calcium/calmodulin-dependent protein kinase II, delta |
| chr3 | 126621977 | 126622229 | 253 | * | NA_peak_15 | 26 | Intron (ENSM | 3 | 126597414 | 126844327 | 246914 | 1 | 108058 | ENSMUST0( | 24563 | ENSMUSG0( | Camk2d | calcium/calmodulin-dependent protein kinase II, delta |
| chr3 | 126865504 | 126865787 | 284 | * | NA_peak_15 | 19 | Distal Interge | 3 | 126805862 | 126842893 | 37032 | 1 | 108058 | ENSMUST0( | 59642 | ENSMUSG0( | Camk2d | calcium/calmodulin-dependent protein kinase II, delta |
| chr3 | 128139720 | 128140128 | 409 | * | NA_peak_15 | 70 | Intron (ENSM | 3 | 128124205 | 128133360 | 9156 | 2 | 402774 | ENSMUST0( | -6360 | ENSMUSG0( | D030025E07 | RIKEN cDNA D030025E07 gene |
| chr3 | 130991803 | 130992064 | 262 | * | NA_peak_15 | 16 | Distal Interge | 3 | 131110471 | 131224356 | 113886 | 1 | 16842 | ENSMUST0( | -118407 | ENSMUSG0( | Lef1 | lymphoid enhancer binding factor 1 |
| chr3 | 132827457 | 132827834 | 378 | * | NA_peak_15 | 46 | Intron (ENSM | 3 | 132786910 | 132838506 | 51597 | 1 | 271981 | ENSMUST0( | 40547 | ENSMUSG0( | Tbck | TBC1 domain containing kinase |
| chr3 | 132990163 | 132990446 | 284 | * | NA_peak_15 | 23 | Intron (ENSM | 3 | 132996111 | 133005677 | 9567 | 2 | 67553 | ENSMUST0( | 15231 | ENSMUSG0( | Gstcd | glutathione S-transferase, C-terminal domain containing |
| chr3 | 133837166 | 133837450 | 285 | * | NA_peak_15 | 23 | Intron (ENSM | 3 | 133488535 | 133545139 | 56605 | 2 | 214133 | ENSMUST0( | -292027 | ENSMUSG0( | Tet2 | tet methylcytosine dioxygenase 2 |
| chr3 | 135521814 | 135521848 | 35 | * | NA_peak_15 | 15 | Promoter (1-: | 3 | 135522935 | 135524559 | 1625 | 1 | 110173 | ENSMUST0( | -1087 | ENSMUSG0( | Manba | mannosidase, beta A, lysosomal |
| chr3 | 135579087 | 135579163 | 77 | * | NA_peak_15 | 29 | Distal Interge | 3 | 135590780 | 135594441 | 3662 | 2 | 18033 | ENSMUST0( | 15278 | ENSMUSG0( | Nfkb1 | nuclear factor of kappa light polypeptide gene enhancer in B cells 1, p105 |
| chr3 | 135826090 | 135826469 | 380 | * | NA_peak_15 | 23 | Promoter (<= | 3 | 135826090 | 135888386 | 62297 | 1 | 67547 | ENSMUST0( | 0 | ENSMUSG0( | Slc39a8 | solute carrier family 39 (metal ion transporter), member 8 |
| chr3 | 135880534 | 135880730 | 197 | * | NA_peak_15 | 23 | Intron (ENSM | 3 | 135884160 | 135888568 | 4409 | 1 | 67547 | ENSMUST0( | -3430 | ENSMUSG0( | Slc39a8 | solute carrier family 39 (metal ion transporter), member 8 |
| chr3 | 136174645 | 136174881 | 237 | * | NA_peak_15 | 17 | Intron (ENSM | 3 | 136210162 | 136247755 | 37594 | 2 | 242248 | ENSMUST0( | 72874 | ENSMUSG0( | Bank1 | B cell scaffold protein with ankyrin repeats 1 |
| chr3 | 136955815 | 136956198 | 384 | * | NA_peak_15 | 65 | Distal Interge | 3 | 136903149 | 136905659 | 2511 | 1 | 19055 | ENSMUST0( | 52666 | ENSMUSG0( | Ppp3ca | protein phosphatase 3, catalytic subunit, alpha isoform |
| chr3 | 137670384 | 137670760 | 377 | * | NA_peak_15 | 43 | Distal Interge | 3 | 137624037 | 137626174 | 2138 | 1 | 73284 | ENSMUST0( | 46347 | ENSMUSG0( | Ddit4l | DNA-damage-inducible transcript 4-like |
| chr3 | 137911248 | 137911545 | 298 | * | NA_peak_15 | 19 | 3' UTR | 3 | 137918506 | 137928219 | 9714 | 1 | 56692 | ENSMUST0( | -6961 | ENSMUSG0( | Lamtor3 | late endosomal/lysosomal adaptor, MAPK and MTOR activator 3 |
| chr3 | 138493673 | 138494081 | 409 | * | NA_peak_15 | 23 | Distal Interge | 3 | 138458956 | 138489515 | 30560 | 2 | 75624 | ENSMUST0( | -4158 | ENSMUSG0( | Metap1 | methionyl aminopeptidase 1 |
| chr3 | 138645391 | 138645587 | 197 | * | NA_peak_15 | 30 | Distal Interge | 3 | 138711535 | 138726984 | 15450 | 2 | 74635 | ENSMUST0( | 81397 | ENSMUSG0( | 4930425O10 | RIKEN cDNA 4930425O10 gene |
| chr3 | 138656682 | 138656707 | 26 | * | NA_peak_15 | 26 | Distal Interge | 3 | 138711535 | 138726984 | 15450 | 2 | 74635 | ENSMUST0( | 70277 | ENSMUSG0( | 4930425O10 | RIKEN cDNA 4930425O10 gene |
| chr3 | 138742469 | 138742579 | 111 | * | NA_peak_15 | 16 | Promoter (<= | 3 | 138742505 | 138896861 | 154357 | 1 | 56224 | ENSMUST0( | 0 | ENSMUSG0( | Tspan5 | tetraspanin 5 |
| chr3 | 140538236 | 140538608 | 373 | * | NA_peak_15 | 30 | Distal Interge | 3 | 140028249 | 140028370 | 122 | 1 | 102465198 | ENSMUST0( | 509987 | ENSMUSG0( | Mir7j | microRNA 7j |
| chr3 | 142265813 | 142266131 | 319 | * | NA_peak_15 | 26 | Intron (ENSM | 3 | 142249368 | 142303979 | 54612 | 2 | 56376 | ENSMUST0( | 37848 | ENSMUSG0( | Pdlim5 | PDZ and LIM domain 5 |
| chr3 | 142320892 | 142321342 | 451 | * | NA_peak_15 | 22 | Intron (ENSM | 3 | 142249427 | 142304442 | 55016 | 2 | 56376 | ENSMUST0( | -16450 | ENSMUSG0( | Pdlim5 | PDZ and LIM domain 5 |
| chr3 | 142355210 | 142355624 | 415 | * | NA_peak_15 | 23 | Intron (ENSM | 3 | 142242680 | 142391808 | 149129 | 2 | 56376 | ENSMUST0( | 36184 | ENSMUSG0( | Pdlim5 | PDZ and LIM domain 5 |
| chr3 | 142454810 | 142455072 | 263 | * | NA_peak_15 | 35 | Distal Interge | 3 | 142493978 | 142511104 | 17127 | 1 | 229898 | ENSMUST0( | -38906 | ENSMUSG0( | Gbp5 | guanylate binding protein 5 |
| chr3 | 143724145 | 143724618 | 474 | * | NA_peak_15 | 19 | Distal Interge | 3 | 143694189 | 143712210 | 18022 | 2 | 621809 | ENSMUST0( | -11935 | ENSMUSG0( | Gm6260 | predicted gene 6260 |
| chr3 | 144205343 | 144205900 | 558 | * | NA_peak_15 | 14 | Promoter (<= | 3 | 144188683 | 144205220 | 16538 | 2 | 16911 | ENSMUST0( | -123 | ENSMUSG0( | Lmo4 | LIM domain only 4 |
| chr3 | 144214046 | 144214278 | 233 | * | NA_peak_15 | 16 | Distal Interge | 3 | 144188683 | 144205220 | 16538 | 2 | 16911 | ENSMUST0( | -8826 | ENSMUSG0( | Lmo4 | LIM domain only 4 |
| chr3 | 144739030 | 144739321 | 292 | * | NA_peak_15 | 21 | Intron (ENSM | 3 | 144688844 | 144720335 | 31492 | 2 | 54673 | ENSMUST0( | -18695 | ENSMUSG0( | Sh3glb1 | SH3-domain GRB2-like B1 (endophilin) |
| chr3 | 144829657 | 144829939 | 283 | * | NA_peak_15 | 49 | Intron (ENSM | 3 | 144796559 | 144819494 | 22936 | 2 | 80797 | ENSMUST0( | -10163 | ENSMUSG0( | Clca3a2 | chloride channel accessory 3A2 |
| chr3 | 144869578 | 144869649 | 72 | * | NA_peak_15 | 23 | Intron (ENSM | 3 | 144861826 | 144889745 | 27920 | 2 | 622139 | ENSMUST0( | 20096 | ENSMUSG0( | Clca4c-ps | chloride channel accessory 4C, pseudogene |
| chr3 | 144900289 | 144900359 | 71 | * | NA_peak_15 | 26 | Distal Interge | 3 | 144879322 | 144896011 | 16690 | 2 | 622139 | ENSMUST0( | -4278 | ENSMUSG0( | Clca4c-ps | chloride channel accessory 4C, pseudogene |
| chr3 | 144940043 | 144940362 | 320 | * | NA_peak_15 | 20 | Distal Interge | 3 | 144910921 | 144932529 | 21609 | 2 | 99709 | ENSMUST0( | -7514 | ENSMUSG0( | Clca4b | chloride channel accessory 4B |
| chr3 | 144942228 | 144942406 | 179 | * | NA_peak_15 | 23 | Distal Interge | 3 | 144910921 | 144932529 | 21609 | 2 | 99709 | ENSMUST0( | -9699 | ENSMUSG0( | Clca4b | chloride channel accessory 4B |
| chr3 | 145016182 | 145016466 | 285 | * | NA_peak_15 | 29 | Promoter (2-: | 3 | 145013961 | 145018915 | 4955 | 2 | 23844 | ENSMUST0( | 2449 | ENSMUSG0( | Clca1 | chloride channel accessory 1 |
| chr3 | 145067959 | 145068199 | 241 | * | NA_peak_15 | 23 | Distal Interge | 3 | 145070263 | 145099104 | 28842 | 2 | 229933 | ENSMUST0( | 30905 | ENSMUSG0( | Clca2 | chloride channel accessory 2 |
| chr3 | 145216622 | 145216942 | 321 | * | NA_peak_15 | 26 | Distal Interge | 3 | 145149018 | 145153626 | 4609 | 1 | 52184 | ENSMUST0( | 67604 | ENSMUSG0( | Odf2l | outer dense fiber of sperm tails 2-like |
| chr3 | 145455391 | 145455678 | 288 | * | NA_peak_15 | 39 | Intron (ENSM | 3 | 145515368 | 145538623 | 23256 | 1 | 71355 | ENSMUST0( | -59690 | ENSMUSG0( | Col24a1 | collagen, type XXIV, alpha 1 |
| chr3 | 145667012 | 145667172 | 161 | * | NA_peak_15 | 32 | Intron (ENSM | 3 | 145646976 | 145649981 | 3006 | 2 | 16007 | ENSMUST0( | -17031 | ENSMUSG0( | Ccn1 | cellular communication network factor 1 |
| chr3 | 145758727 | 145758994 | 268 | * | NA_peak_15 | 46 | Promoter (<= | 3 | 145758675 | 145894277 | 135603 | 1 | 69219 | ENSMUST0( | 52 | ENSMUSG0( | Ddah1 | dimethylarginine dimethylaminohydrolase 1 |
| chr3 | 145761875 | 145762027 | 153 | * | NA_peak_15 | 25 | Promoter (1-: | 3 | 145760748 | 145893266 | 132519 | 1 | 69219 | ENSMUST0( | 1127 | ENSMUSG0( | Ddah1 | dimethylarginine dimethylaminohydrolase 1 |
| chr3 | 146685056 | 146685349 | 294 | * | NA_peak_15 | 18 | Promoter (<= | 3 | 146680450 | 146685471 | 5022 | 2 | 75015 | ENSMUST0( | 122 | ENSMUSG0( | Samd13 | sterile alpha motif domain containing 13 |
| chr3 | 146691381 | 146691808 | 428 | * | NA_peak_15 | 20 | Distal Interge | 3 | 146645195 | 146685592 | 40398 | 2 | 75015 | ENSMUST0( | -5789 | ENSMUSG0( | Samd13 | sterile alpha motif domain containing 13 |
| chr3 | 147112173 | 147112763 | 591 | * | NA_peak_15 | 58 | Intron (ENSM | 3 | 146943920 | 146981920 | 38001 | 1 | 70892 | ENSMUST0( | 168253 | ENSMUSG0( | Ttll7 | tubulin tyrosine ligase-like family, member 7 |
| chr3 | 147229543 | 147229933 | 391 | * | NA_peak_15 | 17 | Distal Interge | 3 | 146943920 | 146981920 | 38001 | 1 | 70892 | ENSMUST0( | 285623 | ENSMUSG0( | Ttll7 | tubulin tyrosine ligase-like family, member 7 |
| chr3 | 147732827 | 147733122 | 296 | * | NA_peak_15 | 13 | Distal Interge | 3 | 146943920 | 146981920 | 38001 | 1 | 70892 | ENSMUST0( | 788907 | ENSMUSG0( | Ttll7 | tubulin tyrosine ligase-like family, member 7 |
| chr3 | 155149387 | 155149454 | 68 | * | NA_peak_15 | 17 | Intron (ENSM | 3 | 155093588 | 155137748 | 44161 | 1 | 74435 | ENSMUST0( | 55799 | ENSMUSG0( | Lrriq3 | leucine-rich repeats and IQ motif containing 3 |
| chr3 | 155767329 | 155767667 | 339 | * | NA_peak_15 | 43 | Distal Interge | 3 | 155976354 | 156008592 | 32239 | 1 | 329780 | ENSMUST0( | -208687 | ENSMUSG0( | 9330178D15 | RIKEN cDNA 9330178D15 gene |
| chr3 | 155772044 | 155772454 | 411 | * | NA_peak_15 | 26 | Exon (ENSM | 3 | 155976354 | 156008592 | 32239 | 1 | 329780 | ENSMUST0( | -203900 | ENSMUSG0( | 9330178D15 | RIKEN cDNA 9330178D15 gene |
| chr3 | 156164991 | 156165242 | 252 | * | NA_peak_15 | 22 | Distal Interge | 3 | 155976354 | 156008592 | 32239 | 1 | 329780 | ENSMUST0( | 188637 | ENSMUSG0( | 9330178D15 | RIKEN cDNA 9330178D15 gene |
| chr3 | 157646987 | 157647352 | 366 | * | NA_peak_15 | 60 | Distal Interge | 3 | 157566900 | 157644756 | 77857 | 1 | 19218 | ENSMUST0( | 80087 | ENSMUSG0( | Ptger3 | prostaglandin E receptor 3 (subtype EP3) |
| chr3 | 157946819 | 157947162 | 344 | * | NA_peak_15 | 31 | Promoter (<= | 3 | 157947239 | 158007428 | 60190 | 1 | 433667 | ENSMUST0( | -77 | ENSMUSG0( | Ankrd13c | ankyrin repeat domain 13c |
| chr3 | 157947569 | 157947742 | 174 | * | NA_peak_15 | 31 | Promoter (<= | 3 | 157947466 | 158006837 | 59372 | 1 | 433667 | ENSMUST0( | 103 | ENSMUSG0( | Ankrd13c | ankyrin repeat domain 13c |
| chr3 | 158113371 | 158113670 | 300 | * | NA_peak_15 | 22 | Intron (ENSM | 3 | 158062679 | 158064056 | 1378 | 1 | 67144 | ENSMUST0( | 50692 | ENSMUSG0( | Lrrc40 | leucine rich repeat containing 40 |
| chr3 | 158117362 | 158117634 | 273 | * | NA_peak_15 | 22 | Intron (ENSM | 3 | 158062679 | 158064056 | 1378 | 1 | 67144 | ENSMUST0( | 54683 | ENSMUSG0( | Lrrc40 | leucine rich repeat containing 40 |
| chr3 | 158574019 | 158574329 | 311 | * | NA_peak_15 | 14 | Distal Interge | 3 | 158084439 | 158562221 | 477783 | 2 | 242274 | ENSMUST0( | -11798 | ENSMUSG0( | Lrrc7 | leucine rich repeat containing 7 |
| chr3 | 159385483 | 159385825 | 343 | * | NA_peak_15 | 13 | Distal Interge | 3 | 159495433 | 159529955 | 34523 | 1 | 76131 | ENSMUST0( | -109608 | ENSMUSG0( | Depdc1a | DEP domain containing 1a |
| chr3 | 159539417 | 159539505 | 89 | * | NA_peak_15 | 23 | Distal Interge | 3 | 159495458 | 159529459 | 34002 | 1 | 76131 | ENSMUST0( | 43959 | ENSMUSG0( | Depdc1a | DEP domain containing 1a |
| chr3 | 159553122 | 159553731 | 610 | * | NA_peak_15 | 26 | Distal Interge | 3 | 159599175 | 159625321 | 26147 | 1 | 19892 | ENSMUST0( | -45444 | ENSMUSG0( | Rpe65 | retinal pigment epithelium 65 |
| chr3 | 159684712 | 159684712 | 1 | * | NA_peak_15 | 27 | Distal Interge | 3 | 159599233 | 159624799 | 25567 | 1 | 19892 | ENSMUST0( | 85479 | ENSMUSG0( | Rpe65 | retinal pigment epithelium 65 |
| chr3 | 159697046 | 159697555 | 510 | * | NA_peak_15 | 42 | Distal Interge | 3 | 159599233 | 159624799 | 25567 | 1 | 19892 | ENSMUST0( | 97813 | ENSMUSG0( | Rpe65 | retinal pigment epithelium 65 |
| chr3 | 159749449 | 159749461 | 13 | * | NA_peak_15 | 30 | Distal Interge | 3 | 159839672 | 159936023 | 96352 | 1 | 68151 | ENSMUST0( | -90211 | ENSMUSG0( | Wls | wntless WNT ligand secretion mediator |
| chr4 | 5821548 | 5821741 | 194 | * | NA_peak_16 | 28 | Intron (ENSM | 4 | 5798084 | 5798531 | 448 | 1 | 242297 | ENSMUST0( | 23464 | ENSMUSG0( | Fam110b | family with sequence similarity 110, member B |
| chr4 | 6607396 | 6607682 | 287 | * | NA_peak_16 | 19 | Distal Interge | 4 | 6688046 | 6689351 | 1306 | 2 | 252838 | ENSMUST0( | 81669 | ENSMUSG0( | Tox | thymocyte selection-associated high mobility group box |
| chr4 | 12133812 | 12134018 | 207 | * | NA_peak_16 | 18 | Distal Interge | 4 | 12140264 | 12144688 | 4425 | 1 | 72397 | ENSMUST0( | -6246 | ENSMUSG0( | Rbm12b1 | RNA binding motif protein 12 B1 |
| chr4 | 12136740 | 12137364 | 625 | * | NA_peak_16 | 30 | Promoter (2-: | 4 | 12140264 | 12144688 | 4425 | 1 | 72397 | ENSMUST0( | -2900 | ENSMUSG0( | Rbm12b1 | RNA binding motif protein 12 B1 |
| chr4 | 12848107 | 12848466 | 360 | * | NA_peak_16 | 188 | Distal Interge | 4 | 12906838 | 12980873 | 74036 | 1 | 208820 | ENSMUST0( | -58372 | ENSMUSG0( | Triqk | triple QxxK/R motif containing |
| chr4 | 13294388 | 13294459 | 72 | * | NA_peak_16 | 23 | Distal Interge | 4 | 12917967 | 12981070 | 63104 | 1 | 208820 | ENSMUST0( | 376421 | ENSMUSG0( | Triqk | triple QxxK/R motif containing |
| chr4 | 15913158 | 15913460 | 303 | * | NA_peak_16 | 45 | Distal Interge | 4 | 15885457 | 15896075 | 10619 | 1 | 12307 | ENSMUST0( | 27701 | ENSMUSG0( | Calb1 | calbindin 1 |
| chr4 | 15915735 | 15916022 | 288 | * | NA_peak_16 | 44 | Distal Interge | 4 | 15942900 | 15942991 | 92 | 2 | 102466644 | ENSMUST0( | 26969 | ENSMUSG0( | Mir6400 | microRNA 6400 |
| chr4 | 20422443 | 20422808 | 366 | * | NA_peak_16 | 22 | Intron (ENSM | 4 | 20118874 | 20778567 | 659694 | 2 | 269513 | ENSMUST0( | 355759 | ENSMUSG0( | Nkain3 | Na+/K+ transporting ATPase interacting 3 |
| chr4 | 20944337 | 20944575 | 239 | * | NA_peak_16 | 17 | Distal Interge | 4 | 20243842 | 20778866 | 535025 | 2 | 269513 | ENSMUST0( | -165471 | ENSMUSG0( | Nkain3 | Na+/K+ transporting ATPase interacting 3 |
| chr4 | 22540952 | 22541097 | 146 | * | NA_peak_16 | 15 | Distal Interge | 4 | 22490548 | 22493126 | 2579 | 1 | 105500046 | ENSMUST0( | 50404 | ENSMUSG0( | Gm30731 | predicted gene, 30731 |
| chr4 | 23546529 | 23546922 | 394 | * | NA_peak_16 | 39 | Distal Interge | 4 | 24496451 | 24581338 | 84888 | 1 | 212377 | ENSMUST0( | -949529 | ENSMUSG0( | Mms22l | MMS22-like, DNA repair protein |
| chr4 | 24496282 | 24496576 | 295 | * | NA_peak_16 | 35 | Promoter (<= | 4 | 24496451 | 24581338 | 84888 | 1 | 212377 | ENSMUST0( | 0 | ENSMUSG0( | Mms22l | MMS22-like, DNA repair protein |
| chr4 | 28339559 | 28340078 | 520 | * | NA_peak_16 | 19 | Distal Interge | 4 | 28813131 | 28947047 | 133917 | 1 | 13841 | ENSMUST0( | -473053 | ENSMUSG0( | Epha7 | Eph receptor A7 |
| chr4 | 34540990 | 34541171 | 182 | * | NA_peak_16 | 22 | Distal Interge | 4 | 34550937 | 34566908 | 15972 | 1 | 433693 | ENSMUST0( | -9766 | ENSMUSG0( | Akirin2 | akirin 2 |
| chr4 | 37344169 | 37344600 | 432 | * | NA_peak_16 | 30 | Distal Interge | 4 | 37249403 | 37332515 | 83113 | 1 | 102636890 | ENSMUST0( | 94766 | ENSMUSG0( | Gm12374 | predicted gene 12374 |
| chr4 | 37920197 | 37920519 | 323 | * | NA_peak_16 | 13 | Distal Interge | 4 | 37249403 | 37332515 | 83113 | 1 | 102636890 | ENSMUST0( | 670794 | ENSMUSG0( | Gm12374 | predicted gene 12374 |
| chr4 | 38320314 | 38320656 | 343 | * | NA_peak_16 | 26 | Distal Interge | 4 | 38571667 | 38855483 | 283817 | 2 | 102636968 | ENSMUST0( | 534827 | ENSMUSG0( | Gm12381 | predicted gene 12381 |
| chr4 | 41455447 | 41455746 | 300 | * | NA_peak_16 | 21 | Intron (ENSM | 4 | 41390745 | 41464840 | 74096 | 2 | 109242 | ENSMUST0( | 9094 | ENSMUSG0( | Kif24 | kinesin family member 24 |
| chr4 | 43578721 | 43578751 | 31 | * | NA_peak_16 | 30 | Promoter (<= | 4 | 43578735 | 43587485 | 8751 | 1 | 242406 | ENSMUST0( | 0 | ENSMUSG0( | Rgp1 | RAB6A GEF compex partner 1 |
| chr4 | 43814932 | 43815171 | 240 | * | NA_peak_16 | 25 | Distal Interge | 4 | 43820335 | 43821433 | 1099 | 2 | 29846 | ENSMUST0( | 6262 | ENSMUSG0( | Olfr156 | olfactory receptor 156 |
| chr4 | 45452368 | 45452742 | 375 | * | NA_peak_16 | 20 | Intron (ENSM | 4 | 45423278 | 45482802 | 59525 | 2 | 230126 | ENSMUST0( | 30060 | ENSMUSG0( | Shb | src homology 2 domain-containing transforming protein B |
| chr4 | 45896707 | 45896810 | 104 | * | NA_peak_16 | 13 | Intron (ENSM | 4 | 45890852 | 45950767 | 59916 | 1 | 381522 | ENSMUST0( | 5855 | ENSMUSG0( | Ccdc180 | coiled-coil domain containing 180 |
| chr4 | 46649825 | 46650192 | 368 | * | NA_peak_16 | 26 | Promoter (<= | 4 | 46604390 | 46650209 | 45820 | 2 | 381605 | ENSMUST0( | 17 | ENSMUSG0( | Tbc1d2 | TBC1 domain family, member 2 |
| chr4 | 48798175 | 48798351 | 177 | * | NA_peak_16 | 23 | Distal Interge | 4 | 48663514 | 48673502 | 9989 | 1 | 68016 | ENSMUST0( | 134661 | ENSMUSG0( | Cavin4 | caveolae associated 4 |
| chr4 | 49330514 | 49330861 | 348 | * | NA_peak_16 | 39 | Intron (ENSM | 4 | 49379920 | 49383576 | 3657 | 2 | 209186 | ENSMUST0( | 52715 | ENSMUSG0( | Acnat2 | acyl-coenzyme A amino acid N-acyltransferase 2 |
| chr4 | 50529873 | 50530237 | 365 | * | NA_peak_16 | 108 | Distal Interge | 4 | 49769635 | 49845744 | 76110 | 2 | 242443 | ENSMUST0( | -684129 | ENSMUSG0( | Grin3a | glutamate receptor ionotropic, NMDA3A |
| chr4 | 55350425 | 55350662 | 238 | * | NA_peak_16 | 17 | Promoter (<= | 4 | 55350090 | 55381116 | 31027 | 1 | 19359 | ENSMUST0( | 335 | ENSMUSG0( | Rad23b | RAD23 homolog B, nucleotide excision repair protein |
| chr4 | 57497937 | 57498128 | 192 | * | NA_peak_16 | 37 | Intron (ENSM | 4 | 57434247 | 57712016 | 277770 | 1 | 242481 | ENSMUST0( | 63690 | NA | NA | NA |
| chr4 | 57659089 | 57659151 | 63 | * | NA_peak_16 | 44 | Intron (ENSM | 4 | 57637975 | 57709489 | 71515 | 1 | 242481 | ENSMUST0( | 21114 | NA | NA | NA |
| chr4 | 57664215 | 57664599 | 385 | * | NA_peak_16 | 22 | Intron (ENSM | 4 | 57637975 | 57709489 | 71515 | 1 | 242481 | ENSMUST0( | 26240 | NA | NA | NA |
| chr4 | 58388715 | 58389112 | 398 | * | NA_peak_16 | 41 | Distal Interge | 4 | 58441012 | 58441098 | 87 | 2 | 100526502 | ENSMUST0( | 51986 | ENSMUSG0( | Mir3095 | microRNA 3095 |
| chr4 | 58408119 | 58408385 | 267 | * | NA_peak_16 | 30 | Distal Interge | 4 | 58441012 | 58441098 | 87 | 2 | 100526502 | ENSMUST0( | 32713 | ENSMUSG0( | Mir3095 | microRNA 3095 |
| chr4 | 59343980 | 59344324 | 345 | * | NA_peak_16 | 34 | Intron (ENSM | 4 | 59329210 | 59333022 | 3813 | 2 | 634731 | ENSMUST0( | -10958 | ENSMUSG0( | Susd1 | sushi domain containing 1 |
| chr4 | 62448167 | 62448564 | 398 | * | NA_peak_16 | 30 | Distal Interge | 4 | 62457541 | 62459273 | 1733 | 2 | 71354 | ENSMUST0( | 10709 | ENSMUSG0( | Wdr31 | WD repeat domain 31 |
| chr4 | 62931203 | 62931590 | 388 | * | NA_peak_16 | 29 | Distal Interge | 4 | 62965573 | 63115748 | 150176 | 1 | 72701 | ENSMUST0( | -33983 | ENSMUSG0( | Zfp618 | zinc finger protein 618 |
| chr4 | 64416294 | 64416655 | 362 | * | NA_peak_16 | 26 | Distal Interge | 4 | 64254053 | 64276603 | 22551 | 2 | 102633436 | ENSMUST0( | -139691 | ENSMUSG0( | Gm11217 | predicted gene 11217 |
| chr4 | 64709540 | 64709848 | 309 | * | NA_peak_16 | 23 | Distal Interge | 4 | 65124174 | 65357509 | 233336 | 1 | 18491 | ENSMUST0( | -414326 | ENSMUSG0( | Pappa | pregnancy-associated plasma protein A |
| chr4 | 65996931 | 65997060 | 130 | * | NA_peak_16 | 37 | Intron (ENSM | 4 | 65605249 | 65613756 | 8508 | 1 | 69807 | ENSMUST0( | 391682 | ENSMUSG0( | Trim32 | tripartite motif-containing 32 |
| chr4 | 67940481 | 67940792 | 312 | * | NA_peak_16 | 60 | Distal Interge | 4 | 68312657 | 68321961 | 9305 | 2 | 102633975 | ENSMUST0( | 381169 | ENSMUSG0( | Gm12911 | predicted gene 12911 |
| chr4 | 68042535 | 68042904 | 370 | * | NA_peak_16 | 35 | Intron (ENSM | 4 | 68312657 | 68321961 | 9305 | 2 | 102633975 | ENSMUST0( | 279057 | ENSMUSG0( | Gm12911 | predicted gene 12911 |
| chr4 | 69711727 | 69712113 | 387 | * | NA_peak_16 | 40 | Distal Interge | 4 | 70242471 | 70243852 | 1382 | 2 | 214444 | ENSMUST0( | 531739 | ENSMUSG0( | Cdk5rap2 | CDK5 regulatory subunit associated protein 2 |
| chr4 | 70868602 | 70868848 | 247 | * | NA_peak_16 | 44 | Distal Interge | 4 | 70461826 | 70534995 | 73170 | 2 | 230316 | ENSMUST0( | -333607 | ENSMUSG0( | Megf9 | multiple EGF-like-domains 9 |
| chr4 | 74055445 | 74055787 | 343 | * | NA_peak_16 | 22 | Intron (ENSM | 4 | 74013444 | 74189928 | 176485 | 1 | 242506 | ENSMUST0( | 42001 | ENSMUSG0( | Frmd3 | FERM domain containing 3 |
| chr4 | 75353794 | 75354074 | 281 | * | NA_peak_16 | 30 | Distal Interge | 4 | 75277354 | 75278305 | 952 | 2 | 66928 | ENSMUST0( | -75489 | ENSMUSG0( | Dmac1 | distal membrane arm assembly complex 1 |
| chr4 | 75509984 | 75510217 | 234 | * | NA_peak_16 | 21 | Distal Interge | 4 | 75277354 | 75278305 | 952 | 2 | 66928 | ENSMUST0( | -231679 | ENSMUSG0( | Dmac1 | distal membrane arm assembly complex 1 |
| chr4 | 75925598 | 75925915 | 318 | * | NA_peak_16 | 44 | Distal Interge | 4 | 76039337 | 76084457 | 45121 | 2 | 19266 | ENSMUST0( | 158542 | ENSMUSG0( | Ptprd | protein tyrosine phosphatase, receptor type, D |
| chr4 | 82193145 | 82193515 | 371 | * | NA_peak_16 | 65 | Distal Interge | 4 | 82065957 | 82102673 | 36717 | 1 | 545641 | ENSMUST0( | 127188 | ENSMUSG0( | Gm5860 | predicted gene 5860 |
| chr4 | 85894707 | 85895009 | 303 | * | NA_peak_16 | 26 | Intron (ENSM | 4 | 86053915 | 86087961 | 34047 | 1 | 77739 | ENSMUST0( | -158906 | ENSMUSG0( | Adamtsl1 | ADAMTS-like 1 |
| chr4 | 87619743 | 87619881 | 139 | * | NA_peak_16 | 17 | Distal Interge | 4 | 87773953 | 87806294 | 32342 | 2 | 70122 | ENSMUST0( | 186413 | ENSMUSG0( | Mllt3 | myeloid/lymphoid or mixed-lineage leukemia; translocated to, 3 |
| chr4 | 87678778 | 87679190 | 413 | * | NA_peak_16 | 47 | Distal Interge | 4 | 87773953 | 87806294 | 32342 | 2 | 70122 | ENSMUST0( | 127104 | ENSMUSG0( | Mllt3 | myeloid/lymphoid or mixed-lineage leukemia; translocated to, 3 |
| chr4 | 88056333 | 88056513 | 181 | * | NA_peak_16 | 26 | Distal Interge | 4 | 87769925 | 88033364 | 263440 | 2 | 70122 | ENSMUST0( | -22969 | ENSMUSG0( | Mllt3 | myeloid/lymphoid or mixed-lineage leukemia; translocated to, 3 |
| chr4 | 88286060 | 88286080 | 21 | * | NA_peak_16 | 34 | Promoter (1-: | 4 | 88287635 | 88303630 | 15996 | 1 | 230393 | ENSMUST0( | -1555 | ENSMUSG0( | Focad | focadhesin |
| chr4 | 88773622 | 88774031 | 410 | * | NA_peak_16 | 40 | Promoter (<= | 4 | 88773834 | 88774801 | 968 | 1 | 545647 | ENSMUST0( | 0 | ENSMUSG0( | Gm13289 | predicted gene 13289 |
| chr4 | 88774560 | 88774838 | 279 | * | NA_peak_16 | 19 | Promoter (<= | 4 | 88773834 | 88774801 | 968 | 1 | 545647 | ENSMUST0( | 726 | ENSMUSG0( | Gm13289 | predicted gene 13289 |
| chr4 | 88777449 | 88777768 | 320 | * | NA_peak_16 | 36 | Promoter (<= | 4 | 88776922 | 88777470 | 549 | 1 | 545654 | ENSMUST0( | 527 | ENSMUSG0( | Gm13285 | predicted gene 13285 |
| chr4 | 88779690 | 88780162 | 473 | * | NA_peak_16 | 32 | Promoter (<= | 4 | 88779850 | 88780660 | 811 | 1 | 545648 | ENSMUST0( | 0 | ENSMUSG0( | Gm13272 | predicted gene 13272 |
| chr4 | 88781338 | 88781837 | 500 | * | NA_peak_16 | 107 | Promoter (<= | 4 | 88782131 | 88783592 | 1462 | 1 | 319146 | ENSMUST0( | -294 | ENSMUSG0( | Ifnz | interferon zeta |
| chr4 | 88785785 | 88786122 | 338 | * | NA_peak_16 | 44 | Promoter (<= | 4 | 88785709 | 88786257 | 549 | 1 | 545649 | ENSMUST0( | 76 | ENSMUSG0( | Gm13276 | predicted gene 13276 |
| chr4 | 88792316 | 88792731 | 416 | * | NA_peak_16 | 32 | Promoter (<= | 4 | 88791550 | 88792098 | 549 | 1 | 545654 | ENSMUST0( | 766 | ENSMUSG0( | Gm13285 | predicted gene 13285 |
| chr4 | 88793347 | 88793812 | 466 | * | NA_peak_16 | 30 | Promoter (<= | 4 | 88794487 | 88795035 | 549 | 1 | 545654 | ENSMUST0( | -675 | ENSMUSG0( | Gm13285 | predicted gene 13285 |
| chr4 | 88794692 | 88795057 | 366 | * | NA_peak_16 | 57 | Promoter (<= | 4 | 88794487 | 88795035 | 549 | 1 | 545654 | ENSMUST0( | 205 | ENSMUSG0( | Gm13285 | predicted gene 13285 |
| chr4 | 88799866 | 88800179 | 314 | * | NA_peak_16 | 66 | Promoter (<= | 4 | 88799665 | 88801126 | 1462 | 1 | 545652 | ENSMUST0( | 201 | ENSMUSG0( | Gm13275 | predicted gene 13275 |
| chr4 | 88802943 | 88803612 | 670 | * | NA_peak_16 | 57 | Promoter (<= | 4 | 88803254 | 88803802 | 549 | 1 | 545654 | ENSMUST0( | 0 | ENSMUSG0( | Gm13285 | predicted gene 13285 |
| chr4 | 88805359 | 88805730 | 372 | * | NA_peak_16 | 19 | Promoter (<= | 4 | 88805531 | 88808380 | 2850 | 1 | 668208 | ENSMUST0( | 0 | ENSMUSG0( | Gm13288 | predicted gene 13288 |
| chr4 | 89227023 | 89227315 | 293 | * | NA_peak_16 | 23 | Distal Interge | 4 | 89176374 | 89179474 | 3101 | 1 | 66902 | ENSMUST0( | 50649 | ENSMUSG0( | Mtap | methylthioadenosine phosphorylase |
| chr4 | 89310621 | 89311067 | 447 | * | NA_peak_16 | 35 | Promoter (<= | 4 | 89306299 | 89311039 | 4741 | 2 | 12579 | ENSMUST0( | 0 | ENSMUSG0( | Cdkn2b | cyclin dependent kinase inhibitor 2B |
| chr4 | 95511014 | 95511510 | 497 | * | NA_peak_16 | 46 | Distal Interge | 4 | 95557507 | 95926939 | 369433 | 1 | 75578 | ENSMUST0( | -45997 | ENSMUSG0( | Fggy | FGGY carbohydrate kinase domain containing |
| chr4 | 100560887 | 100561334 | 448 | * | NA_peak_16 | 17 | Distal Interge | 4 | 100478867 | 100550145 | 71279 | 1 | 381534 | ENSMUST0( | 82020 | ENSMUSG0( | Ube2u | ubiquitin-conjugating enzyme E2U (putative) |
| chr4 | 103273614 | 103274198 | 585 | * | NA_peak_16 | 20 | Intron (ENSM | 4 | 103230445 | 103290863 | 60419 | 2 | 70941 | ENSMUST0( | 16665 | ENSMUSG0( | 4921539E11 | RIKEN cDNA 4921539E11 gene |
| chr4 | 105740406 | 105740713 | 308 | * | NA_peak_16 | 70 | Distal Interge | 4 | 106241216 | 106290026 | 48811 | 1 | 105244641 | ENSMUST0( | -500503 | NA | Gm12724 | predicted gene 12724 |
| chr4 | 107922999 | 107923184 | 186 | * | NA_peak_16 | 19 | Promoter (<= | 4 | 107907865 | 107923447 | 15583 | 2 | 12896 | ENSMUST0( | 263 | ENSMUSG0( | Cpt2 | carnitine palmitoyltransferase 2 |
| chr4 | 108165486 | 108165616 | 131 | * | NA_peak_16 | 26 | Promoter (<= | 4 | 108165488 | 108179306 | 13819 | 1 | 52430 | ENSMUST0( | 0 | ENSMUSG0( | Echdc2 | enoyl Coenzyme A hydratase domain containing 2 |
| chr4 | 108848420 | 108848784 | 365 | * | NA_peak_16 | 26 | Promoter (<= | 4 | 108847785 | 108849413 | 1629 | 1 | 100087 | ENSMUST0( | 635 | ENSMUSG0( | Kti12 | KTI12 homolog, chromatin associated |
| chr4 | 109657084 | 109657336 | 253 | * | NA_peak_16 | 15 | Exon (ENSM | 4 | 109660876 | 109665372 | 4497 | 2 | 12580 | ENSMUST0( | 8036 | ENSMUSG0( | Cdkn2c | cyclin dependent kinase inhibitor 2C |
| chr4 | 109665920 | 109666220 | 301 | * | NA_peak_16 | 35 | Promoter (<= | 4 | 109660876 | 109665372 | 4497 | 2 | 12580 | ENSMUST0( | -548 | ENSMUSG0( | Cdkn2c | cyclin dependent kinase inhibitor 2C |
| chr4 | 110188474 | 110188671 | 198 | * | NA_peak_16 | 20 | Distal Interge | 4 | 110206359 | 110223148 | 16790 | 2 | 15572 | ENSMUST0( | 34477 | ENSMUSG0( | Elavl4 | ELAV like RNA binding protein 4 |
| chr4 | 110962647 | 110963008 | 362 | * | NA_peak_16 | 23 | Intron (ENSM | 4 | 111118841 | 111259039 | 140199 | 1 | 78933 | ENSMUST0( | -155833 | ENSMUSG0( | Agbl4 | ATP/GTP binding protein-like 4 |
| chr4 | 111572571 | 111572771 | 201 | * | NA_peak_16 | 16 | Intron (ENSM | 4 | 111566725 | 111657507 | 90783 | 1 | 78933 | ENSMUST0( | 5846 | ENSMUSG0( | Agbl4 | ATP/GTP binding protein-like 4 |
| chr4 | 115670652 | 115670802 | 151 | * | NA_peak_16 | 30 | Distal Interge | 4 | 115624725 | 115647723 | 22999 | 2 | 13120 | ENSMUST0( | -22929 | ENSMUSG0( | Cyp4b1 | cytochrome P450, family 4, subfamily b, polypeptide 1 |
| chr4 | 116276205 | 116276593 | 389 | * | NA_peak_16 | 20 | Intron (ENSM | 4 | 116308008 | 116309205 | 1198 | 2 | 17776 | ENSMUST0( | 32612 | ENSMUSG0( | Mast2 | microtubule associated serine/threonine kinase 2 |
| chr4 | 116685363 | 116685845 | 483 | * | NA_peak_16 | 23 | Promoter (<= | 4 | 116685544 | 116700822 | 15279 | 1 | 18477 | ENSMUST0( | 0 | ENSMUSG0( | Prdx1 | peroxiredoxin 1 |
| chr4 | 116826893 | 116827064 | 172 | * | NA_peak_16 | 23 | Intron (ENSM | 4 | 116819904 | 116821707 | 1804 | 2 | 242642 | ENSMUST0( | -5186 | ENSMUSG0( | Hpdl | 4-hydroxyphenylpyruvate dioxygenase-like |
| chr4 | 117971434 | 117971644 | 211 | * | NA_peak_16 | 13 | Promoter (2-: | 4 | 117940446 | 117968448 | 28003 | 2 | 20441 | ENSMUST0( | -2986 | ENSMUSG0( | St3gal3 | ST3 beta-galactoside alpha-2,3-sialyltransferase 3 |
| chr4 | 118179580 | 118179701 | 122 | * | NA_peak_16 | 20 | Promoter (<= | 4 | 118136959 | 118179654 | 42696 | 2 | 230674 | ENSMUST0( | 0 | ENSMUSG0( | Kdm4a | lysine (K)-specific demethylase 4A |
| chr4 | 123325565 | 123325716 | 152 | * | NA_peak_16 | 39 | Promoter (1-: | 4 | 123321875 | 123323634 | 1760 | 2 | 64059 | ENSMUST0( | -1931 | ENSMUSG0( | Oxct2a | 3-oxoacid CoA transferase 2A |
| chr4 | 126306824 | 126306872 | 49 | * | NA_peak_16 | 24 | Intron (ENSM | 4 | 126300530 | 126309785 | 9256 | 1 | 329941 | ENSMUST0( | 6294 | ENSMUSG0( | Col8a2 | collagen, type VIII, alpha 2 |
| chr4 | 126452505 | 126452684 | 180 | * | NA_peak_16 | 26 | Intron (ENSM | 4 | 126439503 | 126459207 | 19705 | 2 | 236511 | ENSMUST0( | 6523 | ENSMUSG0( | Ago1 | argonaute RISC catalytic subunit 1 |
| chr4 | 127010693 | 127010970 | 278 | * | NA_peak_16 | 40 | Distal Interge | 4 | 127021324 | 127031236 | 9913 | 1 | 71514 | ENSMUST0( | -10354 | ENSMUSG0( | Sfpq | splicing factor proline/glutamine rich (polypyrimidine tract binding protein associated) |
| chr4 | 128883498 | 128883833 | 336 | * | NA_peak_16 | 44 | Promoter (<= | 4 | 128883580 | 128903565 | 19986 | 1 | 67525 | ENSMUST0( | 0 | ENSMUSG0( | Trim62 | tripartite motif-containing 62 |
| chr4 | 132809446 | 132809725 | 280 | * | NA_peak_16 | 17 | Distal Interge | 4 | 132781843 | 132796387 | 14545 | 2 | 230787 | ENSMUST0( | -13059 | ENSMUSG0( | Themis2 | thymocyte selection associated family member 2 |
| chr4 | 133025399 | 133025734 | 336 | * | NA_peak_16 | 30 | Intron (ENSM | 4 | 133012204 | 133077645 | 65442 | 1 | 230793 | ENSMUST0( | 13195 | ENSMUSG0( | Ahdc1 | AT hook, DNA binding motif, containing 1 |
| chr4 | 133038473 | 133038920 | 448 | * | NA_peak_16 | 18 | Promoter (<= | 4 | 133039493 | 133077242 | 37750 | 1 | 230793 | ENSMUST0( | -573 | ENSMUSG0( | Ahdc1 | AT hook, DNA binding motif, containing 1 |
| chr4 | 133240951 | 133241519 | 569 | * | NA_peak_16 | 19 | Promoter (<= | 4 | 133240818 | 133252929 | 12112 | 1 | 53608 | ENSMUST0( | 133 | ENSMUSG0( | Map3k6 | mitogen-activated protein kinase kinase kinase 6 |
| chr4 | 133574689 | 133574960 | 272 | * | NA_peak_16 | 14 | Promoter (<= | 4 | 133574745 | 133584243 | 9499 | 1 | 242691 | ENSMUST0( | 0 | ENSMUSG0( | Gpatch3 | G patch domain containing 3 |
| chr4 | 133752968 | 133753310 | 343 | * | NA_peak_16 | 24 | Promoter (<= | 4 | 133679008 | 133753611 | 74604 | 2 | 93760 | ENSMUST0( | 301 | ENSMUSG0( | Arid1a | AT rich interactive domain 1A (SWI-like) |
| chr4 | 134018167 | 134018607 | 441 | * | NA_peak_16 | 22 | Promoter (<= | 4 | 134007948 | 134018382 | 10435 | 2 | 83557 | ENSMUST0( | 0 | ENSMUSG0( | Lin28a | lin-28 homolog A (C. elegans) |
| chr4 | 135285737 | 135285957 | 221 | * | NA_peak_16 | 30 | Distal Interge | 4 | 135223464 | 135272814 | 49351 | 2 | 29876 | ENSMUST0( | -12923 | ENSMUSG0( | Clic4 | chloride intracellular channel 4 (mitochondrial) |
| chr4 | 136135148 | 136135424 | 277 | * | NA_peak_16 | 24 | Distal Interge | 4 | 136143497 | 136145392 | 1896 | 1 | 15903 | ENSMUST0( | -8073 | ENSMUSG0( | Id3 | inhibitor of DNA binding 3 |
| chr4 | 137994185 | 137994509 | 325 | * | NA_peak_16 | 23 | Promoter (<= | 4 | 137993513 | 138206154 | 212642 | 1 | 230861 | ENSMUST0( | 672 | ENSMUSG0( | Eif4g3 | eukaryotic translation initiation factor 4 gamma, 3 |
| chr4 | 138394917 | 138395234 | 318 | * | NA_peak_16 | 19 | Promoter (1-: | 4 | 138394092 | 138396528 | 2437 | 2 | 625638 | ENSMUST0( | 1294 | ENSMUSG0( | Fam43b | family with sequence similarity 43, member B |
| chr4 | 140695506 | 140695527 | 22 | * | NA_peak_16 | 16 | Distal Interge | 4 | 140700541 | 140723220 | 22680 | 1 | 108911 | ENSMUST0( | -5014 | ENSMUSG0( | Rcc2 | regulator of chromosome condensation 2 |
| chr4 | 140962603 | 140962874 | 272 | * | NA_peak_16 | 28 | Promoter (1-: | 4 | 140961403 | 140974297 | 12895 | 1 | 67680 | ENSMUST0( | 1200 | ENSMUSG0( | Sdhb | succinate dehydrogenase complex, subunit B, iron sulfur (Ip) |
| chr4 | 141139469 | 141139729 | 261 | * | NA_peak_16 | 23 | Promoter (<= | 4 | 141113001 | 141139727 | 26727 | 2 | 213491 | ENSMUST0( | 0 | ENSMUSG0( | Szrd1 | SUZ RNA binding domain containing 1 |
| chr4 | 141210102 | 141210492 | 391 | * | NA_peak_16 | 13 | Distal Interge | 4 | 141213956 | 141220148 | 6193 | 1 | 76166 | ENSMUST0( | -3464 | ENSMUSG0( | Cplane2 | ciliogenesis and planar polarity effector 2 |
| chr4 | 141722726 | 141723055 | 330 | * | NA_peak_16 | 30 | Promoter (<= | 4 | 141677549 | 141723419 | 45871 | 2 | 68817 | ENSMUST0( | 364 | ENSMUSG0( | Ddi2 | DNA-damage inducible protein 2 |
| chr4 | 143138031 | 143138190 | 160 | * | NA_peak_16 | 30 | Promoter (1-: | 4 | 143136100 | 143136162 | 63 | 2 | 102465618 | ENSMUST0( | -1869 | ENSMUSG0( | Mir7021 | microRNA 7021 |

| | | | | | | | | | | | | | | | | | | |
|---|---|---|---|---|---|---|---|---|---|---|---|---|---|---|---|---|---|---|
| chr4 | 144089822 | 144090192 | 371 | * | NA_peak_16 | 37 | Distal Interge | 4 | 144106474 | 144110101 | 3628 | 1 | 100041077 | ENSMUST0 | -16282 | ENSMUSG0 | Gm13102 | predicted gene 13102 |
| chr4 | 145429905 | 145430097 | 193 | * | NA_peak_16 | 28 | Intron (ENSM | 4 | 145464204 | 145465487 | 1284 | 1 | 545700 | ENSMUST0 | -34107 | ENSMUSG0 | Smarca5-ps | SWI/SNF related, matrix associated, actin depenent ragulator of chromatin, subfamily a, member 5, pseudogene |
| chr4 | 145440684 | 145441023 | 340 | * | NA_peak_16 | 32 | Intron (ENSM | 4 | 145464204 | 145465487 | 1284 | 1 | 545700 | ENSMUST0 | -23181 | ENSMUSG0 | Smarca5-ps | SWI/SNF related, matrix associated, actin depenent ragulator of chromatin, subfamily a, member 5, pseudogene |
| chr4 | 145494207 | 145494438 | 232 | * | NA_peak_16 | 16 | Distal Interge | 4 | 145510759 | 145539188 | 28430 | 1 | 101056073 | ENSMUST0 | -16321 | ENSMUSG0 | Zfp990 | zinc finger protein 990 |
| chr4 | 145594009 | 145595197 | 1189 | * | NA_peak_16 | 30 | Intron (ENSM | 4 | 145585203 | 145625341 | 40139 | 1 | 433801 | ENSMUST0 | 8806 | ENSMUSG0 | Zfp268 | zinc finger protein 268 |
| chr4 | 145690302 | 145690926 | 625 | * | NA_peak_16 | 62 | Exon (ENSM | 4 | 145696161 | 145702640 | 6480 | 1 | 100041379 | ENSMUST0 | -5235 | ENSMUSG0 | Zfp980 | zinc finger protein 980 |
| chr4 | 145704818 | 145705031 | 214 | * | NA_peak_16 | 23 | Distal Interge | 4 | 145696161 | 145702640 | 6480 | 1 | 100041379 | ENSMUST0 | 8657 | ENSMUSG0 | Zfp980 | zinc finger protein 980 |
| chr4 | 145714052 | 145714448 | 397 | * | NA_peak_16 | 37 | Distal Interge | 4 | 145696161 | 145702640 | 6480 | 1 | 100041379 | ENSMUST0 | 17891 | ENSMUSG0 | Zfp980 | zinc finger protein 980 |
| chr4 | 145717403 | 145717564 | 162 | * | NA_peak_16 | 32 | Distal Interge | 4 | 145696161 | 145702640 | 6480 | 1 | 100041379 | ENSMUST0 | 21242 | ENSMUSG0 | Zfp980 | zinc finger protein 980 |
| chr4 | 145768251 | 145768737 | 487 | * | NA_peak_16 | 24 | Distal Interge | 4 | 145825237 | 145829942 | 4706 | 1 | 70005 | ENSMUST0 | -56500 | NA | Znf41-ps | ZNF41, pseudogene |
| chr4 | 145774455 | 145774856 | 402 | * | NA_peak_16 | 24 | Exon (ENSM | 4 | 145825237 | 145829942 | 4706 | 1 | 70005 | ENSMUST0 | -50381 | NA | Znf41-ps | ZNF41, pseudogene |
| chr4 | 145780020 | 145780365 | 346 | * | NA_peak_16 | 82 | Distal Interge | 4 | 145825237 | 145829942 | 4706 | 1 | 70005 | ENSMUST0 | -44872 | NA | Znf41-ps | ZNF41, pseudogene |
| chr4 | 145804484 | 145804522 | 39 | * | NA_peak_16 | 34 | Distal Interge | 4 | 145825237 | 145829942 | 4706 | 1 | 70005 | ENSMUST0 | -20715 | NA | Znf41-ps | ZNF41, pseudogene |
| chr4 | 145805029 | 145805196 | 168 | * | NA_peak_16 | 34 | Distal Interge | 4 | 145825237 | 145829942 | 4706 | 1 | 70005 | ENSMUST0 | -20041 | NA | Znf41-ps | ZNF41, pseudogene |
| chr4 | 145846718 | 145847382 | 665 | * | NA_peak_16 | 24 | Distal Interge | 4 | 145837219 | 145837998 | 780 | 1 | 236069 | ENSMUST0 | 9499 | ENSMUSG0 | Gm13238 | carnitine deficiency-associated gene expressed in ventricle 3 pseudogene |
| chr4 | 145848038 | 145848282 | 245 | * | NA_peak_16 | 24 | Distal Interge | 4 | 145837219 | 145837998 | 780 | 1 | 236069 | ENSMUST0 | 10819 | ENSMUSG0 | Gm13238 | carnitine deficiency-associated gene expressed in ventricle 3 pseudogene |
| chr4 | 145862138 | 145862679 | 542 | * | NA_peak_16 | 24 | Distal Interge | 4 | 145837219 | 145837998 | 780 | 1 | 236069 | ENSMUST0 | 24919 | ENSMUSG0 | Gm13238 | carnitine deficiency-associated gene expressed in ventricle 3 pseudogene |
| chr4 | 145866175 | 145866523 | 349 | * | NA_peak_16 | 22 | Distal Interge | 4 | 145837219 | 145837998 | 780 | 1 | 236069 | ENSMUST0 | 28956 | ENSMUSG0 | Gm13238 | carnitine deficiency-associated gene expressed in ventricle 3 pseudogene |
| chr4 | 145881228 | 145881314 | 87 | * | NA_peak_16 | 29 | Intron (ENSM | 4 | 145837219 | 145837998 | 780 | 1 | 236069 | ENSMUST0 | 44009 | ENSMUSG0 | Gm13238 | carnitine deficiency-associated gene expressed in ventricle 3 pseudogene |
| chr4 | 145887572 | 145887894 | 323 | * | NA_peak_16 | 28 | Intron (ENSM | 4 | 145837219 | 145837998 | 780 | 1 | 236069 | ENSMUST0 | 50353 | ENSMUSG0 | Gm13238 | carnitine deficiency-associated gene expressed in ventricle 3 pseudogene |
| chr4 | 146054446 | 146054936 | 491 | * | NA_peak_16 | 14 | Distal Interge | 4 | 146067752 | 146070928 | 3177 | 1 | 627585 | ENSMUST0 | -12816 | NA | Gm13034 | SWI/SNF related, matrix associated, actin dependent regulator of chromatin, subfamily a, member 5 pseudogene |
| chr4 | 146096354 | 146096572 | 219 | * | NA_peak_16 | 49 | Promoter (<= | 4 | 146097298 | 146098597 | 1300 | 1 | 626316 | ENSMUST0 | -726 | ENSMUSG0 | Zfp987 | zinc finger protein 987 |
| chr4 | 146153001 | 146153407 | 407 | * | NA_peak_16 | 23 | Distal Interge | 4 | 146156824 | 146158927 | 2104 | 1 | 667666 | ENSMUST0 | -3417 | ENSMUSG0 | Zfp600 | zinc finger protein 600 |
| chr4 | 146178007 | 146178345 | 339 | * | NA_peak_16 | 14 | Exon (ENSM | 4 | 146161909 | 146195169 | 33261 | 1 | 667666 | ENSMUST0 | 16098 | ENSMUSG0 | Zfp600 | zinc finger protein 600 |
| chr4 | 146189417 | 146190151 | 735 | * | NA_peak_16 | 25 | Intron (ENSM | 4 | 146161909 | 146195169 | 33261 | 1 | 667666 | ENSMUST0 | 27508 | ENSMUSG0 | Zfp600 | zinc finger protein 600 |
| chr4 | 146193060 | 146193275 | 216 | * | NA_peak_16 | 15 | Intron (ENSM | 4 | 146161909 | 146195169 | 33261 | 1 | 667666 | ENSMUST0 | 31151 | ENSMUSG0 | Zfp600 | zinc finger protein 600 |
| chr4 | 146244079 | 146244388 | 310 | * | NA_peak_16 | 32 | Distal Interge | 4 | 146252173 | 146272975 | 20803 | 1 | 666532 | ENSMUST0 | -7785 | ENSMUSG0 | Zfp991 | zinc finger protein 991 |
| chr4 | 146427376 | 146427694 | 319 | * | NA_peak_16 | 24 | Distal Interge | 4 | 146449023 | 146470292 | 21270 | 1 | 433791 | ENSMUST0 | -21329 | ENSMUSG0 | Zfp992 | zinc finger protein 992 |
| chr4 | 146463534 | 146463570 | 37 | * | NA_peak_16 | 19 | Exon (ENSM | 4 | 146449023 | 146470292 | 21270 | 1 | 433791 | ENSMUST0 | 14511 | ENSMUSG0 | Zfp992 | zinc finger protein 992 |
| chr4 | 146478668 | 146478807 | 140 | * | NA_peak_16 | 52 | Distal Interge | 4 | 146502027 | 146539392 | 37366 | 1 | 100041433 | ENSMUST0 | -23220 | ENSMUSG0 | Zfp981 | zinc finger protein 981 |
| chr4 | 146524317 | 146524485 | 169 | * | NA_peak_16 | 18 | Intron (ENSM | 4 | 146514920 | 146539395 | 24476 | 1 | 100041433 | ENSMUST0 | 9397 | ENSMUSG0 | Zfp981 | zinc finger protein 981 |
| chr4 | 146567541 | 146567965 | 425 | * | NA_peak_16 | 34 | Distal Interge | 4 | 146519039 | 146546873 | 27835 | 2 | 67524 | ENSMUST0 | -20668 | ENSMUSG0 | 1700095A21 | RIKEN cDNA 1700095A21 gene |
| chr4 | 146578589 | 146579083 | 495 | * | NA_peak_16 | 22 | Distal Interge | 4 | 146519039 | 146546873 | 27835 | 2 | 67524 | ENSMUST0 | -31716 | ENSMUSG0 | 1700095A21 | RIKEN cDNA 1700095A21 gene |
| chr4 | 146601208 | 146601462 | 255 | * | NA_peak_16 | 22 | Distal Interge | 4 | 146519039 | 146546873 | 27835 | 2 | 67524 | ENSMUST0 | -54335 | ENSMUSG0 | 1700095A21 | RIKEN cDNA 1700095A21 gene |
| chr4 | 146630091 | 146630564 | 474 | * | NA_peak_16 | 26 | Intron (ENSM | 4 | 146519039 | 146546873 | 27835 | 2 | 67524 | ENSMUST0 | -83218 | ENSMUSG0 | 1700095A21 | RIKEN cDNA 1700095A21 gene |
| chr4 | 146710677 | 146711669 | 993 | * | NA_peak_16 | 51 | Distal Interge | 4 | 146519039 | 146546873 | 27835 | 2 | 67524 | ENSMUST0 | -163804 | ENSMUSG0 | 1700095A21 | RIKEN cDNA 1700095A21 gene |
| chr4 | 146739334 | 146739909 | 576 | * | NA_peak_16 | 40 | Distal Interge | 4 | 146519039 | 146546873 | 27835 | 2 | 67524 | ENSMUST0 | -192461 | ENSMUSG0 | 1700095A21 | RIKEN cDNA 1700095A21 gene |
| chr4 | 146742161 | 146742554 | 394 | * | NA_peak_16 | 17 | Distal Interge | 4 | 146519039 | 146546873 | 27835 | 2 | 67524 | ENSMUST0 | -195288 | ENSMUSG0 | 1700095A21 | RIKEN cDNA 1700095A21 gene |
| chr4 | 146893687 | 146894314 | 628 | * | NA_peak_16 | 13 | Exon (ENSM | 4 | 147021850 | 147060794 | 38945 | 1 | 100043034 | ENSMUST0 | -127536 | ENSMUSG0 | Rex2 | reduced expression 2 |
| chr4 | 147022785 | 147023092 | 308 | * | NA_peak_16 | 60 | Promoter (<= | 4 | 147021878 | 147023811 | 1934 | 1 | 100043034 | ENSMUST0 | 907 | ENSMUSG0 | Rex2 | reduced expression 2 |
| chr4 | 147114143 | 147114648 | 506 | * | NA_peak_16 | 17 | Distal Interge | 4 | 147063366 | 147068436 | 5071 | 2 | 71270 | ENSMUST0 | -45707 | ENSMUSG0 | 4933438K21 | RIKEN cDNA 4933438K21 gene |
| chr4 | 147115359 | 147115870 | 512 | * | NA_peak_16 | 22 | Distal Interge | 4 | 147063366 | 147068436 | 5071 | 2 | 71270 | ENSMUST0 | -46923 | ENSMUSG0 | 4933438K21 | RIKEN cDNA 4933438K21 gene |
| chr4 | 147117789 | 147117993 | 205 | * | NA_peak_16 | 19 | Distal Interge | 4 | 147063366 | 147068436 | 5071 | 2 | 71270 | ENSMUST0 | -49353 | ENSMUSG0 | 4933438K21 | RIKEN cDNA 4933438K21 gene |
| chr4 | 147124429 | 147124991 | 563 | * | NA_peak_16 | 32 | Distal Interge | 4 | 147063366 | 147068436 | 5071 | 2 | 71270 | ENSMUST0 | -55993 | ENSMUSG0 | 4933438K21 | RIKEN cDNA 4933438K21 gene |
| chr4 | 147126171 | 147126552 | 382 | * | NA_peak_16 | 33 | Exon (ENSM | 4 | 147063366 | 147068436 | 5071 | 2 | 71270 | ENSMUST0 | -57735 | ENSMUSG0 | 4933438K21 | RIKEN cDNA 4933438K21 gene |
| chr4 | 147134905 | 147135369 | 465 | * | NA_peak_16 | 30 | Intron (ENSM | 4 | 147063366 | 147068436 | 5071 | 2 | 71270 | ENSMUST0 | -66469 | ENSMUSG0 | 4933438K21 | RIKEN cDNA 4933438K21 gene |
| chr4 | 147140676 | 147140946 | 271 | * | NA_peak_16 | 14 | Intron (ENSM | 4 | 147063366 | 147068436 | 5071 | 2 | 71270 | ENSMUST0 | -72240 | ENSMUSG0 | 4933438K21 | RIKEN cDNA 4933438K21 gene |
| chr4 | 147176398 | 147176714 | 317 | * | NA_peak_16 | 37 | Intron (ENSM | 4 | 147063366 | 147068436 | 5071 | 2 | 71270 | ENSMUST0 | -107962 | ENSMUSG0 | 4933438K21 | RIKEN cDNA 4933438K21 gene |
| chr4 | 147180907 | 147181536 | 630 | * | NA_peak_17 | 30 | Distal Interge | 4 | 147063366 | 147068436 | 5071 | 2 | 71270 | ENSMUST0 | -112471 | ENSMUSG0 | 4933438K21 | RIKEN cDNA 4933438K21 gene |
| chr4 | 147232370 | 147232808 | 439 | * | NA_peak_17 | 23 | Exon (ENSM | 4 | 147063366 | 147068436 | 5071 | 2 | 71270 | ENSMUST0 | -163934 | ENSMUSG0 | 4933438K21 | RIKEN cDNA 4933438K21 gene |
| chr4 | 147236696 | 147237096 | 401 | * | NA_peak_17 | 13 | Intron (ENSM | 4 | 147063366 | 147068436 | 5071 | 2 | 71270 | ENSMUST0 | -168260 | ENSMUSG0 | 4933438K21 | RIKEN cDNA 4933438K21 gene |
| chr4 | 147241560 | 147241908 | 349 | * | NA_peak_17 | 20 | Intron (ENSM | 4 | 147063366 | 147068436 | 5071 | 2 | 71270 | ENSMUST0 | -173124 | ENSMUSG0 | 4933438K21 | RIKEN cDNA 4933438K21 gene |
| chr4 | 147243019 | 147243377 | 359 | * | NA_peak_17 | 18 | Intron (ENSM | 4 | 147063366 | 147068436 | 5071 | 2 | 71270 | ENSMUST0 | -174583 | ENSMUSG0 | 4933438K21 | RIKEN cDNA 4933438K21 gene |
| chr4 | 147300202 | 147300535 | 334 | * | NA_peak_17 | 20 | Distal Interge | 4 | 147492417 | 147513486 | 21070 | 1 | 195531 | ENSMUST0 | -191882 | ENSMUSG0 | Zfp982 | zinc finger protein 982 |
| chr4 | 147312752 | 147313107 | 356 | * | NA_peak_17 | 19 | Intron (ENSM | 4 | 147492417 | 147513486 | 21070 | 1 | 195531 | ENSMUST0 | -179310 | ENSMUSG0 | Zfp982 | zinc finger protein 982 |
| chr4 | 147321975 | 147322390 | 416 | * | NA_peak_17 | 22 | Intron (ENSM | 4 | 147492417 | 147513486 | 21070 | 1 | 195531 | ENSMUST0 | -170027 | ENSMUSG0 | Zfp982 | zinc finger protein 982 |
| chr4 | 147339802 | 147340504 | 703 | * | NA_peak_17 | 24 | Distal Interge | 4 | 147492417 | 147513486 | 21070 | 1 | 195531 | ENSMUST0 | -151913 | ENSMUSG0 | Zfp982 | zinc finger protein 982 |
| chr4 | 147365021 | 147365366 | 346 | * | NA_peak_17 | 15 | Intron (ENSM | 4 | 147492417 | 147513486 | 21070 | 1 | 195531 | ENSMUST0 | -127051 | ENSMUSG0 | Zfp982 | zinc finger protein 982 |
| chr4 | 147382848 | 147383077 | 230 | * | NA_peak_17 | 20 | Intron (ENSM | 4 | 147492417 | 147513486 | 21070 | 1 | 195531 | ENSMUST0 | -109340 | ENSMUSG0 | Zfp982 | zinc finger protein 982 |
| chr4 | 147398766 | 147398880 | 115 | * | NA_peak_17 | 33 | Distal Interge | 4 | 147492417 | 147513486 | 21070 | 1 | 195531 | ENSMUST0 | -93537 | ENSMUSG0 | Zfp982 | zinc finger protein 982 |
| chr4 | 147427948 | 147428533 | 586 | * | NA_peak_17 | 30 | Distal Interge | 4 | 147492417 | 147513486 | 21070 | 1 | 195531 | ENSMUST0 | -63884 | ENSMUSG0 | Zfp982 | zinc finger protein 982 |
| chr4 | 147510057 | 147510380 | 324 | * | NA_peak_17 | 37 | Promoter (2- | 4 | 147507600 | 147513420 | 5821 | 1 | 195531 | ENSMUST0 | 2457 | ENSMUSG0 | Zfp982 | zinc finger protein 982 |
| chr4 | 147527846 | 147528260 | 415 | * | NA_peak_17 | 26 | Exon (ENSM | 4 | 147507600 | 147513420 | 5821 | 1 | 195531 | ENSMUST0 | 20246 | ENSMUSG0 | Zfp982 | zinc finger protein 982 |
| chr4 | 147624031 | 147624360 | 330 | * | NA_peak_17 | 15 | Exon (ENSM | 4 | 147613911 | 147642496 | 28586 | 2 | 112422 | ENSMUST0 | 18136 | ENSMUSG0 | Zfp979 | zinc finger protein 979 |
| chr4 | 147643825 | 147644170 | 346 | * | NA_peak_17 | 63 | Promoter (1- | 4 | 147611937 | 147642513 | 30577 | 2 | 112422 | ENSMUST0 | -1312 | ENSMUSG0 | Zfp979 | zinc finger protein 979 |
| chr4 | 147655254 | 147655569 | 316 | * | NA_peak_17 | 28 | Distal Interge | 4 | 147611937 | 147642513 | 30577 | 2 | 112422 | ENSMUST0 | -12741 | ENSMUSG0 | Zfp979 | zinc finger protein 979 |
| chr4 | 147718414 | 147718920 | 507 | * | NA_peak_17 | 26 | Distal Interge | 4 | 147673502 | 147702553 | 29052 | 2 | 100503584 | ENSMUST0 | -15861 | ENSMUSG0 | Zfp534 | zinc finger protein 534 |
| chr4 | 147723435 | 147723698 | 264 | * | NA_peak_17 | 36 | Distal Interge | 4 | 147673502 | 147702553 | 29052 | 2 | 100503584 | ENSMUST0 | -20882 | ENSMUSG0 | Zfp534 | zinc finger protein 534 |
| chr4 | 147800721 | 147801089 | 369 | * | NA_peak_17 | 15 | Intron (ENSM | 4 | 147753974 | 147809788 | 55815 | 2 | 100041677 | ENSMUST0 | 8699 | ENSMUSG0 | Zfp984 | zinc finger protein 984 |
| chr4 | 149007866 | 149008141 | 276 | * | NA_peak_17 | 15 | Intron (ENSM | 4 | 148947492 | 148948914 | 1423 | 2 | 108168987 | ENSMUST0 | -58952 | NA | Gm13205 | predicted gene 13205 |
| chr4 | 149306993 | 149307415 | 423 | * | NA_peak_17 | 35 | Promoter (<= | 4 | 149180236 | 149307506 | 127271 | 2 | 16561 | ENSMUST0 | 91 | ENSMUSG0 | Kif1b | kinesin family member 1B |
| chr4 | 150066810 | 150067026 | 217 | * | NA_peak_17 | 28 | Promoter (1- | 4 | 150068454 | 150068555 | 102 | 1 | 723848 | ENSMUST0 | -1428 | ENSMUSG0 | Mir34a | microRNA 34a |
| chr4 | 152009205 | 152009709 | 505 | * | NA_peak_17 | 26 | Promoter (<= | 4 | 152008803 | 152017680 | 8878 | 1 | 242785 | ENSMUST0 | 402 | ENSMUSG0 | Klhl21 | kelch-like 21 |
| chr4 | 152042497 | 152043091 | 595 | * | NA_peak_17 | 17 | Intron (ENSM | 4 | 152039363 | 152060042 | 20680 | 1 | 74035 | ENSMUST0 | 3134 | ENSMUSG0 | Nol9 | nucleolar protein 9 |
| chr4 | 155682446 | 155682803 | 358 | * | NA_peak_17 | 26 | Distal Interge | 4 | 155694342 | 155696481 | 2140 | 1 | 230991 | ENSMUST0 | -11539 | ENSMUSG0 | Fndc10 | fibronectin type III domain containing 10 |
| chr4 | 155901786 | 155902041 | 256 | * | NA_peak_17 | 18 | Promoter (<= | 4 | 155901668 | 155902770 | 1103 | 1 | 140500 | ENSMUST0 | 118 | ENSMUSG0 | Acap3 | ArfGAP with coiled-coil, ankyrin repeat and PH domains 3 |
| chr4 | 155943774 | 155944103 | 330 | * | NA_peak_17 | 24 | Promoter (<= | 4 | 155943831 | 155957711 | 13881 | 1 | 140499 | ENSMUST0 | 0 | ENSMUSG0 | Ube2j2 | ubiquitin-conjugating enzyme E2J 2 |
| chr4 | 156323665 | 156324190 | 526 | * | NA_peak_17 | 30 | Distal Interge | 4 | 156331103 | 156339496 | 8394 | 1 | 22314 | ENSMUST0 | -6913 | ENSMUSG0 | Vmn2r129 | vomeronasal 2, receptor 129 |
| chr4 | 156330413 | 156330892 | 480 | * | NA_peak_17 | 14 | Promoter (<= | 4 | 156331103 | 156339496 | 8394 | 1 | 22314 | ENSMUST0 | -211 | ENSMUSG0 | Vmn2r129 | vomeronasal 2, receptor 129 |
| chr5 | 3185368 | 3185680 | 313 | * | NA_peak_17 | 19 | Distal Interge | 5 | 3341485 | 3344453 | 2969 | 1 | 12571 | ENSMUST0 | -155805 | ENSMUSG0 | Cdk6 | cyclin-dependent kinase 6 |
| chr5 | 3764839 | 3765613 | 775 | * | NA_peak_17 | 35 | Intron (ENSM | 5 | 3693880 | 3795490 | 101611 | 2 | 70797 | ENSMUST0 | 29877 | ENSMUSG0 | Ankib1 | ankyrin repeat and IBR domain containing 1 |
| chr5 | 3828611 | 3828941 | 331 | * | NA_peak_17 | 13 | Exon (ENSM | 5 | 3812778 | 3818569 | 5792 | 1 | 79264 | ENSMUST0 | 15833 | ENSMUSG0 | Krit1 | KRIT1, ankyrin repeat containing |
| chr5 | 4300660 | 4300899 | 240 | * | NA_peak_17 | 19 | Distal Interge | 5 | 4192367 | 4197651 | 5285 | 1 | 208595 | ENSMUST0 | 108293 | ENSMUSG0 | Mterf1b | mitochondrial transcription termination factor 1b |
| chr5 | 4387229 | 4387922 | 694 | * | NA_peak_17 | 35 | Distal Interge | 5 | 4192367 | 4197651 | 5285 | 1 | 208595 | ENSMUST0 | 194862 | ENSMUSG0 | Mterf1b | mitochondrial transcription termination factor 1b |
| chr5 | 4516514 | 4516809 | 296 | * | NA_peak_17 | 38 | Distal Interge | 5 | 4753839 | 4758035 | 4197 | 2 | 14362 | ENSMUST0 | 241226 | ENSMUSG0 | Fzd1 | frizzled class receptor 1 |
| chr5 | 5662188 | 5662344 | 157 | * | NA_peak_17 | 30 | Promoter (<= | 5 | 5649808 | 5663292 | 13485 | 2 | 207686 | ENSMUST0 | 948 | ENSMUSG0 | Cfap69 | cilia and flagella associated protein 69 |
| chr5 | 5820717 | 5820782 | 66 | * | NA_peak_17 | 38 | Distal Interge | 5 | 5736317 | 5749326 | 13010 | 2 | 70358 | ENSMUST0 | -71391 | ENSMUSG0 | Steap1 | six transmembrane epithelial antigen of the prostate 1 |
| chr5 | 6142066 | 6142411 | 346 | * | NA_peak_17 | 54 | Distal Interge | 5 | 5736317 | 5749326 | 13010 | 2 | 70358 | ENSMUST0 | -392740 | ENSMUSG0 | Steap1 | six transmembrane epithelial antigen of the prostate 1 |
| chr5 | 6562993 | 6563242 | 250 | * | NA_peak_17 | 46 | Distal Interge | 5 | 6769030 | 6876523 | 107494 | 2 | 207618 | ENSMUST0 | 313281 | ENSMUSG0 | Zfp804b | zinc finger protein 804B |
| chr5 | 6740209 | 6740606 | 398 | * | NA_peak_17 | 49 | Distal Interge | 5 | 6769030 | 6876523 | 107494 | 2 | 207618 | ENSMUST0 | 135917 | ENSMUSG0 | Zfp804b | zinc finger protein 804B |
| chr5 | 8130248 | 8130396 | 149 | * | NA_peak_17 | 14 | Exon (ENSM | 5 | 8077839 | 8121095 | 43257 | 2 | 11496 | ENSMUST0 | -9153 | ENSMUSG0 | Adam22 | a disintegrin and metallopeptidase domain 22 |
| chr5 | 8154569 | 8154647 | 79 | * | NA_peak_17 | 16 | Intron (ENSM | 5 | 8077839 | 8121095 | 43257 | 2 | 11496 | ENSMUST0 | -33474 | ENSMUSG0 | Adam22 | a disintegrin and metallopeptidase domain 22 |
| chr5 | 8645205 | 8645492 | 288 | * | NA_peak_17 | 26 | Distal Interge | 5 | 8660077 | 8748575 | 88499 | 1 | 18671 | ENSMUST0 | -14585 | ENSMUSG0 | Abcb1a | ATP-binding cassette, sub-family B (MDR/TAP), member 1A |
| chr5 | 10570929 | 10570939 | 11 | * | NA_peak_17 | 22 | Distal Interge | 5 | 10865029 | 10870808 | 5780 | 1 | 623849 | ENSMUST0 | -294090 | ENSMUSG0 | Gm6455 | predicted gene 6455 |
| chr5 | 13857465 | 13857704 | 240 | * | NA_peak_17 | 22 | Distal Interge | 5 | 13791619 | 13796820 | 5202 | 1 | 71026 | ENSMUST0 | 65846 | ENSMUSG0 | Speer3 | spermatogenesis associated glutamate (E)-rich protein 3 |
| chr5 | 14908265 | 14908689 | 425 | * | NA_peak_17 | 39 | Distal Interge | 5 | 14913397 | 14914853 | 1457 | 2 | 381714 | ENSMUST0 | 6164 | ENSMUSG0 | Gm9758 | predicted gene 9758 |
| chr5 | 14918854 | 14919220 | 367 | * | NA_peak_17 | 14 | Distal Interge | 5 | 14910122 | 14914899 | 4778 | 2 | 381714 | ENSMUST0 | -3955 | ENSMUSG0 | Gm9758 | predicted gene 9758 |
| chr5 | 14933457 | 14934355 | 899 | * | NA_peak_17 | 49 | 3' UTR | 5 | 14933221 | 14938429 | 5209 | 2 | 624245 | ENSMUST0 | 4074 | ENSMUSG0 | Speer4e | spermatogenesis associated glutamate (E)-rich protein 4e |
| chr5 | 14955204 | 14955578 | 375 | * | NA_peak_17 | 43 | Distal Interge | 5 | 14933221 | 14938429 | 5209 | 2 | 624245 | ENSMUST0 | -16775 | ENSMUSG0 | Speer4e | spermatogenesis associated glutamate (E)-rich protein 4e |
| chr5 | 14990659 | 14991158 | 500 | * | NA_peak_17 | 16 | Distal Interge | 5 | 14974113 | 14978935 | 4823 | 2 | 100503549 | ENSMUST0 | -11724 | ENSMUSG0 | Gm10354 | predicted gene 10354 |
| chr5 | 15027703 | 15028084 | 382 | * | NA_peak_17 | 29 | Distal Interge | 5 | 15028950 | 15032998 | 4049 | 2 | 66773 | ENSMUST0 | 4914 | ENSMUSG0 | Gm17019 | predicted gene 17019 |
| chr5 | 15471795 | 15472537 | 743 | * | NA_peak_17 | 41 | 3' UTR | 5 | 15524456 | 15529222 | 4767 | 2 | 100861755 | ENSMUST0 | 56685 | ENSMUSG0 | Gm21190 | predicted gene, 21190 |
| chr5 | 15479986 | 15480477 | 492 | * | NA_peak_17 | 28 | Distal Interge | 5 | 15524456 | 15529222 | 4767 | 2 | 100861755 | ENSMUST0 | 48745 | ENSMUSG0 | Gm21190 | predicted gene, 21190 |
| chr5 | 15487841 | 15488406 | 566 | * | NA_peak_17 | 56 | Distal Interge | 5 | 15524456 | 15529222 | 4767 | 2 | 100861755 | ENSMUST0 | 40816 | ENSMUSG0 | Gm21190 | predicted gene, 21190 |
| chr5 | 15512073 | 15512599 | 527 | * | NA_peak_17 | 15 | Distal Interge | 5 | 15524456 | 15529222 | 4767 | 2 | 100861755 | ENSMUST0 | 16623 | ENSMUSG0 | Gm21190 | predicted gene, 21190 |
| chr5 | 15524429 | 15524693 | 265 | * | NA_peak_17 | 68 | 3' UTR | 5 | 15524456 | 15529222 | 4767 | 2 | 100861755 | ENSMUST0 | 4529 | ENSMUSG0 | Gm21190 | predicted gene, 21190 |
| chr5 | 15589344 | 15589707 | 364 | * | NA_peak_17 | 23 | Intron (ENSM | 5 | 15577333 | 15582078 | 4746 | 1 | 105242399 | ENSMUST0 | 12011 | ENSMUSG0 | Gm21083 | predicted gene, 21083 |
| chr5 | 15597756 | 15598300 | 545 | * | NA_peak_17 | 73 | Intron (ENSM | 5 | 15577333 | 15582078 | 4746 | 1 | 105242399 | ENSMUST0 | 20423 | ENSMUSG0 | Gm21083 | predicted gene, 21083 |
| chr5 | 15601125 | 15601740 | 616 | * | NA_peak_17 | 47 | Intron (ENSM | 5 | 15619064 | 15623864 | 4801 | 1 | 360220 | ENSMUST0 | -17324 | ENSMUSG0 | Speer4d | spermatogenesis associated glutamate (E)-rich protein 4D |
| chr5 | 15609105 | 15609371 | 267 | * | NA_peak_17 | 17 | Intron (ENSM | 5 | 15619064 | 15623864 | 4801 | 1 | 360220 | ENSMUST0 | -9693 | ENSMUSG0 | Speer4d | spermatogenesis associated glutamate (E)-rich protein 4D |
| chr5 | 15661587 | 15661983 | 397 | * | NA_peak_17 | 22 | Distal Interge | 5 | 15652286 | 15657059 | 4774 | 2 | 381715 | ENSMUST0 | -4528 | ENSMUSG0 | 4930572O03 | spermatogenesis associated glutamate (E)-rich protein pseudogene |
| chr5 | 15677830 | 15678092 | 263 | * | NA_peak_17 | 60 | Promoter (2- | 5 | 15680710 | 15714596 | 33887 | 1 | 75858 | ENSMUST0 | -2618 | ENSMUSG0 | Speer4cos | spermatogenesis associated glutamate (E)-rich protein 4C, opposite strand transcript |
| chr5 | 15689755 | 15689964 | 210 | * | NA_peak_17 | 25 | Intron (ENSM | 5 | 15680710 | 15714596 | 33887 | 1 | 75858 | ENSMUST0 | 9045 | ENSMUSG0 | Speer4cos | spermatogenesis associated glutamate (E)-rich protein 4C, opposite strand transcript |
| chr5 | 15697872 | 15698206 | 335 | * | NA_peak_17 | 31 | Intron (ENSM | 5 | 15709501 | 15714236 | 4736 | 2 | 100502967 | ENSMUST0 | 16030 | ENSMUSG0 | Speer4c | spermatogenesis associated glutamate (E)-rich protein 4C |
| chr5 | 15715268 | 15715605 | 338 | * | NA_peak_17 | 25 | Promoter (<= | 5 | 15711557 | 15714271 | 2715 | 2 | 100502967 | ENSMUST0 | -997 | ENSMUSG0 | Speer4c | spermatogenesis associated glutamate (E)-rich protein 4C |
| chr5 | 15808641 | 15808975 | 335 | * | NA_peak_17 | 20 | Distal Interge | 5 | 15863928 | 15883766 | 19839 | 2 | 75058 | ENSMUST0 | 74791 | ENSMUSG0 | 4930519H02 | RIKEN cDNA 4930519H02 gene |
| chr5 | 17794821 | 17795144 | 324 | * | NA_peak_17 | 20 | Intron (ENSM | 5 | 17782016 | 17835696 | 53681 | 2 | 12491 | ENSMUST0 | 40552 | ENSMUSG0 | Cd36 | CD36 molecule |
| chr5 | 18512573 | 18512894 | 322 | * | NA_peak_17 | 13 | Distal Interge | 5 | 18265135 | 18360355 | 95221 | 2 | 14677 | ENSMUST0 | -152218 | ENSMUSG0 | Gnai1 | guanine nucleotide binding protein (G protein), alpha inhibiting 1 |
| chr5 | 18855721 | 18855757 | 37 | * | NA_peak_17 | 23 | Distal Interge | 5 | 19203293 | 19226168 | 22876 | 2 | 70832 | ENSMUST0 | 370411 | ENSMUSG0 | 4921504A21 | RIKEN cDNA 4921504A21 gene |
| chr5 | 18856178 | 18856235 | 58 | * | NA_peak_17 | 23 | Distal Interge | 5 | 19203293 | 19226168 | 22876 | 2 | 70832 | ENSMUST0 | 369933 | ENSMUSG0 | 4921504A21 | RIKEN cDNA 4921504A21 gene |
| chr5 | 19477154 | 19478324 | 1171 | * | NA_peak_17 | 48 | Intron (ENSM | 5 | 19227055 | 20704773 | 1477719 | 1 | 50791 | ENSMUST0 | 250099 | ENSMUSG0 | Magi2 | membrane associated guanylate kinase, WW and PDZ domain containing 2 |
| chr5 | 19977381 | 19978031 | 651 | * | NA_peak_17 | 75 | Intron (ENSM | 5 | 19907961 | 20195087 | 287127 | 1 | 50791 | ENSMUST0 | 69420 | ENSMUSG0 | Magi2 | membrane associated guanylate kinase, WW and PDZ domain containing 2 |
| chr5 | 20365470 | 20365862 | 393 | * | NA_peak_17 | 26 | Intron (ENSM | 5 | 20228186 | 20651728 | 423543 | 1 | 50791 | ENSMUST0 | 137284 | ENSMUSG0 | Magi2 | membrane associated guanylate kinase, WW and PDZ domain containing 2 |
| chr5 | 22215317 | 22215532 | 216 | * | NA_peak_17 | 19 | Intron (ENSM | 5 | 22060313 | 22134669 | 74357 | 2 | 19699 | ENSMUST0 | -80648 | ENSMUSG0 | Reln | reelin |
| chr5 | 24429400 | 24429830 | 431 | * | NA_peak_17 | 21 | Promoter (<= | 5 | 24429018 | 24429779 | 762 | 1 | 20535 | ENSMUST0 | 382 | ENSMUSG0 | Slc4a2 | solute carrier family 4 (anion exchanger), member 2 |
| chr5 | 24850605 | 24850917 | 313 | * | NA_peak_17 | 15 | Distal Interge | 5 | 24802823 | 24842624 | 39802 | 2 | 19744 | ENSMUST0 | -7981 | ENSMUSG0 | Rheb | Ras homolog enriched in brain |
| chr5 | 25132004 | 25132631 | 628 | * | NA_peak_17 | 28 | Distal Interge | 5 | 25100983 | 25103007 | 2025 | 1 | 67261 | ENSMUST0 | 31021 | ENSMUSG0 | 2900005J15 | RIKEN cDNA 2900005J15 gene |
| chr5 | 25135819 | 25136323 | 505 | * | NA_peak_17 | 15 | Distal Interge | 5 | 25100983 | 25103007 | 2025 | 1 | 67261 | ENSMUST0 | 34836 | ENSMUSG0 | 2900005J15 | RIKEN cDNA 2900005J15 gene |
| chr5 | 25980135 | 25980547 | 413 | * | NA_peak_17 | 21 | Distal Interge | 5 | 26000172 | 26003597 | 3426 | 2 | 100038949 | ENSMUST0 | 23050 | ENSMUSG0 | Gm1979 | predicted gene 1979 |
| chr5 | 26007225 | 26007297 | 73 | * | NA_peak_17 | 27 | Promoter (2- | 5 | 25999022 | 26004798 | 5777 | 2 | 100038949 | ENSMUST0 | -2427 | ENSMUSG0 | Gm1979 | predicted gene 1979 |
| chr5 | 27229528 | 27229659 | 132 | * | NA_peak_17 | 30 | Intron (ENSM | 5 | 27261975 | 27399011 | 137037 | 1 | 13483 | ENSMUST0 | -32316 | ENSMUSG0 | Dpp6 | dipeptidylpeptidase 6 |
| chr5 | 29178069 | 29178090 | 22 | * | NA_peak_17 | 14 | Distal Interge | 5 | 29195992 | 29203135 | 7144 | 1 | 56874 | ENSMUST0 | -17902 | ENSMUSG0 | Rnf32 | ring finger protein 32 |
| chr5 | 30174304 | 30174641 | 338 | * | NA_peak_17 | 19 | Promoter (<= | 5 | 30174958 | 30179094 | 4137 | 1 | 231086 | ENSMUST0 | -317 | ENSMUSG0 | Hadhb | hydroxyacyl-CoA dehydrogenase trifunctional multienzyme complex subunit beta |
| chr5 | 30777125 | 30777146 | 22 | * | NA_peak_17 | 30 | Promoter (<= | 5 | 30778130 | 30786934 | 8805 | 1 | 65254 | ENSMUST0 | -984 | ENSMUSG0 | Dpysl5 | dihydropyrimidinase-like 5 |
| chr5 | 31436344 | 31436385 | 42 | * | NA_peak_17 | 27 | Distal Interge | 5 | 31452431 | 31466657 | 14227 | 1 | 269639 | ENSMUST0 | -16046 | ENSMUSG0 | Zfp512 | zinc finger protein 512 |
| chr5 | 31586652 | 31587005 | 354 | * | NA_peak_17 | 17 | Distal Interge | 5 | 31596938 | 31701972 | 105035 | 1 | 107976 | ENSMUST0 | -9933 | ENSMUSG0 | Babam2 | BRISC and BRCA1 A complex member 2 |
| chr5 | 31623710 | 31624008 | 299 | * | NA_peak_17 | 17 | Intron (ENSM | 5 | 31614152 | 31622572 | 8421 | 1 | 66845 | ENSMUST0 | 9558 | ENSMUSG0 | Mrpl33 | mitochondrial ribosomal protein L33 |
| chr5 | 32014918 | 32015480 | 563 | * | NA_peak_17 | 26 | Intron (ENSM | 5 | 32045978 | 32084753 | 38776 | 1 | 107976 | ENSMUST0 | -30498 | ENSMUSG0 | Babam2 | BRISC and BRCA1 A complex member 2 |
| chr5 | 32038021 | 32038183 | 163 | * | NA_peak_17 | 22 | Intron (ENSM | 5 | 32045978 | 32084753 | 38776 | 1 | 107976 | ENSMUST0 | -7795 | ENSMUSG0 | Babam2 | BRISC and BRCA1 A complex member 2 |
| chr5 | 32466157 | 32466502 | 346 | * | NA_peak_17 | 35 | Intron (ENSM | 5 | 32459222 | 32491039 | 31818 | 1 | 19046 | ENSMUST0 | 6935 | ENSMUSG0 | Ppp1cb | protein phosphatase 1 catalytic subunit beta |
| chr5 | 32900474 | 32900475 | 2 | * | NA_peak_17 | 30 | Intron (ENSM | 5 | 32904287 | 32917988 | 13702 | 1 | 277854 | ENSMUST0 | -3812 | ENSMUSG0 | Depdc5 | DEP domain containing 5 |
| chr5 | 32940955 | 32941265 | 311 | * | NA_peak_17 | 26 | Promoter (2- | 5 | 32943818 | 32945032 | 1215 | 1 | 277854 | ENSMUST0 | -2553 | ENSMUSG0 | Depdc5 | DEP domain containing 5 |
| chr5 | 33331676 | 33331860 | 185 | * | NA_peak_17 | 13 | Distal Interge | 5 | 33335509 | 33373296 | 37788 | 1 | 59003 | ENSMUST0 | -3649 | ENSMUSG0 | Maea | macrophage erythroblast attacher |
| chr5 | 33433699 | 33433785 | 87 | * | NA_peak_17 | 21 | Promoter (<= | 5 | 33430734 | 33433976 | 3243 | 2 | 672284 | ENSMUST0 | 191 | ENSMUSG0 | Nkx1-1 | NK1 homeobox 1 |
| chr5 | 33444318 | 33444354 | 37 | * | NA_peak_17 | 22 | Distal Interge | 5 | 33430734 | 33433976 | 3243 | 2 | 672284 | ENSMUST0 | -10342 | ENSMUSG0 | Nkx1-1 | NK1 homeobox 1 |
| chr5 | 33651973 | 33652256 | 284 | * | NA_peak_17 | 22 | Promoter (<= | 5 | 33645629 | 33652296 | 6668 | 2 | 20492 | ENSMUST0 | 40 | ENSMUSG0 | Slbp | stem-loop binding protein |
| chr5 | 33657518 | 33657593 | 76 | * | NA_peak_17 | 22 | Promoter (<= | 5 | 33655558 | 33657856 | 2299 | 2 | 68366 | ENSMUST0 | 263 | ENSMUSG0 | Tmem129 | transmembrane protein 129 |
| chr5 | 33689297 | 33689381 | 85 | * | NA_peak_17 | 24 | Distal Interge | 5 | 33668724 | 33678985 | 10262 | 1 | 21335 | ENSMUST0 | 20573 | ENSMUSG0 | Tacc3 | transforming, acidic coiled-coil containing protein 3 |
| chr5 | 33692566 | 33692905 | 340 | * | NA_peak_17 | 29 | Distal Interge | 5 | 33668724 | 33678985 | 10262 | 1 | 21335 | ENSMUST0 | 23842 | ENSMUSG0 | Tacc3 | transforming, acidic coiled-coil containing protein 3 |
| chr5 | 33822566 | 33822583 | 18 | * | NA_peak_17 | 23 | Promoter (1- | 5 | 33820829 | 33846182 | 25354 | 1 | 107823 | ENSMUST0 | 1737 | ENSMUSG0 | Nsd2 | nuclear receptor binding SET domain protein 2 |
| chr5 | 33935819 | 33936131 | 313 | * | NA_peak_17 | 20 | Promoter (<= | 5 | 33919125 | 33936250 | 17126 | 2 | 24116 | ENSMUST0 | 119 | ENSMUSG0 | Nelfa | negative elongation factor complex member A, Whsc2 |
| chr5 | 34411186 | 34411385 | 200 | * | NA_peak_17 | 19 | Intron (ENSM | 5 | 34426337 | 34485747 | 59411 | 1 | 231128 | ENSMUST0 | -14952 | ENSMUSG0 | Fam193a | family with sequence homology 193, member A |
| chr5 | 34725904 | 34726464 | 561 | * | NA_peak_17 | 21 | Intron (ENSM | 5 | 34732853 | 34732972 | 120 | 2 | 100316674 | ENSMUST0 | 6508 | ENSMUSG0 | Mir467g | microRNA 467g |
| chr5 | 35257387 | 35257595 | 209 | * | NA_peak_17 | 19 | Distal Interge | 5 | 35278319 | 35281763 | 3445 | 1 | 11553 | ENSMUST0 | -20724 | ENSMUSG0 | Adra2c | adrenergic receptor, alpha 2c |
| chr5 | 35677855 | 35678017 | 163 | * | NA_peak_17 | 19 | Promoter (1- | 5 | 35653017 | 35679094 | 26078 | 2 | 78558 | ENSMUST0 | 1077 | ENSMUSG0 | Htra3 | HtrA serine peptidase 3 |
| chr5 | 36713166 | 36713357 | 192 | * | NA_peak_17 | 13 | Distal Interge | 5 | 36671399 | 36696024 | 24626 | 2 | 320661 | ENSMUST0 | -17142 | ENSMUSG0 | D5Ertd579e | DNA segment, Chr 5, ERATO Doi 579, expressed |
| chr5 | 36796255 | 36796571 | 317 | * | NA_peak_17 | 41 | Promoter (<= | 5 | 36794867 | 36796772 | 1906 | 2 | 67568 | ENSMUST0 | 201 | ENSMUSG0 | Mrfap1 | Morf4 family associated protein 1 |
| chr5 | 36989349 | 36989602 | 254 | * | NA_peak_17 | 16 | Promoter (<= | 5 | 36966104 | 36989205 | 23102 | 2 | 22393 | ENSMUST0 | -144 | ENSMUSG0 | Wfs1 | wolframin ER transmembrane glycoprotein |
| chr5 | 37036574 | 37036929 | 356 | * | NA_peak_17 | 21 | Intron (ENSM | 5 | 37029887 | 37091242 | 61356 | 1 | 76071 | ENSMUST0 | 6687 | ENSMUSG0 | Jakmip1 | janus kinase and microtubule interacting protein 1 |
| chr5 | 37163290 | 37163300 | 11 | * | NA_peak_17 | 14 | Exon (ENSM | 5 | 37175490 | 37229547 | 54058 | 1 | 381634 | ENSMUST0 | -12190 | ENSMUSG0 | Gm1043 | predicted gene 1043 |
| chr5 | 37818549 | 37818788 | 240 | * | NA_peak_17 | 19 | Distal Interge | 5 | 37820485 | 37824583 | 4099 | 2 | 17701 | ENSMUST0 | 5795 | ENSMUSG0 | Msx1 | msh homeobox 1 |
| chr5 | 37988498 | 37988525 | 28 | * | NA_peak_17 | 35 | Distal Interge | 5 | 38038742 | 38137765 | 99024 | 1 | 71116 | ENSMUST0 | -50217 | ENSMUSG0 | Stx18 | syntaxin 18 |
| chr5 | 38406786 | 38406904 | 119 | * | NA_peak_17 | 15 | Intron (ENSM | 5 | 38415440 | 38440829 | 25390 | 2 | 117591 | ENSMUST0 | 33925 | ENSMUSG0 | Slc2a9 | solute carrier family 2 (facilitated glucose transporter), member 9 |
| chr5 | 40548629 | 40548939 | 311 | * | NA_peak_17 | 53 | Distal Interge | 5 | 39613935 | 39755475 | 141541 | 2 | 15476 | ENSMUST0 | -793154 | ENSMUSG0 | Hs3st1 | heparan sulfate (glucosamine) 3-O-sulfotransferase 1 |
| chr5 | 42136733 | 42137162 | 430 | * | NA_peak_17 | 23 | Intron (ENSM | 5 | 42067960 | 42216798 | 148839 | 1 | 433882 | ENSMUST0 | 68773 | ENSMUSG0 | Gm16223 | predicted gene 16223 |
| chr5 | 42654090 | 42654229 | 140 | * | NA_peak_17 | 22 | Distal Interge | 5 | 43233170 | 43289723 | 56554 | 1 | 231207 | ENSMUST0 | -578941 | ENSMUSG0 | Cpeb2 | cytoplasmic polyadenylation element binding protein 2 |
| chr5 | 43097065 | 43097121 | 57 | * | NA_peak_17 | 14 | Distal Interge | 5 | 43233170 | 43289723 | 56554 | 1 | 231207 | ENSMUST0 | -136049 | ENSMUSG0 | Cpeb2 | cytoplasmic polyadenylation element binding protein 2 |
| chr5 | 43233583 | 43233903 | 321 | * | NA_peak_17 | 23 | Promoter (<= | 5 | 43233927 | 43237919 | 3993 | 1 | 231207 | ENSMUST0 | -24 | ENSMUSG0 | Cpeb2 | cytoplasmic polyadenylation element binding protein 2 |
| chr5 | 43412843 | 43413016 | 174 | * | NA_peak_17 | 30 | Distal Interge | 5 | 43515538 | 43618803 | 103266 | 1 | 109323 | ENSMUST0 | -102522 | ENSMUSG0 | C1qtnf7 | C1q and tumor necrosis factor related protein 7 |
| chr5 | 45245380 | 45245591 | 212 | * | NA_peak_17 | 19 | Exon (ENSM | 5 | 45110629 | 45175693 | 65065 | 2 | 52401 | ENSMUST0 | -69687 | ENSMUSG0 | D5Ertd615e | DNA segment, Chr 5, ERATO Doi 615, expressed |
| chr5 | 45604326 | 45604446 | 121 | * | NA_peak_17 | 19 | Intron (ENSM | 5 | 45638699 | 45643768 | 5070 | 1 | 319251 | ENSMUST0 | -34253 | ENSMUSG0 | 9630001P10 | RIKEN cDNA 9630001P10 gene |
| chr5 | 45639493 | 45639674 | 182 | * | NA_peak_17 | 19 | Promoter (<= | 5 | 45529705 | 45639614 | 109910 | 2 | 58227 | ENSMUST0 | 0 | ENSMUSG0 | Fam184b | family with sequence similarity 184, member B |
| chr5 | 46400759 | 46401180 | 422 | * | NA_peak_17 | 28 | Distal Interge | 5 | 45926818 | 45932409 | 5592 | 2 | 78106 | ENSMUST0 | -468350 | ENSMUSG0 | 4930405L22 | RIKEN cDNA 4930405L22 gene |
| chr5 | 46464379 | 46464544 | 166 | * | NA_peak_17 | 27 | Distal Interge | 5 | 45926818 | 45932409 | 5592 | 2 | 78106 | ENSMUST0 | -531970 | ENSMUSG0 | 4930405L22 | RIKEN cDNA 4930405L22 gene |
| chr5 | 47163017 | 47163368 | 352 | * | NA_peak_17 | 31 | Distal Interge | 5 | 47983138 | 48307733 | 324596 | 1 | 20563 | ENSMUST0 | -819770 | ENSMUSG0 | Slit2 | slit guidance ligand 2 |
| chr5 | 49195980 | 49196282 | 303 | * | NA_peak_17 | 16 | Intron (ENSM | 5 | 48389512 | 49285659 | 896148 | 2 | 80334 | ENSMUST0 | 89377 | ENSMUSG0 | Kcnip4 | Kv channel interacting protein 4 |
| chr5 | 49752153 | 49752476 | 324 | * | NA_peak_17 | 28 | Distal Interge | 5 | 49966110 | 49972075 | 5966 | 2 | 70693 | ENSMUST0 | 219599 | ENSMUSG0 | Adgra3 | adhesion G protein-coupled receptor A3 |
| chr5 | 51276811 | 51276904 | 94 | * | NA_peak_17 | 23 | Distal Interge | 5 | 51457653 | 51553871 | 96219 | 2 | 19017 | ENSMUST0 | 276967 | ENSMUSG0 | Ppargc1a | peroxisome proliferative activated receptor, gamma, coactivator 1 alpha |
| chr5 | 51583563 | 51583876 | 314 | * | NA_peak_17 | 23 | Distal Interge | 5 | 51495780 | 51567726 | 71947 | 2 | 19017 | ENSMUST0 | -15837 | ENSMUSG0 | Ppargc1a | peroxisome proliferative activated receptor, gamma, coactivator 1 alpha |
| chr5 | 53049153 | 53049196 | 44 | * | NA_peak_17 | 48 | Promoter (<= | 5 | 53049352 | 53071664 | 22313 | 1 | 20531 | ENSMUST0 | -156 | ENSMUSG0 | Slc34a2 | solute carrier family 34 (sodium phosphate), member 2 |
| chr5 | 53460559 | 53460764 | 206 | * | NA_peak_17 | 23 | Distal Interge | 5 | 53466152 | 53649390 | 183239 | 1 | 19664 | ENSMUST0 | -5388 | ENSMUSG0 | Rbpj | recombination signal binding protein for immunoglobulin kappa J region |
| chr5 | 53999328 | 53999561 | 234 | * | NA_peak_17 | 32 | Promoter (<= | 5 | 53998565 | 54121057 | 122493 | 1 | 116873 | ENSMUST0 | 763 | ENSMUSG0 | Stim2 | stromal interaction molecule 2 |
| chr5 | 54003533 | 54003907 | 375 | * | NA_peak_17 | 23 | Intron (ENSM | 5 | 53998565 | 54121057 | 122493 | 1 | 116873 | ENSMUST0 | 4968 | ENSMUSG0 | Stim2 | stromal interaction molecule 2 |
| chr5 | 54994526 | 54994836 | 311 | * | NA_peak_17 | 22 | Distal Interge | 5 | 54349991 | 54358543 | 8553 | 1 | 330086 | ENSMUST0 | 644535 | ENSMUSG0 | Gm10440 | predicted gene 10440 |
| chr5 | 55652801 | 55653210 | 410 | * | NA_peak_17 | 40 | Exon (ENSM | 5 | 54349991 | 54358543 | 8553 | 1 | 330086 | ENSMUST0 | 1302810 | ENSMUSG0 | Gm10440 | predicted gene 10440 |
| chr5 | 55656480 | 55656947 | 468 | * | NA_peak_17 | 26 | Distal Interge | 5 | 54349991 | 54358543 | 8553 | 1 | 330086 | ENSMUST0 | 1306489 | ENSMUSG0 | Gm10440 | predicted gene 10440 |
| chr5 | 56472904 | 56473842 | 939 | * | NA_peak_17 | 49 | Intron (ENSM | 5 | 57570082 | 57717920 | 147839 | 2 | 319216 | ENSMUST0 | 1244078 | ENSMUSG0 | 4932441J04 | RIKEN cDNA 4932441J04 gene |
| chr5 | 57032204 | 57032526 | 323 | * | NA_peak_17 | 39 | Distal Interge | 5 | 57570082 | 57717920 | 147839 | 2 | 319216 | ENSMUST0 | 685394 | ENSMUSG0 | 4932441J04 | RIKEN cDNA 4932441J04 gene |
| chr5 | 57715692 | 57716069 | 378 | * | NA_peak_17 | 28 | Promoter (1- | 5 | 57570082 | 57717920 | 147839 | 2 | 319216 | ENSMUST0 | 1851 | ENSMUSG0 | 4932441J04 | RIKEN cDNA 4932441J04 gene |

| | | | | | | | | | | | | | | | | | |
|---|---|---|---|---|---|---|---|---|---|---|---|---|---|---|---|---|---|
| chr5 | 59311417 | 59311726 | 310 | * | NA_peak_17 | 49 | Distal Interge | 5 | 59963124 | 59985998 | 22875 | 2 | 71057 | ENSMUST00 | 674272 | ENSMUSG00 | 4933402J10R | RIKEN cDNA 4933402J10 gene |
| chr5 | 60757416 | 60757469 | 54 | * | NA_peak_17 | 39 | Distal Interge | 5 | 59963119 | 59986004 | 22886 | 2 | 71057 | ENSMUST00 | -771412 | ENSMUSG00 | 4933402J10R | RIKEN cDNA 4933402J10 gene |
| chr5 | 61216296 | 61216410 | 115 | * | NA_peak_17 | 44 | Distal Interge | 5 | 61808816 | 61811163 | 2348 | 1 | 14380 | ENSMUST00 | -592406 | ENSMUSG00 | G6pd2 | glucose-6-phosphate dehydrogenase 2 |
| chr5 | 62209365 | 62209788 | 424 | * | NA_peak_17 | 26 | Distal Interge | 5 | 61808816 | 61811163 | 2348 | 1 | 14380 | ENSMUST00 | 400549 | ENSMUSG00 | G6pd2 | glucose-6-phosphate dehydrogenase 2 |
| chr5 | 63645318 | 63645598 | 281 | * | NA_peak_17 | 23 | Distal Interge | 5 | 63649102 | 63793387 | 144286 | 1 | 319807 | ENSMUST00 | -3504 | ENSMUSG00 | Nwd2 | NACHT and WD repeat domain containing 2 |
| chr5 | 64061039 | 64061423 | 385 | * | NA_peak_17 | 19 | Distal Interge | 5 | 64044978 | 64050228 | 5251 | 1 | 74757 | ENSMUST00 | 16061 | ENSMUSG00 | 5830416I19R | RIKEN cDNA 5830416I19 gene |
| chr5 | 64230276 | 64230668 | 393 | * | NA_peak_17 | 20 | Promoter (<= | 5 | 64230399 | 64351486 | 121088 | 1 | 57915 | ENSMUST00 | 0 | ENSMUSG00 | Tbc1d1 | TBC1 domain family, member 1 |
| chr5 | 64251211 | 64251396 | 186 | * | NA_peak_17 | 22 | Intron (ENSM | 5 | 64230399 | 64351486 | 121088 | 1 | 57915 | ENSMUST00 | 20812 | ENSMUSG00 | Tbc1d1 | TBC1 domain family, member 1 |
| chr5 | 64316791 | 64317165 | 375 | * | NA_peak_17 | 27 | Intron (ENSM | 5 | 64298530 | 64351481 | 52952 | 1 | 57915 | ENSMUST00 | 18261 | ENSMUSG00 | Tbc1d1 | TBC1 domain family, member 1 |
| chr5 | 65086522 | 65086835 | 314 | * | NA_peak_17 | 20 | Intron (ENSM | 5 | 65062536 | 65091584 | 29049 | 2 | 243025 | ENSMUST00 | 4749 | ENSMUSG00 | Tmem156 | transmembrane protein 156 |
| chr5 | 65687252 | 65687641 | 390 | * | NA_peak_17 | 23 | Intron (ENSM | 5 | 65665953 | 65696865 | 30913 | 2 | 71521 | ENSMUST00 | 9224 | ENSMUSG00 | Pds5a | PDS5 cohesin associated factor A |
| chr5 | 66503706 | 66503947 | 242 | * | NA_peak_18 | 27 | Intron (ENSM | 5 | 66451995 | 66514830 | 62836 | 2 | 11787 | ENSMUST00 | 10883 | ENSMUSG00 | Apbb2 | amyloid beta (A4) precursor protein-binding, family B, member 2 |
| chr5 | 67915108 | 67915538 | 431 | * | NA_peak_18 | 93 | Distal Interge | 5 | 67941669 | 67977070 | 35402 | 1 | 330097 | ENSMUST00 | -26131 | ENSMUSG00 | Gm5108 | predicted gene 5108 |
| chr5 | 68437065 | 68437395 | 331 | * | NA_peak_18 | 21 | Distal Interge | 5 | 68031835 | 68166398 | 134564 | 1 | 433899 | ENSMUST00 | 405230 | ENSMUSG00 | Grxcr1 | glutaredoxin, cysteine rich 1 |
| chr5 | 69119436 | 69119732 | 297 | * | NA_peak_18 | 29 | Intron (ENSM | 5 | 69296209 | 69341677 | 45469 | 2 | 243043 | ENSMUST00 | 221945 | ENSMUSG00 | Kctd8 | potassium channel tetramerisation domain containing 8 |
| chr5 | 70648480 | 70648841 | 362 | * | NA_peak_18 | 22 | Distal Interge | 5 | 70751775 | 70842204 | 90430 | 2 | 14405 | ENSMUST00 | 193363 | ENSMUSG00 | Gabrg1 | gamma-aminobutyric acid (GABA) A receptor, subunit gamma 1 |
| chr5 | 72074273 | 72074304 | 32 | * | NA_peak_18 | 26 | Intron (ENSM | 5 | 72159011 | 72161138 | 2128 | 2 | 27784 | ENSMUST00 | 86834 | ENSMUSG00 | Commd8 | COMM domain containing 8 |
| chr5 | 73066649 | 73066802 | 154 | * | NA_peak_18 | 19 | Promoter (1- | 5 | 73064356 | 73065233 | 878 | 2 | 72313 | ENSMUST00 | -1416 | ENSMUSG00 | Fryl | FRY like transcription coactivator |
| chr5 | 74494103 | 74494392 | 290 | * | NA_peak_18 | 44 | Intron (ENSM | 5 | 74458373 | 74479126 | 20754 | 2 | 212986 | ENSMUST00 | -14977 | ENSMUSG00 | Scfd2 | Sec1 family domain containing 2 |
| chr5 | 76303776 | 76303910 | 135 | * | NA_peak_18 | 36 | Promoter (<= | 5 | 76212177 | 76304548 | 92372 | 2 | 12753 | ENSMUST00 | 638 | ENSMUSG00 | Clock | circadian locomotor output cycles kaput |
| chr5 | 77416328 | 77416635 | 308 | * | NA_peak_18 | 27 | Distal Interge | 5 | 77349240 | 77408041 | 58802 | 2 | 29817 | ENSMUST00 | -8287 | ENSMUSG00 | Igfbp7 | insulin-like growth factor binding protein 7 |
| chr5 | 77433909 | 77434044 | 136 | * | NA_peak_18 | 34 | Exon (ENSM | 5 | 77349240 | 77408041 | 58802 | 2 | 29817 | ENSMUST00 | -25868 | ENSMUSG00 | Igfbp7 | insulin-like growth factor binding protein 7 |
| chr5 | 77964782 | 77965153 | 372 | * | NA_peak_18 | 39 | Distal Interge | 5 | 77465488 | 77468851 | 3364 | 1 | 69370 | ENSMUST00 | 499294 | ENSMUSG00 | 1700017L05R | RIKEN cDNA 1700017L05 gene |
| chr5 | 78087404 | 78088138 | 735 | * | NA_peak_18 | 35 | Distal Interge | 5 | 77465488 | 77468851 | 3364 | 1 | 69370 | ENSMUST00 | 621916 | ENSMUSG00 | 1700017L05R | RIKEN cDNA 1700017L05 gene |
| chr5 | 78157931 | 78157966 | 36 | * | NA_peak_18 | 19 | Distal Interge | 5 | 77465488 | 77468851 | 3364 | 1 | 69370 | ENSMUST00 | 692443 | ENSMUSG00 | 1700017L05R | RIKEN cDNA 1700017L05 gene |
| chr5 | 79251165 | 79251491 | 327 | * | NA_peak_18 | 27 | Distal Interge | 5 | 81020138 | 81388784 | 368647 | 1 | 319387 | ENSMUST00 | -1768647 | ENSMUSG00 | Adgrl3 | adhesion G protein-coupled receptor L3 |
| chr5 | 79419207 | 79419488 | 282 | * | NA_peak_18 | 40 | Distal Interge | 5 | 81020138 | 81388784 | 368647 | 1 | 319387 | ENSMUST00 | -1600650 | ENSMUSG00 | Adgrl3 | adhesion G protein-coupled receptor L3 |
| chr5 | 79654432 | 79654753 | 322 | * | NA_peak_18 | 19 | Distal Interge | 5 | 81020138 | 81388784 | 368647 | 1 | 319387 | ENSMUST00 | -1365385 | ENSMUSG00 | Adgrl3 | adhesion G protein-coupled receptor L3 |
| chr5 | 81453990 | 81454268 | 279 | * | NA_peak_18 | 35 | Intron (ENSM | 5 | 81310025 | 81796634 | 486610 | 1 | 319387 | ENSMUST00 | 143965 | ENSMUSG00 | Adgrl3 | adhesion G protein-coupled receptor L3 |
| chr5 | 82653862 | 82654158 | 297 | * | NA_peak_18 | 26 | Distal Interge | 5 | 82798944 | 82799065 | 122 | 2 | 100316803 | ENSMUST00 | 144907 | ENSMUSG00 | Mir1187 | microRNA 1187 |
| chr5 | 83139016 | 83139287 | 272 | * | NA_peak_18 | 26 | Distal Interge | 5 | 83289996 | 83354956 | 64961 | 2 | 243078 | ENSMUST00 | 215669 | ENSMUSG00 | Tecrl | trans-2,3-enoyl-CoA reductase-like |
| chr5 | 83451379 | 83451754 | 376 | * | NA_peak_18 | 39 | Distal Interge | 5 | 83278145 | 83355195 | 77051 | 2 | 243078 | ENSMUST00 | -96184 | ENSMUSG00 | Tecrl | trans-2,3-enoyl-CoA reductase-like |
| chr5 | 83452138 | 83452382 | 245 | * | NA_peak_18 | 39 | Distal Interge | 5 | 83278145 | 83355195 | 77051 | 2 | 243078 | ENSMUST00 | -96943 | ENSMUSG00 | Tecrl | trans-2,3-enoyl-CoA reductase-like |
| chr5 | 84376109 | 84376301 | 193 | * | NA_peak_18 | 31 | Intron (ENSM | 5 | 84057791 | 84416806 | 359016 | 2 | 13839 | ENSMUST00 | 40505 | ENSMUSG00 | Epha5 | Eph receptor A5 |
| chr5 | 85169289 | 85169601 | 313 | * | NA_peak_18 | 53 | Distal Interge | 5 | 84330775 | 84417382 | 86608 | 2 | 13839 | ENSMUST00 | -751907 | ENSMUSG00 | Epha5 | Eph receptor A5 |
| chr5 | 85265060 | 85265373 | 314 | * | NA_peak_18 | 37 | Distal Interge | 5 | 86057116 | 86065522 | 8407 | 2 | 12617 | ENSMUST00 | 800149 | ENSMUSG00 | Cenpc1 | centromere protein C1 |
| chr5 | 86365743 | 86366036 | 294 | * | NA_peak_18 | 15 | Intron (ENSM | 5 | 86302217 | 86373420 | 71204 | 2 | 231382 | ENSMUST00 | 7384 | ENSMUSG00 | Tmprss11d | transmembrane protease, serine 11d |
| chr5 | 87099839 | 87100138 | 300 | * | NA_peak_18 | 23 | Intron (ENSM | 5 | 87065927 | 87092555 | 26629 | 2 | 231396 | ENSMUST00 | -7284 | ENSMUSG00 | Ugt2b36 | UDP glucuronosyltransferase 2 family, polypeptide B36 |
| chr5 | 87263111 | 87263391 | 281 | * | NA_peak_18 | 26 | Distal Interge | 5 | 87240493 | 87254804 | 14312 | 2 | 112417 | ENSMUST00 | -8307 | ENSMUSG00 | Ugt2b37 | UDP glucuronosyltransferase 2 family, polypeptide B37 |
| chr5 | 88163740 | 88164479 | 740 | * | NA_peak_18 | 34 | Distal Interge | 5 | 88086556 | 88109053 | 22498 | 1 | 20600 | ENSMUST00 | 77184 | ENSMUSG00 | Smr2 | submaxillary gland androgen regulated protein 2 |
| chr5 | 88165232 | 88166119 | 888 | * | NA_peak_18 | 13 | Distal Interge | 5 | 88086556 | 88109053 | 22498 | 1 | 20600 | ENSMUST00 | 78676 | ENSMUSG00 | Smr2 | submaxillary gland androgen regulated protein 2 |
| chr5 | 88166486 | 88166725 | 240 | * | NA_peak_18 | 31 | Distal Interge | 5 | 88086556 | 88109053 | 22498 | 1 | 20600 | ENSMUST00 | 79930 | ENSMUSG00 | Smr2 | submaxillary gland androgen regulated protein 2 |
| chr5 | 89204050 | 89204469 | 420 | * | NA_peak_18 | 13 | Intron (ENSM | 5 | 89028092 | 89122453 | 94362 | 1 | 54403 | ENSMUST00 | 175958 | ENSMUSG00 | Slc4a4 | solute carrier family 4 (anion exchanger), member 4 |
| chr5 | 89292270 | 89292659 | 390 | * | NA_peak_18 | 35 | Distal Interge | 5 | 89421885 | 89435628 | 13744 | 2 | 14473 | ENSMUST00 | 142969 | ENSMUSG00 | Gc | vitamin D binding protein |
| chr5 | 90698134 | 90698761 | 628 | * | NA_peak_18 | 44 | Distal Interge | 5 | 90603831 | 90640657 | 36827 | 2 | 73246 | ENSMUST00 | -57477 | ENSMUSG00 | Rassf6 | Ras association (RalGDS/AF-6) domain family member 6 |
| chr5 | 90713279 | 90713653 | 375 | * | NA_peak_18 | 13 | Distal Interge | 5 | 90759360 | 90761624 | 2265 | 1 | 20311 | ENSMUST00 | -45707 | ENSMUSG00 | Cxcl5 | chemokine (C-X-C motif) ligand 5 |
| chr5 | 90741588 | 90741865 | 278 | * | NA_peak_18 | 23 | Distal Interge | 5 | 90759360 | 90761624 | 2265 | 1 | 20311 | ENSMUST00 | -17495 | ENSMUSG00 | Cxcl5 | chemokine (C-X-C motif) ligand 5 |
| chr5 | 90884382 | 90884494 | 113 | * | NA_peak_18 | 38 | Distal Interge | 5 | 90891241 | 90893115 | 1875 | 1 | 14825 | ENSMUST00 | -6747 | ENSMUSG00 | Cxcl1 | chemokine (C-X-C motif) ligand 1 |
| chr5 | 91021722 | 91021738 | 17 | * | NA_peak_18 | 19 | Distal Interge | 5 | 91027464 | 91034213 | 6750 | 1 | 71920 | ENSMUST00 | -5726 | ENSMUSG00 | Epgn | epithelial mitogen |
| chr5 | 91685149 | 91685451 | 303 | * | NA_peak_18 | 22 | Distal Interge | 5 | 91766242 | 91767300 | 1059 | 2 | 73452 | ENSMUST00 | 81849 | ENSMUSG00 | 1700063O14 | RIKEN cDNA 1700063O14 gene |
| chr5 | 92060986 | 92061276 | 291 | * | NA_peak_18 | 23 | Intron (ENSM | 5 | 92057821 | 92067523 | 9703 | 2 | 23881 | ENSMUST00 | 6247 | ENSMUSG00 | G3bp2 | GTPase activating protein (SH3 domain) binding protein 2 |
| chr5 | 92555188 | 92555513 | 326 | * | NA_peak_18 | 31 | Promoter (<= | 5 | 92555102 | 92590660 | 35559 | 1 | 384198 | ENSMUST00 | 86 | ENSMUSG00 | Fam47e | family with sequence similarity 47, member E |
| chr5 | 92615729 | 92616444 | 716 | * | NA_peak_18 | 21 | Intron (ENSM | 5 | 92607954 | 92612503 | 4550 | 2 | 320696 | ENSMUST00 | -3226 | ENSMUSG00 | Ccdc158 | coiled-coil domain containing 158 |
| chr5 | 92806841 | 92807152 | 312 | * | NA_peak_18 | 17 | Promoter (2- | 5 | 92809378 | 92965318 | 155941 | 1 | 27428 | ENSMUST00 | -2226 | ENSMUSG00 | Shroom3 | shroom family member 3 |
| chr5 | 93044848 | 93045119 | 272 | * | NA_peak_18 | 26 | Promoter (<= | 5 | 93041123 | 93045022 | 3900 | 2 | 78088 | ENSMUST00 | 0 | ENSMUSG00 | Sowahb | sosondowah ankyrin repeat domain family member B |
| chr5 | 93049750 | 93049845 | 96 | * | NA_peak_18 | 23 | Distal Interge | 5 | 93041123 | 93045022 | 3900 | 2 | 78088 | ENSMUST00 | -4728 | ENSMUSG00 | Sowahb | sosondowah ankyrin repeat domain family member B |
| chr5 | 93101302 | 93101452 | 151 | * | NA_peak_18 | 23 | Intron (ENSM | 5 | 93093571 | 93176077 | 82507 | 1 | 52398 | ENSMUST00 | 7731 | ENSMUSG00 | Septin11 | septin 11 |
| chr5 | 93807747 | 93808107 | 361 | * | NA_peak_18 | 118 | Distal Interge | 5 | 93634993 | 93671551 | 36559 | 2 | 381654 | ENSMUST00 | -136196 | ENSMUSG00 | Pramel34 | PRAME like 34 |
| chr5 | 94338294 | 94338575 | 282 | * | NA_peak_18 | 20 | Promoter (1- | 5 | 94339749 | 94385010 | 45262 | 1 | 100554 | ENSMUST00 | -1174 | ENSMUSG00 | AA792892 | expressed sequence AA792892 |
| chr5 | 94472018 | 94472097 | 80 | * | NA_peak_18 | 24 | Distal Interge | 5 | 94526287 | 94546423 | 20137 | 1 | 100041102 | ENSMUST00 | -54190 | ENSMUSG00 | Pramel42 | PRAME like 42 |
| chr5 | 94512714 | 94513023 | 310 | * | NA_peak_18 | 23 | Distal Interge | 5 | 94526287 | 94546423 | 20137 | 1 | 100041102 | ENSMUST00 | -13264 | ENSMUSG00 | Pramel42 | PRAME like 42 |
| chr5 | 94543921 | 94544226 | 306 | * | NA_peak_18 | 24 | Intron (ENSM | 5 | 94534849 | 94537928 | 3080 | 1 | 100041102 | ENSMUST00 | 9072 | ENSMUSG00 | Pramel42 | PRAME like 42 |
| chr5 | 94561547 | 94561865 | 319 | * | NA_peak_18 | 26 | Distal Interge | 5 | 94534849 | 94537928 | 3080 | 1 | 100041102 | ENSMUST00 | 26698 | ENSMUSG00 | Pramel42 | PRAME like 42 |
| chr5 | 94597619 | 94597962 | 344 | * | NA_peak_18 | 23 | Distal Interge | 5 | 94534849 | 94537928 | 3080 | 1 | 100041102 | ENSMUST00 | 62770 | ENSMUSG00 | Pramel42 | PRAME like 42 |
| chr5 | 94602766 | 94603097 | 332 | * | NA_peak_18 | 20 | Distal Interge | 5 | 94534849 | 94537928 | 3080 | 1 | 100041102 | ENSMUST00 | 67917 | ENSMUSG00 | Pramel42 | PRAME like 42 |
| chr5 | 94603528 | 94603711 | 184 | * | NA_peak_18 | 20 | Distal Interge | 5 | 94534849 | 94537928 | 3080 | 1 | 100041102 | ENSMUST00 | 68679 | ENSMUSG00 | Pramel42 | PRAME like 42 |
| chr5 | 94638360 | 94638494 | 135 | * | NA_peak_18 | 13 | Distal Interge | 5 | 94534849 | 94537928 | 3080 | 1 | 100041102 | ENSMUST00 | 103511 | ENSMUSG00 | Pramel42 | PRAME like 42 |
| chr5 | 94716694 | 94717040 | 347 | * | NA_peak_18 | 26 | Distal Interge | 5 | 94865436 | 94885388 | 19953 | 1 | 100041253 | ENSMUST00 | -148396 | ENSMUSG00 | Gm16513 | predicted gene, Gm16513 |
| chr5 | 94724775 | 94724943 | 169 | * | NA_peak_18 | 16 | Distal Interge | 5 | 94865436 | 94885388 | 19953 | 1 | 100041253 | ENSMUST00 | -140493 | ENSMUSG00 | Gm16513 | predicted gene, Gm16513 |
| chr5 | 94755700 | 94756060 | 361 | * | NA_peak_18 | 22 | Distal Interge | 5 | 94865436 | 94885388 | 19953 | 1 | 100041253 | ENSMUST00 | -109376 | ENSMUSG00 | Gm16513 | predicted gene, Gm16513 |
| chr5 | 94830665 | 94830947 | 283 | * | NA_peak_18 | 15 | Distal Interge | 5 | 94865436 | 94885388 | 19953 | 1 | 100041253 | ENSMUST00 | -34489 | ENSMUSG00 | Gm16513 | predicted gene, Gm16513 |
| chr5 | 94858797 | 94859207 | 411 | * | NA_peak_18 | 15 | Distal Interge | 5 | 94865436 | 94885388 | 19953 | 1 | 100041253 | ENSMUST00 | -6229 | ENSMUSG00 | Gm16513 | predicted gene, Gm16513 |
| chr5 | 94896608 | 94897251 | 644 | * | NA_peak_18 | 54 | Distal Interge | 5 | 94873797 | 94876892 | 3096 | 1 | 545762 | ENSMUST00 | 22811 | ENSMUSG00 | Gm16367 | predicted gene 16367 |
| chr5 | 94898373 | 94898537 | 165 | * | NA_peak_18 | 22 | Distal Interge | 5 | 94873797 | 94876892 | 3096 | 1 | 545762 | ENSMUST00 | 24576 | ENSMUSG00 | Gm16367 | predicted gene 16367 |
| chr5 | 94945954 | 94945966 | 13 | * | NA_peak_18 | 13 | Distal Interge | 5 | 94951537 | 94954134 | 2598 | 2 | 331195 | ENSMUST00 | 8168 | ENSMUSG00 | Pramel39 | PRAME like 39 |
| chr5 | 95000468 | 95000855 | 388 | * | NA_peak_18 | 56 | Intron (ENSM | 5 | 94988626 | 95026536 | 37911 | 1 | 622894 | ENSMUST00 | 11842 | ENSMUSG00 | Gm6367 | predicted gene 6367 |
| chr5 | 95107917 | 95108317 | 401 | * | NA_peak_18 | 35 | Intron (ENSM | 5 | 95182624 | 95194966 | 12343 | 1 | 100041253 | ENSMUST00 | -74307 | ENSMUSG00 | Gm16513 | predicted gene, Gm16513 |
| chr5 | 95162472 | 95162910 | 439 | * | NA_peak_18 | 24 | Distal Interge | 5 | 95182624 | 95194966 | 12343 | 1 | 100041253 | ENSMUST00 | -19714 | ENSMUSG00 | Gm16513 | predicted gene, Gm16513 |
| chr5 | 95165131 | 95165387 | 257 | * | NA_peak_18 | 16 | Distal Interge | 5 | 95182624 | 95194966 | 12343 | 1 | 100041253 | ENSMUST00 | -17237 | ENSMUSG00 | Gm16513 | predicted gene, Gm16513 |
| chr5 | 95317525 | 95318007 | 483 | * | NA_peak_18 | 19 | Intron (ENSM | 5 | 95305966 | 95343860 | 37895 | 1 | 100041296 | ENSMUST00 | 11559 | ENSMUSG00 | Pramel47 | PRAME like 47 |
| chr5 | 95394330 | 95394670 | 341 | * | NA_peak_18 | 19 | Distal Interge | 5 | 95340209 | 95342587 | 2379 | 1 | 100041296 | ENSMUST00 | 54121 | ENSMUSG00 | Pramel47 | PRAME like 47 |
| chr5 | 95418470 | 95418529 | 60 | * | NA_peak_18 | 17 | Exon (ENSM | 5 | 95456806 | 95485564 | 28759 | 1 | 320549 | ENSMUST00 | -38277 | ENSMUSG00 | Pramel48 | PRAME like 48 |
| chr5 | 95756463 | 95756886 | 424 | * | NA_peak_18 | 15 | Distal Interge | 5 | 95775990 | 95804708 | 28719 | 1 | 665943 | ENSMUST00 | -19104 | ENSMUSG00 | E330014E10 | RIKEN cDNA E330014E10 gene |
| chr5 | 96121271 | 96121541 | 271 | * | NA_peak_18 | 18 | Intron (ENSM | 5 | 96105610 | 96129028 | 23419 | 2 | 231464 | ENSMUST00 | 7487 | ENSMUSG00 | Cnot6l | CCR4-NOT transcription complex, subunit 6-like |
| chr5 | 96273558 | 96274220 | 663 | * | NA_peak_18 | 61 | Distal Interge | 5 | 96210144 | 96239684 | 29541 | 1 | 94061 | ENSMUST00 | 63414 | ENSMUSG00 | Mrpl1 | mitochondrial ribosomal protein L1 |
| chr5 | 96812715 | 96813081 | 367 | * | NA_peak_18 | 15 | Promoter (1- | 5 | 96811066 | 96834906 | 23841 | 1 | 11745 | ENSMUST00 | 1649 | ENSMUSG00 | Anxa3 | annexin A3 |
| chr5 | 99252541 | 99252916 | 376 | * | NA_peak_18 | 26 | Promoter (<= | 5 | 99217426 | 99252927 | 35502 | 2 | 320292 | ENSMUST00 | 11 | ENSMUSG00 | Rasgef1b | RasGEF domain family, member 1B |
| chr5 | 99928388 | 99928596 | 209 | * | NA_peak_18 | 16 | 3' UTR | 5 | 99961168 | 99964750 | 3583 | 2 | 11991 | ENSMUST00 | 36154 | ENSMUSG00 | Hnrnpd | heterogeneous nuclear ribonucleoprotein D |
| chr5 | 100503962 | 100504179 | 218 | * | NA_peak_18 | 35 | Distal Interge | 5 | 100442200 | 100500639 | 58440 | 2 | 231506 | ENSMUST00 | -3323 | ENSMUSG00 | Lin54 | lin-54 homolog (C. elegans) |
| chr5 | 100862719 | 100863139 | 421 | * | NA_peak_18 | 23 | Intron (ENSM | 5 | 100846417 | 100846850 | 434 | 1 | 231510 | ENSMUST00 | 16302 | ENSMUSG00 | Gpat3 | glycerol-3-phosphate acyltransferase 3 |
| chr5 | 102196005 | 102196394 | 390 | * | NA_peak_18 | 37 | Distal Interge | 5 | 101929306 | 102069921 | 140616 | 2 | 72145 | ENSMUST00 | -126084 | ENSMUSG00 | Wdfy3 | WD repeat and FYVE domain containing 3 |
| chr5 | 103329462 | 103329759 | 298 | * | NA_peak_18 | 23 | Distal Interge | 5 | 103346594 | 103348299 | 1706 | 1 | 75519 | ENSMUST00 | -16835 | ENSMUSG00 | 1700021F02R | RIKEN cDNA 1700021F02 gene |
| chr5 | 103687293 | 103687495 | 203 | * | NA_peak_18 | 30 | Distal Interge | 5 | 103692374 | 103816428 | 124055 | 1 | 17355 | ENSMUST00 | -4879 | ENSMUSG00 | Aff1 | AF4/FMR2 family, member 1 |
| chr5 | 103692590 | 103692736 | 147 | * | NA_peak_18 | 27 | Promoter (<= | 5 | 103692374 | 103816428 | 124055 | 1 | 17355 | ENSMUST00 | 216 | ENSMUSG00 | Aff1 | AF4/FMR2 family, member 1 |
| chr5 | 104135618 | 104135886 | 269 | * | NA_peak_18 | 20 | Distal Interge | 5 | 104079111 | 104113733 | 34623 | 2 | 13602 | ENSMUST00 | -21885 | ENSMUSG00 | Sparcl1 | SPARC-like 1 |
| chr5 | 104839007 | 104839311 | 305 | * | NA_peak_18 | 19 | Intron (ENSM | 5 | 104814343 | 104828957 | 14615 | 2 | 626391 | ENSMUST00 | -10050 | ENSMUSG00 | Zfp951 | zinc finger protein 951 |
| chr5 | 105366573 | 105366969 | 397 | * | NA_peak_18 | 32 | Distal Interge | 5 | 105323042 | 105346472 | 23431 | 2 | 634650 | ENSMUST00 | -20101 | ENSMUSG00 | Gbp11 | guanylate binding protein 11 |
| chr5 | 105700253 | 105700383 | 131 | * | NA_peak_18 | 18 | Promoter (<= | 5 | 105699969 | 105815215 | 115247 | 1 | 231549 | ENSMUST00 | 284 | ENSMUSG00 | Lrrc8d | leucine rich repeat containing 8D |
| chr5 | 106624646 | 106624952 | 307 | * | NA_peak_18 | 19 | Promoter (<= | 5 | 106617073 | 106624390 | 7318 | 2 | 52397 | ENSMUST00 | -256 | ENSMUSG00 | Zfp644 | zinc finger protein 644 |
| chr5 | 107576520 | 107577025 | 506 | * | NA_peak_18 | 15 | Intron (ENSM | 5 | 107597373 | 107642922 | 45550 | 1 | 231571 | ENSMUST00 | -20348 | ENSMUSG00 | Rpap2 | RNA polymerase II associated protein 2 |
| chr5 | 107579149 | 107580610 | 1462 | * | NA_peak_18 | 79 | 3' UTR | 5 | 107597373 | 107642922 | 45550 | 1 | 231571 | ENSMUST00 | -16763 | ENSMUSG00 | Rpap2 | RNA polymerase II associated protein 2 |
| chr5 | 107869102 | 107869394 | 293 | * | NA_peak_18 | 58 | Promoter (<= | 5 | 107788287 | 107869168 | 80882 | 2 | 14020 | ENSMUST00 | 0 | ENSMUSG00 | Evi5 | ecotropic viral integration site 5 |
| chr5 | 108003267 | 108003595 | 329 | * | NA_peak_18 | 19 | Distal Interge | 5 | 107908053 | 107987085 | 79033 | 2 | 67266 | ENSMUST00 | -16182 | ENSMUSG00 | Dipk1a | divergent protein kinase domain 1A |
| chr5 | 108492102 | 108492424 | 323 | * | NA_peak_18 | 20 | Intron (ENSM | 5 | 108486220 | 108496290 | 10071 | 1 | 69587 | ENSMUST00 | 5882 | ENSMUSG00 | Pcgf3 | polycomb group ring finger 3 |
| chr5 | 108549373 | 108549709 | 337 | * | NA_peak_18 | 44 | Promoter (<= | 5 | 108520259 | 108550000 | 29742 | 2 | 12889 | ENSMUST00 | 291 | ENSMUSG00 | Cplx1 | complexin 1 |
| chr5 | 113538855 | 113539423 | 569 | * | NA_peak_18 | 23 | Intron (ENSM | 5 | 113570919 | 113589718 | 18800 | 1 | 320916 | ENSMUST00 | -31496 | ENSMUSG00 | Wscd2 | WSC domain containing 2 |
| chr5 | 113771191 | 113771433 | 243 | * | NA_peak_18 | 36 | Promoter (<= | 5 | 113742448 | 113771638 | 29191 | 2 | 53890 | ENSMUST00 | 205 | ENSMUSG00 | Sart3 | squamous cell carcinoma antigen recognized by T cells 3 |
| chr5 | 113983600 | 113984043 | 444 | * | NA_peak_18 | 27 | Intron (ENSM | 5 | 113942192 | 113989743 | 47552 | 2 | 231637 | ENSMUST00 | 5700 | ENSMUSG00 | Ssh1 | slingshot protein phosphatase 1 |
| chr5 | 114042515 | 114042882 | 368 | * | NA_peak_18 | 17 | Promoter (<= | 5 | 114034951 | 114042881 | 7931 | 2 | 68666 | ENSMUST00 | 0 | ENSMUSG00 | Svop | SV2 related protein |
| chr5 | 114273605 | 114274218 | 614 | * | NA_peak_18 | 32 | Promoter (<= | 5 | 114254164 | 114273807 | 19644 | 2 | 116810 | ENSMUST00 | 0 | ENSMUSG00 | Foxn4 | forkhead box N4 |
| chr5 | 114727628 | 114728026 | 399 | * | NA_peak_18 | 17 | 3' UTR | 5 | 114733216 | 114734166 | 951 | 2 | 26431 | ENSMUST00 | 6140 | ENSMUSG00 | Git2 | GIT ArfGAP 2 |
| chr5 | 114823158 | 114823567 | 410 | * | NA_peak_18 | 28 | Promoter (<= | 5 | 114821935 | 114823468 | 1534 | 2 | 330173 | ENSMUST00 | 0 | ENSMUSG00 | 2610524H06 | RIKEN cDNA 2610524H06 gene |
| chr5 | 114897841 | 114898082 | 242 | * | NA_peak_18 | 23 | Promoter (<= | 5 | 114897742 | 114912226 | 14485 | 1 | 23962 | ENSMUST00 | 99 | ENSMUSG00 | Oasl2 | 2'-5' oligoadenylate synthetase-like 2 |
| chr5 | 114992452 | 114992546 | 95 | * | NA_peak_18 | 22 | Intron (ENSM | 5 | 115011137 | 115098790 | 87654 | 1 | 74585 | ENSMUST00 | -18591 | ENSMUSG00 | Sppl3 | signal peptide peptidase 3 |
| chr5 | 115158128 | 115158661 | 534 | * | NA_peak_18 | 39 | Promoter (<= | 5 | 115142981 | 115158179 | 15199 | 2 | 109154 | ENSMUST00 | 0 | ENSMUSG00 | Mlec | malectin |
| chr5 | 115565006 | 115565536 | 531 | * | NA_peak_18 | 14 | Promoter (<= | 5 | 115565254 | 115622654 | 57401 | 1 | 231659 | ENSMUST00 | 0 | ENSMUSG00 | Gcn1 | GCN1 activator of EIF2AK4 |
| chr5 | 119726611 | 119726959 | 349 | * | NA_peak_18 | 16 | Distal Interge | 5 | 119685095 | 119691218 | 6124 | 2 | 433944 | ENSMUST00 | -35393 | ENSMUSG00 | Tbx3os2 | T-box 3, opposite strand 2 |
| chr5 | 121831376 | 121831584 | 209 | * | NA_peak_18 | 16 | Intron (ENSM | 5 | 121817010 | 121836104 | 19095 | 2 | 16923 | ENSMUST00 | 4520 | ENSMUSG00 | Sh2b3 | SH2B adaptor protein 3 |
| chr5 | 121925054 | 121925335 | 282 | * | NA_peak_18 | 26 | Promoter (<= | 5 | 121860308 | 121924921 | 64614 | 2 | 13048 | ENSMUST00 | -133 | ENSMUSG00 | Cux2 | cut-like homeobox 2 |
| chr5 | 122850254 | 122850639 | 386 | * | NA_peak_18 | 40 | Promoter (<= | 5 | 122850157 | 122864295 | 14139 | 1 | 80751 | ENSMUST00 | 97 | ENSMUSG00 | Rnf34 | ring finger protein 34 |
| chr5 | 123015400 | 123015735 | 336 | * | NA_peak_18 | 34 | Promoter (<= | 5 | 123015338 | 123029490 | 14153 | 1 | 109305 | ENSMUST00 | 62 | ENSMUSG00 | Orai1 | ORAI calcium release-activated calcium modulator 1 |
| chr5 | 123114003 | 123114142 | 140 | * | NA_peak_18 | 24 | 3' UTR | 5 | 123099901 | 123115819 | 15919 | 1 | 330189 | ENSMUST00 | 14102 | ENSMUSG00 | Tmem120b | transmembrane protein 120B |
| chr5 | 123138746 | 123139092 | 347 | * | NA_peak_18 | 14 | Exon (ENSM | 5 | 123142193 | 123149981 | 7789 | 1 | 208043 | ENSMUST00 | -3101 | ENSMUSG00 | Setd1b | SET domain containing 1B |
| chr5 | 123314258 | 123314619 | 362 | * | NA_peak_18 | 23 | Intron (ENSM | 5 | 123302516 | 123326649 | 24134 | 1 | 269701 | ENSMUST00 | 11742 | ENSMUSG00 | Wdr66 | WD repeat domain 66 |
| chr5 | 123394880 | 123395264 | 385 | * | NA_peak_18 | 36 | Promoter (<= | 5 | 123394800 | 123457932 | 63133 | 1 | 208104 | ENSMUST00 | 80 | ENSMUSG00 | Mlxip | MLX interacting protein |
| chr5 | 124031993 | 124032376 | 384 | * | NA_peak_18 | 13 | Promoter (<= | 5 | 124004641 | 124032270 | 27630 | 2 | 330192 | ENSMUST00 | 0 | ENSMUSG00 | Vps37b | vacuolar protein sorting 37B |
| chr5 | 124540699 | 124541552 | 854 | * | NA_peak_18 | 48 | Promoter (<= | 5 | 124540955 | 124549192 | 8238 | 1 | 56334 | ENSMUST00 | 0 | ENSMUSG00 | Tmed2 | transmembrane p24 trafficking protein 2 |
| chr5 | 124551780 | 124551926 | 147 | * | NA_peak_18 | 26 | Promoter (<= | 5 | 124552864 | 124569660 | 16797 | 1 | 67848 | ENSMUST00 | -938 | ENSMUSG00 | Ddx55 | DEAD box helicase 55 |
| chr5 | 125389428 | 125389977 | 550 | * | NA_peak_18 | 16 | Promoter (<= | 5 | 125385965 | 125390014 | 4050 | 2 | 22190 | ENSMUST00 | 37 | ENSMUSG00 | Ubc | ubiquitin C |
| chr5 | 125399940 | 125400299 | 360 | * | NA_peak_18 | 22 | Distal Interge | 5 | 125385977 | 125390202 | 4226 | 2 | 22190 | ENSMUST00 | -9738 | ENSMUSG00 | Ubc | ubiquitin C |
| chr5 | 128724954 | 128725257 | 304 | * | NA_peak_18 | 22 | Intron (ENSM | 5 | 128736174 | 128755474 | 19301 | 1 | 57749 | ENSMUST00 | -10917 | ENSMUSG00 | Piwil1 | piwi-like RNA-mediated gene silencing 1 |
| chr5 | 129501068 | 129501302 | 235 | * | NA_peak_18 | 34 | Promoter (<= | 5 | 129501221 | 129571384 | 70164 | 1 | 231769 | ENSMUST00 | 0 | ENSMUSG00 | Sfswap | splicing factor SWAP |
| chr5 | 129778976 | 129779412 | 437 | * | NA_peak_18 | 13 | Promoter (<= | 5 | 129765558 | 129779173 | 13616 | 2 | 100678 | ENSMUST00 | 0 | ENSMUSG00 | Psph | phosphoserine phosphatase |
| chr5 | 130194737 | 130194972 | 236 | * | NA_peak_18 | 14 | Intron (ENSM | 5 | 130187220 | 130191321 | 4102 | 1 | 56715 | ENSMUST00 | 7517 | ENSMUSG00 | Rabgef1 | RAB guanine nucleotide exchange factor (GEF) 1 |
| chr5 | 130222438 | 130222944 | 507 | * | NA_peak_18 | 22 | Promoter (<= | 5 | 130222335 | 130243666 | 21332 | 1 | 71667 | ENSMUST00 | 103 | ENSMUSG00 | Tmem248 | transmembrane protein 248 |
| chr5 | 135715597 | 135716176 | 580 | * | NA_peak_18 | 16 | Promoter (<= | 5 | 135716005 | 135731878 | 15874 | 1 | 18984 | ENSMUST00 | 0 | ENSMUSG00 | Por | cytochrome p450 oxidoreductase |
| chr5 | 135965583 | 135965846 | 264 | * | NA_peak_18 | 18 | Promoter (<= | 5 | 135963212 | 135965874 | 2663 | 2 | 109267 | ENSMUST00 | 28 | ENSMUSG00 | Ssc4d | scavenger receptor cysteine rich family, 4 domains |
| chr5 | 136085093 | 136085677 | 585 | * | NA_peak_18 | 30 | Promoter (<= | 5 | 136085070 | 136091264 | 6195 | 1 | 54153 | ENSMUST00 | 23 | ENSMUSG00 | Rasa4 | RAS p21 protein activator 4 |
| chr5 | 136274860 | 136275244 | 385 | * | NA_peak_18 | 93 | Exon (ENSM | 5 | 136274493 | 136302521 | 28029 | 2 | 13047 | ENSMUST00 | 27277 | ENSMUSG00 | Cux1 | cut-like homeobox 1 |
| chr5 | 136565328 | 136565590 | 263 | * | NA_peak_18 | 14 | Promoter (<= | 5 | 136274486 | 136565432 | 290947 | 2 | 13047 | ENSMUST00 | 0 | ENSMUSG00 | Cux1 | cut-like homeobox 1 |
| chr5 | 137078568 | 137079266 | 699 | * | NA_peak_18 | 35 | Distal Interge | 5 | 137070177 | 137072268 | 2092 | 2 | 18787 | ENSMUST00 | -6300 | ENSMUSG00 | Serpine1 | serine (or cysteine) peptidase inhibitor, clade E, member 1 |
| chr5 | 137189211 | 137189480 | 270 | * | NA_peak_18 | 26 | Distal Interge | 5 | 137154029 | 137166070 | 12042 | 2 | 666339 | ENSMUST00 | -23141 | ENSMUSG00 | Muc3 | mucin 3, intestinal |
| chr5 | 138154111 | 138154318 | 208 | * | NA_peak_18 | 30 | Promoter (1- | 5 | 138144667 | 138152899 | 8233 | 2 | 56314 | ENSMUST00 | -1212 | ENSMUSG00 | Zfp113 | zinc finger protein 113 |
| chr5 | 138239161 | 138239232 | 72 | * | NA_peak_18 | 13 | Intron (ENSM | 5 | 138229822 | 138253363 | 23542 | 1 | 381680 | ENSMUST00 | 9339 | ENSMUSG00 | Nxpe5 | neurexophilin and PC-esterase domain family, member 5 |
| chr5 | 139516294 | 139516336 | 43 | * | NA_peak_18 | 14 | Distal Interge | 5 | 139523749 | 139541339 | 17591 | 1 | 74973 | ENSMUST00 | -7413 | ENSMUSG00 | 4930500L23R | RIKEN cDNA 4930500L23 gene |
| chr5 | 139540410 | 139540612 | 203 | * | NA_peak_18 | 24 | Promoter (2- | 5 | 139543494 | 139548179 | 4686 | 1 | 22255 | ENSMUST00 | -2882 | ENSMUSG00 | Uncx | UNC homeobox |
| chr5 | 139908949 | 139909280 | 332 | * | NA_peak_18 | 22 | Promoter (<= | 5 | 139908474 | 139971050 | 62577 | 1 | 243312 | ENSMUST00 | 475 | ENSMUSG00 | Elfn1 | leucine rich repeat and fibronectin type III, extracellular 1 |
| chr5 | 140533482 | 140533842 | 361 | * | NA_peak_19 | 83 | Distal Interge | 5 | 140505575 | 140524153 | 18579 | 1 | 59031 | ENSMUST00 | 27907 | ENSMUSG00 | Chst12 | carbohydrate sulfotransferase 12 |
| chr5 | 140668064 | 140668237 | 174 | * | NA_peak_19 | 25 | Promoter (1- | 5 | 140666158 | 140669359 | 3202 | 2 | 74239 | ENSMUST00 | 1122 | ENSMUSG00 | Iqce | IQ motif containing E |
| chr5 | 141926440 | 141926742 | 303 | * | NA_peak_19 | 43 | Intron (ENSM | 5 | 141856787 | 142213194 | 356408 | 1 | 330222 | ENSMUST00 | 69653 | ENSMUSG00 | Sdk1 | sidekick cell adhesion molecule 1 |
| chr5 | 142814939 | 142815286 | 348 | * | NA_peak_19 | 14 | Promoter (2- | 5 | 142793005 | 142817380 | 24376 | 2 | 231861 | ENSMUST00 | 2094 | ENSMUSG00 | Tnrc18 | trinucleotide repeat containing 18 |
| chr5 | 142834726 | 142834762 | 37 | * | NA_peak_19 | 30 | Distal Interge | 5 | 142754789 | 142817662 | 62874 | 2 | 231861 | ENSMUST00 | -17064 | ENSMUSG00 | Tnrc18 | trinucleotide repeat containing 18 |
| chr5 | 142943253 | 142943296 | 44 | * | NA_peak_19 | 34 | Distal Interge | 5 | 142960343 | 142973185 | 12843 | 1 | 14086 | ENSMUST00 | -17047 | ENSMUSG00 | Fscn1 | fascin actin-bundling protein 1 |
| chr5 | 143454119 | 143454533 | 415 | * | NA_peak_19 | 19 | Distal Interge | 5 | 143464584 | 143505942 | 41359 | 1 | 231871 | ENSMUST00 | -10051 | ENSMUSG00 | Daglb | diacylglycerol lipase, beta |
| chr5 | 143487749 | 143487940 | 192 | * | NA_peak_19 | 26 | Promoter (1- | 5 | 143486678 | 143489994 | 3317 | 1 | 231871 | ENSMUST00 | 1071 | ENSMUSG00 | Daglb | diacylglycerol lipase, beta |
| chr5 | 143789971 | 143790399 | 429 | * | NA_peak_19 | 44 | Distal Interge | 5 | 143817788 | 143904251 | 86464 | 1 | 15467 | ENSMUST00 | -27389 | ENSMUSG00 | Eif2ak1 | eukaryotic translation initiation factor 2 alpha kinase 1 |
| chr5 | 143974469 | 143974660 | 192 | * | NA_peak_19 | 13 | Intron (ENSM | 5 | 143991475 | 143992584 | 1110 | 2 | 231874 | ENSMUST00 | 17924 | ENSMUSG00 | Ccz1 | CCZ1 vacuolar protein trafficking and biogenesis associated |
| chr5 | 146484751 | 146484889 | 139 | * | NA_peak_19 | 21 | Promoter (2- | 5 | 146481965 | 146484704 | 2740 | 1 | 623198 | ENSMUST00 | 2786 | ENSMUSG00 | Gm6408 | predicted gene 6408 |
| chr5 | 148802775 | 148803101 | 327 | * | NA_peak_19 | 23 | Distal Interge | 5 | 148777963 | 148783080 | 5118 | 1 | 78015 | ENSMUST00 | 24812 | ENSMUSG00 | 4930505K14R | RIKEN cDNA 4930505K14 gene |
| chr5 | 149051342 | 149051722 | 381 | * | NA_peak_19 | 30 | Promoter (<= | 5 | 149048645 | 149051616 | 2972 | 2 | 15289 | ENSMUST00 | 0 | ENSMUSG00 | Hmgb1 | high mobility group box 1 |
| chr5 | 150594115 | 150594434 | 320 | * | NA_peak_19 | 28 | Promoter (<= | 5 | 150572665 | 150594462 | 21798 | 2 | 100637 | ENSMUST00 | 28 | ENSMUSG00 | N4bp2l1 | NEDD4 binding protein 2-like 1 |
| chr5 | 151228561 | 151228936 | 376 | * | NA_peak_19 | 23 | Intron (ENSM | 5 | 151063546 | 151233774 | 170229 | 2 | 243362 | ENSMUST00 | 4838 | ENSMUSG00 | Stard13 | StAR-related lipid transfer (START) domain containing 13 |
| chr5 | 151352094 | 151352325 | 232 | * | NA_peak_19 | 19 | Intron (ENSM | 5 | 151368675 | 151382183 | 13509 | 1 | 625175 | ENSMUST00 | -16350 | ENSMUSG00 | 1700028E10R | RIKEN cDNA 1700028E10 gene |
| chr6 | 3160352 | 3160644 | 293 | * | NA_peak_19 | 13 | Distal Interge | 6 | 3288519 | 3317019 | 28501 | 1 | 667335 | ENSMUST00 | -127875 | ENSMUSG00 | Gm8579 | spermatogenesis associated glutamate (E)-rich protein 4a pseudogene |
| chr6 | 3184103 | 3184261 | 159 | * | NA_peak_19 | 39 | Distal Interge | 6 | 3288519 | 3317019 | 28501 | 1 | 667335 | ENSMUST00 | -104258 | ENSMUSG00 | Gm8579 | spermatogenesis associated glutamate (E)-rich protein 4a pseudogene |
| chr6 | 4282821 | 4283145 | 325 | * | NA_peak_19 | 35 | Distal Interge | 6 | 4080318 | 4086972 | 6655 | 2 | 12068 | ENSMUST00 | -195849 | ENSMUSG00 | Bet1 | Bet1 golgi vesicular membrane trafficking protein |
| chr6 | 5484374 | 5484412 | 39 | * | NA_peak_19 | 17 | 3' UTR | 6 | 5491591 | 5496261 | 4671 | 2 | 27273 | ENSMUST00 | 11849 | ENSMUSG00 | Pdk4 | pyruvate dehydrogenase kinase, isoenzyme 4 |
| chr6 | 7105538 | 7105843 | 306 | * | NA_peak_19 | 55 | Distal Interge | 6 | 6956007 | 6965658 | 9652 | 1 | 71238 | ENSMUST00 | 149531 | ENSMUSG00 | Sdhaf3 | succinate dehydrogenase complex assembly factor 3 |
| chr6 | 7483909 | 7484118 | 210 | * | NA_peak_19 | 20 | Distal Interge | 6 | 7554879 | 7562977 | 8099 | 1 | 21333 | ENSMUST00 | -70761 | ENSMUSG00 | Tac1 | tachykinin 1 |
| chr6 | 8928705 | 8928772 | 68 | * | NA_peak_19 | 15 | Distal Interge | 6 | 8948431 | 8952430 | 4000 | 1 | 18231 | ENSMUST00 | -19659 | ENSMUSG00 | Nxph1 | neurexophilin 1 |
| chr6 | 12622194 | 12622592 | 399 | * | NA_peak_19 | 128 | Intron (ENSM | 6 | 12465280 | 12555113 | 89834 | 2 | 330267 | ENSMUST00 | -67081 | ENSMUSG00 | Thsd7a | thrombospondin, type I, domain containing 7A |
| chr6 | 14878581 | 14878859 | 279 | * | NA_peak_19 | 65 | Distal Interge | 6 | 14901349 | 15441977 | 540629 | 1 | 114142 | ENSMUST00 | -22490 | ENSMUSG00 | Foxp2 | forkhead box P2 |
| chr6 | 17006026 | 17006191 | 166 | * | NA_peak_19 | 19 | Distal Interge | 6 | 17065149 | 17105823 | 40675 | 1 | 21753 | ENSMUST00 | -58958 | ENSMUSG00 | Tes | testin LIM domain protein |
| chr6 | 17464342 | 17464579 | 238 | * | NA_peak_19 | 41 | Promoter (<= | 6 | 17464107 | 17541412 | 77306 | 1 | 17295 | ENSMUST00 | 235 | ENSMUSG00 | Met | met proto-oncogene |
| chr6 | 19071657 | 19072325 | 669 | * | NA_peak_19 | 56 | Distal Interge | 6 | 18866350 | 18879582 | 13233 | 1 | 75196 | ENSMUST00 | 205307 | ENSMUSG00 | Ankrd7 | ankyrin repeat domain 7 |
| chr6 | 19148564 | 19148774 | 211 | * | NA_peak_19 | 44 | Distal Interge | 6 | 19247899 | 19278843 | 30945 | 2 | 74725 | ENSMUST00 | 130069 | ENSMUSG00 | 4930524B17R | RIKEN cDNA 4930524B17 gene |
| chr6 | 20701878 | 20702200 | 323 | * | NA_peak_19 | 29 | Distal Interge | 6 | 21215503 | 21729805 | 514303 | 1 | 16508 | ENSMUST00 | -513303 | ENSMUSG00 | Kcnd2 | potassium voltage-gated channel, Shal-related family, member 2 |
| chr6 | 23480228 | 23480524 | 297 | * | NA_peak_19 | 44 | Intron (ENSM | 6 | 23263508 | 23587424 | 323917 | 2 | 320405 | ENSMUST00 | 106900 | ENSMUSG00 | Cadps2 | Ca2+-dependent activator protein for secretion 2 |
| chr6 | 23672996 | 23673319 | 324 | * | NA_peak_19 | 54 | Intron (ENSM | 6 | 23653898 | 23655136 | 1239 | 2 | 71300 | ENSMUST00 | -17860 | ENSMUSG00 | Rnf148 | ring finger protein 148 |
| chr6 | 24281184 | 24281513 | 330 | * | NA_peak_19 | 29 | Distal Interge | 6 | 24088283 | 24168092 | 79810 | 2 | 55961 | ENSMUST00 | -113092 | ENSMUSG00 | Slc13a1 | solute carrier family 13 (sodium/sulfate symporters), member 1 |
| chr6 | 24494612 | 24494753 | 142 | * | NA_peak_19 | 54 | Intron (ENSM | 6 | 24444865 | 24515067 | 70203 | 2 | 214704 | ENSMUST00 | 20314 | ENSMUSG00 | Iqub | IQ motif and ubiquitin domain containing |
| chr6 | 25556280 | 25556605 | 326 | * | NA_peak_19 | 49 | Distal Interge | 6 | 25665878 | 25689792 | 23915 | 2 | 14763 | ENSMUST00 | 133187 | ENSMUSG00 | Gpr37 | G protein-coupled receptor 37 |
| chr6 | 27639035 | 27639251 | 217 | * | NA_peak_19 | 65 | Intron (ENSM | 6 | 27936655 | 27936750 | 96 | 2 | 735266 | ENSMUST00 | 297499 | ENSMUSG00 | Mir592 | microRNA 592 |
| chr6 | 29348055 | 29348363 | 309 | * | NA_peak_19 | 18 | Promoter (<= | 6 | 29348069 | 29374503 | 26435 | 1 | 12321 | ENSMUST00 | 0 | ENSMUSG00 | Calu | calumenin |
| chr6 | 29396638 | 29396983 | 346 | * | NA_peak_19 | 22 | Promoter (<= | 6 | 29396643 | 29426948 | 30306 | 1 | 232664 | ENSMUST00 | 0 | ENSMUSG00 | Ccdc136 | coiled-coil domain containing 136 |
| chr6 | 29917246 | 29917619 | 374 | * | NA_peak_19 | 24 | Promoter (<= | 6 | 29917013 | 29959681 | 42669 | 1 | 320609 | ENSMUST00 | 233 | ENSMUSG00 | Strip2 | striatin interacting protein 2 |
| chr6 | 31217908 | 31218346 | 439 | * | NA_peak_19 | 20 | Promoter (<= | 6 | 31165498 | 31218222 | 52725 | 2 | 232685 | ENSMUST00 | 0 | ENSMUSG00 | Lncpint | long non-protein coding RNA, Trp53 induced transcript |
| chr6 | 34312842 | 34313176 | 335 | * | NA_peak_19 | 19 | Intron (ENSM | 6 | 34309793 | 34317430 | 7638 | 2 | 626009 | ENSMUST00 | 4254 | NA | Gm6644 | Akr1b3 pseudogene |
| chr6 | 34430730 | 34431055 | 326 | * | NA_peak_19 | 24 | Intron (ENSM | 6 | 34412334 | 34423142 | 10809 | 1 | 11997 | ENSMUST00 | 18396 | ENSMUSG00 | Akr1b7 | aldo-keto reductase family 1, member B7 |

| | | | | | | | | | | | | | | | | | | |
|---|---|---|---|---|---|---|---|---|---|---|---|---|---|---|---|---|---|---|
| chr6 | 34721833 | 34722135 | 303 | * | NA_peak_19 | 22 | Intron (ENSM | 6 | 34709634 | 34773694 | 64061 | 1 | 109624 | ENSMUST0 | 12199 | ENSMUSG0 | Cald1 | caldesmon 1 |
| chr6 | 35578156 | 35578451 | 296 | * | NA_peak_19 | 44 | Distal Interge | 6 | 35508906 | 35539888 | 30983 | 2 | 14489 | ENSMUST0 | -38268 | ENSMUSG0 | Mtpn | myotrophin |
| chr6 | 36085459 | 36085726 | 268 | * | NA_peak_19 | 17 | Distal Interge | 6 | 36209620 | 36387851 | 178232 | 2 | 319472 | ENSMUST0 | 302125 | ENSMUSG0 | 9330158H04 | RIKEN cDNA 9330158H04 gene |
| chr6 | 36709822 | 36710109 | 288 | * | NA_peak_19 | 13 | Distal Interge | 6 | 36739459 | 36810179 | 70721 | 2 | 19242 | ENSMUST0 | 100070 | ENSMUSG0 | Ptn | pleiotrophin |
| chr6 | 39118609 | 39118766 | 158 | * | NA_peak_19 | 13 | Promoter (<= | 6 | 39118473 | 39148312 | 29840 | 1 | 75379 | ENSMUST0 | 136 | ENSMUSG0 | 4930599N23 | RIKEN cDNA 4930599N23 gene |
| chr6 | 41879703 | 41880155 | 453 | * | NA_peak_19 | 15 | Distal Interge | 6 | 41863340 | 41864289 | 950 | 1 | 84543 | ENSMUST0 | 16363 | ENSMUSG0 | Sval2 | seminal vesicle antigen-like 2 |
| chr6 | 42157442 | 42157879 | 438 | * | NA_peak_19 | 50 | Distal Interge | 6 | 42140936 | 42141895 | 960 | 1 | 353148 | ENSMUST0 | 16506 | ENSMUSG0 | Tas2r139 | taste receptor, type 2, member 139 |
| chr6 | 46532117 | 46532417 | 301 | * | NA_peak_19 | 26 | Intron (ENSM | 6 | 46170215 | 46435576 | 265362 | 1 | 66797 | ENSMUST0 | 361902 | ENSMUSG0 | Cntnap2 | contactin associated protein-like 2 |
| chr6 | 46582317 | 46582347 | 31 | * | NA_peak_19 | 23 | Intron (ENSM | 6 | 46170215 | 46435576 | 265362 | 1 | 66797 | ENSMUST0 | 412102 | ENSMUSG0 | Cntnap2 | contactin associated protein-like 2 |
| chr6 | 47653766 | 47654585 | 820 | * | NA_peak_19 | 34 | Distal Interge | 6 | 47552386 | 47595341 | 42956 | 2 | 14056 | ENSMUST0 | -58425 | ENSMUSG0 | Ezh2 | enhancer of zeste 2 polycomb repressive complex 2 subunit |
| chr6 | 47740537 | 47740886 | 350 | * | NA_peak_19 | 70 | Distal Interge | 6 | 47803576 | 47803652 | 77 | 2 | 735289 | ENSMUST0 | 62766 | ENSMUSG0 | Mir704 | microRNA 704 |
| chr6 | 47744861 | 47745404 | 544 | * | NA_peak_19 | 51 | Distal Interge | 6 | 47803576 | 47803652 | 77 | 2 | 735289 | ENSMUST0 | 58248 | ENSMUSG0 | Mir704 | microRNA 704 |
| chr6 | 47765946 | 47766255 | 310 | * | NA_peak_19 | 55 | Distal Interge | 6 | 47803576 | 47803652 | 77 | 2 | 735289 | ENSMUST0 | 37397 | ENSMUSG0 | Mir704 | microRNA 704 |
| chr6 | 47943212 | 47943722 | 511 | * | NA_peak_19 | 23 | Promoter (<= | 6 | 47943250 | 47953574 | 10325 | 1 | 232785 | ENSMUST0 | 0 | ENSMUSG0 | Zfp783 | zinc finger protein 783 |
| chr6 | 49616594 | 49616833 | 240 | * | NA_peak_19 | 19 | Distal Interge | 6 | 49438564 | 49469500 | 30937 | 1 | 77485 | ENSMUST0 | 178030 | ENSMUSG0 | Stk31 | serine threonine kinase 31 |
| chr6 | 49617160 | 49617597 | 438 | * | NA_peak_19 | 34 | Distal Interge | 6 | 49438564 | 49469500 | 30937 | 1 | 77485 | ENSMUST0 | 178596 | ENSMUSG0 | Stk31 | serine threonine kinase 31 |
| chr6 | 49891709 | 49891798 | 90 | * | NA_peak_19 | 19 | Distal Interge | 6 | 49822710 | 49829507 | 6798 | 1 | 109648 | ENSMUST0 | 68999 | ENSMUSG0 | Npy | neuropeptide Y |
| chr6 | 50367254 | 50367599 | 346 | * | NA_peak_19 | 26 | Promoter (2- | 6 | 50295687 | 50370215 | 74529 | 2 | 71720 | ENSMUST0 | 2616 | ENSMUSG0 | Osbpl3 | oxysterol binding protein-like 3 |
| chr6 | 57292581 | 57292918 | 338 | * | NA_peak_19 | 18 | Distal Interge | 6 | 57322724 | 57323635 | 912 | 2 | 171202 | ENSMUST0 | 30717 | ENSMUSG0 | Vmn1r16 | vomeronasal 1 receptor 16 |
| chr6 | 57486658 | 57487150 | 493 | * | NA_peak_19 | 26 | Promoter (1- | 6 | 57485121 | 57626030 | 140910 | 1 | 67138 | ENSMUST0 | 1537 | ENSMUSG0 | Herc6 | hect domain and RLD 6 |
| chr6 | 58589117 | 58589419 | 303 | * | NA_peak_19 | 24 | Intron (ENSM | 6 | 58584527 | 58586560 | 2034 | 1 | 26357 | ENSMUST0 | 4590 | ENSMUSG0 | Abcg2 | ATP binding cassette subfamily G member 2 (Junior blood group) |
| chr6 | 58600264 | 58600547 | 284 | * | NA_peak_19 | 54 | Intron (ENSM | 6 | 58596671 | 58695676 | 99006 | 1 | 26357 | ENSMUST0 | 3593 | ENSMUSG0 | Abcg2 | ATP binding cassette subfamily G member 2 (Junior blood group) |
| chr6 | 62039613 | 62040045 | 433 | * | NA_peak_19 | 40 | Intron (ENSM | 6 | 61956689 | 62382865 | 426177 | 1 | 232035 | ENSMUST0 | 82924 | ENSMUSG0 | Ccser1 | coiled-coil serine rich 1 |
| chr6 | 62882733 | 62883093 | 361 | * | NA_peak_19 | 30 | Distal Interge | 6 | 63255876 | 63314071 | 58196 | 1 | 14804 | ENSMUST0 | -372783 | ENSMUSG0 | Grid2 | glutamate receptor, ionotropic, delta 2 |
| chr6 | 64788913 | 64789219 | 307 | * | NA_peak_19 | 23 | Distal Interge | 6 | 64729125 | 64731245 | 2121 | 1 | 11921 | ENSMUST0 | 59788 | ENSMUSG0 | Atoh1 | atonal bHLH transcription factor 1 |
| chr6 | 66462337 | 66462560 | 224 | * | NA_peak_19 | 22 | Intron (ENSM | 6 | 66535390 | 66547220 | 11831 | 1 | 56150 | ENSMUST0 | -72830 | ENSMUSG0 | Mad2l1 | MAD2 mitotic arrest deficient-like 1 |
| chr6 | 67267210 | 67267687 | 478 | * | NA_peak_19 | 22 | Promoter (<= | 6 | 67267257 | 67272855 | 5599 | 1 | 66870 | ENSMUST0 | 0 | ENSMUSG0 | Serbp1 | serpine1 mRNA binding protein 1 |
| chr6 | 67683337 | 67683684 | 348 | * | NA_peak_19 | 15 | Distal Interge | 6 | 67565191 | 67566446 | 1256 | 2 | 75070 | ENSMUST0 | -116891 | NA | 4930515G16 | myc induced nuclear antigen pseudogene |
| chr6 | 67690178 | 67690596 | 419 | * | NA_peak_19 | 23 | Distal Interge | 6 | 67565191 | 67566446 | 1256 | 2 | 75070 | ENSMUST0 | -123732 | NA | 4930515G16 | myc induced nuclear antigen pseudogene |
| chr6 | 67694303 | 67694510 | 208 | * | NA_peak_19 | 19 | Distal Interge | 6 | 67565191 | 67566446 | 1256 | 2 | 75070 | ENSMUST0 | -127857 | NA | 4930515G16 | myc induced nuclear antigen pseudogene |
| chr6 | 71831352 | 71831488 | 137 | * | NA_peak_19 | 17 | Promoter (<= | 6 | 71831379 | 71855186 | 23808 | 1 | 76614 | ENSMUST0 | 0 | ENSMUSG0 | Immt | inner membrane protein, mitochondrial |
| chr6 | 78155860 | 78156231 | 372 | * | NA_peak_19 | 75 | Distal Interge | 6 | 76881637 | 77979699 | 1098063 | 2 | 12386 | ENSMUST0 | -176161 | ENSMUSG0 | Ctnna2 | catenin (cadherin associated protein), alpha 2 |
| chr6 | 79375576 | 79375946 | 371 | * | NA_peak_19 | 15 | Distal Interge | 6 | 79032044 | 79035869 | 3826 | 1 | 100504701 | ENSMUST0 | 343532 | ENSMUSG0 | Gm20362 | predicted gene, 20362 |
| chr6 | 88077761 | 88077874 | 114 | * | NA_peak_19 | 19 | Distal Interge | 6 | 88084482 | 88105304 | 20823 | 1 | 103963 | ENSMUST0 | -6608 | ENSMUSG0 | Rpn1 | ribophorin I |
| chr6 | 88572626 | 88572938 | 313 | * | NA_peak_19 | 16 | Intron (ENSM | 6 | 88545120 | 88618501 | 73382 | 2 | 74589 | ENSMUST0 | 45563 | ENSMUSG0 | Kbtbd12 | kelch repeat and BTB (POZ) domain containing 12 |
| chr6 | 95214666 | 95215009 | 344 | * | NA_peak_19 | 51 | Distal Interge | 6 | 95117984 | 95121797 | 3814 | 1 | 243574 | ENSMUST0 | 96682 | ENSMUSG0 | Kbtbd8 | kelch repeat and BTB (POZ) domain containing 8 |
| chr6 | 96963408 | 96963755 | 348 | * | NA_peak_19 | 34 | Intron (ENSM | 6 | 96831203 | 97060413 | 229211 | 2 | 320701 | ENSMUST0 | 96658 | ENSMUSG0 | Tafa4 | TAFA chemokine like family member 4 |
| chr6 | 96970751 | 96971150 | 400 | * | NA_peak_19 | 42 | Intron (ENSM | 6 | 96831203 | 97060413 | 229211 | 2 | 320701 | ENSMUST0 | 89263 | ENSMUSG0 | Tafa4 | TAFA chemokine like family member 4 |
| chr6 | 107332362 | 107332705 | 344 | * | NA_peak_19 | 54 | Distal Interge | 6 | 107529768 | 107570214 | 40447 | 1 | 16979 | ENSMUST0 | -197063 | ENSMUSG0 | Lrrn1 | leucine rich repeat protein 1, neuronal |
| chr6 | 110849116 | 110849476 | 361 | * | NA_peak_19 | 49 | Intron (ENSM | 6 | 110645582 | 111567230 | 921649 | 1 | 108073 | ENSMUST0 | 203534 | ENSMUSG0 | Grm7 | glutamate receptor, metabotropic 7 |
| chr6 | 113494836 | 113495179 | 344 | * | NA_peak_19 | 15 | Promoter (2- | 6 | 113492019 | 113493335 | 1317 | 1 | 171508 | ENSMUST0 | 2817 | ENSMUSG0 | Creld1 | cysteine-rich with EGF-like domains 1 |
| chr6 | 113697384 | 113697755 | 372 | * | NA_peak_19 | 19 | Promoter (<= | 6 | 113697564 | 113710949 | 13386 | 1 | 381801 | ENSMUST0 | 0 | ENSMUSG0 | Tatdn2 | TatD DNase domain containing 2 |
| chr6 | 116548824 | 116549272 | 449 | * | NA_peak_19 | 68 | Distal Interge | 6 | 116556405 | 116557412 | 1008 | 1 | 258754 | ENSMUST0 | -7133 | ENSMUSG0 | Olfr214 | olfactory receptor 214 |
| chr6 | 118209770 | 118210072 | 303 | * | NA_peak_19 | 54 | Distal Interge | 6 | 118154566 | 118197718 | 43153 | 2 | 19713 | ENSMUST0 | -12052 | ENSMUSG0 | Ret | ret proto-oncogene |
| chr6 | 120037094 | 120037481 | 388 | * | NA_peak_19 | 23 | Promoter (<= | 6 | 119924621 | 120037659 | 113039 | 2 | 232341 | ENSMUST0 | 178 | ENSMUSG0 | Wnk1 | WNK lysine deficient protein kinase 1 |
| chr6 | 120134324 | 120134588 | 265 | * | NA_peak_19 | 33 | Intron (ENSM | 6 | 120093350 | 120200339 | 106990 | 1 | 29862 | ENSMUST0 | 40974 | ENSMUSG0 | Ninj2 | ninjurin 2 |
| chr6 | 122915618 | 122915690 | 73 | * | NA_peak_19 | 20 | Distal Interge | 6 | 122921848 | 122934619 | 12772 | 1 | 269799 | ENSMUST0 | -6158 | ENSMUSG0 | Clec4a1 | C-type lectin domain family 4, member a1 |
| chr6 | 126456299 | 126456398 | 100 | * | NA_peak_19 | 23 | Distal Interge | 6 | 126532551 | 126535412 | 2862 | 2 | 16493 | ENSMUST0 | 79014 | ENSMUSG0 | Kcna5 | potassium voltage-gated channel, shaker-related subfamily, member 5 |
| chr6 | 128681130 | 128681881 | 752 | * | NA_peak_19 | 53 | Distal Interge | 6 | 128662391 | 128674969 | 12579 | 1 | 94071 | ENSMUST0 | 18739 | ENSMUSG0 | Clec2h | C-type lectin domain family 2, member h |
| chr6 | 132876675 | 132876832 | 158 | * | NA_peak_19 | 26 | Distal Interge | 6 | 132868008 | 132869009 | 1002 | 1 | 387344 | ENSMUST0 | 8667 | ENSMUSG0 | Tas2r110 | taste receptor, type 2, member 110 |
| chr6 | 132967134 | 132967448 | 315 | * | NA_peak_19 | 23 | Distal Interge | 6 | 132956884 | 132957919 | 1036 | 2 | 387356 | ENSMUST0 | -9215 | ENSMUSG0 | Tas2r131 | taste receptor, type 2, member 131 |
| chr6 | 149503561 | 149503787 | 227 | * | NA_peak_19 | 28 | Intron (ENSM | 6 | 149513830 | 149519313 | 5484 | 1 | 12121 | ENSMUST0 | -10043 | ENSMUSG0 | Bicd1 | BICD cargo adaptor 1 |
| chr7 | 5206393 | 5206827 | 435 | * | NA_peak_19 | 263 | Distal Interge | 7 | 5194916 | 5196747 | 1832 | 2 | 81015 | ENSMUST0 | -9646 | ENSMUSG0 | Vmn1r56 | vomeronasal 1 receptor 56 |
| chr7 | 7364805 | 7365096 | 292 | * | NA_peak_19 | 26 | Intron (ENSM | 7 | 7334149 | 7374573 | 40425 | 2 | 22306 | ENSMUST0 | 9477 | ENSMUSG0 | Vmn2r30 | vomeronasal 2, receptor 30 |
| chr7 | 7407362 | 7407840 | 479 | * | NA_peak_19 | 66 | Distal Interge | 7 | 7378251 | 7399703 | 21453 | 2 | 100042591 | ENSMUST0 | -7659 | ENSMUSG0 | Vmn2r31 | vomeronasal 2, receptor 31 |
| chr7 | 7472793 | 7473385 | 593 | * | NA_peak_19 | 20 | Intron (ENSM | 7 | 7462096 | 7480018 | 17923 | 2 | 22311 | ENSMUST0 | 6633 | ENSMUSG0 | Vmn2r32 | vomeronasal 2, receptor 32 |
| chr7 | 7508067 | 7508520 | 454 | * | NA_peak_19 | 20 | Distal Interge | 7 | 7462096 | 7480018 | 17923 | 2 | 22311 | ENSMUST0 | -28049 | ENSMUSG0 | Vmn2r32 | vomeronasal 2, receptor 32 |
| chr7 | 7515768 | 7516124 | 357 | * | NA_peak_19 | 33 | Distal Interge | 7 | 7462096 | 7480018 | 17923 | 2 | 22311 | ENSMUST0 | -35750 | ENSMUSG0 | Vmn2r32 | vomeronasal 2, receptor 32 |
| chr7 | 7527429 | 7527497 | 69 | * | NA_peak_19 | 30 | Distal Interge | 7 | 7550967 | 7566786 | 15820 | 2 | 624512 | ENSMUST0 | 39289 | ENSMUSG0 | Vmn2r33 | vomeronasal 2, receptor 33 |
| chr7 | 7546243 | 7546640 | 398 | * | NA_peak_19 | 60 | Distal Interge | 7 | 7550967 | 7566786 | 15820 | 2 | 624512 | ENSMUST0 | 20146 | ENSMUSG0 | Vmn2r33 | vomeronasal 2, receptor 33 |
| chr7 | 7569462 | 7569625 | 164 | * | NA_peak_19 | 19 | Promoter (2- | 7 | 7550967 | 7566786 | 15820 | 2 | 624512 | ENSMUST0 | -2676 | ENSMUSG0 | Vmn2r33 | vomeronasal 2, receptor 33 |
| chr7 | 8077478 | 8077674 | 197 | * | NA_peak_19 | 29 | Distal Interge | 7 | 8086387 | 8092838 | 6452 | 2 | 72716 | ENSMUST0 | 15164 | ENSMUSG0 | Gm45351 | predicted gene 45351 |
| chr7 | 8101255 | 8101577 | 323 | * | NA_peak_19 | 20 | Distal Interge | 7 | 8086326 | 8093041 | 6716 | 2 | 72716 | ENSMUST0 | -8214 | ENSMUSG0 | Gm45351 | predicted gene 45351 |
| chr7 | 8233951 | 8234283 | 333 | * | NA_peak_19 | 31 | Distal Interge | 7 | 8244348 | 8260599 | 16252 | 2 | 381838 | ENSMUST0 | 26316 | ENSMUSG0 | Vmn2r43 | vomeronasal 2, receptor 43 |
| chr7 | 8238572 | 8238906 | 335 | * | NA_peak_19 | 13 | Distal Interge | 7 | 8244348 | 8260599 | 16252 | 2 | 381838 | ENSMUST0 | 21693 | ENSMUSG0 | Vmn2r43 | vomeronasal 2, receptor 43 |
| chr7 | 8244829 | 8245198 | 370 | * | NA_peak_19 | 16 | Exon (ENSM | 7 | 8244348 | 8260599 | 16252 | 2 | 381838 | ENSMUST0 | 15401 | ENSMUSG0 | Vmn2r43 | vomeronasal 2, receptor 43 |
| chr7 | 8385377 | 8385565 | 189 | * | NA_peak_19 | 23 | Promoter (2- | 7 | 8366164 | 8383333 | 17170 | 2 | 434113 | ENSMUST0 | -2044 | ENSMUSG0 | Vmn2r44 | vomeronasal 2, receptor 44 |
| chr7 | 8868276 | 8868572 | 297 | * | NA_peak_19 | 34 | Distal Interge | 7 | 8855847 | 8862851 | 7005 | 2 | 72716 | ENSMUST0 | -5425 | ENSMUSG0 | Gm45351 | predicted gene 45351 |
| chr7 | 9004209 | 9004473 | 265 | * | NA_peak_19 | 39 | Distal Interge | 7 | 9013806 | 9030798 | 16993 | 2 | 545909 | ENSMUST0 | 26325 | ENSMUSG0 | Vmn2r39 | vomeronasal 2, receptor 39 |
| chr7 | 10036930 | 10037253 | 324 | * | NA_peak_19 | 26 | 3' UTR | 7 | 10035836 | 10053296 | 17461 | 2 | 434117 | ENSMUST0 | 16043 | ENSMUSG0 | Vmn2r50 | vomeronasal 2, receptor 50 |
| chr7 | 14843872 | 14844103 | 232 | * | NA_peak_19 | 19 | Distal Interge | 7 | 14659745 | 14664574 | 4830 | 1 | 194588 | ENSMUST0 | 184127 | ENSMUSG0 | Obox7 | oocyte specific homeobox 7 |
| chr7 | 15198144 | 15198716 | 573 | * | NA_peak_19 | 42 | Distal Interge | 7 | 15388721 | 15398545 | 9825 | 1 | 246792 | ENSMUST0 | -190005 | ENSMUSG0 | Obox2 | oocyte specific homeobox 2 |
| chr7 | 15226115 | 15226803 | 689 | * | NA_peak_19 | 35 | Distal Interge | 7 | 15388721 | 15398545 | 9825 | 1 | 246792 | ENSMUST0 | -161918 | ENSMUSG0 | Obox2 | oocyte specific homeobox 2 |
| chr7 | 15229878 | 15230050 | 173 | * | NA_peak_19 | 14 | Exon (ENSM | 7 | 15388721 | 15398545 | 9825 | 1 | 246792 | ENSMUST0 | -158671 | ENSMUSG0 | Obox2 | oocyte specific homeobox 2 |
| chr7 | 15258180 | 15258854 | 675 | * | NA_peak_19 | 28 | Distal Interge | 7 | 15388721 | 15398545 | 9825 | 1 | 246792 | ENSMUST0 | -129867 | ENSMUSG0 | Obox2 | oocyte specific homeobox 2 |
| chr7 | 15438749 | 15439290 | 542 | * | NA_peak_19 | 112 | Intron (ENSM | 7 | 15390176 | 15398129 | 7954 | 1 | 246792 | ENSMUST0 | 48573 | ENSMUSG0 | Obox2 | oocyte specific homeobox 2 |
| chr7 | 15481121 | 15481493 | 373 | * | NA_peak_19 | 40 | Exon (ENSM | 7 | 15512340 | 15556446 | 44107 | 1 | 71468 | ENSMUST0 | -30847 | ENSMUSG0 | Obox1 | oocyte specific homeobox 1 |
| chr7 | 15482824 | 15483161 | 338 | * | NA_peak_19 | 35 | Intron (ENSM | 7 | 15512340 | 15556446 | 44107 | 1 | 71468 | ENSMUST0 | -29179 | ENSMUSG0 | Obox1 | oocyte specific homeobox 1 |
| chr7 | 16299745 | 16300030 | 286 | * | NA_peak_19 | 16 | Distal Interge | 7 | 16308393 | 16312414 | 4022 | 1 | 170770 | ENSMUST0 | -8363 | ENSMUSG0 | Bbc3 | BCL2 binding component 3 |
| chr7 | 27015346 | 27015737 | 392 | * | NA_peak_19 | 17 | Distal Interge | 7 | 27029081 | 27037375 | 8295 | 1 | 13085 | ENSMUST0 | -13344 | ENSMUSG0 | Cyp2a12 | cytochrome P450, family 2, subfamily a, polypeptide 12 |
| chr7 | 38686317 | 38686627 | 311 | * | NA_peak_19 | 46 | Distal Interge | 7 | 38791661 | 38860532 | 68872 | 1 | 105247250 | ENSMUST0 | -105034 | ENSMUSG0 | 1700010N08 | RIKEN cDNA 1700010N08 gene |
| chr7 | 38836214 | 38836524 | 311 | * | NA_peak_19 | 23 | Intron (ENSM | 7 | 38791661 | 38860532 | 68872 | 1 | 105247250 | ENSMUST0 | 44553 | ENSMUSG0 | 1700010N08 | RIKEN cDNA 1700010N08 gene |
| chr7 | 41199923 | 41200351 | 429 | * | NA_peak_19 | 13 | Intron (ENSM | 7 | 41153841 | 41289889 | 136049 | 1 | 434172 | ENSMUST0 | 46082 | ENSMUSG0 | Gm5592 | predicted gene 5592 |
| chr7 | 41322932 | 41323364 | 433 | * | NA_peak_19 | 42 | Distal Interge | 7 | 41284327 | 41290183 | 5857 | 1 | 434172 | ENSMUST0 | 38605 | ENSMUSG0 | Gm5592 | predicted gene 5592 |
| chr7 | 43132150 | 43132351 | 202 | * | NA_peak_19 | 19 | Intron (ENSM | 7 | 43114229 | 43189654 | 75426 | 1 | 668620 | ENSMUST0 | 17921 | ENSMUSG0 | Zfp936 | zinc finger protein 936 |
| chr7 | 48371470 | 48372009 | 540 | * | NA_peak_19 | 14 | Distal Interge | 7 | 48388526 | 48389648 | 1123 | 1 | 404240 | ENSMUST0 | -16517 | ENSMUSG0 | Mrgprb8 | MAS-related GPR, member B8 |
| chr7 | 50956046 | 50956369 | 324 | * | NA_peak_19 | 33 | Distal Interge | 7 | 50599190 | 50600530 | 1341 | 1 | 243996 | ENSMUST0 | 356856 | ENSMUSG0 | 4933405O20 | RIKEN cDNA 4933405O20 gene |
| chr7 | 50963027 | 50963274 | 248 | * | NA_peak_19 | 29 | Distal Interge | 7 | 50599190 | 50600530 | 1341 | 1 | 243996 | ENSMUST0 | 363837 | ENSMUSG0 | 4933405O20 | RIKEN cDNA 4933405O20 gene |
| chr7 | 52514915 | 52515148 | 234 | * | NA_peak_19 | 35 | Distal Interge | 7 | 52011683 | 52015728 | 4046 | 2 | 100503036 | ENSMUST0 | -499187 | ENSMUSG0 | 1700015G11 | RIKEN cDNA 1700015G11 gene |
| chr7 | 54925126 | 54925540 | 415 | * | NA_peak_19 | 28 | Intron (ENSM | 7 | 54835615 | 55268885 | 433271 | 1 | 233271 | ENSMUST0 | 89511 | ENSMUSG0 | Luzp2 | leucine zipper protein 2 |
| chr7 | 57371942 | 57372039 | 98 | * | NA_peak_19 | 26 | Intron (ENSM | 7 | 57178382 | 57386871 | 208490 | 2 | 14407 | ENSMUST0 | 14832 | ENSMUSG0 | Gabrg3 | gamma-aminobutyric acid (GABA) A receptor, subunit gamma 3 |
| chr7 | 58240089 | 58240286 | 198 | * | NA_peak_19 | 26 | Distal Interge | 7 | 58658246 | 58829420 | 171175 | 1 | 11982 | ENSMUST0 | -417960 | ENSMUSG0 | Atp10a | ATPase, class V, type 10A |
| chr7 | 59296928 | 59297054 | 127 | * | NA_peak_19 | 44 | Promoter (2- | 7 | 59300042 | 59306729 | 6688 | 1 | 22215 | ENSMUST0 | -2988 | ENSMUSG0 | Ube3a | ubiquitin protein ligase E3A |
| chr7 | 59577538 | 59577855 | 318 | * | NA_peak_19 | 27 | Distal Interge | 7 | 59675988 | 59676081 | 94 | 2 | 64244 | ENSMUST0 | 98226 | ENSMUSG0 | Snord116l1 | small nucleolar RNA, C/D box 116-like 1 |
| chr7 | 59742468 | 59742770 | 303 | * | NA_peak_19 | 74 | Intron (ENSM | 7 | 59751489 | 59751582 | 94 | 2 | 64243 | ENSMUST0 | 8812 | ENSMUSG0 | Gm26504 | predicted gene, 26504 |
| chr7 | 59760033 | 59760374 | 342 | * | NA_peak_19 | 71 | Intron (ENSM | 7 | 59751489 | 59751582 | 94 | 2 | 64243 | ENSMUST0 | -8451 | ENSMUSG0 | Gm26504 | predicted gene, 26504 |
| chr7 | 59862764 | 59863115 | 352 | * | NA_peak_19 | 27 | Exon (ENSM | 7 | 59619158 | 59874749 | 255592 | 2 | 52480 | ENSMUST0 | 11634 | NA | Snhg14 | small nucleolar RNA host gene 14 |
| chr7 | 60680374 | 60680932 | 559 | * | NA_peak_19 | 13 | Distal Interge | 7 | 59982502 | 60140219 | 157718 | 2 | 20646 | ENSMUST0 | -540155 | ENSMUSG0 | Snrpn | small nuclear ribonucleoprotein N |
| chr7 | 63374078 | 63374488 | 411 | * | NA_peak_19 | 120 | Intron (ENSM | 7 | 63344690 | 63444529 | 99840 | 2 | 77631 | ENSMUST0 | 70041 | ENSMUSG0 | 4930554H23 | RIKEN cDNA 4930554H23 gene |
| chr7 | 74801504 | 74801665 | 162 | * | NA_peak_19 | 20 | Distal Interge | 7 | 75030194 | 75058162 | 27969 | 1 | 75205 | ENSMUST0 | -228529 | ENSMUSG0 | 4930533N22 | RIKEN cDNA 4930533N22 gene |
| chr7 | 77652896 | 77653176 | 281 | * | NA_peak_19 | 22 | Distal Interge | 7 | 78247388 | 78261887 | 14500 | 2 | 18213 | ENSMUST0 | 608711 | ENSMUSG0 | Ntrk3 | neurotrophic tyrosine kinase, receptor, type 3 |
| chr7 | 77654226 | 77654831 | 606 | * | NA_peak_19 | 26 | Distal Interge | 7 | 78247388 | 78261887 | 14500 | 2 | 18213 | ENSMUST0 | 607056 | ENSMUSG0 | Ntrk3 | neurotrophic tyrosine kinase, receptor, type 3 |
| chr7 | 82469566 | 82469695 | 130 | * | NA_peak_19 | 43 | Intron (ENSM | 7 | 82408474 | 82465844 | 57371 | 1 | 269959 | ENSMUST0 | 61092 | ENSMUSG0 | Adamtsl3 | ADAMTS-like 3 |
| chr7 | 85100119 | 85100578 | 460 | * | NA_peak_19 | 36 | Distal Interge | 7 | 85126967 | 85155998 | 29032 | 2 | 620672 | ENSMUST0 | 55420 | ENSMUSG0 | Vmn2r67 | vomeronasal 2, receptor 67 |
| chr7 | 87512755 | 87513059 | 305 | * | NA_peak_19 | 404 | Distal Interge | 7 | 87424771 | 87493512 | 68742 | 2 | 22173 | ENSMUST0 | -19243 | ENSMUSG0 | Tyr | tyrosinase |
| chr7 | 87904830 | 87905159 | 330 | * | NA_peak_19 | 35 | Intron (ENSM | 7 | 87603089 | 87809620 | 206532 | 1 | 108071 | ENSMUST0 | 301741 | ENSMUSG0 | Grm5 | glutamate receptor, metabotropic 5 |
| chr7 | 96196356 | 96196813 | 458 | * | NA_peak_19 | 18 | Intron (ENSM | 7 | 96209663 | 96437138 | 227476 | 1 | 23966 | ENSMUST0 | -12850 | ENSMUSG0 | Tenm4 | teneurin transmembrane protein 4 |
| chr7 | 97116333 | 97116672 | 340 | * | NA_peak_19 | 30 | Intron (ENSM | 7 | 97081776 | 97305036 | 223261 | 1 | 14389 | ENSMUST0 | 34557 | ENSMUSG0 | Gab2 | growth factor receptor bound protein 2-associated protein 2 |
| chr7 | 99182705 | 99183105 | 401 | * | NA_peak_19 | 30 | Promoter (<= | 7 | 99153658 | 99182719 | 29062 | 2 | 67800 | ENSMUST0 | 0 | ENSMUSG0 | Dgat2 | diacylglycerol O-acyltransferase 2 |
| chr7 | 99268439 | 99268761 | 323 | * | NA_peak_19 | 26 | Promoter (<= | 7 | 99268406 | 99337137 | 68732 | 1 | 17760 | ENSMUST0 | 33 | ENSMUSG0 | Map6 | microtubule-associated protein 6 |
| chr7 | 103104497 | 103104888 | 392 | * | NA_peak_19 | 27 | Distal Interge | 7 | 103097720 | 103098749 | 1030 | 1 | 259091 | ENSMUST0 | 6777 | ENSMUSG0 | Olfr585 | olfactory receptor 585 |
| chr7 | 103921751 | 103922134 | 384 | * | NA_peak_19 | 30 | Distal Interge | 7 | 103937382 | 103938335 | 954 | 1 | 259123 | ENSMUST0 | -15248 | ENSMUSG0 | Olfr632 | olfactory receptor 632 |
| chr7 | 104033329 | 104033755 | 427 | * | NA_peak_19 | 17 | Intron (ENSM | 7 | 104039798 | 104040736 | 939 | 1 | 259075 | ENSMUST0 | -6043 | ENSMUSG0 | Olfr641 | olfactory receptor 641 |
| chr7 | 104122778 | 104122938 | 161 | * | NA_peak_19 | 23 | Intron (ENSM | 7 | 104127912 | 104129826 | 1915 | 2 | 70980 | ENSMUST0 | 6888 | ENSMUSG0 | Ubqln5 | ubiquilin 5 |
| chr7 | 108430107 | 108430699 | 593 | * | NA_peak_19 | 19 | Distal Interge | 7 | 108422573 | 108423517 | 945 | 1 | 258733 | ENSMUST0 | 7534 | ENSMUSG0 | Olfr497 | olfactory receptor 497 |
| chr7 | 113381902 | 113382276 | 375 | * | NA_peak_19 | 29 | Distal Interge | 7 | 113385580 | 113388570 | 2991 | 2 | 19226 | ENSMUST0 | 6294 | ENSMUSG0 | Pth | parathyroid hormone |
| chr7 | 118070650 | 118071116 | 467 | * | NA_peak_19 | 44 | Distal Interge | 7 | 118109663 | 118115228 | 5566 | 2 | 267019 | ENSMUST0 | 44112 | ENSMUSG0 | Rps15a | ribosomal protein S15A |
| chr7 | 122542038 | 122542341 | 304 | * | NA_peak_20 | 18 | Intron (ENSM | 7 | 122599982 | 122628078 | 28097 | 1 | 18751 | ENSMUST0 | -57641 | ENSMUSG0 | Prkcb | protein kinase C, beta |
| chr7 | 123767421 | 123767607 | 187 | * | NA_peak_20 | 30 | Distal Interge | 7 | 123982869 | 124398989 | 416121 | 1 | 628779 | ENSMUST0 | -215262 | ENSMUSG0 | Hs3st4 | heparan sulfate (glucosamine) 3-O-sulfotransferase 4 |
| chr7 | 125010993 | 125011444 | 452 | * | NA_peak_20 | 75 | Distal Interge | 7 | 124625670 | 124708935 | 83266 | 2 | 73076 | ENSMUST0 | -302058 | ENSMUSG0 | 3100003L05 | RIKEN cDNA 3100003L05 gene |
| chr7 | 125532923 | 125533033 | 111 | * | NA_peak_20 | 13 | Distal Interge | 7 | 125552120 | 125579474 | 27355 | 1 | 16190 | ENSMUST0 | -19087 | ENSMUSG0 | Il4ra | interleukin 4 receptor, alpha |
| chr7 | 127475326 | 127475735 | 410 | * | NA_peak_20 | 44 | Promoter (<= | 7 | 127475501 | 127476757 | 1257 | 1 | 233895 | ENSMUST0 | 0 | ENSMUSG0 | Prr14 | proline rich 14 |
| chr7 | 131488924 | 131489048 | 125 | * | NA_peak_20 | 19 | Intron (ENSM | 7 | 131542867 | 131544925 | 2059 | 1 | 15373 | ENSMUST0 | -53819 | ENSMUSG0 | Hmx3 | H6 homeobox 3 |
| chr7 | 140617338 | 140617435 | 98 | * | NA_peak_20 | 35 | 3' UTR | 7 | 140601340 | 140611124 | 9785 | 1 | 18345 | ENSMUST0 | 15998 | ENSMUSG0 | Olfr46 | olfactory receptor 46 |
| chr7 | 143658458 | 143659067 | 610 | * | NA_peak_20 | 35 | Intron (ENSM | 7 | 143636725 | 143649661 | 12937 | 2 | 79202 | ENSMUST0 | -8797 | ENSMUSG0 | Tnfrsf22 | tumor necrosis factor receptor superfamily, member 22 |
| chr7 | 144726395 | 144726668 | 274 | * | NA_peak_20 | 31 | Intron (ENSM | 7 | 144588549 | 144738543 | 149995 | 2 | 101772 | ENSMUST0 | 11875 | ENSMUSG0 | Ano1 | anoctamin 1, calcium activated chloride channel |
| chr8 | 3286296 | 3286351 | 56 | * | NA_peak_20 | 29 | Distal Interge | 8 | 3150922 | 3279617 | 128696 | 2 | 16337 | ENSMUST0 | -6679 | ENSMUSG0 | Insr | insulin receptor |
| chr8 | 3313145 | 3313502 | 358 | * | NA_peak_20 | 32 | Distal Interge | 8 | 3150922 | 3279617 | 128696 | 2 | 16337 | ENSMUST0 | -33528 | ENSMUSG0 | Insr | insulin receptor |
| chr8 | 4178981 | 4179027 | 47 | * | NA_peak_20 | 15 | Intron (ENSM | 8 | 4187188 | 4188029 | 842 | 1 | 213027 | ENSMUST0 | -8161 | ENSMUSG0 | Evi5l | ecotropic viral integration site 5 like |
| chr8 | 4393285 | 4393504 | 220 | * | NA_peak_20 | 26 | Intron (ENSM | 8 | 4349588 | 4360020 | 10433 | 1 | 20300 | ENSMUST0 | 43697 | ENSMUSG0 | Ccl25 | chemokine (C-C motif) ligand 25 |
| chr8 | 4571510 | 4571943 | 434 | * | NA_peak_20 | 16 | Distal Interge | 8 | 4613167 | 4630231 | 17065 | 1 | 233987 | ENSMUST0 | -41224 | ENSMUSG0 | Zfp958 | zinc finger protein 958 |
| chr8 | 5506327 | 5506408 | 82 | * | NA_peak_20 | 17 | Distal Interge | 8 | 5083219 | 5105351 | 22133 | 2 | 20494 | ENSMUST0 | -400976 | ENSMUSG0 | Slc10a2 | solute carrier family 10, member 2 |
| chr8 | 6054623 | 6054960 | 338 | * | NA_peak_20 | 30 | Distal Interge | 8 | 5083219 | 5105351 | 22133 | 2 | 20494 | ENSMUST0 | -949272 | ENSMUSG0 | Slc10a2 | solute carrier family 10, member 2 |
| chr8 | 7130491 | 7130821 | 331 | * | NA_peak_20 | 44 | Distal Interge | 8 | 8620597 | 8639363 | 18767 | 2 | 13642 | ENSMUST0 | 1508542 | ENSMUSG0 | Efnb2 | ephrin B2 |
| chr8 | 7148722 | 7149002 | 281 | * | NA_peak_20 | 25 | Distal Interge | 8 | 8620597 | 8639363 | 18767 | 2 | 13642 | ENSMUST0 | 1490361 | ENSMUSG0 | Efnb2 | ephrin B2 |
| chr8 | 7278534 | 7278994 | 461 | * | NA_peak_20 | 30 | Distal Interge | 8 | 8620597 | 8639363 | 18767 | 2 | 13642 | ENSMUST0 | 1360369 | ENSMUSG0 | Efnb2 | ephrin B2 |
| chr8 | 7391061 | 7391428 | 368 | * | NA_peak_20 | 23 | Intron (ENSM | 8 | 8620597 | 8639363 | 18767 | 2 | 13642 | ENSMUST0 | 1247935 | ENSMUSG0 | Efnb2 | ephrin B2 |
| chr8 | 7542174 | 7542440 | 267 | * | NA_peak_20 | 37 | Intron (ENSM | 8 | 8620597 | 8639363 | 18767 | 2 | 13642 | ENSMUST0 | 1096923 | ENSMUSG0 | Efnb2 | ephrin B2 |
| chr8 | 9514714 | 9514962 | 249 | * | NA_peak_20 | 26 | Intron (ENSM | 8 | 9207238 | 9323711 | 116474 | 2 | 270028 | ENSMUST0 | -191003 | ENSMUSG0 | Fam155a | family with sequence similarity 155, member A |
| chr8 | 10221281 | 10221743 | 463 | * | NA_peak_20 | 23 | Intron (ENSM | 8 | 10249288 | 10369447 | 120160 | 1 | 244281 | ENSMUST0 | -27545 | ENSMUSG0 | Myo16 | myosin XVI |
| chr8 | 10511023 | 10511261 | 239 | * | NA_peak_20 | 39 | Intron (ENSM | 8 | 10272572 | 10634741 | 362170 | 1 | 244281 | ENSMUST0 | 238451 | ENSMUSG0 | Myo16 | myosin XVI |
| chr8 | 11550483 | 11550732 | 250 | * | NA_peak_20 | 31 | Promoter (<= | 8 | 11526024 | 11550692 | 24669 | 2 | 71941 | ENSMUST0 | 0 | ENSMUSG0 | Cars2 | cysteinyl-tRNA synthetase 2 (mitochondrial)(putative) |
| chr8 | 11634996 | 11635131 | 136 | * | NA_peak_20 | 25 | Promoter (<= | 8 | 11614963 | 11635036 | 20074 | 2 | 102334 | ENSMUST0 | 0 | ENSMUSG0 | Ankrd10 | ankyrin repeat domain 10 |
| chr8 | 11648926 | 11649264 | 339 | * | NA_peak_20 | 20 | Distal Interge | 8 | 11655478 | 11660735 | 5258 | 2 | 76413 | ENSMUST0 | 11471 | ENSMUSG0 | 1700016D06 | RIKEN cDNA 1700016D06 gene |
| chr8 | 11877195 | 11877318 | 124 | * | NA_peak_20 | 19 | Intron (ENSM | 8 | 11842061 | 11855757 | 13697 | 1 | 75528 | ENSMUST0 | 35134 | ENSMUSG0 | Tex29 | testis expressed 29 |
| chr8 | 12109977 | 12110167 | 191 | * | NA_peak_20 | 26 | Distal Interge | 8 | 11842061 | 11855757 | 13697 | 1 | 75528 | ENSMUST0 | 267916 | ENSMUSG0 | Tex29 | testis expressed 29 |
| chr8 | 13838966 | 13839127 | 162 | * | NA_peak_20 | 39 | Promoter (<= | 8 | 13835233 | 13838389 | 3157 | 2 | 231201 | ENSMUST0 | -577 | ENSMUSG0 | AF366264 | cDNA sequence AF366264 |
| chr8 | 14658888 | 14659066 | 179 | * | NA_peak_20 | 22 | Intron (ENSM | 8 | 14742753 | 14760902 | 18150 | 1 | 244310 | ENSMUST0 | -83687 | ENSMUSG0 | Dlgap2 | DLG associated protein 2 |
| chr8 | 16902084 | 16902425 | 342 | * | NA_peak_20 | 19 | Intron (ENSM | 8 | 15892537 | 17535281 | 1642745 | 2 | 94109 | ENSMUST0 | 632856 | ENSMUSG0 | Csmd1 | CUB and Sushi multiple domains 1 |
| chr8 | 18042802 | 18043173 | 372 | * | NA_peak_20 | 22 | Distal Interge | 8 | 16344863 | 17535586 | 1190724 | 2 | 94109 | ENSMUST0 | -507216 | ENSMUSG0 | Csmd1 | CUB and Sushi multiple domains 1 |
| chr8 | 19076579 | 19076785 | 207 | * | NA_peak_20 | 44 | Distal Interge | 8 | 19052826 | 19064810 | 11985 | 2 | 360214 | ENSMUST0 | -11769 | ENSMUSG0 | Defb39 | defensin beta 39 |
| chr8 | 19705173 | 19705480 | 308 | * | NA_peak_20 | 51 | Distal Interge | 8 | 19729576 | 19753527 | 23952 | 1 | 626415 | ENSMUST0 | -24096 | ENSMUSG0 | 4930467E23 | RIKEN cDNA 4930467E23 gene |
| chr8 | 19721388 | 19721831 | 444 | * | NA_peak_20 | 20 | Distal Interge | 8 | 19729576 | 19753527 | 23952 | 1 | 626415 | ENSMUST0 | -7745 | ENSMUSG0 | 4930467E23 | RIKEN cDNA 4930467E23 gene |
| chr8 | 19727769 | 19728215 | 447 | * | NA_peak_20 | 27 | Promoter (1- | 8 | 19729576 | 19753527 | 23952 | 1 | 626415 | ENSMUST0 | -1361 | ENSMUSG0 | 4930467E23 | RIKEN cDNA 4930467E23 gene |
| chr8 | 19773734 | 19774171 | 438 | * | NA_peak_20 | 83 | Distal Interge | 8 | 19729576 | 19753527 | 23952 | 1 | 626415 | ENSMUST0 | 44158 | ENSMUSG0 | 4930467E23 | RIKEN cDNA 4930467E23 gene |
| chr8 | 19801895 | 19802205 | 311 | * | NA_peak_20 | 48 | Distal Interge | 8 | 19729576 | 19753527 | 23952 | 1 | 626415 | ENSMUST0 | 72319 | ENSMUSG0 | 4930467E23 | RIKEN cDNA 4930467E23 gene |
| chr8 | 19933796 | 19934523 | 728 | * | NA_peak_20 | 44 | Distal Interge | 8 | 19729576 | 19753527 | 23952 | 1 | 626415 | ENSMUST0 | 204220 | ENSMUSG0 | 4930467E23 | RIKEN cDNA 4930467E23 gene |
| chr8 | 19966738 | 19967459 | 722 | * | NA_peak_20 | 104 | Distal Interge | 8 | 19729576 | 19753527 | 23952 | 1 | 626415 | ENSMUST0 | 237162 | ENSMUSG0 | 4930467E23 | RIKEN cDNA 4930467E23 gene |
| chr8 | 19967910 | 19968113 | 204 | * | NA_peak_20 | 30 | Distal Interge | 8 | 19729576 | 19753527 | 23952 | 1 | 626415 | ENSMUST0 | 238334 | ENSMUSG0 | 4930467E23 | RIKEN cDNA 4930467E23 gene |
| chr8 | 20042143 | 20042330 | 188 | * | NA_peak_20 | 20 | Distal Interge | 8 | 20262613 | 20283266 | 20654 | 2 | 547150 | ENSMUST0 | 240936 | ENSMUSG0 | 6820431F20 | cadherin 11 pseudogene |
| chr8 | 20047816 | 20048618 | 803 | * | NA_peak_20 | 28 | Distal Interge | 8 | 20262613 | 20283266 | 20654 | 2 | 547150 | ENSMUST0 | 234648 | ENSMUSG0 | 6820431F20 | cadherin 11 pseudogene |
| chr8 | 20049068 | 20049688 | 621 | * | NA_peak_20 | 59 | Distal Interge | 8 | 20262613 | 20283266 | 20654 | 2 | 547150 | ENSMUST0 | 233578 | ENSMUSG0 | 6820431F20 | cadherin 11 pseudogene |
| chr8 | 20087393 | 20087760 | 368 | * | NA_peak_20 | 15 | Distal Interge | 8 | 20262613 | 20283266 | 20654 | 2 | 547150 | ENSMUST0 | 195506 | ENSMUSG0 | 6820431F20 | cadherin 11 pseudogene |
| chr8 | 20090851 | 20091204 | 354 | * | NA_peak_20 | 29 | Distal Interge | 8 | 20262613 | 20283266 | 20654 | 2 | 547150 | ENSMUST0 | 192062 | ENSMUSG0 | 6820431F20 | cadherin 11 pseudogene |
| chr8 | 20099906 | 20100358 | 453 | * | NA_peak_20 | 14 | Distal Interge | 8 | 20262613 | 20283266 | 20654 | 2 | 547150 | ENSMUST0 | 182908 | ENSMUSG0 | 6820431F20 | cadherin 11 pseudogene |
| chr8 | 20102272 | 20102654 | 383 | * | NA_peak_20 | 13 | Distal Interge | 8 | 20262613 | 20283266 | 20654 | 2 | 547150 | ENSMUST0 | 180612 | ENSMUSG0 | 6820431F20 | cadherin 11 pseudogene |
| chr8 | 20107277 | 20107621 | 345 | * | NA_peak_20 | 51 | Distal Interge | 8 | 20262613 | 20283266 | 20654 | 2 | 547150 | ENSMUST0 | 175645 | ENSMUSG0 | 6820431F20 | cadherin 11 pseudogene |
| chr8 | 20107967 | 20108924 | 958 | * | NA_peak_20 | 148 | Distal Interge | 8 | 20262613 | 20283266 | 20654 | 2 | 547150 | ENSMUST0 | 174342 | ENSMUSG0 | 6820431F20 | cadherin 11 pseudogene |
| chr8 | 20119062 | 20119563 | 502 | * | NA_peak_20 | 40 | Distal Interge | 8 | 20262613 | 20283266 | 20654 | 2 | 547150 | ENSMUST0 | 163703 | ENSMUSG0 | 6820431F20 | cadherin 11 pseudogene |
| chr8 | 20131682 | 20132312 | 631 | * | NA_peak_20 | 95 | Distal Interge | 8 | 20262613 | 20283266 | 20654 | 2 | 547150 | ENSMUST0 | 150954 | ENSMUSG0 | 6820431F20 | cadherin 11 pseudogene |
| chr8 | 20139051 | 20139359 | 309 | * | NA_peak_20 | 29 | Intron (ENSM | 8 | 20262613 | 20283266 | 20654 | 2 | 547150 | ENSMUST0 | 143907 | ENSMUSG0 | 6820431F20 | cadherin 11 pseudogene |
| chr8 | 20283853 | 20284179 | 327 | * | NA_peak_20 | 26 | Promoter (<= | 8 | 20262613 | 20283266 | 20654 | 2 | 547150 | ENSMUST0 | -587 | ENSMUSG0 | 6820431F20 | cadherin 11 pseudogene |
| chr8 | 20288562 | 20288664 | 103 | * | NA_peak_20 | 24 | Intron (ENSM | 8 | 20262613 | 20283266 | 20654 | 2 | 547150 | ENSMUST0 | -5296 | ENSMUSG0 | 6820431F20 | cadherin 11 pseudogene |
| chr8 | 20292898 | 20293281 | 384 | * | NA_peak_20 | 50 | Intron (ENSM | 8 | 20268286 | 20297432 | 29147 | 2 | 547150 | ENSMUST0 | 4151 | ENSMUSG0 | 6820431F20 | cadherin 11 pseudogene |
| chr8 | 20311542 | 20312053 | 512 | * | NA_peak_20 | 19 | Distal Interge | 8 | 20268286 | 20297432 | 29147 | 2 | 547150 | ENSMUST0 | -14110 | ENSMUSG0 | 6820431F20 | cadherin 11 pseudogene |
| chr8 | 20323495 | 20323800 | 306 | * | NA_peak_20 | 33 | Distal Interge | 8 | 20268286 | 20297432 | 29147 | 2 | 547150 | ENSMUST0 | -26063 | ENSMUSG0 | 6820431F20 | cadherin 11 pseudogene |
| chr8 | 20334064 | 20334546 | 483 | * | NA_peak_20 | 42 | Distal Interge | 8 | 20337664 | 20363210 | 25547 | 2 | 100040599 | ENSMUST0 | 28664 | ENSMUSG0 | Gm15319 | predicted gene 15319 |
| chr8 | 20345834 | 20346439 | 606 | * | NA_peak_20 | 15 | Intron (ENSM | 8 | 20337664 | 20363210 | 25547 | 2 | 100040599 | ENSMUST0 | 16771 | ENSMUSG0 | Gm15319 | predicted gene 15319 |
| chr8 | 20348907 | 20349412 | 506 | * | NA_peak_20 | 25 | Intron (ENSM | 8 | 20337664 | 20363210 | 25547 | 2 | 100040599 | ENSMUST0 | 13798 | ENSMUSG0 | Gm15319 | predicted gene 15319 |
| chr8 | 20351331 | 20351713 | 383 | * | NA_peak_20 | 22 | Intron (ENSM | 8 | 20337664 | 20363210 | 25547 | 2 | 100040599 | ENSMUST0 | 11497 | ENSMUSG0 | Gm15319 | predicted gene 15319 |
| chr8 | 20371856 | 20372177 | 322 | * | NA_peak_20 | 125 | Distal Interge | 8 | 20337664 | 20363210 | 25547 | 2 | 100040599 | ENSMUST0 | -8646 | ENSMUSG0 | Gm15319 | predicted gene 15319 |
| chr8 | 20394665 | 20395013 | 349 | * | NA_peak_20 | 22 | Intron (ENSM | 8 | 20395392 | 20410565 | 15174 | 2 | 381598 | ENSMUST0 | 15552 | ENSMUSG0 | 2610005L07 | cadherin 11 pseudogene |
| chr8 | 20407505 | 20408215 | 711 | * | NA_peak_20 | 29 | Promoter (2- | 8 | 20395392 | 20410565 | 15174 | 2 | 381598 | ENSMUST0 | 2350 | ENSMUSG0 | 2610005L07 | cadherin 11 pseudogene |
| chr8 | 20435824 | 20436249 | 426 | * | NA_peak_20 | 36 | Distal Interge | 8 | 20385782 | 20424814 | 39033 | 2 | 381598 | ENSMUST0 | -11010 | ENSMUSG0 | 2610005L07 | cadherin 11 pseudogene |
| chr8 | 20441919 | 20442263 | 345 | * | NA_peak_20 | 41 | Distal Interge | 8 | 20385782 | 20424814 | 39033 | 2 | 381598 | ENSMUST0 | -17105 | ENSMUSG0 | 2610005L07 | cadherin 11 pseudogene |
| chr8 | 20493725 | 20494251 | 527 | * | NA_peak_20 | 51 | Distal Interge | 8 | 20385782 | 20424814 | 39033 | 2 | 381598 | ENSMUST0 | -68911 | ENSMUSG0 | 2610005L07 | cadherin 11 pseudogene |
| chr8 | 20520363 | 20520687 | 325 | * | NA_peak_20 | 123 | Distal Interge | 8 | 20614919 | 20640703 | 25785 | 1 | 100861668 | ENSMUST0 | -94232 | ENSMUSG0 | Gm21119 | predicted gene, 21119 |
| chr8 | 20530092 | 20530435 | 344 | * | NA_peak_20 | 43 | Distal Interge | 8 | 20614919 | 20640703 | 25785 | 1 | 100861668 | ENSMUST0 | -84484 | ENSMUSG0 | Gm21119 | predicted gene, 21119 |
| chr8 | 20537091 | 20537360 | 270 | * | NA_peak_20 | 48 | Distal Interge | 8 | 20614919 | 20640703 | 25785 | 1 | 100861668 | ENSMUST0 | -77559 | ENSMUSG0 | Gm21119 | predicted gene, 21119 |

| | | | | | | | | | | | | | | | | | | |
|---|---|---|---|---|---|---|---|---|---|---|---|---|---|---|---|---|---|---|
| chr8 | 20539747 | 20540120 | 374 | * | NA_peak_20 | 28 | Distal Interge | 8 | 20614919 | 20640703 | 25785 | 1 | 100861668 | ENSMUST0( | -74799 | ENSMUSG0( | Gm21119 | predicted gene, 21119 |
| chr8 | 20570743 | 20571247 | 505 | * | NA_peak_20 | 36 | Intron (ENSM | 8 | 20614919 | 20640703 | 25785 | 1 | 100861668 | ENSMUST0( | -43672 | ENSMUSG0( | Gm21119 | predicted gene, 21119 |
| chr8 | 20578083 | 20578453 | 371 | * | NA_peak_20 | 53 | Intron (ENSM | 8 | 20614919 | 20640703 | 25785 | 1 | 100861668 | ENSMUST0( | -36466 | ENSMUSG0( | Gm21119 | predicted gene, 21119 |
| chr8 | 20608366 | 20608931 | 566 | * | NA_peak_20 | 36 | Distal Interge | 8 | 20614919 | 20640703 | 25785 | 1 | 100861668 | ENSMUST0( | -5988 | ENSMUSG0( | Gm21119 | predicted gene, 21119 |
| chr8 | 20624143 | 20624457 | 315 | * | NA_peak_20 | 39 | Intron (ENSM | 8 | 20614919 | 20640703 | 25785 | 1 | 100861668 | ENSMUST0( | 9224 | ENSMUSG0( | Gm21119 | predicted gene, 21119 |
| chr8 | 20631865 | 20632228 | 364 | * | NA_peak_20 | 18 | Intron (ENSM | 8 | 20614919 | 20640703 | 25785 | 1 | 100861668 | ENSMUST0( | 16946 | ENSMUSG0( | Gm21119 | predicted gene, 21119 |
| chr8 | 20642611 | 20643092 | 482 | * | NA_peak_20 | 67 | Distal Interge | 8 | 20614919 | 20640703 | 25785 | 1 | 100861668 | ENSMUST0( | 27692 | ENSMUSG0( | Gm21119 | predicted gene, 21119 |
| chr8 | 20652356 | 20652696 | 341 | * | NA_peak_20 | 126 | Distal Interge | 8 | 20614919 | 20640703 | 25785 | 1 | 100861668 | ENSMUST0( | 37437 | ENSMUSG0( | Gm21119 | predicted gene, 21119 |
| chr8 | 20668421 | 20668813 | 393 | * | NA_peak_20 | 59 | Distal Interge | 8 | 20614919 | 20640703 | 25785 | 1 | 100861668 | ENSMUST0( | 53502 | ENSMUSG0( | Gm21119 | predicted gene, 21119 |
| chr8 | 20673962 | 20674221 | 260 | * | NA_peak_20 | 126 | Distal Interge | 8 | 20614919 | 20640703 | 25785 | 1 | 100861668 | ENSMUST0( | 59043 | ENSMUSG0( | Gm21119 | predicted gene, 21119 |
| chr8 | 20690462 | 20690869 | 408 | * | NA_peak_20 | 48 | Distal Interge | 8 | 20614919 | 20640703 | 25785 | 1 | 100861668 | ENSMUST0( | 75543 | ENSMUSG0( | Gm21119 | predicted gene, 21119 |
| chr8 | 20813639 | 20813802 | 164 | * | NA_peak_20 | 25 | Distal Interge | 8 | 20892669 | 20897723 | 5055 | 1 | 654453 | ENSMUST0( | -78867 | ENSMUSG0( | Defb33 | defensin beta 33 |
| chr8 | 20814095 | 20814095 | 1 | * | NA_peak_20 | 25 | Distal Interge | 8 | 20892669 | 20897723 | 5055 | 1 | 654453 | ENSMUST0( | -78574 | ENSMUSG0( | Defb33 | defensin beta 33 |
| chr8 | 20831605 | 20831931 | 327 | * | NA_peak_20 | 57 | Intron (ENSM | 8 | 20892669 | 20897723 | 5055 | 1 | 654453 | ENSMUST0( | -60738 | ENSMUSG0( | Defb33 | defensin beta 33 |
| chr8 | 21274478 | 21274849 | 372 | * | NA_peak_20 | 16 | Distal Interge | 8 | 21287409 | 21288377 | 969 | 1 | 13237 | ENSMUST0( | -12560 | ENSMUSG0( | Defa3 | defensin, alpha, 3 |
| chr8 | 22593999 | 22594166 | 168 | * | NA_peak_20 | 17 | Promoter (<= | 8 | 22577075 | 22593813 | 16739 | 2 | 22335 | ENSMUST0( | -186 | ENSMUSG0( | Vdac3 | voltage-dependent anion channel 3 |
| chr8 | 22819716 | 22819906 | 191 | * | NA_peak_20 | 49 | Distal Interge | 8 | 22792287 | 22805622 | 13336 | 2 | 64933 | ENSMUST0( | -14094 | ENSMUSG0( | Ap3m2 | adaptor-related protein complex 3, mu 2 subunit |
| chr8 | 23300583 | 23300694 | 112 | * | NA_peak_20 | 44 | Distal Interge | 8 | 23241346 | 23257074 | 15729 | 2 | 57437 | ENSMUST0( | -43509 | ENSMUSG0( | Golga7 | golgi autoantigen, golgin subfamily a, 7 |
| chr8 | 23641829 | 23642121 | 293 | * | NA_peak_20 | 23 | Intron (ENSM | 8 | 23636019 | 23902111 | 266093 | 1 | 320158 | ENSMUST0( | 5810 | ENSMUSG0( | Zmat4 | zinc finger, matrin type 4 |
| chr8 | 24895748 | 24895969 | 222 | * | NA_peak_20 | 38 | Intron (ENSM | 8 | 24836146 | 24884595 | 48450 | 2 | 353188 | ENSMUST0( | -11153 | ENSMUSG0( | Adam32 | a disintegrin and metallopeptidase domain 32 |
| chr8 | 25071760 | 25072297 | 538 | * | NA_peak_20 | 16 | Exon (ENSM | 8 | 25062433 | 25091346 | 28914 | 2 | 83436 | ENSMUST0( | 19049 | ENSMUSG0( | Plekha2 | pleckstrin homology domain-containing, family A (phosphoinositide binding specific) member 2 |
| chr8 | 25212495 | 25212788 | 294 | * | NA_peak_20 | 26 | Intron (ENSM | 8 | 25154552 | 25201542 | 46991 | 2 | 320165 | ENSMUST0( | -10953 | ENSMUSG0( | Tacc1 | transforming, acidic coiled-coil containing protein 1 |
| chr8 | 25519391 | 25519675 | 285 | * | NA_peak_20 | 22 | Promoter (<= | 8 | 25519344 | 25558227 | 38884 | 1 | 14182 | ENSMUST0( | 47 | ENSMUSG0( | Fgfr1 | fibroblast growth factor receptor 1 |
| chr8 | 25606703 | 25607139 | 437 | * | NA_peak_20 | 39 | Intron (ENSM | 8 | 25602319 | 25663599 | 61281 | 1 | 234135 | ENSMUST0( | 4384 | ENSMUSG0( | Nsd3 | nuclear receptor binding SET domain protein 3 |
| chr8 | 25712567 | 25712927 | 361 | * | NA_peak_20 | 28 | Intron (ENSM | 8 | 25720037 | 25724887 | 4851 | 1 | 71910 | ENSMUST0( | -7110 | ENSMUSG0( | Plpp5 | phospholipid phosphatase 5 |
| chr8 | 27066987 | 27067188 | 202 | * | NA_peak_20 | 30 | Distal Interge | 8 | 27050774 | 27052341 | 1568 | 1 | 114863 | ENSMUST0( | 16213 | ENSMUSG0( | Plpbp | pyridoxal phosphate binding protein |
| chr8 | 27304396 | 27304408 | 13 | * | NA_peak_20 | 30 | Distal Interge | 8 | 27296055 | 27296801 | 747 | 1 | 732482 | ENSMUST0( | 8341 | NA | Gm9731 | Fam58a pseudogene |
| chr8 | 28021018 | 28021464 | 447 | * | NA_peak_20 | 30 | Distal Interge | 8 | 28082694 | 28084266 | 1573 | 2 | 666422 | ENSMUST0( | 62802 | ENSMUSG0( | Gm8096 | 3-phosphoglycerate dehydrogenase pseudogene |
| chr8 | 28535761 | 28536080 | 320 | * | NA_peak_20 | 43 | Distal Interge | 8 | 28083330 | 28084920 | 1591 | 2 | 666422 | ENSMUST0( | -450841 | ENSMUSG0( | Gm8096 | 3-phosphoglycerate dehydrogenase pseudogene |
| chr8 | 28905657 | 28905876 | 220 | * | NA_peak_20 | 35 | Intron (ENSM | 8 | 28652425 | 29219427 | 567003 | 2 | 210801 | ENSMUST0( | 313551 | ENSMUSG0( | Unc5d | unc-5 netrin receptor D |
| chr8 | 29358885 | 29359141 | 257 | * | NA_peak_20 | 30 | Distal Interge | 8 | 28646717 | 29219636 | 572920 | 2 | 210801 | ENSMUST0( | -139249 | ENSMUSG0( | Unc5d | unc-5 netrin receptor D |
| chr8 | 31415775 | 31416223 | 449 | * | NA_peak_20 | 19 | Distal Interge | 8 | 31378813 | 31382439 | 3627 | 1 | 74517 | ENSMUST0( | 36962 | ENSMUSG0( | 7420700N18 | RIKEN cDNA 7420700N18 gene |
| chr8 | 31706974 | 31707283 | 310 | * | NA_peak_20 | 19 | Distal Interge | 8 | 31826288 | 31849375 | 23088 | 2 | 211323 | ENSMUST0( | 142092 | ENSMUSG0( | Nrg1 | neuregulin 1 |
| chr8 | 33311836 | 33312738 | 903 | * | NA_peak_20 | 16 | Intron (ENSM | 8 | 33235684 | 33330550 | 94867 | 2 | 22427 | ENSMUST0( | 17812 | ENSMUSG0( | Wrn | Werner syndrome RecQ like helicase |
| chr8 | 33370520 | 33370940 | 421 | * | NA_peak_20 | 17 | Intron (ENSM | 8 | 33235682 | 33385472 | 149791 | 2 | 22427 | ENSMUST0( | 14532 | ENSMUSG0( | Wrn | Werner syndrome RecQ like helicase |
| chr8 | 33385298 | 33385660 | 363 | * | NA_peak_20 | 22 | Promoter (<= | 8 | 33234384 | 33385527 | 151144 | 2 | 22427 | ENSMUST0( | 0 | ENSMUSG0( | Wrn | Werner syndrome RecQ like helicase |
| chr8 | 33458188 | 33458520 | 333 | * | NA_peak_20 | 30 | Distal Interge | 8 | 33484196 | 33485167 | 972 | 1 | 628431 | ENSMUST0( | -25676 | NA | Hmgb1-rs17 | high mobility group box 1, related sequence 17 |
| chr8 | 33662701 | 33662799 | 99 | * | NA_peak_20 | 30 | Intron (ENSM | 8 | 33669002 | 33685652 | 16651 | 1 | 14782 | ENSMUST0( | -6203 | ENSMUSG0( | Gsr | glutathione reductase |
| chr8 | 34895375 | 34895776 | 402 | * | NA_peak_20 | 13 | Intron (ENSM | 8 | 34831571 | 34890151 | 58581 | 2 | 21951 | ENSMUST0( | -5224 | ENSMUSG0( | Tnks | tankyrase, TRF1-interacting ankyrin-related ADP-ribose polymerase |
| chr8 | 35085393 | 35085544 | 152 | * | NA_peak_20 | 30 | Distal Interge | 8 | 34826460 | 34965690 | 139231 | 2 | 21951 | ENSMUST0( | -119703 | ENSMUSG0( | Tnks | tankyrase, TRF1-interacting ankyrin-related ADP-ribose polymerase |
| chr8 | 35605879 | 35606359 | 481 | * | NA_peak_20 | 23 | Promoter (<= | 8 | 35607277 | 35677020 | 69744 | 1 | 52065 | ENSMUST0( | -918 | ENSMUSG0( | Mfhas1 | malignant fibrous histiocytoma amplified sequence 1 |
| chr8 | 35724699 | 35725056 | 358 | * | NA_peak_20 | 22 | Intron (ENSM | 8 | 35765790 | 35818047 | 52258 | 1 | 100502846 | ENSMUST0( | -40734 | ENSMUSG0( | Gm19410 | predicted gene, 19410 |
| chr8 | 36511976 | 36512234 | 259 | * | NA_peak_20 | 23 | 3' UTR | 8 | 36505489 | 36512593 | 7105 | 1 | 319582 | ENSMUST0( | 6487 | ENSMUSG0( | Trmt9b | tRNA methyltransferase 9B |
| chr8 | 36739712 | 36740066 | 355 | * | NA_peak_20 | 19 | Intron (ENSM | 8 | 36567751 | 36733054 | 165304 | 2 | 50768 | ENSMUST0( | -6658 | ENSMUSG0( | Dlc1 | deleted in liver cancer 1 |
| chr8 | 36774140 | 36774442 | 303 | * | NA_peak_20 | 35 | Intron (ENSM | 8 | 36567751 | 36733054 | 165304 | 2 | 50768 | ENSMUST0( | -41086 | ENSMUSG0( | Dlc1 | deleted in liver cancer 1 |
| chr8 | 36862506 | 36862642 | 137 | * | NA_peak_20 | 23 | Intron (ENSM | 8 | 36849126 | 36952357 | 103232 | 2 | 50768 | ENSMUST0( | 89715 | ENSMUSG0( | Dlc1 | deleted in liver cancer 1 |
| chr8 | 36873755 | 36873987 | 233 | * | NA_peak_20 | 22 | Intron (ENSM | 8 | 36849126 | 36952357 | 103232 | 2 | 50768 | ENSMUST0( | 78370 | ENSMUSG0( | Dlc1 | deleted in liver cancer 1 |
| chr8 | 38676520 | 38676776 | 257 | * | NA_peak_20 | 24 | Distal Interge | 8 | 37522554 | 38661508 | 1138955 | 2 | 244431 | ENSMUST0( | -15012 | ENSMUSG0( | Sgcz | sarcoglycan zeta |
| chr8 | 40551502 | 40551780 | 279 | * | NA_peak_20 | 16 | 3' UTR | 8 | 40543424 | 40546543 | 3120 | 1 | 52348 | ENSMUST0( | 8078 | ENSMUSG0( | Vps37a | vacuolar protein sorting 37A |
| chr8 | 40634491 | 40634759 | 269 | * | NA_peak_20 | 24 | Promoter (<= | 8 | 40564031 | 40634759 | 70729 | 2 | 54384 | ENSMUST0( | 0 | ENSMUSG0( | Mtmr7 | myotubularin related protein 7 |
| chr8 | 40710466 | 40710695 | 230 | * | NA_peak_20 | 19 | Distal Interge | 8 | 40675077 | 40682199 | 7123 | 1 | 13526 | ENSMUST0( | 35389 | ENSMUSG0( | Adam24 | a disintegrin and metallopeptidase domain 24 (testase 1) |
| chr8 | 40819586 | 40819972 | 387 | * | NA_peak_20 | 18 | Distal Interge | 8 | 40822990 | 40827037 | 4048 | 1 | 546055 | ENSMUST0( | -3018 | ENSMUSG0( | Adam39 | a disintegrin and metallopeptidase domain 39 |
| chr8 | 40869033 | 40869356 | 324 | * | NA_peak_20 | 15 | Intron (ENSM | 8 | 40874948 | 40922070 | 47123 | 1 | 11988 | ENSMUST0( | -5592 | ENSMUSG0( | Slc7a2 | solute carrier family 7 (cationic amino acid transporter, y+ system), member 2 |
| chr8 | 41001596 | 41002132 | 537 | * | NA_peak_20 | 19 | Promoter (1- | 8 | 40998770 | 41000205 | 1436 | 2 | 102103 | ENSMUST0( | -1391 | ENSMUSG0( | Mtus1 | mitochondrial tumor suppressor 1 |
| chr8 | 41233842 | 41234219 | 378 | * | NA_peak_20 | 31 | Distal Interge | 8 | 41239752 | 41275094 | 35343 | 1 | 18536 | ENSMUST0( | -5533 | ENSMUSG0( | Pcm1 | pericentriolar material 1 |
| chr8 | 41239609 | 41239652 | 44 | * | NA_peak_20 | 35 | Promoter (<= | 8 | 41239752 | 41275094 | 35343 | 1 | 18536 | ENSMUST0( | -100 | ENSMUSG0( | Pcm1 | pericentriolar material 1 |
| chr8 | 43791661 | 43791848 | 188 | * | NA_peak_20 | 22 | Distal Interge | 8 | 43650309 | 43710049 | 59741 | 2 | 252866 | ENSMUST0( | -81612 | ENSMUSG0( | Adam34 | a disintegrin and metallopeptidase domain 34 |
| chr8 | 43858183 | 43858649 | 467 | * | NA_peak_20 | 30 | Distal Interge | 8 | 43650309 | 43710049 | 59741 | 2 | 252866 | ENSMUST0( | -148134 | ENSMUSG0( | Adam34 | a disintegrin and metallopeptidase domain 34 |
| chr8 | 45164001 | 45164060 | 60 | * | NA_peak_20 | 23 | Promoter (1- | 8 | 45160494 | 45165219 | 4726 | 2 | 622523 | ENSMUST0( | 1159 | ENSMUSG0( | Gm6329 | predicted gene 6329 |
| chr8 | 45180262 | 45180852 | 591 | * | NA_peak_20 | 37 | Intron (ENSM | 8 | 45160494 | 45165219 | 4726 | 2 | 622523 | ENSMUST0( | -15043 | ENSMUSG0( | Gm6329 | predicted gene 6329 |
| chr8 | 45685825 | 45686129 | 305 | * | NA_peak_20 | 22 | Promoter (2- | 8 | 45688333 | 45759141 | 70809 | 1 | 234214 | ENSMUST0( | -2204 | ENSMUSG0( | Sorbs2 | sorbin and SH3 domain containing 2 |
| chr8 | 45923701 | 45924026 | 326 | * | NA_peak_20 | 26 | Distal Interge | 8 | 45915032 | 45919238 | 4207 | 1 | 53318 | ENSMUST0( | 8669 | ENSMUSG0( | Pdlim3 | PDZ and LIM domain 3 |
| chr8 | 46226211 | 46226534 | 324 | * | NA_peak_20 | 14 | Distal Interge | 8 | 46206797 | 46211284 | 4488 | 2 | 11739 | ENSMUST0( | -14927 | ENSMUSG0( | Slc25a4 | solute carrier family 25 (mitochondrial carrier, adenine nucleotide translocator), member 4 |
| chr8 | 46292911 | 46293249 | 339 | * | NA_peak_20 | 25 | Promoter (<= | 8 | 46292444 | 46293335 | 892 | 2 | 234219 | ENSMUST0( | 86 | ENSMUSG0( | Helt | helt bHLH transcription factor |
| chr8 | 46487999 | 46488348 | 350 | * | NA_peak_20 | 33 | Promoter (2- | 8 | 46490966 | 46535581 | 44616 | 1 | 14081 | ENSMUST0( | -2618 | ENSMUSG0( | Acsl1 | acyl-CoA synthetase long-chain family member 1 |
| chr8 | 46960641 | 46961208 | 568 | * | NA_peak_20 | 31 | Distal Interge | 8 | 46986887 | 47096762 | 109876 | 1 | 320981 | ENSMUST0( | -25679 | ENSMUSG0( | Enpp6 | ectonucleotide pyrophosphatase/phosphodiesterase 6 |
| chr8 | 47146940 | 47147485 | 546 | * | NA_peak_20 | 27 | Intron (ENSM | 8 | 47082321 | 47093255 | 10935 | 1 | 320981 | ENSMUST0( | 64619 | ENSMUSG0( | Enpp6 | ectonucleotide pyrophosphatase/phosphodiesterase 6 |
| chr8 | 47475803 | 47476297 | 495 | * | NA_peak_20 | 25 | Distal Interge | 8 | 47504245 | 47511348 | 7104 | 2 | 320714 | ENSMUST0( | 35051 | ENSMUSG0( | Trappc11 | trafficking protein particle complex 11 |
| chr8 | 47509067 | 47509394 | 328 | * | NA_peak_20 | 24 | Promoter (1- | 8 | 47504245 | 47511348 | 7104 | 2 | 320714 | ENSMUST0( | 1954 | ENSMUSG0( | Trappc11 | trafficking protein particle complex 11 |
| chr8 | 47533736 | 47534153 | 418 | * | NA_peak_20 | 24 | Promoter (<= | 8 | 47533686 | 47544143 | 10458 | 1 | 192174 | ENSMUST0( | 50 | ENSMUSG0( | Rwdd4a | RWD domain containing 4A |
| chr8 | 47713886 | 47713959 | 74 | * | NA_peak_20 | 19 | Promoter (<= | 8 | 47709344 | 47713932 | 4589 | 2 | 70925 | ENSMUST0( | 0 | ENSMUSG0( | Cdkn2aip | CDKN2A interacting protein |
| chr8 | 47754334 | 47754870 | 537 | * | NA_peak_20 | 36 | Distal Interge | 8 | 47709344 | 47713932 | 4589 | 2 | 70925 | ENSMUST0( | -40402 | ENSMUSG0( | Cdkn2aip | CDKN2A interacting protein |
| chr8 | 49325044 | 49325489 | 446 | * | NA_peak_20 | 35 | Distal Interge | 8 | 48646261 | 48843951 | 197691 | 2 | 23965 | ENSMUST0( | -481093 | ENSMUSG0( | Tenm3 | teneurin transmembrane protein 3 |
| chr8 | 51734526 | 51735283 | 758 | * | NA_peak_20 | 35 | Distal Interge | 8 | 52356611 | 52403339 | 46729 | 1 | 330767 | ENSMUST0( | -621328 | ENSMUSG0( | 1190028D05 | RIKEN cDNA 1190028D05 gene |
| chr8 | 53117530 | 53118067 | 538 | * | NA_peak_20 | 39 | Distal Interge | 8 | 53511727 | 53523334 | 11608 | 1 | 11593 | ENSMUST0( | -393660 | ENSMUSG0( | Aga | aspartylglucosaminidase |
| chr8 | 54709380 | 54709900 | 521 | * | NA_peak_20 | 50 | Promoter (2- | 8 | 54696269 | 54706981 | 10713 | 2 | 244484 | ENSMUST0( | -2399 | ENSMUSG0( | Wdr17 | WD repeat domain 17 |
| chr8 | 54868488 | 54868555 | 68 | * | NA_peak_20 | 40 | Intron (ENSM | 8 | 54693231 | 54887150 | 193920 | 2 | 244484 | ENSMUST0( | 18595 | ENSMUSG0( | Wdr17 | WD repeat domain 17 |
| chr8 | 55118972 | 55119330 | 359 | * | NA_peak_20 | 26 | Distal Interge | 8 | 54954843 | 55060871 | 106029 | 1 | 234267 | ENSMUST0( | 164129 | ENSMUSG0( | Gpm6a | glycoprotein m6a |
| chr8 | 55123087 | 55123442 | 356 | * | NA_peak_20 | 22 | Distal Interge | 8 | 54954843 | 55060871 | 106029 | 1 | 234267 | ENSMUST0( | 168244 | ENSMUSG0( | Gpm6a | glycoprotein m6a |
| chr8 | 55736215 | 55736512 | 298 | * | NA_peak_20 | 30 | Distal Interge | 8 | 55870912 | 55906948 | 36037 | 2 | 244486 | ENSMUST0( | 170436 | ENSMUSG0( | Adam29 | a disintegrin and metallopeptidase domain 29 |
| chr8 | 57099312 | 57099651 | 340 | * | NA_peak_21 | 17 | Distal Interge | 8 | 57304271 | 57305358 | 1088 | 2 | 102636514 | ENSMUST0( | 205707 | NA | Hand2os1 | Hand2, opposite strand 1 |
| chr8 | 57216565 | 57216860 | 296 | * | NA_peak_21 | 26 | Distal Interge | 8 | 57304271 | 57305358 | 1088 | 2 | 102636514 | ENSMUST0( | 88498 | NA | Hand2os1 | Hand2, opposite strand 1 |
| chr8 | 57498712 | 57498771 | 60 | * | NA_peak_21 | 19 | Intron (ENSM | 8 | 57505815 | 57506731 | 917 | 1 | 78353 | ENSMUST0( | -7044 | ENSMUSG0( | 2500002B13 | RIKEN cDNA 2500002B13 gene |
| chr8 | 57598884 | 57599281 | 398 | * | NA_peak_21 | 26 | Intron (ENSM | 8 | 57651757 | 57662548 | 10792 | 1 | 100502619 | ENSMUST0( | -52476 | NA | AW046200 | expressed sequence AW046200 |
| chr8 | 59677797 | 59678078 | 282 | * | NA_peak_21 | 22 | Distal Interge | 8 | 59825649 | 59844723 | 19075 | 1 | 75017 | ENSMUST0( | -147571 | ENSMUSG0( | 4930470O06 | RIKEN cDNA 4930470O06 gene |
| chr8 | 60400062 | 60400277 | 216 | * | NA_peak_21 | 22 | Distal Interge | 8 | 60505932 | 60545677 | 39746 | 1 | 23923 | ENSMUST0( | -105655 | ENSMUSG0( | Aadat | aminoadipate aminotransferase |
| chr8 | 60738340 | 60738653 | 314 | * | NA_peak_21 | 24 | Distal Interge | 8 | 60655540 | 60672362 | 16823 | 1 | 71306 | ENSMUST0( | 82800 | ENSMUSG0( | Mfap3l | microfibrillar-associated protein 3-like |
| chr8 | 61083935 | 61083953 | 19 | * | NA_peak_21 | 44 | Intron (ENSM | 8 | 61049835 | 61089455 | 39621 | 1 | 18004 | ENSMUST0( | 34100 | ENSMUSG0( | Nek1 | NIMA (never in mitosis gene a)-related expressed kinase 1 |
| chr8 | 61145082 | 61145204 | 123 | * | NA_peak_21 | 13 | Intron (ENSM | 8 | 61223872 | 61396071 | 172200 | 1 | 59009 | ENSMUST0( | -78668 | ENSMUSG0( | Sh3rf1 | SH3 domain containing ring finger 1 |
| chr8 | 61224659 | 61224802 | 144 | * | NA_peak_21 | 14 | Promoter (<= | 8 | 61224244 | 61386641 | 162398 | 1 | 59009 | ENSMUST0( | 415 | ENSMUSG0( | Sh3rf1 | SH3 domain containing ring finger 1 |
| chr8 | 62102348 | 62102681 | 334 | * | NA_peak_21 | 34 | Intron (ENSM | 8 | 62057042 | 62123116 | 66075 | 2 | 26359 | ENSMUST0( | 20435 | ENSMUSG0( | Anxa10 | annexin A10 |
| chr8 | 63674495 | 63674840 | 346 | * | NA_peak_21 | 48 | Distal Interge | 8 | 63924694 | 63952170 | 27477 | 2 | 244495 | ENSMUST0( | 277330 | ENSMUSG0( | Sgo2b | shugoshin 2B |
| chr8 | 63817310 | 63817771 | 462 | * | NA_peak_21 | 19 | Distal Interge | 8 | 63924694 | 63952170 | 27477 | 2 | 244495 | ENSMUST0( | 134399 | ENSMUSG0( | Sgo2b | shugoshin 2B |
| chr8 | 63909845 | 63910149 | 305 | * | NA_peak_21 | 44 | Distal Interge | 8 | 63924694 | 63952170 | 27477 | 2 | 244495 | ENSMUST0( | 42021 | ENSMUSG0( | Sgo2b | shugoshin 2B |
| chr8 | 64343732 | 64343901 | 170 | * | NA_peak_21 | 16 | Distal Interge | 8 | 64084326 | 64206008 | 121683 | 2 | 21892 | ENSMUST0( | -137724 | ENSMUSG0( | Tll1 | tolloid-like |
| chr8 | 65185711 | 65186016 | 306 | * | NA_peak_21 | 26 | Distal Interge | 8 | 65085613 | 65129563 | 43951 | 2 | 407795 | ENSMUST0( | -56148 | ENSMUSG0( | Smim31 | small integral membrane protein 31 |
| chr8 | 66001604 | 66001920 | 317 | * | NA_peak_21 | 39 | Intron (ENSM | 8 | 65967157 | 66471625 | 504469 | 1 | 72925 | ENSMUST0( | 34447 | ENSMUSG0( | Marchf1 | membrane associated ring-CH-type finger 1 |
| chr8 | 66657038 | 66657307 | 270 | * | NA_peak_21 | 20 | Distal Interge | 8 | 66680569 | 66688047 | 7479 | 2 | 18168 | ENSMUST0( | 30740 | ENSMUSG0( | Npy5r | neuropeptide Y receptor Y5 |
| chr8 | 67186834 | 67187129 | 296 | * | NA_peak_21 | 22 | Distal Interge | 8 | 67490758 | 67492104 | 1347 | 1 | 17960 | ENSMUST0( | -303629 | ENSMUSG0( | Nat1 | N-acetyl transferase 1 |
| chr8 | 67429945 | 67430110 | 166 | * | NA_peak_21 | 22 | Distal Interge | 8 | 67490758 | 67492104 | 1347 | 1 | 17960 | ENSMUST0( | -60648 | ENSMUSG0( | Nat1 | N-acetyl transferase 1 |
| chr8 | 68328047 | 68328096 | 50 | * | NA_peak_21 | 23 | Intron (ENSM | 8 | 68368434 | 68372293 | 3860 | 1 | 73609 | ENSMUST0( | -40338 | ENSMUSG0( | 1700125H03 | RIKEN cDNA 1700125H03 gene |
| chr8 | 68407360 | 68407686 | 327 | * | NA_peak_21 | 27 | Intron (ENSM | 8 | 68368466 | 68375710 | 7245 | 1 | 73609 | ENSMUST0( | 38894 | ENSMUSG0( | 1700125H03 | RIKEN cDNA 1700125H03 gene |
| chr8 | 69385587 | 69385987 | 401 | * | NA_peak_21 | 23 | Intron (ENSM | 8 | 69366510 | 69395522 | 29013 | 2 | 378466 | ENSMUST0( | 9535 | ENSMUSG0( | Gm10033 | predicted gene 10033 |
| chr8 | 69625176 | 69625538 | 363 | * | NA_peak_21 | 21 | Promoter (<= | 8 | 69618869 | 69625545 | 6677 | 2 | 234362 | ENSMUST0( | 7 | ENSMUSG0( | Zfp868 | zinc finger protein 868 |
| chr8 | 69664703 | 69664997 | 295 | * | NA_peak_21 | 46 | 3' UTR | 8 | 69659263 | 69664170 | 4908 | 1 | 636741 | ENSMUST0( | 5440 | ENSMUSG0( | Zfp964 | zinc finger protein 964 |
| chr8 | 69791176 | 69791445 | 270 | * | NA_peak_21 | 30 | Promoter (<= | 8 | 69791250 | 69807745 | 16496 | 1 | 170759 | ENSMUST0( | 0 | ENSMUSG0( | Atp13a1 | ATPase type 13A1 |
| chr8 | 70283394 | 70283582 | 189 | * | NA_peak_21 | 19 | Promoter (<= | 8 | 70283034 | 70286302 | 3269 | 1 | 26558 | ENSMUST0( | 360 | ENSMUSG0( | Homer3 | homer scaffolding protein 3 |
| chr8 | 70332191 | 70332323 | 133 | * | NA_peak_21 | 19 | Promoter (2- | 8 | 70329817 | 70331587 | 1771 | 1 | 14559 | ENSMUST0( | 2374 | ENSMUSG0( | Gdf1 | growth differentiation factor 1 |
| chr8 | 70353250 | 70353642 | 393 | * | NA_peak_21 | 25 | Promoter (<= | 8 | 70331525 | 70353278 | 21754 | 2 | 19704 | ENSMUST0( | 0 | ENSMUSG0( | Upf1 | UPF1 regulator of nonsense transcripts homolog (yeast) |
| chr8 | 70375315 | 70375520 | 206 | * | NA_peak_21 | 33 | Promoter (<= | 8 | 70375038 | 70375753 | 716 | 1 | 12845 | ENSMUST0( | 277 | ENSMUSG0( | Comp | cartilage oligomeric matrix protein |
| chr8 | 70527456 | 70527948 | 493 | * | NA_peak_21 | 34 | Promoter (<= | 8 | 70527724 | 70535328 | 7605 | 1 | 14232 | ENSMUST0( | 0 | ENSMUSG0( | Fkbp8 | FK506 binding protein 8 |
| chr8 | 70730922 | 70731245 | 324 | * | NA_peak_21 | 13 | Promoter (<= | 8 | 70730112 | 70745893 | 15782 | 1 | 110385 | ENSMUST0( | 810 | ENSMUSG0( | Pde4c | phosphodiesterase 4C, cAMP specific |
| chr8 | 70758010 | 70758115 | 106 | * | NA_peak_21 | 15 | Promoter (1- | 8 | 70756457 | 70757700 | 1244 | 1 | 19339 | ENSMUST0( | 1553 | ENSMUSG0( | Rab3a | RAB3A, member RAS oncogene family |
| chr8 | 70794343 | 70794674 | 332 | * | NA_peak_21 | 20 | Promoter (1- | 8 | 70779204 | 70792437 | 13234 | 2 | 546071 | ENSMUST0( | -1906 | ENSMUSG0( | Mast3 | microtubule associated serine/threonine kinase 3 |
| chr8 | 70873301 | 70873843 | 543 | * | NA_peak_21 | 16 | Promoter (<= | 8 | 70869988 | 70873383 | 3396 | 2 | 234388 | ENSMUST0( | 0 | ENSMUSG0( | Ccdc124 | coiled-coil domain containing 124 |
| chr8 | 71289996 | 71290164 | 169 | * | NA_peak_21 | 22 | Intron (ENSM | 8 | 71251125 | 71272934 | 21810 | 2 | 76478 | ENSMUST0( | -17062 | ENSMUSG0( | Haus8 | 4HAUS augmin-like complex, subunit 8 |
| chr8 | 71340577 | 71341258 | 682 | * | NA_peak_21 | 33 | Exon (ENSM | 8 | 71333817 | 71339008 | 5192 | 1 | 17925 | ENSMUST0( | 6760 | ENSMUSG0( | Myo9b | myosin IXb |
| chr8 | 71476175 | 71476394 | 220 | * | NA_peak_21 | 16 | 3' UTR | 8 | 71470411 | 71476098 | 5688 | 1 | 66498 | ENSMUST0( | 5764 | ENSMUSG0( | Dda1 | DET1 and DDB1 associated 1 |
| chr8 | 71679063 | 71679406 | 344 | * | NA_peak_21 | 29 | Promoter (<= | 8 | 71678335 | 71690224 | 11890 | 1 | 16453 | ENSMUST0( | 728 | ENSMUSG0( | Jak3 | Janus kinase 3 |
| chr8 | 71802916 | 71803562 | 647 | * | NA_peak_21 | 30 | Distal Interge | 8 | 71708394 | 71725716 | 17323 | 2 | 74015 | ENSMUST0( | -77200 | ENSMUSG0( | Fcho1 | FCH domain only 1 |
| chr8 | 71808858 | 71809276 | 419 | * | NA_peak_21 | 26 | Distal Interge | 8 | 71882019 | 71895727 | 13709 | 1 | 236193 | ENSMUST0( | -72743 | ENSMUSG0( | Zfp709 | zinc finger protein 709 |
| chr8 | 71835259 | 71835568 | 310 | * | NA_peak_21 | 36 | Intron (ENSM | 8 | 71882019 | 71895727 | 13709 | 1 | 236193 | ENSMUST0( | -46451 | ENSMUSG0( | Zfp709 | zinc finger protein 709 |
| chr8 | 73012808 | 73012886 | 79 | * | NA_peak_21 | 36 | Intron (ENSM | 8 | 73016512 | 73016591 | 80 | 1 | 100628574 | ENSMUST0( | -3626 | ENSMUSG0( | Mir28b | microRNA 28b |
| chr8 | 73276405 | 73276605 | 201 | * | NA_peak_21 | 15 | Intron (ENSM | 8 | 72814599 | 73352556 | 537958 | 2 | 16795 | ENSMUST0( | 75951 | ENSMUSG0( | Large1 | LARGE xylosyl- and glucuronyltransferase 1 |
| chr8 | 73560072 | 73560456 | 385 | * | NA_peak_21 | 32 | Distal Interge | 8 | 72814599 | 73353540 | 538942 | 2 | 16795 | ENSMUST0( | -206532 | ENSMUSG0( | Large1 | LARGE xylosyl- and glucuronyltransferase 1 |
| chr8 | 74729038 | 74729383 | 346 | * | NA_peak_21 | 30 | Intron (ENSM | 8 | 74873173 | 74893506 | 20334 | 1 | 71597 | ENSMUST0( | -143790 | ENSMUSG0( | Isx | intestine specific homeobox |
| chr8 | 74749692 | 74749963 | 272 | * | NA_peak_21 | 26 | Intron (ENSM | 8 | 74873173 | 74893506 | 20334 | 1 | 71597 | ENSMUST0( | -123210 | ENSMUSG0( | Isx | intestine specific homeobox |
| chr8 | 74798932 | 74799198 | 267 | * | NA_peak_21 | 16 | Intron (ENSM | 8 | 74873173 | 74893506 | 20334 | 1 | 71597 | ENSMUST0( | -73975 | ENSMUSG0( | Isx | intestine specific homeobox |
| chr8 | 74980829 | 74980836 | 8 | * | NA_peak_21 | 22 | Distal Interge | 8 | 74993356 | 75031972 | 38617 | 1 | 70823 | ENSMUST0( | -12520 | ENSMUSG0( | Hmgxb4 | HMG box domain containing 4 |
| chr8 | 75020131 | 75020275 | 145 | * | NA_peak_21 | 13 | Promoter (2- | 8 | 75022855 | 75024076 | 1222 | 1 | 70823 | ENSMUST0( | -2580 | ENSMUSG0( | Hmgxb4 | HMG box domain containing 4 |
| chr8 | 75137694 | 75138034 | 341 | * | NA_peak_21 | 18 | Distal Interge | 8 | 75123067 | 75125228 | 2162 | 1 | 17218 | ENSMUST0( | 14627 | ENSMUSG0( | Mcm5 | minichromosome maintenance complex component 5 |
| chr8 | 77416559 | 77416860 | 302 | * | NA_peak_21 | 18 | Intron (ENSM | 8 | 77517056 | 77518579 | 1524 | 1 | 70345 | ENSMUST0( | -100196 | ENSMUSG0( | 0610038B21 | RIKEN cDNA 0610038B21 gene |
| chr8 | 78267942 | 78268229 | 288 | * | NA_peak_21 | 23 | Intron (ENSM | 8 | 78251491 | 78276829 | 25339 | 1 | 73301 | ENSMUST0( | 16451 | ENSMUSG0( | Ttc29 | tetratricopeptide repeat domain 29 |
| chr8 | 79055532 | 79055853 | 322 | * | NA_peak_21 | 17 | Intron (ENSM | 8 | 79028587 | 79190856 | 162270 | 1 | 622675 | ENSMUST0( | 26945 | ENSMUSG0( | Zfp827 | zinc finger protein 827 |
| chr8 | 79711545 | 79711700 | 156 | * | NA_peak_21 | 33 | Promoter (<= | 8 | 79686206 | 79711670 | 25465 | 2 | 24015 | ENSMUST0( | 0 | ENSMUSG0( | Abce1 | ATP-binding cassette, sub-family E (OABP), member 1 |
| chr8 | 79715970 | 79716109 | 140 | * | NA_peak_21 | 44 | Intron (ENSM | 8 | 79711851 | 79779059 | 67209 | 1 | 68999 | ENSMUST0( | 4119 | ENSMUSG0( | Anapc10 | anaphase promoting complex subunit 10 |
| chr8 | 80627623 | 80628051 | 429 | * | NA_peak_21 | 22 | Intron (ENSM | 8 | 80611080 | 80695356 | 84277 | 1 | 333315 | ENSMUST0( | 16543 | ENSMUSG0( | Frem3 | Fras1 related extracellular matrix protein 3 |
| chr8 | 80644549 | 80644887 | 339 | * | NA_peak_21 | 43 | Intron (ENSM | 8 | 80611080 | 80695356 | 84277 | 1 | 333315 | ENSMUST0( | 33469 | ENSMUSG0( | Frem3 | Fras1 related extracellular matrix protein 3 |
| chr8 | 80750657 | 80750864 | 208 | * | NA_peak_21 | 23 | Distal Interge | 8 | 80698507 | 80739497 | 40991 | 2 | 93762 | ENSMUST0( | -11160 | ENSMUSG0( | Smarca5 | SWI/SNF related, matrix associated, actin dependent regulator of chromatin, subfamily a, member 5 |
| chr8 | 80929641 | 80930007 | 367 | * | NA_peak_21 | 26 | Distal Interge | 8 | 80764438 | 80880519 | 116082 | 2 | 14388 | ENSMUST0( | -49122 | ENSMUSG0( | Gab1 | growth factor receptor bound protein 2-associated protein 1 |
| chr8 | 81681318 | 81681431 | 114 | * | NA_peak_21 | 20 | Intron (ENSM | 8 | 81715200 | 82122559 | 407360 | 1 | 234515 | ENSMUST0( | -33769 | ENSMUSG0( | Inpp4b | inositol polyphosphate-4-phosphatase, type II |
| chr8 | 81915233 | 81915638 | 406 | * | NA_peak_21 | 15 | Intron (ENSM | 8 | 81890319 | 81902942 | 12624 | 1 | 234515 | ENSMUST0( | 24914 | ENSMUSG0( | Inpp4b | inositol polyphosphate-4-phosphatase, type II |
| chr8 | 83363392 | 83363687 | 296 | * | NA_peak_21 | 19 | Intron (ENSM | 8 | 83358026 | 83382320 | 24295 | 1 | 67555 | ENSMUST0( | 5366 | ENSMUSG0( | Mgat4d | MGAT4 family, member C |
| chr8 | 83413776 | 83413824 | 49 | * | NA_peak_21 | 17 | Intron (ENSM | 8 | 83389878 | 83426831 | 36954 | 1 | 12745 | ENSMUST0( | 23898 | ENSMUSG0( | Clgn | calmegin |
| chr8 | 83548187 | 83548423 | 237 | * | NA_peak_21 | 32 | Exon (ENSM | 8 | 83541093 | 83542188 | 1096 | 1 | 258267 | ENSMUST0( | 7094 | ENSMUSG0( | Olfr370 | olfactory receptor 370 |
| chr8 | 83980869 | 83980888 | 20 | * | NA_peak_21 | 15 | Intron (ENSM | 8 | 83976883 | 83995594 | 18712 | 1 | 18747 | ENSMUST0( | 3986 | ENSMUSG0( | Prkaca | protein kinase, cAMP dependent, catalytic, alpha |
| chr8 | 83986768 | 83987193 | 426 | * | NA_peak_21 | 25 | Exon (ENSM | 8 | 83976883 | 83995594 | 18712 | 1 | 18747 | ENSMUST0( | 9885 | ENSMUSG0( | Prkaca | protein kinase, cAMP dependent, catalytic, alpha |
| chr8 | 84067122 | 84067344 | 223 | * | NA_peak_21 | 14 | Promoter (<= | 8 | 84066836 | 84096992 | 30157 | 1 | 19724 | ENSMUST0( | 286 | ENSMUSG0( | Rfx1 | regulatory factor X, 1 (influences HLA class II expression) |
| chr8 | 84168901 | 84169354 | 454 | * | NA_peak_21 | 21 | Intron (ENSM | 8 | 84173733 | 84176552 | 2820 | 2 | 244551 | ENSMUST0( | 7198 | ENSMUSG0( | Nanos3 | nanos C2HC-type zinc finger 3 |
| chr8 | 84197573 | 84197673 | 101 | * | NA_peak_21 | 28 | Exon (ENSM | 8 | 84208518 | 84208592 | 75 | 1 | 387216 | ENSMUST0( | -10845 | ENSMUSG0( | Mir23a | microRNA 23a |
| chr8 | 84351035 | 84351172 | 138 | * | NA_peak_21 | 17 | Intron (ENSM | 8 | 84338639 | 84640246 | 301608 | 1 | 12286 | ENSMUST0( | 12396 | ENSMUSG0( | Cacna1a | calcium channel, voltage-dependent, P/Q type, alpha 1A subunit |
| chr8 | 84393723 | 84394092 | 370 | * | NA_peak_21 | 22 | Intron (ENSM | 8 | 84388440 | 84519039 | 130600 | 1 | 12286 | ENSMUST0( | 5283 | ENSMUSG0( | Cacna1a | calcium channel, voltage-dependent, P/Q type, alpha 1A subunit |
| chr8 | 84655517 | 84656096 | 580 | * | NA_peak_21 | 30 | Distal Interge | 8 | 84661331 | 84662854 | 1524 | 2 | 15936 | ENSMUST0( | 6758 | ENSMUSG0( | Ier2 | immediate early response 2 |
| chr8 | 84856803 | 84857066 | 264 | * | NA_peak_21 | 22 | Promoter (<= | 8 | 84856989 | 84869257 | 12269 | 1 | 66590 | ENSMUST0( | 0 | ENSMUSG0( | Farsa | phenylalanyl-tRNA synthetase, alpha subunit |
| chr8 | 84903233 | 84903830 | 598 | * | NA_peak_21 | 39 | Promoter (1- | 8 | 84901928 | 84905291 | 3364 | 1 | 16596 | ENSMUST0( | 1305 | ENSMUSG0( | Klf1 | Kruppel-like factor 1 (erythroid) |
| chr8 | 84944227 | 84944574 | 348 | * | NA_peak_21 | 16 | Promoter (2- | 8 | 84946991 | 84956603 | 9613 | 1 | 234542 | ENSMUST0( | -2417 | ENSMUSG0( | Rtbdn | retbindin |
| chr8 | 84945363 | 84945750 | 388 | * | NA_peak_21 | 25 | Promoter (1- | 8 | 84946991 | 84956603 | 9613 | 1 | 234542 | ENSMUST0( | -1241 | ENSMUSG0( | Rtbdn | retbindin |
| chr8 | 84957871 | 84958277 | 407 | * | NA_peak_21 | 28 | Promoter (<= | 8 | 84957678 | 84958116 | 439 | 2 | 69724 | ENSMUST0( | 0 | ENSMUSG0( | Rnaseh2a | ribonuclease H2, large subunit |
| chr8 | 84976004 | 84976316 | 313 | * | NA_peak_21 | 30 | Promoter (2- | 8 | 84976910 | 84978718 | 1809 | 2 | 16477 | ENSMUST0( | 2402 | ENSMUSG0( | Junb | jun B proto-oncogene |
| chr8 | 85191700 | 85192003 | 304 | * | NA_peak_21 | 23 | Distal Interge | 8 | 85230497 | 85231435 | 939 | 1 | 258858 | ENSMUST0( | -38494 | ENSMUSG0( | Olfr371 | olfactory receptor 371 |
| chr8 | 85258057 | 85258099 | 43 | * | NA_peak_21 | 44 | Distal Interge | 8 | 85230497 | 85231435 | 939 | 1 | 258858 | ENSMUST0( | 27560 | ENSMUSG0( | Olfr371 | olfactory receptor 371 |
| chr8 | 85976020 | 85976034 | 15 | * | NA_peak_21 | 22 | Intron (ENSM | 8 | 86017432 | 86030391 | 12960 | 1 | 102093 | ENSMUST0( | -41398 | ENSMUSG0( | Phkb | phosphorylase kinase beta |
| chr8 | 86159817 | 86160131 | 315 | * | NA_peak_21 | 22 | Distal Interge | 8 | 86035646 | 86060641 | 24996 | 1 | 102093 | ENSMUST0( | 124171 | ENSMUSG0( | Phkb | phosphorylase kinase beta |
| chr8 | 87962452 | 87962454 | 3 | * | NA_peak_21 | 18 | Promoter (2- | 8 | 87661810 | 87959595 | 297786 | 2 | 94187 | ENSMUST0( | -2857 | ENSMUSG0( | Zfp423 | zinc finger protein 423 |
| chr8 | 88138241 | 88138633 | 393 | * | NA_peak_21 | 30 | Promoter (<= | 8 | 88138206 | 88138274 | 69 | 1 | 102465649 | ENSMUST0( | 35 | ENSMUSG0( | Mir7071 | microRNA 7071 |
| chr8 | 88268174 | 88268202 | 29 | * | NA_peak_21 | 23 | Distal Interge | 8 | 88272403 | 88329039 | 56637 | 1 | 11513 | ENSMUST0( | -4201 | ENSMUSG0( | Adcy7 | adenylate cyclase 7 |
| chr8 | 89041853 | 89042199 | 347 | * | NA_peak_21 | 13 | Promoter (1- | 8 | 89027235 | 89044162 | 16928 | 2 | 58198 | ENSMUST0( | 1963 | ENSMUSG0( | Sall1 | spalt like transcription factor 1 |
| chr8 | 90858715 | 90858785 | 71 | * | NA_peak_21 | 33 | Intron (ENSM | 8 | 90849667 | 90872090 | 22424 | 1 | 109151 | ENSMUST0( | 9048 | ENSMUSG0( | Chd9 | chromodomain helicase DNA binding protein 9 |
| chr8 | 90859074 | 90859157 | 84 | * | NA_peak_21 | 33 | Intron (ENSM | 8 | 90849667 | 90872090 | 22424 | 1 | 109151 | ENSMUST0( | 9407 | ENSMUSG0( | Chd9 | chromodomain helicase DNA binding protein 9 |
| chr8 | 90960311 | 90960746 | 436 | * | NA_peak_21 | 26 | Intron (ENSM | 8 | 90955435 | 90990632 | 35198 | 1 | 109151 | ENSMUST0( | 4876 | ENSMUSG0( | Chd9 | chromodomain helicase DNA binding protein 9 |
| chr8 | 91087153 | 91087455 | 303 | * | NA_peak_21 | 20 | Exon (ENSM | 8 | 91070158 | 91123842 | 53685 | 1 | 19651 | ENSMUST0( | 16995 | ENSMUSG0( | Rbl2 | RB transcriptional corepressor like 2 |
| chr8 | 92341127 | 92341197 | 71 | * | NA_peak_21 | 18 | Intron (ENSM | 8 | 92326033 | 92346870 | 20838 | 2 | 71296 | ENSMUST0( | 5673 | ENSMUSG0( | Crnde | colorectal neoplasia differentially expressed (non-protein coding) |
| chr8 | 92366529 | 92366579 | 51 | * | NA_peak_21 | 18 | Intron (ENSM | 8 | 92357749 | 92361456 | 3708 | 1 | 54352 | ENSMUST0( | 8780 | ENSMUSG0( | Irx5 | Iroquois homeobox 5 |
| chr8 | 93505825 | 93506147 | 323 | * | NA_peak_21 | 35 | Promoter (<= | 8 | 93499213 | 93506768 | 7556 | 2 | 67935 | ENSMUST0( | 621 | ENSMUSG0( | Ces5a | carboxylesterase 5A |
| chr8 | 94038235 | 94038326 | 92 | * | NA_peak_21 | 16 | Promoter (<= | 8 | 94037296 | 94056111 | 18816 | 1 | 270086 | ENSMUST0( | 939 | ENSMUSG0( | Ogfod1 | 2-oxoglutarate and iron-dependent oxygenase domain containing 1 |
| chr8 | 94097931 | 94098275 | 345 | * | NA_peak_21 | 20 | Promoter (<= | 8 | 94067954 | 94098928 | 30975 | 2 | 67378 | ENSMUST0( | 653 | ENSMUSG0( | Bbs2 | Bardet-Biedl syndrome 2 (human) |
| chr8 | 94173172 | 94173334 | 163 | * | NA_peak_21 | 22 | Promoter (<= | 8 | 94172860 | 94173564 | 705 | 1 | 17750 | ENSMUST0( | 312 | ENSMUSG0( | Mt2 | metallothionein 2 |
| chr8 | 94463152 | 94463905 | 754 | * | NA_peak_21 | 71 | Intron (ENSM | 8 | 94470010 | 94506941 | 36932 | 1 | 434341 | ENSMUST0( | -6105 | ENSMUSG0( | Nlrc5 | NLR family, CARD domain containing 5 |
| chr8 | 94603648 | 94603797 | 150 | * | NA_peak_21 | 16 | Promoter (1- | 8 | 94602066 | 94660275 | 58210 | 1 | 67610 | ENSMUST0( | 1582 | ENSMUSG0( | Rspry1 | ring finger and SPRY domain containing 1 |
| chr8 | 94604992 | 94605182 | 191 | * | NA_peak_21 | 16 | Promoter (2- | 8 | 94602066 | 94660275 | 58210 | 1 | 67610 | ENSMUST0( | 2926 | ENSMUSG0( | Rspry1 | ring finger and SPRY domain containing 1 |
| chr8 | 94608536 | 94608860 | 325 | * | NA_peak_21 | 39 | Intron (ENSM | 8 | 94602066 | 94660275 | 58210 | 1 | 67610 | ENSMUST0( | 6470 | ENSMUSG0( | Rspry1 | ring finger and SPRY domain containing 1 |
| chr8 | 94651319 | 94651379 | 61 | * | NA_peak_21 | 13 | Intron (ENSM | 8 | 94666600 | 94674425 | 7826 | 1 | 107566 | ENSMUST0( | -15221 | ENSMUSG0( | Arl2bp | ADP-ribosylation factor-like 2 binding protein |
| chr8 | 94653301 | 94653653 | 353 | * | NA_peak_21 | 22 | Exon (ENSM | 8 | 94666600 | 94674425 | 7826 | 1 | 107566 | ENSMUST0( | -12947 | ENSMUSG0( | Arl2bp | ADP-ribosylation factor-like 2 binding protein |
| chr8 | 94696004 | 94696398 | 395 | * | NA_peak_21 | 34 | Promoter (<= | 8 | 94674894 | 94696278 | 21385 | 2 | 67801 | ENSMUST0( | 0 | ENSMUSG0( | Pllp | plasma membrane proteolipid |
| chr8 | 94917450 | 94918047 | 598 | * | NA_peak_21 | 29 | Promoter (<= | 8 | 94913565 | 94917521 | 3957 | 2 | 234582 | ENSMUST0( | 0 | ENSMUSG0( | Ccdc102a | coiled-coil domain containing 102A |
| chr8 | 95332238 | 95332342 | 105 | * | NA_peak_21 | 28 | Promoter (<= | 8 | 95332284 | 95347507 | 15224 | 1 | 101985 | ENSMUST0( | 0 | ENSMUSG0( | Usb1 | U6 snRNA biogenesis 1 |
| chr8 | 96419256 | 96419585 | 330 | * | NA_peak_21 | 23 | Distal Interge | 8 | 95864134 | 95888547 | 24414 | 2 | 14719 | ENSMUST0( | -530709 | ENSMUSG0( | Got2 | glutamatic-oxaloacetic transaminase 2, mitochondrial |
| chr8 | 97115012 | 97115264 | 253 | * | NA_peak_21 | 63 | Distal Interge | 8 | 95864134 | 95888547 | 24414 | 2 | 14719 | ENSMUST0( | -1226465 | ENSMUSG0( | Got2 | glutamatic-oxaloacetic transaminase 2, mitochondrial |
| chr8 | 99663140 | 99663180 | 41 | * | NA_peak_21 | 16 | Intron (ENSM | 8 | 99416674 | 99733438 | 316765 | 1 | 234624 | ENSMUST0( | 246466 | ENSMUSG0( | A330008L17 | RIKEN cDNA A330008L17 gene |

| | | | | | | | | | | | | | | | | | | |
|---|---|---|---|---|---|---|---|---|---|---|---|---|---|---|---|---|---|---|
| chr8 | 100279897 | 100280198 | 302 | * | NA_peak_21 | 38 | Distal Interge | 8 | 99416674 | 99733438 | 316765 | 1 | 234624 | ENSMUST0( | 863223 | ENSMUSG0( | A330008L17I | RIKEN cDNA A330008L17 gene |
| chr8 | 102952251 | 102952562 | 312 | * | NA_peak_21 | 23 | Distal Interge | 8 | 102661487 | 102785642 | 124156 | 2 | 12552 | ENSMUST0( | -166609 | ENSMUSG0( | Cdh11 | cadherin 11 |
| chr8 | 104533972 | 104534414 | 443 | * | NA_peak_21 | 34 | Promoter (<= | 8 | 104522905 | 104534608 | 11704 | 2 | 234664 | ENSMUST0( | 194 | ENSMUSG0( | Nae1 | NEDD8 activating enzyme E1 subunit 1 |
| chr8 | 104649424 | 104650226 | 803 | * | NA_peak_21 | 31 | Intron (ENSM | 8 | 104639838 | 104641965 | 2128 | 2 | 68523 | ENSMUST0( | -7459 | ENSMUSG0( | Ciao2b | cytosolic iron-sulfur assembly component 2B |
| chr8 | 105156168 | 105156451 | 284 | * | NA_peak_21 | 21 | Distal Interge | 8 | 105170674 | 105217989 | 47316 | 1 | 12400 | ENSMUST0( | -14223 | ENSMUSG0( | Cbfb | core binding factor beta |
| chr8 | 105171521 | 105171867 | 347 | * | NA_peak_21 | 38 | Promoter (<= | 8 | 105171564 | 105194535 | 22972 | 1 | 12400 | ENSMUST0( | 0 | ENSMUSG0( | Cbfb | core binding factor beta |
| chr8 | 105229883 | 105229989 | 107 | * | NA_peak_21 | 25 | Promoter (<= | 8 | 105230968 | 105240039 | 9072 | 1 | 234678 | ENSMUST0( | -979 | ENSMUSG0( | D230025D16 | RIKEN cDNA D230025D16 gene |
| chr8 | 105237625 | 105238168 | 544 | * | NA_peak_21 | 29 | Intron (ENSM | 8 | 105230968 | 105240039 | 9072 | 1 | 234678 | ENSMUST0( | 6657 | ENSMUSG0( | D230025D16 | RIKEN cDNA D230025D16 gene |
| chr8 | 105254055 | 105254253 | 199 | * | NA_peak_21 | 20 | Promoter (<= | 8 | 105252639 | 105255151 | 2513 | 2 | 97440 | ENSMUST0( | 898 | ENSMUSG0( | B3gnt9 | UDP-GlcNAc:betaGal beta-1,3-N-acetylglucosaminyltransferase 9 |
| chr8 | 105397746 | 105397881 | 136 | * | NA_peak_21 | 21 | Intron (ENSM | 8 | 105382942 | 105385417 | 2476 | 2 | 279499 | ENSMUST0( | -12329 | ENSMUSG0( | Kctd19 | potassium channel tetramerisation domain containing 19 |
| chr8 | 105632501 | 105632731 | 231 | * | NA_peak_21 | 15 | Intron (ENSM | 8 | 105636568 | 105681647 | 45080 | 1 | 13018 | ENSMUST0( | -3837 | ENSMUSG0( | Ctcf | CCCTC-binding factor |
| chr8 | 106242816 | 106243093 | 278 | * | NA_peak_22 | 29 | Exon (ENSM | 8 | 106237761 | 106244723 | 6963 | 1 | 214572 | ENSMUST0( | 5055 | ENSMUSG0( | Prmt7 | protein arginine N-methyltransferase 7 |
| chr8 | 106406606 | 106406937 | 332 | * | NA_peak_22 | 22 | Distal Interge | 8 | 106415327 | 106426598 | 11272 | 1 | 22751 | ENSMUST0( | -8390 | ENSMUSG0( | Zfp90 | zinc finger protein 90 |
| chr8 | 106683122 | 106683642 | 521 | * | NA_peak_22 | 29 | Promoter (<= | 8 | 106683146 | 106706262 | 23117 | 1 | 272538 | ENSMUST0( | 0 | ENSMUSG0( | Tango6 | transport and golgi organization 6 |
| chr8 | 106785715 | 106786042 | 328 | * | NA_peak_22 | 17 | Intron (ENSM | 8 | 106870242 | 106882902 | 12661 | 1 | 15118 | ENSMUST0( | -84200 | ENSMUSG0( | Has3 | hyaluronan synthase 3 |
| chr8 | 106925666 | 106925863 | 198 | * | NA_peak_22 | 25 | Distal Interge | 8 | 106935750 | 107019714 | 83965 | 1 | 20650 | ENSMUST0( | -9887 | ENSMUSG0( | Sntb2 | syntrophin, basic 2 |
| chr8 | 106926296 | 106926488 | 193 | * | NA_peak_22 | 25 | Distal Interge | 8 | 106935750 | 107019714 | 83965 | 1 | 20650 | ENSMUST0( | -9262 | ENSMUSG0( | Sntb2 | syntrophin, basic 2 |
| chr8 | 106932821 | 106933050 | 230 | * | NA_peak_22 | 35 | Promoter (2- | 8 | 106935750 | 107019714 | 83965 | 1 | 20650 | ENSMUST0( | -2700 | ENSMUSG0( | Sntb2 | syntrophin, basic 2 |
| chr8 | 106935972 | 106936327 | 356 | * | NA_peak_22 | 30 | Promoter (<= | 8 | 106936149 | 106981744 | 45596 | 1 | 20650 | ENSMUST0( | 0 | ENSMUSG0( | Sntb2 | syntrophin, basic 2 |
| chr8 | 106937230 | 106937537 | 308 | * | NA_peak_22 | 19 | Promoter (1- | 8 | 106936149 | 106981744 | 45596 | 1 | 20650 | ENSMUST0( | 1081 | ENSMUSG0( | Sntb2 | syntrophin, basic 2 |
| chr8 | 107037712 | 107038003 | 292 | * | NA_peak_22 | 21 | Promoter (1- | 8 | 107039019 | 107042687 | 3669 | 1 | 116733 | ENSMUST0( | -1016 | ENSMUSG0( | Vps4a | vacuolar protein sorting 4A |
| chr8 | 107056317 | 107056701 | 385 | * | NA_peak_22 | 35 | Promoter (<= | 8 | 107046289 | 107056689 | 10401 | 2 | 97484 | ENSMUST0( | 0 | ENSMUSG0( | Cog8 | component of oligomeric golgi complex 8 |
| chr8 | 107057011 | 107057064 | 54 | * | NA_peak_22 | 16 | Promoter (<= | 8 | 107056906 | 107058188 | 1283 | 1 | 66164 | ENSMUST0( | 105 | ENSMUSG0( | Nip7 | NIP7, nucleolar pre-rRNA processing protein |
| chr8 | 107330777 | 107331133 | 357 | * | NA_peak_22 | 30 | Intron (ENSM | 8 | 107338508 | 107355740 | 17233 | 1 | 54446 | ENSMUST0( | -7375 | ENSMUSG0( | Nfat5 | nuclear factor of activated T cells 5 |
| chr8 | 107390138 | 107390437 | 300 | * | NA_peak_22 | 19 | Intron (ENSM | 8 | 107388225 | 107403206 | 14982 | 2 | 18104 | ENSMUST0( | 12769 | ENSMUSG0( | Nqo1 | NAD(P)H dehydrogenase, quinone 1 |
| chr8 | 107463177 | 107463517 | 341 | * | NA_peak_22 | 35 | Intron (ENSM | 8 | 107483332 | 107546152 | 62821 | 1 | 66894 | ENSMUST0( | -19815 | ENSMUSG0( | Wwp2 | WW domain containing E3 ubiquitin protein ligase 2 |
| chr8 | 107744087 | 107744512 | 426 | * | NA_peak_22 | 13 | Distal Interge | 8 | 107580381 | 107588464 | 8084 | 2 | 17463 | ENSMUST0( | -155623 | ENSMUSG0( | Psmd7 | proteasome (prosome, macropain) 26S subunit, non-ATPase, 7 |
| chr8 | 108436555 | 108436855 | 301 | * | NA_peak_22 | 28 | Intron (ENSM | 8 | 108553252 | 108584595 | 31344 | 2 | 109215 | ENSMUST0( | 147740 | ENSMUSG0( | Lncbate1 | brown adipose tissue enriched long non-coding RNA 1 |
| chr8 | 108663458 | 108663662 | 205 | * | NA_peak_22 | 23 | Intron (ENSM | 8 | 108703100 | 108800270 | 97171 | 1 | 11906 | ENSMUST0( | -39438 | ENSMUSG0( | Zfhx3 | zinc finger homeobox 3 |
| chr8 | 108695828 | 108696549 | 722 | * | NA_peak_22 | 24 | Intron (ENSM | 8 | 108703100 | 108800270 | 97171 | 1 | 11906 | ENSMUST0( | -6551 | ENSMUSG0( | Zfhx3 | zinc finger homeobox 3 |
| chr8 | 108852731 | 108853119 | 389 | * | NA_peak_22 | 22 | Intron (ENSM | 8 | 108936860 | 108936938 | 79 | 1 | 100526511 | ENSMUST0( | -83741 | ENSMUSG0( | Mir3108 | microRNA 3108 |
| chr8 | 109130583 | 109130948 | 366 | * | NA_peak_22 | 45 | Intron (ENSM | 8 | 109075245 | 109228058 | 152814 | 1 | 628686 | ENSMUST0( | 55338 | ENSMUSG0( | D030068K23 | RIKEN cDNA D030068K23 gene |
| chr8 | 109573009 | 109573275 | 267 | * | NA_peak_22 | 15 | 3' UTR | 8 | 109568888 | 109573523 | 4636 | 1 | 234723 | ENSMUST0( | 4121 | ENSMUSG0( | Txnl4b | thioredoxin-like 4B |
| chr8 | 109656464 | 109656832 | 369 | * | NA_peak_22 | 14 | Intron (ENSM | 8 | 109647485 | 109674386 | 26902 | 1 | 244646 | ENSMUST0( | 8979 | ENSMUSG0( | Pkd1l3 | polycystic kidney disease 1 like 3 |
| chr8 | 109678998 | 109679418 | 421 | * | NA_peak_22 | 21 | Promoter (1- | 8 | 109671773 | 109677594 | 5822 | 2 | 71955 | ENSMUST0( | -1404 | ENSMUSG0( | Ist1 | increased sodium tolerance 1 homolog (yeast) |
| chr8 | 109759296 | 109759670 | 375 | * | NA_peak_22 | 17 | Distal Interge | 8 | 109778554 | 109864204 | 85651 | 1 | 11765 | ENSMUST0( | -18884 | ENSMUSG0( | Ap1g1 | adaptor protein complex AP-1, gamma 1 subunit |
| chr8 | 109778566 | 109779339 | 774 | * | NA_peak_22 | 45 | Promoter (<= | 8 | 109778897 | 109805668 | 26772 | 1 | 11765 | ENSMUST0( | 0 | ENSMUSG0( | Ap1g1 | adaptor protein complex AP-1, gamma 1 subunit |
| chr8 | 109816002 | 109816345 | 344 | * | NA_peak_22 | 21 | Intron (ENSM | 8 | 109839295 | 109839379 | 85 | 1 | 100217432 | ENSMUST0( | -22950 | NA | Snord71 | small nucleolar RNA, C/D box 71 |
| chr8 | 109816641 | 109816658 | 18 | * | NA_peak_22 | 21 | Intron (ENSM | 8 | 109839295 | 109839379 | 85 | 1 | 100217432 | ENSMUST0( | -22637 | NA | Snord71 | small nucleolar RNA, C/D box 71 |
| chr8 | 109865907 | 109866032 | 126 | * | NA_peak_22 | 19 | Promoter (2- | 8 | 109868542 | 109904971 | 36430 | 1 | 244650 | ENSMUST0( | -2510 | ENSMUSG0( | Phlpp2 | PH domain and leucine rich repeat protein phosphatase 2 |
| chr8 | 109936868 | 109937277 | 410 | * | NA_peak_22 | 15 | Exon (ENSM | 8 | 109959625 | 109962053 | 2429 | 2 | 73608 | ENSMUST0( | 24776 | ENSMUSG0( | Marveld3 | MARVEL (membrane-associating) domain containing 3 |
| chr8 | 111103679 | 111104040 | 362 | * | NA_peak_22 | 39 | Exon (ENSM | 8 | 111094655 | 111135147 | 40493 | 1 | 319518 | ENSMUST0( | 9024 | ENSMUSG0( | Pdpr | pyruvate dehydrogenase phosphatase regulatory subunit |
| chr8 | 111267644 | 111267934 | 291 | * | NA_peak_22 | 13 | Distal Interge | 8 | 111157898 | 111259216 | 101319 | 2 | 20340 | ENSMUST0( | -8428 | ENSMUSG0( | Glg1 | golgi apparatus protein 1 |
| chr8 | 111474697 | 111474989 | 293 | * | NA_peak_22 | 29 | Intron (ENSM | 8 | 111450208 | 111521987 | 71780 | 2 | 319481 | ENSMUST0( | 46998 | ENSMUSG0( | Wdr59 | WD repeat domain 59 |
| chr8 | 111613611 | 111613767 | 157 | * | NA_peak_22 | 30 | Intron (ENSM | 8 | 111609296 | 111621775 | 12480 | 1 | 170737 | ENSMUST0( | 4315 | ENSMUSG0( | Znrf1 | zinc and ring finger 1 |
| chr8 | 111650763 | 111650937 | 175 | * | NA_peak_22 | 17 | Intron (ENSM | 8 | 111643537 | 111670682 | 27146 | 1 | 22640 | ENSMUST0( | 7226 | ENSMUSG0( | Zfp1 | zinc finger protein 1 |
| chr8 | 111685761 | 111685811 | 51 | * | NA_peak_22 | 13 | Distal Interge | 8 | 111689601 | 111689674 | 74 | 2 | 102466993 | ENSMUST0( | 3863 | ENSMUSG0( | Mir7076 | microRNA 7076 |
| chr8 | 111872757 | 111872898 | 142 | * | NA_peak_22 | 25 | Intron (ENSM | 8 | 111863038 | 111876750 | 13713 | 2 | 66817 | ENSMUST0( | 3852 | ENSMUSG0( | Tmem170 | transmembrane protein 170 |
| chr8 | 111897545 | 111897902 | 358 | * | NA_peak_22 | 18 | Intron (ENSM | 8 | 111889136 | 111910447 | 21312 | 2 | 56773 | ENSMUST0( | 12545 | ENSMUSG0( | Chst5 | carbohydrate (N-acetylglucosamine 6-O) sulfotransferase 5 |
| chr8 | 111913066 | 111913331 | 266 | * | NA_peak_22 | 17 | Promoter (2- | 8 | 111889136 | 111910447 | 21312 | 2 | 56773 | ENSMUST0( | -2619 | ENSMUSG0( | Chst5 | carbohydrate (N-acetylglucosamine 6-O) sulfotransferase 5 |
| chr8 | 113282696 | 113282802 | 107 | * | NA_peak_22 | 20 | Distal Interge | 8 | 113067262 | 113093334 | 26073 | 1 | 791378 | ENSMUST0( | 215434 | ENSMUSG0( | Gm10280 | predicted gene 10280 |
| chr8 | 113285251 | 113285558 | 308 | * | NA_peak_22 | 23 | Distal Interge | 8 | 113067262 | 113093334 | 26073 | 1 | 791378 | ENSMUST0( | 217989 | ENSMUSG0( | Gm10280 | predicted gene 10280 |
| chr8 | 114622213 | 114622536 | 324 | * | NA_peak_22 | 18 | Intron (ENSM | 8 | 114608436 | 114706324 | 97889 | 1 | 80707 | ENSMUST0( | 13777 | ENSMUSG0( | Wwox | WW domain-containing oxidoreductase |
| chr8 | 115016022 | 115016376 | 355 | * | NA_peak_22 | 20 | Intron (ENSM | 8 | 114712167 | 115352143 | 639977 | 1 | 80707 | ENSMUST0( | 303855 | ENSMUSG0( | Wwox | WW domain-containing oxidoreductase |
| chr8 | 115214771 | 115215164 | 394 | * | NA_peak_22 | 19 | Intron (ENSM | 8 | 115701435 | 115707794 | 6360 | 2 | 17132 | ENSMUST0( | 492630 | ENSMUSG0( | Maf | avian musculoaponeurotic fibrosarcoma oncogene homolog |
| chr8 | 115624805 | 115625039 | 235 | * | NA_peak_22 | 24 | Distal Interge | 8 | 115701435 | 115707794 | 6360 | 2 | 17132 | ENSMUST0( | 82755 | ENSMUSG0( | Maf | avian musculoaponeurotic fibrosarcoma oncogene homolog |
| chr8 | 115706774 | 115707494 | 721 | * | NA_peak_22 | 18 | Promoter (<= | 8 | 115701435 | 115707794 | 6360 | 2 | 17132 | ENSMUST0( | 300 | ENSMUSG0( | Maf | avian musculoaponeurotic fibrosarcoma oncogene homolog |
| chr8 | 115708131 | 115708489 | 359 | * | NA_peak_22 | 22 | Promoter (<= | 8 | 115701435 | 115707794 | 6360 | 2 | 17132 | ENSMUST0( | -337 | ENSMUSG0( | Maf | avian musculoaponeurotic fibrosarcoma oncogene homolog |
| chr8 | 116505478 | 116505826 | 349 | * | NA_peak_22 | 30 | Promoter (<= | 8 | 116505015 | 116515915 | 10901 | 1 | 75465 | ENSMUST0( | 463 | ENSMUSG0( | Dynlrb2 | dynein light chain roadblock-type 2 |
| chr8 | 116978764 | 116979232 | 469 | * | NA_peak_22 | 13 | Promoter (<= | 8 | 116969594 | 116978943 | 9350 | 2 | 69528 | ENSMUST0( | 0 | ENSMUSG0( | 1700030J22I | RIKEN cDNA 1700030J22 gene |
| chr8 | 117979051 | 117979374 | 324 | * | NA_peak_22 | 23 | Distal Interge | 8 | 117791645 | 117801943 | 10299 | 2 | 68533 | ENSMUST0( | -177108 | ENSMUSG0( | Mphosph6 | M phase phosphoprotein 6 |
| chr8 | 119794056 | 119794361 | 306 | * | NA_peak_22 | 28 | Distal Interge | 8 | 119760076 | 119778416 | 18341 | 2 | 74347 | ENSMUST0( | -15640 | ENSMUSG0( | Meak7 | MTOR associated protein , eak-7 homolog |
| chr8 | 119840739 | 119840981 | 243 | * | NA_peak_22 | 40 | Promoter (<= | 8 | 119809222 | 119840544 | 31323 | 2 | 72042 | ENSMUST0( | -195 | ENSMUSG0( | Cotl1 | coactosin-like 1 (Dictyostelium) |
| chr8 | 119894174 | 119894389 | 216 | * | NA_peak_22 | 20 | Distal Interge | 8 | 119910360 | 119957555 | 47196 | 1 | 22224 | ENSMUST0( | -15971 | ENSMUSG0( | Usp10 | ubiquitin specific peptidase 10 |
| chr8 | 120605793 | 120606225 | 433 | * | NA_peak_22 | 29 | Intron (ENSM | 8 | 120578633 | 120589304 | 10672 | 2 | 272551 | ENSMUST0( | -16489 | ENSMUSG0( | Gins2 | GINS complex subunit 2 (Psf2 homolog) |
| chr8 | 121590260 | 121590763 | 504 | * | NA_peak_22 | 18 | Promoter (<= | 8 | 121590361 | 121598760 | 8400 | 1 | 67443 | ENSMUST0( | 0 | ENSMUSG0( | Map1lc3b | microtubule-associated protein 1 light chain 3 beta |
| chr8 | 122444703 | 122444864 | 162 | * | NA_peak_22 | 14 | Promoter (1- | 8 | 122433601 | 122443422 | 9822 | 2 | 192156 | ENSMUST0( | -1281 | ENSMUSG0( | Mvd | mevalonate (diphospho) decarboxylase |
| chr8 | 122558496 | 122558830 | 335 | * | NA_peak_22 | 42 | Intron (ENSM | 8 | 122481698 | 122551329 | 69632 | 2 | 234839 | ENSMUST0( | -7167 | ENSMUSG0( | Piezo1 | piezo-type mechanosensitive ion channel component 1 |
| chr8 | 122567748 | 122567935 | 188 | * | NA_peak_22 | 33 | Promoter (<= | 8 | 122568015 | 122573128 | 5114 | 1 | 67177 | ENSMUST0( | -80 | ENSMUSG0( | Cdt1 | chromatin licensing and DNA replication factor 1 |
| chr8 | 123027073 | 123027258 | 186 | * | NA_peak_22 | 30 | Intron (ENSM | 8 | 122883822 | 123042020 | 158199 | 2 | 77087 | ENSMUST0( | 14762 | ENSMUSG0( | Ankrd11 | ankyrin repeat domain 11 |
| chr8 | 123420928 | 123420985 | 58 | * | NA_peak_22 | 13 | Promoter (2- | 8 | 123423527 | 123452793 | 29267 | 1 | 23854 | ENSMUST0( | -2542 | ENSMUSG0( | Def8 | differentially expressed in FDCP 8 |
| chr8 | 123538817 | 123539175 | 359 | * | NA_peak_22 | 26 | Promoter (<= | 8 | 123538786 | 123538904 | 119 | 2 | 100861480 | ENSMUST0( | 0 | ENSMUSG0( | Gm25018 | predicted gene, 25018 |
| chr8 | 123587500 | 123587816 | 317 | * | NA_peak_22 | 23 | Intron (ENSM | 8 | 123565969 | 123566087 | 119 | 2 | 100861494 | ENSMUST0( | -21413 | ENSMUSG0( | Gm22109 | predicted gene, 22109 |
| chr8 | 123601672 | 123601986 | 315 | * | NA_peak_22 | 26 | Exon (ENSM | 8 | 123565969 | 123566087 | 119 | 2 | 100861494 | ENSMUST0( | -35585 | ENSMUSG0( | Gm22109 | predicted gene, 22109 |
| chr8 | 123655519 | 123655831 | 313 | * | NA_peak_22 | 18 | Promoter (1- | 8 | 123653929 | 123663884 | 9956 | 1 | 69581 | ENSMUST0( | 1590 | ENSMUSG0( | Rhou | ras homolog family member U |
| chr8 | 123816532 | 123816711 | 180 | * | NA_peak_22 | 17 | Intron (ENSM | 8 | 123806068 | 123834485 | 28418 | 1 | 19341 | ENSMUST0( | 10464 | ENSMUSG0( | Rab4a | RAB4A, member RAS oncogene family |
| chr8 | 123991596 | 123991869 | 274 | * | NA_peak_22 | 16 | Intron (ENSM | 8 | 123952459 | 123983122 | 30664 | 2 | 56199 | ENSMUST0( | -8474 | ENSMUSG0( | Abcb10 | ATP-binding cassette, sub-family B (MDR/TAP), member 10 |
| chr8 | 125420671 | 125420937 | 267 | * | NA_peak_22 | 22 | Intron (ENSM | 8 | 125418063 | 125492710 | 74648 | 2 | 244668 | ENSMUST0( | 71773 | ENSMUSG0( | Sipa1l2 | signal-induced proliferation-associated 1 like 2 |
| chr8 | 127064292 | 127064468 | 177 | * | NA_peak_22 | 16 | Promoter (<= | 8 | 127064295 | 127610805 | 546511 | 1 | 93742 | ENSMUST0( | 0 | ENSMUSG0( | Pard3 | par-3 family cell polarity regulator |
| chr8 | 127237954 | 127238078 | 125 | * | NA_peak_22 | 15 | Intron (ENSM | 8 | 127299585 | 127402577 | 102993 | 1 | 93742 | ENSMUST0( | -61507 | ENSMUSG0( | Pard3 | par-3 family cell polarity regulator |
| chr8 | 128090920 | 128091320 | 401 | * | NA_peak_22 | 16 | Distal Interge | 8 | 128278148 | 128278225 | 78 | 2 | 102465212 | ENSMUST0( | 186905 | ENSMUSG0( | Mir21c | microRNA 21c |
| chr8 | 128784611 | 128784888 | 278 | * | NA_peak_22 | 16 | Intron (ENSM | 8 | 128724333 | 128732389 | 8057 | 1 | 16412 | ENSMUST0( | 60278 | ENSMUSG0( | Itgb1 | integrin beta 1 (fibronectin receptor beta) |
| chr8 | 128939405 | 128939779 | 375 | * | NA_peak_22 | 16 | Intron (ENSM | 8 | 128980101 | 129065488 | 85388 | 2 | 74703 | ENSMUST0( | 125709 | ENSMUSG0( | Ccdc7a | coiled-coil domain containing 7A |
| chr8 | 128949259 | 128949635 | 377 | * | NA_peak_22 | 35 | Intron (ENSM | 8 | 128980101 | 129065488 | 85388 | 2 | 74703 | ENSMUST0( | 115853 | ENSMUSG0( | Ccdc7a | coiled-coil domain containing 7A |
| chr9 | 3002005 | 3003652 | 1648 | * | NA_peak_22 | 42 | Exon (ENSM | 9 | 3038669 | 3038743 | 75 | 2 | 100628572 | ENSMUST0( | 35091 | ENSMUSG0( | Mir101c | microRNA 101c |
| chr9 | 3006336 | 3007251 | 916 | * | NA_peak_22 | 75 | Exon (ENSM | 9 | 3038669 | 3038743 | 75 | 2 | 100628572 | ENSMUST0( | 31492 | ENSMUSG0( | Mir101c | microRNA 101c |
| chr9 | 3009381 | 3009892 | 512 | * | NA_peak_22 | 60 | Distal Interge | 9 | 3038669 | 3038743 | 75 | 2 | 100628572 | ENSMUST0( | 28851 | ENSMUSG0( | Mir101c | microRNA 101c |
| chr9 | 3012774 | 3013323 | 550 | * | NA_peak_22 | 33 | Exon (ENSM | 9 | 3038669 | 3038743 | 75 | 2 | 100628572 | ENSMUST0( | 25420 | ENSMUSG0( | Mir101c | microRNA 101c |
| chr9 | 3019858 | 3020925 | 1068 | * | NA_peak_22 | 18 | Exon (ENSM | 9 | 3038669 | 3038743 | 75 | 2 | 100628572 | ENSMUST0( | 17818 | ENSMUSG0( | Mir101c | microRNA 101c |
| chr9 | 3024073 | 3024560 | 488 | * | NA_peak_22 | 115 | Exon (ENSM | 9 | 3038669 | 3038743 | 75 | 2 | 100628572 | ENSMUST0( | 14183 | ENSMUSG0( | Mir101c | microRNA 101c |
| chr9 | 3027518 | 3028187 | 670 | * | NA_peak_22 | 57 | Exon (ENSM | 9 | 3038669 | 3038743 | 75 | 2 | 100628572 | ENSMUST0( | 10556 | ENSMUSG0( | Mir101c | microRNA 101c |
| chr9 | 3029412 | 3029844 | 433 | * | NA_peak_22 | 87 | Exon (ENSM | 9 | 3038669 | 3038743 | 75 | 2 | 100628572 | ENSMUST0( | 8899 | ENSMUSG0( | Mir101c | microRNA 101c |
| chr9 | 3030553 | 3030907 | 355 | * | NA_peak_22 | 78 | Exon (ENSM | 9 | 3038669 | 3038743 | 75 | 2 | 100628572 | ENSMUST0( | 7836 | ENSMUSG0( | Mir101c | microRNA 101c |
| chr9 | 3031685 | 3032208 | 524 | * | NA_peak_22 | 25 | Exon (ENSM | 9 | 3038669 | 3038743 | 75 | 2 | 100628572 | ENSMUST0( | 6535 | ENSMUSG0( | Mir101c | microRNA 101c |
| chr9 | 3035712 | 3036290 | 579 | * | NA_peak_22 | 64 | Promoter (2- | 9 | 3038669 | 3038743 | 75 | 2 | 100628572 | ENSMUST0( | 2453 | ENSMUSG0( | Mir101c | microRNA 101c |
| chr9 | 3305110 | 3305454 | 345 | * | NA_peak_22 | 14 | Distal Interge | 9 | 3335140 | 3386175 | 51036 | 1 | 67667 | ENSMUST0( | -29686 | ENSMUSG0( | Alkbh8 | alkB homolog 8, tRNA methyltransferase |
| chr9 | 3311600 | 3312272 | 673 | * | NA_peak_22 | 36 | Distal Interge | 9 | 3335140 | 3386175 | 51036 | 1 | 67667 | ENSMUST0( | -22868 | ENSMUSG0( | Alkbh8 | alkB homolog 8, tRNA methyltransferase |
| chr9 | 3659570 | 3659912 | 343 | * | NA_peak_22 | 50 | Intron (ENSM | 9 | 3532888 | 3761177 | 228290 | 1 | 234889 | ENSMUST0( | 126682 | ENSMUSG0( | Gucy1a2 | guanylate cyclase 1, soluble, alpha 2 |
| chr9 | 4138709 | 4138976 | 268 | * | NA_peak_22 | 49 | Distal Interge | 9 | 4294793 | 4309467 | 14675 | 2 | 67618 | ENSMUST0( | 170491 | ENSMUSG0( | Aasdhppt | aminoadipate-semialdehyde dehydrogenase-phosphopantetheinyl transferase |
| chr9 | 11725439 | 11725740 | 302 | * | NA_peak_22 | 26 | Distal Interge | 9 | 9660969 | 10904775 | 1243807 | 2 | 244682 | ENSMUST0( | -820664 | ENSMUSG0( | Cntn5 | contactin 5 |
| chr9 | 18142482 | 18142812 | 331 | * | NA_peak_22 | 35 | Distal Interge | 9 | 18292125 | 18317442 | 25318 | 1 | 66917 | ENSMUST0( | -149313 | ENSMUSG0( | Chordc1 | cysteine and histidine-rich domain (CHORD)-containing, zinc-binding protein 1 |
| chr9 | 18981339 | 18981632 | 294 | * | NA_peak_22 | 54 | Distal Interge | 9 | 18987990 | 18988928 | 939 | 1 | 258074 | ENSMUST0( | -6358 | ENSMUSG0( | Olfr834 | olfactory receptor 834 |
| chr9 | 21521005 | 21521162 | 158 | * | NA_peak_22 | 26 | Distal Interge | 9 | 21526176 | 21545333 | 19158 | 1 | 382062 | ENSMUST0( | -5014 | ENSMUSG0( | AB124611 | cDNA sequence AB124611 |
| chr9 | 21521611 | 21521773 | 163 | * | NA_peak_22 | 19 | Distal Interge | 9 | 21526176 | 21545333 | 19158 | 1 | 382062 | ENSMUST0( | -4403 | ENSMUSG0( | AB124611 | cDNA sequence AB124611 |
| chr9 | 23744217 | 23744503 | 287 | * | NA_peak_22 | 39 | Distal Interge | 9 | 24097996 | 24289892 | 191897 | 1 | 319239 | ENSMUST0( | -353493 | ENSMUSG0( | Npsr1 | neuropeptide S receptor 1 |
| chr9 | 26403893 | 26404165 | 273 | * | NA_peak_22 | 48 | Distal Interge | 9 | 26733728 | 26761350 | 27623 | 1 | 76898 | ENSMUST0( | -329563 | ENSMUSG0( | B3gat1 | beta-1,3-glucuronyltransferase 1 (glucuronosyltransferase P) |
| chr9 | 37882261 | 37882529 | 269 | * | NA_peak_22 | 26 | Distal Interge | 9 | 37896724 | 37903519 | 6796 | 1 | 258310 | ENSMUST0( | -14195 | ENSMUSG0( | Olfr145 | olfactory receptor 145 |
| chr9 | 56223568 | 56223945 | 378 | * | NA_peak_22 | 51 | Exon (ENSM | 9 | 56201126 | 56244065 | 42940 | 2 | 244895 | ENSMUST0( | 20120 | ENSMUSG0( | Peak1 | pseudopodium-enriched atypical kinase 1 |
| chr9 | 59691634 | 59691966 | 333 | * | NA_peak_22 | 43 | Intron (ENSM | 9 | 59680144 | 59713923 | 33780 | 1 | 546134 | ENSMUST0( | 11490 | ENSMUSG0( | Gramd2 | GRAM domain containing 2 |
| chr9 | 59704155 | 59704164 | 10 | * | NA_peak_22 | 23 | Intron (ENSM | 9 | 59707637 | 59718874 | 11238 | 1 | 546134 | ENSMUST0( | -3473 | ENSMUSG0( | Gramd2 | GRAM domain containing 2 |
| chr9 | 59751129 | 59751473 | 345 | * | NA_peak_22 | 34 | Promoter (<= | 9 | 59750896 | 59928866 | 177971 | 1 | 270163 | ENSMUST0( | 233 | ENSMUSG0( | Myo9a | myosin IXa |
| chr9 | 61896314 | 61896338 | 25 | * | NA_peak_22 | 22 | Distal Interge | 9 | 61913284 | 61914542 | 1259 | 2 | 56040 | ENSMUST0( | 18204 | ENSMUSG0( | Rplp1 | ribosomal protein, large, P1 |
| chr9 | 62379998 | 62380281 | 284 | * | NA_peak_22 | 25 | Distal Interge | 9 | 62373439 | 62377781 | 4343 | 1 | 11737 | ENSMUST0( | 6559 | ENSMUSG0( | Anp32a | acidic (leucine-rich) nuclear phosphoprotein 32 family, member A |
| chr9 | 62505714 | 62505828 | 115 | * | NA_peak_22 | 22 | Intron (ENSM | 9 | 62427936 | 62510498 | 82563 | 2 | 235431 | ENSMUST0( | 4670 | ENSMUSG0( | Coro2b | coronin, actin binding protein, 2B |
| chr9 | 62512998 | 62513208 | 211 | * | NA_peak_22 | 22 | Promoter (2- | 9 | 62427936 | 62510498 | 82563 | 2 | 235431 | ENSMUST0( | -2500 | ENSMUSG0( | Coro2b | coronin, actin binding protein, 2B |
| chr9 | 64101054 | 64101082 | 29 | * | NA_peak_22 | 18 | Distal Interge | 9 | 64117147 | 64138118 | 20972 | 1 | 235435 | ENSMUST0( | -16065 | ENSMUSG0( | Lctl | lactase-like |
| chr9 | 65215328 | 65215660 | 333 | * | NA_peak_22 | 42 | Promoter (<= | 9 | 65215145 | 65239224 | 24080 | 1 | 214424 | ENSMUST0( | 183 | ENSMUSG0( | Parp16 | poly (ADP-ribose) polymerase family, member 16 |
| chr9 | 65470521 | 65470570 | 50 | * | NA_peak_22 | 13 | Promoter (1- | 9 | 65468789 | 65474284 | 5496 | 1 | 27965 | ENSMUST0( | 1732 | ENSMUSG0( | Spg21 | SPG21, maspardin |
| chr9 | 66946210 | 66946407 | 198 | * | NA_peak_22 | 44 | Promoter (<= | 9 | 66946355 | 66949516 | 3162 | 1 | 67941 | ENSMUST0( | 0 | ENSMUSG0( | Rps27l | ribosomal protein S27-like |
| chr9 | 67018930 | 67019110 | 181 | * | NA_peak_22 | 22 | Distal Interge | 9 | 67029920 | 67032825 | 2906 | 2 | 22003 | ENSMUST0( | 13715 | ENSMUSG0( | Tpm1 | tropomyosin 1, alpha |
| chr9 | 67049506 | 67049533 | 28 | * | NA_peak_22 | 19 | Promoter (<= | 9 | 67022590 | 67049406 | 26817 | 2 | 22003 | ENSMUST0( | -100 | ENSMUSG0( | Tpm1 | tropomyosin 1, alpha |
| chr9 | 67763274 | 67763440 | 167 | * | NA_peak_22 | 19 | Distal Interge | 9 | 67758713 | 67760230 | 1518 | 2 | 102633151 | ENSMUST0( | -3044 | ENSMUSG0( | M5C1000I18I | RIKEN cDNA M5C1000I18 gene |
| chr9 | 69453556 | 69453879 | 324 | * | NA_peak_22 | 24 | Promoter (<= | 9 | 69453620 | 69491795 | 38176 | 1 | 12306 | ENSMUST0( | 0 | ENSMUSG0( | Anxa2 | annexin A2 |
| chr9 | 70319277 | 70319630 | 354 | * | NA_peak_22 | 30 | Intron (ENSM | 9 | 70405694 | 70405753 | 60 | 2 | 100885839 | ENSMUST0( | 86123 | ENSMUSG0( | Mir5626 | microRNA 5626 |
| chr9 | 71469414 | 71469715 | 302 | * | NA_peak_22 | 23 | Distal Interge | 9 | 71478471 | 71484958 | 6488 | 2 | 28015 | ENSMUST0( | 15243 | ENSMUSG0( | Polr2m | polymerase (RNA) II (DNA directed) polypeptide M |
| chr9 | 74953341 | 74953505 | 165 | * | NA_peak_22 | 31 | Promoter (<= | 9 | 74953072 | 74953990 | 919 | 1 | 235493 | ENSMUST0( | 269 | ENSMUSG0( | Fam214a | family with sequence similarity 214, member A |
| chr9 | 75070939 | 75071564 | 626 | * | NA_peak_22 | 21 | Promoter (<= | 9 | 75071015 | 75117308 | 46294 | 1 | 17918 | ENSMUST0( | 0 | ENSMUSG0( | Myo5a | myosin VA |
| chr9 | 78252046 | 78252367 | 322 | * | NA_peak_22 | 18 | Distal Interge | 9 | 78257129 | 78269166 | 12038 | 1 | 100042295 | ENSMUST0( | -4762 | ENSMUSG0( | Gm3776 | predicted gene 3776 |
| chr9 | 79260583 | 79260874 | 292 | * | NA_peak_22 | 39 | Intron (ENSM | 9 | 79444037 | 79519302 | 75266 | 2 | 100038575 | ENSMUST0( | 258428 | ENSMUSG0( | Gm10635 | predicted gene 10635 |
| chr9 | 79906340 | 79906836 | 497 | * | NA_peak_22 | 59 | Intron (ENSM | 9 | 79820661 | 79962033 | 141373 | 2 | 70598 | ENSMUST0( | 55197 | ENSMUSG0( | Filip1 | filamin A interacting protein 1 |
| chr9 | 82352801 | 82353013 | 213 | * | NA_peak_22 | 23 | Distal Interge | 9 | 82829540 | 82847687 | 18148 | 1 | 65099 | ENSMUST0( | -476527 | ENSMUSG0( | Irak1bp1 | interleukin-1 receptor-associated kinase 1 binding protein 1 |
| chr9 | 82568169 | 82568473 | 305 | * | NA_peak_22 | 37 | Distal Interge | 9 | 82829540 | 82847687 | 18148 | 1 | 65099 | ENSMUST0( | -261067 | ENSMUSG0( | Irak1bp1 | interleukin-1 receptor-associated kinase 1 binding protein 1 |
| chr9 | 82974196 | 82974530 | 335 | * | NA_peak_22 | 23 | Promoter (<= | 9 | 82927158 | 82974268 | 47111 | 2 | 83946 | ENSMUST0( | 0 | ENSMUSG0( | Phip | pleckstrin homology domain interacting protein |
| chr9 | 83944878 | 83944910 | 33 | * | NA_peak_22 | 35 | Intron (ENSM | 9 | 83948781 | 84124240 | 175460 | 1 | 12040 | ENSMUST0( | -3871 | ENSMUSG0( | Bckdhb | branched chain ketoacid dehydrogenase E1, beta polypeptide |
| chr9 | 84888042 | 84888444 | 403 | * | NA_peak_22 | 49 | Distal Interge | 9 | 84774512 | 84826183 | 51672 | 1 | 75253 | ENSMUST0( | 113530 | ENSMUSG0( | 4930554C24 | RIKEN cDNA 4930554C24 gene |
| chr9 | 86459206 | 86459604 | 399 | * | NA_peak_22 | 30 | Intron (ENSM | 9 | 86307306 | 86464950 | 157645 | 2 | 70348 | ENSMUST0( | 5346 | ENSMUSG0( | Ube2cbp | ubiquitin-conjugating enzyme E2C binding protein |
| chr9 | 86459919 | 86460380 | 462 | * | NA_peak_22 | 30 | Intron (ENSM | 9 | 86307306 | 86464950 | 157645 | 2 | 70348 | ENSMUST0( | 4570 | ENSMUSG0( | Ube2cbp | ubiquitin-conjugating enzyme E2C binding protein |
| chr9 | 88481380 | 88482309 | 930 | * | NA_peak_22 | 31 | Promoter (<= | 9 | 88449731 | 88482248 | 32518 | 2 | 56403 | ENSMUST0( | 0 | ENSMUSG0( | Syncrip | synaptotagmin binding, cytoplasmic RNA interacting protein |
| chr9 | 88807938 | 88808251 | 314 | * | NA_peak_22 | 29 | Distal Interge | 9 | 88839164 | 88848190 | 9027 | 1 | 666731 | ENSMUST0( | -30913 | ENSMUSG0( | Trim43c | tripartite motif-containing 43C |
| chr9 | 91263866 | 91264109 | 244 | * | NA_peak_22 | 20 | Distal Interge | 9 | 91362413 | 91378915 | 16503 | 1 | 22774 | ENSMUST0( | -98304 | ENSMUSG0( | Zic4 | zinc finger protein of the cerebellum 4 |
| chr9 | 91647711 | 91647950 | 240 | * | NA_peak_22 | 26 | Distal Interge | 9 | 91382410 | 91388044 | 5635 | 1 | 22774 | ENSMUST0( | 265301 | ENSMUSG0( | Zic4 | zinc finger protein of the cerebellum 4 |
| chr9 | 91861696 | 91862017 | 322 | * | NA_peak_22 | 26 | Distal Interge | 9 | 92192936 | 92209772 | 16837 | 1 | 100504689 | ENSMUST0( | -330919 | ENSMUSG0( | Plscr5 | phospholipid scramblase family, member 5 |
| chr9 | 93033858 | 93034393 | 536 | * | NA_peak_22 | 30 | Distal Interge | 9 | 92979995 | 93005739 | 25745 | 2 | 78349 | ENSMUST0( | -28119 | ENSMUSG0( | 1700034K08I | RIKEN cDNA 1700034K08 gene |
| chr9 | 94641383 | 94641733 | 351 | * | NA_peak_22 | 20 | Distal Interge | 9 | 94669909 | 95230445 | 560537 | 1 | 331004 | ENSMUST0( | -28176 | ENSMUSG0( | Slc9a9 | solute carrier family 9 (sodium/hydrogen exchanger), member 9 |
| chr9 | 94821891 | 94822267 | 377 | * | NA_peak_22 | 48 | Intron (ENSM | 9 | 94751829 | 94751890 | 62 | 1 | 102465763 | ENSMUST0( | 70062 | ENSMUSG0( | Mir7656 | microRNA 7656 |
| chr9 | 95296267 | 95296616 | 350 | * | NA_peak_22 | 64 | Distal Interge | 9 | 95399292 | 95406722 | 7431 | 2 | 54371 | ENSMUST0( | 110106 | ENSMUSG0( | Chst2 | carbohydrate sulfotransferase 2 |
| chr9 | 95406925 | 95407241 | 317 | * | NA_peak_22 | 23 | Promoter (<= | 9 | 95399292 | 95406722 | 7431 | 2 | 54371 | ENSMUST0( | -203 | ENSMUSG0( | Chst2 | carbohydrate sulfotransferase 2 |
| chr9 | 96485076 | 96485335 | 260 | * | NA_peak_22 | 26 | Distal Interge | 9 | 96470937 | 96478675 | 7739 | 2 | 19823 | ENSMUST0( | -6401 | ENSMUSG0( | Rnf7 | ring finger protein 7 |
| chr9 | 98252063 | 98252159 | 97 | * | NA_peak_22 | 54 | Distal Interge | 9 | 98287435 | 98399516 | 112082 | 1 | 74080 | ENSMUST0( | -35276 | ENSMUSG0( | Nmnat3 | nicotinamide nucleotide adenylyltransferase 3 |
| chr9 | 101133110 | 101133407 | 298 | * | NA_peak_22 | 30 | Intron (ENSM | 9 | 101107883 | 101125125 | 17243 | 2 | 235542 | ENSMUST0( | -7985 | ENSMUSG0( | Ppp2r3a | protein phosphatase 2, regulatory subunit B'', alpha |
| chr9 | 104223586 | 104223975 | 390 | * | NA_peak_23 | 22 | Intron (ENSM | 9 | 104228354 | 104228993 | 640 | 2 | 235567 | ENSMUST0( | 5018 | ENSMUSG0( | Dnajc13 | DnaJ heat shock protein family (Hsp40) member C13 |
| chr9 | 106161468 | 106161556 | 89 | * | NA_peak_23 | 17 | Distal Interge | 9 | 106152869 | 106158138 | 5270 | 2 | 235582 | ENSMUST0( | -3330 | ENSMUSG0( | Glyctk | glycerate kinase |
| chr9 | 107698780 | 107699013 | 234 | * | NA_peak_23 | 13 | Intron (ENSM | 9 | 107687275 | 107692714 | 5440 | 2 | 20350 | ENSMUST0( | -6066 | ENSMUSG0( | Sema3f | sema domain, immunoglobulin domain (Ig), short basic domain, secreted, (semaphorin) 3F |
| chr9 | 108046597 | 108046830 | 234 | * | NA_peak_23 | 27 | Promoter (1- | 9 | 108044863 | 108045539 | 677 | 1 | 27399 | ENSMUST0( | 1734 | ENSMUSG0( | Ip6k1 | inositol hexaphosphate kinase 1 |
| chr9 | 108160501 | 108160761 | 261 | * | NA_peak_23 | 40 | Exon (ENSM | 9 | 108137328 | 108137715 | 388 | 1 | 76873 | ENSMUST0( | 23173 | ENSMUSG0( | 4930447F24I | RIKEN cDNA 4930447F24 gene |
| chr9 | 108306055 | 108306236 | 182 | * | NA_peak_23 | 30 | Promoter (<= | 9 | 108303654 | 108306160 | 2507 | 2 | 102791 | ENSMUST0( | 0 | ENSMUSG0( | Tcta | T cell leukemia translocation altered gene |
| chr9 | 108578471 | 108578883 | 413 | * | NA_peak_23 | 29 | Promoter (<= | 9 | 108572311 | 108578739 | 6429 | 2 | 83669 | ENSMUST0( | 0 | ENSMUSG0( | Wdr6 | WD repeat domain 6 |
| chr9 | 108648965 | 108649312 | 348 | * | NA_peak_23 | 26 | Promoter (<= | 9 | 108615768 | 108649318 | 33551 | 2 | 23807 | ENSMUST0( | 6 | ENSMUSG0( | Arih2 | ariadne RBR E3 ubiquitin protein ligase 2 |
| chr9 | 109372627 | 109372894 | 268 | * | NA_peak_23 | 32 | Distal Interge | 9 | 109378400 | 109404296 | 25897 | 2 | 382156 | ENSMUST0( | 31402 | ENSMUSG0( | Fbxw22 | F-box and WD-40 domain protein 22 |
| chr9 | 110742271 | 110742547 | 277 | * | NA_peak_23 | 19 | Promoter (<= | 9 | 110722142 | 110742719 | 20578 | 2 | 19228 | ENSMUST0( | 172 | ENSMUSG0( | Pth1r | parathyroid hormone 1 receptor |
| chr9 | 114145810 | 114146120 | 311 | * | NA_peak_23 | 23 | Intron (ENSM | 9 | 114057140 | 114098728 | 41589 | 1 | 382111 | ENSMUST0( | 88670 | ENSMUSG0( | Susd5 | sushi domain containing 5 |
| chr9 | 114781221 | 114781639 | 419 | * | NA_peak_23 | 21 | Promoter (<= | 9 | 114756839 | 114781832 | 24994 | 2 | 102545 | ENSMUST0( | 193 | ENSMUSG0( | Cmtm7 | CKLF-like MARVEL transmembrane domain containing 7 |
| chr9 | 114973786 | 114974053 | 268 | * | NA_peak_23 | 23 | Distal Interge | 9 | 114978569 | 115232223 | 253655 | 1 | 74486 | ENSMUST0( | -4516 | ENSMUSG0( | Osbpl10 | oxysterol binding protein-like 10 |
| chr9 | 115310399 | 115310798 | 400 | * | NA_peak_23 | 28 | Promoter (<= | 9 | 115242581 | 115310421 | 67841 | 2 | 68292 | ENSMUST0( | 0 | ENSMUSG0( | Stt3b | STT3, subunit of the oligosaccharyltransferase complex, homolog B (S. cerevisiae) |
| chr9 | 116980084 | 116980162 | 79 | * | NA_peak_23 | 23 | Intron (ENSM | 9 | 116582822 | 116822819 | 239998 | 2 | 207181 | ENSMUST0( | -157265 | ENSMUSG0( | Rbms3 | RNA binding motif, single stranded interacting protein |
| chr9 | 116986128 | 116986466 | 339 | * | NA_peak_23 | 35 | Intron (ENSM | 9 | 116582822 | 116822819 | 239998 | 2 | 207181 | ENSMUST0( | -163309 | ENSMUSG0( | Rbms3 | RNA binding motif, single stranded interacting protein |
| chr9 | 118040358 | 118040814 | 457 | * | NA_peak_23 | 23 | Promoter (<= | 9 | 118040499 | 118063839 | 23341 | 1 | 27215 | ENSMUST0( | 0 | ENSMUSG0( | Azi2 | 5-azacytidine induced gene 2 |
| chr9 | 119444993 | 119445290 | 298 | * | NA_peak_23 | 20 | Promoter (<= | 9 | 119444987 | 119464239 | 19253 | 1 | 208194 | ENSMUST0( | 6 | ENSMUSG0( | Exog | endo/exonuclease (5'-3'), endonuclease G-like |
| chr9 | 119868351 | 119868562 | 212 | * | NA_peak_23 | 24 | Distal Interge | 9 | 119894878 | 119926586 | 31709 | 1 | 67561 | ENSMUST0( | -26316 | ENSMUSG0( | Wdr48 | WD repeat domain 48 |
| chr9 | 122299884 | 122300280 | 397 | * | NA_peak_23 | 54 | Intron (ENSM | 9 | 122175874 | 122294423 | 118550 | 2 | 102566 | ENSMUST0( | -5461 | ENSMUSG0( | Ano10 | anoctamin 10 |
| chr9 | 123988837 | 123989283 | 447 | * | NA_peak_23 | 19 | Distal Interge | 9 | 123962124 | 123968692 | 6569 | 2 | 12768 | ENSMUST0( | -20145 | ENSMUSG0( | Ccr1 | chemokine (C-C motif) receptor 1 |
| chrX | 3083804 | 3084037 | 234 | * | NA_peak_23 | 23 | Distal Interge | 20 | 3076875 | 3078817 | 1943 | 1 | 100861966 | ENSMUST0( | 6929 | ENSMUSG0( | Btbd35f23 | BTB domain containing 35, family member 23 |
| chrX | 3084886 | 3085401 | 516 | * | NA_peak_23 | 24 | Distal Interge | 20 | 3076875 | 3078817 | 1943 | 1 | 100861966 | ENSMUST0( | 8011 | ENSMUSG0( | Btbd35f23 | BTB domain containing 35, family member 23 |
| chrX | 3087417 | 3087693 | 277 | * | NA_peak_23 | 39 | Distal Interge | 20 | 3076875 | 3078817 | 1943 | 1 | 100861966 | ENSMUST0( | 10542 | ENSMUSG0( | Btbd35f23 | BTB domain containing 35, family member 23 |
| chrX | 3225842 | 3225905 | 64 | * | NA_peak_23 | 17 | Intron (ENSM | 20 | 3076875 | 3078817 | 1943 | 1 | 100861966 | ENSMUST0( | 148967 | ENSMUSG0( | Btbd35f23 | BTB domain containing 35, family member 23 |
| chrX | 3232163 | 3232707 | 545 | * | NA_peak_23 | 49 | Distal Interge | 20 | 3076875 | 3078817 | 1943 | 1 | 100861966 | ENSMUST0( | 155288 | ENSMUSG0( | Btbd35f23 | BTB domain containing 35, family member 23 |
| chrX | 3238814 | 3239130 | 317 | * | NA_peak_23 | 28 | Distal Interge | 20 | 3076875 | 3078817 | 1943 | 1 | 100861966 | ENSMUST0( | 161939 | ENSMUSG0( | Btbd35f23 | BTB domain containing 35, family member 23 |
| chrX | 3242678 | 3243356 | 679 | * | NA_peak_23 | 24 | Distal Interge | 20 | 3076875 | 3078817 | 1943 | 1 | 100861966 | ENSMUST0( | 165803 | ENSMUSG0( | Btbd35f23 | BTB domain containing 35, family member 23 |
| chrX | 3431352 | 3431993 | 642 | * | NA_peak_23 | 38 | Distal Interge | 20 | 3441731 | 3443690 | 1960 | 2 | 630022 | ENSMUST0( | 11697 | ENSMUSG0( | Btbd35f18 | BTB domain containing 35, family member 18 |
| chrX | 3764716 | 3765037 | 322 | * | NA_peak_23 | 44 | Distal Interge | 20 | 3750934 | 3752885 | 1952 | 2 | 668960 | ENSMUST0( | -11831 | ENSMUSG0( | Btbd35f10 | BTB domain containing 35, family member 10 |
| chrX | 3775224 | 3775585 | 362 | * | NA_peak_23 | 57 | Distal Interge | 20 | 3750934 | 3752885 | 1952 | 2 | 668960 | ENSMUST0( | -22339 | ENSMUSG0( | Btbd35f10 | BTB domain containing 35, family member 10 |
| chrX | 4117947 | 4118322 | 376 | * | NA_peak_23 | 26 | Exon (ENSM | 20 | 4196576 | 4198535 | 1960 | 1 | 627264 | ENSMUST0( | -78254 | ENSMUSG0( | Btbd35f28 | BTB domain containing 35, family member 28 |
| chrX | 4135180 | 4135382 | 203 | * | NA_peak_23 | 17 | Distal Interge | 20 | 4196576 | 4198535 | 1960 | 1 | 627264 | ENSMUST0( | -61194 | ENSMUSG0( | Btbd35f28 | BTB domain containing 35, family member 28 |
| chrX | 4137078 | 4137650 | 573 | * | NA_peak_23 | 20 | Distal Interge | 20 | 4196576 | 4198535 | 1960 | 1 | 627264 | ENSMUST0( | -58926 | ENSMUSG0( | Btbd35f28 | BTB domain containing 35, family member 28 |
| chrX | 4142922 | 4143421 | 500 | * | NA_peak_23 | 37 | Distal Interge | 20 | 4196576 | 4198535 | 1960 | 1 | 627264 | ENSMUST0( | -53155 | ENSMUSG0( | Btbd35f28 | BTB domain containing 35, family member 28 |
| chrX | 4353742 | 4354118 | 377 | * | NA_peak_23 | 13 | Distal Interge | 20 | 4370636 | 4372595 | 1960 | 1 | 668964 | ENSMUST0( | -16518 | ENSMUSG0( | Btbd35f17 | BTB domain containing 35, family member 17 |
| chrX | 5081306 | 5081833 | 528 | * | NA_peak_23 | 53 | Distal Interge | 20 | 4952135 | 4954077 | 1943 | 1 | 100042275 | ENSMUST0( | 129171 | ENSMUSG0( | Btbd35f27 | BTB domain containing 35, family member 27 |
| chrX | 5089307 | 5089488 | 182 | * | NA_peak_23 | 35 | Distal Interge | 20 | 4952135 | 4954077 | 1943 | 1 | 100042275 | ENSMUST0( | 137172 | ENSMUSG0( | Btbd35f27 | BTB domain containing 35, family member 27 |
| chrX | 5103099 | 5103189 | 91 | * | NA_peak_23 | 37 | Distal Interge | 20 | 4952135 | 4954077 | 1943 | 1 | 100042275 | ENSMUST0( | 150964 | ENSMUSG0( | Btbd35f27 | BTB domain containing 35, family member 27 |

| | | | | | | | | | | | | | | | | | | |
|---|---|---|---|---|---|---|---|---|---|---|---|---|---|---|---|---|---|---|
| chrX | 5300088 | 5300455 | 368 | * | NA_peak_23 | 44 | Distal Interge | 20 | 5466898 | 5469265 | 2368 | 2 | 17870 | ENSMUST0( | 168810 | ENSMUSG0( | Mycs | myc-like oncogene, s-myc protein |
| chrX | 5315651 | 5315966 | 316 | * | NA_peak_23 | 31 | Distal Interge | 20 | 5466898 | 5469265 | 2368 | 2 | 17870 | ENSMUST0( | 153299 | ENSMUSG0( | Mycs | myc-like oncogene, s-myc protein |
| chrX | 5322076 | 5322419 | 344 | * | NA_peak_23 | 20 | Distal Interge | 20 | 5466898 | 5469265 | 2368 | 2 | 17870 | ENSMUST0( | 146846 | ENSMUSG0( | Mycs | myc-like oncogene, s-myc protein |
| chrX | 7770469 | 7770888 | 420 | * | NA_peak_23 | 18 | Promoter (1-: | 20 | 7769430 | 7771318 | 1889 | 1 | 209446 | ENSMUST0( | 1039 | ENSMUSG0( | Tfe3 | transcription factor E3 |
| chrX | 8519124 | 8519429 | 306 | * | NA_peak_23 | 31 | Intron (ENSM | 20 | 8514052 | 8524954 | 10903 | 2 | 546263 | ENSMUST0( | 5525 | ENSMUSG0( | Gm14459 | predicted gene 14459 |
| chrX | 15966071 | 15966246 | 176 | * | NA_peak_23 | 27 | Distal Interge | 20 | 16522869 | 16523689 | 821 | 1 | 66742 | ENSMUST0( | -556623 | ENSMUSG0( | Cypt1 | cysteine-rich perinuclear theca 1 |
| chrX | 17386337 | 17386627 | 291 | * | NA_peak_23 | 43 | Distal Interge | 20 | 17132049 | 17319368 | 187320 | 2 | 74405 | ENSMUST0( | -66969 | ENSMUSG0( | Efhc2 | EF-hand domain (C-terminal) containing 2 |
| chrX | 19651921 | 19652116 | 196 | * | NA_peak_23 | 44 | Distal Interge | 20 | 19479420 | 19481319 | 1900 | 2 | 278255 | ENSMUST0( | -170602 | ENSMUSG0( | Tex13c3 | TEX13 family member C3 |
| chrX | 20036573 | 20036840 | 268 | * | NA_peak_23 | 30 | Distal Interge | 20 | 20059560 | 20097521 | 37962 | 1 | 60322 | ENSMUST0( | -22720 | ENSMUSG0( | Chst7 | carbohydrate (N-acetylglucosamino) sulfotransferase 7 |
| chrX | 20476174 | 20476289 | 116 | * | NA_peak_23 | 15 | Intron (ENSM | 20 | 20460527 | 20517896 | 57370 | 1 | 382207 | ENSMUST0( | 15647 | ENSMUSG0( | Jade3 | jade family PHD finger 3 |
| chrX | 20767316 | 20767802 | 487 | * | NA_peak_23 | 23 | Distal Interge | 20 | 20797814 | 20860497 | 62684 | 1 | 11836 | ENSMUST0( | -30012 | ENSMUSG0( | Araf | Araf proto-oncogene, serine/threonine kinase |
| chrX | 31183591 | 31183888 | 298 | * | NA_peak_23 | 22 | Distal Interge | 20 | 31336401 | 31338101 | 1701 | 1 | 668960 | ENSMUST0( | -152513 | ENSMUSG0( | Btbd35f10 | BTB domain containing 35, family member 10 |
| chrX | 31187208 | 31187352 | 145 | * | NA_peak_23 | 15 | Distal Interge | 20 | 31336401 | 31338101 | 1701 | 1 | 668960 | ENSMUST0( | -149049 | ENSMUSG0( | Btbd35f10 | BTB domain containing 35, family member 10 |
| chrX | 31200045 | 31200339 | 295 | * | NA_peak_23 | 18 | Distal Interge | 20 | 31336401 | 31338101 | 1701 | 1 | 668960 | ENSMUST0( | -136062 | ENSMUSG0( | Btbd35f10 | BTB domain containing 35, family member 10 |
| chrX | 31219857 | 31220204 | 348 | * | NA_peak_23 | 20 | Distal Interge | 20 | 31336401 | 31338101 | 1701 | 1 | 668960 | ENSMUST0( | -116197 | ENSMUSG0( | Btbd35f10 | BTB domain containing 35, family member 10 |
| chrX | 31228449 | 31228655 | 207 | * | NA_peak_23 | 14 | Distal Interge | 20 | 31336401 | 31338101 | 1701 | 1 | 668960 | ENSMUST0( | -107746 | ENSMUSG0( | Btbd35f10 | BTB domain containing 35, family member 10 |
| chrX | 31229996 | 31230338 | 343 | * | NA_peak_23 | 42 | Distal Interge | 20 | 31336401 | 31338101 | 1701 | 1 | 668960 | ENSMUST0( | -106063 | ENSMUSG0( | Btbd35f10 | BTB domain containing 35, family member 10 |
| chrX | 31267765 | 31268056 | 292 | * | NA_peak_23 | 19 | Distal Interge | 20 | 31336401 | 31338101 | 1701 | 1 | 668960 | ENSMUST0( | -68345 | ENSMUSG0( | Btbd35f10 | BTB domain containing 35, family member 10 |
| chrX | 31625246 | 31625527 | 282 | * | NA_peak_23 | 44 | Distal Interge | 20 | 31382186 | 31383886 | 1701 | 2 | 668963 | ENSMUST0( | -241360 | ENSMUSG0( | Btbd35f20 | BTB domain containing 35, family member 20 |
| chrX | 31703488 | 31703782 | 295 | * | NA_peak_23 | 65 | Distal Interge | 20 | 31962044 | 31962821 | 778 | 1 | 74851 | ENSMUST0( | -258262 | ENSMUSG0( | Spin2-ps1 | spindlin family, member 2, pseudogene 1 |
| chrX | 32146778 | 32147012 | 235 | * | NA_peak_23 | 23 | Distal Interge | 20 | 32278435 | 32280341 | 1907 | 1 | 100040482 | ENSMUST0( | -131423 | ENSMUSG0( | Btbd35f2 | BTB domain containing 35, family member 2 |
| chrX | 32257473 | 32257742 | 270 | * | NA_peak_23 | 24 | Distal Interge | 20 | 32278435 | 32280341 | 1907 | 1 | 100040482 | ENSMUST0( | -20693 | ENSMUSG0( | Btbd35f2 | BTB domain containing 35, family member 2 |
| chrX | 32306268 | 32306774 | 507 | * | NA_peak_23 | 22 | Distal Interge | 20 | 32278435 | 32280341 | 1907 | 1 | 100040482 | ENSMUST0( | 27833 | ENSMUSG0( | Btbd35f2 | BTB domain containing 35, family member 2 |
| chrX | 32638282 | 32638608 | 327 | * | NA_peak_23 | 24 | Distal Interge | 20 | 32650716 | 32651426 | 711 | 1 | 100862314 | ENSMUST0( | -12108 | ENSMUSG0( | Gm21637 | predicted gene, 21637 |
| chrX | 32698150 | 32698194 | 45 | * | NA_peak_23 | 19 | Distal Interge | 20 | 32726109 | 32728051 | 1943 | 1 | 100862329 | ENSMUST0( | -27915 | ENSMUSG0( | Btbd35f21 | BTB domain containing 35, family member 21 |
| chrX | 32943669 | 32943981 | 313 | * | NA_peak_23 | 26 | Distal Interge | 20 | 32973938 | 32975845 | 1908 | 1 | 100040533 | ENSMUST0( | -29957 | ENSMUSG0( | Btbd35f5 | BTB domain containing 35, family member 5 |
| chrX | 33089739 | 33090056 | 318 | * | NA_peak_23 | 46 | Distal Interge | 20 | 32973938 | 32975845 | 1908 | 1 | 100040533 | ENSMUST0( | 115801 | ENSMUSG0( | Btbd35f5 | BTB domain containing 35, family member 5 |
| chrX | 33110145 | 33110453 | 309 | * | NA_peak_23 | 25 | Distal Interge | 20 | 32973938 | 32975845 | 1908 | 1 | 100040533 | ENSMUST0( | 136207 | ENSMUSG0( | Btbd35f5 | BTB domain containing 35, family member 5 |
| chrX | 33249981 | 33250235 | 255 | * | NA_peak_23 | 26 | Distal Interge | 20 | 33313379 | 33315286 | 1908 | 1 | 100040606 | ENSMUST0( | -63144 | ENSMUSG0( | Btbd35f12 | BTB domain containing 35, family member 12 |
| chrX | 33353277 | 33353597 | 321 | * | NA_peak_23 | 40 | Distal Interge | 20 | 33313379 | 33315286 | 1908 | 1 | 100040606 | ENSMUST0( | 39898 | ENSMUSG0( | Btbd35f12 | BTB domain containing 35, family member 12 |
| chrX | 33557513 | 33557982 | 470 | * | NA_peak_23 | 70 | Distal Interge | 20 | 33575398 | 33577314 | 1917 | 2 | 668960 | ENSMUST0( | 19332 | ENSMUSG0( | Btbd35f10 | BTB domain containing 35, family member 10 |
| chrX | 33930147 | 33930478 | 332 | * | NA_peak_23 | 21 | Distal Interge | 20 | 33700649 | 33702349 | 1701 | 2 | 100040732 | ENSMUST0( | -227798 | ENSMUSG0( | Btbd35f14 | BTB domain containing 35, family member 14 |
| chrX | 33936200 | 33936447 | 248 | * | NA_peak_23 | 16 | Distal Interge | 20 | 34159636 | 34161564 | 1929 | 2 | 668960 | ENSMUST0( | 225117 | ENSMUSG0( | Btbd35f10 | BTB domain containing 35, family member 10 |
| chrX | 33987905 | 33988612 | 708 | * | NA_peak_23 | 30 | Distal Interge | 20 | 34159636 | 34161564 | 1929 | 2 | 668960 | ENSMUST0( | 172952 | ENSMUSG0( | Btbd35f10 | BTB domain containing 35, family member 10 |
| chrX | 33995652 | 33996038 | 387 | * | NA_peak_23 | 29 | Distal Interge | 20 | 34159636 | 34161564 | 1929 | 2 | 668960 | ENSMUST0( | 165526 | ENSMUSG0( | Btbd35f10 | BTB domain containing 35, family member 10 |
| chrX | 34018059 | 34018704 | 646 | * | NA_peak_23 | 24 | Distal Interge | 20 | 34159636 | 34161564 | 1929 | 2 | 668960 | ENSMUST0( | 142860 | ENSMUSG0( | Btbd35f10 | BTB domain containing 35, family member 10 |
| chrX | 34210471 | 34210943 | 473 | * | NA_peak_23 | 34 | Distal Interge | 20 | 34159636 | 34161564 | 1929 | 2 | 668960 | ENSMUST0( | -48907 | ENSMUSG0( | Btbd35f10 | BTB domain containing 35, family member 10 |
| chrX | 34229440 | 34229555 | 116 | * | NA_peak_23 | 22 | Distal Interge | 20 | 34159636 | 34161564 | 1929 | 2 | 668960 | ENSMUST0( | -67876 | ENSMUSG0( | Btbd35f10 | BTB domain containing 35, family member 10 |
| chrX | 34325274 | 34326127 | 854 | * | NA_peak_23 | 22 | Distal Interge | 20 | 34159636 | 34161564 | 1929 | 2 | 668960 | ENSMUST0( | -163710 | ENSMUSG0( | Btbd35f10 | BTB domain containing 35, family member 10 |
| chrX | 34331929 | 34332245 | 317 | * | NA_peak_23 | 37 | Distal Interge | 20 | 34159636 | 34161564 | 1929 | 2 | 668960 | ENSMUST0( | -170365 | ENSMUSG0( | Btbd35f10 | BTB domain containing 35, family member 10 |
| chrX | 34464091 | 34464381 | 291 | * | NA_peak_23 | 21 | Distal Interge | 20 | 34159636 | 34161564 | 1929 | 2 | 668960 | ENSMUST0( | -302527 | ENSMUSG0( | Btbd35f10 | BTB domain containing 35, family member 10 |
| chrX | 34468220 | 34468588 | 369 | * | NA_peak_23 | 35 | Distal Interge | 20 | 34159636 | 34161564 | 1929 | 2 | 668960 | ENSMUST0( | -306656 | ENSMUSG0( | Btbd35f10 | BTB domain containing 35, family member 10 |
| chrX | 35827993 | 35828018 | 26 | * | NA_peak_23 | 39 | Distal Interge | 20 | 35888832 | 36076562 | 187731 | 1 | 75974 | ENSMUST0( | -60814 | ENSMUSG0( | Dock11 | dedicator of cytokinesis 11 |
| chrX | 39665694 | 39665979 | 286 | * | NA_peak_23 | 76 | Distal Interge | 20 | 39862919 | 39863604 | 686 | 2 | 100504400 | ENSMUST0( | 197625 | ENSMUSG0( | Cypt14 | cysteine-rich perinuclear theca 14 |
| chrX | 40835095 | 40835142 | 48 | * | NA_peak_23 | 35 | Distal Interge | 20 | 41400854 | 41613099 | 212246 | 1 | 53623 | ENSMUST0( | -565712 | ENSMUSG0( | Gria3 | glutamate receptor, ionotropic, AMPA3 (alpha 3) |
| chrX | 47135700 | 47136221 | 522 | * | NA_peak_23 | 26 | Distal Interge | 20 | 47841930 | 47892396 | 50467 | 2 | 93761 | ENSMUST0( | 756175 | ENSMUSG0( | Smarca1 | SWI/SNF related, matrix associated, actin dependent regulator of chromatin, subfamily a, member 1 |
| chrX | 48703861 | 48704056 | 196 | * | NA_peak_23 | 30 | Promoter (2-: | 20 | 48706180 | 48709066 | 2887 | 1 | 209003 | ENSMUST0( | -2124 | ENSMUSG0( | Rbmx2 | RNA binding motif protein, X-linked 2 |
| chrX | 56885338 | 56885595 | 258 | * | NA_peak_23 | 13 | Intron (ENSM | 20 | 56894372 | 57001558 | 107187 | 1 | 236798 | ENSMUST0( | -8777 | ENSMUSG0( | Adgrg4 | adhesion G protein-coupled receptor G4 |
| chrX | 60621821 | 60622036 | 216 | * | NA_peak_23 | 62 | Intron (ENSM | 20 | 60235461 | 60592698 | 357238 | 2 | 320940 | ENSMUST0( | -29123 | ENSMUSG0( | Atp11c | ATPase, class VI, type 11C |
| chrX | 61257677 | 61258005 | 329 | * | NA_peak_23 | 21 | Intron (ENSM | 20 | 61288188 | 61292484 | 4297 | 1 | 245440 | ENSMUST0( | -30183 | ENSMUSG0( | Gm4988 | predicted gene 4988 |
| chrX | 64888000 | 64889087 | 1088 | * | NA_peak_23 | 35 | Distal Interge | 20 | 65047644 | 65049014 | 1371 | 1 | 70062 | ENSMUST0( | -158557 | ENSMUSG0( | Ctag2 | cancer/testis antigen 2 |
| chrX | 65089760 | 65090091 | 332 | * | NA_peak_23 | 29 | Distal Interge | 20 | 65047644 | 65049014 | 1371 | 1 | 70062 | ENSMUST0( | 42116 | ENSMUSG0( | Ctag2 | cancer/testis antigen 2 |
| chrX | 65091600 | 65091886 | 287 | * | NA_peak_23 | 20 | Distal Interge | 20 | 65047644 | 65049014 | 1371 | 1 | 70062 | ENSMUST0( | 43956 | ENSMUSG0( | Ctag2 | cancer/testis antigen 2 |
| chrX | 69936375 | 69936626 | 252 | * | NA_peak_23 | 39 | Intron (ENSM | 20 | 69945300 | 69945978 | 679 | 1 | 75509 | ENSMUST0( | -8674 | ENSMUSG0( | 1700020N15 | RIKEN cDNA 1700020N15 gene |
| chrX | 72129494 | 72129732 | 239 | * | NA_peak_23 | 44 | Distal Interge | 20 | 72222017 | 72222964 | 948 | 1 | 17140 | ENSMUST0( | -92285 | ENSMUSG0( | Magea4 | MAGE family member A4 |
| chrX | 74539632 | 74539999 | 368 | * | NA_peak_23 | 23 | Distal Interge | 20 | 74594328 | 74595275 | 948 | 1 | 258393 | ENSMUST0( | -54329 | ENSMUSG0( | Olfr1325 | olfactory receptor 1325 |
| chrX | 75198794 | 75199068 | 275 | * | NA_peak_23 | 30 | Intron (ENSM | 20 | 75149121 | 75163770 | 14650 | 2 | 434800 | ENSMUST0( | -35024 | ENSMUSG0( | Smim9 | small integral membrane protein 9 |
| chrX | 75199577 | 75199685 | 109 | * | NA_peak_23 | 30 | Intron (ENSM | 20 | 75149121 | 75163770 | 14650 | 2 | 434800 | ENSMUST0( | -35807 | ENSMUSG0( | Smim9 | small integral membrane protein 9 |
| chrX | 76148098 | 76148433 | 336 | * | NA_peak_23 | 39 | Distal Interge | 20 | 76264304 | 76267738 | 3435 | 1 | 238829 | ENSMUST0( | -115871 | ENSMUSG0( | Cldn34b3 | claudin 34B3 |
| chrX | 78077833 | 78078095 | 263 | * | NA_peak_23 | 23 | Distal Interge | 20 | 78085505 | 78091374 | 5870 | 2 | 18249 | ENSMUST0( | 13279 | ENSMUSG0( | Obp1a | odorant binding protein IA |
| chrX | 78709914 | 78710204 | 291 | * | NA_peak_23 | 16 | Distal Interge | 20 | 78737763 | 78739401 | 1639 | 1 | 70864 | ENSMUST0( | -27559 | ENSMUSG0( | Fam47c | family with sequence similarity 47, member C |
| chrX | 82660584 | 82661107 | 524 | * | NA_peak_23 | 93 | Distal Interge | 20 | 82948870 | 83492739 | 543870 | 1 | 13405 | ENSMUST0( | -287763 | ENSMUSG0( | Dmd | dystrophin, muscular dystrophy |
| chrX | 84154608 | 84155015 | 408 | * | NA_peak_23 | 80 | Intron (ENSM | 20 | 84076649 | 84309894 | 233246 | 1 | 13405 | ENSMUST0( | 77959 | ENSMUSG0( | Dmd | dystrophin, muscular dystrophy |
| chrX | 86897359 | 86897746 | 388 | * | NA_peak_23 | 127 | Intron (ENSM | 20 | 86250256 | 86305093 | 54838 | 2 | 434903 | ENSMUST0( | -592266 | ENSMUSG0( | Mageb4 | MAGE family member B4 |
| chrX | 89018184 | 89018490 | 307 | * | NA_peak_23 | 30 | Distal Interge | 20 | 88759471 | 88760312 | 842 | 2 | 100316809 | ENSMUST0( | -257872 | ENSMUSG0( | Mir1906-1 | microRNA 1906-1 |
| chrX | 89495836 | 89496167 | 332 | * | NA_peak_23 | 17 | Distal Interge | 20 | 89403848 | 89409689 | 5842 | 2 | 18630 | ENSMUST0( | -86147 | ENSMUSG0( | Dcaf8l | DDB1 and CUL4 associated factor 8 like |
| chrX | 91839312 | 91840238 | 927 | * | NA_peak_23 | 44 | Distal Interge | 20 | 91779544 | 91783018 | 3475 | 2 | 74271 | ENSMUST0( | -56294 | ENSMUSG0( | Mageb5 | MAGE family member B5 |
| chrX | 92802359 | 92802726 | 368 | * | NA_peak_23 | 20 | Distal Interge | 20 | 92118879 | 92599572 | 480694 | 2 | 215641 | ENSMUST0( | -202787 | ENSMUSG0( | Mageb18 | MAGE family member B18 |
| chrX | 92911883 | 92912203 | 321 | * | NA_peak_23 | 44 | Distal Interge | 20 | 93156154 | 93156873 | 720 | 1 | 69357 | ENSMUST0( | -243951 | ENSMUSG0( | BC061195 | cDNA sequence BC061195 |
| chrX | 94877883 | 94878251 | 369 | * | NA_peak_23 | 144 | Distal Interge | 20 | 94795563 | 94798183 | 2621 | 2 | 668171 | ENSMUST0( | -79700 | NA | Zxda | zinc finger, X-linked, duplicated A |
| chrX | 94878960 | 94879256 | 297 | * | NA_peak_23 | 30 | Distal Interge | 20 | 94795563 | 94798183 | 2621 | 2 | 668171 | ENSMUST0( | -80777 | NA | Zxda | zinc finger, X-linked, duplicated A |
| chrX | 94882071 | 94882816 | 746 | * | NA_peak_23 | 120 | Distal Interge | 20 | 94795563 | 94798183 | 2621 | 2 | 668171 | ENSMUST0( | -83888 | NA | Zxda | zinc finger, X-linked, duplicated A |
| chrX | 94883207 | 94883820 | 614 | * | NA_peak_23 | 40 | Distal Interge | 20 | 94795563 | 94798183 | 2621 | 2 | 668171 | ENSMUST0( | -85024 | NA | Zxda | zinc finger, X-linked, duplicated A |
| chrX | 94885584 | 94886154 | 571 | * | NA_peak_23 | 57 | Distal Interge | 20 | 94795563 | 94798183 | 2621 | 2 | 668171 | ENSMUST0( | -87401 | NA | Zxda | zinc finger, X-linked, duplicated A |
| chrX | 94886707 | 94887629 | 923 | * | NA_peak_23 | 80 | Distal Interge | 20 | 94795563 | 94798183 | 2621 | 2 | 668171 | ENSMUST0( | -88524 | NA | Zxda | zinc finger, X-linked, duplicated A |
| chrX | 94890646 | 94891107 | 462 | * | NA_peak_23 | 33 | Distal Interge | 20 | 94795563 | 94798183 | 2621 | 2 | 668171 | ENSMUST0( | -92463 | NA | Zxda | zinc finger, X-linked, duplicated A |
| chrX | 94892285 | 94892649 | 365 | * | NA_peak_23 | 28 | Distal Interge | 20 | 94795563 | 94798183 | 2621 | 2 | 668171 | ENSMUST0( | -94102 | NA | Zxda | zinc finger, X-linked, duplicated A |
| chrX | 94893544 | 94893869 | 326 | * | NA_peak_23 | 53 | Distal Interge | 20 | 94795563 | 94798183 | 2621 | 2 | 668171 | ENSMUST0( | -95361 | NA | Zxda | zinc finger, X-linked, duplicated A |
| chrX | 94901238 | 94902631 | 1394 | * | NA_peak_23 | 83 | Distal Interge | 20 | 94795563 | 94798183 | 2621 | 2 | 668171 | ENSMUST0( | -103055 | NA | Zxda | zinc finger, X-linked, duplicated A |
| chrX | 94903232 | 94903727 | 496 | * | NA_peak_23 | 49 | Distal Interge | 20 | 94795563 | 94798183 | 2621 | 2 | 668171 | ENSMUST0( | -105049 | NA | Zxda | zinc finger, X-linked, duplicated A |
| chrX | 94905456 | 94906011 | 556 | * | NA_peak_23 | 58 | Distal Interge | 20 | 94795563 | 94798183 | 2621 | 2 | 668171 | ENSMUST0( | -107273 | NA | Zxda | zinc finger, X-linked, duplicated A |
| chrX | 94910222 | 94910646 | 425 | * | NA_peak_23 | 33 | Distal Interge | 20 | 94795563 | 94798183 | 2621 | 2 | 668171 | ENSMUST0( | -112039 | NA | Zxda | zinc finger, X-linked, duplicated A |
| chrX | 94911314 | 94911811 | 498 | * | NA_peak_23 | 109 | Distal Interge | 20 | 94795563 | 94798183 | 2621 | 2 | 668171 | ENSMUST0( | -113131 | NA | Zxda | zinc finger, X-linked, duplicated A |
| chrX | 94913636 | 94913929 | 294 | * | NA_peak_23 | 131 | Distal Interge | 20 | 95022510 | 95026682 | 4173 | 2 | 270624 | ENSMUST0( | 112753 | ENSMUSG0( | Spin4 | spindlin family, member 4 |
| chrX | 94914252 | 94915723 | 1472 | * | NA_peak_23 | 131 | Distal Interge | 20 | 95022510 | 95026682 | 4173 | 2 | 270624 | ENSMUST0( | 110959 | ENSMUSG0( | Spin4 | spindlin family, member 4 |
| chrX | 94968842 | 94969565 | 724 | * | NA_peak_23 | 40 | Distal Interge | 20 | 95022510 | 95026682 | 4173 | 2 | 270624 | ENSMUST0( | 57117 | ENSMUSG0( | Spin4 | spindlin family, member 4 |
| chrX | 94970341 | 94970868 | 528 | * | NA_peak_23 | 34 | Distal Interge | 20 | 95022510 | 95026682 | 4173 | 2 | 270624 | ENSMUST0( | 55814 | ENSMUSG0( | Spin4 | spindlin family, member 4 |
| chrX | 94979600 | 94979838 | 239 | * | NA_peak_23 | 26 | Distal Interge | 20 | 95022510 | 95026682 | 4173 | 2 | 270624 | ENSMUST0( | 46844 | ENSMUSG0( | Spin4 | spindlin family, member 4 |
| chrX | 95745177 | 95745548 | 372 | * | NA_peak_23 | 92 | Intron (ENSM | 20 | 95732586 | 95733265 | 680 | 2 | 76386 | ENSMUST0( | -11912 | ENSMUSG0( | 1700010D01 | RIKEN cDNA 1700010D01 gene |
| chrX | 97487888 | 97488272 | 385 | * | NA_peak_23 | 19 | Distal Interge | 20 | 97337791 | 97377216 | 39426 | 2 | 245527 | ENSMUST0( | -110672 | ENSMUSG0( | Eda2r | ectodysplasin A2 receptor |
| chrX | 97966216 | 97966432 | 217 | * | NA_peak_23 | 32 | Distal Interge | 20 | 98148769 | 98323215 | 174447 | 1 | 11835 | ENSMUST0( | -182337 | ENSMUSG0( | Ar | androgen receptor |
| chrX | 100007004 | 100007404 | 401 | * | NA_peak_23 | 56 | Intron (ENSM | 20 | 99975781 | 100397156 | 421376 | 1 | 13607 | ENSMUST0( | 31223 | ENSMUSG0( | Eda | ectodysplasin-A |
| chrX | 100864600 | 100865300 | 701 | * | NA_peak_23 | 31 | Intron (ENSM | 20 | 100808000 | 100814830 | 6831 | 1 | 53310 | ENSMUST0( | 56600 | ENSMUSG0( | Dlg3 | discs large MAGUK scaffold protein 3 |
| chrX | 104627649 | 104627939 | 291 | * | NA_peak_23 | 17 | Intron (ENSM | 20 | 104536969 | 104671064 | 134096 | 2 | 108672 | ENSMUST0( | 43125 | ENSMUSG0( | Zdhhc15 | zinc finger, DHHC domain containing 15 |
| chrX | 105021850 | 105022220 | 371 | * | NA_peak_23 | 44 | Distal Interge | 20 | 105044001 | 105070035 | 26035 | 2 | 71445 | ENSMUST0( | 47815 | ENSMUSG0( | 5530601H04 | RIKEN cDNA 5530601H04 gene |
| chrX | 105615962 | 105616360 | 399 | * | NA_peak_23 | 35 | Distal Interge | 20 | 105499772 | 105500639 | 868 | 1 | 245566 | ENSMUST0( | 116190 | ENSMUSG0( | Cypt2 | cysteine-rich perinuclear theca 2 |
| chrX | 110865753 | 110865958 | 206 | * | NA_peak_23 | 29 | Distal Interge | 20 | 110814280 | 110817207 | 2928 | 1 | 18994 | ENSMUST0( | 51473 | ENSMUSG0( | Pou3f4 | POU domain, class 3, transcription factor 4 |
| chrX | 114140616 | 114141201 | 586 | * | NA_peak_23 | 36 | Distal Interge | 20 | 114474333 | 114560829 | 86497 | 1 | 237010 | ENSMUST0( | -333132 | ENSMUSG0( | Klhl4 | kelch-like 4 |
| chrX | 115386535 | 115386849 | 315 | * | NA_peak_23 | 35 | Distal Interge | 20 | 114980970 | 114981317 | 348 | 2 | 664969 | ENSMUST0( | -405218 | ENSMUSG0( | Gm7429 | predicted pseudogene 7429 |
| chrX | 120977823 | 120978042 | 220 | * | NA_peak_23 | 38 | Distal Interge | 20 | 120401364 | 120906344 | 504981 | 1 | 245578 | ENSMUST0( | 576459 | ENSMUSG0( | Pcdh11x | protocadherin 11 X-linked |
| chrX | 121585488 | 121585883 | 396 | * | NA_peak_23 | 27 | Distal Interge | 20 | 122394565 | 122397401 | 2837 | 2 | 54561 | ENSMUST0( | 811518 | ENSMUSG0( | Nap1l3 | nucleosome assembly protein 1-like 3 |
| chrX | 123038427 | 123038930 | 504 | * | NA_peak_23 | 52 | Distal Interge | 20 | 123103495 | 123145651 | 42157 | 1 | 73061 | ENSMUST0( | -64565 | ENSMUSG0( | Cldn34c1 | claudin 34C1 |
| chrX | 123043813 | 123044233 | 421 | * | NA_peak_23 | 44 | Distal Interge | 20 | 123103495 | 123145651 | 42157 | 1 | 73061 | ENSMUST0( | -59262 | ENSMUSG0( | Cldn34c1 | claudin 34C1 |
| chrX | 123044937 | 123045815 | 879 | * | NA_peak_23 | 92 | Distal Interge | 20 | 123103495 | 123145651 | 42157 | 1 | 73061 | ENSMUST0( | -57680 | ENSMUSG0( | Cldn34c1 | claudin 34C1 |
| chrX | 123075889 | 123076108 | 220 | * | NA_peak_23 | 114 | Distal Interge | 20 | 123103495 | 123145651 | 42157 | 1 | 73061 | ENSMUST0( | -27387 | ENSMUSG0( | Cldn34c1 | claudin 34C1 |
| chrX | 123076456 | 123076503 | 48 | * | NA_peak_23 | 30 | Distal Interge | 20 | 123103495 | 123145651 | 42157 | 1 | 73061 | ENSMUST0( | -26992 | ENSMUSG0( | Cldn34c1 | claudin 34C1 |
| chrX | 124217568 | 124218190 | 623 | * | NA_peak_23 | 35 | Distal Interge | 20 | 124124961 | 124135952 | 10992 | 2 | 100038941 | ENSMUST0( | -81616 | ENSMUSG0( | Vmn2r121 | vomeronasal 2, receptor 121 |
| chrX | 124227712 | 124228204 | 493 | * | NA_peak_23 | 65 | Distal Interge | 20 | 124124961 | 124135952 | 10992 | 2 | 100038941 | ENSMUST0( | -91760 | ENSMUSG0( | Vmn2r121 | vomeronasal 2, receptor 121 |
| chrX | 127728775 | 127728874 | 100 | * | NA_peak_23 | 22 | Intron (ENSM | 20 | 127721182 | 127736554 | 15373 | 2 | 73934 | ENSMUST0( | 7680 | ENSMUSG0( | Cldn34c4 | claudin 34C4 |
| chrX | 127959523 | 127959863 | 341 | * | NA_peak_23 | 39 | Distal Interge | 20 | 127721182 | 127736554 | 15373 | 2 | 73934 | ENSMUST0( | -222969 | ENSMUSG0( | Cldn34c4 | claudin 34C4 |
| chrX | 135298906 | 135299205 | 300 | * | NA_peak_23 | 26 | Distal Interge | 20 | 135302307 | 135312652 | 10346 | 1 | 83565 | ENSMUST0( | -3102 | ENSMUSG0( | Pramel3 | PRAME like 3 |
| chrX | 154505708 | 154506090 | 383 | * | NA_peak_23 | 70 | Distal Interge | 20 | 154482002 | 154484682 | 2681 | 1 | 75185 | ENSMUST0( | 23706 | ENSMUSG0( | Samt4 | spermatogenesis associated multipass transmembrane protein 4 |
| chrX | 156582078 | 156582450 | 373 | * | NA_peak_23 | 27 | Distal Interge | 20 | 157162075 | 157415286 | 253212 | 2 | 18675 | ENSMUST0( | 832836 | ENSMUSG0( | Phex | phosphate regulating endopeptidase homolog, X-linked |
| chrX | 158146873 | 158147094 | 222 | * | NA_peak_23 | 23 | Distal Interge | 20 | 157832743 | 158043294 | 210552 | 2 | 245684 | ENSMUST0( | -103579 | ENSMUSG0( | Cnksr2 | connector enhancer of kinase suppressor of Ras 2 |
| chrX | 158872107 | 158872228 | 122 | * | NA_peak_23 | 24 | Distal Interge | 20 | 159210307 | 159364502 | 154196 | 1 | 110651 | ENSMUST0( | -338079 | ENSMUSG0( | Rps6ka3 | ribosomal protein S6 kinase polypeptide 3 |
| chrX | 161711901 | 161712526 | 626 | * | NA_peak_23 | 23 | Distal Interge | 20 | 161717069 | 161779494 | 62426 | 1 | 24004 | ENSMUST0( | -4543 | ENSMUSG0( | Rai2 | retinoic acid induced 2 |
| chrX | 165468893 | 165469035 | 143 | * | NA_peak_23 | 23 | Distal Interge | 20 | 165129017 | 165327393 | 198377 | 2 | 237213 | ENSMUST0( | -141500 | ENSMUSG0( | Glra2 | glycine receptor, alpha 2 subunit |
| chrY | 90728709 | 90729286 | 578 | * | NA_peak_23 | 25 | Distal Interge | 21 | 90752427 | 90755467 | 3041 | 2 | 654820 | ENSMUST0( | 26181 | ENSMUSG0( | G530011O06 | RIKEN cDNA G530011O06 gene |
| chrY | 90804506 | 90804783 | 278 | * | NA_peak_23 | 30 | Intron (ENSM | 21 | 90790451 | 90816465 | 26015 | 1 | 170942 | ENSMUST0( | 14055 | ENSMUSG0( | Erdr1 | erythroid differentiation regulator 1 |
